# Global transcriptome analysis reveals partial estrogen-like effects of karanjin in MCF-7 breast cancer cells

**DOI:** 10.1101/2021.10.28.466373

**Authors:** Gaurav Bhatt, Akshita Gupta, Latha Rangan, Anil Mukund Limaye

## Abstract

Karanjin, an abundantly occurring furanoflavonoid in edible and non-edible legumes, exerts diverse biological effects *in vivo*, and *in vitro*. Its potential as an anticancer agent is also gaining traction following recent demonstrations of its anti-proliferative, cell cycle inhibitory, and pro-apoptotic effects. However, the universality of its anticancer potential is yet to be scrutinized, particularly so because flavonoids can act as selective estrogen receptor modulators (SERMs). Even the genomic correlates of its biological activities are yet to be examined in hormone responsive cells. This paper presents the early and direct transcriptomic footprint of 10 μM karanjin in MCF-7 breast cancer cells, using next generation sequencing technology (RNA-seq). We show that karanjin-modulated gene-expression repertoire is enriched in several hallmark gene sets, which include early estrogen-response, and G2/M checkpoint genes. Genes modulated by karanjin overlapped with those modulated by 1 nM 17β-estradiol (E2), or 1 μM tamoxifen. Karanjin altered the expression of selected estrogen-regulated genes in a cell-type, and concentration dependent manner. It downmodulated the expression of ERα protein in MCF-7 cells. Furthermore, ERα knockdown negatively impacted karanjin’s ability to modulate the expression of selected E2 target genes. Our data suggest that karanjin exerts its effects on ERα-positive breast cancer cells, at least in part, via ERα. The apparent SERM-like effects of karanjin pose a caveat to the anticancer potential of karanjin. In-depth studies on cell-type and concentration-dependent effects of karanjin may bring out its true potential in endocrine therapies.

## 1. Introduction

Karanjin is a furanoflavonoid occurring widely in Leguminosae. It produces a myriad of biological effects *in vitro* or *in vivo*, such as anti-proliferative, glucose-uptake-promoting, anti-hyperglycemic, anti-inflammatory, and anti-ulcer (Singh et al., 2021). The cell-cycle inhibitory, and pro-apoptotic effects of karanjin *in vitro* have fuelled speculations about its anticancer properties (Guo et al., 2015; Roy et al., 2019). Recently, Roy and co-workers (Roy et al., 2019) reiterated karanjin-mediated G2/M arrest, and apoptosis in HeLa cells. Furthermore, they demonstrated karanjin induced DNA damage, and P53 expression, associated with lowered reactive oxygen species (ROS), and restricted nuclear translocation of NF-κB via cytoplasmic I-κB induction (Roy et al., 2019). The fact that karanjin exerts a much lower growth inhibitory effect on normal mouse embryonic fibroblasts (Roy et al., 2019) augurs well with its anticancer potential. All cancer cell lines tested so far are growth-inhibited by karanjin with varying IC_50_ values.

Flavonoids can induce cell cycle arrest and apoptosis, thereby inhibiting proliferation of cells *in vitro* (Fotsis et al., 1997; Haddad et al., 2006; Memariani et al., 2021; Park et al., 2012; S. Singh et al., 2021; Xu et al., 2013; Zava and Duwe, 1997). Given the flavonoid structure of karanjin, its negative impact on cell proliferation is not surprising. However, flavonoids also bind ERα (Choi et al., 2008; Hong et al., 2015), modulate its transactivation function (Choi et al., 2008), and differentially effects cell proliferation and gene expression in a concentration and cell context dependent manner (Constantinou et al., 1998; Fioravanti et al., 1998; Hsu et al., 1999; Lavigne et al., 2008; Maggiolini et al., 2001; Yang et al., 2007). Thus, universality of the anti-tumor efficacy of karanjin cannot be concluded merely on the basis of its antiproliferative effects demonstrated in a few cell lines. The flavonoid structure of karanjin presents a caveat to its potential in anticancer therapy, lest it be counterproductive in hormone dependent tumors. Arguably, the molecular phenotype of cells determines their response to karanjin in terms of alteration in growth, as well as global gene expression. The latter is crucial to the understanding of the biological effects of karanjin. However, the impact of karanjin on global gene expression, as yet, remains unaddressed.

In this study we used next generation sequencing technology to assess the transcriptomic response of MCF-7 breast cancer cells treated with 10 μM karanjin for 24 h. Among the diverse repertoire of gene sets regulated by karanjin, we found G2/M-checkpoint, and estrogen-response-early genes. 10 µM karanjin modulated G2/M checkpoint genes in a manner that is consistent with cell cycle progression, rather than cell cycle arrest. Modulation of estrogen-response-early genes, and few well-established estrogen-regulated genes in cell-type- and concentration-dependent manner, suggests partial estrogen-like effect of karanjin on gene expression in ER-positive breast cancer cells.

## 2. Materials and Methods

### 2.1. Chemicals, reagents, and plasticware

Karanjin was purchased from Yucca Enterprises (Batch No. Yucca/KG/2019/04/21, Mumbai, India). The purity and identity of the compound was independently verified by HPLC, HRMS, and NMR. The data matched with the previously isolated karanjin from *Pongamia pinnata* (L.) Pierre seeds as described earlier (Singh et al., 2016) (Supplementary data 1). 17β-estradiol (E2, Cat. No. E8875) was purchased from Sigma-Aldrich (MO, USA). Dulbecco’s Modified Eagle Medium (DMEM) with (Cat. No. AT007) or without phenol red (Cat. No. AT187), Roswell Park Memorial Institute (RPMI)-1640 with (Cat. No. AT162) or without phenol red (Cat. No. AT171), Dulbecco’s Phosphate Buffered Saline (DPBS, Cat. No. TS1006), trypsin-EDTA (Cat. No. TCL014), antibiotic solution (Cat. No. A001), fetal bovine serum (FBS, Cat. No. RM10432), and charcoal-stripped FBS (cs-FBS, Cat. No. RM10416), were purchased from HiMedia (Mumbai, India). PowerUp SYBR Green PCR Master Mix (Cat. No. A25743), High-Capacity cDNA Reverse Transcription Kit (Cat. No. 4368814), ERα siRNA (Cat No.4392420), scrambled siRNA (Cat. No. AM4611), and Lipofectamine RNAiMAX (Cat. No. 13778-075) were from Invitrogen (CA, USA). ERα antibody (Cat. No. 8644S), β-actin (Cat. No. 4970T) and anti-rabbit immunoglobin G (Cat. No. 7074S) were from Cell Signalling Technology (Massachusetts, USA). Polyclonal histone H3 antibody (Cat. No. BB-AB0055) was purchased from BioBharati LifeScience Pvt. Ltd. (Kolkata, India). All other chemicals and buffers were purchased from Merck (Mumbai, India), Sigma (St Louis, MO, USA), or SRL (Mumbai, India). All cell culture plasticware was purchased from Eppendorf (Hamburg, Germany).

### 2.2. Cell culture

MCF-7 and T47D cells were obtained from National Centre for Cell Science (NCCS) (Pune, India). MCF-7 or T47D cells were routinely cultured and maintained under standard conditions of 37°C and 5% CO_2_ in phenol red-containing DMEM or RPMI-1640, respectively, which were supplemented with 10% heat-inactivated FBS, 100 units/mL penicillin, and 100 μg/mL streptomycin (M1 medium).

### 2.3. Treatment

Cells were seeded in M1 medium. When 60% confluent, the cells were shifted to phenol red-free DMEM or RPMI-1640, supplemented with 10% heat-inactivated cs-FBS, 100 units/mL penicillin, and 100 μg/mL streptomycin (M2 medium) for 24 h. Thereafter, the cells were treated with vehicle (0.1% DMSO), 10 nM E2, or indicated concentrations of karanjin in M2 medium for indicated periods of time.

### 2.4. Cell viability

40,000 MCF-7 cells were seeded in 35 mm dishes with M1 medium. After 36 h, cells were washed twice with DPBS and incubated in M2 medium for 3 h. The cells were treated with vehicle (DMSO), 10 nM E2, or the indicated concentrations of karanjin in M2 medium for 0, 24, or 120 h. Thereafter, the cells were washed with DPBS, trypsinised, and viable cells were counted on the basis of trypan blue dye exclusion (Strober, 2001). 10 nM E2 treatment was used as a standard reference. The 0 h treatment group provided the starting viable counts. In case of longer (120 h) treatment durations, the treatment medium was replenished every 48 h.

### 2.5. Transcriptome profiling

Total RNA quality was assessed using Bioanalyzer 2100 (Agilent Technologies, CA, US). RNA samples which were used for library preparation using TruSeq RNA Kit (Illumina, USA) had a concentration of 1 µg/ml and RIN value above 9.5. cDNAs reverse transcribed from mRNA were amplified, and purified using AMPureXP beads (Beckman Coulter, USA). The double-stranded cDNAs were end-repaired, polyadenylated, and ligated with adapter sequences followed by size selection for approximately 250-450 bp using AMPure XP beads. Uracil-containing strands were degraded by treatment with USER Enzyme (New England Biolabs, USA). Sequencing was performed using Illumina Novaseq 6000 Platform in paired-end format with the Phred score of 64. Approximately 40 million reads per sample were sequenced. The FASTQ files obtained after sequencing were subjected to QC using the FASTQC tool (Andrews et al., 2015). Trimming to remove the adapter sequences and short reads was performed using Trimmomatic (Bolger et al., 2014). The trimmed reads files were aligned using STAR aligner (Dobin et al., 2013). BAM files generated were used to obtain the read counts using the featureCounts tool (Liao et al., 2014). The read count files were merged and processed further using the DESeq2 package in R (Love et al., 2014). The merged read counts file was subjected to count normalization to obtain the normalized counts. Quality control (QC) was performed on the normalized counts using unsupervised clustering analysis, and data visualization in the form of principal component analysis (PCA) plots and correlation heatmaps. Following QC, the normalized counts were fitted on the negative binomial model. After estimation of dispersion values, the normalized counts were subjected to statistical analyses to obtain differentially expressed genes. The differentially modulated genes were identified by applying Wald statistic with α equal to 0.05, followed by FDR correction with a 5% cut-off. RNA-seq data generated in this study has been submitted to NCBI GEO (GSE183913).

### 2.6. Gene set enrichment analysis

The hallmark gene sets enriched upon karanjin simulation were identified by enrichment analysis using fGSEA package in R with FDR correction of 25%. (Sergushichev, 2016). Enrichment plots, and normalized enrichment scores (NES) plots were generated using additional R packages.

### 2.7. Identification of genes regulated by estrogen, tamoxifen and karanjin

Curated gene expression dataset corresponding to 1 nM E2 or 1 μM tamoxifen (GSE117942) treatment in MCF-7 cells for 24 hours was obtained from GEO (Guan et al., 2019). The read count data were analysed using the DESeq2 package in R to identify genes regulated by 1 nM estrogen or 1 μM tamoxifen. The differentially modulated genes were identified by applying Wald statistic with α equal to 0.05, followed by FDR correction with a 5% cut-off. They were then compared to the karanjin regulated genes (this study) to generate sets of overlapping genes depicted in the form of a Venn diagram.

### 2.8. Total RNA isolation and cDNA synthesis

Total RNA was isolated using RNA extraction reagent prepared in-house. The RNA integrity was checked by agarose gel electrophoresis and quantified using BioSpectrometer® (Eppendorf, Hamburg, Germany). Typically, 2 µg of total RNA was reverse transcribed using High-Capacity cDNA Reverse Transcription kit as per the manufacturer’s instructions.

### 2.9. qRT-PCR

Typically, 2 µl of 1:10 diluted cDNA was used as template for PCR reactions with gene-specific primers (Supplementary data 2). Cyclophilin A served as an internal control. Real-time PCRs were carried out in Agilent AriaMx Real-time PCR System (Agilent technologies, CA, US). The comparative ΔΔCt method (Livak and Schmittgen, 2001) was used for relative quantification of gene expression.

### 2.10. Western blotting

Total protein was isolated from the phenolic fraction of the RNA extraction reagent. Alternatively, total protein was extracted using the Laemmli sample buffer. Protein was quantified by Lowry (Lowry et al., 1951) or TCA (Choi et al., 1993) method, respectively. Protein was fractionated on 10% SDS-PAGE gel, and transferred to a nitrocellulose membrane. The blots were probed for with ERα, β-actin or histone (H3)-specific antibodies followed by HRP-conjugated anti-rabbit secondary antibody. Chemiluminescence signals obtained with Clarity Western ECL Substrate (Bio-Rad, California, US, Cat. No. 170-5060) were captured with ChemiDoc XRS+ system (Bio-Rad, California, US). Histone H3 or β-actin served as an internal control (Supplementary data 3).

### 2.11. siRNA transfection

MCF-7 cells were seeded in six-well plates. Cells were transfected with siRNA for 24 h using Lipofectamine RNAiMAX transfection reagent according to manufacturer’s instructions. Each well received 25 pmol of scrambled (control) or ERα-specific siRNA.

### 2.12. Statistical analysis

Two-group data were analysed by one-tailed t-tests. Multiple group data were analysed by one-way ANOVA followed by multiple comparison tests with TukeyHSD. To study the effect of karanjin concentration on viable cells after 24, or 120 h, and the effect of ERα knockdown on karanjin-modulated gene expression, the data were analysed by 2 × 2 factorial ANOVA. All statistical analyses were performed using the R statistical package. Figure-wise raw data, and results of statistical analyses are provided as Supplementary data 4.

## 3. Results

### 3.1. Determining the optimal concentration of karanjin

A study of the genome-wide transcriptomic effects of karanjin necessitated the identification of an optimal concentration, and duration of treatment, which captured direct and early responses, free from strong mitotic or toxic effects. The short- and long-term effects of varying concentrations of karanjin on MCF-7 cell viability were analysed. 10 nM E2, which induces proliferation of MCF-7 cells (Lippman et al., 1976), served as a reference. MCF-7 cells were treated with 0 to 50 μM concentrations of karanjin and viable cells were recorded after 0 or 24 h. We applied 2 × 2 factorial ANOVA to study the difference in viable cells after 0 and 24 h (short-term) of treatment as a function of karanjin concentration. There was no interaction between concentration and time (p ≈ 1). The effect of time was significant. The viable cell counts were significantly higher after 24 h of treatment (p < 0.0001). There was no effect of concentration. The viable cell counts after 24 h of treatment with all concentrations of karanjin (0-50 μM) were not significantly different (Fig. 1A). We concluded that karanjin had no significant impact on MCF-7 cell viability over a period of 24 h, although proliferation was evident, comparable to that induced by E2 (Fig. 1A).

**Fig. 1.**
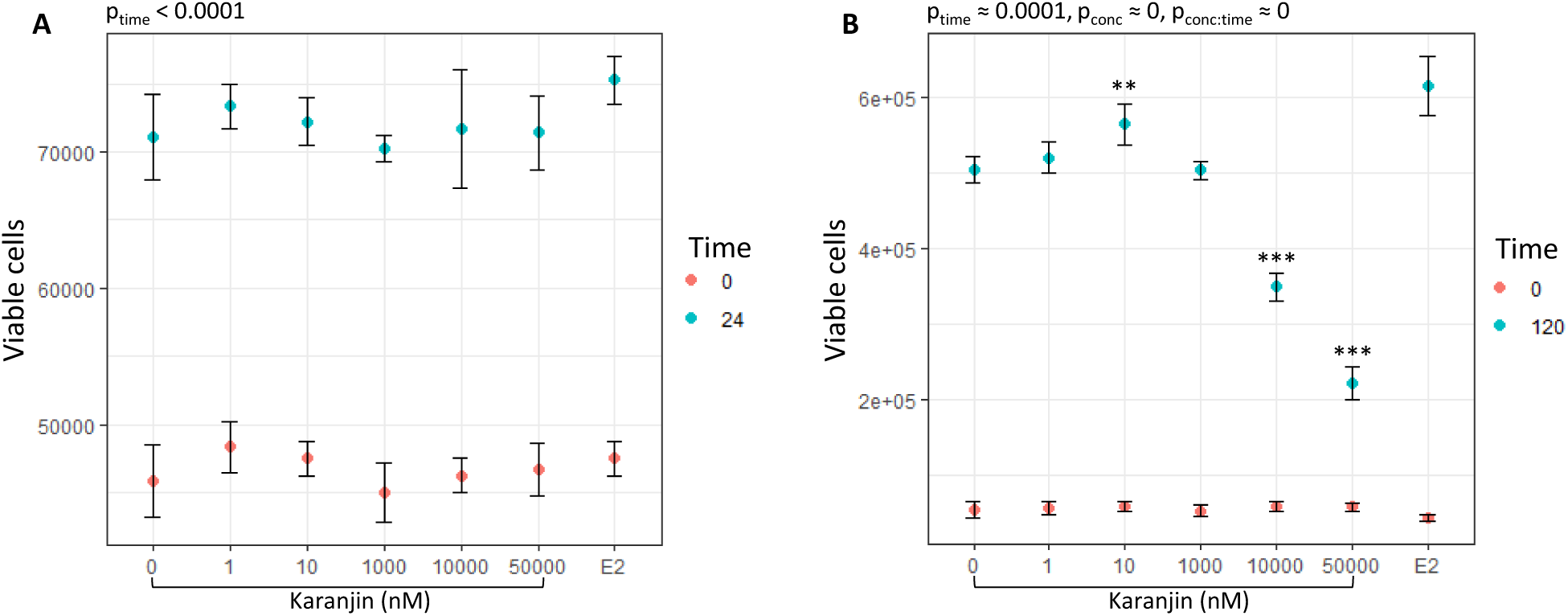
Concentration dependent effect of short- (A) or long-term (B) stimulation with karanjin on MCF-7 cell viability. 4 × 10^4^ MCF-7 cells were seeded in 35 mm dishes and grown in M1 medium for a period of 36 h. The cells were then shifted to M2 medium and incubated for 3 h. Thereafter, the cells were treated with vehicle, or indicated concentrations of karanjin. After 24 (A) or 120 h (B) the viable cells were counted (blue dots) based on trypan blue dye exclusion using a hemocytometer. Cells treated for 0 h provided the initial viable counts for each treatment group (red dots). In the 120-h experiment (B), the treatment medium was replenished every 48 h. Each dot represents mean viable count ± sd (n = 3). The data were analysed by 2 × 2 factorial ANOVA to test whether the increase in viable count after 24 (A) or 120 h (B) was dependent on concentration of karanjin. 10 nM E2 treatment was used as a reference, and was not a part of the statistical analysis. Asterisks represent significant result with respect to control for 120 h data. (**p < 0.01, ***p < 0.001). p_time_, and p_conc_ are p values for main effects of time, and concentration respectively. p_conc:time_ is the p value for the interaction.

A similar experiment designed to analyse the viable cells after treatment with karanjin for 0 and 120 h (long-term) revealed a significant interaction between time and concentration. Although the viable cell counts increased at all concentrations of karanjin as revealed by significant main effects of time (p ≈ 0), the number of viable cells after 120 h of treatment depended on the concentration of karanjin (Fig. 1B, p value for interaction term ≈ 0). 10 nM karanjin treatment yielded significantly higher viable cell count (p = 0.002), whereas 10 or 50 μM karanjin yielded significantly lower viable cell counts with respect to control (p ≈ 0). Thus, 10 μM karanjin had no effect on MCF-7 cell viability over a period of 24 h, and had an intermediate effect on viability over a period of 120 h. Based on these results, we studied the genome-wide alteration in MCF-7 cell transcriptome following short-term (24 h) treatment with 10 μM karanjin.

### 3.2. RNA-seq analysis

Total RNA isolated from MCF-7 cells treated with DMSO (control) or 10 μM karanjin were subjected to next generation sequencing (RNA-seq) to identify karanjin-modulated genes. The RIN values of RNA samples were greater than 9.5 (Supplementary data 5). Sequencing libraries prepared using the RNA samples were of acceptable quality; all having Q30 values greater than 95 (Supplementary data 5). Paired-end reads were generated, and the quality of reads before and after trimming were assessed using FastQC. Less than 1% data was lost due to trimming (Supplementary data 5). The sequences were aligned with the human genome build GRCh38 using STAR, and feature read counts files (Supplementary data 6) were analysed using DESeq2. Unsupervised clustering analysis of read count data from three control (G1, G2, and G3) and three karanjin treated (K1, K2, and K3) samples revealed that K1 was an outlier (Supplementary data 7). Hence, K1 was omitted from differential gene expression analysis.

### 3.3. Karanjin regulated genes

We identified karanjin-modulated genes by interrogating the read count data with DESeq2. The volcano plot in Fig. 2A, shows that 736 genes (blue dots) were modulated by karanjin, based on a threshold of 0 for LFC, and p < 0.05 for Wald statistic and FDR (Supplementary Data 8). 362 genes were repressed, and 374 genes were induced by karanjin as illustrated by Fig. 2B. Among the top 25 induced (Table 1) or repressed genes (Table 2) were those that encode proteins with diverse functions. These include receptors, signal transducers, enzymes, enzyme inhibitors, transcription factors, and non-coding RNAs. Selected genes from Table 1, and 2 were validated by qRT-PCR. As shown in Fig. 2C, the significant downmodulation of *B3GALT5* (n = 3, p = 0.009), *BRINP2* (n = 4, p = 0.002), *CHST1* (n = 3, p = 0.002), and *CSTA* (n = 3, p = 0.002), and significant upregulation of *CYPIA1* (n = 3, p = 0.002), *CYPIB1* (n = 4, p < 0.0001), *MRVI1* (n = 4, p < 0.001), and *CREG2* (n = 4, p < 0.0001) following 10 μM karanjin treatment is consistent with the DESeq2 results.

**Table 1.**
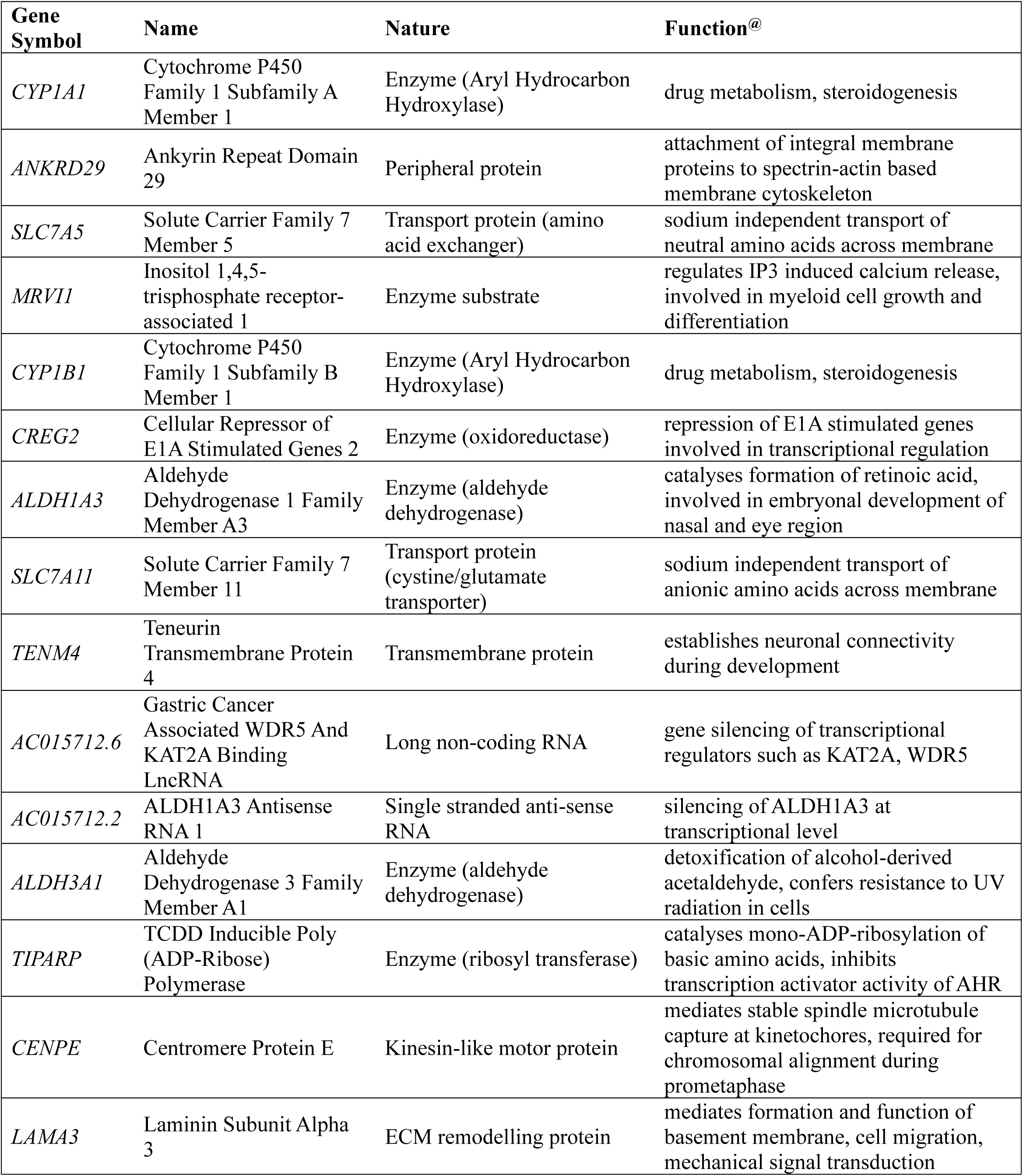
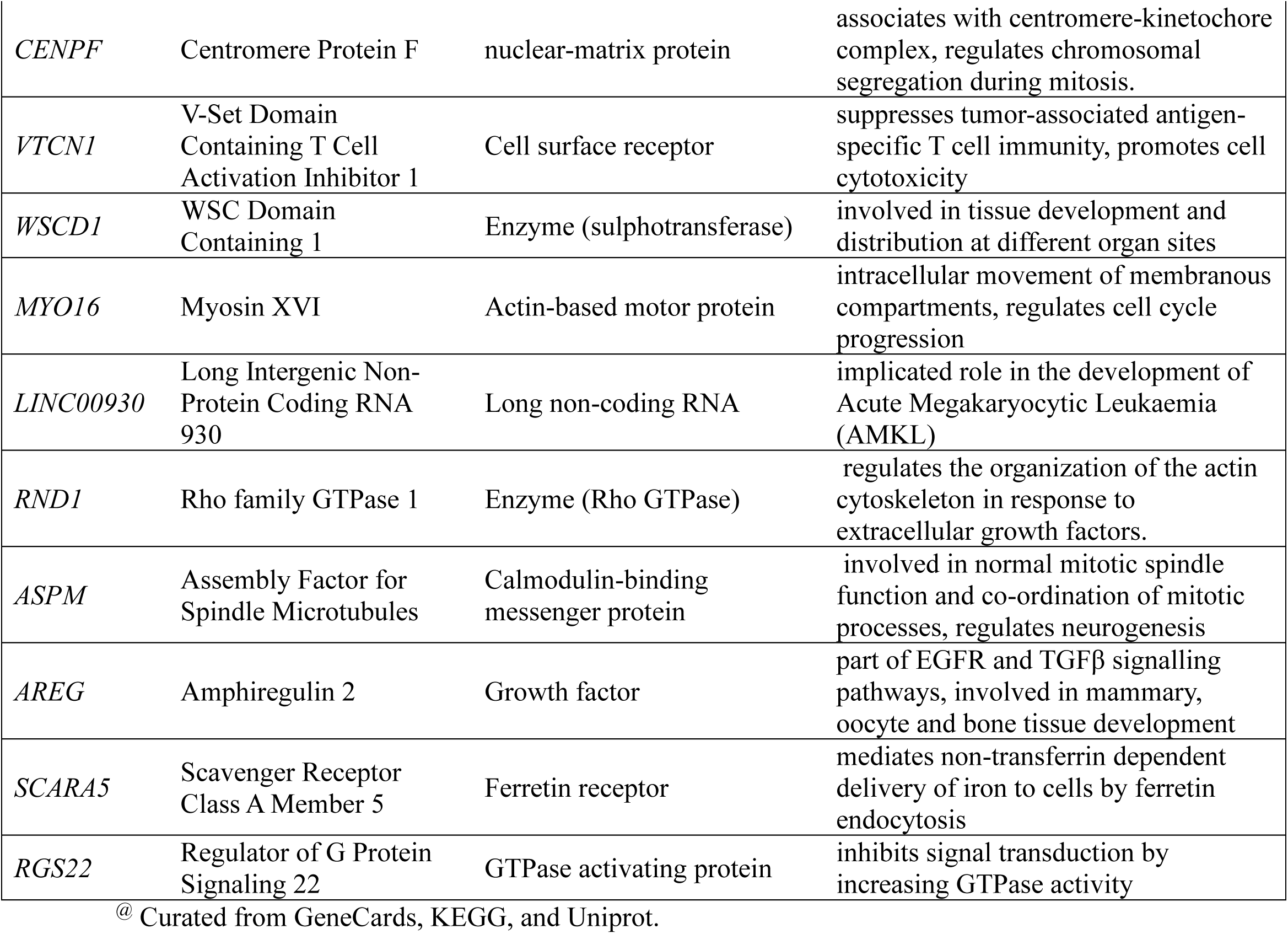
Top 25 up-regulated genes

**Table 2:** Top 25 down-regulated genes.

| Gene Symbol | Name | Nature | Function<sup>@</sup> |
| --- | --- | --- | --- |
| *BRINP2* | Bone Morphogenetic Protein/Retinoic Acid Inducible Neural-Specific 2 | Regulatory protein | cell cycle, positive regulation of neuron differentiation, negative regulation of mitotic cell cycle |
| *B3GALT5-AS1* | B3GALT5 Antisense RNA 1 | Long non-coding RNA | implicated role in development of Rectum Adenocarcinoma |
| *B3GALT5* | Beta-1,3-galactosyltransferase 5 | Enzyme (Transferase) | glycosphingolipid biosynthesis; protein glycosylation; acetylgalactosaminyltransferase activity |
| *CHST1* | Keratan sulphate 6-sulfotransferase 1 | Enzyme (Sulfotransferase) | glycan biosynthesis; sulfotransferase activity and keratan sulfotransferase activity |
| *CSTA* | Cystatin A | Cysteine Protease Inhibitor | epidermal development and maintenance; cell-cell adhesion; keratinocyte differentiation |
| *ADAM2* | ADAM Metallopeptidase Domain 2 | Enzyme (Peptidase)/membrane-anchored proteins | cell-cell and cell-matrix interactions, including fertilization, muscle development, and neurogenesis. |
| *SLIT1* | Slit Guidance Ligand 1 | Glycosaminoglycan binding protein | calcium ion binding; axon guidance; negative chemotaxis |
| *GHR* | Growth hormone receptor | Cytokine receptor | PI3K-AKT, JAK-STAT signaling pathway, growth hormone synthesis, secretion and action |
| *CXCR4* | C-X-C chemokine receptor type 4 | Chemokine receptor | regulates calcium, chemokine signaling pathway |
| *RET* | Proto-oncogene tyrosine-protein kinase Ret | Enzyme (Tyrosine protein kinase) | regulates ERK signaling, PI3K signaling, calcium signaling pathway, involved in central carbon metabolism in cancer |
| *RASSF4* | Ras association domain family member 4 | KRAS effector protein | Regulates hippo signaling pathway, involved in tumor suppression |
| *SIM1* | SIM BHLH transcription factor 1 | Transcription factor | Genetic information processing; DNA-binding transcription factor activity and obsolete signal transducer activity |
| *NR4A3* | Nuclear receptor subfamily 4 group A member 3 | Nuclear receptor | Genetic information processing; signaling and cellular processes; transcriptional mis-regulation in cancer |
| *NELL2* | Neural EGFL like 2 | Laminin domain-containing proteins | Signaling and cellular processes; plays a role in neural cell growth and differentiation as well as in oncogenesis |
| *DIO1* | Iodothyronine deiodinase 1 | Enzyme (Oxidoreductase) | Thyroid hormone metabolism and signaling pathway; cellular amino acid metabolic process |
| *CLEC3A* | C-type lectin domain family 3 member A | Carbohydrate-binding protein | promotes cell adhesion to laminin-332 and fibronectin |
| *FUT2* | Fucosyltransferase 2 | Enzyme (Transferases) | membrane trafficking and genetic information processing |
| *HOXB2* | Homeobox B2 | Transcription factor | involved in development and part of developmental regulatory system that provides cells with specific positional identities on the anterior-posterior axis. |
| *C10orf10* | DEPP1 autophagy regulator | Transcription factor activator | critical modulator of FOXO3-induced autophagy via increased cellular ROS. |
| *PLA2G10* | Secretory phospholipase A2 | Hydrolases (phospholipase A2) | phospholipid remodelling; maturation and activation of innate immune cells |
| *KCNJ8* | Potassium inwardly rectifying channel subfamily J member 8 | Integral membrane protein | role in cGMP-PKG signaling pathway, potassium transport inside the cell |
| *ADGRA2* | Adhesion G protein-coupled receptor A2 | Enzyme (Peptidase) and inhibitor; G protein-coupled receptor | metabolism; signaling and cellular processes |
| *IL1R1* | Interleukin 1 receptor type 1 | Cytokine receptor | signal transduction; NF-kappa B signaling pathway; Cytokine-cytokine receptor interaction |
| *SLC35C1* | Solute carrier family 35 (GDP-fucose transporter), member C1 | Transporters (Solute carrier family) | signaling and cellular processes; N-Glycan biosynthesis |
| *RGS11* | Regulator of G Protein Signaling 11 | GTPase-activating protein | involved in cAMP Pathway and G<sub>ai</sub> signaling |

<sup>@</sup> Curated from GeneCards, KEGG, and Uniprot.

**Fig. 2.**
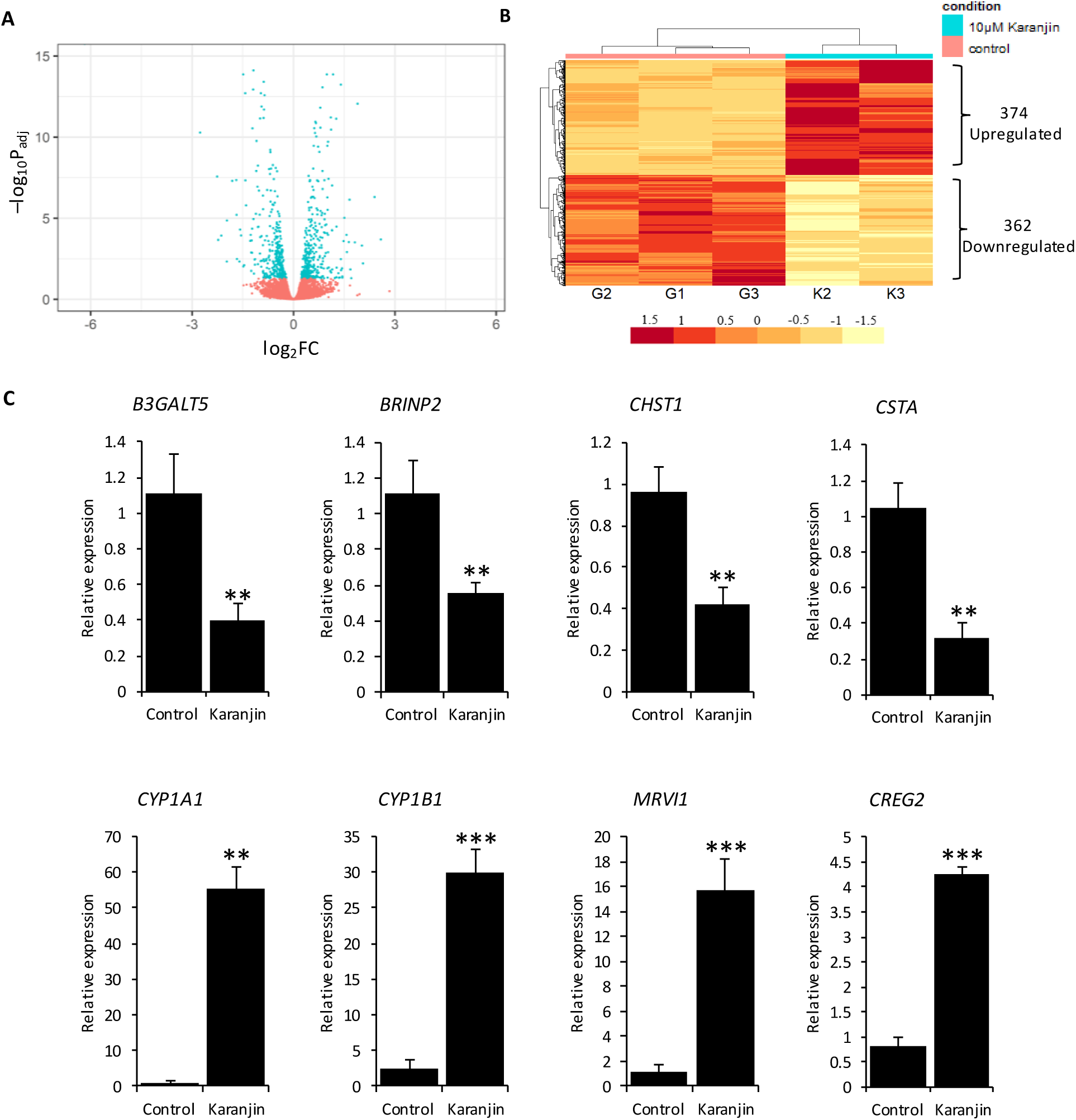
Summary of the RNA-seq data. A. Volcano plot. Each point represents a gene, which is plotted according to its –log_10_P_adj_ and log_2_FC. The blue dots represent genes (n = 736) that are significantly modulated by 10 µM karanjin. The red dots represent non-significant genes. B. Expression heatmap of 736 significantly modulated genes. G1, G2 and G3 are control samples. K2 and K3 are karanjin treated samples. The color for each gene on the heatmap represents its log-normalized count. C. Validation. Total RNA isolated from MCF-7 cells treated with vehicle or karanjin (10 μM) for a period of 24 h was subjected to qRT-PCR analysis of the indicated genes chosen from Table 1, and 2. The ΔΔCt method was used for analysing the qRT-PCR data. The experiment had three replicated dishes of control or karanjin treated cells. The expression in one control sample was set to 1 and those determined for the others were expressed relative to control. Bars represent mean relative expression ± sd (n = 3). For each gene the data were analysed by one-tailed t-tests. Asterisks represent significant result. (**p < 0.01, ***p < 0.001).

### 3.4. Karanjin modulates G2/M checkpoint, and estrogen-response-early genes

The diversity of genes regulated by karanjin motivated the mining of enriched gene-sets using fGSEA package. The karanjin-modulated genes were enriched in several hallmark genesets (Fig 3A), which include G2/M checkpoint, and estrogen-response-early genes. The enrichment plots for G2/M checkpoint, and estrogen-response-early genes are shown in Fig 3B, and 3C, respectively. The leading-edge genes in the G2/M checkpoint set were *SLC7A5, CENPF, CDC25B, UBE2C, MYC, CENPE, KPNA2, AURKA, CKS2, ATRX, CENPA, UBE2S, CCNA2, TPX2, CCNB2, PLK1, HMMR, PTTG1* and *BUB1*. The leading-edge genes in estrogen-response-early set were *SLC7A5, STC2, TIPARP, TSKU, AREG, CD44* and *TFF1* (also known as *pS2*). The manner in which karanjin regulated the leading-edge genes within the two-hallmark gene-sets are shown in Fig. 3D and 3E, respectively.

**Fig. 3.**
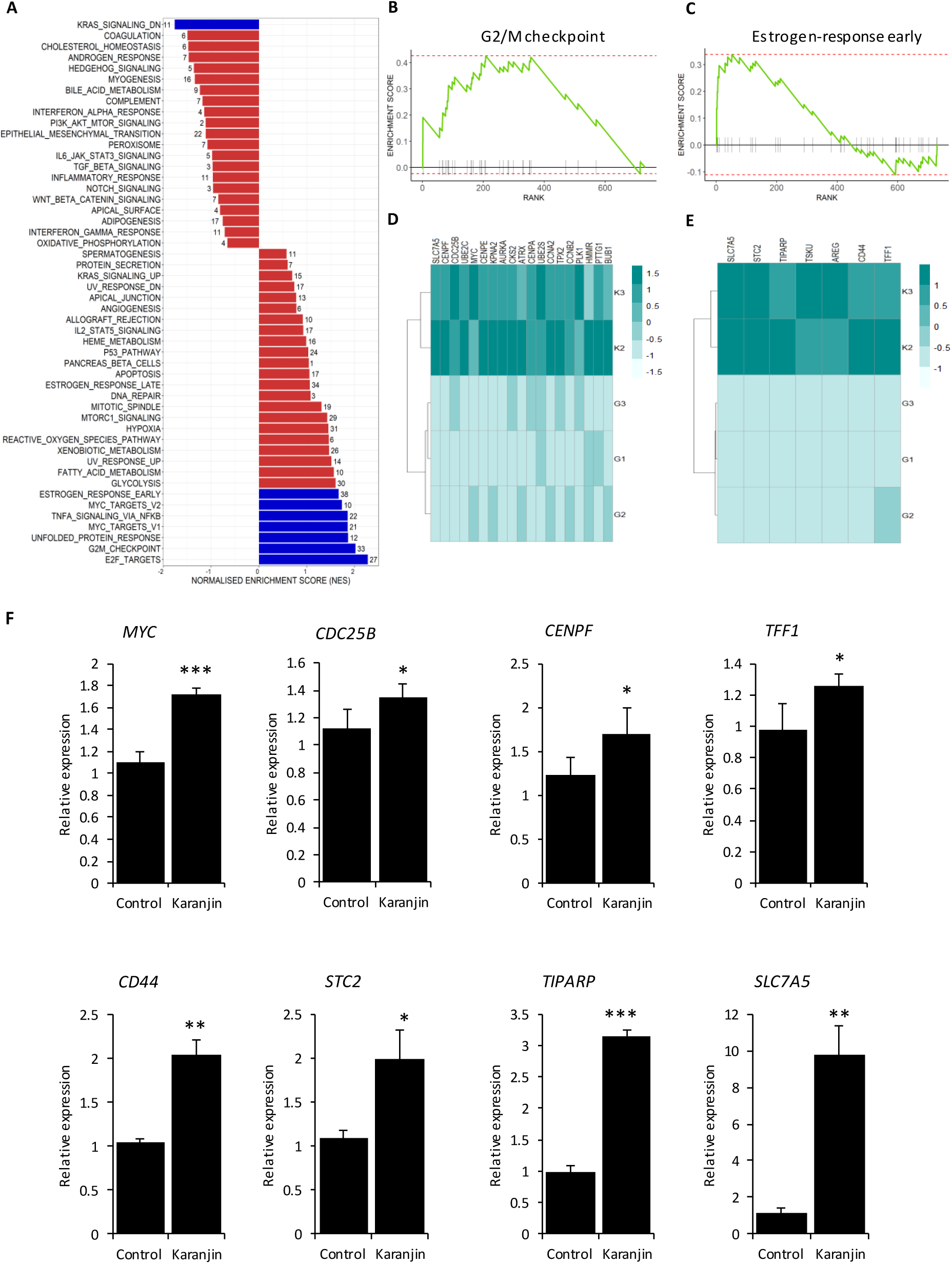
Gene set enrichment analysis. A. Bar chart showing the fGSEA results. The fGSEA package in R was used to analyse the RNA-seq data to identify the enriched hallmark gene sets in MCF-7 cells following 10μM karanjin treatment. Bars represent NES. Blue bars correspond to the significantly enriched gene sets based on a FDR cut-off 25%. The number of leading-edge genes in each gene set is indicated beside the bars. B, C. Enrichment plots for G2/M checkpoint, and estrogen-response-early genes, respectively. D, E. The pattern of expression of leading-edge genes are shown as heatmaps below the respective enrichment plots for G2/M checkpoint, and estrogen-response-early genes. G1, G2, and G3 are controls. K2, and K3 are karanjin treated samples. The color on the heatmap represents the expression in terms of log-normalized counts for genes (rows) across samples (columns). F. Validation. Total RNA isolated from control and karanjin treated cells were subjected to qRT-PCR analysis of the G2/M checkpoint, and estrogen-response-early genes. The ΔΔCt method was used for analysing the qPCR data. The experiment had three replicate dishes of control or karanjin treated (10 μM) cells. The expression in one control sample was set to 1, relative to which the expression in other samples were expressed. Bars represent mean relative expression ± sd (n = 3). For each gene the data were analysed by one-tailed t-tests. Asterisks represent significant result. (*p < 0.05, **p < 0.01, ***p < 0.001).

Modulation of selected leading-edge genes from both hallmark gene-sets was validated by qRT-PCR (Fig. 3F). Among the G2/M checkpoint genes, we confirmed the induction of *MYC* (n = 3, p < 0.001), *CDC25B* (n = 3, p = 0.0042), and *CENPF* (n = 3, p = 0.046). Among estrogen-response-early genes, we confirmed the induction of *TFF1* (n = 4, p = 0.019), *CD44* (n = 3, p = 0.003), *STC2* (n = 3, p = 0.017), and *TIPARP* (n = 3, p < 0.001). *SLC7A5*, which belonged to both hallmark gene-sets, was significantly induced (n = 3, p = 0.004).

### 3.5. Karanjin-modulated genes overlap with E2- or tamoxifen-modulated genes

The identification and validation of estrogen-response-early genes instigated us to examine the similarity or differences between genomic effects of karanjin, E2, and tamoxifen, which is a SERM. Re-analysis of RNA-seq data (GSE117942) of MCF-7 cells treated with 1 nM E2, or 1 μM tamoxifen (Guan et al., 2019) yielded two overlapping sets of 3727 estrogen-regulated, and 435 tamoxifen-regulated genes. Karanjin-modulated genes were matched with those regulated by estrogen or tamoxifen. 419 genes were regulated by karanjin and estrogen, whereas, 94 genes were regulated by karanjin and tamoxifen. 72 genes were regulated by estrogen, tamoxifen and karanjin (Fig 4A). Heatmaps in Fig. 4B and 4C show that karanjin had similar, or opposite effects on estrogen or tamoxifen regulated genes, respectively. For instance, *STC2, TIPARP, SLC7A5*, and *AREG* were upregulated, whereas *CSTA, ADAMTS19, and NCOA3* were downregulated by karanjin and estrogen treatment. *CYP1A1, BMF, and RET* were regulated by karanjin, and estrogen in opposite directions. Karanjin also induced similar or opposite effects on genes regulated by tamoxifen. *SLC7A5, FOSL2*, and *PHLDA1* genes were upregulated, whereas *IGFBP4, CHRD*, and *PCDH7* were downregulated by both karanjin and tamoxifen. *GREB1, TFF1*, and *CYP1B1* were regulated by both, albeit in opposite directions (Supplementary data 9).

**Fig. 4.**
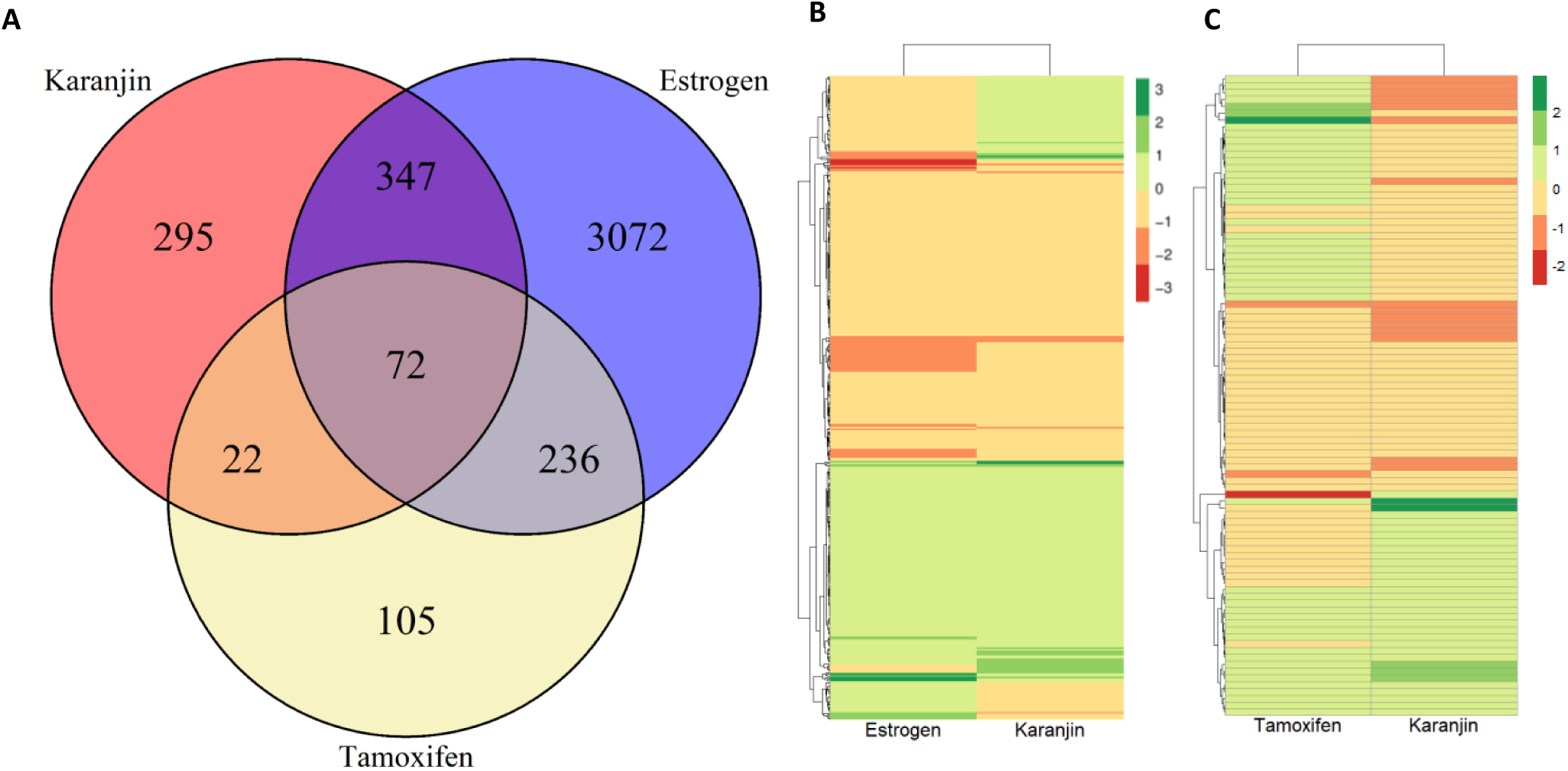
Overlapping genomic targets of karanjin, estrogen and tamoxifen. A. Venn diagram showing the number of genes modulated by karanjin (10 µM), estrogen (1 nM), or tamoxifen (1 µM) in MCF-7 cells. The karanjin modulated genes were from this study. Estrogen or tamoxifen modulated genes were obtained by analyzing the GSE117942 dataset as described in materials and methods. Values in the areas of intersection are the number of genes commonly regulated by two or more compounds. B. and C. Heatmaps showing the patterns of gene modulation by karanjin compared with estrogen (n = 419), or tamoxifen (n = 94), respectively. Colors represent log_2_FC expression compared to their respective controls. For better visualization of expression patterns across tamoxifen and estrogen modulated genes, *CYP1A1* has been omitted during generation of the heatmap due to its high fold change.

### 3.6. Concentration- and cell type-dependent effect of karanjin on gene expression

The effects of varying concentrations of karanjin on the expression of few estrogen-response-early genes was also addressed in two ER-positive cell lines, namely MCF-7 and T47D. The experimental design included cells treated with 10 nM E2 as a reference compound. E2 induces TFF1 expression in ER-positive breast cancer cells (Brown et al., 1984). As expected, 10 nM E2 induced *TFF1* mRNA in both the cell lines (Fig. 5A, B; n = 3, p < 0.001). 10 nM karanjin did not affect *TFF1* mRNA expression in MCF-7 cells. In our qRT-PCR based validation of RNA-seq data, we had confirmed the significant induction (∼1.2 fold) of *TFF1* mRNA in MCF-7 cells by 10 μM karanjin (Fig. 3F). The 1.2-fold induction of *TFF1* mRNA by 10 μM karanjin in Fig. 5A (left panel) is consistent with RNA-seq data, although one-way ANOVA followed by TukeyHSD does not show significant result. 50 μM karanjin significantly induced *TFF1* mRNA expression in MCF-7 (Fig. 5A, left panel, n = 3, p < 0.001). None of the concentrations of karanjin modulated the expression of TFF1 in T47D cells (Fig. 5A, right panel). 10 nM E2 did not affect *TIPARP* mRNA expression in MCF-7 cells. In agreement with RNA-seq data, 10 μM karanjin induced *TIPARP* mRNA expression (Fig. 5B, left panel; n = 3, p < 0.001). In contrast, 10 nM karanjin repressed its expression (Fig. 5B, left panel; n = 3, p < 0.01). In T47D cells, 10 nM E2 induced the expression of *TIPARP* mRNA (Fig. 5B, right panel; n = 3, p < 0.01). Karanjin did not modulate the expression of *TIPARP* mRNA in T47D cells. STC2 mRNA, another estrogen-response-early gene, was induced by 10 nM E2 in both cell lines (Fig. 5C, n = 3, p < 0.001 for MCF-7 cells, p < 0.05 for T47D). In MCF-7 cells *STC2* mRNA was induced by karanjin only at a concentration of 10 µM (Fig. 5C, left panel, n = 3, p < 0.01). 10 nM and 50 μM karanjin induced *STC2* mRNA expression more than two-fold (Fig. 5C, left panel). However, the results were not statistically significant when analysed by ANOVA followed by TukeyHSD (p = 0.07 for 10 nM, and p = 0.09 for 50 μM). None of the concentrations of karanjin significantly modulated the expression of *STC2* mRNA in T47D cells (Fig. 5C, right panel). *SLC7A5* mRNA expression was significantly induced by 10 nM E2 in T47D cells (Fig. 5D, right panel, n = 3, p < 0.001), but not in MCF-7 cell (Fig 5D, left panel). There was significant induction of *SLC7A5* mRNA expression by 10 and 50 µM karanjin (Fig. 5D, left panel, n = 3, p < 0.001 for 10 μM, and p < 0.01 for 50 μM). None of the concentrations of karanjin modulated its expression in T47D cells. *CD44* was not modulated by 10 nM E2 in both cell lines (Fig. 5E). Karanjin induced its expression only in MCF-7 cells at a concentration of 10 µM (Fig. 5E, left panel, n = 3, p < 0.01). Besides, estrogen-response-early genes, we also studied the effect of karanjin on the expression of *CSTA*, a known estrogen suppressed gene (John Mary et al., 2020). As expected, 10 nM E2 suppressed *CSTA* mRNA expression in MCF-7 cells (Fig. 5F, left panel, n = 3, p < 0.001). A significant and progressive dose-dependent suppression of *CSTA* mRNA by karanjin was observed in MCF-7 cells (Fig. 5F, n = 3, p < 0.001). In T47D cells, 10 nM E2 did not affect *CSTA* mRNA expression, which is consistent with the previous findings (John Mary et al., 2020). 10 nM and 10 μM karanjin did not modulate *CSTA* mRNA expression in T47D cells. However, in sharp contrast to MCF-7 cells, 50 μM karanjin significantly induced *CSTA* mRNA in T47D cells (Fig. 5F, right panel, p < 0.001). These data demonstrate that karanjin partially mimics estrogen-like effects on gene expression in a concentration- or cell type-dependent manner.

**Fig. 5.**
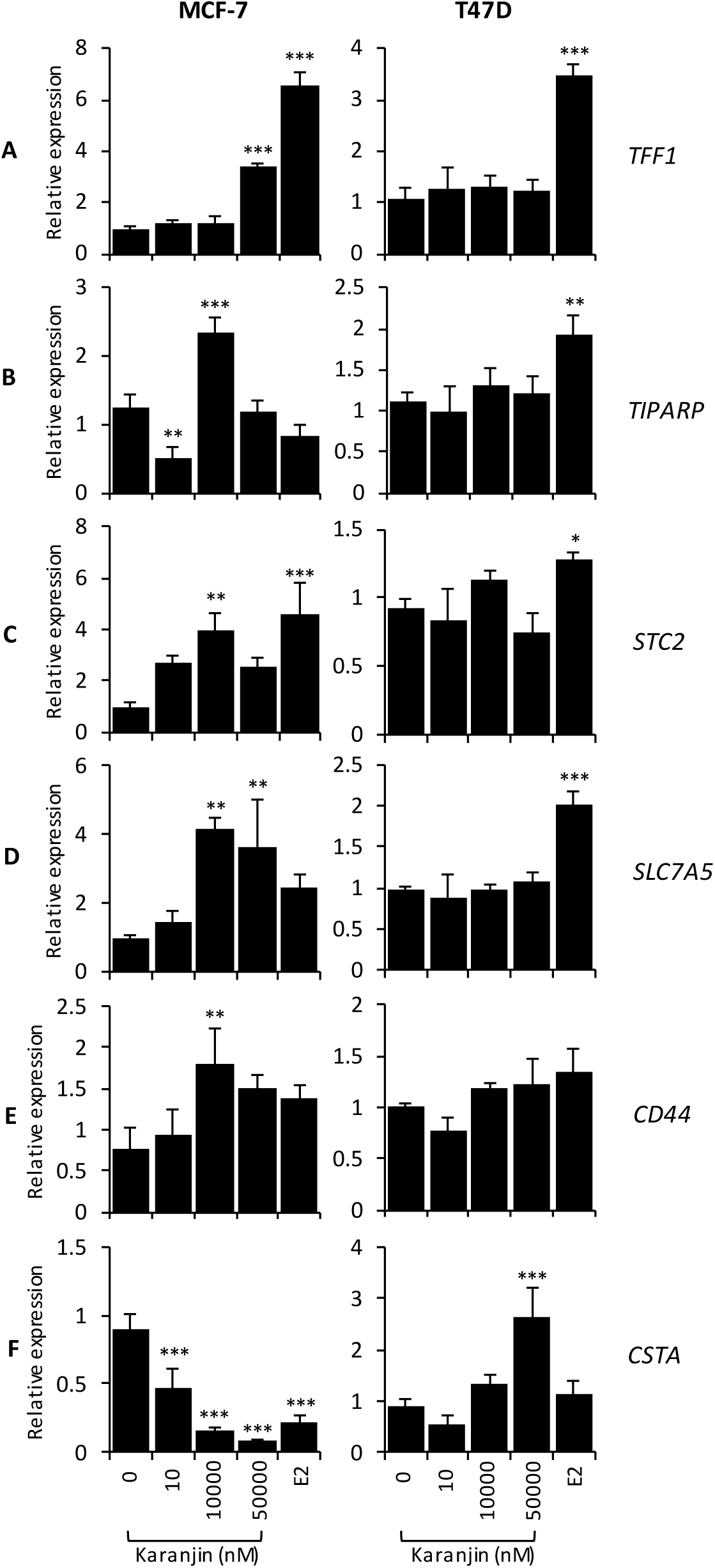
Concentration dependent effect of karanjin on gene expression in MCF-7 and T47D cells. Cells were treated with indicated concentrations of karanjin or 10 nM E2 for 24 h. Thereafter, total RNA was isolated and subjected to qRT-PCR analysis of the indicated genes using the ΔΔCt method. The experiment was done with three replicate dishes of control or karanjin treated cells. The expression in each of the samples was expressed relative to one of the controls, which was set to 1. Bars represent mean relative expression ± sd (n = 3). For each gene the data were analysed by one-way ANOVA followed by TukeyHSD. Significantly different means with respect to control (0 µM karanjin) are indicated by asterisks. (*p < 0.05, **p < 0.01, ***p < 0.001)

### 3.7. Karanjin modulates gene expression via ERα in a gene-dependent manner

Modulation of estrogen-response-early genes led us to hypothesize, the involvement of ERα, at least in part. Post E2-mediated activation, ERα is typically turned over by proteasomal degradation (Reid et al., 2003). There was a significant decrease in ERα protein in MCF-7 cells after treatment with 10 μM karanjin for 24 h, which further decreased with time (Fig. 6A). Thus, post karanjin stimulation, the fate of ERα is similar to that brought about by estrogen.

**Fig. 6.**
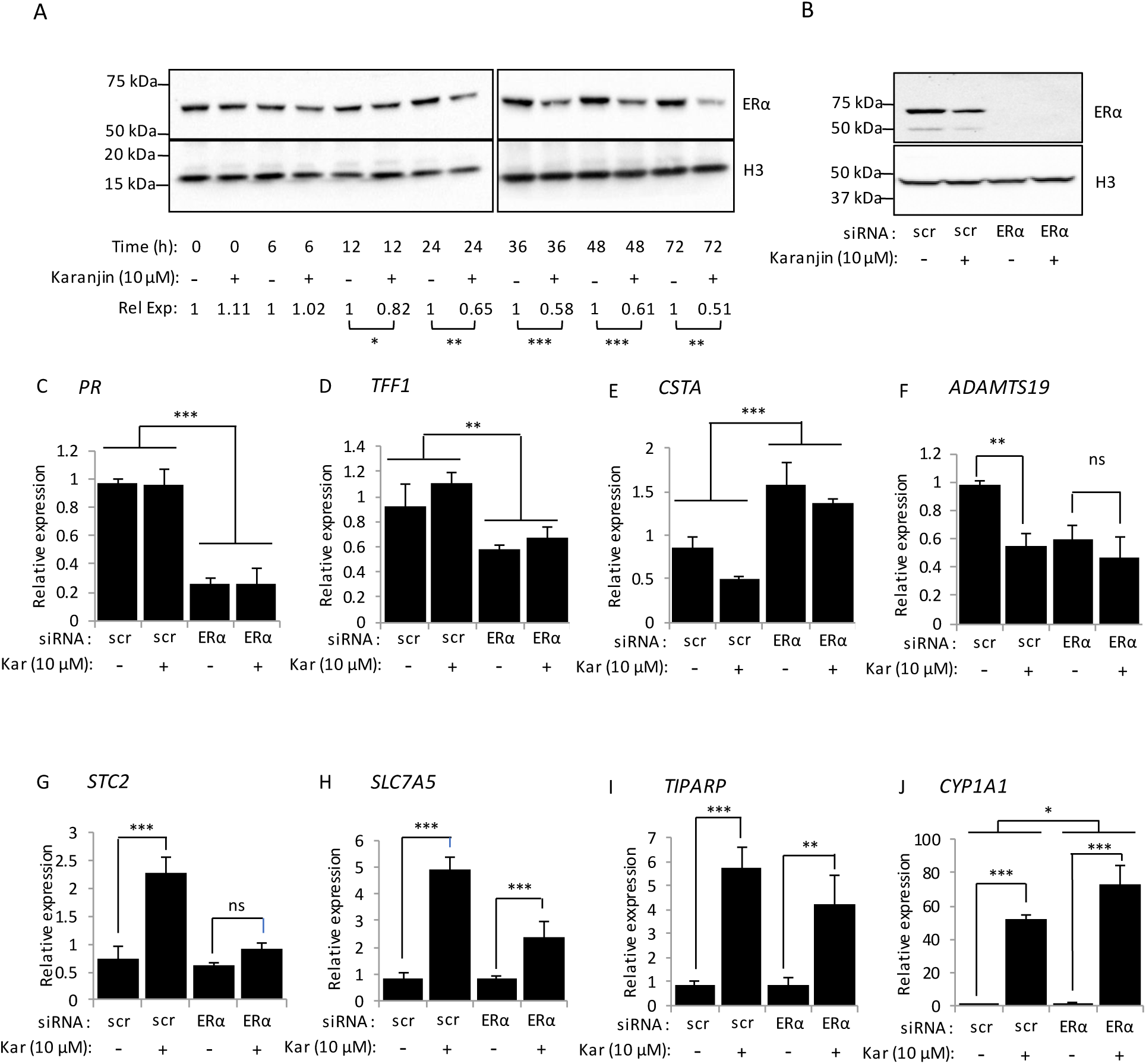
Differential involvement of ERα in karanjin-mediated modulation of gene expression in MCF-7 cells. A. Karanjin-mediated reduction in ERα protein expression in MCF-7 cells. Cells were treated with 10 µM karanjin for the indicated periods of time. At each time point, the untreated cells (0 µM karanjin) served as control. Total protein extracted from Laemmli buffer was subjected to western blotting analysis using the ERα specific antibody. Histone H3 served as an internal control, which was probed with H3-specific antibody. Chemiluminescence signals were imaged in ChemiDoc XRS+ system. The images were processed in ImageJ. The background subtracted band intensity for ERα normalized against that obtained for Histone-H3 served as a measure of ERα protein expression. For each time point ERα expression in control sample was set to 1 and that obtained for karanjin treated sample was expressed relative to control. Mean relative expression (Rel Exp) of ERα ± sd (n = 5) for each time-point is show below. For each time point the data were analysed by a one-tailed t-test. Significant results are indicated by asterisks. (*p < 0.05, **p < 0.01, ***p < 0.001). B. ERα knockdown. MCF-7 cells pre-treated with scrambled (scr) or ERα-specific siRNA were treated with vehicle or 10 µM karanjin (Kar) for a period of 24 h. Total protein was extracted from the phenolic fraction of RNA extraction reagent, and subjected to western blot analysis using ERα and β-actin specific antibodies. β-actin served as an internal control. C-J. Total RNA was extracted from MCF-7 cells, which were treated as in B. The expression of the indicated genes was analysed by qRT-PCR. The experiment was done with three replicate dishes for each experimental group. The expression in one control sample (scr + 0 µM karanjin) was set to 1, and those determined for the others were expressed relative to control. Bars represent mean relative expression ± sd (n = 3). For each gene the data were analysed by 2 × 2 factorial ANOVA.

To gather more evidences for the involvement of ERα, we examined the effect of ERα knockdown on karanjin-mediated modulation of gene expression. qRT-PCR was applied to analyse the expression of selected karanjin-modulated genes in MCF-7 cells, which were treated with vehicle or 10 μM karanjin after prior treatment with scrambled or ERα-specific siRNA. As shown in Fig. 6B, ERα protein was undetectable in cells treated with ERα-specific siRNA. Gene expression data were analysed by 2 × 2 factorial ANOVA to test the effect of karanjin treatment as a function of ERα status. We first analysed the expression levels of *PR* and *TFF1* mRNAs. These genes are ERα-dependent classical estrogen-induced genes, whose expression are significantly reduced upon ERα knockdown (Brown et al., 1984; John Mary et al., 2017; Kumar et al., 2021; Nardulli et al., 1988). As expected, a significant main effect of ERα was observed on *PR* mRNA levels in MCF-7 cells (Fig. 6C, p ≈ 0). However, there was no evidence for an interaction between karanjin and ERα (p = 0.98). Irrespective of the ERα status, karanjin did not modulate the levels of *PR* mRNA. This result is not surprising, since our RNA-seq experiment did not reveal *PR* mRNA modulation by karanjin. A similar result was obtained for *TFF1*. There was no interaction between karanjin and ERα (p = 0.485), but there was a significant main effect of ERα (Fig. 6D, p < 0.001). Karanjin induced *TFF1* mRNA levels by 1.2-fold in cells treated with scrambled siRNA. However, this induction was not significant after multiple comparison, although the induction is comparable to that significant induction inferred from the RNA-seq data (Fig. 3F). In case of *CSTA* mRNA, there was no evidence for the interaction between karanjin and ERα (Fig. 6E, p = 0.37). However, there were significant main effects of karanjin treatment (p = 0.007), and ERα (p ≈ 0). In contrast, analysis of *ADAMTS19* revealed that there was a significant interaction between karanjin and ERα (Fig. 6F, p = 0.02). Karanjin treatment significantly reduced *ADAMTS19* mRNA in scrambled siRNA treated cells, but not in ERα siRNA treated cells. We also analysed the mRNA expression of other estrogen-response early genes, such as *STC2, SLC7A5*, and *TIPARP*. For *STC2* and *SLC7A5*, a significant interaction between karanjin and ERα (Fig. 6G, H; p < 0.001) was found; the induction in mRNA expression by karanjin being greater in cells treated with scrambled siRNA compared to ERα-specific siRNA. In case of *TIPARP* only the main effect of karanjin was significant (Fig. 6I, p < 0.001). We also analysed *CYP1A1* mRNA expression, which does not fall under the estrogen-response-early gene set. There was a modest but significant interaction between karanjin and ERα (Fig. 6J, p = 0.042), with significant main effects of karanjin (p < 0.001) and ERα (p = 0.03).

## 4. Discussion

Karanjin is a bioactive compound, with anti-hyperglycaemic (Tamrakar et al., 2008), anti-inflammatory (Bose et al., 2014), anti-ulcer (Vismaya et al., 2011), and anti-colitis (Patel and Trivedi, 2017) effects *in vivo*, and anti-proliferative (Guo et al., 2015; Raghav et al., 2019; Roy et al., 2019), cell cycle inhibitory (Guo et al., 2015; Roy et al., 2019), ROS limiting (Roy et al., 2019), and glucose-uptake inducing effects *in vitro* (Jaiswal et al., 2011). Its molecular effects include inhibition of TNFα production, modulation of NF-κB activity (Bose et al., 2014), GLUT-4 translocation, protein phosphorylation, protein kinase activation (Jaiswal et al., 2011), enzyme inhibition (Joshi et al., 2018), and interference with transporters (Michaelis et al., 2014). However, the genomic correlates of these effects were unknown. This study explored the genome-wide alteration of gene expression at the mRNA level in MCF-7 cells, using next generation sequencing. The transcriptomic- or gene-modulatory footprint of karanjin encompasses a multitude of cellular processes, such as metabolism (glycolysis, fatty acid, and xenobiotic), signalling (TNFα, KRAS), ROS-scavenging, unfolded protein response, modulation of transcription factor targets (E2F and MYC), and hormonal response (androgen, and estrogen response early).

Various research groups have previously demonstrated *in vitro* effects of karanjin over a wide concentration range. We found no evidence for any short-term (24 h) effect of karanjin (0-50 µM) on MCF-7 viable cell count. However, long-term (120 h) effect was significantly concentration-dependent, although the viable cell count increased at all concentrations of karanjin. Viable count was significantly lower at higher concentrations (10 and 50 μM), whereas it was, modestly, but significantly higher at lower concentration (10 nM). These data suggest that the anti-proliferative effect of karanjin may not be universal, but rather be cell- and concentration-dependent. This presents a caveat to the anticancer potential of karanjin, and underscore the importance of dosage for various healthcare applications suggested in the literature.

The RNA-seq data, from the 24 h experiment, capture the early effects of karanjin. Since the viability of MCF-7 cells treated for 24 h with 10 µM karanjin is as good as vehicle, the changes in gene expression reflected in RNA-seq data should be free from those associated with proliferation or toxicity. In this context, the enrichment of G2/M checkpoint genes among those modulated by karanjin is worthy of attention, as it generates contradictions. The enrichment of G2/M checkpoint genes appears to contradict the observation that none of the concentrations of karanjin resulted in significantly different viable counts compared to vehicle treated control, in 24 h. Furthermore, 10 μM karanjin modulated the G2/M checkpoint genes in a manner that is consistent with cell cycle progression, rather than cell cycle arrest. For example, *SLC7A5*, a sodium-independent transporter is one of the karanjin induced genes. It is an amino acid exchanger that maintains intracellular levels of leucine, an established master regulator of the mTORC1 pathway, and overtly expressed in a variety of cancers (El Ansari et al., 2018). *MYC* is a proto-oncogene that encodes a nuclear phosphoprotein. The regulation of MYC expression in cells is closely linked with cell proliferation. Elevated expression of MYC promotes the activation of cyclins and CDKs, and impairs the functionality of cell cycle inhibitors (Miller et al., 2012). CDC25B is a MYC target (Zörnig and Evan, 1996). It facilitates entry of cells into mitosis by dephosphorylating cell dependent kinase-CDC2 (Lammer et al., 1998). Induction of the selected G2/M checkpoint genes by 10 μM karanjin contradicts the observation that over a period of 120 h, 10 μM karanjin treatment results in significantly lesser viable counts compared to control. Furthermore, it contradicts other reports of cell cycle arrest and apoptosis (Guo et al., 2015; Roy et al., 2019).

Here we present possible explanations behind the contradictions. Although 10 μM karanjin treatment for 24 or 120 h produces similar, or significantly lower viable counts, respectively, compared to control, proliferation of cells is evident from a significant (1.55- fold and 5.9-fold, respectively) increase compared to baseline (0 h) viable count. The cells could not have proliferated without cell cycle progression. In the event of cell cycle arrest followed by apoptosis, the viable counts are expected to go below the baseline. Thus, it is possible, that at 10 μM, karanjin delays cell cycle progression rather than induce arrest. The delay could be due to other yet unknown effects of karanjin. It is to be noted that even at 50 μM, there is proliferation of cells, although the final viable count after 120 h is even lower than that resulting from 10 μM. On the other hand, 10 nM karanjin results in significantly more viable count compared to control. The concentration dependent effect indicates that there is more than one receptor for karanjin in MCF-7 cells. The high affinity receptors could be responsible for proliferative actions of karanjin at lower concentrations. Whereas, low affinity receptors could be responsible for cell cycle arrest at high concentrations.

In some studies, conclusions about anti-proliferative effects of karanjin were based on MTT assays, which are end-point assays. This obscures the initial viability at the start of the experiment, which makes it difficult to conclude whether the treatment caused cell death or cell cycle arrest. This distinction is possible with trypan blue dye exclusion assay. Furthermore, mechanisms of cell cycle arrest, and pro-apoptotic effects of karanjin, were shown in HeLa, Hep G2, A549, and HL-60, but not in MCF-7 cells (Guo et al., 2015; Roy et al., 2019). Thus, the apparent contradictions could be due to the intrinsic properties of MCF-7 cells.

Enrichment of estrogen-response-early genes and overlap of karanjin modulated genes with those regulated by 1 nM E2 or 1 μM tamoxifen suggest partial estrogen like effects of karanjin. Enhanced ERα protein turnover following karanjin treatment, and the negative impact of ERα knockdown on karanjin-mediated alteration of gene expression provide enticing evidences in favour of ERα mediated actions of karanjin. It remains to be seen whether karanjin directly binds ERα. However, it is possible, given the flavonoid structure of SERMs (Rosenberg Zand et al., 2000). They have cell-type and concentration dependent effects on proliferation (Wang et al., 1996) and gene expression in hormone responsive cells (Diel et al., 2001; Lavigne et al., 2008). It is possible that karanjin may have SERM-like activity since our study shows that karanjin not only enhances ERα protein turnover, but also exerts concentration-, gene-, and cell-type-dependent effect on gene expression. This selectivity can potentially be attributed to the differential recruitment of co-activators or co-repressors on karanjin-ER-complex at target gene promoters. ERα plays a central role in estrogen-mediated proliferation of breast cancer cells. The partial estrogen-like or SERM-like activity of karanjin could underlie the concentration dependent effect on MCF-7 cell proliferation. The compromised proliferation of MCF-7 cells treated with 10 μM karanjin in the face of induced expression of estrogen-response-early genes could be due to the additional effects of karanjin mediated via non-ERα targets. Thus, the present study provides a valuable insight into the hitherto unexplored effects of karanjin on endocrine responsive cells. The caveat exposed by the data with respect to the anticancer potential of karanjin cannot be overlooked.

In summary, karanjin affects proliferation of MCF-7 cells in a concentration dependent manner. 10 μM karanjin induces genome-wide alteration in MCF-7 cell transcriptome, which likely impacts diverse cellular processes including G2/M transition and early estrogen responses. It modulates gene expression in the ERα-positive cell lines in a manner that suggests SERM-like activity. The findings call for an in-depth investigation into karanjin’s effects on breast cancer cells and the mechanisms, particularly the involvement of ERα, before considering its usage in breast cancer therapy. This study, while exposing a caveat to the anticancer potential of karanjin, will inspire further investigations into the possible use of karanjin or its derivatives in endocrine therapies.

## Supporting information

All supplementary material

## CRediT authorship contribution statement

**Gaurav Bhatt:** Data curation, Investigation. **Akshita:** Data curation, In-silico investigation. **Latha Rangan:** Conceptualization, Supervision. **Anil Mukund Limaye:** Conceptualization, Supervision, Validation, Writing - review & editing.

## Acknowledgments

The work was funded by a core research grant from the Department of Science and Technology, Govt. of India (CRG/2020/002109). Infrastructural support from the Department of Biosciences and Bioengineering, IIT Guwahati. is acknowledged. GB, and AG acknowledge the fellowship from IIT Guwahati. Technical support received from Mr. Debojit Bhattacharjee regarding HRMS is acknowledged. The authors thank Prathibha Ranganathan for critical reading of the manuscript.

## References

Andrews, S., Krueger, F., Seconds-Pichon, A., Biggins, F., Wingett, S., 2015. FastQC. A quality control tool for high throughput sequence data. Babraham Bioinformatics http://www.bioinformatics.babraham.ac.uk/projects/fastqc/,2010 (accessed 1.25.21).

Bolger, A.M., Lohse, M., Usadel, B., 2014. Trimmomatic: a flexible trimmer for Illumina sequence data. Bioinformatics 30, 2114–20. https://doi.org/10.1093/bioinformatics/btu170

Bose, M., Chakraborty, M., Bhattacharya, S., Bhattacharjee, P., Mandal, S., Kar, M., Mishra, R., n.d. Suppression of NF-κB p65 nuclear translocation and tumor necrosis factor-α by Pongamia pinnata seed extract in adjuvant-induced arthritis. J. Immunotoxicol. 11, 222–30. https://doi.org/10.3109/1547691X.2013.824931

Bose, M., Chakraborty, M., Bhattacharya, S., Mukherjee, D., Mandal, S., Mishra, R., 2014. Prevention of arthritis markers in experimental animal and inflammation signalling in macrophage by Karanjin isolated from Pongamia pinnata seed extract. Phytother. Res. 28, 1188–95. https://doi.org/10.1002/ptr.5113

Brown, A.M., Jeltsch, J.M., Roberts, M., Chambon, P., 1984. Activation of pS2 gene transcription is a primary response to estrogen in the human breast cancer cell line MCF-7. Proc. Natl. Acad. Sci. U. S. A. 81, 6344–8. https://doi.org/10.1073/pnas.81.20.6344

Choi, S.Y., Ha, T.Y., Ahn, J.Y., Kim, S.R., Kang, K.S., Hwang, I.K., Kim, S., 2008. Estrogenic activities of isoflavones and flavones and their structure-activity relationships. Planta Med. 74, 25–32. https://doi.org/10.1055/s-2007-993760

Choi, W.S., Chung, K.J., Chang, M.S., Chun, J.K., Lee, H.W., Hong, S.Y., 1993. A turbidimetric determination of protein by trichloroacetic acid. Arch. Pharm. Res. 16, 57–61. https://doi.org/10.1007/BF02974129

Constantinou, A.I., Krygier, A.E., Mehta, R.R., 1998. Genistein induces maturation of cultured human breast cancer cells and prevents tumor growth in nude mice. Am. J. Clin. Nutr. 68, 1426S–1430S. https://doi.org/10.1093/ajcn/68.6.1426S

Diel, P., Olff, S., Schmidt, S., Michna, H., 2001. Molecular identification of potential selective estrogen receptor modulator (SERM) like properties of phytoestrogens in the human breast cancer cell line MCF-7. Planta Med. 67, 510–4. https://doi.org/10.1055/s-2001-16474

Dobin, A., Davis, C.A., Schlesinger, F., Drenkow, J., Zaleski, C., Jha, S., Batut, P., Chaisson, M., Gingeras, T.R., 2013. STAR: ultrafast universal RNA-seq aligner. Bioinformatics 29, 15–21. https://doi.org/10.1093/bioinformatics/bts635

El Ansari, R., Craze, M.L., Miligy, I., Diez-Rodriguez, M., Nolan, C.C., Ellis, I.O., Rakha, E.A., Green, A.R., 2018. The amino acid transporter SLC7A5 confers a poor prognosis in the highly proliferative breast cancer subtypes and is a key therapeutic target in luminal B tumours. Breast Cancer Res. 20, 21. https://doi.org/10.1186/s13058-018-0946-6

Fioravanti, L., Cappelletti, V., Miodini, P., Ronchi, E., Brivio, M., Di Fronzo, G., 1998. Genistein in the control of breast cancer cell growth: insights into the mechanism of action in vitro. Cancer Lett. 130, 143–52. https://doi.org/10.1016/s0304-3835(98)00130-x

Fotsis, T., Pepper, M.S., Aktas, E., Breit, S., Rasku, S., Adlercreutz, H., Wähälä, K., Montesano, R., Schweigerer, L., 1997. Flavonoids, dietary-derived inhibitors of cell proliferation and in vitro angiogenesis. Cancer Res. 57, 2916–21.

Guan, J., Zhou, W., Hafner, M., Blake, R.A., Chalouni, C., Chen, I.P., De Bruyn, T., Giltnane, J.M., Hartman, S.J., Heidersbach, A., Houtman, R., Ingalla, E., Kategaya, L., Kleinheinz, T., Li, J., Martin, S.E., Modrusan, Z., Nannini, M., Oeh, J., Ubhayakar, S., Wang, X., Wertz, I.E., Young, A., Yu, M., Sampath, D., Hager, J.H., Friedman, L.S., Daemen, A., Metcalfe, C., 2019. Therapeutic Ligands Antagonize Estrogen Receptor Function by Impairing Its Mobility. Cell 178, 949-963.e18. https://doi.org/10.1016/j.cell.2019.06.026

Guo, J.R., Chen, Q.Q., Wai-Kei Lam, C., Zhang, W., 2015. Effects of karanjin on cell cycle arrest and apoptosis in human A549, HepG2 and HL-60 cancer cells. Biol. Res. 48. https://doi.org/10.1186/s40659-015-0031-x

Haddad, A.Q., Venkateswaran, V., Viswanathan, L., Teahan, S.J., Fleshner, N.E., Klotz, L.H., 2006. Novel antiproliferative flavonoids induce cell cycle arrest in human prostate cancer cell lines. Prostate Cancer Prostatic Dis. 9, 68–76. https://doi.org/10.1038/sj.pcan.4500845

Hong, H., Branham, W.S., Ng, H.W., Moland, C.L., Dial, S.L., Fang, H., Perkins, R., Sheehan, D., Tong, W., 2015. Human sex hormone-binding globulin binding affinities of 125 structurally diverse chemicals and comparison with their binding to androgen receptor, estrogen receptor, and α-fetoprotein. Toxicol. Sci. 143, 333–48. https://doi.org/10.1093/toxsci/kfu231

Hsu, J.T., Hung, H.C., Chen, C.J., Hsu, W.L., Ying, C., 1999. Effects of the dietary phytoestrogen biochanin A on cell growth in the mammary carcinoma cell line MCF-7. J. Nutr. Biochem. 10, 510–7. https://doi.org/10.1016/s0955-2863(99)00037-6

Jaiswal, N., Yadav, P.P., Maurya, R., Srivastava, A.K., Tamrakar, A.K., 2011. Karanjin from Pongamia pinnata induces GLUT4 translocation in skeletal muscle cells in a phosphatidylinositol-3-kinase-independent manner. Eur. J. Pharmacol. 670, 22–8. https://doi.org/10.1016/j.ejphar.2011.08.049

John Mary, D.J.S., Manjegowda, M.C., Kumar, A., Dutta, S., Limaye, A.M., 2017. The role of cystatin A in breast cancer and its functional link with ERα. J. Genet. Genomics 44, 593–597. https://doi.org/10.1016/j.jgg.2017.10.001

John Mary, D.J.S., Sikarwar, G., Kumar, A., Limaye, A.M., 2020. Interplay of ERα binding and DNA methylation in the intron-2 determines the expression and estrogen regulation of cystatin A in breast cancer cells. Mol. Cell. Endocrinol. 504, 110701. https://doi.org/10.1016/j.mce.2020.110701

Joshi, P., Sonawane, V.R., Williams, I.S., McCann, G.J.P., Gatchie, L., Sharma, R., Satti, N., Chaudhuri, B., Bharate, S.B., 2018. Identification of karanjin isolated from the Indian beech tree as a potent CYP1 enzyme inhibitor with cellular efficacy: Via screening of a natural product repository. Medchemcomm 9, 371–382. https://doi.org/10.1039/c7md00388a

Kumar, A., Dhillon, A., Manjegowda, M.C., Singh, N., Mary, D.J.S.J., Kumar, S., Modi, D., Limaye, A.M., 2021. Estrogen suppresses HOXB2 expression via ERα in breast cancer cells. Gene 794, 145746. https://doi.org/10.1016/j.gene.2021.145746

Lammer, C., Wagerer, S., Saffrich, R., Mertens, D., Ansorge, W., Hoffmann, I., 1998. The cdc25B phosphatase is essential for the G2/M phase transition in human cells. J. Cell Sci. 111, 2445–2453. https://doi.org/10.1242/jcs.111.16.2445

Lavigne, J.A., Takahashi, Y., Chandramouli, G.V.R., Liu, H., Perkins, S.N., Hursting, S.D., Wang, T.T.Y., 2008. Concentration-dependent effects of genistein on global gene expression in MCF-7 breast cancer cells: an oligo microarray study. Breast Cancer Res. Treat. 110, 85–98. https://doi.org/10.1007/s10549-007-9705-6

Liao, Y., Smyth, G.K., Shi, W., 2014. featureCounts: an efficient general purpose program for assigning sequence reads to genomic features. Bioinformatics 30, 923–30. https://doi.org/10.1093/bioinformatics/btt656

Lippman, M., Bolan, G., Huff, K., 1976. The effects of estrogens and antiestrogens on hormone-responsive human breast cancer in long-term tissue culture. Cancer Res. 36, 4595–601.

Livak, K.J., Schmittgen, T.D., 2001. Analysis of relative gene expression data using real-time quantitative PCR and the 2(-Delta Delta C(T)) Method. Methods 25, 402–8. https://doi.org/10.1006/meth.2001.1262

Love, M.I., Huber, W., Anders, S., 2014. Moderated estimation of fold change and dispersion for RNA-seq data with DESeq2. Genome Biol. 15, 550. https://doi.org/10.1186/s13059-014-0550-8

Lowry, O.H., Rosebrough, N.J., Farr, A.L., Randall, R.J., 1951. Protein measurement with the Folin phenol reagent. J. Biol. Chem. 193, 265–275. https://doi.org/10.1016/s0021-9258(19)52451-6

Maggiolini, M., Bonofiglio, D., Marsico, S., Panno, M.L., Cenni, B., Picard, D., Andò, S., 2001. Estrogen receptor alpha mediates the proliferative but not the cytotoxic dose-dependent effects of two major phytoestrogens on human breast cancer cells. Mol. Pharmacol. 60, 595–602.

Memariani, Z., Abbas, S.Q., ul Hassan, S.S., Ahmadi, A., Chabra, A., 2021. Naringin and naringenin as anticancer agents and adjuvants in cancer combination therapy: Efficacy and molecular mechanisms of action, a comprehensive narrative review. Pharmacol. Res. 171, 105264. https://doi.org/10.1016/j.phrs.2020.105264

Michaelis, M., Rothweiler, F., Nerreter, T., Sharifi, M., Ghafourian, T., Cinatl, J., 2014. Karanjin interferes with ABCB1, ABCC1, and ABCG2. J. Pharm. Pharm. Sci. 17, 92–105. https://doi.org/10.18433/j3bw2s

Miller, D.M., Thomas, S.D., Islam, A., Muench, D., Sedoris, K., 2012. c-Myc and cancer metabolism. Clin. Cancer Res. 18, 5546–5553. https://doi.org/10.1158/1078-0432.CCR-12-0977

Nardulli, A.M., Greene, G.L., O’Malley, B.W., Katzenellenbogen, B.S., 1988. Regulation of progesterone receptor messenger ribonucleic acid and protein levels in MCF-7 cells by estradiol: analysis of estrogen’s effect on progesterone receptor synthesis and degradation. Endocrinology 122, 935–44. https://doi.org/10.1210/endo-122-3-935

Park, K. Il, Park, H.S., Nagappan, A., Hong, G.E., Lee, D.H., Kang, S.R., Kim, J.A., Zhang, J., Kim, E.H., Lee, W.S., Shin, S.C., Hah, Y.S., Kim, G.S., 2012. Induction of the cell cycle arrest and apoptosis by flavonoids isolated from Korean Citrus aurantium L. in non-small-cell lung cancer cells. Food Chem. 135, 2728–35. https://doi.org/10.1016/j.foodchem.2012.06.097

Patel, P.P., Trivedi, N.D., 2017. Effect of karanjin on 2,4,6-trinitrobenzenesulfonic acid-induced colitis in Balb/c mice. Indian J. Pharmacol. 49, 161–167. https://doi.org/10.4103/ijp.IJP_234_15

Raghav, D., Mahanty, S., Rathinasamy, K., 2019. Biochemical and toxicological investigation of karanjin, a bio-pesticide isolated from Pongamia seed oil. Pestic. Biochem. Physiol. 157, 108–121. https://doi.org/10.1016/j.pestbp.2019.03.011

Reid, G., Hübner, M.R., Métivier, R., Brand, H., Denger, S., Manu, D., Beaudouin, J., Ellenberg, J., Gannon, F., 2003. Cyclic, proteasome-mediated turnover of unliganded and liganded ERα on responsive promoters is an integral feature of estrogen signaling. Mol. Cell 11, 695–707. https://doi.org/10.1016/S1097-2765(03)00090-X

Rosenberg Zand, R.S., Jenkins, D.J.A., Diamandis, E.P., 2000. Steroid hormone activity of flavonoids and related compounds. Breast Cancer Res. Treat. 62, 35–49. https://doi.org/10.1023/A:1006422302173

Roy, R., Pal, D., Sur, S., Mandal, S., Saha, P., Panda, C.K., 2019. Pongapin and Karanjin, furanoflavanoids of Pongamia pinnata, induce G2/M arrest and apoptosis in cervical cancer cells by differential reactive oxygen species modulation, DNA damage, and nuclear factor kappa-light-chain-enhancer of activated B cell signal. Phyther. Res. 33, 1084–1094. https://doi.org/10.1002/ptr.6302

Sergushichev, A.A., 2016. An algorithm for fast preranked gene set enrichment analysis using cumulative statistic calculation. bioRxiv 060012.

Singh, A., Bhatt, G., Gujre, N., Mitra, S., Swaminathan, R., Limaye, A.M., Rangan, L., 2021. Karanjin. Phytochemistry 183, 112641. https://doi.org/10.1016/j.phytochem.2020.112641

Singh, A., Jahan, I., Sharma, M., Rangan, L., Khare, A., Panda, A., 2016. Structural Characterization, In Silico Studies and In Vitro Antibacterial Evaluation of a Furanoflavonoid from Karanj. Planta Medica Lett. 3, e91–e95. https://doi.org/10.1055/s-0042-105159

Singh, S., Meena, A., Luqman, S., 2021. Baicalin mediated regulation of key signaling pathways in cancer. Pharmacol. Res. 164, 105387. https://doi.org/10.1016/j.phrs.2020.105387

Strober, W., 2001. Trypan blue exclusion test of cell viability. Curr. Protoc. Immunol. Appendix 3, Appendix 3B. https://doi.org/10.1002/0471142735.ima03bs21

Tamrakar, A.K., Yadav, P.P., Tiwari, P., Maurya, R., Srivastava, A.K., 2008. Identification of pongamol and karanjin as lead compounds with antihyperglycemic activity from Pongamia pinnata fruits. J. Ethnopharmacol. 118, 435–9. https://doi.org/10.1016/j.jep.2008.05.008

Vismaya Belagihally, S.M., Rajashekhar, S., Jayaram, V.B., Dharmesh, S.M., Thirumakudalu, S.K.C., 2011. Gastroprotective Properties of Karanjin from Karanja (Pongamia pinnata) Seeds; Role as Antioxidant and H, K-ATPase Inhibitor. Evid. Based. Complement. Alternat. Med. 2011, 747246. https://doi.org/10.1093/ecam/neq027

Wang, T.T.Y., Sathyamoorthy, N., Phang, J.M., 1996. Molecular effects of genistein on estrogen receptor mediated pathways. Carcinogenesis 17, 271–275. https://doi.org/10.1093/carcin/17.2.271

Xu, R., Zhang, Y., Ye, X., Xue, S., Shi, J., Pan, J., Chen, Q., 2013. Inhibition effects and induction of apoptosis of flavonoids on the prostate cancer cell line PC-3 in vitro. Food Chem. 138, 48–53. https://doi.org/10.1016/j.foodchem.2012.09.102

Yang, S., Zhou, Q., Yang, X., 2007. Caspase-3 status is a determinant of the differential responses to genistein between MDA-MB-231 and MCF-7 breast cancer cells. Biochim. Biophys. Acta 1773, 903–11. https://doi.org/10.1016/j.bbamcr.2007.03.021

Zava, D.T., Duwe, G., 1997. Estrogenic and antiproliferative properties of genistein and other flavonoids in human breast cancer cells in vitro. Nutr. Cancer 27, 31–40. https://doi.org/10.1080/01635589709514498

Zörnig, M., Evan, G.I., 1996. Cell cycle: On target with Myc. Curr. Biol. 6, 1553–1556. https://doi.org/10.1016/S0960-9822(02)70769-0

