## Supplementary material for "Global transcriptome analysis reveals partial estrogen-like effects of karanjin in MCF-7 breast cancer cells": All supplementary material: Supplementary data 1.pdf

### Certificate of analysis for Karanjin from Yucca Enterprises

Yucca Enterprises

**YUCCA ENTERPRISES**
**A-246, Antop Hill warehousing Co.,**
**Barkat Ali Naka, VIT Marg,**
**WADALA (E), MUMBAI 400 037.**
****

***Certificate of Analysis***

**KARANGIN**

**Batch** : Yucca/KG/2019/04/21 **Date** : 22.04.2019

**Chemical Name** : 3-Methoxy-2-phenyl-4H-furo(2,3-h)(1)benzopyran-4-one; Karanjin
**Chemical Formula** : $C_{18}H_{12}O_4$
**Chemical Family** : Flavonoid
**Molecular Weight** : 292.29
**CAS Number** : 521-88-0
**Description** : White powder with a faint yellow cast

**Molecular structure** :

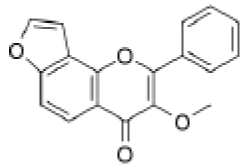

**Intended use** : For laboratory use only
**Solubility** : Soluble in ethyl and methyl alcohol, benzene and petroleum ether.
**Storage condition** : +2°C to +8°C, Protect from light

**Analytical test** :

| S. No. | Test | Result |
|---|---|---|
| 1 | Test for identity (by TLC) | Complies |
| 2 | Purity test (by HPLC) | ≥ 90 % |

**For Yucca Enterprises**

YUCCA ENTERPRISES

### Characterization of karanjin

Karanjin isolated earlier from *P. pinnata* seed oil, and standard purchased from Yucca Enterprises was subjected to HRMS, HPLC and NMR for validation.

### High-resolution mass spectrometry (HRMS)

About, 1 mg of karanjin was dissolved in 1 mL of HPLC grade acetonitrile and filtered. The samples were injected via auto-sampler to record the mass spectra using standard protocol. Mass spectrum was recorded on an Agilent Technologies 6520 Accurate Mass Q-TOF LC/MS (Agilent, CA, USA) in ESI +ive ion mode with a flow rate of 0.2 mL/min over a total run time of 30 min. HRMS data for karanjin isolated in-house and that procured from Yucca Enterprises are shown in Fig. S1A and Fig. S1B.

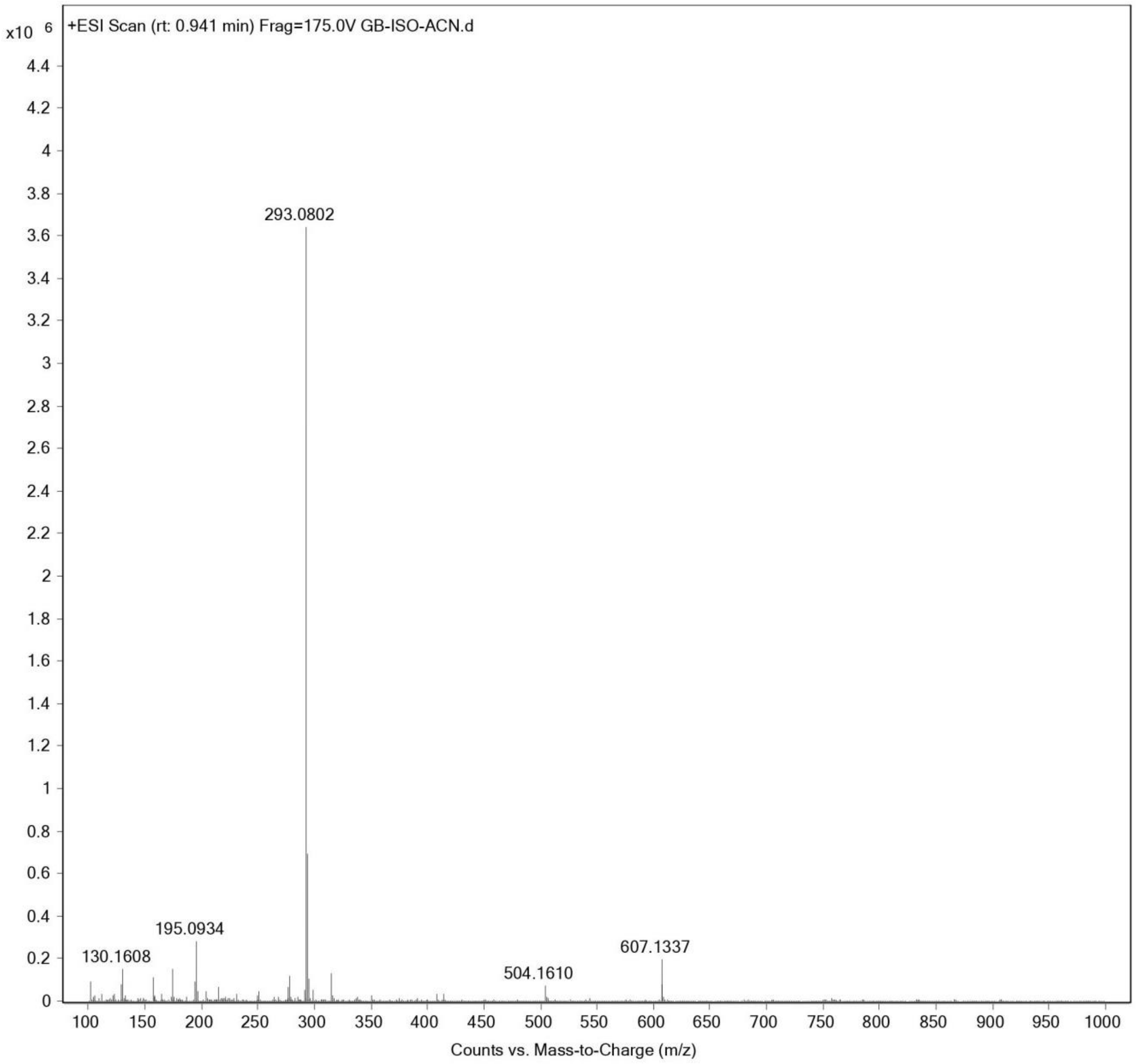


**Fig. S1A.** Mass spectrum of isolated karanjin. The *m/z* of the compound in the positive mode $[M + H]^+$ is 293.0802.

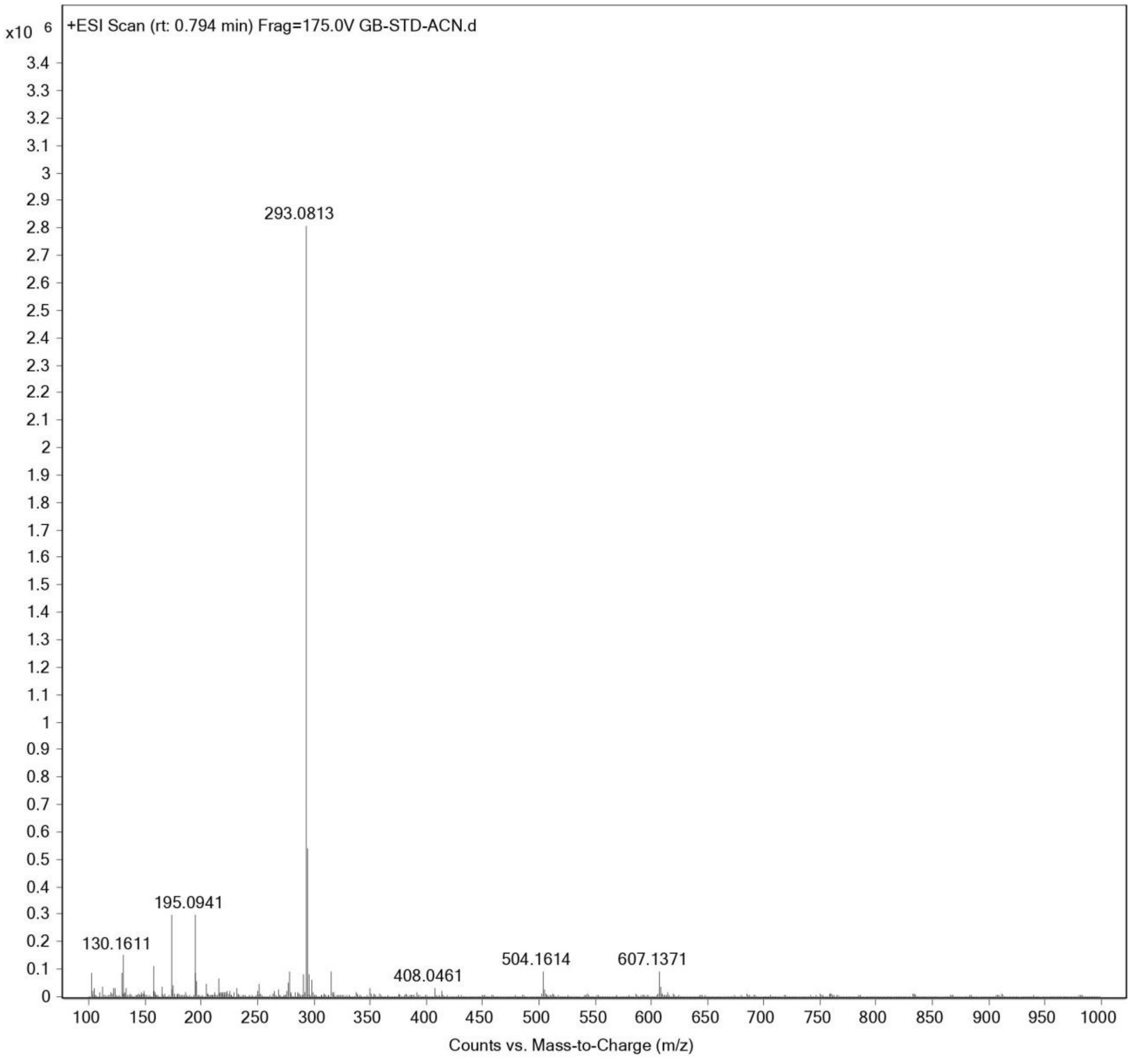


**Fig. S1B.** Mass spectrum of karanjin from Yucca enterprise. The *m/z* of the compound in the positive mode $[M + H]^+$ is 293.0813.

**Purification and detection by HPLC**

To assess the purity of karanjin, high performance liquid chromatography analysis was performed using analytical HPLC system (Shimdazu, Japan) with a degassing unit (DGU 20ASR) and autosampler (SIL 20AHT) and liquid chromatogram (LC 20AD) system. The chromatographic separation was performed using reverse-phase C-18 column (Luna ® , 5μM, 100 Å, 250×4.6 mm ). Karanjin isolated in-house or standard (Yucca enterprises, India) stock of 1 mg/mL was prepared in methanol:water:acetic acid (85:13.5:1.5) filtered through a 0.45 μm syringe filter. 10 μL of filtered samples were then injected and analysed in isocratic mode methanol:water:acetic acid (85:13.5:1.5) for 20 mins at a flow rate of 1.0 mL/min. Karanjin was detected using a UV/Vis detector (SPD-20A) at a set wavelength of 264 nm. HPLC data for karanjin isolated in-house and standard procured from Yucca Enterprises are shown in Fig. S2A and Fig. S2B.

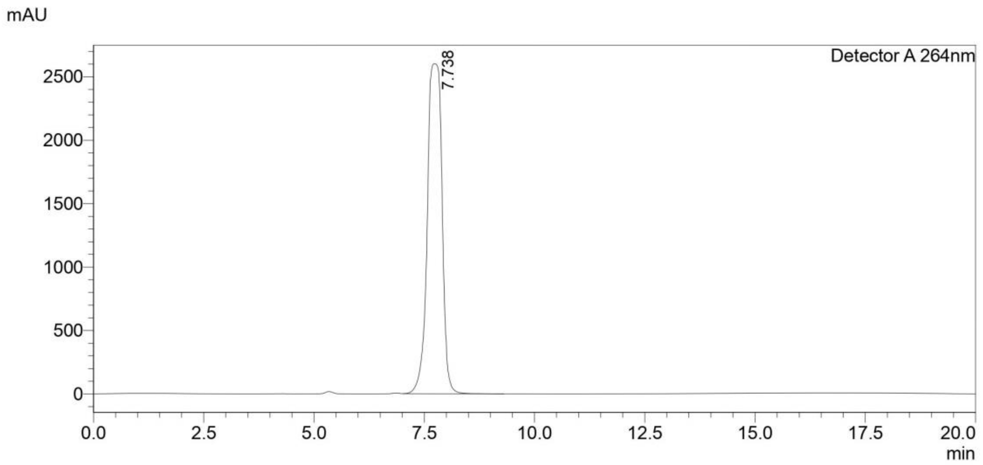


**Fig. S2A** HPLC chromatogram of karanjin purified from *P. pinnata*. The compound showed a retention time of 7.738 min and greater than 98% purity.

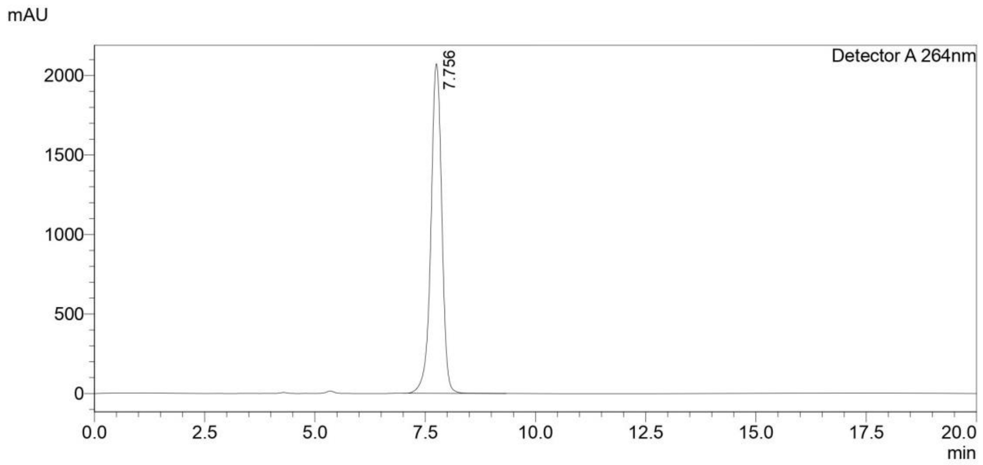


**Fig. S2B.** HPLC chromatogram of karanjin standard from Yucca Enterprises. The compound showed a retention time of 7.756 min, and greater than 98% purity.

### Nuclear magnetic resonance

One-dimension spectral analysis was carried out by dissolving 10 mg of isolated and standard karanjin from Yucca Enterprises in 600 μL $CDCl_3$. $^1$H NMR spectra at 600 MHz and $^{13}$C spectra at 150 MHz were acquired on a Bruker Ascend ™ 600 (Bruker BioSpin AG, Switzerland) with a central peak of the $CDCl_3$ triplet ($\delta$ 77.04 ppm) as the internal standard along with tetramethylsilane (TMS) as an internal reference. $^1$H and $^{13}$C spectral chemical shifts and coupling constants are expressed in $\delta$ and Hz, respectively. NMR was controlled by the software TopSpin 2.1. NMR data for karanjin isolated in-house and standard procured from Yucca Enterprises are shown in Fig. S3A, B, C and D.

A.

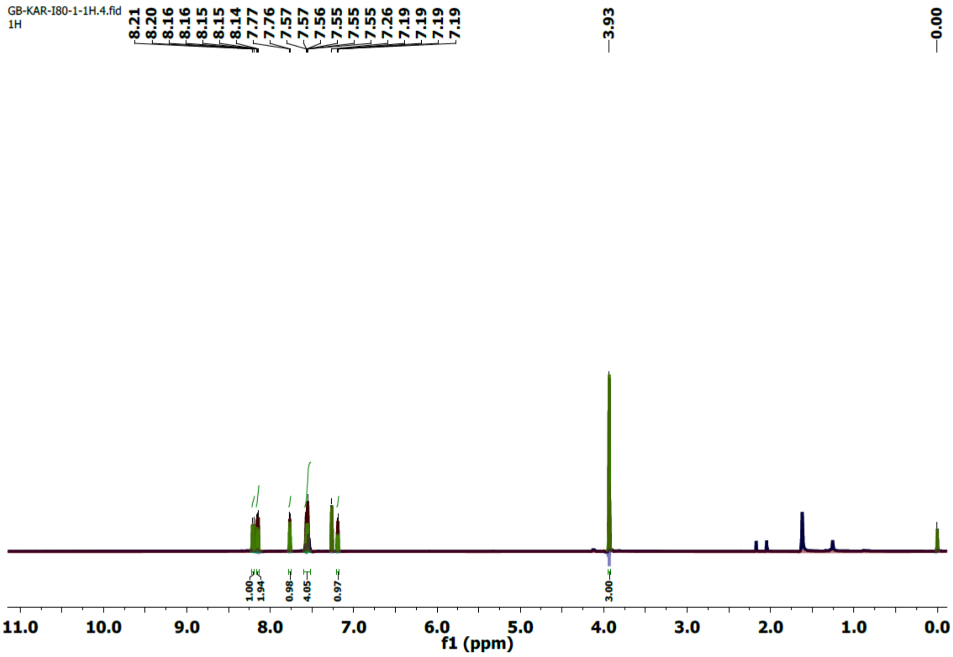
GB-KAR-I80-1-1H.4.fid
1H
8.21
8.20
8.16
8.16
8.15
8.15
8.14
7.77
7.76
7.57
7.57
7.56
7.55
7.55
7.55
7.26
7.19
7.19
7.19
7.19
3.93
0.00
1.00
1.94
0.98
4.05
0.97
3.00
11.0 10.0 9.0 8.0 7.0 6.0 5.0 4.0 3.0 2.0 1.0 0.0
f1 (ppm)

B.

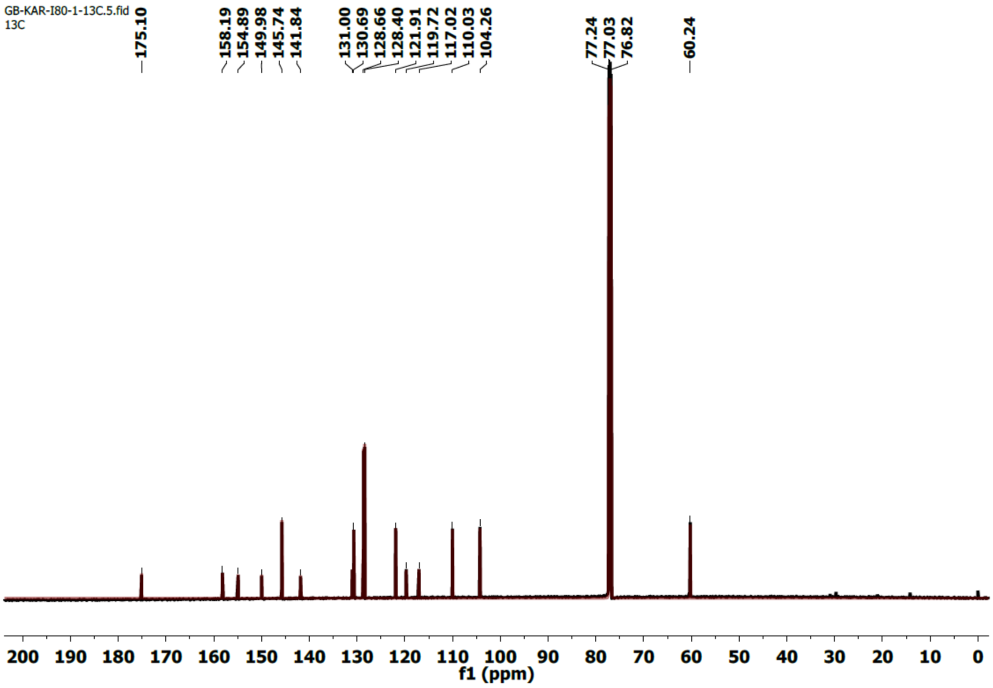
GB-KAR-I80-1-13C.5.fid
13C
175.10
158.19
154.89
149.98
145.74
141.84
131.00
130.69
128.66
128.40
121.91
119.72
117.02
110.03
104.26
77.24
77.03
76.82
60.24
200 190 180 170 160 150 140 130 120 110 100 90 80 70 60 50 40 30 20 10 0
f1 (ppm)

**C.**

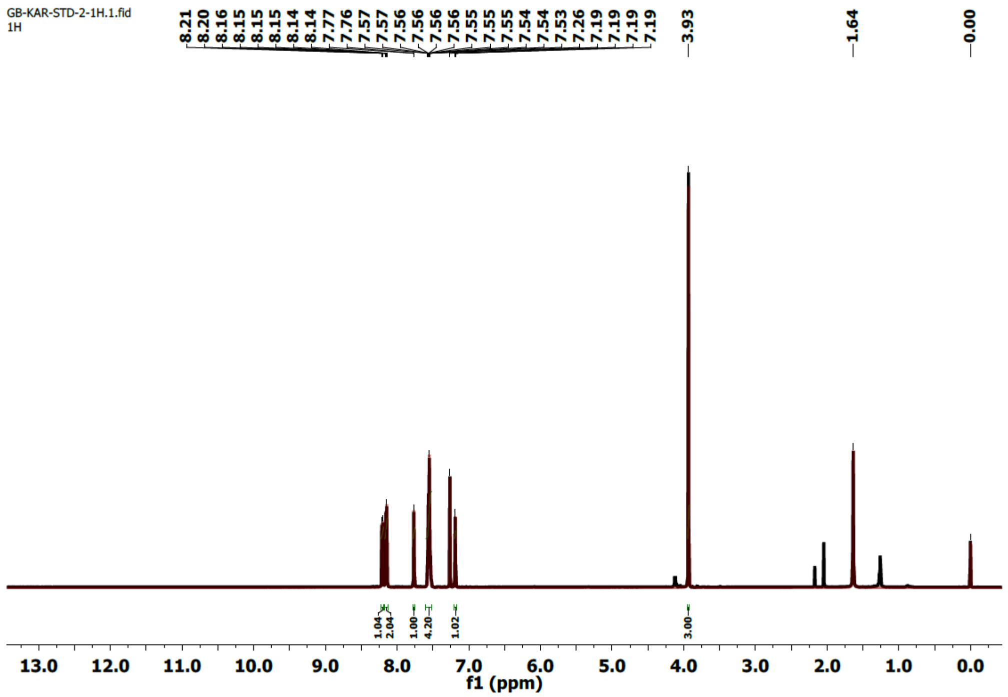


**D.**

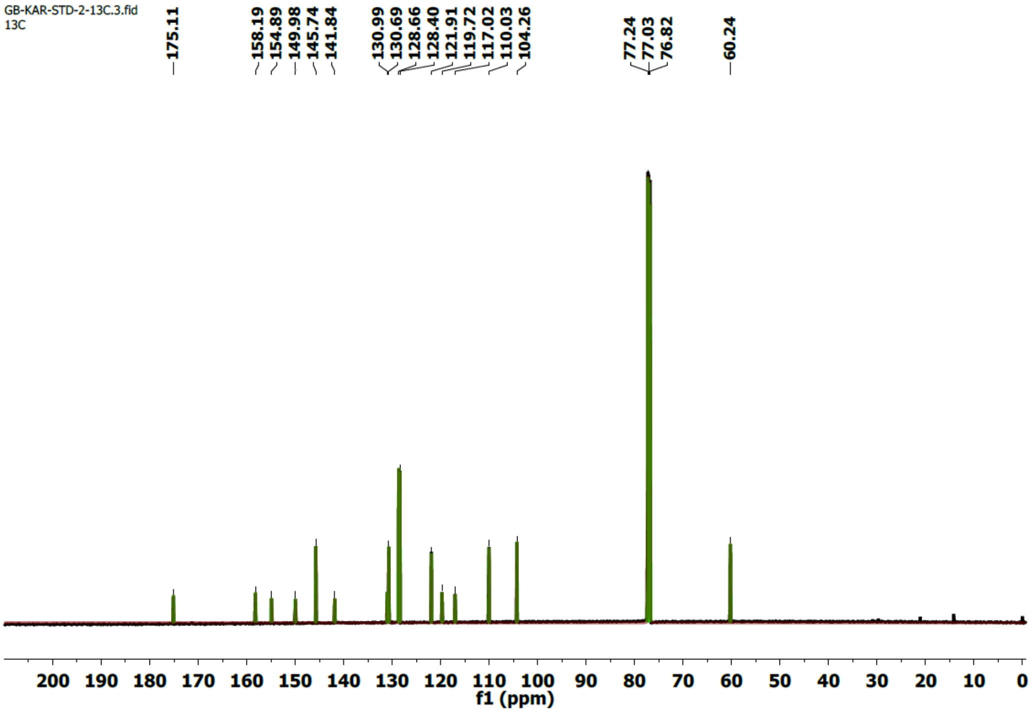


**Fig. S3.** $^{1}$H NMR (A, C), $^{13}$C NMR spectra (B, D) for karanjin isolated in-house (A, B) and standard purchased from Yucca Enterprises (C, D).
