## Supplementary material for "Global transcriptome analysis reveals partial estrogen-like effects of karanjin in MCF-7 breast cancer cells": All supplementary material: Supplementary data 2.pdf

**Supplementary data 2:** List of primers.

| Primer name | Sequence | Amplicon length (bp) | Annealing temperature (°C) |
|---|---|---|---|
| CSTA-F | ATCTGAGGCCAAACCCGCC | 275 | 60 |
| CSTA-R | AGCCCGTCAGCTCGTCATC | | |
| CycA-F | GGGCCGCGTCTCCTTTGAGC | 158 | 60 |
| CycA-R | GGCGTGTGAAGTCACCACCC | | |
| BRINP2-F | GACTGGCTGCTCACAGACC | 126 | 60 |
| BRINP2-R | CACCTTCCAACGGGCAAAC | | |
| B3GALT5-F | CCTCTTGGCATTTACACTGTGG | 115 | 60 |
| B3GALT5-R | CCCCCAGAACCAGAAGGC | | |
| TFF1-F | GGGTCCCCTGGTGCTTCTAT | 140 | 60 |
| TFF1-R | AGCCGAGCTCTGGGACTAA | | |
| CHST1-F | GCCCTTTCGACCTGGAGG | 134 | 60 |
| CHST1-R | CAAGGGGTGAGGTCAAAGAGG | | |
| SLC7A5-F | GTGGACTTCGGGAACTATCACC | 126 | 60 |
| SLC7A5-R | GGACCCCACGAAGAAGAGC | | |
| MRVI1-F | CACCGGGAAACCTACCAGAAG | 130 | 60 |
| MRVI1-R | CTTCCGTTGCTTTCGACATGC | | |
| CYP1B1-F | GCCACTATCACTGACATCTTCG | 129 | 60 |
| CYP1B1-R | CACGACCTGATCCAATTCTGC | | |
| CREG2-F | CAGATGATCGCAGTGTCTCCA | 153 | 60 |
| CREG2-R | GCCTCCATACCATTTCTGAAGC | | |
| TIPARP-F | TCATTGGCAGATCAAAAGGACAAC | 160 | 60 |
| TIPARP-R | CACGTTCATGGCATTCAAATCTGC | | |
| STC2-F | ATGCTACCTCAAGCACGACC | 129 | 60 |

| | | | |
|---|---|---|---|
| STC2-R | CAGGTCAGCAGCAAGTTCAC | | |
| CD44-F | CAAGTTTTGGTGGCACGCAG | 135 | 60 |
| CD44-R | GTCCGAGAGATGCTGTAGCG | | |
| CYP1A1-F | ACCTTTGAGAAGGGCCACATCCG | 154 | 60 |
| CYP1A1-R | TGACTGTGTCAAACCCAGCTCCAAAG | | |
| CDC25B- F | CTGTAGCCTGGACAAGAGAGTC | 112 | 60 |
| CDC25B- R | GTAGTCGTTGACAGCACGGT | | |
| CENPF-F | GGCTGCACAGAAGTTAGCG | 143 | 60 |
| CENPF-R | GGAGGATGGTGCCTGAATCTAC | | |
| PR-F | CGCGCTCTACCCTGCACTC | 121 | 60 |
| PR-R | TGAATCCGGCCTCAGGTAGTT | | |
| ADAMTS19-F | CGGAGTGTGGTCTGAGTGT | 111 | 60 |
| ADAMTS19-R | GCCTCCCTGGGTCTGGTAG | | |
| MYC-F | CCGTCCTCGGATTCTCTGCT | 231 | 60 |
| MYC-R | TGGGCTGTGAGGAGGTTTGC | | |

F and R indicate sense and antisense primers respectively.
