## Supplementary material for "Global transcriptome analysis reveals partial estrogen-like effects of karanjin in MCF-7 breast cancer cells": All supplementary material: Supplementary data 3.pdf

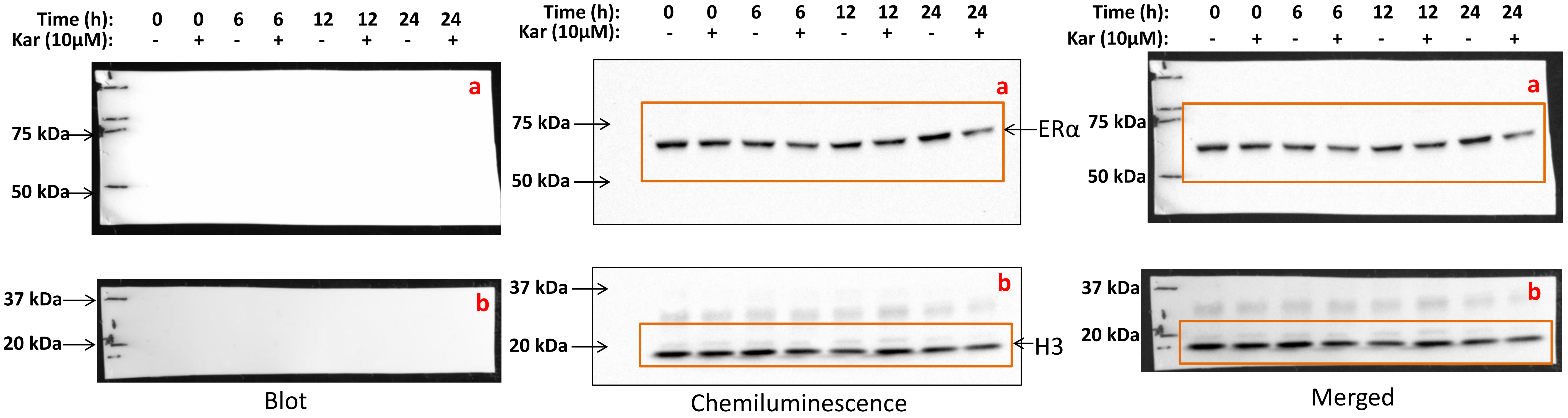


**Supplementary data 3A. Karanjin mediated reduction in ERα protein expression in MCF-7 cells.** Cells were treated with 10 µM karanjin (kar 10 µM) for the indicated periods of time. At each time point, the untreated cells (0 µM karanjin) served as control. Total protein extracted with Laemmli buffer was subjected to western blotting analysis using the ERα specific antibody. Post transfer, the blot was cut horizontally between the 50 kDa and 37 kDa marker to probe for ERα (blot a) followed by histone (H3) (blot b) as described. The specific band for ERα is indicated by the arrow. The portions of the Chemiluminescence image within the red colored rectangles were shown in Fig. 6A and comprise of data from 0 hour to 24 hours. Data shown here is 1 representative replicate , out of 5 replicates.

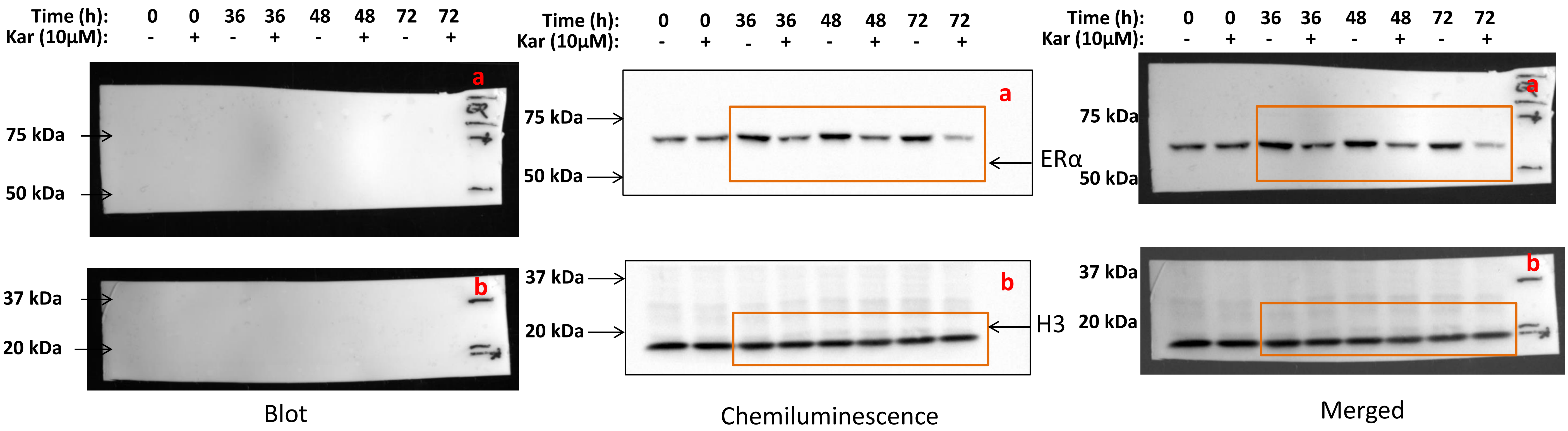


**Supplementary data 3B. Karanjin mediated reduction in ERα protein expression in MCF-7 cells.** Cells were treated with 10 µM karanjin (kar 10 µM) for the indicated periods of time. At each time point, the untreated cells (0 µM karanjin) served as control. Total protein extracted with Laemmli buffer was subjected to western blotting analysis using the ERα specific antibody. Post transfer, the blot was cut horizontally between the 50 kDa and 37 kDa marker to probe for ERα (blot a) followed by histone (H3) (blot b) as described. The specific band for ERα is indicated by the arrow. The portions of the chemiluminescence image within the red colored rectangles were shown in Fig. 6A and comprise of data from 36 hours to 72 hours. Data shown here is 1 representative replicate , out of 5 replicates.

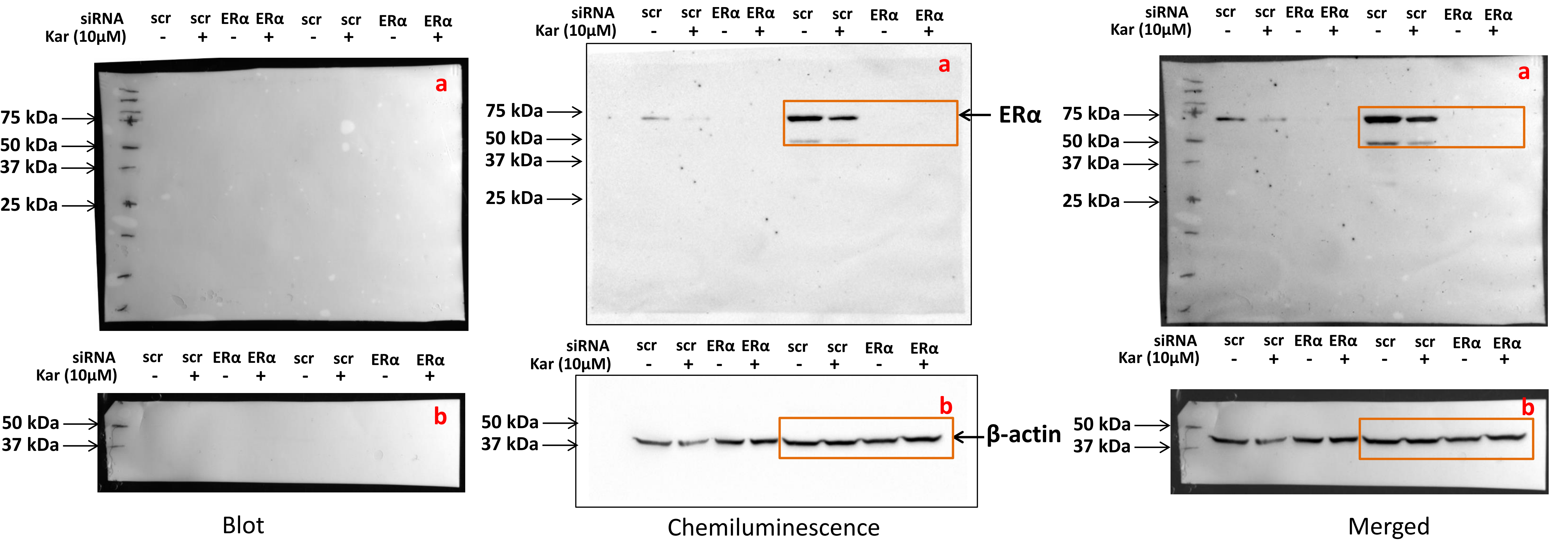


**Supplementary data 3C. Effect of ERα knockdown.** MCF-7 pre-treated with scrambled (scr) or ERα-specific siRNA were treated with vehicle or 10 µM karanjin (Kar 10µM) for a period of 24 h. Total protein, isolated from the phenolic fraction of RNA extraction reagent, was separated by 10 % SDS-PAGE and transferred to nitrocellulose membrane. The transferred, full blot was probed for ERα (blot a). The specific band for ERα is indicated by the arrow. To probe β-actin (blot b), the full blot was cut horizontally at two positions; between 50 kDa & 75 kDa, and between 25 kDa & 37 kDa markers. The portions of the chemiluminescence data within the red-colored rectangles were shown in Fig. 6B. Shown are the 2 representative replicates out of 3.
