## Supplementary material for "Global transcriptome analysis reveals partial estrogen-like effects of karanjin in MCF-7 breast cancer cells": All supplementary material: Supplementary data 4.pdf

Figure 1A

```
> Data
   Time Conc_K    Count
1     0      0 48750.00
2     0      0 45000.00
3     0      0 43750.00
4    24      0 73333.33
5    24      0 72500.00
6    24      0 67500.00
7     0      1 46250.00
8     0      1 50000.00
9     0      1 48750.00
10   24      1 75000.00
11   24      1 71666.67
12   24      1 73333.33
13    0     10 48750.00
14    0     10 47500.00
15    0     10 46250.00
16   24     10 74166.67
17   24     10 70833.33
18   24     10 71666.67
19    0   1000 43750.00
20    0   1000 47500.00
21    0   1000 43750.00
22   24   1000 69166.67
23   24   1000 70833.33
24   24   1000 70833.33
25    0  10000 45000.00
26    0  10000 47500.00
27    0  10000 46250.00
28   24  10000 69166.67
29   24  10000 69166.67
30   24  10000 76666.67
31    0  50000 45000.00
32    0  50000 48750.00
33    0  50000 46250.00
34   24  50000 68333.33
35   24  50000 73333.33
36   24  50000 72500.00
>
> Data$Conc_K <- as.factor(Data$Conc_K)
> Data$Time <- as.factor(Data$Time)
>
> anova2 <- aov(formula = Count ~ Time + Conc_K + Time:Conc_K, data = Data)
> summary(anova2)
            Df    Sum Sq   Mean Sq  F value Pr(>F)
Time         1 5.656e+09 5.656e+09 1052.863 <2e-16 ***
Conc_K       5 3.679e+07 7.358e+06    1.370  0.271
Time:Conc_K  5 6.800e+05 1.360e+05    0.025  1.000
Residuals   24 1.289e+08 5.372e+06
---
Signif. codes:  0 ‘***’ 0.001 ‘**’ 0.01 ‘*’ 0.05 ‘.’ 0.1 ‘ ’ 1
> TukeyHSD(anova2)
```

```
  Tukey multiple comparisons of means
    95% family-wise confidence level

Fit: aov(formula = Count ~ Time + Conc_K + Time:Conc_K, data = Data)

$Time
         diff      lwr      upr p adj
24-0 25069.44 23474.86 26664.03     0

$Conc_K
                  diff       lwr       upr     p adj
1-0          2361.11111 -1776.496 6498.7185 0.5056372
10-0         1388.88889 -2748.718 5526.4963 0.9003172
1000-0       -833.33333 -4970.941 3304.2740 0.9881995
10000-0       486.11111 -3651.496 4623.7185 0.9990592
50000-0       555.55555 -3582.052 4693.1629 0.9982138
10-1         -972.22222 -5109.830 3165.3852 0.9766531
1000-1      -3194.44445 -7332.052  943.1629 0.2003362
10000-1     -1875.00000 -6012.607 2262.6074 0.7259817
50000-1     -1805.55556 -5943.163 2332.0518 0.7553601
1000-10     -2222.22222 -6359.830 1915.3852 0.5690027
10000-10     -902.77778 -5040.385 3234.8296 0.9831283
50000-10     -833.33334 -4970.941 3304.2740 0.9881995
10000-1000   1319.44445 -2818.163 5457.0518 0.9179623
50000-1000   1388.88889 -2748.718 5526.4963 0.9003172
50000-10000    69.44444 -4068.163 4207.0518 0.9999999

$`Time:Conc_K`
                        diff        lwr        upr     p adj
24:0-0:0          25277.7778  18454.147  32101.409 0.0000000
0:1-0:0            2500.0000  -4323.631   9323.631 0.9677688
24:1-0:0          27500.0000  20676.369  34323.631 0.0000000
0:10-0:0           1666.6667  -5156.964   8490.298 0.9987303
24:10-0:0         26388.8889  19565.258  33212.520 0.0000000
0:1000-0:0         -833.3333  -7656.964   5990.298 0.9999986
24:1000-0:0       24444.4444  17620.813  31268.076 0.0000000
0:10000-0:0         416.6667  -6406.964   7240.298 1.0000000
24:10000-0:0      25833.3333  19009.702  32656.964 0.0000000
0:50000-0:0         833.3333  -5990.298   7656.964 0.9999986
24:50000-0:0      25555.5556  18731.924  32379.187 0.0000000
0:1-24:0         -22777.7778 -29601.409 -15954.147 0.0000000
24:1-24:0          2222.2222  -4601.409   9045.853 0.9862708
0:10-24:0        -23611.1111 -30434.742 -16787.480 0.0000000
24:10-24:0         1111.1111  -5712.520   7934.742 0.9999730
0:1000-24:0      -26111.1111 -32934.742 -19287.480 0.0000000
24:1000-24:0       -833.3333  -7656.964   5990.298 0.9999986
0:10000-24:0     -24861.1111 -31684.742 -18037.480 0.0000000
24:10000-24:0       555.5556  -6268.076   7379.187 1.0000000
0:50000-24:0     -24444.4444 -31268.076 -17620.813 0.0000000
24:50000-24:0       277.7778  -6545.853   7101.409 1.0000000
24:1-0:1          25000.0000  18176.369  31823.631 0.0000000
0:10-0:1           -833.3333  -7656.964   5990.298 0.9999986
24:10-0:1         23888.8889  17065.258  30712.520 0.0000000
0:1000-0:1        -3333.3333 -10156.964   3490.298 0.8225284
```

```
24:1000-0:1        21944.4444  15120.813  28768.076 0.0000000
0:10000-0:1        -2083.3333  -8906.964   4740.298 0.9916750
24:10000-0:1       23333.3333  16509.702  30156.964 0.0000000
0:50000-0:1        -1666.6667  -8490.298   5156.964 0.9987303
24:50000-0:1       23055.5556  16231.924  29879.187 0.0000000
0:10-24:1         -25833.3333 -32656.964 -19009.702 0.0000000
24:10-24:1         -1111.1111  -7934.742   5712.520 0.9999730
0:1000-24:1       -28333.3333 -35156.964 -21509.702 0.0000000
24:1000-24:1       -3055.5556  -9879.187   3768.076 0.8873175
0:10000-24:1      -27083.3333 -33906.964 -20259.702 0.0000000
24:10000-24:1      -1666.6667  -8490.298   5156.964 0.9987303
0:50000-24:1      -26666.6667 -33490.298 -19843.036 0.0000000
24:50000-24:1      -1944.4444  -8768.076   4879.187 0.9952347
24:10-0:10         24722.2222  17898.591  31545.853 0.0000000
0:1000-0:10        -2500.0000  -9323.631   4323.631 0.9677688
24:1000-0:10       22777.7778  15954.147  29601.409 0.0000000
0:10000-0:10       -1250.0000  -8073.631   5573.631 0.9999137
24:10000-0:10      24166.6667  17343.036  30990.298 0.0000000
0:50000-0:10        -833.3333  -7656.964   5990.298 0.9999986
24:50000-0:10      23888.8889  17065.258  30712.520 0.0000000
0:1000-24:10      -27222.2222 -34045.853 -20398.591 0.0000000
24:1000-24:10      -1944.4444  -8768.076   4879.187 0.9952347
0:10000-24:10     -25972.2222 -32795.853 -19148.591 0.0000000
24:10000-24:10      -555.5556  -7379.187   6268.076 1.0000000
0:50000-24:10     -25555.5556 -32379.187 -18731.924 0.0000000
24:50000-24:10      -833.3333  -7656.964   5990.298 0.9999986
24:1000-0:1000     25277.7778  18454.147  32101.409 0.0000000
0:10000-0:1000      1250.0000  -5573.631   8073.631 0.9999137
24:10000-0:1000    26666.6667  19843.036  33490.298 0.0000000
0:50000-0:1000      1666.6667  -5156.964   8490.298 0.9987303
24:50000-0:1000    26388.8889  19565.258  33212.520 0.0000000
0:10000-24:1000   -24027.7778 -30851.409 -17204.147 0.0000000
24:10000-24:1000    1388.8889  -5434.742   8212.520 0.9997625
0:50000-24:1000   -23611.1111 -30434.742 -16787.480 0.0000000
24:50000-24:1000    1111.1111  -5712.520   7934.742 0.9999730
24:10000-0:10000   25416.6667  18593.036  32240.298 0.0000000
0:50000-0:10000      416.6667  -6406.964   7240.298 1.0000000
24:50000-0:10000   25138.8889  18315.258  31962.520 0.0000000
0:50000-24:10000  -25000.0000 -31823.631 -18176.369 0.0000000
24:50000-24:10000   -277.7778  -7101.409   6545.853 1.0000000
24:50000-0:50000   24722.2222  17898.591  31545.853 0.0000000

>
```

Figure 1B

```
> Data
   Time Conc_K    Count
1     0      0  67500.0
2     0      0  46250.0
3     0      0  50000.0
4   120      0 516666.7
5   120      0 508333.3
6   120      0 483333.3
```

```
7     0      1  50000.0
8     0      1  66250.0
9     0      1  55000.0
10  120      1 516666.7
11  120      1 500000.0
12  120      1 541666.7
13    0     10  67500.0
14    0     10  56250.0
15    0     10  55000.0
16  120     10 583333.3
17  120     10 533333.3
18  120     10 575000.0
19    0   1000  61250.0
20    0   1000  52500.0
21    0   1000  47500.0
22  120   1000 500000.0
23  120   1000 516666.7
24  120   1000 491666.7
25    0  10000  63750.0
26    0  10000  52500.0
27    0  10000  61250.0
28  120  10000 327916.7
29  120  10000 355833.3
30  120  10000 363750.0
31    0  50000  63750.0
32    0  50000  55000.0
33    0  50000  56250.0
34  120  50000 225000.0
35  120  50000 200000.0
36  120  50000 241666.7
>
> Data$Conc_K <- as.factor(Data$Conc_K)
> Data$Time <- as.factor(Data$Time)
>
> anova2 <- aov(formula = Count ~ Time + Conc_K + Time:Conc_K, data = Data)
> summary(anova2)
            Df    Sum Sq   Mean Sq F value   Pr(>F)
Time         1 1.343e+12 1.343e+12  5827.5  < 2e-16 ***
Conc_K       5 1.262e+11 2.524e+10   109.5 1.02e-15 ***
Time:Conc_K  5 1.293e+11 2.587e+10   112.2 7.76e-16 ***
Residuals   24 5.531e+09 2.305e+08
---
Signif. codes:  0 ‘***’ 0.001 ‘**’ 0.01 ‘*’ 0.05 ‘.’ 0.1 ‘ ’ 1
> TukeyHSD(anova2)
  Tukey multiple comparisons of means
    95% family-wise confidence level

Fit: aov(formula = Count ~ Time + Conc_K + Time:Conc_K, data = Data)

$Time
          diff      lwr      upr p adj
120-0 386296.3 375852.3 396740.3     0

$Conc_K
```

| | diff | lwr | upr | p adj |
|---|---|---|---|---|
| 1-0 | 9583.3334 | -17516.752 | 36683.418 | 0.8792133 |
| 10-0 | 33055.5556 | 5955.470 | 60155.641 | 0.0106904 |
| 1000-0 | -416.6667 | -27516.752 | 26683.418 | 1.0000000 |
| 10000-0 | -74513.8889 | -101613.974 | -47413.804 | 0.0000001 |
| 50000-0 | -138402.7778 | -165502.863 | -111302.693 | 0.0000000 |
| 10-1 | 23472.2222 | -3627.863 | 50572.307 | 0.1169885 |
| 1000-1 | -10000.0000 | -37100.085 | 17100.085 | 0.8593603 |
| 10000-1 | -84097.2222 | -111197.307 | -56997.137 | 0.0000000 |
| 50000-1 | -147986.1111 | -175086.196 | -120886.026 | 0.0000000 |
| 1000-10 | -33472.2222 | -60572.307 | -6372.137 | 0.0095518 |
| 10000-10 | -107569.4444 | -134669.529 | -80469.359 | 0.0000000 |
| 50000-10 | -171458.3333 | -198558.418 | -144358.248 | 0.0000000 |
| 10000-1000 | -74097.2222 | -101197.307 | -46997.137 | 0.0000002 |
| 50000-1000 | -137986.1111 | -165086.196 | -110886.026 | 0.0000000 |
| 50000-10000 | -63888.8889 | -90988.974 | -36788.804 | 0.0000022 |

$`Time:Conc_K`

| | diff | lwr | upr | p adj |
|---|---|---|---|---|
| 120:0-0:0 | 4.481944e+05 | 403501.7109 | 492887.18 | 0.0000000 |
| 0:1-0:0 | 2.500000e+03 | -42192.7335 | 47192.73 | 1.0000000 |
| 120:1-0:0 | 4.648611e+05 | 420168.3776 | 509553.84 | 0.0000000 |
| 0:10-0:0 | 5.000000e+03 | -39692.7335 | 49692.73 | 0.9999994 |
| 120:10-0:0 | 5.093056e+05 | 464612.8220 | 553998.29 | 0.0000000 |
| 0:1000-0:0 | -8.333333e+02 | -45526.0669 | 43859.40 | 1.0000000 |
| 120:1000-0:0 | 4.481944e+05 | 403501.7109 | 492887.18 | 0.0000000 |
| 0:10000-0:0 | 4.583333e+03 | -40109.4002 | 49276.07 | 0.9999998 |
| 120:10000-0:0 | 2.945833e+05 | 249890.5998 | 339276.07 | 0.0000000 |
| 0:50000-0:0 | 3.750000e+03 | -40942.7335 | 48442.73 | 1.0000000 |
| 120:50000-0:0 | 1.676389e+05 | 122946.1554 | 212331.62 | 0.0000000 |
| 0:1-120:0 | -4.456944e+05 | -490387.1780 | -401001.71 | 0.0000000 |
| 120:1-120:0 | 1.666667e+04 | -28026.0668 | 61359.40 | 0.9636195 |
| 0:10-120:0 | -4.431944e+05 | -487887.1780 | -398501.71 | 0.0000000 |
| 120:10-120:0 | 6.111111e+04 | 16418.3776 | 105803.84 | 0.0023233 |
| 0:1000-120:0 | -4.490278e+05 | -493720.5113 | -404335.04 | 0.0000000 |
| 120:1000-120:0 | 3.333343e-05 | -44692.7335 | 44692.73 | 1.0000000 |
| 0:10000-120:0 | -4.436111e+05 | -488303.8446 | -398918.38 | 0.0000000 |
| 120:10000-120:0 | -1.536111e+05 | -198303.8446 | -108918.38 | 0.0000000 |
| 0:50000-120:0 | -4.444444e+05 | -489137.1780 | -399751.71 | 0.0000000 |
| 120:50000-120:0 | -2.805556e+05 | -325248.2891 | -235862.82 | 0.0000000 |
| 120:1-0:1 | 4.623611e+05 | 417668.3776 | 507053.84 | 0.0000000 |
| 0:10-0:1 | 2.500000e+03 | -42192.7335 | 47192.73 | 1.0000000 |
| 120:10-0:1 | 5.068056e+05 | 462112.8220 | 551498.29 | 0.0000000 |
| 0:1000-0:1 | -3.333333e+03 | -48026.0669 | 41359.40 | 1.0000000 |
| 120:1000-0:1 | 4.456944e+05 | 401001.7109 | 490387.18 | 0.0000000 |
| 0:10000-0:1 | 2.083333e+03 | -42609.4002 | 46776.07 | 1.0000000 |
| 120:10000-0:1 | 2.920833e+05 | 247390.5998 | 336776.07 | 0.0000000 |
| 0:50000-0:1 | 1.250000e+03 | -43442.7335 | 45942.73 | 1.0000000 |
| 120:50000-0:1 | 1.651389e+05 | 120446.1554 | 209831.62 | 0.0000000 |
| 0:10-120:1 | -4.598611e+05 | -504553.8447 | -415168.38 | 0.0000000 |
| 120:10-120:1 | 4.444444e+04 | -248.2891 | 89137.18 | 0.0522004 |
| 0:1000-120:1 | -4.656944e+05 | -510387.1780 | -421001.71 | 0.0000000 |
| 120:1000-120:1 | -1.666667e+04 | -61359.4002 | 28026.07 | 0.9636195 |
| 0:10000-120:1 | -4.602778e+05 | -504970.5113 | -415585.04 | 0.0000000 |

```
120:10000-120:1     -1.702778e+05 -214970.5113 -125585.04 0.0000000
0:50000-120:1       -4.611111e+05 -505803.8447 -416418.38 0.0000000
120:50000-120:1     -2.972222e+05 -341914.9558 -252529.49 0.0000000
120:10-0:10          5.043056e+05  459612.8220  548998.29 0.0000000
0:1000-0:10         -5.833333e+03  -50526.0669   38859.40 0.9999972
120:1000-0:10        4.431944e+05  398501.7109  487887.18 0.0000000
0:10000-0:10        -4.166667e+02  -45109.4002   44276.07 1.0000000
120:10000-0:10       2.895833e+05  244890.5998  334276.07 0.0000000
0:50000-0:10        -1.250000e+03  -45942.7335   43442.73 1.0000000
120:50000-0:10       1.626389e+05  117946.1554  207331.62 0.0000000
0:1000-120:10       -5.101389e+05 -554831.6224 -465446.16 0.0000000
120:1000-120:10     -6.111111e+04 -105803.8446  -16418.38 0.0023233
0:10000-120:10      -5.047222e+05 -549414.9557 -460029.49 0.0000000
120:10000-120:10    -2.147222e+05 -259414.9557 -170029.49 0.0000000
0:50000-120:10      -5.055556e+05 -550248.2891 -460862.82 0.0000000
120:50000-120:10    -3.416667e+05 -386359.4002 -296973.93 0.0000000
120:1000-0:1000      4.490278e+05  404335.0443  493720.51 0.0000000
0:10000-0:1000       5.416667e+03  -39276.0669   50109.40 0.9999987
120:10000-0:1000     2.954167e+05  250723.9331  340109.40 0.0000000
0:50000-0:1000       4.583333e+03  -40109.4002   49276.07 0.9999998
120:50000-0:1000     1.684722e+05  123779.4887  213164.96 0.0000000
0:10000-120:1000    -4.436111e+05 -488303.8447 -398918.38 0.0000000
120:10000-120:1000  -1.536111e+05 -198303.8447 -108918.38 0.0000000
0:50000-120:1000    -4.444444e+05 -489137.1780 -399751.71 0.0000000
120:50000-120:1000  -2.805556e+05 -325248.2891 -235862.82 0.0000000
120:10000-0:10000    2.900000e+05  245307.2665  334692.73 0.0000000
0:50000-0:10000     -8.333333e+02  -45526.0669   43859.40 1.0000000
120:50000-0:10000    1.630556e+05  118362.8220  207748.29 0.0000000
0:50000-120:10000   -2.908333e+05 -335526.0669 -246140.60 0.0000000
120:50000-120:10000 -1.269444e+05 -171637.1780  -82251.71 0.0000000
120:50000-0:50000    1.638889e+05  119196.1554  208581.62 0.0000000

>
```

Figure 2C

B3GALT5

```
> Data <- read.table(file.choose(), header=T, sep="\t")
> Data
    Control  Karanjin
1 1.0000000 0.3815648
2 0.9726549 0.4976948
3 1.3628877 0.3253355
> summary(Data)
    Control          Karanjin
 Min.   :0.9727   Min.   :0.3253
 1st Qu.:0.9863   1st Qu.:0.3535
 Median :1.0000   Median :0.3816
 Mean   :1.1118   Mean   :0.4015
 3rd Qu.:1.1814   3rd Qu.:0.4396
 Max.   :1.3629   Max.   :0.4977
> t.test(Data$Control, Data$Karanjin, alternative="greater", var.equal=FALSE)
```

```
        Welch Two Sample t-test

data:  Data$Control and Data$Karanjin
t = 5.2375, df = 2.6344, p-value = 0.009303
alternative hypothesis: true difference in means is greater than 0
95 percent confidence interval:
 0.3722609       Inf
sample estimates:
mean of x mean of y
1.1118475 0.4015317

>
```

BRINP2

```
> Data <- read.table(file.choose(), header=T, sep="\t")
> Data
    Control  Karanjin
1 1.0000000 0.5743492
2 1.3597424 0.5743492
3 1.1486984 0.4741230
4 0.9637071 0.6070974
> summary(Data)
    Control          Karanjin
 Min.   :0.9637   Min.   :0.4741
 1st Qu.:0.9909   1st Qu.:0.5493
 Median :1.0743   Median :0.5743
 Mean   :1.1180   Mean   :0.5575
 3rd Qu.:1.2015   3rd Qu.:0.5825
 Max.   :1.3597   Max.   :0.6071
> t.test(Data$Control, Data$Karanjin, alternative="greater", var.equal=FALSE)

        Welch Two Sample t-test

data:  Data$Control and Data$Karanjin
t = 5.9338, df = 3.6101, p-value = 0.002779
alternative hypothesis: true difference in means is greater than 0
95 percent confidence interval:
 0.3527505       Inf
sample estimates:
mean of x mean of y
1.1180370 0.5574797

>
```

CHST1

```
> Data <- read.table(file.choose(), header=T, sep="\t")
> Data
    Control  Karanjin
1 1.0000000 0.3643339
2 0.8274056 0.5152460
3 1.0545786 0.3745768
> summary(Data)
```

```
    Control          Karanjin
 Min.   :0.8274   Min.   :0.3643
 1st Qu.:0.9137   1st Qu.:0.3695
 Median :1.0000   Median :0.3746
 Mean   :0.9607   Mean   :0.4181
 3rd Qu.:1.0273   3rd Qu.:0.4449
 Max.   :1.0546   Max.   :0.5152
> t.test(Data$Control, Data$Karanjin, alternative="greater", var.equal=FALSE)

        Welch Two Sample t-test

data:  Data$Control and Data$Karanjin
t = 6.4588, df = 3.6108, p-value = 0.00209
alternative hypothesis: true difference in means is greater than 0
95 percent confidence interval:
 0.3578169       Inf
sample estimates:
mean of x mean of y
0.9606614 0.4180522

>
```

CSTA

```
> Data <- read.table(file.choose(), header=T, sep="\t")
> Data
    Control  Karanjin
1 1.0000000 0.3719894
2 1.2085971 0.3601492
3 0.9244497 0.2327199
> summary(Data)
    Control          Karanjin
 Min.   :0.9244   Min.   :0.2327
 1st Qu.:0.9622   1st Qu.:0.2964
 Median :1.0000   Median :0.3601
 Mean   :1.0443   Mean   :0.3216
 3rd Qu.:1.1043   3rd Qu.:0.3661
 Max.   :1.2086   Max.   :0.3720
> t.test(Data$Control, Data$Karanjin, alternative="greater", var.equal=FALSE)

        Welch Two Sample t-test

data:  Data$Control and Data$Karanjin
t = 7.5319, df = 3.0235, p-value = 0.002361
alternative hypothesis: true difference in means is greater than 0
95 percent confidence interval:
 0.4976255       Inf
sample estimates:
mean of x mean of y
1.0443489 0.3216195

>
```

CYP1A1

```
> Data <- read.table(file.choose(), header=T, sep="\t")
> Data
    Control Karanjin
1 1.0000000 48.84029
2 0.6070974 61.10986
3 1.3692001 56.36262
> summary(Data)
    Control          Karanjin    
 Min.   :0.6071   Min.   :48.84  
 1st Qu.:0.8035   1st Qu.:52.60  
 Median :1.0000   Median :56.36  
 Mean   :0.9921   Mean   :55.44  
 3rd Qu.:1.1846   3rd Qu.:58.74  
 Max.   :1.3692   Max.   :61.11  
> t.test(Data$Karanjin, Data$Control, alternative="greater", var.equal=FALSE)

        Welch Two Sample t-test

data:  Data$Karanjin and Data$Control
t = 15.214, df = 2.0152, p-value = 0.00208
alternative hypothesis: true difference in means is greater than 0
95 percent confidence interval:
 44.04904      Inf
sample estimates:
 mean of x  mean of y 
55.4375911  0.9920992 

>
```

CYP1B1

```
> Data <- read.table(file.choose(), header=T, sep="\t")
> Data
   Control Karanjin
1 1.000000 32.44670
2 2.614738 25.28132
3 4.131503 29.65082
4 1.972465 32.07402
> summary(Data)
    Control         Karanjin    
 Min.   :1.000   Min.   :25.28  
 1st Qu.:1.729   1st Qu.:28.56  
 Median :2.294   Median :30.86  
 Mean   :2.430   Mean   :29.86  
 3rd Qu.:2.994   3rd Qu.:32.17  
 Max.   :4.132   Max.   :32.45  
> t.test(Data$Karanjin, Data$Control, alternative="greater", var.equal=FALSE)

        Welch Two Sample t-test

data:  Data$Karanjin and Data$Control
t = 15.46, df = 3.9305, p-value = 5.739e-05
alternative hypothesis: true difference in means is greater than 0
```

```
95 percent confidence interval:
 23.63136      Inf
sample estimates:
mean of x mean of y
29.863216  2.429677

>
```

MRVI1

```
> Data <- read.table(file.choose(), header=T, sep="\t")
> Data
    Control Karanjin
1 1.0000000 12.99604
2 1.3883135 14.72300
3 1.8192368 18.63574
4 0.6227248 16.60255
> summary(Data)
    Control          Karanjin
 Min.   :0.6227   Min.   :13.00
 1st Qu.:0.9057   1st Qu.:14.29
 Median :1.1942   Median :15.66
 Mean   :1.2076   Mean   :15.74
 3rd Qu.:1.4960   3rd Qu.:17.11
 Max.   :1.8192   Max.   :18.64
> t.test(Data$Karanjin, Data$Control, alternative="greater", var.equal=FALSE)

        Welch Two Sample t-test

data:  Data$Karanjin and Data$Control
t = 11.708, df = 3.268, p-value = 0.0004425
alternative hypothesis: true difference in means is greater than 0
95 percent confidence interval:
 11.70605      Inf
sample estimates:
mean of x mean of y
15.739333  1.207569

>
```

CREG2

```
> Data <- read.table(file.choose(), header=T, sep="\t")
> Data
    Control Karanjin
1 1.0000000 4.084049
2 0.8408964 4.160240
3 0.8160145 4.448544
4 0.5547847 4.267329
> summary(Data)
    Control          Karanjin
 Min.   :0.5548   Min.   :4.084
 1st Qu.:0.7507   1st Qu.:4.141
 Median :0.8285   Median :4.214
```

```
 Mean   :0.8029   Mean   :4.240  
 3rd Qu.:0.8807   3rd Qu.:4.313  
 Max.   :1.0000   Max.   :4.449  
> t.test(Data$Karanjin, Data$Control, alternative="greater", var.equal=FALSE)

        Welch Two Sample t-test

data:  Data$Karanjin and Data$Control
t = 28.305, df = 5.8624, p-value = 8.517e-08
alternative hypothesis: true difference in means is greater than 0
95 percent confidence interval:
 3.200159      Inf
sample estimates:
mean of x mean of y 
4.2400404 0.8029239 

>
```

Figure 3F

MYC

```
> Data <- read.table(file.choose(), header=T, sep="\t")
> Data
   Control Karanjin
1 1.000000 1.765406
2 1.086735 1.650992
3 1.200249 1.749165
> summary(Data)
    Control         Karanjin    
 Min.   :1.000   Min.   :1.651  
 1st Qu.:1.043   1st Qu.:1.700  
 Median :1.087   Median :1.749  
 Mean   :1.096   Mean   :1.722  
 3rd Qu.:1.143   3rd Qu.:1.757  
 Max.   :1.200   Max.   :1.765  
> t.test(Data$Karanjin, Data$Control, alternative="greater", var.equal=FALSE)

        Welch Two Sample t-test

data:  Data$Karanjin and Data$Control
t = 9.1939, df = 3.3282, p-value = 0.0008856
alternative hypothesis: true difference in means is greater than 0
95 percent confidence interval:
 0.4721496       Inf
sample estimates:
mean of x mean of y 
 1.721855  1.095661 

>
```

CDC25B

```
> Data <- read.table(file.choose(), header=T, sep="\t")
```

```
> Data
   Control Karanjin
1 1.000000 1.242575
2 1.274561 1.417485
3 1.079228 1.394744
> summary(Data)
    Control         Karanjin    
 Min.   :1.000   Min.   :1.243  
 1st Qu.:1.040   1st Qu.:1.319  
 Median :1.079   Median :1.395  
 Mean   :1.118   Mean   :1.352  
 3rd Qu.:1.177   3rd Qu.:1.406  
 Max.   :1.275   Max.   :1.417  
> t.test(Data$Karanjin, Data$Control, alternative="greater", var.equal=FALSE)

        Welch Two Sample t-test

data:  Data$Karanjin and Data$Control
t = 2.3761, df = 3.5033, p-value = 0.04275
alternative hypothesis: true difference in means is greater than 0
95 percent confidence interval:
 0.01518186        Inf
sample estimates:
mean of x mean of y 
 1.351601  1.117930 

>
```

CENPF

```
> Data <- read.table(file.choose(), header=T, sep="\t")
> Data
   Control Karanjin
1 1.000000 1.372367
2 1.319508 1.918528
3 1.385109 1.827663
> summary(Data)
    Control         Karanjin    
 Min.   :1.000   Min.   :1.372  
 1st Qu.:1.160   1st Qu.:1.600  
 Median :1.320   Median :1.828  
 Mean   :1.235   Mean   :1.706  
 3rd Qu.:1.352   3rd Qu.:1.873  
 Max.   :1.385   Max.   :1.919  
> t.test(Data$Karanjin, Data$Control, alternative="greater", var.equal=FALSE)

        Welch Two Sample t-test

data:  Data$Karanjin and Data$Control
t = 2.2809, df = 3.5916, p-value = 0.04613
alternative hypothesis: true difference in means is greater than 0
95 percent confidence interval:
 0.01601452        Inf
sample estimates:
```

```
mean of x mean of y
 1.706186  1.234872

>
```

TFF1

```
> Data <- read.table(file.choose(), header=T, sep="\t")
> Data
    Control Karanjin
1 1.0000000 1.359742
2 0.9930925 1.245450
3 0.7465617 1.175548
4 1.1620456 1.236847
> summary(Data)
    Control          Karanjin
 Min.   :0.7466   Min.   :1.176
 1st Qu.:0.9315   1st Qu.:1.222
 Median :0.9965   Median :1.241
 Mean   :0.9754   Mean   :1.254
 3rd Qu.:1.0405   3rd Qu.:1.274
 Max.   :1.1620   Max.   :1.360
> t.test(Data$Karanjin, Data$Control, alternative="greater", var.equal=FALSE)

        Welch Two Sample t-test

data:  Data$Karanjin and Data$Control
t = 2.9707, df = 4.1586, p-value = 0.01958
alternative hypothesis: true difference in means is greater than 0
95 percent confidence interval:
 0.08094774        Inf
sample estimates:
mean of x mean of y
1.2543966 0.9754249

>
```

CD44

```
> Data <- read.table(file.choose(), header=T, sep="\t")
> Data
   Control Karanjin
1 1.000000 1.892115
2 1.054579 2.214018
3 1.076738 2.027919
> summary(Data)
    Control         Karanjin
 Min.   :1.000   Min.   :1.892
 1st Qu.:1.027   1st Qu.:1.960
 Median :1.055   Median :2.028
 Mean   :1.044   Mean   :2.045
 3rd Qu.:1.066   3rd Qu.:2.121
 Max.   :1.077   Max.   :2.214
> t.test(Data$Karanjin, Data$Control, alternative="greater", var.equal=FALSE)
```

```
        Welch Two Sample t-test

data:  Data$Karanjin and Data$Control
t = 10.421, df = 2.238, p-value = 0.003052
alternative hypothesis: true difference in means is greater than 0
95 percent confidence interval:
 0.7397735       Inf
sample estimates:
mean of x mean of y
 2.044684  1.043772

>
```

STC2

```
> Data <- read.table(file.choose(), header=T, sep="\t")
> Data
   Control Karanjin
1 1.000000 1.609560
2 1.069300 2.255322
3 1.178267 2.094588
> summary(Data)
    Control         Karanjin
 Min.   :1.000   Min.   :1.610
 1st Qu.:1.035   1st Qu.:1.852
 Median :1.069   Median :2.095
 Mean   :1.083   Mean   :1.986
 3rd Qu.:1.124   3rd Qu.:2.175
 Max.   :1.178   Max.   :2.255
> t.test(Data$Karanjin, Data$Control, alternative="greater", var.equal=FALSE)

        Welch Two Sample t-test

data:  Data$Karanjin and Data$Control
t = 4.4994, df = 2.2844, p-value = 0.01787
alternative hypothesis: true difference in means is greater than 0
95 percent confidence interval:
 0.3642131       Inf
sample estimates:
mean of x mean of y
 1.986490  1.082522

>
```

TIPARP

```
> Data <- read.table(file.choose(), header=T, sep="\t")
> Data
   Control Karanjin
1 1.000000 3.073750
2 0.941696 3.087987
3 1.000000 3.256525
> summary(Data)
```

```
    Control          Karanjin
 Min.   :0.9417   Min.   :3.074
 1st Qu.:0.9708   1st Qu.:3.081
 Median :1.0000   Median :3.088
 Mean   :0.9806   Mean   :3.139
 3rd Qu.:1.0000   3rd Qu.:3.172
 Max.   :1.0000   Max.   :3.257
> t.test(Data$Karanjin, Data$Control, alternative="greater", var.equal=FALSE)

        Welch Two Sample t-test

data:  Data$Karanjin and Data$Control
t = 34.916, df = 2.4333, p-value = 0.0001199
alternative hypothesis: true difference in means is greater than 0
95 percent confidence interval:
 1.998436      Inf
sample estimates:
mean of x mean of y
3.1394208 0.9805653

>
```

SLC7A5

```
> Data <- read.table(file.choose(), header=T, sep="\t")
> Data
    Control  Karanjin
1 1.0000000  8.378353
2 1.4674724  9.382680
3 0.9012505 11.498178
> summary(Data)
    Control          Karanjin
 Min.   :0.9013   Min.   : 8.378
 1st Qu.:0.9506   1st Qu.: 8.881
 Median :1.0000   Median : 9.383
 Mean   :1.1229   Mean   : 9.753
 3rd Qu.:1.2337   3rd Qu.:10.440
 Max.   :1.4675   Max.   :11.498
> t.test(Data$Karanjin, Data$Control, alternative="greater", var.equal=FALSE)

        Welch Two Sample t-test

data:  Data$Karanjin and Data$Control
t = 9.2213, df = 2.1441, p-value = 0.004612
alternative hypothesis: true difference in means is greater than 0
95 percent confidence interval:
 6.018581      Inf
sample estimates:
mean of x mean of y
 9.753070  1.122908

>
```

Figure 5 (MCF-7)

TFF1

```
> Data <- read.table(file.choose(), header=TRUE, sep="\t")
> Data
         C      K10   K10000   K50000       E2
1 1.000000 1.325619 0.952638 3.363586 7.110741
2 1.002313 1.099362 1.151355 3.538979 6.147501
3 0.856584 1.057018 1.512219 3.363586 6.408559
> boxplot(Data)
> summary(Data)
       C                K10             K10000           K50000            E2

 Min.   :0.8566   Min.   :1.057   Min.   :0.9526   Min.   :3.364   Min.   :6.148

 1st Qu.:0.9283   1st Qu.:1.078   1st Qu.:1.0520   1st Qu.:3.364   1st Qu.:6.278

 Median :1.0000   Median :1.099   Median :1.1514   Median :3.364   Median :6.409

 Mean   :0.9530   Mean   :1.161   Mean   :1.2054   Mean   :3.422   Mean   :6.556

 3rd Qu.:1.0012   3rd Qu.:1.212   3rd Qu.:1.3318   3rd Qu.:3.451   3rd Qu.:6.760

 Max.   :1.0023   Max.   :1.326   Max.   :1.5122   Max.   :3.539   Max.   :7.111

> Stacked_Data <- stack(Data)
> Stacked_Data
     values    ind
1  1.000000      C
2  1.002313      C
3  0.856584      C
4  1.325619    K10
5  1.099362    K10
6  1.057018    K10
7  0.952638 K10000
8  1.151355 K10000
9  1.512219 K10000
10 3.363586 K50000
11 3.538979 K50000
12 3.363586 K50000
13 7.110741     E2
14 6.147501     E2
15 6.408559     E2
> Anova_Result <- aov(values~ind, data=Stacked_Data)
> summary(Anova_Result)
            Df Sum Sq Mean Sq F value  Pr(>F)
ind          4  69.10  17.276   235.5 7.6e-10 ***
Residuals   10   0.73   0.073
---
Signif. codes:  0 ‘***’ 0.001 ‘**’ 0.01 ‘*’ 0.05 ‘.’ 0.1 ‘ ’ 1
> TukeyHSD(Anova_Result)
  Tukey multiple comparisons of means
    95% family-wise confidence level
```

```
Fit: aov(formula = values ~ ind, data = Stacked_Data)

$ind
                    diff        lwr       upr     p adj
K10-C         0.20770081 -0.5200434 0.9354450 0.8752233
K10000-C      0.25243829 -0.4753059 0.9801825 0.7821236
K50000-C      2.46908449  1.7413403 3.1968287 0.0000045
E2-C          5.60263467  4.8748905 6.3303789 0.0000000
K10000-K10    0.04473748 -0.6830067 0.7724817 0.9995565
K50000-K10    2.26138368  1.5336395 2.9891279 0.0000100
E2-K10        5.39493387  4.6671897 6.1226781 0.0000000
K50000-K10000 2.21664620  1.4889020 2.9443904 0.0000120
E2-K10000     5.35019639  4.6224522 6.0779406 0.0000000
E2-K50000     3.13355018  2.4058060 3.8612944 0.0000005

>
```

TIPARP

```
> Data <- read.table(file.choose(), header=TRUE, sep="\t")
> Data
         C       K10   K10000   K50000        E2
1 1.000000 0.5439955 2.321408 1.201636 0.6191382
2 1.305860 0.3272201 2.569819 1.330222 0.9648211
3 1.396356 0.6514230 2.136131 1.010451 0.8960596
> boxplot(Data)
> summary(Data)
       C               K10              K10000           K50000             E2

 Min.   :1.000   Min.   :0.3272   Min.   :2.136   Min.   :1.010   Min.   :0.6191

 1st Qu.:1.153   1st Qu.:0.4356   1st Qu.:2.229   1st Qu.:1.106   1st Qu.:0.7576

 Median :1.306   Median :0.5440   Median :2.321   Median :1.202   Median :0.8961

 Mean   :1.234   Mean   :0.5075   Mean   :2.342   Mean   :1.181   Mean   :0.8267

 3rd Qu.:1.351   3rd Qu.:0.5977   3rd Qu.:2.446   3rd Qu.:1.266   3rd Qu.:0.9304

 Max.   :1.396   Max.   :0.6514   Max.   :2.570   Max.   :1.330   Max.   :0.9648

> Stacked_Data <- stack(Data)
> Stacked_Data
      values    ind
1  1.0000000      C
2  1.3058598      C
3  1.3963559      C
4  0.5439955    K10
5  0.3272201    K10
6  0.6514230    K10
7  2.3214078 K10000
8  2.5698189 K10000
9  2.1361308 K10000
10 1.2016361 K50000
```

```
11 1.3302217 K50000
12 1.0104514 K50000
13 0.6191382     E2
14 0.9648211     E2
15 0.8960596     E2
> Anova_Result <- aov(values~ind, data=Stacked_Data)
> summary(Anova_Result)
            Df Sum Sq Mean Sq F value   Pr(>F)
ind          4  5.772  1.4429   40.73 3.69e-06 ***
Residuals   10  0.354  0.0354
---
Signif. codes:  0 ‘***’ 0.001 ‘**’ 0.01 ‘*’ 0.05 ‘.’ 0.1 ‘ ’ 1
> TukeyHSD(Anova_Result)
  Tukey multiple comparisons of means
    95% family-wise confidence level

Fit: aov(formula = values ~ ind, data = Stacked_Data)

$ind
                     diff        lwr         upr     p adj
K10-C         -0.72652566 -1.2323130 -0.22073832 0.0056265
K10000-C       1.10838061  0.6025933  1.61416796 0.0002167
K50000-C      -0.05330217 -0.5590895  0.45248518 0.9963598
E2-C          -0.40739892 -0.9131863  0.09838843 0.1332094
K10000-K10     1.83490627  1.3291189  2.34069362 0.0000024
K50000-K10     0.67322349  0.1674361  1.17901084 0.0093946
E2-K10         0.31912674 -0.1866606  0.82491408 0.3002920
K50000-K10000 -1.16168278 -1.6674701 -0.65589544 0.0001456
E2-K10000     -1.51577953 -2.0215669 -1.00999219 0.0000139
E2-K50000     -0.35409675 -0.8598841  0.15169059 0.2203272

>
```

STC2

```
> Data <- read.table(file.choose(), header=TRUE, sep="\t")
> Data
          C      K10   K10000   K50000        E
1 1.0000000 2.996614 3.279176 2.163449 3.152872
2 0.8029229 2.351096 4.121968 2.861292 5.253698
3 1.1947151 2.706947 4.594793 2.663519 5.327037
> boxplot(Data)
> summary(Data)
       C               K10            K10000          K50000            E

 Min.   :0.8029   Min.   :2.351   Min.   :3.279   Min.   :2.163   Min.   :3.153

 1st Qu.:0.9015   1st Qu.:2.529   1st Qu.:3.701   1st Qu.:2.413   1st Qu.:4.203

 Median :1.0000   Median :2.707   Median :4.122   Median :2.664   Median :5.254

 Mean   :0.9992   Mean   :2.685   Mean   :3.999   Mean   :2.563   Mean   :4.578

 3rd Qu.:1.0974   3rd Qu.:2.852   3rd Qu.:4.358   3rd Qu.:2.762   3rd Qu.:5.290
```

```
 Max.   :1.1947   Max.   :2.997   Max.   :4.595   Max.   :2.861   Max.   :5.327

> Stacked_Data <- stack(Data)
> Stacked_Data
      values    ind
1  1.0000000      C
2  0.8029229      C
3  1.1947151      C
4  2.9966142    K10
5  2.3510958    K10
6  2.7069470    K10
7  3.2791760 K10000
8  4.1219681 K10000
9  4.5947934 K10000
10 2.1634493 K50000
11 2.8612919 K50000
12 2.6635186 K50000
13 3.1528721      E
14 5.2536982      E
15 5.3270371      E
> Anova_Result <- aov(values~ind, data=Stacked_Data)
> summary(Anova_Result)
            Df Sum Sq Mean Sq F value   Pr(>F)    
ind          4 23.323   5.831   13.01 0.000565 ***
Residuals   10  4.481   0.448                     
---
Signif. codes:  0 '***' 0.001 '**' 0.01 '*' 0.05 '.' 0.1 ' ' 1
> TukeyHSD(Anova_Result)
  Tukey multiple comparisons of means
    95% family-wise confidence level

Fit: aov(formula = values ~ ind, data = Stacked_Data)

$ind
                    diff        lwr       upr     p adj
K10-C          1.6856730 -0.1131982 3.4845442 0.0689076
K10000-C       2.9994332  1.2005620 4.7983044 0.0019194
K50000-C       1.5635406 -0.2353306 3.3624118 0.0970938
E-C            3.5786565  1.7797853 5.3775277 0.0004819
K10000-K10     1.3137602 -0.4851110 3.1126314 0.1913320
K50000-K10    -0.1221324 -1.9210036 1.6767388 0.9993439
E-K10          1.8929835  0.0941123 3.6918547 0.0382684
K50000-K10000 -1.4358926 -3.2347638 0.3629786 0.1380306
E-K10000       0.5792233 -1.2196479 2.3780945 0.8223956
E-K50000       2.0151159  0.2162447 3.8139871 0.0270624

>
```

SLC7A5

```
> Data <- read.table(file.choose(), header=TRUE, sep="\t")
> Data
         C      K10   K10000   K50000        E
```

```
1 1.0000000 1.295342 4.297011 4.901874 2.094588
2 0.7773664 1.844632 4.367073 2.075319 2.867910
3 1.0424658 1.203025 3.749418 3.881641 2.411616
> boxplot(Data)
> summary(Data)
       C                K10             K10000          K50000             E

 Min.   :0.7774   Min.   :1.203   Min.   :3.749   Min.   :2.075   Min.   :2.095

 1st Qu.:0.8887   1st Qu.:1.249   1st Qu.:4.023   1st Qu.:2.978   1st Qu.:2.253

 Median :1.0000   Median :1.295   Median :4.297   Median :3.882   Median :2.412

 Mean   :0.9399   Mean   :1.448   Mean   :4.138   Mean   :3.620   Mean   :2.458

 3rd Qu.:1.0212   3rd Qu.:1.570   3rd Qu.:4.332   3rd Qu.:4.392   3rd Qu.:2.640

 Max.   :1.0425   Max.   :1.845   Max.   :4.367   Max.   :4.902   Max.   :2.868

> Stacked_Data <- stack(Data)
> Stacked_Data
      values    ind
1  1.0000000      C
2  0.7773664      C
3  1.0424658      C
4  1.2953423    K10
5  1.8446324    K10
6  1.2030250    K10
7  4.2970106 K10000
8  4.3670731 K10000
9  3.7494180 K10000
10 4.9018738 K50000
11 2.0753193 K50000
12 3.8816409 K50000
13 2.0945882      E
14 2.8679105      E
15 2.4116157      E
> Anova_Result <- aov(values~ind, data=Stacked_Data)
> summary(Anova_Result)
            Df Sum Sq Mean Sq F value   Pr(>F)
ind          4  22.43   5.608   11.42 0.000953 ***
Residuals   10   4.91   0.491
---
Signif. codes:  0 ‘***’ 0.001 ‘**’ 0.01 ‘*’ 0.05 ‘.’ 0.1 ‘ ’ 1
> TukeyHSD(Anova_Result)
  Tukey multiple comparisons of means
    95% family-wise confidence level

Fit: aov(formula = values ~ ind, data = Stacked_Data)

$ind
                    diff        lwr       upr     p adj
K10-C          0.5077225 -1.3751827 2.3906277 0.8952160
K10000-C       3.1978898  1.3149846 5.0807950 0.0016704
```

```
K50000-C       2.6796673  0.7967621 4.5625725 0.0059974
E-C            1.5180941 -0.3648111 3.4009993 0.1327029
K10000-K10     2.6901673  0.8072621 4.5730725 0.0058385
K50000-K10     2.1719448  0.2890396 4.0548500 0.0228485
E-K10          1.0103716 -0.8725336 2.8932768 0.4409239
K50000-K10000 -0.5182225 -2.4011278 1.3646827 0.8883434
E-K10000      -1.6797957 -3.5627010 0.2031095 0.0865122
E-K50000      -1.1615732 -3.0444784 0.7213320 0.3189555

>
```

CD44

```
> Data <- read.table(file.choose(), header=TRUE, sep="\t")
> Data
          C       K10   K10000   K50000        E
1 1.0000000 0.8197940 2.000000 1.554733 1.362888
2 0.5058097 1.2953423 1.316463 1.341022 1.547565
3 0.7937005 0.6941567 2.118926 1.645280 1.251218
> boxplot(Data)
> summary(Data)
       C                K10              K10000          K50000              E

 Min.   :0.5058   Min.   :0.6942   Min.   :1.316   Min.   :1.341   Min.   :1.251

 1st Qu.:0.6498   1st Qu.:0.7570   1st Qu.:1.658   1st Qu.:1.448   1st Qu.:1.307

 Median :0.7937   Median :0.8198   Median :2.000   Median :1.555   Median :1.363

 Mean   :0.7665   Mean   :0.9364   Mean   :1.812   Mean   :1.514   Mean   :1.387

 3rd Qu.:0.8969   3rd Qu.:1.0576   3rd Qu.:2.059   3rd Qu.:1.600   3rd Qu.:1.455

 Max.   :1.0000   Max.   :1.2953   Max.   :2.119   Max.   :1.645   Max.   :1.548

> Stacked_Data <- stack(Data)
> Stacked_Data
      values    ind
1  1.0000000      C
2  0.5058097      C
3  0.7937005      C
4  0.8197940    K10
5  1.2953423    K10
6  0.6941567    K10
7  2.0000000 K10000
8  1.3164627 K10000
9  2.1189262 K10000
10 1.5547328 K50000
11 1.3410224 K50000
12 1.6452802 K50000
13 1.3628877      E
14 1.5475650      E
15 1.2512181      E
> Anova_Result <- aov(values~ind, data=Stacked_Data)
```

```
> summary(Anova_Result)
            Df Sum Sq Mean Sq F value  Pr(>F)
ind          4 2.1917  0.5479   6.909 0.00619 **
Residuals   10 0.7931  0.0793
---
Signif. codes:  0 ‘***’ 0.001 ‘**’ 0.01 ‘*’ 0.05 ‘.’ 0.1 ‘ ’ 1
> TukeyHSD(Anova_Result)
  Tukey multiple comparisons of means
    95% family-wise confidence level

Fit: aov(formula = values ~ ind, data = Stacked_Data)

$ind
                    diff          lwr       upr     p adj
K10-C          0.1699276 -0.586810616 0.9266658 0.9421427
K10000-C       1.0452929  0.288554695 1.8020311 0.0073448
K50000-C       0.7471751 -0.009563127 1.5039133 0.0533333
E-C            0.6207202 -0.136018005 1.3774584 0.1238612
K10000-K10     0.8753653  0.118627118 1.6321035 0.0224744
K50000-K10     0.5772475 -0.179490703 1.3339857 0.1639017
E-K10          0.4507926 -0.305945581 1.2075308 0.3486776
K50000-K10000 -0.2981178 -1.054856014 0.4586204 0.6992258
E-K10000      -0.4245727 -1.181310892 0.3321655 0.4011585
E-K50000      -0.1264549 -0.883193070 0.6302833 0.9794650

>
```

CSTA

```
> Data <- read.table(file.choose(), header=TRUE, sep="\t")
> Data
          C       K10    K10000      K50000        E2
1 1.0000000 0.3824474 0.1393386 0.07802066 0.2354240
2 0.8029229 0.6299605 0.1364710 0.09192918 0.2570285
3 0.9159453 0.4070656 0.1813271 0.08114616 0.1676278
> boxplot(Data)
> summary(Data)
       C               K10              K10000           K50000              E2

 Min.   :0.8029   Min.   :0.3824   Min.   :0.1365   Min.   :0.07802   Min.
:0.1676
 1st Qu.:0.8594   1st Qu.:0.3948   1st Qu.:0.1379   1st Qu.:0.07958   1st
Qu.:0.2015
 Median :0.9159   Median :0.4071   Median :0.1393   Median :0.08115   Median
:0.2354
 Mean   :0.9063   Mean   :0.4732   Mean   :0.1524   Mean   :0.08370   Mean
:0.2200
 3rd Qu.:0.9580   3rd Qu.:0.5185   3rd Qu.:0.1603   3rd Qu.:0.08654   3rd
Qu.:0.2462
 Max.   :1.0000   Max.   :0.6300   Max.   :0.1813   Max.   :0.09193   Max.
:0.2570
> Stacked_Data <- stack(Data)
> Stacked_Data
       values    ind
```

```
1  1.00000000      C
2  0.80292288      C
3  0.91594529      C
4  0.38244742    K10
5  0.62996052    K10
6  0.40706563    K10
7  0.13933858 K10000
8  0.13647103 K10000
9  0.18132713 K10000
10 0.07802066 K50000
11 0.09192918 K50000
12 0.08114616 K50000
13 0.23542400     E2
14 0.25702846     E2
15 0.16762780     E2
> Anova_Result <- aov(values~ind, data=Stacked_Data)
> summary(Anova_Result)
            Df Sum Sq Mean Sq F value   Pr(>F)
ind          4 1.3501  0.3375   54.03 9.77e-07 ***
Residuals   10 0.0625  0.0062
---
Signif. codes:  0 ‘***’ 0.001 ‘**’ 0.01 ‘*’ 0.05 ‘.’ 0.1 ‘ ’ 1
> TukeyHSD(Anova_Result)
  Tukey multiple comparisons of means
    95% family-wise confidence level

Fit: aov(formula = values ~ ind, data = Stacked_Data)

$ind
                     diff         lwr         upr     p adj
K10-C         -0.43313153 -0.64550649 -0.22075657 0.0003935
K10000-C      -0.75391048 -0.96628543 -0.54153552 0.0000029
K50000-C      -0.82259072 -1.03496568 -0.61021577 0.0000013
E2-C          -0.68626264 -0.89863759 -0.47388768 0.0000070
K10000-K10    -0.32077895 -0.53315390 -0.10840399 0.0039567
K50000-K10    -0.38945919 -0.60183415 -0.17708424 0.0009237
E2-K10        -0.25313111 -0.46550606 -0.04075615 0.0188118
K50000-K10000 -0.06868025 -0.28105520  0.14369471 0.8202161
E2-K10000      0.06764784 -0.14472711  0.28002280 0.8277438
E2-K50000      0.13632809 -0.07604687  0.34870304 0.2862871

>
```

Figure 5 (T47D)

TFF1

```
> Data <- read.table(file.choose(), header=TRUE, sep="\t")
> Data
         C      K10   K10000   K50000       E2
1 1.000000 0.917004 1.359742 1.296840 3.702073
2 0.938438 1.121166 1.487958 1.038859 3.478182
3 1.308880 1.723092 1.047294 1.399586 3.298172
```

```
> boxplot(Data)
> summary(Data)
       C                K10             K10000           K50000             E2

 Min.   :0.9384   Min.   :0.917   Min.   :1.047   Min.   :1.039   Min.   :3.298

 1st Qu.:0.9692   1st Qu.:1.019   1st Qu.:1.204   1st Qu.:1.168   1st Qu.:3.388

 Median :1.0000   Median :1.121   Median :1.360   Median :1.297   Median :3.478

 Mean   :1.0824   Mean   :1.254   Mean   :1.298   Mean   :1.245   Mean   :3.493

 3rd Qu.:1.1544   3rd Qu.:1.422   3rd Qu.:1.424   3rd Qu.:1.348   3rd Qu.:3.590

 Max.   :1.3089   Max.   :1.723   Max.   :1.488   Max.   :1.400   Max.   :3.702

> Stacked_Data <- stack(Data)
> Stacked_Data
     values    ind
1  1.000000      C
2  0.938438      C
3  1.308880      C
4  0.917004    K10
5  1.121166    K10
6  1.723092    K10
7  1.359742 K10000
8  1.487958 K10000
9  1.047294 K10000
10 1.296840 K50000
11 1.038859 K50000
12 1.399586 K50000
13 3.702073     E2
14 3.478182     E2
15 3.298172     E2
> Anova_Result <- aov(values~ind, data=Stacked_Data)
> summary(Anova_Result)
            Df Sum Sq Mean Sq F value   Pr(>F)
ind          4 12.479  3.1198   45.62 2.17e-06 ***
Residuals   10  0.684  0.0684
---
Signif. codes:  0 ‘***’ 0.001 ‘**’ 0.01 ‘*’ 0.05 ‘.’ 0.1 ‘ ’ 1
> TukeyHSD(Anova_Result)
  Tukey multiple comparisons of means
    95% family-wise confidence level

Fit: aov(formula = values ~ ind, data = Stacked_Data)

$ind
                      diff        lwr       upr     p adj
K10-C          0.171314664 -0.5313566 0.8739860 0.9240275
K10000-C       0.215891854 -0.4867795 0.9185632 0.8446938
K50000-C       0.162655358 -0.5400160 0.8653267 0.9359561
E2-C           2.410369457  1.7076981 3.1130408 0.0000040
K10000-K10     0.044577190 -0.6580941 0.7472485 0.9994978
```

```
K50000-K10    -0.008659305 -0.7113306 0.6940120 0.9999993
E2-K10         2.239054793  1.5363835 2.9417261 0.0000080
K50000-K10000 -0.053236495 -0.7559078 0.6494348 0.9989907
E2-K10000      2.194477603  1.4918063 2.8971489 0.0000096
E2-K50000      2.247714098  1.5450428 2.9503854 0.0000077

>
```

TIPARP

```
> Data <- read.table(file.choose(), header=TRUE, sep="\t")
> Data
         C       K10   K10000   K50000        E
1 1.000000 1.3379276 1.298339 1.049717 2.143547
2 1.239708 0.8546072 1.540430 1.112136 1.697408
3 1.096825 0.7439788 1.119872 1.460707 1.963372
> boxplot(Data)
> summary(Data)
       C              K10              K10000           K50000            E

 Min.   :1.000   Min.   :0.7440   Min.   :1.120   Min.   :1.050   Min.   :1.697

 1st Qu.:1.048   1st Qu.:0.7993   1st Qu.:1.209   1st Qu.:1.081   1st Qu.:1.830

 Median :1.097   Median :0.8546   Median :1.298   Median :1.112   Median :1.963

 Mean   :1.112   Mean   :0.9788   Mean   :1.320   Mean   :1.208   Mean   :1.935

 3rd Qu.:1.168   3rd Qu.:1.0963   3rd Qu.:1.419   3rd Qu.:1.286   3rd Qu.:2.053

 Max.   :1.240   Max.   :1.3379   Max.   :1.540   Max.   :1.461   Max.   :2.144

> Stacked_Data <- stack(Data)
> Stacked_Data
      values    ind
1  1.0000000      C
2  1.2397077      C
3  1.0968250      C
4  1.3379276    K10
5  0.8546072    K10
6  0.7439788    K10
7  1.2983386 K10000
8  1.5404302 K10000
9  1.1198716 K10000
10 1.0497167 K50000
11 1.1121361 K50000
12 1.4607068 K50000
13 2.1435469      E
14 1.6974079      E
15 1.9633717      E
> Anova_Result <- aov(values~ind, data=Stacked_Data)
> summary(Anova_Result)
            Df Sum Sq Mean Sq F value  Pr(>F)
ind          4 1.6492  0.4123   7.981 0.00371 **
```

```
Residuals   10 0.5166  0.0517
---
Signif. codes:  0 ‘***’ 0.001 ‘**’ 0.01 ‘*’ 0.05 ‘.’ 0.1 ‘ ’ 1
> TukeyHSD(Anova_Result)
  Tukey multiple comparisons of means
    95% family-wise confidence level

Fit: aov(formula = values ~ ind, data = Stacked_Data)

$ind
                    diff          lwr       upr     p adj
K10-C         -0.13333973 -0.744087823 0.4774084 0.9473867
K10000-C       0.20736924 -0.403378847 0.8181173 0.7943233
K50000-C       0.09534231 -0.515405780 0.7060904 0.9839847
E-C            0.82259797  0.211849874 1.4333461 0.0086914
K10000-K10     0.34070898 -0.270039116 0.9514571 0.4062412
K50000-K10     0.22868204 -0.382066049 0.8394301 0.7344325
E-K10          0.95593770  0.345189605 1.5666858 0.0030630
K50000-K10000 -0.11202693 -0.722775025 0.4987212 0.9712953
E-K10000       0.61522872  0.004480629 1.2259768 0.0481606
E-K50000       0.72725565  0.116507562 1.3380037 0.0189209

>
```

STC2

```
> Data <- read.table(file.choose(), header=TRUE, sep="\t")
> Data
          C       K10   K10000    K50000       E2
1 1.0000000 0.9930925 1.162046 0.6099093 1.211393
2 0.8745827 0.5770094 1.189207 0.7827734 1.319508
3 0.8888427 0.9351912 1.057018 0.8685415 1.292353
> boxplot(Data)
> summary(Data)
       C                K10              K10000          K50000             E2

 Min.   :0.8746   Min.   :0.5770   Min.   :1.057   Min.   :0.6099   Min.
:1.211
 1st Qu.:0.8817   1st Qu.:0.7561   1st Qu.:1.110   1st Qu.:0.6963   1st
Qu.:1.252
 Median :0.8888   Median :0.9352   Median :1.162   Median :0.7828   Median
:1.292
 Mean   :0.9211   Mean   :0.8351   Mean   :1.136   Mean   :0.7537   Mean
:1.274
 3rd Qu.:0.9444   3rd Qu.:0.9641   3rd Qu.:1.176   3rd Qu.:0.8257   3rd
Qu.:1.306
 Max.   :1.0000   Max.   :0.9931   Max.   :1.189   Max.   :0.8685   Max.
:1.320
> Stacked_Data <- stack(Data)
> Stacked_Data
      values    ind
1  1.0000000      C
2  0.8745827      C
3  0.8888427      C
```

```
4  0.9930925    K10
5  0.5770094    K10
6  0.9351912    K10
7  1.1620456 K10000
8  1.1892071 K10000
9  1.0570180 K10000
10 0.6099093 K50000
11 0.7827734 K50000
12 0.8685415 K50000
13 1.2113927     E2
14 1.3195079     E2
15 1.2923528     E2
> Anova_Result <- aov(values~ind, data=Stacked_Data)
> summary(Anova_Result)
            Df Sum Sq Mean Sq F value  Pr(>F)
ind          4 0.5598 0.13996    8.65 0.00276 **
Residuals   10 0.1618 0.01618
---
Signif. codes:  0 ‘***’ 0.001 ‘**’ 0.01 ‘*’ 0.05 ‘.’ 0.1 ‘ ’ 1
> TukeyHSD(Anova_Result)
  Tukey multiple comparisons of means
    95% family-wise confidence level

Fit: aov(formula = values ~ ind, data = Stacked_Data)

$ind
                     diff         lwr         upr     p adj
K10-C         -0.08604408 -0.42785670  0.25576855 0.9157348
K10000-C       0.21494846 -0.12686416  0.55676109 0.3030321
K50000-C      -0.16740038 -0.50921300  0.17441225 0.5224162
E2-C           0.35327604  0.01146342  0.69508867 0.0421242
K10000-K10     0.30099254 -0.04082009  0.64280517 0.0916929
K50000-K10    -0.08135630 -0.42316893  0.26045633 0.9297799
E2-K10         0.43932012  0.09750749  0.78113275 0.0117821
K50000-K10000 -0.38234884 -0.72416147 -0.04053621 0.0272865
E2-K10000      0.13832758 -0.20348505  0.48014021 0.6795220
E2-K50000      0.52067642  0.17886379  0.86248905 0.0037248

>
```

SLC7A5

```
> Data <- read.table(file.choose(), header=TRUE, sep="\t")
> Data
          C       K10    K10000   K50000        E
1 1.0000000 1.0448772 0.9794203 1.004632 2.173470
2 0.9351912 0.5383688 1.0424658 1.047294 1.836128
3 1.0116194 1.0376597 0.9308797 1.211393 2.018570
> boxplot(Data)
> summary(Data)
       C                K10              K10000           K50000            E

 Min.   :0.9352   Min.   :0.5384   Min.   :0.9309   Min.   :1.005   Min.
:1.836
```

```
 1st Qu.:0.9676   1st Qu.:0.7880   1st Qu.:0.9552   1st Qu.:1.026   1st
Qu.:1.927
 Median :1.0000   Median :1.0377   Median :0.9794   Median :1.047   Median
:2.019
 Mean   :0.9823   Mean   :0.8736   Mean   :0.9843   Mean   :1.088   Mean
:2.009
 3rd Qu.:1.0058   3rd Qu.:1.0413   3rd Qu.:1.0109   3rd Qu.:1.129   3rd
Qu.:2.096
 Max.   :1.0116   Max.   :1.0449   Max.   :1.0425   Max.   :1.211   Max.
:2.173
> Stacked_Data <- stack(Data)
> Stacked_Data
      values    ind
1  1.0000000      C
2  0.9351912      C
3  1.0116194      C
4  1.0448772    K10
5  0.5383688    K10
6  1.0376597    K10
7  0.9794203 K10000
8  1.0424658 K10000
9  0.9308797 K10000
10 1.0046317 K50000
11 1.0472941 K50000
12 1.2113927 K50000
13 2.1734697      E
14 1.8361280      E
15 2.0185696      E
> Anova_Result <- aov(values~ind, data=Stacked_Data)
> summary(Anova_Result)
            Df Sum Sq Mean Sq F value   Pr(>F)    
ind          4 2.6022  0.6505    25.1 3.38e-05 ***
Residuals   10 0.2591  0.0259                     
---
Signif. codes:  0 ‘***’ 0.001 ‘**’ 0.01 ‘*’ 0.05 ‘.’ 0.1 ‘ ’ 1
> TukeyHSD(Anova_Result)
  Tukey multiple comparisons of means
    95% family-wise confidence level

Fit: aov(formula = values ~ ind, data = Stacked_Data)

$ind
                      diff        lwr       upr     p adj
K10-C         -0.108635031 -0.5412117 0.3239417 0.9163733
K10000-C       0.001985029 -0.4305917 0.4345617 1.0000000
K50000-C       0.105502615 -0.3270741 0.5380793 0.9239375
E-C            1.027118893  0.5945422 1.4596956 0.0001095
K10000-K10     0.110620060 -0.3219566 0.5431967 0.9113710
K50000-K10     0.214137646 -0.2184390 0.6467143 0.5129338
E-K10          1.135753924  0.7031772 1.5683306 0.0000456
K50000-K10000  0.103517586 -0.3290591 0.5360943 0.9285207
E-K10000       1.025133864  0.5925572 1.4577106 0.0001114
E-K50000       0.921616278  0.4890396 1.3541930 0.0002743
```

```
>

CD44

> Data <- read.table(file.choose(), header=TRUE, sep="\t")
> Data
         C       K10   K10000   K50000        E
1 1.000000 0.9244497 1.208597 1.156688 1.594753
2 1.002313 0.6476711 1.228303 1.044877 1.191958
3 1.032876 0.7269863 1.117287 1.494849 1.254112
> boxplot(Data)
> summary(Data)
       C              K10              K10000          K50000            E

 Min.   :1.000   Min.   :0.6477   Min.   :1.117   Min.   :1.045   Min.   :1.192

 1st Qu.:1.001   1st Qu.:0.6873   1st Qu.:1.163   1st Qu.:1.101   1st Qu.:1.223

 Median :1.002   Median :0.7270   Median :1.209   Median :1.157   Median :1.254

 Mean   :1.012   Mean   :0.7664   Mean   :1.185   Mean   :1.232   Mean   :1.347

 3rd Qu.:1.018   3rd Qu.:0.8257   3rd Qu.:1.218   3rd Qu.:1.326   3rd Qu.:1.424

 Max.   :1.033   Max.   :0.9244   Max.   :1.228   Max.   :1.495   Max.   :1.595

> Stacked_Data <- stack(Data)
> Stacked_Data
      values    ind
1  1.0000000      C
2  1.0023132      C
3  1.0328757      C
4  0.9244497    K10
5  0.6476711    K10
6  0.7269863    K10
7  1.2085971 K10000
8  1.2283031 K10000
9  1.1172871 K10000
10 1.1566882 K50000
11 1.0448772 K50000
12 1.4948492 K50000
13 1.5947534      E
14 1.1919579      E
15 1.2541124      E
> Anova_Result <- aov(values~ind, data=Stacked_Data)
> summary(Anova_Result)
            Df Sum Sq Mean Sq F value  Pr(>F)
ind          4 0.6131 0.15328   6.079 0.00955 **
Residuals   10 0.2521 0.02521
---
Signif. codes:  0 ‘***’ 0.001 ‘**’ 0.01 ‘*’ 0.05 ‘.’ 0.1 ‘ ’ 1
> TukeyHSD(Anova_Result)
  Tukey multiple comparisons of means
    95% family-wise confidence level
```

```
Fit: aov(formula = values ~ ind, data = Stacked_Data)

$ind
                    diff          lwr       upr     p adj
K10-C         -0.24536061 -0.672055694 0.1813345 0.3794325
K10000-C       0.17299949 -0.253695595 0.5996946 0.6781348
K50000-C       0.22040857 -0.206286514 0.6471037 0.4750365
E-C            0.33521162 -0.091483466 0.7619067 0.1467947
K10000-K10     0.41836010 -0.008334984 0.8450552 0.0552441
K50000-K10     0.46576918  0.039074097 0.8924643 0.0313159
E-K10          0.58057223  0.153877145 1.0072673 0.0081250
K50000-K10000  0.04740908 -0.379286003 0.4741042 0.9955405
E-K10000       0.16221213 -0.264482954 0.5889072 0.7241978
E-K50000       0.11480305 -0.311892035 0.5414981 0.8959380

>
```

CSTA

```
> Data <- read.table(file.choose(), header=TRUE, sep="\t")
> Data
          C       K10   K10000   K50000        E2
1 1.0000000 0.4763190 1.359742 1.922966 0.8908987
2 0.9330330 0.4569157 1.487958 3.010493 1.4306459
3 0.7791646 0.7236346 1.198863 2.921414 1.0304920
> boxplot(Data)
> summary(Data)
       C                K10              K10000           K50000             E2

 Min.   :0.7792   Min.   :0.4569   Min.   :1.199   Min.   :1.923   Min.
:0.8909
 1st Qu.:0.8561   1st Qu.:0.4666   1st Qu.:1.279   1st Qu.:2.422   1st
Qu.:0.9607
 Median :0.9330   Median :0.4763   Median :1.360   Median :2.921   Median
:1.0305
 Mean   :0.9041   Mean   :0.5523   Mean   :1.349   Mean   :2.618   Mean
:1.1173
 3rd Qu.:0.9665   3rd Qu.:0.6000   3rd Qu.:1.424   3rd Qu.:2.966   3rd
Qu.:1.2306
 Max.   :1.0000   Max.   :0.7236   Max.   :1.488   Max.   :3.010   Max.
:1.4306
> Stacked_Data <- stack(Data)
> Stacked_Data
      values    ind
1  1.0000000      C
2  0.9330330      C
3  0.7791646      C
4  0.4763190    K10
5  0.4569157    K10
6  0.7236346    K10
7  1.3597424 K10000
8  1.4879575 K10000
9  1.1988629 K10000
```

```
10 1.9229661 K50000
11 3.0104935 K50000
12 2.9214137 K50000
13 0.8908987     E2
14 1.4306459     E2
15 1.0304920     E2
> Anova_Result <- aov(values~ind, data=Stacked_Data)
> summary(Anova_Result)
            Df Sum Sq Mean Sq F value   Pr(>F)    
ind          4  7.467  1.8669   18.71 0.000123 ***
Residuals   10  0.998  0.0998                     
---
Signif. codes:  0 ‘***’ 0.001 ‘**’ 0.01 ‘*’ 0.05 ‘.’ 0.1 ‘ ’ 1
> TukeyHSD(Anova_Result)
  Tukey multiple comparisons of means
    95% family-wise confidence level

Fit: aov(formula = values ~ ind, data = Stacked_Data)

$ind
                    diff         lwr        upr     p adj
K10-C         -0.3517761 -1.20067839  0.4971262 0.6615794
K10000-C       0.4447884 -0.40411391  1.2936907 0.4622877
K50000-C       1.7142252  0.86532293  2.5631275 0.0004267
E2-C           0.2132797 -0.63562261  1.0621820 0.9162585
K10000-K10     0.7965645 -0.05233783  1.6454668 0.0684630
K50000-K10     2.0660013  1.21709901  2.9149036 0.0000885
E2-K10         0.5650558 -0.28384653  1.4139581 0.2577337
K50000-K10000  1.2694368  0.42053453  2.1183391 0.0042480
E2-K10000     -0.2315087 -1.08041101  0.6173936 0.8914643
E2-K50000     -1.5009455 -2.34984785 -0.6520432 0.0012277

>
```

Figure 6A

```
> Data <- read.table(file.choose(), header=T, sep="\t")
> Data
  C0        K0 C6        K6 C12       K12 C24       K24 C36       K36 C48
K48 C72       K72
1  1 1.1551301  1 1.0495125   1 0.7270701   1 0.7239968   1 0.7559323   1
0.7462179   1 0.7073575
2  1 1.3046432  1 1.2647325   1 0.8052722   1 0.5113620   1 0.4869743   1
0.5901115   1 0.4597401
3  1 0.8253401  1 0.9823255   1 1.1105916   1 0.7970731   1 0.6234904   1
0.6997466   1 0.6982828
4  1 0.9772993  1 1.0395061   1 0.6942474   1 0.5805090   1 0.5589845   1
0.5227752   1 0.2844275
5  1 1.3093908  1 0.7759095   1 0.7559991   1 0.6255584   1 0.4978086   1
0.4738820   1 0.4122345
>
```

0h

```
> t.test(Data$K0, Data$C0, alternative = "less", var.equal=FALSE)

        Welch Two Sample t-test

data:  Data$K0 and Data$C0
t = 1.2115, df = 4, p-value = 0.8538
alternative hypothesis: true difference in means is less than 0
95 percent confidence interval:
      -Inf 0.3156058
sample estimates:
mean of x mean of y 
 1.114361  1.000000 
>
```

6 h

```
> t.test(Data$K6, Data$C6, alternative = "less", var.equal=FALSE)

        Welch Two Sample t-test

data:  Data$K6 and Data$C6
t = 0.28663, df = 4, p-value = 0.6057
alternative hypothesis: true difference in means is less than 0
95 percent confidence interval:
      -Inf 0.1889791
sample estimates:
mean of x mean of y 
 1.022397  1.000000 
>
```

12 h

```
> t.test(Data$K12, Data$C12, alternative = "less", var.equal=FALSE)

        Welch Two Sample t-test

data:  Data$K12 and Data$C12
t = -2.4107, df = 4, p-value = 0.03675
alternative hypothesis: true difference in means is less than 0
95 percent confidence interval:
        -Inf -0.02098094
sample estimates:
mean of x mean of y 
0.8186361 1.0000000 
>
```

24 h

```
> t.test(Data$K24, Data$C24, alternative = "less", var.equal=FALSE)

        Welch Two Sample t-test

data:  Data$K24 and Data$C24
t = -6.9274, df = 4, p-value = 0.00114
```

```
alternative hypothesis: true difference in means is less than 0
95 percent confidence interval:
       -Inf -0.2438832
sample estimates:
mean of x mean of y
0.6476998 1.0000000
>

36 h

> t.test(Data$K36, Data$C36, alternative = "less", var.equal=FALSE)

        Welch Two Sample t-test

data:  Data$K36 and Data$C36
t = -8.4272, df = 4, p-value = 0.0005429
alternative hypothesis: true difference in means is less than 0
95 percent confidence interval:
       -Inf -0.3102867
sample estimates:
mean of x mean of y
 0.584638  1.000000
>

48 h

> t.test(Data$K48, Data$C48, alternative = "less", var.equal=FALSE)

        Welch Two Sample t-test

data:  Data$K48 and Data$C48
t = -7.6372, df = 4, p-value = 0.0007894
alternative hypothesis: true difference in means is less than 0
95 percent confidence interval:
      -Inf -0.283625
sample estimates:
mean of x mean of y
0.6065466 1.0000000
>

74 h

> t.test(Data$K72, Data$C72, alternative = "less", var.equal=FALSE)

        Welch Two Sample t-test

data:  Data$K72 and Data$C72
t = -5.884, df = 4, p-value = 0.002085
alternative hypothesis: true difference in means is less than 0
95 percent confidence interval:
      -Inf -0.310932
sample estimates:
mean of x mean of y
0.5124085 1.0000000
```

```
>
```

Figure 6C

PR

```
> Data <- read.table(file.choose(), header=TRUE, sep="\t")
> Data
     T   S       exp
1  Veh scr 1.0000000
2  Veh scr 0.9265881
3  Veh scr 0.9704102
4  Kar scr 0.9930925
5  Kar scr 0.8487040
6  Kar scr 1.0472941
7  Veh  ER 0.3084981
8  Veh  ER 0.2097389
9  Veh  ER 0.2448551
10 Kar  ER 0.1604282
11 Kar  ER 0.2232399
12 Kar  ER 0.3789291
> summary(Data)
      T                  S                  exp
 Length:12          Length:12          Min.   :0.1604
 Class :character   Class :character   1st Qu.:0.2395
 Mode  :character   Mode  :character   Median :0.6138
                                       Mean   :0.6093
                                       3rd Qu.:0.9761
                                       Max.   :1.0473
> Data$T <- as.factor(Data$T)
> Data$S <- as.factor(Data$S)
> str(Data)
'data.frame':   12 obs. of  3 variables:
 $ T  : Factor w/ 2 levels "Kar","Veh": 2 2 2 1 1 1 2 2 2 1 ...
 $ S  : Factor w/ 2 levels "ER","scr": 2 2 2 2 2 2 1 1 1 1 ...
 $ exp: num  1 0.927 0.97 0.993 0.849 ...
>
> #without interaction term
> anova1 <- aov(formula = exp ~ T + S, data = Data)
> summary(anova1)
            Df Sum Sq Mean Sq F value   Pr(>F)
T            1 0.0000   0.000   0.001    0.976
S            1 1.5126   1.513 251.497 6.95e-08 ***
Residuals    9 0.0541   0.006
---
Signif. codes:  0 ‘***’ 0.001 ‘**’ 0.01 ‘*’ 0.05 ‘.’ 0.1 ‘ ’ 1
> TukeyHSD(anova1)
  Tukey multiple comparisons of means
    95% family-wise confidence level

Fit: aov(formula = exp ~ T + S, data = Data)

$T
```

```
               diff         lwr       upr     p adj
Veh-Kar 0.001400421 -0.09988707 0.1026879 0.9757312

$S
            diff       lwr       upr p adj
scr-ER 0.7100666 0.6087791 0.8113541 1e-07

>
> #with interaction terms
> anova2 <- aov(formula = exp ~ S + T + S:T, data = Data)
> summary(anova2)
            Df Sum Sq Mean Sq F value   Pr(>F)
S            1 1.5126  1.5126 223.571 3.95e-07 ***
T            1 0.0000  0.0000   0.001    0.977
S:T          1 0.0000  0.0000   0.001    0.980
Residuals    8 0.0541  0.0068
---
Signif. codes:  0 '***' 0.001 '**' 0.01 '*' 0.05 '.' 0.1 ' ' 1
> TukeyHSD(anova2)
  Tukey multiple comparisons of means
    95% family-wise confidence level

Fit: aov(formula = exp ~ S + T + S:T, data = Data)

$S
            diff       lwr       upr p adj
scr-ER 0.7100666 0.6005573 0.8195759 4e-07

$T
               diff        lwr       upr     p adj
Veh-Kar 0.001400421 -0.1081089 0.1109097 0.9771966

$`S:T`
                        diff        lwr        upr     p adj
scr:Kar-ER:Kar   0.708831111  0.4937635  0.9238987 0.0000265
ER:Veh-ER:Kar    0.000164941 -0.2149027  0.2152325 1.0000000
scr:Veh-ER:Kar   0.711467012  0.4963994  0.9265346 0.0000257
ER:Veh-scr:Kar  -0.708666170 -0.9237338 -0.4935986 0.0000265
scr:Veh-scr:Kar  0.002635901 -0.2124317  0.2177035 0.9999763
scr:Veh-ER:Veh   0.711302071  0.4962345  0.9263697 0.0000258

>
```

TFF1

```
> Data <- read.table(file.choose(), header=TRUE, sep="\t")
> Data
     T   S       exp
1  veh scr 1.0000000
2  veh scr 0.7186361
3  veh scr 1.0497167
4  kar scr 1.0139595
5  kar scr 1.0892487
6  kar scr 1.2002487
```

```
7  veh  ER 0.5471469
8  veh  ER 0.6042985
9  veh  ER 0.6056964
10 kar  ER 0.5783441
11 kar  ER 0.6973718
12 kar  ER 0.7371346
> summary(Data)
      T                  S                  exp        
 Length:12          Length:12          Min.   :0.5471  
 Class :character   Class :character   1st Qu.:0.6053  
 Mode  :character   Mode  :character   Median :0.7279  
                                       Mean   :0.8202  
                                       3rd Qu.:1.0229  
                                       Max.   :1.2002  
> Data$T <- as.factor(Data$T)
> Data$S <- as.factor(Data$S)
> str(Data)
'data.frame':   12 obs. of  3 variables:
 $ T  : Factor w/ 2 levels "kar","veh": 2 2 2 1 1 1 2 2 2 1 ...
 $ S  : Factor w/ 2 levels "ER","scr": 2 2 2 2 2 2 1 1 1 1 ...
 $ exp: num  1 0.719 1.05 1.014 1.089 ...
> 
> #without interaction term
> anova1 <- aov(formula = exp ~ T + S, data = Data)
> summary(anova1)
            Df Sum Sq Mean Sq F value   Pr(>F)    
T            1 0.0521  0.0521   4.523 0.062358 .  
S            1 0.4415  0.4415  38.317 0.000161 ***
Residuals    9 0.1037  0.0115                     
---
Signif. codes:  0 ‘***’ 0.001 ‘**’ 0.01 ‘*’ 0.05 ‘.’ 0.1 ‘ ’ 1
> TukeyHSD(anova1)
  Tukey multiple comparisons of means
    95% family-wise confidence level

Fit: aov(formula = exp ~ T + S, data = Data)

$T
              diff        lwr         upr     p adj
veh-kar -0.1318021 -0.2720008 0.008396579 0.0623575

$S
            diff       lwr       upr     p adj
scr-ER 0.3836362 0.2434375 0.5238349 0.0001608

> 
> #with interaction terms
> anova2 <- aov(formula = exp ~ S + T + S:T, data = Data)
> summary(anova2)
            Df Sum Sq Mean Sq F value   Pr(>F)    
S            1 0.4415  0.4415  36.339 0.000313 ***
T            1 0.0521  0.0521   4.289 0.072111 .  
S:T          1 0.0065  0.0065   0.535 0.485227    
Residuals    8 0.0972  0.0122                     
```

```
---
Signif. codes:  0 ‘***’ 0.001 ‘**’ 0.01 ‘*’ 0.05 ‘.’ 0.1 ‘ ’ 1
> TukeyHSD(anova2)
  Tukey multiple comparisons of means
    95% family-wise confidence level

Fit: aov(formula = exp ~ S + T + S:T, data = Data)

$S
            diff       lwr       upr     p adj
scr-ER 0.3836362 0.2368819 0.5303905 0.0003134

$T
              diff        lwr        upr     p adj
veh-kar -0.1318021 -0.2785564 0.01495215 0.0721107

$`S:T`
                       diff         lwr        upr     p adj
scr:kar-ER:kar   0.43020209  0.14198829  0.7184159 0.0060656
ER:veh-ER:kar   -0.08523626 -0.37345006  0.2029775 0.7815758
scr:veh-ER:kar   0.25183409 -0.03637971  0.5400479 0.0884612
ER:veh-scr:kar  -0.51543835 -0.80365215 -0.2272246 0.0019702
scr:veh-scr:kar -0.17836800 -0.46658180  0.1098458 0.2703079
scr:veh-ER:veh   0.33707035  0.04885655  0.6252841 0.0235169

>
```

CSTA

```
> Data <- read.table(file.choose(), header=TRUE, sep="\t")
> Data
     T   S       exp
1  veh scr 1.0000000
2  veh scr 0.7791646
3  veh scr 0.8216903
4  kar scr 0.5011566
5  kar scr 0.5309569
6  kar scr 0.4818536
7  veh  ER 1.6132835
8  veh  ER 1.3256194
9  veh  ER 1.8234450
10 kar  ER 1.4044449
11 kar  ER 1.4028233
12 kar  ER 1.3286858
> summary(Data)
      T                  S                  exp
 Length:12          Length:12          Min.   :0.4819
 Class :character   Class :character   1st Qu.:0.7171
 Mode  :character   Mode  :character   Median :1.1628
                                       Mean   :1.0844
                                       3rd Qu.:1.4032
                                       Max.   :1.8234
> Data$T <- as.factor(Data$T)
> Data$S <- as.factor(Data$S)
```

```
> str(Data)
'data.frame':   12 obs. of  3 variables:
 $ T  : Factor w/ 2 levels "kar","veh": 2 2 2 1 1 1 2 2 2 1 ...
 $ S  : Factor w/ 2 levels "ER","scr": 2 2 2 2 2 2 1 1 1 1 ...
 $ exp: num  1 0.779 0.822 0.501 0.531 ...
>
> #without interaction term
> anova1 <- aov(formula = exp ~ T + S, data = Data)
> summary(anova1)
            Df Sum Sq Mean Sq F value   Pr(>F)
T            1 0.2446  0.2446   12.58  0.00625 **
S            1 1.9068  1.9068   98.06 3.88e-06 ***
Residuals    9 0.1750  0.0194
---
Signif. codes:  0 ‘***’ 0.001 ‘**’ 0.01 ‘*’ 0.05 ‘.’ 0.1 ‘ ’ 1
> TukeyHSD(anova1)
  Tukey multiple comparisons of means
    95% family-wise confidence level

Fit: aov(formula = exp ~ T + S, data = Data)

$T
            diff       lwr       upr     p adj
veh-kar 0.285547 0.1034184 0.4676755 0.0062483

$S
             diff        lwr        upr   p adj
scr-ER -0.7972467 -0.9793752 -0.6151181 3.9e-06

>
> #with interaction terms
> anova2 <- aov(formula = exp ~ S + T + S:T, data = Data)
> summary(anova2)
            Df Sum Sq Mean Sq F value   Pr(>F)
S            1 1.9068  1.9068  96.950 9.53e-06 ***
T            1 0.2446  0.2446  12.437  0.00777 **
S:T          1 0.0177  0.0177   0.898  0.37094
Residuals    8 0.1573  0.0197
---
Signif. codes:  0 ‘***’ 0.001 ‘**’ 0.01 ‘*’ 0.05 ‘.’ 0.1 ‘ ’ 1
> TukeyHSD(anova2)
  Tukey multiple comparisons of means
    95% family-wise confidence level

Fit: aov(formula = exp ~ S + T + S:T, data = Data)

$S
             diff        lwr        upr   p adj
scr-ER -0.7972467 -0.9839614 -0.6105319 9.5e-06

$T
            diff       lwr       upr     p adj
veh-kar 0.285547 0.0988322 0.4722617 0.0077703
```

```
$`S:T`
                        diff          lwr        upr     p adj
scr:kar-ER:kar  -0.8739957 -1.240688671 -0.5073027 0.0002809
ER:veh-ER:kar    0.2087980 -0.157895039  0.5754910 0.3300540
scr:veh-ER:kar  -0.5116997 -0.878392720 -0.1450067 0.0089965
ER:veh-scr:kar   1.0827936  0.716100622  1.4494866 0.0000598
scr:veh-scr:kar  0.3622960 -0.004397059  0.7289890 0.0527773
scr:veh-ER:veh  -0.7204977 -1.087190691 -0.3538047 0.0010610

>
```

ADAMTS19

```
> Data <- read.table(file.choose(), header=TRUE, sep="\t")
> Data
     T   S       exp
1  veh scr 1.0000000
2  veh scr 0.9373545
3  veh scr 0.9953897
4  kar scr 0.5946036
5  kar scr 0.4622248
6  kar scr 0.6029039
7  veh  ER 0.7103819
8  veh  ER 0.5248583
9  veh  ER 0.5433674
10 kar  ER 0.3763117
11 kar  ER 0.4014614
12 kar  ER 0.6299605
> summary(Data)
      T                  S                  exp
 Length:12          Length:12          Min.   :0.3763
 Class :character   Class :character   1st Qu.:0.5092
 Mode  :character   Mode  :character   Median :0.5988
                                       Mean   :0.6482
                                       3rd Qu.:0.7671
                                       Max.   :1.0000
> Data$T <- as.factor(Data$T)
> Data$S <- as.factor(Data$S)
> str(Data)
'data.frame':   12 obs. of  3 variables:
 $ T  : Factor w/ 2 levels "kar","veh": 2 2 2 1 1 1 2 2 2 1 ...
 $ S  : Factor w/ 2 levels "ER","scr": 2 2 2 2 2 2 1 1 1 1 ...
 $ exp: num  1 0.937 0.995 0.595 0.462 ...
>
> #without interaction term
> anova1 <- aov(formula = exp ~ T + S, data = Data)
> summary(anova1)
            Df Sum Sq Mean Sq F value  Pr(>F)
T            1 0.2252 0.22520   14.21 0.00442 **
S            1 0.1648 0.16477   10.39 0.01042 *
Residuals    9 0.1427 0.01585
---
Signif. codes:  0 ‘***’ 0.001 ‘**’ 0.01 ‘*’ 0.05 ‘.’ 0.1 ‘ ’ 1
> TukeyHSD(anova1)
```

```
  Tukey multiple comparisons of means
    95% family-wise confidence level

Fit: aov(formula = exp ~ T + S, data = Data)

$T
            diff       lwr       upr     p adj
veh-kar 0.273981 0.1095434 0.4384186 0.0044223

$S
            diff        lwr       upr     p adj
scr-ER 0.2343559 0.06991825 0.3987935 0.0104221

>
> #with interaction terms
> anova2 <- aov(formula = exp ~ S + T + S:T, data = Data)
> summary(anova2)
            Df  Sum Sq Mean Sq F value  Pr(>F)
S            1 0.16477 0.16477  17.612 0.00301 **
T            1 0.22520 0.22520  24.071 0.00118 **
S:T          1 0.06782 0.06782   7.249 0.02739 *
Residuals    8 0.07484 0.00936
---
Signif. codes:  0 ‘***’ 0.001 ‘**’ 0.01 ‘*’ 0.05 ‘.’ 0.1 ‘ ’ 1
> TukeyHSD(anova2)
  Tukey multiple comparisons of means
    95% family-wise confidence level

Fit: aov(formula = exp ~ S + T + S:T, data = Data)

$S
            diff       lwr       upr     p adj
scr-ER 0.2343559 0.1055796 0.3631321 0.0030104

$T
            diff       lwr       upr     p adj
veh-kar 0.273981 0.1452047 0.4027572 0.0011845

$`S:T`
                      diff        lwr       upr     p adj
scr:kar-ER:kar  0.08399955 -0.1689068 0.3369059 0.7194734
ER:veh-ER:kar   0.12362466 -0.1292817 0.3765310 0.4469501
scr:veh-ER:kar  0.50833684  0.2554305 0.7612432 0.0009110
ER:veh-scr:kar  0.03962511 -0.2132812 0.2925315 0.9563958
scr:veh-scr:kar 0.42433729  0.1714309 0.6772436 0.0029606
scr:veh-ER:veh  0.38471218  0.1318058 0.6376185 0.0054157

>
```

CYP1A1

```
> Data <- read.table(file.choose(), header=TRUE, sep="\t")
> Data
     T   S        exp
```

```
1  veh scr  1.0000000
2  veh scr  0.7992212
3  veh scr  0.9138314
4  kar scr 55.4583709
5  kar scr 52.2249270
6  kar scr 48.1122823
7  veh  ER  1.8488993
8  veh  ER  1.2893703
9  veh  ER  1.5052467
10 kar  ER 87.2248113
11 kar  ER 71.5063768
12 kar  ER 59.3016360
> summary(Data)
      T                   S                  exp
 Length:12          Length:12          Min.   : 0.7992
 Class :character   Class :character   1st Qu.: 1.2170
 Mode  :character   Mode  :character   Median :24.9806
                                       Mean   :31.7654
                                       3rd Qu.:56.4192
                                       Max.   :87.2248
> Data$T <- as.factor(Data$T)
> Data$S <- as.factor(Data$S)
> str(Data)
'data.frame':   12 obs. of  3 variables:
 $ T  : Factor w/ 2 levels "kar","veh": 2 2 2 1 1 1 2 2 2 1 ...
 $ S  : Factor w/ 2 levels "ER","scr": 2 2 2 2 2 2 1 1 1 1 ...
 $ exp: num  1 0.799 0.914 55.458 52.225 ...
>
> #without interaction term
> anova1 <- aov(formula = exp ~ T + S, data = Data)
> summary(anova1)
            Df Sum Sq Mean Sq F value   Pr(>F)
T            1  11192   11192 139.457 8.83e-07 ***
S            1    343     343   4.276   0.0686 .
Residuals    9    722      80
---
Signif. codes:  0 ‘***’ 0.001 ‘**’ 0.01 ‘*’ 0.05 ‘.’ 0.1 ‘ ’ 1
> TukeyHSD(anova1)
  Tukey multiple comparisons of means
    95% family-wise confidence level

Fit: aov(formula = exp ~ T + S, data = Data)

$T
             diff       lwr       upr p adj
veh-kar -61.07864 -72.77882 -49.37846 9e-07

$S
            diff      lwr      upr    p adj
scr-ER -10.69462 -22.3948 1.005561 0.068615

>
> #with interaction terms
> anova2 <- aov(formula = exp ~ S + T + S:T, data = Data)
```

```
> summary(anova2)
            Df Sum Sq Mean Sq F value   Pr(>F)    
S            1    343     343   6.548   0.0337 *  
T            1  11192   11192 213.584 4.71e-07 ***
S:T          1    303     303   5.784   0.0428 *  
Residuals    8    419      52                     
---
Signif. codes:  0 ‘***’ 0.001 ‘**’ 0.01 ‘*’ 0.05 ‘.’ 0.1 ‘ ’ 1
> TukeyHSD(anova2)
  Tukey multiple comparisons of means
    95% family-wise confidence level

Fit: aov(formula = exp ~ S + T + S:T, data = Data)

$S
            diff       lwr       upr     p adj
scr-ER -10.69462 -20.33214 -1.057097 0.0337025

$T
             diff       lwr       upr p adj
veh-kar -61.07864 -70.71616 -51.44112 5e-07

$`S:T`
                       diff       lwr       upr     p adj
scr:kar-ER:kar  -20.7457479 -39.67308  -1.81842 0.0325182
ER:veh-ER:kar   -71.1297693 -90.05710 -52.20244 0.0000099
scr:veh-ER:kar  -71.7732572 -90.70058 -52.84593 0.0000092
ER:veh-scr:kar  -50.3840213 -69.31135 -31.45669 0.0001275
scr:veh-scr:kar -51.0275092 -69.95484 -32.10018 0.0001163
scr:veh-ER:veh   -0.6434879 -19.57082  18.28384 0.9994976

>
```

SLC7A5

```
> Data <- read.table(file.choose(), header=TRUE, sep="\t")
> Data
     T   S       exp
1  veh scr 1.0000000
2  veh scr 0.6567123
3  veh scr 0.8908987
4  kar scr 5.1218559
5  kar scr 4.4382779
6  kar scr 5.2174083
7  veh  ER 0.7038468
8  veh  ER 0.9482460
9  veh  ER 0.7827734
10 kar  ER 2.6329254
11 kar  ER 1.7694897
12 kar  ER 2.7830497
> summary(Data)
```

```
      T                  S                  exp
 Length:12          Length:12          Min.   :0.6567
 Class :character   Class :character   1st Qu.:0.8639
 Mode  :character   Mode  :character   Median :1.3847
                                       Mean   :2.2455
                                       3rd Qu.:3.1969
                                       Max.   :5.2174
> Data$T <- as.factor(Data$T)
> Data$S <- as.factor(Data$S)
> str(Data)
'data.frame':   12 obs. of  3 variables:
 $ T  : Factor w/ 2 levels "kar","veh": 2 2 2 1 1 1 2 2 2 1 ...
 $ S  : Factor w/ 2 levels "ER","scr": 2 2 2 2 2 2 1 1 1 1 ...
 $ exp: num  1 0.657 0.891 5.122 4.438 ...
>
> #without interaction term
> anova1 <- aov(formula = exp ~ T + S, data = Data)
> summary(anova1)
            Df Sum Sq Mean Sq F value   Pr(>F)
T            1 24.028  24.028  37.847 0.000168 ***
S            1  4.947   4.947   7.792 0.021005 *
Residuals    9  5.714   0.635
---
Signif. codes:  0 ‘***’ 0.001 ‘**’ 0.01 ‘*’ 0.05 ‘.’ 0.1 ‘ ’ 1
> TukeyHSD(anova1)
  Tukey multiple comparisons of means
    95% family-wise confidence level

Fit: aov(formula = exp ~ T + S, data = Data)

$T
             diff       lwr       upr     p adj
veh-kar -2.830088 -3.870749 -1.789428 0.0001683

$S
           diff       lwr      upr     p adj
scr-ER 1.284137 0.2434763 2.324798 0.0210051

>
> #with interaction terms
> anova2 <- aov(formula = exp ~ S + T + S:T, data = Data)
> summary(anova2)
            Df Sum Sq Mean Sq F value   Pr(>F)
S            1  4.947   4.947   37.61 0.000279 ***
T            1 24.028  24.028  182.68 8.62e-07 ***
S:T          1  4.662   4.662   35.44 0.000341 ***
Residuals    8  1.052   0.132
---
Signif. codes:  0 ‘***’ 0.001 ‘**’ 0.01 ‘*’ 0.05 ‘.’ 0.1 ‘ ’ 1
> TukeyHSD(anova2)
  Tukey multiple comparisons of means
    95% family-wise confidence level

Fit: aov(formula = exp ~ S + T + S:T, data = Data)
```

```
$S
           diff       lwr      upr     p adj
scr-ER 1.284137 0.8012851 1.766989 0.0002792

$T
             diff      lwr       upr p adj
veh-kar -2.830088 -3.31294 -2.347236 9e-07

$`S:T`
                       diff        lwr        upr     p adj
scr:kar-ER:kar   2.53069243  1.5824095  3.4789754 0.0001252
ER:veh-ER:kar   -1.58353285 -2.5318158 -0.6352499 0.0030503
scr:veh-ER:kar  -1.54595126 -2.4942342 -0.5976683 0.0035446
ER:veh-scr:kar  -4.11422528 -5.0625082 -3.1659423 0.0000033
scr:veh-scr:kar -4.07664369 -5.0249266 -3.1283608 0.0000036
scr:veh-ER:veh   0.03758159 -0.9107013  0.9858645 0.9992060

>
```

TIPARP

```
> Data <- read.table(file.choose(), header=TRUE, sep="\t")
> Data
     T   S       exp
1  Veh scr 1.0000000
2  Veh scr 0.8970954
3  Veh scr 0.7006018
4  Kar scr 6.7739625
5  Kar scr 5.2294770
6  Kar scr 5.1813690
7  Veh  ER 1.1905817
8  Veh  ER 0.5717012
9  Veh  ER 0.7405488
10 Kar  ER 3.7321320
11 Kar  ER 3.3558231
12 Kar  ER 5.5918797
> summary(Data)
      T                  S                  exp
 Length:12          Length:12          Min.   :0.5717
 Class :character   Class :character   1st Qu.:0.8580
 Mode  :character   Mode  :character   Median :2.2732
                                       Mean   :2.9138
                                       3rd Qu.:5.1934
                                       Max.   :6.7740
> Data$T <- as.factor(Data$T)
> Data$S <- as.factor(Data$S)
> str(Data)
'data.frame':   12 obs. of  3 variables:
 $ T  : Factor w/ 2 levels "Kar","Veh": 2 2 2 1 1 1 2 2 2 1 ...
 $ S  : Factor w/ 2 levels "ER","scr": 2 2 2 2 2 2 1 1 1 1 ...
 $ exp: num  1 0.897 0.701 6.774 5.229 ...
>
> #without interaction term
```

```
> anova1 <- aov(formula = exp ~ T + S, data = Data)
> summary(anova1)
            Df Sum Sq Mean Sq F value   Pr(>F)    
T            1  51.11   51.11  72.094 1.37e-05 ***
S            1   1.76    1.76   2.487    0.149    
Residuals    9   6.38    0.71                     
---
Signif. codes:  0 ‘***’ 0.001 ‘**’ 0.01 ‘*’ 0.05 ‘.’ 0.1 ‘ ’ 1
> TukeyHSD(anova1)
  Tukey multiple comparisons of means
    95% family-wise confidence level

Fit: aov(formula = exp ~ T + S, data = Data)

$T
             diff       lwr       upr    p adj
Veh-Kar -4.127352 -5.226979 -3.027726 1.37e-05

$S
            diff        lwr      upr     p adj
scr-ER 0.7666399 -0.3329869 1.866267 0.1492194

> 
> #with interaction terms
> anova2 <- aov(formula = exp ~ S + T + S:T, data = Data)
> summary(anova2)
            Df Sum Sq Mean Sq F value   Pr(>F)    
S            1   1.76    1.76   2.964    0.123    
T            1  51.11   51.11  85.908 1.49e-05 ***
S:T          1   1.62    1.62   2.724    0.137    
Residuals    8   4.76    0.59                     
---
Signif. codes:  0 ‘***’ 0.001 ‘**’ 0.01 ‘*’ 0.05 ‘.’ 0.1 ‘ ’ 1
> TukeyHSD(anova2)
  Tukey multiple comparisons of means
    95% family-wise confidence level

Fit: aov(formula = exp ~ S + T + S:T, data = Data)

$S
            diff        lwr     upr     p adj
scr-ER 0.7666399 -0.2602304 1.79351 0.1234429

$T
             diff       lwr       upr    p adj
Veh-Kar -4.127352 -5.154223 -3.100482 1.49e-05

$`S:T`
                        diff       lwr       upr     p adj
scr:Kar-ER:Kar   1.50165790 -0.515034  3.518350 0.1577251
ER:Veh-ER:Kar   -3.39233435 -5.409026 -1.375642 0.0029134
scr:Veh-ER:Kar  -3.36071252 -5.377404 -1.344021 0.0030902
ER:Veh-scr:Kar  -4.89399225 -6.910684 -2.877300 0.0002473
scr:Veh-scr:Kar -4.86237042 -6.879062 -2.845679 0.0002589
```

```
scr:Veh-ER:Veh   0.03162183 -1.985070  2.048314 0.9999505

>

STC2

> Data <- read.table(file.choose(), header=TRUE, sep="\t")
> Data
     T   S       exp
1  veh scr 1.0000000
2  veh scr 0.5946036
3  veh scr 0.5946036
4  kar scr 2.5432383
5  kar scr 2.3240912
6  kar scr 1.9724654
7  veh  ER 0.5509526
8  veh  ER 0.6402320
9  veh  ER 0.6372803
10 kar  ER 0.9416960
11 kar  ER 0.8293195
12 kar  ER 1.0092848
> summary(Data)
      T                  S                  exp
 Length:12          Length:12          Min.   :0.5510
 Class :character   Class :character   1st Qu.:0.6266
 Mode  :character   Mode  :character   Median :0.8855
                                       Mean   :1.1365
                                       3rd Qu.:1.2501
                                       Max.   :2.5432
> Data$T <- as.factor(Data$T)
> Data$S <- as.factor(Data$S)
> str(Data)
'data.frame':   12 obs. of  3 variables:
 $ T  : Factor w/ 2 levels "kar","veh": 2 2 2 1 1 1 2 2 2 1 ...
 $ S  : Factor w/ 2 levels "ER","scr": 2 2 2 2 2 2 1 1 1 1 ...
 $ exp: num  1 0.595 0.595 2.543 2.324 ...
>
> #without interaction term
> anova1 <- aov(formula = exp ~ T + S, data = Data)
> summary(anova1)
            Df Sum Sq Mean Sq F value Pr(>F)
T            1  2.616  2.6156   16.38 0.0029 **
S            1  1.628  1.6282   10.20 0.0109 *
Residuals    9  1.437  0.1597
---
Signif. codes:  0 ‘***’ 0.001 ‘**’ 0.01 ‘*’ 0.05 ‘.’ 0.1 ‘ ’ 1
> TukeyHSD(anova1)
  Tukey multiple comparisons of means
    95% family-wise confidence level

Fit: aov(formula = exp ~ T + S, data = Data)

$T
              diff       lwr        upr     p adj
```

```
veh-kar -0.9337372 -1.455637 -0.4118378 0.0028968

$S
            diff       lwr      upr     p adj
scr-ER 0.7367061 0.2148067 1.258606 0.0109496

>
> #with interaction terms
> anova2 <- aov(formula = exp ~ S + T + S:T, data = Data)
> summary(anova2)
            Df Sum Sq Mean Sq F value   Pr(>F)
S            1 1.6282  1.6282   43.85 0.000166 ***
T            1 2.6156  2.6156   70.44 3.09e-05 ***
S:T          1 1.1401  1.1401   30.70 0.000547 ***
Residuals    8 0.2971  0.0371
---
Signif. codes:  0 ‘***’ 0.001 ‘**’ 0.01 ‘*’ 0.05 ‘.’ 0.1 ‘ ’ 1
> TukeyHSD(anova2)
  Tukey multiple comparisons of means
    95% family-wise confidence level

Fit: aov(formula = exp ~ S + T + S:T, data = Data)

$S
            diff       lwr       upr     p adj
scr-ER 0.7367061 0.4801568 0.9932555 0.0001656

$T
              diff       lwr        upr    p adj
veh-kar -0.9337372 -1.190287 -0.6771879 3.09e-05

$`S:T`
                     diff        lwr        upr     p adj
scr:kar-ER:kar   1.3531649  0.8493223  1.8570074 0.0001196
ER:veh-ER:kar   -0.3172785 -0.8211211  0.1865641 0.2584424
scr:veh-ER:kar  -0.1970311 -0.7008736  0.3068115 0.6143320
ER:veh-scr:kar  -1.6704434 -2.1742859 -1.1666008 0.0000253
scr:veh-scr:kar -1.5501959 -2.0540385 -1.0463534 0.0000442
scr:veh-ER:veh   0.1202474 -0.3835951  0.6240900 0.8682250

>
```
