## Supplementary material for "Global transcriptome analysis reveals partial estrogen-like effects of karanjin in MCF-7 breast cancer cells": All supplementary material: Supplementary data 5.pdf

**Table 1:** Quality measures of RNA samples used for RNA-seq

| Sr.No | Sample Name | Conc (ng/µl) | Final Volume (µl) | Total Amount (µg) | RIN Value | rRNA Ratio |
|---|---|---|---|---|---|---|
| 1 | G1 | 381.066 | 19 | 7.24 | 9.6 | 1.9 |
| 2 | G2 | 311.152 | 21 | 6.534 | 9.9 | 2.2 |
| 3 | G3 | 511.198 | 19 | 9.713 | 9.8 | 2.4 |
| 4 | K1 | 438.895 | 19 | 8.339 | 9.9 | 2.1 |
| 5 | K2 | 383.067 | 21 | 8.044 | 9.8 | 2.2 |
| 6 | K3 | 421.955 | 21 | 8.861 | 10 | 2.1 |

G1, G2, G3- three biological replicates of total RNA from MCF-7 cells treated with vehicle (DMSO)
K1, K2, K3- three biological replicates of total RNA from MCF-7 cells treated with 10 µM karanjin.

**Table 2:** Sequencing read statistics of raw data using Illumina Novaseq 6000.

| Sr.No | Sample Name | Total read bases[@] | Total reads[@] | GC(%) | AT(%) | Q20(%) | Q30(%) |
|---|---|---|---|---|---|---|---|
| 1 | G1 | 3845557830 | 38074830 | 50.26 | 49.74 | 98.39 | 95.34 |
| 2 | G2 | 3741880522 | 37048322 | 49.97 | 50.03 | 98.3 | 95.12 |
| 3 | G3 | 4551822752 | 45067552 | 50.35 | 49.65 | 98.44 | 95.45 |
| 4 | K1 | 3896588484 | 38580084 | 51.16 | 48.84 | 98.5 | 95.64 |
| 5 | K2 | 4301862094 | 42592694 | 49.84 | 50.16 | 98.51 | 95.55 |
| 6 | K3 | 4175723396 | 41343796 | 50.19 | 49.81 | 98.37 | 95.29 |

[@]paired-end reads
G1, G2, G3- correspond to samples from MCF-7 cells treated with vehicle (DMSO)
K1, K2, K3- correspond to samples from MCF-7 cells treated with 10 µMkaranjin.

**Table 3:** Read statistics before and after trimming.

| Sr.No | Sample | File Names (paired-end) | Total Reads | | % data lost |
|---|---|---|---|---|---|
| | | | Before trimming | After trimming | |
| 1 | G1 | G1_1 | 19037415 | 18876043 | 0.8477 |
| 2 | | G1_2 | 19037415 | 18876043 | |
| 3 | G2 | G2_1 | 18524161 | 18384899 | 0.7518 |
| 4 | | G2_2 | 18524161 | 18384899 | |
| 5 | G3 | G3_1 | 22533776 | 22366336 | 0.7431 |
| 6 | | G3_2 | 22533776 | 22366336 | |
| 7 | K1 | K1_1 | 19290042 | 19149767 | 0.7272 |
| 8 | | K1_2 | 19290042 | 19149767 | |
| 9 | K2 | K2_1 | 21296347 | 21128011 | 0.7904 |
| 10 | | K2_2 | 21296347 | 21128011 | |
| 11 | K3 | K3_1 | 20671898 | 20501108 | 0.8262 |
| 12 | | K3_2 | 20671898 | 20501108 | |

G1, G2, G3- correspond to samples from MCF-7 cells treated with vehicle (DMSO)
K1, K2, K3- correspond to samples from MCF-7 cells treated with 10 µM karanjin.
