## Supplementary material for "Global transcriptome analysis reveals partial estrogen-like effects of karanjin in MCF-7 breast cancer cells": All supplementary material: Supplementary data 6.pdf

| Gene ID | G1 | G2 | G3 | K1 | K2 | K3 |
|---|---|---|---|---|---|---|
| ENSG00000223972 | 0 | 0 | 0 | 1 | 0 | 0 |
| ENSG00000227232 | 34 | 27 | 42 | 34 | 27 | 28 |
| ENSG00000278267 | 21 | 4 | 6 | 6 | 1 | 6 |
| ENSG00000243485 | 0 | 0 | 0 | 0 | 0 | 0 |
| ENSG00000284332 | 0 | 0 | 0 | 0 | 0 | 0 |
| ENSG00000237613 | 0 | 0 | 0 | 0 | 0 | 0 |
| ENSG00000268020 | 0 | 0 | 0 | 0 | 0 | 0 |
| ENSG00000240361 | 0 | 0 | 0 | 0 | 0 | 0 |
| ENSG00000186092 | 0 | 0 | 0 | 0 | 0 | 0 |
| ENSG00000238009 | 0 | 0 | 0 | 2 | 1 | 0 |
| ENSG00000239945 | 0 | 0 | 0 | 0 | 0 | 0 |
| ENSG00000233750 | 6 | 6 | 8 | 3 | 4 | 4 |
| ENSG00000268903 | 156 | 73 | 128 | 78 | 90 | 86 |
| ENSG00000269981 | 72 | 38 | 63 | 48 | 45 | 56 |
| ENSG00000239906 | 2 | 0 | 0 | 0 | 0 | 0 |
| ENSG00000241860 | 8 | 3 | 12 | 2 | 3 | 10 |
| ENSG00000222623 | 0 | 0 | 0 | 0 | 0 | 0 |
| ENSG00000241599 | 0 | 0 | 0 | 0 | 0 | 0 |
| ENSG00000279928 | 0 | 0 | 0 | 0 | 0 | 0 |
| ENSG00000279457 | 128 | 71 | 84 | 84 | 74 | 78 |
| ENSG00000273874 | 1 | 0 | 0 | 0 | 1 | 0 |
| ENSG00000228463 | 22 | 11 | 12 | 8 | 10 | 8 |
| ENSG00000236679 | 1 | 0 | 0 | 0 | 0 | 0 |
| ENSG00000236601 | 34 | 14 | 28 | 31 | 31 | 19 |
| ENSG00000237094 | 29 | 15 | 37 | 9 | 14 | 14 |
| ENSG00000269732 | 0 | 0 | 0 | 0 | 0 | 0 |
| ENSG00000284733 | 0 | 0 | 0 | 0 | 0 | 0 |
| ENSG00000233653 | 0 | 0 | 0 | 0 | 0 | 0 |
| ENSG00000250575 | 1 | 0 | 0 | 0 | 0 | 0 |
| ENSG00000278757 | 1 | 0 | 2 | 0 | 1 | 0 |
| ENSG00000230021 | 26 | 3 | 10 | 3 | 9 | 4 |
| ENSG00000235146 | 0 | 0 | 0 | 0 | 1 | 0 |
| ENSG00000225972 | 4 | 2 | 1 | 2 | 0 | 2 |
| ENSG00000225630 | 2663 | 1401 | 1411 | 1354 | 1392 | 1429 |
| ENSG00000237973 | 1836 | 880 | 1213 | 976 | 1163 | 1340 |
| ENSG00000278791 | 0 | 0 | 0 | 0 | 0 | 0 |
| ENSG00000229344 | 16 | 10 | 23 | 15 | 23 | 16 |
| ENSG00000240409 | 4 | 2 | 0 | 1 | 0 | 0 |
| ENSG00000248527 | 20979 | 10983 | 11517 | 11053 | 10493 | 10692 |
| ENSG00000198744 | 29 | 9 | 23 | 14 | 14 | 16 |
| ENSG00000268663 | 0 | 0 | 0 | 0 | 0 | 0 |
| ENSG00000284662 | 0 | 0 | 0 | 0 | 0 | 0 |
| ENSG00000229376 | 6 | 3 | 1 | 0 | 1 | 4 |
| ENSG00000228327 | 60 | 26 | 46 | 17 | 42 | 34 |
| ENSG00000223181 | 0 | 0 | 0 | 0 | 0 | 0 |
| ENSG00000229905 | 1 | 0 | 0 | 0 | 0 | 0 |
| ENSG00000237491 | 36 | 7 | 21 | 8 | 24 | 11 |
| ENSG00000285268 | 0 | 0 | 0 | 0 | 0 | 0 |
| ENSG00000230092 | 16 | 5 | 3 | 3 | 5 | 3 |

| | | | | | | |
|---|---|---|---|---|---|---|
| ENSG00000177757 | 3 | 3 | 7 | 4 | 2 | 6 |
| ENSG00000228794 | 252 | 114 | 143 | 117 | 133 | 144 |
| ENSG00000225880 | 12 | 5 | 14 | 11 | 12 | 8 |
| ENSG00000230368 | 0 | 0 | 0 | 0 | 0 | 0 |
| ENSG00000234711 | 2 | 0 | 0 | 0 | 0 | 1 |
| ENSG00000283040 | 0 | 0 | 0 | 0 | 0 | 0 |
| ENSG00000272438 | 17 | 3 | 5 | 1 | 4 | 7 |
| ENSG00000230699 | 60 | 16 | 21 | 23 | 21 | 26 |
| ENSG00000241180 | 0 | 0 | 0 | 0 | 0 | 0 |
| ENSG00000223764 | 2156 | 855 | 1144 | 913 | 932 | 842 |
| ENSG00000187634 | 1483 | 611 | 792 | 749 | 672 | 620 |
| ENSG00000188976 | 4196 | 1914 | 2530 | 2922 | 2459 | 2459 |
| ENSG00000187961 | 574 | 270 | 378 | 334 | 356 | 329 |
| ENSG00000187583 | 331 | 133 | 189 | 150 | 139 | 169 |
| ENSG00000187642 | 93 | 57 | 86 | 58 | 37 | 58 |
| ENSG00000272512 | 135 | 45 | 52 | 58 | 40 | 45 |
| ENSG00000188290 | 956 | 442 | 504 | 564 | 450 | 515 |
| ENSG00000187608 | 1622 | 766 | 954 | 1074 | 888 | 810 |
| ENSG00000231702 | 1 | 0 | 2 | 0 | 1 | 3 |
| ENSG00000224969 | 27 | 11 | 25 | 14 | 8 | 15 |
| ENSG00000188157 | 13441 | 6108 | 8565 | 7491 | 7072 | 6914 |
| ENSG00000242590 | 0 | 0 | 0 | 0 | 0 | 0 |
| ENSG00000217801 | 457 | 184 | 353 | 290 | 276 | 293 |
| ENSG00000273443 | 23 | 7 | 6 | 16 | 18 | 15 |
| ENSG00000237330 | 1072 | 565 | 601 | 813 | 794 | 726 |
| ENSG00000131591 | 841 | 365 | 558 | 536 | 453 | 507 |
| ENSG00000285812 | 16 | 10 | 9 | 1 | 16 | 8 |
| ENSG00000223823 | 12 | 5 | 8 | 3 | 8 | 1 |
| ENSG00000207730 | 3 | 3 | 8 | 2 | 7 | 2 |
| ENSG00000207607 | 4 | 1 | 1 | 1 | 6 | 5 |
| ENSG00000198976 | 8 | 3 | 10 | 11 | 10 | 10 |
| ENSG00000272141 | 111 | 52 | 77 | 65 | 42 | 57 |
| ENSG00000205231 | 0 | 5 | 0 | 0 | 1 | 0 |
| ENSG00000162571 | 8 | 6 | 8 | 7 | 2 | 5 |
| ENSG00000186891 | 209 | 113 | 119 | 120 | 86 | 114 |
| ENSG00000186827 | 0 | 0 | 0 | 2 | 0 | 1 |
| ENSG00000078808 | 5438 | 2464 | 3362 | 3176 | 2672 | 2761 |
| ENSG00000176022 | 1057 | 520 | 625 | 712 | 589 | 681 |
| ENSG00000184163 | 26 | 7 | 21 | 19 | 16 | 16 |
| ENSG00000260179 | 5 | 3 | 5 | 5 | 5 | 3 |
| ENSG00000160087 | 1721 | 828 | 1099 | 1140 | 968 | 1106 |
| ENSG00000230415 | 16 | 7 | 7 | 5 | 8 | 3 |
| ENSG00000162572 | 140 | 39 | 81 | 59 | 61 | 71 |
| ENSG00000131584 | 2388 | 877 | 1411 | 1067 | 1055 | 1007 |
| ENSG00000278073 | 0 | 0 | 0 | 0 | 0 | 0 |
| ENSG00000169972 | 479 | 300 | 350 | 412 | 371 | 342 |
| ENSG00000127054 | 2820 | 1370 | 1766 | 1614 | 1496 | 1621 |
| ENSG00000283712 | 0 | 0 | 0 | 0 | 0 | 0 |
| ENSG00000240731 | 102 | 51 | 85 | 49 | 57 | 83 |
| ENSG00000224051 | 859 | 371 | 502 | 594 | 515 | 482 |

| | | | | | | |
|---|---|---|---|---|---|---|
| ENSG00000169962 | 34 | 7 | 17 | 5 | 9 | 17 |
| ENSG00000107404 | 2208 | 1093 | 1393 | 1566 | 1437 | 1419 |
| ENSG00000284372 | 0 | 0 | 0 | 0 | 0 | 0 |
| ENSG00000162576 | 68 | 20 | 43 | 26 | 42 | 44 |
| ENSG00000175756 | 3658 | 1722 | 2017 | 2642 | 2050 | 2025 |
| ENSG00000223663 | 0 | 0 | 2 | 0 | 0 | 0 |
| ENSG00000221978 | 7114 | 2886 | 4741 | 2811 | 3501 | 3529 |
| ENSG00000224870 | 903 | 383 | 506 | 504 | 444 | 480 |
| ENSG00000242485 | 1990 | 1085 | 1289 | 1377 | 1103 | 1208 |
| ENSG00000264293 | 0 | 0 | 0 | 0 | 0 | 0 |
| ENSG00000272455 | 193 | 98 | 120 | 121 | 105 | 88 |
| ENSG00000235098 | 12 | 2 | 3 | 3 | 5 | 2 |
| ENSG00000225905 | 0 | 0 | 5 | 1 | 1 | 1 |
| ENSG00000205116 | 0 | 0 | 0 | 0 | 0 | 0 |
| ENSG00000225285 | 8 | 5 | 8 | 4 | 3 | 3 |
| ENSG00000179403 | 1123 | 537 | 837 | 657 | 529 | 582 |
| ENSG00000215915 | 19 | 11 | 23 | 6 | 12 | 5 |
| ENSG00000160072 | 596 | 268 | 378 | 389 | 381 | 396 |
| ENSG00000284740 | 11 | 1 | 7 | 3 | 1 | 3 |
| ENSG00000197785 | 972 | 533 | 636 | 680 | 612 | 645 |
| ENSG00000205090 | 6 | 6 | 6 | 4 | 8 | 9 |
| ENSG00000160075 | 3700 | 1760 | 2290 | 2447 | 2083 | 2079 |
| ENSG00000215014 | 20 | 9 | 11 | 7 | 10 | 7 |
| ENSG00000274481 | 0 | 0 | 0 | 0 | 0 | 0 |
| ENSG00000228594 | 1288 | 600 | 823 | 991 | 629 | 813 |
| ENSG00000272106 | 268 | 154 | 212 | 159 | 151 | 160 |
| ENSG00000197530 | 858 | 407 | 491 | 536 | 427 | 415 |
| ENSG00000189409 | 7 | 2 | 1 | 0 | 4 | 1 |
| ENSG00000248333 | 1236 | 559 | 736 | 634 | 674 | 662 |
| ENSG00000272004 | 0 | 0 | 5 | 0 | 0 | 0 |
| ENSG00000189339 | 2369 | 1166 | 1575 | 1076 | 1219 | 1348 |
| ENSG00000269737 | 34 | 14 | 18 | 8 | 18 | 15 |
| ENSG00000215914 | 9 | 1 | 3 | 2 | 0 | 1 |
| ENSG00000008128 | 59 | 45 | 69 | 48 | 77 | 54 |
| ENSG00000268575 | 125 | 100 | 122 | 60 | 93 | 98 |
| ENSG00000227775 | 18 | 9 | 12 | 8 | 11 | 13 |
| ENSG00000215790 | 695 | 333 | 491 | 328 | 376 | 388 |
| ENSG00000008130 | 3974 | 1931 | 2411 | 2744 | 2985 | 2910 |
| ENSG00000078369 | 14458 | 7392 | 9262 | 7379 | 8399 | 8361 |
| ENSG00000231050 | 3 | 4 | 2 | 2 | 3 | 2 |
| ENSG00000169885 | 38 | 24 | 40 | 18 | 27 | 44 |
| ENSG00000178821 | 110 | 44 | 64 | 98 | 64 | 75 |
| ENSG00000142609 | 16 | 7 | 11 | 8 | 5 | 5 |
| ENSG00000233542 | 0 | 0 | 0 | 0 | 0 | 0 |
| ENSG00000187730 | 52 | 18 | 40 | 34 | 28 | 22 |
| ENSG00000226969 | 1 | 0 | 1 | 2 | 1 | 2 |
| ENSG00000067606 | 949 | 529 | 688 | 589 | 603 | 582 |
| ENSG00000271806 | 6 | 0 | 1 | 2 | 1 | 1 |
| ENSG00000182873 | 94 | 36 | 48 | 50 | 54 | 46 |
| ENSG00000162585 | 964 | 515 | 630 | 683 | 569 | 565 |

| | | | | | | |
|---|---|---|---|---|---|---|
| ENSG00000234396 | 0 | 0 | 0 | 0 | 0 | 0 |
| ENSG00000157933 | 1649 | 709 | 921 | 811 | 801 | 896 |
| ENSG00000116151 | 267 | 135 | 154 | 107 | 120 | 125 |
| ENSG00000272161 | 0 | 1 | 0 | 1 | 0 | 0 |
| ENSG00000269896 | 81 | 32 | 38 | 36 | 26 | 35 |
| ENSG00000272420 | 15 | 5 | 11 | 12 | 3 | 10 |
| ENSG00000157916 | 4191 | 1918 | 2564 | 2246 | 2321 | 2226 |
| ENSG00000157911 | 758 | 428 | 467 | 524 | 443 | 507 |
| ENSG00000149527 | 57 | 14 | 26 | 24 | 17 | 22 |
| ENSG00000224387 | 0 | 0 | 2 | 0 | 0 | 0 |
| ENSG00000229393 | 2 | 0 | 1 | 0 | 0 | 0 |
| ENSG00000157881 | 768 | 392 | 539 | 517 | 510 | 496 |
| ENSG00000197921 | 0 | 0 | 0 | 0 | 0 | 1 |
| ENSG00000272449 | 239 | 109 | 136 | 187 | 186 | 164 |
| ENSG00000238164 | 0 | 0 | 1 | 3 | 2 | 1 |
| ENSG00000157873 | 0 | 1 | 0 | 0 | 1 | 1 |
| ENSG00000225931 | 0 | 0 | 0 | 1 | 2 | 1 |
| ENSG00000228037 | 6 | 2 | 2 | 1 | 1 | 2 |
| ENSG00000157870 | 1555 | 704 | 1068 | 970 | 733 | 926 |
| ENSG00000142606 | 120 | 50 | 81 | 50 | 49 | 66 |
| ENSG00000237058 | 0 | 0 | 0 | 0 | 0 | 0 |
| ENSG00000215912 | 19 | 9 | 18 | 5 | 5 | 12 |
| ENSG00000285945 | 0 | 0 | 0 | 0 | 0 | 0 |
| ENSG00000283259 | 0 | 0 | 0 | 0 | 0 | 0 |
| ENSG00000233234 | 0 | 0 | 0 | 0 | 0 | 0 |
| ENSG00000231630 | 0 | 0 | 0 | 0 | 0 | 0 |
| ENSG00000284745 | 0 | 0 | 0 | 0 | 0 | 0 |
| ENSG00000169717 | 0 | 0 | 0 | 0 | 0 | 0 |
| ENSG00000177133 | 34 | 20 | 21 | 10 | 4 | 12 |
| ENSG00000142611 | 154 | 51 | 104 | 49 | 53 | 72 |
| ENSG00000283572 | 0 | 0 | 0 | 0 | 0 | 1 |
| ENSG00000226286 | 0 | 4 | 3 | 3 | 1 | 6 |
| ENSG00000279839 | 0 | 0 | 0 | 0 | 0 | 0 |
| ENSG00000272235 | 0 | 0 | 0 | 0 | 0 | 0 |
| ENSG00000130762 | 2861 | 1331 | 1913 | 1715 | 1623 | 1678 |
| ENSG00000272088 | 6 | 3 | 5 | 2 | 3 | 7 |
| ENSG00000162591 | 783 | 336 | 498 | 325 | 341 | 365 |
| ENSG00000207776 | 0 | 0 | 0 | 0 | 0 | 0 |
| ENSG00000238260 | 2 | 3 | 12 | 12 | 10 | 4 |
| ENSG00000158109 | 2230 | 1106 | 1273 | 1260 | 1178 | 1290 |
| ENSG00000116213 | 1091 | 440 | 595 | 633 | 625 | 592 |
| ENSG00000078900 | 211 | 114 | 141 | 87 | 88 | 99 |
| ENSG00000227589 | 0 | 0 | 0 | 0 | 0 | 0 |
| ENSG00000235131 | 0 | 0 | 0 | 0 | 0 | 0 |
| ENSG00000227372 | 0 | 0 | 0 | 0 | 0 | 0 |
| ENSG00000276189 | 0 | 0 | 0 | 0 | 0 | 0 |
| ENSG00000162592 | 4 | 5 | 5 | 6 | 1 | 6 |
| ENSG00000235169 | 113 | 44 | 58 | 46 | 44 | 56 |
| ENSG00000130764 | 2143 | 1069 | 1474 | 1339 | 1216 | 1204 |
| ENSG00000266075 | 10 | 4 | 4 | 2 | 8 | 3 |

| | | | | | | |
|---|---|---|---|---|---|---|
| ENSG00000272153 | 36 | 22 | 19 | 17 | 25 | 21 |
| ENSG00000116198 | 1482 | 726 | 898 | 676 | 845 | 828 |
| ENSG00000169598 | 319 | 137 | 186 | 143 | 168 | 157 |
| ENSG00000198912 | 523 | 246 | 295 | 225 | 352 | 252 |
| ENSG00000236423 | 27 | 10 | 8 | 14 | 9 | 11 |
| ENSG00000233304 | 0 | 0 | 0 | 0 | 0 | 0 |
| ENSG00000226374 | 0 | 0 | 0 | 0 | 0 | 0 |
| ENSG00000284668 | 0 | 0 | 0 | 0 | 0 | 0 |
| ENSG00000284703 | 0 | 0 | 0 | 0 | 0 | 0 |
| ENSG00000229280 | 0 | 0 | 0 | 0 | 0 | 0 |
| ENSG00000235054 | 0 | 0 | 0 | 0 | 0 | 0 |
| ENSG00000284694 | 0 | 0 | 0 | 0 | 0 | 0 |
| ENSG00000227169 | 0 | 0 | 0 | 0 | 0 | 0 |
| ENSG00000232596 | 0 | 0 | 0 | 0 | 0 | 0 |
| ENSG00000196581 | 11 | 9 | 5 | 2 | 5 | 4 |
| ENSG00000284739 | 0 | 0 | 0 | 0 | 0 | 0 |
| ENSG00000284674 | 0 | 0 | 0 | 0 | 0 | 0 |
| ENSG00000231510 | 0 | 0 | 0 | 0 | 0 | 0 |
| ENSG00000284616 | 0 | 0 | 0 | 0 | 0 | 0 |
| ENSG00000284692 | 0 | 0 | 0 | 0 | 0 | 0 |
| ENSG00000284666 | 0 | 0 | 0 | 0 | 0 | 0 |
| ENSG00000260972 | 0 | 0 | 0 | 0 | 0 | 0 |
| ENSG00000283356 | 0 | 0 | 0 | 0 | 0 | 0 |
| ENSG00000236948 | 0 | 0 | 0 | 0 | 0 | 0 |
| ENSG00000264341 | 0 | 0 | 0 | 0 | 0 | 0 |
| ENSG00000264101 | 0 | 0 | 0 | 0 | 0 | 0 |
| ENSG00000131697 | 603 | 227 | 360 | 292 | 294 | 319 |
| ENSG00000069424 | 787 | 386 | 468 | 442 | 504 | 467 |
| ENSG00000116254 | 217 | 84 | 131 | 97 | 81 | 81 |
| ENSG00000285629 | 0 | 0 | 0 | 0 | 1 | 0 |
| ENSG00000116251 | 2785 | 1583 | 1842 | 2007 | 2122 | 2050 |
| ENSG00000226944 | 18 | 15 | 18 | 13 | 8 | 9 |
| ENSG00000158286 | 538 | 237 | 348 | 166 | 194 | 213 |
| ENSG00000116237 | 4186 | 2163 | 2687 | 2328 | 2203 | 2499 |
| ENSG00000225077 | 26 | 26 | 27 | 16 | 20 | 34 |
| ENSG00000173673 | 0 | 0 | 0 | 0 | 0 | 0 |
| ENSG00000158292 | 668 | 295 | 360 | 413 | 371 | 321 |
| ENSG00000097021 | 1761 | 808 | 1061 | 1225 | 979 | 1006 |
| ENSG00000271746 | 0 | 0 | 0 | 0 | 0 | 0 |
| ENSG00000069812 | 128 | 83 | 92 | 81 | 108 | 65 |
| ENSG00000187017 | 1596 | 716 | 1045 | 1333 | 1247 | 1159 |
| ENSG00000265392 | 0 | 0 | 0 | 0 | 1 | 0 |
| ENSG00000231868 | 12 | 3 | 12 | 11 | 8 | 13 |
| ENSG00000215788 | 314 | 136 | 244 | 170 | 168 | 204 |
| ENSG00000171680 | 466 | 182 | 314 | 219 | 255 | 239 |
| ENSG00000162408 | 758 | 446 | 471 | 386 | 418 | 481 |
| ENSG00000253022 | 1 | 1 | 0 | 0 | 0 | 0 |
| ENSG00000229519 | 2 | 0 | 5 | 1 | 3 | 3 |
| ENSG00000173662 | 2 | 0 | 4 | 1 | 4 | 1 |
| ENSG00000204859 | 672 | 299 | 368 | 368 | 325 | 317 |

| | | | | | | |
|---|---|---|---|---|---|---|
| ENSG00000162413 | 1362 | 599 | 784 | 708 | 709 | 711 |
| ENSG00000116273 | 1401 | 649 | 877 | 709 | 662 | 736 |
| ENSG00000041988 | 438 | 207 | 223 | 231 | 245 | 198 |
| ENSG00000007923 | 1368 | 608 | 800 | 741 | 776 | 902 |
| ENSG00000228750 | 0 | 0 | 0 | 0 | 0 | 0 |
| ENSG00000284744 | 0 | 0 | 0 | 0 | 0 | 0 |
| ENSG00000237436 | 26 | 9 | 9 | 14 | 15 | 15 |
| ENSG00000171735 | 1076 | 587 | 684 | 634 | 732 | 660 |
| ENSG00000227950 | 0 | 0 | 0 | 0 | 0 | 0 |
| ENSG00000237365 | 0 | 0 | 0 | 0 | 0 | 0 |
| ENSG00000207056 | 0 | 0 | 0 | 0 | 0 | 0 |
| ENSG00000237402 | 0 | 0 | 0 | 0 | 0 | 1 |
| ENSG00000237728 | 0 | 0 | 0 | 0 | 0 | 0 |
| ENSG00000225126 | 0 | 0 | 0 | 0 | 1 | 0 |
| ENSG00000270171 | 4 | 5 | 7 | 3 | 2 | 1 |
| ENSG00000270035 | 12 | 6 | 18 | 6 | 6 | 8 |
| ENSG00000269978 | 36 | 14 | 22 | 14 | 28 | 16 |
| ENSG00000049245 | 3204 | 1603 | 1948 | 1529 | 1829 | 1731 |
| ENSG00000269925 | 18 | 5 | 6 | 4 | 6 | 3 |
| ENSG00000049246 | 321 | 177 | 256 | 100 | 141 | 157 |
| ENSG00000236266 | 0 | 2 | 0 | 0 | 0 | 1 |
| ENSG00000049247 | 0 | 0 | 0 | 0 | 0 | 0 |
| ENSG00000049249 | 0 | 0 | 1 | 0 | 1 | 1 |
| ENSG00000270330 | 0 | 0 | 0 | 0 | 0 | 0 |
| ENSG00000116288 | 6300 | 3366 | 3963 | 4091 | 3766 | 3830 |
| ENSG00000200344 | 2 | 1 | 0 | 1 | 1 | 2 |
| ENSG00000284747 | 12 | 2 | 6 | 6 | 8 | 8 |
| ENSG00000284716 | 2 | 0 | 1 | 0 | 0 | 1 |
| ENSG00000116285 | 1317 | 582 | 713 | 484 | 709 | 641 |
| ENSG00000238290 | 1 | 0 | 0 | 2 | 0 | 0 |
| ENSG00000232848 | 0 | 0 | 0 | 0 | 0 | 0 |
| ENSG00000229305 | 0 | 0 | 0 | 0 | 0 | 0 |
| ENSG00000227634 | 0 | 0 | 0 | 0 | 0 | 0 |
| ENSG00000200975 | 0 | 0 | 0 | 0 | 0 | 0 |
| ENSG00000233645 | 0 | 0 | 0 | 0 | 0 | 0 |
| ENSG00000251977 | 0 | 0 | 0 | 0 | 0 | 0 |
| ENSG00000162426 | 6 | 4 | 10 | 4 | 1 | 1 |
| ENSG00000252254 | 0 | 0 | 0 | 0 | 0 | 0 |
| ENSG00000142599 | 2129 | 902 | 1347 | 811 | 912 | 1028 |
| ENSG00000232912 | 4 | 0 | 0 | 0 | 0 | 1 |
| ENSG00000221083 | 0 | 0 | 0 | 0 | 0 | 0 |
| ENSG00000270282 | 0 | 0 | 0 | 0 | 0 | 0 |
| ENSG00000234619 | 2 | 0 | 0 | 0 | 1 | 0 |
| ENSG00000224315 | 0 | 0 | 1 | 0 | 0 | 0 |
| ENSG00000222511 | 0 | 0 | 0 | 0 | 0 | 0 |
| ENSG00000226545 | 0 | 0 | 3 | 0 | 0 | 0 |
| ENSG00000228423 | 0 | 0 | 0 | 0 | 0 | 0 |
| ENSG00000229716 | 0 | 0 | 0 | 0 | 0 | 0 |
| ENSG00000074800 | 60244 | 29483 | 39562 | 36163 | 31019 | 34175 |
| ENSG00000274258 | 0 | 0 | 0 | 0 | 0 | 2 |

| | | | | | | |
|---|---|---|---|---|---|---|
| ENSG00000230679 | 14 | 6 | 10 | 5 | 12 | 5 |
| ENSG00000201725 | 0 | 1 | 2 | 0 | 0 | 0 |
| ENSG00000238249 | 2 | 2 | 1 | 0 | 0 | 0 |
| ENSG00000232208 | 0 | 0 | 0 | 0 | 0 | 0 |
| ENSG00000232663 | 0 | 0 | 0 | 0 | 0 | 0 |
| ENSG00000131686 | 2 | 0 | 3 | 0 | 0 | 0 |
| ENSG00000265141 | 0 | 0 | 0 | 0 | 0 | 0 |
| ENSG00000197241 | 2 | 0 | 0 | 1 | 0 | 0 |
| ENSG00000142583 | 7 | 3 | 4 | 0 | 6 | 4 |
| ENSG00000252404 | 0 | 0 | 0 | 0 | 0 | 0 |
| ENSG00000180758 | 1757 | 882 | 1107 | 1082 | 1212 | 1300 |
| ENSG00000228526 | 492 | 233 | 362 | 225 | 311 | 278 |
| ENSG00000284357 | 0 | 0 | 0 | 0 | 0 | 0 |
| ENSG00000234546 | 80 | 24 | 33 | 42 | 38 | 28 |
| ENSG00000049239 | 1940 | 863 | 1067 | 893 | 981 | 1002 |
| ENSG00000171621 | 582 | 288 | 326 | 344 | 356 | 349 |
| ENSG00000284652 | 2 | 0 | 1 | 1 | 0 | 1 |
| ENSG00000284693 | 14 | 1 | 18 | 8 | 4 | 8 |
| ENSG00000252956 | 0 | 0 | 0 | 0 | 0 | 0 |
| ENSG00000235263 | 0 | 0 | 0 | 1 | 0 | 0 |
| ENSG00000171612 | 337 | 144 | 221 | 201 | 200 | 234 |
| ENSG00000231181 | 5 | 1 | 1 | 1 | 1 | 1 |
| ENSG00000188807 | 521 | 273 | 334 | 321 | 285 | 364 |
| ENSG00000171608 | 61 | 39 | 47 | 49 | 52 | 54 |
| ENSG00000179840 | 0 | 0 | 0 | 0 | 0 | 0 |
| ENSG00000233268 | 0 | 0 | 0 | 0 | 0 | 0 |
| ENSG00000231789 | 4 | 5 | 5 | 7 | 12 | 8 |
| ENSG00000171603 | 7042 | 3141 | 4307 | 3013 | 3123 | 3182 |
| ENSG00000271264 | 0 | 0 | 0 | 0 | 0 | 0 |
| ENSG00000280113 | 0 | 1 | 0 | 0 | 0 | 2 |
| ENSG00000178585 | 1262 | 702 | 809 | 927 | 833 | 779 |
| ENSG00000223989 | 0 | 0 | 0 | 0 | 0 | 0 |
| ENSG00000285701 | 0 | 1 | 1 | 0 | 1 | 0 |
| ENSG00000162441 | 744 | 407 | 480 | 374 | 477 | 438 |
| ENSG00000228150 | 18 | 13 | 16 | 8 | 9 | 10 |
| ENSG00000173614 | 555 | 293 | 325 | 272 | 357 | 335 |
| ENSG00000202415 | 5 | 1 | 0 | 1 | 1 | 2 |
| ENSG00000283611 | 0 | 1 | 1 | 0 | 0 | 0 |
| ENSG00000265521 | 2 | 0 | 0 | 0 | 0 | 0 |
| ENSG00000241326 | 0 | 0 | 0 | 0 | 0 | 0 |
| ENSG00000162444 | 82 | 49 | 44 | 85 | 77 | 66 |
| ENSG00000130939 | 2971 | 1297 | 1732 | 1164 | 1523 | 1527 |
| ENSG00000224340 | 0 | 1 | 0 | 0 | 0 | 0 |
| ENSG00000233623 | 0 | 0 | 1 | 1 | 1 | 1 |
| ENSG00000201746 | 1 | 0 | 2 | 0 | 0 | 0 |
| ENSG00000054523 | 2723 | 1221 | 1648 | 691 | 1405 | 1267 |
| ENSG00000264881 | 0 | 0 | 0 | 0 | 0 | 0 |
| ENSG00000199562 | 0 | 0 | 0 | 0 | 1 | 0 |
| ENSG00000264501 | 0 | 0 | 0 | 0 | 0 | 0 |
| ENSG00000284735 | 0 | 9 | 8 | 3 | 8 | 3 |

| | | | | | | |
|---|---|---|---|---|---|---|
| ENSG00000284642 | 15 | 5 | 9 | 7 | 8 | 11 |
| ENSG00000142657 | 13984 | 6576 | 8526 | 8760 | 8069 | 9037 |
| ENSG00000271989 | 0 | 1 | 1 | 0 | 1 | 1 |
| ENSG00000251503 | 0 | 1 | 2 | 2 | 3 | 0 |
| ENSG00000175279 | 63 | 22 | 30 | 38 | 39 | 37 |
| ENSG00000241563 | 0 | 3 | 2 | 0 | 6 | 0 |
| ENSG00000160049 | 1820 | 1019 | 1203 | 1180 | 1268 | 1399 |
| ENSG00000203469 | 0 | 0 | 0 | 0 | 0 | 0 |
| ENSG00000142655 | 799 | 358 | 537 | 464 | 354 | 453 |
| ENSG00000243267 | 0 | 2 | 1 | 1 | 0 | 1 |
| ENSG00000130940 | 731 | 283 | 401 | 222 | 265 | 259 |
| ENSG00000272078 | 5 | 8 | 7 | 5 | 8 | 8 |
| ENSG00000225158 | 0 | 0 | 0 | 0 | 0 | 0 |
| ENSG00000175262 | 0 | 1 | 2 | 2 | 0 | 1 |
| ENSG00000215785 | 0 | 0 | 0 | 1 | 0 | 1 |
| ENSG00000120948 | 2836 | 1629 | 1835 | 1381 | 1995 | 1734 |
| ENSG00000277726 | 0 | 1 | 0 | 0 | 0 | 1 |
| ENSG00000009724 | 21 | 13 | 22 | 9 | 12 | 15 |
| ENSG00000271895 | 25 | 9 | 19 | 6 | 12 | 13 |
| ENSG00000116649 | 1495 | 852 | 920 | 1446 | 1011 | 1118 |
| ENSG00000171824 | 1644 | 912 | 1162 | 738 | 1111 | 972 |
| ENSG00000226849 | 5 | 3 | 7 | 3 | 3 | 5 |
| ENSG00000230337 | 9 | 12 | 3 | 6 | 8 | 11 |
| ENSG00000198793 | 3332 | 1393 | 1954 | 1292 | 1720 | 1870 |
| ENSG00000225602 | 0 | 0 | 0 | 0 | 0 | 0 |
| ENSG00000253086 | 0 | 0 | 0 | 0 | 0 | 0 |
| ENSG00000171819 | 0 | 1 | 0 | 0 | 0 | 0 |
| ENSG00000207451 | 0 | 0 | 0 | 0 | 0 | 0 |
| ENSG00000238173 | 0 | 0 | 0 | 1 | 0 | 0 |
| ENSG00000120942 | 769 | 347 | 409 | 518 | 415 | 489 |
| ENSG00000238199 | 3 | 0 | 0 | 0 | 0 | 1 |
| ENSG00000284646 | 0 | 0 | 0 | 0 | 0 | 0 |
| ENSG00000224338 | 0 | 0 | 0 | 0 | 0 | 0 |
| ENSG00000204624 | 57 | 34 | 70 | 30 | 23 | 17 |
| ENSG00000285833 | 0 | 0 | 1 | 0 | 0 | 0 |
| ENSG00000235643 | 0 | 1 | 0 | 0 | 0 | 0 |
| ENSG00000284708 | 2 | 0 | 0 | 0 | 0 | 0 |
| ENSG00000116661 | 312 | 143 | 186 | 142 | 170 | 118 |
| ENSG00000132879 | 758 | 327 | 454 | 425 | 410 | 385 |
| ENSG00000116663 | 397 | 197 | 238 | 224 | 241 | 215 |
| ENSG00000116670 | 746 | 398 | 492 | 540 | 470 | 420 |
| ENSG00000162490 | 6 | 2 | 4 | 3 | 6 | 3 |
| ENSG00000177674 | 930 | 428 | 604 | 613 | 497 | 538 |
| ENSG00000215910 | 0 | 0 | 0 | 0 | 0 | 0 |
| ENSG00000177553 | 0 | 0 | 0 | 0 | 0 | 0 |
| ENSG00000177000 | 902 | 573 | 602 | 365 | 396 | 444 |
| ENSG00000011021 | 551 | 306 | 429 | 231 | 297 | 352 |
| ENSG00000276470 | 0 | 0 | 0 | 0 | 0 | 0 |
| ENSG00000278852 | 0 | 0 | 0 | 0 | 0 | 0 |
| ENSG00000175206 | 0 | 1 | 5 | 1 | 1 | 1 |

| | | | | | | |
|---|---|---|---|---|---|---|
| ENSG00000275915 | 0 | 0 | 0 | 0 | 0 | 0 |
| ENSG00000120937 | 0 | 0 | 0 | 0 | 0 | 0 |
| ENSG00000224904 | 0 | 0 | 0 | 0 | 0 | 0 |
| ENSG00000285646 | 10 | 3 | 6 | 1 | 5 | 3 |
| ENSG00000199347 | 0 | 0 | 0 | 0 | 0 | 0 |
| ENSG00000201801 | 0 | 0 | 0 | 0 | 0 | 0 |
| ENSG00000116685 | 2379 | 1180 | 1406 | 1495 | 1328 | 1347 |
| ENSG00000083444 | 2027 | 957 | 1096 | 1238 | 1016 | 1084 |
| ENSG00000116688 | 3406 | 1652 | 2045 | 1823 | 1887 | 1943 |
| ENSG00000270914 | 0 | 0 | 0 | 0 | 0 | 0 |
| ENSG00000116691 | 551 | 276 | 331 | 419 | 338 | 349 |
| ENSG00000202461 | 0 | 1 | 3 | 0 | 0 | 2 |
| ENSG00000276869 | 0 | 0 | 0 | 0 | 0 | 0 |
| ENSG00000243539 | 0 | 0 | 0 | 0 | 0 | 0 |
| ENSG00000120949 | 0 | 0 | 0 | 0 | 0 | 1 |
| ENSG00000201135 | 0 | 0 | 0 | 0 | 0 | 0 |
| ENSG00000227875 | 0 | 0 | 0 | 0 | 0 | 0 |
| ENSG00000285604 | 0 | 0 | 0 | 0 | 0 | 0 |
| ENSG00000283789 | 0 | 0 | 0 | 0 | 0 | 0 |
| ENSG00000028137 | 0 | 0 | 0 | 0 | 0 | 0 |
| ENSG00000263676 | 0 | 0 | 0 | 0 | 0 | 0 |
| ENSG00000225196 | 0 | 0 | 0 | 0 | 0 | 0 |
| ENSG00000252969 | 0 | 0 | 0 | 0 | 0 | 0 |
| ENSG00000048707 | 1729 | 842 | 902 | 370 | 995 | 869 |
| ENSG00000239149 | 4 | 2 | 0 | 0 | 0 | 0 |
| ENSG00000229484 | 0 | 0 | 0 | 0 | 0 | 0 |
| ENSG00000162496 | 336 | 158 | 171 | 98 | 111 | 134 |
| ENSG00000221340 | 2 | 0 | 0 | 1 | 0 | 0 |
| ENSG00000276830 | 0 | 1 | 0 | 0 | 0 | 0 |
| ENSG00000272482 | 0 | 2 | 1 | 0 | 0 | 1 |
| ENSG00000204518 | 0 | 0 | 0 | 0 | 0 | 0 |
| ENSG00000226166 | 0 | 0 | 0 | 0 | 0 | 0 |
| ENSG00000188984 | 0 | 0 | 0 | 0 | 0 | 0 |
| ENSG00000157330 | 0 | 0 | 0 | 0 | 0 | 0 |
| ENSG00000116726 | 0 | 0 | 0 | 0 | 0 | 0 |
| ENSG00000116721 | 0 | 0 | 0 | 0 | 0 | 0 |
| ENSG00000228338 | 0 | 0 | 0 | 0 | 0 | 0 |
| ENSG00000239810 | 0 | 0 | 0 | 0 | 0 | 0 |
| ENSG00000231103 | 0 | 0 | 0 | 0 | 0 | 0 |
| ENSG00000179172 | 15 | 0 | 1 | 0 | 0 | 0 |
| ENSG00000120952 | 0 | 0 | 0 | 0 | 0 | 0 |
| ENSG00000243073 | 0 | 0 | 0 | 0 | 0 | 0 |
| ENSG00000187545 | 0 | 0 | 0 | 0 | 0 | 0 |
| ENSG00000204510 | 0 | 0 | 0 | 0 | 0 | 0 |
| ENSG00000207434 | 0 | 0 | 0 | 0 | 0 | 0 |
| ENSG00000234064 | 0 | 0 | 0 | 0 | 0 | 0 |
| ENSG00000232423 | 0 | 0 | 0 | 0 | 0 | 0 |
| ENSG00000274178 | 0 | 0 | 0 | 0 | 0 | 0 |
| ENSG00000274764 | 0 | 0 | 0 | 0 | 0 | 0 |
| ENSG00000277058 | 0 | 0 | 0 | 0 | 1 | 0 |

| | | | | | | |
|---|---|---|---|---|---|---|
| ENSG00000229571 | 0 | 0 | 0 | 0 | 0 | 0 |
| ENSG00000277862 | 0 | 0 | 0 | 0 | 0 | 0 |
| ENSG00000236179 | 0 | 0 | 0 | 0 | 0 | 0 |
| ENSG00000275774 | 9 | 0 | 0 | 0 | 0 | 0 |
| ENSG00000229978 | 0 | 0 | 0 | 0 | 0 | 0 |
| ENSG00000233336 | 0 | 0 | 0 | 0 | 0 | 0 |
| ENSG00000280267 | 0 | 0 | 0 | 0 | 0 | 0 |
| ENSG00000179412 | 4 | 0 | 0 | 0 | 0 | 0 |
| ENSG00000204505 | 0 | 0 | 0 | 0 | 0 | 0 |
| ENSG00000279169 | 0 | 0 | 0 | 0 | 0 | 0 |
| ENSG00000279804 | 0 | 0 | 0 | 0 | 0 | 0 |
| ENSG00000250894 | 0 | 0 | 0 | 0 | 0 | 0 |
| ENSG00000270601 | 0 | 0 | 0 | 0 | 0 | 0 |
| ENSG00000231219 | 0 | 0 | 0 | 0 | 0 | 0 |
| ENSG00000207511 | 0 | 0 | 0 | 0 | 0 | 0 |
| ENSG00000182330 | 0 | 0 | 0 | 0 | 0 | 0 |
| ENSG00000237700 | 0 | 0 | 0 | 0 | 0 | 0 |
| ENSG00000204501 | 0 | 0 | 0 | 0 | 0 | 0 |
| ENSG00000228229 | 0 | 0 | 0 | 0 | 0 | 0 |
| ENSG00000204481 | 0 | 0 | 0 | 0 | 0 | 0 |
| ENSG00000204480 | 0 | 0 | 0 | 0 | 0 | 0 |
| ENSG00000204479 | 0 | 0 | 0 | 0 | 0 | 0 |
| ENSG00000204478 | 0 | 0 | 0 | 0 | 0 | 0 |
| ENSG00000162494 | 0 | 0 | 0 | 0 | 0 | 0 |
| ENSG00000259961 | 0 | 0 | 0 | 0 | 0 | 0 |
| ENSG00000215909 | 0 | 0 | 0 | 0 | 0 | 0 |
| ENSG00000162493 | 0 | 1 | 1 | 0 | 1 | 0 |
| ENSG00000222952 | 0 | 0 | 0 | 0 | 0 | 0 |
| ENSG00000237445 | 0 | 0 | 0 | 0 | 0 | 0 |
| ENSG00000253085 | 0 | 0 | 0 | 0 | 1 | 0 |
| ENSG00000116731 | 1262 | 480 | 728 | 434 | 584 | 596 |
| ENSG00000189337 | 812 | 371 | 469 | 340 | 382 | 409 |
| ENSG00000252151 | 0 | 0 | 0 | 0 | 0 | 0 |
| ENSG00000231606 | 0 | 0 | 0 | 0 | 0 | 0 |
| ENSG00000234593 | 0 | 0 | 0 | 0 | 0 | 0 |
| ENSG00000235829 | 0 | 0 | 0 | 0 | 0 | 0 |
| ENSG00000175147 | 39 | 26 | 33 | 3 | 14 | 27 |
| ENSG00000171729 | 549 | 237 | 284 | 273 | 231 | 285 |
| ENSG00000204464 | 0 | 0 | 0 | 0 | 0 | 0 |
| ENSG00000215720 | 0 | 0 | 0 | 0 | 1 | 0 |
| ENSG00000278480 | 0 | 1 | 0 | 0 | 0 | 0 |
| ENSG00000142621 | 26 | 3 | 2 | 2 | 3 | 1 |
| ENSG00000233485 | 0 | 0 | 0 | 0 | 0 | 0 |
| ENSG00000236045 | 0 | 0 | 0 | 0 | 0 | 0 |
| ENSG00000228140 | 4 | 3 | 4 | 0 | 2 | 0 |
| ENSG00000142634 | 1290 | 711 | 890 | 923 | 809 | 822 |
| ENSG00000162438 | 2 | 0 | 1 | 2 | 2 | 2 |
| ENSG00000142615 | 0 | 0 | 0 | 0 | 0 | 0 |
| ENSG00000215704 | 0 | 0 | 0 | 2 | 1 | 2 |
| ENSG00000132906 | 591 | 274 | 328 | 312 | 318 | 331 |

| | | | | | | |
|---|---|---|---|---|---|---|
| ENSG00000116138 | 1063 | 465 | 584 | 328 | 500 | 482 |
| ENSG00000251866 | 0 | 0 | 0 | 0 | 0 | 0 |
| ENSG00000272510 | 0 | 0 | 0 | 0 | 0 | 0 |
| ENSG00000116771 | 289 | 128 | 173 | 186 | 176 | 156 |
| ENSG00000237301 | 8 | 14 | 6 | 8 | 8 | 10 |
| ENSG00000235084 | 8 | 8 | 6 | 8 | 9 | 5 |
| ENSG00000252417 | 0 | 0 | 0 | 0 | 0 | 0 |
| ENSG00000236500 | 0 | 0 | 0 | 0 | 0 | 0 |
| ENSG00000197312 | 1853 | 901 | 947 | 426 | 974 | 818 |
| ENSG00000215695 | 0 | 0 | 0 | 0 | 0 | 0 |
| ENSG00000271742 | 0 | 0 | 0 | 0 | 1 | 0 |
| ENSG00000116786 | 2076 | 913 | 1191 | 1275 | 1099 | 1088 |
| ENSG00000237938 | 6 | 4 | 5 | 2 | 4 | 2 |
| ENSG00000162461 | 32 | 15 | 17 | 20 | 19 | 14 |
| ENSG00000224459 | 0 | 0 | 0 | 0 | 0 | 0 |
| ENSG00000162460 | 0 | 1 | 0 | 0 | 0 | 0 |
| ENSG00000162458 | 827 | 443 | 480 | 435 | 459 | 394 |
| ENSG00000224321 | 2 | 1 | 0 | 0 | 2 | 0 |
| ENSG00000233954 | 614 | 369 | 415 | 440 | 446 | 404 |
| ENSG00000178715 | 6 | 2 | 0 | 0 | 0 | 2 |
| ENSG00000179743 | 265 | 155 | 212 | 140 | 135 | 160 |
| ENSG00000065526 | 1904 | 800 | 1072 | 576 | 953 | 948 |
| ENSG00000270620 | 0 | 0 | 0 | 0 | 0 | 0 |
| ENSG00000116809 | 985 | 425 | 543 | 513 | 512 | 470 |
| ENSG00000234607 | 1 | 0 | 0 | 0 | 0 | 0 |
| ENSG00000231353 | 0 | 0 | 0 | 0 | 0 | 0 |
| ENSG00000275503 | 0 | 0 | 0 | 0 | 0 | 0 |
| ENSG00000224966 | 0 | 0 | 0 | 0 | 0 | 0 |
| ENSG00000183888 | 3 | 1 | 2 | 2 | 0 | 0 |
| ENSG00000233078 | 0 | 0 | 0 | 0 | 0 | 0 |
| ENSG00000173641 | 0 | 0 | 3 | 0 | 0 | 0 |
| ENSG00000186510 | 2 | 1 | 0 | 0 | 1 | 1 |
| ENSG00000232456 | 3 | 0 | 3 | 5 | 3 | 4 |
| ENSG00000184908 | 0 | 0 | 0 | 0 | 0 | 0 |
| ENSG00000185519 | 99 | 39 | 47 | 95 | 65 | 47 |
| ENSG00000142627 | 1256 | 682 | 756 | 766 | 716 | 750 |
| ENSG00000227959 | 16 | 4 | 5 | 5 | 8 | 10 |
| ENSG00000224621 | 7 | 1 | 3 | 1 | 2 | 0 |
| ENSG00000234166 | 0 | 0 | 0 | 0 | 0 | 0 |
| ENSG00000142632 | 679 | 297 | 429 | 373 | 317 | 371 |
| ENSG00000237276 | 22 | 21 | 10 | 14 | 7 | 8 |
| ENSG00000132881 | 241 | 85 | 113 | 104 | 102 | 100 |
| ENSG00000233929 | 0 | 0 | 0 | 0 | 0 | 0 |
| ENSG00000037637 | 1239 | 505 | 798 | 566 | 581 | 694 |
| ENSG00000055070 | 3911 | 1987 | 2534 | 2316 | 2337 | 2516 |
| ENSG00000226457 | 0 | 0 | 0 | 1 | 1 | 0 |
| ENSG00000187144 | 0 | 1 | 1 | 2 | 1 | 0 |
| ENSG00000157191 | 1933 | 962 | 1155 | 1235 | 1140 | 1094 |
| ENSG00000226029 | 57 | 20 | 37 | 16 | 23 | 20 |
| ENSG00000080947 | 128 | 71 | 73 | 40 | 68 | 66 |

| | | | | | | |
|---|---|---|---|---|---|---|
| ENSG00000206652 | 0 | 0 | 0 | 0 | 0 | 0 |
| ENSG00000261135 | 4 | 8 | 9 | 7 | 1 | 9 |
| ENSG00000285853 | 1 | 0 | 1 | 0 | 0 | 1 |
| ENSG00000224174 | 0 | 0 | 0 | 0 | 0 | 0 |
| ENSG00000233421 | 0 | 0 | 0 | 0 | 0 | 0 |
| ENSG00000277344 | 0 | 0 | 0 | 0 | 0 | 0 |
| ENSG00000225710 | 1 | 0 | 0 | 0 | 0 | 0 |
| ENSG00000219481 | 1903 | 918 | 1221 | 1509 | 2155 | 2038 |
| ENSG00000271732 | 15 | 3 | 5 | 15 | 16 | 20 |
| ENSG00000215908 | 713 | 397 | 489 | 336 | 449 | 458 |
| ENSG00000283773 | 0 | 0 | 0 | 0 | 0 | 0 |
| ENSG00000186301 | 123 | 43 | 77 | 43 | 50 | 66 |
| ENSG00000282143 | 0 | 0 | 0 | 0 | 0 | 0 |
| ENSG00000207513 | 0 | 0 | 0 | 0 | 0 | 0 |
| ENSG00000280114 | 4 | 4 | 4 | 4 | 2 | 0 |
| ENSG00000236698 | 2 | 1 | 9 | 3 | 1 | 3 |
| ENSG00000268869 | 11 | 0 | 4 | 2 | 6 | 2 |
| ENSG00000279151 | 0 | 0 | 0 | 0 | 0 | 0 |
| ENSG00000282740 | 0 | 0 | 0 | 0 | 0 | 0 |
| ENSG00000058453 | 665 | 257 | 409 | 376 | 352 | 428 |
| ENSG00000227684 | 0 | 0 | 0 | 0 | 0 | 0 |
| ENSG00000207389 | 0 | 0 | 0 | 0 | 0 | 0 |
| ENSG00000186715 | 49 | 17 | 25 | 11 | 13 | 12 |
| ENSG00000223643 | 0 | 0 | 0 | 0 | 0 | 0 |
| ENSG00000230239 | 0 | 0 | 0 | 0 | 0 | 0 |
| ENSG00000263811 | 0 | 0 | 0 | 0 | 0 | 0 |
| ENSG00000228549 | 0 | 1 | 0 | 0 | 1 | 0 |
| ENSG00000277234 | 0 | 0 | 0 | 0 | 0 | 0 |
| ENSG00000238142 | 0 | 0 | 0 | 0 | 0 | 0 |
| ENSG00000235241 | 0 | 0 | 0 | 0 | 0 | 0 |
| ENSG00000207005 | 0 | 0 | 0 | 1 | 1 | 0 |
| ENSG00000272426 | 6 | 0 | 2 | 1 | 1 | 1 |
| ENSG00000117122 | 90 | 51 | 60 | 41 | 29 | 49 |
| ENSG00000226526 | 0 | 0 | 0 | 0 | 0 | 0 |
| ENSG00000159363 | 2006 | 959 | 1289 | 1438 | 1279 | 1234 |
| ENSG00000117118 | 2522 | 1295 | 1479 | 1503 | 1440 | 1479 |
| ENSG00000117115 | 201 | 90 | 93 | 88 | 97 | 111 |
| ENSG00000204362 | 0 | 0 | 0 | 0 | 0 | 0 |
| ENSG00000282843 | 0 | 0 | 0 | 0 | 0 | 0 |
| ENSG00000142623 | 0 | 1 | 2 | 11 | 5 | 2 |
| ENSG00000142619 | 21 | 22 | 23 | 37 | 39 | 38 |
| ENSG00000266634 | 0 | 0 | 0 | 0 | 0 | 0 |
| ENSG00000159339 | 2 | 1 | 2 | 13 | 12 | 1 |
| ENSG00000276747 | 5 | 1 | 4 | 2 | 2 | 4 |
| ENSG00000227751 | 0 | 0 | 0 | 0 | 0 | 0 |
| ENSG00000179051 | 10185 | 5177 | 6744 | 6355 | 6947 | 7647 |
| ENSG00000074964 | 2116 | 951 | 1411 | 1392 | 1473 | 1383 |
| ENSG00000236648 | 0 | 0 | 0 | 0 | 0 | 0 |
| ENSG00000117148 | 0 | 0 | 6 | 5 | 1 | 4 |
| ENSG00000284653 | 0 | 0 | 0 | 0 | 0 | 0 |

| | | | | | | |
|---|---|---|---|---|---|---|
| ENSG00000261781 | 0 | 0 | 0 | 0 | 0 | 0 |
| ENSG00000117154 | 1 | 0 | 0 | 1 | 1 | 0 |
| ENSG00000280222 | 0 | 0 | 0 | 0 | 0 | 0 |
| ENSG00000230035 | 0 | 0 | 0 | 0 | 0 | 0 |
| ENSG00000225387 | 0 | 0 | 0 | 0 | 0 | 0 |
| ENSG00000179023 | 104 | 67 | 64 | 52 | 43 | 40 |
| ENSG00000235282 | 0 | 0 | 0 | 0 | 0 | 0 |
| ENSG00000225478 | 0 | 0 | 0 | 0 | 0 | 0 |
| ENSG00000009709 | 0 | 0 | 2 | 1 | 1 | 0 |
| ENSG00000179002 | 0 | 0 | 0 | 0 | 0 | 0 |
| ENSG00000255275 | 0 | 0 | 0 | 0 | 0 | 0 |
| ENSG00000159423 | 1735 | 846 | 1014 | 906 | 728 | 816 |
| ENSG00000265606 | 0 | 0 | 0 | 0 | 0 | 0 |
| ENSG00000221662 | 0 | 0 | 0 | 0 | 0 | 0 |
| ENSG00000169991 | 827 | 390 | 402 | 507 | 605 | 532 |
| ENSG00000272084 | 18 | 10 | 9 | 7 | 11 | 3 |
| ENSG00000127481 | 5912 | 2512 | 3472 | 1791 | 3070 | 3182 |
| ENSG00000230424 | 10 | 5 | 2 | 12 | 7 | 2 |
| ENSG00000127463 | 2564 | 1311 | 1537 | 1341 | 1352 | 1493 |
| ENSG00000053372 | 1532 | 841 | 953 | 1211 | 1097 | 1132 |
| ENSG00000211454 | 10 | 5 | 11 | 4 | 6 | 8 |
| ENSG00000162482 | 26 | 4 | 20 | 6 | 12 | 16 |
| ENSG00000270728 | 43 | 34 | 26 | 32 | 23 | 25 |
| ENSG00000053371 | 2319 | 1071 | 1393 | 1559 | 1191 | 1258 |
| ENSG00000200403 | 0 | 1 | 1 | 2 | 0 | 0 |
| ENSG00000040487 | 497 | 226 | 300 | 254 | 262 | 261 |
| ENSG00000077549 | 6882 | 3483 | 4134 | 3930 | 3951 | 3626 |
| ENSG00000240490 | 0 | 0 | 0 | 0 | 0 | 0 |
| ENSG00000173436 | 421 | 278 | 322 | 284 | 279 | 289 |
| ENSG00000201609 | 0 | 0 | 0 | 0 | 0 | 0 |
| ENSG00000235185 | 0 | 0 | 0 | 0 | 0 | 0 |
| ENSG00000158747 | 606 | 327 | 332 | 515 | 368 | 260 |
| ENSG00000270136 | 0 | 0 | 0 | 0 | 0 | 0 |
| ENSG00000226396 | 8 | 9 | 9 | 4 | 3 | 6 |
| ENSG00000158748 | 3 | 4 | 0 | 1 | 2 | 2 |
| ENSG00000162542 | 466 | 216 | 272 | 288 | 262 | 302 |
| ENSG00000178828 | 0 | 0 | 0 | 0 | 0 | 1 |
| ENSG00000235434 | 0 | 0 | 0 | 0 | 0 | 0 |
| ENSG00000169914 | 523 | 259 | 322 | 157 | 282 | 306 |
| ENSG00000188784 | 0 | 0 | 0 | 0 | 0 | 0 |
| ENSG00000242688 | 0 | 0 | 0 | 0 | 0 | 0 |
| ENSG00000188257 | 18 | 6 | 7 | 4 | 4 | 2 |
| ENSG00000127472 | 0 | 0 | 1 | 0 | 0 | 0 |
| ENSG00000117215 | 0 | 0 | 0 | 0 | 0 | 0 |
| ENSG00000158786 | 6 | 1 | 2 | 2 | 1 | 1 |
| ENSG00000227066 | 0 | 0 | 1 | 0 | 0 | 0 |
| ENSG00000187980 | 0 | 0 | 0 | 0 | 0 | 0 |
| ENSG00000225986 | 0 | 0 | 0 | 0 | 0 | 0 |
| ENSG00000162543 | 0 | 0 | 0 | 0 | 0 | 0 |
| ENSG00000228105 | 0 | 0 | 0 | 0 | 0 | 0 |

| | | | | | | |
|---|---|---|---|---|---|---|
| ENSG00000284710 | 0 | 0 | 2 | 0 | 0 | 0 |
| ENSG00000158816 | 0 | 1 | 0 | 0 | 0 | 0 |
| ENSG00000226664 | 0 | 0 | 0 | 0 | 0 | 0 |
| ENSG00000236963 | 0 | 0 | 0 | 0 | 0 | 0 |
| ENSG00000226487 | 0 | 0 | 0 | 0 | 0 | 0 |
| ENSG00000284641 | 0 | 0 | 0 | 0 | 0 | 0 |
| ENSG00000284743 | 0 | 0 | 0 | 0 | 0 | 0 |
| ENSG00000162545 | 3436 | 1579 | 1921 | 1568 | 1388 | 1526 |
| ENSG00000090432 | 1081 | 430 | 645 | 649 | 557 | 646 |
| ENSG00000215520 | 0 | 0 | 0 | 0 | 0 | 0 |
| ENSG00000183114 | 0 | 0 | 0 | 0 | 0 | 0 |
| ENSG00000158825 | 0 | 0 | 2 | 1 | 0 | 1 |
| ENSG00000158828 | 687 | 451 | 487 | 508 | 559 | 445 |
| ENSG00000284005 | 0 | 0 | 0 | 0 | 0 | 0 |
| ENSG00000117242 | 91 | 82 | 81 | 87 | 84 | 61 |
| ENSG00000244038 | 9723 | 4967 | 5909 | 5456 | 5576 | 5356 |
| ENSG00000117245 | 11 | 6 | 5 | 4 | 8 | 4 |
| ENSG00000235432 | 0 | 0 | 0 | 0 | 0 | 0 |
| ENSG00000189410 | 15 | 8 | 17 | 21 | 21 | 21 |
| ENSG00000233069 | 0 | 0 | 0 | 0 | 1 | 0 |
| ENSG00000127483 | 4800 | 2233 | 3063 | 1691 | 2474 | 2645 |
| ENSG00000075151 | 3195 | 1545 | 1753 | 931 | 2066 | 1524 |
| ENSG00000251914 | 0 | 0 | 0 | 0 | 0 | 0 |
| ENSG00000221808 | 0 | 0 | 0 | 0 | 0 | 0 |
| ENSG00000233072 | 0 | 0 | 0 | 0 | 0 | 0 |
| ENSG00000236073 | 0 | 0 | 0 | 0 | 0 | 0 |
| ENSG00000235112 | 0 | 0 | 0 | 0 | 0 | 0 |
| ENSG00000117298 | 3719 | 1500 | 2285 | 1404 | 1304 | 1556 |
| ENSG00000236936 | 0 | 0 | 0 | 0 | 0 | 0 |
| ENSG00000231105 | 0 | 1 | 0 | 1 | 0 | 0 |
| ENSG00000236216 | 0 | 0 | 0 | 0 | 0 | 0 |
| ENSG00000228823 | 0 | 0 | 0 | 0 | 0 | 0 |
| ENSG00000236009 | 0 | 0 | 0 | 0 | 0 | 0 |
| ENSG00000227001 | 16 | 10 | 10 | 9 | 11 | 13 |
| ENSG00000187952 | 10 | 2 | 5 | 3 | 2 | 2 |
| ENSG00000186543 | 0 | 0 | 0 | 0 | 0 | 0 |
| ENSG00000142794 | 700 | 331 | 429 | 298 | 348 | 392 |
| ENSG00000176378 | 0 | 0 | 0 | 0 | 0 | 0 |
| ENSG00000162551 | 0 | 1 | 0 | 0 | 0 | 0 |
| ENSG00000233431 | 0 | 0 | 0 | 1 | 0 | 0 |
| ENSG00000076864 | 1705 | 757 | 1121 | 1481 | 1398 | 1538 |
| ENSG00000090686 | 1652 | 792 | 872 | 488 | 1061 | 812 |
| ENSG00000231978 | 0 | 0 | 0 | 0 | 0 | 0 |
| ENSG00000187942 | 0 | 0 | 1 | 1 | 0 | 1 |
| ENSG00000142798 | 924 | 360 | 645 | 434 | 477 | 467 |
| ENSG00000232037 | 0 | 0 | 1 | 2 | 0 | 0 |
| ENSG00000283234 | 1 | 1 | 1 | 1 | 0 | 0 |
| ENSG00000219073 | 0 | 0 | 0 | 0 | 0 | 0 |
| ENSG00000285959 | 0 | 0 | 0 | 0 | 0 | 0 |
| ENSG00000273610 | 0 | 0 | 0 | 0 | 0 | 0 |

| | | | | | | |
|---|---|---|---|---|---|---|
| ENSG00000201919 | 0 | 0 | 0 | 0 | 0 | 0 |
| ENSG00000142789 | 0 | 0 | 0 | 0 | 0 | 0 |
| ENSG00000278124 | 0 | 0 | 0 | 0 | 0 | 0 |
| ENSG00000201273 | 0 | 0 | 0 | 0 | 0 | 0 |
| ENSG00000228397 | 0 | 2 | 0 | 0 | 0 | 0 |
| ENSG00000218510 | 68 | 52 | 47 | 27 | 46 | 38 |
| ENSG00000285721 | 16 | 5 | 6 | 4 | 1 | 2 |
| ENSG00000285752 | 20 | 9 | 16 | 14 | 12 | 8 |
| ENSG00000285794 | 0 | 0 | 0 | 0 | 0 | 0 |
| ENSG00000070831 | 21 | 30 | 25 | 16 | 19 | 27 |
| ENSG00000230068 | 0 | 5 | 1 | 0 | 0 | 0 |
| ENSG00000271428 | 1 | 0 | 1 | 0 | 0 | 0 |
| ENSG00000162552 | 239 | 105 | 119 | 98 | 85 | 89 |
| ENSG00000285873 | 17 | 7 | 13 | 13 | 5 | 6 |
| ENSG00000266564 | 0 | 0 | 0 | 0 | 0 | 0 |
| ENSG00000234397 | 0 | 0 | 0 | 0 | 0 | 0 |
| ENSG00000279625 | 0 | 0 | 0 | 0 | 0 | 0 |
| ENSG00000184677 | 1495 | 665 | 796 | 542 | 736 | 751 |
| ENSG00000237200 | 6 | 2 | 4 | 0 | 4 | 5 |
| ENSG00000070886 | 0 | 2 | 1 | 4 | 0 | 4 |
| ENSG00000276835 | 0 | 0 | 0 | 0 | 0 | 0 |
| ENSG00000173372 | 0 | 0 | 0 | 0 | 0 | 0 |
| ENSG00000159189 | 0 | 0 | 0 | 0 | 0 | 0 |
| ENSG00000173369 | 0 | 0 | 0 | 0 | 0 | 0 |
| ENSG00000133216 | 337 | 163 | 206 | 183 | 146 | 150 |
| ENSG00000265422 | 0 | 0 | 0 | 0 | 0 | 0 |
| ENSG00000225952 | 0 | 0 | 0 | 0 | 0 | 0 |
| ENSG00000264014 | 0 | 0 | 0 | 0 | 0 | 0 |
| ENSG00000215906 | 0 | 0 | 0 | 0 | 0 | 0 |
| ENSG00000227868 | 2 | 3 | 0 | 2 | 1 | 0 |
| ENSG00000004487 | 3810 | 1859 | 2404 | 1946 | 2348 | 2540 |
| ENSG00000240553 | 0 | 0 | 0 | 0 | 0 | 0 |
| ENSG00000263793 | 0 | 0 | 0 | 0 | 0 | 0 |
| ENSG00000266802 | 0 | 0 | 0 | 0 | 0 | 0 |
| ENSG00000169641 | 1210 | 520 | 701 | 434 | 560 | 656 |
| ENSG00000229010 | 0 | 1 | 0 | 0 | 0 | 0 |
| ENSG00000206935 | 0 | 0 | 0 | 0 | 0 | 1 |
| ENSG00000252578 | 0 | 0 | 0 | 0 | 0 | 0 |
| ENSG00000179546 | 79 | 53 | 32 | 21 | 41 | 43 |
| ENSG00000215381 | 0 | 0 | 0 | 0 | 0 | 0 |
| ENSG00000261326 | 124 | 48 | 42 | 16 | 73 | 41 |
| ENSG00000284726 | 6 | 3 | 2 | 2 | 8 | 1 |
| ENSG00000125944 | 4858 | 2450 | 2952 | 2146 | 3228 | 2987 |
| ENSG00000125945 | 772 | 347 | 446 | 327 | 374 | 403 |
| ENSG00000249087 | 64 | 30 | 40 | 33 | 49 | 35 |
| ENSG00000201405 | 0 | 0 | 0 | 0 | 0 | 0 |
| ENSG00000271420 | 0 | 0 | 0 | 1 | 0 | 0 |
| ENSG00000204219 | 67 | 43 | 42 | 37 | 54 | 28 |
| ENSG00000232482 | 4 | 0 | 0 | 0 | 0 | 0 |
| ENSG00000088280 | 1347 | 569 | 820 | 626 | 594 | 562 |

| | | | | | | |
|---|---|---|---|---|---|---|
| ENSG00000007968 | 294 | 150 | 181 | 91 | 123 | 146 |
| ENSG00000235052 | 33 | 9 | 19 | 10 | 19 | 21 |
| ENSG00000117318 | 7907 | 3648 | 4330 | 5831 | 5025 | 4716 |
| ENSG00000285802 | 3 | 8 | 5 | 3 | 7 | 9 |
| ENSG00000197880 | 0 | 0 | 0 | 0 | 1 | 0 |
| ENSG00000229239 | 0 | 2 | 1 | 0 | 0 | 0 |
| ENSG00000142676 | 22950 | 12234 | 13761 | 14805 | 14099 | 13727 |
| ENSG00000011007 | 2089 | 1107 | 1358 | 960 | 1123 | 1352 |
| ENSG00000236810 | 16 | 7 | 11 | 15 | 17 | 7 |
| ENSG00000057757 | 1248 | 592 | 668 | 566 | 595 | 634 |
| ENSG00000011009 | 3402 | 1688 | 1942 | 2150 | 1813 | 1842 |
| ENSG00000117308 | 1148 | 592 | 644 | 758 | 704 | 642 |
| ENSG00000117305 | 1686 | 802 | 1027 | 907 | 853 | 859 |
| ENSG00000179163 | 2260 | 1164 | 1459 | 1030 | 1142 | 1114 |
| ENSG00000188822 | 2 | 1 | 3 | 0 | 1 | 0 |
| ENSG00000229106 | 0 | 0 | 0 | 2 | 0 | 0 |
| ENSG00000232557 | 0 | 0 | 0 | 0 | 0 | 0 |
| ENSG00000264926 | 0 | 0 | 0 | 0 | 0 | 0 |
| ENSG00000230228 | 0 | 1 | 0 | 0 | 1 | 0 |
| ENSG00000189266 | 3540 | 1830 | 2302 | 1420 | 1735 | 1647 |
| ENSG00000188529 | 4415 | 2197 | 2754 | 1497 | 2188 | 2033 |
| ENSG00000215905 | 0 | 0 | 0 | 0 | 0 | 0 |
| ENSG00000225315 | 4 | 3 | 6 | 2 | 5 | 1 |
| ENSG00000142661 | 2 | 1 | 2 | 2 | 1 | 2 |
| ENSG00000230703 | 0 | 0 | 0 | 0 | 0 | 0 |
| ENSG00000142677 | 0 | 0 | 0 | 0 | 0 | 0 |
| ENSG00000185436 | 36 | 21 | 30 | 26 | 15 | 11 |
| ENSG00000230023 | 0 | 0 | 1 | 1 | 0 | 0 |
| ENSG00000232298 | 2 | 1 | 0 | 2 | 0 | 1 |
| ENSG00000158055 | 433 | 236 | 283 | 248 | 261 | 296 |
| ENSG00000001460 | 261 | 123 | 128 | 142 | 138 | 130 |
| ENSG00000001461 | 1193 | 495 | 732 | 426 | 638 | 593 |
| ENSG00000117602 | 701 | 353 | 404 | 299 | 378 | 390 |
| ENSG00000264443 | 0 | 0 | 0 | 0 | 0 | 0 |
| ENSG00000184454 | 10 | 7 | 5 | 2 | 0 | 2 |
| ENSG00000227312 | 0 | 0 | 0 | 0 | 0 | 0 |
| ENSG00000239106 | 0 | 0 | 0 | 0 | 0 | 0 |
| ENSG00000133226 | 2864 | 1434 | 1516 | 928 | 1350 | 1374 |
| ENSG00000284699 | 0 | 0 | 0 | 0 | 0 | 0 |
| ENSG00000169504 | 4094 | 2044 | 2295 | 1491 | 2287 | 1969 |
| ENSG00000238482 | 0 | 0 | 0 | 0 | 0 | 0 |
| ENSG00000020633 | 0 | 0 | 1 | 0 | 0 | 1 |
| ENSG00000278034 | 0 | 0 | 0 | 0 | 0 | 0 |
| ENSG00000229162 | 0 | 0 | 0 | 0 | 0 | 0 |
| ENSG00000261025 | 0 | 0 | 0 | 0 | 0 | 0 |
| ENSG00000264371 | 0 | 0 | 0 | 0 | 0 | 0 |
| ENSG00000285823 | 0 | 0 | 0 | 0 | 0 | 0 |
| ENSG00000233755 | 0 | 0 | 0 | 0 | 0 | 0 |
| ENSG00000233419 | 0 | 0 | 0 | 0 | 0 | 0 |
| ENSG00000231953 | 0 | 0 | 0 | 0 | 0 | 0 |

| | | | | | | |
|---|---|---|---|---|---|---|
| ENSG00000117614 | 929 | 548 | 573 | 421 | 667 | 498 |
| ENSG00000284602 | 30 | 7 | 15 | 6 | 4 | 9 |
| ENSG00000284657 | 5 | 4 | 3 | 2 | 3 | 1 |
| ENSG00000117616 | 2277 | 981 | 1368 | 741 | 1204 | 873 |
| ENSG00000272432 | 10 | 5 | 2 | 3 | 3 | 6 |
| ENSG00000261349 | 12 | 4 | 7 | 4 | 7 | 6 |
| ENSG00000187010 | 9 | 9 | 7 | 3 | 7 | 6 |
| ENSG00000224183 | 5 | 1 | 1 | 2 | 0 | 1 |
| ENSG00000259984 | 7 | 0 | 3 | 0 | 1 | 1 |
| ENSG00000183726 | 3245 | 1523 | 1903 | 1604 | 1838 | 1864 |
| ENSG00000188672 | 12 | 8 | 5 | 3 | 4 | 6 |
| ENSG00000238084 | 0 | 1 | 1 | 1 | 0 | 0 |
| ENSG00000204178 | 1419 | 599 | 753 | 662 | 738 | 758 |
| ENSG00000157978 | 1818 | 842 | 1045 | 886 | 829 | 801 |
| ENSG00000225643 | 4 | 6 | 1 | 0 | 4 | 2 |
| ENSG00000117643 | 442 | 182 | 251 | 174 | 153 | 170 |
| ENSG00000233478 | 2 | 1 | 0 | 1 | 1 | 0 |
| ENSG00000162430 | 1829 | 999 | 1318 | 1179 | 1050 | 1205 |
| ENSG00000255054 | 0 | 0 | 0 | 0 | 0 | 0 |
| ENSG00000228172 | 25 | 17 | 21 | 24 | 16 | 21 |
| ENSG00000117640 | 1099 | 559 | 749 | 576 | 633 | 704 |
| ENSG00000272478 | 0 | 0 | 0 | 0 | 0 | 0 |
| ENSG00000127423 | 197 | 120 | 135 | 134 | 149 | 129 |
| ENSG00000236528 | 4 | 3 | 2 | 3 | 2 | 3 |
| ENSG00000182749 | 138 | 86 | 110 | 113 | 119 | 122 |
| ENSG00000207302 | 0 | 0 | 0 | 0 | 1 | 0 |
| ENSG00000117632 | 3724 | 1898 | 2272 | 2310 | 2440 | 2371 |
| ENSG00000229786 | 0 | 1 | 0 | 0 | 0 | 0 |
| ENSG00000223624 | 0 | 0 | 0 | 0 | 0 | 0 |
| ENSG00000283938 | 0 | 0 | 0 | 0 | 0 | 0 |
| ENSG00000158006 | 1212 | 536 | 752 | 540 | 632 | 655 |
| ENSG00000207237 | 0 | 0 | 0 | 0 | 0 | 0 |
| ENSG00000252190 | 0 | 0 | 0 | 0 | 0 | 0 |
| ENSG00000252691 | 0 | 0 | 0 | 0 | 0 | 0 |
| ENSG00000158008 | 4 | 1 | 2 | 3 | 5 | 1 |
| ENSG00000158014 | 0 | 0 | 0 | 0 | 0 | 0 |
| ENSG00000284309 | 0 | 0 | 0 | 0 | 0 | 0 |
| ENSG00000158022 | 2 | 0 | 0 | 0 | 1 | 0 |
| ENSG00000175087 | 565 | 235 | 295 | 190 | 273 | 256 |
| ENSG00000197245 | 4 | 4 | 2 | 0 | 0 | 2 |
| ENSG00000282872 | 0 | 0 | 0 | 0 | 0 | 0 |
| ENSG00000236782 | 0 | 1 | 4 | 8 | 12 | 2 |
| ENSG00000142684 | 0 | 0 | 0 | 0 | 0 | 0 |
| ENSG00000142675 | 792 | 372 | 500 | 414 | 446 | 412 |
| ENSG00000188782 | 0 | 1 | 0 | 1 | 1 | 0 |
| ENSG00000236155 | 0 | 0 | 0 | 0 | 0 | 0 |
| ENSG00000278572 | 0 | 0 | 0 | 0 | 0 | 0 |
| ENSG00000130695 | 967 | 435 | 598 | 477 | 572 | 566 |
| ENSG00000270733 | 2 | 1 | 0 | 0 | 0 | 1 |
| ENSG00000142669 | 2083 | 1084 | 1368 | 1453 | 1276 | 1289 |

| | | | | | | |
|---|---|---|---|---|---|---|
| ENSG00000158062 | 490 | 214 | 292 | 208 | 188 | 234 |
| ENSG00000169442 | 10 | 4 | 10 | 4 | 0 | 7 |
| ENSG00000176092 | 74 | 34 | 43 | 51 | 49 | 43 |
| ENSG00000225854 | 0 | 0 | 0 | 0 | 0 | 1 |
| ENSG00000176083 | 0 | 0 | 0 | 0 | 0 | 0 |
| ENSG00000131914 | 2 | 0 | 0 | 0 | 0 | 0 |
| ENSG00000117682 | 1291 | 599 | 792 | 571 | 648 | 715 |
| ENSG00000223583 | 2 | 0 | 2 | 1 | 0 | 0 |
| ENSG00000225891 | 3 | 4 | 7 | 0 | 4 | 4 |
| ENSG00000198830 | 11446 | 5247 | 6885 | 5468 | 5624 | 5790 |
| ENSG00000224129 | 0 | 0 | 0 | 0 | 0 | 0 |
| ENSG00000117676 | 1161 | 593 | 714 | 764 | 789 | 657 |
| ENSG00000238705 | 0 | 0 | 0 | 0 | 0 | 0 |
| ENSG00000243905 | 0 | 0 | 0 | 0 | 0 | 0 |
| ENSG00000252802 | 0 | 0 | 0 | 0 | 0 | 0 |
| ENSG00000252429 | 0 | 0 | 0 | 0 | 0 | 0 |
| ENSG00000229247 | 0 | 0 | 0 | 0 | 0 | 0 |
| ENSG00000238316 | 0 | 0 | 0 | 0 | 0 | 0 |
| ENSG00000235069 | 0 | 0 | 0 | 0 | 0 | 0 |
| ENSG00000260063 | 24 | 8 | 27 | 14 | 14 | 24 |
| ENSG00000117713 | 3681 | 1490 | 2198 | 1363 | 1579 | 1920 |
| ENSG00000060642 | 1131 | 509 | 688 | 570 | 501 | 554 |
| ENSG00000241188 | 0 | 0 | 0 | 0 | 0 | 0 |
| ENSG00000204160 | 1825 | 769 | 977 | 1088 | 880 | 998 |
| ENSG00000175793 | 1979 | 950 | 1202 | 1319 | 1083 | 1055 |
| ENSG00000142751 | 1361 | 725 | 885 | 785 | 783 | 767 |
| ENSG00000226698 | 0 | 0 | 0 | 0 | 0 | 0 |
| ENSG00000198746 | 395 | 175 | 246 | 259 | 237 | 204 |
| ENSG00000090273 | 2492 | 1327 | 1443 | 1732 | 1684 | 1617 |
| ENSG00000131910 | 0 | 1 | 0 | 0 | 1 | 3 |
| ENSG00000175707 | 764 | 362 | 448 | 501 | 362 | 393 |
| ENSG00000172746 | 0 | 0 | 0 | 2 | 1 | 0 |
| ENSG00000224064 | 0 | 0 | 1 | 0 | 1 | 0 |
| ENSG00000225294 | 10 | 0 | 1 | 1 | 1 | 2 |
| ENSG00000253368 | 244 | 90 | 107 | 120 | 105 | 119 |
| ENSG00000158246 | 173 | 94 | 113 | 88 | 102 | 88 |
| ENSG00000090020 | 6015 | 2620 | 3239 | 3370 | 3523 | 3350 |
| ENSG00000229985 | 0 | 0 | 0 | 0 | 0 | 0 |
| ENSG00000214973 | 0 | 0 | 0 | 0 | 0 | 3 |
| ENSG00000225159 | 12 | 4 | 7 | 3 | 5 | 4 |
| ENSG00000225990 | 0 | 0 | 0 | 0 | 0 | 0 |
| ENSG00000224311 | 0 | 0 | 0 | 0 | 0 | 0 |
| ENSG00000142784 | 2241 | 1040 | 1369 | 1236 | 1227 | 1211 |
| ENSG00000252098 | 0 | 0 | 0 | 0 | 0 | 0 |
| ENSG00000243659 | 0 | 1 | 0 | 0 | 2 | 1 |
| ENSG00000186501 | 1172 | 566 | 698 | 726 | 632 | 666 |
| ENSG00000206888 | 0 | 0 | 1 | 0 | 0 | 2 |
| ENSG00000241547 | 8 | 2 | 1 | 2 | 1 | 2 |
| ENSG00000142765 | 2098 | 914 | 1331 | 1195 | 914 | 1031 |
| ENSG00000142733 | 790 | 384 | 488 | 432 | 389 | 414 |

| | | | | | | |
|---|---|---|---|---|---|---|
| ENSG00000142748 | 8 | 4 | 7 | 3 | 3 | 0 |
| ENSG00000174950 | 8 | 12 | 7 | 9 | 9 | 8 |
| ENSG00000231207 | 0 | 0 | 0 | 0 | 0 | 0 |
| ENSG00000181773 | 2 | 5 | 3 | 9 | 7 | 5 |
| ENSG00000158195 | 4182 | 1993 | 2534 | 2220 | 2468 | 2620 |
| ENSG00000241169 | 0 | 0 | 0 | 0 | 0 | 0 |
| ENSG00000237429 | 0 | 0 | 0 | 1 | 1 | 2 |
| ENSG00000126705 | 1174 | 513 | 663 | 724 | 748 | 778 |
| ENSG00000000938 | 0 | 0 | 0 | 0 | 0 | 0 |
| ENSG00000235912 | 4 | 1 | 0 | 0 | 3 | 2 |
| ENSG00000233975 | 0 | 0 | 0 | 0 | 0 | 0 |
| ENSG00000126709 | 788 | 412 | 458 | 389 | 403 | 331 |
| ENSG00000225886 | 0 | 1 | 1 | 1 | 1 | 0 |
| ENSG00000206767 | 0 | 0 | 0 | 0 | 0 | 0 |
| ENSG00000226662 | 0 | 0 | 0 | 0 | 0 | 0 |
| ENSG00000207095 | 0 | 0 | 0 | 0 | 0 | 0 |
| ENSG00000009780 | 326 | 165 | 184 | 131 | 192 | 153 |
| ENSG00000231344 | 2 | 2 | 1 | 3 | 1 | 2 |
| ENSG00000230833 | 0 | 0 | 0 | 0 | 0 | 0 |
| ENSG00000117758 | 2273 | 1036 | 1300 | 992 | 1174 | 1175 |
| ENSG00000269971 | 0 | 2 | 0 | 0 | 0 | 0 |
| ENSG00000270031 | 2 | 3 | 1 | 2 | 1 | 0 |
| ENSG00000223062 | 0 | 0 | 0 | 0 | 0 | 0 |
| ENSG00000117751 | 1144 | 631 | 752 | 613 | 736 | 735 |
| ENSG00000252947 | 0 | 0 | 0 | 0 | 1 | 0 |
| ENSG00000130775 | 4 | 8 | 3 | 2 | 5 | 4 |
| ENSG00000117748 | 1872 | 954 | 1120 | 998 | 1000 | 1088 |
| ENSG00000130768 | 74 | 31 | 48 | 52 | 74 | 83 |
| ENSG00000227050 | 3 | 0 | 0 | 0 | 2 | 1 |
| ENSG00000158156 | 379 | 184 | 210 | 284 | 265 | 266 |
| ENSG00000158161 | 636 | 320 | 387 | 243 | 364 | 310 |
| ENSG00000238231 | 1 | 0 | 1 | 0 | 0 | 0 |
| ENSG00000240750 | 1 | 1 | 0 | 0 | 1 | 0 |
| ENSG00000228589 | 10 | 5 | 6 | 7 | 5 | 7 |
| ENSG00000228943 | 0 | 0 | 0 | 0 | 0 | 0 |
| ENSG00000214812 | 0 | 1 | 0 | 0 | 0 | 0 |
| ENSG00000253005 | 0 | 0 | 0 | 0 | 0 | 0 |
| ENSG00000169403 | 75 | 64 | 58 | 59 | 76 | 78 |
| ENSG00000126698 | 3647 | 1884 | 2124 | 2066 | 2227 | 2106 |
| ENSG00000130770 | 4028 | 1903 | 2571 | 2249 | 2120 | 1994 |
| ENSG00000270605 | 118 | 54 | 86 | 42 | 49 | 49 |
| ENSG00000271398 | 1 | 0 | 1 | 0 | 1 | 1 |
| ENSG00000130766 | 1213 | 527 | 714 | 704 | 598 | 622 |
| ENSG00000130772 | 840 | 355 | 417 | 464 | 430 | 407 |
| ENSG00000204138 | 1146 | 598 | 717 | 481 | 621 | 597 |
| ENSG00000206779 | 0 | 0 | 0 | 1 | 0 | 0 |
| ENSG00000229820 | 0 | 1 | 0 | 0 | 0 | 0 |
| ENSG00000221216 | 0 | 0 | 0 | 0 | 0 | 0 |
| ENSG00000180198 | 5781 | 3004 | 3474 | 3031 | 3310 | 3307 |
| ENSG00000242125 | 1556 | 808 | 928 | 968 | 1209 | 1135 |

| | | | | | | |
|---|---|---|---|---|---|---|
| ENSG00000274266 | 0 | 0 | 0 | 0 | 0 | 0 |
| ENSG00000200087 | 16 | 16 | 9 | 5 | 16 | 14 |
| ENSG00000231468 | 0 | 4 | 3 | 3 | 0 | 1 |
| ENSG00000279443 | 156 | 105 | 115 | 62 | 87 | 96 |
| ENSG00000180098 | 806 | 440 | 503 | 428 | 444 | 484 |
| ENSG00000197989 | 663 | 350 | 424 | 436 | 450 | 449 |
| ENSG00000221539 | 7 | 1 | 6 | 2 | 5 | 7 |
| ENSG00000278274 | 0 | 0 | 0 | 0 | 0 | 0 |
| ENSG00000273544 | 0 | 0 | 0 | 0 | 0 | 0 |
| ENSG00000280498 | 0 | 0 | 0 | 0 | 0 | 0 |
| ENSG00000274582 | 0 | 0 | 0 | 0 | 0 | 0 |
| ENSG00000120656 | 883 | 471 | 527 | 549 | 545 | 497 |
| ENSG00000188060 | 28 | 16 | 13 | 9 | 8 | 13 |
| ENSG00000229388 | 16 | 8 | 12 | 13 | 10 | 4 |
| ENSG00000270103 | 0 | 0 | 0 | 0 | 0 | 0 |
| ENSG00000274978 | 0 | 0 | 0 | 0 | 0 | 0 |
| ENSG00000162419 | 443 | 268 | 263 | 193 | 271 | 287 |
| ENSG00000252777 | 0 | 0 | 2 | 1 | 0 | 0 |
| ENSG00000198492 | 2859 | 1320 | 1600 | 1281 | 1559 | 1655 |
| ENSG00000116329 | 4 | 0 | 3 | 0 | 3 | 2 |
| ENSG00000233427 | 0 | 1 | 0 | 1 | 1 | 1 |
| ENSG00000199756 | 0 | 0 | 0 | 0 | 0 | 0 |
| ENSG00000159023 | 1669 | 800 | 1062 | 541 | 843 | 756 |
| ENSG00000225616 | 0 | 0 | 0 | 0 | 0 | 1 |
| ENSG00000206704 | 0 | 0 | 0 | 0 | 0 | 0 |
| ENSG00000277958 | 0 | 0 | 0 | 0 | 0 | 0 |
| ENSG00000253304 | 3 | 1 | 1 | 2 | 1 | 2 |
| ENSG00000225750 | 0 | 1 | 0 | 0 | 0 | 0 |
| ENSG00000116350 | 2158 | 956 | 1344 | 942 | 1191 | 1195 |
| ENSG00000277916 | 0 | 0 | 0 | 0 | 0 | 0 |
| ENSG00000116353 | 582 | 236 | 275 | 366 | 298 | 325 |
| ENSG00000237934 | 4 | 2 | 3 | 0 | 2 | 4 |
| ENSG00000060656 | 2894 | 1291 | 1683 | 1385 | 1472 | 1387 |
| ENSG00000230523 | 0 | 2 | 0 | 0 | 0 | 1 |
| ENSG00000225011 | 0 | 0 | 0 | 0 | 0 | 0 |
| ENSG00000228176 | 0 | 0 | 0 | 0 | 0 | 0 |
| ENSG00000284676 | 0 | 0 | 0 | 0 | 0 | 0 |
| ENSG00000270927 | 0 | 0 | 0 | 0 | 0 | 0 |
| ENSG00000233399 | 0 | 0 | 0 | 0 | 0 | 3 |
| ENSG00000233372 | 0 | 0 | 0 | 0 | 0 | 0 |
| ENSG00000231251 | 0 | 0 | 0 | 0 | 0 | 0 |
| ENSG00000236335 | 0 | 0 | 0 | 0 | 0 | 0 |
| ENSG00000231949 | 0 | 0 | 0 | 0 | 0 | 0 |
| ENSG00000162510 | 10 | 0 | 4 | 4 | 6 | 4 |
| ENSG00000186056 | 71 | 58 | 68 | 39 | 48 | 56 |
| ENSG00000162511 | 37 | 15 | 10 | 20 | 33 | 14 |
| ENSG00000264773 | 0 | 0 | 0 | 0 | 0 | 0 |
| ENSG00000229607 | 0 | 0 | 1 | 0 | 5 | 5 |
| ENSG00000223382 | 10 | 6 | 7 | 8 | 17 | 15 |
| ENSG00000222784 | 0 | 0 | 0 | 0 | 0 | 0 |

| | | | | | | |
|---|---|---|---|---|---|---|
| ENSG00000235143 | 0 | 0 | 0 | 0 | 0 | 1 |
| ENSG00000162512 | 1803 | 890 | 1052 | 1062 | 1095 | 980 |
| ENSG00000266210 | 0 | 0 | 0 | 0 | 0 | 0 |
| ENSG00000134644 | 3295 | 1547 | 1949 | 1222 | 1728 | 1759 |
| ENSG00000237329 | 0 | 0 | 1 | 0 | 0 | 0 |
| ENSG00000232768 | 0 | 0 | 0 | 0 | 0 | 0 |
| ENSG00000215900 | 0 | 0 | 0 | 0 | 0 | 0 |
| ENSG00000225142 | 0 | 0 | 0 | 0 | 0 | 0 |
| ENSG00000084628 | 125 | 59 | 54 | 154 | 83 | 123 |
| ENSG00000060688 | 1178 | 583 | 601 | 681 | 708 | 649 |
| ENSG00000229447 | 1 | 1 | 5 | 2 | 3 | 4 |
| ENSG00000121766 | 879 | 495 | 606 | 596 | 611 | 582 |
| ENSG00000229044 | 0 | 0 | 0 | 0 | 1 | 0 |
| ENSG00000121769 | 19 | 6 | 10 | 4 | 8 | 8 |
| ENSG00000168528 | 318 | 153 | 203 | 186 | 149 | 131 |
| ENSG00000215899 | 0 | 0 | 0 | 0 | 0 | 0 |
| ENSG00000206981 | 0 | 0 | 0 | 0 | 0 | 0 |
| ENSG00000260386 | 3 | 0 | 0 | 0 | 0 | 0 |
| ENSG00000265961 | 0 | 0 | 0 | 0 | 0 | 0 |
| ENSG00000284543 | 2 | 1 | 0 | 0 | 0 | 0 |
| ENSG00000229167 | 0 | 0 | 0 | 0 | 0 | 0 |
| ENSG00000142910 | 34 | 21 | 26 | 10 | 10 | 12 |
| ENSG00000121764 | 5 | 0 | 0 | 1 | 0 | 1 |
| ENSG00000162517 | 2332 | 1124 | 1480 | 1472 | 1284 | 1321 |
| ENSG00000235790 | 2 | 0 | 5 | 0 | 0 | 0 |
| ENSG00000264078 | 1 | 1 | 1 | 1 | 0 | 2 |
| ENSG00000084636 | 65 | 20 | 43 | 27 | 18 | 24 |
| ENSG00000121753 | 124 | 51 | 71 | 44 | 57 | 47 |
| ENSG00000266580 | 0 | 0 | 0 | 0 | 0 | 0 |
| ENSG00000254545 | 0 | 0 | 0 | 0 | 0 | 0 |
| ENSG00000134668 | 46 | 20 | 21 | 20 | 31 | 26 |
| ENSG00000203620 | 0 | 0 | 0 | 0 | 0 | 0 |
| ENSG00000269967 | 0 | 1 | 3 | 1 | 0 | 1 |
| ENSG00000184007 | 5727 | 2593 | 3040 | 2504 | 2922 | 2881 |
| ENSG00000228634 | 4 | 3 | 1 | 0 | 0 | 3 |
| ENSG00000284702 | 2 | 0 | 0 | 0 | 0 | 0 |
| ENSG00000121774 | 7080 | 3414 | 4402 | 3469 | 4002 | 4132 |
| ENSG00000203325 | 1 | 1 | 1 | 0 | 1 | 0 |
| ENSG00000121775 | 300 | 188 | 200 | 199 | 181 | 148 |
| ENSG00000266203 | 0 | 0 | 0 | 0 | 0 | 0 |
| ENSG00000276493 | 0 | 0 | 0 | 0 | 0 | 0 |
| ENSG00000025800 | 4015 | 1844 | 2421 | 1517 | 2206 | 2044 |
| ENSG00000250135 | 0 | 0 | 0 | 0 | 0 | 0 |
| ENSG00000084652 | 2811 | 1276 | 1795 | 1433 | 1495 | 1659 |
| ENSG00000160050 | 18 | 14 | 30 | 16 | 16 | 16 |
| ENSG00000224066 | 3 | 0 | 4 | 0 | 1 | 1 |
| ENSG00000160051 | 154 | 72 | 99 | 87 | 83 | 99 |
| ENSG00000222046 | 6 | 5 | 3 | 4 | 4 | 1 |
| ENSG00000160055 | 204 | 116 | 144 | 127 | 152 | 137 |
| ENSG00000084623 | 6669 | 3379 | 4018 | 3634 | 4204 | 3710 |

| | | | | | | |
|---|---|---|---|---|---|---|
| ENSG00000220785 | 56 | 20 | 35 | 18 | 16 | 30 |
| ENSG00000183615 | 22 | 14 | 14 | 9 | 22 | 19 |
| ENSG00000182866 | 0 | 0 | 0 | 0 | 0 | 0 |
| ENSG00000277617 | 0 | 0 | 0 | 0 | 0 | 0 |
| ENSG00000252450 | 0 | 0 | 0 | 0 | 0 | 0 |
| ENSG00000116478 | 6741 | 2878 | 4156 | 2860 | 3309 | 3532 |
| ENSG00000175130 | 9943 | 5104 | 5917 | 6050 | 5556 | 5472 |
| ENSG00000233775 | 2 | 0 | 1 | 0 | 3 | 2 |
| ENSG00000162526 | 17 | 7 | 20 | 0 | 10 | 5 |
| ENSG00000225828 | 113 | 55 | 68 | 58 | 60 | 70 |
| ENSG00000160058 | 3883 | 1868 | 2575 | 1924 | 2088 | 2205 |
| ENSG00000224333 | 0 | 2 | 0 | 0 | 0 | 0 |
| ENSG00000270850 | 0 | 0 | 0 | 0 | 0 | 0 |
| ENSG00000229259 | 0 | 0 | 0 | 0 | 0 | 0 |
| ENSG00000263529 | 0 | 0 | 0 | 0 | 0 | 0 |
| ENSG00000254553 | 0 | 0 | 0 | 0 | 0 | 0 |
| ENSG00000273274 | 1 | 5 | 7 | 3 | 5 | 3 |
| ENSG00000160062 | 85 | 72 | 84 | 37 | 73 | 63 |
| ENSG00000176261 | 746 | 338 | 467 | 381 | 483 | 437 |
| ENSG00000200591 | 0 | 0 | 1 | 0 | 0 | 0 |
| ENSG00000162521 | 4502 | 2091 | 2894 | 1917 | 2311 | 2449 |
| ENSG00000162520 | 2 | 4 | 3 | 2 | 6 | 8 |
| ENSG00000224409 | 4 | 10 | 6 | 8 | 3 | 1 |
| ENSG00000162522 | 11856 | 5237 | 7061 | 5644 | 5873 | 5933 |
| ENSG00000134684 | 2767 | 1371 | 1678 | 2017 | 1781 | 1834 |
| ENSG00000116497 | 1159 | 516 | 684 | 401 | 518 | 476 |
| ENSG00000160097 | 28 | 11 | 16 | 8 | 11 | 17 |
| ENSG00000121905 | 23 | 5 | 22 | 7 | 8 | 9 |
| ENSG00000121900 | 1976 | 1047 | 1275 | 1265 | 1054 | 1119 |
| ENSG00000116514 | 500 | 237 | 334 | 292 | 268 | 278 |
| ENSG00000278966 | 0 | 0 | 0 | 0 | 0 | 0 |
| ENSG00000217644 | 0 | 1 | 0 | 0 | 0 | 1 |
| ENSG00000239670 | 2 | 0 | 0 | 0 | 0 | 0 |
| ENSG00000236065 | 1 | 0 | 0 | 0 | 0 | 0 |
| ENSG00000004455 | 3050 | 1530 | 2034 | 1873 | 1833 | 1882 |
| ENSG00000276645 | 1 | 0 | 0 | 0 | 0 | 0 |
| ENSG00000142920 | 54 | 39 | 38 | 26 | 34 | 26 |
| ENSG00000278997 | 32 | 29 | 24 | 19 | 27 | 23 |
| ENSG00000116525 | 856 | 452 | 539 | 451 | 418 | 412 |
| ENSG00000279179 | 23 | 11 | 23 | 9 | 6 | 23 |
| ENSG00000284721 | 0 | 0 | 0 | 0 | 0 | 0 |
| ENSG00000160094 | 1304 | 581 | 817 | 609 | 609 | 676 |
| ENSG00000270115 | 10 | 12 | 9 | 4 | 2 | 2 |
| ENSG00000184389 | 0 | 0 | 0 | 0 | 0 | 0 |
| ENSG00000225313 | 34 | 13 | 27 | 10 | 23 | 11 |
| ENSG00000134686 | 1385 | 665 | 872 | 860 | 818 | 757 |
| ENSG00000284154 | 3 | 1 | 2 | 1 | 1 | 1 |
| ENSG00000222112 | 0 | 0 | 1 | 4 | 0 | 1 |
| ENSG00000233246 | 7 | 3 | 3 | 4 | 6 | 1 |
| ENSG00000234512 | 0 | 0 | 0 | 0 | 0 | 0 |

| | | | | | | |
|---|---|---|---|---|---|---|
| ENSG00000121903 | 191 | 81 | 91 | 71 | 71 | 70 |
| ENSG00000235907 | 0 | 0 | 0 | 0 | 0 | 0 |
| ENSG00000121904 | 0 | 1 | 2 | 0 | 0 | 0 |
| ENSG00000227000 | 0 | 0 | 0 | 0 | 0 | 0 |
| ENSG00000176256 | 0 | 0 | 0 | 0 | 0 | 0 |
| ENSG00000231163 | 0 | 0 | 0 | 0 | 0 | 0 |
| ENSG00000201148 | 0 | 0 | 0 | 0 | 0 | 0 |
| ENSG00000142698 | 0 | 0 | 0 | 0 | 0 | 0 |
| ENSG00000270241 | 0 | 0 | 0 | 0 | 0 | 0 |
| ENSG00000207941 | 0 | 0 | 0 | 0 | 0 | 0 |
| ENSG00000163866 | 879 | 430 | 533 | 583 | 479 | 565 |
| ENSG00000189280 | 2 | 3 | 5 | 2 | 6 | 3 |
| ENSG00000189433 | 0 | 0 | 1 | 3 | 2 | 2 |
| ENSG00000255811 | 0 | 0 | 0 | 0 | 0 | 0 |
| ENSG00000188910 | 58 | 41 | 31 | 49 | 42 | 49 |
| ENSG00000187513 | 0 | 0 | 0 | 0 | 0 | 0 |
| ENSG00000230163 | 38 | 19 | 27 | 21 | 16 | 20 |
| ENSG00000116544 | 52 | 33 | 40 | 32 | 32 | 24 |
| ENSG00000284773 | 14 | 15 | 10 | 11 | 11 | 10 |
| ENSG00000241014 | 8 | 4 | 4 | 7 | 9 | 1 |
| ENSG00000271741 | 0 | 1 | 0 | 0 | 0 | 0 |
| ENSG00000243749 | 0 | 7 | 12 | 7 | 3 | 6 |
| ENSG00000163867 | 269 | 109 | 154 | 82 | 139 | 105 |
| ENSG00000197056 | 223 | 136 | 121 | 75 | 225 | 112 |
| ENSG00000233940 | 0 | 0 | 0 | 0 | 0 | 0 |
| ENSG00000228348 | 0 | 0 | 0 | 0 | 1 | 0 |
| ENSG00000284640 | 0 | 0 | 1 | 2 | 0 | 1 |
| ENSG00000116560 | 8052 | 3688 | 4695 | 3966 | 4569 | 4742 |
| ENSG00000201868 | 0 | 0 | 0 | 0 | 0 | 0 |
| ENSG00000146463 | 1281 | 674 | 634 | 366 | 962 | 647 |
| ENSG00000240374 | 0 | 0 | 0 | 0 | 0 | 0 |
| ENSG00000201542 | 0 | 0 | 0 | 0 | 0 | 0 |
| ENSG00000229994 | 42 | 35 | 24 | 25 | 35 | 37 |
| ENSG00000227409 | 2 | 0 | 0 | 0 | 0 | 0 |
| ENSG00000278702 | 0 | 0 | 0 | 1 | 0 | 1 |
| ENSG00000142687 | 3019 | 1445 | 1908 | 1232 | 1384 | 1496 |
| ENSG00000236274 | 0 | 0 | 0 | 0 | 0 | 0 |
| ENSG00000020129 | 2433 | 1093 | 1481 | 1265 | 1129 | 1266 |
| ENSG00000239636 | 0 | 2 | 3 | 2 | 0 | 1 |
| ENSG00000116819 | 10 | 6 | 8 | 6 | 3 | 5 |
| ENSG00000126067 | 4219 | 2258 | 2706 | 2714 | 2747 | 2773 |
| ENSG00000142686 | 601 | 296 | 327 | 361 | 358 | 334 |
| ENSG00000092853 | 191 | 128 | 121 | 57 | 98 | 94 |
| ENSG00000232335 | 0 | 1 | 0 | 0 | 0 | 0 |
| ENSG00000134698 | 635 | 313 | 457 | 202 | 346 | 383 |
| ENSG00000092847 | 1834 | 803 | 1015 | 754 | 1008 | 1001 |
| ENSG00000271164 | 0 | 0 | 0 | 0 | 0 | 0 |
| ENSG00000271914 | 0 | 0 | 0 | 0 | 0 | 0 |
| ENSG00000126070 | 565 | 261 | 265 | 156 | 333 | 262 |
| ENSG00000271554 | 1 | 1 | 0 | 0 | 0 | 0 |

| | | | | | | |
|---|---|---|---|---|---|---|
| ENSG00000232862 | 0 | 0 | 0 | 0 | 1 | 0 |
| ENSG00000092850 | 0 | 0 | 0 | 0 | 3 | 0 |
| ENSG00000116863 | 867 | 379 | 512 | 497 | 434 | 456 |
| ENSG00000171812 | 0 | 0 | 0 | 0 | 0 | 0 |
| ENSG00000054116 | 2147 | 1144 | 1272 | 1517 | 1194 | 1348 |
| ENSG00000116871 | 1791 | 902 | 1017 | 1265 | 1164 | 1022 |
| ENSG00000264592 | 0 | 0 | 0 | 0 | 0 | 0 |
| ENSG00000054118 | 6449 | 2840 | 3807 | 2388 | 3103 | 3381 |
| ENSG00000231849 | 0 | 0 | 1 | 0 | 0 | 0 |
| ENSG00000214193 | 302 | 148 | 197 | 117 | 158 | 159 |
| ENSG00000142694 | 144 | 95 | 106 | 145 | 139 | 131 |
| ENSG00000116883 | 89 | 41 | 79 | 53 | 67 | 59 |
| ENSG00000196182 | 1848 | 932 | 1130 | 1137 | 1061 | 1076 |
| ENSG00000181817 | 843 | 407 | 479 | 648 | 504 | 511 |
| ENSG00000222821 | 0 | 0 | 0 | 0 | 0 | 0 |
| ENSG00000116885 | 335 | 167 | 179 | 155 | 175 | 167 |
| ENSG00000201448 | 4 | 1 | 1 | 0 | 3 | 2 |
| ENSG00000116898 | 2012 | 1005 | 1170 | 1268 | 1141 | 1169 |
| ENSG00000119535 | 14 | 11 | 5 | 1 | 7 | 5 |
| ENSG00000230214 | 0 | 0 | 0 | 0 | 0 | 0 |
| ENSG00000284705 | 0 | 0 | 0 | 0 | 0 | 0 |
| ENSG00000284720 | 0 | 0 | 0 | 0 | 0 | 0 |
| ENSG00000234481 | 0 | 0 | 0 | 0 | 0 | 0 |
| ENSG00000163873 | 4 | 2 | 5 | 0 | 1 | 0 |
| ENSG00000284650 | 0 | 0 | 0 | 0 | 0 | 0 |
| ENSG00000264698 | 0 | 0 | 0 | 0 | 0 | 0 |
| ENSG00000207328 | 0 | 0 | 0 | 0 | 0 | 0 |
| ENSG00000223944 | 0 | 0 | 0 | 0 | 0 | 0 |
| ENSG00000252368 | 0 | 0 | 0 | 0 | 0 | 0 |
| ENSG00000230451 | 0 | 0 | 0 | 0 | 0 | 0 |
| ENSG00000233621 | 333 | 203 | 240 | 213 | 167 | 183 |
| ENSG00000163874 | 1492 | 745 | 952 | 745 | 808 | 824 |
| ENSG00000283724 | 0 | 0 | 0 | 0 | 0 | 0 |
| ENSG00000163875 | 2322 | 1204 | 1606 | 1168 | 1205 | 1307 |
| ENSG00000263675 | 5 | 3 | 10 | 3 | 4 | 1 |
| ENSG00000163877 | 977 | 529 | 684 | 491 | 559 | 598 |
| ENSG00000232273 | 0 | 0 | 0 | 0 | 0 | 0 |
| ENSG00000237749 | 0 | 0 | 0 | 1 | 0 | 1 |
| ENSG00000163879 | 0 | 1 | 3 | 1 | 2 | 1 |
| ENSG00000134697 | 1200 | 707 | 806 | 612 | 897 | 797 |
| ENSG00000284748 | 0 | 0 | 0 | 0 | 0 | 0 |
| ENSG00000169218 | 2 | 0 | 0 | 1 | 0 | 2 |
| ENSG00000116922 | 721 | 328 | 446 | 310 | 427 | 404 |
| ENSG00000134690 | 666 | 296 | 414 | 436 | 483 | 504 |
| ENSG00000183317 | 371 | 174 | 218 | 178 | 196 | 192 |
| ENSG00000200421 | 0 | 0 | 0 | 0 | 0 | 0 |
| ENSG00000214141 | 6 | 0 | 0 | 0 | 0 | 0 |
| ENSG00000227416 | 0 | 0 | 2 | 0 | 0 | 0 |
| ENSG00000185090 | 664 | 387 | 463 | 487 | 452 | 521 |
| ENSG00000233728 | 0 | 0 | 0 | 0 | 0 | 0 |

| | | | | | | |
|---|---|---|---|---|---|---|
| ENSG00000196449 | 679 | 346 | 423 | 469 | 456 | 476 |
| ENSG00000197982 | 1248 | 641 | 715 | 1011 | 760 | 713 |
| ENSG00000188786 | 897 | 441 | 538 | 327 | 443 | 440 |
| ENSG00000235673 | 0 | 0 | 0 | 0 | 0 | 0 |
| ENSG00000204084 | 422 | 205 | 252 | 172 | 234 | 216 |
| ENSG00000230955 | 0 | 0 | 0 | 0 | 0 | 0 |
| ENSG00000252448 | 1 | 1 | 1 | 1 | 0 | 0 |
| ENSG00000222282 | 0 | 0 | 0 | 0 | 0 | 0 |
| ENSG00000277168 | 0 | 0 | 0 | 0 | 0 | 0 |
| ENSG00000183431 | 2352 | 1209 | 1410 | 1149 | 1411 | 1351 |
| ENSG00000212541 | 1 | 2 | 1 | 0 | 0 | 1 |
| ENSG00000183386 | 70 | 41 | 45 | 58 | 50 | 41 |
| ENSG00000183520 | 771 | 417 | 467 | 363 | 605 | 476 |
| ENSG00000185668 | 0 | 1 | 0 | 1 | 1 | 0 |
| ENSG00000224592 | 1 | 0 | 0 | 0 | 0 | 0 |
| ENSG00000223589 | 0 | 0 | 0 | 0 | 0 | 0 |
| ENSG00000265596 | 0 | 0 | 0 | 0 | 0 | 0 |
| ENSG00000284623 | 0 | 0 | 0 | 0 | 0 | 0 |
| ENSG00000237290 | 0 | 0 | 0 | 0 | 0 | 0 |
| ENSG00000275350 | 0 | 0 | 0 | 0 | 0 | 0 |
| ENSG00000200796 | 0 | 0 | 0 | 0 | 0 | 0 |
| ENSG00000238063 | 0 | 0 | 0 | 0 | 0 | 0 |
| ENSG00000215895 | 2 | 0 | 0 | 0 | 0 | 1 |
| ENSG00000284632 | 0 | 0 | 0 | 0 | 0 | 0 |
| ENSG00000116954 | 552 | 244 | 285 | 242 | 284 | 295 |
| ENSG00000273637 | 0 | 0 | 0 | 0 | 0 | 0 |
| ENSG00000228436 | 13 | 7 | 12 | 9 | 6 | 11 |
| ENSG00000214114 | 941 | 505 | 621 | 508 | 577 | 580 |
| ENSG00000274944 | 0 | 0 | 0 | 0 | 1 | 1 |
| ENSG00000131233 | 1 | 1 | 1 | 0 | 0 | 2 |
| ENSG00000158315 | 8 | 1 | 8 | 2 | 5 | 1 |
| ENSG00000199963 | 0 | 0 | 0 | 0 | 0 | 0 |
| ENSG00000207466 | 0 | 0 | 0 | 0 | 0 | 0 |
| ENSG00000235549 | 0 | 0 | 0 | 0 | 0 | 0 |
| ENSG00000174574 | 1992 | 916 | 1076 | 907 | 1156 | 1187 |
| ENSG00000168653 | 3364 | 1712 | 1889 | 2276 | 2135 | 1897 |
| ENSG00000127603 | 1625 | 848 | 999 | 476 | 1070 | 987 |
| ENSG00000206654 | 0 | 0 | 0 | 0 | 0 | 0 |
| ENSG00000222378 | 0 | 0 | 0 | 0 | 0 | 0 |
| ENSG00000270040 | 0 | 0 | 0 | 1 | 0 | 0 |
| ENSG00000226438 | 0 | 1 | 0 | 0 | 0 | 0 |
| ENSG00000217897 | 0 | 0 | 0 | 0 | 0 | 0 |
| ENSG00000183682 | 6 | 5 | 8 | 7 | 2 | 7 |
| ENSG00000237624 | 0 | 0 | 0 | 0 | 0 | 0 |
| ENSG00000243970 | 18 | 15 | 24 | 15 | 21 | 14 |
| ENSG00000090621 | 3541 | 1940 | 2358 | 2297 | 2489 | 2445 |
| ENSG00000228060 | 10 | 6 | 7 | 8 | 11 | 5 |
| ENSG00000201457 | 1 | 4 | 0 | 1 | 3 | 2 |
| ENSG00000163909 | 36 | 20 | 26 | 18 | 34 | 23 |
| ENSG00000225903 | 0 | 0 | 0 | 0 | 0 | 0 |

| | | | | | | |
|---|---|---|---|---|---|---|
| ENSG00000116981 | 0 | 0 | 0 | 0 | 0 | 0 |
| ENSG00000116983 | 10 | 2 | 1 | 2 | 0 | 2 |
| ENSG00000084072 | 1635 | 811 | 969 | 946 | 905 | 882 |
| ENSG00000225333 | 0 | 0 | 0 | 0 | 0 | 0 |
| ENSG00000252413 | 0 | 0 | 0 | 0 | 0 | 0 |
| ENSG00000116985 | 208 | 139 | 177 | 102 | 117 | 119 |
| ENSG00000198754 | 2 | 2 | 4 | 5 | 4 | 4 |
| ENSG00000285905 | 2 | 0 | 1 | 0 | 0 | 0 |
| ENSG00000261798 | 7 | 3 | 5 | 6 | 5 | 7 |
| ENSG00000229213 | 0 | 0 | 0 | 0 | 0 | 0 |
| ENSG00000284719 | 3 | 6 | 1 | 6 | 2 | 5 |
| ENSG00000284678 | 0 | 0 | 1 | 0 | 0 | 0 |
| ENSG00000043514 | 575 | 270 | 352 | 278 | 353 | 324 |
| ENSG00000202222 | 0 | 0 | 0 | 0 | 0 | 0 |
| ENSG00000116990 | 343 | 169 | 203 | 126 | 135 | 168 |
| ENSG00000236546 | 0 | 1 | 0 | 0 | 1 | 2 |
| ENSG00000207356 | 0 | 0 | 0 | 0 | 0 | 0 |
| ENSG00000168389 | 26 | 18 | 16 | 23 | 22 | 12 |
| ENSG00000228477 | 2 | 4 | 0 | 4 | 3 | 2 |
| ENSG00000131236 | 4647 | 2476 | 2920 | 2238 | 2894 | 2728 |
| ENSG00000131238 | 6457 | 3242 | 3948 | 2490 | 2921 | 2904 |
| ENSG00000226745 | 0 | 0 | 0 | 0 | 0 | 0 |
| ENSG00000117000 | 388 | 230 | 140 | 74 | 342 | 173 |
| ENSG00000207508 | 0 | 0 | 0 | 0 | 0 | 0 |
| ENSG00000188800 | 0 | 0 | 0 | 1 | 0 | 0 |
| ENSG00000259943 | 324 | 134 | 201 | 129 | 151 | 126 |
| ENSG00000084073 | 2373 | 1232 | 1440 | 1021 | 1378 | 1311 |
| ENSG00000231296 | 1 | 0 | 4 | 0 | 0 | 1 |
| ENSG00000049089 | 708 | 325 | 459 | 234 | 189 | 256 |
| ENSG00000227311 | 0 | 0 | 0 | 0 | 0 | 0 |
| ENSG00000084070 | 1756 | 696 | 978 | 813 | 786 | 894 |
| ENSG00000213172 | 0 | 0 | 0 | 0 | 0 | 0 |
| ENSG00000284677 | 0 | 0 | 0 | 0 | 0 | 1 |
| ENSG00000187801 | 60 | 28 | 29 | 21 | 46 | 30 |
| ENSG00000260920 | 53 | 42 | 36 | 23 | 44 | 30 |
| ENSG00000279667 | 7 | 1 | 3 | 0 | 6 | 0 |
| ENSG00000187815 | 155 | 62 | 78 | 69 | 108 | 74 |
| ENSG00000238287 | 9 | 5 | 8 | 6 | 11 | 7 |
| ENSG00000164002 | 140 | 81 | 99 | 92 | 104 | 117 |
| ENSG00000227278 | 0 | 0 | 0 | 0 | 0 | 0 |
| ENSG00000238186 | 8 | 4 | 4 | 4 | 2 | 3 |
| ENSG00000117010 | 53 | 32 | 34 | 28 | 29 | 23 |
| ENSG00000236505 | 0 | 0 | 0 | 0 | 0 | 0 |
| ENSG00000233910 | 0 | 0 | 0 | 0 | 0 | 0 |
| ENSG00000117016 | 692 | 298 | 369 | 236 | 292 | 282 |
| ENSG00000237899 | 13 | 3 | 9 | 2 | 7 | 7 |
| ENSG00000272145 | 52 | 34 | 53 | 11 | 36 | 26 |
| ENSG00000066136 | 1329 | 643 | 848 | 585 | 589 | 657 |
| ENSG00000198974 | 0 | 0 | 0 | 0 | 0 | 0 |
| ENSG00000207962 | 0 | 0 | 0 | 0 | 0 | 0 |

| | | | | | | |
|---|---|---|---|---|---|---|
| ENSG00000117013 | 48 | 23 | 31 | 30 | 36 | 42 |
| ENSG00000264582 | 0 | 0 | 0 | 0 | 0 | 1 |
| ENSG00000179862 | 242 | 125 | 176 | 190 | 139 | 152 |
| ENSG00000229528 | 0 | 0 | 0 | 0 | 0 | 0 |
| ENSG00000228940 | 0 | 0 | 0 | 0 | 0 | 0 |
| ENSG00000233118 | 0 | 0 | 0 | 0 | 0 | 0 |
| ENSG00000171793 | 1567 | 722 | 853 | 828 | 966 | 1092 |
| ENSG00000281207 | 6 | 5 | 11 | 3 | 9 | 9 |
| ENSG00000171790 | 18 | 5 | 5 | 0 | 5 | 6 |
| ENSG00000010803 | 775 | 374 | 455 | 379 | 492 | 478 |
| ENSG00000215893 | 0 | 0 | 0 | 0 | 0 | 0 |
| ENSG00000235358 | 0 | 2 | 0 | 2 | 0 | 3 |
| ENSG00000213987 | 0 | 0 | 0 | 0 | 0 | 0 |
| ENSG00000204060 | 374 | 183 | 193 | 259 | 232 | 179 |
| ENSG00000229901 | 0 | 2 | 2 | 3 | 4 | 0 |
| ENSG00000252563 | 0 | 0 | 0 | 0 | 0 | 0 |
| ENSG00000127129 | 50 | 39 | 47 | 39 | 56 | 33 |
| ENSG00000127124 | 574 | 239 | 301 | 181 | 204 | 211 |
| ENSG00000230881 | 0 | 0 | 1 | 1 | 1 | 0 |
| ENSG00000230638 | 19 | 14 | 14 | 3 | 12 | 8 |
| ENSG00000284895 | 0 | 0 | 0 | 0 | 0 | 0 |
| ENSG00000227538 | 0 | 0 | 0 | 0 | 0 | 0 |
| ENSG00000228776 | 0 | 0 | 0 | 0 | 0 | 0 |
| ENSG00000044012 | 0 | 0 | 0 | 0 | 0 | 0 |
| ENSG00000197273 | 0 | 0 | 0 | 0 | 0 | 0 |
| ENSG00000198815 | 1865 | 899 | 1109 | 578 | 910 | 915 |
| ENSG00000227527 | 9 | 3 | 11 | 3 | 3 | 3 |
| ENSG00000177181 | 146 | 58 | 93 | 46 | 58 | 53 |
| ENSG00000235002 | 0 | 0 | 0 | 0 | 0 | 0 |
| ENSG00000066185 | 28 | 8 | 15 | 6 | 15 | 13 |
| ENSG00000127125 | 2482 | 1254 | 1511 | 1346 | 1305 | 1374 |
| ENSG00000186409 | 51 | 36 | 41 | 14 | 38 | 21 |
| ENSG00000230116 | 0 | 0 | 0 | 0 | 0 | 0 |
| ENSG00000236876 | 0 | 1 | 2 | 2 | 0 | 0 |
| ENSG00000200254 | 0 | 0 | 0 | 0 | 0 | 0 |
| ENSG00000236180 | 0 | 0 | 0 | 0 | 0 | 0 |
| ENSG00000171960 | 475 | 287 | 340 | 383 | 320 | 339 |
| ENSG00000285728 | 23 | 16 | 18 | 15 | 19 | 26 |
| ENSG00000234917 | 0 | 1 | 4 | 0 | 4 | 3 |
| ENSG00000065978 | 11201 | 5600 | 6180 | 6978 | 7159 | 7005 |
| ENSG00000164007 | 0 | 0 | 0 | 0 | 0 | 0 |
| ENSG00000117385 | 748 | 331 | 433 | 393 | 415 | 358 |
| ENSG00000164008 | 93 | 25 | 66 | 34 | 54 | 65 |
| ENSG00000283580 | 0 | 0 | 0 | 1 | 1 | 0 |
| ENSG00000228452 | 0 | 0 | 0 | 0 | 0 | 0 |
| ENSG00000274386 | 16 | 3 | 8 | 6 | 15 | 9 |
| ENSG00000177868 | 286 | 174 | 174 | 163 | 188 | 152 |
| ENSG00000164010 | 630 | 318 | 432 | 278 | 328 | 319 |
| ENSG00000228192 | 7 | 10 | 9 | 8 | 6 | 5 |
| ENSG00000164011 | 332 | 200 | 163 | 197 | 178 | 175 |

| | | | | | | |
|---|---|---|---|---|---|---|
| ENSG00000233708 | 0 | 0 | 0 | 0 | 0 | 1 |
| ENSG00000237090 | 0 | 0 | 0 | 0 | 0 | 0 |
| ENSG00000225099 | 0 | 2 | 0 | 0 | 0 | 0 |
| ENSG00000117394 | 6225 | 2810 | 3742 | 3212 | 2667 | 3022 |
| ENSG00000284138 | 1 | 2 | 3 | 0 | 0 | 1 |
| ENSG00000227533 | 24 | 17 | 19 | 18 | 16 | 19 |
| ENSG00000283973 | 2 | 0 | 1 | 0 | 0 | 1 |
| ENSG00000207256 | 0 | 0 | 1 | 0 | 0 | 0 |
| ENSG00000283136 | 0 | 0 | 0 | 0 | 0 | 0 |
| ENSG00000277513 | 0 | 0 | 0 | 0 | 0 | 0 |
| ENSG00000186973 | 67 | 20 | 36 | 16 | 24 | 36 |
| ENSG00000117395 | 2213 | 1245 | 1455 | 1591 | 1561 | 1677 |
| ENSG00000284410 | 0 | 0 | 0 | 0 | 0 | 0 |
| ENSG00000243710 | 13 | 13 | 10 | 1 | 6 | 4 |
| ENSG00000199240 | 0 | 0 | 0 | 0 | 0 | 0 |
| ENSG00000179178 | 735 | 339 | 424 | 485 | 379 | 402 |
| ENSG00000253313 | 651 | 300 | 412 | 324 | 360 | 360 |
| ENSG00000066056 | 2 | 0 | 0 | 0 | 0 | 1 |
| ENSG00000117400 | 1 | 4 | 6 | 2 | 1 | 5 |
| ENSG00000234694 | 26 | 17 | 18 | 21 | 23 | 13 |
| ENSG00000117399 | 1342 | 620 | 730 | 1028 | 792 | 918 |
| ENSG00000066322 | 4491 | 2442 | 2773 | 3001 | 2648 | 2793 |
| ENSG00000283836 | 0 | 0 | 0 | 0 | 0 | 0 |
| ENSG00000159479 | 1050 | 509 | 669 | 574 | 602 | 614 |
| ENSG00000229431 | 20 | 3 | 7 | 1 | 5 | 4 |
| ENSG00000198198 | 906 | 391 | 664 | 338 | 426 | 482 |
| ENSG00000229372 | 0 | 0 | 0 | 0 | 0 | 0 |
| ENSG00000274975 | 0 | 0 | 0 | 0 | 0 | 0 |
| ENSG00000178922 | 0 | 0 | 0 | 0 | 0 | 0 |
| ENSG00000229348 | 2 | 0 | 0 | 0 | 0 | 2 |
| ENSG00000142949 | 19971 | 8519 | 12287 | 7783 | 9057 | 9390 |
| ENSG00000284989 | 0 | 0 | 0 | 0 | 1 | 0 |
| ENSG00000066135 | 1347 | 646 | 894 | 714 | 823 | 798 |
| ENSG00000236200 | 43 | 25 | 17 | 23 | 29 | 26 |
| ENSG00000126091 | 22 | 17 | 22 | 11 | 18 | 19 |
| ENSG00000229444 | 0 | 0 | 0 | 0 | 0 | 0 |
| ENSG00000283414 | 0 | 0 | 0 | 0 | 0 | 0 |
| ENSG00000233674 | 0 | 0 | 0 | 0 | 0 | 0 |
| ENSG00000283477 | 0 | 0 | 0 | 0 | 0 | 0 |
| ENSG00000224751 | 0 | 0 | 0 | 0 | 0 | 0 |
| ENSG00000117407 | 42 | 28 | 19 | 32 | 19 | 21 |
| ENSG00000237950 | 18 | 3 | 6 | 13 | 6 | 9 |
| ENSG00000117408 | 1457 | 643 | 858 | 848 | 745 | 902 |
| ENSG00000285649 | 1 | 4 | 1 | 0 | 1 | 0 |
| ENSG00000132768 | 1069 | 482 | 701 | 778 | 660 | 754 |
| ENSG00000117410 | 4435 | 2077 | 2609 | 2744 | 2168 | 2247 |
| ENSG00000117411 | 1112 | 567 | 670 | 893 | 674 | 737 |
| ENSG00000159214 | 329 | 167 | 212 | 182 | 167 | 161 |
| ENSG00000196517 | 483 | 229 | 314 | 267 | 251 | 255 |
| ENSG00000230615 | 77 | 48 | 54 | 43 | 53 | 59 |

| | | | | | | |
|---|---|---|---|---|---|---|
| ENSG00000227163 | 0 | 0 | 0 | 0 | 0 | 0 |
| ENSG00000271647 | 0 | 0 | 0 | 0 | 0 | 0 |
| ENSG00000239560 | 0 | 0 | 0 | 0 | 0 | 0 |
| ENSG00000171872 | 0 | 0 | 0 | 0 | 0 | 0 |
| ENSG00000233514 | 2 | 1 | 0 | 0 | 0 | 0 |
| ENSG00000283039 | 0 | 0 | 0 | 0 | 0 | 0 |
| ENSG00000226804 | 0 | 0 | 0 | 0 | 0 | 0 |
| ENSG00000200171 | 0 | 0 | 0 | 0 | 0 | 0 |
| ENSG00000238037 | 0 | 0 | 0 | 0 | 0 | 0 |
| ENSG00000227994 | 0 | 0 | 0 | 0 | 0 | 0 |
| ENSG00000271329 | 0 | 0 | 0 | 0 | 0 | 1 |
| ENSG00000178028 | 1366 | 603 | 812 | 645 | 619 | 652 |
| ENSG00000117419 | 1365 | 675 | 790 | 953 | 820 | 835 |
| ENSG00000233602 | 0 | 2 | 0 | 0 | 0 | 2 |
| ENSG00000199385 | 0 | 0 | 0 | 0 | 0 | 0 |
| ENSG00000187147 | 1253 | 632 | 841 | 790 | 817 | 839 |
| ENSG00000263381 | 0 | 0 | 0 | 0 | 0 | 0 |
| ENSG00000126106 | 228 | 128 | 200 | 193 | 181 | 177 |
| ENSG00000198520 | 120 | 61 | 89 | 74 | 76 | 62 |
| ENSG00000199377 | 0 | 0 | 0 | 0 | 0 | 2 |
| ENSG00000200169 | 0 | 1 | 0 | 0 | 1 | 0 |
| ENSG00000142945 | 699 | 365 | 461 | 486 | 560 | 536 |
| ENSG00000225721 | 0 | 0 | 1 | 1 | 3 | 0 |
| ENSG00000142937 | 32829 | 16205 | 19880 | 23050 | 19266 | 19630 |
| ENSG00000264294 | 0 | 0 | 0 | 0 | 0 | 0 |
| ENSG00000200913 | 6 | 2 | 2 | 1 | 1 | 5 |
| ENSG00000202031 | 3 | 2 | 3 | 1 | 0 | 2 |
| ENSG00000207421 | 0 | 0 | 0 | 0 | 0 | 0 |
| ENSG00000281859 | 0 | 0 | 0 | 0 | 0 | 0 |
| ENSG00000234093 | 2 | 2 | 0 | 1 | 3 | 1 |
| ENSG00000142959 | 8 | 2 | 8 | 3 | 6 | 8 |
| ENSG00000173846 | 104 | 61 | 73 | 95 | 72 | 69 |
| ENSG00000188396 | 0 | 1 | 2 | 1 | 0 | 0 |
| ENSG00000222009 | 6 | 2 | 2 | 2 | 4 | 4 |
| ENSG00000117425 | 13 | 1 | 5 | 7 | 7 | 3 |
| ENSG00000202444 | 0 | 0 | 0 | 0 | 0 | 0 |
| ENSG00000226499 | 0 | 0 | 0 | 0 | 0 | 1 |
| ENSG00000070785 | 457 | 268 | 302 | 291 | 338 | 363 |
| ENSG00000253039 | 0 | 0 | 0 | 0 | 0 | 0 |
| ENSG00000229815 | 0 | 0 | 0 | 0 | 0 | 0 |
| ENSG00000232369 | 2 | 1 | 0 | 1 | 0 | 2 |
| ENSG00000268949 | 7 | 1 | 5 | 1 | 3 | 2 |
| ENSG00000126107 | 2084 | 902 | 1133 | 1075 | 890 | 973 |
| ENSG00000126088 | 1399 | 633 | 812 | 906 | 710 | 809 |
| ENSG00000162415 | 1006 | 501 | 575 | 320 | 376 | 400 |
| ENSG00000234315 | 0 | 0 | 0 | 0 | 0 | 0 |
| ENSG00000281912 | 27 | 16 | 23 | 17 | 9 | 24 |
| ENSG00000186603 | 242 | 158 | 201 | 151 | 163 | 202 |
| ENSG00000132781 | 491 | 245 | 296 | 311 | 325 | 275 |
| ENSG00000132773 | 355 | 191 | 235 | 193 | 177 | 194 |

| | | | | | | |
|---|---|---|---|---|---|---|
| ENSG00000070759 | 476 | 222 | 266 | 170 | 216 | 207 |
| ENSG00000236872 | 0 | 0 | 0 | 0 | 0 | 0 |
| ENSG00000280670 | 69 | 49 | 58 | 47 | 36 | 57 |
| ENSG00000132763 | 373 | 180 | 226 | 213 | 233 | 249 |
| ENSG00000117450 | 16341 | 9786 | 10906 | 11142 | 11891 | 11380 |
| ENSG00000234379 | 0 | 0 | 0 | 0 | 0 | 0 |
| ENSG00000117448 | 2305 | 1064 | 1319 | 1473 | 1272 | 1309 |
| ENSG00000281133 | 2 | 0 | 1 | 0 | 0 | 0 |
| ENSG00000280836 | 0 | 1 | 1 | 1 | 0 | 0 |
| ENSG00000132780 | 2206 | 1142 | 1251 | 1016 | 1446 | 1351 |
| ENSG00000281112 | 0 | 0 | 0 | 0 | 0 | 0 |
| ENSG00000281825 | 0 | 0 | 1 | 0 | 0 | 1 |
| ENSG00000159588 | 43 | 12 | 21 | 15 | 14 | 17 |
| ENSG00000159592 | 3455 | 1535 | 2119 | 1308 | 1826 | 1854 |
| ENSG00000225447 | 0 | 0 | 2 | 0 | 1 | 0 |
| ENSG00000234329 | 9 | 4 | 2 | 0 | 9 | 0 |
| ENSG00000159596 | 826 | 433 | 515 | 368 | 522 | 462 |
| ENSG00000197429 | 551 | 281 | 315 | 205 | 278 | 243 |
| ENSG00000230896 | 39 | 24 | 36 | 32 | 32 | 21 |
| ENSG00000236111 | 0 | 0 | 0 | 0 | 0 | 0 |
| ENSG00000086015 | 1226 | 581 | 733 | 691 | 704 | 708 |
| ENSG00000232467 | 0 | 0 | 0 | 0 | 0 | 1 |
| ENSG00000117461 | 6680 | 2830 | 3729 | 2657 | 3259 | 3162 |
| ENSG00000278139 | 0 | 0 | 0 | 0 | 0 | 0 |
| ENSG00000226957 | 6 | 1 | 4 | 1 | 2 | 0 |
| ENSG00000233114 | 0 | 1 | 0 | 0 | 0 | 0 |
| ENSG00000227857 | 19 | 17 | 15 | 11 | 13 | 8 |
| ENSG00000117472 | 990 | 461 | 636 | 389 | 316 | 354 |
| ENSG00000250719 | 0 | 0 | 0 | 0 | 0 | 0 |
| ENSG00000085998 | 3247 | 1406 | 1911 | 1793 | 1621 | 1675 |
| ENSG00000171357 | 4 | 3 | 2 | 1 | 3 | 3 |
| ENSG00000085999 | 158 | 88 | 110 | 76 | 103 | 112 |
| ENSG00000132128 | 3761 | 1632 | 2234 | 2141 | 1836 | 1950 |
| ENSG00000173660 | 3373 | 1636 | 1905 | 2168 | 1986 | 1929 |
| ENSG00000117481 | 1046 | 462 | 596 | 491 | 562 | 581 |
| ENSG00000225779 | 0 | 0 | 0 | 0 | 0 | 0 |
| ENSG00000117480 | 1197 | 534 | 731 | 597 | 572 | 605 |
| ENSG00000232022 | 7 | 4 | 1 | 4 | 2 | 7 |
| ENSG00000224863 | 0 | 0 | 0 | 0 | 0 | 0 |
| ENSG00000197587 | 0 | 2 | 2 | 3 | 7 | 4 |
| ENSG00000282881 | 0 | 0 | 0 | 0 | 0 | 0 |
| ENSG00000269956 | 13 | 5 | 4 | 6 | 12 | 8 |
| ENSG00000162456 | 0 | 0 | 0 | 0 | 0 | 0 |
| ENSG00000079277 | 961 | 541 | 701 | 588 | 747 | 646 |
| ENSG00000142961 | 430 | 201 | 240 | 269 | 329 | 289 |
| ENSG00000123472 | 1759 | 822 | 1053 | 882 | 1017 | 966 |
| ENSG00000233647 | 0 | 0 | 1 | 0 | 0 | 5 |
| ENSG00000186118 | 12 | 9 | 10 | 6 | 11 | 8 |
| ENSG00000228237 | 7 | 0 | 3 | 1 | 1 | 3 |
| ENSG00000159658 | 4262 | 2111 | 2595 | 1493 | 2363 | 2191 |

| | | | | | | |
|---|---|---|---|---|---|---|
| ENSG00000236476 | 0 | 0 | 0 | 0 | 0 | 1 |
| ENSG00000142973 | 218 | 84 | 105 | 53 | 60 | 59 |
| ENSG00000154198 | 10 | 6 | 5 | 7 | 8 | 4 |
| ENSG00000238145 | 0 | 0 | 0 | 0 | 0 | 0 |
| ENSG00000187048 | 0 | 0 | 0 | 0 | 0 | 0 |
| ENSG00000259832 | 0 | 0 | 0 | 0 | 0 | 0 |
| ENSG00000261196 | 0 | 0 | 0 | 0 | 0 | 0 |
| ENSG00000261593 | 0 | 0 | 0 | 0 | 0 | 0 |
| ENSG00000261290 | 0 | 0 | 0 | 0 | 0 | 0 |
| ENSG00000186377 | 0 | 2 | 5 | 1 | 2 | 3 |
| ENSG00000186160 | 0 | 0 | 1 | 0 | 1 | 0 |
| ENSG00000225154 | 0 | 0 | 1 | 0 | 1 | 0 |
| ENSG00000225506 | 6 | 6 | 6 | 5 | 7 | 12 |
| ENSG00000162365 | 0 | 0 | 0 | 0 | 0 | 0 |
| ENSG00000225514 | 0 | 0 | 0 | 0 | 0 | 1 |
| ENSG00000224805 | 26 | 15 | 26 | 15 | 9 | 14 |
| ENSG00000162366 | 2 | 0 | 0 | 0 | 0 | 0 |
| ENSG00000162367 | 0 | 0 | 0 | 0 | 0 | 0 |
| ENSG00000226252 | 0 | 0 | 0 | 0 | 0 | 0 |
| ENSG00000123473 | 404 | 208 | 251 | 139 | 309 | 240 |
| ENSG00000162368 | 2999 | 1542 | 1646 | 1318 | 2041 | 1793 |
| ENSG00000225762 | 13 | 10 | 12 | 11 | 12 | 17 |
| ENSG00000186790 | 7 | 0 | 2 | 0 | 1 | 0 |
| ENSG00000237424 | 28 | 16 | 10 | 21 | 18 | 5 |
| ENSG00000186564 | 35 | 13 | 8 | 12 | 7 | 15 |
| ENSG00000271355 | 0 | 0 | 0 | 0 | 0 | 0 |
| ENSG00000229554 | 0 | 0 | 0 | 0 | 0 | 0 |
| ENSG00000227883 | 0 | 0 | 0 | 0 | 0 | 0 |
| ENSG00000227947 | 0 | 0 | 0 | 0 | 0 | 0 |
| ENSG00000269113 | 2 | 2 | 2 | 2 | 0 | 1 |
| ENSG00000223814 | 0 | 0 | 0 | 0 | 0 | 0 |
| ENSG00000225028 | 0 | 0 | 0 | 0 | 0 | 0 |
| ENSG00000226133 | 0 | 0 | 0 | 0 | 0 | 0 |
| ENSG00000231413 | 0 | 0 | 0 | 0 | 0 | 0 |
| ENSG00000237180 | 0 | 0 | 0 | 0 | 0 | 0 |
| ENSG00000242267 | 0 | 0 | 0 | 0 | 0 | 0 |
| ENSG00000279096 | 0 | 0 | 0 | 0 | 0 | 0 |
| ENSG00000223720 | 0 | 0 | 0 | 0 | 0 | 0 |
| ENSG00000117834 | 2 | 0 | 0 | 0 | 0 | 0 |
| ENSG00000272491 | 0 | 0 | 0 | 0 | 0 | 0 |
| ENSG00000279214 | 0 | 0 | 0 | 0 | 0 | 0 |
| ENSG00000132122 | 34 | 24 | 29 | 6 | 25 | 14 |
| ENSG00000224986 | 2 | 0 | 0 | 0 | 0 | 0 |
| ENSG00000206700 | 0 | 0 | 0 | 0 | 0 | 0 |
| ENSG00000235105 | 0 | 0 | 0 | 1 | 2 | 0 |
| ENSG00000223175 | 0 | 0 | 0 | 0 | 0 | 0 |
| ENSG00000232514 | 0 | 0 | 0 | 0 | 0 | 0 |
| ENSG00000186094 | 0 | 2 | 0 | 0 | 0 | 1 |
| ENSG00000162373 | 0 | 0 | 1 | 2 | 3 | 0 |
| ENSG00000229846 | 0 | 0 | 0 | 0 | 0 | 0 |

| | | | | | | |
|---|---|---|---|---|---|---|
| ENSG00000230114 | 0 | 0 | 0 | 0 | 0 | 0 |
| ENSG00000225623 | 0 | 0 | 0 | 0 | 0 | 0 |
| ENSG00000237478 | 0 | 0 | 0 | 0 | 0 | 0 |
| ENSG00000215887 | 0 | 0 | 0 | 0 | 0 | 0 |
| ENSG00000279324 | 0 | 0 | 0 | 0 | 0 | 0 |
| ENSG00000233595 | 0 | 0 | 0 | 0 | 0 | 0 |
| ENSG00000162374 | 0 | 0 | 0 | 0 | 0 | 0 |
| ENSG00000230828 | 0 | 0 | 0 | 0 | 0 | 0 |
| ENSG00000237337 | 0 | 0 | 0 | 0 | 0 | 0 |
| ENSG00000233407 | 0 | 0 | 0 | 0 | 0 | 0 |
| ENSG00000284696 | 0 | 0 | 0 | 0 | 0 | 0 |
| ENSG00000284645 | 0 | 0 | 0 | 0 | 0 | 0 |
| ENSG00000234080 | 0 | 0 | 0 | 0 | 0 | 0 |
| ENSG00000229316 | 0 | 0 | 0 | 0 | 0 | 0 |
| ENSG00000225671 | 0 | 0 | 0 | 0 | 0 | 0 |
| ENSG00000142700 | 0 | 0 | 1 | 0 | 0 | 0 |
| ENSG00000284700 | 0 | 2 | 0 | 0 | 0 | 0 |
| ENSG00000185104 | 1269 | 624 | 739 | 702 | 690 | 701 |
| ENSG00000225767 | 0 | 2 | 0 | 0 | 0 | 0 |
| ENSG00000207194 | 0 | 0 | 0 | 0 | 0 | 0 |
| ENSG00000230585 | 3 | 3 | 0 | 1 | 1 | 0 |
| ENSG00000237130 | 0 | 0 | 0 | 0 | 0 | 0 |
| ENSG00000123080 | 442 | 211 | 263 | 226 | 267 | 269 |
| ENSG00000265538 | 0 | 0 | 0 | 0 | 0 | 0 |
| ENSG00000275816 | 0 | 0 | 0 | 0 | 0 | 0 |
| ENSG00000204006 | 0 | 0 | 0 | 0 | 0 | 0 |
| ENSG00000200262 | 0 | 0 | 0 | 0 | 0 | 0 |
| ENSG00000213478 | 0 | 1 | 0 | 0 | 0 | 0 |
| ENSG00000203356 | 8 | 0 | 0 | 0 | 1 | 3 |
| ENSG00000123091 | 3595 | 1744 | 2099 | 1755 | 2073 | 1894 |
| ENSG00000233406 | 0 | 0 | 1 | 0 | 1 | 1 |
| ENSG00000236434 | 2 | 1 | 2 | 2 | 0 | 0 |
| ENSG00000085831 | 5425 | 2588 | 3226 | 2923 | 3009 | 2930 |
| ENSG00000261664 | 52 | 23 | 27 | 30 | 30 | 22 |
| ENSG00000085832 | 1680 | 931 | 876 | 543 | 1139 | 798 |
| ENSG00000232027 | 8 | 1 | 0 | 1 | 2 | 1 |
| ENSG00000206595 | 0 | 0 | 0 | 0 | 0 | 0 |
| ENSG00000238140 | 0 | 0 | 1 | 0 | 1 | 0 |
| ENSG00000227070 | 32 | 14 | 10 | 3 | 13 | 9 |
| ENSG00000252032 | 0 | 0 | 0 | 0 | 0 | 0 |
| ENSG00000227742 | 28 | 16 | 22 | 15 | 17 | 18 |
| ENSG00000117859 | 1753 | 987 | 1063 | 544 | 1101 | 803 |
| ENSG00000236973 | 0 | 0 | 1 | 1 | 0 | 0 |
| ENSG00000232846 | 0 | 0 | 0 | 0 | 0 | 0 |
| ENSG00000212624 | 0 | 0 | 0 | 0 | 0 | 0 |
| ENSG00000078618 | 4235 | 2217 | 2679 | 1897 | 2851 | 2513 |
| ENSG00000266993 | 9 | 5 | 8 | 2 | 9 | 6 |
| ENSG00000272175 | 3 | 0 | 0 | 0 | 0 | 0 |
| ENSG00000283899 | 0 | 0 | 0 | 0 | 0 | 0 |
| ENSG00000230604 | 0 | 1 | 1 | 0 | 0 | 0 |

| | | | | | | |
|---|---|---|---|---|---|---|
| ENSG00000169213 | 66 | 46 | 47 | 13 | 26 | 14 |
| ENSG00000200839 | 0 | 0 | 0 | 0 | 0 | 0 |
| ENSG00000229032 | 0 | 0 | 0 | 0 | 0 | 0 |
| ENSG00000264613 | 0 | 0 | 0 | 0 | 0 | 0 |
| ENSG00000117862 | 219 | 158 | 187 | 159 | 158 | 154 |
| ENSG00000285839 | 0 | 0 | 1 | 0 | 0 | 1 |
| ENSG00000198841 | 136 | 71 | 70 | 112 | 73 | 117 |
| ENSG00000223390 | 0 | 0 | 1 | 0 | 0 | 2 |
| ENSG00000228369 | 1 | 0 | 0 | 1 | 1 | 3 |
| ENSG00000134717 | 2351 | 1220 | 1325 | 1261 | 1427 | 1357 |
| ENSG00000157077 | 698 | 323 | 408 | 269 | 383 | 324 |
| ENSG00000241745 | 0 | 0 | 0 | 0 | 0 | 0 |
| ENSG00000228407 | 0 | 0 | 3 | 1 | 1 | 3 |
| ENSG00000223429 | 0 | 0 | 0 | 0 | 0 | 0 |
| ENSG00000224255 | 0 | 0 | 0 | 0 | 0 | 0 |
| ENSG00000236004 | 0 | 0 | 0 | 0 | 0 | 0 |
| ENSG00000237663 | 0 | 0 | 0 | 0 | 0 | 0 |
| ENSG00000213735 | 0 | 0 | 0 | 0 | 0 | 0 |
| ENSG00000154222 | 1308 | 612 | 830 | 774 | 812 | 781 |
| ENSG00000272100 | 5 | 2 | 6 | 2 | 3 | 3 |
| ENSG00000224680 | 4 | 2 | 13 | 4 | 6 | 7 |
| ENSG00000085840 | 213 | 92 | 111 | 75 | 102 | 124 |
| ENSG00000134748 | 1350 | 612 | 782 | 658 | 785 | 692 |
| ENSG00000134744 | 98 | 47 | 51 | 17 | 84 | 61 |
| ENSG00000272371 | 0 | 0 | 0 | 0 | 0 | 0 |
| ENSG00000236290 | 0 | 0 | 0 | 0 | 0 | 0 |
| ENSG00000116157 | 0 | 0 | 0 | 0 | 0 | 0 |
| ENSG00000182183 | 0 | 0 | 0 | 0 | 0 | 0 |
| ENSG00000162377 | 784 | 383 | 494 | 351 | 518 | 496 |
| ENSG00000233664 | 0 | 0 | 0 | 0 | 0 | 0 |
| ENSG00000239640 | 0 | 0 | 0 | 0 | 0 | 0 |
| ENSG00000162378 | 1196 | 505 | 662 | 323 | 623 | 484 |
| ENSG00000252018 | 0 | 0 | 0 | 0 | 0 | 0 |
| ENSG00000228929 | 12 | 7 | 6 | 4 | 6 | 3 |
| ENSG00000206627 | 2 | 0 | 0 | 1 | 0 | 0 |
| ENSG00000203995 | 492 | 280 | 304 | 228 | 241 | 272 |
| ENSG00000242391 | 1 | 2 | 4 | 2 | 2 | 5 |
| ENSG00000121310 | 2196 | 948 | 1426 | 974 | 1037 | 1131 |
| ENSG00000230953 | 10 | 14 | 18 | 4 | 14 | 10 |
| ENSG00000231866 | 0 | 0 | 0 | 0 | 0 | 0 |
| ENSG00000116171 | 2614 | 1294 | 1591 | 1151 | 1543 | 1357 |
| ENSG00000264834 | 0 | 0 | 0 | 0 | 0 | 0 |
| ENSG00000266188 | 0 | 0 | 0 | 0 | 0 | 0 |
| ENSG00000265815 | 0 | 0 | 0 | 0 | 0 | 0 |
| ENSG00000237279 | 0 | 0 | 0 | 0 | 0 | 0 |
| ENSG00000236360 | 1 | 0 | 2 | 0 | 0 | 0 |
| ENSG00000226147 | 0 | 0 | 0 | 1 | 1 | 0 |
| ENSG00000174348 | 0 | 0 | 0 | 0 | 0 | 0 |
| ENSG00000232993 | 0 | 0 | 0 | 0 | 0 | 0 |
| ENSG00000227644 | 0 | 0 | 0 | 0 | 0 | 0 |

| | | | | | | |
|---|---|---|---|---|---|---|
| ENSG00000162383 | 0 | 0 | 0 | 1 | 0 | 1 |
| ENSG00000235563 | 0 | 0 | 0 | 0 | 0 | 0 |
| ENSG00000277397 | 0 | 0 | 0 | 0 | 0 | 0 |
| ENSG00000157184 | 773 | 454 | 553 | 464 | 547 | 507 |
| ENSG00000236723 | 1 | 3 | 2 | 1 | 2 | 0 |
| ENSG00000162384 | 930 | 455 | 534 | 524 | 495 | 511 |
| ENSG00000259818 | 4 | 0 | 1 | 1 | 0 | 1 |
| ENSG00000162385 | 740 | 369 | 422 | 440 | 427 | 382 |
| ENSG00000226754 | 64 | 29 | 45 | 29 | 29 | 36 |
| ENSG00000157193 | 1200 | 550 | 773 | 642 | 826 | 931 |
| ENSG00000234578 | 0 | 0 | 0 | 0 | 0 | 0 |
| ENSG00000228838 | 0 | 0 | 0 | 0 | 1 | 0 |
| ENSG00000225030 | 0 | 0 | 0 | 0 | 0 | 1 |
| ENSG00000232762 | 0 | 2 | 0 | 0 | 1 | 1 |
| ENSG00000225675 | 7 | 4 | 3 | 5 | 4 | 11 |
| ENSG00000285954 | 0 | 0 | 0 | 0 | 0 | 0 |
| ENSG00000226938 | 0 | 0 | 0 | 0 | 0 | 0 |
| ENSG00000230138 | 0 | 0 | 0 | 0 | 0 | 0 |
| ENSG00000236253 | 0 | 0 | 0 | 0 | 0 | 0 |
| ENSG00000143006 | 0 | 0 | 0 | 0 | 0 | 0 |
| ENSG00000174332 | 0 | 0 | 0 | 0 | 0 | 0 |
| ENSG00000239007 | 0 | 0 | 0 | 0 | 0 | 0 |
| ENSG00000058804 | 1055 | 510 | 621 | 466 | 701 | 644 |
| ENSG00000201003 | 0 | 0 | 1 | 0 | 0 | 0 |
| ENSG00000229687 | 0 | 0 | 0 | 0 | 0 | 0 |
| ENSG00000058799 | 970 | 479 | 611 | 467 | 480 | 526 |
| ENSG00000211452 | 157 | 55 | 114 | 36 | 21 | 41 |
| ENSG00000081870 | 469 | 257 | 329 | 221 | 211 | 263 |
| ENSG00000116212 | 1002 | 522 | 620 | 590 | 571 | 605 |
| ENSG00000219102 | 14 | 7 | 6 | 1 | 6 | 8 |
| ENSG00000203985 | 40 | 25 | 30 | 11 | 2 | 14 |
| ENSG00000116209 | 7004 | 3471 | 4287 | 3445 | 3625 | 3686 |
| ENSG00000280378 | 18 | 4 | 6 | 3 | 3 | 3 |
| ENSG00000116205 | 519 | 255 | 313 | 217 | 281 | 289 |
| ENSG00000283749 | 0 | 0 | 0 | 0 | 0 | 0 |
| ENSG00000225183 | 0 | 0 | 0 | 0 | 0 | 0 |
| ENSG00000279049 | 0 | 0 | 0 | 0 | 0 | 0 |
| ENSG00000256407 | 0 | 0 | 0 | 0 | 0 | 0 |
| ENSG00000157211 | 0 | 0 | 0 | 0 | 0 | 0 |
| ENSG00000280425 | 2 | 0 | 0 | 0 | 0 | 0 |
| ENSG00000215883 | 216 | 124 | 138 | 98 | 112 | 127 |
| ENSG00000116221 | 3000 | 1530 | 1800 | 2046 | 1724 | 1759 |
| ENSG00000157216 | 1390 | 746 | 852 | 688 | 727 | 698 |
| ENSG00000198711 | 4 | 5 | 6 | 2 | 1 | 8 |
| ENSG00000225632 | 0 | 0 | 0 | 0 | 0 | 0 |
| ENSG00000232245 | 0 | 0 | 0 | 0 | 0 | 0 |
| ENSG00000235741 | 0 | 0 | 0 | 0 | 0 | 0 |
| ENSG00000237173 | 0 | 0 | 0 | 0 | 0 | 0 |
| ENSG00000227453 | 0 | 0 | 0 | 0 | 0 | 0 |
| ENSG00000162390 | 103 | 62 | 91 | 103 | 86 | 86 |

| | | | | | | |
|---|---|---|---|---|---|---|
| ENSG00000162391 | 3 | 1 | 6 | 2 | 2 | 5 |
| ENSG00000230728 | 0 | 0 | 0 | 0 | 0 | 0 |
| ENSG00000184313 | 4 | 0 | 1 | 1 | 1 | 2 |
| ENSG00000271723 | 0 | 0 | 0 | 1 | 1 | 1 |
| ENSG00000243725 | 32 | 20 | 40 | 29 | 30 | 33 |
| ENSG00000162396 | 371 | 163 | 207 | 219 | 205 | 179 |
| ENSG00000006555 | 18 | 13 | 16 | 11 | 16 | 18 |
| ENSG00000237453 | 0 | 0 | 0 | 0 | 0 | 0 |
| ENSG00000162398 | 0 | 1 | 0 | 0 | 0 | 1 |
| ENSG00000116133 | 14700 | 6899 | 8217 | 7214 | 6989 | 6958 |
| ENSG00000242396 | 3 | 0 | 2 | 0 | 3 | 3 |
| ENSG00000233203 | 6 | 4 | 5 | 6 | 8 | 12 |
| ENSG00000284601 | 0 | 0 | 0 | 0 | 0 | 0 |
| ENSG00000143001 | 8 | 4 | 2 | 3 | 8 | 4 |
| ENSG00000233271 | 0 | 0 | 0 | 0 | 0 | 0 |
| ENSG00000162399 | 0 | 1 | 0 | 0 | 1 | 1 |
| ENSG00000169174 | 0 | 0 | 0 | 0 | 0 | 0 |
| ENSG00000162402 | 992 | 527 | 402 | 205 | 862 | 454 |
| ENSG00000231090 | 0 | 0 | 0 | 0 | 0 | 0 |
| ENSG00000231095 | 0 | 0 | 0 | 0 | 1 | 0 |
| ENSG00000265822 | 0 | 0 | 0 | 0 | 0 | 0 |
| ENSG00000234810 | 0 | 0 | 0 | 0 | 0 | 0 |
| ENSG00000231900 | 0 | 0 | 0 | 0 | 0 | 0 |
| ENSG00000232198 | 0 | 0 | 0 | 0 | 0 | 0 |
| ENSG00000199831 | 0 | 0 | 0 | 0 | 0 | 0 |
| ENSG00000252162 | 0 | 0 | 0 | 0 | 0 | 0 |
| ENSG00000272051 | 0 | 0 | 0 | 0 | 0 | 0 |
| ENSG00000233079 | 0 | 0 | 0 | 0 | 0 | 0 |
| ENSG00000230250 | 0 | 0 | 0 | 0 | 0 | 0 |
| ENSG00000227104 | 0 | 0 | 0 | 0 | 0 | 0 |
| ENSG00000235612 | 0 | 0 | 0 | 0 | 0 | 0 |
| ENSG00000260971 | 0 | 0 | 0 | 0 | 0 | 0 |
| ENSG00000284686 | 2 | 0 | 1 | 1 | 0 | 4 |
| ENSG00000229871 | 0 | 0 | 0 | 0 | 0 | 0 |
| ENSG00000223956 | 2 | 1 | 1 | 1 | 1 | 0 |
| ENSG00000162407 | 252 | 119 | 147 | 119 | 149 | 140 |
| ENSG00000236776 | 0 | 0 | 0 | 0 | 0 | 0 |
| ENSG00000224980 | 0 | 0 | 0 | 0 | 0 | 0 |
| ENSG00000225475 | 0 | 0 | 0 | 0 | 1 | 0 |
| ENSG00000162409 | 0 | 1 | 1 | 0 | 0 | 0 |
| ENSG00000187889 | 0 | 0 | 0 | 0 | 0 | 0 |
| ENSG00000229913 | 0 | 0 | 0 | 0 | 0 | 0 |
| ENSG00000157131 | 0 | 0 | 0 | 0 | 0 | 0 |
| ENSG00000021852 | 0 | 0 | 0 | 0 | 0 | 0 |
| ENSG00000236341 | 0 | 0 | 0 | 0 | 0 | 0 |
| ENSG00000173406 | 0 | 0 | 0 | 0 | 0 | 1 |
| ENSG00000227935 | 0 | 0 | 0 | 0 | 0 | 0 |
| ENSG00000236888 | 0 | 0 | 0 | 0 | 0 | 0 |
| ENSG00000226759 | 0 | 0 | 0 | 0 | 0 | 0 |
| ENSG00000229887 | 0 | 0 | 0 | 0 | 0 | 0 |

| | | | | | | |
|---|---|---|---|---|---|---|
| ENSG00000223416 | 0 | 0 | 0 | 0 | 0 | 0 |
| ENSG00000235038 | 0 | 0 | 0 | 0 | 0 | 0 |
| ENSG00000230546 | 0 | 0 | 0 | 0 | 0 | 0 |
| ENSG00000233216 | 0 | 0 | 0 | 0 | 0 | 0 |
| ENSG00000162600 | 417 | 217 | 275 | 190 | 299 | 223 |
| ENSG00000270209 | 0 | 0 | 0 | 0 | 0 | 0 |
| ENSG00000264128 | 0 | 0 | 0 | 0 | 0 | 0 |
| ENSG00000184292 | 15735 | 7457 | 10269 | 10782 | 10200 | 9975 |
| ENSG00000185839 | 6 | 2 | 2 | 4 | 3 | 0 |
| ENSG00000162601 | 734 | 397 | 369 | 119 | 500 | 343 |
| ENSG00000283445 | 0 | 2 | 1 | 0 | 0 | 1 |
| ENSG00000177606 | 2544 | 1105 | 1621 | 1576 | 1537 | 1659 |
| ENSG00000234807 | 68 | 30 | 36 | 34 | 34 | 32 |
| ENSG00000272226 | 4 | 3 | 2 | 2 | 1 | 0 |
| ENSG00000231740 | 0 | 0 | 0 | 0 | 0 | 0 |
| ENSG00000232453 | 10 | 3 | 5 | 4 | 8 | 1 |
| ENSG00000237352 | 0 | 0 | 1 | 1 | 2 | 0 |
| ENSG00000228121 | 2 | 0 | 0 | 0 | 1 | 0 |
| ENSG00000241042 | 0 | 0 | 0 | 0 | 0 | 0 |
| ENSG00000235215 | 0 | 0 | 0 | 0 | 0 | 0 |
| ENSG00000224609 | 1 | 2 | 3 | 1 | 2 | 3 |
| ENSG00000270457 | 0 | 0 | 0 | 0 | 1 | 0 |
| ENSG00000172456 | 403 | 232 | 254 | 242 | 260 | 256 |
| ENSG00000266150 | 0 | 0 | 0 | 0 | 0 | 0 |
| ENSG00000226883 | 2 | 2 | 1 | 0 | 0 | 0 |
| ENSG00000134709 | 649 | 383 | 294 | 153 | 546 | 278 |
| ENSG00000134716 | 337 | 135 | 170 | 108 | 135 | 139 |
| ENSG00000265535 | 0 | 0 | 0 | 0 | 0 | 0 |
| ENSG00000162598 | 0 | 0 | 0 | 0 | 0 | 0 |
| ENSG00000229332 | 0 | 0 | 0 | 0 | 0 | 0 |
| ENSG00000238242 | 0 | 0 | 0 | 0 | 0 | 0 |
| ENSG00000226476 | 30 | 13 | 11 | 4 | 7 | 9 |
| ENSG00000231252 | 0 | 0 | 0 | 0 | 0 | 0 |
| ENSG00000162599 | 12 | 11 | 10 | 7 | 14 | 11 |
| ENSG00000237928 | 0 | 0 | 0 | 0 | 0 | 0 |
| ENSG00000270742 | 0 | 0 | 0 | 0 | 0 | 0 |
| ENSG00000237853 | 0 | 0 | 0 | 0 | 0 | 0 |
| ENSG00000284808 | 0 | 0 | 0 | 0 | 0 | 0 |
| ENSG00000223920 | 0 | 0 | 0 | 1 | 0 | 0 |
| ENSG00000162604 | 661 | 312 | 400 | 329 | 388 | 425 |
| ENSG00000271200 | 0 | 0 | 0 | 0 | 1 | 1 |
| ENSG00000132849 | 2146 | 1216 | 1275 | 585 | 1175 | 979 |
| ENSG00000200575 | 0 | 0 | 0 | 1 | 0 | 0 |
| ENSG00000212360 | 0 | 0 | 0 | 0 | 0 | 0 |
| ENSG00000236646 | 0 | 0 | 0 | 0 | 0 | 0 |
| ENSG00000242860 | 0 | 0 | 3 | 0 | 0 | 0 |
| ENSG00000263908 | 0 | 0 | 0 | 0 | 0 | 0 |
| ENSG00000283690 | 0 | 0 | 0 | 0 | 0 | 0 |
| ENSG00000234204 | 0 | 0 | 0 | 0 | 0 | 0 |
| ENSG00000234088 | 0 | 0 | 0 | 0 | 0 | 0 |

| | | | | | | |
|---|---|---|---|---|---|---|
| ENSG00000240563 | 0 | 0 | 0 | 0 | 0 | 0 |
| ENSG00000237227 | 0 | 0 | 0 | 0 | 0 | 0 |
| ENSG00000200174 | 0 | 0 | 0 | 0 | 0 | 0 |
| ENSG00000132854 | 0 | 0 | 0 | 0 | 0 | 1 |
| ENSG00000201153 | 0 | 0 | 0 | 0 | 0 | 0 |
| ENSG00000162607 | 976 | 499 | 485 | 299 | 693 | 529 |
| ENSG00000116641 | 702 | 439 | 320 | 168 | 595 | 338 |
| ENSG00000270549 | 0 | 0 | 0 | 0 | 0 | 0 |
| ENSG00000132855 | 0 | 0 | 0 | 0 | 0 | 0 |
| ENSG00000278967 | 0 | 0 | 0 | 0 | 0 | 0 |
| ENSG00000213703 | 0 | 0 | 0 | 0 | 0 | 0 |
| ENSG00000235545 | 0 | 4 | 2 | 3 | 0 | 3 |
| ENSG00000125703 | 190 | 134 | 144 | 102 | 164 | 137 |
| ENSG00000234318 | 0 | 0 | 0 | 2 | 3 | 1 |
| ENSG00000237163 | 0 | 0 | 0 | 1 | 0 | 0 |
| ENSG00000229537 | 0 | 0 | 0 | 0 | 0 | 0 |
| ENSG00000227485 | 0 | 0 | 0 | 0 | 0 | 0 |
| ENSG00000229225 | 0 | 0 | 0 | 0 | 0 | 0 |
| ENSG00000203605 | 0 | 0 | 0 | 0 | 0 | 0 |
| ENSG00000224209 | 0 | 0 | 0 | 0 | 1 | 3 |
| ENSG00000252259 | 0 | 0 | 0 | 0 | 0 | 0 |
| ENSG00000228734 | 0 | 0 | 0 | 1 | 0 | 0 |
| ENSG00000223683 | 0 | 0 | 0 | 0 | 0 | 0 |
| ENSG00000230798 | 68 | 35 | 52 | 38 | 37 | 30 |
| ENSG00000187140 | 27 | 21 | 16 | 23 | 17 | 15 |
| ENSG00000275836 | 0 | 0 | 0 | 0 | 0 | 0 |
| ENSG00000252784 | 0 | 0 | 0 | 0 | 0 | 0 |
| ENSG00000236674 | 0 | 0 | 0 | 0 | 0 | 0 |
| ENSG00000088035 | 567 | 268 | 303 | 195 | 295 | 281 |
| ENSG00000142856 | 197 | 84 | 118 | 109 | 160 | 121 |
| ENSG00000203965 | 62 | 42 | 53 | 39 | 68 | 47 |
| ENSG00000264271 | 0 | 0 | 1 | 0 | 0 | 0 |
| ENSG00000251720 | 0 | 0 | 0 | 0 | 0 | 0 |
| ENSG00000116652 | 2 | 7 | 1 | 0 | 2 | 2 |
| ENSG00000276884 | 0 | 0 | 0 | 0 | 0 | 0 |
| ENSG00000278287 | 0 | 0 | 0 | 0 | 0 | 0 |
| ENSG00000277565 | 0 | 0 | 0 | 0 | 0 | 0 |
| ENSG00000275823 | 0 | 0 | 0 | 0 | 0 | 0 |
| ENSG00000277689 | 0 | 0 | 0 | 0 | 0 | 0 |
| ENSG00000274199 | 0 | 0 | 0 | 0 | 0 | 0 |
| ENSG00000079739 | 125 | 41 | 58 | 44 | 54 | 49 |
| ENSG00000244256 | 2 | 0 | 0 | 0 | 0 | 0 |
| ENSG00000185483 | 0 | 3 | 3 | 0 | 4 | 1 |
| ENSG00000177452 | 0 | 0 | 0 | 0 | 0 | 0 |
| ENSG00000228056 | 0 | 0 | 0 | 0 | 0 | 0 |
| ENSG00000207190 | 0 | 0 | 0 | 0 | 0 | 0 |
| ENSG00000200508 | 0 | 0 | 0 | 0 | 0 | 0 |
| ENSG00000223949 | 0 | 0 | 0 | 0 | 0 | 0 |
| ENSG00000284928 | 0 | 0 | 0 | 0 | 0 | 0 |
| ENSG00000177414 | 0 | 0 | 0 | 0 | 0 | 0 |

| | | | | | | |
|---|---|---|---|---|---|---|
| ENSG00000238653 | 0 | 0 | 0 | 0 | 0 | 0 |
| ENSG00000158966 | 108 | 55 | 81 | 28 | 50 | 56 |
| ENSG00000264470 | 0 | 0 | 0 | 0 | 0 | 0 |
| ENSG00000162437 | 574 | 340 | 321 | 249 | 321 | 305 |
| ENSG00000162434 | 6468 | 3403 | 4024 | 2251 | 3247 | 3015 |
| ENSG00000234784 | 0 | 0 | 0 | 0 | 0 | 0 |
| ENSG00000233877 | 0 | 1 | 0 | 0 | 0 | 0 |
| ENSG00000226891 | 2 | 3 | 5 | 0 | 0 | 5 |
| ENSG00000185031 | 0 | 0 | 0 | 0 | 0 | 0 |
| ENSG00000272506 | 0 | 0 | 0 | 0 | 0 | 0 |
| ENSG00000212257 | 0 | 0 | 0 | 0 | 1 | 0 |
| ENSG00000265996 | 1 | 1 | 0 | 0 | 0 | 0 |
| ENSG00000199135 | 0 | 0 | 0 | 0 | 0 | 0 |
| ENSG00000231485 | 0 | 1 | 0 | 1 | 0 | 0 |
| ENSG00000235804 | 0 | 0 | 0 | 0 | 0 | 0 |
| ENSG00000234496 | 0 | 0 | 0 | 0 | 0 | 0 |
| ENSG00000162433 | 344 | 202 | 232 | 166 | 220 | 281 |
| ENSG00000231622 | 0 | 0 | 0 | 0 | 0 | 0 |
| ENSG00000116675 | 352 | 199 | 218 | 133 | 221 | 182 |
| ENSG00000229294 | 0 | 0 | 0 | 0 | 0 | 0 |
| ENSG00000234144 | 0 | 0 | 0 | 0 | 0 | 0 |
| ENSG00000222624 | 0 | 0 | 0 | 0 | 0 | 0 |
| ENSG00000213625 | 2328 | 1115 | 1437 | 1060 | 1372 | 1263 |
| ENSG00000116678 | 54 | 46 | 41 | 24 | 83 | 39 |
| ENSG00000237852 | 3 | 0 | 1 | 1 | 1 | 0 |
| ENSG00000224570 | 0 | 0 | 0 | 0 | 1 | 3 |
| ENSG00000285079 | 0 | 0 | 0 | 0 | 0 | 0 |
| ENSG00000239319 | 0 | 0 | 0 | 0 | 0 | 0 |
| ENSG00000184588 | 0 | 0 | 0 | 0 | 0 | 0 |
| ENSG00000227466 | 0 | 0 | 0 | 0 | 0 | 0 |
| ENSG00000223152 | 0 | 0 | 0 | 0 | 0 | 0 |
| ENSG00000118473 | 0 | 0 | 0 | 0 | 0 | 0 |
| ENSG00000264720 | 0 | 0 | 0 | 0 | 0 | 0 |
| ENSG00000248458 | 0 | 0 | 0 | 0 | 0 | 0 |
| ENSG00000152760 | 0 | 0 | 0 | 0 | 0 | 0 |
| ENSG00000172410 | 0 | 0 | 0 | 0 | 0 | 0 |
| ENSG00000152763 | 76 | 42 | 55 | 35 | 86 | 44 |
| ENSG00000231080 | 0 | 0 | 0 | 0 | 0 | 0 |
| ENSG00000198160 | 557 | 296 | 308 | 165 | 345 | 323 |
| ENSG00000116704 | 1047 | 436 | 606 | 286 | 457 | 445 |
| ENSG00000203963 | 0 | 0 | 0 | 0 | 0 | 0 |
| ENSG00000252433 | 0 | 0 | 0 | 0 | 0 | 0 |
| ENSG00000275678 | 0 | 0 | 0 | 0 | 0 | 0 |
| ENSG00000162594 | 0 | 1 | 0 | 0 | 0 | 0 |
| ENSG00000253074 | 0 | 0 | 0 | 0 | 0 | 0 |
| ENSG00000252116 | 0 | 0 | 0 | 0 | 0 | 0 |
| ENSG00000228523 | 0 | 0 | 0 | 0 | 0 | 0 |
| ENSG00000081985 | 0 | 0 | 1 | 0 | 0 | 0 |
| ENSG00000142864 | 9869 | 5182 | 5929 | 4700 | 6273 | 5882 |
| ENSG00000223263 | 0 | 0 | 1 | 1 | 0 | 0 |

| | | | | | | |
|---|---|---|---|---|---|---|
| ENSG00000235200 | 0 | 0 | 0 | 0 | 0 | 0 |
| ENSG00000207504 | 0 | 0 | 0 | 0 | 0 | 0 |
| ENSG00000238139 | 0 | 0 | 0 | 0 | 0 | 0 |
| ENSG00000242482 | 0 | 0 | 0 | 0 | 0 | 0 |
| ENSG00000232048 | 0 | 0 | 0 | 0 | 0 | 0 |
| ENSG00000116717 | 313 | 125 | 188 | 210 | 198 | 185 |
| ENSG00000172380 | 1555 | 774 | 831 | 571 | 929 | 910 |
| ENSG00000238778 | 0 | 0 | 0 | 0 | 0 | 0 |
| ENSG00000232284 | 2 | 2 | 1 | 2 | 1 | 4 |
| ENSG00000235055 | 0 | 0 | 0 | 0 | 0 | 0 |
| ENSG00000162595 | 0 | 0 | 0 | 0 | 0 | 0 |
| ENSG00000228431 | 0 | 0 | 0 | 0 | 0 | 0 |
| ENSG00000116729 | 14 | 8 | 4 | 6 | 14 | 8 |
| ENSG00000234383 | 2 | 0 | 0 | 0 | 0 | 0 |
| ENSG00000221203 | 0 | 0 | 0 | 0 | 0 | 0 |
| ENSG00000229133 | 0 | 2 | 1 | 1 | 0 | 2 |
| ENSG00000225242 | 0 | 0 | 0 | 0 | 0 | 0 |
| ENSG00000237622 | 0 | 0 | 0 | 0 | 0 | 0 |
| ENSG00000229440 | 0 | 0 | 0 | 0 | 0 | 0 |
| ENSG00000116745 | 0 | 0 | 0 | 0 | 0 | 0 |
| ENSG00000024526 | 150 | 92 | 83 | 59 | 196 | 108 |
| ENSG00000233589 | 3 | 1 | 2 | 0 | 3 | 2 |
| ENSG00000234264 | 2 | 1 | 0 | 2 | 1 | 4 |
| ENSG00000225167 | 0 | 0 | 0 | 0 | 0 | 0 |
| ENSG00000285041 | 0 | 0 | 0 | 0 | 0 | 0 |
| ENSG00000285407 | 0 | 2 | 0 | 0 | 0 | 0 |
| ENSG00000285473 | 0 | 0 | 0 | 0 | 0 | 0 |
| ENSG00000223883 | 0 | 0 | 0 | 0 | 0 | 0 |
| ENSG00000235446 | 0 | 0 | 0 | 0 | 0 | 0 |
| ENSG00000229639 | 0 | 0 | 0 | 0 | 0 | 0 |
| ENSG00000033122 | 10 | 3 | 1 | 0 | 11 | 2 |
| ENSG00000270615 | 0 | 0 | 0 | 0 | 0 | 0 |
| ENSG00000240692 | 0 | 0 | 0 | 0 | 0 | 0 |
| ENSG00000229359 | 0 | 0 | 0 | 0 | 0 | 0 |
| ENSG00000237919 | 0 | 0 | 0 | 0 | 0 | 0 |
| ENSG00000066557 | 318 | 177 | 142 | 86 | 250 | 131 |
| ENSG00000244389 | 0 | 0 | 0 | 0 | 0 | 0 |
| ENSG00000116754 | 2934 | 1427 | 1482 | 825 | 1775 | 1301 |
| ENSG00000228988 | 0 | 1 | 0 | 0 | 0 | 0 |
| ENSG00000118454 | 730 | 369 | 459 | 279 | 441 | 359 |
| ENSG00000197568 | 221 | 99 | 100 | 112 | 103 | 90 |
| ENSG00000226088 | 1 | 2 | 2 | 0 | 2 | 0 |
| ENSG00000116761 | 88 | 39 | 45 | 44 | 33 | 26 |
| ENSG00000278636 | 0 | 0 | 0 | 0 | 0 | 0 |
| ENSG00000271992 | 0 | 2 | 3 | 0 | 0 | 1 |
| ENSG00000233020 | 0 | 0 | 0 | 0 | 0 | 0 |
| ENSG00000237033 | 0 | 0 | 0 | 0 | 0 | 0 |
| ENSG00000229051 | 0 | 0 | 0 | 0 | 0 | 0 |
| ENSG00000226208 | 0 | 0 | 0 | 0 | 0 | 0 |
| ENSG00000050628 | 10 | 4 | 8 | 1 | 3 | 8 |

| | | | | | | |
|---|---|---|---|---|---|---|
| ENSG00000235782 | 0 | 0 | 0 | 0 | 0 | 0 |
| ENSG00000269933 | 0 | 0 | 0 | 0 | 0 | 0 |
| ENSG00000235079 | 0 | 0 | 0 | 0 | 0 | 1 |
| ENSG00000132485 | 1201 | 741 | 709 | 439 | 939 | 680 |
| ENSG00000207721 | 4 | 2 | 3 | 2 | 4 | 1 |
| ENSG00000229956 | 2 | 3 | 2 | 0 | 0 | 2 |
| ENSG00000226324 | 0 | 0 | 0 | 0 | 0 | 0 |
| ENSG00000172260 | 0 | 0 | 0 | 0 | 0 | 0 |
| ENSG00000231985 | 0 | 0 | 0 | 0 | 0 | 0 |
| ENSG00000271618 | 0 | 0 | 0 | 0 | 0 | 0 |
| ENSG00000228853 | 0 | 0 | 0 | 0 | 0 | 0 |
| ENSG00000233994 | 0 | 0 | 0 | 0 | 1 | 1 |
| ENSG00000227207 | 0 | 0 | 0 | 0 | 0 | 0 |
| ENSG00000212366 | 0 | 0 | 0 | 0 | 0 | 0 |
| ENSG00000225087 | 0 | 0 | 0 | 0 | 1 | 0 |
| ENSG00000227016 | 0 | 0 | 0 | 0 | 0 | 0 |
| ENSG00000285808 | 0 | 0 | 0 | 0 | 0 | 0 |
| ENSG00000280317 | 0 | 0 | 0 | 0 | 0 | 0 |
| ENSG00000229636 | 0 | 0 | 0 | 0 | 0 | 0 |
| ENSG00000251825 | 0 | 0 | 0 | 0 | 0 | 0 |
| ENSG00000233973 | 0 | 0 | 0 | 0 | 0 | 0 |
| ENSG00000223479 | 0 | 0 | 0 | 0 | 0 | 0 |
| ENSG00000200601 | 0 | 0 | 0 | 0 | 0 | 0 |
| ENSG00000285778 | 0 | 0 | 0 | 0 | 0 | 0 |
| ENSG00000252005 | 0 | 0 | 0 | 0 | 0 | 0 |
| ENSG00000162620 | 8 | 7 | 4 | 12 | 9 | 6 |
| ENSG00000254685 | 226 | 145 | 147 | 108 | 178 | 130 |
| ENSG00000259030 | 0 | 0 | 0 | 0 | 2 | 2 |
| ENSG00000116783 | 0 | 0 | 0 | 0 | 0 | 0 |
| ENSG00000237324 | 0 | 0 | 0 | 0 | 0 | 0 |
| ENSG00000233894 | 0 | 0 | 0 | 0 | 0 | 0 |
| ENSG00000162621 | 0 | 0 | 0 | 0 | 0 | 0 |
| ENSG00000178965 | 0 | 0 | 0 | 0 | 0 | 0 |
| ENSG00000234497 | 0 | 0 | 0 | 0 | 0 | 0 |
| ENSG00000272864 | 0 | 0 | 0 | 0 | 0 | 0 |
| ENSG00000116791 | 635 | 294 | 358 | 261 | 448 | 338 |
| ENSG00000162623 | 322 | 218 | 168 | 135 | 301 | 190 |
| ENSG00000229943 | 0 | 0 | 0 | 0 | 0 | 0 |
| ENSG00000261213 | 0 | 0 | 0 | 0 | 0 | 0 |
| ENSG00000224127 | 0 | 0 | 0 | 0 | 0 | 0 |
| ENSG00000162624 | 0 | 0 | 0 | 0 | 0 | 0 |
| ENSG00000224149 | 0 | 0 | 0 | 0 | 0 | 0 |
| ENSG00000206999 | 0 | 0 | 0 | 0 | 0 | 0 |
| ENSG00000137968 | 2 | 3 | 7 | 0 | 3 | 3 |
| ENSG00000224493 | 0 | 0 | 0 | 0 | 0 | 0 |
| ENSG00000252338 | 0 | 0 | 0 | 0 | 0 | 0 |
| ENSG00000213579 | 0 | 0 | 0 | 0 | 0 | 0 |
| ENSG00000230863 | 2 | 2 | 0 | 0 | 1 | 0 |
| ENSG00000117054 | 424 | 240 | 243 | 139 | 283 | 233 |
| ENSG00000181227 | 10 | 3 | 1 | 1 | 5 | 2 |

| | | | | | | |
|---|---|---|---|---|---|---|
| ENSG00000137955 | 1023 | 569 | 657 | 608 | 866 | 728 |
| ENSG00000206620 | 0 | 0 | 0 | 0 | 0 | 0 |
| ENSG00000207241 | 0 | 0 | 0 | 0 | 0 | 0 |
| ENSG00000201487 | 0 | 0 | 0 | 0 | 0 | 0 |
| ENSG00000057468 | 0 | 0 | 2 | 0 | 1 | 0 |
| ENSG00000154007 | 0 | 0 | 0 | 0 | 0 | 0 |
| ENSG00000225605 | 0 | 0 | 0 | 0 | 0 | 0 |
| ENSG00000230027 | 0 | 0 | 0 | 0 | 0 | 0 |
| ENSG00000184005 | 0 | 0 | 0 | 0 | 0 | 0 |
| ENSG00000223905 | 0 | 0 | 0 | 0 | 0 | 0 |
| ENSG00000272855 | 0 | 0 | 0 | 0 | 0 | 0 |
| ENSG00000226415 | 13 | 13 | 7 | 10 | 8 | 20 |
| ENSG00000199921 | 0 | 0 | 0 | 0 | 0 | 0 |
| ENSG00000237552 | 0 | 0 | 0 | 0 | 0 | 0 |
| ENSG00000117069 | 2 | 0 | 8 | 1 | 4 | 0 |
| ENSG00000276081 | 0 | 0 | 0 | 0 | 0 | 0 |
| ENSG00000230498 | 0 | 0 | 0 | 0 | 0 | 0 |
| ENSG00000142892 | 493 | 228 | 256 | 183 | 278 | 258 |
| ENSG00000226084 | 44 | 7 | 10 | 9 | 8 | 15 |
| ENSG00000228187 | 0 | 0 | 0 | 0 | 0 | 0 |
| ENSG00000154027 | 0 | 0 | 0 | 0 | 0 | 0 |
| ENSG00000233099 | 0 | 0 | 0 | 0 | 0 | 0 |
| ENSG00000251767 | 0 | 0 | 0 | 0 | 0 | 0 |
| ENSG00000036549 | 812 | 486 | 455 | 236 | 753 | 416 |
| ENSG00000251785 | 0 | 0 | 0 | 0 | 0 | 0 |
| ENSG00000243437 | 0 | 0 | 0 | 0 | 0 | 0 |
| ENSG00000077254 | 1510 | 693 | 966 | 486 | 916 | 694 |
| ENSG00000232672 | 0 | 0 | 0 | 0 | 0 | 0 |
| ENSG00000180488 | 541 | 292 | 295 | 122 | 325 | 244 |
| ENSG00000222849 | 0 | 0 | 0 | 0 | 0 | 0 |
| ENSG00000219201 | 2 | 3 | 3 | 2 | 1 | 0 |
| ENSG00000235613 | 0 | 0 | 0 | 0 | 0 | 0 |
| ENSG00000232015 | 0 | 0 | 0 | 0 | 0 | 0 |
| ENSG00000235927 | 2 | 2 | 0 | 1 | 1 | 0 |
| ENSG00000162614 | 4 | 4 | 1 | 0 | 4 | 3 |
| ENSG00000162613 | 4085 | 2181 | 2568 | 1819 | 2766 | 2588 |
| ENSG00000162616 | 95 | 54 | 58 | 38 | 50 | 46 |
| ENSG00000137960 | 0 | 0 | 0 | 0 | 0 | 0 |
| ENSG00000273338 | 2 | 1 | 1 | 0 | 1 | 0 |
| ENSG00000285928 | 0 | 0 | 2 | 1 | 0 | 0 |
| ENSG00000213561 | 0 | 0 | 0 | 0 | 0 | 0 |
| ENSG00000251958 | 0 | 0 | 0 | 0 | 0 | 0 |
| ENSG00000213560 | 0 | 0 | 0 | 0 | 0 | 0 |
| ENSG00000202263 | 0 | 0 | 0 | 0 | 0 | 0 |
| ENSG00000233333 | 0 | 0 | 0 | 0 | 0 | 0 |
| ENSG00000237413 | 0 | 0 | 0 | 0 | 0 | 0 |
| ENSG00000122420 | 0 | 0 | 0 | 0 | 0 | 0 |
| ENSG00000227556 | 0 | 0 | 0 | 0 | 0 | 0 |
| ENSG00000212308 | 0 | 0 | 0 | 0 | 0 | 0 |
| ENSG00000137959 | 0 | 2 | 1 | 0 | 1 | 0 |

| | | | | | | |
|---|---|---|---|---|---|---|
| ENSG00000137965 | 0 | 0 | 1 | 1 | 0 | 0 |
| ENSG00000238015 | 0 | 0 | 0 | 0 | 0 | 0 |
| ENSG00000235400 | 0 | 0 | 0 | 0 | 0 | 0 |
| ENSG00000162618 | 0 | 0 | 0 | 0 | 0 | 0 |
| ENSG00000230787 | 0 | 0 | 0 | 0 | 0 | 0 |
| ENSG00000235011 | 0 | 0 | 0 | 0 | 0 | 0 |
| ENSG00000282898 | 0 | 0 | 0 | 0 | 0 | 0 |
| ENSG00000285462 | 0 | 0 | 0 | 0 | 0 | 0 |
| ENSG00000232676 | 0 | 0 | 0 | 0 | 0 | 0 |
| ENSG00000285409 | 0 | 0 | 0 | 0 | 0 | 0 |
| ENSG00000235756 | 0 | 0 | 0 | 0 | 0 | 0 |
| ENSG00000260322 | 0 | 0 | 0 | 0 | 0 | 0 |
| ENSG00000277670 | 0 | 0 | 0 | 0 | 0 | 0 |
| ENSG00000236407 | 0 | 0 | 0 | 0 | 0 | 0 |
| ENSG00000225598 | 14 | 12 | 11 | 7 | 12 | 9 |
| ENSG00000213559 | 2 | 0 | 3 | 4 | 2 | 2 |
| ENSG00000227062 | 0 | 0 | 0 | 0 | 0 | 0 |
| ENSG00000234108 | 0 | 0 | 0 | 0 | 0 | 0 |
| ENSG00000224326 | 9 | 1 | 5 | 6 | 9 | 3 |
| ENSG00000234184 | 0 | 0 | 0 | 0 | 0 | 0 |
| ENSG00000227937 | 0 | 0 | 0 | 0 | 0 | 0 |
| ENSG00000229752 | 0 | 2 | 0 | 0 | 0 | 1 |
| ENSG00000235089 | 0 | 4 | 0 | 0 | 0 | 0 |
| ENSG00000285179 | 0 | 0 | 0 | 0 | 0 | 0 |
| ENSG00000223026 | 0 | 0 | 0 | 0 | 0 | 0 |
| ENSG00000117114 | 214 | 71 | 99 | 62 | 114 | 105 |
| ENSG00000224494 | 0 | 0 | 0 | 0 | 0 | 0 |
| ENSG00000227288 | 0 | 1 | 0 | 0 | 0 | 0 |
| ENSG00000227960 | 0 | 0 | 0 | 0 | 0 | 0 |
| ENSG00000234953 | 0 | 0 | 0 | 0 | 0 | 0 |
| ENSG00000242598 | 0 | 0 | 0 | 0 | 0 | 0 |
| ENSG00000236676 | 0 | 0 | 0 | 0 | 1 | 0 |
| ENSG00000234683 | 0 | 0 | 0 | 0 | 0 | 0 |
| ENSG00000215875 | 0 | 0 | 0 | 0 | 0 | 0 |
| ENSG00000233290 | 0 | 0 | 0 | 0 | 1 | 0 |
| ENSG00000230817 | 0 | 0 | 1 | 0 | 0 | 0 |
| ENSG00000236268 | 0 | 0 | 0 | 0 | 0 | 0 |
| ENSG00000285782 | 0 | 0 | 0 | 0 | 0 | 1 |
| ENSG00000231364 | 0 | 0 | 0 | 0 | 0 | 1 |
| ENSG00000233008 | 17 | 10 | 11 | 19 | 15 | 17 |
| ENSG00000237076 | 0 | 0 | 0 | 0 | 0 | 0 |
| ENSG00000137941 | 203 | 72 | 82 | 34 | 109 | 55 |
| ENSG00000233061 | 0 | 0 | 0 | 0 | 0 | 0 |
| ENSG00000229486 | 0 | 0 | 0 | 0 | 0 | 0 |
| ENSG00000285201 | 2 | 0 | 0 | 2 | 4 | 1 |
| ENSG00000271576 | 6 | 3 | 5 | 4 | 1 | 1 |
| ENSG00000142875 | 717 | 364 | 307 | 237 | 442 | 321 |
| ENSG00000213540 | 0 | 0 | 0 | 0 | 0 | 0 |
| ENSG00000226970 | 4 | 1 | 1 | 1 | 2 | 1 |
| ENSG00000199959 | 0 | 0 | 0 | 0 | 0 | 0 |

| | | | | | | |
|---|---|---|---|---|---|---|
| ENSG00000203943 | 50 | 9 | 30 | 8 | 19 | 26 |
| ENSG00000249237 | 0 | 0 | 0 | 0 | 0 | 0 |
| ENSG00000240520 | 12 | 3 | 1 | 1 | 0 | 3 |
| ENSG00000137976 | 10 | 4 | 9 | 1 | 2 | 7 |
| ENSG00000285361 | 0 | 1 | 0 | 1 | 0 | 0 |
| ENSG00000117133 | 1169 | 606 | 737 | 658 | 653 | 624 |
| ENSG00000174021 | 340 | 303 | 360 | 358 | 311 | 327 |
| ENSG00000285851 | 5 | 3 | 5 | 4 | 2 | 1 |
| ENSG00000122432 | 5 | 8 | 7 | 1 | 6 | 3 |
| ENSG00000117151 | 216 | 132 | 180 | 112 | 147 | 126 |
| ENSG00000284882 | 2 | 0 | 2 | 0 | 0 | 1 |
| ENSG00000285374 | 1 | 0 | 0 | 0 | 0 | 0 |
| ENSG00000180869 | 2 | 0 | 3 | 0 | 0 | 2 |
| ENSG00000232622 | 0 | 0 | 0 | 0 | 1 | 0 |
| ENSG00000117155 | 424 | 295 | 283 | 134 | 396 | 213 |
| ENSG00000285325 | 0 | 0 | 0 | 0 | 0 | 0 |
| ENSG00000171517 | 2 | 2 | 0 | 1 | 1 | 0 |
| ENSG00000153898 | 53 | 33 | 49 | 73 | 90 | 85 |
| ENSG00000162643 | 9 | 7 | 9 | 6 | 19 | 5 |
| ENSG00000055732 | 208 | 122 | 157 | 55 | 130 | 144 |
| ENSG00000266110 | 0 | 0 | 0 | 0 | 0 | 0 |
| ENSG00000280099 | 139 | 86 | 74 | 58 | 101 | 80 |
| ENSG00000097096 | 790 | 367 | 459 | 203 | 373 | 374 |
| ENSG00000162642 | 643 | 337 | 389 | 422 | 368 | 428 |
| ENSG00000223254 | 0 | 0 | 0 | 0 | 1 | 0 |
| ENSG00000142867 | 702 | 354 | 417 | 280 | 387 | 342 |
| ENSG00000223653 | 0 | 0 | 1 | 2 | 2 | 4 |
| ENSG00000153904 | 736 | 318 | 380 | 241 | 368 | 292 |
| ENSG00000199459 | 0 | 0 | 0 | 0 | 0 | 0 |
| ENSG00000273264 | 4 | 2 | 0 | 1 | 2 | 2 |
| ENSG00000282057 | 5 | 3 | 3 | 2 | 1 | 0 |
| ENSG00000272691 | 0 | 0 | 1 | 0 | 0 | 0 |
| ENSG00000142871 | 158 | 77 | 100 | 110 | 99 | 112 |
| ENSG00000199934 | 0 | 0 | 0 | 0 | 0 | 0 |
| ENSG00000230285 | 0 | 0 | 0 | 0 | 0 | 0 |
| ENSG00000117174 | 334 | 170 | 163 | 160 | 236 | 154 |
| ENSG00000171502 | 31 | 25 | 36 | 16 | 14 | 18 |
| ENSG00000201620 | 0 | 0 | 0 | 0 | 0 | 0 |
| ENSG00000229505 | 0 | 0 | 0 | 0 | 0 | 0 |
| ENSG00000285210 | 0 | 3 | 1 | 0 | 0 | 0 |
| ENSG00000122417 | 123 | 67 | 57 | 32 | 104 | 63 |
| ENSG00000278281 | 0 | 0 | 0 | 0 | 0 | 0 |
| ENSG00000231349 | 0 | 0 | 0 | 0 | 0 | 0 |
| ENSG00000137975 | 4 | 1 | 2 | 1 | 2 | 0 |
| ENSG00000016490 | 0 | 0 | 0 | 0 | 0 | 0 |
| ENSG00000016602 | 1 | 0 | 1 | 0 | 1 | 0 |
| ENSG00000224466 | 0 | 0 | 0 | 0 | 0 | 0 |
| ENSG00000236915 | 4 | 2 | 1 | 3 | 5 | 5 |
| ENSG00000153923 | 1 | 0 | 0 | 0 | 0 | 0 |
| ENSG00000261737 | 13 | 6 | 9 | 7 | 7 | 1 |

| | | | | | | |
|---|---|---|---|---|---|---|
| ENSG00000097033 | 1212 | 611 | 654 | 584 | 733 | 598 |
| ENSG00000230721 | 0 | 0 | 0 | 0 | 0 | 0 |
| ENSG00000284846 | 0 | 0 | 0 | 0 | 0 | 0 |
| ENSG00000183291 | 2440 | 1211 | 1438 | 1284 | 1402 | 1449 |
| ENSG00000153936 | 1213 | 624 | 678 | 436 | 728 | 601 |
| ENSG00000267734 | 0 | 0 | 0 | 0 | 0 | 1 |
| ENSG00000267561 | 0 | 0 | 0 | 0 | 0 | 0 |
| ENSG00000235251 | 0 | 0 | 0 | 0 | 0 | 0 |
| ENSG00000225568 | 0 | 0 | 0 | 0 | 0 | 0 |
| ENSG00000267272 | 2 | 1 | 1 | 4 | 0 | 3 |
| ENSG00000284240 | 0 | 0 | 0 | 0 | 0 | 0 |
| ENSG00000143013 | 300 | 126 | 184 | 116 | 128 | 153 |
| ENSG00000227290 | 0 | 0 | 0 | 0 | 0 | 0 |
| ENSG00000199318 | 0 | 0 | 0 | 0 | 0 | 0 |
| ENSG00000237505 | 9 | 6 | 9 | 8 | 2 | 6 |
| ENSG00000279778 | 0 | 0 | 0 | 0 | 0 | 0 |
| ENSG00000270507 | 0 | 0 | 0 | 0 | 0 | 0 |
| ENSG00000239504 | 0 | 0 | 0 | 0 | 0 | 0 |
| ENSG00000230053 | 0 | 0 | 0 | 0 | 0 | 0 |
| ENSG00000274321 | 0 | 0 | 0 | 0 | 0 | 0 |
| ENSG00000237959 | 0 | 0 | 0 | 0 | 0 | 0 |
| ENSG00000065243 | 750 | 426 | 363 | 157 | 537 | 318 |
| ENSG00000207234 | 0 | 1 | 0 | 0 | 1 | 0 |
| ENSG00000241975 | 4 | 3 | 3 | 0 | 4 | 3 |
| ENSG00000137947 | 500 | 261 | 357 | 277 | 334 | 302 |
| ENSG00000235308 | 2 | 0 | 0 | 0 | 0 | 1 |
| ENSG00000137944 | 392 | 200 | 270 | 165 | 268 | 194 |
| ENSG00000213516 | 276 | 116 | 155 | 90 | 135 | 142 |
| ENSG00000117226 | 12 | 17 | 15 | 5 | 22 | 7 |
| ENSG00000202385 | 0 | 0 | 0 | 0 | 0 | 0 |
| ENSG00000117228 | 2 | 0 | 0 | 4 | 2 | 1 |
| ENSG00000234518 | 52 | 36 | 43 | 30 | 37 | 49 |
| ENSG00000162645 | 304 | 198 | 202 | 139 | 229 | 191 |
| ENSG00000213512 | 0 | 0 | 0 | 0 | 0 | 0 |
| ENSG00000162654 | 8 | 0 | 5 | 2 | 2 | 2 |
| ENSG00000284734 | 0 | 0 | 0 | 0 | 0 | 0 |
| ENSG00000284637 | 0 | 0 | 0 | 0 | 0 | 0 |
| ENSG00000154451 | 0 | 0 | 0 | 0 | 0 | 2 |
| ENSG00000237568 | 0 | 0 | 0 | 0 | 0 | 0 |
| ENSG00000238081 | 0 | 0 | 0 | 0 | 0 | 0 |
| ENSG00000233235 | 0 | 0 | 0 | 0 | 0 | 0 |
| ENSG00000183347 | 0 | 0 | 0 | 1 | 0 | 0 |
| ENSG00000215874 | 0 | 0 | 0 | 0 | 0 | 0 |
| ENSG00000225492 | 0 | 0 | 0 | 0 | 0 | 0 |
| ENSG00000197147 | 422 | 198 | 257 | 105 | 232 | 210 |
| ENSG00000231999 | 7 | 0 | 0 | 3 | 0 | 0 |
| ENSG00000230735 | 0 | 0 | 0 | 0 | 0 | 0 |
| ENSG00000171488 | 30 | 16 | 15 | 12 | 7 | 4 |
| ENSG00000271949 | 0 | 0 | 0 | 0 | 0 | 0 |
| ENSG00000226394 | 0 | 0 | 0 | 0 | 0 | 0 |

| | | | | | | |
|---|---|---|---|---|---|---|
| ENSG00000231613 | 0 | 0 | 0 | 0 | 0 | 0 |
| ENSG00000272931 | 20 | 14 | 12 | 13 | 13 | 15 |
| ENSG00000171492 | 498 | 291 | 325 | 264 | 347 | 286 |
| ENSG00000272672 | 0 | 0 | 0 | 3 | 0 | 0 |
| ENSG00000252797 | 1 | 0 | 0 | 0 | 0 | 0 |
| ENSG00000228175 | 6 | 8 | 9 | 5 | 9 | 6 |
| ENSG00000162664 | 478 | 284 | 273 | 159 | 367 | 295 |
| ENSG00000212459 | 0 | 0 | 0 | 0 | 0 | 0 |
| ENSG00000229201 | 0 | 0 | 0 | 0 | 0 | 0 |
| ENSG00000199666 | 0 | 0 | 0 | 0 | 0 | 0 |
| ENSG00000143032 | 0 | 0 | 0 | 0 | 0 | 0 |
| ENSG00000233593 | 8 | 3 | 2 | 1 | 2 | 1 |
| ENSG00000285532 | 0 | 0 | 0 | 0 | 0 | 0 |
| ENSG00000225446 | 0 | 0 | 0 | 0 | 0 | 0 |
| ENSG00000272094 | 0 | 0 | 0 | 0 | 0 | 0 |
| ENSG00000232882 | 4 | 4 | 1 | 1 | 2 | 1 |
| ENSG00000122482 | 315 | 163 | 117 | 64 | 325 | 122 |
| ENSG00000232393 | 0 | 0 | 0 | 0 | 0 | 0 |
| ENSG00000162669 | 0 | 0 | 0 | 0 | 0 | 0 |
| ENSG00000206817 | 0 | 0 | 0 | 0 | 0 | 0 |
| ENSG00000215873 | 0 | 0 | 0 | 0 | 0 | 0 |
| ENSG00000097046 | 229 | 148 | 147 | 71 | 147 | 170 |
| ENSG00000241318 | 0 | 0 | 0 | 0 | 0 | 0 |
| ENSG00000229067 | 0 | 0 | 0 | 0 | 0 | 0 |
| ENSG00000203914 | 0 | 0 | 0 | 0 | 1 | 1 |
| ENSG00000069702 | 16 | 12 | 6 | 4 | 7 | 6 |
| ENSG00000239794 | 0 | 0 | 0 | 0 | 0 | 0 |
| ENSG00000137948 | 9 | 5 | 5 | 1 | 1 | 1 |
| ENSG00000172031 | 56 | 27 | 52 | 16 | 22 | 23 |
| ENSG00000233228 | 0 | 0 | 0 | 0 | 0 | 0 |
| ENSG00000230667 | 0 | 1 | 1 | 0 | 1 | 0 |
| ENSG00000189195 | 42 | 16 | 13 | 2 | 50 | 16 |
| ENSG00000224678 | 0 | 1 | 0 | 0 | 0 | 0 |
| ENSG00000233401 | 0 | 0 | 0 | 0 | 0 | 0 |
| ENSG00000273487 | 0 | 0 | 0 | 0 | 0 | 2 |
| ENSG00000226773 | 0 | 0 | 0 | 0 | 0 | 0 |
| ENSG00000203910 | 0 | 0 | 0 | 0 | 0 | 1 |
| ENSG00000233125 | 0 | 0 | 0 | 0 | 0 | 0 |
| ENSG00000174842 | 178 | 88 | 109 | 83 | 94 | 97 |
| ENSG00000122484 | 231 | 123 | 137 | 87 | 172 | 94 |
| ENSG00000242764 | 0 | 0 | 0 | 0 | 0 | 0 |
| ENSG00000162676 | 50 | 24 | 22 | 17 | 12 | 19 |
| ENSG00000067208 | 320 | 151 | 115 | 67 | 189 | 118 |
| ENSG00000223787 | 0 | 0 | 0 | 0 | 0 | 0 |
| ENSG00000229992 | 0 | 0 | 0 | 2 | 0 | 0 |
| ENSG00000201317 | 0 | 0 | 0 | 0 | 0 | 0 |
| ENSG00000225505 | 1 | 0 | 0 | 0 | 0 | 1 |
| ENSG00000234202 | 0 | 0 | 0 | 0 | 0 | 0 |
| ENSG00000223896 | 0 | 0 | 0 | 0 | 0 | 0 |
| ENSG00000122406 | 17757 | 9101 | 10757 | 11139 | 11407 | 10763 |

| | | | | | | |
|---|---|---|---|---|---|---|
| ENSG00000154511 | 160 | 96 | 79 | 77 | 99 | 89 |
| ENSG00000206680 | 0 | 0 | 0 | 0 | 0 | 0 |
| ENSG00000207523 | 0 | 2 | 2 | 1 | 3 | 3 |
| ENSG00000207022 | 0 | 0 | 0 | 0 | 1 | 0 |
| ENSG00000229052 | 0 | 0 | 0 | 0 | 0 | 0 |
| ENSG00000252121 | 0 | 0 | 0 | 0 | 0 | 0 |
| ENSG00000239710 | 0 | 0 | 0 | 0 | 0 | 0 |
| ENSG00000252752 | 0 | 0 | 0 | 0 | 0 | 0 |
| ENSG00000222664 | 0 | 0 | 0 | 0 | 0 | 0 |
| ENSG00000251837 | 0 | 0 | 0 | 0 | 0 | 0 |
| ENSG00000143033 | 493 | 247 | 270 | 178 | 327 | 285 |
| ENSG00000117500 | 882 | 444 | 398 | 267 | 556 | 401 |
| ENSG00000122483 | 20 | 12 | 12 | 9 | 21 | 15 |
| ENSG00000237003 | 0 | 0 | 0 | 0 | 0 | 1 |
| ENSG00000225297 | 0 | 0 | 0 | 0 | 0 | 0 |
| ENSG00000223745 | 299 | 123 | 247 | 114 | 164 | 177 |
| ENSG00000229567 | 0 | 1 | 0 | 0 | 0 | 0 |
| ENSG00000117505 | 1115 | 566 | 589 | 517 | 630 | 569 |
| ENSG00000229635 | 0 | 0 | 2 | 0 | 2 | 1 |
| ENSG00000207379 | 0 | 0 | 0 | 0 | 0 | 0 |
| ENSG00000137942 | 1613 | 874 | 714 | 372 | 1196 | 670 |
| ENSG00000212601 | 0 | 0 | 0 | 0 | 0 | 0 |
| ENSG00000137936 | 786 | 416 | 499 | 382 | 475 | 444 |
| ENSG00000224093 | 5 | 4 | 6 | 2 | 3 | 2 |
| ENSG00000211575 | 0 | 0 | 0 | 0 | 0 | 0 |
| ENSG00000260464 | 10 | 14 | 12 | 10 | 13 | 9 |
| ENSG00000067334 | 304 | 178 | 145 | 84 | 412 | 211 |
| ENSG00000023909 | 572 | 248 | 331 | 243 | 386 | 406 |
| ENSG00000223490 | 0 | 0 | 0 | 0 | 2 | 0 |
| ENSG00000234099 | 0 | 0 | 1 | 0 | 0 | 0 |
| ENSG00000235794 | 0 | 0 | 0 | 0 | 0 | 0 |
| ENSG00000232752 | 0 | 0 | 0 | 0 | 0 | 0 |
| ENSG00000229842 | 0 | 0 | 0 | 0 | 0 | 0 |
| ENSG00000250890 | 0 | 0 | 0 | 0 | 0 | 0 |
| ENSG00000236689 | 0 | 0 | 0 | 0 | 0 | 0 |
| ENSG00000226960 | 0 | 0 | 0 | 0 | 0 | 0 |
| ENSG00000233129 | 0 | 0 | 0 | 0 | 0 | 2 |
| ENSG00000198691 | 16 | 10 | 10 | 11 | 9 | 17 |
| ENSG00000233482 | 0 | 0 | 0 | 0 | 0 | 0 |
| ENSG00000137962 | 274 | 143 | 132 | 57 | 208 | 124 |
| ENSG00000264963 | 0 | 0 | 0 | 0 | 0 | 0 |
| ENSG00000226835 | 0 | 0 | 1 | 0 | 0 | 0 |
| ENSG00000178596 | 0 | 0 | 0 | 0 | 0 | 0 |
| ENSG00000236098 | 0 | 0 | 0 | 0 | 0 | 0 |
| ENSG00000232918 | 0 | 0 | 1 | 0 | 0 | 0 |
| ENSG00000117528 | 906 | 471 | 457 | 278 | 611 | 446 |
| ENSG00000117525 | 52 | 27 | 37 | 32 | 38 | 35 |
| ENSG00000223675 | 0 | 0 | 0 | 0 | 0 | 0 |
| ENSG00000224081 | 5 | 6 | 4 | 5 | 4 | 1 |
| ENSG00000232857 | 0 | 0 | 0 | 0 | 0 | 0 |

| | | | | | | |
|---|---|---|---|---|---|---|
| ENSG00000263526 | 0 | 0 | 0 | 0 | 0 | 0 |
| ENSG00000143036 | 74 | 37 | 44 | 26 | 46 | 42 |
| ENSG00000117519 | 0 | 0 | 2 | 0 | 0 | 1 |
| ENSG00000235501 | 0 | 0 | 0 | 0 | 0 | 0 |
| ENSG00000172339 | 172 | 80 | 101 | 88 | 94 | 100 |
| ENSG00000230427 | 0 | 0 | 0 | 0 | 0 | 0 |
| ENSG00000152078 | 86 | 36 | 31 | 12 | 46 | 44 |
| ENSG00000271092 | 4 | 5 | 3 | 0 | 8 | 2 |
| ENSG00000231992 | 0 | 0 | 0 | 0 | 0 | 0 |
| ENSG00000206679 | 0 | 0 | 0 | 0 | 0 | 0 |
| ENSG00000226026 | 8 | 4 | 0 | 1 | 0 | 1 |
| ENSG00000122481 | 53 | 34 | 25 | 19 | 29 | 16 |
| ENSG00000228852 | 3 | 3 | 2 | 1 | 3 | 0 |
| ENSG00000228504 | 0 | 0 | 0 | 0 | 0 | 0 |
| ENSG00000230434 | 0 | 0 | 0 | 0 | 0 | 0 |
| ENSG00000237954 | 0 | 0 | 0 | 0 | 0 | 0 |
| ENSG00000233907 | 0 | 0 | 0 | 0 | 0 | 0 |
| ENSG00000228971 | 0 | 0 | 0 | 0 | 0 | 0 |
| ENSG00000271252 | 0 | 0 | 0 | 0 | 0 | 0 |
| ENSG00000237435 | 0 | 1 | 3 | 1 | 1 | 0 |
| ENSG00000200800 | 0 | 0 | 0 | 0 | 0 | 1 |
| ENSG00000231987 | 1 | 2 | 0 | 0 | 0 | 0 |
| ENSG00000225923 | 0 | 0 | 0 | 0 | 0 | 0 |
| ENSG00000234422 | 0 | 0 | 0 | 0 | 0 | 0 |
| ENSG00000228502 | 11 | 9 | 8 | 7 | 8 | 6 |
| ENSG00000241992 | 0 | 0 | 0 | 0 | 0 | 0 |
| ENSG00000234226 | 0 | 0 | 0 | 0 | 0 | 0 |
| ENSG00000137970 | 54 | 36 | 34 | 43 | 48 | 38 |
| ENSG00000223229 | 0 | 0 | 0 | 0 | 0 | 0 |
| ENSG00000117569 | 315 | 124 | 191 | 90 | 154 | 177 |
| ENSG00000188641 | 481 | 241 | 339 | 133 | 276 | 189 |
| ENSG00000232878 | 0 | 1 | 0 | 0 | 0 | 0 |
| ENSG00000232542 | 0 | 0 | 0 | 0 | 0 | 0 |
| ENSG00000228057 | 2 | 1 | 4 | 1 | 3 | 1 |
| ENSG00000226128 | 0 | 0 | 0 | 0 | 0 | 0 |
| ENSG00000230718 | 0 | 0 | 0 | 0 | 0 | 0 |
| ENSG00000235777 | 0 | 0 | 0 | 0 | 0 | 0 |
| ENSG00000270911 | 0 | 0 | 0 | 0 | 0 | 0 |
| ENSG00000225206 | 0 | 0 | 0 | 0 | 0 | 0 |
| ENSG00000259946 | 0 | 0 | 0 | 0 | 0 | 0 |
| ENSG00000284247 | 0 | 0 | 0 | 0 | 0 | 0 |
| ENSG00000284202 | 0 | 0 | 0 | 0 | 0 | 0 |
| ENSG00000285922 | 0 | 0 | 0 | 0 | 0 | 0 |
| ENSG00000233557 | 0 | 0 | 0 | 0 | 0 | 0 |
| ENSG00000226053 | 0 | 0 | 0 | 0 | 0 | 0 |
| ENSG00000280040 | 4 | 0 | 0 | 0 | 0 | 3 |
| ENSG00000162627 | 787 | 408 | 498 | 364 | 416 | 413 |
| ENSG00000117598 | 159 | 65 | 70 | 76 | 125 | 85 |
| ENSG00000232825 | 0 | 0 | 0 | 0 | 0 | 0 |
| ENSG00000227034 | 6 | 0 | 1 | 4 | 1 | 0 |

| | | | | | | |
|---|---|---|---|---|---|---|
| ENSG00000117600 | 0 | 0 | 0 | 0 | 0 | 0 |
| ENSG00000233983 | 0 | 0 | 0 | 0 | 0 | 0 |
| ENSG00000224445 | 0 | 0 | 0 | 0 | 0 | 0 |
| ENSG00000099260 | 6 | 3 | 7 | 1 | 0 | 1 |
| ENSG00000223656 | 0 | 0 | 0 | 0 | 0 | 0 |
| ENSG00000156869 | 71 | 29 | 27 | 22 | 30 | 13 |
| ENSG00000201491 | 0 | 0 | 0 | 0 | 0 | 0 |
| ENSG00000202254 | 0 | 0 | 0 | 0 | 0 | 0 |
| ENSG00000231996 | 0 | 0 | 0 | 0 | 0 | 0 |
| ENSG00000162688 | 416 | 200 | 176 | 85 | 292 | 185 |
| ENSG00000228084 | 5 | 7 | 5 | 6 | 5 | 6 |
| ENSG00000117620 | 751 | 333 | 402 | 191 | 453 | 333 |
| ENSG00000283761 | 0 | 1 | 3 | 1 | 2 | 2 |
| ENSG00000212248 | 0 | 0 | 0 | 0 | 0 | 0 |
| ENSG00000202259 | 0 | 0 | 0 | 0 | 0 | 0 |
| ENSG00000156875 | 33 | 16 | 43 | 25 | 24 | 44 |
| ENSG00000241073 | 0 | 0 | 0 | 0 | 0 | 0 |
| ENSG00000156876 | 66 | 52 | 37 | 12 | 67 | 45 |
| ENSG00000226952 | 0 | 0 | 1 | 0 | 0 | 0 |
| ENSG00000122435 | 201 | 103 | 112 | 60 | 124 | 93 |
| ENSG00000122477 | 11 | 5 | 6 | 2 | 8 | 7 |
| ENSG00000137992 | 446 | 229 | 253 | 123 | 267 | 195 |
| ENSG00000271415 | 0 | 0 | 0 | 0 | 0 | 0 |
| ENSG00000225169 | 0 | 0 | 1 | 0 | 1 | 0 |
| ENSG00000285530 | 0 | 0 | 0 | 0 | 0 | 0 |
| ENSG00000230287 | 0 | 0 | 0 | 0 | 0 | 0 |
| ENSG00000224616 | 6 | 0 | 2 | 2 | 0 | 1 |
| ENSG00000137996 | 601 | 287 | 365 | 282 | 410 | 355 |
| ENSG00000207750 | 0 | 0 | 0 | 0 | 0 | 1 |
| ENSG00000215871 | 0 | 0 | 0 | 0 | 0 | 0 |
| ENSG00000223906 | 0 | 0 | 0 | 0 | 0 | 0 |
| ENSG00000079335 | 86 | 51 | 56 | 14 | 49 | 41 |
| ENSG00000234332 | 0 | 0 | 0 | 1 | 0 | 0 |
| ENSG00000228086 | 0 | 0 | 0 | 0 | 0 | 0 |
| ENSG00000181656 | 0 | 0 | 0 | 0 | 0 | 1 |
| ENSG00000213421 | 0 | 0 | 0 | 0 | 0 | 0 |
| ENSG00000230402 | 0 | 0 | 0 | 0 | 0 | 0 |
| ENSG00000285525 | 0 | 0 | 0 | 0 | 0 | 0 |
| ENSG00000224685 | 0 | 0 | 0 | 0 | 0 | 0 |
| ENSG00000162692 | 0 | 0 | 0 | 0 | 0 | 0 |
| ENSG00000162694 | 76 | 59 | 61 | 46 | 61 | 80 |
| ENSG00000273204 | 0 | 0 | 0 | 0 | 0 | 0 |
| ENSG00000162695 | 500 | 226 | 232 | 166 | 306 | 269 |
| ENSG00000230946 | 0 | 0 | 0 | 0 | 0 | 0 |
| ENSG00000117543 | 549 | 284 | 319 | 366 | 380 | 371 |
| ENSG00000235795 | 6 | 1 | 1 | 1 | 5 | 1 |
| ENSG00000233184 | 78 | 57 | 55 | 54 | 77 | 45 |
| ENSG00000252765 | 0 | 0 | 0 | 0 | 0 | 0 |
| ENSG00000271578 | 0 | 0 | 0 | 0 | 0 | 0 |
| ENSG00000225938 | 0 | 0 | 0 | 0 | 0 | 0 |

| | | | | | | |
|---|---|---|---|---|---|---|
| ENSG00000170989 | 0 | 0 | 0 | 0 | 0 | 0 |
| ENSG00000228399 | 14 | 11 | 12 | 9 | 5 | 11 |
| ENSG00000173810 | 0 | 0 | 0 | 0 | 0 | 0 |
| ENSG00000231671 | 0 | 0 | 0 | 0 | 0 | 0 |
| ENSG00000226715 | 0 | 0 | 0 | 0 | 0 | 0 |
| ENSG00000252530 | 0 | 0 | 0 | 0 | 0 | 0 |
| ENSG00000183298 | 0 | 1 | 1 | 1 | 1 | 0 |
| ENSG00000118733 | 0 | 0 | 0 | 0 | 0 | 0 |
| ENSG00000252717 | 0 | 0 | 0 | 0 | 0 | 0 |
| ENSG00000271277 | 0 | 0 | 0 | 0 | 0 | 0 |
| ENSG00000162699 | 0 | 0 | 0 | 0 | 0 | 0 |
| ENSG00000233359 | 0 | 0 | 0 | 0 | 0 | 0 |
| ENSG00000230864 | 0 | 0 | 0 | 0 | 0 | 0 |
| ENSG00000060718 | 0 | 1 | 1 | 0 | 0 | 0 |
| ENSG00000234257 | 0 | 0 | 0 | 0 | 0 | 0 |
| ENSG00000230759 | 0 | 0 | 0 | 0 | 0 | 0 |
| ENSG00000224613 | 0 | 0 | 0 | 0 | 0 | 0 |
| ENSG00000222069 | 0 | 0 | 0 | 0 | 0 | 0 |
| ENSG00000185946 | 175 | 112 | 128 | 68 | 123 | 74 |
| ENSG00000240038 | 6 | 2 | 0 | 1 | 1 | 0 |
| ENSG00000236085 | 0 | 0 | 0 | 0 | 0 | 0 |
| ENSG00000243480 | 0 | 0 | 0 | 0 | 0 | 0 |
| ENSG00000237763 | 0 | 0 | 0 | 0 | 0 | 0 |
| ENSG00000234441 | 0 | 0 | 0 | 0 | 0 | 0 |
| ENSG00000174876 | 0 | 0 | 0 | 0 | 0 | 0 |
| ENSG00000227408 | 0 | 0 | 0 | 0 | 0 | 0 |
| ENSG00000187733 | 0 | 0 | 0 | 0 | 0 | 0 |
| ENSG00000225191 | 0 | 0 | 0 | 0 | 0 | 0 |
| ENSG00000215869 | 6 | 0 | 4 | 2 | 0 | 0 |
| ENSG00000228347 | 0 | 0 | 0 | 0 | 0 | 0 |
| ENSG00000285981 | 0 | 0 | 0 | 0 | 0 | 0 |
| ENSG00000225036 | 0 | 0 | 0 | 0 | 0 | 0 |
| ENSG00000230768 | 0 | 0 | 0 | 0 | 0 | 0 |
| ENSG00000227850 | 0 | 0 | 0 | 0 | 0 | 0 |
| ENSG00000237897 | 0 | 0 | 0 | 0 | 0 | 0 |
| ENSG00000232952 | 0 | 0 | 0 | 1 | 0 | 0 |
| ENSG00000233047 | 0 | 0 | 0 | 0 | 0 | 0 |
| ENSG00000237480 | 0 | 0 | 0 | 0 | 0 | 0 |
| ENSG00000230932 | 0 | 0 | 0 | 0 | 0 | 0 |
| ENSG00000270342 | 0 | 0 | 1 | 0 | 2 | 1 |
| ENSG00000270976 | 0 | 0 | 0 | 0 | 0 | 0 |
| ENSG00000233449 | 0 | 0 | 0 | 0 | 0 | 0 |
| ENSG00000232551 | 0 | 0 | 1 | 0 | 0 | 2 |
| ENSG00000237460 | 6 | 6 | 7 | 2 | 5 | 8 |
| ENSG00000198890 | 8790 | 4189 | 5216 | 4841 | 4839 | 5333 |
| ENSG00000162631 | 2 | 0 | 0 | 0 | 1 | 1 |
| ENSG00000224550 | 0 | 1 | 0 | 2 | 1 | 3 |
| ENSG00000134215 | 1456 | 740 | 853 | 489 | 820 | 754 |
| ENSG00000275455 | 0 | 0 | 0 | 0 | 0 | 0 |
| ENSG00000230489 | 0 | 0 | 0 | 0 | 0 | 0 |

| | | | | | | |
|---|---|---|---|---|---|---|
| ENSG00000226822 | 1 | 0 | 0 | 0 | 0 | 1 |
| ENSG00000085491 | 7296 | 3623 | 4295 | 3277 | 4494 | 4385 |
| ENSG00000260879 | 0 | 0 | 0 | 0 | 0 | 0 |
| ENSG00000280186 | 44 | 15 | 32 | 21 | 19 | 21 |
| ENSG00000196427 | 86 | 39 | 49 | 51 | 53 | 66 |
| ENSG00000238122 | 1 | 0 | 0 | 1 | 0 | 0 |
| ENSG00000241361 | 37 | 17 | 23 | 12 | 39 | 22 |
| ENSG00000243967 | 0 | 2 | 0 | 1 | 1 | 2 |
| ENSG00000238118 | 1 | 2 | 4 | 0 | 1 | 3 |
| ENSG00000224698 | 0 | 0 | 0 | 0 | 0 | 0 |
| ENSG00000186086 | 328 | 189 | 239 | 245 | 258 | 283 |
| ENSG00000283354 | 0 | 0 | 0 | 0 | 0 | 1 |
| ENSG00000227832 | 0 | 0 | 0 | 0 | 0 | 0 |
| ENSG00000226483 | 0 | 3 | 1 | 1 | 2 | 0 |
| ENSG00000162636 | 8215 | 4069 | 4976 | 2417 | 4689 | 4555 |
| ENSG00000162639 | 1719 | 888 | 1038 | 842 | 957 | 975 |
| ENSG00000285923 | 13 | 4 | 10 | 2 | 4 | 8 |
| ENSG00000134186 | 2134 | 1012 | 1247 | 733 | 1356 | 1093 |
| ENSG00000143107 | 6 | 2 | 2 | 1 | 1 | 0 |
| ENSG00000232971 | 0 | 0 | 0 | 0 | 0 | 0 |
| ENSG00000116266 | 1697 | 942 | 879 | 507 | 1378 | 917 |
| ENSG00000228076 | 0 | 0 | 0 | 0 | 0 | 0 |
| ENSG00000162641 | 23 | 16 | 9 | 22 | 30 | 30 |
| ENSG00000274068 | 0 | 0 | 0 | 0 | 0 | 0 |
| ENSG00000203897 | 1 | 2 | 1 | 0 | 2 | 0 |
| ENSG00000121957 | 980 | 562 | 646 | 565 | 836 | 704 |
| ENSG00000121940 | 977 | 552 | 718 | 495 | 748 | 609 |
| ENSG00000085433 | 837 | 393 | 565 | 300 | 453 | 421 |
| ENSG00000237349 | 0 | 0 | 0 | 0 | 0 | 0 |
| ENSG00000244716 | 61 | 28 | 52 | 51 | 35 | 33 |
| ENSG00000228665 | 0 | 0 | 0 | 2 | 0 | 0 |
| ENSG00000228054 | 0 | 0 | 0 | 0 | 0 | 0 |
| ENSG00000197780 | 1536 | 828 | 894 | 694 | 1045 | 828 |
| ENSG00000273382 | 156 | 118 | 127 | 71 | 101 | 99 |
| ENSG00000215717 | 2361 | 1252 | 1508 | 1103 | 1400 | 1347 |
| ENSG00000278249 | 0 | 0 | 0 | 0 | 0 | 0 |
| ENSG00000270066 | 0 | 0 | 0 | 0 | 0 | 0 |
| ENSG00000251484 | 1 | 9 | 1 | 1 | 0 | 0 |
| ENSG00000179902 | 8 | 7 | 6 | 2 | 3 | 3 |
| ENSG00000116299 | 6214 | 2582 | 4020 | 2118 | 1903 | 2209 |
| ENSG00000031698 | 7185 | 3498 | 4039 | 4226 | 4160 | 3878 |
| ENSG00000143126 | 12592 | 5133 | 7759 | 5805 | 5883 | 7085 |
| ENSG00000134222 | 955 | 507 | 613 | 623 | 684 | 611 |
| ENSG00000221986 | 0 | 0 | 0 | 0 | 1 | 0 |
| ENSG00000134243 | 7830 | 3341 | 4664 | 2866 | 3592 | 3889 |
| ENSG00000143106 | 8787 | 4175 | 5179 | 5067 | 5355 | 5041 |
| ENSG00000143028 | 8 | 2 | 0 | 2 | 3 | 3 |
| ENSG00000162650 | 490 | 163 | 235 | 244 | 209 | 223 |
| ENSG00000174151 | 2662 | 1232 | 1599 | 1306 | 1533 | 1606 |
| ENSG00000181754 | 1153 | 541 | 650 | 521 | 550 | 537 |

| | | | | | | |
|---|---|---|---|---|---|---|
| ENSG00000156097 | 8 | 6 | 8 | 3 | 11 | 5 |
| ENSG00000254942 | 0 | 0 | 0 | 0 | 0 | 0 |
| ENSG00000065135 | 5609 | 3023 | 3384 | 2692 | 3656 | 3152 |
| ENSG00000206832 | 3 | 1 | 1 | 0 | 1 | 0 |
| ENSG00000225113 | 0 | 0 | 0 | 0 | 0 | 0 |
| ENSG00000284443 | 0 | 0 | 0 | 0 | 0 | 0 |
| ENSG00000134183 | 0 | 0 | 0 | 0 | 0 | 0 |
| ENSG00000116337 | 1835 | 980 | 1350 | 1461 | 1518 | 1404 |
| ENSG00000228703 | 0 | 0 | 0 | 0 | 0 | 0 |
| ENSG00000235045 | 0 | 0 | 2 | 0 | 0 | 0 |
| ENSG00000168765 | 1288 | 617 | 734 | 995 | 1041 | 920 |
| ENSG00000213366 | 29 | 12 | 22 | 20 | 40 | 29 |
| ENSG00000134184 | 4 | 3 | 2 | 4 | 1 | 2 |
| ENSG00000260246 | 0 | 0 | 0 | 0 | 0 | 0 |
| ENSG00000134201 | 13 | 11 | 4 | 8 | 5 | 15 |
| ENSG00000241720 | 27 | 15 | 17 | 27 | 31 | 29 |
| ENSG00000134202 | 16982 | 9569 | 10811 | 12765 | 12895 | 11825 |
| ENSG00000198758 | 12 | 9 | 8 | 7 | 11 | 12 |
| ENSG00000224927 | 6 | 1 | 0 | 0 | 1 | 3 |
| ENSG00000228420 | 19 | 13 | 9 | 7 | 17 | 11 |
| ENSG00000235005 | 2 | 2 | 1 | 0 | 5 | 2 |
| ENSG00000261055 | 0 | 0 | 1 | 0 | 2 | 0 |
| ENSG00000184371 | 72 | 38 | 56 | 33 | 39 | 47 |
| ENSG00000168710 | 6969 | 3402 | 4370 | 2981 | 3812 | 3934 |
| ENSG00000143093 | 2363 | 1045 | 1358 | 1022 | 1116 | 1294 |
| ENSG00000258634 | 332 | 159 | 223 | 193 | 201 | 235 |
| ENSG00000156150 | 0 | 0 | 0 | 0 | 0 | 0 |
| ENSG00000258673 | 3 | 0 | 1 | 0 | 3 | 2 |
| ENSG00000186150 | 0 | 0 | 1 | 0 | 0 | 0 |
| ENSG00000197106 | 31 | 18 | 15 | 21 | 20 | 23 |
| ENSG00000227091 | 0 | 0 | 1 | 0 | 0 | 0 |
| ENSG00000235526 | 4 | 1 | 3 | 2 | 3 | 3 |
| ENSG00000224965 | 0 | 0 | 1 | 1 | 0 | 1 |
| ENSG00000116396 | 58 | 24 | 44 | 35 | 36 | 31 |
| ENSG00000282852 | 0 | 0 | 0 | 0 | 0 | 0 |
| ENSG00000200536 | 0 | 0 | 0 | 0 | 0 | 0 |
| ENSG00000227963 | 6 | 4 | 9 | 6 | 3 | 10 |
| ENSG00000162775 | 489 | 275 | 355 | 206 | 288 | 283 |
| ENSG00000224699 | 105 | 62 | 91 | 53 | 69 | 71 |
| ENSG00000168679 | 40 | 21 | 31 | 11 | 36 | 25 |
| ENSG00000273373 | 197 | 85 | 141 | 62 | 114 | 65 |
| ENSG00000134248 | 2566 | 1500 | 1774 | 1886 | 1760 | 1806 |
| ENSG00000143125 | 2 | 0 | 0 | 1 | 1 | 0 |
| ENSG00000270380 | 0 | 0 | 0 | 0 | 0 | 0 |
| ENSG00000282887 | 0 | 0 | 0 | 0 | 0 | 0 |
| ENSG00000283999 | 0 | 0 | 0 | 0 | 0 | 0 |
| ENSG00000240194 | 0 | 0 | 0 | 0 | 0 | 0 |
| ENSG00000235407 | 0 | 0 | 0 | 0 | 0 | 0 |
| ENSG00000143105 | 0 | 0 | 0 | 0 | 0 | 0 |
| ENSG00000177301 | 0 | 0 | 0 | 0 | 0 | 0 |

| | | | | | | |
|---|---|---|---|---|---|---|
| ENSG00000259834 | 0 | 0 | 0 | 0 | 0 | 0 |
| ENSG00000177272 | 0 | 1 | 0 | 0 | 0 | 0 |
| ENSG00000199710 | 0 | 0 | 0 | 0 | 0 | 0 |
| ENSG00000237415 | 0 | 0 | 0 | 0 | 0 | 0 |
| ENSG00000232791 | 0 | 0 | 0 | 0 | 0 | 0 |
| ENSG00000143119 | 3 | 2 | 0 | 1 | 0 | 2 |
| ENSG00000261654 | 0 | 6 | 7 | 2 | 4 | 5 |
| ENSG00000232811 | 16 | 3 | 3 | 1 | 11 | 7 |
| ENSG00000121931 | 345 | 195 | 194 | 107 | 251 | 204 |
| ENSG00000273010 | 3 | 4 | 5 | 3 | 4 | 9 |
| ENSG00000225672 | 0 | 0 | 0 | 0 | 0 | 0 |
| ENSG00000252760 | 0 | 0 | 0 | 0 | 0 | 0 |
| ENSG00000156171 | 1129 | 633 | 721 | 580 | 827 | 651 |
| ENSG00000134255 | 855 | 485 | 521 | 432 | 549 | 512 |
| ENSG00000272982 | 0 | 0 | 0 | 0 | 0 | 0 |
| ENSG00000273221 | 0 | 0 | 0 | 0 | 0 | 0 |
| ENSG00000162777 | 2157 | 1045 | 1260 | 1068 | 1158 | 1105 |
| ENSG00000064886 | 20 | 9 | 22 | 16 | 12 | 9 |
| ENSG00000236040 | 0 | 0 | 0 | 0 | 0 | 0 |
| ENSG00000203878 | 0 | 0 | 0 | 0 | 0 | 0 |
| ENSG00000134216 | 0 | 0 | 0 | 0 | 0 | 2 |
| ENSG00000229283 | 0 | 0 | 0 | 0 | 0 | 0 |
| ENSG00000232240 | 0 | 0 | 0 | 0 | 0 | 0 |
| ENSG00000173947 | 82 | 64 | 58 | 29 | 39 | 36 |
| ENSG00000234020 | 0 | 0 | 0 | 0 | 0 | 0 |
| ENSG00000236012 | 0 | 0 | 0 | 0 | 0 | 0 |
| ENSG00000227179 | 0 | 0 | 0 | 0 | 0 | 1 |
| ENSG00000085465 | 72 | 40 | 42 | 66 | 83 | 64 |
| ENSG00000260948 | 0 | 4 | 2 | 1 | 3 | 1 |
| ENSG00000233337 | 2 | 4 | 4 | 1 | 1 | 10 |
| ENSG00000243960 | 2 | 2 | 3 | 1 | 0 | 3 |
| ENSG00000116455 | 919 | 506 | 650 | 692 | 689 | 693 |
| ENSG00000199890 | 2 | 0 | 0 | 0 | 2 | 0 |
| ENSG00000116459 | 6400 | 3517 | 3587 | 3488 | 4150 | 3491 |
| ENSG00000143110 | 3 | 2 | 4 | 7 | 6 | 2 |
| ENSG00000121933 | 2 | 1 | 0 | 0 | 0 | 0 |
| ENSG00000200360 | 0 | 0 | 0 | 0 | 0 | 0 |
| ENSG00000282608 | 0 | 1 | 0 | 1 | 0 | 0 |
| ENSG00000116473 | 1719 | 828 | 909 | 652 | 903 | 862 |
| ENSG00000231346 | 2 | 2 | 1 | 1 | 3 | 0 |
| ENSG00000215867 | 2 | 0 | 0 | 0 | 0 | 0 |
| ENSG00000201028 | 0 | 0 | 0 | 0 | 0 | 0 |
| ENSG00000197852 | 5 | 3 | 2 | 2 | 3 | 4 |
| ENSG00000284755 | 0 | 0 | 0 | 0 | 0 | 0 |
| ENSG00000227811 | 2 | 1 | 1 | 0 | 2 | 1 |
| ENSG00000284830 | 4 | 2 | 3 | 2 | 2 | 3 |
| ENSG00000064703 | 740 | 319 | 426 | 309 | 392 | 385 |
| ENSG00000171385 | 64 | 25 | 28 | 19 | 36 | 45 |
| ENSG00000232558 | 0 | 0 | 0 | 0 | 0 | 0 |
| ENSG00000237556 | 2 | 0 | 0 | 0 | 0 | 1 |

| | | | | | | |
|---|---|---|---|---|---|---|
| ENSG00000231437 | 381 | 183 | 238 | 160 | 215 | 181 |
| ENSG00000231246 | 0 | 0 | 0 | 0 | 0 | 0 |
| ENSG00000234388 | 0 | 0 | 0 | 0 | 0 | 0 |
| ENSG00000238761 | 0 | 0 | 0 | 0 | 0 | 0 |
| ENSG00000143079 | 666 | 377 | 369 | 194 | 419 | 304 |
| ENSG00000283296 | 0 | 0 | 0 | 0 | 0 | 0 |
| ENSG00000134245 | 101 | 47 | 60 | 19 | 38 | 31 |
| ENSG00000273483 | 0 | 1 | 2 | 0 | 0 | 1 |
| ENSG00000007341 | 341 | 155 | 185 | 174 | 191 | 195 |
| ENSG00000116489 | 6002 | 3224 | 3399 | 2409 | 3601 | 2882 |
| ENSG00000235299 | 0 | 0 | 0 | 0 | 0 | 0 |
| ENSG00000252750 | 1 | 0 | 0 | 0 | 0 | 0 |
| ENSG00000155363 | 2690 | 1316 | 1784 | 1257 | 1162 | 1390 |
| ENSG00000225075 | 19 | 5 | 7 | 9 | 5 | 14 |
| ENSG00000155366 | 6376 | 3313 | 4222 | 3679 | 2800 | 3148 |
| ENSG00000271810 | 0 | 1 | 1 | 0 | 0 | 0 |
| ENSG00000155367 | 101 | 27 | 60 | 31 | 33 | 48 |
| ENSG00000261595 | 6 | 0 | 2 | 0 | 1 | 2 |
| ENSG00000184599 | 4 | 0 | 0 | 0 | 3 | 1 |
| ENSG00000234790 | 0 | 0 | 0 | 0 | 0 | 0 |
| ENSG00000215866 | 0 | 0 | 0 | 0 | 0 | 0 |
| ENSG00000224167 | 0 | 0 | 0 | 0 | 0 | 0 |
| ENSG00000228040 | 0 | 0 | 0 | 0 | 0 | 0 |
| ENSG00000155380 | 376 | 160 | 187 | 146 | 198 | 178 |
| ENSG00000229020 | 0 | 0 | 0 | 0 | 0 | 0 |
| ENSG00000226419 | 8 | 3 | 7 | 3 | 5 | 2 |
| ENSG00000238198 | 2 | 0 | 1 | 0 | 2 | 0 |
| ENSG00000198799 | 527 | 251 | 323 | 151 | 265 | 233 |
| ENSG00000237278 | 0 | 0 | 0 | 0 | 0 | 0 |
| ENSG00000233839 | 0 | 0 | 0 | 0 | 0 | 0 |
| ENSG00000236887 | 0 | 0 | 0 | 0 | 0 | 0 |
| ENSG00000081026 | 359 | 185 | 181 | 104 | 248 | 163 |
| ENSG00000232499 | 1 | 1 | 1 | 1 | 0 | 1 |
| ENSG00000238256 | 0 | 0 | 0 | 0 | 0 | 0 |
| ENSG00000116793 | 1336 | 631 | 804 | 672 | 824 | 821 |
| ENSG00000232450 | 43 | 27 | 32 | 25 | 27 | 26 |
| ENSG00000081019 | 1642 | 719 | 855 | 593 | 895 | 760 |
| ENSG00000231128 | 2 | 2 | 3 | 0 | 2 | 1 |
| ENSG00000134242 | 23 | 11 | 8 | 6 | 19 | 4 |
| ENSG00000226167 | 286 | 143 | 198 | 113 | 167 | 168 |
| ENSG00000188761 | 48 | 22 | 27 | 22 | 36 | 30 |
| ENSG00000134262 | 1406 | 714 | 978 | 719 | 888 | 830 |
| ENSG00000118655 | 2421 | 1169 | 1496 | 1224 | 1378 | 1538 |
| ENSG00000235527 | 73 | 29 | 28 | 26 | 21 | 22 |
| ENSG00000163349 | 14402 | 6395 | 7993 | 4441 | 6978 | 6921 |
| ENSG00000116774 | 19 | 14 | 10 | 19 | 11 | 9 |
| ENSG00000270780 | 0 | 0 | 0 | 0 | 0 | 0 |
| ENSG00000282048 | 0 | 2 | 0 | 0 | 0 | 0 |
| ENSG00000134207 | 0 | 0 | 0 | 0 | 0 | 1 |
| ENSG00000232895 | 0 | 0 | 0 | 0 | 0 | 0 |

| | | | | | | |
|---|---|---|---|---|---|---|
| ENSG00000273698 | 2 | 0 | 0 | 0 | 0 | 0 |
| ENSG00000271419 | 0 | 0 | 0 | 0 | 0 | 0 |
| ENSG00000197323 | 7613 | 3942 | 3552 | 1981 | 4745 | 3582 |
| ENSG00000226984 | 0 | 0 | 0 | 0 | 0 | 0 |
| ENSG00000225650 | 0 | 2 | 0 | 0 | 0 | 0 |
| ENSG00000201114 | 0 | 0 | 0 | 0 | 0 | 0 |
| ENSG00000236480 | 4 | 1 | 2 | 2 | 1 | 0 |
| ENSG00000116752 | 5337 | 2950 | 2865 | 2128 | 3775 | 2724 |
| ENSG00000175984 | 214 | 110 | 136 | 61 | 133 | 103 |
| ENSG00000116748 | 34 | 17 | 25 | 13 | 8 | 17 |
| ENSG00000242769 | 0 | 0 | 0 | 0 | 0 | 0 |
| ENSG00000213281 | 12961 | 6262 | 7317 | 4915 | 7787 | 6687 |
| ENSG00000009307 | 95300 | 48407 | 57884 | 41458 | 57560 | 51821 |
| ENSG00000201900 | 4 | 8 | 7 | 0 | 4 | 2 |
| ENSG00000052723 | 4290 | 2020 | 2320 | 1446 | 2614 | 2093 |
| ENSG00000227970 | 0 | 0 | 0 | 0 | 0 | 0 |
| ENSG00000198765 | 0 | 0 | 0 | 0 | 0 | 0 |
| ENSG00000134200 | 0 | 0 | 0 | 0 | 0 | 0 |
| ENSG00000134198 | 0 | 0 | 1 | 1 | 1 | 2 |
| ENSG00000233730 | 0 | 0 | 0 | 0 | 0 | 0 |
| ENSG00000285698 | 0 | 0 | 0 | 0 | 0 | 0 |
| ENSG00000228035 | 0 | 0 | 0 | 0 | 0 | 0 |
| ENSG00000134259 | 0 | 0 | 0 | 0 | 0 | 0 |
| ENSG00000226973 | 0 | 0 | 0 | 0 | 0 | 0 |
| ENSG00000274468 | 0 | 0 | 0 | 0 | 0 | 0 |
| ENSG00000229101 | 0 | 0 | 0 | 0 | 0 | 0 |
| ENSG00000232185 | 0 | 0 | 0 | 0 | 0 | 0 |
| ENSG00000270631 | 0 | 0 | 0 | 0 | 0 | 0 |
| ENSG00000239984 | 0 | 0 | 0 | 0 | 0 | 0 |
| ENSG00000207502 | 0 | 0 | 0 | 0 | 0 | 0 |
| ENSG00000173218 | 4420 | 2023 | 2545 | 2063 | 2568 | 2813 |
| ENSG00000118729 | 4 | 2 | 5 | 2 | 1 | 3 |
| ENSG00000177551 | 0 | 1 | 1 | 1 | 1 | 0 |
| ENSG00000214204 | 0 | 0 | 0 | 0 | 0 | 0 |
| ENSG00000271143 | 0 | 0 | 0 | 0 | 1 | 0 |
| ENSG00000228127 | 4 | 0 | 0 | 0 | 1 | 1 |
| ENSG00000163393 | 368 | 168 | 160 | 94 | 166 | 144 |
| ENSG00000237993 | 0 | 0 | 0 | 0 | 0 | 0 |
| ENSG00000173212 | 2 | 3 | 0 | 0 | 1 | 1 |
| ENSG00000235933 | 0 | 0 | 0 | 0 | 0 | 0 |
| ENSG00000221040 | 0 | 0 | 0 | 0 | 0 | 0 |
| ENSG00000163399 | 26786 | 13854 | 16905 | 14101 | 15776 | 15805 |
| ENSG00000203865 | 218 | 78 | 120 | 143 | 106 | 94 |
| ENSG00000212385 | 8 | 0 | 3 | 3 | 3 | 1 |
| ENSG00000233154 | 53 | 24 | 18 | 15 | 28 | 17 |
| ENSG00000230381 | 0 | 0 | 0 | 0 | 0 | 0 |
| ENSG00000200547 | 2 | 0 | 0 | 0 | 0 | 0 |
| ENSG00000224950 | 71 | 20 | 25 | 19 | 42 | 19 |
| ENSG00000116815 | 596 | 301 | 323 | 266 | 368 | 330 |
| ENSG00000177173 | 60 | 29 | 28 | 18 | 22 | 22 |

| | | | | | | |
|---|---|---|---|---|---|---|
| ENSG00000264419 | 0 | 0 | 0 | 0 | 0 | 0 |
| ENSG00000143061 | 7761 | 3562 | 4409 | 2628 | 3509 | 3411 |
| ENSG00000211543 | 0 | 0 | 0 | 0 | 0 | 0 |
| ENSG00000203864 | 0 | 0 | 0 | 0 | 0 | 0 |
| ENSG00000231072 | 0 | 0 | 1 | 0 | 0 | 0 |
| ENSG00000223489 | 0 | 0 | 0 | 0 | 0 | 0 |
| ENSG00000116824 | 0 | 0 | 0 | 0 | 0 | 0 |
| ENSG00000225079 | 0 | 0 | 0 | 0 | 0 | 0 |
| ENSG00000272715 | 0 | 0 | 1 | 1 | 0 | 0 |
| ENSG00000134247 | 2648 | 1257 | 1741 | 1055 | 1336 | 1377 |
| ENSG00000252510 | 0 | 0 | 0 | 0 | 0 | 0 |
| ENSG00000134256 | 61 | 35 | 40 | 13 | 17 | 28 |
| ENSG00000236137 | 108 | 39 | 58 | 50 | 66 | 62 |
| ENSG00000116830 | 1477 | 637 | 837 | 526 | 945 | 858 |
| ENSG00000215930 | 1 | 0 | 1 | 0 | 0 | 0 |
| ENSG00000134253 | 610 | 259 | 451 | 265 | 328 | 329 |
| ENSG00000228453 | 0 | 0 | 0 | 0 | 0 | 0 |
| ENSG00000134258 | 64 | 40 | 65 | 115 | 157 | 146 |
| ENSG00000277141 | 0 | 0 | 0 | 0 | 0 | 0 |
| ENSG00000235202 | 0 | 0 | 0 | 0 | 0 | 0 |
| ENSG00000271427 | 39 | 13 | 38 | 15 | 18 | 18 |
| ENSG00000198162 | 3478 | 1625 | 1890 | 860 | 1769 | 1507 |
| ENSG00000279513 | 4 | 4 | 2 | 0 | 1 | 2 |
| ENSG00000226755 | 1 | 0 | 1 | 0 | 0 | 0 |
| ENSG00000236866 | 0 | 0 | 0 | 0 | 0 | 1 |
| ENSG00000183508 | 482 | 199 | 248 | 152 | 194 | 159 |
| ENSG00000213262 | 0 | 0 | 0 | 0 | 0 | 0 |
| ENSG00000212266 | 0 | 0 | 0 | 0 | 0 | 0 |
| ENSG00000270719 | 0 | 0 | 0 | 0 | 0 | 0 |
| ENSG00000228217 | 0 | 0 | 1 | 0 | 2 | 0 |
| ENSG00000196505 | 703 | 413 | 366 | 250 | 400 | 345 |
| ENSG00000065183 | 1005 | 533 | 686 | 584 | 729 | 725 |
| ENSG00000155761 | 4 | 3 | 1 | 1 | 1 | 1 |
| ENSG00000222209 | 0 | 0 | 0 | 0 | 0 | 0 |
| ENSG00000226126 | 4 | 0 | 0 | 0 | 0 | 0 |
| ENSG00000229911 | 0 | 0 | 0 | 0 | 0 | 0 |
| ENSG00000092607 | 0 | 0 | 1 | 0 | 0 | 0 |
| ENSG00000226172 | 0 | 0 | 0 | 0 | 0 | 0 |
| ENSG00000239216 | 0 | 0 | 0 | 0 | 0 | 0 |
| ENSG00000116874 | 515 | 279 | 292 | 222 | 338 | 266 |
| ENSG00000224238 | 0 | 0 | 2 | 0 | 0 | 0 |
| ENSG00000223575 | 0 | 0 | 0 | 0 | 0 | 0 |
| ENSG00000236804 | 0 | 0 | 0 | 0 | 0 | 0 |
| ENSG00000231365 | 258 | 123 | 162 | 116 | 139 | 129 |
| ENSG00000227056 | 0 | 1 | 0 | 0 | 0 | 0 |
| ENSG00000227712 | 0 | 0 | 0 | 0 | 0 | 0 |
| ENSG00000232650 | 0 | 0 | 0 | 0 | 0 | 0 |
| ENSG00000194297 | 0 | 0 | 0 | 0 | 0 | 0 |
| ENSG00000116882 | 0 | 0 | 0 | 1 | 0 | 0 |
| ENSG00000230921 | 0 | 0 | 0 | 0 | 0 | 0 |

| | | | | | | |
|---|---|---|---|---|---|---|
| ENSG00000203859 | 1 | 0 | 0 | 0 | 0 | 0 |
| ENSG00000232780 | 0 | 0 | 0 | 0 | 0 | 0 |
| ENSG00000203858 | 0 | 0 | 0 | 0 | 0 | 0 |
| ENSG00000235595 | 0 | 0 | 0 | 0 | 0 | 0 |
| ENSG00000187481 | 0 | 0 | 0 | 0 | 0 | 0 |
| ENSG00000229570 | 0 | 0 | 0 | 0 | 0 | 0 |
| ENSG00000203857 | 0 | 0 | 0 | 0 | 0 | 0 |
| ENSG00000226443 | 0 | 0 | 0 | 1 | 0 | 1 |
| ENSG00000249798 | 0 | 0 | 0 | 0 | 0 | 0 |
| ENSG00000239873 | 0 | 0 | 0 | 0 | 0 | 0 |
| ENSG00000203855 | 0 | 0 | 0 | 0 | 0 | 0 |
| ENSG00000240244 | 0 | 0 | 0 | 0 | 0 | 0 |
| ENSG00000260941 | 4 | 0 | 0 | 0 | 0 | 0 |
| ENSG00000198857 | 2 | 0 | 0 | 0 | 0 | 1 |
| ENSG00000143067 | 173 | 64 | 123 | 65 | 96 | 91 |
| ENSG00000092621 | 264 | 174 | 181 | 282 | 250 | 190 |
| ENSG00000134240 | 8 | 8 | 7 | 5 | 5 | 1 |
| ENSG00000134193 | 468 | 198 | 243 | 232 | 258 | 236 |
| ENSG00000215864 | 1 | 1 | 0 | 0 | 0 | 0 |
| ENSG00000227205 | 0 | 0 | 0 | 0 | 0 | 0 |
| ENSG00000226446 | 0 | 0 | 0 | 0 | 0 | 0 |
| ENSG00000134249 | 0 | 0 | 0 | 0 | 0 | 0 |
| ENSG00000261662 | 10 | 3 | 6 | 5 | 4 | 5 |
| ENSG00000134250 | 4031 | 1764 | 2322 | 1276 | 1651 | 1879 |
| ENSG00000273406 | 0 | 0 | 0 | 0 | 0 | 0 |
| ENSG00000207149 | 0 | 0 | 0 | 0 | 0 | 0 |
| ENSG00000265808 | 5024 | 2636 | 3148 | 2400 | 2865 | 2736 |
| ENSG00000274642 | 0 | 0 | 0 | 2 | 2 | 0 |
| ENSG00000277313 | 0 | 0 | 0 | 0 | 0 | 0 |
| ENSG00000223804 | 1 | 0 | 2 | 0 | 0 | 0 |
| ENSG00000273727 | 0 | 0 | 0 | 0 | 0 | 0 |
| ENSG00000270392 | 11 | 8 | 6 | 1 | 6 | 8 |
| ENSG00000270231 | 365 | 171 | 245 | 140 | 178 | 186 |
| ENSG00000275131 | 257 | 87 | 146 | 71 | 109 | 97 |
| ENSG00000273694 | 0 | 0 | 0 | 0 | 0 | 0 |
| ENSG00000275933 | 0 | 0 | 0 | 0 | 0 | 0 |
| ENSG00000273136 | 280 | 137 | 135 | 88 | 175 | 200 |
| ENSG00000275538 | 0 | 1 | 0 | 0 | 0 | 1 |
| ENSG00000263353 | 0 | 0 | 0 | 1 | 0 | 0 |
| ENSG00000281741 | 1 | 1 | 4 | 0 | 1 | 0 |
| ENSG00000226067 | 7 | 5 | 10 | 13 | 15 | 7 |
| ENSG00000277610 | 0 | 0 | 0 | 0 | 0 | 0 |
| ENSG00000285215 | 0 | 0 | 0 | 0 | 0 | 0 |
| ENSG00000275585 | 0 | 0 | 0 | 0 | 0 | 0 |
| ENSG00000273516 | 0 | 0 | 0 | 0 | 0 | 0 |
| ENSG00000275296 | 0 | 0 | 0 | 0 | 0 | 0 |
| ENSG00000198019 | 0 | 0 | 2 | 0 | 2 | 0 |
| ENSG00000234998 | 0 | 0 | 0 | 0 | 0 | 0 |
| ENSG00000233029 | 6 | 1 | 2 | 1 | 0 | 1 |
| ENSG00000223345 | 0 | 1 | 0 | 0 | 0 | 1 |

| | | | | | | |
|---|---|---|---|---|---|---|
| ENSG00000227193 | 0 | 1 | 1 | 1 | 0 | 0 |
| ENSG00000213244 | 0 | 0 | 0 | 0 | 0 | 0 |
| ENSG00000268993 | 0 | 0 | 0 | 0 | 0 | 0 |
| ENSG00000188610 | 54 | 46 | 36 | 41 | 67 | 48 |
| ENSG00000171943 | 188 | 63 | 107 | 67 | 97 | 74 |
| ENSG00000275129 | 0 | 0 | 0 | 0 | 0 | 0 |
| ENSG00000230806 | 0 | 0 | 1 | 0 | 0 | 1 |
| ENSG00000227082 | 8 | 4 | 5 | 1 | 3 | 1 |
| ENSG00000228826 | 0 | 0 | 0 | 0 | 0 | 0 |
| ENSG00000233432 | 0 | 0 | 0 | 0 | 0 | 0 |
| ENSG00000272583 | 0 | 0 | 0 | 0 | 0 | 0 |
| ENSG00000231752 | 70 | 43 | 31 | 24 | 35 | 30 |
| ENSG00000224857 | 0 | 0 | 0 | 0 | 0 | 0 |
| ENSG00000276118 | 0 | 0 | 0 | 0 | 0 | 0 |
| ENSG00000277095 | 0 | 0 | 0 | 0 | 0 | 0 |
| ENSG00000277702 | 0 | 0 | 0 | 0 | 0 | 0 |
| ENSG00000252830 | 0 | 0 | 0 | 0 | 0 | 0 |
| ENSG00000264145 | 0 | 0 | 0 | 0 | 0 | 0 |
| ENSG00000274927 | 0 | 0 | 0 | 0 | 0 | 0 |
| ENSG00000242943 | 0 | 0 | 0 | 0 | 0 | 0 |
| ENSG00000223495 | 0 | 0 | 0 | 0 | 0 | 0 |
| ENSG00000235887 | 0 | 0 | 0 | 0 | 0 | 0 |
| ENSG00000203825 | 0 | 0 | 0 | 0 | 0 | 0 |
| ENSG00000179571 | 2 | 1 | 0 | 0 | 1 | 0 |
| ENSG00000226028 | 0 | 0 | 0 | 0 | 0 | 0 |
| ENSG00000202167 | 0 | 0 | 0 | 0 | 0 | 0 |
| ENSG00000207349 | 0 | 0 | 0 | 0 | 0 | 0 |
| ENSG00000252826 | 0 | 0 | 0 | 0 | 0 | 0 |
| ENSG00000206737 | 0 | 0 | 0 | 0 | 0 | 0 |
| ENSG00000232721 | 0 | 0 | 0 | 0 | 0 | 0 |
| ENSG00000223779 | 0 | 0 | 0 | 0 | 0 | 0 |
| ENSG00000242352 | 0 | 0 | 0 | 0 | 0 | 0 |
| ENSG00000270962 | 0 | 0 | 0 | 0 | 0 | 0 |
| ENSG00000263590 | 0 | 0 | 0 | 0 | 0 | 0 |
| ENSG00000252105 | 0 | 0 | 0 | 0 | 0 | 0 |
| ENSG00000233430 | 0 | 0 | 0 | 0 | 0 | 0 |
| ENSG00000237503 | 0 | 0 | 0 | 0 | 0 | 0 |
| ENSG00000265531 | 0 | 0 | 0 | 0 | 0 | 0 |
| ENSG00000240929 | 0 | 0 | 0 | 0 | 0 | 0 |
| ENSG00000234571 | 0 | 0 | 0 | 0 | 0 | 0 |
| ENSG00000273213 | 0 | 0 | 0 | 0 | 0 | 0 |
| ENSG00000230186 | 0 | 0 | 0 | 0 | 0 | 0 |
| ENSG00000203818 | 0 | 0 | 0 | 0 | 0 | 0 |
| ENSG00000268074 | 0 | 0 | 0 | 0 | 0 | 0 |
| ENSG00000263513 | 0 | 0 | 1 | 3 | 3 | 4 |
| ENSG00000270872 | 0 | 0 | 0 | 0 | 0 | 0 |
| ENSG00000276674 | 0 | 0 | 0 | 0 | 0 | 0 |
| ENSG00000275075 | 0 | 0 | 0 | 0 | 0 | 0 |
| ENSG00000275767 | 0 | 0 | 0 | 0 | 0 | 0 |
| ENSG00000232527 | 0 | 0 | 0 | 0 | 0 | 0 |

| | | | | | | |
|---|---|---|---|---|---|---|
| ENSG00000276442 | 0 | 0 | 0 | 0 | 0 | 0 |
| ENSG00000175658 | 0 | 0 | 0 | 0 | 0 | 0 |
| ENSG00000276756 | 0 | 0 | 0 | 0 | 0 | 0 |
| ENSG00000271567 | 0 | 0 | 0 | 0 | 0 | 0 |
| ENSG00000271439 | 0 | 0 | 0 | 0 | 0 | 0 |
| ENSG00000207205 | 1 | 0 | 2 | 1 | 1 | 0 |
| ENSG00000237343 | 0 | 0 | 0 | 2 | 0 | 1 |
| ENSG00000233586 | 4 | 3 | 6 | 1 | 3 | 1 |
| ENSG00000266338 | 1578 | 807 | 1161 | 596 | 990 | 983 |
| ENSG00000227212 | 17 | 7 | 14 | 5 | 20 | 10 |
| ENSG00000229002 | 0 | 0 | 0 | 0 | 0 | 0 |
| ENSG00000275291 | 0 | 0 | 0 | 0 | 0 | 0 |
| ENSG00000278099 | 0 | 0 | 0 | 0 | 0 | 0 |
| ENSG00000277918 | 0 | 0 | 0 | 0 | 0 | 0 |
| ENSG00000279782 | 0 | 0 | 0 | 0 | 0 | 0 |
| ENSG00000277199 | 0 | 0 | 0 | 0 | 0 | 0 |
| ENSG00000224363 | 0 | 0 | 0 | 0 | 0 | 0 |
| ENSG00000278273 | 0 | 0 | 0 | 0 | 0 | 0 |
| ENSG00000222854 | 0 | 0 | 0 | 0 | 0 | 0 |
| ENSG00000273825 | 0 | 0 | 0 | 0 | 0 | 0 |
| ENSG00000196369 | 133 | 64 | 76 | 45 | 91 | 90 |
| ENSG00000271644 | 0 | 0 | 0 | 0 | 0 | 0 |
| ENSG00000215784 | 66 | 18 | 26 | 43 | 40 | 39 |
| ENSG00000283752 | 0 | 0 | 0 | 0 | 0 | 1 |
| ENSG00000272419 | 89 | 39 | 47 | 44 | 52 | 42 |
| ENSG00000277678 | 0 | 3 | 0 | 0 | 2 | 0 |
| ENSG00000281571 | 10 | 4 | 11 | 12 | 9 | 5 |
| ENSG00000223612 | 38 | 8 | 14 | 9 | 13 | 20 |
| ENSG00000256374 | 0 | 0 | 0 | 0 | 0 | 0 |
| ENSG00000271291 | 0 | 0 | 0 | 0 | 0 | 0 |
| ENSG00000276216 | 7 | 1 | 6 | 1 | 0 | 0 |
| ENSG00000207501 | 0 | 0 | 0 | 0 | 0 | 0 |
| ENSG00000236140 | 2 | 1 | 4 | 2 | 2 | 1 |
| ENSG00000162825 | 196 | 99 | 79 | 66 | 109 | 80 |
| ENSG00000234367 | 2 | 2 | 3 | 0 | 3 | 1 |
| ENSG00000270384 | 0 | 0 | 0 | 0 | 0 | 0 |
| ENSG00000270722 | 0 | 0 | 0 | 0 | 1 | 0 |
| ENSG00000278182 | 0 | 0 | 0 | 0 | 0 | 0 |
| ENSG00000275064 | 0 | 0 | 0 | 0 | 0 | 0 |
| ENSG00000272150 | 175 | 84 | 96 | 114 | 152 | 135 |
| ENSG00000117262 | 329 | 158 | 231 | 158 | 198 | 197 |
| ENSG00000174827 | 20 | 13 | 9 | 3 | 10 | 8 |
| ENSG00000117281 | 4 | 6 | 17 | 7 | 5 | 4 |
| ENSG00000265491 | 3083 | 1504 | 1900 | 1508 | 1734 | 1744 |
| ENSG00000186141 | 1343 | 707 | 926 | 754 | 841 | 866 |
| ENSG00000186364 | 245 | 112 | 207 | 159 | 141 | 186 |
| ENSG00000131788 | 6044 | 2734 | 3568 | 3149 | 3375 | 3619 |
| ENSG00000278549 | 0 | 1 | 0 | 0 | 0 | 0 |
| ENSG00000198483 | 106 | 50 | 52 | 65 | 70 | 70 |
| ENSG00000143127 | 36 | 11 | 15 | 9 | 24 | 22 |

| | | | | | | |
|---|---|---|---|---|---|---|
| ENSG00000244619 | 1 | 0 | 0 | 0 | 0 | 2 |
| ENSG00000131779 | 2295 | 1284 | 1362 | 1321 | 1317 | 1320 |
| ENSG00000265241 | 4826 | 2650 | 3403 | 3083 | 3198 | 3498 |
| ENSG00000211451 | 80 | 49 | 83 | 59 | 66 | 60 |
| ENSG00000234222 | 21 | 19 | 50 | 14 | 28 | 27 |
| ENSG00000280778 | 0 | 0 | 1 | 0 | 0 | 0 |
| ENSG00000271601 | 11 | 9 | 21 | 6 | 10 | 13 |
| ENSG00000272031 | 202 | 89 | 137 | 116 | 131 | 136 |
| ENSG00000278431 | 0 | 0 | 0 | 0 | 0 | 0 |
| ENSG00000121851 | 529 | 274 | 350 | 358 | 347 | 326 |
| ENSG00000265972 | 13515 | 6243 | 9282 | 5462 | 6518 | 5838 |
| ENSG00000168509 | 0 | 1 | 1 | 0 | 2 | 0 |
| ENSG00000201558 | 0 | 2 | 0 | 1 | 0 | 1 |
| ENSG00000233396 | 135 | 60 | 81 | 62 | 81 | 79 |
| ENSG00000271425 | 184 | 89 | 78 | 38 | 93 | 73 |
| ENSG00000213240 | 0 | 1 | 0 | 0 | 0 | 0 |
| ENSG00000264343 | 16 | 6 | 9 | 4 | 9 | 11 |
| ENSG00000276509 | 0 | 1 | 1 | 0 | 0 | 0 |
| ENSG00000201789 | 0 | 0 | 0 | 0 | 0 | 0 |
| ENSG00000271121 | 0 | 0 | 1 | 0 | 0 | 0 |
| ENSG00000277406 | 52 | 20 | 29 | 17 | 26 | 19 |
| ENSG00000270339 | 0 | 0 | 0 | 0 | 0 | 0 |
| ENSG00000273768 | 4 | 1 | 1 | 0 | 0 | 0 |
| ENSG00000274428 | 0 | 0 | 0 | 0 | 0 | 0 |
| ENSG00000277655 | 0 | 0 | 0 | 0 | 0 | 0 |
| ENSG00000276975 | 0 | 0 | 0 | 0 | 0 | 0 |
| ENSG00000274408 | 0 | 0 | 0 | 0 | 0 | 0 |
| ENSG00000274450 | 0 | 0 | 0 | 0 | 0 | 0 |
| ENSG00000268043 | 889 | 397 | 537 | 296 | 438 | 446 |
| ENSG00000244371 | 0 | 2 | 0 | 0 | 0 | 0 |
| ENSG00000234225 | 0 | 0 | 1 | 0 | 0 | 0 |
| ENSG00000227242 | 0 | 0 | 2 | 0 | 0 | 1 |
| ENSG00000225603 | 0 | 2 | 0 | 0 | 0 | 0 |
| ENSG00000201142 | 0 | 0 | 0 | 0 | 0 | 0 |
| ENSG00000230832 | 6 | 3 | 2 | 4 | 1 | 4 |
| ENSG00000283342 | 2 | 5 | 9 | 3 | 4 | 2 |
| ENSG00000131791 | 2605 | 1199 | 1662 | 1023 | 1351 | 1313 |
| ENSG00000180867 | 307 | 173 | 192 | 129 | 179 | 164 |
| ENSG00000237188 | 2 | 4 | 1 | 0 | 3 | 2 |
| ENSG00000131781 | 212 | 99 | 130 | 70 | 135 | 106 |
| ENSG00000226015 | 35 | 13 | 25 | 18 | 24 | 22 |
| ENSG00000236806 | 0 | 5 | 1 | 0 | 3 | 1 |
| ENSG00000131778 | 3052 | 1482 | 1936 | 1664 | 1856 | 1891 |
| ENSG00000278811 | 39 | 18 | 18 | 9 | 11 | 14 |
| ENSG00000213226 | 0 | 1 | 0 | 0 | 0 | 0 |
| ENSG00000226653 | 0 | 0 | 0 | 0 | 0 | 0 |
| ENSG00000207209 | 0 | 0 | 0 | 0 | 0 | 0 |
| ENSG00000272443 | 2 | 1 | 1 | 0 | 4 | 5 |
| ENSG00000272480 | 0 | 0 | 0 | 0 | 0 | 0 |
| ENSG00000116128 | 1940 | 797 | 1128 | 706 | 833 | 918 |

| | | | | | | |
|---|---|---|---|---|---|---|
| ENSG00000162836 | 1532 | 788 | 896 | 896 | 1001 | 920 |
| ENSG00000277762 | 0 | 0 | 0 | 0 | 0 | 0 |
| ENSG00000227139 | 0 | 0 | 0 | 0 | 0 | 0 |
| ENSG00000265107 | 0 | 0 | 0 | 0 | 0 | 0 |
| ENSG00000274415 | 0 | 0 | 0 | 0 | 0 | 0 |
| ENSG00000234190 | 0 | 0 | 0 | 0 | 0 | 0 |
| ENSG00000223728 | 0 | 0 | 0 | 0 | 0 | 0 |
| ENSG00000121634 | 0 | 0 | 0 | 0 | 0 | 0 |
| ENSG00000188092 | 277 | 151 | 254 | 146 | 199 | 175 |
| ENSG00000215859 | 18 | 6 | 6 | 2 | 13 | 9 |
| ENSG00000273059 | 0 | 0 | 0 | 0 | 0 | 0 |
| ENSG00000206791 | 0 | 0 | 0 | 0 | 1 | 0 |
| ENSG00000274372 | 2 | 2 | 5 | 2 | 4 | 6 |
| ENSG00000206585 | 1 | 1 | 0 | 0 | 0 | 0 |
| ENSG00000229828 | 0 | 0 | 0 | 0 | 0 | 1 |
| ENSG00000263956 | 959 | 408 | 538 | 306 | 382 | 434 |
| ENSG00000225080 | 4 | 2 | 5 | 0 | 3 | 2 |
| ENSG00000198658 | 2 | 1 | 1 | 3 | 1 | 0 |
| ENSG00000227733 | 0 | 0 | 0 | 0 | 0 | 0 |
| ENSG00000232265 | 0 | 0 | 0 | 0 | 0 | 0 |
| ENSG00000238765 | 0 | 0 | 0 | 0 | 0 | 0 |
| ENSG00000224335 | 0 | 0 | 0 | 0 | 0 | 0 |
| ENSG00000227700 | 0 | 0 | 0 | 0 | 0 | 0 |
| ENSG00000199879 | 1 | 0 | 0 | 0 | 3 | 2 |
| ENSG00000234283 | 0 | 2 | 1 | 0 | 0 | 0 |
| ENSG00000228626 | 2 | 7 | 12 | 7 | 11 | 11 |
| ENSG00000274020 | 19 | 16 | 14 | 8 | 16 | 18 |
| ENSG00000238107 | 0 | 0 | 0 | 1 | 0 | 0 |
| ENSG00000224481 | 3 | 1 | 5 | 2 | 0 | 6 |
| ENSG00000235988 | 0 | 0 | 0 | 0 | 0 | 0 |
| ENSG00000252744 | 0 | 0 | 0 | 0 | 0 | 0 |
| ENSG00000283676 | 0 | 0 | 3 | 0 | 0 | 0 |
| ENSG00000202408 | 0 | 2 | 1 | 0 | 0 | 0 |
| ENSG00000272824 | 0 | 0 | 0 | 0 | 0 | 0 |
| ENSG00000207340 | 0 | 0 | 0 | 0 | 0 | 0 |
| ENSG00000252135 | 0 | 0 | 0 | 0 | 1 | 0 |
| ENSG00000278596 | 0 | 0 | 0 | 0 | 0 | 0 |
| ENSG00000238825 | 0 | 0 | 0 | 0 | 0 | 2 |
| ENSG00000231551 | 13 | 7 | 1 | 4 | 14 | 5 |
| ENSG00000201183 | 0 | 0 | 0 | 0 | 0 | 0 |
| ENSG00000225871 | 2 | 0 | 0 | 0 | 1 | 0 |
| ENSG00000285062 | 0 | 0 | 0 | 0 | 0 | 0 |
| ENSG00000280649 | 0 | 2 | 1 | 1 | 1 | 1 |
| ENSG00000236334 | 0 | 0 | 0 | 0 | 1 | 0 |
| ENSG00000274210 | 2 | 2 | 2 | 1 | 1 | 0 |
| ENSG00000270629 | 327 | 160 | 182 | 116 | 209 | 201 |
| ENSG00000252515 | 0 | 0 | 0 | 0 | 0 | 0 |
| ENSG00000177144 | 0 | 0 | 0 | 0 | 0 | 0 |
| ENSG00000275140 | 2 | 0 | 0 | 0 | 0 | 0 |
| ENSG00000271546 | 0 | 0 | 0 | 0 | 0 | 0 |

| | | | | | | |
|---|---|---|---|---|---|---|
| ENSG00000178104 | 2628 | 1211 | 1448 | 1085 | 1348 | 1113 |
| ENSG00000252656 | 0 | 0 | 0 | 0 | 0 | 0 |
| ENSG00000222788 | 1 | 0 | 0 | 0 | 0 | 1 |
| ENSG00000254913 | 0 | 0 | 0 | 0 | 0 | 0 |
| ENSG00000255148 | 0 | 0 | 0 | 0 | 0 | 0 |
| ENSG00000254539 | 0 | 0 | 0 | 1 | 0 | 0 |
| ENSG00000269713 | 1003 | 430 | 621 | 343 | 564 | 519 |
| ENSG00000274019 | 2 | 1 | 0 | 0 | 0 | 0 |
| ENSG00000201699 | 0 | 0 | 0 | 0 | 0 | 0 |
| ENSG00000274265 | 3 | 1 | 2 | 3 | 3 | 1 |
| ENSG00000215861 | 34 | 22 | 21 | 23 | 25 | 15 |
| ENSG00000272755 | 0 | 0 | 0 | 0 | 0 | 1 |
| ENSG00000273591 | 0 | 0 | 0 | 0 | 0 | 0 |
| ENSG00000274423 | 8 | 2 | 3 | 2 | 0 | 3 |
| ENSG00000271383 | 458 | 249 | 280 | 148 | 267 | 244 |
| ENSG00000263464 | 0 | 0 | 0 | 0 | 0 | 0 |
| ENSG00000277147 | 149 | 87 | 109 | 113 | 107 | 109 |
| ENSG00000284964 | 0 | 0 | 0 | 0 | 0 | 0 |
| ENSG00000275557 | 18 | 13 | 15 | 31 | 33 | 12 |
| ENSG00000206828 | 4 | 1 | 4 | 1 | 3 | 0 |
| ENSG00000284842 | 0 | 0 | 0 | 0 | 0 | 0 |
| ENSG00000234232 | 0 | 0 | 0 | 0 | 1 | 0 |
| ENSG00000275229 | 0 | 0 | 0 | 0 | 0 | 0 |
| ENSG00000226500 | 0 | 0 | 0 | 0 | 0 | 0 |
| ENSG00000150337 | 22 | 27 | 15 | 4 | 11 | 9 |
| ENSG00000203814 | 267 | 126 | 163 | 79 | 112 | 129 |
| ENSG00000233030 | 67 | 53 | 41 | 30 | 51 | 27 |
| ENSG00000183598 | 22 | 8 | 13 | 7 | 11 | 13 |
| ENSG00000270882 | 122 | 56 | 60 | 74 | 53 | 52 |
| ENSG00000203811 | 1 | 0 | 0 | 0 | 0 | 0 |
| ENSG00000203812 | 20 | 4 | 13 | 5 | 3 | 6 |
| ENSG00000220323 | 439 | 199 | 306 | 170 | 236 | 225 |
| ENSG00000261716 | 2102 | 820 | 1245 | 810 | 1028 | 1107 |
| ENSG00000272196 | 3 | 2 | 1 | 0 | 2 | 1 |
| ENSG00000203852 | 0 | 0 | 0 | 0 | 0 | 0 |
| ENSG00000270276 | 2 | 0 | 1 | 2 | 0 | 0 |
| ENSG00000264207 | 2 | 0 | 0 | 0 | 0 | 0 |
| ENSG00000184678 | 5567 | 2472 | 3151 | 2422 | 2514 | 2850 |
| ENSG00000184260 | 78 | 31 | 31 | 45 | 37 | 28 |
| ENSG00000184270 | 1 | 1 | 3 | 1 | 6 | 3 |
| ENSG00000178096 | 1271 | 620 | 788 | 794 | 717 | 711 |
| ENSG00000159164 | 9 | 2 | 4 | 1 | 4 | 3 |
| ENSG00000143368 | 5690 | 2778 | 3252 | 3310 | 2836 | 3062 |
| ENSG00000014914 | 931 | 339 | 513 | 409 | 475 | 488 |
| ENSG00000264522 | 2252 | 1079 | 1415 | 844 | 1040 | 1204 |
| ENSG00000285184 | 154 | 78 | 93 | 73 | 94 | 112 |
| ENSG00000223945 | 15 | 4 | 14 | 2 | 5 | 6 |
| ENSG00000136631 | 2703 | 1466 | 1843 | 1196 | 1797 | 1561 |
| ENSG00000023902 | 141 | 90 | 126 | 129 | 128 | 112 |
| ENSG00000285554 | 0 | 0 | 1 | 0 | 0 | 0 |

| | | | | | | |
|---|---|---|---|---|---|---|
| ENSG00000266187 | 0 | 0 | 0 | 0 | 0 | 0 |
| ENSG00000143401 | 1309 | 728 | 793 | 557 | 947 | 858 |
| ENSG00000222222 | 0 | 0 | 1 | 0 | 0 | 1 |
| ENSG00000276110 | 0 | 2 | 0 | 0 | 2 | 1 |
| ENSG00000118298 | 53 | 27 | 34 | 21 | 40 | 44 |
| ENSG00000117362 | 11294 | 5633 | 6882 | 7128 | 6241 | 6284 |
| ENSG00000118292 | 26 | 17 | 27 | 17 | 22 | 18 |
| ENSG00000159208 | 294 | 98 | 141 | 154 | 138 | 148 |
| ENSG00000266472 | 4973 | 2567 | 3178 | 3069 | 3159 | 3025 |
| ENSG00000117360 | 3577 | 1696 | 2095 | 1731 | 2164 | 1839 |
| ENSG00000163125 | 3498 | 1586 | 1837 | 1014 | 1570 | 1685 |
| ENSG00000143374 | 2149 | 1022 | 1292 | 1241 | 1200 | 1222 |
| ENSG00000283933 | 0 | 0 | 0 | 0 | 0 | 0 |
| ENSG00000143369 | 36 | 14 | 27 | 11 | 17 | 30 |
| ENSG00000228126 | 24 | 7 | 3 | 8 | 14 | 9 |
| ENSG00000237781 | 5 | 1 | 8 | 1 | 2 | 3 |
| ENSG00000143382 | 56 | 26 | 39 | 24 | 23 | 24 |
| ENSG00000264553 | 0 | 0 | 1 | 0 | 0 | 1 |
| ENSG00000203804 | 14 | 12 | 2 | 1 | 9 | 5 |
| ENSG00000277452 | 0 | 2 | 0 | 0 | 0 | 1 |
| ENSG00000274963 | 0 | 0 | 0 | 0 | 1 | 0 |
| ENSG00000143384 | 17123 | 8159 | 10407 | 7917 | 9729 | 9363 |
| ENSG00000253047 | 6 | 6 | 4 | 6 | 6 | 4 |
| ENSG00000143420 | 17940 | 9012 | 10940 | 9386 | 9539 | 9622 |
| ENSG00000276103 | 0 | 2 | 1 | 1 | 1 | 0 |
| ENSG00000143457 | 2260 | 1072 | 1354 | 888 | 1073 | 1022 |
| ENSG00000143452 | 0 | 0 | 0 | 0 | 0 | 0 |
| ENSG00000206931 | 0 | 0 | 0 | 0 | 0 | 0 |
| ENSG00000163131 | 26 | 3 | 13 | 7 | 13 | 12 |
| ENSG00000236713 | 0 | 0 | 0 | 0 | 2 | 0 |
| ENSG00000143387 | 283 | 131 | 208 | 115 | 151 | 169 |
| ENSG00000224690 | 4 | 3 | 3 | 1 | 0 | 1 |
| ENSG00000143437 | 3870 | 1649 | 2396 | 1384 | 1718 | 1815 |
| ENSG00000200175 | 0 | 0 | 0 | 0 | 0 | 0 |
| ENSG00000224800 | 2 | 0 | 1 | 1 | 0 | 2 |
| ENSG00000283324 | 1 | 0 | 0 | 0 | 0 | 0 |
| ENSG00000229357 | 0 | 0 | 0 | 0 | 0 | 0 |
| ENSG00000143379 | 2649 | 1278 | 1772 | 1282 | 1478 | 1593 |
| ENSG00000143418 | 25349 | 13112 | 16871 | 9825 | 10444 | 11114 |
| ENSG00000259357 | 0 | 0 | 0 | 0 | 0 | 0 |
| ENSG00000231073 | 18 | 15 | 25 | 6 | 9 | 12 |
| ENSG00000143412 | 4662 | 2361 | 2985 | 1424 | 1425 | 1563 |
| ENSG00000143409 | 2166 | 1142 | 1418 | 894 | 1072 | 1015 |
| ENSG00000143363 | 2653 | 1341 | 1611 | 1199 | 1259 | 1360 |
| ENSG00000200759 | 0 | 0 | 1 | 0 | 0 | 0 |
| ENSG00000163141 | 386 | 174 | 228 | 153 | 144 | 160 |
| ENSG00000143443 | 861 | 370 | 546 | 491 | 433 | 477 |
| ENSG00000197622 | 9320 | 4460 | 5750 | 4064 | 4600 | 4767 |
| ENSG00000213190 | 322 | 139 | 170 | 125 | 149 | 148 |
| ENSG00000143458 | 1000 | 494 | 643 | 366 | 424 | 492 |

| | | | | | | |
|---|---|---|---|---|---|---|
| ENSG00000233717 | 0 | 0 | 0 | 0 | 0 | 0 |
| ENSG00000261168 | 14 | 2 | 5 | 1 | 8 | 1 |
| ENSG00000143434 | 209 | 80 | 135 | 70 | 102 | 72 |
| ENSG00000163154 | 0 | 5 | 4 | 0 | 0 | 0 |
| ENSG00000163156 | 1214 | 606 | 754 | 715 | 652 | 695 |
| ENSG00000163155 | 390 | 182 | 266 | 245 | 248 | 303 |
| ENSG00000163157 | 0 | 0 | 0 | 0 | 0 | 0 |
| ENSG00000163159 | 3687 | 1671 | 2113 | 2258 | 2014 | 2033 |
| ENSG00000143398 | 6397 | 2952 | 3842 | 2532 | 3127 | 3505 |
| ENSG00000159352 | 11090 | 5459 | 6612 | 6789 | 6242 | 6312 |
| ENSG00000232671 | 31 | 33 | 28 | 10 | 13 | 24 |
| ENSG00000143373 | 4820 | 2096 | 2814 | 2453 | 2470 | 2462 |
| ENSG00000143393 | 6212 | 2702 | 3795 | 3208 | 3201 | 3222 |
| ENSG00000265753 | 2 | 4 | 3 | 0 | 1 | 0 |
| ENSG00000273481 | 3 | 1 | 0 | 0 | 1 | 2 |
| ENSG00000143390 | 3308 | 1595 | 2076 | 1331 | 1703 | 1678 |
| ENSG00000224645 | 0 | 0 | 0 | 0 | 0 | 0 |
| ENSG00000237976 | 5 | 5 | 6 | 6 | 3 | 5 |
| ENSG00000143416 | 6152 | 2541 | 3758 | 2246 | 2165 | 2252 |
| ENSG00000159377 | 17980 | 8690 | 10606 | 10944 | 10151 | 9758 |
| ENSG00000143442 | 5785 | 2493 | 3343 | 1911 | 2738 | 2758 |
| ENSG00000238711 | 1 | 2 | 0 | 0 | 0 | 1 |
| ENSG00000143375 | 4556 | 2257 | 2814 | 1872 | 2510 | 2577 |
| ENSG00000252840 | 2 | 0 | 1 | 0 | 1 | 1 |
| ENSG00000143367 | 3886 | 2114 | 2716 | 2258 | 3013 | 2920 |
| ENSG00000232536 | 6 | 5 | 11 | 4 | 3 | 7 |
| ENSG00000207606 | 0 | 2 | 0 | 0 | 0 | 0 |
| ENSG00000223861 | 15 | 7 | 3 | 3 | 10 | 2 |
| ENSG00000143376 | 3498 | 1623 | 1966 | 1099 | 1879 | 1680 |
| ENSG00000250734 | 1 | 2 | 1 | 0 | 0 | 1 |
| ENSG00000206635 | 0 | 1 | 0 | 0 | 0 | 0 |
| ENSG00000227045 | 2 | 0 | 1 | 0 | 2 | 2 |
| ENSG00000159409 | 39 | 15 | 31 | 8 | 9 | 12 |
| ENSG00000178796 | 6 | 5 | 3 | 1 | 1 | 2 |
| ENSG00000206980 | 0 | 0 | 0 | 0 | 0 | 0 |
| ENSG00000269621 | 0 | 0 | 0 | 0 | 0 | 0 |
| ENSG00000236940 | 0 | 0 | 0 | 0 | 0 | 0 |
| ENSG00000143436 | 2360 | 1483 | 1815 | 1726 | 1700 | 1737 |
| ENSG00000143450 | 90 | 55 | 84 | 44 | 59 | 60 |
| ENSG00000249602 | 13 | 19 | 19 | 14 | 16 | 17 |
| ENSG00000232937 | 0 | 0 | 0 | 0 | 0 | 0 |
| ENSG00000268288 | 4 | 3 | 0 | 0 | 4 | 2 |
| ENSG00000182134 | 1946 | 955 | 1175 | 873 | 1099 | 1033 |
| ENSG00000203288 | 258 | 139 | 171 | 140 | 142 | 153 |
| ENSG00000269489 | 17 | 9 | 19 | 8 | 8 | 7 |
| ENSG00000213171 | 4 | 4 | 8 | 5 | 3 | 4 |
| ENSG00000143365 | 503 | 188 | 373 | 256 | 192 | 211 |
| ENSG00000225556 | 74 | 26 | 48 | 37 | 35 | 41 |
| ENSG00000201134 | 12 | 5 | 8 | 3 | 6 | 0 |
| ENSG00000234614 | 173 | 91 | 126 | 70 | 82 | 94 |

| | | | | | | |
|---|---|---|---|---|---|---|
| ENSG00000196407 | 110 | 71 | 130 | 70 | 113 | 91 |
| ENSG00000159445 | 853 | 366 | 460 | 311 | 461 | 451 |
| ENSG00000285651 | 3 | 11 | 1 | 1 | 0 | 1 |
| ENSG00000223810 | 0 | 0 | 0 | 0 | 0 | 0 |
| ENSG00000197747 | 8651 | 4793 | 5023 | 5940 | 5805 | 4735 |
| ENSG00000229021 | 4 | 10 | 14 | 3 | 8 | 9 |
| ENSG00000240667 | 0 | 0 | 0 | 0 | 0 | 0 |
| ENSG00000163191 | 22358 | 11469 | 12530 | 13058 | 12013 | 11104 |
| ENSG00000232678 | 0 | 0 | 0 | 0 | 0 | 0 |
| ENSG00000285818 | 0 | 0 | 0 | 0 | 1 | 0 |
| ENSG00000182898 | 10 | 4 | 2 | 0 | 4 | 4 |
| ENSG00000159450 | 209 | 92 | 150 | 86 | 104 | 114 |
| ENSG00000226716 | 0 | 0 | 1 | 0 | 0 | 0 |
| ENSG00000233792 | 0 | 0 | 0 | 1 | 0 | 1 |
| ENSG00000215853 | 0 | 0 | 0 | 0 | 0 | 0 |
| ENSG00000237975 | 0 | 0 | 1 | 0 | 0 | 0 |
| ENSG00000236427 | 0 | 0 | 0 | 0 | 0 | 0 |
| ENSG00000197915 | 0 | 0 | 0 | 0 | 0 | 0 |
| ENSG00000143631 | 0 | 0 | 0 | 0 | 0 | 0 |
| ENSG00000143520 | 0 | 0 | 0 | 0 | 0 | 0 |
| ENSG00000183586 | 0 | 0 | 0 | 0 | 0 | 0 |
| ENSG00000143536 | 0 | 0 | 1 | 0 | 0 | 0 |
| ENSG00000186207 | 0 | 0 | 0 | 0 | 0 | 0 |
| ENSG00000169509 | 0 | 0 | 0 | 0 | 0 | 0 |
| ENSG00000185966 | 0 | 0 | 0 | 0 | 0 | 0 |
| ENSG00000163202 | 0 | 0 | 0 | 0 | 0 | 0 |
| ENSG00000244057 | 0 | 0 | 0 | 0 | 0 | 0 |
| ENSG00000187238 | 0 | 0 | 0 | 0 | 0 | 0 |
| ENSG00000185962 | 0 | 0 | 0 | 0 | 0 | 0 |
| ENSG00000226947 | 0 | 0 | 0 | 0 | 0 | 0 |
| ENSG00000285753 | 0 | 0 | 0 | 0 | 0 | 0 |
| ENSG00000187223 | 0 | 0 | 0 | 0 | 0 | 0 |
| ENSG00000187180 | 0 | 0 | 0 | 0 | 0 | 0 |
| ENSG00000159455 | 0 | 0 | 0 | 0 | 0 | 0 |
| ENSG00000187173 | 0 | 0 | 0 | 0 | 0 | 0 |
| ENSG00000187170 | 0 | 0 | 0 | 0 | 0 | 0 |
| ENSG00000198854 | 0 | 0 | 0 | 0 | 0 | 0 |
| ENSG00000229713 | 0 | 0 | 0 | 0 | 0 | 0 |
| ENSG00000233819 | 0 | 0 | 0 | 0 | 0 | 0 |
| ENSG00000203786 | 0 | 0 | 0 | 0 | 0 | 0 |
| ENSG00000240386 | 0 | 0 | 0 | 0 | 0 | 0 |
| ENSG00000186226 | 0 | 0 | 0 | 0 | 0 | 0 |
| ENSG00000172155 | 0 | 0 | 0 | 0 | 0 | 0 |
| ENSG00000197084 | 0 | 0 | 0 | 0 | 0 | 0 |
| ENSG00000196734 | 0 | 1 | 0 | 0 | 0 | 0 |
| ENSG00000186844 | 0 | 0 | 0 | 0 | 0 | 0 |
| ENSG00000235942 | 0 | 0 | 0 | 0 | 0 | 0 |
| ENSG00000285946 | 0 | 0 | 0 | 0 | 0 | 0 |
| ENSG00000163206 | 0 | 0 | 0 | 0 | 0 | 0 |
| ENSG00000163207 | 0 | 0 | 1 | 0 | 1 | 0 |

| | | | | | | |
|---|---|---|---|---|---|---|
| ENSG00000224308 | 0 | 0 | 0 | 0 | 1 | 0 |
| ENSG00000283227 | 0 | 0 | 0 | 0 | 0 | 0 |
| ENSG00000184148 | 2 | 0 | 0 | 0 | 0 | 0 |
| ENSG00000169474 | 1 | 0 | 0 | 0 | 0 | 0 |
| ENSG00000163209 | 0 | 0 | 0 | 1 | 1 | 0 |
| ENSG00000252920 | 0 | 0 | 0 | 0 | 0 | 0 |
| ENSG00000234262 | 0 | 0 | 0 | 0 | 0 | 0 |
| ENSG00000169469 | 0 | 0 | 0 | 1 | 1 | 0 |
| ENSG00000163216 | 0 | 0 | 0 | 0 | 0 | 0 |
| ENSG00000241794 | 0 | 0 | 0 | 0 | 0 | 0 |
| ENSG00000196805 | 0 | 0 | 0 | 0 | 0 | 0 |
| ENSG00000203785 | 0 | 0 | 0 | 0 | 0 | 0 |
| ENSG00000244094 | 0 | 0 | 0 | 0 | 0 | 1 |
| ENSG00000229035 | 0 | 0 | 0 | 0 | 0 | 0 |
| ENSG00000159516 | 0 | 0 | 0 | 0 | 0 | 0 |
| ENSG00000229699 | 0 | 0 | 0 | 0 | 0 | 0 |
| ENSG00000203784 | 0 | 0 | 0 | 0 | 0 | 0 |
| ENSG00000203783 | 0 | 0 | 0 | 0 | 0 | 0 |
| ENSG00000230779 | 0 | 0 | 0 | 0 | 0 | 0 |
| ENSG00000203782 | 0 | 0 | 0 | 0 | 0 | 0 |
| ENSG00000159527 | 0 | 0 | 0 | 0 | 0 | 0 |
| ENSG00000163218 | 8 | 0 | 7 | 3 | 7 | 3 |
| ENSG00000207321 | 0 | 0 | 0 | 0 | 0 | 0 |
| ENSG00000163220 | 610 | 465 | 411 | 398 | 517 | 387 |
| ENSG00000163221 | 6 | 3 | 8 | 3 | 4 | 4 |
| ENSG00000237008 | 0 | 0 | 0 | 0 | 0 | 0 |
| ENSG00000143546 | 92 | 78 | 64 | 49 | 83 | 65 |
| ENSG00000224784 | 0 | 0 | 0 | 0 | 0 | 0 |
| ENSG00000184330 | 0 | 0 | 0 | 0 | 0 | 0 |
| ENSG00000203781 | 0 | 0 | 0 | 0 | 0 | 0 |
| ENSG00000197364 | 0 | 0 | 0 | 0 | 0 | 0 |
| ENSG00000143556 | 2 | 1 | 1 | 0 | 3 | 2 |
| ENSG00000263841 | 0 | 0 | 0 | 0 | 0 | 0 |
| ENSG00000238279 | 1 | 0 | 0 | 0 | 1 | 0 |
| ENSG00000197956 | 936 | 452 | 601 | 664 | 515 | 491 |
| ENSG00000196420 | 0 | 0 | 0 | 2 | 0 | 3 |
| ENSG00000196154 | 104 | 61 | 62 | 95 | 69 | 62 |
| ENSG00000188015 | 2 | 2 | 2 | 5 | 4 | 6 |
| ENSG00000196754 | 1 | 2 | 10 | 6 | 2 | 9 |
| ENSG00000285867 | 21 | 12 | 9 | 11 | 9 | 6 |
| ENSG00000188643 | 3557 | 1815 | 2098 | 1870 | 1684 | 1622 |
| ENSG00000189334 | 10402 | 5124 | 6219 | 6306 | 5731 | 5445 |
| ENSG00000189171 | 2271 | 1698 | 2027 | 1972 | 1807 | 1558 |
| ENSG00000271853 | 0 | 12 | 7 | 9 | 12 | 5 |
| ENSG00000160678 | 40 | 19 | 26 | 24 | 15 | 17 |
| ENSG00000272030 | 10 | 1 | 6 | 3 | 1 | 5 |
| ENSG00000160679 | 3665 | 1891 | 2405 | 1783 | 1930 | 2106 |
| ENSG00000143553 | 1430 | 693 | 832 | 802 | 779 | 766 |
| ENSG00000143621 | 8334 | 4141 | 4871 | 4702 | 4895 | 5156 |
| ENSG00000169418 | 40 | 12 | 18 | 47 | 33 | 35 |

| | | | | | | |
|---|---|---|---|---|---|---|
| ENSG00000274940 | 0 | 0 | 0 | 0 | 0 | 0 |
| ENSG00000242565 | 0 | 0 | 0 | 0 | 0 | 0 |
| ENSG00000235015 | 0 | 0 | 0 | 0 | 1 | 0 |
| ENSG00000199565 | 0 | 0 | 0 | 0 | 0 | 0 |
| ENSG00000143624 | 3068 | 1324 | 1927 | 1257 | 1518 | 1613 |
| ENSG00000243613 | 0 | 0 | 0 | 0 | 0 | 0 |
| ENSG00000233222 | 0 | 0 | 0 | 0 | 0 | 0 |
| ENSG00000143554 | 1615 | 710 | 971 | 880 | 765 | 805 |
| ENSG00000207039 | 0 | 0 | 0 | 0 | 0 | 0 |
| ENSG00000143614 | 1455 | 657 | 882 | 499 | 652 | 696 |
| ENSG00000231827 | 9 | 0 | 5 | 0 | 1 | 2 |
| ENSG00000223599 | 2 | 2 | 4 | 0 | 1 | 0 |
| ENSG00000236327 | 0 | 0 | 0 | 0 | 0 | 0 |
| ENSG00000284738 | 14 | 11 | 6 | 9 | 7 | 8 |
| ENSG00000198837 | 2382 | 1102 | 1483 | 1265 | 1186 | 1259 |
| ENSG00000160741 | 2143 | 942 | 1221 | 1113 | 1074 | 1148 |
| ENSG00000143570 | 129 | 38 | 96 | 65 | 72 | 75 |
| ENSG00000285779 | 0 | 0 | 0 | 0 | 0 | 0 |
| ENSG00000276584 | 1 | 0 | 0 | 0 | 0 | 0 |
| ENSG00000282386 | 69 | 48 | 56 | 41 | 48 | 57 |
| ENSG00000273026 | 30 | 30 | 33 | 30 | 29 | 22 |
| ENSG00000143578 | 1071 | 521 | 738 | 572 | 504 | 530 |
| ENSG00000143543 | 597 | 405 | 513 | 606 | 487 | 577 |
| ENSG00000285641 | 0 | 5 | 3 | 3 | 2 | 1 |
| ENSG00000272654 | 65 | 46 | 41 | 42 | 26 | 37 |
| ENSG00000143545 | 668 | 443 | 508 | 440 | 502 | 412 |
| ENSG00000177954 | 19213 | 9436 | 11001 | 12665 | 11063 | 10454 |
| ENSG00000143552 | 638 | 303 | 377 | 351 | 340 | 383 |
| ENSG00000231416 | 11 | 8 | 2 | 1 | 3 | 3 |
| ENSG00000200220 | 0 | 0 | 0 | 0 | 0 | 0 |
| ENSG00000231837 | 0 | 0 | 0 | 0 | 0 | 0 |
| ENSG00000263987 | 0 | 0 | 0 | 0 | 0 | 0 |
| ENSG00000143549 | 12335 | 6208 | 7190 | 6726 | 7128 | 6926 |
| ENSG00000264384 | 0 | 1 | 1 | 0 | 1 | 0 |
| ENSG00000215938 | 0 | 0 | 0 | 0 | 0 | 0 |
| ENSG00000163263 | 0 | 0 | 0 | 0 | 0 | 0 |
| ENSG00000143612 | 10286 | 5015 | 5958 | 6256 | 6095 | 5904 |
| ENSG00000143569 | 6039 | 2719 | 3455 | 2682 | 3025 | 3312 |
| ENSG00000201129 | 0 | 0 | 0 | 0 | 0 | 0 |
| ENSG00000143575 | 3252 | 1614 | 1913 | 2026 | 1791 | 1834 |
| ENSG00000252682 | 0 | 0 | 0 | 1 | 0 | 0 |
| ENSG00000212292 | 0 | 0 | 0 | 0 | 0 | 0 |
| ENSG00000222457 | 0 | 0 | 0 | 0 | 0 | 0 |
| ENSG00000278694 | 0 | 0 | 0 | 0 | 0 | 0 |
| ENSG00000143595 | 2 | 0 | 1 | 0 | 1 | 0 |
| ENSG00000143515 | 3778 | 1622 | 2427 | 1088 | 1319 | 1550 |
| ENSG00000238365 | 2 | 2 | 4 | 0 | 3 | 0 |
| ENSG00000237920 | 0 | 0 | 0 | 0 | 0 | 0 |
| ENSG00000226855 | 2 | 0 | 0 | 3 | 0 | 0 |
| ENSG00000228013 | 10 | 3 | 1 | 1 | 4 | 7 |

| | | | | | | |
|---|---|---|---|---|---|---|
| ENSG00000160712 | 267 | 132 | 157 | 149 | 228 | 169 |
| ENSG00000228264 | 0 | 0 | 0 | 0 | 0 | 0 |
| ENSG00000169291 | 0 | 1 | 0 | 0 | 0 | 0 |
| ENSG00000273110 | 0 | 0 | 0 | 0 | 0 | 0 |
| ENSG00000163239 | 0 | 0 | 0 | 0 | 0 | 0 |
| ENSG00000160714 | 4118 | 2104 | 2607 | 2346 | 2430 | 2405 |
| ENSG00000229780 | 10 | 4 | 3 | 2 | 3 | 0 |
| ENSG00000160716 | 8 | 2 | 0 | 0 | 0 | 6 |
| ENSG00000233875 | 0 | 0 | 0 | 0 | 0 | 0 |
| ENSG00000160710 | 10639 | 4658 | 6361 | 4119 | 5386 | 5614 |
| ENSG00000143603 | 0 | 0 | 0 | 0 | 4 | 0 |
| ENSG00000163344 | 1310 | 572 | 723 | 795 | 677 | 655 |
| ENSG00000270361 | 4 | 2 | 0 | 2 | 4 | 0 |
| ENSG00000163346 | 4111 | 1766 | 2254 | 1896 | 1832 | 1704 |
| ENSG00000268154 | 0 | 0 | 0 | 0 | 0 | 0 |
| ENSG00000163348 | 3417 | 1566 | 2090 | 1788 | 1730 | 1757 |
| ENSG00000271380 | 30 | 15 | 25 | 17 | 15 | 21 |
| ENSG00000160691 | 2996 | 1368 | 1857 | 1682 | 1734 | 1783 |
| ENSG00000173207 | 901 | 438 | 563 | 628 | 651 | 616 |
| ENSG00000264349 | 0 | 0 | 1 | 2 | 1 | 1 |
| ENSG00000160688 | 1617 | 864 | 988 | 1223 | 1121 | 1126 |
| ENSG00000163352 | 12 | 10 | 5 | 4 | 5 | 3 |
| ENSG00000160685 | 4076 | 1784 | 2338 | 1953 | 1917 | 1882 |
| ENSG00000163354 | 110 | 38 | 97 | 51 | 57 | 58 |
| ENSG00000163357 | 39 | 23 | 20 | 17 | 20 | 23 |
| ENSG00000232093 | 41 | 19 | 26 | 25 | 24 | 23 |
| ENSG00000143537 | 6322 | 2963 | 3861 | 4101 | 3322 | 3595 |
| ENSG00000243364 | 506 | 293 | 348 | 383 | 325 | 297 |
| ENSG00000251246 | 0 | 0 | 3 | 0 | 1 | 0 |
| ENSG00000143590 | 314 | 164 | 238 | 185 | 157 | 184 |
| ENSG00000202027 | 0 | 0 | 0 | 0 | 0 | 0 |
| ENSG00000169242 | 2463 | 1182 | 1532 | 1137 | 1071 | 1249 |
| ENSG00000169241 | 4157 | 1918 | 2617 | 1908 | 2012 | 1904 |
| ENSG00000179085 | 836 | 404 | 546 | 572 | 440 | 430 |
| ENSG00000223452 | 0 | 0 | 0 | 0 | 0 | 0 |
| ENSG00000163463 | 234 | 125 | 154 | 119 | 130 | 147 |
| ENSG00000273088 | 0 | 0 | 0 | 0 | 0 | 0 |
| ENSG00000163462 | 59 | 19 | 39 | 29 | 33 | 30 |
| ENSG00000185499 | 1117 | 533 | 903 | 582 | 493 | 589 |
| ENSG00000231064 | 28 | 15 | 32 | 16 | 13 | 16 |
| ENSG00000284586 | 0 | 0 | 0 | 0 | 0 | 0 |
| ENSG00000169231 | 789 | 375 | 597 | 417 | 431 | 465 |
| ENSG00000173171 | 636 | 323 | 430 | 504 | 364 | 433 |
| ENSG00000236263 | 0 | 0 | 0 | 0 | 0 | 0 |
| ENSG00000160766 | 465 | 212 | 256 | 227 | 257 | 249 |
| ENSG00000236675 | 77 | 53 | 72 | 54 | 73 | 63 |
| ENSG00000177628 | 2410 | 1185 | 1439 | 1323 | 1299 | 1373 |
| ENSG00000160767 | 1280 | 559 | 719 | 778 | 704 | 745 |
| ENSG00000116521 | 2961 | 1443 | 1736 | 1891 | 1578 | 1586 |
| ENSG00000176444 | 1835 | 829 | 1034 | 783 | 905 | 942 |

| | | | | | | |
|---|---|---|---|---|---|---|
| ENSG00000143630 | 432 | 178 | 279 | 171 | 253 | 220 |
| ENSG00000143627 | 0 | 0 | 0 | 0 | 0 | 2 |
| ENSG00000160752 | 2589 | 1620 | 2052 | 2197 | 2046 | 2024 |
| ENSG00000225855 | 280 | 77 | 142 | 81 | 112 | 142 |
| ENSG00000160753 | 3366 | 1486 | 2027 | 1807 | 1507 | 1754 |
| ENSG00000116539 | 1267 | 732 | 553 | 246 | 1170 | 708 |
| ENSG00000283701 | 0 | 0 | 0 | 0 | 0 | 0 |
| ENSG00000207134 | 0 | 0 | 0 | 0 | 0 | 0 |
| ENSG00000227773 | 0 | 0 | 0 | 0 | 0 | 0 |
| ENSG00000207144 | 0 | 0 | 0 | 0 | 0 | 0 |
| ENSG00000237872 | 0 | 1 | 0 | 0 | 0 | 0 |
| ENSG00000235919 | 160 | 68 | 90 | 53 | 70 | 82 |
| ENSG00000271267 | 10 | 6 | 5 | 1 | 10 | 5 |
| ENSG00000225082 | 3 | 0 | 1 | 2 | 0 | 4 |
| ENSG00000223503 | 3 | 0 | 1 | 0 | 2 | 1 |
| ENSG00000232519 | 8 | 1 | 0 | 4 | 1 | 2 |
| ENSG00000125459 | 428 | 147 | 275 | 270 | 211 | 247 |
| ENSG00000246203 | 10 | 6 | 5 | 5 | 4 | 2 |
| ENSG00000163374 | 1627 | 849 | 966 | 641 | 890 | 850 |
| ENSG00000132676 | 5720 | 2953 | 3610 | 3013 | 3392 | 3380 |
| ENSG00000227673 | 0 | 0 | 0 | 0 | 0 | 0 |
| ENSG00000203761 | 115 | 45 | 75 | 49 | 43 | 74 |
| ENSG00000116580 | 1576 | 676 | 884 | 508 | 828 | 763 |
| ENSG00000234937 | 3 | 1 | 0 | 0 | 1 | 1 |
| ENSG00000132718 | 2 | 1 | 2 | 0 | 1 | 0 |
| ENSG00000283442 | 0 | 0 | 0 | 0 | 0 | 0 |
| ENSG00000143622 | 1590 | 830 | 942 | 660 | 798 | 767 |
| ENSG00000132680 | 1677 | 713 | 1059 | 612 | 980 | 652 |
| ENSG00000207475 | 0 | 2 | 1 | 1 | 1 | 1 |
| ENSG00000280466 | 0 | 0 | 0 | 0 | 0 | 0 |
| ENSG00000281394 | 0 | 0 | 0 | 0 | 0 | 0 |
| ENSG00000173080 | 0 | 3 | 3 | 2 | 0 | 0 |
| ENSG00000116584 | 1002 | 483 | 662 | 503 | 503 | 460 |
| ENSG00000277817 | 0 | 0 | 0 | 0 | 0 | 0 |
| ENSG00000273002 | 18 | 9 | 19 | 4 | 12 | 3 |
| ENSG00000224276 | 0 | 0 | 0 | 0 | 0 | 0 |
| ENSG00000285677 | 2 | 2 | 0 | 0 | 3 | 2 |
| ENSG00000163479 | 6617 | 3508 | 4113 | 4563 | 3874 | 3986 |
| ENSG00000160803 | 2977 | 1405 | 1649 | 1781 | 1844 | 1808 |
| ENSG00000116586 | 1815 | 940 | 1108 | 1201 | 1054 | 978 |
| ENSG00000132698 | 4969 | 2441 | 2887 | 3009 | 2633 | 2780 |
| ENSG00000254726 | 931 | 417 | 545 | 622 | 621 | 683 |
| ENSG00000160789 | 8208 | 3910 | 4690 | 6167 | 5561 | 4811 |
| ENSG00000196189 | 809 | 385 | 488 | 354 | 333 | 441 |
| ENSG00000252236 | 0 | 0 | 0 | 0 | 0 | 1 |
| ENSG00000160785 | 2261 | 992 | 1306 | 1024 | 1193 | 1211 |
| ENSG00000260238 | 0 | 1 | 1 | 2 | 0 | 2 |
| ENSG00000160783 | 657 | 355 | 429 | 416 | 398 | 426 |
| ENSG00000242252 | 4 | 6 | 5 | 2 | 2 | 6 |
| ENSG00000160781 | 725 | 327 | 519 | 334 | 380 | 374 |

| | | | | | | |
|---|---|---|---|---|---|---|
| ENSG00000198952 | 5625 | 2397 | 3420 | 2940 | 2950 | 3067 |
| ENSG00000163472 | 626 | 327 | 411 | 381 | 350 | 323 |
| ENSG00000198715 | 985 | 498 | 574 | 599 | 531 | 525 |
| ENSG00000189030 | 0 | 0 | 0 | 0 | 0 | 0 |
| ENSG00000163468 | 16032 | 7959 | 9109 | 10356 | 10110 | 9704 |
| ENSG00000163467 | 39 | 23 | 21 | 20 | 21 | 26 |
| ENSG00000132677 | 395 | 179 | 231 | 163 | 160 | 196 |
| ENSG00000237390 | 4 | 1 | 3 | 1 | 2 | 1 |
| ENSG00000125462 | 0 | 2 | 2 | 2 | 1 | 1 |
| ENSG00000207933 | 0 | 0 | 0 | 0 | 0 | 0 |
| ENSG00000236947 | 18 | 4 | 21 | 9 | 8 | 10 |
| ENSG00000116604 | 1692 | 866 | 1147 | 750 | 785 | 894 |
| ENSG00000206651 | 0 | 0 | 0 | 0 | 0 | 0 |
| ENSG00000260460 | 0 | 0 | 0 | 0 | 0 | 0 |
| ENSG00000183856 | 1071 | 451 | 685 | 578 | 651 | 735 |
| ENSG00000187862 | 2 | 0 | 0 | 0 | 0 | 0 |
| ENSG00000163382 | 2323 | 1176 | 1427 | 1751 | 1327 | 1416 |
| ENSG00000160818 | 895 | 473 | 514 | 479 | 560 | 565 |
| ENSG00000272971 | 0 | 0 | 2 | 1 | 0 | 2 |
| ENSG00000132702 | 1 | 1 | 4 | 1 | 2 | 3 |
| ENSG00000272068 | 276 | 125 | 166 | 196 | 227 | 223 |
| ENSG00000132692 | 25 | 7 | 19 | 11 | 19 | 26 |
| ENSG00000272405 | 313 | 121 | 191 | 217 | 245 | 286 |
| ENSG00000229953 | 31 | 19 | 21 | 17 | 37 | 21 |
| ENSG00000132688 | 236 | 102 | 155 | 123 | 114 | 137 |
| ENSG00000237588 | 3 | 5 | 8 | 3 | 1 | 4 |
| ENSG00000285570 | 35 | 11 | 15 | 8 | 15 | 16 |
| ENSG00000143320 | 22764 | 10349 | 14568 | 11901 | 9219 | 10963 |
| ENSG00000223356 | 0 | 0 | 3 | 1 | 0 | 1 |
| ENSG00000143319 | 2578 | 1308 | 1462 | 1297 | 1492 | 1463 |
| ENSG00000143303 | 1053 | 459 | 662 | 640 | 588 | 555 |
| ENSG00000143314 | 2032 | 1018 | 1284 | 1608 | 1311 | 1432 |
| ENSG00000143321 | 17957 | 9013 | 10830 | 11197 | 10880 | 10681 |
| ENSG00000143294 | 3757 | 1544 | 1957 | 1962 | 1740 | 1872 |
| ENSG00000027869 | 0 | 1 | 0 | 0 | 0 | 0 |
| ENSG00000198400 | 0 | 0 | 2 | 0 | 1 | 0 |
| ENSG00000027644 | 0 | 0 | 0 | 0 | 0 | 1 |
| ENSG00000187800 | 0 | 2 | 1 | 1 | 0 | 0 |
| ENSG00000160838 | 3 | 0 | 3 | 2 | 0 | 3 |
| ENSG00000132694 | 2008 | 809 | 1223 | 759 | 908 | 1066 |
| ENSG00000211581 | 0 | 0 | 0 | 0 | 0 | 1 |
| ENSG00000266160 | 0 | 0 | 0 | 0 | 0 | 0 |
| ENSG00000237842 | 0 | 1 | 1 | 0 | 0 | 0 |
| ENSG00000224520 | 46 | 22 | 30 | 21 | 20 | 27 |
| ENSG00000253831 | 0 | 0 | 0 | 0 | 0 | 0 |
| ENSG00000117036 | 1306 | 562 | 777 | 447 | 668 | 658 |
| ENSG00000235700 | 0 | 0 | 1 | 1 | 0 | 0 |
| ENSG00000229961 | 0 | 0 | 0 | 0 | 0 | 0 |
| ENSG00000284592 | 0 | 0 | 0 | 0 | 0 | 0 |
| ENSG00000228239 | 0 | 0 | 0 | 0 | 0 | 0 |

| | | | | | | |
|---|---|---|---|---|---|---|
| ENSG00000271736 | 0 | 0 | 0 | 0 | 0 | 0 |
| ENSG00000237189 | 0 | 0 | 0 | 0 | 0 | 0 |
| ENSG00000143297 | 0 | 0 | 0 | 0 | 0 | 0 |
| ENSG00000163518 | 0 | 0 | 0 | 0 | 0 | 0 |
| ENSG00000236731 | 0 | 0 | 0 | 0 | 0 | 0 |
| ENSG00000233712 | 0 | 0 | 0 | 0 | 0 | 0 |
| ENSG00000160856 | 1 | 0 | 0 | 1 | 0 | 0 |
| ENSG00000227217 | 0 | 0 | 0 | 0 | 0 | 0 |
| ENSG00000231700 | 0 | 0 | 0 | 0 | 0 | 0 |
| ENSG00000232366 | 0 | 0 | 0 | 0 | 0 | 0 |
| ENSG00000132704 | 0 | 0 | 0 | 0 | 0 | 0 |
| ENSG00000163534 | 0 | 0 | 0 | 0 | 0 | 0 |
| ENSG00000073754 | 0 | 0 | 0 | 0 | 0 | 0 |
| ENSG00000227425 | 0 | 0 | 0 | 0 | 0 | 0 |
| ENSG00000236957 | 0 | 0 | 0 | 0 | 0 | 0 |
| ENSG00000183853 | 0 | 0 | 0 | 0 | 0 | 0 |
| ENSG00000226520 | 0 | 0 | 0 | 0 | 0 | 0 |
| ENSG00000231666 | 0 | 0 | 0 | 0 | 0 | 0 |
| ENSG00000227295 | 2 | 2 | 0 | 0 | 0 | 1 |
| ENSG00000158473 | 0 | 0 | 1 | 0 | 0 | 0 |
| ENSG00000229914 | 4 | 4 | 0 | 1 | 2 | 1 |
| ENSG00000176320 | 0 | 1 | 2 | 1 | 0 | 1 |
| ENSG00000158477 | 0 | 0 | 0 | 0 | 0 | 0 |
| ENSG00000230942 | 0 | 0 | 0 | 0 | 1 | 0 |
| ENSG00000158481 | 0 | 0 | 0 | 0 | 0 | 0 |
| ENSG00000158485 | 0 | 0 | 0 | 0 | 0 | 0 |
| ENSG00000158488 | 0 | 1 | 1 | 0 | 0 | 0 |
| ENSG00000186306 | 0 | 0 | 0 | 0 | 1 | 2 |
| ENSG00000180708 | 3 | 4 | 2 | 2 | 3 | 4 |
| ENSG00000203758 | 0 | 0 | 0 | 0 | 0 | 0 |
| ENSG00000236257 | 2 | 1 | 0 | 0 | 0 | 2 |
| ENSG00000173285 | 0 | 0 | 0 | 0 | 0 | 0 |
| ENSG00000198965 | 0 | 0 | 0 | 0 | 0 | 0 |
| ENSG00000236656 | 0 | 0 | 0 | 0 | 0 | 0 |
| ENSG00000198703 | 0 | 0 | 0 | 0 | 0 | 0 |
| ENSG00000227445 | 0 | 0 | 0 | 0 | 0 | 0 |
| ENSG00000231434 | 0 | 0 | 0 | 0 | 0 | 1 |
| ENSG00000197532 | 0 | 0 | 0 | 0 | 0 | 0 |
| ENSG00000186440 | 0 | 0 | 0 | 0 | 0 | 0 |
| ENSG00000279111 | 0 | 0 | 0 | 0 | 0 | 0 |
| ENSG00000198967 | 0 | 0 | 0 | 0 | 0 | 0 |
| ENSG00000163554 | 0 | 0 | 0 | 0 | 0 | 0 |
| ENSG00000231237 | 0 | 0 | 0 | 0 | 0 | 0 |
| ENSG00000196171 | 0 | 0 | 0 | 0 | 0 | 0 |
| ENSG00000203757 | 0 | 0 | 0 | 0 | 0 | 0 |
| ENSG00000180437 | 0 | 0 | 0 | 0 | 0 | 0 |
| ENSG00000226949 | 0 | 0 | 0 | 0 | 0 | 0 |
| ENSG00000197403 | 0 | 0 | 0 | 0 | 0 | 0 |
| ENSG00000180433 | 0 | 0 | 0 | 0 | 0 | 0 |
| ENSG00000188340 | 0 | 0 | 0 | 0 | 0 | 0 |

| | | | | | | |
|---|---|---|---|---|---|---|
| ENSG00000229724 | 0 | 0 | 0 | 0 | 0 | 0 |
| ENSG00000180409 | 0 | 0 | 0 | 0 | 0 | 0 |
| ENSG00000163563 | 0 | 0 | 0 | 0 | 0 | 0 |
| ENSG00000229849 | 0 | 0 | 0 | 0 | 0 | 0 |
| ENSG00000163564 | 0 | 0 | 0 | 0 | 0 | 0 |
| ENSG00000163565 | 0 | 0 | 0 | 0 | 0 | 0 |
| ENSG00000163568 | 0 | 0 | 0 | 0 | 0 | 0 |
| ENSG00000233473 | 0 | 0 | 0 | 0 | 0 | 0 |
| ENSG00000162706 | 0 | 0 | 0 | 0 | 0 | 0 |
| ENSG00000222552 | 0 | 0 | 0 | 0 | 0 | 0 |
| ENSG00000225670 | 0 | 0 | 0 | 0 | 0 | 0 |
| ENSG00000213088 | 0 | 0 | 0 | 0 | 0 | 0 |
| ENSG00000215846 | 0 | 0 | 0 | 0 | 0 | 0 |
| ENSG00000248642 | 0 | 0 | 0 | 0 | 0 | 0 |
| ENSG00000179639 | 0 | 0 | 0 | 0 | 0 | 0 |
| ENSG00000196266 | 0 | 0 | 0 | 0 | 0 | 0 |
| ENSG00000228560 | 0 | 0 | 0 | 0 | 0 | 0 |
| ENSG00000248442 | 0 | 0 | 0 | 0 | 0 | 0 |
| ENSG00000251686 | 0 | 0 | 0 | 0 | 0 | 0 |
| ENSG00000249197 | 0 | 0 | 0 | 0 | 0 | 0 |
| ENSG00000249730 | 0 | 0 | 0 | 0 | 0 | 0 |
| ENSG00000196184 | 0 | 0 | 0 | 0 | 0 | 0 |
| ENSG00000224943 | 0 | 0 | 0 | 0 | 0 | 0 |
| ENSG00000184155 | 0 | 0 | 0 | 0 | 0 | 0 |
| ENSG00000231100 | 0 | 0 | 0 | 0 | 0 | 0 |
| ENSG00000271890 | 0 | 0 | 0 | 0 | 0 | 0 |
| ENSG00000132703 | 0 | 0 | 0 | 0 | 0 | 0 |
| ENSG00000158731 | 0 | 0 | 0 | 0 | 0 | 0 |
| ENSG00000276632 | 0 | 0 | 0 | 0 | 0 | 0 |
| ENSG00000223603 | 0 | 0 | 0 | 0 | 0 | 0 |
| ENSG00000132693 | 0 | 0 | 0 | 0 | 0 | 0 |
| ENSG00000235226 | 0 | 0 | 0 | 0 | 0 | 0 |
| ENSG00000158716 | 534 | 252 | 358 | 459 | 339 | 354 |
| ENSG00000181036 | 0 | 0 | 0 | 0 | 0 | 0 |
| ENSG00000158714 | 0 | 0 | 0 | 0 | 1 | 0 |
| ENSG00000256029 | 8 | 1 | 3 | 1 | 2 | 0 |
| ENSG00000188004 | 0 | 0 | 0 | 0 | 0 | 0 |
| ENSG00000212161 | 0 | 0 | 0 | 0 | 0 | 0 |
| ENSG00000243284 | 0 | 0 | 0 | 0 | 0 | 0 |
| ENSG00000272668 | 24 | 4 | 6 | 3 | 2 | 3 |
| ENSG00000213085 | 51 | 24 | 30 | 15 | 25 | 17 |
| ENSG00000266458 | 0 | 0 | 0 | 0 | 0 | 0 |
| ENSG00000279430 | 0 | 0 | 0 | 1 | 0 | 0 |
| ENSG00000158710 | 12491 | 6545 | 7184 | 8837 | 8223 | 7513 |
| ENSG00000085552 | 1184 | 537 | 657 | 742 | 679 | 651 |
| ENSG00000162723 | 0 | 0 | 0 | 0 | 2 | 0 |
| ENSG00000224259 | 0 | 0 | 0 | 0 | 0 | 0 |
| ENSG00000273933 | 0 | 0 | 0 | 0 | 0 | 0 |
| ENSG00000177807 | 4 | 0 | 4 | 1 | 0 | 0 |
| ENSG00000237409 | 0 | 0 | 0 | 0 | 0 | 1 |

| | | | | | | |
|---|---|---|---|---|---|---|
| ENSG00000143315 | 1134 | 568 | 669 | 376 | 575 | 598 |
| ENSG00000225279 | 0 | 0 | 0 | 0 | 0 | 0 |
| ENSG00000162728 | 2 | 0 | 0 | 0 | 0 | 0 |
| ENSG00000162729 | 626 | 327 | 421 | 435 | 383 | 399 |
| ENSG00000018625 | 0 | 0 | 1 | 1 | 0 | 0 |
| ENSG00000132681 | 0 | 1 | 1 | 0 | 0 | 2 |
| ENSG00000143318 | 5 | 0 | 0 | 0 | 1 | 1 |
| ENSG00000227741 | 8 | 8 | 9 | 1 | 2 | 1 |
| ENSG00000162734 | 2361 | 1161 | 1431 | 1533 | 1508 | 1432 |
| ENSG00000132716 | 1377 | 683 | 971 | 680 | 801 | 789 |
| ENSG00000258465 | 4 | 1 | 1 | 0 | 0 | 0 |
| ENSG00000228606 | 18 | 14 | 17 | 10 | 19 | 24 |
| ENSG00000224261 | 4 | 0 | 2 | 2 | 3 | 1 |
| ENSG00000162735 | 1489 | 840 | 950 | 783 | 917 | 892 |
| ENSG00000122218 | 6208 | 2924 | 3659 | 2178 | 3396 | 2813 |
| ENSG00000235082 | 8 | 2 | 1 | 2 | 1 | 3 |
| ENSG00000202078 | 0 | 0 | 0 | 0 | 0 | 0 |
| ENSG00000162736 | 3183 | 1459 | 1869 | 1265 | 1520 | 1591 |
| ENSG00000171786 | 4 | 1 | 0 | 1 | 0 | 1 |
| ENSG00000201608 | 0 | 0 | 0 | 0 | 0 | 0 |
| ENSG00000162738 | 8 | 1 | 8 | 5 | 9 | 6 |
| ENSG00000162739 | 0 | 0 | 0 | 0 | 0 | 0 |
| ENSG00000234425 | 0 | 0 | 0 | 0 | 0 | 0 |
| ENSG00000066294 | 0 | 0 | 2 | 0 | 0 | 0 |
| ENSG00000117090 | 0 | 0 | 0 | 0 | 0 | 0 |
| ENSG00000235101 | 0 | 0 | 0 | 0 | 0 | 0 |
| ENSG00000228863 | 0 | 0 | 1 | 0 | 0 | 0 |
| ENSG00000117091 | 0 | 0 | 0 | 0 | 0 | 0 |
| ENSG00000026751 | 0 | 0 | 1 | 0 | 0 | 0 |
| ENSG00000275801 | 0 | 0 | 0 | 0 | 0 | 0 |
| ENSG00000274562 | 0 | 0 | 0 | 0 | 0 | 0 |
| ENSG00000122224 | 0 | 0 | 0 | 0 | 0 | 0 |
| ENSG00000122223 | 0 | 0 | 0 | 0 | 0 | 0 |
| ENSG00000226003 | 0 | 0 | 0 | 0 | 0 | 0 |
| ENSG00000179914 | 0 | 0 | 0 | 0 | 0 | 0 |
| ENSG00000213080 | 12 | 10 | 3 | 3 | 6 | 4 |
| ENSG00000232188 | 0 | 0 | 0 | 0 | 1 | 0 |
| ENSG00000198358 | 0 | 0 | 0 | 0 | 0 | 0 |
| ENSG00000233691 | 0 | 0 | 0 | 0 | 0 | 0 |
| ENSG00000158764 | 0 | 0 | 0 | 0 | 0 | 0 |
| ENSG00000158769 | 4371 | 2112 | 2639 | 2092 | 2148 | 2266 |
| ENSG00000270149 | 16 | 10 | 11 | 3 | 4 | 10 |
| ENSG00000232879 | 0 | 3 | 3 | 0 | 0 | 2 |
| ENSG00000215845 | 1867 | 953 | 1240 | 1074 | 986 | 999 |
| ENSG00000158773 | 1522 | 847 | 975 | 904 | 862 | 946 |
| ENSG00000186517 | 75 | 39 | 61 | 30 | 41 | 42 |
| ENSG00000143217 | 1651 | 680 | 859 | 819 | 970 | 883 |
| ENSG00000228917 | 0 | 0 | 1 | 0 | 1 | 2 |
| ENSG00000162755 | 282 | 148 | 201 | 166 | 161 | 177 |
| ENSG00000143256 | 854 | 427 | 466 | 573 | 502 | 528 |

| | | | | | | |
|---|---|---|---|---|---|---|
| ENSG00000158793 | 764 | 384 | 548 | 452 | 490 | 487 |
| ENSG00000158796 | 1260 | 624 | 796 | 697 | 635 | 702 |
| ENSG00000238934 | 2 | 0 | 0 | 0 | 0 | 1 |
| ENSG00000143222 | 2041 | 1075 | 1367 | 1199 | 1250 | 1194 |
| ENSG00000224985 | 0 | 2 | 0 | 4 | 0 | 0 |
| ENSG00000143258 | 1052 | 507 | 635 | 565 | 489 | 560 |
| ENSG00000143224 | 510 | 239 | 367 | 248 | 263 | 239 |
| ENSG00000158850 | 1723 | 829 | 1051 | 1070 | 1114 | 973 |
| ENSG00000158859 | 2 | 2 | 1 | 0 | 2 | 1 |
| ENSG00000158864 | 4296 | 2176 | 2589 | 2325 | 2360 | 2456 |
| ENSG00000158869 | 0 | 2 | 1 | 5 | 0 | 1 |
| ENSG00000158874 | 0 | 0 | 0 | 0 | 0 | 0 |
| ENSG00000158882 | 497 | 229 | 299 | 275 | 270 | 297 |
| ENSG00000284035 | 0 | 0 | 0 | 1 | 2 | 0 |
| ENSG00000143257 | 8 | 6 | 7 | 3 | 5 | 8 |
| ENSG00000248485 | 10 | 12 | 4 | 3 | 14 | 3 |
| ENSG00000158887 | 0 | 1 | 1 | 2 | 2 | 2 |
| ENSG00000143252 | 4218 | 1897 | 2319 | 2199 | 2249 | 2222 |
| ENSG00000188931 | 0 | 0 | 0 | 0 | 0 | 0 |
| ENSG00000236484 | 0 | 0 | 1 | 0 | 0 | 0 |
| ENSG00000283696 | 2 | 1 | 0 | 0 | 2 | 4 |
| ENSG00000206921 | 0 | 0 | 2 | 0 | 0 | 2 |
| ENSG00000283360 | 1 | 0 | 0 | 0 | 1 | 0 |
| ENSG00000215840 | 2 | 1 | 1 | 0 | 0 | 0 |
| ENSG00000277882 | 0 | 0 | 0 | 0 | 0 | 0 |
| ENSG00000283317 | 0 | 0 | 1 | 0 | 1 | 0 |
| ENSG00000143226 | 1 | 7 | 1 | 2 | 2 | 0 |
| ENSG00000273112 | 0 | 0 | 0 | 0 | 0 | 0 |
| ENSG00000173110 | 7 | 10 | 12 | 5 | 6 | 4 |
| ENSG00000224203 | 0 | 0 | 0 | 0 | 0 | 0 |
| ENSG00000203747 | 0 | 0 | 0 | 0 | 0 | 0 |
| ENSG00000224515 | 0 | 0 | 0 | 0 | 0 | 0 |
| ENSG00000244682 | 1 | 0 | 2 | 0 | 1 | 0 |
| ENSG00000225217 | 4 | 0 | 0 | 0 | 0 | 0 |
| ENSG00000231586 | 0 | 0 | 0 | 0 | 0 | 0 |
| ENSG00000162747 | 0 | 0 | 0 | 0 | 0 | 0 |
| ENSG00000072694 | 0 | 0 | 2 | 0 | 2 | 0 |
| ENSG00000234211 | 0 | 0 | 0 | 0 | 0 | 0 |
| ENSG00000213075 | 0 | 0 | 0 | 0 | 0 | 0 |
| ENSG00000199595 | 0 | 0 | 0 | 0 | 0 | 0 |
| ENSG00000132185 | 0 | 1 | 0 | 0 | 0 | 0 |
| ENSG00000162746 | 40 | 6 | 21 | 16 | 28 | 16 |
| ENSG00000241347 | 0 | 0 | 0 | 0 | 0 | 0 |
| ENSG00000081721 | 550 | 341 | 406 | 401 | 379 | 417 |
| ENSG00000226889 | 0 | 0 | 0 | 0 | 0 | 0 |
| ENSG00000118217 | 1366 | 606 | 743 | 481 | 701 | 659 |
| ENSG00000229808 | 6 | 1 | 1 | 0 | 1 | 1 |
| ENSG00000162745 | 12 | 3 | 6 | 2 | 6 | 4 |
| ENSG00000227818 | 0 | 0 | 0 | 0 | 0 | 0 |
| ENSG00000198929 | 203 | 83 | 134 | 50 | 65 | 83 |

| | | | | | | |
|---|---|---|---|---|---|---|
| ENSG00000285636 | 0 | 1 | 0 | 0 | 0 | 0 |
| ENSG00000266144 | 0 | 0 | 0 | 0 | 0 | 0 |
| ENSG00000227094 | 0 | 0 | 0 | 0 | 0 | 0 |
| ENSG00000252262 | 0 | 0 | 0 | 0 | 0 | 0 |
| ENSG00000207729 | 0 | 0 | 0 | 0 | 0 | 0 |
| ENSG00000254706 | 0 | 0 | 0 | 0 | 0 | 0 |
| ENSG00000239887 | 691 | 265 | 439 | 302 | 262 | 284 |
| ENSG00000171722 | 7 | 6 | 10 | 0 | 5 | 2 |
| ENSG00000198574 | 0 | 0 | 0 | 0 | 1 | 0 |
| ENSG00000227243 | 0 | 0 | 0 | 0 | 0 | 0 |
| ENSG00000152332 | 4191 | 1886 | 2171 | 1023 | 2163 | 1886 |
| ENSG00000227941 | 0 | 0 | 0 | 0 | 0 | 0 |
| ENSG00000259788 | 0 | 0 | 0 | 0 | 0 | 0 |
| ENSG00000117143 | 741 | 399 | 473 | 288 | 474 | 367 |
| ENSG00000272574 | 6 | 4 | 3 | 1 | 1 | 1 |
| ENSG00000162733 | 10 | 4 | 6 | 2 | 2 | 1 |
| ENSG00000243173 | 0 | 0 | 0 | 0 | 0 | 0 |
| ENSG00000132196 | 295 | 130 | 158 | 149 | 158 | 167 |
| ENSG00000185860 | 0 | 0 | 0 | 0 | 0 | 0 |
| ENSG00000230739 | 0 | 0 | 0 | 0 | 0 | 0 |
| ENSG00000227667 | 0 | 0 | 0 | 0 | 0 | 0 |
| ENSG00000117152 | 0 | 0 | 0 | 0 | 0 | 0 |
| ENSG00000143248 | 2 | 0 | 0 | 1 | 2 | 2 |
| ENSG00000232892 | 0 | 0 | 0 | 0 | 0 | 0 |
| ENSG00000225755 | 0 | 0 | 0 | 0 | 0 | 0 |
| ENSG00000232995 | 7 | 5 | 12 | 4 | 6 | 4 |
| ENSG00000237756 | 0 | 0 | 0 | 0 | 0 | 0 |
| ENSG00000143228 | 148 | 113 | 91 | 74 | 191 | 124 |
| ENSG00000225122 | 0 | 0 | 0 | 0 | 0 | 0 |
| ENSG00000200327 | 0 | 0 | 0 | 0 | 0 | 0 |
| ENSG00000212527 | 0 | 0 | 0 | 0 | 0 | 0 |
| ENSG00000228289 | 0 | 0 | 0 | 1 | 0 | 0 |
| ENSG00000212538 | 0 | 0 | 0 | 0 | 0 | 0 |
| ENSG00000225719 | 0 | 0 | 0 | 0 | 0 | 0 |
| ENSG00000199849 | 0 | 0 | 0 | 0 | 0 | 0 |
| ENSG00000213070 | 4 | 0 | 0 | 1 | 0 | 0 |
| ENSG00000185630 | 4835 | 2222 | 2959 | 1326 | 1904 | 2073 |
| ENSG00000207082 | 0 | 0 | 0 | 0 | 0 | 0 |
| ENSG00000269887 | 25 | 9 | 10 | 3 | 11 | 9 |
| ENSG00000276359 | 0 | 0 | 0 | 0 | 0 | 0 |
| ENSG00000233693 | 0 | 2 | 1 | 0 | 0 | 0 |
| ENSG00000271917 | 2 | 0 | 0 | 0 | 0 | 0 |
| ENSG00000201987 | 0 | 0 | 0 | 0 | 0 | 0 |
| ENSG00000225487 | 0 | 0 | 0 | 0 | 0 | 0 |
| ENSG00000201270 | 0 | 0 | 0 | 0 | 0 | 0 |
| ENSG00000162761 | 0 | 0 | 0 | 0 | 0 | 0 |
| ENSG00000224702 | 0 | 0 | 0 | 0 | 0 | 0 |
| ENSG00000238022 | 0 | 0 | 0 | 0 | 0 | 0 |
| ENSG00000143171 | 0 | 0 | 0 | 0 | 0 | 0 |
| ENSG00000237463 | 0 | 0 | 0 | 0 | 0 | 0 |

| | | | | | | |
|---|---|---|---|---|---|---|
| ENSG00000230748 | 0 | 0 | 0 | 0 | 0 | 0 |
| ENSG00000162763 | 0 | 0 | 0 | 0 | 0 | 0 |
| ENSG00000237783 | 0 | 0 | 0 | 0 | 0 | 0 |
| ENSG00000236206 | 0 | 0 | 0 | 0 | 0 | 0 |
| ENSG00000143198 | 928 | 475 | 567 | 557 | 535 | 461 |
| ENSG00000143149 | 1768 | 867 | 1119 | 837 | 936 | 922 |
| ENSG00000206990 | 0 | 0 | 0 | 0 | 0 | 0 |
| ENSG00000230175 | 0 | 0 | 0 | 0 | 0 | 0 |
| ENSG00000225272 | 0 | 0 | 0 | 0 | 1 | 0 |
| ENSG00000215838 | 4 | 0 | 2 | 0 | 0 | 0 |
| ENSG00000273365 | 0 | 0 | 0 | 0 | 1 | 0 |
| ENSG00000143183 | 3462 | 1640 | 1961 | 1777 | 1841 | 1789 |
| ENSG00000224358 | 23 | 12 | 15 | 12 | 10 | 10 |
| ENSG00000230659 | 0 | 0 | 0 | 0 | 0 | 0 |
| ENSG00000143179 | 936 | 516 | 606 | 620 | 696 | 718 |
| ENSG00000236364 | 8 | 7 | 6 | 4 | 0 | 2 |
| ENSG00000283936 | 0 | 0 | 0 | 0 | 0 | 0 |
| ENSG00000271527 | 0 | 0 | 0 | 0 | 0 | 0 |
| ENSG00000226292 | 0 | 0 | 0 | 0 | 0 | 0 |
| ENSG00000207341 | 0 | 0 | 0 | 0 | 0 | 0 |
| ENSG00000188859 | 8 | 6 | 10 | 4 | 6 | 2 |
| ENSG00000203307 | 0 | 0 | 0 | 0 | 0 | 0 |
| ENSG00000230898 | 0 | 0 | 1 | 0 | 0 | 0 |
| ENSG00000215952 | 0 | 0 | 0 | 0 | 0 | 0 |
| ENSG00000215835 | 50 | 20 | 38 | 21 | 30 | 26 |
| ENSG00000229588 | 0 | 0 | 0 | 0 | 0 | 0 |
| ENSG00000225325 | 0 | 0 | 0 | 0 | 0 | 0 |
| ENSG00000230231 | 0 | 0 | 0 | 0 | 0 | 0 |
| ENSG00000234142 | 0 | 0 | 0 | 0 | 0 | 0 |
| ENSG00000238087 | 0 | 0 | 0 | 0 | 0 | 0 |
| ENSG00000215834 | 0 | 0 | 0 | 0 | 0 | 0 |
| ENSG00000234984 | 0 | 0 | 0 | 0 | 0 | 0 |
| ENSG00000232341 | 0 | 3 | 0 | 0 | 0 | 0 |
| ENSG00000232148 | 0 | 0 | 0 | 0 | 0 | 0 |
| ENSG00000232223 | 0 | 0 | 0 | 0 | 0 | 0 |
| ENSG00000143157 | 3086 | 1432 | 1934 | 1581 | 1754 | 1917 |
| ENSG00000152382 | 698 | 280 | 381 | 261 | 305 | 372 |
| ENSG00000225171 | 2 | 0 | 1 | 1 | 1 | 4 |
| ENSG00000143195 | 0 | 0 | 0 | 0 | 1 | 0 |
| ENSG00000143194 | 0 | 0 | 0 | 0 | 0 | 0 |
| ENSG00000200036 | 0 | 0 | 0 | 0 | 0 | 0 |
| ENSG00000227907 | 0 | 0 | 0 | 0 | 0 | 0 |
| ENSG00000143167 | 0 | 0 | 0 | 0 | 0 | 1 |
| ENSG00000198842 | 0 | 0 | 0 | 0 | 0 | 0 |
| ENSG00000213068 | 0 | 0 | 0 | 0 | 0 | 0 |
| ENSG00000231605 | 0 | 0 | 0 | 0 | 0 | 0 |
| ENSG00000272205 | 8 | 5 | 2 | 2 | 7 | 1 |
| ENSG00000143190 | 928 | 417 | 444 | 208 | 462 | 386 |
| ENSG00000272033 | 0 | 0 | 0 | 0 | 0 | 1 |
| ENSG00000198821 | 0 | 0 | 1 | 0 | 1 | 0 |

| | | | | | | |
|---|---|---|---|---|---|---|
| ENSG00000233411 | 2 | 3 | 0 | 1 | 0 | 0 |
| ENSG00000273160 | 2 | 0 | 1 | 1 | 0 | 1 |
| ENSG00000227776 | 0 | 0 | 0 | 0 | 0 | 0 |
| ENSG00000143162 | 2387 | 1260 | 1319 | 1177 | 1484 | 1249 |
| ENSG00000231029 | 0 | 0 | 0 | 0 | 0 | 0 |
| ENSG00000241666 | 0 | 0 | 0 | 0 | 0 | 0 |
| ENSG00000198771 | 0 | 0 | 0 | 0 | 0 | 0 |
| ENSG00000197965 | 1890 | 879 | 1028 | 903 | 1067 | 1096 |
| ENSG00000143199 | 2 | 1 | 2 | 0 | 3 | 2 |
| ENSG00000250762 | 0 | 0 | 0 | 0 | 3 | 1 |
| ENSG00000232194 | 0 | 0 | 1 | 0 | 0 | 0 |
| ENSG00000143158 | 921 | 493 | 643 | 537 | 501 | 548 |
| ENSG00000143164 | 1438 | 684 | 897 | 608 | 857 | 742 |
| ENSG00000221545 | 0 | 0 | 0 | 0 | 0 | 0 |
| ENSG00000224837 | 2 | 1 | 1 | 1 | 5 | 0 |
| ENSG00000143147 | 38 | 20 | 18 | 25 | 34 | 20 |
| ENSG00000143155 | 1609 | 791 | 963 | 827 | 1000 | 932 |
| ENSG00000237049 | 0 | 1 | 0 | 0 | 0 | 1 |
| ENSG00000237131 | 0 | 0 | 0 | 0 | 0 | 0 |
| ENSG00000213064 | 1153 | 654 | 664 | 561 | 739 | 633 |
| ENSG00000214262 | 0 | 0 | 0 | 0 | 0 | 0 |
| ENSG00000206880 | 0 | 0 | 0 | 0 | 0 | 0 |
| ENSG00000143178 | 46 | 22 | 38 | 12 | 35 | 28 |
| ENSG00000227722 | 0 | 0 | 0 | 0 | 0 | 0 |
| ENSG00000207974 | 0 | 0 | 0 | 0 | 0 | 0 |
| ENSG00000228697 | 0 | 0 | 0 | 0 | 0 | 0 |
| ENSG00000283255 | 0 | 0 | 0 | 0 | 0 | 0 |
| ENSG00000215833 | 0 | 0 | 0 | 0 | 0 | 0 |
| ENSG00000143185 | 0 | 0 | 0 | 0 | 1 | 0 |
| ENSG00000227777 | 0 | 0 | 0 | 0 | 0 | 0 |
| ENSG00000143184 | 0 | 0 | 0 | 0 | 0 | 0 |
| ENSG00000229757 | 0 | 0 | 0 | 0 | 0 | 1 |
| ENSG00000143196 | 0 | 0 | 0 | 0 | 0 | 0 |
| ENSG00000285622 | 0 | 0 | 0 | 0 | 0 | 0 |
| ENSG00000235736 | 0 | 0 | 0 | 0 | 0 | 1 |
| ENSG00000225826 | 0 | 0 | 0 | 0 | 0 | 0 |
| ENSG00000229612 | 0 | 0 | 0 | 0 | 0 | 0 |
| ENSG00000203601 | 0 | 0 | 0 | 0 | 0 | 0 |
| ENSG00000213063 | 6 | 0 | 0 | 0 | 0 | 2 |
| ENSG00000252987 | 0 | 0 | 0 | 0 | 0 | 0 |
| ENSG00000237707 | 2 | 0 | 1 | 0 | 0 | 0 |
| ENSG00000143153 | 5824 | 2902 | 3830 | 2703 | 3758 | 3737 |
| ENSG00000143156 | 252 | 162 | 153 | 118 | 205 | 152 |
| ENSG00000235575 | 0 | 0 | 1 | 1 | 2 | 0 |
| ENSG00000117475 | 557 | 370 | 353 | 186 | 569 | 315 |
| ENSG00000117477 | 2 | 2 | 3 | 1 | 1 | 0 |
| ENSG00000117479 | 1456 | 744 | 958 | 543 | 792 | 773 |
| ENSG00000234604 | 8 | 3 | 6 | 0 | 2 | 3 |
| ENSG00000213062 | 74 | 30 | 41 | 24 | 36 | 32 |
| ENSG00000198734 | 2 | 1 | 2 | 0 | 4 | 3 |

| | | | | | | |
|---|---|---|---|---|---|---|
| ENSG00000174175 | 0 | 0 | 0 | 0 | 0 | 0 |
| ENSG00000000460 | 449 | 230 | 290 | 168 | 251 | 220 |
| ENSG00000188404 | 217 | 124 | 191 | 46 | 65 | 108 |
| ENSG00000007908 | 18 | 8 | 5 | 1 | 3 | 3 |
| ENSG00000230704 | 0 | 2 | 1 | 0 | 0 | 0 |
| ENSG00000171806 | 391 | 212 | 239 | 163 | 287 | 204 |
| ENSG00000000457 | 653 | 353 | 454 | 256 | 330 | 323 |
| ENSG00000239494 | 7 | 4 | 4 | 1 | 7 | 4 |
| ENSG00000075945 | 587 | 349 | 414 | 278 | 394 | 317 |
| ENSG00000243051 | 0 | 0 | 0 | 0 | 0 | 0 |
| ENSG00000235378 | 0 | 0 | 0 | 0 | 0 | 0 |
| ENSG00000232959 | 0 | 0 | 0 | 0 | 0 | 0 |
| ENSG00000224896 | 0 | 0 | 0 | 0 | 0 | 0 |
| ENSG00000203740 | 2 | 0 | 1 | 1 | 0 | 0 |
| ENSG00000263390 | 0 | 0 | 0 | 0 | 0 | 0 |
| ENSG00000283340 | 0 | 0 | 0 | 0 | 0 | 0 |
| ENSG00000233985 | 0 | 0 | 0 | 0 | 0 | 0 |
| ENSG00000238051 | 0 | 0 | 0 | 0 | 0 | 0 |
| ENSG00000224286 | 0 | 0 | 0 | 0 | 0 | 0 |
| ENSG00000226967 | 0 | 0 | 0 | 0 | 0 | 0 |
| ENSG00000231407 | 0 | 1 | 0 | 3 | 4 | 1 |
| ENSG00000120370 | 312 | 131 | 150 | 77 | 170 | 123 |
| ENSG00000225545 | 0 | 0 | 0 | 0 | 0 | 0 |
| ENSG00000235303 | 0 | 0 | 0 | 0 | 0 | 0 |
| ENSG00000116132 | 0 | 0 | 0 | 0 | 0 | 0 |
| ENSG00000271811 | 0 | 1 | 0 | 0 | 0 | 0 |
| ENSG00000117501 | 0 | 0 | 0 | 0 | 0 | 0 |
| ENSG00000226552 | 0 | 0 | 0 | 0 | 0 | 0 |
| ENSG00000007933 | 0 | 0 | 0 | 0 | 0 | 0 |
| ENSG00000221390 | 0 | 0 | 0 | 0 | 0 | 0 |
| ENSG00000283598 | 0 | 0 | 0 | 0 | 0 | 0 |
| ENSG00000117507 | 0 | 0 | 0 | 0 | 0 | 0 |
| ENSG00000094963 | 2 | 1 | 2 | 1 | 0 | 0 |
| ENSG00000225243 | 0 | 0 | 0 | 0 | 0 | 0 |
| ENSG00000231424 | 0 | 1 | 1 | 0 | 1 | 0 |
| ENSG00000010932 | 11 | 3 | 21 | 3 | 8 | 13 |
| ENSG00000206692 | 0 | 0 | 0 | 0 | 0 | 0 |
| ENSG00000230547 | 6 | 3 | 3 | 0 | 4 | 1 |
| ENSG00000076258 | 177 | 96 | 123 | 82 | 113 | 94 |
| ENSG00000285665 | 6 | 8 | 6 | 5 | 7 | 10 |
| ENSG00000268062 | 0 | 2 | 0 | 1 | 2 | 1 |
| ENSG00000239428 | 0 | 0 | 0 | 0 | 0 | 0 |
| ENSG00000234333 | 0 | 0 | 0 | 0 | 0 | 0 |
| ENSG00000117523 | 3967 | 2396 | 2093 | 887 | 3162 | 2181 |
| ENSG00000283683 | 0 | 0 | 0 | 0 | 0 | 0 |
| ENSG00000034971 | 0 | 0 | 0 | 0 | 0 | 0 |
| ENSG00000233328 | 4 | 1 | 3 | 4 | 3 | 5 |
| ENSG00000230364 | 2 | 0 | 0 | 2 | 0 | 0 |
| ENSG00000117533 | 360 | 183 | 213 | 148 | 250 | 177 |
| ENSG00000232261 | 0 | 0 | 0 | 0 | 0 | 0 |

| | | | | | | |
|---|---|---|---|---|---|---|
| ENSG00000271459 | 0 | 0 | 0 | 0 | 0 | 0 |
| ENSG00000236741 | 0 | 0 | 0 | 0 | 0 | 0 |
| ENSG00000253060 | 0 | 0 | 0 | 0 | 0 | 0 |
| ENSG00000010165 | 2224 | 1127 | 1415 | 1431 | 1412 | 1481 |
| ENSG00000213060 | 0 | 0 | 1 | 0 | 0 | 0 |
| ENSG00000224600 | 0 | 0 | 0 | 0 | 0 | 0 |
| ENSG00000197959 | 38 | 25 | 22 | 13 | 41 | 21 |
| ENSG00000233540 | 2 | 0 | 0 | 1 | 0 | 0 |
| ENSG00000230630 | 0 | 0 | 0 | 0 | 0 | 0 |
| ENSG00000283844 | 0 | 0 | 0 | 0 | 0 | 0 |
| ENSG00000283152 | 0 | 0 | 0 | 0 | 0 | 0 |
| ENSG00000208024 | 0 | 0 | 0 | 0 | 0 | 0 |
| ENSG00000206684 | 0 | 0 | 0 | 0 | 0 | 0 |
| ENSG00000135845 | 1590 | 762 | 817 | 804 | 810 | 849 |
| ENSG00000180999 | 8 | 4 | 5 | 7 | 14 | 14 |
| ENSG00000094975 | 1705 | 869 | 763 | 444 | 1374 | 810 |
| ENSG00000251943 | 0 | 0 | 0 | 0 | 0 | 0 |
| ENSG00000117560 | 0 | 0 | 0 | 0 | 0 | 0 |
| ENSG00000229785 | 0 | 0 | 0 | 0 | 0 | 0 |
| ENSG00000279061 | 0 | 0 | 0 | 0 | 0 | 0 |
| ENSG00000224228 | 0 | 0 | 0 | 0 | 0 | 0 |
| ENSG00000227177 | 0 | 0 | 0 | 0 | 0 | 0 |
| ENSG00000224000 | 0 | 0 | 0 | 0 | 0 | 0 |
| ENSG00000120337 | 0 | 0 | 0 | 0 | 0 | 0 |
| ENSG00000230849 | 0 | 1 | 0 | 0 | 0 | 0 |
| ENSG00000226375 | 1 | 0 | 0 | 0 | 0 | 0 |
| ENSG00000117586 | 18 | 2 | 4 | 2 | 1 | 5 |
| ENSG00000232751 | 0 | 0 | 0 | 0 | 0 | 0 |
| ENSG00000231615 | 0 | 0 | 0 | 0 | 0 | 0 |
| ENSG00000203739 | 25 | 18 | 30 | 27 | 14 | 25 |
| ENSG00000117592 | 8136 | 4151 | 5115 | 4585 | 4916 | 4681 |
| ENSG00000162753 | 0 | 0 | 0 | 0 | 0 | 0 |
| ENSG00000238272 | 0 | 0 | 0 | 0 | 0 | 0 |
| ENSG00000285777 | 0 | 0 | 0 | 0 | 0 | 0 |
| ENSG00000183831 | 2 | 4 | 0 | 0 | 0 | 0 |
| ENSG00000232113 | 0 | 0 | 0 | 0 | 0 | 0 |
| ENSG00000076321 | 824 | 396 | 498 | 289 | 445 | 426 |
| ENSG00000225591 | 0 | 0 | 0 | 0 | 2 | 0 |
| ENSG00000200674 | 0 | 1 | 0 | 0 | 1 | 0 |
| ENSG00000120334 | 378 | 166 | 213 | 130 | 218 | 176 |
| ENSG00000252357 | 0 | 0 | 0 | 0 | 0 | 0 |
| ENSG00000117593 | 2376 | 1257 | 1606 | 1328 | 1707 | 1682 |
| ENSG00000270084 | 26 | 9 | 13 | 10 | 4 | 8 |
| ENSG00000234741 | 8655 | 4420 | 5545 | 6496 | 6312 | 6255 |
| ENSG00000185278 | 661 | 323 | 349 | 164 | 394 | 260 |
| ENSG00000117601 | 2 | 1 | 6 | 3 | 1 | 0 |
| ENSG00000252231 | 0 | 0 | 0 | 0 | 0 | 1 |
| ENSG00000135870 | 1145 | 529 | 536 | 302 | 649 | 467 |
| ENSG00000200755 | 0 | 0 | 0 | 0 | 0 | 0 |
| ENSG00000236535 | 0 | 0 | 2 | 0 | 0 | 0 |

| | | | | | | |
|---|---|---|---|---|---|---|
| ENSG00000224977 | 3 | 0 | 4 | 3 | 3 | 3 |
| ENSG00000225713 | 0 | 0 | 0 | 0 | 0 | 0 |
| ENSG00000227373 | 2 | 2 | 1 | 2 | 1 | 3 |
| ENSG00000152061 | 678 | 355 | 334 | 196 | 438 | 287 |
| ENSG00000237317 | 0 | 0 | 0 | 0 | 0 | 0 |
| ENSG00000203737 | 2 | 0 | 0 | 0 | 2 | 0 |
| ENSG00000223828 | 0 | 0 | 0 | 0 | 0 | 0 |
| ENSG00000231378 | 0 | 1 | 1 | 0 | 0 | 1 |
| ENSG00000237249 | 0 | 2 | 1 | 0 | 0 | 0 |
| ENSG00000223525 | 0 | 0 | 0 | 0 | 0 | 0 |
| ENSG00000235869 | 0 | 0 | 0 | 0 | 0 | 0 |
| ENSG00000229531 | 0 | 0 | 0 | 0 | 0 | 0 |
| ENSG00000252552 | 0 | 1 | 0 | 0 | 1 | 0 |
| ENSG00000116161 | 3219 | 1786 | 2013 | 1997 | 2243 | 2025 |
| ENSG00000120333 | 1464 | 744 | 806 | 788 | 853 | 726 |
| ENSG00000206659 | 0 | 0 | 0 | 0 | 0 | 0 |
| ENSG00000181101 | 0 | 0 | 0 | 0 | 0 | 0 |
| ENSG00000120332 | 0 | 0 | 0 | 0 | 0 | 0 |
| ENSG00000235750 | 426 | 214 | 338 | 250 | 329 | 278 |
| ENSG00000230687 | 0 | 0 | 0 | 0 | 0 | 0 |
| ENSG00000233946 | 0 | 0 | 0 | 0 | 0 | 0 |
| ENSG00000260990 | 0 | 0 | 0 | 0 | 1 | 0 |
| ENSG00000116147 | 0 | 3 | 1 | 1 | 0 | 0 |
| ENSG00000235628 | 0 | 0 | 0 | 0 | 0 | 0 |
| ENSG00000224718 | 0 | 1 | 0 | 0 | 0 | 0 |
| ENSG00000227740 | 2 | 4 | 5 | 1 | 5 | 5 |
| ENSG00000230777 | 1 | 1 | 3 | 0 | 1 | 3 |
| ENSG00000143207 | 2212 | 1057 | 1389 | 1108 | 1288 | 1257 |
| ENSG00000252906 | 0 | 0 | 0 | 0 | 1 | 1 |
| ENSG00000228686 | 0 | 3 | 0 | 2 | 0 | 0 |
| ENSG00000236021 | 2 | 4 | 6 | 2 | 6 | 6 |
| ENSG00000273629 | 1 | 0 | 0 | 0 | 0 | 0 |
| ENSG00000232463 | 0 | 0 | 0 | 0 | 0 | 0 |
| ENSG00000253025 | 0 | 0 | 0 | 0 | 0 | 0 |
| ENSG00000227815 | 0 | 0 | 0 | 3 | 0 | 1 |
| ENSG00000231020 | 0 | 0 | 0 | 0 | 0 | 0 |
| ENSG00000225904 | 0 | 0 | 0 | 0 | 0 | 0 |
| ENSG00000116183 | 0 | 0 | 1 | 0 | 0 | 0 |
| ENSG00000237514 | 0 | 0 | 0 | 0 | 0 | 0 |
| ENSG00000152092 | 0 | 0 | 0 | 0 | 0 | 0 |
| ENSG00000202609 | 0 | 0 | 0 | 0 | 0 | 0 |
| ENSG00000198797 | 145 | 39 | 49 | 27 | 12 | 5 |
| ENSG00000224968 | 0 | 0 | 0 | 0 | 0 | 0 |
| ENSG00000227579 | 2 | 0 | 3 | 3 | 0 | 1 |
| ENSG00000236720 | 0 | 0 | 0 | 0 | 0 | 0 |
| ENSG00000120341 | 0 | 0 | 0 | 0 | 0 | 0 |
| ENSG00000254154 | 4 | 0 | 0 | 0 | 0 | 3 |
| ENSG00000242193 | 93 | 42 | 62 | 32 | 64 | 61 |
| ENSG00000276563 | 0 | 0 | 0 | 0 | 2 | 0 |
| ENSG00000224687 | 18 | 8 | 13 | 6 | 11 | 9 |

| | | | | | | |
|---|---|---|---|---|---|---|
| ENSG00000075391 | 343 | 156 | 193 | 105 | 244 | 176 |
| ENSG00000270575 | 0 | 0 | 0 | 0 | 0 | 0 |
| ENSG00000213058 | 0 | 0 | 0 | 0 | 2 | 0 |
| ENSG00000188585 | 0 | 0 | 0 | 0 | 0 | 0 |
| ENSG00000264809 | 0 | 0 | 0 | 0 | 0 | 0 |
| ENSG00000285910 | 0 | 0 | 0 | 0 | 0 | 0 |
| ENSG00000240021 | 0 | 0 | 0 | 0 | 0 | 0 |
| ENSG00000213057 | 101 | 52 | 62 | 31 | 63 | 51 |
| ENSG00000201347 | 0 | 0 | 0 | 0 | 0 | 0 |
| ENSG00000273384 | 3 | 3 | 3 | 1 | 5 | 6 |
| ENSG00000266417 | 0 | 0 | 0 | 0 | 0 | 0 |
| ENSG00000273062 | 31 | 13 | 15 | 11 | 12 | 23 |
| ENSG00000116191 | 1492 | 858 | 800 | 341 | 927 | 650 |
| ENSG00000228196 | 2 | 1 | 0 | 0 | 2 | 1 |
| ENSG00000201791 | 0 | 0 | 2 | 0 | 0 | 0 |
| ENSG00000116194 | 0 | 0 | 4 | 0 | 0 | 0 |
| ENSG00000116199 | 3095 | 1726 | 1996 | 1400 | 1755 | 1915 |
| ENSG00000232750 | 0 | 0 | 0 | 0 | 1 | 0 |
| ENSG00000186283 | 2866 | 1383 | 1729 | 1856 | 1524 | 1775 |
| ENSG00000143322 | 1184 | 451 | 581 | 348 | 638 | 551 |
| ENSG00000234041 | 0 | 0 | 0 | 0 | 0 | 0 |
| ENSG00000231905 | 2 | 1 | 1 | 0 | 0 | 0 |
| ENSG00000212338 | 0 | 0 | 0 | 0 | 0 | 0 |
| ENSG00000201619 | 0 | 0 | 0 | 0 | 0 | 0 |
| ENSG00000231564 | 0 | 1 | 0 | 0 | 0 | 0 |
| ENSG00000225711 | 0 | 1 | 1 | 0 | 0 | 0 |
| ENSG00000223794 | 0 | 0 | 0 | 0 | 0 | 0 |
| ENSG00000228191 | 0 | 0 | 0 | 0 | 0 | 0 |
| ENSG00000057252 | 943 | 482 | 583 | 398 | 572 | 500 |
| ENSG00000162779 | 7 | 7 | 10 | 3 | 6 | 5 |
| ENSG00000224620 | 0 | 0 | 0 | 0 | 0 | 0 |
| ENSG00000236539 | 0 | 0 | 1 | 0 | 0 | 1 |
| ENSG00000261250 | 0 | 0 | 0 | 0 | 0 | 0 |
| ENSG00000116218 | 0 | 0 | 0 | 0 | 0 | 1 |
| ENSG00000251875 | 0 | 0 | 0 | 0 | 0 | 0 |
| ENSG00000227141 | 12 | 6 | 8 | 4 | 8 | 5 |
| ENSG00000261060 | 2 | 1 | 2 | 1 | 0 | 2 |
| ENSG00000162782 | 451 | 213 | 222 | 161 | 274 | 188 |
| ENSG00000243062 | 0 | 0 | 0 | 0 | 0 | 0 |
| ENSG00000143340 | 0 | 0 | 0 | 0 | 0 | 0 |
| ENSG00000229407 | 0 | 0 | 0 | 1 | 0 | 1 |
| ENSG00000231966 | 0 | 0 | 1 | 0 | 1 | 0 |
| ENSG00000169905 | 1670 | 807 | 917 | 668 | 981 | 939 |
| ENSG00000272906 | 14 | 8 | 14 | 10 | 12 | 7 |
| ENSG00000143337 | 1970 | 968 | 1037 | 631 | 1236 | 878 |
| ENSG00000264916 | 0 | 0 | 0 | 0 | 0 | 0 |
| ENSG00000261831 | 0 | 0 | 0 | 0 | 1 | 0 |
| ENSG00000260360 | 6 | 6 | 4 | 3 | 3 | 3 |
| ENSG00000135837 | 523 | 371 | 253 | 81 | 614 | 300 |
| ENSG00000235145 | 0 | 0 | 0 | 0 | 0 | 1 |

| | | | | | | |
|---|---|---|---|---|---|---|
| ENSG00000223450 | 0 | 0 | 0 | 0 | 0 | 0 |
| ENSG00000261817 | 0 | 0 | 0 | 0 | 0 | 0 |
| ENSG00000116260 | 7452 | 3294 | 4094 | 4639 | 4646 | 4500 |
| ENSG00000121454 | 159 | 86 | 109 | 71 | 119 | 105 |
| ENSG00000230124 | 1922 | 842 | 1087 | 910 | 1123 | 1063 |
| ENSG00000235060 | 0 | 0 | 0 | 0 | 0 | 0 |
| ENSG00000265435 | 0 | 0 | 0 | 0 | 0 | 0 |
| ENSG00000206905 | 0 | 0 | 0 | 0 | 0 | 0 |
| ENSG00000236719 | 0 | 2 | 1 | 0 | 0 | 0 |
| ENSG00000143324 | 2611 | 1200 | 1496 | 726 | 1251 | 1100 |
| ENSG00000283575 | 0 | 0 | 0 | 0 | 0 | 0 |
| ENSG00000225857 | 7 | 0 | 0 | 0 | 1 | 2 |
| ENSG00000135835 | 110 | 51 | 71 | 30 | 48 | 69 |
| ENSG00000243155 | 42 | 21 | 26 | 23 | 18 | 15 |
| ENSG00000232586 | 0 | 1 | 0 | 0 | 2 | 1 |
| ENSG00000251520 | 3 | 2 | 0 | 0 | 2 | 2 |
| ENSG00000270711 | 0 | 0 | 3 | 1 | 1 | 0 |
| ENSG00000135823 | 3625 | 1872 | 2182 | 1813 | 2231 | 2054 |
| ENSG00000153029 | 1516 | 710 | 942 | 588 | 849 | 786 |
| ENSG00000162783 | 1978 | 979 | 1094 | 1431 | 1527 | 1293 |
| ENSG00000237292 | 6 | 1 | 1 | 0 | 1 | 1 |
| ENSG00000225359 | 0 | 0 | 0 | 0 | 0 | 0 |
| ENSG00000179452 | 0 | 0 | 0 | 0 | 0 | 0 |
| ENSG00000198216 | 0 | 0 | 0 | 1 | 0 | 0 |
| ENSG00000273573 | 0 | 0 | 0 | 0 | 0 | 0 |
| ENSG00000252977 | 0 | 0 | 0 | 0 | 0 | 0 |
| ENSG00000272880 | 0 | 0 | 0 | 0 | 0 | 0 |
| ENSG00000222397 | 0 | 0 | 0 | 0 | 0 | 0 |
| ENSG00000179930 | 0 | 0 | 0 | 0 | 1 | 0 |
| ENSG00000224810 | 0 | 0 | 1 | 0 | 0 | 0 |
| ENSG00000225982 | 0 | 0 | 1 | 0 | 1 | 1 |
| ENSG00000232309 | 0 | 0 | 0 | 0 | 0 | 0 |
| ENSG00000228918 | 0 | 0 | 0 | 0 | 0 | 0 |
| ENSG00000232096 | 0 | 0 | 0 | 0 | 0 | 0 |
| ENSG00000206764 | 0 | 0 | 0 | 0 | 0 | 0 |
| ENSG00000228664 | 0 | 0 | 0 | 0 | 0 | 0 |
| ENSG00000231684 | 0 | 2 | 0 | 1 | 0 | 0 |
| ENSG00000135821 | 6954 | 3571 | 4223 | 3217 | 3280 | 3461 |
| ENSG00000203730 | 2 | 0 | 0 | 0 | 1 | 0 |
| ENSG00000203729 | 0 | 0 | 0 | 0 | 0 | 0 |
| ENSG00000121446 | 0 | 1 | 0 | 0 | 0 | 0 |
| ENSG00000270443 | 0 | 0 | 0 | 0 | 0 | 0 |
| ENSG00000233537 | 0 | 0 | 0 | 0 | 0 | 0 |
| ENSG00000135828 | 776 | 359 | 465 | 285 | 382 | 365 |
| ENSG00000276839 | 0 | 1 | 0 | 0 | 0 | 0 |
| ENSG00000143333 | 291 | 144 | 147 | 118 | 133 | 168 |
| ENSG00000261504 | 4 | 2 | 1 | 3 | 1 | 5 |
| ENSG00000135824 | 2 | 0 | 0 | 0 | 1 | 0 |
| ENSG00000227181 | 0 | 0 | 0 | 0 | 0 | 0 |
| ENSG00000271269 | 0 | 0 | 0 | 0 | 0 | 0 |

| | | | | | | |
|---|---|---|---|---|---|---|
| ENSG00000135838 | 31 | 28 | 14 | 20 | 20 | 27 |
| ENSG00000135829 | 8406 | 4108 | 4913 | 3159 | 4772 | 4546 |
| ENSG00000157060 | 0 | 0 | 0 | 0 | 0 | 0 |
| ENSG00000224040 | 10 | 6 | 7 | 5 | 10 | 4 |
| ENSG00000199837 | 0 | 0 | 0 | 0 | 0 | 0 |
| ENSG00000285609 | 0 | 0 | 0 | 0 | 0 | 0 |
| ENSG00000226570 | 0 | 0 | 0 | 0 | 0 | 0 |
| ENSG00000176855 | 37 | 5 | 5 | 15 | 18 | 11 |
| ENSG00000207472 | 0 | 0 | 0 | 0 | 0 | 0 |
| ENSG00000135862 | 5406 | 2469 | 3117 | 1829 | 2452 | 2399 |
| ENSG00000224468 | 4 | 5 | 5 | 4 | 8 | 1 |
| ENSG00000058085 | 239 | 122 | 154 | 181 | 236 | 184 |
| ENSG00000157064 | 54 | 24 | 19 | 18 | 23 | 30 |
| ENSG00000233562 | 0 | 0 | 0 | 0 | 0 | 0 |
| ENSG00000232860 | 33 | 6 | 15 | 10 | 11 | 12 |
| ENSG00000116698 | 4709 | 2137 | 2741 | 1681 | 2532 | 2519 |
| ENSG00000116701 | 12 | 5 | 3 | 3 | 7 | 2 |
| ENSG00000162704 | 6803 | 3241 | 3900 | 3395 | 3915 | 3948 |
| ENSG00000143344 | 60 | 17 | 20 | 27 | 40 | 32 |
| ENSG00000173627 | 0 | 0 | 0 | 0 | 0 | 0 |
| ENSG00000270994 | 0 | 0 | 0 | 0 | 0 | 0 |
| ENSG00000227554 | 0 | 0 | 0 | 0 | 0 | 0 |
| ENSG00000198756 | 8 | 6 | 6 | 5 | 8 | 9 |
| ENSG00000198860 | 1042 | 491 | 581 | 589 | 660 | 624 |
| ENSG00000199840 | 0 | 0 | 0 | 0 | 0 | 0 |
| ENSG00000207080 | 0 | 0 | 0 | 0 | 0 | 0 |
| ENSG00000231791 | 0 | 0 | 0 | 0 | 0 | 0 |
| ENSG00000244568 | 0 | 0 | 0 | 0 | 0 | 0 |
| ENSG00000271387 | 10 | 8 | 10 | 5 | 9 | 12 |
| ENSG00000116667 | 2600 | 1216 | 1685 | 927 | 965 | 1031 |
| ENSG00000230470 | 0 | 0 | 3 | 0 | 0 | 0 |
| ENSG00000232036 | 0 | 0 | 0 | 0 | 1 | 1 |
| ENSG00000285847 | 1 | 0 | 0 | 0 | 0 | 0 |
| ENSG00000116406 | 1385 | 627 | 619 | 316 | 852 | 586 |
| ENSG00000135842 | 6 | 3 | 2 | 1 | 2 | 2 |
| ENSG00000252222 | 0 | 0 | 0 | 0 | 0 | 0 |
| ENSG00000260976 | 0 | 3 | 1 | 0 | 1 | 4 |
| ENSG00000201312 | 0 | 0 | 0 | 0 | 0 | 0 |
| ENSG00000121481 | 546 | 258 | 282 | 221 | 283 | 240 |
| ENSG00000232792 | 2 | 0 | 0 | 0 | 0 | 0 |
| ENSG00000121486 | 891 | 497 | 413 | 308 | 536 | 411 |
| ENSG00000116668 | 116 | 52 | 64 | 40 | 74 | 51 |
| ENSG00000231084 | 14 | 3 | 15 | 10 | 18 | 8 |
| ENSG00000213051 | 0 | 0 | 0 | 0 | 1 | 0 |
| ENSG00000201421 | 0 | 0 | 0 | 0 | 0 | 0 |
| ENSG00000206640 | 0 | 0 | 0 | 0 | 0 | 0 |
| ENSG00000233583 | 0 | 0 | 0 | 0 | 0 | 0 |
| ENSG00000252612 | 0 | 0 | 0 | 1 | 0 | 0 |
| ENSG00000238061 | 1 | 0 | 1 | 0 | 0 | 1 |
| ENSG00000279838 | 4 | 2 | 4 | 1 | 0 | 1 |

| | | | | | | |
|---|---|---|---|---|---|---|
| ENSG00000116679 | 3613 | 1749 | 2092 | 1161 | 1848 | 1663 |
| ENSG00000273004 | 2 | 6 | 6 | 1 | 5 | 4 |
| ENSG00000261024 | 0 | 1 | 0 | 0 | 0 | 0 |
| ENSG00000252407 | 0 | 0 | 0 | 0 | 0 | 0 |
| ENSG00000270920 | 0 | 0 | 0 | 0 | 0 | 0 |
| ENSG00000279401 | 0 | 0 | 0 | 0 | 0 | 0 |
| ENSG00000228309 | 0 | 0 | 0 | 0 | 0 | 0 |
| ENSG00000207108 | 0 | 0 | 0 | 0 | 0 | 0 |
| ENSG00000201596 | 0 | 0 | 0 | 0 | 0 | 0 |
| ENSG00000261729 | 0 | 0 | 0 | 0 | 0 | 0 |
| ENSG00000143341 | 12638 | 5937 | 7373 | 2623 | 5293 | 4558 |
| ENSG00000224691 | 9 | 7 | 8 | 3 | 13 | 3 |
| ENSG00000116690 | 23 | 10 | 20 | 11 | 6 | 15 |
| ENSG00000047410 | 787 | 513 | 562 | 158 | 1130 | 706 |
| ENSG00000202025 | 0 | 0 | 0 | 0 | 0 | 0 |
| ENSG00000157181 | 845 | 444 | 407 | 212 | 701 | 411 |
| ENSG00000262180 | 10 | 4 | 3 | 1 | 15 | 2 |
| ENSG00000229739 | 1 | 0 | 0 | 1 | 0 | 0 |
| ENSG00000116703 | 2 | 0 | 0 | 0 | 0 | 0 |
| ENSG00000273198 | 0 | 0 | 0 | 0 | 0 | 0 |
| ENSG00000228238 | 0 | 0 | 0 | 0 | 0 | 0 |
| ENSG00000233196 | 0 | 0 | 0 | 0 | 0 | 1 |
| ENSG00000073756 | 0 | 0 | 0 | 0 | 0 | 0 |
| ENSG00000273129 | 0 | 0 | 0 | 0 | 0 | 0 |
| ENSG00000116711 | 0 | 0 | 0 | 0 | 0 | 0 |
| ENSG00000230426 | 0 | 0 | 0 | 0 | 0 | 0 |
| ENSG00000271558 | 0 | 0 | 0 | 0 | 0 | 0 |
| ENSG00000225462 | 0 | 1 | 0 | 0 | 0 | 1 |
| ENSG00000274702 | 0 | 0 | 0 | 0 | 0 | 0 |
| ENSG00000236401 | 0 | 0 | 0 | 0 | 0 | 0 |
| ENSG00000223847 | 0 | 0 | 0 | 0 | 0 | 0 |
| ENSG00000285894 | 0 | 0 | 0 | 0 | 0 | 0 |
| ENSG00000150732 | 0 | 0 | 0 | 0 | 0 | 0 |
| ENSG00000222240 | 0 | 0 | 0 | 0 | 0 | 0 |
| ENSG00000261182 | 0 | 0 | 0 | 0 | 0 | 0 |
| ENSG00000224278 | 0 | 0 | 0 | 0 | 0 | 0 |
| ENSG00000225006 | 0 | 0 | 0 | 0 | 0 | 0 |
| ENSG00000235083 | 0 | 0 | 0 | 0 | 0 | 0 |
| ENSG00000231022 | 0 | 0 | 0 | 0 | 0 | 0 |
| ENSG00000237283 | 0 | 0 | 0 | 0 | 0 | 0 |
| ENSG00000238054 | 0 | 0 | 2 | 1 | 0 | 0 |
| ENSG00000226486 | 0 | 0 | 0 | 0 | 0 | 0 |
| ENSG00000271036 | 0 | 0 | 0 | 0 | 0 | 0 |
| ENSG00000226196 | 0 | 0 | 0 | 0 | 0 | 0 |
| ENSG00000252553 | 0 | 0 | 0 | 0 | 0 | 0 |
| ENSG00000232212 | 0 | 0 | 0 | 0 | 0 | 0 |
| ENSG00000238270 | 0 | 0 | 0 | 0 | 0 | 0 |
| ENSG00000230987 | 0 | 0 | 0 | 0 | 0 | 0 |
| ENSG00000162670 | 1111 | 498 | 719 | 325 | 350 | 451 |
| ENSG00000225811 | 0 | 0 | 0 | 0 | 0 | 0 |

| | | | | | | |
|---|---|---|---|---|---|---|
| ENSG00000237457 | 25 | 18 | 29 | 13 | 23 | 26 |
| ENSG00000241505 | 0 | 0 | 0 | 0 | 0 | 0 |
| ENSG00000231175 | 0 | 0 | 0 | 0 | 0 | 0 |
| ENSG00000236025 | 0 | 0 | 0 | 0 | 0 | 0 |
| ENSG00000238108 | 0 | 0 | 0 | 0 | 0 | 0 |
| ENSG00000285638 | 3 | 1 | 2 | 1 | 2 | 0 |
| ENSG00000228020 | 8 | 3 | 3 | 1 | 4 | 0 |
| ENSG00000261642 | 5 | 4 | 0 | 0 | 0 | 0 |
| ENSG00000271187 | 0 | 0 | 0 | 0 | 0 | 0 |
| ENSG00000233882 | 0 | 0 | 0 | 0 | 0 | 0 |
| ENSG00000223344 | 0 | 0 | 0 | 0 | 0 | 0 |
| ENSG00000228215 | 0 | 0 | 0 | 0 | 0 | 0 |
| ENSG00000150681 | 0 | 0 | 0 | 0 | 0 | 0 |
| ENSG00000285280 | 0 | 0 | 0 | 0 | 0 | 0 |
| ENSG00000226723 | 0 | 0 | 0 | 0 | 0 | 1 |
| ENSG00000236792 | 0 | 0 | 0 | 0 | 0 | 0 |
| ENSG00000253148 | 0 | 0 | 0 | 0 | 0 | 0 |
| ENSG00000236069 | 0 | 0 | 0 | 0 | 0 | 0 |
| ENSG00000090104 | 0 | 0 | 0 | 0 | 0 | 1 |
| ENSG00000127074 | 0 | 0 | 0 | 0 | 0 | 0 |
| ENSG00000231767 | 4 | 4 | 8 | 2 | 2 | 5 |
| ENSG00000283795 | 0 | 0 | 0 | 0 | 0 | 0 |
| ENSG00000228687 | 0 | 0 | 0 | 0 | 0 | 0 |
| ENSG00000226814 | 0 | 0 | 0 | 0 | 0 | 0 |
| ENSG00000116741 | 2 | 3 | 5 | 1 | 3 | 1 |
| ENSG00000223075 | 0 | 0 | 0 | 0 | 0 | 0 |
| ENSG00000234473 | 0 | 0 | 0 | 2 | 2 | 0 |
| ENSG00000116750 | 770 | 457 | 399 | 294 | 608 | 430 |
| ENSG00000238754 | 0 | 0 | 0 | 0 | 0 | 0 |
| ENSG00000116747 | 949 | 521 | 462 | 314 | 679 | 473 |
| ENSG00000023572 | 710 | 323 | 473 | 332 | 433 | 386 |
| ENSG00000134371 | 1588 | 866 | 864 | 531 | 1073 | 868 |
| ENSG00000221680 | 0 | 0 | 0 | 0 | 0 | 0 |
| ENSG00000162630 | 0 | 1 | 0 | 0 | 0 | 0 |
| ENSG00000276735 | 0 | 0 | 0 | 0 | 0 | 0 |
| ENSG00000232077 | 2 | 0 | 1 | 0 | 0 | 0 |
| ENSG00000226640 | 0 | 0 | 0 | 0 | 0 | 0 |
| ENSG00000237011 | 0 | 0 | 0 | 0 | 0 | 0 |
| ENSG00000237213 | 0 | 0 | 0 | 0 | 0 | 0 |
| ENSG00000227240 | 0 | 0 | 0 | 0 | 0 | 0 |
| ENSG00000233057 | 1 | 0 | 0 | 3 | 1 | 0 |
| ENSG00000231714 | 0 | 0 | 0 | 0 | 0 | 0 |
| ENSG00000251813 | 0 | 0 | 0 | 0 | 0 | 0 |
| ENSG00000228167 | 0 | 0 | 0 | 0 | 0 | 0 |
| ENSG00000285718 | 0 | 0 | 0 | 0 | 0 | 0 |
| ENSG00000227421 | 0 | 0 | 0 | 0 | 0 | 0 |
| ENSG00000162687 | 0 | 0 | 0 | 0 | 0 | 0 |
| ENSG00000265986 | 0 | 0 | 0 | 0 | 0 | 0 |
| ENSG00000000971 | 0 | 0 | 0 | 0 | 0 | 0 |
| ENSG00000116785 | 0 | 0 | 0 | 0 | 0 | 0 |

| | | | | | | |
|---|---|---|---|---|---|---|
| ENSG00000244414 | 0 | 0 | 0 | 0 | 0 | 0 |
| ENSG00000285986 | 0 | 0 | 0 | 0 | 0 | 0 |
| ENSG00000134365 | 0 | 0 | 0 | 0 | 0 | 0 |
| ENSG00000080910 | 0 | 0 | 0 | 0 | 0 | 0 |
| ENSG00000273844 | 0 | 0 | 0 | 0 | 0 | 0 |
| ENSG00000134389 | 4 | 0 | 0 | 0 | 0 | 0 |
| ENSG00000143278 | 0 | 0 | 0 | 0 | 0 | 0 |
| ENSG00000066279 | 100 | 52 | 64 | 28 | 238 | 95 |
| ENSG00000235748 | 0 | 0 | 1 | 0 | 1 | 0 |
| ENSG00000177888 | 294 | 213 | 139 | 84 | 460 | 153 |
| ENSG00000237861 | 0 | 0 | 0 | 0 | 1 | 0 |
| ENSG00000134376 | 0 | 0 | 0 | 0 | 0 | 0 |
| ENSG00000221879 | 0 | 0 | 0 | 0 | 0 | 0 |
| ENSG00000230260 | 0 | 0 | 0 | 0 | 0 | 0 |
| ENSG00000213047 | 305 | 185 | 140 | 86 | 298 | 127 |
| ENSG00000207139 | 0 | 0 | 0 | 0 | 0 | 0 |
| ENSG00000213046 | 0 | 0 | 0 | 0 | 0 | 1 |
| ENSG00000235582 | 2 | 1 | 1 | 1 | 0 | 0 |
| ENSG00000233040 | 0 | 0 | 0 | 0 | 0 | 0 |
| ENSG00000224901 | 0 | 0 | 1 | 0 | 0 | 0 |
| ENSG00000203724 | 105 | 64 | 73 | 53 | 67 | 45 |
| ENSG00000143355 | 0 | 1 | 1 | 0 | 0 | 0 |
| ENSG00000151414 | 1282 | 701 | 703 | 404 | 817 | 606 |
| ENSG00000232824 | 0 | 0 | 0 | 0 | 0 | 0 |
| ENSG00000151418 | 0 | 0 | 0 | 0 | 0 | 0 |
| ENSG00000223881 | 0 | 0 | 0 | 0 | 0 | 0 |
| ENSG00000081237 | 0 | 0 | 0 | 2 | 1 | 0 |
| ENSG00000261573 | 0 | 0 | 0 | 0 | 0 | 0 |
| ENSG00000236278 | 0 | 0 | 0 | 0 | 0 | 0 |
| ENSG00000229989 | 0 | 4 | 2 | 5 | 6 | 3 |
| ENSG00000207975 | 1 | 0 | 1 | 0 | 1 | 0 |
| ENSG00000207759 | 0 | 0 | 0 | 0 | 0 | 0 |
| ENSG00000227747 | 0 | 0 | 0 | 0 | 0 | 0 |
| ENSG00000225172 | 0 | 0 | 0 | 0 | 0 | 0 |
| ENSG00000233410 | 0 | 0 | 0 | 0 | 0 | 0 |
| ENSG00000235492 | 0 | 0 | 0 | 0 | 0 | 0 |
| ENSG00000231718 | 0 | 0 | 0 | 0 | 0 | 0 |
| ENSG00000228190 | 0 | 0 | 0 | 0 | 0 | 0 |
| ENSG00000236468 | 2 | 1 | 0 | 0 | 0 | 1 |
| ENSG00000231984 | 0 | 0 | 0 | 0 | 0 | 0 |
| ENSG00000228530 | 0 | 0 | 0 | 0 | 0 | 0 |
| ENSG00000200139 | 0 | 0 | 0 | 0 | 0 | 0 |
| ENSG00000229747 | 0 | 0 | 0 | 0 | 0 | 0 |
| ENSG00000202491 | 0 | 0 | 0 | 0 | 0 | 0 |
| ENSG00000202329 | 0 | 0 | 0 | 0 | 0 | 0 |
| ENSG00000116833 | 26 | 11 | 10 | 5 | 12 | 8 |
| ENSG00000252860 | 0 | 0 | 0 | 0 | 0 | 0 |
| ENSG00000229220 | 0 | 0 | 0 | 0 | 0 | 0 |
| ENSG00000233827 | 0 | 0 | 0 | 0 | 0 | 0 |
| ENSG00000203721 | 2 | 4 | 2 | 2 | 6 | 5 |

| | | | | | | |
|---|---|---|---|---|---|---|
| ENSG00000273093 | 0 | 0 | 0 | 0 | 0 | 0 |
| ENSG00000213045 | 0 | 0 | 0 | 0 | 0 | 0 |
| ENSG00000230623 | 0 | 0 | 1 | 0 | 0 | 1 |
| ENSG00000162702 | 1765 | 819 | 896 | 624 | 1001 | 792 |
| ENSG00000282849 | 0 | 0 | 0 | 0 | 0 | 0 |
| ENSG00000118193 | 113 | 74 | 51 | 26 | 209 | 81 |
| ENSG00000118197 | 851 | 419 | 567 | 367 | 521 | 445 |
| ENSG00000260088 | 0 | 3 | 6 | 3 | 4 | 3 |
| ENSG00000118200 | 1562 | 830 | 714 | 354 | 1225 | 799 |
| ENSG00000231333 | 0 | 0 | 1 | 0 | 0 | 0 |
| ENSG00000170128 | 0 | 0 | 0 | 0 | 0 | 0 |
| ENSG00000163362 | 192 | 104 | 99 | 78 | 104 | 65 |
| ENSG00000233217 | 1 | 2 | 0 | 0 | 0 | 0 |
| ENSG00000201032 | 0 | 0 | 0 | 0 | 0 | 0 |
| ENSG00000116852 | 40 | 37 | 19 | 30 | 22 | 35 |
| ENSG00000229191 | 0 | 0 | 0 | 0 | 1 | 0 |
| ENSG00000234132 | 0 | 0 | 0 | 0 | 0 | 0 |
| ENSG00000081248 | 0 | 0 | 0 | 0 | 0 | 0 |
| ENSG00000232237 | 22 | 12 | 28 | 10 | 20 | 8 |
| ENSG00000116857 | 4380 | 2116 | 2670 | 2651 | 2223 | 2323 |
| ENSG00000163395 | 6 | 5 | 3 | 6 | 9 | 3 |
| ENSG00000229821 | 0 | 0 | 0 | 0 | 0 | 0 |
| ENSG00000081277 | 357 | 186 | 235 | 228 | 268 | 246 |
| ENSG00000118194 | 50 | 15 | 25 | 26 | 28 | 15 |
| ENSG00000159166 | 2121 | 1076 | 1384 | 1631 | 1475 | 1587 |
| ENSG00000282221 | 12 | 5 | 11 | 8 | 6 | 4 |
| ENSG00000159173 | 21 | 9 | 22 | 13 | 12 | 15 |
| ENSG00000227048 | 0 | 0 | 0 | 0 | 0 | 0 |
| ENSG00000224818 | 0 | 0 | 0 | 0 | 0 | 0 |
| ENSG00000174307 | 2477 | 1314 | 1585 | 1845 | 1637 | 1431 |
| ENSG00000159176 | 10622 | 5163 | 6880 | 5605 | 5212 | 5744 |
| ENSG00000224536 | 84 | 47 | 61 | 51 | 58 | 51 |
| ENSG00000223396 | 131 | 55 | 75 | 75 | 92 | 73 |
| ENSG00000134369 | 101 | 45 | 65 | 52 | 41 | 69 |
| ENSG00000236390 | 0 | 0 | 0 | 0 | 0 | 0 |
| ENSG00000231871 | 2 | 2 | 5 | 0 | 3 | 0 |
| ENSG00000264802 | 0 | 0 | 0 | 0 | 0 | 0 |
| ENSG00000235121 | 22 | 22 | 18 | 9 | 7 | 12 |
| ENSG00000200942 | 3 | 1 | 2 | 0 | 0 | 0 |
| ENSG00000221028 | 0 | 0 | 0 | 0 | 0 | 0 |
| ENSG00000198700 | 1868 | 924 | 1120 | 760 | 1050 | 1006 |
| ENSG00000277681 | 0 | 0 | 0 | 0 | 1 | 1 |
| ENSG00000198892 | 204 | 96 | 146 | 142 | 141 | 128 |
| ENSG00000223774 | 0 | 0 | 0 | 0 | 1 | 1 |
| ENSG00000163431 | 5 | 3 | 4 | 2 | 3 | 6 |
| ENSG00000134375 | 1360 | 686 | 870 | 806 | 852 | 802 |
| ENSG00000206637 | 0 | 0 | 1 | 0 | 0 | 0 |
| ENSG00000238003 | 0 | 0 | 1 | 0 | 0 | 1 |
| ENSG00000176393 | 2844 | 1441 | 1932 | 1777 | 1638 | 1754 |
| ENSG00000234678 | 213 | 136 | 131 | 77 | 105 | 108 |

| | | | | | | |
|---|---|---|---|---|---|---|
| ENSG00000275207 | 1 | 1 | 0 | 1 | 0 | 0 |
| ENSG00000163435 | 5487 | 2788 | 3713 | 2914 | 2528 | 3333 |
| ENSG00000249007 | 9 | 11 | 7 | 4 | 7 | 16 |
| ENSG00000232296 | 1 | 4 | 3 | 1 | 0 | 3 |
| ENSG00000235811 | 3 | 2 | 6 | 1 | 0 | 5 |
| ENSG00000212512 | 2 | 1 | 0 | 1 | 1 | 0 |
| ENSG00000227109 | 5 | 1 | 1 | 0 | 3 | 1 |
| ENSG00000170075 | 688 | 309 | 486 | 389 | 332 | 512 |
| ENSG00000143862 | 1924 | 815 | 1134 | 1079 | 980 | 1019 |
| ENSG00000143851 | 2 | 2 | 6 | 0 | 1 | 0 |
| ENSG00000243323 | 24 | 18 | 36 | 3 | 19 | 16 |
| ENSG00000133067 | 5 | 2 | 8 | 8 | 7 | 15 |
| ENSG00000077152 | 1017 | 541 | 716 | 653 | 673 | 712 |
| ENSG00000077157 | 995 | 484 | 634 | 267 | 488 | 477 |
| ENSG00000229120 | 0 | 0 | 0 | 0 | 0 | 0 |
| ENSG00000272262 | 0 | 0 | 0 | 0 | 0 | 0 |
| ENSG00000232626 | 0 | 4 | 3 | 1 | 1 | 1 |
| ENSG00000236439 | 10 | 15 | 2 | 5 | 4 | 7 |
| ENSG00000253042 | 0 | 0 | 0 | 0 | 0 | 0 |
| ENSG00000143858 | 24 | 3 | 5 | 4 | 7 | 10 |
| ENSG00000226862 | 5 | 2 | 3 | 2 | 3 | 7 |
| ENSG00000225620 | 0 | 0 | 0 | 0 | 0 | 0 |
| ENSG00000117139 | 10123 | 4645 | 5719 | 3422 | 5091 | 4457 |
| ENSG00000235449 | 2 | 1 | 1 | 0 | 3 | 0 |
| ENSG00000226148 | 0 | 0 | 0 | 1 | 2 | 0 |
| ENSG00000260021 | 0 | 1 | 0 | 0 | 0 | 0 |
| ENSG00000228288 | 174 | 81 | 102 | 85 | 77 | 82 |
| ENSG00000184774 | 38 | 23 | 21 | 12 | 23 | 21 |
| ENSG00000214796 | 772 | 402 | 452 | 388 | 373 | 439 |
| ENSG00000234996 | 74 | 43 | 48 | 32 | 34 | 43 |
| ENSG00000183155 | 1955 | 981 | 1165 | 951 | 1053 | 1057 |
| ENSG00000117153 | 2996 | 1329 | 1776 | 1222 | 1511 | 1510 |
| ENSG00000230280 | 19 | 23 | 27 | 6 | 23 | 14 |
| ENSG00000199471 | 3 | 3 | 5 | 0 | 1 | 0 |
| ENSG00000159346 | 8336 | 4263 | 5082 | 4637 | 4531 | 4631 |
| ENSG00000159348 | 4657 | 2213 | 2861 | 1974 | 2069 | 1933 |
| ENSG00000234761 | 0 | 0 | 0 | 0 | 0 | 0 |
| ENSG00000231547 | 0 | 0 | 0 | 0 | 0 | 0 |
| ENSG00000163444 | 3569 | 1883 | 2105 | 2164 | 2103 | 2127 |
| ENSG00000143847 | 42 | 13 | 29 | 12 | 24 | 29 |
| ENSG00000122180 | 0 | 0 | 1 | 0 | 0 | 0 |
| ENSG00000163485 | 16 | 13 | 19 | 16 | 18 | 15 |
| ENSG00000234775 | 0 | 0 | 0 | 0 | 0 | 0 |
| ENSG00000224671 | 0 | 0 | 0 | 0 | 0 | 0 |
| ENSG00000133055 | 0 | 1 | 6 | 5 | 1 | 0 |
| ENSG00000133048 | 0 | 0 | 0 | 3 | 0 | 1 |
| ENSG00000133063 | 6 | 0 | 0 | 0 | 1 | 0 |
| ENSG00000236523 | 0 | 0 | 0 | 0 | 1 | 0 |
| ENSG00000231507 | 12 | 6 | 4 | 1 | 4 | 4 |
| ENSG00000233791 | 17 | 10 | 9 | 1 | 3 | 4 |

| | | | | | | |
|---|---|---|---|---|---|---|
| ENSG00000159388 | 4236 | 2025 | 2336 | 1439 | 1473 | 1780 |
| ENSG00000202300 | 0 | 0 | 0 | 0 | 0 | 0 |
| ENSG00000122176 | 0 | 0 | 0 | 0 | 0 | 0 |
| ENSG00000229652 | 0 | 0 | 0 | 0 | 0 | 0 |
| ENSG00000271588 | 0 | 0 | 0 | 0 | 0 | 0 |
| ENSG00000188783 | 0 | 0 | 1 | 0 | 0 | 2 |
| ENSG00000188770 | 0 | 0 | 0 | 0 | 0 | 0 |
| ENSG00000058668 | 1028 | 391 | 576 | 263 | 429 | 460 |
| ENSG00000236035 | 0 | 0 | 0 | 0 | 0 | 0 |
| ENSG00000221643 | 0 | 0 | 1 | 0 | 0 | 1 |
| ENSG00000122188 | 0 | 0 | 0 | 0 | 0 | 1 |
| ENSG00000058673 | 15 | 9 | 12 | 5 | 11 | 12 |
| ENSG00000257315 | 23 | 15 | 19 | 3 | 18 | 9 |
| ENSG00000227417 | 0 | 2 | 0 | 0 | 0 | 0 |
| ENSG00000223505 | 0 | 1 | 0 | 0 | 0 | 3 |
| ENSG00000182004 | 3292 | 1695 | 1940 | 2103 | 2140 | 2068 |
| ENSG00000236430 | 0 | 1 | 0 | 0 | 0 | 0 |
| ENSG00000232917 | 0 | 0 | 0 | 0 | 1 | 0 |
| ENSG00000237379 | 0 | 0 | 0 | 0 | 0 | 0 |
| ENSG00000176754 | 0 | 0 | 0 | 1 | 1 | 0 |
| ENSG00000143842 | 2228 | 946 | 1266 | 965 | 1005 | 1021 |
| ENSG00000261065 | 0 | 0 | 0 | 0 | 0 | 0 |
| ENSG00000143845 | 422 | 236 | 337 | 232 | 220 | 223 |
| ENSG00000230550 | 1 | 0 | 0 | 0 | 4 | 1 |
| ENSG00000143839 | 1 | 2 | 0 | 5 | 0 | 0 |
| ENSG00000225522 | 0 | 0 | 0 | 0 | 0 | 0 |
| ENSG00000170498 | 12 | 3 | 6 | 8 | 12 | 3 |
| ENSG00000174567 | 530 | 265 | 324 | 295 | 303 | 297 |
| ENSG00000143850 | 3415 | 1481 | 2277 | 1408 | 1482 | 1700 |
| ENSG00000231691 | 5 | 4 | 9 | 5 | 6 | 4 |
| ENSG00000219133 | 2 | 1 | 0 | 1 | 0 | 1 |
| ENSG00000280924 | 4 | 6 | 3 | 9 | 6 | 4 |
| ENSG00000226330 | 22 | 6 | 16 | 12 | 17 | 12 |
| ENSG00000237848 | 2 | 0 | 0 | 0 | 0 | 1 |
| ENSG00000158615 | 5278 | 2434 | 3485 | 2455 | 3403 | 3573 |
| ENSG00000133056 | 2495 | 1187 | 1475 | 1165 | 1522 | 1470 |
| ENSG00000198625 | 1730 | 829 | 1103 | 550 | 785 | 868 |
| ENSG00000236779 | 2 | 1 | 0 | 0 | 0 | 0 |
| ENSG00000200408 | 0 | 0 | 0 | 0 | 0 | 0 |
| ENSG00000240710 | 2 | 0 | 0 | 0 | 0 | 2 |
| ENSG00000170382 | 86 | 40 | 66 | 48 | 45 | 45 |
| ENSG00000240219 | 11 | 10 | 12 | 4 | 4 | 2 |
| ENSG00000228153 | 0 | 0 | 2 | 0 | 0 | 0 |
| ENSG00000252650 | 0 | 0 | 0 | 0 | 0 | 0 |
| ENSG00000251861 | 0 | 0 | 0 | 0 | 0 | 0 |
| ENSG00000163531 | 10 | 12 | 23 | 2 | 9 | 3 |
| ENSG00000229657 | 0 | 0 | 0 | 0 | 0 | 0 |
| ENSG00000184144 | 0 | 2 | 2 | 0 | 0 | 2 |
| ENSG00000174529 | 65 | 26 | 34 | 35 | 46 | 39 |
| ENSG00000117222 | 1293 | 638 | 836 | 450 | 728 | 681 |

| | | | | | | |
|---|---|---|---|---|---|---|
| ENSG00000271580 | 6 | 5 | 7 | 1 | 7 | 6 |
| ENSG00000236108 | 2 | 0 | 0 | 2 | 0 | 0 |
| ENSG00000133059 | 1782 | 849 | 1162 | 530 | 916 | 855 |
| ENSG00000213041 | 0 | 4 | 0 | 0 | 0 | 0 |
| ENSG00000133069 | 28 | 7 | 10 | 9 | 17 | 15 |
| ENSG00000225063 | 0 | 0 | 0 | 0 | 0 | 0 |
| ENSG00000163545 | 154 | 64 | 81 | 82 | 55 | 53 |
| ENSG00000162873 | 2 | 1 | 0 | 0 | 1 | 1 |
| ENSG00000235363 | 263 | 142 | 186 | 178 | 159 | 153 |
| ENSG00000226235 | 0 | 0 | 1 | 0 | 0 | 0 |
| ENSG00000186007 | 1 | 0 | 0 | 0 | 1 | 0 |
| ENSG00000281406 | 4 | 0 | 0 | 0 | 0 | 2 |
| ENSG00000199059 | 0 | 0 | 0 | 0 | 0 | 0 |
| ENSG00000224717 | 0 | 0 | 0 | 0 | 0 | 0 |
| ENSG00000117266 | 885 | 348 | 520 | 474 | 393 | 530 |
| ENSG00000253097 | 0 | 0 | 0 | 0 | 0 | 0 |
| ENSG00000174514 | 25 | 10 | 15 | 10 | 11 | 10 |
| ENSG00000206762 | 0 | 0 | 1 | 1 | 1 | 0 |
| ENSG00000158711 | 2091 | 1012 | 976 | 529 | 1395 | 996 |
| ENSG00000236942 | 1 | 1 | 0 | 0 | 1 | 0 |
| ENSG00000158715 | 286 | 119 | 151 | 221 | 253 | 233 |
| ENSG00000275213 | 0 | 1 | 0 | 0 | 0 | 0 |
| ENSG00000069275 | 24977 | 12150 | 14721 | 9045 | 13830 | 12631 |
| ENSG00000201944 | 0 | 3 | 0 | 0 | 0 | 0 |
| ENSG00000117280 | 2342 | 960 | 1349 | 643 | 1014 | 1025 |
| ENSG00000285521 | 0 | 0 | 0 | 0 | 0 | 0 |
| ENSG00000133065 | 2156 | 912 | 1299 | 760 | 981 | 1079 |
| ENSG00000162877 | 0 | 5 | 3 | 0 | 0 | 0 |
| ENSG00000174502 | 2 | 0 | 2 | 0 | 0 | 0 |
| ENSG00000227687 | 0 | 0 | 0 | 0 | 0 | 0 |
| ENSG00000276600 | 24 | 16 | 15 | 15 | 21 | 13 |
| ENSG00000196188 | 26 | 21 | 13 | 5 | 24 | 19 |
| ENSG00000285417 | 0 | 0 | 0 | 0 | 0 | 0 |
| ENSG00000263961 | 0 | 0 | 0 | 0 | 0 | 0 |
| ENSG00000252692 | 0 | 0 | 0 | 0 | 0 | 0 |
| ENSG00000198049 | 0 | 0 | 0 | 0 | 0 | 0 |
| ENSG00000226780 | 0 | 0 | 0 | 0 | 0 | 0 |
| ENSG00000225483 | 0 | 0 | 0 | 0 | 0 | 0 |
| ENSG00000236889 | 0 | 0 | 0 | 0 | 0 | 0 |
| ENSG00000196550 | 76 | 50 | 50 | 57 | 57 | 73 |
| ENSG00000266028 | 1637 | 754 | 1012 | 690 | 880 | 912 |
| ENSG00000229509 | 0 | 0 | 0 | 0 | 0 | 0 |
| ENSG00000263528 | 391 | 130 | 191 | 160 | 187 | 181 |
| ENSG00000278465 | 0 | 0 | 0 | 0 | 0 | 0 |
| ENSG00000162888 | 14 | 3 | 2 | 1 | 3 | 4 |
| ENSG00000261000 | 4 | 0 | 0 | 0 | 0 | 0 |
| ENSG00000266094 | 148 | 74 | 101 | 62 | 94 | 110 |
| ENSG00000234981 | 0 | 0 | 2 | 0 | 1 | 0 |
| ENSG00000279946 | 0 | 0 | 0 | 0 | 0 | 0 |
| ENSG00000143486 | 2511 | 1137 | 1553 | 1514 | 1465 | 1485 |

| | | | | | | |
|---|---|---|---|---|---|---|
| ENSG00000237605 | 2 | 4 | 1 | 0 | 3 | 3 |
| ENSG00000143479 | 61 | 26 | 38 | 38 | 30 | 33 |
| ENSG00000162889 | 6348 | 3222 | 3564 | 3470 | 3373 | 3350 |
| ENSG00000224114 | 2 | 1 | 4 | 1 | 1 | 0 |
| ENSG00000199349 | 2 | 0 | 0 | 0 | 0 | 0 |
| ENSG00000136634 | 0 | 0 | 0 | 0 | 0 | 1 |
| ENSG00000142224 | 1 | 0 | 0 | 0 | 0 | 0 |
| ENSG00000162891 | 2 | 0 | 0 | 0 | 0 | 0 |
| ENSG00000162892 | 0 | 1 | 1 | 0 | 1 | 0 |
| ENSG00000162894 | 16 | 9 | 9 | 12 | 3 | 6 |
| ENSG00000271680 | 0 | 0 | 0 | 0 | 0 | 0 |
| ENSG00000226945 | 0 | 0 | 0 | 0 | 1 | 0 |
| ENSG00000162896 | 4 | 0 | 1 | 0 | 0 | 0 |
| ENSG00000162897 | 0 | 0 | 0 | 0 | 0 | 1 |
| ENSG00000182795 | 22 | 9 | 12 | 12 | 17 | 19 |
| ENSG00000123836 | 723 | 351 | 520 | 276 | 398 | 400 |
| ENSG00000180667 | 15 | 7 | 13 | 13 | 9 | 5 |
| ENSG00000123843 | 0 | 0 | 0 | 0 | 0 | 0 |
| ENSG00000123838 | 0 | 0 | 1 | 0 | 0 | 0 |
| ENSG00000275392 | 0 | 0 | 0 | 2 | 0 | 0 |
| ENSG00000226565 | 0 | 0 | 0 | 0 | 0 | 0 |
| ENSG00000224462 | 0 | 0 | 0 | 0 | 0 | 0 |
| ENSG00000243636 | 0 | 0 | 0 | 0 | 0 | 0 |
| ENSG00000232621 | 0 | 0 | 0 | 0 | 0 | 0 |
| ENSG00000237074 | 0 | 0 | 0 | 0 | 0 | 0 |
| ENSG00000196352 | 3326 | 1675 | 2022 | 1372 | 2014 | 1755 |
| ENSG00000283044 | 0 | 0 | 0 | 0 | 0 | 0 |
| ENSG00000117322 | 2 | 4 | 5 | 0 | 1 | 0 |
| ENSG00000203710 | 0 | 0 | 0 | 0 | 1 | 0 |
| ENSG00000236911 | 0 | 0 | 0 | 0 | 0 | 0 |
| ENSG00000197721 | 0 | 1 | 0 | 0 | 2 | 0 |
| ENSG00000244703 | 2 | 0 | 0 | 0 | 0 | 0 |
| ENSG00000226289 | 0 | 0 | 0 | 0 | 0 | 0 |
| ENSG00000274059 | 0 | 0 | 0 | 0 | 0 | 0 |
| ENSG00000285239 | 0 | 0 | 0 | 0 | 0 | 0 |
| ENSG00000117335 | 5912 | 2942 | 3266 | 1953 | 3526 | 2925 |
| ENSG00000234219 | 0 | 0 | 0 | 0 | 0 | 0 |
| ENSG00000203709 | 143 | 82 | 140 | 52 | 110 | 84 |
| ENSG00000284214 | 0 | 0 | 0 | 0 | 0 | 0 |
| ENSG00000284203 | 0 | 0 | 0 | 0 | 0 | 0 |
| ENSG00000174059 | 2 | 0 | 0 | 0 | 0 | 0 |
| ENSG00000285719 | 0 | 0 | 0 | 0 | 0 | 0 |
| ENSG00000284237 | 2 | 2 | 9 | 2 | 2 | 3 |
| ENSG00000076356 | 54 | 19 | 25 | 30 | 34 | 25 |
| ENSG00000226843 | 0 | 0 | 0 | 0 | 0 | 0 |
| ENSG00000261453 | 0 | 0 | 0 | 0 | 0 | 0 |
| ENSG00000259815 | 0 | 0 | 0 | 0 | 0 | 0 |
| ENSG00000227887 | 0 | 0 | 0 | 0 | 0 | 0 |
| ENSG00000232812 | 0 | 0 | 0 | 0 | 0 | 0 |
| ENSG00000236950 | 0 | 0 | 0 | 0 | 0 | 0 |

| | | | | | | |
|---|---|---|---|---|---|---|
| ENSG00000278684 | 0 | 0 | 0 | 0 | 0 | 0 |
| ENSG00000232537 | 0 | 0 | 0 | 0 | 0 | 0 |
| ENSG00000228081 | 0 | 0 | 0 | 0 | 0 | 0 |
| ENSG00000224540 | 0 | 0 | 0 | 0 | 0 | 0 |
| ENSG00000225712 | 0 | 0 | 0 | 1 | 0 | 0 |
| ENSG00000227940 | 0 | 0 | 0 | 0 | 0 | 0 |
| ENSG00000231648 | 0 | 0 | 0 | 0 | 0 | 0 |
| ENSG00000230937 | 8 | 7 | 3 | 2 | 8 | 3 |
| ENSG00000284485 | 0 | 0 | 0 | 0 | 0 | 0 |
| ENSG00000224260 | 0 | 0 | 0 | 0 | 0 | 0 |
| ENSG00000008118 | 8 | 4 | 7 | 1 | 6 | 3 |
| ENSG00000196878 | 314 | 183 | 187 | 263 | 358 | 226 |
| ENSG00000283876 | 0 | 0 | 0 | 0 | 0 | 0 |
| ENSG00000227591 | 0 | 0 | 0 | 0 | 0 | 0 |
| ENSG00000123689 | 0 | 0 | 0 | 0 | 0 | 0 |
| ENSG00000117594 | 0 | 0 | 0 | 0 | 0 | 0 |
| ENSG00000236136 | 0 | 0 | 0 | 0 | 0 | 0 |
| ENSG00000009790 | 0 | 0 | 2 | 2 | 2 | 1 |
| ENSG00000162757 | 136 | 74 | 71 | 73 | 63 | 78 |
| ENSG00000117595 | 1327 | 650 | 867 | 555 | 714 | 627 |
| ENSG00000117597 | 398 | 212 | 255 | 240 | 281 | 291 |
| ENSG00000143469 | 0 | 0 | 0 | 0 | 0 | 0 |
| ENSG00000203706 | 37 | 18 | 30 | 19 | 33 | 23 |
| ENSG00000082497 | 299 | 134 | 187 | 91 | 168 | 132 |
| ENSG00000228110 | 4 | 3 | 1 | 0 | 1 | 0 |
| ENSG00000233455 | 2 | 0 | 1 | 0 | 0 | 0 |
| ENSG00000054392 | 290 | 136 | 165 | 135 | 139 | 142 |
| ENSG00000200972 | 0 | 0 | 0 | 0 | 0 | 0 |
| ENSG00000270936 | 0 | 0 | 0 | 0 | 0 | 0 |
| ENSG00000143473 | 8 | 3 | 1 | 1 | 2 | 2 |
| ENSG00000279333 | 0 | 0 | 0 | 0 | 0 | 0 |
| ENSG00000284299 | 0 | 0 | 0 | 0 | 0 | 0 |
| ENSG00000224668 | 0 | 0 | 0 | 0 | 0 | 0 |
| ENSG00000283952 | 0 | 0 | 0 | 0 | 0 | 0 |
| ENSG00000284376 | 0 | 1 | 0 | 0 | 0 | 0 |
| ENSG00000234233 | 0 | 0 | 0 | 0 | 0 | 0 |
| ENSG00000234004 | 0 | 0 | 1 | 0 | 0 | 2 |
| ENSG00000226986 | 0 | 0 | 0 | 0 | 0 | 2 |
| ENSG00000117625 | 654 | 312 | 358 | 191 | 322 | 247 |
| ENSG00000082512 | 649 | 314 | 353 | 177 | 367 | 232 |
| ENSG00000153363 | 328 | 211 | 243 | 256 | 213 | 217 |
| ENSG00000236809 | 31 | 5 | 20 | 15 | 21 | 14 |
| ENSG00000238137 | 0 | 0 | 0 | 1 | 0 | 1 |
| ENSG00000198570 | 0 | 0 | 0 | 1 | 0 | 0 |
| ENSG00000223649 | 0 | 0 | 0 | 0 | 0 | 0 |
| ENSG00000170385 | 862 | 383 | 545 | 389 | 576 | 506 |
| ENSG00000261314 | 0 | 0 | 0 | 0 | 0 | 0 |
| ENSG00000228792 | 0 | 0 | 0 | 0 | 0 | 0 |
| ENSG00000227764 | 0 | 0 | 0 | 0 | 0 | 0 |
| ENSG00000117650 | 135 | 83 | 104 | 77 | 150 | 121 |

| | | | | | | |
|---|---|---|---|---|---|---|
| ENSG00000224763 | 0 | 0 | 0 | 0 | 0 | 0 |
| ENSG00000231057 | 2 | 0 | 0 | 0 | 0 | 0 |
| ENSG00000226868 | 0 | 0 | 0 | 0 | 0 | 0 |
| ENSG00000123684 | 1732 | 778 | 1030 | 532 | 944 | 870 |
| ENSG00000241395 | 0 | 1 | 0 | 0 | 0 | 0 |
| ENSG00000212205 | 4 | 0 | 0 | 0 | 0 | 0 |
| ENSG00000229258 | 1 | 0 | 0 | 0 | 0 | 0 |
| ENSG00000233626 | 0 | 0 | 0 | 2 | 0 | 0 |
| ENSG00000143493 | 386 | 208 | 175 | 133 | 256 | 170 |
| ENSG00000212187 | 0 | 0 | 0 | 0 | 0 | 0 |
| ENSG00000143476 | 190 | 111 | 126 | 66 | 68 | 99 |
| ENSG00000220749 | 21 | 8 | 12 | 11 | 15 | 14 |
| ENSG00000264358 | 0 | 0 | 0 | 0 | 0 | 0 |
| ENSG00000252879 | 0 | 0 | 0 | 0 | 0 | 0 |
| ENSG00000229983 | 0 | 0 | 0 | 0 | 0 | 0 |
| ENSG00000226251 | 0 | 0 | 0 | 0 | 0 | 0 |
| ENSG00000066027 | 1444 | 678 | 831 | 594 | 730 | 738 |
| ENSG00000230063 | 0 | 0 | 1 | 0 | 0 | 0 |
| ENSG00000234915 | 0 | 0 | 0 | 0 | 0 | 0 |
| ENSG00000225338 | 0 | 2 | 0 | 0 | 0 | 0 |
| ENSG00000201544 | 0 | 0 | 0 | 0 | 0 | 1 |
| ENSG00000229832 | 4 | 1 | 0 | 1 | 0 | 1 |
| ENSG00000065600 | 193 | 96 | 116 | 110 | 119 | 134 |
| ENSG00000117691 | 768 | 403 | 495 | 521 | 475 | 406 |
| ENSG00000224535 | 0 | 0 | 0 | 0 | 1 | 0 |
| ENSG00000228067 | 0 | 0 | 0 | 0 | 0 | 0 |
| ENSG00000260805 | 44 | 26 | 17 | 13 | 28 | 27 |
| ENSG00000162772 | 38 | 25 | 32 | 39 | 26 | 26 |
| ENSG00000235862 | 0 | 0 | 0 | 0 | 0 | 0 |
| ENSG00000162771 | 0 | 0 | 0 | 0 | 0 | 0 |
| ENSG00000226036 | 4 | 0 | 0 | 0 | 0 | 0 |
| ENSG00000237980 | 0 | 0 | 0 | 0 | 0 | 0 |
| ENSG00000123685 | 0 | 0 | 0 | 0 | 0 | 0 |
| ENSG00000117697 | 395 | 202 | 250 | 216 | 216 | 223 |
| ENSG00000203705 | 190 | 92 | 98 | 110 | 110 | 108 |
| ENSG00000236905 | 0 | 0 | 0 | 0 | 0 | 0 |
| ENSG00000185523 | 0 | 3 | 1 | 1 | 3 | 0 |
| ENSG00000198468 | 29 | 18 | 24 | 13 | 21 | 29 |
| ENSG00000235182 | 0 | 0 | 0 | 0 | 0 | 0 |
| ENSG00000236317 | 0 | 0 | 1 | 0 | 0 | 0 |
| ENSG00000162769 | 316 | 168 | 230 | 148 | 241 | 217 |
| ENSG00000282718 | 0 | 0 | 0 | 0 | 0 | 0 |
| ENSG00000143494 | 124 | 48 | 66 | 27 | 47 | 68 |
| ENSG00000174606 | 517 | 234 | 257 | 189 | 271 | 230 |
| ENSG00000136643 | 537 | 241 | 275 | 214 | 312 | 237 |
| ENSG00000228646 | 0 | 0 | 0 | 0 | 0 | 0 |
| ENSG00000225233 | 0 | 0 | 0 | 0 | 0 | 0 |
| ENSG00000228255 | 0 | 0 | 0 | 0 | 0 | 0 |
| ENSG00000230461 | 0 | 0 | 0 | 0 | 0 | 0 |
| ENSG00000281664 | 0 | 0 | 0 | 0 | 0 | 0 |

| | | | | | | |
|---|---|---|---|---|---|---|
| ENSG00000117707 | 0 | 0 | 0 | 0 | 0 | 0 |
| ENSG00000274895 | 0 | 0 | 0 | 0 | 0 | 0 |
| ENSG00000272167 | 0 | 0 | 0 | 0 | 0 | 0 |
| ENSG00000283518 | 0 | 0 | 0 | 0 | 0 | 0 |
| ENSG00000143499 | 592 | 319 | 348 | 309 | 352 | 304 |
| ENSG00000228470 | 0 | 0 | 0 | 1 | 2 | 0 |
| ENSG00000152104 | 322 | 159 | 206 | 88 | 151 | 133 |
| ENSG00000213036 | 2 | 1 | 0 | 2 | 0 | 2 |
| ENSG00000215819 | 0 | 0 | 0 | 0 | 0 | 0 |
| ENSG00000117724 | 48 | 24 | 38 | 7 | 117 | 87 |
| ENSG00000250536 | 0 | 0 | 0 | 0 | 1 | 0 |
| ENSG00000224584 | 0 | 0 | 0 | 0 | 0 | 0 |
| ENSG00000185221 | 0 | 0 | 0 | 0 | 0 | 0 |
| ENSG00000223869 | 0 | 0 | 0 | 0 | 0 | 0 |
| ENSG00000082482 | 144 | 65 | 110 | 115 | 140 | 184 |
| ENSG00000232809 | 0 | 0 | 0 | 0 | 1 | 0 |
| ENSG00000282265 | 0 | 0 | 0 | 0 | 1 | 0 |
| ENSG00000136636 | 3214 | 1678 | 2102 | 1077 | 1855 | 1762 |
| ENSG00000042781 | 2 | 0 | 0 | 0 | 1 | 1 |
| ENSG00000202498 | 0 | 0 | 0 | 0 | 0 | 0 |
| ENSG00000229242 | 0 | 0 | 0 | 0 | 0 | 0 |
| ENSG00000233620 | 0 | 0 | 0 | 0 | 0 | 0 |
| ENSG00000236292 | 0 | 0 | 0 | 0 | 0 | 0 |
| ENSG00000226013 | 0 | 0 | 0 | 0 | 0 | 0 |
| ENSG00000196482 | 20 | 13 | 10 | 11 | 30 | 24 |
| ENSG00000092978 | 1635 | 796 | 1028 | 486 | 721 | 662 |
| ENSG00000162814 | 77 | 55 | 80 | 33 | 37 | 42 |
| ENSG00000234070 | 0 | 0 | 1 | 0 | 0 | 0 |
| ENSG00000228247 | 0 | 0 | 0 | 0 | 0 | 0 |
| ENSG00000231814 | 0 | 0 | 0 | 0 | 0 | 0 |
| ENSG00000230714 | 0 | 0 | 0 | 0 | 0 | 0 |
| ENSG00000232100 | 0 | 0 | 0 | 0 | 0 | 0 |
| ENSG00000201493 | 0 | 0 | 0 | 0 | 0 | 0 |
| ENSG00000067533 | 645 | 394 | 417 | 321 | 544 | 455 |
| ENSG00000229016 | 0 | 0 | 0 | 0 | 0 | 0 |
| ENSG00000223375 | 0 | 0 | 0 | 0 | 0 | 0 |
| ENSG00000232480 | 16 | 11 | 20 | 9 | 15 | 16 |
| ENSG00000092969 | 370 | 221 | 256 | 99 | 120 | 123 |
| ENSG00000281453 | 0 | 0 | 0 | 0 | 0 | 0 |
| ENSG00000228208 | 0 | 1 | 0 | 0 | 0 | 0 |
| ENSG00000212610 | 0 | 0 | 0 | 0 | 0 | 0 |
| ENSG00000226693 | 0 | 0 | 0 | 0 | 0 | 0 |
| ENSG00000225561 | 0 | 0 | 0 | 0 | 0 | 0 |
| ENSG00000228063 | 0 | 0 | 0 | 0 | 0 | 0 |
| ENSG00000143353 | 446 | 203 | 289 | 285 | 292 | 283 |
| ENSG00000225920 | 3 | 1 | 2 | 2 | 4 | 5 |
| ENSG00000223842 | 4 | 1 | 1 | 1 | 1 | 1 |
| ENSG00000277007 | 6 | 0 | 4 | 0 | 14 | 2 |
| ENSG00000228536 | 0 | 0 | 0 | 0 | 0 | 0 |
| ENSG00000238232 | 0 | 0 | 0 | 0 | 0 | 0 |

| | | | | | | |
|---|---|---|---|---|---|---|
| ENSG00000215817 | 2 | 0 | 2 | 1 | 2 | 1 |
| ENSG00000196660 | 0 | 0 | 0 | 0 | 0 | 3 |
| ENSG00000252086 | 0 | 0 | 0 | 0 | 0 | 0 |
| ENSG00000221673 | 0 | 0 | 0 | 0 | 0 | 0 |
| ENSG00000136628 | 3508 | 2101 | 1933 | 1218 | 2812 | 1861 |
| ENSG00000162813 | 1778 | 927 | 1055 | 928 | 1001 | 1001 |
| ENSG00000067704 | 4263 | 2238 | 2685 | 1795 | 2525 | 2309 |
| ENSG00000207590 | 0 | 0 | 0 | 0 | 0 | 0 |
| ENSG00000207624 | 0 | 1 | 0 | 0 | 0 | 0 |
| ENSG00000232134 | 4 | 2 | 1 | 2 | 2 | 1 |
| ENSG00000118873 | 879 | 444 | 474 | 238 | 670 | 402 |
| ENSG00000281696 | 0 | 0 | 0 | 0 | 0 | 0 |
| ENSG00000222370 | 0 | 0 | 0 | 0 | 0 | 0 |
| ENSG00000270585 | 0 | 1 | 1 | 0 | 0 | 0 |
| ENSG00000270708 | 0 | 0 | 0 | 0 | 0 | 0 |
| ENSG00000218283 | 626 | 312 | 368 | 265 | 366 | 302 |
| ENSG00000213033 | 1 | 0 | 0 | 0 | 0 | 1 |
| ENSG00000230318 | 0 | 0 | 0 | 0 | 0 | 0 |
| ENSG00000232883 | 0 | 0 | 0 | 1 | 0 | 0 |
| ENSG00000226927 | 0 | 0 | 0 | 0 | 0 | 0 |
| ENSG00000260505 | 0 | 0 | 0 | 0 | 0 | 0 |
| ENSG00000235163 | 0 | 0 | 0 | 0 | 0 | 0 |
| ENSG00000234863 | 0 | 0 | 0 | 0 | 0 | 0 |
| ENSG00000226226 | 0 | 0 | 0 | 0 | 0 | 0 |
| ENSG00000231529 | 0 | 0 | 0 | 0 | 0 | 0 |
| ENSG00000116141 | 10 | 8 | 3 | 3 | 5 | 0 |
| ENSG00000243872 | 0 | 0 | 0 | 0 | 0 | 0 |
| ENSG00000233012 | 0 | 0 | 0 | 0 | 0 | 0 |
| ENSG00000162817 | 258 | 148 | 145 | 113 | 152 | 106 |
| ENSG00000117791 | 131 | 70 | 88 | 83 | 76 | 64 |
| ENSG00000186205 | 831 | 406 | 395 | 346 | 411 | 367 |
| ENSG00000221571 | 1 | 0 | 0 | 0 | 0 | 0 |
| ENSG00000276642 | 0 | 0 | 0 | 0 | 0 | 0 |
| ENSG00000272823 | 0 | 0 | 0 | 0 | 0 | 0 |
| ENSG00000238078 | 0 | 0 | 0 | 0 | 0 | 0 |
| ENSG00000257551 | 0 | 0 | 0 | 0 | 0 | 0 |
| ENSG00000136630 | 0 | 0 | 0 | 0 | 0 | 0 |
| ENSG00000226211 | 0 | 0 | 0 | 0 | 0 | 0 |
| ENSG00000234754 | 0 | 0 | 0 | 0 | 0 | 0 |
| ENSG00000232436 | 0 | 0 | 0 | 0 | 0 | 0 |
| ENSG00000227585 | 0 | 0 | 0 | 0 | 0 | 0 |
| ENSG00000143507 | 252 | 98 | 87 | 104 | 128 | 99 |
| ENSG00000227925 | 0 | 0 | 0 | 0 | 0 | 0 |
| ENSG00000200033 | 0 | 0 | 0 | 0 | 0 | 0 |
| ENSG00000238042 | 0 | 0 | 0 | 0 | 0 | 0 |
| ENSG00000228437 | 0 | 0 | 0 | 0 | 0 | 0 |
| ENSG00000232679 | 0 | 0 | 0 | 0 | 0 | 0 |
| ENSG00000236230 | 0 | 0 | 0 | 0 | 0 | 0 |
| ENSG00000227613 | 0 | 0 | 0 | 0 | 0 | 0 |
| ENSG00000226643 | 0 | 0 | 0 | 0 | 0 | 0 |

| | | | | | | |
|---|---|---|---|---|---|---|
| ENSG00000234419 | 0 | 0 | 0 | 0 | 0 | 0 |
| ENSG00000276997 | 2 | 7 | 8 | 6 | 3 | 3 |
| ENSG00000222399 | 0 | 0 | 0 | 0 | 0 | 0 |
| ENSG00000267305 | 0 | 0 | 0 | 0 | 0 | 0 |
| ENSG00000143512 | 19 | 14 | 17 | 14 | 31 | 17 |
| ENSG00000143498 | 89 | 42 | 45 | 49 | 66 | 55 |
| ENSG00000225265 | 55 | 37 | 38 | 28 | 40 | 56 |
| ENSG00000154305 | 792 | 511 | 382 | 226 | 697 | 411 |
| ENSG00000229399 | 0 | 0 | 0 | 0 | 1 | 0 |
| ENSG00000272750 | 12 | 2 | 5 | 0 | 5 | 3 |
| ENSG00000186063 | 676 | 362 | 323 | 289 | 459 | 326 |
| ENSG00000162819 | 1306 | 675 | 703 | 338 | 852 | 600 |
| ENSG00000197520 | 2 | 2 | 0 | 0 | 0 | 0 |
| ENSG00000228106 | 72 | 33 | 49 | 47 | 59 | 59 |
| ENSG00000278610 | 0 | 0 | 1 | 0 | 0 | 2 |
| ENSG00000154309 | 183 | 91 | 90 | 58 | 97 | 83 |
| ENSG00000234523 | 0 | 0 | 1 | 0 | 0 | 0 |
| ENSG00000187554 | 162 | 65 | 81 | 62 | 73 | 70 |
| ENSG00000236846 | 0 | 0 | 0 | 0 | 0 | 1 |
| ENSG00000226601 | 0 | 1 | 0 | 0 | 0 | 0 |
| ENSG00000143502 | 757 | 363 | 416 | 354 | 434 | 446 |
| ENSG00000251789 | 0 | 0 | 0 | 0 | 0 | 0 |
| ENSG00000178395 | 2 | 1 | 3 | 2 | 1 | 3 |
| ENSG00000203697 | 371 | 190 | 324 | 147 | 265 | 249 |
| ENSG00000227002 | 0 | 1 | 0 | 0 | 0 | 0 |
| ENSG00000212398 | 0 | 0 | 0 | 0 | 0 | 0 |
| ENSG00000162909 | 965 | 469 | 623 | 606 | 754 | 626 |
| ENSG00000143514 | 743 | 392 | 429 | 288 | 450 | 368 |
| ENSG00000227621 | 2 | 0 | 1 | 0 | 0 | 0 |
| ENSG00000188460 | 0 | 0 | 0 | 0 | 0 | 0 |
| ENSG00000233771 | 0 | 0 | 0 | 0 | 0 | 0 |
| ENSG00000272645 | 245 | 106 | 157 | 90 | 122 | 100 |
| ENSG00000212157 | 0 | 0 | 0 | 0 | 0 | 0 |
| ENSG00000185495 | 67 | 29 | 29 | 22 | 50 | 31 |
| ENSG00000278467 | 3 | 1 | 4 | 1 | 6 | 1 |
| ENSG00000229930 | 2 | 1 | 0 | 3 | 0 | 0 |
| ENSG00000252484 | 0 | 0 | 0 | 0 | 0 | 0 |
| ENSG00000143756 | 1158 | 599 | 624 | 402 | 648 | 541 |
| ENSG00000236773 | 2 | 5 | 2 | 2 | 0 | 2 |
| ENSG00000143753 | 1062 | 465 | 496 | 348 | 508 | 400 |
| ENSG00000201898 | 0 | 0 | 0 | 0 | 0 | 0 |
| ENSG00000232628 | 0 | 2 | 0 | 0 | 0 | 0 |
| ENSG00000237101 | 2 | 2 | 1 | 1 | 0 | 1 |
| ENSG00000143748 | 1069 | 452 | 548 | 370 | 445 | 441 |
| ENSG00000221406 | 0 | 0 | 0 | 1 | 0 | 0 |
| ENSG00000229742 | 0 | 0 | 0 | 0 | 0 | 0 |
| ENSG00000206887 | 0 | 0 | 0 | 1 | 1 | 0 |
| ENSG00000143771 | 1733 | 802 | 899 | 877 | 915 | 796 |
| ENSG00000162923 | 4162 | 1919 | 2426 | 1481 | 2175 | 1980 |
| ENSG00000266618 | 0 | 0 | 0 | 0 | 0 | 0 |

| | | | | | | |
|---|---|---|---|---|---|---|
| ENSG00000143786 | 55 | 30 | 29 | 29 | 24 | 45 |
| ENSG00000229991 | 0 | 0 | 0 | 0 | 0 | 0 |
| ENSG00000233384 | 0 | 1 | 0 | 0 | 0 | 1 |
| ENSG00000234247 | 0 | 0 | 0 | 0 | 0 | 0 |
| ENSG00000229400 | 0 | 0 | 0 | 0 | 0 | 0 |
| ENSG00000225334 | 0 | 0 | 0 | 0 | 0 | 0 |
| ENSG00000185842 | 170 | 81 | 61 | 76 | 125 | 98 |
| ENSG00000143815 | 1024 | 530 | 582 | 479 | 662 | 585 |
| ENSG00000234476 | 0 | 0 | 0 | 0 | 0 | 0 |
| ENSG00000282418 | 0 | 0 | 0 | 0 | 0 | 0 |
| ENSG00000154380 | 1858 | 856 | 1035 | 637 | 1129 | 1013 |
| ENSG00000227496 | 0 | 0 | 0 | 0 | 0 | 0 |
| ENSG00000226349 | 0 | 0 | 0 | 0 | 0 | 0 |
| ENSG00000223306 | 0 | 0 | 0 | 0 | 0 | 0 |
| ENSG00000143742 | 4563 | 2203 | 2499 | 2250 | 2883 | 2252 |
| ENSG00000234112 | 3 | 3 | 0 | 0 | 1 | 3 |
| ENSG00000143819 | 1754 | 727 | 977 | 1001 | 870 | 893 |
| ENSG00000242861 | 117 | 76 | 106 | 77 | 97 | 78 |
| ENSG00000196187 | 2573 | 1203 | 1628 | 1067 | 1359 | 1395 |
| ENSG00000243709 | 3 | 1 | 0 | 1 | 1 | 0 |
| ENSG00000255835 | 0 | 0 | 0 | 0 | 0 | 0 |
| ENSG00000143811 | 826 | 435 | 601 | 565 | 596 | 590 |
| ENSG00000284519 | 0 | 0 | 0 | 0 | 0 | 0 |
| ENSG00000248322 | 0 | 0 | 0 | 0 | 0 | 0 |
| ENSG00000143768 | 0 | 0 | 0 | 0 | 0 | 0 |
| ENSG00000213032 | 0 | 0 | 0 | 0 | 0 | 0 |
| ENSG00000143751 | 710 | 379 | 465 | 320 | 418 | 364 |
| ENSG00000272562 | 2 | 5 | 8 | 3 | 2 | 8 |
| ENSG00000163041 | 3295 | 1614 | 2031 | 1503 | 1874 | 1768 |
| ENSG00000225518 | 54 | 29 | 42 | 32 | 31 | 37 |
| ENSG00000270598 | 2 | 0 | 0 | 0 | 0 | 0 |
| ENSG00000182827 | 1567 | 893 | 1000 | 590 | 1010 | 814 |
| ENSG00000234478 | 3 | 5 | 0 | 2 | 4 | 3 |
| ENSG00000223570 | 0 | 0 | 0 | 1 | 0 | 0 |
| ENSG00000185155 | 0 | 1 | 0 | 0 | 0 | 0 |
| ENSG00000183814 | 190 | 106 | 113 | 79 | 113 | 119 |
| ENSG00000236994 | 0 | 0 | 0 | 0 | 0 | 0 |
| ENSG00000275406 | 0 | 0 | 0 | 0 | 0 | 0 |
| ENSG00000143799 | 5564 | 2608 | 3300 | 2745 | 2944 | 3261 |
| ENSG00000270287 | 0 | 0 | 0 | 0 | 0 | 0 |
| ENSG00000229965 | 0 | 0 | 0 | 0 | 0 | 0 |
| ENSG00000223282 | 0 | 0 | 0 | 0 | 0 | 0 |
| ENSG00000177770 | 0 | 0 | 0 | 0 | 0 | 0 |
| ENSG00000203685 | 54 | 27 | 21 | 24 | 14 | 33 |
| ENSG00000143772 | 598 | 277 | 308 | 301 | 282 | 311 |
| ENSG00000228382 | 0 | 0 | 0 | 0 | 0 | 0 |
| ENSG00000228548 | 0 | 0 | 0 | 0 | 0 | 1 |
| ENSG00000238183 | 0 | 0 | 0 | 0 | 0 | 0 |
| ENSG00000143801 | 736 | 342 | 498 | 438 | 399 | 471 |
| ENSG00000163050 | 950 | 456 | 584 | 526 | 454 | 513 |

| | | | | | | |
|---|---|---|---|---|---|---|
| ENSG00000228525 | 0 | 0 | 0 | 0 | 0 | 0 |
| ENSG00000143776 | 1097 | 500 | 494 | 209 | 704 | 428 |
| ENSG00000233706 | 0 | 0 | 0 | 0 | 0 | 0 |
| ENSG00000228625 | 0 | 0 | 0 | 0 | 0 | 0 |
| ENSG00000228729 | 0 | 0 | 0 | 0 | 0 | 0 |
| ENSG00000236636 | 0 | 0 | 0 | 0 | 0 | 0 |
| ENSG00000234277 | 1 | 0 | 0 | 0 | 0 | 0 |
| ENSG00000232864 | 0 | 0 | 0 | 0 | 0 | 0 |
| ENSG00000242757 | 0 | 0 | 0 | 0 | 0 | 0 |
| ENSG00000224834 | 0 | 0 | 0 | 0 | 0 | 0 |
| ENSG00000237193 | 0 | 0 | 0 | 0 | 0 | 0 |
| ENSG00000278180 | 0 | 0 | 0 | 0 | 0 | 0 |
| ENSG00000237469 | 0 | 0 | 0 | 0 | 0 | 0 |
| ENSG00000235827 | 2 | 0 | 1 | 0 | 0 | 0 |
| ENSG00000227711 | 0 | 1 | 0 | 0 | 0 | 0 |
| ENSG00000202264 | 0 | 0 | 0 | 0 | 0 | 0 |
| ENSG00000181450 | 142 | 68 | 49 | 38 | 127 | 59 |
| ENSG00000225934 | 0 | 1 | 0 | 0 | 0 | 0 |
| ENSG00000215812 | 0 | 0 | 0 | 0 | 0 | 0 |
| ENSG00000143740 | 1077 | 457 | 608 | 560 | 551 | 502 |
| ENSG00000081692 | 222 | 122 | 200 | 216 | 154 | 169 |
| ENSG00000230005 | 0 | 1 | 0 | 1 | 0 | 0 |
| ENSG00000185888 | 0 | 0 | 0 | 0 | 0 | 0 |
| ENSG00000143816 | 254 | 92 | 158 | 134 | 111 | 100 |
| ENSG00000264483 | 0 | 0 | 0 | 0 | 0 | 0 |
| ENSG00000233003 | 0 | 0 | 0 | 0 | 0 | 0 |
| ENSG00000274886 | 0 | 0 | 0 | 0 | 0 | 0 |
| ENSG00000154342 | 10 | 2 | 11 | 4 | 2 | 5 |
| ENSG00000276255 | 10 | 3 | 3 | 4 | 3 | 1 |
| ENSG00000143761 | 14657 | 6776 | 8758 | 8417 | 7411 | 8028 |
| ENSG00000284163 | 0 | 0 | 0 | 0 | 0 | 0 |
| ENSG00000143793 | 1034 | 477 | 614 | 623 | 532 | 518 |
| ENSG00000162910 | 786 | 385 | 496 | 561 | 470 | 440 |
| ENSG00000224566 | 5 | 3 | 5 | 0 | 2 | 0 |
| ENSG00000280157 | 6 | 7 | 7 | 2 | 7 | 5 |
| ENSG00000227625 | 0 | 0 | 0 | 0 | 0 | 0 |
| ENSG00000143774 | 3808 | 1823 | 2209 | 2486 | 1918 | 1888 |
| ENSG00000198835 | 10 | 8 | 12 | 15 | 9 | 3 |
| ENSG00000203684 | 0 | 4 | 2 | 4 | 0 | 5 |
| ENSG00000181873 | 342 | 168 | 238 | 178 | 137 | 182 |
| ENSG00000162913 | 106 | 37 | 44 | 35 | 37 | 41 |
| ENSG00000154358 | 394 | 161 | 294 | 152 | 189 | 209 |
| ENSG00000269890 | 0 | 0 | 0 | 0 | 0 | 0 |
| ENSG00000269934 | 2 | 0 | 0 | 0 | 0 | 0 |
| ENSG00000270110 | 6 | 1 | 3 | 0 | 0 | 0 |
| ENSG00000270104 | 0 | 1 | 0 | 0 | 0 | 1 |
| ENSG00000154370 | 1395 | 667 | 857 | 886 | 757 | 752 |
| ENSG00000270094 | 0 | 0 | 0 | 0 | 0 | 0 |
| ENSG00000284311 | 0 | 0 | 0 | 0 | 0 | 0 |
| ENSG00000231563 | 6 | 3 | 7 | 1 | 8 | 4 |

| | | | | | | |
|---|---|---|---|---|---|---|
| ENSG00000162931 | 28 | 30 | 25 | 11 | 11 | 12 |
| ENSG00000168148 | 4 | 5 | 2 | 0 | 4 | 1 |
| ENSG00000230841 | 0 | 1 | 3 | 0 | 1 | 1 |
| ENSG00000181218 | 1096 | 528 | 661 | 773 | 694 | 668 |
| ENSG00000196890 | 78 | 40 | 43 | 30 | 37 | 41 |
| ENSG00000266174 | 0 | 1 | 4 | 0 | 0 | 0 |
| ENSG00000181201 | 0 | 0 | 0 | 0 | 0 | 0 |
| ENSG00000279306 | 3 | 0 | 0 | 2 | 4 | 2 |
| ENSG00000168159 | 3260 | 1587 | 1939 | 2086 | 1718 | 1758 |
| ENSG00000215811 | 5 | 7 | 5 | 1 | 4 | 2 |
| ENSG00000202056 | 0 | 0 | 0 | 0 | 0 | 0 |
| ENSG00000251823 | 0 | 0 | 0 | 0 | 0 | 0 |
| ENSG00000199352 | 0 | 0 | 0 | 0 | 0 | 0 |
| ENSG00000201588 | 0 | 0 | 0 | 0 | 0 | 0 |
| ENSG00000199337 | 0 | 0 | 0 | 0 | 0 | 0 |
| ENSG00000200381 | 0 | 0 | 0 | 0 | 0 | 0 |
| ENSG00000199396 | 0 | 0 | 0 | 0 | 0 | 0 |
| ENSG00000200624 | 0 | 0 | 0 | 0 | 0 | 0 |
| ENSG00000202521 | 0 | 0 | 0 | 0 | 0 | 0 |
| ENSG00000200343 | 0 | 0 | 0 | 0 | 0 | 0 |
| ENSG00000201321 | 0 | 0 | 0 | 0 | 0 | 0 |
| ENSG00000199910 | 0 | 0 | 0 | 0 | 0 | 0 |
| ENSG00000199334 | 0 | 0 | 0 | 0 | 0 | 0 |
| ENSG00000199270 | 0 | 0 | 0 | 0 | 0 | 0 |
| ENSG00000202526 | 0 | 0 | 0 | 0 | 0 | 0 |
| ENSG00000201355 | 0 | 0 | 0 | 0 | 0 | 0 |
| ENSG00000201925 | 0 | 0 | 0 | 0 | 0 | 0 |
| ENSG00000202257 | 0 | 0 | 0 | 0 | 0 | 0 |
| ENSG00000200370 | 0 | 0 | 0 | 0 | 0 | 0 |
| ENSG00000212237 | 0 | 2 | 1 | 0 | 0 | 1 |
| ENSG00000183929 | 0 | 0 | 0 | 0 | 0 | 1 |
| ENSG00000206878 | 0 | 0 | 0 | 0 | 0 | 1 |
| ENSG00000234975 | 8 | 5 | 6 | 2 | 4 | 9 |
| ENSG00000116574 | 984 | 450 | 586 | 282 | 442 | 351 |
| ENSG00000271475 | 0 | 0 | 0 | 0 | 0 | 0 |
| ENSG00000271399 | 0 | 0 | 0 | 0 | 0 | 0 |
| ENSG00000282111 | 0 | 0 | 0 | 0 | 0 | 0 |
| ENSG00000229840 | 0 | 0 | 0 | 0 | 0 | 0 |
| ENSG00000236975 | 2 | 1 | 3 | 0 | 0 | 1 |
| ENSG00000233920 | 2 | 0 | 0 | 0 | 0 | 0 |
| ENSG00000177800 | 0 | 0 | 0 | 0 | 0 | 0 |
| ENSG00000177788 | 26 | 15 | 14 | 16 | 7 | 13 |
| ENSG00000168118 | 2105 | 1063 | 1264 | 1076 | 1067 | 1094 |
| ENSG00000213029 | 0 | 0 | 0 | 0 | 0 | 0 |
| ENSG00000237481 | 0 | 1 | 1 | 0 | 0 | 1 |
| ENSG00000154429 | 440 | 203 | 214 | 181 | 273 | 210 |
| ENSG00000252506 | 0 | 0 | 0 | 0 | 0 | 0 |
| ENSG00000252051 | 0 | 0 | 0 | 0 | 0 | 0 |
| ENSG00000230331 | 0 | 0 | 0 | 0 | 0 | 0 |
| ENSG00000143632 | 14 | 8 | 3 | 7 | 6 | 1 |

| | | | | | | |
|---|---|---|---|---|---|---|
| ENSG00000069248 | 1504 | 786 | 919 | 610 | 975 | 835 |
| ENSG00000226920 | 0 | 0 | 0 | 0 | 0 | 0 |
| ENSG00000223635 | 0 | 0 | 1 | 1 | 3 | 2 |
| ENSG00000135776 | 608 | 246 | 349 | 255 | 323 | 348 |
| ENSG00000199672 | 1 | 0 | 0 | 0 | 0 | 0 |
| ENSG00000201492 | 0 | 1 | 1 | 0 | 1 | 1 |
| ENSG00000229367 | 0 | 0 | 0 | 1 | 0 | 2 |
| ENSG00000135801 | 847 | 407 | 482 | 401 | 500 | 485 |
| ENSG00000135763 | 456 | 265 | 311 | 305 | 323 | 373 |
| ENSG00000213028 | 0 | 0 | 0 | 0 | 0 | 1 |
| ENSG00000225984 | 0 | 0 | 0 | 0 | 0 | 0 |
| ENSG00000227043 | 0 | 0 | 0 | 0 | 0 | 0 |
| ENSG00000228058 | 0 | 0 | 0 | 0 | 0 | 0 |
| ENSG00000143641 | 1713 | 887 | 1053 | 1136 | 1439 | 1204 |
| ENSG00000227006 | 8 | 3 | 4 | 0 | 2 | 4 |
| ENSG00000224407 | 0 | 0 | 0 | 0 | 0 | 0 |
| ENSG00000177614 | 42 | 31 | 37 | 21 | 19 | 23 |
| ENSG00000282564 | 0 | 0 | 0 | 0 | 0 | 0 |
| ENSG00000232964 | 0 | 0 | 0 | 0 | 0 | 0 |
| ENSG00000235817 | 0 | 0 | 0 | 0 | 0 | 0 |
| ENSG00000135775 | 1455 | 688 | 900 | 736 | 736 | 802 |
| ENSG00000135744 | 0 | 0 | 0 | 3 | 1 | 0 |
| ENSG00000244137 | 23 | 10 | 12 | 3 | 16 | 21 |
| ENSG00000135773 | 828 | 353 | 516 | 371 | 376 | 386 |
| ENSG00000202063 | 0 | 0 | 0 | 0 | 0 | 0 |
| ENSG00000225656 | 0 | 0 | 0 | 0 | 0 | 0 |
| ENSG00000119280 | 1265 | 664 | 690 | 637 | 711 | 720 |
| ENSG00000223393 | 11 | 5 | 4 | 7 | 5 | 9 |
| ENSG00000240502 | 0 | 0 | 0 | 0 | 0 | 0 |
| ENSG00000143643 | 637 | 298 | 382 | 248 | 397 | 360 |
| ENSG00000173409 | 479 | 245 | 279 | 254 | 301 | 300 |
| ENSG00000182118 | 60 | 32 | 41 | 43 | 44 | 43 |
| ENSG00000283799 | 0 | 0 | 0 | 0 | 0 | 0 |
| ENSG00000227934 | 2 | 0 | 1 | 0 | 1 | 0 |
| ENSG00000229595 | 0 | 0 | 0 | 0 | 0 | 0 |
| ENSG00000119283 | 2 | 0 | 2 | 2 | 3 | 4 |
| ENSG00000235710 | 0 | 0 | 0 | 0 | 0 | 0 |
| ENSG00000143633 | 550 | 270 | 346 | 269 | 308 | 321 |
| ENSG00000116906 | 1218 | 574 | 718 | 605 | 695 | 656 |
| ENSG00000222407 | 0 | 0 | 0 | 0 | 0 | 0 |
| ENSG00000116903 | 616 | 288 | 378 | 224 | 324 | 314 |
| ENSG00000010072 | 643 | 238 | 326 | 284 | 366 | 327 |
| ENSG00000135766 | 1044 | 504 | 569 | 470 | 544 | 519 |
| ENSG00000223982 | 0 | 0 | 0 | 0 | 0 | 0 |
| ENSG00000233461 | 64 | 29 | 25 | 45 | 37 | 45 |
| ENSG00000270106 | 0 | 0 | 0 | 1 | 0 | 1 |
| ENSG00000116918 | 523 | 271 | 309 | 156 | 328 | 250 |
| ENSG00000229228 | 0 | 0 | 1 | 0 | 0 | 0 |
| ENSG00000162946 | 30 | 8 | 28 | 25 | 22 | 18 |
| ENSG00000222986 | 0 | 0 | 0 | 0 | 1 | 0 |

| | | | | | | |
|---|---|---|---|---|---|---|
| ENSG00000274121 | 0 | 0 | 0 | 0 | 0 | 0 |
| ENSG00000226758 | 0 | 0 | 0 | 0 | 0 | 0 |
| ENSG00000235152 | 0 | 0 | 0 | 0 | 1 | 0 |
| ENSG00000236372 | 0 | 0 | 0 | 0 | 0 | 0 |
| ENSG00000224436 | 0 | 0 | 0 | 0 | 0 | 0 |
| ENSG00000242794 | 0 | 0 | 0 | 0 | 0 | 0 |
| ENSG00000116991 | 816 | 366 | 522 | 491 | 639 | 601 |
| ENSG00000238382 | 0 | 0 | 0 | 0 | 0 | 0 |
| ENSG00000225919 | 0 | 0 | 0 | 0 | 0 | 0 |
| ENSG00000236497 | 0 | 0 | 0 | 0 | 0 | 0 |
| ENSG00000212916 | 261 | 121 | 194 | 96 | 145 | 144 |
| ENSG00000206835 | 0 | 0 | 0 | 0 | 0 | 0 |
| ENSG00000232175 | 0 | 0 | 0 | 0 | 0 | 0 |
| ENSG00000135778 | 888 | 465 | 543 | 565 | 577 | 561 |
| ENSG00000135749 | 341 | 158 | 239 | 184 | 225 | 217 |
| ENSG00000231940 | 3 | 1 | 2 | 0 | 1 | 2 |
| ENSG00000251508 | 0 | 0 | 0 | 0 | 0 | 0 |
| ENSG00000143674 | 370 | 193 | 222 | 159 | 249 | 197 |
| ENSG00000252501 | 0 | 0 | 0 | 0 | 0 | 0 |
| ENSG00000135750 | 808 | 388 | 445 | 377 | 423 | 404 |
| ENSG00000265744 | 0 | 0 | 0 | 0 | 0 | 1 |
| ENSG00000225486 | 0 | 0 | 0 | 0 | 0 | 0 |
| ENSG00000183780 | 4 | 0 | 3 | 3 | 3 | 4 |
| ENSG00000236101 | 0 | 0 | 0 | 0 | 0 | 0 |
| ENSG00000233332 | 0 | 0 | 0 | 0 | 1 | 0 |
| ENSG00000231272 | 0 | 0 | 0 | 0 | 0 | 0 |
| ENSG00000236244 | 0 | 0 | 0 | 0 | 0 | 0 |
| ENSG00000232166 | 0 | 0 | 0 | 0 | 0 | 0 |
| ENSG00000264377 | 0 | 0 | 0 | 0 | 0 | 0 |
| ENSG00000235605 | 1 | 0 | 0 | 3 | 0 | 0 |
| ENSG00000236358 | 4 | 1 | 1 | 1 | 0 | 2 |
| ENSG00000273367 | 4 | 0 | 0 | 5 | 6 | 1 |
| ENSG00000231663 | 18 | 14 | 18 | 12 | 7 | 6 |
| ENSG00000168275 | 510 | 258 | 319 | 318 | 339 | 349 |
| ENSG00000059588 | 941 | 456 | 602 | 375 | 561 | 545 |
| ENSG00000231768 | 0 | 0 | 3 | 1 | 2 | 0 |
| ENSG00000230404 | 0 | 0 | 0 | 0 | 0 | 0 |
| ENSG00000212144 | 0 | 0 | 0 | 0 | 0 | 0 |
| ENSG00000168264 | 2532 | 1215 | 1613 | 1523 | 1609 | 1817 |
| ENSG00000228830 | 0 | 0 | 0 | 0 | 0 | 0 |
| ENSG00000224939 | 2 | 2 | 5 | 2 | 1 | 1 |
| ENSG00000272362 | 0 | 0 | 0 | 0 | 0 | 0 |
| ENSG00000228044 | 0 | 0 | 0 | 3 | 3 | 0 |
| ENSG00000241475 | 0 | 0 | 0 | 2 | 1 | 0 |
| ENSG00000230628 | 0 | 0 | 0 | 0 | 0 | 0 |
| ENSG00000282097 | 0 | 0 | 0 | 0 | 0 | 0 |
| ENSG00000227630 | 14 | 9 | 13 | 6 | 5 | 7 |
| ENSG00000279261 | 0 | 0 | 0 | 0 | 0 | 0 |
| ENSG00000201638 | 0 | 0 | 0 | 0 | 0 | 0 |
| ENSG00000239690 | 0 | 0 | 0 | 0 | 0 | 0 |

| | | | | | | |
|---|---|---|---|---|---|---|
| ENSG00000237520 | 0 | 1 | 1 | 0 | 2 | 0 |
| ENSG00000238005 | 13 | 8 | 13 | 13 | 8 | 12 |
| ENSG00000258082 | 4 | 0 | 0 | 0 | 0 | 0 |
| ENSG00000280587 | 48 | 30 | 58 | 34 | 23 | 54 |
| ENSG00000233018 | 0 | 0 | 0 | 0 | 0 | 0 |
| ENSG00000273416 | 2 | 0 | 0 | 5 | 1 | 0 |
| ENSG00000173726 | 5209 | 2573 | 3167 | 2658 | 3233 | 3105 |
| ENSG00000207181 | 2 | 4 | 7 | 2 | 9 | 3 |
| ENSG00000188739 | 7 | 15 | 16 | 8 | 11 | 9 |
| ENSG00000054267 | 691 | 430 | 281 | 122 | 579 | 281 |
| ENSG00000263439 | 0 | 0 | 0 | 0 | 0 | 0 |
| ENSG00000236863 | 0 | 1 | 0 | 0 | 0 | 0 |
| ENSG00000152904 | 387 | 257 | 305 | 220 | 320 | 257 |
| ENSG00000285053 | 110 | 62 | 70 | 37 | 93 | 68 |
| ENSG00000227962 | 2 | 0 | 2 | 0 | 0 | 0 |
| ENSG00000230026 | 0 | 0 | 0 | 0 | 1 | 0 |
| ENSG00000285177 | 0 | 0 | 1 | 1 | 0 | 1 |
| ENSG00000284770 | 210 | 131 | 153 | 88 | 128 | 126 |
| ENSG00000229795 | 0 | 0 | 0 | 0 | 0 | 1 |
| ENSG00000162885 | 298 | 150 | 165 | 107 | 164 | 158 |
| ENSG00000238236 | 0 | 0 | 0 | 0 | 0 | 0 |
| ENSG00000235638 | 0 | 0 | 0 | 0 | 0 | 0 |
| ENSG00000231199 | 0 | 0 | 0 | 0 | 0 | 0 |
| ENSG00000226663 | 0 | 0 | 0 | 0 | 0 | 0 |
| ENSG00000231582 | 0 | 0 | 0 | 0 | 0 | 0 |
| ENSG00000232714 | 0 | 0 | 0 | 0 | 0 | 0 |
| ENSG00000235761 | 0 | 0 | 0 | 0 | 0 | 0 |
| ENSG00000168243 | 36 | 26 | 27 | 42 | 43 | 42 |
| ENSG00000270710 | 0 | 0 | 0 | 0 | 1 | 0 |
| ENSG00000227236 | 0 | 0 | 0 | 0 | 0 | 0 |
| ENSG00000143669 | 101 | 73 | 41 | 16 | 153 | 72 |
| ENSG00000235674 | 0 | 0 | 0 | 0 | 0 | 1 |
| ENSG00000229463 | 0 | 1 | 0 | 0 | 0 | 0 |
| ENSG00000222831 | 0 | 0 | 0 | 0 | 0 | 0 |
| ENSG00000206803 | 0 | 0 | 0 | 0 | 0 | 0 |
| ENSG00000237845 | 0 | 0 | 0 | 0 | 0 | 0 |
| ENSG00000229291 | 0 | 0 | 0 | 0 | 0 | 0 |
| ENSG00000116962 | 0 | 0 | 0 | 0 | 0 | 0 |
| ENSG00000252822 | 0 | 0 | 0 | 0 | 0 | 0 |
| ENSG00000235371 | 2 | 0 | 0 | 0 | 0 | 0 |
| ENSG00000077585 | 1485 | 699 | 924 | 713 | 751 | 703 |
| ENSG00000086619 | 335 | 134 | 203 | 116 | 173 | 141 |
| ENSG00000222650 | 0 | 0 | 0 | 0 | 0 | 0 |
| ENSG00000237922 | 0 | 0 | 0 | 1 | 0 | 0 |
| ENSG00000186197 | 100 | 77 | 106 | 89 | 95 | 104 |
| ENSG00000244457 | 0 | 0 | 0 | 0 | 0 | 0 |
| ENSG00000116977 | 2720 | 1387 | 1843 | 781 | 1413 | 1270 |
| ENSG00000223776 | 54 | 29 | 39 | 21 | 29 | 25 |
| ENSG00000273058 | 9 | 3 | 5 | 3 | 3 | 3 |
| ENSG00000230325 | 6 | 6 | 3 | 3 | 9 | 1 |

| | | | | | | |
|---|---|---|---|---|---|---|
| ENSG00000119285 | 791 | 489 | 437 | 275 | 701 | 480 |
| ENSG00000077522 | 0 | 1 | 0 | 0 | 0 | 0 |
| ENSG00000116984 | 1238 | 599 | 783 | 369 | 567 | 694 |
| ENSG00000226498 | 0 | 0 | 0 | 0 | 0 | 0 |
| ENSG00000237991 | 2 | 1 | 1 | 0 | 0 | 1 |
| ENSG00000244020 | 0 | 0 | 0 | 0 | 0 | 0 |
| ENSG00000198626 | 0 | 1 | 1 | 0 | 1 | 0 |
| ENSG00000252396 | 0 | 0 | 0 | 0 | 0 | 0 |
| ENSG00000266262 | 0 | 0 | 0 | 0 | 0 | 0 |
| ENSG00000252290 | 0 | 0 | 0 | 0 | 0 | 0 |
| ENSG00000237250 | 0 | 0 | 0 | 0 | 0 | 0 |
| ENSG00000116996 | 0 | 0 | 0 | 0 | 0 | 0 |
| ENSG00000243781 | 0 | 0 | 0 | 0 | 0 | 0 |
| ENSG00000283174 | 0 | 0 | 0 | 0 | 0 | 0 |
| ENSG00000283166 | 0 | 0 | 0 | 0 | 0 | 0 |
| ENSG00000283377 | 0 | 0 | 0 | 0 | 0 | 0 |
| ENSG00000270188 | 0 | 0 | 0 | 0 | 0 | 0 |
| ENSG00000232154 | 0 | 0 | 0 | 0 | 0 | 0 |
| ENSG00000225723 | 0 | 0 | 0 | 0 | 0 | 0 |
| ENSG00000227860 | 0 | 0 | 0 | 0 | 0 | 0 |
| ENSG00000230019 | 0 | 0 | 0 | 0 | 0 | 0 |
| ENSG00000232989 | 0 | 0 | 0 | 0 | 0 | 0 |
| ENSG00000234464 | 0 | 0 | 0 | 0 | 0 | 0 |
| ENSG00000252371 | 0 | 0 | 0 | 0 | 0 | 0 |
| ENSG00000215808 | 2 | 0 | 4 | 0 | 1 | 2 |
| ENSG00000231877 | 0 | 0 | 0 | 0 | 0 | 0 |
| ENSG00000215149 | 2 | 0 | 0 | 0 | 0 | 0 |
| ENSG00000224783 | 0 | 0 | 0 | 0 | 0 | 0 |
| ENSG00000227854 | 0 | 0 | 0 | 0 | 0 | 0 |
| ENSG00000259776 | 0 | 0 | 0 | 0 | 0 | 0 |
| ENSG00000133019 | 0 | 3 | 0 | 0 | 0 | 0 |
| ENSG00000233355 | 0 | 0 | 0 | 0 | 0 | 0 |
| ENSG00000234601 | 0 | 0 | 0 | 0 | 0 | 0 |
| ENSG00000231979 | 0 | 0 | 0 | 0 | 0 | 0 |
| ENSG00000215805 | 0 | 0 | 0 | 0 | 0 | 0 |
| ENSG00000217327 | 0 | 0 | 0 | 0 | 0 | 0 |
| ENSG00000155816 | 6 | 7 | 8 | 2 | 2 | 5 |
| ENSG00000228818 | 4 | 0 | 2 | 1 | 2 | 3 |
| ENSG00000202041 | 0 | 0 | 0 | 0 | 0 | 0 |
| ENSG00000223694 | 0 | 1 | 0 | 0 | 0 | 0 |
| ENSG00000233735 | 0 | 0 | 0 | 0 | 0 | 0 |
| ENSG00000252317 | 0 | 0 | 0 | 0 | 0 | 0 |
| ENSG00000233519 | 0 | 0 | 0 | 0 | 0 | 0 |
| ENSG00000180875 | 0 | 0 | 0 | 0 | 0 | 0 |
| ENSG00000224359 | 0 | 0 | 0 | 0 | 0 | 0 |
| ENSG00000226014 | 0 | 0 | 0 | 0 | 0 | 0 |
| ENSG00000231440 | 0 | 0 | 0 | 0 | 0 | 0 |
| ENSG00000228844 | 0 | 0 | 0 | 0 | 0 | 0 |
| ENSG00000251750 | 0 | 0 | 0 | 0 | 0 | 0 |
| ENSG00000234872 | 0 | 0 | 0 | 0 | 0 | 0 |

| | | | | | | |
|---|---|---|---|---|---|---|
| ENSG00000199241 | 0 | 0 | 0 | 0 | 0 | 0 |
| ENSG00000230015 | 0 | 0 | 0 | 0 | 0 | 0 |
| ENSG00000226919 | 0 | 0 | 0 | 0 | 0 | 0 |
| ENSG00000223963 | 0 | 0 | 0 | 0 | 0 | 0 |
| ENSG00000182901 | 0 | 0 | 0 | 0 | 0 | 1 |
| ENSG00000215802 | 0 | 0 | 0 | 0 | 0 | 0 |
| ENSG00000233615 | 0 | 0 | 0 | 0 | 0 | 0 |
| ENSG00000238085 | 0 | 0 | 0 | 1 | 0 | 0 |
| ENSG00000265831 | 0 | 0 | 0 | 0 | 0 | 0 |
| ENSG00000224348 | 0 | 0 | 0 | 0 | 0 | 0 |
| ENSG00000225554 | 0 | 0 | 0 | 0 | 1 | 0 |
| ENSG00000091483 | 1597 | 794 | 989 | 852 | 914 | 898 |
| ENSG00000117009 | 17 | 17 | 11 | 7 | 20 | 11 |
| ENSG00000054277 | 158 | 112 | 98 | 97 | 110 | 94 |
| ENSG00000203668 | 288 | 211 | 126 | 74 | 325 | 208 |
| ENSG00000162843 | 3 | 4 | 0 | 1 | 2 | 3 |
| ENSG00000229022 | 0 | 0 | 1 | 0 | 0 | 0 |
| ENSG00000174371 | 196 | 136 | 110 | 60 | 114 | 103 |
| ENSG00000235990 | 0 | 0 | 0 | 0 | 0 | 0 |
| ENSG00000196289 | 0 | 0 | 0 | 0 | 0 | 0 |
| ENSG00000213026 | 0 | 0 | 0 | 0 | 0 | 0 |
| ENSG00000197769 | 2 | 0 | 0 | 1 | 0 | 0 |
| ENSG00000224625 | 0 | 0 | 0 | 0 | 0 | 0 |
| ENSG00000277099 | 0 | 0 | 0 | 0 | 0 | 1 |
| ENSG00000180287 | 0 | 0 | 0 | 0 | 0 | 0 |
| ENSG00000272865 | 0 | 0 | 0 | 0 | 1 | 0 |
| ENSG00000252084 | 0 | 0 | 0 | 0 | 0 | 0 |
| ENSG00000224525 | 0 | 0 | 0 | 0 | 0 | 0 |
| ENSG00000226750 | 0 | 0 | 0 | 0 | 0 | 0 |
| ENSG00000235467 | 0 | 0 | 0 | 0 | 0 | 0 |
| ENSG00000213690 | 0 | 0 | 0 | 0 | 0 | 0 |
| ENSG00000270859 | 0 | 0 | 0 | 0 | 0 | 0 |
| ENSG00000215800 | 0 | 0 | 0 | 0 | 0 | 0 |
| ENSG00000270818 | 0 | 0 | 0 | 0 | 0 | 0 |
| ENSG00000277704 | 0 | 0 | 0 | 0 | 0 | 0 |
| ENSG00000230199 | 0 | 0 | 1 | 0 | 0 | 0 |
| ENSG00000234116 | 0 | 0 | 0 | 0 | 0 | 0 |
| ENSG00000231512 | 0 | 0 | 0 | 0 | 0 | 0 |
| ENSG00000278455 | 0 | 0 | 0 | 0 | 0 | 0 |
| ENSG00000253326 | 0 | 0 | 0 | 0 | 0 | 1 |
| ENSG00000214837 | 26 | 12 | 12 | 10 | 14 | 25 |
| ENSG00000212230 | 0 | 0 | 0 | 0 | 0 | 0 |
| ENSG00000143702 | 253 | 196 | 133 | 63 | 229 | 120 |
| ENSG00000227230 | 0 | 3 | 0 | 0 | 1 | 0 |
| ENSG00000232085 | 0 | 0 | 0 | 0 | 0 | 0 |
| ENSG00000054282 | 287 | 171 | 217 | 102 | 199 | 232 |
| ENSG00000224727 | 1 | 0 | 1 | 0 | 0 | 0 |
| ENSG00000265201 | 2 | 0 | 0 | 0 | 0 | 0 |
| ENSG00000117020 | 0 | 0 | 3 | 1 | 1 | 2 |
| ENSG00000236031 | 0 | 0 | 0 | 0 | 0 | 0 |

| | | | | | | |
|---|---|---|---|---|---|---|
| ENSG00000226766 | 0 | 0 | 0 | 0 | 0 | 0 |
| ENSG00000232184 | 0 | 0 | 0 | 0 | 0 | 0 |
| ENSG00000228939 | 0 | 0 | 0 | 0 | 0 | 0 |
| ENSG00000226828 | 0 | 0 | 0 | 0 | 0 | 0 |
| ENSG00000179456 | 20 | 9 | 8 | 3 | 16 | 6 |
| ENSG00000279774 | 0 | 0 | 0 | 0 | 0 | 0 |
| ENSG00000229960 | 0 | 0 | 0 | 0 | 0 | 0 |
| ENSG00000244066 | 0 | 0 | 0 | 0 | 0 | 0 |
| ENSG00000237759 | 0 | 0 | 0 | 0 | 0 | 0 |
| ENSG00000173728 | 3 | 1 | 0 | 1 | 0 | 2 |
| ENSG00000240963 | 0 | 0 | 0 | 0 | 0 | 0 |
| ENSG00000225401 | 0 | 0 | 0 | 0 | 0 | 0 |
| ENSG00000035687 | 1945 | 1053 | 1256 | 997 | 1262 | 1229 |
| ENSG00000179397 | 5 | 2 | 8 | 3 | 3 | 9 |
| ENSG00000227735 | 0 | 0 | 0 | 0 | 0 | 0 |
| ENSG00000121644 | 984 | 454 | 572 | 465 | 536 | 540 |
| ENSG00000232059 | 1 | 0 | 0 | 0 | 0 | 0 |
| ENSG00000284188 | 0 | 0 | 0 | 0 | 0 | 0 |
| ENSG00000282317 | 0 | 0 | 1 | 0 | 0 | 0 |
| ENSG00000203667 | 53 | 29 | 33 | 23 | 34 | 27 |
| ENSG00000153187 | 15609 | 7805 | 9768 | 6798 | 8973 | 9252 |
| ENSG00000273175 | 0 | 0 | 2 | 0 | 1 | 0 |
| ENSG00000201758 | 0 | 0 | 0 | 0 | 0 | 0 |
| ENSG00000252073 | 0 | 0 | 0 | 0 | 0 | 0 |
| ENSG00000272195 | 5 | 1 | 2 | 2 | 2 | 2 |
| ENSG00000203666 | 158 | 81 | 86 | 66 | 81 | 92 |
| ENSG00000252282 | 0 | 0 | 0 | 0 | 0 | 0 |
| ENSG00000251754 | 0 | 0 | 0 | 0 | 0 | 0 |
| ENSG00000223353 | 0 | 0 | 0 | 0 | 0 | 0 |
| ENSG00000201170 | 0 | 0 | 0 | 0 | 0 | 0 |
| ENSG00000162849 | 6 | 1 | 4 | 5 | 5 | 3 |
| ENSG00000232192 | 0 | 0 | 0 | 0 | 0 | 0 |
| ENSG00000232793 | 0 | 0 | 0 | 0 | 0 | 0 |
| ENSG00000238224 | 0 | 0 | 0 | 0 | 1 | 0 |
| ENSG00000231612 | 3 | 1 | 0 | 3 | 1 | 1 |
| ENSG00000185420 | 754 | 425 | 525 | 440 | 479 | 468 |
| ENSG00000235096 | 0 | 0 | 0 | 0 | 0 | 0 |
| ENSG00000226876 | 0 | 1 | 0 | 0 | 0 | 0 |
| ENSG00000225222 | 0 | 0 | 0 | 0 | 0 | 0 |
| ENSG00000202184 | 0 | 0 | 0 | 0 | 0 | 0 |
| ENSG00000230184 | 0 | 0 | 0 | 0 | 0 | 0 |
| ENSG00000207326 | 0 | 0 | 0 | 0 | 0 | 0 |
| ENSG00000227728 | 0 | 0 | 0 | 0 | 0 | 0 |
| ENSG00000242042 | 0 | 0 | 0 | 0 | 0 | 0 |
| ENSG00000230813 | 0 | 0 | 0 | 0 | 0 | 0 |
| ENSG00000229112 | 0 | 0 | 0 | 0 | 0 | 0 |
| ENSG00000162851 | 406 | 222 | 233 | 216 | 282 | 227 |
| ENSG00000162852 | 775 | 376 | 464 | 272 | 428 | 367 |
| ENSG00000225300 | 2 | 1 | 0 | 0 | 0 | 0 |
| ENSG00000228955 | 1 | 1 | 0 | 0 | 0 | 0 |

| | | | | | | |
|---|---|---|---|---|---|---|
| ENSG00000260698 | 9 | 2 | 4 | 3 | 6 | 4 |
| ENSG00000235021 | 14 | 6 | 14 | 5 | 3 | 11 |
| ENSG00000143653 | 6750 | 3371 | 4192 | 3878 | 4071 | 4090 |
| ENSG00000228879 | 2 | 0 | 0 | 0 | 0 | 0 |
| ENSG00000223519 | 42 | 20 | 29 | 10 | 21 | 23 |
| ENSG00000260855 | 87 | 27 | 83 | 14 | 28 | 61 |
| ENSG00000227953 | 110 | 55 | 88 | 77 | 68 | 76 |
| ENSG00000235386 | 0 | 0 | 0 | 0 | 0 | 0 |
| ENSG00000252011 | 0 | 0 | 0 | 0 | 0 | 0 |
| ENSG00000153207 | 606 | 374 | 252 | 148 | 695 | 325 |
| ENSG00000197472 | 22 | 16 | 9 | 9 | 16 | 14 |
| ENSG00000135747 | 6 | 3 | 1 | 4 | 1 | 2 |
| ENSG00000236775 | 0 | 0 | 0 | 0 | 0 | 0 |
| ENSG00000277462 | 74 | 32 | 27 | 29 | 62 | 38 |
| ENSG00000215796 | 0 | 0 | 1 | 0 | 0 | 0 |
| ENSG00000228982 | 0 | 0 | 0 | 0 | 0 | 0 |
| ENSG00000188295 | 172 | 102 | 109 | 80 | 97 | 101 |
| ENSG00000221953 | 20 | 8 | 9 | 13 | 6 | 12 |
| ENSG00000226164 | 0 | 0 | 0 | 0 | 0 | 0 |
| ENSG00000196418 | 142 | 99 | 91 | 80 | 120 | 106 |
| ENSG00000224014 | 8 | 9 | 8 | 1 | 5 | 10 |
| ENSG00000215795 | 8 | 9 | 14 | 7 | 14 | 15 |
| ENSG00000259865 | 99 | 35 | 47 | 25 | 42 | 46 |
| ENSG00000227671 | 736 | 391 | 412 | 125 | 400 | 372 |
| ENSG00000283881 | 0 | 0 | 0 | 0 | 0 | 0 |
| ENSG00000252516 | 3 | 2 | 1 | 1 | 1 | 2 |
| ENSG00000232347 | 0 | 0 | 0 | 0 | 0 | 0 |
| ENSG00000214144 | 0 | 0 | 0 | 0 | 0 | 0 |
| ENSG00000235818 | 0 | 0 | 0 | 0 | 0 | 0 |
| ENSG00000197617 | 0 | 0 | 0 | 0 | 1 | 0 |
| ENSG00000202079 | 0 | 0 | 0 | 2 | 3 | 0 |
| ENSG00000162714 | 1363 | 671 | 810 | 757 | 892 | 812 |
| ENSG00000162711 | 0 | 0 | 0 | 0 | 0 | 0 |
| ENSG00000177535 | 0 | 0 | 0 | 0 | 0 | 0 |
| ENSG00000203664 | 0 | 0 | 0 | 0 | 0 | 0 |
| ENSG00000169224 | 0 | 0 | 0 | 0 | 0 | 0 |
| ENSG00000196242 | 0 | 0 | 0 | 0 | 0 | 0 |
| ENSG00000284824 | 0 | 0 | 0 | 0 | 0 | 0 |
| ENSG00000236817 | 0 | 0 | 0 | 0 | 0 | 0 |
| ENSG00000177489 | 0 | 0 | 0 | 0 | 0 | 0 |
| ENSG00000177476 | 0 | 0 | 0 | 0 | 0 | 0 |
| ENSG00000283369 | 0 | 0 | 0 | 0 | 0 | 0 |
| ENSG00000198452 | 0 | 0 | 0 | 0 | 0 | 0 |
| ENSG00000235749 | 0 | 0 | 0 | 0 | 0 | 0 |
| ENSG00000230411 | 0 | 0 | 0 | 0 | 0 | 0 |
| ENSG00000197437 | 0 | 0 | 0 | 0 | 0 | 0 |
| ENSG00000169214 | 0 | 0 | 0 | 0 | 0 | 0 |
| ENSG00000239395 | 0 | 0 | 0 | 0 | 0 | 0 |
| ENSG00000241128 | 0 | 0 | 0 | 0 | 0 | 0 |
| ENSG00000153230 | 0 | 0 | 0 | 0 | 0 | 0 |

| | | | | | | |
|---|---|---|---|---|---|---|
| ENSG00000221888 | 0 | 0 | 0 | 0 | 0 | 0 |
| ENSG00000228336 | 0 | 0 | 0 | 0 | 0 | 0 |
| ENSG00000196772 | 0 | 0 | 0 | 0 | 0 | 0 |
| ENSG00000232908 | 0 | 0 | 0 | 0 | 0 | 0 |
| ENSG00000230576 | 0 | 0 | 0 | 0 | 0 | 0 |
| ENSG00000197591 | 0 | 0 | 0 | 0 | 0 | 0 |
| ENSG00000162722 | 0 | 0 | 0 | 0 | 0 | 0 |
| ENSG00000238243 | 0 | 0 | 0 | 0 | 0 | 0 |
| ENSG00000177462 | 0 | 0 | 0 | 0 | 0 | 0 |
| ENSG00000177275 | 0 | 0 | 0 | 0 | 0 | 0 |
| ENSG00000226191 | 0 | 0 | 0 | 0 | 0 | 0 |
| ENSG00000224830 | 0 | 0 | 0 | 0 | 0 | 0 |
| ENSG00000279263 | 0 | 0 | 0 | 0 | 0 | 0 |
| ENSG00000187080 | 0 | 0 | 0 | 0 | 0 | 0 |
| ENSG00000237492 | 0 | 0 | 0 | 0 | 0 | 0 |
| ENSG00000224227 | 0 | 0 | 0 | 0 | 0 | 0 |
| ENSG00000199442 | 0 | 0 | 0 | 0 | 0 | 0 |
| ENSG00000232215 | 0 | 0 | 0 | 0 | 0 | 0 |
| ENSG00000197454 | 0 | 0 | 0 | 0 | 0 | 0 |
| ENSG00000200085 | 0 | 0 | 0 | 0 | 0 | 0 |
| ENSG00000203663 | 0 | 0 | 2 | 0 | 5 | 0 |
| ENSG00000200982 | 0 | 0 | 0 | 0 | 0 | 0 |
| ENSG00000198128 | 0 | 0 | 0 | 0 | 0 | 0 |
| ENSG00000197067 | 0 | 0 | 0 | 0 | 0 | 0 |
| ENSG00000196071 | 0 | 0 | 0 | 0 | 0 | 0 |
| ENSG00000177233 | 0 | 0 | 0 | 0 | 0 | 0 |
| ENSG00000162727 | 0 | 0 | 0 | 0 | 0 | 0 |
| ENSG00000198601 | 0 | 0 | 0 | 0 | 0 | 0 |
| ENSG00000228198 | 0 | 0 | 0 | 0 | 0 | 0 |
| ENSG00000171180 | 0 | 0 | 0 | 0 | 0 | 0 |
| ENSG00000177212 | 0 | 0 | 0 | 0 | 0 | 0 |
| ENSG00000177201 | 0 | 0 | 0 | 0 | 0 | 0 |
| ENSG00000177186 | 0 | 0 | 0 | 0 | 0 | 0 |
| ENSG00000177174 | 0 | 0 | 0 | 0 | 0 | 0 |
| ENSG00000196944 | 0 | 0 | 0 | 0 | 0 | 0 |
| ENSG00000198104 | 0 | 0 | 0 | 0 | 0 | 0 |
| ENSG00000175143 | 0 | 0 | 0 | 0 | 0 | 0 |
| ENSG00000227152 | 0 | 0 | 0 | 0 | 0 | 0 |
| ENSG00000196240 | 0 | 0 | 0 | 0 | 0 | 0 |
| ENSG00000196539 | 0 | 0 | 0 | 0 | 0 | 0 |
| ENSG00000229255 | 0 | 0 | 0 | 0 | 0 | 0 |
| ENSG00000203661 | 0 | 0 | 0 | 0 | 0 | 0 |
| ENSG00000271934 | 0 | 0 | 0 | 0 | 0 | 0 |
| ENSG00000188558 | 0 | 0 | 0 | 0 | 0 | 0 |
| ENSG00000224521 | 0 | 0 | 0 | 0 | 0 | 0 |
| ENSG00000227102 | 0 | 0 | 0 | 0 | 0 | 0 |
| ENSG00000182783 | 0 | 0 | 0 | 0 | 0 | 0 |
| ENSG00000183310 | 0 | 0 | 0 | 0 | 0 | 0 |
| ENSG00000184022 | 0 | 0 | 0 | 0 | 0 | 0 |
| ENSG00000201602 | 0 | 0 | 0 | 0 | 0 | 0 |

| | | | | | | |
|---|---|---|---|---|---|---|
| ENSG00000271701 | 0 | 0 | 0 | 0 | 0 | 0 |
| ENSG00000279301 | 0 | 0 | 0 | 0 | 0 | 0 |
| ENSG00000177151 | 0 | 0 | 0 | 0 | 0 | 0 |
| ENSG00000187701 | 0 | 0 | 0 | 0 | 0 | 0 |
| ENSG00000189181 | 0 | 0 | 0 | 0 | 0 | 0 |
| ENSG00000229703 | 0 | 0 | 0 | 0 | 0 | 0 |
| ENSG00000276520 | 0 | 0 | 0 | 0 | 0 | 0 |
| ENSG00000259823 | 0 | 0 | 0 | 0 | 0 | 0 |
| ENSG00000242529 | 0 | 0 | 0 | 0 | 0 | 0 |
| ENSG00000229663 | 0 | 0 | 0 | 0 | 0 | 0 |
| ENSG00000175137 | 1535 | 699 | 882 | 898 | 753 | 867 |
| ENSG00000264500 | 0 | 0 | 0 | 0 | 0 | 0 |
| ENSG00000171161 | 808 | 389 | 526 | 504 | 432 | 469 |
| ENSG00000171163 | 870 | 398 | 519 | 467 | 458 | 478 |
| ENSG00000227237 | 0 | 2 | 2 | 3 | 1 | 0 |
| ENSG00000185220 | 301 | 153 | 234 | 162 | 186 | 219 |
| ENSG00000200495 | 0 | 2 | 0 | 0 | 0 | 0 |
| ENSG00000233084 | 4 | 2 | 1 | 0 | 0 | 1 |
| ENSG00000184731 | 1581 | 638 | 910 | 573 | 823 | 890 |
| ENSG00000227061 | 2 | 0 | 0 | 0 | 0 | 0 |
| ENSG00000035115 | 3087 | 1582 | 1971 | 1344 | 1872 | 1635 |
| ENSG00000143727 | 3478 | 1753 | 2250 | 2025 | 2090 | 2148 |
| ENSG00000189292 | 0 | 1 | 3 | 0 | 1 | 0 |
| ENSG00000228643 | 2 | 3 | 4 | 1 | 1 | 4 |
| ENSG00000235779 | 0 | 0 | 0 | 0 | 0 | 0 |
| ENSG00000277997 | 0 | 0 | 0 | 0 | 0 | 0 |
| ENSG00000233684 | 0 | 0 | 3 | 1 | 0 | 0 |
| ENSG00000236856 | 0 | 0 | 0 | 1 | 0 | 0 |
| ENSG00000226277 | 0 | 0 | 0 | 0 | 0 | 0 |
| ENSG00000223985 | 0 | 0 | 0 | 0 | 0 | 0 |
| ENSG00000284452 | 0 | 0 | 0 | 0 | 0 | 0 |
| ENSG00000225942 | 0 | 0 | 0 | 0 | 0 | 0 |
| ENSG00000233633 | 40 | 12 | 34 | 17 | 12 | 12 |
| ENSG00000151353 | 1816 | 885 | 1229 | 983 | 978 | 1025 |
| ENSG00000233296 | 4 | 5 | 9 | 5 | 1 | 5 |
| ENSG00000280360 | 3 | 2 | 2 | 2 | 1 | 0 |
| ENSG00000233970 | 0 | 0 | 0 | 0 | 0 | 0 |
| ENSG00000227713 | 0 | 0 | 0 | 0 | 0 | 0 |
| ENSG00000272342 | 4 | 9 | 7 | 1 | 8 | 6 |
| ENSG00000231173 | 1 | 0 | 1 | 3 | 0 | 3 |
| ENSG00000223751 | 11 | 2 | 5 | 3 | 1 | 5 |
| ENSG00000237667 | 0 | 3 | 0 | 2 | 6 | 2 |
| ENSG00000228799 | 0 | 0 | 0 | 0 | 0 | 0 |
| ENSG00000234796 | 0 | 0 | 0 | 0 | 0 | 0 |
| ENSG00000235688 | 0 | 0 | 0 | 0 | 0 | 0 |
| ENSG00000172554 | 0 | 0 | 0 | 0 | 0 | 0 |
| ENSG00000235403 | 0 | 0 | 0 | 0 | 0 | 0 |
| ENSG00000236665 | 0 | 0 | 0 | 0 | 0 | 0 |
| ENSG00000233553 | 0 | 0 | 0 | 0 | 0 | 0 |
| ENSG00000231482 | 2 | 2 | 0 | 0 | 3 | 2 |

| | | | | | | |
|---|---|---|---|---|---|---|
| ENSG00000115705 | 36 | 11 | 26 | 3 | 12 | 8 |
| ENSG00000228613 | 21 | 15 | 15 | 9 | 10 | 6 |
| ENSG00000283766 | 5 | 2 | 1 | 1 | 5 | 3 |
| ENSG00000203635 | 24 | 17 | 13 | 3 | 11 | 16 |
| ENSG00000130508 | 2847 | 1197 | 1791 | 1087 | 1287 | 1498 |
| ENSG00000186487 | 0 | 0 | 0 | 0 | 0 | 0 |
| ENSG00000284600 | 0 | 0 | 0 | 0 | 0 | 0 |
| ENSG00000232057 | 0 | 0 | 0 | 0 | 0 | 0 |
| ENSG00000225619 | 0 | 0 | 0 | 0 | 0 | 0 |
| ENSG00000234929 | 0 | 0 | 0 | 0 | 0 | 0 |
| ENSG00000227364 | 0 | 0 | 0 | 0 | 0 | 0 |
| ENSG00000237720 | 0 | 0 | 0 | 0 | 0 | 0 |
| ENSG00000228391 | 0 | 0 | 0 | 0 | 0 | 0 |
| ENSG00000234423 | 0 | 0 | 0 | 1 | 0 | 0 |
| ENSG00000236760 | 2 | 0 | 0 | 0 | 0 | 0 |
| ENSG00000226649 | 0 | 0 | 0 | 0 | 0 | 2 |
| ENSG00000032389 | 1788 | 849 | 985 | 1097 | 937 | 1041 |
| ENSG00000224885 | 5 | 3 | 1 | 1 | 4 | 2 |
| ENSG00000171853 | 1213 | 567 | 733 | 684 | 626 | 609 |
| ENSG00000225234 | 15 | 9 | 17 | 9 | 8 | 9 |
| ENSG00000271868 | 5 | 7 | 3 | 1 | 0 | 2 |
| ENSG00000182551 | 4390 | 2365 | 2796 | 2445 | 2263 | 2385 |
| ENSG00000235078 | 5 | 5 | 11 | 6 | 5 | 4 |
| ENSG00000242282 | 33 | 11 | 37 | 20 | 21 | 24 |
| ENSG00000255767 | 0 | 0 | 0 | 0 | 0 | 0 |
| ENSG00000171865 | 798 | 499 | 650 | 534 | 608 | 677 |
| ENSG00000234171 | 207 | 114 | 133 | 146 | 125 | 138 |
| ENSG00000171863 | 23893 | 12256 | 14498 | 15529 | 14777 | 14128 |
| ENSG00000118004 | 4 | 1 | 0 | 3 | 0 | 2 |
| ENSG00000237370 | 0 | 0 | 0 | 0 | 0 | 0 |
| ENSG00000188765 | 5 | 1 | 0 | 1 | 1 | 0 |
| ENSG00000151360 | 0 | 0 | 0 | 0 | 1 | 0 |
| ENSG00000232718 | 0 | 0 | 0 | 0 | 0 | 0 |
| ENSG00000224661 | 0 | 0 | 0 | 0 | 0 | 0 |
| ENSG00000214866 | 0 | 0 | 0 | 0 | 0 | 0 |
| ENSG00000237401 | 0 | 0 | 0 | 0 | 0 | 0 |
| ENSG00000229550 | 0 | 0 | 0 | 0 | 0 | 0 |
| ENSG00000231821 | 0 | 0 | 0 | 0 | 0 | 0 |
| ENSG00000231532 | 2 | 0 | 0 | 0 | 0 | 0 |
| ENSG00000252238 | 0 | 0 | 0 | 0 | 0 | 0 |
| ENSG00000207192 | 0 | 0 | 0 | 0 | 0 | 0 |
| ENSG00000228585 | 0 | 0 | 0 | 0 | 0 | 0 |
| ENSG00000232835 | 0 | 0 | 0 | 0 | 0 | 0 |
| ENSG00000224128 | 0 | 0 | 0 | 0 | 0 | 0 |
| ENSG00000230090 | 0 | 0 | 0 | 0 | 0 | 0 |
| ENSG00000176887 | 0 | 0 | 0 | 0 | 0 | 0 |
| ENSG00000242540 | 0 | 0 | 0 | 0 | 0 | 0 |
| ENSG00000236106 | 0 | 0 | 0 | 0 | 0 | 0 |
| ENSG00000237266 | 0 | 0 | 0 | 0 | 0 | 0 |
| ENSG00000232044 | 0 | 0 | 0 | 0 | 0 | 0 |

| | | | | | | |
|---|---|---|---|---|---|---|
| ENSG00000276733 | 0 | 0 | 0 | 0 | 0 | 0 |
| ENSG00000234275 | 0 | 0 | 0 | 0 | 0 | 0 |
| ENSG00000227007 | 0 | 0 | 0 | 0 | 0 | 0 |
| ENSG00000231610 | 0 | 0 | 0 | 0 | 0 | 0 |
| ENSG00000226488 | 0 | 0 | 0 | 0 | 0 | 0 |
| ENSG00000236172 | 0 | 0 | 0 | 0 | 0 | 0 |
| ENSG00000283394 | 0 | 0 | 0 | 0 | 0 | 0 |
| ENSG00000205837 | 0 | 0 | 0 | 0 | 0 | 0 |
| ENSG00000225964 | 0 | 0 | 0 | 0 | 0 | 0 |
| ENSG00000134326 | 8 | 4 | 4 | 8 | 0 | 8 |
| ENSG00000134321 | 1 | 1 | 0 | 1 | 1 | 0 |
| ENSG00000271947 | 0 | 0 | 0 | 0 | 0 | 0 |
| ENSG00000228203 | 0 | 0 | 0 | 0 | 0 | 0 |
| ENSG00000151692 | 154 | 64 | 84 | 58 | 55 | 68 |
| ENSG00000223884 | 2 | 0 | 1 | 0 | 0 | 2 |
| ENSG00000223145 | 0 | 0 | 0 | 0 | 0 | 0 |
| ENSG00000272002 | 0 | 0 | 0 | 0 | 0 | 0 |
| ENSG00000213774 | 0 | 0 | 0 | 0 | 0 | 0 |
| ENSG00000229727 | 0 | 0 | 0 | 0 | 0 | 0 |
| ENSG00000221255 | 0 | 0 | 0 | 0 | 0 | 0 |
| ENSG00000230515 | 0 | 0 | 0 | 0 | 0 | 0 |
| ENSG00000235576 | 0 | 0 | 0 | 0 | 0 | 0 |
| ENSG00000232979 | 0 | 0 | 0 | 0 | 0 | 0 |
| ENSG00000229405 | 0 | 0 | 0 | 0 | 0 | 0 |
| ENSG00000226506 | 0 | 0 | 0 | 0 | 0 | 0 |
| ENSG00000235665 | 0 | 0 | 0 | 0 | 0 | 0 |
| ENSG00000236790 | 0 | 0 | 1 | 0 | 1 | 1 |
| ENSG00000229740 | 0 | 0 | 0 | 0 | 0 | 0 |
| ENSG00000285591 | 0 | 0 | 0 | 0 | 0 | 0 |
| ENSG00000236008 | 38 | 19 | 19 | 6 | 15 | 9 |
| ENSG00000231083 | 0 | 3 | 1 | 0 | 2 | 1 |
| ENSG00000225642 | 0 | 0 | 0 | 0 | 0 | 0 |
| ENSG00000235092 | 234 | 88 | 134 | 62 | 109 | 117 |
| ENSG00000115738 | 6516 | 2815 | 3622 | 3324 | 3523 | 3548 |
| ENSG00000134313 | 1866 | 968 | 969 | 383 | 1138 | 970 |
| ENSG00000143797 | 1711 | 857 | 975 | 601 | 980 | 791 |
| ENSG00000230886 | 0 | 0 | 0 | 0 | 0 | 0 |
| ENSG00000223640 | 0 | 0 | 0 | 0 | 0 | 0 |
| ENSG00000242136 | 9 | 1 | 0 | 2 | 10 | 6 |
| ENSG00000261104 | 16 | 5 | 18 | 17 | 21 | 14 |
| ENSG00000244310 | 0 | 0 | 0 | 0 | 0 | 0 |
| ENSG00000240980 | 0 | 0 | 0 | 0 | 0 | 0 |
| ENSG00000151693 | 1327 | 542 | 790 | 422 | 717 | 670 |
| ENSG00000213772 | 0 | 0 | 0 | 0 | 0 | 0 |
| ENSG00000119185 | 1791 | 846 | 1018 | 1039 | 1066 | 988 |
| ENSG00000119203 | 1426 | 835 | 928 | 661 | 1108 | 872 |
| ENSG00000134330 | 1421 | 746 | 862 | 853 | 876 | 790 |
| ENSG00000151694 | 1042 | 481 | 630 | 380 | 551 | 571 |
| ENSG00000239300 | 0 | 2 | 3 | 0 | 2 | 7 |
| ENSG00000271855 | 17 | 8 | 10 | 6 | 17 | 12 |

| | | | | | | |
|---|---|---|---|---|---|---|
| ENSG00000134308 | 11245 | 5527 | 6438 | 5468 | 6245 | 6210 |
| ENSG00000207086 | 0 | 0 | 0 | 0 | 0 | 0 |
| ENSG00000240687 | 0 | 0 | 0 | 0 | 0 | 0 |
| ENSG00000244260 | 0 | 0 | 0 | 0 | 0 | 0 |
| ENSG00000200034 | 0 | 0 | 0 | 0 | 0 | 0 |
| ENSG00000240960 | 0 | 0 | 0 | 0 | 0 | 0 |
| ENSG00000243491 | 0 | 0 | 0 | 0 | 0 | 0 |
| ENSG00000115750 | 532 | 337 | 316 | 237 | 358 | 307 |
| ENSG00000269973 | 108 | 42 | 36 | 51 | 81 | 53 |
| ENSG00000134317 | 1330 | 641 | 824 | 500 | 739 | 785 |
| ENSG00000188525 | 0 | 1 | 0 | 0 | 0 | 0 |
| ENSG00000260476 | 0 | 0 | 0 | 0 | 0 | 0 |
| ENSG00000260077 | 28 | 13 | 26 | 12 | 18 | 31 |
| ENSG00000172059 | 406 | 246 | 316 | 254 | 299 | 321 |
| ENSG00000271787 | 0 | 0 | 0 | 0 | 0 | 0 |
| ENSG00000205795 | 10 | 5 | 3 | 7 | 9 | 3 |
| ENSG00000233502 | 0 | 0 | 0 | 0 | 0 | 2 |
| ENSG00000212558 | 0 | 0 | 1 | 0 | 0 | 0 |
| ENSG00000171848 | 1417 | 817 | 1085 | 684 | 995 | 1036 |
| ENSG00000284681 | 0 | 1 | 1 | 2 | 1 | 4 |
| ENSG00000163009 | 3 | 5 | 0 | 0 | 0 | 2 |
| ENSG00000206647 | 0 | 0 | 0 | 0 | 0 | 1 |
| ENSG00000265418 | 0 | 0 | 0 | 0 | 0 | 0 |
| ENSG00000264030 | 0 | 0 | 0 | 0 | 0 | 0 |
| ENSG00000285872 | 32 | 13 | 17 | 14 | 20 | 16 |
| ENSG00000206898 | 0 | 0 | 0 | 1 | 0 | 1 |
| ENSG00000115756 | 1243 | 595 | 823 | 895 | 735 | 742 |
| ENSG00000115758 | 1886 | 899 | 1305 | 1073 | 973 | 1154 |
| ENSG00000206633 | 3 | 0 | 2 | 0 | 0 | 0 |
| ENSG00000257135 | 0 | 2 | 0 | 4 | 1 | 1 |
| ENSG00000217258 | 0 | 0 | 0 | 0 | 0 | 0 |
| ENSG00000115761 | 1005 | 579 | 675 | 540 | 783 | 655 |
| ENSG00000234818 | 0 | 0 | 0 | 0 | 0 | 0 |
| ENSG00000243819 | 40 | 25 | 24 | 22 | 21 | 16 |
| ENSG00000277446 | 0 | 0 | 0 | 0 | 0 | 0 |
| ENSG00000236162 | 0 | 0 | 0 | 0 | 0 | 0 |
| ENSG00000143882 | 59 | 18 | 23 | 35 | 29 | 42 |
| ENSG00000239053 | 0 | 0 | 0 | 0 | 0 | 0 |
| ENSG00000272275 | 2 | 3 | 3 | 1 | 3 | 2 |
| ENSG00000143870 | 5340 | 2951 | 3471 | 2770 | 3206 | 3167 |
| ENSG00000238962 | 0 | 0 | 0 | 0 | 0 | 0 |
| ENSG00000232056 | 0 | 0 | 0 | 0 | 0 | 0 |
| ENSG00000270488 | 0 | 0 | 0 | 0 | 0 | 0 |
| ENSG00000271952 | 0 | 0 | 0 | 0 | 0 | 0 |
| ENSG00000162975 | 30 | 17 | 28 | 31 | 25 | 25 |
| ENSG00000226080 | 0 | 0 | 0 | 0 | 0 | 0 |
| ENSG00000145063 | 12 | 6 | 5 | 6 | 3 | 6 |
| ENSG00000150873 | 10 | 4 | 5 | 3 | 7 | 14 |
| ENSG00000162976 | 1905 | 1076 | 1293 | 1051 | 1127 | 1030 |
| ENSG00000134318 | 607 | 311 | 230 | 116 | 610 | 300 |

| | | | | | | |
|---|---|---|---|---|---|---|
| ENSG00000207267 | 0 | 0 | 0 | 0 | 2 | 0 |
| ENSG00000230154 | 0 | 0 | 0 | 0 | 1 | 0 |
| ENSG00000226961 | 0 | 0 | 0 | 0 | 0 | 0 |
| ENSG00000230952 | 0 | 0 | 0 | 0 | 0 | 0 |
| ENSG00000231403 | 0 | 0 | 0 | 0 | 0 | 0 |
| ENSG00000224177 | 0 | 0 | 0 | 0 | 0 | 0 |
| ENSG00000285569 | 0 | 1 | 0 | 1 | 0 | 0 |
| ENSG00000169016 | 939 | 428 | 600 | 479 | 567 | 659 |
| ENSG00000196208 | 3137 | 1535 | 2101 | 1987 | 2200 | 2869 |
| ENSG00000201610 | 0 | 0 | 0 | 0 | 0 | 0 |
| ENSG00000264010 | 1 | 0 | 0 | 0 | 0 | 0 |
| ENSG00000251718 | 0 | 0 | 0 | 0 | 0 | 0 |
| ENSG00000202248 | 0 | 0 | 0 | 0 | 0 | 0 |
| ENSG00000285626 | 0 | 0 | 0 | 0 | 0 | 0 |
| ENSG00000239899 | 1 | 3 | 1 | 0 | 2 | 2 |
| ENSG00000169006 | 2 | 0 | 3 | 0 | 0 | 0 |
| ENSG00000230044 | 4 | 0 | 1 | 0 | 0 | 1 |
| ENSG00000276411 | 0 | 0 | 0 | 0 | 0 | 0 |
| ENSG00000134324 | 1043 | 496 | 678 | 313 | 581 | 505 |
| ENSG00000228496 | 6 | 3 | 0 | 3 | 2 | 0 |
| ENSG00000203643 | 0 | 0 | 0 | 0 | 0 | 0 |
| ENSG00000230790 | 0 | 0 | 0 | 0 | 0 | 0 |
| ENSG00000265056 | 0 | 0 | 0 | 0 | 0 | 0 |
| ENSG00000265172 | 0 | 0 | 0 | 0 | 0 | 0 |
| ENSG00000224184 | 0 | 0 | 0 | 0 | 0 | 0 |
| ENSG00000238503 | 0 | 0 | 0 | 0 | 0 | 0 |
| ENSG00000264089 | 0 | 0 | 0 | 0 | 0 | 0 |
| ENSG00000207183 | 0 | 0 | 0 | 0 | 0 | 0 |
| ENSG00000223360 | 0 | 0 | 0 | 0 | 0 | 0 |
| ENSG00000261117 | 2 | 1 | 3 | 1 | 0 | 6 |
| ENSG00000071575 | 605 | 266 | 488 | 229 | 250 | 413 |
| ENSG00000264370 | 1 | 0 | 0 | 0 | 0 | 0 |
| ENSG00000225649 | 0 | 0 | 0 | 0 | 0 | 0 |
| ENSG00000285876 | 0 | 0 | 0 | 0 | 0 | 0 |
| ENSG00000229370 | 0 | 0 | 0 | 0 | 0 | 0 |
| ENSG00000230448 | 2 | 0 | 0 | 0 | 0 | 0 |
| ENSG00000227718 | 0 | 0 | 0 | 0 | 0 | 0 |
| ENSG00000251859 | 0 | 0 | 0 | 0 | 0 | 0 |
| ENSG00000237261 | 0 | 0 | 0 | 0 | 0 | 0 |
| ENSG00000228505 | 0 | 0 | 0 | 0 | 0 | 0 |
| ENSG00000162981 | 1296 | 503 | 712 | 687 | 773 | 794 |
| ENSG00000235127 | 0 | 0 | 0 | 0 | 0 | 0 |
| ENSG00000151779 | 1106 | 551 | 599 | 496 | 655 | 525 |
| ENSG00000229087 | 0 | 0 | 0 | 0 | 0 | 0 |
| ENSG00000234022 | 0 | 0 | 0 | 0 | 0 | 0 |
| ENSG00000079785 | 3262 | 1800 | 1868 | 1316 | 2353 | 1861 |
| ENSG00000224194 | 0 | 0 | 0 | 0 | 0 | 0 |
| ENSG00000231031 | 0 | 0 | 0 | 0 | 0 | 0 |
| ENSG00000237326 | 0 | 0 | 0 | 0 | 0 | 0 |
| ENSG00000202160 | 0 | 0 | 0 | 0 | 0 | 0 |

| | | | | | | |
|---|---|---|---|---|---|---|
| ENSG00000214657 | 0 | 0 | 0 | 0 | 0 | 0 |
| ENSG00000238371 | 0 | 0 | 0 | 0 | 0 | 0 |
| ENSG00000223850 | 0 | 0 | 0 | 0 | 0 | 0 |
| ENSG00000233718 | 0 | 0 | 0 | 0 | 0 | 1 |
| ENSG00000134323 | 16 | 15 | 14 | 5 | 9 | 4 |
| ENSG00000243541 | 0 | 0 | 0 | 0 | 0 | 0 |
| ENSG00000226764 | 0 | 0 | 0 | 0 | 0 | 0 |
| ENSG00000236289 | 0 | 0 | 0 | 0 | 0 | 0 |
| ENSG00000236989 | 0 | 0 | 0 | 0 | 0 | 0 |
| ENSG00000271315 | 0 | 0 | 0 | 0 | 0 | 0 |
| ENSG00000251704 | 0 | 0 | 0 | 0 | 0 | 0 |
| ENSG00000226041 | 0 | 0 | 0 | 0 | 0 | 0 |
| ENSG00000228876 | 0 | 0 | 0 | 0 | 0 | 0 |
| ENSG00000231266 | 0 | 0 | 0 | 0 | 0 | 0 |
| ENSG00000285354 | 0 | 0 | 0 | 0 | 0 | 0 |
| ENSG00000232444 | 0 | 0 | 0 | 0 | 0 | 0 |
| ENSG00000224400 | 0 | 0 | 0 | 0 | 0 | 0 |
| ENSG00000237633 | 0 | 0 | 0 | 0 | 0 | 0 |
| ENSG00000279663 | 0 | 0 | 0 | 0 | 0 | 0 |
| ENSG00000197872 | 164 | 105 | 149 | 90 | 147 | 94 |
| ENSG00000223536 | 0 | 1 | 0 | 0 | 0 | 0 |
| ENSG00000224604 | 182 | 102 | 195 | 79 | 82 | 158 |
| ENSG00000237670 | 1 | 0 | 4 | 0 | 0 | 1 |
| ENSG00000222842 | 0 | 0 | 0 | 0 | 0 | 0 |
| ENSG00000214843 | 2 | 2 | 0 | 3 | 0 | 1 |
| ENSG00000236348 | 6 | 3 | 6 | 2 | 1 | 3 |
| ENSG00000214842 | 0 | 0 | 0 | 0 | 0 | 0 |
| ENSG00000163032 | 2 | 2 | 4 | 0 | 0 | 0 |
| ENSG00000163029 | 410 | 258 | 163 | 87 | 507 | 237 |
| ENSG00000178295 | 444 | 246 | 200 | 88 | 375 | 214 |
| ENSG00000151379 | 0 | 0 | 0 | 0 | 0 | 0 |
| ENSG00000170745 | 325 | 204 | 176 | 183 | 214 | 205 |
| ENSG00000212455 | 0 | 0 | 0 | 0 | 0 | 0 |
| ENSG00000260331 | 3 | 0 | 0 | 0 | 0 | 0 |
| ENSG00000240857 | 705 | 400 | 438 | 477 | 423 | 449 |
| ENSG00000250741 | 0 | 0 | 0 | 0 | 0 | 0 |
| ENSG00000185013 | 0 | 7 | 9 | 2 | 10 | 3 |
| ENSG00000207170 | 0 | 0 | 0 | 0 | 0 | 0 |
| ENSG00000224626 | 0 | 0 | 2 | 0 | 0 | 0 |
| ENSG00000236204 | 163 | 106 | 131 | 71 | 105 | 107 |
| ENSG00000266738 | 0 | 0 | 0 | 0 | 0 | 0 |
| ENSG00000143867 | 16 | 2 | 1 | 4 | 1 | 5 |
| ENSG00000234597 | 0 | 2 | 0 | 0 | 0 | 0 |
| ENSG00000237992 | 0 | 0 | 1 | 0 | 0 | 1 |
| ENSG00000235911 | 0 | 0 | 0 | 0 | 0 | 0 |
| ENSG00000213403 | 0 | 0 | 0 | 0 | 0 | 0 |
| ENSG00000228784 | 11 | 7 | 8 | 1 | 4 | 7 |
| ENSG00000183891 | 254 | 127 | 199 | 97 | 141 | 167 |
| ENSG00000271991 | 5 | 1 | 5 | 2 | 6 | 2 |
| ENSG00000118965 | 189 | 138 | 165 | 93 | 180 | 185 |

| | | | | | | |
|---|---|---|---|---|---|---|
| ENSG00000227210 | 29 | 11 | 18 | 15 | 29 | 20 |
| ENSG00000132031 | 388 | 209 | 232 | 218 | 199 | 219 |
| ENSG00000068697 | 11848 | 5819 | 6820 | 6366 | 6913 | 6411 |
| ENSG00000223734 | 2 | 1 | 1 | 2 | 0 | 5 |
| ENSG00000234378 | 0 | 3 | 0 | 0 | 2 | 1 |
| ENSG00000213729 | 0 | 0 | 0 | 0 | 0 | 0 |
| ENSG00000266059 | 0 | 0 | 0 | 0 | 0 | 0 |
| ENSG00000200763 | 0 | 0 | 0 | 0 | 0 | 0 |
| ENSG00000115884 | 9943 | 4629 | 6044 | 4703 | 3890 | 4449 |
| ENSG00000235411 | 4 | 0 | 1 | 0 | 0 | 0 |
| ENSG00000238735 | 1 | 0 | 0 | 0 | 0 | 0 |
| ENSG00000055917 | 4470 | 2160 | 2656 | 1380 | 2551 | 2160 |
| ENSG00000232800 | 0 | 0 | 0 | 0 | 0 | 0 |
| ENSG00000212171 | 0 | 0 | 0 | 0 | 0 | 0 |
| ENSG00000143878 | 10199 | 4873 | 6077 | 6105 | 5548 | 6263 |
| ENSG00000228950 | 0 | 3 | 1 | 1 | 1 | 1 |
| ENSG00000200829 | 0 | 0 | 0 | 0 | 0 | 0 |
| ENSG00000227047 | 2 | 0 | 0 | 0 | 0 | 0 |
| ENSG00000279307 | 2 | 0 | 5 | 3 | 2 | 3 |
| ENSG00000118960 | 1611 | 771 | 1036 | 998 | 815 | 865 |
| ENSG00000269976 | 0 | 0 | 3 | 0 | 0 | 3 |
| ENSG00000231948 | 0 | 2 | 2 | 1 | 2 | 2 |
| ENSG00000233416 | 2 | 0 | 0 | 1 | 2 | 0 |
| ENSG00000143869 | 2 | 1 | 2 | 0 | 4 | 1 |
| ENSG00000270100 | 0 | 0 | 0 | 0 | 0 | 0 |
| ENSG00000118961 | 534 | 352 | 306 | 232 | 380 | 319 |
| ENSG00000236436 | 0 | 0 | 0 | 0 | 0 | 0 |
| ENSG00000261012 | 0 | 0 | 0 | 0 | 0 | 0 |
| ENSG00000084674 | 0 | 0 | 0 | 0 | 0 | 0 |
| ENSG00000280390 | 0 | 0 | 0 | 0 | 0 | 0 |
| ENSG00000218819 | 0 | 0 | 0 | 0 | 0 | 0 |
| ENSG00000271629 | 0 | 0 | 0 | 0 | 0 | 0 |
| ENSG00000233005 | 0 | 0 | 0 | 0 | 0 | 0 |
| ENSG00000231204 | 0 | 0 | 0 | 0 | 0 | 0 |
| ENSG00000270654 | 0 | 0 | 0 | 0 | 0 | 0 |
| ENSG00000235537 | 0 | 0 | 0 | 0 | 0 | 0 |
| ENSG00000228538 | 0 | 0 | 0 | 0 | 0 | 0 |
| ENSG00000229621 | 0 | 0 | 0 | 0 | 0 | 0 |
| ENSG00000264192 | 0 | 0 | 0 | 0 | 0 | 0 |
| ENSG00000228999 | 0 | 0 | 0 | 0 | 0 | 0 |
| ENSG00000223691 | 0 | 0 | 0 | 0 | 0 | 0 |
| ENSG00000234189 | 0 | 0 | 0 | 0 | 0 | 0 |
| ENSG00000206882 | 0 | 0 | 0 | 0 | 0 | 0 |
| ENSG00000234207 | 0 | 0 | 0 | 0 | 0 | 0 |
| ENSG00000233587 | 0 | 0 | 0 | 0 | 0 | 0 |
| ENSG00000222616 | 0 | 0 | 0 | 0 | 0 | 0 |
| ENSG00000232451 | 0 | 0 | 0 | 0 | 0 | 0 |
| ENSG00000223530 | 0 | 0 | 0 | 0 | 0 | 0 |
| ENSG00000233714 | 0 | 0 | 0 | 0 | 0 | 0 |
| ENSG00000223634 | 0 | 1 | 0 | 0 | 0 | 0 |

| | | | | | | |
|---|---|---|---|---|---|---|
| ENSG00000235497 | 2 | 0 | 0 | 0 | 0 | 0 |
| ENSG00000119771 | 70 | 27 | 50 | 39 | 44 | 41 |
| ENSG00000224361 | 0 | 0 | 0 | 0 | 0 | 0 |
| ENSG00000279526 | 0 | 0 | 0 | 0 | 0 | 0 |
| ENSG00000283031 | 5 | 4 | 1 | 3 | 2 | 1 |
| ENSG00000119778 | 965 | 413 | 426 | 215 | 605 | 385 |
| ENSG00000224464 | 0 | 0 | 0 | 1 | 0 | 0 |
| ENSG00000238111 | 0 | 0 | 0 | 0 | 0 | 0 |
| ENSG00000173960 | 1029 | 521 | 589 | 374 | 587 | 620 |
| ENSG00000234946 | 0 | 0 | 0 | 0 | 0 | 0 |
| ENSG00000243847 | 0 | 0 | 1 | 0 | 0 | 1 |
| ENSG00000205639 | 2 | 1 | 4 | 1 | 0 | 5 |
| ENSG00000163026 | 362 | 160 | 187 | 171 | 198 | 181 |
| ENSG00000222940 | 0 | 0 | 0 | 0 | 0 | 0 |
| ENSG00000119782 | 15 | 7 | 4 | 6 | 6 | 1 |
| ENSG00000115128 | 3532 | 1775 | 2145 | 2254 | 2214 | 2164 |
| ENSG00000219626 | 61 | 41 | 36 | 29 | 37 | 35 |
| ENSG00000115129 | 274 | 165 | 169 | 238 | 192 | 162 |
| ENSG00000176732 | 36 | 16 | 16 | 14 | 17 | 18 |
| ENSG00000276087 | 0 | 0 | 0 | 0 | 0 | 0 |
| ENSG00000232642 | 0 | 0 | 2 | 0 | 0 | 0 |
| ENSG00000186453 | 0 | 0 | 2 | 0 | 0 | 1 |
| ENSG00000223754 | 0 | 0 | 0 | 0 | 0 | 0 |
| ENSG00000198399 | 1029 | 494 | 512 | 248 | 706 | 434 |
| ENSG00000242628 | 0 | 0 | 0 | 0 | 0 | 1 |
| ENSG00000251805 | 0 | 0 | 0 | 0 | 0 | 0 |
| ENSG00000232963 | 0 | 0 | 0 | 0 | 0 | 0 |
| ENSG00000233747 | 6 | 0 | 1 | 0 | 0 | 0 |
| ENSG00000084676 | 1479 | 717 | 834 | 389 | 873 | 694 |
| ENSG00000202430 | 0 | 0 | 0 | 0 | 0 | 0 |
| ENSG00000206732 | 0 | 0 | 0 | 0 | 0 | 0 |
| ENSG00000184924 | 375 | 236 | 275 | 273 | 234 | 242 |
| ENSG00000138092 | 609 | 326 | 451 | 352 | 365 | 436 |
| ENSG00000138031 | 587 | 308 | 367 | 339 | 379 | 408 |
| ENSG00000271936 | 46 | 21 | 23 | 17 | 18 | 18 |
| ENSG00000115137 | 351 | 160 | 226 | 116 | 162 | 185 |
| ENSG00000237953 | 0 | 0 | 0 | 0 | 0 | 0 |
| ENSG00000202479 | 0 | 0 | 0 | 0 | 0 | 0 |
| ENSG00000224165 | 148 | 81 | 76 | 42 | 52 | 59 |
| ENSG00000207069 | 0 | 0 | 0 | 0 | 0 | 0 |
| ENSG00000084710 | 986 | 433 | 576 | 391 | 538 | 492 |
| ENSG00000276653 | 0 | 0 | 0 | 0 | 0 | 0 |
| ENSG00000229593 | 0 | 0 | 1 | 0 | 0 | 0 |
| ENSG00000115138 | 2 | 0 | 0 | 0 | 0 | 0 |
| ENSG00000230452 | 0 | 0 | 0 | 0 | 0 | 0 |
| ENSG00000119772 | 3190 | 1267 | 1849 | 1480 | 1692 | 1793 |
| ENSG00000221445 | 0 | 0 | 0 | 0 | 0 | 0 |
| ENSG00000235072 | 7 | 3 | 6 | 9 | 10 | 6 |
| ENSG00000138101 | 670 | 310 | 376 | 326 | 318 | 308 |
| ENSG00000224220 | 0 | 0 | 0 | 0 | 0 | 0 |

| | | | | | | |
|---|---|---|---|---|---|---|
| ENSG00000201160 | 0 | 0 | 0 | 0 | 0 | 0 |
| ENSG00000143970 | 1216 | 553 | 597 | 251 | 696 | 601 |
| ENSG00000187536 | 0 | 0 | 0 | 0 | 0 | 0 |
| ENSG00000217643 | 0 | 0 | 0 | 0 | 0 | 0 |
| ENSG00000218682 | 3 | 0 | 0 | 0 | 0 | 0 |
| ENSG00000217718 | 0 | 0 | 0 | 0 | 0 | 0 |
| ENSG00000084731 | 1453 | 628 | 855 | 852 | 1095 | 902 |
| ENSG00000237629 | 0 | 0 | 1 | 0 | 0 | 0 |
| ENSG00000268412 | 2 | 1 | 1 | 0 | 1 | 1 |
| ENSG00000084733 | 6357 | 2993 | 3712 | 3190 | 3798 | 3613 |
| ENSG00000199872 | 0 | 0 | 0 | 0 | 0 | 0 |
| ENSG00000235440 | 0 | 0 | 0 | 0 | 0 | 0 |
| ENSG00000231655 | 0 | 0 | 0 | 0 | 1 | 0 |
| ENSG00000214465 | 1 | 1 | 1 | 0 | 0 | 0 |
| ENSG00000227133 | 0 | 0 | 0 | 0 | 0 | 1 |
| ENSG00000237951 | 0 | 0 | 0 | 0 | 0 | 1 |
| ENSG00000157833 | 6 | 2 | 6 | 4 | 12 | 8 |
| ENSG00000084754 | 8899 | 4335 | 5185 | 4747 | 5074 | 5082 |
| ENSG00000138029 | 4506 | 2312 | 2955 | 2185 | 2541 | 2456 |
| ENSG00000173567 | 10 | 13 | 10 | 6 | 9 | 7 |
| ENSG00000138018 | 1813 | 970 | 1052 | 643 | 1314 | 1080 |
| ENSG00000157856 | 0 | 0 | 0 | 0 | 0 | 0 |
| ENSG00000115155 | 0 | 0 | 1 | 3 | 0 | 0 |
| ENSG00000173557 | 0 | 0 | 1 | 0 | 0 | 1 |
| ENSG00000157884 | 1 | 2 | 0 | 0 | 0 | 0 |
| ENSG00000225378 | 0 | 0 | 0 | 0 | 0 | 0 |
| ENSG00000225024 | 0 | 0 | 0 | 0 | 0 | 0 |
| ENSG00000171303 | 0 | 0 | 1 | 1 | 0 | 0 |
| ENSG00000213699 | 1603 | 738 | 992 | 976 | 885 | 998 |
| ENSG00000115163 | 249 | 127 | 159 | 194 | 227 | 218 |
| ENSG00000145835 | 0 | 0 | 0 | 0 | 0 | 0 |
| ENSG00000157851 | 62 | 30 | 33 | 34 | 19 | 24 |
| ENSG00000230286 | 0 | 2 | 0 | 0 | 0 | 0 |
| ENSG00000084764 | 1106 | 479 | 651 | 622 | 603 | 554 |
| ENSG00000205500 | 0 | 1 | 1 | 1 | 0 | 0 |
| ENSG00000119777 | 2845 | 1350 | 1652 | 1580 | 1475 | 1472 |
| ENSG00000084693 | 1619 | 818 | 1023 | 911 | 988 | 938 |
| ENSG00000231636 | 0 | 0 | 1 | 1 | 2 | 0 |
| ENSG00000272056 | 1 | 0 | 1 | 1 | 1 | 4 |
| ENSG00000229122 | 7 | 3 | 4 | 7 | 8 | 8 |
| ENSG00000272148 | 6 | 6 | 9 | 8 | 5 | 10 |
| ENSG00000228474 | 2717 | 1673 | 1926 | 2123 | 1864 | 1688 |
| ENSG00000138080 | 54 | 36 | 41 | 30 | 44 | 40 |
| ENSG00000138030 | 239 | 127 | 146 | 156 | 112 | 129 |
| ENSG00000138028 | 531 | 287 | 350 | 428 | 343 | 407 |
| ENSG00000143994 | 24 | 8 | 11 | 5 | 9 | 11 |
| ENSG00000138073 | 2740 | 1437 | 1632 | 1794 | 1542 | 1553 |
| ENSG00000186143 | 0 | 0 | 0 | 0 | 0 | 0 |
| ENSG00000163792 | 0 | 0 | 0 | 0 | 0 | 0 |
| ENSG00000236099 | 1 | 0 | 0 | 0 | 0 | 0 |

| | | | | | | |
|---|---|---|---|---|---|---|
| ENSG00000138074 | 5630 | 2913 | 3646 | 3668 | 3558 | 3965 |
| ENSG00000138085 | 3913 | 2042 | 2604 | 2407 | 2197 | 2177 |
| ENSG00000084774 | 3020 | 1192 | 1763 | 1738 | 1571 | 1833 |
| ENSG00000115194 | 132 | 59 | 66 | 71 | 90 | 65 |
| ENSG00000163793 | 0 | 0 | 0 | 0 | 0 | 2 |
| ENSG00000138100 | 22 | 3 | 4 | 3 | 8 | 5 |
| ENSG00000163794 | 44 | 32 | 25 | 35 | 25 | 31 |
| ENSG00000115204 | 1781 | 834 | 961 | 935 | 1001 | 966 |
| ENSG00000115207 | 2541 | 1214 | 1617 | 1341 | 1343 | 1556 |
| ENSG00000234945 | 1 | 7 | 3 | 1 | 2 | 5 |
| ENSG00000234072 | 593 | 262 | 381 | 301 | 305 | 385 |
| ENSG00000115211 | 1698 | 921 | 1104 | 966 | 984 | 931 |
| ENSG00000115234 | 4204 | 1906 | 2440 | 2749 | 2303 | 2429 |
| ENSG00000163795 | 109 | 65 | 76 | 69 | 51 | 46 |
| ENSG00000115241 | 11808 | 5916 | 7176 | 5295 | 4981 | 5130 |
| ENSG00000213453 | 5 | 0 | 3 | 3 | 4 | 7 |
| ENSG00000115216 | 18007 | 9009 | 10186 | 7300 | 7061 | 6843 |
| ENSG00000157992 | 654 | 305 | 386 | 420 | 363 | 420 |
| ENSG00000138002 | 543 | 262 | 395 | 217 | 266 | 287 |
| ENSG00000235267 | 0 | 0 | 3 | 0 | 0 | 0 |
| ENSG00000200003 | 0 | 0 | 0 | 0 | 0 | 0 |
| ENSG00000115226 | 8 | 2 | 2 | 0 | 2 | 0 |
| ENSG00000084734 | 0 | 0 | 0 | 0 | 0 | 0 |
| ENSG00000221843 | 129 | 59 | 59 | 45 | 67 | 72 |
| ENSG00000243943 | 305 | 141 | 180 | 135 | 181 | 173 |
| ENSG00000259080 | 0 | 0 | 0 | 0 | 0 | 0 |
| ENSG00000176714 | 128 | 69 | 94 | 55 | 83 | 61 |
| ENSG00000198522 | 1320 | 727 | 814 | 738 | 831 | 850 |
| ENSG00000206731 | 2 | 2 | 1 | 0 | 2 | 0 |
| ENSG00000119760 | 2350 | 1012 | 1448 | 969 | 1213 | 1199 |
| ENSG00000163798 | 1334 | 699 | 826 | 604 | 792 | 717 |
| ENSG00000205334 | 0 | 0 | 0 | 0 | 0 | 0 |
| ENSG00000243147 | 1600 | 788 | 941 | 1002 | 946 | 1007 |
| ENSG00000171174 | 167 | 97 | 107 | 96 | 63 | 87 |
| ENSG00000158019 | 916 | 450 | 519 | 491 | 470 | 536 |
| ENSG00000270846 | 0 | 0 | 0 | 0 | 0 | 0 |
| ENSG00000265321 | 8 | 12 | 6 | 4 | 10 | 12 |
| ENSG00000270528 | 0 | 0 | 0 | 0 | 1 | 1 |
| ENSG00000277035 | 0 | 0 | 0 | 0 | 0 | 0 |
| ENSG00000223522 | 5 | 1 | 0 | 1 | 0 | 4 |
| ENSG00000225991 | 0 | 0 | 0 | 0 | 0 | 0 |
| ENSG00000229951 | 62 | 45 | 62 | 85 | 122 | 93 |
| ENSG00000075426 | 2287 | 1057 | 1420 | 2039 | 2312 | 2378 |
| ENSG00000270640 | 0 | 0 | 0 | 0 | 2 | 0 |
| ENSG00000270210 | 2 | 0 | 0 | 0 | 4 | 3 |
| ENSG00000227938 | 0 | 0 | 0 | 0 | 0 | 0 |
| ENSG00000163803 | 6 | 1 | 3 | 2 | 0 | 1 |
| ENSG00000242915 | 0 | 0 | 0 | 0 | 0 | 0 |
| ENSG00000230730 | 4 | 0 | 0 | 0 | 0 | 0 |
| ENSG00000222232 | 0 | 0 | 0 | 0 | 0 | 0 |

| | | | | | | |
|---|---|---|---|---|---|---|
| ENSG00000226833 | 16 | 9 | 8 | 5 | 11 | 16 |
| ENSG00000213639 | 3627 | 2053 | 2097 | 1417 | 2428 | 1875 |
| ENSG00000163806 | 6 | 9 | 7 | 6 | 9 | 4 |
| ENSG00000273233 | 0 | 0 | 4 | 1 | 0 | 3 |
| ENSG00000171103 | 420 | 224 | 251 | 242 | 283 | 277 |
| ENSG00000163811 | 1544 | 778 | 915 | 805 | 1046 | 1047 |
| ENSG00000264994 | 0 | 0 | 0 | 0 | 0 | 1 |
| ENSG00000265145 | 0 | 1 | 0 | 0 | 0 | 0 |
| ENSG00000222224 | 0 | 0 | 0 | 0 | 0 | 0 |
| ENSG00000265706 | 2 | 1 | 0 | 0 | 0 | 0 |
| ENSG00000189350 | 11 | 1 | 4 | 0 | 0 | 0 |
| ENSG00000200283 | 0 | 0 | 0 | 0 | 0 | 0 |
| ENSG00000179270 | 0 | 0 | 1 | 0 | 0 | 0 |
| ENSG00000229224 | 0 | 0 | 0 | 0 | 0 | 0 |
| ENSG00000115295 | 419 | 220 | 361 | 122 | 211 | 231 |
| ENSG00000171094 | 0 | 0 | 1 | 2 | 0 | 0 |
| ENSG00000242699 | 0 | 0 | 0 | 0 | 0 | 0 |
| ENSG00000230737 | 0 | 0 | 0 | 0 | 0 | 0 |
| ENSG00000197644 | 0 | 0 | 0 | 0 | 0 | 0 |
| ENSG00000233862 | 14 | 0 | 1 | 0 | 2 | 0 |
| ENSG00000225943 | 0 | 0 | 0 | 0 | 0 | 0 |
| ENSG00000119801 | 3722 | 1902 | 2168 | 1636 | 2134 | 1763 |
| ENSG00000207187 | 0 | 0 | 0 | 0 | 0 | 0 |
| ENSG00000231712 | 0 | 0 | 0 | 0 | 0 | 0 |
| ENSG00000213626 | 2 | 1 | 0 | 0 | 0 | 0 |
| ENSG00000235997 | 0 | 0 | 0 | 0 | 0 | 0 |
| ENSG00000280154 | 0 | 0 | 0 | 0 | 0 | 0 |
| ENSG00000172954 | 519 | 287 | 298 | 178 | 310 | 257 |
| ENSG00000278908 | 4 | 5 | 4 | 3 | 1 | 6 |
| ENSG00000162949 | 1883 | 948 | 1123 | 635 | 818 | 842 |
| ENSG00000230118 | 5 | 8 | 8 | 5 | 8 | 6 |
| ENSG00000285984 | 0 | 0 | 0 | 0 | 0 | 0 |
| ENSG00000158089 | 44 | 26 | 21 | 24 | 15 | 28 |
| ENSG00000234579 | 0 | 0 | 0 | 0 | 0 | 0 |
| ENSG00000214711 | 7 | 1 | 5 | 2 | 3 | 2 |
| ENSG00000201671 | 0 | 0 | 0 | 0 | 0 | 0 |
| ENSG00000013016 | 564 | 246 | 283 | 206 | 238 | 239 |
| ENSG00000213620 | 0 | 0 | 0 | 0 | 0 | 0 |
| ENSG00000158125 | 0 | 0 | 0 | 2 | 0 | 1 |
| ENSG00000277893 | 0 | 0 | 0 | 0 | 0 | 0 |
| ENSG00000228563 | 0 | 0 | 0 | 0 | 0 | 0 |
| ENSG00000270422 | 0 | 0 | 0 | 0 | 0 | 0 |
| ENSG00000223647 | 0 | 0 | 0 | 0 | 0 | 0 |
| ENSG00000237007 | 1 | 1 | 1 | 0 | 0 | 0 |
| ENSG00000242272 | 0 | 0 | 0 | 0 | 0 | 0 |
| ENSG00000273165 | 0 | 0 | 0 | 0 | 0 | 0 |
| ENSG00000162959 | 82 | 42 | 50 | 32 | 54 | 58 |
| ENSG00000162961 | 1945 | 976 | 1279 | 1044 | 1167 | 1057 |
| ENSG00000271228 | 2 | 1 | 2 | 3 | 1 | 0 |
| ENSG00000021574 | 722 | 387 | 351 | 190 | 432 | 296 |

| | | | | | | |
|---|---|---|---|---|---|---|
| ENSG00000272716 | 4 | 7 | 7 | 5 | 7 | 11 |
| ENSG00000152683 | 1389 | 653 | 806 | 487 | 796 | 734 |
| ENSG00000229816 | 8 | 8 | 12 | 2 | 9 | 6 |
| ENSG00000201113 | 0 | 0 | 0 | 0 | 0 | 0 |
| ENSG00000091106 | 8 | 2 | 7 | 6 | 5 | 4 |
| ENSG00000119820 | 1845 | 947 | 1082 | 870 | 1044 | 943 |
| ENSG00000272754 | 0 | 0 | 0 | 0 | 0 | 0 |
| ENSG00000115760 | 1552 | 988 | 779 | 293 | 1711 | 860 |
| ENSG00000230046 | 0 | 1 | 1 | 0 | 0 | 0 |
| ENSG00000276334 | 10 | 8 | 9 | 3 | 10 | 2 |
| ENSG00000276517 | 6 | 8 | 9 | 1 | 10 | 7 |
| ENSG00000207653 | 0 | 0 | 0 | 0 | 0 | 0 |
| ENSG00000274159 | 0 | 0 | 1 | 0 | 0 | 0 |
| ENSG00000279897 | 2 | 1 | 0 | 0 | 0 | 0 |
| ENSG00000279544 | 2 | 2 | 0 | 0 | 0 | 0 |
| ENSG00000018699 | 521 | 276 | 321 | 259 | 379 | 372 |
| ENSG00000265057 | 0 | 0 | 0 | 0 | 0 | 0 |
| ENSG00000230876 | 0 | 0 | 1 | 0 | 0 | 0 |
| ENSG00000236854 | 0 | 1 | 0 | 0 | 0 | 0 |
| ENSG00000206951 | 1 | 0 | 0 | 0 | 0 | 0 |
| ENSG00000049323 | 1201 | 510 | 760 | 388 | 510 | 512 |
| ENSG00000223951 | 0 | 0 | 0 | 0 | 0 | 0 |
| ENSG00000285577 | 0 | 0 | 0 | 0 | 0 | 0 |
| ENSG00000212204 | 0 | 0 | 0 | 0 | 0 | 0 |
| ENSG00000252246 | 0 | 0 | 0 | 0 | 0 | 0 |
| ENSG00000283591 | 0 | 0 | 0 | 0 | 0 | 0 |
| ENSG00000152689 | 6 | 5 | 12 | 9 | 18 | 14 |
| ENSG00000237133 | 0 | 0 | 0 | 0 | 1 | 2 |
| ENSG00000119812 | 2100 | 1019 | 1141 | 887 | 1212 | 1142 |
| ENSG00000238038 | 0 | 0 | 0 | 0 | 0 | 0 |
| ENSG00000187999 | 0 | 0 | 0 | 0 | 0 | 1 |
| ENSG00000228262 | 0 | 0 | 0 | 1 | 0 | 0 |
| ENSG00000239649 | 0 | 0 | 0 | 0 | 0 | 0 |
| ENSG00000235064 | 0 | 0 | 1 | 0 | 1 | 0 |
| ENSG00000237790 | 0 | 0 | 0 | 0 | 0 | 0 |
| ENSG00000285754 | 0 | 0 | 0 | 0 | 0 | 0 |
| ENSG00000272027 | 0 | 0 | 0 | 0 | 0 | 0 |
| ENSG00000232153 | 0 | 0 | 0 | 0 | 0 | 0 |
| ENSG00000226994 | 0 | 0 | 0 | 0 | 0 | 0 |
| ENSG00000264352 | 0 | 0 | 0 | 0 | 0 | 0 |
| ENSG00000237320 | 0 | 0 | 0 | 0 | 0 | 0 |
| ENSG00000196085 | 0 | 0 | 0 | 0 | 0 | 0 |
| ENSG00000265301 | 0 | 0 | 0 | 0 | 0 | 0 |
| ENSG00000207255 | 0 | 0 | 0 | 0 | 0 | 0 |
| ENSG00000229013 | 0 | 0 | 0 | 0 | 0 | 0 |
| ENSG00000234587 | 0 | 0 | 0 | 0 | 0 | 0 |
| ENSG00000233752 | 0 | 0 | 0 | 0 | 0 | 0 |
| ENSG00000260025 | 39 | 11 | 18 | 15 | 12 | 11 |
| ENSG00000150938 | 775 | 393 | 541 | 237 | 348 | 332 |
| ENSG00000273090 | 0 | 0 | 1 | 0 | 0 | 0 |

| | | | | | | |
|---|---|---|---|---|---|---|
| ENSG00000171055 | 1496 | 827 | 870 | 688 | 840 | 767 |
| ENSG00000270677 | 0 | 0 | 0 | 0 | 0 | 0 |
| ENSG00000205221 | 1 | 0 | 0 | 0 | 1 | 1 |
| ENSG00000279519 | 6 | 11 | 10 | 4 | 18 | 6 |
| ENSG00000115808 | 1143 | 601 | 584 | 316 | 764 | 583 |
| ENSG00000252756 | 0 | 1 | 0 | 0 | 0 | 0 |
| ENSG00000008869 | 1472 | 713 | 839 | 361 | 842 | 665 |
| ENSG00000152133 | 382 | 193 | 209 | 114 | 218 | 192 |
| ENSG00000055332 | 2008 | 1071 | 1244 | 680 | 1136 | 1101 |
| ENSG00000224891 | 0 | 0 | 1 | 0 | 0 | 0 |
| ENSG00000138068 | 0 | 1 | 0 | 0 | 0 | 0 |
| ENSG00000233159 | 0 | 0 | 0 | 0 | 0 | 0 |
| ENSG00000218739 | 1149 | 665 | 778 | 660 | 784 | 718 |
| ENSG00000115816 | 416 | 305 | 203 | 116 | 482 | 272 |
| ENSG00000272054 | 5 | 8 | 22 | 8 | 14 | 9 |
| ENSG00000003509 | 607 | 296 | 385 | 307 | 351 | 304 |
| ENSG00000115825 | 226 | 172 | 147 | 77 | 213 | 135 |
| ENSG00000232028 | 0 | 0 | 0 | 0 | 0 | 0 |
| ENSG00000207364 | 0 | 0 | 0 | 0 | 0 | 0 |
| ENSG00000285925 | 0 | 0 | 0 | 0 | 0 | 0 |
| ENSG00000115828 | 28 | 23 | 32 | 22 | 33 | 34 |
| ENSG00000253078 | 0 | 0 | 0 | 0 | 0 | 0 |
| ENSG00000236213 | 0 | 0 | 0 | 0 | 0 | 0 |
| ENSG00000163171 | 11 | 6 | 4 | 5 | 6 | 8 |
| ENSG00000236572 | 0 | 0 | 0 | 0 | 0 | 0 |
| ENSG00000225402 | 0 | 0 | 0 | 0 | 0 | 0 |
| ENSG00000237803 | 0 | 0 | 0 | 0 | 0 | 0 |
| ENSG00000115841 | 58 | 28 | 27 | 31 | 31 | 34 |
| ENSG00000235848 | 1 | 2 | 2 | 3 | 6 | 5 |
| ENSG00000138061 | 8947 | 5058 | 5565 | 22770 | 39676 | 29873 |
| ENSG00000232973 | 103 | 68 | 54 | 115 | 194 | 149 |
| ENSG00000280276 | 2 | 0 | 0 | 0 | 3 | 0 |
| ENSG00000229160 | 0 | 0 | 1 | 1 | 3 | 1 |
| ENSG00000199603 | 0 | 0 | 0 | 0 | 2 | 0 |
| ENSG00000273006 | 0 | 0 | 0 | 0 | 0 | 2 |
| ENSG00000227292 | 0 | 0 | 1 | 0 | 0 | 0 |
| ENSG00000227770 | 0 | 0 | 0 | 0 | 0 | 0 |
| ENSG00000213594 | 0 | 0 | 0 | 0 | 0 | 0 |
| ENSG00000119787 | 2023 | 994 | 1274 | 865 | 1335 | 1158 |
| ENSG00000231367 | 0 | 2 | 1 | 0 | 1 | 2 |
| ENSG00000235009 | 0 | 1 | 0 | 0 | 0 | 0 |
| ENSG00000213553 | 218 | 91 | 98 | 123 | 93 | 97 |
| ENSG00000271443 | 0 | 0 | 0 | 0 | 2 | 0 |
| ENSG00000143889 | 1631 | 840 | 976 | 851 | 1042 | 977 |
| ENSG00000235586 | 2 | 1 | 0 | 1 | 1 | 2 |
| ENSG00000143891 | 156 | 68 | 94 | 57 | 83 | 100 |
| ENSG00000232518 | 0 | 0 | 0 | 0 | 0 | 0 |
| ENSG00000115875 | 3977 | 2143 | 2498 | 2088 | 2428 | 2364 |
| ENSG00000152147 | 580 | 259 | 342 | 406 | 340 | 401 |
| ENSG00000227238 | 0 | 0 | 0 | 0 | 0 | 0 |

| | | | | | | |
|---|---|---|---|---|---|---|
| ENSG00000232114 | 0 | 0 | 0 | 0 | 0 | 0 |
| ENSG00000163214 | 801 | 429 | 475 | 283 | 596 | 431 |
| ENSG00000223922 | 6 | 2 | 2 | 2 | 2 | 2 |
| ENSG00000225284 | 0 | 0 | 0 | 0 | 1 | 0 |
| ENSG00000188010 | 451 | 213 | 242 | 252 | 271 | 241 |
| ENSG00000207295 | 0 | 0 | 0 | 0 | 0 | 0 |
| ENSG00000214694 | 4 | 2 | 2 | 1 | 1 | 0 |
| ENSG00000265905 | 0 | 0 | 0 | 0 | 0 | 0 |
| ENSG00000269210 | 0 | 0 | 0 | 0 | 0 | 0 |
| ENSG00000115904 | 1431 | 759 | 618 | 337 | 1161 | 631 |
| ENSG00000229692 | 48 | 28 | 23 | 20 | 37 | 12 |
| ENSG00000206985 | 0 | 1 | 0 | 0 | 0 | 0 |
| ENSG00000225900 | 0 | 0 | 0 | 0 | 1 | 0 |
| ENSG00000202309 | 0 | 0 | 0 | 0 | 0 | 0 |
| ENSG00000205111 | 2 | 2 | 4 | 1 | 1 | 0 |
| ENSG00000011566 | 1323 | 712 | 703 | 352 | 834 | 610 |
| ENSG00000252473 | 0 | 0 | 3 | 0 | 1 | 2 |
| ENSG00000273035 | 0 | 0 | 0 | 0 | 1 | 1 |
| ENSG00000231312 | 308 | 159 | 168 | 165 | 161 | 162 |
| ENSG00000152154 | 15 | 8 | 9 | 6 | 2 | 7 |
| ENSG00000138050 | 224 | 126 | 153 | 104 | 158 | 139 |
| ENSG00000227028 | 1 | 1 | 1 | 0 | 2 | 1 |
| ENSG00000235653 | 0 | 0 | 0 | 0 | 0 | 0 |
| ENSG00000183023 | 463 | 226 | 268 | 156 | 226 | 186 |
| ENSG00000285898 | 0 | 0 | 0 | 0 | 0 | 0 |
| ENSG00000233128 | 0 | 0 | 0 | 0 | 0 | 0 |
| ENSG00000237442 | 0 | 0 | 1 | 0 | 0 | 0 |
| ENSG00000236090 | 2 | 0 | 0 | 0 | 0 | 0 |
| ENSG00000236533 | 0 | 0 | 0 | 0 | 1 | 0 |
| ENSG00000207218 | 0 | 0 | 0 | 0 | 0 | 0 |
| ENSG00000223245 | 0 | 0 | 0 | 0 | 0 | 0 |
| ENSG00000214691 | 0 | 0 | 0 | 0 | 0 | 0 |
| ENSG00000234362 | 0 | 0 | 0 | 0 | 0 | 0 |
| ENSG00000205086 | 0 | 0 | 0 | 0 | 0 | 0 |
| ENSG00000226398 | 0 | 0 | 0 | 0 | 0 | 0 |
| ENSG00000162878 | 8 | 12 | 1 | 7 | 3 | 9 |
| ENSG00000224875 | 0 | 0 | 0 | 0 | 0 | 0 |
| ENSG00000143924 | 2127 | 1097 | 1332 | 689 | 1171 | 1111 |
| ENSG00000115944 | 5331 | 2641 | 3209 | 3397 | 2962 | 2996 |
| ENSG00000221300 | 0 | 1 | 0 | 0 | 0 | 0 |
| ENSG00000171126 | 2 | 2 | 3 | 6 | 0 | 7 |
| ENSG00000232526 | 0 | 0 | 0 | 0 | 0 | 0 |
| ENSG00000236913 | 0 | 0 | 0 | 0 | 0 | 0 |
| ENSG00000057935 | 2025 | 1055 | 1317 | 918 | 1221 | 1102 |
| ENSG00000273913 | 0 | 0 | 0 | 0 | 0 | 0 |
| ENSG00000215263 | 0 | 0 | 0 | 0 | 0 | 0 |
| ENSG00000226523 | 0 | 1 | 1 | 0 | 0 | 0 |
| ENSG00000200550 | 0 | 0 | 0 | 1 | 1 | 0 |
| ENSG00000162881 | 0 | 0 | 1 | 1 | 0 | 1 |
| ENSG00000162882 | 2 | 0 | 0 | 1 | 1 | 5 |

| | | | | | | |
|---|---|---|---|---|---|---|
| ENSG00000226491 | 0 | 0 | 0 | 0 | 0 | 3 |
| ENSG00000232202 | 0 | 0 | 0 | 0 | 0 | 1 |
| ENSG00000224739 | 0 | 0 | 0 | 0 | 0 | 0 |
| ENSG00000231826 | 0 | 1 | 0 | 1 | 0 | 0 |
| ENSG00000233978 | 0 | 0 | 0 | 0 | 0 | 0 |
| ENSG00000207087 | 0 | 0 | 0 | 0 | 0 | 0 |
| ENSG00000230587 | 0 | 0 | 0 | 0 | 0 | 0 |
| ENSG00000152518 | 3652 | 1849 | 1651 | 1151 | 1289 | 1079 |
| ENSG00000279873 | 23 | 22 | 13 | 7 | 15 | 12 |
| ENSG00000234936 | 16 | 6 | 19 | 6 | 12 | 12 |
| ENSG00000115970 | 1485 | 757 | 755 | 640 | 842 | 670 |
| ENSG00000252804 | 0 | 0 | 0 | 0 | 0 | 0 |
| ENSG00000264071 | 0 | 0 | 0 | 0 | 0 | 0 |
| ENSG00000207411 | 0 | 0 | 0 | 0 | 0 | 0 |
| ENSG00000152527 | 311 | 144 | 114 | 54 | 195 | 97 |
| ENSG00000223658 | 28 | 16 | 23 | 22 | 21 | 12 |
| ENSG00000229695 | 0 | 0 | 0 | 0 | 0 | 0 |
| ENSG00000252490 | 0 | 0 | 0 | 0 | 0 | 0 |
| ENSG00000138036 | 486 | 285 | 296 | 217 | 314 | 258 |
| ENSG00000138075 | 0 | 1 | 2 | 1 | 0 | 0 |
| ENSG00000143921 | 2 | 2 | 0 | 0 | 0 | 0 |
| ENSG00000138095 | 3756 | 2442 | 1963 | 1223 | 3757 | 2230 |
| ENSG00000200924 | 0 | 0 | 2 | 0 | 0 | 0 |
| ENSG00000264047 | 0 | 0 | 1 | 0 | 0 | 0 |
| ENSG00000219391 | 0 | 0 | 0 | 0 | 1 | 0 |
| ENSG00000252599 | 0 | 0 | 0 | 0 | 0 | 0 |
| ENSG00000224637 | 0 | 0 | 1 | 1 | 0 | 0 |
| ENSG00000273106 | 0 | 1 | 0 | 0 | 1 | 0 |
| ENSG00000138032 | 570 | 266 | 360 | 253 | 356 | 356 |
| ENSG00000285542 | 0 | 0 | 0 | 0 | 0 | 0 |
| ENSG00000226972 | 0 | 0 | 0 | 1 | 1 | 0 |
| ENSG00000138079 | 0 | 0 | 1 | 0 | 0 | 1 |
| ENSG00000138078 | 1599 | 811 | 926 | 469 | 1062 | 747 |
| ENSG00000143919 | 285 | 118 | 130 | 117 | 125 | 136 |
| ENSG00000259439 | 2 | 2 | 1 | 1 | 0 | 1 |
| ENSG00000278957 | 0 | 0 | 0 | 0 | 0 | 0 |
| ENSG00000274520 | 0 | 0 | 0 | 0 | 0 | 0 |
| ENSG00000277311 | 0 | 0 | 0 | 0 | 0 | 0 |
| ENSG00000276229 | 0 | 0 | 0 | 0 | 0 | 0 |
| ENSG00000277809 | 0 | 0 | 0 | 0 | 0 | 0 |
| ENSG00000276048 | 0 | 0 | 0 | 0 | 0 | 0 |
| ENSG00000274547 | 0 | 0 | 0 | 0 | 0 | 0 |
| ENSG00000276670 | 0 | 0 | 0 | 0 | 0 | 0 |
| ENSG00000236502 | 0 | 0 | 0 | 0 | 0 | 0 |
| ENSG00000138083 | 0 | 0 | 0 | 0 | 0 | 0 |
| ENSG00000225156 | 0 | 0 | 0 | 0 | 0 | 0 |
| ENSG00000231848 | 2 | 1 | 0 | 1 | 0 | 0 |
| ENSG00000170577 | 79 | 40 | 45 | 32 | 31 | 31 |
| ENSG00000231156 | 0 | 0 | 0 | 0 | 0 | 0 |
| ENSG00000205054 | 0 | 0 | 1 | 0 | 0 | 0 |

| | | | | | | |
|---|---|---|---|---|---|---|
| ENSG00000231054 | 0 | 0 | 0 | 0 | 0 | 0 |
| ENSG00000068784 | 420 | 204 | 249 | 131 | 333 | 265 |
| ENSG00000239396 | 2 | 0 | 0 | 0 | 0 | 0 |
| ENSG00000171132 | 561 | 270 | 332 | 225 | 288 | 349 |
| ENSG00000228481 | 1 | 0 | 0 | 0 | 0 | 0 |
| ENSG00000233829 | 0 | 0 | 0 | 0 | 0 | 0 |
| ENSG00000232696 | 0 | 0 | 0 | 0 | 0 | 0 |
| ENSG00000231336 | 0 | 0 | 0 | 0 | 1 | 0 |
| ENSG00000233335 | 0 | 0 | 0 | 0 | 0 | 0 |
| ENSG00000278764 | 0 | 0 | 0 | 0 | 0 | 0 |
| ENSG00000116016 | 3513 | 2068 | 1724 | 1282 | 1815 | 1509 |
| ENSG00000228100 | 2 | 0 | 0 | 0 | 2 | 0 |
| ENSG00000187600 | 0 | 0 | 0 | 0 | 0 | 0 |
| ENSG00000284608 | 0 | 0 | 0 | 0 | 0 | 0 |
| ENSG00000241791 | 0 | 0 | 0 | 0 | 0 | 0 |
| ENSG00000284701 | 0 | 0 | 0 | 0 | 0 | 0 |
| ENSG00000250565 | 272 | 122 | 151 | 162 | 190 | 161 |
| ENSG00000253515 | 4 | 0 | 1 | 0 | 0 | 0 |
| ENSG00000119729 | 1616 | 820 | 957 | 752 | 1118 | 877 |
| ENSG00000250116 | 13 | 4 | 4 | 4 | 7 | 3 |
| ENSG00000151665 | 376 | 277 | 383 | 321 | 321 | 335 |
| ENSG00000119878 | 622 | 312 | 394 | 367 | 393 | 338 |
| ENSG00000279254 | 0 | 1 | 0 | 0 | 1 | 0 |
| ENSG00000222005 | 4 | 2 | 2 | 1 | 2 | 2 |
| ENSG00000171150 | 468 | 238 | 271 | 162 | 303 | 204 |
| ENSG00000239332 | 4 | 0 | 1 | 3 | 5 | 2 |
| ENSG00000226548 | 0 | 0 | 0 | 0 | 0 | 1 |
| ENSG00000228925 | 9 | 4 | 5 | 1 | 5 | 4 |
| ENSG00000180398 | 3903 | 1944 | 2357 | 1828 | 2393 | 2275 |
| ENSG00000068724 | 1194 | 581 | 738 | 744 | 722 | 681 |
| ENSG00000272814 | 0 | 0 | 0 | 0 | 1 | 0 |
| ENSG00000233845 | 0 | 1 | 0 | 0 | 0 | 1 |
| ENSG00000239605 | 0 | 1 | 1 | 0 | 1 | 1 |
| ENSG00000273269 | 0 | 0 | 0 | 0 | 0 | 0 |
| ENSG00000225187 | 0 | 0 | 2 | 0 | 3 | 0 |
| ENSG00000143933 | 7815 | 4404 | 5548 | 4272 | 6107 | 4234 |
| ENSG00000234690 | 17 | 15 | 24 | 17 | 12 | 16 |
| ENSG00000226087 | 0 | 0 | 1 | 0 | 0 | 0 |
| ENSG00000236824 | 2 | 1 | 4 | 8 | 6 | 1 |
| ENSG00000119888 | 10906 | 5668 | 6567 | 4888 | 6167 | 5274 |
| ENSG00000222685 | 0 | 0 | 0 | 0 | 0 | 1 |
| ENSG00000207923 | 7 | 0 | 3 | 2 | 6 | 2 |
| ENSG00000277396 | 2 | 0 | 1 | 0 | 1 | 2 |
| ENSG00000095002 | 1431 | 787 | 885 | 448 | 967 | 781 |
| ENSG00000184261 | 6 | 9 | 7 | 8 | 7 | 14 |
| ENSG00000235760 | 0 | 4 | 4 | 0 | 1 | 4 |
| ENSG00000230979 | 8 | 2 | 1 | 6 | 1 | 1 |
| ENSG00000116062 | 2556 | 1417 | 1702 | 984 | 1507 | 1413 |
| ENSG00000224443 | 0 | 0 | 0 | 0 | 0 | 0 |
| ENSG00000224058 | 0 | 0 | 0 | 0 | 0 | 0 |

| | | | | | | |
|---|---|---|---|---|---|---|
| ENSG00000283502 | 0 | 1 | 0 | 0 | 0 | 0 |
| ENSG00000138081 | 1014 | 522 | 682 | 371 | 589 | 547 |
| ENSG00000235369 | 5 | 0 | 1 | 1 | 0 | 0 |
| ENSG00000236549 | 0 | 1 | 0 | 0 | 0 | 0 |
| ENSG00000233230 | 22 | 18 | 21 | 9 | 19 | 17 |
| ENSG00000226473 | 0 | 0 | 0 | 0 | 0 | 0 |
| ENSG00000230773 | 0 | 0 | 1 | 0 | 0 | 0 |
| ENSG00000275423 | 0 | 0 | 0 | 0 | 0 | 0 |
| ENSG00000201010 | 0 | 0 | 0 | 0 | 0 | 0 |
| ENSG00000170802 | 410 | 216 | 179 | 80 | 297 | 185 |
| ENSG00000251889 | 0 | 0 | 0 | 0 | 0 | 0 |
| ENSG00000272663 | 9 | 9 | 7 | 0 | 2 | 5 |
| ENSG00000162869 | 894 | 445 | 543 | 351 | 538 | 464 |
| ENSG00000202227 | 2 | 1 | 0 | 0 | 0 | 0 |
| ENSG00000243244 | 433 | 226 | 149 | 122 | 315 | 124 |
| ENSG00000068781 | 0 | 0 | 0 | 0 | 1 | 0 |
| ENSG00000242441 | 0 | 0 | 0 | 0 | 1 | 0 |
| ENSG00000279956 | 0 | 0 | 0 | 0 | 0 | 0 |
| ENSG00000239995 | 0 | 0 | 0 | 0 | 0 | 0 |
| ENSG00000138039 | 2 | 0 | 1 | 0 | 0 | 0 |
| ENSG00000232285 | 0 | 0 | 0 | 0 | 0 | 0 |
| ENSG00000282890 | 0 | 0 | 0 | 0 | 0 | 0 |
| ENSG00000214602 | 0 | 0 | 0 | 0 | 0 | 0 |
| ENSG00000170820 | 0 | 1 | 0 | 0 | 0 | 0 |
| ENSG00000212036 | 0 | 0 | 0 | 0 | 0 | 0 |
| ENSG00000282998 | 0 | 0 | 0 | 0 | 0 | 0 |
| ENSG00000206915 | 0 | 0 | 0 | 0 | 0 | 0 |
| ENSG00000282828 | 0 | 0 | 0 | 0 | 0 | 0 |
| ENSG00000234253 | 0 | 0 | 0 | 0 | 1 | 0 |
| ENSG00000212580 | 0 | 0 | 0 | 0 | 0 | 0 |
| ENSG00000179915 | 0 | 0 | 0 | 0 | 0 | 0 |
| ENSG00000285548 | 0 | 0 | 0 | 0 | 0 | 0 |
| ENSG00000230327 | 0 | 0 | 0 | 0 | 0 | 0 |
| ENSG00000283058 | 0 | 0 | 0 | 0 | 0 | 0 |
| ENSG00000216191 | 0 | 0 | 0 | 0 | 0 | 0 |
| ENSG00000238165 | 0 | 0 | 0 | 0 | 0 | 0 |
| ENSG00000231918 | 0 | 0 | 0 | 0 | 0 | 0 |
| ENSG00000224344 | 0 | 0 | 0 | 0 | 0 | 0 |
| ENSG00000223710 | 0 | 0 | 0 | 0 | 0 | 0 |
| ENSG00000230840 | 0 | 0 | 0 | 0 | 0 | 0 |
| ENSG00000226366 | 0 | 0 | 0 | 0 | 0 | 0 |
| ENSG00000236538 | 0 | 0 | 0 | 0 | 0 | 0 |
| ENSG00000202195 | 0 | 0 | 0 | 0 | 0 | 0 |
| ENSG00000225444 | 0 | 0 | 0 | 0 | 0 | 0 |
| ENSG00000225842 | 0 | 0 | 0 | 0 | 0 | 0 |
| ENSG00000236837 | 0 | 0 | 0 | 0 | 0 | 0 |
| ENSG00000204993 | 0 | 0 | 0 | 0 | 0 | 0 |
| ENSG00000233083 | 0 | 0 | 0 | 0 | 0 | 0 |
| ENSG00000264975 | 0 | 0 | 0 | 0 | 0 | 0 |
| ENSG00000228033 | 0 | 0 | 0 | 0 | 0 | 0 |

| | | | | | | |
|---|---|---|---|---|---|---|
| ENSG00000232604 | 0 | 0 | 0 | 0 | 0 | 0 |
| ENSG00000232668 | 0 | 0 | 0 | 0 | 0 | 0 |
| ENSG00000251942 | 0 | 0 | 0 | 0 | 0 | 0 |
| ENSG00000223897 | 0 | 0 | 0 | 0 | 0 | 0 |
| ENSG00000115239 | 49 | 27 | 29 | 30 | 32 | 31 |
| ENSG00000207456 | 0 | 0 | 0 | 0 | 0 | 0 |
| ENSG00000270898 | 0 | 1 | 0 | 0 | 0 | 0 |
| ENSG00000143942 | 134 | 55 | 53 | 77 | 58 | 85 |
| ENSG00000068912 | 2361 | 1160 | 1574 | 1018 | 1431 | 1429 |
| ENSG00000265452 | 1 | 1 | 4 | 2 | 2 | 4 |
| ENSG00000119737 | 24 | 15 | 10 | 13 | 10 | 10 |
| ENSG00000068878 | 2062 | 1132 | 1195 | 727 | 1745 | 1103 |
| ENSG00000170634 | 296 | 125 | 145 | 179 | 170 | 173 |
| ENSG00000235937 | 0 | 0 | 0 | 0 | 0 | 0 |
| ENSG00000233266 | 1 | 4 | 5 | 5 | 2 | 2 |
| ENSG00000241114 | 0 | 0 | 0 | 0 | 0 | 0 |
| ENSG00000272156 | 0 | 5 | 0 | 10 | 5 | 5 |
| ENSG00000252934 | 0 | 0 | 0 | 0 | 0 | 0 |
| ENSG00000178021 | 0 | 0 | 0 | 0 | 0 | 0 |
| ENSG00000177994 | 0 | 0 | 0 | 1 | 0 | 0 |
| ENSG00000115306 | 3350 | 1600 | 2213 | 1577 | 2344 | 2306 |
| ENSG00000228108 | 1 | 0 | 0 | 0 | 0 | 0 |
| ENSG00000237887 | 0 | 1 | 0 | 0 | 0 | 0 |
| ENSG00000234943 | 3 | 2 | 0 | 0 | 1 | 0 |
| ENSG00000238018 | 11 | 5 | 4 | 5 | 3 | 5 |
| ENSG00000214595 | 130 | 61 | 80 | 32 | 60 | 76 |
| ENSG00000231334 | 0 | 0 | 0 | 0 | 0 | 0 |
| ENSG00000285519 | 0 | 0 | 0 | 0 | 0 | 1 |
| ENSG00000252507 | 0 | 0 | 1 | 0 | 0 | 0 |
| ENSG00000115310 | 5140 | 2508 | 3031 | 2851 | 3555 | 3297 |
| ENSG00000200086 | 0 | 0 | 0 | 0 | 0 | 0 |
| ENSG00000162994 | 60 | 32 | 57 | 20 | 55 | 46 |
| ENSG00000203327 | 2 | 0 | 3 | 4 | 2 | 0 |
| ENSG00000227799 | 54 | 20 | 24 | 25 | 24 | 14 |
| ENSG00000143947 | 23865 | 12427 | 14174 | 15980 | 15553 | 13794 |
| ENSG00000085760 | 1000 | 555 | 648 | 381 | 822 | 614 |
| ENSG00000162997 | 66 | 42 | 68 | 39 | 60 | 43 |
| ENSG00000240401 | 14 | 12 | 11 | 4 | 5 | 9 |
| ENSG00000206964 | 3 | 1 | 0 | 3 | 3 | 1 |
| ENSG00000115355 | 12 | 9 | 6 | 5 | 9 | 4 |
| ENSG00000233594 | 0 | 0 | 0 | 0 | 1 | 3 |
| ENSG00000238619 | 0 | 0 | 0 | 0 | 0 | 0 |
| ENSG00000238493 | 0 | 0 | 0 | 0 | 0 | 0 |
| ENSG00000239189 | 0 | 0 | 0 | 0 | 0 | 0 |
| ENSG00000163001 | 1397 | 680 | 854 | 745 | 834 | 884 |
| ENSG00000275052 | 2826 | 1554 | 1462 | 880 | 2050 | 1279 |
| ENSG00000212175 | 0 | 1 | 0 | 1 | 0 | 0 |
| ENSG00000272606 | 0 | 0 | 0 | 0 | 0 | 0 |
| ENSG00000138035 | 954 | 539 | 603 | 410 | 667 | 667 |
| ENSG00000115380 | 251 | 148 | 151 | 96 | 180 | 126 |

| | | | | | | |
|---|---|---|---|---|---|---|
| ENSG00000202344 | 0 | 0 | 0 | 0 | 0 | 0 |
| ENSG00000272180 | 12 | 5 | 10 | 1 | 5 | 6 |
| ENSG00000226702 | 0 | 0 | 0 | 0 | 0 | 0 |
| ENSG00000207548 | 0 | 0 | 0 | 0 | 0 | 0 |
| ENSG00000207798 | 0 | 0 | 0 | 0 | 0 | 0 |
| ENSG00000211520 | 0 | 0 | 0 | 0 | 0 | 0 |
| ENSG00000229805 | 0 | 0 | 0 | 0 | 0 | 0 |
| ENSG00000271894 | 0 | 0 | 0 | 0 | 0 | 0 |
| ENSG00000233251 | 378 | 189 | 186 | 111 | 265 | 189 |
| ENSG00000055813 | 129 | 71 | 68 | 62 | 75 | 76 |
| ENSG00000199395 | 0 | 0 | 0 | 0 | 0 | 0 |
| ENSG00000231570 | 0 | 0 | 0 | 0 | 0 | 0 |
| ENSG00000271115 | 0 | 0 | 0 | 0 | 0 | 0 |
| ENSG00000285755 | 3 | 2 | 2 | 0 | 3 | 3 |
| ENSG00000270569 | 0 | 1 | 0 | 0 | 0 | 0 |
| ENSG00000212168 | 0 | 0 | 0 | 0 | 0 | 0 |
| ENSG00000271615 | 0 | 0 | 0 | 0 | 0 | 0 |
| ENSG00000028116 | 214 | 126 | 131 | 69 | 230 | 143 |
| ENSG00000115392 | 237 | 170 | 206 | 114 | 271 | 164 |
| ENSG00000273063 | 0 | 0 | 0 | 0 | 0 | 0 |
| ENSG00000233426 | 36 | 9 | 8 | 10 | 16 | 17 |
| ENSG00000225226 | 0 | 0 | 0 | 0 | 0 | 0 |
| ENSG00000233723 | 2 | 0 | 2 | 0 | 0 | 0 |
| ENSG00000231043 | 76 | 51 | 64 | 43 | 80 | 42 |
| ENSG00000285673 | 0 | 0 | 0 | 0 | 0 | 0 |
| ENSG00000222030 | 0 | 0 | 0 | 0 | 0 | 0 |
| ENSG00000271955 | 0 | 0 | 0 | 0 | 0 | 0 |
| ENSG00000233891 | 0 | 0 | 0 | 0 | 1 | 0 |
| ENSG00000231815 | 0 | 0 | 0 | 0 | 0 | 0 |
| ENSG00000270447 | 0 | 0 | 0 | 0 | 0 | 0 |
| ENSG00000200101 | 0 | 0 | 0 | 0 | 0 | 0 |
| ENSG00000252726 | 0 | 0 | 0 | 0 | 0 | 0 |
| ENSG00000285611 | 0 | 0 | 0 | 0 | 0 | 0 |
| ENSG00000228590 | 0 | 0 | 0 | 0 | 0 | 0 |
| ENSG00000200807 | 0 | 0 | 0 | 0 | 0 | 0 |
| ENSG00000266078 | 0 | 0 | 0 | 0 | 0 | 0 |
| ENSG00000119866 | 0 | 0 | 1 | 0 | 0 | 0 |
| ENSG00000233953 | 0 | 0 | 0 | 0 | 0 | 0 |
| ENSG00000242158 | 0 | 0 | 0 | 0 | 0 | 0 |
| ENSG00000271134 | 0 | 0 | 0 | 0 | 0 | 0 |
| ENSG00000213495 | 0 | 0 | 0 | 0 | 0 | 0 |
| ENSG00000252718 | 0 | 0 | 0 | 0 | 0 | 0 |
| ENSG00000271707 | 0 | 0 | 0 | 0 | 1 | 0 |
| ENSG00000115421 | 606 | 299 | 290 | 182 | 348 | 272 |
| ENSG00000228414 | 5 | 0 | 2 | 2 | 1 | 0 |
| ENSG00000241524 | 0 | 0 | 0 | 0 | 0 | 0 |
| ENSG00000237024 | 0 | 0 | 0 | 0 | 0 | 0 |
| ENSG00000162924 | 252 | 152 | 95 | 37 | 168 | 91 |
| ENSG00000201076 | 0 | 0 | 0 | 0 | 0 | 0 |
| ENSG00000267520 | 0 | 0 | 0 | 0 | 0 | 0 |

| | | | | | | |
|---|---|---|---|---|---|---|
| ENSG00000237522 | 7 | 5 | 6 | 2 | 9 | 3 |
| ENSG00000232713 | 6 | 2 | 0 | 2 | 6 | 5 |
| ENSG00000162927 | 171 | 98 | 119 | 70 | 114 | 97 |
| ENSG00000222251 | 0 | 0 | 0 | 0 | 0 | 0 |
| ENSG00000162928 | 1853 | 879 | 1057 | 800 | 1050 | 1068 |
| ENSG00000162929 | 437 | 283 | 257 | 177 | 290 | 284 |
| ENSG00000274769 | 0 | 0 | 0 | 0 | 0 | 2 |
| ENSG00000212978 | 279 | 171 | 155 | 141 | 167 | 161 |
| ENSG00000237651 | 27 | 17 | 10 | 6 | 20 | 14 |
| ENSG00000271889 | 9 | 4 | 4 | 3 | 7 | 3 |
| ENSG00000173209 | 2230 | 921 | 1552 | 878 | 1188 | 1150 |
| ENSG00000115464 | 844 | 551 | 422 | 134 | 1059 | 509 |
| ENSG00000273302 | 2 | 1 | 1 | 1 | 3 | 0 |
| ENSG00000271146 | 0 | 0 | 0 | 0 | 0 | 0 |
| ENSG00000234624 | 1 | 0 | 0 | 0 | 0 | 0 |
| ENSG00000206937 | 0 | 0 | 0 | 0 | 0 | 0 |
| ENSG00000270820 | 73 | 31 | 51 | 20 | 32 | 21 |
| ENSG00000082898 | 7501 | 3704 | 4050 | 2215 | 4515 | 3618 |
| ENSG00000285857 | 4 | 0 | 2 | 0 | 1 | 0 |
| ENSG00000271243 | 0 | 0 | 0 | 0 | 0 | 0 |
| ENSG00000226884 | 0 | 0 | 0 | 0 | 0 | 0 |
| ENSG00000206973 | 0 | 0 | 0 | 0 | 0 | 0 |
| ENSG00000213486 | 0 | 0 | 0 | 0 | 0 | 0 |
| ENSG00000227361 | 0 | 0 | 0 | 0 | 0 | 0 |
| ENSG00000229831 | 0 | 0 | 0 | 0 | 0 | 0 |
| ENSG00000170264 | 164 | 50 | 68 | 21 | 65 | 55 |
| ENSG00000230702 | 0 | 1 | 0 | 0 | 1 | 0 |
| ENSG00000277498 | 0 | 0 | 0 | 0 | 0 | 0 |
| ENSG00000115484 | 8003 | 3901 | 4923 | 5156 | 5286 | 4846 |
| ENSG00000236498 | 2 | 2 | 1 | 4 | 1 | 2 |
| ENSG00000173163 | 565 | 305 | 334 | 315 | 279 | 250 |
| ENSG00000278523 | 0 | 0 | 0 | 0 | 0 | 0 |
| ENSG00000229839 | 0 | 3 | 1 | 3 | 1 | 5 |
| ENSG00000242735 | 0 | 0 | 0 | 0 | 0 | 0 |
| ENSG00000271657 | 0 | 0 | 0 | 0 | 0 | 0 |
| ENSG00000170340 | 903 | 414 | 508 | 460 | 516 | 519 |
| ENSG00000266097 | 0 | 0 | 0 | 0 | 0 | 0 |
| ENSG00000239958 | 0 | 0 | 0 | 0 | 0 | 0 |
| ENSG00000270335 | 0 | 0 | 0 | 0 | 0 | 0 |
| ENSG00000228541 | 0 | 2 | 5 | 0 | 2 | 0 |
| ENSG00000241625 | 0 | 0 | 0 | 0 | 0 | 0 |
| ENSG00000186889 | 48 | 22 | 44 | 22 | 30 | 28 |
| ENSG00000233702 | 0 | 0 | 0 | 0 | 0 | 0 |
| ENSG00000229503 | 0 | 1 | 0 | 1 | 0 | 1 |
| ENSG00000232144 | 0 | 0 | 0 | 0 | 0 | 0 |
| ENSG00000230595 | 0 | 0 | 0 | 0 | 0 | 0 |
| ENSG00000226622 | 0 | 0 | 0 | 0 | 0 | 0 |
| ENSG00000115504 | 399 | 239 | 243 | 134 | 383 | 251 |
| ENSG00000252436 | 0 | 0 | 0 | 0 | 0 | 0 |
| ENSG00000226605 | 0 | 0 | 0 | 0 | 0 | 0 |

| | | | | | | |
|---|---|---|---|---|---|---|
| ENSG00000237162 | 0 | 0 | 0 | 0 | 0 | 0 |
| ENSG00000231609 | 7 | 5 | 5 | 2 | 5 | 6 |
| ENSG00000115507 | 153 | 58 | 78 | 50 | 58 | 100 |
| ENSG00000270437 | 0 | 1 | 0 | 0 | 0 | 0 |
| ENSG00000232403 | 0 | 0 | 0 | 0 | 1 | 0 |
| ENSG00000242412 | 0 | 0 | 0 | 0 | 0 | 0 |
| ENSG00000143951 | 141 | 76 | 105 | 73 | 86 | 89 |
| ENSG00000283636 | 0 | 0 | 0 | 0 | 0 | 0 |
| ENSG00000233716 | 0 | 0 | 0 | 0 | 0 | 0 |
| ENSG00000279485 | 0 | 0 | 0 | 0 | 0 | 0 |
| ENSG00000014641 | 4793 | 2494 | 2797 | 2672 | 2860 | 2728 |
| ENSG00000228305 | 6 | 1 | 3 | 1 | 3 | 3 |
| ENSG00000229920 | 0 | 1 | 0 | 0 | 0 | 0 |
| ENSG00000234488 | 0 | 0 | 0 | 0 | 0 | 1 |
| ENSG00000228872 | 0 | 0 | 0 | 0 | 0 | 0 |
| ENSG00000229221 | 0 | 0 | 0 | 0 | 0 | 0 |
| ENSG00000169764 | 2864 | 1603 | 1763 | 1314 | 2021 | 1504 |
| ENSG00000251775 | 1 | 2 | 2 | 0 | 1 | 1 |
| ENSG00000143952 | 1042 | 534 | 640 | 391 | 736 | 502 |
| ENSG00000228079 | 0 | 0 | 0 | 0 | 0 | 0 |
| ENSG00000197329 | 176 | 85 | 81 | 69 | 111 | 64 |
| ENSG00000225889 | 0 | 0 | 0 | 0 | 0 | 0 |
| ENSG00000230923 | 0 | 0 | 0 | 0 | 0 | 0 |
| ENSG00000270354 | 0 | 0 | 0 | 0 | 0 | 0 |
| ENSG00000238012 | 0 | 0 | 0 | 0 | 0 | 0 |
| ENSG00000238201 | 0 | 0 | 0 | 0 | 0 | 0 |
| ENSG00000264297 | 0 | 0 | 0 | 0 | 0 | 0 |
| ENSG00000283222 | 0 | 0 | 0 | 0 | 0 | 0 |
| ENSG00000236148 | 0 | 0 | 0 | 0 | 0 | 0 |
| ENSG00000223935 | 0 | 0 | 2 | 0 | 0 | 0 |
| ENSG00000237217 | 0 | 0 | 0 | 0 | 0 | 0 |
| ENSG00000119862 | 314 | 136 | 147 | 130 | 128 | 130 |
| ENSG00000223863 | 0 | 0 | 0 | 0 | 0 | 0 |
| ENSG00000260101 | 6 | 9 | 7 | 1 | 4 | 7 |
| ENSG00000119844 | 1692 | 891 | 1038 | 617 | 1040 | 897 |
| ENSG00000283204 | 0 | 0 | 0 | 0 | 0 | 0 |
| ENSG00000252414 | 0 | 0 | 1 | 0 | 0 | 0 |
| ENSG00000233694 | 3 | 0 | 1 | 1 | 1 | 1 |
| ENSG00000179833 | 88 | 41 | 65 | 46 | 78 | 44 |
| ENSG00000226756 | 0 | 0 | 0 | 0 | 0 | 0 |
| ENSG00000227791 | 0 | 0 | 0 | 0 | 0 | 0 |
| ENSG00000275588 | 0 | 0 | 0 | 0 | 0 | 0 |
| ENSG00000199964 | 0 | 0 | 0 | 0 | 0 | 0 |
| ENSG00000234572 | 0 | 0 | 0 | 0 | 0 | 0 |
| ENSG00000237638 | 0 | 0 | 0 | 0 | 0 | 0 |
| ENSG00000244534 | 0 | 0 | 0 | 0 | 0 | 0 |
| ENSG00000227394 | 0 | 0 | 1 | 1 | 2 | 0 |
| ENSG00000115902 | 147 | 77 | 82 | 81 | 66 | 80 |
| ENSG00000252892 | 0 | 1 | 0 | 0 | 0 | 0 |
| ENSG00000232613 | 1 | 0 | 0 | 0 | 1 | 0 |

| | | | | | | |
|---|---|---|---|---|---|---|
| ENSG00000011523 | 549 | 295 | 327 | 265 | 319 | 285 |
| ENSG00000138069 | 5077 | 2507 | 2874 | 1900 | 3084 | 2669 |
| ENSG00000272025 | 0 | 0 | 0 | 0 | 0 | 0 |
| ENSG00000273763 | 29 | 9 | 15 | 12 | 15 | 16 |
| ENSG00000172974 | 55 | 32 | 19 | 35 | 25 | 41 |
| ENSG00000138071 | 10414 | 4975 | 5898 | 3736 | 6052 | 5744 |
| ENSG00000198369 | 1469 | 718 | 788 | 697 | 884 | 871 |
| ENSG00000232693 | 0 | 0 | 0 | 1 | 0 | 0 |
| ENSG00000204929 | 4 | 0 | 0 | 2 | 0 | 0 |
| ENSG00000234255 | 0 | 0 | 0 | 0 | 0 | 0 |
| ENSG00000237979 | 0 | 0 | 0 | 1 | 0 | 1 |
| ENSG00000223739 | 0 | 0 | 0 | 0 | 0 | 0 |
| ENSG00000243029 | 0 | 0 | 0 | 0 | 0 | 0 |
| ENSG00000251900 | 0 | 0 | 0 | 0 | 0 | 0 |
| ENSG00000235725 | 0 | 0 | 0 | 0 | 0 | 0 |
| ENSG00000281920 | 0 | 0 | 0 | 0 | 0 | 0 |
| ENSG00000214533 | 4 | 1 | 2 | 2 | 1 | 0 |
| ENSG00000280257 | 0 | 0 | 0 | 0 | 0 | 0 |
| ENSG00000227293 | 0 | 0 | 0 | 0 | 0 | 0 |
| ENSG00000225815 | 5 | 2 | 0 | 0 | 1 | 1 |
| ENSG00000264525 | 0 | 0 | 0 | 0 | 0 | 0 |
| ENSG00000232164 | 5 | 9 | 7 | 3 | 4 | 6 |
| ENSG00000226819 | 3 | 0 | 2 | 2 | 1 | 5 |
| ENSG00000143995 | 383 | 170 | 206 | 166 | 192 | 221 |
| ENSG00000230749 | 0 | 0 | 1 | 0 | 0 | 0 |
| ENSG00000232046 | 0 | 0 | 0 | 0 | 0 | 0 |
| ENSG00000237179 | 0 | 0 | 0 | 0 | 0 | 0 |
| ENSG00000224071 | 0 | 0 | 0 | 0 | 0 | 0 |
| ENSG00000230355 | 0 | 0 | 0 | 0 | 0 | 0 |
| ENSG00000230525 | 0 | 0 | 0 | 0 | 0 | 0 |
| ENSG00000232688 | 0 | 0 | 0 | 0 | 0 | 0 |
| ENSG00000223859 | 0 | 0 | 0 | 0 | 0 | 0 |
| ENSG00000235885 | 0 | 0 | 0 | 0 | 0 | 0 |
| ENSG00000236780 | 0 | 0 | 0 | 0 | 0 | 1 |
| ENSG00000236605 | 0 | 0 | 0 | 0 | 0 | 0 |
| ENSG00000143971 | 142 | 88 | 75 | 43 | 205 | 94 |
| ENSG00000224173 | 0 | 0 | 0 | 0 | 0 | 0 |
| ENSG00000235495 | 0 | 0 | 0 | 0 | 0 | 0 |
| ENSG00000237013 | 0 | 0 | 0 | 0 | 0 | 0 |
| ENSG00000235968 | 0 | 0 | 0 | 0 | 0 | 0 |
| ENSG00000197223 | 563 | 232 | 340 | 265 | 324 | 336 |
| ENSG00000284932 | 0 | 0 | 0 | 0 | 0 | 0 |
| ENSG00000243667 | 150 | 92 | 107 | 102 | 129 | 103 |
| ENSG00000236259 | 0 | 0 | 0 | 0 | 0 | 0 |
| ENSG00000273398 | 0 | 1 | 0 | 0 | 0 | 0 |
| ENSG00000115946 | 853 | 442 | 529 | 532 | 643 | 644 |
| ENSG00000221823 | 1113 | 614 | 791 | 662 | 740 | 797 |
| ENSG00000273275 | 0 | 0 | 0 | 0 | 0 | 0 |
| ENSG00000273064 | 11 | 1 | 5 | 2 | 8 | 8 |
| ENSG00000119865 | 0 | 0 | 0 | 0 | 0 | 0 |

| | | | | | | |
|---|---|---|---|---|---|---|
| ENSG00000203395 | 0 | 0 | 0 | 0 | 0 | 0 |
| ENSG00000115956 | 0 | 0 | 1 | 0 | 0 | 0 |
| ENSG00000229462 | 0 | 0 | 0 | 0 | 0 | 0 |
| ENSG00000204923 | 53 | 19 | 26 | 8 | 36 | 16 |
| ENSG00000169621 | 4 | 2 | 8 | 0 | 3 | 3 |
| ENSG00000214525 | 0 | 0 | 0 | 0 | 0 | 0 |
| ENSG00000169618 | 8 | 1 | 1 | 0 | 3 | 2 |
| ENSG00000163219 | 0 | 0 | 0 | 0 | 1 | 1 |
| ENSG00000237576 | 0 | 0 | 0 | 0 | 0 | 0 |
| ENSG00000228329 | 0 | 0 | 0 | 0 | 0 | 0 |
| ENSG00000163217 | 0 | 0 | 0 | 0 | 0 | 0 |
| ENSG00000283586 | 0 | 0 | 0 | 0 | 0 | 0 |
| ENSG00000183607 | 0 | 0 | 0 | 0 | 0 | 0 |
| ENSG00000169605 | 0 | 0 | 0 | 0 | 0 | 0 |
| ENSG00000169604 | 450 | 203 | 273 | 193 | 216 | 224 |
| ENSG00000266649 | 0 | 0 | 0 | 0 | 0 | 0 |
| ENSG00000251850 | 0 | 0 | 0 | 0 | 0 | 0 |
| ENSG00000199460 | 0 | 0 | 0 | 0 | 0 | 0 |
| ENSG00000271597 | 0 | 0 | 0 | 0 | 0 | 0 |
| ENSG00000198380 | 2207 | 1092 | 1179 | 659 | 1218 | 970 |
| ENSG00000252250 | 0 | 0 | 0 | 0 | 1 | 0 |
| ENSG00000169599 | 599 | 276 | 355 | 273 | 291 | 319 |
| ENSG00000115977 | 1021 | 526 | 606 | 317 | 543 | 494 |
| ENSG00000244236 | 0 | 0 | 0 | 0 | 0 | 0 |
| ENSG00000207016 | 0 | 0 | 0 | 0 | 0 | 0 |
| ENSG00000232228 | 0 | 0 | 1 | 0 | 0 | 0 |
| ENSG00000229948 | 0 | 0 | 0 | 0 | 0 | 0 |
| ENSG00000196975 | 2363 | 1290 | 1404 | 1251 | 1413 | 1435 |
| ENSG00000231024 | 1 | 2 | 2 | 0 | 4 | 0 |
| ENSG00000087338 | 1246 | 632 | 692 | 357 | 675 | 517 |
| ENSG00000275381 | 2 | 2 | 1 | 0 | 0 | 0 |
| ENSG00000270723 | 2 | 0 | 1 | 0 | 0 | 0 |
| ENSG00000124380 | 802 | 414 | 537 | 450 | 525 | 488 |
| ENSG00000059728 | 970 | 434 | 616 | 352 | 480 | 554 |
| ENSG00000244617 | 8 | 7 | 8 | 5 | 10 | 6 |
| ENSG00000179818 | 482 | 194 | 303 | 181 | 193 | 207 |
| ENSG00000263669 | 0 | 0 | 0 | 0 | 0 | 0 |
| ENSG00000169564 | 3382 | 2047 | 2700 | 2419 | 2203 | 2393 |
| ENSG00000233060 | 0 | 0 | 2 | 0 | 0 | 0 |
| ENSG00000225392 | 0 | 0 | 0 | 0 | 0 | 1 |
| ENSG00000231327 | 6 | 6 | 8 | 15 | 11 | 7 |
| ENSG00000115998 | 370 | 182 | 228 | 152 | 194 | 234 |
| ENSG00000116001 | 1855 | 952 | 1073 | 578 | 1165 | 1019 |
| ENSG00000221238 | 0 | 0 | 0 | 0 | 0 | 0 |
| ENSG00000231414 | 0 | 2 | 1 | 1 | 0 | 1 |
| ENSG00000116005 | 4763 | 2179 | 2807 | 1396 | 2690 | 2146 |
| ENSG00000143977 | 2147 | 1114 | 1287 | 1357 | 1308 | 1328 |
| ENSG00000035141 | 3858 | 2155 | 2593 | 2202 | 2380 | 2490 |
| ENSG00000233849 | 0 | 0 | 0 | 0 | 0 | 0 |
| ENSG00000235289 | 0 | 0 | 0 | 0 | 0 | 0 |

| | | | | | | |
|---|---|---|---|---|---|---|
| ENSG00000229229 | 0 | 0 | 0 | 0 | 0 | 0 |
| ENSG00000163235 | 165 | 90 | 92 | 68 | 113 | 102 |
| ENSG00000224606 | 0 | 0 | 0 | 0 | 0 | 0 |
| ENSG00000075340 | 2 | 0 | 0 | 0 | 0 | 0 |
| ENSG00000235035 | 0 | 0 | 0 | 0 | 0 | 0 |
| ENSG00000183733 | 0 | 0 | 0 | 0 | 0 | 0 |
| ENSG00000228034 | 2 | 2 | 0 | 1 | 0 | 0 |
| ENSG00000152672 | 4 | 0 | 2 | 0 | 0 | 1 |
| ENSG00000231386 | 0 | 0 | 0 | 0 | 0 | 0 |
| ENSG00000116031 | 0 | 1 | 0 | 3 | 1 | 1 |
| ENSG00000237751 | 8 | 9 | 6 | 5 | 3 | 4 |
| ENSG00000277764 | 0 | 0 | 0 | 3 | 0 | 0 |
| ENSG00000116035 | 124 | 68 | 70 | 75 | 71 | 69 |
| ENSG00000116039 | 12 | 12 | 18 | 10 | 9 | 13 |
| ENSG00000258881 | 0 | 2 | 2 | 3 | 0 | 3 |
| ENSG00000239322 | 0 | 0 | 0 | 0 | 0 | 0 |
| ENSG00000226186 | 5 | 2 | 4 | 2 | 3 | 6 |
| ENSG00000241159 | 0 | 0 | 0 | 1 | 0 | 0 |
| ENSG00000144031 | 5 | 2 | 4 | 3 | 2 | 0 |
| ENSG00000144043 | 4940 | 2446 | 2982 | 2723 | 2709 | 2892 |
| ENSG00000228384 | 0 | 0 | 0 | 1 | 1 | 0 |
| ENSG00000236469 | 0 | 0 | 0 | 0 | 0 | 2 |
| ENSG00000205847 | 8 | 4 | 4 | 5 | 5 | 6 |
| ENSG00000234485 | 0 | 0 | 0 | 0 | 0 | 0 |
| ENSG00000234896 | 2 | 0 | 1 | 2 | 2 | 0 |
| ENSG00000124357 | 2354 | 1115 | 1459 | 1296 | 1328 | 1249 |
| ENSG00000272735 | 12 | 8 | 8 | 7 | 8 | 2 |
| ENSG00000124370 | 307 | 139 | 178 | 139 | 154 | 199 |
| ENSG00000124383 | 936 | 548 | 543 | 311 | 736 | 511 |
| ENSG00000233870 | 2 | 0 | 0 | 0 | 0 | 0 |
| ENSG00000124374 | 476 | 216 | 262 | 191 | 279 | 249 |
| ENSG00000075292 | 942 | 627 | 462 | 202 | 1303 | 590 |
| ENSG00000281195 | 4 | 3 | 1 | 3 | 6 | 3 |
| ENSG00000200779 | 0 | 0 | 0 | 0 | 0 | 0 |
| ENSG00000135636 | 6 | 1 | 0 | 4 | 7 | 2 |
| ENSG00000233971 | 6 | 0 | 0 | 0 | 2 | 3 |
| ENSG00000003137 | 10 | 4 | 2 | 7 | 7 | 6 |
| ENSG00000144036 | 1492 | 733 | 815 | 425 | 750 | 643 |
| ENSG00000228568 | 0 | 0 | 0 | 0 | 0 | 0 |
| ENSG00000222536 | 0 | 0 | 0 | 0 | 0 | 0 |
| ENSG00000212378 | 0 | 0 | 0 | 0 | 0 | 0 |
| ENSG00000116096 | 2170 | 1154 | 1380 | 1535 | 1215 | 1238 |
| ENSG00000135638 | 0 | 2 | 3 | 1 | 1 | 0 |
| ENSG00000278060 | 1 | 0 | 0 | 0 | 0 | 0 |
| ENSG00000144040 | 2012 | 874 | 1253 | 776 | 831 | 967 |
| ENSG00000135631 | 479 | 212 | 278 | 262 | 229 | 273 |
| ENSG00000272702 | 8 | 4 | 8 | 10 | 6 | 4 |
| ENSG00000214513 | 0 | 0 | 0 | 0 | 0 | 0 |
| ENSG00000135632 | 989 | 491 | 637 | 704 | 569 | 678 |
| ENSG00000135617 | 634 | 281 | 354 | 337 | 288 | 274 |

| | | | | | | |
|---|---|---|---|---|---|---|
| ENSG00000135624 | 13087 | 6454 | 7775 | 8747 | 8142 | 8189 |
| ENSG00000163013 | 748 | 368 | 450 | 376 | 433 | 441 |
| ENSG00000135625 | 0 | 0 | 1 | 0 | 0 | 2 |
| ENSG00000252988 | 0 | 0 | 0 | 0 | 0 | 0 |
| ENSG00000285068 | 0 | 0 | 0 | 0 | 0 | 0 |
| ENSG00000214484 | 0 | 0 | 0 | 0 | 0 | 0 |
| ENSG00000116127 | 273 | 146 | 111 | 61 | 291 | 132 |
| ENSG00000230002 | 8 | 2 | 4 | 3 | 7 | 9 |
| ENSG00000284902 | 0 | 0 | 0 | 0 | 0 | 0 |
| ENSG00000144035 | 2 | 1 | 1 | 0 | 1 | 1 |
| ENSG00000163016 | 8 | 1 | 0 | 4 | 7 | 2 |
| ENSG00000204872 | 0 | 0 | 1 | 0 | 0 | 0 |
| ENSG00000144034 | 609 | 299 | 336 | 345 | 394 | 371 |
| ENSG00000273245 | 0 | 0 | 0 | 2 | 0 | 0 |
| ENSG00000280037 | 1 | 0 | 1 | 0 | 1 | 1 |
| ENSG00000144048 | 865 | 476 | 570 | 474 | 535 | 544 |
| ENSG00000187833 | 0 | 0 | 0 | 0 | 0 | 0 |
| ENSG00000124356 | 2173 | 1045 | 1243 | 1045 | 1274 | 1215 |
| ENSG00000271536 | 0 | 0 | 0 | 0 | 0 | 0 |
| ENSG00000163017 | 18 | 10 | 5 | 9 | 4 | 6 |
| ENSG00000114956 | 1428 | 882 | 1012 | 962 | 997 | 973 |
| ENSG00000237883 | 40 | 37 | 31 | 37 | 30 | 41 |
| ENSG00000201876 | 0 | 0 | 0 | 0 | 0 | 0 |
| ENSG00000235499 | 34 | 14 | 24 | 13 | 13 | 25 |
| ENSG00000187605 | 3397 | 1285 | 2059 | 1295 | 1648 | 1964 |
| ENSG00000279181 | 0 | 1 | 0 | 0 | 0 | 2 |
| ENSG00000257800 | 53 | 24 | 45 | 18 | 22 | 27 |
| ENSG00000217702 | 5 | 2 | 3 | 1 | 3 | 1 |
| ENSG00000163170 | 878 | 422 | 494 | 558 | 517 | 551 |
| ENSG00000225439 | 8 | 4 | 6 | 7 | 5 | 5 |
| ENSG00000114978 | 6903 | 3082 | 3604 | 2504 | 3535 | 3339 |
| ENSG00000279201 | 0 | 0 | 0 | 0 | 0 | 0 |
| ENSG00000279070 | 0 | 0 | 0 | 0 | 0 | 0 |
| ENSG00000065911 | 2931 | 1755 | 1975 | 1734 | 2015 | 1933 |
| ENSG00000264324 | 0 | 0 | 1 | 1 | 0 | 1 |
| ENSG00000188687 | 7 | 6 | 10 | 2 | 3 | 5 |
| ENSG00000229798 | 0 | 0 | 0 | 0 | 0 | 0 |
| ENSG00000252214 | 0 | 0 | 0 | 0 | 0 | 0 |
| ENSG00000234632 | 0 | 0 | 0 | 0 | 0 | 0 |
| ENSG00000225658 | 0 | 0 | 0 | 0 | 0 | 0 |
| ENSG00000204843 | 4968 | 2425 | 3181 | 2698 | 2690 | 2818 |
| ENSG00000237737 | 3 | 1 | 0 | 0 | 4 | 3 |
| ENSG00000159239 | 15 | 3 | 11 | 7 | 2 | 2 |
| ENSG00000234521 | 5 | 3 | 2 | 3 | 1 | 2 |
| ENSG00000284308 | 73 | 72 | 64 | 65 | 46 | 61 |
| ENSG00000233155 | 7 | 4 | 13 | 14 | 15 | 7 |
| ENSG00000005448 | 1373 | 643 | 783 | 960 | 711 | 700 |
| ENSG00000114993 | 180 | 74 | 89 | 74 | 90 | 96 |
| ENSG00000115274 | 37 | 24 | 37 | 39 | 39 | 25 |
| ENSG00000274049 | 0 | 3 | 0 | 0 | 0 | 2 |

| | | | | | | |
|---|---|---|---|---|---|---|
| ENSG00000239779 | 543 | 264 | 417 | 312 | 290 | 311 |
| ENSG00000115275 | 3530 | 1638 | 2059 | 2048 | 1926 | 1846 |
| ENSG00000204822 | 135 | 78 | 78 | 64 | 84 | 81 |
| ENSG00000135637 | 440 | 193 | 293 | 171 | 272 | 241 |
| ENSG00000115282 | 1654 | 734 | 872 | 642 | 693 | 784 |
| ENSG00000179528 | 69 | 33 | 43 | 40 | 48 | 30 |
| ENSG00000272183 | 12 | 5 | 6 | 5 | 5 | 9 |
| ENSG00000257702 | 82 | 35 | 59 | 37 | 33 | 42 |
| ENSG00000115289 | 701 | 341 | 451 | 390 | 407 | 388 |
| ENSG00000115297 | 0 | 0 | 4 | 2 | 2 | 2 |
| ENSG00000144045 | 43 | 10 | 41 | 11 | 8 | 11 |
| ENSG00000115307 | 4144 | 2150 | 2589 | 2857 | 2625 | 2683 |
| ENSG00000115317 | 602 | 304 | 417 | 376 | 399 | 388 |
| ENSG00000115318 | 69 | 35 | 65 | 55 | 44 | 59 |
| ENSG00000115325 | 135 | 107 | 117 | 169 | 113 | 117 |
| ENSG00000159374 | 2 | 3 | 1 | 2 | 0 | 2 |
| ENSG00000233872 | 0 | 0 | 0 | 0 | 0 | 0 |
| ENSG00000108442 | 0 | 0 | 0 | 0 | 0 | 0 |
| ENSG00000135622 | 395 | 199 | 199 | 234 | 258 | 233 |
| ENSG00000224646 | 1 | 1 | 4 | 1 | 0 | 2 |
| ENSG00000272711 | 50 | 21 | 21 | 24 | 18 | 33 |
| ENSG00000159399 | 2319 | 1080 | 1514 | 717 | 991 | 1158 |
| ENSG00000204792 | 203 | 129 | 105 | 30 | 95 | 57 |
| ENSG00000236209 | 2 | 12 | 1 | 0 | 1 | 2 |
| ENSG00000230836 | 589 | 279 | 276 | 109 | 234 | 143 |
| ENSG00000115350 | 1456 | 724 | 850 | 852 | 763 | 695 |
| ENSG00000115353 | 0 | 5 | 1 | 0 | 0 | 0 |
| ENSG00000263909 | 0 | 0 | 0 | 0 | 0 | 0 |
| ENSG00000270571 | 3 | 1 | 4 | 1 | 4 | 4 |
| ENSG00000236167 | 0 | 0 | 0 | 0 | 0 | 0 |
| ENSG00000115363 | 0 | 0 | 4 | 0 | 0 | 1 |
| ENSG00000237293 | 0 | 0 | 0 | 0 | 0 | 0 |
| ENSG00000238410 | 0 | 0 | 0 | 0 | 0 | 0 |
| ENSG00000231172 | 0 | 0 | 0 | 0 | 0 | 0 |
| ENSG00000230477 | 0 | 0 | 0 | 0 | 0 | 0 |
| ENSG00000221638 | 0 | 0 | 0 | 0 | 0 | 0 |
| ENSG00000115364 | 1777 | 983 | 1084 | 928 | 1152 | 1058 |
| ENSG00000226247 | 0 | 0 | 0 | 0 | 0 | 0 |
| ENSG00000005436 | 260 | 159 | 132 | 121 | 193 | 164 |
| ENSG00000270696 | 308 | 164 | 132 | 116 | 188 | 147 |
| ENSG00000271452 | 11 | 3 | 3 | 0 | 2 | 7 |
| ENSG00000270462 | 2 | 0 | 0 | 0 | 0 | 0 |
| ENSG00000270996 | 23 | 4 | 11 | 8 | 17 | 1 |
| ENSG00000233107 | 0 | 0 | 0 | 0 | 0 | 0 |
| ENSG00000228209 | 0 | 0 | 0 | 0 | 0 | 0 |
| ENSG00000271356 | 0 | 0 | 0 | 0 | 0 | 0 |
| ENSG00000200488 | 0 | 0 | 0 | 0 | 0 | 0 |
| ENSG00000251947 | 0 | 0 | 0 | 0 | 0 | 0 |
| ENSG00000176204 | 0 | 0 | 0 | 0 | 0 | 0 |
| ENSG00000200028 | 0 | 0 | 0 | 0 | 0 | 0 |

| | | | | | | |
|---|---|---|---|---|---|---|
| ENSG00000234653 | 0 | 0 | 0 | 0 | 0 | 0 |
| ENSG00000227088 | 0 | 0 | 1 | 1 | 1 | 0 |
| ENSG00000230968 | 0 | 0 | 0 | 0 | 0 | 0 |
| ENSG00000237077 | 0 | 0 | 0 | 0 | 0 | 0 |
| ENSG00000233605 | 0 | 0 | 0 | 0 | 0 | 0 |
| ENSG00000229494 | 0 | 0 | 0 | 0 | 1 | 0 |
| ENSG00000214429 | 1 | 5 | 5 | 2 | 4 | 5 |
| ENSG00000270470 | 0 | 0 | 0 | 0 | 0 | 0 |
| ENSG00000234877 | 0 | 0 | 0 | 0 | 0 | 0 |
| ENSG00000202423 | 0 | 0 | 0 | 0 | 0 | 0 |
| ENSG00000200048 | 0 | 0 | 0 | 0 | 0 | 0 |
| ENSG00000143954 | 0 | 0 | 0 | 0 | 0 | 0 |
| ENSG00000172023 | 0 | 0 | 0 | 0 | 0 | 0 |
| ENSG00000115386 | 0 | 0 | 0 | 0 | 0 | 0 |
| ENSG00000204787 | 0 | 0 | 0 | 0 | 0 | 0 |
| ENSG00000172016 | 0 | 0 | 0 | 0 | 0 | 0 |
| ENSG00000224879 | 0 | 0 | 0 | 0 | 0 | 0 |
| ENSG00000066032 | 0 | 0 | 0 | 0 | 0 | 0 |
| ENSG00000229385 | 0 | 0 | 0 | 0 | 0 | 0 |
| ENSG00000277171 | 0 | 0 | 0 | 0 | 0 | 0 |
| ENSG00000230237 | 0 | 0 | 0 | 0 | 0 | 0 |
| ENSG00000252031 | 0 | 0 | 0 | 0 | 0 | 0 |
| ENSG00000266436 | 0 | 0 | 0 | 0 | 0 | 0 |
| ENSG00000276917 | 0 | 0 | 0 | 0 | 0 | 0 |
| ENSG00000224731 | 0 | 0 | 0 | 0 | 0 | 0 |
| ENSG00000223427 | 0 | 0 | 0 | 0 | 0 | 0 |
| ENSG00000162951 | 0 | 0 | 0 | 0 | 0 | 0 |
| ENSG00000237031 | 0 | 0 | 0 | 0 | 0 | 0 |
| ENSG00000230975 | 0 | 0 | 0 | 0 | 0 | 0 |
| ENSG00000234429 | 0 | 0 | 0 | 0 | 0 | 0 |
| ENSG00000231722 | 0 | 0 | 0 | 0 | 0 | 0 |
| ENSG00000235519 | 0 | 0 | 0 | 0 | 0 | 0 |
| ENSG00000200890 | 0 | 0 | 0 | 0 | 0 | 0 |
| ENSG00000233444 | 0 | 0 | 0 | 0 | 0 | 0 |
| ENSG00000242118 | 0 | 0 | 0 | 0 | 0 | 0 |
| ENSG00000231781 | 0 | 0 | 0 | 0 | 0 | 0 |
| ENSG00000229560 | 0 | 0 | 0 | 0 | 0 | 0 |
| ENSG00000252897 | 0 | 0 | 0 | 0 | 0 | 0 |
| ENSG00000201311 | 0 | 0 | 0 | 0 | 0 | 0 |
| ENSG00000237498 | 0 | 0 | 0 | 0 | 0 | 0 |
| ENSG00000226860 | 0 | 0 | 0 | 0 | 0 | 0 |
| ENSG00000226700 | 0 | 0 | 0 | 0 | 0 | 0 |
| ENSG00000225045 | 0 | 0 | 0 | 0 | 0 | 0 |
| ENSG00000224695 | 0 | 0 | 0 | 0 | 0 | 0 |
| ENSG00000227789 | 0 | 0 | 0 | 0 | 0 | 0 |
| ENSG00000168129 | 0 | 0 | 0 | 0 | 0 | 0 |
| ENSG00000223977 | 0 | 0 | 0 | 0 | 0 | 0 |
| ENSG00000232548 | 0 | 0 | 0 | 0 | 0 | 0 |
| ENSG00000224627 | 0 | 0 | 0 | 0 | 0 | 0 |
| ENSG00000199295 | 0 | 0 | 0 | 0 | 0 | 0 |

| | | | | | | |
|---|---|---|---|---|---|---|
| ENSG00000228225 | 0 | 0 | 0 | 0 | 0 | 0 |
| ENSG00000228902 | 0 | 0 | 0 | 0 | 0 | 0 |
| ENSG00000182814 | 0 | 0 | 0 | 0 | 1 | 0 |
| ENSG00000228272 | 0 | 0 | 0 | 0 | 0 | 0 |
| ENSG00000163541 | 3011 | 1390 | 1816 | 1638 | 1739 | 1618 |
| ENSG00000115423 | 8 | 1 | 1 | 2 | 4 | 2 |
| ENSG00000259056 | 0 | 0 | 0 | 0 | 0 | 0 |
| ENSG00000235847 | 0 | 0 | 1 | 0 | 0 | 0 |
| ENSG00000186854 | 2 | 2 | 3 | 3 | 3 | 0 |
| ENSG00000276362 | 0 | 0 | 0 | 0 | 0 | 0 |
| ENSG00000213400 | 0 | 0 | 0 | 0 | 0 | 0 |
| ENSG00000034510 | 16597 | 7966 | 10296 | 10447 | 8989 | 8725 |
| ENSG00000213399 | 6 | 1 | 0 | 2 | 1 | 0 |
| ENSG00000176407 | 2832 | 1341 | 1499 | 1553 | 1818 | 1605 |
| ENSG00000260840 | 0 | 0 | 0 | 0 | 0 | 0 |
| ENSG00000235463 | 0 | 0 | 0 | 0 | 0 | 0 |
| ENSG00000225787 | 0 | 0 | 0 | 0 | 0 | 0 |
| ENSG00000152284 | 262 | 144 | 159 | 92 | 109 | 75 |
| ENSG00000231134 | 0 | 0 | 0 | 0 | 0 | 0 |
| ENSG00000200701 | 0 | 0 | 0 | 0 | 0 | 0 |
| ENSG00000230299 | 0 | 0 | 0 | 0 | 0 | 0 |
| ENSG00000246575 | 96 | 47 | 71 | 32 | 62 | 55 |
| ENSG00000152291 | 8245 | 3948 | 5085 | 2537 | 4260 | 3936 |
| ENSG00000239143 | 0 | 0 | 0 | 1 | 0 | 0 |
| ENSG00000270532 | 0 | 0 | 1 | 0 | 0 | 0 |
| ENSG00000042445 | 1760 | 860 | 1099 | 815 | 899 | 975 |
| ENSG00000115459 | 853 | 407 | 498 | 447 | 496 | 439 |
| ENSG00000271014 | 0 | 0 | 1 | 0 | 0 | 0 |
| ENSG00000202382 | 2 | 2 | 0 | 0 | 1 | 1 |
| ENSG00000278590 | 7 | 0 | 1 | 0 | 1 | 2 |
| ENSG00000273196 | 1 | 1 | 0 | 1 | 0 | 0 |
| ENSG00000042493 | 948 | 493 | 627 | 567 | 434 | 495 |
| ENSG00000152292 | 15 | 4 | 4 | 7 | 1 | 7 |
| ENSG00000207207 | 0 | 0 | 1 | 0 | 0 | 0 |
| ENSG00000244399 | 0 | 0 | 0 | 0 | 0 | 0 |
| ENSG00000199936 | 0 | 0 | 0 | 0 | 0 | 0 |
| ENSG00000229648 | 0 | 0 | 0 | 0 | 0 | 0 |
| ENSG00000168906 | 8348 | 3914 | 5624 | 3538 | 4581 | 4923 |
| ENSG00000115486 | 2001 | 891 | 1195 | 853 | 809 | 943 |
| ENSG00000118640 | 7388 | 4027 | 4832 | 4202 | 3764 | 3949 |
| ENSG00000276281 | 0 | 0 | 0 | 0 | 0 | 0 |
| ENSG00000168899 | 491 | 262 | 352 | 293 | 231 | 235 |
| ENSG00000168894 | 3263 | 1685 | 2177 | 1986 | 1689 | 1761 |
| ENSG00000168890 | 1198 | 472 | 768 | 604 | 481 | 576 |
| ENSG00000168883 | 2831 | 1415 | 1758 | 1439 | 1512 | 1609 |
| ENSG00000168887 | 1953 | 984 | 1195 | 939 | 1098 | 1022 |
| ENSG00000168878 | 0 | 1 | 0 | 1 | 0 | 2 |
| ENSG00000115523 | 0 | 0 | 1 | 0 | 0 | 1 |
| ENSG00000203363 | 0 | 0 | 0 | 0 | 0 | 0 |
| ENSG00000199687 | 0 | 0 | 0 | 1 | 0 | 0 |

| | | | | | | |
|---|---|---|---|---|---|---|
| ENSG00000168874 | 259 | 146 | 138 | 519 | 435 | 347 |
| ENSG00000278783 | 0 | 0 | 1 | 1 | 1 | 1 |
| ENSG00000229498 | 2 | 0 | 0 | 1 | 0 | 1 |
| ENSG00000252321 | 0 | 0 | 0 | 0 | 0 | 0 |
| ENSG00000115525 | 2640 | 1169 | 1544 | 1298 | 1564 | 1665 |
| ENSG00000232504 | 15 | 12 | 21 | 8 | 14 | 18 |
| ENSG00000272564 | 2 | 3 | 5 | 5 | 4 | 5 |
| ENSG00000068654 | 2754 | 1306 | 1585 | 1199 | 1398 | 1723 |
| ENSG00000132300 | 2090 | 1044 | 1232 | 827 | 1402 | 1194 |
| ENSG00000208772 | 14 | 11 | 5 | 5 | 13 | 12 |
| ENSG00000132305 | 3252 | 1691 | 1990 | 1572 | 2276 | 1927 |
| ENSG00000200241 | 0 | 1 | 0 | 0 | 0 | 0 |
| ENSG00000265420 | 0 | 0 | 0 | 0 | 0 | 0 |
| ENSG00000273080 | 17 | 14 | 19 | 19 | 17 | 19 |
| ENSG00000132313 | 1575 | 729 | 835 | 746 | 994 | 869 |
| ENSG00000068615 | 123 | 67 | 55 | 30 | 47 | 49 |
| ENSG00000202537 | 0 | 0 | 0 | 0 | 0 | 0 |
| ENSG00000251974 | 0 | 0 | 0 | 0 | 0 | 0 |
| ENSG00000115548 | 1308 | 643 | 880 | 480 | 787 | 701 |
| ENSG00000115561 | 4231 | 1962 | 2463 | 1837 | 2201 | 2390 |
| ENSG00000249884 | 3 | 2 | 3 | 0 | 3 | 0 |
| ENSG00000200563 | 0 | 0 | 0 | 0 | 0 | 0 |
| ENSG00000228363 | 7 | 3 | 10 | 4 | 7 | 3 |
| ENSG00000239305 | 1441 | 749 | 989 | 629 | 799 | 774 |
| ENSG00000153561 | 2320 | 1080 | 1344 | 877 | 1257 | 1277 |
| ENSG00000153563 | 38 | 18 | 20 | 20 | 28 | 33 |
| ENSG00000172116 | 0 | 0 | 0 | 1 | 0 | 0 |
| ENSG00000233673 | 6 | 3 | 2 | 0 | 3 | 1 |
| ENSG00000213605 | 0 | 0 | 0 | 0 | 0 | 0 |
| ENSG00000187627 | 0 | 0 | 1 | 0 | 0 | 0 |
| ENSG00000231351 | 0 | 1 | 0 | 2 | 0 | 0 |
| ENSG00000226114 | 0 | 0 | 0 | 0 | 0 | 0 |
| ENSG00000183281 | 9 | 9 | 12 | 8 | 3 | 7 |
| ENSG00000285793 | 19 | 6 | 5 | 11 | 6 | 14 |
| ENSG00000231259 | 0 | 0 | 0 | 0 | 0 | 0 |
| ENSG00000266931 | 0 | 0 | 0 | 0 | 0 | 0 |
| ENSG00000230395 | 0 | 0 | 0 | 0 | 0 | 0 |
| ENSG00000204745 | 49 | 29 | 39 | 23 | 40 | 53 |
| ENSG00000264793 | 0 | 0 | 0 | 0 | 0 | 0 |
| ENSG00000233671 | 0 | 0 | 0 | 0 | 0 | 0 |
| ENSG00000281477 | 0 | 0 | 0 | 0 | 0 | 0 |
| ENSG00000271097 | 0 | 0 | 0 | 0 | 0 | 0 |
| ENSG00000233999 | 0 | 1 | 0 | 0 | 1 | 0 |
| ENSG00000224881 | 3 | 0 | 0 | 0 | 0 | 1 |
| ENSG00000280721 | 8 | 1 | 0 | 1 | 0 | 0 |
| ENSG00000222041 | 368 | 251 | 270 | 182 | 204 | 176 |
| ENSG00000284879 | 7 | 10 | 11 | 3 | 11 | 7 |
| ENSG00000273445 | 42 | 30 | 22 | 32 | 46 | 28 |
| ENSG00000214354 | 0 | 0 | 0 | 0 | 0 | 0 |
| ENSG00000265507 | 0 | 0 | 0 | 0 | 0 | 0 |

| | | | | | | |
|---|---|---|---|---|---|---|
| ENSG00000225933 | 0 | 1 | 0 | 0 | 0 | 0 |
| ENSG00000234231 | 0 | 0 | 0 | 0 | 0 | 0 |
| ENSG00000125551 | 3 | 2 | 2 | 1 | 3 | 1 |
| ENSG00000185304 | 24 | 9 | 13 | 3 | 7 | 4 |
| ENSG00000229604 | 0 | 2 | 0 | 0 | 0 | 0 |
| ENSG00000225428 | 0 | 0 | 0 | 0 | 0 | 0 |
| ENSG00000233205 | 1 | 0 | 1 | 1 | 0 | 1 |
| ENSG00000275597 | 0 | 0 | 0 | 0 | 0 | 0 |
| ENSG00000222724 | 0 | 0 | 0 | 0 | 0 | 0 |
| ENSG00000172086 | 1434 | 776 | 862 | 673 | 895 | 794 |
| ENSG00000115593 | 0 | 0 | 0 | 0 | 0 | 0 |
| ENSG00000265003 | 0 | 0 | 0 | 0 | 0 | 0 |
| ENSG00000163586 | 0 | 0 | 1 | 0 | 0 | 0 |
| ENSG00000144115 | 1056 | 473 | 719 | 586 | 531 | 577 |
| ENSG00000232508 | 0 | 0 | 0 | 0 | 0 | 0 |
| ENSG00000283262 | 0 | 0 | 0 | 0 | 0 | 0 |
| ENSG00000212133 | 0 | 0 | 0 | 0 | 0 | 0 |
| ENSG00000252395 | 0 | 0 | 0 | 0 | 0 | 0 |
| ENSG00000224291 | 0 | 0 | 0 | 0 | 0 | 0 |
| ENSG00000214336 | 16 | 6 | 3 | 4 | 7 | 6 |
| ENSG00000172073 | 0 | 0 | 0 | 0 | 0 | 0 |
| ENSG00000225420 | 0 | 1 | 2 | 0 | 0 | 0 |
| ENSG00000172071 | 975 | 557 | 636 | 391 | 641 | 569 |
| ENSG00000234028 | 198 | 69 | 132 | 64 | 99 | 84 |
| ENSG00000153574 | 1034 | 505 | 657 | 658 | 675 | 687 |
| ENSG00000230006 | 0 | 0 | 0 | 0 | 0 | 0 |
| ENSG00000240040 | 0 | 0 | 0 | 0 | 0 | 0 |
| ENSG00000283270 | 0 | 0 | 0 | 0 | 0 | 0 |
| ENSG00000265510 | 0 | 0 | 0 | 0 | 0 | 0 |
| ENSG00000211592 | 0 | 0 | 0 | 0 | 0 | 0 |
| ENSG00000211593 | 0 | 0 | 0 | 0 | 0 | 0 |
| ENSG00000211594 | 0 | 0 | 0 | 0 | 0 | 0 |
| ENSG00000211595 | 0 | 0 | 0 | 0 | 0 | 0 |
| ENSG00000211596 | 0 | 0 | 0 | 0 | 0 | 0 |
| ENSG00000211597 | 0 | 0 | 0 | 0 | 0 | 0 |
| ENSG00000211598 | 0 | 0 | 0 | 0 | 0 | 0 |
| ENSG00000211599 | 0 | 0 | 0 | 0 | 0 | 0 |
| ENSG00000197794 | 0 | 0 | 0 | 0 | 0 | 0 |
| ENSG00000225136 | 0 | 0 | 0 | 0 | 0 | 0 |
| ENSG00000253435 | 0 | 0 | 0 | 0 | 0 | 0 |
| ENSG00000243466 | 0 | 0 | 0 | 0 | 0 | 0 |
| ENSG00000239855 | 0 | 0 | 0 | 0 | 0 | 0 |
| ENSG00000243063 | 0 | 0 | 0 | 0 | 0 | 0 |
| ENSG00000240671 | 0 | 0 | 0 | 0 | 0 | 0 |
| ENSG00000241755 | 0 | 0 | 0 | 0 | 0 | 0 |
| ENSG00000253278 | 0 | 0 | 0 | 0 | 0 | 0 |
| ENSG00000241351 | 0 | 0 | 0 | 0 | 0 | 0 |
| ENSG00000243290 | 0 | 0 | 0 | 0 | 0 | 0 |
| ENSG00000253497 | 0 | 0 | 0 | 0 | 0 | 0 |
| ENSG00000253265 | 0 | 0 | 0 | 0 | 0 | 0 |

| | | | | | | |
|---|---|---|---|---|---|---|
| ENSG00000244437 | 0 | 0 | 0 | 0 | 0 | 0 |
| ENSG00000240864 | 0 | 0 | 0 | 0 | 0 | 0 |
| ENSG00000240382 | 0 | 0 | 0 | 0 | 0 | 0 |
| ENSG00000254157 | 0 | 0 | 0 | 0 | 0 | 0 |
| ENSG00000253732 | 0 | 0 | 0 | 0 | 0 | 0 |
| ENSG00000239951 | 0 | 0 | 0 | 0 | 0 | 0 |
| ENSG00000211611 | 0 | 0 | 0 | 0 | 0 | 0 |
| ENSG00000253578 | 0 | 0 | 0 | 0 | 0 | 0 |
| ENSG00000253625 | 0 | 0 | 0 | 0 | 0 | 0 |
| ENSG00000241294 | 0 | 0 | 0 | 0 | 0 | 0 |
| ENSG00000253202 | 0 | 0 | 0 | 0 | 0 | 0 |
| ENSG00000254098 | 0 | 0 | 0 | 0 | 0 | 0 |
| ENSG00000277852 | 0 | 0 | 0 | 0 | 0 | 0 |
| ENSG00000244575 | 0 | 0 | 0 | 0 | 0 | 0 |
| ENSG00000244116 | 0 | 0 | 0 | 0 | 0 | 0 |
| ENSG00000253998 | 0 | 0 | 0 | 0 | 0 | 0 |
| ENSG00000243238 | 0 | 0 | 0 | 0 | 0 | 0 |
| ENSG00000253158 | 0 | 0 | 0 | 0 | 0 | 0 |
| ENSG00000253870 | 0 | 0 | 0 | 0 | 0 | 0 |
| ENSG00000242076 | 0 | 0 | 0 | 0 | 0 | 0 |
| ENSG00000253860 | 0 | 0 | 0 | 0 | 0 | 0 |
| ENSG00000253461 | 0 | 0 | 0 | 0 | 0 | 0 |
| ENSG00000253487 | 0 | 0 | 0 | 0 | 0 | 0 |
| ENSG00000239862 | 0 | 0 | 0 | 0 | 0 | 0 |
| ENSG00000253592 | 0 | 0 | 0 | 0 | 0 | 0 |
| ENSG00000242371 | 0 | 0 | 0 | 0 | 0 | 0 |
| ENSG00000273962 | 0 | 0 | 0 | 0 | 0 | 0 |
| ENSG00000283427 | 0 | 0 | 0 | 0 | 0 | 0 |
| ENSG00000283196 | 215 | 133 | 130 | 142 | 122 | 152 |
| ENSG00000276861 | 0 | 0 | 0 | 0 | 0 | 0 |
| ENSG00000283132 | 0 | 0 | 0 | 0 | 0 | 0 |
| ENSG00000283214 | 0 | 0 | 0 | 0 | 0 | 0 |
| ENSG00000283659 | 0 | 0 | 0 | 0 | 0 | 0 |
| ENSG00000251039 | 0 | 0 | 0 | 0 | 0 | 0 |
| ENSG00000251546 | 0 | 0 | 0 | 0 | 0 | 0 |
| ENSG00000254009 | 0 | 0 | 0 | 0 | 0 | 0 |
| ENSG00000250036 | 0 | 0 | 0 | 0 | 0 | 0 |
| ENSG00000253127 | 0 | 0 | 0 | 0 | 0 | 0 |
| ENSG00000232747 | 0 | 0 | 0 | 0 | 0 | 0 |
| ENSG00000253501 | 0 | 0 | 0 | 0 | 0 | 0 |
| ENSG00000239975 | 0 | 0 | 0 | 0 | 0 | 0 |
| ENSG00000253191 | 0 | 0 | 0 | 0 | 0 | 0 |
| ENSG00000253999 | 0 | 0 | 0 | 0 | 0 | 0 |
| ENSG00000239571 | 0 | 0 | 0 | 0 | 0 | 0 |
| ENSG00000243264 | 0 | 0 | 0 | 0 | 0 | 0 |
| ENSG00000242534 | 0 | 0 | 0 | 0 | 0 | 0 |
| ENSG00000224607 | 0 | 0 | 0 | 0 | 0 | 0 |
| ENSG00000211623 | 0 | 0 | 0 | 0 | 0 | 0 |
| ENSG00000254097 | 0 | 0 | 0 | 0 | 0 | 0 |
| ENSG00000241566 | 0 | 0 | 0 | 0 | 0 | 0 |

| | | | | | | |
|---|---|---|---|---|---|---|
| ENSG00000254345 | 0 | 0 | 0 | 0 | 0 | 0 |
| ENSG00000253365 | 0 | 0 | 0 | 0 | 0 | 0 |
| ENSG00000225523 | 0 | 0 | 0 | 0 | 0 | 0 |
| ENSG00000211625 | 0 | 0 | 0 | 0 | 0 | 0 |
| ENSG00000253765 | 0 | 0 | 0 | 0 | 0 | 0 |
| ENSG00000254220 | 0 | 0 | 0 | 0 | 0 | 0 |
| ENSG00000211626 | 0 | 0 | 0 | 0 | 0 | 0 |
| ENSG00000242766 | 0 | 0 | 0 | 0 | 0 | 0 |
| ENSG00000241244 | 0 | 0 | 0 | 0 | 0 | 0 |
| ENSG00000224041 | 0 | 0 | 0 | 0 | 0 | 0 |
| ENSG00000254292 | 0 | 0 | 0 | 0 | 0 | 0 |
| ENSG00000276566 | 0 | 0 | 0 | 0 | 0 | 0 |
| ENSG00000278857 | 0 | 0 | 0 | 0 | 0 | 0 |
| ENSG00000211632 | 0 | 0 | 0 | 0 | 0 | 0 |
| ENSG00000253906 | 0 | 0 | 0 | 0 | 0 | 0 |
| ENSG00000211633 | 0 | 0 | 0 | 0 | 0 | 0 |
| ENSG00000242580 | 0 | 0 | 0 | 0 | 0 | 0 |
| ENSG00000239819 | 0 | 0 | 0 | 0 | 0 | 0 |
| ENSG00000228325 | 0 | 0 | 0 | 0 | 0 | 0 |
| ENSG00000275490 | 0 | 0 | 0 | 0 | 0 | 0 |
| ENSG00000270999 | 0 | 0 | 0 | 0 | 0 | 0 |
| ENSG00000265897 | 0 | 0 | 0 | 0 | 0 | 0 |
| ENSG00000281904 | 2 | 0 | 0 | 1 | 0 | 0 |
| ENSG00000230964 | 1 | 0 | 0 | 0 | 1 | 2 |
| ENSG00000261600 | 30 | 16 | 10 | 20 | 13 | 4 |
| ENSG00000278131 | 0 | 0 | 0 | 0 | 0 | 0 |
| ENSG00000143429 | 38 | 15 | 25 | 15 | 17 | 16 |
| ENSG00000222835 | 0 | 0 | 0 | 0 | 0 | 0 |
| ENSG00000184303 | 0 | 0 | 0 | 0 | 0 | 0 |
| ENSG00000230572 | 0 | 0 | 0 | 0 | 0 | 0 |
| ENSG00000271627 | 0 | 0 | 0 | 0 | 0 | 0 |
| ENSG00000223703 | 2 | 0 | 0 | 0 | 0 | 1 |
| ENSG00000232531 | 2 | 0 | 0 | 2 | 0 | 1 |
| ENSG00000236969 | 0 | 0 | 0 | 0 | 0 | 0 |
| ENSG00000235235 | 0 | 0 | 0 | 0 | 0 | 0 |
| ENSG00000214330 | 0 | 0 | 0 | 0 | 0 | 0 |
| ENSG00000237085 | 0 | 0 | 0 | 0 | 0 | 0 |
| ENSG00000227497 | 0 | 0 | 0 | 0 | 0 | 0 |
| ENSG00000214329 | 0 | 0 | 0 | 0 | 0 | 0 |
| ENSG00000226481 | 0 | 0 | 0 | 0 | 0 | 0 |
| ENSG00000234898 | 0 | 0 | 0 | 0 | 0 | 0 |
| ENSG00000235147 | 0 | 0 | 0 | 0 | 0 | 0 |
| ENSG00000223816 | 0 | 0 | 0 | 0 | 0 | 0 |
| ENSG00000261522 | 0 | 1 | 1 | 0 | 0 | 0 |
| ENSG00000259916 | 2 | 5 | 5 | 1 | 1 | 3 |
| ENSG00000237383 | 0 | 0 | 0 | 0 | 0 | 0 |
| ENSG00000227265 | 0 | 0 | 0 | 0 | 0 | 0 |
| ENSG00000213355 | 0 | 0 | 0 | 0 | 0 | 0 |
| ENSG00000234837 | 0 | 0 | 0 | 0 | 0 | 0 |
| ENSG00000223564 | 0 | 0 | 0 | 0 | 0 | 0 |

| | | | | | | |
|---|---|---|---|---|---|---|
| ENSG00000232502 | 0 | 0 | 0 | 0 | 0 | 0 |
| ENSG00000235571 | 0 | 0 | 0 | 0 | 0 | 0 |
| ENSG00000229089 | 0 | 0 | 0 | 0 | 0 | 0 |
| ENSG00000212140 | 0 | 0 | 0 | 0 | 0 | 0 |
| ENSG00000278631 | 0 | 0 | 0 | 0 | 0 | 0 |
| ENSG00000259848 | 0 | 0 | 1 | 1 | 0 | 0 |
| ENSG00000163060 | 0 | 0 | 0 | 0 | 0 | 0 |
| ENSG00000237055 | 0 | 0 | 0 | 0 | 0 | 0 |
| ENSG00000232128 | 0 | 0 | 0 | 0 | 0 | 0 |
| ENSG00000233158 | 0 | 0 | 0 | 0 | 0 | 0 |
| ENSG00000231331 | 0 | 0 | 0 | 0 | 0 | 0 |
| ENSG00000232594 | 0 | 0 | 0 | 0 | 0 | 0 |
| ENSG00000233105 | 0 | 0 | 0 | 0 | 0 | 0 |
| ENSG00000224585 | 0 | 0 | 0 | 0 | 0 | 0 |
| ENSG00000233447 | 0 | 0 | 0 | 0 | 0 | 0 |
| ENSG00000235186 | 0 | 0 | 0 | 0 | 0 | 0 |
| ENSG00000277591 | 0 | 0 | 0 | 0 | 0 | 0 |
| ENSG00000233850 | 0 | 0 | 0 | 0 | 0 | 0 |
| ENSG00000172005 | 0 | 0 | 0 | 0 | 1 | 0 |
| ENSG00000231062 | 0 | 0 | 1 | 0 | 1 | 0 |
| ENSG00000144029 | 2471 | 1185 | 1438 | 1419 | 1465 | 1465 |
| ENSG00000144026 | 678 | 330 | 481 | 268 | 442 | 342 |
| ENSG00000275111 | 327 | 147 | 194 | 158 | 165 | 169 |
| ENSG00000275768 | 0 | 0 | 0 | 0 | 0 | 0 |
| ENSG00000233757 | 14 | 8 | 4 | 0 | 9 | 5 |
| ENSG00000155066 | 3309 | 1371 | 2152 | 1485 | 1375 | 1424 |
| ENSG00000115041 | 72 | 29 | 52 | 43 | 38 | 44 |
| ENSG00000235129 | 0 | 0 | 0 | 0 | 0 | 0 |
| ENSG00000115042 | 831 | 359 | 468 | 532 | 441 | 487 |
| ENSG00000248821 | 0 | 0 | 0 | 0 | 0 | 0 |
| ENSG00000233275 | 0 | 0 | 0 | 0 | 0 | 0 |
| ENSG00000227120 | 0 | 0 | 0 | 0 | 0 | 0 |
| ENSG00000204705 | 0 | 0 | 0 | 0 | 0 | 0 |
| ENSG00000277747 | 0 | 0 | 0 | 0 | 0 | 0 |
| ENSG00000234903 | 0 | 0 | 0 | 0 | 0 | 0 |
| ENSG00000144010 | 0 | 0 | 0 | 0 | 0 | 0 |
| ENSG00000238162 | 0 | 0 | 0 | 0 | 0 | 0 |
| ENSG00000223917 | 0 | 0 | 0 | 0 | 0 | 0 |
| ENSG00000237308 | 0 | 0 | 0 | 0 | 0 | 0 |
| ENSG00000224719 | 0 | 0 | 0 | 0 | 0 | 0 |
| ENSG00000226185 | 1 | 0 | 2 | 0 | 0 | 0 |
| ENSG00000272913 | 327 | 194 | 206 | 195 | 209 | 217 |
| ENSG00000229689 | 198 | 98 | 133 | 86 | 125 | 124 |
| ENSG00000236431 | 30 | 17 | 23 | 8 | 19 | 15 |
| ENSG00000273305 | 23 | 26 | 18 | 4 | 20 | 17 |
| ENSG00000233037 | 0 | 0 | 0 | 0 | 1 | 1 |
| ENSG00000168992 | 6 | 6 | 7 | 5 | 7 | 7 |
| ENSG00000232717 | 0 | 0 | 0 | 0 | 0 | 0 |
| ENSG00000274028 | 0 | 0 | 0 | 0 | 0 | 0 |
| ENSG00000230083 | 0 | 0 | 0 | 0 | 0 | 0 |

| | | | | | | |
|---|---|---|---|---|---|---|
| ENSG00000144015 | 0 | 0 | 0 | 0 | 0 | 0 |
| ENSG00000236026 | 0 | 0 | 0 | 0 | 0 | 0 |
| ENSG00000271003 | 0 | 0 | 0 | 0 | 0 | 0 |
| ENSG00000270193 | 0 | 0 | 0 | 0 | 0 | 0 |
| ENSG00000204699 | 0 | 0 | 0 | 0 | 0 | 0 |
| ENSG00000275497 | 0 | 0 | 0 | 0 | 0 | 0 |
| ENSG00000235959 | 0 | 0 | 0 | 0 | 0 | 0 |
| ENSG00000236750 | 0 | 0 | 0 | 0 | 0 | 0 |
| ENSG00000235584 | 52 | 18 | 52 | 18 | 6 | 4 |
| ENSG00000237510 | 0 | 0 | 0 | 0 | 0 | 0 |
| ENSG00000232931 | 113 | 68 | 77 | 34 | 144 | 73 |
| ENSG00000174501 | 23 | 39 | 22 | 20 | 49 | 36 |
| ENSG00000275961 | 0 | 0 | 0 | 0 | 0 | 0 |
| ENSG00000231584 | 496 | 286 | 317 | 350 | 431 | 346 |
| ENSG00000186281 | 8 | 1 | 2 | 5 | 10 | 2 |
| ENSG00000274286 | 41 | 22 | 27 | 24 | 26 | 27 |
| ENSG00000188886 | 111 | 50 | 61 | 198 | 162 | 146 |
| ENSG00000158050 | 1209 | 573 | 607 | 1244 | 1025 | 859 |
| ENSG00000228873 | 50 | 21 | 28 | 33 | 56 | 43 |
| ENSG00000084090 | 7476 | 3719 | 4592 | 4024 | 4280 | 4387 |
| ENSG00000204685 | 122 | 68 | 75 | 73 | 61 | 88 |
| ENSG00000135956 | 5164 | 2642 | 3213 | 2742 | 2595 | 2830 |
| ENSG00000144021 | 4830 | 2248 | 2832 | 2399 | 2655 | 2591 |
| ENSG00000144028 | 12846 | 5829 | 7929 | 5665 | 7222 | 7034 |
| ENSG00000230747 | 0 | 0 | 0 | 0 | 0 | 0 |
| ENSG00000198885 | 6 | 0 | 0 | 0 | 0 | 0 |
| ENSG00000121152 | 439 | 303 | 356 | 268 | 383 | 406 |
| ENSG00000163121 | 8 | 2 | 1 | 6 | 1 | 2 |
| ENSG00000235480 | 0 | 0 | 0 | 0 | 0 | 0 |
| ENSG00000196843 | 115 | 49 | 71 | 79 | 73 | 70 |
| ENSG00000114982 | 2397 | 991 | 1446 | 888 | 1060 | 1175 |
| ENSG00000249715 | 4 | 6 | 1 | 2 | 5 | 4 |
| ENSG00000114988 | 1074 | 512 | 561 | 508 | 527 | 620 |
| ENSG00000158158 | 3420 | 1516 | 2111 | 1274 | 1339 | 1571 |
| ENSG00000264157 | 1 | 0 | 0 | 0 | 0 | 1 |
| ENSG00000273265 | 17 | 11 | 12 | 12 | 13 | 15 |
| ENSG00000168763 | 1684 | 854 | 1174 | 857 | 803 | 988 |
| ENSG00000163126 | 61 | 36 | 49 | 26 | 47 | 44 |
| ENSG00000213337 | 513 | 329 | 433 | 388 | 352 | 341 |
| ENSG00000168758 | 15693 | 6831 | 10291 | 5848 | 5781 | 6937 |
| ENSG00000168754 | 196 | 105 | 159 | 83 | 81 | 87 |
| ENSG00000274206 | 0 | 1 | 1 | 0 | 0 | 0 |
| ENSG00000252845 | 16 | 12 | 11 | 9 | 3 | 5 |
| ENSG00000275094 | 0 | 0 | 0 | 0 | 0 | 0 |
| ENSG00000273634 | 0 | 0 | 0 | 0 | 0 | 0 |
| ENSG00000248134 | 0 | 0 | 0 | 0 | 0 | 0 |
| ENSG00000144188 | 0 | 0 | 0 | 0 | 0 | 0 |
| ENSG00000230343 | 0 | 0 | 0 | 0 | 0 | 0 |
| ENSG00000236847 | 0 | 0 | 0 | 0 | 0 | 0 |
| ENSG00000270748 | 0 | 0 | 0 | 0 | 0 | 0 |

| | | | | | | |
|---|---|---|---|---|---|---|
| ENSG00000271402 | 0 | 0 | 0 | 0 | 0 | 0 |
| ENSG00000204670 | 0 | 0 | 0 | 0 | 0 | 0 |
| ENSG00000188383 | 2 | 1 | 0 | 0 | 2 | 1 |
| ENSG00000144199 | 126 | 76 | 105 | 86 | 77 | 68 |
| ENSG00000279791 | 0 | 0 | 1 | 0 | 1 | 0 |
| ENSG00000275655 | 0 | 0 | 0 | 0 | 0 | 0 |
| ENSG00000135976 | 67 | 43 | 40 | 15 | 59 | 36 |
| ENSG00000277701 | 0 | 0 | 0 | 0 | 0 | 0 |
| ENSG00000237837 | 0 | 0 | 0 | 0 | 0 | 0 |
| ENSG00000271351 | 0 | 0 | 0 | 0 | 0 | 0 |
| ENSG00000276050 | 0 | 0 | 0 | 0 | 0 | 0 |
| ENSG00000271015 | 0 | 0 | 0 | 0 | 0 | 0 |
| ENSG00000270252 | 0 | 0 | 0 | 0 | 0 | 0 |
| ENSG00000271569 | 0 | 0 | 0 | 0 | 0 | 0 |
| ENSG00000270450 | 0 | 0 | 0 | 0 | 0 | 0 |
| ENSG00000278537 | 0 | 0 | 0 | 0 | 0 | 0 |
| ENSG00000270187 | 0 | 0 | 0 | 0 | 0 | 0 |
| ENSG00000230606 | 1 | 2 | 2 | 1 | 0 | 1 |
| ENSG00000278766 | 0 | 0 | 0 | 0 | 1 | 1 |
| ENSG00000196912 | 21 | 9 | 16 | 10 | 21 | 22 |
| ENSG00000235833 | 0 | 0 | 0 | 0 | 0 | 0 |
| ENSG00000228970 | 0 | 0 | 0 | 0 | 0 | 1 |
| ENSG00000135940 | 4686 | 2282 | 2832 | 3290 | 2772 | 2620 |
| ENSG00000115073 | 3258 | 1529 | 2082 | 2045 | 1725 | 1899 |
| ENSG00000228486 | 148 | 73 | 88 | 44 | 83 | 74 |
| ENSG00000201806 | 6 | 2 | 0 | 1 | 0 | 1 |
| ENSG00000115085 | 0 | 0 | 0 | 2 | 0 | 0 |
| ENSG00000075568 | 2185 | 1074 | 1033 | 521 | 1189 | 928 |
| ENSG00000235734 | 14 | 14 | 28 | 12 | 10 | 16 |
| ENSG00000238719 | 0 | 0 | 0 | 0 | 0 | 0 |
| ENSG00000168658 | 14 | 4 | 3 | 17 | 14 | 6 |
| ENSG00000231635 | 0 | 0 | 0 | 0 | 0 | 0 |
| ENSG00000222000 | 0 | 0 | 0 | 0 | 0 | 0 |
| ENSG00000144191 | 0 | 0 | 0 | 0 | 0 | 0 |
| ENSG00000227987 | 0 | 0 | 0 | 0 | 0 | 0 |
| ENSG00000040933 | 1473 | 714 | 903 | 813 | 1225 | 1165 |
| ENSG00000183513 | 1466 | 719 | 860 | 708 | 737 | 744 |
| ENSG00000115446 | 1116 | 541 | 718 | 553 | 647 | 673 |
| ENSG00000071073 | 1630 | 806 | 818 | 362 | 901 | 593 |
| ENSG00000236564 | 0 | 0 | 0 | 0 | 0 | 0 |
| ENSG00000226791 | 0 | 2 | 0 | 0 | 0 | 0 |
| ENSG00000201070 | 0 | 0 | 0 | 0 | 0 | 0 |
| ENSG00000196872 | 888 | 400 | 525 | 411 | 400 | 426 |
| ENSG00000238830 | 0 | 0 | 0 | 0 | 0 | 0 |
| ENSG00000135951 | 184 | 83 | 117 | 56 | 117 | 97 |
| ENSG00000231822 | 3 | 2 | 2 | 1 | 4 | 4 |
| ENSG00000241962 | 4 | 11 | 6 | 1 | 4 | 4 |
| ENSG00000273045 | 310 | 201 | 224 | 160 | 209 | 219 |
| ENSG00000144182 | 110 | 55 | 64 | 62 | 65 | 54 |
| ENSG00000273155 | 0 | 0 | 0 | 0 | 0 | 0 |

| | | | | | | |
|---|---|---|---|---|---|---|
| ENSG00000158411 | 650 | 327 | 392 | 289 | 334 | 383 |
| ENSG00000185414 | 1599 | 845 | 959 | 1010 | 995 | 988 |
| ENSG00000185674 | 1 | 2 | 2 | 2 | 0 | 2 |
| ENSG00000144214 | 20 | 4 | 4 | 10 | 17 | 6 |
| ENSG00000115514 | 842 | 496 | 566 | 479 | 650 | 479 |
| ENSG00000158417 | 1257 | 867 | 502 | 249 | 1920 | 669 |
| ENSG00000135945 | 1118 | 516 | 643 | 333 | 577 | 548 |
| ENSG00000273306 | 17 | 6 | 9 | 0 | 16 | 8 |
| ENSG00000144218 | 123 | 70 | 68 | 25 | 44 | 40 |
| ENSG00000230393 | 2 | 1 | 0 | 1 | 1 | 2 |
| ENSG00000232084 | 0 | 0 | 0 | 0 | 0 | 0 |
| ENSG00000170500 | 2103 | 812 | 1299 | 624 | 971 | 1094 |
| ENSG00000269383 | 0 | 0 | 0 | 0 | 0 | 0 |
| ENSG00000270699 | 0 | 0 | 0 | 0 | 0 | 0 |
| ENSG00000115526 | 128 | 93 | 142 | 115 | 129 | 180 |
| ENSG00000238029 | 0 | 0 | 0 | 0 | 0 | 0 |
| ENSG00000204640 | 0 | 0 | 0 | 0 | 0 | 0 |
| ENSG00000204637 | 0 | 0 | 0 | 0 | 0 | 0 |
| ENSG00000115539 | 1356 | 747 | 834 | 664 | 756 | 790 |
| ENSG00000227259 | 0 | 0 | 0 | 0 | 0 | 0 |
| ENSG00000234988 | 0 | 0 | 0 | 0 | 0 | 0 |
| ENSG00000226034 | 0 | 0 | 0 | 0 | 0 | 0 |
| ENSG00000234169 | 0 | 0 | 0 | 0 | 0 | 0 |
| ENSG00000228488 | 0 | 0 | 0 | 0 | 0 | 0 |
| ENSG00000170485 | 281 | 165 | 175 | 133 | 158 | 198 |
| ENSG00000232034 | 0 | 1 | 0 | 0 | 0 | 2 |
| ENSG00000230140 | 0 | 0 | 0 | 0 | 0 | 0 |
| ENSG00000223947 | 6 | 5 | 3 | 0 | 3 | 2 |
| ENSG00000071082 | 15319 | 7595 | 8859 | 10364 | 9460 | 9042 |
| ENSG00000204634 | 647 | 365 | 469 | 255 | 398 | 429 |
| ENSG00000271532 | 1 | 2 | 0 | 0 | 2 | 3 |
| ENSG00000272902 | 33 | 12 | 21 | 16 | 21 | 12 |
| ENSG00000158435 | 3687 | 1747 | 2076 | 1884 | 1988 | 2045 |
| ENSG00000163162 | 1868 | 914 | 1078 | 1174 | 1410 | 1270 |
| ENSG00000212283 | 6 | 2 | 1 | 2 | 1 | 2 |
| ENSG00000264857 | 0 | 0 | 0 | 0 | 0 | 0 |
| ENSG00000175874 | 73 | 42 | 41 | 181 | 347 | 181 |
| ENSG00000226058 | 0 | 0 | 0 | 1 | 0 | 0 |
| ENSG00000196460 | 2 | 0 | 0 | 7 | 20 | 4 |
| ENSG00000223826 | 0 | 0 | 0 | 0 | 0 | 0 |
| ENSG00000237549 | 0 | 0 | 0 | 0 | 0 | 0 |
| ENSG00000232556 | 0 | 0 | 0 | 0 | 0 | 0 |
| ENSG00000071054 | 3123 | 1429 | 1740 | 1393 | 1851 | 1680 |
| ENSG00000281162 | 0 | 1 | 0 | 0 | 0 | 0 |
| ENSG00000115590 | 0 | 2 | 3 | 1 | 0 | 1 |
| ENSG00000115594 | 1445 | 585 | 854 | 240 | 323 | 337 |
| ENSG00000226925 | 0 | 0 | 0 | 0 | 0 | 0 |
| ENSG00000115598 | 0 | 0 | 0 | 0 | 0 | 0 |
| ENSG00000205716 | 0 | 0 | 0 | 0 | 0 | 0 |
| ENSG00000115602 | 0 | 0 | 0 | 0 | 0 | 1 |

| | | | | | | |
|---|---|---|---|---|---|---|
| ENSG00000115604 | 2 | 0 | 1 | 0 | 0 | 0 |
| ENSG00000236785 | 0 | 0 | 0 | 0 | 0 | 0 |
| ENSG00000115607 | 0 | 0 | 0 | 0 | 0 | 0 |
| ENSG00000264764 | 0 | 0 | 0 | 0 | 0 | 0 |
| ENSG00000236525 | 0 | 0 | 0 | 0 | 0 | 0 |
| ENSG00000234389 | 0 | 0 | 0 | 0 | 0 | 0 |
| ENSG00000180251 | 73 | 30 | 29 | 29 | 27 | 14 |
| ENSG00000115616 | 1948 | 863 | 1061 | 662 | 858 | 831 |
| ENSG00000135953 | 1056 | 475 | 659 | 535 | 647 | 662 |
| ENSG00000170417 | 215 | 77 | 118 | 95 | 122 | 144 |
| ENSG00000277527 | 0 | 0 | 0 | 0 | 0 | 0 |
| ENSG00000224095 | 0 | 0 | 0 | 0 | 0 | 0 |
| ENSG00000227680 | 0 | 0 | 0 | 0 | 0 | 0 |
| ENSG00000229209 | 0 | 0 | 0 | 0 | 0 | 0 |
| ENSG00000227623 | 0 | 0 | 0 | 0 | 0 | 0 |
| ENSG00000236596 | 0 | 0 | 0 | 0 | 0 | 0 |
| ENSG00000221849 | 0 | 0 | 0 | 0 | 0 | 0 |
| ENSG00000231723 | 0 | 0 | 0 | 0 | 0 | 0 |
| ENSG00000232597 | 0 | 0 | 0 | 0 | 0 | 0 |
| ENSG00000236141 | 0 | 0 | 0 | 0 | 0 | 0 |
| ENSG00000256637 | 1 | 0 | 0 | 0 | 0 | 0 |
| ENSG00000273515 | 0 | 0 | 0 | 0 | 0 | 0 |
| ENSG00000228968 | 0 | 0 | 0 | 0 | 0 | 0 |
| ENSG00000233955 | 2 | 1 | 0 | 0 | 0 | 0 |
| ENSG00000227157 | 0 | 0 | 0 | 0 | 0 | 0 |
| ENSG00000225765 | 0 | 0 | 0 | 0 | 0 | 0 |
| ENSG00000235597 | 0 | 0 | 0 | 0 | 0 | 0 |
| ENSG00000234781 | 0 | 0 | 0 | 0 | 0 | 0 |
| ENSG00000231626 | 0 | 0 | 0 | 0 | 0 | 0 |
| ENSG00000236109 | 0 | 0 | 0 | 0 | 0 | 0 |
| ENSG00000230690 | 0 | 0 | 0 | 0 | 0 | 0 |
| ENSG00000225669 | 0 | 0 | 0 | 0 | 0 | 0 |
| ENSG00000228528 | 0 | 0 | 0 | 0 | 0 | 0 |
| ENSG00000279904 | 0 | 0 | 0 | 0 | 0 | 0 |
| ENSG00000207249 | 0 | 0 | 0 | 0 | 0 | 0 |
| ENSG00000234177 | 0 | 0 | 0 | 0 | 0 | 0 |
| ENSG00000213301 | 0 | 0 | 0 | 0 | 0 | 0 |
| ENSG00000233639 | 0 | 0 | 0 | 0 | 0 | 0 |
| ENSG00000269707 | 0 | 0 | 0 | 0 | 0 | 0 |
| ENSG00000198914 | 0 | 0 | 0 | 0 | 0 | 0 |
| ENSG00000239587 | 0 | 0 | 0 | 0 | 0 | 0 |
| ENSG00000229743 | 0 | 0 | 0 | 0 | 0 | 1 |
| ENSG00000224509 | 0 | 0 | 0 | 0 | 0 | 0 |
| ENSG00000135972 | 906 | 524 | 596 | 505 | 681 | 615 |
| ENSG00000231851 | 5 | 3 | 2 | 0 | 5 | 0 |
| ENSG00000226508 | 83 | 29 | 37 | 35 | 34 | 35 |
| ENSG00000135973 | 6 | 2 | 1 | 0 | 1 | 2 |
| ENSG00000272861 | 1 | 4 | 5 | 6 | 0 | 1 |
| ENSG00000135966 | 1700 | 809 | 1030 | 796 | 933 | 972 |
| ENSG00000235319 | 6 | 11 | 5 | 12 | 6 | 5 |

| | | | | | | |
|---|---|---|---|---|---|---|
| ENSG00000272994 | 172 | 84 | 118 | 65 | 103 | 115 |
| ENSG00000135974 | 929 | 454 | 589 | 339 | 487 | 435 |
| ENSG00000115641 | 67 | 58 | 27 | 69 | 75 | 57 |
| ENSG00000238273 | 0 | 0 | 0 | 0 | 1 | 0 |
| ENSG00000071051 | 1073 | 540 | 652 | 607 | 606 | 624 |
| ENSG00000235522 | 2 | 0 | 3 | 2 | 4 | 0 |
| ENSG00000234162 | 0 | 0 | 0 | 0 | 0 | 0 |
| ENSG00000119147 | 0 | 0 | 0 | 0 | 0 | 0 |
| ENSG00000115652 | 1995 | 1001 | 1167 | 1122 | 1218 | 1317 |
| ENSG00000233339 | 0 | 0 | 0 | 0 | 0 | 0 |
| ENSG00000273595 | 0 | 0 | 0 | 0 | 0 | 0 |
| ENSG00000213295 | 0 | 0 | 0 | 0 | 0 | 1 |
| ENSG00000224061 | 0 | 0 | 0 | 0 | 0 | 0 |
| ENSG00000230353 | 0 | 0 | 0 | 0 | 0 | 0 |
| ENSG00000233939 | 0 | 0 | 0 | 0 | 0 | 0 |
| ENSG00000227513 | 0 | 0 | 0 | 0 | 0 | 0 |
| ENSG00000229682 | 0 | 0 | 0 | 0 | 0 | 0 |
| ENSG00000235486 | 0 | 0 | 0 | 0 | 0 | 0 |
| ENSG00000240935 | 0 | 0 | 0 | 0 | 0 | 0 |
| ENSG00000153165 | 2 | 0 | 1 | 3 | 0 | 2 |
| ENSG00000237666 | 0 | 0 | 0 | 0 | 0 | 0 |
| ENSG00000254126 | 0 | 0 | 0 | 0 | 0 | 0 |
| ENSG00000232001 | 0 | 0 | 0 | 0 | 0 | 0 |
| ENSG00000231505 | 0 | 0 | 0 | 0 | 0 | 0 |
| ENSG00000214199 | 6 | 6 | 5 | 10 | 2 | 5 |
| ENSG00000144057 | 0 | 0 | 0 | 0 | 0 | 0 |
| ENSG00000238250 | 0 | 0 | 0 | 0 | 0 | 0 |
| ENSG00000227294 | 0 | 0 | 0 | 0 | 0 | 0 |
| ENSG00000229360 | 0 | 0 | 0 | 0 | 0 | 0 |
| ENSG00000226905 | 0 | 0 | 0 | 0 | 0 | 0 |
| ENSG00000229457 | 0 | 0 | 0 | 0 | 0 | 0 |
| ENSG00000225588 | 0 | 0 | 0 | 0 | 0 | 0 |
| ENSG00000237880 | 0 | 0 | 0 | 0 | 0 | 0 |
| ENSG00000224568 | 0 | 0 | 0 | 0 | 0 | 0 |
| ENSG00000232991 | 0 | 0 | 0 | 0 | 0 | 0 |
| ENSG00000230651 | 0 | 0 | 0 | 1 | 0 | 0 |
| ENSG00000196862 | 1 | 0 | 0 | 0 | 0 | 0 |
| ENSG00000233494 | 0 | 0 | 0 | 0 | 0 | 0 |
| ENSG00000227270 | 0 | 0 | 0 | 0 | 0 | 0 |
| ENSG00000237745 | 1 | 0 | 0 | 0 | 0 | 0 |
| ENSG00000235325 | 0 | 0 | 0 | 0 | 0 | 0 |
| ENSG00000115665 | 0 | 0 | 0 | 0 | 0 | 0 |
| ENSG00000231221 | 0 | 0 | 0 | 0 | 0 | 0 |
| ENSG00000229890 | 0 | 0 | 0 | 0 | 0 | 0 |
| ENSG00000225328 | 0 | 0 | 0 | 0 | 0 | 0 |
| ENSG00000236877 | 0 | 0 | 0 | 0 | 0 | 0 |
| ENSG00000196228 | 0 | 0 | 0 | 0 | 0 | 0 |
| ENSG00000227597 | 0 | 0 | 0 | 0 | 0 | 0 |
| ENSG00000198203 | 0 | 0 | 0 | 0 | 1 | 0 |
| ENSG00000261209 | 0 | 0 | 0 | 0 | 0 | 0 |

| | | | | | | |
|---|---|---|---|---|---|---|
| ENSG00000237223 | 0 | 0 | 0 | 0 | 0 | 0 |
| ENSG00000271070 | 0 | 0 | 0 | 0 | 0 | 0 |
| ENSG00000198075 | 0 | 0 | 1 | 0 | 0 | 0 |
| ENSG00000271284 | 0 | 0 | 0 | 0 | 0 | 0 |
| ENSG00000135968 | 302 | 165 | 134 | 42 | 328 | 172 |
| ENSG00000214184 | 7 | 7 | 8 | 6 | 9 | 8 |
| ENSG00000169756 | 1028 | 496 | 606 | 351 | 623 | 491 |
| ENSG00000228763 | 0 | 0 | 2 | 0 | 0 | 0 |
| ENSG00000233648 | 0 | 0 | 0 | 0 | 0 | 0 |
| ENSG00000235644 | 0 | 0 | 0 | 0 | 0 | 0 |
| ENSG00000153201 | 557 | 362 | 278 | 107 | 1097 | 462 |
| ENSG00000163006 | 145 | 75 | 82 | 49 | 101 | 86 |
| ENSG00000227470 | 0 | 1 | 1 | 0 | 2 | 3 |
| ENSG00000135960 | 0 | 0 | 0 | 0 | 2 | 1 |
| ENSG00000277628 | 0 | 0 | 0 | 0 | 0 | 0 |
| ENSG00000259863 | 0 | 0 | 0 | 0 | 0 | 0 |
| ENSG00000172985 | 0 | 0 | 4 | 0 | 0 | 0 |
| ENSG00000264934 | 0 | 0 | 0 | 0 | 0 | 0 |
| ENSG00000228551 | 0 | 0 | 0 | 0 | 0 | 0 |
| ENSG00000265965 | 0 | 0 | 0 | 0 | 0 | 0 |
| ENSG00000186522 | 428 | 241 | 251 | 123 | 241 | 206 |
| ENSG00000237916 | 3 | 3 | 1 | 0 | 0 | 1 |
| ENSG00000230696 | 0 | 0 | 0 | 0 | 0 | 0 |
| ENSG00000198142 | 446 | 217 | 321 | 174 | 216 | 226 |
| ENSG00000230603 | 0 | 0 | 0 | 0 | 0 | 0 |
| ENSG00000231099 | 0 | 0 | 0 | 0 | 0 | 0 |
| ENSG00000233307 | 0 | 0 | 0 | 0 | 0 | 0 |
| ENSG00000233707 | 0 | 0 | 1 | 0 | 0 | 2 |
| ENSG00000282033 | 0 | 0 | 0 | 0 | 0 | 0 |
| ENSG00000015568 | 0 | 2 | 0 | 0 | 0 | 1 |
| ENSG00000256977 | 0 | 0 | 0 | 0 | 0 | 0 |
| ENSG00000284337 | 0 | 0 | 0 | 0 | 0 | 0 |
| ENSG00000186148 | 0 | 0 | 0 | 0 | 0 | 0 |
| ENSG00000283684 | 0 | 0 | 0 | 0 | 0 | 0 |
| ENSG00000240069 | 0 | 0 | 0 | 0 | 0 | 0 |
| ENSG00000225108 | 0 | 0 | 0 | 0 | 0 | 0 |
| ENSG00000204588 | 138 | 73 | 61 | 52 | 100 | 81 |
| ENSG00000235721 | 33 | 16 | 14 | 8 | 16 | 22 |
| ENSG00000229730 | 0 | 0 | 0 | 0 | 0 | 0 |
| ENSG00000283283 | 5 | 5 | 9 | 2 | 9 | 9 |
| ENSG00000265682 | 0 | 0 | 0 | 0 | 0 | 0 |
| ENSG00000144063 | 104 | 37 | 48 | 62 | 46 | 67 |
| ENSG00000264979 | 0 | 0 | 0 | 0 | 0 | 0 |
| ENSG00000144061 | 122 | 53 | 58 | 45 | 47 | 45 |
| ENSG00000233370 | 0 | 1 | 0 | 0 | 0 | 0 |
| ENSG00000175701 | 509 | 264 | 348 | 384 | 334 | 325 |
| ENSG00000261760 | 0 | 0 | 0 | 0 | 0 | 0 |
| ENSG00000282304 | 0 | 0 | 0 | 0 | 0 | 0 |
| ENSG00000282785 | 0 | 0 | 0 | 0 | 0 | 0 |
| ENSG00000263881 | 0 | 0 | 0 | 0 | 0 | 0 |

| | | | | | | |
|---|---|---|---|---|---|---|
| ENSG00000283167 | 4 | 2 | 1 | 0 | 3 | 0 |
| ENSG00000226991 | 0 | 0 | 0 | 0 | 1 | 0 |
| ENSG00000230650 | 28 | 19 | 15 | 10 | 18 | 20 |
| ENSG00000175772 | 48 | 25 | 40 | 32 | 51 | 36 |
| ENSG00000226065 | 0 | 0 | 0 | 0 | 0 | 0 |
| ENSG00000230596 | 0 | 0 | 0 | 0 | 0 | 1 |
| ENSG00000273471 | 0 | 0 | 0 | 0 | 0 | 0 |
| ENSG00000184115 | 0 | 0 | 0 | 0 | 0 | 0 |
| ENSG00000257207 | 0 | 0 | 0 | 0 | 0 | 0 |
| ENSG00000256671 | 0 | 0 | 0 | 0 | 0 | 0 |
| ENSG00000227574 | 0 | 0 | 0 | 0 | 0 | 0 |
| ENSG00000231583 | 0 | 0 | 0 | 0 | 0 | 0 |
| ENSG00000183054 | 0 | 0 | 0 | 0 | 1 | 0 |
| ENSG00000231536 | 0 | 0 | 0 | 0 | 0 | 0 |
| ENSG00000216867 | 0 | 2 | 0 | 0 | 2 | 1 |
| ENSG00000169679 | 787 | 427 | 471 | 430 | 797 | 539 |
| ENSG00000235881 | 0 | 0 | 0 | 0 | 0 | 0 |
| ENSG00000153093 | 70 | 45 | 56 | 98 | 99 | 124 |
| ENSG00000236330 | 0 | 0 | 0 | 0 | 0 | 1 |
| ENSG00000172965 | 236 | 165 | 222 | 160 | 173 | 164 |
| ENSG00000204581 | 30 | 23 | 28 | 19 | 17 | 16 |
| ENSG00000153094 | 1722 | 846 | 1054 | 754 | 1044 | 1043 |
| ENSG00000230499 | 0 | 0 | 0 | 0 | 0 | 1 |
| ENSG00000227992 | 2 | 0 | 0 | 1 | 0 | 1 |
| ENSG00000271590 | 0 | 0 | 0 | 0 | 2 | 0 |
| ENSG00000229118 | 0 | 0 | 0 | 0 | 0 | 0 |
| ENSG00000270190 | 0 | 0 | 0 | 0 | 0 | 0 |
| ENSG00000226928 | 0 | 0 | 0 | 0 | 0 | 0 |
| ENSG00000266139 | 0 | 0 | 0 | 0 | 0 | 0 |
| ENSG00000279191 | 10 | 6 | 8 | 1 | 3 | 1 |
| ENSG00000223973 | 0 | 0 | 0 | 0 | 0 | 0 |
| ENSG00000285016 | 2 | 0 | 1 | 2 | 0 | 1 |
| ENSG00000280878 | 0 | 0 | 0 | 0 | 0 | 0 |
| ENSG00000240350 | 0 | 0 | 0 | 0 | 0 | 1 |
| ENSG00000224959 | 1 | 0 | 1 | 0 | 0 | 2 |
| ENSG00000271635 | 0 | 0 | 0 | 0 | 0 | 0 |
| ENSG00000225744 | 0 | 0 | 0 | 0 | 0 | 0 |
| ENSG00000153107 | 988 | 534 | 602 | 385 | 711 | 632 |
| ENSG00000237583 | 0 | 0 | 0 | 0 | 0 | 0 |
| ENSG00000227842 | 0 | 0 | 0 | 0 | 0 | 0 |
| ENSG00000228615 | 1 | 0 | 0 | 0 | 0 | 0 |
| ENSG00000266063 | 0 | 0 | 0 | 0 | 0 | 0 |
| ENSG00000230958 | 0 | 0 | 0 | 0 | 0 | 0 |
| ENSG00000236307 | 12 | 7 | 6 | 11 | 8 | 11 |
| ENSG00000153208 | 51 | 25 | 33 | 8 | 22 | 26 |
| ENSG00000240183 | 0 | 0 | 0 | 0 | 0 | 0 |
| ENSG00000225496 | 0 | 0 | 1 | 0 | 0 | 0 |
| ENSG00000232277 | 0 | 0 | 0 | 0 | 0 | 0 |
| ENSG00000153214 | 2270 | 1123 | 1304 | 917 | 1309 | 1177 |
| ENSG00000144152 | 12 | 6 | 6 | 7 | 8 | 7 |

| | | | | | | |
|---|---|---|---|---|---|---|
| ENSG00000278962 | 0 | 0 | 0 | 1 | 0 | 3 |
| ENSG00000144161 | 443 | 202 | 247 | 190 | 324 | 282 |
| ENSG00000188177 | 346 | 229 | 193 | 88 | 199 | 156 |
| ENSG00000235258 | 0 | 0 | 0 | 0 | 0 | 0 |
| ENSG00000236555 | 0 | 0 | 0 | 0 | 0 | 0 |
| ENSG00000169629 | 31 | 19 | 9 | 2 | 33 | 15 |
| ENSG00000283698 | 0 | 0 | 0 | 0 | 0 | 0 |
| ENSG00000114999 | 1456 | 616 | 799 | 657 | 826 | 839 |
| ENSG00000125630 | 1474 | 700 | 888 | 684 | 805 | 873 |
| ENSG00000207383 | 20 | 13 | 12 | 3 | 11 | 7 |
| ENSG00000125611 | 865 | 399 | 567 | 527 | 472 | 451 |
| ENSG00000243389 | 0 | 4 | 2 | 1 | 0 | 1 |
| ENSG00000228251 | 3 | 3 | 2 | 4 | 7 | 7 |
| ENSG00000231747 | 8 | 0 | 4 | 4 | 1 | 2 |
| ENSG00000237753 | 34 | 25 | 24 | 22 | 15 | 28 |
| ENSG00000144136 | 2158 | 1035 | 1296 | 1038 | 1259 | 1270 |
| ENSG00000144130 | 0 | 1 | 3 | 1 | 1 | 3 |
| ENSG00000169607 | 135 | 89 | 104 | 39 | 174 | 125 |
| ENSG00000115008 | 2 | 0 | 1 | 3 | 0 | 1 |
| ENSG00000125538 | 0 | 0 | 0 | 0 | 0 | 2 |
| ENSG00000280228 | 0 | 0 | 0 | 0 | 0 | 1 |
| ENSG00000180152 | 0 | 0 | 0 | 0 | 0 | 1 |
| ENSG00000125571 | 6 | 1 | 2 | 0 | 0 | 0 |
| ENSG00000227368 | 0 | 0 | 0 | 0 | 0 | 1 |
| ENSG00000136688 | 0 | 0 | 0 | 0 | 0 | 2 |
| ENSG00000236124 | 0 | 0 | 0 | 0 | 0 | 0 |
| ENSG00000232090 | 0 | 0 | 0 | 0 | 0 | 0 |
| ENSG00000136694 | 0 | 0 | 0 | 0 | 0 | 0 |
| ENSG00000136696 | 0 | 0 | 0 | 0 | 0 | 0 |
| ENSG00000136695 | 2 | 1 | 7 | 9 | 3 | 6 |
| ENSG00000136697 | 0 | 0 | 0 | 0 | 0 | 0 |
| ENSG00000201805 | 0 | 0 | 0 | 0 | 0 | 0 |
| ENSG00000136689 | 2 | 4 | 3 | 12 | 13 | 10 |
| ENSG00000125637 | 3962 | 1781 | 2227 | 1642 | 1838 | 1739 |
| ENSG00000234174 | 2 | 6 | 3 | 0 | 0 | 3 |
| ENSG00000189223 | 1356 | 655 | 801 | 660 | 783 | 747 |
| ENSG00000125618 | 169 | 61 | 101 | 65 | 83 | 72 |
| ENSG00000231292 | 5 | 1 | 0 | 1 | 0 | 0 |
| ENSG00000234997 | 2 | 1 | 1 | 1 | 2 | 1 |
| ENSG00000272563 | 4 | 0 | 0 | 0 | 0 | 0 |
| ENSG00000136682 | 1046 | 634 | 634 | 416 | 583 | 522 |
| ENSG00000184492 | 42 | 17 | 14 | 13 | 8 | 6 |
| ENSG00000234148 | 0 | 0 | 0 | 0 | 0 | 0 |
| ENSG00000231943 | 0 | 0 | 0 | 0 | 0 | 0 |
| ENSG00000225398 | 0 | 0 | 0 | 0 | 0 | 0 |
| ENSG00000226516 | 0 | 0 | 0 | 0 | 0 | 0 |
| ENSG00000221055 | 0 | 3 | 3 | 2 | 1 | 3 |
| ENSG00000146556 | 138 | 62 | 105 | 84 | 89 | 78 |
| ENSG00000236397 | 41 | 17 | 27 | 16 | 23 | 21 |
| ENSG00000279267 | 4 | 4 | 1 | 1 | 6 | 3 |

| | | | | | | |
|---|---|---|---|---|---|---|
| ENSG00000240356 | 66 | 31 | 51 | 45 | 44 | 36 |
| ENSG00000144134 | 215 | 84 | 132 | 87 | 128 | 112 |
| ENSG00000144158 | 4 | 2 | 2 | 1 | 2 | 6 |
| ENSG00000175509 | 0 | 0 | 0 | 0 | 0 | 1 |
| ENSG00000233479 | 0 | 0 | 0 | 0 | 0 | 0 |
| ENSG00000227359 | 19 | 12 | 31 | 14 | 17 | 13 |
| ENSG00000202427 | 0 | 0 | 0 | 0 | 0 | 0 |
| ENSG00000115084 | 2250 | 1151 | 1365 | 782 | 1175 | 1061 |
| ENSG00000265429 | 1 | 0 | 0 | 0 | 0 | 1 |
| ENSG00000244063 | 0 | 0 | 0 | 1 | 0 | 1 |
| ENSG00000228857 | 0 | 1 | 0 | 1 | 3 | 1 |
| ENSG00000270019 | 10 | 3 | 3 | 2 | 1 | 0 |
| ENSG00000115091 | 6784 | 3675 | 4021 | 2797 | 4151 | 3624 |
| ENSG00000279957 | 0 | 0 | 0 | 0 | 0 | 0 |
| ENSG00000234199 | 0 | 0 | 0 | 0 | 0 | 0 |
| ENSG00000243179 | 0 | 0 | 0 | 0 | 0 | 0 |
| ENSG00000212182 | 0 | 0 | 0 | 0 | 0 | 0 |
| ENSG00000235242 | 0 | 0 | 0 | 0 | 0 | 0 |
| ENSG00000231926 | 0 | 0 | 0 | 0 | 0 | 0 |
| ENSG00000222923 | 0 | 0 | 0 | 0 | 0 | 0 |
| ENSG00000175497 | 0 | 0 | 0 | 0 | 0 | 0 |
| ENSG00000231538 | 0 | 0 | 0 | 0 | 0 | 0 |
| ENSG00000235717 | 0 | 0 | 0 | 0 | 0 | 0 |
| ENSG00000235026 | 0 | 0 | 0 | 0 | 0 | 0 |
| ENSG00000230391 | 0 | 0 | 0 | 0 | 0 | 0 |
| ENSG00000271667 | 0 | 0 | 0 | 0 | 0 | 0 |
| ENSG00000224869 | 0 | 0 | 0 | 0 | 0 | 0 |
| ENSG00000226341 | 0 | 0 | 0 | 0 | 0 | 0 |
| ENSG00000235871 | 0 | 0 | 0 | 0 | 0 | 0 |
| ENSG00000230158 | 0 | 0 | 0 | 0 | 0 | 0 |
| ENSG00000230950 | 0 | 0 | 0 | 0 | 0 | 0 |
| ENSG00000228898 | 0 | 0 | 0 | 0 | 0 | 0 |
| ENSG00000239185 | 0 | 0 | 0 | 0 | 0 | 0 |
| ENSG00000227291 | 0 | 0 | 0 | 0 | 0 | 0 |
| ENSG00000238207 | 0 | 0 | 0 | 0 | 0 | 0 |
| ENSG00000088205 | 2720 | 1446 | 1557 | 1152 | 1798 | 1533 |
| ENSG00000236255 | 4 | 2 | 11 | 4 | 8 | 2 |
| ENSG00000125631 | 0 | 1 | 2 | 0 | 2 | 1 |
| ENSG00000125633 | 1352 | 645 | 766 | 426 | 687 | 625 |
| ENSG00000224967 | 0 | 0 | 2 | 0 | 2 | 0 |
| ENSG00000235066 | 0 | 0 | 1 | 0 | 1 | 0 |
| ENSG00000272895 | 0 | 0 | 0 | 0 | 0 | 0 |
| ENSG00000279227 | 12 | 17 | 8 | 18 | 10 | 14 |
| ENSG00000243510 | 0 | 0 | 0 | 0 | 0 | 0 |
| ENSG00000125629 | 471 | 208 | 233 | 163 | 221 | 226 |
| ENSG00000226856 | 0 | 0 | 0 | 1 | 0 | 0 |
| ENSG00000258910 | 0 | 0 | 0 | 0 | 0 | 0 |
| ENSG00000163064 | 0 | 0 | 0 | 0 | 0 | 0 |
| ENSG00000019169 | 0 | 0 | 0 | 0 | 0 | 0 |
| ENSG00000259094 | 0 | 0 | 0 | 0 | 0 | 0 |

| | | | | | | |
|---|---|---|---|---|---|---|
| ENSG00000144119 | 0 | 0 | 0 | 0 | 0 | 0 |
| ENSG00000264833 | 0 | 0 | 0 | 0 | 0 | 0 |
| ENSG00000115107 | 1047 | 499 | 591 | 622 | 619 | 674 |
| ENSG00000229867 | 16 | 2 | 2 | 4 | 4 | 2 |
| ENSG00000186132 | 380 | 167 | 203 | 194 | 182 | 148 |
| ENSG00000155368 | 9797 | 5147 | 5483 | 5356 | 5300 | 4892 |
| ENSG00000171227 | 180 | 112 | 110 | 91 | 126 | 103 |
| ENSG00000080293 | 3 | 1 | 3 | 0 | 4 | 2 |
| ENSG00000231013 | 0 | 1 | 0 | 0 | 0 | 0 |
| ENSG00000163075 | 8 | 2 | 4 | 4 | 4 | 3 |
| ENSG00000144120 | 358 | 186 | 195 | 220 | 195 | 252 |
| ENSG00000229326 | 0 | 0 | 0 | 0 | 0 | 1 |
| ENSG00000088179 | 534 | 297 | 268 | 188 | 373 | 241 |
| ENSG00000233741 | 0 | 0 | 0 | 0 | 0 | 0 |
| ENSG00000115109 | 5019 | 2671 | 3098 | 1484 | 2980 | 2576 |
| ENSG00000224789 | 0 | 0 | 1 | 1 | 0 | 0 |
| ENSG00000202046 | 0 | 0 | 0 | 0 | 0 | 0 |
| ENSG00000236878 | 2 | 0 | 1 | 0 | 1 | 1 |
| ENSG00000227788 | 0 | 0 | 0 | 0 | 0 | 2 |
| ENSG00000233286 | 0 | 0 | 0 | 0 | 0 | 0 |
| ENSG00000243104 | 0 | 0 | 1 | 0 | 0 | 0 |
| ENSG00000234065 | 2 | 2 | 0 | 0 | 3 | 2 |
| ENSG00000223549 | 0 | 3 | 2 | 0 | 2 | 0 |
| ENSG00000226479 | 1081 | 502 | 622 | 604 | 546 | 679 |
| ENSG00000144118 | 5269 | 2605 | 2934 | 2636 | 3367 | 2981 |
| ENSG00000235840 | 2 | 0 | 2 | 0 | 0 | 0 |
| ENSG00000163083 | 2653 | 1454 | 1632 | 1261 | 1452 | 1349 |
| ENSG00000280409 | 0 | 0 | 0 | 0 | 0 | 0 |
| ENSG00000237614 | 6 | 13 | 6 | 8 | 5 | 8 |
| ENSG00000232140 | 0 | 0 | 0 | 0 | 0 | 0 |
| ENSG00000201006 | 0 | 0 | 0 | 0 | 0 | 0 |
| ENSG00000229774 | 0 | 0 | 0 | 0 | 0 | 0 |
| ENSG00000074047 | 2 | 0 | 0 | 1 | 0 | 0 |
| ENSG00000271709 | 0 | 0 | 0 | 0 | 0 | 0 |
| ENSG00000201584 | 0 | 0 | 0 | 0 | 0 | 0 |
| ENSG00000234455 | 0 | 0 | 0 | 0 | 0 | 0 |
| ENSG00000236145 | 0 | 0 | 0 | 0 | 0 | 0 |
| ENSG00000115112 | 0 | 1 | 2 | 0 | 0 | 2 |
| ENSG00000232723 | 0 | 0 | 0 | 0 | 0 | 0 |
| ENSG00000074054 | 2006 | 884 | 1117 | 531 | 1091 | 1000 |
| ENSG00000265451 | 2 | 0 | 5 | 0 | 1 | 2 |
| ENSG00000264229 | 0 | 0 | 1 | 1 | 2 | 0 |
| ENSG00000222467 | 0 | 0 | 0 | 0 | 0 | 0 |
| ENSG00000236859 | 197 | 111 | 103 | 93 | 114 | 110 |
| ENSG00000230993 | 2 | 0 | 0 | 0 | 0 | 0 |
| ENSG00000279721 | 0 | 0 | 2 | 2 | 0 | 0 |
| ENSG00000213239 | 0 | 0 | 0 | 0 | 0 | 0 |
| ENSG00000155438 | 1054 | 635 | 750 | 742 | 789 | 760 |
| ENSG00000211460 | 3480 | 1648 | 2064 | 1412 | 1896 | 2058 |
| ENSG00000270798 | 0 | 0 | 0 | 0 | 0 | 0 |

| | | | | | | |
|---|---|---|---|---|---|---|
| ENSG00000224655 | 0 | 0 | 0 | 0 | 0 | 0 |
| ENSG00000237856 | 0 | 0 | 0 | 0 | 0 | 0 |
| ENSG00000232740 | 0 | 0 | 0 | 0 | 0 | 0 |
| ENSG00000226708 | 0 | 0 | 0 | 0 | 0 | 0 |
| ENSG00000235640 | 0 | 0 | 0 | 0 | 0 | 0 |
| ENSG00000227738 | 0 | 0 | 0 | 0 | 0 | 0 |
| ENSG00000279874 | 0 | 0 | 0 | 0 | 0 | 0 |
| ENSG00000223118 | 0 | 0 | 0 | 0 | 0 | 0 |
| ENSG00000228400 | 0 | 0 | 0 | 0 | 0 | 0 |
| ENSG00000155052 | 0 | 0 | 0 | 0 | 0 | 0 |
| ENSG00000238046 | 0 | 0 | 0 | 0 | 0 | 0 |
| ENSG00000270815 | 0 | 0 | 0 | 0 | 0 | 0 |
| ENSG00000252674 | 0 | 0 | 0 | 0 | 0 | 0 |
| ENSG00000206726 | 0 | 0 | 0 | 0 | 0 | 0 |
| ENSG00000235491 | 0 | 0 | 0 | 0 | 0 | 0 |
| ENSG00000224410 | 0 | 0 | 0 | 0 | 0 | 0 |
| ENSG00000226813 | 0 | 0 | 0 | 0 | 0 | 0 |
| ENSG00000235774 | 0 | 0 | 0 | 0 | 0 | 0 |
| ENSG00000236655 | 0 | 0 | 0 | 0 | 0 | 0 |
| ENSG00000213236 | 28 | 21 | 13 | 12 | 8 | 17 |
| ENSG00000136732 | 0 | 0 | 0 | 0 | 0 | 0 |
| ENSG00000235128 | 0 | 0 | 0 | 0 | 0 | 0 |
| ENSG00000206963 | 0 | 0 | 0 | 0 | 0 | 0 |
| ENSG00000237524 | 0 | 0 | 0 | 0 | 0 | 0 |
| ENSG00000238788 | 0 | 0 | 0 | 0 | 0 | 0 |
| ENSG00000260634 | 0 | 0 | 0 | 0 | 0 | 0 |
| ENSG00000260163 | 0 | 0 | 0 | 0 | 0 | 0 |
| ENSG00000136717 | 107 | 72 | 40 | 48 | 61 | 42 |
| ENSG00000237630 | 0 | 0 | 0 | 0 | 0 | 0 |
| ENSG00000186684 | 0 | 0 | 0 | 0 | 0 | 0 |
| ENSG00000226789 | 0 | 0 | 0 | 0 | 1 | 0 |
| ENSG00000163161 | 3619 | 1693 | 2279 | 1670 | 2001 | 1956 |
| ENSG00000169967 | 1910 | 944 | 951 | 496 | 1217 | 906 |
| ENSG00000200250 | 0 | 0 | 1 | 0 | 0 | 0 |
| ENSG00000236682 | 17 | 14 | 19 | 12 | 17 | 18 |
| ENSG00000115718 | 4 | 7 | 5 | 3 | 2 | 2 |
| ENSG00000264075 | 0 | 0 | 2 | 0 | 0 | 0 |
| ENSG00000163166 | 1288 | 731 | 800 | 535 | 987 | 828 |
| ENSG00000231731 | 19 | 15 | 5 | 6 | 9 | 8 |
| ENSG00000202429 | 0 | 0 | 0 | 0 | 0 | 0 |
| ENSG00000169994 | 0 | 0 | 0 | 0 | 0 | 0 |
| ENSG00000272789 | 0 | 0 | 0 | 0 | 0 | 0 |
| ENSG00000072163 | 2 | 0 | 1 | 0 | 0 | 1 |
| ENSG00000144230 | 0 | 0 | 2 | 1 | 0 | 0 |
| ENSG00000136709 | 2616 | 1225 | 1565 | 1121 | 1469 | 1364 |
| ENSG00000173349 | 0 | 0 | 0 | 0 | 0 | 0 |
| ENSG00000201470 | 1 | 0 | 0 | 0 | 0 | 0 |
| ENSG00000200390 | 0 | 0 | 0 | 0 | 0 | 0 |
| ENSG00000274010 | 0 | 1 | 2 | 1 | 1 | 0 |
| ENSG00000144231 | 1290 | 645 | 772 | 700 | 797 | 778 |

| | | | | | | |
|---|---|---|---|---|---|---|
| ENSG00000202532 | 0 | 0 | 0 | 0 | 0 | 0 |
| ENSG00000244563 | 0 | 0 | 0 | 0 | 0 | 0 |
| ENSG00000273950 | 0 | 0 | 1 | 0 | 1 | 0 |
| ENSG00000144233 | 2355 | 968 | 1312 | 890 | 1039 | 1186 |
| ENSG00000272667 | 113 | 58 | 65 | 70 | 77 | 53 |
| ENSG00000204399 | 0 | 0 | 0 | 0 | 0 | 0 |
| ENSG00000136715 | 1705 | 798 | 906 | 633 | 840 | 839 |
| ENSG00000234044 | 0 | 0 | 0 | 0 | 0 | 0 |
| ENSG00000200432 | 0 | 0 | 0 | 0 | 0 | 0 |
| ENSG00000136731 | 2813 | 1546 | 1580 | 881 | 1605 | 1483 |
| ENSG00000229173 | 0 | 0 | 0 | 0 | 0 | 0 |
| ENSG00000232101 | 0 | 0 | 0 | 0 | 0 | 0 |
| ENSG00000229758 | 1 | 1 | 1 | 0 | 1 | 0 |
| ENSG00000185631 | 0 | 0 | 0 | 0 | 0 | 0 |
| ENSG00000238180 | 3 | 0 | 0 | 0 | 0 | 0 |
| ENSG00000136720 | 3031 | 1445 | 1910 | 1684 | 1596 | 1868 |
| ENSG00000252742 | 0 | 0 | 1 | 0 | 2 | 0 |
| ENSG00000238379 | 0 | 0 | 0 | 0 | 0 | 0 |
| ENSG00000227653 | 0 | 0 | 0 | 1 | 0 | 1 |
| ENSG00000237532 | 0 | 0 | 0 | 0 | 0 | 0 |
| ENSG00000238277 | 0 | 0 | 0 | 0 | 0 | 0 |
| ENSG00000204460 | 0 | 0 | 0 | 0 | 0 | 0 |
| ENSG00000233428 | 0 | 0 | 0 | 0 | 0 | 0 |
| ENSG00000229536 | 2 | 2 | 1 | 1 | 2 | 4 |
| ENSG00000214100 | 9 | 3 | 5 | 3 | 1 | 3 |
| ENSG00000237574 | 0 | 1 | 0 | 1 | 0 | 0 |
| ENSG00000234559 | 0 | 0 | 0 | 0 | 0 | 0 |
| ENSG00000225449 | 79 | 36 | 47 | 27 | 50 | 40 |
| ENSG00000222014 | 130 | 69 | 81 | 54 | 57 | 59 |
| ENSG00000284619 | 0 | 0 | 0 | 0 | 0 | 1 |
| ENSG00000180178 | 62 | 28 | 36 | 12 | 17 | 24 |
| ENSG00000250207 | 0 | 0 | 0 | 0 | 0 | 0 |
| ENSG00000217289 | 2 | 1 | 2 | 1 | 0 | 0 |
| ENSG00000231147 | 2 | 1 | 1 | 0 | 6 | 0 |
| ENSG00000231240 | 16 | 10 | 9 | 20 | 8 | 19 |
| ENSG00000248313 | 0 | 0 | 0 | 0 | 0 | 0 |
| ENSG00000196604 | 0 | 5 | 1 | 0 | 3 | 0 |
| ENSG00000228471 | 0 | 0 | 0 | 0 | 0 | 0 |
| ENSG00000224087 | 0 | 0 | 0 | 0 | 0 | 0 |
| ENSG00000252223 | 0 | 0 | 0 | 0 | 0 | 0 |
| ENSG00000223760 | 0 | 0 | 0 | 0 | 0 | 0 |
| ENSG00000152076 | 249 | 81 | 139 | 97 | 94 | 98 |
| ENSG00000136699 | 3446 | 1518 | 2048 | 1736 | 1686 | 1852 |
| ENSG00000152082 | 1740 | 1110 | 1436 | 1738 | 1267 | 1254 |
| ENSG00000152086 | 0 | 3 | 1 | 0 | 2 | 0 |
| ENSG00000227632 | 7 | 6 | 7 | 9 | 5 | 3 |
| ENSG00000230756 | 2 | 1 | 1 | 0 | 3 | 2 |
| ENSG00000213225 | 693 | 355 | 490 | 390 | 407 | 422 |
| ENSG00000234398 | 0 | 0 | 0 | 0 | 0 | 0 |
| ENSG00000236595 | 0 | 0 | 0 | 0 | 0 | 0 |

| | | | | | | |
|---|---|---|---|---|---|---|
| ENSG00000237722 | 0 | 0 | 0 | 0 | 0 | 0 |
| ENSG00000231954 | 0 | 0 | 0 | 0 | 0 | 0 |
| ENSG00000236211 | 0 | 0 | 0 | 0 | 0 | 0 |
| ENSG00000230496 | 0 | 0 | 0 | 0 | 0 | 0 |
| ENSG00000232990 | 0 | 0 | 0 | 0 | 0 | 0 |
| ENSG00000249997 | 0 | 0 | 0 | 0 | 0 | 0 |
| ENSG00000283940 | 0 | 0 | 0 | 0 | 0 | 0 |
| ENSG00000284171 | 0 | 0 | 0 | 0 | 0 | 0 |
| ENSG00000230548 | 0 | 0 | 0 | 0 | 0 | 0 |
| ENSG00000234934 | 0 | 0 | 0 | 0 | 0 | 0 |
| ENSG00000248136 | 0 | 0 | 0 | 0 | 0 | 0 |
| ENSG00000233550 | 0 | 0 | 0 | 0 | 0 | 0 |
| ENSG00000226915 | 0 | 0 | 0 | 0 | 0 | 0 |
| ENSG00000271435 | 0 | 0 | 0 | 0 | 0 | 0 |
| ENSG00000224352 | 0 | 0 | 0 | 0 | 0 | 0 |
| ENSG00000136710 | 2267 | 1035 | 1249 | 1082 | 1003 | 1070 |
| ENSG00000136718 | 3651 | 1775 | 2155 | 2280 | 1950 | 2152 |
| ENSG00000072135 | 3106 | 1351 | 2204 | 1570 | 1386 | 1627 |
| ENSG00000178162 | 45 | 10 | 37 | 3 | 10 | 25 |
| ENSG00000230646 | 4 | 1 | 8 | 0 | 1 | 5 |
| ENSG00000239402 | 16 | 9 | 35 | 4 | 5 | 27 |
| ENSG00000196834 | 32 | 18 | 28 | 4 | 8 | 9 |
| ENSG00000251973 | 0 | 0 | 0 | 0 | 0 | 0 |
| ENSG00000225819 | 0 | 0 | 0 | 0 | 0 | 0 |
| ENSG00000274711 | 0 | 0 | 0 | 0 | 0 | 0 |
| ENSG00000152093 | 0 | 0 | 0 | 0 | 0 | 0 |
| ENSG00000225341 | 0 | 0 | 0 | 0 | 0 | 0 |
| ENSG00000184761 | 0 | 0 | 0 | 0 | 0 | 0 |
| ENSG00000183292 | 0 | 0 | 0 | 0 | 0 | 0 |
| ENSG00000232408 | 0 | 0 | 0 | 0 | 0 | 0 |
| ENSG00000136698 | 0 | 0 | 0 | 0 | 0 | 0 |
| ENSG00000277668 | 0 | 0 | 0 | 0 | 0 | 0 |
| ENSG00000222038 | 31 | 2 | 4 | 0 | 0 | 1 |
| ENSG00000251757 | 0 | 0 | 0 | 0 | 0 | 0 |
| ENSG00000214081 | 0 | 4 | 2 | 1 | 1 | 1 |
| ENSG00000282944 | 1 | 2 | 3 | 0 | 1 | 1 |
| ENSG00000240253 | 10 | 3 | 12 | 3 | 5 | 5 |
| ENSG00000223349 | 12 | 0 | 2 | 0 | 2 | 2 |
| ENSG00000173302 | 2 | 0 | 1 | 0 | 0 | 0 |
| ENSG00000229797 | 0 | 0 | 0 | 0 | 0 | 0 |
| ENSG00000178171 | 0 | 0 | 1 | 0 | 0 | 0 |
| ENSG00000232503 | 0 | 0 | 0 | 0 | 0 | 0 |
| ENSG00000233221 | 2 | 0 | 0 | 3 | 2 | 0 |
| ENSG00000136002 | 248 | 105 | 151 | 155 | 148 | 142 |
| ENSG00000281516 | 0 | 0 | 0 | 0 | 0 | 0 |
| ENSG00000281159 | 0 | 0 | 0 | 0 | 0 | 0 |
| ENSG00000284479 | 0 | 0 | 0 | 0 | 0 | 0 |
| ENSG00000152102 | 6337 | 2809 | 3844 | 2591 | 3179 | 3497 |
| ENSG00000115762 | 8223 | 4162 | 5276 | 3227 | 4523 | 4154 |
| ENSG00000230450 | 0 | 0 | 0 | 0 | 0 | 0 |

| | | | | | | |
|---|---|---|---|---|---|---|
| ENSG00000225448 | 0 | 0 | 0 | 0 | 0 | 0 |
| ENSG00000236956 | 0 | 0 | 0 | 0 | 0 | 0 |
| ENSG00000285819 | 0 | 0 | 0 | 0 | 0 | 0 |
| ENSG00000248796 | 0 | 0 | 0 | 0 | 0 | 0 |
| ENSG00000188219 | 28 | 2 | 0 | 1 | 1 | 2 |
| ENSG00000212199 | 0 | 0 | 0 | 0 | 0 | 0 |
| ENSG00000231431 | 3 | 1 | 3 | 0 | 0 | 0 |
| ENSG00000236516 | 0 | 0 | 3 | 1 | 0 | 2 |
| ENSG00000284604 | 0 | 0 | 0 | 0 | 0 | 0 |
| ENSG00000227817 | 0 | 0 | 0 | 0 | 0 | 0 |
| ENSG00000284706 | 0 | 0 | 0 | 0 | 0 | 0 |
| ENSG00000224425 | 0 | 1 | 4 | 0 | 0 | 0 |
| ENSG00000233087 | 6 | 1 | 6 | 4 | 5 | 4 |
| ENSG00000283303 | 0 | 0 | 0 | 0 | 0 | 0 |
| ENSG00000233285 | 0 | 0 | 0 | 0 | 0 | 0 |
| ENSG00000227275 | 0 | 0 | 0 | 0 | 0 | 0 |
| ENSG00000235080 | 0 | 0 | 0 | 0 | 0 | 0 |
| ENSG00000231777 | 0 | 0 | 0 | 0 | 0 | 0 |
| ENSG00000284003 | 0 | 0 | 0 | 0 | 0 | 0 |
| ENSG00000283865 | 0 | 0 | 0 | 0 | 0 | 0 |
| ENSG00000227661 | 0 | 0 | 0 | 0 | 0 | 0 |
| ENSG00000236590 | 0 | 0 | 0 | 0 | 0 | 0 |
| ENSG00000234480 | 0 | 0 | 0 | 0 | 0 | 0 |
| ENSG00000237910 | 0 | 0 | 0 | 0 | 0 | 0 |
| ENSG00000234939 | 0 | 0 | 0 | 0 | 0 | 0 |
| ENSG00000234576 | 0 | 0 | 0 | 0 | 0 | 0 |
| ENSG00000226831 | 0 | 0 | 0 | 0 | 0 | 0 |
| ENSG00000276460 | 0 | 0 | 0 | 0 | 0 | 0 |
| ENSG00000223631 | 0 | 0 | 0 | 0 | 0 | 0 |
| ENSG00000284659 | 4 | 1 | 4 | 1 | 1 | 2 |
| ENSG00000214077 | 28 | 13 | 22 | 12 | 10 | 16 |
| ENSG00000217950 | 6 | 2 | 5 | 8 | 3 | 2 |
| ENSG00000232742 | 2 | 4 | 7 | 7 | 7 | 7 |
| ENSG00000273073 | 15 | 10 | 10 | 8 | 4 | 15 |
| ENSG00000173272 | 2096 | 1129 | 1377 | 1439 | 1209 | 1165 |
| ENSG00000075886 | 33 | 34 | 28 | 14 | 20 | 10 |
| ENSG00000284149 | 0 | 0 | 0 | 0 | 0 | 0 |
| ENSG00000152117 | 947 | 477 | 581 | 394 | 426 | 480 |
| ENSG00000163040 | 762 | 290 | 452 | 368 | 309 | 414 |
| ENSG00000224679 | 5 | 2 | 5 | 0 | 1 | 4 |
| ENSG00000204434 | 340 | 154 | 178 | 44 | 136 | 156 |
| ENSG00000251956 | 0 | 0 | 0 | 0 | 0 | 0 |
| ENSG00000273588 | 39 | 16 | 25 | 5 | 17 | 28 |
| ENSG00000224559 | 509 | 244 | 363 | 94 | 204 | 245 |
| ENSG00000270930 | 19 | 9 | 14 | 2 | 9 | 10 |
| ENSG00000197927 | 447 | 247 | 405 | 185 | 225 | 265 |
| ENSG00000213222 | 185 | 90 | 113 | 30 | 66 | 76 |
| ENSG00000186825 | 0 | 1 | 2 | 0 | 0 | 0 |
| ENSG00000274353 | 0 | 0 | 0 | 1 | 0 | 0 |
| ENSG00000229203 | 2 | 0 | 0 | 0 | 1 | 0 |

| | | | | | | |
|---|---|---|---|---|---|---|
| ENSG00000232760 | 0 | 0 | 0 | 0 | 0 | 0 |
| ENSG00000244337 | 0 | 0 | 0 | 0 | 0 | 0 |
| ENSG00000235615 | 0 | 0 | 0 | 0 | 0 | 0 |
| ENSG00000236485 | 0 | 0 | 0 | 0 | 0 | 0 |
| ENSG00000226886 | 0 | 0 | 0 | 0 | 0 | 0 |
| ENSG00000271179 | 0 | 0 | 0 | 0 | 0 | 0 |
| ENSG00000225594 | 0 | 0 | 0 | 0 | 0 | 0 |
| ENSG00000227745 | 0 | 0 | 0 | 0 | 0 | 0 |
| ENSG00000163046 | 0 | 0 | 0 | 0 | 0 | 0 |
| ENSG00000239108 | 0 | 0 | 0 | 0 | 0 | 0 |
| ENSG00000202174 | 0 | 0 | 0 | 0 | 0 | 0 |
| ENSG00000221288 | 0 | 0 | 0 | 0 | 0 | 0 |
| ENSG00000233786 | 0 | 0 | 0 | 0 | 0 | 0 |
| ENSG00000230803 | 0 | 0 | 0 | 0 | 0 | 0 |
| ENSG00000018607 | 0 | 0 | 0 | 0 | 0 | 0 |
| ENSG00000272769 | 6 | 4 | 4 | 7 | 6 | 6 |
| ENSG00000230992 | 0 | 5 | 1 | 2 | 5 | 3 |
| ENSG00000252705 | 0 | 0 | 0 | 0 | 0 | 0 |
| ENSG00000183840 | 49 | 33 | 40 | 28 | 37 | 29 |
| ENSG00000232349 | 0 | 0 | 0 | 0 | 0 | 0 |
| ENSG00000200718 | 0 | 0 | 0 | 0 | 0 | 0 |
| ENSG00000150551 | 55 | 51 | 23 | 23 | 29 | 19 |
| ENSG00000230065 | 2 | 0 | 0 | 0 | 0 | 0 |
| ENSG00000176771 | 116 | 50 | 72 | 18 | 39 | 37 |
| ENSG00000228721 | 0 | 0 | 0 | 0 | 0 | 0 |
| ENSG00000233729 | 0 | 0 | 0 | 0 | 0 | 0 |
| ENSG00000279512 | 0 | 0 | 1 | 0 | 0 | 0 |
| ENSG00000226953 | 6 | 3 | 4 | 2 | 1 | 1 |
| ENSG00000252688 | 0 | 0 | 0 | 0 | 0 | 0 |
| ENSG00000222068 | 0 | 0 | 0 | 0 | 0 | 0 |
| ENSG00000232474 | 0 | 0 | 0 | 0 | 0 | 0 |
| ENSG00000223430 | 0 | 0 | 0 | 0 | 0 | 0 |
| ENSG00000200708 | 0 | 0 | 0 | 0 | 0 | 0 |
| ENSG00000152127 | 1844 | 796 | 1147 | 795 | 992 | 1073 |
| ENSG00000263813 | 0 | 0 | 0 | 0 | 0 | 0 |
| ENSG00000222921 | 0 | 0 | 0 | 0 | 0 | 0 |
| ENSG00000234060 | 0 | 0 | 0 | 0 | 0 | 0 |
| ENSG00000152128 | 6 | 2 | 2 | 1 | 3 | 2 |
| ENSG00000224043 | 30 | 25 | 32 | 31 | 30 | 23 |
| ENSG00000234682 | 0 | 0 | 0 | 0 | 0 | 0 |
| ENSG00000228098 | 0 | 0 | 0 | 0 | 0 | 0 |
| ENSG00000153086 | 0 | 1 | 1 | 0 | 3 | 1 |
| ENSG00000263783 | 0 | 0 | 0 | 0 | 0 | 0 |
| ENSG00000082258 | 1279 | 583 | 602 | 297 | 742 | 551 |
| ENSG00000176601 | 0 | 0 | 1 | 1 | 0 | 1 |
| ENSG00000115839 | 2089 | 1094 | 1177 | 764 | 1426 | 1117 |
| ENSG00000208308 | 0 | 0 | 0 | 0 | 0 | 0 |
| ENSG00000121988 | 338 | 195 | 196 | 109 | 233 | 159 |
| ENSG00000235651 | 14 | 3 | 0 | 0 | 4 | 2 |
| ENSG00000048991 | 1563 | 807 | 729 | 509 | 951 | 715 |

| | | | | | | |
|---|---|---|---|---|---|---|
| ENSG00000201308 | 0 | 0 | 0 | 0 | 0 | 0 |
| ENSG00000207654 | 0 | 0 | 0 | 0 | 0 | 0 |
| ENSG00000144224 | 5625 | 3062 | 3216 | 1788 | 3818 | 2904 |
| ENSG00000115850 | 2 | 1 | 0 | 0 | 0 | 3 |
| ENSG00000200664 | 0 | 0 | 0 | 0 | 0 | 0 |
| ENSG00000226806 | 0 | 0 | 0 | 0 | 0 | 0 |
| ENSG00000076003 | 2431 | 1302 | 1543 | 1033 | 1220 | 1332 |
| ENSG00000214070 | 0 | 0 | 0 | 1 | 0 | 0 |
| ENSG00000115866 | 2323 | 1171 | 1326 | 1038 | 1370 | 1246 |
| ENSG00000231890 | 34 | 19 | 17 | 22 | 11 | 22 |
| ENSG00000237262 | 0 | 0 | 0 | 0 | 0 | 0 |
| ENSG00000121966 | 262 | 90 | 150 | 62 | 33 | 59 |
| ENSG00000227347 | 3 | 0 | 1 | 0 | 2 | 6 |
| ENSG00000279024 | 2 | 0 | 1 | 0 | 1 | 0 |
| ENSG00000230037 | 0 | 0 | 0 | 0 | 0 | 0 |
| ENSG00000251976 | 0 | 0 | 0 | 0 | 0 | 0 |
| ENSG00000229568 | 0 | 0 | 0 | 0 | 0 | 0 |
| ENSG00000144229 | 0 | 1 | 0 | 1 | 2 | 0 |
| ENSG00000212425 | 0 | 0 | 0 | 0 | 0 | 0 |
| ENSG00000222394 | 0 | 0 | 0 | 0 | 0 | 0 |
| ENSG00000150540 | 333 | 253 | 233 | 202 | 312 | 202 |
| ENSG00000224231 | 0 | 0 | 0 | 0 | 0 | 0 |
| ENSG00000233045 | 1 | 1 | 1 | 1 | 4 | 2 |
| ENSG00000234645 | 0 | 0 | 0 | 0 | 0 | 0 |
| ENSG00000237939 | 0 | 1 | 0 | 0 | 0 | 0 |
| ENSG00000232915 | 0 | 0 | 0 | 0 | 0 | 0 |
| ENSG00000228043 | 13 | 5 | 12 | 5 | 10 | 13 |
| ENSG00000230569 | 0 | 0 | 0 | 0 | 0 | 0 |
| ENSG00000144228 | 1977 | 1148 | 1189 | 586 | 1272 | 892 |
| ENSG00000241772 | 0 | 0 | 0 | 0 | 1 | 0 |
| ENSG00000260059 | 2 | 2 | 3 | 1 | 0 | 1 |
| ENSG00000237772 | 0 | 2 | 2 | 0 | 1 | 0 |
| ENSG00000144227 | 0 | 0 | 0 | 0 | 0 | 0 |
| ENSG00000222826 | 0 | 0 | 0 | 0 | 0 | 0 |
| ENSG00000229104 | 0 | 0 | 0 | 0 | 0 | 0 |
| ENSG00000232992 | 0 | 0 | 0 | 0 | 0 | 0 |
| ENSG00000226939 | 0 | 1 | 0 | 0 | 0 | 0 |
| ENSG00000229131 | 0 | 0 | 0 | 0 | 0 | 0 |
| ENSG00000206901 | 0 | 0 | 0 | 0 | 0 | 0 |
| ENSG00000235232 | 0 | 0 | 0 | 0 | 0 | 0 |
| ENSG00000228374 | 0 | 0 | 0 | 0 | 0 | 0 |
| ENSG00000223554 | 0 | 0 | 0 | 0 | 0 | 0 |
| ENSG00000240977 | 0 | 0 | 0 | 0 | 0 | 0 |
| ENSG00000224028 | 0 | 0 | 0 | 0 | 0 | 0 |
| ENSG00000227149 | 0 | 0 | 0 | 0 | 0 | 0 |
| ENSG00000231567 | 0 | 0 | 0 | 0 | 0 | 0 |
| ENSG00000223921 | 0 | 0 | 0 | 0 | 0 | 0 |
| ENSG00000168702 | 131 | 90 | 84 | 46 | 116 | 80 |
| ENSG00000277306 | 0 | 0 | 0 | 0 | 0 | 0 |
| ENSG00000283506 | 0 | 0 | 0 | 0 | 0 | 0 |

| | | | | | | |
|---|---|---|---|---|---|---|
| ENSG00000236440 | 0 | 0 | 0 | 0 | 0 | 0 |
| ENSG00000238065 | 0 | 0 | 1 | 0 | 1 | 0 |
| ENSG00000201892 | 0 | 0 | 0 | 0 | 0 | 0 |
| ENSG00000252015 | 0 | 0 | 0 | 0 | 0 | 0 |
| ENSG00000233329 | 0 | 0 | 0 | 0 | 0 | 0 |
| ENSG00000244125 | 0 | 0 | 0 | 0 | 0 | 0 |
| ENSG00000228134 | 0 | 0 | 0 | 0 | 0 | 0 |
| ENSG00000229781 | 4 | 1 | 4 | 1 | 3 | 3 |
| ENSG00000115919 | 5914 | 3536 | 3971 | 2214 | 3860 | 3100 |
| ENSG00000270390 | 4 | 1 | 0 | 0 | 1 | 1 |
| ENSG00000075884 | 0 | 0 | 0 | 1 | 0 | 0 |
| ENSG00000224288 | 0 | 0 | 0 | 0 | 0 | 0 |
| ENSG00000226839 | 0 | 0 | 0 | 0 | 0 | 0 |
| ENSG00000224852 | 0 | 0 | 0 | 0 | 0 | 0 |
| ENSG00000232772 | 0 | 0 | 0 | 0 | 0 | 0 |
| ENSG00000236300 | 0 | 0 | 0 | 0 | 0 | 0 |
| ENSG00000227557 | 0 | 0 | 0 | 0 | 0 | 1 |
| ENSG00000225476 | 0 | 0 | 0 | 0 | 0 | 0 |
| ENSG00000284167 | 0 | 0 | 0 | 0 | 0 | 0 |
| ENSG00000231766 | 0 | 0 | 0 | 0 | 0 | 0 |
| ENSG00000257284 | 0 | 0 | 0 | 0 | 0 | 0 |
| ENSG00000228655 | 0 | 0 | 0 | 0 | 0 | 0 |
| ENSG00000257640 | 0 | 0 | 0 | 0 | 0 | 0 |
| ENSG00000258268 | 0 | 0 | 0 | 0 | 0 | 0 |
| ENSG00000283994 | 0 | 0 | 0 | 0 | 0 | 0 |
| ENSG00000231758 | 0 | 0 | 0 | 0 | 0 | 0 |
| ENSG00000257277 | 0 | 0 | 0 | 0 | 0 | 0 |
| ENSG00000257226 | 0 | 0 | 0 | 0 | 0 | 0 |
| ENSG00000236356 | 0 | 0 | 0 | 0 | 0 | 0 |
| ENSG00000200287 | 0 | 0 | 0 | 0 | 0 | 0 |
| ENSG00000232377 | 0 | 0 | 0 | 0 | 0 | 0 |
| ENSG00000121964 | 376 | 177 | 175 | 152 | 212 | 198 |
| ENSG00000169554 | 0 | 0 | 0 | 0 | 0 | 0 |
| ENSG00000283118 | 0 | 0 | 0 | 0 | 0 | 0 |
| ENSG00000279166 | 0 | 0 | 0 | 0 | 0 | 0 |
| ENSG00000238057 | 0 | 0 | 0 | 0 | 0 | 0 |
| ENSG00000275372 | 0 | 0 | 0 | 0 | 0 | 0 |
| ENSG00000273537 | 0 | 0 | 0 | 0 | 0 | 0 |
| ENSG00000277444 | 0 | 0 | 0 | 0 | 0 | 0 |
| ENSG00000232606 | 0 | 0 | 0 | 0 | 0 | 0 |
| ENSG00000226674 | 2 | 1 | 2 | 0 | 1 | 0 |
| ENSG00000234940 | 0 | 0 | 0 | 0 | 0 | 0 |
| ENSG00000276706 | 0 | 0 | 0 | 0 | 0 | 0 |
| ENSG00000235352 | 0 | 0 | 0 | 0 | 0 | 0 |
| ENSG00000268580 | 0 | 0 | 0 | 0 | 0 | 0 |
| ENSG00000235435 | 0 | 0 | 0 | 0 | 0 | 0 |
| ENSG00000214064 | 0 | 0 | 0 | 0 | 0 | 0 |
| ENSG00000225107 | 0 | 0 | 0 | 0 | 0 | 0 |
| ENSG00000226218 | 0 | 0 | 0 | 0 | 0 | 0 |
| ENSG00000234119 | 0 | 0 | 0 | 0 | 0 | 0 |

| | | | | | | |
|---|---|---|---|---|---|---|
| ENSG00000279876 | 0 | 0 | 0 | 0 | 0 | 0 |
| ENSG00000233842 | 0 | 0 | 0 | 0 | 0 | 0 |
| ENSG00000251905 | 0 | 0 | 0 | 0 | 0 | 0 |
| ENSG00000230559 | 0 | 0 | 0 | 0 | 0 | 0 |
| ENSG00000227134 | 0 | 0 | 0 | 0 | 0 | 0 |
| ENSG00000198526 | 0 | 0 | 0 | 0 | 0 | 0 |
| ENSG00000271583 | 0 | 0 | 0 | 0 | 0 | 0 |
| ENSG00000231040 | 0 | 0 | 0 | 0 | 0 | 0 |
| ENSG00000227902 | 0 | 0 | 0 | 0 | 0 | 0 |
| ENSG00000207225 | 0 | 0 | 0 | 0 | 0 | 0 |
| ENSG00000207161 | 0 | 0 | 0 | 0 | 0 | 0 |
| ENSG00000202074 | 0 | 0 | 0 | 0 | 0 | 0 |
| ENSG00000253083 | 0 | 0 | 0 | 0 | 0 | 0 |
| ENSG00000229143 | 0 | 0 | 0 | 0 | 0 | 0 |
| ENSG00000121989 | 308 | 123 | 144 | 90 | 147 | 116 |
| ENSG00000212389 | 0 | 0 | 0 | 0 | 0 | 0 |
| ENSG00000223911 | 0 | 0 | 0 | 0 | 0 | 0 |
| ENSG00000115947 | 969 | 498 | 508 | 415 | 625 | 511 |
| ENSG00000204406 | 792 | 379 | 387 | 200 | 397 | 370 |
| ENSG00000281469 | 4 | 0 | 0 | 0 | 0 | 2 |
| ENSG00000281772 | 0 | 0 | 0 | 0 | 0 | 0 |
| ENSG00000212181 | 0 | 0 | 0 | 0 | 0 | 0 |
| ENSG00000224007 | 0 | 0 | 0 | 0 | 0 | 0 |
| ENSG00000232238 | 1 | 0 | 0 | 0 | 0 | 0 |
| ENSG00000135999 | 755 | 340 | 394 | 220 | 361 | 333 |
| ENSG00000222126 | 0 | 0 | 0 | 0 | 0 | 0 |
| ENSG00000223962 | 0 | 0 | 1 | 0 | 0 | 0 |
| ENSG00000231079 | 0 | 0 | 0 | 0 | 0 | 0 |
| ENSG00000228848 | 0 | 0 | 0 | 0 | 0 | 0 |
| ENSG00000226340 | 0 | 0 | 0 | 0 | 0 | 0 |
| ENSG00000168280 | 565 | 268 | 406 | 223 | 313 | 413 |
| ENSG00000150556 | 607 | 302 | 368 | 478 | 555 | 512 |
| ENSG00000228236 | 2 | 0 | 1 | 0 | 4 | 1 |
| ENSG00000224337 | 0 | 0 | 0 | 0 | 0 | 0 |
| ENSG00000187123 | 521 | 241 | 347 | 251 | 336 | 322 |
| ENSG00000223604 | 0 | 0 | 0 | 0 | 0 | 0 |
| ENSG00000168288 | 3652 | 1896 | 2352 | 1832 | 2205 | 2076 |
| ENSG00000231969 | 0 | 0 | 0 | 1 | 0 | 0 |
| ENSG00000207270 | 0 | 0 | 0 | 0 | 0 | 0 |
| ENSG00000162947 | 0 | 0 | 0 | 0 | 0 | 0 |
| ENSG00000230645 | 0 | 0 | 0 | 0 | 0 | 0 |
| ENSG00000231420 | 0 | 0 | 0 | 0 | 0 | 0 |
| ENSG00000115963 | 1173 | 689 | 632 | 455 | 874 | 725 |
| ENSG00000236049 | 0 | 0 | 0 | 0 | 0 | 0 |
| ENSG00000237220 | 0 | 0 | 0 | 0 | 0 | 0 |
| ENSG00000232359 | 0 | 0 | 0 | 0 | 0 | 0 |
| ENSG00000224048 | 2 | 0 | 1 | 0 | 0 | 0 |
| ENSG00000228064 | 0 | 0 | 0 | 0 | 0 | 0 |
| ENSG00000222031 | 0 | 0 | 0 | 0 | 0 | 0 |
| ENSG00000236885 | 0 | 0 | 0 | 0 | 0 | 0 |

| | | | | | | |
|---|---|---|---|---|---|---|
| ENSG00000213201 | 0 | 0 | 0 | 0 | 0 | 0 |
| ENSG00000184898 | 360 | 145 | 177 | 94 | 148 | 123 |
| ENSG00000123609 | 208 | 123 | 169 | 88 | 159 | 144 |
| ENSG00000123610 | 0 | 0 | 0 | 0 | 0 | 0 |
| ENSG00000264684 | 0 | 0 | 0 | 0 | 0 | 0 |
| ENSG00000283569 | 0 | 0 | 0 | 0 | 0 | 0 |
| ENSG00000242113 | 0 | 0 | 0 | 0 | 0 | 0 |
| ENSG00000080345 | 298 | 203 | 165 | 45 | 489 | 214 |
| ENSG00000183091 | 16 | 12 | 17 | 5 | 22 | 18 |
| ENSG00000162980 | 415 | 216 | 254 | 187 | 373 | 251 |
| ENSG00000283228 | 0 | 0 | 0 | 0 | 0 | 0 |
| ENSG00000182389 | 5 | 6 | 5 | 1 | 3 | 1 |
| ENSG00000225214 | 0 | 0 | 0 | 0 | 0 | 0 |
| ENSG00000115145 | 1385 | 685 | 764 | 378 | 781 | 621 |
| ENSG00000234272 | 1 | 0 | 0 | 1 | 1 | 0 |
| ENSG00000199884 | 0 | 0 | 0 | 0 | 0 | 0 |
| ENSG00000157827 | 348 | 159 | 175 | 108 | 251 | 191 |
| ENSG00000213197 | 0 | 1 | 0 | 0 | 0 | 0 |
| ENSG00000270776 | 0 | 0 | 0 | 0 | 0 | 0 |
| ENSG00000196504 | 2575 | 1495 | 1181 | 615 | 2151 | 1265 |
| ENSG00000177917 | 691 | 327 | 385 | 308 | 304 | 335 |
| ENSG00000234932 | 0 | 0 | 0 | 0 | 0 | 0 |
| ENSG00000226213 | 0 | 0 | 0 | 0 | 0 | 0 |
| ENSG00000214025 | 0 | 0 | 0 | 0 | 0 | 0 |
| ENSG00000224612 | 0 | 0 | 0 | 0 | 0 | 0 |
| ENSG00000226338 | 0 | 0 | 0 | 0 | 0 | 0 |
| ENSG00000227708 | 0 | 0 | 0 | 0 | 0 | 0 |
| ENSG00000214024 | 0 | 0 | 0 | 0 | 0 | 0 |
| ENSG00000242575 | 0 | 0 | 1 | 0 | 3 | 1 |
| ENSG00000227400 | 0 | 0 | 0 | 0 | 0 | 0 |
| ENSG00000200377 | 0 | 0 | 0 | 0 | 0 | 0 |
| ENSG00000177519 | 66 | 18 | 29 | 11 | 11 | 19 |
| ENSG00000236917 | 0 | 0 | 0 | 0 | 0 | 0 |
| ENSG00000144278 | 0 | 0 | 0 | 0 | 1 | 0 |
| ENSG00000230251 | 0 | 0 | 0 | 0 | 0 | 0 |
| ENSG00000224675 | 0 | 0 | 0 | 0 | 0 | 0 |
| ENSG00000238004 | 0 | 0 | 0 | 0 | 0 | 0 |
| ENSG00000223290 | 0 | 0 | 0 | 0 | 0 | 0 |
| ENSG00000235949 | 98 | 66 | 100 | 45 | 56 | 75 |
| ENSG00000162989 | 2565 | 1101 | 1668 | 648 | 978 | 1026 |
| ENSG00000228837 | 0 | 0 | 0 | 0 | 0 | 0 |
| ENSG00000230991 | 0 | 0 | 2 | 0 | 0 | 0 |
| ENSG00000223197 | 0 | 0 | 0 | 0 | 0 | 0 |
| ENSG00000232982 | 0 | 0 | 0 | 0 | 0 | 0 |
| ENSG00000229701 | 0 | 0 | 0 | 0 | 0 | 0 |
| ENSG00000227682 | 0 | 1 | 0 | 0 | 0 | 0 |
| ENSG00000233709 | 0 | 0 | 0 | 0 | 0 | 0 |
| ENSG00000233039 | 0 | 0 | 0 | 0 | 0 | 0 |
| ENSG00000233248 | 0 | 0 | 0 | 0 | 0 | 0 |
| ENSG00000229935 | 0 | 0 | 0 | 0 | 0 | 0 |

| | | | | | | |
|---|---|---|---|---|---|---|
| ENSG00000206718 | 0 | 0 | 0 | 0 | 0 | 0 |
| ENSG00000226383 | 4 | 1 | 0 | 0 | 0 | 0 |
| ENSG00000235548 | 0 | 0 | 0 | 0 | 0 | 0 |
| ENSG00000282440 | 0 | 0 | 0 | 0 | 0 | 0 |
| ENSG00000153234 | 562 | 241 | 370 | 177 | 235 | 301 |
| ENSG00000115159 | 3122 | 1519 | 1921 | 821 | 1829 | 1490 |
| ENSG00000283436 | 0 | 0 | 0 | 0 | 0 | 0 |
| ENSG00000238069 | 0 | 0 | 0 | 0 | 0 | 0 |
| ENSG00000213194 | 0 | 0 | 1 | 0 | 0 | 0 |
| ENSG00000136542 | 0 | 0 | 0 | 1 | 0 | 0 |
| ENSG00000222404 | 0 | 0 | 0 | 0 | 0 | 0 |
| ENSG00000136541 | 0 | 0 | 0 | 0 | 0 | 0 |
| ENSG00000230562 | 14 | 4 | 16 | 5 | 6 | 13 |
| ENSG00000115165 | 0 | 0 | 0 | 0 | 0 | 0 |
| ENSG00000123612 | 0 | 6 | 2 | 0 | 3 | 1 |
| ENSG00000234584 | 2 | 0 | 1 | 0 | 0 | 1 |
| ENSG00000115170 | 670 | 284 | 414 | 260 | 342 | 362 |
| ENSG00000007001 | 5 | 0 | 1 | 1 | 1 | 0 |
| ENSG00000270557 | 13 | 5 | 4 | 5 | 7 | 4 |
| ENSG00000251980 | 0 | 0 | 0 | 0 | 0 | 0 |
| ENSG00000271320 | 2 | 0 | 3 | 1 | 1 | 0 |
| ENSG00000228937 | 0 | 0 | 0 | 0 | 0 | 0 |
| ENSG00000212410 | 0 | 0 | 0 | 0 | 0 | 0 |
| ENSG00000231158 | 0 | 0 | 0 | 0 | 0 | 0 |
| ENSG00000237327 | 0 | 0 | 0 | 0 | 0 | 0 |
| ENSG00000227480 | 0 | 0 | 0 | 0 | 0 | 0 |
| ENSG00000153237 | 32 | 19 | 18 | 14 | 28 | 12 |
| ENSG00000234890 | 0 | 0 | 0 | 0 | 0 | 0 |
| ENSG00000243723 | 0 | 0 | 0 | 0 | 0 | 0 |
| ENSG00000144283 | 4987 | 2524 | 3008 | 1932 | 2695 | 2455 |
| ENSG00000227331 | 0 | 0 | 0 | 0 | 1 | 0 |
| ENSG00000204380 | 21 | 14 | 18 | 6 | 14 | 9 |
| ENSG00000228586 | 0 | 0 | 0 | 0 | 0 | 0 |
| ENSG00000231675 | 0 | 0 | 0 | 0 | 0 | 0 |
| ENSG00000163331 | 0 | 0 | 1 | 1 | 1 | 0 |
| ENSG00000243792 | 0 | 0 | 0 | 0 | 0 | 0 |
| ENSG00000251491 | 0 | 0 | 0 | 0 | 0 | 0 |
| ENSG00000188668 | 0 | 0 | 0 | 0 | 0 | 0 |
| ENSG00000222300 | 0 | 0 | 0 | 0 | 0 | 0 |
| ENSG00000115183 | 1087 | 502 | 594 | 285 | 515 | 478 |
| ENSG00000213189 | 20 | 7 | 4 | 11 | 10 | 8 |
| ENSG00000202029 | 0 | 0 | 0 | 0 | 0 | 0 |
| ENSG00000226878 | 0 | 0 | 0 | 0 | 0 | 0 |
| ENSG00000275141 | 0 | 0 | 0 | 0 | 0 | 0 |
| ENSG00000196151 | 586 | 284 | 454 | 276 | 355 | 374 |
| ENSG00000123636 | 344 | 226 | 195 | 45 | 414 | 187 |
| ENSG00000223642 | 0 | 1 | 0 | 0 | 0 | 0 |
| ENSG00000270893 | 0 | 0 | 1 | 0 | 0 | 0 |
| ENSG00000225369 | 0 | 0 | 0 | 0 | 0 | 0 |
| ENSG00000207117 | 0 | 0 | 0 | 0 | 0 | 0 |

| | | | | | | |
|---|---|---|---|---|---|---|
| ENSG00000224152 | 179 | 81 | 96 | 76 | 102 | 95 |
| ENSG00000226266 | 28 | 10 | 15 | 12 | 10 | 12 |
| ENSG00000230783 | 2 | 0 | 0 | 0 | 0 | 1 |
| ENSG00000136536 | 2017 | 1031 | 998 | 514 | 1347 | 956 |
| ENSG00000241399 | 21 | 19 | 15 | 15 | 13 | 16 |
| ENSG00000248672 | 0 | 0 | 0 | 0 | 0 | 0 |
| ENSG00000054219 | 2 | 0 | 1 | 1 | 0 | 0 |
| ENSG00000227055 | 0 | 0 | 0 | 0 | 0 | 0 |
| ENSG00000153246 | 256 | 139 | 134 | 66 | 135 | 75 |
| ENSG00000115221 | 1268 | 722 | 716 | 339 | 649 | 449 |
| ENSG00000285155 | 1 | 0 | 3 | 0 | 4 | 5 |
| ENSG00000285373 | 2 | 1 | 2 | 0 | 1 | 3 |
| ENSG00000153250 | 1248 | 599 | 734 | 424 | 740 | 593 |
| ENSG00000283734 | 0 | 0 | 0 | 0 | 0 | 0 |
| ENSG00000279844 | 0 | 0 | 0 | 0 | 0 | 0 |
| ENSG00000244372 | 0 | 0 | 0 | 0 | 0 | 0 |
| ENSG00000232337 | 0 | 0 | 0 | 0 | 0 | 0 |
| ENSG00000224467 | 0 | 0 | 0 | 0 | 0 | 0 |
| ENSG00000136560 | 1316 | 748 | 783 | 444 | 851 | 615 |
| ENSG00000235724 | 8 | 2 | 6 | 1 | 8 | 3 |
| ENSG00000227403 | 107 | 39 | 69 | 49 | 56 | 76 |
| ENSG00000225813 | 0 | 0 | 0 | 0 | 0 | 2 |
| ENSG00000115233 | 3725 | 1987 | 2193 | 1777 | 2457 | 2020 |
| ENSG00000232320 | 1 | 0 | 0 | 0 | 0 | 0 |
| ENSG00000271924 | 4 | 1 | 3 | 0 | 1 | 2 |
| ENSG00000136535 | 12 | 0 | 8 | 2 | 4 | 0 |
| ENSG00000224076 | 0 | 0 | 0 | 0 | 0 | 0 |
| ENSG00000251621 | 0 | 0 | 0 | 0 | 0 | 0 |
| ENSG00000144290 | 50 | 14 | 31 | 10 | 21 | 21 |
| ENSG00000280105 | 0 | 0 | 0 | 0 | 0 | 0 |
| ENSG00000226121 | 4 | 1 | 2 | 0 | 3 | 2 |
| ENSG00000238570 | 0 | 0 | 0 | 0 | 0 | 0 |
| ENSG00000226946 | 0 | 0 | 0 | 0 | 0 | 0 |
| ENSG00000235716 | 0 | 0 | 0 | 0 | 0 | 0 |
| ENSG00000234732 | 0 | 0 | 0 | 0 | 0 | 0 |
| ENSG00000197635 | 14 | 6 | 6 | 3 | 12 | 12 |
| ENSG00000230918 | 0 | 0 | 0 | 0 | 0 | 0 |
| ENSG00000231445 | 0 | 0 | 0 | 0 | 0 | 0 |
| ENSG00000224674 | 0 | 0 | 0 | 0 | 0 | 0 |
| ENSG00000233397 | 0 | 0 | 0 | 0 | 0 | 0 |
| ENSG00000115263 | 0 | 0 | 0 | 0 | 0 | 0 |
| ENSG00000236841 | 0 | 0 | 0 | 0 | 0 | 0 |
| ENSG00000078098 | 0 | 0 | 0 | 0 | 0 | 0 |
| ENSG00000115267 | 256 | 96 | 135 | 57 | 117 | 96 |
| ENSG00000115271 | 1121 | 547 | 703 | 518 | 687 | 605 |
| ENSG00000184611 | 6 | 1 | 3 | 0 | 1 | 3 |
| ENSG00000212312 | 0 | 0 | 0 | 0 | 0 | 0 |
| ENSG00000237750 | 0 | 0 | 0 | 0 | 0 | 0 |
| ENSG00000230282 | 0 | 0 | 0 | 0 | 0 | 0 |
| ENSG00000200902 | 0 | 0 | 0 | 0 | 0 | 0 |

| | | | | | | |
|---|---|---|---|---|---|---|
| ENSG00000182263 | 253 | 143 | 117 | 67 | 232 | 116 |
| ENSG00000237844 | 0 | 0 | 0 | 0 | 1 | 0 |
| ENSG00000226847 | 0 | 0 | 0 | 0 | 0 | 0 |
| ENSG00000229641 | 0 | 0 | 0 | 0 | 0 | 0 |
| ENSG00000115290 | 405 | 201 | 240 | 159 | 233 | 191 |
| ENSG00000277998 | 4 | 1 | 3 | 2 | 7 | 1 |
| ENSG00000082438 | 599 | 373 | 321 | 111 | 386 | 221 |
| ENSG00000224331 | 3 | 2 | 2 | 0 | 3 | 0 |
| ENSG00000206869 | 2 | 0 | 1 | 1 | 0 | 0 |
| ENSG00000236283 | 15 | 13 | 8 | 9 | 11 | 6 |
| ENSG00000199508 | 0 | 0 | 0 | 0 | 0 | 0 |
| ENSG00000223318 | 0 | 0 | 0 | 0 | 0 | 0 |
| ENSG00000169507 | 4 | 1 | 2 | 0 | 0 | 1 |
| ENSG00000153253 | 0 | 0 | 0 | 0 | 0 | 0 |
| ENSG00000136531 | 0 | 0 | 0 | 0 | 0 | 0 |
| ENSG00000233204 | 0 | 0 | 0 | 0 | 0 | 0 |
| ENSG00000178662 | 0 | 1 | 0 | 0 | 0 | 0 |
| ENSG00000115339 | 1725 | 840 | 864 | 695 | 1126 | 850 |
| ENSG00000235192 | 6 | 3 | 1 | 4 | 4 | 5 |
| ENSG00000229195 | 0 | 1 | 0 | 0 | 0 | 0 |
| ENSG00000232411 | 0 | 0 | 0 | 0 | 0 | 0 |
| ENSG00000123607 | 367 | 234 | 180 | 102 | 249 | 154 |
| ENSG00000224490 | 0 | 2 | 0 | 1 | 0 | 3 |
| ENSG00000236107 | 0 | 0 | 0 | 0 | 0 | 0 |
| ENSG00000144285 | 0 | 0 | 0 | 0 | 0 | 0 |
| ENSG00000238813 | 0 | 0 | 0 | 0 | 0 | 0 |
| ENSG00000222376 | 0 | 0 | 0 | 0 | 0 | 0 |
| ENSG00000169432 | 0 | 0 | 0 | 0 | 0 | 0 |
| ENSG00000136546 | 0 | 0 | 0 | 0 | 0 | 0 |
| ENSG00000283657 | 0 | 0 | 0 | 0 | 0 | 0 |
| ENSG00000163092 | 0 | 1 | 0 | 0 | 0 | 0 |
| ENSG00000225182 | 0 | 0 | 0 | 0 | 0 | 0 |
| ENSG00000254552 | 0 | 0 | 0 | 0 | 0 | 0 |
| ENSG00000228222 | 0 | 0 | 0 | 0 | 0 | 0 |
| ENSG00000238357 | 0 | 0 | 0 | 0 | 0 | 0 |
| ENSG00000214211 | 0 | 0 | 0 | 1 | 0 | 0 |
| ENSG00000235335 | 0 | 0 | 0 | 0 | 1 | 0 |
| ENSG00000172318 | 0 | 1 | 1 | 0 | 0 | 0 |
| ENSG00000198648 | 1728 | 818 | 1016 | 632 | 832 | 744 |
| ENSG00000238259 | 0 | 0 | 0 | 0 | 0 | 0 |
| ENSG00000241074 | 0 | 0 | 0 | 0 | 0 | 1 |
| ENSG00000172292 | 1124 | 558 | 580 | 280 | 509 | 476 |
| ENSG00000265694 | 0 | 0 | 0 | 0 | 0 | 0 |
| ENSG00000199348 | 0 | 0 | 0 | 0 | 0 | 0 |
| ENSG00000227617 | 0 | 0 | 0 | 0 | 1 | 0 |
| ENSG00000163072 | 5 | 8 | 10 | 6 | 8 | 7 |
| ENSG00000152253 | 224 | 142 | 162 | 124 | 194 | 152 |
| ENSG00000152254 | 0 | 0 | 0 | 0 | 0 | 0 |
| ENSG00000073734 | 0 | 0 | 0 | 0 | 0 | 0 |
| ENSG00000073737 | 0 | 0 | 0 | 0 | 0 | 0 |

| | | | | | | |
|---|---|---|---|---|---|---|
| ENSG00000235321 | 0 | 0 | 0 | 0 | 0 | 0 |
| ENSG00000235995 | 0 | 0 | 0 | 0 | 0 | 0 |
| ENSG00000081479 | 6 | 4 | 8 | 0 | 0 | 3 |
| ENSG00000163093 | 116 | 82 | 70 | 69 | 80 | 79 |
| ENSG00000251569 | 0 | 0 | 0 | 0 | 0 | 0 |
| ENSG00000239474 | 1 | 2 | 2 | 2 | 2 | 0 |
| ENSG00000138399 | 256 | 177 | 195 | 160 | 208 | 210 |
| ENSG00000229827 | 0 | 1 | 0 | 0 | 0 | 0 |
| ENSG00000138398 | 749 | 452 | 295 | 129 | 874 | 364 |
| ENSG00000154479 | 8 | 4 | 4 | 4 | 5 | 4 |
| ENSG00000226072 | 0 | 0 | 0 | 0 | 0 | 0 |
| ENSG00000144362 | 203 | 90 | 126 | 94 | 102 | 104 |
| ENSG00000213160 | 264 | 104 | 135 | 102 | 194 | 145 |
| ENSG00000206961 | 0 | 0 | 0 | 0 | 1 | 0 |
| ENSG00000231550 | 2 | 0 | 1 | 0 | 0 | 0 |
| ENSG00000214369 | 0 | 0 | 0 | 1 | 0 | 0 |
| ENSG00000138385 | 2918 | 1784 | 1819 | 1379 | 2316 | 1830 |
| ENSG00000138382 | 819 | 446 | 515 | 487 | 626 | 525 |
| ENSG00000144357 | 1632 | 827 | 788 | 336 | 1031 | 694 |
| ENSG00000227176 | 1 | 0 | 0 | 2 | 1 | 2 |
| ENSG00000252807 | 0 | 0 | 0 | 0 | 0 | 0 |
| ENSG00000071909 | 8 | 6 | 3 | 0 | 0 | 0 |
| ENSG00000231898 | 0 | 0 | 2 | 0 | 0 | 0 |
| ENSG00000214821 | 0 | 0 | 0 | 0 | 0 | 0 |
| ENSG00000213981 | 0 | 0 | 0 | 0 | 0 | 0 |
| ENSG00000275414 | 0 | 0 | 0 | 0 | 0 | 0 |
| ENSG00000222033 | 4 | 1 | 3 | 0 | 1 | 1 |
| ENSG00000204335 | 2 | 1 | 1 | 0 | 0 | 2 |
| ENSG00000234350 | 5 | 5 | 6 | 3 | 5 | 2 |
| ENSG00000128692 | 14 | 15 | 10 | 11 | 10 | 12 |
| ENSG00000239467 | 51 | 42 | 38 | 44 | 37 | 45 |
| ENSG00000222509 | 0 | 0 | 0 | 0 | 0 | 0 |
| ENSG00000204334 | 31 | 20 | 14 | 10 | 32 | 10 |
| ENSG00000128683 | 33 | 24 | 25 | 54 | 59 | 38 |
| ENSG00000235934 | 0 | 0 | 0 | 0 | 0 | 1 |
| ENSG00000115806 | 5134 | 2374 | 3032 | 2411 | 2622 | 2665 |
| ENSG00000198586 | 1380 | 718 | 733 | 463 | 923 | 770 |
| ENSG00000123600 | 351 | 181 | 191 | 167 | 208 | 191 |
| ENSG00000224553 | 0 | 0 | 0 | 0 | 1 | 0 |
| ENSG00000115827 | 493 | 286 | 291 | 164 | 354 | 292 |
| ENSG00000228507 | 0 | 0 | 0 | 0 | 0 | 0 |
| ENSG00000234061 | 0 | 0 | 1 | 0 | 2 | 0 |
| ENSG00000233762 | 16 | 10 | 13 | 8 | 12 | 13 |
| ENSG00000071967 | 4 | 6 | 1 | 3 | 1 | 4 |
| ENSG00000233934 | 0 | 0 | 0 | 0 | 0 | 0 |
| ENSG00000077380 | 2708 | 1433 | 1542 | 988 | 1806 | 1442 |
| ENSG00000115840 | 385 | 167 | 272 | 184 | 237 | 255 |
| ENSG00000252779 | 0 | 0 | 0 | 0 | 0 | 0 |
| ENSG00000128708 | 1213 | 805 | 888 | 527 | 1150 | 695 |
| ENSG00000172878 | 225 | 151 | 139 | 171 | 257 | 196 |

| | | | | | | |
|---|---|---|---|---|---|---|
| ENSG00000144355 | 258 | 190 | 191 | 241 | 300 | 288 |
| ENSG00000115844 | 76 | 55 | 50 | 105 | 101 | 81 |
| ENSG00000236651 | 2 | 1 | 5 | 4 | 2 | 6 |
| ENSG00000232555 | 0 | 0 | 0 | 0 | 0 | 0 |
| ENSG00000278924 | 0 | 0 | 0 | 0 | 0 | 0 |
| ENSG00000091409 | 261 | 143 | 99 | 79 | 124 | 96 |
| ENSG00000226963 | 0 | 0 | 0 | 0 | 0 | 0 |
| ENSG00000232788 | 0 | 0 | 0 | 0 | 0 | 0 |
| ENSG00000225205 | 4 | 1 | 3 | 0 | 3 | 2 |
| ENSG00000152256 | 266 | 129 | 174 | 89 | 157 | 137 |
| ENSG00000199866 | 0 | 0 | 0 | 0 | 0 | 1 |
| ENSG00000230104 | 0 | 0 | 0 | 0 | 0 | 0 |
| ENSG00000228016 | 0 | 0 | 0 | 1 | 0 | 0 |
| ENSG00000239041 | 0 | 0 | 0 | 0 | 0 | 0 |
| ENSG00000091428 | 14 | 21 | 17 | 11 | 9 | 10 |
| ENSG00000232292 | 0 | 0 | 0 | 0 | 0 | 0 |
| ENSG00000091436 | 1141 | 521 | 587 | 375 | 701 | 543 |
| ENSG00000238133 | 0 | 3 | 4 | 4 | 0 | 4 |
| ENSG00000231735 | 0 | 0 | 0 | 0 | 0 | 0 |
| ENSG00000236391 | 0 | 0 | 0 | 0 | 1 | 0 |
| ENSG00000144354 | 141 | 67 | 84 | 55 | 70 | 81 |
| ENSG00000213148 | 0 | 0 | 0 | 0 | 0 | 0 |
| ENSG00000236832 | 0 | 0 | 0 | 0 | 0 | 0 |
| ENSG00000226131 | 0 | 1 | 0 | 0 | 0 | 0 |
| ENSG00000237016 | 0 | 0 | 0 | 0 | 0 | 0 |
| ENSG00000237617 | 3 | 2 | 0 | 0 | 0 | 1 |
| ENSG00000270460 | 17 | 6 | 20 | 4 | 16 | 13 |
| ENSG00000234415 | 2 | 0 | 0 | 0 | 0 | 0 |
| ENSG00000224638 | 0 | 0 | 0 | 0 | 0 | 0 |
| ENSG00000172845 | 2523 | 1376 | 1514 | 837 | 1583 | 1249 |
| ENSG00000271151 | 46 | 11 | 19 | 10 | 15 | 13 |
| ENSG00000273258 | 0 | 0 | 0 | 0 | 0 | 0 |
| ENSG00000260868 | 0 | 1 | 0 | 1 | 0 | 1 |
| ENSG00000235414 | 0 | 0 | 0 | 0 | 0 | 0 |
| ENSG00000138430 | 4840 | 2325 | 2706 | 2400 | 2977 | 2688 |
| ENSG00000270799 | 0 | 0 | 0 | 0 | 0 | 0 |
| ENSG00000243770 | 0 | 0 | 0 | 0 | 0 | 0 |
| ENSG00000201088 | 0 | 0 | 0 | 0 | 0 | 0 |
| ENSG00000227180 | 0 | 0 | 0 | 0 | 0 | 0 |
| ENSG00000237804 | 0 | 0 | 0 | 0 | 0 | 0 |
| ENSG00000231453 | 2 | 2 | 7 | 0 | 0 | 4 |
| ENSG00000280414 | 2 | 0 | 2 | 0 | 2 | 0 |
| ENSG00000217236 | 10 | 1 | 3 | 6 | 3 | 3 |
| ENSG00000138433 | 677 | 406 | 394 | 236 | 504 | 303 |
| ENSG00000144306 | 658 | 343 | 373 | 225 | 385 | 316 |
| ENSG00000163328 | 207 | 94 | 115 | 58 | 135 | 85 |
| ENSG00000226853 | 18 | 2 | 6 | 7 | 9 | 17 |
| ENSG00000179997 | 0 | 0 | 0 | 0 | 0 | 0 |
| ENSG00000279884 | 0 | 0 | 2 | 0 | 0 | 0 |
| ENSG00000236449 | 9 | 6 | 5 | 4 | 1 | 2 |

| | | | | | | |
|---|---|---|---|---|---|---|
| ENSG00000206965 | 0 | 0 | 1 | 0 | 1 | 1 |
| ENSG00000115935 | 896 | 418 | 613 | 431 | 504 | 609 |
| ENSG00000237798 | 0 | 1 | 0 | 0 | 0 | 1 |
| ENSG00000235655 | 947 | 701 | 837 | 680 | 678 | 703 |
| ENSG00000138435 | 0 | 0 | 0 | 0 | 0 | 0 |
| ENSG00000128656 | 263 | 147 | 141 | 121 | 147 | 180 |
| ENSG00000212167 | 0 | 0 | 0 | 0 | 0 | 0 |
| ENSG00000199827 | 0 | 0 | 0 | 0 | 0 | 0 |
| ENSG00000227187 | 0 | 0 | 0 | 0 | 0 | 0 |
| ENSG00000115966 | 678 | 415 | 376 | 204 | 462 | 337 |
| ENSG00000277948 | 0 | 1 | 0 | 1 | 1 | 2 |
| ENSG00000215973 | 0 | 0 | 0 | 0 | 0 | 0 |
| ENSG00000229750 | 0 | 0 | 1 | 0 | 0 | 0 |
| ENSG00000154518 | 5355 | 2608 | 3055 | 3094 | 3002 | 2778 |
| ENSG00000200121 | 0 | 0 | 0 | 0 | 0 | 0 |
| ENSG00000233131 | 0 | 0 | 0 | 0 | 0 | 0 |
| ENSG00000229066 | 0 | 0 | 0 | 0 | 0 | 0 |
| ENSG00000223976 | 0 | 0 | 0 | 0 | 0 | 0 |
| ENSG00000235047 | 0 | 0 | 0 | 0 | 0 | 0 |
| ENSG00000229779 | 0 | 0 | 0 | 0 | 0 | 0 |
| ENSG00000144320 | 984 | 477 | 467 | 311 | 677 | 482 |
| ENSG00000174279 | 0 | 0 | 0 | 0 | 0 | 0 |
| ENSG00000128714 | 0 | 0 | 0 | 0 | 0 | 0 |
| ENSG00000170178 | 0 | 1 | 3 | 0 | 0 | 1 |
| ENSG00000128713 | 4 | 0 | 3 | 0 | 0 | 0 |
| ENSG00000128710 | 0 | 0 | 0 | 0 | 0 | 0 |
| ENSG00000237380 | 0 | 0 | 0 | 0 | 0 | 0 |
| ENSG00000128709 | 0 | 0 | 0 | 0 | 0 | 0 |
| ENSG00000175879 | 50 | 25 | 26 | 32 | 31 | 20 |
| ENSG00000128652 | 0 | 0 | 1 | 1 | 0 | 0 |
| ENSG00000207744 | 0 | 0 | 0 | 0 | 0 | 0 |
| ENSG00000278500 | 0 | 0 | 0 | 0 | 0 | 0 |
| ENSG00000170166 | 0 | 0 | 0 | 0 | 0 | 0 |
| ENSG00000224189 | 82 | 38 | 59 | 33 | 39 | 25 |
| ENSG00000272729 | 2 | 0 | 0 | 0 | 0 | 0 |
| ENSG00000226363 | 24 | 15 | 11 | 14 | 9 | 8 |
| ENSG00000128645 | 0 | 0 | 2 | 0 | 1 | 0 |
| ENSG00000283880 | 0 | 0 | 0 | 0 | 0 | 0 |
| ENSG00000279160 | 0 | 0 | 0 | 0 | 0 | 0 |
| ENSG00000218175 | 20 | 13 | 10 | 13 | 7 | 5 |
| ENSG00000230384 | 0 | 0 | 0 | 0 | 0 | 0 |
| ENSG00000128654 | 1404 | 642 | 762 | 730 | 832 | 794 |
| ENSG00000237260 | 2 | 0 | 0 | 1 | 1 | 0 |
| ENSG00000229434 | 0 | 0 | 0 | 0 | 0 | 0 |
| ENSG00000236231 | 0 | 0 | 0 | 0 | 0 | 0 |
| ENSG00000283203 | 0 | 0 | 0 | 0 | 0 | 0 |
| ENSG00000272551 | 6 | 0 | 0 | 0 | 0 | 0 |
| ENSG00000163364 | 93 | 45 | 69 | 56 | 64 | 67 |
| ENSG00000224577 | 14 | 12 | 22 | 8 | 13 | 16 |
| ENSG00000252027 | 0 | 0 | 0 | 0 | 0 | 0 |

| | | | | | | |
|---|---|---|---|---|---|---|
| ENSG00000230552 | 2 | 2 | 1 | 0 | 0 | 0 |
| ENSG00000237053 | 0 | 0 | 0 | 2 | 1 | 0 |
| ENSG00000223777 | 0 | 0 | 0 | 0 | 0 | 1 |
| ENSG00000279205 | 4 | 5 | 2 | 3 | 2 | 0 |
| ENSG00000227098 | 0 | 0 | 0 | 1 | 0 | 0 |
| ENSG00000206866 | 0 | 0 | 0 | 0 | 0 | 0 |
| ENSG00000236501 | 0 | 0 | 0 | 0 | 0 | 3 |
| ENSG00000251746 | 0 | 0 | 0 | 0 | 0 | 0 |
| ENSG00000229337 | 0 | 0 | 0 | 0 | 0 | 1 |
| ENSG00000226273 | 0 | 0 | 0 | 0 | 0 | 0 |
| ENSG00000201102 | 0 | 0 | 0 | 0 | 0 | 0 |
| ENSG00000213069 | 0 | 0 | 2 | 0 | 1 | 0 |
| ENSG00000170144 | 9976 | 5412 | 6226 | 3865 | 6632 | 5706 |
| ENSG00000284252 | 0 | 0 | 0 | 0 | 0 | 0 |
| ENSG00000116044 | 1976 | 1225 | 1040 | 932 | 2315 | 1536 |
| ENSG00000225808 | 4 | 7 | 2 | 1 | 0 | 4 |
| ENSG00000265396 | 0 | 0 | 0 | 0 | 0 | 0 |
| ENSG00000222043 | 15 | 10 | 21 | 15 | 11 | 8 |
| ENSG00000213963 | 0 | 0 | 0 | 0 | 1 | 0 |
| ENSG00000271825 | 0 | 0 | 0 | 0 | 0 | 0 |
| ENSG00000271996 | 0 | 0 | 0 | 0 | 0 | 1 |
| ENSG00000278418 | 0 | 0 | 0 | 0 | 0 | 0 |
| ENSG00000280374 | 0 | 0 | 0 | 0 | 0 | 0 |
| ENSG00000237178 | 0 | 0 | 0 | 0 | 0 | 0 |
| ENSG00000238295 | 0 | 0 | 0 | 0 | 0 | 0 |
| ENSG00000018510 | 4 | 0 | 0 | 0 | 0 | 0 |
| ENSG00000196659 | 284 | 151 | 206 | 153 | 126 | 175 |
| ENSG00000237655 | 0 | 0 | 0 | 0 | 0 | 0 |
| ENSG00000197557 | 251 | 117 | 138 | 112 | 173 | 133 |
| ENSG00000128655 | 0 | 1 | 1 | 1 | 1 | 1 |
| ENSG00000284741 | 0 | 0 | 0 | 0 | 0 | 0 |
| ENSG00000229941 | 0 | 0 | 0 | 0 | 0 | 0 |
| ENSG00000236664 | 0 | 0 | 0 | 0 | 0 | 0 |
| ENSG00000213962 | 0 | 0 | 0 | 0 | 0 | 0 |
| ENSG00000229744 | 0 | 0 | 0 | 0 | 0 | 0 |
| ENSG00000206788 | 0 | 0 | 0 | 0 | 0 | 0 |
| ENSG00000249909 | 0 | 0 | 0 | 0 | 0 | 0 |
| ENSG00000227241 | 0 | 0 | 0 | 0 | 0 | 0 |
| ENSG00000155636 | 507 | 282 | 313 | 219 | 276 | 224 |
| ENSG00000223096 | 0 | 0 | 0 | 0 | 0 | 0 |
| ENSG00000079156 | 10 | 25 | 12 | 7 | 15 | 16 |
| ENSG00000223960 | 11 | 3 | 7 | 2 | 8 | 5 |
| ENSG00000180228 | 953 | 431 | 566 | 481 | 631 | 597 |
| ENSG00000204311 | 20 | 20 | 24 | 13 | 27 | 27 |
| ENSG00000238082 | 2 | 4 | 2 | 1 | 0 | 3 |
| ENSG00000079150 | 12 | 8 | 9 | 11 | 11 | 8 |
| ENSG00000116095 | 781 | 364 | 416 | 335 | 432 | 414 |
| ENSG00000237298 | 43 | 24 | 32 | 14 | 28 | 15 |
| ENSG00000155657 | 100 | 73 | 90 | 46 | 101 | 58 |
| ENSG00000270956 | 5 | 3 | 0 | 1 | 4 | 2 |

| | | | | | | |
|---|---|---|---|---|---|---|
| ENSG00000270277 | 3 | 4 | 3 | 1 | 2 | 8 |
| ENSG00000279598 | 0 | 0 | 0 | 0 | 0 | 0 |
| ENSG00000271011 | 0 | 0 | 0 | 0 | 0 | 0 |
| ENSG00000270574 | 1 | 2 | 1 | 0 | 1 | 1 |
| ENSG00000271141 | 0 | 0 | 0 | 0 | 0 | 0 |
| ENSG00000271401 | 0 | 0 | 0 | 0 | 0 | 0 |
| ENSG00000267784 | 0 | 0 | 0 | 0 | 0 | 0 |
| ENSG00000163492 | 0 | 0 | 0 | 0 | 0 | 0 |
| ENSG00000238542 | 0 | 0 | 0 | 0 | 0 | 0 |
| ENSG00000231541 | 0 | 0 | 0 | 0 | 0 | 0 |
| ENSG00000187231 | 860 | 466 | 528 | 275 | 504 | 368 |
| ENSG00000237477 | 0 | 0 | 0 | 0 | 0 | 0 |
| ENSG00000237964 | 0 | 0 | 0 | 0 | 0 | 0 |
| ENSG00000144331 | 8 | 2 | 7 | 2 | 2 | 6 |
| ENSG00000213126 | 0 | 0 | 0 | 0 | 0 | 0 |
| ENSG00000221240 | 0 | 0 | 0 | 0 | 0 | 0 |
| ENSG00000280636 | 0 | 0 | 0 | 0 | 0 | 0 |
| ENSG00000202216 | 0 | 0 | 0 | 0 | 0 | 0 |
| ENSG00000163510 | 288 | 198 | 122 | 73 | 345 | 157 |
| ENSG00000270289 | 0 | 0 | 0 | 0 | 0 | 0 |
| ENSG00000225258 | 0 | 0 | 0 | 0 | 0 | 0 |
| ENSG00000281131 | 0 | 0 | 0 | 0 | 0 | 0 |
| ENSG00000279559 | 0 | 0 | 0 | 0 | 0 | 0 |
| ENSG00000226564 | 8 | 10 | 5 | 2 | 4 | 3 |
| ENSG00000236667 | 0 | 0 | 0 | 0 | 0 | 0 |
| ENSG00000170035 | 1215 | 565 | 742 | 700 | 784 | 751 |
| ENSG00000236153 | 0 | 0 | 0 | 0 | 1 | 0 |
| ENSG00000238171 | 0 | 0 | 0 | 0 | 0 | 0 |
| ENSG00000234663 | 0 | 0 | 0 | 0 | 0 | 0 |
| ENSG00000266705 | 0 | 0 | 0 | 0 | 0 | 0 |
| ENSG00000226681 | 0 | 0 | 0 | 0 | 0 | 0 |
| ENSG00000115232 | 2 | 2 | 0 | 0 | 0 | 0 |
| ENSG00000188452 | 22 | 9 | 15 | 5 | 10 | 6 |
| ENSG00000162992 | 0 | 0 | 0 | 0 | 0 | 0 |
| ENSG00000234595 | 0 | 0 | 0 | 0 | 0 | 0 |
| ENSG00000225570 | 0 | 0 | 0 | 0 | 0 | 0 |
| ENSG00000223873 | 0 | 0 | 0 | 0 | 0 | 0 |
| ENSG00000221023 | 0 | 0 | 0 | 0 | 0 | 0 |
| ENSG00000260742 | 28 | 4 | 7 | 7 | 11 | 8 |
| ENSG00000138434 | 3035 | 1500 | 1747 | 1005 | 1840 | 1421 |
| ENSG00000150722 | 38 | 34 | 17 | 18 | 27 | 24 |
| ENSG00000213958 | 8 | 6 | 6 | 1 | 5 | 0 |
| ENSG00000222418 | 0 | 0 | 0 | 0 | 0 | 0 |
| ENSG00000115252 | 25 | 12 | 23 | 5 | 9 | 1 |
| ENSG00000242121 | 0 | 0 | 0 | 0 | 0 | 0 |
| ENSG00000077232 | 1866 | 1010 | 966 | 595 | 1393 | 1017 |
| ENSG00000232430 | 0 | 0 | 0 | 0 | 0 | 0 |
| ENSG00000212404 | 0 | 0 | 0 | 0 | 0 | 0 |
| ENSG00000162998 | 0 | 4 | 4 | 2 | 3 | 3 |
| ENSG00000207178 | 0 | 0 | 0 | 0 | 0 | 0 |

| | | | | | | |
|---|---|---|---|---|---|---|
| ENSG00000061676 | 5201 | 2845 | 2751 | 1613 | 4192 | 2647 |
| ENSG00000202141 | 0 | 0 | 0 | 0 | 0 | 0 |
| ENSG00000221498 | 0 | 0 | 0 | 0 | 0 | 0 |
| ENSG00000231203 | 26 | 17 | 9 | 14 | 8 | 9 |
| ENSG00000162999 | 47 | 31 | 21 | 17 | 26 | 25 |
| ENSG00000224643 | 0 | 0 | 0 | 0 | 0 | 0 |
| ENSG00000163002 | 328 | 186 | 199 | 146 | 249 | 221 |
| ENSG00000272800 | 10 | 1 | 2 | 7 | 6 | 4 |
| ENSG00000213120 | 0 | 0 | 0 | 0 | 0 | 0 |
| ENSG00000177855 | 16 | 9 | 22 | 12 | 13 | 14 |
| ENSG00000234172 | 0 | 0 | 0 | 0 | 0 | 0 |
| ENSG00000266808 | 0 | 0 | 0 | 0 | 0 | 0 |
| ENSG00000283839 | 18 | 7 | 5 | 3 | 3 | 3 |
| ENSG00000170396 | 48 | 33 | 20 | 9 | 20 | 18 |
| ENSG00000237824 | 0 | 0 | 0 | 0 | 0 | 0 |
| ENSG00000225406 | 0 | 0 | 0 | 0 | 0 | 0 |
| ENSG00000212581 | 0 | 0 | 0 | 0 | 0 | 0 |
| ENSG00000226747 | 0 | 0 | 0 | 0 | 1 | 0 |
| ENSG00000188738 | 14 | 4 | 2 | 3 | 11 | 5 |
| ENSG00000231646 | 0 | 0 | 0 | 0 | 0 | 0 |
| ENSG00000224019 | 0 | 0 | 0 | 0 | 0 | 0 |
| ENSG00000237877 | 11 | 5 | 10 | 0 | 6 | 8 |
| ENSG00000226410 | 0 | 0 | 0 | 0 | 0 | 0 |
| ENSG00000232568 | 0 | 0 | 0 | 0 | 0 | 0 |
| ENSG00000259915 | 0 | 0 | 0 | 0 | 1 | 0 |
| ENSG00000227692 | 3 | 1 | 0 | 0 | 1 | 1 |
| ENSG00000065548 | 1268 | 910 | 635 | 428 | 1288 | 755 |
| ENSG00000224323 | 0 | 0 | 0 | 0 | 0 | 0 |
| ENSG00000138448 | 3680 | 1812 | 1960 | 871 | 2256 | 1510 |
| ENSG00000227227 | 0 | 1 | 0 | 2 | 0 | 0 |
| ENSG00000144369 | 1066 | 490 | 485 | 189 | 539 | 385 |
| ENSG00000163012 | 0 | 0 | 0 | 0 | 0 | 0 |
| ENSG00000226553 | 0 | 0 | 0 | 0 | 0 | 0 |
| ENSG00000224063 | 8 | 2 | 5 | 4 | 11 | 2 |
| ENSG00000222845 | 0 | 0 | 0 | 0 | 0 | 0 |
| ENSG00000199626 | 0 | 0 | 0 | 0 | 0 | 0 |
| ENSG00000064989 | 2 | 1 | 1 | 2 | 1 | 0 |
| ENSG00000231739 | 0 | 0 | 0 | 0 | 0 | 0 |
| ENSG00000003436 | 3197 | 2018 | 2188 | 1302 | 2975 | 2216 |
| ENSG00000231689 | 0 | 0 | 0 | 1 | 0 | 0 |
| ENSG00000235616 | 0 | 0 | 0 | 0 | 0 | 0 |
| ENSG00000213115 | 0 | 0 | 0 | 0 | 0 | 0 |
| ENSG00000200650 | 0 | 0 | 0 | 0 | 0 | 0 |
| ENSG00000144366 | 311 | 171 | 158 | 76 | 172 | 127 |
| ENSG00000207951 | 0 | 1 | 0 | 0 | 0 | 0 |
| ENSG00000223523 | 338 | 267 | 341 | 84 | 138 | 167 |
| ENSG00000174325 | 0 | 0 | 0 | 0 | 1 | 1 |
| ENSG00000168542 | 273 | 110 | 146 | 71 | 93 | 70 |
| ENSG00000221502 | 0 | 0 | 0 | 0 | 0 | 0 |
| ENSG00000283238 | 0 | 0 | 0 | 0 | 0 | 0 |

| | | | | | | |
|---|---|---|---|---|---|---|
| ENSG00000284180 | 0 | 0 | 0 | 0 | 0 | 0 |
| ENSG00000204262 | 267 | 98 | 242 | 88 | 130 | 152 |
| ENSG00000228073 | 0 | 0 | 0 | 0 | 0 | 0 |
| ENSG00000264725 | 0 | 0 | 0 | 0 | 0 | 0 |
| ENSG00000213601 | 0 | 0 | 1 | 1 | 1 | 1 |
| ENSG00000115368 | 1294 | 719 | 765 | 611 | 1039 | 832 |
| ENSG00000233996 | 0 | 0 | 0 | 0 | 0 | 0 |
| ENSG00000138449 | 2 | 0 | 0 | 0 | 0 | 0 |
| ENSG00000276828 | 0 | 0 | 0 | 0 | 0 | 0 |
| ENSG00000138381 | 1052 | 581 | 651 | 504 | 814 | 651 |
| ENSG00000151687 | 61 | 25 | 38 | 30 | 43 | 20 |
| ENSG00000128694 | 203 | 138 | 142 | 117 | 140 | 171 |
| ENSG00000253559 | 12 | 2 | 5 | 3 | 7 | 7 |
| ENSG00000273240 | 4 | 2 | 1 | 0 | 1 | 0 |
| ENSG00000128699 | 1813 | 902 | 1243 | 792 | 997 | 980 |
| ENSG00000064933 | 378 | 211 | 194 | 111 | 273 | 191 |
| ENSG00000187699 | 6 | 6 | 0 | 2 | 9 | 1 |
| ENSG00000223824 | 0 | 0 | 0 | 0 | 0 | 0 |
| ENSG00000204253 | 52 | 32 | 21 | 25 | 40 | 41 |
| ENSG00000213110 | 0 | 0 | 0 | 0 | 0 | 0 |
| ENSG00000138379 | 4 | 0 | 0 | 0 | 0 | 0 |
| ENSG00000198130 | 526 | 321 | 328 | 270 | 362 | 297 |
| ENSG00000151689 | 75 | 36 | 43 | 41 | 59 | 57 |
| ENSG00000151690 | 868 | 453 | 522 | 339 | 532 | 477 |
| ENSG00000272979 | 1 | 0 | 3 | 1 | 2 | 0 |
| ENSG00000189362 | 98 | 64 | 80 | 64 | 60 | 76 |
| ENSG00000233654 | 6 | 3 | 6 | 4 | 5 | 4 |
| ENSG00000251935 | 0 | 0 | 0 | 0 | 0 | 0 |
| ENSG00000284052 | 0 | 3 | 1 | 1 | 3 | 1 |
| ENSG00000138386 | 137 | 89 | 133 | 57 | 121 | 104 |
| ENSG00000228509 | 0 | 0 | 0 | 0 | 1 | 0 |
| ENSG00000235852 | 0 | 0 | 0 | 0 | 0 | 0 |
| ENSG00000115419 | 1122 | 554 | 663 | 443 | 737 | 596 |
| ENSG00000115415 | 4409 | 2241 | 2542 | 1736 | 2511 | 2218 |
| ENSG00000229023 | 0 | 2 | 0 | 0 | 0 | 1 |
| ENSG00000230686 | 0 | 0 | 0 | 0 | 0 | 0 |
| ENSG00000231858 | 3 | 0 | 0 | 2 | 0 | 1 |
| ENSG00000138378 | 48 | 32 | 20 | 17 | 20 | 23 |
| ENSG00000207402 | 0 | 0 | 0 | 0 | 0 | 0 |
| ENSG00000280083 | 2 | 1 | 1 | 0 | 1 | 0 |
| ENSG00000230611 | 0 | 0 | 1 | 0 | 0 | 0 |
| ENSG00000227542 | 27 | 17 | 21 | 10 | 12 | 19 |
| ENSG00000128641 | 10069 | 6077 | 6353 | 3332 | 6183 | 5110 |
| ENSG00000252130 | 0 | 0 | 0 | 0 | 0 | 0 |
| ENSG00000173559 | 483 | 329 | 364 | 372 | 608 | 457 |
| ENSG00000274845 | 0 | 0 | 0 | 0 | 0 | 0 |
| ENSG00000225884 | 0 | 0 | 0 | 0 | 0 | 0 |
| ENSG00000168497 | 2 | 0 | 0 | 0 | 4 | 1 |
| ENSG00000233766 | 0 | 0 | 0 | 0 | 0 | 0 |
| ENSG00000213946 | 0 | 0 | 0 | 0 | 0 | 0 |

| | | | | | | |
|---|---|---|---|---|---|---|
| ENSG00000144339 | 0 | 0 | 0 | 0 | 0 | 0 |
| ENSG00000229395 | 0 | 0 | 0 | 0 | 0 | 0 |
| ENSG00000232227 | 5 | 0 | 2 | 0 | 0 | 1 |
| ENSG00000227418 | 2 | 1 | 6 | 2 | 2 | 2 |
| ENSG00000226242 | 0 | 0 | 0 | 0 | 1 | 0 |
| ENSG00000278406 | 0 | 0 | 0 | 0 | 0 | 0 |
| ENSG00000232658 | 0 | 0 | 0 | 0 | 0 | 0 |
| ENSG00000235218 | 0 | 0 | 0 | 0 | 0 | 0 |
| ENSG00000224219 | 0 | 0 | 0 | 0 | 0 | 0 |
| ENSG00000260142 | 0 | 0 | 0 | 0 | 0 | 0 |
| ENSG00000224670 | 0 | 0 | 0 | 0 | 0 | 0 |
| ENSG00000215043 | 0 | 0 | 0 | 0 | 0 | 0 |
| ENSG00000236695 | 0 | 0 | 0 | 0 | 0 | 0 |
| ENSG00000225539 | 0 | 0 | 0 | 0 | 0 | 0 |
| ENSG00000225394 | 0 | 0 | 0 | 0 | 0 | 0 |
| ENSG00000276123 | 0 | 0 | 0 | 0 | 0 | 0 |
| ENSG00000252517 | 0 | 0 | 0 | 0 | 0 | 0 |
| ENSG00000230173 | 0 | 0 | 0 | 0 | 0 | 0 |
| ENSG00000235056 | 0 | 0 | 0 | 0 | 0 | 0 |
| ENSG00000224099 | 0 | 0 | 0 | 0 | 0 | 0 |
| ENSG00000271893 | 0 | 0 | 0 | 0 | 0 | 0 |
| ENSG00000202206 | 0 | 0 | 0 | 0 | 0 | 0 |
| ENSG00000223466 | 0 | 0 | 0 | 0 | 0 | 0 |
| ENSG00000234919 | 0 | 0 | 0 | 0 | 0 | 0 |
| ENSG00000196950 | 544 | 259 | 289 | 124 | 277 | 234 |
| ENSG00000213107 | 0 | 0 | 0 | 0 | 0 | 0 |
| ENSG00000201813 | 0 | 0 | 0 | 0 | 0 | 0 |
| ENSG00000118997 | 40 | 13 | 13 | 10 | 18 | 19 |
| ENSG00000203435 | 2 | 1 | 5 | 1 | 0 | 2 |
| ENSG00000227878 | 0 | 0 | 0 | 0 | 0 | 0 |
| ENSG00000081320 | 332 | 136 | 165 | 104 | 184 | 169 |
| ENSG00000272211 | 2 | 1 | 3 | 0 | 0 | 0 |
| ENSG00000138411 | 434 | 226 | 284 | 253 | 425 | 390 |
| ENSG00000229056 | 0 | 0 | 2 | 0 | 2 | 0 |
| ENSG00000276200 | 0 | 0 | 0 | 0 | 0 | 0 |
| ENSG00000282836 | 0 | 0 | 0 | 1 | 4 | 3 |
| ENSG00000144395 | 35 | 26 | 38 | 8 | 58 | 47 |
| ENSG00000252923 | 0 | 0 | 0 | 0 | 0 | 0 |
| ENSG00000119041 | 1743 | 950 | 972 | 725 | 1326 | 971 |
| ENSG00000187944 | 0 | 0 | 1 | 0 | 0 | 0 |
| ENSG00000280176 | 0 | 0 | 0 | 0 | 0 | 0 |
| ENSG00000197121 | 95 | 55 | 49 | 18 | 98 | 45 |
| ENSG00000065413 | 230 | 117 | 117 | 52 | 135 | 106 |
| ENSG00000225636 | 2 | 0 | 0 | 0 | 0 | 0 |
| ENSG00000224442 | 0 | 0 | 0 | 0 | 0 | 0 |
| ENSG00000231621 | 0 | 0 | 0 | 0 | 1 | 0 |
| ENSG00000236977 | 0 | 0 | 0 | 0 | 1 | 0 |
| ENSG00000230482 | 0 | 0 | 0 | 0 | 0 | 0 |
| ENSG00000225979 | 0 | 1 | 0 | 0 | 0 | 0 |
| ENSG00000213104 | 0 | 1 | 0 | 3 | 0 | 1 |

| | | | | | | |
|---|---|---|---|---|---|---|
| ENSG00000115524 | 8424 | 4714 | 4834 | 2385 | 6368 | 4461 |
| ENSG00000202434 | 0 | 0 | 0 | 0 | 0 | 0 |
| ENSG00000206836 | 2 | 0 | 0 | 0 | 0 | 0 |
| ENSG00000115520 | 1722 | 781 | 1023 | 576 | 991 | 910 |
| ENSG00000144381 | 13324 | 7194 | 8039 | 8130 | 11171 | 9210 |
| ENSG00000115541 | 889 | 596 | 715 | 898 | 1029 | 859 |
| ENSG00000270757 | 0 | 0 | 0 | 0 | 2 | 0 |
| ENSG00000115540 | 793 | 414 | 430 | 347 | 508 | 401 |
| ENSG00000231699 | 0 | 0 | 0 | 0 | 0 | 1 |
| ENSG00000162944 | 37 | 16 | 7 | 8 | 17 | 13 |
| ENSG00000222017 | 42 | 33 | 27 | 21 | 25 | 30 |
| ENSG00000247626 | 526 | 257 | 322 | 321 | 347 | 379 |
| ENSG00000152430 | 7 | 3 | 6 | 2 | 3 | 3 |
| ENSG00000115896 | 16 | 7 | 3 | 2 | 12 | 3 |
| ENSG00000237035 | 0 | 0 | 0 | 0 | 0 | 0 |
| ENSG00000236653 | 0 | 0 | 0 | 0 | 0 | 0 |
| ENSG00000225421 | 0 | 0 | 0 | 0 | 0 | 0 |
| ENSG00000231557 | 0 | 0 | 0 | 0 | 0 | 0 |
| ENSG00000230407 | 0 | 0 | 0 | 0 | 0 | 0 |
| ENSG00000238698 | 0 | 0 | 0 | 0 | 0 | 0 |
| ENSG00000119042 | 500 | 214 | 325 | 187 | 255 | 274 |
| ENSG00000257045 | 1 | 0 | 0 | 0 | 0 | 0 |
| ENSG00000225953 | 15 | 5 | 5 | 3 | 4 | 4 |
| ENSG00000238217 | 0 | 0 | 0 | 0 | 0 | 0 |
| ENSG00000213938 | 2 | 0 | 0 | 0 | 2 | 1 |
| ENSG00000226124 | 163 | 84 | 85 | 99 | 101 | 107 |
| ENSG00000232732 | 7 | 1 | 1 | 1 | 0 | 0 |
| ENSG00000227004 | 0 | 0 | 0 | 0 | 0 | 0 |
| ENSG00000240302 | 0 | 0 | 0 | 0 | 0 | 0 |
| ENSG00000178074 | 516 | 223 | 215 | 139 | 275 | 213 |
| ENSG00000162971 | 563 | 295 | 359 | 268 | 366 | 292 |
| ENSG00000162972 | 946 | 450 | 541 | 506 | 501 | 512 |
| ENSG00000196141 | 2469 | 1374 | 1372 | 1274 | 1986 | 1585 |
| ENSG00000201649 | 0 | 0 | 0 | 0 | 0 | 0 |
| ENSG00000155729 | 1101 | 495 | 628 | 387 | 565 | 601 |
| ENSG00000163535 | 90 | 67 | 38 | 19 | 127 | 64 |
| ENSG00000138356 | 387 | 197 | 410 | 95 | 121 | 185 |
| ENSG00000244301 | 0 | 0 | 1 | 0 | 0 | 0 |
| ENSG00000284084 | 0 | 0 | 0 | 0 | 0 | 0 |
| ENSG00000237166 | 0 | 2 | 1 | 2 | 4 | 1 |
| ENSG00000243478 | 5 | 1 | 0 | 1 | 2 | 1 |
| ENSG00000201737 | 0 | 0 | 0 | 0 | 0 | 0 |
| ENSG00000230408 | 0 | 2 | 2 | 0 | 0 | 2 |
| ENSG00000082153 | 7131 | 4117 | 4204 | 2754 | 4755 | 3905 |
| ENSG00000207116 | 0 | 0 | 0 | 0 | 0 | 0 |
| ENSG00000224346 | 0 | 0 | 0 | 0 | 0 | 0 |
| ENSG00000223092 | 0 | 0 | 0 | 0 | 0 | 0 |
| ENSG00000013441 | 1499 | 854 | 1011 | 580 | 1086 | 667 |
| ENSG00000252759 | 1 | 5 | 4 | 0 | 0 | 2 |
| ENSG00000240344 | 863 | 426 | 541 | 342 | 462 | 382 |

| | | | | | | |
|---|---|---|---|---|---|---|
| ENSG00000201499 | 5 | 2 | 4 | 1 | 0 | 3 |
| ENSG00000196290 | 1198 | 621 | 779 | 581 | 670 | 710 |
| ENSG00000252916 | 5 | 0 | 3 | 0 | 1 | 0 |
| ENSG00000115942 | 1656 | 809 | 1041 | 842 | 989 | 949 |
| ENSG00000183308 | 0 | 4 | 1 | 0 | 1 | 0 |
| ENSG00000155744 | 600 | 284 | 300 | 145 | 374 | 248 |
| ENSG00000225695 | 0 | 0 | 1 | 0 | 2 | 1 |
| ENSG00000226658 | 0 | 1 | 0 | 0 | 0 | 0 |
| ENSG00000119013 | 1540 | 768 | 857 | 822 | 900 | 787 |
| ENSG00000252148 | 0 | 0 | 1 | 0 | 0 | 1 |
| ENSG00000273975 | 0 | 1 | 0 | 2 | 0 | 0 |
| ENSG00000232719 | 0 | 0 | 0 | 0 | 0 | 0 |
| ENSG00000224672 | 0 | 0 | 0 | 0 | 0 | 0 |
| ENSG00000003402 | 1287 | 644 | 824 | 456 | 687 | 682 |
| ENSG00000232133 | 0 | 1 | 0 | 1 | 1 | 0 |
| ENSG00000226312 | 4 | 4 | 2 | 1 | 1 | 2 |
| ENSG00000238829 | 0 | 0 | 0 | 0 | 0 | 1 |
| ENSG00000235579 | 0 | 0 | 0 | 0 | 0 | 0 |
| ENSG00000234431 | 0 | 0 | 0 | 0 | 0 | 0 |
| ENSG00000003400 | 28 | 21 | 12 | 12 | 11 | 9 |
| ENSG00000227348 | 0 | 0 | 0 | 0 | 0 | 0 |
| ENSG00000225796 | 0 | 0 | 0 | 0 | 0 | 0 |
| ENSG00000235318 | 0 | 0 | 0 | 0 | 0 | 0 |
| ENSG00000064012 | 545 | 245 | 328 | 261 | 357 | 363 |
| ENSG00000155749 | 0 | 2 | 5 | 1 | 0 | 2 |
| ENSG00000115993 | 1854 | 946 | 1200 | 722 | 1054 | 996 |
| ENSG00000082146 | 610 | 256 | 393 | 302 | 347 | 333 |
| ENSG00000213090 | 5 | 1 | 2 | 0 | 3 | 2 |
| ENSG00000202008 | 0 | 0 | 0 | 0 | 0 | 0 |
| ENSG00000155754 | 3 | 0 | 0 | 1 | 1 | 7 |
| ENSG00000234842 | 0 | 0 | 0 | 0 | 0 | 0 |
| ENSG00000224299 | 0 | 0 | 0 | 0 | 0 | 0 |
| ENSG00000227736 | 0 | 0 | 0 | 0 | 0 | 0 |
| ENSG00000224590 | 0 | 0 | 0 | 0 | 0 | 0 |
| ENSG00000234731 | 0 | 0 | 0 | 0 | 0 | 0 |
| ENSG00000249177 | 0 | 0 | 0 | 0 | 0 | 0 |
| ENSG00000227841 | 0 | 0 | 0 | 0 | 0 | 0 |
| ENSG00000155755 | 459 | 197 | 236 | 212 | 296 | 261 |
| ENSG00000241790 | 0 | 0 | 0 | 0 | 0 | 0 |
| ENSG00000082126 | 8 | 0 | 5 | 4 | 0 | 2 |
| ENSG00000222972 | 0 | 0 | 0 | 0 | 0 | 0 |
| ENSG00000003393 | 1310 | 604 | 757 | 501 | 712 | 724 |
| ENSG00000230799 | 0 | 0 | 0 | 0 | 1 | 0 |
| ENSG00000138395 | 0 | 1 | 0 | 0 | 0 | 0 |
| ENSG00000202137 | 0 | 0 | 0 | 0 | 0 | 0 |
| ENSG00000231124 | 0 | 0 | 0 | 0 | 0 | 0 |
| ENSG00000212184 | 0 | 0 | 0 | 0 | 0 | 0 |
| ENSG00000273209 | 10 | 11 | 10 | 14 | 11 | 8 |
| ENSG00000155760 | 793 | 389 | 472 | 460 | 471 | 502 |
| ENSG00000182329 | 23 | 12 | 8 | 19 | 22 | 21 |

| | | | | | | |
|---|---|---|---|---|---|---|
| ENSG00000222035 | 21 | 15 | 25 | 21 | 19 | 39 |
| ENSG00000231903 | 1 | 0 | 0 | 2 | 1 | 1 |
| ENSG00000227890 | 0 | 0 | 0 | 0 | 0 | 0 |
| ENSG00000223825 | 1 | 0 | 0 | 0 | 2 | 0 |
| ENSG00000116030 | 5502 | 2585 | 3139 | 2693 | 3001 | 2876 |
| ENSG00000224463 | 0 | 0 | 0 | 0 | 0 | 0 |
| ENSG00000055044 | 1352 | 844 | 791 | 597 | 1317 | 988 |
| ENSG00000212534 | 0 | 0 | 2 | 0 | 0 | 0 |
| ENSG00000212309 | 0 | 0 | 0 | 0 | 1 | 0 |
| ENSG00000271852 | 0 | 0 | 0 | 0 | 0 | 0 |
| ENSG00000238317 | 0 | 0 | 0 | 1 | 0 | 1 |
| ENSG00000264275 | 0 | 0 | 2 | 0 | 0 | 1 |
| ENSG00000226261 | 0 | 1 | 6 | 0 | 0 | 0 |
| ENSG00000272966 | 4 | 1 | 4 | 0 | 0 | 0 |
| ENSG00000242696 | 0 | 0 | 0 | 1 | 1 | 0 |
| ENSG00000273456 | 101 | 61 | 78 | 59 | 58 | 64 |
| ENSG00000204217 | 1639 | 929 | 866 | 408 | 1011 | 786 |
| ENSG00000270915 | 2 | 0 | 0 | 0 | 0 | 0 |
| ENSG00000236047 | 0 | 0 | 2 | 2 | 0 | 1 |
| ENSG00000223619 | 0 | 0 | 0 | 0 | 0 | 0 |
| ENSG00000233888 | 0 | 0 | 0 | 0 | 0 | 0 |
| ENSG00000235433 | 0 | 0 | 1 | 0 | 0 | 0 |
| ENSG00000237542 | 0 | 0 | 0 | 0 | 0 | 0 |
| ENSG00000233134 | 0 | 0 | 0 | 0 | 0 | 0 |
| ENSG00000241221 | 0 | 0 | 0 | 0 | 0 | 0 |
| ENSG00000227454 | 0 | 0 | 2 | 0 | 0 | 0 |
| ENSG00000233311 | 0 | 0 | 0 | 0 | 0 | 0 |
| ENSG00000138439 | 463 | 248 | 262 | 208 | 281 | 258 |
| ENSG00000163596 | 177 | 110 | 126 | 79 | 119 | 106 |
| ENSG00000240761 | 0 | 0 | 0 | 0 | 0 | 0 |
| ENSG00000233579 | 0 | 1 | 0 | 0 | 0 | 0 |
| ENSG00000138442 | 1160 | 593 | 750 | 716 | 821 | 823 |
| ENSG00000138380 | 364 | 156 | 232 | 72 | 186 | 146 |
| ENSG00000232479 | 2 | 4 | 4 | 3 | 2 | 3 |
| ENSG00000270664 | 0 | 0 | 0 | 0 | 0 | 0 |
| ENSG00000144426 | 234 | 133 | 97 | 23 | 194 | 106 |
| ENSG00000232994 | 0 | 0 | 0 | 0 | 0 | 0 |
| ENSG00000202059 | 0 | 0 | 0 | 0 | 0 | 0 |
| ENSG00000228513 | 0 | 0 | 0 | 0 | 0 | 0 |
| ENSG00000225124 | 4 | 0 | 0 | 0 | 0 | 0 |
| ENSG00000204196 | 0 | 0 | 0 | 0 | 0 | 0 |
| ENSG00000119004 | 427 | 215 | 257 | 174 | 244 | 212 |
| ENSG00000264041 | 0 | 0 | 0 | 0 | 0 | 0 |
| ENSG00000238285 | 0 | 0 | 0 | 0 | 0 | 0 |
| ENSG00000138443 | 2246 | 1090 | 1413 | 911 | 1266 | 1179 |
| ENSG00000256458 | 0 | 0 | 0 | 0 | 0 | 0 |
| ENSG00000173166 | 287 | 110 | 202 | 70 | 160 | 166 |
| ENSG00000233459 | 0 | 0 | 0 | 0 | 0 | 0 |
| ENSG00000233031 | 0 | 0 | 0 | 0 | 0 | 0 |
| ENSG00000178562 | 0 | 2 | 0 | 0 | 2 | 1 |

| | | | | | | |
|---|---|---|---|---|---|---|
| ENSG00000224791 | 4 | 2 | 2 | 2 | 2 | 1 |
| ENSG00000213925 | 0 | 0 | 0 | 0 | 0 | 0 |
| ENSG00000206970 | 0 | 0 | 0 | 0 | 0 | 0 |
| ENSG00000163599 | 0 | 0 | 0 | 0 | 0 | 0 |
| ENSG00000163600 | 0 | 0 | 0 | 0 | 0 | 0 |
| ENSG00000235951 | 0 | 0 | 0 | 0 | 0 | 0 |
| ENSG00000234714 | 0 | 0 | 0 | 0 | 0 | 0 |
| ENSG00000237271 | 0 | 0 | 0 | 0 | 0 | 0 |
| ENSG00000236681 | 0 | 0 | 0 | 0 | 0 | 0 |
| ENSG00000225304 | 0 | 0 | 0 | 0 | 0 | 0 |
| ENSG00000236634 | 0 | 0 | 0 | 0 | 0 | 0 |
| ENSG00000237843 | 0 | 0 | 0 | 0 | 0 | 0 |
| ENSG00000116117 | 283 | 140 | 175 | 82 | 143 | 110 |
| ENSG00000223890 | 0 | 0 | 0 | 0 | 0 | 0 |
| ENSG00000118257 | 28 | 8 | 14 | 1 | 6 | 5 |
| ENSG00000225216 | 0 | 0 | 0 | 0 | 0 | 0 |
| ENSG00000201875 | 0 | 0 | 0 | 0 | 0 | 0 |
| ENSG00000213739 | 0 | 0 | 0 | 0 | 0 | 0 |
| ENSG00000225610 | 0 | 0 | 0 | 0 | 0 | 0 |
| ENSG00000114933 | 359 | 169 | 158 | 69 | 225 | 147 |
| ENSG00000252485 | 0 | 0 | 0 | 0 | 0 | 0 |
| ENSG00000207386 | 0 | 0 | 0 | 0 | 0 | 0 |
| ENSG00000237041 | 0 | 0 | 0 | 0 | 0 | 0 |
| ENSG00000227946 | 74 | 57 | 72 | 54 | 64 | 77 |
| ENSG00000023228 | 4517 | 2105 | 2640 | 1834 | 2475 | 2421 |
| ENSG00000231955 | 0 | 0 | 0 | 0 | 0 | 0 |
| ENSG00000237580 | 0 | 0 | 0 | 0 | 0 | 0 |
| ENSG00000114942 | 11692 | 6106 | 6585 | 7815 | 7207 | 7025 |
| ENSG00000277502 | 0 | 0 | 0 | 0 | 0 | 0 |
| ENSG00000281780 | 1 | 0 | 0 | 0 | 3 | 2 |
| ENSG00000207047 | 0 | 0 | 2 | 1 | 1 | 0 |
| ENSG00000207406 | 0 | 0 | 0 | 0 | 0 | 0 |
| ENSG00000183671 | 10 | 5 | 3 | 1 | 7 | 4 |
| ENSG00000279220 | 0 | 0 | 0 | 0 | 0 | 0 |
| ENSG00000222496 | 0 | 0 | 0 | 0 | 0 | 0 |
| ENSG00000159186 | 0 | 0 | 0 | 0 | 0 | 0 |
| ENSG00000224070 | 0 | 0 | 0 | 0 | 0 | 0 |
| ENSG00000252716 | 0 | 0 | 0 | 0 | 0 | 0 |
| ENSG00000204186 | 68 | 35 | 23 | 7 | 53 | 28 |
| ENSG00000180105 | 0 | 0 | 0 | 0 | 0 | 0 |
| ENSG00000234491 | 0 | 0 | 0 | 0 | 0 | 0 |
| ENSG00000114948 | 19 | 9 | 8 | 16 | 21 | 20 |
| ENSG00000228577 | 0 | 0 | 0 | 0 | 0 | 0 |
| ENSG00000231653 | 0 | 0 | 0 | 0 | 0 | 0 |
| ENSG00000235118 | 0 | 0 | 0 | 0 | 0 | 0 |
| ENSG00000232125 | 0 | 0 | 1 | 0 | 0 | 0 |
| ENSG00000213083 | 0 | 0 | 0 | 0 | 0 | 0 |
| ENSG00000207319 | 0 | 0 | 0 | 0 | 0 | 0 |
| ENSG00000138400 | 75 | 31 | 75 | 37 | 52 | 49 |
| ENSG00000118246 | 697 | 464 | 383 | 294 | 625 | 490 |

| | | | | | | |
|---|---|---|---|---|---|---|
| ENSG00000263468 | 0 | 0 | 0 | 0 | 0 | 0 |
| ENSG00000283469 | 0 | 0 | 0 | 0 | 0 | 0 |
| ENSG00000260171 | 0 | 0 | 0 | 0 | 0 | 0 |
| ENSG00000229321 | 0 | 1 | 0 | 0 | 0 | 0 |
| ENSG00000144410 | 0 | 0 | 1 | 0 | 0 | 0 |
| ENSG00000118263 | 650 | 258 | 354 | 254 | 426 | 370 |
| ENSG00000253008 | 0 | 0 | 0 | 0 | 0 | 0 |
| ENSG00000237892 | 3 | 2 | 3 | 0 | 0 | 0 |
| ENSG00000277590 | 1 | 0 | 2 | 0 | 0 | 0 |
| ENSG00000224342 | 0 | 0 | 1 | 0 | 0 | 0 |
| ENSG00000234902 | 8 | 2 | 3 | 0 | 0 | 4 |
| ENSG00000225916 | 0 | 2 | 0 | 0 | 0 | 0 |
| ENSG00000240440 | 2 | 0 | 1 | 0 | 1 | 1 |
| ENSG00000229647 | 1 | 0 | 0 | 0 | 1 | 0 |
| ENSG00000223725 | 3 | 1 | 3 | 3 | 1 | 3 |
| ENSG00000225064 | 0 | 0 | 0 | 0 | 0 | 1 |
| ENSG00000221628 | 0 | 0 | 0 | 0 | 0 | 0 |
| ENSG00000118260 | 930 | 508 | 507 | 278 | 615 | 442 |
| ENSG00000144401 | 509 | 232 | 337 | 325 | 373 | 326 |
| ENSG00000223433 | 0 | 0 | 0 | 0 | 0 | 0 |
| ENSG00000224137 | 0 | 0 | 0 | 0 | 0 | 0 |
| ENSG00000199251 | 0 | 0 | 0 | 0 | 0 | 0 |
| ENSG00000213082 | 0 | 0 | 0 | 0 | 0 | 0 |
| ENSG00000200764 | 0 | 0 | 0 | 0 | 0 | 0 |
| ENSG00000163249 | 491 | 236 | 308 | 244 | 370 | 346 |
| ENSG00000272851 | 0 | 0 | 0 | 0 | 0 | 0 |
| ENSG00000284079 | 0 | 0 | 0 | 0 | 0 | 0 |
| ENSG00000163251 | 764 | 338 | 374 | 368 | 489 | 440 |
| ENSG00000178385 | 618 | 228 | 316 | 142 | 233 | 199 |
| ENSG00000225111 | 1 | 1 | 0 | 0 | 1 | 0 |
| ENSG00000212246 | 0 | 0 | 0 | 0 | 0 | 0 |
| ENSG00000185078 | 0 | 0 | 0 | 0 | 0 | 0 |
| ENSG00000251941 | 0 | 0 | 0 | 0 | 0 | 0 |
| ENSG00000224839 | 0 | 0 | 0 | 0 | 0 | 0 |
| ENSG00000229150 | 0 | 0 | 0 | 0 | 0 | 0 |
| ENSG00000118231 | 0 | 0 | 0 | 0 | 0 | 0 |
| ENSG00000163254 | 0 | 0 | 0 | 0 | 0 | 0 |
| ENSG00000224529 | 0 | 0 | 0 | 0 | 0 | 0 |
| ENSG00000182187 | 0 | 0 | 0 | 0 | 0 | 0 |
| ENSG00000168582 | 0 | 0 | 0 | 0 | 0 | 0 |
| ENSG00000188674 | 0 | 0 | 0 | 0 | 0 | 0 |
| ENSG00000233869 | 0 | 0 | 0 | 0 | 0 | 0 |
| ENSG00000228111 | 0 | 0 | 0 | 0 | 0 | 0 |
| ENSG00000138413 | 9253 | 4523 | 5707 | 3543 | 4074 | 4405 |
| ENSG00000231908 | 49 | 23 | 31 | 32 | 35 | 28 |
| ENSG00000115020 | 776 | 335 | 326 | 123 | 452 | 319 |
| ENSG00000228380 | 0 | 0 | 0 | 0 | 0 | 0 |
| ENSG00000144407 | 0 | 1 | 0 | 0 | 0 | 1 |
| ENSG00000213081 | 0 | 0 | 0 | 0 | 0 | 0 |
| ENSG00000227430 | 0 | 0 | 0 | 0 | 0 | 0 |

| | | | | | | |
|---|---|---|---|---|---|---|
| ENSG00000231896 | 0 | 0 | 0 | 0 | 0 | 0 |
| ENSG00000232409 | 0 | 0 | 0 | 0 | 0 | 0 |
| ENSG00000202164 | 0 | 0 | 0 | 0 | 0 | 0 |
| ENSG00000235864 | 0 | 0 | 0 | 0 | 0 | 0 |
| ENSG00000232799 | 0 | 0 | 0 | 0 | 0 | 0 |
| ENSG00000240286 | 0 | 0 | 0 | 0 | 0 | 0 |
| ENSG00000229670 | 0 | 2 | 1 | 0 | 0 | 0 |
| ENSG00000078018 | 458 | 213 | 286 | 164 | 290 | 268 |
| ENSG00000222317 | 0 | 0 | 0 | 0 | 0 | 0 |
| ENSG00000144406 | 0 | 0 | 0 | 0 | 0 | 0 |
| ENSG00000237194 | 0 | 0 | 0 | 0 | 0 | 0 |
| ENSG00000197713 | 2385 | 1252 | 1341 | 1148 | 1653 | 1361 |
| ENSG00000144445 | 267 | 134 | 159 | 70 | 153 | 116 |
| ENSG00000272807 | 6 | 7 | 2 | 5 | 11 | 10 |
| ENSG00000229127 | 8 | 2 | 0 | 2 | 1 | 1 |
| ENSG00000231063 | 0 | 0 | 0 | 1 | 0 | 1 |
| ENSG00000279317 | 0 | 1 | 2 | 0 | 0 | 1 |
| ENSG00000115361 | 12 | 5 | 4 | 8 | 4 | 6 |
| ENSG00000199203 | 0 | 0 | 0 | 0 | 0 | 0 |
| ENSG00000168530 | 0 | 1 | 1 | 0 | 1 | 0 |
| ENSG00000234281 | 3 | 0 | 1 | 0 | 2 | 0 |
| ENSG00000115365 | 1905 | 979 | 1109 | 650 | 944 | 973 |
| ENSG00000021826 | 26 | 13 | 26 | 5 | 19 | 23 |
| ENSG00000280837 | 0 | 0 | 0 | 0 | 0 | 0 |
| ENSG00000271074 | 0 | 0 | 0 | 0 | 0 | 0 |
| ENSG00000231017 | 0 | 0 | 0 | 0 | 0 | 0 |
| ENSG00000178568 | 553 | 287 | 262 | 107 | 317 | 245 |
| ENSG00000199585 | 0 | 0 | 0 | 0 | 0 | 0 |
| ENSG00000232631 | 0 | 0 | 0 | 0 | 0 | 0 |
| ENSG00000236007 | 0 | 0 | 0 | 0 | 0 | 1 |
| ENSG00000221782 | 0 | 0 | 0 | 0 | 0 | 0 |
| ENSG00000273118 | 0 | 0 | 1 | 1 | 0 | 0 |
| ENSG00000234308 | 0 | 0 | 0 | 0 | 0 | 0 |
| ENSG00000231254 | 0 | 0 | 0 | 0 | 0 | 0 |
| ENSG00000223373 | 0 | 0 | 0 | 0 | 0 | 0 |
| ENSG00000266354 | 0 | 0 | 0 | 0 | 0 | 0 |
| ENSG00000283637 | 0 | 0 | 0 | 0 | 0 | 0 |
| ENSG00000030419 | 373 | 184 | 172 | 77 | 233 | 168 |
| ENSG00000270659 | 62 | 33 | 50 | 31 | 29 | 49 |
| ENSG00000236193 | 0 | 1 | 0 | 0 | 0 | 0 |
| ENSG00000272519 | 0 | 0 | 0 | 0 | 0 | 0 |
| ENSG00000196096 | 0 | 0 | 0 | 0 | 0 | 0 |
| ENSG00000144451 | 1 | 4 | 5 | 3 | 4 | 1 |
| ENSG00000265734 | 0 | 0 | 0 | 0 | 0 | 0 |
| ENSG00000197585 | 4 | 1 | 2 | 1 | 2 | 1 |
| ENSG00000213917 | 0 | 0 | 1 | 0 | 0 | 0 |
| ENSG00000174453 | 0 | 0 | 0 | 0 | 0 | 0 |
| ENSG00000224257 | 0 | 0 | 0 | 0 | 0 | 0 |
| ENSG00000177261 | 0 | 0 | 0 | 0 | 0 | 0 |
| ENSG00000138376 | 504 | 253 | 331 | 139 | 210 | 233 |

| | | | | | | |
|---|---|---|---|---|---|---|
| ENSG00000229267 | 14 | 4 | 10 | 1 | 5 | 5 |
| ENSG00000207274 | 0 | 0 | 2 | 0 | 0 | 0 |
| ENSG00000230076 | 229 | 84 | 112 | 171 | 119 | 131 |
| ENSG00000144452 | 301 | 152 | 273 | 38 | 132 | 112 |
| ENSG00000227769 | 0 | 2 | 1 | 2 | 1 | 2 |
| ENSG00000237571 | 2 | 0 | 1 | 0 | 0 | 4 |
| ENSG00000138363 | 9037 | 4347 | 5792 | 4153 | 4480 | 4967 |
| ENSG00000115414 | 946 | 450 | 544 | 229 | 319 | 300 |
| ENSG00000230695 | 0 | 0 | 0 | 0 | 0 | 0 |
| ENSG00000225166 | 0 | 0 | 0 | 0 | 0 | 0 |
| ENSG00000228618 | 0 | 0 | 0 | 0 | 0 | 0 |
| ENSG00000234938 | 0 | 0 | 0 | 0 | 0 | 0 |
| ENSG00000237525 | 0 | 0 | 0 | 0 | 0 | 1 |
| ENSG00000227981 | 0 | 0 | 0 | 0 | 0 | 0 |
| ENSG00000235770 | 0 | 8 | 8 | 6 | 11 | 5 |
| ENSG00000230838 | 0 | 0 | 0 | 0 | 0 | 0 |
| ENSG00000216721 | 0 | 0 | 0 | 0 | 0 | 0 |
| ENSG00000227824 | 0 | 0 | 0 | 0 | 0 | 0 |
| ENSG00000226276 | 6 | 9 | 4 | 5 | 3 | 0 |
| ENSG00000118242 | 2787 | 1555 | 1869 | 1487 | 1539 | 1699 |
| ENSG00000115425 | 413 | 201 | 257 | 162 | 236 | 213 |
| ENSG00000163449 | 76 | 33 | 39 | 21 | 25 | 32 |
| ENSG00000079246 | 10615 | 5846 | 6646 | 5139 | 7394 | 6414 |
| ENSG00000232970 | 0 | 0 | 0 | 0 | 0 | 0 |
| ENSG00000236478 | 0 | 1 | 0 | 1 | 2 | 0 |
| ENSG00000279348 | 40 | 17 | 29 | 9 | 20 | 24 |
| ENSG00000260804 | 128 | 68 | 82 | 43 | 93 | 86 |
| ENSG00000144583 | 0 | 1 | 0 | 0 | 0 | 0 |
| ENSG00000231092 | 0 | 0 | 0 | 0 | 0 | 0 |
| ENSG00000233581 | 0 | 0 | 1 | 0 | 0 | 0 |
| ENSG00000207303 | 0 | 0 | 0 | 0 | 0 | 0 |
| ENSG00000235042 | 0 | 0 | 0 | 0 | 0 | 2 |
| ENSG00000138375 | 816 | 359 | 453 | 445 | 426 | 414 |
| ENSG00000232485 | 12 | 13 | 20 | 16 | 14 | 10 |
| ENSG00000241520 | 10 | 3 | 2 | 2 | 4 | 6 |
| ENSG00000197756 | 26792 | 13557 | 16070 | 19157 | 16286 | 15607 |
| ENSG00000224391 | 0 | 0 | 0 | 0 | 0 | 0 |
| ENSG00000225794 | 0 | 0 | 0 | 0 | 0 | 0 |
| ENSG00000235444 | 0 | 0 | 0 | 0 | 0 | 0 |
| ENSG00000115457 | 2490 | 1434 | 1398 | 1533 | 1369 | 1038 |
| ENSG00000115461 | 1395 | 526 | 856 | 276 | 286 | 416 |
| ENSG00000236886 | 0 | 0 | 0 | 0 | 0 | 0 |
| ENSG00000237930 | 0 | 0 | 0 | 0 | 0 | 0 |
| ENSG00000222832 | 0 | 0 | 0 | 0 | 0 | 0 |
| ENSG00000229352 | 0 | 0 | 0 | 0 | 0 | 0 |
| ENSG00000234588 | 0 | 0 | 0 | 0 | 0 | 0 |
| ENSG00000231597 | 0 | 0 | 0 | 0 | 0 | 0 |
| ENSG00000118245 | 0 | 0 | 0 | 0 | 0 | 0 |
| ENSG00000237479 | 0 | 0 | 0 | 0 | 0 | 0 |
| ENSG00000227021 | 0 | 0 | 0 | 0 | 0 | 0 |

| | | | | | | |
|---|---|---|---|---|---|---|
| ENSG00000236295 | 0 | 0 | 0 | 0 | 0 | 0 |
| ENSG00000223874 | 0 | 0 | 0 | 0 | 0 | 0 |
| ENSG00000259855 | 0 | 0 | 0 | 0 | 0 | 0 |
| ENSG00000251982 | 0 | 0 | 0 | 0 | 0 | 0 |
| ENSG00000233143 | 0 | 0 | 0 | 0 | 0 | 0 |
| ENSG00000278004 | 0 | 0 | 0 | 0 | 1 | 0 |
| ENSG00000231672 | 56 | 24 | 27 | 14 | 20 | 20 |
| ENSG00000224414 | 0 | 0 | 0 | 0 | 0 | 0 |
| ENSG00000231035 | 0 | 0 | 0 | 0 | 0 | 0 |
| ENSG00000079308 | 102 | 48 | 81 | 19 | 41 | 31 |
| ENSG00000275458 | 0 | 0 | 0 | 0 | 0 | 0 |
| ENSG00000223923 | 1 | 0 | 0 | 0 | 0 | 0 |
| ENSG00000188282 | 0 | 0 | 0 | 0 | 0 | 0 |
| ENSG00000229754 | 0 | 0 | 0 | 0 | 0 | 0 |
| ENSG00000180871 | 0 | 0 | 0 | 0 | 0 | 1 |
| ENSG00000163464 | 0 | 0 | 0 | 2 | 1 | 3 |
| ENSG00000224159 | 0 | 0 | 0 | 0 | 0 | 0 |
| ENSG00000163466 | 11383 | 5952 | 6905 | 6148 | 7052 | 6652 |
| ENSG00000261338 | 24 | 15 | 17 | 24 | 24 | 12 |
| ENSG00000179921 | 2 | 3 | 3 | 1 | 0 | 3 |
| ENSG00000127837 | 7479 | 3818 | 4177 | 4689 | 4258 | 4178 |
| ENSG00000127838 | 4986 | 2309 | 3116 | 2422 | 2364 | 2327 |
| ENSG00000135926 | 184 | 96 | 114 | 109 | 136 | 120 |
| ENSG00000284173 | 0 | 0 | 0 | 0 | 0 | 0 |
| ENSG00000237281 | 0 | 1 | 0 | 0 | 0 | 0 |
| ENSG00000278524 | 0 | 0 | 0 | 0 | 0 | 0 |
| ENSG00000230580 | 0 | 0 | 0 | 0 | 3 | 1 |
| ENSG00000158428 | 14 | 5 | 5 | 1 | 2 | 0 |
| ENSG00000225062 | 1 | 1 | 0 | 1 | 3 | 1 |
| ENSG00000018280 | 165 | 58 | 68 | 53 | 26 | 61 |
| ENSG00000144579 | 5613 | 2498 | 3287 | 2556 | 2263 | 2468 |
| ENSG00000273361 | 0 | 1 | 1 | 0 | 0 | 0 |
| ENSG00000199121 | 0 | 0 | 0 | 1 | 0 | 0 |
| ENSG00000127831 | 62 | 32 | 30 | 22 | 27 | 16 |
| ENSG00000135913 | 336 | 215 | 203 | 61 | 194 | 188 |
| ENSG00000222714 | 0 | 0 | 0 | 0 | 0 | 0 |
| ENSG00000144580 | 2006 | 1004 | 1260 | 957 | 1263 | 1253 |
| ENSG00000207393 | 0 | 0 | 0 | 0 | 0 | 0 |
| ENSG00000115556 | 99 | 49 | 57 | 47 | 57 | 69 |
| ENSG00000273466 | 1 | 5 | 2 | 4 | 0 | 2 |
| ENSG00000115568 | 1037 | 430 | 605 | 496 | 494 | 571 |
| ENSG00000074582 | 907 | 393 | 548 | 587 | 479 | 563 |
| ENSG00000163481 | 1104 | 504 | 645 | 689 | 598 | 618 |
| ENSG00000163482 | 1450 | 605 | 831 | 625 | 766 | 813 |
| ENSG00000135912 | 750 | 299 | 413 | 337 | 401 | 396 |
| ENSG00000135929 | 2 | 4 | 6 | 2 | 5 | 5 |
| ENSG00000272555 | 0 | 0 | 0 | 0 | 0 | 0 |
| ENSG00000115592 | 0 | 0 | 1 | 0 | 0 | 0 |
| ENSG00000283639 | 0 | 0 | 0 | 0 | 0 | 0 |
| ENSG00000232625 | 0 | 0 | 0 | 0 | 0 | 0 |

| | | | | | | |
|---|---|---|---|---|---|---|
| ENSG00000115596 | 0 | 0 | 0 | 0 | 0 | 0 |
| ENSG00000135925 | 0 | 0 | 0 | 2 | 0 | 1 |
| ENSG00000228135 | 0 | 0 | 0 | 0 | 1 | 0 |
| ENSG00000200029 | 0 | 0 | 0 | 0 | 0 | 0 |
| ENSG00000235024 | 2 | 0 | 0 | 1 | 3 | 0 |
| ENSG00000224928 | 4 | 2 | 2 | 1 | 1 | 2 |
| ENSG00000171450 | 18 | 2 | 8 | 9 | 7 | 7 |
| ENSG00000236445 | 0 | 0 | 0 | 0 | 0 | 0 |
| ENSG00000163497 | 0 | 0 | 0 | 0 | 0 | 0 |
| ENSG00000163499 | 0 | 0 | 0 | 0 | 0 | 3 |
| ENSG00000198973 | 0 | 0 | 0 | 0 | 0 | 0 |
| ENSG00000224090 | 0 | 0 | 0 | 0 | 0 | 1 |
| ENSG00000181378 | 2 | 1 | 3 | 1 | 1 | 0 |
| ENSG00000163501 | 2 | 2 | 0 | 0 | 0 | 0 |
| ENSG00000264755 | 0 | 0 | 0 | 0 | 0 | 0 |
| ENSG00000272644 | 14 | 10 | 6 | 10 | 8 | 8 |
| ENSG00000187736 | 4 | 4 | 4 | 1 | 0 | 2 |
| ENSG00000280537 | 12 | 3 | 10 | 3 | 2 | 4 |
| ENSG00000240317 | 0 | 0 | 0 | 0 | 0 | 0 |
| ENSG00000213901 | 17 | 15 | 6 | 5 | 12 | 6 |
| ENSG00000115649 | 3328 | 1642 | 2052 | 1924 | 1764 | 1823 |
| ENSG00000144567 | 4289 | 2003 | 2615 | 2645 | 2436 | 2359 |
| ENSG00000158552 | 1052 | 521 | 628 | 583 | 501 | 531 |
| ENSG00000115657 | 118 | 81 | 118 | 118 | 110 | 119 |
| ENSG00000284820 | 2 | 5 | 6 | 1 | 3 | 4 |
| ENSG00000198925 | 1336 | 613 | 809 | 673 | 719 | 681 |
| ENSG00000163516 | 2186 | 912 | 1380 | 1046 | 1107 | 1227 |
| ENSG00000163521 | 182 | 111 | 140 | 74 | 106 | 93 |
| ENSG00000115661 | 1310 | 550 | 735 | 751 | 626 | 653 |
| ENSG00000127824 | 161 | 131 | 104 | 170 | 200 | 125 |
| ENSG00000243910 | 2 | 0 | 0 | 0 | 0 | 0 |
| ENSG00000135924 | 2487 | 1172 | 1459 | 1350 | 1267 | 1135 |
| ENSG00000054356 | 2 | 1 | 0 | 0 | 0 | 0 |
| ENSG00000207647 | 0 | 0 | 0 | 0 | 0 | 0 |
| ENSG00000230432 | 0 | 0 | 0 | 0 | 0 | 0 |
| ENSG00000182698 | 0 | 0 | 0 | 0 | 0 | 0 |
| ENSG00000232789 | 0 | 0 | 0 | 0 | 0 | 0 |
| ENSG00000123992 | 3996 | 1950 | 2538 | 2089 | 1995 | 1873 |
| ENSG00000229525 | 33 | 15 | 23 | 17 | 18 | 17 |
| ENSG00000175084 | 0 | 3 | 1 | 4 | 3 | 0 |
| ENSG00000234638 | 0 | 0 | 0 | 0 | 0 | 0 |
| ENSG00000280323 | 0 | 0 | 0 | 0 | 0 | 0 |
| ENSG00000072195 | 744 | 307 | 457 | 231 | 244 | 245 |
| ENSG00000227432 | 0 | 0 | 0 | 2 | 1 | 1 |
| ENSG00000268603 | 1 | 2 | 2 | 4 | 3 | 2 |
| ENSG00000144591 | 1776 | 873 | 1187 | 1067 | 1010 | 966 |
| ENSG00000072182 | 12 | 5 | 6 | 5 | 4 | 3 |
| ENSG00000123989 | 4807 | 2301 | 2840 | 3268 | 2631 | 2699 |
| ENSG00000188760 | 66 | 31 | 33 | 38 | 43 | 44 |
| ENSG00000268896 | 0 | 0 | 0 | 0 | 1 | 0 |

| | | | | | | |
|---|---|---|---|---|---|---|
| ENSG00000265252 | 0 | 0 | 0 | 0 | 0 | 0 |
| ENSG00000124006 | 3948 | 1707 | 2600 | 2122 | 1984 | 2133 |
| ENSG00000269068 | 0 | 0 | 1 | 1 | 0 | 1 |
| ENSG00000123999 | 86 | 46 | 55 | 50 | 53 | 57 |
| ENSG00000222678 | 0 | 0 | 0 | 0 | 0 | 0 |
| ENSG00000144589 | 1428 | 567 | 805 | 736 | 633 | 678 |
| ENSG00000228973 | 0 | 0 | 0 | 1 | 0 | 0 |
| ENSG00000114923 | 277 | 109 | 134 | 146 | 116 | 128 |
| ENSG00000267919 | 0 | 0 | 1 | 3 | 1 | 2 |
| ENSG00000231802 | 1 | 0 | 2 | 0 | 1 | 0 |
| ENSG00000227308 | 2 | 1 | 1 | 1 | 1 | 1 |
| ENSG00000223903 | 0 | 0 | 0 | 0 | 0 | 0 |
| ENSG00000228909 | 0 | 0 | 0 | 0 | 0 | 0 |
| ENSG00000266518 | 0 | 0 | 0 | 0 | 0 | 0 |
| ENSG00000225911 | 0 | 0 | 0 | 0 | 0 | 0 |
| ENSG00000239498 | 7 | 3 | 9 | 5 | 3 | 1 |
| ENSG00000235337 | 1 | 0 | 0 | 0 | 0 | 2 |
| ENSG00000236451 | 2 | 0 | 0 | 1 | 1 | 0 |
| ENSG00000224819 | 0 | 0 | 0 | 0 | 0 | 0 |
| ENSG00000232089 | 0 | 0 | 0 | 0 | 0 | 0 |
| ENSG00000116106 | 1591 | 773 | 935 | 508 | 881 | 790 |
| ENSG00000226726 | 0 | 0 | 0 | 0 | 0 | 0 |
| ENSG00000272944 | 44 | 22 | 34 | 26 | 48 | 34 |
| ENSG00000230406 | 0 | 0 | 0 | 0 | 0 | 1 |
| ENSG00000234446 | 0 | 0 | 0 | 0 | 0 | 0 |
| ENSG00000233868 | 0 | 0 | 0 | 2 | 0 | 1 |
| ENSG00000226666 | 2 | 0 | 2 | 1 | 1 | 2 |
| ENSG00000226331 | 0 | 0 | 0 | 0 | 0 | 0 |
| ENSG00000135903 | 0 | 0 | 2 | 1 | 0 | 1 |
| ENSG00000163081 | 0 | 0 | 0 | 0 | 0 | 0 |
| ENSG00000267034 | 15 | 7 | 13 | 3 | 7 | 8 |
| ENSG00000237732 | 28 | 21 | 29 | 21 | 25 | 28 |
| ENSG00000200822 | 0 | 0 | 0 | 0 | 0 | 0 |
| ENSG00000163082 | 54 | 25 | 35 | 11 | 20 | 38 |
| ENSG00000228670 | 0 | 0 | 0 | 0 | 0 | 0 |
| ENSG00000202016 | 0 | 0 | 0 | 0 | 0 | 0 |
| ENSG00000234285 | 0 | 0 | 0 | 0 | 0 | 0 |
| ENSG00000261428 | 10 | 4 | 4 | 7 | 6 | 7 |
| ENSG00000116120 | 1268 | 719 | 803 | 649 | 966 | 899 |
| ENSG00000124003 | 0 | 0 | 0 | 0 | 0 | 0 |
| ENSG00000233751 | 0 | 0 | 0 | 0 | 0 | 0 |
| ENSG00000203531 | 0 | 0 | 0 | 0 | 0 | 0 |
| ENSG00000235225 | 0 | 0 | 0 | 0 | 0 | 0 |
| ENSG00000123983 | 7811 | 4046 | 4252 | 2410 | 4085 | 3350 |
| ENSG00000224121 | 2 | 3 | 0 | 0 | 0 | 0 |
| ENSG00000234193 | 1 | 2 | 0 | 0 | 1 | 4 |
| ENSG00000278083 | 0 | 1 | 0 | 0 | 0 | 0 |
| ENSG00000152049 | 3 | 0 | 1 | 2 | 3 | 4 |
| ENSG00000243125 | 0 | 0 | 0 | 0 | 0 | 0 |
| ENSG00000233299 | 0 | 0 | 0 | 0 | 0 | 0 |

| | | | | | | |
|---|---|---|---|---|---|---|
| ENSG00000231189 | 0 | 0 | 0 | 0 | 0 | 0 |
| ENSG00000235610 | 0 | 0 | 0 | 0 | 0 | 0 |
| ENSG00000171951 | 0 | 0 | 0 | 0 | 1 | 0 |
| ENSG00000226539 | 0 | 1 | 2 | 2 | 0 | 0 |
| ENSG00000152056 | 188 | 114 | 127 | 68 | 127 | 120 |
| ENSG00000085449 | 1725 | 740 | 994 | 580 | 849 | 806 |
| ENSG00000135900 | 1096 | 498 | 609 | 531 | 649 | 574 |
| ENSG00000228802 | 0 | 0 | 0 | 1 | 0 | 3 |
| ENSG00000135919 | 16 | 14 | 9 | 8 | 13 | 9 |
| ENSG00000224826 | 0 | 0 | 0 | 0 | 0 | 0 |
| ENSG00000228681 | 0 | 0 | 0 | 0 | 0 | 0 |
| ENSG00000124019 | 0 | 0 | 0 | 0 | 1 | 0 |
| ENSG00000274629 | 0 | 0 | 0 | 0 | 0 | 1 |
| ENSG00000036257 | 2246 | 1093 | 1340 | 854 | 1380 | 1093 |
| ENSG00000228446 | 0 | 1 | 2 | 0 | 1 | 0 |
| ENSG00000283428 | 0 | 0 | 0 | 0 | 0 | 0 |
| ENSG00000135905 | 24 | 13 | 16 | 5 | 24 | 8 |
| ENSG00000263828 | 0 | 0 | 0 | 0 | 0 | 0 |
| ENSG00000144460 | 0 | 0 | 0 | 1 | 2 | 2 |
| ENSG00000228206 | 0 | 0 | 0 | 0 | 1 | 1 |
| ENSG00000273301 | 2 | 2 | 4 | 1 | 6 | 5 |
| ENSG00000279015 | 0 | 0 | 0 | 0 | 0 | 2 |
| ENSG00000230621 | 0 | 0 | 0 | 0 | 0 | 0 |
| ENSG00000235070 | 29 | 10 | 11 | 7 | 20 | 10 |
| ENSG00000263363 | 0 | 0 | 0 | 0 | 0 | 0 |
| ENSG00000169047 | 12000 | 5358 | 7825 | 5199 | 6408 | 7243 |
| ENSG00000272622 | 1390 | 777 | 1016 | 606 | 832 | 943 |
| ENSG00000261379 | 699 | 302 | 565 | 194 | 423 | 393 |
| ENSG00000144468 | 722 | 386 | 480 | 314 | 373 | 433 |
| ENSG00000212391 | 0 | 0 | 0 | 0 | 0 | 0 |
| ENSG00000081052 | 4 | 4 | 5 | 4 | 3 | 3 |
| ENSG00000169031 | 10 | 4 | 10 | 4 | 9 | 6 |
| ENSG00000236432 | 15 | 11 | 11 | 12 | 11 | 6 |
| ENSG00000168958 | 1763 | 923 | 1002 | 974 | 1040 | 1006 |
| ENSG00000168955 | 0 | 3 | 1 | 0 | 3 | 1 |
| ENSG00000284629 | 0 | 2 | 1 | 0 | 0 | 0 |
| ENSG00000284092 | 0 | 0 | 0 | 0 | 0 | 0 |
| ENSG00000173744 | 2229 | 1098 | 1199 | 878 | 1312 | 1182 |
| ENSG00000285843 | 0 | 0 | 0 | 0 | 0 | 0 |
| ENSG00000284643 | 0 | 0 | 0 | 0 | 0 | 0 |
| ENSG00000284622 | 0 | 1 | 0 | 0 | 0 | 0 |
| ENSG00000042304 | 0 | 0 | 0 | 0 | 0 | 0 |
| ENSG00000284704 | 0 | 0 | 0 | 0 | 0 | 0 |
| ENSG00000236116 | 0 | 0 | 0 | 0 | 0 | 0 |
| ENSG00000284631 | 0 | 0 | 0 | 0 | 0 | 0 |
| ENSG00000284667 | 0 | 0 | 0 | 0 | 0 | 0 |
| ENSG00000135917 | 2 | 1 | 5 | 1 | 1 | 1 |
| ENSG00000284725 | 0 | 0 | 0 | 0 | 0 | 0 |
| ENSG00000284718 | 0 | 0 | 0 | 0 | 0 | 0 |
| ENSG00000284635 | 0 | 0 | 0 | 0 | 0 | 0 |

| | | | | | | |
|---|---|---|---|---|---|---|
| ENSG00000252346 | 0 | 0 | 0 | 0 | 0 | 0 |
| ENSG00000226718 | 0 | 0 | 0 | 0 | 0 | 0 |
| ENSG00000229172 | 0 | 0 | 0 | 0 | 0 | 0 |
| ENSG00000115009 | 0 | 0 | 1 | 0 | 0 | 0 |
| ENSG00000183514 | 0 | 0 | 0 | 0 | 0 | 0 |
| ENSG00000123977 | 0 | 0 | 0 | 0 | 0 | 0 |
| ENSG00000272237 | 0 | 0 | 0 | 0 | 0 | 0 |
| ENSG00000153820 | 0 | 0 | 1 | 0 | 0 | 0 |
| ENSG00000200281 | 0 | 0 | 0 | 0 | 0 | 0 |
| ENSG00000203387 | 0 | 0 | 0 | 0 | 0 | 0 |
| ENSG00000232023 | 51 | 22 | 37 | 19 | 25 | 25 |
| ENSG00000228226 | 0 | 0 | 0 | 0 | 0 | 0 |
| ENSG00000237322 | 0 | 0 | 0 | 0 | 0 | 0 |
| ENSG00000153823 | 4 | 4 | 1 | 6 | 9 | 0 |
| ENSG00000226297 | 0 | 0 | 0 | 0 | 0 | 0 |
| ENSG00000253006 | 0 | 0 | 0 | 0 | 0 | 0 |
| ENSG00000187957 | 16 | 10 | 8 | 6 | 8 | 14 |
| ENSG00000238782 | 0 | 0 | 0 | 0 | 0 | 0 |
| ENSG00000235993 | 0 | 0 | 0 | 0 | 0 | 0 |
| ENSG00000153827 | 5897 | 2968 | 3213 | 1805 | 3472 | 2662 |
| ENSG00000222344 | 0 | 0 | 0 | 0 | 0 | 0 |
| ENSG00000153832 | 348 | 186 | 193 | 150 | 180 | 150 |
| ENSG00000252333 | 0 | 0 | 0 | 2 | 0 | 0 |
| ENSG00000231534 | 0 | 0 | 0 | 0 | 0 | 0 |
| ENSG00000206725 | 0 | 0 | 0 | 0 | 0 | 0 |
| ENSG00000163053 | 870 | 413 | 560 | 440 | 510 | 509 |
| ENSG00000199400 | 0 | 1 | 0 | 1 | 3 | 2 |
| ENSG00000225963 | 3 | 1 | 0 | 3 | 4 | 0 |
| ENSG00000283451 | 0 | 0 | 0 | 0 | 0 | 0 |
| ENSG00000135899 | 329 | 178 | 207 | 109 | 129 | 114 |
| ENSG00000079263 | 8 | 6 | 6 | 1 | 1 | 0 |
| ENSG00000185404 | 555 | 246 | 352 | 202 | 270 | 280 |
| ENSG00000279964 | 0 | 0 | 0 | 0 | 0 | 0 |
| ENSG00000067066 | 375 | 243 | 299 | 226 | 305 | 301 |
| ENSG00000243650 | 1 | 1 | 3 | 1 | 3 | 4 |
| ENSG00000250011 | 0 | 0 | 0 | 0 | 0 | 1 |
| ENSG00000235419 | 0 | 0 | 0 | 0 | 0 | 0 |
| ENSG00000183022 | 2 | 1 | 0 | 0 | 1 | 0 |
| ENSG00000283202 | 0 | 0 | 0 | 0 | 0 | 0 |
| ENSG00000199791 | 0 | 0 | 0 | 0 | 0 | 0 |
| ENSG00000226125 | 0 | 3 | 2 | 1 | 0 | 2 |
| ENSG00000135932 | 3333 | 1747 | 2031 | 1353 | 2092 | 1761 |
| ENSG00000237758 | 4 | 1 | 0 | 1 | 0 | 2 |
| ENSG00000201044 | 0 | 0 | 0 | 0 | 0 | 0 |
| ENSG00000270891 | 0 | 0 | 0 | 0 | 0 | 0 |
| ENSG00000135916 | 178 | 113 | 84 | 109 | 98 | 65 |
| ENSG00000232520 | 0 | 0 | 0 | 0 | 0 | 0 |
| ENSG00000135898 | 0 | 0 | 0 | 0 | 0 | 0 |
| ENSG00000230385 | 0 | 0 | 0 | 0 | 0 | 0 |
| ENSG00000283164 | 0 | 0 | 0 | 0 | 0 | 0 |

| | | | | | | |
|---|---|---|---|---|---|---|
| ENSG00000235013 | 0 | 0 | 0 | 0 | 0 | 0 |
| ENSG00000238062 | 0 | 0 | 0 | 0 | 1 | 0 |
| ENSG00000173699 | 0 | 0 | 0 | 0 | 0 | 0 |
| ENSG00000204128 | 64 | 30 | 22 | 18 | 15 | 15 |
| ENSG00000261829 | 0 | 2 | 0 | 0 | 1 | 0 |
| ENSG00000173692 | 3697 | 1971 | 2262 | 1689 | 2485 | 2101 |
| ENSG00000135914 | 3 | 2 | 0 | 0 | 2 | 0 |
| ENSG00000201574 | 0 | 0 | 0 | 0 | 0 | 0 |
| ENSG00000135931 | 1321 | 649 | 810 | 596 | 702 | 698 |
| ENSG00000237039 | 43 | 28 | 57 | 53 | 55 | 51 |
| ENSG00000263641 | 0 | 1 | 0 | 0 | 0 | 0 |
| ENSG00000224376 | 13 | 15 | 14 | 16 | 14 | 8 |
| ENSG00000156966 | 11 | 5 | 12 | 5 | 7 | 12 |
| ENSG00000172799 | 0 | 0 | 0 | 0 | 1 | 0 |
| ENSG00000230047 | 0 | 0 | 0 | 1 | 1 | 0 |
| ENSG00000233538 | 1 | 0 | 0 | 0 | 1 | 1 |
| ENSG00000115053 | 15179 | 8167 | 9490 | 7587 | 10134 | 9764 |
| ENSG00000206885 | 4 | 0 | 5 | 4 | 1 | 2 |
| ENSG00000207280 | 4 | 3 | 2 | 4 | 3 | 1 |
| ENSG00000202400 | 0 | 0 | 0 | 0 | 0 | 0 |
| ENSG00000283312 | 10 | 4 | 3 | 3 | 2 | 2 |
| ENSG00000283491 | 10 | 5 | 2 | 6 | 6 | 4 |
| ENSG00000223198 | 0 | 1 | 0 | 0 | 0 | 0 |
| ENSG00000181798 | 20 | 6 | 11 | 3 | 8 | 10 |
| ENSG00000171596 | 0 | 0 | 0 | 0 | 0 | 0 |
| ENSG00000231494 | 0 | 0 | 0 | 0 | 0 | 0 |
| ENSG00000228884 | 0 | 0 | 0 | 0 | 0 | 0 |
| ENSG00000177673 | 0 | 0 | 0 | 0 | 0 | 0 |
| ENSG00000239202 | 0 | 0 | 0 | 0 | 0 | 0 |
| ENSG00000187514 | 34704 | 19392 | 22527 | 19254 | 21156 | 22746 |
| ENSG00000277986 | 0 | 0 | 0 | 0 | 0 | 0 |
| ENSG00000284378 | 0 | 0 | 0 | 0 | 0 | 0 |
| ENSG00000156973 | 843 | 490 | 534 | 498 | 540 | 537 |
| ENSG00000281072 | 0 | 0 | 0 | 0 | 0 | 0 |
| ENSG00000144524 | 1381 | 599 | 775 | 657 | 673 | 735 |
| ENSG00000237641 | 5 | 4 | 6 | 4 | 5 | 3 |
| ENSG00000261096 | 2 | 2 | 6 | 0 | 0 | 2 |
| ENSG00000260622 | 0 | 0 | 0 | 0 | 0 | 0 |
| ENSG00000222246 | 0 | 0 | 0 | 0 | 0 | 0 |
| ENSG00000163273 | 14 | 8 | 6 | 5 | 1 | 3 |
| ENSG00000144535 | 473 | 226 | 319 | 279 | 242 | 314 |
| ENSG00000227033 | 0 | 0 | 0 | 0 | 0 | 0 |
| ENSG00000207626 | 0 | 0 | 0 | 0 | 0 | 0 |
| ENSG00000251485 | 16 | 9 | 6 | 13 | 10 | 13 |
| ENSG00000230122 | 0 | 0 | 0 | 0 | 0 | 2 |
| ENSG00000275157 | 0 | 0 | 0 | 0 | 0 | 0 |
| ENSG00000163283 | 0 | 3 | 0 | 0 | 0 | 4 |
| ENSG00000224516 | 0 | 0 | 0 | 0 | 0 | 0 |
| ENSG00000244280 | 0 | 1 | 0 | 0 | 0 | 0 |
| ENSG00000163286 | 0 | 1 | 0 | 0 | 0 | 1 |

| | | | | | | |
|---|---|---|---|---|---|---|
| ENSG00000204121 | 2 | 2 | 0 | 0 | 1 | 0 |
| ENSG00000237087 | 0 | 0 | 0 | 0 | 0 | 0 |
| ENSG00000223788 | 0 | 0 | 0 | 0 | 0 | 0 |
| ENSG00000163295 | 0 | 0 | 0 | 1 | 0 | 0 |
| ENSG00000171551 | 110 | 41 | 43 | 48 | 33 | 52 |
| ENSG00000237412 | 0 | 3 | 0 | 0 | 1 | 0 |
| ENSG00000135902 | 6 | 4 | 2 | 1 | 4 | 3 |
| ENSG00000196811 | 4 | 3 | 3 | 1 | 7 | 2 |
| ENSG00000221944 | 294 | 149 | 184 | 153 | 174 | 184 |
| ENSG00000135930 | 3376 | 1599 | 2039 | 1878 | 1874 | 1918 |
| ENSG00000284407 | 0 | 0 | 0 | 0 | 0 | 0 |
| ENSG00000237126 | 56 | 36 | 29 | 14 | 30 | 26 |
| ENSG00000115468 | 1606 | 779 | 1022 | 682 | 618 | 759 |
| ENSG00000244384 | 0 | 0 | 0 | 0 | 0 | 0 |
| ENSG00000204120 | 2908 | 1343 | 1851 | 955 | 1492 | 1355 |
| ENSG00000213055 | 0 | 1 | 0 | 0 | 0 | 0 |
| ENSG00000241409 | 0 | 0 | 2 | 0 | 0 | 0 |
| ENSG00000115474 | 9 | 5 | 4 | 0 | 0 | 0 |
| ENSG00000252452 | 0 | 0 | 0 | 0 | 0 | 0 |
| ENSG00000199781 | 0 | 0 | 0 | 0 | 0 | 0 |
| ENSG00000182600 | 54 | 26 | 43 | 29 | 31 | 35 |
| ENSG00000066248 | 107 | 53 | 61 | 41 | 57 | 65 |
| ENSG00000222001 | 0 | 0 | 0 | 0 | 0 | 0 |
| ENSG00000115488 | 0 | 0 | 0 | 0 | 0 | 0 |
| ENSG00000168918 | 4 | 3 | 0 | 0 | 2 | 0 |
| ENSG00000263941 | 0 | 0 | 0 | 0 | 0 | 0 |
| ENSG00000085978 | 1718 | 851 | 979 | 734 | 868 | 835 |
| ENSG00000252010 | 0 | 0 | 0 | 0 | 1 | 2 |
| ENSG00000251791 | 0 | 0 | 3 | 3 | 2 | 2 |
| ENSG00000130561 | 2 | 0 | 0 | 0 | 1 | 0 |
| ENSG00000259793 | 2 | 1 | 0 | 1 | 0 | 3 |
| ENSG00000077044 | 1511 | 664 | 897 | 629 | 729 | 744 |
| ENSG00000201013 | 0 | 0 | 0 | 0 | 0 | 0 |
| ENSG00000085982 | 1653 | 795 | 1038 | 646 | 877 | 919 |
| ENSG00000271639 | 2 | 0 | 0 | 0 | 1 | 1 |
| ENSG00000228949 | 1 | 0 | 3 | 0 | 2 | 3 |
| ENSG00000227846 | 0 | 0 | 0 | 0 | 0 | 0 |
| ENSG00000242366 | 0 | 0 | 0 | 1 | 0 | 1 |
| ENSG00000242515 | 0 | 4 | 0 | 0 | 0 | 0 |
| ENSG00000234143 | 0 | 0 | 0 | 0 | 0 | 0 |
| ENSG00000241119 | 6 | 2 | 0 | 2 | 0 | 0 |
| ENSG00000244122 | 25 | 11 | 17 | 17 | 21 | 31 |
| ENSG00000167165 | 100 | 87 | 108 | 125 | 160 | 166 |
| ENSG00000224814 | 2 | 0 | 1 | 1 | 2 | 0 |
| ENSG00000240224 | 0 | 2 | 0 | 0 | 0 | 0 |
| ENSG00000244474 | 0 | 0 | 1 | 1 | 1 | 1 |
| ENSG00000233445 | 0 | 0 | 0 | 0 | 0 | 0 |
| ENSG00000243135 | 0 | 0 | 0 | 0 | 0 | 1 |
| ENSG00000248114 | 0 | 0 | 0 | 0 | 0 | 0 |
| ENSG00000227802 | 1 | 1 | 1 | 2 | 1 | 1 |

| | | | | | | |
|---|---|---|---|---|---|---|
| ENSG00000228445 | 0 | 0 | 1 | 6 | 4 | 1 |
| ENSG00000248705 | 0 | 1 | 0 | 0 | 0 | 0 |
| ENSG00000178836 | 0 | 0 | 1 | 1 | 0 | 4 |
| ENSG00000241635 | 48 | 20 | 23 | 26 | 38 | 34 |
| ENSG00000185038 | 13 | 9 | 13 | 6 | 13 | 10 |
| ENSG00000123485 | 578 | 309 | 390 | 366 | 414 | 463 |
| ENSG00000224287 | 8 | 10 | 4 | 4 | 11 | 12 |
| ENSG00000279809 | 0 | 0 | 0 | 0 | 0 | 0 |
| ENSG00000144481 | 2 | 0 | 1 | 0 | 0 | 0 |
| ENSG00000237581 | 0 | 0 | 0 | 0 | 0 | 0 |
| ENSG00000072080 | 0 | 0 | 0 | 0 | 0 | 0 |
| ENSG00000279218 | 0 | 0 | 0 | 0 | 0 | 0 |
| ENSG00000226442 | 0 | 0 | 0 | 0 | 0 | 0 |
| ENSG00000261298 | 0 | 0 | 0 | 0 | 0 | 0 |
| ENSG00000237784 | 0 | 0 | 0 | 0 | 0 | 0 |
| ENSG00000228162 | 0 | 0 | 0 | 0 | 0 | 0 |
| ENSG00000231682 | 0 | 0 | 0 | 0 | 0 | 0 |
| ENSG00000232690 | 0 | 0 | 0 | 0 | 0 | 0 |
| ENSG00000188042 | 330 | 156 | 231 | 309 | 354 | 412 |
| ENSG00000280744 | 0 | 0 | 0 | 0 | 0 | 0 |
| ENSG00000278593 | 0 | 0 | 0 | 0 | 0 | 0 |
| ENSG00000235726 | 9 | 8 | 11 | 5 | 9 | 6 |
| ENSG00000130147 | 1962 | 980 | 1280 | 861 | 1030 | 1011 |
| ENSG00000226542 | 0 | 0 | 0 | 0 | 0 | 0 |
| ENSG00000231032 | 0 | 0 | 0 | 0 | 0 | 0 |
| ENSG00000235293 | 0 | 0 | 0 | 0 | 0 | 0 |
| ENSG00000157985 | 2037 | 880 | 1218 | 848 | 951 | 1043 |
| ENSG00000235529 | 8 | 4 | 6 | 10 | 4 | 11 |
| ENSG00000222007 | 0 | 0 | 0 | 0 | 0 | 1 |
| ENSG00000228499 | 0 | 0 | 0 | 0 | 0 | 0 |
| ENSG00000238812 | 0 | 0 | 0 | 0 | 0 | 0 |
| ENSG00000264676 | 0 | 1 | 0 | 0 | 0 | 0 |
| ENSG00000207049 | 0 | 0 | 0 | 0 | 0 | 0 |
| ENSG00000168505 | 12 | 6 | 7 | 6 | 6 | 8 |
| ENSG00000233611 | 3 | 3 | 2 | 0 | 1 | 1 |
| ENSG00000182177 | 4 | 3 | 0 | 1 | 1 | 0 |
| ENSG00000200745 | 0 | 3 | 0 | 0 | 0 | 0 |
| ENSG00000132321 | 9 | 1 | 2 | 2 | 0 | 3 |
| ENSG00000270540 | 2 | 1 | 4 | 0 | 1 | 0 |
| ENSG00000232893 | 4 | 2 | 3 | 3 | 0 | 2 |
| ENSG00000231546 | 0 | 0 | 0 | 0 | 0 | 0 |
| ENSG00000144476 | 5247 | 2514 | 3365 | 1756 | 1733 | 2059 |
| ENSG00000232328 | 11 | 10 | 5 | 7 | 6 | 7 |
| ENSG00000202341 | 0 | 0 | 0 | 0 | 0 | 0 |
| ENSG00000124835 | 0 | 0 | 0 | 0 | 0 | 0 |
| ENSG00000227252 | 18 | 8 | 19 | 9 | 9 | 14 |
| ENSG00000198612 | 1695 | 866 | 1021 | 854 | 950 | 957 |
| ENSG00000224844 | 0 | 0 | 0 | 1 | 0 | 1 |
| ENSG00000224132 | 0 | 0 | 1 | 0 | 0 | 0 |
| ENSG00000163359 | 108 | 48 | 60 | 40 | 48 | 58 |

| | | | | | | |
|---|---|---|---|---|---|---|
| ENSG00000222022 | 4 | 1 | 1 | 3 | 3 | 3 |
| ENSG00000222032 | 0 | 1 | 5 | 0 | 1 | 1 |
| ENSG00000115648 | 24102 | 10472 | 16445 | 9981 | 9820 | 11831 |
| ENSG00000277842 | 0 | 0 | 0 | 0 | 0 | 0 |
| ENSG00000222449 | 0 | 0 | 1 | 0 | 0 | 2 |
| ENSG00000071677 | 1 | 1 | 10 | 7 | 3 | 2 |
| ENSG00000124839 | 2341 | 1055 | 1421 | 1073 | 1185 | 1183 |
| ENSG00000234949 | 211 | 78 | 99 | 81 | 109 | 108 |
| ENSG00000227107 | 2 | 3 | 2 | 0 | 0 | 0 |
| ENSG00000124831 | 2953 | 1477 | 1490 | 637 | 1636 | 1220 |
| ENSG00000177483 | 4 | 1 | 0 | 2 | 2 | 4 |
| ENSG00000132329 | 164 | 68 | 104 | 118 | 88 | 73 |
| ENSG00000263723 | 0 | 0 | 0 | 0 | 0 | 0 |
| ENSG00000184182 | 311 | 148 | 197 | 202 | 202 | 199 |
| ENSG00000258984 | 1 | 3 | 1 | 1 | 1 | 6 |
| ENSG00000251971 | 1 | 0 | 0 | 0 | 0 | 0 |
| ENSG00000132330 | 68 | 26 | 39 | 42 | 32 | 40 |
| ENSG00000144488 | 0 | 1 | 2 | 0 | 0 | 0 |
| ENSG00000168427 | 2 | 0 | 0 | 1 | 2 | 0 |
| ENSG00000279484 | 0 | 0 | 0 | 0 | 0 | 0 |
| ENSG00000178752 | 26 | 14 | 17 | 13 | 21 | 17 |
| ENSG00000132323 | 760 | 333 | 479 | 430 | 428 | 418 |
| ENSG00000186235 | 0 | 0 | 0 | 0 | 0 | 0 |
| ENSG00000225057 | 1 | 2 | 2 | 1 | 3 | 1 |
| ENSG00000144485 | 570 | 245 | 354 | 468 | 390 | 360 |
| ENSG00000132326 | 571 | 289 | 356 | 198 | 256 | 297 |
| ENSG00000283635 | 1 | 1 | 0 | 0 | 2 | 0 |
| ENSG00000232306 | 0 | 0 | 0 | 0 | 0 | 0 |
| ENSG00000204104 | 675 | 324 | 418 | 248 | 387 | 389 |
| ENSG00000202099 | 0 | 0 | 1 | 0 | 0 | 0 |
| ENSG00000065802 | 1241 | 617 | 774 | 563 | 712 | 717 |
| ENSG00000229915 | 0 | 0 | 3 | 1 | 1 | 1 |
| ENSG00000274346 | 0 | 0 | 0 | 0 | 0 | 0 |
| ENSG00000225493 | 0 | 4 | 0 | 0 | 0 | 1 |
| ENSG00000234279 | 0 | 0 | 0 | 0 | 0 | 0 |
| ENSG00000224252 | 0 | 0 | 0 | 0 | 0 | 0 |
| ENSG00000226992 | 0 | 0 | 0 | 0 | 0 | 0 |
| ENSG00000233608 | 0 | 0 | 0 | 0 | 0 | 0 |
| ENSG00000227744 | 0 | 0 | 0 | 0 | 0 | 0 |
| ENSG00000068024 | 355 | 179 | 260 | 207 | 217 | 225 |
| ENSG00000266109 | 0 | 0 | 0 | 0 | 0 | 0 |
| ENSG00000264810 | 0 | 0 | 0 | 0 | 0 | 0 |
| ENSG00000265215 | 0 | 0 | 0 | 0 | 0 | 0 |
| ENSG00000264292 | 0 | 0 | 0 | 0 | 0 | 0 |
| ENSG00000222020 | 16 | 4 | 18 | 12 | 7 | 20 |
| ENSG00000196758 | 0 | 0 | 0 | 0 | 0 | 0 |
| ENSG00000227459 | 0 | 0 | 0 | 0 | 0 | 0 |
| ENSG00000259889 | 0 | 0 | 0 | 0 | 0 | 0 |
| ENSG00000220256 | 0 | 0 | 0 | 0 | 0 | 0 |
| ENSG00000130414 | 4574 | 2095 | 2651 | 2368 | 2364 | 2402 |

| | | | | | | |
|---|---|---|---|---|---|---|
| ENSG00000264279 | 0 | 0 | 1 | 0 | 1 | 0 |
| ENSG00000182083 | 2 | 0 | 0 | 0 | 1 | 0 |
| ENSG00000178586 | 2 | 0 | 0 | 0 | 0 | 0 |
| ENSG00000231278 | 0 | 0 | 0 | 0 | 0 | 0 |
| ENSG00000213048 | 2 | 0 | 0 | 1 | 0 | 0 |
| ENSG00000172428 | 1257 | 601 | 691 | 813 | 683 | 673 |
| ENSG00000178602 | 0 | 0 | 0 | 0 | 0 | 1 |
| ENSG00000227479 | 0 | 0 | 0 | 0 | 0 | 0 |
| ENSG00000276366 | 0 | 0 | 0 | 0 | 0 | 0 |
| ENSG00000063660 | 3873 | 1419 | 2555 | 1788 | 1391 | 1647 |
| ENSG00000218416 | 1928 | 856 | 1267 | 559 | 490 | 652 |
| ENSG00000207611 | 31 | 10 | 23 | 10 | 8 | 16 |
| ENSG00000144504 | 570 | 252 | 371 | 270 | 265 | 336 |
| ENSG00000188542 | 209 | 120 | 138 | 101 | 117 | 139 |
| ENSG00000142327 | 1852 | 939 | 1277 | 1224 | 1016 | 1042 |
| ENSG00000260942 | 126 | 77 | 93 | 53 | 90 | 50 |
| ENSG00000142330 | 690 | 307 | 395 | 402 | 378 | 357 |
| ENSG00000178623 | 66 | 21 | 29 | 13 | 23 | 27 |
| ENSG00000276949 | 0 | 0 | 0 | 0 | 0 | 0 |
| ENSG00000185176 | 0 | 0 | 0 | 0 | 0 | 0 |
| ENSG00000219159 | 0 | 0 | 0 | 0 | 0 | 1 |
| ENSG00000184945 | 0 | 0 | 0 | 0 | 0 | 0 |
| ENSG00000274977 | 0 | 0 | 0 | 0 | 0 | 0 |
| ENSG00000130294 | 842 | 362 | 477 | 440 | 423 | 424 |
| ENSG00000172482 | 3 | 3 | 2 | 2 | 6 | 3 |
| ENSG00000172478 | 340 | 189 | 222 | 110 | 121 | 94 |
| ENSG00000226321 | 8 | 5 | 8 | 4 | 0 | 2 |
| ENSG00000233392 | 0 | 0 | 0 | 0 | 0 | 0 |
| ENSG00000223991 | 0 | 0 | 0 | 0 | 0 | 0 |
| ENSG00000162804 | 646 | 283 | 421 | 215 | 263 | 249 |
| ENSG00000229996 | 0 | 2 | 1 | 0 | 3 | 0 |
| ENSG00000225521 | 2 | 1 | 0 | 1 | 2 | 1 |
| ENSG00000122085 | 2292 | 1031 | 1193 | 1002 | 1135 | 1060 |
| ENSG00000115687 | 1190 | 439 | 613 | 514 | 604 | 659 |
| ENSG00000235861 | 0 | 0 | 0 | 0 | 0 | 0 |
| ENSG00000115685 | 3781 | 1911 | 2335 | 2186 | 2162 | 2118 |
| ENSG00000146205 | 179 | 92 | 96 | 72 | 86 | 96 |
| ENSG00000115677 | 23412 | 11034 | 13951 | 10936 | 12678 | 12004 |
| ENSG00000168385 | 13790 | 7091 | 8671 | 5805 | 8011 | 7561 |
| ENSG00000223374 | 15 | 6 | 11 | 2 | 12 | 6 |
| ENSG00000006607 | 2090 | 948 | 1137 | 732 | 994 | 1107 |
| ENSG00000263752 | 0 | 0 | 0 | 0 | 0 | 0 |
| ENSG00000224232 | 0 | 2 | 1 | 0 | 2 | 1 |
| ENSG00000115694 | 4752 | 2153 | 2767 | 2673 | 2473 | 2498 |
| ENSG00000234235 | 2 | 1 | 3 | 4 | 3 | 2 |
| ENSG00000176720 | 1493 | 717 | 865 | 1072 | 928 | 803 |
| ENSG00000273113 | 37 | 9 | 20 | 23 | 24 | 10 |
| ENSG00000176946 | 1743 | 860 | 1081 | 1288 | 967 | 1068 |
| ENSG00000207186 | 0 | 0 | 0 | 0 | 1 | 1 |
| ENSG00000168397 | 2947 | 1309 | 1741 | 1680 | 1584 | 1644 |

| | | | | | | |
|---|---|---|---|---|---|---|
| ENSG00000168393 | 1593 | 770 | 920 | 1192 | 992 | 961 |
| ENSG00000168395 | 1659 | 871 | 1098 | 1115 | 1078 | 1142 |
| ENSG00000235351 | 17 | 6 | 7 | 7 | 10 | 6 |
| ENSG00000180902 | 1313 | 624 | 921 | 707 | 583 | 703 |
| ENSG00000215692 | 0 | 0 | 2 | 0 | 0 | 1 |
| ENSG00000234793 | 0 | 0 | 0 | 0 | 0 | 0 |
| ENSG00000154252 | 6 | 2 | 1 | 0 | 0 | 4 |
| ENSG00000215023 | 1 | 1 | 2 | 0 | 1 | 0 |
| ENSG00000224272 | 5 | 2 | 5 | 2 | 1 | 0 |
| ENSG00000204099 | 6 | 12 | 8 | 8 | 2 | 4 |
| ENSG00000235151 | 2 | 1 | 4 | 1 | 1 | 0 |
| ENSG00000188389 | 0 | 0 | 1 | 0 | 0 | 3 |
| ENSG00000188011 | 0 | 0 | 0 | 0 | 0 | 0 |
| ENSG00000233806 | 19 | 11 | 12 | 3 | 11 | 10 |
| ENSG00000216921 | 1 | 0 | 1 | 0 | 1 | 1 |
| ENSG00000237940 | 21 | 7 | 15 | 4 | 8 | 5 |
| ENSG00000280119 | 1 | 2 | 0 | 0 | 2 | 1 |
| ENSG00000226423 | 0 | 0 | 0 | 0 | 0 | 0 |
| ENSG00000232002 | 0 | 0 | 0 | 0 | 0 | 0 |
| ENSG00000261186 | 18 | 6 | 11 | 6 | 6 | 7 |
| ENSG00000220804 | 45 | 27 | 42 | 19 | 36 | 40 |
| ENSG00000224160 | 0 | 0 | 0 | 0 | 0 | 0 |
| ENSG00000244528 | 0 | 0 | 0 | 0 | 0 | 0 |
| ENSG00000271153 | 0 | 0 | 0 | 1 | 0 | 1 |
| ENSG00000223587 | 0 | 0 | 0 | 0 | 0 | 0 |
| ENSG00000224918 | 0 | 0 | 0 | 0 | 0 | 0 |
| ENSG00000224318 | 0 | 0 | 0 | 0 | 0 | 0 |
| ENSG00000134121 | 0 | 0 | 0 | 0 | 0 | 0 |
| ENSG00000252017 | 0 | 0 | 0 | 0 | 0 | 0 |
| ENSG00000231660 | 0 | 0 | 0 | 0 | 0 | 0 |
| ENSG00000234661 | 0 | 0 | 0 | 0 | 0 | 0 |
| ENSG00000224957 | 0 | 0 | 0 | 0 | 0 | 0 |
| ENSG00000238075 | 0 | 0 | 0 | 0 | 0 | 0 |
| ENSG00000224239 | 0 | 0 | 0 | 0 | 0 | 0 |
| ENSG00000235158 | 0 | 0 | 0 | 0 | 0 | 0 |
| ENSG00000244169 | 0 | 0 | 0 | 0 | 0 | 0 |
| ENSG00000134115 | 0 | 0 | 0 | 0 | 0 | 0 |
| ENSG00000232563 | 0 | 0 | 0 | 0 | 0 | 0 |
| ENSG00000184423 | 0 | 0 | 0 | 0 | 0 | 0 |
| ENSG00000214074 | 0 | 0 | 0 | 0 | 0 | 0 |
| ENSG00000214073 | 0 | 0 | 0 | 0 | 0 | 0 |
| ENSG00000225044 | 0 | 0 | 0 | 0 | 0 | 0 |
| ENSG00000223040 | 0 | 0 | 0 | 0 | 0 | 0 |
| ENSG00000144619 | 0 | 0 | 1 | 0 | 0 | 0 |
| ENSG00000227588 | 0 | 0 | 0 | 0 | 0 | 0 |
| ENSG00000230398 | 0 | 0 | 0 | 0 | 0 | 0 |
| ENSG00000225310 | 0 | 0 | 0 | 0 | 0 | 0 |
| ENSG00000237990 | 0 | 0 | 0 | 0 | 0 | 0 |
| ENSG00000091181 | 2 | 1 | 2 | 4 | 0 | 0 |
| ENSG00000253049 | 0 | 0 | 0 | 0 | 0 | 0 |

| | | | | | | |
|---|---|---|---|---|---|---|
| ENSG00000072756 | 947 | 505 | 537 | 478 | 720 | 618 |
| ENSG00000113851 | 999 | 526 | 626 | 438 | 638 | 541 |
| ENSG00000271870 | 27 | 24 | 14 | 6 | 30 | 20 |
| ENSG00000223727 | 0 | 0 | 0 | 0 | 0 | 0 |
| ENSG00000144455 | 2552 | 1196 | 1469 | 1320 | 1375 | 1333 |
| ENSG00000175928 | 0 | 0 | 0 | 0 | 0 | 0 |
| ENSG00000229241 | 0 | 0 | 0 | 0 | 1 | 0 |
| ENSG00000170364 | 353 | 199 | 220 | 191 | 229 | 234 |
| ENSG00000234325 | 0 | 0 | 0 | 0 | 0 | 0 |
| ENSG00000231249 | 48 | 19 | 39 | 25 | 41 | 38 |
| ENSG00000150995 | 115 | 54 | 71 | 66 | 135 | 107 |
| ENSG00000235947 | 0 | 0 | 0 | 0 | 1 | 0 |
| ENSG00000275981 | 0 | 0 | 0 | 0 | 0 | 0 |
| ENSG00000235978 | 1 | 0 | 0 | 4 | 10 | 6 |
| ENSG00000235831 | 8 | 8 | 4 | 2 | 11 | 5 |
| ENSG00000134107 | 427 | 210 | 271 | 215 | 257 | 307 |
| ENSG00000230182 | 0 | 0 | 0 | 0 | 0 | 0 |
| ENSG00000224082 | 0 | 0 | 0 | 0 | 0 | 0 |
| ENSG00000207283 | 0 | 0 | 0 | 0 | 0 | 0 |
| ENSG00000237566 | 0 | 0 | 0 | 0 | 0 | 0 |
| ENSG00000134108 | 3994 | 1957 | 2308 | 1830 | 2218 | 2008 |
| ENSG00000233912 | 2 | 4 | 2 | 1 | 0 | 1 |
| ENSG00000268509 | 0 | 0 | 0 | 0 | 0 | 0 |
| ENSG00000134109 | 1786 | 830 | 1206 | 683 | 985 | 1089 |
| ENSG00000265180 | 0 | 0 | 0 | 0 | 0 | 0 |
| ENSG00000241227 | 0 | 0 | 0 | 0 | 0 | 0 |
| ENSG00000230944 | 0 | 0 | 0 | 0 | 0 | 0 |
| ENSG00000229642 | 0 | 0 | 0 | 0 | 0 | 0 |
| ENSG00000226022 | 0 | 0 | 0 | 0 | 0 | 0 |
| ENSG00000232958 | 0 | 0 | 0 | 0 | 0 | 0 |
| ENSG00000189229 | 0 | 0 | 0 | 0 | 0 | 0 |
| ENSG00000226258 | 0 | 0 | 0 | 0 | 0 | 0 |
| ENSG00000223719 | 0 | 0 | 0 | 0 | 0 | 0 |
| ENSG00000196277 | 0 | 0 | 0 | 0 | 0 | 0 |
| ENSG00000236152 | 0 | 0 | 0 | 0 | 0 | 0 |
| ENSG00000237665 | 0 | 0 | 0 | 0 | 0 | 0 |
| ENSG00000236202 | 0 | 0 | 0 | 0 | 0 | 0 |
| ENSG00000270207 | 0 | 0 | 0 | 0 | 0 | 0 |
| ENSG00000227110 | 47 | 31 | 43 | 36 | 32 | 37 |
| ENSG00000228351 | 3 | 0 | 0 | 0 | 0 | 0 |
| ENSG00000253081 | 0 | 0 | 1 | 0 | 1 | 0 |
| ENSG00000231401 | 0 | 2 | 0 | 0 | 0 | 0 |
| ENSG00000071282 | 448 | 243 | 289 | 597 | 633 | 629 |
| ENSG00000224884 | 1 | 2 | 0 | 1 | 0 | 0 |
| ENSG00000125046 | 0 | 1 | 0 | 0 | 1 | 2 |
| ENSG00000215160 | 0 | 0 | 0 | 2 | 0 | 0 |
| ENSG00000182533 | 0 | 0 | 2 | 0 | 2 | 0 |
| ENSG00000180914 | 54 | 20 | 24 | 15 | 23 | 19 |
| ENSG00000070950 | 517 | 293 | 267 | 183 | 326 | 265 |
| ENSG00000281863 | 0 | 0 | 0 | 0 | 0 | 0 |

| | | | | | | |
|---|---|---|---|---|---|---|
| ENSG00000199815 | 0 | 0 | 0 | 0 | 0 | 0 |
| ENSG00000196220 | 240 | 101 | 140 | 152 | 130 | 185 |
| ENSG00000224808 | 0 | 0 | 0 | 0 | 0 | 0 |
| ENSG00000228723 | 0 | 0 | 0 | 0 | 0 | 0 |
| ENSG00000227929 | 0 | 3 | 0 | 0 | 0 | 0 |
| ENSG00000235830 | 0 | 1 | 2 | 0 | 0 | 0 |
| ENSG00000254485 | 4 | 13 | 14 | 6 | 11 | 8 |
| ENSG00000214041 | 0 | 1 | 2 | 0 | 0 | 0 |
| ENSG00000206573 | 664 | 344 | 417 | 317 | 433 | 328 |
| ENSG00000134077 | 1593 | 845 | 960 | 636 | 1015 | 934 |
| ENSG00000168137 | 5216 | 2334 | 2995 | 1656 | 2660 | 2770 |
| ENSG00000156959 | 0 | 0 | 0 | 1 | 1 | 0 |
| ENSG00000235338 | 0 | 0 | 0 | 0 | 0 | 0 |
| ENSG00000163719 | 2738 | 1325 | 1540 | 1570 | 1382 | 1521 |
| ENSG00000144550 | 2 | 1 | 0 | 1 | 0 | 0 |
| ENSG00000156983 | 1410 | 608 | 847 | 718 | 733 | 750 |
| ENSG00000114026 | 734 | 382 | 471 | 413 | 398 | 416 |
| ENSG00000134072 | 56 | 17 | 28 | 23 | 27 | 30 |
| ENSG00000171148 | 4761 | 2333 | 2772 | 3154 | 2628 | 2593 |
| ENSG00000241553 | 4045 | 2066 | 2641 | 2868 | 2488 | 2523 |
| ENSG00000250151 | 0 | 1 | 1 | 1 | 1 | 1 |
| ENSG00000214021 | 648 | 259 | 415 | 239 | 293 | 335 |
| ENSG00000269886 | 2 | 1 | 0 | 0 | 0 | 0 |
| ENSG00000156990 | 1734 | 850 | 1068 | 1195 | 1013 | 987 |
| ENSG00000187288 | 4 | 0 | 4 | 0 | 1 | 5 |
| ENSG00000171135 | 1853 | 911 | 1144 | 1357 | 1181 | 1206 |
| ENSG00000163701 | 62 | 34 | 44 | 29 | 31 | 38 |
| ENSG00000163702 | 1385 | 678 | 823 | 721 | 678 | 633 |
| ENSG00000212327 | 10 | 0 | 7 | 5 | 4 | 4 |
| ENSG00000163703 | 798 | 356 | 485 | 425 | 407 | 434 |
| ENSG00000269894 | 19 | 13 | 16 | 16 | 10 | 12 |
| ENSG00000163704 | 2490 | 1029 | 1551 | 1314 | 1080 | 1373 |
| ENSG00000230082 | 0 | 4 | 3 | 1 | 2 | 7 |
| ENSG00000269982 | 95 | 53 | 67 | 35 | 45 | 62 |
| ENSG00000125037 | 4759 | 2318 | 3009 | 2780 | 3215 | 2932 |
| ENSG00000180385 | 261 | 136 | 181 | 102 | 207 | 205 |
| ENSG00000232901 | 16 | 19 | 12 | 11 | 6 | 11 |
| ENSG00000206567 | 229 | 113 | 172 | 139 | 195 | 165 |
| ENSG00000186162 | 570 | 290 | 334 | 349 | 304 | 350 |
| ENSG00000144554 | 547 | 312 | 330 | 202 | 437 | 325 |
| ENSG00000201182 | 0 | 0 | 0 | 0 | 0 | 2 |
| ENSG00000229217 | 0 | 0 | 0 | 0 | 0 | 0 |
| ENSG00000163705 | 0 | 3 | 2 | 1 | 0 | 0 |
| ENSG00000254999 | 9822 | 4850 | 5830 | 5684 | 5533 | 5257 |
| ENSG00000134086 | 3993 | 2270 | 2521 | 1822 | 2280 | 2448 |
| ENSG00000134070 | 22 | 12 | 16 | 19 | 21 | 26 |
| ENSG00000157014 | 1219 | 541 | 770 | 775 | 734 | 781 |
| ENSG00000272410 | 0 | 0 | 0 | 0 | 0 | 0 |
| ENSG00000231177 | 4 | 3 | 6 | 13 | 2 | 7 |
| ENSG00000157017 | 12 | 2 | 8 | 2 | 5 | 5 |

| | | | | | | |
|---|---|---|---|---|---|---|
| ENSG00000240288 | 8 | 10 | 7 | 5 | 11 | 3 |
| ENSG00000276678 | 0 | 0 | 0 | 0 | 0 | 0 |
| ENSG00000157020 | 5220 | 2353 | 2930 | 3148 | 2874 | 2851 |
| ENSG00000157087 | 0 | 1 | 2 | 0 | 0 | 0 |
| ENSG00000264534 | 0 | 0 | 0 | 0 | 0 | 0 |
| ENSG00000216135 | 0 | 0 | 0 | 0 | 0 | 0 |
| ENSG00000236999 | 0 | 0 | 0 | 0 | 0 | 0 |
| ENSG00000224771 | 0 | 0 | 0 | 0 | 0 | 0 |
| ENSG00000226567 | 0 | 0 | 0 | 0 | 0 | 0 |
| ENSG00000230599 | 0 | 0 | 0 | 0 | 0 | 0 |
| ENSG00000132164 | 2 | 0 | 0 | 0 | 0 | 0 |
| ENSG00000157103 | 0 | 0 | 0 | 0 | 0 | 0 |
| ENSG00000232287 | 0 | 0 | 0 | 0 | 0 | 0 |
| ENSG00000196639 | 0 | 0 | 2 | 0 | 1 | 1 |
| ENSG00000285906 | 0 | 0 | 1 | 0 | 0 | 0 |
| ENSG00000213964 | 0 | 0 | 0 | 0 | 0 | 0 |
| ENSG00000197548 | 1095 | 522 | 633 | 549 | 575 | 514 |
| ENSG00000226621 | 0 | 0 | 0 | 0 | 0 | 0 |
| ENSG00000235240 | 0 | 0 | 0 | 0 | 0 | 0 |
| ENSG00000270531 | 0 | 0 | 0 | 0 | 0 | 0 |
| ENSG00000144560 | 1965 | 812 | 1170 | 780 | 999 | 1007 |
| ENSG00000271716 | 2 | 0 | 0 | 0 | 0 | 0 |
| ENSG00000272483 | 0 | 1 | 0 | 0 | 0 | 0 |
| ENSG00000144559 | 420 | 223 | 278 | 300 | 297 | 330 |
| ENSG00000230342 | 0 | 0 | 0 | 0 | 0 | 0 |
| ENSG00000224263 | 0 | 0 | 0 | 0 | 0 | 0 |
| ENSG00000225275 | 0 | 0 | 0 | 0 | 0 | 0 |
| ENSG00000231897 | 0 | 0 | 0 | 0 | 0 | 0 |
| ENSG00000251038 | 0 | 0 | 0 | 0 | 0 | 0 |
| ENSG00000263988 | 0 | 0 | 0 | 0 | 0 | 0 |
| ENSG00000157152 | 0 | 5 | 2 | 0 | 3 | 2 |
| ENSG00000226642 | 0 | 0 | 0 | 0 | 0 | 0 |
| ENSG00000157150 | 6 | 8 | 7 | 18 | 7 | 12 |
| ENSG00000233026 | 0 | 0 | 0 | 0 | 0 | 0 |
| ENSG00000236062 | 0 | 0 | 0 | 0 | 0 | 0 |
| ENSG00000132170 | 820 | 422 | 418 | 632 | 767 | 703 |
| ENSG00000225026 | 0 | 1 | 0 | 0 | 0 | 0 |
| ENSG00000154743 | 573 | 275 | 373 | 325 | 413 | 437 |
| ENSG00000200114 | 0 | 0 | 0 | 0 | 0 | 0 |
| ENSG00000251774 | 0 | 0 | 0 | 0 | 0 | 1 |
| ENSG00000225526 | 194 | 63 | 94 | 96 | 88 | 93 |
| ENSG00000075975 | 1403 | 722 | 843 | 695 | 827 | 820 |
| ENSG00000274514 | 0 | 1 | 1 | 0 | 2 | 0 |
| ENSG00000132155 | 5364 | 2422 | 3307 | 2486 | 2894 | 2953 |
| ENSG00000233252 | 0 | 0 | 0 | 0 | 0 | 0 |
| ENSG00000088726 | 27 | 26 | 21 | 32 | 25 | 25 |
| ENSG00000213943 | 21 | 15 | 9 | 8 | 12 | 13 |
| ENSG00000144712 | 5 | 4 | 3 | 5 | 3 | 8 |
| ENSG00000272263 | 8 | 2 | 0 | 7 | 2 | 4 |
| ENSG00000144713 | 27506 | 14093 | 16650 | 20800 | 18076 | 17460 |

| | | | | | | |
|---|---|---|---|---|---|---|
| ENSG00000207496 | 0 | 0 | 0 | 0 | 0 | 0 |
| ENSG00000250939 | 0 | 0 | 0 | 0 | 0 | 0 |
| ENSG00000232746 | 0 | 0 | 0 | 0 | 0 | 0 |
| ENSG00000144711 | 2261 | 992 | 1447 | 962 | 994 | 1099 |
| ENSG00000231362 | 0 | 0 | 0 | 0 | 0 | 0 |
| ENSG00000132182 | 5345 | 2209 | 3314 | 2420 | 2585 | 3015 |
| ENSG00000238138 | 0 | 0 | 0 | 0 | 0 | 0 |
| ENSG00000223398 | 0 | 0 | 1 | 0 | 0 | 0 |
| ENSG00000244502 | 1 | 4 | 2 | 4 | 8 | 0 |
| ENSG00000163517 | 1736 | 830 | 1041 | 1045 | 836 | 929 |
| ENSG00000163520 | 12 | 5 | 2 | 7 | 7 | 6 |
| ENSG00000224514 | 0 | 0 | 0 | 0 | 0 | 0 |
| ENSG00000154764 | 0 | 0 | 0 | 0 | 0 | 0 |
| ENSG00000233121 | 6 | 0 | 1 | 1 | 1 | 1 |
| ENSG00000275340 | 0 | 1 | 1 | 0 | 1 | 0 |
| ENSG00000180438 | 4 | 4 | 3 | 1 | 0 | 3 |
| ENSG00000224752 | 0 | 0 | 0 | 0 | 0 | 0 |
| ENSG00000163528 | 873 | 471 | 469 | 508 | 513 | 560 |
| ENSG00000170876 | 3990 | 2092 | 2385 | 2161 | 2294 | 2440 |
| ENSG00000268279 | 14 | 10 | 8 | 14 | 14 | 19 |
| ENSG00000228242 | 56 | 32 | 35 | 9 | 27 | 16 |
| ENSG00000154767 | 1509 | 830 | 1155 | 731 | 873 | 985 |
| ENSG00000170860 | 2083 | 1098 | 1274 | 1298 | 1269 | 1290 |
| ENSG00000255021 | 0 | 0 | 0 | 0 | 0 | 0 |
| ENSG00000251576 | 0 | 0 | 0 | 0 | 1 | 0 |
| ENSG00000199609 | 0 | 0 | 0 | 0 | 0 | 0 |
| ENSG00000131389 | 4399 | 2058 | 2613 | 2038 | 3087 | 2965 |
| ENSG00000144596 | 4 | 2 | 1 | 0 | 2 | 1 |
| ENSG00000207163 | 0 | 0 | 0 | 0 | 0 | 0 |
| ENSG00000234123 | 0 | 0 | 0 | 0 | 0 | 0 |
| ENSG00000283392 | 0 | 0 | 1 | 0 | 2 | 0 |
| ENSG00000235629 | 0 | 1 | 0 | 1 | 0 | 1 |
| ENSG00000154781 | 1081 | 533 | 618 | 494 | 615 | 537 |
| ENSG00000131379 | 3 | 2 | 1 | 1 | 2 | 1 |
| ENSG00000230172 | 0 | 0 | 0 | 0 | 0 | 0 |
| ENSG00000285969 | 8 | 2 | 1 | 2 | 2 | 1 |
| ENSG00000260902 | 0 | 0 | 0 | 0 | 0 | 0 |
| ENSG00000154783 | 29 | 23 | 15 | 29 | 18 | 26 |
| ENSG00000225733 | 8688 | 3722 | 5090 | 3221 | 4212 | 4568 |
| ENSG00000177463 | 2200 | 917 | 1408 | 604 | 1134 | 1127 |
| ENSG00000131368 | 1811 | 1003 | 1283 | 1226 | 1379 | 1289 |
| ENSG00000131381 | 1929 | 882 | 1077 | 775 | 1079 | 1028 |
| ENSG00000228874 | 2 | 1 | 7 | 0 | 2 | 1 |
| ENSG00000227578 | 0 | 1 | 0 | 0 | 0 | 0 |
| ENSG00000230524 | 45 | 16 | 19 | 7 | 17 | 16 |
| ENSG00000131375 | 2218 | 1048 | 1226 | 849 | 1352 | 1062 |
| ENSG00000224660 | 357 | 174 | 226 | 116 | 199 | 235 |
| ENSG00000131370 | 1462 | 780 | 1011 | 657 | 767 | 844 |
| ENSG00000201162 | 0 | 0 | 0 | 0 | 0 | 0 |
| ENSG00000226427 | 0 | 0 | 0 | 0 | 0 | 0 |

| | | | | | | |
|---|---|---|---|---|---|---|
| ENSG00000206562 | 733 | 355 | 470 | 379 | 450 | 459 |
| ENSG00000144597 | 1994 | 998 | 1313 | 939 | 1250 | 1273 |
| ENSG00000206926 | 2 | 1 | 1 | 0 | 0 | 4 |
| ENSG00000249786 | 2 | 0 | 1 | 0 | 2 | 1 |
| ENSG00000206561 | 32 | 13 | 14 | 9 | 18 | 15 |
| ENSG00000263573 | 0 | 0 | 0 | 0 | 0 | 0 |
| ENSG00000241064 | 0 | 0 | 0 | 0 | 0 | 0 |
| ENSG00000131373 | 683 | 376 | 466 | 406 | 513 | 520 |
| ENSG00000169814 | 759 | 343 | 505 | 349 | 412 | 371 |
| ENSG00000206560 | 1908 | 932 | 1112 | 566 | 1069 | 916 |
| ENSG00000264354 | 0 | 0 | 0 | 0 | 0 | 0 |
| ENSG00000279465 | 0 | 0 | 0 | 0 | 0 | 0 |
| ENSG00000270409 | 0 | 1 | 0 | 0 | 0 | 0 |
| ENSG00000263740 | 2 | 3 | 2 | 0 | 0 | 0 |
| ENSG00000207815 | 0 | 0 | 1 | 0 | 0 | 0 |
| ENSG00000224728 | 12 | 4 | 6 | 3 | 11 | 4 |
| ENSG00000131386 | 2 | 1 | 1 | 0 | 1 | 0 |
| ENSG00000285654 | 0 | 0 | 0 | 0 | 0 | 0 |
| ENSG00000154813 | 852 | 460 | 379 | 422 | 601 | 491 |
| ENSG00000154814 | 499 | 257 | 312 | 262 | 293 | 306 |
| ENSG00000270739 | 0 | 0 | 0 | 1 | 0 | 0 |
| ENSG00000131378 | 856 | 414 | 635 | 537 | 497 | 596 |
| ENSG00000271964 | 1 | 2 | 4 | 4 | 2 | 5 |
| ENSG00000272498 | 2 | 1 | 3 | 1 | 0 | 2 |
| ENSG00000272529 | 1 | 2 | 1 | 2 | 5 | 2 |
| ENSG00000269391 | 0 | 0 | 0 | 0 | 0 | 0 |
| ENSG00000233570 | 0 | 0 | 0 | 0 | 0 | 0 |
| ENSG00000092345 | 0 | 0 | 0 | 0 | 0 | 0 |
| ENSG00000229271 | 0 | 0 | 0 | 0 | 0 | 0 |
| ENSG00000224039 | 0 | 0 | 0 | 0 | 0 | 0 |
| ENSG00000154822 | 4 | 0 | 0 | 1 | 0 | 0 |
| ENSG00000283885 | 0 | 0 | 0 | 0 | 0 | 0 |
| ENSG00000226441 | 0 | 0 | 0 | 0 | 0 | 0 |
| ENSG00000285731 | 0 | 0 | 1 | 0 | 0 | 0 |
| ENSG00000131374 | 3733 | 1552 | 2207 | 1273 | 1842 | 1861 |
| ENSG00000231780 | 0 | 0 | 0 | 0 | 0 | 0 |
| ENSG00000227309 | 0 | 0 | 0 | 0 | 0 | 0 |
| ENSG00000271841 | 0 | 0 | 0 | 0 | 0 | 0 |
| ENSG00000233133 | 2 | 0 | 1 | 0 | 0 | 0 |
| ENSG00000213383 | 0 | 0 | 0 | 0 | 0 | 0 |
| ENSG00000214286 | 0 | 0 | 0 | 0 | 0 | 0 |
| ENSG00000226238 | 0 | 0 | 0 | 0 | 0 | 0 |
| ENSG00000274840 | 0 | 0 | 0 | 0 | 0 | 0 |
| ENSG00000182568 | 1 | 0 | 0 | 0 | 0 | 1 |
| ENSG00000272477 | 0 | 0 | 0 | 0 | 0 | 0 |
| ENSG00000228956 | 0 | 0 | 1 | 0 | 1 | 0 |
| ENSG00000236570 | 2 | 0 | 1 | 0 | 1 | 0 |
| ENSG00000251761 | 0 | 0 | 0 | 0 | 0 | 0 |
| ENSG00000183960 | 0 | 2 | 0 | 0 | 0 | 1 |
| ENSG00000271898 | 0 | 0 | 0 | 0 | 0 | 0 |

| | | | | | | |
|---|---|---|---|---|---|---|
| ENSG00000163576 | 36 | 13 | 22 | 24 | 20 | 16 |
| ENSG00000232953 | 2 | 0 | 0 | 0 | 0 | 1 |
| ENSG00000144566 | 3741 | 1881 | 2167 | 1536 | 2155 | 2004 |
| ENSG00000183977 | 16 | 10 | 9 | 2 | 8 | 11 |
| ENSG00000201545 | 0 | 0 | 0 | 0 | 0 | 0 |
| ENSG00000230697 | 0 | 0 | 4 | 0 | 0 | 0 |
| ENSG00000229468 | 2 | 1 | 0 | 0 | 0 | 0 |
| ENSG00000114166 | 1267 | 628 | 723 | 330 | 636 | 500 |
| ENSG00000266745 | 0 | 0 | 0 | 0 | 0 | 0 |
| ENSG00000129810 | 94 | 77 | 90 | 64 | 121 | 94 |
| ENSG00000231304 | 3 | 0 | 2 | 1 | 2 | 2 |
| ENSG00000199577 | 0 | 0 | 0 | 0 | 0 | 0 |
| ENSG00000261734 | 0 | 0 | 0 | 2 | 0 | 0 |
| ENSG00000206807 | 0 | 0 | 0 | 0 | 0 | 0 |
| ENSG00000282987 | 0 | 1 | 0 | 0 | 0 | 0 |
| ENSG00000236380 | 0 | 0 | 0 | 0 | 0 | 0 |
| ENSG00000151789 | 0 | 0 | 0 | 0 | 0 | 0 |
| ENSG00000225542 | 0 | 0 | 0 | 0 | 0 | 0 |
| ENSG00000223351 | 0 | 0 | 0 | 0 | 0 | 0 |
| ENSG00000132967 | 84 | 57 | 52 | 56 | 62 | 65 |
| ENSG00000234723 | 0 | 0 | 0 | 0 | 0 | 0 |
| ENSG00000271235 | 0 | 0 | 0 | 0 | 0 | 0 |
| ENSG00000226183 | 0 | 0 | 0 | 0 | 0 | 0 |
| ENSG00000231915 | 0 | 0 | 0 | 0 | 0 | 0 |
| ENSG00000215016 | 0 | 0 | 0 | 0 | 0 | 1 |
| ENSG00000233153 | 2 | 0 | 0 | 0 | 0 | 0 |
| ENSG00000182247 | 508 | 270 | 274 | 215 | 217 | 199 |
| ENSG00000206728 | 0 | 0 | 0 | 2 | 0 | 1 |
| ENSG00000228171 | 0 | 0 | 0 | 0 | 0 | 0 |
| ENSG00000252562 | 0 | 0 | 0 | 0 | 0 | 0 |
| ENSG00000212269 | 0 | 0 | 0 | 0 | 0 | 0 |
| ENSG00000223791 | 2 | 0 | 0 | 1 | 0 | 0 |
| ENSG00000170142 | 2394 | 1216 | 1442 | 1228 | 1504 | 1452 |
| ENSG00000231395 | 0 | 3 | 0 | 0 | 5 | 0 |
| ENSG00000197885 | 378 | 182 | 212 | 185 | 225 | 235 |
| ENSG00000174748 | 44079 | 21474 | 25597 | 29204 | 25909 | 26093 |
| ENSG00000174738 | 943 | 504 | 571 | 249 | 607 | 538 |
| ENSG00000224074 | 0 | 1 | 0 | 0 | 0 | 0 |
| ENSG00000242109 | 0 | 0 | 0 | 0 | 0 | 0 |
| ENSG00000151090 | 473 | 259 | 233 | 116 | 283 | 230 |
| ENSG00000229756 | 0 | 0 | 0 | 0 | 0 | 0 |
| ENSG00000224822 | 0 | 1 | 0 | 3 | 0 | 0 |
| ENSG00000228791 | 18 | 13 | 7 | 4 | 10 | 6 |
| ENSG00000272554 | 0 | 0 | 0 | 0 | 0 | 0 |
| ENSG00000265028 | 0 | 0 | 0 | 0 | 0 | 0 |
| ENSG00000233544 | 0 | 0 | 0 | 0 | 0 | 0 |
| ENSG00000237838 | 0 | 0 | 0 | 0 | 0 | 0 |
| ENSG00000244404 | 0 | 0 | 0 | 0 | 0 | 0 |
| ENSG00000201713 | 0 | 0 | 0 | 0 | 0 | 0 |
| ENSG00000077092 | 2 | 0 | 3 | 3 | 1 | 2 |

| | | | | | | |
|---|---|---|---|---|---|---|
| ENSG00000213877 | 0 | 0 | 0 | 0 | 0 | 0 |
| ENSG00000201962 | 0 | 0 | 0 | 0 | 0 | 0 |
| ENSG00000229421 | 0 | 0 | 0 | 0 | 0 | 0 |
| ENSG00000272166 | 0 | 0 | 0 | 0 | 0 | 0 |
| ENSG00000077097 | 3692 | 2269 | 1685 | 800 | 3562 | 1604 |
| ENSG00000275227 | 1 | 1 | 2 | 0 | 0 | 1 |
| ENSG00000283971 | 0 | 0 | 1 | 0 | 2 | 0 |
| ENSG00000238225 | 0 | 0 | 0 | 0 | 0 | 0 |
| ENSG00000151092 | 700 | 314 | 361 | 256 | 361 | 300 |
| ENSG00000213872 | 2 | 0 | 3 | 1 | 0 | 1 |
| ENSG00000213871 | 0 | 0 | 0 | 0 | 0 | 1 |
| ENSG00000151093 | 525 | 261 | 325 | 342 | 320 | 340 |
| ENSG00000230891 | 0 | 0 | 0 | 0 | 0 | 0 |
| ENSG00000234158 | 0 | 0 | 0 | 0 | 0 | 0 |
| ENSG00000237603 | 0 | 0 | 0 | 0 | 0 | 0 |
| ENSG00000230465 | 0 | 0 | 0 | 0 | 0 | 0 |
| ENSG00000225386 | 0 | 0 | 0 | 0 | 0 | 0 |
| ENSG00000179796 | 0 | 0 | 0 | 0 | 2 | 0 |
| ENSG00000163491 | 0 | 4 | 5 | 0 | 2 | 2 |
| ENSG00000224237 | 0 | 0 | 0 | 0 | 0 | 0 |
| ENSG00000207251 | 0 | 0 | 0 | 0 | 0 | 0 |
| ENSG00000033867 | 812 | 447 | 399 | 206 | 647 | 456 |
| ENSG00000227523 | 0 | 0 | 1 | 1 | 1 | 0 |
| ENSG00000230807 | 1 | 1 | 0 | 1 | 1 | 0 |
| ENSG00000201033 | 0 | 0 | 0 | 0 | 0 | 0 |
| ENSG00000232167 | 0 | 0 | 0 | 0 | 0 | 0 |
| ENSG00000238912 | 0 | 0 | 0 | 0 | 0 | 0 |
| ENSG00000223968 | 0 | 0 | 0 | 0 | 0 | 0 |
| ENSG00000213846 | 54 | 10 | 28 | 18 | 33 | 24 |
| ENSG00000271943 | 1 | 0 | 1 | 0 | 1 | 0 |
| ENSG00000232766 | 0 | 0 | 0 | 0 | 0 | 0 |
| ENSG00000272282 | 0 | 0 | 0 | 0 | 0 | 0 |
| ENSG00000163508 | 20 | 10 | 6 | 6 | 2 | 11 |
| ENSG00000225548 | 2 | 0 | 0 | 1 | 1 | 0 |
| ENSG00000229243 | 0 | 0 | 0 | 0 | 0 | 0 |
| ENSG00000235493 | 0 | 0 | 0 | 0 | 0 | 0 |
| ENSG00000187118 | 645 | 327 | 439 | 376 | 418 | 411 |
| ENSG00000163512 | 454 | 227 | 286 | 185 | 276 | 216 |
| ENSG00000206559 | 1 | 3 | 6 | 4 | 7 | 4 |
| ENSG00000283563 | 0 | 0 | 0 | 0 | 0 | 0 |
| ENSG00000234048 | 0 | 2 | 0 | 0 | 0 | 0 |
| ENSG00000230480 | 0 | 0 | 0 | 0 | 0 | 0 |
| ENSG00000144642 | 0 | 1 | 1 | 1 | 0 | 0 |
| ENSG00000215004 | 0 | 0 | 0 | 0 | 0 | 0 |
| ENSG00000235904 | 0 | 0 | 0 | 0 | 0 | 0 |
| ENSG00000226216 | 0 | 0 | 0 | 0 | 0 | 0 |
| ENSG00000203506 | 0 | 0 | 0 | 0 | 0 | 0 |
| ENSG00000227795 | 0 | 0 | 0 | 0 | 0 | 0 |
| ENSG00000235593 | 0 | 0 | 0 | 0 | 0 | 0 |
| ENSG00000199927 | 0 | 0 | 0 | 0 | 0 | 0 |

| | | | | | | |
|---|---|---|---|---|---|---|
| ENSG00000281710 | 0 | 0 | 0 | 0 | 0 | 0 |
| ENSG00000227260 | 0 | 0 | 0 | 0 | 0 | 0 |
| ENSG00000227549 | 0 | 0 | 0 | 0 | 0 | 0 |
| ENSG00000163513 | 694 | 346 | 419 | 270 | 440 | 384 |
| ENSG00000224618 | 0 | 0 | 0 | 0 | 0 | 0 |
| ENSG00000261468 | 0 | 0 | 0 | 0 | 0 | 0 |
| ENSG00000144644 | 2 | 0 | 0 | 0 | 0 | 0 |
| ENSG00000265376 | 0 | 0 | 0 | 0 | 0 | 0 |
| ENSG00000213854 | 1 | 0 | 0 | 0 | 0 | 1 |
| ENSG00000222983 | 0 | 0 | 0 | 0 | 0 | 0 |
| ENSG00000230111 | 0 | 0 | 0 | 0 | 0 | 0 |
| ENSG00000227339 | 4 | 0 | 2 | 0 | 0 | 0 |
| ENSG00000163527 | 5628 | 2793 | 3173 | 2041 | 2901 | 2557 |
| ENSG00000144645 | 1034 | 447 | 646 | 486 | 673 | 651 |
| ENSG00000232490 | 3 | 0 | 0 | 0 | 0 | 0 |
| ENSG00000223965 | 0 | 0 | 0 | 0 | 0 | 1 |
| ENSG00000197385 | 114 | 73 | 76 | 17 | 60 | 45 |
| ENSG00000235670 | 4 | 2 | 0 | 4 | 2 | 1 |
| ENSG00000201701 | 0 | 0 | 0 | 0 | 0 | 0 |
| ENSG00000252398 | 0 | 0 | 0 | 0 | 0 | 0 |
| ENSG00000251590 | 0 | 0 | 0 | 0 | 0 | 0 |
| ENSG00000152642 | 4422 | 2151 | 2656 | 1646 | 2320 | 2356 |
| ENSG00000236732 | 5 | 5 | 14 | 2 | 4 | 7 |
| ENSG00000237433 | 0 | 2 | 1 | 2 | 0 | 1 |
| ENSG00000261572 | 16 | 7 | 7 | 6 | 10 | 6 |
| ENSG00000170293 | 169 | 75 | 97 | 93 | 98 | 72 |
| ENSG00000270377 | 0 | 0 | 0 | 0 | 0 | 0 |
| ENSG00000234737 | 22 | 9 | 16 | 7 | 6 | 11 |
| ENSG00000153551 | 6 | 4 | 4 | 5 | 4 | 5 |
| ENSG00000091317 | 3998 | 1906 | 2483 | 1808 | 2297 | 2397 |
| ENSG00000276147 | 0 | 0 | 1 | 0 | 2 | 0 |
| ENSG00000213849 | 0 | 0 | 0 | 0 | 0 | 0 |
| ENSG00000144635 | 1665 | 790 | 962 | 777 | 881 | 791 |
| ENSG00000226955 | 0 | 0 | 1 | 0 | 0 | 0 |
| ENSG00000231552 | 0 | 0 | 0 | 0 | 0 | 0 |
| ENSG00000237676 | 2 | 0 | 0 | 1 | 0 | 0 |
| ENSG00000182973 | 838 | 473 | 533 | 400 | 500 | 498 |
| ENSG00000251224 | 0 | 0 | 0 | 0 | 0 | 0 |
| ENSG00000213842 | 0 | 0 | 1 | 0 | 2 | 1 |
| ENSG00000227337 | 0 | 0 | 0 | 0 | 1 | 0 |
| ENSG00000206557 | 52 | 17 | 14 | 11 | 35 | 24 |
| ENSG00000183813 | 4 | 0 | 1 | 0 | 1 | 1 |
| ENSG00000170266 | 4384 | 2081 | 2696 | 2044 | 2200 | 2189 |
| ENSG00000223899 | 1 | 1 | 1 | 0 | 0 | 2 |
| ENSG00000235424 | 1 | 0 | 0 | 0 | 0 | 0 |
| ENSG00000188167 | 61 | 43 | 44 | 41 | 34 | 39 |
| ENSG00000275090 | 0 | 0 | 0 | 0 | 0 | 0 |
| ENSG00000170275 | 5291 | 2605 | 3216 | 2793 | 2961 | 2897 |
| ENSG00000272149 | 0 | 0 | 0 | 0 | 0 | 0 |
| ENSG00000173705 | 0 | 4 | 3 | 6 | 4 | 1 |

| | | | | | | |
|---|---|---|---|---|---|---|
| ENSG00000230119 | 0 | 0 | 0 | 0 | 0 | 0 |
| ENSG00000153558 | 157 | 69 | 78 | 68 | 90 | 68 |
| ENSG00000153560 | 4593 | 2062 | 2691 | 1736 | 2109 | 2168 |
| ENSG00000252700 | 0 | 0 | 0 | 0 | 0 | 0 |
| ENSG00000202175 | 0 | 0 | 0 | 0 | 0 | 0 |
| ENSG00000163539 | 895 | 512 | 507 | 323 | 679 | 501 |
| ENSG00000238008 | 0 | 0 | 0 | 0 | 0 | 0 |
| ENSG00000228112 | 0 | 0 | 0 | 0 | 0 | 0 |
| ENSG00000227707 | 0 | 0 | 0 | 0 | 0 | 0 |
| ENSG00000271643 | 258 | 115 | 149 | 91 | 154 | 114 |
| ENSG00000170248 | 3729 | 1958 | 1998 | 1212 | 2336 | 1861 |
| ENSG00000226320 | 0 | 0 | 0 | 0 | 0 | 1 |
| ENSG00000236452 | 0 | 0 | 0 | 0 | 0 | 0 |
| ENSG00000227498 | 0 | 0 | 0 | 0 | 0 | 0 |
| ENSG00000235534 | 0 | 0 | 0 | 0 | 0 | 0 |
| ENSG00000228767 | 4 | 0 | 0 | 1 | 1 | 0 |
| ENSG00000212442 | 0 | 0 | 0 | 0 | 0 | 0 |
| ENSG00000172995 | 2 | 2 | 0 | 0 | 2 | 1 |
| ENSG00000230830 | 0 | 0 | 0 | 0 | 0 | 0 |
| ENSG00000207625 | 0 | 0 | 0 | 0 | 0 | 0 |
| ENSG00000226489 | 0 | 0 | 0 | 0 | 0 | 0 |
| ENSG00000236158 | 0 | 0 | 0 | 0 | 0 | 0 |
| ENSG00000144681 | 0 | 0 | 0 | 0 | 0 | 0 |
| ENSG00000231382 | 0 | 0 | 0 | 0 | 0 | 0 |
| ENSG00000201315 | 0 | 0 | 0 | 0 | 0 | 0 |
| ENSG00000163673 | 0 | 0 | 0 | 0 | 0 | 0 |
| ENSG00000230067 | 2 | 0 | 0 | 0 | 2 | 0 |
| ENSG00000279727 | 0 | 0 | 0 | 0 | 0 | 0 |
| ENSG00000281100 | 6 | 2 | 2 | 0 | 2 | 3 |
| ENSG00000168016 | 209 | 118 | 148 | 89 | 117 | 98 |
| ENSG00000234073 | 0 | 0 | 1 | 1 | 0 | 0 |
| ENSG00000210181 | 0 | 0 | 0 | 0 | 0 | 0 |
| ENSG00000272334 | 0 | 3 | 1 | 0 | 2 | 0 |
| ENSG00000236165 | 0 | 1 | 1 | 1 | 1 | 0 |
| ENSG00000178567 | 1416 | 637 | 783 | 431 | 783 | 618 |
| ENSG00000076242 | 1310 | 719 | 887 | 635 | 817 | 718 |
| ENSG00000224858 | 1 | 2 | 5 | 3 | 2 | 3 |
| ENSG00000093167 | 855 | 424 | 540 | 415 | 556 | 444 |
| ENSG00000206645 | 0 | 1 | 0 | 0 | 0 | 0 |
| ENSG00000225558 | 0 | 0 | 0 | 0 | 0 | 0 |
| ENSG00000199594 | 0 | 0 | 0 | 1 | 0 | 0 |
| ENSG00000224080 | 16 | 6 | 7 | 7 | 13 | 7 |
| ENSG00000271993 | 1 | 5 | 0 | 2 | 9 | 5 |
| ENSG00000231449 | 0 | 0 | 1 | 1 | 0 | 0 |
| ENSG00000271653 | 0 | 0 | 0 | 0 | 0 | 0 |
| ENSG00000270194 | 49 | 34 | 25 | 16 | 21 | 31 |
| ENSG00000144674 | 221 | 107 | 155 | 39 | 305 | 270 |
| ENSG00000230409 | 17 | 6 | 9 | 8 | 6 | 6 |
| ENSG00000222208 | 0 | 1 | 0 | 0 | 0 | 2 |
| ENSG00000198590 | 8 | 4 | 13 | 4 | 9 | 8 |

| | | | | | | |
|---|---|---|---|---|---|---|
| ENSG00000144668 | 0 | 0 | 0 | 0 | 0 | 0 |
| ENSG00000239105 | 0 | 0 | 0 | 0 | 0 | 0 |
| ENSG00000235257 | 0 | 0 | 0 | 0 | 0 | 0 |
| ENSG00000235482 | 0 | 0 | 0 | 0 | 0 | 0 |
| ENSG00000236411 | 0 | 0 | 0 | 0 | 0 | 0 |
| ENSG00000144677 | 156 | 85 | 101 | 56 | 61 | 61 |
| ENSG00000199075 | 0 | 0 | 0 | 0 | 0 | 0 |
| ENSG00000136059 | 4 | 3 | 1 | 2 | 3 | 1 |
| ENSG00000187091 | 338 | 177 | 224 | 163 | 169 | 158 |
| ENSG00000200579 | 0 | 0 | 1 | 0 | 1 | 1 |
| ENSG00000008226 | 18 | 6 | 8 | 5 | 14 | 11 |
| ENSG00000231668 | 0 | 0 | 0 | 0 | 0 | 0 |
| ENSG00000060971 | 2094 | 1175 | 1288 | 1478 | 1308 | 1265 |
| ENSG00000201965 | 0 | 0 | 0 | 0 | 0 | 0 |
| ENSG00000172936 | 1922 | 856 | 1113 | 992 | 1055 | 1009 |
| ENSG00000172939 | 2311 | 1070 | 1435 | 966 | 1300 | 1355 |
| ENSG00000172940 | 0 | 1 | 0 | 0 | 0 | 1 |
| ENSG00000237982 | 0 | 0 | 0 | 0 | 0 | 0 |
| ENSG00000144671 | 0 | 0 | 1 | 2 | 0 | 0 |
| ENSG00000199514 | 0 | 0 | 0 | 0 | 0 | 0 |
| ENSG00000230123 | 0 | 0 | 0 | 0 | 0 | 0 |
| ENSG00000093217 | 99 | 55 | 52 | 35 | 54 | 63 |
| ENSG00000229589 | 48 | 28 | 37 | 27 | 33 | 37 |
| ENSG00000114739 | 608 | 272 | 370 | 165 | 285 | 272 |
| ENSG00000157036 | 267 | 117 | 156 | 100 | 134 | 153 |
| ENSG00000207512 | 0 | 0 | 0 | 0 | 0 | 0 |
| ENSG00000230683 | 0 | 0 | 0 | 0 | 0 | 0 |
| ENSG00000232439 | 1 | 3 | 0 | 0 | 0 | 4 |
| ENSG00000183873 | 6 | 3 | 8 | 6 | 1 | 2 |
| ENSG00000185313 | 0 | 0 | 0 | 0 | 0 | 0 |
| ENSG00000231243 | 0 | 0 | 0 | 0 | 0 | 0 |
| ENSG00000168356 | 0 | 0 | 0 | 0 | 0 | 0 |
| ENSG00000206708 | 0 | 0 | 0 | 0 | 0 | 0 |
| ENSG00000114742 | 1114 | 497 | 650 | 408 | 572 | 554 |
| ENSG00000114745 | 2666 | 1187 | 1450 | 1313 | 1291 | 1419 |
| ENSG00000168026 | 84 | 38 | 82 | 41 | 41 | 43 |
| ENSG00000284158 | 0 | 0 | 0 | 0 | 0 | 0 |
| ENSG00000144655 | 619 | 344 | 426 | 307 | 307 | 274 |
| ENSG00000284669 | 12 | 2 | 3 | 2 | 1 | 4 |
| ENSG00000283849 | 4 | 2 | 3 | 0 | 0 | 2 |
| ENSG00000214305 | 0 | 0 | 0 | 0 | 0 | 0 |
| ENSG00000168334 | 0 | 0 | 0 | 0 | 0 | 0 |
| ENSG00000215034 | 0 | 0 | 0 | 0 | 0 | 0 |
| ENSG00000168329 | 8 | 2 | 0 | 1 | 1 | 2 |
| ENSG00000270367 | 0 | 0 | 0 | 0 | 0 | 0 |
| ENSG00000179934 | 0 | 0 | 0 | 0 | 0 | 0 |
| ENSG00000228168 | 2 | 1 | 1 | 0 | 0 | 0 |
| ENSG00000238267 | 0 | 0 | 1 | 0 | 0 | 1 |
| ENSG00000223668 | 1 | 2 | 0 | 1 | 1 | 1 |
| ENSG00000144659 | 2265 | 1087 | 1386 | 1344 | 1261 | 1232 |

| | | | | | | |
|---|---|---|---|---|---|---|
| ENSG00000168028 | 21681 | 10834 | 12234 | 16064 | 13381 | 12644 |
| ENSG00000206760 | 0 | 0 | 0 | 0 | 0 | 0 |
| ENSG00000202363 | 0 | 0 | 0 | 0 | 0 | 0 |
| ENSG00000168314 | 0 | 0 | 0 | 0 | 0 | 1 |
| ENSG00000285885 | 0 | 0 | 0 | 0 | 0 | 0 |
| ENSG00000230757 | 0 | 0 | 0 | 0 | 0 | 0 |
| ENSG00000170011 | 10 | 1 | 4 | 0 | 4 | 1 |
| ENSG00000201570 | 0 | 0 | 0 | 0 | 0 | 0 |
| ENSG00000280739 | 11 | 20 | 12 | 6 | 15 | 16 |
| ENSG00000265327 | 0 | 0 | 1 | 0 | 0 | 0 |
| ENSG00000202517 | 0 | 0 | 0 | 0 | 0 | 0 |
| ENSG00000114784 | 770 | 355 | 417 | 392 | 427 | 442 |
| ENSG00000223797 | 100 | 52 | 79 | 78 | 62 | 69 |
| ENSG00000230274 | 0 | 0 | 0 | 0 | 0 | 0 |
| ENSG00000168032 | 31 | 21 | 4 | 18 | 30 | 17 |
| ENSG00000188846 | 12501 | 6496 | 7265 | 8329 | 8517 | 7867 |
| ENSG00000177873 | 202 | 107 | 146 | 95 | 135 | 127 |
| ENSG00000177842 | 254 | 92 | 143 | 72 | 103 | 137 |
| ENSG00000199906 | 1 | 0 | 1 | 0 | 0 | 1 |
| ENSG00000172888 | 825 | 503 | 514 | 346 | 443 | 446 |
| ENSG00000234067 | 0 | 0 | 0 | 0 | 0 | 0 |
| ENSG00000226845 | 0 | 0 | 0 | 0 | 0 | 0 |
| ENSG00000226302 | 0 | 0 | 0 | 0 | 0 | 0 |
| ENSG00000229072 | 0 | 0 | 0 | 0 | 0 | 0 |
| ENSG00000234287 | 8 | 5 | 9 | 5 | 6 | 9 |
| ENSG00000231873 | 0 | 0 | 0 | 0 | 0 | 0 |
| ENSG00000233096 | 0 | 0 | 0 | 0 | 0 | 0 |
| ENSG00000233919 | 0 | 0 | 0 | 0 | 0 | 0 |
| ENSG00000275256 | 0 | 0 | 0 | 0 | 0 | 0 |
| ENSG00000168036 | 9299 | 4263 | 5576 | 3568 | 5004 | 4618 |
| ENSG00000168038 | 76 | 47 | 45 | 39 | 63 | 47 |
| ENSG00000252409 | 0 | 0 | 0 | 0 | 0 | 0 |
| ENSG00000222942 | 0 | 0 | 0 | 0 | 0 | 0 |
| ENSG00000223513 | 0 | 0 | 0 | 0 | 0 | 0 |
| ENSG00000251986 | 0 | 0 | 0 | 0 | 0 | 0 |
| ENSG00000212145 | 0 | 0 | 0 | 0 | 0 | 0 |
| ENSG00000234270 | 0 | 0 | 0 | 0 | 0 | 0 |
| ENSG00000235339 | 0 | 0 | 0 | 0 | 0 | 0 |
| ENSG00000182606 | 2264 | 1082 | 1479 | 767 | 930 | 966 |
| ENSG00000229339 | 0 | 0 | 0 | 0 | 0 | 0 |
| ENSG00000234562 | 0 | 0 | 0 | 0 | 0 | 0 |
| ENSG00000222872 | 0 | 0 | 1 | 0 | 0 | 0 |
| ENSG00000187094 | 0 | 1 | 1 | 0 | 0 | 0 |
| ENSG00000281160 | 0 | 0 | 0 | 0 | 0 | 0 |
| ENSG00000231280 | 0 | 0 | 0 | 0 | 0 | 0 |
| ENSG00000229572 | 0 | 1 | 0 | 1 | 0 | 0 |
| ENSG00000157093 | 0 | 0 | 0 | 0 | 0 | 0 |
| ENSG00000235787 | 0 | 0 | 0 | 0 | 0 | 0 |
| ENSG00000114812 | 253 | 99 | 162 | 201 | 211 | 227 |
| ENSG00000232354 | 17 | 3 | 5 | 5 | 17 | 16 |

| | | | | | | |
|---|---|---|---|---|---|---|
| ENSG00000093183 | 962 | 478 | 618 | 413 | 549 | 562 |
| ENSG00000008324 | 451 | 202 | 306 | 240 | 246 | 232 |
| ENSG00000114857 | 717 | 462 | 322 | 124 | 712 | 359 |
| ENSG00000230084 | 1 | 0 | 1 | 1 | 1 | 1 |
| ENSG00000114853 | 46 | 24 | 24 | 26 | 24 | 16 |
| ENSG00000240203 | 0 | 0 | 0 | 0 | 0 | 0 |
| ENSG00000157119 | 0 | 0 | 0 | 0 | 0 | 0 |
| ENSG00000010282 | 2 | 0 | 0 | 0 | 0 | 0 |
| ENSG00000285539 | 0 | 0 | 0 | 0 | 0 | 0 |
| ENSG00000230970 | 0 | 0 | 0 | 0 | 0 | 0 |
| ENSG00000280571 | 0 | 0 | 0 | 0 | 0 | 0 |
| ENSG00000244607 | 2 | 0 | 0 | 0 | 0 | 1 |
| ENSG00000173811 | 0 | 0 | 0 | 0 | 4 | 1 |
| ENSG00000225611 | 0 | 0 | 0 | 1 | 0 | 0 |
| ENSG00000181061 | 722 | 428 | 475 | 509 | 557 | 497 |
| ENSG00000235288 | 0 | 0 | 0 | 0 | 0 | 0 |
| ENSG00000144648 | 26 | 9 | 11 | 5 | 7 | 8 |
| ENSG00000273328 | 8 | 5 | 3 | 4 | 3 | 5 |
| ENSG00000273291 | 0 | 0 | 0 | 1 | 0 | 0 |
| ENSG00000240747 | 0 | 0 | 0 | 0 | 0 | 0 |
| ENSG00000180432 | 0 | 1 | 0 | 2 | 1 | 0 |
| ENSG00000236459 | 0 | 3 | 0 | 0 | 0 | 0 |
| ENSG00000182983 | 0 | 1 | 2 | 0 | 2 | 1 |
| ENSG00000206552 | 0 | 0 | 0 | 0 | 0 | 0 |
| ENSG00000144649 | 6 | 4 | 5 | 10 | 9 | 9 |
| ENSG00000232375 | 0 | 0 | 0 | 0 | 0 | 0 |
| ENSG00000144647 | 344 | 207 | 236 | 262 | 265 | 217 |
| ENSG00000227245 | 0 | 0 | 0 | 0 | 0 | 0 |
| ENSG00000285766 | 0 | 1 | 0 | 0 | 0 | 0 |
| ENSG00000163788 | 292 | 127 | 171 | 95 | 148 | 132 |
| ENSG00000241939 | 0 | 0 | 0 | 0 | 0 | 1 |
| ENSG00000234617 | 28 | 13 | 19 | 7 | 19 | 8 |
| ENSG00000160746 | 1086 | 540 | 701 | 568 | 612 | 606 |
| ENSG00000251811 | 0 | 0 | 0 | 0 | 0 | 0 |
| ENSG00000223916 | 0 | 0 | 1 | 0 | 0 | 0 |
| ENSG00000011198 | 686 | 371 | 459 | 337 | 508 | 379 |
| ENSG00000235886 | 0 | 0 | 0 | 0 | 0 | 0 |
| ENSG00000271192 | 0 | 0 | 0 | 0 | 0 | 0 |
| ENSG00000272121 | 0 | 0 | 0 | 0 | 0 | 0 |
| ENSG00000252980 | 0 | 0 | 0 | 0 | 0 | 0 |
| ENSG00000207954 | 0 | 0 | 0 | 0 | 0 | 0 |
| ENSG00000280435 | 0 | 0 | 0 | 0 | 0 | 0 |
| ENSG00000261786 | 0 | 0 | 0 | 0 | 0 | 0 |
| ENSG00000173769 | 0 | 0 | 0 | 0 | 0 | 0 |
| ENSG00000271937 | 0 | 1 | 1 | 0 | 0 | 0 |
| ENSG00000179152 | 695 | 360 | 342 | 229 | 465 | 340 |
| ENSG00000225873 | 0 | 0 | 0 | 0 | 0 | 1 |
| ENSG00000283036 | 0 | 0 | 0 | 1 | 0 | 0 |
| ENSG00000185219 | 624 | 301 | 373 | 205 | 298 | 285 |
| ENSG00000178917 | 78 | 45 | 37 | 18 | 48 | 39 |

| | | | | | | |
|---|---|---|---|---|---|---|
| ENSG00000214825 | 0 | 0 | 0 | 0 | 0 | 0 |
| ENSG00000277509 | 0 | 0 | 0 | 0 | 1 | 0 |
| ENSG00000196345 | 9 | 2 | 5 | 3 | 5 | 5 |
| ENSG00000236869 | 1 | 2 | 3 | 0 | 0 | 2 |
| ENSG00000144792 | 0 | 0 | 0 | 0 | 1 | 0 |
| ENSG00000214820 | 0 | 0 | 2 | 1 | 1 | 0 |
| ENSG00000186448 | 263 | 130 | 132 | 69 | 200 | 125 |
| ENSG00000233509 | 0 | 1 | 0 | 0 | 0 | 0 |
| ENSG00000169981 | 165 | 77 | 95 | 80 | 76 | 69 |
| ENSG00000272077 | 10 | 1 | 2 | 0 | 3 | 7 |
| ENSG00000235805 | 0 | 0 | 0 | 0 | 0 | 0 |
| ENSG00000196653 | 0 | 0 | 0 | 0 | 0 | 0 |
| ENSG00000186446 | 0 | 0 | 1 | 0 | 0 | 0 |
| ENSG00000163807 | 1058 | 442 | 568 | 424 | 534 | 507 |
| ENSG00000163808 | 77 | 52 | 45 | 16 | 119 | 38 |
| ENSG00000169964 | 366 | 167 | 211 | 199 | 200 | 189 |
| ENSG00000284498 | 0 | 0 | 0 | 0 | 0 | 0 |
| ENSG00000163810 | 0 | 0 | 3 | 0 | 0 | 0 |
| ENSG00000235845 | 0 | 0 | 0 | 0 | 0 | 0 |
| ENSG00000163812 | 2138 | 963 | 1200 | 1101 | 1264 | 1149 |
| ENSG00000227123 | 0 | 0 | 1 | 1 | 0 | 0 |
| ENSG00000075914 | 476 | 277 | 340 | 331 | 366 | 393 |
| ENSG00000163815 | 0 | 0 | 1 | 2 | 0 | 0 |
| ENSG00000252410 | 0 | 0 | 0 | 0 | 0 | 0 |
| ENSG00000163814 | 0 | 0 | 0 | 0 | 0 | 0 |
| ENSG00000224094 | 15 | 3 | 5 | 4 | 2 | 4 |
| ENSG00000249992 | 16 | 5 | 8 | 3 | 6 | 8 |
| ENSG00000202268 | 0 | 0 | 0 | 0 | 0 | 0 |
| ENSG00000011376 | 624 | 292 | 387 | 330 | 378 | 364 |
| ENSG00000232455 | 0 | 0 | 0 | 0 | 0 | 0 |
| ENSG00000144791 | 888 | 393 | 583 | 375 | 488 | 475 |
| ENSG00000230530 | 4 | 2 | 3 | 2 | 1 | 1 |
| ENSG00000211456 | 519 | 276 | 328 | 213 | 321 | 266 |
| ENSG00000244357 | 0 | 1 | 0 | 0 | 0 | 0 |
| ENSG00000279017 | 0 | 0 | 0 | 0 | 0 | 0 |
| ENSG00000163817 | 0 | 0 | 0 | 0 | 0 | 0 |
| ENSG00000163818 | 237 | 108 | 159 | 70 | 122 | 89 |
| ENSG00000285788 | 0 | 0 | 0 | 0 | 0 | 0 |
| ENSG00000236930 | 0 | 0 | 0 | 0 | 0 | 0 |
| ENSG00000173585 | 0 | 0 | 1 | 0 | 0 | 0 |
| ENSG00000201635 | 0 | 0 | 0 | 0 | 0 | 0 |
| ENSG00000163820 | 1834 | 780 | 1080 | 699 | 888 | 1032 |
| ENSG00000172215 | 14 | 5 | 6 | 3 | 7 | 1 |
| ENSG00000173578 | 0 | 0 | 0 | 0 | 1 | 0 |
| ENSG00000226933 | 0 | 0 | 0 | 0 | 0 | 0 |
| ENSG00000229515 | 0 | 0 | 0 | 0 | 0 | 0 |
| ENSG00000183625 | 14 | 8 | 7 | 6 | 19 | 12 |
| ENSG00000163823 | 33 | 3 | 28 | 8 | 18 | 12 |
| ENSG00000236736 | 0 | 0 | 0 | 0 | 0 | 0 |
| ENSG00000121807 | 0 | 0 | 0 | 0 | 0 | 0 |

| | | | | | | |
|---|---|---|---|---|---|---|
| ENSG00000223552 | 1 | 0 | 0 | 3 | 2 | 0 |
| ENSG00000160791 | 0 | 0 | 0 | 0 | 0 | 0 |
| ENSG00000121797 | 2 | 1 | 1 | 1 | 0 | 2 |
| ENSG00000283646 | 0 | 0 | 0 | 0 | 0 | 0 |
| ENSG00000012223 | 0 | 4 | 1 | 2 | 1 | 0 |
| ENSG00000163825 | 0 | 0 | 0 | 1 | 1 | 1 |
| ENSG00000163827 | 0 | 3 | 1 | 0 | 1 | 1 |
| ENSG00000268324 | 0 | 0 | 0 | 0 | 0 | 2 |
| ENSG00000241186 | 0 | 0 | 0 | 0 | 0 | 0 |
| ENSG00000283877 | 0 | 0 | 0 | 0 | 0 | 0 |
| ENSG00000283473 | 0 | 0 | 0 | 0 | 0 | 0 |
| ENSG00000251967 | 0 | 0 | 0 | 0 | 0 | 0 |
| ENSG00000178038 | 365 | 137 | 176 | 170 | 200 | 232 |
| ENSG00000181585 | 0 | 0 | 1 | 2 | 1 | 1 |
| ENSG00000206549 | 0 | 0 | 0 | 0 | 0 | 0 |
| ENSG00000283706 | 0 | 0 | 0 | 0 | 0 | 0 |
| ENSG00000261603 | 0 | 0 | 0 | 0 | 0 | 0 |
| ENSG00000188086 | 0 | 0 | 0 | 0 | 0 | 0 |
| ENSG00000284672 | 0 | 0 | 0 | 0 | 0 | 0 |
| ENSG00000238013 | 0 | 0 | 0 | 0 | 0 | 0 |
| ENSG00000226074 | 0 | 0 | 0 | 0 | 0 | 0 |
| ENSG00000178055 | 0 | 0 | 0 | 0 | 0 | 0 |
| ENSG00000160808 | 0 | 0 | 1 | 0 | 0 | 0 |
| ENSG00000160801 | 6 | 1 | 2 | 0 | 1 | 0 |
| ENSG00000273682 | 0 | 0 | 2 | 1 | 1 | 1 |
| ENSG00000160799 | 661 | 360 | 405 | 413 | 336 | 324 |
| ENSG00000160796 | 5205 | 2171 | 2894 | 1922 | 2117 | 2081 |
| ENSG00000236409 | 4 | 4 | 6 | 10 | 11 | 6 |
| ENSG00000181555 | 870 | 446 | 396 | 178 | 706 | 413 |
| ENSG00000275940 | 0 | 0 | 0 | 0 | 0 | 0 |
| ENSG00000227398 | 89 | 42 | 66 | 44 | 41 | 46 |
| ENSG00000088727 | 123 | 70 | 78 | 67 | 72 | 70 |
| ENSG00000239128 | 0 | 0 | 0 | 1 | 0 | 1 |
| ENSG00000114648 | 716 | 250 | 347 | 319 | 341 | 391 |
| ENSG00000280173 | 58 | 23 | 21 | 14 | 22 | 21 |
| ENSG00000260236 | 21 | 15 | 12 | 19 | 13 | 10 |
| ENSG00000076201 | 1221 | 485 | 714 | 632 | 609 | 608 |
| ENSG00000114650 | 2033 | 973 | 1244 | 1297 | 1239 | 1251 |
| ENSG00000276925 | 0 | 1 | 5 | 2 | 0 | 2 |
| ENSG00000163832 | 695 | 377 | 406 | 466 | 389 | 358 |
| ENSG00000271161 | 2 | 1 | 1 | 0 | 3 | 1 |
| ENSG00000244732 | 0 | 0 | 0 | 0 | 0 | 0 |
| ENSG00000114646 | 28 | 32 | 26 | 30 | 29 | 32 |
| ENSG00000173473 | 2738 | 1340 | 1687 | 1228 | 1552 | 1606 |
| ENSG00000214773 | 0 | 0 | 0 | 0 | 0 | 0 |
| ENSG00000235695 | 0 | 0 | 0 | 0 | 0 | 0 |
| ENSG00000132153 | 2435 | 1113 | 1515 | 1462 | 1335 | 1362 |
| ENSG00000274420 | 0 | 0 | 0 | 0 | 0 | 0 |
| ENSG00000221585 | 0 | 0 | 0 | 0 | 0 | 0 |
| ENSG00000047849 | 5997 | 2700 | 3443 | 2891 | 3259 | 3379 |

| | | | | | | |
|---|---|---|---|---|---|---|
| ENSG00000199591 | 0 | 0 | 0 | 0 | 0 | 0 |
| ENSG00000224895 | 0 | 0 | 0 | 0 | 0 | 0 |
| ENSG00000270285 | 0 | 0 | 0 | 0 | 0 | 0 |
| ENSG00000265078 | 0 | 0 | 0 | 0 | 1 | 0 |
| ENSG00000236768 | 0 | 0 | 0 | 0 | 0 | 0 |
| ENSG00000164045 | 262 | 123 | 138 | 123 | 130 | 139 |
| ENSG00000235627 | 0 | 0 | 1 | 0 | 0 | 0 |
| ENSG00000253056 | 0 | 0 | 0 | 0 | 0 | 0 |
| ENSG00000231884 | 0 | 2 | 1 | 0 | 0 | 0 |
| ENSG00000265483 | 0 | 0 | 0 | 0 | 0 | 0 |
| ENSG00000164047 | 10 | 3 | 4 | 1 | 7 | 2 |
| ENSG00000164048 | 214 | 95 | 154 | 120 | 132 | 130 |
| ENSG00000229759 | 2 | 0 | 1 | 1 | 0 | 1 |
| ENSG00000199476 | 0 | 0 | 0 | 0 | 0 | 0 |
| ENSG00000172113 | 708 | 340 | 398 | 246 | 411 | 329 |
| ENSG00000228638 | 0 | 0 | 0 | 0 | 0 | 0 |
| ENSG00000229453 | 4 | 1 | 1 | 2 | 0 | 2 |
| ENSG00000252466 | 0 | 0 | 0 | 0 | 0 | 0 |
| ENSG00000164049 | 11 | 6 | 7 | 2 | 5 | 4 |
| ENSG00000277483 | 0 | 0 | 0 | 0 | 0 | 0 |
| ENSG00000164050 | 2314 | 949 | 1503 | 1033 | 1088 | 1189 |
| ENSG00000164051 | 340 | 199 | 231 | 256 | 176 | 208 |
| ENSG00000232112 | 763 | 616 | 756 | 758 | 721 | 706 |
| ENSG00000244380 | 0 | 0 | 1 | 0 | 0 | 0 |
| ENSG00000164053 | 363 | 223 | 269 | 255 | 211 | 285 |
| ENSG00000213689 | 0 | 0 | 0 | 0 | 0 | 0 |
| ENSG00000164054 | 4910 | 2291 | 2974 | 2966 | 2524 | 2579 |
| ENSG00000114268 | 418 | 186 | 238 | 246 | 207 | 239 |
| ENSG00000278799 | 0 | 0 | 0 | 0 | 0 | 0 |
| ENSG00000145040 | 0 | 0 | 2 | 2 | 0 | 4 |
| ENSG00000114270 | 211 | 59 | 91 | 111 | 65 | 66 |
| ENSG00000284251 | 0 | 0 | 1 | 0 | 0 | 0 |
| ENSG00000010256 | 3698 | 1650 | 2018 | 2406 | 1993 | 1823 |
| ENSG00000183396 | 1 | 3 | 0 | 0 | 0 | 1 |
| ENSG00000225697 | 646 | 307 | 394 | 392 | 333 | 408 |
| ENSG00000284364 | 0 | 0 | 0 | 0 | 0 | 0 |
| ENSG00000008300 | 1180 | 494 | 642 | 519 | 630 | 574 |
| ENSG00000284575 | 0 | 0 | 0 | 0 | 0 | 0 |
| ENSG00000228350 | 11 | 1 | 5 | 1 | 3 | 2 |
| ENSG00000271973 | 3 | 2 | 1 | 3 | 1 | 1 |
| ENSG00000213672 | 670 | 315 | 449 | 391 | 338 | 421 |
| ENSG00000068745 | 2852 | 1264 | 1682 | 1370 | 1436 | 1423 |
| ENSG00000114302 | 2080 | 962 | 1216 | 852 | 1037 | 1102 |
| ENSG00000224424 | 34 | 8 | 18 | 18 | 14 | 14 |
| ENSG00000178537 | 565 | 271 | 336 | 253 | 259 | 289 |
| ENSG00000221883 | 46 | 23 | 27 | 19 | 28 | 23 |
| ENSG00000177479 | 1668 | 859 | 1071 | 836 | 1047 | 1063 |
| ENSG00000235236 | 2 | 1 | 2 | 7 | 1 | 3 |
| ENSG00000223343 | 13 | 5 | 9 | 7 | 10 | 6 |
| ENSG00000273211 | 0 | 0 | 0 | 0 | 0 | 0 |

| | | | | | | |
|---|---|---|---|---|---|---|
| ENSG00000178467 | 2295 | 1077 | 1468 | 1380 | 1231 | 1215 |
| ENSG00000285416 | 35 | 16 | 18 | 12 | 24 | 14 |
| ENSG00000178252 | 6073 | 2687 | 3939 | 2941 | 2728 | 3169 |
| ENSG00000178149 | 671 | 344 | 454 | 360 | 364 | 364 |
| ENSG00000199032 | 0 | 0 | 2 | 0 | 1 | 0 |
| ENSG00000178057 | 560 | 288 | 358 | 365 | 329 | 303 |
| ENSG00000207605 | 0 | 0 | 0 | 0 | 0 | 0 |
| ENSG00000178035 | 5536 | 2878 | 3450 | 4230 | 3663 | 3700 |
| ENSG00000272434 | 8 | 2 | 1 | 3 | 4 | 3 |
| ENSG00000198218 | 2150 | 1048 | 1310 | 968 | 1144 | 1185 |
| ENSG00000241461 | 0 | 0 | 2 | 0 | 0 | 0 |
| ENSG00000172053 | 5608 | 2556 | 3066 | 3435 | 3094 | 3041 |
| ENSG00000284488 | 0 | 0 | 0 | 0 | 0 | 0 |
| ENSG00000172046 | 2107 | 951 | 1245 | 1042 | 1129 | 1164 |
| ENSG00000172037 | 5217 | 2231 | 3179 | 2282 | 2153 | 2325 |
| ENSG00000270441 | 18 | 10 | 21 | 22 | 13 | 17 |
| ENSG00000177352 | 521 | 250 | 314 | 313 | 270 | 304 |
| ENSG00000185909 | 301 | 135 | 168 | 192 | 151 | 167 |
| ENSG00000236980 | 0 | 0 | 2 | 0 | 0 | 0 |
| ENSG00000173421 | 0 | 0 | 1 | 2 | 0 | 0 |
| ENSG00000270538 | 0 | 0 | 0 | 0 | 0 | 0 |
| ENSG00000225399 | 0 | 1 | 2 | 0 | 0 | 0 |
| ENSG00000188315 | 197 | 89 | 135 | 102 | 104 | 110 |
| ENSG00000199546 | 0 | 0 | 0 | 0 | 0 | 0 |
| ENSG00000264633 | 0 | 0 | 0 | 0 | 0 | 0 |
| ENSG00000114316 | 1133 | 565 | 672 | 539 | 568 | 630 |
| ENSG00000233276 | 0 | 2 | 1 | 0 | 0 | 2 |
| ENSG00000067560 | 10373 | 4861 | 5787 | 5968 | 5861 | 5868 |
| ENSG00000235908 | 2 | 3 | 2 | 0 | 1 | 1 |
| ENSG00000145022 | 988 | 515 | 673 | 517 | 598 | 578 |
| ENSG00000145020 | 3 | 4 | 3 | 3 | 0 | 2 |
| ENSG00000283189 | 7 | 3 | 4 | 2 | 1 | 5 |
| ENSG00000145029 | 0 | 0 | 0 | 0 | 0 | 0 |
| ENSG00000201301 | 0 | 0 | 0 | 0 | 0 | 0 |
| ENSG00000173402 | 9143 | 4281 | 5614 | 3974 | 4126 | 4675 |
| ENSG00000226913 | 13 | 7 | 10 | 1 | 4 | 3 |
| ENSG00000164061 | 56 | 30 | 44 | 27 | 41 | 33 |
| ENSG00000235120 | 0 | 0 | 0 | 0 | 0 | 0 |
| ENSG00000164062 | 4056 | 1883 | 2297 | 2748 | 2323 | 2396 |
| ENSG00000173531 | 167 | 72 | 120 | 62 | 76 | 75 |
| ENSG00000259970 | 0 | 0 | 0 | 0 | 0 | 0 |
| ENSG00000164068 | 869 | 418 | 550 | 491 | 481 | 487 |
| ENSG00000176020 | 0 | 0 | 1 | 0 | 0 | 0 |
| ENSG00000173540 | 656 | 324 | 458 | 357 | 356 | 328 |
| ENSG00000176095 | 1368 | 579 | 792 | 782 | 670 | 674 |
| ENSG00000239576 | 0 | 0 | 0 | 0 | 0 | 0 |
| ENSG00000244730 | 0 | 0 | 1 | 0 | 0 | 0 |
| ENSG00000187492 | 0 | 0 | 0 | 0 | 0 | 0 |
| ENSG00000185614 | 4 | 2 | 1 | 5 | 4 | 0 |
| ENSG00000182179 | 8 | 0 | 4 | 3 | 3 | 3 |

| | | | | | | |
|---|---|---|---|---|---|---|
| ENSG00000283726 | 0 | 0 | 0 | 0 | 0 | 0 |
| ENSG00000183763 | 214 | 105 | 89 | 99 | 106 | 90 |
| ENSG00000164076 | 0 | 1 | 1 | 1 | 0 | 4 |
| ENSG00000264706 | 0 | 0 | 0 | 0 | 0 | 0 |
| ENSG00000234667 | 0 | 0 | 0 | 0 | 1 | 0 |
| ENSG00000164078 | 205 | 96 | 133 | 161 | 177 | 135 |
| ENSG00000230698 | 0 | 0 | 0 | 0 | 1 | 0 |
| ENSG00000228008 | 2 | 0 | 4 | 0 | 1 | 2 |
| ENSG00000164077 | 229 | 101 | 141 | 161 | 108 | 152 |
| ENSG00000004534 | 1846 | 754 | 1319 | 712 | 926 | 1101 |
| ENSG00000003756 | 2194 | 1039 | 1510 | 839 | 1061 | 1075 |
| ENSG00000281691 | 3 | 7 | 5 | 2 | 5 | 2 |
| ENSG00000235016 | 23 | 21 | 21 | 13 | 18 | 17 |
| ENSG00000001617 | 3969 | 1812 | 2673 | 2367 | 1979 | 2234 |
| ENSG00000207922 | 0 | 0 | 0 | 0 | 0 | 0 |
| ENSG00000114349 | 6 | 1 | 4 | 6 | 2 | 1 |
| ENSG00000188338 | 18 | 8 | 15 | 2 | 6 | 6 |
| ENSG00000114353 | 6845 | 2949 | 4212 | 3880 | 3306 | 3763 |
| ENSG00000275334 | 0 | 0 | 0 | 0 | 0 | 0 |
| ENSG00000213600 | 0 | 0 | 1 | 0 | 1 | 1 |
| ENSG00000230454 | 53 | 14 | 26 | 16 | 23 | 26 |
| ENSG00000232352 | 14 | 8 | 21 | 6 | 9 | 23 |
| ENSG00000012171 | 921 | 397 | 660 | 508 | 461 | 541 |
| ENSG00000283848 | 0 | 0 | 0 | 0 | 0 | 0 |
| ENSG00000179564 | 0 | 2 | 1 | 0 | 0 | 3 |
| ENSG00000214706 | 1034 | 468 | 633 | 917 | 674 | 713 |
| ENSG00000186792 | 131 | 58 | 50 | 110 | 109 | 92 |
| ENSG00000243477 | 162 | 69 | 131 | 96 | 77 | 134 |
| ENSG00000114378 | 17 | 6 | 5 | 1 | 8 | 6 |
| ENSG00000068001 | 657 | 382 | 448 | 546 | 405 | 438 |
| ENSG00000114383 | 948 | 506 | 614 | 662 | 583 | 619 |
| ENSG00000068028 | 351 | 188 | 232 | 216 | 192 | 176 |
| ENSG00000281358 | 3 | 0 | 1 | 1 | 0 | 3 |
| ENSG00000235058 | 0 | 0 | 0 | 0 | 0 | 0 |
| ENSG00000004838 | 22 | 7 | 17 | 9 | 8 | 6 |
| ENSG00000114388 | 580 | 232 | 351 | 303 | 255 | 309 |
| ENSG00000114395 | 342 | 131 | 190 | 188 | 171 | 167 |
| ENSG00000272104 | 0 | 1 | 0 | 0 | 0 | 0 |
| ENSG00000126062 | 1256 | 720 | 795 | 832 | 682 | 742 |
| ENSG00000007402 | 74 | 27 | 42 | 30 | 32 | 25 |
| ENSG00000271858 | 15 | 10 | 12 | 6 | 12 | 13 |
| ENSG00000202322 | 0 | 0 | 0 | 0 | 0 | 0 |
| ENSG00000088543 | 25 | 13 | 17 | 18 | 18 | 18 |
| ENSG00000114735 | 594 | 287 | 417 | 297 | 298 | 313 |
| ENSG00000280382 | 0 | 0 | 0 | 0 | 0 | 0 |
| ENSG00000114737 | 346 | 192 | 231 | 208 | 238 | 209 |
| ENSG00000114738 | 1224 | 584 | 745 | 837 | 685 | 801 |
| ENSG00000273356 | 7 | 1 | 1 | 3 | 6 | 6 |
| ENSG00000272543 | 0 | 0 | 0 | 0 | 0 | 0 |
| ENSG00000088538 | 54 | 29 | 37 | 15 | 21 | 35 |

| | | | | | | |
|---|---|---|---|---|---|---|
| ENSG00000228337 | 0 | 0 | 0 | 0 | 0 | 0 |
| ENSG00000213587 | 0 | 0 | 0 | 0 | 0 | 0 |
| ENSG00000235278 | 0 | 0 | 0 | 0 | 0 | 0 |
| ENSG00000240959 | 0 | 0 | 0 | 0 | 0 | 0 |
| ENSG00000145050 | 732 | 378 | 518 | 474 | 483 | 453 |
| ENSG00000259956 | 2096 | 924 | 1269 | 1214 | 1187 | 1200 |
| ENSG00000145041 | 803 | 422 | 563 | 353 | 513 | 543 |
| ENSG00000164080 | 583 | 243 | 382 | 244 | 386 | 367 |
| ENSG00000221015 | 0 | 0 | 0 | 0 | 0 | 0 |
| ENSG00000164081 | 1261 | 559 | 803 | 840 | 612 | 715 |
| ENSG00000201595 | 0 | 0 | 2 | 0 | 0 | 0 |
| ENSG00000164082 | 8 | 3 | 4 | 2 | 2 | 8 |
| ENSG00000214686 | 0 | 0 | 0 | 1 | 0 | 0 |
| ENSG00000285749 | 0 | 0 | 0 | 0 | 0 | 0 |
| ENSG00000224792 | 0 | 0 | 0 | 0 | 0 | 0 |
| ENSG00000229972 | 0 | 0 | 0 | 0 | 0 | 0 |
| ENSG00000184345 | 0 | 0 | 0 | 0 | 0 | 0 |
| ENSG00000241789 | 0 | 0 | 0 | 0 | 0 | 0 |
| ENSG00000235455 | 0 | 0 | 0 | 0 | 0 | 0 |
| ENSG00000214681 | 0 | 0 | 0 | 0 | 0 | 0 |
| ENSG00000173389 | 0 | 0 | 0 | 0 | 0 | 0 |
| ENSG00000114767 | 429 | 235 | 276 | 396 | 323 | 341 |
| ENSG00000041880 | 113 | 47 | 67 | 51 | 51 | 64 |
| ENSG00000280422 | 0 | 2 | 0 | 0 | 2 | 2 |
| ENSG00000180929 | 0 | 0 | 0 | 0 | 0 | 0 |
| ENSG00000090097 | 377 | 143 | 192 | 203 | 168 | 179 |
| ENSG00000272762 | 0 | 0 | 0 | 0 | 0 | 0 |
| ENSG00000114779 | 996 | 527 | 630 | 589 | 496 | 565 |
| ENSG00000248487 | 36 | 16 | 18 | 35 | 24 | 30 |
| ENSG00000114786 | 2 | 1 | 0 | 0 | 0 | 0 |
| ENSG00000243989 | 33 | 20 | 22 | 27 | 18 | 24 |
| ENSG00000162244 | 16942 | 8422 | 9801 | 12208 | 9875 | 9440 |
| ENSG00000285685 | 0 | 0 | 0 | 0 | 0 | 0 |
| ENSG00000164086 | 210 | 90 | 101 | 110 | 102 | 117 |
| ENSG00000164087 | 288 | 137 | 194 | 223 | 145 | 196 |
| ENSG00000242849 | 0 | 0 | 0 | 0 | 0 | 0 |
| ENSG00000023330 | 1084 | 471 | 699 | 545 | 614 | 682 |
| ENSG00000280003 | 0 | 1 | 0 | 0 | 0 | 0 |
| ENSG00000239732 | 0 | 0 | 0 | 0 | 0 | 0 |
| ENSG00000173366 | 0 | 0 | 0 | 0 | 0 | 0 |
| ENSG00000247596 | 634 | 390 | 451 | 552 | 476 | 455 |
| ENSG00000243224 | 5 | 7 | 9 | 9 | 8 | 4 |
| ENSG00000164088 | 199 | 89 | 125 | 102 | 99 | 76 |
| ENSG00000164091 | 2421 | 1220 | 1451 | 964 | 1358 | 1379 |
| ENSG00000199150 | 1 | 0 | 0 | 0 | 0 | 0 |
| ENSG00000168237 | 52 | 27 | 51 | 38 | 41 | 48 |
| ENSG00000242797 | 4 | 2 | 0 | 1 | 1 | 1 |
| ENSG00000207926 | 0 | 1 | 0 | 2 | 1 | 1 |
| ENSG00000114841 | 146 | 66 | 83 | 50 | 52 | 75 |
| ENSG00000239557 | 9 | 5 | 4 | 2 | 3 | 0 |

| | | | | | | |
|---|---|---|---|---|---|---|
| ENSG00000163930 | 1646 | 755 | 874 | 927 | 871 | 906 |
| ENSG00000010318 | 85 | 45 | 89 | 56 | 62 | 60 |
| ENSG00000010319 | 12 | 7 | 4 | 5 | 6 | 2 |
| ENSG00000114854 | 17 | 3 | 3 | 3 | 3 | 4 |
| ENSG00000010322 | 1736 | 745 | 998 | 758 | 793 | 803 |
| ENSG00000010327 | 0 | 1 | 1 | 1 | 0 | 1 |
| ENSG00000168268 | 2546 | 1187 | 1405 | 1708 | 1374 | 1454 |
| ENSG00000168273 | 283 | 166 | 211 | 171 | 184 | 193 |
| ENSG00000163939 | 1032 | 524 | 498 | 243 | 655 | 420 |
| ENSG00000252768 | 2 | 0 | 0 | 1 | 0 | 0 |
| ENSG00000221518 | 0 | 0 | 0 | 0 | 0 | 0 |
| ENSG00000163938 | 1524 | 821 | 1014 | 830 | 1320 | 1112 |
| ENSG00000280904 | 0 | 0 | 0 | 0 | 0 | 0 |
| ENSG00000252787 | 0 | 0 | 0 | 0 | 0 | 0 |
| ENSG00000212493 | 2 | 1 | 3 | 0 | 0 | 0 |
| ENSG00000238862 | 0 | 0 | 0 | 0 | 0 | 0 |
| ENSG00000222345 | 8 | 8 | 7 | 1 | 3 | 2 |
| ENSG00000212452 | 4 | 1 | 4 | 2 | 3 | 0 |
| ENSG00000016864 | 904 | 449 | 595 | 417 | 541 | 494 |
| ENSG00000114902 | 1847 | 878 | 1069 | 944 | 972 | 952 |
| ENSG00000114904 | 578 | 279 | 331 | 241 | 310 | 304 |
| ENSG00000275956 | 3 | 2 | 2 | 2 | 0 | 1 |
| ENSG00000270941 | 0 | 0 | 0 | 0 | 1 | 0 |
| ENSG00000271137 | 0 | 0 | 0 | 0 | 0 | 0 |
| ENSG00000055957 | 2 | 0 | 0 | 0 | 0 | 0 |
| ENSG00000162267 | 1 | 0 | 0 | 0 | 0 | 0 |
| ENSG00000055955 | 0 | 0 | 2 | 2 | 0 | 0 |
| ENSG00000243696 | 6 | 2 | 3 | 1 | 2 | 4 |
| ENSG00000239799 | 0 | 1 | 0 | 0 | 0 | 0 |
| ENSG00000272573 | 2 | 1 | 1 | 1 | 0 | 0 |
| ENSG00000248592 | 0 | 4 | 2 | 5 | 6 | 4 |
| ENSG00000213533 | 24 | 15 | 26 | 28 | 21 | 16 |
| ENSG00000275789 | 0 | 0 | 1 | 0 | 0 | 0 |
| ENSG00000279144 | 0 | 0 | 0 | 1 | 2 | 4 |
| ENSG00000163935 | 503 | 247 | 306 | 219 | 283 | 279 |
| ENSG00000272305 | 0 | 3 | 5 | 0 | 6 | 5 |
| ENSG00000280417 | 35 | 13 | 22 | 18 | 24 | 16 |
| ENSG00000242142 | 0 | 0 | 0 | 0 | 0 | 0 |
| ENSG00000163933 | 615 | 295 | 401 | 384 | 354 | 378 |
| ENSG00000163932 | 6922 | 3352 | 4094 | 3093 | 3367 | 3226 |
| ENSG00000163931 | 22527 | 10280 | 13236 | 18917 | 15728 | 16597 |
| ENSG00000272886 | 622 | 246 | 373 | 185 | 309 | 301 |
| ENSG00000274967 | 0 | 1 | 0 | 0 | 1 | 0 |
| ENSG00000157388 | 94 | 35 | 58 | 73 | 86 | 93 |
| ENSG00000207109 | 0 | 0 | 0 | 0 | 0 | 0 |
| ENSG00000212608 | 0 | 0 | 0 | 0 | 0 | 0 |
| ENSG00000227224 | 0 | 0 | 0 | 0 | 0 | 0 |
| ENSG00000251987 | 0 | 0 | 0 | 0 | 0 | 0 |
| ENSG00000271916 | 0 | 3 | 2 | 1 | 0 | 0 |
| ENSG00000016391 | 361 | 209 | 247 | 149 | 190 | 212 |

| | | | | | | |
|---|---|---|---|---|---|---|
| ENSG00000056736 | 107 | 83 | 89 | 49 | 71 | 66 |
| ENSG00000271976 | 42 | 32 | 39 | 33 | 27 | 31 |
| ENSG00000113812 | 653 | 345 | 383 | 262 | 362 | 375 |
| ENSG00000113811 | 517 | 212 | 289 | 296 | 339 | 286 |
| ENSG00000242983 | 0 | 0 | 0 | 0 | 0 | 0 |
| ENSG00000157445 | 0 | 0 | 3 | 0 | 0 | 3 |
| ENSG00000229806 | 0 | 0 | 0 | 0 | 0 | 1 |
| ENSG00000265992 | 0 | 0 | 0 | 0 | 0 | 0 |
| ENSG00000243715 | 0 | 0 | 0 | 0 | 0 | 0 |
| ENSG00000144771 | 0 | 0 | 0 | 0 | 0 | 0 |
| ENSG00000239991 | 0 | 0 | 0 | 0 | 0 | 0 |
| ENSG00000243572 | 0 | 0 | 0 | 0 | 0 | 0 |
| ENSG00000240708 | 0 | 0 | 0 | 0 | 0 | 0 |
| ENSG00000242568 | 0 | 0 | 0 | 0 | 0 | 0 |
| ENSG00000242775 | 0 | 0 | 0 | 0 | 0 | 0 |
| ENSG00000114251 | 11 | 11 | 4 | 6 | 8 | 15 |
| ENSG00000244586 | 2 | 1 | 1 | 0 | 0 | 0 |
| ENSG00000242317 | 0 | 0 | 0 | 0 | 0 | 0 |
| ENSG00000187672 | 120 | 55 | 133 | 82 | 131 | 145 |
| ENSG00000240777 | 0 | 0 | 0 | 0 | 0 | 0 |
| ENSG00000281708 | 0 | 0 | 0 | 0 | 0 | 0 |
| ENSG00000264013 | 0 | 0 | 0 | 0 | 0 | 0 |
| ENSG00000200379 | 0 | 0 | 0 | 0 | 0 | 0 |
| ENSG00000200238 | 0 | 0 | 0 | 0 | 0 | 0 |
| ENSG00000180376 | 106 | 60 | 47 | 34 | 105 | 60 |
| ENSG00000163946 | 1445 | 750 | 703 | 320 | 1019 | 623 |
| ENSG00000163947 | 182 | 124 | 109 | 74 | 123 | 95 |
| ENSG00000240198 | 0 | 0 | 0 | 0 | 0 | 0 |
| ENSG00000186451 | 6 | 1 | 0 | 0 | 1 | 0 |
| ENSG00000272202 | 0 | 0 | 1 | 0 | 0 | 1 |
| ENSG00000144730 | 92 | 34 | 48 | 26 | 68 | 47 |
| ENSG00000229863 | 0 | 0 | 0 | 0 | 0 | 0 |
| ENSG00000163666 | 13 | 2 | 14 | 11 | 7 | 5 |
| ENSG00000157500 | 774 | 395 | 382 | 179 | 427 | 333 |
| ENSG00000239388 | 22 | 7 | 9 | 2 | 4 | 10 |
| ENSG00000174844 | 0 | 1 | 0 | 0 | 3 | 0 |
| ENSG00000212392 | 1 | 0 | 0 | 0 | 0 | 0 |
| ENSG00000206918 | 0 | 0 | 0 | 0 | 0 | 0 |
| ENSG00000239831 | 2 | 0 | 0 | 0 | 0 | 0 |
| ENSG00000237186 | 0 | 0 | 0 | 0 | 0 | 0 |
| ENSG00000202428 | 0 | 0 | 0 | 0 | 0 | 0 |
| ENSG00000206815 | 2 | 0 | 0 | 2 | 0 | 1 |
| ENSG00000174840 | 912 | 404 | 491 | 447 | 482 | 542 |
| ENSG00000168374 | 3613 | 1838 | 2177 | 1782 | 1968 | 1865 |
| ENSG00000272146 | 0 | 1 | 1 | 1 | 2 | 2 |
| ENSG00000210841 | 0 | 0 | 0 | 0 | 0 | 0 |
| ENSG00000174839 | 585 | 300 | 385 | 245 | 282 | 397 |
| ENSG00000239801 | 0 | 0 | 1 | 2 | 2 | 0 |
| ENSG00000241933 | 2 | 0 | 0 | 0 | 2 | 2 |
| ENSG00000163681 | 777 | 469 | 436 | 247 | 552 | 395 |

| | | | | | | |
|---|---|---|---|---|---|---|
| ENSG00000270772 | 0 | 0 | 0 | 0 | 0 | 0 |
| ENSG00000213509 | 0 | 0 | 1 | 0 | 0 | 1 |
| ENSG00000136068 | 9997 | 4069 | 5937 | 3625 | 4124 | 4276 |
| ENSG00000244161 | 40 | 19 | 24 | 7 | 26 | 16 |
| ENSG00000283148 | 2 | 1 | 2 | 0 | 0 | 0 |
| ENSG00000283511 | 17 | 7 | 4 | 4 | 4 | 4 |
| ENSG00000163687 | 0 | 1 | 0 | 0 | 0 | 0 |
| ENSG00000163686 | 114 | 52 | 59 | 64 | 68 | 49 |
| ENSG00000163684 | 391 | 237 | 219 | 187 | 240 | 204 |
| ENSG00000255154 | 0 | 2 | 1 | 1 | 4 | 0 |
| ENSG00000273493 | 0 | 0 | 0 | 0 | 0 | 0 |
| ENSG00000168297 | 260 | 109 | 112 | 94 | 103 | 99 |
| ENSG00000275607 | 0 | 0 | 0 | 0 | 0 | 0 |
| ENSG00000168291 | 1368 | 762 | 916 | 794 | 805 | 837 |
| ENSG00000272182 | 0 | 0 | 0 | 0 | 0 | 2 |
| ENSG00000272360 | 0 | 0 | 0 | 0 | 0 | 0 |
| ENSG00000168301 | 144 | 73 | 79 | 40 | 53 | 36 |
| ENSG00000168306 | 21 | 14 | 11 | 2 | 3 | 2 |
| ENSG00000236722 | 0 | 0 | 0 | 0 | 0 | 0 |
| ENSG00000168309 | 0 | 0 | 1 | 1 | 0 | 0 |
| ENSG00000243384 | 0 | 0 | 0 | 0 | 1 | 0 |
| ENSG00000244383 | 2 | 0 | 0 | 0 | 0 | 1 |
| ENSG00000198643 | 0 | 0 | 0 | 0 | 0 | 0 |
| ENSG00000163689 | 58 | 46 | 51 | 27 | 46 | 32 |
| ENSG00000242428 | 0 | 0 | 0 | 0 | 0 | 0 |
| ENSG00000243295 | 0 | 0 | 0 | 0 | 0 | 0 |
| ENSG00000243903 | 0 | 0 | 0 | 0 | 0 | 0 |
| ENSG00000242545 | 0 | 0 | 0 | 0 | 0 | 0 |
| ENSG00000241804 | 0 | 0 | 0 | 0 | 0 | 0 |
| ENSG00000277592 | 0 | 0 | 0 | 0 | 0 | 0 |
| ENSG00000189283 | 35 | 15 | 26 | 25 | 18 | 31 |
| ENSG00000240097 | 0 | 1 | 0 | 0 | 0 | 0 |
| ENSG00000281426 | 0 | 0 | 0 | 0 | 0 | 0 |
| ENSG00000232407 | 1 | 0 | 0 | 0 | 1 | 0 |
| ENSG00000225673 | 0 | 0 | 0 | 0 | 0 | 0 |
| ENSG00000244183 | 0 | 0 | 0 | 0 | 0 | 0 |
| ENSG00000212211 | 0 | 1 | 0 | 0 | 0 | 0 |
| ENSG00000144724 | 694 | 317 | 366 | 148 | 293 | 243 |
| ENSG00000252420 | 0 | 0 | 0 | 0 | 0 | 0 |
| ENSG00000226360 | 24 | 12 | 11 | 7 | 16 | 22 |
| ENSG00000252184 | 0 | 0 | 0 | 0 | 0 | 0 |
| ENSG00000237456 | 0 | 0 | 0 | 0 | 0 | 0 |
| ENSG00000241472 | 162 | 112 | 132 | 73 | 91 | 89 |
| ENSG00000114405 | 5443 | 2624 | 3361 | 2476 | 2596 | 2433 |
| ENSG00000153266 | 0 | 0 | 0 | 0 | 0 | 0 |
| ENSG00000163618 | 2 | 0 | 0 | 2 | 0 | 1 |
| ENSG00000225662 | 0 | 0 | 0 | 0 | 0 | 0 |
| ENSG00000244632 | 0 | 0 | 0 | 0 | 0 | 0 |
| ENSG00000252449 | 0 | 0 | 0 | 0 | 0 | 0 |
| ENSG00000226082 | 0 | 0 | 0 | 0 | 0 | 0 |

| | | | | | | |
|---|---|---|---|---|---|---|
| ENSG00000244342 | 0 | 1 | 0 | 0 | 0 | 0 |
| ENSG00000242841 | 14 | 5 | 3 | 9 | 4 | 6 |
| ENSG00000240359 | 0 | 0 | 0 | 0 | 0 | 0 |
| ENSG00000163630 | 0 | 0 | 0 | 0 | 0 | 0 |
| ENSG00000241359 | 0 | 0 | 4 | 0 | 0 | 0 |
| ENSG00000252301 | 0 | 0 | 0 | 0 | 0 | 0 |
| ENSG00000188817 | 2 | 2 | 1 | 2 | 1 | 0 |
| ENSG00000224479 | 156 | 88 | 98 | 69 | 104 | 87 |
| ENSG00000163632 | 16 | 17 | 7 | 5 | 10 | 18 |
| ENSG00000163634 | 9313 | 4754 | 5512 | 5056 | 5402 | 5089 |
| ENSG00000240549 | 5 | 6 | 2 | 1 | 4 | 0 |
| ENSG00000285399 | 60 | 35 | 34 | 18 | 45 | 40 |
| ENSG00000285258 | 4 | 6 | 4 | 1 | 1 | 2 |
| ENSG00000163635 | 64 | 32 | 65 | 17 | 41 | 29 |
| ENSG00000280620 | 14 | 6 | 11 | 9 | 6 | 11 |
| ENSG00000239653 | 47 | 29 | 22 | 7 | 25 | 28 |
| ENSG00000271843 | 19 | 14 | 26 | 31 | 31 | 30 |
| ENSG00000163636 | 49894 | 25349 | 30155 | 24316 | 30168 | 26957 |
| ENSG00000243410 | 0 | 0 | 0 | 0 | 0 | 0 |
| ENSG00000272181 | 9 | 3 | 2 | 2 | 4 | 4 |
| ENSG00000241111 | 78 | 30 | 30 | 21 | 30 | 30 |
| ENSG00000200222 | 1 | 0 | 0 | 2 | 2 | 0 |
| ENSG00000189196 | 35 | 19 | 16 | 5 | 11 | 8 |
| ENSG00000163637 | 1228 | 578 | 701 | 594 | 790 | 706 |
| ENSG00000241572 | 0 | 0 | 0 | 0 | 0 | 0 |
| ENSG00000241101 | 5 | 5 | 14 | 4 | 4 | 10 |
| ENSG00000239926 | 0 | 0 | 0 | 1 | 0 | 0 |
| ENSG00000226017 | 6 | 10 | 14 | 6 | 7 | 9 |
| ENSG00000244564 | 9 | 9 | 6 | 8 | 6 | 11 |
| ENSG00000212340 | 0 | 0 | 0 | 0 | 0 | 0 |
| ENSG00000163638 | 6 | 3 | 3 | 7 | 6 | 5 |
| ENSG00000241158 | 0 | 0 | 0 | 0 | 0 | 0 |
| ENSG00000241684 | 0 | 0 | 0 | 0 | 0 | 0 |
| ENSG00000243149 | 0 | 0 | 0 | 0 | 0 | 0 |
| ENSG00000151276 | 1136 | 516 | 672 | 349 | 532 | 527 |
| ENSG00000270059 | 0 | 1 | 4 | 0 | 3 | 5 |
| ENSG00000244226 | 0 | 0 | 0 | 0 | 0 | 0 |
| ENSG00000223409 | 0 | 0 | 0 | 0 | 0 | 0 |
| ENSG00000272610 | 2 | 3 | 7 | 1 | 2 | 2 |
| ENSG00000240175 | 0 | 0 | 0 | 0 | 0 | 0 |
| ENSG00000202071 | 0 | 0 | 0 | 0 | 0 | 0 |
| ENSG00000203647 | 0 | 1 | 0 | 1 | 0 | 0 |
| ENSG00000144741 | 491 | 237 | 294 | 334 | 263 | 278 |
| ENSG00000206759 | 0 | 0 | 0 | 0 | 0 | 0 |
| ENSG00000241587 | 0 | 0 | 1 | 0 | 0 | 0 |
| ENSG00000144749 | 402 | 196 | 252 | 200 | 255 | 290 |
| ENSG00000271174 | 0 | 0 | 0 | 0 | 0 | 0 |
| ENSG00000214560 | 0 | 0 | 0 | 0 | 0 | 0 |
| ENSG00000285738 | 0 | 0 | 0 | 0 | 0 | 1 |
| ENSG00000163376 | 20 | 19 | 19 | 16 | 42 | 30 |

| | | | | | | |
|---|---|---|---|---|---|---|
| ENSG00000283298 | 0 | 0 | 0 | 0 | 0 | 0 |
| ENSG00000241884 | 0 | 0 | 1 | 0 | 0 | 0 |
| ENSG00000172340 | 1769 | 850 | 1107 | 777 | 959 | 984 |
| ENSG00000242510 | 0 | 0 | 0 | 0 | 0 | 0 |
| ENSG00000241316 | 14 | 7 | 3 | 1 | 3 | 6 |
| ENSG00000183662 | 0 | 0 | 0 | 0 | 0 | 0 |
| ENSG00000214552 | 0 | 0 | 0 | 0 | 0 | 1 |
| ENSG00000240568 | 0 | 0 | 0 | 0 | 0 | 0 |
| ENSG00000244640 | 0 | 0 | 0 | 0 | 0 | 0 |
| ENSG00000241506 | 43 | 25 | 19 | 19 | 23 | 20 |
| ENSG00000163377 | 0 | 0 | 0 | 0 | 0 | 0 |
| ENSG00000252828 | 0 | 0 | 0 | 0 | 0 | 0 |
| ENSG00000163378 | 289 | 162 | 148 | 85 | 170 | 144 |
| ENSG00000244513 | 24 | 14 | 17 | 11 | 16 | 17 |
| ENSG00000144747 | 796 | 425 | 353 | 143 | 682 | 319 |
| ENSG00000265355 | 0 | 1 | 0 | 0 | 0 | 0 |
| ENSG00000144744 | 1272 | 688 | 806 | 564 | 741 | 743 |
| ENSG00000144746 | 3298 | 1584 | 1884 | 1633 | 1985 | 1745 |
| ENSG00000163380 | 28 | 3 | 17 | 7 | 12 | 20 |
| ENSG00000114541 | 48 | 19 | 20 | 6 | 21 | 11 |
| ENSG00000239263 | 0 | 0 | 0 | 0 | 0 | 0 |
| ENSG00000240265 | 0 | 0 | 0 | 0 | 0 | 0 |
| ENSG00000240842 | 0 | 0 | 0 | 0 | 0 | 0 |
| ENSG00000241667 | 0 | 0 | 0 | 0 | 0 | 0 |
| ENSG00000187098 | 84 | 56 | 58 | 32 | 48 | 50 |
| ENSG00000241665 | 0 | 0 | 0 | 0 | 0 | 0 |
| ENSG00000240405 | 0 | 0 | 0 | 0 | 1 | 0 |
| ENSG00000271034 | 0 | 0 | 0 | 0 | 0 | 0 |
| ENSG00000242728 | 0 | 0 | 0 | 0 | 0 | 0 |
| ENSG00000242120 | 0 | 0 | 0 | 0 | 0 | 0 |
| ENSG00000240156 | 0 | 0 | 0 | 0 | 0 | 0 |
| ENSG00000243873 | 0 | 0 | 0 | 0 | 0 | 0 |
| ENSG00000206939 | 0 | 0 | 0 | 0 | 0 | 0 |
| ENSG00000114861 | 712 | 286 | 367 | 217 | 300 | 278 |
| ENSG00000285708 | 0 | 0 | 0 | 0 | 0 | 0 |
| ENSG00000244203 | 0 | 1 | 0 | 0 | 0 | 0 |
| ENSG00000221264 | 0 | 0 | 0 | 0 | 0 | 0 |
| ENSG00000274387 | 0 | 0 | 0 | 0 | 0 | 0 |
| ENSG00000242094 | 2 | 3 | 1 | 0 | 2 | 1 |
| ENSG00000277855 | 5 | 6 | 2 | 0 | 7 | 7 |
| ENSG00000270562 | 44 | 15 | 31 | 7 | 20 | 10 |
| ENSG00000163412 | 503 | 301 | 321 | 238 | 315 | 284 |
| ENSG00000170837 | 392 | 157 | 248 | 239 | 235 | 266 |
| ENSG00000163421 | 0 | 0 | 0 | 0 | 0 | 0 |
| ENSG00000239250 | 0 | 0 | 0 | 0 | 0 | 0 |
| ENSG00000244130 | 0 | 0 | 0 | 0 | 0 | 0 |
| ENSG00000241163 | 0 | 0 | 0 | 0 | 0 | 0 |
| ENSG00000273461 | 0 | 0 | 0 | 0 | 0 | 0 |
| ENSG00000276471 | 0 | 0 | 0 | 0 | 0 | 0 |
| ENSG00000239350 | 0 | 0 | 0 | 0 | 0 | 0 |

| | | | | | | |
|---|---|---|---|---|---|---|
| ENSG00000242609 | 0 | 0 | 0 | 0 | 0 | 0 |
| ENSG00000243083 | 0 | 0 | 0 | 0 | 1 | 2 |
| ENSG00000272774 | 0 | 0 | 0 | 0 | 0 | 0 |
| ENSG00000242618 | 0 | 0 | 0 | 0 | 0 | 0 |
| ENSG00000260670 | 0 | 0 | 0 | 0 | 0 | 0 |
| ENSG00000163602 | 1230 | 571 | 701 | 468 | 752 | 626 |
| ENSG00000244345 | 0 | 0 | 0 | 0 | 0 | 0 |
| ENSG00000199469 | 0 | 0 | 0 | 0 | 0 | 0 |
| ENSG00000239568 | 0 | 0 | 0 | 0 | 0 | 0 |
| ENSG00000222838 | 0 | 0 | 0 | 0 | 0 | 0 |
| ENSG00000144736 | 587 | 269 | 293 | 310 | 361 | 324 |
| ENSG00000241537 | 0 | 0 | 0 | 0 | 0 | 0 |
| ENSG00000240135 | 0 | 0 | 1 | 1 | 0 | 0 |
| ENSG00000271266 | 0 | 1 | 0 | 0 | 0 | 0 |
| ENSG00000172986 | 56 | 29 | 48 | 24 | 32 | 29 |
| ENSG00000276890 | 0 | 0 | 0 | 0 | 0 | 0 |
| ENSG00000242960 | 0 | 0 | 1 | 2 | 3 | 3 |
| ENSG00000163605 | 1894 | 1001 | 1023 | 664 | 1215 | 960 |
| ENSG00000238959 | 0 | 0 | 0 | 0 | 0 | 0 |
| ENSG00000255423 | 59 | 17 | 17 | 7 | 13 | 17 |
| ENSG00000252651 | 0 | 0 | 0 | 0 | 0 | 0 |
| ENSG00000223247 | 0 | 0 | 0 | 0 | 0 | 0 |
| ENSG00000182921 | 0 | 0 | 0 | 0 | 1 | 0 |
| ENSG00000252937 | 0 | 0 | 0 | 0 | 0 | 0 |
| ENSG00000121440 | 360 | 175 | 200 | 144 | 217 | 188 |
| ENSG00000239119 | 0 | 0 | 0 | 0 | 0 | 0 |
| ENSG00000239677 | 0 | 0 | 1 | 0 | 0 | 0 |
| ENSG00000278934 | 2 | 3 | 2 | 2 | 0 | 1 |
| ENSG00000242364 | 0 | 0 | 0 | 0 | 0 | 0 |
| ENSG00000242741 | 0 | 4 | 6 | 0 | 3 | 4 |
| ENSG00000278554 | 0 | 0 | 0 | 0 | 0 | 0 |
| ENSG00000239454 | 0 | 0 | 0 | 0 | 1 | 0 |
| ENSG00000223738 | 0 | 0 | 0 | 0 | 0 | 0 |
| ENSG00000213452 | 0 | 0 | 0 | 0 | 0 | 0 |
| ENSG00000113805 | 0 | 0 | 0 | 0 | 1 | 0 |
| ENSG00000241022 | 0 | 0 | 0 | 0 | 0 | 0 |
| ENSG00000242638 | 0 | 0 | 0 | 0 | 0 | 0 |
| ENSG00000266780 | 0 | 0 | 0 | 0 | 0 | 0 |
| ENSG00000213300 | 38 | 18 | 36 | 21 | 27 | 26 |
| ENSG00000228868 | 0 | 0 | 0 | 0 | 0 | 0 |
| ENSG00000243674 | 0 | 0 | 0 | 0 | 0 | 0 |
| ENSG00000179799 | 0 | 0 | 0 | 0 | 0 | 0 |
| ENSG00000230655 | 0 | 0 | 0 | 0 | 0 | 0 |
| ENSG00000244699 | 0 | 0 | 2 | 1 | 1 | 0 |
| ENSG00000240665 | 0 | 0 | 0 | 0 | 0 | 0 |
| ENSG00000242953 | 2 | 0 | 2 | 0 | 0 | 2 |
| ENSG00000238278 | 26 | 26 | 24 | 27 | 25 | 30 |
| ENSG00000244026 | 185 | 105 | 115 | 170 | 128 | 169 |
| ENSG00000272690 | 2 | 0 | 7 | 1 | 4 | 0 |
| ENSG00000239959 | 0 | 0 | 0 | 0 | 2 | 1 |

| | | | | | | |
|---|---|---|---|---|---|---|
| ENSG00000243794 | 0 | 0 | 0 | 0 | 0 | 0 |
| ENSG00000272710 | 0 | 0 | 0 | 0 | 0 | 0 |
| ENSG00000244674 | 0 | 0 | 0 | 0 | 0 | 0 |
| ENSG00000239383 | 0 | 0 | 0 | 0 | 0 | 0 |
| ENSG00000244222 | 0 | 0 | 0 | 0 | 0 | 0 |
| ENSG00000172971 | 0 | 2 | 2 | 0 | 2 | 1 |
| ENSG00000243422 | 6 | 2 | 4 | 1 | 4 | 3 |
| ENSG00000282912 | 0 | 0 | 0 | 0 | 0 | 1 |
| ENSG00000282511 | 0 | 0 | 0 | 0 | 0 | 0 |
| ENSG00000283894 | 0 | 0 | 0 | 0 | 0 | 0 |
| ENSG00000241532 | 6 | 1 | 3 | 0 | 3 | 2 |
| ENSG00000241622 | 0 | 2 | 0 | 5 | 0 | 0 |
| ENSG00000172969 | 0 | 0 | 0 | 0 | 2 | 0 |
| ENSG00000236138 | 10 | 1 | 0 | 2 | 2 | 1 |
| ENSG00000242516 | 197 | 93 | 126 | 116 | 109 | 99 |
| ENSG00000227124 | 2 | 1 | 0 | 1 | 0 | 0 |
| ENSG00000266396 | 0 | 0 | 0 | 0 | 0 | 0 |
| ENSG00000185008 | 0 | 0 | 0 | 0 | 2 | 0 |
| ENSG00000222883 | 0 | 0 | 0 | 0 | 0 | 0 |
| ENSG00000240809 | 4 | 0 | 1 | 2 | 2 | 2 |
| ENSG00000199520 | 0 | 0 | 0 | 0 | 0 | 0 |
| ENSG00000213450 | 0 | 0 | 0 | 0 | 0 | 0 |
| ENSG00000206891 | 0 | 0 | 0 | 0 | 0 | 0 |
| ENSG00000244461 | 0 | 0 | 0 | 0 | 0 | 0 |
| ENSG00000240573 | 0 | 0 | 0 | 0 | 0 | 0 |
| ENSG00000243957 | 0 | 0 | 0 | 0 | 0 | 0 |
| ENSG00000222574 | 0 | 0 | 0 | 0 | 0 | 0 |
| ENSG00000240241 | 0 | 0 | 0 | 0 | 0 | 0 |
| ENSG00000239226 | 0 | 0 | 0 | 0 | 0 | 0 |
| ENSG00000169855 | 410 | 205 | 244 | 79 | 164 | 170 |
| ENSG00000240752 | 0 | 0 | 0 | 0 | 0 | 0 |
| ENSG00000240964 | 0 | 0 | 1 | 0 | 0 | 0 |
| ENSG00000265193 | 0 | 0 | 0 | 0 | 0 | 0 |
| ENSG00000243877 | 1 | 0 | 3 | 0 | 0 | 0 |
| ENSG00000271389 | 0 | 0 | 0 | 0 | 0 | 0 |
| ENSG00000239699 | 0 | 0 | 0 | 0 | 0 | 0 |
| ENSG00000242586 | 0 | 0 | 0 | 0 | 0 | 0 |
| ENSG00000242828 | 0 | 4 | 0 | 0 | 0 | 0 |
| ENSG00000242781 | 0 | 0 | 0 | 0 | 0 | 0 |
| ENSG00000243694 | 0 | 0 | 0 | 0 | 0 | 0 |
| ENSG00000242145 | 0 | 0 | 0 | 0 | 0 | 0 |
| ENSG00000241593 | 0 | 0 | 0 | 0 | 0 | 0 |
| ENSG00000244281 | 0 | 0 | 0 | 0 | 0 | 0 |
| ENSG00000114480 | 644 | 344 | 390 | 229 | 358 | 349 |
| ENSG00000222389 | 0 | 0 | 0 | 0 | 0 | 0 |
| ENSG00000241319 | 0 | 0 | 0 | 0 | 0 | 0 |
| ENSG00000240404 | 0 | 0 | 0 | 0 | 0 | 0 |
| ENSG00000242190 | 0 | 0 | 2 | 0 | 1 | 1 |
| ENSG00000242009 | 0 | 0 | 0 | 0 | 0 | 0 |
| ENSG00000239440 | 0 | 0 | 0 | 0 | 0 | 0 |

| | | | | | | |
|---|---|---|---|---|---|---|
| ENSG00000241367 | 0 | 0 | 0 | 0 | 0 | 0 |
| ENSG00000241095 | 0 | 1 | 0 | 0 | 0 | 2 |
| ENSG00000242195 | 0 | 0 | 1 | 0 | 0 | 0 |
| ENSG00000242641 | 0 | 0 | 0 | 0 | 0 | 0 |
| ENSG00000241776 | 0 | 0 | 0 | 0 | 0 | 0 |
| ENSG00000242339 | 0 | 0 | 0 | 0 | 0 | 0 |
| ENSG00000175161 | 0 | 0 | 0 | 0 | 0 | 0 |
| ENSG00000264084 | 0 | 0 | 0 | 0 | 0 | 0 |
| ENSG00000241648 | 0 | 0 | 0 | 0 | 0 | 0 |
| ENSG00000239519 | 0 | 0 | 0 | 0 | 0 | 0 |
| ENSG00000243915 | 0 | 0 | 0 | 0 | 0 | 0 |
| ENSG00000201410 | 0 | 0 | 0 | 0 | 0 | 0 |
| ENSG00000251773 | 0 | 0 | 0 | 0 | 0 | 0 |
| ENSG00000239767 | 0 | 0 | 0 | 0 | 0 | 0 |
| ENSG00000222934 | 0 | 0 | 0 | 0 | 0 | 0 |
| ENSG00000241328 | 0 | 0 | 0 | 0 | 0 | 0 |
| ENSG00000277746 | 0 | 0 | 0 | 0 | 0 | 0 |
| ENSG00000206538 | 0 | 0 | 0 | 0 | 0 | 0 |
| ENSG00000241293 | 0 | 1 | 0 | 0 | 0 | 0 |
| ENSG00000281392 | 0 | 1 | 0 | 0 | 0 | 0 |
| ENSG00000264119 | 1 | 0 | 2 | 0 | 1 | 1 |
| ENSG00000083937 | 2334 | 1282 | 1456 | 1066 | 1490 | 1356 |
| ENSG00000064835 | 2 | 0 | 0 | 0 | 0 | 0 |
| ENSG00000239829 | 0 | 0 | 0 | 0 | 0 | 0 |
| ENSG00000240902 | 0 | 1 | 2 | 1 | 0 | 1 |
| ENSG00000240255 | 0 | 0 | 0 | 0 | 0 | 0 |
| ENSG00000239572 | 0 | 3 | 3 | 0 | 0 | 1 |
| ENSG00000200703 | 0 | 0 | 0 | 0 | 0 | 0 |
| ENSG00000221059 | 0 | 0 | 0 | 0 | 0 | 0 |
| ENSG00000179097 | 2 | 0 | 0 | 1 | 0 | 1 |
| ENSG00000163320 | 2921 | 1480 | 1660 | 797 | 1744 | 1332 |
| ENSG00000175105 | 385 | 221 | 163 | 83 | 255 | 160 |
| ENSG00000241535 | 1 | 0 | 0 | 0 | 0 | 0 |
| ENSG00000179021 | 942 | 407 | 495 | 398 | 505 | 398 |
| ENSG00000242159 | 2 | 0 | 0 | 0 | 0 | 0 |
| ENSG00000283434 | 0 | 0 | 0 | 0 | 0 | 0 |
| ENSG00000207316 | 0 | 0 | 0 | 0 | 0 | 0 |
| ENSG00000242365 | 0 | 0 | 0 | 0 | 0 | 0 |
| ENSG00000242705 | 0 | 0 | 0 | 0 | 0 | 0 |
| ENSG00000241861 | 0 | 0 | 0 | 0 | 0 | 0 |
| ENSG00000044524 | 12 | 6 | 17 | 2 | 4 | 3 |
| ENSG00000285780 | 0 | 0 | 0 | 0 | 0 | 0 |
| ENSG00000240951 | 0 | 0 | 0 | 0 | 0 | 0 |
| ENSG00000240309 | 0 | 0 | 0 | 0 | 0 | 0 |
| ENSG00000212598 | 0 | 0 | 0 | 0 | 0 | 0 |
| ENSG00000222490 | 0 | 0 | 0 | 0 | 0 | 0 |
| ENSG00000189002 | 0 | 0 | 0 | 0 | 0 | 0 |
| ENSG00000271024 | 0 | 0 | 0 | 0 | 0 | 0 |
| ENSG00000283367 | 0 | 0 | 0 | 0 | 0 | 0 |
| ENSG00000283544 | 2 | 0 | 0 | 0 | 0 | 0 |

| | | | | | | |
|---|---|---|---|---|---|---|
| ENSG00000251727 | 0 | 0 | 0 | 0 | 0 | 0 |
| ENSG00000184500 | 86 | 56 | 58 | 31 | 53 | 65 |
| ENSG00000201339 | 0 | 0 | 0 | 0 | 0 | 0 |
| ENSG00000200366 | 0 | 0 | 0 | 0 | 0 | 0 |
| ENSG00000169379 | 304 | 174 | 158 | 87 | 168 | 130 |
| ENSG00000239614 | 0 | 0 | 0 | 0 | 0 | 0 |
| ENSG00000178750 | 0 | 1 | 0 | 0 | 0 | 1 |
| ENSG00000178700 | 374 | 194 | 247 | 152 | 210 | 242 |
| ENSG00000178694 | 306 | 173 | 191 | 128 | 211 | 186 |
| ENSG00000242457 | 0 | 2 | 2 | 1 | 1 | 0 |
| ENSG00000178660 | 5 | 4 | 3 | 4 | 1 | 3 |
| ENSG00000242911 | 0 | 0 | 0 | 0 | 0 | 0 |
| ENSG00000234629 | 0 | 0 | 1 | 0 | 0 | 0 |
| ENSG00000239589 | 0 | 0 | 0 | 0 | 0 | 0 |
| ENSG00000244048 | 0 | 0 | 0 | 0 | 0 | 0 |
| ENSG00000244681 | 0 | 0 | 0 | 0 | 0 | 0 |
| ENSG00000271671 | 0 | 0 | 0 | 0 | 0 | 0 |
| ENSG00000243547 | 34 | 8 | 11 | 9 | 7 | 11 |
| ENSG00000278103 | 0 | 0 | 0 | 0 | 0 | 0 |
| ENSG00000241815 | 0 | 0 | 0 | 0 | 0 | 0 |
| ENSG00000199286 | 0 | 0 | 0 | 0 | 0 | 0 |
| ENSG00000269028 | 9 | 2 | 9 | 2 | 11 | 10 |
| ENSG00000218754 | 0 | 0 | 0 | 0 | 0 | 0 |
| ENSG00000241018 | 0 | 0 | 0 | 0 | 0 | 0 |
| ENSG00000176243 | 0 | 0 | 0 | 0 | 0 | 0 |
| ENSG00000080224 | 303 | 126 | 182 | 59 | 93 | 85 |
| ENSG00000251172 | 0 | 0 | 0 | 0 | 0 | 0 |
| ENSG00000261292 | 6 | 3 | 5 | 0 | 3 | 2 |
| ENSG00000113966 | 115 | 55 | 56 | 60 | 81 | 49 |
| ENSG00000282527 | 0 | 0 | 0 | 0 | 0 | 0 |
| ENSG00000080200 | 26 | 4 | 5 | 1 | 16 | 5 |
| ENSG00000279658 | 0 | 0 | 0 | 0 | 0 | 0 |
| ENSG00000170854 | 615 | 325 | 376 | 271 | 371 | 345 |
| ENSG00000285614 | 2 | 0 | 0 | 0 | 0 | 1 |
| ENSG00000183185 | 0 | 0 | 0 | 0 | 0 | 0 |
| ENSG00000250750 | 0 | 0 | 0 | 0 | 0 | 0 |
| ENSG00000213439 | 0 | 0 | 0 | 0 | 0 | 0 |
| ENSG00000196578 | 0 | 0 | 0 | 0 | 0 | 0 |
| ENSG00000249225 | 0 | 0 | 0 | 0 | 0 | 0 |
| ENSG00000251090 | 0 | 0 | 0 | 0 | 0 | 0 |
| ENSG00000248365 | 0 | 0 | 0 | 0 | 0 | 0 |
| ENSG00000231192 | 0 | 0 | 0 | 0 | 0 | 0 |
| ENSG00000236032 | 0 | 0 | 0 | 0 | 0 | 0 |
| ENSG00000233412 | 0 | 0 | 0 | 0 | 0 | 0 |
| ENSG00000249321 | 0 | 0 | 0 | 0 | 0 | 0 |
| ENSG00000249975 | 0 | 0 | 0 | 0 | 0 | 0 |
| ENSG00000248507 | 0 | 0 | 0 | 0 | 0 | 0 |
| ENSG00000251088 | 0 | 0 | 0 | 0 | 0 | 0 |
| ENSG00000187900 | 0 | 0 | 0 | 0 | 0 | 0 |
| ENSG00000230301 | 0 | 0 | 0 | 0 | 0 | 0 |

| | | | | | | |
|---|---|---|---|---|---|---|
| ENSG00000197938 | 0 | 0 | 0 | 0 | 0 | 0 |
| ENSG00000232535 | 0 | 0 | 0 | 0 | 0 | 0 |
| ENSG00000196098 | 0 | 0 | 0 | 0 | 0 | 0 |
| ENSG00000206536 | 0 | 0 | 0 | 0 | 0 | 0 |
| ENSG00000244653 | 0 | 0 | 0 | 0 | 0 | 0 |
| ENSG00000232382 | 0 | 0 | 0 | 0 | 0 | 0 |
| ENSG00000231861 | 0 | 0 | 0 | 0 | 0 | 0 |
| ENSG00000080822 | 1006 | 550 | 642 | 473 | 607 | 540 |
| ENSG00000285635 | 0 | 0 | 0 | 0 | 0 | 0 |
| ENSG00000248839 | 0 | 0 | 0 | 0 | 0 | 0 |
| ENSG00000250562 | 0 | 0 | 0 | 0 | 0 | 0 |
| ENSG00000154165 | 0 | 0 | 0 | 0 | 0 | 0 |
| ENSG00000080819 | 329 | 151 | 248 | 173 | 234 | 225 |
| ENSG00000240827 | 0 | 0 | 0 | 0 | 0 | 0 |
| ENSG00000244153 | 4 | 2 | 0 | 1 | 2 | 1 |
| ENSG00000239445 | 0 | 0 | 0 | 0 | 0 | 0 |
| ENSG00000064225 | 0 | 0 | 0 | 2 | 1 | 0 |
| ENSG00000249274 | 0 | 0 | 0 | 0 | 0 | 0 |
| ENSG00000057019 | 1013 | 501 | 599 | 391 | 632 | 530 |
| ENSG00000206712 | 0 | 0 | 1 | 0 | 0 | 0 |
| ENSG00000239462 | 0 | 0 | 0 | 0 | 0 | 0 |
| ENSG00000207331 | 0 | 0 | 0 | 0 | 0 | 0 |
| ENSG00000240476 | 0 | 0 | 0 | 0 | 0 | 0 |
| ENSG00000275108 | 0 | 0 | 0 | 0 | 0 | 0 |
| ENSG00000241305 | 0 | 0 | 0 | 0 | 0 | 0 |
| ENSG00000240572 | 0 | 0 | 0 | 0 | 0 | 0 |
| ENSG00000243296 | 0 | 0 | 0 | 0 | 0 | 0 |
| ENSG00000243089 | 0 | 0 | 0 | 0 | 0 | 0 |
| ENSG00000244464 | 0 | 0 | 0 | 0 | 0 | 0 |
| ENSG00000144810 | 0 | 1 | 0 | 1 | 0 | 0 |
| ENSG00000273374 | 0 | 0 | 3 | 2 | 4 | 2 |
| ENSG00000184220 | 463 | 269 | 263 | 284 | 354 | 324 |
| ENSG00000168386 | 80 | 55 | 52 | 23 | 50 | 39 |
| ENSG00000264897 | 0 | 0 | 0 | 0 | 0 | 0 |
| ENSG00000235156 | 0 | 0 | 0 | 0 | 0 | 0 |
| ENSG00000277000 | 0 | 0 | 0 | 0 | 0 | 0 |
| ENSG00000243974 | 0 | 0 | 1 | 0 | 0 | 0 |
| ENSG00000036054 | 825 | 423 | 497 | 272 | 643 | 426 |
| ENSG00000114021 | 2007 | 1145 | 1347 | 1278 | 1499 | 1461 |
| ENSG00000154174 | 3580 | 1724 | 2261 | 1681 | 2124 | 2082 |
| ENSG00000206535 | 68 | 26 | 37 | 51 | 51 | 55 |
| ENSG00000181458 | 20 | 5 | 8 | 10 | 12 | 8 |
| ENSG00000144820 | 0 | 0 | 0 | 0 | 0 | 0 |
| ENSG00000252989 | 0 | 0 | 0 | 0 | 0 | 0 |
| ENSG00000243495 | 0 | 0 | 0 | 0 | 0 | 0 |
| ENSG00000114354 | 4018 | 1970 | 2310 | 1845 | 2386 | 2176 |
| ENSG00000154175 | 2 | 0 | 1 | 0 | 1 | 0 |
| ENSG00000213431 | 0 | 0 | 0 | 0 | 0 | 0 |
| ENSG00000201642 | 0 | 0 | 0 | 0 | 0 | 0 |
| ENSG00000242573 | 0 | 0 | 0 | 0 | 0 | 0 |

| | | | | | | |
|---|---|---|---|---|---|---|
| ENSG00000081148 | 0 | 0 | 4 | 0 | 2 | 3 |
| ENSG00000138468 | 398 | 197 | 169 | 58 | 339 | 162 |
| ENSG00000241738 | 0 | 0 | 0 | 0 | 0 | 0 |
| ENSG00000232830 | 0 | 0 | 0 | 0 | 0 | 0 |
| ENSG00000282978 | 4 | 1 | 5 | 0 | 0 | 1 |
| ENSG00000201554 | 0 | 0 | 0 | 0 | 0 | 0 |
| ENSG00000175841 | 0 | 0 | 0 | 0 | 0 | 0 |
| ENSG00000174173 | 612 | 370 | 362 | 351 | 517 | 396 |
| ENSG00000081154 | 2617 | 1285 | 1522 | 1174 | 1707 | 1435 |
| ENSG00000242299 | 69 | 42 | 38 | 34 | 34 | 51 |
| ENSG00000243396 | 1 | 0 | 0 | 0 | 0 | 2 |
| ENSG00000252348 | 0 | 0 | 0 | 0 | 0 | 0 |
| ENSG00000212319 | 0 | 0 | 0 | 0 | 0 | 0 |
| ENSG00000201511 | 0 | 0 | 0 | 0 | 0 | 0 |
| ENSG00000066422 | 1165 | 589 | 601 | 268 | 876 | 474 |
| ENSG00000201121 | 0 | 0 | 0 | 0 | 0 | 0 |
| ENSG00000256628 | 77 | 35 | 38 | 38 | 32 | 53 |
| ENSG00000114391 | 15470 | 7896 | 9279 | 10468 | 9607 | 8919 |
| ENSG00000244119 | 20 | 16 | 12 | 15 | 26 | 26 |
| ENSG00000182504 | 302 | 153 | 150 | 56 | 164 | 127 |
| ENSG00000144815 | 768 | 370 | 473 | 273 | 348 | 412 |
| ENSG00000249474 | 0 | 0 | 0 | 0 | 0 | 0 |
| ENSG00000144802 | 997 | 455 | 606 | 283 | 479 | 546 |
| ENSG00000214407 | 0 | 0 | 0 | 0 | 0 | 0 |
| ENSG00000202177 | 0 | 0 | 0 | 0 | 0 | 0 |
| ENSG00000241280 | 1 | 0 | 0 | 0 | 0 | 0 |
| ENSG00000170044 | 0 | 0 | 0 | 0 | 0 | 0 |
| ENSG00000241754 | 2 | 0 | 0 | 0 | 0 | 0 |
| ENSG00000201065 | 0 | 0 | 0 | 0 | 0 | 0 |
| ENSG00000201922 | 0 | 0 | 0 | 0 | 0 | 0 |
| ENSG00000236691 | 0 | 0 | 0 | 0 | 0 | 0 |
| ENSG00000265328 | 0 | 0 | 0 | 0 | 0 | 0 |
| ENSG00000240107 | 0 | 0 | 0 | 0 | 0 | 0 |
| ENSG00000243195 | 0 | 0 | 0 | 0 | 0 | 0 |
| ENSG00000214405 | 0 | 0 | 0 | 0 | 0 | 0 |
| ENSG00000243635 | 0 | 0 | 0 | 0 | 0 | 0 |
| ENSG00000170017 | 3882 | 1955 | 2077 | 862 | 1855 | 1356 |
| ENSG00000114423 | 917 | 422 | 486 | 227 | 400 | 329 |
| ENSG00000244652 | 0 | 0 | 0 | 0 | 0 | 0 |
| ENSG00000200610 | 0 | 0 | 0 | 0 | 0 | 0 |
| ENSG00000200361 | 0 | 0 | 0 | 0 | 0 | 0 |
| ENSG00000242759 | 3 | 2 | 3 | 3 | 3 | 0 |
| ENSG00000239997 | 0 | 0 | 0 | 0 | 0 | 0 |
| ENSG00000241787 | 0 | 0 | 0 | 0 | 0 | 0 |
| ENSG00000239632 | 0 | 0 | 0 | 0 | 0 | 0 |
| ENSG00000242214 | 0 | 0 | 0 | 0 | 0 | 0 |
| ENSG00000242836 | 0 | 0 | 0 | 0 | 0 | 0 |
| ENSG00000249580 | 0 | 0 | 0 | 0 | 0 | 0 |
| ENSG00000243658 | 0 | 0 | 0 | 0 | 0 | 0 |
| ENSG00000240034 | 0 | 0 | 0 | 0 | 0 | 0 |

| | | | | | | |
|---|---|---|---|---|---|---|
| ENSG00000250630 | 0 | 0 | 0 | 0 | 0 | 0 |
| ENSG00000239586 | 0 | 0 | 0 | 0 | 0 | 0 |
| ENSG00000240796 | 0 | 0 | 0 | 0 | 0 | 0 |
| ENSG00000252626 | 0 | 0 | 0 | 0 | 0 | 0 |
| ENSG00000239455 | 0 | 0 | 0 | 0 | 0 | 0 |
| ENSG00000243701 | 142 | 87 | 78 | 72 | 101 | 88 |
| ENSG00000239828 | 0 | 0 | 0 | 0 | 1 | 0 |
| ENSG00000241218 | 0 | 0 | 0 | 0 | 0 | 0 |
| ENSG00000272597 | 0 | 0 | 0 | 0 | 0 | 0 |
| ENSG00000138483 | 0 | 0 | 0 | 0 | 0 | 1 |
| ENSG00000273125 | 3 | 0 | 0 | 0 | 1 | 1 |
| ENSG00000114439 | 884 | 458 | 364 | 138 | 651 | 364 |
| ENSG00000240423 | 10 | 9 | 10 | 9 | 10 | 11 |
| ENSG00000241469 | 0 | 0 | 1 | 1 | 1 | 2 |
| ENSG00000284828 | 8 | 7 | 6 | 2 | 6 | 2 |
| ENSG00000279277 | 16 | 7 | 7 | 1 | 7 | 6 |
| ENSG00000196776 | 4887 | 2631 | 3343 | 2593 | 3160 | 3263 |
| ENSG00000271856 | 0 | 0 | 0 | 0 | 0 | 0 |
| ENSG00000114446 | 927 | 490 | 587 | 513 | 692 | 570 |
| ENSG00000114455 | 0 | 0 | 1 | 1 | 0 | 0 |
| ENSG00000243188 | 0 | 0 | 0 | 0 | 0 | 0 |
| ENSG00000144821 | 4 | 0 | 2 | 0 | 0 | 2 |
| ENSG00000241634 | 6 | 2 | 2 | 0 | 2 | 3 |
| ENSG00000163507 | 295 | 162 | 138 | 83 | 263 | 142 |
| ENSG00000202379 | 0 | 6 | 0 | 0 | 2 | 3 |
| ENSG00000276763 | 7 | 5 | 7 | 4 | 4 | 6 |
| ENSG00000198919 | 470 | 251 | 204 | 65 | 371 | 213 |
| ENSG00000241777 | 0 | 0 | 0 | 0 | 0 | 0 |
| ENSG00000163515 | 0 | 0 | 0 | 0 | 0 | 0 |
| ENSG00000163519 | 0 | 0 | 0 | 0 | 0 | 0 |
| ENSG00000138472 | 0 | 0 | 0 | 0 | 0 | 0 |
| ENSG00000114487 | 0 | 0 | 0 | 0 | 0 | 0 |
| ENSG00000221633 | 0 | 0 | 0 | 0 | 0 | 0 |
| ENSG00000239314 | 0 | 0 | 0 | 0 | 0 | 0 |
| ENSG00000241224 | 0 | 0 | 0 | 0 | 0 | 0 |
| ENSG00000261364 | 0 | 0 | 0 | 0 | 0 | 0 |
| ENSG00000214381 | 0 | 0 | 0 | 0 | 0 | 0 |
| ENSG00000252889 | 0 | 0 | 0 | 0 | 0 | 0 |
| ENSG00000239288 | 0 | 0 | 0 | 0 | 0 | 0 |
| ENSG00000163530 | 0 | 0 | 0 | 0 | 0 | 0 |
| ENSG00000277635 | 0 | 0 | 0 | 0 | 0 | 0 |
| ENSG00000121570 | 0 | 0 | 0 | 0 | 0 | 0 |
| ENSG00000241257 | 0 | 0 | 0 | 0 | 0 | 0 |
| ENSG00000244561 | 0 | 0 | 0 | 0 | 0 | 0 |
| ENSG00000230618 | 0 | 0 | 0 | 0 | 0 | 0 |
| ENSG00000228980 | 0 | 0 | 0 | 0 | 0 | 0 |
| ENSG00000244196 | 0 | 0 | 0 | 0 | 0 | 0 |
| ENSG00000265956 | 0 | 0 | 0 | 0 | 0 | 0 |
| ENSG00000242029 | 0 | 0 | 0 | 0 | 0 | 0 |
| ENSG00000271249 | 0 | 0 | 0 | 0 | 0 | 0 |

| | | | | | | |
|---|---|---|---|---|---|---|
| ENSG00000214380 | 0 | 0 | 0 | 0 | 0 | 0 |
| ENSG00000243945 | 0 | 0 | 0 | 0 | 0 | 0 |
| ENSG00000240895 | 0 | 0 | 0 | 0 | 0 | 0 |
| ENSG00000221206 | 0 | 0 | 0 | 0 | 0 | 0 |
| ENSG00000244722 | 0 | 0 | 0 | 1 | 0 | 1 |
| ENSG00000201426 | 0 | 0 | 0 | 0 | 0 | 0 |
| ENSG00000206532 | 12 | 4 | 4 | 0 | 2 | 1 |
| ENSG00000242242 | 4 | 1 | 4 | 0 | 1 | 3 |
| ENSG00000177707 | 179 | 58 | 94 | 51 | 58 | 52 |
| ENSG00000153283 | 2 | 0 | 2 | 2 | 4 | 2 |
| ENSG00000239311 | 0 | 0 | 0 | 0 | 1 | 0 |
| ENSG00000244142 | 0 | 0 | 0 | 0 | 0 | 0 |
| ENSG00000240787 | 1 | 0 | 0 | 0 | 0 | 0 |
| ENSG00000177494 | 0 | 0 | 0 | 0 | 0 | 0 |
| ENSG00000244720 | 0 | 0 | 0 | 0 | 0 | 0 |
| ENSG00000240891 | 14 | 5 | 7 | 1 | 5 | 3 |
| ENSG00000240766 | 0 | 0 | 0 | 0 | 0 | 0 |
| ENSG00000144824 | 77 | 52 | 48 | 20 | 53 | 52 |
| ENSG00000144827 | 733 | 299 | 402 | 341 | 303 | 360 |
| ENSG00000144834 | 0 | 0 | 0 | 0 | 0 | 0 |
| ENSG00000239439 | 0 | 0 | 0 | 0 | 0 | 0 |
| ENSG00000176040 | 2 | 2 | 11 | 1 | 5 | 2 |
| ENSG00000114529 | 162 | 104 | 131 | 87 | 123 | 86 |
| ENSG00000207940 | 0 | 0 | 1 | 0 | 0 | 0 |
| ENSG00000174500 | 22 | 4 | 11 | 3 | 14 | 9 |
| ENSG00000261488 | 7 | 8 | 6 | 2 | 2 | 1 |
| ENSG00000172139 | 0 | 0 | 0 | 0 | 0 | 0 |
| ENSG00000244144 | 0 | 0 | 0 | 0 | 0 | 0 |
| ENSG00000239482 | 0 | 0 | 0 | 0 | 0 | 0 |
| ENSG00000283669 | 0 | 0 | 0 | 0 | 0 | 0 |
| ENSG00000091972 | 0 | 0 | 0 | 0 | 0 | 0 |
| ENSG00000243081 | 0 | 0 | 0 | 0 | 0 | 0 |
| ENSG00000186265 | 0 | 0 | 0 | 0 | 1 | 1 |
| ENSG00000242525 | 0 | 0 | 0 | 0 | 0 | 0 |
| ENSG00000279349 | 0 | 1 | 0 | 0 | 1 | 1 |
| ENSG00000144848 | 1665 | 829 | 1012 | 827 | 952 | 836 |
| ENSG00000138459 | 753 | 342 | 427 | 321 | 390 | 390 |
| ENSG00000091986 | 16 | 15 | 15 | 10 | 12 | 8 |
| ENSG00000242308 | 0 | 0 | 0 | 0 | 0 | 0 |
| ENSG00000240893 | 4 | 0 | 0 | 0 | 0 | 0 |
| ENSG00000242770 | 0 | 0 | 0 | 1 | 1 | 0 |
| ENSG00000206531 | 0 | 0 | 3 | 0 | 1 | 0 |
| ENSG00000163606 | 18 | 4 | 5 | 0 | 8 | 4 |
| ENSG00000272844 | 11 | 3 | 4 | 5 | 1 | 3 |
| ENSG00000163607 | 229 | 127 | 119 | 99 | 161 | 126 |
| ENSG00000163608 | 1262 | 771 | 945 | 638 | 986 | 739 |
| ENSG00000240057 | 0 | 3 | 1 | 1 | 0 | 2 |
| ENSG00000243483 | 0 | 0 | 0 | 0 | 0 | 0 |
| ENSG00000241219 | 0 | 0 | 0 | 0 | 0 | 0 |
| ENSG00000243795 | 0 | 0 | 0 | 0 | 0 | 0 |

| | | | | | | |
|---|---|---|---|---|---|---|
| ENSG00000144857 | 20 | 11 | 12 | 3 | 24 | 17 |
| ENSG00000206530 | 413 | 221 | 217 | 61 | 319 | 180 |
| ENSG00000240751 | 0 | 3 | 2 | 1 | 0 | 0 |
| ENSG00000285943 | 12 | 14 | 14 | 6 | 7 | 4 |
| ENSG00000243849 | 1 | 1 | 0 | 1 | 0 | 0 |
| ENSG00000275670 | 0 | 0 | 0 | 0 | 0 | 0 |
| ENSG00000163611 | 85 | 42 | 55 | 40 | 57 | 44 |
| ENSG00000072858 | 2908 | 1142 | 1467 | 885 | 1346 | 1142 |
| ENSG00000239453 | 5 | 1 | 2 | 0 | 7 | 2 |
| ENSG00000265253 | 0 | 0 | 0 | 0 | 0 | 0 |
| ENSG00000241529 | 0 | 2 | 0 | 0 | 0 | 0 |
| ENSG00000176542 | 668 | 375 | 370 | 113 | 450 | 346 |
| ENSG00000121579 | 5497 | 2938 | 3074 | 2258 | 3350 | 3001 |
| ENSG00000239280 | 1 | 0 | 1 | 0 | 1 | 0 |
| ENSG00000242659 | 5 | 2 | 5 | 1 | 0 | 1 |
| ENSG00000114573 | 5094 | 2152 | 2805 | 1507 | 2553 | 2176 |
| ENSG00000178075 | 50 | 26 | 21 | 12 | 35 | 27 |
| ENSG00000240776 | 0 | 0 | 1 | 0 | 0 | 1 |
| ENSG00000241889 | 4 | 1 | 0 | 0 | 1 | 3 |
| ENSG00000240375 | 0 | 0 | 0 | 0 | 0 | 0 |
| ENSG00000273394 | 16 | 4 | 3 | 5 | 3 | 2 |
| ENSG00000184307 | 1246 | 548 | 659 | 470 | 746 | 619 |
| ENSG00000163617 | 150 | 93 | 114 | 70 | 169 | 95 |
| ENSG00000270880 | 0 | 0 | 0 | 0 | 0 | 0 |
| ENSG00000285836 | 0 | 0 | 0 | 0 | 2 | 4 |
| ENSG00000151576 | 1024 | 521 | 653 | 462 | 653 | 651 |
| ENSG00000240912 | 0 | 1 | 0 | 1 | 1 | 1 |
| ENSG00000151577 | 0 | 0 | 0 | 0 | 0 | 0 |
| ENSG00000241490 | 0 | 0 | 0 | 0 | 0 | 0 |
| ENSG00000240487 | 0 | 0 | 0 | 0 | 0 | 0 |
| ENSG00000174255 | 0 | 0 | 0 | 0 | 0 | 0 |
| ENSG00000181847 | 0 | 0 | 0 | 0 | 0 | 0 |
| ENSG00000181722 | 3 | 4 | 3 | 2 | 3 | 3 |
| ENSG00000259976 | 0 | 0 | 0 | 0 | 0 | 0 |
| ENSG00000284134 | 0 | 0 | 0 | 0 | 0 | 0 |
| ENSG00000241560 | 0 | 0 | 0 | 0 | 0 | 0 |
| ENSG00000242290 | 0 | 0 | 0 | 0 | 0 | 0 |
| ENSG00000241295 | 0 | 0 | 0 | 0 | 0 | 0 |
| ENSG00000264623 | 0 | 0 | 0 | 0 | 0 | 0 |
| ENSG00000239946 | 0 | 0 | 0 | 0 | 0 | 0 |
| ENSG00000240002 | 0 | 0 | 0 | 0 | 0 | 0 |
| ENSG00000242767 | 0 | 0 | 0 | 0 | 0 | 0 |
| ENSG00000242880 | 0 | 0 | 0 | 0 | 0 | 0 |
| ENSG00000244157 | 0 | 0 | 0 | 0 | 0 | 0 |
| ENSG00000172020 | 0 | 0 | 0 | 0 | 0 | 0 |
| ENSG00000241596 | 0 | 0 | 0 | 0 | 0 | 0 |
| ENSG00000185565 | 2 | 0 | 0 | 2 | 0 | 0 |
| ENSG00000243359 | 0 | 0 | 0 | 0 | 0 | 0 |
| ENSG00000240922 | 0 | 4 | 1 | 1 | 1 | 0 |
| ENSG00000241397 | 0 | 0 | 0 | 0 | 0 | 0 |

| | | | | | | |
|---|---|---|---|---|---|---|
| ENSG00000241156 | 0 | 0 | 0 | 0 | 0 | 0 |
| ENSG00000198406 | 42 | 27 | 27 | 19 | 38 | 23 |
| ENSG00000243197 | 0 | 0 | 0 | 0 | 0 | 0 |
| ENSG00000278072 | 0 | 0 | 0 | 0 | 0 | 0 |
| ENSG00000265433 | 0 | 0 | 0 | 0 | 0 | 0 |
| ENSG00000274732 | 0 | 0 | 0 | 0 | 0 | 0 |
| ENSG00000242385 | 0 | 0 | 0 | 0 | 0 | 0 |
| ENSG00000200372 | 0 | 0 | 0 | 0 | 0 | 0 |
| ENSG00000243014 | 0 | 0 | 0 | 0 | 0 | 0 |
| ENSG00000206889 | 0 | 0 | 0 | 0 | 0 | 0 |
| ENSG00000239268 | 0 | 0 | 0 | 0 | 0 | 0 |
| ENSG00000240393 | 0 | 0 | 0 | 0 | 0 | 0 |
| ENSG00000241213 | 0 | 0 | 0 | 0 | 0 | 0 |
| ENSG00000242816 | 0 | 0 | 0 | 0 | 0 | 0 |
| ENSG00000243276 | 0 | 0 | 0 | 0 | 0 | 0 |
| ENSG00000144847 | 4 | 1 | 1 | 0 | 0 | 0 |
| ENSG00000239877 | 0 | 0 | 0 | 0 | 0 | 0 |
| ENSG00000163424 | 0 | 0 | 0 | 0 | 0 | 0 |
| ENSG00000251012 | 0 | 0 | 0 | 0 | 0 | 0 |
| ENSG00000114638 | 2 | 1 | 2 | 0 | 0 | 4 |
| ENSG00000121578 | 838 | 329 | 492 | 376 | 417 | 425 |
| ENSG00000240254 | 0 | 2 | 0 | 0 | 0 | 0 |
| ENSG00000031081 | 29 | 20 | 19 | 6 | 10 | 12 |
| ENSG00000242829 | 0 | 0 | 0 | 0 | 0 | 0 |
| ENSG00000241155 | 2 | 0 | 1 | 0 | 3 | 1 |
| ENSG00000207310 | 0 | 0 | 0 | 0 | 0 | 0 |
| ENSG00000176142 | 1103 | 498 | 688 | 509 | 588 | 519 |
| ENSG00000163389 | 652 | 348 | 380 | 313 | 362 | 420 |
| ENSG00000272662 | 1 | 1 | 3 | 0 | 1 | 0 |
| ENSG00000113845 | 2570 | 1217 | 1464 | 1397 | 1479 | 1446 |
| ENSG00000121594 | 0 | 0 | 0 | 0 | 0 | 0 |
| ENSG00000272967 | 0 | 0 | 0 | 0 | 0 | 0 |
| ENSG00000144843 | 26 | 18 | 16 | 11 | 18 | 15 |
| ENSG00000144837 | 0 | 0 | 0 | 0 | 0 | 0 |
| ENSG00000242052 | 2 | 0 | 0 | 0 | 0 | 0 |
| ENSG00000121577 | 26 | 5 | 7 | 5 | 14 | 10 |
| ENSG00000138495 | 1366 | 681 | 772 | 947 | 900 | 814 |
| ENSG00000183833 | 21 | 9 | 21 | 1 | 5 | 4 |
| ENSG00000285585 | 0 | 0 | 1 | 0 | 0 | 0 |
| ENSG00000239994 | 0 | 0 | 0 | 0 | 0 | 0 |
| ENSG00000144852 | 4 | 2 | 1 | 3 | 2 | 2 |
| ENSG00000240471 | 2 | 1 | 0 | 0 | 0 | 0 |
| ENSG00000082701 | 3923 | 1761 | 2269 | 1392 | 2001 | 1940 |
| ENSG00000239835 | 0 | 0 | 0 | 0 | 0 | 0 |
| ENSG00000241546 | 0 | 0 | 0 | 0 | 0 | 0 |
| ENSG00000242622 | 66 | 60 | 90 | 35 | 46 | 59 |
| ENSG00000244308 | 0 | 1 | 0 | 1 | 0 | 0 |
| ENSG00000244139 | 0 | 0 | 0 | 0 | 0 | 0 |
| ENSG00000175697 | 2 | 5 | 4 | 6 | 4 | 7 |
| ENSG00000206721 | 0 | 0 | 0 | 0 | 0 | 0 |

| | | | | | | |
|---|---|---|---|---|---|---|
| ENSG00000240882 | 0 | 0 | 4 | 0 | 0 | 0 |
| ENSG00000163428 | 1675 | 801 | 632 | 372 | 1218 | 637 |
| ENSG00000240661 | 7 | 9 | 11 | 3 | 7 | 9 |
| ENSG00000282950 | 7 | 4 | 2 | 1 | 2 | 4 |
| ENSG00000242613 | 0 | 0 | 0 | 0 | 0 | 0 |
| ENSG00000163430 | 8 | 9 | 7 | 11 | 4 | 10 |
| ENSG00000284121 | 0 | 0 | 0 | 0 | 0 | 0 |
| ENSG00000240774 | 0 | 0 | 0 | 0 | 0 | 0 |
| ENSG00000244441 | 0 | 0 | 0 | 0 | 0 | 0 |
| ENSG00000065518 | 3721 | 1836 | 2218 | 2181 | 2111 | 2006 |
| ENSG00000113924 | 2 | 4 | 1 | 4 | 2 | 2 |
| ENSG00000144840 | 1251 | 560 | 734 | 432 | 624 | 583 |
| ENSG00000240452 | 3 | 1 | 0 | 0 | 1 | 2 |
| ENSG00000243257 | 0 | 1 | 0 | 0 | 0 | 0 |
| ENSG00000153767 | 579 | 270 | 377 | 242 | 342 | 334 |
| ENSG00000213371 | 4 | 5 | 6 | 5 | 7 | 4 |
| ENSG00000243593 | 0 | 0 | 0 | 0 | 0 | 0 |
| ENSG00000145087 | 0 | 0 | 0 | 0 | 0 | 0 |
| ENSG00000270782 | 0 | 0 | 0 | 0 | 0 | 0 |
| ENSG00000264477 | 0 | 0 | 0 | 0 | 0 | 0 |
| ENSG00000243813 | 0 | 0 | 0 | 0 | 0 | 0 |
| ENSG00000242103 | 0 | 0 | 0 | 0 | 0 | 0 |
| ENSG00000051341 | 171 | 111 | 133 | 41 | 197 | 115 |
| ENSG00000242445 | 4 | 3 | 4 | 3 | 0 | 0 |
| ENSG00000186103 | 0 | 0 | 0 | 0 | 0 | 0 |
| ENSG00000163833 | 0 | 0 | 0 | 0 | 0 | 0 |
| ENSG00000180353 | 8 | 3 | 2 | 7 | 5 | 3 |
| ENSG00000243544 | 0 | 0 | 0 | 0 | 0 | 0 |
| ENSG00000222057 | 0 | 0 | 0 | 0 | 0 | 0 |
| ENSG00000173230 | 541 | 290 | 330 | 109 | 690 | 472 |
| ENSG00000173226 | 833 | 381 | 498 | 387 | 465 | 494 |
| ENSG00000145088 | 40 | 18 | 29 | 27 | 27 | 25 |
| ENSG00000163406 | 271 | 150 | 150 | 118 | 161 | 164 |
| ENSG00000145103 | 631 | 235 | 373 | 260 | 304 | 315 |
| ENSG00000202388 | 0 | 0 | 0 | 0 | 0 | 0 |
| ENSG00000114013 | 0 | 0 | 0 | 0 | 0 | 1 |
| ENSG00000036828 | 1 | 2 | 3 | 0 | 0 | 0 |
| ENSG00000240236 | 0 | 0 | 0 | 0 | 0 | 0 |
| ENSG00000121552 | 1794 | 724 | 1747 | 251 | 178 | 454 |
| ENSG00000160124 | 412 | 207 | 249 | 239 | 263 | 220 |
| ENSG00000114023 | 1267 | 611 | 979 | 462 | 560 | 645 |
| ENSG00000196981 | 577 | 304 | 363 | 265 | 344 | 265 |
| ENSG00000272758 | 61 | 31 | 38 | 27 | 31 | 31 |
| ENSG00000242531 | 1 | 0 | 4 | 0 | 2 | 0 |
| ENSG00000114030 | 1627 | 874 | 965 | 570 | 1021 | 891 |
| ENSG00000275348 | 0 | 0 | 0 | 0 | 0 | 0 |
| ENSG00000138496 | 739 | 347 | 451 | 203 | 255 | 293 |
| ENSG00000163840 | 724 | 285 | 455 | 250 | 377 | 414 |
| ENSG00000173200 | 0 | 0 | 0 | 2 | 0 | 0 |
| ENSG00000243918 | 0 | 0 | 0 | 0 | 0 | 0 |

| | | | | | | |
|---|---|---|---|---|---|---|
| ENSG00000173193 | 308 | 127 | 176 | 51 | 160 | 122 |
| ENSG00000169087 | 118 | 66 | 92 | 53 | 55 | 70 |
| ENSG00000138463 | 937 | 467 | 587 | 453 | 549 | 552 |
| ENSG00000273033 | 182 | 66 | 73 | 48 | 85 | 66 |
| ENSG00000082684 | 12 | 1 | 4 | 2 | 1 | 1 |
| ENSG00000065485 | 748 | 390 | 539 | 406 | 438 | 437 |
| ENSG00000284421 | 0 | 0 | 0 | 0 | 0 | 0 |
| ENSG00000121542 | 527 | 271 | 277 | 259 | 313 | 271 |
| ENSG00000273454 | 4 | 1 | 0 | 0 | 1 | 2 |
| ENSG00000173175 | 4301 | 2025 | 2686 | 1265 | 1344 | 1717 |
| ENSG00000272678 | 0 | 0 | 0 | 0 | 0 | 0 |
| ENSG00000206527 | 5309 | 2590 | 2905 | 1860 | 3221 | 2635 |
| ENSG00000239523 | 85 | 76 | 97 | 86 | 80 | 66 |
| ENSG00000065534 | 87 | 35 | 61 | 37 | 67 | 60 |
| ENSG00000250174 | 2 | 0 | 0 | 0 | 1 | 0 |
| ENSG00000273123 | 0 | 0 | 0 | 0 | 0 | 0 |
| ENSG00000207002 | 0 | 0 | 0 | 0 | 0 | 0 |
| ENSG00000175455 | 709 | 436 | 423 | 226 | 676 | 459 |
| ENSG00000065371 | 0 | 0 | 1 | 0 | 0 | 0 |
| ENSG00000160145 | 511 | 243 | 355 | 212 | 257 | 256 |
| ENSG00000266383 | 0 | 0 | 0 | 0 | 0 | 0 |
| ENSG00000240622 | 0 | 0 | 0 | 0 | 0 | 0 |
| ENSG00000276656 | 0 | 0 | 0 | 0 | 0 | 0 |
| ENSG00000252751 | 0 | 0 | 0 | 0 | 0 | 0 |
| ENSG00000260391 | 0 | 0 | 0 | 0 | 0 | 0 |
| ENSG00000114491 | 1483 | 684 | 869 | 741 | 789 | 872 |
| ENSG00000265981 | 0 | 0 | 0 | 0 | 1 | 0 |
| ENSG00000242199 | 0 | 2 | 0 | 0 | 1 | 0 |
| ENSG00000082781 | 3608 | 1843 | 2424 | 2162 | 2154 | 2365 |
| ENSG00000244286 | 0 | 0 | 1 | 0 | 0 | 0 |
| ENSG00000280042 | 0 | 4 | 2 | 0 | 2 | 0 |
| ENSG00000276626 | 0 | 0 | 0 | 0 | 0 | 0 |
| ENSG00000243986 | 0 | 0 | 0 | 0 | 0 | 0 |
| ENSG00000173702 | 0 | 1 | 1 | 0 | 0 | 0 |
| ENSG00000173706 | 104 | 44 | 54 | 49 | 47 | 41 |
| ENSG00000252642 | 0 | 0 | 0 | 0 | 0 | 0 |
| ENSG00000221955 | 127 | 63 | 58 | 79 | 80 | 62 |
| ENSG00000199327 | 0 | 0 | 0 | 0 | 0 | 0 |
| ENSG00000264986 | 0 | 0 | 0 | 0 | 0 | 0 |
| ENSG00000163848 | 686 | 421 | 256 | 155 | 644 | 289 |
| ENSG00000229048 | 0 | 0 | 0 | 0 | 0 | 1 |
| ENSG00000243508 | 0 | 0 | 0 | 1 | 0 | 0 |
| ENSG00000252953 | 0 | 0 | 0 | 0 | 0 | 0 |
| ENSG00000114520 | 1539 | 793 | 934 | 741 | 1076 | 880 |
| ENSG00000201800 | 0 | 0 | 0 | 0 | 0 | 0 |
| ENSG00000252965 | 3 | 1 | 5 | 1 | 6 | 1 |
| ENSG00000144909 | 907 | 457 | 481 | 266 | 503 | 419 |
| ENSG00000243072 | 0 | 0 | 0 | 0 | 0 | 0 |
| ENSG00000242001 | 0 | 0 | 0 | 0 | 0 | 0 |
| ENSG00000234458 | 0 | 0 | 2 | 0 | 0 | 0 |

| | | | | | | |
|---|---|---|---|---|---|---|
| ENSG00000243429 | 2 | 0 | 0 | 0 | 0 | 1 |
| ENSG00000228963 | 0 | 0 | 0 | 0 | 0 | 0 |
| ENSG00000239978 | 0 | 0 | 0 | 0 | 0 | 0 |
| ENSG00000179170 | 0 | 0 | 0 | 0 | 0 | 0 |
| ENSG00000239432 | 0 | 0 | 0 | 0 | 0 | 0 |
| ENSG00000284624 | 4 | 2 | 2 | 0 | 0 | 3 |
| ENSG00000284660 | 0 | 0 | 0 | 0 | 0 | 0 |
| ENSG00000272840 | 0 | 0 | 0 | 0 | 0 | 0 |
| ENSG00000239804 | 2 | 2 | 2 | 0 | 0 | 0 |
| ENSG00000221737 | 0 | 0 | 0 | 0 | 1 | 0 |
| ENSG00000242229 | 0 | 0 | 0 | 0 | 0 | 2 |
| ENSG00000241346 | 0 | 0 | 1 | 0 | 0 | 0 |
| ENSG00000241288 | 163 | 96 | 135 | 42 | 94 | 84 |
| ENSG00000241278 | 12 | 8 | 6 | 1 | 7 | 3 |
| ENSG00000248787 | 20 | 9 | 16 | 9 | 7 | 6 |
| ENSG00000171084 | 248 | 143 | 149 | 173 | 135 | 167 |
| ENSG00000189366 | 534 | 321 | 369 | 330 | 255 | 285 |
| ENSG00000241439 | 0 | 0 | 0 | 0 | 1 | 0 |
| ENSG00000114547 | 19 | 13 | 13 | 17 | 11 | 14 |
| ENSG00000114544 | 1856 | 846 | 1149 | 1076 | 1011 | 1058 |
| ENSG00000248607 | 0 | 0 | 1 | 0 | 1 | 0 |
| ENSG00000250012 | 27 | 20 | 22 | 19 | 27 | 19 |
| ENSG00000144908 | 11 | 4 | 6 | 8 | 3 | 6 |
| ENSG00000250218 | 0 | 0 | 2 | 0 | 0 | 0 |
| ENSG00000272020 | 0 | 0 | 0 | 0 | 0 | 0 |
| ENSG00000246022 | 0 | 0 | 0 | 0 | 0 | 0 |
| ENSG00000250934 | 0 | 0 | 0 | 0 | 0 | 0 |
| ENSG00000251448 | 0 | 0 | 0 | 0 | 0 | 0 |
| ENSG00000163884 | 8 | 1 | 2 | 1 | 5 | 3 |
| ENSG00000249833 | 0 | 0 | 0 | 0 | 0 | 0 |
| ENSG00000163885 | 0 | 0 | 4 | 0 | 0 | 0 |
| ENSG00000279328 | 0 | 0 | 2 | 0 | 1 | 3 |
| ENSG00000070476 | 1313 | 528 | 819 | 562 | 679 | 753 |
| ENSG00000159650 | 0 | 0 | 0 | 0 | 0 | 1 |
| ENSG00000180767 | 0 | 0 | 0 | 0 | 0 | 0 |
| ENSG00000180697 | 0 | 0 | 0 | 1 | 0 | 0 |
| ENSG00000201968 | 0 | 0 | 0 | 0 | 0 | 0 |
| ENSG00000197763 | 458 | 236 | 312 | 244 | 307 | 297 |
| ENSG00000248557 | 6 | 1 | 4 | 3 | 1 | 2 |
| ENSG00000198284 | 0 | 0 | 0 | 0 | 0 | 0 |
| ENSG00000248850 | 0 | 0 | 0 | 0 | 0 | 0 |
| ENSG00000159685 | 255 | 123 | 157 | 184 | 176 | 185 |
| ENSG00000249558 | 0 | 0 | 0 | 0 | 0 | 0 |
| ENSG00000280053 | 0 | 0 | 1 | 0 | 2 | 0 |
| ENSG00000114554 | 3992 | 1762 | 2399 | 1668 | 1895 | 2007 |
| ENSG00000214324 | 0 | 0 | 0 | 0 | 0 | 0 |
| ENSG00000212556 | 0 | 0 | 0 | 0 | 0 | 0 |
| ENSG00000243016 | 0 | 0 | 0 | 0 | 0 | 0 |
| ENSG00000201288 | 0 | 0 | 0 | 0 | 0 | 0 |
| ENSG00000239620 | 0 | 0 | 0 | 0 | 0 | 0 |

| | | | | | | |
|---|---|---|---|---|---|---|
| ENSG00000244215 | 0 | 0 | 1 | 0 | 0 | 0 |
| ENSG00000239921 | 0 | 0 | 0 | 0 | 0 | 0 |
| ENSG00000240562 | 0 | 0 | 0 | 0 | 0 | 0 |
| ENSG00000282975 | 0 | 0 | 0 | 0 | 2 | 0 |
| ENSG00000282860 | 0 | 0 | 0 | 0 | 0 | 0 |
| ENSG00000163870 | 1669 | 818 | 964 | 1271 | 1247 | 1039 |
| ENSG00000275067 | 0 | 0 | 0 | 0 | 0 | 1 |
| ENSG00000276083 | 0 | 0 | 0 | 0 | 0 | 0 |
| ENSG00000073111 | 1599 | 819 | 1050 | 889 | 798 | 981 |
| ENSG00000285600 | 4 | 0 | 6 | 0 | 4 | 3 |
| ENSG00000114631 | 1355 | 726 | 779 | 1039 | 908 | 887 |
| ENSG00000114626 | 667 | 346 | 453 | 383 | 393 | 338 |
| ENSG00000074416 | 2539 | 1475 | 1782 | 1641 | 1837 | 1720 |
| ENSG00000187715 | 0 | 1 | 0 | 0 | 1 | 1 |
| ENSG00000201210 | 0 | 0 | 0 | 0 | 0 | 0 |
| ENSG00000058262 | 12335 | 6092 | 7575 | 7365 | 7478 | 7284 |
| ENSG00000175792 | 2318 | 1206 | 1435 | 1836 | 1632 | 1688 |
| ENSG00000222627 | 0 | 0 | 0 | 0 | 0 | 1 |
| ENSG00000239608 | 0 | 0 | 1 | 0 | 0 | 0 |
| ENSG00000212344 | 0 | 0 | 0 | 0 | 0 | 0 |
| ENSG00000132394 | 1046 | 494 | 646 | 676 | 504 | 532 |
| ENSG00000285619 | 0 | 0 | 0 | 0 | 0 | 0 |
| ENSG00000179407 | 0 | 0 | 0 | 0 | 0 | 0 |
| ENSG00000242049 | 0 | 0 | 0 | 0 | 0 | 0 |
| ENSG00000179348 | 2516 | 1116 | 1367 | 1109 | 1057 | 1206 |
| ENSG00000244300 | 18 | 13 | 14 | 5 | 19 | 16 |
| ENSG00000239405 | 1 | 0 | 2 | 0 | 0 | 0 |
| ENSG00000198685 | 0 | 0 | 0 | 0 | 0 | 0 |
| ENSG00000163902 | 10592 | 5122 | 6088 | 6252 | 6056 | 6010 |
| ENSG00000277250 | 0 | 0 | 0 | 0 | 0 | 1 |
| ENSG00000242551 | 2 | 0 | 1 | 0 | 0 | 0 |
| ENSG00000207130 | 0 | 0 | 0 | 0 | 0 | 0 |
| ENSG00000075785 | 11568 | 5653 | 6534 | 5432 | 6487 | 6605 |
| ENSG00000239503 | 1 | 0 | 0 | 0 | 0 | 0 |
| ENSG00000242992 | 0 | 3 | 1 | 1 | 0 | 1 |
| ENSG00000244232 | 0 | 0 | 0 | 0 | 0 | 1 |
| ENSG00000239483 | 0 | 0 | 0 | 0 | 0 | 2 |
| ENSG00000244065 | 0 | 0 | 0 | 0 | 0 | 0 |
| ENSG00000243022 | 0 | 0 | 0 | 0 | 0 | 0 |
| ENSG00000242062 | 0 | 0 | 0 | 0 | 0 | 0 |
| ENSG00000261159 | 18 | 12 | 9 | 9 | 7 | 7 |
| ENSG00000231305 | 12 | 9 | 20 | 10 | 16 | 16 |
| ENSG00000250796 | 3 | 2 | 2 | 0 | 0 | 0 |
| ENSG00000177646 | 2005 | 939 | 1228 | 1136 | 1089 | 1142 |
| ENSG00000114656 | 185 | 76 | 142 | 66 | 72 | 59 |
| ENSG00000279507 | 3 | 0 | 1 | 1 | 1 | 1 |
| ENSG00000114654 | 10 | 9 | 16 | 7 | 6 | 5 |
| ENSG00000249505 | 0 | 0 | 0 | 0 | 0 | 0 |
| ENSG00000169704 | 0 | 0 | 0 | 0 | 0 | 0 |
| ENSG00000172780 | 97 | 59 | 91 | 62 | 72 | 95 |

| | | | | | | |
|---|---|---|---|---|---|---|
| ENSG00000261796 | 0 | 0 | 0 | 0 | 0 | 0 |
| ENSG00000273174 | 5 | 5 | 4 | 6 | 2 | 8 |
| ENSG00000240682 | 577 | 347 | 418 | 320 | 362 | 341 |
| ENSG00000273437 | 2 | 6 | 4 | 3 | 4 | 5 |
| ENSG00000169714 | 10950 | 5647 | 6822 | 6063 | 7096 | 7080 |
| ENSG00000241673 | 0 | 0 | 0 | 0 | 0 | 0 |
| ENSG00000181789 | 10500 | 4828 | 6244 | 5020 | 5524 | 5360 |
| ENSG00000278658 | 0 | 1 | 1 | 0 | 2 | 1 |
| ENSG00000183624 | 3667 | 1749 | 2246 | 1897 | 1882 | 1991 |
| ENSG00000184897 | 1813 | 1060 | 1390 | 1503 | 1192 | 1228 |
| ENSG00000206417 | 97 | 45 | 57 | 29 | 42 | 33 |
| ENSG00000248916 | 1 | 1 | 0 | 0 | 1 | 0 |
| ENSG00000270492 | 0 | 1 | 1 | 1 | 0 | 0 |
| ENSG00000270773 | 0 | 0 | 1 | 0 | 0 | 2 |
| ENSG00000244932 | 40 | 31 | 55 | 14 | 35 | 27 |
| ENSG00000251474 | 491 | 194 | 307 | 134 | 221 | 197 |
| ENSG00000207088 | 6 | 3 | 3 | 0 | 1 | 3 |
| ENSG00000172771 | 22 | 8 | 6 | 7 | 5 | 6 |
| ENSG00000129071 | 1952 | 1177 | 1314 | 776 | 1429 | 1091 |
| ENSG00000163913 | 1636 | 847 | 1182 | 847 | 914 | 957 |
| ENSG00000163914 | 2 | 1 | 1 | 2 | 1 | 1 |
| ENSG00000178804 | 0 | 0 | 3 | 0 | 0 | 0 |
| ENSG00000004399 | 8465 | 4017 | 5098 | 3817 | 4119 | 4219 |
| ENSG00000239437 | 0 | 0 | 0 | 0 | 0 | 0 |
| ENSG00000248659 | 4 | 0 | 0 | 0 | 0 | 2 |
| ENSG00000172765 | 1649 | 833 | 891 | 585 | 855 | 791 |
| ENSG00000202412 | 0 | 1 | 0 | 0 | 0 | 0 |
| ENSG00000283509 | 0 | 0 | 0 | 0 | 0 | 0 |
| ENSG00000203644 | 245 | 118 | 183 | 79 | 143 | 136 |
| ENSG00000271270 | 108 | 48 | 79 | 40 | 61 | 48 |
| ENSG00000250643 | 1 | 0 | 2 | 1 | 0 | 0 |
| ENSG00000170893 | 0 | 0 | 0 | 0 | 0 | 0 |
| ENSG00000248459 | 0 | 0 | 0 | 0 | 0 | 0 |
| ENSG00000284731 | 0 | 1 | 0 | 0 | 0 | 0 |
| ENSG00000249869 | 0 | 0 | 0 | 0 | 0 | 0 |
| ENSG00000180770 | 0 | 0 | 0 | 0 | 0 | 0 |
| ENSG00000251178 | 0 | 0 | 0 | 0 | 0 | 0 |
| ENSG00000251610 | 0 | 0 | 0 | 0 | 0 | 0 |
| ENSG00000251287 | 27 | 17 | 29 | 14 | 15 | 10 |
| ENSG00000248243 | 2 | 1 | 3 | 3 | 4 | 2 |
| ENSG00000253540 | 245 | 172 | 211 | 72 | 88 | 101 |
| ENSG00000249846 | 313 | 153 | 210 | 51 | 56 | 62 |
| ENSG00000248618 | 0 | 0 | 1 | 0 | 0 | 0 |
| ENSG00000250580 | 0 | 0 | 0 | 0 | 0 | 0 |
| ENSG00000228252 | 10 | 6 | 5 | 0 | 1 | 0 |
| ENSG00000249253 | 0 | 0 | 0 | 0 | 0 | 0 |
| ENSG00000172752 | 0 | 0 | 0 | 0 | 0 | 0 |
| ENSG00000206384 | 0 | 0 | 0 | 2 | 2 | 0 |
| ENSG00000196455 | 1170 | 545 | 583 | 377 | 717 | 506 |
| ENSG00000222783 | 0 | 0 | 0 | 0 | 0 | 0 |

| | | | | | | |
|---|---|---|---|---|---|---|
| ENSG00000285631 | 0 | 0 | 0 | 0 | 0 | 0 |
| ENSG00000232829 | 0 | 0 | 0 | 0 | 0 | 0 |
| ENSG00000017260 | 2160 | 1194 | 1222 | 942 | 1580 | 1282 |
| ENSG00000284095 | 0 | 0 | 0 | 0 | 0 | 0 |
| ENSG00000250592 | 0 | 0 | 0 | 0 | 0 | 0 |
| ENSG00000252894 | 0 | 0 | 0 | 0 | 0 | 0 |
| ENSG00000249098 | 0 | 0 | 0 | 0 | 0 | 0 |
| ENSG00000251579 | 0 | 0 | 0 | 0 | 0 | 0 |
| ENSG00000034533 | 250 | 137 | 165 | 112 | 127 | 136 |
| ENSG00000114670 | 344 | 133 | 190 | 133 | 176 | 150 |
| ENSG00000250129 | 0 | 0 | 0 | 0 | 1 | 0 |
| ENSG00000201550 | 0 | 0 | 0 | 0 | 0 | 0 |
| ENSG00000250608 | 1 | 5 | 2 | 2 | 3 | 2 |
| ENSG00000246082 | 2 | 4 | 0 | 1 | 2 | 0 |
| ENSG00000198585 | 2159 | 964 | 1355 | 846 | 1061 | 1059 |
| ENSG00000261167 | 20 | 5 | 6 | 4 | 15 | 12 |
| ENSG00000114686 | 2897 | 1489 | 1770 | 1861 | 1976 | 2050 |
| ENSG00000249020 | 0 | 2 | 0 | 0 | 0 | 2 |
| ENSG00000248468 | 2 | 0 | 0 | 0 | 0 | 0 |
| ENSG00000248530 | 0 | 1 | 0 | 0 | 0 | 0 |
| ENSG00000196353 | 3 | 1 | 5 | 3 | 6 | 9 |
| ENSG00000251447 | 0 | 0 | 0 | 0 | 0 | 0 |
| ENSG00000265859 | 0 | 0 | 0 | 0 | 0 | 0 |
| ENSG00000240854 | 2 | 0 | 3 | 1 | 0 | 0 |
| ENSG00000242899 | 0 | 0 | 0 | 0 | 0 | 0 |
| ENSG00000014257 | 78 | 33 | 49 | 18 | 35 | 13 |
| ENSG00000240890 | 0 | 1 | 0 | 0 | 0 | 0 |
| ENSG00000213264 | 0 | 0 | 0 | 0 | 0 | 0 |
| ENSG00000138246 | 820 | 503 | 392 | 161 | 1018 | 456 |
| ENSG00000240303 | 2 | 6 | 17 | 1 | 3 | 6 |
| ENSG00000274810 | 1 | 1 | 0 | 0 | 0 | 1 |
| ENSG00000129048 | 1 | 1 | 0 | 0 | 1 | 1 |
| ENSG00000251614 | 0 | 0 | 0 | 0 | 0 | 0 |
| ENSG00000081307 | 1874 | 1077 | 1173 | 667 | 1166 | 950 |
| ENSG00000113971 | 190 | 110 | 125 | 33 | 108 | 94 |
| ENSG00000248724 | 0 | 1 | 0 | 0 | 0 | 0 |
| ENSG00000249725 | 0 | 0 | 0 | 0 | 0 | 0 |
| ENSG00000144868 | 0 | 0 | 0 | 0 | 0 | 0 |
| ENSG00000251011 | 0 | 0 | 0 | 0 | 0 | 0 |
| ENSG00000170819 | 0 | 0 | 0 | 6 | 1 | 0 |
| ENSG00000249820 | 0 | 0 | 0 | 0 | 0 | 0 |
| ENSG00000249993 | 0 | 0 | 0 | 0 | 0 | 0 |
| ENSG00000214301 | 0 | 0 | 0 | 0 | 0 | 0 |
| ENSG00000272832 | 2 | 0 | 1 | 0 | 0 | 0 |
| ENSG00000250983 | 0 | 0 | 0 | 0 | 0 | 0 |
| ENSG00000201827 | 0 | 0 | 0 | 0 | 0 | 0 |
| ENSG00000091527 | 5028 | 2408 | 2845 | 2526 | 2799 | 2989 |
| ENSG00000163781 | 809 | 546 | 427 | 223 | 827 | 490 |
| ENSG00000242337 | 0 | 2 | 2 | 1 | 1 | 1 |
| ENSG00000252641 | 1 | 0 | 0 | 1 | 0 | 0 |

| | | | | | | |
|---|---|---|---|---|---|---|
| ENSG00000201394 | 0 | 0 | 0 | 0 | 0 | 0 |
| ENSG00000091513 | 11 | 1 | 4 | 3 | 5 | 3 |
| ENSG00000244062 | 0 | 0 | 0 | 0 | 0 | 0 |
| ENSG00000144867 | 1884 | 918 | 1082 | 1091 | 1070 | 1198 |
| ENSG00000285908 | 0 | 0 | 0 | 0 | 0 | 0 |
| ENSG00000154917 | 518 | 271 | 334 | 226 | 268 | 260 |
| ENSG00000174640 | 10 | 6 | 4 | 0 | 3 | 0 |
| ENSG00000243832 | 0 | 0 | 0 | 0 | 0 | 0 |
| ENSG00000163785 | 2018 | 924 | 1213 | 898 | 1115 | 1085 |
| ENSG00000231845 | 0 | 1 | 0 | 1 | 0 | 0 |
| ENSG00000240006 | 2 | 1 | 4 | 2 | 1 | 1 |
| ENSG00000260633 | 10 | 1 | 5 | 1 | 0 | 1 |
| ENSG00000214289 | 1 | 0 | 1 | 0 | 0 | 1 |
| ENSG00000114019 | 3526 | 1754 | 2399 | 1602 | 1767 | 1969 |
| ENSG00000277723 | 2 | 0 | 0 | 0 | 1 | 0 |
| ENSG00000248377 | 0 | 0 | 0 | 0 | 0 | 0 |
| ENSG00000214288 | 0 | 0 | 0 | 0 | 0 | 0 |
| ENSG00000284561 | 0 | 0 | 0 | 0 | 0 | 0 |
| ENSG00000129055 | 1734 | 975 | 936 | 932 | 1253 | 959 |
| ENSG00000182923 | 294 | 170 | 160 | 89 | 251 | 186 |
| ENSG00000249691 | 0 | 1 | 0 | 0 | 0 | 0 |
| ENSG00000154928 | 0 | 3 | 2 | 1 | 1 | 1 |
| ENSG00000174611 | 2 | 1 | 6 | 3 | 3 | 0 |
| ENSG00000253087 | 0 | 0 | 0 | 0 | 0 | 0 |
| ENSG00000201413 | 0 | 0 | 0 | 0 | 0 | 0 |
| ENSG00000240086 | 58 | 31 | 36 | 23 | 17 | 22 |
| ENSG00000242222 | 0 | 0 | 1 | 0 | 0 | 0 |
| ENSG00000241905 | 0 | 0 | 0 | 0 | 0 | 0 |
| ENSG00000073711 | 229 | 168 | 109 | 67 | 243 | 132 |
| ENSG00000227267 | 0 | 0 | 0 | 0 | 0 | 0 |
| ENSG00000273455 | 2 | 0 | 0 | 0 | 1 | 0 |
| ENSG00000174579 | 966 | 404 | 534 | 351 | 511 | 464 |
| ENSG00000241438 | 0 | 1 | 0 | 0 | 0 | 0 |
| ENSG00000244740 | 0 | 0 | 0 | 0 | 0 | 0 |
| ENSG00000114054 | 3126 | 1572 | 1963 | 1611 | 1672 | 1717 |
| ENSG00000118007 | 704 | 406 | 313 | 186 | 568 | 332 |
| ENSG00000200571 | 0 | 0 | 0 | 0 | 0 | 0 |
| ENSG00000244101 | 0 | 0 | 0 | 0 | 0 | 0 |
| ENSG00000252914 | 0 | 0 | 0 | 0 | 0 | 0 |
| ENSG00000240695 | 4 | 1 | 1 | 0 | 3 | 1 |
| ENSG00000261758 | 2 | 6 | 6 | 5 | 4 | 3 |
| ENSG00000242968 | 2 | 1 | 4 | 0 | 1 | 2 |
| ENSG00000168917 | 3 | 3 | 6 | 4 | 1 | 5 |
| ENSG00000273486 | 64 | 34 | 39 | 37 | 48 | 43 |
| ENSG00000239213 | 71 | 78 | 97 | 67 | 84 | 81 |
| ENSG00000158092 | 572 | 288 | 346 | 231 | 352 | 320 |
| ENSG00000214283 | 0 | 0 | 0 | 0 | 0 | 0 |
| ENSG00000174564 | 20 | 21 | 11 | 12 | 20 | 22 |
| ENSG00000249407 | 0 | 1 | 0 | 0 | 0 | 1 |
| ENSG00000201325 | 0 | 0 | 0 | 0 | 0 | 0 |

| | | | | | | |
|---|---|---|---|---|---|---|
| ENSG00000243886 | 2 | 0 | 1 | 0 | 1 | 0 |
| ENSG00000242070 | 0 | 0 | 0 | 0 | 0 | 0 |
| ENSG00000168875 | 0 | 0 | 0 | 0 | 0 | 0 |
| ENSG00000239513 | 0 | 0 | 0 | 0 | 0 | 0 |
| ENSG00000261146 | 0 | 0 | 0 | 0 | 0 | 0 |
| ENSG00000241478 | 0 | 0 | 2 | 0 | 0 | 0 |
| ENSG00000066405 | 0 | 0 | 0 | 1 | 0 | 0 |
| ENSG00000272609 | 0 | 0 | 0 | 0 | 0 | 0 |
| ENSG00000158163 | 0 | 2 | 1 | 1 | 0 | 0 |
| ENSG00000240668 | 0 | 2 | 2 | 2 | 2 | 0 |
| ENSG00000118017 | 0 | 0 | 0 | 0 | 0 | 0 |
| ENSG00000138231 | 764 | 371 | 460 | 322 | 464 | 447 |
| ENSG00000114098 | 1049 | 570 | 589 | 394 | 796 | 614 |
| ENSG00000181322 | 6 | 6 | 14 | 3 | 4 | 2 |
| ENSG00000158186 | 74 | 37 | 55 | 46 | 46 | 42 |
| ENSG00000158220 | 118 | 47 | 66 | 46 | 66 | 68 |
| ENSG00000280399 | 8 | 2 | 6 | 1 | 2 | 6 |
| ENSG00000114107 | 281 | 247 | 183 | 107 | 323 | 167 |
| ENSG00000158234 | 371 | 192 | 200 | 147 | 253 | 178 |
| ENSG00000174715 | 0 | 0 | 0 | 0 | 0 | 3 |
| ENSG00000051382 | 2144 | 1078 | 1251 | 778 | 1283 | 1075 |
| ENSG00000239207 | 0 | 0 | 1 | 1 | 0 | 0 |
| ENSG00000242321 | 0 | 0 | 0 | 0 | 0 | 0 |
| ENSG00000241429 | 2 | 4 | 1 | 3 | 1 | 2 |
| ENSG00000242244 | 1 | 0 | 1 | 0 | 0 | 0 |
| ENSG00000244578 | 6 | 0 | 0 | 0 | 0 | 0 |
| ENSG00000183770 | 0 | 0 | 0 | 0 | 0 | 0 |
| ENSG00000206262 | 0 | 2 | 1 | 2 | 0 | 3 |
| ENSG00000206260 | 0 | 0 | 0 | 0 | 0 | 0 |
| ENSG00000175110 | 1177 | 627 | 729 | 597 | 714 | 685 |
| ENSG00000184814 | 0 | 0 | 0 | 0 | 0 | 0 |
| ENSG00000233701 | 0 | 0 | 0 | 0 | 0 | 0 |
| ENSG00000251137 | 0 | 0 | 0 | 0 | 0 | 0 |
| ENSG00000232416 | 0 | 0 | 0 | 0 | 0 | 0 |
| ENSG00000281473 | 0 | 0 | 0 | 0 | 0 | 0 |
| ENSG00000276304 | 0 | 0 | 0 | 0 | 0 | 0 |
| ENSG00000272656 | 10 | 1 | 6 | 1 | 1 | 2 |
| ENSG00000184432 | 8314 | 4280 | 4971 | 3303 | 4951 | 4220 |
| ENSG00000283418 | 0 | 6 | 2 | 1 | 2 | 3 |
| ENSG00000248932 | 121 | 67 | 91 | 75 | 75 | 50 |
| ENSG00000114113 | 1 | 2 | 2 | 5 | 1 | 4 |
| ENSG00000248790 | 0 | 0 | 0 | 0 | 0 | 0 |
| ENSG00000178631 | 31 | 14 | 12 | 14 | 8 | 9 |
| ENSG00000114115 | 20 | 8 | 5 | 5 | 4 | 2 |
| ENSG00000163864 | 73 | 30 | 54 | 64 | 57 | 56 |
| ENSG00000214280 | 4 | 4 | 2 | 0 | 0 | 2 |
| ENSG00000201600 | 0 | 0 | 1 | 0 | 0 | 0 |
| ENSG00000244692 | 0 | 0 | 0 | 0 | 0 | 0 |
| ENSG00000261763 | 0 | 0 | 0 | 0 | 0 | 0 |
| ENSG00000250543 | 0 | 0 | 0 | 0 | 0 | 0 |

| | | | | | | |
|---|---|---|---|---|---|---|
| ENSG00000251471 | 0 | 0 | 0 | 0 | 0 | 0 |
| ENSG00000250161 | 0 | 0 | 0 | 0 | 0 | 0 |
| ENSG00000158258 | 1904 | 816 | 938 | 1144 | 1460 | 1517 |
| ENSG00000249290 | 0 | 0 | 0 | 0 | 0 | 0 |
| ENSG00000251058 | 0 | 0 | 1 | 0 | 0 | 0 |
| ENSG00000250433 | 2 | 0 | 0 | 0 | 0 | 1 |
| ENSG00000249305 | 0 | 0 | 0 | 0 | 0 | 0 |
| ENSG00000155890 | 0 | 0 | 0 | 0 | 0 | 0 |
| ENSG00000261826 | 0 | 0 | 0 | 0 | 0 | 0 |
| ENSG00000224085 | 0 | 0 | 0 | 0 | 0 | 0 |
| ENSG00000114120 | 2560 | 1293 | 1200 | 767 | 1783 | 1259 |
| ENSG00000248773 | 5 | 0 | 0 | 1 | 3 | 1 |
| ENSG00000175093 | 52 | 26 | 21 | 35 | 24 | 33 |
| ENSG00000251270 | 0 | 0 | 0 | 0 | 0 | 0 |
| ENSG00000155893 | 397 | 188 | 216 | 151 | 173 | 203 |
| ENSG00000241400 | 0 | 0 | 1 | 0 | 0 | 0 |
| ENSG00000249417 | 0 | 0 | 1 | 0 | 0 | 1 |
| ENSG00000249540 | 0 | 0 | 0 | 0 | 0 | 0 |
| ENSG00000177311 | 446 | 254 | 202 | 110 | 376 | 218 |
| ENSG00000215606 | 0 | 0 | 0 | 0 | 0 | 0 |
| ENSG00000155903 | 270 | 145 | 134 | 60 | 198 | 113 |
| ENSG00000250170 | 0 | 0 | 0 | 0 | 0 | 0 |
| ENSG00000249986 | 0 | 0 | 0 | 0 | 0 | 0 |
| ENSG00000242104 | 0 | 1 | 0 | 0 | 0 | 0 |
| ENSG00000241899 | 0 | 0 | 0 | 0 | 0 | 0 |
| ENSG00000241526 | 0 | 0 | 0 | 0 | 0 | 0 |
| ENSG00000114125 | 2947 | 1574 | 1884 | 1924 | 1792 | 1812 |
| ENSG00000285558 | 0 | 0 | 0 | 0 | 0 | 0 |
| ENSG00000114124 | 0 | 1 | 5 | 0 | 0 | 3 |
| ENSG00000228398 | 0 | 0 | 0 | 0 | 0 | 0 |
| ENSG00000069849 | 2590 | 1344 | 1413 | 1307 | 1570 | 1418 |
| ENSG00000200389 | 0 | 0 | 0 | 0 | 0 | 0 |
| ENSG00000244124 | 1 | 2 | 3 | 3 | 1 | 4 |
| ENSG00000279147 | 2 | 0 | 0 | 0 | 0 | 0 |
| ENSG00000114126 | 1879 | 1015 | 1147 | 669 | 888 | 884 |
| ENSG00000233597 | 0 | 0 | 2 | 0 | 2 | 0 |
| ENSG00000206604 | 0 | 0 | 0 | 0 | 0 | 0 |
| ENSG00000175066 | 881 | 444 | 597 | 277 | 477 | 436 |
| ENSG00000114127 | 757 | 386 | 281 | 74 | 531 | 285 |
| ENSG00000251804 | 0 | 0 | 0 | 0 | 0 | 0 |
| ENSG00000202125 | 0 | 0 | 0 | 0 | 0 | 0 |
| ENSG00000175054 | 404 | 265 | 192 | 101 | 438 | 248 |
| ENSG00000242479 | 0 | 0 | 0 | 0 | 0 | 0 |
| ENSG00000244327 | 0 | 0 | 0 | 0 | 0 | 0 |
| ENSG00000242390 | 1 | 3 | 0 | 0 | 0 | 0 |
| ENSG00000252745 | 0 | 0 | 0 | 0 | 0 | 0 |
| ENSG00000120756 | 1133 | 656 | 606 | 293 | 782 | 516 |
| ENSG00000239641 | 0 | 0 | 0 | 0 | 0 | 0 |
| ENSG00000199319 | 0 | 0 | 0 | 0 | 0 | 0 |
| ENSG00000144935 | 200 | 116 | 137 | 67 | 131 | 103 |

| | | | | | | |
|---|---|---|---|---|---|---|
| ENSG00000251787 | 0 | 1 | 0 | 0 | 0 | 2 |
| ENSG00000163710 | 12 | 8 | 4 | 9 | 5 | 6 |
| ENSG00000240950 | 0 | 1 | 2 | 3 | 2 | 0 |
| ENSG00000243818 | 0 | 0 | 0 | 0 | 0 | 0 |
| ENSG00000188582 | 0 | 0 | 0 | 0 | 0 | 0 |
| ENSG00000241592 | 0 | 0 | 0 | 0 | 0 | 0 |
| ENSG00000241570 | 0 | 0 | 0 | 0 | 0 | 0 |
| ENSG00000163714 | 1592 | 1069 | 724 | 389 | 1598 | 918 |
| ENSG00000268129 | 0 | 0 | 0 | 0 | 0 | 1 |
| ENSG00000243733 | 0 | 0 | 0 | 0 | 0 | 0 |
| ENSG00000175040 | 0 | 0 | 0 | 0 | 0 | 0 |
| ENSG00000241679 | 0 | 0 | 0 | 0 | 0 | 0 |
| ENSG00000244171 | 0 | 1 | 2 | 2 | 1 | 2 |
| ENSG00000181804 | 0 | 0 | 0 | 0 | 0 | 1 |
| ENSG00000240888 | 0 | 0 | 0 | 0 | 0 | 0 |
| ENSG00000240012 | 0 | 0 | 0 | 0 | 0 | 0 |
| ENSG00000244493 | 0 | 0 | 0 | 0 | 0 | 0 |
| ENSG00000243033 | 0 | 0 | 0 | 0 | 0 | 0 |
| ENSG00000243759 | 0 | 2 | 0 | 0 | 0 | 0 |
| ENSG00000248374 | 0 | 0 | 0 | 0 | 0 | 0 |
| ENSG00000235110 | 0 | 0 | 0 | 0 | 0 | 0 |
| ENSG00000181744 | 971 | 433 | 597 | 356 | 431 | 456 |
| ENSG00000222778 | 0 | 0 | 0 | 0 | 0 | 0 |
| ENSG00000280397 | 0 | 0 | 0 | 0 | 0 | 0 |
| ENSG00000244024 | 0 | 0 | 0 | 0 | 0 | 0 |
| ENSG00000241695 | 0 | 0 | 0 | 0 | 0 | 0 |
| ENSG00000239797 | 2 | 0 | 0 | 0 | 0 | 1 |
| ENSG00000244358 | 0 | 0 | 0 | 0 | 0 | 0 |
| ENSG00000240095 | 0 | 0 | 0 | 0 | 0 | 0 |
| ENSG00000261051 | 0 | 2 | 0 | 0 | 1 | 2 |
| ENSG00000243415 | 0 | 0 | 0 | 0 | 0 | 0 |
| ENSG00000240032 | 0 | 0 | 3 | 0 | 1 | 3 |
| ENSG00000152952 | 1275 | 705 | 758 | 390 | 879 | 626 |
| ENSG00000114698 | 6 | 6 | 2 | 1 | 9 | 4 |
| ENSG00000243596 | 0 | 0 | 0 | 0 | 0 | 0 |
| ENSG00000241358 | 0 | 0 | 0 | 0 | 1 | 0 |
| ENSG00000163746 | 0 | 0 | 0 | 0 | 0 | 0 |
| ENSG00000279320 | 0 | 0 | 0 | 0 | 2 | 0 |
| ENSG00000188313 | 406 | 181 | 260 | 173 | 245 | 249 |
| ENSG00000199812 | 0 | 0 | 0 | 0 | 0 | 0 |
| ENSG00000231213 | 0 | 0 | 0 | 0 | 0 | 0 |
| ENSG00000241457 | 0 | 0 | 0 | 0 | 0 | 0 |
| ENSG00000243620 | 0 | 0 | 0 | 0 | 0 | 0 |
| ENSG00000242671 | 0 | 0 | 0 | 0 | 0 | 0 |
| ENSG00000207156 | 0 | 0 | 0 | 0 | 0 | 0 |
| ENSG00000239412 | 0 | 0 | 0 | 0 | 0 | 0 |
| ENSG00000174963 | 429 | 205 | 272 | 140 | 193 | 202 |
| ENSG00000241202 | 0 | 0 | 0 | 0 | 0 | 0 |
| ENSG00000152977 | 770 | 309 | 492 | 380 | 463 | 474 |
| ENSG00000240804 | 0 | 0 | 0 | 0 | 0 | 0 |

| | | | | | | |
|---|---|---|---|---|---|---|
| ENSG00000239661 | 0 | 0 | 0 | 0 | 0 | 0 |
| ENSG00000239922 | 0 | 0 | 0 | 0 | 0 | 0 |
| ENSG00000285798 | 0 | 0 | 0 | 0 | 0 | 0 |
| ENSG00000241131 | 0 | 0 | 0 | 0 | 0 | 0 |
| ENSG00000285660 | 0 | 0 | 0 | 0 | 0 | 0 |
| ENSG00000241792 | 0 | 0 | 0 | 0 | 0 | 0 |
| ENSG00000244650 | 0 | 0 | 0 | 0 | 0 | 0 |
| ENSG00000205412 | 0 | 0 | 0 | 0 | 0 | 0 |
| ENSG00000243347 | 0 | 0 | 0 | 0 | 0 | 0 |
| ENSG00000242440 | 0 | 0 | 0 | 0 | 0 | 0 |
| ENSG00000285557 | 2 | 0 | 0 | 0 | 0 | 1 |
| ENSG00000241993 | 0 | 0 | 0 | 0 | 0 | 0 |
| ENSG00000144891 | 0 | 0 | 0 | 0 | 0 | 0 |
| ENSG00000153002 | 0 | 0 | 2 | 0 | 0 | 0 |
| ENSG00000244089 | 0 | 0 | 0 | 0 | 0 | 0 |
| ENSG00000240521 | 0 | 1 | 0 | 2 | 0 | 0 |
| ENSG00000163751 | 0 | 0 | 0 | 0 | 0 | 0 |
| ENSG00000241627 | 6 | 4 | 2 | 0 | 5 | 2 |
| ENSG00000163754 | 2769 | 1378 | 1609 | 1266 | 1590 | 1499 |
| ENSG00000071794 | 1196 | 701 | 657 | 289 | 823 | 545 |
| ENSG00000239718 | 0 | 0 | 0 | 0 | 0 | 0 |
| ENSG00000201031 | 0 | 0 | 0 | 0 | 0 | 0 |
| ENSG00000240511 | 0 | 0 | 0 | 0 | 0 | 0 |
| ENSG00000163755 | 1722 | 873 | 991 | 464 | 920 | 711 |
| ENSG00000047457 | 13 | 10 | 4 | 4 | 3 | 6 |
| ENSG00000240216 | 49 | 23 | 50 | 10 | 20 | 22 |
| ENSG00000244468 | 79 | 49 | 64 | 11 | 18 | 17 |
| ENSG00000163762 | 14 | 7 | 14 | 3 | 11 | 7 |
| ENSG00000243822 | 0 | 0 | 0 | 0 | 0 | 0 |
| ENSG00000169908 | 20 | 9 | 25 | 8 | 15 | 15 |
| ENSG00000240541 | 2 | 4 | 3 | 0 | 4 | 4 |
| ENSG00000243885 | 3 | 6 | 6 | 4 | 7 | 3 |
| ENSG00000240246 | 0 | 0 | 0 | 0 | 0 | 0 |
| ENSG00000241929 | 0 | 0 | 0 | 0 | 0 | 0 |
| ENSG00000282502 | 0 | 0 | 0 | 0 | 0 | 0 |
| ENSG00000239408 | 0 | 0 | 0 | 0 | 0 | 0 |
| ENSG00000169903 | 2 | 3 | 1 | 2 | 1 | 2 |
| ENSG00000244503 | 0 | 0 | 0 | 0 | 0 | 0 |
| ENSG00000018408 | 465 | 240 | 304 | 165 | 237 | 216 |
| ENSG00000252581 | 0 | 0 | 0 | 0 | 0 | 0 |
| ENSG00000241985 | 0 | 0 | 0 | 1 | 0 | 0 |
| ENSG00000241313 | 26 | 9 | 16 | 8 | 11 | 12 |
| ENSG00000114744 | 1102 | 501 | 565 | 470 | 670 | 588 |
| ENSG00000206199 | 0 | 0 | 1 | 0 | 0 | 1 |
| ENSG00000251854 | 0 | 0 | 0 | 0 | 0 | 0 |
| ENSG00000082996 | 1203 | 642 | 710 | 464 | 810 | 686 |
| ENSG00000252172 | 0 | 0 | 0 | 0 | 0 | 0 |
| ENSG00000239351 | 0 | 0 | 0 | 0 | 0 | 2 |
| ENSG00000070087 | 8620 | 4538 | 5390 | 4463 | 5458 | 5112 |
| ENSG00000242791 | 1 | 1 | 4 | 1 | 2 | 1 |

| | | | | | | |
|---|---|---|---|---|---|---|
| ENSG00000224831 | 23 | 16 | 7 | 16 | 8 | 17 |
| ENSG00000182000 | 2 | 0 | 0 | 1 | 0 | 0 |
| ENSG00000241947 | 0 | 0 | 0 | 0 | 0 | 0 |
| ENSG00000243944 | 0 | 1 | 0 | 0 | 0 | 0 |
| ENSG00000240477 | 0 | 0 | 0 | 0 | 0 | 0 |
| ENSG00000243321 | 0 | 0 | 0 | 0 | 0 | 0 |
| ENSG00000241143 | 0 | 0 | 0 | 0 | 0 | 0 |
| ENSG00000241048 | 0 | 1 | 0 | 1 | 2 | 2 |
| ENSG00000244541 | 49 | 33 | 29 | 18 | 32 | 18 |
| ENSG00000278234 | 1 | 0 | 0 | 0 | 1 | 0 |
| ENSG00000243550 | 6 | 6 | 3 | 11 | 14 | 13 |
| ENSG00000196428 | 1864 | 828 | 1042 | 837 | 1104 | 1079 |
| ENSG00000120742 | 5720 | 2620 | 3300 | 2725 | 3225 | 3016 |
| ENSG00000144895 | 2492 | 1477 | 1558 | 1236 | 2101 | 1549 |
| ENSG00000198843 | 4895 | 2438 | 2776 | 2326 | 2722 | 2652 |
| ENSG00000271711 | 0 | 1 | 0 | 0 | 1 | 0 |
| ENSG00000163645 | 22 | 3 | 10 | 5 | 2 | 5 |
| ENSG00000240137 | 56 | 27 | 50 | 43 | 31 | 36 |
| ENSG00000244668 | 12 | 2 | 4 | 5 | 8 | 5 |
| ENSG00000181788 | 1761 | 847 | 1005 | 926 | 876 | 956 |
| ENSG00000244265 | 4 | 2 | 2 | 1 | 2 | 2 |
| ENSG00000239265 | 44 | 24 | 26 | 13 | 22 | 25 |
| ENSG00000214237 | 0 | 0 | 2 | 0 | 1 | 0 |
| ENSG00000260234 | 0 | 0 | 0 | 0 | 0 | 0 |
| ENSG00000243273 | 0 | 1 | 1 | 1 | 0 | 1 |
| ENSG00000163646 | 0 | 0 | 0 | 0 | 0 | 0 |
| ENSG00000276055 | 0 | 0 | 0 | 0 | 0 | 0 |
| ENSG00000144893 | 358 | 183 | 233 | 95 | 205 | 202 |
| ENSG00000199994 | 0 | 0 | 0 | 0 | 0 | 0 |
| ENSG00000174946 | 0 | 0 | 0 | 0 | 0 | 0 |
| ENSG00000174944 | 1 | 1 | 0 | 0 | 0 | 0 |
| ENSG00000239570 | 0 | 0 | 0 | 0 | 0 | 0 |
| ENSG00000138271 | 41 | 24 | 38 | 33 | 59 | 55 |
| ENSG00000181631 | 0 | 0 | 0 | 0 | 0 | 0 |
| ENSG00000169313 | 0 | 0 | 0 | 0 | 0 | 0 |
| ENSG00000152580 | 2 | 1 | 2 | 0 | 0 | 7 |
| ENSG00000242810 | 0 | 0 | 0 | 0 | 0 | 0 |
| ENSG00000242119 | 0 | 0 | 0 | 0 | 0 | 0 |
| ENSG00000264330 | 0 | 0 | 0 | 0 | 0 | 0 |
| ENSG00000241151 | 0 | 0 | 0 | 0 | 0 | 0 |
| ENSG00000197953 | 0 | 0 | 0 | 0 | 0 | 0 |
| ENSG00000242908 | 0 | 0 | 0 | 0 | 0 | 0 |
| ENSG00000242097 | 0 | 0 | 0 | 0 | 0 | 0 |
| ENSG00000240602 | 0 | 0 | 0 | 0 | 0 | 0 |
| ENSG00000250271 | 0 | 0 | 0 | 0 | 0 | 0 |
| ENSG00000114771 | 0 | 0 | 0 | 1 | 0 | 0 |
| ENSG00000198829 | 0 | 0 | 0 | 0 | 0 | 0 |
| ENSG00000239941 | 0 | 0 | 0 | 0 | 0 | 0 |
| ENSG00000152601 | 1539 | 844 | 843 | 502 | 1325 | 912 |
| ENSG00000229619 | 28 | 18 | 18 | 6 | 19 | 16 |

| | | | | | | |
|---|---|---|---|---|---|---|
| ENSG00000221962 | 0 | 0 | 0 | 0 | 0 | 0 |
| ENSG00000201217 | 0 | 0 | 0 | 0 | 0 | 0 |
| ENSG00000243305 | 0 | 0 | 0 | 0 | 0 | 0 |
| ENSG00000244545 | 0 | 0 | 0 | 0 | 0 | 0 |
| ENSG00000242561 | 0 | 0 | 0 | 0 | 0 | 0 |
| ENSG00000169860 | 2 | 3 | 0 | 4 | 1 | 2 |
| ENSG00000241810 | 0 | 0 | 0 | 0 | 0 | 0 |
| ENSG00000241671 | 0 | 0 | 0 | 0 | 0 | 0 |
| ENSG00000244268 | 3 | 0 | 0 | 2 | 0 | 0 |
| ENSG00000181467 | 1618 | 889 | 946 | 942 | 1009 | 1074 |
| ENSG00000265813 | 0 | 0 | 0 | 0 | 0 | 0 |
| ENSG00000241220 | 0 | 0 | 3 | 0 | 0 | 0 |
| ENSG00000241912 | 0 | 0 | 0 | 0 | 0 | 0 |
| ENSG00000238755 | 0 | 0 | 0 | 0 | 0 | 0 |
| ENSG00000237787 | 0 | 0 | 0 | 0 | 0 | 0 |
| ENSG00000207323 | 0 | 0 | 0 | 0 | 0 | 0 |
| ENSG00000200162 | 0 | 0 | 0 | 0 | 0 | 0 |
| ENSG00000243486 | 0 | 0 | 0 | 0 | 0 | 0 |
| ENSG00000201398 | 0 | 0 | 0 | 0 | 0 | 0 |
| ENSG00000240068 | 0 | 0 | 0 | 0 | 0 | 0 |
| ENSG00000243069 | 7 | 5 | 9 | 4 | 11 | 6 |
| ENSG00000114790 | 790 | 302 | 401 | 177 | 265 | 244 |
| ENSG00000174953 | 1351 | 795 | 647 | 416 | 1192 | 724 |
| ENSG00000271922 | 0 | 0 | 0 | 0 | 0 | 0 |
| ENSG00000279305 | 0 | 0 | 0 | 0 | 0 | 0 |
| ENSG00000174948 | 0 | 0 | 0 | 0 | 0 | 0 |
| ENSG00000240048 | 0 | 0 | 0 | 0 | 0 | 0 |
| ENSG00000243861 | 0 | 0 | 0 | 0 | 0 | 0 |
| ENSG00000243154 | 0 | 0 | 0 | 0 | 0 | 0 |
| ENSG00000242151 | 0 | 0 | 0 | 0 | 0 | 0 |
| ENSG00000240601 | 0 | 0 | 0 | 0 | 0 | 0 |
| ENSG00000243969 | 0 | 0 | 0 | 0 | 0 | 0 |
| ENSG00000243953 | 0 | 0 | 0 | 0 | 0 | 0 |
| ENSG00000196549 | 102 | 53 | 86 | 27 | 58 | 44 |
| ENSG00000240666 | 0 | 0 | 3 | 0 | 1 | 3 |
| ENSG00000241336 | 0 | 0 | 0 | 0 | 0 | 0 |
| ENSG00000240045 | 0 | 0 | 0 | 0 | 0 | 0 |
| ENSG00000242790 | 1 | 0 | 0 | 0 | 0 | 0 |
| ENSG00000239443 | 0 | 0 | 0 | 0 | 0 | 1 |
| ENSG00000114805 | 600 | 295 | 332 | 165 | 339 | 283 |
| ENSG00000248772 | 0 | 0 | 0 | 0 | 0 | 0 |
| ENSG00000243518 | 0 | 0 | 0 | 0 | 0 | 0 |
| ENSG00000242152 | 0 | 0 | 0 | 0 | 0 | 0 |
| ENSG00000272096 | 0 | 0 | 0 | 0 | 0 | 0 |
| ENSG00000239508 | 1 | 1 | 0 | 0 | 0 | 0 |
| ENSG00000242925 | 0 | 0 | 0 | 0 | 0 | 0 |
| ENSG00000239212 | 0 | 0 | 0 | 0 | 0 | 0 |
| ENSG00000214919 | 0 | 1 | 0 | 0 | 4 | 0 |
| ENSG00000241307 | 0 | 0 | 0 | 0 | 0 | 2 |
| ENSG00000174928 | 57 | 24 | 53 | 28 | 48 | 45 |

| | | | | | | |
|---|---|---|---|---|---|---|
| ENSG00000284952 | 0 | 0 | 1 | 2 | 0 | 1 |
| ENSG00000169359 | 494 | 236 | 301 | 296 | 270 | 303 |
| ENSG00000163655 | 3268 | 1631 | 1877 | 1532 | 2008 | 1862 |
| ENSG00000240489 | 37 | 21 | 42 | 33 | 35 | 42 |
| ENSG00000174930 | 0 | 0 | 0 | 0 | 0 | 0 |
| ENSG00000169282 | 10 | 2 | 13 | 4 | 8 | 11 |
| ENSG00000242017 | 0 | 0 | 0 | 1 | 0 | 0 |
| ENSG00000242537 | 0 | 0 | 0 | 0 | 0 | 0 |
| ENSG00000227709 | 0 | 0 | 0 | 0 | 0 | 0 |
| ENSG00000240596 | 0 | 0 | 0 | 0 | 0 | 0 |
| ENSG00000242370 | 0 | 0 | 0 | 0 | 0 | 0 |
| ENSG00000272990 | 0 | 0 | 0 | 2 | 2 | 0 |
| ENSG00000114850 | 3853 | 1825 | 2130 | 2098 | 2620 | 2330 |
| ENSG00000243926 | 85 | 65 | 69 | 99 | 60 | 69 |
| ENSG00000163659 | 573 | 305 | 306 | 975 | 1495 | 1079 |
| ENSG00000174912 | 16 | 11 | 11 | 9 | 23 | 20 |
| ENSG00000240875 | 391 | 200 | 294 | 430 | 567 | 516 |
| ENSG00000230457 | 6 | 5 | 7 | 4 | 5 | 6 |
| ENSG00000197980 | 43 | 24 | 28 | 37 | 54 | 47 |
| ENSG00000241879 | 0 | 0 | 0 | 0 | 0 | 0 |
| ENSG00000239706 | 0 | 0 | 0 | 0 | 0 | 0 |
| ENSG00000222499 | 0 | 0 | 0 | 0 | 0 | 0 |
| ENSG00000243629 | 0 | 0 | 0 | 0 | 0 | 0 |
| ENSG00000241544 | 0 | 0 | 0 | 0 | 0 | 0 |
| ENSG00000241135 | 0 | 0 | 0 | 0 | 0 | 0 |
| ENSG00000163660 | 3004 | 1578 | 1812 | 965 | 1947 | 1592 |
| ENSG00000201778 | 0 | 0 | 0 | 0 | 0 | 0 |
| ENSG00000244515 | 12 | 4 | 6 | 5 | 1 | 4 |
| ENSG00000241770 | 1 | 1 | 1 | 1 | 5 | 2 |
| ENSG00000222675 | 0 | 0 | 0 | 0 | 0 | 0 |
| ENSG00000243176 | 14 | 10 | 7 | 8 | 8 | 6 |
| ENSG00000270981 | 0 | 0 | 0 | 0 | 0 | 0 |
| ENSG00000197415 | 18 | 14 | 25 | 5 | 9 | 12 |
| ENSG00000163661 | 8 | 2 | 3 | 0 | 1 | 0 |
| ENSG00000174899 | 2 | 1 | 1 | 0 | 4 | 2 |
| ENSG00000251751 | 0 | 0 | 0 | 0 | 0 | 0 |
| ENSG00000239716 | 0 | 0 | 0 | 0 | 0 | 0 |
| ENSG00000242536 | 0 | 0 | 0 | 0 | 0 | 0 |
| ENSG00000168779 | 34 | 11 | 29 | 18 | 25 | 27 |
| ENSG00000174891 | 546 | 282 | 326 | 221 | 412 | 307 |
| ENSG00000241723 | 0 | 0 | 0 | 0 | 0 | 0 |
| ENSG00000277143 | 0 | 0 | 0 | 0 | 0 | 0 |
| ENSG00000243314 | 0 | 0 | 0 | 0 | 0 | 0 |
| ENSG00000243150 | 0 | 0 | 0 | 0 | 0 | 0 |
| ENSG00000178053 | 614 | 405 | 423 | 409 | 534 | 441 |
| ENSG00000243025 | 0 | 0 | 0 | 0 | 0 | 0 |
| ENSG00000243165 | 0 | 0 | 0 | 0 | 0 | 0 |
| ENSG00000168827 | 1681 | 903 | 1088 | 665 | 894 | 884 |
| ENSG00000079257 | 13585 | 8839 | 8358 | 9881 | 14810 | 12248 |
| ENSG00000272087 | 12 | 4 | 7 | 1 | 2 | 6 |

| | | | | | | |
|---|---|---|---|---|---|---|
| ENSG00000272440 | 5 | 1 | 1 | 1 | 3 | 2 |
| ENSG00000118849 | 0 | 0 | 1 | 1 | 2 | 2 |
| ENSG00000118855 | 3490 | 1734 | 2216 | 1901 | 2475 | 2556 |
| ENSG00000240207 | 24 | 17 | 22 | 15 | 23 | 21 |
| ENSG00000243675 | 0 | 1 | 0 | 0 | 1 | 1 |
| ENSG00000271778 | 0 | 2 | 1 | 0 | 3 | 4 |
| ENSG00000272247 | 6 | 5 | 6 | 1 | 1 | 3 |
| ENSG00000232374 | 2 | 0 | 1 | 0 | 0 | 0 |
| ENSG00000279311 | 0 | 0 | 0 | 0 | 0 | 0 |
| ENSG00000283154 | 6 | 7 | 7 | 12 | 12 | 7 |
| ENSG00000214216 | 0 | 0 | 0 | 0 | 0 | 0 |
| ENSG00000263634 | 0 | 0 | 0 | 0 | 0 | 0 |
| ENSG00000244249 | 0 | 0 | 0 | 0 | 0 | 0 |
| ENSG00000243116 | 0 | 0 | 0 | 0 | 0 | 0 |
| ENSG00000241211 | 0 | 0 | 0 | 0 | 0 | 0 |
| ENSG00000252763 | 0 | 0 | 0 | 0 | 0 | 0 |
| ENSG00000151967 | 4 | 2 | 3 | 2 | 0 | 3 |
| ENSG00000244040 | 0 | 0 | 0 | 0 | 0 | 0 |
| ENSG00000168811 | 6 | 1 | 2 | 0 | 1 | 2 |
| ENSG00000242107 | 0 | 0 | 0 | 0 | 0 | 0 |
| ENSG00000184100 | 0 | 1 | 2 | 0 | 1 | 1 |
| ENSG00000180044 | 4 | 4 | 5 | 4 | 1 | 6 |
| ENSG00000248710 | 0 | 0 | 0 | 0 | 0 | 0 |
| ENSG00000068885 | 246 | 154 | 124 | 46 | 239 | 109 |
| ENSG00000243687 | 0 | 0 | 0 | 0 | 0 | 0 |
| ENSG00000113810 | 1029 | 580 | 454 | 214 | 995 | 514 |
| ENSG00000207779 | 0 | 0 | 0 | 0 | 0 | 0 |
| ENSG00000198987 | 0 | 0 | 0 | 0 | 0 | 0 |
| ENSG00000213186 | 319 | 151 | 155 | 103 | 198 | 177 |
| ENSG00000244009 | 4 | 0 | 1 | 2 | 0 | 1 |
| ENSG00000238427 | 0 | 0 | 0 | 0 | 0 | 0 |
| ENSG00000186432 | 2905 | 1438 | 1659 | 1073 | 1687 | 1451 |
| ENSG00000238741 | 3 | 3 | 2 | 1 | 0 | 0 |
| ENSG00000229320 | 227 | 101 | 131 | 163 | 131 | 166 |
| ENSG00000242285 | 0 | 0 | 0 | 0 | 0 | 0 |
| ENSG00000179674 | 0 | 2 | 1 | 2 | 5 | 2 |
| ENSG00000207084 | 0 | 0 | 0 | 0 | 0 | 0 |
| ENSG00000260572 | 16 | 7 | 10 | 8 | 5 | 9 |
| ENSG00000163590 | 712 | 308 | 363 | 209 | 383 | 327 |
| ENSG00000169255 | 2522 | 1329 | 1603 | 1329 | 1665 | 1509 |
| ENSG00000169251 | 11203 | 6743 | 6255 | 4290 | 10101 | 6602 |
| ENSG00000271052 | 3 | 0 | 1 | 0 | 4 | 2 |
| ENSG00000240138 | 10 | 1 | 0 | 1 | 5 | 2 |
| ENSG00000243838 | 12 | 5 | 5 | 3 | 7 | 6 |
| ENSG00000196542 | 60883 | 30792 | 38079 | 30570 | 41566 | 37052 |
| ENSG00000240567 | 48 | 30 | 36 | 19 | 37 | 30 |
| ENSG00000269888 | 1 | 0 | 0 | 1 | 0 | 0 |
| ENSG00000234851 | 4 | 6 | 5 | 11 | 20 | 12 |
| ENSG00000182447 | 3 | 0 | 2 | 1 | 0 | 0 |
| ENSG00000240354 | 0 | 0 | 0 | 0 | 0 | 0 |

| | | | | | | |
|---|---|---|---|---|---|---|
| ENSG00000241874 | 0 | 0 | 0 | 0 | 0 | 0 |
| ENSG00000243044 | 0 | 0 | 0 | 0 | 0 | 0 |
| ENSG00000241168 | 2 | 0 | 0 | 0 | 0 | 0 |
| ENSG00000244571 | 0 | 0 | 0 | 0 | 0 | 0 |
| ENSG00000241369 | 0 | 0 | 0 | 0 | 0 | 0 |
| ENSG00000238924 | 0 | 0 | 0 | 0 | 0 | 0 |
| ENSG00000240792 | 0 | 0 | 0 | 0 | 0 | 0 |
| ENSG00000214210 | 0 | 0 | 0 | 0 | 0 | 0 |
| ENSG00000221251 | 0 | 0 | 0 | 0 | 0 | 0 |
| ENSG00000241636 | 0 | 0 | 0 | 0 | 0 | 0 |
| ENSG00000241767 | 1 | 0 | 0 | 0 | 0 | 0 |
| ENSG00000090402 | 4 | 0 | 1 | 0 | 0 | 0 |
| ENSG00000200052 | 0 | 0 | 0 | 0 | 0 | 0 |
| ENSG00000239205 | 0 | 0 | 0 | 0 | 0 | 0 |
| ENSG00000121871 | 0 | 0 | 0 | 0 | 0 | 0 |
| ENSG00000244128 | 0 | 0 | 0 | 0 | 0 | 0 |
| ENSG00000114200 | 2 | 0 | 0 | 0 | 0 | 0 |
| ENSG00000242178 | 0 | 0 | 0 | 0 | 0 | 0 |
| ENSG00000240084 | 0 | 0 | 0 | 0 | 0 | 0 |
| ENSG00000240824 | 0 | 0 | 0 | 0 | 0 | 0 |
| ENSG00000242456 | 0 | 0 | 0 | 0 | 0 | 0 |
| ENSG00000242326 | 0 | 0 | 0 | 0 | 0 | 0 |
| ENSG00000244429 | 0 | 0 | 0 | 0 | 0 | 0 |
| ENSG00000244321 | 0 | 0 | 0 | 0 | 0 | 0 |
| ENSG00000274149 | 0 | 0 | 0 | 0 | 0 | 0 |
| ENSG00000251759 | 0 | 0 | 0 | 0 | 0 | 0 |
| ENSG00000224962 | 0 | 0 | 0 | 0 | 0 | 0 |
| ENSG00000240458 | 0 | 0 | 0 | 0 | 0 | 0 |
| ENSG00000231516 | 0 | 0 | 0 | 0 | 0 | 0 |
| ENSG00000240033 | 0 | 1 | 0 | 0 | 0 | 0 |
| ENSG00000239227 | 0 | 0 | 0 | 0 | 0 | 0 |
| ENSG00000220343 | 0 | 0 | 0 | 0 | 0 | 0 |
| ENSG00000169064 | 0 | 0 | 0 | 0 | 0 | 0 |
| ENSG00000241669 | 0 | 0 | 0 | 0 | 0 | 0 |
| ENSG00000114204 | 0 | 0 | 0 | 0 | 0 | 0 |
| ENSG00000174776 | 0 | 0 | 0 | 0 | 0 | 1 |
| ENSG00000114209 | 1565 | 838 | 788 | 684 | 1061 | 757 |
| ENSG00000241120 | 0 | 2 | 0 | 0 | 1 | 0 |
| ENSG00000163536 | 18 | 12 | 8 | 5 | 3 | 13 |
| ENSG00000244227 | 0 | 0 | 0 | 0 | 0 | 0 |
| ENSG00000244706 | 0 | 0 | 0 | 0 | 0 | 0 |
| ENSG00000239238 | 0 | 0 | 0 | 0 | 0 | 0 |
| ENSG00000242036 | 0 | 0 | 0 | 0 | 0 | 0 |
| ENSG00000241203 | 0 | 0 | 0 | 0 | 0 | 0 |
| ENSG00000173905 | 360 | 151 | 196 | 97 | 196 | 152 |
| ENSG00000206120 | 0 | 0 | 0 | 0 | 0 | 0 |
| ENSG00000241882 | 0 | 0 | 0 | 0 | 0 | 0 |
| ENSG00000222357 | 0 | 0 | 0 | 0 | 0 | 0 |
| ENSG00000207717 | 0 | 0 | 0 | 0 | 0 | 0 |
| ENSG00000226765 | 0 | 0 | 0 | 0 | 0 | 0 |

| | | | | | | |
|---|---|---|---|---|---|---|
| ENSG00000231688 | 0 | 0 | 0 | 0 | 0 | 0 |
| ENSG00000242268 | 0 | 0 | 0 | 0 | 0 | 0 |
| ENSG00000272797 | 0 | 0 | 0 | 0 | 0 | 0 |
| ENSG00000273177 | 0 | 0 | 0 | 0 | 0 | 0 |
| ENSG00000241383 | 0 | 0 | 0 | 0 | 0 | 0 |
| ENSG00000085276 | 332 | 187 | 155 | 54 | 162 | 96 |
| ENSG00000241479 | 4 | 0 | 1 | 0 | 0 | 2 |
| ENSG00000213178 | 18 | 13 | 9 | 11 | 13 | 6 |
| ENSG00000242795 | 0 | 0 | 0 | 0 | 0 | 0 |
| ENSG00000244381 | 0 | 0 | 0 | 2 | 0 | 0 |
| ENSG00000206777 | 0 | 0 | 0 | 0 | 0 | 0 |
| ENSG00000270141 | 1 | 0 | 1 | 0 | 0 | 0 |
| ENSG00000277925 | 0 | 0 | 0 | 0 | 0 | 0 |
| ENSG00000184378 | 137 | 66 | 59 | 75 | 91 | 89 |
| ENSG00000269889 | 2 | 4 | 5 | 4 | 8 | 7 |
| ENSG00000085274 | 632 | 329 | 400 | 277 | 375 | 350 |
| ENSG00000269984 | 39 | 17 | 15 | 5 | 14 | 9 |
| ENSG00000171757 | 2 | 0 | 2 | 1 | 2 | 0 |
| ENSG00000270135 | 0 | 0 | 0 | 0 | 0 | 0 |
| ENSG00000270096 | 0 | 0 | 0 | 0 | 0 | 0 |
| ENSG00000188306 | 7 | 1 | 0 | 1 | 0 | 2 |
| ENSG00000114248 | 0 | 0 | 0 | 1 | 0 | 0 |
| ENSG00000240128 | 0 | 0 | 0 | 0 | 0 | 0 |
| ENSG00000187033 | 0 | 0 | 0 | 0 | 0 | 0 |
| ENSG00000244039 | 0 | 0 | 0 | 0 | 0 | 0 |
| ENSG00000239219 | 0 | 0 | 0 | 0 | 0 | 1 |
| ENSG00000244193 | 0 | 0 | 0 | 0 | 0 | 0 |
| ENSG00000008952 | 2212 | 1224 | 1108 | 684 | 1745 | 1075 |
| ENSG00000240373 | 5 | 3 | 2 | 1 | 1 | 3 |
| ENSG00000173890 | 1994 | 974 | 1093 | 732 | 978 | 929 |
| ENSG00000201342 | 0 | 2 | 0 | 0 | 0 | 0 |
| ENSG00000242583 | 0 | 1 | 0 | 1 | 0 | 1 |
| ENSG00000173889 | 1203 | 706 | 566 | 231 | 721 | 511 |
| ENSG00000268220 | 0 | 0 | 0 | 0 | 0 | 0 |
| ENSG00000199536 | 0 | 1 | 0 | 0 | 1 | 1 |
| ENSG00000163558 | 2323 | 1033 | 1462 | 739 | 1290 | 1124 |
| ENSG00000200118 | 0 | 0 | 0 | 0 | 0 | 0 |
| ENSG00000239628 | 0 | 0 | 0 | 0 | 0 | 0 |
| ENSG00000136603 | 1414 | 727 | 603 | 255 | 948 | 534 |
| ENSG00000242578 | 4 | 1 | 0 | 1 | 2 | 1 |
| ENSG00000013297 | 4 | 2 | 3 | 3 | 1 | 3 |
| ENSG00000285218 | 0 | 0 | 0 | 0 | 0 | 0 |
| ENSG00000283474 | 0 | 0 | 0 | 0 | 0 | 0 |
| ENSG00000285051 | 0 | 0 | 0 | 0 | 0 | 0 |
| ENSG00000013293 | 0 | 0 | 0 | 0 | 0 | 0 |
| ENSG00000241416 | 2 | 1 | 0 | 2 | 3 | 4 |
| ENSG00000213174 | 1 | 0 | 0 | 0 | 0 | 0 |
| ENSG00000244738 | 0 | 0 | 0 | 0 | 0 | 0 |
| ENSG00000239739 | 0 | 0 | 0 | 0 | 0 | 0 |
| ENSG00000163584 | 807 | 465 | 481 | 565 | 584 | 506 |

| | | | | | | |
|---|---|---|---|---|---|---|
| ENSG00000163577 | 200 | 96 | 109 | 71 | 132 | 107 |
| ENSG00000240704 | 0 | 0 | 0 | 0 | 0 | 0 |
| ENSG00000199488 | 0 | 2 | 0 | 0 | 1 | 2 |
| ENSG00000163581 | 0 | 0 | 1 | 0 | 0 | 0 |
| ENSG00000154310 | 45 | 20 | 21 | 14 | 18 | 18 |
| ENSG00000207963 | 0 | 0 | 0 | 0 | 0 | 0 |
| ENSG00000222506 | 0 | 0 | 0 | 0 | 0 | 0 |
| ENSG00000279673 | 4 | 1 | 0 | 2 | 1 | 0 |
| ENSG00000240497 | 0 | 0 | 0 | 0 | 0 | 0 |
| ENSG00000207114 | 0 | 0 | 0 | 0 | 0 | 0 |
| ENSG00000234594 | 0 | 0 | 0 | 0 | 0 | 0 |
| ENSG00000075651 | 321 | 189 | 173 | 141 | 300 | 230 |
| ENSG00000186329 | 0 | 0 | 0 | 0 | 0 | 0 |
| ENSG00000234717 | 0 | 0 | 0 | 0 | 0 | 0 |
| ENSG00000235943 | 0 | 0 | 0 | 0 | 0 | 0 |
| ENSG00000075420 | 631 | 267 | 352 | 183 | 378 | 300 |
| ENSG00000237483 | 0 | 0 | 0 | 0 | 0 | 0 |
| ENSG00000243398 | 0 | 0 | 0 | 0 | 0 | 0 |
| ENSG00000279862 | 2 | 2 | 0 | 0 | 0 | 0 |
| ENSG00000236686 | 0 | 0 | 0 | 0 | 0 | 0 |
| ENSG00000121853 | 0 | 0 | 0 | 0 | 0 | 0 |
| ENSG00000121858 | 102 | 53 | 23 | 15 | 29 | 23 |
| ENSG00000224611 | 0 | 0 | 0 | 0 | 0 | 0 |
| ENSG00000223387 | 1 | 0 | 0 | 0 | 1 | 0 |
| ENSG00000224426 | 0 | 1 | 0 | 0 | 0 | 0 |
| ENSG00000144959 | 767 | 363 | 408 | 248 | 393 | 363 |
| ENSG00000231384 | 0 | 0 | 0 | 0 | 0 | 0 |
| ENSG00000252082 | 0 | 0 | 0 | 0 | 0 | 0 |
| ENSG00000114346 | 752 | 497 | 353 | 202 | 882 | 463 |
| ENSG00000200355 | 0 | 0 | 0 | 0 | 0 | 0 |
| ENSG00000201458 | 0 | 0 | 0 | 0 | 0 | 0 |
| ENSG00000227440 | 10 | 8 | 6 | 4 | 6 | 1 |
| ENSG00000237501 | 0 | 0 | 0 | 0 | 0 | 0 |
| ENSG00000144962 | 0 | 0 | 0 | 0 | 0 | 0 |
| ENSG00000237473 | 0 | 0 | 0 | 0 | 0 | 0 |
| ENSG00000169760 | 0 | 0 | 0 | 0 | 0 | 0 |
| ENSG00000237645 | 0 | 0 | 0 | 0 | 0 | 0 |
| ENSG00000228213 | 0 | 0 | 0 | 0 | 0 | 0 |
| ENSG00000252982 | 0 | 0 | 0 | 0 | 0 | 0 |
| ENSG00000232601 | 0 | 0 | 0 | 0 | 0 | 0 |
| ENSG00000213169 | 0 | 0 | 0 | 0 | 2 | 0 |
| ENSG00000177694 | 76 | 40 | 26 | 37 | 87 | 62 |
| ENSG00000253020 | 0 | 0 | 0 | 0 | 0 | 0 |
| ENSG00000230497 | 0 | 0 | 0 | 0 | 0 | 0 |
| ENSG00000230292 | 0 | 0 | 0 | 0 | 0 | 0 |
| ENSG00000226779 | 2 | 0 | 0 | 0 | 0 | 0 |
| ENSG00000264974 | 0 | 0 | 0 | 0 | 0 | 0 |
| ENSG00000199792 | 0 | 0 | 0 | 0 | 0 | 0 |
| ENSG00000201648 | 0 | 0 | 0 | 0 | 0 | 0 |
| ENSG00000236241 | 2 | 0 | 0 | 0 | 0 | 0 |

| | | | | | | |
|---|---|---|---|---|---|---|
| ENSG00000214192 | 2 | 2 | 1 | 1 | 1 | 1 |
| ENSG00000201452 | 0 | 0 | 0 | 0 | 0 | 0 |
| ENSG00000225552 | 0 | 0 | 0 | 0 | 0 | 0 |
| ENSG00000260217 | 0 | 0 | 0 | 0 | 0 | 0 |
| ENSG00000230362 | 0 | 0 | 0 | 0 | 0 | 0 |
| ENSG00000236357 | 0 | 0 | 0 | 0 | 0 | 0 |
| ENSG00000271413 | 0 | 0 | 0 | 0 | 0 | 0 |
| ENSG00000238026 | 0 | 0 | 0 | 0 | 0 | 0 |
| ENSG00000283333 | 0 | 0 | 0 | 0 | 0 | 0 |
| ENSG00000223715 | 0 | 0 | 0 | 0 | 0 | 0 |
| ENSG00000232461 | 0 | 0 | 0 | 0 | 0 | 0 |
| ENSG00000252302 | 0 | 0 | 0 | 0 | 0 | 0 |
| ENSG00000228308 | 0 | 0 | 0 | 0 | 0 | 0 |
| ENSG00000231888 | 0 | 0 | 0 | 0 | 0 | 0 |
| ENSG00000177565 | 1501 | 833 | 802 | 547 | 941 | 740 |
| ENSG00000231310 | 0 | 0 | 0 | 0 | 0 | 0 |
| ENSG00000201343 | 0 | 0 | 0 | 0 | 0 | 0 |
| ENSG00000200882 | 0 | 0 | 0 | 1 | 0 | 0 |
| ENSG00000203645 | 0 | 0 | 0 | 1 | 0 | 0 |
| ENSG00000236893 | 0 | 0 | 0 | 0 | 0 | 0 |
| ENSG00000226782 | 0 | 0 | 0 | 0 | 0 | 0 |
| ENSG00000228221 | 0 | 3 | 5 | 3 | 1 | 2 |
| ENSG00000252028 | 0 | 0 | 0 | 0 | 0 | 0 |
| ENSG00000236385 | 0 | 0 | 1 | 0 | 0 | 0 |
| ENSG00000200288 | 0 | 1 | 0 | 0 | 0 | 0 |
| ENSG00000228561 | 3 | 2 | 4 | 3 | 5 | 5 |
| ENSG00000277241 | 0 | 1 | 1 | 1 | 0 | 1 |
| ENSG00000231574 | 63 | 34 | 37 | 27 | 36 | 46 |
| ENSG00000225790 | 4 | 2 | 2 | 1 | 1 | 2 |
| ENSG00000270321 | 1 | 0 | 0 | 1 | 0 | 0 |
| ENSG00000199858 | 0 | 0 | 0 | 0 | 0 | 0 |
| ENSG00000223930 | 0 | 0 | 0 | 0 | 0 | 0 |
| ENSG00000197584 | 2 | 0 | 0 | 0 | 1 | 0 |
| ENSG00000275163 | 0 | 0 | 0 | 0 | 0 | 0 |
| ENSG00000223941 | 0 | 0 | 0 | 0 | 0 | 0 |
| ENSG00000252336 | 0 | 0 | 0 | 0 | 0 | 0 |
| ENSG00000271131 | 2 | 0 | 0 | 0 | 0 | 0 |
| ENSG00000237978 | 160 | 78 | 105 | 56 | 82 | 77 |
| ENSG00000172667 | 912 | 500 | 462 | 263 | 555 | 403 |
| ENSG00000200616 | 0 | 0 | 0 | 0 | 0 | 0 |
| ENSG00000229102 | 0 | 0 | 0 | 0 | 0 | 0 |
| ENSG00000121879 | 569 | 283 | 252 | 145 | 457 | 243 |
| ENSG00000201957 | 0 | 0 | 0 | 0 | 0 | 0 |
| ENSG00000171121 | 148 | 65 | 99 | 48 | 75 | 63 |
| ENSG00000240429 | 84 | 55 | 57 | 32 | 44 | 56 |
| ENSG00000121864 | 1115 | 445 | 574 | 440 | 633 | 552 |
| ENSG00000260743 | 2 | 3 | 2 | 3 | 2 | 0 |
| ENSG00000171109 | 1587 | 820 | 926 | 540 | 1237 | 852 |
| ENSG00000114450 | 55 | 40 | 36 | 29 | 61 | 39 |
| ENSG00000242539 | 0 | 0 | 0 | 0 | 0 | 0 |

| | | | | | | |
|---|---|---|---|---|---|---|
| ENSG00000272699 | 0 | 0 | 0 | 0 | 0 | 0 |
| ENSG00000239255 | 0 | 0 | 0 | 0 | 0 | 0 |
| ENSG00000181260 | 0 | 0 | 1 | 0 | 0 | 0 |
| ENSG00000136518 | 1999 | 1255 | 1361 | 1101 | 1341 | 1379 |
| ENSG00000272910 | 1 | 2 | 1 | 1 | 5 | 2 |
| ENSG00000272660 | 2 | 0 | 0 | 2 | 1 | 1 |
| ENSG00000136522 | 958 | 527 | 569 | 633 | 711 | 559 |
| ENSG00000136521 | 3122 | 1451 | 1790 | 1664 | 1727 | 1635 |
| ENSG00000239096 | 0 | 1 | 2 | 0 | 0 | 0 |
| ENSG00000243977 | 0 | 0 | 0 | 2 | 0 | 0 |
| ENSG00000058056 | 1240 | 604 | 820 | 563 | 723 | 739 |
| ENSG00000114757 | 3 | 3 | 5 | 4 | 1 | 2 |
| ENSG00000270178 | 0 | 0 | 0 | 0 | 0 | 0 |
| ENSG00000243799 | 0 | 0 | 0 | 0 | 0 | 0 |
| ENSG00000244302 | 0 | 0 | 0 | 0 | 0 | 0 |
| ENSG00000201822 | 0 | 0 | 0 | 0 | 0 | 0 |
| ENSG00000282327 | 0 | 0 | 0 | 0 | 0 | 0 |
| ENSG00000213158 | 0 | 0 | 0 | 0 | 0 | 0 |
| ENSG00000239880 | 0 | 0 | 0 | 0 | 0 | 0 |
| ENSG00000241696 | 0 | 0 | 0 | 0 | 0 | 0 |
| ENSG00000201810 | 0 | 0 | 0 | 0 | 0 | 0 |
| ENSG00000239774 | 0 | 0 | 0 | 1 | 1 | 0 |
| ENSG00000199986 | 0 | 0 | 0 | 0 | 0 | 0 |
| ENSG00000241946 | 0 | 0 | 0 | 0 | 0 | 0 |
| ENSG00000163728 | 616 | 368 | 402 | 206 | 514 | 316 |
| ENSG00000284862 | 0 | 1 | 0 | 0 | 1 | 0 |
| ENSG00000145075 | 4 | 0 | 1 | 4 | 8 | 4 |
| ENSG00000243187 | 1 | 0 | 1 | 0 | 0 | 0 |
| ENSG00000285336 | 0 | 0 | 0 | 0 | 0 | 0 |
| ENSG00000242811 | 0 | 0 | 0 | 0 | 0 | 0 |
| ENSG00000242068 | 0 | 0 | 0 | 0 | 0 | 0 |
| ENSG00000241473 | 0 | 0 | 0 | 0 | 0 | 0 |
| ENSG00000239516 | 0 | 0 | 0 | 0 | 0 | 0 |
| ENSG00000114416 | 2558 | 1600 | 1594 | 870 | 2257 | 1470 |
| ENSG00000213157 | 0 | 0 | 0 | 0 | 0 | 0 |
| ENSG00000205981 | 822 | 410 | 482 | 434 | 414 | 425 |
| ENSG00000242808 | 4 | 1 | 3 | 2 | 5 | 2 |
| ENSG00000242140 | 0 | 0 | 0 | 0 | 1 | 1 |
| ENSG00000206932 | 4 | 4 | 0 | 1 | 1 | 3 |
| ENSG00000237806 | 0 | 0 | 0 | 0 | 0 | 0 |
| ENSG00000241231 | 0 | 0 | 0 | 0 | 0 | 0 |
| ENSG00000235294 | 0 | 0 | 0 | 0 | 0 | 0 |
| ENSG00000239381 | 0 | 0 | 0 | 0 | 0 | 0 |
| ENSG00000276074 | 0 | 0 | 0 | 0 | 0 | 0 |
| ENSG00000276690 | 0 | 0 | 0 | 0 | 0 | 0 |
| ENSG00000181449 | 266 | 127 | 117 | 81 | 95 | 90 |
| ENSG00000278026 | 0 | 0 | 0 | 0 | 0 | 0 |
| ENSG00000266317 | 0 | 0 | 0 | 0 | 0 | 0 |
| ENSG00000199839 | 0 | 0 | 0 | 0 | 0 | 0 |
| ENSG00000285546 | 0 | 0 | 0 | 0 | 0 | 0 |

| | | | | | | |
|---|---|---|---|---|---|---|
| ENSG00000239805 | 0 | 0 | 0 | 0 | 0 | 0 |
| ENSG00000213155 | 0 | 0 | 0 | 0 | 0 | 0 |
| ENSG00000242512 | 0 | 0 | 0 | 0 | 0 | 0 |
| ENSG00000252257 | 0 | 0 | 0 | 0 | 0 | 0 |
| ENSG00000240478 | 0 | 0 | 0 | 0 | 0 | 0 |
| ENSG00000242012 | 0 | 0 | 0 | 0 | 0 | 0 |
| ENSG00000241098 | 0 | 0 | 0 | 0 | 0 | 0 |
| ENSG00000244199 | 0 | 0 | 0 | 0 | 0 | 0 |
| ENSG00000244247 | 0 | 0 | 0 | 0 | 0 | 0 |
| ENSG00000228766 | 0 | 1 | 1 | 0 | 0 | 1 |
| ENSG00000238837 | 0 | 0 | 0 | 0 | 0 | 0 |
| ENSG00000243378 | 0 | 0 | 0 | 0 | 0 | 0 |
| ENSG00000260377 | 0 | 0 | 0 | 0 | 1 | 0 |
| ENSG00000240063 | 4 | 1 | 1 | 0 | 0 | 0 |
| ENSG00000058063 | 1316 | 745 | 637 | 381 | 948 | 633 |
| ENSG00000270554 | 0 | 0 | 0 | 0 | 0 | 0 |
| ENSG00000273261 | 7 | 2 | 0 | 0 | 0 | 1 |
| ENSG00000043093 | 1282 | 722 | 623 | 426 | 996 | 674 |
| ENSG00000244346 | 0 | 1 | 0 | 0 | 1 | 1 |
| ENSG00000078070 | 1801 | 976 | 1293 | 775 | 1031 | 986 |
| ENSG00000243368 | 16 | 3 | 7 | 5 | 7 | 4 |
| ENSG00000078081 | 18 | 6 | 3 | 7 | 3 | 4 |
| ENSG00000053524 | 30 | 16 | 14 | 9 | 16 | 12 |
| ENSG00000176597 | 133 | 55 | 77 | 39 | 46 | 38 |
| ENSG00000202502 | 0 | 0 | 0 | 0 | 1 | 0 |
| ENSG00000240024 | 240 | 130 | 115 | 118 | 129 | 142 |
| ENSG00000201229 | 0 | 0 | 0 | 0 | 0 | 0 |
| ENSG00000253092 | 3 | 2 | 0 | 1 | 1 | 2 |
| ENSG00000199363 | 0 | 0 | 0 | 0 | 0 | 0 |
| ENSG00000251730 | 0 | 0 | 0 | 0 | 0 | 0 |
| ENSG00000271682 | 0 | 0 | 0 | 0 | 0 | 0 |
| ENSG00000172578 | 0 | 0 | 0 | 0 | 2 | 0 |
| ENSG00000242522 | 0 | 0 | 0 | 0 | 0 | 0 |
| ENSG00000243584 | 0 | 0 | 0 | 0 | 0 | 0 |
| ENSG00000114796 | 1015 | 506 | 556 | 251 | 583 | 356 |
| ENSG00000163872 | 814 | 408 | 533 | 371 | 576 | 455 |
| ENSG00000233885 | 33 | 14 | 9 | 6 | 19 | 17 |
| ENSG00000283765 | 4 | 1 | 0 | 1 | 0 | 0 |
| ENSG00000180834 | 16 | 20 | 9 | 16 | 9 | 18 |
| ENSG00000175193 | 292 | 207 | 273 | 270 | 233 | 240 |
| ENSG00000234371 | 0 | 0 | 1 | 0 | 0 | 1 |
| ENSG00000263932 | 0 | 0 | 0 | 0 | 0 | 0 |
| ENSG00000233441 | 0 | 0 | 0 | 0 | 0 | 0 |
| ENSG00000114770 | 12660 | 5226 | 7882 | 4119 | 5394 | 5626 |
| ENSG00000223882 | 0 | 1 | 2 | 0 | 1 | 1 |
| ENSG00000223529 | 7 | 1 | 5 | 1 | 3 | 1 |
| ENSG00000186090 | 0 | 0 | 0 | 0 | 0 | 0 |
| ENSG00000178084 | 0 | 0 | 0 | 0 | 0 | 0 |
| ENSG00000233576 | 1 | 0 | 0 | 1 | 0 | 0 |
| ENSG00000207351 | 0 | 0 | 0 | 0 | 0 | 0 |

| | | | | | | |
|---|---|---|---|---|---|---|
| ENSG00000238020 | 0 | 0 | 0 | 0 | 0 | 0 |
| ENSG00000186038 | 0 | 0 | 0 | 1 | 0 | 1 |
| ENSG00000205955 | 2 | 2 | 2 | 2 | 0 | 2 |
| ENSG00000273181 | 9 | 10 | 6 | 7 | 8 | 2 |
| ENSG00000272721 | 17 | 8 | 18 | 10 | 7 | 11 |
| ENSG00000145191 | 3040 | 1457 | 1869 | 1548 | 1663 | 1710 |
| ENSG00000228205 | 21 | 9 | 13 | 10 | 9 | 18 |
| ENSG00000161202 | 5551 | 2300 | 3195 | 2342 | 2798 | 2932 |
| ENSG00000161203 | 19219 | 9249 | 11328 | 11626 | 10659 | 10537 |
| ENSG00000161204 | 3689 | 1630 | 2034 | 1997 | 1807 | 1900 |
| ENSG00000145198 | 24 | 7 | 20 | 13 | 12 | 13 |
| ENSG00000221120 | 0 | 0 | 0 | 0 | 0 | 0 |
| ENSG00000214160 | 1884 | 896 | 1059 | 1311 | 1011 | 1116 |
| ENSG00000284753 | 231 | 123 | 155 | 197 | 171 | 159 |
| ENSG00000284917 | 0 | 0 | 0 | 0 | 0 | 0 |
| ENSG00000163888 | 81 | 40 | 58 | 53 | 40 | 54 |
| ENSG00000145194 | 0 | 0 | 0 | 0 | 0 | 0 |
| ENSG00000175166 | 13643 | 6586 | 7958 | 7927 | 8182 | 7969 |
| ENSG00000114867 | 24908 | 11140 | 14977 | 12768 | 13261 | 13981 |
| ENSG00000212158 | 0 | 0 | 0 | 0 | 0 | 0 |
| ENSG00000175182 | 613 | 264 | 350 | 322 | 323 | 303 |
| ENSG00000114859 | 328 | 167 | 197 | 141 | 156 | 153 |
| ENSG00000163882 | 2866 | 1334 | 1695 | 1739 | 1561 | 1558 |
| ENSG00000090534 | 19 | 10 | 3 | 5 | 7 | 6 |
| ENSG00000090539 | 400 | 176 | 239 | 122 | 108 | 85 |
| ENSG00000224153 | 2 | 1 | 0 | 0 | 0 | 0 |
| ENSG00000283625 | 0 | 0 | 0 | 0 | 0 | 0 |
| ENSG00000227509 | 0 | 0 | 0 | 0 | 0 | 0 |
| ENSG00000230215 | 0 | 0 | 0 | 0 | 0 | 0 |
| ENSG00000182580 | 522 | 257 | 326 | 329 | 326 | 305 |
| ENSG00000177383 | 3091 | 1543 | 1867 | 1669 | 1583 | 1802 |
| ENSG00000272970 | 6 | 5 | 0 | 0 | 3 | 4 |
| ENSG00000273403 | 2 | 0 | 2 | 0 | 2 | 1 |
| ENSG00000272922 | 0 | 1 | 0 | 0 | 1 | 0 |
| ENSG00000229433 | 0 | 0 | 0 | 0 | 0 | 0 |
| ENSG00000156931 | 1327 | 779 | 796 | 544 | 1010 | 751 |
| ENSG00000187068 | 62 | 24 | 26 | 12 | 20 | 12 |
| ENSG00000223358 | 0 | 0 | 0 | 0 | 1 | 0 |
| ENSG00000113790 | 227 | 140 | 114 | 77 | 158 | 115 |
| ENSG00000229704 | 0 | 1 | 0 | 0 | 0 | 0 |
| ENSG00000264614 | 0 | 0 | 0 | 0 | 0 | 0 |
| ENSG00000073803 | 1158 | 506 | 650 | 417 | 591 | 565 |
| ENSG00000229638 | 54 | 52 | 49 | 44 | 45 | 56 |
| ENSG00000163900 | 1573 | 785 | 925 | 666 | 828 | 737 |
| ENSG00000163898 | 535 | 281 | 344 | 185 | 279 | 236 |
| ENSG00000163904 | 2017 | 872 | 1181 | 764 | 991 | 1017 |
| ENSG00000236489 | 2 | 1 | 1 | 1 | 1 | 0 |
| ENSG00000073792 | 229 | 117 | 127 | 76 | 92 | 87 |
| ENSG00000163915 | 0 | 0 | 0 | 0 | 0 | 0 |
| ENSG00000265470 | 0 | 0 | 0 | 0 | 0 | 0 |

| | | | | | | |
|---|---|---|---|---|---|---|
| ENSG00000136527 | 4668 | 2232 | 2612 | 2325 | 2722 | 2699 |
| ENSG00000171658 | 0 | 1 | 2 | 0 | 0 | 4 |
| ENSG00000244405 | 48 | 32 | 31 | 31 | 40 | 31 |
| ENSG00000277118 | 0 | 0 | 0 | 0 | 0 | 0 |
| ENSG00000234197 | 0 | 0 | 0 | 0 | 0 | 0 |
| ENSG00000058866 | 4 | 0 | 4 | 1 | 1 | 0 |
| ENSG00000234548 | 0 | 0 | 0 | 0 | 0 | 0 |
| ENSG00000224406 | 0 | 0 | 0 | 0 | 0 | 0 |
| ENSG00000226859 | 0 | 0 | 0 | 0 | 0 | 0 |
| ENSG00000213139 | 30 | 9 | 11 | 22 | 17 | 7 |
| ENSG00000113838 | 428 | 207 | 222 | 175 | 246 | 224 |
| ENSG00000090520 | 1642 | 884 | 1160 | 982 | 1066 | 1051 |
| ENSG00000197099 | 0 | 0 | 1 | 0 | 1 | 0 |
| ENSG00000283149 | 0 | 0 | 0 | 0 | 0 | 0 |
| ENSG00000283390 | 0 | 0 | 0 | 0 | 1 | 0 |
| ENSG00000145192 | 0 | 0 | 0 | 0 | 0 | 0 |
| ENSG00000275696 | 0 | 0 | 0 | 0 | 0 | 0 |
| ENSG00000090512 | 0 | 0 | 0 | 0 | 0 | 0 |
| ENSG00000113905 | 2 | 1 | 0 | 0 | 0 | 0 |
| ENSG00000225175 | 0 | 0 | 0 | 0 | 0 | 0 |
| ENSG00000226724 | 0 | 0 | 0 | 0 | 0 | 0 |
| ENSG00000113889 | 0 | 2 | 0 | 0 | 0 | 0 |
| ENSG00000226652 | 0 | 0 | 0 | 0 | 0 | 1 |
| ENSG00000207505 | 0 | 0 | 0 | 0 | 0 | 0 |
| ENSG00000230951 | 0 | 0 | 0 | 0 | 0 | 0 |
| ENSG00000263826 | 38 | 23 | 21 | 13 | 17 | 22 |
| ENSG00000156976 | 14308 | 7163 | 9175 | 6167 | 8066 | 7349 |
| ENSG00000238942 | 0 | 0 | 0 | 0 | 0 | 0 |
| ENSG00000200418 | 0 | 0 | 0 | 0 | 0 | 0 |
| ENSG00000283958 | 0 | 0 | 0 | 0 | 0 | 0 |
| ENSG00000221420 | 0 | 0 | 0 | 0 | 0 | 0 |
| ENSG00000200320 | 0 | 0 | 0 | 0 | 0 | 0 |
| ENSG00000263776 | 0 | 0 | 0 | 0 | 0 | 0 |
| ENSG00000163918 | 539 | 307 | 335 | 292 | 350 | 352 |
| ENSG00000231724 | 0 | 0 | 0 | 0 | 0 | 0 |
| ENSG00000232233 | 10 | 6 | 5 | 2 | 7 | 4 |
| ENSG00000231982 | 11 | 18 | 11 | 11 | 29 | 17 |
| ENSG00000181092 | 0 | 0 | 0 | 0 | 0 | 1 |
| ENSG00000226482 | 0 | 0 | 0 | 0 | 0 | 0 |
| ENSG00000223803 | 2 | 2 | 4 | 5 | 2 | 7 |
| ENSG00000233565 | 0 | 0 | 0 | 0 | 0 | 0 |
| ENSG00000073849 | 17 | 20 | 18 | 18 | 23 | 12 |
| ENSG00000163923 | 273 | 120 | 134 | 220 | 194 | 198 |
| ENSG00000198491 | 0 | 0 | 0 | 0 | 0 | 0 |
| ENSG00000175077 | 0 | 0 | 0 | 0 | 0 | 0 |
| ENSG00000127241 | 2 | 0 | 3 | 1 | 0 | 0 |
| ENSG00000283175 | 0 | 0 | 0 | 0 | 0 | 0 |
| ENSG00000136514 | 10 | 0 | 0 | 0 | 0 | 1 |
| ENSG00000280270 | 0 | 0 | 0 | 0 | 0 | 0 |
| ENSG00000239093 | 0 | 1 | 2 | 1 | 0 | 1 |

| | | | | | | |
|---|---|---|---|---|---|---|
| ENSG00000228952 | 28 | 15 | 26 | 11 | 19 | 21 |
| ENSG00000157005 | 0 | 0 | 0 | 0 | 0 | 0 |
| ENSG00000198471 | 0 | 0 | 0 | 0 | 0 | 0 |
| ENSG00000228804 | 0 | 0 | 0 | 0 | 0 | 0 |
| ENSG00000113916 | 560 | 233 | 259 | 188 | 220 | 221 |
| ENSG00000285938 | 4 | 1 | 1 | 0 | 1 | 1 |
| ENSG00000224049 | 0 | 0 | 0 | 0 | 0 | 0 |
| ENSG00000236864 | 0 | 0 | 0 | 0 | 0 | 0 |
| ENSG00000224187 | 0 | 0 | 0 | 0 | 0 | 0 |
| ENSG00000225058 | 0 | 0 | 0 | 0 | 0 | 0 |
| ENSG00000236412 | 0 | 0 | 0 | 0 | 0 | 0 |
| ENSG00000234238 | 0 | 0 | 0 | 0 | 0 | 0 |
| ENSG00000270959 | 110 | 62 | 73 | 49 | 58 | 50 |
| ENSG00000145012 | 1831 | 976 | 899 | 491 | 1707 | 1170 |
| ENSG00000279891 | 0 | 0 | 0 | 0 | 0 | 0 |
| ENSG00000224563 | 0 | 0 | 0 | 0 | 0 | 0 |
| ENSG00000207651 | 0 | 0 | 0 | 0 | 0 | 0 |
| ENSG00000234076 | 0 | 0 | 0 | 0 | 1 | 0 |
| ENSG00000188001 | 42 | 28 | 36 | 30 | 27 | 44 |
| ENSG00000236182 | 14 | 2 | 7 | 1 | 3 | 4 |
| ENSG00000230115 | 0 | 0 | 0 | 0 | 0 | 0 |
| ENSG00000073282 | 68 | 37 | 30 | 18 | 32 | 13 |
| ENSG00000227971 | 0 | 0 | 0 | 0 | 0 | 0 |
| ENSG00000216058 | 0 | 0 | 0 | 0 | 0 | 0 |
| ENSG00000090530 | 0 | 1 | 1 | 0 | 0 | 2 |
| ENSG00000230077 | 0 | 0 | 0 | 0 | 0 | 0 |
| ENSG00000225764 | 0 | 0 | 0 | 0 | 0 | 0 |
| ENSG00000212489 | 0 | 0 | 0 | 0 | 0 | 0 |
| ENSG00000238077 | 0 | 0 | 0 | 0 | 0 | 0 |
| ENSG00000163347 | 50 | 36 | 25 | 31 | 52 | 42 |
| ENSG00000113946 | 0 | 0 | 0 | 0 | 0 | 0 |
| ENSG00000198398 | 0 | 0 | 0 | 0 | 0 | 0 |
| ENSG00000196083 | 128 | 64 | 95 | 45 | 104 | 54 |
| ENSG00000231047 | 0 | 0 | 0 | 0 | 0 | 0 |
| ENSG00000223117 | 0 | 0 | 1 | 0 | 0 | 1 |
| ENSG00000273370 | 0 | 0 | 0 | 0 | 0 | 0 |
| ENSG00000228730 | 0 | 0 | 0 | 0 | 0 | 0 |
| ENSG00000232610 | 0 | 0 | 0 | 0 | 0 | 0 |
| ENSG00000214147 | 0 | 0 | 0 | 0 | 0 | 0 |
| ENSG00000205835 | 0 | 0 | 0 | 0 | 0 | 0 |
| ENSG00000188729 | 0 | 0 | 0 | 0 | 0 | 0 |
| ENSG00000233308 | 0 | 0 | 0 | 0 | 0 | 0 |
| ENSG00000188958 | 0 | 1 | 0 | 0 | 0 | 0 |
| ENSG00000152492 | 838 | 445 | 531 | 362 | 507 | 418 |
| ENSG00000223812 | 0 | 1 | 0 | 0 | 0 | 0 |
| ENSG00000253548 | 0 | 0 | 0 | 0 | 0 | 0 |
| ENSG00000201860 | 0 | 0 | 0 | 0 | 0 | 0 |
| ENSG00000237653 | 0 | 0 | 0 | 0 | 0 | 1 |
| ENSG00000232353 | 0 | 0 | 0 | 0 | 1 | 0 |
| ENSG00000222583 | 0 | 0 | 0 | 0 | 0 | 0 |

| | | | | | | |
|---|---|---|---|---|---|---|
| ENSG00000114279 | 167 | 82 | 102 | 70 | 92 | 63 |
| ENSG00000231383 | 0 | 0 | 0 | 0 | 0 | 0 |
| ENSG00000230126 | 10 | 10 | 6 | 9 | 6 | 5 |
| ENSG00000226709 | 3 | 2 | 4 | 0 | 3 | 2 |
| ENSG00000238902 | 0 | 0 | 0 | 0 | 0 | 0 |
| ENSG00000200184 | 0 | 0 | 0 | 0 | 0 | 0 |
| ENSG00000180611 | 1267 | 646 | 824 | 834 | 1019 | 994 |
| ENSG00000232130 | 0 | 0 | 0 | 0 | 0 | 0 |
| ENSG00000236229 | 0 | 0 | 0 | 0 | 0 | 0 |
| ENSG00000127252 | 0 | 0 | 0 | 0 | 0 | 0 |
| ENSG00000187527 | 0 | 0 | 1 | 0 | 0 | 0 |
| ENSG00000236508 | 0 | 0 | 0 | 0 | 0 | 0 |
| ENSG00000127249 | 26 | 11 | 13 | 2 | 4 | 6 |
| ENSG00000225473 | 0 | 0 | 0 | 0 | 0 | 0 |
| ENSG00000236297 | 0 | 0 | 1 | 0 | 3 | 1 |
| ENSG00000198836 | 1286 | 899 | 703 | 388 | 1518 | 782 |
| ENSG00000224855 | 3 | 2 | 1 | 0 | 1 | 2 |
| ENSG00000207370 | 0 | 0 | 0 | 0 | 0 | 1 |
| ENSG00000243991 | 0 | 0 | 0 | 0 | 0 | 0 |
| ENSG00000223986 | 0 | 0 | 0 | 0 | 0 | 0 |
| ENSG00000229155 | 0 | 0 | 0 | 0 | 0 | 1 |
| ENSG00000214146 | 3 | 6 | 4 | 1 | 2 | 3 |
| ENSG00000238043 | 0 | 0 | 1 | 1 | 0 | 1 |
| ENSG00000224403 | 0 | 0 | 0 | 0 | 1 | 0 |
| ENSG00000230102 | 20 | 7 | 16 | 3 | 4 | 2 |
| ENSG00000276407 | 0 | 0 | 0 | 0 | 0 | 0 |
| ENSG00000229964 | 0 | 0 | 0 | 2 | 0 | 0 |
| ENSG00000232874 | 6 | 9 | 0 | 2 | 2 | 3 |
| ENSG00000114315 | 2343 | 1029 | 1212 | 915 | 892 | 1260 |
| ENSG00000242201 | 0 | 0 | 0 | 0 | 0 | 0 |
| ENSG00000225742 | 0 | 1 | 2 | 0 | 1 | 1 |
| ENSG00000238097 | 0 | 0 | 0 | 0 | 0 | 0 |
| ENSG00000228271 | 0 | 0 | 0 | 0 | 0 | 0 |
| ENSG00000214145 | 1 | 0 | 1 | 0 | 2 | 0 |
| ENSG00000178772 | 0 | 1 | 0 | 0 | 0 | 1 |
| ENSG00000172061 | 2 | 2 | 3 | 5 | 3 | 13 |
| ENSG00000178732 | 0 | 0 | 0 | 0 | 0 | 0 |
| ENSG00000133657 | 2080 | 1047 | 1058 | 493 | 1385 | 1014 |
| ENSG00000271483 | 0 | 0 | 0 | 0 | 0 | 0 |
| ENSG00000233058 | 9 | 6 | 5 | 1 | 2 | 4 |
| ENSG00000244675 | 0 | 0 | 0 | 1 | 0 | 0 |
| ENSG00000252053 | 0 | 0 | 0 | 0 | 0 | 0 |
| ENSG00000232873 | 0 | 0 | 0 | 0 | 0 | 0 |
| ENSG00000231770 | 205 | 111 | 157 | 124 | 134 | 156 |
| ENSG00000145014 | 610 | 278 | 275 | 297 | 324 | 309 |
| ENSG00000229334 | 15 | 8 | 10 | 1 | 13 | 14 |
| ENSG00000272707 | 0 | 2 | 2 | 0 | 2 | 2 |
| ENSG00000041802 | 1743 | 899 | 1099 | 893 | 1156 | 1041 |
| ENSG00000185112 | 216 | 82 | 101 | 67 | 69 | 80 |
| ENSG00000233987 | 0 | 0 | 0 | 0 | 0 | 0 |

| | | | | | | |
|---|---|---|---|---|---|---|
| ENSG00000237222 | 0 | 0 | 3 | 0 | 1 | 0 |
| ENSG00000234805 | 0 | 0 | 0 | 0 | 0 | 0 |
| ENSG00000230401 | 0 | 0 | 0 | 0 | 0 | 0 |
| ENSG00000238031 | 0 | 0 | 0 | 0 | 0 | 0 |
| ENSG00000173950 | 852 | 407 | 521 | 450 | 524 | 521 |
| ENSG00000233303 | 0 | 0 | 0 | 0 | 0 | 0 |
| ENSG00000265333 | 0 | 0 | 0 | 0 | 0 | 0 |
| ENSG00000244314 | 0 | 0 | 0 | 0 | 0 | 0 |
| ENSG00000230266 | 0 | 1 | 1 | 0 | 0 | 0 |
| ENSG00000206600 | 0 | 0 | 0 | 0 | 0 | 0 |
| ENSG00000114331 | 1244 | 663 | 686 | 293 | 853 | 627 |
| ENSG00000229325 | 4 | 4 | 2 | 3 | 3 | 3 |
| ENSG00000227433 | 0 | 0 | 0 | 0 | 0 | 0 |
| ENSG00000207368 | 0 | 0 | 0 | 0 | 0 | 0 |
| ENSG00000184203 | 1871 | 1012 | 1166 | 1221 | 1462 | 1297 |
| ENSG00000252620 | 2 | 0 | 3 | 0 | 2 | 2 |
| ENSG00000223711 | 2 | 1 | 0 | 0 | 0 | 0 |
| ENSG00000265882 | 0 | 0 | 0 | 0 | 0 | 0 |
| ENSG00000236844 | 0 | 0 | 0 | 0 | 0 | 1 |
| ENSG00000236425 | 0 | 0 | 0 | 0 | 0 | 0 |
| ENSG00000189058 | 1017 | 507 | 560 | 324 | 408 | 386 |
| ENSG00000224769 | 59 | 22 | 25 | 14 | 22 | 21 |
| ENSG00000282206 | 1 | 3 | 8 | 2 | 4 | 5 |
| ENSG00000271662 | 40 | 22 | 30 | 22 | 19 | 27 |
| ENSG00000229178 | 22 | 10 | 15 | 11 | 12 | 11 |
| ENSG00000242086 | 565 | 344 | 487 | 233 | 331 | 360 |
| ENSG00000215837 | 0 | 0 | 0 | 0 | 0 | 0 |
| ENSG00000207650 | 26 | 9 | 24 | 22 | 30 | 18 |
| ENSG00000283426 | 0 | 0 | 0 | 0 | 0 | 0 |
| ENSG00000176945 | 136 | 88 | 106 | 71 | 74 | 76 |
| ENSG00000145113 | 7 | 1 | 0 | 0 | 1 | 1 |
| ENSG00000223783 | 0 | 0 | 1 | 1 | 1 | 0 |
| ENSG00000235836 | 0 | 0 | 0 | 0 | 0 | 0 |
| ENSG00000061938 | 2925 | 1271 | 1670 | 1429 | 1454 | 1486 |
| ENSG00000276489 | 0 | 0 | 0 | 0 | 0 | 0 |
| ENSG00000224614 | 20 | 5 | 10 | 5 | 14 | 5 |
| ENSG00000226155 | 2 | 2 | 4 | 5 | 0 | 5 |
| ENSG00000260261 | 329 | 141 | 269 | 136 | 178 | 159 |
| ENSG00000239122 | 7 | 2 | 10 | 8 | 3 | 5 |
| ENSG00000231443 | 24 | 25 | 18 | 8 | 21 | 25 |
| ENSG00000185485 | 592 | 247 | 373 | 192 | 255 | 292 |
| ENSG00000231464 | 5 | 2 | 2 | 0 | 1 | 0 |
| ENSG00000072274 | 4025 | 2173 | 2494 | 1698 | 2865 | 2794 |
| ENSG00000228413 | 4 | 0 | 1 | 0 | 2 | 0 |
| ENSG00000252174 | 0 | 0 | 0 | 0 | 1 | 0 |
| ENSG00000224652 | 24 | 21 | 11 | 16 | 10 | 18 |
| ENSG00000163958 | 0 | 2 | 0 | 1 | 0 | 0 |
| ENSG00000222335 | 0 | 0 | 0 | 0 | 0 | 0 |
| ENSG00000163959 | 46 | 45 | 34 | 34 | 40 | 42 |
| ENSG00000161217 | 3732 | 1881 | 2417 | 1693 | 2121 | 2189 |

| | | | | | | |
|---|---|---|---|---|---|---|
| ENSG00000272741 | 0 | 0 | 0 | 0 | 0 | 0 |
| ENSG00000228028 | 4 | 0 | 0 | 0 | 1 | 1 |
| ENSG00000213123 | 152 | 103 | 105 | 105 | 114 | 97 |
| ENSG00000237418 | 8 | 5 | 3 | 1 | 3 | 3 |
| ENSG00000273331 | 0 | 0 | 0 | 0 | 0 | 0 |
| ENSG00000235897 | 15 | 17 | 4 | 1 | 3 | 10 |
| ENSG00000145107 | 0 | 1 | 2 | 0 | 0 | 1 |
| ENSG00000212146 | 0 | 0 | 0 | 0 | 0 | 0 |
| ENSG00000163960 | 1663 | 774 | 840 | 471 | 893 | 720 |
| ENSG00000206644 | 0 | 0 | 0 | 0 | 0 | 0 |
| ENSG00000241868 | 0 | 0 | 0 | 0 | 0 | 1 |
| ENSG00000243339 | 5 | 3 | 4 | 2 | 3 | 3 |
| ENSG00000226345 | 0 | 0 | 2 | 0 | 0 | 0 |
| ENSG00000225822 | 0 | 0 | 0 | 0 | 0 | 0 |
| ENSG00000225240 | 0 | 0 | 0 | 0 | 0 | 0 |
| ENSG00000163961 | 514 | 273 | 350 | 216 | 328 | 319 |
| ENSG00000273013 | 0 | 0 | 1 | 0 | 2 | 1 |
| ENSG00000214097 | 0 | 0 | 0 | 0 | 1 | 0 |
| ENSG00000213358 | 0 | 0 | 0 | 0 | 0 | 0 |
| ENSG00000225770 | 0 | 4 | 1 | 1 | 0 | 1 |
| ENSG00000185798 | 295 | 176 | 224 | 167 | 213 | 167 |
| ENSG00000174013 | 1444 | 653 | 832 | 582 | 853 | 736 |
| ENSG00000232065 | 17 | 10 | 11 | 10 | 7 | 9 |
| ENSG00000174004 | 4 | 4 | 2 | 5 | 1 | 2 |
| ENSG00000163964 | 1627 | 672 | 903 | 658 | 907 | 921 |
| ENSG00000174007 | 143 | 79 | 86 | 60 | 132 | 77 |
| ENSG00000201441 | 3 | 1 | 0 | 1 | 0 | 0 |
| ENSG00000180370 | 10617 | 5160 | 6388 | 3879 | 5606 | 5612 |
| ENSG00000272359 | 1 | 0 | 1 | 0 | 1 | 0 |
| ENSG00000206892 | 2 | 0 | 0 | 0 | 1 | 1 |
| ENSG00000119231 | 1613 | 805 | 1026 | 604 | 921 | 837 |
| ENSG00000230732 | 2 | 12 | 9 | 4 | 7 | 3 |
| ENSG00000114503 | 3693 | 1843 | 2281 | 1532 | 2046 | 2110 |
| ENSG00000225578 | 19 | 10 | 14 | 11 | 10 | 9 |
| ENSG00000270170 | 1059 | 565 | 676 | 790 | 688 | 720 |
| ENSG00000119227 | 411 | 230 | 245 | 230 | 217 | 224 |
| ENSG00000233487 | 11 | 11 | 13 | 9 | 6 | 7 |
| ENSG00000163975 | 148 | 68 | 89 | 77 | 75 | 87 |
| ENSG00000228109 | 83 | 37 | 56 | 47 | 49 | 51 |
| ENSG00000075711 | 2511 | 1234 | 1378 | 787 | 1515 | 1168 |
| ENSG00000265850 | 0 | 0 | 0 | 0 | 0 | 0 |
| ENSG00000227375 | 2 | 4 | 1 | 0 | 0 | 2 |
| ENSG00000235126 | 0 | 0 | 0 | 0 | 0 | 0 |
| ENSG00000229912 | 0 | 2 | 3 | 1 | 1 | 0 |
| ENSG00000237167 | 0 | 0 | 0 | 0 | 1 | 0 |
| ENSG00000272989 | 8 | 3 | 7 | 10 | 7 | 17 |
| ENSG00000161267 | 1259 | 574 | 851 | 710 | 678 | 746 |
| ENSG00000214135 | 775 | 361 | 571 | 287 | 415 | 387 |
| ENSG00000213519 | 1 | 2 | 0 | 0 | 0 | 1 |
| ENSG00000249626 | 103 | 47 | 77 | 42 | 60 | 70 |

| | | | | | | |
|---|---|---|---|---|---|---|
| ENSG00000273308 | 3 | 4 | 7 | 2 | 2 | 1 |
| ENSG00000236833 | 12 | 1 | 1 | 2 | 0 | 0 |
| ENSG00000145016 | 1275 | 573 | 719 | 534 | 670 | 743 |
| ENSG00000284146 | 0 | 0 | 0 | 0 | 0 | 0 |
| ENSG00000122068 | 2603 | 1393 | 1353 | 967 | 1779 | 1453 |
| ENSG00000273375 | 2 | 0 | 0 | 0 | 0 | 0 |
| ENSG00000186001 | 1209 | 582 | 712 | 453 | 654 | 608 |
| ENSG00000234136 | 0 | 1 | 0 | 0 | 0 | 0 |
| ENSG00000234742 | 6 | 3 | 0 | 1 | 6 | 2 |
| ENSG00000114473 | 97 | 71 | 82 | 57 | 64 | 75 |
| ENSG00000199306 | 0 | 0 | 0 | 0 | 0 | 0 |
| ENSG00000182899 | 14598 | 7217 | 8755 | 9683 | 8665 | 8594 |
| ENSG00000185621 | 333 | 130 | 152 | 66 | 201 | 104 |
| ENSG00000201622 | 0 | 0 | 0 | 0 | 2 | 0 |
| ENSG00000232832 | 0 | 0 | 0 | 0 | 0 | 0 |
| ENSG00000226435 | 0 | 0 | 0 | 0 | 0 | 0 |
| ENSG00000212297 | 0 | 0 | 0 | 0 | 0 | 0 |
| ENSG00000232783 | 0 | 0 | 0 | 0 | 0 | 0 |
| ENSG00000213113 | 12 | 0 | 1 | 0 | 2 | 0 |
| ENSG00000236246 | 0 | 0 | 0 | 0 | 0 | 0 |
| ENSG00000236438 | 32 | 12 | 12 | 8 | 15 | 20 |
| ENSG00000222792 | 0 | 0 | 0 | 0 | 0 | 0 |
| ENSG00000230304 | 1 | 0 | 0 | 0 | 0 | 0 |
| ENSG00000226008 | 0 | 0 | 0 | 0 | 0 | 0 |
| ENSG00000248302 | 0 | 0 | 0 | 0 | 0 | 0 |
| ENSG00000272602 | 451 | 241 | 264 | 174 | 258 | 253 |
| ENSG00000250238 | 0 | 0 | 0 | 0 | 0 | 0 |
| ENSG00000250312 | 284 | 139 | 148 | 57 | 158 | 111 |
| ENSG00000275426 | 2 | 2 | 3 | 1 | 5 | 2 |
| ENSG00000250892 | 0 | 0 | 5 | 2 | 1 | 4 |
| ENSG00000198155 | 36 | 21 | 19 | 12 | 14 | 24 |
| ENSG00000251188 | 5 | 5 | 3 | 4 | 4 | 5 |
| ENSG00000251689 | 5 | 3 | 1 | 1 | 1 | 2 |
| ENSG00000186777 | 19 | 22 | 15 | 5 | 13 | 11 |
| ENSG00000251535 | 0 | 0 | 0 | 0 | 1 | 1 |
| ENSG00000131127 | 150 | 69 | 76 | 28 | 118 | 59 |
| ENSG00000248564 | 0 | 0 | 1 | 0 | 0 | 0 |
| ENSG00000250321 | 2 | 1 | 3 | 1 | 1 | 4 |
| ENSG00000207642 | 2 | 0 | 0 | 0 | 0 | 0 |
| ENSG00000281016 | 2 | 2 | 2 | 0 | 3 | 0 |
| ENSG00000249025 | 0 | 0 | 0 | 0 | 0 | 0 |
| ENSG00000272885 | 2 | 6 | 2 | 0 | 3 | 1 |
| ENSG00000251595 | 21 | 12 | 15 | 11 | 12 | 12 |
| ENSG00000182903 | 114 | 73 | 68 | 41 | 175 | 81 |
| ENSG00000174227 | 2185 | 964 | 1294 | 933 | 1087 | 1106 |
| ENSG00000273238 | 2 | 0 | 0 | 0 | 1 | 1 |
| ENSG00000283183 | 0 | 2 | 0 | 0 | 0 | 0 |
| ENSG00000133256 | 139 | 56 | 79 | 59 | 65 | 66 |
| ENSG00000242686 | 10 | 1 | 7 | 4 | 10 | 3 |
| ENSG00000272927 | 14 | 2 | 10 | 6 | 4 | 7 |

| | | | | | | |
|---|---|---|---|---|---|---|
| ENSG00000169020 | 684 | 584 | 626 | 729 | 683 | 609 |
| ENSG00000215375 | 374 | 162 | 254 | 194 | 137 | 202 |
| ENSG00000169026 | 15 | 12 | 14 | 11 | 23 | 8 |
| ENSG00000185619 | 3262 | 1416 | 2009 | 1565 | 1820 | 2028 |
| ENSG00000248416 | 0 | 0 | 2 | 0 | 0 | 1 |
| ENSG00000272588 | 5 | 6 | 11 | 1 | 5 | 11 |
| ENSG00000249592 | 94 | 53 | 53 | 30 | 57 | 58 |
| ENSG00000233799 | 8 | 5 | 4 | 6 | 7 | 3 |
| ENSG00000168993 | 178 | 97 | 110 | 61 | 42 | 83 |
| ENSG00000260262 | 0 | 0 | 1 | 0 | 0 | 0 |
| ENSG00000178950 | 4403 | 2027 | 2674 | 2346 | 2304 | 2330 |
| ENSG00000127419 | 743 | 322 | 483 | 336 | 354 | 365 |
| ENSG00000145214 | 870 | 408 | 557 | 543 | 498 | 487 |
| ENSG00000145217 | 106 | 41 | 84 | 62 | 46 | 79 |
| ENSG00000127415 | 598 | 228 | 402 | 315 | 272 | 306 |
| ENSG00000127418 | 856 | 417 | 504 | 518 | 484 | 451 |
| ENSG00000250259 | 0 | 0 | 0 | 0 | 0 | 0 |
| ENSG00000178222 | 0 | 1 | 0 | 0 | 0 | 1 |
| ENSG00000251639 | 1 | 0 | 0 | 0 | 0 | 0 |
| ENSG00000251652 | 0 | 0 | 0 | 0 | 0 | 0 |
| ENSG00000215367 | 0 | 0 | 1 | 0 | 0 | 0 |
| ENSG00000212458 | 0 | 0 | 0 | 0 | 0 | 0 |
| ENSG00000227189 | 0 | 2 | 1 | 1 | 1 | 1 |
| ENSG00000159674 | 15 | 16 | 20 | 6 | 4 | 11 |
| ENSG00000273179 | 0 | 1 | 1 | 2 | 0 | 6 |
| ENSG00000280927 | 90 | 35 | 43 | 32 | 34 | 45 |
| ENSG00000159692 | 3818 | 1843 | 2688 | 2402 | 2114 | 2385 |
| ENSG00000196810 | 776 | 389 | 428 | 367 | 445 | 455 |
| ENSG00000090316 | 2946 | 1405 | 1791 | 1699 | 1670 | 1719 |
| ENSG00000163945 | 610 | 246 | 432 | 192 | 245 | 368 |
| ENSG00000254094 | 0 | 5 | 0 | 0 | 1 | 0 |
| ENSG00000253399 | 0 | 0 | 0 | 2 | 0 | 0 |
| ENSG00000235608 | 0 | 0 | 0 | 0 | 0 | 0 |
| ENSG00000272783 | 0 | 0 | 0 | 0 | 0 | 0 |
| ENSG00000244459 | 36 | 15 | 13 | 20 | 22 | 15 |
| ENSG00000174137 | 68 | 31 | 34 | 38 | 28 | 45 |
| ENSG00000207009 | 0 | 0 | 0 | 0 | 0 | 0 |
| ENSG00000163950 | 1374 | 667 | 801 | 564 | 643 | 714 |
| ENSG00000270195 | 22 | 11 | 21 | 10 | 15 | 16 |
| ENSG00000168936 | 1947 | 846 | 1178 | 1054 | 1011 | 944 |
| ENSG00000013810 | 1442 | 752 | 956 | 1102 | 1084 | 1003 |
| ENSG00000068078 | 662 | 316 | 379 | 280 | 306 | 287 |
| ENSG00000168924 | 1774 | 830 | 1128 | 929 | 929 | 1078 |
| ENSG00000109685 | 4096 | 1752 | 2407 | 1771 | 2159 | 2318 |
| ENSG00000249784 | 9 | 4 | 2 | 0 | 2 | 4 |
| ENSG00000185049 | 1294 | 662 | 777 | 820 | 660 | 728 |
| ENSG00000284587 | 0 | 0 | 0 | 0 | 0 | 0 |
| ENSG00000243449 | 422 | 176 | 234 | 273 | 269 | 227 |
| ENSG00000185818 | 296 | 149 | 170 | 212 | 159 | 185 |
| ENSG00000130997 | 9 | 18 | 9 | 6 | 9 | 8 |

| | | | | | | |
|---|---|---|---|---|---|---|
| ENSG00000249006 | 0 | 0 | 0 | 0 | 0 | 0 |
| ENSG00000250623 | 0 | 0 | 0 | 0 | 0 | 0 |
| ENSG00000214367 | 476 | 246 | 268 | 156 | 229 | 250 |
| ENSG00000249047 | 0 | 0 | 0 | 0 | 1 | 1 |
| ENSG00000123933 | 4782 | 2226 | 2823 | 2747 | 2439 | 2144 |
| ENSG00000284276 | 0 | 0 | 0 | 0 | 0 | 0 |
| ENSG00000159733 | 456 | 194 | 251 | 237 | 231 | 215 |
| ENSG00000251148 | 2 | 1 | 0 | 0 | 1 | 0 |
| ENSG00000239247 | 4 | 0 | 1 | 0 | 0 | 0 |
| ENSG00000249077 | 2 | 0 | 1 | 0 | 0 | 0 |
| ENSG00000206113 | 20 | 7 | 11 | 9 | 12 | 13 |
| ENSG00000063978 | 3142 | 1657 | 1807 | 1569 | 1854 | 1776 |
| ENSG00000248399 | 0 | 0 | 1 | 1 | 0 | 0 |
| ENSG00000251229 | 1 | 3 | 1 | 0 | 1 | 2 |
| ENSG00000125386 | 911 | 396 | 497 | 356 | 450 | 485 |
| ENSG00000248155 | 0 | 0 | 0 | 0 | 0 | 0 |
| ENSG00000206738 | 0 | 0 | 0 | 0 | 0 | 0 |
| ENSG00000168884 | 1528 | 743 | 976 | 1020 | 884 | 902 |
| ENSG00000087266 | 2387 | 1133 | 1535 | 1181 | 1456 | 1478 |
| ENSG00000087274 | 5626 | 2624 | 3507 | 2595 | 2598 | 2800 |
| ENSG00000109736 | 3255 | 1589 | 2065 | 2005 | 1710 | 1704 |
| ENSG00000249673 | 710 | 335 | 463 | 393 | 377 | 339 |
| ENSG00000087269 | 1378 | 850 | 1020 | 1064 | 1034 | 1147 |
| ENSG00000125388 | 118 | 79 | 84 | 60 | 71 | 67 |
| ENSG00000197386 | 6762 | 2922 | 4258 | 2816 | 3317 | 3688 |
| ENSG00000199335 | 0 | 0 | 0 | 0 | 0 | 0 |
| ENSG00000251075 | 0 | 0 | 1 | 0 | 0 | 1 |
| ENSG00000278778 | 0 | 0 | 0 | 0 | 0 | 0 |
| ENSG00000274693 | 0 | 0 | 0 | 0 | 0 | 0 |
| ENSG00000188981 | 20 | 13 | 17 | 13 | 9 | 16 |
| ENSG00000159788 | 711 | 357 | 505 | 368 | 454 | 438 |
| ENSG00000248840 | 0 | 0 | 0 | 0 | 0 | 0 |
| ENSG00000248843 | 0 | 0 | 0 | 0 | 0 | 0 |
| ENSG00000109758 | 38 | 14 | 15 | 16 | 14 | 16 |
| ENSG00000175920 | 958 | 419 | 607 | 562 | 458 | 471 |
| ENSG00000163956 | 4888 | 2187 | 2839 | 2674 | 2309 | 2354 |
| ENSG00000270090 | 0 | 0 | 0 | 0 | 0 | 0 |
| ENSG00000216560 | 0 | 0 | 0 | 0 | 1 | 1 |
| ENSG00000250681 | 0 | 0 | 1 | 0 | 0 | 0 |
| ENSG00000250632 | 0 | 0 | 0 | 0 | 0 | 0 |
| ENSG00000250986 | 488 | 219 | 229 | 247 | 240 | 251 |
| ENSG00000184160 | 2161 | 1111 | 1306 | 1475 | 1212 | 1257 |
| ENSG00000251166 | 0 | 0 | 0 | 0 | 0 | 0 |
| ENSG00000234386 | 0 | 0 | 0 | 0 | 0 | 0 |
| ENSG00000249522 | 0 | 0 | 0 | 0 | 0 | 0 |
| ENSG00000253917 | 0 | 0 | 1 | 0 | 0 | 0 |
| ENSG00000271977 | 0 | 0 | 0 | 0 | 0 | 0 |
| ENSG00000251271 | 5 | 3 | 4 | 1 | 1 | 4 |
| ENSG00000251669 | 143 | 120 | 133 | 120 | 88 | 112 |
| ENSG00000250476 | 0 | 0 | 0 | 0 | 0 | 0 |

| | | | | | | |
|---|---|---|---|---|---|---|
| ENSG00000284727 | 0 | 0 | 0 | 0 | 0 | 0 |
| ENSG00000243894 | 0 | 1 | 0 | 0 | 0 | 0 |
| ENSG00000250940 | 0 | 0 | 0 | 0 | 0 | 0 |
| ENSG00000248669 | 0 | 0 | 0 | 0 | 0 | 0 |
| ENSG00000198787 | 0 | 0 | 0 | 0 | 0 | 0 |
| ENSG00000250381 | 0 | 0 | 0 | 0 | 0 | 0 |
| ENSG00000250710 | 0 | 0 | 0 | 0 | 0 | 0 |
| ENSG00000249844 | 0 | 0 | 0 | 0 | 0 | 0 |
| ENSG00000163982 | 0 | 0 | 0 | 0 | 0 | 0 |
| ENSG00000132406 | 684 | 301 | 396 | 329 | 363 | 393 |
| ENSG00000145220 | 351 | 163 | 161 | 205 | 280 | 242 |
| ENSG00000168826 | 147 | 62 | 95 | 70 | 74 | 74 |
| ENSG00000248516 | 0 | 0 | 0 | 0 | 0 | 2 |
| ENSG00000168824 | 0 | 0 | 0 | 1 | 1 | 1 |
| ENSG00000168818 | 801 | 350 | 410 | 416 | 401 | 441 |
| ENSG00000248221 | 0 | 0 | 0 | 0 | 0 | 0 |
| ENSG00000237170 | 0 | 0 | 0 | 0 | 0 | 0 |
| ENSG00000247708 | 12 | 12 | 12 | 15 | 5 | 12 |
| ENSG00000280310 | 0 | 1 | 0 | 0 | 0 | 0 |
| ENSG00000273396 | 0 | 0 | 0 | 0 | 0 | 0 |
| ENSG00000163132 | 56 | 25 | 21 | 38 | 18 | 39 |
| ENSG00000251587 | 0 | 0 | 0 | 0 | 0 | 0 |
| ENSG00000200761 | 0 | 0 | 0 | 0 | 0 | 0 |
| ENSG00000170891 | 0 | 0 | 0 | 0 | 0 | 0 |
| ENSG00000152953 | 0 | 0 | 0 | 0 | 0 | 0 |
| ENSG00000252266 | 0 | 0 | 0 | 0 | 0 | 0 |
| ENSG00000201496 | 0 | 0 | 0 | 0 | 0 | 0 |
| ENSG00000082929 | 33 | 30 | 38 | 25 | 25 | 22 |
| ENSG00000173040 | 0 | 0 | 0 | 0 | 0 | 0 |
| ENSG00000072840 | 0 | 0 | 0 | 0 | 0 | 0 |
| ENSG00000072832 | 0 | 0 | 0 | 2 | 0 | 1 |
| ENSG00000181215 | 2 | 0 | 1 | 0 | 0 | 0 |
| ENSG00000263631 | 0 | 0 | 0 | 0 | 0 | 0 |
| ENSG00000152969 | 0 | 0 | 1 | 2 | 0 | 2 |
| ENSG00000284684 | 0 | 0 | 0 | 0 | 0 | 0 |
| ENSG00000249896 | 0 | 2 | 4 | 2 | 3 | 6 |
| ENSG00000251408 | 0 | 0 | 0 | 0 | 0 | 0 |
| ENSG00000109501 | 2100 | 1018 | 1186 | 1399 | 1306 | 1169 |
| ENSG00000074211 | 1799 | 874 | 1091 | 944 | 1023 | 1143 |
| ENSG00000013288 | 2017 | 869 | 1199 | 893 | 883 | 1017 |
| ENSG00000179010 | 18767 | 9090 | 11129 | 8376 | 9234 | 9094 |
| ENSG00000251580 | 60 | 49 | 47 | 41 | 39 | 52 |
| ENSG00000284847 | 2 | 1 | 3 | 0 | 1 | 2 |
| ENSG00000170846 | 1475 | 714 | 841 | 747 | 799 | 789 |
| ENSG00000246526 | 31 | 15 | 28 | 26 | 25 | 19 |
| ENSG00000279859 | 1 | 1 | 4 | 0 | 2 | 1 |
| ENSG00000163993 | 1391 | 723 | 795 | 1107 | 1056 | 866 |
| ENSG00000282742 | 0 | 3 | 0 | 0 | 2 | 0 |
| ENSG00000178988 | 3943 | 2017 | 2452 | 1854 | 2315 | 2156 |
| ENSG00000186222 | 954 | 407 | 492 | 526 | 496 | 504 |

| | | | | | | |
|---|---|---|---|---|---|---|
| ENSG00000170871 | 1956 | 872 | 1025 | 488 | 1111 | 905 |
| ENSG00000132405 | 859 | 420 | 563 | 381 | 558 | 616 |
| ENSG00000187904 | 0 | 2 | 0 | 1 | 0 | 0 |
| ENSG00000202392 | 0 | 1 | 0 | 0 | 0 | 0 |
| ENSG00000245748 | 1 | 1 | 0 | 2 | 0 | 2 |
| ENSG00000173013 | 73 | 34 | 34 | 38 | 31 | 33 |
| ENSG00000173011 | 1114 | 586 | 724 | 524 | 577 | 634 |
| ENSG00000109519 | 1790 | 917 | 1067 | 1066 | 1073 | 1150 |
| ENSG00000245468 | 23 | 16 | 13 | 16 | 10 | 10 |
| ENSG00000200867 | 0 | 0 | 0 | 0 | 0 | 0 |
| ENSG00000184985 | 41 | 15 | 25 | 45 | 31 | 42 |
| ENSG00000264658 | 0 | 0 | 0 | 0 | 0 | 0 |
| ENSG00000178597 | 4 | 1 | 5 | 5 | 3 | 0 |
| ENSG00000266690 | 0 | 0 | 0 | 0 | 0 | 0 |
| ENSG00000272620 | 2 | 3 | 3 | 2 | 2 | 1 |
| ENSG00000196526 | 1351 | 618 | 896 | 580 | 725 | 751 |
| ENSG00000228919 | 0 | 0 | 0 | 0 | 0 | 0 |
| ENSG00000163995 | 148 | 56 | 93 | 84 | 75 | 70 |
| ENSG00000207807 | 0 | 0 | 0 | 0 | 0 | 0 |
| ENSG00000273267 | 2 | 0 | 0 | 0 | 0 | 2 |
| ENSG00000251460 | 0 | 0 | 0 | 0 | 0 | 0 |
| ENSG00000250471 | 2 | 1 | 1 | 0 | 1 | 0 |
| ENSG00000125089 | 0 | 5 | 10 | 1 | 3 | 0 |
| ENSG00000170801 | 0 | 0 | 0 | 0 | 0 | 0 |
| ENSG00000249145 | 0 | 0 | 0 | 0 | 0 | 0 |
| ENSG00000251615 | 115 | 48 | 56 | 34 | 42 | 36 |
| ENSG00000087008 | 911 | 468 | 540 | 446 | 439 | 473 |
| ENSG00000202054 | 0 | 0 | 0 | 0 | 0 | 0 |
| ENSG00000155275 | 351 | 150 | 240 | 157 | 168 | 195 |
| ENSG00000251186 | 0 | 0 | 1 | 0 | 0 | 1 |
| ENSG00000205959 | 0 | 2 | 1 | 0 | 1 | 2 |
| ENSG00000155269 | 11 | 9 | 14 | 12 | 6 | 12 |
| ENSG00000109625 | 9 | 8 | 4 | 3 | 7 | 3 |
| ENSG00000250915 | 0 | 0 | 0 | 0 | 0 | 0 |
| ENSG00000215612 | 0 | 0 | 0 | 0 | 0 | 0 |
| ENSG00000258507 | 0 | 0 | 0 | 0 | 0 | 0 |
| ENSG00000197468 | 0 | 0 | 0 | 0 | 0 | 0 |
| ENSG00000249272 | 1 | 0 | 0 | 0 | 0 | 0 |
| ENSG00000250804 | 0 | 0 | 0 | 0 | 0 | 0 |
| ENSG00000249347 | 0 | 0 | 0 | 0 | 0 | 0 |
| ENSG00000284636 | 0 | 0 | 0 | 0 | 2 | 0 |
| ENSG00000271057 | 0 | 0 | 1 | 0 | 0 | 0 |
| ENSG00000251313 | 0 | 0 | 0 | 0 | 0 | 0 |
| ENSG00000250342 | 1 | 0 | 0 | 0 | 0 | 0 |
| ENSG00000249767 | 2 | 1 | 1 | 0 | 2 | 0 |
| ENSG00000163612 | 0 | 1 | 2 | 1 | 1 | 1 |
| ENSG00000250268 | 0 | 0 | 0 | 0 | 0 | 1 |
| ENSG00000229924 | 0 | 0 | 1 | 0 | 0 | 0 |
| ENSG00000231396 | 0 | 0 | 0 | 0 | 0 | 0 |
| ENSG00000233136 | 0 | 0 | 0 | 0 | 0 | 0 |

| | | | | | | |
|---|---|---|---|---|---|---|
| ENSG00000227551 | 0 | 0 | 0 | 0 | 0 | 0 |
| ENSG00000232399 | 0 | 0 | 0 | 0 | 0 | 0 |
| ENSG00000249482 | 0 | 0 | 1 | 0 | 0 | 0 |
| ENSG00000223569 | 0 | 0 | 0 | 0 | 0 | 0 |
| ENSG00000250231 | 0 | 0 | 0 | 0 | 0 | 0 |
| ENSG00000249104 | 0 | 0 | 0 | 0 | 0 | 0 |
| ENSG00000250844 | 0 | 0 | 0 | 0 | 0 | 0 |
| ENSG00000248920 | 0 | 0 | 0 | 0 | 0 | 0 |
| ENSG00000250745 | 0 | 0 | 0 | 0 | 0 | 0 |
| ENSG00000249811 | 0 | 0 | 0 | 0 | 0 | 0 |
| ENSG00000248933 | 0 | 0 | 0 | 0 | 0 | 0 |
| ENSG00000250913 | 0 | 0 | 0 | 0 | 0 | 0 |
| ENSG00000232264 | 0 | 0 | 0 | 0 | 0 | 0 |
| ENSG00000230430 | 0 | 0 | 0 | 0 | 0 | 0 |
| ENSG00000229579 | 0 | 0 | 0 | 0 | 0 | 0 |
| ENSG00000227140 | 0 | 0 | 0 | 0 | 0 | 0 |
| ENSG00000235780 | 0 | 0 | 0 | 0 | 0 | 0 |
| ENSG00000231051 | 0 | 0 | 0 | 0 | 0 | 0 |
| ENSG00000231637 | 0 | 0 | 0 | 0 | 0 | 0 |
| ENSG00000251694 | 0 | 0 | 0 | 0 | 0 | 0 |
| ENSG00000228856 | 0 | 0 | 0 | 0 | 0 | 0 |
| ENSG00000205946 | 0 | 0 | 0 | 0 | 0 | 0 |
| ENSG00000219492 | 0 | 0 | 0 | 0 | 0 | 0 |
| ENSG00000188438 | 0 | 0 | 0 | 0 | 0 | 0 |
| ENSG00000252307 | 0 | 0 | 0 | 0 | 0 | 0 |
| ENSG00000232773 | 0 | 0 | 0 | 0 | 0 | 0 |
| ENSG00000186146 | 0 | 0 | 0 | 0 | 0 | 0 |
| ENSG00000248364 | 0 | 0 | 0 | 0 | 0 | 0 |
| ENSG00000284689 | 0 | 0 | 0 | 0 | 0 | 0 |
| ENSG00000250884 | 0 | 0 | 0 | 0 | 0 | 0 |
| ENSG00000251624 | 1 | 0 | 0 | 0 | 0 | 0 |
| ENSG00000249866 | 0 | 0 | 0 | 0 | 0 | 0 |
| ENSG00000249443 | 0 | 0 | 0 | 0 | 0 | 0 |
| ENSG00000284648 | 3 | 2 | 1 | 2 | 0 | 0 |
| ENSG00000242034 | 0 | 0 | 0 | 0 | 0 | 0 |
| ENSG00000221275 | 0 | 0 | 0 | 0 | 0 | 0 |
| ENSG00000244004 | 0 | 0 | 0 | 0 | 0 | 0 |
| ENSG00000249799 | 1 | 0 | 0 | 0 | 0 | 0 |
| ENSG00000250942 | 2 | 1 | 0 | 0 | 0 | 0 |
| ENSG00000186234 | 0 | 0 | 1 | 0 | 0 | 1 |
| ENSG00000251087 | 0 | 0 | 0 | 0 | 0 | 0 |
| ENSG00000203436 | 0 | 0 | 0 | 0 | 1 | 0 |
| ENSG00000249563 | 0 | 0 | 0 | 0 | 0 | 0 |
| ENSG00000250425 | 0 | 0 | 0 | 0 | 0 | 0 |
| ENSG00000109667 | 0 | 0 | 0 | 0 | 0 | 0 |
| ENSG00000169676 | 0 | 0 | 0 | 0 | 0 | 0 |
| ENSG00000252002 | 0 | 0 | 0 | 0 | 0 | 0 |
| ENSG00000249219 | 0 | 0 | 0 | 0 | 0 | 0 |
| ENSG00000250413 | 0 | 0 | 0 | 0 | 0 | 0 |
| ENSG00000261490 | 19 | 8 | 12 | 5 | 13 | 8 |

| | | | | | | |
|---|---|---|---|---|---|---|
| ENSG00000071127 | 10652 | 5423 | 6465 | 6212 | 6607 | 6148 |
| ENSG00000264931 | 2 | 1 | 0 | 0 | 0 | 0 |
| ENSG00000223086 | 0 | 0 | 0 | 0 | 0 | 0 |
| ENSG00000250393 | 0 | 0 | 0 | 0 | 0 | 0 |
| ENSG00000251296 | 0 | 0 | 0 | 0 | 0 | 0 |
| ENSG00000248777 | 0 | 0 | 0 | 0 | 0 | 0 |
| ENSG00000271544 | 0 | 0 | 0 | 0 | 0 | 0 |
| ENSG00000250040 | 0 | 0 | 0 | 0 | 0 | 0 |
| ENSG00000250573 | 0 | 0 | 0 | 0 | 0 | 0 |
| ENSG00000250074 | 0 | 0 | 0 | 0 | 0 | 0 |
| ENSG00000250505 | 0 | 0 | 0 | 0 | 0 | 0 |
| ENSG00000251278 | 0 | 0 | 0 | 0 | 0 | 0 |
| ENSG00000251338 | 0 | 0 | 0 | 0 | 0 | 0 |
| ENSG00000250613 | 0 | 0 | 0 | 0 | 0 | 0 |
| ENSG00000178163 | 0 | 0 | 0 | 0 | 0 | 0 |
| ENSG00000248419 | 0 | 0 | 0 | 0 | 0 | 0 |
| ENSG00000109684 | 0 | 0 | 0 | 0 | 0 | 0 |
| ENSG00000249334 | 0 | 0 | 0 | 0 | 0 | 0 |
| ENSG00000283083 | 0 | 0 | 0 | 0 | 2 | 0 |
| ENSG00000207716 | 0 | 0 | 0 | 0 | 0 | 0 |
| ENSG00000250896 | 8 | 4 | 5 | 6 | 10 | 4 |
| ENSG00000002587 | 6 | 11 | 9 | 18 | 10 | 5 |
| ENSG00000222182 | 0 | 0 | 0 | 0 | 0 | 0 |
| ENSG00000251152 | 0 | 0 | 0 | 0 | 0 | 0 |
| ENSG00000249631 | 4 | 0 | 0 | 3 | 3 | 0 |
| ENSG00000248300 | 0 | 0 | 0 | 0 | 0 | 0 |
| ENSG00000248262 | 0 | 0 | 0 | 0 | 0 | 0 |
| ENSG00000251210 | 0 | 0 | 0 | 0 | 0 | 0 |
| ENSG00000212282 | 0 | 0 | 0 | 0 | 0 | 0 |
| ENSG00000249995 | 0 | 0 | 0 | 0 | 0 | 0 |
| ENSG00000249780 | 0 | 0 | 0 | 0 | 0 | 0 |
| ENSG00000250371 | 0 | 0 | 0 | 0 | 0 | 0 |
| ENSG00000250098 | 0 | 0 | 0 | 0 | 0 | 0 |
| ENSG00000283271 | 0 | 0 | 0 | 0 | 0 | 0 |
| ENSG00000205940 | 0 | 2 | 1 | 1 | 3 | 5 |
| ENSG00000157869 | 404 | 159 | 195 | 166 | 208 | 190 |
| ENSG00000281202 | 0 | 0 | 0 | 0 | 0 | 0 |
| ENSG00000109705 | 45 | 22 | 24 | 14 | 19 | 13 |
| ENSG00000246095 | 0 | 0 | 0 | 0 | 0 | 0 |
| ENSG00000038219 | 265 | 169 | 142 | 74 | 326 | 178 |
| ENSG00000266240 | 0 | 0 | 0 | 0 | 0 | 0 |
| ENSG00000249148 | 0 | 0 | 0 | 0 | 0 | 0 |
| ENSG00000242209 | 0 | 0 | 0 | 0 | 0 | 0 |
| ENSG00000250634 | 0 | 0 | 0 | 0 | 0 | 0 |
| ENSG00000250497 | 0 | 0 | 0 | 0 | 0 | 0 |
| ENSG00000224569 | 0 | 0 | 0 | 0 | 0 | 0 |
| ENSG00000248698 | 0 | 0 | 0 | 0 | 0 | 0 |
| ENSG00000251679 | 0 | 0 | 0 | 0 | 0 | 0 |
| ENSG00000249988 | 0 | 0 | 0 | 0 | 0 | 0 |
| ENSG00000251412 | 0 | 0 | 0 | 0 | 0 | 0 |

| | | | | | | |
|---|---|---|---|---|---|---|
| ENSG00000248425 | 0 | 0 | 0 | 0 | 0 | 0 |
| ENSG00000248360 | 4 | 0 | 3 | 0 | 4 | 1 |
| ENSG00000251252 | 0 | 0 | 0 | 0 | 0 | 0 |
| ENSG00000202449 | 0 | 0 | 0 | 0 | 0 | 0 |
| ENSG00000247624 | 1 | 1 | 2 | 1 | 1 | 1 |
| ENSG00000137449 | 555 | 269 | 318 | 182 | 315 | 282 |
| ENSG00000249252 | 0 | 0 | 0 | 1 | 1 | 0 |
| ENSG00000199420 | 0 | 0 | 0 | 0 | 0 | 0 |
| ENSG00000163145 | 0 | 0 | 0 | 0 | 0 | 1 |
| ENSG00000251379 | 0 | 0 | 0 | 0 | 0 | 0 |
| ENSG00000048342 | 221 | 146 | 141 | 108 | 179 | 155 |
| ENSG00000249930 | 1 | 0 | 0 | 0 | 0 | 0 |
| ENSG00000273133 | 0 | 3 | 0 | 2 | 1 | 2 |
| ENSG00000118564 | 1372 | 669 | 727 | 628 | 765 | 698 |
| ENSG00000237765 | 361 | 195 | 279 | 157 | 277 | 215 |
| ENSG00000109743 | 2 | 1 | 2 | 2 | 0 | 6 |
| ENSG00000214846 | 2 | 0 | 1 | 0 | 1 | 0 |
| ENSG00000248188 | 0 | 0 | 0 | 0 | 0 | 0 |
| ENSG00000004468 | 6 | 3 | 4 | 4 | 2 | 2 |
| ENSG00000251644 | 0 | 0 | 0 | 0 | 0 | 0 |
| ENSG00000137440 | 0 | 0 | 0 | 2 | 0 | 1 |
| ENSG00000137441 | 4 | 0 | 0 | 0 | 0 | 1 |
| ENSG00000007062 | 0 | 0 | 0 | 0 | 0 | 0 |
| ENSG00000169762 | 769 | 330 | 427 | 264 | 335 | 387 |
| ENSG00000249234 | 2 | 0 | 0 | 0 | 0 | 0 |
| ENSG00000263327 | 158 | 50 | 67 | 79 | 69 | 60 |
| ENSG00000242358 | 0 | 0 | 0 | 0 | 0 | 0 |
| ENSG00000248851 | 0 | 0 | 0 | 0 | 0 | 0 |
| ENSG00000249506 | 9 | 2 | 3 | 2 | 4 | 3 |
| ENSG00000248138 | 13 | 8 | 14 | 11 | 9 | 6 |
| ENSG00000169744 | 0 | 1 | 1 | 3 | 0 | 3 |
| ENSG00000249998 | 0 | 0 | 0 | 0 | 0 | 0 |
| ENSG00000251212 | 0 | 0 | 0 | 0 | 0 | 0 |
| ENSG00000250819 | 0 | 0 | 0 | 0 | 0 | 0 |
| ENSG00000206780 | 0 | 0 | 0 | 0 | 0 | 0 |
| ENSG00000215278 | 0 | 0 | 0 | 0 | 0 | 0 |
| ENSG00000151552 | 212 | 108 | 133 | 106 | 116 | 108 |
| ENSG00000249581 | 0 | 0 | 0 | 0 | 0 | 0 |
| ENSG00000249488 | 0 | 0 | 0 | 0 | 0 | 0 |
| ENSG00000283219 | 0 | 0 | 0 | 0 | 0 | 0 |
| ENSG00000002549 | 2011 | 960 | 1200 | 892 | 1140 | 1146 |
| ENSG00000249502 | 2 | 3 | 0 | 2 | 1 | 1 |
| ENSG00000118579 | 1994 | 959 | 1178 | 885 | 1061 | 1009 |
| ENSG00000047662 | 8 | 2 | 4 | 2 | 1 | 1 |
| ENSG00000163257 | 1014 | 526 | 646 | 389 | 602 | 609 |
| ENSG00000109805 | 186 | 115 | 106 | 73 | 264 | 145 |
| ENSG00000178177 | 206 | 117 | 101 | 41 | 146 | 99 |
| ENSG00000235413 | 4 | 2 | 4 | 1 | 0 | 0 |
| ENSG00000251048 | 0 | 0 | 0 | 0 | 0 | 0 |
| ENSG00000279918 | 0 | 0 | 0 | 0 | 0 | 0 |

| | | | | | | |
|---|---|---|---|---|---|---|
| ENSG00000248238 | 4 | 1 | 1 | 2 | 3 | 1 |
| ENSG00000251816 | 0 | 0 | 0 | 0 | 0 | 0 |
| ENSG00000279379 | 0 | 0 | 0 | 0 | 0 | 0 |
| ENSG00000248515 | 4 | 3 | 11 | 4 | 4 | 2 |
| ENSG00000249441 | 0 | 0 | 0 | 0 | 0 | 0 |
| ENSG00000241612 | 0 | 0 | 0 | 0 | 1 | 0 |
| ENSG00000250611 | 0 | 0 | 0 | 0 | 0 | 0 |
| ENSG00000145147 | 280 | 134 | 194 | 55 | 55 | 91 |
| ENSG00000248228 | 0 | 0 | 0 | 0 | 0 | 0 |
| ENSG00000207732 | 0 | 0 | 0 | 0 | 0 | 0 |
| ENSG00000163138 | 255 | 125 | 183 | 134 | 185 | 146 |
| ENSG00000185774 | 21 | 6 | 7 | 5 | 12 | 15 |
| ENSG00000272995 | 0 | 0 | 0 | 0 | 0 | 1 |
| ENSG00000250243 | 0 | 0 | 0 | 0 | 0 | 0 |
| ENSG00000283281 | 0 | 0 | 0 | 0 | 0 | 0 |
| ENSG00000250092 | 0 | 0 | 0 | 0 | 0 | 0 |
| ENSG00000242768 | 0 | 0 | 0 | 0 | 0 | 0 |
| ENSG00000248343 | 0 | 0 | 0 | 0 | 0 | 0 |
| ENSG00000280650 | 0 | 0 | 0 | 0 | 0 | 0 |
| ENSG00000250039 | 0 | 0 | 0 | 0 | 1 | 0 |
| ENSG00000239001 | 0 | 0 | 0 | 0 | 0 | 0 |
| ENSG00000152990 | 904 | 509 | 531 | 435 | 628 | 556 |
| ENSG00000244538 | 1 | 0 | 0 | 0 | 1 | 0 |
| ENSG00000251516 | 0 | 0 | 0 | 0 | 0 | 0 |
| ENSG00000249948 | 0 | 0 | 0 | 0 | 0 | 0 |
| ENSG00000237350 | 12 | 5 | 6 | 3 | 8 | 4 |
| ENSG00000248837 | 0 | 0 | 0 | 0 | 0 | 0 |
| ENSG00000249645 | 0 | 0 | 0 | 0 | 0 | 0 |
| ENSG00000249547 | 0 | 0 | 0 | 0 | 0 | 0 |
| ENSG00000251220 | 0 | 0 | 0 | 0 | 0 | 0 |
| ENSG00000250137 | 0 | 0 | 0 | 0 | 0 | 0 |
| ENSG00000251292 | 0 | 0 | 0 | 0 | 0 | 0 |
| ENSG00000109819 | 0 | 0 | 0 | 0 | 0 | 0 |
| ENSG00000249453 | 0 | 0 | 0 | 0 | 0 | 0 |
| ENSG00000109606 | 5450 | 3035 | 3183 | 2030 | 3398 | 2849 |
| ENSG00000207697 | 1 | 0 | 0 | 0 | 3 | 1 |
| ENSG00000243005 | 0 | 0 | 0 | 0 | 0 | 0 |
| ENSG00000249256 | 0 | 0 | 0 | 0 | 0 | 0 |
| ENSG00000261121 | 2 | 0 | 3 | 0 | 0 | 1 |
| ENSG00000251638 | 0 | 0 | 0 | 0 | 0 | 0 |
| ENSG00000248444 | 0 | 0 | 0 | 0 | 0 | 0 |
| ENSG00000109610 | 1 | 1 | 1 | 0 | 1 | 0 |
| ENSG00000181982 | 520 | 224 | 304 | 248 | 308 | 230 |
| ENSG00000153012 | 24 | 4 | 7 | 3 | 8 | 13 |
| ENSG00000109618 | 576 | 284 | 350 | 176 | 340 | 258 |
| ENSG00000281501 | 78 | 56 | 56 | 41 | 56 | 36 |
| ENSG00000281028 | 0 | 0 | 0 | 0 | 0 | 0 |
| ENSG00000271172 | 0 | 0 | 1 | 0 | 0 | 0 |
| ENSG00000038210 | 752 | 369 | 444 | 335 | 409 | 412 |
| ENSG00000278135 | 0 | 0 | 0 | 0 | 0 | 0 |

| | | | | | | |
|---|---|---|---|---|---|---|
| ENSG00000168228 | 202 | 84 | 121 | 73 | 118 | 88 |
| ENSG00000053900 | 721 | 379 | 425 | 281 | 492 | 401 |
| ENSG00000249320 | 0 | 0 | 0 | 0 | 0 | 0 |
| ENSG00000248608 | 0 | 0 | 0 | 0 | 0 | 0 |
| ENSG00000238632 | 0 | 0 | 0 | 0 | 0 | 0 |
| ENSG00000248545 | 0 | 0 | 0 | 0 | 0 | 0 |
| ENSG00000228154 | 0 | 0 | 0 | 0 | 0 | 0 |
| ENSG00000157765 | 0 | 0 | 0 | 0 | 0 | 0 |
| ENSG00000242640 | 0 | 0 | 0 | 0 | 0 | 0 |
| ENSG00000249277 | 0 | 0 | 0 | 0 | 0 | 0 |
| ENSG00000250050 | 0 | 0 | 0 | 0 | 0 | 0 |
| ENSG00000248133 | 0 | 0 | 0 | 0 | 0 | 0 |
| ENSG00000251353 | 0 | 0 | 0 | 0 | 0 | 0 |
| ENSG00000248654 | 0 | 0 | 0 | 0 | 0 | 0 |
| ENSG00000091490 | 248 | 142 | 185 | 192 | 223 | 185 |
| ENSG00000250541 | 0 | 0 | 0 | 0 | 0 | 0 |
| ENSG00000250317 | 552 | 266 | 279 | 319 | 312 | 339 |
| ENSG00000251009 | 0 | 0 | 0 | 0 | 0 | 0 |
| ENSG00000283314 | 0 | 0 | 0 | 0 | 0 | 0 |
| ENSG00000248452 | 0 | 0 | 0 | 0 | 0 | 0 |
| ENSG00000168214 | 1516 | 803 | 708 | 401 | 946 | 738 |
| ENSG00000163394 | 0 | 0 | 0 | 0 | 0 | 0 |
| ENSG00000109680 | 57 | 28 | 44 | 43 | 53 | 39 |
| ENSG00000200999 | 0 | 0 | 0 | 0 | 0 | 0 |
| ENSG00000109689 | 811 | 312 | 454 | 205 | 276 | 404 |
| ENSG00000240005 | 6 | 0 | 2 | 4 | 9 | 3 |
| ENSG00000248340 | 0 | 0 | 0 | 0 | 0 | 0 |
| ENSG00000251080 | 0 | 0 | 0 | 0 | 0 | 0 |
| ENSG00000205830 | 0 | 0 | 0 | 0 | 0 | 0 |
| ENSG00000249699 | 0 | 0 | 0 | 0 | 0 | 0 |
| ENSG00000222206 | 0 | 0 | 0 | 0 | 0 | 0 |
| ENSG00000251325 | 0 | 0 | 0 | 0 | 0 | 0 |
| ENSG00000250858 | 0 | 0 | 0 | 0 | 0 | 0 |
| ENSG00000251410 | 1 | 1 | 0 | 0 | 0 | 0 |
| ENSG00000250038 | 2 | 0 | 0 | 0 | 0 | 0 |
| ENSG00000250064 | 0 | 0 | 0 | 0 | 0 | 2 |
| ENSG00000243548 | 0 | 0 | 0 | 0 | 0 | 0 |
| ENSG00000283275 | 0 | 0 | 0 | 0 | 0 | 0 |
| ENSG00000250927 | 0 | 0 | 0 | 0 | 0 | 0 |
| ENSG00000250190 | 0 | 0 | 0 | 0 | 0 | 0 |
| ENSG00000276507 | 0 | 0 | 0 | 0 | 0 | 0 |
| ENSG00000249228 | 0 | 0 | 0 | 0 | 0 | 0 |
| ENSG00000248176 | 0 | 0 | 0 | 0 | 0 | 0 |
| ENSG00000248338 | 0 | 0 | 0 | 0 | 0 | 0 |
| ENSG00000251373 | 0 | 0 | 0 | 0 | 0 | 0 |
| ENSG00000251113 | 0 | 0 | 0 | 0 | 0 | 0 |
| ENSG00000283666 | 0 | 0 | 0 | 0 | 0 | 0 |
| ENSG00000249542 | 0 | 0 | 0 | 0 | 0 | 0 |
| ENSG00000249564 | 0 | 0 | 0 | 0 | 0 | 0 |
| ENSG00000244031 | 0 | 0 | 0 | 0 | 0 | 0 |

| | | | | | | |
|---|---|---|---|---|---|---|
| ENSG00000248281 | 0 | 0 | 0 | 0 | 0 | 0 |
| ENSG00000169851 | 496 | 189 | 443 | 79 | 101 | 197 |
| ENSG00000249678 | 0 | 0 | 0 | 0 | 0 | 0 |
| ENSG00000250624 | 0 | 0 | 0 | 0 | 0 | 0 |
| ENSG00000251182 | 0 | 0 | 0 | 0 | 0 | 0 |
| ENSG00000251434 | 0 | 3 | 2 | 0 | 1 | 2 |
| ENSG00000249882 | 0 | 0 | 0 | 0 | 0 | 0 |
| ENSG00000251129 | 129 | 60 | 76 | 44 | 74 | 56 |
| ENSG00000251329 | 0 | 0 | 0 | 0 | 0 | 0 |
| ENSG00000250076 | 0 | 0 | 0 | 0 | 0 | 0 |
| ENSG00000249452 | 0 | 0 | 0 | 0 | 0 | 0 |
| ENSG00000249766 | 0 | 0 | 0 | 0 | 0 | 0 |
| ENSG00000248417 | 0 | 0 | 0 | 0 | 0 | 0 |
| ENSG00000250954 | 0 | 1 | 0 | 1 | 0 | 0 |
| ENSG00000250723 | 0 | 0 | 0 | 0 | 0 | 0 |
| ENSG00000239983 | 0 | 1 | 0 | 0 | 0 | 0 |
| ENSG00000248685 | 0 | 0 | 0 | 0 | 0 | 0 |
| ENSG00000250597 | 0 | 0 | 0 | 0 | 0 | 0 |
| ENSG00000275250 | 0 | 0 | 0 | 0 | 0 | 0 |
| ENSG00000238694 | 0 | 0 | 0 | 0 | 0 | 0 |
| ENSG00000250416 | 0 | 0 | 0 | 0 | 0 | 0 |
| ENSG00000206665 | 0 | 0 | 0 | 0 | 0 | 0 |
| ENSG00000047365 | 116 | 65 | 46 | 36 | 158 | 74 |
| ENSG00000248466 | 0 | 0 | 0 | 0 | 0 | 0 |
| ENSG00000247193 | 0 | 0 | 0 | 0 | 0 | 0 |
| ENSG00000197057 | 0 | 0 | 0 | 0 | 0 | 0 |
| ENSG00000251438 | 0 | 0 | 0 | 0 | 0 | 0 |
| ENSG00000283207 | 0 | 0 | 0 | 0 | 0 | 0 |
| ENSG00000248215 | 0 | 0 | 0 | 0 | 0 | 0 |
| ENSG00000251588 | 0 | 0 | 0 | 0 | 0 | 0 |
| ENSG00000261761 | 0 | 0 | 0 | 0 | 0 | 0 |
| ENSG00000247810 | 0 | 0 | 0 | 0 | 0 | 0 |
| ENSG00000264319 | 0 | 0 | 0 | 0 | 0 | 0 |
| ENSG00000174145 | 4 | 0 | 1 | 1 | 3 | 3 |
| ENSG00000154274 | 273 | 220 | 130 | 150 | 225 | 147 |
| ENSG00000248936 | 2 | 0 | 2 | 1 | 3 | 1 |
| ENSG00000181826 | 173 | 64 | 112 | 43 | 74 | 84 |
| ENSG00000207075 | 0 | 0 | 0 | 0 | 0 | 0 |
| ENSG00000239532 | 0 | 0 | 0 | 0 | 0 | 0 |
| ENSG00000169299 | 1609 | 817 | 970 | 709 | 1175 | 952 |
| ENSG00000279386 | 8 | 0 | 3 | 0 | 2 | 2 |
| ENSG00000249863 | 23 | 5 | 7 | 3 | 11 | 11 |
| ENSG00000065882 | 864 | 495 | 538 | 390 | 489 | 466 |
| ENSG00000250254 | 0 | 0 | 0 | 0 | 1 | 0 |
| ENSG00000251332 | 0 | 0 | 0 | 0 | 0 | 0 |
| ENSG00000249649 | 0 | 0 | 0 | 0 | 0 | 0 |
| ENSG00000251642 | 0 | 0 | 0 | 0 | 0 | 0 |
| ENSG00000248227 | 0 | 0 | 0 | 0 | 0 | 0 |
| ENSG00000280015 | 0 | 0 | 0 | 0 | 0 | 0 |
| ENSG00000249534 | 0 | 0 | 1 | 0 | 0 | 0 |

| | | | | | | |
|---|---|---|---|---|---|---|
| ENSG00000249667 | 0 | 0 | 0 | 0 | 0 | 0 |
| ENSG00000251635 | 0 | 0 | 0 | 0 | 0 | 0 |
| ENSG00000231160 | 14 | 9 | 19 | 7 | 2 | 9 |
| ENSG00000271958 | 0 | 0 | 0 | 0 | 0 | 0 |
| ENSG00000109787 | 1594 | 811 | 902 | 450 | 785 | 694 |
| ENSG00000222230 | 0 | 0 | 0 | 0 | 0 | 0 |
| ENSG00000174123 | 0 | 0 | 0 | 0 | 0 | 1 |
| ENSG00000174125 | 0 | 0 | 0 | 0 | 0 | 0 |
| ENSG00000174130 | 0 | 1 | 0 | 1 | 1 | 0 |
| ENSG00000197712 | 721 | 323 | 389 | 326 | 439 | 326 |
| ENSG00000207944 | 0 | 1 | 0 | 0 | 0 | 0 |
| ENSG00000121895 | 3 | 2 | 1 | 16 | 29 | 17 |
| ENSG00000109790 | 787 | 496 | 512 | 504 | 844 | 619 |
| ENSG00000249207 | 3 | 4 | 4 | 3 | 4 | 1 |
| ENSG00000249685 | 3 | 3 | 2 | 1 | 3 | 1 |
| ENSG00000157796 | 506 | 290 | 362 | 163 | 357 | 260 |
| ENSG00000035928 | 1235 | 649 | 613 | 332 | 890 | 552 |
| ENSG00000206675 | 0 | 1 | 0 | 0 | 0 | 0 |
| ENSG00000222592 | 0 | 0 | 0 | 0 | 0 | 0 |
| ENSG00000134962 | 70 | 37 | 34 | 24 | 31 | 32 |
| ENSG00000264621 | 0 | 0 | 0 | 0 | 0 | 0 |
| ENSG00000238797 | 0 | 0 | 0 | 0 | 0 | 0 |
| ENSG00000163682 | 16992 | 8209 | 10138 | 10628 | 10355 | 10346 |
| ENSG00000121897 | 353 | 160 | 195 | 130 | 225 | 179 |
| ENSG00000224097 | 2 | 1 | 3 | 1 | 2 | 1 |
| ENSG00000109814 | 26519 | 11912 | 15164 | 8227 | 12800 | 11625 |
| ENSG00000249348 | 76 | 52 | 74 | 57 | 59 | 49 |
| ENSG00000163683 | 24385 | 11242 | 14820 | 8159 | 11391 | 11045 |
| ENSG00000252796 | 0 | 0 | 0 | 0 | 0 | 0 |
| ENSG00000255458 | 20 | 8 | 11 | 8 | 8 | 9 |
| ENSG00000078140 | 2901 | 1362 | 1633 | 1201 | 1729 | 1625 |
| ENSG00000252975 | 0 | 0 | 1 | 0 | 0 | 0 |
| ENSG00000249019 | 0 | 0 | 0 | 0 | 0 | 0 |
| ENSG00000243260 | 0 | 1 | 0 | 0 | 0 | 1 |
| ENSG00000180610 | 0 | 2 | 6 | 1 | 1 | 0 |
| ENSG00000121892 | 2292 | 1342 | 1077 | 590 | 2171 | 1220 |
| ENSG00000271278 | 0 | 0 | 0 | 0 | 0 | 0 |
| ENSG00000252970 | 0 | 0 | 0 | 1 | 0 | 0 |
| ENSG00000250568 | 0 | 0 | 0 | 0 | 1 | 1 |
| ENSG00000231707 | 4 | 0 | 1 | 0 | 0 | 1 |
| ENSG00000249064 | 0 | 0 | 3 | 2 | 0 | 2 |
| ENSG00000205794 | 0 | 1 | 0 | 1 | 1 | 0 |
| ENSG00000078177 | 147 | 63 | 52 | 22 | 151 | 61 |
| ENSG00000200455 | 0 | 0 | 0 | 0 | 0 | 0 |
| ENSG00000201863 | 0 | 0 | 0 | 0 | 0 | 0 |
| ENSG00000248977 | 0 | 0 | 0 | 0 | 0 | 0 |
| ENSG00000260296 | 6 | 5 | 10 | 3 | 3 | 5 |
| ENSG00000168421 | 21 | 5 | 5 | 15 | 10 | 12 |
| ENSG00000250338 | 1 | 0 | 0 | 0 | 0 | 0 |
| ENSG00000249241 | 0 | 0 | 0 | 0 | 0 | 0 |

| | | | | | | |
|---|---|---|---|---|---|---|
| ENSG00000174343 | 0 | 0 | 0 | 0 | 1 | 0 |
| ENSG00000239010 | 0 | 0 | 0 | 0 | 0 | 0 |
| ENSG00000163694 | 3900 | 1720 | 2343 | 1340 | 1809 | 2026 |
| ENSG00000250893 | 0 | 0 | 3 | 0 | 6 | 1 |
| ENSG00000242197 | 0 | 0 | 0 | 0 | 0 | 0 |
| ENSG00000263642 | 0 | 0 | 0 | 0 | 0 | 0 |
| ENSG00000179299 | 259 | 126 | 167 | 140 | 185 | 163 |
| ENSG00000248780 | 5 | 3 | 4 | 3 | 2 | 3 |
| ENSG00000163697 | 2679 | 1400 | 1340 | 806 | 1426 | 1088 |
| ENSG00000250906 | 4 | 1 | 2 | 0 | 0 | 0 |
| ENSG00000207195 | 1 | 1 | 1 | 0 | 0 | 0 |
| ENSG00000201736 | 0 | 1 | 0 | 0 | 0 | 0 |
| ENSG00000199790 | 0 | 0 | 0 | 0 | 0 | 0 |
| ENSG00000207198 | 0 | 0 | 0 | 0 | 0 | 0 |
| ENSG00000251173 | 0 | 0 | 0 | 0 | 0 | 0 |
| ENSG00000154277 | 64 | 31 | 29 | 30 | 29 | 40 |
| ENSG00000207481 | 0 | 0 | 0 | 0 | 0 | 0 |
| ENSG00000064042 | 216 | 127 | 82 | 71 | 120 | 58 |
| ENSG00000239627 | 0 | 0 | 0 | 0 | 0 | 0 |
| ENSG00000249216 | 0 | 0 | 0 | 0 | 0 | 0 |
| ENSG00000249081 | 0 | 0 | 0 | 0 | 0 | 0 |
| ENSG00000109132 | 0 | 0 | 0 | 0 | 0 | 0 |
| ENSG00000250467 | 0 | 0 | 0 | 0 | 0 | 0 |
| ENSG00000249122 | 0 | 0 | 0 | 0 | 0 | 0 |
| ENSG00000200338 | 0 | 0 | 0 | 0 | 0 | 0 |
| ENSG00000249745 | 0 | 0 | 0 | 0 | 0 | 0 |
| ENSG00000245870 | 0 | 0 | 0 | 0 | 0 | 0 |
| ENSG00000249771 | 0 | 1 | 0 | 0 | 0 | 0 |
| ENSG00000249887 | 0 | 0 | 0 | 0 | 0 | 0 |
| ENSG00000109133 | 1617 | 805 | 801 | 578 | 1014 | 743 |
| ENSG00000182308 | 8 | 0 | 2 | 3 | 0 | 2 |
| ENSG00000272862 | 0 | 0 | 0 | 0 | 1 | 0 |
| ENSG00000014824 | 1603 | 907 | 911 | 622 | 1153 | 947 |
| ENSG00000249212 | 0 | 0 | 0 | 0 | 0 | 0 |
| ENSG00000188848 | 0 | 0 | 0 | 0 | 1 | 0 |
| ENSG00000285454 | 0 | 0 | 0 | 0 | 0 | 0 |
| ENSG00000250781 | 0 | 0 | 0 | 0 | 0 | 0 |
| ENSG00000178343 | 0 | 0 | 0 | 0 | 0 | 0 |
| ENSG00000124406 | 309 | 155 | 146 | 55 | 121 | 96 |
| ENSG00000240167 | 0 | 0 | 0 | 0 | 0 | 0 |
| ENSG00000248283 | 0 | 0 | 3 | 0 | 0 | 0 |
| ENSG00000260519 | 2 | 0 | 0 | 1 | 0 | 2 |
| ENSG00000251517 | 0 | 0 | 0 | 0 | 0 | 0 |
| ENSG00000252595 | 0 | 0 | 0 | 0 | 0 | 0 |
| ENSG00000215203 | 0 | 0 | 0 | 0 | 0 | 0 |
| ENSG00000248939 | 0 | 0 | 0 | 0 | 0 | 0 |
| ENSG00000250657 | 0 | 0 | 0 | 0 | 0 | 0 |
| ENSG00000277096 | 0 | 0 | 0 | 0 | 0 | 0 |
| ENSG00000213851 | 0 | 0 | 0 | 0 | 0 | 0 |
| ENSG00000248143 | 0 | 0 | 0 | 0 | 0 | 0 |

| | | | | | | |
|---|---|---|---|---|---|---|
| ENSG00000251501 | 0 | 0 | 0 | 0 | 0 | 0 |
| ENSG00000239464 | 0 | 0 | 0 | 0 | 0 | 0 |
| ENSG00000239679 | 0 | 0 | 0 | 0 | 0 | 0 |
| ENSG00000124399 | 0 | 2 | 1 | 0 | 0 | 0 |
| ENSG00000250863 | 0 | 0 | 0 | 0 | 0 | 0 |
| ENSG00000251350 | 0 | 0 | 0 | 0 | 0 | 2 |
| ENSG00000183783 | 0 | 0 | 0 | 0 | 0 | 0 |
| ENSG00000251159 | 0 | 0 | 0 | 0 | 0 | 0 |
| ENSG00000177752 | 0 | 0 | 0 | 0 | 0 | 0 |
| ENSG00000151806 | 501 | 247 | 285 | 170 | 297 | 256 |
| ENSG00000163281 | 63 | 32 | 33 | 19 | 41 | 38 |
| ENSG00000273369 | 7 | 3 | 8 | 1 | 8 | 0 |
| ENSG00000272936 | 0 | 0 | 0 | 0 | 0 | 0 |
| ENSG00000249729 | 0 | 0 | 0 | 0 | 0 | 0 |
| ENSG00000248987 | 0 | 0 | 0 | 0 | 0 | 0 |
| ENSG00000275959 | 0 | 0 | 0 | 0 | 0 | 0 |
| ENSG00000248744 | 0 | 0 | 0 | 0 | 0 | 0 |
| ENSG00000248320 | 0 | 0 | 0 | 0 | 0 | 0 |
| ENSG00000224560 | 0 | 0 | 0 | 0 | 0 | 0 |
| ENSG00000200720 | 0 | 0 | 0 | 0 | 0 | 0 |
| ENSG00000222257 | 0 | 0 | 0 | 0 | 0 | 0 |
| ENSG00000163285 | 0 | 0 | 0 | 0 | 0 | 0 |
| ENSG00000260878 | 0 | 0 | 0 | 0 | 0 | 0 |
| ENSG00000151834 | 0 | 0 | 0 | 0 | 0 | 0 |
| ENSG00000249330 | 0 | 0 | 0 | 0 | 0 | 0 |
| ENSG00000252243 | 0 | 0 | 0 | 0 | 0 | 0 |
| ENSG00000249936 | 26 | 15 | 22 | 9 | 6 | 22 |
| ENSG00000170516 | 0 | 0 | 0 | 0 | 0 | 0 |
| ENSG00000109158 | 0 | 0 | 0 | 0 | 0 | 0 |
| ENSG00000163288 | 0 | 0 | 0 | 0 | 0 | 0 |
| ENSG00000260918 | 0 | 0 | 0 | 0 | 0 | 0 |
| ENSG00000169019 | 193 | 91 | 110 | 113 | 195 | 115 |
| ENSG00000282904 | 2 | 0 | 0 | 0 | 0 | 0 |
| ENSG00000145246 | 14 | 2 | 1 | 2 | 3 | 4 |
| ENSG00000242431 | 0 | 0 | 0 | 0 | 0 | 0 |
| ENSG00000248254 | 0 | 0 | 0 | 0 | 0 | 0 |
| ENSG00000145244 | 0 | 0 | 0 | 0 | 0 | 0 |
| ENSG00000277388 | 0 | 0 | 0 | 0 | 0 | 0 |
| ENSG00000242262 | 0 | 2 | 3 | 2 | 3 | 3 |
| ENSG00000282917 | 0 | 0 | 0 | 0 | 0 | 0 |
| ENSG00000259959 | 42 | 29 | 17 | 14 | 23 | 35 |
| ENSG00000170448 | 361 | 199 | 199 | 136 | 249 | 187 |
| ENSG00000163293 | 107 | 53 | 64 | 49 | 91 | 80 |
| ENSG00000198515 | 8 | 6 | 0 | 1 | 6 | 4 |
| ENSG00000074966 | 1 | 2 | 0 | 0 | 0 | 1 |
| ENSG00000200269 | 0 | 0 | 0 | 0 | 0 | 0 |
| ENSG00000283499 | 0 | 0 | 0 | 0 | 0 | 0 |
| ENSG00000135605 | 93 | 53 | 74 | 25 | 66 | 53 |
| ENSG00000202014 | 0 | 0 | 0 | 0 | 0 | 0 |
| ENSG00000109171 | 1294 | 629 | 689 | 548 | 735 | 688 |

| | | | | | | |
|---|---|---|---|---|---|---|
| ENSG00000145248 | 0 | 0 | 0 | 0 | 2 | 2 |
| ENSG00000182223 | 0 | 0 | 0 | 0 | 0 | 0 |
| ENSG00000075539 | 666 | 429 | 312 | 124 | 729 | 341 |
| ENSG00000251722 | 0 | 0 | 0 | 0 | 0 | 0 |
| ENSG00000109180 | 3086 | 1659 | 1954 | 1578 | 1941 | 1826 |
| ENSG00000248256 | 2 | 2 | 0 | 4 | 2 | 2 |
| ENSG00000145247 | 1279 | 669 | 766 | 939 | 954 | 836 |
| ENSG00000252913 | 0 | 0 | 0 | 0 | 0 | 0 |
| ENSG00000251334 | 0 | 0 | 0 | 0 | 0 | 0 |
| ENSG00000109182 | 2 | 0 | 0 | 0 | 2 | 0 |
| ENSG00000225455 | 0 | 0 | 0 | 1 | 0 | 0 |
| ENSG00000249828 | 0 | 0 | 0 | 0 | 0 | 0 |
| ENSG00000249079 | 0 | 0 | 0 | 0 | 0 | 0 |
| ENSG00000280043 | 0 | 0 | 0 | 0 | 0 | 0 |
| ENSG00000250016 | 0 | 0 | 0 | 0 | 0 | 0 |
| ENSG00000237961 | 0 | 0 | 0 | 0 | 0 | 1 |
| ENSG00000268736 | 0 | 0 | 0 | 0 | 0 | 0 |
| ENSG00000248946 | 0 | 0 | 1 | 0 | 0 | 0 |
| ENSG00000248583 | 0 | 0 | 0 | 0 | 0 | 0 |
| ENSG00000251056 | 0 | 0 | 0 | 0 | 0 | 0 |
| ENSG00000250769 | 0 | 0 | 0 | 0 | 0 | 0 |
| ENSG00000248532 | 0 | 0 | 0 | 0 | 0 | 0 |
| ENSG00000269848 | 0 | 0 | 0 | 0 | 0 | 0 |
| ENSG00000268967 | 0 | 0 | 0 | 0 | 0 | 0 |
| ENSG00000268391 | 0 | 0 | 0 | 0 | 0 | 0 |
| ENSG00000232471 | 0 | 0 | 0 | 0 | 0 | 0 |
| ENSG00000251630 | 0 | 0 | 1 | 0 | 1 | 1 |
| ENSG00000251539 | 0 | 0 | 0 | 0 | 0 | 0 |
| ENSG00000250753 | 4 | 0 | 2 | 0 | 0 | 0 |
| ENSG00000249337 | 0 | 0 | 0 | 0 | 0 | 0 |
| ENSG00000109184 | 1020 | 516 | 606 | 316 | 669 | 562 |
| ENSG00000250473 | 0 | 0 | 0 | 0 | 0 | 0 |
| ENSG00000272576 | 4 | 0 | 2 | 0 | 3 | 1 |
| ENSG00000243929 | 0 | 0 | 0 | 0 | 0 | 0 |
| ENSG00000188993 | 0 | 1 | 2 | 0 | 0 | 0 |
| ENSG00000163069 | 464 | 255 | 256 | 182 | 312 | 229 |
| ENSG00000251040 | 0 | 0 | 0 | 0 | 0 | 0 |
| ENSG00000163071 | 111 | 78 | 81 | 57 | 99 | 73 |
| ENSG00000251286 | 0 | 0 | 0 | 0 | 0 | 0 |
| ENSG00000199924 | 0 | 0 | 0 | 0 | 0 | 0 |
| ENSG00000109189 | 961 | 438 | 619 | 312 | 498 | 455 |
| ENSG00000248866 | 56 | 15 | 48 | 30 | 28 | 35 |
| ENSG00000260120 | 0 | 0 | 0 | 0 | 0 | 0 |
| ENSG00000226950 | 1437 | 807 | 891 | 1033 | 1096 | 1033 |
| ENSG00000264585 | 0 | 0 | 0 | 0 | 1 | 0 |
| ENSG00000212588 | 19 | 3 | 10 | 10 | 9 | 9 |
| ENSG00000248375 | 1 | 2 | 0 | 1 | 4 | 0 |
| ENSG00000226887 | 77 | 33 | 30 | 31 | 47 | 41 |
| ENSG00000212490 | 0 | 0 | 0 | 0 | 0 | 0 |
| ENSG00000199901 | 0 | 0 | 0 | 0 | 0 | 0 |

| | | | | | | |
|---|---|---|---|---|---|---|
| ENSG00000250302 | 0 | 0 | 0 | 1 | 0 | 0 |
| ENSG00000128045 | 18 | 15 | 14 | 21 | 26 | 12 |
| ENSG00000184178 | 410 | 208 | 195 | 248 | 256 | 291 |
| ENSG00000248115 | 0 | 0 | 0 | 0 | 0 | 0 |
| ENSG00000207385 | 0 | 0 | 0 | 0 | 0 | 0 |
| ENSG00000145216 | 384 | 286 | 279 | 161 | 276 | 267 |
| ENSG00000282278 | 0 | 0 | 0 | 0 | 0 | 0 |
| ENSG00000072201 | 423 | 178 | 253 | 163 | 211 | 188 |
| ENSG00000250930 | 0 | 0 | 0 | 0 | 0 | 0 |
| ENSG00000173966 | 0 | 0 | 1 | 0 | 0 | 0 |
| ENSG00000248494 | 18 | 8 | 14 | 13 | 14 | 8 |
| ENSG00000249341 | 0 | 0 | 0 | 0 | 0 | 0 |
| ENSG00000249706 | 0 | 0 | 0 | 0 | 0 | 0 |
| ENSG00000250815 | 0 | 0 | 0 | 0 | 0 | 0 |
| ENSG00000229585 | 0 | 0 | 0 | 0 | 0 | 0 |
| ENSG00000272650 | 0 | 0 | 0 | 1 | 0 | 0 |
| ENSG00000109220 | 268 | 116 | 148 | 126 | 130 | 138 |
| ENSG00000269506 | 0 | 0 | 0 | 0 | 0 | 0 |
| ENSG00000251105 | 0 | 0 | 0 | 0 | 0 | 0 |
| ENSG00000250147 | 0 | 0 | 0 | 0 | 0 | 0 |
| ENSG00000180613 | 0 | 0 | 0 | 0 | 0 | 0 |
| ENSG00000242670 | 0 | 0 | 0 | 0 | 1 | 0 |
| ENSG00000134853 | 0 | 0 | 0 | 0 | 4 | 0 |
| ENSG00000248184 | 0 | 0 | 0 | 0 | 0 | 0 |
| ENSG00000248518 | 0 | 0 | 0 | 0 | 0 | 0 |
| ENSG00000251264 | 0 | 0 | 0 | 0 | 0 | 0 |
| ENSG00000250456 | 0 | 0 | 0 | 0 | 0 | 0 |
| ENSG00000157404 | 22 | 11 | 16 | 7 | 14 | 12 |
| ENSG00000249727 | 0 | 0 | 0 | 0 | 0 | 0 |
| ENSG00000251256 | 0 | 0 | 0 | 0 | 0 | 0 |
| ENSG00000248317 | 0 | 0 | 0 | 0 | 0 | 0 |
| ENSG00000244422 | 0 | 0 | 0 | 0 | 0 | 0 |
| ENSG00000252815 | 0 | 0 | 0 | 0 | 0 | 0 |
| ENSG00000250646 | 0 | 0 | 0 | 0 | 0 | 0 |
| ENSG00000244034 | 0 | 0 | 0 | 0 | 0 | 0 |
| ENSG00000128052 | 26 | 4 | 12 | 0 | 2 | 1 |
| ENSG00000239545 | 0 | 0 | 0 | 0 | 0 | 0 |
| ENSG00000250812 | 0 | 0 | 0 | 0 | 0 | 0 |
| ENSG00000206839 | 0 | 0 | 0 | 0 | 0 | 0 |
| ENSG00000250464 | 0 | 0 | 0 | 0 | 0 | 0 |
| ENSG00000128039 | 1395 | 685 | 802 | 704 | 737 | 764 |
| ENSG00000251111 | 0 | 0 | 0 | 0 | 0 | 0 |
| ENSG00000249700 | 37 | 12 | 14 | 15 | 14 | 27 |
| ENSG00000273257 | 0 | 1 | 3 | 2 | 1 | 4 |
| ENSG00000134851 | 2550 | 1306 | 1574 | 1398 | 1617 | 1584 |
| ENSG00000239040 | 5 | 0 | 9 | 0 | 2 | 1 |
| ENSG00000134852 | 615 | 372 | 287 | 172 | 624 | 317 |
| ENSG00000238585 | 0 | 1 | 0 | 0 | 0 | 0 |
| ENSG00000223305 | 2 | 0 | 0 | 2 | 0 | 0 |
| ENSG00000272969 | 3 | 2 | 1 | 0 | 2 | 0 |

| | | | | | | |
|---|---|---|---|---|---|---|
| ENSG00000163440 | 0 | 0 | 0 | 0 | 0 | 0 |
| ENSG00000109255 | 16 | 2 | 1 | 3 | 2 | 5 |
| ENSG00000251923 | 0 | 0 | 0 | 0 | 0 | 0 |
| ENSG00000250821 | 0 | 0 | 0 | 0 | 0 | 0 |
| ENSG00000090989 | 1090 | 604 | 607 | 413 | 823 | 547 |
| ENSG00000202358 | 0 | 0 | 1 | 0 | 5 | 0 |
| ENSG00000174799 | 129 | 72 | 60 | 28 | 123 | 60 |
| ENSG00000109265 | 204 | 104 | 113 | 78 | 138 | 111 |
| ENSG00000200741 | 0 | 0 | 0 | 0 | 1 | 0 |
| ENSG00000252489 | 0 | 0 | 0 | 0 | 0 | 0 |
| ENSG00000250324 | 0 | 0 | 0 | 0 | 0 | 0 |
| ENSG00000157426 | 416 | 193 | 215 | 151 | 238 | 207 |
| ENSG00000242083 | 2 | 0 | 0 | 0 | 3 | 0 |
| ENSG00000269921 | 6 | 4 | 5 | 6 | 6 | 5 |
| ENSG00000128059 | 897 | 450 | 511 | 293 | 552 | 524 |
| ENSG00000270147 | 5 | 0 | 4 | 3 | 8 | 1 |
| ENSG00000283156 | 4 | 4 | 0 | 1 | 1 | 2 |
| ENSG00000128050 | 4872 | 2401 | 2867 | 2421 | 3027 | 3045 |
| ENSG00000174780 | 3394 | 1839 | 1943 | 1419 | 2484 | 1792 |
| ENSG00000196503 | 0 | 0 | 3 | 0 | 0 | 1 |
| ENSG00000249693 | 2 | 0 | 0 | 0 | 1 | 1 |
| ENSG00000248590 | 0 | 0 | 0 | 1 | 0 | 2 |
| ENSG00000171476 | 6 | 9 | 10 | 9 | 11 | 6 |
| ENSG00000280285 | 0 | 0 | 0 | 0 | 0 | 0 |
| ENSG00000241250 | 0 | 0 | 0 | 0 | 0 | 0 |
| ENSG00000227040 | 0 | 0 | 0 | 0 | 1 | 0 |
| ENSG00000240545 | 0 | 0 | 0 | 0 | 0 | 0 |
| ENSG00000265894 | 0 | 0 | 0 | 0 | 0 | 0 |
| ENSG00000128040 | 8 | 4 | 1 | 1 | 1 | 1 |
| ENSG00000278226 | 0 | 0 | 0 | 0 | 0 | 0 |
| ENSG00000251799 | 0 | 0 | 0 | 0 | 0 | 0 |
| ENSG00000084093 | 1976 | 952 | 886 | 418 | 1132 | 861 |
| ENSG00000269949 | 0 | 3 | 1 | 3 | 4 | 1 |
| ENSG00000084092 | 2189 | 1017 | 1287 | 1174 | 1280 | 1152 |
| ENSG00000047315 | 3753 | 1872 | 2118 | 1363 | 2442 | 1940 |
| ENSG00000251703 | 0 | 0 | 1 | 0 | 0 | 0 |
| ENSG00000163453 | 0 | 2 | 2 | 1 | 1 | 3 |
| ENSG00000250384 | 0 | 0 | 0 | 0 | 0 | 0 |
| ENSG00000245067 | 0 | 0 | 0 | 0 | 0 | 0 |
| ENSG00000250192 | 0 | 0 | 0 | 0 | 0 | 0 |
| ENSG00000243920 | 0 | 0 | 0 | 0 | 0 | 0 |
| ENSG00000248505 | 0 | 0 | 0 | 0 | 0 | 0 |
| ENSG00000251459 | 0 | 0 | 0 | 0 | 0 | 0 |
| ENSG00000251049 | 0 | 0 | 0 | 0 | 0 | 0 |
| ENSG00000250333 | 0 | 0 | 0 | 0 | 0 | 0 |
| ENSG00000249831 | 0 | 0 | 0 | 0 | 0 | 0 |
| ENSG00000248660 | 0 | 0 | 0 | 0 | 0 | 0 |
| ENSG00000249105 | 0 | 0 | 0 | 0 | 0 | 0 |
| ENSG00000250340 | 0 | 0 | 0 | 0 | 0 | 0 |
| ENSG00000250078 | 0 | 0 | 0 | 0 | 0 | 0 |

| | | | | | | |
|---|---|---|---|---|---|---|
| ENSG00000251339 | 0 | 0 | 0 | 0 | 0 | 0 |
| ENSG00000251266 | 0 | 0 | 0 | 0 | 0 | 0 |
| ENSG00000249382 | 0 | 0 | 0 | 0 | 0 | 0 |
| ENSG00000250375 | 0 | 0 | 0 | 0 | 0 | 0 |
| ENSG00000249111 | 0 | 0 | 0 | 0 | 0 | 0 |
| ENSG00000199780 | 0 | 0 | 0 | 0 | 0 | 0 |
| ENSG00000249392 | 0 | 0 | 0 | 0 | 0 | 0 |
| ENSG00000249892 | 0 | 0 | 0 | 0 | 0 | 0 |
| ENSG00000201676 | 0 | 0 | 0 | 0 | 0 | 0 |
| ENSG00000201775 | 0 | 0 | 0 | 0 | 0 | 0 |
| ENSG00000283043 | 0 | 0 | 0 | 0 | 0 | 0 |
| ENSG00000249066 | 0 | 0 | 0 | 0 | 0 | 0 |
| ENSG00000248237 | 0 | 0 | 0 | 0 | 0 | 0 |
| ENSG00000265829 | 0 | 0 | 0 | 0 | 0 | 0 |
| ENSG00000240152 | 0 | 0 | 0 | 0 | 0 | 0 |
| ENSG00000150471 | 0 | 0 | 0 | 0 | 1 | 1 |
| ENSG00000241261 | 0 | 0 | 0 | 0 | 0 | 0 |
| ENSG00000205682 | 0 | 0 | 0 | 0 | 0 | 0 |
| ENSG00000226439 | 0 | 0 | 0 | 2 | 2 | 0 |
| ENSG00000202169 | 0 | 0 | 0 | 0 | 0 | 0 |
| ENSG00000248692 | 0 | 0 | 0 | 0 | 0 | 0 |
| ENSG00000225405 | 0 | 0 | 0 | 0 | 4 | 3 |
| ENSG00000241344 | 0 | 0 | 0 | 0 | 0 | 0 |
| ENSG00000250249 | 0 | 0 | 0 | 0 | 0 | 0 |
| ENSG00000248847 | 0 | 0 | 0 | 0 | 0 | 0 |
| ENSG00000248336 | 0 | 0 | 0 | 0 | 1 | 0 |
| ENSG00000180673 | 0 | 0 | 0 | 1 | 2 | 1 |
| ENSG00000251055 | 0 | 0 | 0 | 0 | 0 | 0 |
| ENSG00000231442 | 0 | 0 | 0 | 0 | 0 | 0 |
| ENSG00000250775 | 0 | 0 | 0 | 0 | 0 | 0 |
| ENSG00000205678 | 0 | 0 | 0 | 0 | 0 | 0 |
| ENSG00000252104 | 0 | 0 | 0 | 0 | 0 | 0 |
| ENSG00000250768 | 0 | 0 | 0 | 0 | 0 | 0 |
| ENSG00000251430 | 0 | 0 | 0 | 0 | 0 | 0 |
| ENSG00000251385 | 0 | 0 | 0 | 0 | 0 | 0 |
| ENSG00000249893 | 0 | 0 | 0 | 0 | 0 | 0 |
| ENSG00000250169 | 0 | 0 | 0 | 0 | 0 | 0 |
| ENSG00000248259 | 0 | 0 | 0 | 0 | 0 | 0 |
| ENSG00000249045 | 0 | 0 | 0 | 0 | 0 | 0 |
| ENSG00000248328 | 0 | 0 | 0 | 0 | 0 | 0 |
| ENSG00000244669 | 0 | 0 | 0 | 0 | 0 | 0 |
| ENSG00000250125 | 0 | 0 | 0 | 0 | 0 | 0 |
| ENSG00000248447 | 0 | 0 | 0 | 0 | 0 | 0 |
| ENSG00000248998 | 0 | 0 | 0 | 1 | 0 | 0 |
| ENSG00000249351 | 0 | 0 | 0 | 0 | 0 | 0 |
| ENSG00000285711 | 5 | 2 | 2 | 0 | 0 | 2 |
| ENSG00000145242 | 0 | 0 | 0 | 0 | 0 | 0 |
| ENSG00000243199 | 132 | 71 | 98 | 85 | 102 | 90 |
| ENSG00000250846 | 0 | 0 | 0 | 0 | 0 | 0 |
| ENSG00000248479 | 0 | 0 | 0 | 0 | 0 | 0 |

| | | | | | | |
|---|---|---|---|---|---|---|
| ENSG00000222765 | 0 | 0 | 0 | 0 | 0 | 0 |
| ENSG00000251527 | 0 | 0 | 0 | 0 | 0 | 0 |
| ENSG00000249413 | 0 | 0 | 0 | 0 | 0 | 0 |
| ENSG00000272304 | 1 | 0 | 0 | 0 | 0 | 0 |
| ENSG00000236562 | 0 | 0 | 1 | 1 | 0 | 0 |
| ENSG00000221563 | 0 | 0 | 0 | 0 | 0 | 0 |
| ENSG00000244099 | 0 | 0 | 0 | 0 | 0 | 0 |
| ENSG00000252890 | 0 | 0 | 0 | 0 | 0 | 0 |
| ENSG00000250075 | 4 | 0 | 1 | 0 | 2 | 2 |
| ENSG00000206629 | 0 | 0 | 0 | 0 | 0 | 0 |
| ENSG00000274759 | 0 | 0 | 0 | 0 | 0 | 0 |
| ENSG00000250030 | 0 | 0 | 1 | 0 | 0 | 0 |
| ENSG00000145241 | 294 | 148 | 141 | 53 | 229 | 160 |
| ENSG00000035720 | 0 | 0 | 0 | 0 | 0 | 0 |
| ENSG00000279464 | 7 | 3 | 2 | 1 | 0 | 1 |
| ENSG00000033178 | 1020 | 627 | 566 | 280 | 825 | 495 |
| ENSG00000270257 | 0 | 0 | 0 | 1 | 1 | 0 |
| ENSG00000248049 | 217 | 94 | 124 | 96 | 95 | 101 |
| ENSG00000250656 | 6 | 1 | 1 | 0 | 2 | 2 |
| ENSG00000270228 | 8 | 5 | 3 | 6 | 5 | 6 |
| ENSG00000270292 | 0 | 1 | 0 | 0 | 1 | 0 |
| ENSG00000109163 | 1 | 1 | 2 | 1 | 2 | 1 |
| ENSG00000202374 | 0 | 0 | 1 | 0 | 0 | 0 |
| ENSG00000232398 | 0 | 0 | 0 | 0 | 1 | 0 |
| ENSG00000153802 | 0 | 0 | 1 | 0 | 0 | 0 |
| ENSG00000187054 | 0 | 0 | 0 | 0 | 0 | 0 |
| ENSG00000229009 | 0 | 0 | 0 | 0 | 0 | 0 |
| ENSG00000207428 | 0 | 0 | 0 | 0 | 0 | 0 |
| ENSG00000227725 | 0 | 1 | 0 | 1 | 0 | 0 |
| ENSG00000283131 | 0 | 0 | 0 | 0 | 0 | 0 |
| ENSG00000198092 | 0 | 0 | 0 | 0 | 0 | 0 |
| ENSG00000215127 | 2 | 0 | 0 | 0 | 0 | 0 |
| ENSG00000250426 | 0 | 0 | 0 | 0 | 0 | 0 |
| ENSG00000250026 | 0 | 0 | 0 | 0 | 0 | 0 |
| ENSG00000185873 | 2 | 0 | 0 | 0 | 0 | 0 |
| ENSG00000250642 | 0 | 0 | 0 | 0 | 0 | 0 |
| ENSG00000248639 | 0 | 0 | 0 | 0 | 0 | 0 |
| ENSG00000249531 | 0 | 0 | 0 | 0 | 0 | 0 |
| ENSG00000248122 | 0 | 0 | 0 | 0 | 0 | 0 |
| ENSG00000083896 | 2462 | 1202 | 1417 | 858 | 1528 | 1244 |
| ENSG00000162840 | 0 | 0 | 0 | 0 | 0 | 1 |
| ENSG00000087128 | 2 | 3 | 9 | 4 | 8 | 5 |
| ENSG00000250566 | 0 | 0 | 0 | 0 | 0 | 0 |
| ENSG00000197888 | 0 | 1 | 0 | 0 | 0 | 1 |
| ENSG00000249472 | 0 | 0 | 0 | 0 | 0 | 0 |
| ENSG00000251101 | 0 | 0 | 0 | 0 | 0 | 0 |
| ENSG00000251074 | 0 | 0 | 0 | 0 | 0 | 0 |
| ENSG00000196620 | 36 | 16 | 21 | 6 | 10 | 16 |
| ENSG00000248635 | 10 | 1 | 3 | 0 | 1 | 2 |
| ENSG00000250376 | 2 | 0 | 2 | 0 | 0 | 0 |

| | | | | | | |
|---|---|---|---|---|---|---|
| ENSG00000251236 | 0 | 0 | 0 | 0 | 0 | 0 |
| ENSG00000109181 | 4 | 0 | 0 | 0 | 0 | 1 |
| ENSG00000251424 | 0 | 0 | 0 | 0 | 0 | 0 |
| ENSG00000249235 | 0 | 0 | 0 | 0 | 1 | 0 |
| ENSG00000251529 | 0 | 0 | 0 | 0 | 0 | 0 |
| ENSG00000248547 | 0 | 0 | 0 | 0 | 0 | 0 |
| ENSG00000248613 | 0 | 0 | 0 | 0 | 0 | 0 |
| ENSG00000272626 | 0 | 0 | 0 | 0 | 0 | 0 |
| ENSG00000249985 | 0 | 0 | 0 | 0 | 0 | 0 |
| ENSG00000198277 | 0 | 0 | 0 | 0 | 0 | 0 |
| ENSG00000135220 | 0 | 0 | 0 | 0 | 0 | 0 |
| ENSG00000251498 | 0 | 0 | 0 | 0 | 0 | 0 |
| ENSG00000249735 | 0 | 0 | 0 | 0 | 0 | 0 |
| ENSG00000250277 | 0 | 0 | 0 | 0 | 0 | 0 |
| ENSG00000251685 | 0 | 0 | 0 | 0 | 0 | 0 |
| ENSG00000249890 | 0 | 0 | 0 | 0 | 0 | 0 |
| ENSG00000250919 | 0 | 0 | 0 | 0 | 0 | 0 |
| ENSG00000171234 | 0 | 0 | 0 | 0 | 0 | 0 |
| ENSG00000248763 | 0 | 0 | 0 | 0 | 0 | 0 |
| ENSG00000251284 | 0 | 0 | 0 | 0 | 0 | 0 |
| ENSG00000250100 | 0 | 0 | 0 | 0 | 0 | 0 |
| ENSG00000251427 | 0 | 0 | 0 | 0 | 0 | 0 |
| ENSG00000196472 | 0 | 0 | 0 | 0 | 0 | 0 |
| ENSG00000250696 | 0 | 0 | 0 | 0 | 0 | 0 |
| ENSG00000213759 | 2 | 0 | 0 | 0 | 0 | 0 |
| ENSG00000268803 | 0 | 0 | 0 | 0 | 0 | 0 |
| ENSG00000249763 | 0 | 0 | 0 | 0 | 0 | 0 |
| ENSG00000135226 | 0 | 0 | 0 | 0 | 0 | 0 |
| ENSG00000251691 | 0 | 0 | 0 | 0 | 0 | 0 |
| ENSG00000250612 | 0 | 0 | 0 | 0 | 0 | 0 |
| ENSG00000268209 | 0 | 0 | 0 | 0 | 0 | 0 |
| ENSG00000215110 | 0 | 0 | 0 | 0 | 0 | 0 |
| ENSG00000248824 | 0 | 0 | 0 | 0 | 0 | 0 |
| ENSG00000249956 | 0 | 0 | 0 | 0 | 0 | 0 |
| ENSG00000250828 | 0 | 0 | 0 | 0 | 0 | 0 |
| ENSG00000249686 | 0 | 0 | 0 | 0 | 0 | 0 |
| ENSG00000156096 | 0 | 0 | 0 | 0 | 0 | 0 |
| ENSG00000248886 | 0 | 0 | 0 | 0 | 0 | 0 |
| ENSG00000251177 | 0 | 0 | 0 | 0 | 0 | 0 |
| ENSG00000173610 | 0 | 0 | 0 | 0 | 1 | 0 |
| ENSG00000271271 | 0 | 0 | 0 | 0 | 0 | 0 |
| ENSG00000173597 | 0 | 0 | 0 | 0 | 0 | 0 |
| ENSG00000236959 | 0 | 0 | 0 | 0 | 0 | 0 |
| ENSG00000284695 | 0 | 0 | 0 | 0 | 0 | 0 |
| ENSG00000109193 | 2 | 0 | 0 | 0 | 0 | 0 |
| ENSG00000251489 | 0 | 0 | 0 | 0 | 0 | 0 |
| ENSG00000126545 | 0 | 0 | 0 | 0 | 0 | 0 |
| ENSG00000135222 | 0 | 0 | 0 | 0 | 0 | 0 |
| ENSG00000126549 | 0 | 0 | 0 | 0 | 0 | 0 |
| ENSG00000205649 | 0 | 0 | 0 | 0 | 0 | 0 |

| | | | | | | |
|---|---|---|---|---|---|---|
| ENSG00000126550 | 0 | 0 | 0 | 0 | 0 | 0 |
| ENSG00000234124 | 0 | 0 | 0 | 0 | 0 | 0 |
| ENSG00000187533 | 0 | 0 | 0 | 0 | 0 | 0 |
| ENSG00000109205 | 0 | 0 | 1 | 0 | 0 | 0 |
| ENSG00000181617 | 0 | 0 | 0 | 0 | 0 | 0 |
| ENSG00000171209 | 0 | 0 | 0 | 0 | 0 | 0 |
| ENSG00000145309 | 0 | 0 | 0 | 0 | 0 | 1 |
| ENSG00000109208 | 0 | 0 | 0 | 0 | 0 | 0 |
| ENSG00000171201 | 0 | 0 | 0 | 0 | 0 | 0 |
| ENSG00000171199 | 0 | 0 | 0 | 0 | 0 | 0 |
| ENSG00000171195 | 0 | 0 | 0 | 0 | 0 | 1 |
| ENSG00000187689 | 65 | 37 | 43 | 36 | 39 | 25 |
| ENSG00000178522 | 0 | 0 | 0 | 0 | 0 | 0 |
| ENSG00000132464 | 0 | 0 | 0 | 0 | 0 | 0 |
| ENSG00000132465 | 0 | 0 | 2 | 1 | 1 | 1 |
| ENSG00000132467 | 884 | 455 | 526 | 401 | 587 | 507 |
| ENSG00000207448 | 0 | 0 | 0 | 0 | 1 | 1 |
| ENSG00000207058 | 0 | 0 | 0 | 0 | 0 | 0 |
| ENSG00000272986 | 6 | 2 | 5 | 3 | 0 | 3 |
| ENSG00000018189 | 373 | 198 | 227 | 167 | 220 | 224 |
| ENSG00000132463 | 4324 | 2130 | 2563 | 2253 | 2581 | 2548 |
| ENSG00000283313 | 1 | 0 | 0 | 0 | 0 | 0 |
| ENSG00000207007 | 0 | 0 | 0 | 0 | 0 | 0 |
| ENSG00000173542 | 617 | 298 | 249 | 136 | 469 | 276 |
| ENSG00000156136 | 603 | 296 | 402 | 305 | 380 | 399 |
| ENSG00000080493 | 0 | 0 | 1 | 0 | 2 | 0 |
| ENSG00000249170 | 0 | 0 | 0 | 0 | 0 | 0 |
| ENSG00000270098 | 0 | 0 | 0 | 0 | 0 | 0 |
| ENSG00000145321 | 0 | 0 | 0 | 0 | 0 | 0 |
| ENSG00000222236 | 0 | 0 | 0 | 0 | 0 | 0 |
| ENSG00000248567 | 0 | 0 | 0 | 0 | 0 | 0 |
| ENSG00000056291 | 0 | 1 | 0 | 0 | 0 | 0 |
| ENSG00000156140 | 0 | 0 | 0 | 0 | 0 | 0 |
| ENSG00000250877 | 0 | 0 | 0 | 0 | 0 | 0 |
| ENSG00000270665 | 0 | 0 | 1 | 0 | 0 | 0 |
| ENSG00000252955 | 0 | 0 | 0 | 0 | 0 | 0 |
| ENSG00000210839 | 0 | 0 | 0 | 0 | 0 | 0 |
| ENSG00000163626 | 484 | 269 | 310 | 247 | 285 | 284 |
| ENSG00000132466 | 2521 | 1262 | 1247 | 665 | 1514 | 1219 |
| ENSG00000248641 | 0 | 5 | 0 | 0 | 2 | 0 |
| ENSG00000250220 | 10 | 9 | 10 | 7 | 8 | 3 |
| ENSG00000221639 | 1 | 0 | 0 | 0 | 0 | 0 |
| ENSG00000249976 | 0 | 0 | 0 | 0 | 0 | 0 |
| ENSG00000163631 | 14 | 5 | 0 | 3 | 1 | 0 |
| ENSG00000081051 | 0 | 0 | 0 | 0 | 0 | 0 |
| ENSG00000079557 | 0 | 0 | 0 | 0 | 0 | 0 |
| ENSG00000250436 | 0 | 0 | 0 | 0 | 0 | 0 |
| ENSG00000249970 | 0 | 0 | 0 | 0 | 0 | 0 |
| ENSG00000169435 | 0 | 0 | 0 | 0 | 1 | 1 |
| ENSG00000228277 | 0 | 0 | 0 | 0 | 0 | 0 |

| | | | | | | |
|---|---|---|---|---|---|---|
| ENSG00000169429 | 0 | 1 | 1 | 1 | 4 | 5 |
| ENSG00000249051 | 0 | 0 | 0 | 0 | 0 | 0 |
| ENSG00000124875 | 0 | 0 | 0 | 0 | 0 | 0 |
| ENSG00000250550 | 0 | 0 | 0 | 0 | 0 | 0 |
| ENSG00000109272 | 0 | 0 | 0 | 0 | 0 | 0 |
| ENSG00000163739 | 0 | 0 | 0 | 0 | 0 | 0 |
| ENSG00000249920 | 0 | 0 | 0 | 0 | 0 | 0 |
| ENSG00000250339 | 0 | 0 | 0 | 0 | 0 | 0 |
| ENSG00000163737 | 0 | 0 | 0 | 0 | 0 | 0 |
| ENSG00000163736 | 0 | 0 | 0 | 0 | 0 | 0 |
| ENSG00000163735 | 0 | 0 | 0 | 0 | 1 | 0 |
| ENSG00000244194 | 0 | 0 | 0 | 0 | 0 | 0 |
| ENSG00000163734 | 0 | 0 | 0 | 0 | 0 | 0 |
| ENSG00000248848 | 0 | 0 | 0 | 0 | 0 | 0 |
| ENSG00000251017 | 0 | 0 | 0 | 0 | 0 | 0 |
| ENSG00000081041 | 0 | 0 | 0 | 0 | 0 | 0 |
| ENSG00000163738 | 95 | 58 | 69 | 58 | 62 | 65 |
| ENSG00000269559 | 2 | 2 | 2 | 0 | 3 | 1 |
| ENSG00000182585 | 0 | 0 | 0 | 0 | 0 | 0 |
| ENSG00000124882 | 0 | 0 | 3 | 1 | 3 | 1 |
| ENSG00000250532 | 0 | 0 | 0 | 0 | 0 | 0 |
| ENSG00000109321 | 208 | 127 | 170 | 307 | 374 | 419 |
| ENSG00000249942 | 2 | 1 | 0 | 0 | 0 | 1 |
| ENSG00000174808 | 14 | 6 | 6 | 6 | 5 | 7 |
| ENSG00000250356 | 0 | 0 | 0 | 0 | 0 | 0 |
| ENSG00000169116 | 50 | 27 | 35 | 26 | 33 | 34 |
| ENSG00000249717 | 0 | 0 | 0 | 0 | 0 | 0 |
| ENSG00000248165 | 0 | 0 | 0 | 0 | 0 | 0 |
| ENSG00000260265 | 0 | 0 | 0 | 0 | 0 | 0 |
| ENSG00000250315 | 0 | 0 | 0 | 0 | 0 | 0 |
| ENSG00000250560 | 0 | 0 | 0 | 0 | 0 | 0 |
| ENSG00000251185 | 4 | 0 | 0 | 0 | 0 | 0 |
| ENSG00000251454 | 0 | 0 | 0 | 0 | 0 | 0 |
| ENSG00000251383 | 0 | 0 | 0 | 0 | 0 | 0 |
| ENSG00000248646 | 0 | 0 | 0 | 0 | 0 | 0 |
| ENSG00000250735 | 0 | 0 | 0 | 1 | 0 | 0 |
| ENSG00000163743 | 391 | 208 | 216 | 153 | 217 | 154 |
| ENSG00000174796 | 391 | 244 | 246 | 109 | 261 | 234 |
| ENSG00000174792 | 0 | 0 | 1 | 0 | 4 | 3 |
| ENSG00000138769 | 10 | 4 | 5 | 4 | 5 | 6 |
| ENSG00000138757 | 3588 | 1721 | 2131 | 1360 | 1945 | 2008 |
| ENSG00000201644 | 0 | 0 | 0 | 0 | 0 | 0 |
| ENSG00000138768 | 1144 | 690 | 554 | 316 | 1078 | 606 |
| ENSG00000222644 | 0 | 0 | 0 | 0 | 0 | 0 |
| ENSG00000229717 | 0 | 0 | 0 | 0 | 0 | 0 |
| ENSG00000156194 | 2 | 0 | 0 | 0 | 1 | 0 |
| ENSG00000138744 | 1024 | 542 | 645 | 500 | 547 | 566 |
| ENSG00000198301 | 632 | 422 | 381 | 219 | 675 | 496 |
| ENSG00000245928 | 0 | 0 | 1 | 0 | 0 | 0 |
| ENSG00000138755 | 0 | 0 | 0 | 0 | 0 | 0 |

| | | | | | | |
|---|---|---|---|---|---|---|
| ENSG00000156219 | 1 | 0 | 0 | 0 | 1 | 0 |
| ENSG00000169245 | 2 | 0 | 0 | 0 | 0 | 1 |
| ENSG00000169248 | 0 | 2 | 0 | 0 | 0 | 0 |
| ENSG00000225846 | 0 | 0 | 0 | 0 | 0 | 0 |
| ENSG00000138750 | 854 | 494 | 524 | 323 | 530 | 511 |
| ENSG00000138760 | 5332 | 2646 | 3422 | 2222 | 2949 | 2665 |
| ENSG00000189157 | 75 | 33 | 49 | 32 | 60 | 41 |
| ENSG00000272414 | 0 | 1 | 0 | 1 | 1 | 0 |
| ENSG00000118804 | 136 | 66 | 96 | 96 | 119 | 85 |
| ENSG00000163749 | 4 | 2 | 0 | 4 | 1 | 2 |
| ENSG00000249858 | 0 | 0 | 0 | 0 | 0 | 0 |
| ENSG00000212368 | 0 | 0 | 0 | 0 | 0 | 0 |
| ENSG00000240426 | 0 | 0 | 0 | 0 | 0 | 0 |
| ENSG00000199857 | 0 | 0 | 0 | 0 | 0 | 0 |
| ENSG00000138771 | 2076 | 897 | 1220 | 815 | 907 | 918 |
| ENSG00000249278 | 0 | 0 | 0 | 0 | 0 | 0 |
| ENSG00000207307 | 0 | 0 | 0 | 0 | 0 | 0 |
| ENSG00000263445 | 0 | 0 | 0 | 0 | 0 | 0 |
| ENSG00000283682 | 0 | 0 | 0 | 0 | 0 | 0 |
| ENSG00000233860 | 0 | 0 | 0 | 1 | 0 | 0 |
| ENSG00000224218 | 0 | 0 | 0 | 0 | 0 | 0 |
| ENSG00000270244 | 0 | 0 | 0 | 0 | 0 | 0 |
| ENSG00000242727 | 0 | 0 | 0 | 0 | 0 | 1 |
| ENSG00000186212 | 245 | 147 | 177 | 137 | 136 | 137 |
| ENSG00000138758 | 975 | 509 | 509 | 287 | 553 | 389 |
| ENSG00000234036 | 0 | 0 | 0 | 0 | 2 | 2 |
| ENSG00000118816 | 8885 | 4222 | 5169 | 5049 | 5066 | 5052 |
| ENSG00000214062 | 1 | 2 | 0 | 0 | 0 | 3 |
| ENSG00000271676 | 0 | 0 | 0 | 0 | 0 | 0 |
| ENSG00000201641 | 0 | 0 | 0 | 0 | 0 | 0 |
| ENSG00000138764 | 3761 | 1692 | 2390 | 1603 | 2180 | 2222 |
| ENSG00000270669 | 0 | 0 | 0 | 0 | 0 | 0 |
| ENSG00000250006 | 0 | 0 | 0 | 0 | 0 | 0 |
| ENSG00000248831 | 0 | 0 | 0 | 0 | 0 | 0 |
| ENSG00000249036 | 0 | 0 | 0 | 0 | 0 | 0 |
| ENSG00000156234 | 0 | 0 | 0 | 0 | 0 | 0 |
| ENSG00000221711 | 0 | 0 | 0 | 0 | 0 | 0 |
| ENSG00000138767 | 913 | 377 | 455 | 328 | 463 | 417 |
| ENSG00000169288 | 410 | 220 | 254 | 214 | 292 | 217 |
| ENSG00000250214 | 0 | 0 | 0 | 0 | 0 | 0 |
| ENSG00000248926 | 0 | 0 | 0 | 0 | 1 | 0 |
| ENSG00000248956 | 0 | 0 | 0 | 0 | 0 | 0 |
| ENSG00000248640 | 1 | 0 | 0 | 0 | 0 | 0 |
| ENSG00000248128 | 0 | 0 | 0 | 0 | 0 | 0 |
| ENSG00000249072 | 0 | 0 | 0 | 0 | 0 | 0 |
| ENSG00000138759 | 622 | 268 | 410 | 74 | 153 | 187 |
| ENSG00000249565 | 6 | 3 | 4 | 7 | 1 | 3 |
| ENSG00000248117 | 0 | 0 | 0 | 0 | 0 | 0 |
| ENSG00000138772 | 1035 | 627 | 625 | 502 | 585 | 472 |
| ENSG00000200998 | 0 | 0 | 0 | 0 | 0 | 0 |

| | | | | | | |
|---|---|---|---|---|---|---|
| ENSG00000251442 | 0 | 0 | 0 | 0 | 0 | 0 |
| ENSG00000251019 | 0 | 0 | 0 | 0 | 0 | 0 |
| ENSG00000239793 | 0 | 0 | 0 | 0 | 0 | 0 |
| ENSG00000260278 | 0 | 0 | 0 | 2 | 1 | 0 |
| ENSG00000138756 | 167 | 70 | 85 | 45 | 90 | 74 |
| ENSG00000266270 | 0 | 0 | 0 | 0 | 0 | 0 |
| ENSG00000163291 | 214 | 91 | 87 | 77 | 113 | 87 |
| ENSG00000242175 | 0 | 0 | 0 | 0 | 0 | 0 |
| ENSG00000249307 | 0 | 0 | 0 | 0 | 0 | 0 |
| ENSG00000156269 | 0 | 0 | 0 | 0 | 0 | 0 |
| ENSG00000196475 | 0 | 0 | 0 | 0 | 0 | 0 |
| ENSG00000250334 | 0 | 0 | 0 | 0 | 0 | 0 |
| ENSG00000249646 | 0 | 0 | 0 | 0 | 0 | 0 |
| ENSG00000251399 | 1 | 0 | 0 | 0 | 0 | 0 |
| ENSG00000248408 | 0 | 0 | 0 | 0 | 0 | 0 |
| ENSG00000251321 | 0 | 0 | 0 | 0 | 0 | 0 |
| ENSG00000212620 | 0 | 0 | 0 | 0 | 0 | 0 |
| ENSG00000163297 | 0 | 1 | 1 | 1 | 0 | 0 |
| ENSG00000236530 | 0 | 0 | 0 | 0 | 0 | 0 |
| ENSG00000244002 | 2 | 0 | 0 | 0 | 0 | 0 |
| ENSG00000248719 | 0 | 0 | 0 | 0 | 0 | 0 |
| ENSG00000152784 | 0 | 0 | 0 | 0 | 0 | 0 |
| ENSG00000138675 | 0 | 0 | 0 | 0 | 0 | 0 |
| ENSG00000197826 | 2 | 0 | 0 | 0 | 0 | 0 |
| ENSG00000232327 | 0 | 0 | 0 | 0 | 0 | 0 |
| ENSG00000152785 | 0 | 0 | 0 | 0 | 0 | 0 |
| ENSG00000138669 | 8 | 4 | 7 | 1 | 9 | 6 |
| ENSG00000251059 | 0 | 0 | 0 | 0 | 0 | 0 |
| ENSG00000207065 | 0 | 0 | 0 | 0 | 0 | 0 |
| ENSG00000138670 | 8 | 6 | 5 | 3 | 5 | 7 |
| ENSG00000248282 | 0 | 0 | 0 | 0 | 0 | 0 |
| ENSG00000248448 | 0 | 0 | 0 | 0 | 0 | 0 |
| ENSG00000252762 | 0 | 0 | 0 | 0 | 0 | 0 |
| ENSG00000248543 | 0 | 0 | 0 | 0 | 0 | 0 |
| ENSG00000250826 | 0 | 0 | 0 | 0 | 0 | 0 |
| ENSG00000202485 | 0 | 0 | 0 | 0 | 0 | 0 |
| ENSG00000234568 | 0 | 0 | 0 | 0 | 0 | 0 |
| ENSG00000284516 | 0 | 0 | 0 | 0 | 0 | 0 |
| ENSG00000273156 | 0 | 1 | 0 | 0 | 2 | 0 |
| ENSG00000138668 | 4580 | 2329 | 3104 | 2027 | 2358 | 2651 |
| ENSG00000272677 | 5 | 6 | 3 | 5 | 3 | 5 |
| ENSG00000250325 | 6 | 0 | 0 | 0 | 0 | 1 |
| ENSG00000202440 | 0 | 0 | 0 | 0 | 0 | 0 |
| ENSG00000152795 | 4988 | 2290 | 3271 | 1794 | 2574 | 2821 |
| ENSG00000145293 | 1738 | 846 | 1097 | 890 | 1080 | 1064 |
| ENSG00000249242 | 4462 | 2202 | 3229 | 1248 | 1913 | 1936 |
| ENSG00000214988 | 0 | 13 | 5 | 0 | 1 | 1 |
| ENSG00000227304 | 2 | 2 | 9 | 2 | 4 | 3 |
| ENSG00000249960 | 0 | 0 | 0 | 0 | 0 | 0 |
| ENSG00000248113 | 0 | 0 | 0 | 0 | 1 | 0 |

| | | | | | | |
|---|---|---|---|---|---|---|
| ENSG00000231782 | 0 | 0 | 0 | 0 | 0 | 0 |
| ENSG00000145284 | 314 | 155 | 214 | 154 | 171 | 201 |
| ENSG00000270480 | 1 | 1 | 1 | 0 | 3 | 0 |
| ENSG00000207746 | 0 | 0 | 0 | 0 | 0 | 0 |
| ENSG00000270842 | 0 | 0 | 0 | 0 | 0 | 0 |
| ENSG00000138674 | 4268 | 2248 | 2276 | 1371 | 2440 | 1989 |
| ENSG00000251022 | 337 | 187 | 226 | 136 | 258 | 219 |
| ENSG00000168152 | 52 | 44 | 18 | 17 | 49 | 29 |
| ENSG00000189308 | 468 | 172 | 252 | 120 | 231 | 231 |
| ENSG00000199530 | 0 | 0 | 0 | 0 | 0 | 0 |
| ENSG00000251874 | 0 | 0 | 0 | 0 | 0 | 0 |
| ENSG00000138663 | 1339 | 656 | 707 | 676 | 777 | 681 |
| ENSG00000145287 | 25 | 14 | 10 | 16 | 12 | 16 |
| ENSG00000250057 | 0 | 0 | 0 | 0 | 1 | 4 |
| ENSG00000250677 | 2 | 2 | 1 | 0 | 2 | 2 |
| ENSG00000248401 | 0 | 1 | 1 | 0 | 1 | 1 |
| ENSG00000173085 | 719 | 345 | 411 | 363 | 360 | 396 |
| ENSG00000173083 | 68 | 28 | 34 | 17 | 30 | 15 |
| ENSG00000274154 | 0 | 0 | 0 | 0 | 0 | 0 |
| ENSG00000251647 | 0 | 0 | 0 | 0 | 0 | 0 |
| ENSG00000163312 | 406 | 226 | 242 | 138 | 219 | 202 |
| ENSG00000163319 | 305 | 225 | 283 | 251 | 275 | 239 |
| ENSG00000163322 | 248 | 187 | 160 | 80 | 230 | 142 |
| ENSG00000213608 | 6 | 4 | 1 | 0 | 0 | 1 |
| ENSG00000239525 | 0 | 0 | 0 | 0 | 0 | 0 |
| ENSG00000138678 | 262 | 119 | 131 | 122 | 174 | 167 |
| ENSG00000201633 | 0 | 0 | 0 | 0 | 0 | 0 |
| ENSG00000249171 | 0 | 0 | 0 | 0 | 0 | 0 |
| ENSG00000250546 | 10 | 7 | 2 | 7 | 3 | 1 |
| ENSG00000200108 | 0 | 0 | 1 | 0 | 0 | 0 |
| ENSG00000214980 | 0 | 0 | 0 | 0 | 0 | 0 |
| ENSG00000248749 | 0 | 0 | 0 | 0 | 0 | 0 |
| ENSG00000163623 | 0 | 0 | 0 | 0 | 0 | 0 |
| ENSG00000258013 | 0 | 0 | 0 | 0 | 0 | 1 |
| ENSG00000163624 | 2257 | 1159 | 1384 | 1002 | 1413 | 1211 |
| ENSG00000163625 | 1518 | 758 | 893 | 364 | 877 | 740 |
| ENSG00000239466 | 0 | 0 | 0 | 0 | 0 | 0 |
| ENSG00000251260 | 0 | 0 | 0 | 0 | 0 | 0 |
| ENSG00000252062 | 0 | 0 | 0 | 0 | 0 | 0 |
| ENSG00000180769 | 146 | 78 | 114 | 63 | 95 | 110 |
| ENSG00000201901 | 0 | 0 | 0 | 0 | 0 | 0 |
| ENSG00000248725 | 0 | 0 | 0 | 0 | 0 | 0 |
| ENSG00000138639 | 282 | 109 | 157 | 57 | 103 | 107 |
| ENSG00000266421 | 0 | 0 | 0 | 0 | 0 | 0 |
| ENSG00000109339 | 75 | 36 | 47 | 19 | 66 | 35 |
| ENSG00000250062 | 0 | 1 | 0 | 0 | 0 | 0 |
| ENSG00000252757 | 0 | 0 | 0 | 0 | 0 | 0 |
| ENSG00000266515 | 0 | 0 | 0 | 0 | 0 | 0 |
| ENSG00000163629 | 767 | 410 | 354 | 147 | 476 | 271 |
| ENSG00000145283 | 0 | 0 | 0 | 2 | 0 | 0 |

| | | | | | | |
|---|---|---|---|---|---|---|
| ENSG00000243312 | 0 | 1 | 0 | 0 | 1 | 0 |
| ENSG00000163633 | 1 | 2 | 4 | 7 | 6 | 5 |
| ENSG00000250202 | 0 | 0 | 0 | 0 | 0 | 0 |
| ENSG00000285458 | 1 | 5 | 0 | 1 | 3 | 0 |
| ENSG00000251411 | 2 | 2 | 0 | 1 | 3 | 0 |
| ENSG00000284968 | 15 | 14 | 16 | 10 | 20 | 14 |
| ENSG00000172493 | 1108 | 462 | 633 | 315 | 549 | 508 |
| ENSG00000235043 | 12 | 10 | 5 | 8 | 4 | 3 |
| ENSG00000145332 | 930 | 451 | 491 | 190 | 474 | 344 |
| ENSG00000248196 | 0 | 2 | 0 | 0 | 1 | 1 |
| ENSG00000248180 | 6 | 2 | 0 | 0 | 0 | 1 |
| ENSG00000250572 | 0 | 0 | 0 | 0 | 0 | 0 |
| ENSG00000263981 | 0 | 0 | 0 | 0 | 0 | 0 |
| ENSG00000170509 | 25 | 20 | 33 | 13 | 15 | 13 |
| ENSG00000255723 | 1 | 4 | 0 | 0 | 1 | 0 |
| ENSG00000198189 | 243 | 105 | 133 | 119 | 119 | 107 |
| ENSG00000240966 | 0 | 0 | 0 | 0 | 0 | 0 |
| ENSG00000249262 | 0 | 0 | 0 | 0 | 0 | 0 |
| ENSG00000170502 | 803 | 446 | 473 | 389 | 531 | 430 |
| ENSG00000272856 | 0 | 0 | 0 | 0 | 0 | 0 |
| ENSG00000152583 | 9 | 7 | 10 | 3 | 7 | 5 |
| ENSG00000249001 | 21 | 4 | 13 | 13 | 21 | 10 |
| ENSG00000152591 | 0 | 0 | 0 | 2 | 0 | 2 |
| ENSG00000152592 | 4 | 4 | 4 | 1 | 2 | 2 |
| ENSG00000200024 | 0 | 0 | 0 | 0 | 0 | 0 |
| ENSG00000249973 | 0 | 0 | 0 | 0 | 0 | 0 |
| ENSG00000029559 | 19 | 16 | 15 | 8 | 27 | 24 |
| ENSG00000152595 | 0 | 0 | 1 | 0 | 1 | 0 |
| ENSG00000183199 | 19 | 8 | 19 | 9 | 13 | 9 |
| ENSG00000118785 | 0 | 0 | 0 | 0 | 0 | 0 |
| ENSG00000202000 | 0 | 1 | 0 | 0 | 0 | 0 |
| ENSG00000118762 | 758 | 316 | 445 | 257 | 444 | 410 |
| ENSG00000118777 | 1638 | 982 | 1178 | 1792 | 2597 | 2208 |
| ENSG00000241853 | 0 | 0 | 0 | 0 | 0 | 2 |
| ENSG00000201707 | 0 | 0 | 0 | 0 | 0 | 0 |
| ENSG00000221440 | 0 | 0 | 0 | 0 | 0 | 0 |
| ENSG00000251285 | 0 | 0 | 0 | 0 | 0 | 0 |
| ENSG00000199326 | 0 | 0 | 0 | 0 | 0 | 0 |
| ENSG00000163644 | 447 | 252 | 241 | 152 | 213 | 207 |
| ENSG00000200469 | 0 | 0 | 0 | 0 | 0 | 0 |
| ENSG00000246375 | 4 | 1 | 1 | 2 | 0 | 0 |
| ENSG00000207480 | 0 | 0 | 0 | 0 | 0 | 0 |
| ENSG00000138642 | 134 | 73 | 78 | 71 | 57 | 48 |
| ENSG00000138646 | 20 | 15 | 21 | 15 | 10 | 10 |
| ENSG00000249934 | 0 | 0 | 0 | 0 | 1 | 0 |
| ENSG00000145337 | 1570 | 837 | 966 | 1020 | 996 | 992 |
| ENSG00000255072 | 0 | 0 | 0 | 0 | 0 | 0 |
| ENSG00000285385 | 0 | 0 | 0 | 0 | 0 | 0 |
| ENSG00000285122 | 0 | 0 | 0 | 0 | 0 | 0 |
| ENSG00000249755 | 0 | 1 | 0 | 0 | 0 | 0 |

| | | | | | | |
|---|---|---|---|---|---|---|
| ENSG00000252322 | 0 | 1 | 0 | 0 | 0 | 0 |
| ENSG00000138641 | 1304 | 771 | 836 | 361 | 619 | 523 |
| ENSG00000207524 | 4 | 0 | 1 | 2 | 2 | 4 |
| ENSG00000177432 | 16 | 7 | 6 | 7 | 10 | 5 |
| ENSG00000248019 | 47 | 28 | 37 | 9 | 19 | 22 |
| ENSG00000278151 | 0 | 0 | 0 | 0 | 0 | 0 |
| ENSG00000276003 | 0 | 0 | 1 | 0 | 0 | 0 |
| ENSG00000138640 | 40 | 22 | 28 | 12 | 31 | 23 |
| ENSG00000180346 | 223 | 135 | 125 | 95 | 131 | 118 |
| ENSG00000270720 | 0 | 0 | 1 | 0 | 0 | 1 |
| ENSG00000212226 | 0 | 0 | 0 | 0 | 0 | 0 |
| ENSG00000185477 | 56 | 32 | 27 | 11 | 50 | 39 |
| ENSG00000251095 | 0 | 4 | 7 | 3 | 9 | 3 |
| ENSG00000145335 | 0 | 0 | 0 | 0 | 0 | 0 |
| ENSG00000274238 | 0 | 0 | 0 | 0 | 0 | 0 |
| ENSG00000277695 | 0 | 0 | 0 | 0 | 0 | 0 |
| ENSG00000276542 | 0 | 0 | 0 | 0 | 0 | 0 |
| ENSG00000247775 | 0 | 0 | 0 | 0 | 0 | 0 |
| ENSG00000138722 | 0 | 0 | 0 | 0 | 0 | 1 |
| ENSG00000184305 | 155 | 79 | 82 | 52 | 96 | 74 |
| ENSG00000252087 | 0 | 0 | 0 | 0 | 0 | 1 |
| ENSG00000251401 | 0 | 0 | 0 | 0 | 0 | 0 |
| ENSG00000187653 | 66 | 59 | 55 | 53 | 40 | 56 |
| ENSG00000248984 | 0 | 0 | 0 | 0 | 0 | 0 |
| ENSG00000249049 | 0 | 0 | 0 | 0 | 0 | 2 |
| ENSG00000279013 | 0 | 2 | 0 | 0 | 3 | 2 |
| ENSG00000249891 | 0 | 0 | 0 | 0 | 0 | 0 |
| ENSG00000249052 | 0 | 0 | 0 | 0 | 0 | 0 |
| ENSG00000248817 | 0 | 0 | 0 | 0 | 0 | 0 |
| ENSG00000280056 | 0 | 0 | 0 | 0 | 0 | 0 |
| ENSG00000249152 | 0 | 0 | 0 | 0 | 0 | 0 |
| ENSG00000248511 | 0 | 0 | 0 | 0 | 0 | 0 |
| ENSG00000152208 | 0 | 0 | 1 | 0 | 0 | 0 |
| ENSG00000251449 | 0 | 0 | 0 | 0 | 0 | 0 |
| ENSG00000249239 | 0 | 0 | 0 | 0 | 0 | 0 |
| ENSG00000248627 | 0 | 0 | 0 | 0 | 0 | 0 |
| ENSG00000248750 | 0 | 0 | 0 | 0 | 0 | 0 |
| ENSG00000250908 | 0 | 0 | 0 | 0 | 0 | 0 |
| ENSG00000252342 | 0 | 0 | 0 | 0 | 0 | 0 |
| ENSG00000172238 | 0 | 0 | 0 | 0 | 0 | 0 |
| ENSG00000246541 | 0 | 1 | 0 | 0 | 0 | 0 |
| ENSG00000250897 | 0 | 0 | 0 | 0 | 0 | 0 |
| ENSG00000163104 | 499 | 361 | 235 | 149 | 535 | 304 |
| ENSG00000163106 | 0 | 0 | 0 | 0 | 0 | 0 |
| ENSG00000241103 | 0 | 0 | 0 | 0 | 0 | 0 |
| ENSG00000163110 | 1166 | 645 | 667 | 366 | 743 | 544 |
| ENSG00000249951 | 0 | 0 | 0 | 0 | 0 | 0 |
| ENSG00000249599 | 2 | 0 | 3 | 2 | 3 | 0 |
| ENSG00000138696 | 147 | 50 | 39 | 44 | 66 | 54 |
| ENSG00000182168 | 13 | 5 | 8 | 0 | 0 | 2 |

| | | | | | | |
|---|---|---|---|---|---|---|
| ENSG00000271474 | 4 | 0 | 3 | 1 | 1 | 1 |
| ENSG00000242936 | 0 | 0 | 0 | 0 | 0 | 0 |
| ENSG00000163114 | 0 | 0 | 0 | 0 | 0 | 0 |
| ENSG00000200622 | 0 | 0 | 0 | 0 | 0 | 0 |
| ENSG00000201744 | 0 | 0 | 0 | 0 | 0 | 0 |
| ENSG00000250655 | 0 | 0 | 0 | 0 | 0 | 0 |
| ENSG00000248510 | 0 | 0 | 0 | 0 | 0 | 0 |
| ENSG00000201640 | 0 | 0 | 0 | 0 | 0 | 0 |
| ENSG00000236764 | 0 | 0 | 1 | 0 | 0 | 0 |
| ENSG00000253170 | 0 | 0 | 1 | 0 | 0 | 0 |
| ENSG00000163116 | 4 | 2 | 0 | 1 | 1 | 2 |
| ENSG00000254044 | 1 | 0 | 0 | 0 | 0 | 0 |
| ENSG00000251620 | 0 | 1 | 1 | 0 | 0 | 0 |
| ENSG00000250293 | 0 | 0 | 0 | 0 | 0 | 0 |
| ENSG00000242697 | 6 | 1 | 5 | 8 | 5 | 5 |
| ENSG00000214904 | 0 | 0 | 0 | 0 | 0 | 0 |
| ENSG00000214559 | 0 | 0 | 0 | 2 | 2 | 1 |
| ENSG00000138698 | 1076 | 530 | 576 | 447 | 654 | 573 |
| ENSG00000242318 | 0 | 0 | 0 | 0 | 0 | 0 |
| ENSG00000168785 | 28 | 11 | 25 | 10 | 14 | 29 |
| ENSG00000251523 | 0 | 0 | 0 | 0 | 0 | 0 |
| ENSG00000260641 | 0 | 0 | 1 | 2 | 0 | 0 |
| ENSG00000249764 | 0 | 0 | 0 | 0 | 0 | 0 |
| ENSG00000227118 | 0 | 0 | 0 | 0 | 0 | 0 |
| ENSG00000151247 | 1486 | 739 | 779 | 597 | 873 | 752 |
| ENSG00000249055 | 0 | 1 | 1 | 1 | 1 | 0 |
| ENSG00000263923 | 1 | 0 | 0 | 0 | 0 | 0 |
| ENSG00000214886 | 0 | 0 | 0 | 0 | 0 | 0 |
| ENSG00000252548 | 0 | 0 | 0 | 0 | 0 | 0 |
| ENSG00000251437 | 0 | 0 | 0 | 0 | 0 | 0 |
| ENSG00000164024 | 2614 | 1199 | 1528 | 1002 | 1468 | 1412 |
| ENSG00000265213 | 0 | 0 | 0 | 0 | 0 | 0 |
| ENSG00000250496 | 0 | 0 | 0 | 0 | 0 | 0 |
| ENSG00000272777 | 12 | 0 | 3 | 5 | 5 | 2 |
| ENSG00000197894 | 4006 | 1955 | 2498 | 2027 | 2190 | 2074 |
| ENSG00000246090 | 56 | 16 | 38 | 18 | 38 | 24 |
| ENSG00000198099 | 0 | 0 | 0 | 0 | 1 | 0 |
| ENSG00000249065 | 0 | 0 | 0 | 0 | 0 | 0 |
| ENSG00000172955 | 3 | 0 | 2 | 2 | 3 | 4 |
| ENSG00000187758 | 1 | 0 | 0 | 0 | 1 | 0 |
| ENSG00000196616 | 0 | 2 | 0 | 0 | 0 | 0 |
| ENSG00000248144 | 0 | 0 | 0 | 0 | 0 | 0 |
| ENSG00000196344 | 0 | 0 | 0 | 0 | 0 | 0 |
| ENSG00000250300 | 0 | 0 | 0 | 0 | 0 | 0 |
| ENSG00000138813 | 0 | 0 | 0 | 0 | 0 | 0 |
| ENSG00000145331 | 402 | 183 | 234 | 140 | 208 | 214 |
| ENSG00000138823 | 0 | 0 | 5 | 1 | 0 | 1 |
| ENSG00000248676 | 0 | 0 | 0 | 0 | 0 | 0 |
| ENSG00000248713 | 2 | 0 | 0 | 0 | 0 | 0 |
| ENSG00000070190 | 24 | 6 | 24 | 17 | 50 | 22 |

| | | | | | | |
|---|---|---|---|---|---|---|
| ENSG00000279913 | 0 | 0 | 0 | 0 | 0 | 0 |
| ENSG00000109270 | 1192 | 571 | 642 | 520 | 650 | 537 |
| ENSG00000164031 | 620 | 311 | 248 | 157 | 519 | 283 |
| ENSG00000279098 | 0 | 0 | 0 | 0 | 0 | 0 |
| ENSG00000164032 | 6711 | 3698 | 4387 | 4740 | 4424 | 4347 |
| ENSG00000245322 | 5 | 0 | 1 | 0 | 2 | 0 |
| ENSG00000250403 | 0 | 0 | 0 | 0 | 0 | 0 |
| ENSG00000248820 | 0 | 0 | 0 | 0 | 0 | 0 |
| ENSG00000145358 | 14 | 1 | 7 | 1 | 1 | 1 |
| ENSG00000249710 | 0 | 0 | 0 | 0 | 0 | 0 |
| ENSG00000164035 | 71 | 56 | 54 | 33 | 61 | 56 |
| ENSG00000250223 | 0 | 0 | 1 | 0 | 1 | 0 |
| ENSG00000251219 | 0 | 0 | 0 | 0 | 0 | 0 |
| ENSG00000251636 | 0 | 0 | 0 | 0 | 0 | 0 |
| ENSG00000138814 | 3852 | 1760 | 1690 | 1133 | 1712 | 1350 |
| ENSG00000273882 | 0 | 0 | 0 | 0 | 0 | 0 |
| ENSG00000221265 | 0 | 0 | 0 | 0 | 0 | 0 |
| ENSG00000254531 | 330 | 205 | 136 | 186 | 163 | 133 |
| ENSG00000153064 | 52 | 43 | 26 | 14 | 37 | 23 |
| ENSG00000199529 | 0 | 0 | 0 | 0 | 0 | 0 |
| ENSG00000251149 | 0 | 0 | 0 | 0 | 0 | 0 |
| ENSG00000251309 | 0 | 0 | 0 | 0 | 0 | 0 |
| ENSG00000138821 | 540 | 279 | 233 | 224 | 325 | 231 |
| ENSG00000241144 | 0 | 0 | 0 | 0 | 0 | 0 |
| ENSG00000248161 | 2 | 5 | 4 | 4 | 3 | 4 |
| ENSG00000251572 | 0 | 0 | 0 | 0 | 0 | 0 |
| ENSG00000260651 | 3 | 0 | 1 | 2 | 0 | 1 |
| ENSG00000109320 | 1545 | 716 | 830 | 609 | 734 | 777 |
| ENSG00000109323 | 1241 | 625 | 753 | 516 | 728 | 657 |
| ENSG00000241981 | 10 | 0 | 0 | 0 | 1 | 0 |
| ENSG00000230069 | 3 | 0 | 1 | 1 | 0 | 1 |
| ENSG00000248971 | 20 | 11 | 6 | 3 | 8 | 5 |
| ENSG00000224207 | 0 | 0 | 0 | 0 | 1 | 0 |
| ENSG00000251288 | 0 | 2 | 0 | 0 | 0 | 1 |
| ENSG00000109332 | 6287 | 2977 | 3447 | 2854 | 3595 | 3239 |
| ENSG00000246560 | 11 | 14 | 12 | 12 | 12 | 13 |
| ENSG00000252739 | 0 | 0 | 0 | 0 | 0 | 0 |
| ENSG00000145354 | 769 | 397 | 507 | 473 | 469 | 475 |
| ENSG00000164037 | 8 | 13 | 10 | 7 | 9 | 16 |
| ENSG00000250651 | 0 | 5 | 2 | 1 | 0 | 1 |
| ENSG00000249283 | 0 | 0 | 0 | 0 | 0 | 0 |
| ENSG00000164038 | 142 | 95 | 120 | 123 | 156 | 148 |
| ENSG00000164039 | 157 | 105 | 105 | 94 | 109 | 143 |
| ENSG00000138778 | 20 | 20 | 24 | 10 | 82 | 60 |
| ENSG00000248740 | 0 | 0 | 0 | 1 | 0 | 0 |
| ENSG00000251577 | 4 | 0 | 0 | 0 | 0 | 0 |
| ENSG00000250920 | 0 | 0 | 0 | 0 | 0 | 0 |
| ENSG00000251571 | 0 | 0 | 0 | 0 | 0 | 0 |
| ENSG00000169836 | 0 | 1 | 1 | 1 | 0 | 1 |
| ENSG00000252460 | 0 | 0 | 0 | 0 | 0 | 0 |

| | | | | | | |
|---|---|---|---|---|---|---|
| ENSG00000249614 | 0 | 0 | 0 | 0 | 0 | 0 |
| ENSG00000251170 | 0 | 0 | 0 | 0 | 0 | 0 |
| ENSG00000168772 | 7 | 4 | 1 | 3 | 2 | 4 |
| ENSG00000245384 | 0 | 0 | 0 | 0 | 0 | 0 |
| ENSG00000250670 | 0 | 0 | 0 | 0 | 0 | 0 |
| ENSG00000248242 | 0 | 0 | 0 | 0 | 1 | 0 |
| ENSG00000244470 | 0 | 0 | 0 | 0 | 0 | 0 |
| ENSG00000248373 | 0 | 0 | 0 | 0 | 0 | 0 |
| ENSG00000251906 | 0 | 0 | 0 | 0 | 0 | 0 |
| ENSG00000276992 | 0 | 0 | 0 | 0 | 0 | 0 |
| ENSG00000251473 | 0 | 0 | 0 | 0 | 0 | 0 |
| ENSG00000252136 | 0 | 0 | 0 | 0 | 0 | 0 |
| ENSG00000251259 | 6 | 1 | 6 | 4 | 1 | 4 |
| ENSG00000168769 | 496 | 275 | 198 | 76 | 401 | 247 |
| ENSG00000251586 | 0 | 0 | 0 | 1 | 0 | 0 |
| ENSG00000243383 | 0 | 0 | 0 | 0 | 0 | 0 |
| ENSG00000138777 | 2895 | 1402 | 1754 | 1639 | 1594 | 1732 |
| ENSG00000200917 | 2 | 0 | 1 | 0 | 0 | 3 |
| ENSG00000249264 | 14 | 0 | 6 | 3 | 9 | 6 |
| ENSG00000216425 | 0 | 0 | 0 | 0 | 0 | 0 |
| ENSG00000250922 | 0 | 0 | 0 | 0 | 0 | 0 |
| ENSG00000250522 | 0 | 0 | 0 | 0 | 0 | 2 |
| ENSG00000236699 | 8 | 5 | 8 | 4 | 13 | 6 |
| ENSG00000249885 | 0 | 1 | 0 | 0 | 0 | 0 |
| ENSG00000248778 | 0 | 0 | 0 | 0 | 0 | 0 |
| ENSG00000138785 | 360 | 233 | 287 | 257 | 297 | 294 |
| ENSG00000138780 | 614 | 293 | 344 | 221 | 350 | 309 |
| ENSG00000251175 | 0 | 0 | 3 | 0 | 1 | 0 |
| ENSG00000168743 | 4127 | 1794 | 2427 | 1000 | 1379 | 1362 |
| ENSG00000250740 | 0 | 1 | 0 | 0 | 1 | 0 |
| ENSG00000249635 | 0 | 0 | 0 | 0 | 1 | 0 |
| ENSG00000145348 | 656 | 347 | 316 | 202 | 471 | 300 |
| ENSG00000164022 | 1137 | 656 | 681 | 568 | 919 | 686 |
| ENSG00000250298 | 0 | 0 | 0 | 0 | 0 | 0 |
| ENSG00000249613 | 0 | 0 | 0 | 0 | 0 | 0 |
| ENSG00000248915 | 0 | 0 | 0 | 0 | 0 | 0 |
| ENSG00000155011 | 0 | 1 | 1 | 0 | 0 | 1 |
| ENSG00000213522 | 0 | 0 | 0 | 0 | 0 | 0 |
| ENSG00000251081 | 29 | 6 | 13 | 7 | 10 | 9 |
| ENSG00000252470 | 0 | 0 | 0 | 0 | 0 | 0 |
| ENSG00000138801 | 2303 | 1041 | 1456 | 867 | 1163 | 1229 |
| ENSG00000164023 | 115 | 71 | 93 | 42 | 57 | 42 |
| ENSG00000245293 | 0 | 0 | 0 | 0 | 0 | 0 |
| ENSG00000222691 | 0 | 0 | 0 | 0 | 0 | 0 |
| ENSG00000155016 | 241 | 104 | 148 | 92 | 131 | 103 |
| ENSG00000249604 | 0 | 0 | 0 | 0 | 2 | 0 |
| ENSG00000138796 | 1448 | 622 | 817 | 602 | 633 | 718 |
| ENSG00000138795 | 4 | 5 | 2 | 4 | 3 | 5 |
| ENSG00000232021 | 0 | 0 | 0 | 0 | 0 | 0 |
| ENSG00000239939 | 0 | 0 | 0 | 0 | 0 | 0 |

| | | | | | | |
|---|---|---|---|---|---|---|
| ENSG00000278589 | 0 | 0 | 0 | 0 | 0 | 0 |
| ENSG00000250485 | 0 | 0 | 0 | 0 | 0 | 0 |
| ENSG00000234492 | 5 | 0 | 2 | 0 | 1 | 2 |
| ENSG00000109475 | 8811 | 4434 | 5275 | 5825 | 5333 | 5215 |
| ENSG00000198856 | 2101 | 1026 | 1161 | 1152 | 1313 | 1125 |
| ENSG00000206601 | 0 | 0 | 0 | 0 | 0 | 0 |
| ENSG00000164089 | 0 | 0 | 0 | 1 | 0 | 0 |
| ENSG00000249257 | 0 | 0 | 0 | 0 | 0 | 0 |
| ENSG00000250185 | 0 | 0 | 0 | 0 | 0 | 0 |
| ENSG00000188517 | 0 | 0 | 0 | 0 | 0 | 0 |
| ENSG00000246774 | 0 | 0 | 0 | 0 | 0 | 0 |
| ENSG00000249465 | 0 | 0 | 1 | 0 | 0 | 1 |
| ENSG00000247950 | 17 | 13 | 20 | 9 | 11 | 18 |
| ENSG00000138802 | 924 | 513 | 468 | 253 | 621 | 469 |
| ENSG00000243227 | 0 | 0 | 0 | 0 | 0 | 0 |
| ENSG00000207988 | 0 | 0 | 1 | 0 | 0 | 0 |
| ENSG00000225094 | 4 | 3 | 4 | 4 | 3 | 6 |
| ENSG00000249965 | 0 | 1 | 0 | 0 | 5 | 3 |
| ENSG00000005059 | 464 | 262 | 304 | 250 | 319 | 301 |
| ENSG00000248785 | 0 | 0 | 0 | 1 | 0 | 0 |
| ENSG00000138794 | 896 | 404 | 455 | 313 | 473 | 486 |
| ENSG00000273447 | 9 | 2 | 7 | 2 | 2 | 4 |
| ENSG00000123739 | 1069 | 438 | 583 | 350 | 603 | 502 |
| ENSG00000285330 | 0 | 0 | 0 | 0 | 0 | 0 |
| ENSG00000205403 | 0 | 0 | 0 | 0 | 0 | 0 |
| ENSG00000272795 | 0 | 1 | 2 | 1 | 0 | 0 |
| ENSG00000109534 | 546 | 299 | 360 | 333 | 400 | 368 |
| ENSG00000180245 | 0 | 1 | 0 | 0 | 0 | 2 |
| ENSG00000183423 | 0 | 1 | 1 | 0 | 1 | 0 |
| ENSG00000249681 | 0 | 0 | 0 | 0 | 0 | 0 |
| ENSG00000138798 | 56 | 34 | 58 | 30 | 42 | 42 |
| ENSG00000207260 | 0 | 0 | 0 | 0 | 0 | 1 |
| ENSG00000170522 | 1062 | 567 | 689 | 393 | 652 | 621 |
| ENSG00000263940 | 0 | 0 | 0 | 0 | 0 | 0 |
| ENSG00000248200 | 0 | 0 | 0 | 0 | 0 | 0 |
| ENSG00000166530 | 0 | 0 | 0 | 0 | 0 | 0 |
| ENSG00000250594 | 0 | 0 | 0 | 0 | 0 | 0 |
| ENSG00000212160 | 0 | 0 | 0 | 0 | 0 | 0 |
| ENSG00000250304 | 0 | 1 | 0 | 0 | 0 | 0 |
| ENSG00000138792 | 292 | 133 | 183 | 91 | 181 | 150 |
| ENSG00000251464 | 0 | 0 | 0 | 0 | 0 | 0 |
| ENSG00000251046 | 0 | 1 | 0 | 0 | 0 | 3 |
| ENSG00000199894 | 0 | 0 | 0 | 0 | 0 | 0 |
| ENSG00000250511 | 0 | 0 | 0 | 0 | 0 | 0 |
| ENSG00000250103 | 0 | 0 | 0 | 0 | 0 | 0 |
| ENSG00000164093 | 96 | 50 | 54 | 42 | 54 | 34 |
| ENSG00000249519 | 0 | 0 | 0 | 0 | 0 | 0 |
| ENSG00000215961 | 0 | 0 | 0 | 0 | 0 | 0 |
| ENSG00000251483 | 0 | 0 | 0 | 0 | 0 | 0 |
| ENSG00000251312 | 0 | 0 | 0 | 0 | 0 | 0 |

| | | | | | | |
|---|---|---|---|---|---|---|
| ENSG00000200963 | 0 | 0 | 0 | 0 | 0 | 0 |
| ENSG00000241722 | 0 | 0 | 0 | 0 | 0 | 0 |
| ENSG00000248656 | 0 | 0 | 0 | 0 | 0 | 0 |
| ENSG00000250855 | 0 | 0 | 0 | 0 | 0 | 0 |
| ENSG00000248432 | 0 | 1 | 0 | 0 | 0 | 0 |
| ENSG00000249815 | 0 | 0 | 0 | 0 | 0 | 1 |
| ENSG00000250556 | 0 | 0 | 0 | 0 | 0 | 0 |
| ENSG00000250307 | 0 | 0 | 0 | 0 | 0 | 0 |
| ENSG00000240459 | 0 | 0 | 0 | 0 | 0 | 0 |
| ENSG00000174749 | 909 | 394 | 471 | 365 | 510 | 515 |
| ENSG00000243280 | 0 | 1 | 0 | 0 | 1 | 0 |
| ENSG00000260526 | 34 | 17 | 24 | 12 | 11 | 19 |
| ENSG00000138660 | 869 | 430 | 480 | 319 | 553 | 537 |
| ENSG00000272567 | 6 | 0 | 0 | 1 | 0 | 1 |
| ENSG00000145365 | 394 | 172 | 203 | 205 | 203 | 218 |
| ENSG00000073331 | 378 | 136 | 209 | 143 | 167 | 183 |
| ENSG00000251441 | 0 | 2 | 0 | 0 | 1 | 1 |
| ENSG00000248697 | 0 | 0 | 0 | 0 | 2 | 2 |
| ENSG00000178403 | 2 | 0 | 1 | 0 | 0 | 1 |
| ENSG00000249509 | 2 | 0 | 0 | 0 | 2 | 0 |
| ENSG00000248439 | 0 | 0 | 0 | 0 | 0 | 0 |
| ENSG00000138658 | 87 | 39 | 32 | 18 | 78 | 47 |
| ENSG00000250966 | 4 | 4 | 4 | 1 | 1 | 4 |
| ENSG00000174720 | 300 | 249 | 205 | 106 | 432 | 239 |
| ENSG00000249532 | 4 | 2 | 1 | 0 | 0 | 1 |
| ENSG00000199169 | 0 | 0 | 0 | 0 | 0 | 0 |
| ENSG00000199145 | 0 | 1 | 0 | 0 | 0 | 0 |
| ENSG00000207927 | 0 | 0 | 0 | 0 | 0 | 0 |
| ENSG00000199102 | 0 | 0 | 0 | 0 | 0 | 0 |
| ENSG00000284463 | 0 | 0 | 0 | 0 | 0 | 0 |
| ENSG00000220924 | 0 | 0 | 0 | 0 | 0 | 0 |
| ENSG00000250046 | 0 | 0 | 0 | 0 | 0 | 0 |
| ENSG00000248633 | 0 | 0 | 0 | 0 | 0 | 0 |
| ENSG00000241002 | 0 | 0 | 0 | 0 | 0 | 0 |
| ENSG00000202536 | 0 | 0 | 0 | 0 | 0 | 0 |
| ENSG00000241741 | 14 | 9 | 8 | 13 | 13 | 14 |
| ENSG00000250500 | 0 | 0 | 0 | 0 | 0 | 0 |
| ENSG00000145362 | 4 | 1 | 1 | 0 | 0 | 2 |
| ENSG00000234841 | 0 | 0 | 0 | 0 | 0 | 0 |
| ENSG00000251126 | 0 | 0 | 0 | 0 | 0 | 0 |
| ENSG00000228814 | 0 | 0 | 0 | 0 | 0 | 0 |
| ENSG00000248152 | 0 | 0 | 0 | 0 | 0 | 0 |
| ENSG00000249373 | 0 | 0 | 0 | 0 | 0 | 0 |
| ENSG00000283193 | 0 | 0 | 0 | 0 | 0 | 0 |
| ENSG00000275810 | 0 | 0 | 0 | 0 | 0 | 0 |
| ENSG00000196656 | 22 | 9 | 5 | 6 | 6 | 4 |
| ENSG00000239279 | 0 | 0 | 0 | 0 | 0 | 0 |
| ENSG00000206820 | 0 | 0 | 0 | 0 | 0 | 0 |
| ENSG00000145349 | 399 | 144 | 257 | 123 | 165 | 159 |
| ENSG00000250229 | 0 | 0 | 0 | 0 | 0 | 0 |

| | | | | | | |
|---|---|---|---|---|---|---|
| ENSG00000180801 | 20 | 3 | 10 | 3 | 5 | 4 |
| ENSG00000249304 | 0 | 0 | 0 | 0 | 1 | 0 |
| ENSG00000248716 | 0 | 0 | 0 | 0 | 0 | 0 |
| ENSG00000174607 | 0 | 0 | 0 | 0 | 0 | 0 |
| ENSG00000207931 | 0 | 0 | 0 | 0 | 0 | 0 |
| ENSG00000250215 | 0 | 0 | 0 | 0 | 0 | 0 |
| ENSG00000240637 | 0 | 0 | 0 | 0 | 0 | 0 |
| ENSG00000138653 | 0 | 0 | 0 | 0 | 0 | 0 |
| ENSG00000251308 | 0 | 0 | 0 | 0 | 0 | 0 |
| ENSG00000250693 | 0 | 0 | 0 | 0 | 0 | 0 |
| ENSG00000248488 | 0 | 0 | 0 | 0 | 0 | 0 |
| ENSG00000250363 | 0 | 1 | 0 | 1 | 0 | 0 |
| ENSG00000249259 | 0 | 0 | 0 | 0 | 0 | 0 |
| ENSG00000250442 | 0 | 0 | 0 | 0 | 0 | 0 |
| ENSG00000250627 | 0 | 0 | 0 | 0 | 0 | 0 |
| ENSG00000270394 | 0 | 0 | 0 | 0 | 0 | 0 |
| ENSG00000284253 | 0 | 0 | 0 | 0 | 0 | 0 |
| ENSG00000250272 | 0 | 0 | 0 | 0 | 0 | 0 |
| ENSG00000248139 | 0 | 0 | 0 | 0 | 0 | 0 |
| ENSG00000250791 | 0 | 0 | 0 | 0 | 0 | 0 |
| ENSG00000213493 | 0 | 1 | 0 | 0 | 0 | 0 |
| ENSG00000201752 | 0 | 0 | 0 | 0 | 0 | 0 |
| ENSG00000271053 | 0 | 0 | 0 | 0 | 0 | 0 |
| ENSG00000174599 | 14 | 1 | 8 | 5 | 5 | 4 |
| ENSG00000223605 | 0 | 0 | 0 | 0 | 0 | 0 |
| ENSG00000224932 | 0 | 0 | 0 | 0 | 1 | 0 |
| ENSG00000228358 | 0 | 0 | 0 | 0 | 0 | 0 |
| ENSG00000236922 | 0 | 0 | 0 | 0 | 0 | 0 |
| ENSG00000237684 | 0 | 0 | 0 | 0 | 0 | 0 |
| ENSG00000213492 | 0 | 0 | 0 | 0 | 0 | 0 |
| ENSG00000229565 | 0 | 0 | 0 | 0 | 0 | 0 |
| ENSG00000164100 | 0 | 0 | 0 | 0 | 0 | 0 |
| ENSG00000251463 | 0 | 0 | 0 | 0 | 0 | 1 |
| ENSG00000269893 | 1907 | 956 | 1164 | 1597 | 1345 | 1371 |
| ENSG00000275994 | 0 | 0 | 0 | 0 | 0 | 0 |
| ENSG00000164099 | 6 | 0 | 0 | 3 | 1 | 1 |
| ENSG00000230671 | 0 | 0 | 0 | 0 | 0 | 0 |
| ENSG00000273716 | 0 | 0 | 1 | 0 | 2 | 0 |
| ENSG00000154608 | 4 | 1 | 3 | 1 | 2 | 0 |
| ENSG00000260404 | 177 | 73 | 124 | 46 | 88 | 86 |
| ENSG00000223225 | 0 | 0 | 0 | 0 | 0 | 0 |
| ENSG00000248213 | 13 | 3 | 0 | 1 | 7 | 2 |
| ENSG00000251155 | 0 | 1 | 1 | 0 | 1 | 1 |
| ENSG00000281731 | 3 | 0 | 2 | 0 | 0 | 1 |
| ENSG00000145388 | 728 | 401 | 388 | 226 | 447 | 342 |
| ENSG00000150961 | 1333 | 607 | 739 | 488 | 778 | 667 |
| ENSG00000172403 | 36 | 22 | 18 | 5 | 11 | 5 |
| ENSG00000249002 | 0 | 0 | 0 | 0 | 0 | 0 |
| ENSG00000280551 | 0 | 0 | 0 | 0 | 0 | 0 |
| ENSG00000172399 | 4 | 7 | 5 | 6 | 3 | 5 |

| | | | | | | |
|---|---|---|---|---|---|---|
| ENSG00000178636 | 0 | 0 | 0 | 0 | 0 | 0 |
| ENSG00000145390 | 290 | 185 | 149 | 62 | 358 | 174 |
| ENSG00000164096 | 2691 | 1322 | 1608 | 1210 | 1479 | 1421 |
| ENSG00000145384 | 0 | 0 | 0 | 0 | 0 | 0 |
| ENSG00000250412 | 5 | 3 | 3 | 1 | 4 | 5 |
| ENSG00000201186 | 0 | 0 | 0 | 0 | 0 | 0 |
| ENSG00000249244 | 8 | 3 | 2 | 1 | 7 | 7 |
| ENSG00000283050 | 168 | 66 | 98 | 53 | 79 | 53 |
| ENSG00000260091 | 0 | 0 | 0 | 0 | 0 | 0 |
| ENSG00000223208 | 0 | 0 | 0 | 0 | 0 | 0 |
| ENSG00000245958 | 221 | 147 | 220 | 153 | 178 | 181 |
| ENSG00000250950 | 0 | 0 | 0 | 0 | 0 | 0 |
| ENSG00000138735 | 330 | 215 | 190 | 121 | 249 | 154 |
| ENSG00000221245 | 0 | 0 | 0 | 0 | 0 | 0 |
| ENSG00000250772 | 0 | 0 | 0 | 0 | 0 | 0 |
| ENSG00000250183 | 0 | 0 | 0 | 0 | 0 | 0 |
| ENSG00000250392 | 0 | 0 | 0 | 0 | 0 | 0 |
| ENSG00000200553 | 0 | 0 | 0 | 0 | 0 | 0 |
| ENSG00000253825 | 0 | 0 | 0 | 0 | 0 | 0 |
| ENSG00000164109 | 643 | 349 | 375 | 427 | 429 | 429 |
| ENSG00000250938 | 1 | 0 | 3 | 0 | 0 | 0 |
| ENSG00000250219 | 0 | 0 | 0 | 0 | 0 | 0 |
| ENSG00000225665 | 0 | 0 | 0 | 0 | 0 | 0 |
| ENSG00000249409 | 0 | 0 | 0 | 0 | 0 | 0 |
| ENSG00000138738 | 34 | 8 | 14 | 7 | 15 | 12 |
| ENSG00000212359 | 0 | 0 | 0 | 0 | 0 | 0 |
| ENSG00000251609 | 0 | 1 | 0 | 0 | 0 | 0 |
| ENSG00000173376 | 0 | 0 | 0 | 0 | 0 | 0 |
| ENSG00000223006 | 0 | 0 | 0 | 0 | 0 | 0 |
| ENSG00000250344 | 0 | 0 | 0 | 0 | 0 | 0 |
| ENSG00000271302 | 0 | 0 | 0 | 0 | 0 | 0 |
| ENSG00000050730 | 0 | 0 | 0 | 0 | 0 | 0 |
| ENSG00000252183 | 0 | 0 | 0 | 0 | 0 | 0 |
| ENSG00000207148 | 0 | 0 | 0 | 0 | 0 | 0 |
| ENSG00000186867 | 0 | 0 | 0 | 0 | 0 | 0 |
| ENSG00000213480 | 0 | 0 | 0 | 0 | 0 | 0 |
| ENSG00000224062 | 0 | 0 | 0 | 0 | 0 | 0 |
| ENSG00000164111 | 4363 | 2253 | 2586 | 1973 | 2262 | 2076 |
| ENSG00000164112 | 0 | 0 | 0 | 0 | 0 | 0 |
| ENSG00000226757 | 0 | 0 | 0 | 0 | 0 | 0 |
| ENSG00000123737 | 585 | 272 | 352 | 251 | 379 | 324 |
| ENSG00000145386 | 730 | 373 | 397 | 443 | 568 | 547 |
| ENSG00000138686 | 237 | 108 | 107 | 96 | 122 | 112 |
| ENSG00000261501 | 0 | 0 | 0 | 0 | 0 | 0 |
| ENSG00000138741 | 0 | 0 | 0 | 0 | 0 | 0 |
| ENSG00000237868 | 0 | 0 | 0 | 0 | 0 | 0 |
| ENSG00000241037 | 0 | 0 | 0 | 0 | 0 | 0 |
| ENSG00000138688 | 403 | 248 | 206 | 69 | 585 | 263 |
| ENSG00000164113 | 0 | 0 | 0 | 0 | 0 | 0 |
| ENSG00000109471 | 0 | 0 | 0 | 0 | 0 | 1 |

| | | | | | | |
|---|---|---|---|---|---|---|
| ENSG00000138684 | 0 | 0 | 0 | 0 | 0 | 0 |
| ENSG00000227145 | 0 | 0 | 1 | 0 | 0 | 0 |
| ENSG00000224786 | 0 | 0 | 0 | 1 | 0 | 0 |
| ENSG00000181004 | 113 | 41 | 65 | 54 | 62 | 59 |
| ENSG00000244083 | 0 | 0 | 0 | 0 | 0 | 0 |
| ENSG00000138685 | 0 | 1 | 0 | 1 | 1 | 0 |
| ENSG00000240775 | 0 | 0 | 0 | 0 | 0 | 0 |
| ENSG00000273007 | 2 | 1 | 0 | 1 | 2 | 4 |
| ENSG00000170917 | 72 | 65 | 69 | 39 | 62 | 55 |
| ENSG00000145375 | 287 | 172 | 157 | 104 | 170 | 141 |
| ENSG00000250322 | 0 | 0 | 0 | 0 | 0 | 0 |
| ENSG00000164056 | 142 | 35 | 67 | 39 | 51 | 75 |
| ENSG00000249125 | 0 | 0 | 0 | 0 | 0 | 0 |
| ENSG00000251526 | 4 | 0 | 1 | 0 | 0 | 1 |
| ENSG00000249464 | 2 | 4 | 5 | 2 | 2 | 1 |
| ENSG00000226655 | 0 | 0 | 0 | 0 | 0 | 0 |
| ENSG00000228489 | 0 | 0 | 0 | 0 | 0 | 0 |
| ENSG00000250458 | 0 | 0 | 0 | 0 | 0 | 0 |
| ENSG00000250484 | 0 | 0 | 0 | 0 | 0 | 0 |
| ENSG00000249837 | 0 | 0 | 0 | 0 | 0 | 0 |
| ENSG00000248506 | 0 | 0 | 0 | 0 | 0 | 0 |
| ENSG00000248565 | 0 | 0 | 0 | 0 | 0 | 0 |
| ENSG00000261083 | 0 | 0 | 1 | 0 | 0 | 0 |
| ENSG00000151458 | 4445 | 2298 | 2415 | 673 | 2023 | 1629 |
| ENSG00000196159 | 88 | 40 | 40 | 9 | 30 | 24 |
| ENSG00000201297 | 0 | 0 | 0 | 0 | 0 | 0 |
| ENSG00000283550 | 0 | 0 | 0 | 0 | 0 | 0 |
| ENSG00000250149 | 0 | 0 | 0 | 0 | 0 | 0 |
| ENSG00000251436 | 0 | 0 | 0 | 0 | 0 | 0 |
| ENSG00000249555 | 0 | 0 | 0 | 0 | 0 | 0 |
| ENSG00000251649 | 0 | 0 | 0 | 0 | 0 | 0 |
| ENSG00000251005 | 0 | 0 | 0 | 0 | 0 | 0 |
| ENSG00000223620 | 0 | 0 | 0 | 0 | 0 | 0 |
| ENSG00000199862 | 0 | 0 | 0 | 0 | 0 | 0 |
| ENSG00000250945 | 0 | 0 | 0 | 0 | 0 | 0 |
| ENSG00000248491 | 0 | 0 | 0 | 0 | 0 | 0 |
| ENSG00000248209 | 0 | 0 | 0 | 0 | 0 | 0 |
| ENSG00000249795 | 0 | 0 | 0 | 0 | 0 | 0 |
| ENSG00000164066 | 112 | 67 | 74 | 30 | 79 | 60 |
| ENSG00000151475 | 0 | 0 | 0 | 0 | 0 | 0 |
| ENSG00000164070 | 244 | 139 | 130 | 59 | 177 | 118 |
| ENSG00000282855 | 2 | 0 | 0 | 0 | 1 | 0 |
| ENSG00000244021 | 0 | 0 | 0 | 0 | 2 | 0 |
| ENSG00000261668 | 0 | 1 | 0 | 1 | 0 | 0 |
| ENSG00000142731 | 369 | 210 | 250 | 117 | 284 | 227 |
| ENSG00000251821 | 0 | 1 | 1 | 0 | 3 | 0 |
| ENSG00000164073 | 757 | 427 | 480 | 288 | 429 | 414 |
| ENSG00000164074 | 423 | 208 | 271 | 150 | 259 | 251 |
| ENSG00000138709 | 525 | 259 | 291 | 210 | 342 | 310 |
| ENSG00000241651 | 0 | 0 | 0 | 0 | 1 | 1 |

| | | | | | | |
|---|---|---|---|---|---|---|
| ENSG00000248394 | 0 | 0 | 2 | 0 | 0 | 0 |
| ENSG00000164040 | 2215 | 1051 | 1358 | 1100 | 1278 | 1294 |
| ENSG00000251432 | 59 | 24 | 56 | 25 | 40 | 44 |
| ENSG00000273077 | 2 | 0 | 1 | 0 | 2 | 0 |
| ENSG00000248187 | 0 | 0 | 0 | 0 | 0 | 0 |
| ENSG00000248802 | 0 | 0 | 0 | 0 | 0 | 0 |
| ENSG00000280711 | 0 | 0 | 0 | 0 | 0 | 0 |
| ENSG00000077684 | 402 | 214 | 219 | 154 | 211 | 215 |
| ENSG00000151466 | 95 | 53 | 47 | 44 | 84 | 62 |
| ENSG00000275647 | 0 | 0 | 0 | 0 | 0 | 0 |
| ENSG00000281369 | 0 | 0 | 0 | 1 | 0 | 0 |
| ENSG00000151470 | 204 | 109 | 123 | 104 | 131 | 136 |
| ENSG00000251345 | 0 | 0 | 0 | 0 | 0 | 0 |
| ENSG00000248958 | 0 | 0 | 0 | 0 | 0 | 0 |
| ENSG00000251195 | 0 | 0 | 0 | 0 | 0 | 0 |
| ENSG00000250193 | 0 | 0 | 0 | 0 | 0 | 0 |
| ENSG00000246876 | 0 | 0 | 0 | 0 | 0 | 0 |
| ENSG00000249618 | 0 | 0 | 0 | 0 | 0 | 0 |
| ENSG00000234125 | 0 | 0 | 0 | 0 | 0 | 0 |
| ENSG00000251326 | 0 | 0 | 0 | 0 | 0 | 0 |
| ENSG00000248877 | 0 | 0 | 0 | 0 | 0 | 0 |
| ENSG00000249018 | 0 | 0 | 0 | 0 | 0 | 0 |
| ENSG00000280059 | 0 | 0 | 0 | 0 | 0 | 0 |
| ENSG00000250503 | 0 | 0 | 0 | 0 | 0 | 0 |
| ENSG00000250213 | 0 | 0 | 0 | 0 | 0 | 0 |
| ENSG00000206782 | 0 | 0 | 0 | 0 | 0 | 0 |
| ENSG00000250102 | 0 | 0 | 0 | 0 | 0 | 0 |
| ENSG00000248715 | 0 | 0 | 0 | 0 | 0 | 0 |
| ENSG00000241392 | 0 | 0 | 0 | 0 | 0 | 0 |
| ENSG00000252014 | 0 | 0 | 0 | 0 | 0 | 0 |
| ENSG00000249416 | 0 | 0 | 0 | 0 | 0 | 0 |
| ENSG00000251676 | 0 | 0 | 0 | 0 | 0 | 0 |
| ENSG00000184139 | 0 | 0 | 0 | 0 | 0 | 0 |
| ENSG00000251555 | 2 | 0 | 0 | 0 | 1 | 0 |
| ENSG00000251598 | 0 | 0 | 0 | 0 | 0 | 0 |
| ENSG00000249463 | 0 | 0 | 0 | 0 | 0 | 0 |
| ENSG00000251051 | 0 | 0 | 0 | 0 | 0 | 0 |
| ENSG00000248369 | 0 | 0 | 0 | 0 | 0 | 0 |
| ENSG00000251398 | 0 | 0 | 0 | 0 | 1 | 0 |
| ENSG00000249038 | 0 | 0 | 0 | 0 | 0 | 0 |
| ENSG00000251488 | 7 | 2 | 2 | 0 | 1 | 0 |
| ENSG00000249513 | 5 | 0 | 1 | 0 | 0 | 0 |
| ENSG00000250241 | 345 | 165 | 232 | 90 | 144 | 132 |
| ENSG00000248735 | 4 | 1 | 0 | 1 | 3 | 2 |
| ENSG00000138650 | 14043 | 5703 | 8649 | 3600 | 3949 | 5049 |
| ENSG00000250191 | 0 | 0 | 0 | 0 | 0 | 0 |
| ENSG00000251388 | 1 | 2 | 1 | 0 | 0 | 0 |
| ENSG00000254535 | 321 | 145 | 141 | 93 | 141 | 108 |
| ENSG00000251199 | 0 | 0 | 0 | 0 | 0 | 0 |
| ENSG00000248191 | 0 | 0 | 1 | 0 | 0 | 0 |

| | | | | | | |
|---|---|---|---|---|---|---|
| ENSG00000251291 | 0 | 0 | 0 | 0 | 0 | 1 |
| ENSG00000249000 | 0 | 0 | 0 | 0 | 0 | 0 |
| ENSG00000249847 | 0 | 0 | 0 | 0 | 0 | 0 |
| ENSG00000251228 | 0 | 0 | 0 | 0 | 0 | 0 |
| ENSG00000241411 | 4 | 1 | 0 | 0 | 0 | 0 |
| ENSG00000250144 | 2 | 2 | 1 | 3 | 1 | 4 |
| ENSG00000248434 | 0 | 0 | 0 | 0 | 0 | 0 |
| ENSG00000248456 | 0 | 0 | 0 | 0 | 0 | 0 |
| ENSG00000283432 | 0 | 0 | 0 | 0 | 0 | 0 |
| ENSG00000250834 | 0 | 0 | 0 | 1 | 0 | 0 |
| ENSG00000248104 | 0 | 0 | 0 | 0 | 0 | 0 |
| ENSG00000250865 | 0 | 0 | 0 | 0 | 0 | 0 |
| ENSG00000248330 | 0 | 0 | 0 | 0 | 0 | 0 |
| ENSG00000207322 | 0 | 0 | 0 | 0 | 0 | 0 |
| ENSG00000251492 | 0 | 0 | 0 | 0 | 0 | 0 |
| ENSG00000261166 | 0 | 0 | 0 | 0 | 0 | 0 |
| ENSG00000251567 | 0 | 0 | 0 | 0 | 0 | 0 |
| ENSG00000250665 | 0 | 0 | 0 | 0 | 0 | 0 |
| ENSG00000249311 | 0 | 0 | 0 | 0 | 0 | 0 |
| ENSG00000279749 | 0 | 0 | 0 | 0 | 0 | 0 |
| ENSG00000248869 | 0 | 0 | 0 | 0 | 0 | 0 |
| ENSG00000250341 | 0 | 0 | 0 | 0 | 0 | 0 |
| ENSG00000250813 | 0 | 0 | 0 | 0 | 0 | 0 |
| ENSG00000251440 | 0 | 0 | 0 | 0 | 0 | 0 |
| ENSG00000213448 | 0 | 0 | 0 | 0 | 0 | 0 |
| ENSG00000189184 | 0 | 0 | 0 | 0 | 0 | 0 |
| ENSG00000251632 | 0 | 0 | 0 | 0 | 0 | 0 |
| ENSG00000280352 | 0 | 0 | 0 | 0 | 0 | 0 |
| ENSG00000250126 | 0 | 0 | 0 | 0 | 0 | 0 |
| ENSG00000250777 | 0 | 0 | 0 | 0 | 0 | 0 |
| ENSG00000250034 | 0 | 0 | 0 | 0 | 0 | 0 |
| ENSG00000248307 | 0 | 0 | 0 | 0 | 0 | 0 |
| ENSG00000250033 | 0 | 0 | 0 | 2 | 3 | 1 |
| ENSG00000151012 | 97 | 76 | 59 | 123 | 454 | 257 |
| ENSG00000249091 | 0 | 0 | 0 | 0 | 0 | 0 |
| ENSG00000248397 | 0 | 0 | 0 | 0 | 0 | 0 |
| ENSG00000251372 | 0 | 0 | 0 | 0 | 0 | 0 |
| ENSG00000249381 | 0 | 0 | 0 | 0 | 0 | 0 |
| ENSG00000248305 | 0 | 0 | 0 | 0 | 0 | 0 |
| ENSG00000248795 | 0 | 0 | 0 | 1 | 1 | 0 |
| ENSG00000250977 | 0 | 0 | 0 | 0 | 0 | 0 |
| ENSG00000250501 | 0 | 0 | 0 | 0 | 0 | 0 |
| ENSG00000250195 | 0 | 0 | 0 | 0 | 0 | 0 |
| ENSG00000250577 | 0 | 0 | 0 | 0 | 0 | 0 |
| ENSG00000151014 | 148 | 77 | 83 | 76 | 98 | 104 |
| ENSG00000109381 | 963 | 439 | 576 | 349 | 541 | 518 |
| ENSG00000252503 | 0 | 4 | 1 | 1 | 4 | 0 |
| ENSG00000240723 | 0 | 0 | 0 | 0 | 0 | 0 |
| ENSG00000179967 | 17 | 12 | 14 | 19 | 17 | 13 |
| ENSG00000276396 | 0 | 0 | 0 | 0 | 0 | 0 |

| | | | | | | |
|---|---|---|---|---|---|---|
| ENSG00000265892 | 0 | 0 | 0 | 0 | 0 | 0 |
| ENSG00000207384 | 0 | 0 | 0 | 0 | 0 | 0 |
| ENSG00000252362 | 0 | 0 | 0 | 0 | 0 | 0 |
| ENSG00000206722 | 0 | 0 | 0 | 0 | 0 | 0 |
| ENSG00000137463 | 3 | 1 | 1 | 3 | 0 | 0 |
| ENSG00000109390 | 1493 | 773 | 922 | 827 | 872 | 810 |
| ENSG00000164134 | 1231 | 721 | 619 | 360 | 1045 | 646 |
| ENSG00000200520 | 0 | 0 | 0 | 0 | 0 | 0 |
| ENSG00000273247 | 122 | 59 | 88 | 46 | 71 | 49 |
| ENSG00000248863 | 0 | 0 | 0 | 0 | 0 | 0 |
| ENSG00000239005 | 2 | 0 | 0 | 0 | 4 | 0 |
| ENSG00000172007 | 398 | 187 | 228 | 135 | 206 | 187 |
| ENSG00000145391 | 5505 | 2573 | 3201 | 1803 | 3107 | 2979 |
| ENSG00000249302 | 0 | 1 | 0 | 0 | 0 | 0 |
| ENSG00000272717 | 12 | 7 | 5 | 5 | 11 | 11 |
| ENSG00000261268 | 0 | 0 | 0 | 0 | 0 | 0 |
| ENSG00000085871 | 591 | 289 | 382 | 353 | 322 | 369 |
| ENSG00000201533 | 0 | 0 | 0 | 0 | 0 | 0 |
| ENSG00000178458 | 9 | 1 | 2 | 1 | 0 | 2 |
| ENSG00000196782 | 372 | 150 | 221 | 149 | 171 | 204 |
| ENSG00000252233 | 0 | 0 | 0 | 0 | 0 | 0 |
| ENSG00000250698 | 0 | 0 | 0 | 0 | 0 | 0 |
| ENSG00000153130 | 1096 | 637 | 600 | 544 | 828 | 616 |
| ENSG00000196951 | 25 | 7 | 4 | 10 | 19 | 10 |
| ENSG00000249754 | 0 | 0 | 0 | 0 | 0 | 0 |
| ENSG00000241923 | 4 | 2 | 0 | 1 | 3 | 1 |
| ENSG00000153132 | 24 | 20 | 16 | 7 | 13 | 6 |
| ENSG00000205301 | 0 | 0 | 0 | 0 | 0 | 0 |
| ENSG00000280262 | 0 | 0 | 0 | 0 | 0 | 0 |
| ENSG00000179387 | 668 | 353 | 375 | 256 | 487 | 346 |
| ENSG00000109424 | 0 | 0 | 0 | 1 | 0 | 0 |
| ENSG00000242102 | 0 | 0 | 0 | 0 | 0 | 0 |
| ENSG00000248621 | 0 | 0 | 0 | 0 | 0 | 0 |
| ENSG00000109436 | 13777 | 6184 | 9551 | 3506 | 4762 | 5486 |
| ENSG00000249661 | 131 | 63 | 99 | 25 | 28 | 50 |
| ENSG00000207155 | 0 | 0 | 0 | 0 | 0 | 0 |
| ENSG00000248335 | 0 | 0 | 0 | 0 | 0 | 0 |
| ENSG00000273472 | 341 | 157 | 290 | 105 | 144 | 141 |
| ENSG00000222613 | 0 | 0 | 0 | 0 | 0 | 0 |
| ENSG00000170153 | 228 | 105 | 145 | 91 | 134 | 122 |
| ENSG00000109445 | 718 | 416 | 458 | 423 | 480 | 434 |
| ENSG00000248747 | 0 | 0 | 0 | 0 | 0 | 0 |
| ENSG00000248810 | 0 | 0 | 0 | 0 | 0 | 0 |
| ENSG00000250189 | 0 | 0 | 0 | 0 | 0 | 0 |
| ENSG00000272632 | 0 | 0 | 0 | 0 | 0 | 0 |
| ENSG00000250141 | 0 | 0 | 0 | 0 | 0 | 0 |
| ENSG00000164136 | 6 | 2 | 3 | 3 | 2 | 3 |
| ENSG00000109452 | 1954 | 1211 | 1094 | 713 | 1878 | 1161 |
| ENSG00000242134 | 0 | 0 | 0 | 0 | 0 | 0 |
| ENSG00000250835 | 0 | 0 | 1 | 0 | 0 | 0 |

| | | | | | | |
|---|---|---|---|---|---|---|
| ENSG00000251248 | 0 | 0 | 0 | 0 | 0 | 0 |
| ENSG00000249806 | 0 | 0 | 0 | 1 | 0 | 0 |
| ENSG00000250326 | 6 | 3 | 4 | 6 | 5 | 1 |
| ENSG00000251300 | 0 | 0 | 0 | 0 | 0 | 0 |
| ENSG00000170185 | 1100 | 569 | 585 | 354 | 609 | 549 |
| ENSG00000250969 | 0 | 0 | 0 | 0 | 0 | 0 |
| ENSG00000200049 | 0 | 0 | 0 | 0 | 0 | 0 |
| ENSG00000109458 | 522 | 280 | 290 | 98 | 293 | 215 |
| ENSG00000265623 | 0 | 0 | 0 | 0 | 0 | 0 |
| ENSG00000228981 | 210 | 105 | 129 | 188 | 119 | 127 |
| ENSG00000243175 | 0 | 0 | 0 | 0 | 3 | 0 |
| ENSG00000250540 | 0 | 0 | 0 | 0 | 0 | 0 |
| ENSG00000153147 | 2567 | 1494 | 1420 | 906 | 1880 | 1302 |
| ENSG00000245112 | 2 | 0 | 1 | 0 | 0 | 0 |
| ENSG00000248924 | 0 | 0 | 0 | 0 | 0 | 0 |
| ENSG00000251600 | 2 | 1 | 5 | 2 | 3 | 0 |
| ENSG00000236296 | 5 | 6 | 3 | 3 | 7 | 5 |
| ENSG00000183090 | 0 | 0 | 0 | 0 | 0 | 0 |
| ENSG00000250345 | 0 | 0 | 0 | 0 | 0 | 0 |
| ENSG00000197465 | 0 | 0 | 0 | 0 | 0 | 0 |
| ENSG00000249741 | 0 | 0 | 0 | 0 | 0 | 0 |
| ENSG00000250361 | 0 | 0 | 0 | 0 | 0 | 0 |
| ENSG00000170180 | 0 | 0 | 1 | 0 | 0 | 0 |
| ENSG00000285783 | 1 | 0 | 1 | 0 | 0 | 0 |
| ENSG00000285713 | 0 | 0 | 0 | 0 | 0 | 0 |
| ENSG00000261129 | 0 | 0 | 0 | 0 | 0 | 0 |
| ENSG00000250504 | 0 | 0 | 0 | 1 | 2 | 0 |
| ENSG00000248890 | 0 | 3 | 0 | 0 | 1 | 0 |
| ENSG00000164161 | 0 | 0 | 0 | 0 | 0 | 0 |
| ENSG00000273621 | 0 | 0 | 0 | 0 | 0 | 0 |
| ENSG00000164162 | 137 | 93 | 107 | 80 | 95 | 102 |
| ENSG00000249193 | 0 | 0 | 0 | 1 | 0 | 2 |
| ENSG00000250406 | 0 | 0 | 0 | 0 | 0 | 0 |
| ENSG00000251452 | 0 | 0 | 0 | 0 | 0 | 0 |
| ENSG00000222594 | 1 | 0 | 0 | 0 | 0 | 0 |
| ENSG00000164163 | 1192 | 820 | 784 | 656 | 1245 | 973 |
| ENSG00000164164 | 1264 | 648 | 584 | 316 | 799 | 677 |
| ENSG00000238713 | 0 | 0 | 0 | 0 | 0 | 0 |
| ENSG00000240992 | 0 | 0 | 0 | 0 | 0 | 0 |
| ENSG00000272727 | 0 | 0 | 0 | 0 | 0 | 0 |
| ENSG00000251333 | 2 | 0 | 1 | 0 | 1 | 1 |
| ENSG00000278943 | 4 | 0 | 0 | 1 | 0 | 1 |
| ENSG00000248745 | 0 | 0 | 0 | 0 | 0 | 0 |
| ENSG00000170365 | 248 | 138 | 126 | 120 | 159 | 148 |
| ENSG00000250582 | 0 | 0 | 0 | 0 | 0 | 0 |
| ENSG00000250902 | 0 | 0 | 0 | 0 | 0 | 0 |
| ENSG00000151611 | 122 | 69 | 68 | 68 | 91 | 73 |
| ENSG00000249245 | 0 | 1 | 0 | 0 | 0 | 0 |
| ENSG00000248356 | 3 | 0 | 2 | 0 | 0 | 0 |
| ENSG00000237136 | 0 | 0 | 0 | 0 | 0 | 0 |

| | | | | | | |
|---|---|---|---|---|---|---|
| ENSG00000151612 | 26 | 22 | 15 | 17 | 14 | 9 |
| ENSG00000279481 | 0 | 0 | 0 | 0 | 0 | 0 |
| ENSG00000251687 | 0 | 0 | 0 | 0 | 0 | 0 |
| ENSG00000201314 | 0 | 0 | 0 | 0 | 0 | 0 |
| ENSG00000260303 | 0 | 0 | 0 | 0 | 0 | 0 |
| ENSG00000251024 | 0 | 0 | 0 | 0 | 0 | 0 |
| ENSG00000248809 | 0 | 0 | 0 | 0 | 0 | 0 |
| ENSG00000164167 | 277 | 143 | 194 | 180 | 167 | 157 |
| ENSG00000279845 | 4 | 2 | 3 | 1 | 3 | 0 |
| ENSG00000250673 | 0 | 0 | 0 | 0 | 0 | 1 |
| ENSG00000280250 | 0 | 0 | 0 | 0 | 0 | 0 |
| ENSG00000251010 | 0 | 0 | 0 | 0 | 0 | 0 |
| ENSG00000120519 | 153 | 96 | 94 | 39 | 105 | 101 |
| ENSG00000278438 | 0 | 0 | 0 | 0 | 0 | 0 |
| ENSG00000207182 | 0 | 0 | 0 | 0 | 0 | 0 |
| ENSG00000151615 | 0 | 0 | 0 | 0 | 0 | 0 |
| ENSG00000137473 | 0 | 0 | 0 | 0 | 0 | 0 |
| ENSG00000248764 | 0 | 0 | 0 | 0 | 0 | 0 |
| ENSG00000248346 | 0 | 0 | 0 | 0 | 0 | 0 |
| ENSG00000260090 | 0 | 0 | 0 | 0 | 0 | 0 |
| ENSG00000279376 | 0 | 0 | 0 | 0 | 0 | 0 |
| ENSG00000221369 | 0 | 0 | 0 | 0 | 0 | 0 |
| ENSG00000241281 | 0 | 0 | 0 | 0 | 0 | 0 |
| ENSG00000249004 | 0 | 0 | 0 | 0 | 0 | 0 |
| ENSG00000151617 | 24 | 8 | 22 | 9 | 23 | 14 |
| ENSG00000280219 | 6 | 6 | 11 | 1 | 8 | 8 |
| ENSG00000251254 | 0 | 0 | 0 | 0 | 0 | 0 |
| ENSG00000251688 | 2 | 1 | 1 | 0 | 0 | 0 |
| ENSG00000251298 | 1 | 1 | 1 | 2 | 1 | 3 |
| ENSG00000164168 | 1536 | 781 | 842 | 579 | 919 | 816 |
| ENSG00000164169 | 405 | 212 | 208 | 161 | 248 | 206 |
| ENSG00000071205 | 797 | 351 | 451 | 368 | 429 | 425 |
| ENSG00000252951 | 0 | 0 | 0 | 0 | 1 | 0 |
| ENSG00000264274 | 0 | 0 | 0 | 0 | 0 | 0 |
| ENSG00000240014 | 0 | 0 | 0 | 0 | 0 | 0 |
| ENSG00000270387 | 0 | 0 | 1 | 0 | 0 | 1 |
| ENSG00000151623 | 17 | 1 | 5 | 1 | 2 | 3 |
| ENSG00000250354 | 0 | 0 | 0 | 0 | 0 | 0 |
| ENSG00000251418 | 0 | 0 | 0 | 0 | 0 | 0 |
| ENSG00000201846 | 0 | 0 | 0 | 0 | 0 | 0 |
| ENSG00000250357 | 0 | 0 | 0 | 0 | 0 | 0 |
| ENSG00000225176 | 0 | 0 | 0 | 0 | 0 | 0 |
| ENSG00000279016 | 0 | 0 | 0 | 0 | 0 | 0 |
| ENSG00000249317 | 0 | 0 | 0 | 0 | 0 | 0 |
| ENSG00000239168 | 0 | 0 | 0 | 0 | 0 | 0 |
| ENSG00000249752 | 0 | 0 | 0 | 0 | 0 | 0 |
| ENSG00000250252 | 0 | 0 | 0 | 0 | 0 | 0 |
| ENSG00000248210 | 0 | 0 | 0 | 0 | 0 | 0 |
| ENSG00000234828 | 0 | 0 | 0 | 0 | 0 | 0 |
| ENSG00000250905 | 0 | 0 | 0 | 0 | 0 | 0 |

| | | | | | | |
|---|---|---|---|---|---|---|
| ENSG00000251482 | 0 | 0 | 0 | 0 | 0 | 0 |
| ENSG00000202331 | 0 | 0 | 0 | 0 | 0 | 0 |
| ENSG00000207121 | 0 | 0 | 0 | 0 | 0 | 0 |
| ENSG00000250704 | 0 | 0 | 0 | 0 | 0 | 0 |
| ENSG00000170390 | 2 | 3 | 3 | 1 | 2 | 7 |
| ENSG00000238721 | 0 | 0 | 0 | 0 | 0 | 0 |
| ENSG00000198589 | 1482 | 898 | 763 | 264 | 1293 | 664 |
| ENSG00000248521 | 0 | 0 | 0 | 0 | 0 | 0 |
| ENSG00000249690 | 0 | 0 | 0 | 1 | 0 | 1 |
| ENSG00000181541 | 0 | 0 | 0 | 0 | 1 | 0 |
| ENSG00000183432 | 0 | 2 | 0 | 0 | 0 | 0 |
| ENSG00000252451 | 0 | 0 | 0 | 0 | 0 | 0 |
| ENSG00000235606 | 0 | 0 | 0 | 0 | 1 | 0 |
| ENSG00000145425 | 14198 | 8552 | 10019 | 11884 | 10381 | 9840 |
| ENSG00000201264 | 2 | 0 | 0 | 0 | 2 | 0 |
| ENSG00000109686 | 743 | 351 | 309 | 235 | 403 | 364 |
| ENSG00000208797 | 0 | 0 | 0 | 0 | 0 | 0 |
| ENSG00000270681 | 5 | 9 | 7 | 13 | 11 | 6 |
| ENSG00000249012 | 0 | 0 | 0 | 1 | 0 | 0 |
| ENSG00000189099 | 0 | 0 | 0 | 0 | 0 | 0 |
| ENSG00000250350 | 0 | 0 | 0 | 0 | 0 | 0 |
| ENSG00000252544 | 0 | 0 | 0 | 0 | 0 | 0 |
| ENSG00000270265 | 8 | 5 | 5 | 0 | 1 | 3 |
| ENSG00000251611 | 14 | 4 | 11 | 7 | 5 | 8 |
| ENSG00000164142 | 189 | 60 | 94 | 47 | 54 | 74 |
| ENSG00000253019 | 0 | 0 | 0 | 0 | 0 | 0 |
| ENSG00000251603 | 9 | 8 | 3 | 5 | 7 | 7 |
| ENSG00000278978 | 52 | 31 | 34 | 10 | 13 | 25 |
| ENSG00000059691 | 620 | 250 | 312 | 324 | 286 | 309 |
| ENSG00000251455 | 3 | 0 | 3 | 1 | 2 | 3 |
| ENSG00000249184 | 0 | 0 | 0 | 0 | 0 | 0 |
| ENSG00000249708 | 0 | 0 | 0 | 0 | 0 | 0 |
| ENSG00000249818 | 0 | 0 | 0 | 0 | 0 | 0 |
| ENSG00000248991 | 0 | 0 | 0 | 0 | 0 | 0 |
| ENSG00000251249 | 0 | 0 | 0 | 0 | 0 | 0 |
| ENSG00000253077 | 0 | 0 | 0 | 0 | 0 | 0 |
| ENSG00000245954 | 2 | 0 | 0 | 0 | 0 | 0 |
| ENSG00000250706 | 0 | 0 | 0 | 0 | 0 | 0 |
| ENSG00000251377 | 0 | 0 | 0 | 0 | 0 | 0 |
| ENSG00000244544 | 0 | 0 | 0 | 0 | 0 | 0 |
| ENSG00000270302 | 0 | 0 | 0 | 0 | 0 | 0 |
| ENSG00000270883 | 0 | 0 | 0 | 0 | 1 | 0 |
| ENSG00000109670 | 624 | 275 | 320 | 197 | 321 | 314 |
| ENSG00000270751 | 0 | 0 | 0 | 0 | 0 | 0 |
| ENSG00000264678 | 0 | 0 | 0 | 0 | 0 | 0 |
| ENSG00000268471 | 163 | 66 | 84 | 57 | 91 | 74 |
| ENSG00000283792 | 0 | 0 | 0 | 0 | 0 | 0 |
| ENSG00000243417 | 0 | 0 | 0 | 1 | 3 | 1 |
| ENSG00000240669 | 0 | 0 | 0 | 0 | 0 | 0 |
| ENSG00000170006 | 306 | 155 | 195 | 145 | 268 | 191 |

| | | | | | | |
|---|---|---|---|---|---|---|
| ENSG00000248571 | 0 | 0 | 0 | 0 | 0 | 0 |
| ENSG00000169989 | 12 | 5 | 8 | 5 | 8 | 9 |
| ENSG00000164144 | 1401 | 747 | 800 | 534 | 922 | 746 |
| ENSG00000250980 | 0 | 2 | 0 | 0 | 0 | 0 |
| ENSG00000231841 | 2 | 2 | 1 | 1 | 2 | 3 |
| ENSG00000137460 | 523 | 272 | 321 | 255 | 364 | 359 |
| ENSG00000249022 | 0 | 0 | 0 | 0 | 1 | 0 |
| ENSG00000109654 | 101 | 52 | 80 | 24 | 56 | 41 |
| ENSG00000252030 | 0 | 0 | 0 | 0 | 0 | 0 |
| ENSG00000201786 | 0 | 0 | 0 | 0 | 0 | 0 |
| ENSG00000213406 | 0 | 0 | 0 | 0 | 0 | 0 |
| ENSG00000121211 | 78 | 36 | 31 | 35 | 64 | 47 |
| ENSG00000243883 | 0 | 0 | 0 | 0 | 0 | 0 |
| ENSG00000121210 | 1129 | 544 | 657 | 364 | 565 | 513 |
| ENSG00000250771 | 1 | 1 | 0 | 0 | 0 | 0 |
| ENSG00000248208 | 0 | 0 | 0 | 0 | 0 | 0 |
| ENSG00000137462 | 90 | 40 | 50 | 57 | 94 | 63 |
| ENSG00000145428 | 0 | 0 | 0 | 0 | 0 | 0 |
| ENSG00000249309 | 0 | 0 | 0 | 0 | 0 | 0 |
| ENSG00000145423 | 0 | 0 | 0 | 0 | 0 | 0 |
| ENSG00000280241 | 0 | 0 | 0 | 0 | 0 | 0 |
| ENSG00000197410 | 0 | 0 | 0 | 0 | 0 | 0 |
| ENSG00000278981 | 0 | 0 | 0 | 0 | 0 | 0 |
| ENSG00000280005 | 0 | 0 | 0 | 0 | 0 | 0 |
| ENSG00000249200 | 0 | 0 | 0 | 0 | 0 | 0 |
| ENSG00000250609 | 0 | 0 | 0 | 0 | 0 | 0 |
| ENSG00000171566 | 1247 | 565 | 661 | 620 | 833 | 727 |
| ENSG00000200350 | 0 | 0 | 0 | 0 | 2 | 0 |
| ENSG00000248611 | 0 | 0 | 0 | 0 | 0 | 0 |
| ENSG00000171564 | 4 | 0 | 0 | 0 | 0 | 1 |
| ENSG00000171560 | 0 | 1 | 0 | 0 | 0 | 0 |
| ENSG00000171557 | 0 | 0 | 0 | 0 | 0 | 0 |
| ENSG00000121207 | 2 | 1 | 0 | 2 | 1 | 1 |
| ENSG00000249074 | 0 | 0 | 0 | 0 | 0 | 0 |
| ENSG00000249041 | 0 | 0 | 0 | 0 | 0 | 0 |
| ENSG00000249692 | 0 | 0 | 0 | 0 | 0 | 0 |
| ENSG00000151962 | 0 | 0 | 0 | 0 | 0 | 0 |
| ENSG00000222582 | 0 | 0 | 0 | 0 | 0 | 0 |
| ENSG00000252604 | 0 | 0 | 0 | 0 | 0 | 0 |
| ENSG00000250910 | 0 | 0 | 0 | 0 | 0 | 0 |
| ENSG00000250538 | 0 | 0 | 0 | 0 | 0 | 0 |
| ENSG00000185149 | 0 | 0 | 0 | 0 | 0 | 0 |
| ENSG00000249924 | 0 | 0 | 0 | 0 | 0 | 0 |
| ENSG00000164114 | 133 | 76 | 70 | 21 | 137 | 52 |
| ENSG00000250205 | 0 | 0 | 0 | 0 | 0 | 0 |
| ENSG00000250398 | 0 | 0 | 0 | 0 | 0 | 0 |
| ENSG00000271606 | 0 | 0 | 0 | 0 | 0 | 0 |
| ENSG00000249247 | 0 | 0 | 0 | 0 | 0 | 0 |
| ENSG00000249770 | 0 | 0 | 0 | 0 | 0 | 0 |
| ENSG00000248748 | 0 | 0 | 0 | 0 | 0 | 0 |

| | | | | | | |
|---|---|---|---|---|---|---|
| ENSG00000248548 | 0 | 0 | 0 | 0 | 0 | 0 |
| ENSG00000249127 | 0 | 0 | 0 | 0 | 0 | 0 |
| ENSG00000249698 | 0 | 0 | 0 | 0 | 0 | 0 |
| ENSG00000248420 | 0 | 0 | 0 | 0 | 0 | 0 |
| ENSG00000249386 | 0 | 0 | 0 | 0 | 0 | 0 |
| ENSG00000251223 | 0 | 0 | 0 | 0 | 0 | 0 |
| ENSG00000248907 | 0 | 0 | 0 | 0 | 0 | 0 |
| ENSG00000251407 | 0 | 0 | 0 | 0 | 0 | 0 |
| ENSG00000251244 | 0 | 0 | 0 | 0 | 0 | 0 |
| ENSG00000164116 | 2 | 0 | 0 | 0 | 0 | 0 |
| ENSG00000260244 | 0 | 0 | 0 | 0 | 0 | 0 |
| ENSG00000061918 | 0 | 1 | 1 | 0 | 0 | 1 |
| ENSG00000256394 | 0 | 0 | 0 | 0 | 0 | 0 |
| ENSG00000151790 | 0 | 0 | 1 | 0 | 0 | 0 |
| ENSG00000256043 | 18 | 12 | 10 | 8 | 10 | 7 |
| ENSG00000249013 | 0 | 0 | 0 | 0 | 0 | 0 |
| ENSG00000251511 | 0 | 0 | 0 | 0 | 0 | 0 |
| ENSG00000251283 | 0 | 0 | 0 | 0 | 0 | 0 |
| ENSG00000249479 | 0 | 0 | 0 | 0 | 0 | 0 |
| ENSG00000248912 | 0 | 0 | 0 | 0 | 0 | 0 |
| ENSG00000145431 | 4 | 3 | 1 | 1 | 0 | 1 |
| ENSG00000248629 | 0 | 0 | 0 | 0 | 0 | 0 |
| ENSG00000280284 | 0 | 0 | 0 | 0 | 0 | 0 |
| ENSG00000278880 | 0 | 0 | 0 | 0 | 0 | 0 |
| ENSG00000250622 | 0 | 0 | 0 | 0 | 0 | 0 |
| ENSG00000252350 | 0 | 0 | 0 | 0 | 0 | 0 |
| ENSG00000252524 | 0 | 0 | 0 | 0 | 0 | 0 |
| ENSG00000109738 | 4 | 4 | 1 | 0 | 0 | 0 |
| ENSG00000199859 | 0 | 0 | 0 | 0 | 0 | 0 |
| ENSG00000120251 | 0 | 0 | 0 | 0 | 0 | 0 |
| ENSG00000249819 | 0 | 0 | 0 | 0 | 0 | 0 |
| ENSG00000234111 | 0 | 0 | 0 | 0 | 0 | 0 |
| ENSG00000249275 | 0 | 0 | 0 | 0 | 0 | 0 |
| ENSG00000200521 | 0 | 0 | 0 | 0 | 0 | 0 |
| ENSG00000249627 | 0 | 0 | 0 | 0 | 0 | 0 |
| ENSG00000279703 | 0 | 0 | 0 | 0 | 0 | 0 |
| ENSG00000279994 | 0 | 0 | 0 | 0 | 0 | 0 |
| ENSG00000250475 | 0 | 0 | 0 | 0 | 0 | 0 |
| ENSG00000279139 | 0 | 0 | 0 | 0 | 0 | 0 |
| ENSG00000280140 | 0 | 0 | 0 | 1 | 0 | 0 |
| ENSG00000164125 | 183 | 128 | 95 | 140 | 261 | 153 |
| ENSG00000248429 | 72 | 50 | 32 | 52 | 47 | 52 |
| ENSG00000251073 | 0 | 0 | 0 | 0 | 0 | 0 |
| ENSG00000250604 | 0 | 0 | 0 | 0 | 0 | 0 |
| ENSG00000164124 | 252 | 126 | 157 | 104 | 200 | 142 |
| ENSG00000251429 | 0 | 0 | 0 | 0 | 0 | 0 |
| ENSG00000171509 | 0 | 0 | 0 | 0 | 0 | 0 |
| ENSG00000199634 | 0 | 0 | 0 | 0 | 0 | 0 |
| ENSG00000248387 | 0 | 0 | 0 | 0 | 0 | 0 |
| ENSG00000249901 | 0 | 0 | 0 | 0 | 0 | 0 |

| | | | | | | |
|---|---|---|---|---|---|---|
| ENSG00000205208 | 331 | 184 | 279 | 215 | 204 | 234 |
| ENSG00000171503 | 644 | 332 | 421 | 259 | 376 | 313 |
| ENSG00000271817 | 1 | 0 | 0 | 0 | 1 | 1 |
| ENSG00000171497 | 747 | 340 | 493 | 403 | 508 | 453 |
| ENSG00000052795 | 234 | 100 | 124 | 60 | 100 | 109 |
| ENSG00000206703 | 0 | 0 | 0 | 0 | 0 | 0 |
| ENSG00000164123 | 0 | 0 | 1 | 1 | 0 | 1 |
| ENSG00000274535 | 0 | 0 | 0 | 0 | 0 | 0 |
| ENSG00000248988 | 4 | 3 | 2 | 1 | 2 | 1 |
| ENSG00000249367 | 0 | 0 | 0 | 0 | 0 | 0 |
| ENSG00000206978 | 0 | 0 | 0 | 0 | 0 | 0 |
| ENSG00000109756 | 947 | 448 | 492 | 268 | 564 | 471 |
| ENSG00000264105 | 0 | 0 | 0 | 0 | 0 | 0 |
| ENSG00000283379 | 0 | 0 | 0 | 0 | 0 | 0 |
| ENSG00000248552 | 0 | 0 | 0 | 0 | 0 | 0 |
| ENSG00000250180 | 0 | 0 | 0 | 0 | 0 | 0 |
| ENSG00000250488 | 0 | 0 | 0 | 0 | 0 | 0 |
| ENSG00000250997 | 0 | 0 | 0 | 0 | 0 | 0 |
| ENSG00000249425 | 0 | 1 | 0 | 0 | 0 | 0 |
| ENSG00000242635 | 0 | 0 | 0 | 0 | 0 | 0 |
| ENSG00000249568 | 11 | 5 | 17 | 0 | 4 | 3 |
| ENSG00000168843 | 11 | 9 | 13 | 1 | 1 | 1 |
| ENSG00000249419 | 0 | 0 | 0 | 0 | 0 | 0 |
| ENSG00000251253 | 0 | 0 | 0 | 0 | 0 | 0 |
| ENSG00000248396 | 0 | 0 | 0 | 0 | 0 | 0 |
| ENSG00000248431 | 0 | 0 | 0 | 0 | 0 | 0 |
| ENSG00000251123 | 0 | 0 | 0 | 0 | 0 | 0 |
| ENSG00000283468 | 0 | 0 | 0 | 0 | 0 | 0 |
| ENSG00000250027 | 2 | 2 | 2 | 0 | 0 | 0 |
| ENSG00000145414 | 331 | 162 | 198 | 188 | 229 | 197 |
| ENSG00000250547 | 0 | 0 | 0 | 0 | 0 | 0 |
| ENSG00000164128 | 278 | 128 | 140 | 115 | 144 | 124 |
| ENSG00000164129 | 0 | 2 | 1 | 2 | 6 | 4 |
| ENSG00000227283 | 0 | 0 | 0 | 0 | 0 | 0 |
| ENSG00000151005 | 2 | 2 | 0 | 0 | 0 | 0 |
| ENSG00000198498 | 604 | 371 | 396 | 388 | 458 | 369 |
| ENSG00000145416 | 86 | 60 | 42 | 21 | 82 | 33 |
| ENSG00000273449 | 19 | 5 | 3 | 4 | 7 | 6 |
| ENSG00000253001 | 0 | 0 | 0 | 0 | 0 | 0 |
| ENSG00000177803 | 0 | 0 | 0 | 0 | 0 | 0 |
| ENSG00000250746 | 0 | 1 | 1 | 0 | 0 | 0 |
| ENSG00000248546 | 0 | 1 | 0 | 0 | 0 | 2 |
| ENSG00000251162 | 0 | 0 | 0 | 0 | 0 | 0 |
| ENSG00000223037 | 0 | 0 | 0 | 0 | 0 | 0 |
| ENSG00000248162 | 0 | 0 | 0 | 0 | 0 | 0 |
| ENSG00000248771 | 0 | 0 | 1 | 1 | 0 | 0 |
| ENSG00000201050 | 0 | 0 | 0 | 0 | 0 | 0 |
| ENSG00000248329 | 2 | 1 | 0 | 0 | 0 | 2 |
| ENSG00000250227 | 0 | 0 | 0 | 0 | 0 | 0 |
| ENSG00000244146 | 0 | 0 | 0 | 0 | 0 | 0 |

| | | | | | | |
|---|---|---|---|---|---|---|
| ENSG00000285563 | 0 | 0 | 0 | 0 | 0 | 0 |
| ENSG00000221891 | 0 | 0 | 0 | 0 | 0 | 0 |
| ENSG00000270960 | 0 | 0 | 0 | 0 | 0 | 0 |
| ENSG00000248645 | 0 | 0 | 0 | 0 | 0 | 0 |
| ENSG00000121089 | 16 | 18 | 14 | 7 | 17 | 12 |
| ENSG00000236941 | 0 | 0 | 0 | 0 | 0 | 0 |
| ENSG00000183439 | 0 | 0 | 0 | 0 | 0 | 0 |
| ENSG00000250486 | 46 | 16 | 43 | 24 | 39 | 23 |
| ENSG00000248632 | 2 | 1 | 0 | 0 | 0 | 0 |
| ENSG00000248229 | 0 | 0 | 0 | 0 | 0 | 0 |
| ENSG00000250887 | 0 | 0 | 0 | 0 | 0 | 0 |
| ENSG00000250712 | 0 | 0 | 1 | 0 | 0 | 0 |
| ENSG00000240674 | 0 | 0 | 0 | 1 | 0 | 0 |
| ENSG00000176979 | 0 | 0 | 0 | 0 | 0 | 0 |
| ENSG00000276623 | 0 | 0 | 0 | 0 | 0 | 0 |
| ENSG00000250430 | 0 | 0 | 0 | 0 | 0 | 0 |
| ENSG00000249617 | 0 | 0 | 0 | 0 | 0 | 1 |
| ENSG00000250374 | 2 | 0 | 0 | 0 | 0 | 1 |
| ENSG00000170088 | 1544 | 812 | 938 | 624 | 918 | 896 |
| ENSG00000109466 | 383 | 196 | 212 | 122 | 209 | 178 |
| ENSG00000200974 | 0 | 0 | 0 | 0 | 0 | 0 |
| ENSG00000229894 | 4 | 2 | 1 | 0 | 0 | 0 |
| ENSG00000052802 | 2164 | 1019 | 1033 | 985 | 1339 | 1170 |
| ENSG00000109472 | 322 | 151 | 190 | 160 | 199 | 149 |
| ENSG00000207559 | 0 | 0 | 0 | 0 | 0 | 0 |
| ENSG00000251596 | 2 | 0 | 4 | 0 | 0 | 0 |
| ENSG00000248287 | 0 | 0 | 0 | 0 | 0 | 0 |
| ENSG00000248327 | 0 | 0 | 0 | 0 | 0 | 0 |
| ENSG00000250725 | 0 | 0 | 0 | 0 | 0 | 0 |
| ENSG00000249500 | 0 | 0 | 0 | 0 | 0 | 0 |
| ENSG00000038295 | 0 | 0 | 0 | 0 | 0 | 0 |
| ENSG00000252959 | 0 | 0 | 0 | 0 | 0 | 0 |
| ENSG00000206711 | 0 | 0 | 0 | 0 | 0 | 0 |
| ENSG00000249675 | 0 | 0 | 0 | 0 | 0 | 0 |
| ENSG00000196104 | 0 | 0 | 0 | 0 | 0 | 0 |
| ENSG00000212373 | 0 | 0 | 0 | 0 | 0 | 0 |
| ENSG00000250150 | 0 | 0 | 0 | 0 | 0 | 0 |
| ENSG00000240772 | 0 | 0 | 0 | 0 | 0 | 0 |
| ENSG00000248913 | 0 | 0 | 0 | 0 | 0 | 0 |
| ENSG00000222721 | 0 | 0 | 0 | 0 | 0 | 0 |
| ENSG00000270336 | 0 | 0 | 0 | 0 | 0 | 0 |
| ENSG00000248601 | 0 | 0 | 0 | 0 | 0 | 0 |
| ENSG00000109511 | 0 | 0 | 0 | 0 | 0 | 0 |
| ENSG00000137628 | 6 | 2 | 2 | 0 | 3 | 0 |
| ENSG00000181381 | 20 | 16 | 4 | 11 | 24 | 16 |
| ENSG00000207171 | 0 | 0 | 0 | 0 | 0 | 0 |
| ENSG00000129116 | 619 | 348 | 332 | 281 | 372 | 261 |
| ENSG00000229226 | 0 | 0 | 0 | 0 | 0 | 0 |
| ENSG00000249623 | 0 | 0 | 0 | 0 | 0 | 0 |
| ENSG00000206613 | 0 | 0 | 0 | 0 | 0 | 0 |

| | | | | | | |
|---|---|---|---|---|---|---|
| ENSG00000214676 | 0 | 0 | 0 | 0 | 0 | 0 |
| ENSG00000249609 | 0 | 0 | 0 | 0 | 0 | 0 |
| ENSG00000279384 | 0 | 0 | 1 | 2 | 1 | 1 |
| ENSG00000145439 | 547 | 262 | 373 | 252 | 281 | 350 |
| ENSG00000201176 | 0 | 0 | 0 | 0 | 0 | 0 |
| ENSG00000201818 | 0 | 0 | 0 | 0 | 0 | 0 |
| ENSG00000251445 | 0 | 0 | 0 | 0 | 0 | 0 |
| ENSG00000154447 | 516 | 220 | 279 | 189 | 224 | 256 |
| ENSG00000214671 | 0 | 0 | 2 | 0 | 0 | 1 |
| ENSG00000251171 | 0 | 0 | 0 | 0 | 0 | 0 |
| ENSG00000137601 | 110 | 85 | 51 | 15 | 133 | 71 |
| ENSG00000109572 | 1777 | 817 | 983 | 521 | 983 | 855 |
| ENSG00000056050 | 258 | 158 | 153 | 133 | 183 | 165 |
| ENSG00000229204 | 3 | 0 | 5 | 3 | 5 | 1 |
| ENSG00000248625 | 0 | 0 | 0 | 0 | 0 | 0 |
| ENSG00000248319 | 0 | 0 | 0 | 0 | 0 | 0 |
| ENSG00000251200 | 0 | 0 | 0 | 0 | 0 | 0 |
| ENSG00000249955 | 0 | 0 | 0 | 0 | 0 | 0 |
| ENSG00000198948 | 121 | 45 | 55 | 42 | 83 | 63 |
| ENSG00000109576 | 141 | 74 | 128 | 74 | 108 | 93 |
| ENSG00000248822 | 0 | 0 | 0 | 0 | 0 | 0 |
| ENSG00000248739 | 0 | 0 | 0 | 0 | 0 | 0 |
| ENSG00000250266 | 2 | 0 | 0 | 0 | 1 | 0 |
| ENSG00000251061 | 0 | 0 | 0 | 0 | 0 | 0 |
| ENSG00000250082 | 0 | 0 | 0 | 0 | 0 | 0 |
| ENSG00000181359 | 2 | 0 | 0 | 0 | 0 | 1 |
| ENSG00000250968 | 0 | 0 | 0 | 0 | 0 | 0 |
| ENSG00000249943 | 0 | 0 | 0 | 0 | 0 | 0 |
| ENSG00000251961 | 0 | 0 | 0 | 0 | 0 | 0 |
| ENSG00000248872 | 0 | 0 | 0 | 0 | 0 | 0 |
| ENSG00000249525 | 0 | 0 | 0 | 0 | 0 | 0 |
| ENSG00000248755 | 0 | 0 | 0 | 0 | 0 | 0 |
| ENSG00000278502 | 0 | 0 | 0 | 0 | 0 | 0 |
| ENSG00000248339 | 0 | 0 | 0 | 0 | 0 | 0 |
| ENSG00000251176 | 0 | 0 | 0 | 0 | 0 | 0 |
| ENSG00000249297 | 0 | 0 | 0 | 0 | 0 | 0 |
| ENSG00000174473 | 148 | 56 | 96 | 47 | 35 | 42 |
| ENSG00000248637 | 0 | 0 | 0 | 0 | 0 | 0 |
| ENSG00000245213 | 46 | 18 | 41 | 22 | 39 | 39 |
| ENSG00000241652 | 0 | 0 | 0 | 0 | 0 | 0 |
| ENSG00000109586 | 1388 | 731 | 827 | 407 | 868 | 751 |
| ENSG00000221296 | 0 | 0 | 0 | 0 | 0 | 0 |
| ENSG00000248774 | 5 | 0 | 1 | 1 | 0 | 2 |
| ENSG00000164104 | 3146 | 1601 | 2072 | 1741 | 2108 | 2075 |
| ENSG00000272870 | 27 | 6 | 13 | 6 | 4 | 8 |
| ENSG00000164105 | 386 | 198 | 233 | 223 | 220 | 229 |
| ENSG00000164106 | 4 | 2 | 0 | 1 | 2 | 0 |
| ENSG00000242882 | 0 | 0 | 0 | 0 | 0 | 0 |
| ENSG00000252898 | 0 | 0 | 0 | 0 | 0 | 0 |
| ENSG00000261646 | 0 | 1 | 1 | 0 | 0 | 0 |

| | | | | | | |
|---|---|---|---|---|---|---|
| ENSG00000164107 | 0 | 0 | 0 | 0 | 0 | 0 |
| ENSG00000237125 | 45 | 25 | 34 | 26 | 42 | 27 |
| ENSG00000234801 | 0 | 0 | 0 | 0 | 1 | 2 |
| ENSG00000213370 | 0 | 2 | 0 | 0 | 1 | 0 |
| ENSG00000250043 | 5 | 7 | 4 | 4 | 4 | 7 |
| ENSG00000250708 | 11 | 5 | 13 | 8 | 7 | 8 |
| ENSG00000251216 | 0 | 0 | 0 | 0 | 0 | 0 |
| ENSG00000248174 | 2 | 0 | 1 | 0 | 0 | 0 |
| ENSG00000250957 | 0 | 0 | 0 | 0 | 0 | 0 |
| ENSG00000251213 | 0 | 0 | 0 | 0 | 0 | 0 |
| ENSG00000164117 | 667 | 334 | 375 | 269 | 319 | 341 |
| ENSG00000164118 | 299 | 150 | 146 | 75 | 175 | 118 |
| ENSG00000249875 | 0 | 0 | 0 | 0 | 0 | 0 |
| ENSG00000265846 | 0 | 0 | 0 | 0 | 0 | 0 |
| ENSG00000164120 | 75 | 29 | 34 | 27 | 45 | 56 |
| ENSG00000250596 | 0 | 0 | 0 | 0 | 0 | 0 |
| ENSG00000251584 | 0 | 0 | 0 | 0 | 0 | 0 |
| ENSG00000145451 | 86 | 46 | 56 | 21 | 32 | 37 |
| ENSG00000168594 | 5 | 2 | 4 | 3 | 4 | 1 |
| ENSG00000250200 | 0 | 0 | 0 | 0 | 0 | 0 |
| ENSG00000250431 | 0 | 0 | 0 | 0 | 0 | 1 |
| ENSG00000249411 | 0 | 0 | 0 | 0 | 0 | 0 |
| ENSG00000249284 | 0 | 0 | 0 | 0 | 0 | 0 |
| ENSG00000248551 | 0 | 1 | 0 | 0 | 0 | 0 |
| ENSG00000249945 | 0 | 0 | 0 | 0 | 0 | 0 |
| ENSG00000251174 | 0 | 0 | 0 | 0 | 0 | 0 |
| ENSG00000248574 | 0 | 0 | 0 | 0 | 0 | 0 |
| ENSG00000150625 | 16 | 6 | 5 | 3 | 3 | 4 |
| ENSG00000249106 | 0 | 0 | 0 | 0 | 0 | 0 |
| ENSG00000248249 | 0 | 0 | 0 | 0 | 0 | 0 |
| ENSG00000250372 | 0 | 0 | 0 | 0 | 0 | 0 |
| ENSG00000150627 | 98 | 58 | 45 | 15 | 50 | 49 |
| ENSG00000201516 | 0 | 0 | 0 | 0 | 0 | 0 |
| ENSG00000150628 | 1 | 2 | 1 | 0 | 1 | 2 |
| ENSG00000164122 | 0 | 0 | 0 | 0 | 0 | 1 |
| ENSG00000248980 | 0 | 0 | 0 | 0 | 0 | 1 |
| ENSG00000129128 | 1850 | 912 | 1061 | 800 | 1145 | 1029 |
| ENSG00000248480 | 0 | 0 | 0 | 0 | 0 | 0 |
| ENSG00000248388 | 0 | 0 | 0 | 0 | 0 | 0 |
| ENSG00000150630 | 75 | 33 | 34 | 56 | 42 | 38 |
| ENSG00000249458 | 0 | 0 | 0 | 0 | 0 | 0 |
| ENSG00000250263 | 0 | 0 | 0 | 0 | 0 | 0 |
| ENSG00000222859 | 0 | 0 | 0 | 0 | 0 | 0 |
| ENSG00000249008 | 0 | 0 | 0 | 0 | 0 | 0 |
| ENSG00000109674 | 98 | 41 | 27 | 46 | 58 | 52 |
| ENSG00000249084 | 0 | 0 | 1 | 0 | 0 | 0 |
| ENSG00000038002 | 394 | 194 | 230 | 205 | 240 | 187 |
| ENSG00000250131 | 4 | 1 | 1 | 0 | 2 | 2 |
| ENSG00000285574 | 0 | 0 | 0 | 0 | 0 | 0 |
| ENSG00000201931 | 0 | 0 | 0 | 0 | 0 | 0 |

| | | | | | | |
|---|---|---|---|---|---|---|
| ENSG00000251416 | 0 | 0 | 0 | 0 | 0 | 0 |
| ENSG00000231171 | 0 | 0 | 0 | 0 | 0 | 0 |
| ENSG00000251504 | 0 | 0 | 0 | 0 | 0 | 0 |
| ENSG00000253000 | 0 | 0 | 0 | 0 | 0 | 0 |
| ENSG00000241877 | 0 | 0 | 0 | 0 | 0 | 0 |
| ENSG00000201727 | 0 | 0 | 0 | 0 | 0 | 0 |
| ENSG00000212191 | 0 | 0 | 0 | 0 | 0 | 0 |
| ENSG00000250839 | 0 | 0 | 0 | 0 | 0 | 0 |
| ENSG00000250993 | 0 | 0 | 0 | 0 | 0 | 0 |
| ENSG00000248921 | 0 | 0 | 0 | 0 | 0 | 0 |
| ENSG00000251025 | 0 | 0 | 0 | 0 | 0 | 0 |
| ENSG00000249883 | 0 | 0 | 0 | 0 | 0 | 0 |
| ENSG00000248197 | 0 | 0 | 0 | 0 | 0 | 0 |
| ENSG00000248770 | 0 | 0 | 0 | 0 | 0 | 0 |
| ENSG00000249460 | 0 | 0 | 0 | 0 | 0 | 0 |
| ENSG00000225356 | 0 | 0 | 0 | 0 | 0 | 0 |
| ENSG00000251336 | 0 | 0 | 0 | 0 | 0 | 0 |
| ENSG00000251742 | 0 | 0 | 0 | 0 | 0 | 0 |
| ENSG00000177822 | 0 | 0 | 0 | 0 | 0 | 0 |
| ENSG00000251433 | 0 | 0 | 0 | 0 | 0 | 0 |
| ENSG00000218336 | 50 | 18 | 19 | 14 | 22 | 23 |
| ENSG00000248266 | 0 | 0 | 0 | 0 | 0 | 0 |
| ENSG00000221227 | 0 | 0 | 0 | 0 | 0 | 0 |
| ENSG00000252343 | 0 | 0 | 0 | 0 | 0 | 0 |
| ENSG00000251714 | 0 | 0 | 0 | 0 | 0 | 0 |
| ENSG00000248816 | 0 | 0 | 0 | 0 | 0 | 0 |
| ENSG00000248694 | 0 | 0 | 0 | 0 | 0 | 0 |
| ENSG00000272646 | 0 | 0 | 0 | 0 | 0 | 0 |
| ENSG00000129187 | 935 | 457 | 546 | 585 | 675 | 648 |
| ENSG00000250636 | 0 | 0 | 0 | 0 | 0 | 0 |
| ENSG00000230219 | 0 | 0 | 0 | 0 | 0 | 0 |
| ENSG00000273180 | 0 | 0 | 0 | 0 | 0 | 0 |
| ENSG00000272784 | 2 | 0 | 0 | 1 | 0 | 0 |
| ENSG00000251359 | 8 | 1 | 1 | 4 | 1 | 1 |
| ENSG00000151718 | 266 | 97 | 126 | 58 | 126 | 89 |
| ENSG00000251128 | 0 | 0 | 0 | 0 | 0 | 0 |
| ENSG00000177300 | 0 | 0 | 0 | 0 | 0 | 0 |
| ENSG00000185758 | 0 | 0 | 0 | 0 | 0 | 0 |
| ENSG00000250292 | 0 | 0 | 0 | 0 | 0 | 0 |
| ENSG00000248376 | 0 | 0 | 0 | 0 | 0 | 0 |
| ENSG00000168564 | 191 | 76 | 118 | 68 | 86 | 91 |
| ENSG00000213434 | 0 | 0 | 1 | 0 | 0 | 0 |
| ENSG00000232648 | 3 | 1 | 3 | 3 | 5 | 1 |
| ENSG00000168556 | 134 | 53 | 80 | 48 | 75 | 73 |
| ENSG00000272744 | 2 | 0 | 2 | 0 | 3 | 1 |
| ENSG00000182552 | 182 | 103 | 99 | 109 | 123 | 99 |
| ENSG00000252157 | 0 | 0 | 0 | 0 | 0 | 0 |
| ENSG00000168538 | 417 | 194 | 219 | 123 | 238 | 194 |
| ENSG00000201433 | 0 | 0 | 0 | 0 | 0 | 0 |
| ENSG00000251739 | 1 | 2 | 2 | 0 | 0 | 0 |

| | | | | | | |
|---|---|---|---|---|---|---|
| ENSG00000283383 | 0 | 0 | 0 | 0 | 0 | 0 |
| ENSG00000173320 | 100 | 42 | 36 | 27 | 53 | 37 |
| ENSG00000248206 | 4 | 1 | 0 | 0 | 0 | 4 |
| ENSG00000283355 | 0 | 0 | 3 | 1 | 0 | 1 |
| ENSG00000250726 | 0 | 0 | 0 | 0 | 0 | 0 |
| ENSG00000164303 | 2 | 0 | 0 | 0 | 0 | 0 |
| ENSG00000240159 | 0 | 0 | 0 | 0 | 0 | 0 |
| ENSG00000244512 | 0 | 0 | 0 | 0 | 0 | 0 |
| ENSG00000227765 | 2 | 5 | 1 | 2 | 0 | 2 |
| ENSG00000180712 | 6 | 3 | 3 | 3 | 7 | 5 |
| ENSG00000249096 | 0 | 0 | 0 | 0 | 0 | 0 |
| ENSG00000168310 | 454 | 205 | 258 | 142 | 226 | 240 |
| ENSG00000251878 | 0 | 0 | 0 | 0 | 0 | 0 |
| ENSG00000270426 | 4 | 1 | 2 | 2 | 0 | 0 |
| ENSG00000271646 | 18 | 27 | 25 | 12 | 23 | 15 |
| ENSG00000271538 | 10 | 8 | 5 | 1 | 3 | 3 |
| ENSG00000249056 | 0 | 0 | 0 | 0 | 0 | 0 |
| ENSG00000254233 | 2 | 0 | 0 | 0 | 0 | 0 |
| ENSG00000164305 | 236 | 135 | 164 | 128 | 147 | 131 |
| ENSG00000164306 | 99 | 40 | 49 | 37 | 40 | 46 |
| ENSG00000151725 | 206 | 105 | 106 | 66 | 118 | 102 |
| ENSG00000151726 | 888 | 420 | 512 | 410 | 601 | 523 |
| ENSG00000249138 | 2 | 0 | 0 | 0 | 0 | 0 |
| ENSG00000251139 | 0 | 0 | 0 | 0 | 0 | 0 |
| ENSG00000251230 | 0 | 0 | 0 | 0 | 0 | 0 |
| ENSG00000266698 | 0 | 0 | 0 | 0 | 0 | 0 |
| ENSG00000249173 | 0 | 0 | 0 | 0 | 0 | 0 |
| ENSG00000251147 | 0 | 0 | 0 | 0 | 0 | 0 |
| ENSG00000263458 | 0 | 0 | 0 | 0 | 0 | 0 |
| ENSG00000279460 | 0 | 0 | 0 | 0 | 0 | 0 |
| ENSG00000248517 | 0 | 0 | 0 | 0 | 0 | 0 |
| ENSG00000187821 | 0 | 0 | 0 | 0 | 0 | 0 |
| ENSG00000250754 | 0 | 0 | 0 | 0 | 0 | 0 |
| ENSG00000249520 | 0 | 0 | 0 | 0 | 0 | 0 |
| ENSG00000151729 | 489 | 230 | 292 | 259 | 263 | 270 |
| ENSG00000164323 | 276 | 154 | 153 | 95 | 220 | 152 |
| ENSG00000109762 | 162 | 74 | 111 | 96 | 115 | 116 |
| ENSG00000109771 | 6 | 10 | 8 | 4 | 3 | 5 |
| ENSG00000250410 | 0 | 0 | 1 | 0 | 1 | 3 |
| ENSG00000186352 | 14 | 4 | 7 | 2 | 5 | 3 |
| ENSG00000109775 | 223 | 108 | 136 | 91 | 153 | 116 |
| ENSG00000207231 | 0 | 0 | 0 | 0 | 0 | 0 |
| ENSG00000205129 | 0 | 1 | 0 | 1 | 2 | 1 |
| ENSG00000168491 | 0 | 1 | 1 | 2 | 1 | 0 |
| ENSG00000249679 | 0 | 0 | 0 | 0 | 1 | 0 |
| ENSG00000154553 | 0 | 0 | 0 | 0 | 0 | 1 |
| ENSG00000154556 | 0 | 1 | 0 | 0 | 0 | 0 |
| ENSG00000233110 | 0 | 0 | 0 | 0 | 0 | 0 |
| ENSG00000235902 | 0 | 0 | 0 | 0 | 0 | 0 |
| ENSG00000207497 | 0 | 0 | 0 | 0 | 0 | 0 |

| | | | | | | |
|---|---|---|---|---|---|---|
| ENSG00000285745 | 0 | 0 | 0 | 0 | 0 | 0 |
| ENSG00000235149 | 0 | 0 | 0 | 0 | 0 | 0 |
| ENSG00000222727 | 0 | 0 | 0 | 0 | 0 | 0 |
| ENSG00000164342 | 0 | 2 | 0 | 0 | 0 | 0 |
| ENSG00000109794 | 9 | 8 | 19 | 4 | 6 | 5 |
| ENSG00000253013 | 0 | 0 | 0 | 0 | 0 | 0 |
| ENSG00000250327 | 2 | 0 | 3 | 1 | 2 | 0 |
| ENSG00000251008 | 0 | 0 | 0 | 0 | 0 | 0 |
| ENSG00000279943 | 2 | 0 | 1 | 1 | 0 | 1 |
| ENSG00000145476 | 148 | 55 | 101 | 58 | 85 | 101 |
| ENSG00000164344 | 0 | 0 | 0 | 0 | 0 | 1 |
| ENSG00000088926 | 0 | 0 | 0 | 0 | 0 | 0 |
| ENSG00000251165 | 0 | 0 | 0 | 0 | 0 | 0 |
| ENSG00000213332 | 0 | 0 | 0 | 0 | 1 | 0 |
| ENSG00000272297 | 0 | 0 | 0 | 0 | 0 | 0 |
| ENSG00000212387 | 0 | 0 | 0 | 0 | 0 | 0 |
| ENSG00000168412 | 0 | 0 | 0 | 0 | 1 | 0 |
| ENSG00000083857 | 71 | 53 | 24 | 35 | 115 | 51 |
| ENSG00000250829 | 0 | 0 | 0 | 0 | 0 | 0 |
| ENSG00000272218 | 0 | 0 | 0 | 0 | 0 | 0 |
| ENSG00000249539 | 0 | 0 | 0 | 0 | 0 | 0 |
| ENSG00000249742 | 0 | 0 | 0 | 0 | 0 | 0 |
| ENSG00000250971 | 0 | 0 | 0 | 0 | 0 | 0 |
| ENSG00000249747 | 0 | 0 | 0 | 0 | 0 | 0 |
| ENSG00000250658 | 0 | 0 | 0 | 0 | 0 | 0 |
| ENSG00000250042 | 0 | 0 | 0 | 0 | 0 | 0 |
| ENSG00000250620 | 0 | 0 | 0 | 0 | 0 | 0 |
| ENSG00000251643 | 0 | 0 | 0 | 0 | 0 | 0 |
| ENSG00000251490 | 0 | 0 | 0 | 0 | 0 | 0 |
| ENSG00000250590 | 0 | 0 | 0 | 0 | 0 | 0 |
| ENSG00000249642 | 0 | 0 | 0 | 0 | 0 | 0 |
| ENSG00000249162 | 0 | 0 | 0 | 0 | 0 | 0 |
| ENSG00000213331 | 0 | 0 | 0 | 0 | 0 | 0 |
| ENSG00000179059 | 0 | 0 | 0 | 0 | 0 | 0 |
| ENSG00000179046 | 0 | 0 | 0 | 0 | 0 | 0 |
| ENSG00000241011 | 0 | 0 | 0 | 0 | 0 | 0 |
| ENSG00000201085 | 0 | 0 | 0 | 0 | 0 | 0 |
| ENSG00000184108 | 0 | 0 | 0 | 0 | 0 | 0 |
| ENSG00000247130 | 0 | 0 | 1 | 0 | 0 | 0 |
| ENSG00000248370 | 0 | 0 | 0 | 0 | 0 | 0 |
| ENSG00000242169 | 0 | 0 | 1 | 0 | 0 | 0 |
| ENSG00000248880 | 0 | 0 | 0 | 0 | 0 | 0 |
| ENSG00000249378 | 0 | 0 | 0 | 0 | 0 | 0 |
| ENSG00000250523 | 0 | 0 | 0 | 0 | 0 | 0 |
| ENSG00000275826 | 0 | 0 | 0 | 0 | 0 | 0 |
| ENSG00000252275 | 0 | 0 | 0 | 0 | 0 | 0 |
| ENSG00000180015 | 6 | 3 | 7 | 8 | 8 | 9 |
| ENSG00000278974 | 71 | 30 | 25 | 25 | 39 | 24 |
| ENSG00000249269 | 0 | 0 | 0 | 0 | 0 | 0 |
| ENSG00000251619 | 0 | 0 | 0 | 0 | 0 | 0 |

| | | | | | | |
|---|---|---|---|---|---|---|
| ENSG00000250626 | 0 | 0 | 0 | 0 | 0 | 0 |
| ENSG00000250024 | 0 | 0 | 0 | 0 | 0 | 0 |
| ENSG00000251310 | 0 | 0 | 0 | 0 | 0 | 0 |
| ENSG00000249877 | 0 | 0 | 0 | 0 | 0 | 0 |
| ENSG00000205100 | 0 | 0 | 0 | 0 | 0 | 0 |
| ENSG00000250739 | 0 | 0 | 0 | 0 | 0 | 0 |
| ENSG00000272566 | 0 | 0 | 0 | 0 | 0 | 0 |
| ENSG00000202215 | 0 | 0 | 0 | 0 | 0 | 0 |
| ENSG00000245685 | 4 | 1 | 1 | 2 | 0 | 1 |
| ENSG00000250666 | 0 | 0 | 0 | 0 | 0 | 0 |
| ENSG00000109536 | 245 | 103 | 143 | 95 | 127 | 107 |
| ENSG00000249462 | 0 | 0 | 0 | 0 | 0 | 0 |
| ENSG00000271307 | 0 | 0 | 0 | 0 | 0 | 0 |
| ENSG00000251297 | 8 | 2 | 3 | 3 | 3 | 0 |
| ENSG00000199572 | 0 | 0 | 0 | 0 | 0 | 0 |
| ENSG00000201145 | 0 | 0 | 0 | 0 | 0 | 0 |
| ENSG00000224807 | 0 | 0 | 0 | 0 | 0 | 1 |
| ENSG00000205097 | 0 | 0 | 0 | 0 | 0 | 0 |
| ENSG00000250053 | 0 | 0 | 0 | 0 | 0 | 0 |
| ENSG00000214273 | 0 | 0 | 0 | 0 | 0 | 0 |
| ENSG00000249003 | 0 | 0 | 0 | 0 | 0 | 0 |
| ENSG00000281591 | 0 | 0 | 0 | 0 | 0 | 0 |
| ENSG00000277162 | 0 | 0 | 0 | 0 | 0 | 0 |
| ENSG00000281720 | 0 | 0 | 0 | 0 | 0 | 0 |
| ENSG00000281652 | 0 | 0 | 0 | 0 | 0 | 0 |
| ENSG00000281058 | 0 | 0 | 0 | 0 | 0 | 0 |
| ENSG00000280799 | 0 | 0 | 0 | 0 | 0 | 0 |
| ENSG00000258834 | 0 | 0 | 0 | 0 | 0 | 0 |
| ENSG00000280757 | 0 | 0 | 0 | 0 | 0 | 0 |
| ENSG00000281627 | 0 | 0 | 0 | 0 | 0 | 0 |
| ENSG00000280457 | 0 | 0 | 0 | 0 | 0 | 0 |
| ENSG00000260596 | 0 | 0 | 0 | 0 | 0 | 0 |
| ENSG00000261914 | 0 | 0 | 0 | 0 | 0 | 0 |
| ENSG00000282994 | 0 | 0 | 0 | 0 | 0 | 0 |
| ENSG00000250020 | 0 | 0 | 0 | 0 | 0 | 3 |
| ENSG00000153404 | 8 | 11 | 14 | 8 | 13 | 10 |
| ENSG00000199540 | 0 | 0 | 0 | 0 | 0 | 0 |
| ENSG00000185028 | 2 | 0 | 0 | 0 | 0 | 5 |
| ENSG00000261434 | 0 | 0 | 0 | 0 | 0 | 0 |
| ENSG00000164366 | 792 | 375 | 476 | 438 | 422 | 431 |
| ENSG00000260774 | 29 | 7 | 21 | 9 | 24 | 16 |
| ENSG00000073578 | 7619 | 3811 | 4492 | 4366 | 4413 | 4403 |
| ENSG00000248925 | 4 | 3 | 1 | 3 | 2 | 3 |
| ENSG00000249915 | 4633 | 2489 | 2961 | 2958 | 3126 | 3115 |
| ENSG00000250848 | 2 | 4 | 2 | 1 | 4 | 0 |
| ENSG00000063438 | 1539 | 688 | 961 | 1179 | 1614 | 1590 |
| ENSG00000214278 | 0 | 0 | 0 | 0 | 2 | 2 |
| ENSG00000250645 | 16 | 7 | 7 | 7 | 12 | 4 |
| ENSG00000221990 | 461 | 224 | 280 | 299 | 263 | 289 |
| ENSG00000180104 | 2991 | 1388 | 1824 | 1922 | 1987 | 1874 |

| | | | | | | |
|---|---|---|---|---|---|---|
| ENSG00000188242 | 9 | 1 | 8 | 2 | 4 | 7 |
| ENSG00000066230 | 124 | 75 | 142 | 116 | 158 | 133 |
| ENSG00000225138 | 1101 | 481 | 721 | 803 | 780 | 813 |
| ENSG00000250385 | 0 | 0 | 0 | 1 | 0 | 0 |
| ENSG00000264233 | 0 | 0 | 0 | 0 | 0 | 0 |
| ENSG00000249650 | 2 | 0 | 3 | 1 | 1 | 0 |
| ENSG00000112877 | 118 | 61 | 96 | 75 | 72 | 78 |
| ENSG00000171368 | 61 | 29 | 29 | 34 | 38 | 48 |
| ENSG00000271781 | 63 | 36 | 29 | 41 | 27 | 41 |
| ENSG00000206077 | 18 | 10 | 11 | 13 | 12 | 13 |
| ENSG00000249908 | 0 | 0 | 0 | 0 | 0 | 1 |
| ENSG00000285522 | 0 | 0 | 0 | 0 | 0 | 0 |
| ENSG00000188818 | 193 | 87 | 117 | 90 | 88 | 76 |
| ENSG00000247872 | 2 | 0 | 0 | 0 | 0 | 1 |
| ENSG00000028310 | 1640 | 785 | 941 | 861 | 857 | 923 |
| ENSG00000071539 | 717 | 405 | 427 | 369 | 422 | 533 |
| ENSG00000248949 | 0 | 0 | 0 | 0 | 0 | 0 |
| ENSG00000250173 | 0 | 0 | 0 | 0 | 0 | 0 |
| ENSG00000215246 | 0 | 0 | 1 | 0 | 0 | 0 |
| ENSG00000272347 | 2 | 1 | 2 | 0 | 2 | 3 |
| ENSG00000145506 | 161 | 84 | 96 | 107 | 69 | 92 |
| ENSG00000113504 | 2220 | 1005 | 1509 | 1116 | 1015 | 1224 |
| ENSG00000263834 | 0 | 0 | 0 | 0 | 0 | 1 |
| ENSG00000249201 | 0 | 0 | 0 | 0 | 0 | 2 |
| ENSG00000174358 | 6 | 1 | 1 | 4 | 4 | 0 |
| ENSG00000164363 | 0 | 0 | 0 | 0 | 0 | 0 |
| ENSG00000164362 | 4 | 1 | 4 | 0 | 2 | 4 |
| ENSG00000263670 | 0 | 0 | 0 | 0 | 0 | 0 |
| ENSG00000049656 | 4814 | 2445 | 2922 | 2977 | 2938 | 2871 |
| ENSG00000250584 | 0 | 0 | 0 | 0 | 0 | 1 |
| ENSG00000248951 | 0 | 0 | 0 | 0 | 0 | 0 |
| ENSG00000279908 | 29 | 17 | 14 | 10 | 18 | 9 |
| ENSG00000142319 | 15 | 1 | 9 | 4 | 8 | 7 |
| ENSG00000153395 | 13784 | 6446 | 7676 | 9365 | 9401 | 8773 |
| ENSG00000273742 | 0 | 1 | 0 | 0 | 0 | 0 |
| ENSG00000251532 | 3 | 6 | 4 | 5 | 4 | 4 |
| ENSG00000185986 | 237 | 121 | 154 | 94 | 128 | 149 |
| ENSG00000271119 | 27 | 28 | 36 | 22 | 37 | 31 |
| ENSG00000248126 | 11 | 9 | 19 | 11 | 8 | 9 |
| ENSG00000188002 | 1174 | 522 | 852 | 631 | 669 | 779 |
| ENSG00000232615 | 9 | 4 | 4 | 1 | 4 | 3 |
| ENSG00000263746 | 0 | 0 | 0 | 0 | 0 | 0 |
| ENSG00000260066 | 0 | 0 | 0 | 0 | 0 | 0 |
| ENSG00000171421 | 1219 | 606 | 640 | 674 | 633 | 688 |
| ENSG00000145494 | 2411 | 1181 | 1449 | 1540 | 1407 | 1280 |
| ENSG00000249966 | 0 | 0 | 0 | 0 | 0 | 0 |
| ENSG00000250417 | 0 | 0 | 0 | 0 | 0 | 0 |
| ENSG00000279222 | 0 | 0 | 0 | 0 | 0 | 0 |
| ENSG00000113430 | 428 | 223 | 291 | 260 | 234 | 194 |
| ENSG00000249116 | 49 | 28 | 46 | 53 | 33 | 36 |

| | | | | | | |
|---|---|---|---|---|---|---|
| ENSG00000249326 | 35 | 17 | 24 | 18 | 14 | 15 |
| ENSG00000248994 | 0 | 0 | 0 | 0 | 0 | 1 |
| ENSG00000248597 | 0 | 0 | 0 | 0 | 0 | 0 |
| ENSG00000249731 | 0 | 0 | 0 | 0 | 0 | 0 |
| ENSG00000201026 | 0 | 0 | 0 | 0 | 0 | 0 |
| ENSG00000275733 | 0 | 0 | 0 | 0 | 0 | 0 |
| ENSG00000281560 | 0 | 0 | 0 | 0 | 0 | 0 |
| ENSG00000259757 | 0 | 0 | 0 | 0 | 0 | 0 |
| ENSG00000170561 | 999 | 495 | 601 | 541 | 648 | 567 |
| ENSG00000186493 | 823 | 401 | 513 | 628 | 455 | 471 |
| ENSG00000249541 | 2 | 0 | 0 | 0 | 0 | 0 |
| ENSG00000248137 | 0 | 0 | 0 | 0 | 0 | 0 |
| ENSG00000248736 | 0 | 0 | 0 | 0 | 0 | 0 |
| ENSG00000249808 | 0 | 0 | 0 | 0 | 0 | 0 |
| ENSG00000248118 | 0 | 0 | 0 | 0 | 0 | 0 |
| ENSG00000249830 | 0 | 0 | 0 | 0 | 0 | 0 |
| ENSG00000250716 | 0 | 0 | 0 | 0 | 0 | 0 |
| ENSG00000170549 | 0 | 0 | 0 | 0 | 0 | 0 |
| ENSG00000259603 | 0 | 0 | 0 | 0 | 0 | 0 |
| ENSG00000248962 | 0 | 0 | 0 | 0 | 0 | 0 |
| ENSG00000249994 | 0 | 0 | 0 | 0 | 0 | 0 |
| ENSG00000250921 | 0 | 0 | 0 | 0 | 0 | 0 |
| ENSG00000260763 | 0 | 0 | 0 | 0 | 0 | 0 |
| ENSG00000248973 | 0 | 0 | 0 | 0 | 0 | 0 |
| ENSG00000231291 | 0 | 0 | 0 | 0 | 0 | 0 |
| ENSG00000251371 | 0 | 0 | 0 | 0 | 0 | 0 |
| ENSG00000249521 | 0 | 0 | 0 | 0 | 0 | 0 |
| ENSG00000248311 | 0 | 0 | 0 | 0 | 0 | 0 |
| ENSG00000248467 | 0 | 0 | 0 | 0 | 0 | 0 |
| ENSG00000249941 | 0 | 0 | 0 | 0 | 0 | 0 |
| ENSG00000251506 | 0 | 0 | 0 | 0 | 0 | 0 |
| ENSG00000250481 | 0 | 0 | 0 | 0 | 0 | 0 |
| ENSG00000215231 | 0 | 0 | 0 | 0 | 0 | 0 |
| ENSG00000251426 | 0 | 0 | 0 | 0 | 0 | 0 |
| ENSG00000250529 | 0 | 0 | 0 | 0 | 0 | 0 |
| ENSG00000223007 | 0 | 0 | 0 | 0 | 0 | 0 |
| ENSG00000250579 | 0 | 0 | 0 | 0 | 0 | 0 |
| ENSG00000145536 | 0 | 1 | 0 | 0 | 0 | 0 |
| ENSG00000250866 | 0 | 0 | 0 | 0 | 0 | 0 |
| ENSG00000251568 | 0 | 0 | 0 | 0 | 0 | 0 |
| ENSG00000248729 | 2 | 0 | 0 | 0 | 0 | 0 |
| ENSG00000250808 | 0 | 0 | 0 | 0 | 0 | 0 |
| ENSG00000251476 | 0 | 0 | 0 | 0 | 0 | 0 |
| ENSG00000164151 | 1015 | 544 | 518 | 233 | 804 | 467 |
| ENSG00000261037 | 0 | 0 | 0 | 0 | 0 | 0 |
| ENSG00000259861 | 0 | 0 | 0 | 0 | 0 | 0 |
| ENSG00000278913 | 0 | 0 | 0 | 0 | 0 | 0 |
| ENSG00000250730 | 0 | 0 | 0 | 0 | 0 | 0 |
| ENSG00000250490 | 0 | 0 | 0 | 0 | 0 | 0 |
| ENSG00000133398 | 1041 | 484 | 598 | 559 | 561 | 598 |

| | | | | | | |
|---|---|---|---|---|---|---|
| ENSG00000215218 | 94 | 41 | 60 | 35 | 47 | 53 |
| ENSG00000271230 | 0 | 0 | 0 | 0 | 0 | 0 |
| ENSG00000250056 | 0 | 0 | 0 | 0 | 0 | 0 |
| ENSG00000037474 | 3307 | 1618 | 2016 | 1796 | 2051 | 2155 |
| ENSG00000145545 | 1708 | 845 | 993 | 893 | 966 | 936 |
| ENSG00000248677 | 5 | 2 | 2 | 1 | 1 | 1 |
| ENSG00000112941 | 1470 | 741 | 888 | 755 | 788 | 899 |
| ENSG00000251365 | 8 | 3 | 13 | 5 | 11 | 4 |
| ENSG00000272071 | 0 | 2 | 1 | 0 | 1 | 2 |
| ENSG00000283420 | 0 | 0 | 0 | 0 | 0 | 0 |
| ENSG00000250974 | 0 | 0 | 0 | 0 | 0 | 0 |
| ENSG00000200243 | 0 | 0 | 0 | 0 | 0 | 0 |
| ENSG00000250060 | 0 | 0 | 0 | 0 | 0 | 0 |
| ENSG00000212258 | 0 | 0 | 0 | 0 | 0 | 0 |
| ENSG00000237611 | 0 | 0 | 0 | 0 | 0 | 0 |
| ENSG00000285689 | 0 | 0 | 0 | 0 | 0 | 0 |
| ENSG00000248775 | 0 | 0 | 0 | 0 | 0 | 0 |
| ENSG00000251250 | 0 | 0 | 0 | 0 | 0 | 0 |
| ENSG00000205976 | 0 | 0 | 0 | 0 | 0 | 0 |
| ENSG00000251678 | 0 | 0 | 0 | 0 | 0 | 0 |
| ENSG00000250668 | 0 | 0 | 0 | 0 | 0 | 0 |
| ENSG00000249638 | 0 | 0 | 0 | 0 | 0 | 0 |
| ENSG00000249865 | 0 | 0 | 0 | 0 | 0 | 0 |
| ENSG00000223597 | 0 | 0 | 0 | 0 | 0 | 0 |
| ENSG00000078295 | 0 | 0 | 0 | 0 | 0 | 0 |
| ENSG00000250761 | 0 | 0 | 0 | 0 | 0 | 0 |
| ENSG00000215217 | 14 | 11 | 8 | 9 | 12 | 3 |
| ENSG00000124275 | 723 | 389 | 388 | 317 | 383 | 379 |
| ENSG00000124279 | 291 | 174 | 234 | 163 | 229 | 174 |
| ENSG00000251168 | 2 | 0 | 0 | 0 | 1 | 0 |
| ENSG00000199773 | 0 | 0 | 0 | 0 | 0 | 0 |
| ENSG00000280338 | 0 | 0 | 0 | 0 | 0 | 0 |
| ENSG00000280251 | 0 | 0 | 0 | 0 | 0 | 0 |
| ENSG00000249486 | 0 | 0 | 0 | 0 | 0 | 0 |
| ENSG00000245729 | 0 | 0 | 0 | 0 | 0 | 0 |
| ENSG00000249159 | 2 | 0 | 0 | 1 | 2 | 0 |
| ENSG00000251549 | 0 | 0 | 0 | 0 | 0 | 0 |
| ENSG00000272049 | 3 | 0 | 2 | 0 | 3 | 1 |
| ENSG00000247516 | 72 | 36 | 45 | 32 | 36 | 51 |
| ENSG00000250037 | 0 | 0 | 0 | 0 | 0 | 0 |
| ENSG00000279075 | 1 | 2 | 1 | 0 | 1 | 1 |
| ENSG00000284000 | 0 | 0 | 0 | 0 | 0 | 0 |
| ENSG00000249782 | 0 | 0 | 0 | 0 | 0 | 0 |
| ENSG00000279097 | 0 | 0 | 0 | 0 | 0 | 0 |
| ENSG00000279264 | 0 | 0 | 0 | 0 | 0 | 0 |
| ENSG00000250389 | 0 | 0 | 0 | 0 | 0 | 0 |
| ENSG00000248765 | 0 | 0 | 0 | 0 | 0 | 0 |
| ENSG00000279002 | 0 | 0 | 0 | 0 | 0 | 0 |
| ENSG00000280373 | 0 | 0 | 0 | 0 | 0 | 0 |
| ENSG00000279125 | 0 | 0 | 0 | 0 | 0 | 0 |

| | | | | | | |
|---|---|---|---|---|---|---|
| ENSG00000280438 | 0 | 0 | 0 | 0 | 0 | 0 |
| ENSG00000278877 | 0 | 0 | 0 | 0 | 0 | 0 |
| ENSG00000260981 | 0 | 0 | 0 | 0 | 0 | 0 |
| ENSG00000280185 | 0 | 0 | 0 | 0 | 0 | 0 |
| ENSG00000250198 | 0 | 0 | 0 | 0 | 0 | 0 |
| ENSG00000279635 | 0 | 0 | 0 | 0 | 0 | 0 |
| ENSG00000250619 | 0 | 0 | 0 | 0 | 0 | 0 |
| ENSG00000112902 | 506 | 236 | 257 | 102 | 240 | 203 |
| ENSG00000266415 | 0 | 0 | 0 | 0 | 0 | 0 |
| ENSG00000248537 | 0 | 0 | 0 | 0 | 0 | 0 |
| ENSG00000279698 | 0 | 0 | 0 | 0 | 0 | 0 |
| ENSG00000251370 | 0 | 0 | 0 | 0 | 1 | 0 |
| ENSG00000250001 | 0 | 1 | 0 | 0 | 0 | 0 |
| ENSG00000250786 | 122 | 63 | 77 | 52 | 74 | 56 |
| ENSG00000239112 | 0 | 0 | 3 | 1 | 0 | 0 |
| ENSG00000248525 | 0 | 0 | 0 | 0 | 0 | 0 |
| ENSG00000169777 | 0 | 0 | 0 | 0 | 0 | 0 |
| ENSG00000249781 | 0 | 0 | 0 | 0 | 0 | 0 |
| ENSG00000222054 | 0 | 0 | 0 | 0 | 0 | 0 |
| ENSG00000250814 | 0 | 0 | 0 | 0 | 0 | 0 |
| ENSG00000279011 | 0 | 0 | 0 | 0 | 0 | 0 |
| ENSG00000279755 | 0 | 0 | 0 | 0 | 0 | 0 |
| ENSG00000278874 | 0 | 0 | 0 | 0 | 0 | 0 |
| ENSG00000280016 | 0 | 0 | 0 | 0 | 0 | 0 |
| ENSG00000249807 | 0 | 0 | 0 | 0 | 0 | 0 |
| ENSG00000280449 | 0 | 0 | 0 | 0 | 0 | 0 |
| ENSG00000280368 | 0 | 0 | 0 | 0 | 0 | 0 |
| ENSG00000260515 | 0 | 0 | 0 | 0 | 0 | 0 |
| ENSG00000271998 | 0 | 0 | 0 | 0 | 0 | 0 |
| ENSG00000272417 | 0 | 0 | 1 | 1 | 0 | 0 |
| ENSG00000150756 | 737 | 346 | 388 | 269 | 389 | 370 |
| ENSG00000248968 | 0 | 1 | 1 | 1 | 0 | 1 |
| ENSG00000150753 | 9511 | 5361 | 6349 | 6044 | 7301 | 7353 |
| ENSG00000271980 | 3 | 1 | 0 | 0 | 1 | 2 |
| ENSG00000271715 | 0 | 1 | 0 | 1 | 0 | 0 |
| ENSG00000164237 | 2333 | 1032 | 1264 | 658 | 916 | 1003 |
| ENSG00000252341 | 0 | 0 | 0 | 0 | 0 | 0 |
| ENSG00000252671 | 0 | 0 | 0 | 0 | 0 | 0 |
| ENSG00000280369 | 1 | 0 | 3 | 0 | 0 | 0 |
| ENSG00000259802 | 146 | 65 | 79 | 49 | 62 | 77 |
| ENSG00000145495 | 7062 | 3521 | 4610 | 2567 | 3866 | 3559 |
| ENSG00000250600 | 1 | 1 | 1 | 0 | 0 | 0 |
| ENSG00000145491 | 8 | 7 | 11 | 0 | 2 | 5 |
| ENSG00000277073 | 0 | 0 | 0 | 0 | 0 | 0 |
| ENSG00000246016 | 0 | 0 | 0 | 0 | 2 | 0 |
| ENSG00000244361 | 0 | 0 | 0 | 0 | 0 | 0 |
| ENSG00000249166 | 0 | 0 | 0 | 0 | 0 | 0 |
| ENSG00000249396 | 0 | 1 | 0 | 0 | 0 | 0 |
| ENSG00000249160 | 0 | 0 | 0 | 0 | 0 | 0 |
| ENSG00000280017 | 0 | 0 | 0 | 0 | 0 | 0 |

| | | | | | | |
|---|---|---|---|---|---|---|
| ENSG00000279450 | 0 | 0 | 0 | 0 | 0 | 0 |
| ENSG00000164236 | 24 | 7 | 16 | 5 | 7 | 4 |
| ENSG00000250106 | 0 | 0 | 0 | 0 | 0 | 0 |
| ENSG00000251196 | 0 | 0 | 0 | 0 | 0 | 0 |
| ENSG00000251548 | 0 | 0 | 0 | 0 | 0 | 0 |
| ENSG00000279979 | 0 | 0 | 0 | 0 | 0 | 0 |
| ENSG00000278865 | 0 | 0 | 0 | 0 | 0 | 0 |
| ENSG00000112977 | 3936 | 2079 | 2628 | 2441 | 2119 | 2326 |
| ENSG00000272324 | 1 | 1 | 0 | 0 | 0 | 0 |
| ENSG00000250576 | 0 | 0 | 0 | 0 | 0 | 0 |
| ENSG00000169862 | 3087 | 1331 | 1868 | 941 | 1023 | 1239 |
| ENSG00000207312 | 0 | 0 | 0 | 0 | 0 | 0 |
| ENSG00000212305 | 0 | 0 | 0 | 0 | 0 | 0 |
| ENSG00000248783 | 0 | 0 | 0 | 0 | 0 | 0 |
| ENSG00000248131 | 0 | 0 | 0 | 0 | 0 | 0 |
| ENSG00000251112 | 0 | 0 | 0 | 0 | 0 | 0 |
| ENSG00000249053 | 0 | 0 | 0 | 0 | 0 | 0 |
| ENSG00000248457 | 0 | 0 | 0 | 0 | 0 | 0 |
| ENSG00000248145 | 0 | 0 | 0 | 0 | 0 | 0 |
| ENSG00000242608 | 0 | 0 | 0 | 0 | 0 | 0 |
| ENSG00000248462 | 0 | 0 | 0 | 0 | 0 | 0 |
| ENSG00000243744 | 0 | 0 | 0 | 0 | 0 | 0 |
| ENSG00000039139 | 70 | 50 | 52 | 28 | 74 | 51 |
| ENSG00000251423 | 0 | 0 | 0 | 0 | 0 | 0 |
| ENSG00000038382 | 2693 | 1267 | 1741 | 1082 | 1635 | 1532 |
| ENSG00000145569 | 617 | 273 | 372 | 226 | 316 | 360 |
| ENSG00000248791 | 0 | 1 | 0 | 0 | 0 | 2 |
| ENSG00000250526 | 2 | 1 | 0 | 0 | 0 | 0 |
| ENSG00000250182 | 27 | 9 | 12 | 7 | 15 | 25 |
| ENSG00000261360 | 38 | 21 | 41 | 20 | 22 | 10 |
| ENSG00000154124 | 1135 | 597 | 695 | 509 | 723 | 661 |
| ENSG00000154122 | 189 | 108 | 135 | 71 | 85 | 101 |
| ENSG00000249485 | 34 | 15 | 34 | 14 | 13 | 25 |
| ENSG00000264792 | 0 | 0 | 0 | 0 | 0 | 0 |
| ENSG00000272057 | 0 | 0 | 0 | 0 | 0 | 0 |
| ENSG00000249928 | 0 | 0 | 0 | 0 | 0 | 0 |
| ENSG00000250662 | 0 | 0 | 0 | 0 | 0 | 0 |
| ENSG00000250247 | 0 | 0 | 0 | 0 | 0 | 0 |
| ENSG00000202269 | 0 | 0 | 0 | 0 | 0 | 0 |
| ENSG00000249332 | 0 | 0 | 0 | 0 | 0 | 0 |
| ENSG00000248486 | 0 | 0 | 0 | 0 | 0 | 0 |
| ENSG00000248834 | 0 | 0 | 0 | 0 | 0 | 0 |
| ENSG00000248699 | 0 | 0 | 0 | 0 | 0 | 0 |
| ENSG00000183580 | 0 | 2 | 0 | 0 | 0 | 0 |
| ENSG00000250250 | 0 | 0 | 0 | 0 | 0 | 0 |
| ENSG00000216077 | 0 | 0 | 0 | 0 | 0 | 0 |
| ENSG00000222312 | 0 | 0 | 0 | 0 | 0 | 0 |
| ENSG00000183654 | 0 | 0 | 0 | 0 | 0 | 0 |
| ENSG00000250981 | 0 | 0 | 0 | 0 | 0 | 0 |
| ENSG00000250448 | 0 | 0 | 0 | 0 | 0 | 0 |

| | | | | | | |
|---|---|---|---|---|---|---|
| ENSG00000248403 | 0 | 0 | 0 | 0 | 0 | 0 |
| ENSG00000248150 | 0 | 1 | 1 | 0 | 0 | 0 |
| ENSG00000249737 | 0 | 0 | 0 | 0 | 0 | 0 |
| ENSG00000173545 | 1692 | 848 | 993 | 1111 | 970 | 947 |
| ENSG00000154153 | 327 | 148 | 192 | 89 | 142 | 99 |
| ENSG00000246214 | 1 | 3 | 0 | 0 | 0 | 3 |
| ENSG00000250415 | 0 | 0 | 0 | 0 | 1 | 0 |
| ENSG00000201370 | 0 | 0 | 0 | 0 | 0 | 0 |
| ENSG00000145555 | 292 | 128 | 129 | 117 | 182 | 134 |
| ENSG00000253093 | 0 | 0 | 0 | 0 | 0 | 0 |
| ENSG00000243538 | 0 | 0 | 0 | 0 | 0 | 0 |
| ENSG00000201436 | 0 | 0 | 0 | 0 | 0 | 0 |
| ENSG00000176788 | 11842 | 5919 | 7305 | 5483 | 6356 | 6724 |
| ENSG00000271892 | 0 | 0 | 0 | 0 | 0 | 1 |
| ENSG00000206814 | 0 | 0 | 0 | 0 | 0 | 0 |
| ENSG00000215196 | 33 | 20 | 15 | 5 | 19 | 8 |
| ENSG00000251990 | 0 | 0 | 0 | 0 | 0 | 0 |
| ENSG00000250816 | 0 | 0 | 0 | 0 | 0 | 0 |
| ENSG00000252908 | 0 | 0 | 0 | 0 | 0 | 0 |
| ENSG00000185296 | 14 | 8 | 7 | 12 | 9 | 15 |
| ENSG00000201715 | 0 | 0 | 0 | 0 | 0 | 0 |
| ENSG00000223361 | 2 | 4 | 0 | 2 | 0 | 2 |
| ENSG00000248223 | 0 | 0 | 0 | 0 | 0 | 0 |
| ENSG00000249199 | 11 | 5 | 5 | 10 | 11 | 4 |
| ENSG00000250822 | 2 | 3 | 2 | 0 | 0 | 1 |
| ENSG00000248455 | 0 | 0 | 0 | 0 | 0 | 0 |
| ENSG00000249662 | 0 | 1 | 0 | 0 | 0 | 0 |
| ENSG00000268799 | 0 | 0 | 0 | 0 | 0 | 0 |
| ENSG00000250667 | 0 | 0 | 0 | 0 | 0 | 0 |
| ENSG00000284373 | 0 | 0 | 0 | 0 | 0 | 0 |
| ENSG00000248205 | 0 | 0 | 0 | 0 | 0 | 0 |
| ENSG00000249620 | 0 | 0 | 0 | 0 | 0 | 0 |
| ENSG00000284439 | 0 | 0 | 0 | 0 | 0 | 0 |
| ENSG00000284283 | 0 | 0 | 0 | 0 | 0 | 0 |
| ENSG00000284234 | 0 | 0 | 0 | 0 | 0 | 0 |
| ENSG00000284042 | 0 | 0 | 0 | 0 | 0 | 0 |
| ENSG00000284465 | 0 | 0 | 0 | 0 | 0 | 0 |
| ENSG00000250088 | 0 | 0 | 0 | 0 | 0 | 0 |
| ENSG00000283967 | 0 | 0 | 0 | 0 | 0 | 0 |
| ENSG00000283988 | 0 | 0 | 0 | 0 | 0 | 0 |
| ENSG00000284356 | 0 | 0 | 0 | 0 | 0 | 0 |
| ENSG00000248422 | 0 | 0 | 0 | 0 | 0 | 0 |
| ENSG00000249282 | 0 | 0 | 0 | 0 | 0 | 0 |
| ENSG00000283740 | 0 | 0 | 0 | 0 | 0 | 0 |
| ENSG00000249156 | 0 | 0 | 0 | 0 | 0 | 0 |
| ENSG00000271296 | 0 | 0 | 0 | 0 | 0 | 0 |
| ENSG00000250351 | 0 | 0 | 0 | 0 | 0 | 0 |
| ENSG00000250386 | 0 | 0 | 0 | 0 | 0 | 0 |
| ENSG00000283776 | 0 | 0 | 0 | 0 | 0 | 0 |
| ENSG00000250782 | 0 | 0 | 0 | 0 | 0 | 0 |

| | | | | | | |
|---|---|---|---|---|---|---|
| ENSG00000248471 | 0 | 0 | 0 | 0 | 0 | 0 |
| ENSG00000248160 | 0 | 0 | 0 | 0 | 0 | 0 |
| ENSG00000248542 | 0 | 0 | 0 | 0 | 0 | 0 |
| ENSG00000248337 | 0 | 0 | 0 | 0 | 0 | 0 |
| ENSG00000269466 | 0 | 3 | 1 | 0 | 0 | 0 |
| ENSG00000274832 | 0 | 0 | 0 | 0 | 0 | 0 |
| ENSG00000250715 | 0 | 0 | 0 | 0 | 0 | 0 |
| ENSG00000249937 | 5 | 1 | 0 | 0 | 0 | 4 |
| ENSG00000240785 | 0 | 0 | 0 | 0 | 0 | 0 |
| ENSG00000212278 | 0 | 0 | 0 | 0 | 0 | 0 |
| ENSG00000241552 | 0 | 0 | 0 | 0 | 0 | 0 |
| ENSG00000248693 | 2 | 1 | 4 | 1 | 1 | 0 |
| ENSG00000249191 | 0 | 0 | 0 | 0 | 0 | 0 |
| ENSG00000249927 | 0 | 0 | 0 | 0 | 0 | 0 |
| ENSG00000249285 | 0 | 0 | 0 | 0 | 0 | 0 |
| ENSG00000276764 | 0 | 0 | 0 | 0 | 0 | 0 |
| ENSG00000249174 | 0 | 0 | 0 | 0 | 0 | 0 |
| ENSG00000251487 | 0 | 0 | 1 | 0 | 0 | 0 |
| ENSG00000242654 | 0 | 0 | 0 | 0 | 0 | 0 |
| ENSG00000249557 | 0 | 0 | 0 | 0 | 0 | 0 |
| ENSG00000145526 | 61 | 18 | 54 | 8 | 19 | 20 |
| ENSG00000248766 | 0 | 0 | 0 | 0 | 0 | 0 |
| ENSG00000214132 | 0 | 0 | 0 | 0 | 0 | 0 |
| ENSG00000249854 | 0 | 0 | 0 | 0 | 0 | 0 |
| ENSG00000251629 | 0 | 0 | 0 | 0 | 0 | 0 |
| ENSG00000249490 | 0 | 0 | 0 | 0 | 0 | 0 |
| ENSG00000248308 | 0 | 0 | 0 | 0 | 0 | 0 |
| ENSG00000248286 | 0 | 0 | 0 | 0 | 0 | 0 |
| ENSG00000249359 | 0 | 0 | 0 | 0 | 1 | 0 |
| ENSG00000183666 | 287 | 115 | 153 | 105 | 152 | 138 |
| ENSG00000233974 | 2 | 3 | 4 | 0 | 0 | 3 |
| ENSG00000279107 | 0 | 0 | 0 | 0 | 0 | 0 |
| ENSG00000272130 | 0 | 0 | 0 | 0 | 0 | 0 |
| ENSG00000253766 | 0 | 0 | 0 | 0 | 0 | 0 |
| ENSG00000154162 | 4 | 0 | 10 | 2 | 2 | 3 |
| ENSG00000213430 | 8 | 5 | 6 | 5 | 4 | 9 |
| ENSG00000253104 | 0 | 0 | 0 | 0 | 0 | 0 |
| ENSG00000168967 | 0 | 0 | 0 | 0 | 0 | 0 |
| ENSG00000249206 | 0 | 0 | 0 | 0 | 0 | 0 |
| ENSG00000249404 | 0 | 0 | 0 | 0 | 0 | 0 |
| ENSG00000250319 | 0 | 0 | 0 | 0 | 0 | 0 |
| ENSG00000206912 | 0 | 0 | 0 | 0 | 0 | 0 |
| ENSG00000250332 | 0 | 0 | 2 | 0 | 0 | 1 |
| ENSG00000249289 | 0 | 0 | 0 | 0 | 0 | 0 |
| ENSG00000164256 | 0 | 0 | 5 | 0 | 0 | 0 |
| ENSG00000248591 | 0 | 0 | 0 | 0 | 0 | 0 |
| ENSG00000249607 | 0 | 0 | 0 | 0 | 0 | 0 |
| ENSG00000248874 | 0 | 0 | 0 | 0 | 0 | 0 |
| ENSG00000252083 | 0 | 0 | 0 | 0 | 0 | 0 |
| ENSG00000234865 | 0 | 0 | 0 | 0 | 0 | 0 |

| | | | | | | |
|---|---|---|---|---|---|---|
| ENSG00000200600 | 0 | 0 | 0 | 0 | 0 | 0 |
| ENSG00000226432 | 2 | 1 | 1 | 0 | 0 | 0 |
| ENSG00000250694 | 0 | 0 | 0 | 0 | 0 | 0 |
| ENSG00000277455 | 0 | 0 | 0 | 0 | 0 | 0 |
| ENSG00000040731 | 0 | 0 | 0 | 0 | 1 | 0 |
| ENSG00000251294 | 0 | 0 | 0 | 0 | 0 | 0 |
| ENSG00000249562 | 0 | 0 | 0 | 0 | 0 | 0 |
| ENSG00000250099 | 0 | 0 | 0 | 0 | 0 | 0 |
| ENSG00000248156 | 0 | 0 | 0 | 0 | 0 | 0 |
| ENSG00000223004 | 0 | 0 | 0 | 0 | 0 | 0 |
| ENSG00000248908 | 0 | 0 | 0 | 0 | 0 | 0 |
| ENSG00000250524 | 0 | 0 | 0 | 0 | 0 | 0 |
| ENSG00000199911 | 0 | 0 | 0 | 0 | 0 | 0 |
| ENSG00000251273 | 2 | 4 | 4 | 1 | 1 | 5 |
| ENSG00000245662 | 0 | 0 | 0 | 0 | 0 | 1 |
| ENSG00000285042 | 0 | 0 | 0 | 0 | 0 | 0 |
| ENSG00000251654 | 0 | 0 | 0 | 0 | 0 | 0 |
| ENSG00000260761 | 0 | 0 | 0 | 0 | 0 | 0 |
| ENSG00000279671 | 0 | 0 | 0 | 0 | 0 | 0 |
| ENSG00000248605 | 0 | 0 | 0 | 0 | 0 | 0 |
| ENSG00000279531 | 0 | 0 | 0 | 0 | 0 | 0 |
| ENSG00000201016 | 0 | 0 | 0 | 0 | 0 | 0 |
| ENSG00000279869 | 0 | 0 | 0 | 0 | 0 | 0 |
| ENSG00000251593 | 0 | 0 | 0 | 0 | 0 | 0 |
| ENSG00000222560 | 0 | 0 | 0 | 0 | 0 | 0 |
| ENSG00000249099 | 0 | 0 | 0 | 0 | 0 | 0 |
| ENSG00000230571 | 0 | 0 | 0 | 0 | 0 | 0 |
| ENSG00000248426 | 0 | 0 | 0 | 0 | 0 | 0 |
| ENSG00000250703 | 0 | 0 | 0 | 0 | 0 | 0 |
| ENSG00000251033 | 0 | 0 | 0 | 0 | 0 | 0 |
| ENSG00000249616 | 0 | 0 | 0 | 0 | 0 | 0 |
| ENSG00000113100 | 0 | 0 | 0 | 0 | 0 | 0 |
| ENSG00000207261 | 0 | 0 | 0 | 0 | 0 | 0 |
| ENSG00000248490 | 0 | 0 | 0 | 0 | 0 | 0 |
| ENSG00000250337 | 4 | 0 | 0 | 0 | 0 | 3 |
| ENSG00000250079 | 0 | 0 | 0 | 0 | 0 | 0 |
| ENSG00000249785 | 0 | 0 | 0 | 0 | 0 | 0 |
| ENSG00000252867 | 0 | 0 | 0 | 0 | 0 | 0 |
| ENSG00000250453 | 0 | 0 | 0 | 0 | 0 | 0 |
| ENSG00000270495 | 0 | 0 | 0 | 0 | 0 | 0 |
| ENSG00000271035 | 0 | 0 | 0 | 0 | 0 | 0 |
| ENSG00000162685 | 0 | 0 | 0 | 0 | 0 | 0 |
| ENSG00000226400 | 0 | 0 | 0 | 0 | 0 | 0 |
| ENSG00000277878 | 0 | 0 | 0 | 0 | 0 | 0 |
| ENSG00000249226 | 0 | 0 | 0 | 0 | 0 | 0 |
| ENSG00000252601 | 0 | 0 | 0 | 0 | 0 | 0 |
| ENSG00000248687 | 0 | 0 | 0 | 0 | 0 | 0 |
| ENSG00000270430 | 0 | 0 | 0 | 0 | 0 | 0 |
| ENSG00000259786 | 0 | 0 | 0 | 0 | 0 | 0 |
| ENSG00000248391 | 0 | 0 | 0 | 0 | 0 | 0 |

| | | | | | | |
|---|---|---|---|---|---|---|
| ENSG00000276098 | 0 | 0 | 0 | 0 | 0 | 0 |
| ENSG00000248418 | 0 | 0 | 0 | 0 | 0 | 0 |
| ENSG00000250984 | 0 | 0 | 0 | 0 | 0 | 0 |
| ENSG00000249744 | 0 | 0 | 0 | 0 | 0 | 0 |
| ENSG00000233787 | 0 | 0 | 0 | 0 | 0 | 0 |
| ENSG00000223136 | 0 | 0 | 0 | 0 | 0 | 0 |
| ENSG00000280305 | 0 | 0 | 0 | 0 | 0 | 0 |
| ENSG00000250587 | 0 | 0 | 0 | 0 | 0 | 0 |
| ENSG00000250118 | 0 | 0 | 0 | 0 | 0 | 0 |
| ENSG00000241668 | 0 | 0 | 0 | 0 | 0 | 0 |
| ENSG00000253940 | 0 | 0 | 0 | 0 | 0 | 0 |
| ENSG00000254138 | 0 | 0 | 0 | 0 | 0 | 0 |
| ENSG00000113361 | 0 | 0 | 0 | 0 | 0 | 0 |
| ENSG00000250482 | 0 | 0 | 0 | 0 | 0 | 0 |
| ENSG00000113360 | 1791 | 837 | 973 | 737 | 1006 | 944 |
| ENSG00000206682 | 0 | 0 | 0 | 0 | 0 | 0 |
| ENSG00000082213 | 1095 | 614 | 610 | 407 | 698 | 573 |
| ENSG00000279923 | 0 | 2 | 4 | 1 | 3 | 1 |
| ENSG00000199765 | 0 | 0 | 0 | 0 | 0 | 0 |
| ENSG00000133401 | 16 | 6 | 21 | 10 | 15 | 10 |
| ENSG00000248624 | 0 | 0 | 0 | 0 | 0 | 0 |
| ENSG00000248878 | 0 | 0 | 0 | 0 | 0 | 0 |
| ENSG00000249114 | 0 | 0 | 0 | 0 | 0 | 0 |
| ENSG00000248813 | 0 | 0 | 0 | 0 | 0 | 0 |
| ENSG00000248378 | 0 | 0 | 0 | 0 | 0 | 0 |
| ENSG00000250164 | 0 | 0 | 0 | 0 | 0 | 0 |
| ENSG00000249688 | 0 | 0 | 0 | 0 | 0 | 0 |
| ENSG00000252373 | 0 | 0 | 0 | 0 | 0 | 0 |
| ENSG00000224282 | 0 | 0 | 0 | 0 | 0 | 0 |
| ENSG00000251887 | 0 | 0 | 0 | 0 | 0 | 0 |
| ENSG00000241739 | 0 | 0 | 0 | 0 | 1 | 0 |
| ENSG00000243914 | 0 | 0 | 0 | 0 | 0 | 0 |
| ENSG00000214108 | 0 | 0 | 0 | 0 | 0 | 1 |
| ENSG00000249774 | 2 | 0 | 0 | 0 | 0 | 0 |
| ENSG00000266243 | 0 | 0 | 0 | 0 | 0 | 0 |
| ENSG00000250764 | 0 | 5 | 11 | 1 | 4 | 7 |
| ENSG00000113384 | 4741 | 2144 | 2805 | 1931 | 2263 | 2440 |
| ENSG00000200065 | 2 | 0 | 0 | 0 | 0 | 0 |
| ENSG00000272086 | 82 | 35 | 61 | 46 | 36 | 47 |
| ENSG00000230829 | 0 | 0 | 0 | 0 | 0 | 0 |
| ENSG00000150712 | 1459 | 597 | 809 | 405 | 716 | 674 |
| ENSG00000199731 | 0 | 0 | 0 | 0 | 0 | 0 |
| ENSG00000207052 | 0 | 0 | 0 | 0 | 0 | 1 |
| ENSG00000056097 | 2730 | 1384 | 1367 | 850 | 1629 | 1383 |
| ENSG00000207956 | 0 | 2 | 0 | 0 | 1 | 0 |
| ENSG00000249142 | 0 | 0 | 0 | 0 | 0 | 0 |
| ENSG00000113387 | 3312 | 1695 | 1861 | 1519 | 2115 | 1667 |
| ENSG00000250885 | 0 | 0 | 0 | 0 | 0 | 0 |
| ENSG00000113389 | 36 | 22 | 39 | 15 | 20 | 23 |
| ENSG00000251062 | 0 | 0 | 0 | 0 | 0 | 0 |

| | | | | | | |
|---|---|---|---|---|---|---|
| ENSG00000250697 | 4 | 0 | 1 | 1 | 0 | 1 |
| ENSG00000248279 | 0 | 0 | 0 | 0 | 0 | 0 |
| ENSG00000249102 | 0 | 0 | 0 | 0 | 0 | 0 |
| ENSG00000251281 | 0 | 0 | 0 | 0 | 3 | 0 |
| ENSG00000250672 | 0 | 0 | 0 | 0 | 0 | 0 |
| ENSG00000240376 | 12 | 9 | 3 | 7 | 4 | 7 |
| ENSG00000251443 | 0 | 0 | 0 | 0 | 0 | 0 |
| ENSG00000249572 | 40 | 37 | 27 | 25 | 19 | 26 |
| ENSG00000113407 | 3726 | 1977 | 2254 | 1755 | 2577 | 2320 |
| ENSG00000249360 | 0 | 0 | 0 | 0 | 0 | 0 |
| ENSG00000250583 | 0 | 0 | 0 | 0 | 0 | 0 |
| ENSG00000151388 | 4 | 1 | 4 | 0 | 2 | 1 |
| ENSG00000201623 | 0 | 0 | 0 | 0 | 0 | 0 |
| ENSG00000182631 | 0 | 0 | 0 | 0 | 1 | 0 |
| ENSG00000164175 | 0 | 0 | 0 | 0 | 0 | 0 |
| ENSG00000250914 | 0 | 0 | 0 | 0 | 0 | 0 |
| ENSG00000242110 | 79 | 55 | 68 | 63 | 47 | 60 |
| ENSG00000273294 | 2 | 1 | 2 | 1 | 0 | 0 |
| ENSG00000250767 | 0 | 0 | 0 | 0 | 0 | 1 |
| ENSG00000082196 | 54 | 30 | 42 | 23 | 22 | 32 |
| ENSG00000278900 | 0 | 0 | 0 | 0 | 0 | 0 |
| ENSG00000279995 | 0 | 0 | 0 | 0 | 1 | 0 |
| ENSG00000271771 | 0 | 0 | 0 | 0 | 0 | 0 |
| ENSG00000215158 | 82 | 39 | 38 | 24 | 31 | 37 |
| ENSG00000215156 | 8 | 5 | 1 | 3 | 0 | 1 |
| ENSG00000249380 | 0 | 0 | 0 | 0 | 0 | 0 |
| ENSG00000271874 | 0 | 0 | 0 | 0 | 0 | 0 |
| ENSG00000039560 | 61 | 54 | 23 | 26 | 92 | 49 |
| ENSG00000250234 | 0 | 0 | 0 | 0 | 0 | 0 |
| ENSG00000272323 | 59 | 29 | 42 | 26 | 46 | 31 |
| ENSG00000205838 | 6 | 16 | 13 | 3 | 8 | 14 |
| ENSG00000250140 | 0 | 0 | 0 | 0 | 0 | 0 |
| ENSG00000241829 | 0 | 0 | 0 | 0 | 0 | 0 |
| ENSG00000113456 | 1032 | 523 | 642 | 531 | 667 | 584 |
| ENSG00000113460 | 863 | 430 | 496 | 429 | 577 | 607 |
| ENSG00000168724 | 1412 | 743 | 809 | 522 | 1004 | 791 |
| ENSG00000113492 | 0 | 0 | 0 | 0 | 0 | 0 |
| ENSG00000113494 | 1008 | 645 | 526 | 231 | 942 | 533 |
| ENSG00000250886 | 0 | 1 | 2 | 0 | 0 | 0 |
| ENSG00000201368 | 0 | 0 | 0 | 0 | 0 | 0 |
| ENSG00000152582 | 172 | 83 | 87 | 76 | 92 | 72 |
| ENSG00000238441 | 0 | 0 | 0 | 0 | 0 | 0 |
| ENSG00000248969 | 0 | 0 | 0 | 0 | 0 | 0 |
| ENSG00000168685 | 0 | 0 | 0 | 0 | 0 | 0 |
| ENSG00000279737 | 0 | 0 | 0 | 0 | 0 | 0 |
| ENSG00000248349 | 0 | 0 | 0 | 0 | 0 | 0 |
| ENSG00000152611 | 0 | 0 | 0 | 0 | 0 | 0 |
| ENSG00000249748 | 0 | 0 | 0 | 0 | 0 | 0 |
| ENSG00000145626 | 0 | 0 | 0 | 0 | 0 | 0 |
| ENSG00000168671 | 8 | 2 | 2 | 3 | 0 | 4 |

| | | | | | | |
|---|---|---|---|---|---|---|
| ENSG00000251591 | 0 | 0 | 0 | 0 | 0 | 0 |
| ENSG00000164187 | 675 | 352 | 316 | 142 | 488 | 267 |
| ENSG00000207756 | 0 | 0 | 0 | 0 | 0 | 0 |
| ENSG00000145604 | 413 | 218 | 246 | 185 | 220 | 230 |
| ENSG00000201223 | 0 | 0 | 0 | 0 | 0 | 0 |
| ENSG00000152620 | 624 | 310 | 393 | 230 | 378 | 298 |
| ENSG00000245711 | 6 | 1 | 2 | 2 | 5 | 3 |
| ENSG00000164188 | 0 | 0 | 0 | 0 | 0 | 0 |
| ENSG00000222178 | 0 | 0 | 0 | 0 | 0 | 0 |
| ENSG00000242292 | 0 | 0 | 0 | 0 | 0 | 0 |
| ENSG00000079215 | 2 | 1 | 1 | 1 | 0 | 0 |
| ENSG00000274441 | 0 | 0 | 0 | 0 | 0 | 0 |
| ENSG00000250155 | 0 | 0 | 0 | 0 | 0 | 0 |
| ENSG00000279400 | 0 | 0 | 0 | 0 | 0 | 0 |
| ENSG00000272103 | 2 | 0 | 4 | 0 | 0 | 2 |
| ENSG00000285967 | 662 | 262 | 393 | 230 | 349 | 339 |
| ENSG00000164190 | 492 | 321 | 287 | 99 | 709 | 370 |
| ENSG00000249850 | 2 | 0 | 1 | 0 | 1 | 1 |
| ENSG00000244073 | 0 | 2 | 1 | 2 | 1 | 1 |
| ENSG00000197603 | 188 | 99 | 90 | 42 | 183 | 92 |
| ENSG00000271378 | 0 | 0 | 0 | 0 | 0 | 0 |
| ENSG00000228212 | 1 | 0 | 2 | 0 | 0 | 0 |
| ENSG00000242493 | 0 | 0 | 0 | 0 | 0 | 0 |
| ENSG00000270558 | 13 | 4 | 7 | 6 | 6 | 6 |
| ENSG00000113569 | 1483 | 835 | 961 | 604 | 964 | 847 |
| ENSG00000251880 | 0 | 1 | 1 | 0 | 2 | 2 |
| ENSG00000250455 | 0 | 0 | 0 | 0 | 0 | 0 |
| ENSG00000082068 | 873 | 452 | 499 | 391 | 513 | 472 |
| ENSG00000248216 | 0 | 0 | 0 | 0 | 0 | 0 |
| ENSG00000222743 | 0 | 0 | 0 | 0 | 0 | 0 |
| ENSG00000206743 | 2 | 0 | 0 | 0 | 0 | 0 |
| ENSG00000248587 | 7 | 4 | 3 | 1 | 1 | 0 |
| ENSG00000168621 | 2 | 2 | 4 | 2 | 2 | 5 |
| ENSG00000248539 | 0 | 0 | 0 | 0 | 0 | 0 |
| ENSG00000250032 | 0 | 0 | 0 | 0 | 0 | 0 |
| ENSG00000251518 | 0 | 0 | 0 | 0 | 0 | 0 |
| ENSG00000248533 | 0 | 0 | 0 | 0 | 0 | 0 |
| ENSG00000250003 | 0 | 0 | 0 | 0 | 0 | 0 |
| ENSG00000248461 | 0 | 0 | 0 | 0 | 0 | 0 |
| ENSG00000276332 | 0 | 0 | 0 | 0 | 0 | 0 |
| ENSG00000164318 | 0 | 0 | 0 | 0 | 0 | 0 |
| ENSG00000248730 | 0 | 0 | 0 | 0 | 0 | 0 |
| ENSG00000249071 | 0 | 0 | 0 | 0 | 0 | 0 |
| ENSG00000248572 | 0 | 0 | 0 | 0 | 0 | 0 |
| ENSG00000249491 | 0 | 0 | 0 | 0 | 0 | 0 |
| ENSG00000248234 | 0 | 0 | 0 | 0 | 0 | 0 |
| ENSG00000251257 | 0 | 0 | 0 | 0 | 0 | 0 |
| ENSG00000113594 | 66 | 29 | 21 | 12 | 37 | 20 |
| ENSG00000244968 | 28 | 16 | 26 | 17 | 24 | 15 |
| ENSG00000265304 | 0 | 0 | 0 | 0 | 0 | 0 |

| | | | | | | |
|---|---|---|---|---|---|---|
| ENSG00000251189 | 0 | 0 | 0 | 0 | 0 | 0 |
| ENSG00000249740 | 4 | 0 | 0 | 1 | 1 | 4 |
| ENSG00000248577 | 0 | 0 | 0 | 0 | 0 | 0 |
| ENSG00000250629 | 0 | 0 | 0 | 0 | 0 | 0 |
| ENSG00000145623 | 248 | 101 | 99 | 89 | 166 | 107 |
| ENSG00000164327 | 364 | 280 | 162 | 68 | 432 | 170 |
| ENSG00000249089 | 0 | 0 | 0 | 0 | 0 | 0 |
| ENSG00000082074 | 0 | 0 | 0 | 1 | 0 | 1 |
| ENSG00000251215 | 0 | 1 | 0 | 0 | 0 | 0 |
| ENSG00000113600 | 0 | 0 | 0 | 0 | 0 | 0 |
| ENSG00000153071 | 13 | 21 | 15 | 5 | 7 | 10 |
| ENSG00000271334 | 0 | 0 | 0 | 0 | 0 | 0 |
| ENSG00000251515 | 0 | 0 | 0 | 0 | 0 | 0 |
| ENSG00000250492 | 2 | 3 | 3 | 1 | 0 | 1 |
| ENSG00000248651 | 0 | 0 | 0 | 0 | 0 | 0 |
| ENSG00000250048 | 0 | 0 | 0 | 0 | 0 | 0 |
| ENSG00000249668 | 0 | 0 | 0 | 2 | 1 | 0 |
| ENSG00000250585 | 0 | 0 | 0 | 0 | 0 | 0 |
| ENSG00000199361 | 0 | 0 | 0 | 0 | 0 | 0 |
| ENSG00000283286 | 0 | 0 | 0 | 0 | 0 | 0 |
| ENSG00000285552 | 0 | 0 | 0 | 0 | 0 | 0 |
| ENSG00000285616 | 0 | 0 | 0 | 0 | 0 | 0 |
| ENSG00000113638 | 238 | 114 | 133 | 81 | 151 | 103 |
| ENSG00000199552 | 0 | 0 | 0 | 0 | 0 | 0 |
| ENSG00000171522 | 397 | 189 | 239 | 282 | 316 | 349 |
| ENSG00000132356 | 1955 | 983 | 1228 | 588 | 1072 | 918 |
| ENSG00000212567 | 0 | 0 | 0 | 0 | 0 | 0 |
| ENSG00000145592 | 20718 | 10344 | 11904 | 14107 | 12550 | 11913 |
| ENSG00000212296 | 3 | 2 | 1 | 1 | 0 | 0 |
| ENSG00000132357 | 12 | 5 | 10 | 5 | 8 | 5 |
| ENSG00000112936 | 2 | 0 | 0 | 0 | 0 | 0 |
| ENSG00000253098 | 0 | 0 | 0 | 0 | 0 | 0 |
| ENSG00000248120 | 0 | 0 | 0 | 0 | 0 | 0 |
| ENSG00000171495 | 0 | 0 | 0 | 0 | 0 | 0 |
| ENSG00000039537 | 0 | 0 | 0 | 0 | 0 | 0 |
| ENSG00000182836 | 78 | 45 | 49 | 39 | 118 | 97 |
| ENSG00000251478 | 0 | 0 | 0 | 0 | 0 | 1 |
| ENSG00000083720 | 356 | 207 | 266 | 143 | 197 | 155 |
| ENSG00000248668 | 2 | 2 | 2 | 1 | 1 | 0 |
| ENSG00000240281 | 0 | 0 | 0 | 0 | 0 | 0 |
| ENSG00000205765 | 1044 | 552 | 609 | 303 | 606 | 488 |
| ENSG00000151876 | 364 | 176 | 203 | 158 | 159 | 184 |
| ENSG00000239694 | 0 | 0 | 0 | 0 | 0 | 0 |
| ENSG00000199487 | 0 | 0 | 0 | 0 | 0 | 0 |
| ENSG00000249266 | 0 | 0 | 0 | 0 | 0 | 0 |
| ENSG00000260786 | 0 | 0 | 0 | 0 | 0 | 0 |
| ENSG00000112964 | 194 | 63 | 76 | 19 | 33 | 21 |
| ENSG00000248873 | 2 | 0 | 1 | 0 | 1 | 1 |
| ENSG00000250860 | 0 | 0 | 0 | 0 | 0 | 0 |
| ENSG00000270513 | 0 | 0 | 0 | 0 | 0 | 0 |

| | | | | | | |
|---|---|---|---|---|---|---|
| ENSG00000198865 | 0 | 0 | 0 | 0 | 0 | 0 |
| ENSG00000250722 | 28 | 16 | 23 | 8 | 6 | 11 |
| ENSG00000272234 | 0 | 0 | 0 | 0 | 0 | 0 |
| ENSG00000249092 | 0 | 1 | 0 | 0 | 1 | 0 |
| ENSG00000251163 | 0 | 0 | 0 | 0 | 0 | 0 |
| ENSG00000251656 | 0 | 0 | 0 | 0 | 0 | 0 |
| ENSG00000249516 | 0 | 0 | 0 | 0 | 0 | 0 |
| ENSG00000250711 | 1 | 1 | 0 | 0 | 0 | 0 |
| ENSG00000271788 | 8 | 3 | 7 | 1 | 2 | 4 |
| ENSG00000177738 | 187 | 108 | 178 | 97 | 120 | 124 |
| ENSG00000251131 | 2 | 5 | 0 | 3 | 2 | 3 |
| ENSG00000272382 | 6 | 5 | 4 | 3 | 2 | 6 |
| ENSG00000177721 | 55 | 29 | 42 | 46 | 54 | 33 |
| ENSG00000215068 | 197 | 102 | 109 | 79 | 86 | 108 |
| ENSG00000272144 | 16 | 4 | 6 | 5 | 6 | 0 |
| ENSG00000172262 | 965 | 600 | 616 | 438 | 681 | 573 |
| ENSG00000278487 | 0 | 0 | 0 | 0 | 0 | 0 |
| ENSG00000177453 | 4 | 4 | 8 | 4 | 7 | 5 |
| ENSG00000249779 | 2 | 0 | 0 | 0 | 1 | 1 |
| ENSG00000249888 | 0 | 0 | 0 | 0 | 0 | 1 |
| ENSG00000247763 | 1 | 0 | 3 | 0 | 0 | 1 |
| ENSG00000261604 | 2 | 0 | 1 | 0 | 1 | 0 |
| ENSG00000112972 | 2046 | 1067 | 1282 | 1020 | 1345 | 1327 |
| ENSG00000188850 | 0 | 0 | 0 | 0 | 1 | 0 |
| ENSG00000151882 | 319 | 187 | 239 | 135 | 151 | 193 |
| ENSG00000151881 | 347 | 187 | 228 | 134 | 196 | 192 |
| ENSG00000249492 | 16 | 7 | 11 | 6 | 9 | 5 |
| ENSG00000172244 | 63 | 35 | 51 | 25 | 77 | 41 |
| ENSG00000249855 | 14 | 6 | 9 | 2 | 8 | 9 |
| ENSG00000248554 | 7 | 3 | 5 | 3 | 7 | 6 |
| ENSG00000248240 | 3 | 1 | 1 | 6 | 5 | 3 |
| ENSG00000172239 | 2948 | 1417 | 1707 | 1528 | 1874 | 1749 |
| ENSG00000248092 | 311 | 153 | 214 | 142 | 247 | 218 |
| ENSG00000279557 | 1 | 0 | 4 | 0 | 0 | 0 |
| ENSG00000249286 | 1 | 0 | 0 | 0 | 0 | 1 |
| ENSG00000112992 | 2319 | 1090 | 1378 | 744 | 1134 | 1198 |
| ENSG00000224993 | 4 | 2 | 2 | 1 | 1 | 0 |
| ENSG00000249276 | 0 | 0 | 0 | 0 | 0 | 0 |
| ENSG00000212561 | 0 | 0 | 0 | 0 | 0 | 0 |
| ENSG00000070193 | 0 | 0 | 0 | 0 | 0 | 0 |
| ENSG00000248464 | 0 | 0 | 0 | 0 | 0 | 0 |
| ENSG00000249203 | 0 | 0 | 0 | 0 | 0 | 0 |
| ENSG00000250418 | 0 | 0 | 0 | 0 | 0 | 0 |
| ENSG00000263556 | 0 | 0 | 0 | 0 | 0 | 0 |
| ENSG00000251141 | 0 | 3 | 1 | 0 | 3 | 0 |
| ENSG00000248779 | 0 | 0 | 0 | 0 | 0 | 0 |
| ENSG00000112996 | 1373 | 644 | 755 | 734 | 682 | 749 |
| ENSG00000272335 | 95 | 51 | 46 | 33 | 69 | 46 |
| ENSG00000248148 | 0 | 0 | 0 | 0 | 0 | 0 |
| ENSG00000164588 | 0 | 0 | 0 | 0 | 0 | 0 |

| | | | | | | |
|---|---|---|---|---|---|---|
| ENSG00000250422 | 0 | 0 | 0 | 0 | 0 | 0 |
| ENSG00000250017 | 0 | 0 | 0 | 0 | 0 | 0 |
| ENSG00000250122 | 0 | 2 | 0 | 1 | 0 | 0 |
| ENSG00000170571 | 2830 | 1440 | 1651 | 836 | 1604 | 1287 |
| ENSG00000271455 | 0 | 0 | 0 | 0 | 0 | 0 |
| ENSG00000213956 | 0 | 0 | 0 | 0 | 0 | 1 |
| ENSG00000271752 | 0 | 0 | 1 | 2 | 2 | 0 |
| ENSG00000151883 | 758 | 436 | 483 | 273 | 461 | 377 |
| ENSG00000249405 | 0 | 0 | 0 | 0 | 0 | 0 |
| ENSG00000270830 | 0 | 0 | 0 | 0 | 0 | 0 |
| ENSG00000251601 | 0 | 0 | 0 | 0 | 0 | 0 |
| ENSG00000250360 | 0 | 0 | 0 | 0 | 0 | 0 |
| ENSG00000251573 | 0 | 0 | 0 | 0 | 0 | 0 |
| ENSG00000248918 | 0 | 0 | 0 | 0 | 0 | 0 |
| ENSG00000207060 | 0 | 0 | 0 | 0 | 0 | 0 |
| ENSG00000278734 | 0 | 0 | 0 | 0 | 0 | 0 |
| ENSG00000206606 | 0 | 0 | 0 | 0 | 0 | 0 |
| ENSG00000240627 | 0 | 0 | 0 | 0 | 0 | 0 |
| ENSG00000259663 | 0 | 0 | 0 | 0 | 0 | 0 |
| ENSG00000016082 | 4 | 0 | 0 | 0 | 0 | 0 |
| ENSG00000249338 | 0 | 0 | 0 | 0 | 0 | 0 |
| ENSG00000248891 | 0 | 0 | 0 | 0 | 0 | 0 |
| ENSG00000251125 | 0 | 0 | 0 | 0 | 0 | 0 |
| ENSG00000251873 | 0 | 0 | 0 | 0 | 0 | 0 |
| ENSG00000249270 | 0 | 0 | 0 | 0 | 0 | 0 |
| ENSG00000279470 | 0 | 0 | 0 | 0 | 0 | 0 |
| ENSG00000248789 | 0 | 0 | 0 | 0 | 0 | 0 |
| ENSG00000244621 | 0 | 0 | 0 | 0 | 0 | 0 |
| ENSG00000250827 | 0 | 0 | 0 | 0 | 0 | 0 |
| ENSG00000248898 | 0 | 0 | 0 | 0 | 0 | 0 |
| ENSG00000213949 | 0 | 1 | 0 | 0 | 0 | 1 |
| ENSG00000152684 | 947 | 448 | 552 | 621 | 572 | 594 |
| ENSG00000251237 | 0 | 0 | 0 | 0 | 0 | 0 |
| ENSG00000241809 | 0 | 0 | 0 | 0 | 0 | 0 |
| ENSG00000249899 | 0 | 0 | 0 | 0 | 0 | 0 |
| ENSG00000164171 | 286 | 147 | 141 | 83 | 273 | 151 |
| ENSG00000272123 | 0 | 0 | 2 | 0 | 0 | 0 |
| ENSG00000164172 | 1202 | 614 | 783 | 644 | 749 | 677 |
| ENSG00000247796 | 125 | 69 | 75 | 69 | 79 | 75 |
| ENSG00000213940 | 0 | 0 | 0 | 0 | 0 | 0 |
| ENSG00000285831 | 0 | 0 | 0 | 0 | 0 | 0 |
| ENSG00000240052 | 0 | 0 | 0 | 0 | 0 | 0 |
| ENSG00000134363 | 0 | 1 | 0 | 3 | 1 | 3 |
| ENSG00000164258 | 1113 | 574 | 610 | 602 | 736 | 626 |
| ENSG00000250447 | 0 | 0 | 0 | 0 | 0 | 0 |
| ENSG00000217791 | 0 | 0 | 0 | 0 | 0 | 0 |
| ENSG00000272416 | 2 | 0 | 2 | 0 | 3 | 1 |
| ENSG00000185305 | 531 | 262 | 275 | 247 | 344 | 263 |
| ENSG00000222960 | 0 | 0 | 0 | 0 | 0 | 0 |
| ENSG00000207627 | 0 | 0 | 0 | 0 | 0 | 0 |

| | | | | | | |
|---|---|---|---|---|---|---|
| ENSG00000241230 | 0 | 0 | 0 | 0 | 0 | 0 |
| ENSG00000265421 | 0 | 0 | 0 | 0 | 0 | 0 |
| ENSG00000279638 | 0 | 0 | 0 | 0 | 0 | 0 |
| ENSG00000249069 | 0 | 0 | 0 | 1 | 3 | 0 |
| ENSG00000251597 | 0 | 0 | 0 | 0 | 0 | 0 |
| ENSG00000169271 | 0 | 0 | 0 | 0 | 0 | 0 |
| ENSG00000178996 | 802 | 328 | 428 | 372 | 407 | 409 |
| ENSG00000280159 | 0 | 0 | 0 | 2 | 1 | 2 |
| ENSG00000232517 | 0 | 0 | 0 | 0 | 0 | 0 |
| ENSG00000226810 | 0 | 0 | 0 | 0 | 0 | 0 |
| ENSG00000249843 | 0 | 0 | 0 | 0 | 0 | 0 |
| ENSG00000249016 | 0 | 0 | 0 | 0 | 0 | 0 |
| ENSG00000164283 | 0 | 0 | 0 | 0 | 0 | 0 |
| ENSG00000240535 | 0 | 0 | 0 | 0 | 0 | 0 |
| ENSG00000113088 | 0 | 0 | 0 | 0 | 0 | 0 |
| ENSG00000207229 | 0 | 0 | 0 | 0 | 0 | 0 |
| ENSG00000251376 | 0 | 0 | 0 | 0 | 0 | 0 |
| ENSG00000248029 | 0 | 0 | 0 | 0 | 0 | 0 |
| ENSG00000249454 | 0 | 0 | 0 | 0 | 0 | 0 |
| ENSG00000145649 | 0 | 0 | 0 | 0 | 0 | 0 |
| ENSG00000164287 | 2 | 0 | 0 | 0 | 0 | 1 |
| ENSG00000164294 | 100 | 63 | 64 | 54 | 60 | 60 |
| ENSG00000198983 | 0 | 0 | 0 | 0 | 0 | 0 |
| ENSG00000207728 | 0 | 0 | 0 | 0 | 0 | 0 |
| ENSG00000251856 | 0 | 0 | 0 | 0 | 0 | 0 |
| ENSG00000234602 | 38 | 21 | 20 | 14 | 12 | 11 |
| ENSG00000152669 | 198 | 80 | 93 | 79 | 86 | 72 |
| ENSG00000251307 | 5 | 0 | 1 | 0 | 1 | 2 |
| ENSG00000067248 | 1444 | 851 | 657 | 389 | 998 | 702 |
| ENSG00000039123 | 1054 | 651 | 542 | 289 | 1108 | 572 |
| ENSG00000271410 | 0 | 0 | 0 | 0 | 0 | 0 |
| ENSG00000067113 | 325 | 174 | 182 | 177 | 198 | 166 |
| ENSG00000279883 | 0 | 0 | 0 | 0 | 0 | 0 |
| ENSG00000265135 | 0 | 0 | 0 | 0 | 0 | 0 |
| ENSG00000250853 | 0 | 2 | 0 | 0 | 1 | 0 |
| ENSG00000249410 | 0 | 0 | 0 | 0 | 0 | 0 |
| ENSG00000177058 | 723 | 335 | 401 | 320 | 446 | 367 |
| ENSG00000152670 | 0 | 3 | 1 | 0 | 1 | 5 |
| ENSG00000251368 | 0 | 0 | 0 | 0 | 0 | 0 |
| ENSG00000199322 | 0 | 0 | 0 | 0 | 0 | 0 |
| ENSG00000248854 | 0 | 0 | 0 | 0 | 0 | 0 |
| ENSG00000164509 | 0 | 0 | 0 | 0 | 0 | 0 |
| ENSG00000134352 | 439 | 289 | 202 | 99 | 478 | 185 |
| ENSG00000262211 | 0 | 0 | 0 | 0 | 0 | 0 |
| ENSG00000264281 | 1 | 1 | 3 | 3 | 0 | 2 |
| ENSG00000249236 | 0 | 1 | 0 | 0 | 0 | 0 |
| ENSG00000227908 | 8 | 3 | 3 | 2 | 3 | 1 |
| ENSG00000238326 | 0 | 0 | 0 | 0 | 0 | 0 |
| ENSG00000164512 | 0 | 0 | 0 | 0 | 0 | 0 |
| ENSG00000253032 | 0 | 0 | 0 | 0 | 0 | 0 |

| | | | | | | |
|---|---|---|---|---|---|---|
| ENSG00000243592 | 0 | 0 | 0 | 0 | 0 | 0 |
| ENSG00000223003 | 0 | 0 | 0 | 0 | 0 | 0 |
| ENSG00000251435 | 0 | 0 | 0 | 0 | 0 | 0 |
| ENSG00000212265 | 0 | 0 | 0 | 0 | 0 | 0 |
| ENSG00000248781 | 0 | 0 | 0 | 0 | 0 | 0 |
| ENSG00000210678 | 0 | 0 | 0 | 0 | 0 | 0 |
| ENSG00000249697 | 0 | 0 | 0 | 0 | 0 | 0 |
| ENSG00000250787 | 5 | 1 | 2 | 0 | 1 | 2 |
| ENSG00000248727 | 35 | 28 | 28 | 12 | 20 | 17 |
| ENSG00000226221 | 22 | 15 | 13 | 12 | 17 | 14 |
| ENSG00000225940 | 0 | 0 | 0 | 0 | 0 | 0 |
| ENSG00000234553 | 0 | 0 | 0 | 0 | 0 | 0 |
| ENSG00000228650 | 0 | 0 | 0 | 1 | 0 | 0 |
| ENSG00000095015 | 2107 | 1036 | 1184 | 484 | 957 | 765 |
| ENSG00000237705 | 0 | 1 | 0 | 0 | 0 | 1 |
| ENSG00000225230 | 2 | 1 | 1 | 1 | 1 | 2 |
| ENSG00000155542 | 223 | 145 | 179 | 158 | 139 | 143 |
| ENSG00000155545 | 821 | 463 | 445 | 249 | 572 | 407 |
| ENSG00000271828 | 9 | 4 | 2 | 1 | 10 | 5 |
| ENSG00000235635 | 0 | 0 | 0 | 0 | 0 | 3 |
| ENSG00000233943 | 0 | 0 | 0 | 0 | 0 | 0 |
| ENSG00000248652 | 0 | 0 | 0 | 0 | 0 | 0 |
| ENSG00000250480 | 0 | 0 | 0 | 0 | 0 | 0 |
| ENSG00000207293 | 0 | 0 | 1 | 0 | 0 | 0 |
| ENSG00000062194 | 2122 | 1235 | 1124 | 765 | 1685 | 1118 |
| ENSG00000248586 | 0 | 0 | 2 | 0 | 0 | 0 |
| ENSG00000250961 | 0 | 0 | 0 | 0 | 0 | 0 |
| ENSG00000250308 | 0 | 0 | 0 | 0 | 0 | 0 |
| ENSG00000169067 | 0 | 0 | 0 | 0 | 0 | 0 |
| ENSG00000248294 | 0 | 0 | 0 | 0 | 0 | 0 |
| ENSG00000249436 | 0 | 0 | 0 | 0 | 0 | 0 |
| ENSG00000249198 | 0 | 0 | 0 | 0 | 0 | 0 |
| ENSG00000249584 | 0 | 0 | 0 | 1 | 0 | 0 |
| ENSG00000248132 | 0 | 3 | 0 | 0 | 1 | 2 |
| ENSG00000248271 | 0 | 1 | 2 | 0 | 1 | 0 |
| ENSG00000145632 | 16571 | 7745 | 10909 | 5668 | 6862 | 8242 |
| ENSG00000175857 | 0 | 0 | 0 | 0 | 0 | 0 |
| ENSG00000271257 | 0 | 0 | 0 | 0 | 0 | 0 |
| ENSG00000265699 | 0 | 0 | 0 | 0 | 0 | 0 |
| ENSG00000250127 | 0 | 0 | 0 | 0 | 0 | 0 |
| ENSG00000152932 | 0 | 0 | 0 | 0 | 0 | 0 |
| ENSG00000248475 | 0 | 0 | 0 | 0 | 0 | 0 |
| ENSG00000240412 | 0 | 0 | 0 | 0 | 0 | 0 |
| ENSG00000248733 | 0 | 0 | 0 | 0 | 0 | 0 |
| ENSG00000113448 | 24 | 18 | 15 | 12 | 21 | 8 |
| ENSG00000247345 | 0 | 0 | 0 | 0 | 0 | 0 |
| ENSG00000276214 | 0 | 0 | 0 | 0 | 0 | 0 |
| ENSG00000251306 | 0 | 0 | 0 | 0 | 0 | 0 |
| ENSG00000202601 | 0 | 0 | 0 | 0 | 0 | 0 |
| ENSG00000248935 | 0 | 0 | 0 | 0 | 0 | 0 |

| | | | | | | |
|---|---|---|---|---|---|---|
| ENSG00000202017 | 0 | 0 | 0 | 0 | 0 | 0 |
| ENSG00000213896 | 0 | 0 | 0 | 0 | 0 | 0 |
| ENSG00000214244 | 0 | 0 | 0 | 0 | 0 | 0 |
| ENSG00000273701 | 0 | 0 | 0 | 0 | 0 | 0 |
| ENSG00000152931 | 2 | 0 | 0 | 1 | 0 | 0 |
| ENSG00000275634 | 0 | 0 | 0 | 0 | 0 | 0 |
| ENSG00000276233 | 0 | 0 | 0 | 0 | 0 | 0 |
| ENSG00000035499 | 386 | 185 | 236 | 175 | 305 | 211 |
| ENSG00000251303 | 4 | 1 | 0 | 1 | 0 | 2 |
| ENSG00000250148 | 0 | 3 | 4 | 1 | 3 | 0 |
| ENSG00000164181 | 1636 | 807 | 935 | 510 | 811 | 723 |
| ENSG00000272308 | 4 | 5 | 1 | 0 | 4 | 2 |
| ENSG00000049167 | 378 | 198 | 239 | 204 | 257 | 227 |
| ENSG00000215032 | 3 | 6 | 8 | 3 | 5 | 8 |
| ENSG00000233847 | 1 | 0 | 1 | 1 | 0 | 1 |
| ENSG00000164182 | 558 | 313 | 329 | 444 | 386 | 373 |
| ENSG00000279985 | 0 | 0 | 0 | 0 | 0 | 0 |
| ENSG00000276945 | 0 | 2 | 0 | 1 | 0 | 2 |
| ENSG00000188725 | 1654 | 875 | 1038 | 788 | 980 | 1000 |
| ENSG00000251279 | 0 | 4 | 2 | 0 | 2 | 1 |
| ENSG00000249279 | 0 | 0 | 0 | 0 | 0 | 0 |
| ENSG00000130449 | 801 | 345 | 400 | 300 | 486 | 441 |
| ENSG00000251682 | 0 | 0 | 1 | 1 | 0 | 1 |
| ENSG00000250461 | 6 | 1 | 4 | 1 | 0 | 0 |
| ENSG00000213891 | 0 | 0 | 0 | 0 | 0 | 0 |
| ENSG00000178722 | 2 | 0 | 0 | 0 | 0 | 0 |
| ENSG00000248529 | 0 | 0 | 0 | 0 | 0 | 0 |
| ENSG00000212215 | 0 | 0 | 0 | 0 | 0 | 0 |
| ENSG00000251575 | 0 | 0 | 0 | 0 | 1 | 0 |
| ENSG00000249878 | 0 | 0 | 0 | 0 | 0 | 0 |
| ENSG00000248199 | 0 | 0 | 0 | 0 | 0 | 0 |
| ENSG00000251983 | 0 | 0 | 0 | 0 | 0 | 0 |
| ENSG00000068796 | 974 | 567 | 528 | 358 | 650 | 553 |
| ENSG00000086189 | 1081 | 553 | 603 | 554 | 684 | 737 |
| ENSG00000086200 | 656 | 355 | 387 | 287 | 462 | 343 |
| ENSG00000199279 | 0 | 0 | 0 | 0 | 0 | 0 |
| ENSG00000268942 | 4 | 1 | 0 | 0 | 0 | 0 |
| ENSG00000241991 | 0 | 0 | 0 | 0 | 0 | 0 |
| ENSG00000240436 | 0 | 0 | 0 | 0 | 0 | 0 |
| ENSG00000186105 | 0 | 2 | 0 | 0 | 0 | 0 |
| ENSG00000217416 | 1 | 0 | 1 | 3 | 1 | 0 |
| ENSG00000249579 | 0 | 0 | 0 | 0 | 0 | 0 |
| ENSG00000248185 | 0 | 0 | 0 | 0 | 0 | 0 |
| ENSG00000248285 | 0 | 0 | 0 | 0 | 0 | 0 |
| ENSG00000178394 | 0 | 0 | 0 | 0 | 0 | 0 |
| ENSG00000164197 | 22 | 14 | 18 | 17 | 12 | 16 |
| ENSG00000236068 | 0 | 0 | 0 | 0 | 0 | 0 |
| ENSG00000186479 | 0 | 0 | 0 | 0 | 0 | 0 |
| ENSG00000242547 | 0 | 0 | 0 | 0 | 0 | 0 |
| ENSG00000201869 | 0 | 0 | 0 | 0 | 0 | 0 |

| | | | | | | |
|---|---|---|---|---|---|---|
| ENSG00000224296 | 0 | 0 | 0 | 0 | 0 | 0 |
| ENSG00000145642 | 0 | 0 | 0 | 0 | 0 | 0 |
| ENSG00000153006 | 426 | 194 | 218 | 161 | 262 | 216 |
| ENSG00000153015 | 505 | 267 | 218 | 189 | 394 | 255 |
| ENSG00000272370 | 0 | 2 | 5 | 2 | 1 | 1 |
| ENSG00000272354 | 2 | 0 | 1 | 0 | 0 | 0 |
| ENSG00000207439 | 0 | 0 | 0 | 0 | 0 | 0 |
| ENSG00000049192 | 3 | 6 | 4 | 0 | 6 | 1 |
| ENSG00000248288 | 0 | 0 | 0 | 0 | 0 | 0 |
| ENSG00000250732 | 0 | 0 | 0 | 0 | 0 | 0 |
| ENSG00000205644 | 0 | 0 | 0 | 0 | 0 | 0 |
| ENSG00000250081 | 1 | 0 | 0 | 0 | 0 | 1 |
| ENSG00000123219 | 198 | 124 | 120 | 92 | 165 | 123 |
| ENSG00000113593 | 661 | 326 | 417 | 236 | 501 | 339 |
| ENSG00000113595 | 343 | 201 | 179 | 78 | 313 | 157 |
| ENSG00000207352 | 0 | 0 | 0 | 0 | 0 | 0 |
| ENSG00000113597 | 1048 | 534 | 625 | 348 | 617 | 450 |
| ENSG00000253251 | 89 | 52 | 57 | 34 | 54 | 41 |
| ENSG00000197860 | 143 | 83 | 87 | 66 | 108 | 84 |
| ENSG00000123213 | 1478 | 798 | 875 | 804 | 1069 | 1130 |
| ENSG00000251648 | 0 | 0 | 0 | 0 | 0 | 0 |
| ENSG00000269961 | 28 | 15 | 13 | 12 | 17 | 15 |
| ENSG00000112851 | 2369 | 1122 | 1064 | 505 | 1501 | 839 |
| ENSG00000252904 | 0 | 0 | 0 | 0 | 0 | 0 |
| ENSG00000153914 | 939 | 622 | 472 | 184 | 936 | 508 |
| ENSG00000253744 | 1 | 0 | 0 | 0 | 0 | 1 |
| ENSG00000248846 | 0 | 0 | 0 | 0 | 2 | 1 |
| ENSG00000251391 | 1 | 1 | 0 | 0 | 0 | 0 |
| ENSG00000250313 | 0 | 0 | 0 | 0 | 0 | 0 |
| ENSG00000234838 | 0 | 0 | 0 | 0 | 0 | 0 |
| ENSG00000069020 | 666 | 333 | 558 | 267 | 341 | 458 |
| ENSG00000248803 | 0 | 0 | 0 | 0 | 0 | 0 |
| ENSG00000249057 | 0 | 0 | 0 | 0 | 0 | 0 |
| ENSG00000229666 | 9 | 4 | 13 | 17 | 18 | 16 |
| ENSG00000134061 | 4 | 0 | 1 | 4 | 1 | 2 |
| ENSG00000251206 | 0 | 1 | 2 | 1 | 0 | 3 |
| ENSG00000249364 | 89 | 42 | 41 | 49 | 78 | 64 |
| ENSG00000250978 | 0 | 0 | 0 | 0 | 0 | 0 |
| ENSG00000239870 | 0 | 0 | 0 | 0 | 0 | 0 |
| ENSG00000249736 | 2 | 0 | 0 | 0 | 0 | 0 |
| ENSG00000249238 | 0 | 0 | 0 | 0 | 0 | 0 |
| ENSG00000250669 | 0 | 1 | 1 | 0 | 0 | 0 |
| ENSG00000249721 | 0 | 0 | 0 | 0 | 0 | 0 |
| ENSG00000250421 | 0 | 0 | 0 | 0 | 0 | 0 |
| ENSG00000249894 | 0 | 0 | 0 | 0 | 0 | 0 |
| ENSG00000273860 | 0 | 0 | 0 | 0 | 0 | 0 |
| ENSG00000253801 | 0 | 0 | 0 | 0 | 0 | 0 |
| ENSG00000252108 | 0 | 0 | 0 | 0 | 0 | 0 |
| ENSG00000213864 | 2 | 0 | 0 | 0 | 1 | 0 |
| ENSG00000253787 | 0 | 0 | 0 | 0 | 0 | 0 |

| | | | | | | |
|---|---|---|---|---|---|---|
| ENSG00000145675 | 1767 | 1020 | 947 | 423 | 1148 | 846 |
| ENSG00000251044 | 0 | 0 | 0 | 0 | 0 | 0 |
| ENSG00000248884 | 2 | 0 | 0 | 1 | 1 | 0 |
| ENSG00000249588 | 0 | 0 | 0 | 0 | 0 | 0 |
| ENSG00000248359 | 0 | 0 | 0 | 0 | 0 | 0 |
| ENSG00000249335 | 0 | 0 | 0 | 0 | 0 | 0 |
| ENSG00000250289 | 0 | 0 | 0 | 0 | 0 | 0 |
| ENSG00000212249 | 0 | 0 | 0 | 0 | 0 | 0 |
| ENSG00000250066 | 0 | 0 | 0 | 0 | 0 | 0 |
| ENSG00000276309 | 0 | 0 | 0 | 0 | 0 | 0 |
| ENSG00000249352 | 0 | 1 | 2 | 0 | 0 | 0 |
| ENSG00000250237 | 0 | 3 | 0 | 0 | 1 | 0 |
| ENSG00000249183 | 0 | 0 | 0 | 0 | 0 | 0 |
| ENSG00000145740 | 2468 | 1252 | 1580 | 1079 | 1382 | 1268 |
| ENSG00000248664 | 0 | 3 | 2 | 0 | 1 | 1 |
| ENSG00000220986 | 0 | 0 | 0 | 0 | 0 | 0 |
| ENSG00000280894 | 0 | 0 | 0 | 0 | 0 | 0 |
| ENSG00000134057 | 3520 | 1665 | 2032 | 2340 | 2586 | 2375 |
| ENSG00000280187 | 281 | 152 | 178 | 96 | 152 | 154 |
| ENSG00000153044 | 508 | 237 | 269 | 265 | 285 | 310 |
| ENSG00000134056 | 1082 | 538 | 634 | 543 | 585 | 559 |
| ENSG00000134058 | 2352 | 1164 | 1443 | 1284 | 1455 | 1275 |
| ENSG00000183323 | 1712 | 827 | 1072 | 570 | 743 | 798 |
| ENSG00000213830 | 31 | 19 | 30 | 6 | 19 | 19 |
| ENSG00000215006 | 4 | 2 | 6 | 8 | 4 | 6 |
| ENSG00000271045 | 1 | 1 | 0 | 1 | 0 | 0 |
| ENSG00000085231 | 336 | 215 | 239 | 199 | 329 | 220 |
| ENSG00000273841 | 1716 | 868 | 1048 | 831 | 1234 | 1084 |
| ENSG00000152942 | 1064 | 593 | 636 | 425 | 798 | 625 |
| ENSG00000152939 | 2341 | 1110 | 1395 | 919 | 1154 | 1196 |
| ENSG00000244717 | 0 | 0 | 0 | 0 | 0 | 0 |
| ENSG00000249295 | 1 | 3 | 2 | 1 | 0 | 0 |
| ENSG00000266477 | 0 | 0 | 0 | 0 | 0 | 0 |
| ENSG00000197822 | 1649 | 832 | 995 | 539 | 870 | 788 |
| ENSG00000212260 | 1 | 0 | 0 | 0 | 0 | 1 |
| ENSG00000183474 | 342 | 147 | 183 | 138 | 155 | 151 |
| ENSG00000248477 | 0 | 1 | 1 | 0 | 0 | 0 |
| ENSG00000250138 | 13 | 3 | 8 | 0 | 2 | 7 |
| ENSG00000253203 | 2 | 0 | 0 | 4 | 2 | 2 |
| ENSG00000248769 | 0 | 0 | 0 | 0 | 0 | 0 |
| ENSG00000198237 | 0 | 4 | 1 | 2 | 0 | 2 |
| ENSG00000251158 | 0 | 0 | 0 | 0 | 0 | 0 |
| ENSG00000249230 | 0 | 0 | 0 | 0 | 0 | 0 |
| ENSG00000205572 | 33 | 16 | 24 | 20 | 22 | 22 |
| ENSG00000205571 | 214 | 150 | 157 | 121 | 140 | 161 |
| ENSG00000285151 | 0 | 0 | 0 | 0 | 0 | 0 |
| ENSG00000179978 | 22 | 7 | 15 | 2 | 7 | 10 |
| ENSG00000253492 | 0 | 0 | 0 | 0 | 0 | 0 |
| ENSG00000254701 | 8 | 1 | 11 | 2 | 4 | 3 |
| ENSG00000253816 | 15 | 4 | 7 | 3 | 7 | 13 |

| | | | | | | |
|---|---|---|---|---|---|---|
| ENSG00000226259 | 35 | 10 | 21 | 10 | 15 | 13 |
| ENSG00000269983 | 5 | 1 | 3 | 2 | 1 | 3 |
| ENSG00000250687 | 0 | 0 | 0 | 0 | 0 | 0 |
| ENSG00000253333 | 5 | 0 | 0 | 1 | 0 | 0 |
| ENSG00000196302 | 5 | 0 | 0 | 0 | 0 | 0 |
| ENSG00000253366 | 0 | 0 | 0 | 0 | 0 | 1 |
| ENSG00000254335 | 0 | 0 | 0 | 0 | 0 | 0 |
| ENSG00000172058 | 0 | 0 | 0 | 0 | 0 | 0 |
| ENSG00000172062 | 389 | 193 | 204 | 192 | 191 | 220 |
| ENSG00000285204 | 0 | 0 | 0 | 0 | 0 | 0 |
| ENSG00000249437 | 23 | 14 | 16 | 0 | 9 | 13 |
| ENSG00000145736 | 216 | 106 | 117 | 88 | 119 | 129 |
| ENSG00000230847 | 7 | 5 | 8 | 2 | 3 | 10 |
| ENSG00000251634 | 16 | 12 | 7 | 3 | 6 | 4 |
| ENSG00000253900 | 0 | 0 | 0 | 0 | 0 | 0 |
| ENSG00000215630 | 10 | 2 | 3 | 4 | 4 | 1 |
| ENSG00000235558 | 2 | 1 | 1 | 0 | 0 | 1 |
| ENSG00000250387 | 0 | 0 | 0 | 0 | 0 | 0 |
| ENSG00000253985 | 0 | 0 | 0 | 0 | 0 | 0 |
| ENSG00000169040 | 0 | 0 | 0 | 0 | 0 | 0 |
| ENSG00000249981 | 0 | 0 | 0 | 0 | 0 | 0 |
| ENSG00000145734 | 318 | 192 | 185 | 67 | 439 | 218 |
| ENSG00000253536 | 0 | 0 | 0 | 0 | 0 | 0 |
| ENSG00000247911 | 0 | 1 | 0 | 0 | 0 | 0 |
| ENSG00000131844 | 10873 | 5237 | 6519 | 3958 | 4947 | 5481 |
| ENSG00000164326 | 0 | 0 | 0 | 0 | 0 | 0 |
| ENSG00000278824 | 0 | 0 | 0 | 0 | 0 | 0 |
| ENSG00000285804 | 0 | 0 | 1 | 0 | 1 | 0 |
| ENSG00000131711 | 146 | 71 | 77 | 41 | 79 | 77 |
| ENSG00000264099 | 0 | 0 | 0 | 0 | 0 | 0 |
| ENSG00000113048 | 2389 | 1090 | 1517 | 1243 | 1373 | 1459 |
| ENSG00000244748 | 2 | 1 | 1 | 0 | 0 | 0 |
| ENSG00000049883 | 400 | 218 | 234 | 152 | 290 | 224 |
| ENSG00000244061 | 0 | 0 | 0 | 0 | 0 | 0 |
| ENSG00000251108 | 0 | 0 | 0 | 0 | 0 | 0 |
| ENSG00000261269 | 0 | 0 | 0 | 0 | 0 | 0 |
| ENSG00000178175 | 0 | 0 | 0 | 0 | 0 | 0 |
| ENSG00000248371 | 0 | 0 | 0 | 0 | 0 | 0 |
| ENSG00000251613 | 3 | 0 | 2 | 1 | 2 | 1 |
| ENSG00000226544 | 2 | 2 | 2 | 1 | 2 | 2 |
| ENSG00000223497 | 0 | 0 | 0 | 0 | 0 | 0 |
| ENSG00000206770 | 0 | 0 | 0 | 0 | 0 | 0 |
| ENSG00000249085 | 1 | 1 | 1 | 0 | 3 | 2 |
| ENSG00000083312 | 3310 | 1728 | 1639 | 980 | 2528 | 1481 |
| ENSG00000263593 | 0 | 0 | 0 | 0 | 0 | 0 |
| ENSG00000242602 | 2 | 0 | 0 | 0 | 0 | 0 |
| ENSG00000271926 | 7 | 5 | 0 | 1 | 4 | 5 |
| ENSG00000272081 | 20 | 9 | 10 | 6 | 5 | 5 |
| ENSG00000157107 | 457 | 231 | 220 | 106 | 328 | 193 |
| ENSG00000251467 | 2 | 0 | 0 | 0 | 1 | 0 |

| | | | | | | |
|---|---|---|---|---|---|---|
| ENSG00000248930 | 0 | 0 | 0 | 0 | 0 | 0 |
| ENSG00000157111 | 4 | 3 | 2 | 8 | 5 | 7 |
| ENSG00000251599 | 0 | 0 | 0 | 0 | 0 | 0 |
| ENSG00000250747 | 0 | 0 | 0 | 0 | 0 | 0 |
| ENSG00000164325 | 0 | 0 | 0 | 0 | 0 | 0 |
| ENSG00000249743 | 0 | 0 | 0 | 0 | 0 | 0 |
| ENSG00000251543 | 0 | 0 | 0 | 0 | 0 | 0 |
| ENSG00000249149 | 0 | 0 | 0 | 0 | 0 | 0 |
| ENSG00000251493 | 106 | 71 | 92 | 72 | 71 | 74 |
| ENSG00000247993 | 0 | 0 | 0 | 0 | 0 | 0 |
| ENSG00000251604 | 2 | 0 | 1 | 0 | 0 | 0 |
| ENSG00000251324 | 0 | 0 | 0 | 0 | 0 | 0 |
| ENSG00000272525 | 70 | 17 | 36 | 30 | 26 | 31 |
| ENSG00000145741 | 14232 | 7300 | 8517 | 7780 | 8881 | 8334 |
| ENSG00000255883 | 0 | 0 | 1 | 0 | 0 | 2 |
| ENSG00000164331 | 720 | 333 | 461 | 311 | 435 | 372 |
| ENSG00000164338 | 728 | 322 | 367 | 311 | 447 | 388 |
| ENSG00000214944 | 449 | 180 | 197 | 144 | 204 | 188 |
| ENSG00000249293 | 0 | 0 | 0 | 0 | 0 | 0 |
| ENSG00000184084 | 0 | 0 | 0 | 0 | 0 | 0 |
| ENSG00000238913 | 0 | 0 | 0 | 0 | 0 | 0 |
| ENSG00000259968 | 0 | 0 | 0 | 0 | 0 | 0 |
| ENSG00000248474 | 0 | 0 | 0 | 0 | 0 | 0 |
| ENSG00000248942 | 0 | 0 | 0 | 0 | 0 | 0 |
| ENSG00000244326 | 0 | 0 | 0 | 0 | 0 | 0 |
| ENSG00000251041 | 0 | 0 | 0 | 0 | 0 | 0 |
| ENSG00000249343 | 0 | 0 | 0 | 0 | 0 | 0 |
| ENSG00000250446 | 0 | 0 | 0 | 0 | 0 | 0 |
| ENSG00000248673 | 0 | 0 | 0 | 0 | 0 | 0 |
| ENSG00000171617 | 2132 | 1141 | 1373 | 863 | 1254 | 1126 |
| ENSG00000049860 | 4684 | 2250 | 2718 | 2371 | 2443 | 2445 |
| ENSG00000164347 | 1468 | 742 | 881 | 707 | 923 | 782 |
| ENSG00000207336 | 3 | 1 | 0 | 1 | 2 | 2 |
| ENSG00000164346 | 3182 | 1769 | 1939 | 1963 | 2225 | 1870 |
| ENSG00000198780 | 116 | 60 | 43 | 30 | 69 | 63 |
| ENSG00000199645 | 0 | 0 | 0 | 0 | 0 | 0 |
| ENSG00000250071 | 2 | 0 | 0 | 0 | 1 | 0 |
| ENSG00000271714 | 3 | 4 | 3 | 1 | 4 | 4 |
| ENSG00000212363 | 0 | 0 | 0 | 0 | 0 | 0 |
| ENSG00000249856 | 0 | 0 | 0 | 0 | 0 | 0 |
| ENSG00000196390 | 0 | 0 | 0 | 0 | 0 | 0 |
| ENSG00000239517 | 0 | 0 | 0 | 0 | 0 | 0 |
| ENSG00000182383 | 0 | 0 | 0 | 0 | 0 | 0 |
| ENSG00000249157 | 0 | 0 | 0 | 0 | 0 | 0 |
| ENSG00000176928 | 63 | 33 | 38 | 18 | 14 | 21 |
| ENSG00000250889 | 4 | 0 | 0 | 0 | 5 | 1 |
| ENSG00000145700 | 2 | 4 | 8 | 0 | 9 | 5 |
| ENSG00000251663 | 0 | 0 | 0 | 0 | 0 | 0 |
| ENSG00000247372 | 0 | 0 | 1 | 2 | 2 | 0 |
| ENSG00000113161 | 3135 | 1515 | 2037 | 1569 | 2017 | 1832 |

| | | | | | | |
|---|---|---|---|---|---|---|
| ENSG00000113163 | 1732 | 852 | 853 | 410 | 961 | 782 |
| ENSG00000271815 | 1 | 1 | 4 | 0 | 1 | 0 |
| ENSG00000242198 | 0 | 0 | 0 | 0 | 0 | 0 |
| ENSG00000122008 | 166 | 93 | 83 | 32 | 168 | 74 |
| ENSG00000251724 | 2 | 0 | 0 | 0 | 0 | 0 |
| ENSG00000248881 | 0 | 0 | 0 | 0 | 0 | 0 |
| ENSG00000272040 | 0 | 0 | 1 | 3 | 1 | 2 |
| ENSG00000189045 | 0 | 0 | 2 | 0 | 1 | 2 |
| ENSG00000152359 | 380 | 208 | 248 | 160 | 281 | 241 |
| ENSG00000207333 | 0 | 0 | 0 | 0 | 0 | 0 |
| ENSG00000251078 | 0 | 0 | 0 | 0 | 0 | 0 |
| ENSG00000251419 | 0 | 0 | 0 | 0 | 0 | 0 |
| ENSG00000249301 | 0 | 0 | 0 | 0 | 0 | 0 |
| ENSG00000248684 | 0 | 0 | 0 | 0 | 0 | 0 |
| ENSG00000228367 | 0 | 0 | 0 | 0 | 0 | 0 |
| ENSG00000249777 | 0 | 0 | 0 | 0 | 0 | 0 |
| ENSG00000251342 | 0 | 0 | 0 | 0 | 0 | 0 |
| ENSG00000248127 | 0 | 0 | 0 | 0 | 0 | 0 |
| ENSG00000122012 | 0 | 0 | 1 | 0 | 0 | 0 |
| ENSG00000252833 | 0 | 0 | 0 | 0 | 0 | 0 |
| ENSG00000251668 | 0 | 0 | 0 | 0 | 0 | 0 |
| ENSG00000254893 | 9 | 9 | 4 | 2 | 6 | 8 |
| ENSG00000249014 | 0 | 1 | 1 | 2 | 0 | 2 |
| ENSG00000251107 | 0 | 0 | 0 | 0 | 0 | 0 |
| ENSG00000250348 | 0 | 0 | 0 | 0 | 0 | 0 |
| ENSG00000251235 | 0 | 0 | 0 | 0 | 1 | 1 |
| ENSG00000145703 | 0 | 0 | 0 | 0 | 0 | 0 |
| ENSG00000250574 | 0 | 0 | 0 | 0 | 0 | 0 |
| ENSG00000249713 | 0 | 0 | 0 | 0 | 0 | 0 |
| ENSG00000164220 | 0 | 1 | 0 | 0 | 0 | 0 |
| ENSG00000225407 | 0 | 0 | 0 | 0 | 0 | 0 |
| ENSG00000277188 | 0 | 0 | 0 | 0 | 0 | 0 |
| ENSG00000278237 | 0 | 0 | 0 | 0 | 0 | 0 |
| ENSG00000181104 | 61 | 23 | 24 | 23 | 34 | 10 |
| ENSG00000164251 | 226 | 104 | 133 | 85 | 109 | 101 |
| ENSG00000264391 | 0 | 0 | 0 | 0 | 0 | 0 |
| ENSG00000171643 | 0 | 0 | 0 | 0 | 0 | 0 |
| ENSG00000252684 | 0 | 0 | 0 | 0 | 0 | 0 |
| ENSG00000145708 | 0 | 0 | 0 | 0 | 0 | 0 |
| ENSG00000164252 | 649 | 343 | 413 | 265 | 505 | 366 |
| ENSG00000285000 | 0 | 0 | 0 | 0 | 1 | 0 |
| ENSG00000132846 | 206 | 114 | 136 | 83 | 106 | 102 |
| ENSG00000250615 | 0 | 0 | 0 | 0 | 0 | 0 |
| ENSG00000238961 | 2 | 1 | 0 | 1 | 0 | 0 |
| ENSG00000250802 | 45 | 31 | 38 | 15 | 31 | 33 |
| ENSG00000284762 | 0 | 0 | 0 | 0 | 0 | 0 |
| ENSG00000251566 | 0 | 0 | 0 | 0 | 0 | 0 |
| ENSG00000113231 | 26 | 19 | 11 | 12 | 28 | 15 |
| ENSG00000251400 | 0 | 2 | 1 | 1 | 0 | 4 |
| ENSG00000164253 | 1193 | 552 | 688 | 547 | 693 | 658 |

| | | | | | | |
|---|---|---|---|---|---|---|
| ENSG00000244019 | 0 | 0 | 0 | 0 | 1 | 0 |
| ENSG00000244363 | 2 | 4 | 3 | 3 | 8 | 3 |
| ENSG00000171540 | 6 | 1 | 4 | 0 | 3 | 3 |
| ENSG00000171530 | 4527 | 2231 | 2671 | 2461 | 2571 | 2590 |
| ENSG00000213763 | 2 | 1 | 1 | 1 | 1 | 0 |
| ENSG00000253572 | 0 | 0 | 0 | 0 | 0 | 0 |
| ENSG00000132842 | 2072 | 1062 | 1039 | 682 | 1350 | 941 |
| ENSG00000253558 | 0 | 0 | 0 | 0 | 0 | 0 |
| ENSG00000249372 | 0 | 0 | 0 | 0 | 0 | 0 |
| ENSG00000251605 | 0 | 0 | 0 | 0 | 0 | 0 |
| ENSG00000200331 | 0 | 0 | 0 | 0 | 0 | 0 |
| ENSG00000206714 | 0 | 0 | 0 | 0 | 0 | 0 |
| ENSG00000245556 | 561 | 249 | 332 | 293 | 317 | 299 |
| ENSG00000085365 | 2004 | 1039 | 1078 | 712 | 1203 | 1010 |
| ENSG00000145685 | 1963 | 887 | 1112 | 752 | 1047 | 1022 |
| ENSG00000248909 | 2 | 1 | 0 | 0 | 0 | 0 |
| ENSG00000254170 | 5 | 0 | 0 | 1 | 1 | 0 |
| ENSG00000254310 | 0 | 0 | 0 | 0 | 0 | 0 |
| ENSG00000113273 | 329 | 157 | 181 | 97 | 157 | 163 |
| ENSG00000132837 | 13 | 8 | 13 | 7 | 3 | 5 |
| ENSG00000214890 | 0 | 0 | 0 | 0 | 0 | 0 |
| ENSG00000132840 | 0 | 0 | 0 | 1 | 0 | 0 |
| ENSG00000145692 | 0 | 3 | 0 | 0 | 1 | 0 |
| ENSG00000201955 | 0 | 0 | 0 | 0 | 0 | 0 |
| ENSG00000249829 | 0 | 0 | 0 | 0 | 0 | 0 |
| ENSG00000206592 | 0 | 3 | 2 | 0 | 0 | 0 |
| ENSG00000250330 | 1 | 1 | 1 | 1 | 0 | 0 |
| ENSG00000152409 | 1009 | 467 | 501 | 272 | 585 | 514 |
| ENSG00000213757 | 5 | 1 | 0 | 0 | 1 | 0 |
| ENSG00000152413 | 494 | 241 | 297 | 180 | 294 | 280 |
| ENSG00000213755 | 0 | 0 | 0 | 0 | 0 | 0 |
| ENSG00000243160 | 0 | 0 | 0 | 0 | 0 | 0 |
| ENSG00000207391 | 0 | 0 | 0 | 0 | 0 | 0 |
| ENSG00000277818 | 0 | 0 | 0 | 0 | 0 | 0 |
| ENSG00000164329 | 1070 | 534 | 597 | 278 | 664 | 493 |
| ENSG00000164309 | 162 | 82 | 129 | 22 | 93 | 87 |
| ENSG00000250258 | 6 | 2 | 7 | 11 | 10 | 8 |
| ENSG00000238254 | 0 | 0 | 0 | 0 | 0 | 0 |
| ENSG00000250888 | 0 | 0 | 0 | 0 | 0 | 0 |
| ENSG00000177034 | 1209 | 526 | 656 | 296 | 632 | 609 |
| ENSG00000113296 | 5 | 6 | 5 | 2 | 5 | 0 |
| ENSG00000251050 | 0 | 0 | 0 | 0 | 0 | 0 |
| ENSG00000249778 | 0 | 0 | 0 | 0 | 0 | 0 |
| ENSG00000249825 | 2 | 2 | 3 | 2 | 2 | 4 |
| ENSG00000164300 | 2577 | 1209 | 1662 | 973 | 1533 | 1641 |
| ENSG00000251675 | 2 | 2 | 1 | 0 | 0 | 0 |
| ENSG00000249438 | 1 | 0 | 0 | 0 | 0 | 0 |
| ENSG00000184188 | 0 | 0 | 0 | 0 | 0 | 1 |
| ENSG00000251828 | 0 | 0 | 0 | 0 | 0 | 0 |
| ENSG00000244076 | 0 | 0 | 0 | 0 | 0 | 0 |

| | | | | | | |
|---|---|---|---|---|---|---|
| ENSG00000164299 | 0 | 0 | 0 | 0 | 0 | 0 |
| ENSG00000227087 | 0 | 0 | 2 | 0 | 1 | 0 |
| ENSG00000248569 | 4 | 5 | 7 | 6 | 5 | 4 |
| ENSG00000220157 | 0 | 0 | 0 | 0 | 3 | 1 |
| ENSG00000206774 | 0 | 0 | 0 | 0 | 0 | 0 |
| ENSG00000039319 | 848 | 494 | 402 | 241 | 876 | 411 |
| ENSG00000249042 | 613 | 320 | 336 | 282 | 398 | 303 |
| ENSG00000152380 | 54 | 33 | 39 | 13 | 30 | 35 |
| ENSG00000242706 | 0 | 0 | 0 | 0 | 0 | 0 |
| ENSG00000240003 | 2 | 1 | 2 | 2 | 2 | 2 |
| ENSG00000189127 | 260 | 102 | 103 | 70 | 175 | 103 |
| ENSG00000251366 | 0 | 0 | 0 | 0 | 0 | 0 |
| ENSG00000251221 | 0 | 3 | 0 | 0 | 0 | 1 |
| ENSG00000228716 | 1468 | 729 | 844 | 539 | 771 | 792 |
| ENSG00000249655 | 5 | 3 | 4 | 5 | 1 | 1 |
| ENSG00000271043 | 1 | 0 | 0 | 0 | 0 | 0 |
| ENSG00000113318 | 592 | 278 | 285 | 135 | 342 | 271 |
| ENSG00000248967 | 0 | 0 | 0 | 0 | 1 | 0 |
| ENSG00000244630 | 0 | 0 | 0 | 0 | 0 | 0 |
| ENSG00000251450 | 0 | 0 | 0 | 0 | 0 | 0 |
| ENSG00000113319 | 35 | 12 | 17 | 11 | 14 | 21 |
| ENSG00000251001 | 0 | 0 | 0 | 0 | 0 | 0 |
| ENSG00000249772 | 0 | 0 | 0 | 0 | 0 | 0 |
| ENSG00000247572 | 177 | 111 | 136 | 103 | 118 | 125 |
| ENSG00000131730 | 16 | 7 | 8 | 7 | 6 | 9 |
| ENSG00000248794 | 9 | 4 | 3 | 1 | 4 | 0 |
| ENSG00000131732 | 680 | 369 | 400 | 408 | 438 | 396 |
| ENSG00000172497 | 0 | 1 | 0 | 0 | 1 | 5 |
| ENSG00000145687 | 1153 | 515 | 696 | 526 | 514 | 621 |
| ENSG00000249483 | 0 | 0 | 0 | 0 | 2 | 0 |
| ENSG00000214857 | 1 | 1 | 0 | 0 | 0 | 0 |
| ENSG00000152348 | 445 | 197 | 264 | 182 | 251 | 237 |
| ENSG00000236447 | 0 | 0 | 0 | 1 | 0 | 0 |
| ENSG00000251495 | 5 | 4 | 2 | 1 | 1 | 0 |
| ENSG00000248192 | 0 | 0 | 0 | 0 | 0 | 0 |
| ENSG00000265684 | 0 | 0 | 0 | 0 | 0 | 0 |
| ENSG00000251374 | 0 | 1 | 0 | 0 | 0 | 0 |
| ENSG00000186468 | 5613 | 4611 | 5471 | 6483 | 6602 | 5865 |
| ENSG00000205464 | 100 | 30 | 70 | 46 | 34 | 30 |
| ENSG00000248393 | 0 | 0 | 0 | 0 | 0 | 0 |
| ENSG00000248870 | 0 | 0 | 0 | 0 | 0 | 0 |
| ENSG00000249100 | 0 | 0 | 0 | 0 | 0 | 0 |
| ENSG00000240411 | 0 | 0 | 0 | 0 | 0 | 0 |
| ENSG00000248105 | 0 | 0 | 0 | 0 | 0 | 0 |
| ENSG00000283359 | 0 | 0 | 0 | 0 | 0 | 0 |
| ENSG00000281327 | 0 | 0 | 0 | 0 | 0 | 0 |
| ENSG00000248112 | 0 | 0 | 0 | 0 | 0 | 0 |
| ENSG00000249857 | 0 | 0 | 0 | 0 | 0 | 0 |
| ENSG00000248400 | 0 | 0 | 0 | 0 | 0 | 0 |
| ENSG00000249664 | 4 | 0 | 0 | 0 | 0 | 0 |

| | | | | | | |
|---|---|---|---|---|---|---|
| ENSG00000271862 | 7 | 8 | 9 | 3 | 6 | 6 |
| ENSG00000174695 | 2886 | 1596 | 1594 | 1180 | 2031 | 1520 |
| ENSG00000238835 | 0 | 0 | 0 | 0 | 0 | 0 |
| ENSG00000152422 | 122 | 75 | 73 | 44 | 105 | 63 |
| ENSG00000243385 | 0 | 0 | 0 | 0 | 0 | 0 |
| ENSG00000251552 | 0 | 0 | 0 | 0 | 0 | 0 |
| ENSG00000251395 | 0 | 0 | 0 | 0 | 0 | 0 |
| ENSG00000038427 | 0 | 0 | 0 | 0 | 0 | 0 |
| ENSG00000249835 | 0 | 0 | 0 | 0 | 0 | 0 |
| ENSG00000145681 | 0 | 0 | 1 | 0 | 0 | 0 |
| ENSG00000253073 | 0 | 0 | 0 | 0 | 0 | 0 |
| ENSG00000206744 | 0 | 0 | 0 | 0 | 0 | 0 |
| ENSG00000242858 | 0 | 0 | 0 | 1 | 0 | 0 |
| ENSG00000202157 | 0 | 0 | 0 | 0 | 0 | 0 |
| ENSG00000250013 | 0 | 0 | 0 | 0 | 0 | 0 |
| ENSG00000164176 | 0 | 0 | 0 | 0 | 0 | 0 |
| ENSG00000252861 | 0 | 0 | 0 | 0 | 0 | 0 |
| ENSG00000250320 | 0 | 0 | 0 | 0 | 0 | 0 |
| ENSG00000214853 | 0 | 0 | 0 | 0 | 0 | 0 |
| ENSG00000249349 | 0 | 0 | 0 | 0 | 0 | 0 |
| ENSG00000248674 | 0 | 0 | 0 | 0 | 0 | 0 |
| ENSG00000250253 | 0 | 0 | 0 | 0 | 0 | 0 |
| ENSG00000249792 | 0 | 0 | 0 | 0 | 0 | 0 |
| ENSG00000248170 | 0 | 0 | 0 | 0 | 0 | 0 |
| ENSG00000248667 | 0 | 0 | 0 | 0 | 0 | 0 |
| ENSG00000249023 | 0 | 0 | 0 | 0 | 0 | 0 |
| ENSG00000249842 | 0 | 0 | 0 | 0 | 0 | 0 |
| ENSG00000243859 | 0 | 0 | 0 | 0 | 1 | 1 |
| ENSG00000251066 | 0 | 1 | 0 | 1 | 1 | 1 |
| ENSG00000250874 | 9 | 5 | 7 | 5 | 4 | 7 |
| ENSG00000241965 | 0 | 0 | 0 | 0 | 0 | 0 |
| ENSG00000249153 | 0 | 0 | 0 | 0 | 0 | 0 |
| ENSG00000250359 | 0 | 0 | 0 | 0 | 0 | 0 |
| ENSG00000205449 | 1 | 0 | 1 | 2 | 0 | 3 |
| ENSG00000280009 | 0 | 0 | 0 | 0 | 0 | 0 |
| ENSG00000248195 | 0 | 0 | 0 | 0 | 0 | 0 |
| ENSG00000248363 | 0 | 0 | 0 | 0 | 0 | 0 |
| ENSG00000250124 | 0 | 0 | 0 | 0 | 0 | 0 |
| ENSG00000251585 | 0 | 0 | 0 | 0 | 0 | 0 |
| ENSG00000127184 | 7268 | 3809 | 4450 | 4400 | 4241 | 4082 |
| ENSG00000284078 | 0 | 0 | 0 | 0 | 0 | 0 |
| ENSG00000207013 | 0 | 0 | 0 | 0 | 0 | 0 |
| ENSG00000250544 | 2 | 0 | 0 | 0 | 1 | 0 |
| ENSG00000248701 | 0 | 0 | 0 | 0 | 0 | 0 |
| ENSG00000242477 | 0 | 0 | 0 | 1 | 2 | 1 |
| ENSG00000249061 | 0 | 0 | 0 | 0 | 0 | 0 |
| ENSG00000251951 | 0 | 0 | 0 | 0 | 0 | 0 |
| ENSG00000265919 | 0 | 0 | 0 | 0 | 0 | 0 |
| ENSG00000233828 | 0 | 0 | 0 | 0 | 0 | 0 |
| ENSG00000145715 | 1258 | 632 | 770 | 336 | 820 | 613 |

| | | | | | | |
|---|---|---|---|---|---|---|
| ENSG00000241243 | 0 | 0 | 0 | 0 | 0 | 0 |
| ENSG00000207452 | 0 | 0 | 0 | 0 | 0 | 0 |
| ENSG00000134480 | 826 | 443 | 460 | 416 | 596 | 435 |
| ENSG00000285190 | 0 | 0 | 1 | 0 | 0 | 0 |
| ENSG00000200462 | 0 | 0 | 0 | 0 | 0 | 0 |
| ENSG00000249362 | 0 | 0 | 0 | 0 | 0 | 0 |
| ENSG00000248708 | 0 | 0 | 0 | 0 | 0 | 0 |
| ENSG00000164180 | 882 | 429 | 503 | 299 | 642 | 443 |
| ENSG00000247828 | 179 | 108 | 110 | 50 | 114 | 81 |
| ENSG00000207129 | 4 | 2 | 0 | 0 | 2 | 2 |
| ENSG00000240083 | 0 | 0 | 0 | 0 | 0 | 0 |
| ENSG00000206958 | 0 | 0 | 0 | 0 | 0 | 0 |
| ENSG00000250306 | 0 | 0 | 0 | 0 | 0 | 0 |
| ENSG00000250156 | 0 | 0 | 0 | 0 | 0 | 0 |
| ENSG00000271904 | 0 | 0 | 0 | 0 | 0 | 0 |
| ENSG00000245526 | 129 | 76 | 98 | 35 | 70 | 79 |
| ENSG00000250705 | 0 | 0 | 0 | 0 | 0 | 0 |
| ENSG00000284447 | 0 | 0 | 0 | 0 | 0 | 0 |
| ENSG00000245864 | 0 | 0 | 0 | 0 | 0 | 0 |
| ENSG00000250377 | 0 | 0 | 0 | 0 | 0 | 0 |
| ENSG00000250555 | 0 | 0 | 0 | 0 | 0 | 0 |
| ENSG00000081189 | 2 | 3 | 10 | 3 | 1 | 4 |
| ENSG00000248309 | 0 | 0 | 0 | 0 | 0 | 0 |
| ENSG00000250831 | 0 | 0 | 0 | 0 | 0 | 0 |
| ENSG00000250437 | 0 | 0 | 0 | 0 | 0 | 0 |
| ENSG00000214942 | 0 | 0 | 0 | 0 | 0 | 0 |
| ENSG00000249265 | 0 | 0 | 0 | 0 | 0 | 0 |
| ENSG00000264342 | 0 | 0 | 0 | 0 | 0 | 0 |
| ENSG00000248555 | 0 | 0 | 0 | 0 | 0 | 0 |
| ENSG00000253321 | 0 | 0 | 0 | 1 | 2 | 0 |
| ENSG00000260871 | 0 | 0 | 0 | 0 | 1 | 0 |
| ENSG00000153140 | 666 | 333 | 404 | 271 | 429 | 347 |
| ENSG00000255647 | 5 | 4 | 4 | 2 | 4 | 2 |
| ENSG00000176055 | 302 | 150 | 186 | 144 | 183 | 166 |
| ENSG00000113356 | 105 | 73 | 62 | 63 | 105 | 91 |
| ENSG00000176018 | 744 | 422 | 422 | 179 | 571 | 363 |
| ENSG00000164199 | 364 | 199 | 192 | 138 | 321 | 203 |
| ENSG00000249514 | 0 | 0 | 0 | 0 | 1 | 0 |
| ENSG00000248323 | 12 | 7 | 4 | 8 | 8 | 7 |
| ENSG00000285618 | 0 | 1 | 0 | 0 | 0 | 0 |
| ENSG00000212930 | 0 | 0 | 0 | 0 | 0 | 0 |
| ENSG00000251093 | 0 | 0 | 0 | 0 | 0 | 0 |
| ENSG00000199643 | 0 | 0 | 0 | 0 | 0 | 0 |
| ENSG00000234292 | 0 | 0 | 1 | 0 | 0 | 0 |
| ENSG00000277001 | 0 | 0 | 0 | 0 | 0 | 0 |
| ENSG00000240388 | 0 | 0 | 0 | 0 | 0 | 0 |
| ENSG00000241059 | 0 | 0 | 0 | 0 | 0 | 0 |
| ENSG00000113369 | 971 | 535 | 752 | 303 | 564 | 503 |
| ENSG00000281357 | 64 | 26 | 46 | 25 | 31 | 39 |
| ENSG00000251356 | 2 | 2 | 0 | 1 | 0 | 0 |

| | | | | | | |
|---|---|---|---|---|---|---|
| ENSG00000248826 | 0 | 0 | 0 | 0 | 0 | 0 |
| ENSG00000276426 | 0 | 0 | 0 | 0 | 0 | 0 |
| ENSG00000250049 | 0 | 1 | 0 | 0 | 1 | 0 |
| ENSG00000248864 | 0 | 0 | 0 | 0 | 0 | 0 |
| ENSG00000249776 | 4 | 0 | 0 | 0 | 0 | 0 |
| ENSG00000249169 | 0 | 0 | 0 | 0 | 0 | 0 |
| ENSG00000249984 | 0 | 0 | 0 | 0 | 0 | 0 |
| ENSG00000248588 | 0 | 0 | 0 | 0 | 0 | 0 |
| ENSG00000248950 | 0 | 0 | 0 | 0 | 0 | 0 |
| ENSG00000249958 | 0 | 1 | 0 | 0 | 0 | 0 |
| ENSG00000248528 | 0 | 0 | 0 | 0 | 0 | 0 |
| ENSG00000251361 | 5 | 0 | 0 | 0 | 2 | 0 |
| ENSG00000213730 | 0 | 0 | 0 | 0 | 2 | 0 |
| ENSG00000237187 | 6 | 1 | 1 | 1 | 2 | 1 |
| ENSG00000175745 | 11 | 1 | 5 | 3 | 2 | 5 |
| ENSG00000278905 | 0 | 0 | 0 | 0 | 0 | 0 |
| ENSG00000113391 | 889 | 405 | 468 | 288 | 504 | 368 |
| ENSG00000284336 | 0 | 0 | 0 | 0 | 0 | 0 |
| ENSG00000249353 | 534 | 248 | 276 | 280 | 370 | 301 |
| ENSG00000248483 | 2 | 2 | 0 | 0 | 1 | 0 |
| ENSG00000272406 | 0 | 1 | 0 | 0 | 0 | 0 |
| ENSG00000251023 | 2 | 1 | 0 | 0 | 0 | 0 |
| ENSG00000185261 | 25 | 12 | 15 | 5 | 20 | 13 |
| ENSG00000251544 | 0 | 0 | 0 | 0 | 0 | 0 |
| ENSG00000254132 | 0 | 1 | 0 | 0 | 0 | 0 |
| ENSG00000251340 | 0 | 0 | 0 | 0 | 0 | 0 |
| ENSG00000232578 | 0 | 0 | 1 | 0 | 0 | 0 |
| ENSG00000270133 | 4 | 2 | 3 | 1 | 0 | 3 |
| ENSG00000133302 | 200 | 125 | 112 | 68 | 167 | 112 |
| ENSG00000175471 | 26 | 10 | 12 | 7 | 13 | 5 |
| ENSG00000249175 | 0 | 0 | 0 | 0 | 0 | 0 |
| ENSG00000243806 | 0 | 0 | 0 | 0 | 0 | 0 |
| ENSG00000276514 | 0 | 0 | 0 | 0 | 0 | 0 |
| ENSG00000249545 | 0 | 0 | 0 | 0 | 0 | 0 |
| ENSG00000153347 | 0 | 0 | 0 | 0 | 0 | 1 |
| ENSG00000225165 | 0 | 0 | 0 | 0 | 0 | 0 |
| ENSG00000198677 | 690 | 431 | 273 | 112 | 818 | 384 |
| ENSG00000212259 | 0 | 0 | 0 | 0 | 0 | 0 |
| ENSG00000164291 | 181 | 60 | 64 | 47 | 80 | 58 |
| ENSG00000178015 | 2 | 0 | 5 | 1 | 3 | 2 |
| ENSG00000175449 | 48 | 13 | 21 | 26 | 25 | 21 |
| ENSG00000145757 | 160 | 78 | 79 | 86 | 102 | 77 |
| ENSG00000250240 | 97 | 41 | 50 | 41 | 60 | 34 |
| ENSG00000164292 | 26528 | 11168 | 12565 | 6843 | 13043 | 9130 |
| ENSG00000173221 | 179 | 105 | 135 | 95 | 118 | 112 |
| ENSG00000251348 | 44 | 34 | 20 | 3 | 29 | 12 |
| ENSG00000251000 | 9 | 9 | 3 | 3 | 6 | 6 |
| ENSG00000251409 | 0 | 2 | 0 | 0 | 1 | 0 |
| ENSG00000272021 | 0 | 0 | 0 | 0 | 0 | 1 |
| ENSG00000236882 | 4 | 0 | 5 | 0 | 3 | 0 |

| | | | | | | |
|---|---|---|---|---|---|---|
| ENSG00000250362 | 0 | 0 | 0 | 0 | 0 | 0 |
| ENSG00000118985 | 388 | 198 | 186 | 132 | 265 | 170 |
| ENSG00000251314 | 0 | 0 | 3 | 0 | 2 | 0 |
| ENSG00000250955 | 0 | 0 | 0 | 0 | 0 | 0 |
| ENSG00000213716 | 0 | 0 | 0 | 0 | 0 | 0 |
| ENSG00000250551 | 0 | 0 | 0 | 0 | 0 | 0 |
| ENSG00000207578 | 0 | 0 | 0 | 0 | 0 | 0 |
| ENSG00000206997 | 0 | 0 | 0 | 0 | 0 | 0 |
| ENSG00000249746 | 0 | 0 | 0 | 0 | 0 | 0 |
| ENSG00000250158 | 0 | 0 | 0 | 0 | 0 | 0 |
| ENSG00000175426 | 0 | 0 | 0 | 0 | 1 | 1 |
| ENSG00000153113 | 3079 | 1935 | 2002 | 1291 | 2184 | 1642 |
| ENSG00000249180 | 0 | 0 | 0 | 0 | 0 | 0 |
| ENSG00000164307 | 685 | 352 | 400 | 206 | 300 | 309 |
| ENSG00000248734 | 3 | 1 | 3 | 1 | 0 | 0 |
| ENSG00000272109 | 2 | 1 | 2 | 0 | 0 | 2 |
| ENSG00000247121 | 52 | 19 | 27 | 17 | 26 | 30 |
| ENSG00000164308 | 4 | 6 | 6 | 11 | 5 | 3 |
| ENSG00000113441 | 1636 | 821 | 763 | 302 | 1090 | 682 |
| ENSG00000227836 | 0 | 0 | 1 | 0 | 0 | 1 |
| ENSG00000200884 | 0 | 0 | 0 | 0 | 0 | 0 |
| ENSG00000213701 | 0 | 0 | 0 | 0 | 0 | 1 |
| ENSG00000251513 | 8 | 0 | 3 | 0 | 1 | 0 |
| ENSG00000145721 | 0 | 0 | 0 | 0 | 1 | 0 |
| ENSG00000058729 | 443 | 281 | 277 | 207 | 391 | 289 |
| ENSG00000206698 | 0 | 1 | 0 | 0 | 0 | 0 |
| ENSG00000248758 | 0 | 0 | 0 | 0 | 0 | 0 |
| ENSG00000251054 | 0 | 1 | 1 | 0 | 2 | 2 |
| ENSG00000249761 | 0 | 0 | 0 | 0 | 0 | 0 |
| ENSG00000251389 | 0 | 0 | 0 | 1 | 0 | 1 |
| ENSG00000249977 | 1 | 1 | 0 | 0 | 1 | 0 |
| ENSG00000250331 | 0 | 2 | 0 | 0 | 0 | 0 |
| ENSG00000248931 | 0 | 0 | 1 | 0 | 0 | 0 |
| ENSG00000251193 | 0 | 0 | 0 | 0 | 0 | 0 |
| ENSG00000248202 | 0 | 0 | 2 | 2 | 3 | 0 |
| ENSG00000249101 | 0 | 0 | 0 | 0 | 0 | 0 |
| ENSG00000248901 | 0 | 0 | 0 | 0 | 0 | 0 |
| ENSG00000249135 | 0 | 0 | 0 | 0 | 0 | 0 |
| ENSG00000238000 | 64 | 27 | 34 | 36 | 27 | 28 |
| ENSG00000176183 | 0 | 0 | 0 | 0 | 0 | 0 |
| ENSG00000250221 | 12 | 8 | 7 | 4 | 2 | 3 |
| ENSG00000250830 | 0 | 0 | 0 | 0 | 0 | 0 |
| ENSG00000249990 | 0 | 0 | 0 | 0 | 0 | 0 |
| ENSG00000249448 | 0 | 0 | 0 | 0 | 0 | 0 |
| ENSG00000251102 | 0 | 0 | 0 | 0 | 0 | 0 |
| ENSG00000251553 | 0 | 0 | 0 | 0 | 0 | 0 |
| ENSG00000174136 | 321 | 149 | 194 | 175 | 157 | 196 |
| ENSG00000246763 | 41 | 16 | 20 | 11 | 11 | 10 |
| ENSG00000279232 | 4 | 1 | 1 | 0 | 1 | 1 |
| ENSG00000250162 | 0 | 0 | 0 | 0 | 1 | 0 |

| | | | | | | |
|---|---|---|---|---|---|---|
| ENSG00000153922 | 797 | 414 | 307 | 138 | 664 | 350 |
| ENSG00000222610 | 0 | 0 | 0 | 0 | 0 | 0 |
| ENSG00000248489 | 11 | 13 | 11 | 7 | 2 | 3 |
| ENSG00000200351 | 0 | 0 | 0 | 0 | 0 | 0 |
| ENSG00000241597 | 2 | 1 | 0 | 0 | 0 | 0 |
| ENSG00000249444 | 0 | 0 | 0 | 0 | 0 | 0 |
| ENSG00000206356 | 0 | 0 | 0 | 0 | 0 | 0 |
| ENSG00000227762 | 0 | 0 | 0 | 0 | 0 | 0 |
| ENSG00000249515 | 0 | 0 | 0 | 0 | 0 | 0 |
| ENSG00000248236 | 0 | 0 | 0 | 0 | 0 | 0 |
| ENSG00000248928 | 0 | 0 | 0 | 0 | 0 | 0 |
| ENSG00000250391 | 0 | 0 | 0 | 0 | 0 | 0 |
| ENSG00000249963 | 0 | 0 | 0 | 0 | 0 | 0 |
| ENSG00000199378 | 0 | 0 | 0 | 0 | 0 | 0 |
| ENSG00000270232 | 0 | 0 | 0 | 0 | 0 | 0 |
| ENSG00000270230 | 0 | 0 | 0 | 1 | 0 | 1 |
| ENSG00000271687 | 0 | 0 | 0 | 0 | 1 | 0 |
| ENSG00000270906 | 0 | 0 | 0 | 0 | 0 | 1 |
| ENSG00000270350 | 0 | 0 | 0 | 0 | 0 | 0 |
| ENSG00000271480 | 0 | 0 | 0 | 0 | 0 | 0 |
| ENSG00000270388 | 0 | 0 | 0 | 0 | 0 | 0 |
| ENSG00000270307 | 0 | 0 | 0 | 0 | 0 | 0 |
| ENSG00000270225 | 0 | 0 | 0 | 0 | 0 | 0 |
| ENSG00000271207 | 0 | 0 | 0 | 0 | 0 | 0 |
| ENSG00000274455 | 0 | 0 | 0 | 0 | 0 | 0 |
| ENSG00000207077 | 0 | 0 | 0 | 0 | 0 | 0 |
| ENSG00000275374 | 3 | 0 | 0 | 2 | 0 | 1 |
| ENSG00000273957 | 0 | 0 | 0 | 0 | 0 | 0 |
| ENSG00000249787 | 0 | 0 | 0 | 0 | 0 | 0 |
| ENSG00000247877 | 0 | 3 | 1 | 2 | 1 | 3 |
| ENSG00000174132 | 905 | 416 | 533 | 502 | 510 | 479 |
| ENSG00000250806 | 0 | 0 | 0 | 0 | 0 | 0 |
| ENSG00000207269 | 0 | 0 | 0 | 0 | 0 | 0 |
| ENSG00000113532 | 1442 | 670 | 1174 | 397 | 771 | 923 |
| ENSG00000221263 | 0 | 0 | 0 | 0 | 0 | 0 |
| ENSG00000277506 | 0 | 0 | 0 | 0 | 0 | 0 |
| ENSG00000251261 | 0 | 0 | 0 | 0 | 0 | 0 |
| ENSG00000249495 | 0 | 0 | 0 | 0 | 0 | 0 |
| ENSG00000199786 | 0 | 0 | 0 | 0 | 0 | 0 |
| ENSG00000248261 | 0 | 0 | 0 | 0 | 0 | 0 |
| ENSG00000173930 | 54 | 22 | 35 | 8 | 11 | 14 |
| ENSG00000222987 | 0 | 0 | 0 | 0 | 0 | 0 |
| ENSG00000205359 | 0 | 1 | 0 | 0 | 0 | 0 |
| ENSG00000249017 | 0 | 0 | 0 | 0 | 0 | 0 |
| ENSG00000250958 | 0 | 0 | 0 | 0 | 0 | 0 |
| ENSG00000247402 | 0 | 0 | 0 | 0 | 0 | 0 |
| ENSG00000250682 | 0 | 0 | 0 | 0 | 0 | 0 |
| ENSG00000145730 | 1526 | 713 | 741 | 653 | 979 | 768 |
| ENSG00000175749 | 0 | 0 | 0 | 0 | 0 | 0 |
| ENSG00000145723 | 152 | 81 | 80 | 51 | 93 | 75 |

| | | | | | | |
|---|---|---|---|---|---|---|
| ENSG00000145725 | 631 | 421 | 271 | 146 | 651 | 312 |
| ENSG00000248203 | 0 | 0 | 0 | 0 | 0 | 0 |
| ENSG00000181751 | 941 | 432 | 546 | 399 | 522 | 506 |
| ENSG00000250567 | 0 | 0 | 0 | 0 | 0 | 1 |
| ENSG00000226926 | 2 | 0 | 0 | 2 | 2 | 3 |
| ENSG00000248757 | 0 | 0 | 1 | 0 | 0 | 0 |
| ENSG00000112874 | 790 | 410 | 430 | 297 | 515 | 425 |
| ENSG00000283462 | 0 | 0 | 0 | 1 | 1 | 0 |
| ENSG00000251026 | 2 | 1 | 1 | 0 | 0 | 1 |
| ENSG00000201910 | 0 | 0 | 0 | 0 | 0 | 0 |
| ENSG00000239808 | 0 | 0 | 0 | 0 | 0 | 0 |
| ENSG00000251574 | 59 | 26 | 44 | 24 | 30 | 51 |
| ENSG00000278958 | 0 | 1 | 0 | 0 | 0 | 0 |
| ENSG00000253776 | 0 | 0 | 0 | 0 | 0 | 0 |
| ENSG00000252881 | 0 | 0 | 0 | 1 | 0 | 0 |
| ENSG00000253584 | 0 | 0 | 0 | 0 | 0 | 0 |
| ENSG00000232159 | 0 | 0 | 0 | 0 | 0 | 0 |
| ENSG00000201790 | 0 | 0 | 0 | 0 | 0 | 0 |
| ENSG00000251204 | 0 | 0 | 1 | 0 | 0 | 0 |
| ENSG00000250145 | 0 | 0 | 0 | 0 | 0 | 0 |
| ENSG00000252337 | 0 | 0 | 0 | 0 | 0 | 0 |
| ENSG00000251027 | 0 | 0 | 0 | 0 | 0 | 2 |
| ENSG00000250273 | 0 | 0 | 0 | 0 | 1 | 0 |
| ENSG00000184349 | 276 | 142 | 171 | 143 | 198 | 207 |
| ENSG00000249959 | 0 | 0 | 0 | 0 | 0 | 0 |
| ENSG00000248827 | 0 | 0 | 0 | 0 | 0 | 0 |
| ENSG00000239708 | 0 | 0 | 0 | 0 | 0 | 0 |
| ENSG00000251732 | 0 | 0 | 0 | 0 | 0 | 0 |
| ENSG00000145743 | 1372 | 544 | 803 | 445 | 664 | 642 |
| ENSG00000244245 | 0 | 1 | 3 | 2 | 1 | 1 |
| ENSG00000272523 | 73 | 15 | 38 | 32 | 32 | 35 |
| ENSG00000151422 | 210 | 93 | 75 | 30 | 188 | 86 |
| ENSG00000249921 | 0 | 0 | 0 | 0 | 0 | 0 |
| ENSG00000250383 | 0 | 0 | 0 | 0 | 0 | 0 |
| ENSG00000207404 | 0 | 0 | 0 | 0 | 0 | 0 |
| ENSG00000250441 | 0 | 0 | 0 | 0 | 0 | 0 |
| ENSG00000206593 | 0 | 0 | 0 | 0 | 0 | 0 |
| ENSG00000176857 | 0 | 1 | 0 | 0 | 0 | 0 |
| ENSG00000249476 | 6 | 3 | 8 | 1 | 7 | 8 |
| ENSG00000198961 | 2068 | 1208 | 1121 | 553 | 1586 | 1095 |
| ENSG00000248867 | 0 | 0 | 0 | 0 | 0 | 0 |
| ENSG00000251367 | 0 | 0 | 0 | 0 | 0 | 0 |
| ENSG00000248876 | 0 | 0 | 0 | 0 | 0 | 0 |
| ENSG00000248440 | 0 | 0 | 0 | 0 | 0 | 0 |
| ENSG00000214794 | 2 | 0 | 0 | 0 | 0 | 0 |
| ENSG00000271849 | 5 | 2 | 2 | 5 | 4 | 1 |
| ENSG00000112893 | 2156 | 1207 | 1137 | 605 | 1525 | 1064 |
| ENSG00000202512 | 0 | 0 | 0 | 0 | 0 | 0 |
| ENSG00000249068 | 0 | 0 | 0 | 0 | 0 | 0 |
| ENSG00000254106 | 0 | 0 | 0 | 0 | 1 | 0 |

| | | | | | | |
|---|---|---|---|---|---|---|
| ENSG00000253224 | 0 | 0 | 0 | 0 | 0 | 0 |
| ENSG00000186952 | 3 | 1 | 3 | 4 | 6 | 3 |
| ENSG00000221436 | 0 | 0 | 0 | 0 | 0 | 0 |
| ENSG00000207177 | 0 | 0 | 0 | 0 | 0 | 0 |
| ENSG00000164209 | 1146 | 588 | 644 | 462 | 779 | 644 |
| ENSG00000270779 | 0 | 0 | 0 | 0 | 0 | 0 |
| ENSG00000248966 | 0 | 0 | 0 | 0 | 0 | 0 |
| ENSG00000248428 | 0 | 0 | 0 | 0 | 0 | 0 |
| ENSG00000251627 | 0 | 0 | 0 | 0 | 0 | 0 |
| ENSG00000145777 | 0 | 0 | 0 | 0 | 0 | 0 |
| ENSG00000253613 | 0 | 0 | 0 | 0 | 0 | 0 |
| ENSG00000134987 | 921 | 500 | 538 | 334 | 538 | 511 |
| ENSG00000250728 | 0 | 0 | 0 | 0 | 0 | 0 |
| ENSG00000214784 | 2 | 5 | 1 | 1 | 2 | 3 |
| ENSG00000152495 | 0 | 1 | 0 | 0 | 0 | 0 |
| ENSG00000249318 | 0 | 0 | 0 | 0 | 0 | 0 |
| ENSG00000248268 | 0 | 0 | 0 | 0 | 0 | 0 |
| ENSG00000164211 | 80 | 25 | 38 | 18 | 38 | 38 |
| ENSG00000246859 | 14 | 9 | 12 | 3 | 21 | 6 |
| ENSG00000249619 | 0 | 0 | 0 | 0 | 0 | 0 |
| ENSG00000251099 | 0 | 0 | 0 | 0 | 0 | 0 |
| ENSG00000134986 | 210 | 121 | 137 | 70 | 112 | 106 |
| ENSG00000253057 | 0 | 0 | 0 | 0 | 0 | 0 |
| ENSG00000249439 | 0 | 0 | 0 | 0 | 0 | 0 |
| ENSG00000250095 | 0 | 0 | 0 | 0 | 0 | 0 |
| ENSG00000129595 | 413 | 210 | 217 | 91 | 173 | 148 |
| ENSG00000224032 | 1371 | 679 | 855 | 1063 | 914 | 869 |
| ENSG00000238363 | 9 | 9 | 8 | 8 | 4 | 1 |
| ENSG00000250882 | 0 | 0 | 1 | 0 | 0 | 0 |
| ENSG00000251187 | 0 | 2 | 0 | 0 | 0 | 0 |
| ENSG00000251076 | 0 | 0 | 0 | 0 | 0 | 0 |
| ENSG00000248350 | 0 | 0 | 0 | 0 | 3 | 0 |
| ENSG00000278921 | 60 | 29 | 23 | 19 | 21 | 22 |
| ENSG00000248430 | 0 | 0 | 0 | 0 | 0 | 0 |
| ENSG00000251014 | 0 | 0 | 0 | 0 | 0 | 0 |
| ENSG00000250358 | 8 | 2 | 1 | 1 | 1 | 0 |
| ENSG00000134982 | 279 | 192 | 121 | 36 | 366 | 142 |
| ENSG00000250045 | 0 | 0 | 0 | 0 | 0 | 0 |
| ENSG00000212370 | 0 | 0 | 0 | 0 | 0 | 0 |
| ENSG00000258864 | 0 | 0 | 0 | 0 | 0 | 0 |
| ENSG00000153037 | 482 | 397 | 417 | 358 | 477 | 419 |
| ENSG00000129625 | 7743 | 3961 | 4461 | 3472 | 4146 | 3806 |
| ENSG00000249947 | 3 | 7 | 3 | 4 | 0 | 3 |
| ENSG00000212643 | 0 | 0 | 0 | 0 | 0 | 0 |
| ENSG00000172795 | 1613 | 796 | 753 | 368 | 1026 | 699 |
| ENSG00000171444 | 436 | 206 | 232 | 123 | 224 | 174 |
| ENSG00000232633 | 4 | 0 | 0 | 0 | 1 | 0 |
| ENSG00000212122 | 0 | 0 | 0 | 0 | 0 | 0 |
| ENSG00000251741 | 0 | 0 | 0 | 0 | 0 | 0 |
| ENSG00000047188 | 734 | 380 | 406 | 140 | 483 | 302 |

| | | | | | | |
|---|---|---|---|---|---|---|
| ENSG00000080709 | 22 | 10 | 12 | 16 | 8 | 11 |
| ENSG00000222706 | 0 | 0 | 0 | 0 | 0 | 0 |
| ENSG00000246316 | 0 | 0 | 0 | 0 | 0 | 0 |
| ENSG00000251542 | 0 | 0 | 0 | 0 | 0 | 0 |
| ENSG00000251132 | 0 | 0 | 0 | 0 | 0 | 0 |
| ENSG00000250242 | 0 | 0 | 0 | 0 | 0 | 0 |
| ENSG00000152503 | 137 | 84 | 73 | 34 | 105 | 53 |
| ENSG00000250472 | 0 | 0 | 0 | 0 | 0 | 0 |
| ENSG00000249791 | 0 | 0 | 0 | 0 | 0 | 0 |
| ENSG00000164219 | 798 | 451 | 461 | 371 | 574 | 451 |
| ENSG00000271797 | 8 | 1 | 4 | 3 | 2 | 3 |
| ENSG00000243304 | 10 | 0 | 2 | 4 | 0 | 4 |
| ENSG00000164221 | 108 | 62 | 71 | 42 | 114 | 66 |
| ENSG00000250197 | 0 | 1 | 0 | 0 | 0 | 0 |
| ENSG00000249026 | 0 | 0 | 0 | 0 | 1 | 0 |
| ENSG00000248722 | 0 | 0 | 0 | 0 | 0 | 0 |
| ENSG00000249944 | 0 | 0 | 0 | 0 | 0 | 0 |
| ENSG00000249433 | 0 | 0 | 0 | 0 | 0 | 0 |
| ENSG00000250444 | 2 | 1 | 3 | 1 | 0 | 1 |
| ENSG00000145780 | 498 | 226 | 283 | 158 | 283 | 257 |
| ENSG00000251201 | 0 | 0 | 3 | 0 | 3 | 3 |
| ENSG00000243414 | 0 | 0 | 0 | 0 | 0 | 0 |
| ENSG00000249249 | 6 | 5 | 8 | 1 | 6 | 6 |
| ENSG00000134970 | 2053 | 1020 | 1062 | 888 | 1235 | 1070 |
| ENSG00000249021 | 0 | 0 | 0 | 0 | 0 | 0 |
| ENSG00000248709 | 0 | 0 | 0 | 0 | 0 | 0 |
| ENSG00000251118 | 0 | 1 | 0 | 0 | 0 | 0 |
| ENSG00000222598 | 0 | 0 | 0 | 0 | 0 | 0 |
| ENSG00000129596 | 0 | 4 | 3 | 2 | 6 | 0 |
| ENSG00000145782 | 1877 | 939 | 1182 | 791 | 1117 | 1032 |
| ENSG00000177879 | 2137 | 1035 | 1228 | 1018 | 1200 | 1147 |
| ENSG00000172901 | 15 | 2 | 4 | 3 | 11 | 2 |
| ENSG00000248295 | 0 | 0 | 0 | 0 | 0 | 0 |
| ENSG00000268223 | 0 | 0 | 0 | 0 | 0 | 0 |
| ENSG00000185641 | 28 | 9 | 11 | 24 | 17 | 29 |
| ENSG00000272265 | 0 | 1 | 2 | 1 | 1 | 0 |
| ENSG00000271918 | 67 | 39 | 53 | 41 | 36 | 42 |
| ENSG00000145781 | 739 | 378 | 491 | 373 | 500 | 405 |
| ENSG00000212457 | 0 | 0 | 0 | 0 | 0 | 0 |
| ENSG00000232916 | 0 | 0 | 0 | 0 | 0 | 0 |
| ENSG00000250015 | 2 | 0 | 0 | 0 | 1 | 0 |
| ENSG00000092421 | 194 | 93 | 93 | 65 | 101 | 71 |
| ENSG00000248445 | 3 | 12 | 20 | 8 | 6 | 6 |
| ENSG00000239528 | 2 | 1 | 0 | 0 | 0 | 0 |
| ENSG00000249167 | 0 | 0 | 0 | 0 | 0 | 1 |
| ENSG00000197575 | 0 | 0 | 1 | 1 | 1 | 0 |
| ENSG00000251311 | 0 | 0 | 0 | 0 | 0 | 0 |
| ENSG00000249150 | 0 | 0 | 0 | 0 | 0 | 0 |
| ENSG00000250650 | 0 | 0 | 0 | 0 | 0 | 0 |
| ENSG00000250949 | 0 | 0 | 0 | 0 | 0 | 0 |

| | | | | | | |
|---|---|---|---|---|---|---|
| ENSG00000243758 | 0 | 0 | 0 | 0 | 0 | 0 |
| ENSG00000285809 | 0 | 0 | 0 | 0 | 0 | 0 |
| ENSG00000248663 | 241 | 149 | 158 | 119 | 114 | 133 |
| ENSG00000251477 | 0 | 0 | 0 | 0 | 0 | 0 |
| ENSG00000249582 | 0 | 0 | 0 | 0 | 0 | 0 |
| ENSG00000249797 | 0 | 0 | 8 | 1 | 1 | 0 |
| ENSG00000250427 | 0 | 0 | 1 | 0 | 0 | 0 |
| ENSG00000250891 | 15 | 4 | 2 | 3 | 8 | 4 |
| ENSG00000249551 | 0 | 0 | 0 | 0 | 0 | 0 |
| ENSG00000249128 | 0 | 0 | 0 | 0 | 1 | 0 |
| ENSG00000238904 | 0 | 0 | 0 | 0 | 0 | 0 |
| ENSG00000249426 | 1 | 0 | 1 | 1 | 0 | 0 |
| ENSG00000169570 | 483 | 242 | 345 | 182 | 262 | 274 |
| ENSG00000197744 | 35 | 25 | 18 | 19 | 17 | 29 |
| ENSG00000283498 | 54 | 30 | 28 | 37 | 26 | 37 |
| ENSG00000249494 | 0 | 1 | 0 | 1 | 0 | 2 |
| ENSG00000223179 | 0 | 0 | 0 | 0 | 0 | 0 |
| ENSG00000250678 | 0 | 0 | 0 | 0 | 0 | 0 |
| ENSG00000250928 | 0 | 0 | 0 | 0 | 0 | 0 |
| ENSG00000172869 | 647 | 438 | 306 | 113 | 639 | 314 |
| ENSG00000248885 | 0 | 0 | 0 | 0 | 0 | 0 |
| ENSG00000248979 | 0 | 0 | 0 | 0 | 0 | 0 |
| ENSG00000206786 | 0 | 0 | 0 | 0 | 0 | 0 |
| ENSG00000264536 | 0 | 0 | 0 | 0 | 0 | 0 |
| ENSG00000145779 | 170 | 123 | 91 | 90 | 135 | 82 |
| ENSG00000243333 | 0 | 0 | 0 | 0 | 0 | 0 |
| ENSG00000202092 | 0 | 0 | 0 | 0 | 1 | 0 |
| ENSG00000133835 | 5216 | 2737 | 3279 | 2083 | 3141 | 2864 |
| ENSG00000213663 | 0 | 0 | 0 | 0 | 0 | 0 |
| ENSG00000249919 | 0 | 0 | 0 | 0 | 0 | 0 |
| ENSG00000164334 | 1 | 3 | 2 | 0 | 1 | 0 |
| ENSG00000250895 | 0 | 0 | 0 | 0 | 0 | 0 |
| ENSG00000234259 | 0 | 0 | 1 | 0 | 0 | 1 |
| ENSG00000251293 | 0 | 0 | 0 | 0 | 0 | 0 |
| ENSG00000251975 | 0 | 0 | 0 | 0 | 0 | 0 |
| ENSG00000261036 | 0 | 0 | 0 | 0 | 0 | 0 |
| ENSG00000184838 | 2 | 0 | 1 | 0 | 0 | 0 |
| ENSG00000270741 | 0 | 0 | 0 | 0 | 0 | 0 |
| ENSG00000222609 | 0 | 0 | 0 | 0 | 0 | 0 |
| ENSG00000248927 | 0 | 0 | 0 | 0 | 0 | 0 |
| ENSG00000251135 | 0 | 0 | 0 | 0 | 0 | 0 |
| ENSG00000248853 | 0 | 0 | 0 | 0 | 0 | 0 |
| ENSG00000250847 | 0 | 0 | 0 | 0 | 0 | 0 |
| ENSG00000229855 | 0 | 0 | 0 | 0 | 0 | 0 |
| ENSG00000279860 | 0 | 0 | 0 | 0 | 0 | 0 |
| ENSG00000244088 | 0 | 0 | 0 | 0 | 0 | 0 |
| ENSG00000213661 | 0 | 0 | 0 | 0 | 0 | 0 |
| ENSG00000249600 | 0 | 0 | 0 | 0 | 0 | 0 |
| ENSG00000181867 | 0 | 0 | 0 | 0 | 0 | 0 |
| ENSG00000151304 | 154 | 79 | 60 | 42 | 137 | 70 |

| | | | | | | |
|---|---|---|---|---|---|---|
| ENSG00000113083 | 3 | 2 | 2 | 0 | 1 | 0 |
| ENSG00000247311 | 0 | 0 | 0 | 0 | 0 | 0 |
| ENSG00000164185 | 0 | 0 | 0 | 0 | 0 | 1 |
| ENSG00000250803 | 0 | 0 | 0 | 0 | 0 | 0 |
| ENSG00000249610 | 0 | 0 | 0 | 0 | 1 | 0 |
| ENSG00000272139 | 0 | 0 | 0 | 0 | 0 | 0 |
| ENSG00000064692 | 2 | 1 | 0 | 1 | 3 | 2 |
| ENSG00000249621 | 0 | 0 | 0 | 0 | 0 | 0 |
| ENSG00000249916 | 3 | 3 | 2 | 0 | 2 | 1 |
| ENSG00000250328 | 2 | 0 | 3 | 0 | 1 | 1 |
| ENSG00000251538 | 0 | 0 | 0 | 0 | 0 | 0 |
| ENSG00000240058 | 0 | 0 | 0 | 0 | 0 | 0 |
| ENSG00000205302 | 1588 | 958 | 876 | 501 | 1300 | 926 |
| ENSG00000260686 | 6 | 1 | 3 | 7 | 5 | 1 |
| ENSG00000064652 | 713 | 376 | 455 | 379 | 399 | 347 |
| ENSG00000263432 | 11 | 7 | 13 | 4 | 7 | 3 |
| ENSG00000168938 | 3214 | 2025 | 2320 | 2035 | 1980 | 1897 |
| ENSG00000249996 | 5 | 3 | 2 | 3 | 0 | 1 |
| ENSG00000276047 | 0 | 0 | 0 | 1 | 0 | 0 |
| ENSG00000223652 | 54 | 36 | 51 | 44 | 46 | 33 |
| ENSG00000061455 | 195 | 74 | 87 | 71 | 96 | 78 |
| ENSG00000213655 | 0 | 0 | 0 | 0 | 0 | 0 |
| ENSG00000168944 | 502 | 255 | 254 | 122 | 380 | 254 |
| ENSG00000250539 | 47 | 18 | 17 | 13 | 20 | 11 |
| ENSG00000249400 | 0 | 0 | 0 | 0 | 0 | 0 |
| ENSG00000279772 | 0 | 1 | 0 | 0 | 0 | 0 |
| ENSG00000151292 | 821 | 444 | 398 | 236 | 597 | 425 |
| ENSG00000235275 | 14 | 9 | 7 | 3 | 8 | 5 |
| ENSG00000253807 | 0 | 0 | 0 | 0 | 0 | 0 |
| ENSG00000250862 | 0 | 0 | 0 | 0 | 0 | 0 |
| ENSG00000251421 | 0 | 0 | 0 | 0 | 0 | 0 |
| ENSG00000248600 | 0 | 0 | 0 | 0 | 0 | 0 |
| ENSG00000251214 | 0 | 0 | 0 | 0 | 0 | 0 |
| ENSG00000250438 | 0 | 0 | 0 | 0 | 0 | 0 |
| ENSG00000248296 | 0 | 0 | 2 | 0 | 0 | 0 |
| ENSG00000168916 | 4 | 0 | 1 | 0 | 1 | 0 |
| ENSG00000249112 | 0 | 0 | 0 | 0 | 0 | 0 |
| ENSG00000251456 | 0 | 0 | 0 | 0 | 0 | 0 |
| ENSG00000242814 | 0 | 0 | 0 | 0 | 0 | 0 |
| ENSG00000260192 | 0 | 0 | 0 | 0 | 1 | 0 |
| ENSG00000248443 | 0 | 0 | 0 | 0 | 0 | 0 |
| ENSG00000249261 | 0 | 0 | 0 | 0 | 0 | 0 |
| ENSG00000250194 | 0 | 0 | 0 | 0 | 0 | 0 |
| ENSG00000250721 | 0 | 0 | 0 | 0 | 0 | 0 |
| ENSG00000222107 | 0 | 0 | 0 | 0 | 0 | 0 |
| ENSG00000250402 | 0 | 0 | 0 | 0 | 0 | 0 |
| ENSG00000249950 | 0 | 0 | 0 | 0 | 0 | 0 |
| ENSG00000249904 | 0 | 0 | 0 | 0 | 0 | 0 |
| ENSG00000241888 | 0 | 0 | 0 | 0 | 0 | 0 |
| ENSG00000248752 | 0 | 0 | 0 | 0 | 0 | 0 |

| | | | | | | |
|---|---|---|---|---|---|---|
| ENSG00000248107 | 0 | 0 | 0 | 0 | 0 | 0 |
| ENSG00000155324 | 726 | 431 | 419 | 352 | 476 | 421 |
| ENSG00000250602 | 20 | 6 | 4 | 3 | 9 | 2 |
| ENSG00000279118 | 8 | 8 | 2 | 8 | 5 | 4 |
| ENSG00000164904 | 2729 | 1438 | 1647 | 1142 | 1357 | 1367 |
| ENSG00000202082 | 0 | 0 | 0 | 0 | 0 | 0 |
| ENSG00000164902 | 1436 | 715 | 760 | 481 | 846 | 719 |
| ENSG00000230929 | 4 | 2 | 3 | 1 | 3 | 1 |
| ENSG00000196900 | 0 | 0 | 0 | 0 | 0 | 0 |
| ENSG00000213630 | 0 | 0 | 0 | 0 | 0 | 0 |
| ENSG00000249768 | 0 | 0 | 0 | 0 | 0 | 0 |
| ENSG00000251072 | 5 | 5 | 0 | 0 | 0 | 1 |
| ENSG00000252185 | 0 | 0 | 0 | 0 | 0 | 0 |
| ENSG00000113368 | 3862 | 1855 | 2397 | 1698 | 1989 | 2310 |
| ENSG00000173926 | 100 | 48 | 46 | 53 | 36 | 49 |
| ENSG00000164241 | 223 | 109 | 140 | 103 | 147 | 136 |
| ENSG00000279446 | 0 | 0 | 0 | 0 | 0 | 0 |
| ENSG00000214743 | 0 | 0 | 0 | 0 | 0 | 0 |
| ENSG00000250080 | 0 | 0 | 0 | 0 | 0 | 0 |
| ENSG00000271585 | 0 | 0 | 0 | 0 | 0 | 0 |
| ENSG00000250956 | 0 | 0 | 0 | 0 | 0 | 0 |
| ENSG00000283897 | 61 | 19 | 23 | 23 | 29 | 26 |
| ENSG00000145794 | 16 | 3 | 5 | 2 | 3 | 7 |
| ENSG00000249577 | 0 | 0 | 0 | 0 | 0 | 0 |
| ENSG00000271766 | 0 | 0 | 0 | 0 | 0 | 0 |
| ENSG00000250859 | 10 | 2 | 2 | 3 | 5 | 2 |
| ENSG00000164244 | 3942 | 1833 | 2220 | 1469 | 2039 | 1916 |
| ENSG00000282925 | 0 | 0 | 0 | 0 | 0 | 0 |
| ENSG00000205279 | 0 | 0 | 0 | 0 | 0 | 0 |
| ENSG00000248799 | 0 | 0 | 0 | 0 | 0 | 0 |
| ENSG00000230561 | 46 | 24 | 23 | 14 | 16 | 11 |
| ENSG00000251032 | 0 | 0 | 0 | 0 | 0 | 0 |
| ENSG00000250603 | 0 | 0 | 0 | 0 | 0 | 0 |
| ENSG00000245937 | 1169 | 570 | 751 | 578 | 618 | 574 |
| ENSG00000064651 | 5962 | 2810 | 3471 | 1011 | 2181 | 1888 |
| ENSG00000250329 | 2 | 0 | 1 | 0 | 0 | 1 |
| ENSG00000138829 | 44 | 16 | 39 | 15 | 10 | 14 |
| ENSG00000207290 | 0 | 0 | 0 | 0 | 0 | 0 |
| ENSG00000113396 | 310 | 160 | 196 | 99 | 94 | 128 |
| ENSG00000248634 | 0 | 0 | 0 | 0 | 0 | 0 |
| ENSG00000066583 | 6581 | 2999 | 3604 | 2153 | 2113 | 2410 |
| ENSG00000264563 | 0 | 0 | 0 | 0 | 0 | 0 |
| ENSG00000265691 | 0 | 0 | 0 | 0 | 0 | 0 |
| ENSG00000249421 | 127 | 66 | 90 | 36 | 53 | 54 |
| ENSG00000145808 | 1508 | 685 | 805 | 234 | 355 | 372 |
| ENSG00000251680 | 1 | 0 | 0 | 0 | 0 | 0 |
| ENSG00000186367 | 4 | 4 | 1 | 0 | 0 | 1 |
| ENSG00000198108 | 16 | 6 | 4 | 5 | 6 | 8 |
| ENSG00000221562 | 0 | 0 | 0 | 0 | 0 | 0 |
| ENSG00000199455 | 0 | 0 | 0 | 0 | 0 | 0 |

| | | | | | | |
|---|---|---|---|---|---|---|
| ENSG00000248610 | 0 | 0 | 0 | 0 | 0 | 0 |
| ENSG00000252514 | 0 | 0 | 0 | 0 | 0 | 0 |
| ENSG00000249312 | 2 | 0 | 1 | 0 | 0 | 0 |
| ENSG00000249418 | 0 | 0 | 0 | 0 | 0 | 0 |
| ENSG00000244192 | 0 | 0 | 0 | 0 | 0 | 2 |
| ENSG00000250405 | 0 | 0 | 0 | 0 | 0 | 0 |
| ENSG00000248955 | 0 | 0 | 0 | 0 | 0 | 0 |
| ENSG00000169567 | 5353 | 2617 | 3255 | 3191 | 3312 | 3159 |
| ENSG00000186687 | 397 | 196 | 199 | 129 | 264 | 214 |
| ENSG00000158985 | 2038 | 926 | 1134 | 778 | 1074 | 1035 |
| ENSG00000279370 | 1 | 2 | 0 | 1 | 0 | 0 |
| ENSG00000158987 | 98 | 43 | 39 | 18 | 110 | 46 |
| ENSG00000273217 | 0 | 0 | 0 | 0 | 0 | 0 |
| ENSG00000275939 | 0 | 0 | 0 | 0 | 0 | 0 |
| ENSG00000217128 | 692 | 374 | 420 | 196 | 434 | 302 |
| ENSG00000279584 | 0 | 0 | 2 | 3 | 0 | 7 |
| ENSG00000239642 | 0 | 0 | 0 | 0 | 0 | 0 |
| ENSG00000281938 | 0 | 0 | 0 | 0 | 0 | 0 |
| ENSG00000234758 | 0 | 0 | 0 | 0 | 0 | 0 |
| ENSG00000164398 | 0 | 1 | 0 | 0 | 0 | 0 |
| ENSG00000223548 | 0 | 0 | 0 | 0 | 0 | 0 |
| ENSG00000231585 | 0 | 0 | 0 | 0 | 0 | 0 |
| ENSG00000164399 | 0 | 0 | 0 | 0 | 0 | 0 |
| ENSG00000164400 | 0 | 0 | 0 | 0 | 0 | 0 |
| ENSG00000253067 | 0 | 0 | 0 | 0 | 0 | 0 |
| ENSG00000227616 | 0 | 0 | 0 | 0 | 0 | 0 |
| ENSG00000235775 | 0 | 0 | 0 | 0 | 0 | 0 |
| ENSG00000224015 | 0 | 0 | 0 | 0 | 0 | 0 |
| ENSG00000237714 | 0 | 0 | 0 | 0 | 0 | 0 |
| ENSG00000072682 | 961 | 546 | 515 | 608 | 680 | 514 |
| ENSG00000224431 | 10 | 2 | 8 | 3 | 4 | 10 |
| ENSG00000283678 | 0 | 0 | 0 | 0 | 0 | 0 |
| ENSG00000131435 | 0 | 0 | 0 | 0 | 0 | 0 |
| ENSG00000197208 | 79 | 64 | 55 | 66 | 98 | 98 |
| ENSG00000233006 | 58 | 34 | 28 | 9 | 31 | 27 |
| ENSG00000263597 | 2 | 2 | 1 | 3 | 2 | 7 |
| ENSG00000197375 | 1787 | 928 | 1103 | 828 | 1011 | 1061 |
| ENSG00000197536 | 39 | 24 | 28 | 16 | 37 | 28 |
| ENSG00000283782 | 2 | 0 | 0 | 0 | 3 | 4 |
| ENSG00000238160 | 0 | 0 | 0 | 0 | 0 | 0 |
| ENSG00000202533 | 6 | 0 | 1 | 1 | 1 | 0 |
| ENSG00000234290 | 14 | 12 | 16 | 2 | 13 | 13 |
| ENSG00000125347 | 371 | 203 | 245 | 273 | 269 | 293 |
| ENSG00000113525 | 0 | 0 | 0 | 0 | 0 | 0 |
| ENSG00000113522 | 45 | 30 | 34 | 9 | 45 | 29 |
| ENSG00000223442 | 0 | 5 | 3 | 2 | 7 | 1 |
| ENSG00000169194 | 0 | 3 | 0 | 0 | 0 | 1 |
| ENSG00000113520 | 2 | 1 | 0 | 0 | 0 | 0 |
| ENSG00000230612 | 14 | 9 | 3 | 0 | 5 | 6 |
| ENSG00000131437 | 694 | 380 | 375 | 171 | 531 | 366 |

| | | | | | | |
|---|---|---|---|---|---|---|
| ENSG00000205089 | 12 | 5 | 7 | 9 | 4 | 11 |
| ENSG00000164402 | 4244 | 1933 | 2427 | 1775 | 2169 | 2248 |
| ENSG00000198944 | 12 | 4 | 6 | 1 | 3 | 9 |
| ENSG00000272203 | 0 | 0 | 0 | 0 | 0 | 0 |
| ENSG00000164403 | 193 | 98 | 118 | 102 | 87 | 109 |
| ENSG00000201274 | 0 | 0 | 0 | 0 | 0 | 0 |
| ENSG00000164404 | 45 | 13 | 33 | 21 | 13 | 14 |
| ENSG00000164405 | 3040 | 1955 | 2111 | 2435 | 2208 | 2023 |
| ENSG00000164406 | 71 | 22 | 34 | 31 | 30 | 30 |
| ENSG00000072364 | 4277 | 1982 | 2258 | 1038 | 2324 | 1959 |
| ENSG00000235253 | 1 | 2 | 1 | 0 | 1 | 0 |
| ENSG00000202417 | 0 | 0 | 0 | 0 | 0 | 1 |
| ENSG00000225364 | 0 | 0 | 0 | 0 | 0 | 0 |
| ENSG00000272023 | 4 | 3 | 0 | 0 | 1 | 2 |
| ENSG00000229738 | 0 | 0 | 0 | 0 | 0 | 0 |
| ENSG00000199677 | 0 | 0 | 0 | 0 | 0 | 0 |
| ENSG00000155329 | 458 | 236 | 213 | 201 | 254 | 233 |
| ENSG00000248648 | 0 | 0 | 10 | 0 | 1 | 5 |
| ENSG00000170606 | 4881 | 2530 | 2615 | 2302 | 3370 | 2493 |
| ENSG00000279691 | 2 | 5 | 4 | 3 | 1 | 4 |
| ENSG00000251616 | 0 | 0 | 0 | 1 | 0 | 0 |
| ENSG00000248245 | 0 | 4 | 5 | 2 | 3 | 4 |
| ENSG00000240611 | 0 | 0 | 0 | 0 | 0 | 0 |
| ENSG00000053108 | 207 | 117 | 183 | 120 | 100 | 170 |
| ENSG00000249478 | 2 | 0 | 2 | 0 | 0 | 0 |
| ENSG00000250244 | 8 | 1 | 0 | 2 | 2 | 2 |
| ENSG00000250409 | 0 | 0 | 0 | 0 | 0 | 0 |
| ENSG00000221287 | 0 | 0 | 0 | 0 | 0 | 0 |
| ENSG00000280104 | 0 | 0 | 1 | 0 | 0 | 1 |
| ENSG00000279197 | 0 | 0 | 0 | 0 | 0 | 0 |
| ENSG00000251243 | 0 | 0 | 0 | 0 | 0 | 0 |
| ENSG00000249073 | 0 | 0 | 0 | 0 | 0 | 0 |
| ENSG00000113583 | 3365 | 1530 | 2092 | 1541 | 1803 | 1800 |
| ENSG00000213585 | 10169 | 5249 | 6063 | 5535 | 6515 | 5969 |
| ENSG00000271737 | 41 | 34 | 29 | 29 | 38 | 33 |
| ENSG00000271128 | 0 | 0 | 0 | 0 | 0 | 0 |
| ENSG00000081059 | 20 | 21 | 21 | 5 | 5 | 2 |
| ENSG00000113558 | 7018 | 4089 | 4506 | 4002 | 5012 | 4311 |
| ENSG00000272772 | 0 | 0 | 0 | 0 | 0 | 0 |
| ENSG00000113575 | 5840 | 2920 | 3646 | 2684 | 3536 | 3250 |
| ENSG00000273345 | 2 | 2 | 5 | 0 | 2 | 2 |
| ENSG00000266751 | 0 | 0 | 0 | 0 | 0 | 0 |
| ENSG00000270177 | 47 | 22 | 26 | 24 | 37 | 30 |
| ENSG00000239615 | 0 | 0 | 0 | 0 | 0 | 0 |
| ENSG00000006837 | 32 | 23 | 15 | 15 | 21 | 19 |
| ENSG00000119048 | 1727 | 818 | 1048 | 774 | 1002 | 970 |
| ENSG00000279469 | 8 | 3 | 1 | 1 | 0 | 0 |
| ENSG00000248559 | 6 | 3 | 4 | 4 | 4 | 3 |
| ENSG00000237190 | 1120 | 568 | 675 | 701 | 700 | 638 |
| ENSG00000243648 | 2 | 1 | 0 | 2 | 0 | 3 |

| | | | | | | |
|---|---|---|---|---|---|---|
| ENSG00000250564 | 0 | 1 | 0 | 0 | 0 | 0 |
| ENSG00000250994 | 0 | 1 | 0 | 0 | 0 | 1 |
| ENSG00000207222 | 0 | 0 | 0 | 0 | 0 | 0 |
| ENSG00000251169 | 110 | 45 | 66 | 64 | 54 | 77 |
| ENSG00000240250 | 7 | 2 | 1 | 1 | 5 | 1 |
| ENSG00000280420 | 9 | 2 | 3 | 3 | 3 | 7 |
| ENSG00000043143 | 3209 | 1399 | 1899 | 1562 | 1565 | 1813 |
| ENSG00000152700 | 3021 | 1451 | 1705 | 1267 | 1662 | 1323 |
| ENSG00000207459 | 0 | 0 | 0 | 0 | 0 | 0 |
| ENSG00000113615 | 1491 | 830 | 802 | 438 | 1081 | 768 |
| ENSG00000201298 | 0 | 0 | 0 | 0 | 0 | 0 |
| ENSG00000222266 | 0 | 0 | 1 | 0 | 1 | 0 |
| ENSG00000164615 | 1981 | 967 | 1184 | 1160 | 1127 | 1143 |
| ENSG00000145833 | 2035 | 1210 | 1095 | 474 | 1414 | 1048 |
| ENSG00000241179 | 0 | 0 | 0 | 0 | 0 | 0 |
| ENSG00000181904 | 1625 | 822 | 993 | 519 | 907 | 897 |
| ENSG00000270892 | 0 | 0 | 0 | 0 | 0 | 0 |
| ENSG00000113621 | 1217 | 608 | 784 | 589 | 663 | 634 |
| ENSG00000132570 | 514 | 270 | 377 | 244 | 269 | 292 |
| ENSG00000279799 | 22 | 16 | 10 | 14 | 8 | 9 |
| ENSG00000244921 | 9 | 5 | 7 | 5 | 1 | 8 |
| ENSG00000249119 | 4 | 4 | 8 | 0 | 1 | 4 |
| ENSG00000248923 | 16 | 8 | 14 | 14 | 6 | 16 |
| ENSG00000247627 | 15 | 8 | 13 | 8 | 6 | 10 |
| ENSG00000198868 | 0 | 2 | 0 | 1 | 2 | 1 |
| ENSG00000263963 | 0 | 0 | 0 | 0 | 0 | 0 |
| ENSG00000249192 | 2 | 1 | 2 | 0 | 0 | 0 |
| ENSG00000152705 | 18 | 7 | 12 | 4 | 5 | 7 |
| ENSG00000069011 | 211 | 90 | 92 | 149 | 94 | 98 |
| ENSG00000224186 | 14 | 14 | 17 | 12 | 18 | 17 |
| ENSG00000277619 | 0 | 0 | 0 | 0 | 0 | 0 |
| ENSG00000249082 | 0 | 1 | 0 | 2 | 1 | 0 |
| ENSG00000248753 | 0 | 0 | 0 | 0 | 0 | 0 |
| ENSG00000248482 | 0 | 0 | 0 | 0 | 0 | 0 |
| ENSG00000249647 | 0 | 0 | 0 | 0 | 0 | 0 |
| ENSG00000113648 | 8191 | 4336 | 5445 | 4677 | 4973 | 5299 |
| ENSG00000270021 | 30 | 14 | 24 | 16 | 12 | 21 |
| ENSG00000251380 | 0 | 0 | 0 | 0 | 0 | 0 |
| ENSG00000255833 | 0 | 0 | 0 | 0 | 0 | 0 |
| ENSG00000249639 | 0 | 0 | 0 | 1 | 0 | 0 |
| ENSG00000181965 | 0 | 0 | 0 | 0 | 0 | 0 |
| ENSG00000250167 | 0 | 0 | 0 | 0 | 0 | 0 |
| ENSG00000145824 | 0 | 1 | 0 | 0 | 7 | 4 |
| ENSG00000145832 | 6 | 5 | 0 | 2 | 0 | 2 |
| ENSG00000251045 | 0 | 0 | 0 | 0 | 0 | 0 |
| ENSG00000223539 | 0 | 0 | 0 | 0 | 0 | 0 |
| ENSG00000265924 | 0 | 0 | 0 | 0 | 0 | 0 |
| ENSG00000250378 | 0 | 0 | 0 | 0 | 0 | 0 |
| ENSG00000145839 | 0 | 0 | 0 | 0 | 0 | 0 |
| ENSG00000254239 | 0 | 0 | 0 | 0 | 0 | 0 |

| | | | | | | |
|---|---|---|---|---|---|---|
| ENSG00000253927 | 0 | 0 | 0 | 0 | 0 | 0 |
| ENSG00000145826 | 2 | 0 | 0 | 0 | 0 | 0 |
| ENSG00000164616 | 0 | 0 | 0 | 0 | 0 | 0 |
| ENSG00000120708 | 3149 | 1960 | 1807 | 2298 | 2681 | 2585 |
| ENSG00000270123 | 0 | 0 | 0 | 0 | 0 | 0 |
| ENSG00000278815 | 0 | 0 | 0 | 0 | 0 | 0 |
| ENSG00000164621 | 0 | 1 | 0 | 0 | 0 | 0 |
| ENSG00000277859 | 0 | 0 | 0 | 0 | 0 | 0 |
| ENSG00000113658 | 1541 | 784 | 771 | 313 | 1030 | 651 |
| ENSG00000275646 | 0 | 0 | 0 | 0 | 0 | 0 |
| ENSG00000277524 | 0 | 0 | 0 | 0 | 0 | 0 |
| ENSG00000274934 | 0 | 0 | 0 | 0 | 0 | 0 |
| ENSG00000271824 | 8 | 10 | 1 | 10 | 10 | 13 |
| ENSG00000069018 | 0 | 1 | 0 | 1 | 0 | 0 |
| ENSG00000248211 | 1 | 0 | 2 | 0 | 1 | 1 |
| ENSG00000250947 | 0 | 0 | 0 | 0 | 0 | 0 |
| ENSG00000279240 | 0 | 0 | 0 | 0 | 0 | 0 |
| ENSG00000248351 | 0 | 0 | 0 | 0 | 0 | 0 |
| ENSG00000213568 | 0 | 0 | 0 | 0 | 0 | 0 |
| ENSG00000249803 | 0 | 0 | 0 | 0 | 0 | 0 |
| ENSG00000250284 | 0 | 0 | 0 | 0 | 0 | 0 |
| ENSG00000248616 | 0 | 0 | 0 | 0 | 0 | 0 |
| ENSG00000222285 | 0 | 0 | 0 | 0 | 0 | 0 |
| ENSG00000152377 | 14 | 3 | 7 | 9 | 3 | 10 |
| ENSG00000146021 | 2 | 0 | 4 | 1 | 1 | 1 |
| ENSG00000216009 | 0 | 0 | 0 | 0 | 0 | 0 |
| ENSG00000177733 | 9487 | 4771 | 6039 | 5003 | 5605 | 5731 |
| ENSG00000270697 | 2 | 1 | 1 | 0 | 0 | 0 |
| ENSG00000226306 | 0 | 0 | 0 | 0 | 0 | 0 |
| ENSG00000250159 | 0 | 0 | 0 | 1 | 0 | 0 |
| ENSG00000120729 | 2 | 0 | 2 | 0 | 0 | 0 |
| ENSG00000078795 | 5 | 1 | 0 | 1 | 3 | 1 |
| ENSG00000031003 | 1424 | 823 | 911 | 508 | 975 | 756 |
| ENSG00000246323 | 4 | 4 | 1 | 1 | 2 | 6 |
| ENSG00000250260 | 1 | 0 | 0 | 0 | 1 | 0 |
| ENSG00000222924 | 0 | 0 | 0 | 0 | 0 | 0 |
| ENSG00000061492 | 1 | 0 | 0 | 0 | 0 | 0 |
| ENSG00000112981 | 18 | 6 | 6 | 5 | 17 | 7 |
| ENSG00000212460 | 0 | 0 | 0 | 0 | 0 | 0 |
| ENSG00000199880 | 0 | 0 | 0 | 0 | 0 | 0 |
| ENSG00000112983 | 2151 | 986 | 1366 | 876 | 1287 | 1195 |
| ENSG00000112984 | 1322 | 708 | 757 | 749 | 931 | 855 |
| ENSG00000094880 | 2392 | 1302 | 1484 | 1040 | 1323 | 1221 |
| ENSG00000146013 | 30 | 17 | 12 | 25 | 11 | 15 |
| ENSG00000249851 | 0 | 0 | 0 | 0 | 0 | 0 |
| ENSG00000158402 | 306 | 159 | 191 | 197 | 249 | 202 |
| ENSG00000120709 | 2120 | 1026 | 1253 | 1190 | 1203 | 1208 |
| ENSG00000249971 | 0 | 0 | 0 | 0 | 0 | 0 |
| ENSG00000120733 | 4843 | 2077 | 2896 | 1699 | 2250 | 2377 |
| ENSG00000132563 | 18 | 16 | 31 | 32 | 26 | 20 |

| | | | | | | |
|---|---|---|---|---|---|---|
| ENSG00000271196 | 0 | 0 | 0 | 0 | 0 | 0 |
| ENSG00000120738 | 524 | 179 | 276 | 236 | 161 | 251 |
| ENSG00000241458 | 0 | 1 | 1 | 0 | 0 | 0 |
| ENSG00000120705 | 6600 | 3296 | 3866 | 2929 | 3617 | 3611 |
| ENSG00000113013 | 11716 | 6128 | 7293 | 5527 | 7142 | 6937 |
| ENSG00000222937 | 0 | 0 | 0 | 0 | 0 | 0 |
| ENSG00000206989 | 3 | 2 | 0 | 0 | 0 | 0 |
| ENSG00000044115 | 16495 | 7646 | 9946 | 7833 | 9626 | 9008 |
| ENSG00000232174 | 0 | 0 | 0 | 0 | 0 | 0 |
| ENSG00000238745 | 0 | 0 | 0 | 0 | 0 | 0 |
| ENSG00000253404 | 62 | 48 | 49 | 58 | 64 | 41 |
| ENSG00000146006 | 0 | 0 | 0 | 0 | 3 | 0 |
| ENSG00000120725 | 2388 | 1381 | 1624 | 1482 | 1206 | 1165 |
| ENSG00000201532 | 0 | 0 | 0 | 0 | 0 | 0 |
| ENSG00000285956 | 0 | 0 | 0 | 0 | 0 | 0 |
| ENSG00000249593 | 21 | 7 | 15 | 4 | 9 | 7 |
| ENSG00000242683 | 0 | 0 | 0 | 0 | 0 | 0 |
| ENSG00000252533 | 0 | 0 | 0 | 0 | 0 | 0 |
| ENSG00000280987 | 1 | 2 | 0 | 1 | 3 | 3 |
| ENSG00000281398 | 326 | 187 | 215 | 182 | 300 | 286 |
| ENSG00000252213 | 0 | 0 | 0 | 0 | 0 | 0 |
| ENSG00000200959 | 0 | 0 | 0 | 0 | 1 | 0 |
| ENSG00000015479 | 68 | 44 | 34 | 23 | 41 | 23 |
| ENSG00000199545 | 0 | 2 | 0 | 0 | 1 | 0 |
| ENSG00000253015 | 0 | 0 | 0 | 0 | 0 | 0 |
| ENSG00000120727 | 1584 | 1036 | 1278 | 954 | 1242 | 1048 |
| ENSG00000272742 | 0 | 0 | 0 | 0 | 0 | 0 |
| ENSG00000170482 | 17 | 7 | 17 | 6 | 6 | 11 |
| ENSG00000170476 | 27 | 12 | 23 | 11 | 13 | 14 |
| ENSG00000228672 | 198 | 86 | 112 | 77 | 121 | 90 |
| ENSG00000170469 | 84 | 33 | 49 | 51 | 40 | 61 |
| ENSG00000170464 | 240 | 135 | 167 | 131 | 163 | 160 |
| ENSG00000279686 | 0 | 0 | 0 | 0 | 0 | 0 |
| ENSG00000200378 | 0 | 1 | 0 | 0 | 0 | 0 |
| ENSG00000249751 | 0 | 0 | 0 | 0 | 0 | 0 |
| ENSG00000283288 | 0 | 0 | 0 | 0 | 0 | 0 |
| ENSG00000184584 | 22 | 3 | 14 | 9 | 6 | 13 |
| ENSG00000249689 | 0 | 0 | 0 | 0 | 0 | 0 |
| ENSG00000249215 | 0 | 0 | 0 | 0 | 0 | 0 |
| ENSG00000131508 | 2947 | 1544 | 1851 | 1754 | 1716 | 1811 |
| ENSG00000207020 | 0 | 0 | 0 | 0 | 0 | 0 |
| ENSG00000275506 | 0 | 2 | 1 | 0 | 0 | 0 |
| ENSG00000272255 | 3 | 0 | 0 | 1 | 1 | 1 |
| ENSG00000171604 | 8426 | 4281 | 4834 | 4882 | 4717 | 4680 |
| ENSG00000250635 | 5 | 3 | 6 | 2 | 3 | 2 |
| ENSG00000249526 | 1 | 1 | 1 | 0 | 0 | 2 |
| ENSG00000200756 | 0 | 0 | 0 | 0 | 0 | 0 |
| ENSG00000249131 | 0 | 0 | 0 | 0 | 1 | 0 |
| ENSG00000251387 | 2 | 0 | 4 | 1 | 1 | 1 |
| ENSG00000271792 | 0 | 0 | 0 | 0 | 0 | 0 |

| | | | | | | |
|---|---|---|---|---|---|---|
| ENSG00000146005 | 20 | 15 | 13 | 12 | 8 | 11 |
| ENSG00000158458 | 8 | 0 | 4 | 2 | 0 | 0 |
| ENSG00000250692 | 0 | 0 | 0 | 0 | 0 | 0 |
| ENSG00000207317 | 0 | 0 | 0 | 0 | 0 | 0 |
| ENSG00000245146 | 41 | 35 | 33 | 19 | 26 | 20 |
| ENSG00000185129 | 1191 | 491 | 561 | 426 | 581 | 548 |
| ENSG00000182700 | 68 | 50 | 27 | 10 | 40 | 25 |
| ENSG00000254363 | 29 | 12 | 17 | 7 | 12 | 10 |
| ENSG00000120306 | 1180 | 574 | 649 | 697 | 668 | 561 |
| ENSG00000250069 | 0 | 1 | 1 | 0 | 0 | 0 |
| ENSG00000113068 | 1268 | 755 | 830 | 913 | 824 | 879 |
| ENSG00000113070 | 40 | 14 | 19 | 17 | 39 | 25 |
| ENSG00000253965 | 0 | 0 | 0 | 0 | 0 | 0 |
| ENSG00000113073 | 1 | 4 | 7 | 3 | 3 | 3 |
| ENSG00000249637 | 15 | 24 | 21 | 22 | 13 | 16 |
| ENSG00000222790 | 0 | 0 | 0 | 0 | 0 | 0 |
| ENSG00000131503 | 74 | 63 | 60 | 27 | 65 | 66 |
| ENSG00000254996 | 39 | 44 | 44 | 40 | 63 | 46 |
| ENSG00000200051 | 0 | 0 | 0 | 0 | 0 | 0 |
| ENSG00000213523 | 976 | 564 | 631 | 676 | 579 | 624 |
| ENSG00000243056 | 8 | 2 | 4 | 0 | 0 | 2 |
| ENSG00000113108 | 666 | 249 | 422 | 289 | 260 | 277 |
| ENSG00000283155 | 0 | 0 | 1 | 0 | 0 | 0 |
| ENSG00000283717 | 0 | 0 | 0 | 0 | 0 | 0 |
| ENSG00000176087 | 3532 | 1626 | 2092 | 1990 | 1900 | 2005 |
| ENSG00000283602 | 0 | 0 | 0 | 0 | 1 | 0 |
| ENSG00000200235 | 0 | 0 | 0 | 0 | 0 | 0 |
| ENSG00000251453 | 0 | 0 | 0 | 0 | 0 | 0 |
| ENSG00000240518 | 0 | 0 | 0 | 0 | 0 | 0 |
| ENSG00000170458 | 11 | 11 | 14 | 6 | 0 | 8 |
| ENSG00000131495 | 1434 | 1072 | 1213 | 1277 | 1198 | 1047 |
| ENSG00000113119 | 444 | 245 | 306 | 219 | 230 | 243 |
| ENSG00000113141 | 2839 | 1596 | 1773 | 1140 | 1977 | 1554 |
| ENSG00000284325 | 0 | 0 | 0 | 0 | 0 | 0 |
| ENSG00000120314 | 2291 | 1207 | 1539 | 1333 | 1435 | 1397 |
| ENSG00000256453 | 0 | 0 | 0 | 0 | 0 | 0 |
| ENSG00000170445 | 3470 | 1720 | 1950 | 2161 | 2100 | 2208 |
| ENSG00000112855 | 2490 | 1315 | 1610 | 1550 | 1610 | 1727 |
| ENSG00000146007 | 3611 | 2007 | 2210 | 1849 | 2262 | 2101 |
| ENSG00000199990 | 0 | 1 | 0 | 0 | 0 | 1 |
| ENSG00000202111 | 0 | 1 | 0 | 0 | 0 | 1 |
| ENSG00000202515 | 0 | 1 | 2 | 1 | 1 | 1 |
| ENSG00000250515 | 0 | 0 | 0 | 0 | 0 | 0 |
| ENSG00000248106 | 0 | 0 | 0 | 0 | 0 | 0 |
| ENSG00000204970 | 27 | 25 | 19 | 17 | 29 | 30 |
| ENSG00000279726 | 0 | 0 | 1 | 2 | 0 | 1 |
| ENSG00000204969 | 17 | 7 | 16 | 8 | 10 | 14 |
| ENSG00000255408 | 18 | 7 | 7 | 8 | 16 | 14 |
| ENSG00000204967 | 10 | 3 | 7 | 2 | 2 | 0 |
| ENSG00000204965 | 6 | 2 | 6 | 4 | 9 | 1 |

| | | | | | | |
|---|---|---|---|---|---|---|
| ENSG00000081842 | 200 | 71 | 140 | 113 | 102 | 133 |
| ENSG00000204963 | 75 | 43 | 47 | 49 | 55 | 49 |
| ENSG00000204962 | 9 | 4 | 8 | 8 | 13 | 10 |
| ENSG00000204961 | 13 | 3 | 1 | 4 | 6 | 3 |
| ENSG00000278907 | 0 | 0 | 0 | 0 | 0 | 0 |
| ENSG00000250120 | 71 | 50 | 66 | 36 | 69 | 66 |
| ENSG00000249504 | 0 | 0 | 0 | 0 | 0 | 0 |
| ENSG00000278946 | 0 | 0 | 0 | 0 | 0 | 0 |
| ENSG00000249158 | 651 | 327 | 425 | 270 | 374 | 383 |
| ENSG00000251664 | 120 | 74 | 77 | 52 | 74 | 90 |
| ENSG00000278915 | 0 | 0 | 0 | 0 | 0 | 0 |
| ENSG00000278901 | 2 | 2 | 3 | 0 | 0 | 0 |
| ENSG00000239389 | 54 | 18 | 40 | 13 | 30 | 28 |
| ENSG00000248383 | 75 | 32 | 31 | 17 | 28 | 35 |
| ENSG00000279028 | 0 | 0 | 2 | 0 | 0 | 0 |
| ENSG00000243232 | 153 | 72 | 71 | 56 | 84 | 96 |
| ENSG00000279047 | 8 | 5 | 4 | 4 | 2 | 1 |
| ENSG00000171815 | 8 | 3 | 4 | 4 | 4 | 4 |
| ENSG00000279558 | 4 | 0 | 0 | 0 | 0 | 0 |
| ENSG00000112852 | 232 | 97 | 162 | 136 | 132 | 165 |
| ENSG00000272154 | 1 | 1 | 0 | 0 | 0 | 3 |
| ENSG00000113205 | 292 | 132 | 168 | 113 | 177 | 164 |
| ENSG00000272108 | 8 | 2 | 6 | 5 | 16 | 9 |
| ENSG00000081818 | 72 | 32 | 34 | 22 | 23 | 24 |
| ENSG00000113209 | 4 | 2 | 7 | 5 | 5 | 3 |
| ENSG00000280029 | 1 | 0 | 0 | 0 | 1 | 1 |
| ENSG00000113211 | 17 | 6 | 8 | 3 | 7 | 3 |
| ENSG00000255622 | 8 | 3 | 7 | 1 | 0 | 1 |
| ENSG00000280336 | 0 | 1 | 0 | 0 | 0 | 0 |
| ENSG00000113212 | 19 | 9 | 17 | 8 | 7 | 15 |
| ENSG00000120322 | 110 | 45 | 74 | 59 | 68 | 79 |
| ENSG00000279472 | 0 | 0 | 0 | 0 | 1 | 0 |
| ENSG00000272674 | 303 | 131 | 214 | 146 | 164 | 207 |
| ENSG00000279068 | 4 | 3 | 3 | 6 | 6 | 5 |
| ENSG00000278936 | 0 | 0 | 0 | 0 | 0 | 0 |
| ENSG00000177839 | 80 | 39 | 59 | 52 | 61 | 53 |
| ENSG00000279375 | 5 | 8 | 4 | 4 | 2 | 8 |
| ENSG00000120324 | 80 | 34 | 68 | 29 | 50 | 36 |
| ENSG00000197479 | 20 | 15 | 15 | 7 | 15 | 7 |
| ENSG00000120328 | 28 | 14 | 14 | 1 | 10 | 10 |
| ENSG00000187372 | 84 | 37 | 52 | 21 | 44 | 54 |
| ENSG00000120327 | 179 | 55 | 67 | 45 | 88 | 71 |
| ENSG00000279983 | 0 | 0 | 0 | 0 | 0 | 0 |
| ENSG00000146001 | 4 | 7 | 10 | 3 | 7 | 5 |
| ENSG00000262096 | 2 | 3 | 2 | 1 | 0 | 3 |
| ENSG00000113248 | 25 | 23 | 29 | 7 | 8 | 27 |
| ENSG00000277795 | 1 | 0 | 0 | 0 | 0 | 2 |
| ENSG00000178913 | 6164 | 3032 | 3620 | 3001 | 3306 | 3043 |
| ENSG00000120329 | 2 | 5 | 3 | 2 | 0 | 0 |
| ENSG00000272070 | 93 | 37 | 61 | 27 | 66 | 57 |

| | | | | | | |
|---|---|---|---|---|---|---|
| ENSG00000204956 | 120 | 52 | 75 | 41 | 55 | 66 |
| ENSG00000081853 | 279 | 99 | 155 | 84 | 102 | 127 |
| ENSG00000254245 | 38 | 17 | 13 | 11 | 13 | 16 |
| ENSG00000254221 | 184 | 63 | 98 | 75 | 98 | 87 |
| ENSG00000280026 | 0 | 0 | 0 | 0 | 0 | 0 |
| ENSG00000262576 | 33 | 20 | 17 | 15 | 23 | 23 |
| ENSG00000253910 | 31 | 10 | 13 | 5 | 17 | 7 |
| ENSG00000253485 | 16 | 12 | 4 | 1 | 9 | 12 |
| ENSG00000262209 | 16 | 5 | 5 | 3 | 6 | 9 |
| ENSG00000253731 | 24 | 14 | 15 | 16 | 29 | 26 |
| ENSG00000253537 | 34 | 13 | 20 | 8 | 12 | 18 |
| ENSG00000253953 | 3 | 2 | 1 | 1 | 1 | 6 |
| ENSG00000253767 | 6 | 3 | 3 | 0 | 1 | 1 |
| ENSG00000276547 | 744 | 331 | 486 | 318 | 352 | 420 |
| ENSG00000261934 | 38 | 10 | 8 | 10 | 9 | 12 |
| ENSG00000253305 | 33 | 7 | 11 | 8 | 6 | 12 |
| ENSG00000253846 | 10 | 0 | 4 | 2 | 4 | 3 |
| ENSG00000254122 | 24 | 7 | 9 | 5 | 10 | 12 |
| ENSG00000253873 | 0 | 0 | 1 | 0 | 0 | 3 |
| ENSG00000248449 | 0 | 1 | 0 | 3 | 0 | 0 |
| ENSG00000279855 | 0 | 0 | 0 | 0 | 0 | 0 |
| ENSG00000253159 | 2 | 1 | 4 | 2 | 3 | 2 |
| ENSG00000276545 | 5 | 3 | 6 | 2 | 4 | 1 |
| ENSG00000273557 | 3 | 2 | 0 | 2 | 1 | 0 |
| ENSG00000240184 | 444 | 196 | 341 | 215 | 201 | 279 |
| ENSG00000242020 | 2 | 0 | 0 | 0 | 0 | 0 |
| ENSG00000242419 | 6 | 2 | 1 | 2 | 2 | 2 |
| ENSG00000240764 | 6 | 0 | 2 | 3 | 1 | 4 |
| ENSG00000131504 | 6291 | 2959 | 3819 | 3252 | 3941 | 4075 |
| ENSG00000246422 | 0 | 0 | 0 | 0 | 0 | 0 |
| ENSG00000240974 | 3 | 0 | 0 | 0 | 0 | 0 |
| ENSG00000228737 | 2 | 4 | 4 | 5 | 8 | 6 |
| ENSG00000171720 | 2315 | 1174 | 1462 | 1340 | 1373 | 1371 |
| ENSG00000164620 | 226 | 99 | 147 | 124 | 97 | 119 |
| ENSG00000197948 | 897 | 425 | 638 | 494 | 586 | 528 |
| ENSG00000120318 | 14 | 9 | 7 | 5 | 6 | 8 |
| ENSG00000278925 | 5 | 8 | 9 | 2 | 0 | 5 |
| ENSG00000156453 | 2038 | 911 | 1213 | 903 | 966 | 1004 |
| ENSG00000226040 | 3 | 2 | 0 | 0 | 2 | 1 |
| ENSG00000081791 | 2791 | 1354 | 1667 | 1585 | 1578 | 1562 |
| ENSG00000113555 | 30 | 6 | 14 | 11 | 12 | 12 |
| ENSG00000271871 | 0 | 2 | 2 | 1 | 5 | 0 |
| ENSG00000013561 | 2070 | 993 | 1256 | 974 | 1147 | 1154 |
| ENSG00000254099 | 0 | 0 | 1 | 0 | 0 | 0 |
| ENSG00000113552 | 1972 | 877 | 1130 | 1240 | 1090 | 1151 |
| ENSG00000250347 | 0 | 0 | 0 | 0 | 0 | 0 |
| ENSG00000131507 | 3117 | 1475 | 1890 | 1506 | 1664 | 1702 |
| ENSG00000280047 | 26 | 7 | 8 | 9 | 9 | 11 |
| ENSG00000187678 | 22 | 14 | 22 | 10 | 18 | 15 |
| ENSG00000277968 | 0 | 0 | 0 | 0 | 0 | 0 |

| | | | | | | |
|---|---|---|---|---|---|---|
| ENSG00000274721 | 0 | 0 | 0 | 0 | 0 | 0 |
| ENSG00000231185 | 2 | 1 | 3 | 1 | 5 | 8 |
| ENSG00000225147 | 0 | 0 | 0 | 0 | 0 | 0 |
| ENSG00000113578 | 0 | 0 | 0 | 0 | 0 | 0 |
| ENSG00000261757 | 0 | 0 | 0 | 0 | 0 | 0 |
| ENSG00000236714 | 0 | 0 | 0 | 0 | 0 | 0 |
| ENSG00000145819 | 190 | 81 | 71 | 69 | 68 | 66 |
| ENSG00000226272 | 0 | 0 | 0 | 0 | 0 | 1 |
| ENSG00000285366 | 0 | 0 | 0 | 1 | 0 | 0 |
| ENSG00000230789 | 0 | 0 | 0 | 0 | 0 | 0 |
| ENSG00000113580 | 1562 | 757 | 818 | 649 | 1334 | 1084 |
| ENSG00000231579 | 4 | 0 | 0 | 0 | 0 | 0 |
| ENSG00000279679 | 0 | 0 | 0 | 0 | 1 | 0 |
| ENSG00000279130 | 5 | 2 | 6 | 0 | 1 | 4 |
| ENSG00000249429 | 0 | 0 | 0 | 0 | 0 | 0 |
| ENSG00000253023 | 0 | 0 | 0 | 0 | 0 | 0 |
| ENSG00000251205 | 0 | 0 | 1 | 0 | 0 | 0 |
| ENSG00000249881 | 0 | 0 | 0 | 0 | 0 | 0 |
| ENSG00000266478 | 0 | 0 | 0 | 0 | 0 | 0 |
| ENSG00000225869 | 0 | 0 | 0 | 0 | 0 | 0 |
| ENSG00000158497 | 0 | 0 | 0 | 0 | 0 | 0 |
| ENSG00000239390 | 0 | 0 | 0 | 0 | 0 | 0 |
| ENSG00000145817 | 1632 | 809 | 833 | 547 | 932 | 703 |
| ENSG00000183775 | 0 | 0 | 2 | 0 | 0 | 0 |
| ENSG00000249605 | 0 | 0 | 0 | 0 | 0 | 0 |
| ENSG00000285605 | 0 | 0 | 0 | 0 | 0 | 0 |
| ENSG00000250972 | 0 | 0 | 0 | 0 | 0 | 0 |
| ENSG00000213488 | 0 | 0 | 0 | 0 | 0 | 0 |
| ENSG00000201423 | 0 | 0 | 0 | 0 | 0 | 0 |
| ENSG00000249229 | 0 | 0 | 0 | 0 | 0 | 0 |
| ENSG00000251031 | 0 | 0 | 0 | 0 | 0 | 0 |
| ENSG00000215325 | 0 | 2 | 0 | 1 | 0 | 0 |
| ENSG00000250842 | 0 | 0 | 1 | 0 | 0 | 0 |
| ENSG00000248125 | 0 | 0 | 0 | 0 | 0 | 0 |
| ENSG00000186314 | 118 | 79 | 79 | 87 | 103 | 95 |
| ENSG00000249363 | 4 | 0 | 0 | 1 | 1 | 1 |
| ENSG00000204928 | 0 | 0 | 0 | 1 | 1 | 1 |
| ENSG00000156463 | 55 | 23 | 29 | 66 | 57 | 47 |
| ENSG00000248493 | 0 | 0 | 0 | 0 | 0 | 0 |
| ENSG00000173261 | 12 | 2 | 3 | 3 | 4 | 7 |
| ENSG00000251556 | 3 | 1 | 0 | 0 | 0 | 1 |
| ENSG00000133706 | 2244 | 1158 | 1358 | 1006 | 1727 | 1485 |
| ENSG00000239602 | 2 | 0 | 0 | 0 | 0 | 0 |
| ENSG00000239317 | 0 | 0 | 0 | 0 | 0 | 0 |
| ENSG00000275740 | 0 | 0 | 0 | 0 | 0 | 0 |
| ENSG00000091009 | 560 | 315 | 252 | 151 | 381 | 256 |
| ENSG00000238374 | 0 | 0 | 0 | 0 | 0 | 0 |
| ENSG00000250025 | 0 | 0 | 0 | 0 | 0 | 0 |
| ENSG00000091010 | 6 | 1 | 1 | 7 | 2 | 3 |
| ENSG00000271724 | 0 | 0 | 0 | 0 | 0 | 0 |

| | | | | | | |
|---|---|---|---|---|---|---|
| ENSG00000248842 | 0 | 0 | 0 | 0 | 0 | 0 |
| ENSG00000113649 | 1742 | 977 | 921 | 619 | 1273 | 1035 |
| ENSG00000277568 | 3 | 2 | 1 | 0 | 1 | 0 |
| ENSG00000173250 | 0 | 0 | 2 | 0 | 1 | 0 |
| ENSG00000250407 | 0 | 0 | 0 | 0 | 0 | 0 |
| ENSG00000156475 | 12 | 4 | 6 | 9 | 8 | 7 |
| ENSG00000280646 | 0 | 0 | 0 | 0 | 0 | 0 |
| ENSG00000241291 | 0 | 0 | 0 | 0 | 0 | 0 |
| ENSG00000248568 | 3 | 1 | 2 | 2 | 3 | 0 |
| ENSG00000249553 | 0 | 0 | 0 | 0 | 0 | 0 |
| ENSG00000250343 | 0 | 0 | 0 | 0 | 0 | 0 |
| ENSG00000169302 | 2 | 1 | 1 | 0 | 3 | 0 |
| ENSG00000113657 | 0 | 1 | 1 | 1 | 2 | 3 |
| ENSG00000272239 | 0 | 0 | 0 | 0 | 0 | 0 |
| ENSG00000280780 | 0 | 0 | 0 | 0 | 0 | 0 |
| ENSG00000176049 | 0 | 0 | 0 | 0 | 1 | 0 |
| ENSG00000164266 | 0 | 2 | 1 | 1 | 0 | 1 |
| ENSG00000249518 | 0 | 0 | 0 | 0 | 0 | 0 |
| ENSG00000164265 | 1 | 0 | 0 | 0 | 0 | 0 |
| ENSG00000178776 | 0 | 0 | 1 | 0 | 1 | 2 |
| ENSG00000248362 | 0 | 0 | 0 | 0 | 0 | 0 |
| ENSG00000251320 | 0 | 0 | 0 | 0 | 0 | 0 |
| ENSG00000250346 | 0 | 0 | 1 | 0 | 0 | 0 |
| ENSG00000133710 | 33 | 15 | 23 | 7 | 25 | 16 |
| ENSG00000196800 | 1 | 1 | 1 | 1 | 1 | 10 |
| ENSG00000178172 | 0 | 1 | 0 | 0 | 2 | 0 |
| ENSG00000249503 | 0 | 0 | 0 | 0 | 0 | 0 |
| ENSG00000248109 | 0 | 0 | 0 | 0 | 0 | 0 |
| ENSG00000214510 | 0 | 0 | 1 | 2 | 0 | 1 |
| ENSG00000247199 | 0 | 0 | 1 | 0 | 2 | 0 |
| ENSG00000249834 | 0 | 0 | 0 | 0 | 0 | 0 |
| ENSG00000145879 | 4 | 2 | 0 | 1 | 2 | 0 |
| ENSG00000204909 | 0 | 0 | 0 | 0 | 0 | 1 |
| ENSG00000283653 | 0 | 0 | 0 | 0 | 0 | 0 |
| ENSG00000145868 | 960 | 479 | 572 | 352 | 586 | 461 |
| ENSG00000274799 | 0 | 0 | 0 | 0 | 0 | 0 |
| ENSG00000251330 | 1 | 1 | 1 | 0 | 0 | 0 |
| ENSG00000164270 | 0 | 0 | 0 | 0 | 0 | 0 |
| ENSG00000253297 | 0 | 0 | 0 | 0 | 0 | 0 |
| ENSG00000169252 | 2 | 1 | 1 | 0 | 0 | 2 |
| ENSG00000169247 | 41 | 38 | 53 | 22 | 22 | 51 |
| ENSG00000272411 | 0 | 0 | 0 | 0 | 0 | 0 |
| ENSG00000251842 | 0 | 0 | 0 | 0 | 0 | 0 |
| ENSG00000207714 | 0 | 0 | 0 | 0 | 0 | 0 |
| ENSG00000250072 | 0 | 0 | 0 | 0 | 0 | 0 |
| ENSG00000200161 | 0 | 0 | 0 | 0 | 0 | 0 |
| ENSG00000173210 | 632 | 278 | 407 | 256 | 302 | 292 |
| ENSG00000248647 | 0 | 0 | 0 | 0 | 0 | 1 |
| ENSG00000253406 | 0 | 0 | 0 | 0 | 0 | 0 |
| ENSG00000157510 | 6 | 3 | 4 | 2 | 1 | 5 |

| | | | | | | |
|---|---|---|---|---|---|---|
| ENSG00000285736 | 0 | 1 | 1 | 0 | 0 | 0 |
| ENSG00000164284 | 614 | 350 | 407 | 309 | 418 | 358 |
| ENSG00000253618 | 0 | 0 | 5 | 0 | 1 | 2 |
| ENSG00000145882 | 996 | 465 | 544 | 577 | 533 | 565 |
| ENSG00000127743 | 2 | 0 | 1 | 0 | 0 | 1 |
| ENSG00000253865 | 2 | 0 | 0 | 0 | 1 | 1 |
| ENSG00000249669 | 0 | 0 | 0 | 0 | 0 | 0 |
| ENSG00000275871 | 0 | 0 | 0 | 0 | 0 | 0 |
| ENSG00000284182 | 0 | 0 | 0 | 0 | 0 | 0 |
| ENSG00000276365 | 0 | 0 | 0 | 0 | 0 | 0 |
| ENSG00000113712 | 4939 | 2247 | 3061 | 1742 | 2502 | 2601 |
| ENSG00000230551 | 761 | 379 | 578 | 159 | 460 | 326 |
| ENSG00000241112 | 4 | 4 | 1 | 0 | 0 | 0 |
| ENSG00000183111 | 670 | 250 | 461 | 207 | 213 | 244 |
| ENSG00000222213 | 0 | 0 | 0 | 0 | 0 | 0 |
| ENSG00000221043 | 0 | 0 | 0 | 0 | 0 | 0 |
| ENSG00000242928 | 0 | 0 | 0 | 0 | 0 | 0 |
| ENSG00000155846 | 190 | 64 | 109 | 73 | 105 | 102 |
| ENSG00000199047 | 0 | 0 | 0 | 0 | 0 | 0 |
| ENSG00000132915 | 2 | 1 | 0 | 0 | 1 | 3 |
| ENSG00000251583 | 2 | 0 | 0 | 1 | 1 | 1 |
| ENSG00000200334 | 0 | 0 | 0 | 0 | 0 | 0 |
| ENSG00000155850 | 166 | 101 | 162 | 70 | 199 | 126 |
| ENSG00000164296 | 276 | 109 | 165 | 112 | 151 | 155 |
| ENSG00000113716 | 2232 | 1091 | 1397 | 1263 | 1273 | 1284 |
| ENSG00000248696 | 0 | 1 | 0 | 0 | 0 | 0 |
| ENSG00000241907 | 0 | 4 | 0 | 0 | 0 | 2 |
| ENSG00000182578 | 0 | 3 | 1 | 0 | 0 | 3 |
| ENSG00000214485 | 13 | 13 | 13 | 9 | 17 | 5 |
| ENSG00000199409 | 0 | 0 | 0 | 0 | 0 | 0 |
| ENSG00000113721 | 0 | 0 | 2 | 0 | 0 | 2 |
| ENSG00000113722 | 0 | 0 | 0 | 0 | 1 | 0 |
| ENSG00000011083 | 0 | 0 | 0 | 0 | 0 | 0 |
| ENSG00000070808 | 0 | 0 | 0 | 0 | 0 | 0 |
| ENSG00000183876 | 0 | 0 | 0 | 0 | 0 | 0 |
| ENSG00000070814 | 1844 | 948 | 1099 | 1290 | 1057 | 1281 |
| ENSG00000019582 | 8 | 2 | 4 | 4 | 1 | 2 |
| ENSG00000164587 | 27145 | 14000 | 15857 | 20198 | 16659 | 15863 |
| ENSG00000254333 | 3 | 4 | 2 | 2 | 2 | 0 |
| ENSG00000070614 | 4127 | 1803 | 2360 | 1720 | 1940 | 2239 |
| ENSG00000171992 | 552 | 261 | 278 | 272 | 284 | 227 |
| ENSG00000253852 | 0 | 0 | 0 | 0 | 0 | 0 |
| ENSG00000164591 | 0 | 0 | 0 | 0 | 3 | 1 |
| ENSG00000250309 | 0 | 0 | 0 | 0 | 0 | 0 |
| ENSG00000086589 | 2078 | 939 | 1162 | 936 | 957 | 1074 |
| ENSG00000132912 | 2956 | 1484 | 1938 | 1137 | 1608 | 1451 |
| ENSG00000256235 | 328 | 141 | 194 | 191 | 168 | 166 |
| ENSG00000237693 | 0 | 0 | 2 | 0 | 0 | 0 |
| ENSG00000145908 | 4 | 3 | 3 | 3 | 8 | 3 |
| ENSG00000197083 | 0 | 0 | 0 | 0 | 0 | 0 |

| | | | | | | |
|---|---|---|---|---|---|---|
| ENSG00000274235 | 0 | 0 | 0 | 0 | 0 | 0 |
| ENSG00000277866 | 0 | 0 | 0 | 0 | 0 | 0 |
| ENSG00000211445 | 2728 | 1358 | 1713 | 1511 | 1453 | 1461 |
| ENSG00000145901 | 2288 | 1107 | 1540 | 1618 | 1325 | 1369 |
| ENSG00000197043 | 2237 | 1076 | 1477 | 881 | 737 | 737 |
| ENSG00000254298 | 0 | 0 | 0 | 0 | 0 | 0 |
| ENSG00000285420 | 0 | 0 | 0 | 0 | 0 | 0 |
| ENSG00000198624 | 214 | 111 | 118 | 99 | 105 | 95 |
| ENSG00000196743 | 1686 | 872 | 1014 | 855 | 984 | 928 |
| ENSG00000186334 | 0 | 0 | 0 | 0 | 0 | 0 |
| ENSG00000186335 | 0 | 0 | 0 | 0 | 0 | 0 |
| ENSG00000270904 | 0 | 0 | 0 | 0 | 0 | 0 |
| ENSG00000253897 | 0 | 0 | 0 | 0 | 0 | 0 |
| ENSG00000253472 | 0 | 0 | 0 | 0 | 0 | 0 |
| ENSG00000271494 | 0 | 0 | 0 | 0 | 0 | 0 |
| ENSG00000123643 | 778 | 349 | 489 | 354 | 481 | 450 |
| ENSG00000200227 | 0 | 1 | 0 | 0 | 0 | 1 |
| ENSG00000086570 | 2 | 2 | 2 | 1 | 2 | 1 |
| ENSG00000271795 | 7 | 1 | 3 | 0 | 7 | 4 |
| ENSG00000276622 | 0 | 0 | 0 | 0 | 0 | 0 |
| ENSG00000260581 | 0 | 0 | 0 | 0 | 0 | 0 |
| ENSG00000113140 | 4 | 2 | 3 | 4 | 0 | 0 |
| ENSG00000249035 | 0 | 0 | 0 | 0 | 0 | 0 |
| ENSG00000222102 | 0 | 0 | 0 | 0 | 0 | 0 |
| ENSG00000272112 | 21 | 9 | 25 | 12 | 14 | 12 |
| ENSG00000177556 | 2637 | 1405 | 1662 | 1474 | 1407 | 1297 |
| ENSG00000253921 | 0 | 1 | 0 | 1 | 3 | 3 |
| ENSG00000213433 | 0 | 2 | 1 | 3 | 2 | 2 |
| ENSG00000275765 | 21 | 16 | 12 | 16 | 10 | 9 |
| ENSG00000145907 | 6629 | 3320 | 3636 | 2571 | 3589 | 3255 |
| ENSG00000145888 | 0 | 0 | 0 | 0 | 0 | 0 |
| ENSG00000270978 | 2 | 0 | 0 | 0 | 0 | 0 |
| ENSG00000222308 | 0 | 0 | 0 | 0 | 0 | 0 |
| ENSG00000254226 | 20 | 2 | 3 | 4 | 5 | 2 |
| ENSG00000132911 | 0 | 0 | 0 | 0 | 0 | 0 |
| ENSG00000242080 | 0 | 0 | 0 | 0 | 0 | 0 |
| ENSG00000249484 | 0 | 0 | 0 | 0 | 0 | 0 |
| ENSG00000155511 | 0 | 0 | 0 | 0 | 0 | 0 |
| ENSG00000242976 | 0 | 0 | 0 | 0 | 0 | 0 |
| ENSG00000251183 | 0 | 0 | 0 | 0 | 0 | 0 |
| ENSG00000261382 | 0 | 0 | 0 | 0 | 0 | 0 |
| ENSG00000055147 | 604 | 318 | 338 | 284 | 332 | 318 |
| ENSG00000037749 | 689 | 361 | 441 | 200 | 430 | 314 |
| ENSG00000164574 | 2752 | 1247 | 1833 | 1538 | 1949 | 1868 |
| ENSG00000245275 | 24 | 14 | 12 | 8 | 17 | 12 |
| ENSG00000253119 | 0 | 0 | 0 | 0 | 0 | 0 |
| ENSG00000221430 | 0 | 0 | 0 | 0 | 0 | 0 |
| ENSG00000241963 | 0 | 0 | 0 | 0 | 0 | 0 |
| ENSG00000164576 | 974 | 535 | 635 | 597 | 656 | 710 |
| ENSG00000113196 | 0 | 0 | 0 | 0 | 0 | 0 |

| | | | | | | |
|---|---|---|---|---|---|---|
| ENSG00000254293 | 0 | 0 | 0 | 0 | 0 | 0 |
| ENSG00000253886 | 0 | 0 | 0 | 0 | 0 | 0 |
| ENSG00000253146 | 0 | 0 | 0 | 0 | 0 | 0 |
| ENSG00000264760 | 0 | 0 | 0 | 0 | 0 | 0 |
| ENSG00000221552 | 0 | 0 | 0 | 0 | 0 | 1 |
| ENSG00000274472 | 0 | 0 | 0 | 0 | 1 | 0 |
| ENSG00000155506 | 13677 | 5623 | 8008 | 5196 | 6322 | 7312 |
| ENSG00000272197 | 0 | 0 | 0 | 1 | 0 | 1 |
| ENSG00000241187 | 2 | 0 | 0 | 1 | 0 | 0 |
| ENSG00000170271 | 352 | 145 | 209 | 181 | 199 | 187 |
| ENSG00000277039 | 0 | 0 | 2 | 0 | 0 | 0 |
| ENSG00000263361 | 1 | 0 | 1 | 0 | 0 | 1 |
| ENSG00000155508 | 2833 | 1363 | 1728 | 1243 | 1666 | 1589 |
| ENSG00000082516 | 1633 | 871 | 1047 | 871 | 1021 | 945 |
| ENSG00000082515 | 1485 | 758 | 891 | 929 | 927 | 904 |
| ENSG00000253261 | 0 | 0 | 0 | 0 | 0 | 0 |
| ENSG00000226650 | 0 | 0 | 0 | 0 | 0 | 0 |
| ENSG00000253646 | 0 | 0 | 0 | 0 | 0 | 0 |
| ENSG00000271477 | 0 | 0 | 0 | 0 | 0 | 0 |
| ENSG00000270442 | 0 | 0 | 0 | 0 | 0 | 0 |
| ENSG00000200275 | 0 | 0 | 0 | 0 | 0 | 0 |
| ENSG00000170624 | 0 | 0 | 0 | 0 | 0 | 0 |
| ENSG00000253370 | 0 | 0 | 0 | 0 | 0 | 0 |
| ENSG00000254163 | 0 | 0 | 0 | 0 | 0 | 0 |
| ENSG00000199468 | 0 | 0 | 0 | 0 | 0 | 0 |
| ENSG00000231989 | 6 | 8 | 3 | 6 | 3 | 12 |
| ENSG00000145850 | 0 | 0 | 0 | 0 | 0 | 0 |
| ENSG00000270560 | 0 | 0 | 0 | 0 | 0 | 0 |
| ENSG00000113249 | 0 | 0 | 0 | 0 | 0 | 0 |
| ENSG00000135077 | 0 | 1 | 1 | 0 | 0 | 0 |
| ENSG00000254246 | 0 | 0 | 0 | 0 | 0 | 0 |
| ENSG00000155868 | 527 | 283 | 335 | 226 | 312 | 279 |
| ENSG00000113263 | 0 | 0 | 0 | 0 | 0 | 0 |
| ENSG00000170613 | 0 | 0 | 0 | 0 | 0 | 0 |
| ENSG00000253980 | 0 | 0 | 0 | 0 | 0 | 0 |
| ENSG00000253653 | 1 | 1 | 3 | 2 | 0 | 0 |
| ENSG00000055163 | 1384 | 652 | 766 | 694 | 744 | 754 |
| ENSG00000285868 | 0 | 0 | 0 | 0 | 0 | 0 |
| ENSG00000172568 | 0 | 0 | 0 | 0 | 0 | 0 |
| ENSG00000251405 | 1 | 0 | 0 | 0 | 0 | 0 |
| ENSG00000248544 | 2 | 2 | 0 | 1 | 1 | 1 |
| ENSG00000135074 | 20 | 19 | 16 | 15 | 21 | 19 |
| ENSG00000253155 | 0 | 0 | 0 | 0 | 0 | 0 |
| ENSG00000172548 | 0 | 0 | 0 | 0 | 0 | 0 |
| ENSG00000253519 | 0 | 1 | 0 | 0 | 0 | 0 |
| ENSG00000252068 | 0 | 0 | 0 | 0 | 0 | 0 |
| ENSG00000254336 | 0 | 0 | 0 | 0 | 0 | 0 |
| ENSG00000039600 | 2 | 4 | 2 | 1 | 3 | 3 |
| ENSG00000187658 | 0 | 0 | 0 | 0 | 0 | 0 |
| ENSG00000253798 | 0 | 0 | 0 | 0 | 0 | 0 |

| | | | | | | |
|---|---|---|---|---|---|---|
| ENSG00000113272 | 775 | 473 | 534 | 407 | 540 | 504 |
| ENSG00000155858 | 597 | 319 | 397 | 268 | 331 | 326 |
| ENSG00000253630 | 0 | 0 | 0 | 0 | 0 | 0 |
| ENSG00000113282 | 3846 | 1803 | 2347 | 1358 | 2031 | 1989 |
| ENSG00000206819 | 0 | 0 | 0 | 0 | 0 | 0 |
| ENSG00000244331 | 0 | 0 | 0 | 0 | 0 | 0 |
| ENSG00000222626 | 0 | 0 | 0 | 0 | 0 | 0 |
| ENSG00000253743 | 0 | 0 | 0 | 0 | 0 | 0 |
| ENSG00000213414 | 0 | 0 | 0 | 1 | 1 | 0 |
| ENSG00000253422 | 0 | 0 | 0 | 0 | 0 | 0 |
| ENSG00000253792 | 0 | 0 | 0 | 0 | 0 | 0 |
| ENSG00000253673 | 0 | 0 | 0 | 0 | 0 | 0 |
| ENSG00000254047 | 0 | 0 | 0 | 0 | 0 | 0 |
| ENSG00000253449 | 0 | 0 | 0 | 0 | 0 | 0 |
| ENSG00000253424 | 0 | 0 | 0 | 0 | 0 | 0 |
| ENSG00000253134 | 0 | 0 | 0 | 0 | 0 | 0 |
| ENSG00000276778 | 0 | 0 | 0 | 0 | 0 | 0 |
| ENSG00000254350 | 0 | 0 | 0 | 0 | 0 | 0 |
| ENSG00000272085 | 0 | 0 | 0 | 0 | 0 | 0 |
| ENSG00000254135 | 0 | 0 | 0 | 0 | 0 | 0 |
| ENSG00000164330 | 0 | 0 | 0 | 0 | 0 | 0 |
| ENSG00000253811 | 0 | 0 | 0 | 0 | 0 | 0 |
| ENSG00000253456 | 0 | 0 | 0 | 0 | 0 | 0 |
| ENSG00000245812 | 0 | 0 | 0 | 0 | 0 | 0 |
| ENSG00000279204 | 0 | 0 | 0 | 0 | 0 | 0 |
| ENSG00000145860 | 729 | 345 | 389 | 396 | 508 | 449 |
| ENSG00000253256 | 0 | 0 | 0 | 0 | 0 | 0 |
| ENSG00000253315 | 0 | 0 | 0 | 0 | 0 | 0 |
| ENSG00000252458 | 0 | 0 | 0 | 0 | 0 | 0 |
| ENSG00000164332 | 414 | 238 | 180 | 108 | 334 | 208 |
| ENSG00000249738 | 0 | 0 | 0 | 0 | 0 | 0 |
| ENSG00000113302 | 0 | 0 | 0 | 0 | 0 | 0 |
| ENSG00000221601 | 0 | 0 | 0 | 0 | 0 | 0 |
| ENSG00000253298 | 0 | 0 | 0 | 0 | 0 | 0 |
| ENSG00000283413 | 4 | 2 | 2 | 0 | 2 | 3 |
| ENSG00000253311 | 0 | 0 | 0 | 0 | 0 | 0 |
| ENSG00000170214 | 2 | 0 | 3 | 0 | 0 | 0 |
| ENSG00000248626 | 0 | 1 | 4 | 0 | 1 | 1 |
| ENSG00000199398 | 0 | 0 | 0 | 0 | 0 | 0 |
| ENSG00000113312 | 2099 | 1032 | 1191 | 1186 | 1288 | 1224 |
| ENSG00000223908 | 0 | 0 | 1 | 0 | 0 | 0 |
| ENSG00000170234 | 858 | 406 | 489 | 321 | 491 | 526 |
| ENSG00000170231 | 16 | 6 | 3 | 18 | 13 | 25 |
| ENSG00000247699 | 0 | 0 | 0 | 0 | 0 | 0 |
| ENSG00000253772 | 0 | 0 | 0 | 0 | 0 | 0 |
| ENSG00000135083 | 383 | 210 | 194 | 382 | 448 | 410 |
| ENSG00000254373 | 2 | 1 | 2 | 2 | 1 | 1 |
| ENSG00000270612 | 0 | 0 | 0 | 0 | 0 | 0 |
| ENSG00000145861 | 0 | 1 | 1 | 2 | 3 | 2 |
| ENSG00000221886 | 220 | 119 | 115 | 88 | 155 | 121 |

| | | | | | | |
|---|---|---|---|---|---|---|
| ENSG00000164609 | 1503 | 803 | 979 | 559 | 1035 | 852 |
| ENSG00000164611 | 1059 | 457 | 615 | 811 | 769 | 695 |
| ENSG00000253522 | 0 | 0 | 0 | 0 | 0 | 0 |
| ENSG00000265237 | 0 | 0 | 0 | 0 | 0 | 0 |
| ENSG00000283733 | 0 | 0 | 0 | 0 | 0 | 0 |
| ENSG00000118322 | 0 | 0 | 2 | 0 | 1 | 1 |
| ENSG00000253687 | 0 | 0 | 0 | 0 | 0 | 0 |
| ENSG00000254391 | 0 | 0 | 0 | 0 | 0 | 0 |
| ENSG00000253417 | 0 | 0 | 0 | 0 | 0 | 1 |
| ENSG00000145864 | 0 | 0 | 0 | 0 | 0 | 0 |
| ENSG00000145863 | 0 | 0 | 0 | 0 | 0 | 0 |
| ENSG00000253403 | 0 | 0 | 0 | 0 | 0 | 0 |
| ENSG00000118990 | 0 | 0 | 0 | 0 | 0 | 0 |
| ENSG00000022355 | 0 | 0 | 0 | 0 | 0 | 0 |
| ENSG00000280776 | 2 | 0 | 4 | 0 | 2 | 0 |
| ENSG00000113327 | 3 | 1 | 0 | 0 | 1 | 2 |
| ENSG00000254186 | 0 | 0 | 0 | 0 | 2 | 0 |
| ENSG00000201474 | 0 | 0 | 0 | 0 | 0 | 0 |
| ENSG00000254351 | 0 | 0 | 0 | 0 | 0 | 0 |
| ENSG00000249449 | 0 | 0 | 0 | 0 | 0 | 0 |
| ENSG00000250114 | 0 | 0 | 0 | 0 | 0 | 0 |
| ENSG00000113328 | 2482 | 1332 | 1461 | 1434 | 2049 | 1468 |
| ENSG00000170584 | 1071 | 598 | 653 | 707 | 747 | 737 |
| ENSG00000072571 | 317 | 179 | 158 | 128 | 465 | 236 |
| ENSG00000251018 | 0 | 0 | 4 | 0 | 0 | 0 |
| ENSG00000038274 | 1692 | 798 | 1040 | 834 | 945 | 1004 |
| ENSG00000285590 | 0 | 0 | 0 | 0 | 0 | 0 |
| ENSG00000224012 | 0 | 0 | 0 | 0 | 0 | 0 |
| ENSG00000251998 | 0 | 0 | 0 | 0 | 0 | 0 |
| ENSG00000273569 | 0 | 0 | 0 | 0 | 0 | 0 |
| ENSG00000271679 | 0 | 0 | 0 | 0 | 0 | 0 |
| ENSG00000253331 | 0 | 0 | 0 | 0 | 0 | 0 |
| ENSG00000253538 | 0 | 0 | 0 | 0 | 0 | 0 |
| ENSG00000241956 | 0 | 0 | 0 | 0 | 0 | 0 |
| ENSG00000253236 | 0 | 0 | 0 | 0 | 0 | 0 |
| ENSG00000253600 | 0 | 0 | 0 | 0 | 0 | 0 |
| ENSG00000206845 | 0 | 0 | 0 | 0 | 0 | 0 |
| ENSG00000285549 | 0 | 0 | 0 | 0 | 0 | 0 |
| ENSG00000254066 | 0 | 0 | 0 | 0 | 0 | 0 |
| ENSG00000253693 | 0 | 0 | 0 | 0 | 0 | 0 |
| ENSG00000252794 | 0 | 0 | 0 | 0 | 0 | 0 |
| ENSG00000253512 | 0 | 0 | 0 | 0 | 0 | 0 |
| ENSG00000253946 | 0 | 0 | 0 | 0 | 0 | 0 |
| ENSG00000254252 | 0 | 0 | 0 | 0 | 0 | 0 |
| ENSG00000254171 | 0 | 0 | 0 | 0 | 0 | 0 |
| ENSG00000229119 | 10 | 6 | 7 | 6 | 9 | 7 |
| ENSG00000254130 | 0 | 0 | 0 | 0 | 0 | 0 |
| ENSG00000253713 | 0 | 0 | 0 | 0 | 0 | 0 |
| ENSG00000253469 | 0 | 0 | 0 | 0 | 0 | 0 |
| ENSG00000145934 | 0 | 0 | 1 | 0 | 0 | 0 |

| | | | | | | |
|---|---|---|---|---|---|---|
| ENSG00000254365 | 0 | 0 | 0 | 0 | 0 | 0 |
| ENSG00000254297 | 0 | 0 | 0 | 0 | 0 | 0 |
| ENSG00000253527 | 0 | 0 | 0 | 0 | 0 | 0 |
| ENSG00000254187 | 0 | 0 | 0 | 0 | 0 | 0 |
| ENSG00000253357 | 0 | 0 | 0 | 0 | 0 | 0 |
| ENSG00000253660 | 0 | 0 | 0 | 0 | 0 | 0 |
| ENSG00000253947 | 0 | 0 | 0 | 0 | 0 | 0 |
| ENSG00000253065 | 0 | 0 | 0 | 0 | 0 | 0 |
| ENSG00000253925 | 0 | 0 | 0 | 0 | 0 | 0 |
| ENSG00000279739 | 0 | 0 | 0 | 0 | 0 | 0 |
| ENSG00000253978 | 2 | 3 | 0 | 2 | 0 | 3 |
| ENSG00000113645 | 2790 | 1324 | 1608 | 1280 | 1476 | 1402 |
| ENSG00000113643 | 2754 | 1547 | 1709 | 1244 | 2153 | 1785 |
| ENSG00000188573 | 0 | 2 | 0 | 0 | 0 | 0 |
| ENSG00000120137 | 3312 | 1744 | 1543 | 946 | 2375 | 1528 |
| ENSG00000253861 | 0 | 0 | 0 | 0 | 0 | 0 |
| ENSG00000199035 | 0 | 0 | 0 | 0 | 0 | 0 |
| ENSG00000283612 | 0 | 0 | 0 | 0 | 0 | 0 |
| ENSG00000233913 | 3834 | 1899 | 2224 | 2900 | 2120 | 2053 |
| ENSG00000254192 | 0 | 0 | 0 | 0 | 0 | 0 |
| ENSG00000184347 | 6 | 2 | 1 | 0 | 2 | 0 |
| ENSG00000254042 | 0 | 0 | 0 | 0 | 0 | 0 |
| ENSG00000207739 | 0 | 0 | 0 | 0 | 0 | 0 |
| ENSG00000202345 | 0 | 0 | 0 | 0 | 0 | 0 |
| ENSG00000248965 | 0 | 0 | 0 | 0 | 0 | 0 |
| ENSG00000279752 | 0 | 0 | 0 | 0 | 0 | 0 |
| ENSG00000248222 | 0 | 0 | 0 | 0 | 0 | 0 |
| ENSG00000207619 | 0 | 0 | 0 | 0 | 0 | 0 |
| ENSG00000206614 | 0 | 0 | 0 | 0 | 0 | 0 |
| ENSG00000040275 | 483 | 265 | 242 | 217 | 431 | 266 |
| ENSG00000134516 | 6 | 2 | 3 | 3 | 6 | 9 |
| ENSG00000253269 | 0 | 0 | 0 | 0 | 0 | 0 |
| ENSG00000204767 | 0 | 0 | 0 | 0 | 0 | 0 |
| ENSG00000263831 | 0 | 0 | 0 | 0 | 0 | 0 |
| ENSG00000168269 | 22 | 4 | 11 | 7 | 5 | 8 |
| ENSG00000253219 | 2 | 2 | 1 | 3 | 0 | 0 |
| ENSG00000249601 | 0 | 0 | 0 | 0 | 0 | 1 |
| ENSG00000234511 | 0 | 0 | 0 | 0 | 0 | 2 |
| ENSG00000043462 | 0 | 0 | 0 | 0 | 0 | 1 |
| ENSG00000250274 | 0 | 0 | 0 | 0 | 0 | 0 |
| ENSG00000235172 | 0 | 0 | 0 | 0 | 0 | 0 |
| ENSG00000182132 | 2 | 0 | 0 | 0 | 0 | 1 |
| ENSG00000145936 | 6 | 1 | 5 | 0 | 3 | 1 |
| ENSG00000253647 | 0 | 0 | 0 | 0 | 0 | 0 |
| ENSG00000253858 | 0 | 0 | 0 | 0 | 0 | 0 |
| ENSG00000253591 | 2 | 0 | 0 | 0 | 1 | 1 |
| ENSG00000253348 | 6 | 1 | 1 | 0 | 0 | 0 |
| ENSG00000094755 | 26 | 11 | 10 | 5 | 8 | 8 |
| ENSG00000204764 | 426 | 249 | 242 | 122 | 312 | 227 |
| ENSG00000244501 | 0 | 0 | 0 | 0 | 0 | 0 |

| | | | | | | |
|---|---|---|---|---|---|---|
| ENSG00000253966 | 7 | 1 | 4 | 4 | 1 | 3 |
| ENSG00000253213 | 0 | 0 | 1 | 1 | 0 | 0 |
| ENSG00000275038 | 0 | 0 | 0 | 0 | 1 | 1 |
| ENSG00000164438 | 0 | 0 | 0 | 0 | 0 | 0 |
| ENSG00000213393 | 0 | 0 | 0 | 0 | 0 | 0 |
| ENSG00000265740 | 0 | 0 | 0 | 0 | 0 | 0 |
| ENSG00000254256 | 0 | 0 | 0 | 0 | 0 | 0 |
| ENSG00000206909 | 0 | 0 | 0 | 0 | 0 | 0 |
| ENSG00000254012 | 0 | 1 | 0 | 0 | 1 | 0 |
| ENSG00000264249 | 0 | 0 | 0 | 0 | 0 | 0 |
| ENSG00000181163 | 29498 | 16025 | 17348 | 17605 | 21826 | 18346 |
| ENSG00000156427 | 42 | 15 | 32 | 13 | 7 | 17 |
| ENSG00000285914 | 0 | 0 | 0 | 0 | 0 | 0 |
| ENSG00000253110 | 0 | 0 | 0 | 0 | 0 | 0 |
| ENSG00000185662 | 0 | 1 | 1 | 1 | 0 | 2 |
| ENSG00000072803 | 4452 | 2090 | 2534 | 1893 | 2386 | 2347 |
| ENSG00000072786 | 2898 | 1435 | 1769 | 1638 | 1805 | 1666 |
| ENSG00000214360 | 0 | 1 | 1 | 0 | 0 | 0 |
| ENSG00000168246 | 1158 | 539 | 689 | 560 | 627 | 647 |
| ENSG00000212529 | 0 | 0 | 0 | 0 | 0 | 0 |
| ENSG00000253218 | 0 | 0 | 0 | 0 | 0 | 0 |
| ENSG00000174705 | 320 | 163 | 214 | 178 | 206 | 217 |
| ENSG00000254299 | 0 | 0 | 0 | 0 | 0 | 0 |
| ENSG00000280318 | 0 | 0 | 0 | 0 | 0 | 0 |
| ENSG00000214357 | 749 | 426 | 553 | 460 | 540 | 541 |
| ENSG00000253683 | 11 | 7 | 4 | 4 | 2 | 4 |
| ENSG00000265160 | 0 | 0 | 0 | 0 | 0 | 0 |
| ENSG00000253445 | 0 | 0 | 0 | 0 | 0 | 0 |
| ENSG00000253295 | 2 | 5 | 1 | 3 | 0 | 2 |
| ENSG00000213386 | 0 | 1 | 1 | 0 | 2 | 3 |
| ENSG00000253736 | 10 | 0 | 2 | 1 | 1 | 1 |
| ENSG00000120129 | 414 | 196 | 239 | 264 | 232 | 274 |
| ENSG00000206741 | 0 | 0 | 0 | 0 | 0 | 0 |
| ENSG00000270243 | 0 | 0 | 0 | 0 | 0 | 0 |
| ENSG00000253628 | 0 | 0 | 0 | 0 | 0 | 0 |
| ENSG00000113719 | 7830 | 3488 | 4742 | 3460 | 3394 | 3896 |
| ENSG00000204758 | 20 | 11 | 8 | 7 | 9 | 9 |
| ENSG00000254295 | 0 | 0 | 0 | 0 | 0 | 0 |
| ENSG00000037241 | 1499 | 783 | 932 | 946 | 872 | 855 |
| ENSG00000253785 | 0 | 3 | 0 | 0 | 1 | 0 |
| ENSG00000113732 | 9086 | 4617 | 5584 | 5243 | 5290 | 4896 |
| ENSG00000254328 | 0 | 1 | 0 | 0 | 0 | 0 |
| ENSG00000212402 | 0 | 0 | 1 | 1 | 0 | 0 |
| ENSG00000207210 | 0 | 0 | 0 | 0 | 0 | 0 |
| ENSG00000164463 | 363 | 171 | 150 | 64 | 228 | 133 |
| ENSG00000253439 | 0 | 0 | 0 | 0 | 0 | 0 |
| ENSG00000253172 | 0 | 0 | 0 | 1 | 0 | 0 |
| ENSG00000113734 | 615 | 314 | 373 | 376 | 346 | 343 |
| ENSG00000254200 | 0 | 0 | 0 | 0 | 0 | 1 |
| ENSG00000183072 | 0 | 0 | 0 | 0 | 0 | 0 |

| | | | | | | |
|---|---|---|---|---|---|---|
| ENSG00000201277 | 0 | 0 | 0 | 0 | 0 | 0 |
| ENSG00000252169 | 0 | 0 | 0 | 0 | 0 | 0 |
| ENSG00000113739 | 7914 | 3759 | 4550 | 9503 | 9047 | 9273 |
| ENSG00000274994 | 0 | 0 | 0 | 0 | 0 | 0 |
| ENSG00000199219 | 0 | 0 | 0 | 0 | 0 | 0 |
| ENSG00000253968 | 0 | 1 | 0 | 0 | 0 | 0 |
| ENSG00000253141 | 0 | 1 | 0 | 0 | 0 | 0 |
| ENSG00000253768 | 45 | 31 | 58 | 90 | 84 | 73 |
| ENSG00000254164 | 15 | 14 | 10 | 16 | 9 | 10 |
| ENSG00000253955 | 8 | 7 | 10 | 4 | 12 | 14 |
| ENSG00000145919 | 2728 | 1446 | 1628 | 2038 | 1899 | 1911 |
| ENSG00000253959 | 0 | 1 | 0 | 1 | 0 | 1 |
| ENSG00000253428 | 0 | 0 | 0 | 0 | 0 | 0 |
| ENSG00000253686 | 4 | 1 | 1 | 0 | 3 | 0 |
| ENSG00000254211 | 0 | 0 | 1 | 1 | 2 | 1 |
| ENSG00000113742 | 695 | 330 | 379 | 207 | 448 | 307 |
| ENSG00000185056 | 0 | 0 | 1 | 0 | 2 | 0 |
| ENSG00000253109 | 0 | 0 | 0 | 0 | 0 | 0 |
| ENSG00000170091 | 2 | 1 | 3 | 0 | 1 | 0 |
| ENSG00000253244 | 0 | 0 | 0 | 0 | 0 | 0 |
| ENSG00000253447 | 0 | 0 | 0 | 0 | 0 | 0 |
| ENSG00000254011 | 0 | 0 | 0 | 0 | 0 | 0 |
| ENSG00000249722 | 0 | 0 | 0 | 0 | 0 | 0 |
| ENSG00000249306 | 8 | 3 | 0 | 2 | 1 | 0 |
| ENSG00000213376 | 0 | 0 | 0 | 0 | 2 | 0 |
| ENSG00000249031 | 0 | 0 | 0 | 0 | 0 | 1 |
| ENSG00000250256 | 11 | 3 | 10 | 4 | 2 | 1 |
| ENSG00000248929 | 0 | 0 | 1 | 0 | 0 | 0 |
| ENSG00000120149 | 2414 | 1073 | 1507 | 1097 | 1084 | 1253 |
| ENSG00000251670 | 0 | 0 | 1 | 1 | 0 | 1 |
| ENSG00000266890 | 0 | 0 | 2 | 0 | 1 | 0 |
| ENSG00000251144 | 0 | 1 | 2 | 0 | 0 | 0 |
| ENSG00000204754 | 0 | 0 | 0 | 0 | 0 | 0 |
| ENSG00000251122 | 0 | 0 | 0 | 0 | 0 | 0 |
| ENSG00000248293 | 0 | 0 | 0 | 0 | 0 | 0 |
| ENSG00000249403 | 0 | 0 | 0 | 0 | 0 | 0 |
| ENSG00000248355 | 0 | 0 | 0 | 0 | 0 | 0 |
| ENSG00000184845 | 205 | 115 | 178 | 97 | 124 | 190 |
| ENSG00000164466 | 5014 | 2363 | 3108 | 1931 | 2673 | 2638 |
| ENSG00000270839 | 0 | 0 | 0 | 1 | 1 | 0 |
| ENSG00000222111 | 0 | 0 | 0 | 0 | 0 | 0 |
| ENSG00000113749 | 22 | 5 | 23 | 11 | 11 | 16 |
| ENSG00000200648 | 0 | 0 | 0 | 0 | 0 | 0 |
| ENSG00000145920 | 0 | 0 | 0 | 0 | 0 | 0 |
| ENSG00000280068 | 2 | 1 | 4 | 0 | 0 | 1 |
| ENSG00000250820 | 0 | 0 | 0 | 0 | 0 | 0 |
| ENSG00000051596 | 1326 | 665 | 887 | 805 | 822 | 784 |
| ENSG00000250801 | 0 | 0 | 0 | 0 | 0 | 0 |
| ENSG00000272494 | 0 | 0 | 0 | 0 | 0 | 0 |
| ENSG00000250992 | 0 | 0 | 0 | 0 | 0 | 0 |

| | | | | | | |
|---|---|---|---|---|---|---|
| ENSG00000248469 | 0 | 0 | 0 | 0 | 0 | 0 |
| ENSG00000182230 | 0 | 0 | 0 | 0 | 0 | 0 |
| ENSG00000285476 | 0 | 0 | 0 | 0 | 0 | 0 |
| ENSG00000251623 | 0 | 0 | 0 | 0 | 0 | 0 |
| ENSG00000251458 | 0 | 0 | 0 | 0 | 0 | 0 |
| ENSG00000248596 | 3 | 0 | 2 | 0 | 1 | 1 |
| ENSG00000283559 | 0 | 0 | 0 | 0 | 0 | 0 |
| ENSG00000283235 | 5 | 5 | 0 | 1 | 4 | 2 |
| ENSG00000270989 | 0 | 0 | 0 | 0 | 0 | 0 |
| ENSG00000170085 | 523 | 240 | 285 | 204 | 282 | 273 |
| ENSG00000251667 | 2 | 2 | 2 | 1 | 1 | 0 |
| ENSG00000122203 | 6913 | 3541 | 4294 | 3283 | 3924 | 3980 |
| ENSG00000250909 | 3 | 4 | 6 | 3 | 8 | 11 |
| ENSG00000251414 | 19 | 15 | 16 | 3 | 13 | 9 |
| ENSG00000175414 | 9 | 7 | 6 | 3 | 7 | 7 |
| ENSG00000221464 | 0 | 0 | 0 | 0 | 0 | 0 |
| ENSG00000048162 | 882 | 566 | 586 | 829 | 698 | 698 |
| ENSG00000146066 | 1570 | 965 | 1241 | 1291 | 1051 | 1145 |
| ENSG00000175416 | 3775 | 1825 | 1952 | 2441 | 1974 | 1791 |
| ENSG00000280139 | 0 | 0 | 0 | 0 | 0 | 0 |
| ENSG00000113194 | 5295 | 2342 | 3054 | 2072 | 2558 | 2726 |
| ENSG00000146083 | 2796 | 1249 | 1722 | 1520 | 1625 | 1718 |
| ENSG00000074276 | 50 | 39 | 55 | 39 | 37 | 58 |
| ENSG00000243959 | 0 | 0 | 0 | 0 | 0 | 0 |
| ENSG00000207420 | 0 | 0 | 0 | 0 | 0 | 0 |
| ENSG00000169258 | 494 | 252 | 314 | 368 | 261 | 333 |
| ENSG00000074317 | 4 | 0 | 2 | 1 | 1 | 1 |
| ENSG00000266329 | 0 | 0 | 0 | 0 | 0 | 0 |
| ENSG00000175766 | 2 | 0 | 1 | 0 | 3 | 0 |
| ENSG00000048140 | 3640 | 1756 | 2060 | 2451 | 2156 | 2185 |
| ENSG00000248484 | 0 | 0 | 0 | 0 | 0 | 0 |
| ENSG00000251446 | 0 | 0 | 0 | 0 | 0 | 0 |
| ENSG00000248859 | 0 | 0 | 0 | 0 | 0 | 0 |
| ENSG00000113763 | 115 | 45 | 52 | 66 | 43 | 43 |
| ENSG00000160883 | 0 | 1 | 0 | 2 | 2 | 1 |
| ENSG00000087206 | 1178 | 629 | 767 | 627 | 749 | 726 |
| ENSG00000113761 | 1423 | 598 | 805 | 612 | 634 | 672 |
| ENSG00000251666 | 3 | 6 | 11 | 2 | 2 | 8 |
| ENSG00000160867 | 546 | 215 | 283 | 297 | 227 | 191 |
| ENSG00000165671 | 4261 | 2025 | 2463 | 1192 | 2443 | 2209 |
| ENSG00000240729 | 0 | 1 | 1 | 0 | 1 | 1 |
| ENSG00000270706 | 0 | 1 | 0 | 0 | 0 | 0 |
| ENSG00000169228 | 416 | 278 | 371 | 318 | 304 | 268 |
| ENSG00000213347 | 499 | 251 | 355 | 386 | 382 | 302 |
| ENSG00000169230 | 2413 | 1475 | 1688 | 2113 | 1842 | 1842 |
| ENSG00000169223 | 7717 | 3850 | 4713 | 5194 | 4422 | 4378 |
| ENSG00000239393 | 2 | 1 | 0 | 0 | 0 | 0 |
| ENSG00000169220 | 80 | 35 | 45 | 60 | 55 | 57 |
| ENSG00000131183 | 0 | 3 | 2 | 4 | 5 | 5 |
| ENSG00000196570 | 0 | 1 | 1 | 0 | 0 | 0 |

| | | | | | | |
|---|---|---|---|---|---|---|
| ENSG00000131187 | 224 | 137 | 206 | 259 | 218 | 217 |
| ENSG00000198055 | 1969 | 1019 | 1284 | 1411 | 1173 | 1272 |
| ENSG00000246334 | 37 | 17 | 21 | 19 | 21 | 13 |
| ENSG00000131188 | 804 | 333 | 452 | 514 | 407 | 445 |
| ENSG00000113758 | 4125 | 1869 | 2277 | 2564 | 2294 | 2174 |
| ENSG00000279821 | 26 | 9 | 4 | 19 | 18 | 9 |
| ENSG00000196923 | 2184 | 1028 | 1322 | 1678 | 1468 | 1349 |
| ENSG00000248996 | 5 | 2 | 4 | 1 | 2 | 2 |
| ENSG00000146094 | 254 | 78 | 110 | 117 | 96 | 123 |
| ENSG00000183258 | 3995 | 1897 | 2286 | 2382 | 2180 | 2253 |
| ENSG00000146067 | 1434 | 613 | 918 | 722 | 752 | 774 |
| ENSG00000272459 | 11 | 4 | 6 | 0 | 3 | 2 |
| ENSG00000184840 | 7755 | 3792 | 4335 | 4946 | 4537 | 4250 |
| ENSG00000027847 | 836 | 381 | 513 | 576 | 482 | 528 |
| ENSG00000247679 | 70 | 68 | 51 | 39 | 51 | 53 |
| ENSG00000246596 | 24 | 5 | 15 | 4 | 11 | 12 |
| ENSG00000249849 | 0 | 0 | 0 | 0 | 0 | 0 |
| ENSG00000170074 | 2 | 0 | 0 | 0 | 0 | 2 |
| ENSG00000249109 | 0 | 0 | 0 | 0 | 0 | 0 |
| ENSG00000248943 | 0 | 0 | 0 | 0 | 0 | 0 |
| ENSG00000170089 | 928 | 484 | 569 | 633 | 597 | 632 |
| ENSG00000214351 | 0 | 0 | 0 | 0 | 0 | 0 |
| ENSG00000249186 | 0 | 0 | 0 | 0 | 0 | 0 |
| ENSG00000250101 | 17 | 8 | 9 | 7 | 13 | 10 |
| ENSG00000249684 | 85 | 51 | 46 | 44 | 38 | 37 |
| ENSG00000280207 | 3 | 0 | 1 | 0 | 0 | 1 |
| ENSG00000249129 | 41 | 21 | 30 | 11 | 24 | 21 |
| ENSG00000175325 | 0 | 0 | 1 | 0 | 0 | 1 |
| ENSG00000204677 | 21 | 7 | 15 | 4 | 11 | 17 |
| ENSG00000218227 | 40 | 16 | 29 | 28 | 48 | 29 |
| ENSG00000145911 | 5224 | 2938 | 3172 | 3238 | 4005 | 3227 |
| ENSG00000145916 | 7058 | 3584 | 4333 | 3966 | 4325 | 4395 |
| ENSG00000145912 | 3465 | 2242 | 2733 | 3869 | 3102 | 3371 |
| ENSG00000251211 | 2 | 0 | 2 | 3 | 1 | 4 |
| ENSG00000244234 | 0 | 2 | 0 | 0 | 1 | 0 |
| ENSG00000197451 | 9081 | 4697 | 5574 | 5831 | 5926 | 6545 |
| ENSG00000175309 | 1644 | 769 | 1042 | 691 | 752 | 899 |
| ENSG00000050767 | 62 | 20 | 43 | 19 | 15 | 27 |
| ENSG00000271681 | 0 | 0 | 0 | 0 | 0 | 0 |
| ENSG00000253698 | 0 | 0 | 0 | 0 | 0 | 0 |
| ENSG00000242341 | 0 | 0 | 0 | 0 | 0 | 0 |
| ENSG00000245688 | 1 | 0 | 0 | 0 | 0 | 0 |
| ENSG00000113240 | 601 | 315 | 323 | 134 | 378 | 221 |
| ENSG00000252464 | 0 | 1 | 4 | 0 | 6 | 2 |
| ENSG00000285891 | 0 | 0 | 0 | 0 | 0 | 0 |
| ENSG00000285978 | 0 | 0 | 1 | 1 | 0 | 0 |
| ENSG00000169131 | 515 | 249 | 292 | 204 | 373 | 289 |
| ENSG00000250420 | 0 | 0 | 4 | 1 | 1 | 0 |
| ENSG00000274813 | 0 | 0 | 0 | 0 | 0 | 0 |
| ENSG00000253725 | 0 | 0 | 0 | 0 | 0 | 0 |

| | | | | | | |
|---|---|---|---|---|---|---|
| ENSG00000278441 | 0 | 0 | 0 | 0 | 0 | 0 |
| ENSG00000178338 | 343 | 157 | 178 | 84 | 215 | 171 |
| ENSG00000206624 | 0 | 0 | 0 | 0 | 0 | 0 |
| ENSG00000198939 | 28 | 16 | 13 | 12 | 14 | 18 |
| ENSG00000213328 | 0 | 0 | 0 | 0 | 2 | 0 |
| ENSG00000253869 | 0 | 0 | 0 | 1 | 0 | 1 |
| ENSG00000272431 | 0 | 0 | 0 | 0 | 0 | 0 |
| ENSG00000178187 | 0 | 0 | 0 | 0 | 0 | 0 |
| ENSG00000254035 | 0 | 0 | 1 | 0 | 0 | 0 |
| ENSG00000113262 | 0 | 0 | 0 | 0 | 0 | 0 |
| ENSG00000254158 | 0 | 0 | 0 | 0 | 0 | 0 |
| ENSG00000234284 | 139 | 87 | 70 | 35 | 94 | 46 |
| ENSG00000177932 | 223 | 149 | 154 | 74 | 168 | 96 |
| ENSG00000253144 | 0 | 0 | 0 | 0 | 0 | 0 |
| ENSG00000087116 | 0 | 0 | 0 | 0 | 2 | 1 |
| ENSG00000253979 | 0 | 0 | 0 | 0 | 0 | 0 |
| ENSG00000253163 | 0 | 0 | 0 | 0 | 0 | 0 |
| ENSG00000253617 | 0 | 0 | 0 | 0 | 0 | 0 |
| ENSG00000250674 | 0 | 0 | 0 | 0 | 0 | 0 |
| ENSG00000274716 | 0 | 0 | 0 | 0 | 0 | 0 |
| ENSG00000253652 | 0 | 0 | 0 | 0 | 2 | 0 |
| ENSG00000251545 | 0 | 0 | 0 | 0 | 0 | 0 |
| ENSG00000253520 | 2 | 0 | 2 | 1 | 0 | 2 |
| ENSG00000176783 | 2294 | 1142 | 1406 | 1220 | 1384 | 1279 |
| ENSG00000249140 | 0 | 3 | 1 | 2 | 0 | 0 |
| ENSG00000244945 | 14 | 9 | 10 | 12 | 15 | 23 |
| ENSG00000169045 | 13698 | 6270 | 8275 | 4330 | 7583 | 7528 |
| ENSG00000273549 | 0 | 2 | 0 | 0 | 0 | 0 |
| ENSG00000204661 | 0 | 2 | 4 | 4 | 1 | 6 |
| ENSG00000228259 | 0 | 0 | 1 | 0 | 1 | 1 |
| ENSG00000250999 | 12 | 5 | 11 | 7 | 6 | 9 |
| ENSG00000204659 | 9 | 7 | 5 | 5 | 5 | 4 |
| ENSG00000127022 | 25503 | 12231 | 14688 | 10725 | 14149 | 13905 |
| ENSG00000225051 | 26 | 11 | 14 | 9 | 12 | 13 |
| ENSG00000161021 | 3997 | 1637 | 2199 | 1625 | 1853 | 2103 |
| ENSG00000213316 | 192 | 49 | 74 | 54 | 51 | 68 |
| ENSG00000161013 | 6564 | 3158 | 4123 | 4064 | 3752 | 3941 |
| ENSG00000221394 | 0 | 0 | 0 | 0 | 0 | 0 |
| ENSG00000161011 | 10792 | 5393 | 6946 | 7134 | 6721 | 6783 |
| ENSG00000161010 | 1259 | 549 | 778 | 456 | 607 | 676 |
| ENSG00000222448 | 3 | 4 | 2 | 4 | 4 | 2 |
| ENSG00000252202 | 6 | 1 | 4 | 3 | 5 | 3 |
| ENSG00000245317 | 92 | 52 | 86 | 42 | 51 | 50 |
| ENSG00000197226 | 10527 | 4288 | 6045 | 5540 | 5242 | 5708 |
| ENSG00000113269 | 4136 | 1975 | 2558 | 2123 | 2082 | 2166 |
| ENSG00000242979 | 0 | 0 | 0 | 0 | 0 | 0 |
| ENSG00000249876 | 0 | 0 | 0 | 0 | 0 | 0 |
| ENSG00000249412 | 0 | 2 | 1 | 0 | 0 | 2 |
| ENSG00000285865 | 2 | 2 | 1 | 1 | 4 | 3 |
| ENSG00000198995 | 0 | 0 | 0 | 0 | 0 | 0 |

| | | | | | | |
|---|---|---|---|---|---|---|
| ENSG00000278965 | 0 | 0 | 0 | 0 | 0 | 0 |
| ENSG00000253480 | 0 | 0 | 0 | 0 | 0 | 0 |
| ENSG00000146090 | 10 | 3 | 1 | 5 | 7 | 5 |
| ENSG00000229721 | 0 | 1 | 0 | 0 | 0 | 0 |
| ENSG00000050748 | 3416 | 1721 | 2163 | 1609 | 2067 | 1970 |
| ENSG00000253908 | 12 | 7 | 8 | 2 | 5 | 4 |
| ENSG00000248367 | 53 | 30 | 37 | 31 | 34 | 29 |
| ENSG00000131459 | 184 | 75 | 117 | 136 | 130 | 107 |
| ENSG00000250509 | 0 | 0 | 0 | 0 | 0 | 0 |
| ENSG00000200555 | 0 | 0 | 0 | 0 | 0 | 0 |
| ENSG00000113300 | 3140 | 1619 | 1910 | 1009 | 1844 | 1572 |
| ENSG00000250749 | 0 | 1 | 1 | 0 | 0 | 0 |
| ENSG00000161055 | 0 | 0 | 0 | 1 | 0 | 0 |
| ENSG00000037280 | 23 | 21 | 16 | 9 | 15 | 12 |
| ENSG00000251352 | 0 | 0 | 0 | 0 | 0 | 0 |
| ENSG00000248717 | 0 | 0 | 0 | 0 | 0 | 0 |
| ENSG00000236791 | 0 | 0 | 0 | 0 | 0 | 0 |
| ENSG00000199892 | 0 | 0 | 0 | 0 | 0 | 0 |
| ENSG00000174339 | 0 | 0 | 0 | 0 | 0 | 0 |
| ENSG00000131446 | 2847 | 1226 | 1619 | 1668 | 1576 | 1539 |
| ENSG00000280161 | 35 | 18 | 16 | 21 | 15 | 11 |
| ENSG00000278970 | 869 | 462 | 598 | 566 | 525 | 473 |
| ENSG00000245060 | 718 | 357 | 425 | 408 | 430 | 394 |
| ENSG00000196670 | 1141 | 589 | 740 | 443 | 835 | 657 |
| ENSG00000113303 | 2 | 1 | 1 | 1 | 2 | 1 |
| ENSG00000222852 | 0 | 0 | 0 | 0 | 0 | 0 |
| ENSG00000243664 | 0 | 0 | 0 | 0 | 0 | 0 |
| ENSG00000249287 | 0 | 0 | 0 | 0 | 0 | 0 |
| ENSG00000248761 | 0 | 0 | 0 | 0 | 0 | 0 |
| ENSG00000231228 | 0 | 0 | 0 | 0 | 0 | 0 |
| ENSG00000168903 | 0 | 0 | 0 | 0 | 0 | 1 |
| ENSG00000206686 | 0 | 0 | 0 | 0 | 0 | 0 |
| ENSG00000165810 | 110 | 62 | 67 | 51 | 60 | 62 |
| ENSG00000277912 | 0 | 0 | 0 | 0 | 0 | 0 |
| ENSG00000241870 | 0 | 0 | 0 | 0 | 0 | 0 |
| ENSG00000214295 | 0 | 0 | 2 | 0 | 0 | 0 |
| ENSG00000248560 | 0 | 0 | 0 | 0 | 0 | 0 |
| ENSG00000185372 | 0 | 0 | 0 | 0 | 0 | 0 |
| ENSG00000182613 | 0 | 0 | 0 | 0 | 0 | 0 |
| ENSG00000248473 | 4 | 0 | 0 | 1 | 0 | 0 |
| ENSG00000250222 | 22 | 14 | 19 | 17 | 17 | 17 |
| ENSG00000146054 | 138 | 68 | 86 | 83 | 91 | 60 |
| ENSG00000250900 | 8 | 4 | 2 | 4 | 3 | 5 |
| ENSG00000248514 | 0 | 0 | 0 | 1 | 0 | 1 |
| ENSG00000146063 | 2345 | 1067 | 1501 | 1243 | 1199 | 1267 |
| ENSG00000264732 | 0 | 0 | 0 | 0 | 0 | 0 |
| ENSG00000247049 | 0 | 0 | 0 | 0 | 0 | 0 |
| ENSG00000204628 | 52272 | 26685 | 31128 | 36408 | 32126 | 31481 |
| ENSG00000272296 | 0 | 0 | 0 | 0 | 0 | 0 |
| ENSG00000264549 | 0 | 0 | 0 | 0 | 0 | 0 |

| | | | | | | |
|---|---|---|---|---|---|---|
| ENSG00000233937 | 192 | 97 | 119 | 107 | 116 | 116 |
| ENSG00000183718 | 485 | 223 | 295 | 150 | 252 | 230 |
| ENSG00000248275 | 327 | 150 | 246 | 168 | 197 | 203 |
| ENSG00000248103 | 3 | 1 | 0 | 0 | 1 | 0 |
| ENSG00000274525 | 4 | 1 | 6 | 1 | 4 | 2 |
| ENSG00000222533 | 1 | 0 | 0 | 0 | 1 | 2 |
| ENSG00000238035 | 2 | 3 | 10 | 3 | 5 | 5 |
| ENSG00000274553 | 0 | 0 | 0 | 0 | 0 | 0 |
| ENSG00000250765 | 0 | 0 | 0 | 0 | 0 | 0 |
| ENSG00000230178 | 0 | 0 | 0 | 0 | 0 | 0 |
| ENSG00000251035 | 0 | 0 | 0 | 0 | 0 | 0 |
| ENSG00000213285 | 0 | 0 | 0 | 0 | 0 | 0 |
| ENSG00000271530 | 0 | 0 | 0 | 0 | 0 | 0 |
| ENSG00000220212 | 0 | 0 | 0 | 0 | 0 | 0 |
| ENSG00000217929 | 0 | 0 | 0 | 1 | 0 | 0 |
| ENSG00000218577 | 0 | 0 | 0 | 0 | 0 | 0 |
| ENSG00000263667 | 0 | 0 | 0 | 0 | 0 | 0 |
| ENSG00000285578 | 0 | 0 | 0 | 0 | 0 | 0 |
| ENSG00000205653 | 0 | 0 | 0 | 0 | 0 | 0 |
| ENSG00000112679 | 907 | 460 | 574 | 477 | 453 | 484 |
| ENSG00000137265 | 2 | 1 | 2 | 0 | 0 | 0 |
| ENSG00000112685 | 1546 | 767 | 993 | 639 | 933 | 828 |
| ENSG00000230433 | 0 | 0 | 0 | 2 | 0 | 0 |
| ENSG00000188996 | 0 | 1 | 1 | 1 | 2 | 0 |
| ENSG00000219712 | 0 | 0 | 1 | 0 | 0 | 0 |
| ENSG00000272463 | 0 | 0 | 0 | 0 | 0 | 0 |
| ENSG00000271727 | 0 | 3 | 0 | 2 | 1 | 0 |
| ENSG00000271911 | 1 | 0 | 0 | 0 | 0 | 0 |
| ENSG00000272485 | 0 | 0 | 0 | 0 | 0 | 0 |
| ENSG00000276687 | 0 | 0 | 0 | 0 | 0 | 0 |
| ENSG00000278343 | 0 | 0 | 0 | 0 | 0 | 0 |
| ENSG00000270937 | 0 | 0 | 0 | 0 | 0 | 0 |
| ENSG00000176515 | 0 | 0 | 0 | 0 | 0 | 0 |
| ENSG00000164379 | 0 | 0 | 0 | 0 | 2 | 3 |
| ENSG00000281809 | 0 | 0 | 0 | 0 | 0 | 0 |
| ENSG00000261730 | 0 | 0 | 0 | 0 | 0 | 0 |
| ENSG00000137273 | 0 | 0 | 0 | 0 | 0 | 0 |
| ENSG00000275859 | 0 | 0 | 0 | 0 | 0 | 0 |
| ENSG00000243439 | 0 | 0 | 0 | 0 | 0 | 0 |
| ENSG00000218027 | 0 | 2 | 0 | 0 | 0 | 0 |
| ENSG00000272279 | 0 | 0 | 0 | 1 | 3 | 0 |
| ENSG00000280916 | 0 | 0 | 1 | 2 | 0 | 0 |
| ENSG00000054598 | 246 | 116 | 142 | 130 | 134 | 123 |
| ENSG00000112699 | 647 | 361 | 415 | 467 | 406 | 354 |
| ENSG00000285603 | 0 | 0 | 0 | 0 | 0 | 0 |
| ENSG00000250903 | 115 | 70 | 109 | 88 | 101 | 73 |
| ENSG00000236086 | 0 | 0 | 0 | 0 | 2 | 0 |
| ENSG00000272465 | 4 | 2 | 7 | 4 | 3 | 5 |
| ENSG00000280423 | 0 | 0 | 2 | 0 | 0 | 0 |
| ENSG00000228718 | 0 | 0 | 0 | 0 | 0 | 0 |

| | | | | | | |
|---|---|---|---|---|---|---|
| ENSG00000164385 | 0 | 3 | 2 | 2 | 1 | 1 |
| ENSG00000145949 | 38 | 26 | 22 | 13 | 14 | 17 |
| ENSG00000124535 | 3222 | 1531 | 1887 | 1762 | 1820 | 1867 |
| ENSG00000021355 | 1886 | 952 | 1230 | 899 | 1055 | 1006 |
| ENSG00000266750 | 0 | 0 | 0 | 0 | 0 | 0 |
| ENSG00000230438 | 2 | 0 | 3 | 2 | 1 | 3 |
| ENSG00000170542 | 47 | 23 | 28 | 15 | 33 | 11 |
| ENSG00000232999 | 0 | 0 | 0 | 0 | 0 | 0 |
| ENSG00000228365 | 0 | 0 | 0 | 0 | 0 | 0 |
| ENSG00000124570 | 2832 | 1406 | 1666 | 1559 | 1544 | 1510 |
| ENSG00000244041 | 198 | 77 | 128 | 95 | 87 | 86 |
| ENSG00000124588 | 1726 | 910 | 964 | 994 | 1133 | 1076 |
| ENSG00000224846 | 2 | 0 | 2 | 0 | 1 | 1 |
| ENSG00000271361 | 0 | 0 | 0 | 0 | 0 | 0 |
| ENSG00000274855 | 0 | 0 | 0 | 0 | 0 | 0 |
| ENSG00000232654 | 0 | 0 | 0 | 0 | 0 | 0 |
| ENSG00000217707 | 0 | 0 | 0 | 0 | 0 | 0 |
| ENSG00000229282 | 0 | 1 | 0 | 0 | 0 | 1 |
| ENSG00000137275 | 1561 | 763 | 1056 | 710 | 849 | 842 |
| ENSG00000272277 | 61 | 37 | 64 | 62 | 70 | 47 |
| ENSG00000222248 | 0 | 1 | 1 | 0 | 0 | 0 |
| ENSG00000137274 | 543 | 273 | 332 | 310 | 304 | 335 |
| ENSG00000228170 | 0 | 0 | 0 | 0 | 0 | 0 |
| ENSG00000137267 | 261 | 147 | 144 | 198 | 174 | 166 |
| ENSG00000216819 | 1 | 0 | 0 | 0 | 0 | 0 |
| ENSG00000230269 | 0 | 0 | 0 | 1 | 0 | 0 |
| ENSG00000137285 | 18 | 10 | 11 | 8 | 14 | 15 |
| ENSG00000180822 | 582 | 283 | 346 | 338 | 338 | 339 |
| ENSG00000137266 | 913 | 405 | 555 | 360 | 446 | 438 |
| ENSG00000272320 | 6 | 7 | 5 | 1 | 5 | 3 |
| ENSG00000228793 | 0 | 0 | 0 | 0 | 0 | 0 |
| ENSG00000270477 | 0 | 0 | 0 | 0 | 0 | 0 |
| ENSG00000168994 | 30 | 24 | 22 | 18 | 11 | 19 |
| ENSG00000270504 | 2 | 1 | 3 | 2 | 3 | 3 |
| ENSG00000219992 | 1 | 0 | 0 | 0 | 0 | 0 |
| ENSG00000238158 | 0 | 0 | 0 | 0 | 0 | 0 |
| ENSG00000233068 | 0 | 0 | 0 | 0 | 0 | 0 |
| ENSG00000145945 | 482 | 249 | 317 | 295 | 269 | 257 |
| ENSG00000260604 | 62 | 15 | 10 | 13 | 19 | 10 |
| ENSG00000218472 | 0 | 0 | 0 | 0 | 0 | 0 |
| ENSG00000217135 | 0 | 0 | 0 | 0 | 0 | 0 |
| ENSG00000217566 | 0 | 0 | 0 | 0 | 0 | 2 |
| ENSG00000216657 | 0 | 4 | 1 | 0 | 0 | 1 |
| ENSG00000279926 | 27 | 4 | 2 | 4 | 10 | 9 |
| ENSG00000272248 | 2 | 1 | 3 | 3 | 2 | 2 |
| ENSG00000230648 | 1 | 2 | 0 | 2 | 0 | 1 |
| ENSG00000112739 | 2220 | 1228 | 1130 | 494 | 1572 | 969 |
| ENSG00000145975 | 2 | 2 | 0 | 0 | 0 | 1 |
| ENSG00000185689 | 0 | 3 | 1 | 0 | 3 | 2 |
| ENSG00000198721 | 348 | 234 | 246 | 206 | 280 | 250 |

| | | | | | | |
|---|---|---|---|---|---|---|
| ENSG00000254821 | 0 | 0 | 0 | 0 | 0 | 0 |
| ENSG00000234817 | 8 | 2 | 3 | 1 | 2 | 3 |
| ENSG00000237716 | 0 | 0 | 0 | 0 | 0 | 0 |
| ENSG00000216307 | 0 | 0 | 0 | 0 | 0 | 0 |
| ENSG00000285424 | 0 | 1 | 0 | 0 | 2 | 0 |
| ENSG00000231811 | 0 | 2 | 3 | 0 | 2 | 2 |
| ENSG00000201185 | 0 | 0 | 0 | 0 | 0 | 0 |
| ENSG00000285694 | 0 | 0 | 0 | 1 | 0 | 0 |
| ENSG00000260239 | 51 | 22 | 35 | 11 | 19 | 27 |
| ENSG00000270695 | 0 | 0 | 0 | 0 | 0 | 0 |
| ENSG00000260673 | 0 | 0 | 0 | 0 | 0 | 0 |
| ENSG00000205444 | 2 | 1 | 1 | 0 | 1 | 1 |
| ENSG00000217385 | 0 | 0 | 1 | 0 | 0 | 0 |
| ENSG00000153046 | 2197 | 968 | 1402 | 956 | 1053 | 1147 |
| ENSG00000284823 | 0 | 0 | 0 | 0 | 0 | 0 |
| ENSG00000236336 | 2 | 0 | 0 | 1 | 0 | 0 |
| ENSG00000217030 | 0 | 0 | 0 | 0 | 0 | 0 |
| ENSG00000124787 | 433 | 221 | 218 | 254 | 283 | 283 |
| ENSG00000272142 | 59 | 40 | 39 | 41 | 37 | 41 |
| ENSG00000271978 | 6 | 1 | 9 | 2 | 4 | 6 |
| ENSG00000220685 | 6 | 0 | 0 | 0 | 2 | 1 |
| ENSG00000219607 | 6 | 2 | 3 | 2 | 3 | 1 |
| ENSG00000214113 | 314 | 188 | 188 | 199 | 204 | 200 |
| ENSG00000265083 | 0 | 0 | 0 | 0 | 0 | 0 |
| ENSG00000145982 | 398 | 156 | 247 | 224 | 203 | 215 |
| ENSG00000220446 | 0 | 0 | 0 | 0 | 0 | 0 |
| ENSG00000269985 | 0 | 0 | 0 | 0 | 0 | 0 |
| ENSG00000218574 | 0 | 0 | 0 | 0 | 0 | 1 |
| ENSG00000270174 | 0 | 3 | 0 | 0 | 0 | 0 |
| ENSG00000217239 | 2 | 0 | 0 | 0 | 0 | 1 |
| ENSG00000216360 | 0 | 0 | 0 | 2 | 1 | 1 |
| ENSG00000233064 | 0 | 0 | 1 | 0 | 0 | 0 |
| ENSG00000239472 | 0 | 1 | 0 | 0 | 0 | 0 |
| ENSG00000215057 | 0 | 0 | 0 | 0 | 0 | 0 |
| ENSG00000124785 | 0 | 0 | 0 | 0 | 0 | 0 |
| ENSG00000124491 | 0 | 0 | 0 | 0 | 0 | 0 |
| ENSG00000263572 | 0 | 0 | 0 | 0 | 0 | 0 |
| ENSG00000283616 | 0 | 0 | 0 | 0 | 0 | 0 |
| ENSG00000216863 | 0 | 0 | 0 | 0 | 0 | 0 |
| ENSG00000241216 | 0 | 0 | 0 | 0 | 0 | 0 |
| ENSG00000285717 | 0 | 0 | 0 | 0 | 0 | 0 |
| ENSG00000218868 | 0 | 0 | 0 | 0 | 0 | 0 |
| ENSG00000112799 | 0 | 0 | 0 | 0 | 0 | 0 |
| ENSG00000261211 | 147 | 52 | 69 | 53 | 101 | 57 |
| ENSG00000279368 | 71 | 22 | 45 | 22 | 41 | 24 |
| ENSG00000226281 | 1 | 0 | 6 | 2 | 1 | 3 |
| ENSG00000224532 | 0 | 0 | 0 | 0 | 0 | 0 |
| ENSG00000219986 | 0 | 0 | 0 | 0 | 0 | 0 |
| ENSG00000223342 | 0 | 0 | 0 | 0 | 0 | 0 |
| ENSG00000240936 | 0 | 0 | 0 | 0 | 0 | 0 |

| | | | | | | |
|---|---|---|---|---|---|---|
| ENSG00000203498 | 0 | 0 | 0 | 0 | 0 | 0 |
| ENSG00000124782 | 1789 | 747 | 1058 | 646 | 817 | 848 |
| ENSG00000225092 | 2 | 0 | 4 | 0 | 1 | 2 |
| ENSG00000201483 | 0 | 0 | 0 | 0 | 0 | 0 |
| ENSG00000285771 | 2 | 0 | 0 | 0 | 0 | 0 |
| ENSG00000124783 | 4161 | 2305 | 2163 | 1702 | 2959 | 2272 |
| ENSG00000238221 | 1 | 0 | 0 | 0 | 2 | 0 |
| ENSG00000164304 | 4 | 0 | 1 | 0 | 2 | 0 |
| ENSG00000219375 | 0 | 0 | 0 | 0 | 0 | 0 |
| ENSG00000220472 | 6 | 9 | 6 | 9 | 10 | 9 |
| ENSG00000230570 | 0 | 0 | 0 | 0 | 0 | 0 |
| ENSG00000124784 | 417 | 253 | 265 | 256 | 332 | 328 |
| ENSG00000233503 | 0 | 0 | 1 | 0 | 0 | 2 |
| ENSG00000219993 | 0 | 0 | 0 | 0 | 0 | 0 |
| ENSG00000234115 | 4 | 4 | 2 | 0 | 1 | 2 |
| ENSG00000261189 | 74 | 60 | 57 | 83 | 79 | 82 |
| ENSG00000096696 | 10103 | 5017 | 5225 | 2968 | 7914 | 5920 |
| ENSG00000168566 | 708 | 288 | 404 | 213 | 409 | 372 |
| ENSG00000236512 | 2 | 0 | 0 | 0 | 0 | 0 |
| ENSG00000153162 | 0 | 0 | 0 | 2 | 1 | 3 |
| ENSG00000239264 | 63 | 46 | 39 | 41 | 57 | 93 |
| ENSG00000259040 | 0 | 0 | 0 | 0 | 0 | 0 |
| ENSG00000217746 | 0 | 0 | 0 | 0 | 0 | 0 |
| ENSG00000219294 | 2 | 0 | 0 | 0 | 0 | 0 |
| ENSG00000188428 | 595 | 299 | 350 | 207 | 349 | 262 |
| ENSG00000265818 | 0 | 0 | 0 | 0 | 0 | 0 |
| ENSG00000124802 | 286 | 170 | 190 | 217 | 222 | 232 |
| ENSG00000232234 | 2 | 1 | 2 | 0 | 1 | 2 |
| ENSG00000285216 | 0 | 0 | 0 | 0 | 0 | 0 |
| ENSG00000234763 | 0 | 0 | 0 | 0 | 0 | 0 |
| ENSG00000124786 | 692 | 375 | 428 | 305 | 469 | 377 |
| ENSG00000285219 | 98 | 59 | 85 | 52 | 42 | 66 |
| ENSG00000276019 | 0 | 0 | 0 | 0 | 0 | 0 |
| ENSG00000225775 | 0 | 0 | 0 | 0 | 0 | 0 |
| ENSG00000181355 | 2 | 0 | 1 | 0 | 0 | 0 |
| ENSG00000216368 | 0 | 0 | 0 | 0 | 0 | 1 |
| ENSG00000229765 | 2 | 0 | 0 | 0 | 0 | 0 |
| ENSG00000221583 | 0 | 0 | 0 | 0 | 0 | 0 |
| ENSG00000216781 | 0 | 0 | 0 | 0 | 0 | 0 |
| ENSG00000137203 | 738 | 336 | 458 | 619 | 614 | 675 |
| ENSG00000285278 | 31 | 30 | 29 | 14 | 20 | 21 |
| ENSG00000229950 | 20 | 11 | 15 | 14 | 15 | 17 |
| ENSG00000228478 | 0 | 0 | 0 | 0 | 0 | 0 |
| ENSG00000285759 | 0 | 0 | 0 | 0 | 1 | 1 |
| ENSG00000183674 | 0 | 0 | 0 | 0 | 0 | 0 |
| ENSG00000229401 | 0 | 0 | 0 | 0 | 0 | 0 |
| ENSG00000263705 | 0 | 0 | 0 | 0 | 0 | 0 |
| ENSG00000238076 | 0 | 1 | 0 | 0 | 0 | 0 |
| ENSG00000231056 | 0 | 0 | 0 | 0 | 0 | 0 |
| ENSG00000111846 | 33 | 22 | 29 | 9 | 13 | 13 |

| | | | | | | |
|---|---|---|---|---|---|---|
| ENSG00000237685 | 7 | 4 | 6 | 1 | 1 | 4 |
| ENSG00000216359 | 3 | 2 | 0 | 2 | 0 | 1 |
| ENSG00000217181 | 2 | 0 | 0 | 0 | 0 | 0 |
| ENSG00000205318 | 0 | 0 | 0 | 0 | 0 | 0 |
| ENSG00000285763 | 27 | 29 | 18 | 13 | 19 | 13 |
| ENSG00000137434 | 81 | 28 | 45 | 49 | 41 | 41 |
| ENSG00000201048 | 0 | 0 | 0 | 0 | 0 | 0 |
| ENSG00000111845 | 582 | 300 | 296 | 288 | 377 | 326 |
| ENSG00000111843 | 3966 | 1950 | 2334 | 2443 | 2190 | 2262 |
| ENSG00000272097 | 53 | 12 | 23 | 19 | 27 | 20 |
| ENSG00000137210 | 2692 | 1477 | 1661 | 1852 | 1669 | 1728 |
| ENSG00000272162 | 0 | 0 | 0 | 0 | 0 | 0 |
| ENSG00000222383 | 1 | 0 | 4 | 1 | 1 | 0 |
| ENSG00000111837 | 18 | 13 | 15 | 8 | 8 | 9 |
| ENSG00000124827 | 2 | 1 | 1 | 0 | 0 | 0 |
| ENSG00000235051 | 0 | 0 | 0 | 1 | 0 | 0 |
| ENSG00000153157 | 4 | 3 | 2 | 0 | 3 | 4 |
| ENSG00000197977 | 433 | 198 | 251 | 184 | 229 | 226 |
| ENSG00000274541 | 0 | 0 | 0 | 0 | 0 | 0 |
| ENSG00000230314 | 11 | 2 | 6 | 4 | 7 | 6 |
| ENSG00000224531 | 595 | 295 | 353 | 223 | 349 | 351 |
| ENSG00000244476 | 2 | 0 | 1 | 0 | 0 | 0 |
| ENSG00000271245 | 0 | 0 | 0 | 0 | 0 | 0 |
| ENSG00000247925 | 2 | 0 | 2 | 0 | 1 | 1 |
| ENSG00000111859 | 1803 | 811 | 1220 | 1070 | 1329 | 1623 |
| ENSG00000233656 | 0 | 0 | 1 | 1 | 0 | 0 |
| ENSG00000242753 | 3 | 1 | 1 | 5 | 4 | 2 |
| ENSG00000202313 | 0 | 0 | 0 | 0 | 0 | 0 |
| ENSG00000218020 | 2 | 4 | 0 | 1 | 3 | 3 |
| ENSG00000205269 | 408 | 199 | 191 | 129 | 269 | 187 |
| ENSG00000271897 | 0 | 0 | 0 | 0 | 0 | 0 |
| ENSG00000277473 | 0 | 0 | 0 | 0 | 0 | 0 |
| ENSG00000207419 | 0 | 0 | 0 | 0 | 0 | 0 |
| ENSG00000111863 | 6 | 1 | 0 | 0 | 2 | 0 |
| ENSG00000234427 | 0 | 0 | 0 | 0 | 0 | 0 |
| ENSG00000215124 | 0 | 0 | 0 | 0 | 0 | 0 |
| ENSG00000225102 | 0 | 0 | 0 | 0 | 0 | 0 |
| ENSG00000229896 | 0 | 2 | 0 | 1 | 0 | 0 |
| ENSG00000095951 | 275 | 135 | 129 | 61 | 258 | 156 |
| ENSG00000078401 | 243 | 110 | 261 | 139 | 147 | 283 |
| ENSG00000218893 | 0 | 0 | 0 | 0 | 0 | 0 |
| ENSG00000223321 | 0 | 0 | 0 | 0 | 0 | 0 |
| ENSG00000212802 | 84 | 43 | 56 | 63 | 46 | 50 |
| ENSG00000234015 | 0 | 0 | 0 | 0 | 0 | 0 |
| ENSG00000112137 | 7 | 5 | 11 | 4 | 10 | 7 |
| ENSG00000206702 | 0 | 0 | 0 | 0 | 0 | 0 |
| ENSG00000215022 | 21 | 7 | 7 | 1 | 10 | 7 |
| ENSG00000145979 | 1103 | 584 | 735 | 672 | 781 | 698 |
| ENSG00000272379 | 0 | 0 | 0 | 0 | 0 | 0 |
| ENSG00000145990 | 314 | 136 | 147 | 106 | 142 | 152 |

| | | | | | | |
|---|---|---|---|---|---|---|
| ENSG00000237786 | 0 | 0 | 0 | 0 | 1 | 1 |
| ENSG00000218265 | 0 | 0 | 0 | 0 | 1 | 1 |
| ENSG00000202351 | 0 | 0 | 0 | 0 | 0 | 0 |
| ENSG00000124523 | 735 | 352 | 463 | 364 | 435 | 432 |
| ENSG00000261071 | 0 | 0 | 0 | 0 | 0 | 0 |
| ENSG00000225921 | 2195 | 1115 | 1266 | 1303 | 1324 | 1308 |
| ENSG00000010017 | 2317 | 1180 | 1512 | 1028 | 1395 | 1364 |
| ENSG00000050393 | 1640 | 823 | 929 | 880 | 1063 | 1010 |
| ENSG00000272209 | 16 | 6 | 7 | 8 | 15 | 9 |
| ENSG00000180537 | 6 | 3 | 4 | 5 | 7 | 2 |
| ENSG00000220868 | 0 | 0 | 0 | 0 | 0 | 0 |
| ENSG00000233981 | 0 | 0 | 0 | 0 | 0 | 0 |
| ENSG00000112149 | 60 | 37 | 29 | 28 | 39 | 27 |
| ENSG00000285639 | 0 | 0 | 0 | 0 | 0 | 0 |
| ENSG00000238987 | 0 | 0 | 0 | 0 | 0 | 0 |
| ENSG00000230631 | 0 | 0 | 0 | 0 | 0 | 0 |
| ENSG00000226673 | 0 | 0 | 0 | 0 | 0 | 0 |
| ENSG00000237346 | 0 | 0 | 0 | 0 | 1 | 1 |
| ENSG00000234540 | 0 | 0 | 0 | 0 | 0 | 0 |
| ENSG00000229646 | 1 | 0 | 0 | 0 | 1 | 2 |
| ENSG00000206960 | 0 | 0 | 0 | 0 | 0 | 0 |
| ENSG00000234261 | 77 | 49 | 38 | 45 | 40 | 53 |
| ENSG00000216754 | 0 | 0 | 0 | 0 | 1 | 0 |
| ENSG00000242989 | 0 | 0 | 0 | 0 | 0 | 0 |
| ENSG00000271888 | 61 | 37 | 41 | 43 | 46 | 53 |
| ENSG00000008083 | 4112 | 1786 | 2348 | 1729 | 2058 | 2161 |
| ENSG00000235488 | 11 | 2 | 8 | 4 | 5 | 7 |
| ENSG00000201367 | 0 | 0 | 0 | 0 | 0 | 0 |
| ENSG00000201519 | 0 | 0 | 0 | 0 | 0 | 0 |
| ENSG00000277411 | 4 | 0 | 2 | 0 | 0 | 0 |
| ENSG00000047579 | 712 | 367 | 443 | 425 | 409 | 354 |
| ENSG00000214027 | 0 | 1 | 2 | 1 | 0 | 0 |
| ENSG00000232947 | 0 | 0 | 0 | 0 | 0 | 0 |
| ENSG00000274212 | 0 | 0 | 0 | 0 | 0 | 0 |
| ENSG00000217585 | 0 | 0 | 0 | 0 | 0 | 0 |
| ENSG00000007944 | 2790 | 1290 | 1557 | 1145 | 1549 | 1357 |
| ENSG00000263712 | 0 | 0 | 0 | 0 | 2 | 1 |
| ENSG00000218073 | 2 | 0 | 1 | 0 | 0 | 0 |
| ENSG00000217078 | 0 | 1 | 3 | 0 | 2 | 1 |
| ENSG00000216364 | 0 | 0 | 2 | 0 | 0 | 0 |
| ENSG00000251831 | 0 | 0 | 0 | 0 | 0 | 0 |
| ENSG00000207193 | 0 | 0 | 0 | 0 | 0 | 0 |
| ENSG00000137198 | 184 | 120 | 124 | 89 | 90 | 91 |
| ENSG00000282024 | 4 | 0 | 2 | 0 | 0 | 2 |
| ENSG00000124788 | 1821 | 794 | 1098 | 818 | 1194 | 1101 |
| ENSG00000229931 | 79 | 25 | 40 | 40 | 44 | 49 |
| ENSG00000272341 | 153 | 72 | 82 | 65 | 148 | 93 |
| ENSG00000230873 | 0 | 0 | 0 | 2 | 2 | 1 |
| ENSG00000112183 | 16 | 4 | 8 | 13 | 9 | 11 |
| ENSG00000219314 | 0 | 0 | 0 | 0 | 0 | 0 |

| | | | | | | |
|---|---|---|---|---|---|---|
| ENSG00000112186 | 487 | 218 | 276 | 180 | 232 | 210 |
| ENSG00000184612 | 2 | 4 | 1 | 0 | 0 | 1 |
| ENSG00000218359 | 0 | 0 | 0 | 2 | 1 | 0 |
| ENSG00000137414 | 1817 | 869 | 1076 | 718 | 994 | 841 |
| ENSG00000124789 | 2555 | 1303 | 1496 | 787 | 1483 | 1373 |
| ENSG00000206881 | 0 | 0 | 0 | 0 | 0 | 0 |
| ENSG00000272269 | 18 | 3 | 4 | 7 | 7 | 5 |
| ENSG00000201415 | 0 | 0 | 0 | 0 | 0 | 0 |
| ENSG00000201442 | 0 | 0 | 0 | 0 | 0 | 0 |
| ENSG00000137177 | 1727 | 694 | 982 | 513 | 799 | 754 |
| ENSG00000220920 | 8 | 8 | 2 | 2 | 2 | 4 |
| ENSG00000187566 | 184 | 85 | 146 | 116 | 94 | 112 |
| ENSG00000137364 | 1033 | 450 | 537 | 380 | 538 | 491 |
| ENSG00000165097 | 1220 | 600 | 726 | 475 | 724 | 682 |
| ENSG00000124795 | 2250 | 1322 | 1012 | 627 | 1775 | 1184 |
| ENSG00000199715 | 2 | 1 | 0 | 0 | 0 | 0 |
| ENSG00000278102 | 0 | 0 | 0 | 0 | 0 | 0 |
| ENSG00000200681 | 0 | 0 | 0 | 0 | 0 | 0 |
| ENSG00000217414 | 0 | 1 | 2 | 0 | 0 | 0 |
| ENSG00000218976 | 0 | 0 | 1 | 0 | 0 | 0 |
| ENSG00000137393 | 610 | 316 | 419 | 161 | 241 | 234 |
| ENSG00000283408 | 0 | 0 | 0 | 0 | 0 | 0 |
| ENSG00000207775 | 0 | 0 | 0 | 0 | 0 | 0 |
| ENSG00000217379 | 0 | 0 | 0 | 0 | 0 | 0 |
| ENSG00000231662 | 0 | 0 | 0 | 4 | 0 | 0 |
| ENSG00000228412 | 206 | 110 | 135 | 101 | 121 | 97 |
| ENSG00000218643 | 0 | 0 | 1 | 0 | 0 | 0 |
| ENSG00000201523 | 0 | 0 | 0 | 0 | 0 | 0 |
| ENSG00000214012 | 6 | 8 | 7 | 5 | 1 | 3 |
| ENSG00000217314 | 0 | 0 | 0 | 0 | 0 | 0 |
| ENSG00000200957 | 0 | 0 | 0 | 0 | 0 | 0 |
| ENSG00000172201 | 507 | 237 | 275 | 314 | 290 | 288 |
| ENSG00000229700 | 0 | 0 | 1 | 1 | 0 | 2 |
| ENSG00000172197 | 721 | 269 | 403 | 210 | 199 | 242 |
| ENSG00000227803 | 0 | 2 | 0 | 0 | 0 | 0 |
| ENSG00000112242 | 1297 | 570 | 835 | 549 | 714 | 744 |
| ENSG00000240869 | 0 | 2 | 0 | 0 | 0 | 0 |
| ENSG00000224707 | 0 | 0 | 0 | 0 | 0 | 2 |
| ENSG00000222431 | 0 | 0 | 0 | 0 | 0 | 0 |
| ENSG00000201683 | 0 | 0 | 0 | 0 | 0 | 0 |
| ENSG00000145996 | 454 | 195 | 245 | 146 | 259 | 213 |
| ENSG00000225832 | 0 | 0 | 0 | 0 | 0 | 0 |
| ENSG00000233848 | 0 | 0 | 0 | 0 | 0 | 1 |
| ENSG00000252046 | 0 | 0 | 0 | 0 | 0 | 0 |
| ENSG00000227089 | 0 | 0 | 0 | 0 | 0 | 0 |
| ENSG00000280989 | 0 | 0 | 0 | 0 | 0 | 0 |
| ENSG00000231754 | 0 | 3 | 4 | 0 | 2 | 0 |
| ENSG00000283480 | 4 | 2 | 1 | 2 | 0 | 3 |
| ENSG00000124766 | 6044 | 2917 | 3657 | 3200 | 3930 | 3873 |
| ENSG00000220771 | 0 | 1 | 2 | 2 | 3 | 0 |

| | | | | | | |
|---|---|---|---|---|---|---|
| ENSG00000272168 | 2 | 5 | 6 | 4 | 5 | 7 |
| ENSG00000280443 | 0 | 0 | 0 | 0 | 0 | 0 |
| ENSG00000222515 | 0 | 0 | 0 | 0 | 0 | 0 |
| ENSG00000260455 | 0 | 0 | 0 | 0 | 0 | 0 |
| ENSG00000219404 | 0 | 0 | 0 | 0 | 0 | 0 |
| ENSG00000261568 | 0 | 0 | 0 | 0 | 0 | 0 |
| ENSG00000172179 | 0 | 0 | 0 | 0 | 0 | 0 |
| ENSG00000112273 | 0 | 0 | 0 | 1 | 0 | 0 |
| ENSG00000228772 | 0 | 0 | 0 | 0 | 0 | 0 |
| ENSG00000233358 | 0 | 0 | 0 | 0 | 0 | 0 |
| ENSG00000218476 | 0 | 0 | 0 | 0 | 0 | 0 |
| ENSG00000207394 | 0 | 0 | 0 | 0 | 0 | 0 |
| ENSG00000235743 | 0 | 0 | 0 | 0 | 0 | 0 |
| ENSG00000219453 | 0 | 0 | 0 | 0 | 0 | 0 |
| ENSG00000278128 | 0 | 0 | 0 | 0 | 0 | 0 |
| ENSG00000218512 | 0 | 0 | 0 | 1 | 0 | 0 |
| ENSG00000220748 | 0 | 0 | 0 | 0 | 0 | 0 |
| ENSG00000219582 | 0 | 0 | 0 | 0 | 0 | 0 |
| ENSG00000152954 | 0 | 0 | 0 | 0 | 0 | 1 |
| ENSG00000251830 | 0 | 0 | 0 | 0 | 0 | 0 |
| ENSG00000146038 | 1195 | 525 | 662 | 313 | 445 | 425 |
| ENSG00000146049 | 1 | 3 | 2 | 2 | 3 | 2 |
| ENSG00000202050 | 0 | 0 | 0 | 0 | 0 | 0 |
| ENSG00000124532 | 1440 | 725 | 818 | 738 | 854 | 850 |
| ENSG00000112293 | 119 | 40 | 48 | 39 | 50 | 40 |
| ENSG00000112294 | 3596 | 1594 | 2095 | 1215 | 1604 | 1705 |
| ENSG00000137261 | 109 | 74 | 76 | 81 | 103 | 78 |
| ENSG00000218186 | 0 | 0 | 1 | 1 | 0 | 0 |
| ENSG00000111802 | 1874 | 960 | 1061 | 810 | 1098 | 963 |
| ENSG00000112304 | 885 | 549 | 648 | 613 | 664 | 608 |
| ENSG00000272345 | 12 | 12 | 5 | 4 | 6 | 8 |
| ENSG00000112308 | 5933 | 2844 | 3208 | 2349 | 3361 | 3025 |
| ENSG00000272402 | 18 | 9 | 8 | 4 | 9 | 5 |
| ENSG00000224164 | 0 | 0 | 0 | 0 | 0 | 0 |
| ENSG00000218806 | 0 | 0 | 0 | 0 | 0 | 0 |
| ENSG00000112312 | 554 | 341 | 357 | 318 | 403 | 377 |
| ENSG00000260286 | 0 | 0 | 0 | 0 | 0 | 0 |
| ENSG00000282804 | 0 | 0 | 0 | 0 | 0 | 0 |
| ENSG00000111913 | 0 | 3 | 0 | 0 | 0 | 0 |
| ENSG00000217083 | 0 | 0 | 0 | 0 | 0 | 0 |
| ENSG00000214975 | 58 | 38 | 39 | 33 | 34 | 27 |
| ENSG00000220517 | 0 | 1 | 0 | 0 | 0 | 0 |
| ENSG00000244618 | 0 | 0 | 0 | 0 | 0 | 0 |
| ENSG00000229313 | 0 | 0 | 0 | 0 | 0 | 0 |
| ENSG00000223623 | 0 | 0 | 0 | 0 | 0 | 0 |
| ENSG00000168405 | 260 | 165 | 204 | 91 | 133 | 151 |
| ENSG00000285801 | 0 | 0 | 1 | 1 | 0 | 0 |
| ENSG00000219682 | 2 | 3 | 1 | 1 | 0 | 0 |
| ENSG00000262400 | 0 | 0 | 0 | 0 | 0 | 0 |
| ENSG00000217805 | 0 | 0 | 0 | 0 | 0 | 0 |

| | | | | | | |
|---|---|---|---|---|---|---|
| ENSG00000219681 | 0 | 0 | 0 | 0 | 0 | 0 |
| ENSG00000271108 | 0 | 0 | 0 | 0 | 0 | 0 |
| ENSG00000216718 | 0 | 0 | 0 | 1 | 0 | 0 |
| ENSG00000213972 | 1 | 0 | 0 | 2 | 0 | 1 |
| ENSG00000079691 | 1632 | 739 | 993 | 514 | 817 | 868 |
| ENSG00000207286 | 0 | 0 | 0 | 0 | 0 | 0 |
| ENSG00000207490 | 0 | 0 | 0 | 0 | 0 | 0 |
| ENSG00000079689 | 2 | 1 | 0 | 0 | 0 | 0 |
| ENSG00000217408 | 0 | 0 | 0 | 0 | 0 | 0 |
| ENSG00000164508 | 0 | 0 | 0 | 0 | 0 | 0 |
| ENSG00000146047 | 0 | 0 | 0 | 0 | 0 | 0 |
| ENSG00000178762 | 0 | 0 | 0 | 0 | 0 | 0 |
| ENSG00000216436 | 0 | 0 | 0 | 0 | 0 | 0 |
| ENSG00000146039 | 0 | 0 | 0 | 0 | 0 | 0 |
| ENSG00000124568 | 0 | 0 | 0 | 0 | 0 | 0 |
| ENSG00000124564 | 0 | 0 | 0 | 0 | 0 | 0 |
| ENSG00000242387 | 0 | 0 | 0 | 0 | 0 | 0 |
| ENSG00000277410 | 0 | 0 | 0 | 0 | 0 | 0 |
| ENSG00000112337 | 0 | 0 | 0 | 0 | 0 | 0 |
| ENSG00000112343 | 237 | 126 | 167 | 97 | 169 | 169 |
| ENSG00000272558 | 0 | 0 | 0 | 0 | 0 | 0 |
| ENSG00000272462 | 31 | 20 | 34 | 15 | 29 | 24 |
| ENSG00000272810 | 3 | 2 | 2 | 0 | 0 | 0 |
| ENSG00000224447 | 0 | 0 | 0 | 0 | 0 | 0 |
| ENSG00000124610 | 0 | 0 | 1 | 1 | 0 | 2 |
| ENSG00000275714 | 25 | 21 | 20 | 14 | 14 | 19 |
| ENSG00000278637 | 4 | 0 | 2 | 1 | 0 | 2 |
| ENSG00000278705 | 1 | 5 | 0 | 0 | 1 | 0 |
| ENSG00000274267 | 36 | 20 | 20 | 17 | 14 | 24 |
| ENSG00000278463 | 0 | 4 | 3 | 5 | 0 | 0 |
| ENSG00000276410 | 5 | 1 | 2 | 0 | 2 | 1 |
| ENSG00000234816 | 2 | 2 | 0 | 1 | 5 | 3 |
| ENSG00000278272 | 10 | 2 | 6 | 3 | 6 | 3 |
| ENSG00000187837 | 1518 | 724 | 977 | 913 | 824 | 808 |
| ENSG00000010704 | 169 | 106 | 97 | 82 | 103 | 68 |
| ENSG00000197061 | 8 | 3 | 3 | 2 | 1 | 2 |
| ENSG00000187475 | 4 | 2 | 5 | 1 | 2 | 2 |
| ENSG00000180596 | 525 | 216 | 269 | 208 | 256 | 208 |
| ENSG00000180573 | 4204 | 1760 | 2317 | 2111 | 1929 | 2019 |
| ENSG00000168298 | 2 | 3 | 1 | 0 | 1 | 6 |
| ENSG00000158373 | 2369 | 1182 | 1450 | 1280 | 1256 | 1168 |
| ENSG00000283064 | 20 | 12 | 20 | 13 | 9 | 11 |
| ENSG00000217159 | 5 | 0 | 3 | 2 | 0 | 2 |
| ENSG00000274290 | 98 | 33 | 49 | 51 | 39 | 49 |
| ENSG00000277157 | 27 | 7 | 15 | 14 | 10 | 7 |
| ENSG00000216331 | 1 | 6 | 4 | 5 | 1 | 3 |
| ENSG00000282988 | 47 | 38 | 41 | 32 | 22 | 30 |
| ENSG00000197409 | 0 | 0 | 0 | 0 | 0 | 0 |
| ENSG00000196866 | 0 | 0 | 0 | 0 | 0 | 0 |
| ENSG00000277224 | 94 | 38 | 48 | 17 | 26 | 30 |

| | | | | | | |
|---|---|---|---|---|---|---|
| ENSG00000217275 | 54 | 28 | 32 | 16 | 29 | 35 |
| ENSG00000276966 | 160 | 70 | 89 | 47 | 67 | 76 |
| ENSG00000273802 | 946 | 316 | 530 | 314 | 428 | 412 |
| ENSG00000277075 | 423 | 170 | 218 | 221 | 170 | 177 |
| ENSG00000277720 | 0 | 0 | 1 | 0 | 0 | 2 |
| ENSG00000274750 | 921 | 351 | 606 | 326 | 457 | 403 |
| ENSG00000218281 | 14 | 2 | 11 | 4 | 6 | 4 |
| ENSG00000124575 | 49 | 16 | 20 | 11 | 16 | 18 |
| ENSG00000274618 | 0 | 0 | 1 | 1 | 0 | 0 |
| ENSG00000275663 | 2 | 0 | 1 | 2 | 1 | 0 |
| ENSG00000277775 | 23 | 8 | 13 | 14 | 4 | 15 |
| ENSG00000275713 | 368 | 139 | 193 | 112 | 190 | 158 |
| ENSG00000273983 | 269 | 119 | 153 | 110 | 126 | 130 |
| ENSG00000218690 | 18 | 8 | 18 | 6 | 7 | 10 |
| ENSG00000278588 | 9 | 2 | 1 | 3 | 3 | 3 |
| ENSG00000158406 | 1272 | 558 | 783 | 480 | 559 | 595 |
| ENSG00000220875 | 0 | 1 | 0 | 0 | 0 | 2 |
| ENSG00000186470 | 2 | 4 | 2 | 3 | 6 | 3 |
| ENSG00000124508 | 342 | 201 | 225 | 166 | 229 | 216 |
| ENSG00000026950 | 18 | 4 | 3 | 5 | 6 | 9 |
| ENSG00000124549 | 48 | 41 | 42 | 32 | 44 | 39 |
| ENSG00000111801 | 27 | 4 | 9 | 5 | 16 | 11 |
| ENSG00000112763 | 932 | 410 | 612 | 391 | 534 | 523 |
| ENSG00000284688 | 0 | 0 | 0 | 0 | 0 | 0 |
| ENSG00000124557 | 0 | 0 | 0 | 0 | 0 | 0 |
| ENSG00000199289 | 0 | 0 | 0 | 0 | 0 | 0 |
| ENSG00000228223 | 258 | 109 | 133 | 89 | 132 | 141 |
| ENSG00000182952 | 1758 | 847 | 1155 | 962 | 1068 | 1074 |
| ENSG00000284607 | 6 | 1 | 0 | 2 | 1 | 3 |
| ENSG00000146109 | 1603 | 809 | 903 | 1004 | 834 | 914 |
| ENSG00000275846 | 0 | 3 | 1 | 0 | 0 | 0 |
| ENSG00000271071 | 2 | 0 | 0 | 0 | 0 | 0 |
| ENSG00000181315 | 396 | 168 | 240 | 147 | 264 | 166 |
| ENSG00000285571 | 5 | 1 | 3 | 4 | 6 | 5 |
| ENSG00000261584 | 0 | 0 | 0 | 0 | 2 | 1 |
| ENSG00000241549 | 25 | 21 | 11 | 18 | 7 | 12 |
| ENSG00000243307 | 0 | 0 | 0 | 0 | 0 | 0 |
| ENSG00000224843 | 17 | 6 | 6 | 13 | 4 | 11 |
| ENSG00000272312 | 0 | 0 | 0 | 0 | 0 | 0 |
| ENSG00000220581 | 0 | 0 | 0 | 0 | 0 | 0 |
| ENSG00000216762 | 0 | 0 | 0 | 0 | 0 | 0 |
| ENSG00000219770 | 0 | 0 | 0 | 0 | 0 | 0 |
| ENSG00000222800 | 0 | 0 | 0 | 2 | 0 | 0 |
| ENSG00000272468 | 36 | 10 | 16 | 16 | 22 | 25 |
| ENSG00000124635 | 192 | 87 | 108 | 98 | 114 | 79 |
| ENSG00000196787 | 596 | 258 | 347 | 199 | 249 | 258 |
| ENSG00000276180 | 1242 | 718 | 917 | 828 | 803 | 757 |
| ENSG00000197903 | 2905 | 1422 | 1794 | 1527 | 1555 | 1515 |
| ENSG00000274997 | 6 | 4 | 2 | 1 | 1 | 1 |
| ENSG00000265565 | 0 | 0 | 0 | 0 | 1 | 1 |

| | | | | | | |
|---|---|---|---|---|---|---|
| ENSG00000174572 | 2 | 1 | 0 | 0 | 0 | 0 |
| ENSG00000112812 | 338 | 166 | 170 | 148 | 135 | 117 |
| ENSG00000270935 | 0 | 0 | 0 | 0 | 0 | 0 |
| ENSG00000158553 | 4 | 1 | 0 | 0 | 1 | 0 |
| ENSG00000220758 | 0 | 0 | 0 | 0 | 0 | 0 |
| ENSG00000204789 | 34 | 18 | 16 | 7 | 33 | 19 |
| ENSG00000124613 | 125 | 69 | 91 | 47 | 95 | 71 |
| ENSG00000271755 | 12 | 3 | 4 | 2 | 4 | 2 |
| ENSG00000237154 | 0 | 0 | 0 | 0 | 0 | 0 |
| ENSG00000096654 | 203 | 132 | 124 | 79 | 169 | 134 |
| ENSG00000285849 | 2 | 0 | 2 | 0 | 0 | 1 |
| ENSG00000285703 | 1 | 1 | 0 | 0 | 2 | 2 |
| ENSG00000278332 | 0 | 0 | 0 | 0 | 0 | 0 |
| ENSG00000270666 | 0 | 0 | 0 | 0 | 0 | 0 |
| ENSG00000218347 | 0 | 0 | 0 | 0 | 0 | 0 |
| ENSG00000219738 | 0 | 0 | 0 | 0 | 0 | 0 |
| ENSG00000206671 | 0 | 0 | 0 | 0 | 0 | 0 |
| ENSG00000216676 | 1 | 0 | 0 | 0 | 0 | 0 |
| ENSG00000281706 | 48 | 33 | 34 | 25 | 35 | 40 |
| ENSG00000216915 | 6 | 4 | 8 | 3 | 8 | 7 |
| ENSG00000218069 | 0 | 0 | 0 | 0 | 1 | 0 |
| ENSG00000217862 | 4 | 3 | 5 | 3 | 5 | 4 |
| ENSG00000185130 | 23 | 4 | 10 | 11 | 10 | 13 |
| ENSG00000196747 | 233 | 103 | 158 | 103 | 94 | 123 |
| ENSG00000278828 | 1087 | 479 | 642 | 530 | 456 | 497 |
| ENSG00000276368 | 6 | 5 | 5 | 1 | 4 | 5 |
| ENSG00000273703 | 1 | 1 | 1 | 1 | 0 | 0 |
| ENSG00000197238 | 5 | 2 | 5 | 4 | 2 | 2 |
| ENSG00000273542 | 31 | 19 | 34 | 22 | 22 | 23 |
| ENSG00000233822 | 240 | 107 | 158 | 127 | 126 | 152 |
| ENSG00000275221 | 0 | 0 | 0 | 0 | 0 | 0 |
| ENSG00000217646 | 6 | 3 | 4 | 2 | 2 | 0 |
| ENSG00000276903 | 23 | 4 | 6 | 5 | 7 | 10 |
| ENSG00000184357 | 17 | 6 | 6 | 8 | 3 | 12 |
| ENSG00000275379 | 0 | 0 | 1 | 0 | 0 | 0 |
| ENSG00000275126 | 0 | 0 | 0 | 0 | 0 | 0 |
| ENSG00000197153 | 32 | 11 | 16 | 6 | 15 | 13 |
| ENSG00000278677 | 2 | 1 | 3 | 1 | 2 | 2 |
| ENSG00000274641 | 36 | 16 | 16 | 10 | 18 | 15 |
| ENSG00000238610 | 1 | 0 | 0 | 0 | 0 | 0 |
| ENSG00000168131 | 0 | 0 | 0 | 0 | 0 | 0 |
| ENSG00000168126 | 0 | 0 | 0 | 1 | 0 | 2 |
| ENSG00000124657 | 28 | 11 | 14 | 6 | 11 | 10 |
| ENSG00000214374 | 0 | 0 | 0 | 0 | 0 | 0 |
| ENSG00000216629 | 0 | 0 | 0 | 0 | 0 | 0 |
| ENSG00000217539 | 0 | 0 | 0 | 0 | 0 | 0 |
| ENSG00000199851 | 0 | 0 | 0 | 0 | 0 | 0 |
| ENSG00000217315 | 0 | 0 | 2 | 0 | 0 | 0 |
| ENSG00000187763 | 0 | 0 | 0 | 0 | 0 | 0 |
| ENSG00000182477 | 2 | 0 | 0 | 0 | 0 | 1 |

| | | | | | | |
|---|---|---|---|---|---|---|
| ENSG00000220721 | 0 | 0 | 0 | 0 | 0 | 2 |
| ENSG00000272009 | 41 | 15 | 31 | 21 | 28 | 18 |
| ENSG00000197279 | 80 | 46 | 80 | 42 | 79 | 53 |
| ENSG00000219891 | 10 | 9 | 6 | 5 | 9 | 6 |
| ENSG00000219392 | 0 | 1 | 0 | 2 | 1 | 0 |
| ENSG00000269293 | 48 | 19 | 40 | 36 | 23 | 21 |
| ENSG00000196812 | 337 | 179 | 215 | 188 | 230 | 210 |
| ENSG00000261839 | 0 | 0 | 4 | 1 | 1 | 1 |
| ENSG00000198315 | 1045 | 450 | 568 | 256 | 561 | 472 |
| ENSG00000226314 | 4 | 1 | 3 | 0 | 2 | 2 |
| ENSG00000280107 | 0 | 0 | 0 | 0 | 2 | 0 |
| ENSG00000216901 | 0 | 0 | 0 | 0 | 0 | 0 |
| ENSG00000218016 | 0 | 0 | 0 | 0 | 0 | 0 |
| ENSG00000176933 | 65 | 28 | 55 | 26 | 20 | 31 |
| ENSG00000137185 | 592 | 268 | 374 | 314 | 341 | 365 |
| ENSG00000187626 | 355 | 190 | 223 | 196 | 176 | 211 |
| ENSG00000189134 | 0 | 0 | 0 | 0 | 0 | 0 |
| ENSG00000197062 | 358 | 189 | 211 | 171 | 199 | 182 |
| ENSG00000276302 | 0 | 0 | 0 | 0 | 0 | 1 |
| ENSG00000137338 | 57 | 38 | 54 | 34 | 41 | 44 |
| ENSG00000273712 | 0 | 0 | 0 | 0 | 0 | 0 |
| ENSG00000270326 | 0 | 0 | 1 | 1 | 1 | 0 |
| ENSG00000235109 | 205 | 132 | 154 | 78 | 128 | 143 |
| ENSG00000189298 | 229 | 96 | 129 | 97 | 97 | 95 |
| ENSG00000158691 | 87 | 36 | 47 | 35 | 43 | 57 |
| ENSG00000251877 | 0 | 0 | 0 | 0 | 0 | 0 |
| ENSG00000187987 | 0 | 0 | 0 | 0 | 0 | 0 |
| ENSG00000231162 | 0 | 0 | 0 | 0 | 0 | 1 |
| ENSG00000219262 | 0 | 0 | 0 | 0 | 0 | 0 |
| ENSG00000198704 | 0 | 0 | 0 | 0 | 0 | 0 |
| ENSG00000224586 | 0 | 0 | 0 | 0 | 0 | 0 |
| ENSG00000232040 | 160 | 61 | 80 | 32 | 93 | 87 |
| ENSG00000246350 | 8 | 2 | 1 | 2 | 4 | 4 |
| ENSG00000271440 | 0 | 0 | 0 | 0 | 0 | 0 |
| ENSG00000235570 | 0 | 0 | 0 | 0 | 0 | 0 |
| ENSG00000237425 | 0 | 0 | 0 | 0 | 0 | 0 |
| ENSG00000235559 | 0 | 0 | 1 | 1 | 0 | 0 |
| ENSG00000225173 | 0 | 5 | 2 | 2 | 3 | 3 |
| ENSG00000225595 | 1 | 0 | 0 | 0 | 0 | 0 |
| ENSG00000213916 | 0 | 0 | 0 | 0 | 0 | 0 |
| ENSG00000233366 | 0 | 0 | 0 | 0 | 0 | 0 |
| ENSG00000224157 | 10 | 2 | 4 | 6 | 4 | 7 |
| ENSG00000204713 | 3283 | 1642 | 1980 | 1954 | 2034 | 1873 |
| ENSG00000212240 | 2 | 3 | 0 | 1 | 0 | 0 |
| ENSG00000204709 | 0 | 0 | 0 | 0 | 0 | 0 |
| ENSG00000228666 | 5 | 3 | 2 | 0 | 2 | 2 |
| ENSG00000263426 | 0 | 0 | 0 | 0 | 0 | 0 |
| ENSG00000227214 | 2 | 4 | 9 | 2 | 3 | 8 |
| ENSG00000244349 | 3 | 4 | 2 | 3 | 1 | 2 |
| ENSG00000197935 | 137 | 60 | 73 | 63 | 75 | 57 |

| | | | | | | |
|---|---|---|---|---|---|---|
| ENSG00000223677 | 0 | 0 | 0 | 0 | 0 | 0 |
| ENSG00000277661 | 0 | 0 | 0 | 0 | 0 | 0 |
| ENSG00000204704 | 0 | 0 | 0 | 0 | 0 | 0 |
| ENSG00000236909 | 0 | 0 | 0 | 0 | 0 | 0 |
| ENSG00000228364 | 0 | 0 | 0 | 0 | 0 | 0 |
| ENSG00000204703 | 0 | 0 | 0 | 0 | 0 | 0 |
| ENSG00000204702 | 0 | 0 | 0 | 0 | 0 | 0 |
| ENSG00000204701 | 0 | 0 | 0 | 0 | 0 | 0 |
| ENSG00000227206 | 0 | 0 | 0 | 0 | 0 | 0 |
| ENSG00000203492 | 0 | 0 | 0 | 0 | 0 | 0 |
| ENSG00000284656 | 0 | 0 | 0 | 0 | 0 | 0 |
| ENSG00000204700 | 0 | 0 | 0 | 0 | 0 | 0 |
| ENSG00000224233 | 0 | 0 | 0 | 0 | 0 | 0 |
| ENSG00000230598 | 0 | 0 | 0 | 0 | 0 | 0 |
| ENSG00000197171 | 0 | 0 | 0 | 0 | 0 | 0 |
| ENSG00000213911 | 0 | 0 | 0 | 0 | 0 | 0 |
| ENSG00000204697 | 0 | 0 | 0 | 0 | 0 | 0 |
| ENSG00000242524 | 0 | 0 | 0 | 0 | 0 | 0 |
| ENSG00000204695 | 0 | 0 | 0 | 0 | 0 | 0 |
| ENSG00000230056 | 0 | 0 | 0 | 0 | 0 | 0 |
| ENSG00000243729 | 0 | 0 | 0 | 0 | 0 | 0 |
| ENSG00000112462 | 0 | 0 | 0 | 0 | 0 | 0 |
| ENSG00000280236 | 0 | 0 | 0 | 0 | 0 | 0 |
| ENSG00000251608 | 0 | 0 | 0 | 0 | 0 | 0 |
| ENSG00000204694 | 0 | 0 | 0 | 0 | 0 | 0 |
| ENSG00000206474 | 0 | 0 | 0 | 0 | 0 | 0 |
| ENSG00000277881 | 0 | 0 | 0 | 0 | 0 | 0 |
| ENSG00000204688 | 0 | 0 | 0 | 0 | 0 | 0 |
| ENSG00000225797 | 0 | 0 | 0 | 0 | 0 | 0 |
| ENSG00000230164 | 0 | 0 | 0 | 0 | 0 | 0 |
| ENSG00000204687 | 0 | 0 | 0 | 0 | 0 | 0 |
| ENSG00000213900 | 0 | 0 | 0 | 0 | 1 | 0 |
| ENSG00000229274 | 0 | 0 | 0 | 0 | 0 | 0 |
| ENSG00000224582 | 0 | 0 | 0 | 0 | 0 | 0 |
| ENSG00000229281 | 0 | 0 | 0 | 0 | 0 | 0 |
| ENSG00000237988 | 24 | 8 | 17 | 14 | 16 | 15 |
| ENSG00000213886 | 0 | 0 | 0 | 0 | 0 | 0 |
| ENSG00000204681 | 315 | 115 | 186 | 129 | 145 | 162 |
| ENSG00000232173 | 0 | 0 | 0 | 0 | 0 | 0 |
| ENSG00000227609 | 0 | 0 | 0 | 0 | 0 | 0 |
| ENSG00000201330 | 0 | 0 | 0 | 0 | 0 | 0 |
| ENSG00000231301 | 0 | 0 | 0 | 0 | 0 | 0 |
| ENSG00000204657 | 0 | 0 | 0 | 0 | 0 | 0 |
| ENSG00000235238 | 6 | 0 | 4 | 4 | 2 | 2 |
| ENSG00000204655 | 0 | 0 | 0 | 0 | 0 | 0 |
| ENSG00000204644 | 0 | 1 | 0 | 1 | 0 | 0 |
| ENSG00000233916 | 0 | 0 | 0 | 0 | 0 | 0 |
| ENSG00000204642 | 149 | 79 | 82 | 77 | 80 | 56 |
| ENSG00000225864 | 0 | 0 | 0 | 0 | 0 | 0 |
| ENSG00000214922 | 58 | 32 | 37 | 26 | 36 | 37 |

| | | | | | | |
|---|---|---|---|---|---|---|
| ENSG00000239257 | 0 | 0 | 0 | 0 | 0 | 0 |
| ENSG00000273340 | 26 | 13 | 16 | 9 | 10 | 9 |
| ENSG00000227758 | 0 | 0 | 0 | 0 | 0 | 0 |
| ENSG00000199290 | 0 | 0 | 0 | 0 | 0 | 0 |
| ENSG00000235821 | 0 | 0 | 0 | 0 | 0 | 1 |
| ENSG00000272236 | 0 | 0 | 0 | 0 | 0 | 0 |
| ENSG00000285761 | 0 | 1 | 1 | 0 | 0 | 0 |
| ENSG00000270896 | 0 | 0 | 0 | 0 | 0 | 0 |
| ENSG00000181126 | 1 | 0 | 1 | 0 | 1 | 3 |
| ENSG00000176998 | 0 | 1 | 0 | 3 | 3 | 4 |
| ENSG00000261548 | 0 | 0 | 0 | 0 | 0 | 0 |
| ENSG00000213880 | 0 | 0 | 0 | 0 | 0 | 0 |
| ENSG00000237042 | 0 | 0 | 0 | 0 | 0 | 0 |
| ENSG00000204632 | 21 | 4 | 2 | 1 | 2 | 5 |
| ENSG00000229142 | 0 | 0 | 0 | 0 | 0 | 0 |
| ENSG00000285799 | 12 | 7 | 22 | 2 | 17 | 5 |
| ENSG00000281831 | 0 | 0 | 6 | 0 | 1 | 0 |
| ENSG00000230521 | 5 | 2 | 7 | 3 | 6 | 5 |
| ENSG00000206341 | 51 | 29 | 40 | 32 | 39 | 34 |
| ENSG00000231130 | 0 | 0 | 0 | 0 | 1 | 0 |
| ENSG00000233677 | 0 | 0 | 0 | 0 | 0 | 0 |
| ENSG00000235963 | 0 | 0 | 0 | 0 | 0 | 0 |
| ENSG00000227262 | 0 | 2 | 0 | 0 | 1 | 0 |
| ENSG00000230795 | 2 | 4 | 2 | 4 | 4 | 5 |
| ENSG00000228078 | 1 | 0 | 0 | 0 | 0 | 0 |
| ENSG00000206503 | 1754 | 1059 | 1153 | 1381 | 1134 | 1089 |
| ENSG00000227766 | 0 | 2 | 0 | 0 | 0 | 0 |
| ENSG00000235290 | 5 | 0 | 1 | 0 | 3 | 1 |
| ENSG00000229390 | 0 | 0 | 1 | 1 | 0 | 0 |
| ENSG00000204625 | 0 | 0 | 0 | 4 | 1 | 3 |
| ENSG00000238024 | 2 | 0 | 0 | 4 | 1 | 0 |
| ENSG00000224312 | 0 | 1 | 2 | 0 | 0 | 2 |
| ENSG00000204623 | 220 | 106 | 122 | 69 | 92 | 104 |
| ENSG00000204622 | 2 | 3 | 2 | 0 | 5 | 5 |
| ENSG00000237669 | 2 | 4 | 5 | 6 | 2 | 2 |
| ENSG00000232757 | 3 | 0 | 0 | 0 | 1 | 1 |
| ENSG00000278104 | 0 | 0 | 0 | 0 | 0 | 0 |
| ENSG00000275856 | 0 | 0 | 0 | 0 | 0 | 0 |
| ENSG00000066379 | 379 | 189 | 246 | 302 | 241 | 261 |
| ENSG00000278773 | 0 | 0 | 0 | 0 | 0 | 0 |
| ENSG00000204619 | 2458 | 1107 | 1410 | 1344 | 1334 | 1372 |
| ENSG00000204618 | 162 | 92 | 87 | 92 | 83 | 83 |
| ENSG00000204616 | 21 | 14 | 17 | 14 | 27 | 19 |
| ENSG00000231226 | 2 | 1 | 1 | 1 | 4 | 3 |
| ENSG00000252228 | 0 | 0 | 0 | 0 | 0 | 0 |
| ENSG00000204614 | 0 | 0 | 0 | 0 | 0 | 1 |
| ENSG00000204613 | 0 | 0 | 0 | 0 | 0 | 0 |
| ENSG00000204610 | 0 | 0 | 0 | 0 | 1 | 2 |
| ENSG00000234127 | 3046 | 1381 | 1821 | 1591 | 1510 | 1619 |
| ENSG00000233892 | 11 | 9 | 10 | 6 | 9 | 5 |

| | | | | | | |
|---|---|---|---|---|---|---|
| ENSG00000270604 | 12 | 8 | 3 | 6 | 4 | 8 |
| ENSG00000236475 | 0 | 0 | 1 | 0 | 0 | 2 |
| ENSG00000243753 | 78 | 23 | 43 | 28 | 40 | 28 |
| ENSG00000280128 | 210 | 99 | 90 | 53 | 158 | 79 |
| ENSG00000231074 | 554 | 314 | 390 | 293 | 404 | 410 |
| ENSG00000204599 | 504 | 231 | 301 | 234 | 296 | 285 |
| ENSG00000248167 | 0 | 0 | 0 | 1 | 0 | 0 |
| ENSG00000241370 | 94 | 50 | 69 | 52 | 72 | 71 |
| ENSG00000224372 | 0 | 0 | 0 | 0 | 0 | 0 |
| ENSG00000224486 | 0 | 0 | 0 | 0 | 0 | 0 |
| ENSG00000236405 | 0 | 0 | 0 | 0 | 0 | 0 |
| ENSG00000226577 | 0 | 0 | 0 | 0 | 0 | 0 |
| ENSG00000229068 | 0 | 0 | 0 | 0 | 1 | 0 |
| ENSG00000224936 | 0 | 0 | 0 | 0 | 0 | 0 |
| ENSG00000236603 | 6 | 4 | 6 | 7 | 2 | 12 |
| ENSG00000204592 | 7052 | 3564 | 4285 | 4467 | 4485 | 3849 |
| ENSG00000235781 | 12 | 10 | 8 | 1 | 7 | 4 |
| ENSG00000204590 | 3193 | 1508 | 1982 | 1933 | 1866 | 1674 |
| ENSG00000204576 | 760 | 392 | 443 | 360 | 417 | 439 |
| ENSG00000204574 | 4508 | 2161 | 2613 | 2425 | 2595 | 2541 |
| ENSG00000216101 | 0 | 0 | 0 | 0 | 0 | 0 |
| ENSG00000204569 | 3648 | 1602 | 2323 | 1937 | 1988 | 2368 |
| ENSG00000204568 | 2527 | 1261 | 1616 | 1427 | 1499 | 1485 |
| ENSG00000137343 | 508 | 205 | 331 | 289 | 252 | 332 |
| ENSG00000228415 | 0 | 3 | 2 | 1 | 3 | 3 |
| ENSG00000204564 | 734 | 317 | 435 | 407 | 382 | 410 |
| ENSG00000204560 | 2660 | 1297 | 1721 | 1389 | 1522 | 1470 |
| ENSG00000146112 | 1045 | 526 | 625 | 693 | 605 | 591 |
| ENSG00000137404 | 557 | 241 | 265 | 276 | 258 | 294 |
| ENSG00000230449 | 0 | 1 | 0 | 0 | 0 | 0 |
| ENSG00000137337 | 1147 | 525 | 737 | 540 | 626 | 742 |
| ENSG00000224328 | 0 | 0 | 0 | 0 | 0 | 0 |
| ENSG00000196230 | 17279 | 10518 | 11352 | 14014 | 14404 | 13953 |
| ENSG00000272540 | 3 | 5 | 11 | 10 | 4 | 8 |
| ENSG00000137312 | 7240 | 3299 | 4320 | 3408 | 2810 | 3258 |
| ENSG00000201988 | 2 | 2 | 0 | 0 | 0 | 1 |
| ENSG00000272273 | 1 | 7 | 2 | 0 | 4 | 3 |
| ENSG00000137331 | 3377 | 1588 | 2025 | 3449 | 3238 | 3188 |
| ENSG00000263608 | 0 | 0 | 0 | 0 | 0 | 0 |
| ENSG00000228022 | 4 | 3 | 5 | 3 | 2 | 0 |
| ENSG00000214894 | 141 | 81 | 76 | 93 | 115 | 101 |
| ENSG00000237923 | 0 | 1 | 0 | 0 | 0 | 0 |
| ENSG00000202241 | 0 | 0 | 0 | 0 | 0 | 0 |
| ENSG00000237775 | 0 | 0 | 1 | 0 | 1 | 0 |
| ENSG00000204580 | 20789 | 9274 | 12868 | 11774 | 11251 | 11453 |
| ENSG00000284370 | 0 | 0 | 0 | 0 | 0 | 0 |
| ENSG00000213780 | 13 | 7 | 13 | 15 | 14 | 15 |
| ENSG00000137411 | 1345 | 600 | 855 | 819 | 734 | 751 |
| ENSG00000196260 | 0 | 0 | 0 | 0 | 0 | 0 |
| ENSG00000252761 | 0 | 0 | 0 | 0 | 0 | 0 |

| | | | | | | |
|---|---|---|---|---|---|---|
| ENSG00000168631 | 0 | 0 | 0 | 0 | 0 | 0 |
| ENSG00000233529 | 0 | 0 | 0 | 0 | 0 | 0 |
| ENSG00000275906 | 0 | 0 | 0 | 0 | 0 | 0 |
| ENSG00000204544 | 0 | 0 | 0 | 0 | 0 | 0 |
| ENSG00000261272 | 0 | 0 | 0 | 0 | 0 | 0 |
| ENSG00000228789 | 0 | 0 | 0 | 0 | 0 | 0 |
| ENSG00000222895 | 0 | 0 | 0 | 0 | 0 | 0 |
| ENSG00000204542 | 0 | 1 | 0 | 0 | 0 | 1 |
| ENSG00000204540 | 10 | 11 | 9 | 7 | 17 | 13 |
| ENSG00000204539 | 1 | 1 | 0 | 1 | 2 | 1 |
| ENSG00000204538 | 1 | 3 | 0 | 0 | 2 | 2 |
| ENSG00000238211 | 0 | 0 | 0 | 0 | 0 | 0 |
| ENSG00000204536 | 606 | 295 | 358 | 349 | 333 | 363 |
| ENSG00000137310 | 612 | 383 | 477 | 324 | 370 | 422 |
| ENSG00000204531 | 10 | 2 | 5 | 5 | 3 | 12 |
| ENSG00000204528 | 4 | 1 | 0 | 0 | 0 | 1 |
| ENSG00000272501 | 40 | 26 | 36 | 25 | 25 | 30 |
| ENSG00000206344 | 17 | 12 | 7 | 10 | 10 | 8 |
| ENSG00000271821 | 0 | 0 | 0 | 0 | 0 | 0 |
| ENSG00000255726 | 0 | 0 | 0 | 0 | 0 | 0 |
| ENSG00000255899 | 0 | 0 | 0 | 0 | 0 | 0 |
| ENSG00000204525 | 4262 | 2183 | 2797 | 2514 | 2301 | 2365 |
| ENSG00000234745 | 1038 | 559 | 713 | 678 | 525 | 537 |
| ENSG00000214892 | 2 | 0 | 0 | 0 | 1 | 1 |
| ENSG00000227939 | 5 | 3 | 6 | 5 | 4 | 9 |
| ENSG00000231402 | 2 | 2 | 0 | 0 | 0 | 2 |
| ENSG00000256166 | 11 | 17 | 11 | 9 | 9 | 13 |
| ENSG00000229836 | 26 | 9 | 14 | 16 | 7 | 12 |
| ENSG00000277402 | 0 | 0 | 0 | 0 | 0 | 0 |
| ENSG00000271581 | 1 | 0 | 0 | 0 | 0 | 0 |
| ENSG00000228432 | 0 | 0 | 0 | 0 | 0 | 0 |
| ENSG00000285647 | 0 | 0 | 0 | 0 | 0 | 0 |
| ENSG00000201658 | 0 | 0 | 0 | 0 | 0 | 0 |
| ENSG00000230994 | 0 | 0 | 0 | 0 | 0 | 0 |
| ENSG00000223702 | 0 | 0 | 0 | 0 | 0 | 0 |
| ENSG00000225851 | 0 | 0 | 0 | 0 | 0 | 0 |
| ENSG00000272221 | 27 | 18 | 18 | 20 | 20 | 13 |
| ENSG00000204520 | 1565 | 759 | 951 | 1085 | 964 | 849 |
| ENSG00000206337 | 2 | 8 | 3 | 0 | 7 | 3 |
| ENSG00000199332 | 3 | 1 | 4 | 0 | 2 | 0 |
| ENSG00000230174 | 0 | 0 | 0 | 0 | 0 | 0 |
| ENSG00000233902 | 0 | 0 | 0 | 0 | 0 | 0 |
| ENSG00000204516 | 1329 | 590 | 738 | 563 | 679 | 646 |
| ENSG00000201680 | 0 | 0 | 0 | 0 | 0 | 0 |
| ENSG00000256851 | 0 | 0 | 0 | 0 | 0 | 0 |
| ENSG00000219797 | 0 | 0 | 0 | 0 | 0 | 0 |
| ENSG00000225499 | 0 | 1 | 0 | 0 | 0 | 0 |
| ENSG00000204511 | 6 | 0 | 1 | 0 | 0 | 0 |
| ENSG00000198563 | 3768 | 1657 | 2565 | 1494 | 2180 | 2168 |
| ENSG00000254870 | 0 | 0 | 0 | 0 | 0 | 0 |

| | | | | | | |
|---|---|---|---|---|---|---|
| ENSG00000201785 | 2 | 4 | 0 | 0 | 0 | 0 |
| ENSG00000265236 | 0 | 0 | 0 | 0 | 0 | 0 |
| ENSG00000234006 | 0 | 0 | 1 | 0 | 0 | 0 |
| ENSG00000213760 | 22 | 11 | 11 | 11 | 6 | 11 |
| ENSG00000204498 | 642 | 333 | 424 | 484 | 381 | 357 |
| ENSG00000226979 | 0 | 0 | 3 | 0 | 1 | 0 |
| ENSG00000232810 | 14 | 11 | 10 | 23 | 14 | 26 |
| ENSG00000227507 | 34 | 26 | 22 | 32 | 29 | 35 |
| ENSG00000204482 | 0 | 3 | 1 | 6 | 1 | 3 |
| ENSG00000204475 | 2 | 1 | 0 | 1 | 0 | 0 |
| ENSG00000230622 | 4 | 1 | 0 | 2 | 2 | 1 |
| ENSG00000204472 | 0 | 0 | 0 | 0 | 0 | 1 |
| ENSG00000204469 | 12172 | 5178 | 6980 | 6439 | 6012 | 6313 |
| ENSG00000200816 | 2 | 1 | 0 | 0 | 0 | 0 |
| ENSG00000274494 | 0 | 0 | 0 | 1 | 0 | 0 |
| ENSG00000204463 | 9026 | 3982 | 5914 | 5094 | 4331 | 4690 |
| ENSG00000204444 | 70 | 43 | 50 | 46 | 39 | 41 |
| ENSG00000204439 | 991 | 521 | 607 | 636 | 568 | 572 |
| ENSG00000227198 | 0 | 0 | 0 | 0 | 0 | 0 |
| ENSG00000204438 | 397 | 213 | 261 | 306 | 259 | 238 |
| ENSG00000201207 | 2 | 4 | 3 | 0 | 4 | 2 |
| ENSG00000204435 | 613 | 440 | 519 | 569 | 509 | 506 |
| ENSG00000263020 | 0 | 0 | 0 | 0 | 0 | 0 |
| ENSG00000240053 | 110 | 32 | 45 | 45 | 38 | 69 |
| ENSG00000204428 | 135 | 68 | 88 | 77 | 61 | 60 |
| ENSG00000204427 | 121 | 67 | 107 | 90 | 90 | 68 |
| ENSG00000204422 | 0 | 0 | 0 | 0 | 0 | 0 |
| ENSG00000266776 | 0 | 0 | 0 | 0 | 0 | 0 |
| ENSG00000204424 | 0 | 0 | 0 | 0 | 0 | 0 |
| ENSG00000250641 | 0 | 0 | 0 | 0 | 0 | 0 |
| ENSG00000255552 | 0 | 0 | 0 | 0 | 0 | 0 |
| ENSG00000244355 | 0 | 0 | 0 | 0 | 0 | 0 |
| ENSG00000204420 | 15 | 6 | 10 | 4 | 9 | 11 |
| ENSG00000204421 | 27 | 12 | 18 | 19 | 13 | 10 |
| ENSG00000213722 | 3776 | 1611 | 2095 | 1878 | 1724 | 1741 |
| ENSG00000213719 | 18424 | 9267 | 10848 | 12365 | 11278 | 10309 |
| ENSG00000204410 | 104 | 49 | 85 | 31 | 76 | 62 |
| ENSG00000255152 | 0 | 0 | 2 | 1 | 0 | 0 |
| ENSG00000252743 | 1 | 7 | 2 | 2 | 3 | 2 |
| ENSG00000228727 | 12 | 1 | 5 | 7 | 1 | 7 |
| ENSG00000235663 | 5 | 2 | 6 | 3 | 6 | 5 |
| ENSG00000204396 | 233 | 113 | 180 | 186 | 175 | 213 |
| ENSG00000204394 | 4078 | 1894 | 2495 | 3226 | 2465 | 2981 |
| ENSG00000201555 | 5 | 2 | 5 | 6 | 2 | 1 |
| ENSG00000204392 | 1197 | 568 | 722 | 838 | 727 | 692 |
| ENSG00000204390 | 66 | 45 | 46 | 35 | 25 | 33 |
| ENSG00000204389 | 3246 | 1453 | 1780 | 2035 | 1713 | 1676 |
| ENSG00000204388 | 2651 | 1232 | 1535 | 1726 | 1510 | 1570 |
| ENSG00000285565 | 0 | 0 | 0 | 0 | 0 | 2 |
| ENSG00000204387 | 1618 | 1107 | 1449 | 1520 | 1530 | 1564 |

| | | | | | | |
|---|---|---|---|---|---|---|
| ENSG00000201823 | 2 | 0 | 0 | 0 | 1 | 0 |
| ENSG00000201754 | 0 | 0 | 0 | 0 | 0 | 0 |
| ENSG00000204386 | 4468 | 1998 | 2591 | 2440 | 2187 | 2064 |
| ENSG00000204385 | 28 | 8 | 17 | 8 | 15 | 16 |
| ENSG00000237080 | 10 | 4 | 10 | 3 | 4 | 5 |
| ENSG00000204371 | 2597 | 1199 | 1646 | 1537 | 1417 | 1390 |
| ENSG00000166278 | 13 | 9 | 16 | 20 | 25 | 19 |
| ENSG00000204366 | 301 | 164 | 198 | 187 | 162 | 217 |
| ENSG00000244255 | 0 | 0 | 0 | 0 | 0 | 0 |
| ENSG00000281756 | 0 | 0 | 0 | 0 | 1 | 2 |
| ENSG00000243649 | 52 | 30 | 57 | 24 | 28 | 40 |
| ENSG00000204356 | 2549 | 1210 | 1450 | 1620 | 1532 | 1491 |
| ENSG00000284446 | 0 | 0 | 0 | 0 | 0 | 0 |
| ENSG00000204351 | 2249 | 854 | 1273 | 1119 | 1002 | 1134 |
| ENSG00000204348 | 743 | 334 | 418 | 422 | 396 | 430 |
| ENSG00000204344 | 655 | 293 | 472 | 365 | 324 | 394 |
| ENSG00000244731 | 26 | 10 | 12 | 6 | 5 | 9 |
| ENSG00000233627 | 0 | 0 | 0 | 0 | 0 | 0 |
| ENSG00000204338 | 11 | 3 | 1 | 1 | 4 | 2 |
| ENSG00000248290 | 0 | 0 | 0 | 0 | 0 | 0 |
| ENSG00000250535 | 2 | 9 | 3 | 4 | 5 | 5 |
| ENSG00000224389 | 2 | 3 | 4 | 3 | 5 | 4 |
| ENSG00000229776 | 0 | 0 | 0 | 0 | 0 | 0 |
| ENSG00000231852 | 10 | 3 | 2 | 5 | 2 | 4 |
| ENSG00000168477 | 72 | 32 | 32 | 41 | 39 | 43 |
| ENSG00000252512 | 0 | 0 | 0 | 0 | 0 | 0 |
| ENSG00000284829 | 0 | 0 | 0 | 0 | 0 | 0 |
| ENSG00000213676 | 3823 | 1942 | 2646 | 1917 | 2102 | 2046 |
| ENSG00000204315 | 338 | 208 | 223 | 256 | 173 | 182 |
| ENSG00000204314 | 87 | 29 | 37 | 44 | 23 | 26 |
| ENSG00000285085 | 0 | 0 | 0 | 0 | 2 | 2 |
| ENSG00000284954 | 7 | 13 | 10 | 13 | 17 | 12 |
| ENSG00000221988 | 301 | 176 | 238 | 202 | 183 | 203 |
| ENSG00000258388 | 18 | 13 | 10 | 14 | 15 | 9 |
| ENSG00000241404 | 41 | 11 | 30 | 29 | 14 | 21 |
| ENSG00000204310 | 1107 | 591 | 798 | 734 | 673 | 711 |
| ENSG00000284469 | 0 | 0 | 3 | 2 | 0 | 0 |
| ENSG00000204308 | 2402 | 1186 | 1392 | 1440 | 1285 | 1262 |
| ENSG00000277264 | 0 | 0 | 0 | 0 | 0 | 0 |
| ENSG00000204305 | 204 | 70 | 103 | 74 | 77 | 110 |
| ENSG00000273333 | 0 | 0 | 0 | 0 | 0 | 0 |
| ENSG00000204304 | 2884 | 1254 | 1779 | 1322 | 1392 | 1518 |
| ENSG00000213654 | 30 | 9 | 16 | 19 | 22 | 19 |
| ENSG00000204301 | 15 | 12 | 12 | 10 | 6 | 5 |
| ENSG00000225914 | 4 | 0 | 0 | 0 | 0 | 0 |
| ENSG00000204296 | 0 | 0 | 0 | 0 | 0 | 0 |
| ENSG00000237285 | 0 | 0 | 0 | 1 | 0 | 0 |
| ENSG00000223335 | 0 | 0 | 0 | 0 | 0 | 0 |
| ENSG00000204290 | 0 | 0 | 0 | 0 | 0 | 0 |
| ENSG00000204287 | 4 | 3 | 6 | 0 | 1 | 1 |

| | | | | | | |
|---|---|---|---|---|---|---|
| ENSG00000196301 | 0 | 0 | 0 | 4 | 0 | 1 |
| ENSG00000198502 | 70 | 49 | 41 | 52 | 37 | 40 |
| ENSG00000251916 | 0 | 0 | 0 | 0 | 0 | 0 |
| ENSG00000229391 | 13 | 1 | 2 | 4 | 3 | 1 |
| ENSG00000196126 | 123 | 81 | 83 | 77 | 78 | 63 |
| ENSG00000196735 | 6 | 6 | 1 | 3 | 1 | 1 |
| ENSG00000179344 | 417 | 297 | 293 | 268 | 284 | 213 |
| ENSG00000223534 | 1 | 0 | 0 | 3 | 2 | 3 |
| ENSG00000235040 | 0 | 0 | 0 | 0 | 1 | 0 |
| ENSG00000232080 | 0 | 0 | 0 | 0 | 0 | 0 |
| ENSG00000226030 | 0 | 0 | 0 | 0 | 0 | 0 |
| ENSG00000237541 | 0 | 0 | 0 | 0 | 0 | 0 |
| ENSG00000263649 | 0 | 0 | 0 | 0 | 0 | 0 |
| ENSG00000232629 | 0 | 0 | 0 | 1 | 0 | 0 |
| ENSG00000241106 | 0 | 0 | 1 | 0 | 0 | 1 |
| ENSG00000250264 | 0 | 0 | 0 | 0 | 0 | 0 |
| ENSG00000204267 | 138 | 89 | 81 | 93 | 101 | 103 |
| ENSG00000204264 | 9 | 5 | 3 | 5 | 6 | 0 |
| ENSG00000204261 | 0 | 0 | 1 | 1 | 0 | 1 |
| ENSG00000240065 | 14 | 17 | 12 | 9 | 22 | 14 |
| ENSG00000168394 | 750 | 381 | 449 | 474 | 532 | 435 |
| ENSG00000234515 | 0 | 0 | 1 | 0 | 0 | 0 |
| ENSG00000235301 | 0 | 0 | 0 | 0 | 3 | 1 |
| ENSG00000242574 | 2 | 3 | 1 | 2 | 0 | 3 |
| ENSG00000248993 | 0 | 0 | 0 | 0 | 0 | 0 |
| ENSG00000204257 | 188 | 90 | 111 | 70 | 96 | 84 |
| ENSG00000204256 | 11236 | 5177 | 6822 | 5369 | 5881 | 6125 |
| ENSG00000223837 | 0 | 0 | 0 | 0 | 0 | 0 |
| ENSG00000263756 | 0 | 0 | 0 | 0 | 0 | 0 |
| ENSG00000204252 | 0 | 0 | 0 | 0 | 1 | 2 |
| ENSG00000231389 | 4 | 4 | 1 | 2 | 0 | 1 |
| ENSG00000223865 | 0 | 6 | 3 | 1 | 3 | 2 |
| ENSG00000224796 | 0 | 0 | 0 | 0 | 0 | 0 |
| ENSG00000231461 | 0 | 0 | 0 | 0 | 0 | 0 |
| ENSG00000228688 | 0 | 0 | 0 | 0 | 0 | 0 |
| ENSG00000224557 | 0 | 0 | 0 | 0 | 0 | 0 |
| ENSG00000237398 | 0 | 0 | 0 | 0 | 0 | 0 |
| ENSG00000230313 | 0 | 0 | 0 | 0 | 0 | 0 |
| ENSG00000204248 | 408 | 140 | 244 | 122 | 165 | 163 |
| ENSG00000204231 | 1918 | 830 | 1110 | 947 | 947 | 912 |
| ENSG00000202441 | 14 | 8 | 7 | 4 | 4 | 4 |
| ENSG00000112473 | 6539 | 2967 | 3679 | 3299 | 3379 | 3235 |
| ENSG00000204228 | 226 | 124 | 171 | 124 | 96 | 136 |
| ENSG00000199036 | 0 | 0 | 1 | 0 | 3 | 1 |
| ENSG00000204227 | 1711 | 730 | 1034 | 912 | 817 | 889 |
| ENSG00000225463 | 0 | 0 | 0 | 0 | 0 | 0 |
| ENSG00000223457 | 0 | 0 | 0 | 0 | 0 | 0 |
| ENSG00000272217 | 2 | 2 | 4 | 2 | 3 | 3 |
| ENSG00000232940 | 7 | 5 | 8 | 2 | 7 | 4 |
| ENSG00000223501 | 2919 | 1397 | 1919 | 1298 | 1379 | 1602 |

| | | | | | | |
|---|---|---|---|---|---|---|
| ENSG00000231500 | 42018 | 21168 | 24391 | 29242 | 25176 | 24897 |
| ENSG00000235863 | 452 | 184 | 280 | 220 | 181 | 188 |
| ENSG00000227057 | 2676 | 1346 | 1622 | 1759 | 1582 | 1661 |
| ENSG00000284517 | 0 | 0 | 0 | 0 | 0 | 0 |
| ENSG00000204220 | 1367 | 706 | 871 | 936 | 855 | 807 |
| ENSG00000284064 | 0 | 0 | 0 | 0 | 0 | 0 |
| ENSG00000237441 | 6329 | 2557 | 3950 | 3269 | 2929 | 3233 |
| ENSG00000231925 | 5246 | 2373 | 3309 | 2573 | 2465 | 2621 |
| ENSG00000236104 | 1334 | 592 | 747 | 657 | 639 | 672 |
| ENSG00000204209 | 2199 | 973 | 1287 | 1330 | 1221 | 1336 |
| ENSG00000285064 | 0 | 0 | 0 | 0 | 0 | 0 |
| ENSG00000229596 | 0 | 0 | 1 | 0 | 0 | 0 |
| ENSG00000228285 | 2 | 0 | 0 | 0 | 2 | 1 |
| ENSG00000225644 | 0 | 0 | 0 | 0 | 0 | 0 |
| ENSG00000237649 | 1232 | 525 | 724 | 726 | 778 | 846 |
| ENSG00000204194 | 0 | 0 | 0 | 1 | 0 | 0 |
| ENSG00000112511 | 1522 | 640 | 898 | 779 | 701 | 696 |
| ENSG00000112514 | 4151 | 2018 | 2609 | 2895 | 2417 | 2406 |
| ENSG00000197283 | 887 | 373 | 569 | 371 | 380 | 584 |
| ENSG00000274259 | 8 | 2 | 3 | 6 | 3 | 3 |
| ENSG00000284256 | 1 | 0 | 1 | 0 | 0 | 0 |
| ENSG00000213588 | 0 | 0 | 0 | 1 | 1 | 1 |
| ENSG00000204188 | 1 | 0 | 1 | 1 | 0 | 1 |
| ENSG00000242014 | 0 | 0 | 0 | 0 | 0 | 0 |
| ENSG00000276746 | 0 | 0 | 0 | 0 | 0 | 1 |
| ENSG00000030110 | 3210 | 1722 | 2076 | 1933 | 1777 | 1720 |
| ENSG00000197251 | 0 | 0 | 0 | 0 | 0 | 0 |
| ENSG00000096433 | 2840 | 1223 | 1775 | 1276 | 1513 | 1626 |
| ENSG00000137288 | 1589 | 802 | 1010 | 1171 | 1000 | 954 |
| ENSG00000266509 | 0 | 0 | 0 | 0 | 0 | 0 |
| ENSG00000161896 | 2 | 2 | 0 | 2 | 1 | 0 |
| ENSG00000161904 | 2668 | 1310 | 1588 | 1571 | 1532 | 1475 |
| ENSG00000096395 | 3 | 0 | 0 | 1 | 0 | 0 |
| ENSG00000249346 | 18 | 7 | 8 | 10 | 11 | 12 |
| ENSG00000271362 | 0 | 0 | 0 | 0 | 0 | 0 |
| ENSG00000233183 | 10 | 3 | 5 | 2 | 5 | 5 |
| ENSG00000276824 | 0 | 0 | 0 | 0 | 0 | 0 |
| ENSG00000221697 | 0 | 0 | 0 | 0 | 0 | 0 |
| ENSG00000124493 | 549 | 291 | 388 | 289 | 258 | 251 |
| ENSG00000271231 | 0 | 1 | 0 | 0 | 0 | 0 |
| ENSG00000214810 | 0 | 0 | 0 | 0 | 0 | 0 |
| ENSG00000137309 | 5127 | 2503 | 2913 | 3639 | 2906 | 2940 |
| ENSG00000276404 | 1 | 3 | 5 | 0 | 1 | 4 |
| ENSG00000186577 | 818 | 349 | 482 | 448 | 384 | 379 |
| ENSG00000225339 | 42 | 28 | 33 | 17 | 20 | 15 |
| ENSG00000220583 | 0 | 1 | 0 | 1 | 2 | 0 |
| ENSG00000272325 | 144 | 94 | 93 | 100 | 82 | 91 |
| ENSG00000270800 | 0 | 0 | 0 | 0 | 4 | 0 |
| ENSG00000124614 | 127 | 90 | 112 | 153 | 103 | 98 |
| ENSG00000124507 | 824 | 358 | 481 | 328 | 339 | 311 |

| | | | | | | |
|---|---|---|---|---|---|---|
| ENSG00000124664 | 5545 | 2400 | 3029 | 2564 | 1965 | 2003 |
| ENSG00000196114 | 0 | 0 | 0 | 1 | 0 | 0 |
| ENSG00000196821 | 4764 | 2186 | 2882 | 2675 | 2685 | 2682 |
| ENSG00000216636 | 0 | 0 | 0 | 0 | 0 | 0 |
| ENSG00000265123 | 0 | 0 | 0 | 0 | 0 | 1 |
| ENSG00000220643 | 0 | 0 | 1 | 0 | 0 | 0 |
| ENSG00000272288 | 68 | 40 | 49 | 44 | 26 | 37 |
| ENSG00000186328 | 0 | 0 | 0 | 0 | 0 | 0 |
| ENSG00000217130 | 9 | 3 | 3 | 3 | 2 | 5 |
| ENSG00000124562 | 3282 | 1575 | 1908 | 2134 | 2017 | 1789 |
| ENSG00000065060 | 1629 | 699 | 968 | 560 | 899 | 789 |
| ENSG00000206717 | 0 | 0 | 0 | 0 | 0 | 0 |
| ENSG00000252106 | 0 | 0 | 0 | 0 | 0 | 0 |
| ENSG00000064995 | 1912 | 1084 | 1305 | 1171 | 1349 | 1331 |
| ENSG00000064999 | 1994 | 936 | 1263 | 868 | 1162 | 1180 |
| ENSG00000187762 | 0 | 0 | 0 | 0 | 0 | 1 |
| ENSG00000273870 | 1 | 0 | 0 | 0 | 0 | 0 |
| ENSG00000124678 | 0 | 0 | 0 | 0 | 0 | 0 |
| ENSG00000146197 | 84 | 34 | 47 | 21 | 27 | 15 |
| ENSG00000272374 | 34 | 5 | 16 | 11 | 13 | 5 |
| ENSG00000065029 | 1494 | 645 | 913 | 728 | 715 | 774 |
| ENSG00000217004 | 2 | 1 | 0 | 0 | 2 | 1 |
| ENSG00000023892 | 308 | 121 | 199 | 166 | 135 | 132 |
| ENSG00000112033 | 1209 | 575 | 751 | 600 | 593 | 673 |
| ENSG00000227154 | 0 | 0 | 0 | 0 | 0 | 0 |
| ENSG00000112039 | 285 | 149 | 183 | 151 | 137 | 140 |
| ENSG00000198755 | 22122 | 11193 | 13414 | 14886 | 13561 | 12934 |
| ENSG00000276712 | 5 | 3 | 4 | 2 | 8 | 4 |
| ENSG00000007866 | 1972 | 966 | 1163 | 870 | 980 | 1034 |
| ENSG00000112041 | 26 | 13 | 17 | 6 | 16 | 7 |
| ENSG00000228559 | 5 | 3 | 3 | 3 | 3 | 2 |
| ENSG00000219023 | 2 | 2 | 2 | 3 | 5 | 4 |
| ENSG00000096060 | 691 | 337 | 364 | 276 | 382 | 333 |
| ENSG00000218749 | 0 | 0 | 0 | 0 | 0 | 0 |
| ENSG00000212579 | 0 | 0 | 0 | 0 | 0 | 0 |
| ENSG00000265527 | 0 | 0 | 0 | 0 | 0 | 0 |
| ENSG00000232909 | 2 | 4 | 8 | 1 | 2 | 6 |
| ENSG00000157343 | 22 | 5 | 15 | 5 | 9 | 8 |
| ENSG00000220734 | 0 | 0 | 0 | 0 | 0 | 0 |
| ENSG00000196748 | 5 | 0 | 3 | 1 | 6 | 0 |
| ENSG00000204140 | 0 | 0 | 0 | 0 | 0 | 0 |
| ENSG00000137392 | 0 | 0 | 0 | 0 | 0 | 0 |
| ENSG00000197753 | 0 | 0 | 0 | 0 | 0 | 0 |
| ENSG00000096063 | 2229 | 1205 | 1489 | 1053 | 1563 | 1433 |
| ENSG00000112053 | 0 | 0 | 2 | 0 | 2 | 2 |
| ENSG00000271304 | 0 | 0 | 0 | 0 | 1 | 0 |
| ENSG00000112062 | 3577 | 1708 | 2184 | 1535 | 2009 | 1992 |
| ENSG00000237719 | 6 | 2 | 5 | 0 | 3 | 2 |
| ENSG00000156711 | 725 | 367 | 452 | 355 | 400 | 385 |
| ENSG00000246982 | 49 | 24 | 36 | 11 | 22 | 24 |

| | | | | | | |
|---|---|---|---|---|---|---|
| ENSG00000096070 | 2913 | 1208 | 1696 | 1253 | 1463 | 1580 |
| ENSG00000180316 | 6 | 2 | 2 | 4 | 3 | 6 |
| ENSG00000189325 | 4 | 3 | 6 | 9 | 7 | 8 |
| ENSG00000010030 | 0 | 1 | 3 | 3 | 1 | 5 |
| ENSG00000224666 | 1 | 0 | 0 | 0 | 0 | 1 |
| ENSG00000179165 | 2 | 3 | 1 | 1 | 1 | 1 |
| ENSG00000112078 | 3274 | 1380 | 1880 | 1141 | 1939 | 1624 |
| ENSG00000240733 | 1 | 1 | 0 | 0 | 0 | 0 |
| ENSG00000112079 | 5539 | 2839 | 3155 | 2119 | 3020 | 2615 |
| ENSG00000239964 | 0 | 0 | 0 | 0 | 0 | 0 |
| ENSG00000112081 | 8363 | 4057 | 4875 | 4274 | 4710 | 4515 |
| ENSG00000263894 | 0 | 0 | 0 | 0 | 0 | 0 |
| ENSG00000238554 | 0 | 0 | 0 | 0 | 0 | 1 |
| ENSG00000251864 | 0 | 0 | 0 | 0 | 0 | 0 |
| ENSG00000281450 | 0 | 0 | 0 | 0 | 0 | 0 |
| ENSG00000213500 | 0 | 0 | 0 | 0 | 0 | 0 |
| ENSG00000124762 | 14811 | 7184 | 7999 | 9443 | 9014 | 7702 |
| ENSG00000285244 | 5 | 2 | 6 | 1 | 1 | 0 |
| ENSG00000255587 | 4 | 2 | 1 | 3 | 2 | 0 |
| ENSG00000220349 | 0 | 0 | 0 | 0 | 0 | 0 |
| ENSG00000124772 | 0 | 1 | 0 | 3 | 1 | 0 |
| ENSG00000285888 | 0 | 0 | 0 | 0 | 0 | 0 |
| ENSG00000137168 | 1620 | 843 | 1019 | 1065 | 1112 | 1169 |
| ENSG00000198663 | 5141 | 2147 | 3060 | 2023 | 2430 | 2647 |
| ENSG00000232598 | 0 | 0 | 0 | 0 | 0 | 0 |
| ENSG00000164530 | 0 | 0 | 0 | 0 | 1 | 3 |
| ENSG00000137409 | 7829 | 3690 | 4630 | 4775 | 4315 | 4491 |
| ENSG00000146192 | 6 | 4 | 0 | 7 | 9 | 6 |
| ENSG00000226976 | 49 | 22 | 26 | 23 | 17 | 30 |
| ENSG00000216412 | 0 | 0 | 0 | 0 | 0 | 1 |
| ENSG00000137193 | 314 | 168 | 207 | 287 | 282 | 227 |
| ENSG00000172738 | 7 | 4 | 1 | 4 | 5 | 6 |
| ENSG00000065491 | 847 | 398 | 536 | 439 | 445 | 489 |
| ENSG00000275550 | 0 | 3 | 1 | 0 | 1 | 2 |
| ENSG00000112130 | 624 | 326 | 365 | 337 | 427 | 389 |
| ENSG00000239953 | 0 | 0 | 0 | 1 | 0 | 0 |
| ENSG00000137200 | 2979 | 1399 | 1818 | 1460 | 1596 | 1661 |
| ENSG00000198937 | 503 | 252 | 278 | 303 | 319 | 267 |
| ENSG00000204110 | 2 | 1 | 3 | 0 | 0 | 1 |
| ENSG00000229559 | 0 | 0 | 0 | 0 | 0 | 0 |
| ENSG00000227920 | 0 | 0 | 0 | 0 | 0 | 0 |
| ENSG00000263926 | 0 | 0 | 0 | 0 | 0 | 0 |
| ENSG00000279942 | 2 | 0 | 0 | 0 | 0 | 0 |
| ENSG00000112139 | 19 | 6 | 10 | 1 | 5 | 1 |
| ENSG00000225945 | 9 | 1 | 3 | 2 | 3 | 2 |
| ENSG00000156639 | 3509 | 1669 | 2198 | 1692 | 1738 | 1878 |
| ENSG00000243313 | 0 | 0 | 0 | 0 | 0 | 0 |
| ENSG00000200597 | 0 | 1 | 0 | 0 | 0 | 0 |
| ENSG00000218521 | 0 | 0 | 0 | 0 | 0 | 0 |
| ENSG00000183826 | 986 | 396 | 568 | 361 | 414 | 454 |

| | | | | | | |
|---|---|---|---|---|---|---|
| ENSG00000200706 | 0 | 0 | 0 | 0 | 0 | 0 |
| ENSG00000226533 | 0 | 0 | 0 | 0 | 0 | 0 |
| ENSG00000199938 | 0 | 0 | 0 | 0 | 0 | 0 |
| ENSG00000217227 | 0 | 0 | 0 | 0 | 0 | 0 |
| ENSG00000124767 | 8350 | 4107 | 4876 | 4535 | 5039 | 4946 |
| ENSG00000236355 | 0 | 0 | 0 | 0 | 0 | 0 |
| ENSG00000124721 | 2 | 1 | 3 | 0 | 2 | 0 |
| ENSG00000244297 | 0 | 0 | 0 | 0 | 0 | 0 |
| ENSG00000219273 | 0 | 0 | 0 | 0 | 0 | 0 |
| ENSG00000212586 | 0 | 0 | 0 | 0 | 0 | 0 |
| ENSG00000231150 | 0 | 0 | 0 | 0 | 0 | 0 |
| ENSG00000220076 | 0 | 0 | 0 | 0 | 0 | 0 |
| ENSG00000220556 | 0 | 0 | 0 | 0 | 0 | 0 |
| ENSG00000112164 | 0 | 1 | 1 | 0 | 0 | 0 |
| ENSG00000112167 | 765 | 375 | 435 | 358 | 421 | 409 |
| ENSG00000217165 | 254 | 104 | 141 | 77 | 165 | 113 |
| ENSG00000164626 | 103 | 60 | 77 | 68 | 50 | 66 |
| ENSG00000124780 | 0 | 0 | 0 | 0 | 0 | 0 |
| ENSG00000095981 | 0 | 0 | 0 | 0 | 0 | 0 |
| ENSG00000164627 | 18 | 12 | 10 | 11 | 9 | 6 |
| ENSG00000218986 | 0 | 0 | 0 | 0 | 0 | 0 |
| ENSG00000231276 | 0 | 0 | 0 | 0 | 0 | 0 |
| ENSG00000252990 | 0 | 0 | 0 | 0 | 0 | 0 |
| ENSG00000146122 | 14 | 1 | 4 | 0 | 4 | 3 |
| ENSG00000281969 | 0 | 0 | 0 | 0 | 0 | 0 |
| ENSG00000235033 | 0 | 1 | 0 | 0 | 0 | 0 |
| ENSG00000124615 | 813 | 372 | 438 | 298 | 254 | 330 |
| ENSG00000220586 | 0 | 0 | 1 | 0 | 0 | 0 |
| ENSG00000180211 | 2 | 0 | 0 | 3 | 1 | 2 |
| ENSG00000227131 | 0 | 0 | 0 | 0 | 0 | 0 |
| ENSG00000204091 | 0 | 1 | 0 | 0 | 0 | 0 |
| ENSG00000204092 | 0 | 0 | 0 | 0 | 0 | 0 |
| ENSG00000226070 | 0 | 0 | 0 | 0 | 0 | 0 |
| ENSG00000156564 | 74 | 20 | 55 | 22 | 28 | 28 |
| ENSG00000252767 | 2 | 2 | 1 | 0 | 3 | 1 |
| ENSG00000236075 | 0 | 0 | 0 | 0 | 0 | 0 |
| ENSG00000226454 | 0 | 1 | 0 | 0 | 0 | 0 |
| ENSG00000237947 | 0 | 1 | 0 | 1 | 0 | 0 |
| ENSG00000124602 | 24 | 11 | 20 | 13 | 7 | 9 |
| ENSG00000124596 | 1245 | 694 | 868 | 637 | 840 | 735 |
| ENSG00000112212 | 1 | 0 | 0 | 1 | 0 | 0 |
| ENSG00000124701 | 0 | 0 | 0 | 0 | 2 | 1 |
| ENSG00000001167 | 1775 | 811 | 1057 | 691 | 884 | 1005 |
| ENSG00000161912 | 75 | 46 | 68 | 42 | 42 | 58 |
| ENSG00000161911 | 0 | 0 | 0 | 1 | 0 | 0 |
| ENSG00000095970 | 0 | 0 | 0 | 0 | 0 | 0 |
| ENSG00000112195 | 0 | 0 | 0 | 0 | 0 | 0 |
| ENSG00000184106 | 0 | 0 | 0 | 0 | 4 | 2 |
| ENSG00000188056 | 2 | 0 | 0 | 0 | 0 | 0 |
| ENSG00000212176 | 0 | 0 | 0 | 0 | 0 | 0 |

| | | | | | | |
|---|---|---|---|---|---|---|
| ENSG00000225690 | 0 | 0 | 0 | 0 | 0 | 0 |
| ENSG00000274256 | 0 | 0 | 0 | 0 | 0 | 0 |
| ENSG00000278745 | 0 | 0 | 0 | 0 | 1 | 0 |
| ENSG00000124731 | 0 | 0 | 0 | 0 | 0 | 0 |
| ENSG00000218809 | 0 | 0 | 0 | 0 | 0 | 0 |
| ENSG00000206745 | 0 | 0 | 0 | 0 | 0 | 0 |
| ENSG00000213461 | 0 | 0 | 0 | 0 | 0 | 0 |
| ENSG00000096264 | 0 | 0 | 0 | 0 | 0 | 0 |
| ENSG00000279284 | 0 | 0 | 0 | 0 | 0 | 0 |
| ENSG00000280371 | 0 | 0 | 0 | 0 | 0 | 0 |
| ENSG00000234753 | 15 | 8 | 14 | 11 | 9 | 8 |
| ENSG00000226917 | 0 | 2 | 0 | 0 | 0 | 0 |
| ENSG00000137166 | 3137 | 1368 | 1861 | 1365 | 1265 | 1211 |
| ENSG00000266494 | 0 | 0 | 0 | 0 | 0 | 1 |
| ENSG00000112559 | 0 | 0 | 0 | 0 | 0 | 0 |
| ENSG00000230913 | 0 | 0 | 0 | 0 | 0 | 2 |
| ENSG00000112561 | 192 | 77 | 101 | 65 | 62 | 56 |
| ENSG00000231102 | 0 | 0 | 0 | 0 | 0 | 0 |
| ENSG00000096088 | 0 | 0 | 1 | 0 | 0 | 0 |
| ENSG00000269387 | 0 | 0 | 0 | 0 | 0 | 0 |
| ENSG00000137218 | 98 | 52 | 77 | 59 | 74 | 45 |
| ENSG00000124593 | 97 | 46 | 100 | 40 | 55 | 65 |
| ENSG00000278224 | 3 | 2 | 7 | 3 | 2 | 3 |
| ENSG00000214736 | 0 | 0 | 0 | 0 | 0 | 0 |
| ENSG00000164663 | 323 | 132 | 217 | 83 | 158 | 156 |
| ENSG00000268745 | 0 | 0 | 0 | 0 | 0 | 0 |
| ENSG00000206977 | 0 | 0 | 0 | 0 | 0 | 0 |
| ENSG00000227516 | 0 | 0 | 0 | 0 | 0 | 0 |
| ENSG00000124641 | 1316 | 662 | 779 | 622 | 729 | 711 |
| ENSG00000207371 | 0 | 0 | 0 | 0 | 0 | 0 |
| ENSG00000112578 | 901 | 476 | 549 | 725 | 582 | 631 |
| ENSG00000112576 | 1580 | 846 | 1034 | 1006 | 911 | 897 |
| ENSG00000206875 | 0 | 0 | 2 | 0 | 0 | 0 |
| ENSG00000223946 | 0 | 0 | 0 | 0 | 0 | 0 |
| ENSG00000137413 | 975 | 577 | 617 | 407 | 469 | 512 |
| ENSG00000261068 | 66 | 27 | 22 | 23 | 28 | 18 |
| ENSG00000188112 | 857 | 414 | 567 | 519 | 430 | 519 |
| ENSG00000048545 | 0 | 0 | 0 | 0 | 1 | 0 |
| ENSG00000214732 | 0 | 0 | 0 | 0 | 0 | 0 |
| ENSG00000112599 | 23 | 7 | 8 | 10 | 12 | 10 |
| ENSG00000048544 | 1983 | 955 | 1072 | 785 | 1141 | 948 |
| ENSG00000124496 | 1411 | 559 | 774 | 358 | 363 | 418 |
| ENSG00000237107 | 0 | 0 | 0 | 0 | 0 | 0 |
| ENSG00000024048 | 1691 | 920 | 828 | 443 | 1086 | 658 |
| ENSG00000206848 | 0 | 2 | 2 | 0 | 1 | 0 |
| ENSG00000112619 | 4 | 2 | 1 | 1 | 0 | 3 |
| ENSG00000213435 | 0 | 0 | 0 | 0 | 0 | 0 |
| ENSG00000124659 | 1300 | 560 | 724 | 701 | 656 | 645 |
| ENSG00000112624 | 847 | 409 | 554 | 262 | 361 | 452 |
| ENSG00000146223 | 7626 | 3822 | 4501 | 3077 | 4788 | 4421 |

| | | | | | | |
|---|---|---|---|---|---|---|
| ENSG00000221821 | 368 | 180 | 210 | 200 | 173 | 199 |
| ENSG00000171611 | 0 | 0 | 0 | 0 | 0 | 0 |
| ENSG00000137161 | 1823 | 911 | 1168 | 1133 | 957 | 1075 |
| ENSG00000231113 | 31 | 17 | 28 | 20 | 19 | 17 |
| ENSG00000181524 | 29 | 12 | 11 | 20 | 14 | 20 |
| ENSG00000124713 | 16 | 9 | 10 | 10 | 11 | 16 |
| ENSG00000124587 | 642 | 314 | 426 | 384 | 378 | 358 |
| ENSG00000112640 | 1306 | 671 | 842 | 697 | 798 | 735 |
| ENSG00000124733 | 2729 | 1580 | 1840 | 1935 | 1807 | 1782 |
| ENSG00000124702 | 5293 | 2576 | 3344 | 3405 | 2949 | 3150 |
| ENSG00000124541 | 2013 | 1133 | 1255 | 1114 | 1169 | 1182 |
| ENSG00000272223 | 10 | 9 | 10 | 6 | 2 | 3 |
| ENSG00000240625 | 11 | 0 | 1 | 1 | 0 | 0 |
| ENSG00000044090 | 3386 | 1519 | 2107 | 1822 | 1695 | 1894 |
| ENSG00000137171 | 1119 | 570 | 763 | 624 | 556 | 596 |
| ENSG00000112651 | 1045 | 518 | 672 | 726 | 625 | 591 |
| ENSG00000272170 | 5 | 0 | 4 | 1 | 2 | 2 |
| ENSG00000112655 | 1153 | 535 | 692 | 595 | 581 | 659 |
| ENSG00000112658 | 1867 | 913 | 1128 | 1166 | 1192 | 1098 |
| ENSG00000112659 | 1455 | 624 | 960 | 637 | 721 | 828 |
| ENSG00000245261 | 0 | 0 | 0 | 0 | 0 | 0 |
| ENSG00000112667 | 1649 | 794 | 1019 | 1411 | 1047 | 1128 |
| ENSG00000146216 | 28 | 7 | 13 | 4 | 6 | 10 |
| ENSG00000137204 | 0 | 0 | 0 | 0 | 0 | 0 |
| ENSG00000146215 | 0 | 0 | 0 | 0 | 0 | 0 |
| ENSG00000171467 | 1152 | 540 | 708 | 438 | 603 | 624 |
| ENSG00000219222 | 0 | 0 | 0 | 0 | 0 | 0 |
| ENSG00000220614 | 0 | 0 | 1 | 0 | 0 | 0 |
| ENSG00000179157 | 3 | 0 | 2 | 0 | 0 | 0 |
| ENSG00000226558 | 0 | 0 | 0 | 0 | 0 | 0 |
| ENSG00000124574 | 1419 | 624 | 873 | 735 | 764 | 820 |
| ENSG00000276770 | 0 | 0 | 0 | 0 | 0 | 0 |
| ENSG00000171462 | 43 | 26 | 32 | 34 | 30 | 29 |
| ENSG00000207076 | 0 | 0 | 0 | 0 | 0 | 0 |
| ENSG00000137221 | 856 | 345 | 569 | 479 | 431 | 523 |
| ENSG00000204052 | 113 | 73 | 80 | 92 | 67 | 71 |
| ENSG00000171453 | 917 | 469 | 561 | 617 | 637 | 653 |
| ENSG00000137207 | 5122 | 2664 | 3250 | 3114 | 2621 | 2771 |
| ENSG00000271754 | 2 | 2 | 1 | 4 | 2 | 3 |
| ENSG00000124571 | 2949 | 1500 | 1852 | 1549 | 1841 | 1851 |
| ENSG00000219470 | 0 | 10 | 3 | 3 | 10 | 6 |
| ENSG00000252218 | 2 | 0 | 0 | 1 | 0 | 0 |
| ENSG00000170734 | 463 | 252 | 318 | 193 | 275 | 279 |
| ENSG00000203362 | 27 | 18 | 17 | 14 | 14 | 16 |
| ENSG00000172432 | 2223 | 1126 | 1511 | 1154 | 1458 | 1366 |
| ENSG00000124688 | 1230 | 637 | 659 | 668 | 672 | 635 |
| ENSG00000172426 | 14 | 14 | 6 | 11 | 12 | 9 |
| ENSG00000096080 | 758 | 433 | 501 | 637 | 510 | 421 |
| ENSG00000218107 | 0 | 0 | 0 | 0 | 0 | 0 |
| ENSG00000236961 | 0 | 0 | 0 | 0 | 0 | 0 |

| | | | | | | |
|---|---|---|---|---|---|---|
| ENSG00000112715 | 1763 | 809 | 1056 | 902 | 1081 | 1064 |
| ENSG00000272114 | 0 | 0 | 0 | 0 | 0 | 0 |
| ENSG00000283573 | 0 | 0 | 0 | 0 | 0 | 0 |
| ENSG00000283215 | 0 | 0 | 0 | 0 | 0 | 0 |
| ENSG00000223469 | 0 | 0 | 0 | 0 | 0 | 0 |
| ENSG00000237686 | 0 | 2 | 1 | 1 | 0 | 0 |
| ENSG00000181577 | 0 | 0 | 0 | 0 | 0 | 0 |
| ENSG00000231881 | 0 | 1 | 0 | 1 | 1 | 0 |
| ENSG00000183239 | 0 | 0 | 0 | 0 | 2 | 0 |
| ENSG00000180992 | 1898 | 997 | 1203 | 1423 | 1151 | 1131 |
| ENSG00000137216 | 4847 | 2026 | 2833 | 2098 | 2163 | 2078 |
| ENSG00000137225 | 6 | 2 | 4 | 3 | 6 | 1 |
| ENSG00000265439 | 0 | 0 | 0 | 0 | 0 | 0 |
| ENSG00000262179 | 4 | 2 | 0 | 0 | 0 | 2 |
| ENSG00000112759 | 2079 | 1058 | 1071 | 1048 | 1159 | 1012 |
| ENSG00000096384 | 57485 | 28235 | 33730 | 31535 | 36642 | 36981 |
| ENSG00000157593 | 5252 | 2690 | 3360 | 2967 | 2797 | 2743 |
| ENSG00000284427 | 0 | 0 | 0 | 0 | 0 | 0 |
| ENSG00000146232 | 412 | 174 | 236 | 310 | 249 | 256 |
| ENSG00000178233 | 6 | 1 | 1 | 0 | 0 | 0 |
| ENSG00000272442 | 1 | 2 | 2 | 0 | 1 | 3 |
| ENSG00000146221 | 0 | 1 | 3 | 1 | 2 | 2 |
| ENSG00000124608 | 674 | 351 | 406 | 367 | 363 | 352 |
| ENSG00000249481 | 0 | 0 | 0 | 0 | 0 | 0 |
| ENSG00000096401 | 1894 | 995 | 978 | 725 | 1298 | 911 |
| ENSG00000266619 | 0 | 0 | 0 | 0 | 0 | 0 |
| ENSG00000237530 | 0 | 0 | 0 | 0 | 0 | 0 |
| ENSG00000279076 | 0 | 0 | 0 | 0 | 0 | 0 |
| ENSG00000196284 | 140 | 100 | 82 | 109 | 126 | 128 |
| ENSG00000270900 | 0 | 0 | 0 | 0 | 0 | 0 |
| ENSG00000276156 | 0 | 0 | 0 | 0 | 0 | 0 |
| ENSG00000277769 | 0 | 0 | 0 | 0 | 0 | 0 |
| ENSG00000219384 | 0 | 0 | 0 | 0 | 0 | 0 |
| ENSG00000207769 | 0 | 0 | 0 | 0 | 0 | 0 |
| ENSG00000124813 | 305 | 162 | 184 | 324 | 433 | 496 |
| ENSG00000271857 | 12 | 10 | 6 | 22 | 39 | 27 |
| ENSG00000261080 | 0 | 0 | 0 | 0 | 1 | 0 |
| ENSG00000112782 | 5 | 6 | 3 | 3 | 5 | 1 |
| ENSG00000212468 | 0 | 0 | 0 | 0 | 0 | 0 |
| ENSG00000231769 | 14 | 3 | 10 | 2 | 7 | 4 |
| ENSG00000001561 | 783 | 390 | 416 | 235 | 457 | 367 |
| ENSG00000112796 | 584 | 298 | 377 | 192 | 397 | 350 |
| ENSG00000229349 | 0 | 0 | 1 | 0 | 0 | 1 |
| ENSG00000172348 | 2 | 2 | 3 | 1 | 1 | 1 |
| ENSG00000226541 | 0 | 0 | 0 | 0 | 0 | 0 |
| ENSG00000236466 | 0 | 0 | 0 | 0 | 0 | 0 |
| ENSG00000146233 | 0 | 0 | 0 | 0 | 0 | 1 |
| ENSG00000153291 | 15 | 5 | 6 | 7 | 7 | 7 |
| ENSG00000242973 | 0 | 0 | 0 | 0 | 0 | 0 |
| ENSG00000180113 | 0 | 0 | 0 | 0 | 0 | 0 |

| | | | | | | |
|---|---|---|---|---|---|---|
| ENSG00000146070 | 0 | 0 | 0 | 0 | 0 | 0 |
| ENSG00000230062 | 0 | 0 | 0 | 0 | 0 | 0 |
| ENSG00000236164 | 0 | 0 | 0 | 0 | 0 | 0 |
| ENSG00000112818 | 0 | 0 | 0 | 0 | 0 | 0 |
| ENSG00000069122 | 2 | 0 | 0 | 0 | 0 | 0 |
| ENSG00000225730 | 0 | 0 | 0 | 0 | 0 | 0 |
| ENSG00000153292 | 2 | 2 | 1 | 4 | 5 | 4 |
| ENSG00000146072 | 2256 | 1213 | 1517 | 1444 | 1574 | 1669 |
| ENSG00000271328 | 0 | 0 | 0 | 0 | 0 | 0 |
| ENSG00000270761 | 53 | 17 | 24 | 25 | 17 | 23 |
| ENSG00000198087 | 1419 | 845 | 754 | 340 | 1176 | 660 |
| ENSG00000199762 | 0 | 0 | 0 | 0 | 0 | 0 |
| ENSG00000216813 | 0 | 0 | 0 | 0 | 0 | 0 |
| ENSG00000164393 | 0 | 0 | 0 | 0 | 1 | 0 |
| ENSG00000153294 | 145 | 81 | 119 | 123 | 187 | 183 |
| ENSG00000252711 | 3 | 0 | 0 | 0 | 0 | 0 |
| ENSG00000226594 | 0 | 0 | 0 | 0 | 0 | 0 |
| ENSG00000216616 | 0 | 0 | 0 | 0 | 0 | 0 |
| ENSG00000124818 | 0 | 0 | 0 | 0 | 0 | 0 |
| ENSG00000200340 | 0 | 0 | 0 | 0 | 0 | 0 |
| ENSG00000244694 | 4 | 0 | 0 | 0 | 0 | 1 |
| ENSG00000226068 | 0 | 0 | 0 | 0 | 0 | 0 |
| ENSG00000216835 | 0 | 0 | 0 | 0 | 0 | 0 |
| ENSG00000278736 | 0 | 0 | 0 | 0 | 0 | 0 |
| ENSG00000271162 | 0 | 0 | 0 | 0 | 0 | 0 |
| ENSG00000218337 | 0 | 0 | 0 | 0 | 0 | 0 |
| ENSG00000274867 | 0 | 0 | 0 | 0 | 0 | 0 |
| ENSG00000217631 | 0 | 0 | 0 | 0 | 0 | 0 |
| ENSG00000252457 | 0 | 0 | 0 | 0 | 0 | 0 |
| ENSG00000277043 | 0 | 0 | 0 | 0 | 0 | 0 |
| ENSG00000146085 | 1189 | 571 | 717 | 397 | 651 | 617 |
| ENSG00000031691 | 136 | 72 | 63 | 50 | 103 | 73 |
| ENSG00000203972 | 0 | 0 | 0 | 0 | 0 | 0 |
| ENSG00000197261 | 910 | 349 | 636 | 381 | 420 | 505 |
| ENSG00000216998 | 0 | 0 | 0 | 0 | 0 | 0 |
| ENSG00000112077 | 0 | 0 | 0 | 0 | 2 | 0 |
| ENSG00000124490 | 1 | 0 | 0 | 0 | 2 | 1 |
| ENSG00000279127 | 7 | 2 | 2 | 1 | 2 | 1 |
| ENSG00000096006 | 46 | 35 | 47 | 18 | 18 | 36 |
| ENSG00000170950 | 4 | 4 | 2 | 1 | 6 | 5 |
| ENSG00000235122 | 0 | 0 | 0 | 0 | 0 | 2 |
| ENSG00000226707 | 0 | 0 | 0 | 0 | 0 | 0 |
| ENSG00000124812 | 2 | 1 | 3 | 2 | 2 | 3 |
| ENSG00000214643 | 0 | 0 | 0 | 0 | 0 | 0 |
| ENSG00000177684 | 0 | 0 | 0 | 0 | 0 | 0 |
| ENSG00000214642 | 0 | 0 | 0 | 0 | 0 | 0 |
| ENSG00000203970 | 0 | 0 | 0 | 0 | 0 | 0 |
| ENSG00000180872 | 0 | 0 | 0 | 0 | 0 | 0 |
| ENSG00000226733 | 0 | 0 | 0 | 0 | 0 | 0 |
| ENSG00000233470 | 0 | 0 | 0 | 0 | 0 | 0 |

| | | | | | | |
|---|---|---|---|---|---|---|
| ENSG00000230472 | 0 | 0 | 0 | 0 | 0 | 0 |
| ENSG00000008197 | 0 | 0 | 0 | 0 | 0 | 0 |
| ENSG00000008196 | 0 | 0 | 0 | 0 | 0 | 0 |
| ENSG00000060303 | 0 | 0 | 0 | 0 | 0 | 0 |
| ENSG00000216913 | 0 | 0 | 0 | 0 | 0 | 0 |
| ENSG00000230204 | 2 | 0 | 0 | 0 | 0 | 0 |
| ENSG00000270306 | 0 | 0 | 0 | 0 | 0 | 0 |
| ENSG00000232702 | 0 | 0 | 0 | 0 | 0 | 0 |
| ENSG00000212532 | 0 | 0 | 0 | 0 | 0 | 0 |
| ENSG00000214641 | 0 | 0 | 0 | 0 | 0 | 0 |
| ENSG00000228689 | 0 | 0 | 0 | 0 | 0 | 0 |
| ENSG00000170927 | 0 | 0 | 1 | 0 | 0 | 0 |
| ENSG00000240641 | 0 | 0 | 0 | 0 | 0 | 0 |
| ENSG00000207604 | 0 | 0 | 0 | 0 | 0 | 0 |
| ENSG00000225613 | 0 | 0 | 0 | 0 | 0 | 0 |
| ENSG00000199080 | 0 | 0 | 0 | 0 | 0 | 0 |
| ENSG00000112115 | 0 | 0 | 0 | 0 | 0 | 0 |
| ENSG00000112116 | 0 | 0 | 0 | 0 | 0 | 0 |
| ENSG00000218363 | 0 | 0 | 0 | 0 | 0 | 0 |
| ENSG00000112118 | 3273 | 1730 | 2233 | 1566 | 1708 | 2052 |
| ENSG00000170915 | 162 | 60 | 74 | 81 | 78 | 94 |
| ENSG00000096093 | 916 | 447 | 580 | 329 | 560 | 422 |
| ENSG00000065308 | 2605 | 1149 | 1579 | 820 | 1159 | 1124 |
| ENSG00000225791 | 334 | 177 | 224 | 181 | 179 | 217 |
| ENSG00000270382 | 0 | 0 | 0 | 0 | 0 | 0 |
| ENSG00000216775 | 202 | 93 | 118 | 84 | 115 | 89 |
| ENSG00000096092 | 1009 | 490 | 558 | 528 | 556 | 547 |
| ENSG00000220377 | 0 | 0 | 0 | 0 | 0 | 0 |
| ENSG00000271227 | 0 | 0 | 0 | 0 | 0 | 0 |
| ENSG00000237674 | 0 | 0 | 0 | 0 | 0 | 0 |
| ENSG00000244067 | 4 | 2 | 1 | 1 | 1 | 1 |
| ENSG00000217228 | 0 | 0 | 0 | 0 | 0 | 0 |
| ENSG00000243955 | 2 | 3 | 6 | 2 | 1 | 1 |
| ENSG00000223622 | 0 | 0 | 0 | 0 | 0 | 0 |
| ENSG00000182793 | 0 | 0 | 0 | 0 | 0 | 0 |
| ENSG00000230375 | 0 | 0 | 0 | 0 | 0 | 0 |
| ENSG00000230845 | 2 | 0 | 1 | 1 | 0 | 1 |
| ENSG00000174156 | 0 | 1 | 0 | 2 | 0 | 0 |
| ENSG00000243236 | 0 | 0 | 0 | 0 | 0 | 0 |
| ENSG00000170899 | 68 | 23 | 29 | 40 | 36 | 51 |
| ENSG00000202198 | 0 | 0 | 0 | 0 | 0 | 0 |
| ENSG00000283293 | 0 | 0 | 0 | 0 | 0 | 0 |
| ENSG00000112144 | 1286 | 596 | 698 | 350 | 745 | 654 |
| ENSG00000112146 | 2525 | 1158 | 1531 | 1033 | 1282 | 1297 |
| ENSG00000242865 | 0 | 0 | 0 | 0 | 0 | 0 |
| ENSG00000261745 | 0 | 0 | 0 | 0 | 0 | 0 |
| ENSG00000137270 | 0 | 0 | 0 | 0 | 0 | 0 |
| ENSG00000253026 | 0 | 0 | 0 | 0 | 0 | 0 |
| ENSG00000219074 | 0 | 0 | 0 | 0 | 0 | 0 |
| ENSG00000218713 | 0 | 0 | 0 | 0 | 0 | 0 |

| | | | | | | |
|---|---|---|---|---|---|---|
| ENSG00000206908 | 0 | 0 | 0 | 0 | 0 | 0 |
| ENSG00000219163 | 0 | 0 | 0 | 0 | 0 | 0 |
| ENSG00000012660 | 1899 | 978 | 1057 | 818 | 1063 | 1110 |
| ENSG00000264056 | 0 | 0 | 0 | 0 | 0 | 0 |
| ENSG00000217527 | 0 | 0 | 1 | 0 | 0 | 0 |
| ENSG00000271367 | 1 | 0 | 0 | 0 | 0 | 0 |
| ENSG00000219863 | 0 | 0 | 1 | 0 | 1 | 0 |
| ENSG00000218049 | 0 | 0 | 0 | 0 | 0 | 0 |
| ENSG00000218261 | 0 | 0 | 0 | 0 | 0 | 0 |
| ENSG00000271338 | 0 | 0 | 0 | 0 | 0 | 0 |
| ENSG00000253010 | 0 | 0 | 0 | 0 | 0 | 0 |
| ENSG00000271373 | 0 | 0 | 0 | 0 | 0 | 0 |
| ENSG00000001084 | 1762 | 959 | 1181 | 953 | 1253 | 1245 |
| ENSG00000249379 | 2 | 0 | 3 | 1 | 4 | 1 |
| ENSG00000231683 | 0 | 0 | 0 | 0 | 0 | 0 |
| ENSG00000235899 | 2 | 0 | 0 | 0 | 0 | 0 |
| ENSG00000124743 | 18 | 26 | 19 | 12 | 35 | 21 |
| ENSG00000227885 | 41 | 24 | 38 | 26 | 39 | 28 |
| ENSG00000137269 | 1476 | 785 | 857 | 597 | 854 | 779 |
| ENSG00000271218 | 6 | 5 | 1 | 3 | 6 | 5 |
| ENSG00000146147 | 0 | 1 | 1 | 0 | 0 | 0 |
| ENSG00000236740 | 0 | 0 | 0 | 0 | 0 | 0 |
| ENSG00000235050 | 0 | 0 | 0 | 0 | 0 | 0 |
| ENSG00000236996 | 0 | 0 | 0 | 0 | 0 | 0 |
| ENSG00000218143 | 0 | 0 | 0 | 0 | 0 | 0 |
| ENSG00000137251 | 0 | 0 | 0 | 0 | 0 | 0 |
| ENSG00000228231 | 0 | 0 | 0 | 0 | 0 | 0 |
| ENSG00000213335 | 0 | 0 | 0 | 0 | 0 | 0 |
| ENSG00000216347 | 1 | 0 | 0 | 0 | 0 | 0 |
| ENSG00000220773 | 0 | 0 | 1 | 0 | 2 | 2 |
| ENSG00000220635 | 0 | 0 | 0 | 0 | 0 | 0 |
| ENSG00000251946 | 0 | 0 | 0 | 0 | 0 | 0 |
| ENSG00000224984 | 757 | 411 | 522 | 195 | 288 | 328 |
| ENSG00000168143 | 2125 | 1133 | 1193 | 297 | 1047 | 758 |
| ENSG00000261116 | 1698 | 936 | 805 | 400 | 932 | 622 |
| ENSG00000137252 | 0 | 3 | 0 | 0 | 0 | 0 |
| ENSG00000187871 | 0 | 0 | 0 | 0 | 3 | 0 |
| ENSG00000146151 | 222 | 98 | 182 | 78 | 90 | 140 |
| ENSG00000220725 | 0 | 0 | 0 | 0 | 0 | 0 |
| ENSG00000112175 | 55 | 22 | 40 | 15 | 31 | 34 |
| ENSG00000214604 | 0 | 0 | 0 | 0 | 0 | 0 |
| ENSG00000124749 | 40 | 16 | 17 | 10 | 10 | 9 |
| ENSG00000219095 | 0 | 0 | 0 | 0 | 0 | 0 |
| ENSG00000227602 | 0 | 0 | 0 | 0 | 0 | 0 |
| ENSG00000220666 | 0 | 0 | 1 | 0 | 0 | 0 |
| ENSG00000151914 | 654 | 390 | 446 | 142 | 623 | 466 |
| ENSG00000231441 | 25 | 14 | 19 | 10 | 10 | 7 |
| ENSG00000216854 | 2 | 0 | 0 | 0 | 0 | 0 |
| ENSG00000200224 | 0 | 0 | 0 | 0 | 0 | 0 |
| ENSG00000151917 | 110 | 66 | 72 | 54 | 90 | 72 |

| | | | | | | |
|---|---|---|---|---|---|---|
| ENSG00000219790 | 0 | 0 | 0 | 0 | 0 | 0 |
| ENSG00000218980 | 0 | 1 | 1 | 1 | 1 | 2 |
| ENSG00000276576 | 0 | 0 | 0 | 0 | 0 | 0 |
| ENSG00000168116 | 396 | 201 | 220 | 143 | 289 | 201 |
| ENSG00000112200 | 752 | 426 | 405 | 239 | 485 | 411 |
| ENSG00000226803 | 8 | 3 | 1 | 0 | 5 | 2 |
| ENSG00000112208 | 91 | 55 | 67 | 58 | 63 | 57 |
| ENSG00000112210 | 272 | 173 | 208 | 137 | 193 | 178 |
| ENSG00000146143 | 600 | 299 | 348 | 249 | 382 | 312 |
| ENSG00000212017 | 0 | 0 | 0 | 0 | 0 | 0 |
| ENSG00000266579 | 1 | 0 | 1 | 0 | 0 | 0 |
| ENSG00000272541 | 0 | 0 | 0 | 0 | 0 | 0 |
| ENSG00000271761 | 0 | 0 | 0 | 0 | 0 | 0 |
| ENSG00000272316 | 22 | 7 | 3 | 3 | 5 | 6 |
| ENSG00000239650 | 0 | 0 | 0 | 0 | 0 | 0 |
| ENSG00000283352 | 0 | 0 | 0 | 0 | 0 | 0 |
| ENSG00000218189 | 0 | 0 | 0 | 0 | 0 | 0 |
| ENSG00000215190 | 261 | 159 | 179 | 137 | 176 | 182 |
| ENSG00000225096 | 0 | 1 | 1 | 0 | 0 | 0 |
| ENSG00000214563 | 0 | 0 | 0 | 0 | 0 | 0 |
| ENSG00000216687 | 0 | 0 | 0 | 0 | 0 | 0 |
| ENSG00000214561 | 0 | 0 | 0 | 0 | 0 | 0 |
| ENSG00000275046 | 0 | 0 | 0 | 0 | 0 | 0 |
| ENSG00000283453 | 0 | 1 | 0 | 0 | 0 | 0 |
| ENSG00000277172 | 0 | 0 | 0 | 0 | 0 | 0 |
| ENSG00000275773 | 0 | 0 | 0 | 0 | 0 | 0 |
| ENSG00000274844 | 0 | 0 | 0 | 0 | 0 | 0 |
| ENSG00000276290 | 0 | 0 | 0 | 0 | 0 | 0 |
| ENSG00000220030 | 0 | 0 | 0 | 0 | 0 | 0 |
| ENSG00000255633 | 0 | 0 | 0 | 0 | 0 | 0 |
| ENSG00000250686 | 0 | 0 | 0 | 0 | 0 | 0 |
| ENSG00000112232 | 0 | 0 | 0 | 0 | 0 | 0 |
| ENSG00000216859 | 0 | 0 | 0 | 0 | 0 | 0 |
| ENSG00000217067 | 0 | 0 | 0 | 0 | 0 | 0 |
| ENSG00000262566 | 0 | 0 | 0 | 0 | 0 | 0 |
| ENSG00000217477 | 0 | 0 | 0 | 0 | 0 | 0 |
| ENSG00000198225 | 18 | 5 | 13 | 11 | 13 | 9 |
| ENSG00000219940 | 0 | 0 | 0 | 0 | 0 | 0 |
| ENSG00000218520 | 0 | 0 | 0 | 0 | 0 | 0 |
| ENSG00000146166 | 0 | 0 | 0 | 0 | 0 | 0 |
| ENSG00000218617 | 0 | 0 | 0 | 0 | 0 | 0 |
| ENSG00000218274 | 0 | 0 | 0 | 0 | 0 | 0 |
| ENSG00000218048 | 0 | 0 | 0 | 0 | 0 | 0 |
| ENSG00000225121 | 0 | 0 | 0 | 0 | 0 | 0 |
| ENSG00000112245 | 17 | 12 | 4 | 7 | 10 | 7 |
| ENSG00000213312 | 0 | 0 | 0 | 0 | 0 | 0 |
| ENSG00000266680 | 59 | 37 | 41 | 26 | 37 | 43 |
| ENSG00000285976 | 99 | 52 | 70 | 48 | 64 | 66 |
| ENSG00000262651 | 0 | 0 | 0 | 0 | 0 | 0 |
| ENSG00000213309 | 0 | 0 | 0 | 0 | 1 | 0 |

| | | | | | | |
|---|---|---|---|---|---|---|
| ENSG00000118482 | 475 | 362 | 242 | 100 | 722 | 299 |
| ENSG00000188107 | 5 | 2 | 3 | 3 | 3 | 2 |
| ENSG00000218813 | 0 | 0 | 0 | 0 | 0 | 0 |
| ENSG00000236345 | 1 | 0 | 0 | 1 | 0 | 0 |
| ENSG00000219653 | 0 | 0 | 0 | 0 | 1 | 0 |
| ENSG00000232120 | 0 | 2 | 0 | 0 | 1 | 0 |
| ENSG00000217811 | 0 | 0 | 0 | 0 | 0 | 0 |
| ENSG00000243828 | 0 | 0 | 0 | 0 | 0 | 0 |
| ENSG00000214558 | 1 | 0 | 2 | 0 | 4 | 3 |
| ENSG00000220483 | 0 | 0 | 0 | 0 | 0 | 0 |
| ENSG00000233859 | 2 | 0 | 1 | 3 | 1 | 0 |
| ENSG00000218890 | 0 | 0 | 0 | 0 | 0 | 0 |
| ENSG00000270521 | 0 | 0 | 0 | 0 | 0 | 0 |
| ENSG00000238949 | 0 | 0 | 0 | 0 | 0 | 0 |
| ENSG00000212229 | 0 | 0 | 0 | 0 | 0 | 0 |
| ENSG00000271111 | 0 | 0 | 0 | 0 | 0 | 0 |
| ENSG00000251965 | 0 | 0 | 0 | 0 | 0 | 0 |
| ENSG00000201015 | 0 | 0 | 0 | 0 | 0 | 0 |
| ENSG00000206672 | 0 | 0 | 0 | 0 | 0 | 0 |
| ENSG00000227706 | 0 | 0 | 0 | 0 | 0 | 0 |
| ENSG00000270509 | 0 | 0 | 0 | 0 | 0 | 0 |
| ENSG00000285838 | 0 | 0 | 0 | 0 | 0 | 0 |
| ENSG00000223504 | 0 | 0 | 0 | 0 | 0 | 0 |
| ENSG00000226497 | 0 | 0 | 0 | 0 | 0 | 0 |
| ENSG00000218834 | 0 | 0 | 0 | 0 | 0 | 0 |
| ENSG00000230910 | 2 | 0 | 0 | 0 | 1 | 0 |
| ENSG00000135298 | 0 | 0 | 0 | 0 | 1 | 0 |
| ENSG00000168216 | 1548 | 715 | 905 | 633 | 833 | 755 |
| ENSG00000219085 | 0 | 1 | 0 | 0 | 0 | 0 |
| ENSG00000219881 | 0 | 0 | 1 | 0 | 0 | 0 |
| ENSG00000082293 | 0 | 0 | 0 | 0 | 0 | 0 |
| ENSG00000216365 | 0 | 0 | 0 | 0 | 0 | 0 |
| ENSG00000112280 | 0 | 0 | 0 | 0 | 0 | 0 |
| ENSG00000253809 | 0 | 0 | 0 | 0 | 0 | 0 |
| ENSG00000237643 | 0 | 0 | 0 | 0 | 0 | 0 |
| ENSG00000224349 | 0 | 0 | 0 | 0 | 0 | 0 |
| ENSG00000082269 | 7 | 5 | 10 | 1 | 1 | 9 |
| ENSG00000238386 | 0 | 0 | 0 | 0 | 0 | 0 |
| ENSG00000154079 | 435 | 263 | 289 | 296 | 296 | 278 |
| ENSG00000271967 | 2 | 0 | 1 | 2 | 1 | 0 |
| ENSG00000232389 | 0 | 0 | 0 | 1 | 0 | 1 |
| ENSG00000218870 | 0 | 0 | 0 | 0 | 0 | 0 |
| ENSG00000112305 | 412 | 228 | 279 | 180 | 274 | 199 |
| ENSG00000262803 | 0 | 0 | 1 | 0 | 0 | 0 |
| ENSG00000217767 | 0 | 0 | 1 | 0 | 1 | 0 |
| ENSG00000112309 | 6 | 8 | 10 | 2 | 6 | 2 |
| ENSG00000219669 | 0 | 0 | 0 | 0 | 0 | 0 |
| ENSG00000221345 | 0 | 0 | 0 | 0 | 0 | 0 |
| ENSG00000218350 | 1 | 0 | 3 | 0 | 2 | 0 |
| ENSG00000207180 | 0 | 0 | 0 | 0 | 0 | 0 |

| | | | | | | |
|---|---|---|---|---|---|---|
| ENSG00000232295 | 3 | 1 | 1 | 0 | 3 | 0 |
| ENSG00000276127 | 0 | 0 | 0 | 0 | 0 | 0 |
| ENSG00000119900 | 751 | 324 | 365 | 204 | 317 | 260 |
| ENSG00000269966 | 0 | 0 | 0 | 0 | 0 | 0 |
| ENSG00000230597 | 0 | 0 | 0 | 0 | 0 | 0 |
| ENSG00000233237 | 8 | 0 | 2 | 1 | 1 | 3 |
| ENSG00000279312 | 6 | 0 | 2 | 1 | 0 | 1 |
| ENSG00000199094 | 0 | 0 | 0 | 0 | 0 | 0 |
| ENSG00000279289 | 2 | 0 | 0 | 0 | 0 | 0 |
| ENSG00000207827 | 0 | 0 | 0 | 0 | 0 | 0 |
| ENSG00000223765 | 0 | 0 | 0 | 0 | 0 | 0 |
| ENSG00000285963 | 0 | 0 | 0 | 0 | 0 | 0 |
| ENSG00000219575 | 0 | 0 | 0 | 0 | 0 | 0 |
| ENSG00000218014 | 6 | 7 | 4 | 3 | 3 | 1 |
| ENSG00000202069 | 0 | 0 | 0 | 0 | 0 | 0 |
| ENSG00000079841 | 6 | 3 | 1 | 1 | 2 | 1 |
| ENSG00000218732 | 0 | 0 | 0 | 0 | 0 | 0 |
| ENSG00000217483 | 0 | 0 | 0 | 0 | 0 | 0 |
| ENSG00000181514 | 0 | 0 | 0 | 0 | 0 | 0 |
| ENSG00000285844 | 0 | 0 | 0 | 0 | 0 | 0 |
| ENSG00000185760 | 0 | 0 | 0 | 0 | 0 | 0 |
| ENSG00000233844 | 0 | 0 | 0 | 0 | 0 | 0 |
| ENSG00000266180 | 0 | 0 | 0 | 0 | 0 | 0 |
| ENSG00000232422 | 0 | 0 | 0 | 0 | 0 | 0 |
| ENSG00000220515 | 0 | 0 | 0 | 0 | 0 | 0 |
| ENSG00000229154 | 0 | 0 | 0 | 0 | 0 | 0 |
| ENSG00000234025 | 0 | 0 | 0 | 0 | 0 | 0 |
| ENSG00000251360 | 0 | 0 | 0 | 0 | 0 | 0 |
| ENSG00000243501 | 0 | 0 | 0 | 0 | 0 | 0 |
| ENSG00000256980 | 0 | 0 | 0 | 1 | 0 | 2 |
| ENSG00000135314 | 241 | 145 | 172 | 121 | 154 | 166 |
| ENSG00000229852 | 98 | 33 | 65 | 34 | 46 | 55 |
| ENSG00000239626 | 1 | 1 | 1 | 1 | 0 | 1 |
| ENSG00000234882 | 11 | 7 | 9 | 3 | 6 | 3 |
| ENSG00000223389 | 0 | 0 | 0 | 0 | 0 | 0 |
| ENSG00000226840 | 0 | 0 | 1 | 0 | 0 | 0 |
| ENSG00000203909 | 2 | 0 | 0 | 0 | 2 | 0 |
| ENSG00000203908 | 0 | 0 | 1 | 0 | 0 | 0 |
| ENSG00000203907 | 45 | 23 | 26 | 37 | 32 | 31 |
| ENSG00000231332 | 0 | 1 | 0 | 0 | 1 | 0 |
| ENSG00000235174 | 81 | 37 | 36 | 36 | 41 | 49 |
| ENSG00000224221 | 13 | 2 | 5 | 1 | 4 | 1 |
| ENSG00000080007 | 147 | 102 | 108 | 86 | 166 | 120 |
| ENSG00000164430 | 331 | 153 | 167 | 180 | 191 | 198 |
| ENSG00000135297 | 1057 | 541 | 617 | 407 | 654 | 582 |
| ENSG00000207023 | 0 | 0 | 0 | 0 | 0 | 0 |
| ENSG00000271986 | 0 | 0 | 1 | 0 | 0 | 1 |
| ENSG00000223821 | 0 | 0 | 1 | 0 | 0 | 2 |
| ENSG00000156508 | 213352 | 110285 | 127622 | 128462 | 147412 | 132528 |
| ENSG00000229862 | 2 | 2 | 1 | 0 | 4 | 1 |

| | | | | | | |
|---|---|---|---|---|---|---|
| ENSG00000223967 | 0 | 0 | 0 | 0 | 0 | 0 |
| ENSG00000119899 | 980 | 517 | 592 | 506 | 607 | 639 |
| ENSG00000218459 | 0 | 1 | 0 | 0 | 0 | 0 |
| ENSG00000231652 | 6 | 0 | 2 | 2 | 1 | 2 |
| ENSG00000156535 | 200 | 100 | 116 | 83 | 169 | 116 |
| ENSG00000279790 | 0 | 0 | 0 | 0 | 0 | 0 |
| ENSG00000217268 | 0 | 0 | 0 | 0 | 0 | 0 |
| ENSG00000223786 | 0 | 0 | 0 | 0 | 0 | 0 |
| ENSG00000217178 | 0 | 0 | 0 | 0 | 0 | 0 |
| ENSG00000272243 | 0 | 0 | 0 | 0 | 0 | 0 |
| ENSG00000218483 | 0 | 0 | 0 | 0 | 0 | 0 |
| ENSG00000224583 | 0 | 0 | 0 | 0 | 0 | 0 |
| ENSG00000219736 | 0 | 0 | 0 | 0 | 0 | 0 |
| ENSG00000111799 | 94 | 41 | 60 | 15 | 23 | 25 |
| ENSG00000112695 | 5185 | 2610 | 3226 | 3606 | 3127 | 2924 |
| ENSG00000112697 | 5550 | 2526 | 3255 | 1737 | 3059 | 2524 |
| ENSG00000225793 | 13 | 7 | 10 | 11 | 7 | 8 |
| ENSG00000118407 | 0 | 4 | 0 | 0 | 4 | 2 |
| ENSG00000203489 | 0 | 0 | 0 | 0 | 0 | 0 |
| ENSG00000237174 | 3 | 2 | 4 | 2 | 1 | 1 |
| ENSG00000221332 | 0 | 0 | 0 | 0 | 1 | 0 |
| ENSG00000263533 | 0 | 0 | 0 | 0 | 0 | 0 |
| ENSG00000238156 | 0 | 0 | 0 | 0 | 0 | 0 |
| ENSG00000217680 | 0 | 0 | 0 | 0 | 0 | 0 |
| ENSG00000201291 | 0 | 0 | 0 | 0 | 0 | 0 |
| ENSG00000219722 | 0 | 0 | 0 | 0 | 0 | 0 |
| ENSG00000220730 | 0 | 0 | 0 | 0 | 0 | 0 |
| ENSG00000252137 | 0 | 0 | 0 | 0 | 0 | 0 |
| ENSG00000112701 | 2119 | 1056 | 1181 | 722 | 1421 | 1109 |
| ENSG00000217488 | 0 | 0 | 0 | 0 | 0 | 0 |
| ENSG00000252498 | 0 | 1 | 3 | 0 | 6 | 3 |
| ENSG00000200091 | 0 | 0 | 0 | 0 | 1 | 0 |
| ENSG00000285401 | 0 | 0 | 0 | 0 | 0 | 0 |
| ENSG00000196586 | 4969 | 2540 | 2290 | 1064 | 3284 | 1911 |
| ENSG00000252156 | 0 | 0 | 0 | 0 | 0 | 0 |
| ENSG00000223169 | 0 | 0 | 0 | 0 | 0 | 0 |
| ENSG00000112706 | 16 | 2 | 2 | 1 | 11 | 5 |
| ENSG00000200040 | 0 | 0 | 0 | 0 | 0 | 0 |
| ENSG00000222652 | 0 | 0 | 0 | 0 | 0 | 0 |
| ENSG00000238697 | 0 | 0 | 0 | 0 | 0 | 0 |
| ENSG00000236635 | 0 | 0 | 0 | 0 | 0 | 0 |
| ENSG00000272445 | 0 | 0 | 0 | 0 | 0 | 0 |
| ENSG00000280511 | 0 | 0 | 0 | 0 | 0 | 0 |
| ENSG00000231762 | 0 | 0 | 0 | 0 | 0 | 0 |
| ENSG00000271945 | 0 | 0 | 0 | 0 | 0 | 0 |
| ENSG00000276064 | 0 | 0 | 0 | 0 | 0 | 0 |
| ENSG00000135312 | 0 | 0 | 0 | 0 | 0 | 0 |
| ENSG00000219253 | 0 | 0 | 0 | 0 | 0 | 0 |
| ENSG00000269964 | 0 | 0 | 0 | 0 | 1 | 0 |
| ENSG00000230309 | 0 | 0 | 0 | 0 | 0 | 0 |

| | | | | | | |
|---|---|---|---|---|---|---|
| ENSG00000229495 | 0 | 0 | 0 | 0 | 1 | 0 |
| ENSG00000146243 | 56 | 26 | 37 | 25 | 42 | 26 |
| ENSG00000146247 | 388 | 260 | 219 | 68 | 600 | 286 |
| ENSG00000217512 | 0 | 0 | 0 | 0 | 0 | 0 |
| ENSG00000118418 | 1194 | 680 | 936 | 569 | 729 | 583 |
| ENSG00000270362 | 52 | 32 | 20 | 23 | 33 | 24 |
| ENSG00000220154 | 0 | 0 | 0 | 0 | 0 | 0 |
| ENSG00000201464 | 0 | 0 | 0 | 0 | 0 | 0 |
| ENSG00000217786 | 0 | 0 | 0 | 0 | 0 | 0 |
| ENSG00000218029 | 0 | 0 | 0 | 0 | 0 | 0 |
| ENSG00000262048 | 0 | 0 | 0 | 0 | 0 | 0 |
| ENSG00000231533 | 0 | 0 | 1 | 0 | 0 | 0 |
| ENSG00000218748 | 0 | 0 | 0 | 0 | 0 | 0 |
| ENSG00000135338 | 109 | 71 | 43 | 18 | 79 | 46 |
| ENSG00000261970 | 0 | 0 | 0 | 1 | 0 | 1 |
| ENSG00000272137 | 0 | 1 | 0 | 0 | 2 | 1 |
| ENSG00000279659 | 3 | 6 | 1 | 0 | 3 | 0 |
| ENSG00000198478 | 1559 | 779 | 904 | 645 | 864 | 770 |
| ENSG00000235357 | 0 | 0 | 0 | 0 | 0 | 0 |
| ENSG00000220918 | 0 | 0 | 0 | 0 | 0 | 0 |
| ENSG00000285586 | 0 | 0 | 0 | 0 | 0 | 0 |
| ENSG00000233835 | 0 | 0 | 0 | 0 | 0 | 0 |
| ENSG00000118402 | 158 | 70 | 83 | 55 | 69 | 71 |
| ENSG00000218582 | 3 | 0 | 0 | 0 | 0 | 0 |
| ENSG00000220311 | 0 | 0 | 0 | 0 | 0 | 0 |
| ENSG00000112742 | 210 | 133 | 100 | 78 | 280 | 169 |
| ENSG00000218418 | 0 | 1 | 2 | 1 | 5 | 0 |
| ENSG00000217377 | 0 | 0 | 0 | 0 | 0 | 0 |
| ENSG00000083123 | 798 | 419 | 524 | 276 | 440 | 404 |
| ENSG00000272129 | 45 | 22 | 16 | 15 | 29 | 28 |
| ENSG00000219547 | 0 | 0 | 0 | 0 | 0 | 0 |
| ENSG00000279022 | 17 | 13 | 4 | 3 | 6 | 15 |
| ENSG00000233967 | 50 | 23 | 17 | 2 | 14 | 14 |
| ENSG00000260645 | 22 | 4 | 12 | 9 | 7 | 15 |
| ENSG00000219361 | 2 | 1 | 0 | 0 | 0 | 0 |
| ENSG00000181705 | 0 | 0 | 0 | 0 | 0 | 0 |
| ENSG00000216352 | 0 | 0 | 0 | 0 | 0 | 0 |
| ENSG00000112773 | 2600 | 1204 | 1667 | 696 | 1009 | 1005 |
| ENSG00000226089 | 0 | 0 | 0 | 0 | 0 | 0 |
| ENSG00000277797 | 0 | 0 | 0 | 0 | 0 | 0 |
| ENSG00000212336 | 0 | 0 | 0 | 0 | 0 | 0 |
| ENSG00000232031 | 0 | 0 | 0 | 0 | 0 | 0 |
| ENSG00000206886 | 2 | 4 | 4 | 0 | 1 | 1 |
| ENSG00000219702 | 3 | 3 | 4 | 4 | 5 | 1 |
| ENSG00000224995 | 4 | 0 | 1 | 1 | 3 | 2 |
| ENSG00000226453 | 42 | 34 | 46 | 29 | 36 | 44 |
| ENSG00000260574 | 0 | 0 | 0 | 0 | 0 | 0 |
| ENSG00000005700 | 1686 | 972 | 793 | 338 | 1382 | 775 |
| ENSG00000223044 | 1 | 0 | 2 | 0 | 1 | 0 |
| ENSG00000220537 | 0 | 0 | 0 | 0 | 0 | 0 |

| | | | | | | |
|---|---|---|---|---|---|---|
| ENSG00000146242 | 5706 | 2578 | 3345 | 2801 | 3089 | 3170 |
| ENSG00000118420 | 177 | 107 | 127 | 112 | 130 | 108 |
| ENSG00000220091 | 0 | 0 | 0 | 0 | 0 | 0 |
| ENSG00000227215 | 0 | 0 | 0 | 0 | 0 | 0 |
| ENSG00000083097 | 415 | 236 | 194 | 83 | 380 | 164 |
| ENSG00000013375 | 1287 | 732 | 782 | 432 | 932 | 642 |
| ENSG00000013392 | 663 | 322 | 403 | 249 | 314 | 279 |
| ENSG00000065833 | 2606 | 1270 | 1362 | 1490 | 2010 | 1798 |
| ENSG00000219951 | 0 | 0 | 3 | 0 | 1 | 0 |
| ENSG00000146250 | 16 | 5 | 8 | 9 | 7 | 11 |
| ENSG00000065609 | 2 | 1 | 1 | 0 | 3 | 1 |
| ENSG00000237874 | 0 | 0 | 0 | 0 | 0 | 0 |
| ENSG00000203877 | 0 | 0 | 0 | 0 | 0 | 0 |
| ENSG00000065615 | 641 | 342 | 385 | 261 | 389 | 297 |
| ENSG00000228679 | 0 | 0 | 0 | 0 | 0 | 0 |
| ENSG00000230852 | 0 | 0 | 0 | 0 | 0 | 2 |
| ENSG00000135324 | 0 | 0 | 2 | 0 | 0 | 0 |
| ENSG00000135315 | 28 | 25 | 16 | 7 | 43 | 33 |
| ENSG00000231776 | 4 | 0 | 3 | 0 | 0 | 0 |
| ENSG00000226309 | 0 | 0 | 0 | 0 | 0 | 0 |
| ENSG00000112837 | 0 | 0 | 0 | 0 | 0 | 0 |
| ENSG00000228290 | 0 | 0 | 0 | 0 | 0 | 0 |
| ENSG00000219604 | 0 | 0 | 0 | 0 | 0 | 0 |
| ENSG00000216523 | 0 | 0 | 0 | 0 | 0 | 0 |
| ENSG00000220291 | 0 | 0 | 0 | 0 | 0 | 0 |
| ENSG00000216902 | 0 | 0 | 0 | 0 | 0 | 0 |
| ENSG00000217060 | 0 | 0 | 0 | 0 | 1 | 0 |
| ENSG00000213211 | 2 | 1 | 1 | 2 | 0 | 2 |
| ENSG00000234155 | 0 | 0 | 0 | 0 | 0 | 0 |
| ENSG00000216439 | 0 | 0 | 0 | 0 | 0 | 0 |
| ENSG00000214460 | 0 | 1 | 1 | 0 | 1 | 0 |
| ENSG00000135318 | 205 | 116 | 136 | 50 | 50 | 65 |
| ENSG00000280232 | 2 | 0 | 1 | 0 | 0 | 0 |
| ENSG00000135317 | 3321 | 1713 | 2069 | 1436 | 2004 | 1778 |
| ENSG00000271793 | 0 | 0 | 0 | 0 | 0 | 0 |
| ENSG00000135316 | 7447 | 4041 | 4419 | 3341 | 4940 | 4438 |
| ENSG00000220563 | 1 | 1 | 2 | 4 | 2 | 0 |
| ENSG00000203875 | 2494 | 1283 | 1390 | 1524 | 1800 | 1535 |
| ENSG00000275072 | 0 | 0 | 0 | 0 | 0 | 0 |
| ENSG00000217334 | 0 | 0 | 0 | 0 | 0 | 0 |
| ENSG00000198738 | 0 | 0 | 1 | 0 | 0 | 0 |
| ENSG00000222686 | 0 | 0 | 0 | 0 | 0 | 0 |
| ENSG00000200427 | 0 | 0 | 0 | 0 | 0 | 0 |
| ENSG00000218766 | 0 | 0 | 0 | 0 | 0 | 0 |
| ENSG00000202103 | 0 | 0 | 0 | 0 | 0 | 0 |
| ENSG00000220069 | 0 | 0 | 0 | 0 | 0 | 0 |
| ENSG00000217272 | 0 | 0 | 0 | 0 | 0 | 0 |
| ENSG00000220831 | 0 | 0 | 0 | 0 | 0 | 0 |
| ENSG00000219500 | 0 | 0 | 0 | 0 | 0 | 0 |
| ENSG00000243124 | 0 | 0 | 0 | 0 | 0 | 0 |

| | | | | | | |
|---|---|---|---|---|---|---|
| ENSG00000218561 | 0 | 0 | 0 | 0 | 0 | 0 |
| ENSG00000219190 | 0 | 0 | 0 | 0 | 0 | 0 |
| ENSG00000217769 | 2 | 0 | 0 | 0 | 0 | 0 |
| ENSG00000220240 | 0 | 0 | 0 | 0 | 0 | 0 |
| ENSG00000168830 | 0 | 0 | 3 | 2 | 0 | 0 |
| ENSG00000214457 | 1 | 0 | 0 | 0 | 0 | 0 |
| ENSG00000217169 | 0 | 0 | 0 | 0 | 0 | 0 |
| ENSG00000217776 | 0 | 0 | 0 | 0 | 0 | 0 |
| ENSG00000252697 | 0 | 0 | 0 | 0 | 0 | 0 |
| ENSG00000135346 | 10 | 2 | 5 | 7 | 1 | 3 |
| ENSG00000215477 | 0 | 0 | 0 | 0 | 0 | 0 |
| ENSG00000272008 | 7 | 11 | 7 | 3 | 8 | 7 |
| ENSG00000188994 | 199 | 96 | 84 | 26 | 282 | 137 |
| ENSG00000218313 | 0 | 0 | 3 | 0 | 0 | 0 |
| ENSG00000164411 | 50 | 25 | 22 | 19 | 35 | 21 |
| ENSG00000216990 | 0 | 0 | 0 | 0 | 0 | 1 |
| ENSG00000111850 | 257 | 133 | 146 | 144 | 173 | 146 |
| ENSG00000203872 | 11 | 10 | 7 | 2 | 12 | 11 |
| ENSG00000279616 | 0 | 0 | 1 | 0 | 0 | 0 |
| ENSG00000272514 | 18 | 8 | 6 | 8 | 7 | 10 |
| ENSG00000213204 | 16 | 18 | 27 | 2 | 14 | 12 |
| ENSG00000216324 | 2 | 0 | 2 | 0 | 1 | 0 |
| ENSG00000218793 | 0 | 3 | 1 | 0 | 0 | 0 |
| ENSG00000218698 | 0 | 0 | 0 | 0 | 0 | 0 |
| ENSG00000164414 | 918 | 575 | 678 | 405 | 509 | 470 |
| ENSG00000206715 | 0 | 0 | 0 | 0 | 0 | 1 |
| ENSG00000146282 | 2119 | 1090 | 1346 | 1059 | 1225 | 1349 |
| ENSG00000135336 | 538 | 349 | 286 | 208 | 520 | 310 |
| ENSG00000135334 | 2085 | 1017 | 1433 | 1152 | 1201 | 1216 |
| ENSG00000238628 | 0 | 0 | 0 | 0 | 0 | 0 |
| ENSG00000207131 | 0 | 0 | 0 | 0 | 0 | 0 |
| ENSG00000240997 | 0 | 0 | 0 | 0 | 0 | 0 |
| ENSG00000271114 | 0 | 0 | 0 | 0 | 0 | 0 |
| ENSG00000118434 | 0 | 0 | 0 | 0 | 0 | 0 |
| ENSG00000118432 | 6 | 4 | 4 | 0 | 0 | 4 |
| ENSG00000279565 | 0 | 0 | 0 | 0 | 0 | 0 |
| ENSG00000234426 | 0 | 0 | 0 | 0 | 0 | 0 |
| ENSG00000220131 | 0 | 0 | 0 | 0 | 0 | 0 |
| ENSG00000220267 | 0 | 0 | 0 | 0 | 0 | 0 |
| ENSG00000111880 | 1157 | 615 | 666 | 513 | 729 | 650 |
| ENSG00000222145 | 0 | 0 | 0 | 0 | 0 | 0 |
| ENSG00000280277 | 0 | 0 | 0 | 0 | 1 | 1 |
| ENSG00000220130 | 0 | 0 | 0 | 0 | 0 | 0 |
| ENSG00000220960 | 0 | 0 | 0 | 0 | 0 | 0 |
| ENSG00000223001 | 0 | 0 | 0 | 0 | 0 | 0 |
| ENSG00000271931 | 7 | 0 | 2 | 0 | 1 | 2 |
| ENSG00000146278 | 2039 | 975 | 1210 | 1106 | 1084 | 967 |
| ENSG00000264017 | 2 | 1 | 0 | 0 | 0 | 0 |
| ENSG00000154548 | 3 | 5 | 1 | 0 | 0 | 3 |
| ENSG00000146281 | 598 | 259 | 294 | 207 | 348 | 328 |

| | | | | | | |
|---|---|---|---|---|---|---|
| ENSG00000146276 | 0 | 0 | 0 | 0 | 0 | 1 |
| ENSG00000111886 | 4 | 6 | 1 | 1 | 3 | 3 |
| ENSG00000220418 | 0 | 0 | 0 | 0 | 0 | 0 |
| ENSG00000198833 | 2914 | 1526 | 1737 | 1330 | 1752 | 1582 |
| ENSG00000025039 | 237 | 124 | 174 | 134 | 131 | 158 |
| ENSG00000218965 | 0 | 0 | 0 | 0 | 0 | 0 |
| ENSG00000135299 | 141 | 57 | 86 | 49 | 61 | 66 |
| ENSG00000239316 | 2 | 0 | 0 | 0 | 0 | 0 |
| ENSG00000237027 | 0 | 0 | 0 | 0 | 0 | 0 |
| ENSG00000083099 | 2058 | 1054 | 1183 | 708 | 1195 | 1046 |
| ENSG00000112159 | 1797 | 812 | 1077 | 395 | 1058 | 969 |
| ENSG00000228124 | 1 | 1 | 0 | 1 | 1 | 1 |
| ENSG00000220871 | 0 | 0 | 0 | 0 | 0 | 0 |
| ENSG00000217653 | 0 | 0 | 1 | 0 | 0 | 1 |
| ENSG00000118412 | 184 | 118 | 103 | 26 | 224 | 118 |
| ENSG00000199630 | 0 | 0 | 0 | 0 | 0 | 0 |
| ENSG00000218300 | 0 | 0 | 0 | 0 | 0 | 0 |
| ENSG00000219240 | 0 | 0 | 0 | 0 | 0 | 0 |
| ENSG00000135355 | 0 | 0 | 0 | 0 | 0 | 0 |
| ENSG00000252891 | 0 | 0 | 0 | 0 | 0 | 0 |
| ENSG00000112182 | 0 | 0 | 3 | 2 | 0 | 4 |
| ENSG00000226455 | 0 | 0 | 0 | 0 | 0 | 0 |
| ENSG00000222078 | 0 | 0 | 0 | 0 | 0 | 0 |
| ENSG00000218730 | 0 | 0 | 0 | 0 | 0 | 0 |
| ENSG00000266760 | 0 | 0 | 0 | 0 | 0 | 0 |
| ENSG00000260271 | 0 | 0 | 0 | 0 | 0 | 0 |
| ENSG00000135341 | 1674 | 926 | 858 | 564 | 1212 | 842 |
| ENSG00000224605 | 0 | 0 | 0 | 0 | 0 | 0 |
| ENSG00000263734 | 0 | 0 | 0 | 0 | 0 | 0 |
| ENSG00000224944 | 0 | 0 | 0 | 0 | 0 | 0 |
| ENSG00000262095 | 0 | 0 | 0 | 0 | 0 | 0 |
| ENSG00000219087 | 0 | 0 | 0 | 0 | 0 | 0 |
| ENSG00000220340 | 0 | 0 | 0 | 0 | 0 | 0 |
| ENSG00000240961 | 0 | 0 | 0 | 0 | 0 | 0 |
| ENSG00000219867 | 0 | 0 | 0 | 0 | 0 | 0 |
| ENSG00000285961 | 0 | 0 | 0 | 0 | 0 | 0 |
| ENSG00000220553 | 0 | 0 | 0 | 0 | 0 | 0 |
| ENSG00000200492 | 0 | 0 | 0 | 0 | 0 | 0 |
| ENSG00000261038 | 0 | 0 | 0 | 0 | 0 | 0 |
| ENSG00000220908 | 0 | 0 | 0 | 0 | 0 | 0 |
| ENSG00000221455 | 0 | 0 | 0 | 0 | 0 | 0 |
| ENSG00000219387 | 0 | 0 | 0 | 0 | 0 | 0 |
| ENSG00000235099 | 0 | 0 | 0 | 0 | 0 | 0 |
| ENSG00000213187 | 0 | 0 | 0 | 0 | 0 | 0 |
| ENSG00000135333 | 817 | 412 | 420 | 129 | 330 | 280 |
| ENSG00000280451 | 0 | 0 | 0 | 0 | 0 | 0 |
| ENSG00000252249 | 0 | 0 | 0 | 0 | 0 | 0 |
| ENSG00000229600 | 0 | 0 | 0 | 0 | 0 | 0 |
| ENSG00000231143 | 0 | 0 | 0 | 0 | 0 | 0 |
| ENSG00000220370 | 0 | 0 | 0 | 0 | 0 | 0 |

| | | | | | | |
|---|---|---|---|---|---|---|
| ENSG00000279403 | 0 | 0 | 0 | 0 | 0 | 0 |
| ENSG00000270484 | 0 | 0 | 0 | 0 | 0 | 0 |
| ENSG00000216853 | 0 | 0 | 0 | 0 | 0 | 0 |
| ENSG00000219627 | 0 | 0 | 0 | 0 | 0 | 0 |
| ENSG00000261366 | 4 | 2 | 1 | 3 | 1 | 0 |
| ENSG00000172469 | 198 | 109 | 83 | 42 | 139 | 83 |
| ENSG00000219549 | 0 | 0 | 0 | 0 | 0 | 0 |
| ENSG00000219941 | 0 | 0 | 0 | 0 | 0 | 0 |
| ENSG00000172461 | 334 | 234 | 133 | 56 | 322 | 169 |
| ENSG00000242560 | 0 | 0 | 0 | 0 | 0 | 0 |
| ENSG00000233797 | 2 | 2 | 3 | 1 | 3 | 4 |
| ENSG00000014123 | 1244 | 681 | 723 | 334 | 927 | 602 |
| ENSG00000112214 | 0 | 0 | 0 | 0 | 0 | 0 |
| ENSG00000213183 | 0 | 0 | 0 | 0 | 0 | 0 |
| ENSG00000207309 | 0 | 0 | 0 | 0 | 0 | 0 |
| ENSG00000219139 | 0 | 0 | 0 | 0 | 0 | 0 |
| ENSG00000217331 | 0 | 0 | 0 | 0 | 0 | 0 |
| ENSG00000216518 | 0 | 0 | 0 | 0 | 0 | 0 |
| ENSG00000224384 | 0 | 0 | 0 | 0 | 0 | 0 |
| ENSG00000112218 | 2 | 0 | 2 | 2 | 0 | 1 |
| ENSG00000123545 | 276 | 140 | 164 | 212 | 207 | 209 |
| ENSG00000263479 | 0 | 0 | 0 | 0 | 0 | 0 |
| ENSG00000186231 | 0 | 1 | 0 | 2 | 0 | 1 |
| ENSG00000146263 | 436 | 217 | 201 | 120 | 308 | 214 |
| ENSG00000271860 | 0 | 1 | 0 | 0 | 0 | 3 |
| ENSG00000226207 | 0 | 0 | 0 | 0 | 0 | 0 |
| ENSG00000238367 | 0 | 0 | 0 | 0 | 0 | 0 |
| ENSG00000220505 | 0 | 0 | 0 | 1 | 0 | 0 |
| ENSG00000236920 | 0 | 0 | 0 | 0 | 0 | 0 |
| ENSG00000187472 | 0 | 0 | 0 | 0 | 0 | 0 |
| ENSG00000283010 | 0 | 0 | 0 | 0 | 0 | 0 |
| ENSG00000184486 | 0 | 1 | 0 | 1 | 0 | 1 |
| ENSG00000112234 | 515 | 236 | 268 | 182 | 323 | 236 |
| ENSG00000271042 | 0 | 0 | 0 | 0 | 0 | 0 |
| ENSG00000265226 | 0 | 0 | 0 | 0 | 0 | 0 |
| ENSG00000232699 | 2 | 0 | 0 | 0 | 0 | 0 |
| ENSG00000146267 | 92 | 42 | 61 | 45 | 72 | 62 |
| ENSG00000132423 | 301 | 142 | 213 | 187 | 210 | 206 |
| ENSG00000132424 | 2757 | 1553 | 1643 | 543 | 2175 | 1426 |
| ENSG00000228506 | 121 | 56 | 88 | 51 | 83 | 81 |
| ENSG00000123552 | 255 | 133 | 113 | 70 | 239 | 128 |
| ENSG00000228439 | 143 | 93 | 108 | 76 | 101 | 92 |
| ENSG00000274603 | 0 | 0 | 0 | 0 | 0 | 0 |
| ENSG00000112237 | 2130 | 1072 | 1211 | 1074 | 1366 | 1162 |
| ENSG00000272017 | 0 | 1 | 1 | 0 | 1 | 0 |
| ENSG00000219755 | 20 | 7 | 12 | 8 | 5 | 10 |
| ENSG00000279170 | 4 | 1 | 2 | 0 | 1 | 1 |
| ENSG00000112238 | 0 | 0 | 0 | 0 | 0 | 0 |
| ENSG00000199303 | 0 | 0 | 0 | 0 | 0 | 0 |
| ENSG00000152034 | 2 | 0 | 0 | 0 | 0 | 0 |

| | | | | | | |
|---|---|---|---|---|---|---|
| ENSG00000229315 | 0 | 0 | 0 | 0 | 0 | 0 |
| ENSG00000219257 | 0 | 0 | 0 | 0 | 0 | 0 |
| ENSG00000275716 | 0 | 0 | 0 | 0 | 0 | 0 |
| ENSG00000213177 | 2 | 0 | 0 | 0 | 0 | 0 |
| ENSG00000216378 | 0 | 0 | 0 | 0 | 0 | 0 |
| ENSG00000112246 | 60 | 30 | 37 | 2 | 7 | 18 |
| ENSG00000228082 | 0 | 0 | 0 | 0 | 0 | 0 |
| ENSG00000112249 | 1461 | 916 | 869 | 335 | 1280 | 721 |
| ENSG00000220695 | 0 | 0 | 0 | 0 | 0 | 0 |
| ENSG00000260000 | 101 | 58 | 65 | 42 | 44 | 49 |
| ENSG00000270987 | 2 | 0 | 1 | 3 | 2 | 3 |
| ENSG00000164418 | 68 | 44 | 51 | 12 | 32 | 35 |
| ENSG00000218475 | 0 | 0 | 0 | 0 | 0 | 0 |
| ENSG00000218536 | 0 | 0 | 0 | 0 | 0 | 0 |
| ENSG00000219757 | 0 | 0 | 0 | 0 | 0 | 0 |
| ENSG00000217120 | 0 | 0 | 0 | 0 | 0 | 0 |
| ENSG00000202283 | 0 | 0 | 0 | 0 | 0 | 0 |
| ENSG00000216817 | 0 | 0 | 0 | 0 | 0 | 0 |
| ENSG00000214428 | 0 | 0 | 0 | 0 | 0 | 0 |
| ENSG00000218173 | 0 | 0 | 0 | 0 | 0 | 0 |
| ENSG00000085382 | 270 | 137 | 196 | 109 | 172 | 175 |
| ENSG00000252944 | 0 | 0 | 0 | 0 | 0 | 0 |
| ENSG00000227535 | 0 | 0 | 0 | 0 | 0 | 0 |
| ENSG00000271099 | 0 | 0 | 0 | 0 | 0 | 0 |
| ENSG00000203809 | 0 | 0 | 0 | 0 | 0 | 0 |
| ENSG00000187772 | 0 | 0 | 0 | 0 | 0 | 0 |
| ENSG00000212147 | 0 | 0 | 0 | 0 | 0 | 0 |
| ENSG00000112276 | 6 | 9 | 5 | 9 | 5 | 6 |
| ENSG00000203808 | 0 | 0 | 0 | 0 | 0 | 0 |
| ENSG00000132429 | 0 | 1 | 3 | 3 | 2 | 2 |
| ENSG00000272102 | 0 | 1 | 1 | 1 | 3 | 0 |
| ENSG00000085377 | 1678 | 872 | 1008 | 1041 | 1077 | 1008 |
| ENSG00000231628 | 5 | 4 | 4 | 5 | 10 | 6 |
| ENSG00000216639 | 0 | 0 | 0 | 0 | 0 | 1 |
| ENSG00000219902 | 0 | 1 | 3 | 0 | 0 | 0 |
| ENSG00000237567 | 0 | 0 | 0 | 0 | 0 | 0 |
| ENSG00000219088 | 0 | 0 | 0 | 0 | 0 | 0 |
| ENSG00000284999 | 0 | 0 | 0 | 0 | 0 | 0 |
| ENSG00000200198 | 0 | 0 | 0 | 0 | 0 | 0 |
| ENSG00000057663 | 1205 | 596 | 659 | 546 | 778 | 644 |
| ENSG00000057657 | 0 | 3 | 1 | 1 | 5 | 1 |
| ENSG00000269919 | 0 | 0 | 0 | 0 | 0 | 0 |
| ENSG00000273744 | 0 | 0 | 0 | 0 | 0 | 0 |
| ENSG00000244710 | 0 | 0 | 0 | 0 | 0 | 0 |
| ENSG00000252444 | 0 | 0 | 0 | 0 | 0 | 0 |
| ENSG00000233941 | 0 | 1 | 0 | 0 | 3 | 0 |
| ENSG00000112297 | 3667 | 1553 | 2129 | 1259 | 2306 | 1994 |
| ENSG00000207499 | 0 | 0 | 0 | 0 | 0 | 0 |
| ENSG00000202386 | 0 | 0 | 0 | 0 | 0 | 0 |
| ENSG00000282408 | 0 | 0 | 0 | 0 | 0 | 0 |

| | | | | | | |
|---|---|---|---|---|---|---|
| ENSG00000200314 | 0 | 0 | 0 | 0 | 0 | 0 |
| ENSG00000130347 | 220 | 151 | 181 | 145 | 154 | 138 |
| ENSG00000200295 | 0 | 0 | 0 | 0 | 0 | 0 |
| ENSG00000130348 | 794 | 437 | 492 | 314 | 510 | 424 |
| ENSG00000216977 | 2 | 0 | 0 | 0 | 0 | 0 |
| ENSG00000229654 | 0 | 0 | 0 | 0 | 0 | 0 |
| ENSG00000235142 | 44 | 17 | 36 | 13 | 10 | 13 |
| ENSG00000202285 | 0 | 0 | 0 | 0 | 0 | 0 |
| ENSG00000207577 | 0 | 0 | 0 | 0 | 0 | 0 |
| ENSG00000272398 | 47516 | 22152 | 31398 | 18765 | 23449 | 24172 |
| ENSG00000130349 | 471 | 260 | 303 | 238 | 347 | 278 |
| ENSG00000178409 | 322 | 177 | 221 | 140 | 142 | 208 |
| ENSG00000252125 | 0 | 0 | 0 | 0 | 0 | 0 |
| ENSG00000164494 | 942 | 457 | 540 | 384 | 445 | 516 |
| ENSG00000276620 | 0 | 0 | 0 | 0 | 0 | 0 |
| ENSG00000220237 | 1 | 2 | 3 | 0 | 0 | 1 |
| ENSG00000112320 | 107 | 41 | 46 | 38 | 49 | 37 |
| ENSG00000234206 | 0 | 0 | 0 | 1 | 0 | 1 |
| ENSG00000280135 | 0 | 0 | 0 | 0 | 0 | 0 |
| ENSG00000146285 | 0 | 0 | 0 | 0 | 0 | 0 |
| ENSG00000025796 | 2277 | 1278 | 1257 | 800 | 1691 | 1260 |
| ENSG00000238420 | 0 | 0 | 1 | 0 | 0 | 0 |
| ENSG00000232778 | 0 | 0 | 0 | 0 | 0 | 0 |
| ENSG00000272476 | 20 | 14 | 14 | 11 | 5 | 10 |
| ENSG00000224987 | 0 | 0 | 0 | 0 | 0 | 0 |
| ENSG00000253090 | 0 | 0 | 0 | 0 | 0 | 0 |
| ENSG00000231112 | 0 | 0 | 0 | 0 | 0 | 0 |
| ENSG00000225093 | 4 | 2 | 4 | 2 | 1 | 0 |
| ENSG00000271734 | 2 | 2 | 2 | 2 | 0 | 1 |
| ENSG00000081087 | 1165 | 495 | 624 | 457 | 710 | 564 |
| ENSG00000238490 | 0 | 0 | 0 | 0 | 0 | 0 |
| ENSG00000225174 | 0 | 0 | 0 | 0 | 0 | 0 |
| ENSG00000112333 | 0 | 0 | 0 | 0 | 1 | 0 |
| ENSG00000279398 | 0 | 0 | 0 | 0 | 0 | 0 |
| ENSG00000112335 | 5167 | 2571 | 2818 | 2656 | 2862 | 2734 |
| ENSG00000233511 | 0 | 0 | 0 | 0 | 0 | 0 |
| ENSG00000212525 | 0 | 0 | 0 | 0 | 0 | 0 |
| ENSG00000230248 | 0 | 0 | 0 | 0 | 0 | 0 |
| ENSG00000206974 | 0 | 0 | 0 | 0 | 0 | 0 |
| ENSG00000135537 | 156 | 108 | 114 | 85 | 110 | 103 |
| ENSG00000238055 | 2 | 0 | 0 | 0 | 0 | 0 |
| ENSG00000279498 | 0 | 0 | 0 | 0 | 0 | 0 |
| ENSG00000238079 | 0 | 0 | 0 | 0 | 0 | 0 |
| ENSG00000227072 | 0 | 0 | 0 | 0 | 0 | 0 |
| ENSG00000200799 | 0 | 0 | 0 | 0 | 0 | 0 |
| ENSG00000271608 | 0 | 0 | 0 | 0 | 0 | 0 |
| ENSG00000118689 | 3096 | 1338 | 1732 | 925 | 1617 | 1316 |
| ENSG00000228976 | 0 | 1 | 0 | 0 | 0 | 0 |
| ENSG00000203801 | 4 | 4 | 4 | 3 | 3 | 1 |
| ENSG00000219565 | 0 | 0 | 0 | 0 | 0 | 2 |

| | | | | | | |
|---|---|---|---|---|---|---|
| ENSG00000231559 | 2 | 0 | 0 | 0 | 0 | 0 |
| ENSG00000118690 | 97 | 32 | 52 | 52 | 60 | 50 |
| ENSG00000218872 | 0 | 0 | 0 | 0 | 0 | 0 |
| ENSG00000228834 | 2 | 1 | 0 | 1 | 0 | 1 |
| ENSG00000230290 | 0 | 0 | 1 | 0 | 0 | 0 |
| ENSG00000080546 | 921 | 438 | 489 | 388 | 564 | 470 |
| ENSG00000271730 | 0 | 2 | 2 | 1 | 3 | 1 |
| ENSG00000199944 | 0 | 0 | 0 | 0 | 2 | 0 |
| ENSG00000183137 | 209 | 82 | 98 | 53 | 169 | 102 |
| ENSG00000203799 | 711 | 335 | 565 | 214 | 338 | 307 |
| ENSG00000233908 | 1 | 2 | 0 | 0 | 0 | 1 |
| ENSG00000219700 | 6 | 3 | 8 | 1 | 5 | 5 |
| ENSG00000201023 | 0 | 0 | 1 | 0 | 0 | 0 |
| ENSG00000218632 | 4 | 1 | 5 | 0 | 1 | 2 |
| ENSG00000135535 | 10606 | 5460 | 6078 | 3788 | 5756 | 5006 |
| ENSG00000260273 | 14 | 4 | 8 | 6 | 9 | 6 |
| ENSG00000185250 | 69 | 23 | 44 | 32 | 39 | 45 |
| ENSG00000135587 | 506 | 220 | 300 | 320 | 297 | 262 |
| ENSG00000135596 | 982 | 415 | 578 | 419 | 496 | 463 |
| ENSG00000112365 | 482 | 251 | 353 | 222 | 285 | 309 |
| ENSG00000223537 | 0 | 0 | 1 | 0 | 2 | 2 |
| ENSG00000155085 | 213 | 148 | 165 | 98 | 164 | 139 |
| ENSG00000112367 | 1046 | 541 | 630 | 445 | 711 | 602 |
| ENSG00000232311 | 7 | 2 | 1 | 1 | 3 | 1 |
| ENSG00000146360 | 1 | 0 | 0 | 1 | 0 | 0 |
| ENSG00000112290 | 487 | 244 | 282 | 206 | 242 | 262 |
| ENSG00000168438 | 1100 | 547 | 712 | 388 | 664 | 585 |
| ENSG00000053328 | 0 | 0 | 0 | 0 | 0 | 0 |
| ENSG00000203797 | 2 | 0 | 0 | 0 | 0 | 0 |
| ENSG00000004809 | 0 | 0 | 0 | 0 | 1 | 0 |
| ENSG00000241162 | 0 | 0 | 0 | 0 | 0 | 0 |
| ENSG00000260188 | 0 | 0 | 0 | 0 | 0 | 0 |
| ENSG00000219559 | 0 | 0 | 0 | 1 | 0 | 0 |
| ENSG00000270934 | 0 | 0 | 0 | 0 | 0 | 0 |
| ENSG00000155111 | 2030 | 938 | 1137 | 672 | 1145 | 984 |
| ENSG00000213150 | 0 | 0 | 0 | 0 | 0 | 0 |
| ENSG00000217041 | 0 | 0 | 0 | 0 | 0 | 0 |
| ENSG00000219150 | 0 | 0 | 0 | 0 | 0 | 0 |
| ENSG00000200522 | 0 | 0 | 0 | 0 | 0 | 0 |
| ENSG00000212587 | 0 | 0 | 0 | 0 | 0 | 0 |
| ENSG00000199360 | 0 | 0 | 0 | 0 | 0 | 0 |
| ENSG00000213149 | 3 | 1 | 1 | 1 | 1 | 1 |
| ENSG00000271607 | 0 | 0 | 0 | 0 | 3 | 0 |
| ENSG00000123505 | 1599 | 941 | 808 | 664 | 1288 | 937 |
| ENSG00000219329 | 0 | 0 | 0 | 0 | 0 | 0 |
| ENSG00000155115 | 1027 | 596 | 713 | 553 | 704 | 626 |
| ENSG00000197498 | 808 | 443 | 498 | 483 | 645 | 522 |
| ENSG00000218803 | 0 | 0 | 0 | 1 | 0 | 0 |
| ENSG00000112394 | 27 | 19 | 27 | 25 | 17 | 23 |
| ENSG00000200926 | 0 | 0 | 0 | 0 | 0 | 0 |

| | | | | | | |
|---|---|---|---|---|---|---|
| ENSG00000231046 | 0 | 0 | 0 | 0 | 0 | 0 |
| ENSG00000173214 | 338 | 266 | 184 | 91 | 132 | 138 |
| ENSG00000230177 | 36 | 34 | 25 | 31 | 34 | 35 |
| ENSG00000271789 | 2 | 1 | 1 | 5 | 7 | 1 |
| ENSG00000009413 | 458 | 262 | 232 | 72 | 609 | 297 |
| ENSG00000272356 | 1 | 19 | 8 | 1 | 10 | 9 |
| ENSG00000220392 | 0 | 0 | 0 | 0 | 0 | 0 |
| ENSG00000229276 | 0 | 1 | 0 | 0 | 0 | 0 |
| ENSG00000218676 | 0 | 1 | 1 | 0 | 0 | 0 |
| ENSG00000231889 | 28 | 23 | 30 | 21 | 24 | 33 |
| ENSG00000220506 | 2 | 2 | 2 | 0 | 1 | 2 |
| ENSG00000056972 | 1698 | 850 | 987 | 976 | 997 | 953 |
| ENSG00000255389 | 17 | 4 | 5 | 9 | 5 | 3 |
| ENSG00000010810 | 62 | 16 | 30 | 37 | 27 | 24 |
| ENSG00000227012 | 0 | 0 | 0 | 0 | 0 | 0 |
| ENSG00000112761 | 0 | 0 | 0 | 0 | 0 | 0 |
| ENSG00000074935 | 174 | 83 | 117 | 79 | 127 | 106 |
| ENSG00000203778 | 18 | 9 | 2 | 17 | 5 | 8 |
| ENSG00000112769 | 0 | 0 | 0 | 0 | 0 | 0 |
| ENSG00000237234 | 0 | 0 | 0 | 0 | 0 | 0 |
| ENSG00000207044 | 0 | 0 | 0 | 0 | 0 | 0 |
| ENSG00000270661 | 0 | 0 | 1 | 0 | 0 | 0 |
| ENSG00000271208 | 0 | 0 | 0 | 0 | 0 | 0 |
| ENSG00000226440 | 0 | 0 | 0 | 0 | 0 | 0 |
| ENSG00000281613 | 0 | 0 | 0 | 0 | 0 | 0 |
| ENSG00000216663 | 0 | 0 | 0 | 0 | 0 | 0 |
| ENSG00000251258 | 0 | 2 | 0 | 0 | 0 | 0 |
| ENSG00000219773 | 0 | 0 | 0 | 1 | 1 | 1 |
| ENSG00000215807 | 6 | 1 | 1 | 0 | 1 | 1 |
| ENSG00000217770 | 0 | 0 | 0 | 0 | 0 | 0 |
| ENSG00000232299 | 0 | 0 | 0 | 0 | 0 | 0 |
| ENSG00000225580 | 0 | 1 | 0 | 1 | 0 | 1 |
| ENSG00000271498 | 0 | 0 | 0 | 0 | 0 | 1 |
| ENSG00000201386 | 0 | 0 | 0 | 0 | 0 | 0 |
| ENSG00000218490 | 0 | 0 | 0 | 0 | 0 | 0 |
| ENSG00000216904 | 0 | 0 | 0 | 0 | 0 | 0 |
| ENSG00000223811 | 0 | 0 | 0 | 0 | 0 | 0 |
| ENSG00000232316 | 6 | 8 | 4 | 1 | 2 | 3 |
| ENSG00000236347 | 27 | 22 | 25 | 11 | 31 | 23 |
| ENSG00000218208 | 2 | 0 | 0 | 0 | 0 | 2 |
| ENSG00000230943 | 2 | 0 | 3 | 0 | 0 | 0 |
| ENSG00000231912 | 0 | 0 | 0 | 0 | 0 | 0 |
| ENSG00000219758 | 0 | 0 | 0 | 0 | 0 | 0 |
| ENSG00000277443 | 2473 | 1108 | 1412 | 909 | 921 | 1072 |
| ENSG00000227502 | 2 | 0 | 0 | 0 | 0 | 0 |
| ENSG00000175967 | 0 | 0 | 0 | 0 | 3 | 0 |
| ENSG00000196591 | 3931 | 2099 | 2174 | 1879 | 2560 | 2184 |
| ENSG00000228624 | 6 | 5 | 4 | 3 | 8 | 5 |
| ENSG00000216723 | 0 | 0 | 0 | 0 | 0 | 0 |
| ENSG00000249853 | 34 | 19 | 8 | 9 | 13 | 5 |

| | | | | | | |
|---|---|---|---|---|---|---|
| ENSG00000216471 | 0 | 0 | 0 | 0 | 0 | 0 |
| ENSG00000212333 | 0 | 0 | 0 | 0 | 0 | 0 |
| ENSG00000218089 | 0 | 0 | 0 | 0 | 0 | 0 |
| ENSG00000232395 | 0 | 0 | 0 | 0 | 0 | 0 |
| ENSG00000226079 | 0 | 0 | 0 | 0 | 0 | 0 |
| ENSG00000251882 | 0 | 0 | 0 | 0 | 0 | 0 |
| ENSG00000219619 | 0 | 0 | 0 | 0 | 0 | 0 |
| ENSG00000228777 | 0 | 0 | 0 | 0 | 0 | 0 |
| ENSG00000111816 | 203 | 92 | 103 | 58 | 168 | 112 |
| ENSG00000186743 | 0 | 1 | 0 | 0 | 1 | 0 |
| ENSG00000178425 | 1148 | 615 | 657 | 445 | 661 | 577 |
| ENSG00000123500 | 0 | 1 | 0 | 0 | 1 | 0 |
| ENSG00000218428 | 0 | 0 | 0 | 0 | 0 | 0 |
| ENSG00000236326 | 0 | 0 | 0 | 0 | 0 | 0 |
| ENSG00000187189 | 2533 | 1106 | 1409 | 903 | 1158 | 1144 |
| ENSG00000111817 | 234 | 107 | 107 | 73 | 120 | 110 |
| ENSG00000233558 | 0 | 0 | 0 | 0 | 0 | 0 |
| ENSG00000189241 | 4909 | 2536 | 3243 | 2220 | 2828 | 2730 |
| ENSG00000285446 | 0 | 0 | 0 | 0 | 0 | 0 |
| ENSG00000217241 | 2 | 5 | 2 | 2 | 2 | 3 |
| ENSG00000218153 | 2 | 0 | 0 | 1 | 0 | 0 |
| ENSG00000244158 | 0 | 0 | 0 | 0 | 0 | 0 |
| ENSG00000188820 | 0 | 2 | 0 | 0 | 1 | 1 |
| ENSG00000234117 | 0 | 0 | 0 | 0 | 0 | 0 |
| ENSG00000173626 | 0 | 1 | 0 | 0 | 0 | 0 |
| ENSG00000178033 | 0 | 0 | 0 | 0 | 0 | 0 |
| ENSG00000164451 | 0 | 0 | 0 | 0 | 0 | 0 |
| ENSG00000111832 | 1502 | 715 | 834 | 907 | 885 | 775 |
| ENSG00000111834 | 5 | 4 | 6 | 5 | 1 | 6 |
| ENSG00000153975 | 496 | 283 | 345 | 241 | 396 | 347 |
| ENSG00000196911 | 232 | 131 | 103 | 57 | 139 | 87 |
| ENSG00000183807 | 0 | 0 | 0 | 0 | 0 | 0 |
| ENSG00000173612 | 0 | 0 | 0 | 0 | 0 | 0 |
| ENSG00000185002 | 0 | 0 | 0 | 0 | 0 | 0 |
| ENSG00000217089 | 0 | 0 | 0 | 0 | 0 | 0 |
| ENSG00000200248 | 0 | 0 | 0 | 0 | 0 | 0 |
| ENSG00000170162 | 0 | 0 | 0 | 0 | 0 | 0 |
| ENSG00000047936 | 0 | 0 | 0 | 0 | 0 | 0 |
| ENSG00000251814 | 0 | 0 | 0 | 0 | 0 | 0 |
| ENSG00000222713 | 0 | 0 | 0 | 0 | 0 | 0 |
| ENSG00000282218 | 0 | 0 | 0 | 0 | 0 | 0 |
| ENSG00000219703 | 0 | 0 | 0 | 0 | 0 | 0 |
| ENSG00000226181 | 0 | 0 | 0 | 0 | 0 | 0 |
| ENSG00000164465 | 673 | 323 | 437 | 271 | 279 | 345 |
| ENSG00000207461 | 0 | 0 | 0 | 0 | 0 | 0 |
| ENSG00000047932 | 1176 | 613 | 623 | 349 | 762 | 578 |
| ENSG00000218233 | 0 | 0 | 0 | 0 | 0 | 0 |
| ENSG00000153989 | 2043 | 908 | 1268 | 892 | 1072 | 1088 |
| ENSG00000196376 | 0 | 0 | 3 | 1 | 1 | 1 |
| ENSG00000230202 | 48 | 23 | 17 | 12 | 18 | 11 |

| | | | | | | |
|---|---|---|---|---|---|---|
| ENSG00000216809 | 0 | 2 | 0 | 0 | 0 | 0 |
| ENSG00000111860 | 68 | 41 | 31 | 19 | 45 | 23 |
| ENSG00000169075 | 0 | 1 | 0 | 0 | 0 | 0 |
| ENSG00000198523 | 0 | 0 | 0 | 0 | 0 | 0 |
| ENSG00000217330 | 0 | 0 | 0 | 0 | 0 | 0 |
| ENSG00000225549 | 0 | 0 | 0 | 0 | 0 | 0 |
| ENSG00000111877 | 742 | 331 | 463 | 249 | 394 | 330 |
| ENSG00000111875 | 1041 | 564 | 649 | 530 | 655 | 606 |
| ENSG00000253194 | 18 | 13 | 7 | 3 | 5 | 4 |
| ENSG00000111879 | 40 | 24 | 27 | 14 | 26 | 13 |
| ENSG00000222649 | 0 | 0 | 0 | 0 | 0 | 0 |
| ENSG00000207982 | 0 | 0 | 0 | 0 | 0 | 0 |
| ENSG00000220139 | 0 | 0 | 0 | 0 | 0 | 0 |
| ENSG00000111885 | 2910 | 1447 | 1787 | 736 | 1069 | 1106 |
| ENSG00000216316 | 0 | 0 | 2 | 0 | 1 | 0 |
| ENSG00000200732 | 0 | 0 | 0 | 0 | 0 | 0 |
| ENSG00000265725 | 0 | 0 | 0 | 0 | 0 | 0 |
| ENSG00000206857 | 0 | 0 | 0 | 0 | 0 | 0 |
| ENSG00000200310 | 0 | 0 | 0 | 0 | 0 | 0 |
| ENSG00000216710 | 0 | 0 | 0 | 0 | 0 | 0 |
| ENSG00000146350 | 111 | 27 | 38 | 19 | 63 | 45 |
| ENSG00000206598 | 0 | 0 | 0 | 0 | 0 | 0 |
| ENSG00000207403 | 0 | 0 | 0 | 0 | 0 | 0 |
| ENSG00000220447 | 0 | 0 | 0 | 0 | 0 | 0 |
| ENSG00000272714 | 0 | 0 | 0 | 0 | 0 | 0 |
| ENSG00000152661 | 0 | 4 | 3 | 1 | 0 | 2 |
| ENSG00000199458 | 0 | 0 | 0 | 0 | 0 | 0 |
| ENSG00000219784 | 0 | 0 | 0 | 0 | 0 | 0 |
| ENSG00000201379 | 0 | 0 | 0 | 0 | 0 | 0 |
| ENSG00000222659 | 0 | 0 | 0 | 0 | 0 | 0 |
| ENSG00000220326 | 0 | 0 | 0 | 0 | 0 | 0 |
| ENSG00000218180 | 0 | 1 | 0 | 0 | 0 | 0 |
| ENSG00000219135 | 0 | 0 | 0 | 0 | 0 | 0 |
| ENSG00000217139 | 0 | 0 | 0 | 0 | 0 | 0 |
| ENSG00000220184 | 0 | 0 | 0 | 0 | 0 | 0 |
| ENSG00000199932 | 0 | 0 | 0 | 0 | 0 | 0 |
| ENSG00000025156 | 436 | 308 | 299 | 165 | 329 | 265 |
| ENSG00000279453 | 28 | 28 | 27 | 13 | 32 | 23 |
| ENSG00000111897 | 4094 | 2149 | 2053 | 973 | 2486 | 1809 |
| ENSG00000275339 | 0 | 2 | 2 | 0 | 0 | 0 |
| ENSG00000135549 | 479 | 316 | 300 | 341 | 335 | 333 |
| ENSG00000279114 | 9 | 3 | 3 | 2 | 7 | 4 |
| ENSG00000277408 | 0 | 0 | 0 | 0 | 1 | 0 |
| ENSG00000272472 | 0 | 0 | 0 | 0 | 0 | 0 |
| ENSG00000240606 | 0 | 0 | 0 | 0 | 0 | 0 |
| ENSG00000164434 | 0 | 0 | 0 | 0 | 0 | 0 |
| ENSG00000172594 | 409 | 249 | 269 | 153 | 265 | 183 |
| ENSG00000232362 | 0 | 0 | 0 | 0 | 0 | 0 |
| ENSG00000285652 | 0 | 0 | 0 | 0 | 0 | 0 |
| ENSG00000146352 | 0 | 0 | 0 | 0 | 1 | 0 |

| | | | | | | |
|---|---|---|---|---|---|---|
| ENSG00000186439 | 0 | 1 | 2 | 2 | 0 | 1 |
| ENSG00000235535 | 0 | 0 | 0 | 0 | 1 | 0 |
| ENSG00000285691 | 0 | 0 | 0 | 0 | 0 | 0 |
| ENSG00000285941 | 0 | 0 | 0 | 0 | 0 | 0 |
| ENSG00000188580 | 2 | 0 | 0 | 0 | 0 | 0 |
| ENSG00000237321 | 0 | 0 | 0 | 0 | 0 | 0 |
| ENSG00000236548 | 0 | 0 | 0 | 0 | 0 | 0 |
| ENSG00000146373 | 0 | 0 | 1 | 0 | 0 | 0 |
| ENSG00000111907 | 5770 | 3306 | 3653 | 3723 | 3606 | 3849 |
| ENSG00000111906 | 2201 | 1035 | 1272 | 1280 | 1313 | 1202 |
| ENSG00000226409 | 0 | 0 | 0 | 0 | 0 | 0 |
| ENSG00000237742 | 10 | 3 | 4 | 2 | 0 | 4 |
| ENSG00000224506 | 0 | 0 | 0 | 0 | 0 | 0 |
| ENSG00000135547 | 188 | 69 | 56 | 62 | 61 | 60 |
| ENSG00000111912 | 483 | 234 | 237 | 104 | 210 | 145 |
| ENSG00000232131 | 0 | 2 | 1 | 1 | 0 | 0 |
| ENSG00000222445 | 0 | 0 | 0 | 0 | 0 | 0 |
| ENSG00000111911 | 1760 | 778 | 1072 | 652 | 1045 | 929 |
| ENSG00000251920 | 0 | 0 | 0 | 0 | 0 | 0 |
| ENSG00000066651 | 248 | 178 | 188 | 113 | 254 | 140 |
| ENSG00000213133 | 0 | 0 | 0 | 0 | 0 | 0 |
| ENSG00000203760 | 357 | 193 | 208 | 222 | 190 | 228 |
| ENSG00000207632 | 0 | 0 | 0 | 0 | 0 | 0 |
| ENSG00000201613 | 0 | 0 | 0 | 0 | 0 | 0 |
| ENSG00000220548 | 0 | 0 | 0 | 0 | 0 | 0 |
| ENSG00000275953 | 0 | 0 | 0 | 0 | 0 | 0 |
| ENSG00000217325 | 2 | 1 | 2 | 2 | 0 | 2 |
| ENSG00000217404 | 0 | 0 | 0 | 0 | 0 | 0 |
| ENSG00000146374 | 0 | 0 | 0 | 0 | 0 | 0 |
| ENSG00000118518 | 914 | 471 | 589 | 355 | 619 | 422 |
| ENSG00000093144 | 1075 | 667 | 620 | 471 | 849 | 567 |
| ENSG00000251768 | 0 | 0 | 0 | 0 | 0 | 0 |
| ENSG00000213131 | 9 | 5 | 8 | 0 | 4 | 4 |
| ENSG00000220744 | 4 | 3 | 2 | 0 | 0 | 2 |
| ENSG00000220522 | 0 | 0 | 0 | 0 | 0 | 2 |
| ENSG00000220694 | 0 | 0 | 0 | 0 | 0 | 0 |
| ENSG00000255330 | 0 | 0 | 0 | 0 | 0 | 0 |
| ENSG00000189367 | 0 | 0 | 0 | 0 | 0 | 0 |
| ENSG00000214338 | 0 | 2 | 1 | 2 | 3 | 2 |
| ENSG00000184530 | 3 | 3 | 6 | 4 | 4 | 5 |
| ENSG00000219284 | 0 | 0 | 0 | 0 | 0 | 1 |
| ENSG00000218187 | 0 | 0 | 0 | 0 | 0 | 0 |
| ENSG00000260212 | 0 | 0 | 0 | 0 | 0 | 0 |
| ENSG00000219699 | 0 | 0 | 0 | 0 | 0 | 0 |
| ENSG00000281248 | 0 | 0 | 0 | 0 | 0 | 0 |
| ENSG00000172673 | 0 | 0 | 0 | 0 | 0 | 0 |
| ENSG00000274493 | 0 | 0 | 0 | 0 | 0 | 0 |
| ENSG00000152894 | 4828 | 2648 | 2757 | 1305 | 2830 | 2294 |
| ENSG00000227945 | 2 | 0 | 0 | 1 | 0 | 0 |
| ENSG00000224733 | 0 | 0 | 0 | 0 | 0 | 0 |

| | | | | | | |
|---|---|---|---|---|---|---|
| ENSG00000213130 | 0 | 0 | 1 | 0 | 0 | 0 |
| ENSG00000206982 | 0 | 0 | 0 | 0 | 0 | 0 |
| ENSG00000220110 | 0 | 0 | 0 | 0 | 0 | 0 |
| ENSG00000202200 | 0 | 0 | 0 | 0 | 0 | 0 |
| ENSG00000196569 | 0 | 1 | 0 | 0 | 2 | 0 |
| ENSG00000220884 | 0 | 0 | 1 | 1 | 0 | 0 |
| ENSG00000219642 | 1 | 3 | 0 | 1 | 3 | 0 |
| ENSG00000252554 | 0 | 0 | 0 | 0 | 0 | 0 |
| ENSG00000233351 | 0 | 0 | 0 | 0 | 1 | 0 |
| ENSG00000226149 | 0 | 1 | 0 | 0 | 0 | 0 |
| ENSG00000146376 | 710 | 369 | 358 | 147 | 345 | 262 |
| ENSG00000218689 | 0 | 0 | 0 | 0 | 0 | 0 |
| ENSG00000219666 | 0 | 0 | 0 | 0 | 0 | 0 |
| ENSG00000218725 | 0 | 0 | 0 | 0 | 0 | 0 |
| ENSG00000219302 | 0 | 0 | 0 | 0 | 0 | 0 |
| ENSG00000203756 | 0 | 0 | 0 | 0 | 0 | 0 |
| ENSG00000198945 | 81 | 39 | 46 | 34 | 66 | 59 |
| ENSG00000227678 | 0 | 0 | 0 | 0 | 0 | 0 |
| ENSG00000164483 | 0 | 1 | 1 | 0 | 0 | 1 |
| ENSG00000164484 | 0 | 0 | 2 | 0 | 0 | 0 |
| ENSG00000202438 | 0 | 0 | 0 | 0 | 0 | 0 |
| ENSG00000229923 | 0 | 0 | 0 | 0 | 0 | 0 |
| ENSG00000256162 | 0 | 0 | 1 | 0 | 0 | 0 |
| ENSG00000079819 | 1337 | 639 | 821 | 336 | 515 | 590 |
| ENSG00000118507 | 66 | 27 | 40 | 21 | 26 | 35 |
| ENSG00000218857 | 0 | 0 | 0 | 0 | 0 | 0 |
| ENSG00000219776 | 0 | 0 | 0 | 0 | 0 | 0 |
| ENSG00000118520 | 8 | 5 | 2 | 1 | 1 | 0 |
| ENSG00000112282 | 1581 | 810 | 924 | 419 | 966 | 684 |
| ENSG00000154269 | 8 | 6 | 13 | 3 | 4 | 5 |
| ENSG00000252560 | 0 | 0 | 0 | 0 | 0 | 0 |
| ENSG00000180658 | 0 | 0 | 0 | 0 | 0 | 0 |
| ENSG00000236761 | 1 | 1 | 4 | 0 | 0 | 0 |
| ENSG00000216917 | 0 | 0 | 0 | 0 | 0 | 0 |
| ENSG00000266807 | 0 | 0 | 0 | 0 | 0 | 0 |
| ENSG00000197594 | 2491 | 1229 | 1531 | 689 | 1344 | 1125 |
| ENSG00000219532 | 2 | 0 | 0 | 0 | 0 | 0 |
| ENSG00000200895 | 0 | 0 | 0 | 0 | 0 | 0 |
| ENSG00000237115 | 14 | 6 | 13 | 11 | 5 | 7 |
| ENSG00000236673 | 0 | 0 | 0 | 2 | 0 | 0 |
| ENSG00000280058 | 0 | 0 | 0 | 0 | 0 | 0 |
| ENSG00000118523 | 14 | 15 | 24 | 7 | 10 | 13 |
| ENSG00000227220 | 0 | 0 | 0 | 0 | 0 | 0 |
| ENSG00000236166 | 0 | 1 | 0 | 0 | 1 | 0 |
| ENSG00000265669 | 0 | 0 | 0 | 0 | 0 | 0 |
| ENSG00000280155 | 0 | 0 | 0 | 0 | 0 | 0 |
| ENSG00000228495 | 0 | 0 | 0 | 0 | 0 | 0 |
| ENSG00000223912 | 0 | 0 | 0 | 0 | 0 | 0 |
| ENSG00000079931 | 0 | 1 | 0 | 1 | 0 | 0 |
| ENSG00000079950 | 1587 | 807 | 980 | 667 | 902 | 869 |

| | | | | | | |
|---|---|---|---|---|---|---|
| ENSG00000225343 | 0 | 0 | 0 | 0 | 0 | 0 |
| ENSG00000237110 | 0 | 0 | 0 | 0 | 0 | 0 |
| ENSG00000146385 | 0 | 0 | 0 | 0 | 0 | 0 |
| ENSG00000235998 | 0 | 0 | 0 | 0 | 0 | 0 |
| ENSG00000146383 | 0 | 0 | 0 | 0 | 0 | 0 |
| ENSG00000135569 | 0 | 0 | 0 | 0 | 0 | 0 |
| ENSG00000237449 | 0 | 0 | 0 | 0 | 0 | 0 |
| ENSG00000179073 | 0 | 0 | 0 | 0 | 0 | 0 |
| ENSG00000146378 | 0 | 0 | 0 | 0 | 0 | 0 |
| ENSG00000146399 | 0 | 0 | 0 | 0 | 0 | 0 |
| ENSG00000218194 | 0 | 0 | 0 | 0 | 0 | 0 |
| ENSG00000112299 | 0 | 1 | 0 | 0 | 0 | 0 |
| ENSG00000218991 | 0 | 0 | 0 | 0 | 0 | 0 |
| ENSG00000093134 | 0 | 0 | 1 | 0 | 0 | 0 |
| ENSG00000112303 | 0 | 0 | 0 | 0 | 0 | 0 |
| ENSG00000234484 | 0 | 0 | 0 | 0 | 0 | 0 |
| ENSG00000271488 | 0 | 0 | 0 | 0 | 0 | 0 |
| ENSG00000146409 | 4 | 3 | 0 | 2 | 3 | 5 |
| ENSG00000112306 | 17157 | 8487 | 9663 | 12705 | 11288 | 10355 |
| ENSG00000206754 | 4 | 5 | 5 | 5 | 5 | 2 |
| ENSG00000221500 | 16 | 2 | 6 | 6 | 1 | 2 |
| ENSG00000200534 | 28 | 14 | 16 | 13 | 18 | 15 |
| ENSG00000220557 | 0 | 0 | 0 | 0 | 0 | 0 |
| ENSG00000279960 | 0 | 0 | 0 | 0 | 0 | 0 |
| ENSG00000213122 | 0 | 0 | 0 | 0 | 0 | 0 |
| ENSG00000231023 | 0 | 0 | 0 | 0 | 0 | 0 |
| ENSG00000220113 | 0 | 0 | 0 | 0 | 0 | 0 |
| ENSG00000112319 | 8 | 0 | 0 | 2 | 1 | 1 |
| ENSG00000223542 | 0 | 0 | 0 | 0 | 0 | 0 |
| ENSG00000234567 | 0 | 0 | 0 | 0 | 0 | 0 |
| ENSG00000227954 | 0 | 0 | 0 | 0 | 0 | 0 |
| ENSG00000238059 | 0 | 0 | 0 | 0 | 0 | 0 |
| ENSG00000272428 | 0 | 0 | 0 | 0 | 0 | 0 |
| ENSG00000218213 | 0 | 0 | 0 | 0 | 0 | 0 |
| ENSG00000223586 | 0 | 0 | 0 | 0 | 0 | 0 |
| ENSG00000118526 | 0 | 0 | 0 | 0 | 0 | 0 |
| ENSG00000028839 | 546 | 283 | 286 | 266 | 347 | 268 |
| ENSG00000146411 | 86 | 58 | 45 | 39 | 92 | 52 |
| ENSG00000278744 | 0 | 0 | 0 | 0 | 0 | 0 |
| ENSG00000216753 | 0 | 0 | 0 | 0 | 0 | 0 |
| ENSG00000266875 | 0 | 0 | 0 | 0 | 0 | 0 |
| ENSG00000118515 | 118 | 68 | 75 | 72 | 53 | 60 |
| ENSG00000276703 | 0 | 0 | 0 | 0 | 0 | 0 |
| ENSG00000200058 | 0 | 0 | 0 | 0 | 0 | 0 |
| ENSG00000201309 | 0 | 0 | 0 | 0 | 0 | 0 |
| ENSG00000220378 | 0 | 0 | 1 | 0 | 1 | 0 |
| ENSG00000213118 | 0 | 0 | 0 | 0 | 0 | 0 |
| ENSG00000231971 | 0 | 0 | 0 | 0 | 0 | 0 |
| ENSG00000236700 | 0 | 0 | 0 | 0 | 0 | 0 |
| ENSG00000232310 | 0 | 0 | 0 | 0 | 0 | 0 |

| | | | | | | |
|---|---|---|---|---|---|---|
| ENSG00000218772 | 2 | 0 | 0 | 0 | 0 | 0 |
| ENSG00000229722 | 0 | 0 | 0 | 0 | 0 | 0 |
| ENSG00000227723 | 0 | 0 | 0 | 0 | 0 | 0 |
| ENSG00000236389 | 0 | 0 | 0 | 0 | 0 | 0 |
| ENSG00000213117 | 0 | 0 | 0 | 0 | 0 | 0 |
| ENSG00000118514 | 0 | 2 | 1 | 2 | 1 | 0 |
| ENSG00000240056 | 0 | 0 | 0 | 0 | 0 | 0 |
| ENSG00000112339 | 1438 | 818 | 766 | 551 | 994 | 737 |
| ENSG00000283409 | 1 | 0 | 0 | 0 | 2 | 1 |
| ENSG00000232876 | 4 | 0 | 3 | 0 | 0 | 1 |
| ENSG00000118513 | 632 | 334 | 356 | 227 | 271 | 262 |
| ENSG00000236703 | 0 | 0 | 1 | 0 | 1 | 0 |
| ENSG00000207689 | 0 | 0 | 0 | 0 | 0 | 0 |
| ENSG00000224374 | 0 | 0 | 0 | 0 | 0 | 0 |
| ENSG00000135541 | 357 | 227 | 195 | 56 | 360 | 173 |
| ENSG00000234084 | 10 | 13 | 7 | 1 | 6 | 4 |
| ENSG00000201012 | 0 | 0 | 0 | 0 | 0 | 0 |
| ENSG00000231028 | 119 | 41 | 52 | 32 | 54 | 46 |
| ENSG00000226540 | 0 | 1 | 0 | 0 | 0 | 0 |
| ENSG00000233534 | 0 | 0 | 0 | 0 | 1 | 0 |
| ENSG00000217482 | 2 | 0 | 0 | 0 | 1 | 0 |
| ENSG00000236378 | 0 | 0 | 0 | 0 | 0 | 0 |
| ENSG00000171408 | 17 | 10 | 9 | 6 | 11 | 10 |
| ENSG00000237596 | 0 | 2 | 1 | 0 | 0 | 3 |
| ENSG00000213111 | 0 | 0 | 0 | 0 | 0 | 0 |
| ENSG00000277973 | 0 | 0 | 0 | 0 | 0 | 0 |
| ENSG00000146410 | 120 | 48 | 95 | 77 | 87 | 95 |
| ENSG00000029363 | 2517 | 1607 | 1264 | 633 | 2407 | 1423 |
| ENSG00000216519 | 0 | 0 | 0 | 0 | 0 | 0 |
| ENSG00000260418 | 15 | 7 | 7 | 10 | 8 | 16 |
| ENSG00000135525 | 2748 | 1333 | 1546 | 1182 | 1410 | 1344 |
| ENSG00000220660 | 0 | 3 | 0 | 0 | 0 | 0 |
| ENSG00000216613 | 2 | 0 | 0 | 0 | 0 | 0 |
| ENSG00000218424 | 0 | 0 | 0 | 0 | 0 | 0 |
| ENSG00000271765 | 0 | 1 | 0 | 0 | 0 | 0 |
| ENSG00000272189 | 107 | 45 | 58 | 75 | 79 | 84 |
| ENSG00000197442 | 569 | 279 | 287 | 205 | 307 | 257 |
| ENSG00000234263 | 0 | 2 | 3 | 3 | 2 | 0 |
| ENSG00000212242 | 0 | 0 | 0 | 2 | 1 | 3 |
| ENSG00000112357 | 693 | 276 | 354 | 325 | 387 | 351 |
| ENSG00000201807 | 0 | 0 | 0 | 0 | 0 | 0 |
| ENSG00000213109 | 0 | 0 | 0 | 0 | 0 | 0 |
| ENSG00000182747 | 2 | 0 | 2 | 1 | 0 | 0 |
| ENSG00000207247 | 0 | 0 | 0 | 0 | 0 | 0 |
| ENSG00000230350 | 0 | 0 | 0 | 0 | 0 | 0 |
| ENSG00000225391 | 0 | 0 | 0 | 0 | 0 | 0 |
| ENSG00000235399 | 0 | 0 | 0 | 0 | 0 | 0 |
| ENSG00000016402 | 4 | 1 | 0 | 3 | 1 | 2 |
| ENSG00000164485 | 0 | 0 | 0 | 0 | 1 | 0 |
| ENSG00000027697 | 1463 | 673 | 728 | 717 | 1045 | 866 |

| | | | | | | |
|---|---|---|---|---|---|---|
| ENSG00000177468 | 0 | 0 | 0 | 0 | 0 | 0 |
| ENSG00000213108 | 0 | 0 | 0 | 0 | 0 | 0 |
| ENSG00000230533 | 4 | 4 | 0 | 1 | 2 | 2 |
| ENSG00000283265 | 0 | 0 | 0 | 0 | 0 | 0 |
| ENSG00000220412 | 0 | 0 | 0 | 0 | 0 | 0 |
| ENSG00000234956 | 0 | 0 | 0 | 0 | 0 | 0 |
| ENSG00000207300 | 0 | 0 | 0 | 0 | 0 | 0 |
| ENSG00000237499 | 6 | 2 | 0 | 2 | 5 | 5 |
| ENSG00000118503 | 158 | 88 | 120 | 103 | 157 | 151 |
| ENSG00000226004 | 0 | 0 | 0 | 0 | 0 | 0 |
| ENSG00000229922 | 0 | 0 | 0 | 0 | 0 | 0 |
| ENSG00000219463 | 0 | 0 | 0 | 0 | 0 | 0 |
| ENSG00000112378 | 12539 | 6079 | 7519 | 6930 | 7722 | 7496 |
| ENSG00000112379 | 5319 | 2084 | 2886 | 1071 | 2000 | 2021 |
| ENSG00000254440 | 1 | 4 | 0 | 0 | 0 | 5 |
| ENSG00000262543 | 0 | 1 | 0 | 0 | 0 | 0 |
| ENSG00000218499 | 0 | 0 | 0 | 0 | 0 | 0 |
| ENSG00000215878 | 0 | 1 | 1 | 0 | 2 | 1 |
| ENSG00000051620 | 3663 | 1759 | 2161 | 1848 | 2004 | 1912 |
| ENSG00000135540 | 287 | 158 | 179 | 140 | 169 | 168 |
| ENSG00000266555 | 0 | 1 | 0 | 0 | 0 | 0 |
| ENSG00000274594 | 0 | 0 | 0 | 0 | 0 | 1 |
| ENSG00000216802 | 0 | 0 | 1 | 0 | 2 | 0 |
| ENSG00000225177 | 108 | 37 | 70 | 82 | 67 | 55 |
| ENSG00000279968 | 8 | 4 | 9 | 5 | 6 | 3 |
| ENSG00000024862 | 398 | 228 | 258 | 182 | 259 | 192 |
| ENSG00000203734 | 1 | 2 | 4 | 0 | 0 | 5 |
| ENSG00000225415 | 2 | 0 | 0 | 1 | 0 | 1 |
| ENSG00000220600 | 0 | 0 | 0 | 0 | 0 | 0 |
| ENSG00000135597 | 1327 | 700 | 866 | 692 | 848 | 812 |
| ENSG00000146386 | 1951 | 1013 | 1060 | 1239 | 1412 | 1317 |
| ENSG00000112406 | 666 | 331 | 382 | 290 | 433 | 397 |
| ENSG00000231329 | 0 | 1 | 0 | 0 | 0 | 2 |
| ENSG00000272446 | 0 | 0 | 0 | 0 | 0 | 0 |
| ENSG00000164440 | 25 | 24 | 8 | 14 | 31 | 27 |
| ENSG00000226571 | 0 | 0 | 1 | 1 | 1 | 0 |
| ENSG00000218565 | 0 | 0 | 0 | 0 | 0 | 0 |
| ENSG00000164442 | 2124 | 950 | 1387 | 1122 | 1124 | 1309 |
| ENSG00000238099 | 0 | 0 | 0 | 0 | 0 | 0 |
| ENSG00000219806 | 0 | 0 | 0 | 0 | 0 | 0 |
| ENSG00000205695 | 0 | 0 | 0 | 0 | 0 | 0 |
| ENSG00000231426 | 0 | 0 | 0 | 0 | 0 | 0 |
| ENSG00000225148 | 0 | 0 | 0 | 0 | 0 | 0 |
| ENSG00000236013 | 2 | 0 | 1 | 1 | 0 | 0 |
| ENSG00000252107 | 0 | 0 | 0 | 0 | 0 | 0 |
| ENSG00000263514 | 0 | 0 | 0 | 0 | 0 | 0 |
| ENSG00000285875 | 0 | 0 | 0 | 0 | 0 | 0 |
| ENSG00000264390 | 0 | 0 | 0 | 0 | 0 | 0 |
| ENSG00000217684 | 0 | 0 | 0 | 0 | 0 | 0 |
| ENSG00000234147 | 0 | 0 | 0 | 0 | 0 | 0 |

| | | | | | | |
|---|---|---|---|---|---|---|
| ENSG00000216548 | 0 | 0 | 0 | 0 | 0 | 0 |
| ENSG00000275138 | 0 | 0 | 0 | 0 | 0 | 0 |
| ENSG00000259828 | 0 | 0 | 0 | 0 | 0 | 0 |
| ENSG00000222764 | 0 | 0 | 0 | 0 | 0 | 0 |
| ENSG00000218351 | 0 | 0 | 0 | 0 | 0 | 0 |
| ENSG00000135577 | 0 | 0 | 0 | 0 | 0 | 0 |
| ENSG00000270983 | 0 | 0 | 0 | 0 | 0 | 0 |
| ENSG00000236822 | 0 | 0 | 0 | 0 | 0 | 0 |
| ENSG00000203733 | 0 | 0 | 0 | 0 | 0 | 0 |
| ENSG00000009844 | 2172 | 1102 | 1170 | 990 | 1343 | 1186 |
| ENSG00000237494 | 0 | 2 | 0 | 0 | 0 | 1 |
| ENSG00000112414 | 2636 | 1346 | 1323 | 479 | 1403 | 913 |
| ENSG00000236366 | 0 | 3 | 0 | 2 | 0 | 0 |
| ENSG00000233138 | 0 | 0 | 0 | 0 | 0 | 0 |
| ENSG00000010818 | 467 | 245 | 240 | 112 | 277 | 212 |
| ENSG00000237851 | 0 | 0 | 0 | 0 | 0 | 0 |
| ENSG00000232618 | 3 | 2 | 0 | 0 | 1 | 2 |
| ENSG00000229017 | 0 | 0 | 1 | 1 | 0 | 0 |
| ENSG00000227192 | 6 | 1 | 1 | 1 | 3 | 1 |
| ENSG00000146416 | 985 | 502 | 613 | 471 | 555 | 501 |
| ENSG00000225752 | 0 | 0 | 0 | 0 | 0 | 0 |
| ENSG00000217495 | 0 | 0 | 0 | 0 | 0 | 0 |
| ENSG00000216642 | 0 | 0 | 0 | 0 | 0 | 0 |
| ENSG00000217648 | 46 | 19 | 28 | 14 | 17 | 13 |
| ENSG00000270655 | 0 | 0 | 0 | 0 | 0 | 0 |
| ENSG00000189007 | 153 | 76 | 90 | 76 | 133 | 140 |
| ENSG00000218896 | 0 | 0 | 1 | 0 | 0 | 0 |
| ENSG00000223203 | 0 | 0 | 0 | 1 | 1 | 0 |
| ENSG00000034693 | 420 | 242 | 238 | 187 | 287 | 258 |
| ENSG00000278206 | 15 | 5 | 8 | 2 | 1 | 3 |
| ENSG00000229036 | 109 | 49 | 56 | 49 | 71 | 75 |
| ENSG00000001036 | 3154 | 1732 | 2189 | 2092 | 2191 | 2161 |
| ENSG00000112419 | 1149 | 561 | 521 | 236 | 752 | 540 |
| ENSG00000235740 | 0 | 1 | 0 | 0 | 0 | 0 |
| ENSG00000257065 | 0 | 1 | 0 | 0 | 2 | 0 |
| ENSG00000135521 | 350 | 255 | 242 | 170 | 374 | 298 |
| ENSG00000280148 | 0 | 0 | 0 | 0 | 0 | 0 |
| ENSG00000270890 | 13 | 8 | 5 | 2 | 15 | 9 |
| ENSG00000118491 | 0 | 0 | 0 | 0 | 0 | 0 |
| ENSG00000118495 | 111 | 50 | 71 | 49 | 57 | 59 |
| ENSG00000283122 | 0 | 0 | 2 | 0 | 0 | 0 |
| ENSG00000276680 | 0 | 0 | 0 | 0 | 0 | 0 |
| ENSG00000217231 | 0 | 0 | 0 | 0 | 0 | 0 |
| ENSG00000169976 | 3102 | 1559 | 1782 | 2303 | 1882 | 1847 |
| ENSG00000219297 | 0 | 0 | 0 | 0 | 0 | 0 |
| ENSG00000135604 | 6 | 7 | 6 | 7 | 4 | 10 |
| ENSG00000217027 | 4 | 4 | 1 | 1 | 2 | 2 |
| ENSG00000216475 | 0 | 0 | 0 | 0 | 0 | 0 |
| ENSG00000152818 | 385 | 290 | 188 | 62 | 458 | 195 |
| ENSG00000225311 | 0 | 0 | 0 | 0 | 0 | 0 |

| | | | | | | |
|---|---|---|---|---|---|---|
| ENSG00000219409 | 0 | 0 | 0 | 0 | 1 | 0 |
| ENSG00000217195 | 0 | 0 | 0 | 0 | 0 | 0 |
| ENSG00000220739 | 0 | 0 | 0 | 0 | 0 | 0 |
| ENSG00000285598 | 0 | 0 | 0 | 0 | 0 | 0 |
| ENSG00000112425 | 240 | 108 | 133 | 108 | 112 | 147 |
| ENSG00000201119 | 0 | 0 | 0 | 0 | 0 | 0 |
| ENSG00000217612 | 0 | 0 | 0 | 0 | 0 | 0 |
| ENSG00000270638 | 15 | 5 | 8 | 5 | 7 | 4 |
| ENSG00000270828 | 0 | 0 | 0 | 0 | 0 | 0 |
| ENSG00000280184 | 2 | 2 | 0 | 1 | 4 | 2 |
| ENSG00000118496 | 301 | 145 | 137 | 81 | 239 | 156 |
| ENSG00000235652 | 10 | 10 | 10 | 1 | 7 | 9 |
| ENSG00000146414 | 309 | 151 | 108 | 39 | 185 | 115 |
| ENSG00000152822 | 0 | 3 | 1 | 0 | 0 | 1 |
| ENSG00000187103 | 0 | 0 | 0 | 0 | 0 | 0 |
| ENSG00000222971 | 0 | 0 | 0 | 0 | 0 | 0 |
| ENSG00000216811 | 0 | 0 | 0 | 0 | 0 | 0 |
| ENSG00000118508 | 707 | 337 | 382 | 502 | 515 | 384 |
| ENSG00000237468 | 0 | 0 | 0 | 0 | 0 | 0 |
| ENSG00000118492 | 0 | 0 | 1 | 0 | 0 | 0 |
| ENSG00000273719 | 0 | 0 | 0 | 0 | 0 | 0 |
| ENSG00000207431 | 0 | 0 | 0 | 0 | 0 | 0 |
| ENSG00000228283 | 0 | 1 | 1 | 0 | 2 | 0 |
| ENSG00000233452 | 36 | 12 | 20 | 14 | 33 | 22 |
| ENSG00000227748 | 0 | 0 | 0 | 0 | 0 | 0 |
| ENSG00000272397 | 0 | 0 | 0 | 0 | 0 | 0 |
| ENSG00000196634 | 0 | 0 | 0 | 0 | 0 | 0 |
| ENSG00000164506 | 1325 | 674 | 731 | 435 | 853 | 600 |
| ENSG00000273997 | 0 | 0 | 0 | 0 | 0 | 0 |
| ENSG00000220494 | 0 | 0 | 0 | 0 | 0 | 0 |
| ENSG00000203727 | 0 | 0 | 0 | 0 | 0 | 0 |
| ENSG00000227681 | 0 | 0 | 0 | 0 | 0 | 0 |
| ENSG00000226249 | 0 | 0 | 0 | 0 | 0 | 0 |
| ENSG00000234675 | 0 | 0 | 0 | 0 | 0 | 0 |
| ENSG00000230205 | 0 | 0 | 0 | 0 | 0 | 0 |
| ENSG00000224658 | 0 | 0 | 0 | 0 | 0 | 0 |
| ENSG00000111961 | 1973 | 853 | 1152 | 529 | 740 | 746 |
| ENSG00000207345 | 0 | 0 | 0 | 0 | 0 | 0 |
| ENSG00000215223 | 2 | 0 | 0 | 0 | 1 | 0 |
| ENSG00000217824 | 0 | 0 | 0 | 0 | 0 | 0 |
| ENSG00000231368 | 0 | 0 | 0 | 0 | 0 | 0 |
| ENSG00000111962 | 215 | 115 | 127 | 81 | 117 | 84 |
| ENSG00000227660 | 0 | 0 | 0 | 0 | 0 | 0 |
| ENSG00000236591 | 1 | 3 | 7 | 2 | 6 | 4 |
| ENSG00000219487 | 0 | 0 | 1 | 0 | 0 | 1 |
| ENSG00000228408 | 0 | 0 | 0 | 1 | 0 | 0 |
| ENSG00000055208 | 2814 | 1516 | 1457 | 859 | 1775 | 1220 |
| ENSG00000225924 | 0 | 0 | 0 | 0 | 0 | 0 |
| ENSG00000283608 | 0 | 0 | 0 | 0 | 0 | 0 |
| ENSG00000241391 | 0 | 0 | 0 | 0 | 0 | 0 |

| | | | | | | |
|---|---|---|---|---|---|---|
| ENSG00000216265 | 0 | 0 | 0 | 0 | 0 | 0 |
| ENSG00000177688 | 2 | 0 | 0 | 0 | 0 | 0 |
| ENSG00000178199 | 13 | 4 | 6 | 2 | 5 | 7 |
| ENSG00000219553 | 0 | 0 | 1 | 0 | 2 | 0 |
| ENSG00000131013 | 901 | 492 | 475 | 341 | 540 | 450 |
| ENSG00000252244 | 1 | 0 | 3 | 0 | 0 | 2 |
| ENSG00000235168 | 0 | 0 | 0 | 0 | 0 | 0 |
| ENSG00000055211 | 2000 | 1164 | 1353 | 775 | 1390 | 1181 |
| ENSG00000281021 | 2 | 6 | 6 | 3 | 1 | 1 |
| ENSG00000233330 | 12 | 2 | 10 | 4 | 7 | 11 |
| ENSG00000220848 | 79 | 33 | 47 | 48 | 43 | 57 |
| ENSG00000202343 | 1 | 3 | 8 | 3 | 6 | 4 |
| ENSG00000186625 | 572 | 294 | 391 | 283 | 342 | 336 |
| ENSG00000207292 | 0 | 0 | 0 | 0 | 0 | 0 |
| ENSG00000131023 | 1164 | 569 | 589 | 306 | 705 | 626 |
| ENSG00000278899 | 48 | 21 | 28 | 27 | 26 | 18 |
| ENSG00000120253 | 1058 | 547 | 599 | 433 | 692 | 526 |
| ENSG00000120265 | 2893 | 1658 | 1796 | 1580 | 1740 | 1655 |
| ENSG00000219433 | 5 | 3 | 1 | 2 | 3 | 1 |
| ENSG00000231760 | 12 | 5 | 6 | 5 | 5 | 1 |
| ENSG00000285889 | 0 | 0 | 0 | 0 | 0 | 0 |
| ENSG00000285991 | 4 | 2 | 1 | 2 | 2 | 2 |
| ENSG00000120256 | 101 | 43 | 55 | 34 | 58 | 50 |
| ENSG00000273132 | 0 | 0 | 0 | 0 | 1 | 0 |
| ENSG00000268592 | 32 | 11 | 21 | 8 | 19 | 19 |
| ENSG00000217733 | 0 | 0 | 0 | 0 | 0 | 1 |
| ENSG00000164520 | 0 | 1 | 1 | 1 | 0 | 1 |
| ENSG00000223701 | 2 | 3 | 1 | 1 | 1 | 1 |
| ENSG00000216906 | 21 | 9 | 16 | 3 | 18 | 16 |
| ENSG00000203722 | 20 | 19 | 15 | 10 | 11 | 8 |
| ENSG00000216621 | 5 | 4 | 1 | 0 | 1 | 6 |
| ENSG00000131015 | 475 | 256 | 277 | 231 | 265 | 255 |
| ENSG00000215094 | 0 | 0 | 0 | 0 | 0 | 0 |
| ENSG00000111981 | 100 | 48 | 60 | 42 | 41 | 62 |
| ENSG00000219298 | 0 | 1 | 0 | 0 | 0 | 0 |
| ENSG00000231120 | 4 | 3 | 2 | 1 | 2 | 2 |
| ENSG00000218358 | 40 | 12 | 18 | 21 | 17 | 9 |
| ENSG00000155918 | 1 | 1 | 1 | 0 | 1 | 0 |
| ENSG00000216444 | 0 | 0 | 0 | 0 | 0 | 0 |
| ENSG00000235972 | 0 | 0 | 0 | 0 | 1 | 0 |
| ENSG00000213091 | 0 | 0 | 0 | 0 | 0 | 0 |
| ENSG00000131019 | 36 | 25 | 31 | 13 | 28 | 18 |
| ENSG00000198729 | 38 | 22 | 17 | 16 | 26 | 20 |
| ENSG00000238594 | 0 | 0 | 0 | 0 | 0 | 0 |
| ENSG00000201628 | 0 | 0 | 0 | 0 | 0 | 0 |
| ENSG00000009765 | 4 | 5 | 2 | 2 | 1 | 1 |
| ENSG00000220598 | 0 | 0 | 0 | 0 | 0 | 0 |
| ENSG00000219622 | 0 | 0 | 0 | 0 | 0 | 0 |
| ENSG00000120278 | 52 | 27 | 28 | 27 | 51 | 35 |
| ENSG00000232891 | 0 | 0 | 0 | 0 | 0 | 0 |

| | | | | | | |
|---|---|---|---|---|---|---|
| ENSG00000224029 | 0 | 0 | 0 | 0 | 0 | 0 |
| ENSG00000226599 | 0 | 0 | 0 | 0 | 0 | 0 |
| ENSG00000213089 | 0 | 2 | 0 | 0 | 0 | 0 |
| ENSG00000120254 | 822 | 488 | 545 | 613 | 634 | 637 |
| ENSG00000218996 | 1 | 0 | 1 | 1 | 2 | 0 |
| ENSG00000202119 | 0 | 0 | 0 | 0 | 0 | 0 |
| ENSG00000232290 | 0 | 0 | 0 | 0 | 0 | 0 |
| ENSG00000223598 | 0 | 0 | 0 | 0 | 0 | 0 |
| ENSG00000219747 | 8 | 5 | 8 | 4 | 4 | 6 |
| ENSG00000231883 | 0 | 0 | 0 | 0 | 0 | 0 |
| ENSG00000238616 | 0 | 0 | 0 | 0 | 0 | 0 |
| ENSG00000213087 | 2 | 1 | 0 | 2 | 0 | 0 |
| ENSG00000131016 | 22 | 5 | 3 | 7 | 12 | 4 |
| ENSG00000252431 | 0 | 0 | 0 | 0 | 0 | 0 |
| ENSG00000252487 | 0 | 0 | 0 | 0 | 0 | 0 |
| ENSG00000223039 | 0 | 0 | 0 | 0 | 0 | 0 |
| ENSG00000181472 | 1026 | 499 | 553 | 371 | 474 | 551 |
| ENSG00000252615 | 0 | 0 | 0 | 0 | 0 | 0 |
| ENSG00000155906 | 2850 | 1327 | 1549 | 921 | 1366 | 1149 |
| ENSG00000219395 | 32 | 31 | 15 | 14 | 13 | 12 |
| ENSG00000146476 | 10057 | 5170 | 5387 | 3017 | 4892 | 4048 |
| ENSG00000120262 | 1879 | 898 | 1186 | 491 | 979 | 848 |
| ENSG00000207422 | 365 | 160 | 285 | 81 | 139 | 169 |
| ENSG00000091831 | 9017 | 3759 | 4884 | 1994 | 2405 | 2787 |
| ENSG00000233823 | 8 | 2 | 3 | 2 | 2 | 4 |
| ENSG00000284615 | 0 | 0 | 0 | 0 | 0 | 0 |
| ENSG00000131018 | 131 | 85 | 94 | 26 | 76 | 75 |
| ENSG00000234577 | 0 | 0 | 0 | 0 | 0 | 0 |
| ENSG00000226193 | 0 | 0 | 0 | 0 | 0 | 0 |
| ENSG00000202044 | 0 | 0 | 0 | 0 | 0 | 0 |
| ENSG00000233464 | 0 | 0 | 0 | 0 | 0 | 0 |
| ENSG00000120279 | 0 | 0 | 0 | 0 | 0 | 0 |
| ENSG00000218792 | 0 | 0 | 0 | 0 | 0 | 0 |
| ENSG00000146469 | 0 | 0 | 0 | 0 | 0 | 0 |
| ENSG00000224893 | 0 | 0 | 0 | 0 | 0 | 0 |
| ENSG00000217372 | 0 | 0 | 0 | 0 | 0 | 0 |
| ENSG00000220745 | 0 | 0 | 0 | 0 | 0 | 0 |
| ENSG00000112029 | 443 | 212 | 257 | 200 | 237 | 256 |
| ENSG00000227627 | 1 | 1 | 0 | 1 | 4 | 2 |
| ENSG00000112031 | 411 | 232 | 274 | 200 | 266 | 208 |
| ENSG00000091844 | 338 | 135 | 178 | 127 | 195 | 129 |
| ENSG00000201939 | 0 | 0 | 0 | 0 | 0 | 0 |
| ENSG00000213121 | 0 | 0 | 0 | 0 | 0 | 0 |
| ENSG00000218426 | 236 | 110 | 105 | 114 | 119 | 117 |
| ENSG00000237312 | 0 | 0 | 0 | 0 | 0 | 0 |
| ENSG00000222806 | 0 | 0 | 0 | 0 | 0 | 0 |
| ENSG00000217160 | 0 | 0 | 0 | 0 | 0 | 0 |
| ENSG00000242246 | 0 | 0 | 0 | 0 | 0 | 0 |
| ENSG00000218776 | 0 | 0 | 0 | 0 | 0 | 0 |
| ENSG00000217044 | 0 | 0 | 0 | 0 | 0 | 0 |

| | | | | | | |
|---|---|---|---|---|---|---|
| ENSG00000220733 | 0 | 0 | 0 | 0 | 0 | 0 |
| ENSG00000220514 | 0 | 0 | 0 | 0 | 0 | 0 |
| ENSG00000216713 | 0 | 0 | 0 | 0 | 0 | 0 |
| ENSG00000199246 | 0 | 0 | 0 | 0 | 0 | 0 |
| ENSG00000217085 | 0 | 0 | 0 | 0 | 0 | 0 |
| ENSG00000112038 | 0 | 0 | 0 | 0 | 0 | 0 |
| ENSG00000074706 | 6 | 3 | 4 | 6 | 7 | 0 |
| ENSG00000237593 | 0 | 0 | 0 | 0 | 0 | 0 |
| ENSG00000153721 | 195 | 114 | 105 | 114 | 136 | 110 |
| ENSG00000275419 | 0 | 0 | 0 | 0 | 0 | 0 |
| ENSG00000220181 | 0 | 0 | 0 | 0 | 0 | 0 |
| ENSG00000219146 | 0 | 0 | 0 | 0 | 0 | 0 |
| ENSG00000231316 | 0 | 0 | 0 | 0 | 0 | 0 |
| ENSG00000205625 | 0 | 0 | 0 | 0 | 0 | 0 |
| ENSG00000213079 | 1414 | 641 | 807 | 412 | 725 | 741 |
| ENSG00000200594 | 1 | 0 | 0 | 0 | 0 | 0 |
| ENSG00000146426 | 161 | 61 | 82 | 59 | 100 | 79 |
| ENSG00000218757 | 0 | 1 | 0 | 0 | 0 | 0 |
| ENSG00000264814 | 0 | 0 | 0 | 0 | 0 | 0 |
| ENSG00000238963 | 0 | 1 | 0 | 0 | 0 | 0 |
| ENSG00000274451 | 0 | 0 | 0 | 0 | 0 | 0 |
| ENSG00000235381 | 22 | 8 | 12 | 5 | 11 | 11 |
| ENSG00000029639 | 222 | 120 | 147 | 107 | 114 | 101 |
| ENSG00000171217 | 6 | 1 | 6 | 0 | 1 | 0 |
| ENSG00000232529 | 0 | 0 | 0 | 0 | 0 | 0 |
| ENSG00000074771 | 0 | 0 | 0 | 0 | 0 | 0 |
| ENSG00000238789 | 0 | 0 | 0 | 0 | 0 | 0 |
| ENSG00000221456 | 0 | 0 | 0 | 0 | 0 | 0 |
| ENSG00000285797 | 0 | 0 | 0 | 0 | 0 | 0 |
| ENSG00000212295 | 0 | 0 | 0 | 0 | 0 | 0 |
| ENSG00000227360 | 0 | 0 | 0 | 0 | 0 | 0 |
| ENSG00000220347 | 0 | 0 | 0 | 0 | 0 | 0 |
| ENSG00000271265 | 2 | 1 | 5 | 0 | 1 | 1 |
| ENSG00000049618 | 1144 | 527 | 648 | 397 | 542 | 535 |
| ENSG00000271551 | 5 | 2 | 4 | 3 | 1 | 1 |
| ENSG00000271899 | 0 | 0 | 0 | 0 | 0 | 0 |
| ENSG00000270487 | 0 | 0 | 0 | 0 | 0 | 0 |
| ENSG00000218596 | 0 | 0 | 0 | 0 | 0 | 0 |
| ENSG00000233044 | 0 | 0 | 0 | 0 | 0 | 0 |
| ENSG00000215712 | 272 | 174 | 176 | 169 | 200 | 194 |
| ENSG00000217783 | 0 | 0 | 0 | 0 | 0 | 0 |
| ENSG00000271040 | 6 | 1 | 1 | 7 | 2 | 5 |
| ENSG00000213078 | 8 | 4 | 2 | 1 | 1 | 5 |
| ENSG00000218631 | 24 | 6 | 11 | 9 | 5 | 5 |
| ENSG00000175048 | 204 | 115 | 102 | 114 | 118 | 123 |
| ENSG00000266617 | 0 | 0 | 0 | 0 | 0 | 0 |
| ENSG00000130340 | 1083 | 541 | 729 | 577 | 671 | 672 |
| ENSG00000234361 | 0 | 0 | 0 | 1 | 0 | 0 |
| ENSG00000229502 | 0 | 0 | 0 | 0 | 0 | 0 |
| ENSG00000236324 | 0 | 0 | 0 | 0 | 0 | 0 |

| | | | | | | |
|---|---|---|---|---|---|---|
| ENSG00000220867 | 0 | 0 | 0 | 0 | 0 | 0 |
| ENSG00000252658 | 0 | 0 | 0 | 0 | 0 | 0 |
| ENSG00000078269 | 1363 | 663 | 794 | 481 | 712 | 683 |
| ENSG00000285642 | 0 | 0 | 0 | 0 | 0 | 0 |
| ENSG00000233496 | 0 | 0 | 0 | 0 | 0 | 0 |
| ENSG00000122335 | 248 | 89 | 146 | 96 | 135 | 113 |
| ENSG00000272047 | 489 | 242 | 290 | 280 | 283 | 253 |
| ENSG00000130338 | 1059 | 489 | 636 | 323 | 462 | 536 |
| ENSG00000188451 | 0 | 1 | 2 | 0 | 0 | 0 |
| ENSG00000274023 | 0 | 0 | 0 | 0 | 0 | 0 |
| ENSG00000238019 | 0 | 0 | 0 | 0 | 0 | 0 |
| ENSG00000243373 | 0 | 0 | 0 | 0 | 0 | 0 |
| ENSG00000219188 | 0 | 0 | 0 | 0 | 0 | 0 |
| ENSG00000146433 | 1570 | 684 | 892 | 459 | 806 | 697 |
| ENSG00000218226 | 41 | 32 | 46 | 25 | 44 | 38 |
| ENSG00000278571 | 0 | 0 | 0 | 0 | 0 | 0 |
| ENSG00000146425 | 2351 | 1165 | 1382 | 1218 | 1289 | 1290 |
| ENSG00000164674 | 69 | 36 | 63 | 29 | 36 | 36 |
| ENSG00000219249 | 0 | 0 | 0 | 0 | 0 | 0 |
| ENSG00000265558 | 0 | 0 | 0 | 0 | 0 | 0 |
| ENSG00000092820 | 9066 | 4457 | 6018 | 4632 | 4878 | 5285 |
| ENSG00000233893 | 1 | 1 | 2 | 0 | 1 | 0 |
| ENSG00000243775 | 0 | 0 | 0 | 0 | 0 | 0 |
| ENSG00000285159 | 2 | 1 | 2 | 2 | 5 | 4 |
| ENSG00000203711 | 47 | 25 | 28 | 29 | 25 | 26 |
| ENSG00000213076 | 0 | 0 | 0 | 0 | 0 | 0 |
| ENSG00000223191 | 0 | 0 | 0 | 0 | 0 | 0 |
| ENSG00000130363 | 556 | 222 | 300 | 197 | 322 | 243 |
| ENSG00000271913 | 6 | 4 | 5 | 5 | 8 | 6 |
| ENSG00000164691 | 0 | 2 | 0 | 0 | 0 | 1 |
| ENSG00000226032 | 0 | 0 | 0 | 0 | 0 | 0 |
| ENSG00000285492 | 2 | 0 | 0 | 0 | 0 | 0 |
| ENSG00000224478 | 0 | 0 | 0 | 0 | 0 | 0 |
| ENSG00000233682 | 0 | 0 | 0 | 0 | 0 | 0 |
| ENSG00000164694 | 0 | 0 | 0 | 0 | 0 | 0 |
| ENSG00000235086 | 0 | 0 | 0 | 0 | 0 | 0 |
| ENSG00000231178 | 0 | 0 | 0 | 0 | 0 | 0 |
| ENSG00000234777 | 0 | 0 | 0 | 0 | 0 | 0 |
| ENSG00000216480 | 0 | 0 | 1 | 0 | 1 | 0 |
| ENSG00000276413 | 0 | 1 | 0 | 0 | 0 | 2 |
| ENSG00000237927 | 0 | 0 | 5 | 3 | 1 | 0 |
| ENSG00000112096 | 1114 | 477 | 606 | 480 | 689 | 614 |
| ENSG00000285441 | 0 | 1 | 1 | 0 | 0 | 0 |
| ENSG00000220305 | 4 | 2 | 0 | 2 | 4 | 1 |
| ENSG00000251988 | 2 | 2 | 1 | 2 | 0 | 1 |
| ENSG00000146457 | 1179 | 626 | 737 | 526 | 744 | 647 |
| ENSG00000285427 | 0 | 1 | 0 | 0 | 0 | 0 |
| ENSG00000120437 | 940 | 532 | 601 | 583 | 592 | 619 |
| ENSG00000120438 | 4113 | 2191 | 2570 | 2060 | 2847 | 2527 |
| ENSG00000207392 | 3 | 0 | 1 | 1 | 1 | 0 |

| | | | | | | |
|---|---|---|---|---|---|---|
| ENSG00000206910 | 0 | 0 | 0 | 0 | 0 | 0 |
| ENSG00000112110 | 1163 | 607 | 681 | 737 | 704 | 641 |
| ENSG00000146453 | 0 | 1 | 1 | 0 | 0 | 0 |
| ENSG00000236823 | 0 | 0 | 0 | 0 | 0 | 0 |
| ENSG00000130368 | 0 | 0 | 0 | 0 | 0 | 1 |
| ENSG00000197081 | 5432 | 2244 | 3340 | 2035 | 2878 | 2781 |
| ENSG00000268257 | 0 | 0 | 0 | 0 | 0 | 0 |
| ENSG00000213073 | 65 | 30 | 50 | 18 | 36 | 34 |
| ENSG00000175003 | 6 | 1 | 1 | 0 | 1 | 1 |
| ENSG00000216516 | 0 | 0 | 0 | 0 | 0 | 0 |
| ENSG00000112499 | 0 | 0 | 0 | 0 | 0 | 0 |
| ENSG00000230234 | 0 | 0 | 0 | 0 | 0 | 0 |
| ENSG00000146477 | 0 | 0 | 0 | 0 | 0 | 0 |
| ENSG00000213071 | 0 | 0 | 0 | 0 | 0 | 2 |
| ENSG00000198670 | 0 | 2 | 1 | 0 | 0 | 1 |
| ENSG00000243831 | 0 | 1 | 2 | 1 | 1 | 0 |
| ENSG00000224477 | 0 | 0 | 0 | 0 | 0 | 0 |
| ENSG00000122194 | 0 | 0 | 0 | 0 | 0 | 0 |
| ENSG00000276960 | 0 | 0 | 0 | 0 | 0 | 0 |
| ENSG00000274903 | 0 | 0 | 0 | 0 | 0 | 0 |
| ENSG00000224371 | 0 | 2 | 0 | 0 | 0 | 0 |
| ENSG00000220913 | 0 | 0 | 0 | 0 | 0 | 0 |
| ENSG00000233342 | 0 | 0 | 0 | 0 | 0 | 0 |
| ENSG00000231863 | 0 | 0 | 0 | 0 | 0 | 0 |
| ENSG00000272841 | 23 | 13 | 16 | 15 | 23 | 9 |
| ENSG00000085511 | 393 | 216 | 244 | 172 | 272 | 222 |
| ENSG00000026652 | 12 | 3 | 1 | 2 | 3 | 4 |
| ENSG00000279355 | 0 | 0 | 0 | 0 | 0 | 0 |
| ENSG00000185345 | 24 | 7 | 11 | 10 | 12 | 8 |
| ENSG00000227913 | 0 | 0 | 0 | 0 | 0 | 0 |
| ENSG00000112530 | 9 | 4 | 6 | 6 | 4 | 2 |
| ENSG00000217557 | 0 | 0 | 0 | 0 | 0 | 0 |
| ENSG00000285610 | 0 | 0 | 0 | 0 | 0 | 0 |
| ENSG00000225437 | 0 | 0 | 0 | 0 | 0 | 0 |
| ENSG00000285553 | 0 | 0 | 0 | 0 | 0 | 0 |
| ENSG00000225683 | 0 | 0 | 0 | 0 | 0 | 0 |
| ENSG00000281692 | 0 | 0 | 0 | 1 | 0 | 0 |
| ENSG00000285564 | 0 | 0 | 0 | 0 | 0 | 0 |
| ENSG00000285726 | 0 | 0 | 0 | 0 | 0 | 0 |
| ENSG00000217514 | 0 | 0 | 0 | 0 | 0 | 0 |
| ENSG00000270419 | 0 | 6 | 3 | 4 | 3 | 1 |
| ENSG00000112531 | 956 | 493 | 542 | 322 | 499 | 450 |
| ENSG00000228692 | 0 | 0 | 0 | 0 | 0 | 0 |
| ENSG00000235538 | 1 | 1 | 1 | 1 | 0 | 0 |
| ENSG00000285858 | 0 | 0 | 0 | 0 | 0 | 0 |
| ENSG00000266128 | 0 | 0 | 0 | 0 | 0 | 0 |
| ENSG00000260422 | 0 | 0 | 0 | 0 | 0 | 0 |
| ENSG00000230627 | 0 | 0 | 0 | 0 | 0 | 0 |
| ENSG00000226739 | 0 | 0 | 0 | 0 | 0 | 0 |
| ENSG00000227455 | 0 | 0 | 0 | 0 | 0 | 0 |

| | | | | | | |
|---|---|---|---|---|---|---|
| ENSG00000217878 | 0 | 0 | 0 | 0 | 0 | 0 |
| ENSG00000112539 | 0 | 0 | 0 | 0 | 0 | 0 |
| ENSG00000112541 | 0 | 0 | 0 | 0 | 1 | 0 |
| ENSG00000199523 | 0 | 0 | 1 | 0 | 0 | 0 |
| ENSG00000206681 | 0 | 0 | 0 | 0 | 0 | 0 |
| ENSG00000236627 | 0 | 0 | 0 | 0 | 0 | 0 |
| ENSG00000223942 | 0 | 0 | 0 | 0 | 0 | 0 |
| ENSG00000281832 | 0 | 0 | 0 | 0 | 0 | 0 |
| ENSG00000216624 | 0 | 1 | 1 | 1 | 0 | 1 |
| ENSG00000231297 | 0 | 0 | 0 | 0 | 0 | 0 |
| ENSG00000276643 | 0 | 0 | 0 | 0 | 0 | 0 |
| ENSG00000164458 | 0 | 0 | 0 | 0 | 0 | 0 |
| ENSG00000201292 | 0 | 0 | 0 | 0 | 0 | 0 |
| ENSG00000280991 | 0 | 0 | 0 | 0 | 0 | 0 |
| ENSG00000233365 | 0 | 0 | 0 | 0 | 0 | 0 |
| ENSG00000213536 | 0 | 0 | 0 | 0 | 0 | 0 |
| ENSG00000280850 | 0 | 0 | 0 | 0 | 0 | 0 |
| ENSG00000176381 | 2 | 0 | 1 | 4 | 0 | 0 |
| ENSG00000198818 | 998 | 459 | 568 | 504 | 564 | 462 |
| ENSG00000233231 | 10 | 5 | 5 | 0 | 8 | 6 |
| ENSG00000060762 | 393 | 200 | 237 | 240 | 235 | 234 |
| ENSG00000261420 | 4 | 2 | 5 | 2 | 4 | 3 |
| ENSG00000280057 | 6 | 1 | 0 | 3 | 1 | 0 |
| ENSG00000071242 | 499 | 231 | 308 | 246 | 288 | 371 |
| ENSG00000232082 | 0 | 1 | 2 | 1 | 0 | 0 |
| ENSG00000222958 | 0 | 0 | 0 | 0 | 0 | 0 |
| ENSG00000235272 | 2 | 0 | 1 | 1 | 0 | 0 |
| ENSG00000235008 | 0 | 0 | 0 | 0 | 0 | 0 |
| ENSG00000249141 | 1 | 3 | 3 | 4 | 1 | 2 |
| ENSG00000231654 | 0 | 0 | 0 | 0 | 0 | 0 |
| ENSG00000026297 | 1610 | 799 | 1164 | 950 | 936 | 929 |
| ENSG00000227598 | 5 | 3 | 1 | 1 | 1 | 2 |
| ENSG00000285730 | 2 | 1 | 0 | 0 | 0 | 0 |
| ENSG00000265828 | 0 | 0 | 0 | 0 | 0 | 1 |
| ENSG00000213066 | 198 | 118 | 119 | 90 | 149 | 126 |
| ENSG00000272980 | 0 | 1 | 3 | 1 | 2 | 0 |
| ENSG00000112486 | 0 | 0 | 0 | 0 | 0 | 0 |
| ENSG00000284825 | 0 | 0 | 0 | 0 | 0 | 0 |
| ENSG00000284914 | 0 | 0 | 0 | 0 | 0 | 0 |
| ENSG00000166984 | 0 | 0 | 0 | 0 | 0 | 0 |
| ENSG00000120436 | 0 | 0 | 0 | 0 | 0 | 0 |
| ENSG00000216966 | 0 | 0 | 0 | 0 | 0 | 0 |
| ENSG00000280707 | 0 | 0 | 0 | 0 | 0 | 0 |
| ENSG00000231720 | 0 | 0 | 0 | 0 | 0 | 0 |
| ENSG00000228648 | 0 | 0 | 0 | 0 | 0 | 0 |
| ENSG00000112494 | 0 | 0 | 0 | 0 | 0 | 0 |
| ENSG00000120440 | 0 | 0 | 0 | 0 | 0 | 0 |
| ENSG00000217447 | 0 | 0 | 0 | 0 | 0 | 0 |
| ENSG00000203690 | 0 | 0 | 0 | 0 | 0 | 0 |
| ENSG00000272549 | 0 | 0 | 0 | 0 | 0 | 0 |

| | | | | | | |
|---|---|---|---|---|---|---|
| ENSG00000203688 | 0 | 0 | 0 | 0 | 0 | 0 |
| ENSG00000213065 | 0 | 1 | 1 | 0 | 0 | 0 |
| ENSG00000146521 | 0 | 1 | 0 | 0 | 1 | 0 |
| ENSG00000269155 | 0 | 0 | 0 | 0 | 1 | 0 |
| ENSG00000285212 | 0 | 3 | 1 | 1 | 0 | 0 |
| ENSG00000198221 | 7 | 3 | 8 | 6 | 5 | 4 |
| ENSG00000130396 | 2242 | 1012 | 1463 | 768 | 1132 | 1160 |
| ENSG00000235994 | 0 | 0 | 0 | 0 | 0 | 0 |
| ENSG00000229921 | 0 | 0 | 0 | 0 | 0 | 0 |
| ENSG00000125337 | 0 | 0 | 3 | 1 | 0 | 0 |
| ENSG00000153303 | 0 | 0 | 0 | 0 | 0 | 0 |
| ENSG00000234768 | 0 | 0 | 0 | 0 | 0 | 0 |
| ENSG00000237987 | 0 | 0 | 0 | 0 | 0 | 0 |
| ENSG00000228361 | 0 | 0 | 0 | 0 | 0 | 0 |
| ENSG00000234647 | 0 | 0 | 0 | 0 | 0 | 0 |
| ENSG00000230711 | 0 | 0 | 0 | 0 | 0 | 0 |
| ENSG00000164488 | 0 | 0 | 0 | 0 | 0 | 0 |
| ENSG00000224417 | 0 | 0 | 0 | 0 | 0 | 0 |
| ENSG00000260771 | 0 | 0 | 0 | 0 | 0 | 0 |
| ENSG00000112562 | 24 | 10 | 18 | 1 | 5 | 9 |
| ENSG00000235815 | 0 | 0 | 0 | 0 | 0 | 0 |
| ENSG00000280665 | 0 | 0 | 0 | 0 | 0 | 0 |
| ENSG00000225879 | 0 | 0 | 0 | 0 | 0 | 0 |
| ENSG00000285917 | 0 | 0 | 0 | 0 | 0 | 0 |
| ENSG00000279810 | 0 | 0 | 0 | 0 | 0 | 0 |
| ENSG00000229720 | 0 | 0 | 0 | 0 | 0 | 0 |
| ENSG00000234519 | 0 | 0 | 0 | 0 | 0 | 0 |
| ENSG00000223485 | 0 | 0 | 1 | 1 | 3 | 1 |
| ENSG00000261039 | 0 | 0 | 0 | 0 | 0 | 0 |
| ENSG00000226445 | 14 | 14 | 16 | 14 | 18 | 18 |
| ENSG00000186340 | 54 | 22 | 25 | 16 | 25 | 32 |
| ENSG00000233085 | 0 | 0 | 0 | 0 | 0 | 0 |
| ENSG00000224999 | 0 | 0 | 0 | 0 | 0 | 0 |
| ENSG00000226194 | 0 | 0 | 0 | 0 | 0 | 0 |
| ENSG00000285887 | 0 | 0 | 0 | 0 | 0 | 0 |
| ENSG00000285733 | 0 | 0 | 0 | 0 | 0 | 0 |
| ENSG00000272848 | 8 | 3 | 4 | 0 | 2 | 4 |
| ENSG00000184465 | 662 | 251 | 418 | 209 | 350 | 344 |
| ENSG00000185127 | 1375 | 729 | 834 | 708 | 923 | 831 |
| ENSG00000130024 | 1881 | 1081 | 1311 | 937 | 1239 | 1173 |
| ENSG00000232640 | 19 | 15 | 12 | 22 | 24 | 20 |
| ENSG00000227704 | 0 | 0 | 0 | 0 | 0 | 0 |
| ENSG00000184786 | 94 | 51 | 78 | 38 | 68 | 38 |
| ENSG00000130023 | 834 | 416 | 546 | 333 | 445 | 470 |
| ENSG00000266896 | 32 | 16 | 20 | 11 | 18 | 15 |
| ENSG00000229214 | 0 | 4 | 3 | 8 | 10 | 8 |
| ENSG00000231690 | 0 | 0 | 2 | 0 | 2 | 0 |
| ENSG00000236173 | 0 | 0 | 0 | 0 | 1 | 0 |
| ENSG00000232197 | 4 | 0 | 0 | 4 | 1 | 0 |
| ENSG00000273100 | 0 | 1 | 0 | 0 | 3 | 0 |

| | | | | | | |
|---|---|---|---|---|---|---|
| ENSG00000218716 | 0 | 0 | 0 | 0 | 0 | 0 |
| ENSG00000230960 | 0 | 0 | 0 | 0 | 0 | 0 |
| ENSG00000271820 | 0 | 0 | 0 | 0 | 0 | 1 |
| ENSG00000227508 | 0 | 2 | 2 | 1 | 2 | 2 |
| ENSG00000198719 | 1075 | 456 | 596 | 667 | 776 | 729 |
| ENSG00000112584 | 1911 | 884 | 1084 | 807 | 1119 | 1076 |
| ENSG00000271234 | 3 | 1 | 1 | 0 | 2 | 0 |
| ENSG00000266245 | 0 | 1 | 0 | 0 | 0 | 0 |
| ENSG00000261003 | 0 | 0 | 0 | 0 | 1 | 0 |
| ENSG00000008018 | 7704 | 3884 | 4728 | 4560 | 4650 | 4471 |
| ENSG00000112592 | 633 | 319 | 433 | 284 | 402 | 386 |
| ENSG00000071994 | 3411 | 1704 | 2053 | 2214 | 2093 | 2158 |
| ENSG00000217874 | 0 | 0 | 0 | 0 | 0 | 0 |
| ENSG00000273640 | 0 | 0 | 0 | 0 | 0 | 0 |
| ENSG00000234986 | 0 | 0 | 0 | 0 | 0 | 0 |
| ENSG00000213863 | 0 | 2 | 1 | 0 | 1 | 0 |
| ENSG00000230423 | 4 | 4 | 3 | 0 | 1 | 3 |
| ENSG00000225532 | 0 | 1 | 0 | 2 | 1 | 0 |
| ENSG00000219814 | 0 | 0 | 0 | 0 | 0 | 0 |
| ENSG00000283061 | 0 | 0 | 0 | 0 | 0 | 0 |
| ENSG00000282572 | 12 | 2 | 4 | 1 | 1 | 6 |
| ENSG00000273644 | 0 | 0 | 0 | 0 | 0 | 0 |
| ENSG00000232325 | 0 | 0 | 0 | 0 | 0 | 0 |
| ENSG00000242611 | 0 | 0 | 0 | 1 | 0 | 0 |
| ENSG00000242474 | 47 | 24 | 32 | 23 | 26 | 36 |
| ENSG00000240859 | 32 | 5 | 8 | 9 | 4 | 13 |
| ENSG00000261795 | 0 | 0 | 0 | 1 | 0 | 0 |
| ENSG00000239715 | 0 | 0 | 0 | 0 | 0 | 0 |
| ENSG00000240093 | 0 | 0 | 0 | 0 | 1 | 0 |
| ENSG00000177706 | 805 | 380 | 473 | 503 | 460 | 466 |
| ENSG00000249852 | 10 | 2 | 3 | 1 | 5 | 3 |
| ENSG00000248767 | 0 | 0 | 0 | 0 | 2 | 0 |
| ENSG00000249574 | 0 | 0 | 0 | 0 | 0 | 0 |
| ENSG00000234471 | 0 | 0 | 0 | 0 | 0 | 0 |
| ENSG00000197461 | 643 | 290 | 399 | 361 | 303 | 340 |
| ENSG00000223855 | 460 | 151 | 332 | 177 | 200 | 194 |
| ENSG00000188191 | 1279 | 638 | 809 | 943 | 780 | 772 |
| ENSG00000229380 | 3 | 0 | 0 | 1 | 0 | 0 |
| ENSG00000237181 | 0 | 0 | 0 | 0 | 0 | 0 |
| ENSG00000164818 | 2484 | 1247 | 1563 | 1407 | 1268 | 1481 |
| ENSG00000164828 | 5852 | 2660 | 3756 | 2530 | 3225 | 3299 |
| ENSG00000239857 | 132 | 84 | 157 | 100 | 124 | 113 |
| ENSG00000273151 | 432 | 151 | 245 | 106 | 159 | 148 |
| ENSG00000105963 | 679 | 341 | 388 | 575 | 410 | 378 |
| ENSG00000240230 | 1075 | 555 | 649 | 558 | 522 | 589 |
| ENSG00000073067 | 66 | 28 | 32 | 21 | 23 | 17 |
| ENSG00000146540 | 5646 | 2758 | 3398 | 3659 | 2822 | 2785 |
| ENSG00000199023 | 0 | 4 | 2 | 5 | 2 | 3 |
| ENSG00000225146 | 11 | 5 | 8 | 5 | 4 | 6 |
| ENSG00000164849 | 19 | 14 | 14 | 12 | 10 | 10 |

| | | | | | | |
|---|---|---|---|---|---|---|
| ENSG00000257607 | 0 | 0 | 0 | 0 | 0 | 0 |
| ENSG00000224079 | 0 | 0 | 2 | 1 | 1 | 2 |
| ENSG00000226291 | 1 | 0 | 1 | 0 | 1 | 0 |
| ENSG00000164850 | 561 | 266 | 349 | 165 | 146 | 145 |
| ENSG00000178381 | 935 | 522 | 555 | 526 | 580 | 522 |
| ENSG00000229043 | 608 | 291 | 422 | 291 | 342 | 387 |
| ENSG00000164853 | 0 | 0 | 0 | 0 | 0 | 0 |
| ENSG00000233082 | 3 | 0 | 1 | 0 | 0 | 0 |
| ENSG00000164877 | 2280 | 884 | 1434 | 1191 | 1096 | 1241 |
| ENSG00000225981 | 1 | 0 | 1 | 0 | 0 | 0 |
| ENSG00000273230 | 402 | 201 | 262 | 271 | 255 | 241 |
| ENSG00000164880 | 5097 | 2273 | 3001 | 2833 | 2791 | 2780 |
| ENSG00000231927 | 2 | 0 | 3 | 1 | 1 | 1 |
| ENSG00000198517 | 1144 | 571 | 907 | 752 | 743 | 707 |
| ENSG00000164855 | 3262 | 1484 | 2012 | 1451 | 1462 | 1452 |
| ENSG00000157778 | 1391 | 761 | 908 | 958 | 784 | 819 |
| ENSG00000230487 | 2541 | 1114 | 1494 | 1172 | 1224 | 1147 |
| ENSG00000230444 | 12 | 4 | 12 | 4 | 5 | 7 |
| ENSG00000231476 | 0 | 0 | 0 | 0 | 0 | 0 |
| ENSG00000225968 | 186 | 85 | 108 | 74 | 84 | 67 |
| ENSG00000234977 | 0 | 0 | 0 | 0 | 0 | 0 |
| ENSG00000205971 | 0 | 0 | 0 | 0 | 0 | 0 |
| ENSG00000236081 | 2 | 0 | 0 | 0 | 0 | 0 |
| ENSG00000002822 | 825 | 408 | 527 | 653 | 537 | 508 |
| ENSG00000176349 | 0 | 0 | 0 | 0 | 1 | 0 |
| ENSG00000265089 | 0 | 0 | 0 | 0 | 0 | 0 |
| ENSG00000122687 | 1882 | 867 | 989 | 999 | 950 | 1053 |
| ENSG00000106268 | 589 | 328 | 305 | 434 | 362 | 277 |
| ENSG00000106266 | 1537 | 689 | 914 | 931 | 768 | 804 |
| ENSG00000284047 | 0 | 0 | 0 | 0 | 0 | 0 |
| ENSG00000271098 | 0 | 0 | 0 | 0 | 0 | 0 |
| ENSG00000106263 | 12594 | 6501 | 7832 | 8519 | 8060 | 8407 |
| ENSG00000230914 | 0 | 1 | 0 | 1 | 0 | 0 |
| ENSG00000271163 | 0 | 0 | 0 | 0 | 0 | 0 |
| ENSG00000136213 | 858 | 434 | 509 | 438 | 445 | 427 |
| ENSG00000275572 | 0 | 1 | 0 | 2 | 0 | 0 |
| ENSG00000106003 | 1895 | 876 | 1136 | 673 | 791 | 580 |
| ENSG00000264357 | 0 | 1 | 0 | 0 | 0 | 0 |
| ENSG00000106009 | 3638 | 1565 | 2073 | 2159 | 1873 | 1726 |
| ENSG00000106012 | 3477 | 1510 | 2049 | 1513 | 1712 | 1738 |
| ENSG00000136295 | 2888 | 1198 | 1589 | 1663 | 1673 | 1643 |
| ENSG00000174945 | 208 | 131 | 116 | 189 | 177 | 175 |
| ENSG00000146535 | 2558 | 1235 | 1550 | 1426 | 1413 | 1451 |
| ENSG00000231357 | 0 | 0 | 0 | 0 | 3 | 0 |
| ENSG00000198286 | 0 | 0 | 0 | 0 | 0 | 0 |
| ENSG00000237286 | 0 | 0 | 0 | 0 | 0 | 0 |
| ENSG00000201794 | 0 | 0 | 0 | 0 | 0 | 0 |
| ENSG00000228334 | 0 | 0 | 0 | 0 | 0 | 0 |
| ENSG00000217455 | 2 | 0 | 0 | 0 | 0 | 0 |
| ENSG00000231892 | 0 | 0 | 1 | 1 | 2 | 0 |

| | | | | | | |
|---|---|---|---|---|---|---|
| ENSG00000236708 | 4 | 2 | 6 | 2 | 2 | 4 |
| ENSG00000146555 | 829 | 400 | 581 | 416 | 480 | 589 |
| ENSG00000224593 | 0 | 0 | 0 | 0 | 0 | 0 |
| ENSG00000236510 | 0 | 0 | 0 | 0 | 0 | 0 |
| ENSG00000283991 | 0 | 0 | 0 | 0 | 0 | 0 |
| ENSG00000261809 | 0 | 0 | 0 | 0 | 0 | 0 |
| ENSG00000164916 | 1735 | 958 | 1134 | 859 | 985 | 985 |
| ENSG00000242802 | 2419 | 1066 | 1555 | 1393 | 1364 | 1361 |
| ENSG00000264474 | 0 | 0 | 0 | 0 | 0 | 0 |
| ENSG00000157927 | 98 | 38 | 68 | 37 | 40 | 41 |
| ENSG00000206895 | 0 | 0 | 0 | 0 | 0 | 0 |
| ENSG00000218823 | 0 | 0 | 0 | 0 | 0 | 0 |
| ENSG00000213935 | 0 | 0 | 1 | 0 | 1 | 0 |
| ENSG00000136297 | 0 | 0 | 0 | 0 | 0 | 0 |
| ENSG00000196204 | 450 | 232 | 277 | 286 | 301 | 277 |
| ENSG00000272968 | 0 | 0 | 1 | 1 | 1 | 0 |
| ENSG00000270822 | 0 | 0 | 1 | 1 | 0 | 4 |
| ENSG00000275328 | 5 | 5 | 2 | 3 | 3 | 3 |
| ENSG00000146587 | 898 | 478 | 351 | 192 | 686 | 364 |
| ENSG00000273313 | 1 | 0 | 1 | 0 | 0 | 0 |
| ENSG00000272595 | 0 | 0 | 0 | 0 | 0 | 1 |
| ENSG00000159904 | 15 | 5 | 9 | 5 | 5 | 5 |
| ENSG00000202039 | 0 | 0 | 0 | 0 | 0 | 0 |
| ENSG00000157954 | 4420 | 2051 | 2541 | 2441 | 2436 | 2385 |
| ENSG00000164638 | 577 | 286 | 391 | 377 | 373 | 351 |
| ENSG00000182095 | 5344 | 2135 | 3527 | 2325 | 2212 | 2677 |
| ENSG00000241269 | 3 | 6 | 5 | 2 | 3 | 6 |
| ENSG00000188365 | 4 | 0 | 3 | 2 | 1 | 0 |
| ENSG00000272953 | 26 | 18 | 26 | 10 | 26 | 18 |
| ENSG00000234432 | 43 | 18 | 26 | 17 | 20 | 24 |
| ENSG00000273084 | 33 | 15 | 15 | 17 | 8 | 11 |
| ENSG00000155034 | 945 | 421 | 628 | 389 | 435 | 462 |
| ENSG00000230733 | 1539 | 767 | 979 | 834 | 826 | 752 |
| ENSG00000207973 | 0 | 2 | 0 | 0 | 2 | 1 |
| ENSG00000075624 | 99362 | 52199 | 63252 | 68334 | 64217 | 59126 |
| ENSG00000272719 | 0 | 1 | 1 | 3 | 1 | 0 |
| ENSG00000228974 | 0 | 3 | 0 | 1 | 2 | 8 |
| ENSG00000075618 | 432 | 218 | 261 | 304 | 231 | 230 |
| ENSG00000011275 | 2331 | 1003 | 1368 | 1120 | 1326 | 1329 |
| ENSG00000237738 | 0 | 0 | 0 | 1 | 2 | 0 |
| ENSG00000283419 | 0 | 0 | 0 | 0 | 0 | 0 |
| ENSG00000235944 | 129 | 76 | 101 | 102 | 101 | 109 |
| ENSG00000277265 | 0 | 1 | 0 | 0 | 0 | 0 |
| ENSG00000122543 | 5 | 4 | 3 | 2 | 3 | 3 |
| ENSG00000122674 | 370 | 183 | 268 | 195 | 247 | 223 |
| ENSG00000155026 | 1 | 1 | 0 | 0 | 0 | 0 |
| ENSG00000122512 | 895 | 465 | 556 | 351 | 487 | 557 |
| ENSG00000201990 | 0 | 1 | 0 | 0 | 0 | 0 |
| ENSG00000106305 | 709 | 383 | 441 | 547 | 441 | 532 |
| ENSG00000207217 | 0 | 1 | 0 | 1 | 0 | 1 |

| | | | | | | |
|---|---|---|---|---|---|---|
| ENSG00000086232 | 9129 | 4926 | 5648 | 3866 | 5576 | 4899 |
| ENSG00000157999 | 48 | 16 | 26 | 14 | 28 | 27 |
| ENSG00000240718 | 0 | 2 | 0 | 0 | 0 | 0 |
| ENSG00000252929 | 0 | 0 | 0 | 0 | 0 | 1 |
| ENSG00000231704 | 0 | 0 | 5 | 6 | 0 | 3 |
| ENSG00000106346 | 1017 | 484 | 636 | 507 | 545 | 626 |
| ENSG00000008256 | 1057 | 510 | 604 | 474 | 611 | 593 |
| ENSG00000202273 | 0 | 0 | 0 | 0 | 0 | 0 |
| ENSG00000178397 | 964 | 467 | 560 | 583 | 648 | 538 |
| ENSG00000274319 | 0 | 0 | 0 | 0 | 0 | 0 |
| ENSG00000136238 | 15123 | 7611 | 9052 | 7134 | 8427 | 8379 |
| ENSG00000164535 | 1389 | 751 | 821 | 1030 | 830 | 865 |
| ENSG00000136240 | 9323 | 4785 | 5731 | 5109 | 5546 | 5396 |
| ENSG00000231359 | 11 | 8 | 21 | 4 | 4 | 10 |
| ENSG00000215045 | 86 | 39 | 40 | 18 | 37 | 36 |
| ENSG00000136247 | 1840 | 999 | 1207 | 1156 | 1191 | 1136 |
| ENSG00000232581 | 0 | 0 | 0 | 0 | 0 | 0 |
| ENSG00000146576 | 1295 | 601 | 717 | 750 | 602 | 740 |
| ENSG00000236609 | 40 | 19 | 23 | 34 | 17 | 25 |
| ENSG00000205903 | 2782 | 1149 | 1526 | 1507 | 1509 | 1425 |
| ENSG00000228010 | 16 | 9 | 7 | 2 | 2 | 4 |
| ENSG00000198580 | 12 | 2 | 4 | 2 | 3 | 1 |
| ENSG00000164631 | 1093 | 667 | 470 | 288 | 916 | 550 |
| ENSG00000187953 | 359 | 202 | 264 | 155 | 199 | 207 |
| ENSG00000277327 | 0 | 0 | 0 | 0 | 0 | 0 |
| ENSG00000169402 | 2 | 0 | 4 | 1 | 5 | 2 |
| ENSG00000146574 | 1179 | 627 | 749 | 603 | 787 | 737 |
| ENSG00000273252 | 5 | 2 | 3 | 3 | 0 | 3 |
| ENSG00000155070 | 0 | 0 | 0 | 0 | 0 | 0 |
| ENSG00000205897 | 0 | 0 | 0 | 0 | 0 | 0 |
| ENSG00000250561 | 0 | 0 | 0 | 0 | 0 | 0 |
| ENSG00000243531 | 0 | 0 | 0 | 0 | 0 | 0 |
| ENSG00000226943 | 2 | 0 | 1 | 1 | 0 | 1 |
| ENSG00000241539 | 0 | 0 | 0 | 0 | 0 | 0 |
| ENSG00000242731 | 2 | 0 | 0 | 0 | 0 | 0 |
| ENSG00000266287 | 0 | 0 | 0 | 0 | 0 | 0 |
| ENSG00000239696 | 0 | 0 | 0 | 0 | 0 | 0 |
| ENSG00000201218 | 0 | 0 | 0 | 0 | 0 | 0 |
| ENSG00000106392 | 452 | 199 | 218 | 167 | 279 | 213 |
| ENSG00000244167 | 0 | 0 | 0 | 0 | 0 | 0 |
| ENSG00000230825 | 0 | 0 | 1 | 0 | 2 | 0 |
| ENSG00000236712 | 0 | 0 | 0 | 0 | 0 | 0 |
| ENSG00000215018 | 81 | 35 | 78 | 25 | 37 | 23 |
| ENSG00000272732 | 71 | 32 | 51 | 24 | 26 | 30 |
| ENSG00000272894 | 152 | 71 | 111 | 33 | 55 | 64 |
| ENSG00000164654 | 1077 | 513 | 558 | 381 | 627 | 460 |
| ENSG00000272745 | 2 | 0 | 0 | 0 | 0 | 0 |
| ENSG00000106399 | 1176 | 580 | 690 | 698 | 652 | 633 |
| ENSG00000219545 | 456 | 230 | 273 | 174 | 277 | 212 |
| ENSG00000283549 | 0 | 0 | 0 | 0 | 0 | 0 |

| | | | | | | |
|---|---|---|---|---|---|---|
| ENSG00000270314 | 0 | 0 | 0 | 0 | 1 | 0 |
| ENSG00000234718 | 0 | 1 | 0 | 0 | 0 | 0 |
| ENSG00000201747 | 0 | 0 | 0 | 0 | 0 | 0 |
| ENSG00000214998 | 0 | 0 | 2 | 0 | 0 | 0 |
| ENSG00000234141 | 10 | 1 | 2 | 1 | 1 | 0 |
| ENSG00000233108 | 0 | 1 | 1 | 0 | 3 | 1 |
| ENSG00000106415 | 949 | 514 | 548 | 301 | 625 | 485 |
| ENSG00000233264 | 0 | 3 | 1 | 0 | 1 | 0 |
| ENSG00000003147 | 2094 | 1183 | 1508 | 1121 | 1370 | 1314 |
| ENSG00000227719 | 60 | 21 | 55 | 16 | 37 | 24 |
| ENSG00000244239 | 0 | 0 | 0 | 0 | 0 | 0 |
| ENSG00000229970 | 16 | 11 | 8 | 2 | 3 | 9 |
| ENSG00000272328 | 0 | 2 | 0 | 1 | 1 | 1 |
| ENSG00000122584 | 22 | 10 | 13 | 11 | 8 | 7 |
| ENSG00000236748 | 0 | 0 | 0 | 0 | 0 | 0 |
| ENSG00000278254 | 0 | 2 | 0 | 0 | 0 | 0 |
| ENSG00000271526 | 0 | 0 | 0 | 0 | 0 | 0 |
| ENSG00000278446 | 0 | 0 | 0 | 0 | 0 | 0 |
| ENSG00000233876 | 0 | 0 | 0 | 0 | 0 | 0 |
| ENSG00000271077 | 0 | 0 | 0 | 0 | 0 | 0 |
| ENSG00000197320 | 0 | 0 | 1 | 0 | 0 | 1 |
| ENSG00000234710 | 0 | 0 | 0 | 0 | 0 | 0 |
| ENSG00000270611 | 0 | 0 | 0 | 0 | 0 | 0 |
| ENSG00000235431 | 0 | 0 | 0 | 0 | 0 | 0 |
| ENSG00000212422 | 0 | 0 | 0 | 0 | 0 | 0 |
| ENSG00000283117 | 0 | 0 | 0 | 0 | 0 | 0 |
| ENSG00000229091 | 0 | 0 | 0 | 0 | 0 | 0 |
| ENSG00000234356 | 0 | 0 | 0 | 0 | 0 | 0 |
| ENSG00000236414 | 0 | 0 | 0 | 0 | 0 | 0 |
| ENSG00000201774 | 0 | 0 | 0 | 0 | 0 | 0 |
| ENSG00000189043 | 5752 | 2812 | 3654 | 3298 | 3068 | 3147 |
| ENSG00000271185 | 1 | 1 | 0 | 0 | 2 | 1 |
| ENSG00000106443 | 966 | 556 | 506 | 274 | 671 | 498 |
| ENSG00000230370 | 1 | 1 | 0 | 0 | 0 | 0 |
| ENSG00000230333 | 4 | 1 | 3 | 1 | 3 | 0 |
| ENSG00000226827 | 0 | 0 | 0 | 0 | 0 | 0 |
| ENSG00000005108 | 2 | 0 | 0 | 0 | 0 | 0 |
| ENSG00000230435 | 0 | 0 | 0 | 0 | 1 | 1 |
| ENSG00000235920 | 0 | 0 | 0 | 0 | 0 | 0 |
| ENSG00000106460 | 933 | 408 | 399 | 279 | 633 | 386 |
| ENSG00000146530 | 12 | 5 | 2 | 2 | 2 | 1 |
| ENSG00000274970 | 0 | 0 | 0 | 0 | 0 | 0 |
| ENSG00000226690 | 0 | 0 | 0 | 0 | 1 | 0 |
| ENSG00000236048 | 0 | 0 | 0 | 0 | 0 | 1 |
| ENSG00000203523 | 3 | 0 | 2 | 2 | 1 | 1 |
| ENSG00000224667 | 0 | 0 | 0 | 0 | 0 | 1 |
| ENSG00000225606 | 1 | 4 | 1 | 0 | 1 | 1 |
| ENSG00000006747 | 325 | 157 | 222 | 94 | 145 | 156 |
| ENSG00000229233 | 0 | 0 | 0 | 0 | 0 | 0 |
| ENSG00000271253 | 0 | 2 | 0 | 0 | 0 | 0 |

| | | | | | | |
|---|---|---|---|---|---|---|
| ENSG00000122644 | 976 | 481 | 556 | 423 | 560 | 464 |
| ENSG00000199470 | 0 | 0 | 0 | 0 | 0 | 0 |
| ENSG00000229618 | 2 | 0 | 1 | 1 | 1 | 0 |
| ENSG00000222974 | 0 | 0 | 0 | 0 | 0 | 0 |
| ENSG00000230879 | 0 | 0 | 0 | 0 | 0 | 0 |
| ENSG00000237713 | 0 | 0 | 0 | 0 | 0 | 0 |
| ENSG00000224330 | 0 | 0 | 0 | 0 | 0 | 0 |
| ENSG00000213875 | 0 | 0 | 0 | 0 | 0 | 0 |
| ENSG00000006468 | 0 | 0 | 0 | 0 | 0 | 0 |
| ENSG00000252374 | 0 | 0 | 0 | 0 | 0 | 0 |
| ENSG00000223445 | 0 | 0 | 0 | 0 | 0 | 0 |
| ENSG00000136267 | 2 | 0 | 1 | 1 | 0 | 0 |
| ENSG00000233167 | 0 | 0 | 0 | 0 | 0 | 0 |
| ENSG00000226323 | 0 | 0 | 0 | 0 | 0 | 0 |
| ENSG00000225816 | 0 | 0 | 0 | 0 | 0 | 0 |
| ENSG00000227489 | 0 | 0 | 0 | 0 | 0 | 0 |
| ENSG00000187546 | 0 | 0 | 0 | 0 | 0 | 0 |
| ENSG00000106511 | 0 | 0 | 0 | 0 | 0 | 0 |
| ENSG00000237070 | 0 | 0 | 0 | 0 | 0 | 0 |
| ENSG00000229108 | 0 | 0 | 0 | 0 | 0 | 0 |
| ENSG00000225974 | 0 | 0 | 0 | 0 | 0 | 0 |
| ENSG00000235828 | 0 | 0 | 1 | 0 | 0 | 0 |
| ENSG00000229379 | 0 | 0 | 0 | 0 | 0 | 0 |
| ENSG00000214960 | 78 | 36 | 26 | 24 | 22 | 39 |
| ENSG00000224683 | 0 | 0 | 0 | 0 | 0 | 0 |
| ENSG00000229688 | 2 | 3 | 0 | 0 | 1 | 0 |
| ENSG00000171243 | 0 | 2 | 3 | 0 | 0 | 0 |
| ENSG00000272537 | 0 | 0 | 0 | 0 | 0 | 1 |
| ENSG00000205858 | 0 | 0 | 0 | 0 | 0 | 0 |
| ENSG00000224280 | 0 | 0 | 1 | 0 | 0 | 1 |
| ENSG00000272361 | 0 | 3 | 1 | 1 | 1 | 0 |
| ENSG00000106524 | 493 | 317 | 323 | 209 | 363 | 310 |
| ENSG00000136261 | 2548 | 1500 | 1755 | 1571 | 1964 | 1918 |
| ENSG00000235837 | 4 | 1 | 1 | 0 | 0 | 4 |
| ENSG00000106537 | 6427 | 3009 | 3623 | 2665 | 3088 | 3073 |
| ENSG00000106541 | 7954 | 4137 | 4538 | 4557 | 5135 | 4451 |
| ENSG00000270593 | 0 | 0 | 0 | 0 | 0 | 0 |
| ENSG00000173467 | 520 | 259 | 362 | 179 | 203 | 214 |
| ENSG00000232400 | 0 | 0 | 2 | 0 | 0 | 0 |
| ENSG00000227965 | 2 | 0 | 0 | 0 | 0 | 0 |
| ENSG00000106546 | 884 | 459 | 476 | 258 | 476 | 381 |
| ENSG00000223867 | 0 | 0 | 0 | 0 | 0 | 0 |
| ENSG00000237773 | 17 | 3 | 6 | 6 | 2 | 4 |
| ENSG00000232841 | 0 | 0 | 0 | 0 | 0 | 0 |
| ENSG00000278569 | 0 | 0 | 0 | 0 | 0 | 0 |
| ENSG00000283321 | 0 | 0 | 0 | 0 | 0 | 0 |
| ENSG00000199473 | 0 | 0 | 0 | 0 | 0 | 0 |
| ENSG00000236318 | 0 | 1 | 0 | 0 | 0 | 1 |
| ENSG00000236039 | 12 | 5 | 20 | 4 | 5 | 6 |
| ENSG00000228675 | 0 | 0 | 0 | 0 | 0 | 0 |

| | | | | | | |
|---|---|---|---|---|---|---|
| ENSG00000071189 | 1132 | 642 | 530 | 214 | 808 | 521 |
| ENSG00000279048 | 48 | 19 | 24 | 14 | 15 | 9 |
| ENSG00000236195 | 0 | 0 | 0 | 0 | 0 | 0 |
| ENSG00000280340 | 0 | 0 | 0 | 0 | 0 | 0 |
| ENSG00000229937 | 0 | 0 | 0 | 0 | 0 | 0 |
| ENSG00000048052 | 24 | 6 | 11 | 9 | 7 | 2 |
| ENSG00000221393 | 0 | 0 | 0 | 0 | 0 | 0 |
| ENSG00000225000 | 0 | 0 | 0 | 0 | 0 | 0 |
| ENSG00000222164 | 0 | 0 | 0 | 0 | 0 | 0 |
| ENSG00000226522 | 0 | 0 | 0 | 0 | 0 | 0 |
| ENSG00000232492 | 0 | 0 | 0 | 0 | 0 | 0 |
| ENSG00000122691 | 0 | 0 | 0 | 0 | 0 | 0 |
| ENSG00000232821 | 0 | 0 | 0 | 0 | 0 | 0 |
| ENSG00000236536 | 0 | 0 | 0 | 0 | 0 | 0 |
| ENSG00000229533 | 0 | 0 | 0 | 0 | 0 | 0 |
| ENSG00000146618 | 0 | 0 | 0 | 0 | 0 | 0 |
| ENSG00000223838 | 0 | 0 | 0 | 0 | 0 | 0 |
| ENSG00000105849 | 365 | 224 | 218 | 122 | 258 | 238 |
| ENSG00000265932 | 0 | 0 | 0 | 0 | 0 | 0 |
| ENSG00000173452 | 0 | 0 | 0 | 0 | 0 | 0 |
| ENSG00000237921 | 0 | 0 | 0 | 0 | 0 | 0 |
| ENSG00000243004 | 1 | 2 | 3 | 4 | 2 | 2 |
| ENSG00000213860 | 10 | 7 | 8 | 7 | 5 | 1 |
| ENSG00000267055 | 0 | 0 | 0 | 0 | 0 | 0 |
| ENSG00000183742 | 359 | 240 | 239 | 114 | 267 | 189 |
| ENSG00000228598 | 0 | 0 | 0 | 0 | 0 | 0 |
| ENSG00000233834 | 41 | 19 | 16 | 31 | 20 | 20 |
| ENSG00000226097 | 0 | 0 | 0 | 0 | 0 | 0 |
| ENSG00000271133 | 21 | 3 | 9 | 7 | 15 | 9 |
| ENSG00000105855 | 34 | 27 | 13 | 12 | 20 | 8 |
| ENSG00000200753 | 0 | 0 | 0 | 0 | 0 | 0 |
| ENSG00000179460 | 0 | 0 | 0 | 0 | 0 | 0 |
| ENSG00000231525 | 0 | 0 | 0 | 0 | 0 | 0 |
| ENSG00000004846 | 0 | 0 | 0 | 0 | 0 | 0 |
| ENSG00000223866 | 0 | 0 | 0 | 0 | 0 | 0 |
| ENSG00000164651 | 0 | 0 | 0 | 0 | 0 | 0 |
| ENSG00000219451 | 4 | 2 | 2 | 2 | 0 | 2 |
| ENSG00000232790 | 0 | 0 | 0 | 0 | 0 | 0 |
| ENSG00000234839 | 0 | 0 | 0 | 0 | 0 | 0 |
| ENSG00000285592 | 0 | 0 | 0 | 0 | 0 | 0 |
| ENSG00000243633 | 0 | 0 | 0 | 0 | 0 | 0 |
| ENSG00000225308 | 0 | 0 | 0 | 0 | 0 | 0 |
| ENSG00000195024 | 0 | 0 | 0 | 0 | 0 | 0 |
| ENSG00000229949 | 0 | 0 | 0 | 0 | 0 | 0 |
| ENSG00000105866 | 140 | 42 | 67 | 17 | 58 | 45 |
| ENSG00000221783 | 0 | 0 | 0 | 0 | 0 | 0 |
| ENSG00000105877 | 4 | 6 | 6 | 3 | 7 | 8 |
| ENSG00000164649 | 487 | 253 | 313 | 223 | 283 | 316 |
| ENSG00000136237 | 126 | 85 | 71 | 49 | 73 | 70 |
| ENSG00000251993 | 0 | 0 | 0 | 0 | 0 | 0 |

| | | | | | | |
|---|---|---|---|---|---|---|
| ENSG00000105889 | 0 | 0 | 0 | 0 | 0 | 0 |
| ENSG00000233476 | 49 | 33 | 38 | 38 | 38 | 38 |
| ENSG00000232759 | 0 | 0 | 0 | 0 | 2 | 0 |
| ENSG00000232949 | 0 | 0 | 0 | 0 | 0 | 0 |
| ENSG00000179428 | 0 | 0 | 1 | 1 | 0 | 1 |
| ENSG00000136244 | 3 | 0 | 0 | 6 | 5 | 7 |
| ENSG00000228803 | 0 | 0 | 0 | 0 | 0 | 0 |
| ENSG00000234513 | 0 | 0 | 1 | 0 | 0 | 0 |
| ENSG00000196683 | 3344 | 1601 | 1921 | 2120 | 1848 | 1787 |
| ENSG00000228649 | 3 | 1 | 6 | 4 | 3 | 4 |
| ENSG00000221740 | 0 | 0 | 0 | 0 | 0 | 0 |
| ENSG00000226329 | 0 | 0 | 0 | 0 | 0 | 0 |
| ENSG00000226279 | 0 | 0 | 0 | 0 | 0 | 0 |
| ENSG00000122591 | 64 | 32 | 35 | 27 | 58 | 35 |
| ENSG00000230658 | 8 | 9 | 4 | 2 | 2 | 5 |
| ENSG00000122550 | 671 | 355 | 362 | 266 | 417 | 318 |
| ENSG00000230042 | 6 | 1 | 2 | 0 | 2 | 1 |
| ENSG00000136243 | 351 | 204 | 245 | 159 | 187 | 209 |
| ENSG00000285926 | 0 | 0 | 0 | 0 | 0 | 0 |
| ENSG00000226816 | 0 | 0 | 0 | 0 | 0 | 1 |
| ENSG00000136235 | 144 | 79 | 91 | 30 | 40 | 46 |
| ENSG00000156928 | 529 | 268 | 275 | 351 | 320 | 361 |
| ENSG00000136231 | 0 | 0 | 0 | 0 | 0 | 0 |
| ENSG00000232627 | 0 | 0 | 0 | 0 | 0 | 0 |
| ENSG00000212264 | 0 | 0 | 0 | 0 | 0 | 0 |
| ENSG00000252590 | 0 | 0 | 0 | 0 | 0 | 0 |
| ENSG00000236654 | 0 | 0 | 0 | 0 | 0 | 0 |
| ENSG00000232818 | 2 | 6 | 3 | 1 | 3 | 1 |
| ENSG00000200847 | 0 | 0 | 0 | 0 | 0 | 0 |
| ENSG00000164548 | 2166 | 1123 | 1229 | 1008 | 1250 | 1313 |
| ENSG00000224785 | 0 | 0 | 0 | 0 | 0 | 1 |
| ENSG00000232553 | 2 | 0 | 0 | 1 | 1 | 1 |
| ENSG00000169193 | 220 | 106 | 144 | 129 | 156 | 105 |
| ENSG00000227436 | 0 | 1 | 1 | 0 | 0 | 0 |
| ENSG00000228953 | 0 | 0 | 0 | 0 | 0 | 0 |
| ENSG00000188732 | 190 | 104 | 160 | 108 | 158 | 135 |
| ENSG00000234286 | 4 | 4 | 2 | 6 | 4 | 4 |
| ENSG00000196335 | 0 | 5 | 9 | 2 | 7 | 6 |
| ENSG00000234800 | 0 | 0 | 0 | 0 | 0 | 1 |
| ENSG00000231328 | 0 | 0 | 0 | 0 | 1 | 0 |
| ENSG00000227930 | 0 | 0 | 0 | 0 | 0 | 0 |
| ENSG00000252041 | 0 | 0 | 0 | 0 | 0 | 0 |
| ENSG00000228944 | 0 | 0 | 0 | 0 | 0 | 0 |
| ENSG00000122585 | 0 | 0 | 0 | 0 | 0 | 0 |
| ENSG00000206877 | 0 | 0 | 0 | 0 | 0 | 0 |
| ENSG00000105926 | 94 | 55 | 58 | 56 | 65 | 54 |
| ENSG00000271096 | 0 | 0 | 0 | 0 | 0 | 0 |
| ENSG00000271330 | 0 | 0 | 0 | 0 | 0 | 0 |
| ENSG00000105928 | 4 | 0 | 2 | 0 | 0 | 0 |
| ENSG00000070882 | 10 | 0 | 3 | 0 | 2 | 3 |

| | | | | | | |
|---|---|---|---|---|---|---|
| ENSG00000234999 | 0 | 0 | 0 | 0 | 0 | 0 |
| ENSG00000172115 | 3261 | 1632 | 1812 | 1957 | 2175 | 1921 |
| ENSG00000153790 | 0 | 0 | 0 | 0 | 1 | 0 |
| ENSG00000234406 | 0 | 0 | 0 | 0 | 0 | 0 |
| ENSG00000213798 | 1 | 0 | 0 | 0 | 0 | 0 |
| ENSG00000105954 | 0 | 0 | 0 | 0 | 0 | 0 |
| ENSG00000285716 | 0 | 0 | 0 | 0 | 0 | 0 |
| ENSG00000202233 | 0 | 0 | 0 | 0 | 0 | 0 |
| ENSG00000234922 | 0 | 0 | 0 | 0 | 0 | 0 |
| ENSG00000233824 | 0 | 0 | 0 | 0 | 0 | 0 |
| ENSG00000260951 | 0 | 0 | 0 | 0 | 0 | 0 |
| ENSG00000285960 | 0 | 0 | 0 | 0 | 0 | 0 |
| ENSG00000227386 | 0 | 0 | 0 | 0 | 0 | 0 |
| ENSG00000223561 | 0 | 0 | 0 | 0 | 0 | 0 |
| ENSG00000239622 | 0 | 0 | 0 | 0 | 1 | 0 |
| ENSG00000270933 | 10 | 4 | 3 | 3 | 4 | 1 |
| ENSG00000199085 | 0 | 0 | 0 | 0 | 0 | 0 |
| ENSG00000050344 | 447 | 223 | 266 | 253 | 374 | 313 |
| ENSG00000122566 | 27568 | 14910 | 17266 | 10741 | 19241 | 16388 |
| ENSG00000122565 | 8299 | 4401 | 5016 | 4521 | 5977 | 5226 |
| ENSG00000285860 | 0 | 0 | 0 | 0 | 1 | 0 |
| ENSG00000232383 | 0 | 0 | 0 | 0 | 0 | 0 |
| ENSG00000086300 | 334 | 200 | 182 | 125 | 247 | 168 |
| ENSG00000225792 | 2 | 3 | 8 | 4 | 4 | 7 |
| ENSG00000214870 | 27 | 7 | 15 | 11 | 6 | 8 |
| ENSG00000122548 | 0 | 0 | 0 | 0 | 0 | 0 |
| ENSG00000280255 | 0 | 0 | 1 | 0 | 0 | 0 |
| ENSG00000233760 | 0 | 0 | 0 | 0 | 0 | 0 |
| ENSG00000222004 | 0 | 0 | 0 | 0 | 0 | 0 |
| ENSG00000005020 | 1124 | 535 | 643 | 282 | 611 | 492 |
| ENSG00000213787 | 0 | 0 | 0 | 0 | 0 | 0 |
| ENSG00000226059 | 0 | 0 | 0 | 0 | 0 | 0 |
| ENSG00000214869 | 0 | 1 | 0 | 0 | 0 | 0 |
| ENSG00000213786 | 0 | 1 | 0 | 1 | 0 | 0 |
| ENSG00000105991 | 13 | 13 | 14 | 13 | 14 | 6 |
| ENSG00000233429 | 47 | 61 | 78 | 60 | 61 | 70 |
| ENSG00000276528 | 0 | 0 | 0 | 0 | 0 | 0 |
| ENSG00000276771 | 0 | 0 | 0 | 0 | 0 | 0 |
| ENSG00000274864 | 0 | 0 | 0 | 0 | 0 | 0 |
| ENSG00000274396 | 0 | 0 | 0 | 0 | 0 | 0 |
| ENSG00000277694 | 0 | 0 | 0 | 0 | 0 | 0 |
| ENSG00000105996 | 6 | 0 | 4 | 2 | 0 | 2 |
| ENSG00000105997 | 19 | 7 | 15 | 7 | 6 | 10 |
| ENSG00000253552 | 20 | 8 | 9 | 12 | 8 | 10 |
| ENSG00000197576 | 11 | 19 | 26 | 16 | 10 | 16 |
| ENSG00000273433 | 0 | 0 | 0 | 0 | 0 | 0 |
| ENSG00000254369 | 10 | 6 | 1 | 5 | 4 | 16 |
| ENSG00000106004 | 166 | 97 | 104 | 83 | 84 | 118 |
| ENSG00000106006 | 18 | 9 | 4 | 6 | 6 | 6 |
| ENSG00000272801 | 0 | 0 | 0 | 0 | 0 | 0 |

| | | | | | | |
|---|---|---|---|---|---|---|
| ENSG00000122592 | 20 | 11 | 28 | 7 | 10 | 6 |
| ENSG00000270182 | 2 | 3 | 0 | 0 | 0 | 2 |
| ENSG00000078399 | 1 | 1 | 2 | 0 | 1 | 1 |
| ENSG00000257184 | 0 | 0 | 0 | 0 | 0 | 0 |
| ENSG00000253187 | 0 | 0 | 0 | 0 | 1 | 1 |
| ENSG00000283745 | 0 | 0 | 0 | 0 | 0 | 0 |
| ENSG00000253293 | 116 | 55 | 91 | 78 | 63 | 88 |
| ENSG00000005073 | 12 | 2 | 2 | 6 | 7 | 6 |
| ENSG00000240990 | 16 | 13 | 7 | 7 | 16 | 8 |
| ENSG00000273961 | 0 | 0 | 0 | 0 | 0 | 0 |
| ENSG00000278334 | 0 | 0 | 0 | 0 | 0 | 0 |
| ENSG00000276496 | 0 | 0 | 0 | 0 | 0 | 0 |
| ENSG00000253508 | 0 | 2 | 0 | 1 | 1 | 1 |
| ENSG00000277966 | 0 | 0 | 0 | 0 | 0 | 0 |
| ENSG00000278592 | 0 | 0 | 0 | 0 | 0 | 0 |
| ENSG00000278020 | 0 | 0 | 0 | 0 | 0 | 0 |
| ENSG00000106031 | 22 | 11 | 10 | 12 | 8 | 7 |
| ENSG00000243766 | 0 | 1 | 0 | 0 | 1 | 0 |
| ENSG00000276609 | 0 | 0 | 0 | 0 | 0 | 0 |
| ENSG00000278708 | 0 | 0 | 0 | 0 | 0 | 0 |
| ENSG00000277469 | 0 | 0 | 0 | 0 | 0 | 0 |
| ENSG00000277553 | 0 | 0 | 0 | 0 | 0 | 0 |
| ENSG00000253308 | 2 | 0 | 0 | 0 | 0 | 0 |
| ENSG00000253405 | 0 | 0 | 0 | 0 | 1 | 3 |
| ENSG00000106038 | 7 | 3 | 3 | 1 | 4 | 1 |
| ENSG00000213783 | 1 | 0 | 0 | 0 | 0 | 0 |
| ENSG00000236569 | 0 | 0 | 0 | 0 | 0 | 0 |
| ENSG00000224322 | 0 | 0 | 0 | 0 | 0 | 0 |
| ENSG00000279655 | 0 | 0 | 0 | 0 | 0 | 0 |
| ENSG00000279429 | 0 | 0 | 0 | 0 | 0 | 0 |
| ENSG00000233830 | 26 | 7 | 9 | 10 | 11 | 13 |
| ENSG00000213781 | 0 | 0 | 0 | 0 | 0 | 0 |
| ENSG00000235574 | 0 | 0 | 0 | 0 | 0 | 0 |
| ENSG00000106049 | 2152 | 1135 | 1361 | 913 | 1087 | 1125 |
| ENSG00000229893 | 0 | 0 | 0 | 0 | 1 | 2 |
| ENSG00000106052 | 2810 | 1665 | 1489 | 839 | 2452 | 1527 |
| ENSG00000153814 | 58 | 24 | 24 | 22 | 24 | 20 |
| ENSG00000206623 | 0 | 0 | 0 | 0 | 0 | 0 |
| ENSG00000234336 | 0 | 0 | 1 | 0 | 0 | 0 |
| ENSG00000174495 | 0 | 0 | 0 | 0 | 0 | 0 |
| ENSG00000146592 | 4 | 0 | 0 | 0 | 0 | 0 |
| ENSG00000225286 | 0 | 0 | 0 | 0 | 0 | 0 |
| ENSG00000255690 | 151 | 57 | 85 | 91 | 76 | 76 |
| ENSG00000228421 | 5 | 2 | 1 | 2 | 5 | 2 |
| ENSG00000272568 | 8 | 4 | 9 | 1 | 10 | 5 |
| ENSG00000233517 | 0 | 0 | 1 | 0 | 1 | 1 |
| ENSG00000106066 | 0 | 0 | 0 | 0 | 0 | 0 |
| ENSG00000229452 | 0 | 0 | 0 | 0 | 0 | 0 |
| ENSG00000285412 | 0 | 0 | 0 | 0 | 0 | 0 |
| ENSG00000285162 | 0 | 0 | 0 | 1 | 0 | 0 |

| | | | | | | |
|---|---|---|---|---|---|---|
| ENSG00000106069 | 1 | 0 | 0 | 0 | 0 | 0 |
| ENSG00000285081 | 0 | 0 | 0 | 0 | 0 | 0 |
| ENSG00000235669 | 0 | 0 | 0 | 0 | 0 | 0 |
| ENSG00000237065 | 0 | 0 | 0 | 0 | 0 | 0 |
| ENSG00000223813 | 60 | 43 | 40 | 30 | 32 | 32 |
| ENSG00000270196 | 0 | 0 | 0 | 0 | 0 | 0 |
| ENSG00000176532 | 699 | 297 | 375 | 355 | 342 | 388 |
| ENSG00000225264 | 7 | 5 | 4 | 6 | 7 | 6 |
| ENSG00000223437 | 0 | 0 | 0 | 0 | 0 | 0 |
| ENSG00000227855 | 32 | 27 | 19 | 7 | 24 | 21 |
| ENSG00000212024 | 0 | 0 | 0 | 0 | 0 | 0 |
| ENSG00000122574 | 4 | 1 | 2 | 2 | 0 | 0 |
| ENSG00000238906 | 0 | 0 | 0 | 0 | 0 | 0 |
| ENSG00000136193 | 9486 | 4572 | 6060 | 3627 | 4712 | 5040 |
| ENSG00000227014 | 55 | 38 | 23 | 24 | 38 | 23 |
| ENSG00000106080 | 224 | 98 | 112 | 77 | 123 | 80 |
| ENSG00000106086 | 814 | 371 | 431 | 264 | 503 | 420 |
| ENSG00000231519 | 0 | 2 | 1 | 1 | 1 | 0 |
| ENSG00000283618 | 0 | 0 | 0 | 0 | 0 | 0 |
| ENSG00000180354 | 505 | 216 | 336 | 222 | 240 | 234 |
| ENSG00000230751 | 2 | 1 | 0 | 0 | 0 | 0 |
| ENSG00000251660 | 0 | 0 | 0 | 0 | 0 | 0 |
| ENSG00000225907 | 0 | 0 | 0 | 0 | 0 | 0 |
| ENSG00000227017 | 0 | 0 | 1 | 0 | 0 | 0 |
| ENSG00000180233 | 452 | 201 | 204 | 207 | 249 | 226 |
| ENSG00000207771 | 0 | 0 | 0 | 0 | 0 | 0 |
| ENSG00000283237 | 0 | 0 | 0 | 0 | 0 | 0 |
| ENSG00000281593 | 5 | 3 | 4 | 2 | 2 | 2 |
| ENSG00000235859 | 84 | 39 | 55 | 34 | 55 | 50 |
| ENSG00000281404 | 17 | 20 | 23 | 10 | 24 | 24 |
| ENSG00000106100 | 118 | 72 | 92 | 66 | 70 | 85 |
| ENSG00000272638 | 0 | 0 | 0 | 0 | 0 | 0 |
| ENSG00000006625 | 1590 | 973 | 1209 | 1143 | 1210 | 1112 |
| ENSG00000281039 | 0 | 0 | 0 | 0 | 0 | 1 |
| ENSG00000196295 | 665 | 317 | 431 | 216 | 341 | 295 |
| ENSG00000244480 | 11 | 6 | 7 | 2 | 6 | 6 |
| ENSG00000264520 | 1 | 1 | 2 | 0 | 3 | 1 |
| ENSG00000226874 | 2 | 1 | 2 | 0 | 4 | 6 |
| ENSG00000263683 | 0 | 0 | 0 | 0 | 0 | 0 |
| ENSG00000106105 | 3262 | 1746 | 1818 | 1734 | 2208 | 1943 |
| ENSG00000106113 | 1 | 0 | 0 | 0 | 1 | 1 |
| ENSG00000241644 | 2 | 2 | 1 | 0 | 0 | 0 |
| ENSG00000223419 | 0 | 0 | 0 | 0 | 0 | 0 |
| ENSG00000254959 | 0 | 0 | 0 | 0 | 0 | 0 |
| ENSG00000106125 | 20 | 7 | 4 | 7 | 9 | 7 |
| ENSG00000229263 | 0 | 0 | 1 | 0 | 0 | 0 |
| ENSG00000250424 | 0 | 0 | 0 | 0 | 0 | 0 |
| ENSG00000240583 | 10 | 3 | 1 | 2 | 3 | 6 |
| ENSG00000106128 | 0 | 0 | 0 | 0 | 0 | 0 |
| ENSG00000078549 | 0 | 0 | 0 | 0 | 0 | 0 |

| | | | | | | |
|---|---|---|---|---|---|---|
| ENSG00000232887 | 0 | 0 | 0 | 0 | 0 | 0 |
| ENSG00000271027 | 0 | 0 | 0 | 0 | 0 | 0 |
| ENSG00000164600 | 0 | 0 | 0 | 0 | 0 | 0 |
| ENSG00000231418 | 0 | 0 | 0 | 0 | 0 | 0 |
| ENSG00000180347 | 19 | 5 | 13 | 6 | 8 | 10 |
| ENSG00000106341 | 0 | 0 | 0 | 0 | 0 | 0 |
| ENSG00000154678 | 0 | 0 | 0 | 0 | 1 | 0 |
| ENSG00000236460 | 0 | 0 | 0 | 0 | 0 | 0 |
| ENSG00000284839 | 0 | 0 | 0 | 0 | 1 | 0 |
| ENSG00000226468 | 14 | 5 | 3 | 3 | 6 | 5 |
| ENSG00000226421 | 20 | 2 | 5 | 4 | 7 | 7 |
| ENSG00000106355 | 1783 | 892 | 1042 | 1023 | 1068 | 1039 |
| ENSG00000105778 | 5049 | 2395 | 3400 | 1916 | 2677 | 2691 |
| ENSG00000229358 | 105 | 69 | 70 | 48 | 64 | 66 |
| ENSG00000237004 | 24 | 29 | 30 | 21 | 32 | 20 |
| ENSG00000283461 | 0 | 0 | 0 | 0 | 0 | 0 |
| ENSG00000207573 | 0 | 0 | 0 | 0 | 0 | 0 |
| ENSG00000273014 | 26 | 16 | 15 | 12 | 17 | 21 |
| ENSG00000281332 | 65 | 27 | 47 | 32 | 27 | 46 |
| ENSG00000250756 | 0 | 0 | 0 | 0 | 0 | 0 |
| ENSG00000251999 | 0 | 0 | 0 | 0 | 0 | 0 |
| ENSG00000231952 | 6 | 7 | 8 | 9 | 7 | 6 |
| ENSG00000272905 | 5 | 2 | 1 | 2 | 3 | 2 |
| ENSG00000170852 | 2132 | 984 | 1247 | 962 | 1339 | 1224 |
| ENSG00000205763 | 115 | 51 | 80 | 78 | 69 | 87 |
| ENSG00000224573 | 0 | 0 | 0 | 0 | 1 | 0 |
| ENSG00000122642 | 3116 | 1380 | 2013 | 1630 | 1767 | 1892 |
| ENSG00000252821 | 0 | 0 | 0 | 1 | 1 | 0 |
| ENSG00000122643 | 1242 | 581 | 693 | 585 | 778 | 772 |
| ENSG00000229054 | 0 | 0 | 0 | 0 | 0 | 0 |
| ENSG00000241420 | 0 | 0 | 0 | 0 | 0 | 0 |
| ENSG00000164610 | 541 | 274 | 346 | 357 | 322 | 323 |
| ENSG00000228015 | 0 | 0 | 0 | 0 | 0 | 0 |
| ENSG00000122507 | 478 | 207 | 258 | 154 | 286 | 226 |
| ENSG00000222741 | 0 | 0 | 0 | 0 | 0 | 0 |
| ENSG00000224946 | 0 | 0 | 0 | 0 | 0 | 0 |
| ENSG00000238090 | 0 | 0 | 0 | 0 | 0 | 0 |
| ENSG00000173862 | 0 | 1 | 0 | 0 | 1 | 0 |
| ENSG00000227481 | 0 | 0 | 0 | 0 | 0 | 0 |
| ENSG00000236494 | 0 | 0 | 0 | 0 | 1 | 1 |
| ENSG00000233219 | 0 | 0 | 0 | 0 | 0 | 0 |
| ENSG00000164619 | 35 | 5 | 12 | 7 | 7 | 11 |
| ENSG00000236212 | 6 | 0 | 2 | 1 | 0 | 5 |
| ENSG00000202431 | 5 | 6 | 3 | 2 | 4 | 5 |
| ENSG00000197085 | 0 | 0 | 0 | 0 | 0 | 1 |
| ENSG00000187258 | 0 | 0 | 0 | 0 | 0 | 0 |
| ENSG00000235379 | 0 | 0 | 0 | 0 | 0 | 0 |
| ENSG00000230356 | 0 | 0 | 0 | 0 | 0 | 0 |
| ENSG00000242653 | 0 | 0 | 0 | 0 | 0 | 0 |
| ENSG00000251802 | 0 | 0 | 0 | 0 | 0 | 0 |

| | | | | | | |
|---|---|---|---|---|---|---|
| ENSG00000173852 | 1592 | 775 | 896 | 706 | 931 | 872 |
| ENSG00000221669 | 0 | 0 | 0 | 0 | 1 | 0 |
| ENSG00000189212 | 534 | 247 | 314 | 137 | 200 | 318 |
| ENSG00000236592 | 0 | 0 | 0 | 0 | 0 | 0 |
| ENSG00000164532 | 0 | 0 | 0 | 0 | 0 | 0 |
| ENSG00000226063 | 0 | 0 | 0 | 0 | 0 | 0 |
| ENSG00000235464 | 0 | 0 | 0 | 0 | 0 | 0 |
| ENSG00000122557 | 986 | 469 | 605 | 385 | 584 | 583 |
| ENSG00000271122 | 512 | 234 | 298 | 182 | 271 | 257 |
| ENSG00000227544 | 8 | 2 | 2 | 0 | 0 | 3 |
| ENSG00000228878 | 30 | 23 | 29 | 15 | 23 | 27 |
| ENSG00000232736 | 0 | 0 | 0 | 0 | 0 | 0 |
| ENSG00000122545 | 3334 | 2050 | 1773 | 1018 | 2782 | 1665 |
| ENSG00000274328 | 0 | 0 | 0 | 0 | 0 | 0 |
| ENSG00000200446 | 0 | 0 | 0 | 0 | 0 | 0 |
| ENSG00000273580 | 0 | 0 | 0 | 0 | 0 | 0 |
| ENSG00000226102 | 1 | 0 | 1 | 0 | 0 | 0 |
| ENSG00000270889 | 0 | 0 | 0 | 0 | 0 | 0 |
| ENSG00000232930 | 0 | 0 | 0 | 0 | 0 | 0 |
| ENSG00000285573 | 0 | 0 | 0 | 0 | 0 | 0 |
| ENSG00000205745 | 0 | 0 | 0 | 0 | 0 | 0 |
| ENSG00000261184 | 0 | 0 | 0 | 0 | 0 | 0 |
| ENSG00000228815 | 0 | 0 | 0 | 0 | 0 | 0 |
| ENSG00000271117 | 0 | 0 | 0 | 0 | 0 | 0 |
| ENSG00000122547 | 146 | 58 | 86 | 67 | 63 | 66 |
| ENSG00000228770 | 0 | 0 | 0 | 0 | 0 | 0 |
| ENSG00000229679 | 0 | 0 | 0 | 0 | 0 | 0 |
| ENSG00000234457 | 0 | 0 | 0 | 0 | 0 | 0 |
| ENSG00000260070 | 0 | 0 | 0 | 0 | 0 | 0 |
| ENSG00000164542 | 352 | 176 | 195 | 139 | 206 | 179 |
| ENSG00000201432 | 1 | 0 | 0 | 0 | 0 | 0 |
| ENSG00000011426 | 761 | 539 | 398 | 260 | 934 | 473 |
| ENSG00000237400 | 0 | 0 | 0 | 0 | 0 | 0 |
| ENSG00000136250 | 0 | 0 | 0 | 0 | 0 | 0 |
| ENSG00000230539 | 0 | 0 | 0 | 0 | 0 | 0 |
| ENSG00000272984 | 0 | 0 | 0 | 0 | 0 | 0 |
| ENSG00000229424 | 0 | 0 | 0 | 0 | 0 | 0 |
| ENSG00000235728 | 0 | 0 | 0 | 0 | 0 | 0 |
| ENSG00000225739 | 0 | 0 | 0 | 0 | 0 | 0 |
| ENSG00000230831 | 0 | 0 | 0 | 0 | 0 | 0 |
| ENSG00000155849 | 0 | 2 | 0 | 0 | 0 | 1 |
| ENSG00000221325 | 0 | 0 | 0 | 0 | 0 | 0 |
| ENSG00000224101 | 0 | 0 | 0 | 0 | 0 | 0 |
| ENSG00000224706 | 0 | 0 | 0 | 0 | 0 | 0 |
| ENSG00000222869 | 0 | 0 | 0 | 0 | 0 | 0 |
| ENSG00000213744 | 0 | 0 | 0 | 0 | 0 | 0 |
| ENSG00000200113 | 0 | 0 | 0 | 0 | 0 | 0 |
| ENSG00000201566 | 0 | 0 | 0 | 0 | 0 | 0 |
| ENSG00000178503 | 0 | 0 | 0 | 0 | 0 | 0 |
| ENSG00000187037 | 0 | 0 | 0 | 0 | 0 | 0 |

| | | | | | | |
|---|---|---|---|---|---|---|
| ENSG00000086289 | 66 | 43 | 32 | 36 | 35 | 45 |
| ENSG00000283444 | 0 | 0 | 0 | 0 | 0 | 0 |
| ENSG00000225314 | 0 | 0 | 0 | 0 | 0 | 0 |
| ENSG00000086288 | 0 | 0 | 0 | 0 | 0 | 0 |
| ENSG00000106483 | 0 | 0 | 0 | 0 | 0 | 1 |
| ENSG00000010270 | 942 | 456 | 584 | 516 | 637 | 597 |
| ENSG00000227191 | 7 | 7 | 7 | 1 | 4 | 2 |
| ENSG00000211687 | 0 | 0 | 0 | 0 | 0 | 0 |
| ENSG00000211688 | 0 | 0 | 0 | 0 | 0 | 1 |
| ENSG00000211689 | 287 | 159 | 244 | 80 | 67 | 129 |
| ENSG00000211690 | 0 | 0 | 0 | 0 | 0 | 0 |
| ENSG00000211691 | 0 | 0 | 0 | 0 | 0 | 0 |
| ENSG00000211692 | 0 | 0 | 0 | 0 | 0 | 0 |
| ENSG00000211693 | 0 | 0 | 0 | 0 | 0 | 0 |
| ENSG00000231202 | 0 | 0 | 0 | 0 | 0 | 0 |
| ENSG00000211694 | 0 | 0 | 0 | 0 | 0 | 0 |
| ENSG00000231916 | 7 | 4 | 2 | 0 | 2 | 2 |
| ENSG00000211695 | 36 | 23 | 38 | 5 | 13 | 15 |
| ENSG00000225992 | 0 | 0 | 0 | 0 | 0 | 0 |
| ENSG00000272908 | 0 | 0 | 1 | 0 | 0 | 1 |
| ENSG00000211696 | 0 | 0 | 0 | 0 | 0 | 0 |
| ENSG00000249978 | 0 | 0 | 1 | 2 | 0 | 0 |
| ENSG00000226212 | 0 | 0 | 0 | 0 | 0 | 0 |
| ENSG00000281103 | 4 | 1 | 2 | 2 | 1 | 3 |
| ENSG00000228668 | 2 | 2 | 0 | 0 | 0 | 0 |
| ENSG00000211697 | 0 | 1 | 0 | 0 | 0 | 0 |
| ENSG00000211698 | 0 | 0 | 0 | 0 | 1 | 0 |
| ENSG00000211699 | 0 | 0 | 0 | 0 | 0 | 0 |
| ENSG00000233306 | 2 | 0 | 0 | 0 | 0 | 1 |
| ENSG00000211701 | 0 | 0 | 0 | 0 | 0 | 0 |
| ENSG00000078053 | 2 | 3 | 3 | 2 | 2 | 1 |
| ENSG00000241756 | 0 | 0 | 0 | 0 | 0 | 0 |
| ENSG00000227404 | 0 | 1 | 1 | 1 | 0 | 1 |
| ENSG00000164556 | 0 | 0 | 0 | 0 | 0 | 0 |
| ENSG00000006715 | 2046 | 970 | 1276 | 748 | 1199 | 937 |
| ENSG00000106536 | 0 | 0 | 0 | 0 | 0 | 0 |
| ENSG00000233854 | 0 | 0 | 0 | 0 | 0 | 0 |
| ENSG00000206947 | 0 | 0 | 0 | 0 | 0 | 0 |
| ENSG00000224122 | 0 | 0 | 0 | 0 | 0 | 0 |
| ENSG00000227172 | 2 | 1 | 2 | 1 | 1 | 0 |
| ENSG00000241127 | 225 | 121 | 144 | 131 | 171 | 170 |
| ENSG00000236015 | 15 | 10 | 6 | 4 | 18 | 3 |
| ENSG00000231951 | 0 | 0 | 0 | 0 | 0 | 0 |
| ENSG00000106540 | 0 | 0 | 0 | 0 | 0 | 0 |
| ENSG00000006451 | 2262 | 1164 | 1310 | 1201 | 1367 | 1247 |
| ENSG00000228554 | 2 | 0 | 4 | 0 | 3 | 0 |
| ENSG00000188185 | 383 | 153 | 222 | 178 | 175 | 208 |
| ENSG00000252480 | 0 | 0 | 0 | 0 | 0 | 0 |
| ENSG00000270797 | 0 | 0 | 0 | 0 | 0 | 0 |
| ENSG00000226220 | 5 | 4 | 4 | 1 | 2 | 5 |

| | | | | | | |
|---|---|---|---|---|---|---|
| ENSG00000237251 | 0 | 0 | 2 | 0 | 0 | 0 |
| ENSG00000231198 | 0 | 0 | 0 | 0 | 0 | 0 |
| ENSG00000233225 | 35 | 21 | 26 | 13 | 22 | 27 |
| ENSG00000175741 | 18 | 15 | 17 | 19 | 19 | 9 |
| ENSG00000242855 | 0 | 0 | 0 | 0 | 0 | 0 |
| ENSG00000259826 | 20 | 14 | 23 | 16 | 16 | 11 |
| ENSG00000065883 | 2692 | 1183 | 1475 | 847 | 1188 | 1199 |
| ENSG00000273444 | 0 | 0 | 0 | 0 | 0 | 0 |
| ENSG00000270577 | 3 | 0 | 0 | 0 | 0 | 0 |
| ENSG00000168303 | 1155 | 542 | 654 | 740 | 777 | 733 |
| ENSG00000199273 | 0 | 0 | 0 | 0 | 0 | 0 |
| ENSG00000175600 | 0 | 0 | 1 | 3 | 2 | 4 |
| ENSG00000248711 | 0 | 0 | 0 | 0 | 0 | 0 |
| ENSG00000203446 | 0 | 0 | 0 | 0 | 0 | 0 |
| ENSG00000227658 | 0 | 0 | 0 | 0 | 0 | 0 |
| ENSG00000232458 | 0 | 0 | 0 | 0 | 0 | 0 |
| ENSG00000224017 | 0 | 0 | 0 | 0 | 0 | 0 |
| ENSG00000122641 | 6 | 2 | 1 | 0 | 1 | 3 |
| ENSG00000224116 | 8 | 1 | 0 | 0 | 0 | 0 |
| ENSG00000106571 | 466 | 254 | 316 | 217 | 235 | 245 |
| ENSG00000213721 | 0 | 0 | 1 | 0 | 0 | 0 |
| ENSG00000238284 | 0 | 0 | 0 | 0 | 0 | 0 |
| ENSG00000229161 | 0 | 0 | 0 | 0 | 0 | 0 |
| ENSG00000231539 | 0 | 0 | 0 | 0 | 0 | 0 |
| ENSG00000261019 | 10 | 6 | 2 | 3 | 6 | 3 |
| ENSG00000136197 | 23 | 8 | 6 | 10 | 11 | 7 |
| ENSG00000256646 | 0 | 0 | 0 | 0 | 0 | 1 |
| ENSG00000106588 | 157 | 96 | 122 | 116 | 102 | 98 |
| ENSG00000234983 | 0 | 0 | 0 | 0 | 0 | 0 |
| ENSG00000106591 | 1580 | 812 | 974 | 961 | 920 | 919 |
| ENSG00000232006 | 4 | 0 | 0 | 0 | 0 | 0 |
| ENSG00000201933 | 0 | 0 | 0 | 0 | 0 | 0 |
| ENSG00000002746 | 2 | 0 | 0 | 3 | 7 | 3 |
| ENSG00000181211 | 0 | 0 | 0 | 0 | 0 | 0 |
| ENSG00000264069 | 0 | 0 | 0 | 0 | 0 | 0 |
| ENSG00000252983 | 0 | 0 | 0 | 0 | 0 | 0 |
| ENSG00000223258 | 0 | 0 | 0 | 0 | 0 | 0 |
| ENSG00000228680 | 0 | 0 | 0 | 0 | 0 | 0 |
| ENSG00000213717 | 0 | 0 | 0 | 0 | 0 | 1 |
| ENSG00000231638 | 0 | 0 | 0 | 0 | 0 | 0 |
| ENSG00000164543 | 423 | 206 | 219 | 185 | 271 | 220 |
| ENSG00000106603 | 1543 | 841 | 1021 | 824 | 989 | 945 |
| ENSG00000106605 | 2752 | 1386 | 1621 | 1620 | 1609 | 1533 |
| ENSG00000229497 | 0 | 0 | 1 | 0 | 0 | 0 |
| ENSG00000062582 | 20 | 25 | 18 | 35 | 23 | 27 |
| ENSG00000270617 | 0 | 0 | 0 | 1 | 1 | 0 |
| ENSG00000106608 | 1721 | 710 | 986 | 982 | 917 | 1025 |
| ENSG00000237972 | 4 | 2 | 1 | 0 | 1 | 0 |
| ENSG00000078967 | 217 | 142 | 131 | 159 | 158 | 189 |
| ENSG00000214783 | 31 | 19 | 26 | 18 | 35 | 29 |

| | | | | | | |
|---|---|---|---|---|---|---|
| ENSG00000273432 | 5 | 2 | 6 | 6 | 4 | 2 |
| ENSG00000241057 | 1 | 3 | 0 | 8 | 2 | 6 |
| ENSG00000239556 | 2 | 3 | 9 | 6 | 6 | 7 |
| ENSG00000136206 | 8 | 11 | 9 | 6 | 3 | 9 |
| ENSG00000228434 | 13 | 5 | 8 | 4 | 1 | 7 |
| ENSG00000285596 | 14 | 18 | 8 | 8 | 3 | 5 |
| ENSG00000272655 | 81 | 40 | 43 | 44 | 49 | 36 |
| ENSG00000228903 | 29 | 23 | 24 | 21 | 33 | 10 |
| ENSG00000235314 | 50 | 21 | 22 | 36 | 25 | 28 |
| ENSG00000136279 | 3880 | 1853 | 2457 | 2112 | 1998 | 2044 |
| ENSG00000284067 | 0 | 0 | 0 | 0 | 0 | 0 |
| ENSG00000164708 | 0 | 0 | 0 | 0 | 0 | 0 |
| ENSG00000239775 | 0 | 0 | 0 | 0 | 0 | 0 |
| ENSG00000122678 | 828 | 373 | 456 | 437 | 421 | 455 |
| ENSG00000283969 | 0 | 0 | 0 | 0 | 0 | 0 |
| ENSG00000106624 | 54 | 27 | 40 | 26 | 13 | 21 |
| ENSG00000284412 | 0 | 0 | 0 | 0 | 0 | 0 |
| ENSG00000106628 | 2893 | 1356 | 1634 | 2213 | 1691 | 1922 |
| ENSG00000252848 | 0 | 0 | 0 | 0 | 0 | 0 |
| ENSG00000106631 | 0 | 1 | 0 | 0 | 0 | 0 |
| ENSG00000106633 | 2 | 2 | 0 | 1 | 0 | 1 |
| ENSG00000106636 | 4708 | 2211 | 2690 | 2818 | 2808 | 2645 |
| ENSG00000058404 | 661 | 302 | 439 | 294 | 310 | 316 |
| ENSG00000279853 | 0 | 0 | 1 | 0 | 0 | 0 |
| ENSG00000015676 | 2911 | 1416 | 1873 | 1757 | 1574 | 1650 |
| ENSG00000146677 | 34 | 19 | 16 | 22 | 25 | 23 |
| ENSG00000200456 | 0 | 0 | 0 | 0 | 0 | 0 |
| ENSG00000225443 | 0 | 0 | 0 | 0 | 0 | 0 |
| ENSG00000015520 | 70 | 23 | 55 | 56 | 49 | 60 |
| ENSG00000136271 | 3168 | 1634 | 1897 | 2210 | 1854 | 1938 |
| ENSG00000158604 | 3752 | 1945 | 2363 | 1787 | 2030 | 1977 |
| ENSG00000271586 | 0 | 0 | 0 | 0 | 0 | 0 |
| ENSG00000105953 | 4694 | 2104 | 2870 | 2624 | 2396 | 2539 |
| ENSG00000122515 | 2946 | 1285 | 1720 | 1584 | 1538 | 1663 |
| ENSG00000228596 | 0 | 0 | 0 | 0 | 1 | 0 |
| ENSG00000196262 | 33713 | 17361 | 20346 | 21333 | 20329 | 19773 |
| ENSG00000105968 | 4175 | 2008 | 2676 | 1891 | 2160 | 2239 |
| ENSG00000234183 | 0 | 0 | 1 | 2 | 0 | 0 |
| ENSG00000146676 | 3366 | 1772 | 2013 | 1418 | 1844 | 1779 |
| ENSG00000284261 | 0 | 0 | 0 | 0 | 0 | 0 |
| ENSG00000272768 | 8 | 1 | 5 | 1 | 2 | 2 |
| ENSG00000230160 | 0 | 0 | 0 | 0 | 1 | 0 |
| ENSG00000260997 | 0 | 0 | 0 | 0 | 0 | 0 |
| ENSG00000136286 | 2 | 2 | 1 | 0 | 0 | 1 |
| ENSG00000232956 | 775 | 418 | 456 | 557 | 514 | 503 |
| ENSG00000277184 | 0 | 0 | 0 | 0 | 0 | 0 |
| ENSG00000136280 | 1265 | 639 | 864 | 902 | 846 | 755 |
| ENSG00000136274 | 0 | 0 | 1 | 1 | 0 | 0 |
| ENSG00000136270 | 3187 | 1544 | 1869 | 2303 | 1890 | 2034 |
| ENSG00000206838 | 10 | 1 | 1 | 3 | 3 | 3 |

| | | | | | | |
|---|---|---|---|---|---|---|
| ENSG00000201772 | 11 | 5 | 17 | 10 | 4 | 8 |
| ENSG00000200656 | 0 | 0 | 0 | 0 | 0 | 0 |
| ENSG00000122679 | 143 | 58 | 78 | 80 | 46 | 70 |
| ENSG00000275295 | 0 | 0 | 0 | 0 | 0 | 0 |
| ENSG00000234359 | 0 | 0 | 0 | 0 | 0 | 0 |
| ENSG00000225537 | 0 | 1 | 0 | 1 | 0 | 1 |
| ENSG00000226999 | 0 | 0 | 0 | 0 | 0 | 0 |
| ENSG00000164742 | 4109 | 1711 | 2302 | 1643 | 1731 | 1911 |
| ENSG00000214765 | 538 | 205 | 282 | 205 | 248 | 258 |
| ENSG00000272556 | 179 | 65 | 114 | 67 | 76 | 85 |
| ENSG00000212450 | 0 | 0 | 0 | 0 | 0 | 0 |
| ENSG00000235955 | 2 | 0 | 1 | 0 | 0 | 1 |
| ENSG00000226838 | 0 | 0 | 0 | 0 | 0 | 0 |
| ENSG00000251378 | 0 | 0 | 0 | 0 | 0 | 0 |
| ENSG00000202350 | 0 | 0 | 0 | 0 | 0 | 0 |
| ENSG00000283247 | 0 | 0 | 0 | 1 | 0 | 0 |
| ENSG00000146678 | 0 | 0 | 1 | 0 | 0 | 0 |
| ENSG00000146674 | 1329 | 667 | 576 | 958 | 898 | 843 |
| ENSG00000237471 | 0 | 0 | 0 | 0 | 0 | 0 |
| ENSG00000229628 | 0 | 0 | 0 | 0 | 0 | 1 |
| ENSG00000233952 | 0 | 0 | 0 | 0 | 0 | 0 |
| ENSG00000221971 | 0 | 0 | 0 | 0 | 0 | 0 |
| ENSG00000224547 | 0 | 0 | 0 | 0 | 0 | 0 |
| ENSG00000229459 | 0 | 0 | 0 | 0 | 0 | 0 |
| ENSG00000237760 | 0 | 0 | 0 | 0 | 0 | 0 |
| ENSG00000240355 | 0 | 0 | 0 | 0 | 0 | 0 |
| ENSG00000230680 | 0 | 0 | 0 | 0 | 0 | 0 |
| ENSG00000224519 | 0 | 0 | 0 | 0 | 0 | 0 |
| ENSG00000233539 | 0 | 2 | 1 | 1 | 0 | 0 |
| ENSG00000242948 | 0 | 0 | 0 | 0 | 0 | 0 |
| ENSG00000232072 | 0 | 0 | 0 | 0 | 0 | 0 |
| ENSG00000223829 | 0 | 0 | 0 | 0 | 0 | 0 |
| ENSG00000229192 | 0 | 0 | 0 | 0 | 0 | 0 |
| ENSG00000242552 | 0 | 0 | 0 | 0 | 0 | 0 |
| ENSG00000214754 | 0 | 0 | 0 | 0 | 0 | 0 |
| ENSG00000136205 | 1837 | 888 | 1055 | 918 | 1160 | 1086 |
| ENSG00000236078 | 3 | 3 | 0 | 1 | 1 | 5 |
| ENSG00000221845 | 2 | 0 | 0 | 0 | 0 | 4 |
| ENSG00000158683 | 0 | 2 | 3 | 0 | 0 | 0 |
| ENSG00000146666 | 0 | 0 | 1 | 0 | 0 | 0 |
| ENSG00000136275 | 0 | 0 | 0 | 0 | 0 | 0 |
| ENSG00000225507 | 25 | 13 | 14 | 13 | 13 | 11 |
| ENSG00000136273 | 368 | 206 | 214 | 163 | 252 | 222 |
| ENSG00000164744 | 0 | 0 | 0 | 0 | 0 | 0 |
| ENSG00000164746 | 0 | 0 | 1 | 0 | 0 | 3 |
| ENSG00000183696 | 50 | 27 | 32 | 38 | 48 | 34 |
| ENSG00000179869 | 0 | 0 | 0 | 0 | 0 | 0 |
| ENSG00000228173 | 0 | 0 | 0 | 0 | 0 | 0 |
| ENSG00000285536 | 0 | 0 | 0 | 0 | 0 | 0 |
| ENSG00000225705 | 0 | 0 | 0 | 0 | 0 | 0 |

| | | | | | | |
|---|---|---|---|---|---|---|
| ENSG00000279578 | 0 | 0 | 0 | 0 | 0 | 0 |
| ENSG00000229165 | 0 | 0 | 0 | 0 | 0 | 0 |
| ENSG00000218305 | 0 | 0 | 0 | 0 | 0 | 0 |
| ENSG00000234686 | 0 | 0 | 0 | 0 | 0 | 0 |
| ENSG00000270763 | 0 | 0 | 0 | 0 | 0 | 0 |
| ENSG00000279104 | 0 | 0 | 0 | 0 | 0 | 0 |
| ENSG00000188730 | 0 | 0 | 0 | 0 | 0 | 0 |
| ENSG00000042813 | 0 | 0 | 0 | 1 | 0 | 4 |
| ENSG00000223893 | 0 | 0 | 0 | 0 | 0 | 0 |
| ENSG00000164500 | 0 | 0 | 0 | 0 | 0 | 0 |
| ENSG00000228005 | 0 | 0 | 0 | 0 | 0 | 0 |
| ENSG00000231681 | 0 | 0 | 0 | 0 | 0 | 0 |
| ENSG00000235620 | 0 | 0 | 0 | 0 | 0 | 0 |
| ENSG00000185811 | 0 | 0 | 1 | 0 | 0 | 0 |
| ENSG00000285165 | 0 | 0 | 0 | 0 | 0 | 0 |
| ENSG00000200815 | 0 | 0 | 0 | 0 | 0 | 0 |
| ENSG00000132436 | 409 | 239 | 293 | 143 | 276 | 268 |
| ENSG00000132437 | 3 | 1 | 1 | 1 | 0 | 0 |
| ENSG00000226122 | 0 | 0 | 0 | 0 | 0 | 0 |
| ENSG00000106070 | 277 | 122 | 127 | 90 | 121 | 98 |
| ENSG00000236046 | 0 | 0 | 0 | 0 | 0 | 0 |
| ENSG00000228085 | 0 | 0 | 0 | 0 | 0 | 0 |
| ENSG00000228204 | 3 | 3 | 4 | 0 | 3 | 3 |
| ENSG00000201133 | 0 | 0 | 0 | 0 | 0 | 0 |
| ENSG00000106078 | 1556 | 701 | 923 | 603 | 741 | 794 |
| ENSG00000231490 | 0 | 0 | 0 | 0 | 0 | 0 |
| ENSG00000227233 | 0 | 0 | 0 | 0 | 0 | 0 |
| ENSG00000228897 | 0 | 0 | 0 | 0 | 0 | 0 |
| ENSG00000227080 | 0 | 0 | 0 | 0 | 0 | 0 |
| ENSG00000285741 | 0 | 0 | 0 | 0 | 0 | 0 |
| ENSG00000229478 | 0 | 0 | 0 | 0 | 0 | 0 |
| ENSG00000229403 | 0 | 0 | 0 | 0 | 0 | 0 |
| ENSG00000242650 | 0 | 0 | 0 | 0 | 0 | 0 |
| ENSG00000233960 | 0 | 0 | 0 | 0 | 0 | 0 |
| ENSG00000253028 | 0 | 0 | 0 | 0 | 0 | 0 |
| ENSG00000234304 | 0 | 0 | 0 | 0 | 0 | 0 |
| ENSG00000223970 | 0 | 0 | 0 | 0 | 0 | 0 |
| ENSG00000221900 | 0 | 0 | 0 | 0 | 0 | 0 |
| ENSG00000227344 | 0 | 1 | 0 | 0 | 0 | 0 |
| ENSG00000271194 | 0 | 0 | 0 | 0 | 0 | 0 |
| ENSG00000199629 | 0 | 0 | 0 | 0 | 0 | 0 |
| ENSG00000222154 | 0 | 0 | 0 | 0 | 0 | 0 |
| ENSG00000234105 | 0 | 0 | 0 | 0 | 0 | 0 |
| ENSG00000228627 | 0 | 0 | 0 | 0 | 0 | 0 |
| ENSG00000205628 | 5 | 2 | 4 | 1 | 2 | 2 |
| ENSG00000222355 | 0 | 0 | 0 | 0 | 0 | 0 |
| ENSG00000271306 | 0 | 0 | 0 | 0 | 2 | 0 |
| ENSG00000232418 | 0 | 0 | 0 | 0 | 0 | 0 |
| ENSG00000235454 | 0 | 0 | 0 | 0 | 0 | 0 |
| ENSG00000238131 | 0 | 0 | 0 | 0 | 0 | 0 |

| | | | | | | |
|---|---|---|---|---|---|---|
| ENSG00000280920 | 0 | 0 | 0 | 0 | 0 | 0 |
| ENSG00000231427 | 0 | 0 | 0 | 0 | 0 | 0 |
| ENSG00000213673 | 2 | 0 | 0 | 0 | 0 | 1 |
| ENSG00000170419 | 4 | 0 | 0 | 2 | 1 | 2 |
| ENSG00000224223 | 0 | 0 | 0 | 0 | 0 | 0 |
| ENSG00000228735 | 0 | 0 | 0 | 0 | 0 | 0 |
| ENSG00000200471 | 0 | 0 | 0 | 0 | 0 | 0 |
| ENSG00000232612 | 0 | 0 | 0 | 0 | 0 | 0 |
| ENSG00000132432 | 1975 | 954 | 1150 | 1252 | 1127 | 1026 |
| ENSG00000234707 | 4 | 0 | 2 | 0 | 3 | 3 |
| ENSG00000218586 | 0 | 0 | 0 | 0 | 0 | 0 |
| ENSG00000146648 | 92 | 34 | 49 | 20 | 54 | 28 |
| ENSG00000224057 | 0 | 0 | 0 | 0 | 0 | 0 |
| ENSG00000280890 | 0 | 0 | 1 | 1 | 3 | 0 |
| ENSG00000237210 | 0 | 0 | 0 | 0 | 0 | 0 |
| ENSG00000230564 | 0 | 0 | 0 | 0 | 0 | 0 |
| ENSG00000230936 | 0 | 0 | 0 | 0 | 0 | 1 |
| ENSG00000132434 | 1149 | 544 | 725 | 507 | 562 | 525 |
| ENSG00000154978 | 1543 | 820 | 1008 | 975 | 944 | 1011 |
| ENSG00000223475 | 0 | 0 | 0 | 0 | 0 | 0 |
| ENSG00000233977 | 0 | 1 | 0 | 0 | 0 | 0 |
| ENSG00000231394 | 0 | 1 | 0 | 0 | 0 | 0 |
| ENSG00000234844 | 0 | 0 | 0 | 0 | 0 | 0 |
| ENSG00000235207 | 0 | 0 | 0 | 0 | 0 | 0 |
| ENSG00000231317 | 0 | 0 | 0 | 0 | 0 | 0 |
| ENSG00000176826 | 6 | 0 | 6 | 1 | 1 | 0 |
| ENSG00000252857 | 0 | 0 | 0 | 0 | 0 | 0 |
| ENSG00000275853 | 0 | 0 | 0 | 0 | 0 | 0 |
| ENSG00000224971 | 0 | 0 | 0 | 0 | 0 | 0 |
| ENSG00000237799 | 0 | 0 | 0 | 0 | 0 | 0 |
| ENSG00000275875 | 0 | 0 | 0 | 0 | 0 | 0 |
| ENSG00000273720 | 0 | 0 | 0 | 0 | 0 | 0 |
| ENSG00000226278 | 0 | 0 | 0 | 0 | 0 | 0 |
| ENSG00000206729 | 0 | 0 | 0 | 0 | 0 | 0 |
| ENSG00000154997 | 0 | 1 | 0 | 0 | 0 | 0 |
| ENSG00000235738 | 0 | 0 | 0 | 0 | 0 | 0 |
| ENSG00000225270 | 0 | 0 | 0 | 0 | 0 | 0 |
| ENSG00000229762 | 0 | 0 | 0 | 0 | 0 | 0 |
| ENSG00000225018 | 0 | 0 | 0 | 0 | 0 | 0 |
| ENSG00000227499 | 0 | 0 | 0 | 0 | 0 | 0 |
| ENSG00000249773 | 2 | 0 | 4 | 0 | 3 | 2 |
| ENSG00000178665 | 64 | 69 | 61 | 30 | 52 | 57 |
| ENSG00000146729 | 1322 | 690 | 890 | 578 | 801 | 735 |
| ENSG00000239789 | 140 | 84 | 104 | 95 | 98 | 111 |
| ENSG00000146733 | 674 | 307 | 326 | 333 | 374 | 385 |
| ENSG00000146731 | 6057 | 3094 | 3433 | 2984 | 3756 | 3540 |
| ENSG00000206603 | 3 | 1 | 0 | 0 | 0 | 0 |
| ENSG00000207168 | 0 | 1 | 2 | 0 | 1 | 0 |
| ENSG00000129103 | 5193 | 2786 | 3588 | 2800 | 2737 | 2803 |
| ENSG00000164776 | 17 | 15 | 25 | 18 | 13 | 14 |

| | | | | | | |
|---|---|---|---|---|---|---|
| ENSG00000106153 | 8722 | 4305 | 5143 | 5815 | 5205 | 5141 |
| ENSG00000185290 | 0 | 0 | 0 | 0 | 0 | 0 |
| ENSG00000230191 | 0 | 0 | 0 | 0 | 0 | 0 |
| ENSG00000233028 | 6 | 2 | 2 | 0 | 4 | 2 |
| ENSG00000285670 | 0 | 0 | 0 | 0 | 0 | 0 |
| ENSG00000227491 | 0 | 0 | 0 | 0 | 0 | 0 |
| ENSG00000223559 | 26 | 9 | 8 | 8 | 7 | 10 |
| ENSG00000224155 | 2 | 0 | 1 | 1 | 0 | 0 |
| ENSG00000203462 | 0 | 0 | 0 | 0 | 1 | 0 |
| ENSG00000202296 | 0 | 0 | 0 | 0 | 0 | 0 |
| ENSG00000231299 | 0 | 0 | 0 | 0 | 0 | 0 |
| ENSG00000225371 | 4 | 1 | 1 | 1 | 2 | 1 |
| ENSG00000224370 | 0 | 0 | 0 | 0 | 0 | 0 |
| ENSG00000223330 | 0 | 0 | 0 | 0 | 0 | 0 |
| ENSG00000232944 | 0 | 0 | 0 | 0 | 0 | 0 |
| ENSG00000237268 | 0 | 0 | 0 | 0 | 0 | 0 |
| ENSG00000278205 | 0 | 0 | 0 | 0 | 0 | 0 |
| ENSG00000225488 | 0 | 0 | 0 | 0 | 0 | 0 |
| ENSG00000226485 | 0 | 0 | 0 | 0 | 0 | 0 |
| ENSG00000261275 | 0 | 0 | 0 | 0 | 0 | 0 |
| ENSG00000231504 | 0 | 0 | 0 | 0 | 0 | 0 |
| ENSG00000223740 | 0 | 0 | 0 | 0 | 0 | 0 |
| ENSG00000233288 | 2 | 0 | 0 | 0 | 0 | 0 |
| ENSG00000224947 | 0 | 0 | 0 | 0 | 0 | 0 |
| ENSG00000213650 | 2 | 0 | 1 | 1 | 1 | 1 |
| ENSG00000234716 | 0 | 0 | 0 | 0 | 0 | 0 |
| ENSG00000213067 | 0 | 2 | 0 | 0 | 1 | 0 |
| ENSG00000271047 | 0 | 0 | 0 | 0 | 0 | 0 |
| ENSG00000229909 | 0 | 0 | 0 | 0 | 0 | 0 |
| ENSG00000228303 | 0 | 0 | 0 | 0 | 0 | 0 |
| ENSG00000277174 | 0 | 0 | 0 | 0 | 0 | 0 |
| ENSG00000282879 | 0 | 0 | 0 | 0 | 0 | 0 |
| ENSG00000277253 | 0 | 0 | 0 | 0 | 0 | 0 |
| ENSG00000279072 | 3 | 0 | 0 | 1 | 1 | 0 |
| ENSG00000233437 | 0 | 0 | 0 | 0 | 0 | 0 |
| ENSG00000225244 | 0 | 0 | 0 | 0 | 0 | 0 |
| ENSG00000264426 | 0 | 0 | 0 | 0 | 0 | 0 |
| ENSG00000189166 | 2 | 0 | 0 | 0 | 0 | 0 |
| ENSG00000214668 | 0 | 0 | 0 | 0 | 0 | 0 |
| ENSG00000229508 | 0 | 0 | 0 | 0 | 0 | 0 |
| ENSG00000275061 | 0 | 0 | 0 | 0 | 0 | 0 |
| ENSG00000185177 | 0 | 0 | 0 | 2 | 2 | 1 |
| ENSG00000235095 | 0 | 0 | 0 | 0 | 0 | 0 |
| ENSG00000283431 | 1 | 0 | 1 | 0 | 0 | 0 |
| ENSG00000284474 | 0 | 0 | 0 | 0 | 0 | 0 |
| ENSG00000283547 | 0 | 0 | 0 | 0 | 0 | 0 |
| ENSG00000231537 | 0 | 0 | 0 | 0 | 0 | 0 |
| ENSG00000283230 | 0 | 0 | 0 | 0 | 0 | 0 |
| ENSG00000283481 | 0 | 0 | 0 | 0 | 0 | 0 |
| ENSG00000225295 | 0 | 0 | 0 | 0 | 0 | 0 |

| | | | | | | |
|---|---|---|---|---|---|---|
| ENSG00000228440 | 0 | 0 | 0 | 0 | 0 | 0 |
| ENSG00000283311 | 0 | 0 | 0 | 0 | 0 | 0 |
| ENSG00000224880 | 0 | 0 | 0 | 0 | 0 | 0 |
| ENSG00000224723 | 0 | 0 | 0 | 0 | 0 | 0 |
| ENSG00000230764 | 0 | 0 | 0 | 0 | 0 | 0 |
| ENSG00000237309 | 0 | 0 | 0 | 0 | 0 | 0 |
| ENSG00000236692 | 0 | 0 | 0 | 0 | 0 | 0 |
| ENSG00000284291 | 0 | 0 | 0 | 0 | 0 | 0 |
| ENSG00000284098 | 0 | 0 | 0 | 0 | 0 | 0 |
| ENSG00000283626 | 0 | 0 | 0 | 0 | 0 | 0 |
| ENSG00000228851 | 0 | 0 | 0 | 0 | 0 | 0 |
| ENSG00000284558 | 0 | 0 | 0 | 0 | 0 | 0 |
| ENSG00000284221 | 0 | 0 | 0 | 0 | 0 | 0 |
| ENSG00000227584 | 0 | 0 | 0 | 0 | 0 | 0 |
| ENSG00000230531 | 0 | 0 | 0 | 0 | 0 | 0 |
| ENSG00000284572 | 0 | 0 | 0 | 0 | 0 | 0 |
| ENSG00000234671 | 0 | 0 | 0 | 0 | 0 | 0 |
| ENSG00000169325 | 0 | 0 | 0 | 0 | 0 | 0 |
| ENSG00000234089 | 0 | 0 | 0 | 0 | 0 | 0 |
| ENSG00000238431 | 0 | 0 | 0 | 0 | 0 | 0 |
| ENSG00000274299 | 0 | 0 | 0 | 0 | 0 | 0 |
| ENSG00000280225 | 0 | 0 | 0 | 0 | 0 | 0 |
| ENSG00000260653 | 0 | 0 | 0 | 0 | 0 | 0 |
| ENSG00000266168 | 0 | 0 | 0 | 0 | 0 | 0 |
| ENSG00000227182 | 0 | 0 | 0 | 0 | 0 | 0 |
| ENSG00000235368 | 0 | 0 | 0 | 0 | 0 | 0 |
| ENSG00000182111 | 13 | 5 | 4 | 1 | 5 | 11 |
| ENSG00000270957 | 0 | 0 | 0 | 0 | 0 | 0 |
| ENSG00000233601 | 0 | 0 | 0 | 0 | 0 | 0 |
| ENSG00000243981 | 0 | 0 | 0 | 0 | 0 | 0 |
| ENSG00000270749 | 0 | 0 | 0 | 0 | 0 | 0 |
| ENSG00000229627 | 0 | 0 | 0 | 0 | 0 | 0 |
| ENSG00000230796 | 0 | 0 | 0 | 0 | 0 | 0 |
| ENSG00000237639 | 0 | 0 | 0 | 0 | 0 | 0 |
| ENSG00000233962 | 0 | 0 | 0 | 0 | 0 | 0 |
| ENSG00000236261 | 0 | 0 | 0 | 0 | 0 | 0 |
| ENSG00000234085 | 0 | 0 | 0 | 0 | 0 | 0 |
| ENSG00000227015 | 0 | 0 | 0 | 0 | 0 | 0 |
| ENSG00000232161 | 0 | 0 | 0 | 0 | 0 | 0 |
| ENSG00000230271 | 0 | 0 | 0 | 0 | 0 | 0 |
| ENSG00000250923 | 0 | 0 | 0 | 0 | 0 | 0 |
| ENSG00000224484 | 0 | 0 | 0 | 0 | 0 | 0 |
| ENSG00000236907 | 0 | 0 | 0 | 0 | 0 | 0 |
| ENSG00000250618 | 0 | 0 | 0 | 0 | 0 | 0 |
| ENSG00000224653 | 0 | 0 | 0 | 0 | 0 | 0 |
| ENSG00000223836 | 0 | 0 | 0 | 0 | 0 | 0 |
| ENSG00000237236 | 0 | 0 | 0 | 0 | 0 | 0 |
| ENSG00000231232 | 0 | 0 | 0 | 0 | 0 | 0 |
| ENSG00000231484 | 0 | 0 | 0 | 0 | 0 | 0 |
| ENSG00000271696 | 0 | 0 | 0 | 0 | 0 | 0 |

| | | | | | | |
|---|---|---|---|---|---|---|
| ENSG00000233918 | 0 | 0 | 0 | 0 | 0 | 0 |
| ENSG00000207443 | 0 | 0 | 0 | 0 | 0 | 0 |
| ENSG00000227148 | 0 | 0 | 0 | 0 | 0 | 0 |
| ENSG00000233454 | 0 | 0 | 0 | 0 | 0 | 0 |
| ENSG00000233782 | 0 | 0 | 0 | 0 | 0 | 0 |
| ENSG00000234312 | 0 | 0 | 0 | 0 | 0 | 0 |
| ENSG00000231523 | 0 | 0 | 0 | 0 | 0 | 0 |
| ENSG00000199231 | 0 | 0 | 0 | 0 | 0 | 0 |
| ENSG00000237572 | 0 | 0 | 0 | 0 | 0 | 0 |
| ENSG00000226075 | 0 | 0 | 0 | 0 | 0 | 0 |
| ENSG00000244550 | 0 | 0 | 0 | 0 | 0 | 0 |
| ENSG00000240651 | 0 | 0 | 0 | 0 | 0 | 0 |
| ENSG00000236638 | 0 | 0 | 0 | 0 | 0 | 0 |
| ENSG00000185037 | 0 | 0 | 0 | 0 | 0 | 0 |
| ENSG00000226587 | 0 | 0 | 0 | 0 | 0 | 0 |
| ENSG00000227298 | 0 | 0 | 0 | 0 | 0 | 0 |
| ENSG00000232165 | 0 | 0 | 0 | 0 | 0 | 0 |
| ENSG00000230000 | 0 | 0 | 0 | 0 | 0 | 0 |
| ENSG00000227545 | 0 | 0 | 0 | 0 | 0 | 0 |
| ENSG00000227923 | 0 | 0 | 0 | 0 | 0 | 0 |
| ENSG00000229301 | 0 | 0 | 0 | 0 | 0 | 0 |
| ENSG00000236284 | 0 | 0 | 0 | 0 | 0 | 0 |
| ENSG00000230361 | 0 | 0 | 0 | 0 | 0 | 0 |
| ENSG00000230600 | 0 | 0 | 0 | 0 | 0 | 0 |
| ENSG00000232817 | 0 | 0 | 0 | 0 | 0 | 0 |
| ENSG00000223889 | 0 | 0 | 0 | 0 | 0 | 0 |
| ENSG00000227305 | 0 | 0 | 0 | 0 | 0 | 0 |
| ENSG00000234387 | 0 | 0 | 0 | 0 | 0 | 0 |
| ENSG00000268181 | 0 | 0 | 0 | 0 | 0 | 0 |
| ENSG00000234467 | 0 | 0 | 0 | 0 | 0 | 0 |
| ENSG00000227426 | 0 | 0 | 0 | 0 | 0 | 0 |
| ENSG00000224368 | 0 | 0 | 0 | 0 | 0 | 0 |
| ENSG00000197990 | 0 | 0 | 0 | 0 | 0 | 0 |
| ENSG00000230132 | 0 | 0 | 0 | 0 | 0 | 0 |
| ENSG00000164740 | 0 | 0 | 0 | 0 | 0 | 0 |
| ENSG00000228645 | 0 | 0 | 0 | 0 | 0 | 0 |
| ENSG00000214660 | 0 | 0 | 0 | 0 | 0 | 0 |
| ENSG00000223566 | 0 | 0 | 0 | 0 | 0 | 0 |
| ENSG00000265214 | 0 | 0 | 0 | 0 | 0 | 0 |
| ENSG00000226401 | 0 | 0 | 0 | 0 | 0 | 0 |
| ENSG00000226411 | 0 | 0 | 0 | 0 | 0 | 0 |
| ENSG00000271466 | 0 | 0 | 0 | 0 | 0 | 0 |
| ENSG00000227397 | 0 | 0 | 0 | 0 | 0 | 0 |
| ENSG00000236574 | 0 | 0 | 0 | 0 | 0 | 0 |
| ENSG00000225451 | 0 | 0 | 0 | 0 | 0 | 0 |
| ENSG00000236299 | 0 | 0 | 0 | 0 | 0 | 0 |
| ENSG00000235720 | 0 | 0 | 0 | 0 | 0 | 1 |
| ENSG00000226581 | 0 | 0 | 2 | 1 | 2 | 1 |
| ENSG00000226342 | 7 | 5 | 1 | 0 | 2 | 0 |
| ENSG00000227910 | 39 | 23 | 28 | 16 | 17 | 15 |

| | | | | | | |
|---|---|---|---|---|---|---|
| ENSG00000213645 | 0 | 0 | 0 | 0 | 0 | 0 |
| ENSG00000233173 | 0 | 0 | 0 | 0 | 0 | 0 |
| ENSG00000231264 | 0 | 0 | 0 | 0 | 0 | 0 |
| ENSG00000284534 | 0 | 0 | 0 | 0 | 0 | 0 |
| ENSG00000228660 | 0 | 1 | 0 | 0 | 1 | 0 |
| ENSG00000230639 | 0 | 0 | 0 | 0 | 0 | 0 |
| ENSG00000176232 | 0 | 0 | 0 | 0 | 0 | 0 |
| ENSG00000278577 | 0 | 0 | 0 | 0 | 0 | 0 |
| ENSG00000241149 | 0 | 0 | 0 | 0 | 0 | 0 |
| ENSG00000228564 | 0 | 0 | 0 | 0 | 0 | 0 |
| ENSG00000285544 | 0 | 0 | 0 | 0 | 0 | 0 |
| ENSG00000214652 | 3 | 1 | 1 | 3 | 0 | 0 |
| ENSG00000223558 | 0 | 0 | 0 | 0 | 0 | 0 |
| ENSG00000224458 | 0 | 0 | 0 | 0 | 0 | 0 |
| ENSG00000248704 | 0 | 0 | 0 | 0 | 0 | 0 |
| ENSG00000248315 | 0 | 0 | 0 | 0 | 0 | 0 |
| ENSG00000230052 | 0 | 0 | 0 | 0 | 0 | 0 |
| ENSG00000233385 | 0 | 0 | 0 | 0 | 0 | 0 |
| ENSG00000227035 | 0 | 0 | 0 | 0 | 0 | 0 |
| ENSG00000234234 | 0 | 0 | 0 | 0 | 0 | 0 |
| ENSG00000234700 | 0 | 0 | 0 | 0 | 0 | 0 |
| ENSG00000230252 | 0 | 0 | 0 | 0 | 0 | 0 |
| ENSG00000235646 | 0 | 0 | 0 | 0 | 0 | 0 |
| ENSG00000229881 | 0 | 0 | 0 | 0 | 0 | 0 |
| ENSG00000236976 | 2 | 0 | 1 | 0 | 0 | 0 |
| ENSG00000213643 | 0 | 0 | 0 | 0 | 0 | 0 |
| ENSG00000235227 | 0 | 0 | 0 | 0 | 0 | 0 |
| ENSG00000213644 | 0 | 0 | 0 | 0 | 0 | 0 |
| ENSG00000223614 | 0 | 0 | 0 | 0 | 0 | 0 |
| ENSG00000197123 | 0 | 1 | 0 | 0 | 0 | 0 |
| ENSG00000234444 | 575 | 246 | 294 | 169 | 284 | 264 |
| ENSG00000227986 | 19 | 14 | 9 | 10 | 20 | 14 |
| ENSG00000276650 | 0 | 0 | 0 | 0 | 0 | 0 |
| ENSG00000232727 | 4 | 1 | 1 | 0 | 0 | 0 |
| ENSG00000223845 | 0 | 0 | 0 | 0 | 0 | 0 |
| ENSG00000228014 | 0 | 0 | 0 | 0 | 0 | 0 |
| ENSG00000228653 | 6 | 3 | 2 | 5 | 3 | 2 |
| ENSG00000173041 | 100 | 87 | 56 | 30 | 136 | 83 |
| ENSG00000223974 | 0 | 0 | 0 | 0 | 0 | 0 |
| ENSG00000235349 | 0 | 0 | 0 | 0 | 0 | 0 |
| ENSG00000213642 | 0 | 0 | 0 | 0 | 0 | 0 |
| ENSG00000224669 | 0 | 0 | 0 | 0 | 1 | 0 |
| ENSG00000276475 | 0 | 0 | 0 | 0 | 0 | 0 |
| ENSG00000196247 | 113 | 78 | 75 | 37 | 113 | 79 |
| ENSG00000271550 | 17 | 5 | 14 | 6 | 8 | 9 |
| ENSG00000275667 | 0 | 0 | 0 | 0 | 0 | 0 |
| ENSG00000224172 | 4 | 0 | 2 | 3 | 4 | 0 |
| ENSG00000277206 | 0 | 0 | 0 | 0 | 0 | 0 |
| ENSG00000197008 | 222 | 149 | 111 | 77 | 179 | 129 |
| ENSG00000234338 | 22 | 13 | 12 | 1 | 9 | 9 |

| | | | | | | |
|---|---|---|---|---|---|---|
| ENSG00000182722 | 0 | 0 | 0 | 0 | 0 | 0 |
| ENSG00000213640 | 0 | 0 | 0 | 0 | 0 | 0 |
| ENSG00000198039 | 113 | 68 | 58 | 32 | 71 | 49 |
| ENSG00000189316 | 0 | 0 | 0 | 0 | 0 | 0 |
| ENSG00000223476 | 2 | 1 | 1 | 4 | 2 | 2 |
| ENSG00000270948 | 2 | 3 | 2 | 0 | 0 | 0 |
| ENSG00000152926 | 460 | 307 | 265 | 117 | 408 | 235 |
| ENSG00000213462 | 827 | 350 | 472 | 340 | 534 | 454 |
| ENSG00000201165 | 0 | 0 | 0 | 0 | 0 | 0 |
| ENSG00000234585 | 174 | 77 | 132 | 69 | 99 | 69 |
| ENSG00000239985 | 0 | 0 | 0 | 0 | 0 | 0 |
| ENSG00000207344 | 0 | 0 | 0 | 1 | 0 | 0 |
| ENSG00000207062 | 1 | 1 | 3 | 2 | 2 | 0 |
| ENSG00000227113 | 4 | 3 | 10 | 4 | 1 | 3 |
| ENSG00000226002 | 11 | 4 | 7 | 1 | 5 | 6 |
| ENSG00000164669 | 2 | 2 | 5 | 3 | 8 | 6 |
| ENSG00000237026 | 0 | 0 | 1 | 0 | 0 | 1 |
| ENSG00000229064 | 0 | 0 | 0 | 0 | 0 | 0 |
| ENSG00000282381 | 0 | 0 | 0 | 0 | 1 | 0 |
| ENSG00000277544 | 0 | 0 | 0 | 0 | 0 | 0 |
| ENSG00000231556 | 0 | 0 | 0 | 0 | 0 | 0 |
| ENSG00000232906 | 0 | 0 | 1 | 0 | 0 | 1 |
| ENSG00000146757 | 51 | 46 | 42 | 19 | 84 | 36 |
| ENSG00000238124 | 0 | 0 | 0 | 0 | 0 | 0 |
| ENSG00000226767 | 0 | 0 | 0 | 0 | 0 | 0 |
| ENSG00000235421 | 0 | 0 | 0 | 0 | 0 | 0 |
| ENSG00000230386 | 0 | 0 | 0 | 0 | 0 | 0 |
| ENSG00000272693 | 28 | 17 | 34 | 19 | 28 | 14 |
| ENSG00000273024 | 21 | 17 | 23 | 5 | 19 | 15 |
| ENSG00000228409 | 151 | 84 | 128 | 80 | 82 | 107 |
| ENSG00000206634 | 2 | 3 | 2 | 2 | 5 | 2 |
| ENSG00000206785 | 4 | 3 | 1 | 4 | 0 | 3 |
| ENSG00000275833 | 8 | 8 | 8 | 0 | 8 | 4 |
| ENSG00000224316 | 9 | 12 | 5 | 4 | 4 | 0 |
| ENSG00000200670 | 0 | 0 | 0 | 0 | 0 | 0 |
| ENSG00000236529 | 5 | 1 | 2 | 2 | 3 | 3 |
| ENSG00000201560 | 0 | 0 | 0 | 0 | 0 | 0 |
| ENSG00000196715 | 2094 | 969 | 1232 | 842 | 1242 | 1048 |
| ENSG00000169919 | 3538 | 1689 | 2271 | 1749 | 1542 | 1721 |
| ENSG00000224365 | 1 | 0 | 0 | 0 | 0 | 0 |
| ENSG00000229886 | 0 | 0 | 0 | 0 | 0 | 0 |
| ENSG00000205596 | 0 | 0 | 1 | 0 | 0 | 0 |
| ENSG00000126522 | 1081 | 597 | 733 | 631 | 524 | 545 |
| ENSG00000249319 | 0 | 1 | 1 | 1 | 1 | 0 |
| ENSG00000241258 | 2042 | 1161 | 1215 | 975 | 1211 | 1142 |
| ENSG00000234185 | 0 | 4 | 1 | 0 | 2 | 0 |
| ENSG00000169902 | 333 | 168 | 223 | 165 | 203 | 180 |
| ENSG00000252126 | 0 | 0 | 0 | 0 | 0 | 0 |
| ENSG00000179406 | 423 | 171 | 276 | 119 | 189 | 208 |
| ENSG00000251451 | 0 | 0 | 0 | 0 | 0 | 0 |

| | | | | | | |
|---|---|---|---|---|---|---|
| ENSG00000231234 | 0 | 0 | 0 | 0 | 0 | 0 |
| ENSG00000230189 | 0 | 1 | 1 | 0 | 1 | 1 |
| ENSG00000236928 | 0 | 0 | 0 | 0 | 0 | 2 |
| ENSG00000279785 | 0 | 1 | 0 | 1 | 0 | 0 |
| ENSG00000244510 | 0 | 0 | 0 | 0 | 0 | 0 |
| ENSG00000244657 | 0 | 0 | 0 | 0 | 0 | 0 |
| ENSG00000237310 | 75 | 32 | 40 | 25 | 31 | 27 |
| ENSG00000179342 | 0 | 1 | 1 | 0 | 0 | 0 |
| ENSG00000234500 | 16 | 6 | 10 | 5 | 4 | 5 |
| ENSG00000229180 | 511 | 294 | 343 | 273 | 367 | 359 |
| ENSG00000275400 | 0 | 0 | 0 | 0 | 0 | 0 |
| ENSG00000232491 | 0 | 0 | 0 | 0 | 0 | 0 |
| ENSG00000177418 | 0 | 0 | 0 | 0 | 0 | 0 |
| ENSG00000225573 | 8 | 3 | 4 | 5 | 4 | 6 |
| ENSG00000243335 | 300 | 123 | 170 | 117 | 166 | 134 |
| ENSG00000284461 | 3 | 1 | 3 | 0 | 0 | 0 |
| ENSG00000226824 | 41 | 27 | 41 | 18 | 32 | 31 |
| ENSG00000154710 | 147 | 109 | 130 | 111 | 121 | 91 |
| ENSG00000272831 | 1 | 1 | 3 | 2 | 1 | 1 |
| ENSG00000271064 | 2 | 4 | 3 | 3 | 2 | 1 |
| ENSG00000230583 | 31 | 6 | 11 | 6 | 8 | 10 |
| ENSG00000232546 | 20 | 8 | 13 | 3 | 16 | 8 |
| ENSG00000230295 | 5 | 10 | 21 | 5 | 15 | 12 |
| ENSG00000212568 | 0 | 0 | 0 | 0 | 0 | 0 |
| ENSG00000273142 | 309 | 130 | 207 | 108 | 156 | 152 |
| ENSG00000179131 | 0 | 0 | 0 | 0 | 0 | 0 |
| ENSG00000274135 | 0 | 0 | 0 | 0 | 0 | 0 |
| ENSG00000106609 | 4001 | 1759 | 2268 | 1835 | 2095 | 2306 |
| ENSG00000275878 | 0 | 0 | 0 | 0 | 0 | 0 |
| ENSG00000126524 | 2659 | 1483 | 1589 | 1280 | 1651 | 1614 |
| ENSG00000198874 | 862 | 417 | 526 | 407 | 475 | 499 |
| ENSG00000233383 | 0 | 0 | 0 | 0 | 0 | 0 |
| ENSG00000266525 | 0 | 0 | 0 | 0 | 0 | 0 |
| ENSG00000230358 | 3 | 7 | 5 | 2 | 2 | 4 |
| ENSG00000067601 | 91 | 59 | 78 | 61 | 50 | 45 |
| ENSG00000201565 | 0 | 0 | 0 | 0 | 0 | 0 |
| ENSG00000106610 | 572 | 311 | 362 | 318 | 308 | 328 |
| ENSG00000228019 | 4 | 2 | 4 | 0 | 2 | 0 |
| ENSG00000273448 | 20 | 4 | 13 | 5 | 6 | 11 |
| ENSG00000226392 | 0 | 0 | 0 | 0 | 0 | 0 |
| ENSG00000237466 | 0 | 0 | 0 | 0 | 0 | 0 |
| ENSG00000233423 | 0 | 0 | 0 | 0 | 0 | 0 |
| ENSG00000223948 | 0 | 0 | 0 | 0 | 0 | 0 |
| ENSG00000225209 | 0 | 0 | 0 | 0 | 0 | 0 |
| ENSG00000236531 | 0 | 0 | 0 | 0 | 0 | 0 |
| ENSG00000232975 | 0 | 0 | 0 | 0 | 0 | 0 |
| ENSG00000226829 | 0 | 0 | 0 | 0 | 0 | 0 |
| ENSG00000237754 | 0 | 0 | 0 | 0 | 0 | 0 |
| ENSG00000225959 | 0 | 0 | 0 | 0 | 0 | 0 |
| ENSG00000228429 | 0 | 0 | 0 | 0 | 0 | 0 |

| | | | | | | |
|---|---|---|---|---|---|---|
| ENSG00000222428 | 0 | 0 | 0 | 0 | 0 | 0 |
| ENSG00000233689 | 0 | 0 | 0 | 0 | 0 | 0 |
| ENSG00000200189 | 0 | 0 | 0 | 0 | 0 | 0 |
| ENSG00000236839 | 0 | 0 | 0 | 0 | 0 | 0 |
| ENSG00000225718 | 0 | 0 | 0 | 0 | 0 | 0 |
| ENSG00000230654 | 0 | 0 | 0 | 0 | 0 | 0 |
| ENSG00000229031 | 0 | 0 | 0 | 0 | 0 | 0 |
| ENSG00000252423 | 0 | 0 | 0 | 0 | 0 | 0 |
| ENSG00000200857 | 0 | 0 | 0 | 0 | 0 | 0 |
| ENSG00000234215 | 0 | 0 | 0 | 0 | 0 | 0 |
| ENSG00000158321 | 48 | 13 | 32 | 7 | 9 | 7 |
| ENSG00000236978 | 0 | 0 | 0 | 0 | 0 | 0 |
| ENSG00000185274 | 0 | 0 | 0 | 0 | 0 | 0 |
| ENSG00000265878 | 0 | 0 | 0 | 0 | 0 | 0 |
| ENSG00000283278 | 0 | 0 | 0 | 0 | 0 | 0 |
| ENSG00000199940 | 0 | 0 | 0 | 0 | 0 | 0 |
| ENSG00000183166 | 0 | 0 | 0 | 0 | 0 | 1 |
| ENSG00000201014 | 0 | 0 | 0 | 0 | 0 | 0 |
| ENSG00000235581 | 0 | 0 | 0 | 0 | 0 | 0 |
| ENSG00000228769 | 0 | 0 | 0 | 0 | 0 | 0 |
| ENSG00000277149 | 69 | 40 | 35 | 14 | 34 | 32 |
| ENSG00000264494 | 0 | 0 | 0 | 0 | 0 | 0 |
| ENSG00000270694 | 0 | 0 | 0 | 0 | 0 | 0 |
| ENSG00000270555 | 0 | 0 | 0 | 0 | 0 | 0 |
| ENSG00000266569 | 0 | 0 | 0 | 0 | 1 | 0 |
| ENSG00000225648 | 716 | 425 | 463 | 389 | 452 | 435 |
| ENSG00000274656 | 0 | 0 | 0 | 0 | 0 | 0 |
| ENSG00000179994 | 0 | 0 | 0 | 0 | 0 | 0 |
| ENSG00000196313 | 1563 | 650 | 936 | 745 | 795 | 902 |
| ENSG00000272843 | 2 | 1 | 3 | 0 | 3 | 1 |
| ENSG00000106133 | 18 | 13 | 12 | 13 | 12 | 13 |
| ENSG00000285886 | 2 | 1 | 3 | 1 | 4 | 3 |
| ENSG00000155428 | 4 | 3 | 0 | 2 | 0 | 0 |
| ENSG00000174353 | 107 | 69 | 81 | 60 | 63 | 81 |
| ENSG00000205584 | 1 | 0 | 0 | 0 | 0 | 1 |
| ENSG00000276178 | 0 | 0 | 0 | 0 | 0 | 0 |
| ENSG00000229018 | 8 | 0 | 5 | 3 | 2 | 2 |
| ENSG00000201282 | 0 | 0 | 0 | 0 | 0 | 0 |
| ENSG00000273520 | 0 | 0 | 0 | 0 | 0 | 0 |
| ENSG00000273897 | 0 | 0 | 0 | 0 | 0 | 0 |
| ENSG00000273927 | 0 | 0 | 0 | 0 | 0 | 0 |
| ENSG00000285702 | 0 | 0 | 0 | 0 | 0 | 0 |
| ENSG00000275976 | 0 | 0 | 0 | 0 | 0 | 0 |
| ENSG00000277125 | 0 | 0 | 0 | 0 | 0 | 0 |
| ENSG00000274984 | 0 | 0 | 0 | 0 | 0 | 0 |
| ENSG00000262461 | 0 | 0 | 0 | 0 | 0 | 0 |
| ENSG00000174384 | 7 | 0 | 2 | 2 | 2 | 1 |
| ENSG00000199913 | 0 | 0 | 0 | 0 | 0 | 0 |
| ENSG00000274570 | 2 | 0 | 1 | 0 | 0 | 1 |
| ENSG00000233369 | 1198 | 615 | 745 | 481 | 568 | 580 |

| | | | | | | |
|---|---|---|---|---|---|---|
| ENSG00000233523 | 4 | 0 | 0 | 3 | 0 | 0 |
| ENSG00000182487 | 0 | 0 | 1 | 0 | 0 | 0 |
| ENSG00000214544 | 4 | 6 | 3 | 4 | 3 | 7 |
| ENSG00000205578 | 12 | 3 | 3 | 0 | 0 | 0 |
| ENSG00000130305 | 1329 | 591 | 732 | 911 | 796 | 793 |
| ENSG00000146755 | 8 | 0 | 0 | 1 | 0 | 1 |
| ENSG00000225005 | 2 | 0 | 0 | 0 | 0 | 1 |
| ENSG00000077800 | 4 | 0 | 3 | 0 | 1 | 0 |
| ENSG00000206709 | 0 | 0 | 0 | 0 | 0 | 0 |
| ENSG00000202035 | 0 | 0 | 0 | 0 | 0 | 0 |
| ENSG00000188763 | 8 | 6 | 6 | 7 | 12 | 6 |
| ENSG00000009954 | 3681 | 1801 | 1934 | 1096 | 2133 | 1828 |
| ENSG00000252713 | 0 | 0 | 0 | 0 | 0 | 0 |
| ENSG00000106635 | 1719 | 817 | 1040 | 947 | 871 | 967 |
| ENSG00000106638 | 1644 | 769 | 865 | 971 | 853 | 926 |
| ENSG00000009950 | 210 | 63 | 126 | 66 | 89 | 92 |
| ENSG00000274080 | 0 | 1 | 2 | 2 | 2 | 0 |
| ENSG00000176428 | 212 | 97 | 120 | 129 | 97 | 86 |
| ENSG00000176410 | 631 | 360 | 473 | 415 | 416 | 397 |
| ENSG00000071462 | 2778 | 1470 | 1622 | 1606 | 1870 | 1647 |
| ENSG00000106089 | 359 | 199 | 244 | 241 | 214 | 217 |
| ENSG00000265724 | 0 | 3 | 0 | 0 | 1 | 1 |
| ENSG00000241709 | 0 | 2 | 0 | 1 | 0 | 1 |
| ENSG00000225969 | 20 | 21 | 24 | 22 | 9 | 15 |
| ENSG00000106077 | 3664 | 1853 | 2449 | 1684 | 1655 | 1755 |
| ENSG00000165215 | 9149 | 3858 | 5641 | 5663 | 3783 | 4556 |
| ENSG00000189143 | 15315 | 6999 | 9612 | 9493 | 7753 | 8223 |
| ENSG00000165171 | 96 | 61 | 68 | 43 | 43 | 50 |
| ENSG00000175877 | 13 | 7 | 14 | 12 | 5 | 7 |
| ENSG00000261467 | 0 | 0 | 0 | 0 | 0 | 0 |
| ENSG00000049540 | 0 | 0 | 1 | 0 | 0 | 0 |
| ENSG00000232415 | 0 | 2 | 0 | 0 | 0 | 0 |
| ENSG00000106683 | 1489 | 731 | 850 | 1206 | 1088 | 962 |
| ENSG00000106682 | 13373 | 6514 | 7701 | 6760 | 7507 | 7626 |
| ENSG00000207741 | 9 | 5 | 8 | 1 | 6 | 7 |
| ENSG00000086730 | 29 | 8 | 6 | 11 | 5 | 14 |
| ENSG00000049541 | 1126 | 636 | 752 | 661 | 678 | 616 |
| ENSG00000106665 | 152 | 61 | 82 | 87 | 89 | 76 |
| ENSG00000006704 | 1322 | 686 | 852 | 763 | 762 | 732 |
| ENSG00000238391 | 0 | 0 | 0 | 0 | 0 | 0 |
| ENSG00000273069 | 0 | 0 | 2 | 0 | 1 | 0 |
| ENSG00000279005 | 0 | 0 | 0 | 0 | 0 | 0 |
| ENSG00000263001 | 2411 | 1344 | 1848 | 953 | 1486 | 1444 |
| ENSG00000232729 | 2 | 2 | 6 | 0 | 2 | 1 |
| ENSG00000262648 | 0 | 0 | 0 | 0 | 0 | 0 |
| ENSG00000158517 | 1 | 0 | 0 | 1 | 0 | 0 |
| ENSG00000196275 | 136 | 57 | 80 | 59 | 69 | 61 |
| ENSG00000277072 | 272 | 184 | 192 | 102 | 116 | 146 |
| ENSG00000123965 | 11 | 11 | 15 | 14 | 12 | 13 |
| ENSG00000239069 | 0 | 0 | 0 | 0 | 0 | 0 |

| | | | | | | |
|---|---|---|---|---|---|---|
| ENSG00000184616 | 34 | 19 | 17 | 17 | 16 | 21 |
| ENSG00000274070 | 197 | 90 | 101 | 71 | 84 | 65 |
| ENSG00000274523 | 2820 | 1422 | 1741 | 1766 | 1508 | 1644 |
| ENSG00000277127 | 0 | 0 | 0 | 0 | 0 | 0 |
| ENSG00000174428 | 337 | 158 | 192 | 120 | 164 | 157 |
| ENSG00000165178 | 0 | 0 | 0 | 0 | 0 | 2 |
| ENSG00000277053 | 254 | 116 | 146 | 98 | 146 | 105 |
| ENSG00000276592 | 0 | 0 | 0 | 0 | 0 | 0 |
| ENSG00000277675 | 0 | 0 | 0 | 0 | 0 | 0 |
| ENSG00000263081 | 0 | 0 | 0 | 0 | 0 | 0 |
| ENSG00000275121 | 0 | 0 | 0 | 0 | 0 | 0 |
| ENSG00000274959 | 0 | 0 | 0 | 0 | 0 | 0 |
| ENSG00000276840 | 4 | 7 | 4 | 1 | 0 | 1 |
| ENSG00000267828 | 0 | 0 | 0 | 0 | 0 | 0 |
| ENSG00000278807 | 0 | 0 | 0 | 0 | 0 | 0 |
| ENSG00000273598 | 2 | 0 | 0 | 0 | 0 | 0 |
| ENSG00000202021 | 0 | 0 | 0 | 0 | 0 | 0 |
| ENSG00000278080 | 0 | 0 | 0 | 0 | 0 | 0 |
| ENSG00000278416 | 15 | 7 | 14 | 8 | 5 | 7 |
| ENSG00000199870 | 0 | 0 | 0 | 0 | 0 | 0 |
| ENSG00000205583 | 12 | 11 | 11 | 4 | 8 | 14 |
| ENSG00000275930 | 1 | 0 | 0 | 0 | 1 | 0 |
| ENSG00000146722 | 0 | 1 | 0 | 0 | 1 | 0 |
| ENSG00000178809 | 10 | 3 | 6 | 6 | 1 | 4 |
| ENSG00000223705 | 741 | 299 | 471 | 333 | 384 | 398 |
| ENSG00000272391 | 2145 | 964 | 1255 | 1011 | 1052 | 1178 |
| ENSG00000242073 | 0 | 0 | 0 | 0 | 0 | 0 |
| ENSG00000170092 | 25 | 23 | 27 | 11 | 17 | 10 |
| ENSG00000127957 | 183 | 79 | 102 | 55 | 98 | 85 |
| ENSG00000200874 | 0 | 0 | 0 | 0 | 0 | 0 |
| ENSG00000127946 | 2465 | 1093 | 1488 | 817 | 1290 | 1167 |
| ENSG00000279996 | 0 | 0 | 0 | 0 | 0 | 0 |
| ENSG00000006606 | 12 | 5 | 4 | 5 | 10 | 4 |
| ENSG00000106178 | 0 | 0 | 0 | 0 | 0 | 0 |
| ENSG00000219039 | 0 | 0 | 0 | 3 | 0 | 0 |
| ENSG00000005486 | 3181 | 1372 | 1893 | 1931 | 1523 | 1716 |
| ENSG00000127948 | 5210 | 2321 | 3049 | 3005 | 2427 | 2787 |
| ENSG00000284334 | 0 | 0 | 0 | 0 | 0 | 0 |
| ENSG00000235090 | 2 | 1 | 0 | 1 | 2 | 2 |
| ENSG00000201643 | 1 | 0 | 0 | 0 | 0 | 0 |
| ENSG00000189077 | 1241 | 571 | 768 | 706 | 662 | 638 |
| ENSG00000127952 | 1403 | 722 | 853 | 990 | 869 | 882 |
| ENSG00000280388 | 8 | 7 | 9 | 10 | 7 | 6 |
| ENSG00000146701 | 7606 | 3582 | 4658 | 4917 | 4300 | 4656 |
| ENSG00000230882 | 11 | 11 | 22 | 20 | 20 | 19 |
| ENSG00000251798 | 0 | 0 | 0 | 0 | 0 | 0 |
| ENSG00000227038 | 8 | 1 | 3 | 6 | 3 | 4 |
| ENSG00000224273 | 4 | 0 | 3 | 1 | 0 | 0 |
| ENSG00000213549 | 0 | 0 | 0 | 0 | 1 | 0 |
| ENSG00000228829 | 1 | 0 | 0 | 0 | 0 | 0 |

| | | | | | | |
|---|---|---|---|---|---|---|
| ENSG00000177679 | 577 | 274 | 348 | 357 | 302 | 256 |
| ENSG00000106211 | 45436 | 20253 | 24850 | 29947 | 23570 | 23393 |
| ENSG00000170027 | 10146 | 4776 | 6035 | 4946 | 5886 | 6211 |
| ENSG00000213548 | 0 | 0 | 0 | 0 | 0 | 0 |
| ENSG00000146700 | 47 | 17 | 27 | 17 | 30 | 26 |
| ENSG00000188372 | 428 | 259 | 312 | 447 | 338 | 348 |
| ENSG00000091073 | 757 | 319 | 394 | 407 | 339 | 334 |
| ENSG00000233980 | 0 | 0 | 0 | 0 | 0 | 1 |
| ENSG00000250614 | 2 | 0 | 1 | 0 | 0 | 2 |
| ENSG00000243566 | 2597 | 1221 | 1291 | 1154 | 1032 | 840 |
| ENSG00000250778 | 0 | 0 | 0 | 0 | 0 | 0 |
| ENSG00000238560 | 0 | 0 | 0 | 0 | 0 | 0 |
| ENSG00000230305 | 0 | 0 | 0 | 0 | 0 | 0 |
| ENSG00000185040 | 3 | 4 | 3 | 1 | 1 | 3 |
| ENSG00000205485 | 86 | 50 | 61 | 49 | 59 | 59 |
| ENSG00000146707 | 65 | 38 | 32 | 43 | 25 | 49 |
| ENSG00000214243 | 7 | 2 | 3 | 1 | 1 | 1 |
| ENSG00000285666 | 0 | 0 | 0 | 0 | 0 | 0 |
| ENSG00000231183 | 0 | 0 | 0 | 0 | 0 | 0 |
| ENSG00000265479 | 8 | 7 | 1 | 12 | 7 | 11 |
| ENSG00000283398 | 0 | 1 | 0 | 1 | 0 | 0 |
| ENSG00000231087 | 1 | 0 | 0 | 0 | 0 | 0 |
| ENSG00000236280 | 0 | 0 | 0 | 0 | 0 | 0 |
| ENSG00000225703 | 0 | 0 | 0 | 1 | 0 | 0 |
| ENSG00000186704 | 6 | 2 | 5 | 3 | 3 | 2 |
| ENSG00000242435 | 3 | 0 | 0 | 0 | 0 | 0 |
| ENSG00000241350 | 0 | 0 | 0 | 0 | 0 | 0 |
| ENSG00000201885 | 0 | 0 | 0 | 0 | 0 | 0 |
| ENSG00000186645 | 0 | 0 | 0 | 0 | 0 | 0 |
| ENSG00000233448 | 0 | 0 | 1 | 0 | 1 | 0 |
| ENSG00000201959 | 0 | 0 | 0 | 0 | 0 | 0 |
| ENSG00000259628 | 0 | 0 | 0 | 0 | 0 | 0 |
| ENSG00000205482 | 7 | 9 | 16 | 7 | 6 | 10 |
| ENSG00000225726 | 0 | 1 | 1 | 0 | 1 | 0 |
| ENSG00000214439 | 37 | 19 | 27 | 7 | 10 | 10 |
| ENSG00000213542 | 2 | 3 | 4 | 3 | 3 | 9 |
| ENSG00000135205 | 98 | 51 | 58 | 26 | 81 | 49 |
| ENSG00000127951 | 0 | 1 | 0 | 0 | 0 | 0 |
| ENSG00000250990 | 2 | 2 | 0 | 0 | 3 | 2 |
| ENSG00000186088 | 293 | 148 | 205 | 114 | 157 | 132 |
| ENSG00000273341 | 0 | 0 | 0 | 0 | 1 | 0 |
| ENSG00000237620 | 0 | 0 | 0 | 0 | 0 | 0 |
| ENSG00000127947 | 3138 | 1583 | 1736 | 989 | 1858 | 1398 |
| ENSG00000214293 | 309 | 157 | 201 | 182 | 187 | 185 |
| ENSG00000187257 | 941 | 469 | 554 | 339 | 521 | 448 |
| ENSG00000135211 | 795 | 368 | 485 | 468 | 522 | 429 |
| ENSG00000006576 | 1425 | 876 | 734 | 386 | 959 | 692 |
| ENSG00000222432 | 0 | 0 | 0 | 0 | 0 | 0 |
| ENSG00000232756 | 0 | 0 | 0 | 2 | 3 | 0 |
| ENSG00000187391 | 82 | 34 | 40 | 44 | 38 | 34 |

| | | | | | | |
|---|---|---|---|---|---|---|
| ENSG00000281008 | 0 | 0 | 0 | 0 | 0 | 0 |
| ENSG00000280958 | 0 | 0 | 0 | 0 | 0 | 0 |
| ENSG00000231322 | 0 | 0 | 0 | 0 | 0 | 0 |
| ENSG00000285892 | 0 | 0 | 0 | 0 | 0 | 0 |
| ENSG00000226230 | 0 | 0 | 0 | 0 | 0 | 0 |
| ENSG00000251276 | 0 | 0 | 0 | 0 | 0 | 0 |
| ENSG00000226978 | 0 | 0 | 0 | 0 | 0 | 0 |
| ENSG00000212545 | 0 | 0 | 0 | 0 | 0 | 0 |
| ENSG00000229110 | 0 | 0 | 0 | 0 | 0 | 0 |
| ENSG00000281120 | 0 | 0 | 0 | 0 | 0 | 0 |
| ENSG00000212482 | 0 | 0 | 0 | 0 | 0 | 0 |
| ENSG00000222024 | 0 | 0 | 0 | 0 | 0 | 0 |
| ENSG00000234456 | 0 | 0 | 0 | 0 | 0 | 0 |
| ENSG00000226285 | 0 | 0 | 0 | 0 | 0 | 0 |
| ENSG00000200786 | 0 | 0 | 0 | 0 | 0 | 0 |
| ENSG00000127955 | 40 | 17 | 28 | 10 | 14 | 14 |
| ENSG00000206818 | 0 | 0 | 0 | 0 | 0 | 0 |
| ENSG00000230853 | 0 | 0 | 0 | 0 | 0 | 0 |
| ENSG00000244392 | 0 | 0 | 0 | 0 | 0 | 0 |
| ENSG00000234223 | 0 | 0 | 0 | 0 | 0 | 0 |
| ENSG00000232667 | 0 | 0 | 0 | 0 | 0 | 0 |
| ENSG00000240347 | 0 | 0 | 0 | 0 | 0 | 0 |
| ENSG00000135218 | 94 | 36 | 48 | 22 | 44 | 31 |
| ENSG00000223550 | 0 | 0 | 0 | 0 | 0 | 0 |
| ENSG00000214415 | 0 | 0 | 0 | 0 | 0 | 0 |
| ENSG00000229436 | 0 | 0 | 0 | 0 | 0 | 0 |
| ENSG00000075223 | 9981 | 4872 | 5251 | 2715 | 6110 | 4486 |
| ENSG00000279326 | 0 | 0 | 0 | 0 | 0 | 0 |
| ENSG00000237896 | 0 | 0 | 0 | 0 | 1 | 0 |
| ENSG00000226671 | 0 | 0 | 0 | 0 | 0 | 0 |
| ENSG00000223514 | 0 | 0 | 0 | 0 | 0 | 0 |
| ENSG00000224134 | 0 | 0 | 0 | 0 | 0 | 0 |
| ENSG00000232968 | 0 | 0 | 0 | 0 | 0 | 0 |
| ENSG00000233491 | 0 | 0 | 0 | 0 | 0 | 0 |
| ENSG00000230196 | 0 | 0 | 0 | 0 | 0 | 0 |
| ENSG00000019991 | 0 | 0 | 0 | 0 | 1 | 0 |
| ENSG00000153956 | 0 | 1 | 0 | 0 | 0 | 0 |
| ENSG00000223770 | 0 | 0 | 0 | 0 | 0 | 0 |
| ENSG00000213530 | 0 | 0 | 0 | 0 | 0 | 0 |
| ENSG00000228711 | 0 | 0 | 0 | 0 | 0 | 0 |
| ENSG00000186472 | 80 | 62 | 49 | 11 | 71 | 47 |
| ENSG00000202281 | 0 | 0 | 0 | 0 | 0 | 0 |
| ENSG00000235503 | 0 | 0 | 0 | 0 | 0 | 0 |
| ENSG00000170381 | 185 | 66 | 82 | 52 | 106 | 61 |
| ENSG00000285398 | 0 | 0 | 0 | 0 | 0 | 0 |
| ENSG00000230617 | 0 | 0 | 0 | 0 | 0 | 0 |
| ENSG00000230244 | 0 | 0 | 0 | 0 | 0 | 0 |
| ENSG00000075213 | 0 | 1 | 0 | 1 | 0 | 0 |
| ENSG00000227982 | 0 | 0 | 0 | 0 | 0 | 0 |
| ENSG00000235139 | 0 | 0 | 0 | 0 | 0 | 0 |

| | | | | | | |
|---|---|---|---|---|---|---|
| ENSG00000235243 | 0 | 0 | 0 | 0 | 0 | 0 |
| ENSG00000232605 | 0 | 0 | 0 | 0 | 0 | 0 |
| ENSG00000280325 | 0 | 0 | 0 | 0 | 1 | 0 |
| ENSG00000232019 | 0 | 0 | 0 | 0 | 0 | 0 |
| ENSG00000229251 | 0 | 0 | 0 | 1 | 0 | 2 |
| ENSG00000153993 | 114 | 65 | 62 | 14 | 74 | 59 |
| ENSG00000224059 | 0 | 0 | 0 | 0 | 0 | 0 |
| ENSG00000230614 | 0 | 0 | 0 | 0 | 0 | 0 |
| ENSG00000225128 | 0 | 0 | 0 | 0 | 0 | 0 |
| ENSG00000227785 | 0 | 0 | 0 | 0 | 0 | 0 |
| ENSG00000236610 | 0 | 0 | 0 | 0 | 0 | 0 |
| ENSG00000198822 | 0 | 0 | 0 | 0 | 0 | 3 |
| ENSG00000231255 | 0 | 0 | 0 | 0 | 0 | 0 |
| ENSG00000233073 | 2 | 0 | 0 | 0 | 0 | 0 |
| ENSG00000164659 | 975 | 495 | 578 | 275 | 429 | 425 |
| ENSG00000261462 | 0 | 0 | 0 | 0 | 0 | 1 |
| ENSG00000224046 | 42 | 14 | 23 | 16 | 18 | 23 |
| ENSG00000135164 | 1302 | 663 | 703 | 325 | 945 | 639 |
| ENSG00000278236 | 0 | 0 | 0 | 0 | 2 | 0 |
| ENSG00000135185 | 884 | 396 | 513 | 518 | 488 | 484 |
| ENSG00000200397 | 0 | 0 | 0 | 0 | 0 | 0 |
| ENSG00000182165 | 215 | 222 | 233 | 286 | 266 | 205 |
| ENSG00000273623 | 0 | 0 | 0 | 0 | 0 | 0 |
| ENSG00000276185 | 0 | 0 | 0 | 0 | 0 | 0 |
| ENSG00000005469 | 671 | 348 | 435 | 218 | 518 | 338 |
| ENSG00000005471 | 48 | 25 | 33 | 25 | 24 | 35 |
| ENSG00000085563 | 2 | 1 | 3 | 0 | 1 | 2 |
| ENSG00000231956 | 0 | 0 | 0 | 0 | 0 | 0 |
| ENSG00000105784 | 212 | 88 | 85 | 48 | 78 | 68 |
| ENSG00000075303 | 1355 | 685 | 746 | 434 | 697 | 631 |
| ENSG00000006634 | 526 | 307 | 286 | 234 | 432 | 319 |
| ENSG00000008277 | 427 | 195 | 248 | 116 | 225 | 163 |
| ENSG00000075142 | 1625 | 856 | 1113 | 922 | 945 | 935 |
| ENSG00000254003 | 0 | 0 | 3 | 0 | 0 | 1 |
| ENSG00000228113 | 4 | 0 | 0 | 0 | 0 | 0 |
| ENSG00000127954 | 13 | 8 | 15 | 5 | 2 | 4 |
| ENSG00000225701 | 0 | 0 | 0 | 0 | 0 | 0 |
| ENSG00000207094 | 0 | 0 | 0 | 0 | 0 | 0 |
| ENSG00000270812 | 0 | 0 | 0 | 0 | 0 | 0 |
| ENSG00000235639 | 0 | 0 | 0 | 0 | 0 | 0 |
| ENSG00000228928 | 0 | 0 | 0 | 0 | 0 | 0 |
| ENSG00000231763 | 0 | 0 | 0 | 0 | 0 | 0 |
| ENSG00000237709 | 0 | 0 | 0 | 0 | 0 | 0 |
| ENSG00000233420 | 0 | 0 | 0 | 0 | 0 | 0 |
| ENSG00000227979 | 0 | 0 | 0 | 0 | 0 | 0 |
| ENSG00000182348 | 12 | 5 | 11 | 1 | 6 | 3 |
| ENSG00000164645 | 0 | 1 | 1 | 0 | 0 | 0 |
| ENSG00000227863 | 0 | 0 | 0 | 0 | 0 | 0 |
| ENSG00000239075 | 0 | 0 | 0 | 0 | 0 | 0 |
| ENSG00000227646 | 0 | 0 | 0 | 0 | 0 | 0 |

| | | | | | | |
|---|---|---|---|---|---|---|
| ENSG00000238358 | 0 | 0 | 0 | 0 | 0 | 0 |
| ENSG00000235436 | 2 | 5 | 4 | 0 | 6 | 2 |
| ENSG00000164647 | 8 | 7 | 5 | 4 | 3 | 14 |
| ENSG00000157214 | 10 | 3 | 9 | 5 | 10 | 8 |
| ENSG00000105792 | 95 | 42 | 37 | 17 | 58 | 37 |
| ENSG00000207488 | 0 | 0 | 0 | 0 | 0 | 0 |
| ENSG00000234459 | 2 | 2 | 0 | 1 | 1 | 2 |
| ENSG00000225498 | 0 | 0 | 0 | 0 | 0 | 0 |
| ENSG00000283267 | 0 | 0 | 0 | 0 | 0 | 0 |
| ENSG00000105793 | 582 | 311 | 339 | 230 | 366 | 254 |
| ENSG00000280440 | 0 | 0 | 1 | 0 | 0 | 1 |
| ENSG00000157224 | 864 | 427 | 516 | 432 | 570 | 492 |
| ENSG00000273299 | 0 | 0 | 0 | 0 | 0 | 0 |
| ENSG00000058091 | 453 | 204 | 217 | 93 | 144 | 110 |
| ENSG00000251154 | 0 | 0 | 0 | 0 | 0 | 0 |
| ENSG00000223969 | 4 | 1 | 0 | 0 | 0 | 0 |
| ENSG00000223584 | 0 | 1 | 0 | 0 | 1 | 1 |
| ENSG00000226254 | 0 | 0 | 0 | 0 | 0 | 0 |
| ENSG00000157240 | 906 | 411 | 530 | 353 | 424 | 383 |
| ENSG00000243144 | 1 | 0 | 0 | 0 | 0 | 0 |
| ENSG00000225843 | 0 | 0 | 0 | 0 | 0 | 0 |
| ENSG00000235450 | 0 | 0 | 0 | 0 | 0 | 0 |
| ENSG00000223665 | 0 | 0 | 0 | 0 | 0 | 0 |
| ENSG00000127989 | 387 | 206 | 201 | 151 | 287 | 225 |
| ENSG00000127914 | 564 | 341 | 429 | 76 | 471 | 414 |
| ENSG00000001630 | 183 | 95 | 141 | 89 | 118 | 113 |
| ENSG00000285772 | 0 | 0 | 0 | 0 | 0 | 0 |
| ENSG00000285953 | 0 | 0 | 0 | 0 | 0 | 0 |
| ENSG00000188693 | 9 | 10 | 12 | 3 | 12 | 7 |
| ENSG00000240720 | 3 | 1 | 0 | 0 | 1 | 0 |
| ENSG00000001631 | 174 | 119 | 141 | 52 | 163 | 105 |
| ENSG00000243107 | 0 | 5 | 2 | 0 | 2 | 4 |
| ENSG00000200769 | 0 | 1 | 4 | 1 | 0 | 1 |
| ENSG00000221520 | 2 | 0 | 1 | 0 | 2 | 0 |
| ENSG00000001629 | 3032 | 1618 | 1677 | 848 | 1934 | 1428 |
| ENSG00000230927 | 0 | 0 | 1 | 0 | 0 | 0 |
| ENSG00000270453 | 0 | 0 | 0 | 0 | 0 | 0 |
| ENSG00000157259 | 2352 | 1265 | 1539 | 845 | 1563 | 1314 |
| ENSG00000244055 | 7 | 9 | 2 | 4 | 5 | 6 |
| ENSG00000242950 | 6 | 2 | 1 | 0 | 0 | 2 |
| ENSG00000127980 | 1206 | 605 | 795 | 378 | 719 | 669 |
| ENSG00000127993 | 685 | 271 | 306 | 251 | 316 | 347 |
| ENSG00000278819 | 0 | 1 | 0 | 1 | 0 | 0 |
| ENSG00000234545 | 723 | 375 | 365 | 308 | 388 | 395 |
| ENSG00000105810 | 1246 | 643 | 650 | 254 | 726 | 554 |
| ENSG00000206763 | 0 | 0 | 0 | 0 | 0 | 0 |
| ENSG00000237819 | 0 | 0 | 0 | 0 | 0 | 0 |
| ENSG00000266794 | 0 | 0 | 0 | 0 | 0 | 0 |
| ENSG00000205413 | 8 | 4 | 3 | 1 | 6 | 5 |
| ENSG00000177409 | 0 | 0 | 0 | 0 | 0 | 1 |

| | | | | | | |
|---|---|---|---|---|---|---|
| ENSG00000188175 | 0 | 1 | 1 | 3 | 0 | 0 |
| ENSG00000004766 | 326 | 225 | 204 | 104 | 373 | 198 |
| ENSG00000004948 | 68 | 37 | 33 | 29 | 38 | 29 |
| ENSG00000208014 | 0 | 0 | 0 | 0 | 0 | 0 |
| ENSG00000207656 | 0 | 0 | 0 | 0 | 0 | 0 |
| ENSG00000127928 | 95 | 49 | 49 | 40 | 71 | 52 |
| ENSG00000237729 | 6 | 3 | 0 | 2 | 0 | 4 |
| ENSG00000265423 | 0 | 0 | 0 | 0 | 0 | 0 |
| ENSG00000225898 | 0 | 0 | 0 | 0 | 0 | 0 |
| ENSG00000213483 | 0 | 0 | 0 | 0 | 0 | 1 |
| ENSG00000105825 | 38 | 28 | 11 | 23 | 25 | 23 |
| ENSG00000234695 | 6 | 6 | 9 | 5 | 6 | 7 |
| ENSG00000278388 | 0 | 0 | 0 | 0 | 0 | 0 |
| ENSG00000127920 | 0 | 0 | 0 | 0 | 0 | 0 |
| ENSG00000278959 | 0 | 0 | 0 | 0 | 0 | 0 |
| ENSG00000105829 | 536 | 242 | 329 | 257 | 289 | 291 |
| ENSG00000236861 | 3 | 2 | 2 | 0 | 0 | 0 |
| ENSG00000236453 | 0 | 0 | 0 | 0 | 0 | 0 |
| ENSG00000236938 | 0 | 0 | 0 | 0 | 0 | 0 |
| ENSG00000285090 | 0 | 0 | 0 | 0 | 0 | 0 |
| ENSG00000285964 | 0 | 0 | 0 | 0 | 0 | 0 |
| ENSG00000164692 | 0 | 0 | 0 | 1 | 0 | 0 |
| ENSG00000222870 | 0 | 0 | 0 | 0 | 0 | 0 |
| ENSG00000127995 | 915 | 461 | 525 | 279 | 526 | 413 |
| ENSG00000127990 | 28 | 5 | 11 | 3 | 18 | 15 |
| ENSG00000242265 | 0 | 1 | 2 | 1 | 0 | 0 |
| ENSG00000232385 | 0 | 0 | 0 | 0 | 0 | 0 |
| ENSG00000239030 | 0 | 0 | 0 | 0 | 0 | 0 |
| ENSG00000214342 | 0 | 0 | 0 | 0 | 0 | 0 |
| ENSG00000230682 | 0 | 0 | 0 | 0 | 0 | 0 |
| ENSG00000222220 | 0 | 0 | 0 | 0 | 0 | 0 |
| ENSG00000219807 | 0 | 0 | 0 | 0 | 0 | 0 |
| ENSG00000158528 | 144 | 65 | 60 | 22 | 107 | 43 |
| ENSG00000271159 | 0 | 0 | 0 | 0 | 0 | 0 |
| ENSG00000231153 | 0 | 0 | 0 | 0 | 0 | 0 |
| ENSG00000236197 | 0 | 0 | 0 | 0 | 0 | 0 |
| ENSG00000201607 | 0 | 0 | 0 | 0 | 0 | 0 |
| ENSG00000005421 | 0 | 0 | 0 | 0 | 0 | 0 |
| ENSG00000237160 | 0 | 0 | 0 | 0 | 0 | 0 |
| ENSG00000228751 | 0 | 0 | 0 | 0 | 0 | 0 |
| ENSG00000105852 | 0 | 0 | 0 | 0 | 0 | 0 |
| ENSG00000105854 | 401 | 272 | 226 | 218 | 313 | 222 |
| ENSG00000273138 | 0 | 0 | 0 | 0 | 0 | 0 |
| ENSG00000233942 | 0 | 0 | 0 | 0 | 0 | 0 |
| ENSG00000005981 | 0 | 0 | 0 | 0 | 0 | 0 |
| ENSG00000279525 | 0 | 0 | 0 | 0 | 0 | 0 |
| ENSG00000004799 | 194 | 69 | 82 | 65 | 111 | 64 |
| ENSG00000231170 | 4 | 1 | 2 | 2 | 3 | 2 |
| ENSG00000158560 | 2 | 5 | 4 | 0 | 1 | 1 |
| ENSG00000279067 | 0 | 0 | 0 | 0 | 0 | 0 |

| | | | | | | |
|---|---|---|---|---|---|---|
| ENSG00000004864 | 1679 | 844 | 991 | 870 | 1096 | 1026 |
| ENSG00000208025 | 0 | 0 | 1 | 0 | 0 | 0 |
| ENSG00000237551 | 0 | 0 | 0 | 0 | 0 | 0 |
| ENSG00000275834 | 0 | 0 | 0 | 0 | 0 | 0 |
| ENSG00000207045 | 0 | 0 | 0 | 0 | 0 | 0 |
| ENSG00000207115 | 0 | 0 | 0 | 0 | 0 | 0 |
| ENSG00000252872 | 0 | 0 | 0 | 0 | 0 | 0 |
| ENSG00000127922 | 2014 | 1059 | 1202 | 1170 | 1545 | 1192 |
| ENSG00000227284 | 0 | 0 | 0 | 0 | 0 | 0 |
| ENSG00000244318 | 0 | 0 | 0 | 0 | 0 | 0 |
| ENSG00000231764 | 0 | 1 | 0 | 0 | 1 | 0 |
| ENSG00000274409 | 0 | 0 | 0 | 0 | 0 | 0 |
| ENSG00000277124 | 0 | 0 | 0 | 0 | 0 | 0 |
| ENSG00000006377 | 0 | 0 | 0 | 0 | 0 | 0 |
| ENSG00000274709 | 0 | 0 | 0 | 0 | 0 | 0 |
| ENSG00000105880 | 0 | 1 | 1 | 1 | 1 | 1 |
| ENSG00000196636 | 0 | 0 | 0 | 0 | 0 | 0 |
| ENSG00000224551 | 0 | 0 | 0 | 0 | 0 | 0 |
| ENSG00000224475 | 0 | 0 | 0 | 0 | 0 | 0 |
| ENSG00000226046 | 0 | 0 | 0 | 0 | 0 | 0 |
| ENSG00000199475 | 0 | 0 | 0 | 0 | 0 | 0 |
| ENSG00000006128 | 0 | 0 | 0 | 0 | 0 | 0 |
| ENSG00000284707 | 4 | 2 | 8 | 10 | 3 | 5 |
| ENSG00000070669 | 29 | 8 | 13 | 12 | 7 | 14 |
| ENSG00000226744 | 0 | 0 | 0 | 1 | 0 | 1 |
| ENSG00000231812 | 0 | 0 | 0 | 0 | 0 | 0 |
| ENSG00000237818 | 6 | 0 | 0 | 0 | 0 | 0 |
| ENSG00000231859 | 2 | 0 | 0 | 0 | 0 | 0 |
| ENSG00000232032 | 2 | 0 | 0 | 0 | 0 | 0 |
| ENSG00000284627 | 0 | 0 | 0 | 0 | 0 | 0 |
| ENSG00000232097 | 0 | 0 | 1 | 0 | 0 | 0 |
| ENSG00000238228 | 6 | 1 | 3 | 2 | 2 | 1 |
| ENSG00000266318 | 0 | 0 | 0 | 0 | 0 | 0 |
| ENSG00000266668 | 1 | 0 | 0 | 0 | 0 | 0 |
| ENSG00000183444 | 5 | 3 | 5 | 5 | 9 | 9 |
| ENSG00000285725 | 23 | 21 | 44 | 35 | 36 | 37 |
| ENSG00000243554 | 8 | 16 | 14 | 15 | 13 | 19 |
| ENSG00000135175 | 0 | 0 | 0 | 0 | 0 | 0 |
| ENSG00000277946 | 0 | 0 | 0 | 0 | 0 | 0 |
| ENSG00000164715 | 2505 | 1055 | 1535 | 916 | 1282 | 1431 |
| ENSG00000180535 | 22 | 7 | 12 | 1 | 3 | 2 |
| ENSG00000205356 | 1320 | 542 | 749 | 561 | 598 | 637 |
| ENSG00000164713 | 992 | 465 | 605 | 613 | 591 | 498 |
| ENSG00000006453 | 985 | 543 | 554 | 638 | 787 | 693 |
| ENSG00000272950 | 2 | 0 | 0 | 0 | 1 | 1 |
| ENSG00000214389 | 108 | 28 | 35 | 51 | 53 | 32 |
| ENSG00000213455 | 0 | 0 | 0 | 0 | 0 | 0 |
| ENSG00000223402 | 0 | 0 | 0 | 0 | 0 | 0 |
| ENSG00000106236 | 0 | 0 | 0 | 0 | 0 | 0 |
| ENSG00000207204 | 0 | 0 | 0 | 0 | 0 | 0 |

| | | | | | | |
|---|---|---|---|---|---|---|
| ENSG00000166448 | 27 | 6 | 13 | 14 | 16 | 14 |
| ENSG00000196367 | 5417 | 2195 | 3231 | 1898 | 2790 | 2845 |
| ENSG00000266019 | 0 | 0 | 0 | 0 | 0 | 0 |
| ENSG00000238109 | 0 | 1 | 1 | 0 | 3 | 0 |
| ENSG00000198742 | 2292 | 1275 | 1603 | 1239 | 1619 | 1511 |
| ENSG00000284840 | 0 | 0 | 0 | 0 | 0 | 0 |
| ENSG00000185467 | 31 | 15 | 11 | 14 | 12 | 14 |
| ENSG00000002079 | 21 | 15 | 22 | 8 | 20 | 13 |
| ENSG00000284523 | 35 | 13 | 22 | 10 | 18 | 18 |
| ENSG00000284292 | 1 | 1 | 1 | 1 | 1 | 3 |
| ENSG00000241685 | 1247 | 932 | 1006 | 1114 | 1043 | 1022 |
| ENSG00000130429 | 397 | 281 | 345 | 328 | 316 | 272 |
| ENSG00000106244 | 5887 | 3029 | 3680 | 3680 | 3758 | 3759 |
| ENSG00000106245 | 1398 | 850 | 1055 | 1024 | 1109 | 1090 |
| ENSG00000106246 | 165 | 149 | 163 | 160 | 162 | 149 |
| ENSG00000248919 | 0 | 2 | 2 | 2 | 1 | 1 |
| ENSG00000160917 | 1733 | 833 | 989 | 1075 | 952 | 1088 |
| ENSG00000228335 | 13 | 4 | 12 | 6 | 7 | 8 |
| ENSG00000241468 | 3039 | 2003 | 2365 | 2530 | 2353 | 2193 |
| ENSG00000198556 | 761 | 401 | 475 | 318 | 440 | 398 |
| ENSG00000160908 | 764 | 397 | 454 | 348 | 431 | 430 |
| ENSG00000196652 | 734 | 318 | 446 | 362 | 430 | 398 |
| ENSG00000221909 | 362 | 189 | 181 | 113 | 199 | 137 |
| ENSG00000197343 | 0 | 0 | 0 | 1 | 0 | 0 |
| ENSG00000272647 | 0 | 0 | 0 | 0 | 0 | 0 |
| ENSG00000244219 | 0 | 0 | 0 | 0 | 0 | 0 |
| ENSG00000197037 | 907 | 395 | 478 | 405 | 425 | 492 |
| ENSG00000224448 | 4 | 2 | 2 | 0 | 3 | 0 |
| ENSG00000106258 | 20 | 9 | 15 | 13 | 12 | 10 |
| ENSG00000160870 | 1 | 0 | 5 | 0 | 0 | 1 |
| ENSG00000282277 | 0 | 0 | 0 | 0 | 0 | 0 |
| ENSG00000282301 | 0 | 0 | 0 | 0 | 0 | 0 |
| ENSG00000160868 | 2 | 0 | 0 | 1 | 1 | 1 |
| ENSG00000273407 | 0 | 0 | 0 | 0 | 1 | 0 |
| ENSG00000261511 | 0 | 0 | 0 | 0 | 0 | 0 |
| ENSG00000021461 | 2 | 0 | 0 | 0 | 0 | 2 |
| ENSG00000260445 | 0 | 0 | 0 | 0 | 0 | 0 |
| ENSG00000260524 | 0 | 0 | 0 | 0 | 0 | 0 |
| ENSG00000244623 | 0 | 0 | 0 | 0 | 0 | 0 |
| ENSG00000146833 | 1967 | 966 | 1098 | 899 | 1129 | 1068 |
| ENSG00000176402 | 10 | 3 | 2 | 8 | 4 | 8 |
| ENSG00000237640 | 0 | 0 | 0 | 1 | 0 | 0 |
| ENSG00000199711 | 0 | 0 | 0 | 0 | 0 | 0 |
| ENSG00000222966 | 0 | 0 | 0 | 0 | 0 | 0 |
| ENSG00000160862 | 91 | 38 | 68 | 70 | 39 | 67 |
| ENSG00000214313 | 10 | 4 | 3 | 4 | 7 | 3 |
| ENSG00000235713 | 2 | 0 | 0 | 0 | 0 | 0 |
| ENSG00000106261 | 5475 | 2779 | 2896 | 1265 | 3043 | 2400 |
| ENSG00000166529 | 216 | 129 | 119 | 113 | 154 | 158 |
| ENSG00000166526 | 2286 | 1171 | 1537 | 1222 | 1346 | 1446 |

| | | | | | | |
|---|---|---|---|---|---|---|
| ENSG00000168090 | 5659 | 2813 | 3278 | 3230 | 3018 | 3047 |
| ENSG00000166508 | 6223 | 3304 | 3980 | 3846 | 3377 | 4004 |
| ENSG00000207547 | 42 | 24 | 32 | 34 | 42 | 22 |
| ENSG00000207757 | 5 | 3 | 5 | 2 | 0 | 3 |
| ENSG00000208036 | 5 | 2 | 1 | 1 | 2 | 2 |
| ENSG00000221838 | 699 | 350 | 417 | 392 | 373 | 338 |
| ENSG00000106290 | 1741 | 778 | 978 | 918 | 831 | 959 |
| ENSG00000242798 | 28 | 9 | 26 | 5 | 19 | 14 |
| ENSG00000166997 | 373 | 182 | 187 | 181 | 179 | 206 |
| ENSG00000214309 | 48 | 28 | 38 | 25 | 32 | 35 |
| ENSG00000235077 | 1 | 0 | 0 | 0 | 0 | 1 |
| ENSG00000214142 | 0 | 0 | 0 | 0 | 0 | 0 |
| ENSG00000188186 | 2972 | 1417 | 1726 | 2112 | 1659 | 1647 |
| ENSG00000146826 | 1645 | 805 | 1139 | 971 | 1001 | 1023 |
| ENSG00000284518 | 0 | 0 | 0 | 0 | 0 | 0 |
| ENSG00000197093 | 29 | 16 | 17 | 15 | 18 | 17 |
| ENSG00000213420 | 62 | 33 | 57 | 45 | 60 | 56 |
| ENSG00000066923 | 302 | 177 | 280 | 107 | 202 | 175 |
| ENSG00000239521 | 1032 | 485 | 741 | 451 | 630 | 575 |
| ENSG00000213413 | 35 | 17 | 39 | 9 | 24 | 18 |
| ENSG00000214300 | 256 | 130 | 144 | 177 | 133 | 181 |
| ENSG00000078319 | 532 | 297 | 383 | 317 | 337 | 379 |
| ENSG00000201913 | 0 | 0 | 0 | 0 | 0 | 0 |
| ENSG00000242294 | 166 | 90 | 150 | 107 | 134 | 159 |
| ENSG00000272752 | 5 | 12 | 10 | 15 | 18 | 15 |
| ENSG00000121716 | 167 | 68 | 110 | 84 | 82 | 100 |
| ENSG00000235333 | 0 | 0 | 0 | 0 | 0 | 0 |
| ENSG00000284012 | 0 | 0 | 0 | 0 | 0 | 0 |
| ENSG00000085514 | 11 | 5 | 7 | 6 | 8 | 3 |
| ENSG00000078487 | 270 | 136 | 175 | 181 | 150 | 142 |
| ENSG00000146834 | 3274 | 1652 | 2055 | 1939 | 1705 | 1805 |
| ENSG00000241357 | 0 | 0 | 0 | 0 | 0 | 2 |
| ENSG00000160813 | 591 | 402 | 477 | 509 | 434 | 396 |
| ENSG00000240211 | 6 | 7 | 11 | 5 | 10 | 7 |
| ENSG00000185955 | 11 | 5 | 9 | 9 | 9 | 8 |
| ENSG00000166925 | 2799 | 1309 | 1716 | 1702 | 1440 | 1395 |
| ENSG00000166924 | 16 | 12 | 13 | 6 | 11 | 6 |
| ENSG00000225807 | 0 | 0 | 0 | 0 | 0 | 0 |
| ENSG00000242101 | 0 | 0 | 0 | 0 | 0 | 0 |
| ENSG00000106351 | 930 | 430 | 566 | 455 | 428 | 442 |
| ENSG00000184414 | 0 | 0 | 1 | 0 | 1 | 0 |
| ENSG00000205307 | 1 | 1 | 0 | 0 | 0 | 2 |
| ENSG00000274272 | 77 | 45 | 84 | 82 | 50 | 62 |
| ENSG00000077454 | 861 | 370 | 510 | 431 | 417 | 438 |
| ENSG00000106336 | 13 | 10 | 14 | 12 | 16 | 17 |
| ENSG00000224729 | 2 | 1 | 2 | 2 | 3 | 4 |
| ENSG00000106333 | 391 | 184 | 239 | 192 | 168 | 204 |
| ENSG00000106330 | 1266 | 492 | 778 | 756 | 540 | 649 |
| ENSG00000106327 | 78 | 21 | 31 | 44 | 38 | 38 |
| ENSG00000077080 | 2 | 2 | 2 | 0 | 2 | 3 |

| | | | | | | |
|---|---|---|---|---|---|---|
| ENSG00000172354 | 14415 | 6988 | 8965 | 9662 | 7753 | 7467 |
| ENSG00000146830 | 4556 | 1937 | 2734 | 1840 | 2126 | 2247 |
| ENSG00000172336 | 1757 | 911 | 1006 | 1088 | 934 | 891 |
| ENSG00000130427 | 2 | 0 | 1 | 0 | 1 | 1 |
| ENSG00000146839 | 0 | 1 | 0 | 0 | 2 | 0 |
| ENSG00000196411 | 5556 | 2573 | 3257 | 3037 | 2717 | 2712 |
| ENSG00000263672 | 0 | 0 | 1 | 0 | 1 | 0 |
| ENSG00000146828 | 3705 | 1660 | 2281 | 2024 | 2091 | 2074 |
| ENSG00000236305 | 25 | 13 | 10 | 20 | 25 | 22 |
| ENSG00000087077 | 4112 | 1905 | 2574 | 2850 | 2282 | 2778 |
| ENSG00000283821 | 0 | 0 | 0 | 0 | 0 | 0 |
| ENSG00000087087 | 4188 | 1751 | 2608 | 2304 | 2098 | 2480 |
| ENSG00000176125 | 181 | 72 | 98 | 94 | 68 | 85 |
| ENSG00000087085 | 273 | 158 | 183 | 89 | 100 | 94 |
| ENSG00000239825 | 0 | 0 | 0 | 0 | 0 | 0 |
| ENSG00000235926 | 0 | 0 | 0 | 0 | 0 | 0 |
| ENSG00000169894 | 99 | 61 | 102 | 52 | 53 | 58 |
| ENSG00000274993 | 0 | 0 | 0 | 1 | 0 | 0 |
| ENSG00000205277 | 0 | 0 | 0 | 2 | 1 | 0 |
| ENSG00000227053 | 1 | 0 | 0 | 1 | 1 | 2 |
| ENSG00000169876 | 0 | 0 | 0 | 0 | 0 | 0 |
| ENSG00000222636 | 0 | 0 | 0 | 0 | 0 | 0 |
| ENSG00000169871 | 4445 | 1937 | 2638 | 1548 | 2149 | 2108 |
| ENSG00000106366 | 2 | 0 | 1 | 2 | 1 | 0 |
| ENSG00000106367 | 9031 | 4524 | 5650 | 6608 | 6188 | 5541 |
| ENSG00000264425 | 24 | 10 | 19 | 21 | 21 | 12 |
| ENSG00000128564 | 294 | 142 | 183 | 268 | 211 | 198 |
| ENSG00000167011 | 2 | 1 | 1 | 5 | 3 | 4 |
| ENSG00000106384 | 0 | 0 | 0 | 0 | 1 | 0 |
| ENSG00000205267 | 0 | 0 | 0 | 1 | 2 | 0 |
| ENSG00000213394 | 1 | 0 | 1 | 1 | 0 | 1 |
| ENSG00000106397 | 4651 | 2191 | 2854 | 3043 | 2570 | 2743 |
| ENSG00000106400 | 4037 | 1954 | 2276 | 2777 | 2215 | 2155 |
| ENSG00000106404 | 379 | 130 | 214 | 225 | 176 | 199 |
| ENSG00000214253 | 5289 | 2617 | 3282 | 3474 | 2921 | 3036 |
| ENSG00000207454 | 0 | 0 | 0 | 0 | 0 | 0 |
| ENSG00000278683 | 0 | 0 | 0 | 2 | 0 | 0 |
| ENSG00000214252 | 0 | 0 | 0 | 0 | 0 | 0 |
| ENSG00000232301 | 0 | 0 | 0 | 0 | 0 | 0 |
| ENSG00000232445 | 76 | 45 | 44 | 76 | 81 | 77 |
| ENSG00000128581 | 1821 | 860 | 1126 | 914 | 1024 | 941 |
| ENSG00000160963 | 2 | 0 | 2 | 0 | 0 | 2 |
| ENSG00000233683 | 0 | 0 | 0 | 0 | 0 | 0 |
| ENSG00000233123 | 0 | 0 | 0 | 0 | 0 | 0 |
| ENSG00000106436 | 0 | 0 | 0 | 0 | 0 | 0 |
| ENSG00000257923 | 7210 | 3138 | 4511 | 3721 | 3961 | 4305 |
| ENSG00000259294 | 2 | 2 | 1 | 0 | 0 | 0 |
| ENSG00000272219 | 0 | 4 | 0 | 2 | 0 | 0 |
| ENSG00000261535 | 1 | 1 | 1 | 0 | 0 | 1 |
| ENSG00000280404 | 0 | 2 | 0 | 0 | 0 | 0 |

| | | | | | | |
|---|---|---|---|---|---|---|
| ENSG00000280004 | 0 | 0 | 0 | 0 | 1 | 0 |
| ENSG00000279724 | 0 | 0 | 0 | 0 | 0 | 0 |
| ENSG00000252824 | 0 | 0 | 0 | 0 | 0 | 0 |
| ENSG00000278586 | 0 | 0 | 0 | 0 | 0 | 0 |
| ENSG00000160999 | 119 | 92 | 98 | 108 | 100 | 116 |
| ENSG00000264675 | 0 | 1 | 0 | 0 | 0 | 1 |
| ENSG00000170409 | 6 | 5 | 3 | 0 | 1 | 4 |
| ENSG00000200552 | 0 | 0 | 0 | 0 | 0 | 0 |
| ENSG00000228546 | 0 | 1 | 0 | 1 | 0 | 0 |
| ENSG00000260097 | 20 | 39 | 22 | 11 | 19 | 8 |
| ENSG00000128563 | 1858 | 938 | 1165 | 1085 | 1055 | 992 |
| ENSG00000239969 | 3 | 6 | 1 | 3 | 4 | 4 |
| ENSG00000239486 | 6 | 8 | 6 | 0 | 0 | 2 |
| ENSG00000221510 | 0 | 0 | 0 | 1 | 0 | 0 |
| ENSG00000239480 | 1 | 3 | 3 | 4 | 0 | 5 |
| ENSG00000160991 | 2444 | 1262 | 1457 | 1321 | 1674 | 1394 |
| ENSG00000160993 | 859 | 425 | 476 | 546 | 457 | 436 |
| ENSG00000161036 | 1640 | 768 | 940 | 1054 | 898 | 869 |
| ENSG00000284176 | 0 | 0 | 0 | 0 | 0 | 0 |
| ENSG00000284596 | 0 | 0 | 0 | 0 | 0 | 0 |
| ENSG00000005075 | 2801 | 1302 | 1529 | 1716 | 1528 | 1440 |
| ENSG00000170667 | 8 | 1 | 0 | 5 | 0 | 3 |
| ENSG00000272949 | 3 | 0 | 0 | 0 | 0 | 0 |
| ENSG00000168255 | 152 | 125 | 132 | 108 | 103 | 88 |
| ENSG00000284981 | 1 | 1 | 0 | 0 | 0 | 0 |
| ENSG00000270249 | 0 | 0 | 0 | 0 | 0 | 0 |
| ENSG00000205238 | 48 | 37 | 35 | 12 | 17 | 11 |
| ENSG00000285437 | 68 | 60 | 32 | 22 | 32 | 26 |
| ENSG00000105808 | 45 | 17 | 35 | 31 | 35 | 21 |
| ENSG00000279168 | 17 | 5 | 6 | 6 | 3 | 6 |
| ENSG00000205236 | 0 | 0 | 1 | 0 | 1 | 0 |
| ENSG00000267368 | 0 | 0 | 0 | 0 | 0 | 0 |
| ENSG00000267645 | 0 | 1 | 0 | 0 | 0 | 0 |
| ENSG00000173678 | 5 | 3 | 4 | 1 | 2 | 2 |
| ENSG00000228049 | 13 | 23 | 15 | 7 | 10 | 8 |
| ENSG00000233297 | 14 | 6 | 6 | 3 | 6 | 7 |
| ENSG00000222011 | 525 | 316 | 345 | 250 | 298 | 311 |
| ENSG00000213385 | 28 | 8 | 15 | 9 | 12 | 25 |
| ENSG00000161040 | 10 | 4 | 10 | 2 | 13 | 9 |
| ENSG00000252643 | 2 | 0 | 0 | 0 | 2 | 0 |
| ENSG00000238324 | 2 | 0 | 0 | 0 | 0 | 0 |
| ENSG00000128606 | 11 | 3 | 6 | 2 | 13 | 14 |
| ENSG00000230257 | 0 | 0 | 0 | 0 | 0 | 0 |
| ENSG00000236226 | 0 | 0 | 0 | 0 | 0 | 0 |
| ENSG00000170632 | 2133 | 1042 | 1265 | 1098 | 1254 | 1175 |
| ENSG00000233025 | 12 | 7 | 7 | 4 | 7 | 10 |
| ENSG00000161048 | 837 | 370 | 468 | 252 | 424 | 364 |
| ENSG00000224415 | 2 | 3 | 0 | 0 | 0 | 1 |
| ENSG00000236848 | 2 | 1 | 0 | 0 | 4 | 3 |
| ENSG00000279482 | 0 | 2 | 0 | 0 | 0 | 0 |

| | | | | | | |
|---|---|---|---|---|---|---|
| ENSG00000170629 | 4 | 8 | 8 | 3 | 5 | 9 |
| ENSG00000237632 | 1 | 1 | 0 | 0 | 0 | 0 |
| ENSG00000105819 | 3570 | 1824 | 2360 | 1375 | 2218 | 2101 |
| ENSG00000105821 | 643 | 458 | 317 | 249 | 828 | 407 |
| ENSG00000161057 | 6146 | 3255 | 3861 | 3306 | 4002 | 3673 |
| ENSG00000235354 | 0 | 2 | 4 | 0 | 4 | 2 |
| ENSG00000170615 | 6 | 3 | 2 | 0 | 7 | 4 |
| ENSG00000251852 | 0 | 0 | 0 | 0 | 0 | 0 |
| ENSG00000252047 | 0 | 0 | 0 | 0 | 0 | 0 |
| ENSG00000234715 | 2 | 0 | 0 | 0 | 0 | 0 |
| ENSG00000189056 | 0 | 0 | 0 | 0 | 0 | 0 |
| ENSG00000222386 | 0 | 0 | 0 | 0 | 0 | 0 |
| ENSG00000164815 | 1675 | 850 | 1034 | 775 | 890 | 844 |
| ENSG00000187416 | 4 | 0 | 0 | 0 | 0 | 1 |
| ENSG00000197258 | 91 | 74 | 73 | 75 | 65 | 82 |
| ENSG00000226869 | 0 | 0 | 0 | 0 | 0 | 0 |
| ENSG00000237606 | 0 | 0 | 1 | 0 | 0 | 0 |
| ENSG00000225329 | 20 | 10 | 14 | 5 | 9 | 11 |
| ENSG00000243352 | 0 | 0 | 0 | 0 | 0 | 0 |
| ENSG00000237513 | 4 | 1 | 3 | 0 | 0 | 2 |
| ENSG00000228393 | 130 | 64 | 85 | 31 | 66 | 46 |
| ENSG00000239569 | 108 | 48 | 63 | 62 | 64 | 57 |
| ENSG00000005483 | 1351 | 744 | 650 | 273 | 932 | 578 |
| ENSG00000272918 | 14 | 5 | 15 | 2 | 4 | 8 |
| ENSG00000135250 | 3401 | 1671 | 2048 | 1138 | 1791 | 1582 |
| ENSG00000270764 | 0 | 0 | 0 | 0 | 0 | 0 |
| ENSG00000271482 | 0 | 0 | 0 | 0 | 0 | 0 |
| ENSG00000213361 | 0 | 0 | 0 | 0 | 0 | 0 |
| ENSG00000244490 | 2 | 2 | 1 | 0 | 0 | 0 |
| ENSG00000242154 | 5 | 5 | 5 | 2 | 2 | 3 |
| ENSG00000201179 | 0 | 0 | 0 | 0 | 0 | 0 |
| ENSG00000091127 | 1128 | 545 | 637 | 514 | 668 | 765 |
| ENSG00000223886 | 6 | 5 | 1 | 4 | 2 | 1 |
| ENSG00000135249 | 902 | 530 | 533 | 360 | 717 | 510 |
| ENSG00000185055 | 6 | 1 | 8 | 3 | 12 | 6 |
| ENSG00000272604 | 38 | 18 | 23 | 19 | 22 | 20 |
| ENSG00000231167 | 4 | 4 | 6 | 8 | 7 | 9 |
| ENSG00000146776 | 345 | 164 | 202 | 133 | 165 | 169 |
| ENSG00000226624 | 0 | 0 | 0 | 0 | 1 | 0 |
| ENSG00000128536 | 22 | 21 | 18 | 10 | 27 | 17 |
| ENSG00000280439 | 0 | 0 | 0 | 0 | 0 | 0 |
| ENSG00000008282 | 6218 | 3026 | 3578 | 2689 | 3411 | 3376 |
| ENSG00000242562 | 8 | 2 | 5 | 2 | 4 | 1 |
| ENSG00000201796 | 0 | 0 | 0 | 0 | 0 | 0 |
| ENSG00000105835 | 2769 | 1535 | 1786 | 1203 | 1884 | 1565 |
| ENSG00000273320 | 1 | 1 | 1 | 1 | 5 | 7 |
| ENSG00000243438 | 2 | 0 | 0 | 0 | 0 | 0 |
| ENSG00000243797 | 0 | 0 | 0 | 0 | 0 | 0 |
| ENSG00000267052 | 0 | 0 | 0 | 0 | 0 | 0 |
| ENSG00000253276 | 438 | 182 | 188 | 219 | 260 | 208 |

| | | | | | | |
|---|---|---|---|---|---|---|
| ENSG00000228742 | 0 | 0 | 0 | 0 | 0 | 0 |
| ENSG00000251978 | 0 | 0 | 0 | 0 | 0 | 0 |
| ENSG00000105851 | 0 | 1 | 0 | 0 | 0 | 0 |
| ENSG00000005249 | 376 | 195 | 191 | 154 | 183 | 174 |
| ENSG00000243193 | 0 | 1 | 0 | 0 | 0 | 0 |
| ENSG00000105856 | 2079 | 1107 | 1288 | 777 | 1077 | 902 |
| ENSG00000272072 | 7 | 4 | 6 | 0 | 2 | 5 |
| ENSG00000164597 | 2122 | 1036 | 1095 | 768 | 1211 | 982 |
| ENSG00000228341 | 0 | 0 | 0 | 0 | 0 | 0 |
| ENSG00000172209 | 0 | 1 | 0 | 0 | 2 | 0 |
| ENSG00000105865 | 581 | 304 | 339 | 263 | 367 | 340 |
| ENSG00000272854 | 0 | 0 | 4 | 1 | 5 | 1 |
| ENSG00000075790 | 1197 | 635 | 607 | 499 | 791 | 555 |
| ENSG00000238832 | 3 | 0 | 0 | 0 | 3 | 0 |
| ENSG00000250474 | 2 | 1 | 11 | 7 | 10 | 3 |
| ENSG00000225935 | 0 | 0 | 0 | 0 | 0 | 0 |
| ENSG00000233705 | 0 | 1 | 0 | 0 | 0 | 0 |
| ENSG00000091137 | 6 | 1 | 3 | 1 | 2 | 1 |
| ENSG00000241764 | 73 | 51 | 74 | 64 | 55 | 63 |
| ENSG00000105879 | 1269 | 630 | 730 | 598 | 757 | 747 |
| ENSG00000091138 | 0 | 1 | 0 | 0 | 0 | 0 |
| ENSG00000229626 | 0 | 0 | 0 | 0 | 0 | 0 |
| ENSG00000091140 | 2746 | 1594 | 1627 | 957 | 2252 | 1669 |
| ENSG00000091136 | 65 | 26 | 34 | 17 | 25 | 14 |
| ENSG00000273055 | 0 | 0 | 0 | 0 | 0 | 0 |
| ENSG00000238297 | 0 | 0 | 0 | 0 | 0 | 0 |
| ENSG00000091128 | 2 | 1 | 0 | 1 | 0 | 3 |
| ENSG00000091129 | 6570 | 3580 | 3370 | 1609 | 4149 | 2762 |
| ENSG00000135241 | 1213 | 643 | 671 | 398 | 835 | 586 |
| ENSG00000231006 | 0 | 3 | 0 | 3 | 0 | 0 |
| ENSG00000177683 | 802 | 416 | 462 | 289 | 450 | 433 |
| ENSG00000128590 | 893 | 431 | 513 | 354 | 486 | 460 |
| ENSG00000225647 | 0 | 0 | 0 | 0 | 0 | 1 |
| ENSG00000270425 | 0 | 0 | 0 | 0 | 0 | 0 |
| ENSG00000205174 | 0 | 0 | 0 | 0 | 1 | 0 |
| ENSG00000279043 | 0 | 0 | 0 | 0 | 0 | 1 |
| ENSG00000229603 | 0 | 0 | 0 | 0 | 0 | 0 |
| ENSG00000227968 | 0 | 0 | 0 | 0 | 0 | 0 |
| ENSG00000230941 | 0 | 0 | 0 | 0 | 0 | 0 |
| ENSG00000234273 | 0 | 0 | 0 | 0 | 0 | 0 |
| ENSG00000230192 | 0 | 0 | 0 | 0 | 0 | 0 |
| ENSG00000271368 | 0 | 0 | 0 | 0 | 0 | 1 |
| ENSG00000237064 | 0 | 0 | 0 | 0 | 0 | 0 |
| ENSG00000232199 | 0 | 0 | 0 | 0 | 0 | 0 |
| ENSG00000278424 | 0 | 0 | 0 | 0 | 0 | 0 |
| ENSG00000226965 | 9 | 0 | 4 | 5 | 5 | 5 |
| ENSG00000184903 | 117 | 96 | 106 | 97 | 99 | 83 |
| ENSG00000228540 | 0 | 0 | 1 | 0 | 0 | 0 |
| ENSG00000173114 | 2 | 0 | 0 | 0 | 0 | 2 |
| ENSG00000243621 | 0 | 0 | 0 | 0 | 0 | 0 |

| | | | | | | |
|---|---|---|---|---|---|---|
| ENSG00000227948 | 0 | 0 | 0 | 0 | 0 | 0 |
| ENSG00000128512 | 58 | 31 | 17 | 14 | 24 | 18 |
| ENSG00000225572 | 0 | 0 | 0 | 0 | 0 | 0 |
| ENSG00000222346 | 0 | 0 | 0 | 0 | 0 | 0 |
| ENSG00000234358 | 0 | 0 | 0 | 0 | 0 | 0 |
| ENSG00000198839 | 740 | 438 | 338 | 303 | 555 | 412 |
| ENSG00000202406 | 0 | 0 | 0 | 0 | 0 | 0 |
| ENSG00000226851 | 4 | 2 | 2 | 0 | 0 | 0 |
| ENSG00000230999 | 2 | 0 | 0 | 0 | 0 | 0 |
| ENSG00000232922 | 0 | 0 | 0 | 0 | 0 | 0 |
| ENSG00000236232 | 0 | 0 | 0 | 0 | 0 | 0 |
| ENSG00000006652 | 1400 | 658 | 833 | 779 | 965 | 979 |
| ENSG00000180019 | 0 | 1 | 0 | 1 | 1 | 0 |
| ENSG00000181016 | 48 | 28 | 33 | 23 | 34 | 25 |
| ENSG00000182965 | 0 | 0 | 0 | 0 | 0 | 0 |
| ENSG00000223646 | 0 | 0 | 0 | 0 | 0 | 0 |
| ENSG00000225358 | 0 | 0 | 0 | 0 | 0 | 0 |
| ENSG00000146802 | 985 | 401 | 522 | 379 | 640 | 510 |
| ENSG00000164603 | 467 | 208 | 242 | 164 | 279 | 243 |
| ENSG00000234520 | 0 | 0 | 0 | 0 | 0 | 0 |
| ENSG00000279288 | 0 | 0 | 0 | 0 | 0 | 0 |
| ENSG00000164604 | 0 | 0 | 0 | 0 | 0 | 0 |
| ENSG00000225457 | 0 | 0 | 0 | 0 | 0 | 0 |
| ENSG00000214194 | 517 | 233 | 356 | 346 | 387 | 387 |
| ENSG00000270997 | 0 | 0 | 0 | 0 | 0 | 0 |
| ENSG00000278894 | 0 | 0 | 0 | 0 | 0 | 0 |
| ENSG00000154415 | 2 | 0 | 0 | 2 | 1 | 0 |
| ENSG00000128573 | 0 | 0 | 0 | 1 | 1 | 0 |
| ENSG00000236580 | 0 | 0 | 0 | 0 | 0 | 0 |
| ENSG00000224595 | 0 | 0 | 0 | 0 | 0 | 0 |
| ENSG00000282859 | 0 | 0 | 0 | 0 | 0 | 0 |
| ENSG00000202334 | 0 | 0 | 0 | 0 | 0 | 0 |
| ENSG00000272230 | 0 | 0 | 0 | 0 | 0 | 0 |
| ENSG00000135272 | 0 | 0 | 0 | 0 | 1 | 0 |
| ENSG00000233607 | 0 | 0 | 0 | 0 | 0 | 0 |
| ENSG00000225535 | 0 | 0 | 0 | 0 | 0 | 0 |
| ENSG00000226962 | 0 | 0 | 0 | 0 | 0 | 0 |
| ENSG00000199764 | 0 | 0 | 0 | 0 | 0 | 0 |
| ENSG00000244565 | 0 | 0 | 0 | 0 | 0 | 0 |
| ENSG00000202377 | 0 | 0 | 0 | 0 | 0 | 0 |
| ENSG00000242072 | 0 | 0 | 0 | 0 | 0 | 0 |
| ENSG00000240973 | 0 | 0 | 0 | 0 | 0 | 0 |
| ENSG00000243345 | 0 | 0 | 0 | 0 | 0 | 0 |
| ENSG00000199224 | 0 | 0 | 0 | 0 | 0 | 0 |
| ENSG00000105967 | 0 | 0 | 0 | 0 | 0 | 0 |
| ENSG00000279086 | 0 | 0 | 1 | 0 | 0 | 1 |
| ENSG00000135269 | 1043 | 532 | 642 | 479 | 658 | 596 |
| ENSG00000243243 | 0 | 2 | 3 | 0 | 2 | 4 |
| ENSG00000237813 | 3 | 5 | 1 | 4 | 8 | 3 |
| ENSG00000237870 | 0 | 0 | 0 | 0 | 0 | 0 |

| | | | | | | |
|---|---|---|---|---|---|---|
| ENSG00000105971 | 532 | 272 | 339 | 332 | 421 | 380 |
| ENSG00000213167 | 0 | 0 | 0 | 0 | 1 | 0 |
| ENSG00000105974 | 392 | 254 | 210 | 220 | 235 | 201 |
| ENSG00000235427 | 0 | 0 | 0 | 0 | 0 | 0 |
| ENSG00000243220 | 0 | 0 | 0 | 0 | 0 | 0 |
| ENSG00000231210 | 0 | 1 | 0 | 0 | 0 | 0 |
| ENSG00000105976 | 831 | 380 | 494 | 257 | 351 | 330 |
| ENSG00000198898 | 2356 | 1340 | 1357 | 979 | 1701 | 1122 |
| ENSG00000235945 | 0 | 0 | 0 | 0 | 0 | 0 |
| ENSG00000252115 | 0 | 0 | 0 | 0 | 0 | 0 |
| ENSG00000222150 | 0 | 0 | 0 | 0 | 0 | 0 |
| ENSG00000227199 | 25 | 7 | 21 | 11 | 6 | 12 |
| ENSG00000275359 | 0 | 0 | 0 | 0 | 0 | 0 |
| ENSG00000277583 | 0 | 0 | 0 | 0 | 0 | 0 |
| ENSG00000004866 | 425 | 223 | 294 | 252 | 273 | 234 |
| ENSG00000214188 | 0 | 1 | 2 | 1 | 0 | 0 |
| ENSG00000273596 | 0 | 0 | 0 | 0 | 0 | 0 |
| ENSG00000273571 | 0 | 0 | 0 | 0 | 0 | 0 |
| ENSG00000276100 | 0 | 0 | 0 | 0 | 0 | 0 |
| ENSG00000274606 | 0 | 0 | 0 | 0 | 0 | 0 |
| ENSG00000228368 | 0 | 0 | 0 | 0 | 0 | 0 |
| ENSG00000213050 | 0 | 0 | 0 | 0 | 0 | 0 |
| ENSG00000283923 | 0 | 0 | 0 | 0 | 0 | 0 |
| ENSG00000226367 | 0 | 0 | 1 | 0 | 0 | 1 |
| ENSG00000274344 | 0 | 0 | 0 | 0 | 0 | 0 |
| ENSG00000275054 | 0 | 0 | 0 | 0 | 0 | 0 |
| ENSG00000227532 | 0 | 0 | 0 | 0 | 1 | 0 |
| ENSG00000278526 | 0 | 0 | 0 | 0 | 0 | 0 |
| ENSG00000278482 | 0 | 0 | 0 | 0 | 0 | 0 |
| ENSG00000230785 | 0 | 0 | 0 | 0 | 0 | 0 |
| ENSG00000224042 | 0 | 0 | 0 | 0 | 0 | 0 |
| ENSG00000230605 | 0 | 0 | 0 | 0 | 0 | 0 |
| ENSG00000105989 | 0 | 0 | 0 | 0 | 0 | 0 |
| ENSG00000238202 | 0 | 0 | 0 | 0 | 0 | 0 |
| ENSG00000154438 | 2 | 0 | 0 | 0 | 8 | 1 |
| ENSG00000214684 | 0 | 0 | 0 | 0 | 0 | 0 |
| ENSG00000001626 | 0 | 0 | 1 | 0 | 0 | 1 |
| ENSG00000237974 | 0 | 0 | 0 | 0 | 0 | 0 |
| ENSG00000232661 | 0 | 0 | 0 | 0 | 0 | 0 |
| ENSG00000234001 | 0 | 0 | 0 | 0 | 0 | 0 |
| ENSG00000083622 | 0 | 0 | 0 | 0 | 0 | 0 |
| ENSG00000077063 | 0 | 0 | 0 | 0 | 3 | 0 |
| ENSG00000224136 | 0 | 0 | 0 | 0 | 0 | 0 |
| ENSG00000234826 | 0 | 0 | 0 | 0 | 0 | 0 |
| ENSG00000128534 | 850 | 396 | 478 | 436 | 501 | 482 |
| ENSG00000106013 | 0 | 0 | 0 | 0 | 1 | 0 |
| ENSG00000229982 | 0 | 0 | 0 | 0 | 0 | 0 |
| ENSG00000230520 | 0 | 0 | 0 | 0 | 0 | 0 |
| ENSG00000270516 | 0 | 0 | 0 | 0 | 0 | 0 |
| ENSG00000225546 | 0 | 0 | 0 | 1 | 0 | 0 |

| | | | | | | |
|---|---|---|---|---|---|---|
| ENSG00000271739 | 0 | 0 | 0 | 0 | 0 | 0 |
| ENSG00000233417 | 0 | 0 | 0 | 0 | 0 | 0 |
| ENSG00000233969 | 0 | 0 | 0 | 0 | 0 | 0 |
| ENSG00000184408 | 0 | 0 | 0 | 0 | 0 | 0 |
| ENSG00000252023 | 0 | 0 | 0 | 0 | 0 | 0 |
| ENSG00000231295 | 0 | 0 | 0 | 0 | 0 | 0 |
| ENSG00000106025 | 110 | 59 | 63 | 33 | 47 | 43 |
| ENSG00000071243 | 363 | 179 | 202 | 148 | 177 | 172 |
| ENSG00000202225 | 0 | 0 | 0 | 0 | 0 | 0 |
| ENSG00000106034 | 0 | 0 | 0 | 0 | 0 | 1 |
| ENSG00000234927 | 0 | 0 | 0 | 0 | 0 | 0 |
| ENSG00000212628 | 0 | 0 | 0 | 0 | 0 | 0 |
| ENSG00000207090 | 0 | 0 | 0 | 0 | 0 | 0 |
| ENSG00000002745 | 6 | 0 | 2 | 0 | 0 | 0 |
| ENSG00000196937 | 793 | 439 | 573 | 361 | 473 | 493 |
| ENSG00000213310 | 0 | 0 | 0 | 0 | 0 | 0 |
| ENSG00000227371 | 0 | 0 | 0 | 0 | 0 | 0 |
| ENSG00000234985 | 0 | 1 | 0 | 0 | 0 | 0 |
| ENSG00000227743 | 0 | 0 | 0 | 0 | 0 | 0 |
| ENSG00000252704 | 0 | 0 | 0 | 0 | 0 | 0 |
| ENSG00000233272 | 0 | 0 | 0 | 0 | 0 | 0 |
| ENSG00000106278 | 0 | 0 | 0 | 0 | 0 | 0 |
| ENSG00000008311 | 37 | 26 | 16 | 4 | 11 | 10 |
| ENSG00000252003 | 0 | 0 | 0 | 0 | 0 | 0 |
| ENSG00000226636 | 0 | 0 | 0 | 0 | 0 | 0 |
| ENSG00000234418 | 0 | 0 | 0 | 0 | 0 | 0 |
| ENSG00000213302 | 2 | 0 | 0 | 0 | 0 | 0 |
| ENSG00000128610 | 2 | 0 | 0 | 0 | 0 | 0 |
| ENSG00000230316 | 0 | 1 | 6 | 0 | 3 | 0 |
| ENSG00000081803 | 314 | 125 | 145 | 60 | 136 | 73 |
| ENSG00000240499 | 0 | 0 | 1 | 0 | 0 | 0 |
| ENSG00000225795 | 0 | 0 | 0 | 0 | 0 | 0 |
| ENSG00000235459 | 0 | 0 | 0 | 0 | 0 | 0 |
| ENSG00000188050 | 4 | 0 | 0 | 3 | 1 | 0 |
| ENSG00000235631 | 0 | 4 | 1 | 2 | 0 | 0 |
| ENSG00000227573 | 0 | 0 | 0 | 0 | 0 | 1 |
| ENSG00000128519 | 0 | 0 | 0 | 0 | 0 | 0 |
| ENSG00000226680 | 0 | 0 | 0 | 0 | 0 | 0 |
| ENSG00000081800 | 0 | 0 | 0 | 0 | 0 | 0 |
| ENSG00000223595 | 0 | 0 | 0 | 0 | 0 | 0 |
| ENSG00000164675 | 19 | 6 | 1 | 0 | 6 | 6 |
| ENSG00000232524 | 0 | 2 | 2 | 0 | 0 | 0 |
| ENSG00000207338 | 0 | 0 | 0 | 0 | 0 | 0 |
| ENSG00000128609 | 1720 | 899 | 987 | 812 | 1110 | 894 |
| ENSG00000146809 | 0 | 0 | 0 | 0 | 0 | 0 |
| ENSG00000230442 | 0 | 0 | 0 | 0 | 0 | 0 |
| ENSG00000170807 | 0 | 0 | 0 | 0 | 0 | 0 |
| ENSG00000106299 | 3327 | 1520 | 1788 | 1058 | 1515 | 1530 |
| ENSG00000272686 | 262 | 138 | 131 | 122 | 148 | 143 |
| ENSG00000201104 | 0 | 0 | 0 | 0 | 0 | 0 |

| | | | | | | |
|---|---|---|---|---|---|---|
| ENSG00000228211 | 0 | 0 | 0 | 0 | 0 | 0 |
| ENSG00000106302 | 2 | 0 | 1 | 0 | 0 | 1 |
| ENSG00000106304 | 29 | 12 | 8 | 10 | 12 | 13 |
| ENSG00000225583 | 0 | 0 | 0 | 0 | 0 | 0 |
| ENSG00000241345 | 0 | 0 | 0 | 0 | 0 | 0 |
| ENSG00000234224 | 8 | 3 | 11 | 6 | 1 | 6 |
| ENSG00000242593 | 0 | 0 | 0 | 0 | 0 | 0 |
| ENSG00000243574 | 0 | 0 | 0 | 0 | 0 | 0 |
| ENSG00000241324 | 0 | 0 | 0 | 0 | 0 | 0 |
| ENSG00000230268 | 0 | 0 | 0 | 0 | 0 | 0 |
| ENSG00000213296 | 0 | 0 | 0 | 0 | 0 | 0 |
| ENSG00000207214 | 0 | 0 | 0 | 0 | 0 | 0 |
| ENSG00000279419 | 4 | 0 | 0 | 0 | 0 | 1 |
| ENSG00000170775 | 239 | 96 | 113 | 63 | 80 | 81 |
| ENSG00000275356 | 0 | 0 | 0 | 0 | 0 | 0 |
| ENSG00000128513 | 578 | 331 | 301 | 183 | 459 | 339 |
| ENSG00000224897 | 132 | 45 | 56 | 32 | 59 | 62 |
| ENSG00000186676 | 124 | 46 | 60 | 97 | 78 | 82 |
| ENSG00000224899 | 65 | 41 | 61 | 49 | 50 | 45 |
| ENSG00000205898 | 0 | 0 | 0 | 0 | 0 | 0 |
| ENSG00000237764 | 1 | 0 | 3 | 0 | 3 | 1 |
| ENSG00000219445 | 0 | 0 | 0 | 0 | 0 | 0 |
| ENSG00000213291 | 0 | 0 | 0 | 0 | 0 | 1 |
| ENSG00000234071 | 0 | 0 | 0 | 0 | 0 | 0 |
| ENSG00000227847 | 0 | 0 | 0 | 0 | 0 | 0 |
| ENSG00000227869 | 0 | 0 | 0 | 0 | 0 | 0 |
| ENSG00000197462 | 0 | 0 | 0 | 0 | 0 | 0 |
| ENSG00000241921 | 0 | 0 | 0 | 0 | 0 | 0 |
| ENSG00000179603 | 0 | 0 | 0 | 0 | 0 | 0 |
| ENSG00000227249 | 0 | 0 | 0 | 0 | 0 | 0 |
| ENSG00000224981 | 0 | 0 | 0 | 0 | 0 | 0 |
| ENSG00000230820 | 0 | 0 | 0 | 0 | 0 | 0 |
| ENSG00000207692 | 0 | 0 | 0 | 0 | 0 | 0 |
| ENSG00000236340 | 0 | 0 | 0 | 0 | 0 | 0 |
| ENSG00000236105 | 0 | 0 | 1 | 0 | 0 | 0 |
| ENSG00000048405 | 466 | 231 | 229 | 173 | 339 | 264 |
| ENSG00000279265 | 0 | 1 | 4 | 0 | 2 | 3 |
| ENSG00000224138 | 3 | 5 | 0 | 1 | 17 | 3 |
| ENSG00000280347 | 17 | 18 | 12 | 5 | 21 | 18 |
| ENSG00000226770 | 0 | 0 | 1 | 1 | 0 | 0 |
| ENSG00000179562 | 1231 | 585 | 780 | 635 | 717 | 663 |
| ENSG00000004059 | 5013 | 2646 | 3026 | 3422 | 2946 | 2702 |
| ENSG00000106328 | 1 | 0 | 2 | 0 | 3 | 0 |
| ENSG00000106331 | 0 | 0 | 0 | 0 | 0 | 0 |
| ENSG00000240790 | 0 | 0 | 0 | 0 | 0 | 0 |
| ENSG00000197157 | 6697 | 3142 | 4008 | 3962 | 3660 | 3683 |
| ENSG00000240579 | 0 | 0 | 0 | 0 | 0 | 0 |
| ENSG00000279078 | 0 | 2 | 0 | 0 | 1 | 0 |
| ENSG00000128594 | 15 | 15 | 8 | 10 | 12 | 20 |
| ENSG00000207588 | 0 | 0 | 0 | 0 | 2 | 0 |

| | | | | | | |
|---|---|---|---|---|---|---|
| ENSG00000207705 | 0 | 0 | 0 | 0 | 0 | 0 |
| ENSG00000174697 | 0 | 0 | 0 | 0 | 0 | 0 |
| ENSG00000272915 | 0 | 0 | 2 | 1 | 1 | 0 |
| ENSG00000106344 | 1355 | 621 | 769 | 615 | 761 | 744 |
| ENSG00000242261 | 0 | 0 | 0 | 0 | 0 | 0 |
| ENSG00000238750 | 0 | 0 | 0 | 0 | 0 | 0 |
| ENSG00000224940 | 92 | 35 | 68 | 31 | 43 | 39 |
| ENSG00000106348 | 3927 | 1715 | 2293 | 2111 | 1813 | 1818 |
| ENSG00000228700 | 2 | 4 | 5 | 4 | 3 | 2 |
| ENSG00000238590 | 0 | 0 | 0 | 0 | 0 | 0 |
| ENSG00000273184 | 0 | 0 | 1 | 0 | 0 | 0 |
| ENSG00000135245 | 3011 | 1700 | 2101 | 1217 | 1346 | 1502 |
| ENSG00000240758 | 9 | 10 | 12 | 12 | 14 | 5 |
| ENSG00000272601 | 0 | 0 | 0 | 0 | 0 | 0 |
| ENSG00000165055 | 710 | 412 | 411 | 379 | 491 | 483 |
| ENSG00000276988 | 2 | 1 | 2 | 1 | 1 | 2 |
| ENSG00000273270 | 1021 | 551 | 692 | 558 | 566 | 605 |
| ENSG00000280828 | 15 | 8 | 14 | 1 | 6 | 10 |
| ENSG00000213280 | 5 | 1 | 0 | 0 | 1 | 2 |
| ENSG00000242588 | 89 | 68 | 99 | 49 | 67 | 99 |
| ENSG00000281896 | 0 | 4 | 1 | 0 | 1 | 0 |
| ENSG00000223189 | 0 | 0 | 0 | 0 | 0 | 0 |
| ENSG00000243302 | 255 | 134 | 221 | 106 | 134 | 185 |
| ENSG00000230715 | 255 | 75 | 177 | 89 | 85 | 116 |
| ENSG00000229413 | 30 | 13 | 25 | 16 | 17 | 12 |
| ENSG00000243679 | 312 | 113 | 222 | 112 | 126 | 161 |
| ENSG00000281490 | 565 | 254 | 365 | 182 | 292 | 303 |
| ENSG00000271553 | 12 | 9 | 15 | 8 | 12 | 18 |
| ENSG00000241493 | 0 | 1 | 0 | 2 | 1 | 2 |
| ENSG00000205085 | 57 | 20 | 48 | 12 | 20 | 24 |
| ENSG00000271344 | 12 | 3 | 9 | 5 | 4 | 7 |
| ENSG00000239722 | 0 | 0 | 0 | 0 | 0 | 0 |
| ENSG00000201041 | 0 | 0 | 0 | 0 | 0 | 0 |
| ENSG00000252866 | 0 | 0 | 1 | 0 | 0 | 0 |
| ENSG00000135248 | 4 | 2 | 3 | 2 | 1 | 1 |
| ENSG00000128595 | 5376 | 2708 | 3255 | 2435 | 3222 | 2913 |
| ENSG00000244218 | 6 | 4 | 0 | 2 | 4 | 2 |
| ENSG00000128617 | 8 | 4 | 7 | 9 | 9 | 11 |
| ENSG00000128596 | 75 | 56 | 60 | 29 | 57 | 49 |
| ENSG00000128591 | 4 | 0 | 3 | 0 | 4 | 0 |
| ENSG00000242902 | 0 | 0 | 0 | 0 | 0 | 0 |
| ENSG00000135253 | 5 | 0 | 8 | 5 | 2 | 3 |
| ENSG00000128524 | 2863 | 1648 | 1896 | 1936 | 1837 | 1806 |
| ENSG00000272899 | 70 | 40 | 45 | 53 | 40 | 43 |
| ENSG00000224163 | 0 | 0 | 0 | 0 | 0 | 0 |
| ENSG00000128604 | 632 | 316 | 358 | 323 | 310 | 334 |
| ENSG00000275106 | 7 | 3 | 5 | 1 | 1 | 3 |
| ENSG00000064419 | 3425 | 1873 | 2100 | 1469 | 2086 | 1893 |
| ENSG00000242241 | 0 | 0 | 0 | 0 | 0 | 0 |
| ENSG00000244556 | 7 | 6 | 6 | 3 | 5 | 10 |

| | | | | | | |
|---|---|---|---|---|---|---|
| ENSG00000230359 | 64 | 33 | 35 | 30 | 30 | 46 |
| ENSG00000242162 | 0 | 0 | 0 | 0 | 0 | 0 |
| ENSG00000241102 | 0 | 0 | 0 | 0 | 0 | 0 |
| ENSG00000241573 | 0 | 0 | 0 | 0 | 0 | 0 |
| ENSG00000240416 | 0 | 0 | 0 | 0 | 0 | 0 |
| ENSG00000230626 | 4 | 2 | 1 | 5 | 0 | 4 |
| ENSG00000158457 | 317 | 158 | 174 | 242 | 205 | 189 |
| ENSG00000200629 | 0 | 0 | 0 | 0 | 0 | 0 |
| ENSG00000128602 | 4 | 0 | 0 | 3 | 3 | 1 |
| ENSG00000243230 | 2 | 0 | 0 | 0 | 0 | 0 |
| ENSG00000158467 | 1256 | 625 | 885 | 520 | 687 | 699 |
| ENSG00000238072 | 33 | 26 | 39 | 40 | 24 | 43 |
| ENSG00000128578 | 48 | 27 | 32 | 20 | 20 | 37 |
| ENSG00000239919 | 0 | 0 | 0 | 0 | 0 | 0 |
| ENSG00000199846 | 0 | 0 | 1 | 0 | 0 | 0 |
| ENSG00000240204 | 128 | 65 | 99 | 164 | 164 | 160 |
| ENSG00000273329 | 172 | 102 | 127 | 77 | 119 | 108 |
| ENSG00000106459 | 557 | 269 | 360 | 308 | 300 | 332 |
| ENSG00000212238 | 0 | 0 | 0 | 0 | 0 | 0 |
| ENSG00000207990 | 0 | 0 | 0 | 0 | 0 | 0 |
| ENSG00000199158 | 0 | 0 | 0 | 0 | 0 | 0 |
| ENSG00000207691 | 0 | 0 | 0 | 0 | 0 | 0 |
| ENSG00000242078 | 4 | 5 | 5 | 2 | 3 | 0 |
| ENSG00000186591 | 6117 | 2628 | 3609 | 2194 | 3049 | 2871 |
| ENSG00000244036 | 0 | 0 | 0 | 0 | 0 | 0 |
| ENSG00000199516 | 0 | 0 | 0 | 0 | 0 | 0 |
| ENSG00000091732 | 434 | 189 | 268 | 234 | 236 | 245 |
| ENSG00000201109 | 0 | 0 | 0 | 0 | 0 | 0 |
| ENSG00000128607 | 2754 | 1200 | 1534 | 1032 | 1346 | 1386 |
| ENSG00000229196 | 0 | 0 | 0 | 0 | 0 | 0 |
| ENSG00000146842 | 598 | 262 | 336 | 192 | 315 | 334 |
| ENSG00000240571 | 5 | 4 | 1 | 2 | 7 | 2 |
| ENSG00000165120 | 0 | 0 | 0 | 0 | 0 | 0 |
| ENSG00000270992 | 1 | 0 | 0 | 0 | 0 | 0 |
| ENSG00000158516 | 0 | 0 | 1 | 0 | 0 | 0 |
| ENSG00000128510 | 0 | 0 | 1 | 0 | 1 | 1 |
| ENSG00000158525 | 0 | 0 | 0 | 1 | 0 | 0 |
| ENSG00000091704 | 0 | 0 | 0 | 0 | 0 | 0 |
| ENSG00000106477 | 488 | 224 | 311 | 223 | 299 | 285 |
| ENSG00000259920 | 4 | 2 | 1 | 0 | 2 | 1 |
| ENSG00000272701 | 1 | 1 | 1 | 2 | 2 | 5 |
| ENSG00000106484 | 1597 | 832 | 1027 | 1057 | 1163 | 1149 |
| ENSG00000277269 | 0 | 0 | 0 | 0 | 0 | 0 |
| ENSG00000278055 | 0 | 0 | 0 | 0 | 0 | 0 |
| ENSG00000274500 | 0 | 0 | 0 | 0 | 0 | 0 |
| ENSG00000270823 | 3 | 1 | 0 | 1 | 0 | 0 |
| ENSG00000199043 | 0 | 0 | 0 | 0 | 0 | 0 |
| ENSG00000158623 | 2131 | 1203 | 1502 | 960 | 1254 | 1215 |
| ENSG00000270953 | 13 | 11 | 9 | 1 | 7 | 14 |
| ENSG00000239021 | 0 | 0 | 0 | 0 | 0 | 0 |

| | | | | | | |
|---|---|---|---|---|---|---|
| ENSG00000226045 | 2 | 0 | 0 | 0 | 0 | 0 |
| ENSG00000213265 | 0 | 2 | 0 | 0 | 0 | 1 |
| ENSG00000266265 | 4 | 1 | 0 | 1 | 1 | 0 |
| ENSG00000271522 | 0 | 0 | 0 | 0 | 0 | 0 |
| ENSG00000285106 | 3 | 3 | 5 | 1 | 4 | 1 |
| ENSG00000229858 | 0 | 0 | 0 | 0 | 0 | 0 |
| ENSG00000240680 | 0 | 0 | 0 | 0 | 0 | 0 |
| ENSG00000232716 | 0 | 0 | 0 | 0 | 0 | 0 |
| ENSG00000233559 | 2 | 0 | 0 | 1 | 0 | 0 |
| ENSG00000284032 | 0 | 0 | 0 | 0 | 0 | 0 |
| ENSG00000226380 | 14 | 7 | 5 | 2 | 7 | 7 |
| ENSG00000283797 | 0 | 0 | 0 | 0 | 0 | 0 |
| ENSG00000271204 | 0 | 0 | 0 | 0 | 0 | 1 |
| ENSG00000273319 | 0 | 0 | 0 | 0 | 0 | 0 |
| ENSG00000231721 | 66 | 32 | 50 | 17 | 27 | 33 |
| ENSG00000199627 | 0 | 0 | 0 | 0 | 0 | 1 |
| ENSG00000128585 | 771 | 427 | 380 | 261 | 590 | 360 |
| ENSG00000233287 | 0 | 0 | 0 | 0 | 0 | 0 |
| ENSG00000236753 | 30 | 28 | 25 | 26 | 25 | 25 |
| ENSG00000273489 | 0 | 0 | 0 | 0 | 0 | 0 |
| ENSG00000128567 | 1805 | 912 | 1090 | 976 | 1203 | 1151 |
| ENSG00000224545 | 0 | 0 | 0 | 0 | 0 | 0 |
| ENSG00000213261 | 8 | 6 | 3 | 6 | 4 | 7 |
| ENSG00000236238 | 0 | 0 | 0 | 0 | 0 | 0 |
| ENSG00000236395 | 0 | 0 | 0 | 0 | 0 | 0 |
| ENSG00000236281 | 0 | 0 | 1 | 0 | 0 | 0 |
| ENSG00000233263 | 0 | 0 | 0 | 0 | 0 | 0 |
| ENSG00000236386 | 0 | 0 | 0 | 0 | 0 | 0 |
| ENSG00000224865 | 0 | 0 | 0 | 0 | 0 | 0 |
| ENSG00000229532 | 0 | 0 | 0 | 0 | 0 | 0 |
| ENSG00000221866 | 32 | 11 | 24 | 4 | 6 | 10 |
| ENSG00000225144 | 0 | 0 | 0 | 0 | 0 | 0 |
| ENSG00000223436 | 0 | 0 | 0 | 0 | 0 | 0 |
| ENSG00000183470 | 0 | 0 | 0 | 0 | 0 | 0 |
| ENSG00000201009 | 0 | 0 | 0 | 0 | 0 | 0 |
| ENSG00000225881 | 0 | 0 | 0 | 0 | 0 | 0 |
| ENSG00000106554 | 2273 | 1109 | 1291 | 1254 | 1305 | 1268 |
| ENSG00000227197 | 0 | 0 | 0 | 0 | 0 | 0 |
| ENSG00000283041 | 299 | 201 | 161 | 162 | 122 | 170 |
| ENSG00000284135 | 0 | 0 | 0 | 0 | 0 | 0 |
| ENSG00000272393 | 0 | 0 | 0 | 0 | 0 | 0 |
| ENSG00000237821 | 2 | 1 | 0 | 1 | 0 | 0 |
| ENSG00000131558 | 2859 | 1311 | 1659 | 1179 | 1421 | 1416 |
| ENSG00000276137 | 0 | 0 | 0 | 0 | 2 | 0 |
| ENSG00000235429 | 0 | 0 | 0 | 0 | 0 | 0 |
| ENSG00000235967 | 0 | 0 | 0 | 0 | 0 | 0 |
| ENSG00000226205 | 0 | 0 | 0 | 0 | 0 | 0 |
| ENSG00000155530 | 13 | 4 | 2 | 2 | 2 | 5 |
| ENSG00000231098 | 0 | 0 | 0 | 0 | 2 | 0 |
| ENSG00000205060 | 1173 | 502 | 709 | 476 | 571 | 603 |

| | | | | | | |
|---|---|---|---|---|---|---|
| ENSG00000229177 | 0 | 0 | 0 | 0 | 0 | 0 |
| ENSG00000273297 | 11 | 12 | 20 | 18 | 25 | 20 |
| ENSG00000085662 | 0 | 2 | 0 | 8 | 8 | 7 |
| ENSG00000198074 | 22 | 5 | 6 | 17 | 15 | 19 |
| ENSG00000227471 | 26 | 9 | 15 | 17 | 21 | 30 |
| ENSG00000172331 | 337 | 217 | 236 | 201 | 251 | 227 |
| ENSG00000224375 | 0 | 0 | 0 | 0 | 0 | 0 |
| ENSG00000230101 | 0 | 0 | 0 | 0 | 0 | 0 |
| ENSG00000122786 | 4 | 3 | 2 | 0 | 1 | 4 |
| ENSG00000146856 | 101 | 76 | 69 | 45 | 58 | 48 |
| ENSG00000122783 | 954 | 526 | 682 | 587 | 520 | 591 |
| ENSG00000223500 | 0 | 0 | 0 | 0 | 0 | 0 |
| ENSG00000146859 | 658 | 336 | 455 | 314 | 305 | 326 |
| ENSG00000272941 | 11 | 17 | 12 | 6 | 14 | 9 |
| ENSG00000105875 | 564 | 300 | 334 | 263 | 274 | 253 |
| ENSG00000231794 | 1 | 2 | 1 | 0 | 0 | 0 |
| ENSG00000277138 | 2 | 0 | 0 | 0 | 0 | 0 |
| ENSG00000146857 | 0 | 0 | 0 | 0 | 0 | 0 |
| ENSG00000276067 | 0 | 0 | 1 | 0 | 0 | 0 |
| ENSG00000080802 | 843 | 439 | 479 | 305 | 501 | 371 |
| ENSG00000227073 | 2 | 0 | 0 | 0 | 3 | 0 |
| ENSG00000155561 | 2380 | 1252 | 1309 | 741 | 1732 | 1291 |
| ENSG00000271414 | 0 | 0 | 0 | 0 | 0 | 0 |
| ENSG00000223718 | 0 | 0 | 0 | 0 | 1 | 1 |
| ENSG00000243317 | 3344 | 1562 | 1929 | 1694 | 1898 | 1889 |
| ENSG00000212303 | 0 | 0 | 0 | 0 | 0 | 0 |
| ENSG00000164707 | 16 | 2 | 5 | 3 | 6 | 2 |
| ENSG00000273219 | 0 | 0 | 0 | 0 | 0 | 0 |
| ENSG00000189320 | 2 | 1 | 0 | 0 | 0 | 0 |
| ENSG00000224746 | 10 | 10 | 12 | 7 | 6 | 7 |
| ENSG00000105887 | 7967 | 3904 | 4595 | 3347 | 4581 | 4444 |
| ENSG00000267697 | 0 | 0 | 0 | 0 | 0 | 0 |
| ENSG00000199700 | 0 | 0 | 0 | 0 | 0 | 0 |
| ENSG00000230649 | 0 | 0 | 0 | 0 | 0 | 0 |
| ENSG00000232053 | 0 | 0 | 0 | 0 | 1 | 1 |
| ENSG00000206705 | 0 | 0 | 0 | 0 | 0 | 0 |
| ENSG00000236503 | 0 | 0 | 0 | 0 | 0 | 0 |
| ENSG00000226066 | 0 | 0 | 0 | 0 | 0 | 0 |
| ENSG00000234352 | 0 | 0 | 0 | 0 | 0 | 0 |
| ENSG00000231244 | 0 | 0 | 0 | 0 | 0 | 0 |
| ENSG00000181072 | 0 | 0 | 0 | 0 | 0 | 0 |
| ENSG00000207597 | 0 | 0 | 0 | 0 | 0 | 0 |
| ENSG00000223657 | 0 | 0 | 0 | 0 | 1 | 0 |
| ENSG00000224469 | 0 | 0 | 0 | 0 | 0 | 0 |
| ENSG00000105894 | 0 | 0 | 1 | 0 | 0 | 0 |
| ENSG00000202023 | 0 | 0 | 0 | 0 | 0 | 0 |
| ENSG00000231114 | 0 | 0 | 0 | 0 | 0 | 0 |
| ENSG00000228031 | 0 | 0 | 0 | 0 | 0 | 0 |
| ENSG00000157680 | 2 | 0 | 0 | 0 | 1 | 0 |
| ENSG00000279483 | 40 | 27 | 38 | 46 | 29 | 41 |

| | | | | | | |
|---|---|---|---|---|---|---|
| ENSG00000230383 | 12 | 4 | 3 | 4 | 3 | 4 |
| ENSG00000182158 | 1802 | 789 | 1104 | 509 | 767 | 813 |
| ENSG00000237243 | 0 | 0 | 0 | 1 | 0 | 0 |
| ENSG00000122787 | 2 | 0 | 0 | 2 | 2 | 0 |
| ENSG00000234639 | 0 | 1 | 0 | 0 | 0 | 0 |
| ENSG00000231923 | 0 | 0 | 0 | 0 | 0 | 0 |
| ENSG00000231931 | 0 | 0 | 0 | 0 | 0 | 0 |
| ENSG00000222514 | 0 | 0 | 0 | 0 | 0 | 0 |
| ENSG00000233951 | 0 | 0 | 0 | 0 | 0 | 0 |
| ENSG00000284028 | 0 | 0 | 0 | 0 | 0 | 0 |
| ENSG00000225559 | 0 | 0 | 0 | 0 | 0 | 0 |
| ENSG00000252820 | 0 | 0 | 0 | 0 | 0 | 0 |
| ENSG00000201465 | 0 | 0 | 0 | 0 | 0 | 0 |
| ENSG00000213238 | 0 | 0 | 0 | 0 | 0 | 0 |
| ENSG00000213237 | 0 | 0 | 0 | 1 | 0 | 2 |
| ENSG00000214815 | 0 | 0 | 0 | 0 | 0 | 0 |
| ENSG00000122779 | 1989 | 1143 | 1083 | 513 | 1160 | 746 |
| ENSG00000234995 | 0 | 0 | 0 | 0 | 0 | 0 |
| ENSG00000157703 | 9 | 11 | 10 | 6 | 7 | 9 |
| ENSG00000252188 | 0 | 0 | 0 | 0 | 0 | 0 |
| ENSG00000234658 | 0 | 0 | 0 | 0 | 0 | 0 |
| ENSG00000243099 | 0 | 0 | 0 | 0 | 0 | 0 |
| ENSG00000223721 | 0 | 0 | 0 | 0 | 0 | 0 |
| ENSG00000105929 | 166 | 108 | 127 | 51 | 83 | 64 |
| ENSG00000214128 | 4 | 1 | 0 | 0 | 0 | 0 |
| ENSG00000122778 | 937 | 379 | 531 | 414 | 593 | 537 |
| ENSG00000199646 | 2 | 0 | 0 | 0 | 1 | 0 |
| ENSG00000146858 | 44 | 22 | 27 | 19 | 32 | 25 |
| ENSG00000105939 | 1492 | 790 | 815 | 507 | 897 | 746 |
| ENSG00000229677 | 2 | 0 | 0 | 0 | 0 | 0 |
| ENSG00000105948 | 275 | 151 | 180 | 122 | 163 | 151 |
| ENSG00000239254 | 0 | 2 | 0 | 0 | 1 | 1 |
| ENSG00000271537 | 0 | 1 | 0 | 0 | 0 | 0 |
| ENSG00000228360 | 2 | 0 | 0 | 1 | 1 | 0 |
| ENSG00000271404 | 0 | 0 | 0 | 0 | 1 | 0 |
| ENSG00000157741 | 655 | 386 | 358 | 143 | 525 | 293 |
| ENSG00000206843 | 0 | 0 | 0 | 0 | 0 | 0 |
| ENSG00000164898 | 67 | 42 | 70 | 68 | 56 | 67 |
| ENSG00000146963 | 1076 | 532 | 684 | 502 | 600 | 581 |
| ENSG00000269955 | 0 | 0 | 0 | 0 | 0 | 0 |
| ENSG00000273391 | 29 | 11 | 27 | 22 | 15 | 12 |
| ENSG00000252332 | 0 | 0 | 2 | 0 | 0 | 0 |
| ENSG00000188883 | 296 | 116 | 141 | 144 | 117 | 110 |
| ENSG00000253183 | 0 | 0 | 0 | 0 | 0 | 0 |
| ENSG00000236279 | 0 | 0 | 0 | 0 | 0 | 0 |
| ENSG00000254270 | 0 | 1 | 0 | 0 | 0 | 0 |
| ENSG00000064393 | 1521 | 566 | 1043 | 457 | 652 | 819 |
| ENSG00000059377 | 6 | 3 | 6 | 1 | 7 | 3 |
| ENSG00000059378 | 246 | 127 | 215 | 142 | 131 | 170 |
| ENSG00000006459 | 609 | 305 | 325 | 125 | 309 | 216 |

| | | | | | | |
|---|---|---|---|---|---|---|
| ENSG00000251728 | 3 | 1 | 1 | 1 | 0 | 0 |
| ENSG00000260231 | 228 | 98 | 137 | 101 | 88 | 91 |
| ENSG00000199971 | 0 | 0 | 0 | 0 | 0 | 0 |
| ENSG00000199283 | 0 | 0 | 0 | 0 | 0 | 0 |
| ENSG00000270512 | 0 | 0 | 0 | 0 | 0 | 0 |
| ENSG00000241661 | 2 | 0 | 0 | 0 | 0 | 0 |
| ENSG00000157800 | 1210 | 597 | 804 | 496 | 733 | 666 |
| ENSG00000223113 | 0 | 0 | 1 | 0 | 0 | 0 |
| ENSG00000202472 | 0 | 0 | 0 | 0 | 0 | 0 |
| ENSG00000146955 | 27 | 14 | 21 | 8 | 18 | 14 |
| ENSG00000103200 | 0 | 0 | 0 | 0 | 0 | 0 |
| ENSG00000133606 | 6019 | 2790 | 3531 | 2741 | 3268 | 3188 |
| ENSG00000146966 | 0 | 0 | 0 | 0 | 0 | 1 |
| ENSG00000201354 | 0 | 0 | 0 | 0 | 0 | 0 |
| ENSG00000285904 | 18 | 9 | 11 | 2 | 4 | 3 |
| ENSG00000240173 | 0 | 0 | 0 | 0 | 0 | 0 |
| ENSG00000133597 | 1000 | 576 | 692 | 726 | 570 | 643 |
| ENSG00000090266 | 2257 | 1424 | 1676 | 1816 | 1575 | 1483 |
| ENSG00000240889 | 25 | 9 | 26 | 17 | 11 | 26 |
| ENSG00000157764 | 750 | 395 | 405 | 226 | 358 | 327 |
| ENSG00000271932 | 0 | 0 | 0 | 0 | 0 | 0 |
| ENSG00000271611 | 0 | 0 | 0 | 0 | 0 | 0 |
| ENSG00000242703 | 0 | 0 | 0 | 0 | 0 | 0 |
| ENSG00000090263 | 1467 | 773 | 943 | 914 | 897 | 902 |
| ENSG00000223212 | 0 | 0 | 0 | 0 | 0 | 0 |
| ENSG00000261115 | 1240 | 534 | 709 | 466 | 616 | 641 |
| ENSG00000261778 | 0 | 0 | 0 | 0 | 0 | 0 |
| ENSG00000261624 | 0 | 0 | 0 | 0 | 0 | 0 |
| ENSG00000285841 | 0 | 0 | 0 | 0 | 0 | 0 |
| ENSG00000204990 | 0 | 0 | 1 | 0 | 0 | 0 |
| ENSG00000261629 | 0 | 0 | 1 | 0 | 0 | 0 |
| ENSG00000261797 | 0 | 0 | 0 | 0 | 0 | 0 |
| ENSG00000261570 | 0 | 0 | 0 | 0 | 0 | 0 |
| ENSG00000006530 | 1079 | 535 | 612 | 512 | 640 | 645 |
| ENSG00000244701 | 5 | 4 | 12 | 4 | 4 | 6 |
| ENSG00000257093 | 2448 | 1073 | 1606 | 828 | 1313 | 1249 |
| ENSG00000270157 | 84 | 41 | 66 | 24 | 46 | 48 |
| ENSG00000228775 | 92 | 59 | 47 | 41 | 47 | 43 |
| ENSG00000214102 | 6 | 3 | 5 | 0 | 1 | 1 |
| ENSG00000212153 | 0 | 0 | 0 | 0 | 0 | 0 |
| ENSG00000106028 | 2901 | 1501 | 1733 | 1742 | 1845 | 1797 |
| ENSG00000127362 | 0 | 2 | 0 | 1 | 0 | 0 |
| ENSG00000127364 | 2 | 1 | 3 | 3 | 8 | 3 |
| ENSG00000270923 | 1 | 0 | 1 | 1 | 1 | 1 |
| ENSG00000127366 | 2 | 1 | 1 | 2 | 6 | 7 |
| ENSG00000241926 | 0 | 0 | 0 | 0 | 0 | 0 |
| ENSG00000224354 | 0 | 0 | 0 | 0 | 0 | 0 |
| ENSG00000239648 | 0 | 0 | 0 | 0 | 0 | 0 |
| ENSG00000240677 | 0 | 0 | 0 | 0 | 0 | 0 |
| ENSG00000165076 | 1 | 0 | 0 | 0 | 0 | 0 |

| | | | | | | |
|---|---|---|---|---|---|---|
| ENSG00000240031 | 0 | 0 | 0 | 0 | 0 | 0 |
| ENSG00000237621 | 0 | 0 | 0 | 0 | 0 | 0 |
| ENSG00000257335 | 0 | 0 | 1 | 0 | 0 | 1 |
| ENSG00000244292 | 0 | 0 | 0 | 0 | 0 | 0 |
| ENSG00000258083 | 0 | 0 | 0 | 0 | 0 | 0 |
| ENSG00000258227 | 0 | 0 | 0 | 0 | 0 | 0 |
| ENSG00000257138 | 0 | 0 | 0 | 0 | 0 | 0 |
| ENSG00000257743 | 0 | 0 | 0 | 0 | 0 | 0 |
| ENSG00000240268 | 0 | 0 | 0 | 0 | 0 | 0 |
| ENSG00000258223 | 0 | 0 | 0 | 0 | 0 | 0 |
| ENSG00000186163 | 0 | 0 | 0 | 0 | 0 | 0 |
| ENSG00000241881 | 0 | 0 | 0 | 0 | 0 | 0 |
| ENSG00000277083 | 0 | 0 | 0 | 0 | 0 | 0 |
| ENSG00000242736 | 0 | 0 | 0 | 0 | 0 | 0 |
| ENSG00000226660 | 0 | 0 | 0 | 0 | 0 | 0 |
| ENSG00000237702 | 0 | 0 | 0 | 0 | 0 | 0 |
| ENSG00000211710 | 0 | 0 | 0 | 0 | 0 | 0 |
| ENSG00000211734 | 0 | 0 | 0 | 0 | 0 | 0 |
| ENSG00000211706 | 1 | 1 | 0 | 0 | 0 | 1 |
| ENSG00000211707 | 0 | 0 | 0 | 0 | 0 | 0 |
| ENSG00000211745 | 0 | 0 | 0 | 0 | 0 | 0 |
| ENSG00000283063 | 0 | 0 | 0 | 0 | 0 | 0 |
| ENSG00000282939 | 0 | 0 | 0 | 0 | 0 | 0 |
| ENSG00000243889 | 0 | 0 | 0 | 0 | 0 | 0 |
| ENSG00000242771 | 0 | 0 | 0 | 0 | 0 | 0 |
| ENSG00000211713 | 0 | 0 | 0 | 0 | 0 | 0 |
| ENSG00000211714 | 0 | 0 | 0 | 0 | 0 | 0 |
| ENSG00000239944 | 0 | 0 | 0 | 0 | 0 | 0 |
| ENSG00000211715 | 0 | 0 | 0 | 0 | 0 | 0 |
| ENSG00000211716 | 0 | 0 | 0 | 0 | 0 | 0 |
| ENSG00000211717 | 0 | 0 | 0 | 0 | 0 | 0 |
| ENSG00000211720 | 0 | 0 | 0 | 0 | 0 | 0 |
| ENSG00000244661 | 0 | 0 | 0 | 0 | 0 | 0 |
| ENSG00000229769 | 0 | 0 | 0 | 0 | 0 | 0 |
| ENSG00000241657 | 0 | 0 | 0 | 0 | 0 | 0 |
| ENSG00000211739 | 0 | 0 | 0 | 0 | 0 | 0 |
| ENSG00000211721 | 0 | 1 | 0 | 0 | 0 | 0 |
| ENSG00000253409 | 0 | 0 | 0 | 0 | 0 | 0 |
| ENSG00000230099 | 0 | 0 | 0 | 0 | 0 | 0 |
| ENSG00000211724 | 0 | 0 | 0 | 0 | 0 | 0 |
| ENSG00000227550 | 0 | 0 | 0 | 0 | 0 | 0 |
| ENSG00000211725 | 0 | 0 | 0 | 0 | 0 | 0 |
| ENSG00000253188 | 4 | 1 | 0 | 3 | 0 | 1 |
| ENSG00000211727 | 0 | 0 | 0 | 0 | 0 | 0 |
| ENSG00000211728 | 0 | 0 | 0 | 0 | 0 | 0 |
| ENSG00000253534 | 1 | 0 | 0 | 1 | 0 | 0 |
| ENSG00000253291 | 0 | 0 | 0 | 0 | 0 | 0 |
| ENSG00000211731 | 0 | 0 | 0 | 0 | 0 | 0 |
| ENSG00000278030 | 0 | 0 | 0 | 0 | 0 | 0 |
| ENSG00000276405 | 0 | 0 | 0 | 0 | 0 | 0 |

| | | | | | | |
|---|---|---|---|---|---|---|
| ENSG00000275791 | 0 | 0 | 0 | 0 | 0 | 0 |
| ENSG00000276597 | 0 | 0 | 0 | 0 | 0 | 0 |
| ENSG00000274752 | 0 | 0 | 0 | 0 | 0 | 0 |
| ENSG00000276953 | 0 | 0 | 0 | 0 | 0 | 0 |
| ENSG00000275158 | 0 | 0 | 0 | 0 | 0 | 0 |
| ENSG00000275743 | 0 | 0 | 0 | 0 | 0 | 0 |
| ENSG00000276819 | 0 | 0 | 0 | 0 | 0 | 0 |
| ENSG00000275243 | 0 | 0 | 0 | 0 | 0 | 0 |
| ENSG00000277880 | 0 | 0 | 0 | 0 | 0 | 0 |
| ENSG00000276557 | 0 | 0 | 0 | 0 | 0 | 0 |
| ENSG00000211746 | 0 | 0 | 0 | 0 | 0 | 0 |
| ENSG00000211747 | 0 | 0 | 0 | 0 | 0 | 0 |
| ENSG00000251578 | 0 | 0 | 0 | 0 | 0 | 0 |
| ENSG00000240578 | 0 | 0 | 0 | 0 | 0 | 0 |
| ENSG00000211749 | 0 | 0 | 0 | 0 | 0 | 0 |
| ENSG00000211750 | 0 | 0 | 0 | 0 | 0 | 0 |
| ENSG00000270672 | 0 | 0 | 0 | 0 | 0 | 0 |
| ENSG00000282499 | 0 | 0 | 0 | 0 | 0 | 0 |
| ENSG00000239992 | 0 | 0 | 0 | 0 | 0 | 0 |
| ENSG00000249912 | 0 | 0 | 0 | 0 | 0 | 0 |
| ENSG00000241911 | 0 | 0 | 0 | 0 | 0 | 0 |
| ENSG00000211752 | 0 | 0 | 0 | 0 | 0 | 0 |
| ENSG00000211753 | 0 | 0 | 0 | 0 | 0 | 0 |
| ENSG00000244273 | 0 | 0 | 0 | 0 | 0 | 0 |
| ENSG00000232869 | 0 | 0 | 0 | 0 | 0 | 0 |
| ENSG00000204983 | 0 | 0 | 0 | 1 | 0 | 0 |
| ENSG00000275896 | 0 | 1 | 0 | 0 | 0 | 0 |
| ENSG00000250591 | 0 | 0 | 0 | 0 | 0 | 0 |
| ENSG00000243661 | 0 | 0 | 0 | 0 | 0 | 0 |
| ENSG00000282431 | 0 | 0 | 0 | 0 | 0 | 0 |
| ENSG00000282320 | 0 | 0 | 0 | 0 | 0 | 0 |
| ENSG00000282420 | 0 | 0 | 0 | 0 | 0 | 0 |
| ENSG00000282133 | 0 | 0 | 0 | 0 | 0 | 0 |
| ENSG00000281958 | 0 | 0 | 0 | 0 | 0 | 0 |
| ENSG00000282173 | 0 | 0 | 0 | 0 | 0 | 0 |
| ENSG00000282780 | 0 | 0 | 0 | 0 | 0 | 0 |
| ENSG00000211751 | 0 | 0 | 0 | 0 | 0 | 0 |
| ENSG00000211764 | 0 | 0 | 0 | 0 | 0 | 0 |
| ENSG00000211765 | 0 | 0 | 0 | 0 | 0 | 0 |
| ENSG00000211766 | 0 | 0 | 0 | 0 | 0 | 0 |
| ENSG00000211767 | 0 | 0 | 0 | 0 | 0 | 0 |
| ENSG00000211768 | 0 | 0 | 0 | 0 | 0 | 0 |
| ENSG00000211769 | 0 | 0 | 0 | 0 | 0 | 0 |
| ENSG00000211770 | 0 | 0 | 0 | 0 | 0 | 0 |
| ENSG00000211771 | 0 | 0 | 0 | 0 | 0 | 0 |
| ENSG00000211772 | 0 | 0 | 0 | 0 | 0 | 0 |
| ENSG00000237254 | 0 | 0 | 0 | 2 | 0 | 0 |
| ENSG00000106123 | 0 | 3 | 1 | 5 | 2 | 0 |
| ENSG00000165125 | 10 | 2 | 8 | 0 | 2 | 1 |
| ENSG00000224970 | 0 | 0 | 0 | 0 | 0 | 0 |

| | | | | | | |
|---|---|---|---|---|---|---|
| ENSG00000127412 | 0 | 0 | 0 | 0 | 0 | 0 |
| ENSG00000165131 | 0 | 0 | 0 | 0 | 0 | 0 |
| ENSG00000197993 | 0 | 2 | 0 | 1 | 1 | 1 |
| ENSG00000179468 | 0 | 0 | 0 | 0 | 0 | 0 |
| ENSG00000239293 | 0 | 0 | 0 | 0 | 0 | 0 |
| ENSG00000225781 | 0 | 0 | 0 | 0 | 0 | 0 |
| ENSG00000179420 | 0 | 0 | 0 | 0 | 0 | 0 |
| ENSG00000159763 | 0 | 0 | 0 | 0 | 0 | 0 |
| ENSG00000236398 | 0 | 0 | 0 | 0 | 0 | 0 |
| ENSG00000268170 | 0 | 0 | 0 | 0 | 0 | 0 |
| ENSG00000221937 | 0 | 0 | 0 | 0 | 0 | 0 |
| ENSG00000279223 | 0 | 0 | 0 | 0 | 0 | 0 |
| ENSG00000197448 | 3020 | 1510 | 2005 | 1782 | 1792 | 1740 |
| ENSG00000231840 | 0 | 0 | 1 | 0 | 0 | 1 |
| ENSG00000178826 | 19 | 11 | 18 | 16 | 11 | 11 |
| ENSG00000106144 | 1374 | 663 | 818 | 569 | 744 | 724 |
| ENSG00000239419 | 8 | 1 | 3 | 0 | 3 | 2 |
| ENSG00000240322 | 1 | 0 | 1 | 1 | 0 | 3 |
| ENSG00000231531 | 0 | 0 | 0 | 0 | 0 | 0 |
| ENSG00000188037 | 0 | 0 | 0 | 0 | 0 | 0 |
| ENSG00000159784 | 6 | 3 | 4 | 7 | 10 | 6 |
| ENSG00000272619 | 0 | 0 | 0 | 0 | 0 | 0 |
| ENSG00000232533 | 166 | 103 | 113 | 221 | 149 | 120 |
| ENSG00000159840 | 2282 | 1138 | 1299 | 1807 | 1552 | 1290 |
| ENSG00000278449 | 0 | 0 | 0 | 0 | 0 | 0 |
| ENSG00000146904 | 1396 | 616 | 746 | 745 | 731 | 668 |
| ENSG00000229153 | 1 | 0 | 0 | 0 | 1 | 1 |
| ENSG00000234066 | 0 | 0 | 0 | 0 | 0 | 0 |
| ENSG00000185899 | 0 | 0 | 0 | 0 | 0 | 0 |
| ENSG00000221855 | 0 | 0 | 0 | 0 | 0 | 0 |
| ENSG00000236853 | 0 | 0 | 0 | 0 | 0 | 0 |
| ENSG00000176510 | 0 | 0 | 0 | 0 | 0 | 0 |
| ENSG00000229977 | 0 | 0 | 0 | 0 | 0 | 0 |
| ENSG00000229877 | 0 | 0 | 0 | 0 | 0 | 0 |
| ENSG00000271079 | 2 | 2 | 0 | 0 | 0 | 3 |
| ENSG00000252037 | 0 | 0 | 0 | 0 | 0 | 0 |
| ENSG00000230556 | 0 | 0 | 0 | 0 | 0 | 0 |
| ENSG00000223459 | 105 | 43 | 56 | 11 | 45 | 42 |
| ENSG00000283537 | 37 | 17 | 25 | 16 | 22 | 20 |
| ENSG00000170379 | 242 | 109 | 164 | 116 | 123 | 120 |
| ENSG00000283528 | 0 | 0 | 0 | 0 | 0 | 0 |
| ENSG00000252523 | 0 | 0 | 0 | 0 | 0 | 0 |
| ENSG00000271321 | 0 | 0 | 0 | 0 | 0 | 0 |
| ENSG00000253882 | 10 | 8 | 9 | 5 | 12 | 13 |
| ENSG00000241136 | 0 | 0 | 0 | 0 | 0 | 0 |
| ENSG00000232145 | 0 | 0 | 0 | 0 | 0 | 0 |
| ENSG00000159860 | 4 | 1 | 1 | 3 | 3 | 0 |
| ENSG00000198420 | 1425 | 634 | 802 | 363 | 633 | 605 |
| ENSG00000221910 | 0 | 0 | 0 | 0 | 0 | 0 |
| ENSG00000213215 | 0 | 0 | 0 | 0 | 0 | 0 |

| | | | | | | |
|---|---|---|---|---|---|---|
| ENSG00000270729 | 0 | 0 | 0 | 0 | 0 | 0 |
| ENSG00000273377 | 0 | 0 | 0 | 0 | 0 | 0 |
| ENSG00000242280 | 0 | 0 | 0 | 0 | 0 | 0 |
| ENSG00000221813 | 0 | 0 | 0 | 0 | 0 | 0 |
| ENSG00000221836 | 0 | 0 | 0 | 0 | 1 | 0 |
| ENSG00000221933 | 0 | 0 | 0 | 0 | 0 | 0 |
| ENSG00000239967 | 0 | 0 | 0 | 0 | 0 | 0 |
| ENSG00000221858 | 0 | 0 | 0 | 0 | 0 | 0 |
| ENSG00000221989 | 0 | 0 | 0 | 0 | 0 | 0 |
| ENSG00000239981 | 0 | 0 | 0 | 0 | 0 | 0 |
| ENSG00000221938 | 0 | 0 | 0 | 0 | 0 | 0 |
| ENSG00000273234 | 0 | 2 | 0 | 0 | 1 | 0 |
| ENSG00000183122 | 0 | 0 | 0 | 0 | 0 | 0 |
| ENSG00000240621 | 0 | 0 | 0 | 1 | 0 | 0 |
| ENSG00000225932 | 8 | 4 | 8 | 7 | 8 | 9 |
| ENSG00000213214 | 274 | 112 | 166 | 129 | 161 | 149 |
| ENSG00000244198 | 18 | 19 | 14 | 15 | 17 | 21 |
| ENSG00000212807 | 0 | 0 | 2 | 0 | 0 | 0 |
| ENSG00000170356 | 12 | 1 | 2 | 11 | 3 | 4 |
| ENSG00000284644 | 1 | 0 | 0 | 0 | 0 | 1 |
| ENSG00000243896 | 4 | 1 | 1 | 3 | 0 | 1 |
| ENSG00000244693 | 16 | 0 | 2 | 3 | 6 | 1 |
| ENSG00000204959 | 143 | 45 | 66 | 42 | 69 | 71 |
| ENSG00000228960 | 10 | 2 | 16 | 11 | 11 | 12 |
| ENSG00000244479 | 14 | 6 | 12 | 6 | 14 | 11 |
| ENSG00000221970 | 2 | 0 | 0 | 1 | 3 | 3 |
| ENSG00000050327 | 515 | 268 | 353 | 314 | 328 | 331 |
| ENSG00000106410 | 0 | 0 | 0 | 0 | 0 | 0 |
| ENSG00000242978 | 0 | 0 | 0 | 0 | 0 | 0 |
| ENSG00000221507 | 0 | 1 | 0 | 0 | 0 | 0 |
| ENSG00000196511 | 304 | 164 | 170 | 114 | 159 | 134 |
| ENSG00000243746 | 2 | 2 | 1 | 2 | 1 | 1 |
| ENSG00000200673 | 0 | 0 | 0 | 0 | 0 | 0 |
| ENSG00000236343 | 0 | 0 | 0 | 0 | 0 | 0 |
| ENSG00000214035 | 0 | 0 | 0 | 0 | 0 | 0 |
| ENSG00000225918 | 2 | 0 | 0 | 0 | 0 | 0 |
| ENSG00000226592 | 4 | 0 | 3 | 0 | 1 | 0 |
| ENSG00000231839 | 0 | 0 | 0 | 0 | 0 | 0 |
| ENSG00000230746 | 0 | 0 | 0 | 0 | 0 | 0 |
| ENSG00000280144 | 0 | 0 | 0 | 0 | 0 | 0 |
| ENSG00000174469 | 8 | 4 | 2 | 3 | 3 | 2 |
| ENSG00000274127 | 0 | 0 | 0 | 0 | 0 | 0 |
| ENSG00000274118 | 0 | 0 | 0 | 0 | 0 | 0 |
| ENSG00000236795 | 0 | 0 | 0 | 0 | 0 | 0 |
| ENSG00000226857 | 0 | 0 | 0 | 0 | 0 | 0 |
| ENSG00000232391 | 0 | 1 | 0 | 0 | 0 | 0 |
| ENSG00000221442 | 0 | 0 | 0 | 0 | 0 | 0 |
| ENSG00000230190 | 0 | 0 | 0 | 0 | 0 | 0 |
| ENSG00000270634 | 0 | 0 | 0 | 0 | 0 | 0 |
| ENSG00000207318 | 0 | 0 | 0 | 0 | 0 | 0 |

| | | | | | | |
|---|---|---|---|---|---|---|
| ENSG00000251936 | 0 | 0 | 0 | 0 | 0 | 0 |
| ENSG00000242266 | 0 | 0 | 0 | 0 | 0 | 0 |
| ENSG00000199370 | 0 | 0 | 0 | 0 | 0 | 0 |
| ENSG00000243374 | 0 | 0 | 0 | 0 | 0 | 0 |
| ENSG00000274133 | 0 | 0 | 0 | 0 | 0 | 0 |
| ENSG00000283504 | 0 | 0 | 0 | 0 | 0 | 0 |
| ENSG00000283648 | 0 | 0 | 0 | 0 | 0 | 0 |
| ENSG00000213210 | 0 | 0 | 0 | 0 | 0 | 0 |
| ENSG00000170279 | 0 | 0 | 0 | 0 | 0 | 0 |
| ENSG00000213209 | 0 | 0 | 0 | 0 | 0 | 0 |
| ENSG00000270200 | 0 | 0 | 0 | 0 | 0 | 0 |
| ENSG00000273314 | 86 | 58 | 72 | 70 | 58 | 49 |
| ENSG00000055130 | 2409 | 1234 | 1374 | 1044 | 1391 | 1257 |
| ENSG00000106462 | 1373 | 696 | 906 | 486 | 764 | 805 |
| ENSG00000251712 | 6 | 0 | 0 | 0 | 1 | 0 |
| ENSG00000275586 | 0 | 1 | 0 | 0 | 0 | 0 |
| ENSG00000239468 | 0 | 0 | 2 | 0 | 0 | 0 |
| ENSG00000252316 | 0 | 0 | 0 | 0 | 0 | 0 |
| ENSG00000202354 | 0 | 0 | 0 | 0 | 0 | 0 |
| ENSG00000201098 | 2 | 0 | 0 | 0 | 0 | 0 |
| ENSG00000281189 | 47 | 21 | 34 | 16 | 24 | 24 |
| ENSG00000155660 | 7324 | 3602 | 4453 | 3754 | 4201 | 4040 |
| ENSG00000202528 | 0 | 0 | 0 | 0 | 0 | 0 |
| ENSG00000234565 | 0 | 0 | 0 | 0 | 0 | 0 |
| ENSG00000197362 | 500 | 235 | 320 | 238 | 293 | 295 |
| ENSG00000204947 | 327 | 156 | 209 | 117 | 156 | 126 |
| ENSG00000240877 | 2 | 3 | 4 | 4 | 3 | 1 |
| ENSG00000197024 | 934 | 427 | 483 | 424 | 489 | 479 |
| ENSG00000239719 | 10 | 3 | 3 | 2 | 3 | 2 |
| ENSG00000170265 | 1937 | 866 | 1032 | 1095 | 1023 | 966 |
| ENSG00000170260 | 617 | 280 | 444 | 359 | 337 | 358 |
| ENSG00000204946 | 818 | 416 | 587 | 452 | 459 | 496 |
| ENSG00000271664 | 14 | 6 | 11 | 4 | 7 | 0 |
| ENSG00000244560 | 95 | 47 | 74 | 54 | 45 | 71 |
| ENSG00000231397 | 0 | 0 | 0 | 0 | 0 | 0 |
| ENSG00000215311 | 0 | 0 | 0 | 0 | 0 | 0 |
| ENSG00000228151 | 0 | 1 | 0 | 0 | 1 | 0 |
| ENSG00000261842 | 2 | 0 | 0 | 1 | 0 | 0 |
| ENSG00000196453 | 555 | 254 | 329 | 378 | 324 | 345 |
| ENSG00000181220 | 1046 | 530 | 677 | 608 | 612 | 602 |
| ENSG00000133624 | 499 | 165 | 258 | 146 | 242 | 193 |
| ENSG00000133619 | 219 | 77 | 104 | 86 | 116 | 109 |
| ENSG00000181444 | 2442 | 1058 | 1362 | 1131 | 946 | 1108 |
| ENSG00000197558 | 724 | 227 | 417 | 171 | 210 | 248 |
| ENSG00000106479 | 1087 | 444 | 670 | 428 | 521 | 529 |
| ENSG00000280149 | 10 | 4 | 15 | 10 | 13 | 12 |
| ENSG00000273419 | 10 | 2 | 11 | 8 | 7 | 3 |
| ENSG00000204934 | 97 | 45 | 52 | 34 | 48 | 49 |
| ENSG00000171130 | 2497 | 1171 | 1569 | 1706 | 1419 | 1722 |
| ENSG00000273293 | 0 | 0 | 0 | 0 | 0 | 0 |

| | | | | | | |
|---|---|---|---|---|---|---|
| ENSG00000279536 | 0 | 3 | 0 | 0 | 0 | 0 |
| ENSG00000273011 | 0 | 0 | 0 | 0 | 0 | 0 |
| ENSG00000224016 | 0 | 1 | 2 | 1 | 0 | 0 |
| ENSG00000260555 | 0 | 1 | 0 | 2 | 3 | 0 |
| ENSG00000241449 | 0 | 0 | 0 | 0 | 1 | 0 |
| ENSG00000276538 | 0 | 0 | 0 | 0 | 0 | 0 |
| ENSG00000106526 | 105 | 63 | 82 | 56 | 63 | 53 |
| ENSG00000127399 | 381 | 193 | 290 | 318 | 256 | 250 |
| ENSG00000188707 | 322 | 137 | 188 | 166 | 195 | 225 |
| ENSG00000261305 | 26 | 10 | 19 | 16 | 4 | 10 |
| ENSG00000106538 | 5 | 3 | 5 | 6 | 6 | 8 |
| ENSG00000240449 | 57 | 26 | 54 | 26 | 47 | 36 |
| ENSG00000214022 | 4225 | 2027 | 2524 | 2632 | 2718 | 2705 |
| ENSG00000284041 | 0 | 0 | 0 | 0 | 0 | 1 |
| ENSG00000284048 | 0 | 0 | 0 | 0 | 0 | 0 |
| ENSG00000196456 | 63 | 36 | 34 | 51 | 29 | 29 |
| ENSG00000239377 | 0 | 0 | 0 | 0 | 0 | 0 |
| ENSG00000284691 | 94 | 54 | 55 | 58 | 51 | 61 |
| ENSG00000242258 | 0 | 0 | 0 | 0 | 0 | 0 |
| ENSG00000171115 | 0 | 0 | 0 | 0 | 0 | 0 |
| ENSG00000271582 | 0 | 0 | 0 | 0 | 0 | 0 |
| ENSG00000213205 | 0 | 0 | 0 | 0 | 0 | 0 |
| ENSG00000179144 | 0 | 0 | 0 | 0 | 0 | 0 |
| ENSG00000243853 | 0 | 0 | 0 | 0 | 0 | 0 |
| ENSG00000133574 | 0 | 0 | 0 | 0 | 0 | 0 |
| ENSG00000271568 | 0 | 0 | 0 | 0 | 0 | 0 |
| ENSG00000232361 | 0 | 0 | 0 | 0 | 0 | 0 |
| ENSG00000133561 | 0 | 0 | 0 | 0 | 0 | 0 |
| ENSG00000270990 | 0 | 0 | 0 | 0 | 0 | 0 |
| ENSG00000106560 | 0 | 1 | 2 | 0 | 2 | 0 |
| ENSG00000213203 | 0 | 0 | 0 | 0 | 0 | 0 |
| ENSG00000281887 | 0 | 0 | 0 | 0 | 0 | 0 |
| ENSG00000196329 | 0 | 0 | 0 | 0 | 0 | 0 |
| ENSG00000177590 | 0 | 0 | 0 | 0 | 0 | 0 |
| ENSG00000241134 | 0 | 0 | 0 | 0 | 0 | 0 |
| ENSG00000106565 | 2 | 2 | 1 | 0 | 0 | 2 |
| ENSG00000002933 | 0 | 0 | 0 | 0 | 0 | 0 |
| ENSG00000002726 | 10 | 3 | 4 | 1 | 1 | 4 |
| ENSG00000055118 | 208 | 111 | 111 | 111 | 122 | 121 |
| ENSG00000164867 | 5 | 1 | 5 | 2 | 7 | 1 |
| ENSG00000181652 | 56 | 23 | 34 | 17 | 20 | 18 |
| ENSG00000197150 | 1294 | 627 | 748 | 706 | 569 | 641 |
| ENSG00000243433 | 12 | 2 | 0 | 1 | 1 | 3 |
| ENSG00000213199 | 126 | 48 | 71 | 63 | 64 | 64 |
| ENSG00000164885 | 1449 | 656 | 775 | 868 | 784 | 737 |
| ENSG00000164889 | 7237 | 3492 | 4489 | 4377 | 4235 | 4153 |
| ENSG00000244151 | 14 | 5 | 7 | 6 | 9 | 11 |
| ENSG00000164896 | 2969 | 1416 | 1803 | 1758 | 1558 | 1467 |
| ENSG00000164897 | 2492 | 1076 | 1330 | 1559 | 1301 | 1310 |
| ENSG00000133612 | 3306 | 1536 | 1924 | 1794 | 1734 | 1742 |

| | | | | | | |
|---|---|---|---|---|---|---|
| ENSG00000164900 | 0 | 0 | 1 | 2 | 0 | 1 |
| ENSG00000146926 | 0 | 0 | 0 | 0 | 1 | 0 |
| ENSG00000278685 | 0 | 0 | 0 | 0 | 0 | 0 |
| ENSG00000285292 | 0 | 0 | 0 | 0 | 0 | 0 |
| ENSG00000285480 | 0 | 0 | 2 | 0 | 0 | 2 |
| ENSG00000033050 | 577 | 339 | 413 | 458 | 409 | 453 |
| ENSG00000033100 | 2411 | 1208 | 1617 | 1461 | 1555 | 1423 |
| ENSG00000284191 | 0 | 0 | 0 | 0 | 0 | 0 |
| ENSG00000082014 | 398 | 196 | 275 | 217 | 176 | 215 |
| ENSG00000272661 | 0 | 0 | 0 | 0 | 0 | 0 |
| ENSG00000243018 | 0 | 0 | 2 | 0 | 0 | 0 |
| ENSG00000013374 | 1591 | 811 | 951 | 730 | 927 | 836 |
| ENSG00000187260 | 10 | 3 | 8 | 2 | 4 | 7 |
| ENSG00000243836 | 1 | 0 | 0 | 0 | 0 | 0 |
| ENSG00000241456 | 0 | 0 | 0 | 0 | 0 | 0 |
| ENSG00000127377 | 0 | 0 | 2 | 2 | 0 | 0 |
| ENSG00000265810 | 0 | 0 | 0 | 0 | 0 | 0 |
| ENSG00000241959 | 0 | 0 | 0 | 0 | 0 | 0 |
| ENSG00000106615 | 1956 | 1028 | 1106 | 1112 | 1275 | 1281 |
| ENSG00000240132 | 0 | 5 | 0 | 1 | 0 | 0 |
| ENSG00000106617 | 609 | 340 | 381 | 360 | 390 | 370 |
| ENSG00000242048 | 0 | 0 | 0 | 0 | 0 | 0 |
| ENSG00000239911 | 20 | 5 | 4 | 7 | 6 | 8 |
| ENSG00000212219 | 0 | 0 | 0 | 0 | 0 | 0 |
| ENSG00000222941 | 0 | 0 | 0 | 0 | 0 | 0 |
| ENSG00000227562 | 0 | 0 | 0 | 0 | 0 | 0 |
| ENSG00000106648 | 0 | 0 | 0 | 0 | 0 | 0 |
| ENSG00000276213 | 0 | 0 | 0 | 0 | 0 | 0 |
| ENSG00000178234 | 1576 | 845 | 1034 | 713 | 943 | 894 |
| ENSG00000229591 | 6 | 1 | 5 | 2 | 1 | 2 |
| ENSG00000213188 | 0 | 0 | 2 | 1 | 2 | 3 |
| ENSG00000055609 | 1547 | 827 | 705 | 306 | 1045 | 766 |
| ENSG00000270405 | 0 | 0 | 0 | 0 | 0 | 0 |
| ENSG00000204894 | 0 | 0 | 0 | 0 | 0 | 0 |
| ENSG00000241007 | 0 | 0 | 1 | 0 | 0 | 0 |
| ENSG00000253088 | 0 | 0 | 0 | 0 | 0 | 0 |
| ENSG00000241735 | 4 | 0 | 0 | 0 | 2 | 0 |
| ENSG00000020219 | 2 | 1 | 0 | 0 | 1 | 0 |
| ENSG00000261455 | 388 | 188 | 209 | 190 | 173 | 161 |
| ENSG00000231139 | 0 | 0 | 0 | 0 | 0 | 0 |
| ENSG00000199404 | 0 | 0 | 0 | 0 | 0 | 0 |
| ENSG00000285912 | 0 | 0 | 0 | 0 | 0 | 0 |
| ENSG00000225416 | 0 | 0 | 0 | 0 | 0 | 0 |
| ENSG00000212590 | 0 | 0 | 0 | 0 | 0 | 0 |
| ENSG00000196584 | 237 | 168 | 171 | 95 | 153 | 144 |
| ENSG00000214003 | 0 | 1 | 0 | 0 | 1 | 0 |
| ENSG00000244328 | 0 | 0 | 3 | 0 | 0 | 0 |
| ENSG00000133627 | 411 | 191 | 208 | 202 | 214 | 231 |
| ENSG00000224104 | 0 | 0 | 0 | 0 | 0 | 0 |
| ENSG00000234722 | 2 | 0 | 1 | 0 | 1 | 0 |

| | | | | | | |
|---|---|---|---|---|---|---|
| ENSG00000285524 | 0 | 0 | 0 | 0 | 0 | 0 |
| ENSG00000233489 | 0 | 0 | 0 | 0 | 0 | 0 |
| ENSG00000130226 | 0 | 0 | 0 | 0 | 1 | 0 |
| ENSG00000226509 | 0 | 0 | 0 | 0 | 0 | 0 |
| ENSG00000270114 | 0 | 0 | 0 | 0 | 0 | 0 |
| ENSG00000203335 | 0 | 0 | 0 | 0 | 0 | 0 |
| ENSG00000233363 | 0 | 0 | 0 | 0 | 0 | 0 |
| ENSG00000236408 | 0 | 0 | 0 | 0 | 0 | 0 |
| ENSG00000214106 | 270 | 138 | 141 | 51 | 138 | 93 |
| ENSG00000157212 | 825 | 430 | 486 | 300 | 487 | 433 |
| ENSG00000272760 | 0 | 0 | 0 | 0 | 0 | 0 |
| ENSG00000273183 | 6 | 2 | 3 | 2 | 3 | 3 |
| ENSG00000273344 | 497 | 250 | 291 | 251 | 335 | 288 |
| ENSG00000220575 | 0 | 0 | 0 | 0 | 0 | 0 |
| ENSG00000157219 | 0 | 0 | 0 | 0 | 0 | 0 |
| ENSG00000283339 | 0 | 0 | 0 | 0 | 0 | 0 |
| ENSG00000274637 | 0 | 0 | 0 | 0 | 0 | 0 |
| ENSG00000228806 | 0 | 0 | 0 | 0 | 0 | 0 |
| ENSG00000201066 | 0 | 0 | 0 | 0 | 0 | 0 |
| ENSG00000217825 | 0 | 0 | 0 | 0 | 0 | 0 |
| ENSG00000228521 | 0 | 0 | 3 | 0 | 1 | 0 |
| ENSG00000278998 | 0 | 0 | 0 | 1 | 2 | 0 |
| ENSG00000225612 | 0 | 0 | 0 | 0 | 0 | 0 |
| ENSG00000273117 | 298 | 119 | 157 | 174 | 158 | 158 |
| ENSG00000186480 | 6114 | 2679 | 3426 | 2705 | 2896 | 2967 |
| ENSG00000204960 | 0 | 0 | 0 | 0 | 0 | 0 |
| ENSG00000218672 | 0 | 0 | 0 | 0 | 0 | 0 |
| ENSG00000260426 | 0 | 0 | 0 | 0 | 0 | 0 |
| ENSG00000236544 | 0 | 0 | 1 | 0 | 0 | 2 |
| ENSG00000164778 | 107 | 83 | 48 | 73 | 73 | 65 |
| ENSG00000146910 | 0 | 0 | 0 | 0 | 0 | 0 |
| ENSG00000283128 | 7 | 1 | 7 | 2 | 6 | 4 |
| ENSG00000227365 | 0 | 0 | 0 | 0 | 0 | 0 |
| ENSG00000216895 | 91 | 43 | 55 | 36 | 61 | 45 |
| ENSG00000184863 | 1788 | 775 | 1046 | 625 | 861 | 903 |
| ENSG00000164690 | 424 | 209 | 221 | 208 | 213 | 189 |
| ENSG00000204876 | 0 | 0 | 0 | 0 | 0 | 0 |
| ENSG00000207061 | 0 | 0 | 0 | 0 | 0 | 0 |
| ENSG00000270584 | 0 | 0 | 0 | 0 | 0 | 0 |
| ENSG00000228569 | 4 | 1 | 2 | 1 | 0 | 1 |
| ENSG00000233878 | 12 | 2 | 2 | 2 | 2 | 2 |
| ENSG00000182648 | 631 | 304 | 373 | 373 | 388 | 339 |
| ENSG00000279418 | 0 | 2 | 1 | 1 | 0 | 0 |
| ENSG00000234450 | 2 | 1 | 0 | 0 | 4 | 2 |
| ENSG00000105982 | 115 | 92 | 79 | 68 | 57 | 58 |
| ENSG00000224903 | 0 | 1 | 1 | 1 | 3 | 2 |
| ENSG00000105983 | 2908 | 1354 | 1677 | 1220 | 1510 | 1398 |
| ENSG00000206938 | 0 | 0 | 0 | 0 | 0 | 0 |
| ENSG00000230033 | 0 | 0 | 0 | 0 | 0 | 0 |
| ENSG00000146909 | 977 | 500 | 620 | 464 | 598 | 690 |

| | | | | | | |
|---|---|---|---|---|---|---|
| ENSG00000130675 | 315 | 127 | 174 | 161 | 156 | 138 |
| ENSG00000235029 | 13 | 4 | 10 | 10 | 8 | 6 |
| ENSG00000243479 | 153 | 65 | 65 | 94 | 79 | 84 |
| ENSG00000236102 | 0 | 0 | 0 | 0 | 0 | 0 |
| ENSG00000229365 | 0 | 0 | 0 | 0 | 0 | 0 |
| ENSG00000269446 | 0 | 0 | 0 | 0 | 0 | 0 |
| ENSG00000009335 | 5612 | 2687 | 3376 | 2456 | 3080 | 3256 |
| ENSG00000225224 | 2 | 2 | 0 | 0 | 1 | 1 |
| ENSG00000236018 | 0 | 0 | 1 | 1 | 1 | 2 |
| ENSG00000224629 | 8 | 2 | 2 | 0 | 4 | 3 |
| ENSG00000229660 | 0 | 0 | 1 | 0 | 0 | 0 |
| ENSG00000105993 | 6364 | 2997 | 3580 | 2771 | 3171 | 3415 |
| ENSG00000234210 | 6 | 2 | 1 | 0 | 0 | 0 |
| ENSG00000223872 | 0 | 0 | 0 | 0 | 0 | 0 |
| ENSG00000233191 | 0 | 2 | 0 | 0 | 0 | 0 |
| ENSG00000280266 | 2 | 0 | 0 | 0 | 0 | 0 |
| ENSG00000155093 | 194 | 86 | 132 | 76 | 67 | 98 |
| ENSG00000207960 | 0 | 0 | 0 | 0 | 0 | 0 |
| ENSG00000222012 | 0 | 0 | 0 | 0 | 0 | 0 |
| ENSG00000231515 | 0 | 0 | 0 | 0 | 0 | 0 |
| ENSG00000233038 | 4 | 0 | 2 | 0 | 1 | 0 |
| ENSG00000272839 | 0 | 0 | 0 | 0 | 0 | 0 |
| ENSG00000285821 | 0 | 0 | 0 | 0 | 0 | 0 |
| ENSG00000231980 | 0 | 1 | 0 | 0 | 0 | 1 |
| ENSG00000207637 | 0 | 0 | 0 | 0 | 0 | 0 |
| ENSG00000225365 | 0 | 2 | 5 | 0 | 2 | 3 |
| ENSG00000232715 | 0 | 1 | 1 | 0 | 0 | 0 |
| ENSG00000265598 | 0 | 0 | 1 | 0 | 0 | 0 |
| ENSG00000224324 | 0 | 0 | 0 | 0 | 0 | 0 |
| ENSG00000146918 | 2026 | 1104 | 1406 | 833 | 1543 | 1290 |
| ENSG00000228030 | 0 | 0 | 0 | 0 | 0 | 0 |
| ENSG00000117868 | 7190 | 3281 | 4260 | 2445 | 3503 | 3412 |
| ENSG00000126870 | 724 | 351 | 452 | 282 | 410 | 337 |
| ENSG00000231419 | 53 | 35 | 57 | 22 | 39 | 43 |
| ENSG00000106018 | 116 | 62 | 95 | 73 | 66 | 86 |
| ENSG00000229435 | 0 | 0 | 0 | 0 | 0 | 0 |
| ENSG00000228572 | 0 | 0 | 0 | 0 | 0 | 0 |
| ENSG00000182378 | 0 | 0 | 0 | 0 | 0 | 0 |
| ENSG00000178605 | 0 | 0 | 0 | 0 | 0 | 0 |
| ENSG00000226179 | 0 | 0 | 0 | 0 | 0 | 0 |
| ENSG00000167393 | 0 | 0 | 0 | 0 | 0 | 0 |
| ENSG00000281849 | 0 | 0 | 0 | 0 | 0 | 0 |
| ENSG00000275287 | 0 | 0 | 0 | 0 | 0 | 0 |
| ENSG00000280767 | 0 | 0 | 0 | 0 | 0 | 0 |
| ENSG00000234958 | 0 | 0 | 0 | 0 | 0 | 0 |
| ENSG00000229232 | 0 | 0 | 0 | 0 | 0 | 0 |
| ENSG00000185960 | 0 | 0 | 0 | 0 | 0 | 0 |
| ENSG00000237531 | 0 | 0 | 0 | 0 | 0 | 0 |
| ENSG00000225661 | 0 | 0 | 0 | 0 | 0 | 0 |
| ENSG00000205755 | 0 | 0 | 0 | 0 | 0 | 0 |

| | | | | | | |
|---|---|---|---|---|---|---|
| ENSG00000198223 | 0 | 0 | 0 | 0 | 0 | 0 |
| ENSG00000265658 | 0 | 0 | 0 | 0 | 0 | 0 |
| ENSG00000223274 | 0 | 0 | 0 | 0 | 0 | 0 |
| ENSG00000185291 | 0 | 0 | 0 | 0 | 0 | 0 |
| ENSG00000169100 | 0 | 0 | 0 | 0 | 0 | 0 |
| ENSG00000236871 | 0 | 0 | 0 | 0 | 0 | 0 |
| ENSG00000236017 | 0 | 0 | 0 | 0 | 0 | 0 |
| ENSG00000169093 | 0 | 0 | 0 | 0 | 0 | 0 |
| ENSG00000182162 | 0 | 0 | 0 | 0 | 0 | 0 |
| ENSG00000197976 | 0 | 0 | 0 | 0 | 0 | 0 |
| ENSG00000196433 | 0 | 0 | 0 | 0 | 0 | 0 |
| ENSG00000223511 | 0 | 0 | 0 | 0 | 0 | 0 |
| ENSG00000234622 | 0 | 0 | 0 | 0 | 0 | 0 |
| ENSG00000169084 | 0 | 0 | 0 | 0 | 0 | 0 |
| ENSG00000223571 | 0 | 0 | 0 | 0 | 0 | 0 |
| ENSG00000214717 | 0 | 0 | 0 | 0 | 0 | 0 |
| ENSG00000277120 | 0 | 0 | 0 | 0 | 0 | 0 |
| ENSG00000223773 | 0 | 0 | 0 | 0 | 0 | 0 |
| ENSG00000230542 | 0 | 0 | 0 | 0 | 0 | 0 |
| ENSG00000002586 | 0 | 0 | 0 | 0 | 0 | 0 |
| ENSG00000124343 | 47 | 20 | 15 | 8 | 28 | 22 |
| ENSG00000056998 | 478 | 234 | 304 | 232 | 309 | 281 |
| ENSG00000235483 | 0 | 0 | 0 | 0 | 0 | 0 |
| ENSG00000006756 | 3727 | 1826 | 2627 | 1873 | 1916 | 2239 |
| ENSG00000229851 | 0 | 0 | 0 | 0 | 0 | 0 |
| ENSG00000157399 | 1 | 0 | 1 | 3 | 0 | 0 |
| ENSG00000205667 | 0 | 0 | 1 | 2 | 2 | 2 |
| ENSG00000062096 | 0 | 0 | 0 | 0 | 0 | 0 |
| ENSG00000239228 | 0 | 0 | 0 | 0 | 0 | 0 |
| ENSG00000230249 | 0 | 0 | 0 | 0 | 0 | 0 |
| ENSG00000228459 | 0 | 0 | 0 | 0 | 0 | 0 |
| ENSG00000101825 | 0 | 0 | 0 | 0 | 0 | 0 |
| ENSG00000213487 | 0 | 0 | 0 | 0 | 0 | 0 |
| ENSG00000212214 | 0 | 0 | 0 | 0 | 0 | 0 |
| ENSG00000207435 | 0 | 0 | 0 | 0 | 0 | 0 |
| ENSG00000183943 | 1066 | 540 | 670 | 453 | 623 | 674 |
| ENSG00000207332 | 0 | 0 | 0 | 0 | 0 | 0 |
| ENSG00000207320 | 0 | 0 | 0 | 0 | 0 | 0 |
| ENSG00000236188 | 1 | 0 | 0 | 0 | 0 | 0 |
| ENSG00000182888 | 0 | 0 | 0 | 0 | 0 | 0 |
| ENSG00000285756 | 467 | 284 | 338 | 318 | 342 | 343 |
| ENSG00000225722 | 0 | 0 | 0 | 0 | 0 | 0 |
| ENSG00000236513 | 0 | 0 | 0 | 0 | 0 | 0 |
| ENSG00000235039 | 0 | 0 | 0 | 0 | 0 | 0 |
| ENSG00000226313 | 0 | 0 | 0 | 0 | 0 | 0 |
| ENSG00000238193 | 0 | 0 | 0 | 0 | 0 | 0 |
| ENSG00000201660 | 0 | 0 | 0 | 0 | 0 | 0 |
| ENSG00000236120 | 0 | 0 | 0 | 0 | 0 | 0 |
| ENSG00000146938 | 2 | 0 | 0 | 0 | 0 | 0 |
| ENSG00000252291 | 0 | 0 | 0 | 0 | 0 | 0 |

| | | | | | | |
|---|---|---|---|---|---|---|
| ENSG00000265284 | 0 | 0 | 0 | 0 | 0 | 0 |
| ENSG00000237730 | 0 | 0 | 1 | 1 | 0 | 1 |
| ENSG00000169059 | 0 | 0 | 0 | 0 | 0 | 0 |
| ENSG00000231217 | 2 | 0 | 0 | 0 | 0 | 0 |
| ENSG00000237490 | 0 | 0 | 0 | 0 | 0 | 0 |
| ENSG00000130021 | 1465 | 751 | 907 | 771 | 845 | 828 |
| ENSG00000228550 | 0 | 0 | 0 | 0 | 0 | 0 |
| ENSG00000264268 | 0 | 0 | 1 | 0 | 0 | 1 |
| ENSG00000101846 | 426 | 258 | 259 | 130 | 214 | 219 |
| ENSG00000279682 | 0 | 0 | 0 | 0 | 0 | 0 |
| ENSG00000182583 | 6 | 4 | 6 | 0 | 3 | 6 |
| ENSG00000274923 | 0 | 0 | 0 | 0 | 0 | 0 |
| ENSG00000006757 | 146 | 78 | 95 | 86 | 84 | 95 |
| ENSG00000230394 | 0 | 0 | 0 | 0 | 0 | 0 |
| ENSG00000285679 | 0 | 1 | 0 | 0 | 0 | 1 |
| ENSG00000207628 | 0 | 0 | 0 | 0 | 0 | 0 |
| ENSG00000177504 | 0 | 0 | 0 | 0 | 0 | 1 |
| ENSG00000205642 | 0 | 0 | 0 | 0 | 0 | 0 |
| ENSG00000011201 | 94 | 45 | 35 | 15 | 41 | 35 |
| ENSG00000225574 | 0 | 0 | 0 | 0 | 0 | 0 |
| ENSG00000183304 | 0 | 0 | 0 | 0 | 0 | 0 |
| ENSG00000231619 | 0 | 0 | 0 | 0 | 0 | 0 |
| ENSG00000229012 | 0 | 0 | 0 | 0 | 0 | 0 |
| ENSG00000285896 | 0 | 0 | 0 | 0 | 0 | 0 |
| ENSG00000225271 | 0 | 0 | 0 | 0 | 0 | 0 |
| ENSG00000177138 | 0 | 0 | 0 | 0 | 0 | 0 |
| ENSG00000200852 | 0 | 0 | 0 | 0 | 0 | 0 |
| ENSG00000219870 | 0 | 0 | 0 | 0 | 0 | 0 |
| ENSG00000228543 | 0 | 0 | 0 | 0 | 0 | 0 |
| ENSG00000232111 | 0 | 0 | 0 | 0 | 0 | 0 |
| ENSG00000282306 | 0 | 0 | 0 | 0 | 0 | 0 |
| ENSG00000282501 | 0 | 0 | 0 | 0 | 0 | 0 |
| ENSG00000282358 | 0 | 0 | 0 | 3 | 0 | 0 |
| ENSG00000282668 | 0 | 0 | 0 | 0 | 0 | 0 |
| ENSG00000282299 | 0 | 0 | 0 | 0 | 0 | 0 |
| ENSG00000282697 | 0 | 0 | 0 | 0 | 0 | 0 |
| ENSG00000282222 | 0 | 0 | 0 | 0 | 0 | 0 |
| ENSG00000281955 | 2 | 0 | 0 | 0 | 0 | 0 |
| ENSG00000281941 | 0 | 0 | 0 | 0 | 0 | 0 |
| ENSG00000282300 | 0 | 0 | 0 | 0 | 0 | 0 |
| ENSG00000101849 | 3732 | 1693 | 2236 | 1586 | 1763 | 2047 |
| ENSG00000101850 | 300 | 177 | 192 | 196 | 155 | 167 |
| ENSG00000146950 | 1137 | 486 | 657 | 490 | 503 | 577 |
| ENSG00000215319 | 0 | 1 | 0 | 0 | 0 | 0 |
| ENSG00000230326 | 0 | 0 | 0 | 0 | 0 | 0 |
| ENSG00000206844 | 0 | 0 | 0 | 0 | 0 | 0 |
| ENSG00000234469 | 2 | 0 | 2 | 1 | 0 | 2 |
| ENSG00000047644 | 2750 | 1417 | 1696 | 1096 | 1467 | 1565 |
| ENSG00000225076 | 0 | 0 | 2 | 0 | 1 | 0 |
| ENSG00000202469 | 0 | 0 | 0 | 0 | 0 | 0 |

| | | | | | | |
|---|---|---|---|---|---|---|
| ENSG00000073464 | 24 | 21 | 25 | 19 | 14 | 16 |
| ENSG00000227042 | 0 | 0 | 0 | 0 | 0 | 0 |
| ENSG00000101871 | 2 | 1 | 6 | 5 | 4 | 1 |
| ENSG00000207202 | 0 | 0 | 0 | 0 | 0 | 0 |
| ENSG00000234129 | 10 | 0 | 10 | 4 | 10 | 6 |
| ENSG00000004961 | 1299 | 588 | 702 | 653 | 740 | 731 |
| ENSG00000207151 | 0 | 0 | 0 | 0 | 0 | 3 |
| ENSG00000047648 | 0 | 0 | 0 | 0 | 0 | 0 |
| ENSG00000278218 | 0 | 0 | 0 | 0 | 0 | 0 |
| ENSG00000275194 | 0 | 0 | 0 | 0 | 0 | 0 |
| ENSG00000125363 | 0 | 0 | 0 | 0 | 0 | 0 |
| ENSG00000263652 | 0 | 0 | 0 | 0 | 0 | 0 |
| ENSG00000232392 | 0 | 0 | 0 | 0 | 0 | 0 |
| ENSG00000282840 | 0 | 0 | 1 | 0 | 0 | 2 |
| ENSG00000223418 | 0 | 0 | 0 | 0 | 0 | 0 |
| ENSG00000005302 | 683 | 336 | 419 | 318 | 416 | 378 |
| ENSG00000206792 | 0 | 0 | 0 | 0 | 0 | 0 |
| ENSG00000169933 | 0 | 0 | 0 | 0 | 0 | 1 |
| ENSG00000206795 | 0 | 0 | 0 | 0 | 0 | 0 |
| ENSG00000223487 | 0 | 0 | 0 | 0 | 0 | 0 |
| ENSG00000231211 | 0 | 0 | 0 | 0 | 0 | 0 |
| ENSG00000273800 | 0 | 0 | 0 | 0 | 0 | 0 |
| ENSG00000271814 | 0 | 0 | 0 | 0 | 0 | 0 |
| ENSG00000101911 | 865 | 397 | 502 | 377 | 562 | 498 |
| ENSG00000229083 | 0 | 0 | 1 | 1 | 0 | 0 |
| ENSG00000230691 | 0 | 0 | 0 | 0 | 0 | 0 |
| ENSG00000196664 | 2 | 0 | 0 | 0 | 0 | 0 |
| ENSG00000233338 | 0 | 1 | 0 | 1 | 1 | 0 |
| ENSG00000101916 | 0 | 0 | 0 | 0 | 0 | 0 |
| ENSG00000205542 | 53689 | 26359 | 36616 | 28461 | 27238 | 30346 |
| ENSG00000187268 | 0 | 0 | 0 | 0 | 0 | 0 |
| ENSG00000261030 | 0 | 0 | 0 | 0 | 0 | 0 |
| ENSG00000235385 | 4 | 0 | 5 | 0 | 0 | 0 |
| ENSG00000231216 | 0 | 0 | 0 | 0 | 0 | 0 |
| ENSG00000123594 | 0 | 0 | 0 | 0 | 0 | 0 |
| ENSG00000235592 | 0 | 0 | 0 | 0 | 0 | 0 |
| ENSG00000226985 | 0 | 0 | 0 | 0 | 0 | 0 |
| ENSG00000197582 | 0 | 0 | 0 | 0 | 0 | 0 |
| ENSG00000198759 | 0 | 0 | 0 | 0 | 0 | 0 |
| ENSG00000233357 | 0 | 0 | 0 | 0 | 0 | 0 |
| ENSG00000283493 | 0 | 0 | 0 | 0 | 0 | 0 |
| ENSG00000199622 | 0 | 0 | 0 | 0 | 0 | 0 |
| ENSG00000176896 | 176 | 84 | 90 | 69 | 85 | 59 |
| ENSG00000123595 | 1695 | 790 | 965 | 1027 | 1204 | 1203 |
| ENSG00000196459 | 784 | 339 | 586 | 322 | 466 | 480 |
| ENSG00000046651 | 1113 | 589 | 636 | 303 | 811 | 604 |
| ENSG00000046653 | 30 | 11 | 14 | 4 | 23 | 15 |
| ENSG00000233535 | 0 | 0 | 0 | 0 | 0 | 0 |
| ENSG00000212663 | 0 | 1 | 0 | 0 | 0 | 0 |
| ENSG00000046647 | 608 | 309 | 353 | 286 | 306 | 335 |

| | | | | | | |
|---|---|---|---|---|---|---|
| ENSG00000233247 | 0 | 0 | 0 | 1 | 0 | 0 |
| ENSG00000101958 | 0 | 0 | 0 | 0 | 0 | 0 |
| ENSG00000231066 | 0 | 0 | 2 | 2 | 1 | 1 |
| ENSG00000181544 | 20 | 20 | 26 | 24 | 36 | 24 |
| ENSG00000130150 | 319 | 197 | 152 | 76 | 296 | 134 |
| ENSG00000235929 | 0 | 0 | 0 | 0 | 0 | 0 |
| ENSG00000236929 | 0 | 0 | 0 | 0 | 0 | 0 |
| ENSG00000102048 | 107 | 67 | 94 | 53 | 73 | 66 |
| ENSG00000165192 | 1 | 0 | 0 | 0 | 0 | 0 |
| ENSG00000165195 | 316 | 193 | 194 | 130 | 259 | 208 |
| ENSG00000165197 | 2 | 0 | 1 | 1 | 1 | 0 |
| ENSG00000087842 | 1338 | 624 | 794 | 778 | 784 | 818 |
| ENSG00000102010 | 0 | 0 | 0 | 0 | 0 | 0 |
| ENSG00000130234 | 0 | 0 | 1 | 0 | 0 | 0 |
| ENSG00000285602 | 0 | 2 | 5 | 2 | 1 | 1 |
| ENSG00000225833 | 0 | 0 | 1 | 2 | 2 | 0 |
| ENSG00000147003 | 24 | 11 | 8 | 8 | 10 | 11 |
| ENSG00000237539 | 0 | 0 | 0 | 1 | 0 | 0 |
| ENSG00000186312 | 105 | 66 | 64 | 75 | 78 | 65 |
| ENSG00000169239 | 484 | 265 | 264 | 139 | 217 | 166 |
| ENSG00000279413 | 16 | 10 | 12 | 12 | 12 | 7 |
| ENSG00000200620 | 0 | 1 | 0 | 0 | 0 | 0 |
| ENSG00000281371 | 0 | 0 | 0 | 0 | 0 | 0 |
| ENSG00000169249 | 418 | 234 | 280 | 187 | 300 | 275 |
| ENSG00000182287 | 327 | 163 | 148 | 130 | 183 | 158 |
| ENSG00000200566 | 0 | 0 | 0 | 0 | 0 | 0 |
| ENSG00000226536 | 0 | 0 | 0 | 0 | 0 | 0 |
| ENSG00000126010 | 2 | 2 | 0 | 0 | 1 | 0 |
| ENSG00000238178 | 0 | 1 | 0 | 1 | 1 | 0 |
| ENSG00000182798 | 1 | 0 | 0 | 0 | 0 | 0 |
| ENSG00000215313 | 0 | 0 | 0 | 0 | 0 | 0 |
| ENSG00000235071 | 0 | 0 | 0 | 0 | 0 | 0 |
| ENSG00000239333 | 0 | 0 | 0 | 0 | 0 | 0 |
| ENSG00000047230 | 1745 | 919 | 1100 | 864 | 1086 | 1078 |
| ENSG00000265144 | 0 | 1 | 0 | 0 | 1 | 1 |
| ENSG00000169906 | 0 | 0 | 0 | 1 | 0 | 0 |
| ENSG00000169895 | 4585 | 2562 | 2563 | 1912 | 2780 | 2394 |
| ENSG00000238709 | 0 | 0 | 0 | 0 | 0 | 0 |
| ENSG00000086712 | 1303 | 675 | 735 | 496 | 850 | 746 |
| ENSG00000230180 | 0 | 0 | 0 | 0 | 0 | 0 |
| ENSG00000102054 | 8020 | 3976 | 5091 | 3662 | 4677 | 4513 |
| ENSG00000222736 | 0 | 0 | 0 | 0 | 0 | 0 |
| ENSG00000223498 | 0 | 0 | 0 | 0 | 0 | 0 |
| ENSG00000199410 | 0 | 0 | 0 | 0 | 0 | 0 |
| ENSG00000169891 | 542 | 260 | 363 | 192 | 284 | 264 |
| ENSG00000201467 | 0 | 0 | 0 | 0 | 0 | 0 |
| ENSG00000214322 | 0 | 0 | 0 | 0 | 0 | 0 |
| ENSG00000214321 | 4 | 0 | 0 | 0 | 0 | 0 |
| ENSG00000238764 | 0 | 0 | 0 | 0 | 0 | 0 |
| ENSG00000188158 | 86 | 60 | 52 | 25 | 76 | 53 |

| | | | | | | |
|---|---|---|---|---|---|---|
| ENSG00000265465 | 0 | 0 | 0 | 0 | 1 | 0 |
| ENSG00000235834 | 0 | 0 | 0 | 0 | 0 | 0 |
| ENSG00000230020 | 0 | 0 | 0 | 1 | 0 | 0 |
| ENSG00000226153 | 0 | 0 | 0 | 0 | 0 | 0 |
| ENSG00000047634 | 153 | 83 | 110 | 60 | 89 | 109 |
| ENSG00000227873 | 0 | 0 | 0 | 0 | 0 | 0 |
| ENSG00000131831 | 14 | 8 | 18 | 16 | 22 | 13 |
| ENSG00000276467 | 0 | 0 | 0 | 0 | 0 | 0 |
| ENSG00000225066 | 0 | 0 | 0 | 0 | 0 | 0 |
| ENSG00000225882 | 17 | 7 | 8 | 7 | 15 | 18 |
| ENSG00000177324 | 0 | 1 | 0 | 0 | 0 | 0 |
| ENSG00000102098 | 63 | 31 | 43 | 31 | 52 | 32 |
| ENSG00000235211 | 0 | 0 | 0 | 0 | 0 | 0 |
| ENSG00000207025 | 0 | 0 | 0 | 0 | 0 | 0 |
| ENSG00000008086 | 361 | 184 | 216 | 65 | 193 | 150 |
| ENSG00000236187 | 0 | 0 | 0 | 0 | 0 | 0 |
| ENSG00000102104 | 2 | 0 | 0 | 0 | 0 | 0 |
| ENSG00000086717 | 10 | 2 | 6 | 4 | 13 | 2 |
| ENSG00000237221 | 0 | 0 | 0 | 0 | 0 | 0 |
| ENSG00000196564 | 0 | 0 | 0 | 0 | 0 | 0 |
| ENSG00000224224 | 0 | 0 | 0 | 0 | 0 | 0 |
| ENSG00000237836 | 0 | 4 | 3 | 5 | 6 | 5 |
| ENSG00000044446 | 1041 | 480 | 713 | 443 | 536 | 597 |
| ENSG00000266710 | 0 | 0 | 0 | 1 | 0 | 0 |
| ENSG00000173698 | 31 | 16 | 23 | 10 | 16 | 18 |
| ENSG00000131828 | 3311 | 1623 | 1966 | 1699 | 1939 | 1932 |
| ENSG00000180815 | 1 | 2 | 0 | 3 | 3 | 3 |
| ENSG00000202144 | 0 | 0 | 0 | 0 | 0 | 0 |
| ENSG00000275800 | 0 | 0 | 0 | 0 | 0 | 0 |
| ENSG00000147010 | 514 | 282 | 283 | 260 | 351 | 306 |
| ENSG00000173681 | 295 | 177 | 139 | 90 | 175 | 166 |
| ENSG00000206663 | 0 | 0 | 0 | 0 | 0 | 0 |
| ENSG00000184368 | 28 | 4 | 2 | 2 | 4 | 5 |
| ENSG00000264566 | 0 | 0 | 0 | 0 | 0 | 0 |
| ENSG00000173674 | 2023 | 949 | 1101 | 911 | 1267 | 1159 |
| ENSG00000201882 | 0 | 0 | 0 | 0 | 0 | 0 |
| ENSG00000201592 | 0 | 0 | 0 | 0 | 0 | 1 |
| ENSG00000225037 | 2 | 0 | 4 | 1 | 1 | 1 |
| ENSG00000177189 | 978 | 520 | 562 | 275 | 703 | 525 |
| ENSG00000283631 | 0 | 0 | 0 | 0 | 0 | 0 |
| ENSG00000283400 | 0 | 0 | 0 | 0 | 0 | 0 |
| ENSG00000252978 | 0 | 0 | 0 | 0 | 0 | 0 |
| ENSG00000283380 | 0 | 0 | 0 | 0 | 0 | 0 |
| ENSG00000283546 | 0 | 0 | 0 | 0 | 0 | 0 |
| ENSG00000206716 | 0 | 0 | 0 | 0 | 0 | 0 |
| ENSG00000149970 | 0 | 0 | 0 | 0 | 0 | 0 |
| ENSG00000231913 | 0 | 0 | 0 | 0 | 0 | 0 |
| ENSG00000185915 | 0 | 0 | 0 | 0 | 0 | 0 |
| ENSG00000091482 | 2 | 0 | 0 | 0 | 0 | 0 |
| ENSG00000012174 | 1100 | 550 | 596 | 291 | 589 | 542 |

| | | | | | | |
|---|---|---|---|---|---|---|
| ENSG00000275681 | 0 | 0 | 0 | 0 | 0 | 0 |
| ENSG00000230797 | 38 | 20 | 27 | 14 | 13 | 12 |
| ENSG00000206639 | 0 | 0 | 0 | 0 | 0 | 0 |
| ENSG00000271286 | 0 | 0 | 0 | 0 | 0 | 0 |
| ENSG00000102172 | 4032 | 2076 | 2621 | 2042 | 2377 | 2406 |
| ENSG00000102174 | 83 | 55 | 50 | 48 | 50 | 48 |
| ENSG00000224204 | 0 | 0 | 0 | 0 | 0 | 0 |
| ENSG00000233067 | 0 | 0 | 0 | 0 | 0 | 0 |
| ENSG00000175809 | 0 | 0 | 0 | 0 | 0 | 0 |
| ENSG00000231846 | 0 | 0 | 0 | 0 | 0 | 0 |
| ENSG00000224396 | 0 | 0 | 0 | 0 | 0 | 0 |
| ENSG00000201095 | 0 | 0 | 0 | 0 | 0 | 0 |
| ENSG00000231011 | 0 | 0 | 0 | 0 | 0 | 0 |
| ENSG00000237019 | 0 | 0 | 0 | 0 | 0 | 0 |
| ENSG00000184735 | 2 | 1 | 0 | 0 | 1 | 2 |
| ENSG00000174028 | 63 | 41 | 28 | 22 | 43 | 34 |
| ENSG00000228610 | 0 | 0 | 0 | 0 | 0 | 0 |
| ENSG00000165186 | 0 | 0 | 0 | 0 | 0 | 0 |
| ENSG00000226774 | 0 | 0 | 0 | 0 | 0 | 0 |
| ENSG00000123131 | 2582 | 1304 | 1433 | 1766 | 1657 | 1498 |
| ENSG00000123130 | 862 | 424 | 477 | 467 | 583 | 565 |
| ENSG00000233785 | 2 | 1 | 2 | 1 | 0 | 0 |
| ENSG00000202272 | 0 | 0 | 0 | 0 | 0 | 0 |
| ENSG00000130066 | 3573 | 1905 | 2461 | 1402 | 1960 | 1859 |
| ENSG00000184831 | 369 | 168 | 239 | 231 | 217 | 196 |
| ENSG00000238103 | 0 | 3 | 3 | 4 | 2 | 2 |
| ENSG00000165182 | 8 | 2 | 2 | 1 | 0 | 0 |
| ENSG00000174010 | 126 | 67 | 79 | 54 | 71 | 62 |
| ENSG00000130741 | 5868 | 2900 | 3706 | 2707 | 3812 | 3583 |
| ENSG00000201407 | 0 | 0 | 0 | 0 | 0 | 0 |
| ENSG00000234230 | 1 | 0 | 0 | 1 | 5 | 1 |
| ENSG00000005889 | 807 | 393 | 439 | 206 | 474 | 366 |
| ENSG00000223611 | 0 | 0 | 0 | 0 | 0 | 0 |
| ENSG00000223731 | 0 | 0 | 0 | 0 | 0 | 0 |
| ENSG00000225071 | 0 | 0 | 1 | 1 | 0 | 0 |
| ENSG00000067992 | 1033 | 556 | 600 | 439 | 572 | 535 |
| ENSG00000214124 | 0 | 0 | 0 | 0 | 0 | 0 |
| ENSG00000260822 | 699 | 317 | 305 | 174 | 412 | 311 |
| ENSG00000102230 | 0 | 0 | 0 | 1 | 0 | 0 |
| ENSG00000226222 | 0 | 0 | 0 | 0 | 0 | 0 |
| ENSG00000236836 | 0 | 0 | 0 | 0 | 0 | 0 |
| ENSG00000236487 | 0 | 0 | 0 | 0 | 0 | 0 |
| ENSG00000101868 | 454 | 249 | 198 | 123 | 313 | 218 |
| ENSG00000251869 | 0 | 0 | 1 | 0 | 0 | 0 |
| ENSG00000232472 | 50 | 26 | 34 | 31 | 31 | 33 |
| ENSG00000004848 | 60 | 29 | 47 | 38 | 40 | 18 |
| ENSG00000214111 | 0 | 3 | 0 | 0 | 0 | 1 |
| ENSG00000240439 | 0 | 0 | 0 | 0 | 0 | 0 |
| ENSG00000176812 | 0 | 0 | 0 | 0 | 0 | 0 |
| ENSG00000229310 | 0 | 0 | 0 | 0 | 0 | 0 |

| | | | | | | |
|---|---|---|---|---|---|---|
| ENSG00000228519 | 2 | 2 | 1 | 4 | 4 | 2 |
| ENSG00000233403 | 0 | 0 | 0 | 0 | 0 | 0 |
| ENSG00000176774 | 0 | 0 | 0 | 0 | 0 | 0 |
| ENSG00000232446 | 0 | 0 | 0 | 0 | 0 | 0 |
| ENSG00000232030 | 0 | 0 | 0 | 0 | 0 | 0 |
| ENSG00000227511 | 0 | 0 | 0 | 0 | 0 | 0 |
| ENSG00000176746 | 2 | 2 | 0 | 0 | 3 | 0 |
| ENSG00000232644 | 0 | 0 | 0 | 0 | 0 | 0 |
| ENSG00000188408 | 0 | 0 | 0 | 0 | 0 | 0 |
| ENSG00000237171 | 0 | 0 | 0 | 0 | 0 | 0 |
| ENSG00000228464 | 0 | 0 | 0 | 0 | 0 | 0 |
| ENSG00000231388 | 0 | 0 | 0 | 0 | 0 | 0 |
| ENSG00000230265 | 0 | 0 | 0 | 0 | 0 | 0 |
| ENSG00000259849 | 0 | 0 | 0 | 0 | 0 | 0 |
| ENSG00000236160 | 0 | 0 | 0 | 0 | 0 | 0 |
| ENSG00000216938 | 0 | 0 | 0 | 0 | 0 | 0 |
| ENSG00000236683 | 2 | 0 | 0 | 0 | 0 | 1 |
| ENSG00000252491 | 0 | 0 | 0 | 0 | 0 | 0 |
| ENSG00000242021 | 0 | 0 | 0 | 0 | 0 | 0 |
| ENSG00000228933 | 0 | 0 | 0 | 0 | 0 | 0 |
| ENSG00000233807 | 0 | 0 | 0 | 0 | 0 | 0 |
| ENSG00000252486 | 0 | 0 | 0 | 0 | 0 | 0 |
| ENSG00000235642 | 0 | 0 | 0 | 0 | 0 | 0 |
| ENSG00000224960 | 0 | 0 | 0 | 0 | 0 | 0 |
| ENSG00000223391 | 0 | 0 | 0 | 0 | 0 | 0 |
| ENSG00000238247 | 2 | 0 | 0 | 0 | 0 | 0 |
| ENSG00000189186 | 2 | 1 | 0 | 2 | 1 | 2 |
| ENSG00000177689 | 0 | 0 | 0 | 0 | 0 | 0 |
| ENSG00000229731 | 0 | 0 | 0 | 0 | 0 | 0 |
| ENSG00000226199 | 0 | 0 | 0 | 0 | 0 | 0 |
| ENSG00000232834 | 0 | 0 | 0 | 0 | 0 | 0 |
| ENSG00000228345 | 0 | 0 | 0 | 0 | 0 | 0 |
| ENSG00000233243 | 0 | 0 | 0 | 0 | 0 | 0 |
| ENSG00000224589 | 0 | 0 | 0 | 0 | 0 | 0 |
| ENSG00000226372 | 0 | 0 | 0 | 1 | 0 | 2 |
| ENSG00000277569 | 0 | 0 | 0 | 0 | 0 | 0 |
| ENSG00000223742 | 0 | 0 | 0 | 0 | 0 | 0 |
| ENSG00000169306 | 0 | 0 | 0 | 0 | 0 | 0 |
| ENSG00000226436 | 0 | 0 | 0 | 0 | 0 | 0 |
| ENSG00000201666 | 0 | 0 | 0 | 0 | 0 | 0 |
| ENSG00000237994 | 0 | 0 | 0 | 0 | 0 | 0 |
| ENSG00000201356 | 0 | 0 | 0 | 0 | 0 | 0 |
| ENSG00000227393 | 0 | 0 | 0 | 0 | 0 | 0 |
| ENSG00000264090 | 0 | 0 | 0 | 0 | 0 | 0 |
| ENSG00000227592 | 0 | 0 | 0 | 0 | 0 | 0 |
| ENSG00000270794 | 0 | 0 | 0 | 0 | 0 | 0 |
| ENSG00000099399 | 0 | 0 | 0 | 0 | 0 | 0 |
| ENSG00000198798 | 0 | 0 | 0 | 0 | 0 | 0 |
| ENSG00000120289 | 0 | 0 | 0 | 0 | 0 | 0 |
| ENSG00000214107 | 2 | 0 | 0 | 0 | 0 | 0 |

| | | | | | | |
|---|---|---|---|---|---|---|
| ENSG00000276940 | 0 | 0 | 0 | 0 | 0 | 0 |
| ENSG00000169297 | 0 | 0 | 0 | 0 | 0 | 0 |
| ENSG00000120280 | 0 | 0 | 0 | 0 | 0 | 0 |
| ENSG00000178556 | 0 | 0 | 0 | 0 | 0 | 0 |
| ENSG00000232368 | 0 | 0 | 1 | 0 | 1 | 0 |
| ENSG00000198814 | 105 | 55 | 78 | 39 | 75 | 70 |
| ENSG00000229331 | 0 | 0 | 0 | 0 | 0 | 0 |
| ENSG00000241886 | 0 | 0 | 0 | 1 | 0 | 1 |
| ENSG00000243055 | 0 | 1 | 0 | 0 | 0 | 1 |
| ENSG00000277876 | 0 | 0 | 0 | 0 | 0 | 0 |
| ENSG00000157625 | 1136 | 570 | 575 | 298 | 723 | 571 |
| ENSG00000231542 | 0 | 0 | 0 | 0 | 0 | 0 |
| ENSG00000235512 | 0 | 0 | 0 | 0 | 0 | 0 |
| ENSG00000132446 | 0 | 0 | 0 | 0 | 0 | 0 |
| ENSG00000198947 | 0 | 0 | 0 | 0 | 1 | 0 |
| ENSG00000252903 | 0 | 0 | 0 | 0 | 0 | 0 |
| ENSG00000285900 | 0 | 0 | 0 | 0 | 0 | 0 |
| ENSG00000222922 | 0 | 0 | 0 | 0 | 0 | 0 |
| ENSG00000236285 | 2 | 0 | 0 | 0 | 0 | 0 |
| ENSG00000263600 | 0 | 0 | 0 | 0 | 0 | 0 |
| ENSG00000221348 | 0 | 0 | 0 | 0 | 0 | 0 |
| ENSG00000236828 | 0 | 0 | 0 | 0 | 0 | 0 |
| ENSG00000226781 | 3 | 0 | 4 | 2 | 8 | 3 |
| ENSG00000215310 | 0 | 0 | 0 | 0 | 0 | 0 |
| ENSG00000233928 | 2 | 2 | 1 | 0 | 1 | 0 |
| ENSG00000185448 | 0 | 0 | 0 | 0 | 0 | 0 |
| ENSG00000185638 | 0 | 0 | 0 | 0 | 0 | 0 |
| ENSG00000233571 | 0 | 0 | 0 | 0 | 0 | 0 |
| ENSG00000236072 | 0 | 0 | 0 | 0 | 0 | 0 |
| ENSG00000271136 | 0 | 0 | 0 | 0 | 0 | 0 |
| ENSG00000147027 | 0 | 2 | 0 | 1 | 0 | 2 |
| ENSG00000225384 | 0 | 0 | 0 | 0 | 0 | 0 |
| ENSG00000273704 | 0 | 0 | 0 | 0 | 0 | 0 |
| ENSG00000189132 | 0 | 0 | 0 | 0 | 0 | 0 |
| ENSG00000229347 | 0 | 0 | 0 | 0 | 0 | 0 |
| ENSG00000252411 | 0 | 0 | 0 | 0 | 0 | 0 |
| ENSG00000230227 | 0 | 0 | 0 | 0 | 0 | 0 |
| ENSG00000233663 | 0 | 0 | 0 | 0 | 0 | 0 |
| ENSG00000271088 | 0 | 0 | 0 | 0 | 0 | 0 |
| ENSG00000233880 | 0 | 0 | 0 | 0 | 0 | 0 |
| ENSG00000189023 | 0 | 0 | 0 | 0 | 1 | 0 |
| ENSG00000271519 | 0 | 0 | 0 | 0 | 0 | 0 |
| ENSG00000165164 | 0 | 0 | 0 | 0 | 0 | 1 |
| ENSG00000206733 | 0 | 0 | 0 | 0 | 0 | 0 |
| ENSG00000229384 | 0 | 0 | 0 | 0 | 0 | 0 |
| ENSG00000226484 | 0 | 0 | 0 | 0 | 0 | 0 |
| ENSG00000233921 | 0 | 0 | 0 | 0 | 0 | 0 |
| ENSG00000249467 | 0 | 0 | 0 | 0 | 0 | 0 |
| ENSG00000275314 | 0 | 0 | 0 | 0 | 0 | 0 |
| ENSG00000235510 | 0 | 0 | 0 | 0 | 0 | 0 |

| | | | | | | |
|---|---|---|---|---|---|---|
| ENSG00000223723 | 0 | 0 | 0 | 0 | 1 | 0 |
| ENSG00000231069 | 0 | 0 | 0 | 0 | 0 | 0 |
| ENSG00000198173 | 0 | 0 | 0 | 0 | 0 | 0 |
| ENSG00000243048 | 0 | 0 | 0 | 0 | 0 | 0 |
| ENSG00000214071 | 0 | 0 | 0 | 0 | 0 | 0 |
| ENSG00000233810 | 0 | 0 | 0 | 0 | 0 | 0 |
| ENSG00000130962 | 198 | 116 | 150 | 88 | 149 | 115 |
| ENSG00000250349 | 0 | 0 | 0 | 0 | 0 | 0 |
| ENSG00000241607 | 0 | 0 | 0 | 0 | 0 | 0 |
| ENSG00000219186 | 0 | 1 | 0 | 0 | 1 | 0 |
| ENSG00000234355 | 0 | 0 | 0 | 0 | 0 | 0 |
| ENSG00000174678 | 0 | 0 | 0 | 0 | 0 | 0 |
| ENSG00000147036 | 2 | 1 | 0 | 0 | 0 | 0 |
| ENSG00000047597 | 28 | 21 | 19 | 12 | 16 | 21 |
| ENSG00000285944 | 0 | 0 | 0 | 0 | 0 | 0 |
| ENSG00000165168 | 4 | 2 | 2 | 2 | 3 | 1 |
| ENSG00000165169 | 802 | 457 | 410 | 367 | 573 | 438 |
| ENSG00000206983 | 0 | 0 | 0 | 0 | 0 | 0 |
| ENSG00000259977 | 20 | 5 | 10 | 8 | 3 | 4 |
| ENSG00000224339 | 0 | 0 | 0 | 0 | 0 | 0 |
| ENSG00000187516 | 0 | 0 | 0 | 0 | 0 | 0 |
| ENSG00000229674 | 2 | 0 | 1 | 1 | 0 | 0 |
| ENSG00000147041 | 1138 | 567 | 638 | 334 | 458 | 414 |
| ENSG00000221466 | 0 | 0 | 0 | 0 | 0 | 0 |
| ENSG00000101955 | 38 | 24 | 23 | 5 | 18 | 10 |
| ENSG00000226679 | 0 | 0 | 0 | 0 | 0 | 0 |
| ENSG00000156313 | 126 | 71 | 111 | 49 | 70 | 74 |
| ENSG00000252050 | 1 | 0 | 0 | 0 | 1 | 0 |
| ENSG00000036473 | 0 | 0 | 0 | 0 | 0 | 0 |
| ENSG00000227988 | 0 | 0 | 0 | 0 | 0 | 0 |
| ENSG00000232370 | 0 | 0 | 0 | 0 | 0 | 0 |
| ENSG00000229735 | 0 | 0 | 0 | 0 | 0 | 0 |
| ENSG00000233007 | 0 | 0 | 0 | 0 | 0 | 0 |
| ENSG00000156298 | 0 | 0 | 1 | 0 | 0 | 0 |
| ENSG00000238606 | 0 | 0 | 0 | 0 | 0 | 0 |
| ENSG00000235806 | 2 | 0 | 0 | 2 | 0 | 1 |
| ENSG00000165175 | 3517 | 1633 | 1909 | 1400 | 1443 | 1454 |
| ENSG00000238123 | 28 | 17 | 19 | 8 | 12 | 20 |
| ENSG00000207122 | 0 | 0 | 0 | 0 | 0 | 0 |
| ENSG00000224610 | 0 | 0 | 0 | 0 | 0 | 0 |
| ENSG00000235304 | 0 | 0 | 0 | 0 | 0 | 0 |
| ENSG00000236747 | 0 | 0 | 0 | 0 | 0 | 0 |
| ENSG00000234191 | 0 | 0 | 0 | 0 | 0 | 0 |
| ENSG00000263730 | 0 | 0 | 0 | 0 | 0 | 0 |
| ENSG00000264618 | 0 | 0 | 0 | 0 | 0 | 0 |
| ENSG00000228232 | 84 | 37 | 64 | 49 | 48 | 57 |
| ENSG00000270759 | 0 | 0 | 0 | 0 | 0 | 0 |
| ENSG00000261435 | 0 | 0 | 0 | 1 | 0 | 0 |
| ENSG00000263972 | 0 | 0 | 0 | 0 | 0 | 0 |
| ENSG00000214047 | 0 | 0 | 0 | 0 | 0 | 0 |

| | | | | | | |
|---|---|---|---|---|---|---|
| ENSG00000285709 | 0 | 1 | 0 | 1 | 0 | 0 |
| ENSG00000223486 | 0 | 0 | 0 | 1 | 0 | 1 |
| ENSG00000183337 | 1642 | 688 | 1054 | 642 | 811 | 876 |
| ENSG00000236393 | 0 | 0 | 0 | 1 | 0 | 1 |
| ENSG00000214045 | 0 | 0 | 0 | 0 | 0 | 0 |
| ENSG00000206677 | 0 | 0 | 0 | 0 | 0 | 0 |
| ENSG00000235685 | 0 | 0 | 0 | 0 | 0 | 0 |
| ENSG00000238730 | 0 | 0 | 0 | 0 | 0 | 0 |
| ENSG00000182220 | 4746 | 2385 | 2591 | 1950 | 2694 | 2318 |
| ENSG00000238205 | 0 | 1 | 0 | 0 | 0 | 0 |
| ENSG00000185753 | 960 | 424 | 551 | 317 | 492 | 461 |
| ENSG00000180182 | 1685 | 867 | 942 | 544 | 996 | 827 |
| ENSG00000234636 | 4 | 1 | 6 | 4 | 4 | 6 |
| ENSG00000270791 | 0 | 0 | 0 | 0 | 0 | 0 |
| ENSG00000214031 | 0 | 0 | 0 | 0 | 0 | 0 |
| ENSG00000237661 | 0 | 0 | 0 | 0 | 0 | 0 |
| ENSG00000238222 | 0 | 1 | 0 | 0 | 0 | 0 |
| ENSG00000232801 | 0 | 0 | 0 | 0 | 1 | 0 |
| ENSG00000189145 | 0 | 0 | 0 | 0 | 0 | 0 |
| ENSG00000216866 | 14 | 5 | 6 | 11 | 5 | 7 |
| ENSG00000237931 | 0 | 0 | 1 | 0 | 0 | 0 |
| ENSG00000124486 | 2909 | 1700 | 1335 | 690 | 2643 | 1369 |
| ENSG00000223714 | 8 | 15 | 3 | 3 | 4 | 3 |
| ENSG00000199564 | 0 | 0 | 0 | 0 | 0 | 0 |
| ENSG00000238426 | 0 | 0 | 2 | 0 | 0 | 0 |
| ENSG00000226641 | 0 | 0 | 0 | 0 | 0 | 0 |
| ENSG00000215301 | 10453 | 5510 | 6088 | 3087 | 7133 | 5830 |
| ENSG00000264573 | 0 | 2 | 0 | 0 | 0 | 0 |
| ENSG00000188937 | 0 | 0 | 0 | 0 | 0 | 0 |
| ENSG00000225238 | 0 | 0 | 0 | 0 | 0 | 0 |
| ENSG00000147044 | 1505 | 733 | 927 | 582 | 925 | 850 |
| ENSG00000233033 | 0 | 0 | 1 | 0 | 0 | 0 |
| ENSG00000212207 | 0 | 0 | 0 | 0 | 0 | 0 |
| ENSG00000217624 | 6 | 1 | 3 | 2 | 0 | 1 |
| ENSG00000242559 | 0 | 0 | 0 | 0 | 0 | 0 |
| ENSG00000171659 | 0 | 0 | 0 | 0 | 1 | 0 |
| ENSG00000171657 | 4 | 0 | 1 | 0 | 0 | 1 |
| ENSG00000229822 | 0 | 0 | 0 | 0 | 0 | 0 |
| ENSG00000237875 | 0 | 0 | 0 | 0 | 0 | 0 |
| ENSG00000251807 | 0 | 0 | 0 | 0 | 0 | 0 |
| ENSG00000233103 | 0 | 0 | 0 | 0 | 0 | 0 |
| ENSG00000228847 | 0 | 1 | 1 | 0 | 0 | 0 |
| ENSG00000212560 | 0 | 0 | 0 | 0 | 0 | 0 |
| ENSG00000228780 | 0 | 0 | 0 | 0 | 0 | 0 |
| ENSG00000206896 | 0 | 0 | 0 | 0 | 0 | 0 |
| ENSG00000231772 | 0 | 0 | 0 | 0 | 0 | 0 |
| ENSG00000283737 | 0 | 0 | 0 | 0 | 0 | 0 |
| ENSG00000102055 | 0 | 0 | 0 | 0 | 0 | 0 |
| ENSG00000270280 | 0 | 0 | 0 | 0 | 0 | 0 |
| ENSG00000224294 | 0 | 0 | 0 | 0 | 0 | 0 |

| | | | | | | |
|---|---|---|---|---|---|---|
| ENSG00000237206 | 0 | 0 | 0 | 0 | 0 | 0 |
| ENSG00000285899 | 0 | 0 | 0 | 0 | 0 | 0 |
| ENSG00000231750 | 0 | 0 | 0 | 0 | 0 | 0 |
| ENSG00000189221 | 149 | 81 | 94 | 82 | 89 | 86 |
| ENSG00000069535 | 26 | 11 | 20 | 7 | 11 | 10 |
| ENSG00000124479 | 0 | 0 | 0 | 0 | 0 | 0 |
| ENSG00000236276 | 0 | 0 | 0 | 0 | 0 | 0 |
| ENSG00000214019 | 0 | 2 | 0 | 0 | 0 | 0 |
| ENSG00000183690 | 0 | 1 | 1 | 0 | 3 | 2 |
| ENSG00000198414 | 0 | 0 | 0 | 0 | 0 | 0 |
| ENSG00000214018 | 0 | 0 | 0 | 1 | 1 | 1 |
| ENSG00000200702 | 0 | 0 | 0 | 0 | 0 | 0 |
| ENSG00000224054 | 0 | 0 | 0 | 0 | 0 | 0 |
| ENSG00000215035 | 0 | 0 | 0 | 0 | 0 | 0 |
| ENSG00000069509 | 705 | 317 | 403 | 320 | 400 | 377 |
| ENSG00000233543 | 0 | 0 | 1 | 0 | 0 | 0 |
| ENSG00000229168 | 0 | 0 | 0 | 0 | 0 | 0 |
| ENSG00000214016 | 4 | 1 | 2 | 2 | 1 | 1 |
| ENSG00000242065 | 0 | 0 | 0 | 0 | 0 | 0 |
| ENSG00000189037 | 0 | 0 | 0 | 0 | 0 | 0 |
| ENSG00000147050 | 1017 | 476 | 601 | 329 | 503 | 437 |
| ENSG00000147113 | 0 | 0 | 0 | 0 | 0 | 0 |
| ENSG00000229491 | 2 | 0 | 1 | 0 | 0 | 0 |
| ENSG00000237782 | 0 | 0 | 0 | 0 | 0 | 0 |
| ENSG00000229563 | 0 | 1 | 0 | 0 | 1 | 1 |
| ENSG00000214282 | 2 | 1 | 1 | 0 | 2 | 0 |
| ENSG00000204915 | 0 | 0 | 0 | 0 | 0 | 0 |
| ENSG00000270069 | 0 | 2 | 1 | 1 | 0 | 2 |
| ENSG00000207870 | 0 | 0 | 0 | 0 | 0 | 0 |
| ENSG00000207725 | 0 | 0 | 0 | 0 | 0 | 0 |
| ENSG00000269902 | 0 | 0 | 0 | 0 | 0 | 0 |
| ENSG00000231566 | 0 | 0 | 1 | 0 | 1 | 0 |
| ENSG00000212347 | 0 | 0 | 0 | 0 | 0 | 0 |
| ENSG00000215286 | 0 | 0 | 0 | 0 | 0 | 0 |
| ENSG00000234613 | 0 | 0 | 0 | 0 | 0 | 0 |
| ENSG00000224701 | 0 | 0 | 0 | 0 | 0 | 0 |
| ENSG00000213100 | 0 | 2 | 1 | 1 | 0 | 1 |
| ENSG00000214009 | 0 | 0 | 0 | 0 | 0 | 0 |
| ENSG00000278838 | 0 | 0 | 0 | 0 | 0 | 0 |
| ENSG00000229145 | 0 | 0 | 0 | 0 | 0 | 0 |
| ENSG00000236751 | 1 | 0 | 3 | 1 | 0 | 1 |
| ENSG00000231235 | 0 | 0 | 0 | 0 | 0 | 0 |
| ENSG00000235587 | 38 | 22 | 16 | 21 | 23 | 19 |
| ENSG00000147121 | 308 | 158 | 159 | 137 | 204 | 199 |
| ENSG00000224072 | 6 | 0 | 1 | 0 | 1 | 1 |
| ENSG00000251192 | 124 | 51 | 85 | 52 | 68 | 77 |
| ENSG00000199226 | 0 | 0 | 0 | 0 | 0 | 0 |
| ENSG00000230844 | 120 | 76 | 83 | 56 | 93 | 89 |
| ENSG00000277721 | 0 | 0 | 0 | 3 | 1 | 0 |
| ENSG00000270499 | 4 | 2 | 1 | 0 | 1 | 0 |

| | | | | | | |
|---|---|---|---|---|---|---|
| ENSG00000147119 | 130 | 64 | 71 | 83 | 66 | 69 |
| ENSG00000065923 | 4558 | 2135 | 3076 | 1705 | 2382 | 2494 |
| ENSG00000213997 | 28 | 17 | 25 | 8 | 19 | 33 |
| ENSG00000226048 | 0 | 0 | 0 | 1 | 0 | 1 |
| ENSG00000215284 | 2 | 3 | 2 | 0 | 1 | 0 |
| ENSG00000102218 | 318 | 194 | 246 | 161 | 222 | 198 |
| ENSG00000204904 | 12 | 4 | 3 | 2 | 3 | 5 |
| ENSG00000102221 | 859 | 421 | 530 | 316 | 437 | 445 |
| ENSG00000130988 | 4 | 1 | 0 | 0 | 2 | 1 |
| ENSG00000206685 | 0 | 0 | 0 | 0 | 0 | 1 |
| ENSG00000201659 | 1 | 1 | 0 | 1 | 0 | 0 |
| ENSG00000147123 | 3791 | 1773 | 2227 | 2491 | 1979 | 1982 |
| ENSG00000182872 | 2588 | 1307 | 1597 | 1500 | 1448 | 1420 |
| ENSG00000237351 | 0 | 2 | 1 | 1 | 3 | 1 |
| ENSG00000130985 | 17948 | 8133 | 10502 | 9048 | 8957 | 8850 |
| ENSG00000224975 | 42 | 25 | 39 | 36 | 34 | 27 |
| ENSG00000102225 | 4588 | 2220 | 2743 | 2438 | 2258 | 2367 |
| ENSG00000102226 | 3133 | 1217 | 1862 | 1714 | 1372 | 1705 |
| ENSG00000237623 | 0 | 0 | 0 | 0 | 0 | 0 |
| ENSG00000266158 | 0 | 0 | 0 | 0 | 0 | 0 |
| ENSG00000203402 | 0 | 0 | 0 | 0 | 0 | 0 |
| ENSG00000147117 | 4 | 0 | 4 | 2 | 2 | 1 |
| ENSG00000221459 | 0 | 0 | 0 | 0 | 0 | 0 |
| ENSG00000232009 | 0 | 0 | 0 | 0 | 0 | 0 |
| ENSG00000226556 | 0 | 0 | 0 | 0 | 0 | 0 |
| ENSG00000147124 | 506 | 246 | 276 | 130 | 275 | 217 |
| ENSG00000277605 | 0 | 0 | 0 | 0 | 0 | 0 |
| ENSG00000196741 | 135 | 48 | 88 | 29 | 73 | 46 |
| ENSG00000235636 | 31 | 7 | 10 | 6 | 15 | 11 |
| ENSG00000078061 | 2485 | 1111 | 1575 | 1458 | 1259 | 1363 |
| ENSG00000008056 | 4 | 0 | 4 | 5 | 5 | 0 |
| ENSG00000102265 | 1760 | 814 | 1002 | 1129 | 951 | 920 |
| ENSG00000263858 | 0 | 0 | 1 | 0 | 0 | 0 |
| ENSG00000283743 | 0 | 0 | 0 | 0 | 0 | 0 |
| ENSG00000126759 | 8 | 7 | 11 | 6 | 6 | 8 |
| ENSG00000126767 | 1479 | 643 | 863 | 848 | 794 | 866 |
| ENSG00000126756 | 825 | 509 | 595 | 663 | 576 | 537 |
| ENSG00000267064 | 4 | 5 | 5 | 6 | 2 | 2 |
| ENSG00000187893 | 0 | 0 | 0 | 0 | 0 | 0 |
| ENSG00000224656 | 0 | 0 | 0 | 0 | 0 | 0 |
| ENSG00000229968 | 0 | 0 | 0 | 0 | 0 | 0 |
| ENSG00000188459 | 4 | 1 | 0 | 0 | 0 | 2 |
| ENSG00000197779 | 189 | 108 | 90 | 51 | 116 | 117 |
| ENSG00000224401 | 2 | 2 | 3 | 0 | 2 | 2 |
| ENSG00000147118 | 236 | 124 | 133 | 85 | 186 | 108 |
| ENSG00000221994 | 366 | 180 | 222 | 174 | 217 | 206 |
| ENSG00000171489 | 0 | 0 | 0 | 0 | 0 | 0 |
| ENSG00000277541 | 2 | 0 | 0 | 0 | 0 | 1 |
| ENSG00000171483 | 0 | 0 | 0 | 0 | 0 | 0 |
| ENSG00000171478 | 0 | 0 | 0 | 0 | 0 | 0 |

| | | | | | | |
|---|---|---|---|---|---|---|
| ENSG00000230241 | 0 | 0 | 0 | 0 | 0 | 0 |
| ENSG00000233139 | 0 | 0 | 0 | 0 | 0 | 0 |
| ENSG00000201517 | 0 | 0 | 0 | 0 | 0 | 0 |
| ENSG00000225198 | 0 | 0 | 0 | 0 | 0 | 0 |
| ENSG00000238032 | 0 | 0 | 0 | 0 | 0 | 0 |
| ENSG00000165583 | 0 | 0 | 0 | 0 | 0 | 0 |
| ENSG00000234780 | 0 | 0 | 0 | 0 | 0 | 0 |
| ENSG00000252856 | 0 | 0 | 0 | 0 | 0 | 0 |
| ENSG00000204368 | 0 | 0 | 0 | 0 | 0 | 0 |
| ENSG00000231568 | 0 | 0 | 0 | 0 | 0 | 0 |
| ENSG00000229662 | 0 | 0 | 0 | 0 | 0 | 0 |
| ENSG00000126752 | 0 | 0 | 0 | 0 | 0 | 0 |
| ENSG00000232813 | 0 | 0 | 0 | 0 | 0 | 0 |
| ENSG00000241207 | 0 | 0 | 0 | 0 | 0 | 0 |
| ENSG00000237345 | 0 | 0 | 0 | 0 | 0 | 0 |
| ENSG00000204648 | 0 | 0 | 0 | 0 | 0 | 0 |
| ENSG00000234391 | 0 | 0 | 0 | 0 | 0 | 0 |
| ENSG00000227302 | 0 | 0 | 0 | 0 | 0 | 0 |
| ENSG00000230100 | 0 | 0 | 0 | 0 | 0 | 0 |
| ENSG00000270747 | 0 | 0 | 0 | 0 | 0 | 0 |
| ENSG00000165584 | 0 | 0 | 0 | 0 | 0 | 0 |
| ENSG00000225055 | 0 | 0 | 0 | 0 | 0 | 0 |
| ENSG00000234448 | 0 | 0 | 0 | 0 | 0 | 0 |
| ENSG00000268009 | 0 | 0 | 0 | 0 | 0 | 0 |
| ENSG00000269791 | 0 | 0 | 0 | 0 | 0 | 0 |
| ENSG00000231603 | 0 | 0 | 0 | 0 | 0 | 0 |
| ENSG00000231489 | 0 | 0 | 0 | 0 | 0 | 0 |
| ENSG00000235350 | 3 | 4 | 1 | 0 | 2 | 4 |
| ENSG00000226971 | 0 | 0 | 0 | 0 | 0 | 0 |
| ENSG00000017483 | 0 | 1 | 3 | 3 | 4 | 1 |
| ENSG00000068438 | 1006 | 543 | 627 | 599 | 572 | 656 |
| ENSG00000224292 | 2 | 4 | 5 | 3 | 0 | 1 |
| ENSG00000102312 | 255 | 154 | 155 | 147 | 143 | 145 |
| ENSG00000147155 | 1898 | 971 | 1183 | 1393 | 1142 | 1186 |
| ENSG00000276823 | 0 | 0 | 1 | 0 | 0 | 0 |
| ENSG00000068354 | 707 | 296 | 383 | 407 | 363 | 377 |
| ENSG00000204620 | 21 | 8 | 10 | 7 | 17 | 15 |
| ENSG00000102317 | 6507 | 3457 | 4098 | 2846 | 3819 | 4050 |
| ENSG00000228343 | 85 | 53 | 65 | 66 | 63 | 66 |
| ENSG00000279528 | 37 | 39 | 46 | 45 | 51 | 39 |
| ENSG00000220125 | 0 | 0 | 0 | 0 | 0 | 0 |
| ENSG00000101940 | 1436 | 644 | 851 | 777 | 664 | 720 |
| ENSG00000231725 | 0 | 0 | 0 | 0 | 0 | 0 |
| ENSG00000015285 | 116 | 54 | 102 | 77 | 64 | 60 |
| ENSG00000101945 | 381 | 197 | 234 | 259 | 192 | 231 |
| ENSG00000232828 | 0 | 1 | 0 | 0 | 0 | 2 |
| ENSG00000206723 | 0 | 0 | 0 | 0 | 0 | 0 |
| ENSG00000171433 | 21 | 9 | 7 | 5 | 11 | 6 |
| ENSG00000233585 | 8 | 4 | 3 | 6 | 4 | 6 |
| ENSG00000207367 | 1 | 0 | 2 | 1 | 1 | 0 |

| | | | | | | |
|---|---|---|---|---|---|---|
| ENSG00000102145 | 2 | 0 | 3 | 1 | 0 | 2 |
| ENSG00000094631 | 1118 | 522 | 714 | 585 | 629 | 618 |
| ENSG00000187682 | 2 | 0 | 1 | 1 | 0 | 1 |
| ENSG00000102109 | 2001 | 766 | 1082 | 1363 | 960 | 1029 |
| ENSG00000126768 | 1986 | 958 | 1132 | 1299 | 1137 | 1121 |
| ENSG00000102103 | 1559 | 779 | 985 | 1109 | 931 | 936 |
| ENSG00000102100 | 1850 | 832 | 1073 | 953 | 1001 | 1047 |
| ENSG00000102096 | 458 | 198 | 242 | 256 | 208 | 247 |
| ENSG00000068308 | 2300 | 1055 | 1421 | 1183 | 1138 | 1202 |
| ENSG00000279155 | 4 | 5 | 5 | 1 | 5 | 4 |
| ENSG00000223309 | 1 | 1 | 0 | 3 | 1 | 1 |
| ENSG00000102057 | 115 | 46 | 63 | 50 | 49 | 59 |
| ENSG00000068400 | 2528 | 1053 | 1473 | 1240 | 1243 | 1286 |
| ENSG00000068323 | 2233 | 943 | 1282 | 1262 | 1194 | 1202 |
| ENSG00000147144 | 1271 | 609 | 812 | 593 | 630 | 657 |
| ENSG00000243279 | 322 | 147 | 205 | 214 | 162 | 180 |
| ENSG00000196998 | 1515 | 833 | 993 | 818 | 834 | 860 |
| ENSG00000206936 | 7 | 2 | 13 | 12 | 6 | 4 |
| ENSG00000068394 | 1186 | 610 | 764 | 696 | 709 | 689 |
| ENSG00000265033 | 0 | 0 | 0 | 1 | 0 | 0 |
| ENSG00000280116 | 0 | 0 | 0 | 0 | 0 | 1 |
| ENSG00000269313 | 133 | 61 | 79 | 59 | 56 | 38 |
| ENSG00000279422 | 0 | 1 | 3 | 0 | 0 | 0 |
| ENSG00000102007 | 1949 | 974 | 1122 | 1257 | 1186 | 1145 |
| ENSG00000012211 | 352 | 181 | 211 | 256 | 257 | 236 |
| ENSG00000102003 | 46 | 39 | 23 | 15 | 13 | 37 |
| ENSG00000237341 | 0 | 0 | 0 | 0 | 0 | 0 |
| ENSG00000102001 | 0 | 0 | 0 | 0 | 0 | 0 |
| ENSG00000230216 | 6 | 2 | 3 | 1 | 1 | 1 |
| ENSG00000101997 | 1161 | 468 | 671 | 733 | 511 | 560 |
| ENSG00000049768 | 11 | 13 | 11 | 8 | 10 | 3 |
| ENSG00000049769 | 606 | 252 | 338 | 332 | 273 | 272 |
| ENSG00000270012 | 78 | 26 | 72 | 33 | 40 | 33 |
| ENSG00000215274 | 0 | 1 | 0 | 0 | 0 | 0 |
| ENSG00000224659 | 0 | 0 | 0 | 0 | 0 | 0 |
| ENSG00000274274 | 0 | 0 | 0 | 0 | 0 | 0 |
| ENSG00000275113 | 0 | 0 | 0 | 0 | 0 | 0 |
| ENSG00000236737 | 0 | 0 | 0 | 0 | 0 | 0 |
| ENSG00000237671 | 0 | 0 | 0 | 0 | 0 | 0 |
| ENSG00000227488 | 0 | 0 | 0 | 0 | 0 | 0 |
| ENSG00000236362 | 0 | 0 | 0 | 0 | 0 | 0 |
| ENSG00000216649 | 0 | 0 | 0 | 0 | 0 | 0 |
| ENSG00000215269 | 0 | 0 | 0 | 0 | 0 | 0 |
| ENSG00000224902 | 0 | 0 | 0 | 0 | 0 | 0 |
| ENSG00000205777 | 0 | 0 | 0 | 0 | 0 | 0 |
| ENSG00000189064 | 0 | 0 | 0 | 1 | 0 | 0 |
| ENSG00000213856 | 4 | 5 | 3 | 1 | 0 | 0 |
| ENSG00000227391 | 0 | 0 | 0 | 0 | 0 | 0 |
| ENSG00000068985 | 0 | 0 | 0 | 0 | 0 | 0 |
| ENSG00000101951 | 0 | 0 | 0 | 0 | 0 | 0 |

| | | | | | | |
|---|---|---|---|---|---|---|
| ENSG00000234390 | 79 | 35 | 46 | 32 | 38 | 47 |
| ENSG00000273820 | 240 | 113 | 134 | 131 | 154 | 133 |
| ENSG00000220132 | 0 | 0 | 0 | 0 | 0 | 0 |
| ENSG00000171365 | 338 | 157 | 162 | 94 | 195 | 159 |
| ENSG00000201341 | 0 | 0 | 0 | 0 | 0 | 0 |
| ENSG00000207758 | 0 | 0 | 0 | 0 | 0 | 0 |
| ENSG00000207768 | 0 | 0 | 0 | 0 | 0 | 0 |
| ENSG00000207785 | 0 | 0 | 0 | 0 | 0 | 0 |
| ENSG00000208015 | 0 | 0 | 0 | 0 | 0 | 0 |
| ENSG00000211538 | 0 | 0 | 0 | 0 | 0 | 0 |
| ENSG00000239057 | 0 | 0 | 0 | 0 | 0 | 0 |
| ENSG00000207970 | 0 | 0 | 0 | 0 | 0 | 0 |
| ENSG00000272080 | 0 | 0 | 0 | 0 | 0 | 0 |
| ENSG00000181977 | 0 | 0 | 0 | 0 | 0 | 0 |
| ENSG00000202495 | 0 | 0 | 0 | 0 | 0 | 0 |
| ENSG00000147081 | 0 | 0 | 0 | 0 | 0 | 0 |
| ENSG00000147082 | 0 | 2 | 4 | 1 | 4 | 1 |
| ENSG00000274588 | 0 | 0 | 0 | 0 | 0 | 0 |
| ENSG00000158352 | 12 | 5 | 1 | 5 | 2 | 3 |
| ENSG00000222303 | 0 | 0 | 0 | 0 | 0 | 0 |
| ENSG00000226881 | 0 | 0 | 0 | 0 | 0 | 0 |
| ENSG00000130385 | 0 | 0 | 0 | 0 | 0 | 0 |
| ENSG00000238190 | 0 | 0 | 1 | 0 | 0 | 0 |
| ENSG00000227493 | 0 | 0 | 0 | 0 | 0 | 0 |
| ENSG00000230926 | 0 | 0 | 0 | 0 | 0 | 0 |
| ENSG00000230317 | 0 | 0 | 0 | 0 | 0 | 0 |
| ENSG00000237926 | 0 | 0 | 0 | 0 | 0 | 0 |
| ENSG00000122824 | 4 | 0 | 0 | 1 | 0 | 0 |
| ENSG00000229151 | 0 | 0 | 0 | 0 | 0 | 0 |
| ENSG00000226530 | 0 | 0 | 0 | 0 | 0 | 0 |
| ENSG00000187690 | 0 | 0 | 0 | 0 | 0 | 0 |
| ENSG00000196368 | 4 | 3 | 1 | 2 | 3 | 1 |
| ENSG00000234766 | 0 | 0 | 0 | 0 | 0 | 0 |
| ENSG00000224109 | 0 | 0 | 0 | 0 | 0 | 0 |
| ENSG00000283093 | 0 | 0 | 0 | 0 | 0 | 0 |
| ENSG00000223591 | 0 | 0 | 0 | 0 | 0 | 0 |
| ENSG00000189369 | 0 | 0 | 0 | 0 | 0 | 0 |
| ENSG00000179222 | 6552 | 2964 | 3638 | 2982 | 3215 | 2993 |
| ENSG00000236576 | 0 | 0 | 0 | 0 | 0 | 0 |
| ENSG00000252377 | 0 | 0 | 0 | 0 | 0 | 0 |
| ENSG00000228827 | 0 | 0 | 0 | 0 | 0 | 0 |
| ENSG00000232951 | 0 | 0 | 1 | 0 | 0 | 0 |
| ENSG00000237528 | 0 | 0 | 0 | 0 | 0 | 0 |
| ENSG00000232562 | 0 | 0 | 0 | 0 | 0 | 0 |
| ENSG00000182776 | 0 | 0 | 0 | 0 | 0 | 0 |
| ENSG00000279750 | 0 | 0 | 0 | 0 | 0 | 0 |
| ENSG00000187243 | 0 | 0 | 2 | 1 | 0 | 0 |
| ENSG00000221705 | 0 | 0 | 0 | 0 | 0 | 0 |
| ENSG00000154545 | 0 | 0 | 1 | 0 | 0 | 0 |
| ENSG00000221475 | 0 | 0 | 0 | 0 | 0 | 0 |

| | | | | | | |
|---|---|---|---|---|---|---|
| ENSG00000179028 | 0 | 0 | 0 | 0 | 0 | 0 |
| ENSG00000276474 | 0 | 0 | 0 | 0 | 0 | 0 |
| ENSG00000275335 | 0 | 0 | 0 | 0 | 0 | 0 |
| ENSG00000155622 | 0 | 0 | 0 | 0 | 0 | 0 |
| ENSG00000225957 | 0 | 0 | 0 | 0 | 0 | 0 |
| ENSG00000276897 | 0 | 0 | 0 | 0 | 0 | 0 |
| ENSG00000224556 | 0 | 0 | 0 | 0 | 0 | 0 |
| ENSG00000250084 | 0 | 0 | 0 | 0 | 0 | 0 |
| ENSG00000278358 | 0 | 0 | 0 | 0 | 0 | 0 |
| ENSG00000278160 | 0 | 0 | 0 | 0 | 0 | 0 |
| ENSG00000275643 | 0 | 0 | 0 | 0 | 0 | 0 |
| ENSG00000204379 | 0 | 0 | 0 | 0 | 0 | 0 |
| ENSG00000223395 | 0 | 0 | 0 | 0 | 0 | 0 |
| ENSG00000204382 | 0 | 0 | 0 | 0 | 0 | 0 |
| ENSG00000277289 | 0 | 0 | 0 | 0 | 0 | 0 |
| ENSG00000227058 | 0 | 0 | 0 | 0 | 0 | 0 |
| ENSG00000231593 | 0 | 0 | 0 | 0 | 0 | 0 |
| ENSG00000228771 | 0 | 0 | 0 | 0 | 0 | 0 |
| ENSG00000185319 | 0 | 0 | 0 | 0 | 0 | 0 |
| ENSG00000197185 | 0 | 0 | 0 | 0 | 0 | 0 |
| ENSG00000234792 | 0 | 0 | 0 | 0 | 0 | 0 |
| ENSG00000157965 | 0 | 0 | 0 | 0 | 0 | 0 |
| ENSG00000187754 | 0 | 0 | 0 | 0 | 0 | 0 |
| ENSG00000228354 | 0 | 0 | 0 | 0 | 0 | 0 |
| ENSG00000222608 | 0 | 0 | 0 | 0 | 0 | 0 |
| ENSG00000225508 | 0 | 0 | 0 | 0 | 0 | 0 |
| ENSG00000237275 | 0 | 0 | 0 | 0 | 0 | 0 |
| ENSG00000241476 | 0 | 0 | 0 | 0 | 0 | 0 |
| ENSG00000223958 | 0 | 0 | 0 | 0 | 0 | 0 |
| ENSG00000225397 | 0 | 0 | 0 | 0 | 0 | 0 |
| ENSG00000229826 | 0 | 0 | 0 | 0 | 0 | 0 |
| ENSG00000268447 | 0 | 0 | 0 | 0 | 0 | 0 |
| ENSG00000269281 | 0 | 0 | 0 | 0 | 0 | 0 |
| ENSG00000226867 | 0 | 0 | 0 | 0 | 0 | 0 |
| ENSG00000204363 | 0 | 0 | 0 | 0 | 0 | 0 |
| ENSG00000229885 | 0 | 0 | 0 | 0 | 0 | 0 |
| ENSG00000171405 | 0 | 0 | 0 | 0 | 0 | 0 |
| ENSG00000196395 | 0 | 0 | 0 | 0 | 0 | 0 |
| ENSG00000224781 | 2 | 0 | 1 | 5 | 0 | 1 |
| ENSG00000171402 | 0 | 0 | 0 | 0 | 0 | 0 |
| ENSG00000179304 | 0 | 0 | 1 | 0 | 0 | 0 |
| ENSG00000268350 | 22 | 5 | 13 | 3 | 6 | 11 |
| ENSG00000184194 | 65 | 30 | 47 | 30 | 25 | 22 |
| ENSG00000270497 | 0 | 1 | 1 | 0 | 0 | 0 |
| ENSG00000184205 | 935 | 380 | 559 | 404 | 455 | 441 |
| ENSG00000234019 | 0 | 0 | 0 | 0 | 0 | 0 |
| ENSG00000232593 | 205 | 85 | 104 | 55 | 125 | 115 |
| ENSG00000235224 | 0 | 0 | 0 | 0 | 0 | 0 |
| ENSG00000231340 | 0 | 1 | 0 | 0 | 1 | 1 |
| ENSG00000227329 | 3 | 0 | 2 | 0 | 0 | 3 |

| | | | | | | |
|---|---|---|---|---|---|---|
| ENSG00000126012 | 5536 | 2203 | 3268 | 2448 | 2715 | 2807 |
| ENSG00000276575 | 1 | 0 | 0 | 0 | 0 | 0 |
| ENSG00000277474 | 0 | 0 | 0 | 0 | 0 | 0 |
| ENSG00000235262 | 0 | 0 | 0 | 0 | 0 | 0 |
| ENSG00000124313 | 1017 | 451 | 685 | 555 | 628 | 588 |
| ENSG00000226299 | 0 | 0 | 0 | 0 | 0 | 0 |
| ENSG00000207408 | 0 | 0 | 0 | 0 | 0 | 0 |
| ENSG00000236571 | 0 | 0 | 0 | 0 | 0 | 0 |
| ENSG00000234413 | 0 | 0 | 0 | 0 | 0 | 0 |
| ENSG00000072501 | 3546 | 1689 | 2197 | 1186 | 1936 | 1927 |
| ENSG00000278204 | 0 | 0 | 0 | 0 | 0 | 0 |
| ENSG00000158423 | 111 | 45 | 68 | 39 | 41 | 44 |
| ENSG00000072506 | 3843 | 1904 | 2153 | 2507 | 2206 | 2106 |
| ENSG00000233250 | 4 | 3 | 2 | 4 | 3 | 1 |
| ENSG00000235416 | 0 | 0 | 0 | 0 | 0 | 0 |
| ENSG00000199422 | 0 | 0 | 0 | 0 | 0 | 0 |
| ENSG00000086758 | 10730 | 4507 | 6086 | 3420 | 5337 | 5398 |
| ENSG00000271886 | 0 | 0 | 0 | 0 | 0 | 0 |
| ENSG00000208012 | 0 | 0 | 0 | 0 | 0 | 0 |
| ENSG00000224735 | 0 | 0 | 0 | 0 | 0 | 0 |
| ENSG00000226110 | 0 | 0 | 0 | 0 | 0 | 0 |
| ENSG00000229037 | 1 | 0 | 0 | 0 | 0 | 0 |
| ENSG00000235916 | 0 | 0 | 0 | 1 | 0 | 0 |
| ENSG00000201618 | 0 | 0 | 0 | 0 | 0 | 0 |
| ENSG00000172943 | 2072 | 899 | 1281 | 686 | 963 | 1020 |
| ENSG00000184083 | 1373 | 641 | 889 | 520 | 702 | 715 |
| ENSG00000224949 | 0 | 0 | 0 | 0 | 0 | 0 |
| ENSG00000196632 | 8 | 2 | 0 | 1 | 3 | 1 |
| ENSG00000234566 | 0 | 0 | 0 | 0 | 0 | 0 |
| ENSG00000207104 | 0 | 0 | 0 | 0 | 0 | 0 |
| ENSG00000158526 | 1818 | 909 | 1036 | 1020 | 1021 | 982 |
| ENSG00000102302 | 212 | 136 | 142 | 145 | 157 | 151 |
| ENSG00000130119 | 746 | 609 | 511 | 294 | 452 | 434 |
| ENSG00000215197 | 2 | 0 | 0 | 0 | 0 | 0 |
| ENSG00000102313 | 13 | 7 | 17 | 5 | 5 | 7 |
| ENSG00000102316 | 9574 | 4431 | 5863 | 4148 | 4008 | 3938 |
| ENSG00000221716 | 11 | 2 | 17 | 6 | 6 | 5 |
| ENSG00000275387 | 0 | 0 | 0 | 0 | 0 | 0 |
| ENSG00000067445 | 14 | 2 | 12 | 2 | 7 | 11 |
| ENSG00000221750 | 0 | 0 | 0 | 0 | 0 | 0 |
| ENSG00000158571 | 24 | 17 | 26 | 4 | 7 | 7 |
| ENSG00000169188 | 1778 | 867 | 968 | 919 | 931 | 943 |
| ENSG00000158578 | 0 | 0 | 2 | 0 | 2 | 1 |
| ENSG00000278283 | 0 | 0 | 0 | 0 | 0 | 0 |
| ENSG00000234466 | 0 | 0 | 0 | 0 | 0 | 0 |
| ENSG00000238269 | 0 | 0 | 0 | 0 | 0 | 0 |
| ENSG00000234068 | 0 | 0 | 0 | 0 | 0 | 0 |
| ENSG00000278319 | 0 | 0 | 0 | 0 | 0 | 0 |
| ENSG00000276929 | 0 | 0 | 0 | 0 | 0 | 0 |
| ENSG00000182518 | 370 | 176 | 223 | 211 | 219 | 230 |

| | | | | | | |
|---|---|---|---|---|---|---|
| ENSG00000186678 | 0 | 0 | 0 | 0 | 0 | 0 |
| ENSG00000229760 | 0 | 0 | 0 | 0 | 0 | 0 |
| ENSG00000231947 | 0 | 0 | 0 | 0 | 1 | 0 |
| ENSG00000238047 | 0 | 0 | 0 | 0 | 0 | 0 |
| ENSG00000256045 | 2 | 2 | 1 | 1 | 2 | 1 |
| ENSG00000158639 | 0 | 0 | 0 | 0 | 0 | 0 |
| ENSG00000204279 | 0 | 0 | 0 | 0 | 0 | 0 |
| ENSG00000232765 | 0 | 0 | 0 | 0 | 0 | 0 |
| ENSG00000283334 | 0 | 0 | 0 | 0 | 0 | 0 |
| ENSG00000266328 | 0 | 0 | 0 | 0 | 0 | 0 |
| ENSG00000187601 | 394 | 189 | 248 | 222 | 237 | 233 |
| ENSG00000247746 | 78 | 29 | 43 | 25 | 30 | 28 |
| ENSG00000277516 | 0 | 0 | 0 | 0 | 0 | 0 |
| ENSG00000200635 | 0 | 0 | 0 | 0 | 0 | 0 |
| ENSG00000235136 | 0 | 0 | 0 | 0 | 0 | 0 |
| ENSG00000189299 | 0 | 0 | 0 | 0 | 0 | 0 |
| ENSG00000229594 | 0 | 0 | 0 | 0 | 0 | 0 |
| ENSG00000169164 | 0 | 0 | 0 | 0 | 0 | 0 |
| ENSG00000083750 | 443 | 205 | 328 | 190 | 253 | 250 |
| ENSG00000227486 | 28 | 17 | 25 | 8 | 16 | 20 |
| ENSG00000214745 | 0 | 0 | 2 | 0 | 0 | 1 |
| ENSG00000102349 | 119 | 89 | 104 | 36 | 73 | 52 |
| ENSG00000234135 | 0 | 1 | 0 | 0 | 0 | 0 |
| ENSG00000188021 | 1114 | 561 | 678 | 482 | 591 | 534 |
| ENSG00000230105 | 0 | 1 | 0 | 1 | 0 | 2 |
| ENSG00000204272 | 1089 | 564 | 679 | 626 | 675 | 632 |
| ENSG00000237748 | 4 | 1 | 0 | 1 | 0 | 1 |
| ENSG00000204271 | 1204 | 548 | 734 | 402 | 616 | 530 |
| ENSG00000224799 | 0 | 0 | 0 | 0 | 0 | 0 |
| ENSG00000237708 | 8 | 1 | 0 | 1 | 3 | 2 |
| ENSG00000186787 | 201 | 103 | 173 | 134 | 153 | 119 |
| ENSG00000226310 | 16 | 6 | 1 | 4 | 4 | 10 |
| ENSG00000147059 | 4 | 1 | 1 | 0 | 1 | 0 |
| ENSG00000231765 | 0 | 0 | 0 | 1 | 0 | 0 |
| ENSG00000165591 | 308 | 116 | 128 | 108 | 100 | 105 |
| ENSG00000231831 | 6 | 3 | 4 | 1 | 1 | 0 |
| ENSG00000198455 | 573 | 241 | 282 | 178 | 316 | 240 |
| ENSG00000215174 | 0 | 1 | 0 | 2 | 1 | 1 |
| ENSG00000227427 | 0 | 0 | 0 | 0 | 0 | 0 |
| ENSG00000198205 | 92 | 51 | 58 | 40 | 57 | 62 |
| ENSG00000228004 | 0 | 0 | 0 | 0 | 0 | 0 |
| ENSG00000235124 | 2 | 1 | 0 | 0 | 0 | 0 |
| ENSG00000236852 | 0 | 1 | 0 | 0 | 0 | 0 |
| ENSG00000235170 | 0 | 0 | 0 | 0 | 0 | 0 |
| ENSG00000228182 | 0 | 0 | 0 | 0 | 0 | 0 |
| ENSG00000234570 | 0 | 0 | 0 | 0 | 0 | 0 |
| ENSG00000277499 | 0 | 0 | 0 | 0 | 0 | 0 |
| ENSG00000230823 | 2 | 0 | 0 | 0 | 0 | 0 |
| ENSG00000235437 | 612 | 298 | 404 | 254 | 320 | 327 |
| ENSG00000186767 | 240 | 93 | 171 | 67 | 132 | 125 |

| | | | | | | |
|---|---|---|---|---|---|---|
| ENSG00000233661 | 0 | 0 | 0 | 0 | 1 | 0 |
| ENSG00000131089 | 850 | 385 | 491 | 399 | 397 | 476 |
| ENSG00000231729 | 0 | 0 | 0 | 0 | 0 | 0 |
| ENSG00000236813 | 0 | 0 | 0 | 0 | 0 | 0 |
| ENSG00000222532 | 0 | 0 | 0 | 0 | 0 | 0 |
| ENSG00000232794 | 0 | 1 | 0 | 0 | 0 | 0 |
| ENSG00000226010 | 0 | 0 | 0 | 1 | 1 | 0 |
| ENSG00000184675 | 228 | 110 | 144 | 116 | 185 | 169 |
| ENSG00000244006 | 0 | 0 | 0 | 0 | 2 | 0 |
| ENSG00000198881 | 0 | 0 | 0 | 0 | 0 | 0 |
| ENSG00000102043 | 0 | 0 | 0 | 0 | 3 | 2 |
| ENSG00000213754 | 0 | 0 | 0 | 0 | 0 | 0 |
| ENSG00000283349 | 0 | 0 | 0 | 0 | 0 | 0 |
| ENSG00000230889 | 0 | 0 | 0 | 0 | 0 | 0 |
| ENSG00000227261 | 0 | 0 | 0 | 0 | 0 | 0 |
| ENSG00000226502 | 0 | 0 | 0 | 0 | 0 | 0 |
| ENSG00000225814 | 0 | 0 | 0 | 0 | 0 | 0 |
| ENSG00000236982 | 0 | 0 | 0 | 0 | 0 | 0 |
| ENSG00000126970 | 4 | 1 | 2 | 4 | 3 | 2 |
| ENSG00000237971 | 0 | 0 | 0 | 0 | 0 | 0 |
| ENSG00000225569 | 0 | 0 | 0 | 0 | 0 | 0 |
| ENSG00000231857 | 0 | 0 | 0 | 0 | 0 | 0 |
| ENSG00000237963 | 0 | 0 | 0 | 0 | 0 | 0 |
| ENSG00000102053 | 8 | 2 | 2 | 0 | 6 | 2 |
| ENSG00000228158 | 176 | 53 | 125 | 75 | 91 | 104 |
| ENSG00000227534 | 0 | 0 | 0 | 0 | 0 | 0 |
| ENSG00000001497 | 1632 | 849 | 970 | 1091 | 974 | 1140 |
| ENSG00000227942 | 0 | 0 | 0 | 0 | 0 | 0 |
| ENSG00000147065 | 2 | 0 | 1 | 1 | 0 | 0 |
| ENSG00000231809 | 0 | 0 | 0 | 0 | 0 | 0 |
| ENSG00000276689 | 0 | 0 | 0 | 0 | 0 | 0 |
| ENSG00000233833 | 0 | 0 | 0 | 0 | 0 | 0 |
| ENSG00000230039 | 2 | 0 | 0 | 0 | 0 | 0 |
| ENSG00000217835 | 0 | 5 | 0 | 1 | 0 | 2 |
| ENSG00000237311 | 0 | 0 | 0 | 0 | 0 | 0 |
| ENSG00000274536 | 0 | 0 | 0 | 0 | 0 | 0 |
| ENSG00000284567 | 0 | 0 | 0 | 0 | 0 | 0 |
| ENSG00000155659 | 0 | 0 | 0 | 0 | 0 | 0 |
| ENSG00000215168 | 0 | 0 | 0 | 0 | 0 | 0 |
| ENSG00000237294 | 0 | 0 | 0 | 0 | 0 | 0 |
| ENSG00000089472 | 0 | 1 | 0 | 0 | 1 | 0 |
| ENSG00000231356 | 0 | 0 | 0 | 0 | 0 | 0 |
| ENSG00000226820 | 0 | 0 | 0 | 0 | 0 | 0 |
| ENSG00000235892 | 0 | 0 | 0 | 0 | 0 | 0 |
| ENSG00000131080 | 139 | 56 | 60 | 53 | 60 | 53 |
| ENSG00000222667 | 0 | 0 | 0 | 0 | 0 | 0 |
| ENSG00000226280 | 0 | 0 | 0 | 0 | 0 | 0 |
| ENSG00000233710 | 0 | 0 | 0 | 0 | 0 | 0 |
| ENSG00000169083 | 1814 | 755 | 965 | 429 | 632 | 659 |
| ENSG00000234201 | 0 | 0 | 0 | 0 | 0 | 0 |

| | | | | | | |
|---|---|---|---|---|---|---|
| ENSG00000260118 | 1 | 1 | 0 | 0 | 0 | 0 |
| ENSG00000079482 | 478 | 236 | 273 | 145 | 235 | 237 |
| ENSG00000228328 | 0 | 1 | 1 | 0 | 2 | 0 |
| ENSG00000252145 | 0 | 0 | 0 | 0 | 0 | 0 |
| ENSG00000228125 | 0 | 0 | 0 | 0 | 0 | 0 |
| ENSG00000206646 | 0 | 0 | 0 | 0 | 0 | 0 |
| ENSG00000225925 | 0 | 0 | 0 | 0 | 0 | 0 |
| ENSG00000181704 | 2601 | 1227 | 1547 | 1050 | 1400 | 1314 |
| ENSG00000206747 | 0 | 0 | 0 | 0 | 0 | 0 |
| ENSG00000236735 | 0 | 1 | 0 | 1 | 0 | 0 |
| ENSG00000234463 | 0 | 0 | 0 | 0 | 0 | 0 |
| ENSG00000130052 | 27 | 12 | 10 | 18 | 8 | 13 |
| ENSG00000232267 | 0 | 0 | 0 | 0 | 0 | 0 |
| ENSG00000213740 | 2 | 2 | 1 | 1 | 2 | 2 |
| ENSG00000090776 | 322 | 136 | 164 | 175 | 148 | 145 |
| ENSG00000181191 | 574 | 290 | 386 | 312 | 358 | 291 |
| ENSG00000172186 | 0 | 0 | 0 | 0 | 4 | 0 |
| ENSG00000215162 | 0 | 0 | 0 | 0 | 0 | 0 |
| ENSG00000231706 | 0 | 0 | 0 | 0 | 0 | 0 |
| ENSG00000130054 | 354 | 138 | 202 | 212 | 189 | 172 |
| ENSG00000228160 | 0 | 0 | 0 | 0 | 0 | 0 |
| ENSG00000158813 | 142 | 67 | 94 | 60 | 68 | 67 |
| ENSG00000272351 | 0 | 0 | 0 | 0 | 0 | 0 |
| ENSG00000285776 | 0 | 0 | 0 | 0 | 0 | 0 |
| ENSG00000212434 | 0 | 0 | 0 | 0 | 0 | 0 |
| ENSG00000233229 | 2 | 0 | 0 | 0 | 1 | 2 |
| ENSG00000211991 | 0 | 0 | 0 | 0 | 0 | 0 |
| ENSG00000147160 | 0 | 0 | 0 | 0 | 0 | 0 |
| ENSG00000189401 | 0 | 0 | 0 | 0 | 0 | 0 |
| ENSG00000226579 | 0 | 0 | 0 | 0 | 0 | 0 |
| ENSG00000235382 | 0 | 0 | 0 | 0 | 0 | 0 |
| ENSG00000089289 | 1937 | 1160 | 1390 | 1365 | 1549 | 1318 |
| ENSG00000220925 | 0 | 1 | 0 | 0 | 0 | 0 |
| ENSG00000203588 | 2 | 0 | 7 | 3 | 7 | 1 |
| ENSG00000184210 | 2 | 1 | 0 | 0 | 0 | 0 |
| ENSG00000242657 | 0 | 0 | 0 | 0 | 0 | 0 |
| ENSG00000204195 | 0 | 0 | 0 | 0 | 1 | 0 |
| ENSG00000200473 | 0 | 0 | 0 | 0 | 0 | 0 |
| ENSG00000186912 | 0 | 0 | 0 | 0 | 0 | 0 |
| ENSG00000120500 | 7 | 5 | 5 | 1 | 7 | 1 |
| ENSG00000147127 | 13 | 1 | 5 | 1 | 4 | 5 |
| ENSG00000120509 | 1838 | 888 | 1046 | 972 | 909 | 923 |
| ENSG00000090889 | 708 | 390 | 404 | 346 | 522 | 409 |
| ENSG00000231053 | 0 | 0 | 0 | 1 | 0 | 0 |
| ENSG00000201377 | 0 | 0 | 0 | 0 | 0 | 0 |
| ENSG00000130055 | 69 | 25 | 27 | 13 | 15 | 18 |
| ENSG00000284391 | 11 | 0 | 2 | 0 | 0 | 1 |
| ENSG00000082458 | 3401 | 1590 | 2050 | 1344 | 1634 | 1696 |
| ENSG00000200431 | 0 | 0 | 0 | 0 | 0 | 0 |
| ENSG00000231651 | 6 | 4 | 3 | 7 | 6 | 2 |

| | | | | | | |
|---|---|---|---|---|---|---|
| ENSG00000120498 | 4 | 2 | 1 | 3 | 2 | 0 |
| ENSG00000230187 | 0 | 0 | 0 | 0 | 0 | 0 |
| ENSG00000277492 | 0 | 0 | 0 | 0 | 0 | 0 |
| ENSG00000280704 | 0 | 0 | 0 | 0 | 0 | 0 |
| ENSG00000213747 | 0 | 0 | 0 | 0 | 0 | 0 |
| ENSG00000165349 | 0 | 0 | 0 | 0 | 0 | 0 |
| ENSG00000230629 | 43 | 22 | 20 | 32 | 36 | 37 |
| ENSG00000229324 | 2 | 0 | 0 | 0 | 0 | 0 |
| ENSG00000212605 | 0 | 0 | 0 | 0 | 0 | 0 |
| ENSG00000227536 | 20 | 8 | 7 | 2 | 6 | 9 |
| ENSG00000147164 | 3036 | 1472 | 1884 | 1511 | 1563 | 1718 |
| ENSG00000184481 | 425 | 180 | 257 | 231 | 235 | 265 |
| ENSG00000204165 | 0 | 0 | 0 | 0 | 0 | 1 |
| ENSG00000285171 | 0 | 0 | 0 | 0 | 0 | 0 |
| ENSG00000147168 | 0 | 0 | 0 | 1 | 0 | 0 |
| ENSG00000184634 | 1933 | 776 | 1049 | 753 | 899 | 950 |
| ENSG00000196338 | 28 | 6 | 20 | 20 | 15 | 19 |
| ENSG00000228427 | 0 | 0 | 0 | 0 | 0 | 0 |
| ENSG00000169562 | 0 | 2 | 0 | 0 | 0 | 0 |
| ENSG00000147130 | 2710 | 1119 | 1521 | 1101 | 1161 | 1286 |
| ENSG00000147140 | 16044 | 7502 | 9398 | 7186 | 9100 | 8730 |
| ENSG00000147166 | 12 | 4 | 14 | 4 | 7 | 6 |
| ENSG00000224208 | 0 | 0 | 0 | 0 | 1 | 0 |
| ENSG00000213706 | 0 | 0 | 0 | 0 | 0 | 0 |
| ENSG00000147133 | 1333 | 579 | 895 | 287 | 675 | 583 |
| ENSG00000275975 | 0 | 0 | 0 | 0 | 0 | 0 |
| ENSG00000229601 | 0 | 3 | 1 | 0 | 0 | 0 |
| ENSG00000213018 | 2 | 0 | 2 | 0 | 4 | 1 |
| ENSG00000199751 | 0 | 0 | 0 | 0 | 0 | 0 |
| ENSG00000243468 | 0 | 0 | 0 | 0 | 0 | 0 |
| ENSG00000215120 | 0 | 0 | 1 | 0 | 0 | 0 |
| ENSG00000147162 | 7416 | 3467 | 4873 | 2260 | 3904 | 3568 |
| ENSG00000147174 | 62 | 23 | 30 | 17 | 12 | 23 |
| ENSG00000186810 | 0 | 0 | 0 | 0 | 0 | 1 |
| ENSG00000237265 | 0 | 0 | 0 | 0 | 0 | 0 |
| ENSG00000283599 | 0 | 0 | 0 | 0 | 0 | 0 |
| ENSG00000230934 | 0 | 0 | 0 | 0 | 0 | 0 |
| ENSG00000281852 | 0 | 0 | 0 | 0 | 0 | 0 |
| ENSG00000215115 | 0 | 0 | 0 | 0 | 0 | 0 |
| ENSG00000226870 | 0 | 0 | 0 | 0 | 0 | 0 |
| ENSG00000237182 | 0 | 0 | 0 | 0 | 0 | 0 |
| ENSG00000237717 | 0 | 0 | 0 | 0 | 0 | 0 |
| ENSG00000234442 | 0 | 0 | 0 | 0 | 0 | 0 |
| ENSG00000215113 | 0 | 0 | 0 | 0 | 0 | 0 |
| ENSG00000224617 | 0 | 0 | 0 | 0 | 0 | 0 |
| ENSG00000229030 | 0 | 0 | 0 | 0 | 0 | 0 |
| ENSG00000283178 | 0 | 0 | 0 | 0 | 0 | 0 |
| ENSG00000204131 | 27 | 20 | 14 | 14 | 22 | 8 |
| ENSG00000201392 | 0 | 0 | 0 | 0 | 0 | 0 |
| ENSG00000196933 | 0 | 0 | 0 | 1 | 0 | 0 |

| | | | | | | |
|---|---|---|---|---|---|---|
| ENSG00000242732 | 168 | 85 | 111 | 85 | 132 | 107 |
| ENSG00000225471 | 0 | 0 | 0 | 0 | 0 | 0 |
| ENSG00000102309 | 702 | 359 | 386 | 428 | 423 | 392 |
| ENSG00000239577 | 0 | 0 | 0 | 1 | 0 | 0 |
| ENSG00000186871 | 112 | 78 | 100 | 62 | 120 | 105 |
| ENSG00000198034 | 31631 | 16678 | 19290 | 20890 | 20604 | 19555 |
| ENSG00000238926 | 0 | 0 | 0 | 0 | 0 | 0 |
| ENSG00000125931 | 0 | 0 | 1 | 0 | 0 | 0 |
| ENSG00000285547 | 0 | 0 | 0 | 0 | 0 | 0 |
| ENSG00000147099 | 187 | 89 | 157 | 112 | 141 | 122 |
| ENSG00000222810 | 0 | 0 | 0 | 0 | 0 | 0 |
| ENSG00000067177 | 1040 | 513 | 646 | 466 | 662 | 608 |
| ENSG00000225858 | 0 | 0 | 0 | 0 | 0 | 0 |
| ENSG00000231944 | 0 | 0 | 0 | 0 | 0 | 0 |
| ENSG00000273788 | 0 | 0 | 0 | 0 | 0 | 0 |
| ENSG00000213698 | 0 | 0 | 0 | 0 | 0 | 0 |
| ENSG00000201271 | 0 | 0 | 0 | 0 | 0 | 0 |
| ENSG00000184911 | 0 | 0 | 0 | 1 | 1 | 0 |
| ENSG00000269911 | 0 | 0 | 0 | 0 | 0 | 0 |
| ENSG00000268994 | 0 | 0 | 0 | 0 | 0 | 0 |
| ENSG00000225396 | 0 | 0 | 0 | 0 | 0 | 0 |
| ENSG00000269502 | 0 | 0 | 0 | 0 | 0 | 0 |
| ENSG00000283594 | 0 | 0 | 0 | 0 | 0 | 0 |
| ENSG00000275520 | 0 | 0 | 0 | 0 | 0 | 0 |
| ENSG00000226725 | 0 | 0 | 0 | 0 | 0 | 0 |
| ENSG00000184388 | 2 | 1 | 0 | 0 | 0 | 0 |
| ENSG00000186288 | 0 | 0 | 0 | 0 | 0 | 0 |
| ENSG00000231963 | 0 | 0 | 0 | 0 | 0 | 0 |
| ENSG00000204118 | 42 | 42 | 60 | 29 | 43 | 35 |
| ENSG00000186462 | 324 | 137 | 204 | 96 | 136 | 145 |
| ENSG00000230311 | 4 | 1 | 3 | 0 | 1 | 0 |
| ENSG00000237014 | 0 | 0 | 0 | 0 | 0 | 0 |
| ENSG00000225664 | 0 | 0 | 0 | 0 | 0 | 0 |
| ENSG00000207486 | 0 | 0 | 0 | 0 | 0 | 0 |
| ENSG00000131264 | 0 | 0 | 0 | 0 | 0 | 0 |
| ENSG00000233162 | 0 | 0 | 0 | 0 | 0 | 0 |
| ENSG00000269904 | 26 | 15 | 10 | 8 | 9 | 14 |
| ENSG00000237475 | 0 | 0 | 0 | 0 | 0 | 0 |
| ENSG00000204116 | 279 | 141 | 112 | 74 | 174 | 109 |
| ENSG00000230146 | 2 | 0 | 1 | 0 | 0 | 0 |
| ENSG00000270641 | 0 | 2 | 0 | 0 | 0 | 0 |
| ENSG00000229807 | 27 | 8 | 26 | 2 | 10 | 11 |
| ENSG00000274655 | 0 | 0 | 0 | 0 | 0 | 0 |
| ENSG00000277577 | 0 | 0 | 0 | 0 | 0 | 0 |
| ENSG00000278039 | 0 | 0 | 0 | 0 | 0 | 0 |
| ENSG00000182707 | 0 | 0 | 0 | 0 | 0 | 0 |
| ENSG00000225470 | 16 | 38 | 41 | 32 | 60 | 49 |
| ENSG00000274430 | 0 | 0 | 0 | 0 | 0 | 0 |
| ENSG00000276784 | 0 | 0 | 0 | 0 | 0 | 0 |
| ENSG00000230590 | 1535 | 747 | 914 | 293 | 861 | 811 |

| | | | | | | |
|---|---|---|---|---|---|---|
| ENSG00000228906 | 15 | 3 | 1 | 3 | 1 | 7 |
| ENSG00000271199 | 0 | 0 | 0 | 0 | 1 | 0 |
| ENSG00000233984 | 2 | 2 | 4 | 0 | 0 | 1 |
| ENSG00000203397 | 0 | 0 | 0 | 0 | 0 | 0 |
| ENSG00000234969 | 5 | 1 | 1 | 0 | 2 | 1 |
| ENSG00000223650 | 2 | 0 | 0 | 1 | 0 | 0 |
| ENSG00000232928 | 0 | 0 | 1 | 2 | 0 | 0 |
| ENSG00000280375 | 0 | 1 | 1 | 0 | 1 | 0 |
| ENSG00000230986 | 0 | 1 | 0 | 0 | 0 | 0 |
| ENSG00000224430 | 6 | 0 | 2 | 0 | 2 | 1 |
| ENSG00000231549 | 0 | 0 | 1 | 2 | 0 | 1 |
| ENSG00000204113 | 0 | 0 | 0 | 0 | 0 | 0 |
| ENSG00000228492 | 82 | 30 | 65 | 13 | 44 | 39 |
| ENSG00000271533 | 134 | 64 | 124 | 12 | 87 | 66 |
| ENSG00000202566 | 5 | 2 | 2 | 0 | 2 | 2 |
| ENSG00000212027 | 0 | 0 | 0 | 0 | 0 | 0 |
| ENSG00000283534 | 0 | 0 | 0 | 0 | 0 | 0 |
| ENSG00000265727 | 5 | 4 | 1 | 0 | 2 | 0 |
| ENSG00000278421 | 0 | 0 | 0 | 0 | 0 | 0 |
| ENSG00000273974 | 0 | 0 | 0 | 0 | 0 | 0 |
| ENSG00000275254 | 0 | 0 | 0 | 0 | 0 | 0 |
| ENSG00000275625 | 0 | 0 | 0 | 0 | 0 | 0 |
| ENSG00000207820 | 0 | 0 | 1 | 0 | 0 | 0 |
| ENSG00000199168 | 0 | 0 | 0 | 0 | 0 | 0 |
| ENSG00000277922 | 0 | 0 | 0 | 0 | 0 | 0 |
| ENSG00000187969 | 0 | 0 | 0 | 0 | 0 | 0 |
| ENSG00000226515 | 0 | 0 | 0 | 0 | 0 | 0 |
| ENSG00000213269 | 0 | 0 | 0 | 0 | 0 | 0 |
| ENSG00000239745 | 0 | 0 | 0 | 0 | 0 | 0 |
| ENSG00000230665 | 0 | 0 | 0 | 2 | 3 | 0 |
| ENSG00000213695 | 0 | 1 | 1 | 1 | 0 | 1 |
| ENSG00000147100 | 630 | 292 | 390 | 186 | 235 | 220 |
| ENSG00000231206 | 0 | 0 | 0 | 0 | 0 | 0 |
| ENSG00000230673 | 3 | 4 | 1 | 0 | 1 | 2 |
| ENSG00000131263 | 1230 | 627 | 573 | 285 | 827 | 517 |
| ENSG00000199885 | 0 | 0 | 0 | 0 | 0 | 0 |
| ENSG00000050030 | 2 | 1 | 4 | 0 | 1 | 2 |
| ENSG00000271589 | 0 | 0 | 0 | 0 | 0 | 0 |
| ENSG00000131269 | 506 | 225 | 295 | 186 | 288 | 235 |
| ENSG00000229661 | 0 | 0 | 0 | 0 | 0 | 0 |
| ENSG00000199601 | 0 | 0 | 0 | 0 | 0 | 0 |
| ENSG00000094841 | 346 | 151 | 212 | 149 | 179 | 209 |
| ENSG00000232186 | 0 | 3 | 7 | 0 | 1 | 3 |
| ENSG00000102383 | 2 | 0 | 2 | 0 | 1 | 1 |
| ENSG00000233254 | 0 | 0 | 0 | 2 | 0 | 0 |
| ENSG00000237089 | 0 | 0 | 0 | 0 | 0 | 0 |
| ENSG00000224775 | 0 | 0 | 0 | 0 | 0 | 1 |
| ENSG00000215105 | 53 | 42 | 39 | 13 | 62 | 36 |
| ENSG00000186675 | 0 | 0 | 0 | 0 | 0 | 0 |
| ENSG00000233063 | 0 | 0 | 0 | 0 | 0 | 0 |

| | | | | | | |
|---|---|---|---|---|---|---|
| ENSG00000226753 | 0 | 0 | 0 | 0 | 0 | 0 |
| ENSG00000226854 | 0 | 0 | 0 | 0 | 0 | 0 |
| ENSG00000232332 | 0 | 0 | 0 | 0 | 0 | 0 |
| ENSG00000102390 | 704 | 419 | 409 | 384 | 520 | 405 |
| ENSG00000237252 | 0 | 0 | 0 | 0 | 0 | 0 |
| ENSG00000201826 | 0 | 0 | 0 | 0 | 0 | 0 |
| ENSG00000274398 | 0 | 0 | 0 | 0 | 0 | 0 |
| ENSG00000213684 | 0 | 0 | 0 | 0 | 0 | 0 |
| ENSG00000198934 | 26 | 8 | 10 | 15 | 10 | 21 |
| ENSG00000223259 | 0 | 0 | 0 | 0 | 0 | 0 |
| ENSG00000235447 | 0 | 0 | 0 | 0 | 0 | 0 |
| ENSG00000280870 | 2 | 0 | 0 | 0 | 0 | 0 |
| ENSG00000283242 | 0 | 0 | 0 | 0 | 0 | 0 |
| ENSG00000207995 | 0 | 0 | 0 | 0 | 0 | 0 |
| ENSG00000201447 | 0 | 0 | 0 | 0 | 0 | 0 |
| ENSG00000234404 | 0 | 0 | 0 | 0 | 0 | 0 |
| ENSG00000196468 | 0 | 0 | 0 | 0 | 0 | 0 |
| ENSG00000085224 | 364 | 205 | 196 | 51 | 469 | 308 |
| ENSG00000237106 | 0 | 0 | 0 | 0 | 0 | 0 |
| ENSG00000102158 | 2481 | 1110 | 1393 | 870 | 1274 | 1137 |
| ENSG00000200906 | 0 | 1 | 0 | 0 | 0 | 1 |
| ENSG00000263410 | 0 | 0 | 0 | 0 | 0 | 0 |
| ENSG00000131174 | 2482 | 1215 | 1149 | 1416 | 1284 | 1028 |
| ENSG00000165240 | 641 | 350 | 358 | 137 | 391 | 328 |
| ENSG00000102144 | 13802 | 6788 | 8117 | 7533 | 8144 | 8191 |
| ENSG00000248503 | 5 | 2 | 3 | 3 | 4 | 4 |
| ENSG00000226784 | 0 | 0 | 0 | 0 | 0 | 0 |
| ENSG00000187325 | 1432 | 854 | 862 | 536 | 915 | 751 |
| ENSG00000223592 | 0 | 0 | 0 | 0 | 0 | 0 |
| ENSG00000173198 | 2 | 0 | 0 | 0 | 1 | 0 |
| ENSG00000224755 | 0 | 0 | 0 | 0 | 0 | 0 |
| ENSG00000235061 | 0 | 0 | 0 | 0 | 0 | 0 |
| ENSG00000179300 | 0 | 0 | 3 | 0 | 0 | 1 |
| ENSG00000271433 | 0 | 0 | 0 | 0 | 0 | 0 |
| ENSG00000147145 | 0 | 0 | 1 | 0 | 0 | 0 |
| ENSG00000227645 | 0 | 0 | 0 | 0 | 0 | 0 |
| ENSG00000283622 | 0 | 0 | 0 | 0 | 0 | 0 |
| ENSG00000078589 | 2 | 1 | 4 | 2 | 5 | 5 |
| ENSG00000232168 | 0 | 0 | 0 | 0 | 0 | 0 |
| ENSG00000147138 | 0 | 0 | 0 | 0 | 0 | 0 |
| ENSG00000271005 | 0 | 0 | 0 | 0 | 0 | 0 |
| ENSG00000225025 | 0 | 0 | 0 | 0 | 0 | 0 |
| ENSG00000078596 | 2 | 0 | 1 | 0 | 1 | 0 |
| ENSG00000270268 | 0 | 0 | 0 | 0 | 0 | 0 |
| ENSG00000122145 | 0 | 0 | 0 | 0 | 0 | 0 |
| ENSG00000278530 | 0 | 2 | 4 | 0 | 0 | 0 |
| ENSG00000174016 | 0 | 1 | 1 | 0 | 0 | 0 |
| ENSG00000230247 | 0 | 0 | 0 | 0 | 0 | 0 |
| ENSG00000228051 | 0 | 0 | 0 | 0 | 1 | 0 |
| ENSG00000228612 | 0 | 0 | 0 | 1 | 0 | 0 |

| | | | | | | |
|---|---|---|---|---|---|---|
| ENSG00000165288 | 174 | 101 | 91 | 19 | 171 | 93 |
| ENSG00000225715 | 0 | 0 | 0 | 0 | 0 | 0 |
| ENSG00000239008 | 0 | 0 | 0 | 0 | 0 | 0 |
| ENSG00000212623 | 0 | 0 | 0 | 0 | 0 | 0 |
| ENSG00000073905 | 8 | 2 | 2 | 1 | 5 | 6 |
| ENSG00000212546 | 0 | 0 | 0 | 0 | 0 | 0 |
| ENSG00000284618 | 0 | 0 | 0 | 0 | 0 | 0 |
| ENSG00000198157 | 59 | 34 | 30 | 7 | 48 | 22 |
| ENSG00000131171 | 7031 | 3589 | 3872 | 2029 | 3311 | 2596 |
| ENSG00000231875 | 0 | 1 | 1 | 0 | 0 | 0 |
| ENSG00000224523 | 1 | 2 | 1 | 0 | 1 | 0 |
| ENSG00000233446 | 0 | 0 | 0 | 0 | 0 | 0 |
| ENSG00000206826 | 0 | 0 | 0 | 0 | 0 | 0 |
| ENSG00000228041 | 0 | 0 | 0 | 0 | 0 | 0 |
| ENSG00000213653 | 0 | 0 | 0 | 0 | 0 | 0 |
| ENSG00000202183 | 0 | 0 | 0 | 0 | 0 | 0 |
| ENSG00000236365 | 0 | 0 | 0 | 0 | 0 | 0 |
| ENSG00000232735 | 0 | 0 | 0 | 0 | 0 | 0 |
| ENSG00000279437 | 0 | 0 | 0 | 0 | 0 | 0 |
| ENSG00000196767 | 0 | 0 | 0 | 0 | 0 | 0 |
| ENSG00000236870 | 0 | 0 | 0 | 0 | 0 | 0 |
| ENSG00000215102 | 4 | 0 | 0 | 0 | 2 | 0 |
| ENSG00000183035 | 0 | 0 | 0 | 0 | 0 | 0 |
| ENSG00000072133 | 227 | 142 | 123 | 77 | 143 | 85 |
| ENSG00000221494 | 0 | 0 | 0 | 0 | 0 | 0 |
| ENSG00000165259 | 14 | 4 | 7 | 3 | 10 | 7 |
| ENSG00000233484 | 0 | 0 | 0 | 0 | 0 | 0 |
| ENSG00000237211 | 0 | 0 | 0 | 0 | 0 | 0 |
| ENSG00000229547 | 0 | 0 | 0 | 0 | 0 | 0 |
| ENSG00000230781 | 0 | 0 | 0 | 0 | 0 | 0 |
| ENSG00000155008 | 847 | 458 | 432 | 257 | 499 | 397 |
| ENSG00000184788 | 2 | 0 | 1 | 0 | 0 | 0 |
| ENSG00000283446 | 0 | 0 | 0 | 0 | 0 | 0 |
| ENSG00000276391 | 0 | 0 | 0 | 0 | 0 | 0 |
| ENSG00000235461 | 0 | 0 | 0 | 0 | 0 | 0 |
| ENSG00000147180 | 34 | 26 | 14 | 9 | 18 | 11 |
| ENSG00000124429 | 404 | 261 | 259 | 115 | 339 | 224 |
| ENSG00000283435 | 0 | 0 | 0 | 0 | 0 | 0 |
| ENSG00000223707 | 0 | 0 | 0 | 0 | 0 | 0 |
| ENSG00000276892 | 0 | 0 | 0 | 0 | 0 | 0 |
| ENSG00000188419 | 492 | 229 | 211 | 99 | 341 | 190 |
| ENSG00000199051 | 0 | 0 | 0 | 0 | 0 | 0 |
| ENSG00000199567 | 0 | 0 | 0 | 0 | 0 | 0 |
| ENSG00000234039 | 0 | 0 | 0 | 0 | 0 | 0 |
| ENSG00000225536 | 0 | 0 | 2 | 1 | 3 | 1 |
| ENSG00000223574 | 0 | 0 | 0 | 0 | 0 | 0 |
| ENSG00000126733 | 0 | 0 | 0 | 0 | 0 | 0 |
| ENSG00000215093 | 0 | 1 | 0 | 2 | 0 | 1 |
| ENSG00000223772 | 0 | 0 | 0 | 0 | 0 | 0 |
| ENSG00000277717 | 0 | 0 | 0 | 0 | 0 | 0 |

| | | | | | | |
|---|---|---|---|---|---|---|
| ENSG00000231478 | 0 | 0 | 0 | 0 | 0 | 0 |
| ENSG00000271290 | 0 | 0 | 0 | 0 | 0 | 0 |
| ENSG00000236190 | 0 | 0 | 0 | 0 | 0 | 0 |
| ENSG00000271457 | 0 | 0 | 0 | 0 | 0 | 0 |
| ENSG00000200422 | 0 | 0 | 0 | 0 | 0 | 0 |
| ENSG00000230478 | 0 | 0 | 0 | 0 | 0 | 0 |
| ENSG00000102271 | 8 | 2 | 2 | 0 | 2 | 1 |
| ENSG00000237506 | 12 | 8 | 3 | 6 | 4 | 2 |
| ENSG00000282914 | 0 | 0 | 0 | 0 | 0 | 0 |
| ENSG00000271205 | 0 | 0 | 0 | 0 | 0 | 0 |
| ENSG00000270446 | 0 | 0 | 0 | 0 | 0 | 0 |
| ENSG00000271056 | 0 | 0 | 0 | 0 | 0 | 0 |
| ENSG00000230706 | 0 | 0 | 0 | 0 | 0 | 0 |
| ENSG00000230956 | 0 | 0 | 0 | 0 | 0 | 0 |
| ENSG00000252016 | 0 | 0 | 0 | 0 | 0 | 0 |
| ENSG00000271693 | 0 | 0 | 0 | 0 | 0 | 0 |
| ENSG00000147183 | 0 | 0 | 0 | 0 | 0 | 0 |
| ENSG00000271157 | 0 | 0 | 0 | 0 | 0 | 0 |
| ENSG00000232576 | 0 | 0 | 0 | 0 | 0 | 0 |
| ENSG00000270745 | 0 | 0 | 0 | 0 | 0 | 0 |
| ENSG00000224065 | 0 | 0 | 0 | 0 | 0 | 0 |
| ENSG00000225012 | 0 | 0 | 0 | 0 | 0 | 0 |
| ENSG00000153779 | 2 | 1 | 4 | 1 | 3 | 3 |
| ENSG00000227581 | 0 | 0 | 0 | 0 | 0 | 0 |
| ENSG00000226081 | 0 | 0 | 1 | 0 | 0 | 0 |
| ENSG00000274022 | 0 | 0 | 0 | 0 | 0 | 0 |
| ENSG00000227434 | 0 | 0 | 0 | 0 | 0 | 0 |
| ENSG00000229667 | 0 | 0 | 0 | 0 | 0 | 0 |
| ENSG00000207515 | 0 | 0 | 0 | 0 | 0 | 0 |
| ENSG00000206062 | 0 | 0 | 0 | 0 | 0 | 0 |
| ENSG00000234161 | 0 | 0 | 0 | 0 | 0 | 0 |
| ENSG00000236413 | 0 | 0 | 0 | 0 | 0 | 0 |
| ENSG00000174740 | 0 | 0 | 1 | 0 | 0 | 1 |
| ENSG00000230275 | 0 | 0 | 0 | 0 | 0 | 1 |
| ENSG00000102290 | 0 | 0 | 0 | 0 | 0 | 0 |
| ENSG00000222440 | 0 | 0 | 0 | 0 | 0 | 0 |
| ENSG00000234702 | 0 | 0 | 0 | 0 | 0 | 0 |
| ENSG00000229132 | 14 | 10 | 8 | 11 | 8 | 8 |
| ENSG00000215089 | 34 | 10 | 14 | 18 | 20 | 12 |
| ENSG00000230543 | 0 | 0 | 0 | 0 | 0 | 0 |
| ENSG00000225912 | 2 | 1 | 0 | 2 | 0 | 1 |
| ENSG00000285445 | 0 | 0 | 0 | 0 | 0 | 0 |
| ENSG00000237873 | 0 | 0 | 0 | 0 | 0 | 0 |
| ENSG00000234130 | 0 | 1 | 0 | 0 | 1 | 0 |
| ENSG00000234322 | 0 | 2 | 0 | 0 | 0 | 1 |
| ENSG00000237381 | 0 | 0 | 0 | 0 | 0 | 0 |
| ENSG00000228301 | 0 | 0 | 0 | 0 | 0 | 0 |
| ENSG00000186310 | 0 | 0 | 0 | 1 | 0 | 0 |
| ENSG00000179083 | 10 | 14 | 9 | 10 | 20 | 21 |
| ENSG00000270826 | 0 | 0 | 0 | 0 | 0 | 0 |

| | | | | | | |
|---|---|---|---|---|---|---|
| ENSG00000212495 | 0 | 0 | 0 | 0 | 0 | 0 |
| ENSG00000202410 | 0 | 0 | 0 | 0 | 0 | 0 |
| ENSG00000227521 | 0 | 0 | 0 | 0 | 0 | 0 |
| ENSG00000232293 | 0 | 0 | 0 | 0 | 0 | 0 |
| ENSG00000231001 | 0 | 0 | 0 | 0 | 2 | 1 |
| ENSG00000221187 | 0 | 0 | 0 | 0 | 0 | 0 |
| ENSG00000271094 | 0 | 0 | 0 | 0 | 0 | 0 |
| ENSG00000223467 | 0 | 0 | 0 | 0 | 0 | 0 |
| ENSG00000201567 | 0 | 0 | 0 | 0 | 0 | 0 |
| ENSG00000215085 | 0 | 0 | 0 | 0 | 0 | 0 |
| ENSG00000236185 | 0 | 0 | 0 | 0 | 0 | 0 |
| ENSG00000226307 | 0 | 0 | 0 | 0 | 0 | 0 |
| ENSG00000235425 | 0 | 0 | 0 | 0 | 0 | 0 |
| ENSG00000270583 | 0 | 0 | 0 | 0 | 0 | 0 |
| ENSG00000232842 | 0 | 0 | 1 | 1 | 0 | 0 |
| ENSG00000242998 | 0 | 0 | 0 | 0 | 0 | 0 |
| ENSG00000203262 | 2 | 0 | 0 | 0 | 0 | 0 |
| ENSG00000232069 | 0 | 0 | 0 | 0 | 0 | 0 |
| ENSG00000271209 | 0 | 0 | 0 | 0 | 0 | 0 |
| ENSG00000223260 | 0 | 0 | 0 | 0 | 0 | 0 |
| ENSG00000229693 | 0 | 0 | 0 | 0 | 0 | 0 |
| ENSG00000147202 | 504 | 336 | 208 | 110 | 396 | 209 |
| ENSG00000207426 | 0 | 0 | 0 | 0 | 0 | 0 |
| ENSG00000204086 | 4 | 4 | 2 | 1 | 4 | 1 |
| ENSG00000214628 | 0 | 0 | 0 | 0 | 0 | 0 |
| ENSG00000277900 | 0 | 0 | 0 | 0 | 0 | 0 |
| ENSG00000265260 | 0 | 0 | 0 | 0 | 0 | 0 |
| ENSG00000236256 | 0 | 0 | 0 | 0 | 0 | 1 |
| ENSG00000233887 | 0 | 0 | 0 | 0 | 0 | 0 |
| ENSG00000237752 | 0 | 0 | 0 | 0 | 0 | 0 |
| ENSG00000234889 | 0 | 0 | 0 | 0 | 0 | 0 |
| ENSG00000236243 | 0 | 0 | 0 | 0 | 0 | 0 |
| ENSG00000271322 | 0 | 0 | 0 | 0 | 0 | 0 |
| ENSG00000213704 | 0 | 0 | 0 | 0 | 0 | 0 |
| ENSG00000226837 | 0 | 0 | 0 | 0 | 0 | 0 |
| ENSG00000281566 | 0 | 0 | 2 | 0 | 0 | 0 |
| ENSG00000270308 | 0 | 0 | 0 | 0 | 0 | 0 |
| ENSG00000215070 | 0 | 0 | 0 | 0 | 0 | 0 |
| ENSG00000229946 | 0 | 0 | 0 | 0 | 0 | 0 |
| ENSG00000252296 | 0 | 0 | 0 | 0 | 0 | 0 |
| ENSG00000236988 | 0 | 0 | 0 | 0 | 0 | 0 |
| ENSG00000230592 | 0 | 1 | 1 | 0 | 0 | 1 |
| ENSG00000165194 | 20 | 7 | 12 | 0 | 2 | 3 |
| ENSG00000000005 | 0 | 0 | 0 | 0 | 0 | 0 |
| ENSG00000000003 | 1044 | 528 | 600 | 525 | 611 | 552 |
| ENSG00000102359 | 5 | 2 | 1 | 0 | 1 | 0 |
| ENSG00000102362 | 263 | 143 | 130 | 97 | 103 | 102 |
| ENSG00000223819 | 0 | 0 | 0 | 0 | 0 | 0 |
| ENSG00000238116 | 0 | 0 | 0 | 0 | 0 | 0 |
| ENSG00000275320 | 0 | 0 | 1 | 1 | 1 | 0 |

| | | | | | | |
|---|---|---|---|---|---|---|
| ENSG00000101811 | 1322 | 663 | 750 | 684 | 713 | 729 |
| ENSG00000202231 | 2 | 0 | 0 | 0 | 0 | 0 |
| ENSG00000007952 | 4 | 7 | 9 | 8 | 18 | 8 |
| ENSG00000239345 | 4 | 0 | 0 | 0 | 1 | 0 |
| ENSG00000233680 | 6 | 2 | 1 | 1 | 0 | 2 |
| ENSG00000179031 | 2 | 0 | 4 | 1 | 0 | 0 |
| ENSG00000182489 | 45 | 11 | 14 | 25 | 30 | 15 |
| ENSG00000182574 | 1 | 2 | 1 | 0 | 0 | 0 |
| ENSG00000174225 | 28 | 7 | 14 | 4 | 3 | 14 |
| ENSG00000188917 | 2078 | 1013 | 1265 | 1015 | 1167 | 1241 |
| ENSG00000225839 | 0 | 2 | 2 | 0 | 3 | 3 |
| ENSG00000126950 | 0 | 0 | 0 | 0 | 0 | 0 |
| ENSG00000102384 | 286 | 161 | 198 | 155 | 202 | 171 |
| ENSG00000231643 | 0 | 0 | 0 | 0 | 0 | 0 |
| ENSG00000102385 | 8 | 6 | 3 | 3 | 6 | 8 |
| ENSG00000102387 | 12 | 6 | 5 | 3 | 7 | 6 |
| ENSG00000202024 | 0 | 0 | 0 | 0 | 0 | 0 |
| ENSG00000270813 | 10 | 3 | 7 | 5 | 3 | 2 |
| ENSG00000183911 | 2 | 1 | 6 | 0 | 0 | 3 |
| ENSG00000126953 | 515 | 339 | 331 | 339 | 328 | 395 |
| ENSG00000010671 | 0 | 1 | 0 | 4 | 3 | 3 |
| ENSG00000241343 | 386 | 184 | 238 | 199 | 239 | 272 |
| ENSG00000257529 | 0 | 0 | 0 | 0 | 1 | 0 |
| ENSG00000102393 | 1973 | 948 | 1088 | 1084 | 1169 | 1132 |
| ENSG00000126945 | 3235 | 1952 | 2012 | 1198 | 2232 | 1806 |
| ENSG00000196440 | 36 | 16 | 14 | 15 | 10 | 20 |
| ENSG00000234557 | 0 | 0 | 0 | 0 | 0 | 0 |
| ENSG00000237682 | 2 | 0 | 0 | 0 | 0 | 0 |
| ENSG00000126947 | 1230 | 575 | 603 | 501 | 573 | 562 |
| ENSG00000204072 | 2 | 0 | 0 | 0 | 0 | 0 |
| ENSG00000198960 | 993 | 467 | 587 | 533 | 591 | 566 |
| ENSG00000102401 | 2905 | 1400 | 1736 | 1195 | 1542 | 1474 |
| ENSG00000228275 | 0 | 1 | 0 | 0 | 0 | 0 |
| ENSG00000261101 | 2 | 2 | 5 | 4 | 1 | 9 |
| ENSG00000207291 | 0 | 0 | 2 | 0 | 2 | 1 |
| ENSG00000184867 | 232 | 107 | 169 | 75 | 101 | 147 |
| ENSG00000223753 | 2 | 0 | 0 | 0 | 0 | 1 |
| ENSG00000212584 | 0 | 0 | 0 | 0 | 0 | 0 |
| ENSG00000229090 | 0 | 0 | 0 | 0 | 0 | 0 |
| ENSG00000178146 | 2 | 2 | 0 | 3 | 3 | 1 |
| ENSG00000126952 | 0 | 0 | 0 | 0 | 0 | 1 |
| ENSG00000166432 | 190 | 143 | 112 | 28 | 161 | 122 |
| ENSG00000235196 | 0 | 0 | 0 | 0 | 0 | 0 |
| ENSG00000252810 | 0 | 0 | 0 | 0 | 0 | 0 |
| ENSG00000235976 | 0 | 0 | 0 | 0 | 0 | 0 |
| ENSG00000226119 | 0 | 0 | 0 | 0 | 0 | 0 |
| ENSG00000234901 | 0 | 0 | 0 | 0 | 0 | 0 |
| ENSG00000184905 | 0 | 0 | 0 | 0 | 0 | 1 |
| ENSG00000204071 | 0 | 0 | 0 | 0 | 0 | 0 |
| ENSG00000184515 | 10 | 5 | 8 | 4 | 7 | 5 |

| | | | | | | |
|---|---|---|---|---|---|---|
| ENSG00000251525 | 0 | 0 | 0 | 0 | 0 | 0 |
| ENSG00000268235 | 0 | 0 | 0 | 0 | 0 | 0 |
| ENSG00000269405 | 0 | 0 | 0 | 0 | 0 | 0 |
| ENSG00000273746 | 0 | 0 | 0 | 0 | 0 | 0 |
| ENSG00000284800 | 0 | 0 | 0 | 0 | 0 | 0 |
| ENSG00000269437 | 0 | 0 | 0 | 0 | 0 | 0 |
| ENSG00000231590 | 0 | 0 | 0 | 0 | 0 | 0 |
| ENSG00000215029 | 0 | 0 | 0 | 0 | 0 | 0 |
| ENSG00000158164 | 12 | 2 | 1 | 0 | 2 | 1 |
| ENSG00000176318 | 0 | 0 | 0 | 0 | 0 | 0 |
| ENSG00000196970 | 0 | 0 | 0 | 0 | 0 | 0 |
| ENSG00000125962 | 403 | 195 | 254 | 180 | 210 | 271 |
| ENSG00000271147 | 162 | 80 | 134 | 83 | 88 | 110 |
| ENSG00000270050 | 0 | 0 | 1 | 0 | 0 | 0 |
| ENSG00000227211 | 0 | 2 | 0 | 0 | 0 | 0 |
| ENSG00000198932 | 79 | 32 | 47 | 27 | 38 | 46 |
| ENSG00000252294 | 0 | 1 | 0 | 0 | 0 | 0 |
| ENSG00000230320 | 0 | 0 | 0 | 0 | 0 | 0 |
| ENSG00000158301 | 552 | 251 | 283 | 181 | 252 | 264 |
| ENSG00000198908 | 504 | 190 | 295 | 177 | 235 | 234 |
| ENSG00000223546 | 87 | 40 | 68 | 58 | 78 | 70 |
| ENSG00000232282 | 0 | 1 | 0 | 0 | 0 | 2 |
| ENSG00000229954 | 0 | 0 | 0 | 0 | 0 | 0 |
| ENSG00000237300 | 0 | 0 | 0 | 0 | 0 | 0 |
| ENSG00000237628 | 0 | 0 | 0 | 0 | 0 | 0 |
| ENSG00000228316 | 0 | 0 | 0 | 0 | 0 | 0 |
| ENSG00000224471 | 0 | 0 | 1 | 0 | 0 | 0 |
| ENSG00000226056 | 2 | 1 | 0 | 0 | 0 | 0 |
| ENSG00000234886 | 0 | 0 | 0 | 0 | 1 | 1 |
| ENSG00000227649 | 0 | 0 | 0 | 0 | 0 | 0 |
| ENSG00000229794 | 0 | 0 | 0 | 0 | 0 | 1 |
| ENSG00000239407 | 17 | 11 | 11 | 5 | 13 | 14 |
| ENSG00000102128 | 3 | 0 | 0 | 0 | 0 | 0 |
| ENSG00000234050 | 0 | 0 | 0 | 0 | 0 | 0 |
| ENSG00000224632 | 0 | 0 | 0 | 0 | 0 | 0 |
| ENSG00000133169 | 0 | 0 | 0 | 1 | 0 | 0 |
| ENSG00000147206 | 0 | 1 | 0 | 0 | 0 | 0 |
| ENSG00000226241 | 0 | 0 | 0 | 0 | 2 | 1 |
| ENSG00000233071 | 0 | 0 | 0 | 0 | 0 | 0 |
| ENSG00000102409 | 426 | 230 | 243 | 150 | 200 | 210 |
| ENSG00000180964 | 42 | 21 | 42 | 23 | 34 | 30 |
| ENSG00000224692 | 0 | 0 | 0 | 0 | 0 | 0 |
| ENSG00000204065 | 0 | 0 | 0 | 0 | 0 | 0 |
| ENSG00000133134 | 188 | 95 | 121 | 110 | 139 | 106 |
| ENSG00000182916 | 0 | 0 | 0 | 0 | 0 | 0 |
| ENSG00000238049 | 0 | 0 | 0 | 0 | 0 | 0 |
| ENSG00000185222 | 0 | 0 | 0 | 0 | 0 | 0 |
| ENSG00000166681 | 4958 | 2447 | 3260 | 2739 | 2677 | 2672 |
| ENSG00000281091 | 2 | 2 | 1 | 4 | 4 | 1 |
| ENSG00000234405 | 0 | 1 | 3 | 1 | 0 | 1 |

| | | | | | | |
|---|---|---|---|---|---|---|
| ENSG00000172476 | 21 | 6 | 3 | 4 | 8 | 7 |
| ENSG00000133142 | 5771 | 2929 | 3534 | 2765 | 3500 | 3255 |
| ENSG00000279894 | 9 | 4 | 2 | 4 | 9 | 3 |
| ENSG00000196507 | 1686 | 813 | 1070 | 826 | 903 | 857 |
| ENSG00000224031 | 4 | 0 | 1 | 1 | 0 | 1 |
| ENSG00000172465 | 1262 | 651 | 817 | 700 | 735 | 751 |
| ENSG00000180284 | 0 | 0 | 0 | 0 | 1 | 0 |
| ENSG00000123562 | 12053 | 5785 | 7199 | 5643 | 7044 | 6329 |
| ENSG00000231154 | 32 | 20 | 25 | 10 | 29 | 19 |
| ENSG00000188828 | 0 | 1 | 1 | 0 | 0 | 0 |
| ENSG00000179363 | 9 | 3 | 4 | 4 | 0 | 2 |
| ENSG00000123560 | 0 | 0 | 5 | 1 | 2 | 0 |
| ENSG00000123570 | 249 | 132 | 128 | 87 | 152 | 113 |
| ENSG00000222419 | 0 | 0 | 0 | 0 | 0 | 0 |
| ENSG00000231787 | 4 | 3 | 0 | 2 | 3 | 1 |
| ENSG00000276298 | 0 | 1 | 2 | 0 | 2 | 1 |
| ENSG00000234345 | 0 | 3 | 1 | 1 | 2 | 1 |
| ENSG00000231728 | 11 | 10 | 1 | 5 | 6 | 3 |
| ENSG00000158427 | 24 | 14 | 19 | 16 | 22 | 19 |
| ENSG00000230312 | 0 | 2 | 3 | 0 | 0 | 0 |
| ENSG00000227067 | 0 | 0 | 0 | 0 | 0 | 0 |
| ENSG00000123569 | 3 | 1 | 3 | 1 | 3 | 1 |
| ENSG00000101812 | 0 | 2 | 2 | 2 | 4 | 3 |
| ENSG00000235946 | 2 | 0 | 5 | 2 | 0 | 0 |
| ENSG00000269226 | 10 | 7 | 5 | 3 | 4 | 3 |
| ENSG00000269743 | 134 | 59 | 84 | 54 | 63 | 61 |
| ENSG00000166707 | 24 | 9 | 13 | 16 | 20 | 15 |
| ENSG00000123575 | 3766 | 1778 | 2162 | 1173 | 2020 | 1843 |
| ENSG00000123576 | 0 | 0 | 0 | 0 | 0 | 0 |
| ENSG00000189108 | 13 | 7 | 8 | 5 | 0 | 5 |
| ENSG00000232665 | 0 | 0 | 0 | 0 | 0 | 0 |
| ENSG00000223733 | 0 | 0 | 0 | 0 | 0 | 0 |
| ENSG00000268629 | 0 | 0 | 0 | 0 | 0 | 0 |
| ENSG00000181819 | 0 | 0 | 1 | 0 | 0 | 0 |
| ENSG00000206864 | 0 | 0 | 0 | 0 | 0 | 0 |
| ENSG00000123572 | 66 | 52 | 50 | 17 | 94 | 49 |
| ENSG00000123561 | 0 | 0 | 0 | 0 | 0 | 0 |
| ENSG00000230788 | 0 | 0 | 0 | 0 | 0 | 0 |
| ENSG00000157502 | 17 | 4 | 4 | 3 | 11 | 1 |
| ENSG00000256709 | 0 | 0 | 0 | 0 | 0 | 0 |
| ENSG00000202407 | 0 | 0 | 0 | 0 | 0 | 0 |
| ENSG00000224816 | 0 | 0 | 0 | 0 | 0 | 0 |
| ENSG00000227657 | 0 | 0 | 0 | 0 | 0 | 0 |
| ENSG00000277673 | 0 | 0 | 0 | 0 | 0 | 0 |
| ENSG00000147231 | 0 | 0 | 0 | 0 | 0 | 0 |
| ENSG00000263515 | 0 | 0 | 0 | 0 | 0 | 0 |
| ENSG00000133135 | 4 | 0 | 1 | 0 | 0 | 1 |
| ENSG00000133138 | 36 | 29 | 11 | 3 | 47 | 11 |
| ENSG00000133131 | 632 | 332 | 319 | 279 | 360 | 318 |
| ENSG00000147223 | 0 | 0 | 0 | 0 | 1 | 0 |

| | | | | | | |
|---|---|---|---|---|---|---|
| ENSG00000165376 | 0 | 1 | 0 | 0 | 0 | 0 |
| ENSG00000231144 | 0 | 0 | 0 | 0 | 0 | 0 |
| ENSG00000089682 | 283 | 168 | 171 | 97 | 229 | 155 |
| ENSG00000198088 | 32 | 14 | 20 | 14 | 16 | 18 |
| ENSG00000213569 | 0 | 0 | 0 | 0 | 2 | 0 |
| ENSG00000199832 | 0 | 0 | 0 | 0 | 0 | 0 |
| ENSG00000080572 | 0 | 0 | 0 | 0 | 0 | 1 |
| ENSG00000204053 | 0 | 0 | 0 | 0 | 0 | 0 |
| ENSG00000215007 | 4 | 1 | 1 | 0 | 1 | 1 |
| ENSG00000214720 | 2 | 0 | 0 | 0 | 0 | 0 |
| ENSG00000227610 | 0 | 0 | 0 | 0 | 0 | 0 |
| ENSG00000147234 | 8 | 2 | 4 | 4 | 10 | 6 |
| ENSG00000147224 | 1511 | 771 | 1046 | 1145 | 1363 | 1326 |
| ENSG00000157514 | 6789 | 3275 | 3826 | 2542 | 2601 | 2564 |
| ENSG00000170935 | 12 | 3 | 7 | 3 | 7 | 3 |
| ENSG00000080561 | 1239 | 619 | 720 | 399 | 562 | 510 |
| ENSG00000236064 | 0 | 0 | 1 | 0 | 0 | 0 |
| ENSG00000207363 | 0 | 0 | 0 | 0 | 0 | 1 |
| ENSG00000225728 | 0 | 0 | 0 | 0 | 0 | 0 |
| ENSG00000277883 | 2 | 1 | 0 | 0 | 0 | 1 |
| ENSG00000170925 | 0 | 0 | 0 | 0 | 0 | 1 |
| ENSG00000101842 | 10 | 3 | 4 | 1 | 2 | 3 |
| ENSG00000101843 | 2523 | 1132 | 1398 | 1238 | 1362 | 1347 |
| ENSG00000101844 | 801 | 391 | 443 | 321 | 456 | 428 |
| ENSG00000197565 | 20 | 8 | 7 | 2 | 3 | 7 |
| ENSG00000189372 | 0 | 0 | 0 | 0 | 0 | 0 |
| ENSG00000188153 | 707 | 300 | 559 | 225 | 269 | 382 |
| ENSG00000207146 | 0 | 0 | 1 | 0 | 0 | 0 |
| ENSG00000261409 | 2 | 0 | 0 | 0 | 0 | 1 |
| ENSG00000260548 | 0 | 0 | 0 | 0 | 0 | 0 |
| ENSG00000280322 | 0 | 0 | 0 | 0 | 0 | 0 |
| ENSG00000133124 | 0 | 0 | 1 | 0 | 0 | 0 |
| ENSG00000237863 | 0 | 0 | 0 | 1 | 1 | 0 |
| ENSG00000201443 | 0 | 0 | 0 | 0 | 0 | 0 |
| ENSG00000275110 | 0 | 0 | 0 | 0 | 0 | 1 |
| ENSG00000215000 | 0 | 0 | 0 | 0 | 0 | 0 |
| ENSG00000101890 | 0 | 0 | 0 | 0 | 0 | 0 |
| ENSG00000101888 | 436 | 237 | 236 | 172 | 263 | 230 |
| ENSG00000233874 | 0 | 0 | 0 | 0 | 0 | 1 |
| ENSG00000176076 | 0 | 0 | 0 | 0 | 0 | 0 |
| ENSG00000068366 | 12 | 15 | 13 | 6 | 21 | 16 |
| ENSG00000237319 | 0 | 0 | 0 | 0 | 0 | 0 |
| ENSG00000252069 | 0 | 0 | 0 | 0 | 0 | 0 |
| ENSG00000225381 | 0 | 0 | 0 | 0 | 0 | 0 |
| ENSG00000157600 | 2454 | 1067 | 1099 | 1079 | 1136 | 1198 |
| ENSG00000208013 | 0 | 0 | 0 | 0 | 0 | 0 |
| ENSG00000265584 | 0 | 0 | 0 | 0 | 0 | 0 |
| ENSG00000101935 | 1551 | 721 | 829 | 490 | 824 | 711 |
| ENSG00000208883 | 0 | 0 | 0 | 0 | 0 | 0 |
| ENSG00000224142 | 0 | 0 | 0 | 0 | 0 | 0 |

| | | | | | | |
|---|---|---|---|---|---|---|
| ENSG00000133136 | 0 | 0 | 1 | 0 | 1 | 0 |
| ENSG00000243978 | 0 | 1 | 0 | 0 | 0 | 0 |
| ENSG00000225366 | 0 | 0 | 0 | 0 | 0 | 0 |
| ENSG00000228042 | 0 | 0 | 0 | 0 | 0 | 0 |
| ENSG00000101938 | 0 | 0 | 0 | 0 | 0 | 0 |
| ENSG00000077264 | 0 | 0 | 0 | 0 | 0 | 1 |
| ENSG00000226112 | 0 | 0 | 0 | 0 | 0 | 0 |
| ENSG00000234349 | 0 | 0 | 0 | 0 | 0 | 0 |
| ENSG00000077274 | 2 | 0 | 1 | 0 | 0 | 0 |
| ENSG00000077279 | 0 | 0 | 0 | 0 | 0 | 1 |
| ENSG00000251954 | 0 | 0 | 0 | 0 | 0 | 0 |
| ENSG00000223904 | 0 | 0 | 0 | 0 | 0 | 1 |
| ENSG00000260802 | 0 | 0 | 0 | 0 | 0 | 0 |
| ENSG00000229939 | 84 | 45 | 58 | 49 | 56 | 68 |
| ENSG00000225031 | 18 | 13 | 12 | 13 | 19 | 24 |
| ENSG00000236189 | 0 | 0 | 0 | 0 | 0 | 0 |
| ENSG00000101901 | 1084 | 515 | 649 | 497 | 667 | 596 |
| ENSG00000201420 | 0 | 1 | 0 | 0 | 0 | 0 |
| ENSG00000229487 | 6 | 2 | 0 | 0 | 0 | 0 |
| ENSG00000072315 | 0 | 0 | 0 | 0 | 0 | 0 |
| ENSG00000204025 | 0 | 0 | 0 | 0 | 0 | 0 |
| ENSG00000271421 | 0 | 0 | 0 | 0 | 0 | 0 |
| ENSG00000271464 | 0 | 0 | 0 | 0 | 0 | 0 |
| ENSG00000271400 | 0 | 0 | 0 | 0 | 0 | 0 |
| ENSG00000187823 | 0 | 0 | 0 | 0 | 0 | 1 |
| ENSG00000182508 | 0 | 0 | 0 | 0 | 0 | 0 |
| ENSG00000200436 | 0 | 0 | 0 | 0 | 0 | 0 |
| ENSG00000213652 | 0 | 0 | 0 | 0 | 0 | 0 |
| ENSG00000126016 | 646 | 325 | 385 | 227 | 336 | 314 |
| ENSG00000283775 | 0 | 0 | 0 | 0 | 0 | 0 |
| ENSG00000243011 | 0 | 0 | 0 | 0 | 0 | 0 |
| ENSG00000252930 | 0 | 0 | 0 | 0 | 0 | 0 |
| ENSG00000235329 | 0 | 0 | 0 | 0 | 0 | 0 |
| ENSG00000220541 | 0 | 0 | 0 | 0 | 0 | 0 |
| ENSG00000241743 | 0 | 0 | 0 | 0 | 0 | 0 |
| ENSG00000270524 | 0 | 0 | 0 | 0 | 0 | 0 |
| ENSG00000201674 | 0 | 0 | 0 | 0 | 0 | 0 |
| ENSG00000207220 | 0 | 0 | 0 | 0 | 0 | 0 |
| ENSG00000240327 | 0 | 0 | 0 | 0 | 0 | 0 |
| ENSG00000212611 | 0 | 0 | 0 | 0 | 0 | 0 |
| ENSG00000230502 | 0 | 0 | 0 | 0 | 0 | 0 |
| ENSG00000271686 | 0 | 0 | 0 | 0 | 0 | 0 |
| ENSG00000226547 | 0 | 1 | 0 | 0 | 0 | 0 |
| ENSG00000147246 | 20 | 3 | 8 | 3 | 1 | 9 |
| ENSG00000224664 | 0 | 0 | 0 | 0 | 0 | 0 |
| ENSG00000208839 | 0 | 0 | 0 | 0 | 0 | 0 |
| ENSG00000212100 | 0 | 0 | 0 | 0 | 0 | 0 |
| ENSG00000222321 | 0 | 0 | 0 | 0 | 0 | 0 |
| ENSG00000276158 | 0 | 0 | 0 | 0 | 0 | 0 |
| ENSG00000221710 | 0 | 0 | 0 | 0 | 0 | 0 |

| | | | | | | |
|---|---|---|---|---|---|---|
| ENSG00000222715 | 0 | 0 | 0 | 0 | 0 | 0 |
| ENSG00000224773 | 0 | 0 | 0 | 2 | 1 | 0 |
| ENSG00000199001 | 0 | 0 | 0 | 0 | 0 | 0 |
| ENSG00000270413 | 0 | 0 | 0 | 0 | 0 | 0 |
| ENSG00000123496 | 0 | 0 | 0 | 0 | 0 | 0 |
| ENSG00000222122 | 0 | 0 | 0 | 0 | 0 | 0 |
| ENSG00000238223 | 0 | 0 | 0 | 0 | 0 | 0 |
| ENSG00000236080 | 0 | 0 | 0 | 0 | 0 | 0 |
| ENSG00000130224 | 0 | 0 | 0 | 0 | 0 | 0 |
| ENSG00000271907 | 0 | 0 | 0 | 0 | 0 | 0 |
| ENSG00000212241 | 0 | 0 | 0 | 0 | 0 | 0 |
| ENSG00000175718 | 0 | 0 | 0 | 0 | 0 | 0 |
| ENSG00000235656 | 0 | 0 | 0 | 0 | 0 | 0 |
| ENSG00000102021 | 0 | 0 | 0 | 0 | 0 | 0 |
| ENSG00000274374 | 0 | 0 | 0 | 0 | 0 | 0 |
| ENSG00000271021 | 0 | 0 | 0 | 0 | 0 | 0 |
| ENSG00000271826 | 0 | 0 | 0 | 0 | 0 | 0 |
| ENSG00000252441 | 0 | 0 | 0 | 0 | 0 | 0 |
| ENSG00000102024 | 4 | 0 | 1 | 1 | 4 | 1 |
| ENSG00000270431 | 0 | 0 | 0 | 0 | 0 | 0 |
| ENSG00000227881 | 0 | 0 | 0 | 0 | 0 | 0 |
| ENSG00000234785 | 0 | 0 | 1 | 1 | 0 | 1 |
| ENSG00000228532 | 0 | 3 | 6 | 4 | 2 | 7 |
| ENSG00000229335 | 0 | 0 | 0 | 0 | 0 | 0 |
| ENSG00000235244 | 12 | 5 | 17 | 8 | 7 | 8 |
| ENSG00000266828 | 0 | 0 | 0 | 0 | 0 | 0 |
| ENSG00000231371 | 0 | 0 | 0 | 0 | 0 | 0 |
| ENSG00000234558 | 4 | 0 | 0 | 1 | 0 | 1 |
| ENSG00000180772 | 0 | 1 | 0 | 4 | 1 | 2 |
| ENSG00000207033 | 0 | 0 | 0 | 0 | 0 | 0 |
| ENSG00000268104 | 6909 | 3575 | 3994 | 2194 | 3114 | 2896 |
| ENSG00000204019 | 8 | 3 | 4 | 4 | 4 | 3 |
| ENSG00000236077 | 0 | 0 | 1 | 0 | 0 | 0 |
| ENSG00000237860 | 0 | 0 | 0 | 0 | 0 | 0 |
| ENSG00000230159 | 0 | 0 | 0 | 0 | 0 | 0 |
| ENSG00000224217 | 0 | 0 | 0 | 0 | 0 | 0 |
| ENSG00000206752 | 0 | 0 | 0 | 0 | 0 | 0 |
| ENSG00000213526 | 4 | 0 | 1 | 0 | 0 | 0 |
| ENSG00000238066 | 0 | 0 | 0 | 0 | 0 | 0 |
| ENSG00000224314 | 0 | 0 | 0 | 0 | 0 | 0 |
| ENSG00000279587 | 0 | 2 | 0 | 1 | 0 | 0 |
| ENSG00000003096 | 372 | 170 | 145 | 134 | 162 | 120 |
| ENSG00000200291 | 0 | 0 | 0 | 0 | 0 | 0 |
| ENSG00000237458 | 0 | 0 | 0 | 1 | 0 | 0 |
| ENSG00000234524 | 0 | 0 | 0 | 0 | 0 | 0 |
| ENSG00000270619 | 0 | 0 | 0 | 0 | 0 | 0 |
| ENSG00000131725 | 826 | 380 | 456 | 234 | 516 | 395 |
| ENSG00000221463 | 0 | 0 | 0 | 0 | 0 | 0 |
| ENSG00000147251 | 60 | 53 | 18 | 11 | 69 | 25 |
| ENSG00000206862 | 0 | 0 | 0 | 0 | 0 | 0 |

| | | | | | | |
|---|---|---|---|---|---|---|
| ENSG00000230399 | 0 | 0 | 0 | 0 | 0 | 0 |
| ENSG00000131724 | 4677 | 2317 | 2791 | 1950 | 2709 | 2513 |
| ENSG00000239182 | 0 | 0 | 0 | 0 | 0 | 0 |
| ENSG00000250995 | 22 | 12 | 27 | 8 | 18 | 19 |
| ENSG00000174460 | 6 | 0 | 3 | 1 | 0 | 2 |
| ENSG00000203650 | 0 | 0 | 0 | 0 | 0 | 0 |
| ENSG00000224341 | 0 | 0 | 0 | 0 | 0 | 0 |
| ENSG00000231270 | 0 | 0 | 0 | 0 | 0 | 0 |
| ENSG00000175556 | 4 | 1 | 2 | 7 | 3 | 3 |
| ENSG00000277535 | 0 | 0 | 0 | 0 | 0 | 0 |
| ENSG00000250423 | 0 | 0 | 0 | 0 | 0 | 0 |
| ENSG00000227864 | 0 | 0 | 0 | 0 | 0 | 0 |
| ENSG00000230170 | 0 | 0 | 0 | 0 | 0 | 0 |
| ENSG00000270625 | 0 | 0 | 1 | 0 | 0 | 0 |
| ENSG00000101856 | 3805 | 1756 | 2070 | 1696 | 2070 | 1948 |
| ENSG00000214992 | 0 | 0 | 0 | 0 | 0 | 1 |
| ENSG00000230392 | 0 | 0 | 0 | 1 | 0 | 0 |
| ENSG00000226661 | 2 | 0 | 0 | 0 | 0 | 0 |
| ENSG00000227814 | 0 | 0 | 0 | 0 | 0 | 0 |
| ENSG00000077713 | 2351 | 1125 | 1360 | 990 | 1247 | 1293 |
| ENSG00000237903 | 6 | 2 | 1 | 0 | 1 | 2 |
| ENSG00000207175 | 0 | 1 | 0 | 1 | 1 | 2 |
| ENSG00000266420 | 0 | 1 | 1 | 1 | 2 | 1 |
| ENSG00000252147 | 1 | 3 | 4 | 2 | 6 | 8 |
| ENSG00000224281 | 87 | 53 | 74 | 37 | 59 | 62 |
| ENSG00000005022 | 18627 | 10023 | 11614 | 13283 | 12032 | 12028 |
| ENSG00000018610 | 1525 | 807 | 886 | 930 | 883 | 919 |
| ENSG00000077721 | 4244 | 2094 | 2529 | 2098 | 2390 | 2279 |
| ENSG00000186416 | 631 | 321 | 395 | 379 | 451 | 407 |
| ENSG00000125354 | 30 | 8 | 20 | 15 | 19 | 8 |
| ENSG00000211578 | 0 | 0 | 0 | 0 | 0 | 0 |
| ENSG00000279601 | 0 | 1 | 0 | 0 | 0 | 0 |
| ENSG00000187808 | 15 | 7 | 11 | 9 | 7 | 5 |
| ENSG00000198918 | 6837 | 3371 | 4128 | 4454 | 4608 | 4074 |
| ENSG00000206622 | 1 | 1 | 2 | 1 | 1 | 0 |
| ENSG00000125351 | 547 | 303 | 287 | 129 | 440 | 314 |
| ENSG00000125352 | 793 | 425 | 468 | 478 | 469 | 428 |
| ENSG00000125356 | 3672 | 1851 | 2190 | 2177 | 1874 | 1844 |
| ENSG00000186471 | 0 | 0 | 0 | 0 | 0 | 0 |
| ENSG00000101882 | 667 | 427 | 478 | 404 | 474 | 427 |
| ENSG00000282933 | 2 | 1 | 1 | 1 | 0 | 0 |
| ENSG00000234493 | 0 | 0 | 0 | 0 | 0 | 0 |
| ENSG00000282850 | 0 | 0 | 0 | 0 | 0 | 0 |
| ENSG00000258545 | 4 | 0 | 0 | 4 | 2 | 5 |
| ENSG00000203989 | 0 | 0 | 0 | 0 | 0 | 0 |
| ENSG00000101883 | 0 | 1 | 0 | 0 | 0 | 0 |
| ENSG00000228139 | 0 | 1 | 0 | 1 | 0 | 0 |
| ENSG00000233382 | 41 | 38 | 46 | 27 | 37 | 27 |
| ENSG00000131721 | 0 | 0 | 0 | 0 | 0 | 0 |
| ENSG00000227541 | 0 | 0 | 0 | 0 | 0 | 0 |

| | | | | | | |
|---|---|---|---|---|---|---|
| ENSG00000237757 | 3 | 2 | 3 | 1 | 2 | 0 |
| ENSG00000177485 | 1596 | 850 | 817 | 441 | 1081 | 755 |
| ENSG00000125355 | 0 | 0 | 0 | 0 | 0 | 0 |
| ENSG00000232042 | 0 | 0 | 0 | 0 | 0 | 0 |
| ENSG00000101892 | 0 | 0 | 0 | 0 | 0 | 0 |
| ENSG00000005893 | 7022 | 3538 | 4218 | 2728 | 3941 | 3342 |
| ENSG00000251707 | 0 | 0 | 0 | 0 | 0 | 0 |
| ENSG00000158290 | 2120 | 1159 | 1110 | 551 | 1504 | 972 |
| ENSG00000232119 | 2096 | 1071 | 1226 | 1188 | 1274 | 1207 |
| ENSG00000171155 | 1617 | 884 | 970 | 897 | 915 | 880 |
| ENSG00000278317 | 0 | 0 | 0 | 0 | 0 | 0 |
| ENSG00000181511 | 0 | 0 | 0 | 1 | 1 | 0 |
| ENSG00000237828 | 0 | 1 | 1 | 1 | 0 | 0 |
| ENSG00000236446 | 0 | 0 | 0 | 0 | 0 | 0 |
| ENSG00000271346 | 0 | 0 | 0 | 0 | 0 | 0 |
| ENSG00000278646 | 0 | 0 | 0 | 0 | 0 | 0 |
| ENSG00000226685 | 0 | 0 | 0 | 0 | 0 | 0 |
| ENSG00000270678 | 0 | 0 | 0 | 0 | 0 | 0 |
| ENSG00000226929 | 0 | 0 | 0 | 0 | 0 | 0 |
| ENSG00000270493 | 0 | 0 | 0 | 0 | 0 | 0 |
| ENSG00000224089 | 0 | 0 | 0 | 0 | 0 | 0 |
| ENSG00000271154 | 0 | 0 | 0 | 0 | 0 | 0 |
| ENSG00000226600 | 0 | 0 | 0 | 0 | 0 | 0 |
| ENSG00000271041 | 0 | 0 | 0 | 0 | 0 | 0 |
| ENSG00000230347 | 0 | 0 | 0 | 0 | 0 | 0 |
| ENSG00000270397 | 0 | 0 | 0 | 0 | 0 | 0 |
| ENSG00000228517 | 0 | 0 | 0 | 0 | 0 | 0 |
| ENSG00000271502 | 0 | 0 | 0 | 0 | 0 | 0 |
| ENSG00000226023 | 0 | 0 | 0 | 0 | 0 | 0 |
| ENSG00000271211 | 0 | 0 | 0 | 0 | 0 | 0 |
| ENSG00000237957 | 0 | 0 | 0 | 0 | 0 | 0 |
| ENSG00000270736 | 0 | 0 | 0 | 0 | 0 | 0 |
| ENSG00000230594 | 0 | 0 | 0 | 0 | 0 | 0 |
| ENSG00000270771 | 0 | 0 | 0 | 0 | 0 | 0 |
| ENSG00000236126 | 0 | 0 | 0 | 0 | 0 | 0 |
| ENSG00000270646 | 0 | 0 | 0 | 0 | 0 | 0 |
| ENSG00000242362 | 0 | 0 | 0 | 0 | 0 | 0 |
| ENSG00000270456 | 0 | 0 | 0 | 0 | 0 | 0 |
| ENSG00000236371 | 0 | 0 | 0 | 0 | 0 | 0 |
| ENSG00000270251 | 0 | 0 | 0 | 0 | 0 | 0 |
| ENSG00000233433 | 0 | 0 | 0 | 0 | 0 | 0 |
| ENSG00000182890 | 204 | 90 | 84 | 61 | 65 | 63 |
| ENSG00000234176 | 0 | 0 | 1 | 1 | 0 | 1 |
| ENSG00000265456 | 0 | 0 | 0 | 0 | 0 | 0 |
| ENSG00000213498 | 0 | 0 | 0 | 0 | 0 | 0 |
| ENSG00000260585 | 0 | 0 | 0 | 0 | 0 | 0 |
| ENSG00000237891 | 0 | 0 | 0 | 0 | 0 | 0 |
| ENSG00000280142 | 0 | 0 | 0 | 0 | 0 | 0 |
| ENSG00000232496 | 0 | 0 | 0 | 0 | 1 | 1 |
| ENSG00000212321 | 0 | 0 | 0 | 0 | 0 | 0 |

| | | | | | | |
|---|---|---|---|---|---|---|
| ENSG00000224236 | 0 | 0 | 0 | 0 | 1 | 0 |
| ENSG00000125675 | 0 | 0 | 0 | 0 | 0 | 1 |
| ENSG00000179101 | 2 | 1 | 1 | 0 | 0 | 0 |
| ENSG00000229733 | 0 | 0 | 0 | 0 | 0 | 0 |
| ENSG00000228466 | 0 | 0 | 0 | 0 | 0 | 0 |
| ENSG00000125676 | 1590 | 1128 | 731 | 290 | 2065 | 781 |
| ENSG00000232149 | 6 | 0 | 2 | 1 | 1 | 1 |
| ENSG00000230376 | 0 | 1 | 1 | 0 | 0 | 1 |
| ENSG00000252178 | 0 | 0 | 0 | 0 | 0 | 0 |
| ENSG00000226902 | 0 | 0 | 0 | 0 | 0 | 0 |
| ENSG00000264933 | 0 | 0 | 0 | 0 | 0 | 0 |
| ENSG00000199705 | 1 | 0 | 0 | 0 | 0 | 0 |
| ENSG00000101966 | 3896 | 1914 | 2088 | 1169 | 2174 | 1986 |
| ENSG00000237331 | 0 | 0 | 0 | 0 | 0 | 0 |
| ENSG00000252627 | 0 | 2 | 0 | 0 | 0 | 0 |
| ENSG00000237025 | 2 | 2 | 0 | 0 | 1 | 1 |
| ENSG00000232412 | 4 | 9 | 3 | 3 | 3 | 4 |
| ENSG00000101972 | 1475 | 1124 | 692 | 291 | 2021 | 845 |
| ENSG00000237426 | 0 | 0 | 0 | 0 | 0 | 0 |
| ENSG00000252693 | 0 | 0 | 0 | 0 | 0 | 0 |
| ENSG00000234564 | 0 | 0 | 0 | 0 | 0 | 0 |
| ENSG00000183918 | 0 | 0 | 0 | 0 | 0 | 0 |
| ENSG00000282419 | 0 | 0 | 0 | 0 | 0 | 0 |
| ENSG00000232054 | 0 | 0 | 0 | 0 | 0 | 0 |
| ENSG00000226532 | 0 | 0 | 0 | 0 | 0 | 0 |
| ENSG00000009694 | 0 | 0 | 0 | 0 | 0 | 0 |
| ENSG00000236880 | 0 | 0 | 0 | 0 | 0 | 0 |
| ENSG00000232599 | 0 | 0 | 0 | 0 | 0 | 0 |
| ENSG00000282815 | 0 | 0 | 0 | 1 | 1 | 0 |
| ENSG00000213489 | 0 | 0 | 0 | 0 | 0 | 0 |
| ENSG00000280223 | 0 | 0 | 0 | 0 | 0 | 0 |
| ENSG00000237208 | 14 | 5 | 4 | 6 | 10 | 4 |
| ENSG00000198354 | 2 | 5 | 6 | 6 | 2 | 5 |
| ENSG00000230916 | 0 | 1 | 2 | 1 | 0 | 2 |
| ENSG00000231501 | 0 | 0 | 0 | 0 | 0 | 0 |
| ENSG00000232177 | 2 | 0 | 0 | 0 | 0 | 0 |
| ENSG00000274901 | 0 | 0 | 0 | 0 | 0 | 0 |
| ENSG00000198889 | 0 | 0 | 0 | 0 | 0 | 0 |
| ENSG00000226414 | 0 | 0 | 0 | 0 | 0 | 0 |
| ENSG00000183631 | 0 | 0 | 0 | 0 | 0 | 0 |
| ENSG00000230538 | 0 | 0 | 0 | 0 | 0 | 0 |
| ENSG00000280275 | 0 | 0 | 0 | 0 | 0 | 0 |
| ENSG00000274631 | 0 | 0 | 0 | 0 | 0 | 0 |
| ENSG00000260683 | 0 | 0 | 0 | 0 | 0 | 0 |
| ENSG00000235274 | 0 | 0 | 0 | 0 | 0 | 0 |
| ENSG00000224548 | 0 | 0 | 0 | 0 | 0 | 0 |
| ENSG00000123165 | 0 | 0 | 0 | 0 | 0 | 0 |
| ENSG00000236042 | 0 | 0 | 0 | 0 | 0 | 0 |
| ENSG00000235334 | 0 | 0 | 0 | 0 | 0 | 0 |
| ENSG00000235239 | 0 | 0 | 0 | 0 | 0 | 0 |

| | | | | | | |
|---|---|---|---|---|---|---|
| ENSG00000235735 | 0 | 0 | 0 | 0 | 0 | 0 |
| ENSG00000225689 | 2 | 0 | 1 | 0 | 0 | 0 |
| ENSG00000226802 | 0 | 0 | 0 | 0 | 0 | 0 |
| ENSG00000218728 | 0 | 0 | 0 | 0 | 0 | 0 |
| ENSG00000223995 | 0 | 0 | 0 | 0 | 0 | 0 |
| ENSG00000234592 | 0 | 0 | 0 | 0 | 0 | 0 |
| ENSG00000226339 | 0 | 0 | 0 | 0 | 0 | 0 |
| ENSG00000201518 | 0 | 0 | 0 | 0 | 0 | 0 |
| ENSG00000102038 | 2 | 0 | 0 | 0 | 1 | 1 |
| ENSG00000252170 | 0 | 0 | 0 | 0 | 0 | 0 |
| ENSG00000122126 | 2218 | 1048 | 1422 | 853 | 1361 | 1258 |
| ENSG00000171388 | 4 | 0 | 0 | 0 | 4 | 0 |
| ENSG00000122121 | 0 | 0 | 0 | 0 | 0 | 0 |
| ENSG00000122122 | 0 | 0 | 0 | 1 | 0 | 1 |
| ENSG00000240143 | 0 | 0 | 2 | 3 | 0 | 0 |
| ENSG00000275070 | 2 | 3 | 0 | 0 | 1 | 3 |
| ENSG00000188706 | 3149 | 1509 | 1929 | 2062 | 1956 | 2246 |
| ENSG00000235189 | 0 | 1 | 0 | 0 | 0 | 0 |
| ENSG00000238040 | 0 | 0 | 0 | 0 | 0 | 0 |
| ENSG00000156697 | 1689 | 785 | 1008 | 978 | 968 | 1026 |
| ENSG00000085185 | 787 | 334 | 412 | 318 | 344 | 427 |
| ENSG00000102034 | 3068 | 1484 | 1853 | 2126 | 2327 | 2330 |
| ENSG00000156709 | 3919 | 1905 | 2291 | 1934 | 2239 | 2150 |
| ENSG00000134594 | 5 | 3 | 4 | 3 | 5 | 3 |
| ENSG00000056277 | 238 | 125 | 106 | 56 | 179 | 118 |
| ENSG00000202304 | 0 | 0 | 0 | 0 | 0 | 0 |
| ENSG00000225046 | 0 | 0 | 0 | 0 | 0 | 0 |
| ENSG00000102078 | 560 | 282 | 334 | 274 | 307 | 320 |
| ENSG00000147262 | 0 | 0 | 0 | 0 | 0 | 0 |
| ENSG00000134597 | 1632 | 766 | 939 | 887 | 934 | 932 |
| ENSG00000274775 | 0 | 0 | 0 | 0 | 0 | 2 |
| ENSG00000221930 | 2 | 3 | 5 | 1 | 1 | 1 |
| ENSG00000271677 | 0 | 0 | 0 | 0 | 0 | 0 |
| ENSG00000165675 | 540 | 325 | 324 | 236 | 380 | 293 |
| ENSG00000238188 | 1 | 0 | 0 | 0 | 0 | 0 |
| ENSG00000228659 | 0 | 0 | 0 | 0 | 0 | 0 |
| ENSG00000147256 | 50 | 28 | 15 | 48 | 38 | 31 |
| ENSG00000241198 | 0 | 0 | 0 | 0 | 0 | 0 |
| ENSG00000237650 | 0 | 0 | 0 | 0 | 0 | 0 |
| ENSG00000276205 | 0 | 0 | 0 | 0 | 0 | 0 |
| ENSG00000147255 | 700 | 289 | 347 | 463 | 438 | 533 |
| ENSG00000185903 | 0 | 0 | 0 | 0 | 1 | 0 |
| ENSG00000229082 | 0 | 0 | 0 | 0 | 0 | 0 |
| ENSG00000242610 | 0 | 0 | 0 | 0 | 0 | 0 |
| ENSG00000234288 | 0 | 0 | 0 | 0 | 0 | 0 |
| ENSG00000230973 | 0 | 0 | 0 | 0 | 0 | 0 |
| ENSG00000228450 | 0 | 0 | 2 | 3 | 1 | 0 |
| ENSG00000171054 | 0 | 1 | 0 | 0 | 0 | 0 |
| ENSG00000237225 | 0 | 0 | 0 | 0 | 0 | 0 |
| ENSG00000232787 | 0 | 0 | 0 | 1 | 0 | 0 |

| | | | | | | |
|---|---|---|---|---|---|---|
| ENSG00000235730 | 4 | 0 | 1 | 4 | 2 | 7 |
| ENSG00000227183 | 0 | 1 | 1 | 0 | 2 | 2 |
| ENSG00000213468 | 123 | 61 | 71 | 57 | 101 | 86 |
| ENSG00000241418 | 0 | 0 | 0 | 0 | 0 | 0 |
| ENSG00000228089 | 0 | 1 | 0 | 1 | 0 | 0 |
| ENSG00000235190 | 0 | 0 | 0 | 0 | 0 | 0 |
| ENSG00000200587 | 0 | 0 | 0 | 0 | 0 | 0 |
| ENSG00000134602 | 1552 | 787 | 806 | 554 | 961 | 768 |
| ENSG00000165694 | 2 | 0 | 2 | 1 | 0 | 2 |
| ENSG00000228819 | 0 | 0 | 0 | 0 | 0 | 0 |
| ENSG00000123728 | 2488 | 1292 | 1595 | 1056 | 1284 | 1377 |
| ENSG00000232160 | 145 | 66 | 81 | 53 | 77 | 89 |
| ENSG00000076770 | 388 | 256 | 156 | 69 | 323 | 155 |
| ENSG00000206900 | 0 | 0 | 0 | 0 | 0 | 0 |
| ENSG00000171004 | 140 | 108 | 107 | 106 | 170 | 138 |
| ENSG00000235849 | 0 | 0 | 0 | 0 | 0 | 0 |
| ENSG00000227064 | 0 | 0 | 0 | 0 | 0 | 0 |
| ENSG00000134588 | 1 | 0 | 0 | 1 | 0 | 0 |
| ENSG00000225949 | 0 | 0 | 0 | 0 | 0 | 0 |
| ENSG00000207100 | 0 | 0 | 0 | 0 | 0 | 0 |
| ENSG00000183434 | 0 | 0 | 0 | 0 | 0 | 0 |
| ENSG00000202199 | 0 | 0 | 0 | 0 | 0 | 0 |
| ENSG00000076716 | 582 | 273 | 387 | 175 | 214 | 241 |
| ENSG00000206765 | 0 | 0 | 0 | 0 | 0 | 0 |
| ENSG00000147257 | 18 | 14 | 11 | 16 | 10 | 15 |
| ENSG00000227008 | 10 | 8 | 8 | 6 | 11 | 8 |
| ENSG00000233260 | 0 | 0 | 0 | 0 | 0 | 0 |
| ENSG00000232068 | 0 | 0 | 0 | 0 | 0 | 0 |
| ENSG00000200572 | 0 | 0 | 0 | 0 | 0 | 0 |
| ENSG00000224908 | 0 | 0 | 0 | 0 | 0 | 0 |
| ENSG00000283638 | 0 | 0 | 0 | 0 | 0 | 0 |
| ENSG00000284499 | 0 | 0 | 0 | 0 | 0 | 0 |
| ENSG00000284538 | 0 | 0 | 0 | 0 | 0 | 0 |
| ENSG00000284107 | 0 | 0 | 0 | 0 | 0 | 0 |
| ENSG00000284043 | 0 | 0 | 0 | 0 | 0 | 0 |
| ENSG00000283931 | 0 | 0 | 0 | 0 | 0 | 0 |
| ENSG00000284157 | 0 | 0 | 0 | 0 | 0 | 0 |
| ENSG00000232664 | 0 | 0 | 0 | 0 | 0 | 0 |
| ENSG00000203952 | 10 | 6 | 1 | 2 | 8 | 6 |
| ENSG00000214961 | 0 | 0 | 0 | 0 | 0 | 0 |
| ENSG00000156531 | 1491 | 773 | 829 | 406 | 983 | 767 |
| ENSG00000165704 | 1081 | 585 | 695 | 735 | 813 | 750 |
| ENSG00000226107 | 7 | 2 | 1 | 1 | 3 | 3 |
| ENSG00000216001 | 1 | 0 | 0 | 0 | 0 | 0 |
| ENSG00000199132 | 1 | 0 | 0 | 0 | 0 | 0 |
| ENSG00000207755 | 0 | 0 | 0 | 0 | 0 | 0 |
| ENSG00000207784 | 0 | 0 | 0 | 0 | 0 | 0 |
| ENSG00000223749 | 40 | 16 | 19 | 15 | 21 | 26 |
| ENSG00000208005 | 1 | 1 | 0 | 1 | 0 | 0 |
| ENSG00000284231 | 0 | 0 | 0 | 0 | 0 | 0 |

| | | | | | | |
|---|---|---|---|---|---|---|
| ENSG00000227060 | 0 | 1 | 0 | 2 | 5 | 0 |
| ENSG00000170965 | 4 | 1 | 3 | 6 | 4 | 6 |
| ENSG00000235813 | 0 | 0 | 0 | 0 | 0 | 0 |
| ENSG00000236509 | 0 | 0 | 0 | 0 | 0 | 0 |
| ENSG00000237432 | 0 | 0 | 0 | 0 | 0 | 0 |
| ENSG00000199920 | 0 | 0 | 0 | 0 | 0 | 0 |
| ENSG00000156504 | 2245 | 1108 | 1430 | 966 | 1362 | 1342 |
| ENSG00000156500 | 129 | 42 | 64 | 56 | 53 | 80 |
| ENSG00000280249 | 0 | 0 | 0 | 1 | 0 | 0 |
| ENSG00000101928 | 1059 | 455 | 596 | 443 | 594 | 490 |
| ENSG00000207081 | 0 | 0 | 0 | 0 | 0 | 0 |
| ENSG00000236091 | 2 | 0 | 1 | 0 | 0 | 0 |
| ENSG00000231929 | 0 | 0 | 0 | 0 | 0 | 0 |
| ENSG00000184785 | 0 | 0 | 0 | 0 | 0 | 0 |
| ENSG00000212747 | 1652 | 784 | 967 | 933 | 813 | 851 |
| ENSG00000134590 | 2868 | 1412 | 1711 | 1730 | 1560 | 1503 |
| ENSG00000203950 | 2945 | 1382 | 1741 | 1732 | 1605 | 1622 |
| ENSG00000228372 | 0 | 0 | 0 | 3 | 0 | 0 |
| ENSG00000234228 | 0 | 0 | 0 | 0 | 0 | 0 |
| ENSG00000196972 | 6 | 1 | 3 | 2 | 1 | 3 |
| ENSG00000224107 | 0 | 0 | 0 | 0 | 0 | 0 |
| ENSG00000223809 | 0 | 0 | 0 | 0 | 0 | 0 |
| ENSG00000232582 | 0 | 0 | 0 | 0 | 0 | 0 |
| ENSG00000169551 | 0 | 0 | 0 | 0 | 0 | 0 |
| ENSG00000236491 | 0 | 0 | 1 | 1 | 6 | 2 |
| ENSG00000228922 | 0 | 0 | 0 | 0 | 0 | 0 |
| ENSG00000269833 | 0 | 0 | 0 | 0 | 0 | 0 |
| ENSG00000186376 | 1292 | 561 | 805 | 411 | 581 | 614 |
| ENSG00000238210 | 0 | 0 | 0 | 0 | 0 | 0 |
| ENSG00000274024 | 4 | 0 | 2 | 0 | 1 | 4 |
| ENSG00000283644 | 0 | 0 | 1 | 0 | 0 | 1 |
| ENSG00000284994 | 0 | 2 | 1 | 2 | 1 | 0 |
| ENSG00000173275 | 456 | 210 | 281 | 147 | 255 | 202 |
| ENSG00000229015 | 0 | 0 | 0 | 0 | 0 | 0 |
| ENSG00000178947 | 36 | 25 | 45 | 25 | 34 | 28 |
| ENSG00000201440 | 0 | 0 | 0 | 0 | 1 | 0 |
| ENSG00000225769 | 0 | 0 | 0 | 0 | 0 | 0 |
| ENSG00000273988 | 0 | 0 | 0 | 0 | 0 | 0 |
| ENSG00000271158 | 0 | 0 | 0 | 0 | 0 | 0 |
| ENSG00000225235 | 5 | 3 | 2 | 2 | 0 | 2 |
| ENSG00000165359 | 543 | 283 | 329 | 171 | 266 | 228 |
| ENSG00000237833 | 4 | 0 | 0 | 0 | 0 | 0 |
| ENSG00000224539 | 0 | 0 | 0 | 0 | 0 | 0 |
| ENSG00000277486 | 0 | 0 | 0 | 0 | 0 | 0 |
| ENSG00000198022 | 0 | 0 | 0 | 0 | 0 | 0 |
| ENSG00000274601 | 0 | 0 | 0 | 0 | 0 | 0 |
| ENSG00000268940 | 0 | 0 | 0 | 0 | 1 | 0 |
| ENSG00000233861 | 0 | 0 | 0 | 0 | 0 | 0 |
| ENSG00000238200 | 0 | 0 | 0 | 0 | 0 | 0 |
| ENSG00000269096 | 2 | 0 | 0 | 0 | 0 | 0 |

| | | | | | | |
|---|---|---|---|---|---|---|
| ENSG00000228836 | 0 | 0 | 0 | 0 | 0 | 0 |
| ENSG00000278289 | 0 | 0 | 0 | 0 | 0 | 0 |
| ENSG00000271449 | 0 | 0 | 0 | 0 | 0 | 0 |
| ENSG00000273696 | 0 | 0 | 0 | 0 | 0 | 0 |
| ENSG00000278085 | 0 | 0 | 0 | 0 | 0 | 0 |
| ENSG00000270946 | 0 | 0 | 0 | 0 | 0 | 0 |
| ENSG00000269586 | 0 | 0 | 0 | 0 | 0 | 0 |
| ENSG00000181433 | 0 | 0 | 0 | 0 | 0 | 0 |
| ENSG00000230162 | 0 | 0 | 1 | 0 | 0 | 0 |
| ENSG00000169446 | 1618 | 836 | 931 | 711 | 1001 | 966 |
| ENSG00000198689 | 804 | 389 | 489 | 281 | 397 | 412 |
| ENSG00000219930 | 0 | 0 | 0 | 0 | 0 | 0 |
| ENSG00000231513 | 0 | 0 | 0 | 0 | 0 | 0 |
| ENSG00000239080 | 0 | 0 | 0 | 0 | 0 | 0 |
| ENSG00000022267 | 426 | 214 | 218 | 216 | 214 | 246 |
| ENSG00000129680 | 432 | 189 | 197 | 136 | 367 | 239 |
| ENSG00000230079 | 0 | 0 | 0 | 0 | 0 | 0 |
| ENSG00000156920 | 0 | 0 | 0 | 0 | 1 | 0 |
| ENSG00000102239 | 0 | 0 | 0 | 0 | 0 | 0 |
| ENSG00000102241 | 3251 | 1695 | 2045 | 1304 | 1952 | 1868 |
| ENSG00000102243 | 8 | 6 | 2 | 5 | 2 | 1 |
| ENSG00000216060 | 0 | 0 | 0 | 0 | 0 | 0 |
| ENSG00000199424 | 0 | 0 | 0 | 0 | 0 | 0 |
| ENSG00000252320 | 0 | 0 | 0 | 0 | 0 | 0 |
| ENSG00000233093 | 0 | 0 | 0 | 0 | 0 | 0 |
| ENSG00000102245 | 0 | 0 | 0 | 0 | 0 | 0 |
| ENSG00000129675 | 20 | 15 | 6 | 0 | 3 | 7 |
| ENSG00000212510 | 0 | 0 | 0 | 0 | 0 | 0 |
| ENSG00000271339 | 0 | 0 | 0 | 0 | 0 | 0 |
| ENSG00000231164 | 0 | 0 | 0 | 0 | 0 | 0 |
| ENSG00000225125 | 7 | 3 | 5 | 2 | 5 | 3 |
| ENSG00000202279 | 0 | 0 | 0 | 0 | 1 | 0 |
| ENSG00000232611 | 42 | 10 | 14 | 11 | 15 | 24 |
| ENSG00000225400 | 3 | 3 | 4 | 1 | 6 | 2 |
| ENSG00000147274 | 8985 | 4429 | 5372 | 4360 | 5103 | 5473 |
| ENSG00000206979 | 0 | 0 | 0 | 0 | 0 | 0 |
| ENSG00000234062 | 64 | 31 | 42 | 43 | 42 | 42 |
| ENSG00000232379 | 0 | 0 | 0 | 0 | 0 | 0 |
| ENSG00000237307 | 6 | 5 | 3 | 0 | 0 | 2 |
| ENSG00000165370 | 0 | 0 | 0 | 0 | 0 | 0 |
| ENSG00000214748 | 0 | 0 | 0 | 0 | 0 | 0 |
| ENSG00000237584 | 0 | 0 | 0 | 0 | 0 | 0 |
| ENSG00000224765 | 0 | 2 | 4 | 0 | 1 | 2 |
| ENSG00000156925 | 52 | 17 | 26 | 6 | 10 | 15 |
| ENSG00000283692 | 0 | 0 | 1 | 0 | 0 | 0 |
| ENSG00000239579 | 0 | 0 | 0 | 0 | 0 | 0 |
| ENSG00000229562 | 0 | 0 | 0 | 0 | 0 | 0 |
| ENSG00000222114 | 0 | 0 | 0 | 0 | 0 | 0 |
| ENSG00000252163 | 0 | 0 | 0 | 0 | 0 | 0 |
| ENSG00000183981 | 0 | 0 | 0 | 0 | 0 | 0 |

| | | | | | | |
|---|---|---|---|---|---|---|
| ENSG00000214930 | 0 | 0 | 0 | 0 | 0 | 0 |
| ENSG00000285989 | 0 | 0 | 0 | 0 | 0 | 0 |
| ENSG00000129682 | 523 | 259 | 332 | 171 | 164 | 224 |
| ENSG00000232183 | 0 | 0 | 0 | 0 | 0 | 0 |
| ENSG00000207800 | 0 | 0 | 0 | 0 | 0 | 0 |
| ENSG00000226031 | 0 | 0 | 1 | 0 | 0 | 0 |
| ENSG00000235975 | 0 | 0 | 0 | 0 | 0 | 0 |
| ENSG00000173954 | 0 | 0 | 0 | 0 | 0 | 0 |
| ENSG00000232549 | 0 | 0 | 0 | 0 | 0 | 0 |
| ENSG00000101981 | 0 | 0 | 0 | 0 | 0 | 0 |
| ENSG00000101977 | 4 | 0 | 0 | 0 | 2 | 3 |
| ENSG00000238485 | 0 | 0 | 0 | 0 | 0 | 0 |
| ENSG00000101974 | 742 | 444 | 330 | 196 | 538 | 359 |
| ENSG00000221375 | 0 | 0 | 0 | 0 | 0 | 0 |
| ENSG00000252719 | 0 | 0 | 0 | 0 | 0 | 0 |
| ENSG00000207633 | 0 | 0 | 0 | 0 | 0 | 0 |
| ENSG00000203933 | 0 | 0 | 0 | 0 | 0 | 0 |
| ENSG00000233145 | 0 | 0 | 0 | 0 | 0 | 0 |
| ENSG00000214653 | 3 | 0 | 1 | 0 | 2 | 1 |
| ENSG00000241248 | 0 | 0 | 0 | 0 | 0 | 0 |
| ENSG00000230707 | 0 | 0 | 0 | 1 | 4 | 0 |
| ENSG00000231110 | 0 | 0 | 0 | 0 | 0 | 0 |
| ENSG00000224942 | 0 | 0 | 0 | 0 | 0 | 0 |
| ENSG00000275268 | 0 | 0 | 0 | 0 | 0 | 0 |
| ENSG00000225758 | 0 | 0 | 0 | 0 | 0 | 0 |
| ENSG00000134595 | 405 | 164 | 243 | 210 | 170 | 176 |
| ENSG00000230668 | 1 | 0 | 2 | 0 | 0 | 0 |
| ENSG00000203930 | 8 | 9 | 10 | 0 | 6 | 4 |
| ENSG00000184258 | 0 | 0 | 0 | 0 | 0 | 0 |
| ENSG00000221081 | 0 | 0 | 0 | 0 | 0 | 0 |
| ENSG00000229269 | 0 | 0 | 0 | 0 | 0 | 0 |
| ENSG00000227234 | 4 | 0 | 0 | 0 | 0 | 0 |
| ENSG00000207041 | 0 | 0 | 0 | 0 | 1 | 4 |
| ENSG00000214925 | 0 | 0 | 0 | 1 | 0 | 0 |
| ENSG00000261212 | 4 | 4 | 1 | 1 | 0 | 0 |
| ENSG00000182195 | 4939 | 2481 | 3121 | 2580 | 2306 | 2561 |
| ENSG00000198573 | 0 | 0 | 0 | 0 | 0 | 0 |
| ENSG00000213423 | 0 | 0 | 0 | 0 | 0 | 1 |
| ENSG00000227779 | 0 | 0 | 0 | 0 | 0 | 0 |
| ENSG00000277215 | 10 | 2 | 1 | 0 | 2 | 2 |
| ENSG00000198021 | 0 | 0 | 0 | 0 | 0 | 0 |
| ENSG00000203926 | 0 | 0 | 0 | 0 | 0 | 0 |
| ENSG00000223438 | 0 | 0 | 0 | 0 | 0 | 0 |
| ENSG00000196406 | 0 | 0 | 0 | 0 | 0 | 0 |
| ENSG00000165509 | 2 | 3 | 2 | 0 | 0 | 3 |
| ENSG00000155495 | 0 | 0 | 1 | 0 | 1 | 0 |
| ENSG00000236205 | 0 | 0 | 0 | 0 | 0 | 0 |
| ENSG00000225469 | 0 | 0 | 0 | 0 | 0 | 0 |
| ENSG00000250625 | 0 | 0 | 0 | 0 | 0 | 0 |
| ENSG00000046774 | 0 | 0 | 0 | 0 | 0 | 0 |

| | | | | | | |
|---|---|---|---|---|---|---|
| ENSG00000275508 | 0 | 0 | 0 | 0 | 0 | 0 |
| ENSG00000201000 | 0 | 0 | 0 | 0 | 0 | 0 |
| ENSG00000189326 | 0 | 0 | 0 | 0 | 0 | 0 |
| ENSG00000202473 | 0 | 0 | 0 | 0 | 0 | 0 |
| ENSG00000199878 | 0 | 0 | 0 | 0 | 0 | 0 |
| ENSG00000227303 | 0 | 0 | 0 | 0 | 0 | 0 |
| ENSG00000226064 | 0 | 0 | 0 | 0 | 0 | 0 |
| ENSG00000235539 | 0 | 0 | 0 | 0 | 0 | 0 |
| ENSG00000230130 | 0 | 0 | 0 | 0 | 0 | 0 |
| ENSG00000189252 | 0 | 0 | 0 | 0 | 0 | 0 |
| ENSG00000179542 | 38 | 13 | 15 | 5 | 13 | 20 |
| ENSG00000213924 | 0 | 0 | 0 | 0 | 0 | 0 |
| ENSG00000268988 | 0 | 0 | 0 | 0 | 0 | 0 |
| ENSG00000225970 | 0 | 0 | 0 | 0 | 0 | 0 |
| ENSG00000235729 | 0 | 0 | 0 | 0 | 0 | 0 |
| ENSG00000276380 | 0 | 0 | 0 | 0 | 0 | 0 |
| ENSG00000224630 | 0 | 0 | 1 | 0 | 0 | 0 |
| ENSG00000235598 | 0 | 0 | 0 | 0 | 0 | 0 |
| ENSG00000215054 | 0 | 0 | 0 | 0 | 0 | 0 |
| ENSG00000201912 | 0 | 0 | 0 | 0 | 0 | 0 |
| ENSG00000233548 | 0 | 0 | 0 | 0 | 0 | 0 |
| ENSG00000234712 | 0 | 0 | 0 | 0 | 0 | 0 |
| ENSG00000203923 | 0 | 0 | 0 | 0 | 0 | 0 |
| ENSG00000231093 | 0 | 0 | 0 | 0 | 0 | 0 |
| ENSG00000270709 | 0 | 0 | 0 | 0 | 0 | 0 |
| ENSG00000279585 | 0 | 0 | 0 | 0 | 0 | 0 |
| ENSG00000185985 | 0 | 0 | 0 | 0 | 0 | 0 |
| ENSG00000252219 | 0 | 0 | 0 | 0 | 0 | 0 |
| ENSG00000216075 | 0 | 0 | 0 | 0 | 0 | 0 |
| ENSG00000216005 | 0 | 0 | 0 | 0 | 0 | 0 |
| ENSG00000215943 | 0 | 0 | 0 | 0 | 0 | 0 |
| ENSG00000216098 | 0 | 0 | 0 | 0 | 0 | 0 |
| ENSG00000216064 | 0 | 0 | 0 | 0 | 0 | 0 |
| ENSG00000216056 | 0 | 0 | 0 | 0 | 0 | 0 |
| ENSG00000201594 | 0 | 0 | 0 | 0 | 0 | 0 |
| ENSG00000234748 | 0 | 0 | 0 | 0 | 0 | 0 |
| ENSG00000226388 | 0 | 0 | 0 | 0 | 0 | 0 |
| ENSG00000231686 | 0 | 0 | 0 | 0 | 0 | 2 |
| ENSG00000235699 | 0 | 0 | 0 | 0 | 0 | 0 |
| ENSG00000224440 | 0 | 0 | 0 | 0 | 0 | 0 |
| ENSG00000225660 | 0 | 0 | 0 | 0 | 0 | 0 |
| ENSG00000271660 | 0 | 0 | 0 | 0 | 0 | 0 |
| ENSG00000284377 | 3 | 0 | 0 | 0 | 0 | 1 |
| ENSG00000216171 | 0 | 0 | 0 | 0 | 0 | 0 |
| ENSG00000207871 | 0 | 0 | 0 | 0 | 0 | 0 |
| ENSG00000207873 | 0 | 0 | 0 | 0 | 0 | 0 |
| ENSG00000207984 | 0 | 0 | 0 | 0 | 0 | 0 |
| ENSG00000207731 | 0 | 0 | 0 | 0 | 0 | 0 |
| ENSG00000207969 | 0 | 0 | 0 | 0 | 0 | 0 |
| ENSG00000207589 | 0 | 0 | 0 | 0 | 0 | 0 |

| | | | | | | |
|---|---|---|---|---|---|---|
| ENSG00000252583 | 0 | 0 | 0 | 0 | 0 | 0 |
| ENSG00000212013 | 0 | 0 | 0 | 0 | 0 | 0 |
| ENSG00000212014 | 0 | 0 | 0 | 0 | 0 | 0 |
| ENSG00000208000 | 0 | 0 | 0 | 0 | 0 | 0 |
| ENSG00000207641 | 0 | 0 | 0 | 0 | 0 | 0 |
| ENSG00000207868 | 0 | 0 | 0 | 0 | 0 | 0 |
| ENSG00000207866 | 0 | 0 | 0 | 0 | 0 | 0 |
| ENSG00000207867 | 0 | 0 | 0 | 0 | 0 | 0 |
| ENSG00000274579 | 0 | 1 | 0 | 0 | 0 | 0 |
| ENSG00000228855 | 0 | 0 | 0 | 0 | 0 | 0 |
| ENSG00000227505 | 0 | 0 | 0 | 0 | 0 | 0 |
| ENSG00000271175 | 0 | 0 | 0 | 0 | 0 | 0 |
| ENSG00000202051 | 0 | 0 | 0 | 0 | 0 | 0 |
| ENSG00000227083 | 0 | 0 | 1 | 0 | 2 | 0 |
| ENSG00000268066 | 2 | 0 | 2 | 0 | 3 | 0 |
| ENSG00000274086 | 0 | 0 | 0 | 0 | 0 | 0 |
| ENSG00000102081 | 1964 | 920 | 1143 | 647 | 1066 | 992 |
| ENSG00000236337 | 7 | 5 | 11 | 1 | 2 | 3 |
| ENSG00000176988 | 0 | 0 | 0 | 0 | 0 | 0 |
| ENSG00000201285 | 0 | 0 | 0 | 0 | 0 | 0 |
| ENSG00000219507 | 3 | 5 | 3 | 6 | 7 | 2 |
| ENSG00000233467 | 2 | 0 | 0 | 0 | 0 | 0 |
| ENSG00000176970 | 0 | 0 | 0 | 0 | 0 | 0 |
| ENSG00000237741 | 0 | 0 | 0 | 0 | 0 | 0 |
| ENSG00000155966 | 18 | 4 | 7 | 1 | 0 | 2 |
| ENSG00000223516 | 0 | 0 | 0 | 0 | 0 | 0 |
| ENSG00000222313 | 0 | 0 | 0 | 0 | 0 | 0 |
| ENSG00000242186 | 0 | 0 | 0 | 0 | 0 | 0 |
| ENSG00000010404 | 2655 | 1336 | 1769 | 1241 | 1982 | 1735 |
| ENSG00000241489 | 18 | 28 | 36 | 7 | 23 | 24 |
| ENSG00000238039 | 4 | 6 | 8 | 5 | 3 | 2 |
| ENSG00000176289 | 2 | 2 | 6 | 2 | 1 | 5 |
| ENSG00000241769 | 49 | 26 | 42 | 13 | 21 | 21 |
| ENSG00000197620 | 933 | 417 | 555 | 517 | 527 | 506 |
| ENSG00000283697 | 0 | 0 | 0 | 7 | 0 | 2 |
| ENSG00000267978 | 0 | 0 | 0 | 0 | 0 | 0 |
| ENSG00000268738 | 0 | 0 | 0 | 0 | 0 | 0 |
| ENSG00000269556 | 621 | 288 | 374 | 346 | 383 | 365 |
| ENSG00000229322 | 0 | 0 | 0 | 0 | 0 | 0 |
| ENSG00000185247 | 2 | 0 | 0 | 0 | 0 | 0 |
| ENSG00000228833 | 0 | 0 | 0 | 0 | 0 | 0 |
| ENSG00000223377 | 0 | 0 | 0 | 0 | 0 | 0 |
| ENSG00000231694 | 0 | 0 | 0 | 0 | 0 | 0 |
| ENSG00000235254 | 0 | 0 | 0 | 1 | 1 | 1 |
| ENSG00000171116 | 4 | 7 | 1 | 4 | 5 | 2 |
| ENSG00000123584 | 0 | 0 | 0 | 0 | 0 | 0 |
| ENSG00000224732 | 0 | 0 | 0 | 0 | 0 | 0 |
| ENSG00000229829 | 0 | 0 | 0 | 0 | 0 | 0 |
| ENSG00000280752 | 0 | 0 | 0 | 0 | 0 | 0 |
| ENSG00000230899 | 0 | 1 | 1 | 0 | 0 | 0 |

| | | | | | | |
|---|---|---|---|---|---|---|
| ENSG00000156009 | 0 | 0 | 1 | 0 | 1 | 0 |
| ENSG00000197021 | 1558 | 775 | 992 | 906 | 826 | 895 |
| ENSG00000283463 | 0 | 0 | 0 | 4 | 0 | 3 |
| ENSG00000235703 | 57 | 35 | 43 | 17 | 18 | 40 |
| ENSG00000183171 | 2 | 4 | 4 | 0 | 1 | 0 |
| ENSG00000252454 | 0 | 0 | 0 | 0 | 0 | 0 |
| ENSG00000234825 | 6 | 1 | 2 | 1 | 1 | 0 |
| ENSG00000013619 | 154 | 63 | 110 | 42 | 69 | 56 |
| ENSG00000171100 | 603 | 328 | 386 | 171 | 366 | 319 |
| ENSG00000063601 | 2250 | 1091 | 1384 | 916 | 1171 | 1144 |
| ENSG00000102181 | 1213 | 622 | 786 | 658 | 647 | 655 |
| ENSG00000228728 | 0 | 1 | 4 | 0 | 5 | 1 |
| ENSG00000230153 | 0 | 0 | 0 | 0 | 0 | 0 |
| ENSG00000222796 | 0 | 0 | 0 | 0 | 0 | 0 |
| ENSG00000029993 | 5150 | 2648 | 3203 | 2534 | 3239 | 3005 |
| ENSG00000230508 | 0 | 0 | 0 | 0 | 0 | 0 |
| ENSG00000270916 | 0 | 0 | 0 | 0 | 0 | 0 |
| ENSG00000212595 | 0 | 0 | 0 | 0 | 0 | 0 |
| ENSG00000265789 | 0 | 0 | 0 | 0 | 0 | 0 |
| ENSG00000234696 | 0 | 0 | 0 | 0 | 0 | 0 |
| ENSG00000102195 | 0 | 0 | 0 | 0 | 1 | 0 |
| ENSG00000269993 | 0 | 0 | 0 | 0 | 0 | 0 |
| ENSG00000228717 | 0 | 0 | 0 | 0 | 0 | 0 |
| ENSG00000278724 | 0 | 0 | 0 | 0 | 0 | 0 |
| ENSG00000160131 | 2039 | 983 | 1069 | 895 | 1109 | 1069 |
| ENSG00000270052 | 2 | 1 | 0 | 0 | 2 | 2 |
| ENSG00000214915 | 0 | 0 | 0 | 1 | 0 | 0 |
| ENSG00000166049 | 0 | 0 | 0 | 1 | 0 | 0 |
| ENSG00000239429 | 0 | 0 | 0 | 0 | 0 | 0 |
| ENSG00000130032 | 0 | 0 | 0 | 0 | 0 | 0 |
| ENSG00000147378 | 0 | 0 | 0 | 0 | 0 | 0 |
| ENSG00000183862 | 0 | 0 | 0 | 0 | 0 | 0 |
| ENSG00000229967 | 0 | 0 | 0 | 0 | 0 | 0 |
| ENSG00000147381 | 0 | 0 | 0 | 0 | 0 | 0 |
| ENSG00000102287 | 0 | 0 | 0 | 0 | 0 | 0 |
| ENSG00000284363 | 0 | 0 | 0 | 0 | 0 | 0 |
| ENSG00000283751 | 0 | 0 | 0 | 0 | 0 | 0 |
| ENSG00000228965 | 0 | 0 | 0 | 0 | 0 | 0 |
| ENSG00000242520 | 0 | 0 | 0 | 0 | 0 | 0 |
| ENSG00000266560 | 0 | 0 | 0 | 0 | 0 | 0 |
| ENSG00000124260 | 0 | 0 | 0 | 0 | 0 | 0 |
| ENSG00000231937 | 0 | 0 | 0 | 0 | 0 | 0 |
| ENSG00000011677 | 0 | 1 | 0 | 0 | 0 | 0 |
| ENSG00000252205 | 0 | 0 | 0 | 0 | 0 | 0 |
| ENSG00000207957 | 0 | 0 | 0 | 0 | 0 | 0 |
| ENSG00000211583 | 0 | 0 | 0 | 0 | 0 | 0 |
| ENSG00000207818 | 0 | 0 | 0 | 0 | 0 | 0 |
| ENSG00000223940 | 2 | 2 | 2 | 3 | 2 | 4 |
| ENSG00000268089 | 0 | 0 | 0 | 0 | 0 | 0 |
| ENSG00000269475 | 0 | 0 | 0 | 0 | 0 | 0 |

| | | | | | | |
|---|---|---|---|---|---|---|
| ENSG00000221867 | 0 | 0 | 0 | 0 | 0 | 0 |
| ENSG00000268902 | 0 | 0 | 0 | 0 | 0 | 0 |
| ENSG00000183305 | 0 | 0 | 0 | 0 | 0 | 0 |
| ENSG00000198930 | 0 | 0 | 0 | 0 | 0 | 0 |
| ENSG00000213401 | 0 | 0 | 0 | 0 | 0 | 0 |
| ENSG00000242599 | 0 | 0 | 0 | 0 | 0 | 0 |
| ENSG00000231447 | 0 | 0 | 0 | 0 | 0 | 0 |
| ENSG00000268606 | 0 | 0 | 0 | 0 | 0 | 0 |
| ENSG00000268916 | 0 | 0 | 0 | 0 | 0 | 0 |
| ENSG00000197172 | 0 | 0 | 0 | 0 | 0 | 0 |
| ENSG00000147400 | 2470 | 1291 | 1433 | 1307 | 1570 | 1310 |
| ENSG00000147383 | 2223 | 1146 | 1301 | 1477 | 1313 | 1294 |
| ENSG00000147394 | 2333 | 1117 | 1518 | 1314 | 1436 | 1542 |
| ENSG00000198883 | 0 | 0 | 0 | 0 | 0 | 0 |
| ENSG00000234200 | 0 | 0 | 0 | 0 | 0 | 0 |
| ENSG00000229979 | 0 | 0 | 0 | 0 | 0 | 0 |
| ENSG00000183837 | 0 | 0 | 0 | 0 | 0 | 1 |
| ENSG00000235961 | 10 | 2 | 3 | 2 | 4 | 3 |
| ENSG00000268883 | 2 | 0 | 1 | 0 | 2 | 1 |
| ENSG00000224931 | 0 | 0 | 0 | 0 | 0 | 0 |
| ENSG00000198681 | 0 | 0 | 0 | 0 | 0 | 0 |
| ENSG00000235219 | 0 | 0 | 0 | 0 | 0 | 0 |
| ENSG00000273877 | 0 | 0 | 0 | 0 | 0 | 0 |
| ENSG00000231600 | 0 | 0 | 0 | 0 | 0 | 0 |
| ENSG00000278803 | 0 | 0 | 0 | 0 | 0 | 0 |
| ENSG00000225110 | 0 | 0 | 0 | 0 | 0 | 0 |
| ENSG00000063587 | 879 | 378 | 547 | 378 | 448 | 457 |
| ENSG00000232249 | 0 | 0 | 0 | 0 | 0 | 0 |
| ENSG00000214897 | 4 | 0 | 0 | 0 | 1 | 0 |
| ENSG00000265170 | 0 | 0 | 0 | 0 | 0 | 0 |
| ENSG00000189420 | 27 | 8 | 6 | 8 | 4 | 9 |
| ENSG00000183479 | 51 | 29 | 44 | 46 | 33 | 35 |
| ENSG00000213397 | 31 | 14 | 14 | 18 | 17 | 16 |
| ENSG00000224963 | 0 | 0 | 0 | 0 | 0 | 0 |
| ENSG00000182492 | 2 | 2 | 0 | 0 | 0 | 0 |
| ENSG00000067842 | 0 | 0 | 0 | 0 | 0 | 0 |
| ENSG00000262919 | 939 | 415 | 521 | 694 | 518 | 578 |
| ENSG00000260081 | 152 | 82 | 72 | 98 | 79 | 65 |
| ENSG00000217889 | 23 | 6 | 4 | 5 | 6 | 9 |
| ENSG00000130829 | 66 | 34 | 47 | 32 | 17 | 25 |
| ENSG00000237793 | 0 | 0 | 0 | 0 | 0 | 0 |
| ENSG00000244335 | 0 | 0 | 0 | 0 | 0 | 0 |
| ENSG00000130822 | 15 | 5 | 15 | 7 | 5 | 3 |
| ENSG00000130821 | 774 | 451 | 508 | 528 | 507 | 423 |
| ENSG00000185825 | 11870 | 6071 | 7061 | 8215 | 7057 | 6783 |
| ENSG00000101986 | 676 | 300 | 433 | 390 | 325 | 322 |
| ENSG00000232725 | 7 | 7 | 11 | 12 | 5 | 6 |
| ENSG00000198753 | 531 | 212 | 311 | 221 | 260 | 258 |
| ENSG00000184343 | 71 | 30 | 47 | 23 | 24 | 25 |
| ENSG00000067829 | 3876 | 1924 | 2499 | 2395 | 1952 | 2200 |

| | | | | | | |
|---|---|---|---|---|---|---|
| ENSG00000180879 | 7126 | 3693 | 4294 | 4829 | 3907 | 3844 |
| ENSG00000067840 | 389 | 205 | 241 | 270 | 226 | 238 |
| ENSG00000184844 | 0 | 0 | 1 | 0 | 0 | 0 |
| ENSG00000198910 | 6518 | 3179 | 3401 | 3028 | 3237 | 3153 |
| ENSG00000284987 | 0 | 0 | 0 | 0 | 0 | 0 |
| ENSG00000273769 | 4 | 4 | 2 | 3 | 2 | 2 |
| ENSG00000126895 | 10 | 7 | 4 | 4 | 8 | 1 |
| ENSG00000089820 | 441 | 226 | 297 | 238 | 201 | 226 |
| ENSG00000102030 | 1373 | 807 | 966 | 1247 | 986 | 1009 |
| ENSG00000102032 | 8 | 7 | 6 | 5 | 6 | 5 |
| ENSG00000172534 | 3699 | 1493 | 2346 | 1681 | 1614 | 2062 |
| ENSG00000235802 | 0 | 0 | 1 | 1 | 1 | 0 |
| ENSG00000177854 | 777 | 350 | 464 | 510 | 483 | 413 |
| ENSG00000265176 | 0 | 0 | 0 | 0 | 0 | 0 |
| ENSG00000283694 | 0 | 0 | 0 | 0 | 0 | 0 |
| ENSG00000184216 | 7729 | 4024 | 4989 | 6096 | 5713 | 5460 |
| ENSG00000284286 | 0 | 0 | 0 | 0 | 0 | 0 |
| ENSG00000169057 | 3580 | 1531 | 2104 | 1520 | 1746 | 1801 |
| ENSG00000102076 | 0 | 0 | 0 | 0 | 0 | 0 |
| ENSG00000277008 | 0 | 0 | 0 | 0 | 0 | 0 |
| ENSG00000268221 | 0 | 0 | 0 | 0 | 0 | 0 |
| ENSG00000274962 | 0 | 0 | 0 | 0 | 0 | 0 |
| ENSG00000166160 | 0 | 0 | 0 | 0 | 0 | 0 |
| ENSG00000277009 | 0 | 0 | 0 | 0 | 0 | 0 |
| ENSG00000269433 | 0 | 0 | 0 | 0 | 0 | 0 |
| ENSG00000278057 | 0 | 0 | 0 | 0 | 0 | 0 |
| ENSG00000007350 | 0 | 0 | 1 | 0 | 0 | 0 |
| ENSG00000231830 | 1 | 2 | 2 | 0 | 0 | 1 |
| ENSG00000196924 | 55350 | 25544 | 33629 | 36040 | 36447 | 34003 |
| ENSG00000285018 | 6 | 14 | 4 | 4 | 15 | 9 |
| ENSG00000102119 | 2987 | 1497 | 1902 | 1818 | 1540 | 1637 |
| ENSG00000147403 | 47427 | 26314 | 32326 | 39275 | 31660 | 31103 |
| ENSG00000280195 | 91 | 52 | 70 | 63 | 57 | 56 |
| ENSG00000207165 | 0 | 0 | 0 | 0 | 0 | 0 |
| ENSG00000013563 | 730 | 447 | 539 | 540 | 512 | 522 |
| ENSG00000102125 | 906 | 414 | 628 | 560 | 537 | 591 |
| ENSG00000197180 | 273 | 127 | 182 | 154 | 145 | 148 |
| ENSG00000071553 | 13732 | 6255 | 8340 | 7046 | 6342 | 6714 |
| ENSG00000203879 | 9293 | 4213 | 5875 | 5087 | 4718 | 5039 |
| ENSG00000071859 | 3094 | 1595 | 1836 | 1776 | 1789 | 1605 |
| ENSG00000276350 | 1 | 1 | 0 | 0 | 0 | 0 |
| ENSG00000130827 | 4153 | 1561 | 2532 | 1672 | 1792 | 1908 |
| ENSG00000196976 | 1014 | 548 | 690 | 734 | 620 | 597 |
| ENSG00000102178 | 2675 | 1275 | 1589 | 1668 | 1461 | 1534 |
| ENSG00000126903 | 1339 | 593 | 773 | 736 | 658 | 678 |
| ENSG00000264484 | 0 | 0 | 0 | 0 | 0 | 1 |
| ENSG00000071889 | 1944 | 847 | 1167 | 1298 | 1005 | 1089 |
| ENSG00000261773 | 2 | 5 | 2 | 6 | 1 | 9 |
| ENSG00000160211 | 44570 | 20293 | 27756 | 36274 | 28471 | 31779 |
| ENSG00000269335 | 320 | 184 | 223 | 218 | 198 | 182 |

| | | | | | | |
|---|---|---|---|---|---|---|
| ENSG00000279245 | 0 | 0 | 0 | 0 | 0 | 0 |
| ENSG00000273041 | 0 | 0 | 0 | 0 | 1 | 0 |
| ENSG00000268651 | 0 | 0 | 0 | 0 | 0 | 0 |
| ENSG00000232155 | 0 | 0 | 0 | 0 | 0 | 0 |
| ENSG00000184033 | 0 | 0 | 0 | 0 | 0 | 0 |
| ENSG00000273773 | 0 | 0 | 0 | 0 | 0 | 0 |
| ENSG00000213338 | 0 | 0 | 0 | 0 | 0 | 0 |
| ENSG00000272681 | 0 | 0 | 0 | 0 | 0 | 0 |
| ENSG00000275882 | 0 | 1 | 2 | 3 | 2 | 0 |
| ENSG00000126890 | 0 | 0 | 0 | 0 | 0 | 0 |
| ENSG00000226141 | 0 | 0 | 0 | 0 | 0 | 0 |
| ENSG00000237619 | 0 | 0 | 0 | 0 | 0 | 0 |
| ENSG00000160219 | 0 | 1 | 1 | 0 | 0 | 0 |
| ENSG00000130826 | 2699 | 1499 | 1664 | 1767 | 1923 | 2035 |
| ENSG00000206948 | 1 | 0 | 0 | 0 | 0 | 0 |
| ENSG00000284450 | 0 | 0 | 0 | 0 | 0 | 0 |
| ENSG00000206693 | 0 | 0 | 0 | 0 | 0 | 0 |
| ENSG00000130830 | 0 | 2 | 0 | 0 | 0 | 0 |
| ENSG00000229237 | 2 | 2 | 3 | 0 | 1 | 1 |
| ENSG00000203870 | 0 | 0 | 0 | 0 | 1 | 0 |
| ENSG00000185010 | 102 | 55 | 72 | 32 | 33 | 52 |
| ENSG00000274183 | 0 | 0 | 0 | 0 | 0 | 0 |
| ENSG00000277203 | 1000 | 493 | 600 | 612 | 505 | 547 |
| ENSG00000221533 | 0 | 0 | 0 | 0 | 0 | 0 |
| ENSG00000224069 | 0 | 0 | 0 | 0 | 0 | 0 |
| ENSG00000237859 | 0 | 0 | 0 | 0 | 0 | 0 |
| ENSG00000233688 | 0 | 0 | 0 | 0 | 0 | 0 |
| ENSG00000165775 | 1885 | 1017 | 1066 | 1059 | 1078 | 993 |
| ENSG00000214827 | 59 | 41 | 61 | 32 | 51 | 35 |
| ENSG00000182712 | 0 | 0 | 0 | 0 | 0 | 0 |
| ENSG00000185515 | 1129 | 645 | 540 | 291 | 828 | 510 |
| ENSG00000235991 | 0 | 0 | 0 | 0 | 0 | 1 |
| ENSG00000155959 | 2352 | 1138 | 1348 | 1146 | 1341 | 1274 |
| ENSG00000224978 | 0 | 0 | 0 | 0 | 0 | 0 |
| ENSG00000155961 | 280 | 87 | 155 | 72 | 83 | 87 |
| ENSG00000155962 | 28 | 9 | 15 | 7 | 22 | 8 |
| ENSG00000225008 | 0 | 0 | 1 | 0 | 0 | 0 |
| ENSG00000228986 | 2 | 0 | 0 | 0 | 0 | 4 |
| ENSG00000230491 | 0 | 0 | 0 | 0 | 0 | 0 |
| ENSG00000224216 | 6 | 2 | 1 | 0 | 2 | 0 |
| ENSG00000230578 | 0 | 1 | 0 | 0 | 0 | 0 |
| ENSG00000277858 | 0 | 0 | 0 | 0 | 0 | 0 |
| ENSG00000274791 | 3 | 0 | 0 | 0 | 0 | 0 |
| ENSG00000221190 | 0 | 0 | 0 | 0 | 0 | 0 |
| ENSG00000277150 | 2 | 0 | 2 | 1 | 0 | 1 |
| ENSG00000221603 | 0 | 0 | 0 | 0 | 0 | 0 |
| ENSG00000277745 | 0 | 0 | 0 | 0 | 1 | 0 |
| ENSG00000224533 | 16 | 2 | 5 | 1 | 0 | 6 |
| ENSG00000225393 | 0 | 0 | 0 | 0 | 0 | 0 |
| ENSG00000185973 | 456 | 198 | 257 | 188 | 223 | 202 |

| | | | | | | |
|---|---|---|---|---|---|---|
| ENSG00000168939 | 3 | 0 | 2 | 0 | 3 | 2 |
| ENSG00000237801 | 0 | 0 | 0 | 0 | 0 | 0 |
| ENSG00000237040 | 0 | 0 | 0 | 0 | 0 | 0 |
| ENSG00000124333 | 0 | 0 | 0 | 0 | 0 | 0 |
| ENSG00000228410 | 0 | 0 | 0 | 0 | 0 | 0 |
| ENSG00000223484 | 0 | 0 | 0 | 0 | 0 | 0 |
| ENSG00000124334 | 0 | 0 | 0 | 0 | 0 | 0 |
| ENSG00000270726 | 0 | 0 | 0 | 0 | 0 | 0 |
| ENSG00000185203 | 0 | 0 | 0 | 0 | 0 | 0 |
| ENSG00000182484 | 0 | 0 | 0 | 0 | 0 | 0 |
| ENSG00000227159 | 0 | 0 | 0 | 0 | 0 | 0 |
| ENSG00000253620 | 0 | 0 | 0 | 0 | 0 | 0 |
| ENSG00000253896 | 0 | 0 | 0 | 0 | 0 | 0 |
| ENSG00000254193 | 1 | 0 | 0 | 0 | 0 | 0 |
| ENSG00000253612 | 0 | 0 | 0 | 0 | 0 | 0 |
| ENSG00000176269 | 0 | 0 | 1 | 0 | 1 | 0 |
| ENSG00000255464 | 2 | 1 | 1 | 1 | 0 | 1 |
| ENSG00000253166 | 0 | 1 | 0 | 0 | 0 | 1 |
| ENSG00000223508 | 51 | 27 | 38 | 29 | 39 | 26 |
| ENSG00000172748 | 275 | 155 | 201 | 76 | 181 | 172 |
| ENSG00000272812 | 17 | 12 | 10 | 7 | 12 | 11 |
| ENSG00000272240 | 31 | 13 | 35 | 7 | 19 | 8 |
| ENSG00000273402 | 19 | 5 | 11 | 1 | 8 | 5 |
| ENSG00000254104 | 0 | 0 | 0 | 0 | 0 | 0 |
| ENSG00000249868 | 0 | 0 | 0 | 0 | 0 | 0 |
| ENSG00000182366 | 0 | 0 | 0 | 0 | 0 | 1 |
| ENSG00000147364 | 1243 | 604 | 736 | 573 | 671 | 649 |
| ENSG00000272293 | 0 | 0 | 0 | 0 | 0 | 0 |
| ENSG00000279852 | 0 | 2 | 0 | 0 | 2 | 1 |
| ENSG00000180190 | 6 | 3 | 4 | 2 | 1 | 2 |
| ENSG00000104714 | 591 | 276 | 331 | 280 | 305 | 362 |
| ENSG00000254207 | 5 | 0 | 3 | 2 | 2 | 1 |
| ENSG00000198010 | 6 | 1 | 6 | 2 | 5 | 4 |
| ENSG00000282375 | 0 | 0 | 0 | 0 | 0 | 0 |
| ENSG00000282759 | 0 | 0 | 0 | 0 | 0 | 0 |
| ENSG00000282692 | 0 | 0 | 0 | 0 | 0 | 0 |
| ENSG00000254160 | 0 | 0 | 0 | 0 | 0 | 0 |
| ENSG00000275427 | 0 | 0 | 0 | 0 | 0 | 0 |
| ENSG00000260721 | 0 | 0 | 0 | 0 | 0 | 0 |
| ENSG00000282160 | 0 | 0 | 0 | 0 | 0 | 0 |
| ENSG00000281961 | 0 | 0 | 0 | 0 | 0 | 0 |
| ENSG00000253267 | 0 | 0 | 0 | 0 | 0 | 0 |
| ENSG00000283576 | 0 | 0 | 0 | 0 | 0 | 0 |
| ENSG00000254265 | 0 | 0 | 0 | 0 | 0 | 0 |
| ENSG00000182372 | 2634 | 1341 | 1568 | 1261 | 1491 | 1528 |
| ENSG00000282021 | 162 | 94 | 140 | 84 | 117 | 98 |
| ENSG00000253982 | 401 | 164 | 280 | 190 | 216 | 192 |
| ENSG00000283239 | 0 | 0 | 0 | 0 | 0 | 0 |
| ENSG00000283858 | 0 | 0 | 0 | 0 | 0 | 0 |
| ENSG00000207826 | 0 | 0 | 0 | 0 | 0 | 0 |

| | | | | | | |
|---|---|---|---|---|---|---|
| ENSG00000104728 | 28 | 28 | 22 | 15 | 30 | 8 |
| ENSG00000285957 | 0 | 0 | 1 | 0 | 0 | 0 |
| ENSG00000253764 | 0 | 3 | 0 | 0 | 3 | 2 |
| ENSG00000253696 | 0 | 0 | 0 | 0 | 3 | 1 |
| ENSG00000176595 | 238 | 117 | 148 | 146 | 164 | 175 |
| ENSG00000270988 | 0 | 0 | 0 | 1 | 0 | 0 |
| ENSG00000253805 | 0 | 0 | 0 | 0 | 0 | 0 |
| ENSG00000036448 | 2 | 0 | 0 | 0 | 0 | 1 |
| ENSG00000273593 | 0 | 0 | 0 | 0 | 0 | 0 |
| ENSG00000253444 | 0 | 0 | 0 | 0 | 0 | 0 |
| ENSG00000282686 | 0 | 0 | 0 | 0 | 0 | 0 |
| ENSG00000282142 | 0 | 0 | 0 | 0 | 0 | 0 |
| ENSG00000253855 | 0 | 0 | 0 | 0 | 1 | 0 |
| ENSG00000277526 | 0 | 0 | 0 | 0 | 0 | 0 |
| ENSG00000254319 | 0 | 0 | 0 | 0 | 0 | 0 |
| ENSG00000253853 | 0 | 0 | 0 | 0 | 0 | 0 |
| ENSG00000183117 | 2 | 0 | 0 | 0 | 0 | 0 |
| ENSG00000253162 | 0 | 0 | 0 | 0 | 0 | 0 |
| ENSG00000285861 | 0 | 0 | 0 | 0 | 0 | 0 |
| ENSG00000222546 | 0 | 0 | 0 | 0 | 0 | 0 |
| ENSG00000285977 | 0 | 0 | 0 | 0 | 0 | 0 |
| ENSG00000254244 | 0 | 0 | 0 | 1 | 0 | 1 |
| ENSG00000253262 | 0 | 0 | 0 | 0 | 0 | 0 |
| ENSG00000273876 | 0 | 0 | 0 | 0 | 0 | 0 |
| ENSG00000206661 | 0 | 0 | 0 | 0 | 0 | 0 |
| ENSG00000242079 | 0 | 0 | 0 | 0 | 0 | 0 |
| ENSG00000278256 | 0 | 0 | 0 | 0 | 0 | 0 |
| ENSG00000271248 | 0 | 0 | 0 | 0 | 0 | 0 |
| ENSG00000240486 | 0 | 0 | 0 | 0 | 0 | 0 |
| ENSG00000253510 | 0 | 0 | 0 | 0 | 0 | 0 |
| ENSG00000254007 | 0 | 0 | 0 | 0 | 0 | 0 |
| ENSG00000275461 | 0 | 0 | 0 | 0 | 0 | 0 |
| ENSG00000253571 | 0 | 0 | 0 | 0 | 0 | 0 |
| ENSG00000222589 | 0 | 0 | 0 | 0 | 0 | 0 |
| ENSG00000253189 | 0 | 0 | 0 | 0 | 0 | 0 |
| ENSG00000253880 | 2 | 0 | 0 | 0 | 0 | 0 |
| ENSG00000246089 | 340 | 156 | 212 | 199 | 205 | 216 |
| ENSG00000147316 | 552 | 257 | 303 | 229 | 328 | 315 |
| ENSG00000253550 | 0 | 1 | 0 | 0 | 0 | 0 |
| ENSG00000091879 | 2 | 2 | 1 | 4 | 3 | 2 |
| ENSG00000271743 | 0 | 4 | 3 | 1 | 3 | 3 |
| ENSG00000249898 | 34 | 21 | 31 | 8 | 18 | 20 |
| ENSG00000276580 | 0 | 0 | 0 | 0 | 1 | 0 |
| ENSG00000155189 | 1464 | 693 | 897 | 692 | 888 | 873 |
| ENSG00000266038 | 0 | 0 | 0 | 0 | 0 | 0 |
| ENSG00000283386 | 0 | 0 | 0 | 0 | 0 | 0 |
| ENSG00000275591 | 0 | 0 | 0 | 0 | 0 | 0 |
| ENSG00000245857 | 0 | 0 | 0 | 0 | 0 | 0 |
| ENSG00000164825 | 0 | 2 | 3 | 3 | 0 | 1 |
| ENSG00000250752 | 0 | 0 | 0 | 0 | 0 | 0 |

| | | | | | | |
|---|---|---|---|---|---|---|
| ENSG00000164822 | 0 | 0 | 0 | 0 | 0 | 0 |
| ENSG00000254625 | 0 | 0 | 0 | 0 | 0 | 0 |
| ENSG00000164821 | 0 | 0 | 0 | 0 | 0 | 0 |
| ENSG00000223629 | 0 | 0 | 0 | 0 | 0 | 0 |
| ENSG00000233238 | 0 | 0 | 0 | 0 | 0 | 0 |
| ENSG00000233531 | 0 | 0 | 0 | 0 | 0 | 0 |
| ENSG00000206047 | 0 | 0 | 0 | 0 | 0 | 0 |
| ENSG00000215378 | 0 | 0 | 0 | 0 | 0 | 0 |
| ENSG00000240247 | 0 | 0 | 0 | 0 | 0 | 0 |
| ENSG00000232039 | 0 | 0 | 0 | 0 | 0 | 0 |
| ENSG00000239839 | 0 | 0 | 0 | 0 | 0 | 0 |
| ENSG00000234178 | 0 | 0 | 0 | 0 | 0 | 0 |
| ENSG00000206042 | 0 | 0 | 0 | 0 | 0 | 0 |
| ENSG00000164816 | 0 | 0 | 0 | 0 | 0 | 0 |
| ENSG00000255273 | 0 | 0 | 0 | 0 | 0 | 0 |
| ENSG00000284620 | 0 | 1 | 0 | 0 | 0 | 0 |
| ENSG00000234120 | 0 | 0 | 0 | 0 | 0 | 0 |
| ENSG00000228744 | 0 | 0 | 0 | 0 | 1 | 0 |
| ENSG00000254683 | 0 | 0 | 0 | 0 | 0 | 0 |
| ENSG00000230106 | 0 | 0 | 0 | 0 | 0 | 0 |
| ENSG00000214268 | 0 | 0 | 0 | 0 | 0 | 0 |
| ENSG00000231930 | 0 | 0 | 0 | 0 | 0 | 0 |
| ENSG00000284614 | 0 | 0 | 0 | 0 | 0 | 0 |
| ENSG00000284728 | 0 | 0 | 0 | 0 | 0 | 0 |
| ENSG00000255025 | 0 | 0 | 0 | 0 | 0 | 0 |
| ENSG00000177306 | 0 | 0 | 0 | 0 | 0 | 0 |
| ENSG00000230045 | 0 | 0 | 0 | 0 | 0 | 0 |
| ENSG00000233132 | 0 | 0 | 0 | 0 | 0 | 0 |
| ENSG00000249005 | 0 | 0 | 0 | 0 | 0 | 0 |
| ENSG00000223885 | 0 | 0 | 0 | 0 | 0 | 0 |
| ENSG00000215373 | 0 | 0 | 0 | 0 | 0 | 0 |
| ENSG00000233295 | 0 | 0 | 0 | 0 | 0 | 0 |
| ENSG00000215374 | 11 | 8 | 11 | 11 | 16 | 19 |
| ENSG00000206034 | 0 | 0 | 0 | 0 | 0 | 0 |
| ENSG00000230549 | 0 | 0 | 0 | 0 | 0 | 0 |
| ENSG00000236125 | 0 | 0 | 0 | 0 | 0 | 0 |
| ENSG00000235778 | 0 | 0 | 0 | 0 | 0 | 0 |
| ENSG00000215372 | 0 | 0 | 0 | 0 | 0 | 0 |
| ENSG00000215371 | 0 | 0 | 0 | 0 | 0 | 0 |
| ENSG00000177257 | 0 | 0 | 0 | 0 | 0 | 0 |
| ENSG00000255128 | 0 | 0 | 0 | 0 | 0 | 0 |
| ENSG00000177243 | 0 | 0 | 0 | 0 | 0 | 0 |
| ENSG00000164871 | 0 | 0 | 0 | 0 | 0 | 0 |
| ENSG00000177023 | 0 | 0 | 0 | 0 | 0 | 0 |
| ENSG00000187082 | 0 | 0 | 0 | 0 | 0 | 0 |
| ENSG00000186599 | 0 | 0 | 0 | 0 | 0 | 0 |
| ENSG00000198129 | 0 | 0 | 0 | 0 | 0 | 0 |
| ENSG00000285659 | 0 | 0 | 0 | 0 | 0 | 0 |
| ENSG00000255251 | 0 | 0 | 0 | 0 | 0 | 0 |
| ENSG00000285927 | 0 | 0 | 0 | 0 | 0 | 0 |

| | | | | | | |
|---|---|---|---|---|---|---|
| ENSG00000248944 | 0 | 0 | 0 | 0 | 0 | 0 |
| ENSG00000285975 | 0 | 0 | 0 | 0 | 0 | 0 |
| ENSG00000234749 | 0 | 0 | 0 | 0 | 0 | 0 |
| ENSG00000285687 | 0 | 0 | 0 | 0 | 0 | 0 |
| ENSG00000285765 | 0 | 0 | 0 | 0 | 0 | 0 |
| ENSG00000233176 | 0 | 0 | 0 | 0 | 0 | 0 |
| ENSG00000254796 | 0 | 0 | 0 | 0 | 0 | 0 |
| ENSG00000284617 | 0 | 1 | 0 | 0 | 0 | 1 |
| ENSG00000284655 | 0 | 0 | 0 | 0 | 0 | 0 |
| ENSG00000284698 | 0 | 0 | 0 | 0 | 0 | 0 |
| ENSG00000254889 | 0 | 0 | 0 | 0 | 0 | 0 |
| ENSG00000254715 | 0 | 0 | 0 | 0 | 0 | 0 |
| ENSG00000285814 | 0 | 0 | 0 | 0 | 0 | 0 |
| ENSG00000285913 | 0 | 0 | 0 | 0 | 0 | 0 |
| ENSG00000285620 | 0 | 0 | 0 | 0 | 0 | 0 |
| ENSG00000285937 | 0 | 0 | 0 | 0 | 0 | 0 |
| ENSG00000285720 | 0 | 0 | 0 | 0 | 0 | 0 |
| ENSG00000285657 | 0 | 0 | 0 | 0 | 0 | 0 |
| ENSG00000285607 | 0 | 0 | 0 | 0 | 0 | 0 |
| ENSG00000285950 | 0 | 0 | 0 | 0 | 0 | 0 |
| ENSG00000285862 | 0 | 0 | 0 | 0 | 0 | 0 |
| ENSG00000255378 | 0 | 0 | 0 | 0 | 0 | 0 |
| ENSG00000254776 | 0 | 0 | 0 | 0 | 0 | 0 |
| ENSG00000186572 | 0 | 0 | 0 | 0 | 0 | 0 |
| ENSG00000186562 | 0 | 0 | 0 | 0 | 0 | 0 |
| ENSG00000186579 | 0 | 0 | 0 | 0 | 0 | 0 |
| ENSG00000176782 | 0 | 0 | 0 | 0 | 0 | 0 |
| ENSG00000178287 | 0 | 0 | 0 | 0 | 0 | 0 |
| ENSG00000176797 | 0 | 0 | 0 | 0 | 0 | 0 |
| ENSG00000254543 | 0 | 0 | 0 | 0 | 0 | 0 |
| ENSG00000171711 | 2 | 3 | 2 | 1 | 1 | 4 |
| ENSG00000215356 | 0 | 0 | 0 | 0 | 0 | 0 |
| ENSG00000229907 | 0 | 0 | 0 | 0 | 0 | 0 |
| ENSG00000225725 | 0 | 0 | 0 | 0 | 1 | 0 |
| ENSG00000255211 | 0 | 0 | 0 | 0 | 0 | 0 |
| ENSG00000237038 | 0 | 0 | 0 | 0 | 0 | 0 |
| ENSG00000225327 | 0 | 0 | 0 | 0 | 0 | 0 |
| ENSG00000205989 | 0 | 0 | 0 | 0 | 0 | 0 |
| ENSG00000233115 | 0 | 0 | 0 | 0 | 0 | 0 |
| ENSG00000215354 | 0 | 0 | 0 | 0 | 0 | 0 |
| ENSG00000254229 | 0 | 0 | 0 | 0 | 0 | 0 |
| ENSG00000227013 | 0 | 0 | 0 | 0 | 0 | 0 |
| ENSG00000253881 | 0 | 0 | 0 | 0 | 0 | 0 |
| ENSG00000284603 | 0 | 0 | 0 | 0 | 0 | 0 |
| ENSG00000284606 | 0 | 3 | 2 | 1 | 5 | 3 |
| ENSG00000284661 | 0 | 0 | 0 | 0 | 0 | 0 |
| ENSG00000255459 | 0 | 0 | 0 | 0 | 0 | 0 |
| ENSG00000221305 | 0 | 0 | 0 | 0 | 0 | 0 |
| ENSG00000244427 | 0 | 0 | 0 | 0 | 0 | 0 |
| ENSG00000254311 | 0 | 0 | 0 | 0 | 0 | 0 |

| | | | | | | |
|---|---|---|---|---|---|---|
| ENSG00000249188 | 0 | 0 | 0 | 0 | 0 | 0 |
| ENSG00000253893 | 2 | 0 | 0 | 2 | 3 | 0 |
| ENSG00000268955 | 0 | 1 | 1 | 0 | 1 | 0 |
| ENSG00000173295 | 178 | 76 | 121 | 101 | 110 | 153 |
| ENSG00000253981 | 28 | 15 | 16 | 16 | 15 | 14 |
| ENSG00000275342 | 634 | 295 | 392 | 369 | 363 | 353 |
| ENSG00000253505 | 0 | 0 | 0 | 0 | 0 | 0 |
| ENSG00000254153 | 33 | 17 | 33 | 12 | 22 | 22 |
| ENSG00000253343 | 0 | 0 | 0 | 0 | 0 | 0 |
| ENSG00000253774 | 0 | 0 | 0 | 0 | 0 | 0 |
| ENSG00000253130 | 2 | 3 | 0 | 0 | 1 | 0 |
| ENSG00000263616 | 0 | 0 | 0 | 0 | 0 | 0 |
| ENSG00000253958 | 258 | 130 | 157 | 153 | 150 | 147 |
| ENSG00000284717 | 0 | 0 | 0 | 0 | 1 | 0 |
| ENSG00000254367 | 0 | 0 | 0 | 0 | 0 | 0 |
| ENSG00000147324 | 1359 | 623 | 799 | 656 | 712 | 779 |
| ENSG00000200713 | 0 | 0 | 0 | 0 | 0 | 0 |
| ENSG00000270966 | 0 | 2 | 0 | 0 | 1 | 2 |
| ENSG00000233609 | 8 | 3 | 0 | 0 | 5 | 1 |
| ENSG00000207244 | 0 | 0 | 0 | 0 | 0 | 0 |
| ENSG00000104626 | 976 | 546 | 560 | 369 | 561 | 522 |
| ENSG00000263407 | 0 | 0 | 0 | 0 | 0 | 0 |
| ENSG00000239078 | 0 | 0 | 0 | 0 | 0 | 0 |
| ENSG00000197334 | 0 | 0 | 0 | 0 | 0 | 0 |
| ENSG00000173281 | 985 | 450 | 599 | 262 | 379 | 375 |
| ENSG00000254340 | 1 | 2 | 3 | 1 | 1 | 2 |
| ENSG00000248538 | 16 | 13 | 10 | 9 | 6 | 9 |
| ENSG00000279949 | 4 | 1 | 2 | 2 | 1 | 5 |
| ENSG00000253252 | 0 | 0 | 0 | 0 | 0 | 0 |
| ENSG00000254091 | 0 | 0 | 0 | 0 | 0 | 0 |
| ENSG00000253887 | 0 | 0 | 0 | 0 | 0 | 0 |
| ENSG00000207415 | 0 | 0 | 0 | 0 | 0 | 0 |
| ENSG00000253735 | 0 | 0 | 0 | 0 | 0 | 0 |
| ENSG00000201815 | 0 | 0 | 0 | 0 | 0 | 0 |
| ENSG00000254237 | 0 | 0 | 0 | 0 | 0 | 0 |
| ENSG00000249050 | 0 | 0 | 0 | 0 | 0 | 0 |
| ENSG00000272267 | 18 | 7 | 9 | 7 | 8 | 10 |
| ENSG00000173273 | 1837 | 946 | 1121 | 619 | 1116 | 965 |
| ENSG00000253697 | 0 | 0 | 0 | 0 | 0 | 0 |
| ENSG00000207701 | 0 | 0 | 0 | 0 | 0 | 0 |
| ENSG00000253230 | 0 | 0 | 0 | 0 | 0 | 0 |
| ENSG00000284321 | 0 | 0 | 0 | 0 | 0 | 0 |
| ENSG00000260093 | 6 | 1 | 2 | 0 | 1 | 1 |
| ENSG00000175806 | 404 | 217 | 271 | 249 | 225 | 220 |
| ENSG00000285675 | 0 | 0 | 0 | 0 | 0 | 0 |
| ENSG00000285606 | 0 | 0 | 0 | 0 | 0 | 0 |
| ENSG00000249848 | 0 | 0 | 0 | 0 | 0 | 0 |
| ENSG00000261451 | 2 | 3 | 0 | 0 | 0 | 0 |
| ENSG00000253641 | 0 | 2 | 0 | 2 | 0 | 0 |
| ENSG00000285237 | 0 | 0 | 0 | 0 | 0 | 0 |

| | | | | | | |
|---|---|---|---|---|---|---|
| ENSG00000207128 | 0 | 0 | 0 | 0 | 0 | 0 |
| ENSG00000253678 | 0 | 0 | 0 | 0 | 0 | 0 |
| ENSG00000253649 | 0 | 0 | 1 | 0 | 1 | 2 |
| ENSG00000272505 | 12 | 5 | 4 | 2 | 7 | 5 |
| ENSG00000184647 | 0 | 0 | 0 | 0 | 0 | 0 |
| ENSG00000183638 | 4 | 0 | 0 | 2 | 1 | 3 |
| ENSG00000263762 | 0 | 0 | 0 | 0 | 0 | 0 |
| ENSG00000171060 | 0 | 0 | 0 | 0 | 0 | 0 |
| ENSG00000212433 | 0 | 0 | 0 | 0 | 0 | 0 |
| ENSG00000171056 | 0 | 5 | 2 | 8 | 1 | 3 |
| ENSG00000258724 | 0 | 0 | 0 | 0 | 0 | 0 |
| ENSG00000248896 | 1 | 0 | 1 | 2 | 1 | 0 |
| ENSG00000254093 | 326 | 182 | 185 | 194 | 221 | 200 |
| ENSG00000283210 | 0 | 0 | 0 | 0 | 0 | 0 |
| ENSG00000253695 | 0 | 0 | 1 | 2 | 0 | 0 |
| ENSG00000280294 | 2 | 3 | 0 | 1 | 3 | 1 |
| ENSG00000171044 | 34 | 11 | 20 | 11 | 12 | 13 |
| ENSG00000207600 | 0 | 0 | 0 | 0 | 0 | 0 |
| ENSG00000254839 | 0 | 0 | 0 | 0 | 0 | 1 |
| ENSG00000269918 | 29 | 5 | 14 | 10 | 12 | 13 |
| ENSG00000255310 | 57 | 22 | 14 | 10 | 23 | 23 |
| ENSG00000254936 | 0 | 0 | 0 | 0 | 1 | 2 |
| ENSG00000254556 | 2 | 0 | 0 | 0 | 0 | 0 |
| ENSG00000231965 | 0 | 0 | 0 | 0 | 0 | 0 |
| ENSG00000270076 | 6 | 2 | 2 | 4 | 5 | 2 |
| ENSG00000236827 | 0 | 0 | 0 | 0 | 0 | 0 |
| ENSG00000239315 | 0 | 0 | 0 | 0 | 0 | 0 |
| ENSG00000249316 | 0 | 0 | 0 | 0 | 0 | 0 |
| ENSG00000280273 | 10 | 0 | 3 | 3 | 3 | 4 |
| ENSG00000104643 | 759 | 348 | 399 | 262 | 424 | 332 |
| ENSG00000246477 | 2 | 8 | 5 | 1 | 4 | 2 |
| ENSG00000177710 | 8 | 2 | 1 | 1 | 1 | 2 |
| ENSG00000154316 | 145 | 64 | 86 | 73 | 101 | 118 |
| ENSG00000255020 | 284 | 118 | 175 | 232 | 204 | 202 |
| ENSG00000184608 | 5 | 1 | 1 | 1 | 2 | 0 |
| ENSG00000242483 | 0 | 0 | 0 | 0 | 0 | 0 |
| ENSG00000199368 | 0 | 0 | 0 | 0 | 0 | 0 |
| ENSG00000154319 | 225 | 118 | 148 | 98 | 127 | 168 |
| ENSG00000284957 | 0 | 0 | 0 | 0 | 0 | 0 |
| ENSG00000136573 | 1 | 0 | 1 | 0 | 1 | 0 |
| ENSG00000269954 | 0 | 0 | 1 | 0 | 0 | 2 |
| ENSG00000170983 | 0 | 0 | 0 | 0 | 0 | 0 |
| ENSG00000136574 | 0 | 2 | 1 | 0 | 0 | 1 |
| ENSG00000255394 | 0 | 0 | 0 | 0 | 0 | 0 |
| ENSG00000154328 | 445 | 247 | 278 | 327 | 342 | 341 |
| ENSG00000227203 | 0 | 2 | 0 | 0 | 0 | 0 |
| ENSG00000079459 | 7399 | 3456 | 4330 | 4109 | 4267 | 4087 |
| ENSG00000255046 | 22 | 14 | 8 | 4 | 8 | 12 |
| ENSG00000164733 | 9758 | 4609 | 6202 | 4942 | 4741 | 5225 |
| ENSG00000269899 | 3 | 2 | 4 | 4 | 1 | 4 |

| | | | | | | |
|---|---|---|---|---|---|---|
| ENSG00000255144 | 0 | 0 | 0 | 0 | 0 | 0 |
| ENSG00000254948 | 0 | 1 | 0 | 0 | 0 | 0 |
| ENSG00000206014 | 0 | 1 | 0 | 0 | 0 | 0 |
| ENSG00000205884 | 0 | 0 | 0 | 0 | 0 | 0 |
| ENSG00000205883 | 0 | 0 | 0 | 0 | 0 | 0 |
| ENSG00000205882 | 0 | 0 | 0 | 0 | 0 | 1 |
| ENSG00000255174 | 0 | 0 | 0 | 0 | 0 | 0 |
| ENSG00000254507 | 1 | 0 | 0 | 0 | 0 | 0 |
| ENSG00000284610 | 12 | 5 | 2 | 1 | 2 | 5 |
| ENSG00000255016 | 0 | 0 | 0 | 0 | 0 | 0 |
| ENSG00000254817 | 0 | 0 | 0 | 0 | 0 | 0 |
| ENSG00000254700 | 0 | 0 | 0 | 0 | 0 | 0 |
| ENSG00000233050 | 0 | 0 | 0 | 0 | 0 | 0 |
| ENSG00000252029 | 0 | 0 | 0 | 0 | 0 | 0 |
| ENSG00000255544 | 0 | 0 | 0 | 0 | 0 | 0 |
| ENSG00000215343 | 0 | 0 | 0 | 0 | 0 | 0 |
| ENSG00000255052 | 11 | 6 | 13 | 6 | 10 | 17 |
| ENSG00000254923 | 0 | 0 | 0 | 0 | 0 | 0 |
| ENSG00000226430 | 0 | 0 | 0 | 0 | 0 | 0 |
| ENSG00000223443 | 0 | 0 | 4 | 0 | 0 | 0 |
| ENSG00000254866 | 12 | 7 | 9 | 2 | 7 | 5 |
| ENSG00000205879 | 0 | 1 | 2 | 0 | 2 | 1 |
| ENSG00000249889 | 2 | 1 | 0 | 1 | 0 | 0 |
| ENSG00000186523 | 99 | 72 | 80 | 76 | 67 | 61 |
| ENSG00000255495 | 2 | 1 | 2 | 3 | 6 | 7 |
| ENSG00000254527 | 6 | 2 | 2 | 0 | 2 | 0 |
| ENSG00000237215 | 0 | 0 | 0 | 0 | 0 | 0 |
| ENSG00000232948 | 0 | 0 | 0 | 0 | 0 | 0 |
| ENSG00000252535 | 0 | 0 | 0 | 0 | 0 | 0 |
| ENSG00000254623 | 0 | 0 | 0 | 0 | 0 | 0 |
| ENSG00000215339 | 0 | 0 | 0 | 0 | 0 | 0 |
| ENSG00000227888 | 6 | 2 | 4 | 1 | 1 | 5 |
| ENSG00000254423 | 0 | 0 | 0 | 0 | 0 | 0 |
| ENSG00000255556 | 0 | 0 | 1 | 0 | 0 | 0 |
| ENSG00000242296 | 0 | 0 | 1 | 0 | 0 | 0 |
| ENSG00000270074 | 8 | 1 | 9 | 3 | 4 | 1 |
| ENSG00000251402 | 6 | 0 | 0 | 2 | 1 | 3 |
| ENSG00000250794 | 9 | 2 | 7 | 6 | 1 | 4 |
| ENSG00000145002 | 10 | 4 | 6 | 6 | 4 | 2 |
| ENSG00000255549 | 0 | 0 | 0 | 0 | 0 | 0 |
| ENSG00000283674 | 19 | 10 | 27 | 13 | 14 | 14 |
| ENSG00000270154 | 2 | 5 | 0 | 0 | 1 | 0 |
| ENSG00000242607 | 13 | 1 | 2 | 0 | 0 | 2 |
| ENSG00000255122 | 1 | 2 | 1 | 0 | 0 | 0 |
| ENSG00000284746 | 0 | 0 | 0 | 0 | 0 | 0 |
| ENSG00000284663 | 0 | 0 | 0 | 0 | 0 | 0 |
| ENSG00000244289 | 12 | 0 | 0 | 0 | 0 | 0 |
| ENSG00000255253 | 0 | 0 | 0 | 0 | 0 | 0 |
| ENSG00000284724 | 0 | 0 | 0 | 0 | 0 | 0 |
| ENSG00000284613 | 0 | 0 | 0 | 0 | 0 | 0 |

| | | | | | | |
|---|---|---|---|---|---|---|
| ENSG00000254581 | 0 | 0 | 0 | 0 | 0 | 0 |
| ENSG00000177400 | 0 | 0 | 0 | 0 | 0 | 0 |
| ENSG00000254917 | 0 | 0 | 0 | 0 | 0 | 0 |
| ENSG00000254724 | 0 | 0 | 0 | 0 | 0 | 0 |
| ENSG00000264512 | 0 | 0 | 0 | 0 | 0 | 0 |
| ENSG00000154359 | 864 | 460 | 496 | 390 | 557 | 564 |
| ENSG00000266206 | 0 | 0 | 0 | 0 | 0 | 0 |
| ENSG00000283523 | 0 | 0 | 0 | 0 | 0 | 0 |
| ENSG00000254813 | 0 | 0 | 0 | 0 | 0 | 0 |
| ENSG00000255494 | 0 | 0 | 0 | 0 | 0 | 0 |
| ENSG00000250305 | 8 | 3 | 3 | 4 | 3 | 1 |
| ENSG00000251468 | 4 | 0 | 0 | 0 | 0 | 0 |
| ENSG00000206996 | 0 | 0 | 0 | 0 | 0 | 0 |
| ENSG00000164741 | 184 | 106 | 134 | 44 | 64 | 72 |
| ENSG00000200630 | 0 | 0 | 0 | 0 | 0 | 0 |
| ENSG00000253932 | 0 | 0 | 0 | 0 | 0 | 0 |
| ENSG00000253257 | 0 | 0 | 0 | 0 | 0 | 0 |
| ENSG00000252179 | 0 | 0 | 0 | 0 | 0 | 0 |
| ENSG00000164743 | 10 | 2 | 3 | 4 | 5 | 6 |
| ENSG00000254491 | 0 | 0 | 0 | 0 | 0 | 0 |
| ENSG00000255457 | 0 | 0 | 0 | 0 | 0 | 0 |
| ENSG00000255413 | 0 | 0 | 0 | 0 | 0 | 0 |
| ENSG00000254753 | 0 | 0 | 0 | 0 | 0 | 0 |
| ENSG00000254425 | 0 | 0 | 0 | 0 | 0 | 0 |
| ENSG00000255437 | 0 | 0 | 0 | 0 | 0 | 0 |
| ENSG00000185053 | 0 | 0 | 0 | 0 | 0 | 0 |
| ENSG00000254575 | 0 | 0 | 0 | 0 | 0 | 0 |
| ENSG00000214366 | 0 | 0 | 0 | 0 | 0 | 0 |
| ENSG00000252264 | 0 | 0 | 0 | 0 | 0 | 0 |
| ENSG00000206950 | 0 | 0 | 0 | 0 | 0 | 0 |
| ENSG00000252035 | 0 | 0 | 0 | 0 | 0 | 0 |
| ENSG00000255365 | 0 | 0 | 0 | 0 | 0 | 0 |
| ENSG00000199127 | 0 | 0 | 0 | 0 | 0 | 0 |
| ENSG00000255076 | 0 | 0 | 0 | 0 | 0 | 0 |
| ENSG00000249368 | 0 | 0 | 0 | 0 | 0 | 0 |
| ENSG00000104723 | 2616 | 1315 | 1563 | 1318 | 1519 | 1480 |
| ENSG00000248423 | 0 | 0 | 0 | 0 | 0 | 0 |
| ENSG00000250483 | 0 | 0 | 0 | 0 | 0 | 0 |
| ENSG00000226836 | 0 | 0 | 0 | 0 | 0 | 0 |
| ENSG00000253648 | 0 | 0 | 0 | 0 | 0 | 0 |
| ENSG00000038945 | 0 | 0 | 0 | 0 | 0 | 0 |
| ENSG00000253951 | 0 | 0 | 0 | 0 | 0 | 0 |
| ENSG00000253934 | 0 | 0 | 0 | 0 | 0 | 0 |
| ENSG00000253496 | 0 | 0 | 0 | 0 | 0 | 0 |
| ENSG00000253184 | 0 | 0 | 0 | 0 | 0 | 0 |
| ENSG00000264092 | 0 | 0 | 0 | 0 | 0 | 0 |
| ENSG00000078579 | 0 | 0 | 0 | 0 | 0 | 0 |
| ENSG00000155970 | 0 | 0 | 0 | 0 | 0 | 0 |
| ENSG00000249258 | 0 | 0 | 0 | 0 | 0 | 0 |
| ENSG00000244113 | 0 | 0 | 0 | 0 | 0 | 0 |

| | | | | | | |
|---|---|---|---|---|---|---|
| ENSG00000104219 | 234 | 118 | 102 | 91 | 128 | 92 |
| ENSG00000198791 | 3959 | 1994 | 2254 | 1836 | 2514 | 2164 |
| ENSG00000155975 | 1748 | 841 | 1030 | 732 | 1007 | 884 |
| ENSG00000003987 | 116 | 66 | 67 | 32 | 63 | 76 |
| ENSG00000214607 | 0 | 0 | 0 | 0 | 0 | 0 |
| ENSG00000003989 | 29005 | 12268 | 22862 | 8159 | 13537 | 15519 |
| ENSG00000279271 | 2 | 1 | 1 | 0 | 4 | 1 |
| ENSG00000280453 | 0 | 0 | 0 | 0 | 0 | 0 |
| ENSG00000279903 | 5 | 0 | 1 | 1 | 0 | 0 |
| ENSG00000279932 | 1 | 3 | 1 | 1 | 1 | 0 |
| ENSG00000104213 | 858 | 437 | 596 | 373 | 386 | 431 |
| ENSG00000277466 | 0 | 0 | 0 | 0 | 0 | 0 |
| ENSG00000129422 | 1199 | 679 | 627 | 330 | 767 | 593 |
| ENSG00000265520 | 0 | 0 | 1 | 0 | 0 | 0 |
| ENSG00000253168 | 0 | 0 | 0 | 1 | 0 | 0 |
| ENSG00000212280 | 0 | 0 | 0 | 0 | 0 | 0 |
| ENSG00000253944 | 5 | 4 | 8 | 2 | 1 | 2 |
| ENSG00000253671 | 58 | 22 | 41 | 18 | 22 | 13 |
| ENSG00000104760 | 0 | 0 | 0 | 0 | 1 | 0 |
| ENSG00000253215 | 0 | 1 | 0 | 0 | 0 | 0 |
| ENSG00000254054 | 0 | 0 | 0 | 0 | 0 | 0 |
| ENSG00000078674 | 896 | 598 | 435 | 196 | 1285 | 566 |
| ENSG00000104763 | 4862 | 2581 | 2866 | 2034 | 2811 | 2532 |
| ENSG00000245281 | 11 | 7 | 16 | 19 | 3 | 12 |
| ENSG00000253384 | 2 | 1 | 0 | 0 | 1 | 2 |
| ENSG00000171428 | 535 | 275 | 317 | 222 | 277 | 258 |
| ENSG00000254015 | 0 | 0 | 0 | 0 | 0 | 0 |
| ENSG00000253937 | 0 | 0 | 0 | 0 | 0 | 0 |
| ENSG00000244593 | 0 | 1 | 0 | 0 | 2 | 0 |
| ENSG00000271654 | 0 | 0 | 0 | 0 | 0 | 0 |
| ENSG00000285624 | 0 | 0 | 0 | 0 | 0 | 0 |
| ENSG00000156006 | 1 | 4 | 4 | 2 | 6 | 3 |
| ENSG00000156011 | 2677 | 1289 | 1338 | 598 | 1806 | 1283 |
| ENSG00000187229 | 0 | 0 | 0 | 0 | 0 | 0 |
| ENSG00000244018 | 0 | 1 | 0 | 0 | 0 | 0 |
| ENSG00000283402 | 0 | 1 | 0 | 0 | 0 | 1 |
| ENSG00000278886 | 0 | 0 | 0 | 0 | 0 | 0 |
| ENSG00000201157 | 0 | 0 | 0 | 0 | 0 | 0 |
| ENSG00000253335 | 0 | 1 | 0 | 0 | 0 | 0 |
| ENSG00000270255 | 0 | 1 | 0 | 1 | 0 | 0 |
| ENSG00000253557 | 0 | 0 | 0 | 0 | 0 | 0 |
| ENSG00000254242 | 0 | 0 | 0 | 0 | 0 | 0 |
| ENSG00000250316 | 0 | 0 | 0 | 0 | 0 | 0 |
| ENSG00000254201 | 0 | 0 | 0 | 0 | 0 | 0 |
| ENSG00000253280 | 0 | 0 | 0 | 0 | 0 | 0 |
| ENSG00000279919 | 0 | 0 | 0 | 0 | 0 | 0 |
| ENSG00000104611 | 1123 | 661 | 704 | 482 | 645 | 598 |
| ENSG00000279858 | 0 | 0 | 0 | 0 | 0 | 0 |
| ENSG00000147408 | 159 | 111 | 109 | 101 | 175 | 122 |
| ENSG00000253270 | 0 | 0 | 0 | 0 | 0 | 0 |

| | | | | | | |
|---|---|---|---|---|---|---|
| ENSG00000104613 | 2348 | 1098 | 1315 | 1187 | 1353 | 1387 |
| ENSG00000175445 | 0 | 0 | 0 | 0 | 0 | 0 |
| ENSG00000253775 | 0 | 0 | 0 | 0 | 0 | 0 |
| ENSG00000242709 | 0 | 0 | 0 | 0 | 0 | 0 |
| ENSG00000036565 | 0 | 0 | 0 | 0 | 0 | 0 |
| ENSG00000147416 | 2734 | 1181 | 1518 | 1250 | 1479 | 1375 |
| ENSG00000222267 | 0 | 0 | 0 | 0 | 0 | 0 |
| ENSG00000061337 | 0 | 0 | 0 | 0 | 0 | 0 |
| ENSG00000253733 | 0 | 0 | 0 | 0 | 0 | 0 |
| ENSG00000274467 | 0 | 0 | 0 | 0 | 0 | 0 |
| ENSG00000253164 | 0 | 0 | 0 | 0 | 0 | 0 |
| ENSG00000253178 | 0 | 0 | 0 | 0 | 0 | 0 |
| ENSG00000253300 | 0 | 0 | 0 | 0 | 0 | 0 |
| ENSG00000253866 | 0 | 0 | 0 | 0 | 0 | 0 |
| ENSG00000254260 | 0 | 0 | 0 | 0 | 0 | 0 |
| ENSG00000254092 | 0 | 0 | 0 | 0 | 0 | 0 |
| ENSG00000253147 | 0 | 0 | 0 | 0 | 0 | 0 |
| ENSG00000254026 | 0 | 0 | 0 | 0 | 0 | 0 |
| ENSG00000253199 | 0 | 0 | 0 | 0 | 0 | 0 |
| ENSG00000254040 | 0 | 0 | 0 | 0 | 0 | 0 |
| ENSG00000280215 | 0 | 0 | 0 | 0 | 0 | 0 |
| ENSG00000254334 | 0 | 0 | 0 | 0 | 0 | 0 |
| ENSG00000168546 | 2 | 0 | 0 | 0 | 0 | 0 |
| ENSG00000273484 | 0 | 0 | 0 | 0 | 0 | 0 |
| ENSG00000147443 | 0 | 0 | 1 | 0 | 0 | 0 |
| ENSG00000130227 | 3608 | 1765 | 2073 | 1474 | 2095 | 1860 |
| ENSG00000158806 | 2 | 0 | 1 | 1 | 1 | 3 |
| ENSG00000158815 | 12 | 2 | 1 | 2 | 3 | 0 |
| ENSG00000158856 | 748 | 372 | 458 | 401 | 391 | 337 |
| ENSG00000158863 | 2301 | 1029 | 1258 | 1284 | 1146 | 1162 |
| ENSG00000275074 | 497 | 347 | 315 | 439 | 344 | 328 |
| ENSG00000168453 | 754 | 375 | 415 | 701 | 713 | 550 |
| ENSG00000168476 | 1628 | 760 | 886 | 1129 | 938 | 960 |
| ENSG00000168481 | 4 | 1 | 1 | 5 | 2 | 3 |
| ENSG00000168484 | 0 | 0 | 0 | 0 | 0 | 0 |
| ENSG00000168487 | 918 | 435 | 492 | 557 | 539 | 447 |
| ENSG00000177725 | 0 | 1 | 0 | 0 | 0 | 1 |
| ENSG00000285881 | 0 | 1 | 0 | 0 | 0 | 0 |
| ENSG00000168490 | 0 | 0 | 0 | 1 | 0 | 0 |
| ENSG00000208037 | 2 | 1 | 6 | 2 | 0 | 2 |
| ENSG00000168495 | 1386 | 631 | 757 | 770 | 756 | 814 |
| ENSG00000254064 | 18 | 5 | 8 | 4 | 7 | 8 |
| ENSG00000197181 | 2 | 0 | 0 | 0 | 0 | 1 |
| ENSG00000104635 | 1269 | 588 | 747 | 633 | 672 | 758 |
| ENSG00000241026 | 0 | 0 | 0 | 0 | 1 | 0 |
| ENSG00000201761 | 0 | 0 | 0 | 0 | 0 | 0 |
| ENSG00000120910 | 398 | 187 | 273 | 210 | 201 | 241 |
| ENSG00000254272 | 0 | 1 | 2 | 0 | 0 | 1 |
| ENSG00000251034 | 6 | 1 | 11 | 3 | 14 | 6 |
| ENSG00000120896 | 914 | 416 | 506 | 440 | 394 | 408 |

| | | | | | | |
|---|---|---|---|---|---|---|
| ENSG00000254230 | 0 | 0 | 0 | 0 | 0 | 0 |
| ENSG00000120913 | 303 | 168 | 210 | 162 | 181 | 147 |
| ENSG00000248235 | 0 | 0 | 0 | 0 | 0 | 0 |
| ENSG00000241852 | 165 | 85 | 118 | 121 | 105 | 88 |
| ENSG00000158941 | 3115 | 1570 | 2085 | 1939 | 1718 | 1874 |
| ENSG00000253200 | 45 | 25 | 17 | 15 | 22 | 29 |
| ENSG00000147439 | 497 | 261 | 307 | 305 | 290 | 291 |
| ENSG00000261026 | 2 | 1 | 0 | 0 | 0 | 0 |
| ENSG00000179388 | 42 | 17 | 19 | 28 | 21 | 10 |
| ENSG00000253125 | 36 | 19 | 18 | 33 | 17 | 23 |
| ENSG00000134020 | 0 | 3 | 1 | 1 | 1 | 0 |
| ENSG00000248738 | 0 | 0 | 0 | 0 | 0 | 0 |
| ENSG00000253142 | 0 | 0 | 0 | 0 | 0 | 0 |
| ENSG00000240116 | 0 | 0 | 0 | 0 | 0 | 0 |
| ENSG00000245025 | 4 | 2 | 3 | 0 | 2 | 5 |
| ENSG00000008853 | 710 | 321 | 441 | 468 | 406 | 395 |
| ENSG00000120889 | 1664 | 850 | 1107 | 848 | 1152 | 1076 |
| ENSG00000246130 | 0 | 4 | 0 | 0 | 1 | 0 |
| ENSG00000253616 | 2 | 0 | 2 | 9 | 5 | 5 |
| ENSG00000284948 | 0 | 4 | 4 | 2 | 2 | 2 |
| ENSG00000284956 | 0 | 0 | 0 | 0 | 0 | 0 |
| ENSG00000173535 | 77 | 60 | 76 | 38 | 58 | 71 |
| ENSG00000173530 | 0 | 0 | 3 | 0 | 0 | 0 |
| ENSG00000253930 | 11 | 8 | 8 | 10 | 15 | 22 |
| ENSG00000104689 | 361 | 184 | 212 | 157 | 216 | 219 |
| ENSG00000243265 | 2 | 0 | 1 | 0 | 0 | 0 |
| ENSG00000246582 | 86 | 56 | 72 | 48 | 80 | 63 |
| ENSG00000250714 | 0 | 0 | 0 | 0 | 0 | 0 |
| ENSG00000147457 | 2047 | 1016 | 1223 | 1320 | 1175 | 1151 |
| ENSG00000104679 | 868 | 450 | 524 | 681 | 543 | 482 |
| ENSG00000134013 | 706 | 452 | 363 | 547 | 509 | 374 |
| ENSG00000253837 | 2 | 1 | 1 | 1 | 3 | 0 |
| ENSG00000197217 | 1901 | 852 | 1206 | 696 | 1005 | 971 |
| ENSG00000253390 | 0 | 0 | 3 | 5 | 3 | 0 |
| ENSG00000253483 | 0 | 0 | 0 | 0 | 1 | 0 |
| ENSG00000253986 | 0 | 0 | 2 | 1 | 0 | 1 |
| ENSG00000147454 | 1147 | 512 | 721 | 479 | 630 | 631 |
| ENSG00000252067 | 0 | 0 | 0 | 0 | 0 | 0 |
| ENSG00000253488 | 0 | 0 | 0 | 0 | 0 | 0 |
| ENSG00000167034 | 105 | 64 | 89 | 48 | 74 | 55 |
| ENSG00000180053 | 0 | 0 | 0 | 0 | 0 | 0 |
| ENSG00000253471 | 0 | 0 | 0 | 0 | 0 | 0 |
| ENSG00000254002 | 0 | 0 | 0 | 0 | 0 | 0 |
| ENSG00000271118 | 0 | 0 | 0 | 0 | 0 | 0 |
| ENSG00000207027 | 0 | 0 | 0 | 0 | 0 | 0 |
| ENSG00000159167 | 1562 | 700 | 875 | 868 | 1172 | 1277 |
| ENSG00000207201 | 0 | 0 | 0 | 0 | 0 | 0 |
| ENSG00000253891 | 0 | 0 | 0 | 0 | 0 | 0 |
| ENSG00000042980 | 2 | 0 | 5 | 1 | 2 | 0 |
| ENSG00000253535 | 0 | 0 | 0 | 0 | 0 | 0 |

| | | | | | | |
|---|---|---|---|---|---|---|
| ENSG00000134028 | 0 | 0 | 0 | 0 | 0 | 0 |
| ENSG00000069206 | 0 | 0 | 0 | 0 | 0 | 0 |
| ENSG00000253643 | 0 | 0 | 0 | 0 | 0 | 0 |
| ENSG00000272163 | 0 | 0 | 0 | 0 | 0 | 0 |
| ENSG00000104722 | 0 | 0 | 0 | 0 | 0 | 0 |
| ENSG00000277586 | 0 | 0 | 0 | 0 | 0 | 0 |
| ENSG00000283327 | 0 | 0 | 0 | 0 | 0 | 0 |
| ENSG00000272157 | 0 | 0 | 0 | 0 | 0 | 0 |
| ENSG00000253832 | 0 | 0 | 0 | 0 | 0 | 0 |
| ENSG00000253670 | 0 | 0 | 0 | 0 | 0 | 0 |
| ENSG00000147459 | 1299 | 559 | 727 | 352 | 638 | 572 |
| ENSG00000284858 | 0 | 1 | 0 | 0 | 0 | 0 |
| ENSG00000274988 | 0 | 0 | 0 | 0 | 0 | 0 |
| ENSG00000147437 | 37 | 23 | 13 | 12 | 17 | 7 |
| ENSG00000253476 | 4 | 3 | 3 | 2 | 5 | 1 |
| ENSG00000104756 | 1473 | 644 | 854 | 540 | 858 | 775 |
| ENSG00000184661 | 290 | 175 | 150 | 128 | 239 | 176 |
| ENSG00000253100 | 0 | 0 | 0 | 0 | 0 | 0 |
| ENSG00000253342 | 0 | 0 | 0 | 0 | 0 | 0 |
| ENSG00000285890 | 0 | 0 | 0 | 0 | 0 | 0 |
| ENSG00000254178 | 0 | 0 | 0 | 0 | 0 | 0 |
| ENSG00000221818 | 2 | 0 | 1 | 0 | 0 | 0 |
| ENSG00000199620 | 0 | 0 | 0 | 0 | 0 | 0 |
| ENSG00000249213 | 0 | 0 | 0 | 0 | 1 | 0 |
| ENSG00000221914 | 2167 | 1031 | 1247 | 1062 | 1215 | 1175 |
| ENSG00000228451 | 45 | 19 | 17 | 12 | 12 | 18 |
| ENSG00000104765 | 2467 | 1271 | 1297 | 798 | 1481 | 1048 |
| ENSG00000253430 | 0 | 0 | 0 | 0 | 0 | 0 |
| ENSG00000251203 | 2 | 0 | 0 | 0 | 0 | 0 |
| ENSG00000254362 | 1 | 0 | 0 | 1 | 0 | 1 |
| ENSG00000240694 | 2 | 3 | 0 | 0 | 3 | 2 |
| ENSG00000092964 | 759 | 318 | 400 | 353 | 369 | 382 |
| ENSG00000253208 | 0 | 0 | 0 | 0 | 0 | 0 |
| ENSG00000120907 | 0 | 0 | 0 | 0 | 0 | 0 |
| ENSG00000240519 | 0 | 0 | 0 | 0 | 0 | 0 |
| ENSG00000253813 | 0 | 2 | 1 | 0 | 0 | 0 |
| ENSG00000275115 | 0 | 0 | 0 | 0 | 0 | 0 |
| ENSG00000215288 | 0 | 0 | 0 | 0 | 0 | 0 |
| ENSG00000221616 | 0 | 0 | 0 | 0 | 0 | 0 |
| ENSG00000253899 | 0 | 0 | 0 | 0 | 0 | 0 |
| ENSG00000253903 | 0 | 0 | 0 | 0 | 0 | 0 |
| ENSG00000253888 | 0 | 0 | 0 | 0 | 0 | 0 |
| ENSG00000015592 | 0 | 0 | 2 | 1 | 0 | 0 |
| ENSG00000104228 | 605 | 301 | 369 | 364 | 402 | 386 |
| ENSG00000120899 | 455 | 183 | 249 | 178 | 202 | 200 |
| ENSG00000273836 | 0 | 0 | 0 | 0 | 0 | 0 |
| ENSG00000120903 | 0 | 0 | 0 | 0 | 0 | 0 |
| ENSG00000120915 | 414 | 177 | 245 | 187 | 156 | 199 |
| ENSG00000234770 | 0 | 0 | 2 | 0 | 0 | 0 |
| ENSG00000120885 | 20267 | 8634 | 13225 | 8483 | 6768 | 8638 |

| | | | | | | |
|---|---|---|---|---|---|---|
| ENSG00000284280 | 0 | 0 | 0 | 0 | 0 | 0 |
| ENSG00000168077 | 787 | 359 | 498 | 328 | 294 | 347 |
| ENSG00000222862 | 0 | 0 | 0 | 0 | 0 | 0 |
| ENSG00000253560 | 0 | 0 | 0 | 0 | 0 | 0 |
| ENSG00000279302 | 2 | 0 | 2 | 6 | 1 | 3 |
| ENSG00000265075 | 0 | 0 | 0 | 0 | 0 | 0 |
| ENSG00000284351 | 0 | 0 | 0 | 0 | 0 | 0 |
| ENSG00000147419 | 1865 | 996 | 1117 | 884 | 1171 | 1138 |
| ENSG00000253875 | 0 | 0 | 0 | 0 | 0 | 0 |
| ENSG00000171320 | 194 | 141 | 126 | 57 | 146 | 123 |
| ENSG00000202242 | 0 | 0 | 0 | 0 | 0 | 0 |
| ENSG00000168078 | 529 | 292 | 362 | 258 | 454 | 397 |
| ENSG00000253176 | 0 | 0 | 0 | 0 | 0 | 0 |
| ENSG00000168079 | 24 | 16 | 13 | 34 | 44 | 39 |
| ENSG00000265847 | 0 | 0 | 0 | 0 | 0 | 0 |
| ENSG00000253290 | 0 | 0 | 0 | 0 | 0 | 0 |
| ENSG00000253615 | 0 | 0 | 0 | 0 | 0 | 0 |
| ENSG00000253397 | 0 | 0 | 0 | 0 | 0 | 0 |
| ENSG00000189233 | 2 | 2 | 3 | 2 | 4 | 0 |
| ENSG00000134014 | 1387 | 660 | 841 | 651 | 703 | 785 |
| ENSG00000250516 | 0 | 1 | 0 | 0 | 0 | 0 |
| ENSG00000253907 | 0 | 0 | 0 | 0 | 0 | 0 |
| ENSG00000253690 | 0 | 0 | 1 | 0 | 0 | 0 |
| ENSG00000239198 | 2 | 1 | 0 | 0 | 1 | 0 |
| ENSG00000168081 | 0 | 0 | 2 | 0 | 0 | 0 |
| ENSG00000186918 | 1101 | 574 | 718 | 630 | 636 | 649 |
| ENSG00000214050 | 123 | 81 | 74 | 79 | 79 | 67 |
| ENSG00000254370 | 5 | 0 | 4 | 1 | 3 | 6 |
| ENSG00000207361 | 0 | 0 | 1 | 0 | 2 | 0 |
| ENSG00000253567 | 4 | 1 | 0 | 1 | 1 | 0 |
| ENSG00000104290 | 578 | 316 | 248 | 204 | 381 | 323 |
| ENSG00000265251 | 0 | 0 | 0 | 0 | 0 | 0 |
| ENSG00000202411 | 0 | 0 | 0 | 0 | 0 | 0 |
| ENSG00000012232 | 1090 | 481 | 660 | 470 | 516 | 606 |
| ENSG00000246339 | 45 | 20 | 43 | 23 | 33 | 32 |
| ENSG00000104299 | 562 | 310 | 386 | 322 | 371 | 370 |
| ENSG00000254034 | 4 | 2 | 2 | 0 | 1 | 5 |
| ENSG00000242958 | 0 | 1 | 0 | 0 | 0 | 1 |
| ENSG00000147421 | 472 | 210 | 271 | 153 | 249 | 227 |
| ENSG00000273710 | 0 | 0 | 0 | 0 | 0 | 0 |
| ENSG00000259196 | 0 | 0 | 0 | 0 | 0 | 0 |
| ENSG00000200719 | 0 | 0 | 0 | 0 | 0 | 0 |
| ENSG00000259366 | 29 | 17 | 25 | 11 | 16 | 16 |
| ENSG00000259607 | 0 | 0 | 0 | 0 | 0 | 0 |
| ENSG00000197892 | 1167 | 489 | 704 | 346 | 479 | 559 |
| ENSG00000254129 | 0 | 0 | 0 | 0 | 0 | 0 |
| ENSG00000253770 | 0 | 0 | 0 | 0 | 0 | 0 |
| ENSG00000120875 | 1753 | 768 | 1232 | 1045 | 995 | 1346 |
| ENSG00000285601 | 2 | 4 | 6 | 1 | 4 | 6 |
| ENSG00000260253 | 6 | 1 | 1 | 6 | 1 | 4 |

| | | | | | | |
|---|---|---|---|---|---|---|
| ENSG00000253632 | 0 | 0 | 3 | 0 | 1 | 0 |
| ENSG00000253182 | 0 | 0 | 0 | 0 | 1 | 1 |
| ENSG00000280147 | 0 | 0 | 0 | 0 | 0 | 0 |
| ENSG00000243844 | 0 | 0 | 0 | 0 | 0 | 0 |
| ENSG00000251191 | 22 | 9 | 12 | 17 | 16 | 16 |
| ENSG00000253490 | 2 | 0 | 0 | 0 | 0 | 0 |
| ENSG00000248964 | 0 | 1 | 4 | 0 | 0 | 0 |
| ENSG00000254086 | 0 | 0 | 0 | 0 | 0 | 0 |
| ENSG00000253604 | 0 | 0 | 0 | 0 | 0 | 0 |
| ENSG00000253279 | 0 | 0 | 0 | 0 | 0 | 0 |
| ENSG00000264788 | 0 | 0 | 0 | 0 | 0 | 0 |
| ENSG00000254013 | 0 | 1 | 0 | 0 | 1 | 1 |
| ENSG00000239175 | 0 | 0 | 0 | 0 | 0 | 0 |
| ENSG00000133872 | 3676 | 1815 | 2141 | 1671 | 2082 | 1977 |
| ENSG00000272256 | 1 | 0 | 1 | 1 | 0 | 1 |
| ENSG00000104660 | 1191 | 590 | 749 | 596 | 732 | 708 |
| ENSG00000241511 | 2 | 4 | 0 | 1 | 0 | 1 |
| ENSG00000177669 | 3 | 2 | 0 | 4 | 1 | 4 |
| ENSG00000271869 | 18 | 15 | 21 | 11 | 20 | 14 |
| ENSG00000104671 | 423 | 268 | 337 | 252 | 348 | 312 |
| ENSG00000253708 | 0 | 0 | 0 | 0 | 0 | 0 |
| ENSG00000285669 | 18 | 17 | 10 | 12 | 10 | 15 |
| ENSG00000272375 | 0 | 0 | 3 | 4 | 3 | 3 |
| ENSG00000248159 | 0 | 0 | 0 | 0 | 1 | 0 |
| ENSG00000254316 | 0 | 0 | 0 | 0 | 0 | 0 |
| ENSG00000253346 | 0 | 0 | 0 | 0 | 0 | 0 |
| ENSG00000254152 | 0 | 0 | 1 | 0 | 0 | 0 |
| ENSG00000127589 | 18 | 6 | 5 | 6 | 4 | 7 |
| ENSG00000254109 | 60 | 35 | 34 | 23 | 23 | 16 |
| ENSG00000157110 | 1359 | 686 | 779 | 469 | 507 | 566 |
| ENSG00000279041 | 123 | 72 | 73 | 29 | 44 | 42 |
| ENSG00000197265 | 911 | 424 | 494 | 518 | 552 | 508 |
| ENSG00000253112 | 0 | 2 | 0 | 0 | 0 | 0 |
| ENSG00000253457 | 0 | 2 | 1 | 0 | 1 | 1 |
| ENSG00000254172 | 0 | 0 | 0 | 0 | 0 | 0 |
| ENSG00000104687 | 4621 | 2324 | 2785 | 2202 | 2726 | 2792 |
| ENSG00000104691 | 280 | 151 | 191 | 101 | 144 | 154 |
| ENSG00000271081 | 0 | 0 | 0 | 0 | 0 | 0 |
| ENSG00000104695 | 1674 | 794 | 918 | 780 | 900 | 939 |
| ENSG00000133863 | 0 | 0 | 0 | 0 | 0 | 0 |
| ENSG00000253204 | 2 | 0 | 0 | 0 | 0 | 0 |
| ENSG00000253793 | 0 | 0 | 0 | 0 | 0 | 0 |
| ENSG00000172733 | 0 | 0 | 1 | 0 | 0 | 0 |
| ENSG00000165392 | 193 | 161 | 148 | 63 | 218 | 120 |
| ENSG00000253579 | 0 | 0 | 0 | 0 | 0 | 0 |
| ENSG00000253961 | 1 | 1 | 1 | 0 | 0 | 0 |
| ENSG00000229211 | 0 | 0 | 0 | 0 | 0 | 0 |
| ENSG00000253377 | 0 | 0 | 0 | 0 | 0 | 0 |
| ENSG00000254095 | 0 | 0 | 0 | 0 | 0 | 0 |
| ENSG00000253275 | 0 | 0 | 0 | 0 | 0 | 0 |

| | | | | | | |
|---|---|---|---|---|---|---|
| ENSG00000222468 | 0 | 0 | 0 | 0 | 0 | 0 |
| ENSG00000157168 | 0 | 0 | 0 | 0 | 0 | 0 |
| ENSG00000253974 | 0 | 0 | 0 | 0 | 0 | 0 |
| ENSG00000252261 | 0 | 0 | 0 | 0 | 0 | 0 |
| ENSG00000200246 | 0 | 0 | 0 | 0 | 0 | 0 |
| ENSG00000254049 | 0 | 0 | 0 | 0 | 0 | 0 |
| ENSG00000212407 | 0 | 0 | 0 | 0 | 0 | 0 |
| ENSG00000247134 | 0 | 0 | 0 | 0 | 1 | 1 |
| ENSG00000254346 | 0 | 0 | 0 | 0 | 0 | 0 |
| ENSG00000254090 | 0 | 0 | 0 | 0 | 0 | 0 |
| ENSG00000253393 | 0 | 0 | 0 | 0 | 0 | 0 |
| ENSG00000253993 | 0 | 0 | 0 | 0 | 1 | 0 |
| ENSG00000252735 | 0 | 0 | 0 | 0 | 0 | 0 |
| ENSG00000272338 | 12 | 5 | 5 | 5 | 2 | 2 |
| ENSG00000172728 | 123 | 70 | 97 | 58 | 61 | 64 |
| ENSG00000254061 | 0 | 0 | 0 | 0 | 0 | 0 |
| ENSG00000129696 | 295 | 166 | 213 | 164 | 187 | 184 |
| ENSG00000198042 | 457 | 254 | 288 | 213 | 308 | 300 |
| ENSG00000239039 | 0 | 0 | 0 | 0 | 0 | 0 |
| ENSG00000253232 | 0 | 0 | 0 | 0 | 0 | 0 |
| ENSG00000252505 | 0 | 0 | 0 | 0 | 0 | 0 |
| ENSG00000253995 | 0 | 0 | 0 | 0 | 0 | 0 |
| ENSG00000133874 | 80 | 38 | 46 | 66 | 50 | 41 |
| ENSG00000240189 | 0 | 0 | 0 | 0 | 0 | 0 |
| ENSG00000133878 | 0 | 0 | 0 | 0 | 0 | 0 |
| ENSG00000253642 | 0 | 0 | 0 | 0 | 0 | 0 |
| ENSG00000253684 | 0 | 0 | 0 | 0 | 0 | 0 |
| ENSG00000263794 | 0 | 0 | 0 | 0 | 0 | 0 |
| ENSG00000253569 | 0 | 0 | 0 | 0 | 0 | 0 |
| ENSG00000240738 | 0 | 0 | 0 | 1 | 0 | 2 |
| ENSG00000188512 | 0 | 0 | 0 | 1 | 0 | 1 |
| ENSG00000254194 | 0 | 0 | 0 | 0 | 0 | 0 |
| ENSG00000254302 | 0 | 0 | 0 | 0 | 0 | 0 |
| ENSG00000253108 | 0 | 0 | 0 | 0 | 0 | 0 |
| ENSG00000254033 | 0 | 0 | 0 | 0 | 0 | 0 |
| ENSG00000215264 | 0 | 0 | 0 | 0 | 0 | 0 |
| ENSG00000254344 | 0 | 0 | 0 | 0 | 0 | 0 |
| ENSG00000243181 | 0 | 1 | 0 | 0 | 1 | 1 |
| ENSG00000156687 | 0 | 0 | 0 | 0 | 0 | 0 |
| ENSG00000279099 | 0 | 0 | 0 | 0 | 0 | 0 |
| ENSG00000232024 | 29 | 14 | 19 | 18 | 11 | 18 |
| ENSG00000253452 | 0 | 0 | 0 | 0 | 0 | 0 |
| ENSG00000253524 | 0 | 0 | 0 | 0 | 0 | 0 |
| ENSG00000254309 | 0 | 0 | 0 | 0 | 0 | 0 |
| ENSG00000253504 | 0 | 0 | 0 | 0 | 0 | 0 |
| ENSG00000254384 | 0 | 0 | 0 | 0 | 0 | 0 |
| ENSG00000253685 | 0 | 0 | 0 | 0 | 0 | 0 |
| ENSG00000253610 | 0 | 0 | 0 | 0 | 0 | 0 |
| ENSG00000206871 | 0 | 0 | 0 | 0 | 0 | 0 |
| ENSG00000254320 | 0 | 0 | 0 | 0 | 0 | 0 |

| | | | | | | |
|---|---|---|---|---|---|---|
| ENSG00000253363 | 0 | 0 | 0 | 0 | 0 | 0 |
| ENSG00000244708 | 0 | 0 | 0 | 0 | 0 | 0 |
| ENSG00000199985 | 0 | 0 | 0 | 0 | 0 | 0 |
| ENSG00000215262 | 2 | 0 | 1 | 1 | 6 | 1 |
| ENSG00000274489 | 0 | 0 | 0 | 0 | 0 | 0 |
| ENSG00000219491 | 0 | 0 | 0 | 0 | 0 | 0 |
| ENSG00000243503 | 0 | 0 | 0 | 0 | 0 | 0 |
| ENSG00000254038 | 3 | 1 | 0 | 3 | 4 | 2 |
| ENSG00000253650 | 0 | 0 | 0 | 0 | 0 | 0 |
| ENSG00000253123 | 0 | 1 | 0 | 0 | 0 | 0 |
| ENSG00000253746 | 0 | 0 | 0 | 0 | 0 | 0 |
| ENSG00000253161 | 5 | 1 | 1 | 0 | 1 | 2 |
| ENSG00000253388 | 0 | 0 | 0 | 0 | 0 | 0 |
| ENSG00000253414 | 20 | 5 | 2 | 4 | 3 | 4 |
| ENSG00000254111 | 0 | 0 | 3 | 0 | 0 | 0 |
| ENSG00000254290 | 2 | 1 | 3 | 2 | 4 | 2 |
| ENSG00000254306 | 0 | 0 | 0 | 0 | 0 | 0 |
| ENSG00000253344 | 0 | 0 | 0 | 0 | 0 | 0 |
| ENSG00000223215 | 0 | 0 | 0 | 0 | 0 | 0 |
| ENSG00000183779 | 2597 | 1215 | 1585 | 1641 | 1370 | 1728 |
| ENSG00000253181 | 0 | 0 | 1 | 0 | 0 | 0 |
| ENSG00000183154 | 4 | 0 | 1 | 5 | 3 | 0 |
| ENSG00000147475 | 2329 | 1001 | 1414 | 885 | 1254 | 1256 |
| ENSG00000233170 | 12 | 4 | 3 | 1 | 3 | 2 |
| ENSG00000147471 | 876 | 449 | 543 | 395 | 549 | 497 |
| ENSG00000020181 | 73 | 27 | 43 | 21 | 15 | 14 |
| ENSG00000104221 | 285 | 121 | 155 | 147 | 130 | 129 |
| ENSG00000156675 | 2440 | 1051 | 1584 | 700 | 1088 | 1241 |
| ENSG00000280064 | 0 | 0 | 0 | 0 | 0 | 0 |
| ENSG00000241032 | 0 | 0 | 0 | 0 | 0 | 1 |
| ENSG00000169154 | 0 | 0 | 0 | 0 | 0 | 0 |
| ENSG00000285880 | 0 | 0 | 0 | 0 | 0 | 0 |
| ENSG00000188778 | 0 | 0 | 0 | 0 | 0 | 0 |
| ENSG00000270865 | 0 | 0 | 0 | 0 | 0 | 0 |
| ENSG00000187840 | 1664 | 853 | 988 | 897 | 709 | 737 |
| ENSG00000260949 | 1 | 2 | 4 | 3 | 1 | 2 |
| ENSG00000272128 | 3 | 0 | 0 | 0 | 0 | 1 |
| ENSG00000129691 | 1449 | 676 | 863 | 548 | 678 | 694 |
| ENSG00000253437 | 0 | 0 | 0 | 0 | 0 | 0 |
| ENSG00000240015 | 1 | 0 | 0 | 0 | 0 | 0 |
| ENSG00000147465 | 0 | 0 | 2 | 0 | 1 | 5 |
| ENSG00000253356 | 0 | 0 | 0 | 0 | 0 | 0 |
| ENSG00000175324 | 487 | 243 | 326 | 287 | 320 | 295 |
| ENSG00000253739 | 2 | 0 | 0 | 0 | 0 | 0 |
| ENSG00000156735 | 594 | 275 | 297 | 220 | 283 | 246 |
| ENSG00000248585 | 0 | 0 | 0 | 0 | 0 | 0 |
| ENSG00000285632 | 7 | 0 | 6 | 5 | 3 | 2 |
| ENSG00000085788 | 866 | 469 | 591 | 320 | 519 | 503 |
| ENSG00000147535 | 728 | 332 | 456 | 327 | 431 | 379 |
| ENSG00000147548 | 1192 | 627 | 612 | 395 | 763 | 646 |

| | | | | | | |
|---|---|---|---|---|---|---|
| ENSG00000255487 | 0 | 0 | 0 | 0 | 0 | 0 |
| ENSG00000254898 | 0 | 0 | 0 | 0 | 0 | 0 |
| ENSG00000272092 | 24 | 9 | 14 | 7 | 7 | 9 |
| ENSG00000165046 | 61 | 25 | 50 | 32 | 44 | 38 |
| ENSG00000077782 | 554 | 210 | 217 | 182 | 197 | 194 |
| ENSG00000272159 | 1 | 0 | 0 | 0 | 0 | 0 |
| ENSG00000255201 | 5 | 1 | 4 | 1 | 0 | 1 |
| ENSG00000239218 | 6 | 0 | 1 | 1 | 0 | 3 |
| ENSG00000196166 | 0 | 0 | 2 | 0 | 1 | 0 |
| ENSG00000253361 | 40 | 13 | 10 | 3 | 9 | 1 |
| ENSG00000253570 | 8 | 9 | 2 | 9 | 15 | 13 |
| ENSG00000254321 | 0 | 0 | 0 | 0 | 0 | 0 |
| ENSG00000147526 | 1512 | 724 | 841 | 337 | 703 | 460 |
| ENSG00000207258 | 0 | 0 | 0 | 0 | 0 | 0 |
| ENSG00000253586 | 0 | 0 | 0 | 0 | 0 | 0 |
| ENSG00000253829 | 2 | 0 | 0 | 0 | 0 | 1 |
| ENSG00000169499 | 788 | 416 | 525 | 376 | 484 | 513 |
| ENSG00000253645 | 0 | 0 | 0 | 0 | 0 | 0 |
| ENSG00000169495 | 5 | 2 | 7 | 5 | 8 | 5 |
| ENSG00000169490 | 907 | 421 | 473 | 485 | 502 | 493 |
| ENSG00000168615 | 2886 | 1468 | 1482 | 691 | 1668 | 1147 |
| ENSG00000207199 | 0 | 0 | 0 | 0 | 0 | 0 |
| ENSG00000197140 | 95 | 48 | 55 | 38 | 50 | 45 |
| ENSG00000243095 | 0 | 1 | 0 | 0 | 0 | 3 |
| ENSG00000253160 | 0 | 0 | 0 | 0 | 0 | 0 |
| ENSG00000196115 | 0 | 0 | 0 | 0 | 0 | 0 |
| ENSG00000197475 | 0 | 0 | 2 | 0 | 0 | 0 |
| ENSG00000254067 | 0 | 0 | 0 | 0 | 0 | 0 |
| ENSG00000168619 | 2 | 0 | 1 | 0 | 1 | 0 |
| ENSG00000104755 | 50 | 21 | 34 | 10 | 6 | 9 |
| ENSG00000253332 | 0 | 0 | 0 | 0 | 0 | 0 |
| ENSG00000253233 | 0 | 0 | 0 | 0 | 0 | 0 |
| ENSG00000131203 | 0 | 0 | 0 | 0 | 0 | 0 |
| ENSG00000253939 | 0 | 0 | 0 | 0 | 0 | 0 |
| ENSG00000253838 | 0 | 0 | 0 | 0 | 0 | 0 |
| ENSG00000254287 | 0 | 0 | 0 | 0 | 0 | 0 |
| ENSG00000188676 | 0 | 1 | 0 | 0 | 0 | 0 |
| ENSG00000253790 | 0 | 0 | 0 | 0 | 0 | 0 |
| ENSG00000253381 | 0 | 0 | 0 | 0 | 0 | 0 |
| ENSG00000253665 | 0 | 0 | 0 | 0 | 0 | 0 |
| ENSG00000176907 | 28 | 29 | 15 | 13 | 17 | 7 |
| ENSG00000254143 | 0 | 1 | 3 | 3 | 2 | 1 |
| ENSG00000253802 | 25 | 15 | 11 | 17 | 28 | 20 |
| ENSG00000253354 | 0 | 0 | 0 | 0 | 1 | 0 |
| ENSG00000272479 | 0 | 0 | 0 | 0 | 0 | 0 |
| ENSG00000165061 | 10 | 11 | 13 | 15 | 13 | 20 |
| ENSG00000254383 | 0 | 0 | 0 | 0 | 0 | 0 |
| ENSG00000206867 | 0 | 0 | 0 | 0 | 0 | 0 |
| ENSG00000104332 | 0 | 0 | 0 | 0 | 0 | 0 |
| ENSG00000263372 | 0 | 0 | 0 | 0 | 0 | 0 |

| | | | | | | |
|---|---|---|---|---|---|---|
| ENSG00000239281 | 0 | 0 | 0 | 0 | 0 | 0 |
| ENSG00000253509 | 0 | 0 | 0 | 0 | 0 | 0 |
| ENSG00000206852 | 0 | 0 | 0 | 0 | 0 | 0 |
| ENSG00000238936 | 0 | 0 | 0 | 0 | 0 | 0 |
| ENSG00000253396 | 0 | 0 | 0 | 0 | 0 | 0 |
| ENSG00000147533 | 1322 | 662 | 793 | 656 | 802 | 731 |
| ENSG00000253943 | 0 | 0 | 0 | 0 | 0 | 2 |
| ENSG00000147536 | 97 | 89 | 74 | 92 | 82 | 101 |
| ENSG00000253133 | 0 | 1 | 0 | 1 | 0 | 1 |
| ENSG00000253174 | 0 | 0 | 0 | 0 | 0 | 0 |
| ENSG00000158669 | 1214 | 632 | 724 | 720 | 704 | 779 |
| ENSG00000264578 | 0 | 0 | 0 | 0 | 0 | 0 |
| ENSG00000165066 | 0 | 0 | 0 | 0 | 0 | 0 |
| ENSG00000029534 | 12 | 14 | 12 | 16 | 14 | 8 |
| ENSG00000274705 | 0 | 0 | 0 | 0 | 0 | 0 |
| ENSG00000283450 | 0 | 0 | 0 | 0 | 0 | 0 |
| ENSG00000253389 | 0 | 0 | 0 | 0 | 0 | 0 |
| ENSG00000207101 | 0 | 0 | 0 | 0 | 0 | 0 |
| ENSG00000260588 | 40 | 16 | 31 | 28 | 16 | 25 |
| ENSG00000241834 | 0 | 0 | 0 | 0 | 0 | 0 |
| ENSG00000083168 | 930 | 479 | 591 | 268 | 507 | 469 |
| ENSG00000238966 | 0 | 0 | 1 | 0 | 0 | 0 |
| ENSG00000244235 | 0 | 0 | 0 | 0 | 0 | 0 |
| ENSG00000253135 | 0 | 0 | 0 | 0 | 0 | 0 |
| ENSG00000271938 | 0 | 0 | 0 | 0 | 0 | 0 |
| ENSG00000261449 | 0 | 0 | 0 | 0 | 0 | 0 |
| ENSG00000070718 | 450 | 258 | 287 | 191 | 259 | 245 |
| ENSG00000104368 | 6 | 1 | 0 | 0 | 0 | 0 |
| ENSG00000253408 | 0 | 0 | 0 | 0 | 0 | 0 |
| ENSG00000104365 | 1005 | 489 | 709 | 459 | 602 | 614 |
| ENSG00000070501 | 473 | 215 | 245 | 246 | 265 | 253 |
| ENSG00000240395 | 4 | 3 | 0 | 0 | 0 | 1 |
| ENSG00000104371 | 0 | 0 | 0 | 0 | 0 | 0 |
| ENSG00000078668 | 2412 | 1211 | 1459 | 1353 | 1504 | 1453 |
| ENSG00000168575 | 1560 | 798 | 1014 | 732 | 839 | 924 |
| ENSG00000254131 | 0 | 0 | 0 | 0 | 0 | 3 |
| ENSG00000254165 | 4 | 5 | 2 | 0 | 0 | 2 |
| ENSG00000176209 | 603 | 333 | 379 | 376 | 356 | 337 |
| ENSG00000147432 | 0 | 0 | 0 | 0 | 0 | 0 |
| ENSG00000255101 | 0 | 0 | 0 | 0 | 0 | 0 |
| ENSG00000255361 | 0 | 0 | 0 | 0 | 1 | 0 |
| ENSG00000147434 | 0 | 0 | 0 | 0 | 0 | 0 |
| ENSG00000131931 | 309 | 134 | 129 | 111 | 145 | 149 |
| ENSG00000120925 | 334 | 186 | 265 | 170 | 224 | 206 |
| ENSG00000242719 | 0 | 0 | 0 | 0 | 0 | 0 |
| ENSG00000284062 | 3 | 1 | 0 | 1 | 1 | 0 |
| ENSG00000168172 | 497 | 242 | 234 | 109 | 253 | 180 |
| ENSG00000202514 | 0 | 0 | 0 | 0 | 0 | 0 |
| ENSG00000254673 | 0 | 0 | 0 | 0 | 0 | 0 |
| ENSG00000168522 | 900 | 429 | 531 | 412 | 638 | 472 |

| | | | | | | |
|---|---|---|---|---|---|---|
| ENSG00000200731 | 0 | 0 | 0 | 0 | 0 | 0 |
| ENSG00000185900 | 108 | 74 | 67 | 29 | 25 | 56 |
| ENSG00000254198 | 36 | 23 | 31 | 26 | 23 | 26 |
| ENSG00000165102 | 934 | 459 | 568 | 427 | 494 | 504 |
| ENSG00000253611 | 0 | 0 | 0 | 0 | 0 | 0 |
| ENSG00000253884 | 0 | 0 | 0 | 0 | 0 | 0 |
| ENSG00000255166 | 0 | 0 | 0 | 0 | 0 | 0 |
| ENSG00000253312 | 0 | 0 | 0 | 0 | 0 | 0 |
| ENSG00000253186 | 0 | 0 | 0 | 0 | 0 | 0 |
| ENSG00000253606 | 0 | 0 | 0 | 0 | 0 | 1 |
| ENSG00000234713 | 0 | 0 | 0 | 0 | 0 | 0 |
| ENSG00000253707 | 0 | 0 | 0 | 0 | 0 | 0 |
| ENSG00000188877 | 0 | 0 | 0 | 0 | 0 | 0 |
| ENSG00000238509 | 0 | 0 | 0 | 0 | 0 | 0 |
| ENSG00000255497 | 0 | 0 | 0 | 0 | 0 | 0 |
| ENSG00000254342 | 0 | 0 | 0 | 0 | 0 | 0 |
| ENSG00000201329 | 0 | 0 | 0 | 0 | 0 | 0 |
| ENSG00000250637 | 0 | 0 | 0 | 0 | 0 | 0 |
| ENSG00000254069 | 0 | 0 | 0 | 0 | 0 | 0 |
| ENSG00000253319 | 0 | 0 | 0 | 0 | 0 | 0 |
| ENSG00000253418 | 0 | 0 | 0 | 0 | 0 | 0 |
| ENSG00000254145 | 0 | 0 | 0 | 0 | 0 | 0 |
| ENSG00000253748 | 0 | 0 | 0 | 0 | 0 | 0 |
| ENSG00000253195 | 0 | 0 | 0 | 0 | 0 | 0 |
| ENSG00000253845 | 0 | 0 | 0 | 0 | 0 | 0 |
| ENSG00000253198 | 0 | 0 | 0 | 0 | 0 | 0 |
| ENSG00000276685 | 0 | 0 | 0 | 0 | 0 | 0 |
| ENSG00000253425 | 0 | 0 | 0 | 0 | 0 | 0 |
| ENSG00000248498 | 4 | 2 | 3 | 5 | 10 | 4 |
| ENSG00000254070 | 0 | 0 | 0 | 0 | 0 | 0 |
| ENSG00000251470 | 0 | 0 | 0 | 0 | 0 | 0 |
| ENSG00000228984 | 0 | 0 | 0 | 0 | 0 | 0 |
| ENSG00000250857 | 0 | 0 | 0 | 0 | 0 | 0 |
| ENSG00000253314 | 0 | 0 | 0 | 0 | 0 | 0 |
| ENSG00000253828 | 0 | 0 | 0 | 0 | 0 | 0 |
| ENSG00000282321 | 0 | 0 | 0 | 0 | 0 | 0 |
| ENSG00000252594 | 0 | 0 | 0 | 0 | 0 | 0 |
| ENSG00000254118 | 0 | 0 | 0 | 0 | 0 | 0 |
| ENSG00000254156 | 0 | 0 | 0 | 0 | 0 | 0 |
| ENSG00000253803 | 0 | 0 | 0 | 0 | 0 | 0 |
| ENSG00000253745 | 0 | 0 | 0 | 0 | 0 | 0 |
| ENSG00000253782 | 0 | 0 | 0 | 0 | 0 | 0 |
| ENSG00000215179 | 0 | 0 | 0 | 0 | 1 | 0 |
| ENSG00000222099 | 0 | 0 | 0 | 0 | 0 | 0 |
| ENSG00000248347 | 5 | 7 | 5 | 5 | 5 | 1 |
| ENSG00000285992 | 3 | 0 | 2 | 2 | 0 | 1 |
| ENSG00000200986 | 0 | 0 | 0 | 0 | 0 | 0 |
| ENSG00000188873 | 0 | 0 | 0 | 3 | 1 | 0 |
| ENSG00000248531 | 0 | 0 | 0 | 0 | 0 | 0 |
| ENSG00000254348 | 9 | 1 | 3 | 0 | 1 | 2 |

| | | | | | | |
|---|---|---|---|---|---|---|
| ENSG00000255366 | 18 | 10 | 13 | 6 | 5 | 6 |
| ENSG00000253502 | 0 | 0 | 0 | 0 | 0 | 0 |
| ENSG00000253817 | 0 | 0 | 1 | 0 | 0 | 0 |
| ENSG00000215177 | 0 | 0 | 0 | 0 | 0 | 0 |
| ENSG00000164808 | 2216 | 1041 | 1418 | 912 | 1221 | 1250 |
| ENSG00000253330 | 2 | 4 | 5 | 1 | 6 | 8 |
| ENSG00000269924 | 0 | 0 | 2 | 1 | 2 | 1 |
| ENSG00000251354 | 0 | 1 | 0 | 0 | 0 | 0 |
| ENSG00000254065 | 0 | 1 | 0 | 0 | 2 | 0 |
| ENSG00000207369 | 0 | 0 | 0 | 0 | 0 | 0 |
| ENSG00000221869 | 326 | 126 | 170 | 295 | 281 | 255 |
| ENSG00000253729 | 4978 | 2419 | 2504 | 1266 | 3641 | 2799 |
| ENSG00000207450 | 0 | 0 | 0 | 0 | 0 | 0 |
| ENSG00000104738 | 4898 | 2603 | 2933 | 2681 | 3157 | 3388 |
| ENSG00000222522 | 0 | 0 | 0 | 0 | 0 | 1 |
| ENSG00000169139 | 2250 | 1194 | 1210 | 1042 | 1570 | 1284 |
| ENSG00000253843 | 0 | 0 | 0 | 0 | 0 | 0 |
| ENSG00000253654 | 0 | 0 | 0 | 0 | 0 | 0 |
| ENSG00000252710 | 0 | 0 | 0 | 0 | 0 | 0 |
| ENSG00000277418 | 0 | 0 | 0 | 0 | 0 | 0 |
| ENSG00000224594 | 0 | 0 | 0 | 0 | 0 | 0 |
| ENSG00000279881 | 0 | 0 | 0 | 0 | 0 | 0 |
| ENSG00000253140 | 0 | 0 | 0 | 1 | 0 | 0 |
| ENSG00000253688 | 0 | 0 | 0 | 0 | 0 | 0 |
| ENSG00000253608 | 0 | 0 | 0 | 0 | 0 | 0 |
| ENSG00000233858 | 0 | 0 | 0 | 0 | 0 | 0 |
| ENSG00000253702 | 0 | 0 | 0 | 0 | 0 | 0 |
| ENSG00000253455 | 2 | 0 | 3 | 2 | 5 | 2 |
| ENSG00000253892 | 0 | 0 | 0 | 0 | 0 | 1 |
| ENSG00000034239 | 0 | 0 | 0 | 0 | 0 | 0 |
| ENSG00000233109 | 0 | 0 | 0 | 0 | 0 | 0 |
| ENSG00000019549 | 38 | 12 | 6 | 24 | 14 | 16 |
| ENSG00000168333 | 0 | 0 | 0 | 0 | 0 | 0 |
| ENSG00000253474 | 0 | 0 | 0 | 0 | 0 | 0 |
| ENSG00000199640 | 0 | 0 | 0 | 0 | 0 | 0 |
| ENSG00000254268 | 0 | 0 | 0 | 0 | 0 | 0 |
| ENSG00000253380 | 0 | 0 | 0 | 0 | 0 | 0 |
| ENSG00000253206 | 0 | 0 | 0 | 0 | 0 | 0 |
| ENSG00000253810 | 0 | 0 | 0 | 0 | 0 | 0 |
| ENSG00000253484 | 0 | 0 | 0 | 0 | 0 | 0 |
| ENSG00000271148 | 0 | 0 | 0 | 0 | 0 | 0 |
| ENSG00000147481 | 0 | 0 | 0 | 0 | 0 | 0 |
| ENSG00000253849 | 0 | 0 | 0 | 0 | 0 | 0 |
| ENSG00000254392 | 0 | 0 | 0 | 0 | 0 | 0 |
| ENSG00000254358 | 0 | 0 | 0 | 0 | 0 | 0 |
| ENSG00000253340 | 0 | 0 | 0 | 0 | 0 | 0 |
| ENSG00000201316 | 0 | 0 | 0 | 0 | 0 | 0 |
| ENSG00000253664 | 0 | 0 | 0 | 0 | 0 | 0 |
| ENSG00000147485 | 6 | 1 | 3 | 0 | 2 | 5 |
| ENSG00000254025 | 0 | 0 | 0 | 1 | 0 | 0 |

| | | | | | | |
|---|---|---|---|---|---|---|
| ENSG00000254225 | 0 | 0 | 0 | 0 | 0 | 0 |
| ENSG00000272076 | 0 | 0 | 0 | 0 | 0 | 1 |
| ENSG00000168300 | 1204 | 592 | 665 | 349 | 739 | 573 |
| ENSG00000253475 | 0 | 1 | 0 | 0 | 0 | 0 |
| ENSG00000228801 | 109 | 40 | 71 | 56 | 62 | 66 |
| ENSG00000272024 | 12 | 4 | 4 | 4 | 6 | 2 |
| ENSG00000253844 | 2 | 0 | 5 | 1 | 1 | 0 |
| ENSG00000254136 | 0 | 0 | 0 | 0 | 0 | 0 |
| ENSG00000147488 | 0 | 0 | 0 | 0 | 0 | 0 |
| ENSG00000254314 | 0 | 0 | 0 | 0 | 0 | 0 |
| ENSG00000253551 | 0 | 0 | 0 | 0 | 0 | 0 |
| ENSG00000253924 | 0 | 0 | 0 | 0 | 0 | 0 |
| ENSG00000240534 | 0 | 0 | 0 | 0 | 0 | 0 |
| ENSG00000196711 | 0 | 1 | 0 | 0 | 0 | 0 |
| ENSG00000023287 | 519 | 395 | 223 | 85 | 776 | 307 |
| ENSG00000271555 | 0 | 0 | 1 | 1 | 0 | 1 |
| ENSG00000183729 | 0 | 0 | 0 | 0 | 0 | 0 |
| ENSG00000248911 | 0 | 0 | 0 | 0 | 0 | 0 |
| ENSG00000250494 | 0 | 0 | 0 | 0 | 0 | 0 |
| ENSG00000254687 | 0 | 0 | 0 | 0 | 0 | 0 |
| ENSG00000082556 | 0 | 0 | 0 | 0 | 0 | 0 |
| ENSG00000260484 | 0 | 0 | 0 | 0 | 0 | 0 |
| ENSG00000274295 | 0 | 0 | 0 | 0 | 0 | 0 |
| ENSG00000253369 | 0 | 0 | 0 | 0 | 0 | 0 |
| ENSG00000254204 | 0 | 0 | 0 | 0 | 0 | 0 |
| ENSG00000237807 | 24 | 19 | 7 | 2 | 6 | 6 |
| ENSG00000240919 | 0 | 0 | 0 | 0 | 0 | 0 |
| ENSG00000206144 | 0 | 0 | 0 | 0 | 0 | 0 |
| ENSG00000253668 | 0 | 0 | 0 | 0 | 0 | 0 |
| ENSG00000047249 | 2167 | 975 | 1364 | 1051 | 1237 | 1225 |
| ENSG00000147509 | 9 | 3 | 1 | 4 | 10 | 8 |
| ENSG00000272457 | 0 | 0 | 0 | 0 | 0 | 1 |
| ENSG00000244159 | 0 | 0 | 0 | 0 | 0 | 0 |
| ENSG00000200528 | 0 | 0 | 0 | 0 | 0 | 0 |
| ENSG00000187735 | 2624 | 1306 | 1447 | 1170 | 1431 | 1325 |
| ENSG00000253667 | 2 | 1 | 5 | 3 | 2 | 3 |
| ENSG00000260955 | 6 | 3 | 0 | 1 | 0 | 2 |
| ENSG00000120992 | 2581 | 1203 | 1495 | 1152 | 1351 | 1403 |
| ENSG00000254274 | 0 | 1 | 0 | 0 | 1 | 0 |
| ENSG00000276353 | 0 | 0 | 0 | 0 | 0 | 0 |
| ENSG00000137547 | 1417 | 768 | 847 | 959 | 908 | 942 |
| ENSG00000251835 | 0 | 0 | 0 | 0 | 0 | 0 |
| ENSG00000270980 | 0 | 0 | 0 | 0 | 0 | 0 |
| ENSG00000253840 | 0 | 0 | 0 | 0 | 0 | 0 |
| ENSG00000199212 | 0 | 0 | 0 | 0 | 0 | 0 |
| ENSG00000276605 | 0 | 0 | 0 | 0 | 0 | 0 |
| ENSG00000244107 | 0 | 0 | 0 | 0 | 0 | 0 |
| ENSG00000164736 | 0 | 0 | 0 | 0 | 0 | 0 |
| ENSG00000274310 | 0 | 0 | 0 | 0 | 0 | 0 |
| ENSG00000270275 | 0 | 0 | 0 | 0 | 0 | 0 |

| | | | | | | |
|---|---|---|---|---|---|---|
| ENSG00000104237 | 2 | 0 | 0 | 0 | 0 | 0 |
| ENSG00000226098 | 2 | 0 | 1 | 0 | 0 | 0 |
| ENSG00000253114 | 0 | 0 | 0 | 0 | 0 | 0 |
| ENSG00000206579 | 0 | 0 | 0 | 0 | 0 | 0 |
| ENSG00000253976 | 0 | 0 | 0 | 0 | 0 | 0 |
| ENSG00000253857 | 0 | 0 | 0 | 0 | 0 | 0 |
| ENSG00000248522 | 0 | 0 | 0 | 0 | 0 | 0 |
| ENSG00000254357 | 0 | 0 | 0 | 0 | 0 | 0 |
| ENSG00000167904 | 616 | 277 | 357 | 250 | 343 | 343 |
| ENSG00000222955 | 0 | 0 | 0 | 0 | 0 | 0 |
| ENSG00000137574 | 649 | 334 | 410 | 253 | 374 | 323 |
| ENSG00000254087 | 474 | 214 | 283 | 180 | 247 | 185 |
| ENSG00000254325 | 0 | 0 | 0 | 0 | 0 | 0 |
| ENSG00000199405 | 0 | 0 | 0 | 0 | 0 | 0 |
| ENSG00000240905 | 0 | 0 | 0 | 0 | 0 | 0 |
| ENSG00000241997 | 0 | 0 | 0 | 0 | 0 | 0 |
| ENSG00000236814 | 6 | 6 | 5 | 4 | 4 | 2 |
| ENSG00000008988 | 19416 | 9738 | 11330 | 13126 | 11671 | 11040 |
| ENSG00000238650 | 0 | 0 | 0 | 0 | 0 | 0 |
| ENSG00000253603 | 0 | 0 | 0 | 0 | 0 | 0 |
| ENSG00000248578 | 4 | 0 | 2 | 1 | 0 | 0 |
| ENSG00000172680 | 0 | 0 | 0 | 0 | 0 | 0 |
| ENSG00000221093 | 0 | 0 | 0 | 0 | 0 | 0 |
| ENSG00000181690 | 4 | 0 | 6 | 2 | 4 | 4 |
| ENSG00000170791 | 488 | 261 | 307 | 310 | 259 | 329 |
| ENSG00000272343 | 0 | 0 | 0 | 1 | 0 | 0 |
| ENSG00000254216 | 0 | 0 | 0 | 0 | 0 | 0 |
| ENSG00000170786 | 57 | 46 | 46 | 85 | 86 | 64 |
| ENSG00000253542 | 0 | 0 | 0 | 0 | 0 | 0 |
| ENSG00000181195 | 0 | 0 | 0 | 0 | 0 | 0 |
| ENSG00000254254 | 0 | 0 | 0 | 0 | 0 | 0 |
| ENSG00000253541 | 0 | 0 | 0 | 0 | 0 | 0 |
| ENSG00000246430 | 0 | 0 | 0 | 0 | 0 | 0 |
| ENSG00000253139 | 0 | 0 | 0 | 0 | 0 | 0 |
| ENSG00000241431 | 6 | 2 | 0 | 1 | 0 | 2 |
| ENSG00000254055 | 0 | 0 | 0 | 0 | 0 | 0 |
| ENSG00000270328 | 0 | 0 | 0 | 0 | 0 | 0 |
| ENSG00000206975 | 0 | 0 | 0 | 0 | 0 | 0 |
| ENSG00000104331 | 2629 | 1250 | 1586 | 1017 | 1454 | 1429 |
| ENSG00000222205 | 0 | 0 | 0 | 0 | 0 | 0 |
| ENSG00000253301 | 0 | 0 | 1 | 0 | 0 | 0 |
| ENSG00000251142 | 0 | 0 | 0 | 0 | 0 | 0 |
| ENSG00000253614 | 0 | 0 | 0 | 0 | 0 | 0 |
| ENSG00000215117 | 0 | 0 | 0 | 0 | 0 | 0 |
| ENSG00000202265 | 0 | 0 | 0 | 0 | 0 | 0 |
| ENSG00000253871 | 0 | 0 | 0 | 0 | 0 | 0 |
| ENSG00000243050 | 0 | 0 | 0 | 0 | 0 | 0 |
| ENSG00000253821 | 2 | 0 | 1 | 0 | 1 | 0 |
| ENSG00000253322 | 0 | 0 | 0 | 0 | 0 | 0 |
| ENSG00000254139 | 0 | 0 | 0 | 0 | 0 | 0 |

| | | | | | | |
|---|---|---|---|---|---|---|
| ENSG00000205293 | 0 | 0 | 0 | 0 | 1 | 0 |
| ENSG00000253376 | 0 | 0 | 0 | 0 | 0 | 0 |
| ENSG00000169122 | 360 | 166 | 220 | 255 | 259 | 267 |
| ENSG00000253523 | 0 | 0 | 0 | 0 | 0 | 0 |
| ENSG00000253116 | 0 | 3 | 1 | 0 | 4 | 2 |
| ENSG00000242595 | 0 | 0 | 1 | 0 | 0 | 0 |
| ENSG00000253281 | 0 | 0 | 0 | 0 | 0 | 0 |
| ENSG00000215114 | 1389 | 658 | 712 | 469 | 788 | 704 |
| ENSG00000250031 | 2 | 1 | 2 | 0 | 3 | 1 |
| ENSG00000167910 | 0 | 0 | 0 | 0 | 2 | 0 |
| ENSG00000254103 | 0 | 0 | 0 | 0 | 0 | 0 |
| ENSG00000137575 | 3120 | 1880 | 1694 | 1296 | 2559 | 1670 |
| ENSG00000035681 | 1622 | 844 | 1087 | 815 | 1001 | 1001 |
| ENSG00000242970 | 0 | 0 | 1 | 3 | 0 | 0 |
| ENSG00000198846 | 0 | 0 | 0 | 0 | 0 | 0 |
| ENSG00000201231 | 0 | 0 | 0 | 0 | 0 | 0 |
| ENSG00000254048 | 0 | 0 | 0 | 0 | 0 | 0 |
| ENSG00000167912 | 0 | 0 | 0 | 0 | 0 | 0 |
| ENSG00000206853 | 0 | 0 | 0 | 0 | 0 | 0 |
| ENSG00000201763 | 0 | 0 | 0 | 0 | 0 | 0 |
| ENSG00000254150 | 0 | 0 | 0 | 0 | 0 | 0 |
| ENSG00000253260 | 0 | 0 | 0 | 0 | 0 | 0 |
| ENSG00000253879 | 0 | 0 | 0 | 0 | 0 | 0 |
| ENSG00000253413 | 0 | 0 | 0 | 0 | 0 | 0 |
| ENSG00000253723 | 0 | 0 | 0 | 0 | 0 | 0 |
| ENSG00000278428 | 0 | 0 | 0 | 0 | 0 | 0 |
| ENSG00000254775 | 0 | 0 | 0 | 0 | 0 | 0 |
| ENSG00000254954 | 0 | 0 | 0 | 0 | 0 | 0 |
| ENSG00000178538 | 154 | 105 | 144 | 164 | 184 | 202 |
| ENSG00000251396 | 5 | 5 | 6 | 4 | 2 | 2 |
| ENSG00000226301 | 0 | 0 | 0 | 0 | 0 | 0 |
| ENSG00000104388 | 3944 | 1970 | 2337 | 1931 | 2319 | 2200 |
| ENSG00000255321 | 0 | 1 | 0 | 0 | 0 | 0 |
| ENSG00000228862 | 0 | 1 | 0 | 4 | 6 | 0 |
| ENSG00000255289 | 0 | 0 | 0 | 0 | 0 | 0 |
| ENSG00000171316 | 1080 | 541 | 563 | 268 | 702 | 588 |
| ENSG00000254432 | 1 | 0 | 0 | 0 | 0 | 0 |
| ENSG00000271377 | 0 | 0 | 0 | 0 | 0 | 0 |
| ENSG00000254777 | 0 | 0 | 1 | 0 | 0 | 0 |
| ENSG00000255397 | 4 | 0 | 3 | 3 | 2 | 0 |
| ENSG00000254802 | 0 | 0 | 0 | 0 | 0 | 0 |
| ENSG00000177182 | 0 | 0 | 0 | 0 | 0 | 0 |
| ENSG00000213881 | 4 | 11 | 7 | 10 | 6 | 15 |
| ENSG00000254222 | 0 | 0 | 0 | 0 | 0 | 0 |
| ENSG00000253711 | 0 | 0 | 0 | 0 | 0 | 0 |
| ENSG00000198363 | 1572 | 820 | 823 | 685 | 1141 | 938 |
| ENSG00000254285 | 246 | 153 | 161 | 124 | 100 | 122 |
| ENSG00000222898 | 0 | 0 | 1 | 0 | 1 | 1 |
| ENSG00000264408 | 0 | 0 | 0 | 0 | 0 | 0 |
| ENSG00000251127 | 0 | 0 | 0 | 0 | 0 | 0 |

| | | | | | | |
|---|---|---|---|---|---|---|
| ENSG00000254119 | 0 | 0 | 0 | 0 | 1 | 1 |
| ENSG00000253834 | 0 | 0 | 0 | 0 | 0 | 0 |
| ENSG00000254120 | 0 | 0 | 0 | 0 | 0 | 0 |
| ENSG00000253544 | 0 | 0 | 0 | 0 | 0 | 0 |
| ENSG00000253789 | 0 | 0 | 0 | 0 | 0 | 0 |
| ENSG00000237810 | 0 | 0 | 0 | 0 | 2 | 1 |
| ENSG00000285971 | 0 | 0 | 0 | 0 | 0 | 0 |
| ENSG00000185942 | 0 | 0 | 0 | 0 | 0 | 0 |
| ENSG00000254050 | 0 | 0 | 0 | 0 | 0 | 0 |
| ENSG00000253975 | 0 | 0 | 0 | 0 | 0 | 0 |
| ENSG00000253517 | 0 | 0 | 0 | 0 | 0 | 0 |
| ENSG00000253431 | 0 | 0 | 1 | 1 | 0 | 0 |
| ENSG00000274956 | 0 | 0 | 0 | 0 | 0 | 0 |
| ENSG00000240915 | 0 | 0 | 0 | 0 | 0 | 1 |
| ENSG00000137563 | 728 | 366 | 385 | 387 | 502 | 406 |
| ENSG00000253121 | 0 | 0 | 0 | 0 | 0 | 1 |
| ENSG00000137561 | 2 | 3 | 2 | 1 | 3 | 1 |
| ENSG00000253555 | 0 | 0 | 0 | 0 | 0 | 0 |
| ENSG00000270673 | 56 | 42 | 45 | 34 | 37 | 48 |
| ENSG00000185728 | 2113 | 995 | 1063 | 543 | 1150 | 945 |
| ENSG00000261542 | 10 | 2 | 0 | 2 | 12 | 2 |
| ENSG00000241964 | 0 | 0 | 0 | 0 | 0 | 0 |
| ENSG00000253205 | 0 | 0 | 0 | 0 | 0 | 0 |
| ENSG00000215096 | 0 | 0 | 0 | 0 | 0 | 0 |
| ENSG00000253894 | 0 | 0 | 0 | 0 | 0 | 0 |
| ENSG00000253583 | 0 | 0 | 0 | 0 | 0 | 0 |
| ENSG00000252358 | 0 | 0 | 0 | 0 | 0 | 0 |
| ENSG00000253554 | 0 | 0 | 0 | 0 | 0 | 1 |
| ENSG00000253762 | 0 | 0 | 0 | 0 | 0 | 0 |
| ENSG00000253734 | 0 | 0 | 0 | 0 | 0 | 0 |
| ENSG00000253231 | 0 | 0 | 0 | 0 | 0 | 0 |
| ENSG00000254377 | 0 | 0 | 0 | 0 | 0 | 0 |
| ENSG00000207816 | 0 | 0 | 0 | 0 | 0 | 0 |
| ENSG00000253750 | 0 | 0 | 0 | 0 | 0 | 0 |
| ENSG00000253800 | 0 | 0 | 0 | 0 | 0 | 0 |
| ENSG00000254102 | 0 | 0 | 0 | 0 | 0 | 0 |
| ENSG00000180828 | 0 | 0 | 0 | 0 | 0 | 0 |
| ENSG00000172817 | 0 | 0 | 0 | 0 | 0 | 0 |
| ENSG00000254330 | 0 | 0 | 0 | 0 | 0 | 0 |
| ENSG00000254006 | 0 | 0 | 0 | 0 | 0 | 0 |
| ENSG00000253493 | 0 | 0 | 0 | 0 | 0 | 0 |
| ENSG00000239261 | 0 | 0 | 0 | 0 | 0 | 0 |
| ENSG00000280725 | 0 | 0 | 0 | 0 | 0 | 0 |
| ENSG00000213873 | 0 | 0 | 0 | 0 | 0 | 0 |
| ENSG00000254081 | 0 | 0 | 0 | 0 | 0 | 0 |
| ENSG00000272010 | 2 | 2 | 2 | 0 | 0 | 1 |
| ENSG00000200714 | 2 | 0 | 2 | 2 | 2 | 3 |
| ENSG00000104442 | 1546 | 778 | 995 | 732 | 958 | 900 |
| ENSG00000066855 | 1866 | 864 | 1093 | 879 | 1015 | 952 |
| ENSG00000272155 | 6 | 1 | 2 | 6 | 2 | 6 |

| | | | | | | |
|---|---|---|---|---|---|---|
| ENSG00000205268 | 1128 | 633 | 703 | 459 | 660 | 545 |
| ENSG00000272192 | 2 | 0 | 1 | 1 | 3 | 0 |
| ENSG00000147570 | 0 | 1 | 0 | 1 | 0 | 1 |
| ENSG00000253190 | 0 | 0 | 0 | 0 | 0 | 0 |
| ENSG00000147573 | 0 | 0 | 0 | 1 | 1 | 0 |
| ENSG00000147571 | 0 | 0 | 2 | 1 | 0 | 1 |
| ENSG00000253138 | 0 | 0 | 0 | 0 | 0 | 1 |
| ENSG00000246145 | 2 | 2 | 4 | 2 | 7 | 4 |
| ENSG00000179041 | 897 | 511 | 574 | 718 | 679 | 725 |
| ENSG00000285655 | 0 | 0 | 0 | 0 | 0 | 0 |
| ENSG00000285791 | 0 | 2 | 2 | 3 | 3 | 2 |
| ENSG00000147576 | 57 | 27 | 53 | 26 | 25 | 21 |
| ENSG00000253623 | 0 | 0 | 0 | 0 | 0 | 0 |
| ENSG00000169085 | 1 | 3 | 1 | 1 | 0 | 0 |
| ENSG00000206949 | 0 | 0 | 0 | 0 | 0 | 0 |
| ENSG00000185697 | 168 | 72 | 69 | 59 | 118 | 82 |
| ENSG00000175073 | 648 | 358 | 380 | 235 | 412 | 317 |
| ENSG00000201365 | 0 | 0 | 0 | 0 | 0 | 0 |
| ENSG00000104205 | 235 | 93 | 133 | 44 | 74 | 76 |
| ENSG00000213865 | 120 | 61 | 75 | 66 | 86 | 62 |
| ENSG00000213005 | 0 | 0 | 0 | 0 | 0 | 0 |
| ENSG00000178460 | 94 | 47 | 46 | 32 | 42 | 41 |
| ENSG00000245910 | 982 | 591 | 693 | 871 | 739 | 762 |
| ENSG00000254341 | 0 | 0 | 0 | 0 | 0 | 0 |
| ENSG00000261787 | 114 | 36 | 35 | 37 | 51 | 43 |
| ENSG00000178125 | 8 | 0 | 2 | 2 | 2 | 1 |
| ENSG00000253223 | 0 | 2 | 0 | 0 | 0 | 0 |
| ENSG00000121022 | 2051 | 1189 | 1574 | 1100 | 1424 | 1360 |
| ENSG00000104218 | 373 | 204 | 185 | 82 | 214 | 161 |
| ENSG00000252637 | 0 | 0 | 0 | 0 | 0 | 0 |
| ENSG00000250869 | 0 | 0 | 0 | 0 | 0 | 0 |
| ENSG00000066777 | 1213 | 797 | 609 | 210 | 1077 | 647 |
| ENSG00000271966 | 37 | 26 | 43 | 21 | 26 | 29 |
| ENSG00000241961 | 0 | 0 | 0 | 0 | 0 | 0 |
| ENSG00000165078 | 12 | 10 | 4 | 1 | 4 | 2 |
| ENSG00000254253 | 0 | 0 | 0 | 0 | 0 | 0 |
| ENSG00000255130 | 0 | 0 | 0 | 0 | 0 | 0 |
| ENSG00000255531 | 0 | 0 | 0 | 0 | 0 | 0 |
| ENSG00000046889 | 0 | 0 | 0 | 0 | 0 | 0 |
| ENSG00000255206 | 0 | 0 | 0 | 0 | 0 | 0 |
| ENSG00000254051 | 0 | 0 | 0 | 0 | 0 | 0 |
| ENSG00000252210 | 0 | 0 | 0 | 0 | 0 | 0 |
| ENSG00000253964 | 0 | 0 | 0 | 0 | 0 | 0 |
| ENSG00000248801 | 0 | 0 | 0 | 0 | 0 | 0 |
| ENSG00000165084 | 0 | 0 | 0 | 0 | 0 | 0 |
| ENSG00000236191 | 0 | 0 | 0 | 0 | 0 | 0 |
| ENSG00000239184 | 0 | 0 | 0 | 0 | 0 | 0 |
| ENSG00000254337 | 60 | 34 | 42 | 57 | 48 | 54 |
| ENSG00000253658 | 0 | 0 | 0 | 0 | 0 | 0 |
| ENSG00000238808 | 0 | 0 | 0 | 0 | 0 | 0 |

| | | | | | | |
|---|---|---|---|---|---|---|
| ENSG00000254332 | 4 | 8 | 9 | 10 | 8 | 4 |
| ENSG00000253373 | 0 | 0 | 0 | 0 | 0 | 0 |
| ENSG00000253479 | 0 | 0 | 0 | 1 | 0 | 0 |
| ENSG00000253216 | 0 | 0 | 0 | 0 | 0 | 0 |
| ENSG00000137573 | 99 | 68 | 78 | 55 | 92 | 72 |
| ENSG00000137571 | 0 | 1 | 0 | 2 | 3 | 1 |
| ENSG00000222889 | 0 | 0 | 0 | 0 | 0 | 0 |
| ENSG00000254557 | 0 | 0 | 0 | 0 | 0 | 0 |
| ENSG00000271113 | 0 | 0 | 0 | 0 | 0 | 0 |
| ENSG00000246528 | 2 | 1 | 0 | 0 | 1 | 0 |
| ENSG00000277740 | 0 | 0 | 0 | 0 | 0 | 0 |
| ENSG00000255107 | 0 | 0 | 0 | 0 | 0 | 0 |
| ENSG00000239580 | 0 | 0 | 0 | 0 | 0 | 0 |
| ENSG00000223293 | 0 | 0 | 0 | 0 | 0 | 0 |
| ENSG00000280393 | 0 | 0 | 0 | 0 | 0 | 0 |
| ENSG00000247570 | 0 | 0 | 0 | 0 | 1 | 0 |
| ENSG00000253721 | 0 | 0 | 0 | 0 | 1 | 1 |
| ENSG00000147596 | 0 | 0 | 1 | 0 | 0 | 0 |
| ENSG00000212473 | 0 | 0 | 0 | 0 | 0 | 0 |
| ENSG00000253173 | 2 | 0 | 1 | 2 | 0 | 0 |
| ENSG00000140396 | 1670 | 876 | 831 | 259 | 995 | 709 |
| ENSG00000213003 | 0 | 1 | 0 | 0 | 0 | 0 |
| ENSG00000207036 | 0 | 0 | 0 | 0 | 0 | 1 |
| ENSG00000241204 | 0 | 0 | 0 | 0 | 0 | 0 |
| ENSG00000253143 | 0 | 0 | 0 | 0 | 0 | 0 |
| ENSG00000253967 | 0 | 0 | 0 | 0 | 0 | 0 |
| ENSG00000275128 | 0 | 0 | 0 | 0 | 0 | 0 |
| ENSG00000241105 | 0 | 0 | 0 | 0 | 0 | 0 |
| ENSG00000067167 | 4617 | 2208 | 2459 | 2062 | 2621 | 2342 |
| ENSG00000246366 | 4 | 3 | 3 | 1 | 6 | 1 |
| ENSG00000147592 | 473 | 280 | 348 | 305 | 410 | 352 |
| ENSG00000243532 | 0 | 0 | 0 | 0 | 0 | 0 |
| ENSG00000221947 | 37 | 31 | 36 | 23 | 20 | 27 |
| ENSG00000285579 | 0 | 0 | 0 | 0 | 1 | 0 |
| ENSG00000223220 | 0 | 0 | 0 | 0 | 2 | 1 |
| ENSG00000254031 | 0 | 0 | 0 | 0 | 0 | 0 |
| ENSG00000104313 | 2 | 0 | 0 | 0 | 1 | 1 |
| ENSG00000253124 | 0 | 0 | 0 | 0 | 0 | 0 |
| ENSG00000254277 | 0 | 0 | 0 | 0 | 0 | 0 |
| ENSG00000253287 | 0 | 0 | 0 | 0 | 0 | 0 |
| ENSG00000200191 | 0 | 0 | 0 | 0 | 0 | 0 |
| ENSG00000235531 | 0 | 0 | 0 | 0 | 0 | 0 |
| ENSG00000178860 | 0 | 0 | 0 | 0 | 0 | 0 |
| ENSG00000241942 | 0 | 0 | 0 | 0 | 0 | 0 |
| ENSG00000104321 | 0 | 0 | 0 | 0 | 0 | 0 |
| ENSG00000253317 | 0 | 0 | 0 | 0 | 0 | 0 |
| ENSG00000250979 | 0 | 0 | 0 | 0 | 0 | 0 |
| ENSG00000252509 | 0 | 0 | 0 | 0 | 0 | 0 |
| ENSG00000212989 | 0 | 0 | 0 | 0 | 0 | 0 |
| ENSG00000254347 | 0 | 0 | 0 | 0 | 0 | 0 |

| | | | | | | |
|---|---|---|---|---|---|---|
| ENSG00000182674 | 0 | 0 | 0 | 0 | 0 | 0 |
| ENSG00000253226 | 0 | 0 | 0 | 0 | 0 | 0 |
| ENSG00000278193 | 0 | 0 | 0 | 0 | 0 | 0 |
| ENSG00000253726 | 0 | 0 | 0 | 0 | 0 | 0 |
| ENSG00000253784 | 0 | 0 | 0 | 0 | 0 | 0 |
| ENSG00000260838 | 0 | 0 | 0 | 0 | 0 | 0 |
| ENSG00000147601 | 695 | 359 | 356 | 297 | 485 | 363 |
| ENSG00000222488 | 1 | 0 | 0 | 0 | 0 | 0 |
| ENSG00000272254 | 0 | 0 | 1 | 0 | 0 | 1 |
| ENSG00000253636 | 7 | 7 | 4 | 2 | 5 | 3 |
| ENSG00000164764 | 18 | 5 | 6 | 4 | 10 | 7 |
| ENSG00000237061 | 0 | 0 | 0 | 0 | 0 | 0 |
| ENSG00000253564 | 0 | 0 | 0 | 0 | 0 | 0 |
| ENSG00000274443 | 0 | 0 | 0 | 0 | 0 | 0 |
| ENSG00000254213 | 0 | 0 | 0 | 0 | 0 | 0 |
| ENSG00000147604 | 25002 | 14232 | 15352 | 16452 | 19007 | 14770 |
| ENSG00000121039 | 629 | 316 | 449 | 352 | 469 | 478 |
| ENSG00000250295 | 5 | 3 | 5 | 0 | 5 | 7 |
| ENSG00000253339 | 0 | 0 | 0 | 0 | 0 | 0 |
| ENSG00000253235 | 0 | 0 | 0 | 0 | 0 | 0 |
| ENSG00000253302 | 7 | 1 | 1 | 1 | 0 | 0 |
| ENSG00000040341 | 1621 | 813 | 886 | 571 | 1032 | 690 |
| ENSG00000254273 | 0 | 0 | 0 | 0 | 0 | 0 |
| ENSG00000253383 | 0 | 0 | 0 | 0 | 0 | 0 |
| ENSG00000254105 | 0 | 0 | 0 | 0 | 0 | 0 |
| ENSG00000254538 | 1 | 1 | 2 | 0 | 0 | 0 |
| ENSG00000258677 | 0 | 0 | 0 | 0 | 0 | 0 |
| ENSG00000104343 | 1017 | 559 | 546 | 452 | 685 | 546 |
| ENSG00000253273 | 0 | 0 | 0 | 0 | 0 | 0 |
| ENSG00000272469 | 0 | 0 | 0 | 0 | 0 | 0 |
| ENSG00000251633 | 0 | 0 | 1 | 0 | 0 | 0 |
| ENSG00000154582 | 1884 | 962 | 1137 | 1035 | 1298 | 1154 |
| ENSG00000175606 | 885 | 476 | 472 | 510 | 491 | 491 |
| ENSG00000244295 | 0 | 0 | 0 | 0 | 0 | 0 |
| ENSG00000253833 | 2 | 0 | 0 | 1 | 3 | 1 |
| ENSG00000154589 | 1 | 0 | 0 | 0 | 1 | 1 |
| ENSG00000212348 | 0 | 0 | 0 | 0 | 0 | 0 |
| ENSG00000254288 | 0 | 0 | 0 | 0 | 0 | 0 |
| ENSG00000240047 | 0 | 0 | 0 | 0 | 0 | 0 |
| ENSG00000212485 | 0 | 0 | 0 | 0 | 0 | 0 |
| ENSG00000253115 | 0 | 0 | 0 | 0 | 0 | 0 |
| ENSG00000253983 | 0 | 0 | 0 | 0 | 0 | 0 |
| ENSG00000104369 | 286 | 127 | 173 | 113 | 151 | 148 |
| ENSG00000104381 | 329 | 184 | 233 | 145 | 215 | 225 |
| ENSG00000253596 | 0 | 0 | 0 | 0 | 0 | 0 |
| ENSG00000207417 | 0 | 0 | 0 | 0 | 0 | 0 |
| ENSG00000266502 | 0 | 0 | 0 | 0 | 0 | 0 |
| ENSG00000283505 | 0 | 0 | 0 | 0 | 0 | 0 |
| ENSG00000254349 | 0 | 0 | 0 | 0 | 0 | 0 |
| ENSG00000253341 | 24 | 8 | 16 | 15 | 12 | 46 |

| | | | | | | |
|---|---|---|---|---|---|---|
| ENSG00000254080 | 0 | 0 | 0 | 0 | 0 | 0 |
| ENSG00000283188 | 0 | 0 | 0 | 0 | 0 | 0 |
| ENSG00000253706 | 0 | 0 | 1 | 0 | 0 | 1 |
| ENSG00000253777 | 0 | 0 | 0 | 0 | 0 | 0 |
| ENSG00000137558 | 2 | 1 | 0 | 0 | 0 | 0 |
| ENSG00000254043 | 0 | 0 | 0 | 0 | 0 | 0 |
| ENSG00000121005 | 0 | 0 | 0 | 0 | 0 | 0 |
| ENSG00000254238 | 0 | 0 | 0 | 0 | 0 | 0 |
| ENSG00000253712 | 0 | 2 | 0 | 0 | 1 | 0 |
| ENSG00000249395 | 8 | 1 | 3 | 3 | 3 | 5 |
| ENSG00000253229 | 0 | 0 | 0 | 0 | 0 | 0 |
| ENSG00000254023 | 0 | 0 | 0 | 0 | 0 | 0 |
| ENSG00000164749 | 12 | 1 | 5 | 0 | 4 | 4 |
| ENSG00000222231 | 0 | 0 | 0 | 0 | 0 | 0 |
| ENSG00000270866 | 0 | 0 | 0 | 0 | 0 | 0 |
| ENSG00000271167 | 0 | 0 | 0 | 0 | 0 | 0 |
| ENSG00000254300 | 0 | 0 | 0 | 0 | 0 | 0 |
| ENSG00000253661 | 0 | 0 | 0 | 0 | 0 | 0 |
| ENSG00000253681 | 0 | 0 | 0 | 0 | 0 | 0 |
| ENSG00000254305 | 0 | 0 | 0 | 0 | 0 | 0 |
| ENSG00000091656 | 0 | 0 | 0 | 0 | 0 | 0 |
| ENSG00000276673 | 0 | 0 | 0 | 0 | 0 | 0 |
| ENSG00000253416 | 0 | 0 | 0 | 0 | 0 | 0 |
| ENSG00000266712 | 0 | 0 | 0 | 0 | 0 | 1 |
| ENSG00000164751 | 1684 | 815 | 993 | 836 | 991 | 917 |
| ENSG00000253952 | 0 | 0 | 0 | 0 | 0 | 0 |
| ENSG00000254366 | 0 | 0 | 0 | 0 | 0 | 0 |
| ENSG00000254001 | 0 | 0 | 0 | 0 | 0 | 0 |
| ENSG00000252935 | 0 | 0 | 0 | 0 | 0 | 0 |
| ENSG00000254266 | 6 | 0 | 1 | 0 | 3 | 2 |
| ENSG00000171033 | 805 | 393 | 450 | 247 | 368 | 311 |
| ENSG00000260398 | 4 | 4 | 4 | 2 | 2 | 0 |
| ENSG00000104427 | 374 | 158 | 150 | 87 | 229 | 167 |
| ENSG00000104432 | 1 | 0 | 1 | 1 | 1 | 0 |
| ENSG00000254352 | 6 | 1 | 2 | 0 | 3 | 1 |
| ENSG00000253919 | 12 | 10 | 5 | 3 | 4 | 3 |
| ENSG00000285744 | 2 | 0 | 0 | 0 | 2 | 1 |
| ENSG00000261618 | 2 | 1 | 0 | 0 | 0 | 0 |
| ENSG00000253659 | 0 | 0 | 0 | 0 | 0 | 0 |
| ENSG00000253391 | 0 | 0 | 0 | 0 | 0 | 0 |
| ENSG00000240540 | 0 | 0 | 0 | 1 | 0 | 1 |
| ENSG00000104435 | 0 | 0 | 0 | 0 | 0 | 0 |
| ENSG00000164683 | 70 | 42 | 38 | 65 | 76 | 70 |
| ENSG00000272138 | 20 | 14 | 16 | 26 | 16 | 21 |
| ENSG00000249328 | 2 | 2 | 0 | 0 | 0 | 1 |
| ENSG00000238884 | 0 | 0 | 0 | 0 | 0 | 0 |
| ENSG00000285758 | 0 | 0 | 2 | 0 | 0 | 0 |
| ENSG00000147586 | 165 | 162 | 185 | 142 | 189 | 201 |
| ENSG00000276418 | 0 | 0 | 2 | 0 | 1 | 1 |
| ENSG00000272518 | 27 | 12 | 12 | 17 | 10 | 16 |

| | | | | | | |
|---|---|---|---|---|---|---|
| ENSG00000272264 | 16 | 7 | 12 | 0 | 6 | 6 |
| ENSG00000076554 | 3105 | 1721 | 1860 | 1300 | 1534 | 1530 |
| ENSG00000254205 | 0 | 0 | 0 | 0 | 0 | 0 |
| ENSG00000241550 | 0 | 1 | 0 | 0 | 0 | 0 |
| ENSG00000265588 | 0 | 0 | 0 | 0 | 0 | 0 |
| ENSG00000241746 | 0 | 0 | 0 | 0 | 1 | 1 |
| ENSG00000253238 | 6 | 1 | 6 | 1 | 1 | 2 |
| ENSG00000213791 | 0 | 0 | 0 | 1 | 0 | 0 |
| ENSG00000206649 | 0 | 0 | 0 | 0 | 0 | 0 |
| ENSG00000253237 | 0 | 0 | 0 | 0 | 0 | 0 |
| ENSG00000252884 | 0 | 0 | 0 | 0 | 0 | 0 |
| ENSG00000251867 | 57 | 15 | 47 | 20 | 34 | 28 |
| ENSG00000277604 | 0 | 0 | 0 | 0 | 0 | 0 |
| ENSG00000205189 | 1034 | 503 | 631 | 278 | 547 | 422 |
| ENSG00000254162 | 6 | 4 | 7 | 0 | 3 | 6 |
| ENSG00000260317 | 18 | 7 | 7 | 5 | 7 | 16 |
| ENSG00000188856 | 532 | 278 | 273 | 360 | 313 | 300 |
| ENSG00000252057 | 0 | 0 | 0 | 0 | 0 | 2 |
| ENSG00000253760 | 0 | 0 | 0 | 0 | 0 | 0 |
| ENSG00000223327 | 0 | 0 | 0 | 0 | 0 | 0 |
| ENSG00000254181 | 0 | 0 | 0 | 0 | 0 | 0 |
| ENSG00000243424 | 0 | 0 | 0 | 0 | 0 | 0 |
| ENSG00000164684 | 1059 | 478 | 514 | 213 | 469 | 375 |
| ENSG00000254331 | 0 | 0 | 0 | 0 | 1 | 0 |
| ENSG00000212429 | 0 | 0 | 0 | 0 | 0 | 0 |
| ENSG00000243951 | 0 | 0 | 0 | 0 | 0 | 0 |
| ENSG00000253516 | 0 | 0 | 1 | 3 | 0 | 1 |
| ENSG00000076641 | 44 | 17 | 29 | 7 | 11 | 10 |
| ENSG00000254060 | 0 | 0 | 0 | 0 | 0 | 0 |
| ENSG00000254177 | 0 | 0 | 0 | 0 | 0 | 0 |
| ENSG00000253214 | 0 | 0 | 0 | 0 | 0 | 0 |
| ENSG00000254014 | 0 | 0 | 0 | 0 | 0 | 0 |
| ENSG00000253677 | 0 | 0 | 1 | 0 | 1 | 0 |
| ENSG00000272425 | 0 | 0 | 0 | 1 | 0 | 2 |
| ENSG00000254027 | 0 | 1 | 1 | 1 | 0 | 2 |
| ENSG00000164687 | 19 | 13 | 12 | 20 | 26 | 17 |
| ENSG00000253859 | 0 | 0 | 0 | 0 | 0 | 0 |
| ENSG00000147588 | 0 | 0 | 0 | 0 | 0 | 0 |
| ENSG00000205186 | 0 | 0 | 0 | 0 | 0 | 0 |
| ENSG00000170323 | 0 | 0 | 0 | 0 | 0 | 0 |
| ENSG00000253374 | 0 | 0 | 0 | 0 | 0 | 0 |
| ENSG00000237264 | 3 | 1 | 2 | 2 | 2 | 3 |
| ENSG00000197416 | 0 | 0 | 0 | 0 | 0 | 0 |
| ENSG00000251521 | 0 | 0 | 0 | 0 | 0 | 0 |
| ENSG00000254151 | 0 | 0 | 0 | 0 | 0 | 0 |
| ENSG00000240427 | 0 | 0 | 0 | 0 | 0 | 0 |
| ENSG00000205184 | 0 | 0 | 0 | 0 | 0 | 0 |
| ENSG00000133731 | 259 | 128 | 149 | 115 | 185 | 145 |
| ENSG00000253598 | 25 | 3 | 7 | 5 | 10 | 4 |
| ENSG00000280193 | 9 | 11 | 4 | 5 | 9 | 11 |

| | | | | | | |
|---|---|---|---|---|---|---|
| ENSG00000104231 | 1334 | 787 | 815 | 685 | 1074 | 858 |
| ENSG00000164695 | 1002 | 506 | 582 | 406 | 600 | 554 |
| ENSG00000253334 | 0 | 0 | 0 | 0 | 0 | 0 |
| ENSG00000104497 | 548 | 294 | 291 | 129 | 308 | 230 |
| ENSG00000231942 | 2 | 0 | 1 | 0 | 1 | 1 |
| ENSG00000254689 | 2 | 0 | 0 | 0 | 1 | 3 |
| ENSG00000254123 | 0 | 0 | 0 | 0 | 0 | 0 |
| ENSG00000206228 | 1 | 0 | 2 | 0 | 3 | 0 |
| ENSG00000253503 | 0 | 2 | 0 | 0 | 0 | 0 |
| ENSG00000253836 | 0 | 0 | 0 | 0 | 0 | 0 |
| ENSG00000254394 | 0 | 0 | 0 | 0 | 0 | 0 |
| ENSG00000253581 | 0 | 0 | 0 | 0 | 0 | 0 |
| ENSG00000254189 | 0 | 0 | 0 | 0 | 0 | 0 |
| ENSG00000253423 | 0 | 0 | 0 | 0 | 0 | 0 |
| ENSG00000274116 | 0 | 0 | 0 | 0 | 1 | 0 |
| ENSG00000253898 | 2 | 0 | 0 | 0 | 0 | 0 |
| ENSG00000254202 | 0 | 0 | 0 | 0 | 0 | 0 |
| ENSG00000254195 | 0 | 0 | 0 | 0 | 0 | 0 |
| ENSG00000253362 | 0 | 0 | 0 | 0 | 0 | 0 |
| ENSG00000184672 | 0 | 0 | 0 | 0 | 0 | 0 |
| ENSG00000206701 | 0 | 0 | 0 | 0 | 0 | 0 |
| ENSG00000253137 | 0 | 0 | 0 | 0 | 0 | 0 |
| ENSG00000248978 | 0 | 0 | 0 | 0 | 0 | 0 |
| ENSG00000254212 | 0 | 0 | 0 | 0 | 0 | 0 |
| ENSG00000203413 | 0 | 0 | 0 | 0 | 0 | 0 |
| ENSG00000253655 | 0 | 0 | 0 | 0 | 0 | 0 |
| ENSG00000253283 | 0 | 0 | 1 | 0 | 2 | 1 |
| ENSG00000133739 | 50 | 38 | 30 | 11 | 65 | 33 |
| ENSG00000260493 | 7 | 7 | 9 | 3 | 7 | 4 |
| ENSG00000133740 | 673 | 335 | 453 | 405 | 495 | 441 |
| ENSG00000254208 | 4 | 2 | 3 | 1 | 3 | 1 |
| ENSG00000176731 | 618 | 381 | 436 | 394 | 493 | 451 |
| ENSG00000185015 | 32 | 8 | 18 | 12 | 10 | 8 |
| ENSG00000258256 | 0 | 0 | 0 | 1 | 0 | 1 |
| ENSG00000257962 | 0 | 0 | 0 | 0 | 0 | 0 |
| ENSG00000133742 | 0 | 0 | 0 | 0 | 0 | 0 |
| ENSG00000164879 | 2 | 0 | 0 | 0 | 0 | 0 |
| ENSG00000253549 | 59 | 28 | 29 | 20 | 24 | 16 |
| ENSG00000104267 | 1995 | 763 | 885 | 511 | 537 | 609 |
| ENSG00000254380 | 0 | 0 | 0 | 0 | 0 | 0 |
| ENSG00000253680 | 0 | 0 | 0 | 0 | 0 | 0 |
| ENSG00000270971 | 0 | 0 | 0 | 0 | 0 | 0 |
| ENSG00000273567 | 0 | 0 | 0 | 0 | 0 | 0 |
| ENSG00000205176 | 0 | 0 | 0 | 0 | 0 | 0 |
| ENSG00000276006 | 0 | 0 | 0 | 0 | 0 | 0 |
| ENSG00000275928 | 0 | 0 | 0 | 0 | 0 | 0 |
| ENSG00000276540 | 0 | 0 | 0 | 0 | 0 | 0 |
| ENSG00000271381 | 0 | 0 | 0 | 0 | 0 | 0 |
| ENSG00000276998 | 0 | 0 | 0 | 0 | 0 | 0 |
| ENSG00000274343 | 0 | 0 | 0 | 0 | 0 | 0 |

| | | | | | | |
|---|---|---|---|---|---|---|
| ENSG00000270416 | 0 | 0 | 0 | 0 | 0 | 0 |
| ENSG00000277436 | 0 | 0 | 0 | 0 | 0 | 0 |
| ENSG00000277422 | 0 | 0 | 0 | 0 | 0 | 0 |
| ENSG00000253154 | 122 | 50 | 53 | 53 | 53 | 42 |
| ENSG00000285768 | 2 | 0 | 0 | 0 | 3 | 0 |
| ENSG00000147614 | 6 | 3 | 4 | 4 | 4 | 6 |
| ENSG00000227753 | 0 | 0 | 0 | 0 | 0 | 0 |
| ENSG00000243761 | 0 | 0 | 0 | 0 | 0 | 0 |
| ENSG00000147613 | 0 | 0 | 1 | 0 | 0 | 0 |
| ENSG00000253675 | 0 | 0 | 0 | 0 | 0 | 0 |
| ENSG00000253699 | 1 | 4 | 0 | 0 | 4 | 0 |
| ENSG00000164893 | 25 | 22 | 27 | 11 | 10 | 13 |
| ENSG00000254231 | 777 | 400 | 499 | 129 | 183 | 212 |
| ENSG00000123124 | 10533 | 5550 | 6185 | 2264 | 4793 | 3951 |
| ENSG00000176623 | 3189 | 1443 | 1819 | 1477 | 1816 | 1660 |
| ENSG00000250569 | 47 | 26 | 19 | 13 | 14 | 22 |
| ENSG00000254088 | 6 | 3 | 3 | 2 | 6 | 4 |
| ENSG00000085719 | 11355 | 5673 | 6554 | 4187 | 6481 | 5513 |
| ENSG00000170289 | 2 | 2 | 2 | 1 | 4 | 3 |
| ENSG00000253292 | 0 | 0 | 0 | 0 | 0 | 0 |
| ENSG00000205333 | 0 | 0 | 0 | 0 | 0 | 0 |
| ENSG00000233915 | 0 | 0 | 0 | 0 | 0 | 0 |
| ENSG00000254115 | 0 | 0 | 2 | 0 | 0 | 0 |
| ENSG00000253778 | 0 | 0 | 0 | 0 | 0 | 0 |
| ENSG00000253568 | 0 | 0 | 0 | 0 | 0 | 0 |
| ENSG00000176571 | 0 | 0 | 0 | 0 | 2 | 0 |
| ENSG00000253500 | 0 | 0 | 0 | 1 | 0 | 0 |
| ENSG00000253401 | 0 | 0 | 0 | 0 | 0 | 0 |
| ENSG00000251563 | 0 | 0 | 0 | 0 | 0 | 0 |
| ENSG00000254376 | 0 | 0 | 0 | 0 | 0 | 0 |
| ENSG00000273978 | 0 | 0 | 0 | 0 | 0 | 0 |
| ENSG00000176566 | 0 | 0 | 0 | 0 | 0 | 0 |
| ENSG00000253171 | 0 | 1 | 0 | 0 | 0 | 0 |
| ENSG00000156103 | 544 | 267 | 256 | 60 | 193 | 118 |
| ENSG00000253553 | 8 | 7 | 4 | 1 | 1 | 1 |
| ENSG00000251904 | 4 | 0 | 0 | 0 | 0 | 0 |
| ENSG00000250962 | 0 | 0 | 0 | 0 | 0 | 0 |
| ENSG00000271156 | 0 | 0 | 0 | 0 | 1 | 0 |
| ENSG00000253621 | 0 | 0 | 0 | 0 | 0 | 0 |
| ENSG00000248231 | 0 | 0 | 0 | 0 | 0 | 0 |
| ENSG00000251136 | 335 | 146 | 190 | 151 | 230 | 202 |
| ENSG00000104312 | 1692 | 919 | 1051 | 1033 | 1172 | 966 |
| ENSG00000253349 | 0 | 0 | 0 | 0 | 0 | 0 |
| ENSG00000275103 | 0 | 1 | 0 | 0 | 0 | 0 |
| ENSG00000207359 | 13 | 4 | 8 | 4 | 8 | 9 |
| ENSG00000164823 | 2786 | 1330 | 1572 | 1080 | 1614 | 1520 |
| ENSG00000104320 | 2881 | 1605 | 1500 | 943 | 2058 | 1421 |
| ENSG00000104325 | 4354 | 2384 | 2922 | 2370 | 2684 | 2621 |
| ENSG00000104327 | 8 | 0 | 2 | 0 | 0 | 0 |
| ENSG00000253394 | 3 | 11 | 15 | 9 | 11 | 8 |

| | | | | | | |
|---|---|---|---|---|---|---|
| ENSG00000199354 | 2 | 0 | 0 | 0 | 0 | 0 |
| ENSG00000254180 | 2 | 2 | 4 | 2 | 1 | 1 |
| ENSG00000253799 | 2 | 1 | 1 | 0 | 1 | 2 |
| ENSG00000180694 | 9019 | 4348 | 4921 | 3319 | 3607 | 3776 |
| ENSG00000246792 | 50 | 26 | 38 | 14 | 29 | 31 |
| ENSG00000254182 | 0 | 0 | 0 | 0 | 1 | 0 |
| ENSG00000123119 | 786 | 432 | 297 | 190 | 528 | 274 |
| ENSG00000254251 | 1109 | 594 | 601 | 499 | 505 | 488 |
| ENSG00000253250 | 3 | 4 | 3 | 1 | 5 | 3 |
| ENSG00000155099 | 136 | 95 | 104 | 96 | 112 | 105 |
| ENSG00000253358 | 0 | 0 | 0 | 0 | 0 | 0 |
| ENSG00000254219 | 0 | 0 | 0 | 0 | 1 | 0 |
| ENSG00000253738 | 686 | 390 | 367 | 319 | 593 | 378 |
| ENSG00000155100 | 606 | 323 | 288 | 175 | 433 | 278 |
| ENSG00000214954 | 8 | 7 | 8 | 9 | 22 | 7 |
| ENSG00000253525 | 0 | 0 | 0 | 0 | 0 | 0 |
| ENSG00000266194 | 0 | 0 | 0 | 0 | 0 | 0 |
| ENSG00000147606 | 3 | 1 | 0 | 0 | 1 | 0 |
| ENSG00000253901 | 0 | 1 | 0 | 0 | 0 | 1 |
| ENSG00000200151 | 0 | 0 | 0 | 0 | 0 | 0 |
| ENSG00000271202 | 0 | 0 | 0 | 0 | 0 | 0 |
| ENSG00000229182 | 0 | 0 | 0 | 0 | 0 | 0 |
| ENSG00000079102 | 10 | 5 | 14 | 2 | 9 | 5 |
| ENSG00000253576 | 0 | 0 | 0 | 0 | 0 | 0 |
| ENSG00000233778 | 0 | 0 | 0 | 0 | 0 | 0 |
| ENSG00000253118 | 0 | 0 | 0 | 0 | 0 | 0 |
| ENSG00000253682 | 0 | 0 | 0 | 0 | 0 | 0 |
| ENSG00000253749 | 0 | 0 | 0 | 0 | 0 | 0 |
| ENSG00000253634 | 0 | 0 | 0 | 0 | 0 | 0 |
| ENSG00000253177 | 0 | 0 | 0 | 0 | 0 | 0 |
| ENSG00000248858 | 22 | 5 | 8 | 5 | 13 | 6 |
| ENSG00000254020 | 0 | 0 | 0 | 0 | 0 | 0 |
| ENSG00000253577 | 0 | 0 | 0 | 0 | 0 | 0 |
| ENSG00000253197 | 1 | 2 | 1 | 0 | 0 | 1 |
| ENSG00000205133 | 3208 | 1616 | 1826 | 1127 | 2140 | 1644 |
| ENSG00000251347 | 0 | 0 | 0 | 0 | 0 | 0 |
| ENSG00000277063 | 0 | 0 | 0 | 0 | 0 | 0 |
| ENSG00000279847 | 0 | 0 | 0 | 0 | 0 | 3 |
| ENSG00000246662 | 4 | 3 | 1 | 2 | 1 | 0 |
| ENSG00000253960 | 0 | 0 | 0 | 0 | 0 | 0 |
| ENSG00000268080 | 1 | 0 | 0 | 1 | 0 | 0 |
| ENSG00000254089 | 0 | 0 | 0 | 0 | 0 | 0 |
| ENSG00000222416 | 0 | 0 | 0 | 0 | 0 | 0 |
| ENSG00000253781 | 0 | 0 | 0 | 0 | 0 | 0 |
| ENSG00000188343 | 350 | 177 | 208 | 125 | 252 | 162 |
| ENSG00000271971 | 39 | 21 | 18 | 8 | 41 | 27 |
| ENSG00000253854 | 42 | 22 | 14 | 8 | 24 | 10 |
| ENSG00000183808 | 771 | 342 | 502 | 295 | 432 | 400 |
| ENSG00000253722 | 0 | 0 | 0 | 0 | 0 | 0 |
| ENSG00000279331 | 16 | 17 | 8 | 5 | 12 | 7 |

| | | | | | | |
|---|---|---|---|---|---|---|
| ENSG00000253848 | 0 | 1 | 0 | 1 | 5 | 0 |
| ENSG00000164953 | 301 | 142 | 141 | 83 | 180 | 119 |
| ENSG00000254057 | 0 | 0 | 0 | 0 | 0 | 0 |
| ENSG00000254387 | 0 | 1 | 0 | 0 | 0 | 0 |
| ENSG00000164951 | 915 | 389 | 531 | 325 | 484 | 508 |
| ENSG00000264448 | 1 | 1 | 6 | 2 | 0 | 1 |
| ENSG00000284288 | 0 | 0 | 0 | 0 | 0 | 0 |
| ENSG00000251234 | 0 | 0 | 0 | 0 | 0 | 0 |
| ENSG00000240509 | 2 | 1 | 1 | 5 | 2 | 4 |
| ENSG00000275386 | 0 | 0 | 0 | 0 | 0 | 0 |
| ENSG00000253585 | 0 | 0 | 0 | 0 | 0 | 0 |
| ENSG00000079112 | 4 | 0 | 0 | 0 | 2 | 0 |
| ENSG00000249510 | 0 | 0 | 0 | 0 | 0 | 0 |
| ENSG00000164949 | 10 | 5 | 8 | 12 | 9 | 6 |
| ENSG00000244217 | 0 | 0 | 0 | 0 | 0 | 0 |
| ENSG00000197275 | 103 | 52 | 61 | 30 | 47 | 53 |
| ENSG00000265817 | 12 | 6 | 10 | 0 | 11 | 6 |
| ENSG00000280123 | 67 | 43 | 39 | 22 | 40 | 34 |
| ENSG00000164944 | 5828 | 2669 | 3087 | 1246 | 2917 | 2097 |
| ENSG00000254315 | 1 | 0 | 0 | 0 | 0 | 1 |
| ENSG00000253704 | 29 | 21 | 23 | 15 | 23 | 16 |
| ENSG00000253175 | 2 | 0 | 0 | 1 | 3 | 3 |
| ENSG00000254190 | 0 | 0 | 0 | 1 | 0 | 0 |
| ENSG00000254283 | 2 | 0 | 0 | 0 | 0 | 0 |
| ENSG00000261437 | 248 | 86 | 142 | 96 | 103 | 118 |
| ENSG00000104413 | 6666 | 3367 | 4053 | 2720 | 3902 | 3506 |
| ENSG00000274701 | 2 | 0 | 0 | 0 | 0 | 0 |
| ENSG00000241003 | 0 | 0 | 0 | 0 | 0 | 0 |
| ENSG00000253429 | 0 | 0 | 0 | 0 | 0 | 0 |
| ENSG00000199701 | 0 | 0 | 0 | 0 | 0 | 0 |
| ENSG00000156162 | 2175 | 1105 | 1189 | 725 | 1361 | 1111 |
| ENSG00000254307 | 0 | 0 | 0 | 0 | 0 | 0 |
| ENSG00000164941 | 2123 | 1134 | 1178 | 852 | 1611 | 1106 |
| ENSG00000175305 | 279 | 225 | 197 | 96 | 164 | 133 |
| ENSG00000272509 | 9 | 5 | 3 | 2 | 3 | 3 |
| ENSG00000243979 | 0 | 0 | 0 | 0 | 0 | 3 |
| ENSG00000156170 | 1174 | 604 | 663 | 522 | 519 | 558 |
| ENSG00000164938 | 8316 | 3818 | 4491 | 2221 | 3335 | 2963 |
| ENSG00000253878 | 37 | 9 | 24 | 10 | 16 | 16 |
| ENSG00000253528 | 1 | 1 | 0 | 0 | 0 | 1 |
| ENSG00000200525 | 0 | 0 | 1 | 1 | 1 | 0 |
| ENSG00000245080 | 36 | 23 | 21 | 16 | 22 | 18 |
| ENSG00000254248 | 4 | 2 | 4 | 1 | 2 | 0 |
| ENSG00000284147 | 0 | 0 | 0 | 0 | 0 | 0 |
| ENSG00000283522 | 0 | 0 | 0 | 0 | 0 | 0 |
| ENSG00000175895 | 3928 | 1847 | 2176 | 1214 | 1602 | 1531 |
| ENSG00000253773 | 18 | 4 | 6 | 6 | 4 | 3 |
| ENSG00000253351 | 0 | 0 | 0 | 0 | 0 | 0 |
| ENSG00000156172 | 227 | 102 | 150 | 95 | 122 | 115 |
| ENSG00000253945 | 0 | 0 | 0 | 0 | 0 | 0 |

| | | | | | | |
|---|---|---|---|---|---|---|
| ENSG00000253872 | 2 | 0 | 3 | 0 | 1 | 0 |
| ENSG00000202112 | 0 | 0 | 0 | 0 | 0 | 0 |
| ENSG00000254079 | 0 | 0 | 0 | 0 | 0 | 0 |
| ENSG00000248762 | 0 | 0 | 0 | 0 | 0 | 0 |
| ENSG00000183663 | 0 | 0 | 0 | 0 | 0 | 0 |
| ENSG00000271349 | 0 | 0 | 0 | 0 | 0 | 0 |
| ENSG00000270131 | 0 | 0 | 0 | 0 | 0 | 0 |
| ENSG00000156466 | 0 | 0 | 0 | 0 | 0 | 0 |
| ENSG00000156467 | 6858 | 3641 | 4252 | 3958 | 4668 | 3866 |
| ENSG00000254224 | 0 | 0 | 0 | 1 | 1 | 0 |
| ENSG00000156469 | 623 | 310 | 317 | 309 | 389 | 362 |
| ENSG00000156471 | 4055 | 1923 | 2287 | 2268 | 2297 | 2435 |
| ENSG00000199732 | 0 | 0 | 0 | 0 | 0 | 0 |
| ENSG00000253105 | 0 | 0 | 0 | 0 | 0 | 0 |
| ENSG00000202095 | 0 | 0 | 0 | 0 | 0 | 0 |
| ENSG00000169439 | 116 | 72 | 77 | 46 | 65 | 44 |
| ENSG00000104324 | 78 | 45 | 60 | 44 | 54 | 64 |
| ENSG00000272249 | 0 | 0 | 0 | 0 | 0 | 0 |
| ENSG00000270077 | 0 | 0 | 0 | 0 | 0 | 0 |
| ENSG00000260640 | 0 | 0 | 0 | 0 | 0 | 0 |
| ENSG00000180543 | 3413 | 1590 | 2047 | 1508 | 1894 | 1903 |
| ENSG00000207215 | 0 | 0 | 0 | 0 | 0 | 0 |
| ENSG00000213750 | 2 | 0 | 0 | 0 | 0 | 0 |
| ENSG00000147649 | 6959 | 3571 | 4057 | 2687 | 3946 | 3714 |
| ENSG00000202399 | 4 | 1 | 1 | 2 | 0 | 5 |
| ENSG00000104341 | 7298 | 3966 | 4403 | 4589 | 4727 | 4417 |
| ENSG00000270682 | 0 | 0 | 0 | 0 | 0 | 0 |
| ENSG00000270861 | 0 | 0 | 0 | 0 | 0 | 0 |
| ENSG00000253639 | 0 | 0 | 0 | 0 | 0 | 0 |
| ENSG00000243504 | 4 | 0 | 2 | 1 | 1 | 1 |
| ENSG00000132561 | 641 | 283 | 309 | 210 | 344 | 258 |
| ENSG00000156482 | 16309 | 11544 | 13749 | 15244 | 14064 | 13565 |
| ENSG00000245970 | 107 | 61 | 72 | 58 | 93 | 69 |
| ENSG00000207067 | 0 | 0 | 0 | 0 | 0 | 0 |
| ENSG00000252755 | 4 | 3 | 4 | 1 | 3 | 2 |
| ENSG00000177459 | 28 | 4 | 14 | 10 | 5 | 5 |
| ENSG00000132541 | 1463 | 786 | 921 | 830 | 962 | 835 |
| ENSG00000104356 | 1097 | 526 | 596 | 563 | 573 | 634 |
| ENSG00000104361 | 2451 | 1249 | 1303 | 709 | 1635 | 1081 |
| ENSG00000252558 | 0 | 0 | 0 | 0 | 0 | 0 |
| ENSG00000104375 | 1429 | 834 | 923 | 609 | 864 | 749 |
| ENSG00000278914 | 1 | 2 | 1 | 0 | 0 | 0 |
| ENSG00000253468 | 0 | 1 | 1 | 0 | 0 | 1 |
| ENSG00000156486 | 8 | 6 | 2 | 1 | 1 | 0 |
| ENSG00000272321 | 0 | 0 | 0 | 0 | 0 | 0 |
| ENSG00000207378 | 0 | 0 | 0 | 0 | 0 | 0 |
| ENSG00000253911 | 0 | 0 | 0 | 0 | 0 | 0 |
| ENSG00000254112 | 0 | 1 | 0 | 0 | 0 | 2 |
| ENSG00000240870 | 0 | 0 | 0 | 1 | 0 | 0 |
| ENSG00000253415 | 0 | 0 | 0 | 0 | 0 | 0 |

| | | | | | | |
|---|---|---|---|---|---|---|
| ENSG00000254312 | 0 | 0 | 0 | 0 | 0 | 0 |
| ENSG00000223281 | 0 | 0 | 0 | 0 | 0 | 0 |
| ENSG00000271930 | 15 | 2 | 8 | 6 | 4 | 5 |
| ENSG00000164920 | 1663 | 789 | 985 | 669 | 584 | 721 |
| ENSG00000229625 | 0 | 1 | 0 | 0 | 0 | 0 |
| ENSG00000253948 | 60 | 23 | 32 | 26 | 44 | 30 |
| ENSG00000253180 | 8 | 6 | 8 | 0 | 6 | 2 |
| ENSG00000132549 | 1663 | 821 | 772 | 277 | 1156 | 677 |
| ENSG00000253562 | 0 | 1 | 1 | 0 | 0 | 0 |
| ENSG00000253532 | 0 | 0 | 0 | 0 | 0 | 0 |
| ENSG00000236583 | 0 | 0 | 1 | 0 | 0 | 0 |
| ENSG00000237704 | 0 | 0 | 1 | 0 | 0 | 0 |
| ENSG00000225017 | 0 | 3 | 0 | 0 | 0 | 0 |
| ENSG00000206699 | 0 | 0 | 0 | 0 | 0 | 0 |
| ENSG00000207804 | 0 | 0 | 0 | 0 | 0 | 0 |
| ENSG00000216069 | 0 | 0 | 0 | 0 | 0 | 0 |
| ENSG00000243254 | 0 | 0 | 0 | 0 | 0 | 0 |
| ENSG00000235683 | 0 | 0 | 0 | 0 | 0 | 0 |
| ENSG00000253539 | 2 | 0 | 1 | 1 | 1 | 0 |
| ENSG00000253912 | 0 | 0 | 0 | 0 | 0 | 0 |
| ENSG00000164919 | 14849 | 7175 | 9578 | 10182 | 8946 | 10855 |
| ENSG00000132554 | 44 | 15 | 23 | 34 | 39 | 67 |
| ENSG00000212414 | 0 | 0 | 0 | 0 | 0 | 0 |
| ENSG00000221753 | 0 | 0 | 0 | 0 | 0 | 0 |
| ENSG00000156509 | 6 | 13 | 16 | 6 | 16 | 20 |
| ENSG00000147669 | 2984 | 1504 | 1852 | 1737 | 1793 | 1782 |
| ENSG00000104450 | 909 | 547 | 543 | 422 | 615 | 525 |
| ENSG00000202310 | 0 | 0 | 2 | 0 | 0 | 1 |
| ENSG00000253299 | 0 | 1 | 0 | 0 | 0 | 0 |
| ENSG00000199667 | 1 | 0 | 0 | 0 | 0 | 1 |
| ENSG00000034677 | 2001 | 1089 | 1171 | 684 | 1334 | 1012 |
| ENSG00000253217 | 0 | 0 | 0 | 0 | 0 | 0 |
| ENSG00000253824 | 0 | 1 | 1 | 0 | 1 | 0 |
| ENSG00000265599 | 0 | 0 | 0 | 0 | 0 | 0 |
| ENSG00000253842 | 0 | 0 | 0 | 1 | 1 | 0 |
| ENSG00000253666 | 1 | 1 | 1 | 0 | 1 | 1 |
| ENSG00000254364 | 0 | 0 | 0 | 2 | 5 | 1 |
| ENSG00000186106 | 679 | 363 | 400 | 289 | 411 | 341 |
| ENSG00000251013 | 4 | 0 | 2 | 0 | 1 | 5 |
| ENSG00000174226 | 0 | 0 | 0 | 1 | 1 | 0 |
| ENSG00000253740 | 0 | 1 | 0 | 1 | 0 | 0 |
| ENSG00000252764 | 0 | 0 | 0 | 0 | 0 | 0 |
| ENSG00000070756 | 67800 | 34631 | 39115 | 36927 | 42070 | 39776 |
| ENSG00000284570 | 0 | 0 | 0 | 0 | 0 | 0 |
| ENSG00000221527 | 0 | 0 | 0 | 0 | 0 | 0 |
| ENSG00000202001 | 0 | 0 | 0 | 0 | 0 | 0 |
| ENSG00000242399 | 0 | 0 | 0 | 0 | 0 | 0 |
| ENSG00000222795 | 0 | 0 | 0 | 0 | 0 | 0 |
| ENSG00000212994 | 2 | 0 | 1 | 0 | 0 | 0 |
| ENSG00000275041 | 0 | 0 | 0 | 0 | 0 | 0 |

| | | | | | | |
|---|---|---|---|---|---|---|
| ENSG00000260368 | 8 | 4 | 5 | 10 | 15 | 13 |
| ENSG00000164924 | 52988 | 26812 | 32313 | 24685 | 31634 | 28847 |
| ENSG00000242315 | 0 | 0 | 0 | 0 | 0 | 0 |
| ENSG00000248599 | 2 | 0 | 0 | 0 | 0 | 0 |
| ENSG00000253395 | 38 | 31 | 45 | 28 | 35 | 37 |
| ENSG00000253355 | 4 | 3 | 2 | 1 | 1 | 3 |
| ENSG00000202360 | 0 | 1 | 0 | 1 | 0 | 0 |
| ENSG00000253737 | 4 | 0 | 1 | 2 | 5 | 1 |
| ENSG00000261087 | 49 | 23 | 19 | 15 | 38 | 18 |
| ENSG00000120963 | 2989 | 1450 | 1764 | 1591 | 1822 | 1775 |
| ENSG00000271882 | 2 | 3 | 3 | 2 | 9 | 2 |
| ENSG00000253942 | 0 | 0 | 1 | 0 | 0 | 0 |
| ENSG00000253679 | 0 | 0 | 0 | 0 | 0 | 0 |
| ENSG00000253282 | 0 | 0 | 0 | 1 | 1 | 0 |
| ENSG00000239115 | 0 | 0 | 0 | 0 | 0 | 0 |
| ENSG00000271560 | 0 | 0 | 0 | 0 | 0 | 0 |
| ENSG00000228224 | 4 | 0 | 1 | 2 | 2 | 1 |
| ENSG00000254063 | 0 | 0 | 0 | 0 | 1 | 0 |
| ENSG00000253644 | 0 | 0 | 0 | 0 | 0 | 0 |
| ENSG00000239211 | 0 | 0 | 0 | 0 | 0 | 0 |
| ENSG00000254084 | 0 | 1 | 1 | 1 | 1 | 0 |
| ENSG00000254024 | 65 | 37 | 46 | 34 | 33 | 44 |
| ENSG00000083307 | 5203 | 2302 | 3176 | 2058 | 2496 | 2715 |
| ENSG00000253991 | 0 | 2 | 0 | 0 | 0 | 1 |
| ENSG00000253153 | 0 | 0 | 0 | 0 | 0 | 0 |
| ENSG00000104490 | 107 | 64 | 82 | 47 | 62 | 76 |
| ENSG00000253629 | 0 | 0 | 0 | 0 | 0 | 0 |
| ENSG00000238372 | 0 | 0 | 0 | 0 | 0 | 0 |
| ENSG00000282989 | 0 | 0 | 0 | 0 | 0 | 0 |
| ENSG00000266756 | 0 | 0 | 0 | 0 | 0 | 0 |
| ENSG00000254492 | 0 | 0 | 0 | 0 | 0 | 0 |
| ENSG00000048392 | 2763 | 1510 | 1577 | 844 | 2030 | 1487 |
| ENSG00000246263 | 148 | 67 | 102 | 98 | 113 | 108 |
| ENSG00000253328 | 8 | 0 | 4 | 1 | 1 | 4 |
| ENSG00000104517 | 6072 | 3475 | 3378 | 1294 | 4250 | 2838 |
| ENSG00000272037 | 8 | 7 | 10 | 4 | 25 | 4 |
| ENSG00000253923 | 1 | 3 | 3 | 0 | 1 | 2 |
| ENSG00000221387 | 0 | 0 | 0 | 0 | 3 | 0 |
| ENSG00000271612 | 2 | 0 | 0 | 0 | 0 | 0 |
| ENSG00000240724 | 0 | 0 | 0 | 0 | 0 | 0 |
| ENSG00000253633 | 2 | 0 | 0 | 0 | 0 | 0 |
| ENSG00000252593 | 0 | 0 | 0 | 0 | 0 | 0 |
| ENSG00000155087 | 0 | 0 | 0 | 0 | 0 | 0 |
| ENSG00000253382 | 0 | 0 | 0 | 0 | 0 | 0 |
| ENSG00000155090 | 1887 | 938 | 971 | 800 | 1067 | 992 |
| ENSG00000283959 | 70 | 26 | 31 | 30 | 29 | 33 |
| ENSG00000250684 | 0 | 0 | 0 | 0 | 0 | 1 |
| ENSG00000253669 | 105 | 44 | 38 | 25 | 47 | 47 |
| ENSG00000155096 | 9149 | 4428 | 5205 | 3928 | 5501 | 5233 |
| ENSG00000253385 | 0 | 1 | 0 | 0 | 0 | 0 |

| | | | | | | |
|---|---|---|---|---|---|---|
| ENSG00000253320 | 144 | 99 | 109 | 104 | 119 | 114 |
| ENSG00000253263 | 4 | 0 | 3 | 3 | 4 | 1 |
| ENSG00000244052 | 0 | 0 | 0 | 0 | 2 | 0 |
| ENSG00000201216 | 0 | 0 | 0 | 0 | 0 | 0 |
| ENSG00000253663 | 0 | 0 | 0 | 0 | 0 | 0 |
| ENSG00000254236 | 0 | 1 | 0 | 0 | 0 | 0 |
| ENSG00000155097 | 3582 | 1806 | 2075 | 1376 | 2476 | 1860 |
| ENSG00000253795 | 0 | 0 | 0 | 0 | 0 | 0 |
| ENSG00000253556 | 0 | 0 | 0 | 0 | 0 | 0 |
| ENSG00000253977 | 0 | 0 | 0 | 0 | 0 | 0 |
| ENSG00000250929 | 0 | 0 | 0 | 0 | 0 | 0 |
| ENSG00000236939 | 2 | 3 | 1 | 2 | 3 | 0 |
| ENSG00000164929 | 20 | 5 | 13 | 1 | 5 | 9 |
| ENSG00000265657 | 0 | 0 | 0 | 0 | 0 | 0 |
| ENSG00000247081 | 148 | 106 | 117 | 111 | 125 | 108 |
| ENSG00000253851 | 0 | 0 | 0 | 0 | 0 | 0 |
| ENSG00000164930 | 3327 | 1556 | 1941 | 1153 | 1821 | 1633 |
| ENSG00000164932 | 32 | 5 | 8 | 14 | 15 | 15 |
| ENSG00000266289 | 2 | 0 | 0 | 0 | 1 | 0 |
| ENSG00000207399 | 0 | 0 | 1 | 0 | 0 | 0 |
| ENSG00000164933 | 4 | 11 | 4 | 3 | 7 | 12 |
| ENSG00000285982 | 0 | 0 | 0 | 0 | 0 | 0 |
| ENSG00000164934 | 2759 | 1607 | 1697 | 1596 | 2245 | 1837 |
| ENSG00000261670 | 0 | 0 | 1 | 0 | 0 | 0 |
| ENSG00000271830 | 4 | 2 | 2 | 4 | 2 | 1 |
| ENSG00000253477 | 4 | 2 | 0 | 2 | 3 | 1 |
| ENSG00000176406 | 9 | 11 | 2 | 6 | 15 | 2 |
| ENSG00000255105 | 0 | 0 | 0 | 0 | 0 | 0 |
| ENSG00000254644 | 0 | 0 | 0 | 0 | 0 | 0 |
| ENSG00000239344 | 0 | 0 | 0 | 0 | 1 | 0 |
| ENSG00000147647 | 0 | 0 | 0 | 0 | 0 | 0 |
| ENSG00000164935 | 0 | 0 | 0 | 0 | 0 | 0 |
| ENSG00000179577 | 0 | 0 | 0 | 0 | 0 | 0 |
| ENSG00000279138 | 0 | 0 | 0 | 0 | 0 | 0 |
| ENSG00000208032 | 0 | 0 | 0 | 0 | 0 | 0 |
| ENSG00000147650 | 265 | 105 | 169 | 69 | 136 | 125 |
| ENSG00000253454 | 0 | 0 | 0 | 0 | 0 | 0 |
| ENSG00000169946 | 0 | 0 | 0 | 1 | 1 | 1 |
| ENSG00000283157 | 0 | 0 | 0 | 0 | 0 | 0 |
| ENSG00000243939 | 0 | 0 | 0 | 0 | 0 | 0 |
| ENSG00000253997 | 0 | 0 | 0 | 0 | 0 | 0 |
| ENSG00000240271 | 0 | 0 | 0 | 0 | 0 | 0 |
| ENSG00000254041 | 0 | 0 | 0 | 0 | 0 | 0 |
| ENSG00000253318 | 0 | 0 | 0 | 0 | 0 | 0 |
| ENSG00000243171 | 0 | 0 | 0 | 0 | 0 | 0 |
| ENSG00000251003 | 4 | 2 | 1 | 0 | 6 | 3 |
| ENSG00000253420 | 0 | 0 | 0 | 0 | 0 | 0 |
| ENSG00000254141 | 0 | 0 | 0 | 0 | 0 | 0 |
| ENSG00000243601 | 0 | 0 | 0 | 0 | 0 | 0 |
| ENSG00000253526 | 0 | 0 | 0 | 0 | 0 | 0 |

| | | | | | | |
|---|---|---|---|---|---|---|
| ENSG00000201627 | 0 | 0 | 0 | 0 | 0 | 0 |
| ENSG00000213708 | 0 | 0 | 0 | 0 | 0 | 0 |
| ENSG00000254021 | 0 | 0 | 0 | 0 | 0 | 0 |
| ENSG00000254615 | 0 | 0 | 0 | 0 | 0 | 1 |
| ENSG00000164830 | 1501 | 887 | 810 | 481 | 1253 | 780 |
| ENSG00000253582 | 3 | 0 | 5 | 9 | 6 | 4 |
| ENSG00000253676 | 6 | 6 | 3 | 1 | 2 | 3 |
| ENSG00000251892 | 0 | 1 | 0 | 0 | 0 | 0 |
| ENSG00000174429 | 0 | 1 | 0 | 0 | 0 | 0 |
| ENSG00000275905 | 0 | 0 | 0 | 0 | 0 | 0 |
| ENSG00000254146 | 0 | 0 | 0 | 0 | 0 | 0 |
| ENSG00000154188 | 0 | 0 | 1 | 0 | 0 | 0 |
| ENSG00000248838 | 0 | 0 | 0 | 0 | 0 | 0 |
| ENSG00000200806 | 0 | 0 | 0 | 0 | 0 | 0 |
| ENSG00000147655 | 0 | 0 | 0 | 0 | 0 | 0 |
| ENSG00000253228 | 0 | 0 | 0 | 0 | 0 | 0 |
| ENSG00000230948 | 0 | 2 | 0 | 0 | 0 | 0 |
| ENSG00000104408 | 11198 | 6931 | 7688 | 6359 | 10627 | 6663 |
| ENSG00000253754 | 2 | 0 | 4 | 1 | 0 | 0 |
| ENSG00000243287 | 0 | 0 | 0 | 0 | 0 | 0 |
| ENSG00000104412 | 830 | 513 | 481 | 285 | 767 | 481 |
| ENSG00000253657 | 0 | 0 | 0 | 0 | 0 | 0 |
| ENSG00000253949 | 0 | 0 | 0 | 0 | 0 | 0 |
| ENSG00000164841 | 0 | 0 | 0 | 0 | 0 | 0 |
| ENSG00000253580 | 0 | 0 | 0 | 0 | 0 | 0 |
| ENSG00000253796 | 1 | 0 | 0 | 0 | 0 | 0 |
| ENSG00000174417 | 0 | 0 | 0 | 0 | 0 | 0 |
| ENSG00000120526 | 1606 | 854 | 867 | 841 | 1177 | 944 |
| ENSG00000250267 | 0 | 0 | 0 | 0 | 0 | 0 |
| ENSG00000120533 | 2643 | 1321 | 1537 | 1425 | 1649 | 1441 |
| ENSG00000205038 | 12 | 8 | 5 | 1 | 4 | 4 |
| ENSG00000254185 | 0 | 1 | 0 | 0 | 0 | 0 |
| ENSG00000147654 | 1188 | 610 | 712 | 471 | 676 | 598 |
| ENSG00000147642 | 56 | 35 | 44 | 32 | 33 | 36 |
| ENSG00000248050 | 4 | 0 | 1 | 0 | 0 | 2 |
| ENSG00000255099 | 0 | 0 | 0 | 0 | 0 | 0 |
| ENSG00000283555 | 0 | 0 | 0 | 0 | 0 | 0 |
| ENSG00000241385 | 0 | 0 | 0 | 0 | 0 | 0 |
| ENSG00000164794 | 0 | 3 | 1 | 2 | 2 | 1 |
| ENSG00000255402 | 0 | 0 | 0 | 0 | 0 | 0 |
| ENSG00000240590 | 0 | 0 | 0 | 0 | 0 | 0 |
| ENSG00000253956 | 0 | 0 | 0 | 0 | 0 | 0 |
| ENSG00000253107 | 0 | 0 | 0 | 0 | 0 | 0 |
| ENSG00000253122 | 0 | 0 | 0 | 0 | 0 | 0 |
| ENSG00000253717 | 0 | 0 | 0 | 0 | 0 | 0 |
| ENSG00000253994 | 0 | 0 | 0 | 0 | 0 | 0 |
| ENSG00000254241 | 0 | 0 | 0 | 0 | 0 | 0 |
| ENSG00000253877 | 0 | 0 | 0 | 0 | 0 | 0 |
| ENSG00000253103 | 0 | 0 | 0 | 0 | 0 | 0 |
| ENSG00000253489 | 0 | 0 | 0 | 0 | 0 | 0 |

| | | | | | | |
|---|---|---|---|---|---|---|
| ENSG00000254124 | 0 | 0 | 0 | 0 | 0 | 0 |
| ENSG00000253434 | 0 | 0 | 0 | 0 | 0 | 0 |
| ENSG00000255033 | 0 | 0 | 0 | 0 | 0 | 0 |
| ENSG00000253156 | 0 | 0 | 0 | 0 | 0 | 0 |
| ENSG00000222146 | 0 | 0 | 0 | 0 | 0 | 0 |
| ENSG00000164796 | 32 | 18 | 21 | 8 | 6 | 17 |
| ENSG00000230169 | 0 | 0 | 0 | 0 | 0 | 0 |
| ENSG00000238399 | 0 | 0 | 0 | 0 | 0 | 0 |
| ENSG00000227514 | 0 | 0 | 0 | 0 | 0 | 0 |
| ENSG00000253207 | 2 | 0 | 0 | 0 | 0 | 1 |
| ENSG00000206719 | 0 | 0 | 0 | 0 | 0 | 0 |
| ENSG00000254339 | 0 | 0 | 0 | 0 | 0 | 0 |
| ENSG00000253499 | 0 | 0 | 0 | 0 | 0 | 0 |
| ENSG00000253756 | 0 | 0 | 0 | 0 | 0 | 1 |
| ENSG00000104447 | 3767 | 1731 | 1857 | 855 | 2025 | 1526 |
| ENSG00000227170 | 1 | 0 | 0 | 0 | 0 | 0 |
| ENSG00000249917 | 8 | 4 | 6 | 0 | 5 | 4 |
| ENSG00000199450 | 0 | 0 | 0 | 0 | 0 | 0 |
| ENSG00000253672 | 0 | 0 | 0 | 0 | 0 | 0 |
| ENSG00000147677 | 8262 | 4426 | 4917 | 4056 | 5685 | 4846 |
| ENSG00000147679 | 873 | 394 | 396 | 264 | 540 | 454 |
| ENSG00000164754 | 6456 | 3675 | 3829 | 2215 | 4692 | 3881 |
| ENSG00000253327 | 4 | 3 | 3 | 2 | 4 | 2 |
| ENSG00000283956 | 0 | 0 | 0 | 0 | 0 | 0 |
| ENSG00000205002 | 20 | 16 | 27 | 25 | 22 | 26 |
| ENSG00000164756 | 0 | 0 | 1 | 0 | 0 | 0 |
| ENSG00000244033 | 0 | 0 | 0 | 0 | 0 | 0 |
| ENSG00000253271 | 0 | 0 | 0 | 0 | 0 | 0 |
| ENSG00000253622 | 0 | 0 | 0 | 0 | 0 | 0 |
| ENSG00000240151 | 0 | 0 | 0 | 0 | 0 | 0 |
| ENSG00000252852 | 0 | 0 | 0 | 0 | 0 | 0 |
| ENSG00000164758 | 298 | 130 | 207 | 213 | 194 | 157 |
| ENSG00000239917 | 0 | 0 | 0 | 0 | 0 | 0 |
| ENSG00000182197 | 721 | 305 | 436 | 360 | 389 | 414 |
| ENSG00000177570 | 908 | 440 | 533 | 296 | 534 | 558 |
| ENSG00000225885 | 1 | 0 | 0 | 0 | 1 | 0 |
| ENSG00000281641 | 16 | 13 | 12 | 13 | 14 | 14 |
| ENSG00000242206 | 0 | 0 | 0 | 0 | 0 | 0 |
| ENSG00000164761 | 68 | 27 | 74 | 53 | 72 | 68 |
| ENSG00000207334 | 0 | 0 | 0 | 0 | 0 | 0 |
| ENSG00000254247 | 0 | 0 | 0 | 0 | 0 | 0 |
| ENSG00000184374 | 2 | 2 | 3 | 3 | 6 | 1 |
| ENSG00000254278 | 0 | 0 | 0 | 1 | 0 | 0 |
| ENSG00000147676 | 11159 | 6511 | 7338 | 5011 | 7531 | 6358 |
| ENSG00000253972 | 37 | 17 | 32 | 9 | 10 | 17 |
| ENSG00000276479 | 0 | 0 | 0 | 0 | 0 | 0 |
| ENSG00000206776 | 0 | 0 | 0 | 0 | 0 | 0 |
| ENSG00000136999 | 20 | 11 | 10 | 9 | 12 | 5 |
| ENSG00000253398 | 0 | 1 | 0 | 0 | 2 | 0 |
| ENSG00000254037 | 0 | 0 | 0 | 0 | 1 | 1 |

| | | | | | | |
|---|---|---|---|---|---|---|
| ENSG00000136960 | 2 | 0 | 0 | 1 | 0 | 0 |
| ENSG00000253495 | 0 | 0 | 0 | 0 | 0 | 0 |
| ENSG00000201896 | 0 | 0 | 0 | 0 | 0 | 0 |
| ENSG00000064313 | 1399 | 722 | 682 | 338 | 1008 | 666 |
| ENSG00000136982 | 363 | 190 | 186 | 153 | 206 | 215 |
| ENSG00000279347 | 4 | 0 | 4 | 3 | 1 | 5 |
| ENSG00000244642 | 0 | 1 | 0 | 0 | 0 | 0 |
| ENSG00000245330 | 2 | 3 | 6 | 1 | 3 | 5 |
| ENSG00000155792 | 488 | 306 | 333 | 243 | 256 | 291 |
| ENSG00000202356 | 0 | 0 | 0 | 0 | 0 | 0 |
| ENSG00000254343 | 42 | 26 | 22 | 10 | 35 | 29 |
| ENSG00000187955 | 13 | 2 | 8 | 2 | 4 | 2 |
| ENSG00000172172 | 1854 | 875 | 1029 | 1060 | 1106 | 1050 |
| ENSG00000172167 | 153 | 82 | 105 | 65 | 102 | 86 |
| ENSG00000172164 | 2 | 6 | 5 | 3 | 3 | 5 |
| ENSG00000250290 | 0 | 0 | 0 | 0 | 0 | 0 |
| ENSG00000248318 | 0 | 0 | 0 | 0 | 0 | 0 |
| ENSG00000272502 | 0 | 0 | 0 | 0 | 0 | 0 |
| ENSG00000253619 | 0 | 0 | 0 | 0 | 0 | 0 |
| ENSG00000271219 | 0 | 0 | 0 | 0 | 0 | 0 |
| ENSG00000239872 | 0 | 0 | 0 | 0 | 0 | 0 |
| ENSG00000170961 | 0 | 0 | 0 | 0 | 0 | 0 |
| ENSG00000248690 | 0 | 0 | 0 | 0 | 0 | 0 |
| ENSG00000254209 | 0 | 0 | 0 | 0 | 0 | 0 |
| ENSG00000254303 | 0 | 0 | 0 | 0 | 0 | 0 |
| ENSG00000253814 | 0 | 0 | 0 | 0 | 0 | 0 |
| ENSG00000255364 | 0 | 0 | 0 | 0 | 0 | 0 |
| ENSG00000255325 | 0 | 0 | 0 | 0 | 0 | 0 |
| ENSG00000253819 | 0 | 0 | 0 | 0 | 0 | 0 |
| ENSG00000238901 | 0 | 0 | 0 | 0 | 0 | 0 |
| ENSG00000263443 | 0 | 0 | 0 | 0 | 0 | 0 |
| ENSG00000235165 | 0 | 0 | 0 | 0 | 0 | 0 |
| ENSG00000272384 | 80 | 55 | 67 | 42 | 45 | 60 |
| ENSG00000272043 | 40 | 18 | 18 | 12 | 29 | 20 |
| ENSG00000178764 | 1871 | 838 | 1120 | 735 | 749 | 951 |
| ENSG00000253372 | 4 | 3 | 1 | 0 | 0 | 1 |
| ENSG00000259631 | 1 | 0 | 0 | 1 | 0 | 0 |
| ENSG00000253607 | 0 | 0 | 0 | 0 | 0 | 0 |
| ENSG00000136986 | 6675 | 3390 | 4202 | 3384 | 3848 | 4100 |
| ENSG00000156787 | 221 | 128 | 143 | 93 | 219 | 149 |
| ENSG00000253463 | 0 | 1 | 0 | 0 | 0 | 0 |
| ENSG00000147689 | 427 | 203 | 258 | 132 | 213 | 177 |
| ENSG00000221461 | 0 | 0 | 0 | 0 | 0 | 0 |
| ENSG00000253258 | 3 | 1 | 2 | 1 | 3 | 4 |
| ENSG00000253410 | 0 | 0 | 1 | 0 | 0 | 0 |
| ENSG00000204949 | 18 | 8 | 11 | 5 | 7 | 7 |
| ENSG00000266324 | 0 | 0 | 0 | 0 | 0 | 0 |
| ENSG00000189376 | 211 | 187 | 176 | 188 | 204 | 171 |
| ENSG00000259305 | 24 | 19 | 17 | 16 | 17 | 12 |
| ENSG00000243498 | 3 | 3 | 0 | 0 | 0 | 3 |

| | | | | | | |
|---|---|---|---|---|---|---|
| ENSG00000165156 | 1594 | 935 | 752 | 391 | 1245 | 727 |
| ENSG00000222607 | 0 | 0 | 0 | 0 | 0 | 0 |
| ENSG00000253336 | 2 | 1 | 1 | 0 | 0 | 1 |
| ENSG00000156802 | 1397 | 959 | 654 | 309 | 1366 | 781 |
| ENSG00000252297 | 0 | 0 | 0 | 0 | 1 | 3 |
| ENSG00000207704 | 0 | 0 | 0 | 0 | 0 | 0 |
| ENSG00000283172 | 0 | 0 | 0 | 0 | 0 | 0 |
| ENSG00000254388 | 0 | 3 | 1 | 1 | 0 | 0 |
| ENSG00000254294 | 2 | 1 | 2 | 4 | 1 | 0 |
| ENSG00000156795 | 1074 | 493 | 590 | 612 | 705 | 692 |
| ENSG00000278374 | 1 | 4 | 9 | 1 | 1 | 1 |
| ENSG00000156804 | 390 | 188 | 249 | 151 | 203 | 171 |
| ENSG00000254113 | 0 | 0 | 0 | 0 | 0 | 0 |
| ENSG00000251840 | 0 | 0 | 0 | 0 | 0 | 0 |
| ENSG00000253286 | 0 | 0 | 0 | 0 | 0 | 0 |
| ENSG00000175946 | 0 | 0 | 0 | 0 | 0 | 0 |
| ENSG00000104537 | 0 | 2 | 0 | 0 | 0 | 0 |
| ENSG00000176853 | 4072 | 2164 | 2394 | 1106 | 2685 | 2239 |
| ENSG00000214814 | 18 | 7 | 11 | 4 | 5 | 7 |
| ENSG00000254000 | 0 | 0 | 0 | 0 | 0 | 0 |
| ENSG00000181171 | 0 | 0 | 0 | 0 | 0 | 0 |
| ENSG00000254249 | 0 | 0 | 0 | 0 | 0 | 0 |
| ENSG00000253868 | 0 | 0 | 0 | 0 | 0 | 1 |
| ENSG00000250727 | 0 | 0 | 0 | 1 | 0 | 0 |
| ENSG00000253885 | 0 | 0 | 1 | 0 | 0 | 0 |
| ENSG00000214803 | 4 | 0 | 0 | 0 | 1 | 1 |
| ENSG00000206724 | 0 | 0 | 0 | 0 | 0 | 0 |
| ENSG00000253227 | 6 | 0 | 0 | 5 | 3 | 2 |
| ENSG00000237586 | 0 | 0 | 0 | 0 | 2 | 0 |
| ENSG00000164983 | 1155 | 616 | 710 | 545 | 693 | 640 |
| ENSG00000183665 | 1503 | 672 | 861 | 836 | 822 | 828 |
| ENSG00000245149 | 228 | 103 | 153 | 74 | 143 | 84 |
| ENSG00000170881 | 3502 | 1648 | 2086 | 1609 | 1956 | 1869 |
| ENSG00000147687 | 1590 | 970 | 1043 | 839 | 1309 | 1095 |
| ENSG00000253106 | 141 | 74 | 80 | 42 | 97 | 55 |
| ENSG00000284087 | 0 | 0 | 0 | 0 | 0 | 0 |
| ENSG00000147684 | 4677 | 2738 | 2927 | 3465 | 3216 | 3046 |
| ENSG00000170873 | 1084 | 469 | 534 | 358 | 482 | 460 |
| ENSG00000255491 | 2 | 3 | 0 | 3 | 0 | 2 |
| ENSG00000263735 | 0 | 0 | 0 | 0 | 0 | 0 |
| ENSG00000283609 | 0 | 0 | 0 | 0 | 0 | 0 |
| ENSG00000249816 | 0 | 0 | 0 | 1 | 0 | 0 |
| ENSG00000250428 | 0 | 0 | 0 | 0 | 0 | 0 |
| ENSG00000255313 | 0 | 0 | 0 | 0 | 0 | 0 |
| ENSG00000180938 | 566 | 276 | 395 | 226 | 321 | 242 |
| ENSG00000253513 | 0 | 0 | 0 | 0 | 1 | 0 |
| ENSG00000104549 | 8092 | 3708 | 4536 | 3668 | 4082 | 4391 |
| ENSG00000164961 | 3741 | 2062 | 2175 | 1200 | 2504 | 1819 |
| ENSG00000253167 | 7 | 9 | 13 | 5 | 17 | 8 |
| ENSG00000156831 | 1551 | 831 | 975 | 941 | 1003 | 855 |

| | | | | | | |
|---|---|---|---|---|---|---|
| ENSG00000242170 | 0 | 1 | 0 | 0 | 0 | 0 |
| ENSG00000254431 | 0 | 0 | 1 | 1 | 1 | 0 |
| ENSG00000173334 | 971 | 519 | 606 | 502 | 560 | 576 |
| ENSG00000253111 | 0 | 0 | 0 | 1 | 0 | 0 |
| ENSG00000253470 | 0 | 0 | 0 | 0 | 0 | 0 |
| ENSG00000206695 | 0 | 0 | 0 | 0 | 0 | 0 |
| ENSG00000245164 | 0 | 0 | 0 | 0 | 0 | 0 |
| ENSG00000253651 | 0 | 0 | 0 | 0 | 0 | 0 |
| ENSG00000254227 | 0 | 0 | 0 | 0 | 0 | 0 |
| ENSG00000250563 | 0 | 0 | 0 | 0 | 0 | 0 |
| ENSG00000253841 | 0 | 0 | 0 | 0 | 0 | 0 |
| ENSG00000248720 | 0 | 0 | 0 | 0 | 0 | 0 |
| ENSG00000244791 | 3 | 0 | 0 | 2 | 4 | 1 |
| ENSG00000253427 | 0 | 0 | 0 | 0 | 0 | 0 |
| ENSG00000253220 | 0 | 0 | 0 | 0 | 2 | 1 |
| ENSG00000207138 | 0 | 0 | 0 | 0 | 0 | 1 |
| ENSG00000253530 | 0 | 0 | 0 | 0 | 0 | 0 |
| ENSG00000168672 | 7759 | 3321 | 4421 | 2831 | 4412 | 4356 |
| ENSG00000253438 | 138 | 68 | 131 | 73 | 87 | 120 |
| ENSG00000253543 | 0 | 1 | 0 | 0 | 0 | 0 |
| ENSG00000212451 | 0 | 0 | 0 | 0 | 0 | 0 |
| ENSG00000253573 | 0 | 0 | 0 | 0 | 0 | 1 |
| ENSG00000254166 | 0 | 1 | 1 | 1 | 0 | 1 |
| ENSG00000282961 | 0 | 0 | 0 | 1 | 0 | 0 |
| ENSG00000224722 | 0 | 0 | 0 | 0 | 0 | 0 |
| ENSG00000246228 | 1 | 4 | 1 | 0 | 1 | 1 |
| ENSG00000271509 | 0 | 0 | 0 | 0 | 0 | 0 |
| ENSG00000212993 | 10 | 5 | 4 | 1 | 1 | 2 |
| ENSG00000280997 | 0 | 0 | 0 | 0 | 0 | 0 |
| ENSG00000249375 | 13 | 10 | 8 | 8 | 12 | 8 |
| ENSG00000136997 | 1305 | 747 | 823 | 1386 | 1205 | 1898 |
| ENSG00000249859 | 631 | 388 | 435 | 454 | 493 | 678 |
| ENSG00000276443 | 0 | 0 | 0 | 0 | 0 | 0 |
| ENSG00000283710 | 0 | 0 | 0 | 0 | 0 | 0 |
| ENSG00000278324 | 0 | 0 | 0 | 0 | 0 | 0 |
| ENSG00000280055 | 2 | 6 | 0 | 1 | 5 | 9 |
| ENSG00000221771 | 0 | 0 | 0 | 0 | 0 | 1 |
| ENSG00000207110 | 12 | 12 | 16 | 11 | 17 | 22 |
| ENSG00000283200 | 5 | 0 | 2 | 0 | 5 | 2 |
| ENSG00000278275 | 0 | 1 | 1 | 0 | 0 | 0 |
| ENSG00000222501 | 3 | 2 | 0 | 0 | 0 | 1 |
| ENSG00000221176 | 0 | 0 | 0 | 0 | 0 | 2 |
| ENSG00000221261 | 0 | 0 | 0 | 0 | 0 | 2 |
| ENSG00000201782 | 0 | 0 | 0 | 0 | 0 | 0 |
| ENSG00000254275 | 0 | 4 | 3 | 3 | 1 | 4 |
| ENSG00000229140 | 8 | 4 | 2 | 1 | 3 | 4 |
| ENSG00000253225 | 0 | 0 | 0 | 0 | 0 | 0 |
| ENSG00000222755 | 0 | 0 | 0 | 0 | 0 | 0 |
| ENSG00000253926 | 0 | 0 | 0 | 0 | 0 | 0 |
| ENSG00000263763 | 0 | 0 | 0 | 0 | 0 | 0 |

| | | | | | | |
|---|---|---|---|---|---|---|
| ENSG00000285108 | 0 | 1 | 0 | 1 | 1 | 0 |
| ENSG00000224110 | 0 | 0 | 0 | 0 | 0 | 0 |
| ENSG00000147697 | 0 | 2 | 1 | 0 | 2 | 0 |
| ENSG00000254317 | 8 | 4 | 4 | 1 | 2 | 5 |
| ENSG00000153310 | 3721 | 1839 | 2221 | 1607 | 2099 | 2046 |
| ENSG00000243402 | 0 | 3 | 0 | 0 | 2 | 0 |
| ENSG00000200075 | 0 | 2 | 0 | 0 | 0 | 0 |
| ENSG00000254263 | 3 | 0 | 2 | 0 | 1 | 1 |
| ENSG00000253720 | 0 | 1 | 0 | 2 | 0 | 0 |
| ENSG00000253043 | 0 | 0 | 0 | 0 | 0 | 1 |
| ENSG00000264653 | 0 | 0 | 0 | 0 | 0 | 0 |
| ENSG00000153317 | 1180 | 595 | 643 | 430 | 686 | 599 |
| ENSG00000200304 | 0 | 0 | 0 | 0 | 0 | 0 |
| ENSG00000280543 | 0 | 2 | 1 | 0 | 0 | 0 |
| ENSG00000212342 | 0 | 0 | 0 | 0 | 0 | 0 |
| ENSG00000253656 | 0 | 0 | 0 | 0 | 0 | 0 |
| ENSG00000155897 | 0 | 0 | 0 | 0 | 0 | 0 |
| ENSG00000253259 | 0 | 0 | 0 | 0 | 0 | 0 |
| ENSG00000253992 | 0 | 0 | 0 | 0 | 0 | 0 |
| ENSG00000253407 | 0 | 0 | 0 | 0 | 0 | 0 |
| ENSG00000253507 | 0 | 0 | 0 | 0 | 0 | 0 |
| ENSG00000252158 | 0 | 0 | 0 | 0 | 0 | 0 |
| ENSG00000254255 | 0 | 0 | 0 | 0 | 0 | 0 |
| ENSG00000132294 | 2723 | 1290 | 1519 | 730 | 1547 | 1307 |
| ENSG00000253117 | 0 | 0 | 0 | 0 | 0 | 0 |
| ENSG00000258417 | 0 | 0 | 0 | 0 | 0 | 0 |
| ENSG00000132297 | 0 | 0 | 0 | 0 | 0 | 0 |
| ENSG00000184156 | 0 | 0 | 0 | 0 | 0 | 0 |
| ENSG00000253521 | 0 | 0 | 0 | 0 | 0 | 0 |
| ENSG00000129295 | 12 | 12 | 4 | 7 | 7 | 6 |
| ENSG00000165071 | 0 | 0 | 0 | 0 | 0 | 0 |
| ENSG00000129292 | 1282 | 655 | 675 | 379 | 884 | 629 |
| ENSG00000270137 | 2 | 1 | 0 | 0 | 0 | 0 |
| ENSG00000223697 | 4 | 4 | 3 | 1 | 3 | 5 |
| ENSG00000042832 | 2 | 1 | 2 | 0 | 1 | 4 |
| ENSG00000240237 | 0 | 0 | 0 | 0 | 0 | 0 |
| ENSG00000155926 | 0 | 0 | 0 | 0 | 0 | 0 |
| ENSG00000276961 | 0 | 0 | 0 | 0 | 0 | 0 |
| ENSG00000104415 | 0 | 0 | 0 | 0 | 0 | 0 |
| ENSG00000104419 | 2388 | 965 | 1437 | 571 | 548 | 696 |
| ENSG00000243537 | 6 | 0 | 2 | 0 | 0 | 0 |
| ENSG00000253638 | 0 | 0 | 0 | 0 | 0 | 0 |
| ENSG00000225259 | 0 | 1 | 1 | 0 | 2 | 1 |
| ENSG00000008513 | 5028 | 2497 | 3309 | 3713 | 4744 | 5603 |
| ENSG00000283176 | 0 | 0 | 0 | 0 | 0 | 1 |
| ENSG00000283197 | 0 | 0 | 0 | 0 | 0 | 0 |
| ENSG00000261220 | 55 | 40 | 43 | 51 | 55 | 79 |
| ENSG00000212273 | 0 | 0 | 0 | 0 | 0 | 0 |
| ENSG00000276193 | 0 | 0 | 0 | 0 | 0 | 0 |
| ENSG00000253970 | 0 | 0 | 0 | 0 | 0 | 0 |

| | | | | | | |
|---|---|---|---|---|---|---|
| ENSG00000253916 | 0 | 0 | 0 | 0 | 0 | 0 |
| ENSG00000253561 | 0 | 0 | 0 | 1 | 0 | 0 |
| ENSG00000253165 | 0 | 0 | 0 | 0 | 0 | 0 |
| ENSG00000254313 | 2 | 0 | 1 | 0 | 0 | 0 |
| ENSG00000253593 | 0 | 0 | 0 | 4 | 4 | 9 |
| ENSG00000277576 | 0 | 0 | 0 | 0 | 0 | 0 |
| ENSG00000271240 | 0 | 0 | 0 | 0 | 0 | 0 |
| ENSG00000253627 | 0 | 0 | 0 | 0 | 0 | 0 |
| ENSG00000066827 | 391 | 215 | 263 | 212 | 229 | 242 |
| ENSG00000276140 | 0 | 0 | 0 | 0 | 0 | 0 |
| ENSG00000248492 | 0 | 0 | 0 | 0 | 0 | 0 |
| ENSG00000277732 | 0 | 0 | 0 | 0 | 0 | 0 |
| ENSG00000278454 | 0 | 0 | 0 | 0 | 0 | 0 |
| ENSG00000271048 | 0 | 0 | 0 | 0 | 0 | 0 |
| ENSG00000272456 | 0 | 0 | 0 | 0 | 0 | 0 |
| ENSG00000278898 | 0 | 0 | 0 | 0 | 0 | 0 |
| ENSG00000279518 | 6 | 2 | 1 | 0 | 8 | 6 |
| ENSG00000259820 | 102 | 61 | 60 | 14 | 92 | 32 |
| ENSG00000207582 | 0 | 0 | 0 | 0 | 0 | 0 |
| ENSG00000199153 | 0 | 0 | 0 | 0 | 0 | 0 |
| ENSG00000254028 | 7 | 2 | 5 | 1 | 1 | 2 |
| ENSG00000253433 | 10 | 4 | 5 | 2 | 1 | 1 |
| ENSG00000251218 | 0 | 0 | 0 | 0 | 0 | 0 |
| ENSG00000244413 | 0 | 0 | 0 | 0 | 0 | 0 |
| ENSG00000254083 | 0 | 0 | 0 | 0 | 0 | 0 |
| ENSG00000254372 | 0 | 0 | 0 | 0 | 0 | 0 |
| ENSG00000131773 | 6 | 3 | 3 | 6 | 4 | 5 |
| ENSG00000253915 | 0 | 0 | 0 | 0 | 0 | 0 |
| ENSG00000199652 | 0 | 0 | 0 | 0 | 0 | 0 |
| ENSG00000254101 | 0 | 0 | 0 | 1 | 0 | 0 |
| ENSG00000285683 | 0 | 0 | 0 | 0 | 0 | 0 |
| ENSG00000206678 | 0 | 0 | 0 | 0 | 0 | 0 |
| ENSG00000274572 | 0 | 0 | 0 | 0 | 0 | 0 |
| ENSG00000285817 | 0 | 0 | 0 | 0 | 0 | 0 |
| ENSG00000254076 | 0 | 0 | 0 | 0 | 0 | 0 |
| ENSG00000253288 | 0 | 0 | 0 | 0 | 0 | 0 |
| ENSG00000254361 | 0 | 0 | 0 | 0 | 0 | 0 |
| ENSG00000253988 | 0 | 0 | 0 | 0 | 0 | 0 |
| ENSG00000147724 | 0 | 0 | 0 | 0 | 0 | 0 |
| ENSG00000169436 | 0 | 0 | 0 | 0 | 0 | 0 |
| ENSG00000253432 | 0 | 0 | 0 | 0 | 0 | 0 |
| ENSG00000251744 | 0 | 0 | 0 | 0 | 0 | 0 |
| ENSG00000253783 | 0 | 0 | 0 | 0 | 0 | 0 |
| ENSG00000254258 | 0 | 0 | 0 | 0 | 0 | 0 |
| ENSG00000275078 | 0 | 0 | 0 | 0 | 0 | 0 |
| ENSG00000169427 | 0 | 0 | 1 | 0 | 0 | 1 |
| ENSG00000167632 | 1378 | 608 | 774 | 691 | 618 | 619 |
| ENSG00000253574 | 0 | 0 | 0 | 0 | 1 | 0 |
| ENSG00000250733 | 0 | 0 | 0 | 0 | 0 | 0 |
| ENSG00000274064 | 0 | 0 | 0 | 0 | 1 | 0 |

| | | | | | | |
|---|---|---|---|---|---|---|
| ENSG00000282164 | 50 | 17 | 18 | 12 | 19 | 13 |
| ENSG00000259891 | 101 | 34 | 57 | 42 | 50 | 49 |
| ENSG00000104472 | 1362 | 710 | 874 | 706 | 745 | 810 |
| ENSG00000123908 | 4682 | 1952 | 2711 | 1594 | 2337 | 2408 |
| ENSG00000279766 | 6 | 0 | 4 | 3 | 2 | 7 |
| ENSG00000280326 | 0 | 0 | 0 | 0 | 0 | 0 |
| ENSG00000280303 | 105 | 44 | 50 | 46 | 72 | 51 |
| ENSG00000169398 | 4795 | 2217 | 2795 | 1881 | 2764 | 2445 |
| ENSG00000254324 | 0 | 0 | 0 | 0 | 0 | 0 |
| ENSG00000279786 | 4 | 1 | 4 | 0 | 2 | 0 |
| ENSG00000252864 | 0 | 0 | 0 | 0 | 0 | 0 |
| ENSG00000253266 | 27 | 9 | 11 | 7 | 16 | 14 |
| ENSG00000105339 | 515 | 257 | 334 | 338 | 346 | 307 |
| ENSG00000253210 | 67 | 37 | 52 | 31 | 36 | 40 |
| ENSG00000022567 | 598 | 227 | 363 | 311 | 275 | 352 |
| ENSG00000280035 | 0 | 2 | 0 | 0 | 0 | 0 |
| ENSG00000253307 | 0 | 0 | 0 | 1 | 0 | 0 |
| ENSG00000254019 | 0 | 0 | 0 | 0 | 0 | 0 |
| ENSG00000254197 | 0 | 0 | 0 | 0 | 0 | 0 |
| ENSG00000254291 | 0 | 0 | 0 | 0 | 0 | 0 |
| ENSG00000253595 | 0 | 0 | 1 | 0 | 0 | 0 |
| ENSG00000261655 | 0 | 0 | 2 | 3 | 1 | 1 |
| ENSG00000204882 | 0 | 0 | 0 | 1 | 1 | 0 |
| ENSG00000244998 | 0 | 1 | 0 | 0 | 0 | 1 |
| ENSG00000184489 | 408 | 198 | 247 | 260 | 208 | 224 |
| ENSG00000226807 | 2 | 0 | 0 | 0 | 0 | 0 |
| ENSG00000271959 | 0 | 0 | 1 | 0 | 5 | 1 |
| ENSG00000238854 | 0 | 0 | 0 | 0 | 0 | 0 |
| ENSG00000253605 | 0 | 2 | 0 | 0 | 1 | 2 |
| ENSG00000226490 | 0 | 0 | 0 | 0 | 0 | 0 |
| ENSG00000221768 | 0 | 0 | 0 | 0 | 0 | 0 |
| ENSG00000254183 | 0 | 0 | 0 | 0 | 0 | 0 |
| ENSG00000261710 | 0 | 0 | 0 | 0 | 0 | 0 |
| ENSG00000265247 | 0 | 0 | 0 | 0 | 0 | 0 |
| ENSG00000254008 | 0 | 0 | 0 | 0 | 0 | 0 |
| ENSG00000171045 | 333 | 163 | 202 | 183 | 144 | 168 |
| ENSG00000253602 | 0 | 0 | 0 | 0 | 0 | 0 |
| ENSG00000261693 | 43 | 19 | 33 | 26 | 11 | 20 |
| ENSG00000181790 | 133 | 48 | 76 | 41 | 51 | 55 |
| ENSG00000232722 | 0 | 0 | 0 | 0 | 0 | 0 |
| ENSG00000198576 | 14 | 14 | 27 | 14 | 10 | 21 |
| ENSG00000261044 | 0 | 0 | 0 | 0 | 0 | 0 |
| ENSG00000277332 | 0 | 0 | 0 | 0 | 0 | 0 |
| ENSG00000234616 | 2447 | 1170 | 1540 | 977 | 1152 | 1195 |
| ENSG00000167653 | 1620 | 791 | 952 | 589 | 441 | 485 |
| ENSG00000160886 | 8 | 2 | 1 | 1 | 2 | 2 |
| ENSG00000253741 | 77 | 56 | 69 | 24 | 31 | 40 |
| ENSG00000130193 | 8062 | 3755 | 4760 | 5146 | 4102 | 4374 |
| ENSG00000253806 | 0 | 0 | 0 | 0 | 0 | 0 |
| ENSG00000126233 | 11 | 7 | 7 | 11 | 12 | 5 |

| | | | | | | |
|---|---|---|---|---|---|---|
| ENSG00000197353 | 18 | 5 | 9 | 2 | 4 | 3 |
| ENSG00000253196 | 3 | 0 | 1 | 0 | 1 | 1 |
| ENSG00000284505 | 0 | 0 | 0 | 0 | 0 | 0 |
| ENSG00000283992 | 0 | 0 | 0 | 0 | 0 | 0 |
| ENSG00000180155 | 0 | 0 | 0 | 0 | 0 | 0 |
| ENSG00000167656 | 0 | 2 | 5 | 10 | 10 | 2 |
| ENSG00000253715 | 0 | 0 | 0 | 0 | 0 | 0 |
| ENSG00000283686 | 1 | 0 | 0 | 0 | 0 | 0 |
| ENSG00000253728 | 0 | 0 | 0 | 0 | 0 | 0 |
| ENSG00000104499 | 0 | 0 | 0 | 0 | 0 | 0 |
| ENSG00000253421 | 0 | 0 | 0 | 0 | 0 | 0 |
| ENSG00000160882 | 0 | 0 | 0 | 0 | 0 | 0 |
| ENSG00000179142 | 0 | 0 | 0 | 0 | 0 | 0 |
| ENSG00000247317 | 59 | 28 | 22 | 30 | 20 | 37 |
| ENSG00000253971 | 0 | 1 | 1 | 0 | 3 | 2 |
| ENSG00000160932 | 9597 | 4743 | 5697 | 5759 | 4791 | 4543 |
| ENSG00000177335 | 4 | 2 | 1 | 2 | 0 | 1 |
| ENSG00000250115 | 0 | 0 | 0 | 0 | 0 | 0 |
| ENSG00000243328 | 0 | 0 | 0 | 0 | 0 | 0 |
| ENSG00000261667 | 0 | 0 | 0 | 0 | 0 | 0 |
| ENSG00000176956 | 0 | 0 | 1 | 0 | 0 | 0 |
| ENSG00000277494 | 0 | 0 | 0 | 0 | 0 | 0 |
| ENSG00000181638 | 330 | 144 | 189 | 182 | 182 | 182 |
| ENSG00000264668 | 0 | 0 | 0 | 0 | 2 | 2 |
| ENSG00000250571 | 618 | 333 | 423 | 389 | 325 | 335 |
| ENSG00000253716 | 247 | 134 | 177 | 138 | 158 | 141 |
| ENSG00000185730 | 430 | 220 | 306 | 254 | 269 | 306 |
| ENSG00000272172 | 45 | 21 | 36 | 31 | 14 | 17 |
| ENSG00000184428 | 1741 | 785 | 991 | 1073 | 1015 | 1038 |
| ENSG00000212221 | 0 | 0 | 0 | 0 | 0 | 0 |
| ENSG00000254389 | 354 | 156 | 175 | 138 | 156 | 167 |
| ENSG00000158106 | 4076 | 1756 | 2586 | 2163 | 2015 | 2166 |
| ENSG00000253931 | 19 | 7 | 18 | 6 | 7 | 9 |
| ENSG00000254338 | 22 | 5 | 14 | 5 | 3 | 5 |
| ENSG00000182759 | 260 | 111 | 156 | 121 | 113 | 97 |
| ENSG00000014164 | 1011 | 430 | 536 | 609 | 536 | 492 |
| ENSG00000254859 | 0 | 1 | 1 | 2 | 0 | 4 |
| ENSG00000275558 | 0 | 0 | 0 | 0 | 0 | 0 |
| ENSG00000254144 | 0 | 0 | 1 | 0 | 0 | 0 |
| ENSG00000104518 | 5 | 0 | 6 | 4 | 1 | 7 |
| ENSG00000204839 | 422 | 141 | 253 | 241 | 218 | 233 |
| ENSG00000255050 | 51 | 23 | 22 | 35 | 28 | 40 |
| ENSG00000147813 | 992 | 510 | 714 | 618 | 550 | 568 |
| ENSG00000279605 | 23 | 18 | 20 | 26 | 18 | 27 |
| ENSG00000254741 | 0 | 0 | 0 | 0 | 0 | 0 |
| ENSG00000104529 | 6354 | 3541 | 4631 | 5063 | 4582 | 4487 |
| ENSG00000179886 | 689 | 334 | 403 | 419 | 376 | 363 |
| ENSG00000104524 | 853 | 445 | 517 | 574 | 475 | 577 |
| ENSG00000104522 | 2859 | 1291 | 1639 | 1826 | 1530 | 1440 |
| ENSG00000254812 | 27 | 13 | 17 | 7 | 14 | 6 |

| | | | | | | |
|---|---|---|---|---|---|---|
| ENSG00000183309 | 2078 | 874 | 1181 | 757 | 1051 | 946 |
| ENSG00000181135 | 982 | 467 | 651 | 461 | 491 | 507 |
| ENSG00000181097 | 49 | 23 | 28 | 14 | 32 | 17 |
| ENSG00000255181 | 10 | 2 | 1 | 1 | 3 | 4 |
| ENSG00000254574 | 2 | 2 | 0 | 0 | 1 | 0 |
| ENSG00000181085 | 263 | 112 | 217 | 83 | 109 | 138 |
| ENSG00000254548 | 10 | 6 | 10 | 5 | 4 | 7 |
| ENSG00000180921 | 8168 | 3554 | 4791 | 4706 | 4143 | 4520 |
| ENSG00000265660 | 30 | 19 | 48 | 29 | 39 | 35 |
| ENSG00000203499 | 3742 | 2130 | 2617 | 2081 | 2277 | 2476 |
| ENSG00000254973 | 123 | 69 | 75 | 45 | 61 | 69 |
| ENSG00000180900 | 6514 | 2894 | 3927 | 4202 | 3401 | 3598 |
| ENSG00000284224 | 0 | 0 | 0 | 0 | 0 | 0 |
| ENSG00000179950 | 6309 | 3033 | 3787 | 4105 | 3301 | 3370 |
| ENSG00000255343 | 0 | 0 | 1 | 0 | 0 | 1 |
| ENSG00000185189 | 917 | 360 | 535 | 379 | 404 | 433 |
| ENSG00000284142 | 0 | 0 | 0 | 0 | 0 | 0 |
| ENSG00000285599 | 0 | 0 | 0 | 0 | 0 | 0 |
| ENSG00000261150 | 1618 | 611 | 1064 | 804 | 878 | 694 |
| ENSG00000178209 | 16733 | 6249 | 10293 | 8604 | 8186 | 9057 |
| ENSG00000207574 | 0 | 0 | 0 | 0 | 0 | 0 |
| ENSG00000178685 | 70 | 32 | 52 | 26 | 32 | 28 |
| ENSG00000178719 | 8234 | 3899 | 5235 | 5453 | 4161 | 4383 |
| ENSG00000186583 | 4 | 2 | 3 | 2 | 2 | 1 |
| ENSG00000204791 | 1 | 3 | 3 | 1 | 0 | 1 |
| ENSG00000178814 | 333 | 177 | 259 | 179 | 191 | 155 |
| ENSG00000277888 | 0 | 0 | 0 | 0 | 0 | 0 |
| ENSG00000255224 | 96 | 25 | 50 | 58 | 47 | 34 |
| ENSG00000178896 | 704 | 415 | 476 | 614 | 486 | 464 |
| ENSG00000284200 | 0 | 0 | 0 | 0 | 0 | 0 |
| ENSG00000197858 | 5304 | 2418 | 3281 | 3213 | 2551 | 2705 |
| ENSG00000179091 | 6871 | 3172 | 3789 | 4602 | 3708 | 3720 |
| ENSG00000179526 | 1812 | 870 | 1082 | 1159 | 979 | 939 |
| ENSG00000179632 | 4821 | 2518 | 3002 | 3244 | 2884 | 2689 |
| ENSG00000179698 | 183 | 73 | 110 | 113 | 74 | 130 |
| ENSG00000235173 | 1378 | 608 | 735 | 836 | 567 | 719 |
| ENSG00000227473 | 62 | 37 | 59 | 29 | 56 | 50 |
| ENSG00000179832 | 2308 | 968 | 1461 | 1318 | 1301 | 1376 |
| ENSG00000261236 | 2096 | 998 | 1373 | 1616 | 1267 | 1385 |
| ENSG00000284054 | 0 | 0 | 0 | 0 | 0 | 0 |
| ENSG00000260428 | 80 | 40 | 57 | 51 | 53 | 53 |
| ENSG00000185122 | 2870 | 1404 | 1833 | 1941 | 1853 | 1713 |
| ENSG00000185000 | 1054 | 485 | 641 | 625 | 528 | 537 |
| ENSG00000254690 | 0 | 0 | 0 | 0 | 0 | 0 |
| ENSG00000284229 | 0 | 0 | 0 | 0 | 0 | 0 |
| ENSG00000261678 | 8 | 4 | 7 | 4 | 4 | 5 |
| ENSG00000261587 | 0 | 4 | 1 | 2 | 1 | 2 |
| ENSG00000271698 | 0 | 1 | 2 | 0 | 1 | 0 |
| ENSG00000185803 | 2500 | 1124 | 1402 | 1947 | 1441 | 1480 |
| ENSG00000182325 | 717 | 338 | 480 | 421 | 363 | 416 |

| | | | | | | |
|---|---|---|---|---|---|---|
| ENSG00000173137 | 769 | 327 | 467 | 401 | 347 | 328 |
| ENSG00000071894 | 5589 | 2429 | 3344 | 3145 | 2885 | 3103 |
| ENSG00000284310 | 0 | 0 | 0 | 0 | 0 | 0 |
| ENSG00000284139 | 0 | 0 | 0 | 0 | 0 | 0 |
| ENSG00000283770 | 0 | 0 | 0 | 0 | 0 | 0 |
| ENSG00000272115 | 20 | 3 | 11 | 3 | 4 | 5 |
| ENSG00000147804 | 645 | 290 | 356 | 460 | 402 | 400 |
| ENSG00000160948 | 3796 | 1835 | 2517 | 2364 | 1850 | 2004 |
| ENSG00000160949 | 1315 | 572 | 773 | 619 | 652 | 669 |
| ENSG00000276598 | 0 | 0 | 0 | 0 | 0 | 0 |
| ENSG00000232600 | 16 | 9 | 10 | 6 | 11 | 9 |
| ENSG00000187954 | 2899 | 1357 | 1885 | 1837 | 1637 | 1573 |
| ENSG00000254578 | 0 | 0 | 0 | 0 | 0 | 0 |
| ENSG00000167702 | 2151 | 796 | 1439 | 999 | 1087 | 1194 |
| ENSG00000160973 | 12 | 4 | 2 | 4 | 3 | 3 |
| ENSG00000160972 | 1274 | 590 | 751 | 877 | 796 | 823 |
| ENSG00000255182 | 60 | 37 | 57 | 33 | 37 | 67 |
| ENSG00000167701 | 81 | 22 | 48 | 18 | 29 | 33 |
| ENSG00000167700 | 1512 | 658 | 834 | 1053 | 818 | 813 |
| ENSG00000160957 | 1367 | 632 | 848 | 804 | 698 | 769 |
| ENSG00000265393 | 0 | 0 | 0 | 0 | 0 | 0 |
| ENSG00000160959 | 1155 | 547 | 756 | 689 | 635 | 637 |
| ENSG00000254402 | 0 | 2 | 0 | 0 | 0 | 1 |
| ENSG00000213563 | 1214 | 604 | 820 | 849 | 751 | 718 |
| ENSG00000147799 | 968 | 485 | 627 | 609 | 491 | 537 |
| ENSG00000255456 | 0 | 0 | 0 | 1 | 0 | 0 |
| ENSG00000274772 | 0 | 0 | 0 | 0 | 0 | 0 |
| ENSG00000254533 | 0 | 0 | 0 | 0 | 0 | 0 |
| ENSG00000255085 | 0 | 0 | 0 | 2 | 0 | 0 |
| ENSG00000263612 | 4 | 2 | 2 | 1 | 3 | 1 |
| ENSG00000198169 | 1118 | 485 | 698 | 450 | 538 | 521 |
| ENSG00000196378 | 186 | 103 | 132 | 92 | 131 | 103 |
| ENSG00000244307 | 0 | 1 | 2 | 0 | 2 | 1 |
| ENSG00000161016 | 65528 | 32710 | 37561 | 51088 | 39432 | 38601 |
| ENSG00000283764 | 0 | 0 | 0 | 0 | 0 | 0 |
| ENSG00000197363 | 801 | 340 | 483 | 398 | 438 | 400 |
| ENSG00000147789 | 889 | 478 | 673 | 456 | 516 | 583 |
| ENSG00000170619 | 1278 | 568 | 804 | 807 | 682 | 730 |
| ENSG00000196150 | 589 | 272 | 330 | 245 | 355 | 259 |
| ENSG00000170631 | 570 | 276 | 322 | 250 | 285 | 278 |
| ENSG00000196922 | 791 | 450 | 425 | 245 | 619 | 407 |
| ENSG00000255164 | 0 | 1 | 0 | 1 | 0 | 0 |
| ENSG00000254618 | 0 | 0 | 0 | 0 | 0 | 0 |
| ENSG00000255559 | 31 | 13 | 10 | 9 | 12 | 11 |
| ENSG00000255530 | 0 | 0 | 0 | 0 | 0 | 0 |
| ENSG00000255000 | 0 | 0 | 0 | 0 | 0 | 0 |
| ENSG00000182307 | 2327 | 1137 | 1399 | 1407 | 1225 | 1448 |
| ENSG00000236875 | 0 | 0 | 0 | 0 | 1 | 0 |
| ENSG00000181404 | 81 | 36 | 53 | 40 | 30 | 25 |
| ENSG00000227518 | 2 | 1 | 0 | 1 | 3 | 0 |

| | | | | | | |
|---|---|---|---|---|---|---|
| ENSG00000283921 | 0 | 0 | 0 | 0 | 0 | 0 |
| ENSG00000218839 | 0 | 0 | 0 | 0 | 0 | 0 |
| ENSG00000284986 | 0 | 0 | 0 | 0 | 0 | 0 |
| ENSG00000277631 | 0 | 0 | 0 | 0 | 0 | 0 |
| ENSG00000227917 | 0 | 0 | 0 | 0 | 1 | 0 |
| ENSG00000231808 | 0 | 0 | 0 | 0 | 0 | 0 |
| ENSG00000170122 | 49 | 12 | 34 | 13 | 18 | 17 |
| ENSG00000172785 | 1325 | 765 | 738 | 450 | 803 | 739 |
| ENSG00000183784 | 187 | 109 | 110 | 72 | 86 | 113 |
| ENSG00000107099 | 2470 | 1128 | 1490 | 581 | 954 | 1059 |
| ENSG00000235880 | 0 | 1 | 0 | 0 | 0 | 0 |
| ENSG00000227155 | 24 | 21 | 15 | 6 | 4 | 12 |
| ENSG00000107104 | 518 | 278 | 319 | 239 | 311 | 292 |
| ENSG00000235330 | 14 | 6 | 9 | 7 | 9 | 12 |
| ENSG00000228115 | 0 | 0 | 2 | 1 | 0 | 0 |
| ENSG00000226403 | 0 | 0 | 0 | 0 | 0 | 0 |
| ENSG00000202172 | 0 | 0 | 0 | 0 | 0 | 0 |
| ENSG00000229875 | 0 | 0 | 0 | 0 | 0 | 0 |
| ENSG00000227914 | 0 | 0 | 0 | 0 | 1 | 0 |
| ENSG00000137090 | 0 | 0 | 0 | 0 | 0 | 0 |
| ENSG00000064218 | 0 | 0 | 0 | 0 | 0 | 0 |
| ENSG00000252991 | 0 | 0 | 0 | 0 | 0 | 0 |
| ENSG00000228783 | 0 | 0 | 0 | 0 | 0 | 0 |
| ENSG00000281769 | 0 | 0 | 0 | 0 | 0 | 0 |
| ENSG00000173253 | 12 | 7 | 8 | 6 | 1 | 11 |
| ENSG00000224836 | 0 | 0 | 0 | 0 | 0 | 0 |
| ENSG00000236594 | 0 | 0 | 0 | 0 | 0 | 0 |
| ENSG00000233575 | 0 | 0 | 0 | 0 | 0 | 0 |
| ENSG00000202383 | 0 | 0 | 0 | 0 | 0 | 0 |
| ENSG00000224092 | 0 | 1 | 0 | 0 | 0 | 0 |
| ENSG00000080503 | 2207 | 977 | 1261 | 738 | 1006 | 990 |
| ENSG00000236199 | 0 | 0 | 0 | 0 | 0 | 0 |
| ENSG00000222973 | 0 | 0 | 0 | 0 | 0 | 0 |
| ENSG00000264615 | 0 | 0 | 0 | 0 | 0 | 0 |
| ENSG00000236404 | 17 | 16 | 41 | 11 | 11 | 8 |
| ENSG00000231052 | 0 | 0 | 0 | 0 | 0 | 0 |
| ENSG00000147852 | 1262 | 568 | 818 | 359 | 381 | 512 |
| ENSG00000168263 | 2 | 0 | 3 | 1 | 1 | 1 |
| ENSG00000080608 | 529 | 374 | 330 | 238 | 600 | 429 |
| ENSG00000236496 | 26 | 17 | 11 | 14 | 15 | 22 |
| ENSG00000229184 | 0 | 0 | 0 | 0 | 0 | 1 |
| ENSG00000227835 | 0 | 0 | 0 | 0 | 0 | 0 |
| ENSG00000236511 | 0 | 0 | 0 | 0 | 0 | 0 |
| ENSG00000080298 | 347 | 143 | 172 | 86 | 153 | 139 |
| ENSG00000215297 | 0 | 0 | 0 | 0 | 0 | 0 |
| ENSG00000232104 | 19 | 15 | 12 | 3 | 20 | 14 |
| ENSG00000237359 | 0 | 0 | 1 | 0 | 0 | 0 |
| ENSG00000226669 | 7 | 0 | 0 | 1 | 1 | 0 |
| ENSG00000107249 | 24 | 17 | 11 | 18 | 20 | 23 |
| ENSG00000236724 | 0 | 0 | 0 | 0 | 0 | 0 |

| | | | | | | |
|---|---|---|---|---|---|---|
| ENSG00000237009 | 0 | 0 | 0 | 0 | 0 | 0 |
| ENSG00000230001 | 0 | 0 | 0 | 0 | 0 | 0 |
| ENSG00000228322 | 0 | 0 | 0 | 0 | 0 | 1 |
| ENSG00000200941 | 0 | 0 | 0 | 0 | 0 | 0 |
| ENSG00000106688 | 8 | 6 | 7 | 9 | 1 | 8 |
| ENSG00000106686 | 32 | 14 | 17 | 8 | 14 | 16 |
| ENSG00000225893 | 0 | 0 | 0 | 0 | 0 | 1 |
| ENSG00000205808 | 1481 | 764 | 927 | 640 | 909 | 895 |
| ENSG00000273061 | 39 | 17 | 36 | 26 | 19 | 13 |
| ENSG00000106993 | 746 | 390 | 417 | 335 | 488 | 402 |
| ENSG00000147853 | 3895 | 1871 | 2171 | 1768 | 2386 | 2174 |
| ENSG00000270639 | 0 | 0 | 0 | 0 | 0 | 0 |
| ENSG00000220105 | 2 | 0 | 0 | 2 | 0 | 0 |
| ENSG00000120158 | 1054 | 552 | 613 | 602 | 573 | 599 |
| ENSG00000233186 | 0 | 1 | 0 | 0 | 0 | 0 |
| ENSG00000199065 | 0 | 0 | 0 | 0 | 0 | 0 |
| ENSG00000228165 | 0 | 0 | 0 | 0 | 0 | 0 |
| ENSG00000220937 | 0 | 0 | 0 | 0 | 0 | 0 |
| ENSG00000096968 | 234 | 142 | 148 | 64 | 185 | 99 |
| ENSG00000234534 | 0 | 0 | 0 | 0 | 0 | 0 |
| ENSG00000182347 | 2 | 1 | 0 | 0 | 0 | 0 |
| ENSG00000227730 | 0 | 0 | 0 | 0 | 0 | 0 |
| ENSG00000228166 | 0 | 2 | 0 | 0 | 0 | 0 |
| ENSG00000224083 | 0 | 0 | 0 | 0 | 1 | 0 |
| ENSG00000235917 | 0 | 0 | 0 | 0 | 0 | 0 |
| ENSG00000228097 | 2 | 1 | 0 | 0 | 0 | 0 |
| ENSG00000237711 | 0 | 0 | 0 | 0 | 0 | 0 |
| ENSG00000236254 | 0 | 0 | 0 | 0 | 0 | 0 |
| ENSG00000230225 | 0 | 0 | 0 | 0 | 0 | 0 |
| ENSG00000236567 | 0 | 1 | 0 | 0 | 0 | 0 |
| ENSG00000254017 | 0 | 0 | 1 | 0 | 0 | 0 |
| ENSG00000120210 | 0 | 0 | 0 | 1 | 0 | 1 |
| ENSG00000237553 | 0 | 0 | 0 | 0 | 0 | 0 |
| ENSG00000120211 | 0 | 1 | 0 | 0 | 0 | 0 |
| ENSG00000107014 | 179 | 84 | 86 | 127 | 119 | 91 |
| ENSG00000214195 | 0 | 1 | 0 | 0 | 0 | 0 |
| ENSG00000107018 | 29 | 7 | 18 | 16 | 11 | 12 |
| ENSG00000272866 | 4 | 0 | 0 | 0 | 0 | 0 |
| ENSG00000231967 | 1 | 0 | 1 | 1 | 0 | 1 |
| ENSG00000107020 | 633 | 334 | 393 | 318 | 371 | 318 |
| ENSG00000214190 | 2 | 0 | 1 | 0 | 0 | 0 |
| ENSG00000120217 | 2 | 5 | 1 | 2 | 0 | 1 |
| ENSG00000197646 | 0 | 1 | 0 | 0 | 0 | 0 |
| ENSG00000231509 | 0 | 0 | 0 | 0 | 0 | 0 |
| ENSG00000107036 | 1194 | 643 | 711 | 477 | 742 | 607 |
| ENSG00000225408 | 0 | 1 | 2 | 0 | 1 | 2 |
| ENSG00000099219 | 12880 | 6107 | 7588 | 4013 | 5588 | 5428 |
| ENSG00000223621 | 0 | 2 | 2 | 1 | 3 | 0 |
| ENSG00000183354 | 452 | 240 | 169 | 64 | 355 | 203 |
| ENSG00000120215 | 20 | 7 | 6 | 2 | 7 | 2 |

| | | | | | | |
|---|---|---|---|---|---|---|
| ENSG00000263575 | 0 | 0 | 0 | 0 | 0 | 0 |
| ENSG00000137040 | 689 | 398 | 350 | 247 | 684 | 401 |
| ENSG00000228739 | 0 | 0 | 0 | 0 | 0 | 0 |
| ENSG00000237274 | 0 | 0 | 0 | 0 | 0 | 0 |
| ENSG00000137033 | 0 | 0 | 1 | 0 | 1 | 2 |
| ENSG00000228132 | 0 | 0 | 0 | 0 | 0 | 0 |
| ENSG00000233846 | 2 | 1 | 4 | 1 | 2 | 1 |
| ENSG00000170777 | 0 | 0 | 0 | 0 | 0 | 0 |
| ENSG00000147854 | 1359 | 657 | 694 | 435 | 829 | 634 |
| ENSG00000233367 | 2 | 0 | 0 | 0 | 0 | 0 |
| ENSG00000178445 | 55 | 40 | 42 | 17 | 21 | 25 |
| ENSG00000264530 | 0 | 0 | 0 | 0 | 0 | 0 |
| ENSG00000237158 | 0 | 0 | 0 | 0 | 0 | 0 |
| ENSG00000206147 | 4 | 0 | 0 | 0 | 0 | 0 |
| ENSG00000236924 | 35 | 27 | 22 | 39 | 39 | 28 |
| ENSG00000266863 | 0 | 0 | 0 | 0 | 0 | 0 |
| ENSG00000229057 | 4 | 2 | 1 | 3 | 2 | 0 |
| ENSG00000231381 | 10 | 4 | 6 | 6 | 5 | 9 |
| ENSG00000232946 | 6 | 3 | 1 | 1 | 5 | 5 |
| ENSG00000229611 | 1 | 3 | 2 | 2 | 4 | 0 |
| ENSG00000225489 | 152 | 84 | 128 | 54 | 74 | 91 |
| ENSG00000107077 | 1235 | 522 | 738 | 407 | 624 | 607 |
| ENSG00000231108 | 3 | 0 | 0 | 0 | 0 | 0 |
| ENSG00000256968 | 2 | 6 | 1 | 3 | 1 | 4 |
| ENSG00000230581 | 2 | 0 | 3 | 1 | 3 | 0 |
| ENSG00000224972 | 0 | 0 | 0 | 0 | 0 | 0 |
| ENSG00000285688 | 4 | 3 | 0 | 0 | 0 | 3 |
| ENSG00000230207 | 12 | 3 | 2 | 5 | 2 | 9 |
| ENSG00000178654 | 1 | 0 | 0 | 0 | 0 | 0 |
| ENSG00000273056 | 0 | 0 | 0 | 0 | 0 | 0 |
| ENSG00000137038 | 1399 | 681 | 789 | 990 | 859 | 742 |
| ENSG00000226376 | 0 | 0 | 0 | 0 | 0 | 0 |
| ENSG00000231902 | 0 | 0 | 0 | 0 | 0 | 0 |
| ENSG00000153707 | 0 | 0 | 0 | 0 | 0 | 0 |
| ENSG00000235389 | 0 | 0 | 0 | 0 | 0 | 0 |
| ENSG00000212695 | 0 | 0 | 0 | 0 | 0 | 0 |
| ENSG00000239102 | 0 | 0 | 0 | 0 | 0 | 0 |
| ENSG00000225706 | 0 | 0 | 0 | 0 | 0 | 0 |
| ENSG00000234021 | 0 | 0 | 0 | 0 | 0 | 0 |
| ENSG00000212829 | 0 | 0 | 0 | 0 | 0 | 0 |
| ENSG00000265735 | 0 | 0 | 1 | 0 | 0 | 0 |
| ENSG00000230920 | 0 | 0 | 0 | 0 | 0 | 0 |
| ENSG00000234360 | 0 | 0 | 0 | 0 | 0 | 0 |
| ENSG00000226717 | 0 | 0 | 0 | 0 | 0 | 0 |
| ENSG00000232035 | 0 | 0 | 0 | 0 | 0 | 0 |
| ENSG00000227138 | 0 | 0 | 0 | 0 | 0 | 0 |
| ENSG00000230589 | 0 | 0 | 0 | 0 | 0 | 0 |
| ENSG00000270372 | 0 | 0 | 1 | 1 | 2 | 3 |
| ENSG00000234146 | 0 | 0 | 0 | 0 | 0 | 0 |
| ENSG00000230365 | 0 | 0 | 0 | 0 | 0 | 0 |

| | | | | | | |
|---|---|---|---|---|---|---|
| ENSG00000275130 | 0 | 0 | 0 | 0 | 0 | 0 |
| ENSG00000285784 | 0 | 0 | 0 | 0 | 0 | 0 |
| ENSG00000224935 | 0 | 0 | 0 | 0 | 0 | 0 |
| ENSG00000231491 | 0 | 0 | 0 | 0 | 0 | 0 |
| ENSG00000222581 | 0 | 0 | 0 | 0 | 0 | 0 |
| ENSG00000107165 | 803 | 416 | 484 | 291 | 420 | 349 |
| ENSG00000235448 | 0 | 0 | 3 | 0 | 0 | 1 |
| ENSG00000243115 | 0 | 0 | 0 | 0 | 0 | 0 |
| ENSG00000153714 | 1507 | 728 | 918 | 700 | 764 | 746 |
| ENSG00000222658 | 0 | 0 | 0 | 0 | 0 | 0 |
| ENSG00000285637 | 3 | 1 | 3 | 0 | 3 | 1 |
| ENSG00000214121 | 4 | 2 | 9 | 0 | 5 | 5 |
| ENSG00000234586 | 0 | 0 | 0 | 0 | 0 | 0 |
| ENSG00000107186 | 654 | 339 | 318 | 121 | 392 | 243 |
| ENSG00000234740 | 1 | 0 | 2 | 0 | 1 | 1 |
| ENSG00000223672 | 0 | 0 | 0 | 0 | 0 | 1 |
| ENSG00000270547 | 1 | 1 | 2 | 0 | 1 | 0 |
| ENSG00000226197 | 0 | 0 | 0 | 0 | 0 | 0 |
| ENSG00000205636 | 0 | 0 | 0 | 0 | 0 | 0 |
| ENSG00000229268 | 0 | 0 | 0 | 0 | 0 | 0 |
| ENSG00000235533 | 0 | 0 | 0 | 0 | 0 | 0 |
| ENSG00000223568 | 0 | 0 | 0 | 0 | 0 | 0 |
| ENSG00000213954 | 0 | 0 | 0 | 0 | 0 | 0 |
| ENSG00000147862 | 1149 | 490 | 558 | 275 | 481 | 396 |
| ENSG00000234982 | 0 | 0 | 0 | 0 | 0 | 0 |
| ENSG00000225472 | 0 | 0 | 0 | 0 | 0 | 0 |
| ENSG00000237137 | 0 | 0 | 0 | 0 | 0 | 0 |
| ENSG00000273399 | 0 | 0 | 0 | 0 | 0 | 0 |
| ENSG00000272871 | 4 | 5 | 7 | 2 | 5 | 9 |
| ENSG00000229029 | 2 | 0 | 0 | 0 | 1 | 0 |
| ENSG00000175893 | 893 | 451 | 462 | 271 | 692 | 497 |
| ENSG00000147869 | 0 | 0 | 0 | 0 | 0 | 0 |
| ENSG00000164946 | 12 | 3 | 6 | 0 | 2 | 1 |
| ENSG00000283647 | 0 | 0 | 0 | 0 | 0 | 0 |
| ENSG00000199814 | 0 | 0 | 0 | 0 | 0 | 0 |
| ENSG00000214110 | 44 | 24 | 18 | 18 | 19 | 28 |
| ENSG00000232000 | 0 | 0 | 1 | 0 | 0 | 0 |
| ENSG00000215237 | 0 | 0 | 0 | 0 | 0 | 0 |
| ENSG00000234297 | 0 | 0 | 0 | 0 | 0 | 0 |
| ENSG00000251960 | 0 | 0 | 0 | 0 | 0 | 0 |
| ENSG00000155158 | 1378 | 676 | 845 | 483 | 884 | 697 |
| ENSG00000215236 | 0 | 0 | 0 | 0 | 0 | 0 |
| ENSG00000164975 | 1466 | 792 | 927 | 596 | 997 | 809 |
| ENSG00000251834 | 0 | 0 | 0 | 0 | 0 | 0 |
| ENSG00000164985 | 1162 | 726 | 745 | 500 | 950 | 649 |
| ENSG00000278754 | 0 | 0 | 0 | 0 | 0 | 0 |
| ENSG00000240613 | 0 | 0 | 0 | 0 | 0 | 0 |
| ENSG00000213362 | 1 | 0 | 0 | 2 | 0 | 1 |
| ENSG00000207433 | 0 | 0 | 0 | 0 | 0 | 0 |
| ENSG00000164989 | 57 | 40 | 32 | 14 | 31 | 23 |

| | | | | | | |
|---|---|---|---|---|---|---|
| ENSG00000231376 | 0 | 0 | 0 | 0 | 0 | 0 |
| ENSG00000207360 | 0 | 0 | 0 | 0 | 0 | 0 |
| ENSG00000205549 | 0 | 0 | 0 | 0 | 0 | 0 |
| ENSG00000173068 | 0 | 0 | 0 | 0 | 0 | 0 |
| ENSG00000231756 | 0 | 0 | 0 | 0 | 0 | 0 |
| ENSG00000230694 | 0 | 0 | 0 | 0 | 0 | 0 |
| ENSG00000234779 | 0 | 0 | 0 | 0 | 0 | 0 |
| ENSG00000233551 | 0 | 0 | 0 | 0 | 0 | 0 |
| ENSG00000237153 | 0 | 0 | 0 | 0 | 0 | 0 |
| ENSG00000285169 | 0 | 0 | 0 | 0 | 0 | 0 |
| ENSG00000241152 | 0 | 0 | 0 | 0 | 0 | 0 |
| ENSG00000226470 | 0 | 0 | 0 | 0 | 0 | 0 |
| ENSG00000237083 | 0 | 0 | 0 | 0 | 0 | 0 |
| ENSG00000044459 | 2 | 4 | 1 | 1 | 3 | 0 |
| ENSG00000224488 | 0 | 0 | 0 | 0 | 0 | 0 |
| ENSG00000107295 | 0 | 0 | 0 | 0 | 0 | 0 |
| ENSG00000228810 | 0 | 0 | 0 | 0 | 0 | 0 |
| ENSG00000178031 | 58 | 19 | 28 | 22 | 25 | 30 |
| ENSG00000227167 | 0 | 0 | 0 | 0 | 0 | 0 |
| ENSG00000264638 | 0 | 0 | 0 | 0 | 0 | 0 |
| ENSG00000252960 | 0 | 0 | 0 | 0 | 0 | 0 |
| ENSG00000236857 | 0 | 0 | 0 | 0 | 0 | 0 |
| ENSG00000155875 | 10 | 2 | 3 | 4 | 0 | 2 |
| ENSG00000236680 | 0 | 0 | 2 | 0 | 0 | 1 |
| ENSG00000155876 | 2864 | 1339 | 1570 | 1501 | 1545 | 1366 |
| ENSG00000147874 | 495 | 328 | 215 | 124 | 431 | 227 |
| ENSG00000251733 | 0 | 0 | 2 | 0 | 0 | 0 |
| ENSG00000252943 | 0 | 0 | 0 | 0 | 0 | 0 |
| ENSG00000201118 | 0 | 0 | 0 | 0 | 5 | 0 |
| ENSG00000147872 | 350 | 173 | 203 | 135 | 163 | 128 |
| ENSG00000232176 | 54 | 2 | 5 | 1 | 2 | 1 |
| ENSG00000137145 | 1210 | 739 | 481 | 177 | 893 | 459 |
| ENSG00000232978 | 0 | 0 | 0 | 0 | 0 | 0 |
| ENSG00000272842 | 4 | 4 | 2 | 0 | 4 | 2 |
| ENSG00000273226 | 12 | 6 | 3 | 0 | 4 | 9 |
| ENSG00000137154 | 43085 | 22381 | 25104 | 27152 | 26818 | 24596 |
| ENSG00000234853 | 0 | 1 | 0 | 0 | 0 | 0 |
| ENSG00000285911 | 5 | 5 | 9 | 7 | 5 | 9 |
| ENSG00000177076 | 684 | 316 | 409 | 148 | 303 | 305 |
| ENSG00000260912 | 914 | 420 | 486 | 334 | 419 | 379 |
| ENSG00000231909 | 0 | 0 | 0 | 0 | 0 | 1 |
| ENSG00000155886 | 18 | 16 | 16 | 5 | 18 | 14 |
| ENSG00000232158 | 0 | 0 | 0 | 0 | 0 | 0 |
| ENSG00000276667 | 0 | 0 | 0 | 0 | 0 | 0 |
| ENSG00000261402 | 0 | 0 | 0 | 0 | 0 | 0 |
| ENSG00000270712 | 0 | 0 | 0 | 0 | 0 | 0 |
| ENSG00000277711 | 0 | 0 | 0 | 0 | 0 | 0 |
| ENSG00000237434 | 0 | 0 | 0 | 0 | 0 | 0 |
| ENSG00000171843 | 912 | 488 | 452 | 321 | 588 | 381 |
| ENSG00000263790 | 0 | 0 | 0 | 0 | 0 | 0 |

| | | | | | | |
|---|---|---|---|---|---|---|
| ENSG00000222202 | 0 | 0 | 0 | 0 | 0 | 0 |
| ENSG00000264941 | 0 | 0 | 0 | 0 | 0 | 0 |
| ENSG00000237088 | 0 | 0 | 0 | 0 | 0 | 0 |
| ENSG00000188352 | 1757 | 903 | 996 | 753 | 1054 | 927 |
| ENSG00000227071 | 2 | 3 | 0 | 0 | 3 | 0 |
| ENSG00000207609 | 1 | 1 | 0 | 0 | 0 | 0 |
| ENSG00000202189 | 0 | 0 | 0 | 0 | 0 | 0 |
| ENSG00000188921 | 5 | 9 | 7 | 8 | 10 | 9 |
| ENSG00000230208 | 0 | 0 | 0 | 0 | 0 | 0 |
| ENSG00000171855 | 0 | 0 | 0 | 0 | 0 | 0 |
| ENSG00000225027 | 0 | 0 | 0 | 0 | 0 | 0 |
| ENSG00000275980 | 0 | 0 | 0 | 0 | 0 | 0 |
| ENSG00000177047 | 0 | 0 | 0 | 0 | 0 | 0 |
| ENSG00000199635 | 0 | 0 | 0 | 0 | 0 | 0 |
| ENSG00000137080 | 0 | 0 | 0 | 0 | 0 | 0 |
| ENSG00000232281 | 0 | 0 | 0 | 0 | 0 | 0 |
| ENSG00000236637 | 0 | 0 | 0 | 0 | 0 | 0 |
| ENSG00000226597 | 0 | 0 | 0 | 0 | 0 | 0 |
| ENSG00000214042 | 0 | 0 | 0 | 0 | 0 | 0 |
| ENSG00000186803 | 0 | 0 | 0 | 0 | 0 | 0 |
| ENSG00000223684 | 0 | 0 | 0 | 0 | 0 | 0 |
| ENSG00000147885 | 0 | 0 | 0 | 0 | 0 | 0 |
| ENSG00000234829 | 0 | 0 | 0 | 0 | 0 | 0 |
| ENSG00000232138 | 0 | 0 | 0 | 0 | 0 | 0 |
| ENSG00000228083 | 0 | 0 | 0 | 0 | 0 | 0 |
| ENSG00000224416 | 0 | 0 | 0 | 0 | 0 | 0 |
| ENSG00000147873 | 0 | 0 | 0 | 0 | 0 | 0 |
| ENSG00000226393 | 0 | 1 | 0 | 0 | 0 | 1 |
| ENSG00000198642 | 1357 | 716 | 723 | 543 | 754 | 692 |
| ENSG00000120235 | 0 | 0 | 0 | 0 | 0 | 0 |
| ENSG00000233816 | 0 | 0 | 0 | 0 | 0 | 0 |
| ENSG00000188379 | 0 | 0 | 0 | 0 | 0 | 0 |
| ENSG00000261696 | 0 | 0 | 0 | 0 | 0 | 0 |
| ENSG00000231195 | 0 | 0 | 0 | 0 | 0 | 0 |
| ENSG00000235108 | 0 | 0 | 0 | 0 | 0 | 0 |
| ENSG00000120242 | 0 | 0 | 0 | 0 | 0 | 0 |
| ENSG00000237691 | 0 | 0 | 0 | 0 | 0 | 0 |
| ENSG00000197919 | 0 | 0 | 0 | 0 | 0 | 0 |
| ENSG00000238271 | 0 | 0 | 0 | 0 | 0 | 0 |
| ENSG00000171889 | 0 | 0 | 0 | 0 | 0 | 0 |
| ENSG00000184995 | 0 | 0 | 0 | 0 | 0 | 0 |
| ENSG00000199177 | 0 | 0 | 0 | 0 | 0 | 0 |
| ENSG00000264379 | 0 | 0 | 0 | 0 | 0 | 0 |
| ENSG00000227443 | 0 | 0 | 0 | 0 | 0 | 0 |
| ENSG00000229835 | 3 | 6 | 11 | 3 | 9 | 2 |
| ENSG00000244230 | 4 | 1 | 0 | 0 | 0 | 0 |
| ENSG00000099810 | 18 | 9 | 21 | 7 | 14 | 10 |
| ENSG00000264545 | 0 | 1 | 0 | 0 | 0 | 0 |
| ENSG00000229298 | 21 | 12 | 22 | 9 | 15 | 8 |
| ENSG00000265194 | 0 | 0 | 0 | 0 | 0 | 0 |

| | | | | | | |
|---|---|---|---|---|---|---|
| ENSG00000279670 | 0 | 0 | 0 | 0 | 0 | 0 |
| ENSG00000264801 | 0 | 0 | 0 | 0 | 0 | 0 |
| ENSG00000224854 | 0 | 0 | 0 | 0 | 0 | 0 |
| ENSG00000147889 | 71 | 49 | 74 | 41 | 31 | 37 |
| ENSG00000240498 | 491 | 264 | 337 | 122 | 304 | 253 |
| ENSG00000266446 | 1 | 2 | 0 | 0 | 2 | 0 |
| ENSG00000147883 | 1639 | 823 | 1036 | 583 | 712 | 654 |
| ENSG00000215221 | 10 | 6 | 8 | 2 | 5 | 2 |
| ENSG00000274055 | 0 | 0 | 0 | 0 | 0 | 0 |
| ENSG00000277653 | 0 | 0 | 0 | 0 | 0 | 0 |
| ENSG00000236921 | 4 | 3 | 4 | 3 | 3 | 3 |
| ENSG00000176399 | 501 | 239 | 246 | 156 | 173 | 166 |
| ENSG00000234840 | 25 | 14 | 7 | 4 | 24 | 5 |
| ENSG00000284418 | 1 | 0 | 0 | 1 | 0 | 0 |
| ENSG00000236739 | 0 | 0 | 1 | 0 | 0 | 2 |
| ENSG00000224549 | 0 | 1 | 0 | 1 | 3 | 1 |
| ENSG00000283982 | 11 | 6 | 9 | 2 | 7 | 7 |
| ENSG00000227607 | 0 | 0 | 0 | 0 | 0 | 0 |
| ENSG00000233906 | 0 | 0 | 0 | 0 | 0 | 0 |
| ENSG00000226626 | 0 | 0 | 0 | 0 | 0 | 0 |
| ENSG00000107105 | 0 | 0 | 1 | 1 | 0 | 0 |
| ENSG00000231460 | 0 | 0 | 0 | 0 | 0 | 0 |
| ENSG00000237414 | 0 | 0 | 0 | 0 | 0 | 0 |
| ENSG00000277608 | 0 | 0 | 0 | 0 | 0 | 0 |
| ENSG00000205442 | 0 | 0 | 0 | 0 | 0 | 0 |
| ENSG00000269957 | 0 | 0 | 0 | 0 | 0 | 0 |
| ENSG00000223088 | 0 | 0 | 0 | 0 | 0 | 0 |
| ENSG00000222693 | 0 | 0 | 0 | 0 | 0 | 0 |
| ENSG00000198680 | 815 | 396 | 419 | 410 | 394 | 402 |
| ENSG00000236306 | 0 | 0 | 0 | 0 | 0 | 0 |
| ENSG00000270683 | 0 | 0 | 0 | 0 | 0 | 0 |
| ENSG00000276759 | 0 | 0 | 0 | 0 | 0 | 0 |
| ENSG00000279256 | 0 | 0 | 0 | 0 | 0 | 0 |
| ENSG00000254261 | 0 | 0 | 0 | 0 | 0 | 0 |
| ENSG00000252580 | 0 | 0 | 0 | 0 | 0 | 0 |
| ENSG00000253400 | 0 | 0 | 0 | 0 | 0 | 0 |
| ENSG00000120159 | 943 | 435 | 571 | 501 | 537 | 512 |
| ENSG00000232466 | 0 | 0 | 0 | 0 | 0 | 1 |
| ENSG00000240306 | 0 | 0 | 0 | 0 | 0 | 0 |
| ENSG00000137055 | 1933 | 979 | 1083 | 894 | 1160 | 1118 |
| ENSG00000096872 | 262 | 169 | 135 | 90 | 229 | 121 |
| ENSG00000234676 | 0 | 0 | 2 | 1 | 4 | 1 |
| ENSG00000184434 | 0 | 0 | 1 | 0 | 3 | 0 |
| ENSG00000254396 | 0 | 0 | 2 | 0 | 0 | 0 |
| ENSG00000120156 | 0 | 0 | 1 | 0 | 0 | 0 |
| ENSG00000223162 | 0 | 0 | 0 | 0 | 0 | 0 |
| ENSG00000232867 | 0 | 0 | 0 | 0 | 0 | 0 |
| ENSG00000283945 | 0 | 0 | 0 | 0 | 0 | 0 |
| ENSG00000120160 | 0 | 0 | 0 | 0 | 0 | 0 |
| ENSG00000120162 | 6 | 1 | 0 | 0 | 1 | 2 |

| | | | | | | |
|---|---|---|---|---|---|---|
| ENSG00000237734 | 0 | 0 | 0 | 0 | 0 | 0 |
| ENSG00000147896 | 0 | 0 | 0 | 0 | 0 | 0 |
| ENSG00000285103 | 0 | 2 | 0 | 0 | 0 | 0 |
| ENSG00000147894 | 108 | 74 | 68 | 42 | 51 | 54 |
| ENSG00000215441 | 0 | 0 | 0 | 0 | 0 | 0 |
| ENSG00000260390 | 0 | 0 | 0 | 0 | 0 | 0 |
| ENSG00000260412 | 0 | 0 | 0 | 0 | 0 | 0 |
| ENSG00000174482 | 0 | 0 | 0 | 0 | 0 | 0 |
| ENSG00000229060 | 0 | 0 | 0 | 0 | 0 | 0 |
| ENSG00000224955 | 0 | 0 | 0 | 0 | 0 | 0 |
| ENSG00000215966 | 0 | 0 | 0 | 0 | 0 | 0 |
| ENSG00000215939 | 0 | 0 | 0 | 0 | 0 | 0 |
| ENSG00000230276 | 0 | 0 | 0 | 0 | 0 | 0 |
| ENSG00000277646 | 0 | 0 | 0 | 0 | 0 | 0 |
| ENSG00000229834 | 0 | 0 | 0 | 0 | 0 | 0 |
| ENSG00000230097 | 0 | 0 | 0 | 0 | 0 | 0 |
| ENSG00000280683 | 0 | 0 | 0 | 0 | 0 | 0 |
| ENSG00000232060 | 0 | 0 | 3 | 0 | 0 | 1 |
| ENSG00000239792 | 0 | 0 | 0 | 0 | 0 | 0 |
| ENSG00000215210 | 6 | 11 | 2 | 2 | 2 | 5 |
| ENSG00000235626 | 0 | 0 | 0 | 0 | 0 | 0 |
| ENSG00000226246 | 0 | 0 | 0 | 0 | 0 | 0 |
| ENSG00000233278 | 0 | 0 | 0 | 0 | 0 | 0 |
| ENSG00000214020 | 0 | 0 | 0 | 0 | 0 | 0 |
| ENSG00000228557 | 0 | 0 | 0 | 0 | 0 | 0 |
| ENSG00000223421 | 0 | 0 | 0 | 0 | 0 | 0 |
| ENSG00000232203 | 2 | 0 | 0 | 0 | 0 | 1 |
| ENSG00000260720 | 0 | 0 | 0 | 0 | 0 | 0 |
| ENSG00000226998 | 0 | 0 | 0 | 0 | 0 | 0 |
| ENSG00000229080 | 0 | 0 | 0 | 0 | 0 | 0 |
| ENSG00000235148 | 0 | 0 | 0 | 0 | 0 | 0 |
| ENSG00000252313 | 0 | 0 | 0 | 0 | 0 | 0 |
| ENSG00000225347 | 0 | 0 | 0 | 0 | 0 | 0 |
| ENSG00000122729 | 1323 | 678 | 752 | 571 | 687 | 677 |
| ENSG00000107201 | 151 | 104 | 93 | 42 | 103 | 90 |
| ENSG00000197579 | 828 | 427 | 521 | 302 | 529 | 436 |
| ENSG00000235453 | 89 | 57 | 76 | 75 | 53 | 79 |
| ENSG00000165264 | 1160 | 591 | 723 | 805 | 703 | 734 |
| ENSG00000232303 | 9 | 2 | 5 | 8 | 2 | 5 |
| ENSG00000122728 | 0 | 0 | 1 | 0 | 0 | 0 |
| ENSG00000223440 | 0 | 0 | 0 | 0 | 0 | 0 |
| ENSG00000230516 | 0 | 0 | 0 | 0 | 0 | 1 |
| ENSG00000238001 | 0 | 0 | 0 | 0 | 1 | 0 |
| ENSG00000230867 | 0 | 0 | 0 | 0 | 0 | 0 |
| ENSG00000188133 | 0 | 0 | 0 | 0 | 0 | 0 |
| ENSG00000272113 | 0 | 0 | 0 | 0 | 0 | 0 |
| ENSG00000236796 | 0 | 0 | 0 | 0 | 0 | 0 |
| ENSG00000231193 | 0 | 0 | 0 | 1 | 0 | 0 |
| ENSG00000223807 | 0 | 0 | 2 | 0 | 0 | 0 |
| ENSG00000237437 | 0 | 2 | 10 | 5 | 7 | 8 |

| | | | | | | |
|---|---|---|---|---|---|---|
| ENSG00000137074 | 1070 | 592 | 725 | 648 | 676 | 667 |
| ENSG00000236184 | 18 | 5 | 4 | 2 | 5 | 0 |
| ENSG00000225693 | 0 | 1 | 1 | 0 | 2 | 3 |
| ENSG00000086061 | 5234 | 2826 | 2964 | 2962 | 3873 | 3086 |
| ENSG00000122692 | 2112 | 1113 | 1275 | 986 | 1283 | 1253 |
| ENSG00000086062 | 5834 | 2801 | 3624 | 2697 | 3001 | 3177 |
| ENSG00000233554 | 18 | 7 | 4 | 10 | 3 | 7 |
| ENSG00000252224 | 0 | 0 | 0 | 0 | 0 | 0 |
| ENSG00000122711 | 3 | 1 | 5 | 6 | 7 | 5 |
| ENSG00000107262 | 4218 | 1987 | 2983 | 2975 | 2430 | 3380 |
| ENSG00000086065 | 2965 | 1536 | 1888 | 1590 | 1848 | 1654 |
| ENSG00000086102 | 1526 | 691 | 963 | 556 | 833 | 762 |
| ENSG00000200544 | 0 | 0 | 0 | 0 | 0 | 0 |
| ENSG00000165269 | 6 | 6 | 2 | 2 | 7 | 3 |
| ENSG00000223678 | 0 | 0 | 0 | 0 | 0 | 1 |
| ENSG00000165272 | 3474 | 1808 | 2442 | 2553 | 2585 | 2544 |
| ENSG00000270542 | 0 | 0 | 0 | 0 | 0 | 0 |
| ENSG00000165271 | 1984 | 830 | 1139 | 1264 | 1058 | 1281 |
| ENSG00000275651 | 0 | 0 | 0 | 0 | 0 | 0 |
| ENSG00000226823 | 0 | 2 | 5 | 3 | 0 | 4 |
| ENSG00000228072 | 0 | 0 | 0 | 0 | 0 | 0 |
| ENSG00000230453 | 73 | 33 | 25 | 29 | 47 | 29 |
| ENSG00000225123 | 0 | 2 | 0 | 0 | 0 | 0 |
| ENSG00000234373 | 1 | 4 | 2 | 4 | 3 | 5 |
| ENSG00000226562 | 9 | 1 | 2 | 3 | 9 | 5 |
| ENSG00000204982 | 0 | 0 | 0 | 0 | 0 | 0 |
| ENSG00000205274 | 2 | 0 | 0 | 0 | 0 | 0 |
| ENSG00000231991 | 38 | 17 | 14 | 16 | 23 | 18 |
| ENSG00000183938 | 0 | 0 | 0 | 0 | 0 | 0 |
| ENSG00000234122 | 0 | 0 | 0 | 0 | 0 | 0 |
| ENSG00000229063 | 0 | 0 | 0 | 0 | 0 | 0 |
| ENSG00000215206 | 0 | 0 | 0 | 0 | 0 | 0 |
| ENSG00000240215 | 0 | 0 | 0 | 0 | 0 | 0 |
| ENSG00000237984 | 8 | 4 | 1 | 1 | 1 | 2 |
| ENSG00000281128 | 0 | 0 | 0 | 0 | 0 | 0 |
| ENSG00000231165 | 0 | 0 | 0 | 0 | 0 | 0 |
| ENSG00000260947 | 0 | 0 | 0 | 1 | 0 | 0 |
| ENSG00000227301 | 0 | 0 | 0 | 0 | 0 | 0 |
| ENSG00000233776 | 2 | 1 | 0 | 0 | 0 | 0 |
| ENSG00000010438 | 0 | 0 | 0 | 0 | 1 | 0 |
| ENSG00000235481 | 4 | 1 | 1 | 2 | 1 | 0 |
| ENSG00000223394 | 0 | 0 | 0 | 0 | 0 | 0 |
| ENSG00000107341 | 2519 | 1204 | 1536 | 1415 | 1402 | 1529 |
| ENSG00000200834 | 0 | 0 | 0 | 1 | 0 | 0 |
| ENSG00000251748 | 1 | 0 | 0 | 0 | 0 | 0 |
| ENSG00000137073 | 1655 | 840 | 944 | 689 | 848 | 972 |
| ENSG00000238300 | 0 | 0 | 0 | 0 | 0 | 0 |
| ENSG00000238886 | 0 | 0 | 0 | 0 | 0 | 0 |
| ENSG00000226801 | 2 | 1 | 0 | 0 | 0 | 0 |
| ENSG00000222259 | 0 | 1 | 0 | 0 | 0 | 0 |

| | | | | | | |
|---|---|---|---|---|---|---|
| ENSG00000228352 | 8 | 7 | 2 | 3 | 2 | 3 |
| ENSG00000198876 | 3308 | 1479 | 1935 | 1342 | 1715 | 1753 |
| ENSG00000224802 | 0 | 0 | 0 | 0 | 1 | 0 |
| ENSG00000239392 | 0 | 0 | 0 | 0 | 0 | 0 |
| ENSG00000165006 | 1889 | 967 | 1176 | 990 | 1135 | 999 |
| ENSG00000227383 | 0 | 0 | 0 | 0 | 0 | 0 |
| ENSG00000279336 | 0 | 0 | 0 | 0 | 0 | 0 |
| ENSG00000252164 | 0 | 0 | 0 | 0 | 0 | 0 |
| ENSG00000233668 | 0 | 0 | 0 | 0 | 0 | 0 |
| ENSG00000186638 | 127 | 74 | 97 | 62 | 78 | 90 |
| ENSG00000222426 | 0 | 0 | 0 | 0 | 0 | 0 |
| ENSG00000229207 | 2 | 2 | 0 | 1 | 0 | 2 |
| ENSG00000164978 | 958 | 434 | 535 | 556 | 512 | 489 |
| ENSG00000164976 | 2220 | 1057 | 1272 | 794 | 860 | 869 |
| ENSG00000164972 | 1 | 2 | 1 | 1 | 1 | 6 |
| ENSG00000207032 | 0 | 0 | 1 | 0 | 2 | 0 |
| ENSG00000164970 | 817 | 373 | 502 | 491 | 592 | 598 |
| ENSG00000122735 | 2 | 4 | 5 | 2 | 4 | 3 |
| ENSG00000251697 | 0 | 0 | 0 | 0 | 0 | 0 |
| ENSG00000168913 | 5 | 8 | 5 | 6 | 7 | 9 |
| ENSG00000230729 | 0 | 0 | 0 | 0 | 0 | 0 |
| ENSG00000232983 | 0 | 0 | 0 | 0 | 0 | 0 |
| ENSG00000122756 | 4 | 0 | 0 | 1 | 2 | 2 |
| ENSG00000237159 | 0 | 0 | 0 | 0 | 0 | 0 |
| ENSG00000164967 | 646 | 366 | 383 | 411 | 348 | 360 |
| ENSG00000137100 | 2000 | 921 | 1036 | 1112 | 1016 | 1010 |
| ENSG00000205143 | 3 | 1 | 3 | 3 | 2 | 2 |
| ENSG00000147955 | 1007 | 525 | 674 | 632 | 572 | 643 |
| ENSG00000213930 | 291 | 155 | 208 | 165 | 161 | 197 |
| ENSG00000258728 | 0 | 0 | 0 | 0 | 0 | 0 |
| ENSG00000137070 | 32 | 26 | 30 | 15 | 21 | 27 |
| ENSG00000213927 | 0 | 0 | 0 | 0 | 0 | 0 |
| ENSG00000261215 | 2 | 0 | 4 | 1 | 0 | 0 |
| ENSG00000187186 | 1 | 3 | 1 | 8 | 1 | 2 |
| ENSG00000230074 | 43 | 23 | 34 | 15 | 28 | 22 |
| ENSG00000172724 | 0 | 0 | 0 | 0 | 1 | 0 |
| ENSG00000137077 | 0 | 0 | 0 | 0 | 0 | 0 |
| ENSG00000205108 | 2 | 0 | 2 | 0 | 0 | 0 |
| ENSG00000257198 | 0 | 0 | 0 | 0 | 0 | 0 |
| ENSG00000187791 | 0 | 1 | 0 | 0 | 0 | 0 |
| ENSG00000178723 | 0 | 0 | 0 | 0 | 0 | 0 |
| ENSG00000215199 | 2 | 1 | 1 | 0 | 0 | 1 |
| ENSG00000122733 | 8 | 0 | 1 | 4 | 1 | 0 |
| ENSG00000281491 | 0 | 0 | 0 | 0 | 0 | 0 |
| ENSG00000137094 | 216 | 116 | 136 | 95 | 96 | 89 |
| ENSG00000273597 | 0 | 1 | 0 | 0 | 0 | 0 |
| ENSG00000279489 | 0 | 0 | 0 | 0 | 0 | 0 |
| ENSG00000233148 | 0 | 0 | 0 | 0 | 0 | 0 |
| ENSG00000279608 | 0 | 1 | 0 | 0 | 0 | 0 |
| ENSG00000174038 | 0 | 1 | 0 | 0 | 0 | 0 |

| | | | | | | |
|---|---|---|---|---|---|---|
| ENSG00000241568 | 0 | 0 | 0 | 0 | 0 | 0 |
| ENSG00000165280 | 21376 | 10038 | 12912 | 10925 | 11936 | 11721 |
| ENSG00000221829 | 1098 | 569 | 733 | 733 | 667 | 636 |
| ENSG00000165282 | 2356 | 1021 | 1406 | 1065 | 968 | 1120 |
| ENSG00000234181 | 0 | 0 | 1 | 0 | 2 | 0 |
| ENSG00000165283 | 3773 | 1864 | 2240 | 2697 | 2145 | 2237 |
| ENSG00000005238 | 1311 | 560 | 777 | 596 | 667 | 565 |
| ENSG00000215198 | 0 | 1 | 0 | 0 | 0 | 2 |
| ENSG00000198722 | 2698 | 1146 | 1718 | 908 | 1021 | 1268 |
| ENSG00000279083 | 0 | 0 | 0 | 0 | 1 | 0 |
| ENSG00000179766 | 2 | 1 | 3 | 1 | 2 | 1 |
| ENSG00000244213 | 0 | 0 | 0 | 0 | 0 | 0 |
| ENSG00000198853 | 301 | 134 | 187 | 133 | 129 | 140 |
| ENSG00000235619 | 0 | 2 | 0 | 0 | 0 | 0 |
| ENSG00000215187 | 2 | 1 | 7 | 6 | 11 | 10 |
| ENSG00000227852 | 1 | 1 | 1 | 1 | 4 | 2 |
| ENSG00000107140 | 830 | 384 | 526 | 457 | 417 | 406 |
| ENSG00000263892 | 0 | 0 | 0 | 0 | 0 | 0 |
| ENSG00000137101 | 35 | 12 | 20 | 14 | 22 | 17 |
| ENSG00000231393 | 1 | 0 | 0 | 0 | 1 | 0 |
| ENSG00000227933 | 0 | 0 | 0 | 0 | 0 | 0 |
| ENSG00000137078 | 0 | 0 | 0 | 0 | 0 | 0 |
| ENSG00000269900 | 0 | 0 | 0 | 0 | 0 | 0 |
| ENSG00000277027 | 0 | 0 | 0 | 0 | 0 | 0 |
| ENSG00000159884 | 61 | 36 | 48 | 60 | 43 | 48 |
| ENSG00000137135 | 214 | 118 | 140 | 89 | 124 | 122 |
| ENSG00000243136 | 0 | 0 | 0 | 0 | 0 | 0 |
| ENSG00000107159 | 2 | 1 | 2 | 2 | 0 | 3 |
| ENSG00000198467 | 178 | 86 | 123 | 131 | 117 | 108 |
| ENSG00000137076 | 4512 | 1994 | 2676 | 2245 | 2356 | 2314 |
| ENSG00000284195 | 0 | 0 | 0 | 0 | 0 | 0 |
| ENSG00000107175 | 1566 | 795 | 927 | 971 | 842 | 887 |
| ENSG00000273945 | 0 | 0 | 0 | 0 | 0 | 0 |
| ENSG00000070610 | 1828 | 787 | 1150 | 1074 | 1001 | 1139 |
| ENSG00000107185 | 1698 | 854 | 1147 | 898 | 966 | 1048 |
| ENSG00000215183 | 0 | 0 | 0 | 0 | 0 | 0 |
| ENSG00000228843 | 0 | 0 | 0 | 0 | 0 | 0 |
| ENSG00000227388 | 47 | 11 | 20 | 12 | 26 | 25 |
| ENSG00000159899 | 268 | 106 | 168 | 114 | 141 | 167 |
| ENSG00000137098 | 21 | 6 | 11 | 3 | 5 | 5 |
| ENSG00000285645 | 0 | 0 | 0 | 0 | 0 | 0 |
| ENSG00000137133 | 39 | 32 | 39 | 35 | 32 | 42 |
| ENSG00000137103 | 516 | 209 | 323 | 231 | 190 | 250 |
| ENSG00000204930 | 0 | 0 | 2 | 0 | 3 | 0 |
| ENSG00000236083 | 0 | 0 | 0 | 0 | 0 | 0 |
| ENSG00000168828 | 0 | 0 | 0 | 0 | 0 | 0 |
| ENSG00000233659 | 0 | 0 | 0 | 0 | 0 | 0 |
| ENSG00000196196 | 6 | 5 | 3 | 0 | 1 | 0 |
| ENSG00000235387 | 0 | 0 | 0 | 0 | 0 | 0 |
| ENSG00000225639 | 0 | 0 | 0 | 0 | 0 | 0 |

| | | | | | | |
|---|---|---|---|---|---|---|
| ENSG00000278889 | 0 | 0 | 0 | 0 | 0 | 0 |
| ENSG00000228307 | 0 | 0 | 0 | 0 | 0 | 0 |
| ENSG00000213866 | 78 | 51 | 43 | 40 | 53 | 47 |
| ENSG00000179443 | 0 | 0 | 0 | 0 | 0 | 0 |
| ENSG00000243641 | 0 | 0 | 0 | 0 | 0 | 0 |
| ENSG00000236110 | 0 | 0 | 0 | 0 | 0 | 0 |
| ENSG00000122707 | 28 | 18 | 26 | 14 | 39 | 17 |
| ENSG00000276730 | 0 | 0 | 0 | 0 | 0 | 0 |
| ENSG00000122694 | 112 | 92 | 91 | 79 | 81 | 71 |
| ENSG00000185972 | 0 | 0 | 1 | 0 | 0 | 0 |
| ENSG00000122705 | 4206 | 2047 | 2476 | 2583 | 2337 | 2390 |
| ENSG00000159921 | 616 | 319 | 371 | 239 | 339 | 288 |
| ENSG00000222760 | 1 | 0 | 1 | 1 | 0 | 0 |
| ENSG00000215283 | 14 | 1 | 5 | 2 | 4 | 0 |
| ENSG00000137075 | 763 | 381 | 409 | 221 | 442 | 387 |
| ENSG00000165304 | 752 | 393 | 433 | 354 | 456 | 484 |
| ENSG00000266255 | 0 | 0 | 0 | 0 | 0 | 0 |
| ENSG00000196092 | 20 | 8 | 19 | 15 | 7 | 16 |
| ENSG00000258885 | 0 | 0 | 0 | 0 | 0 | 0 |
| ENSG00000264922 | 0 | 0 | 0 | 0 | 0 | 0 |
| ENSG00000265368 | 0 | 0 | 0 | 0 | 0 | 0 |
| ENSG00000250850 | 0 | 0 | 0 | 0 | 0 | 0 |
| ENSG00000125385 | 0 | 0 | 0 | 0 | 0 | 0 |
| ENSG00000232920 | 0 | 0 | 0 | 0 | 2 | 0 |
| ENSG00000260100 | 0 | 3 | 1 | 1 | 1 | 0 |
| ENSG00000281649 | 2799 | 1242 | 1688 | 835 | 1391 | 1477 |
| ENSG00000233242 | 0 | 0 | 0 | 4 | 1 | 1 |
| ENSG00000147905 | 475 | 257 | 241 | 186 | 324 | 244 |
| ENSG00000200502 | 2 | 2 | 1 | 0 | 0 | 3 |
| ENSG00000228292 | 0 | 0 | 0 | 0 | 0 | 0 |
| ENSG00000206784 | 0 | 0 | 0 | 0 | 0 | 0 |
| ENSG00000252723 | 0 | 0 | 0 | 0 | 0 | 0 |
| ENSG00000224648 | 0 | 0 | 0 | 1 | 0 | 0 |
| ENSG00000137106 | 2779 | 1439 | 1900 | 1969 | 1704 | 1722 |
| ENSG00000236156 | 0 | 0 | 0 | 0 | 0 | 0 |
| ENSG00000168795 | 1347 | 621 | 777 | 536 | 637 | 694 |
| ENSG00000187988 | 0 | 0 | 0 | 0 | 0 | 0 |
| ENSG00000137054 | 582 | 308 | 316 | 299 | 415 | 373 |
| ENSG00000230188 | 0 | 0 | 0 | 0 | 0 | 1 |
| ENSG00000234160 | 40 | 18 | 20 | 32 | 22 | 34 |
| ENSG00000147912 | 109 | 61 | 65 | 84 | 70 | 71 |
| ENSG00000256966 | 0 | 0 | 3 | 4 | 2 | 0 |
| ENSG00000175768 | 103 | 103 | 115 | 150 | 165 | 138 |
| ENSG00000255872 | 0 | 1 | 1 | 2 | 0 | 0 |
| ENSG00000233111 | 0 | 1 | 0 | 0 | 1 | 0 |
| ENSG00000070601 | 6 | 3 | 2 | 4 | 4 | 0 |
| ENSG00000201287 | 0 | 0 | 0 | 0 | 0 | 0 |
| ENSG00000165275 | 258 | 100 | 141 | 89 | 122 | 115 |
| ENSG00000107371 | 556 | 336 | 380 | 388 | 423 | 380 |
| ENSG00000122741 | 1171 | 519 | 656 | 389 | 656 | 539 |

| | | | | | | |
|---|---|---|---|---|---|---|
| ENSG00000232454 | 2 | 2 | 0 | 0 | 0 | 0 |
| ENSG00000226055 | 5 | 4 | 0 | 5 | 3 | 4 |
| ENSG00000122696 | 345 | 212 | 255 | 193 | 225 | 230 |
| ENSG00000213839 | 104 | 46 | 46 | 63 | 58 | 54 |
| ENSG00000107338 | 462 | 188 | 295 | 325 | 305 | 302 |
| ENSG00000251745 | 0 | 0 | 0 | 0 | 0 | 0 |
| ENSG00000200026 | 0 | 0 | 0 | 0 | 0 | 0 |
| ENSG00000223716 | 0 | 0 | 0 | 0 | 0 | 0 |
| ENSG00000137124 | 678 | 272 | 349 | 402 | 386 | 431 |
| ENSG00000137142 | 40 | 16 | 18 | 29 | 17 | 12 |
| ENSG00000283162 | 28 | 16 | 19 | 12 | 24 | 18 |
| ENSG00000225254 | 0 | 0 | 2 | 1 | 0 | 0 |
| ENSG00000215165 | 0 | 0 | 0 | 0 | 2 | 1 |
| ENSG00000229511 | 0 | 0 | 0 | 2 | 0 | 0 |
| ENSG00000237487 | 12 | 0 | 3 | 1 | 4 | 3 |
| ENSG00000243085 | 0 | 1 | 0 | 0 | 0 | 0 |
| ENSG00000176007 | 0 | 0 | 0 | 0 | 0 | 0 |
| ENSG00000273036 | 0 | 0 | 0 | 0 | 0 | 0 |
| ENSG00000283486 | 52 | 10 | 35 | 7 | 27 | 27 |
| ENSG00000272934 | 0 | 0 | 0 | 0 | 0 | 0 |
| ENSG00000272904 | 0 | 0 | 0 | 0 | 0 | 0 |
| ENSG00000250989 | 0 | 0 | 0 | 0 | 0 | 0 |
| ENSG00000225345 | 20 | 10 | 14 | 8 | 8 | 12 |
| ENSG00000180071 | 204 | 105 | 149 | 55 | 138 | 127 |
| ENSG00000204860 | 361 | 148 | 227 | 154 | 191 | 174 |
| ENSG00000237557 | 2 | 0 | 0 | 0 | 0 | 0 |
| ENSG00000284612 | 0 | 0 | 1 | 0 | 0 | 0 |
| ENSG00000252725 | 0 | 0 | 0 | 0 | 0 | 0 |
| ENSG00000283329 | 0 | 0 | 0 | 0 | 0 | 0 |
| ENSG00000228467 | 0 | 0 | 0 | 0 | 0 | 0 |
| ENSG00000234680 | 0 | 0 | 0 | 0 | 0 | 0 |
| ENSG00000106714 | 0 | 0 | 0 | 0 | 0 | 3 |
| ENSG00000226625 | 0 | 0 | 0 | 0 | 0 | 0 |
| ENSG00000271454 | 0 | 0 | 0 | 0 | 0 | 0 |
| ENSG00000243695 | 0 | 0 | 0 | 0 | 0 | 0 |
| ENSG00000278825 | 0 | 0 | 0 | 0 | 0 | 0 |
| ENSG00000204849 | 0 | 0 | 0 | 0 | 0 | 0 |
| ENSG00000277024 | 0 | 0 | 0 | 0 | 0 | 0 |
| ENSG00000215112 | 0 | 0 | 0 | 0 | 0 | 0 |
| ENSG00000224038 | 0 | 0 | 0 | 0 | 0 | 0 |
| ENSG00000231140 | 0 | 0 | 0 | 0 | 0 | 0 |
| ENSG00000198416 | 28 | 7 | 16 | 7 | 18 | 8 |
| ENSG00000229707 | 0 | 0 | 0 | 0 | 0 | 0 |
| ENSG00000269692 | 0 | 0 | 0 | 0 | 0 | 1 |
| ENSG00000273994 | 0 | 0 | 0 | 0 | 0 | 0 |
| ENSG00000234720 | 0 | 0 | 0 | 0 | 0 | 0 |
| ENSG00000226108 | 0 | 0 | 0 | 0 | 0 | 0 |
| ENSG00000232239 | 0 | 0 | 0 | 0 | 0 | 0 |
| ENSG00000278175 | 108 | 57 | 79 | 81 | 78 | 69 |
| ENSG00000229273 | 0 | 0 | 0 | 0 | 0 | 0 |

| | | | | | | |
|---|---|---|---|---|---|---|
| ENSG00000204837 | 0 | 0 | 0 | 0 | 0 | 0 |
| ENSG00000283886 | 0 | 4 | 6 | 2 | 1 | 2 |
| ENSG00000276692 | 0 | 0 | 0 | 0 | 0 | 0 |
| ENSG00000277774 | 0 | 0 | 0 | 0 | 0 | 0 |
| ENSG00000252724 | 0 | 0 | 0 | 0 | 0 | 0 |
| ENSG00000232798 | 0 | 0 | 0 | 0 | 0 | 0 |
| ENSG00000284116 | 0 | 2 | 0 | 0 | 1 | 0 |
| ENSG00000184961 | 0 | 0 | 0 | 0 | 0 | 0 |
| ENSG00000237846 | 0 | 2 | 0 | 0 | 0 | 1 |
| ENSG00000230804 | 0 | 0 | 0 | 0 | 0 | 0 |
| ENSG00000228136 | 0 | 0 | 0 | 0 | 0 | 0 |
| ENSG00000236966 | 0 | 0 | 0 | 0 | 0 | 0 |
| ENSG00000283211 | 0 | 0 | 0 | 0 | 0 | 0 |
| ENSG00000283246 | 0 | 0 | 0 | 0 | 0 | 0 |
| ENSG00000183148 | 0 | 0 | 0 | 0 | 0 | 0 |
| ENSG00000212441 | 0 | 0 | 0 | 0 | 0 | 0 |
| ENSG00000240240 | 0 | 0 | 0 | 0 | 0 | 0 |
| ENSG00000234812 | 0 | 0 | 0 | 0 | 0 | 0 |
| ENSG00000231390 | 0 | 0 | 0 | 0 | 0 | 0 |
| ENSG00000223839 | 0 | 0 | 0 | 0 | 0 | 0 |
| ENSG00000224681 | 0 | 0 | 0 | 0 | 0 | 0 |
| ENSG00000227069 | 0 | 0 | 0 | 0 | 0 | 0 |
| ENSG00000232433 | 0 | 0 | 0 | 0 | 0 | 0 |
| ENSG00000274157 | 0 | 0 | 0 | 0 | 0 | 0 |
| ENSG00000278122 | 0 | 0 | 0 | 0 | 0 | 0 |
| ENSG00000274067 | 0 | 0 | 0 | 0 | 0 | 0 |
| ENSG00000184906 | 0 | 0 | 0 | 0 | 0 | 0 |
| ENSG00000276500 | 0 | 0 | 0 | 0 | 0 | 0 |
| ENSG00000276266 | 0 | 0 | 0 | 0 | 0 | 0 |
| ENSG00000225380 | 0 | 0 | 0 | 0 | 0 | 0 |
| ENSG00000276963 | 0 | 0 | 0 | 0 | 0 | 1 |
| ENSG00000278331 | 0 | 0 | 0 | 0 | 0 | 0 |
| ENSG00000275390 | 0 | 0 | 0 | 0 | 0 | 0 |
| ENSG00000276868 | 0 | 0 | 0 | 0 | 0 | 0 |
| ENSG00000275377 | 0 | 0 | 0 | 0 | 0 | 0 |
| ENSG00000276291 | 158 | 88 | 108 | 54 | 109 | 73 |
| ENSG00000277778 | 1 | 1 | 2 | 0 | 1 | 1 |
| ENSG00000280077 | 76 | 41 | 40 | 16 | 57 | 23 |
| ENSG00000279456 | 0 | 0 | 1 | 0 | 1 | 0 |
| ENSG00000225655 | 0 | 0 | 0 | 0 | 0 | 0 |
| ENSG00000276462 | 0 | 0 | 1 | 0 | 0 | 0 |
| ENSG00000273514 | 0 | 0 | 0 | 0 | 0 | 1 |
| ENSG00000215126 | 90 | 61 | 84 | 61 | 96 | 90 |
| ENSG00000276029 | 0 | 0 | 0 | 0 | 0 | 0 |
| ENSG00000277049 | 5 | 1 | 0 | 0 | 0 | 0 |
| ENSG00000279561 | 143 | 61 | 89 | 48 | 73 | 62 |
| ENSG00000275450 | 4 | 5 | 13 | 1 | 5 | 3 |
| ENSG00000278647 | 0 | 0 | 0 | 0 | 0 | 0 |
| ENSG00000274680 | 0 | 0 | 0 | 0 | 0 | 0 |
| ENSG00000276502 | 0 | 0 | 0 | 0 | 0 | 0 |

| | | | | | | |
|---|---|---|---|---|---|---|
| ENSG00000276832 | 2 | 2 | 1 | 1 | 0 | 2 |
| ENSG00000275160 | 23 | 12 | 20 | 21 | 25 | 19 |
| ENSG00000277340 | 0 | 0 | 0 | 0 | 0 | 0 |
| ENSG00000276094 | 0 | 0 | 0 | 0 | 0 | 0 |
| ENSG00000276562 | 0 | 0 | 0 | 0 | 0 | 0 |
| ENSG00000275493 | 0 | 0 | 0 | 0 | 0 | 0 |
| ENSG00000278077 | 0 | 0 | 0 | 0 | 0 | 0 |
| ENSG00000278054 | 0 | 0 | 0 | 0 | 0 | 0 |
| ENSG00000276135 | 0 | 0 | 0 | 0 | 0 | 0 |
| ENSG00000276412 | 0 | 0 | 0 | 0 | 0 | 0 |
| ENSG00000277979 | 0 | 0 | 0 | 0 | 0 | 0 |
| ENSG00000275239 | 0 | 0 | 0 | 0 | 0 | 0 |
| ENSG00000278028 | 0 | 0 | 0 | 0 | 0 | 0 |
| ENSG00000274688 | 0 | 0 | 0 | 0 | 0 | 0 |
| ENSG00000278001 | 0 | 0 | 0 | 0 | 0 | 0 |
| ENSG00000275297 | 0 | 0 | 0 | 0 | 0 | 0 |
| ENSG00000154529 | 0 | 0 | 0 | 0 | 0 | 0 |
| ENSG00000237207 | 0 | 0 | 0 | 0 | 0 | 0 |
| ENSG00000270909 | 0 | 0 | 0 | 0 | 0 | 0 |
| ENSG00000273717 | 0 | 0 | 0 | 0 | 0 | 0 |
| ENSG00000185775 | 0 | 0 | 0 | 0 | 0 | 0 |
| ENSG00000239986 | 0 | 0 | 0 | 0 | 0 | 0 |
| ENSG00000277260 | 0 | 0 | 0 | 0 | 0 | 0 |
| ENSG00000276277 | 0 | 0 | 0 | 0 | 0 | 0 |
| ENSG00000275649 | 0 | 0 | 0 | 0 | 0 | 0 |
| ENSG00000268234 | 0 | 0 | 1 | 0 | 0 | 0 |
| ENSG00000277213 | 0 | 0 | 0 | 0 | 0 | 0 |
| ENSG00000234861 | 0 | 0 | 0 | 0 | 0 | 0 |
| ENSG00000227353 | 0 | 0 | 0 | 0 | 0 | 0 |
| ENSG00000227924 | 0 | 0 | 0 | 0 | 0 | 0 |
| ENSG00000237357 | 2 | 3 | 6 | 0 | 0 | 1 |
| ENSG00000212952 | 0 | 0 | 0 | 0 | 0 | 0 |
| ENSG00000229311 | 0 | 0 | 0 | 0 | 0 | 0 |
| ENSG00000204816 | 0 | 0 | 0 | 0 | 0 | 0 |
| ENSG00000230245 | 0 | 0 | 0 | 0 | 0 | 0 |
| ENSG00000216829 | 0 | 1 | 0 | 0 | 0 | 0 |
| ENSG00000233424 | 0 | 0 | 0 | 0 | 0 | 0 |
| ENSG00000204818 | 0 | 0 | 0 | 0 | 0 | 0 |
| ENSG00000224537 | 0 | 0 | 0 | 0 | 0 | 0 |
| ENSG00000236816 | 0 | 0 | 0 | 0 | 0 | 0 |
| ENSG00000238529 | 0 | 0 | 0 | 0 | 0 | 0 |
| ENSG00000170165 | 0 | 0 | 0 | 0 | 0 | 0 |
| ENSG00000273900 | 0 | 0 | 0 | 0 | 0 | 0 |
| ENSG00000275144 | 0 | 0 | 0 | 0 | 0 | 0 |
| ENSG00000230635 | 0 | 0 | 0 | 0 | 0 | 0 |
| ENSG00000229079 | 0 | 0 | 0 | 0 | 0 | 0 |
| ENSG00000213816 | 0 | 0 | 0 | 1 | 0 | 0 |
| ENSG00000276591 | 0 | 0 | 0 | 0 | 0 | 0 |
| ENSG00000277697 | 0 | 0 | 0 | 0 | 0 | 0 |
| ENSG00000276128 | 0 | 0 | 0 | 0 | 0 | 0 |

| | | | | | | |
|---|---|---|---|---|---|---|
| ENSG00000277737 | 0 | 0 | 0 | 0 | 0 | 0 |
| ENSG00000277598 | 0 | 0 | 0 | 0 | 0 | 0 |
| ENSG00000276581 | 0 | 0 | 0 | 0 | 0 | 0 |
| ENSG00000275628 | 0 | 0 | 0 | 0 | 0 | 0 |
| ENSG00000274516 | 0 | 0 | 0 | 0 | 0 | 0 |
| ENSG00000276422 | 0 | 0 | 0 | 0 | 0 | 0 |
| ENSG00000278485 | 0 | 0 | 0 | 0 | 0 | 0 |
| ENSG00000283541 | 0 | 0 | 0 | 0 | 0 | 0 |
| ENSG00000276040 | 0 | 0 | 0 | 0 | 0 | 0 |
| ENSG00000277412 | 0 | 0 | 0 | 0 | 0 | 0 |
| ENSG00000276149 | 0 | 0 | 0 | 0 | 0 | 0 |
| ENSG00000274583 | 0 | 0 | 0 | 0 | 0 | 0 |
| ENSG00000275676 | 0 | 0 | 0 | 0 | 0 | 0 |
| ENSG00000283378 | 0 | 0 | 0 | 0 | 0 | 0 |
| ENSG00000276522 | 0 | 0 | 0 | 0 | 0 | 0 |
| ENSG00000277109 | 0 | 0 | 0 | 0 | 0 | 0 |
| ENSG00000231248 | 0 | 0 | 0 | 0 | 0 | 0 |
| ENSG00000238933 | 0 | 0 | 0 | 0 | 0 | 0 |
| ENSG00000204814 | 0 | 0 | 0 | 0 | 0 | 0 |
| ENSG00000231454 | 0 | 0 | 0 | 0 | 0 | 0 |
| ENSG00000273940 | 0 | 0 | 0 | 0 | 0 | 0 |
| ENSG00000154537 | 0 | 0 | 0 | 0 | 0 | 0 |
| ENSG00000277624 | 0 | 0 | 0 | 0 | 0 | 0 |
| ENSG00000170152 | 0 | 0 | 0 | 0 | 0 | 0 |
| ENSG00000204805 | 0 | 0 | 0 | 0 | 0 | 0 |
| ENSG00000235659 | 0 | 0 | 0 | 0 | 0 | 0 |
| ENSG00000223379 | 0 | 0 | 0 | 0 | 0 | 0 |
| ENSG00000238749 | 0 | 0 | 0 | 0 | 0 | 0 |
| ENSG00000229697 | 0 | 0 | 0 | 0 | 0 | 0 |
| ENSG00000276795 | 0 | 0 | 0 | 0 | 0 | 0 |
| ENSG00000276845 | 0 | 0 | 0 | 0 | 0 | 0 |
| ENSG00000231527 | 0 | 0 | 0 | 0 | 0 | 0 |
| ENSG00000284471 | 0 | 0 | 0 | 0 | 0 | 0 |
| ENSG00000231838 | 0 | 0 | 0 | 0 | 0 | 0 |
| ENSG00000231995 | 3 | 1 | 3 | 2 | 3 | 1 |
| ENSG00000231212 | 0 | 0 | 0 | 0 | 0 | 0 |
| ENSG00000236450 | 0 | 0 | 0 | 0 | 0 | 0 |
| ENSG00000252133 | 0 | 0 | 0 | 0 | 0 | 0 |
| ENSG00000275230 | 0 | 0 | 0 | 0 | 0 | 0 |
| ENSG00000204802 | 2 | 0 | 0 | 3 | 1 | 3 |
| ENSG00000227449 | 3 | 1 | 8 | 2 | 4 | 2 |
| ENSG00000233651 | 0 | 0 | 0 | 0 | 0 | 0 |
| ENSG00000226429 | 0 | 0 | 0 | 0 | 0 | 0 |
| ENSG00000224603 | 0 | 0 | 0 | 0 | 0 | 0 |
| ENSG00000204801 | 0 | 0 | 0 | 0 | 0 | 0 |
| ENSG00000232827 | 0 | 0 | 0 | 0 | 0 | 0 |
| ENSG00000226007 | 22 | 5 | 17 | 6 | 9 | 10 |
| ENSG00000212951 | 0 | 0 | 0 | 0 | 0 | 0 |
| ENSG00000278213 | 0 | 0 | 0 | 0 | 0 | 0 |
| ENSG00000234217 | 0 | 0 | 0 | 0 | 0 | 0 |

| | | | | | | |
|---|---|---|---|---|---|---|
| ENSG00000236825 | 0 | 0 | 0 | 0 | 0 | 0 |
| ENSG00000234451 | 0 | 0 | 0 | 0 | 0 | 0 |
| ENSG00000241571 | 0 | 0 | 0 | 0 | 0 | 0 |
| ENSG00000234276 | 0 | 0 | 0 | 0 | 0 | 0 |
| ENSG00000235256 | 0 | 0 | 0 | 0 | 0 | 0 |
| ENSG00000238113 | 81 | 32 | 62 | 37 | 75 | 62 |
| ENSG00000202474 | 23 | 8 | 10 | 8 | 20 | 15 |
| ENSG00000229422 | 140 | 60 | 62 | 34 | 43 | 50 |
| ENSG00000184523 | 1 | 4 | 2 | 8 | 4 | 6 |
| ENSG00000237451 | 8 | 3 | 4 | 8 | 1 | 4 |
| ENSG00000234665 | 0 | 2 | 0 | 6 | 2 | 0 |
| ENSG00000238245 | 0 | 0 | 0 | 1 | 0 | 1 |
| ENSG00000227582 | 6 | 2 | 6 | 4 | 4 | 3 |
| ENSG00000219693 | 1 | 0 | 1 | 0 | 1 | 0 |
| ENSG00000170161 | 94 | 27 | 66 | 45 | 55 | 44 |
| ENSG00000273509 | 4 | 1 | 1 | 0 | 0 | 0 |
| ENSG00000244081 | 0 | 0 | 0 | 0 | 0 | 0 |
| ENSG00000198312 | 0 | 0 | 0 | 0 | 0 | 0 |
| ENSG00000182021 | 0 | 0 | 0 | 0 | 0 | 0 |
| ENSG00000176115 | 0 | 0 | 0 | 0 | 0 | 0 |
| ENSG00000237238 | 4 | 3 | 4 | 0 | 3 | 4 |
| ENSG00000186466 | 2 | 0 | 2 | 0 | 0 | 0 |
| ENSG00000228522 | 0 | 1 | 0 | 0 | 0 | 0 |
| ENSG00000233555 | 0 | 0 | 0 | 0 | 0 | 0 |
| ENSG00000234042 | 2 | 1 | 0 | 0 | 0 | 0 |
| ENSG00000268951 | 0 | 0 | 0 | 0 | 0 | 0 |
| ENSG00000224267 | 0 | 0 | 0 | 0 | 0 | 0 |
| ENSG00000225360 | 0 | 0 | 0 | 0 | 0 | 0 |
| ENSG00000228656 | 0 | 0 | 0 | 0 | 0 | 1 |
| ENSG00000226020 | 0 | 0 | 0 | 0 | 0 | 1 |
| ENSG00000239684 | 0 | 1 | 0 | 0 | 0 | 0 |
| ENSG00000225411 | 6 | 3 | 3 | 0 | 3 | 3 |
| ENSG00000207277 | 1 | 0 | 0 | 0 | 0 | 0 |
| ENSG00000232815 | 106 | 61 | 84 | 64 | 96 | 55 |
| ENSG00000266017 | 14 | 9 | 12 | 19 | 10 | 6 |
| ENSG00000215548 | 24 | 24 | 22 | 8 | 27 | 29 |
| ENSG00000204790 | 2 | 1 | 4 | 1 | 1 | 3 |
| ENSG00000275068 | 0 | 0 | 0 | 0 | 0 | 0 |
| ENSG00000285580 | 0 | 0 | 0 | 0 | 0 | 0 |
| ENSG00000172014 | 0 | 0 | 1 | 0 | 0 | 0 |
| ENSG00000238551 | 0 | 0 | 0 | 0 | 0 | 0 |
| ENSG00000224185 | 0 | 0 | 0 | 0 | 0 | 0 |
| ENSG00000229156 | 0 | 0 | 0 | 0 | 0 | 0 |
| ENSG00000224106 | 0 | 0 | 0 | 0 | 0 | 0 |
| ENSG00000224762 | 0 | 0 | 0 | 0 | 0 | 0 |
| ENSG00000235832 | 0 | 0 | 0 | 0 | 0 | 0 |
| ENSG00000235310 | 0 | 0 | 0 | 0 | 0 | 0 |
| ENSG00000277869 | 0 | 0 | 0 | 0 | 0 | 0 |
| ENSG00000181997 | 0 | 0 | 1 | 0 | 0 | 0 |
| ENSG00000176134 | 0 | 0 | 0 | 0 | 0 | 0 |

| | | | | | | |
|---|---|---|---|---|---|---|
| ENSG00000225883 | 2 | 0 | 0 | 0 | 0 | 0 |
| ENSG00000235932 | 0 | 0 | 0 | 0 | 0 | 0 |
| ENSG00000156755 | 0 | 0 | 0 | 0 | 0 | 0 |
| ENSG00000271225 | 0 | 0 | 0 | 0 | 0 | 0 |
| ENSG00000197550 | 0 | 0 | 0 | 0 | 0 | 0 |
| ENSG00000223602 | 0 | 0 | 0 | 0 | 1 | 0 |
| ENSG00000206804 | 0 | 0 | 0 | 0 | 0 | 0 |
| ENSG00000224599 | 2 | 0 | 0 | 0 | 0 | 0 |
| ENSG00000236252 | 0 | 0 | 0 | 0 | 0 | 0 |
| ENSG00000267085 | 0 | 0 | 0 | 0 | 0 | 0 |
| ENSG00000204780 | 0 | 0 | 0 | 0 | 0 | 0 |
| ENSG00000204779 | 2 | 2 | 1 | 0 | 0 | 0 |
| ENSG00000204778 | 34 | 18 | 16 | 6 | 17 | 10 |
| ENSG00000276723 | 0 | 0 | 0 | 0 | 0 | 0 |
| ENSG00000280286 | 1 | 1 | 1 | 0 | 2 | 3 |
| ENSG00000147996 | 443 | 269 | 275 | 191 | 287 | 247 |
| ENSG00000184659 | 0 | 1 | 2 | 2 | 1 | 0 |
| ENSG00000204776 | 0 | 0 | 0 | 0 | 0 | 0 |
| ENSG00000237904 | 0 | 0 | 0 | 0 | 0 | 0 |
| ENSG00000235893 | 0 | 0 | 0 | 0 | 0 | 0 |
| ENSG00000231701 | 0 | 0 | 0 | 0 | 0 | 0 |
| ENSG00000279955 | 0 | 0 | 0 | 0 | 0 | 0 |
| ENSG00000279688 | 0 | 0 | 0 | 0 | 0 | 0 |
| ENSG00000274893 | 0 | 0 | 0 | 0 | 0 | 0 |
| ENSG00000273668 | 0 | 0 | 0 | 0 | 0 | 0 |
| ENSG00000275026 | 2 | 0 | 0 | 0 | 0 | 0 |
| ENSG00000278068 | 0 | 0 | 0 | 0 | 0 | 0 |
| ENSG00000278720 | 0 | 0 | 0 | 0 | 0 | 0 |
| ENSG00000274628 | 0 | 0 | 0 | 0 | 0 | 0 |
| ENSG00000229146 | 0 | 0 | 0 | 0 | 0 | 0 |
| ENSG00000276257 | 0 | 0 | 0 | 0 | 0 | 0 |
| ENSG00000276203 | 0 | 0 | 0 | 0 | 0 | 0 |
| ENSG00000275030 | 0 | 0 | 0 | 0 | 0 | 0 |
| ENSG00000278849 | 0 | 0 | 0 | 0 | 0 | 0 |
| ENSG00000284611 | 0 | 0 | 0 | 0 | 0 | 0 |
| ENSG00000284658 | 0 | 0 | 0 | 0 | 0 | 0 |
| ENSG00000233765 | 2 | 4 | 8 | 5 | 4 | 0 |
| ENSG00000274181 | 0 | 0 | 0 | 0 | 0 | 0 |
| ENSG00000276894 | 0 | 0 | 0 | 0 | 0 | 0 |
| ENSG00000277297 | 0 | 0 | 0 | 0 | 0 | 0 |
| ENSG00000273849 | 0 | 0 | 0 | 0 | 0 | 0 |
| ENSG00000276457 | 0 | 0 | 0 | 0 | 0 | 0 |
| ENSG00000274349 | 43 | 28 | 9 | 16 | 21 | 14 |
| ENSG00000277490 | 0 | 0 | 0 | 0 | 0 | 0 |
| ENSG00000277350 | 0 | 0 | 0 | 0 | 0 | 0 |
| ENSG00000276039 | 0 | 0 | 0 | 0 | 0 | 0 |
| ENSG00000274355 | 0 | 0 | 0 | 0 | 0 | 0 |
| ENSG00000276348 | 0 | 0 | 0 | 0 | 0 | 0 |
| ENSG00000274820 | 0 | 0 | 0 | 0 | 0 | 0 |
| ENSG00000275969 | 0 | 0 | 0 | 0 | 0 | 0 |

| | | | | | | |
|---|---|---|---|---|---|---|
| ENSG00000276386 | 0 | 0 | 0 | 0 | 0 | 0 |
| ENSG00000274852 | 0 | 0 | 0 | 0 | 0 | 0 |
| ENSG00000277575 | 0 | 0 | 0 | 0 | 0 | 0 |
| ENSG00000273999 | 0 | 0 | 0 | 0 | 0 | 0 |
| ENSG00000276204 | 0 | 0 | 0 | 0 | 0 | 0 |
| ENSG00000276898 | 0 | 0 | 0 | 0 | 0 | 0 |
| ENSG00000275904 | 0 | 0 | 0 | 0 | 0 | 0 |
| ENSG00000277208 | 0 | 0 | 0 | 0 | 0 | 0 |
| ENSG00000275614 | 0 | 0 | 0 | 0 | 0 | 0 |
| ENSG00000274026 | 0 | 0 | 0 | 0 | 0 | 0 |
| ENSG00000278735 | 0 | 0 | 0 | 0 | 0 | 0 |
| ENSG00000274100 | 0 | 0 | 0 | 0 | 0 | 0 |
| ENSG00000278134 | 0 | 0 | 0 | 0 | 0 | 0 |
| ENSG00000276032 | 0 | 0 | 0 | 0 | 0 | 0 |
| ENSG00000278763 | 0 | 0 | 0 | 0 | 0 | 0 |
| ENSG00000274911 | 0 | 0 | 0 | 0 | 0 | 0 |
| ENSG00000274098 | 0 | 0 | 0 | 0 | 0 | 0 |
| ENSG00000278010 | 0 | 0 | 0 | 0 | 0 | 0 |
| ENSG00000277618 | 0 | 0 | 0 | 0 | 0 | 0 |
| ENSG00000260691 | 0 | 0 | 0 | 0 | 0 | 0 |
| ENSG00000278551 | 0 | 0 | 0 | 0 | 0 | 0 |
| ENSG00000284737 | 0 | 0 | 0 | 0 | 0 | 0 |
| ENSG00000226904 | 0 | 1 | 0 | 0 | 0 | 0 |
| ENSG00000196873 | 151 | 97 | 94 | 66 | 99 | 89 |
| ENSG00000279706 | 0 | 0 | 0 | 0 | 0 | 0 |
| ENSG00000187559 | 2 | 0 | 0 | 0 | 0 | 5 |
| ENSG00000234394 | 0 | 0 | 0 | 0 | 0 | 0 |
| ENSG00000225337 | 0 | 0 | 0 | 0 | 0 | 0 |
| ENSG00000154330 | 246 | 113 | 192 | 100 | 110 | 112 |
| ENSG00000224958 | 0 | 1 | 1 | 0 | 1 | 0 |
| ENSG00000233178 | 0 | 0 | 1 | 0 | 0 | 0 |
| ENSG00000229019 | 0 | 0 | 0 | 0 | 0 | 0 |
| ENSG00000181778 | 3 | 0 | 0 | 0 | 0 | 0 |
| ENSG00000226337 | 0 | 0 | 0 | 0 | 0 | 0 |
| ENSG00000234506 | 0 | 0 | 0 | 0 | 0 | 0 |
| ENSG00000224025 | 0 | 0 | 0 | 0 | 0 | 0 |
| ENSG00000107242 | 46 | 40 | 61 | 33 | 50 | 43 |
| ENSG00000187866 | 524 | 244 | 296 | 224 | 288 | 264 |
| ENSG00000236998 | 0 | 0 | 0 | 0 | 0 | 2 |
| ENSG00000236733 | 0 | 0 | 0 | 0 | 0 | 0 |
| ENSG00000207000 | 0 | 0 | 0 | 0 | 0 | 0 |
| ENSG00000165059 | 0 | 0 | 0 | 0 | 0 | 0 |
| ENSG00000165060 | 140 | 109 | 120 | 102 | 106 | 121 |
| ENSG00000285130 | 0 | 0 | 0 | 0 | 0 | 0 |
| ENSG00000270236 | 0 | 0 | 0 | 0 | 0 | 2 |
| ENSG00000119139 | 76 | 33 | 50 | 23 | 44 | 42 |
| ENSG00000278910 | 0 | 0 | 0 | 0 | 0 | 0 |
| ENSG00000135063 | 558 | 255 | 313 | 253 | 297 | 262 |
| ENSG00000107282 | 181 | 77 | 98 | 61 | 79 | 90 |
| ENSG00000243888 | 0 | 0 | 0 | 0 | 0 | 0 |

| | | | | | | |
|---|---|---|---|---|---|---|
| ENSG00000229312 | 131 | 63 | 76 | 83 | 80 | 65 |
| ENSG00000261447 | 10 | 6 | 7 | 3 | 4 | 3 |
| ENSG00000188647 | 512 | 289 | 249 | 125 | 435 | 252 |
| ENSG00000225626 | 0 | 0 | 0 | 0 | 0 | 0 |
| ENSG00000204711 | 0 | 0 | 0 | 0 | 0 | 0 |
| ENSG00000241174 | 0 | 0 | 0 | 0 | 0 | 0 |
| ENSG00000204706 | 0 | 3 | 3 | 2 | 2 | 3 |
| ENSG00000165072 | 16 | 3 | 5 | 2 | 4 | 5 |
| ENSG00000222465 | 0 | 0 | 0 | 0 | 0 | 0 |
| ENSG00000268364 | 22 | 14 | 5 | 8 | 8 | 6 |
| ENSG00000236801 | 8 | 3 | 2 | 4 | 4 | 7 |
| ENSG00000198887 | 525 | 279 | 201 | 97 | 480 | 214 |
| ENSG00000274421 | 0 | 0 | 0 | 0 | 0 | 0 |
| ENSG00000239180 | 0 | 0 | 0 | 0 | 0 | 0 |
| ENSG00000119138 | 1162 | 509 | 716 | 354 | 512 | 544 |
| ENSG00000083067 | 0 | 1 | 0 | 0 | 0 | 1 |
| ENSG00000272232 | 0 | 0 | 0 | 0 | 0 | 0 |
| ENSG00000232086 | 0 | 0 | 0 | 0 | 0 | 0 |
| ENSG00000223966 | 0 | 0 | 0 | 0 | 0 | 0 |
| ENSG00000207935 | 0 | 0 | 0 | 0 | 0 | 0 |
| ENSG00000224437 | 2 | 0 | 1 | 1 | 0 | 0 |
| ENSG00000229814 | 0 | 0 | 1 | 0 | 0 | 0 |
| ENSG00000135048 | 1914 | 974 | 1244 | 610 | 1011 | 894 |
| ENSG00000271166 | 0 | 0 | 0 | 0 | 0 | 0 |
| ENSG00000107362 | 522 | 223 | 285 | 207 | 266 | 259 |
| ENSG00000200922 | 0 | 0 | 0 | 0 | 0 | 0 |
| ENSG00000155621 | 491 | 204 | 242 | 193 | 268 | 260 |
| ENSG00000236060 | 8 | 3 | 1 | 5 | 2 | 3 |
| ENSG00000204669 | 0 | 0 | 0 | 0 | 0 | 0 |
| ENSG00000224820 | 0 | 0 | 0 | 0 | 0 | 0 |
| ENSG00000119125 | 2 | 0 | 0 | 0 | 0 | 1 |
| ENSG00000235523 | 0 | 0 | 0 | 0 | 0 | 0 |
| ENSG00000233933 | 0 | 0 | 0 | 0 | 0 | 0 |
| ENSG00000252428 | 0 | 0 | 0 | 0 | 0 | 0 |
| ENSG00000225434 | 0 | 0 | 0 | 0 | 0 | 0 |
| ENSG00000107372 | 2864 | 1317 | 1703 | 1310 | 1697 | 1653 |
| ENSG00000230003 | 0 | 0 | 0 | 0 | 0 | 0 |
| ENSG00000225947 | 0 | 0 | 0 | 0 | 0 | 0 |
| ENSG00000165091 | 2 | 0 | 1 | 2 | 0 | 1 |
| ENSG00000236862 | 0 | 0 | 0 | 0 | 0 | 0 |
| ENSG00000229753 | 0 | 0 | 0 | 0 | 0 | 0 |
| ENSG00000236849 | 0 | 0 | 0 | 0 | 0 | 1 |
| ENSG00000165092 | 0 | 0 | 0 | 0 | 0 | 0 |
| ENSG00000228024 | 0 | 0 | 0 | 0 | 0 | 0 |
| ENSG00000135046 | 70 | 43 | 49 | 34 | 66 | 32 |
| ENSG00000221844 | 0 | 0 | 0 | 0 | 0 | 0 |
| ENSG00000232590 | 0 | 0 | 0 | 0 | 0 | 0 |
| ENSG00000236701 | 0 | 0 | 0 | 0 | 0 | 0 |
| ENSG00000212454 | 0 | 0 | 0 | 0 | 0 | 0 |
| ENSG00000223368 | 0 | 0 | 0 | 0 | 0 | 0 |

| | | | | | | |
|---|---|---|---|---|---|---|
| ENSG00000227809 | 0 | 0 | 0 | 0 | 0 | 0 |
| ENSG00000224825 | 0 | 0 | 0 | 0 | 0 | 0 |
| ENSG00000198963 | 0 | 0 | 0 | 0 | 0 | 0 |
| ENSG00000119121 | 2 | 2 | 1 | 3 | 1 | 4 |
| ENSG00000202217 | 0 | 0 | 0 | 0 | 0 | 0 |
| ENSG00000270954 | 0 | 0 | 0 | 0 | 0 | 0 |
| ENSG00000252758 | 0 | 0 | 0 | 0 | 0 | 0 |
| ENSG00000224297 | 0 | 0 | 0 | 0 | 0 | 0 |
| ENSG00000135045 | 325 | 147 | 179 | 227 | 181 | 197 |
| ENSG00000203321 | 3 | 1 | 0 | 1 | 3 | 0 |
| ENSG00000156017 | 619 | 280 | 319 | 177 | 305 | 291 |
| ENSG00000229587 | 2 | 1 | 1 | 1 | 0 | 1 |
| ENSG00000278040 | 0 | 0 | 0 | 2 | 0 | 0 |
| ENSG00000106733 | 529 | 270 | 316 | 265 | 351 | 284 |
| ENSG00000134996 | 1222 | 678 | 715 | 612 | 732 | 688 |
| ENSG00000200041 | 0 | 0 | 0 | 0 | 0 | 0 |
| ENSG00000212316 | 0 | 0 | 0 | 0 | 0 | 0 |
| ENSG00000235707 | 0 | 0 | 0 | 0 | 0 | 0 |
| ENSG00000236376 | 0 | 1 | 1 | 4 | 4 | 0 |
| ENSG00000234644 | 0 | 0 | 0 | 0 | 0 | 0 |
| ENSG00000099139 | 0 | 0 | 0 | 0 | 2 | 0 |
| ENSG00000231137 | 0 | 0 | 0 | 0 | 0 | 0 |
| ENSG00000135002 | 938 | 417 | 524 | 425 | 516 | 533 |
| ENSG00000234618 | 12 | 6 | 7 | 6 | 7 | 12 |
| ENSG00000187210 | 1110 | 479 | 629 | 358 | 561 | 503 |
| ENSG00000233814 | 0 | 0 | 0 | 0 | 0 | 0 |
| ENSG00000219149 | 0 | 0 | 0 | 0 | 0 | 0 |
| ENSG00000106772 | 8 | 1 | 1 | 1 | 3 | 3 |
| ENSG00000225937 | 0 | 0 | 0 | 0 | 0 | 0 |
| ENSG00000274326 | 0 | 0 | 0 | 0 | 0 | 0 |
| ENSG00000277320 | 0 | 0 | 0 | 0 | 0 | 0 |
| ENSG00000230335 | 0 | 0 | 0 | 0 | 0 | 0 |
| ENSG00000236604 | 0 | 0 | 0 | 0 | 0 | 0 |
| ENSG00000229857 | 0 | 0 | 0 | 0 | 0 | 0 |
| ENSG00000204612 | 0 | 0 | 0 | 0 | 0 | 0 |
| ENSG00000232851 | 0 | 0 | 0 | 0 | 0 | 0 |
| ENSG00000233526 | 0 | 0 | 0 | 0 | 0 | 0 |
| ENSG00000232998 | 0 | 0 | 1 | 0 | 0 | 0 |
| ENSG00000197969 | 324 | 200 | 190 | 51 | 449 | 215 |
| ENSG00000156049 | 425 | 213 | 261 | 170 | 152 | 173 |
| ENSG00000231373 | 0 | 0 | 0 | 0 | 0 | 0 |
| ENSG00000228248 | 0 | 0 | 0 | 0 | 0 | 0 |
| ENSG00000201711 | 0 | 0 | 0 | 0 | 0 | 0 |
| ENSG00000156052 | 2385 | 1197 | 1271 | 828 | 1439 | 1116 |
| ENSG00000233457 | 0 | 0 | 0 | 0 | 0 | 0 |
| ENSG00000222452 | 0 | 0 | 0 | 0 | 0 | 0 |
| ENSG00000233947 | 0 | 0 | 0 | 0 | 0 | 0 |
| ENSG00000228781 | 0 | 0 | 0 | 0 | 0 | 0 |
| ENSG00000234819 | 0 | 0 | 0 | 2 | 1 | 0 |
| ENSG00000148019 | 758 | 352 | 383 | 296 | 535 | 401 |

| | | | | | | |
|---|---|---|---|---|---|---|
| ENSG00000135069 | 0 | 1 | 0 | 0 | 2 | 0 |
| ENSG00000270627 | 0 | 0 | 0 | 0 | 0 | 0 |
| ENSG00000237257 | 0 | 0 | 0 | 0 | 0 | 0 |
| ENSG00000226234 | 0 | 0 | 0 | 1 | 1 | 1 |
| ENSG00000260995 | 0 | 0 | 0 | 0 | 0 | 0 |
| ENSG00000186940 | 8 | 1 | 3 | 3 | 1 | 0 |
| ENSG00000233086 | 0 | 0 | 0 | 0 | 0 | 0 |
| ENSG00000106829 | 597 | 288 | 374 | 207 | 271 | 325 |
| ENSG00000240979 | 0 | 0 | 0 | 0 | 0 | 0 |
| ENSG00000270691 | 0 | 0 | 0 | 0 | 0 | 0 |
| ENSG00000238002 | 0 | 0 | 0 | 0 | 0 | 0 |
| ENSG00000232494 | 0 | 0 | 0 | 0 | 0 | 0 |
| ENSG00000260564 | 0 | 0 | 0 | 0 | 0 | 0 |
| ENSG00000279715 | 0 | 0 | 0 | 0 | 0 | 0 |
| ENSG00000280366 | 0 | 0 | 0 | 0 | 0 | 0 |
| ENSG00000230945 | 0 | 0 | 0 | 0 | 0 | 0 |
| ENSG00000236521 | 0 | 0 | 0 | 0 | 0 | 0 |
| ENSG00000232981 | 0 | 0 | 0 | 0 | 0 | 0 |
| ENSG00000225901 | 0 | 0 | 0 | 0 | 0 | 0 |
| ENSG00000234637 | 0 | 0 | 0 | 0 | 0 | 0 |
| ENSG00000226798 | 2 | 4 | 10 | 3 | 7 | 9 |
| ENSG00000228375 | 0 | 0 | 0 | 0 | 3 | 0 |
| ENSG00000237072 | 0 | 0 | 0 | 0 | 0 | 0 |
| ENSG00000196781 | 7963 | 3614 | 5034 | 3640 | 3695 | 4236 |
| ENSG00000233926 | 0 | 2 | 4 | 2 | 3 | 2 |
| ENSG00000207206 | 0 | 0 | 0 | 0 | 0 | 0 |
| ENSG00000252167 | 0 | 0 | 0 | 0 | 0 | 0 |
| ENSG00000200969 | 0 | 0 | 0 | 0 | 0 | 0 |
| ENSG00000240632 | 0 | 0 | 0 | 0 | 0 | 0 |
| ENSG00000230846 | 0 | 0 | 0 | 0 | 0 | 0 |
| ENSG00000189357 | 0 | 0 | 0 | 0 | 0 | 0 |
| ENSG00000267559 | 0 | 0 | 0 | 0 | 0 | 0 |
| ENSG00000186788 | 0 | 0 | 0 | 0 | 0 | 0 |
| ENSG00000237770 | 0 | 0 | 0 | 0 | 0 | 0 |
| ENSG00000228430 | 1 | 1 | 2 | 1 | 2 | 3 |
| ENSG00000214929 | 0 | 0 | 0 | 0 | 0 | 0 |
| ENSG00000231649 | 0 | 0 | 0 | 0 | 0 | 0 |
| ENSG00000230360 | 0 | 0 | 0 | 0 | 0 | 0 |
| ENSG00000235377 | 0 | 0 | 0 | 0 | 0 | 0 |
| ENSG00000225085 | 0 | 0 | 0 | 0 | 0 | 0 |
| ENSG00000228046 | 0 | 0 | 0 | 0 | 0 | 0 |
| ENSG00000228123 | 0 | 0 | 0 | 0 | 0 | 0 |
| ENSG00000232749 | 0 | 0 | 0 | 0 | 0 | 0 |
| ENSG00000278988 | 9 | 12 | 3 | 0 | 6 | 5 |
| ENSG00000233309 | 30 | 14 | 8 | 10 | 10 | 12 |
| ENSG00000165105 | 2254 | 1044 | 1305 | 522 | 1042 | 878 |
| ENSG00000251776 | 0 | 0 | 0 | 0 | 0 | 0 |
| ENSG00000273806 | 1 | 0 | 0 | 0 | 0 | 0 |
| ENSG00000229109 | 7 | 1 | 6 | 0 | 4 | 4 |
| ENSG00000172159 | 174 | 103 | 83 | 67 | 124 | 106 |

| | | | | | | |
|---|---|---|---|---|---|---|
| ENSG00000237529 | 6 | 3 | 6 | 1 | 4 | 2 |
| ENSG00000251752 | 1 | 1 | 0 | 0 | 2 | 1 |
| ENSG00000199483 | 0 | 0 | 0 | 0 | 0 | 0 |
| ENSG00000148057 | 230 | 128 | 168 | 159 | 135 | 154 |
| ENSG00000135018 | 6343 | 3072 | 3922 | 3118 | 4134 | 3601 |
| ENSG00000254473 | 67 | 33 | 57 | 46 | 38 | 41 |
| ENSG00000165113 | 246 | 145 | 147 | 88 | 149 | 89 |
| ENSG00000226877 | 4 | 0 | 0 | 0 | 0 | 1 |
| ENSG00000231616 | 6 | 2 | 5 | 2 | 6 | 3 |
| ENSG00000165115 | 140 | 95 | 92 | 47 | 115 | 91 |
| ENSG00000165118 | 1072 | 555 | 654 | 480 | 708 | 689 |
| ENSG00000165119 | 29416 | 15083 | 18705 | 13616 | 16445 | 17247 |
| ENSG00000284179 | 0 | 0 | 0 | 0 | 0 | 0 |
| ENSG00000235298 | 8 | 9 | 7 | 2 | 5 | 1 |
| ENSG00000178966 | 448 | 232 | 242 | 133 | 287 | 226 |
| ENSG00000202523 | 0 | 0 | 0 | 0 | 0 | 0 |
| ENSG00000227463 | 0 | 0 | 0 | 0 | 1 | 0 |
| ENSG00000270166 | 0 | 0 | 0 | 0 | 0 | 0 |
| ENSG00000197506 | 0 | 0 | 3 | 0 | 0 | 0 |
| ENSG00000233262 | 0 | 0 | 0 | 0 | 0 | 0 |
| ENSG00000285987 | 0 | 0 | 0 | 0 | 0 | 0 |
| ENSG00000228095 | 0 | 0 | 0 | 0 | 0 | 0 |
| ENSG00000148053 | 0 | 2 | 0 | 0 | 0 | 0 |
| ENSG00000226590 | 0 | 0 | 0 | 0 | 0 | 0 |
| ENSG00000285556 | 0 | 0 | 0 | 0 | 0 | 0 |
| ENSG00000285634 | 0 | 0 | 0 | 0 | 0 | 0 |
| ENSG00000227412 | 0 | 0 | 0 | 0 | 0 | 0 |
| ENSG00000135049 | 366 | 166 | 186 | 86 | 313 | 184 |
| ENSG00000234424 | 0 | 0 | 0 | 0 | 0 | 0 |
| ENSG00000229454 | 0 | 0 | 0 | 0 | 0 | 0 |
| ENSG00000230303 | 0 | 0 | 1 | 1 | 2 | 0 |
| ENSG00000165121 | 62 | 41 | 62 | 41 | 63 | 56 |
| ENSG00000213711 | 0 | 0 | 0 | 0 | 0 | 0 |
| ENSG00000231741 | 0 | 0 | 0 | 0 | 0 | 2 |
| ENSG00000135040 | 1611 | 889 | 1004 | 621 | 1042 | 862 |
| ENSG00000135052 | 5616 | 2528 | 2971 | 2546 | 2938 | 2702 |
| ENSG00000233713 | 0 | 0 | 0 | 0 | 0 | 0 |
| ENSG00000187753 | 6 | 3 | 1 | 2 | 0 | 1 |
| ENSG00000270688 | 0 | 1 | 0 | 0 | 0 | 0 |
| ENSG00000223012 | 0 | 0 | 0 | 0 | 1 | 0 |
| ENSG00000135070 | 2088 | 952 | 1172 | 1076 | 1187 | 1186 |
| ENSG00000083223 | 938 | 582 | 457 | 263 | 883 | 522 |
| ENSG00000214908 | 0 | 0 | 2 | 0 | 0 | 0 |
| ENSG00000235819 | 0 | 0 | 0 | 0 | 0 | 0 |
| ENSG00000222293 | 0 | 0 | 0 | 0 | 0 | 0 |
| ENSG00000232211 | 0 | 0 | 0 | 0 | 0 | 0 |
| ENSG00000285686 | 0 | 0 | 0 | 0 | 0 | 0 |
| ENSG00000180447 | 2 | 0 | 0 | 0 | 0 | 0 |
| ENSG00000226237 | 0 | 1 | 0 | 0 | 0 | 0 |
| ENSG00000269994 | 71 | 23 | 30 | 18 | 23 | 25 |

| | | | | | | |
|---|---|---|---|---|---|---|
| ENSG00000231007 | 29 | 12 | 29 | 40 | 31 | 27 |
| ENSG00000224887 | 3 | 0 | 0 | 0 | 0 | 0 |
| ENSG00000204446 | 0 | 0 | 0 | 0 | 0 | 0 |
| ENSG00000212421 | 0 | 0 | 0 | 0 | 0 | 0 |
| ENSG00000228373 | 0 | 0 | 0 | 0 | 0 | 0 |
| ENSG00000196730 | 74 | 35 | 45 | 32 | 46 | 49 |
| ENSG00000219928 | 0 | 2 | 1 | 7 | 2 | 3 |
| ENSG00000236709 | 0 | 0 | 0 | 0 | 0 | 0 |
| ENSG00000135047 | 6345 | 3179 | 4090 | 2976 | 3512 | 3369 |
| ENSG00000237747 | 0 | 0 | 0 | 0 | 0 | 0 |
| ENSG00000188029 | 2 | 0 | 0 | 0 | 0 | 0 |
| ENSG00000276478 | 0 | 0 | 0 | 0 | 0 | 0 |
| ENSG00000237743 | 0 | 0 | 0 | 0 | 0 | 0 |
| ENSG00000232743 | 0 | 0 | 0 | 0 | 0 | 0 |
| ENSG00000277357 | 0 | 0 | 0 | 0 | 0 | 0 |
| ENSG00000234575 | 0 | 0 | 0 | 0 | 0 | 0 |
| ENSG00000233884 | 0 | 0 | 0 | 0 | 0 | 0 |
| ENSG00000214888 | 0 | 0 | 0 | 0 | 0 | 0 |
| ENSG00000234460 | 0 | 0 | 1 | 0 | 0 | 0 |
| ENSG00000177992 | 0 | 0 | 0 | 0 | 0 | 0 |
| ENSG00000230246 | 0 | 0 | 0 | 0 | 0 | 1 |
| ENSG00000238110 | 0 | 0 | 0 | 0 | 0 | 0 |
| ENSG00000283205 | 0 | 0 | 0 | 0 | 0 | 0 |
| ENSG00000156345 | 722 | 302 | 429 | 384 | 369 | 470 |
| ENSG00000271923 | 0 | 0 | 0 | 0 | 0 | 0 |
| ENSG00000217716 | 10 | 7 | 7 | 4 | 3 | 2 |
| ENSG00000225385 | 4 | 3 | 5 | 4 | 6 | 3 |
| ENSG00000177910 | 0 | 0 | 0 | 0 | 0 | 0 |
| ENSG00000276535 | 0 | 0 | 0 | 0 | 0 | 0 |
| ENSG00000279154 | 0 | 0 | 0 | 0 | 0 | 0 |
| ENSG00000204429 | 0 | 0 | 0 | 0 | 0 | 0 |
| ENSG00000229288 | 0 | 0 | 0 | 0 | 0 | 0 |
| ENSG00000226327 | 0 | 0 | 0 | 0 | 0 | 0 |
| ENSG00000106723 | 4543 | 2186 | 2799 | 1528 | 2386 | 2416 |
| ENSG00000275017 | 2 | 2 | 1 | 1 | 0 | 3 |
| ENSG00000227638 | 3 | 7 | 2 | 1 | 1 | 3 |
| ENSG00000231804 | 0 | 0 | 1 | 0 | 2 | 0 |
| ENSG00000130045 | 227 | 115 | 127 | 172 | 158 | 145 |
| ENSG00000228189 | 0 | 1 | 0 | 0 | 0 | 0 |
| ENSG00000265873 | 0 | 0 | 0 | 0 | 0 | 0 |
| ENSG00000225856 | 0 | 0 | 0 | 0 | 0 | 0 |
| ENSG00000228658 | 0 | 0 | 0 | 0 | 0 | 0 |
| ENSG00000213694 | 1829 | 822 | 1027 | 900 | 1436 | 1186 |
| ENSG00000148082 | 35 | 7 | 14 | 18 | 13 | 11 |
| ENSG00000224945 | 9 | 8 | 18 | 14 | 8 | 8 |
| ENSG00000123975 | 1247 | 625 | 804 | 1098 | 1035 | 938 |
| ENSG00000265112 | 0 | 2 | 2 | 1 | 0 | 1 |
| ENSG00000187742 | 2952 | 1344 | 1826 | 1165 | 1454 | 1527 |
| ENSG00000187764 | 882 | 369 | 499 | 295 | 419 | 391 |
| ENSG00000231799 | 69 | 43 | 51 | 25 | 45 | 40 |

| | | | | | | |
|---|---|---|---|---|---|---|
| ENSG00000225460 | 0 | 0 | 0 | 0 | 0 | 0 |
| ENSG00000285907 | 65 | 23 | 26 | 31 | 23 | 32 |
| ENSG00000130222 | 258 | 149 | 166 | 178 | 166 | 146 |
| ENSG00000237372 | 2 | 0 | 1 | 4 | 0 | 0 |
| ENSG00000226206 | 0 | 0 | 0 | 0 | 0 | 0 |
| ENSG00000236507 | 0 | 0 | 0 | 0 | 0 | 0 |
| ENSG00000237626 | 0 | 0 | 0 | 0 | 0 | 0 |
| ENSG00000227555 | 0 | 0 | 0 | 0 | 0 | 0 |
| ENSG00000263967 | 0 | 0 | 0 | 0 | 0 | 0 |
| ENSG00000234910 | 0 | 0 | 0 | 0 | 0 | 0 |
| ENSG00000278382 | 0 | 0 | 0 | 0 | 0 | 0 |
| ENSG00000279503 | 0 | 0 | 0 | 0 | 0 | 0 |
| ENSG00000185074 | 0 | 0 | 0 | 0 | 0 | 1 |
| ENSG00000225072 | 0 | 0 | 0 | 0 | 0 | 0 |
| ENSG00000231107 | 49 | 23 | 31 | 11 | 20 | 26 |
| ENSG00000229613 | 0 | 1 | 0 | 0 | 0 | 1 |
| ENSG00000165023 | 23 | 6 | 20 | 4 | 2 | 1 |
| ENSG00000236316 | 0 | 0 | 0 | 0 | 0 | 0 |
| ENSG00000270432 | 0 | 0 | 0 | 0 | 0 | 0 |
| ENSG00000165025 | 767 | 373 | 444 | 357 | 416 | 418 |
| ENSG00000228216 | 0 | 0 | 0 | 2 | 1 | 0 |
| ENSG00000260454 | 0 | 0 | 0 | 0 | 0 | 0 |
| ENSG00000230537 | 0 | 1 | 2 | 0 | 0 | 0 |
| ENSG00000229694 | 0 | 0 | 0 | 0 | 1 | 0 |
| ENSG00000237422 | 0 | 0 | 0 | 0 | 0 | 0 |
| ENSG00000235641 | 0 | 0 | 0 | 0 | 0 | 0 |
| ENSG00000230147 | 0 | 0 | 0 | 0 | 0 | 0 |
| ENSG00000273381 | 2 | 7 | 7 | 0 | 8 | 5 |
| ENSG00000148090 | 373 | 178 | 210 | 140 | 189 | 175 |
| ENSG00000165030 | 284 | 179 | 222 | 152 | 178 | 203 |
| ENSG00000233081 | 6 | 2 | 4 | 2 | 4 | 6 |
| ENSG00000169071 | 6 | 0 | 1 | 0 | 2 | 1 |
| ENSG00000228595 | 0 | 0 | 0 | 0 | 0 | 0 |
| ENSG00000266855 | 0 | 0 | 0 | 0 | 0 | 0 |
| ENSG00000283351 | 0 | 0 | 0 | 0 | 0 | 0 |
| ENSG00000090054 | 4784 | 2536 | 3041 | 2531 | 2856 | 2811 |
| ENSG00000227399 | 0 | 0 | 0 | 0 | 0 | 0 |
| ENSG00000237306 | 0 | 1 | 0 | 0 | 0 | 0 |
| ENSG00000275756 | 0 | 2 | 0 | 3 | 0 | 4 |
| ENSG00000232179 | 0 | 1 | 0 | 0 | 0 | 0 |
| ENSG00000228069 | 4 | 3 | 1 | 0 | 3 | 1 |
| ENSG00000238288 | 0 | 0 | 0 | 0 | 1 | 0 |
| ENSG00000240791 | 0 | 0 | 0 | 0 | 1 | 0 |
| ENSG00000227321 | 2 | 2 | 1 | 0 | 2 | 0 |
| ENSG00000234537 | 12 | 2 | 15 | 11 | 15 | 2 |
| ENSG00000225511 | 38 | 27 | 35 | 37 | 57 | 40 |
| ENSG00000236717 | 1 | 0 | 0 | 3 | 0 | 0 |
| ENSG00000224809 | 0 | 0 | 0 | 0 | 0 | 0 |
| ENSG00000276717 | 0 | 0 | 0 | 0 | 0 | 0 |
| ENSG00000236115 | 0 | 0 | 0 | 1 | 0 | 1 |

| | | | | | | |
|---|---|---|---|---|---|---|
| ENSG00000196305 | 5055 | 2619 | 3228 | 2383 | 3404 | 3123 |
| ENSG00000281156 | 1 | 0 | 0 | 0 | 0 | 0 |
| ENSG00000239183 | 0 | 0 | 0 | 0 | 0 | 0 |
| ENSG00000198000 | 631 | 397 | 285 | 190 | 673 | 324 |
| ENSG00000188312 | 169 | 94 | 119 | 84 | 114 | 95 |
| ENSG00000229530 | 0 | 0 | 0 | 0 | 0 | 0 |
| ENSG00000106809 | 0 | 1 | 0 | 0 | 0 | 0 |
| ENSG00000127083 | 0 | 0 | 0 | 0 | 0 | 0 |
| ENSG00000106819 | 0 | 0 | 1 | 0 | 0 | 1 |
| ENSG00000106823 | 0 | 0 | 0 | 0 | 0 | 0 |
| ENSG00000264158 | 0 | 0 | 0 | 0 | 0 | 0 |
| ENSG00000225551 | 0 | 0 | 0 | 0 | 0 | 0 |
| ENSG00000285650 | 4 | 6 | 0 | 1 | 2 | 0 |
| ENSG00000127080 | 621 | 262 | 325 | 269 | 317 | 328 |
| ENSG00000213669 | 4 | 0 | 0 | 0 | 0 | 0 |
| ENSG00000185963 | 2143 | 998 | 1243 | 1082 | 1234 | 1300 |
| ENSG00000187984 | 4 | 5 | 4 | 5 | 6 | 1 |
| ENSG00000252772 | 0 | 0 | 0 | 0 | 0 | 0 |
| ENSG00000226721 | 0 | 0 | 0 | 0 | 0 | 0 |
| ENSG00000275318 | 5 | 9 | 9 | 2 | 8 | 7 |
| ENSG00000127081 | 148 | 104 | 97 | 60 | 133 | 74 |
| ENSG00000226668 | 0 | 0 | 0 | 1 | 0 | 0 |
| ENSG00000232843 | 0 | 0 | 0 | 0 | 0 | 0 |
| ENSG00000188101 | 0 | 0 | 0 | 0 | 0 | 0 |
| ENSG00000127084 | 221 | 124 | 139 | 109 | 100 | 99 |
| ENSG00000157303 | 42 | 26 | 38 | 29 | 20 | 15 |
| ENSG00000223446 | 0 | 0 | 0 | 0 | 0 | 0 |
| ENSG00000165233 | 1181 | 624 | 720 | 831 | 748 | 677 |
| ENSG00000131669 | 7279 | 3486 | 4046 | 4308 | 3445 | 3413 |
| ENSG00000203364 | 2 | 4 | 8 | 1 | 8 | 3 |
| ENSG00000165238 | 488 | 238 | 270 | 248 | 251 | 263 |
| ENSG00000204352 | 0 | 3 | 1 | 0 | 2 | 1 |
| ENSG00000200570 | 0 | 0 | 0 | 0 | 0 | 0 |
| ENSG00000201074 | 0 | 0 | 0 | 0 | 0 | 0 |
| ENSG00000237385 | 0 | 0 | 0 | 0 | 0 | 0 |
| ENSG00000188938 | 5175 | 2450 | 3038 | 2447 | 2634 | 2509 |
| ENSG00000048828 | 12036 | 5710 | 7022 | 5091 | 6628 | 6497 |
| ENSG00000197724 | 4555 | 1908 | 2628 | 1892 | 2203 | 2317 |
| ENSG00000283146 | 0 | 0 | 0 | 0 | 0 | 0 |
| ENSG00000265347 | 0 | 0 | 0 | 0 | 0 | 0 |
| ENSG00000231990 | 0 | 0 | 0 | 0 | 0 | 0 |
| ENSG00000131668 | 12 | 5 | 8 | 2 | 5 | 2 |
| ENSG00000235601 | 0 | 0 | 2 | 0 | 0 | 2 |
| ENSG00000158079 | 613 | 291 | 339 | 255 | 375 | 319 |
| ENSG00000227666 | 2 | 1 | 4 | 0 | 0 | 6 |
| ENSG00000269946 | 9 | 11 | 8 | 6 | 9 | 6 |
| ENSG00000199165 | 0 | 5 | 0 | 1 | 0 | 2 |
| ENSG00000199072 | 0 | 0 | 0 | 0 | 1 | 0 |
| ENSG00000269929 | 9 | 5 | 10 | 1 | 17 | 11 |
| ENSG00000230262 | 8 | 3 | 2 | 1 | 3 | 2 |

| | | | | | | |
|---|---|---|---|---|---|---|
| ENSG00000199133 | 11 | 8 | 15 | 4 | 7 | 13 |
| ENSG00000175787 | 180 | 96 | 159 | 79 | 129 | 122 |
| ENSG00000237679 | 0 | 1 | 0 | 0 | 0 | 0 |
| ENSG00000130950 | 2 | 0 | 0 | 1 | 1 | 1 |
| ENSG00000232063 | 9 | 2 | 3 | 2 | 0 | 0 |
| ENSG00000224815 | 0 | 1 | 0 | 0 | 0 | 0 |
| ENSG00000282886 | 2 | 0 | 1 | 1 | 1 | 0 |
| ENSG00000148110 | 4260 | 1888 | 2373 | 1792 | 2236 | 2236 |
| ENSG00000278486 | 2 | 2 | 5 | 0 | 0 | 0 |
| ENSG00000224245 | 1 | 3 | 1 | 0 | 0 | 4 |
| ENSG00000202445 | 0 | 0 | 0 | 0 | 0 | 0 |
| ENSG00000231806 | 336 | 164 | 190 | 149 | 199 | 169 |
| ENSG00000130957 | 12 | 7 | 15 | 2 | 9 | 6 |
| ENSG00000165140 | 13533 | 6891 | 9157 | 7729 | 6816 | 7693 |
| ENSG00000148120 | 1129 | 656 | 758 | 795 | 917 | 750 |
| ENSG00000252153 | 0 | 0 | 0 | 0 | 1 | 0 |
| ENSG00000230815 | 0 | 0 | 0 | 0 | 0 | 0 |
| ENSG00000236095 | 0 | 1 | 1 | 0 | 0 | 0 |
| ENSG00000224764 | 1 | 3 | 3 | 1 | 0 | 0 |
| ENSG00000285807 | 0 | 0 | 0 | 0 | 0 | 0 |
| ENSG00000274115 | 0 | 0 | 0 | 0 | 0 | 1 |
| ENSG00000207563 | 4 | 2 | 1 | 2 | 4 | 4 |
| ENSG00000207864 | 21 | 3 | 13 | 8 | 13 | 15 |
| ENSG00000207617 | 0 | 0 | 0 | 0 | 0 | 0 |
| ENSG00000284459 | 0 | 0 | 0 | 0 | 0 | 0 |
| ENSG00000158169 | 718 | 332 | 415 | 375 | 454 | 404 |
| ENSG00000229065 | 1 | 0 | 1 | 0 | 1 | 1 |
| ENSG00000223849 | 0 | 2 | 1 | 0 | 0 | 1 |
| ENSG00000251884 | 0 | 0 | 0 | 0 | 0 | 0 |
| ENSG00000230221 | 0 | 0 | 1 | 0 | 0 | 0 |
| ENSG00000271384 | 8 | 5 | 3 | 4 | 5 | 5 |
| ENSG00000213761 | 0 | 1 | 0 | 0 | 3 | 2 |
| ENSG00000185920 | 143 | 78 | 62 | 42 | 63 | 44 |
| ENSG00000271314 | 0 | 0 | 1 | 0 | 0 | 0 |
| ENSG00000271155 | 0 | 0 | 0 | 0 | 0 | 1 |
| ENSG00000271659 | 2 | 1 | 3 | 3 | 5 | 1 |
| ENSG00000229345 | 0 | 0 | 0 | 0 | 0 | 0 |
| ENSG00000228142 | 0 | 0 | 0 | 0 | 0 | 0 |
| ENSG00000227358 | 0 | 0 | 0 | 0 | 0 | 0 |
| ENSG00000175611 | 343 | 166 | 200 | 148 | 169 | 192 |
| ENSG00000268926 | 0 | 0 | 0 | 0 | 0 | 0 |
| ENSG00000252847 | 0 | 0 | 0 | 0 | 0 | 0 |
| ENSG00000182150 | 360 | 252 | 176 | 61 | 317 | 189 |
| ENSG00000199202 | 0 | 1 | 0 | 0 | 0 | 0 |
| ENSG00000237631 | 0 | 0 | 0 | 0 | 0 | 0 |
| ENSG00000225194 | 5 | 0 | 0 | 0 | 1 | 1 |
| ENSG00000237212 | 0 | 0 | 0 | 0 | 0 | 0 |
| ENSG00000201938 | 0 | 0 | 0 | 0 | 0 | 0 |
| ENSG00000275465 | 0 | 0 | 0 | 0 | 0 | 0 |
| ENSG00000224546 | 284 | 108 | 136 | 124 | 159 | 117 |

| | | | | | | |
|---|---|---|---|---|---|---|
| ENSG00000130948 | 0 | 0 | 0 | 0 | 0 | 0 |
| ENSG00000285269 | 0 | 0 | 1 | 0 | 0 | 1 |
| ENSG00000232283 | 0 | 0 | 0 | 0 | 0 | 0 |
| ENSG00000207276 | 0 | 0 | 0 | 0 | 0 | 0 |
| ENSG00000130958 | 412 | 294 | 346 | 311 | 386 | 365 |
| ENSG00000165244 | 484 | 225 | 299 | 172 | 194 | 267 |
| ENSG00000238251 | 8 | 1 | 3 | 2 | 4 | 2 |
| ENSG00000130956 | 691 | 351 | 418 | 487 | 535 | 550 |
| ENSG00000081377 | 454 | 192 | 249 | 216 | 367 | 314 |
| ENSG00000158122 | 509 | 274 | 295 | 348 | 373 | 379 |
| ENSG00000224848 | 15 | 8 | 6 | 6 | 5 | 4 |
| ENSG00000233820 | 4 | 4 | 5 | 3 | 1 | 4 |
| ENSG00000081386 | 781 | 406 | 453 | 265 | 478 | 366 |
| ENSG00000196597 | 331 | 159 | 213 | 90 | 194 | 143 |
| ENSG00000196312 | 1640 | 871 | 1030 | 637 | 936 | 897 |
| ENSG00000269907 | 4 | 8 | 7 | 0 | 3 | 0 |
| ENSG00000227257 | 0 | 0 | 0 | 0 | 0 | 0 |
| ENSG00000188152 | 24 | 27 | 17 | 27 | 29 | 34 |
| ENSG00000236745 | 1 | 0 | 0 | 0 | 1 | 0 |
| ENSG00000136943 | 326 | 157 | 165 | 117 | 141 | 105 |
| ENSG00000197214 | 0 | 0 | 0 | 0 | 0 | 0 |
| ENSG00000228376 | 0 | 0 | 1 | 0 | 0 | 1 |
| ENSG00000228038 | 0 | 0 | 0 | 0 | 0 | 1 |
| ENSG00000213612 | 4 | 2 | 7 | 0 | 1 | 3 |
| ENSG00000277103 | 1 | 0 | 2 | 0 | 1 | 2 |
| ENSG00000159712 | 8 | 1 | 5 | 5 | 3 | 3 |
| ENSG00000252721 | 0 | 0 | 0 | 0 | 0 | 0 |
| ENSG00000242375 | 0 | 3 | 8 | 2 | 4 | 6 |
| ENSG00000188801 | 0 | 0 | 0 | 0 | 0 | 0 |
| ENSG00000203279 | 5 | 1 | 3 | 2 | 7 | 3 |
| ENSG00000235494 | 2 | 0 | 0 | 0 | 0 | 0 |
| ENSG00000254876 | 35 | 22 | 23 | 26 | 17 | 14 |
| ENSG00000255036 | 1 | 0 | 2 | 2 | 4 | 3 |
| ENSG00000254483 | 0 | 0 | 0 | 0 | 0 | 0 |
| ENSG00000254633 | 0 | 0 | 0 | 0 | 0 | 0 |
| ENSG00000197816 | 24 | 13 | 31 | 12 | 19 | 20 |
| ENSG00000221269 | 0 | 0 | 0 | 1 | 2 | 0 |
| ENSG00000228957 | 0 | 0 | 0 | 0 | 0 | 0 |
| ENSG00000196116 | 1225 | 635 | 771 | 505 | 742 | 640 |
| ENSG00000136842 | 46 | 31 | 30 | 19 | 28 | 32 |
| ENSG00000228174 | 0 | 0 | 0 | 0 | 0 | 0 |
| ENSG00000136925 | 928 | 416 | 569 | 348 | 422 | 443 |
| ENSG00000136937 | 2654 | 1549 | 1507 | 1104 | 1979 | 1511 |
| ENSG00000231521 | 0 | 0 | 0 | 0 | 0 | 0 |
| ENSG00000136936 | 560 | 332 | 353 | 248 | 333 | 348 |
| ENSG00000214417 | 4 | 1 | 0 | 1 | 0 | 1 |
| ENSG00000236130 | 2 | 0 | 2 | 2 | 1 | 3 |
| ENSG00000260677 | 4 | 2 | 0 | 2 | 0 | 1 |
| ENSG00000178919 | 0 | 0 | 0 | 0 | 0 | 0 |
| ENSG00000136932 | 666 | 318 | 423 | 318 | 368 | 368 |

| | | | | | | |
|---|---|---|---|---|---|---|
| ENSG00000201830 | 2 | 0 | 0 | 0 | 0 | 2 |
| ENSG00000136929 | 2 | 0 | 0 | 0 | 0 | 1 |
| ENSG00000136938 | 8333 | 4299 | 5149 | 4867 | 5233 | 5078 |
| ENSG00000236896 | 1 | 2 | 3 | 1 | 0 | 0 |
| ENSG00000212521 | 0 | 0 | 0 | 0 | 0 | 0 |
| ENSG00000095380 | 3839 | 2331 | 2551 | 2191 | 1874 | 2002 |
| ENSG00000106785 | 1551 | 796 | 865 | 634 | 783 | 838 |
| ENSG00000274011 | 5 | 0 | 2 | 1 | 0 | 1 |
| ENSG00000106789 | 4010 | 1912 | 2206 | 957 | 1653 | 1390 |
| ENSG00000095383 | 2183 | 1016 | 1237 | 982 | 846 | 995 |
| ENSG00000278412 | 0 | 0 | 0 | 0 | 0 | 0 |
| ENSG00000224001 | 0 | 1 | 1 | 0 | 0 | 0 |
| ENSG00000136928 | 101 | 55 | 54 | 18 | 25 | 25 |
| ENSG00000229897 | 0 | 0 | 0 | 0 | 0 | 0 |
| ENSG00000165138 | 1418 | 682 | 899 | 704 | 727 | 784 |
| ENSG00000285706 | 0 | 0 | 0 | 0 | 0 | 0 |
| ENSG00000119514 | 132 | 68 | 107 | 75 | 91 | 114 |
| ENSG00000267026 | 0 | 0 | 2 | 0 | 0 | 0 |
| ENSG00000229338 | 0 | 0 | 0 | 0 | 0 | 0 |
| ENSG00000204291 | 0 | 0 | 2 | 0 | 0 | 1 |
| ENSG00000270412 | 0 | 0 | 0 | 0 | 0 | 0 |
| ENSG00000106799 | 1197 | 606 | 687 | 389 | 731 | 605 |
| ENSG00000252942 | 0 | 0 | 0 | 0 | 0 | 0 |
| ENSG00000227269 | 0 | 0 | 0 | 0 | 0 | 0 |
| ENSG00000240235 | 0 | 0 | 0 | 0 | 0 | 0 |
| ENSG00000119523 | 2259 | 1067 | 1417 | 1161 | 1292 | 1248 |
| ENSG00000106803 | 4367 | 2322 | 2512 | 2767 | 2472 | 2413 |
| ENSG00000222337 | 0 | 0 | 0 | 0 | 0 | 0 |
| ENSG00000255815 | 1 | 2 | 1 | 0 | 2 | 1 |
| ENSG00000271086 | 0 | 2 | 3 | 0 | 2 | 0 |
| ENSG00000277828 | 0 | 0 | 0 | 0 | 2 | 0 |
| ENSG00000277465 | 0 | 0 | 0 | 0 | 0 | 0 |
| ENSG00000234860 | 0 | 0 | 0 | 0 | 0 | 0 |
| ENSG00000237461 | 7 | 6 | 8 | 5 | 2 | 3 |
| ENSG00000119508 | 89 | 27 | 45 | 6 | 14 | 16 |
| ENSG00000255145 | 36 | 25 | 18 | 22 | 14 | 16 |
| ENSG00000136874 | 1123 | 592 | 696 | 334 | 677 | 527 |
| ENSG00000254571 | 0 | 0 | 0 | 0 | 0 | 1 |
| ENSG00000023318 | 2634 | 1370 | 1574 | 1169 | 1635 | 1359 |
| ENSG00000234709 | 7 | 1 | 2 | 4 | 4 | 1 |
| ENSG00000119509 | 647 | 278 | 383 | 256 | 347 | 331 |
| ENSG00000239908 | 1 | 1 | 2 | 1 | 1 | 1 |
| ENSG00000238172 | 0 | 2 | 0 | 0 | 0 | 0 |
| ENSG00000270995 | 0 | 0 | 0 | 0 | 0 | 0 |
| ENSG00000231697 | 0 | 0 | 0 | 0 | 0 | 0 |
| ENSG00000136891 | 1626 | 771 | 875 | 888 | 1035 | 1028 |
| ENSG00000066697 | 961 | 495 | 563 | 685 | 711 | 638 |
| ENSG00000251349 | 0 | 0 | 0 | 0 | 0 | 1 |
| ENSG00000241697 | 0 | 0 | 0 | 2 | 0 | 0 |
| ENSG00000170681 | 6 | 10 | 2 | 2 | 6 | 3 |

| | | | | | | |
|---|---|---|---|---|---|---|
| ENSG00000200966 | 0 | 0 | 0 | 0 | 0 | 0 |
| ENSG00000237999 | 0 | 0 | 0 | 0 | 0 | 1 |
| ENSG00000230161 | 0 | 0 | 0 | 0 | 0 | 0 |
| ENSG00000229541 | 0 | 0 | 1 | 0 | 0 | 0 |
| ENSG00000148123 | 0 | 1 | 1 | 1 | 0 | 0 |
| ENSG00000238143 | 0 | 0 | 0 | 0 | 0 | 0 |
| ENSG00000253041 | 0 | 0 | 0 | 0 | 0 | 0 |
| ENSG00000227411 | 0 | 0 | 0 | 0 | 0 | 0 |
| ENSG00000236763 | 0 | 0 | 0 | 0 | 0 | 0 |
| ENSG00000230328 | 0 | 0 | 0 | 0 | 0 | 0 |
| ENSG00000136881 | 0 | 1 | 0 | 0 | 1 | 0 |
| ENSG00000234502 | 0 | 0 | 0 | 0 | 0 | 0 |
| ENSG00000136897 | 1516 | 783 | 863 | 772 | 948 | 773 |
| ENSG00000136870 | 583 | 279 | 351 | 202 | 363 | 308 |
| ENSG00000136872 | 0 | 1 | 0 | 1 | 0 | 0 |
| ENSG00000270982 | 0 | 0 | 0 | 0 | 0 | 0 |
| ENSG00000225376 | 0 | 0 | 0 | 0 | 0 | 0 |
| ENSG00000165152 | 0 | 0 | 0 | 0 | 0 | 0 |
| ENSG00000233055 | 0 | 0 | 0 | 0 | 0 | 0 |
| ENSG00000155827 | 1765 | 949 | 954 | 660 | 1143 | 890 |
| ENSG00000198785 | 2 | 2 | 3 | 1 | 1 | 1 |
| ENSG00000188386 | 0 | 0 | 0 | 0 | 0 | 0 |
| ENSG00000230302 | 0 | 0 | 0 | 0 | 0 | 0 |
| ENSG00000230418 | 0 | 0 | 0 | 0 | 0 | 0 |
| ENSG00000238882 | 0 | 0 | 0 | 0 | 0 | 0 |
| ENSG00000235901 | 0 | 0 | 0 | 0 | 0 | 0 |
| ENSG00000204250 | 0 | 0 | 0 | 0 | 0 | 0 |
| ENSG00000232242 | 0 | 0 | 0 | 0 | 0 | 0 |
| ENSG00000234269 | 0 | 0 | 0 | 0 | 0 | 0 |
| ENSG00000155833 | 0 | 0 | 0 | 0 | 0 | 0 |
| ENSG00000229830 | 0 | 0 | 0 | 0 | 0 | 0 |
| ENSG00000225564 | 0 | 0 | 0 | 0 | 0 | 0 |
| ENSG00000226566 | 0 | 0 | 0 | 0 | 0 | 0 |
| ENSG00000253031 | 0 | 0 | 0 | 0 | 0 | 0 |
| ENSG00000285483 | 0 | 0 | 0 | 0 | 0 | 0 |
| ENSG00000270332 | 7 | 4 | 2 | 3 | 5 | 4 |
| ENSG00000136824 | 383 | 220 | 138 | 70 | 480 | 205 |
| ENSG00000242111 | 0 | 0 | 0 | 0 | 0 | 0 |
| ENSG00000203396 | 0 | 0 | 0 | 0 | 0 | 0 |
| ENSG00000186881 | 0 | 0 | 0 | 0 | 0 | 0 |
| ENSG00000283001 | 0 | 0 | 0 | 0 | 0 | 0 |
| ENSG00000148136 | 0 | 0 | 0 | 0 | 0 | 0 |
| ENSG00000204246 | 0 | 0 | 0 | 0 | 0 | 0 |
| ENSG00000186943 | 0 | 0 | 0 | 0 | 0 | 0 |
| ENSG00000237443 | 0 | 0 | 0 | 0 | 0 | 0 |
| ENSG00000277556 | 0 | 0 | 0 | 0 | 2 | 0 |
| ENSG00000276119 | 0 | 1 | 0 | 0 | 0 | 0 |
| ENSG00000136839 | 0 | 0 | 0 | 0 | 0 | 0 |
| ENSG00000273423 | 0 | 0 | 0 | 0 | 0 | 0 |
| ENSG00000235248 | 0 | 0 | 0 | 0 | 0 | 0 |

| | | | | | | |
|---|---|---|---|---|---|---|
| ENSG00000179055 | 0 | 0 | 0 | 0 | 0 | 0 |
| ENSG00000188712 | 0 | 0 | 0 | 0 | 0 | 0 |
| ENSG00000136783 | 1602 | 705 | 938 | 758 | 827 | 804 |
| ENSG00000165028 | 121 | 83 | 59 | 42 | 55 | 69 |
| ENSG00000165029 | 2955 | 1415 | 1477 | 547 | 1287 | 984 |
| ENSG00000226334 | 36 | 18 | 26 | 11 | 7 | 17 |
| ENSG00000230013 | 7 | 1 | 2 | 6 | 2 | 2 |
| ENSG00000201583 | 0 | 0 | 0 | 0 | 0 | 0 |
| ENSG00000070214 | 10219 | 5181 | 5812 | 4911 | 7114 | 6182 |
| ENSG00000106701 | 170 | 68 | 46 | 34 | 83 | 66 |
| ENSG00000229419 | 10 | 10 | 5 | 1 | 12 | 6 |
| ENSG00000228317 | 0 | 0 | 0 | 0 | 0 | 1 |
| ENSG00000106692 | 863 | 514 | 389 | 220 | 736 | 394 |
| ENSG00000279609 | 0 | 0 | 0 | 0 | 0 | 0 |
| ENSG00000186051 | 2 | 0 | 3 | 0 | 0 | 0 |
| ENSG00000095209 | 908 | 431 | 482 | 445 | 537 | 534 |
| ENSG00000228366 | 0 | 0 | 1 | 0 | 0 | 0 |
| ENSG00000232486 | 0 | 1 | 0 | 0 | 0 | 0 |
| ENSG00000228948 | 0 | 0 | 0 | 0 | 0 | 0 |
| ENSG00000234323 | 2 | 0 | 0 | 0 | 0 | 1 |
| ENSG00000225002 | 0 | 0 | 0 | 0 | 0 | 0 |
| ENSG00000229297 | 0 | 0 | 0 | 0 | 0 | 0 |
| ENSG00000277889 | 0 | 0 | 0 | 0 | 0 | 0 |
| ENSG00000237062 | 0 | 0 | 0 | 0 | 0 | 0 |
| ENSG00000234229 | 0 | 0 | 0 | 0 | 0 | 0 |
| ENSG00000200131 | 0 | 0 | 0 | 0 | 0 | 0 |
| ENSG00000228143 | 0 | 0 | 0 | 0 | 0 | 0 |
| ENSG00000212276 | 0 | 0 | 0 | 0 | 0 | 0 |
| ENSG00000148143 | 36 | 12 | 19 | 15 | 26 | 28 |
| ENSG00000230782 | 0 | 0 | 0 | 0 | 0 | 0 |
| ENSG00000231562 | 0 | 0 | 0 | 0 | 0 | 0 |
| ENSG00000223615 | 0 | 0 | 0 | 0 | 0 | 0 |
| ENSG00000225531 | 0 | 1 | 0 | 0 | 0 | 0 |
| ENSG00000226047 | 0 | 0 | 0 | 0 | 0 | 0 |
| ENSG00000226535 | 0 | 0 | 0 | 0 | 0 | 0 |
| ENSG00000119318 | 13417 | 6539 | 8106 | 6363 | 7948 | 7636 |
| ENSG00000226825 | 0 | 0 | 1 | 0 | 0 | 0 |
| ENSG00000213558 | 0 | 0 | 0 | 0 | 0 | 0 |
| ENSG00000199905 | 0 | 0 | 0 | 0 | 0 | 0 |
| ENSG00000136826 | 1785 | 800 | 1114 | 929 | 969 | 1075 |
| ENSG00000236005 | 0 | 0 | 0 | 0 | 0 | 0 |
| ENSG00000227350 | 0 | 0 | 0 | 0 | 0 | 0 |
| ENSG00000202308 | 0 | 0 | 0 | 0 | 0 | 0 |
| ENSG00000222558 | 0 | 0 | 0 | 0 | 0 | 0 |
| ENSG00000244104 | 2 | 0 | 1 | 0 | 0 | 0 |
| ENSG00000231293 | 0 | 0 | 0 | 0 | 0 | 0 |
| ENSG00000233727 | 0 | 0 | 0 | 0 | 0 | 0 |
| ENSG00000222459 | 0 | 0 | 0 | 0 | 0 | 0 |
| ENSG00000236783 | 0 | 0 | 0 | 0 | 0 | 0 |
| ENSG00000213557 | 0 | 0 | 0 | 0 | 0 | 0 |

| | | | | | | |
|---|---|---|---|---|---|---|
| ENSG00000237360 | 0 | 0 | 0 | 0 | 0 | 0 |
| ENSG00000230030 | 0 | 0 | 0 | 0 | 0 | 0 |
| ENSG00000169253 | 0 | 0 | 0 | 0 | 0 | 0 |
| ENSG00000231678 | 0 | 0 | 0 | 0 | 0 | 0 |
| ENSG00000231810 | 0 | 0 | 0 | 0 | 0 | 0 |
| ENSG00000213556 | 0 | 0 | 0 | 0 | 0 | 0 |
| ENSG00000148156 | 4 | 0 | 2 | 4 | 2 | 0 |
| ENSG00000187003 | 0 | 0 | 0 | 0 | 0 | 0 |
| ENSG00000070061 | 3651 | 1659 | 2160 | 1438 | 2015 | 1985 |
| ENSG00000119328 | 1304 | 683 | 736 | 559 | 843 | 669 |
| ENSG00000119326 | 407 | 259 | 250 | 163 | 264 | 221 |
| ENSG00000106771 | 5162 | 2283 | 3097 | 1674 | 2701 | 2618 |
| ENSG00000199331 | 2 | 0 | 2 | 0 | 1 | 0 |
| ENSG00000207698 | 0 | 0 | 0 | 0 | 0 | 0 |
| ENSG00000260230 | 0 | 0 | 0 | 0 | 0 | 2 |
| ENSG00000095203 | 2118 | 1045 | 1296 | 913 | 1065 | 1145 |
| ENSG00000200106 | 0 | 0 | 0 | 0 | 0 | 0 |
| ENSG00000237124 | 0 | 0 | 0 | 0 | 0 | 0 |
| ENSG00000207329 | 0 | 0 | 0 | 0 | 0 | 0 |
| ENSG00000070159 | 2993 | 1480 | 1874 | 1081 | 1690 | 1596 |
| ENSG00000263861 | 0 | 0 | 0 | 0 | 0 | 0 |
| ENSG00000213539 | 0 | 0 | 1 | 0 | 0 | 1 |
| ENSG00000243444 | 8 | 2 | 3 | 1 | 5 | 1 |
| ENSG00000232939 | 0 | 0 | 1 | 0 | 0 | 0 |
| ENSG00000157654 | 0 | 0 | 0 | 0 | 1 | 0 |
| ENSG00000228053 | 0 | 0 | 0 | 4 | 0 | 0 |
| ENSG00000241978 | 0 | 0 | 0 | 0 | 0 | 0 |
| ENSG00000188959 | 58 | 31 | 28 | 25 | 22 | 25 |
| ENSG00000136810 | 8987 | 4729 | 5511 | 6515 | 6308 | 5919 |
| ENSG00000204193 | 0 | 0 | 0 | 0 | 0 | 0 |
| ENSG00000165124 | 4 | 0 | 0 | 1 | 1 | 0 |
| ENSG00000206658 | 0 | 0 | 0 | 0 | 0 | 0 |
| ENSG00000232896 | 0 | 0 | 0 | 0 | 0 | 0 |
| ENSG00000030304 | 0 | 1 | 0 | 0 | 0 | 0 |
| ENSG00000223918 | 0 | 0 | 0 | 0 | 0 | 0 |
| ENSG00000198121 | 0 | 0 | 0 | 0 | 1 | 0 |
| ENSG00000206923 | 0 | 0 | 0 | 0 | 0 | 0 |
| ENSG00000207401 | 0 | 0 | 0 | 0 | 0 | 0 |
| ENSG00000212409 | 0 | 0 | 0 | 0 | 0 | 0 |
| ENSG00000227531 | 0 | 0 | 0 | 0 | 0 | 0 |
| ENSG00000274263 | 0 | 0 | 0 | 0 | 0 | 0 |
| ENSG00000171133 | 0 | 0 | 0 | 0 | 0 | 0 |
| ENSG00000136813 | 5842 | 2886 | 3596 | 2243 | 3473 | 3155 |
| ENSG00000223238 | 0 | 0 | 0 | 0 | 0 | 0 |
| ENSG00000173258 | 42 | 21 | 34 | 10 | 30 | 27 |
| ENSG00000106853 | 1339 | 735 | 910 | 795 | 1154 | 1091 |
| ENSG00000224644 | 1 | 0 | 0 | 0 | 0 | 0 |
| ENSG00000204173 | 8 | 7 | 6 | 13 | 7 | 10 |
| ENSG00000059769 | 345 | 164 | 216 | 165 | 226 | 223 |
| ENSG00000244115 | 0 | 0 | 0 | 0 | 0 | 0 |

| | | | | | | |
|---|---|---|---|---|---|---|
| ENSG00000242616 | 29 | 38 | 36 | 26 | 37 | 32 |
| ENSG00000165181 | 4 | 8 | 1 | 4 | 2 | 2 |
| ENSG00000224067 | 0 | 0 | 7 | 1 | 1 | 2 |
| ENSG00000206907 | 0 | 0 | 0 | 0 | 0 | 0 |
| ENSG00000148154 | 4610 | 2097 | 2974 | 2114 | 2600 | 3026 |
| ENSG00000266315 | 0 | 0 | 0 | 0 | 0 | 1 |
| ENSG00000222356 | 0 | 0 | 0 | 0 | 0 | 0 |
| ENSG00000259953 | 54 | 21 | 25 | 38 | 44 | 28 |
| ENSG00000106868 | 645 | 372 | 342 | 282 | 421 | 322 |
| ENSG00000222327 | 0 | 0 | 0 | 0 | 0 | 0 |
| ENSG00000200261 | 0 | 0 | 0 | 0 | 1 | 0 |
| ENSG00000225513 | 4 | 0 | 1 | 0 | 2 | 0 |
| ENSG00000119314 | 6340 | 3188 | 3160 | 1788 | 4163 | 3105 |
| ENSG00000242256 | 0 | 0 | 0 | 0 | 0 | 0 |
| ENSG00000199525 | 0 | 0 | 0 | 0 | 0 | 0 |
| ENSG00000243911 | 0 | 0 | 0 | 0 | 0 | 0 |
| ENSG00000230081 | 0 | 0 | 0 | 0 | 0 | 0 |
| ENSG00000213529 | 7 | 0 | 4 | 0 | 0 | 0 |
| ENSG00000119471 | 1842 | 963 | 1058 | 761 | 1180 | 1008 |
| ENSG00000230185 | 15 | 18 | 12 | 7 | 20 | 4 |
| ENSG00000165185 | 621 | 339 | 349 | 265 | 372 | 326 |
| ENSG00000250068 | 1 | 1 | 1 | 2 | 1 | 3 |
| ENSG00000148153 | 2158 | 1051 | 1251 | 926 | 1206 | 1097 |
| ENSG00000226609 | 2 | 2 | 4 | 2 | 3 | 3 |
| ENSG00000148158 | 1602 | 795 | 1046 | 539 | 858 | 1015 |
| ENSG00000119457 | 20 | 1 | 4 | 4 | 2 | 3 |
| ENSG00000271631 | 6 | 1 | 6 | 0 | 7 | 2 |
| ENSG00000228623 | 0 | 0 | 0 | 0 | 1 | 0 |
| ENSG00000237906 | 0 | 0 | 0 | 0 | 0 | 0 |
| ENSG00000285447 | 0 | 0 | 0 | 0 | 0 | 0 |
| ENSG00000136866 | 0 | 1 | 0 | 0 | 0 | 0 |
| ENSG00000285115 | 0 | 0 | 0 | 0 | 0 | 0 |
| ENSG00000284763 | 0 | 0 | 0 | 0 | 0 | 0 |
| ENSG00000284962 | 0 | 0 | 0 | 0 | 0 | 0 |
| ENSG00000285076 | 0 | 0 | 0 | 0 | 0 | 0 |
| ENSG00000284867 | 0 | 0 | 0 | 0 | 0 | 0 |
| ENSG00000285370 | 0 | 0 | 0 | 0 | 0 | 0 |
| ENSG00000225684 | 4 | 7 | 4 | 3 | 4 | 7 |
| ENSG00000231528 | 100 | 60 | 71 | 38 | 50 | 50 |
| ENSG00000136867 | 68 | 46 | 62 | 31 | 42 | 34 |
| ENSG00000119321 | 3050 | 1481 | 1842 | 1334 | 1554 | 1617 |
| ENSG00000136868 | 2494 | 1174 | 1588 | 877 | 1499 | 1290 |
| ENSG00000176386 | 396 | 199 | 240 | 227 | 260 | 251 |
| ENSG00000136875 | 2315 | 1178 | 1386 | 1060 | 1440 | 1274 |
| ENSG00000165188 | 0 | 2 | 1 | 1 | 1 | 0 |
| ENSG00000148225 | 290 | 125 | 177 | 138 | 182 | 158 |
| ENSG00000119411 | 2322 | 1163 | 1390 | 1696 | 1482 | 1505 |
| ENSG00000119431 | 3195 | 1552 | 1870 | 1897 | 1792 | 1672 |
| ENSG00000148218 | 5571 | 2433 | 3419 | 2242 | 2518 | 2687 |
| ENSG00000148229 | 5034 | 2567 | 3090 | 2730 | 2839 | 3110 |

| | | | | | | |
|---|---|---|---|---|---|---|
| ENSG00000157653 | 38 | 21 | 25 | 17 | 26 | 25 |
| ENSG00000138835 | 1401 | 690 | 748 | 717 | 729 | 819 |
| ENSG00000237073 | 3 | 7 | 4 | 1 | 1 | 4 |
| ENSG00000282897 | 0 | 0 | 0 | 0 | 0 | 1 |
| ENSG00000233817 | 0 | 2 | 0 | 0 | 1 | 1 |
| ENSG00000227482 | 2 | 2 | 2 | 4 | 4 | 6 |
| ENSG00000214669 | 0 | 0 | 0 | 0 | 0 | 0 |
| ENSG00000232905 | 0 | 0 | 0 | 0 | 0 | 0 |
| ENSG00000157657 | 1259 | 557 | 749 | 424 | 634 | 683 |
| ENSG00000106927 | 4 | 14 | 2 | 7 | 9 | 6 |
| ENSG00000136883 | 276 | 123 | 148 | 99 | 96 | 96 |
| ENSG00000196739 | 240 | 102 | 109 | 75 | 93 | 114 |
| ENSG00000207726 | 0 | 0 | 0 | 0 | 0 | 0 |
| ENSG00000229314 | 6 | 0 | 0 | 0 | 0 | 1 |
| ENSG00000228278 | 0 | 0 | 0 | 0 | 1 | 0 |
| ENSG00000106948 | 325 | 186 | 233 | 214 | 274 | 244 |
| ENSG00000235119 | 2 | 0 | 0 | 0 | 0 | 0 |
| ENSG00000095397 | 971 | 440 | 551 | 474 | 564 | 501 |
| ENSG00000271214 | 0 | 0 | 0 | 0 | 0 | 0 |
| ENSG00000235797 | 0 | 0 | 0 | 0 | 0 | 0 |
| ENSG00000136888 | 8809 | 4260 | 5235 | 5030 | 5360 | 5030 |
| ENSG00000157693 | 1285 | 647 | 807 | 613 | 717 | 749 |
| ENSG00000201749 | 0 | 0 | 0 | 0 | 0 | 0 |
| ENSG00000230054 | 0 | 0 | 0 | 0 | 0 | 0 |
| ENSG00000230601 | 0 | 0 | 2 | 0 | 0 | 0 |
| ENSG00000230284 | 0 | 0 | 0 | 0 | 0 | 0 |
| ENSG00000274919 | 0 | 0 | 0 | 0 | 0 | 0 |
| ENSG00000181634 | 6 | 7 | 7 | 5 | 6 | 2 |
| ENSG00000229817 | 0 | 0 | 0 | 0 | 0 | 0 |
| ENSG00000173077 | 0 | 0 | 0 | 0 | 0 | 0 |
| ENSG00000106952 | 0 | 0 | 0 | 0 | 0 | 0 |
| ENSG00000041982 | 0 | 0 | 0 | 0 | 0 | 1 |
| ENSG00000236461 | 0 | 0 | 0 | 0 | 0 | 0 |
| ENSG00000234692 | 0 | 0 | 0 | 0 | 0 | 0 |
| ENSG00000228714 | 0 | 3 | 1 | 3 | 0 | 1 |
| ENSG00000274755 | 0 | 0 | 0 | 0 | 0 | 0 |
| ENSG00000204148 | 0 | 0 | 0 | 0 | 1 | 0 |
| ENSG00000228707 | 0 | 0 | 0 | 0 | 0 | 0 |
| ENSG00000182752 | 1 | 1 | 2 | 0 | 3 | 1 |
| ENSG00000226604 | 0 | 0 | 0 | 0 | 0 | 0 |
| ENSG00000244757 | 0 | 0 | 0 | 0 | 1 | 0 |
| ENSG00000256040 | 0 | 0 | 0 | 0 | 0 | 0 |
| ENSG00000148219 | 334 | 165 | 230 | 144 | 137 | 129 |
| ENSG00000229105 | 0 | 0 | 0 | 0 | 0 | 0 |
| ENSG00000230894 | 0 | 1 | 1 | 0 | 0 | 1 |
| ENSG00000119401 | 1216 | 583 | 662 | 615 | 677 | 618 |
| ENSG00000285820 | 0 | 0 | 1 | 0 | 1 | 1 |
| ENSG00000230734 | 0 | 0 | 0 | 0 | 0 | 0 |
| ENSG00000207268 | 0 | 0 | 0 | 0 | 0 | 0 |
| ENSG00000201140 | 0 | 0 | 0 | 0 | 0 | 0 |

| | | | | | | |
|---|---|---|---|---|---|---|
| ENSG00000222413 | 0 | 1 | 0 | 0 | 0 | 0 |
| ENSG00000233516 | 0 | 0 | 0 | 0 | 0 | 0 |
| ENSG00000233569 | 0 | 0 | 0 | 0 | 0 | 0 |
| ENSG00000235133 | 0 | 0 | 0 | 0 | 0 | 0 |
| ENSG00000285082 | 0 | 0 | 0 | 0 | 0 | 0 |
| ENSG00000136869 | 0 | 0 | 0 | 0 | 0 | 0 |
| ENSG00000201444 | 0 | 0 | 0 | 0 | 0 | 0 |
| ENSG00000284977 | 0 | 0 | 0 | 0 | 0 | 0 |
| ENSG00000231901 | 0 | 0 | 0 | 0 | 0 | 0 |
| ENSG00000234782 | 12 | 3 | 6 | 12 | 7 | 10 |
| ENSG00000225050 | 0 | 0 | 0 | 0 | 0 | 0 |
| ENSG00000235201 | 0 | 0 | 0 | 0 | 0 | 0 |
| ENSG00000231166 | 0 | 0 | 0 | 0 | 0 | 0 |
| ENSG00000078725 | 0 | 0 | 0 | 0 | 0 | 0 |
| ENSG00000260970 | 0 | 0 | 0 | 0 | 0 | 0 |
| ENSG00000261432 | 0 | 0 | 0 | 0 | 0 | 0 |
| ENSG00000225960 | 0 | 0 | 0 | 0 | 0 | 0 |
| ENSG00000207814 | 0 | 0 | 0 | 0 | 0 | 0 |
| ENSG00000136861 | 1607 | 759 | 934 | 662 | 1045 | 828 |
| ENSG00000106780 | 1019 | 448 | 527 | 391 | 667 | 560 |
| ENSG00000238181 | 0 | 0 | 0 | 0 | 0 | 0 |
| ENSG00000119402 | 4210 | 2008 | 2591 | 1620 | 2083 | 2127 |
| ENSG00000214654 | 183 | 98 | 99 | 72 | 89 | 120 |
| ENSG00000270917 | 15 | 1 | 4 | 4 | 8 | 7 |
| ENSG00000095261 | 3716 | 1790 | 2105 | 1253 | 1935 | 1788 |
| ENSG00000226752 | 1787 | 767 | 1075 | 548 | 865 | 776 |
| ENSG00000119403 | 1147 | 640 | 746 | 794 | 689 | 776 |
| ENSG00000056558 | 119 | 48 | 73 | 48 | 60 | 57 |
| ENSG00000106804 | 185 | 139 | 101 | 57 | 130 | 86 |
| ENSG00000119397 | 91 | 62 | 73 | 31 | 143 | 78 |
| ENSG00000119396 | 7283 | 3413 | 4000 | 3123 | 3744 | 3656 |
| ENSG00000243738 | 3 | 1 | 3 | 1 | 0 | 1 |
| ENSG00000148180 | 12111 | 5872 | 6700 | 6481 | 6184 | 5312 |
| ENSG00000214651 | 0 | 0 | 1 | 0 | 0 | 0 |
| ENSG00000239593 | 0 | 1 | 0 | 1 | 0 | 0 |
| ENSG00000235865 | 9 | 14 | 12 | 1 | 7 | 2 |
| ENSG00000148175 | 2272 | 1260 | 1448 | 881 | 1211 | 1096 |
| ENSG00000227355 | 8 | 1 | 1 | 3 | 2 | 2 |
| ENSG00000204136 | 4 | 5 | 0 | 0 | 0 | 0 |
| ENSG00000240299 | 0 | 0 | 0 | 0 | 0 | 0 |
| ENSG00000213467 | 0 | 1 | 0 | 1 | 1 | 2 |
| ENSG00000136848 | 1558 | 715 | 927 | 833 | 835 | 851 |
| ENSG00000229854 | 0 | 0 | 0 | 0 | 0 | 0 |
| ENSG00000261534 | 47 | 28 | 29 | 20 | 18 | 23 |
| ENSG00000175764 | 128 | 50 | 83 | 72 | 73 | 65 |
| ENSG00000237548 | 0 | 0 | 0 | 1 | 1 | 1 |
| ENSG00000266583 | 0 | 0 | 0 | 0 | 0 | 0 |
| ENSG00000119421 | 2371 | 1194 | 1429 | 1479 | 1372 | 1418 |
| ENSG00000185681 | 0 | 0 | 0 | 0 | 0 | 1 |
| ENSG00000106852 | 83 | 38 | 54 | 45 | 37 | 54 |

| | | | | | | |
|---|---|---|---|---|---|---|
| ENSG00000119446 | 1296 | 641 | 704 | 520 | 738 | 625 |
| ENSG00000148187 | 1554 | 855 | 997 | 863 | 1026 | 959 |
| ENSG00000095303 | 12 | 2 | 1 | 2 | 4 | 6 |
| ENSG00000269970 | 1 | 0 | 0 | 0 | 1 | 0 |
| ENSG00000233616 | 0 | 0 | 0 | 0 | 0 | 0 |
| ENSG00000229245 | 0 | 0 | 0 | 0 | 0 | 0 |
| ENSG00000231465 | 2 | 1 | 1 | 0 | 2 | 3 |
| ENSG00000136834 | 0 | 1 | 2 | 1 | 0 | 1 |
| ENSG00000197233 | 4 | 0 | 0 | 0 | 0 | 0 |
| ENSG00000239590 | 2 | 0 | 1 | 0 | 0 | 0 |
| ENSG00000171505 | 0 | 0 | 0 | 0 | 0 | 0 |
| ENSG00000171501 | 0 | 0 | 0 | 0 | 0 | 0 |
| ENSG00000171496 | 0 | 0 | 0 | 0 | 0 | 0 |
| ENSG00000228914 | 2 | 1 | 1 | 0 | 0 | 0 |
| ENSG00000234156 | 0 | 0 | 0 | 0 | 1 | 0 |
| ENSG00000165202 | 2 | 0 | 0 | 0 | 0 | 0 |
| ENSG00000280094 | 0 | 0 | 0 | 0 | 0 | 0 |
| ENSG00000173679 | 0 | 0 | 0 | 0 | 0 | 0 |
| ENSG00000171481 | 0 | 0 | 0 | 0 | 0 | 0 |
| ENSG00000226783 | 0 | 0 | 0 | 0 | 0 | 0 |
| ENSG00000136939 | 0 | 0 | 0 | 0 | 0 | 0 |
| ENSG00000171459 | 0 | 0 | 0 | 0 | 0 | 0 |
| ENSG00000232387 | 6 | 0 | 2 | 1 | 1 | 1 |
| ENSG00000148215 | 0 | 0 | 0 | 0 | 0 | 0 |
| ENSG00000136940 | 1901 | 867 | 1180 | 792 | 995 | 983 |
| ENSG00000165204 | 0 | 0 | 0 | 0 | 0 | 0 |
| ENSG00000233425 | 0 | 1 | 1 | 0 | 0 | 0 |
| ENSG00000056586 | 3043 | 1631 | 1577 | 882 | 2323 | 1444 |
| ENSG00000212447 | 1 | 0 | 0 | 0 | 0 | 0 |
| ENSG00000186130 | 752 | 369 | 333 | 257 | 507 | 329 |
| ENSG00000171448 | 1042 | 505 | 596 | 412 | 569 | 504 |
| ENSG00000261094 | 155 | 62 | 85 | 36 | 89 | 69 |
| ENSG00000011454 | 6001 | 2728 | 3380 | 1973 | 3244 | 2704 |
| ENSG00000222351 | 2 | 0 | 0 | 0 | 0 | 0 |
| ENSG00000213216 | 0 | 0 | 0 | 0 | 2 | 0 |
| ENSG00000188394 | 0 | 0 | 0 | 0 | 0 | 0 |
| ENSG00000225078 | 0 | 0 | 0 | 1 | 0 | 1 |
| ENSG00000236901 | 424 | 194 | 274 | 162 | 282 | 263 |
| ENSG00000165209 | 3131 | 1539 | 1749 | 1249 | 2095 | 1952 |
| ENSG00000283941 | 0 | 0 | 0 | 0 | 0 | 0 |
| ENSG00000148204 | 6 | 1 | 1 | 4 | 4 | 4 |
| ENSG00000119522 | 887 | 402 | 564 | 560 | 582 | 530 |
| ENSG00000207991 | 0 | 0 | 1 | 0 | 0 | 0 |
| ENSG00000274932 | 0 | 0 | 0 | 0 | 0 | 0 |
| ENSG00000237336 | 0 | 0 | 0 | 0 | 0 | 0 |
| ENSG00000230826 | 0 | 0 | 1 | 0 | 0 | 0 |
| ENSG00000229115 | 0 | 0 | 0 | 0 | 0 | 0 |
| ENSG00000234881 | 0 | 0 | 0 | 0 | 0 | 0 |
| ENSG00000106689 | 132 | 67 | 89 | 78 | 52 | 65 |
| ENSG00000236668 | 2 | 3 | 4 | 0 | 2 | 0 |

| | | | | | | |
|---|---|---|---|---|---|---|
| ENSG00000231227 | 0 | 2 | 0 | 0 | 3 | 0 |
| ENSG00000234921 | 0 | 0 | 1 | 1 | 2 | 0 |
| ENSG00000119408 | 1317 | 575 | 700 | 747 | 770 | 808 |
| ENSG00000235204 | 0 | 2 | 1 | 0 | 5 | 3 |
| ENSG00000227200 | 2 | 0 | 0 | 0 | 0 | 0 |
| ENSG00000136930 | 7283 | 3897 | 4487 | 4987 | 4642 | 4342 |
| ENSG00000180264 | 12 | 7 | 6 | 6 | 5 | 6 |
| ENSG00000136931 | 4 | 0 | 1 | 0 | 0 | 0 |
| ENSG00000231149 | 0 | 0 | 0 | 0 | 0 | 0 |
| ENSG00000148200 | 256 | 136 | 164 | 110 | 150 | 126 |
| ENSG00000231744 | 0 | 0 | 0 | 0 | 0 | 0 |
| ENSG00000241246 | 0 | 0 | 0 | 0 | 0 | 0 |
| ENSG00000224020 | 0 | 0 | 0 | 0 | 0 | 0 |
| ENSG00000207595 | 0 | 0 | 0 | 0 | 0 | 0 |
| ENSG00000207737 | 0 | 0 | 0 | 0 | 0 | 0 |
| ENSG00000236643 | 12 | 6 | 7 | 2 | 2 | 5 |
| ENSG00000185585 | 114 | 37 | 53 | 38 | 50 | 40 |
| ENSG00000136918 | 0 | 2 | 0 | 1 | 0 | 0 |
| ENSG00000136942 | 21155 | 10874 | 12381 | 15391 | 12375 | 12321 |
| ENSG00000136950 | 3394 | 1614 | 1896 | 1913 | 2038 | 1972 |
| ENSG00000136935 | 2058 | 898 | 1322 | 766 | 1029 | 1086 |
| ENSG00000199313 | 1 | 0 | 1 | 1 | 1 | 2 |
| ENSG00000173611 | 561 | 282 | 260 | 147 | 281 | 191 |
| ENSG00000177736 | 0 | 0 | 0 | 0 | 0 | 1 |
| ENSG00000119414 | 5084 | 2499 | 2885 | 2119 | 2935 | 2507 |
| ENSG00000232630 | 73 | 32 | 41 | 12 | 35 | 30 |
| ENSG00000235332 | 0 | 3 | 1 | 0 | 0 | 1 |
| ENSG00000136933 | 877 | 404 | 481 | 649 | 532 | 548 |
| ENSG00000044574 | 14713 | 7273 | 8759 | 6654 | 7884 | 7390 |
| ENSG00000239705 | 2 | 0 | 0 | 4 | 0 | 0 |
| ENSG00000165219 | 3349 | 1565 | 1882 | 954 | 1777 | 1397 |
| ENSG00000214593 | 0 | 0 | 1 | 0 | 0 | 0 |
| ENSG00000200554 | 0 | 0 | 0 | 0 | 0 | 0 |
| ENSG00000227845 | 0 | 0 | 0 | 0 | 2 | 1 |
| ENSG00000274510 | 0 | 0 | 0 | 0 | 0 | 0 |
| ENSG00000119487 | 4702 | 2294 | 2742 | 2750 | 2993 | 2907 |
| ENSG00000243845 | 0 | 0 | 0 | 0 | 0 | 2 |
| ENSG00000227068 | 0 | 0 | 0 | 0 | 0 | 0 |
| ENSG00000214761 | 0 | 0 | 1 | 0 | 1 | 0 |
| ENSG00000229582 | 0 | 1 | 1 | 3 | 0 | 0 |
| ENSG00000167081 | 979 | 451 | 610 | 467 | 451 | 521 |
| ENSG00000232413 | 0 | 0 | 0 | 0 | 0 | 0 |
| ENSG00000228392 | 0 | 0 | 0 | 0 | 0 | 0 |
| ENSG00000196814 | 654 | 336 | 342 | 283 | 282 | 284 |
| ENSG00000253079 | 0 | 0 | 0 | 0 | 0 | 0 |
| ENSG00000252985 | 0 | 0 | 1 | 0 | 0 | 1 |
| ENSG00000233721 | 0 | 0 | 0 | 0 | 0 | 0 |
| ENSG00000279137 | 0 | 0 | 0 | 0 | 0 | 0 |
| ENSG00000226078 | 0 | 0 | 0 | 0 | 0 | 0 |
| ENSG00000224842 | 41 | 22 | 30 | 22 | 25 | 13 |

| | | | | | | |
|---|---|---|---|---|---|---|
| ENSG00000136944 | 1764 | 738 | 1019 | 749 | 710 | 783 |
| ENSG00000238010 | 0 | 0 | 0 | 0 | 0 | 0 |
| ENSG00000169155 | 820 | 373 | 524 | 268 | 424 | 455 |
| ENSG00000177125 | 832 | 387 | 504 | 308 | 456 | 419 |
| ENSG00000136828 | 1414 | 605 | 827 | 563 | 700 | 630 |
| ENSG00000136859 | 0 | 0 | 0 | 0 | 2 | 1 |
| ENSG00000228487 | 12 | 7 | 8 | 9 | 9 | 12 |
| ENSG00000136895 | 259 | 141 | 145 | 101 | 114 | 163 |
| ENSG00000229369 | 0 | 0 | 0 | 0 | 0 | 0 |
| ENSG00000271833 | 25 | 15 | 19 | 7 | 13 | 10 |
| ENSG00000136856 | 1347 | 611 | 735 | 696 | 594 | 602 |
| ENSG00000196152 | 227 | 125 | 125 | 90 | 114 | 109 |
| ENSG00000197958 | 45660 | 22791 | 25809 | 32895 | 28488 | 27394 |
| ENSG00000201302 | 4 | 2 | 15 | 6 | 7 | 5 |
| ENSG00000148356 | 1839 | 882 | 1198 | 1011 | 972 | 892 |
| ENSG00000200788 | 3 | 2 | 2 | 2 | 1 | 1 |
| ENSG00000136830 | 12216 | 6012 | 7146 | 6413 | 6260 | 6017 |
| ENSG00000136854 | 1248 | 589 | 786 | 557 | 579 | 589 |
| ENSG00000279571 | 0 | 0 | 0 | 0 | 0 | 0 |
| ENSG00000187024 | 7 | 5 | 5 | 2 | 2 | 1 |
| ENSG00000283874 | 0 | 0 | 0 | 0 | 0 | 0 |
| ENSG00000160401 | 137 | 71 | 115 | 73 | 90 | 96 |
| ENSG00000167094 | 0 | 0 | 1 | 5 | 2 | 0 |
| ENSG00000160404 | 1034 | 479 | 603 | 590 | 485 | 481 |
| ENSG00000095370 | 2 | 2 | 4 | 1 | 1 | 3 |
| ENSG00000136807 | 2726 | 1428 | 1607 | 1698 | 1485 | 1503 |
| ENSG00000283863 | 0 | 0 | 0 | 0 | 0 | 0 |
| ENSG00000284547 | 0 | 0 | 0 | 0 | 0 | 0 |
| ENSG00000136877 | 2231 | 1100 | 1369 | 1408 | 1190 | 1197 |
| ENSG00000106991 | 45 | 18 | 38 | 22 | 24 | 32 |
| ENSG00000225032 | 27 | 16 | 25 | 15 | 17 | 23 |
| ENSG00000222421 | 0 | 0 | 0 | 0 | 0 | 0 |
| ENSG00000222455 | 0 | 0 | 0 | 0 | 0 | 0 |
| ENSG00000106992 | 166 | 82 | 92 | 124 | 108 | 105 |
| ENSG00000257524 | 4 | 4 | 5 | 8 | 10 | 4 |
| ENSG00000263979 | 0 | 0 | 0 | 0 | 0 | 0 |
| ENSG00000160408 | 7 | 5 | 7 | 4 | 2 | 5 |
| ENSG00000136840 | 342 | 139 | 239 | 173 | 127 | 128 |
| ENSG00000167103 | 81 | 61 | 70 | 50 | 51 | 50 |
| ENSG00000227218 | 772 | 370 | 576 | 544 | 472 | 715 |
| ENSG00000136908 | 2608 | 1226 | 1714 | 1653 | 1325 | 1593 |
| ENSG00000167106 | 14157 | 6674 | 8632 | 7950 | 7304 | 8369 |
| ENSG00000230848 | 0 | 0 | 0 | 0 | 0 | 0 |
| ENSG00000171169 | 412 | 172 | 210 | 248 | 227 | 216 |
| ENSG00000148339 | 859 | 425 | 606 | 532 | 467 | 473 |
| ENSG00000230536 | 0 | 1 | 0 | 0 | 0 | 0 |
| ENSG00000234771 | 321 | 132 | 187 | 139 | 195 | 185 |
| ENSG00000148334 | 2685 | 1203 | 1656 | 1799 | 1502 | 1601 |
| ENSG00000232850 | 36 | 11 | 7 | 14 | 12 | 8 |
| ENSG00000148346 | 156 | 101 | 134 | 64 | 71 | 120 |

| | | | | | | |
|---|---|---|---|---|---|---|
| ENSG00000171159 | 2563 | 1156 | 1518 | 1752 | 1386 | 1383 |
| ENSG00000148337 | 5406 | 2402 | 3242 | 3225 | 3041 | 3208 |
| ENSG00000106976 | 1162 | 507 | 814 | 464 | 563 | 642 |
| ENSG00000207581 | 0 | 0 | 0 | 1 | 0 | 0 |
| ENSG00000264823 | 0 | 0 | 0 | 0 | 0 | 0 |
| ENSG00000167110 | 8205 | 3592 | 4914 | 3174 | 4276 | 4107 |
| ENSG00000175854 | 936 | 476 | 523 | 588 | 521 | 488 |
| ENSG00000167112 | 4016 | 2017 | 2285 | 2311 | 2247 | 2320 |
| ENSG00000272696 | 17 | 11 | 9 | 12 | 6 | 8 |
| ENSG00000167113 | 2492 | 1136 | 1525 | 1312 | 1264 | 1305 |
| ENSG00000167114 | 2650 | 1264 | 1604 | 1725 | 1353 | 1500 |
| ENSG00000223551 | 65 | 25 | 45 | 35 | 35 | 50 |
| ENSG00000167118 | 3493 | 1596 | 2155 | 2328 | 1901 | 1867 |
| ENSG00000272593 | 0 | 0 | 0 | 0 | 0 | 0 |
| ENSG00000207955 | 4 | 0 | 4 | 1 | 2 | 0 |
| ENSG00000284185 | 0 | 0 | 0 | 0 | 0 | 0 |
| ENSG00000283335 | 0 | 0 | 0 | 0 | 0 | 0 |
| ENSG00000167123 | 4322 | 2062 | 2713 | 2563 | 2125 | 2102 |
| ENSG00000273186 | 27 | 16 | 28 | 17 | 15 | 18 |
| ENSG00000228510 | 0 | 4 | 1 | 2 | 0 | 1 |
| ENSG00000136811 | 2822 | 1410 | 1691 | 1436 | 1664 | 1737 |
| ENSG00000225951 | 3 | 3 | 5 | 4 | 3 | 1 |
| ENSG00000119392 | 3356 | 1695 | 2123 | 1799 | 1996 | 1970 |
| ENSG00000228395 | 45 | 20 | 21 | 15 | 19 | 23 |
| ENSG00000238406 | 1 | 0 | 3 | 0 | 0 | 1 |
| ENSG00000197694 | 7768 | 3322 | 4758 | 3373 | 3823 | 4080 |
| ENSG00000280474 | 0 | 0 | 0 | 0 | 0 | 0 |
| ENSG00000280758 | 6 | 10 | 12 | 4 | 3 | 14 |
| ENSG00000119333 | 7096 | 3542 | 4445 | 4359 | 3456 | 3420 |
| ENSG00000227759 | 1 | 0 | 0 | 0 | 0 | 1 |
| ENSG00000234705 | 24 | 15 | 8 | 8 | 14 | 17 |
| ENSG00000119335 | 20339 | 9967 | 12444 | 10635 | 11465 | 12014 |
| ENSG00000160447 | 278 | 129 | 192 | 178 | 184 | 150 |
| ENSG00000264438 | 0 | 0 | 0 | 0 | 0 | 0 |
| ENSG00000160446 | 3268 | 1676 | 2079 | 2288 | 1842 | 1752 |
| ENSG00000223478 | 309 | 161 | 211 | 202 | 199 | 167 |
| ENSG00000160445 | 4013 | 1852 | 2404 | 2032 | 1950 | 1977 |
| ENSG00000203286 | 7 | 11 | 6 | 0 | 1 | 3 |
| ENSG00000107021 | 2370 | 1142 | 1474 | 1195 | 1239 | 1359 |
| ENSG00000167136 | 370 | 235 | 248 | 322 | 236 | 272 |
| ENSG00000198917 | 2082 | 1083 | 1349 | 1225 | 1231 | 1183 |
| ENSG00000171097 | 193 | 92 | 143 | 111 | 100 | 91 |
| ENSG00000136802 | 2327 | 1013 | 1381 | 1164 | 1246 | 1236 |
| ENSG00000175287 | 1 | 0 | 1 | 0 | 1 | 2 |
| ENSG00000251184 | 0 | 0 | 0 | 0 | 0 | 0 |
| ENSG00000175283 | 1454 | 673 | 855 | 837 | 769 | 798 |
| ENSG00000095319 | 3090 | 1325 | 1840 | 1381 | 1666 | 1796 |
| ENSG00000267834 | 14 | 12 | 14 | 9 | 10 | 9 |
| ENSG00000148341 | 17445 | 8049 | 11214 | 7959 | 8102 | 8116 |
| ENSG00000148343 | 1764 | 754 | 1012 | 953 | 878 | 919 |

| | | | | | | |
|---|---|---|---|---|---|---|
| ENSG00000167130 | 2542 | 1142 | 1494 | 1350 | 1181 | 1383 |
| ENSG00000095321 | 76 | 41 | 47 | 31 | 23 | 25 |
| ENSG00000234055 | 1 | 0 | 0 | 0 | 0 | 0 |
| ENSG00000119383 | 13915 | 6665 | 8610 | 8284 | 6646 | 7134 |
| ENSG00000268707 | 7 | 1 | 3 | 1 | 5 | 6 |
| ENSG00000188483 | 2422 | 1193 | 1477 | 1817 | 1440 | 1704 |
| ENSG00000268050 | 0 | 0 | 0 | 0 | 0 | 0 |
| ENSG00000204055 | 14 | 6 | 11 | 3 | 8 | 5 |
| ENSG00000235007 | 0 | 0 | 0 | 0 | 1 | 2 |
| ENSG00000241084 | 0 | 0 | 0 | 0 | 0 | 0 |
| ENSG00000224307 | 0 | 0 | 0 | 0 | 0 | 1 |
| ENSG00000179082 | 190 | 107 | 139 | 40 | 69 | 63 |
| ENSG00000268615 | 26 | 15 | 39 | 12 | 13 | 17 |
| ENSG00000233901 | 291 | 170 | 223 | 125 | 158 | 140 |
| ENSG00000242281 | 0 | 0 | 0 | 0 | 0 | 0 |
| ENSG00000236146 | 2 | 0 | 1 | 1 | 1 | 3 |
| ENSG00000230676 | 1 | 1 | 0 | 1 | 0 | 3 |
| ENSG00000226355 | 0 | 0 | 0 | 0 | 0 | 1 |
| ENSG00000261334 | 8 | 15 | 7 | 4 | 4 | 5 |
| ENSG00000204054 | 4720 | 2492 | 3345 | 2017 | 2043 | 2587 |
| ENSG00000227619 | 10 | 3 | 13 | 13 | 23 | 8 |
| ENSG00000148335 | 1008 | 562 | 630 | 799 | 680 | 695 |
| ENSG00000179058 | 3 | 11 | 11 | 8 | 16 | 10 |
| ENSG00000148331 | 1613 | 813 | 1072 | 1105 | 990 | 1033 |
| ENSG00000234789 | 0 | 3 | 3 | 5 | 10 | 3 |
| ENSG00000167157 | 31 | 16 | 12 | 26 | 36 | 18 |
| ENSG00000236024 | 0 | 0 | 0 | 0 | 0 | 0 |
| ENSG00000148344 | 80 | 68 | 41 | 79 | 98 | 88 |
| ENSG00000233909 | 0 | 0 | 0 | 0 | 0 | 0 |
| ENSG00000136816 | 1839 | 990 | 1239 | 1026 | 1129 | 1177 |
| ENSG00000136827 | 3880 | 1885 | 2276 | 2160 | 2149 | 2369 |
| ENSG00000136819 | 2702 | 1409 | 1559 | 1316 | 1779 | 1652 |
| ENSG00000136878 | 2072 | 830 | 1189 | 1020 | 1002 | 1054 |
| ENSG00000276124 | 0 | 0 | 0 | 0 | 0 | 0 |
| ENSG00000224988 | 0 | 2 | 2 | 1 | 1 | 0 |
| ENSG00000187239 | 2601 | 1162 | 1696 | 1055 | 1310 | 1362 |
| ENSG00000230684 | 0 | 0 | 0 | 0 | 0 | 0 |
| ENSG00000270755 | 7 | 4 | 6 | 6 | 0 | 3 |
| ENSG00000148358 | 8885 | 3992 | 5587 | 3704 | 4560 | 4695 |
| ENSG00000223188 | 0 | 0 | 1 | 1 | 0 | 0 |
| ENSG00000196979 | 0 | 0 | 0 | 0 | 1 | 0 |
| ENSG00000107130 | 2415 | 1056 | 1286 | 1690 | 1473 | 1525 |
| ENSG00000148357 | 10 | 2 | 0 | 1 | 0 | 4 |
| ENSG00000264169 | 0 | 0 | 0 | 0 | 0 | 0 |
| ENSG00000130707 | 940 | 547 | 638 | 677 | 679 | 609 |
| ENSG00000107164 | 5102 | 2339 | 2939 | 1954 | 2588 | 2720 |
| ENSG00000277713 | 0 | 0 | 0 | 1 | 0 | 0 |
| ENSG00000232172 | 0 | 0 | 0 | 0 | 0 | 0 |
| ENSG00000130711 | 44 | 17 | 26 | 36 | 23 | 39 |
| ENSG00000130713 | 1412 | 804 | 887 | 797 | 919 | 995 |

| | | | | | | |
|---|---|---|---|---|---|---|
| ENSG00000097007 | 3034 | 1398 | 1861 | 1324 | 1387 | 1588 |
| ENSG00000224797 | 0 | 0 | 0 | 0 | 0 | 0 |
| ENSG00000188710 | 26 | 15 | 25 | 13 | 17 | 15 |
| ENSG00000130720 | 1919 | 860 | 1173 | 1021 | 876 | 885 |
| ENSG00000236658 | 6 | 4 | 5 | 2 | 2 | 6 |
| ENSG00000050555 | 0 | 0 | 1 | 2 | 1 | 2 |
| ENSG00000252582 | 0 | 0 | 0 | 0 | 0 | 0 |
| ENSG00000126878 | 8192 | 3802 | 4904 | 3601 | 3374 | 3754 |
| ENSG00000126883 | 2160 | 1025 | 1254 | 793 | 1034 | 1242 |
| ENSG00000252622 | 0 | 0 | 0 | 0 | 0 | 0 |
| ENSG00000236986 | 3 | 0 | 1 | 1 | 2 | 0 |
| ENSG00000246851 | 0 | 3 | 1 | 0 | 2 | 0 |
| ENSG00000226987 | 0 | 0 | 0 | 0 | 0 | 0 |
| ENSG00000126882 | 365 | 153 | 191 | 200 | 198 | 183 |
| ENSG00000160539 | 0 | 1 | 7 | 3 | 3 | 2 |
| ENSG00000285287 | 12 | 11 | 14 | 3 | 6 | 6 |
| ENSG00000130723 | 11567 | 4739 | 7007 | 4708 | 5891 | 6508 |
| ENSG00000235284 | 0 | 1 | 1 | 1 | 2 | 0 |
| ENSG00000231587 | 13 | 4 | 8 | 4 | 4 | 8 |
| ENSG00000176868 | 0 | 0 | 0 | 0 | 0 | 0 |
| ENSG00000130714 | 3486 | 1640 | 2079 | 1932 | 1862 | 1784 |
| ENSG00000230289 | 1 | 6 | 3 | 3 | 2 | 6 |
| ENSG00000130717 | 3970 | 1846 | 2397 | 2298 | 2064 | 2042 |
| ENSG00000283526 | 626 | 370 | 436 | 277 | 345 | 350 |
| ENSG00000107263 | 3797 | 1884 | 2430 | 2052 | 2204 | 2429 |
| ENSG00000240853 | 0 | 0 | 0 | 0 | 0 | 1 |
| ENSG00000231018 | 0 | 0 | 0 | 0 | 0 | 0 |
| ENSG00000212395 | 0 | 0 | 0 | 0 | 0 | 0 |
| ENSG00000236687 | 0 | 0 | 0 | 0 | 0 | 0 |
| ENSG00000160563 | 740 | 402 | 477 | 544 | 464 | 469 |
| ENSG00000196358 | 4 | 0 | 1 | 3 | 5 | 4 |
| ENSG00000107290 | 816 | 466 | 366 | 180 | 818 | 430 |
| ENSG00000125482 | 554 | 284 | 381 | 179 | 336 | 302 |
| ENSG00000188523 | 4 | 0 | 0 | 2 | 2 | 0 |
| ENSG00000252521 | 0 | 0 | 0 | 0 | 0 | 0 |
| ENSG00000125492 | 0 | 1 | 0 | 1 | 0 | 0 |
| ENSG00000125485 | 839 | 403 | 493 | 455 | 498 | 537 |
| ENSG00000125484 | 2405 | 1057 | 1377 | 845 | 1310 | 1305 |
| ENSG00000165695 | 98 | 38 | 55 | 50 | 39 | 44 |
| ENSG00000224992 | 0 | 0 | 0 | 0 | 0 | 0 |
| ENSG00000165698 | 286 | 145 | 197 | 172 | 180 | 135 |
| ENSG00000165699 | 2056 | 848 | 1179 | 573 | 956 | 963 |
| ENSG00000165702 | 2 | 0 | 0 | 1 | 2 | 3 |
| ENSG00000263816 | 0 | 0 | 0 | 0 | 0 | 0 |
| ENSG00000234907 | 0 | 0 | 0 | 0 | 0 | 0 |
| ENSG00000238558 | 0 | 0 | 0 | 0 | 0 | 0 |
| ENSG00000196205 | 806 | 426 | 464 | 440 | 526 | 492 |
| ENSG00000148308 | 3451 | 1593 | 2050 | 2283 | 1972 | 2027 |
| ENSG00000273932 | 0 | 0 | 1 | 0 | 0 | 0 |
| ENSG00000170835 | 56 | 23 | 31 | 16 | 17 | 26 |

| | | | | | | |
|---|---|---|---|---|---|---|
| ENSG00000170827 | 2 | 1 | 1 | 1 | 0 | 0 |
| ENSG00000160271 | 398 | 260 | 292 | 268 | 209 | 245 |
| ENSG00000285245 | 0 | 0 | 0 | 0 | 0 | 0 |
| ENSG00000148288 | 0 | 0 | 0 | 1 | 0 | 0 |
| ENSG00000171102 | 0 | 0 | 0 | 0 | 0 | 0 |
| ENSG00000119440 | 0 | 0 | 0 | 0 | 0 | 0 |
| ENSG00000175164 | 16 | 5 | 14 | 10 | 16 | 8 |
| ENSG00000201843 | 0 | 0 | 0 | 0 | 0 | 0 |
| ENSG00000204031 | 0 | 0 | 0 | 0 | 0 | 0 |
| ENSG00000148296 | 2684 | 1272 | 1439 | 1657 | 1427 | 1551 |
| ENSG00000201451 | 2 | 0 | 5 | 1 | 2 | 0 |
| ENSG00000148297 | 2660 | 1214 | 1519 | 1483 | 1426 | 1554 |
| ENSG00000230064 | 0 | 0 | 0 | 0 | 0 | 0 |
| ENSG00000148303 | 68466 | 34688 | 40365 | 47028 | 40955 | 41157 |
| ENSG00000206611 | 0 | 0 | 0 | 0 | 0 | 0 |
| ENSG00000200831 | 0 | 0 | 0 | 0 | 0 | 0 |
| ENSG00000199744 | 0 | 0 | 0 | 0 | 0 | 0 |
| ENSG00000252542 | 9 | 0 | 1 | 2 | 1 | 4 |
| ENSG00000148290 | 2397 | 1167 | 1448 | 1527 | 1346 | 1311 |
| ENSG00000148291 | 795 | 399 | 510 | 600 | 485 | 497 |
| ENSG00000148248 | 15024 | 6923 | 8707 | 8205 | 7724 | 7965 |
| ENSG00000198870 | 58 | 33 | 37 | 23 | 31 | 35 |
| ENSG00000148300 | 1926 | 920 | 1155 | 1249 | 1132 | 1142 |
| ENSG00000160323 | 179 | 65 | 110 | 54 | 53 | 92 |
| ENSG00000160325 | 1894 | 815 | 1171 | 1023 | 872 | 883 |
| ENSG00000160326 | 974 | 486 | 457 | 555 | 651 | 470 |
| ENSG00000187616 | 0 | 0 | 0 | 0 | 0 | 0 |
| ENSG00000197859 | 0 | 0 | 0 | 0 | 0 | 0 |
| ENSG00000196990 | 0 | 0 | 0 | 0 | 0 | 0 |
| ENSG00000261018 | 0 | 0 | 0 | 0 | 0 | 0 |
| ENSG00000123454 | 1 | 0 | 0 | 4 | 0 | 0 |
| ENSG00000225756 | 2 | 1 | 1 | 0 | 0 | 0 |
| ENSG00000123453 | 18 | 10 | 6 | 4 | 5 | 4 |
| ENSG00000160293 | 4107 | 1996 | 2441 | 1893 | 2013 | 2029 |
| ENSG00000235106 | 1151 | 501 | 681 | 582 | 563 | 614 |
| ENSG00000169925 | 2312 | 997 | 1385 | 944 | 1027 | 1158 |
| ENSG00000235138 | 18 | 8 | 21 | 7 | 9 | 7 |
| ENSG00000271525 | 2 | 0 | 0 | 0 | 0 | 0 |
| ENSG00000273249 | 13 | 8 | 5 | 5 | 5 | 5 |
| ENSG00000196363 | 3619 | 1796 | 2245 | 2080 | 2014 | 2301 |
| ENSG00000221676 | 0 | 1 | 1 | 0 | 0 | 0 |
| ENSG00000273473 | 42 | 17 | 20 | 22 | 19 | 18 |
| ENSG00000223729 | 2 | 4 | 1 | 1 | 1 | 0 |
| ENSG00000279929 | 0 | 0 | 0 | 0 | 0 | 0 |
| ENSG00000186350 | 12347 | 5484 | 7587 | 6586 | 6673 | 7254 |
| ENSG00000263897 | 0 | 1 | 0 | 0 | 1 | 1 |
| ENSG00000223795 | 2 | 0 | 0 | 0 | 0 | 0 |
| ENSG00000228877 | 0 | 0 | 0 | 0 | 1 | 1 |
| ENSG00000227150 | 0 | 0 | 0 | 0 | 0 | 0 |
| ENSG00000130635 | 1268 | 609 | 695 | 585 | 595 | 564 |

| | | | | | | |
|---|---|---|---|---|---|---|
| ENSG00000204011 | 0 | 1 | 1 | 0 | 0 | 2 |
| ENSG00000264744 | 0 | 0 | 0 | 0 | 0 | 0 |
| ENSG00000265872 | 0 | 0 | 0 | 0 | 0 | 0 |
| ENSG00000265848 | 0 | 0 | 0 | 0 | 0 | 0 |
| ENSG00000264163 | 0 | 0 | 0 | 0 | 0 | 0 |
| ENSG00000264136 | 0 | 0 | 0 | 0 | 0 | 0 |
| ENSG00000266827 | 0 | 0 | 0 | 0 | 0 | 0 |
| ENSG00000266514 | 0 | 0 | 0 | 0 | 0 | 0 |
| ENSG00000232355 | 0 | 0 | 0 | 0 | 0 | 0 |
| ENSG00000160339 | 0 | 0 | 0 | 0 | 0 | 0 |
| ENSG00000085265 | 0 | 0 | 0 | 0 | 0 | 0 |
| ENSG00000236403 | 0 | 0 | 0 | 0 | 0 | 0 |
| ENSG00000279935 | 0 | 0 | 0 | 0 | 0 | 0 |
| ENSG00000130558 | 1041 | 446 | 575 | 622 | 553 | 596 |
| ENSG00000227958 | 0 | 0 | 0 | 0 | 0 | 0 |
| ENSG00000233936 | 0 | 0 | 0 | 0 | 0 | 0 |
| ENSG00000178243 | 0 | 0 | 0 | 0 | 0 | 0 |
| ENSG00000235572 | 0 | 0 | 1 | 0 | 0 | 0 |
| ENSG00000224682 | 2 | 4 | 1 | 0 | 0 | 1 |
| ENSG00000225361 | 456 | 198 | 227 | 141 | 179 | 183 |
| ENSG00000196422 | 2425 | 1091 | 1526 | 1360 | 1350 | 1450 |
| ENSG00000160345 | 614 | 331 | 402 | 325 | 309 | 316 |
| ENSG00000122140 | 1946 | 1106 | 1421 | 1740 | 1276 | 1567 |
| ENSG00000226706 | 9 | 7 | 7 | 8 | 13 | 6 |
| ENSG00000160349 | 0 | 0 | 0 | 0 | 0 | 0 |
| ENSG00000122136 | 0 | 0 | 0 | 0 | 0 | 0 |
| ENSG00000122133 | 2 | 0 | 3 | 0 | 0 | 1 |
| ENSG00000237339 | 0 | 0 | 0 | 0 | 0 | 0 |
| ENSG00000224045 | 0 | 0 | 0 | 0 | 0 | 0 |
| ENSG00000236543 | 0 | 0 | 0 | 0 | 0 | 0 |
| ENSG00000204007 | 0 | 0 | 0 | 0 | 0 | 0 |
| ENSG00000148386 | 0 | 0 | 0 | 0 | 0 | 0 |
| ENSG00000165643 | 0 | 0 | 0 | 0 | 0 | 0 |
| ENSG00000107147 | 10 | 1 | 3 | 1 | 0 | 2 |
| ENSG00000130559 | 2278 | 1018 | 1440 | 1133 | 1219 | 1413 |
| ENSG00000238058 | 67 | 35 | 22 | 22 | 45 | 26 |
| ENSG00000130560 | 2164 | 963 | 1206 | 1297 | 1110 | 1155 |
| ENSG00000148411 | 3101 | 1415 | 1684 | 1389 | 1486 | 1458 |
| ENSG00000260193 | 52 | 32 | 37 | 47 | 36 | 30 |
| ENSG00000238227 | 1999 | 855 | 1127 | 1341 | 1123 | 1099 |
| ENSG00000275329 | 63 | 19 | 33 | 21 | 28 | 15 |
| ENSG00000107187 | 0 | 3 | 0 | 2 | 0 | 0 |
| ENSG00000165661 | 1105 | 504 | 664 | 532 | 595 | 662 |
| ENSG00000260220 | 24 | 20 | 23 | 24 | 15 | 13 |
| ENSG00000279813 | 0 | 0 | 0 | 0 | 0 | 0 |
| ENSG00000283769 | 0 | 0 | 0 | 0 | 0 | 0 |
| ENSG00000262075 | 0 | 0 | 4 | 3 | 1 | 2 |
| ENSG00000160360 | 951 | 429 | 642 | 625 | 520 | 562 |
| ENSG00000213221 | 6 | 6 | 1 | 5 | 2 | 2 |
| ENSG00000187796 | 116 | 52 | 75 | 68 | 47 | 74 |

| | | | | | | |
|---|---|---|---|---|---|---|
| ENSG00000165684 | 935 | 396 | 535 | 540 | 537 | 568 |
| ENSG00000165689 | 1793 | 899 | 1077 | 1177 | 1106 | 1196 |
| ENSG00000165688 | 3478 | 1652 | 1996 | 2221 | 1924 | 2044 |
| ENSG00000148384 | 939 | 433 | 550 | 455 | 478 | 484 |
| ENSG00000148396 | 9183 | 3564 | 5741 | 3855 | 3872 | 4691 |
| ENSG00000196366 | 91 | 34 | 70 | 38 | 54 | 35 |
| ENSG00000148400 | 812 | 317 | 517 | 346 | 369 | 412 |
| ENSG00000263403 | 0 | 0 | 0 | 0 | 0 | 0 |
| ENSG00000227512 | 0 | 0 | 0 | 0 | 1 | 0 |
| ENSG00000265181 | 0 | 0 | 0 | 0 | 0 | 0 |
| ENSG00000237886 | 54 | 38 | 46 | 44 | 38 | 46 |
| ENSG00000279141 | 2 | 1 | 0 | 0 | 0 | 0 |
| ENSG00000228401 | 0 | 0 | 0 | 0 | 0 | 0 |
| ENSG00000172889 | 2364 | 1083 | 1414 | 1486 | 1151 | 1095 |
| ENSG00000199161 | 0 | 0 | 0 | 0 | 0 | 0 |
| ENSG00000169692 | 3294 | 1578 | 2021 | 2207 | 1829 | 1780 |
| ENSG00000165716 | 256 | 119 | 179 | 180 | 138 | 165 |
| ENSG00000233016 | 5592 | 2705 | 3454 | 5007 | 4285 | 3928 |
| ENSG00000280496 | 0 | 0 | 0 | 0 | 0 | 0 |
| ENSG00000276161 | 0 | 0 | 0 | 0 | 0 | 0 |
| ENSG00000274998 | 0 | 0 | 0 | 0 | 0 | 0 |
| ENSG00000281808 | 0 | 0 | 0 | 0 | 0 | 0 |
| ENSG00000274356 | 10 | 9 | 11 | 5 | 13 | 9 |
| ENSG00000204003 | 0 | 0 | 0 | 0 | 0 | 0 |
| ENSG00000187922 | 0 | 0 | 0 | 0 | 0 | 0 |
| ENSG00000267206 | 0 | 0 | 0 | 0 | 0 | 0 |
| ENSG00000278756 | 0 | 0 | 0 | 0 | 0 | 0 |
| ENSG00000204001 | 0 | 0 | 0 | 0 | 0 | 0 |
| ENSG00000177984 | 0 | 0 | 0 | 0 | 0 | 0 |
| ENSG00000224662 | 0 | 0 | 0 | 0 | 0 | 0 |
| ENSG00000244187 | 783 | 503 | 694 | 617 | 508 | 485 |
| ENSG00000272896 | 0 | 3 | 0 | 5 | 0 | 0 |
| ENSG00000213213 | 92 | 47 | 66 | 60 | 59 | 54 |
| ENSG00000273066 | 149 | 58 | 97 | 68 | 59 | 69 |
| ENSG00000272679 | 0 | 0 | 0 | 0 | 0 | 0 |
| ENSG00000228544 | 92 | 42 | 86 | 29 | 37 | 44 |
| ENSG00000196642 | 7265 | 3342 | 4530 | 4778 | 4057 | 4112 |
| ENSG00000213212 | 25 | 14 | 4 | 6 | 7 | 6 |
| ENSG00000265806 | 38 | 25 | 32 | 21 | 30 | 32 |
| ENSG00000232434 | 801 | 314 | 467 | 425 | 355 | 421 |
| ENSG00000054148 | 3617 | 1678 | 2305 | 2522 | 1852 | 1971 |
| ENSG00000177943 | 497 | 165 | 284 | 183 | 185 | 235 |
| ENSG00000107223 | 9425 | 4959 | 5754 | 6028 | 4648 | 4919 |
| ENSG00000127191 | 1506 | 694 | 817 | 874 | 751 | 697 |
| ENSG00000266507 | 0 | 0 | 0 | 0 | 0 | 1 |
| ENSG00000213590 | 2 | 3 | 2 | 2 | 1 | 3 |
| ENSG00000260190 | 6 | 1 | 0 | 6 | 2 | 5 |
| ENSG00000159069 | 6049 | 2781 | 3533 | 4040 | 3329 | 3179 |
| ENSG00000176919 | 34 | 15 | 17 | 13 | 22 | 20 |
| ENSG00000184925 | 27 | 11 | 25 | 14 | 14 | 4 |

| | | | | | | |
|---|---|---|---|---|---|---|
| ENSG00000198454 | 0 | 0 | 0 | 0 | 0 | 0 |
| ENSG00000107317 | 0 | 0 | 1 | 0 | 1 | 0 |
| ENSG00000284341 | 0 | 0 | 0 | 0 | 0 | 0 |
| ENSG00000214402 | 0 | 0 | 0 | 0 | 0 | 0 |
| ENSG00000148362 | 2003 | 959 | 1435 | 1473 | 1210 | 1185 |
| ENSG00000169583 | 3688 | 1719 | 1933 | 1830 | 1511 | 1296 |
| ENSG00000107331 | 16695 | 6603 | 10518 | 6781 | 6801 | 7638 |
| ENSG00000180539 | 86 | 20 | 48 | 26 | 23 | 26 |
| ENSG00000180549 | 4 | 4 | 7 | 0 | 3 | 2 |
| ENSG00000279073 | 0 | 0 | 0 | 0 | 0 | 0 |
| ENSG00000107281 | 7094 | 3495 | 5010 | 4752 | 3866 | 3967 |
| ENSG00000054179 | 461 | 189 | 246 | 259 | 202 | 186 |
| ENSG00000236394 | 5 | 0 | 4 | 0 | 3 | 1 |
| ENSG00000229257 | 1 | 0 | 0 | 0 | 0 | 0 |
| ENSG00000186193 | 2085 | 1050 | 1283 | 1638 | 1356 | 1415 |
| ENSG00000231864 | 0 | 0 | 0 | 0 | 0 | 0 |
| ENSG00000197355 | 1546 | 661 | 964 | 770 | 642 | 762 |
| ENSG00000268996 | 390 | 204 | 248 | 209 | 188 | 210 |
| ENSG00000177239 | 5075 | 2328 | 3120 | 2953 | 2540 | 2547 |
| ENSG00000199411 | 0 | 0 | 1 | 0 | 0 | 0 |
| ENSG00000176978 | 4552 | 2185 | 2945 | 2893 | 2409 | 2575 |
| ENSG00000176884 | 552 | 245 | 383 | 315 | 316 | 412 |
| ENSG00000184709 | 45 | 30 | 32 | 49 | 33 | 26 |
| ENSG00000261793 | 0 | 0 | 0 | 0 | 0 | 0 |
| ENSG00000263697 | 0 | 0 | 0 | 0 | 0 | 0 |
| ENSG00000185863 | 4 | 3 | 3 | 5 | 8 | 4 |
| ENSG00000176248 | 2715 | 1189 | 1581 | 1499 | 1465 | 1478 |
| ENSG00000176101 | 2509 | 1150 | 1479 | 1772 | 1441 | 1402 |
| ENSG00000176058 | 2298 | 1038 | 1335 | 1849 | 1502 | 1487 |
| ENSG00000187713 | 1880 | 928 | 1033 | 1200 | 1095 | 1146 |
| ENSG00000188566 | 784 | 329 | 475 | 401 | 421 | 476 |
| ENSG00000284976 | 1275 | 558 | 716 | 985 | 818 | 710 |
| ENSG00000212864 | 581 | 325 | 425 | 603 | 431 | 441 |
| ENSG00000197191 | 569 | 271 | 357 | 584 | 656 | 590 |
| ENSG00000233198 | 254 | 97 | 180 | 297 | 272 | 326 |
| ENSG00000198569 | 141 | 65 | 102 | 149 | 160 | 159 |
| ENSG00000188229 | 12839 | 7345 | 8134 | 11900 | 11178 | 9990 |
| ENSG00000188163 | 26 | 0 | 8 | 6 | 6 | 7 |
| ENSG00000275549 | 55 | 31 | 42 | 21 | 23 | 18 |
| ENSG00000197768 | 32 | 14 | 33 | 17 | 19 | 25 |
| ENSG00000188986 | 3387 | 1722 | 2056 | 2340 | 1975 | 1929 |
| ENSG00000198113 | 194 | 52 | 104 | 94 | 92 | 119 |
| ENSG00000270259 | 0 | 0 | 0 | 0 | 0 | 0 |
| ENSG00000260996 | 2 | 0 | 4 | 0 | 0 | 3 |
| ENSG00000198435 | 321 | 174 | 190 | 210 | 204 | 200 |
| ENSG00000187609 | 864 | 447 | 536 | 520 | 460 | 475 |
| ENSG00000188747 | 1002 | 519 | 772 | 582 | 541 | 613 |
| ENSG00000188833 | 366 | 136 | 229 | 117 | 142 | 135 |
| ENSG00000165802 | 3740 | 1934 | 2391 | 3163 | 2843 | 2766 |
| ENSG00000284317 | 0 | 0 | 0 | 0 | 0 | 0 |

| | | | | | | |
|---|---|---|---|---|---|---|
| ENSG00000130653 | 712 | 272 | 417 | 187 | 188 | 230 |
| ENSG00000182154 | 6070 | 2771 | 3440 | 3983 | 3263 | 3121 |
| ENSG00000148399 | 1735 | 847 | 1065 | 950 | 936 | 1021 |
| ENSG00000165724 | 1559 | 760 | 912 | 960 | 944 | 868 |
| ENSG00000197070 | 6669 | 3295 | 4414 | 3908 | 3483 | 3353 |
| ENSG00000283411 | 0 | 0 | 0 | 0 | 0 | 0 |
| ENSG00000203993 | 564 | 239 | 307 | 296 | 257 | 291 |
| ENSG00000181090 | 4357 | 2032 | 2691 | 1999 | 2111 | 2331 |
| ENSG00000274841 | 0 | 1 | 0 | 1 | 0 | 0 |
| ENSG00000233998 | 0 | 0 | 2 | 1 | 0 | 0 |
| ENSG00000255585 | 21 | 14 | 11 | 8 | 11 | 13 |
| ENSG00000207693 | 2 | 0 | 1 | 0 | 0 | 0 |
| ENSG00000203987 | 0 | 0 | 0 | 0 | 0 | 0 |
| ENSG00000148408 | 4 | 4 | 4 | 2 | 1 | 1 |
| ENSG00000276388 | 0 | 0 | 0 | 0 | 0 | 0 |
| ENSG00000229926 | 0 | 0 | 0 | 0 | 0 | 0 |
| ENSG00000159247 | 160 | 102 | 135 | 84 | 91 | 121 |
| ENSG00000278130 | 0 | 0 | 0 | 0 | 1 | 0 |
| ENSG00000237419 | 0 | 0 | 0 | 0 | 0 | 0 |
| ENSG00000233013 | 8 | 2 | 0 | 0 | 0 | 2 |
| ENSG00000223256 | 0 | 0 | 0 | 0 | 0 | 0 |
| ENSG00000250210 | 0 | 0 | 0 | 0 | 0 | 0 |
| ENSG00000253826 | 0 | 0 | 0 | 0 | 0 | 0 |
| ENSG00000224777 | 0 | 0 | 0 | 0 | 0 | 0 |
| ENSG00000254468 | 0 | 0 | 1 | 0 | 0 | 0 |
| ENSG00000270921 | 0 | 0 | 1 | 0 | 1 | 0 |
| ENSG00000230724 | 22 | 12 | 14 | 8 | 7 | 13 |
| ENSG00000255229 | 0 | 0 | 0 | 0 | 0 | 0 |
| ENSG00000222225 | 0 | 0 | 0 | 0 | 0 | 0 |
| ENSG00000177951 | 1023 | 500 | 617 | 624 | 545 | 546 |
| ENSG00000188076 | 0 | 0 | 0 | 0 | 0 | 0 |
| ENSG00000177947 | 0 | 1 | 0 | 0 | 0 | 0 |
| ENSG00000254559 | 0 | 0 | 0 | 0 | 0 | 0 |
| ENSG00000177963 | 2546 | 1225 | 1579 | 1454 | 1392 | 1415 |
| ENSG00000283920 | 0 | 0 | 0 | 0 | 0 | 0 |
| ENSG00000142082 | 467 | 212 | 300 | 227 | 245 | 242 |
| ENSG00000185627 | 3005 | 1592 | 1728 | 1796 | 1772 | 1693 |
| ENSG00000277290 | 9 | 8 | 9 | 7 | 6 | 8 |
| ENSG00000174885 | 0 | 2 | 2 | 2 | 2 | 2 |
| ENSG00000255026 | 6 | 2 | 7 | 3 | 4 | 5 |
| ENSG00000142102 | 394 | 154 | 216 | 196 | 182 | 204 |
| ENSG00000206013 | 0 | 0 | 0 | 0 | 0 | 0 |
| ENSG00000255533 | 0 | 0 | 0 | 0 | 0 | 0 |
| ENSG00000185201 | 867 | 422 | 565 | 450 | 411 | 504 |
| ENSG00000254910 | 10 | 1 | 10 | 5 | 4 | 4 |
| ENSG00000185885 | 143 | 80 | 103 | 96 | 84 | 79 |
| ENSG00000251661 | 48 | 12 | 28 | 8 | 21 | 21 |
| ENSG00000142089 | 2541 | 1069 | 1488 | 1204 | 902 | 1176 |
| ENSG00000255089 | 0 | 0 | 0 | 0 | 0 | 0 |
| ENSG00000270105 | 0 | 1 | 2 | 0 | 0 | 1 |

| | | | | | | |
|---|---|---|---|---|---|---|
| ENSG00000270030 | 0 | 0 | 0 | 0 | 0 | 0 |
| ENSG00000255328 | 2 | 0 | 0 | 0 | 4 | 1 |
| ENSG00000270972 | 0 | 0 | 0 | 0 | 0 | 0 |
| ENSG00000182272 | 1129 | 521 | 694 | 667 | 626 | 630 |
| ENSG00000184363 | 3280 | 1543 | 1832 | 1985 | 1731 | 1775 |
| ENSG00000185187 | 1815 | 798 | 1118 | 1009 | 850 | 878 |
| ENSG00000185101 | 825 | 298 | 468 | 359 | 329 | 373 |
| ENSG00000243562 | 0 | 0 | 0 | 0 | 0 | 0 |
| ENSG00000174915 | 897 | 355 | 530 | 537 | 473 | 507 |
| ENSG00000255237 | 4 | 1 | 7 | 4 | 1 | 2 |
| ENSG00000023191 | 3581 | 1604 | 2174 | 2347 | 1824 | 1968 |
| ENSG00000254739 | 11 | 6 | 6 | 1 | 4 | 2 |
| ENSG00000174775 | 1079 | 539 | 607 | 681 | 574 | 579 |
| ENSG00000161328 | 349 | 167 | 239 | 161 | 159 | 169 |
| ENSG00000185522 | 146 | 71 | 140 | 77 | 55 | 95 |
| ENSG00000254815 | 64 | 42 | 69 | 40 | 43 | 46 |
| ENSG00000099849 | 1186 | 564 | 748 | 714 | 569 | 587 |
| ENSG00000247095 | 77 | 28 | 51 | 30 | 25 | 32 |
| ENSG00000199038 | 1 | 0 | 0 | 1 | 0 | 0 |
| ENSG00000070047 | 1330 | 523 | 774 | 731 | 702 | 738 |
| ENSG00000185507 | 249 | 126 | 162 | 132 | 123 | 104 |
| ENSG00000099834 | 3 | 3 | 2 | 0 | 0 | 0 |
| ENSG00000070031 | 5 | 2 | 0 | 0 | 0 | 0 |
| ENSG00000069696 | 5 | 3 | 5 | 0 | 0 | 2 |
| ENSG00000177030 | 900 | 410 | 537 | 553 | 430 | 517 |
| ENSG00000255158 | 2 | 3 | 5 | 0 | 3 | 1 |
| ENSG00000177106 | 4169 | 1977 | 2604 | 2373 | 2312 | 2292 |
| ENSG00000177042 | 518 | 292 | 320 | 254 | 257 | 257 |
| ENSG00000269915 | 0 | 0 | 1 | 0 | 0 | 2 |
| ENSG00000177156 | 7114 | 3722 | 4282 | 4964 | 4671 | 4591 |
| ENSG00000177225 | 1308 | 661 | 856 | 757 | 684 | 789 |
| ENSG00000255284 | 114 | 59 | 83 | 53 | 78 | 56 |
| ENSG00000279672 | 26 | 10 | 22 | 12 | 14 | 11 |
| ENSG00000255142 | 8 | 1 | 8 | 1 | 16 | 6 |
| ENSG00000184524 | 12 | 3 | 2 | 0 | 4 | 5 |
| ENSG00000177542 | 621 | 310 | 385 | 393 | 334 | 393 |
| ENSG00000274897 | 32 | 16 | 20 | 17 | 10 | 16 |
| ENSG00000177595 | 454 | 181 | 234 | 198 | 172 | 197 |
| ENSG00000177600 | 8950 | 4334 | 5009 | 6516 | 5316 | 5045 |
| ENSG00000199785 | 0 | 0 | 0 | 0 | 0 | 0 |
| ENSG00000177666 | 1229 | 662 | 893 | 726 | 635 | 674 |
| ENSG00000255108 | 2 | 0 | 2 | 1 | 0 | 0 |
| ENSG00000177685 | 432 | 199 | 264 | 215 | 214 | 235 |
| ENSG00000177697 | 3850 | 2010 | 2365 | 2503 | 2089 | 1992 |
| ENSG00000177700 | 1681 | 850 | 984 | 1062 | 894 | 817 |
| ENSG00000214063 | 553 | 231 | 306 | 278 | 189 | 237 |
| ENSG00000250397 | 31 | 13 | 12 | 15 | 16 | 14 |
| ENSG00000177830 | 1250 | 573 | 709 | 860 | 648 | 704 |
| ENSG00000183020 | 1336 | 657 | 754 | 773 | 719 | 679 |
| ENSG00000222561 | 0 | 0 | 0 | 0 | 0 | 0 |

| | | | | | | |
|---|---|---|---|---|---|---|
| ENSG00000184956 | 6 | 4 | 1 | 3 | 2 | 4 |
| ENSG00000254872 | 11 | 1 | 6 | 2 | 2 | 5 |
| ENSG00000198788 | 2 | 4 | 4 | 4 | 1 | 1 |
| ENSG00000215182 | 15 | 8 | 14 | 12 | 13 | 10 |
| ENSG00000117983 | 312 | 64 | 237 | 149 | 111 | 231 |
| ENSG00000255177 | 0 | 0 | 0 | 0 | 0 | 0 |
| ENSG00000275669 | 0 | 0 | 0 | 0 | 0 | 0 |
| ENSG00000078902 | 1257 | 600 | 693 | 749 | 656 | 641 |
| ENSG00000255153 | 108 | 52 | 74 | 49 | 50 | 51 |
| ENSG00000285985 | 0 | 1 | 0 | 0 | 0 | 0 |
| ENSG00000174672 | 198 | 74 | 129 | 106 | 126 | 99 |
| ENSG00000285581 | 0 | 0 | 0 | 1 | 0 | 2 |
| ENSG00000182208 | 392 | 162 | 221 | 256 | 224 | 213 |
| ENSG00000184545 | 846 | 400 | 548 | 542 | 434 | 506 |
| ENSG00000233930 | 297 | 163 | 233 | 207 | 226 | 228 |
| ENSG00000205869 | 147 | 52 | 62 | 60 | 65 | 67 |
| ENSG00000205867 | 1 | 0 | 5 | 1 | 2 | 3 |
| ENSG00000196224 | 0 | 1 | 0 | 0 | 0 | 0 |
| ENSG00000241598 | 0 | 0 | 0 | 0 | 0 | 0 |
| ENSG00000185940 | 0 | 0 | 0 | 0 | 0 | 0 |
| ENSG00000231487 | 0 | 0 | 0 | 0 | 0 | 0 |
| ENSG00000205866 | 0 | 0 | 0 | 0 | 0 | 0 |
| ENSG00000205865 | 0 | 0 | 0 | 0 | 0 | 0 |
| ENSG00000227306 | 0 | 0 | 0 | 0 | 0 | 0 |
| ENSG00000205864 | 0 | 0 | 0 | 0 | 0 | 1 |
| ENSG00000244242 | 179 | 80 | 126 | 141 | 107 | 108 |
| ENSG00000250644 | 22 | 16 | 37 | 20 | 25 | 24 |
| ENSG00000117984 | 15783 | 8600 | 10313 | 12981 | 9620 | 9414 |
| ENSG00000235027 | 107 | 57 | 67 | 65 | 75 | 72 |
| ENSG00000229512 | 23 | 11 | 24 | 25 | 23 | 15 |
| ENSG00000230834 | 0 | 1 | 5 | 0 | 1 | 5 |
| ENSG00000230980 | 0 | 0 | 0 | 0 | 0 | 0 |
| ENSG00000230080 | 0 | 0 | 0 | 0 | 0 | 0 |
| ENSG00000149043 | 12 | 12 | 8 | 14 | 9 | 13 |
| ENSG00000130598 | 6 | 0 | 1 | 3 | 0 | 4 |
| ENSG00000130592 | 2 | 1 | 0 | 0 | 0 | 0 |
| ENSG00000264493 | 0 | 0 | 0 | 0 | 0 | 0 |
| ENSG00000249086 | 0 | 0 | 0 | 0 | 0 | 0 |
| ENSG00000284594 | 0 | 0 | 0 | 0 | 0 | 0 |
| ENSG00000283787 | 0 | 0 | 0 | 0 | 0 | 0 |
| ENSG00000229671 | 0 | 0 | 0 | 0 | 0 | 0 |
| ENSG00000130595 | 0 | 0 | 1 | 0 | 1 | 0 |
| ENSG00000214026 | 303 | 155 | 217 | 248 | 201 | 194 |
| ENSG00000226416 | 4 | 0 | 1 | 0 | 0 | 1 |
| ENSG00000232987 | 0 | 0 | 0 | 0 | 0 | 0 |
| ENSG00000130600 | 19 | 6 | 18 | 2 | 5 | 8 |
| ENSG00000275266 | 0 | 0 | 0 | 0 | 0 | 0 |
| ENSG00000284010 | 0 | 0 | 0 | 0 | 0 | 0 |
| ENSG00000274866 | 0 | 0 | 0 | 0 | 0 | 0 |
| ENSG00000167244 | 0 | 0 | 0 | 0 | 0 | 0 |

| | | | | | | |
|---|---|---|---|---|---|---|
| ENSG00000284779 | 0 | 0 | 0 | 0 | 0 | 0 |
| ENSG00000240801 | 0 | 0 | 0 | 0 | 0 | 0 |
| ENSG00000129965 | 0 | 0 | 0 | 0 | 0 | 0 |
| ENSG00000207805 | 0 | 0 | 0 | 0 | 0 | 0 |
| ENSG00000099869 | 0 | 0 | 0 | 1 | 2 | 1 |
| ENSG00000254647 | 2 | 0 | 0 | 0 | 0 | 0 |
| ENSG00000180176 | 91 | 67 | 55 | 59 | 65 | 65 |
| ENSG00000265258 | 0 | 0 | 0 | 0 | 0 | 0 |
| ENSG00000183734 | 460 | 205 | 240 | 285 | 222 | 238 |
| ENSG00000110665 | 4 | 1 | 0 | 0 | 0 | 1 |
| ENSG00000064201 | 19 | 6 | 16 | 5 | 10 | 13 |
| ENSG00000207308 | 0 | 0 | 0 | 0 | 0 | 1 |
| ENSG00000238184 | 6 | 4 | 8 | 2 | 2 | 5 |
| ENSG00000236264 | 0 | 2 | 0 | 1 | 0 | 3 |
| ENSG00000199550 | 0 | 2 | 0 | 0 | 0 | 0 |
| ENSG00000110651 | 6168 | 3221 | 3932 | 4482 | 3789 | 3687 |
| ENSG00000184281 | 720 | 354 | 438 | 598 | 441 | 397 |
| ENSG00000230483 | 0 | 0 | 0 | 0 | 0 | 0 |
| ENSG00000070985 | 0 | 0 | 1 | 0 | 2 | 0 |
| ENSG00000053918 | 4 | 4 | 4 | 0 | 11 | 6 |
| ENSG00000269821 | 82 | 50 | 49 | 16 | 55 | 54 |
| ENSG00000275666 | 0 | 0 | 0 | 0 | 0 | 0 |
| ENSG00000276105 | 0 | 0 | 0 | 0 | 0 | 0 |
| ENSG00000276494 | 0 | 0 | 0 | 0 | 0 | 0 |
| ENSG00000276015 | 0 | 0 | 0 | 0 | 0 | 0 |
| ENSG00000236041 | 0 | 0 | 0 | 0 | 0 | 0 |
| ENSG00000229414 | 0 | 0 | 0 | 0 | 0 | 0 |
| ENSG00000237941 | 0 | 0 | 0 | 0 | 0 | 0 |
| ENSG00000274740 | 0 | 0 | 0 | 0 | 0 | 0 |
| ENSG00000129757 | 8 | 8 | 20 | 3 | 7 | 2 |
| ENSG00000254827 | 19 | 4 | 10 | 7 | 10 | 6 |
| ENSG00000110628 | 767 | 379 | 474 | 518 | 408 | 457 |
| ENSG00000181649 | 943 | 395 | 549 | 650 | 469 | 482 |
| ENSG00000205531 | 3962 | 1911 | 2357 | 1969 | 2288 | 2258 |
| ENSG00000207008 | 0 | 0 | 0 | 0 | 0 | 0 |
| ENSG00000183562 | 5 | 2 | 4 | 2 | 4 | 1 |
| ENSG00000110619 | 1269 | 753 | 896 | 839 | 828 | 837 |
| ENSG00000247473 | 0 | 3 | 0 | 0 | 0 | 0 |
| ENSG00000201616 | 0 | 1 | 0 | 0 | 0 | 0 |
| ENSG00000285644 | 26 | 20 | 21 | 14 | 20 | 14 |
| ENSG00000236710 | 2 | 0 | 0 | 0 | 2 | 1 |
| ENSG00000021762 | 456 | 204 | 235 | 250 | 224 | 216 |
| ENSG00000234791 | 0 | 0 | 0 | 0 | 0 | 0 |
| ENSG00000182170 | 0 | 0 | 0 | 0 | 0 | 0 |
| ENSG00000236301 | 1 | 0 | 0 | 0 | 0 | 0 |
| ENSG00000224513 | 0 | 0 | 0 | 0 | 1 | 0 |
| ENSG00000184350 | 0 | 0 | 0 | 0 | 0 | 0 |
| ENSG00000236987 | 0 | 0 | 0 | 0 | 0 | 0 |
| ENSG00000276951 | 0 | 0 | 0 | 0 | 0 | 0 |
| ENSG00000232390 | 0 | 0 | 0 | 0 | 0 | 0 |

| | | | | | | |
|---|---|---|---|---|---|---|
| ENSG00000273813 | 0 | 0 | 0 | 0 | 0 | 0 |
| ENSG00000254592 | 0 | 0 | 0 | 0 | 0 | 0 |
| ENSG00000005801 | 350 | 224 | 189 | 103 | 332 | 217 |
| ENSG00000223756 | 0 | 0 | 0 | 0 | 1 | 0 |
| ENSG00000189398 | 0 | 0 | 0 | 0 | 0 | 0 |
| ENSG00000166492 | 0 | 0 | 6 | 1 | 2 | 1 |
| ENSG00000284605 | 0 | 0 | 1 | 1 | 0 | 1 |
| ENSG00000254757 | 0 | 0 | 1 | 0 | 0 | 0 |
| ENSG00000255367 | 7 | 3 | 5 | 1 | 6 | 6 |
| ENSG00000255522 | 0 | 0 | 0 | 0 | 0 | 0 |
| ENSG00000237226 | 0 | 0 | 0 | 0 | 0 | 0 |
| ENSG00000223688 | 0 | 0 | 0 | 0 | 0 | 0 |
| ENSG00000284639 | 0 | 0 | 0 | 0 | 0 | 0 |
| ENSG00000230851 | 0 | 0 | 0 | 0 | 0 | 0 |
| ENSG00000238161 | 0 | 0 | 0 | 0 | 0 | 0 |
| ENSG00000182048 | 0 | 0 | 0 | 0 | 0 | 0 |
| ENSG00000167311 | 4 | 0 | 0 | 1 | 0 | 0 |
| ENSG00000129744 | 2 | 0 | 0 | 0 | 0 | 0 |
| ENSG00000201279 | 0 | 0 | 0 | 0 | 0 | 0 |
| ENSG00000200201 | 1 | 0 | 0 | 0 | 0 | 2 |
| ENSG00000129749 | 24 | 12 | 10 | 6 | 6 | 5 |
| ENSG00000110713 | 1561 | 628 | 870 | 587 | 828 | 877 |
| ENSG00000251934 | 0 | 0 | 0 | 0 | 0 | 0 |
| ENSG00000238304 | 0 | 0 | 0 | 0 | 0 | 0 |
| ENSG00000148985 | 700 | 353 | 453 | 390 | 386 | 430 |
| ENSG00000177105 | 560 | 316 | 299 | 361 | 265 | 289 |
| ENSG00000228661 | 7 | 2 | 5 | 6 | 4 | 1 |
| ENSG00000167323 | 848 | 357 | 520 | 458 | 406 | 468 |
| ENSG00000229368 | 4 | 3 | 5 | 6 | 0 | 5 |
| ENSG00000284525 | 0 | 0 | 0 | 0 | 0 | 0 |
| ENSG00000230593 | 0 | 0 | 0 | 0 | 0 | 0 |
| ENSG00000206976 | 0 | 0 | 0 | 0 | 0 | 0 |
| ENSG00000255232 | 0 | 0 | 0 | 0 | 0 | 0 |
| ENSG00000255141 | 0 | 0 | 0 | 0 | 0 | 0 |
| ENSG00000224555 | 0 | 0 | 0 | 0 | 0 | 0 |
| ENSG00000240385 | 0 | 0 | 0 | 0 | 0 | 0 |
| ENSG00000167325 | 1901 | 1036 | 1211 | 827 | 1153 | 1087 |
| ENSG00000255276 | 0 | 0 | 0 | 0 | 0 | 0 |
| ENSG00000231548 | 4 | 2 | 0 | 0 | 3 | 1 |
| ENSG00000254480 | 10 | 3 | 6 | 6 | 5 | 6 |
| ENSG00000255387 | 0 | 0 | 0 | 0 | 0 | 0 |
| ENSG00000284018 | 0 | 0 | 0 | 0 | 0 | 0 |
| ENSG00000284306 | 0 | 0 | 0 | 0 | 1 | 0 |
| ENSG00000236175 | 0 | 0 | 0 | 0 | 0 | 0 |
| ENSG00000283873 | 0 | 0 | 0 | 0 | 0 | 0 |
| ENSG00000284546 | 0 | 0 | 0 | 0 | 0 | 0 |
| ENSG00000284438 | 0 | 0 | 0 | 0 | 0 | 0 |
| ENSG00000221996 | 0 | 0 | 0 | 0 | 0 | 0 |
| ENSG00000175800 | 0 | 0 | 0 | 0 | 0 | 0 |
| ENSG00000132109 | 151 | 75 | 91 | 82 | 66 | 48 |

| | | | | | | |
|---|---|---|---|---|---|---|
| ENSG00000225101 | 0 | 0 | 0 | 0 | 0 | 0 |
| ENSG00000237272 | 0 | 0 | 0 | 0 | 0 | 0 |
| ENSG00000171999 | 0 | 0 | 0 | 0 | 0 | 0 |
| ENSG00000181963 | 0 | 0 | 0 | 0 | 0 | 0 |
| ENSG00000196778 | 0 | 0 | 0 | 0 | 0 | 0 |
| ENSG00000226616 | 0 | 0 | 0 | 0 | 0 | 0 |
| ENSG00000197790 | 0 | 0 | 0 | 0 | 0 | 0 |
| ENSG00000171987 | 0 | 0 | 0 | 0 | 0 | 0 |
| ENSG00000226288 | 0 | 0 | 0 | 0 | 0 | 0 |
| ENSG00000232268 | 0 | 0 | 0 | 0 | 0 | 0 |
| ENSG00000167333 | 66 | 36 | 62 | 35 | 44 | 37 |
| ENSG00000197428 | 0 | 0 | 0 | 0 | 0 | 0 |
| ENSG00000180785 | 0 | 0 | 0 | 0 | 0 | 0 |
| ENSG00000180723 | 0 | 0 | 0 | 0 | 0 | 0 |
| ENSG00000167332 | 0 | 0 | 0 | 0 | 0 | 0 |
| ENSG00000197674 | 0 | 0 | 0 | 0 | 0 | 0 |
| ENSG00000167346 | 0 | 0 | 0 | 0 | 0 | 0 |
| ENSG00000272634 | 0 | 0 | 0 | 0 | 0 | 0 |
| ENSG00000272664 | 0 | 0 | 0 | 0 | 0 | 0 |
| ENSG00000271602 | 6 | 3 | 1 | 0 | 0 | 1 |
| ENSG00000272559 | 0 | 0 | 0 | 0 | 0 | 0 |
| ENSG00000273051 | 0 | 0 | 0 | 0 | 0 | 0 |
| ENSG00000201980 | 0 | 0 | 0 | 0 | 0 | 0 |
| ENSG00000280021 | 0 | 0 | 0 | 0 | 0 | 0 |
| ENSG00000176951 | 0 | 0 | 0 | 0 | 0 | 0 |
| ENSG00000231070 | 0 | 0 | 0 | 0 | 0 | 0 |
| ENSG00000279270 | 0 | 0 | 0 | 0 | 0 | 0 |
| ENSG00000176925 | 0 | 0 | 0 | 0 | 0 | 0 |
| ENSG00000197984 | 0 | 0 | 0 | 0 | 0 | 0 |
| ENSG00000176922 | 0 | 0 | 0 | 0 | 0 | 0 |
| ENSG00000176904 | 0 | 0 | 0 | 0 | 0 | 0 |
| ENSG00000198217 | 0 | 0 | 0 | 0 | 0 | 0 |
| ENSG00000176900 | 0 | 0 | 0 | 0 | 0 | 0 |
| ENSG00000197023 | 0 | 0 | 0 | 0 | 0 | 0 |
| ENSG00000176895 | 0 | 0 | 0 | 0 | 0 | 0 |
| ENSG00000176893 | 0 | 0 | 0 | 0 | 0 | 0 |
| ENSG00000278870 | 0 | 0 | 0 | 0 | 0 | 0 |
| ENSG00000227023 | 0 | 0 | 0 | 0 | 0 | 0 |
| ENSG00000205497 | 0 | 0 | 0 | 0 | 0 | 0 |
| ENSG00000205496 | 0 | 0 | 0 | 0 | 0 | 0 |
| ENSG00000225003 | 0 | 0 | 0 | 0 | 0 | 0 |
| ENSG00000176798 | 0 | 0 | 0 | 0 | 0 | 0 |
| ENSG00000224300 | 0 | 0 | 0 | 0 | 0 | 0 |
| ENSG00000230261 | 0 | 0 | 0 | 0 | 0 | 0 |
| ENSG00000205495 | 0 | 0 | 0 | 0 | 0 | 0 |
| ENSG00000176787 | 0 | 0 | 0 | 0 | 0 | 0 |
| ENSG00000273085 | 0 | 0 | 0 | 0 | 0 | 0 |
| ENSG00000237354 | 0 | 0 | 0 | 0 | 0 | 0 |
| ENSG00000226157 | 0 | 0 | 0 | 0 | 0 | 0 |
| ENSG00000203560 | 0 | 0 | 0 | 0 | 0 | 0 |

| | | | | | | |
|---|---|---|---|---|---|---|
| ENSG00000236897 | 0 | 0 | 0 | 0 | 0 | 0 |
| ENSG00000205494 | 0 | 0 | 0 | 0 | 0 | 0 |
| ENSG00000171944 | 0 | 0 | 0 | 0 | 0 | 0 |
| ENSG00000182070 | 0 | 0 | 0 | 0 | 0 | 0 |
| ENSG00000176752 | 0 | 0 | 0 | 0 | 0 | 0 |
| ENSG00000176748 | 0 | 0 | 0 | 0 | 0 | 0 |
| ENSG00000176742 | 0 | 0 | 0 | 0 | 0 | 0 |
| ENSG00000224091 | 0 | 0 | 0 | 0 | 0 | 0 |
| ENSG00000285498 | 0 | 0 | 0 | 0 | 0 | 0 |
| ENSG00000221031 | 0 | 0 | 0 | 0 | 0 | 0 |
| ENSG00000244734 | 0 | 0 | 0 | 0 | 0 | 0 |
| ENSG00000223609 | 0 | 0 | 0 | 0 | 0 | 0 |
| ENSG00000229988 | 0 | 0 | 0 | 0 | 0 | 0 |
| ENSG00000260629 | 0 | 0 | 0 | 0 | 0 | 0 |
| ENSG00000213934 | 0 | 0 | 0 | 0 | 0 | 0 |
| ENSG00000284931 | 0 | 0 | 0 | 0 | 0 | 0 |
| ENSG00000196565 | 0 | 0 | 0 | 0 | 0 | 0 |
| ENSG00000239920 | 0 | 0 | 0 | 0 | 0 | 0 |
| ENSG00000213931 | 0 | 0 | 0 | 0 | 0 | 0 |
| ENSG00000229093 | 0 | 0 | 0 | 0 | 0 | 0 |
| ENSG00000183251 | 0 | 0 | 0 | 0 | 0 | 0 |
| ENSG00000167355 | 0 | 0 | 0 | 0 | 0 | 0 |
| ENSG00000223735 | 0 | 0 | 0 | 0 | 0 | 0 |
| ENSG00000279012 | 0 | 0 | 0 | 0 | 0 | 0 |
| ENSG00000236359 | 0 | 0 | 0 | 0 | 0 | 0 |
| ENSG00000176239 | 0 | 0 | 0 | 0 | 0 | 0 |
| ENSG00000230087 | 0 | 0 | 0 | 0 | 0 | 0 |
| ENSG00000236248 | 0 | 0 | 0 | 0 | 0 | 0 |
| ENSG00000184698 | 0 | 0 | 0 | 0 | 0 | 0 |
| ENSG00000184321 | 0 | 0 | 0 | 0 | 0 | 0 |
| ENSG00000167360 | 0 | 0 | 0 | 0 | 0 | 0 |
| ENSG00000248424 | 0 | 0 | 0 | 0 | 0 | 0 |
| ENSG00000167359 | 0 | 0 | 0 | 0 | 0 | 0 |
| ENSG00000187918 | 0 | 0 | 0 | 0 | 0 | 0 |
| ENSG00000230484 | 0 | 0 | 0 | 0 | 0 | 0 |
| ENSG00000181609 | 0 | 0 | 0 | 0 | 0 | 0 |
| ENSG00000175520 | 0 | 0 | 0 | 0 | 0 | 0 |
| ENSG00000175518 | 0 | 0 | 0 | 0 | 1 | 0 |
| ENSG00000224295 | 0 | 0 | 0 | 0 | 0 | 0 |
| ENSG00000249633 | 0 | 0 | 0 | 0 | 0 | 0 |
| ENSG00000181616 | 0 | 0 | 0 | 0 | 0 | 0 |
| ENSG00000248553 | 0 | 0 | 0 | 0 | 0 | 0 |
| ENSG00000231049 | 0 | 0 | 0 | 0 | 0 | 0 |
| ENSG00000233646 | 0 | 0 | 0 | 0 | 0 | 0 |
| ENSG00000229534 | 0 | 0 | 0 | 0 | 1 | 0 |
| ENSG00000187747 | 0 | 0 | 0 | 0 | 0 | 0 |
| ENSG00000121236 | 4 | 1 | 0 | 0 | 2 | 4 |
| ENSG00000258588 | 0 | 0 | 0 | 0 | 0 | 0 |
| ENSG00000258659 | 0 | 0 | 0 | 0 | 0 | 1 |
| ENSG00000132256 | 274 | 122 | 157 | 113 | 165 | 157 |

| | | | | | | |
|---|---|---|---|---|---|---|
| ENSG00000132274 | 6 | 2 | 1 | 0 | 1 | 0 |
| ENSG00000232381 | 0 | 0 | 0 | 0 | 0 | 0 |
| ENSG00000181109 | 0 | 0 | 0 | 0 | 0 | 0 |
| ENSG00000181023 | 0 | 0 | 0 | 0 | 0 | 0 |
| ENSG00000181074 | 0 | 0 | 0 | 0 | 0 | 0 |
| ENSG00000181017 | 0 | 0 | 0 | 0 | 0 | 0 |
| ENSG00000181009 | 0 | 0 | 0 | 0 | 0 | 0 |
| ENSG00000181001 | 0 | 0 | 0 | 0 | 0 | 0 |
| ENSG00000236965 | 0 | 0 | 0 | 0 | 0 | 0 |
| ENSG00000180988 | 0 | 0 | 0 | 0 | 0 | 0 |
| ENSG00000205409 | 0 | 0 | 0 | 0 | 0 | 0 |
| ENSG00000183269 | 0 | 0 | 0 | 0 | 0 | 0 |
| ENSG00000233563 | 0 | 0 | 0 | 0 | 0 | 0 |
| ENSG00000180974 | 0 | 0 | 0 | 0 | 0 | 0 |
| ENSG00000277932 | 0 | 0 | 0 | 0 | 0 | 0 |
| ENSG00000277514 | 0 | 0 | 0 | 0 | 0 | 0 |
| ENSG00000184478 | 0 | 0 | 0 | 0 | 0 | 0 |
| ENSG00000188691 | 0 | 0 | 0 | 0 | 0 | 0 |
| ENSG00000183313 | 0 | 0 | 0 | 0 | 0 | 0 |
| ENSG00000254831 | 0 | 0 | 0 | 0 | 0 | 0 |
| ENSG00000183389 | 0 | 0 | 0 | 0 | 0 | 0 |
| ENSG00000202147 | 0 | 0 | 0 | 0 | 0 | 0 |
| ENSG00000180934 | 0 | 0 | 0 | 0 | 0 | 0 |
| ENSG00000255481 | 0 | 0 | 0 | 0 | 0 | 0 |
| ENSG00000262980 | 0 | 0 | 0 | 0 | 0 | 0 |
| ENSG00000234895 | 0 | 0 | 0 | 0 | 0 | 0 |
| ENSG00000180919 | 0 | 0 | 0 | 0 | 0 | 0 |
| ENSG00000254444 | 0 | 0 | 0 | 0 | 0 | 0 |
| ENSG00000180913 | 0 | 0 | 0 | 0 | 0 | 0 |
| ENSG00000180909 | 0 | 0 | 0 | 0 | 0 | 0 |
| ENSG00000255307 | 0 | 0 | 0 | 0 | 0 | 0 |
| ENSG00000175485 | 0 | 1 | 0 | 0 | 0 | 0 |
| ENSG00000267940 | 10 | 12 | 15 | 9 | 16 | 6 |
| ENSG00000180878 | 3 | 0 | 3 | 2 | 1 | 1 |
| ENSG00000051009 | 653 | 313 | 432 | 359 | 377 | 411 |
| ENSG00000132259 | 2 | 2 | 2 | 2 | 2 | 1 |
| ENSG00000110148 | 0 | 0 | 0 | 0 | 0 | 0 |
| ENSG00000270435 | 0 | 0 | 0 | 0 | 0 | 0 |
| ENSG00000170955 | 0 | 0 | 0 | 0 | 0 | 0 |
| ENSG00000282556 | 0 | 0 | 0 | 0 | 0 | 0 |
| ENSG00000254443 | 0 | 0 | 0 | 0 | 0 | 0 |
| ENSG00000166311 | 553 | 271 | 334 | 279 | 282 | 259 |
| ENSG00000166313 | 4 | 0 | 2 | 0 | 2 | 2 |
| ENSG00000110169 | 16 | 15 | 28 | 9 | 12 | 7 |
| ENSG00000110171 | 428 | 185 | 227 | 161 | 178 | 187 |
| ENSG00000132254 | 2395 | 1138 | 1505 | 1348 | 1223 | 1190 |
| ENSG00000132286 | 804 | 344 | 458 | 426 | 427 | 433 |
| ENSG00000283977 | 0 | 0 | 0 | 0 | 0 | 0 |
| ENSG00000254595 | 6 | 5 | 1 | 0 | 3 | 6 |
| ENSG00000179532 | 58 | 23 | 30 | 14 | 27 | 26 |

| | | | | | | |
|---|---|---|---|---|---|---|
| ENSG00000132275 | 466 | 190 | 234 | 227 | 214 | 236 |
| ENSG00000254400 | 0 | 0 | 0 | 0 | 0 | 0 |
| ENSG00000166333 | 13 | 38 | 39 | 41 | 37 | 32 |
| ENSG00000166337 | 117 | 34 | 47 | 51 | 74 | 83 |
| ENSG00000254641 | 0 | 0 | 0 | 0 | 0 | 0 |
| ENSG00000285338 | 0 | 0 | 0 | 0 | 0 | 0 |
| ENSG00000166340 | 1394 | 599 | 765 | 579 | 659 | 724 |
| ENSG00000255680 | 0 | 0 | 0 | 0 | 0 | 0 |
| ENSG00000166341 | 23 | 12 | 16 | 7 | 7 | 13 |
| ENSG00000255390 | 0 | 0 | 0 | 0 | 0 | 0 |
| ENSG00000255410 | 0 | 0 | 0 | 1 | 0 | 1 |
| ENSG00000158042 | 910 | 514 | 529 | 686 | 573 | 617 |
| ENSG00000241678 | 0 | 0 | 0 | 0 | 0 | 0 |
| ENSG00000254838 | 0 | 0 | 0 | 0 | 0 | 0 |
| ENSG00000254781 | 0 | 0 | 0 | 0 | 0 | 0 |
| ENSG00000188124 | 0 | 0 | 0 | 1 | 0 | 0 |
| ENSG00000279486 | 0 | 0 | 0 | 0 | 0 | 0 |
| ENSG00000184933 | 0 | 0 | 0 | 0 | 0 | 0 |
| ENSG00000283415 | 0 | 0 | 0 | 0 | 0 | 0 |
| ENSG00000254712 | 0 | 0 | 0 | 0 | 0 | 0 |
| ENSG00000166363 | 0 | 0 | 0 | 0 | 0 | 0 |
| ENSG00000170790 | 0 | 0 | 0 | 0 | 0 | 0 |
| ENSG00000170782 | 0 | 0 | 0 | 0 | 0 | 0 |
| ENSG00000166368 | 0 | 0 | 0 | 0 | 0 | 0 |
| ENSG00000178358 | 0 | 0 | 0 | 0 | 0 | 0 |
| ENSG00000149054 | 0 | 0 | 0 | 0 | 0 | 0 |
| ENSG00000149050 | 0 | 0 | 0 | 0 | 0 | 0 |
| ENSG00000158077 | 0 | 0 | 0 | 0 | 0 | 0 |
| ENSG00000170748 | 0 | 0 | 0 | 0 | 0 | 0 |
| ENSG00000255179 | 0 | 0 | 0 | 0 | 0 | 0 |
| ENSG00000221703 | 0 | 0 | 0 | 0 | 0 | 0 |
| ENSG00000170743 | 2 | 0 | 0 | 0 | 0 | 0 |
| ENSG00000251364 | 29 | 13 | 16 | 3 | 15 | 16 |
| ENSG00000183801 | 2 | 0 | 0 | 0 | 1 | 0 |
| ENSG00000166387 | 584 | 257 | 381 | 174 | 249 | 251 |
| ENSG00000254864 | 12 | 6 | 9 | 3 | 4 | 9 |
| ENSG00000166394 | 8 | 6 | 2 | 6 | 8 | 4 |
| ENSG00000183378 | 0 | 0 | 0 | 0 | 0 | 0 |
| ENSG00000275612 | 0 | 0 | 0 | 0 | 0 | 0 |
| ENSG00000276668 | 0 | 0 | 0 | 0 | 0 | 0 |
| ENSG00000271758 | 0 | 0 | 0 | 0 | 0 | 0 |
| ENSG00000254707 | 0 | 0 | 0 | 0 | 0 | 0 |
| ENSG00000176716 | 0 | 0 | 0 | 0 | 0 | 0 |
| ENSG00000227892 | 0 | 0 | 0 | 0 | 0 | 0 |
| ENSG00000254951 | 0 | 0 | 0 | 0 | 0 | 0 |
| ENSG00000166408 | 0 | 0 | 0 | 0 | 0 | 0 |
| ENSG00000241008 | 0 | 0 | 0 | 0 | 0 | 0 |
| ENSG00000183303 | 0 | 0 | 0 | 0 | 0 | 0 |
| ENSG00000182334 | 0 | 0 | 0 | 0 | 0 | 0 |
| ENSG00000170688 | 0 | 0 | 0 | 0 | 0 | 0 |

| | | | | | | |
|---|---|---|---|---|---|---|
| ENSG00000252769 | 0 | 0 | 0 | 0 | 0 | 0 |
| ENSG00000279000 | 0 | 0 | 0 | 0 | 0 | 0 |
| ENSG00000170683 | 0 | 0 | 0 | 0 | 0 | 0 |
| ENSG00000182261 | 0 | 0 | 0 | 0 | 0 | 0 |
| ENSG00000175390 | 2624 | 1127 | 1325 | 1579 | 1311 | 1346 |
| ENSG00000254642 | 0 | 0 | 0 | 0 | 0 | 0 |
| ENSG00000255420 | 0 | 0 | 0 | 0 | 0 | 0 |
| ENSG00000166402 | 12 | 5 | 7 | 4 | 3 | 11 |
| ENSG00000254921 | 0 | 0 | 0 | 0 | 0 | 0 |
| ENSG00000248332 | 0 | 0 | 0 | 0 | 0 | 0 |
| ENSG00000279391 | 0 | 0 | 0 | 0 | 0 | 0 |
| ENSG00000166405 | 0 | 0 | 0 | 0 | 0 | 0 |
| ENSG00000246820 | 0 | 0 | 0 | 0 | 0 | 0 |
| ENSG00000166407 | 4 | 2 | 1 | 2 | 7 | 0 |
| ENSG00000130413 | 0 | 0 | 0 | 0 | 0 | 0 |
| ENSG00000252778 | 0 | 0 | 0 | 0 | 0 | 0 |
| ENSG00000166436 | 822 | 356 | 497 | 256 | 347 | 446 |
| ENSG00000254900 | 1 | 2 | 1 | 0 | 2 | 0 |
| ENSG00000166441 | 13439 | 6990 | 8207 | 10124 | 8436 | 7933 |
| ENSG00000200983 | 0 | 0 | 0 | 0 | 0 | 0 |
| ENSG00000212607 | 5 | 3 | 4 | 7 | 1 | 5 |
| ENSG00000166444 | 43 | 24 | 23 | 28 | 25 | 25 |
| ENSG00000254665 | 0 | 0 | 0 | 0 | 0 | 0 |
| ENSG00000255159 | 0 | 0 | 0 | 0 | 0 | 0 |
| ENSG00000252905 | 0 | 0 | 0 | 0 | 0 | 0 |
| ENSG00000202276 | 0 | 0 | 0 | 0 | 0 | 0 |
| ENSG00000166452 | 559 | 242 | 339 | 246 | 258 | 318 |
| ENSG00000176029 | 2 | 0 | 0 | 0 | 1 | 0 |
| ENSG00000176009 | 0 | 0 | 0 | 0 | 0 | 0 |
| ENSG00000223080 | 0 | 0 | 0 | 0 | 0 | 0 |
| ENSG00000175348 | 1378 | 715 | 817 | 810 | 864 | 849 |
| ENSG00000254860 | 23 | 14 | 12 | 10 | 15 | 31 |
| ENSG00000175352 | 0 | 0 | 0 | 0 | 0 | 0 |
| ENSG00000253973 | 0 | 0 | 0 | 0 | 0 | 0 |
| ENSG00000175356 | 2451 | 1234 | 2110 | 1893 | 2070 | 2514 |
| ENSG00000264984 | 0 | 0 | 1 | 0 | 0 | 0 |
| ENSG00000213538 | 5 | 3 | 3 | 2 | 5 | 6 |
| ENSG00000184014 | 1013 | 459 | 635 | 480 | 614 | 646 |
| ENSG00000255097 | 0 | 0 | 0 | 0 | 0 | 0 |
| ENSG00000273725 | 0 | 0 | 0 | 0 | 0 | 0 |
| ENSG00000166471 | 1234 | 577 | 769 | 496 | 723 | 604 |
| ENSG00000254884 | 0 | 0 | 0 | 0 | 0 | 1 |
| ENSG00000205339 | 4325 | 2310 | 2138 | 1563 | 3630 | 2328 |
| ENSG00000201998 | 7 | 1 | 0 | 0 | 0 | 0 |
| ENSG00000254397 | 0 | 0 | 0 | 0 | 0 | 0 |
| ENSG00000268403 | 41 | 20 | 36 | 16 | 33 | 12 |
| ENSG00000166478 | 323 | 156 | 147 | 100 | 197 | 172 |
| ENSG00000166483 | 752 | 353 | 444 | 366 | 526 | 451 |
| ENSG00000238387 | 0 | 0 | 0 | 0 | 0 | 0 |
| ENSG00000243964 | 1 | 1 | 6 | 1 | 0 | 0 |

| | | | | | | |
|---|---|---|---|---|---|---|
| ENSG00000255463 | 0 | 0 | 0 | 0 | 0 | 0 |
| ENSG00000239470 | 0 | 1 | 1 | 1 | 1 | 2 |
| ENSG00000133789 | 567 | 279 | 293 | 196 | 369 | 282 |
| ENSG00000201564 | 0 | 0 | 0 | 0 | 0 | 0 |
| ENSG00000245522 | 18 | 9 | 4 | 6 | 6 | 9 |
| ENSG00000246273 | 70 | 37 | 38 | 36 | 48 | 48 |
| ENSG00000133812 | 504 | 309 | 267 | 144 | 372 | 271 |
| ENSG00000255476 | 0 | 0 | 0 | 0 | 0 | 0 |
| ENSG00000252568 | 0 | 0 | 0 | 0 | 0 | 0 |
| ENSG00000254765 | 0 | 0 | 0 | 0 | 0 | 0 |
| ENSG00000254865 | 0 | 2 | 0 | 0 | 0 | 0 |
| ENSG00000254719 | 0 | 0 | 0 | 0 | 0 | 0 |
| ENSG00000254554 | 4 | 3 | 2 | 2 | 1 | 2 |
| ENSG00000148926 | 56 | 12 | 18 | 16 | 26 | 42 |
| ENSG00000133805 | 62 | 38 | 47 | 61 | 55 | 71 |
| ENSG00000221574 | 0 | 0 | 0 | 0 | 0 | 0 |
| ENSG00000255823 | 3 | 0 | 0 | 1 | 1 | 3 |
| ENSG00000283813 | 0 | 0 | 0 | 0 | 0 | 0 |
| ENSG00000110315 | 794 | 393 | 504 | 378 | 665 | 554 |
| ENSG00000177112 | 1 | 1 | 5 | 4 | 0 | 8 |
| ENSG00000133800 | 0 | 0 | 0 | 0 | 0 | 0 |
| ENSG00000072952 | 16 | 9 | 11 | 33 | 73 | 63 |
| ENSG00000206858 | 0 | 0 | 0 | 0 | 0 | 0 |
| ENSG00000198730 | 965 | 523 | 528 | 325 | 827 | 575 |
| ENSG00000110321 | 16994 | 8439 | 9572 | 6966 | 10500 | 9051 |
| ENSG00000238622 | 0 | 0 | 0 | 0 | 0 | 0 |
| ENSG00000246308 | 9 | 5 | 0 | 2 | 6 | 3 |
| ENSG00000236287 | 774 | 346 | 469 | 287 | 441 | 404 |
| ENSG00000247271 | 132 | 76 | 105 | 62 | 113 | 88 |
| ENSG00000250041 | 0 | 0 | 0 | 0 | 0 | 0 |
| ENSG00000254401 | 0 | 0 | 0 | 0 | 0 | 0 |
| ENSG00000254957 | 0 | 0 | 0 | 0 | 0 | 0 |
| ENSG00000270897 | 0 | 0 | 0 | 0 | 0 | 0 |
| ENSG00000255123 | 0 | 0 | 0 | 0 | 0 | 0 |
| ENSG00000255260 | 0 | 0 | 0 | 0 | 0 | 0 |
| ENSG00000110328 | 107 | 62 | 44 | 59 | 88 | 63 |
| ENSG00000254598 | 1 | 6 | 0 | 3 | 5 | 3 |
| ENSG00000255351 | 0 | 0 | 0 | 0 | 0 | 0 |
| ENSG00000255462 | 0 | 0 | 0 | 0 | 0 | 0 |
| ENSG00000266645 | 0 | 0 | 0 | 0 | 0 | 0 |
| ENSG00000280288 | 0 | 0 | 0 | 0 | 0 | 0 |
| ENSG00000255309 | 0 | 0 | 0 | 0 | 0 | 0 |
| ENSG00000278529 | 0 | 0 | 0 | 0 | 0 | 0 |
| ENSG00000170242 | 777 | 459 | 321 | 193 | 684 | 351 |
| ENSG00000255492 | 0 | 0 | 0 | 0 | 0 | 0 |
| ENSG00000050165 | 0 | 0 | 0 | 0 | 1 | 0 |
| ENSG00000254486 | 0 | 0 | 0 | 0 | 0 | 0 |
| ENSG00000254991 | 0 | 0 | 0 | 0 | 0 | 0 |
| ENSG00000255400 | 0 | 0 | 0 | 0 | 0 | 0 |
| ENSG00000133816 | 403 | 188 | 284 | 123 | 141 | 198 |

| | | | | | | |
|---|---|---|---|---|---|---|
| ENSG00000275373 | 0 | 0 | 0 | 0 | 0 | 0 |
| ENSG00000254680 | 0 | 0 | 0 | 0 | 0 | 0 |
| ENSG00000254983 | 0 | 0 | 0 | 0 | 0 | 0 |
| ENSG00000197702 | 1419 | 679 | 836 | 564 | 686 | 629 |
| ENSG00000254847 | 0 | 0 | 0 | 0 | 0 | 1 |
| ENSG00000187079 | 1187 | 588 | 558 | 298 | 824 | 528 |
| ENSG00000254688 | 0 | 0 | 0 | 0 | 0 | 0 |
| ENSG00000255067 | 0 | 0 | 0 | 0 | 0 | 0 |
| ENSG00000252329 | 0 | 0 | 0 | 0 | 0 | 0 |
| ENSG00000203258 | 0 | 0 | 0 | 0 | 0 | 0 |
| ENSG00000251381 | 0 | 0 | 0 | 0 | 0 | 0 |
| ENSG00000254670 | 0 | 2 | 0 | 1 | 0 | 0 |
| ENSG00000189431 | 6 | 8 | 3 | 1 | 1 | 4 |
| ENSG00000255558 | 0 | 0 | 0 | 0 | 0 | 0 |
| ENSG00000255018 | 0 | 0 | 0 | 0 | 0 | 0 |
| ENSG00000133794 | 297 | 163 | 193 | 137 | 173 | 149 |
| ENSG00000222162 | 1 | 0 | 0 | 0 | 0 | 0 |
| ENSG00000148925 | 1231 | 603 | 670 | 530 | 682 | 607 |
| ENSG00000254583 | 0 | 0 | 0 | 0 | 0 | 0 |
| ENSG00000255401 | 0 | 0 | 0 | 0 | 0 | 0 |
| ENSG00000152266 | 0 | 0 | 0 | 0 | 0 | 0 |
| ENSG00000228901 | 2 | 0 | 0 | 0 | 0 | 0 |
| ENSG00000197601 | 1657 | 983 | 766 | 338 | 1025 | 739 |
| ENSG00000254791 | 0 | 0 | 0 | 0 | 0 | 3 |
| ENSG00000240454 | 0 | 0 | 0 | 0 | 0 | 0 |
| ENSG00000200685 | 0 | 0 | 0 | 0 | 0 | 0 |
| ENSG00000255075 | 0 | 0 | 0 | 0 | 0 | 0 |
| ENSG00000255088 | 0 | 0 | 0 | 0 | 0 | 0 |
| ENSG00000255411 | 0 | 0 | 0 | 0 | 0 | 0 |
| ENSG00000254927 | 0 | 0 | 0 | 0 | 0 | 0 |
| ENSG00000254930 | 0 | 0 | 0 | 0 | 0 | 0 |
| ENSG00000201856 | 0 | 0 | 0 | 0 | 0 | 0 |
| ENSG00000254438 | 0 | 0 | 0 | 0 | 0 | 0 |
| ENSG00000262655 | 10 | 3 | 5 | 4 | 3 | 4 |
| ENSG00000212365 | 0 | 0 | 0 | 0 | 0 | 0 |
| ENSG00000254418 | 0 | 0 | 0 | 0 | 1 | 0 |
| ENSG00000133818 | 514 | 233 | 304 | 304 | 300 | 266 |
| ENSG00000255074 | 0 | 0 | 0 | 0 | 0 | 0 |
| ENSG00000129083 | 3471 | 1930 | 1752 | 1133 | 2632 | 1652 |
| ENSG00000251991 | 3 | 1 | 0 | 1 | 2 | 0 |
| ENSG00000256206 | 0 | 0 | 0 | 0 | 0 | 0 |
| ENSG00000129084 | 777 | 503 | 689 | 609 | 674 | 664 |
| ENSG00000152270 | 476 | 296 | 278 | 181 | 323 | 238 |
| ENSG00000255040 | 0 | 0 | 0 | 0 | 0 | 0 |
| ENSG00000186104 | 352 | 200 | 213 | 149 | 234 | 183 |
| ENSG00000175868 | 0 | 0 | 0 | 0 | 0 | 0 |
| ENSG00000253179 | 0 | 0 | 0 | 0 | 0 | 0 |
| ENSG00000110680 | 0 | 0 | 0 | 0 | 0 | 0 |
| ENSG00000250640 | 0 | 0 | 0 | 0 | 0 | 0 |
| ENSG00000188487 | 0 | 0 | 0 | 0 | 0 | 1 |

| | | | | | | |
|---|---|---|---|---|---|---|
| ENSG00000253072 | 0 | 0 | 0 | 0 | 0 | 0 |
| ENSG00000254946 | 0 | 0 | 0 | 0 | 0 | 0 |
| ENSG00000254789 | 0 | 0 | 0 | 0 | 0 | 0 |
| ENSG00000254695 | 0 | 0 | 0 | 0 | 0 | 0 |
| ENSG00000254645 | 0 | 0 | 0 | 0 | 0 | 0 |
| ENSG00000254661 | 0 | 0 | 0 | 0 | 0 | 0 |
| ENSG00000110693 | 35 | 11 | 12 | 1 | 13 | 6 |
| ENSG00000284094 | 0 | 0 | 0 | 0 | 0 | 0 |
| ENSG00000254878 | 0 | 0 | 0 | 0 | 0 | 0 |
| ENSG00000213785 | 0 | 0 | 0 | 0 | 0 | 0 |
| ENSG00000241943 | 0 | 0 | 0 | 0 | 0 | 0 |
| ENSG00000110696 | 3669 | 1791 | 1998 | 1603 | 2077 | 1894 |
| ENSG00000166689 | 2036 | 975 | 1310 | 739 | 886 | 902 |
| ENSG00000199883 | 0 | 0 | 0 | 0 | 0 | 0 |
| ENSG00000244398 | 137 | 92 | 81 | 85 | 82 | 95 |
| ENSG00000184669 | 163 | 97 | 100 | 101 | 99 | 101 |
| ENSG00000110700 | 9440 | 4449 | 5403 | 6324 | 5782 | 5402 |
| ENSG00000272034 | 6 | 10 | 19 | 5 | 13 | 7 |
| ENSG00000201403 | 0 | 0 | 0 | 0 | 0 | 0 |
| ENSG00000011405 | 619 | 365 | 253 | 106 | 630 | 297 |
| ENSG00000201586 | 0 | 0 | 0 | 0 | 0 | 0 |
| ENSG00000240808 | 0 | 0 | 0 | 0 | 0 | 1 |
| ENSG00000213779 | 0 | 0 | 0 | 0 | 0 | 2 |
| ENSG00000070081 | 1410 | 900 | 863 | 452 | 1037 | 678 |
| ENSG00000197149 | 212 | 101 | 141 | 170 | 149 | 103 |
| ENSG00000285545 | 0 | 0 | 1 | 0 | 0 | 0 |
| ENSG00000188211 | 22 | 7 | 7 | 1 | 7 | 7 |
| ENSG00000260196 | 10 | 3 | 6 | 1 | 6 | 2 |
| ENSG00000187486 | 313 | 102 | 170 | 164 | 127 | 118 |
| ENSG00000006071 | 108 | 42 | 72 | 56 | 58 | 48 |
| ENSG00000255035 | 2 | 0 | 1 | 0 | 0 | 0 |
| ENSG00000006611 | 0 | 3 | 2 | 3 | 0 | 0 |
| ENSG00000188162 | 0 | 2 | 0 | 0 | 0 | 0 |
| ENSG00000254586 | 0 | 0 | 0 | 0 | 0 | 0 |
| ENSG00000255335 | 0 | 0 | 0 | 0 | 0 | 0 |
| ENSG00000129152 | 0 | 0 | 0 | 0 | 0 | 0 |
| ENSG00000129159 | 7 | 3 | 2 | 0 | 5 | 1 |
| ENSG00000129158 | 264 | 141 | 189 | 174 | 172 | 147 |
| ENSG00000255448 | 0 | 0 | 0 | 0 | 0 | 0 |
| ENSG00000129167 | 2 | 1 | 1 | 1 | 0 | 2 |
| ENSG00000166788 | 305 | 162 | 177 | 177 | 233 | 186 |
| ENSG00000255254 | 0 | 0 | 0 | 0 | 0 | 0 |
| ENSG00000166787 | 0 | 0 | 0 | 0 | 0 | 0 |
| ENSG00000179826 | 0 | 0 | 0 | 0 | 0 | 0 |
| ENSG00000255470 | 0 | 0 | 0 | 0 | 0 | 0 |
| ENSG00000285562 | 0 | 0 | 0 | 0 | 0 | 0 |
| ENSG00000254857 | 0 | 0 | 0 | 0 | 0 | 0 |
| ENSG00000179817 | 0 | 0 | 0 | 0 | 0 | 0 |
| ENSG00000255138 | 0 | 0 | 0 | 0 | 0 | 0 |
| ENSG00000254546 | 0 | 0 | 0 | 0 | 0 | 0 |

| | | | | | | |
|---|---|---|---|---|---|---|
| ENSG00000189332 | 0 | 0 | 0 | 0 | 0 | 0 |
| ENSG00000148965 | 0 | 0 | 0 | 0 | 0 | 0 |
| ENSG00000255071 | 0 | 0 | 0 | 0 | 0 | 0 |
| ENSG00000134339 | 1 | 0 | 1 | 1 | 2 | 3 |
| ENSG00000200336 | 0 | 0 | 0 | 0 | 0 | 0 |
| ENSG00000212789 | 1 | 1 | 4 | 0 | 0 | 1 |
| ENSG00000173432 | 16 | 6 | 2 | 3 | 3 | 5 |
| ENSG00000201695 | 0 | 0 | 0 | 0 | 0 | 0 |
| ENSG00000110756 | 754 | 320 | 405 | 272 | 458 | 345 |
| ENSG00000110768 | 1618 | 879 | 1079 | 792 | 978 | 954 |
| ENSG00000264603 | 0 | 0 | 0 | 0 | 0 | 0 |
| ENSG00000134333 | 11442 | 5819 | 6417 | 7315 | 8791 | 7757 |
| ENSG00000256006 | 4 | 2 | 2 | 4 | 1 | 2 |
| ENSG00000166796 | 0 | 0 | 0 | 1 | 3 | 1 |
| ENSG00000256734 | 0 | 0 | 0 | 0 | 0 | 0 |
| ENSG00000166800 | 5 | 3 | 4 | 1 | 0 | 1 |
| ENSG00000074319 | 4153 | 1992 | 2435 | 1961 | 2326 | 2268 |
| ENSG00000256464 | 1 | 1 | 3 | 2 | 5 | 3 |
| ENSG00000256588 | 0 | 0 | 0 | 1 | 0 | 0 |
| ENSG00000256361 | 1 | 0 | 0 | 0 | 0 | 0 |
| ENSG00000151116 | 1171 | 508 | 634 | 335 | 679 | 533 |
| ENSG00000274589 | 2 | 0 | 0 | 0 | 0 | 0 |
| ENSG00000247595 | 54 | 32 | 37 | 25 | 29 | 29 |
| ENSG00000256282 | 3 | 2 | 3 | 2 | 3 | 3 |
| ENSG00000279837 | 8 | 7 | 4 | 8 | 7 | 3 |
| ENSG00000179119 | 1224 | 553 | 652 | 327 | 727 | 505 |
| ENSG00000257043 | 0 | 0 | 2 | 0 | 0 | 0 |
| ENSG00000151117 | 194 | 114 | 110 | 61 | 102 | 98 |
| ENSG00000179057 | 17 | 14 | 12 | 12 | 10 | 6 |
| ENSG00000254966 | 6 | 7 | 7 | 4 | 11 | 6 |
| ENSG00000110786 | 0 | 0 | 0 | 0 | 0 | 0 |
| ENSG00000276789 | 0 | 0 | 0 | 0 | 0 | 0 |
| ENSG00000254529 | 0 | 0 | 0 | 0 | 0 | 0 |
| ENSG00000255511 | 0 | 0 | 0 | 0 | 0 | 0 |
| ENSG00000254541 | 0 | 0 | 0 | 0 | 0 | 0 |
| ENSG00000255536 | 0 | 0 | 0 | 0 | 0 | 0 |
| ENSG00000170255 | 0 | 0 | 0 | 0 | 0 | 0 |
| ENSG00000255244 | 0 | 0 | 0 | 0 | 0 | 0 |
| ENSG00000254738 | 0 | 0 | 0 | 0 | 0 | 0 |
| ENSG00000254720 | 0 | 0 | 0 | 0 | 0 | 0 |
| ENSG00000255424 | 0 | 0 | 0 | 0 | 0 | 0 |
| ENSG00000183695 | 0 | 0 | 0 | 0 | 0 | 0 |
| ENSG00000177054 | 744 | 348 | 441 | 331 | 379 | 392 |
| ENSG00000129170 | 0 | 0 | 0 | 0 | 0 | 0 |
| ENSG00000255308 | 3 | 0 | 1 | 0 | 2 | 2 |
| ENSG00000129173 | 299 | 183 | 217 | 88 | 145 | 174 |
| ENSG00000254544 | 0 | 0 | 0 | 0 | 0 | 0 |
| ENSG00000255160 | 3 | 0 | 0 | 0 | 1 | 0 |
| ENSG00000166833 | 23 | 20 | 30 | 11 | 32 | 17 |
| ENSG00000255270 | 0 | 0 | 0 | 0 | 0 | 0 |

| | | | | | | |
|---|---|---|---|---|---|---|
| ENSG00000200687 | 0 | 0 | 0 | 0 | 0 | 0 |
| ENSG00000255043 | 0 | 0 | 0 | 0 | 0 | 0 |
| ENSG00000254622 | 0 | 0 | 0 | 0 | 0 | 0 |
| ENSG00000265210 | 0 | 0 | 0 | 0 | 0 | 0 |
| ENSG00000207407 | 0 | 0 | 0 | 0 | 0 | 0 |
| ENSG00000270607 | 0 | 0 | 0 | 0 | 0 | 0 |
| ENSG00000264309 | 0 | 0 | 0 | 0 | 0 | 0 |
| ENSG00000254542 | 0 | 0 | 0 | 0 | 0 | 0 |
| ENSG00000254453 | 0 | 0 | 0 | 0 | 0 | 0 |
| ENSG00000254894 | 0 | 0 | 0 | 0 | 0 | 0 |
| ENSG00000279266 | 0 | 0 | 0 | 0 | 0 | 0 |
| ENSG00000109851 | 0 | 0 | 0 | 0 | 0 | 0 |
| ENSG00000109854 | 3200 | 1638 | 1852 | 2087 | 2041 | 1928 |
| ENSG00000185238 | 559 | 228 | 341 | 290 | 384 | 399 |
| ENSG00000255291 | 0 | 0 | 0 | 0 | 0 | 0 |
| ENSG00000165970 | 0 | 0 | 0 | 0 | 0 | 0 |
| ENSG00000165973 | 0 | 0 | 0 | 0 | 0 | 0 |
| ENSG00000254906 | 0 | 0 | 0 | 0 | 0 | 0 |
| ENSG00000201059 | 0 | 0 | 0 | 0 | 0 | 0 |
| ENSG00000254816 | 0 | 0 | 0 | 0 | 0 | 0 |
| ENSG00000255167 | 0 | 0 | 0 | 0 | 0 | 0 |
| ENSG00000225477 | 0 | 0 | 0 | 0 | 0 | 0 |
| ENSG00000252816 | 0 | 0 | 0 | 0 | 0 | 0 |
| ENSG00000171714 | 0 | 0 | 0 | 0 | 0 | 0 |
| ENSG00000254820 | 0 | 0 | 0 | 0 | 0 | 0 |
| ENSG00000255372 | 0 | 0 | 0 | 0 | 0 | 0 |
| ENSG00000254768 | 0 | 0 | 0 | 0 | 0 | 0 |
| ENSG00000091664 | 0 | 0 | 0 | 0 | 0 | 0 |
| ENSG00000254540 | 0 | 0 | 0 | 0 | 0 | 0 |
| ENSG00000255323 | 0 | 0 | 0 | 0 | 0 | 0 |
| ENSG00000255357 | 0 | 0 | 0 | 0 | 0 | 0 |
| ENSG00000183161 | 708 | 358 | 463 | 367 | 435 | 398 |
| ENSG00000148935 | 11 | 6 | 5 | 8 | 6 | 4 |
| ENSG00000222427 | 0 | 0 | 0 | 0 | 0 | 0 |
| ENSG00000198168 | 1148 | 581 | 657 | 565 | 730 | 688 |
| ENSG00000246225 | 5 | 9 | 3 | 5 | 6 | 6 |
| ENSG00000255359 | 4 | 1 | 0 | 0 | 0 | 0 |
| ENSG00000255418 | 2 | 0 | 2 | 0 | 1 | 2 |
| ENSG00000271491 | 0 | 0 | 0 | 0 | 0 | 0 |
| ENSG00000255293 | 0 | 0 | 0 | 0 | 0 | 0 |
| ENSG00000275868 | 0 | 0 | 0 | 0 | 0 | 0 |
| ENSG00000254564 | 0 | 0 | 0 | 0 | 0 | 0 |
| ENSG00000254465 | 0 | 0 | 0 | 0 | 0 | 0 |
| ENSG00000240881 | 0 | 0 | 0 | 0 | 0 | 0 |
| ENSG00000255193 | 0 | 0 | 0 | 0 | 0 | 0 |
| ENSG00000254861 | 0 | 0 | 0 | 0 | 0 | 0 |
| ENSG00000252519 | 0 | 0 | 0 | 0 | 0 | 0 |
| ENSG00000277803 | 0 | 0 | 0 | 0 | 0 | 0 |
| ENSG00000254594 | 0 | 0 | 0 | 0 | 0 | 0 |
| ENSG00000207252 | 0 | 0 | 0 | 0 | 0 | 0 |

| | | | | | | |
|---|---|---|---|---|---|---|
| ENSG00000187398 | 6 | 6 | 6 | 0 | 3 | 2 |
| ENSG00000255489 | 0 | 0 | 0 | 0 | 0 | 0 |
| ENSG00000255368 | 0 | 0 | 0 | 0 | 0 | 0 |
| ENSG00000240298 | 0 | 0 | 0 | 0 | 0 | 0 |
| ENSG00000255454 | 0 | 0 | 0 | 0 | 0 | 0 |
| ENSG00000254456 | 0 | 0 | 0 | 0 | 0 | 1 |
| ENSG00000254754 | 0 | 0 | 0 | 0 | 0 | 0 |
| ENSG00000255086 | 0 | 0 | 0 | 0 | 0 | 0 |
| ENSG00000255243 | 0 | 0 | 0 | 0 | 0 | 0 |
| ENSG00000254819 | 0 | 0 | 0 | 0 | 0 | 0 |
| ENSG00000134343 | 0 | 0 | 0 | 0 | 1 | 0 |
| ENSG00000255505 | 0 | 0 | 0 | 0 | 0 | 0 |
| ENSG00000169550 | 0 | 0 | 0 | 0 | 0 | 0 |
| ENSG00000148942 | 0 | 0 | 0 | 0 | 0 | 0 |
| ENSG00000176971 | 0 | 0 | 0 | 0 | 0 | 0 |
| ENSG00000129151 | 0 | 1 | 0 | 0 | 0 | 0 |
| ENSG00000254560 | 2 | 0 | 0 | 0 | 0 | 0 |
| ENSG00000109881 | 305 | 186 | 165 | 165 | 218 | 149 |
| ENSG00000205213 | 875 | 393 | 463 | 300 | 428 | 445 |
| ENSG00000254862 | 0 | 0 | 0 | 0 | 1 | 0 |
| ENSG00000242527 | 2 | 1 | 0 | 0 | 0 | 0 |
| ENSG00000148943 | 658 | 343 | 375 | 257 | 388 | 299 |
| ENSG00000245573 | 27 | 18 | 25 | 8 | 17 | 15 |
| ENSG00000278483 | 0 | 1 | 0 | 1 | 0 | 0 |
| ENSG00000212289 | 0 | 0 | 0 | 0 | 0 | 0 |
| ENSG00000240036 | 7 | 2 | 1 | 3 | 2 | 2 |
| ENSG00000254934 | 0 | 0 | 0 | 0 | 0 | 0 |
| ENSG00000176697 | 2 | 0 | 1 | 0 | 1 | 0 |
| ENSG00000255496 | 0 | 0 | 0 | 0 | 0 | 0 |
| ENSG00000177447 | 0 | 0 | 1 | 0 | 0 | 0 |
| ENSG00000224411 | 2 | 3 | 1 | 1 | 4 | 2 |
| ENSG00000255094 | 2 | 0 | 0 | 0 | 0 | 0 |
| ENSG00000121621 | 45 | 38 | 16 | 15 | 56 | 30 |
| ENSG00000207874 | 0 | 0 | 0 | 0 | 0 | 0 |
| ENSG00000169519 | 464 | 208 | 251 | 145 | 239 | 204 |
| ENSG00000271332 | 0 | 0 | 0 | 0 | 0 | 0 |
| ENSG00000222385 | 0 | 0 | 0 | 0 | 0 | 0 |
| ENSG00000238149 | 0 | 0 | 0 | 0 | 0 | 0 |
| ENSG00000273912 | 0 | 0 | 0 | 0 | 0 | 0 |
| ENSG00000254606 | 2 | 0 | 1 | 0 | 0 | 1 |
| ENSG00000248990 | 0 | 0 | 0 | 0 | 0 | 0 |
| ENSG00000249867 | 0 | 0 | 0 | 0 | 0 | 0 |
| ENSG00000272882 | 0 | 0 | 0 | 0 | 0 | 0 |
| ENSG00000254526 | 0 | 0 | 0 | 0 | 0 | 0 |
| ENSG00000255450 | 0 | 0 | 0 | 0 | 0 | 0 |
| ENSG00000254530 | 0 | 0 | 0 | 0 | 0 | 0 |
| ENSG00000255281 | 0 | 0 | 0 | 0 | 0 | 0 |
| ENSG00000255227 | 0 | 0 | 0 | 0 | 0 | 0 |
| ENSG00000255029 | 0 | 0 | 0 | 0 | 0 | 0 |
| ENSG00000255117 | 0 | 0 | 0 | 0 | 0 | 0 |

| | | | | | | |
|---|---|---|---|---|---|---|
| ENSG00000255047 | 0 | 0 | 0 | 0 | 0 | 0 |
| ENSG00000254734 | 0 | 0 | 0 | 1 | 0 | 0 |
| ENSG00000243505 | 0 | 0 | 0 | 0 | 0 | 0 |
| ENSG00000240554 | 0 | 0 | 0 | 1 | 1 | 0 |
| ENSG00000261340 | 0 | 0 | 0 | 0 | 0 | 0 |
| ENSG00000182255 | 0 | 0 | 0 | 0 | 0 | 0 |
| ENSG00000254532 | 0 | 0 | 0 | 1 | 0 | 0 |
| ENSG00000255288 | 0 | 0 | 0 | 0 | 0 | 0 |
| ENSG00000131808 | 0 | 0 | 0 | 0 | 0 | 0 |
| ENSG00000152219 | 416 | 241 | 219 | 195 | 301 | 202 |
| ENSG00000242353 | 0 | 0 | 0 | 0 | 0 | 0 |
| ENSG00000066382 | 87 | 45 | 30 | 36 | 34 | 20 |
| ENSG00000255480 | 0 | 0 | 0 | 0 | 0 | 0 |
| ENSG00000254489 | 0 | 0 | 0 | 0 | 1 | 0 |
| ENSG00000170959 | 0 | 0 | 0 | 0 | 2 | 1 |
| ENSG00000255370 | 0 | 0 | 0 | 0 | 0 | 0 |
| ENSG00000255525 | 0 | 0 | 0 | 0 | 0 | 0 |
| ENSG00000170946 | 282 | 116 | 161 | 100 | 160 | 126 |
| ENSG00000148950 | 320 | 139 | 157 | 182 | 150 | 151 |
| ENSG00000109911 | 555 | 270 | 354 | 311 | 322 | 313 |
| ENSG00000228061 | 0 | 0 | 0 | 0 | 0 | 0 |
| ENSG00000007372 | 0 | 1 | 0 | 0 | 0 | 0 |
| ENSG00000281880 | 0 | 0 | 0 | 0 | 0 | 0 |
| ENSG00000285283 | 0 | 0 | 0 | 0 | 0 | 0 |
| ENSG00000254584 | 0 | 0 | 0 | 0 | 0 | 0 |
| ENSG00000254836 | 0 | 0 | 0 | 0 | 0 | 0 |
| ENSG00000254627 | 0 | 0 | 0 | 0 | 0 | 0 |
| ENSG00000212551 | 0 | 0 | 0 | 0 | 0 | 0 |
| ENSG00000255490 | 0 | 0 | 0 | 0 | 0 | 0 |
| ENSG00000049449 | 62 | 43 | 61 | 60 | 67 | 72 |
| ENSG00000255252 | 0 | 0 | 0 | 0 | 0 | 0 |
| ENSG00000227160 | 0 | 0 | 0 | 0 | 0 | 0 |
| ENSG00000255375 | 0 | 0 | 0 | 0 | 0 | 0 |
| ENSG00000184937 | 0 | 0 | 0 | 0 | 0 | 0 |
| ENSG00000183242 | 0 | 0 | 0 | 0 | 0 | 0 |
| ENSG00000278822 | 0 | 0 | 0 | 0 | 0 | 0 |
| ENSG00000273677 | 0 | 0 | 0 | 0 | 0 | 0 |
| ENSG00000278045 | 0 | 0 | 0 | 0 | 0 | 0 |
| ENSG00000276530 | 0 | 0 | 0 | 0 | 0 | 0 |
| ENSG00000273908 | 0 | 0 | 0 | 0 | 0 | 0 |
| ENSG00000277119 | 0 | 0 | 0 | 0 | 0 | 0 |
| ENSG00000149100 | 2550 | 1527 | 1573 | 1432 | 2074 | 1656 |
| ENSG00000270903 | 7 | 3 | 0 | 0 | 1 | 0 |
| ENSG00000186714 | 2 | 2 | 5 | 1 | 2 | 2 |
| ENSG00000244535 | 0 | 0 | 0 | 0 | 0 | 0 |
| ENSG00000135378 | 340 | 194 | 191 | 102 | 177 | 171 |
| ENSG00000060749 | 778 | 399 | 335 | 211 | 715 | 378 |
| ENSG00000200615 | 0 | 0 | 0 | 0 | 0 | 0 |
| ENSG00000121690 | 8 | 5 | 3 | 3 | 2 | 0 |
| ENSG00000176148 | 359 | 235 | 247 | 147 | 204 | 183 |

| | | | | | | |
|---|---|---|---|---|---|---|
| ENSG00000213713 | 22 | 22 | 22 | 11 | 30 | 26 |
| ENSG00000280798 | 183 | 81 | 126 | 74 | 91 | 84 |
| ENSG00000176102 | 573 | 390 | 417 | 257 | 472 | 419 |
| ENSG00000247151 | 0 | 0 | 1 | 2 | 1 | 0 |
| ENSG00000206808 | 0 | 0 | 0 | 0 | 0 | 0 |
| ENSG00000241950 | 0 | 0 | 0 | 0 | 0 | 0 |
| ENSG00000200485 | 0 | 0 | 0 | 0 | 0 | 0 |
| ENSG00000241255 | 0 | 2 | 2 | 1 | 2 | 1 |
| ENSG00000110422 | 1147 | 551 | 553 | 242 | 740 | 494 |
| ENSG00000110427 | 0 | 1 | 0 | 1 | 1 | 1 |
| ENSG00000254537 | 0 | 0 | 0 | 0 | 0 | 0 |
| ENSG00000255207 | 0 | 0 | 0 | 0 | 0 | 0 |
| ENSG00000255202 | 0 | 0 | 0 | 0 | 0 | 0 |
| ENSG00000284969 | 0 | 0 | 0 | 0 | 0 | 0 |
| ENSG00000205177 | 0 | 0 | 0 | 0 | 0 | 0 |
| ENSG00000085063 | 2443 | 1297 | 1398 | 1186 | 1599 | 1319 |
| ENSG00000110429 | 1187 | 601 | 663 | 417 | 670 | 591 |
| ENSG00000254508 | 0 | 0 | 0 | 0 | 0 | 0 |
| ENSG00000255272 | 0 | 0 | 0 | 0 | 0 | 0 |
| ENSG00000255161 | 0 | 0 | 0 | 0 | 0 | 0 |
| ENSG00000255169 | 0 | 0 | 0 | 0 | 0 | 0 |
| ENSG00000254619 | 0 | 0 | 0 | 0 | 0 | 0 |
| ENSG00000255133 | 0 | 0 | 0 | 0 | 0 | 0 |
| ENSG00000135363 | 32 | 25 | 11 | 11 | 11 | 19 |
| ENSG00000184566 | 0 | 0 | 0 | 0 | 0 | 0 |
| ENSG00000135387 | 10900 | 5507 | 6302 | 4424 | 5866 | 5139 |
| ENSG00000135372 | 2397 | 1071 | 1259 | 1149 | 1219 | 1365 |
| ENSG00000166016 | 704 | 311 | 437 | 282 | 328 | 347 |
| ENSG00000201867 | 0 | 0 | 0 | 0 | 0 | 0 |
| ENSG00000254708 | 2 | 2 | 0 | 0 | 0 | 0 |
| ENSG00000220204 | 0 | 0 | 0 | 0 | 0 | 0 |
| ENSG00000216708 | 0 | 0 | 0 | 0 | 0 | 0 |
| ENSG00000121691 | 1395 | 557 | 793 | 569 | 684 | 727 |
| ENSG00000135374 | 120 | 58 | 64 | 56 | 65 | 52 |
| ENSG00000255271 | 0 | 0 | 0 | 0 | 0 | 0 |
| ENSG00000255427 | 2 | 0 | 0 | 0 | 0 | 0 |
| ENSG00000135373 | 16 | 11 | 8 | 5 | 9 | 10 |
| ENSG00000271369 | 0 | 0 | 0 | 0 | 0 | 0 |
| ENSG00000270491 | 0 | 0 | 0 | 0 | 0 | 0 |
| ENSG00000149089 | 644 | 313 | 371 | 357 | 389 | 365 |
| ENSG00000110435 | 1025 | 505 | 604 | 383 | 600 | 509 |
| ENSG00000251862 | 0 | 0 | 0 | 0 | 0 | 0 |
| ENSG00000255521 | 2 | 0 | 0 | 0 | 0 | 0 |
| ENSG00000026508 | 739 | 326 | 430 | 505 | 778 | 688 |
| ENSG00000255443 | 4 | 0 | 0 | 3 | 1 | 0 |
| ENSG00000251194 | 2 | 0 | 0 | 0 | 1 | 0 |
| ENSG00000110436 | 143 | 76 | 65 | 29 | 86 | 45 |
| ENSG00000255004 | 0 | 0 | 0 | 0 | 0 | 0 |
| ENSG00000215380 | 0 | 0 | 0 | 0 | 0 | 0 |
| ENSG00000255542 | 0 | 0 | 0 | 0 | 0 | 0 |

| | | | | | | |
|---|---|---|---|---|---|---|
| ENSG00000149090 | 6 | 0 | 0 | 0 | 0 | 1 |
| ENSG00000254669 | 0 | 0 | 0 | 0 | 0 | 0 |
| ENSG00000179431 | 70 | 23 | 28 | 53 | 36 | 60 |
| ENSG00000254686 | 0 | 0 | 0 | 0 | 0 | 0 |
| ENSG00000166326 | 3258 | 1345 | 1895 | 1323 | 1524 | 1743 |
| ENSG00000285705 | 0 | 0 | 0 | 0 | 0 | 0 |
| ENSG00000186082 | 0 | 0 | 0 | 0 | 0 | 0 |
| ENSG00000285740 | 0 | 0 | 0 | 0 | 0 | 0 |
| ENSG00000254919 | 0 | 0 | 0 | 0 | 0 | 0 |
| ENSG00000179241 | 287 | 144 | 127 | 150 | 182 | 173 |
| ENSG00000254672 | 2 | 0 | 0 | 0 | 0 | 0 |
| ENSG00000255256 | 0 | 0 | 0 | 0 | 0 | 0 |
| ENSG00000241228 | 0 | 0 | 0 | 0 | 0 | 0 |
| ENSG00000263389 | 0 | 0 | 0 | 0 | 0 | 0 |
| ENSG00000280321 | 0 | 0 | 1 | 0 | 0 | 0 |
| ENSG00000110442 | 1004 | 489 | 546 | 579 | 611 | 596 |
| ENSG00000135362 | 6 | 2 | 1 | 2 | 2 | 1 |
| ENSG00000254566 | 0 | 0 | 1 | 0 | 0 | 0 |
| ENSG00000255186 | 0 | 0 | 0 | 0 | 0 | 0 |
| ENSG00000255060 | 0 | 0 | 0 | 0 | 0 | 0 |
| ENSG00000175104 | 335 | 137 | 179 | 127 | 159 | 170 |
| ENSG00000280331 | 0 | 0 | 0 | 0 | 0 | 0 |
| ENSG00000166349 | 4 | 3 | 1 | 2 | 2 | 5 |
| ENSG00000175097 | 0 | 0 | 0 | 0 | 0 | 0 |
| ENSG00000166352 | 105 | 50 | 69 | 66 | 73 | 52 |
| ENSG00000254498 | 0 | 0 | 0 | 0 | 0 | 0 |
| ENSG00000251838 | 0 | 0 | 0 | 0 | 0 | 0 |
| ENSG00000242834 | 0 | 0 | 0 | 0 | 0 | 0 |
| ENSG00000254516 | 0 | 0 | 0 | 0 | 0 | 0 |
| ENSG00000279004 | 0 | 0 | 0 | 0 | 0 | 0 |
| ENSG00000254784 | 0 | 0 | 0 | 0 | 0 | 0 |
| ENSG00000255477 | 2 | 0 | 0 | 0 | 0 | 0 |
| ENSG00000255175 | 0 | 0 | 0 | 0 | 0 | 0 |
| ENSG00000254562 | 0 | 0 | 0 | 0 | 0 | 0 |
| ENSG00000285751 | 0 | 0 | 0 | 0 | 0 | 0 |
| ENSG00000240975 | 0 | 0 | 0 | 0 | 0 | 0 |
| ENSG00000201591 | 0 | 0 | 0 | 0 | 0 | 0 |
| ENSG00000270588 | 0 | 0 | 0 | 0 | 0 | 0 |
| ENSG00000255347 | 0 | 0 | 0 | 0 | 0 | 0 |
| ENSG00000279675 | 0 | 0 | 0 | 0 | 0 | 0 |
| ENSG00000148948 | 0 | 0 | 1 | 0 | 0 | 0 |
| ENSG00000206806 | 0 | 0 | 0 | 0 | 0 | 0 |
| ENSG00000254579 | 0 | 0 | 0 | 0 | 0 | 0 |
| ENSG00000252922 | 0 | 0 | 0 | 0 | 0 | 0 |
| ENSG00000255132 | 0 | 0 | 0 | 0 | 0 | 0 |
| ENSG00000255388 | 0 | 0 | 0 | 0 | 0 | 0 |
| ENSG00000242729 | 0 | 0 | 0 | 0 | 0 | 0 |
| ENSG00000255563 | 0 | 0 | 0 | 0 | 0 | 0 |
| ENSG00000255171 | 0 | 0 | 0 | 0 | 0 | 0 |
| ENSG00000254668 | 0 | 0 | 0 | 0 | 0 | 0 |

| | | | | | | |
|---|---|---|---|---|---|---|
| ENSG00000255279 | 0 | 0 | 0 | 0 | 0 | 0 |
| ENSG00000254822 | 0 | 0 | 0 | 0 | 0 | 0 |
| ENSG00000255300 | 0 | 0 | 0 | 0 | 0 | 0 |
| ENSG00000255109 | 0 | 0 | 0 | 0 | 0 | 0 |
| ENSG00000254914 | 0 | 0 | 0 | 0 | 0 | 0 |
| ENSG00000271028 | 0 | 0 | 0 | 0 | 0 | 0 |
| ENSG00000254725 | 0 | 0 | 0 | 0 | 0 | 0 |
| ENSG00000251557 | 0 | 0 | 0 | 0 | 0 | 0 |
| ENSG00000166181 | 2470 | 1455 | 1590 | 1058 | 1688 | 1507 |
| ENSG00000254907 | 5 | 4 | 2 | 2 | 3 | 2 |
| ENSG00000252652 | 0 | 1 | 0 | 0 | 0 | 0 |
| ENSG00000052841 | 1632 | 765 | 953 | 687 | 1070 | 909 |
| ENSG00000255340 | 0 | 0 | 1 | 0 | 0 | 0 |
| ENSG00000254577 | 0 | 1 | 0 | 0 | 0 | 0 |
| ENSG00000252355 | 0 | 0 | 1 | 0 | 0 | 0 |
| ENSG00000254463 | 0 | 0 | 0 | 0 | 2 | 1 |
| ENSG00000213697 | 0 | 0 | 0 | 0 | 0 | 0 |
| ENSG00000283217 | 0 | 0 | 0 | 0 | 0 | 0 |
| ENSG00000211568 | 0 | 0 | 0 | 0 | 0 | 0 |
| ENSG00000235661 | 0 | 0 | 0 | 0 | 0 | 0 |
| ENSG00000283341 | 19 | 12 | 19 | 11 | 12 | 2 |
| ENSG00000199077 | 0 | 0 | 0 | 0 | 0 | 0 |
| ENSG00000149084 | 275 | 194 | 215 | 163 | 234 | 203 |
| ENSG00000243721 | 0 | 0 | 0 | 0 | 0 | 0 |
| ENSG00000283393 | 0 | 0 | 0 | 0 | 0 | 0 |
| ENSG00000254512 | 0 | 0 | 0 | 0 | 0 | 0 |
| ENSG00000283483 | 0 | 0 | 0 | 0 | 0 | 0 |
| ENSG00000246250 | 2 | 0 | 3 | 0 | 1 | 3 |
| ENSG00000283375 | 0 | 0 | 0 | 0 | 0 | 0 |
| ENSG00000166199 | 322 | 167 | 210 | 185 | 217 | 216 |
| ENSG00000213693 | 33 | 9 | 21 | 10 | 22 | 29 |
| ENSG00000244926 | 9 | 5 | 2 | 1 | 6 | 0 |
| ENSG00000254409 | 3 | 2 | 0 | 1 | 0 | 3 |
| ENSG00000187479 | 2 | 0 | 1 | 2 | 3 | 3 |
| ENSG00000244953 | 0 | 0 | 0 | 0 | 0 | 0 |
| ENSG00000205126 | 0 | 0 | 0 | 0 | 0 | 0 |
| ENSG00000110455 | 582 | 242 | 411 | 219 | 267 | 300 |
| ENSG00000255165 | 1 | 1 | 3 | 0 | 3 | 3 |
| ENSG00000151348 | 1631 | 765 | 945 | 833 | 838 | 846 |
| ENSG00000052850 | 4 | 3 | 0 | 1 | 4 | 3 |
| ENSG00000255451 | 0 | 0 | 0 | 0 | 0 | 0 |
| ENSG00000255106 | 0 | 0 | 0 | 0 | 0 | 0 |
| ENSG00000085117 | 67 | 36 | 46 | 21 | 28 | 22 |
| ENSG00000254693 | 0 | 0 | 0 | 0 | 0 | 0 |
| ENSG00000255092 | 0 | 0 | 0 | 0 | 0 | 0 |
| ENSG00000244267 | 0 | 0 | 0 | 0 | 0 | 0 |
| ENSG00000275725 | 0 | 0 | 0 | 0 | 0 | 0 |
| ENSG00000255079 | 0 | 0 | 0 | 0 | 0 | 0 |
| ENSG00000255032 | 0 | 0 | 0 | 0 | 0 | 0 |
| ENSG00000157570 | 7 | 3 | 2 | 1 | 3 | 5 |

| | | | | | | |
|---|---|---|---|---|---|---|
| ENSG00000175274 | 3693 | 1663 | 2410 | 1677 | 1445 | 1726 |
| ENSG00000254654 | 0 | 0 | 0 | 0 | 0 | 0 |
| ENSG00000019485 | 147 | 87 | 121 | 111 | 92 | 121 |
| ENSG00000254664 | 0 | 0 | 0 | 0 | 0 | 0 |
| ENSG00000019505 | 4 | 1 | 3 | 1 | 2 | 6 |
| ENSG00000254497 | 0 | 0 | 0 | 0 | 0 | 0 |
| ENSG00000254427 | 12 | 6 | 2 | 2 | 4 | 9 |
| ENSG00000255267 | 0 | 0 | 0 | 0 | 0 | 0 |
| ENSG00000255041 | 0 | 0 | 0 | 0 | 0 | 0 |
| ENSG00000254651 | 0 | 0 | 0 | 0 | 0 | 0 |
| ENSG00000254746 | 0 | 0 | 0 | 0 | 0 | 0 |
| ENSG00000255519 | 0 | 0 | 0 | 0 | 0 | 0 |
| ENSG00000254514 | 0 | 0 | 0 | 0 | 0 | 0 |
| ENSG00000175264 | 106 | 41 | 54 | 21 | 9 | 19 |
| ENSG00000255091 | 1 | 0 | 1 | 0 | 0 | 0 |
| ENSG00000276639 | 0 | 0 | 0 | 0 | 0 | 0 |
| ENSG00000254519 | 0 | 0 | 0 | 0 | 0 | 0 |
| ENSG00000255426 | 0 | 0 | 0 | 0 | 0 | 0 |
| ENSG00000255226 | 0 | 1 | 0 | 0 | 0 | 0 |
| ENSG00000205106 | 8 | 7 | 7 | 4 | 3 | 6 |
| ENSG00000181830 | 6278 | 2702 | 3755 | 1520 | 1252 | 1644 |
| ENSG00000255447 | 15 | 1 | 4 | 2 | 4 | 2 |
| ENSG00000121671 | 2987 | 1252 | 1758 | 741 | 834 | 878 |
| ENSG00000121653 | 884 | 415 | 540 | 522 | 409 | 426 |
| ENSG00000255498 | 0 | 0 | 0 | 0 | 0 | 0 |
| ENSG00000234776 | 10 | 5 | 4 | 10 | 17 | 11 |
| ENSG00000121680 | 1305 | 604 | 792 | 914 | 656 | 718 |
| ENSG00000165905 | 181 | 100 | 130 | 95 | 86 | 101 |
| ENSG00000135365 | 797 | 383 | 462 | 266 | 464 | 455 |
| ENSG00000254653 | 0 | 0 | 0 | 0 | 0 | 0 |
| ENSG00000255314 | 0 | 1 | 0 | 1 | 0 | 3 |
| ENSG00000254871 | 0 | 0 | 0 | 0 | 1 | 0 |
| ENSG00000255269 | 0 | 0 | 0 | 0 | 0 | 0 |
| ENSG00000254639 | 1 | 0 | 0 | 0 | 0 | 0 |
| ENSG00000255007 | 0 | 0 | 2 | 0 | 0 | 0 |
| ENSG00000157613 | 14 | 3 | 10 | 6 | 2 | 5 |
| ENSG00000149091 | 3473 | 1577 | 1970 | 2057 | 1841 | 1820 |
| ENSG00000264102 | 0 | 0 | 2 | 0 | 0 | 0 |
| ENSG00000110492 | 2439 | 1133 | 1680 | 1260 | 862 | 1124 |
| ENSG00000180720 | 0 | 0 | 0 | 2 | 3 | 1 |
| ENSG00000110497 | 1493 | 658 | 866 | 788 | 787 | 815 |
| ENSG00000244313 | 59 | 42 | 31 | 55 | 43 | 39 |
| ENSG00000265014 | 0 | 0 | 0 | 0 | 0 | 0 |
| ENSG00000283497 | 0 | 0 | 0 | 0 | 0 | 0 |
| ENSG00000285658 | 0 | 2 | 2 | 0 | 0 | 0 |
| ENSG00000180423 | 172 | 64 | 115 | 75 | 80 | 99 |
| ENSG00000175224 | 3183 | 1522 | 1941 | 1639 | 1689 | 1788 |
| ENSG00000175220 | 3118 | 1491 | 1961 | 1525 | 1402 | 1530 |
| ENSG00000175213 | 683 | 272 | 390 | 442 | 311 | 403 |
| ENSG00000180210 | 0 | 1 | 1 | 0 | 0 | 0 |

| | | | | | | |
|---|---|---|---|---|---|---|
| ENSG00000175216 | 2488 | 1574 | 1293 | 732 | 2705 | 1563 |
| ENSG00000263540 | 0 | 0 | 0 | 0 | 0 | 0 |
| ENSG00000252427 | 0 | 0 | 0 | 0 | 0 | 0 |
| ENSG00000212135 | 2 | 1 | 0 | 2 | 0 | 0 |
| ENSG00000247675 | 4 | 4 | 5 | 5 | 9 | 1 |
| ENSG00000134569 | 28 | 8 | 14 | 7 | 11 | 11 |
| ENSG00000149179 | 1116 | 491 | 577 | 627 | 581 | 584 |
| ENSG00000271350 | 0 | 1 | 0 | 0 | 0 | 0 |
| ENSG00000255520 | 0 | 0 | 2 | 0 | 0 | 1 |
| ENSG00000149182 | 5361 | 2464 | 3288 | 2757 | 2540 | 2718 |
| ENSG00000270060 | 20 | 14 | 17 | 10 | 24 | 17 |
| ENSG00000165912 | 2443 | 1157 | 1395 | 1636 | 1469 | 1315 |
| ENSG00000275208 | 0 | 0 | 0 | 0 | 0 | 0 |
| ENSG00000243802 | 3 | 1 | 0 | 0 | 1 | 3 |
| ENSG00000134574 | 1693 | 804 | 991 | 1014 | 1003 | 1071 |
| ENSG00000256897 | 2 | 4 | 5 | 2 | 1 | 2 |
| ENSG00000134575 | 1367 | 651 | 905 | 660 | 623 | 651 |
| ENSG00000025434 | 345 | 151 | 166 | 213 | 163 | 160 |
| ENSG00000110514 | 4060 | 1837 | 2495 | 1730 | 2143 | 2193 |
| ENSG00000256746 | 1 | 0 | 0 | 1 | 1 | 1 |
| ENSG00000283338 | 0 | 3 | 0 | 1 | 0 | 0 |
| ENSG00000134571 | 0 | 0 | 0 | 0 | 0 | 0 |
| ENSG00000066336 | 0 | 0 | 0 | 0 | 0 | 0 |
| ENSG00000255197 | 0 | 1 | 3 | 0 | 0 | 0 |
| ENSG00000264583 | 0 | 0 | 0 | 0 | 0 | 0 |
| ENSG00000165915 | 1350 | 581 | 810 | 760 | 745 | 636 |
| ENSG00000165916 | 5870 | 3050 | 3470 | 3857 | 3711 | 3429 |
| ENSG00000165917 | 8 | 1 | 7 | 4 | 5 | 2 |
| ENSG00000149187 | 6490 | 2736 | 3723 | 2325 | 3080 | 3302 |
| ENSG00000270072 | 0 | 1 | 0 | 0 | 0 | 0 |
| ENSG00000265559 | 0 | 0 | 0 | 0 | 0 | 1 |
| ENSG00000213619 | 2884 | 1472 | 1739 | 1734 | 1699 | 1654 |
| ENSG00000110536 | 337 | 276 | 295 | 323 | 320 | 394 |
| ENSG00000123444 | 360 | 188 | 296 | 178 | 193 | 225 |
| ENSG00000200376 | 0 | 2 | 0 | 0 | 2 | 0 |
| ENSG00000231880 | 0 | 0 | 0 | 0 | 0 | 0 |
| ENSG00000196666 | 0 | 0 | 0 | 1 | 0 | 0 |
| ENSG00000172247 | 4 | 1 | 3 | 1 | 1 | 3 |
| ENSG00000109919 | 5300 | 2628 | 3281 | 3035 | 3018 | 3090 |
| ENSG00000165923 | 113 | 56 | 88 | 48 | 73 | 69 |
| ENSG00000109920 | 2272 | 1078 | 1359 | 535 | 1204 | 1107 |
| ENSG00000200090 | 7 | 8 | 8 | 4 | 4 | 4 |
| ENSG00000030066 | 1872 | 910 | 1044 | 552 | 1118 | 845 |
| ENSG00000271543 | 0 | 1 | 0 | 0 | 0 | 0 |
| ENSG00000252941 | 0 | 0 | 0 | 0 | 0 | 0 |
| ENSG00000252874 | 0 | 0 | 4 | 1 | 2 | 1 |
| ENSG00000271557 | 0 | 2 | 0 | 0 | 0 | 1 |
| ENSG00000254780 | 0 | 1 | 0 | 1 | 0 | 0 |
| ENSG00000149177 | 2069 | 895 | 1314 | 858 | 1292 | 1219 |
| ENSG00000254879 | 0 | 1 | 0 | 0 | 0 | 0 |

| | | | | | | |
|---|---|---|---|---|---|---|
| ENSG00000263693 | 0 | 0 | 0 | 0 | 0 | 0 |
| ENSG00000175619 | 0 | 0 | 0 | 0 | 0 | 0 |
| ENSG00000255514 | 0 | 0 | 0 | 0 | 0 | 0 |
| ENSG00000172208 | 0 | 0 | 0 | 0 | 0 | 0 |
| ENSG00000176567 | 0 | 0 | 0 | 0 | 0 | 0 |
| ENSG00000176555 | 0 | 0 | 0 | 0 | 0 | 0 |
| ENSG00000176547 | 0 | 0 | 0 | 0 | 0 | 0 |
| ENSG00000197161 | 0 | 0 | 0 | 0 | 0 | 0 |
| ENSG00000176540 | 0 | 0 | 0 | 0 | 0 | 0 |
| ENSG00000182565 | 0 | 0 | 0 | 0 | 0 | 0 |
| ENSG00000184789 | 0 | 0 | 0 | 0 | 0 | 0 |
| ENSG00000254925 | 0 | 0 | 0 | 0 | 0 | 0 |
| ENSG00000255215 | 0 | 0 | 0 | 0 | 0 | 0 |
| ENSG00000237388 | 0 | 0 | 0 | 0 | 0 | 0 |
| ENSG00000255113 | 0 | 0 | 0 | 0 | 0 | 0 |
| ENSG00000255304 | 0 | 0 | 0 | 0 | 0 | 0 |
| ENSG00000254832 | 0 | 0 | 0 | 0 | 0 | 0 |
| ENSG00000255534 | 0 | 0 | 0 | 0 | 0 | 0 |
| ENSG00000213607 | 0 | 0 | 0 | 0 | 0 | 0 |
| ENSG00000255297 | 0 | 0 | 0 | 0 | 0 | 0 |
| ENSG00000254674 | 0 | 0 | 0 | 0 | 0 | 0 |
| ENSG00000255053 | 0 | 0 | 0 | 0 | 0 | 0 |
| ENSG00000255551 | 0 | 0 | 0 | 0 | 0 | 0 |
| ENSG00000254728 | 0 | 0 | 0 | 0 | 0 | 0 |
| ENSG00000254920 | 0 | 0 | 0 | 0 | 0 | 0 |
| ENSG00000255527 | 0 | 0 | 0 | 0 | 0 | 0 |
| ENSG00000249910 | 0 | 0 | 0 | 0 | 0 | 0 |
| ENSG00000254517 | 0 | 0 | 0 | 0 | 0 | 0 |
| ENSG00000220948 | 0 | 0 | 0 | 0 | 0 | 0 |
| ENSG00000254764 | 0 | 0 | 0 | 0 | 0 | 0 |
| ENSG00000205044 | 0 | 0 | 0 | 0 | 0 | 0 |
| ENSG00000182053 | 0 | 0 | 0 | 0 | 0 | 0 |
| ENSG00000214891 | 0 | 0 | 0 | 0 | 0 | 0 |
| ENSG00000276637 | 0 | 0 | 0 | 0 | 0 | 0 |
| ENSG00000254801 | 0 | 0 | 0 | 0 | 0 | 0 |
| ENSG00000229361 | 0 | 0 | 0 | 0 | 0 | 0 |
| ENSG00000255452 | 0 | 0 | 0 | 0 | 0 | 0 |
| ENSG00000254412 | 0 | 0 | 0 | 0 | 0 | 0 |
| ENSG00000255338 | 0 | 0 | 0 | 0 | 0 | 0 |
| ENSG00000254993 | 0 | 0 | 0 | 0 | 0 | 0 |
| ENSG00000254567 | 0 | 0 | 0 | 0 | 0 | 0 |
| ENSG00000086205 | 0 | 0 | 0 | 0 | 0 | 0 |
| ENSG00000255532 | 0 | 0 | 0 | 0 | 0 | 0 |
| ENSG00000123447 | 0 | 0 | 0 | 0 | 0 | 0 |
| ENSG00000254496 | 0 | 0 | 0 | 0 | 0 | 0 |
| ENSG00000226268 | 0 | 0 | 1 | 0 | 2 | 0 |
| ENSG00000205035 | 6 | 1 | 0 | 0 | 0 | 0 |
| ENSG00000254487 | 0 | 0 | 0 | 0 | 0 | 0 |
| ENSG00000255268 | 0 | 0 | 0 | 0 | 0 | 0 |
| ENSG00000219061 | 0 | 0 | 0 | 0 | 0 | 0 |

| | | | | | | |
|---|---|---|---|---|---|---|
| ENSG00000265973 | 0 | 0 | 0 | 0 | 0 | 0 |
| ENSG00000254714 | 0 | 0 | 0 | 0 | 0 | 0 |
| ENSG00000255550 | 0 | 0 | 0 | 0 | 0 | 0 |
| ENSG00000255214 | 0 | 0 | 0 | 0 | 0 | 0 |
| ENSG00000255190 | 0 | 0 | 0 | 0 | 0 | 0 |
| ENSG00000254800 | 0 | 0 | 0 | 0 | 0 | 0 |
| ENSG00000255111 | 0 | 0 | 0 | 0 | 0 | 0 |
| ENSG00000254408 | 0 | 0 | 0 | 0 | 0 | 0 |
| ENSG00000254472 | 0 | 0 | 0 | 0 | 0 | 0 |
| ENSG00000254637 | 0 | 0 | 0 | 0 | 0 | 0 |
| ENSG00000254940 | 0 | 0 | 0 | 0 | 0 | 0 |
| ENSG00000254624 | 0 | 0 | 0 | 0 | 0 | 0 |
| ENSG00000258817 | 0 | 0 | 0 | 0 | 0 | 0 |
| ENSG00000221954 | 0 | 0 | 0 | 0 | 0 | 0 |
| ENSG00000283605 | 0 | 0 | 0 | 0 | 0 | 0 |
| ENSG00000260811 | 0 | 0 | 0 | 0 | 0 | 0 |
| ENSG00000260595 | 0 | 0 | 0 | 0 | 0 | 0 |
| ENSG00000255199 | 0 | 0 | 0 | 0 | 0 | 0 |
| ENSG00000255442 | 0 | 1 | 0 | 0 | 0 | 0 |
| ENSG00000236919 | 0 | 0 | 0 | 0 | 0 | 0 |
| ENSG00000255500 | 0 | 0 | 0 | 1 | 0 | 0 |
| ENSG00000255042 | 0 | 1 | 0 | 0 | 0 | 1 |
| ENSG00000255001 | 0 | 0 | 0 | 0 | 0 | 0 |
| ENSG00000254518 | 0 | 0 | 0 | 0 | 0 | 0 |
| ENSG00000254840 | 0 | 0 | 0 | 0 | 0 | 0 |
| ENSG00000214883 | 0 | 0 | 0 | 0 | 0 | 0 |
| ENSG00000237610 | 0 | 0 | 0 | 0 | 0 | 0 |
| ENSG00000185926 | 0 | 0 | 0 | 0 | 0 | 0 |
| ENSG00000255312 | 0 | 0 | 0 | 0 | 0 | 0 |
| ENSG00000255560 | 0 | 0 | 0 | 0 | 0 | 0 |
| ENSG00000254769 | 0 | 0 | 0 | 0 | 0 | 0 |
| ENSG00000254464 | 0 | 0 | 0 | 0 | 0 | 0 |
| ENSG00000227547 | 0 | 0 | 0 | 0 | 0 | 0 |
| ENSG00000225997 | 0 | 0 | 0 | 0 | 0 | 0 |
| ENSG00000255349 | 0 | 0 | 0 | 0 | 0 | 0 |
| ENSG00000221840 | 0 | 0 | 0 | 0 | 0 | 0 |
| ENSG00000255078 | 0 | 0 | 0 | 0 | 0 | 0 |
| ENSG00000150244 | 0 | 0 | 0 | 0 | 0 | 0 |
| ENSG00000255543 | 0 | 0 | 0 | 0 | 0 | 0 |
| ENSG00000255283 | 0 | 0 | 0 | 0 | 0 | 0 |
| ENSG00000255110 | 0 | 0 | 0 | 0 | 0 | 0 |
| ENSG00000166007 | 0 | 0 | 0 | 0 | 0 | 0 |
| ENSG00000254828 | 0 | 0 | 0 | 0 | 0 | 0 |
| ENSG00000254663 | 0 | 0 | 0 | 0 | 0 | 0 |
| ENSG00000254723 | 0 | 0 | 0 | 0 | 0 | 0 |
| ENSG00000181961 | 0 | 0 | 0 | 0 | 0 | 0 |
| ENSG00000181958 | 0 | 0 | 0 | 0 | 0 | 0 |
| ENSG00000254891 | 0 | 0 | 0 | 0 | 0 | 0 |
| ENSG00000255077 | 0 | 0 | 0 | 0 | 0 | 0 |
| ENSG00000255493 | 0 | 0 | 0 | 0 | 0 | 0 |

| | | | | | | |
|---|---|---|---|---|---|---|
| ENSG00000255196 | 0 | 0 | 0 | 0 | 0 | 0 |
| ENSG00000181950 | 0 | 0 | 0 | 0 | 0 | 0 |
| ENSG00000255294 | 0 | 0 | 0 | 0 | 0 | 0 |
| ENSG00000181943 | 0 | 0 | 0 | 0 | 0 | 0 |
| ENSG00000254576 | 0 | 0 | 0 | 0 | 0 | 0 |
| ENSG00000233301 | 0 | 0 | 0 | 0 | 0 | 0 |
| ENSG00000181939 | 0 | 0 | 0 | 0 | 0 | 0 |
| ENSG00000279514 | 0 | 0 | 0 | 0 | 0 | 0 |
| ENSG00000172188 | 0 | 0 | 0 | 0 | 0 | 0 |
| ENSG00000181927 | 0 | 0 | 0 | 0 | 0 | 0 |
| ENSG00000174982 | 0 | 0 | 0 | 0 | 0 | 0 |
| ENSG00000181903 | 0 | 0 | 0 | 0 | 0 | 0 |
| ENSG00000254841 | 0 | 0 | 0 | 0 | 0 | 0 |
| ENSG00000188439 | 0 | 0 | 0 | 0 | 0 | 0 |
| ENSG00000254804 | 0 | 0 | 0 | 0 | 0 | 0 |
| ENSG00000254457 | 0 | 0 | 0 | 0 | 0 | 0 |
| ENSG00000186886 | 0 | 0 | 0 | 0 | 0 | 0 |
| ENSG00000181837 | 0 | 0 | 0 | 0 | 0 | 0 |
| ENSG00000279761 | 0 | 0 | 0 | 0 | 0 | 0 |
| ENSG00000255019 | 0 | 0 | 0 | 0 | 0 | 0 |
| ENSG00000186113 | 0 | 0 | 0 | 0 | 0 | 0 |
| ENSG00000279395 | 0 | 0 | 0 | 0 | 0 | 0 |
| ENSG00000186119 | 0 | 0 | 0 | 0 | 0 | 0 |
| ENSG00000205030 | 0 | 0 | 0 | 0 | 0 | 0 |
| ENSG00000205029 | 0 | 0 | 0 | 0 | 0 | 0 |
| ENSG00000186124 | 0 | 0 | 0 | 0 | 0 | 0 |
| ENSG00000255204 | 0 | 0 | 0 | 0 | 0 | 0 |
| ENSG00000255499 | 0 | 0 | 0 | 0 | 0 | 0 |
| ENSG00000254547 | 0 | 0 | 0 | 0 | 0 | 0 |
| ENSG00000124900 | 0 | 0 | 0 | 0 | 0 | 0 |
| ENSG00000227806 | 0 | 0 | 0 | 0 | 0 | 0 |
| ENSG00000187612 | 0 | 0 | 0 | 0 | 0 | 0 |
| ENSG00000167825 | 0 | 0 | 0 | 0 | 0 | 0 |
| ENSG00000255416 | 0 | 0 | 0 | 0 | 0 | 0 |
| ENSG00000254807 | 0 | 0 | 0 | 0 | 0 | 0 |
| ENSG00000174970 | 0 | 0 | 0 | 0 | 0 | 0 |
| ENSG00000214880 | 0 | 0 | 0 | 0 | 0 | 0 |
| ENSG00000149133 | 0 | 0 | 0 | 0 | 0 | 0 |
| ENSG00000182365 | 0 | 0 | 0 | 0 | 0 | 0 |
| ENSG00000181785 | 0 | 1 | 0 | 0 | 0 | 0 |
| ENSG00000184741 | 0 | 0 | 0 | 0 | 0 | 0 |
| ENSG00000181780 | 0 | 0 | 0 | 0 | 0 | 0 |
| ENSG00000225538 | 0 | 0 | 0 | 0 | 0 | 0 |
| ENSG00000172154 | 0 | 0 | 0 | 0 | 0 | 0 |
| ENSG00000254524 | 0 | 0 | 0 | 0 | 0 | 0 |
| ENSG00000181767 | 0 | 0 | 0 | 0 | 1 | 0 |
| ENSG00000255140 | 0 | 0 | 0 | 0 | 0 | 0 |
| ENSG00000181761 | 0 | 0 | 0 | 0 | 0 | 0 |
| ENSG00000254848 | 0 | 0 | 0 | 0 | 0 | 0 |
| ENSG00000167822 | 0 | 0 | 0 | 0 | 0 | 0 |

| | | | | | | |
|---|---|---|---|---|---|---|
| ENSG00000254947 | 0 | 0 | 0 | 0 | 0 | 0 |
| ENSG00000181752 | 0 | 0 | 0 | 0 | 0 | 0 |
| ENSG00000255217 | 0 | 0 | 0 | 0 | 0 | 0 |
| ENSG00000174957 | 0 | 0 | 0 | 0 | 0 | 0 |
| ENSG00000230213 | 0 | 0 | 0 | 0 | 0 | 0 |
| ENSG00000213604 | 0 | 0 | 0 | 0 | 0 | 0 |
| ENSG00000254658 | 0 | 0 | 0 | 0 | 0 | 0 |
| ENSG00000181718 | 0 | 0 | 0 | 0 | 0 | 0 |
| ENSG00000172489 | 0 | 0 | 0 | 0 | 0 | 0 |
| ENSG00000181698 | 0 | 0 | 0 | 0 | 0 | 0 |
| ENSG00000181693 | 0 | 0 | 0 | 0 | 0 | 0 |
| ENSG00000255461 | 0 | 0 | 0 | 0 | 0 | 0 |
| ENSG00000280314 | 0 | 0 | 0 | 0 | 0 | 0 |
| ENSG00000254722 | 0 | 0 | 0 | 0 | 0 | 0 |
| ENSG00000255134 | 0 | 0 | 0 | 0 | 0 | 0 |
| ENSG00000150261 | 0 | 0 | 0 | 0 | 0 | 0 |
| ENSG00000172487 | 0 | 0 | 0 | 0 | 0 | 0 |
| ENSG00000242208 | 0 | 0 | 0 | 0 | 0 | 0 |
| ENSG00000172199 | 0 | 0 | 0 | 0 | 0 | 0 |
| ENSG00000254903 | 0 | 0 | 0 | 0 | 0 | 0 |
| ENSG00000255466 | 0 | 0 | 0 | 0 | 0 | 0 |
| ENSG00000272987 | 0 | 0 | 0 | 0 | 0 | 0 |
| ENSG00000279961 | 0 | 0 | 0 | 0 | 0 | 0 |
| ENSG00000230416 | 0 | 0 | 0 | 0 | 0 | 0 |
| ENSG00000284732 | 0 | 0 | 0 | 0 | 0 | 0 |
| ENSG00000150269 | 0 | 0 | 0 | 0 | 0 | 0 |
| ENSG00000174937 | 0 | 0 | 0 | 0 | 0 | 0 |
| ENSG00000254752 | 0 | 0 | 0 | 0 | 0 | 0 |
| ENSG00000181371 | 0 | 0 | 0 | 0 | 0 | 0 |
| ENSG00000254411 | 0 | 0 | 0 | 0 | 0 | 0 |
| ENSG00000254490 | 0 | 0 | 0 | 0 | 0 | 0 |
| ENSG00000254660 | 0 | 0 | 0 | 0 | 0 | 0 |
| ENSG00000254603 | 0 | 0 | 0 | 0 | 0 | 0 |
| ENSG00000185701 | 0 | 0 | 1 | 0 | 0 | 0 |
| ENSG00000255223 | 2 | 0 | 1 | 1 | 1 | 0 |
| ENSG00000254834 | 0 | 0 | 0 | 0 | 0 | 0 |
| ENSG00000166693 | 0 | 0 | 0 | 0 | 0 | 0 |
| ENSG00000255012 | 0 | 0 | 0 | 0 | 0 | 0 |
| ENSG00000255172 | 0 | 0 | 0 | 0 | 0 | 0 |
| ENSG00000197866 | 0 | 0 | 0 | 0 | 0 | 0 |
| ENSG00000254795 | 0 | 0 | 0 | 0 | 0 | 0 |
| ENSG00000172464 | 0 | 0 | 0 | 0 | 0 | 0 |
| ENSG00000172459 | 0 | 0 | 0 | 0 | 0 | 0 |
| ENSG00000232511 | 0 | 0 | 0 | 0 | 0 | 0 |
| ENSG00000238041 | 2 | 0 | 0 | 0 | 0 | 0 |
| ENSG00000174914 | 0 | 0 | 0 | 0 | 0 | 0 |
| ENSG00000254953 | 0 | 0 | 0 | 0 | 0 | 0 |
| ENSG00000181325 | 0 | 0 | 0 | 0 | 0 | 0 |
| ENSG00000172457 | 0 | 0 | 0 | 0 | 0 | 0 |
| ENSG00000274770 | 0 | 0 | 0 | 0 | 0 | 0 |

| | | | | | | |
|---|---|---|---|---|---|---|
| ENSG00000254825 | 0 | 0 | 0 | 0 | 0 | 0 |
| ENSG00000181296 | 0 | 0 | 0 | 0 | 0 | 0 |
| ENSG00000255485 | 0 | 0 | 0 | 0 | 0 | 0 |
| ENSG00000205025 | 0 | 0 | 0 | 0 | 0 | 0 |
| ENSG00000241356 | 0 | 0 | 0 | 0 | 0 | 0 |
| ENSG00000255433 | 0 | 0 | 0 | 0 | 0 | 0 |
| ENSG00000274150 | 0 | 0 | 0 | 0 | 0 | 0 |
| ENSG00000181282 | 0 | 0 | 0 | 0 | 0 | 0 |
| ENSG00000181273 | 0 | 0 | 0 | 0 | 0 | 0 |
| ENSG00000255083 | 0 | 0 | 0 | 0 | 0 | 0 |
| ENSG00000273228 | 0 | 0 | 0 | 0 | 0 | 0 |
| ENSG00000283307 | 0 | 0 | 0 | 0 | 0 | 0 |
| ENSG00000272676 | 0 | 0 | 0 | 0 | 0 | 0 |
| ENSG00000273255 | 0 | 0 | 0 | 0 | 0 | 0 |
| ENSG00000183908 | 0 | 0 | 0 | 0 | 0 | 0 |
| ENSG00000134817 | 0 | 0 | 0 | 0 | 0 | 0 |
| ENSG00000149115 | 3732 | 1622 | 2273 | 1852 | 1890 | 1826 |
| ENSG00000254662 | 0 | 0 | 1 | 0 | 0 | 1 |
| ENSG00000149136 | 2929 | 1532 | 1847 | 1757 | 1903 | 1873 |
| ENSG00000109991 | 0 | 0 | 0 | 0 | 0 | 0 |
| ENSG00000156575 | 0 | 0 | 0 | 0 | 0 | 0 |
| ENSG00000186652 | 0 | 0 | 0 | 0 | 0 | 0 |
| ENSG00000254979 | 0 | 0 | 0 | 0 | 0 | 0 |
| ENSG00000134802 | 0 | 0 | 1 | 1 | 0 | 0 |
| ENSG00000252070 | 0 | 0 | 0 | 0 | 0 | 0 |
| ENSG00000222998 | 0 | 0 | 0 | 0 | 0 | 0 |
| ENSG00000186907 | 7 | 1 | 4 | 3 | 5 | 1 |
| ENSG00000255301 | 0 | 0 | 0 | 0 | 0 | 0 |
| ENSG00000149150 | 36 | 25 | 22 | 15 | 11 | 12 |
| ENSG00000265566 | 0 | 1 | 1 | 2 | 2 | 0 |
| ENSG00000134809 | 809 | 356 | 425 | 589 | 409 | 494 |
| ENSG00000214872 | 0 | 0 | 0 | 0 | 0 | 0 |
| ENSG00000156587 | 140 | 68 | 90 | 75 | 52 | 73 |
| ENSG00000240371 | 4 | 1 | 0 | 0 | 1 | 1 |
| ENSG00000149131 | 8 | 3 | 2 | 1 | 1 | 0 |
| ENSG00000254602 | 0 | 0 | 0 | 0 | 0 | 0 |
| ENSG00000208009 | 0 | 0 | 0 | 0 | 0 | 0 |
| ENSG00000166793 | 0 | 0 | 0 | 0 | 0 | 0 |
| ENSG00000172409 | 332 | 151 | 201 | 199 | 204 | 180 |
| ENSG00000156599 | 2702 | 1324 | 1624 | 1437 | 1469 | 1661 |
| ENSG00000156603 | 436 | 202 | 238 | 248 | 209 | 239 |
| ENSG00000213593 | 1746 | 845 | 1022 | 1046 | 941 | 998 |
| ENSG00000254462 | 0 | 0 | 0 | 0 | 0 | 0 |
| ENSG00000213592 | 0 | 2 | 2 | 0 | 0 | 0 |
| ENSG00000211450 | 1482 | 906 | 1109 | 1135 | 977 | 1008 |
| ENSG00000254732 | 6 | 3 | 4 | 0 | 3 | 1 |
| ENSG00000233436 | 0 | 4 | 1 | 0 | 0 | 1 |
| ENSG00000198561 | 10923 | 4990 | 6953 | 3616 | 5120 | 5358 |
| ENSG00000255303 | 2 | 1 | 1 | 1 | 3 | 3 |
| ENSG00000255266 | 0 | 0 | 0 | 0 | 0 | 0 |

| | | | | | | |
|---|---|---|---|---|---|---|
| ENSG00000180714 | 0 | 0 | 0 | 0 | 0 | 0 |
| ENSG00000254749 | 0 | 0 | 0 | 0 | 0 | 0 |
| ENSG00000254601 | 0 | 0 | 0 | 0 | 0 | 0 |
| ENSG00000186509 | 0 | 0 | 0 | 0 | 0 | 0 |
| ENSG00000200817 | 0 | 0 | 0 | 0 | 0 | 0 |
| ENSG00000279051 | 0 | 0 | 0 | 0 | 0 | 0 |
| ENSG00000236541 | 0 | 0 | 0 | 0 | 0 | 0 |
| ENSG00000255146 | 0 | 0 | 0 | 0 | 0 | 0 |
| ENSG00000254853 | 0 | 0 | 0 | 0 | 0 | 0 |
| ENSG00000255131 | 0 | 0 | 0 | 0 | 0 | 0 |
| ENSG00000272685 | 0 | 0 | 0 | 0 | 0 | 0 |
| ENSG00000172377 | 0 | 0 | 0 | 0 | 0 | 0 |
| ENSG00000186508 | 0 | 0 | 0 | 0 | 0 | 0 |
| ENSG00000272569 | 0 | 0 | 0 | 0 | 0 | 0 |
| ENSG00000186513 | 0 | 0 | 0 | 0 | 0 | 0 |
| ENSG00000197887 | 0 | 0 | 0 | 0 | 0 | 0 |
| ENSG00000280204 | 0 | 0 | 0 | 0 | 0 | 0 |
| ENSG00000180475 | 0 | 0 | 0 | 0 | 0 | 0 |
| ENSG00000254589 | 0 | 0 | 0 | 0 | 0 | 0 |
| ENSG00000172772 | 0 | 0 | 0 | 0 | 0 | 0 |
| ENSG00000272900 | 0 | 0 | 0 | 0 | 0 | 0 |
| ENSG00000255218 | 0 | 0 | 0 | 0 | 0 | 0 |
| ENSG00000255431 | 0 | 0 | 0 | 0 | 0 | 0 |
| ENSG00000255096 | 0 | 0 | 0 | 0 | 0 | 0 |
| ENSG00000255333 | 0 | 0 | 0 | 0 | 0 | 0 |
| ENSG00000197786 | 0 | 0 | 0 | 0 | 0 | 0 |
| ENSG00000233499 | 0 | 0 | 0 | 0 | 0 | 0 |
| ENSG00000273408 | 0 | 0 | 0 | 0 | 0 | 0 |
| ENSG00000172769 | 0 | 0 | 0 | 0 | 0 | 0 |
| ENSG00000172365 | 0 | 0 | 0 | 0 | 0 | 0 |
| ENSG00000172362 | 0 | 0 | 0 | 0 | 0 | 0 |
| ENSG00000285895 | 0 | 0 | 0 | 0 | 0 | 0 |
| ENSG00000255299 | 0 | 0 | 0 | 0 | 0 | 0 |
| ENSG00000198283 | 0 | 0 | 0 | 0 | 0 | 0 |
| ENSG00000110031 | 105 | 40 | 50 | 46 | 50 | 54 |
| ENSG00000186660 | 2186 | 949 | 1276 | 926 | 1097 | 1173 |
| ENSG00000255073 | 0 | 0 | 0 | 0 | 0 | 0 |
| ENSG00000269570 | 8 | 1 | 9 | 0 | 3 | 1 |
| ENSG00000242689 | 8 | 0 | 3 | 2 | 4 | 3 |
| ENSG00000280010 | 0 | 0 | 1 | 1 | 3 | 0 |
| ENSG00000149124 | 0 | 0 | 0 | 0 | 0 | 0 |
| ENSG00000197254 | 0 | 0 | 0 | 0 | 0 | 0 |
| ENSG00000254926 | 0 | 0 | 0 | 0 | 0 | 0 |
| ENSG00000255265 | 0 | 0 | 0 | 0 | 0 | 0 |
| ENSG00000156689 | 0 | 0 | 0 | 0 | 0 | 0 |
| ENSG00000254717 | 0 | 0 | 0 | 0 | 0 | 0 |
| ENSG00000166840 | 0 | 0 | 0 | 0 | 0 | 0 |
| ENSG00000255523 | 0 | 0 | 0 | 0 | 0 | 0 |
| ENSG00000255240 | 0 | 0 | 0 | 0 | 0 | 0 |
| ENSG00000254877 | 0 | 0 | 0 | 0 | 0 | 0 |

| | | | | | | |
|---|---|---|---|---|---|---|
| ENSG00000255189 | 0 | 0 | 0 | 0 | 0 | 0 |
| ENSG00000254786 | 0 | 0 | 0 | 0 | 0 | 0 |
| ENSG00000254399 | 0 | 0 | 0 | 0 | 0 | 0 |
| ENSG00000255151 | 0 | 0 | 0 | 0 | 2 | 0 |
| ENSG00000189057 | 136 | 74 | 49 | 8 | 73 | 51 |
| ENSG00000245571 | 153 | 89 | 113 | 75 | 97 | 86 |
| ENSG00000255381 | 0 | 0 | 1 | 0 | 0 | 2 |
| ENSG00000166801 | 1045 | 499 | 682 | 253 | 460 | 415 |
| ENSG00000110042 | 377 | 155 | 233 | 211 | 211 | 220 |
| ENSG00000197629 | 0 | 2 | 0 | 0 | 0 | 0 |
| ENSG00000263999 | 0 | 0 | 0 | 0 | 0 | 0 |
| ENSG00000225802 | 0 | 0 | 0 | 0 | 0 | 0 |
| ENSG00000254799 | 0 | 0 | 0 | 0 | 0 | 0 |
| ENSG00000214797 | 0 | 0 | 0 | 0 | 0 | 0 |
| ENSG00000254704 | 0 | 0 | 0 | 0 | 0 | 0 |
| ENSG00000263944 | 0 | 0 | 0 | 0 | 0 | 0 |
| ENSG00000235678 | 0 | 0 | 0 | 0 | 0 | 0 |
| ENSG00000255386 | 0 | 0 | 0 | 0 | 0 | 0 |
| ENSG00000176495 | 0 | 0 | 0 | 0 | 0 | 0 |
| ENSG00000198261 | 0 | 0 | 0 | 0 | 0 | 0 |
| ENSG00000172324 | 0 | 0 | 0 | 0 | 0 | 0 |
| ENSG00000172320 | 0 | 0 | 0 | 0 | 0 | 0 |
| ENSG00000166884 | 0 | 0 | 0 | 0 | 0 | 0 |
| ENSG00000254466 | 0 | 0 | 0 | 0 | 0 | 0 |
| ENSG00000204989 | 0 | 0 | 0 | 0 | 0 | 0 |
| ENSG00000176200 | 0 | 0 | 0 | 0 | 0 | 0 |
| ENSG00000172742 | 0 | 0 | 0 | 0 | 0 | 0 |
| ENSG00000254770 | 0 | 0 | 0 | 0 | 0 | 0 |
| ENSG00000255008 | 0 | 0 | 0 | 0 | 0 | 1 |
| ENSG00000110048 | 2011 | 974 | 1241 | 801 | 1044 | 1066 |
| ENSG00000264559 | 0 | 1 | 0 | 0 | 0 | 0 |
| ENSG00000255139 | 8 | 5 | 4 | 3 | 4 | 5 |
| ENSG00000166889 | 885 | 493 | 601 | 401 | 515 | 514 |
| ENSG00000255355 | 0 | 0 | 2 | 0 | 2 | 1 |
| ENSG00000223223 | 0 | 0 | 0 | 0 | 0 | 0 |
| ENSG00000172289 | 0 | 0 | 0 | 0 | 0 | 0 |
| ENSG00000166900 | 1914 | 935 | 1227 | 971 | 1248 | 1201 |
| ENSG00000254403 | 0 | 0 | 0 | 0 | 0 | 0 |
| ENSG00000254743 | 0 | 0 | 0 | 0 | 0 | 0 |
| ENSG00000255538 | 0 | 0 | 1 | 0 | 0 | 0 |
| ENSG00000254477 | 2 | 1 | 1 | 4 | 2 | 2 |
| ENSG00000234964 | 0 | 0 | 0 | 0 | 0 | 0 |
| ENSG00000166902 | 1100 | 510 | 623 | 693 | 701 | 689 |
| ENSG00000134812 | 0 | 0 | 0 | 0 | 0 | 0 |
| ENSG00000134827 | 0 | 0 | 0 | 0 | 0 | 0 |
| ENSG00000285231 | 2 | 1 | 0 | 0 | 1 | 0 |
| ENSG00000270929 | 0 | 0 | 0 | 0 | 0 | 0 |
| ENSG00000284873 | 0 | 0 | 0 | 0 | 0 | 0 |
| ENSG00000214788 | 0 | 0 | 0 | 0 | 0 | 0 |
| ENSG00000285010 | 0 | 0 | 0 | 0 | 0 | 0 |

| | | | | | | |
|---|---|---|---|---|---|---|
| ENSG00000255393 | 0 | 0 | 0 | 0 | 0 | 0 |
| ENSG00000149507 | 0 | 0 | 0 | 0 | 0 | 0 |
| ENSG00000149516 | 0 | 0 | 0 | 0 | 0 | 0 |
| ENSG00000255331 | 0 | 0 | 0 | 0 | 0 | 0 |
| ENSG00000149534 | 0 | 0 | 0 | 0 | 0 | 0 |
| ENSG00000254952 | 0 | 0 | 0 | 0 | 0 | 0 |
| ENSG00000110077 | 0 | 0 | 0 | 0 | 0 | 0 |
| ENSG00000110079 | 0 | 0 | 0 | 0 | 0 | 0 |
| ENSG00000214787 | 0 | 0 | 0 | 0 | 0 | 0 |
| ENSG00000275344 | 0 | 0 | 0 | 0 | 0 | 0 |
| ENSG00000166926 | 0 | 0 | 0 | 0 | 0 | 0 |
| ENSG00000166927 | 0 | 0 | 0 | 0 | 0 | 0 |
| ENSG00000166928 | 0 | 0 | 0 | 0 | 0 | 0 |
| ENSG00000166930 | 0 | 0 | 0 | 0 | 0 | 0 |
| ENSG00000156738 | 0 | 0 | 0 | 0 | 0 | 0 |
| ENSG00000071203 | 0 | 0 | 0 | 0 | 0 | 0 |
| ENSG00000204979 | 0 | 0 | 0 | 0 | 0 | 0 |
| ENSG00000283601 | 0 | 0 | 0 | 0 | 0 | 0 |
| ENSG00000181995 | 0 | 0 | 0 | 0 | 0 | 0 |
| ENSG00000166959 | 0 | 0 | 0 | 0 | 0 | 0 |
| ENSG00000214782 | 0 | 0 | 0 | 0 | 0 | 0 |
| ENSG00000166961 | 0 | 0 | 0 | 0 | 0 | 0 |
| ENSG00000172689 | 0 | 0 | 0 | 0 | 0 | 0 |
| ENSG00000255845 | 0 | 0 | 0 | 0 | 0 | 0 |
| ENSG00000255959 | 0 | 1 | 0 | 1 | 3 | 0 |
| ENSG00000256813 | 0 | 0 | 0 | 0 | 0 | 0 |
| ENSG00000110104 | 599 | 368 | 467 | 581 | 476 | 483 |
| ENSG00000183134 | 0 | 0 | 2 | 0 | 0 | 1 |
| ENSG00000149506 | 0 | 0 | 0 | 0 | 0 | 0 |
| ENSG00000110107 | 4255 | 2134 | 2727 | 2629 | 2350 | 2752 |
| ENSG00000257052 | 8 | 7 | 3 | 4 | 6 | 4 |
| ENSG00000256944 | 0 | 1 | 0 | 1 | 0 | 0 |
| ENSG00000110108 | 2492 | 1171 | 1459 | 1492 | 1299 | 1489 |
| ENSG00000256196 | 6 | 1 | 4 | 2 | 4 | 8 |
| ENSG00000006118 | 6404 | 3077 | 4002 | 3978 | 3651 | 3534 |
| ENSG00000110446 | 20 | 7 | 7 | 10 | 8 | 5 |
| ENSG00000013725 | 2 | 3 | 1 | 5 | 2 | 1 |
| ENSG00000207153 | 0 | 0 | 0 | 0 | 0 | 0 |
| ENSG00000256733 | 0 | 0 | 1 | 0 | 0 | 0 |
| ENSG00000110448 | 34 | 10 | 32 | 6 | 9 | 9 |
| ENSG00000167987 | 1796 | 810 | 1090 | 825 | 792 | 821 |
| ENSG00000279549 | 10 | 2 | 4 | 5 | 3 | 4 |
| ENSG00000229859 | 0 | 0 | 0 | 0 | 0 | 0 |
| ENSG00000229183 | 0 | 0 | 0 | 0 | 0 | 0 |
| ENSG00000256220 | 0 | 0 | 0 | 0 | 0 | 0 |
| ENSG00000256713 | 2 | 0 | 0 | 0 | 0 | 0 |
| ENSG00000167992 | 12 | 8 | 6 | 8 | 10 | 5 |
| ENSG00000167986 | 11791 | 5645 | 7010 | 5884 | 6392 | 6901 |
| ENSG00000149476 | 2033 | 976 | 1316 | 1289 | 1028 | 1291 |
| ENSG00000162144 | 911 | 520 | 638 | 600 | 532 | 605 |

| | | | | | | |
|---|---|---|---|---|---|---|
| ENSG00000279246 | 1 | 0 | 1 | 1 | 1 | 1 |
| ENSG00000149483 | 831 | 360 | 519 | 478 | 425 | 513 |
| ENSG00000187049 | 545 | 269 | 324 | 314 | 279 | 314 |
| ENSG00000149532 | 2602 | 1229 | 1652 | 1216 | 1325 | 1543 |
| ENSG00000279632 | 0 | 2 | 0 | 0 | 0 | 1 |
| ENSG00000256591 | 7 | 4 | 9 | 3 | 1 | 9 |
| ENSG00000167985 | 312 | 170 | 240 | 243 | 204 | 184 |
| ENSG00000240823 | 0 | 1 | 3 | 2 | 0 | 1 |
| ENSG00000162148 | 32 | 7 | 23 | 17 | 15 | 16 |
| ENSG00000255931 | 0 | 0 | 2 | 0 | 0 | 0 |
| ENSG00000266006 | 0 | 0 | 0 | 0 | 0 | 0 |
| ENSG00000204950 | 426 | 207 | 258 | 221 | 171 | 185 |
| ENSG00000011347 | 3235 | 1424 | 1979 | 1402 | 1488 | 1514 |
| ENSG00000279878 | 14 | 13 | 22 | 6 | 8 | 11 |
| ENSG00000256443 | 42 | 12 | 27 | 9 | 10 | 13 |
| ENSG00000250230 | 7 | 3 | 9 | 3 | 0 | 8 |
| ENSG00000243742 | 43 | 18 | 21 | 21 | 19 | 24 |
| ENSG00000255947 | 0 | 0 | 0 | 0 | 0 | 0 |
| ENSG00000134780 | 476 | 227 | 291 | 201 | 191 | 262 |
| ENSG00000124915 | 0 | 1 | 0 | 0 | 0 | 0 |
| ENSG00000124920 | 44 | 22 | 35 | 15 | 26 | 16 |
| ENSG00000134825 | 1246 | 752 | 956 | 1049 | 945 | 863 |
| ENSG00000284108 | 0 | 0 | 0 | 0 | 0 | 0 |
| ENSG00000168496 | 1318 | 801 | 903 | 770 | 780 | 860 |
| ENSG00000134824 | 358 | 194 | 221 | 221 | 217 | 234 |
| ENSG00000149485 | 2498 | 1221 | 1608 | 1219 | 1572 | 1683 |
| ENSG00000284416 | 0 | 0 | 0 | 0 | 0 | 0 |
| ENSG00000221968 | 714 | 362 | 524 | 493 | 547 | 452 |
| ENSG00000277892 | 0 | 0 | 0 | 0 | 0 | 0 |
| ENSG00000167994 | 138 | 58 | 65 | 71 | 68 | 67 |
| ENSG00000200898 | 0 | 0 | 0 | 0 | 0 | 0 |
| ENSG00000167995 | 30 | 13 | 27 | 13 | 8 | 9 |
| ENSG00000167996 | 7710 | 4401 | 5644 | 7530 | 6852 | 7163 |
| ENSG00000279491 | 8 | 11 | 14 | 17 | 14 | 10 |
| ENSG00000244176 | 0 | 3 | 1 | 4 | 5 | 0 |
| ENSG00000254424 | 0 | 0 | 0 | 0 | 0 | 0 |
| ENSG00000255553 | 0 | 0 | 0 | 0 | 0 | 0 |
| ENSG00000285656 | 0 | 0 | 0 | 0 | 0 | 0 |
| ENSG00000149503 | 821 | 413 | 573 | 466 | 547 | 598 |
| ENSG00000236607 | 0 | 0 | 0 | 0 | 0 | 0 |
| ENSG00000168515 | 0 | 0 | 0 | 0 | 0 | 0 |
| ENSG00000124939 | 0 | 0 | 0 | 0 | 0 | 0 |
| ENSG00000254404 | 0 | 0 | 0 | 0 | 0 | 0 |
| ENSG00000124935 | 0 | 0 | 0 | 0 | 0 | 0 |
| ENSG00000110484 | 0 | 0 | 1 | 0 | 0 | 0 |
| ENSG00000197745 | 0 | 0 | 0 | 0 | 0 | 0 |
| ENSG00000255213 | 2 | 1 | 1 | 0 | 0 | 0 |
| ENSG00000255118 | 0 | 0 | 0 | 0 | 0 | 0 |
| ENSG00000162174 | 236 | 124 | 139 | 128 | 138 | 156 |
| ENSG00000254454 | 0 | 0 | 1 | 0 | 0 | 0 |

| | | | | | | |
|---|---|---|---|---|---|---|
| ENSG00000255126 | 0 | 0 | 0 | 0 | 0 | 0 |
| ENSG00000149021 | 0 | 0 | 0 | 0 | 1 | 0 |
| ENSG00000255446 | 0 | 0 | 0 | 0 | 0 | 0 |
| ENSG00000124942 | 16430 | 7191 | 10176 | 3951 | 8211 | 6809 |
| ENSG00000250659 | 3 | 1 | 0 | 0 | 0 | 1 |
| ENSG00000257058 | 2 | 0 | 2 | 2 | 1 | 2 |
| ENSG00000254772 | 0 | 0 | 0 | 0 | 0 | 0 |
| ENSG00000255508 | 187 | 69 | 94 | 76 | 79 | 63 |
| ENSG00000276102 | 0 | 0 | 0 | 0 | 0 | 0 |
| ENSG00000149016 | 420 | 226 | 293 | 292 | 265 | 231 |
| ENSG00000149480 | 4043 | 1998 | 2376 | 2213 | 2287 | 2314 |
| ENSG00000149499 | 1632 | 815 | 1130 | 997 | 907 | 985 |
| ENSG00000254964 | 1 | 1 | 1 | 0 | 1 | 0 |
| ENSG00000149489 | 104 | 48 | 91 | 46 | 67 | 46 |
| ENSG00000149541 | 1601 | 796 | 988 | 1143 | 1005 | 960 |
| ENSG00000089597 | 13416 | 6305 | 8108 | 6299 | 6549 | 7082 |
| ENSG00000185085 | 1016 | 435 | 650 | 693 | 490 | 554 |
| ENSG00000255432 | 0 | 0 | 0 | 0 | 0 | 0 |
| ENSG00000278615 | 11 | 13 | 4 | 6 | 7 | 8 |
| ENSG00000162194 | 121 | 89 | 109 | 88 | 109 | 126 |
| ENSG00000214756 | 199 | 99 | 135 | 147 | 133 | 153 |
| ENSG00000206597 | 0 | 0 | 0 | 0 | 0 | 0 |
| ENSG00000204922 | 294 | 204 | 277 | 340 | 308 | 249 |
| ENSG00000162191 | 3010 | 1514 | 1939 | 2145 | 1781 | 1627 |
| ENSG00000177363 | 5 | 2 | 1 | 9 | 4 | 2 |
| ENSG00000234857 | 0 | 0 | 1 | 0 | 1 | 0 |
| ENSG00000168000 | 118 | 75 | 109 | 87 | 82 | 85 |
| ENSG00000162188 | 1 | 2 | 5 | 0 | 5 | 5 |
| ENSG00000214753 | 1511 | 756 | 982 | 888 | 991 | 1012 |
| ENSG00000162222 | 769 | 387 | 404 | 383 | 442 | 419 |
| ENSG00000185670 | 123 | 101 | 82 | 53 | 66 | 61 |
| ENSG00000168002 | 2574 | 1269 | 1486 | 1772 | 1639 | 1624 |
| ENSG00000267811 | 9 | 4 | 3 | 4 | 6 | 5 |
| ENSG00000162227 | 491 | 217 | 304 | 294 | 241 | 264 |
| ENSG00000168569 | 709 | 456 | 540 | 499 | 380 | 428 |
| ENSG00000269176 | 33 | 30 | 38 | 21 | 15 | 28 |
| ENSG00000185475 | 663 | 438 | 523 | 552 | 512 | 502 |
| ENSG00000284535 | 0 | 0 | 0 | 0 | 0 | 0 |
| ENSG00000162231 | 1980 | 962 | 1274 | 1051 | 998 | 1108 |
| ENSG00000274066 | 1 | 0 | 0 | 2 | 0 | 0 |
| ENSG00000162236 | 1334 | 571 | 774 | 687 | 649 | 564 |
| ENSG00000269463 | 14 | 1 | 2 | 2 | 1 | 1 |
| ENSG00000252361 | 2 | 5 | 1 | 3 | 4 | 2 |
| ENSG00000256690 | 4 | 5 | 3 | 4 | 3 | 4 |
| ENSG00000283268 | 0 | 0 | 0 | 0 | 3 | 0 |
| ENSG00000133316 | 884 | 413 | 524 | 699 | 544 | 593 |
| ENSG00000222328 | 0 | 0 | 0 | 0 | 0 | 0 |
| ENSG00000255717 | 2097 | 1181 | 1455 | 1400 | 1592 | 1434 |
| ENSG00000277194 | 0 | 0 | 0 | 0 | 0 | 0 |
| ENSG00000277846 | 1 | 0 | 0 | 0 | 1 | 1 |

| | | | | | | |
|---|---|---|---|---|---|---|
| ENSG00000278527 | 0 | 0 | 0 | 0 | 0 | 0 |
| ENSG00000274544 | 0 | 0 | 0 | 0 | 0 | 0 |
| ENSG00000275996 | 0 | 0 | 0 | 0 | 0 | 0 |
| ENSG00000276788 | 0 | 0 | 0 | 0 | 0 | 0 |
| ENSG00000275043 | 0 | 0 | 0 | 0 | 0 | 0 |
| ENSG00000168003 | 4298 | 2095 | 2454 | 5426 | 4782 | 4304 |
| ENSG00000168539 | 4 | 1 | 1 | 3 | 4 | 3 |
| ENSG00000257002 | 2 | 0 | 7 | 5 | 1 | 2 |
| ENSG00000241082 | 0 | 0 | 0 | 0 | 0 | 0 |
| ENSG00000197901 | 0 | 0 | 0 | 0 | 0 | 0 |
| ENSG00000285388 | 0 | 0 | 0 | 0 | 0 | 0 |
| ENSG00000149452 | 0 | 0 | 0 | 0 | 0 | 0 |
| ENSG00000256041 | 0 | 0 | 0 | 0 | 0 | 0 |
| ENSG00000253547 | 0 | 0 | 0 | 0 | 0 | 0 |
| ENSG00000197658 | 0 | 0 | 0 | 0 | 0 | 0 |
| ENSG00000239924 | 0 | 0 | 0 | 0 | 0 | 0 |
| ENSG00000184999 | 0 | 0 | 0 | 0 | 0 | 0 |
| ENSG00000196600 | 0 | 0 | 0 | 0 | 0 | 0 |
| ENSG00000256863 | 0 | 0 | 0 | 0 | 0 | 0 |
| ENSG00000256847 | 0 | 0 | 0 | 0 | 0 | 0 |
| ENSG00000256181 | 0 | 0 | 0 | 0 | 0 | 0 |
| ENSG00000149742 | 0 | 0 | 0 | 0 | 1 | 0 |
| ENSG00000168004 | 0 | 0 | 0 | 0 | 0 | 0 |
| ENSG00000275598 | 0 | 0 | 0 | 0 | 0 | 0 |
| ENSG00000133317 | 0 | 0 | 0 | 0 | 0 | 0 |
| ENSG00000133321 | 16 | 14 | 9 | 4 | 4 | 4 |
| ENSG00000133328 | 40 | 17 | 38 | 13 | 11 | 9 |
| ENSG00000176485 | 1352 | 572 | 827 | 556 | 550 | 625 |
| ENSG00000203520 | 0 | 0 | 0 | 0 | 0 | 0 |
| ENSG00000184743 | 2582 | 1258 | 1659 | 1005 | 1434 | 1285 |
| ENSG00000270269 | 0 | 1 | 0 | 0 | 0 | 0 |
| ENSG00000256789 | 0 | 0 | 1 | 0 | 0 | 0 |
| ENSG00000133318 | 6653 | 2798 | 3937 | 3111 | 3341 | 3460 |
| ENSG00000271100 | 2 | 1 | 0 | 0 | 0 | 0 |
| ENSG00000188070 | 381 | 168 | 220 | 211 | 198 | 192 |
| ENSG00000264519 | 0 | 0 | 0 | 0 | 0 | 0 |
| ENSG00000168005 | 1023 | 457 | 560 | 604 | 580 | 606 |
| ENSG00000256280 | 2 | 2 | 0 | 1 | 2 | 0 |
| ENSG00000072518 | 2811 | 1310 | 1792 | 1505 | 1449 | 1648 |
| ENSG00000202089 | 0 | 0 | 0 | 0 | 0 | 0 |
| ENSG00000167771 | 74 | 19 | 42 | 37 | 35 | 30 |
| ENSG00000110583 | 1437 | 738 | 889 | 639 | 839 | 815 |
| ENSG00000207200 | 1 | 0 | 1 | 0 | 0 | 0 |
| ENSG00000176340 | 3161 | 2083 | 2490 | 2708 | 2364 | 2303 |
| ENSG00000256100 | 0 | 0 | 0 | 0 | 0 | 0 |
| ENSG00000167770 | 3760 | 1674 | 2314 | 2365 | 1903 | 2197 |
| ENSG00000133315 | 1198 | 551 | 751 | 721 | 558 | 598 |
| ENSG00000256824 | 0 | 0 | 0 | 0 | 0 | 0 |
| ENSG00000256481 | 1 | 1 | 0 | 0 | 0 | 0 |
| ENSG00000126500 | 28 | 6 | 7 | 7 | 13 | 11 |

| | | | | | | |
|---|---|---|---|---|---|---|
| ENSG00000256341 | 0 | 0 | 2 | 0 | 0 | 0 |
| ENSG00000168439 | 7355 | 3594 | 4250 | 4370 | 4475 | 4255 |
| ENSG00000149781 | 26 | 3 | 23 | 11 | 20 | 16 |
| ENSG00000149743 | 900 | 451 | 580 | 575 | 491 | 507 |
| ENSG00000149761 | 932 | 497 | 617 | 661 | 567 | 579 |
| ENSG00000256116 | 0 | 1 | 2 | 1 | 0 | 0 |
| ENSG00000110011 | 800 | 389 | 515 | 521 | 420 | 447 |
| ENSG00000173511 | 651 | 324 | 375 | 409 | 376 | 306 |
| ENSG00000173486 | 1383 | 681 | 830 | 803 | 759 | 723 |
| ENSG00000173457 | 3214 | 1662 | 1905 | 2885 | 2340 | 2266 |
| ENSG00000256940 | 23 | 25 | 42 | 53 | 46 | 61 |
| ENSG00000257086 | 22 | 11 | 19 | 14 | 20 | 13 |
| ENSG00000149782 | 2692 | 1300 | 1585 | 1454 | 1435 | 1343 |
| ENSG00000002330 | 646 | 438 | 588 | 627 | 487 | 506 |
| ENSG00000173264 | 1522 | 661 | 912 | 834 | 630 | 737 |
| ENSG00000182450 | 0 | 0 | 0 | 0 | 0 | 0 |
| ENSG00000257069 | 0 | 0 | 0 | 2 | 1 | 0 |
| ENSG00000207024 | 0 | 0 | 0 | 0 | 0 | 0 |
| ENSG00000219435 | 0 | 0 | 0 | 1 | 0 | 0 |
| ENSG00000173153 | 1503 | 788 | 937 | 1102 | 1086 | 955 |
| ENSG00000173113 | 4013 | 2227 | 2618 | 2781 | 2442 | 2384 |
| ENSG00000126432 | 6681 | 3263 | 3907 | 4272 | 3649 | 3427 |
| ENSG00000236935 | 2 | 0 | 0 | 0 | 0 | 0 |
| ENSG00000168071 | 68 | 18 | 45 | 23 | 32 | 34 |
| ENSG00000278359 | 0 | 0 | 0 | 0 | 0 | 0 |
| ENSG00000162302 | 1255 | 575 | 736 | 845 | 696 | 704 |
| ENSG00000221273 | 0 | 0 | 0 | 0 | 0 | 0 |
| ENSG00000231680 | 0 | 0 | 0 | 0 | 0 | 0 |
| ENSG00000231492 | 0 | 0 | 0 | 0 | 0 | 0 |
| ENSG00000181908 | 2 | 2 | 3 | 1 | 1 | 1 |
| ENSG00000232500 | 0 | 0 | 0 | 0 | 0 | 0 |
| ENSG00000237363 | 0 | 0 | 0 | 0 | 0 | 0 |
| ENSG00000168065 | 0 | 0 | 0 | 0 | 0 | 0 |
| ENSG00000197891 | 0 | 0 | 0 | 0 | 0 | 0 |
| ENSG00000110076 | 0 | 0 | 3 | 0 | 0 | 5 |
| ENSG00000237410 | 0 | 0 | 0 | 0 | 0 | 0 |
| ENSG00000068831 | 22 | 5 | 7 | 4 | 5 | 4 |
| ENSG00000068976 | 20 | 13 | 22 | 15 | 17 | 24 |
| ENSG00000168066 | 4980 | 2423 | 3311 | 2941 | 2593 | 3155 |
| ENSG00000269038 | 5 | 7 | 3 | 4 | 6 | 2 |
| ENSG00000168067 | 537 | 253 | 329 | 259 | 248 | 223 |
| ENSG00000133895 | 2128 | 979 | 1235 | 1147 | 1021 | 1191 |
| ENSG00000171219 | 2883 | 1424 | 1830 | 1420 | 1674 | 1529 |
| ENSG00000110047 | 1717 | 879 | 1033 | 1072 | 964 | 970 |
| ENSG00000230835 | 0 | 0 | 1 | 0 | 0 | 0 |
| ENSG00000229719 | 1 | 2 | 4 | 0 | 1 | 1 |
| ENSG00000283926 | 0 | 0 | 0 | 0 | 0 | 0 |
| ENSG00000284155 | 0 | 0 | 0 | 0 | 0 | 0 |
| ENSG00000110046 | 1580 | 696 | 1095 | 816 | 767 | 875 |
| ENSG00000274986 | 2 | 0 | 0 | 0 | 0 | 0 |

| | | | | | | |
|---|---|---|---|---|---|---|
| ENSG00000274314 | 0 | 0 | 0 | 0 | 0 | 0 |
| ENSG00000068971 | 551 | 217 | 317 | 258 | 257 | 275 |
| ENSG00000149735 | 8 | 3 | 0 | 0 | 2 | 0 |
| ENSG00000168070 | 0 | 0 | 0 | 0 | 0 | 0 |
| ENSG00000168062 | 14 | 3 | 4 | 2 | 2 | 8 |
| ENSG00000213465 | 658 | 425 | 444 | 569 | 459 | 475 |
| ENSG00000273003 | 0 | 0 | 0 | 0 | 0 | 0 |
| ENSG00000278851 | 0 | 0 | 0 | 0 | 0 | 0 |
| ENSG00000270775 | 0 | 0 | 0 | 0 | 0 | 0 |
| ENSG00000110025 | 22 | 13 | 15 | 11 | 24 | 16 |
| ENSG00000168061 | 427 | 248 | 245 | 358 | 274 | 296 |
| ENSG00000168060 | 0 | 0 | 4 | 0 | 1 | 0 |
| ENSG00000146670 | 1027 | 547 | 581 | 589 | 594 | 657 |
| ENSG00000162300 | 234 | 129 | 137 | 158 | 149 | 136 |
| ENSG00000187066 | 19 | 4 | 10 | 4 | 4 | 4 |
| ENSG00000149823 | 2309 | 1080 | 1596 | 1533 | 1278 | 1390 |
| ENSG00000254501 | 0 | 0 | 0 | 0 | 0 | 0 |
| ENSG00000149809 | 7025 | 3071 | 4078 | 3308 | 2726 | 2949 |
| ENSG00000174276 | 580 | 276 | 411 | 382 | 333 | 305 |
| ENSG00000255173 | 0 | 0 | 0 | 0 | 0 | 0 |
| ENSG00000278952 | 222 | 110 | 139 | 127 | 147 | 122 |
| ENSG00000149806 | 6420 | 3871 | 4643 | 5726 | 4589 | 4400 |
| ENSG00000162298 | 1865 | 796 | 1080 | 898 | 927 | 923 |
| ENSG00000149792 | 3501 | 1703 | 2240 | 1878 | 1867 | 1945 |
| ENSG00000284489 | 0 | 0 | 0 | 0 | 0 | 0 |
| ENSG00000254455 | 0 | 0 | 0 | 0 | 0 | 0 |
| ENSG00000222477 | 0 | 0 | 0 | 0 | 0 | 0 |
| ENSG00000255058 | 3 | 0 | 0 | 0 | 0 | 0 |
| ENSG00000204710 | 0 | 1 | 2 | 0 | 0 | 0 |
| ENSG00000255200 | 44 | 24 | 24 | 21 | 35 | 24 |
| ENSG00000254614 | 76 | 62 | 85 | 67 | 76 | 48 |
| ENSG00000014216 | 10487 | 4738 | 6280 | 5798 | 4973 | 5215 |
| ENSG00000197847 | 40 | 17 | 22 | 15 | 25 | 29 |
| ENSG00000014138 | 104 | 87 | 96 | 75 | 83 | 92 |
| ENSG00000285816 | 2 | 0 | 2 | 0 | 3 | 0 |
| ENSG00000149798 | 0 | 0 | 0 | 0 | 0 | 1 |
| ENSG00000133884 | 2339 | 1020 | 1249 | 1142 | 1191 | 1139 |
| ENSG00000173825 | 34 | 30 | 39 | 35 | 26 | 24 |
| ENSG00000255478 | 0 | 0 | 0 | 0 | 1 | 0 |
| ENSG00000162241 | 106 | 51 | 80 | 50 | 70 | 65 |
| ENSG00000126391 | 1506 | 794 | 957 | 902 | 991 | 918 |
| ENSG00000245532 | 9108 | 3215 | 6285 | 3079 | 3897 | 4867 |
| ENSG00000278144 | 0 | 0 | 0 | 0 | 0 | 0 |
| ENSG00000278050 | 0 | 0 | 0 | 0 | 0 | 0 |
| ENSG00000277599 | 0 | 0 | 0 | 0 | 0 | 0 |
| ENSG00000283791 | 0 | 0 | 0 | 0 | 0 | 0 |
| ENSG00000275948 | 0 | 0 | 0 | 0 | 0 | 0 |
| ENSG00000173727 | 238 | 111 | 125 | 122 | 144 | 103 |
| ENSG00000279243 | 3 | 1 | 2 | 2 | 5 | 0 |
| ENSG00000251562 | 11105 | 5940 | 6216 | 1987 | 7410 | 5276 |

| | | | | | | |
|---|---|---|---|---|---|---|
| ENSG00000270117 | 0 | 0 | 0 | 0 | 0 | 0 |
| ENSG00000278217 | 0 | 0 | 0 | 0 | 0 | 0 |
| ENSG00000274072 | 0 | 0 | 0 | 0 | 0 | 0 |
| ENSG00000270526 | 0 | 0 | 0 | 0 | 0 | 0 |
| ENSG00000142186 | 4761 | 2198 | 2759 | 2759 | 2525 | 2442 |
| ENSG00000168056 | 2483 | 1062 | 1513 | 1094 | 860 | 1026 |
| ENSG00000279093 | 0 | 0 | 0 | 0 | 0 | 0 |
| ENSG00000260233 | 91 | 57 | 69 | 42 | 46 | 48 |
| ENSG00000173465 | 831 | 416 | 498 | 629 | 451 | 486 |
| ENSG00000176973 | 300 | 229 | 275 | 309 | 261 | 240 |
| ENSG00000173442 | 2381 | 1112 | 1461 | 1083 | 1253 | 1231 |
| ENSG00000173338 | 32 | 19 | 6 | 26 | 18 | 7 |
| ENSG00000173327 | 3469 | 1696 | 2048 | 2288 | 1972 | 1901 |
| ENSG00000197136 | 5443 | 2269 | 3155 | 2919 | 2739 | 2945 |
| ENSG00000266041 | 0 | 0 | 0 | 0 | 0 | 0 |
| ENSG00000213445 | 415 | 190 | 294 | 271 | 210 | 216 |
| ENSG00000265874 | 4 | 0 | 0 | 1 | 0 | 0 |
| ENSG00000173039 | 2957 | 1448 | 1827 | 1678 | 1670 | 1705 |
| ENSG00000285533 | 195 | 109 | 166 | 137 | 138 | 134 |
| ENSG00000239356 | 0 | 0 | 0 | 0 | 0 | 0 |
| ENSG00000172977 | 706 | 381 | 497 | 417 | 392 | 387 |
| ENSG00000172922 | 1034 | 498 | 640 | 628 | 530 | 520 |
| ENSG00000214659 | 2 | 3 | 1 | 1 | 1 | 0 |
| ENSG00000255557 | 0 | 2 | 1 | 1 | 1 | 1 |
| ENSG00000254470 | 1227 | 632 | 728 | 679 | 672 | 683 |
| ENSG00000172818 | 497 | 260 | 374 | 342 | 360 | 344 |
| ENSG00000255120 | 13 | 4 | 4 | 7 | 13 | 1 |
| ENSG00000255404 | 0 | 0 | 0 | 0 | 0 | 0 |
| ENSG00000172757 | 32669 | 15617 | 19560 | 23649 | 18622 | 20316 |
| ENSG00000172803 | 17 | 10 | 14 | 10 | 11 | 8 |
| ENSG00000172732 | 903 | 436 | 529 | 489 | 555 | 457 |
| ENSG00000172638 | 57 | 24 | 30 | 28 | 18 | 19 |
| ENSG00000172543 | 2 | 0 | 2 | 0 | 0 | 2 |
| ENSG00000172500 | 2759 | 1323 | 1589 | 1957 | 1510 | 1626 |
| ENSG00000175602 | 818 | 323 | 407 | 669 | 491 | 549 |
| ENSG00000175592 | 9 | 4 | 5 | 14 | 9 | 8 |
| ENSG00000175573 | 939 | 512 | 674 | 618 | 602 | 501 |
| ENSG00000175550 | 2654 | 1385 | 1570 | 1871 | 1682 | 1485 |
| ENSG00000175513 | 0 | 0 | 0 | 0 | 0 | 0 |
| ENSG00000175467 | 2341 | 1182 | 1359 | 1418 | 1402 | 1312 |
| ENSG00000254867 | 0 | 1 | 0 | 0 | 1 | 0 |
| ENSG00000175376 | 1017 | 564 | 675 | 516 | 653 | 640 |
| ENSG00000175334 | 5045 | 2400 | 2969 | 3261 | 2800 | 2859 |
| ENSG00000175315 | 4 | 3 | 2 | 3 | 1 | 0 |
| ENSG00000175294 | 4 | 2 | 1 | 0 | 0 | 2 |
| ENSG00000175229 | 0 | 0 | 0 | 0 | 0 | 0 |
| ENSG00000087365 | 9475 | 4651 | 5626 | 5385 | 5220 | 5263 |
| ENSG00000255038 | 17 | 10 | 14 | 9 | 7 | 4 |
| ENSG00000175115 | 6713 | 3193 | 4521 | 3112 | 2964 | 3305 |
| ENSG00000255320 | 63 | 37 | 61 | 40 | 38 | 42 |

| | | | | | | |
|---|---|---|---|---|---|---|
| ENSG00000174996 | 2410 | 1120 | 1705 | 1411 | 1364 | 1320 |
| ENSG00000254461 | 43 | 22 | 24 | 5 | 12 | 27 |
| ENSG00000254855 | 5 | 2 | 1 | 1 | 0 | 3 |
| ENSG00000254762 | 1 | 0 | 2 | 3 | 1 | 0 |
| ENSG00000174903 | 6371 | 3767 | 4811 | 4382 | 3727 | 3724 |
| ENSG00000245156 | 8 | 20 | 21 | 8 | 19 | 15 |
| ENSG00000254452 | 0 | 0 | 0 | 0 | 0 | 0 |
| ENSG00000174871 | 108 | 53 | 82 | 47 | 55 | 43 |
| ENSG00000174851 | 2040 | 1065 | 1277 | 1679 | 1214 | 1365 |
| ENSG00000179292 | 34 | 3 | 22 | 12 | 13 | 17 |
| ENSG00000254458 | 0 | 0 | 0 | 0 | 0 | 0 |
| ENSG00000174807 | 0 | 0 | 0 | 0 | 0 | 0 |
| ENSG00000174791 | 139 | 58 | 103 | 156 | 131 | 137 |
| ENSG00000254756 | 0 | 1 | 3 | 0 | 3 | 2 |
| ENSG00000174744 | 1906 | 908 | 1113 | 1429 | 1142 | 1084 |
| ENSG00000174684 | 766 | 378 | 425 | 424 | 384 | 421 |
| ENSG00000255468 | 96 | 46 | 63 | 46 | 55 | 60 |
| ENSG00000174669 | 1699 | 826 | 1026 | 1004 | 913 | 1031 |
| ENSG00000276918 | 0 | 0 | 0 | 0 | 0 | 0 |
| ENSG00000254736 | 0 | 0 | 0 | 0 | 0 | 0 |
| ENSG00000202317 | 0 | 0 | 0 | 0 | 0 | 0 |
| ENSG00000254510 | 0 | 0 | 0 | 0 | 0 | 0 |
| ENSG00000174576 | 0 | 0 | 0 | 0 | 0 | 0 |
| ENSG00000201733 | 0 | 0 | 0 | 0 | 0 | 0 |
| ENSG00000174547 | 1963 | 960 | 1162 | 1313 | 1123 | 1150 |
| ENSG00000254596 | 0 | 0 | 0 | 0 | 0 | 0 |
| ENSG00000174516 | 460 | 233 | 272 | 299 | 284 | 270 |
| ENSG00000255517 | 89 | 71 | 98 | 93 | 114 | 94 |
| ENSG00000254986 | 4425 | 1961 | 2465 | 2862 | 2374 | 2390 |
| ENSG00000256349 | 0 | 0 | 0 | 0 | 0 | 0 |
| ENSG00000174483 | 32 | 14 | 26 | 15 | 22 | 18 |
| ENSG00000174165 | 1689 | 787 | 1079 | 969 | 937 | 946 |
| ENSG00000248746 | 1 | 0 | 1 | 1 | 4 | 1 |
| ENSG00000250105 | 0 | 0 | 0 | 0 | 0 | 0 |
| ENSG00000174080 | 981 | 432 | 636 | 542 | 415 | 471 |
| ENSG00000182791 | 68 | 35 | 40 | 35 | 19 | 21 |
| ENSG00000173992 | 1000 | 512 | 722 | 653 | 607 | 574 |
| ENSG00000199325 | 4 | 0 | 1 | 0 | 2 | 1 |
| ENSG00000239306 | 1720 | 792 | 1067 | 1190 | 925 | 1211 |
| ENSG00000248643 | 53 | 28 | 41 | 21 | 18 | 19 |
| ENSG00000173933 | 1044 | 513 | 761 | 413 | 593 | 549 |
| ENSG00000173914 | 1058 | 520 | 712 | 536 | 599 | 572 |
| ENSG00000173898 | 3362 | 1488 | 1953 | 1664 | 1736 | 1711 |
| ENSG00000239553 | 1 | 0 | 0 | 0 | 1 | 0 |
| ENSG00000173715 | 1296 | 637 | 853 | 684 | 750 | 693 |
| ENSG00000213409 | 3 | 0 | 0 | 0 | 1 | 1 |
| ENSG00000254984 | 4 | 0 | 0 | 0 | 1 | 0 |
| ENSG00000173653 | 1163 | 474 | 630 | 738 | 604 | 658 |
| ENSG00000173599 | 1036 | 480 | 635 | 635 | 598 | 594 |
| ENSG00000173621 | 468 | 252 | 277 | 440 | 319 | 322 |

| | | | | | | |
|---|---|---|---|---|---|---|
| ENSG00000239099 | 0 | 0 | 0 | 0 | 0 | 0 |
| ENSG00000266423 | 0 | 0 | 0 | 0 | 0 | 0 |
| ENSG00000173237 | 0 | 2 | 0 | 0 | 1 | 2 |
| ENSG00000173227 | 1590 | 767 | 815 | 1289 | 1276 | 1012 |
| ENSG00000283326 | 0 | 0 | 0 | 0 | 0 | 0 |
| ENSG00000173156 | 3938 | 1930 | 2035 | 3273 | 2601 | 2209 |
| ENSG00000173120 | 6888 | 2791 | 3897 | 2784 | 3210 | 3479 |
| ENSG00000179038 | 0 | 2 | 0 | 0 | 0 | 0 |
| ENSG00000173020 | 5680 | 2477 | 3336 | 3108 | 2860 | 2749 |
| ENSG00000172932 | 2173 | 978 | 1313 | 1055 | 1072 | 1162 |
| ENSG00000172830 | 5369 | 2363 | 3186 | 2488 | 2254 | 2489 |
| ENSG00000172613 | 861 | 386 | 533 | 435 | 445 | 499 |
| ENSG00000175482 | 618 | 401 | 592 | 517 | 472 | 477 |
| ENSG00000256514 | 0 | 0 | 0 | 0 | 3 | 0 |
| ENSG00000274251 | 5 | 5 | 6 | 0 | 2 | 2 |
| ENSG00000201684 | 1 | 1 | 4 | 2 | 3 | 3 |
| ENSG00000175505 | 272 | 117 | 186 | 166 | 154 | 180 |
| ENSG00000275484 | 15 | 14 | 15 | 7 | 10 | 10 |
| ENSG00000253024 | 0 | 0 | 2 | 0 | 0 | 0 |
| ENSG00000172531 | 10503 | 5534 | 6882 | 7107 | 6004 | 5910 |
| ENSG00000175463 | 61 | 21 | 23 | 13 | 24 | 29 |
| ENSG00000172508 | 109 | 50 | 65 | 44 | 32 | 33 |
| ENSG00000175634 | 1921 | 1087 | 1263 | 1315 | 1188 | 1138 |
| ENSG00000255949 | 0 | 0 | 0 | 0 | 0 | 0 |
| ENSG00000213402 | 0 | 0 | 0 | 0 | 0 | 0 |
| ENSG00000172725 | 8393 | 4010 | 5082 | 4887 | 4388 | 4034 |
| ENSG00000175514 | 0 | 0 | 1 | 1 | 0 | 1 |
| ENSG00000175544 | 0 | 0 | 1 | 1 | 1 | 4 |
| ENSG00000172663 | 1534 | 664 | 971 | 1025 | 798 | 828 |
| ENSG00000110711 | 2065 | 986 | 1294 | 1360 | 1031 | 1166 |
| ENSG00000276769 | 0 | 0 | 0 | 0 | 0 | 0 |
| ENSG00000110697 | 1289 | 551 | 798 | 790 | 725 | 692 |
| ENSG00000167797 | 1646 | 703 | 962 | 1026 | 782 | 871 |
| ENSG00000167791 | 0 | 0 | 0 | 0 | 0 | 0 |
| ENSG00000283257 | 0 | 0 | 0 | 0 | 0 | 0 |
| ENSG00000084207 | 0 | 0 | 0 | 0 | 1 | 0 |
| ENSG00000184224 | 0 | 0 | 1 | 0 | 1 | 1 |
| ENSG00000255119 | 0 | 0 | 0 | 0 | 0 | 0 |
| ENSG00000167792 | 4462 | 2181 | 2637 | 2912 | 2427 | 2587 |
| ENSG00000231793 | 8 | 3 | 1 | 5 | 4 | 2 |
| ENSG00000167799 | 160 | 103 | 126 | 150 | 134 | 131 |
| ENSG00000167800 | 1 | 4 | 1 | 0 | 1 | 2 |
| ENSG00000132744 | 0 | 0 | 0 | 0 | 0 | 0 |
| ENSG00000227834 | 0 | 0 | 0 | 0 | 0 | 0 |
| ENSG00000132746 | 748 | 269 | 443 | 510 | 495 | 531 |
| ENSG00000239559 | 8 | 7 | 1 | 8 | 4 | 7 |
| ENSG00000184795 | 3 | 0 | 0 | 0 | 0 | 1 |
| ENSG00000235855 | 0 | 0 | 0 | 0 | 0 | 0 |
| ENSG00000254447 | 0 | 0 | 0 | 0 | 0 | 0 |
| ENSG00000285537 | 0 | 0 | 0 | 0 | 0 | 0 |

| | | | | | | |
|---|---|---|---|---|---|---|
| ENSG00000254883 | 0 | 0 | 0 | 0 | 0 | 0 |
| ENSG00000255147 | 0 | 0 | 0 | 0 | 0 | 0 |
| ENSG00000227620 | 0 | 2 | 2 | 1 | 1 | 3 |
| ENSG00000160172 | 265 | 114 | 141 | 122 | 132 | 128 |
| ENSG00000273819 | 0 | 1 | 1 | 0 | 0 | 0 |
| ENSG00000254792 | 0 | 0 | 0 | 0 | 0 | 0 |
| ENSG00000251637 | 0 | 0 | 0 | 0 | 0 | 0 |
| ENSG00000206587 | 0 | 0 | 0 | 0 | 0 | 0 |
| ENSG00000243672 | 0 | 0 | 0 | 0 | 0 | 0 |
| ENSG00000254610 | 9 | 1 | 1 | 1 | 1 | 3 |
| ENSG00000254850 | 0 | 0 | 0 | 0 | 0 | 0 |
| ENSG00000255230 | 0 | 0 | 0 | 0 | 0 | 0 |
| ENSG00000255554 | 0 | 0 | 0 | 0 | 0 | 0 |
| ENSG00000110057 | 1714 | 753 | 1006 | 1074 | 945 | 966 |
| ENSG00000006534 | 513 | 293 | 314 | 343 | 354 | 288 |
| ENSG00000255306 | 12 | 3 | 2 | 4 | 5 | 9 |
| ENSG00000110717 | 2187 | 991 | 1284 | 1437 | 1220 | 1172 |
| ENSG00000284293 | 0 | 0 | 0 | 0 | 0 | 0 |
| ENSG00000284435 | 0 | 0 | 0 | 0 | 0 | 0 |
| ENSG00000110719 | 216 | 84 | 142 | 131 | 130 | 149 |
| ENSG00000275022 | 0 | 0 | 0 | 0 | 0 | 0 |
| ENSG00000255031 | 5 | 4 | 4 | 4 | 8 | 4 |
| ENSG00000110721 | 922 | 508 | 570 | 460 | 577 | 493 |
| ENSG00000255236 | 18 | 8 | 10 | 5 | 8 | 6 |
| ENSG00000110066 | 1952 | 974 | 1143 | 623 | 1114 | 1014 |
| ENSG00000171067 | 815 | 432 | 487 | 635 | 630 | 629 |
| ENSG00000162337 | 3496 | 1582 | 2061 | 2534 | 2740 | 2582 |
| ENSG00000110075 | 4400 | 2233 | 2683 | 1677 | 2742 | 2351 |
| ENSG00000254856 | 0 | 0 | 1 | 0 | 0 | 0 |
| ENSG00000260808 | 3 | 7 | 7 | 2 | 1 | 6 |
| ENSG00000069482 | 78 | 35 | 40 | 72 | 65 | 44 |
| ENSG00000132749 | 939 | 431 | 538 | 451 | 456 | 426 |
| ENSG00000110090 | 8745 | 4201 | 5457 | 3565 | 4133 | 4281 |
| ENSG00000250508 | 0 | 1 | 2 | 1 | 3 | 2 |
| ENSG00000197345 | 1221 | 660 | 698 | 860 | 756 | 785 |
| ENSG00000132740 | 741 | 306 | 447 | 414 | 396 | 444 |
| ENSG00000255741 | 6 | 4 | 7 | 7 | 9 | 6 |
| ENSG00000172938 | 0 | 0 | 0 | 0 | 0 | 0 |
| ENSG00000261625 | 0 | 0 | 0 | 0 | 0 | 0 |
| ENSG00000261276 | 0 | 0 | 0 | 0 | 0 | 0 |
| ENSG00000172935 | 0 | 0 | 0 | 0 | 1 | 2 |
| ENSG00000256508 | 0 | 0 | 0 | 0 | 0 | 0 |
| ENSG00000162341 | 953 | 403 | 527 | 457 | 424 | 437 |
| ENSG00000265539 | 2 | 2 | 1 | 1 | 1 | 0 |
| ENSG00000260895 | 0 | 1 | 0 | 1 | 2 | 0 |
| ENSG00000261070 | 0 | 0 | 1 | 0 | 0 | 0 |
| ENSG00000284713 | 6 | 6 | 5 | 0 | 0 | 1 |
| ENSG00000172927 | 23 | 8 | 14 | 12 | 17 | 15 |
| ENSG00000213275 | 0 | 0 | 0 | 0 | 0 | 0 |
| ENSG00000260877 | 0 | 0 | 1 | 1 | 0 | 0 |

| | | | | | | |
|---|---|---|---|---|---|---|
| ENSG00000259854 | 0 | 0 | 0 | 0 | 0 | 0 |
| ENSG00000255980 | 2 | 1 | 0 | 0 | 0 | 0 |
| ENSG00000255606 | 2 | 0 | 0 | 1 | 0 | 0 |
| ENSG00000261347 | 0 | 0 | 0 | 0 | 0 | 0 |
| ENSG00000255774 | 463 | 250 | 211 | 318 | 278 | 218 |
| ENSG00000285094 | 420 | 215 | 270 | 186 | 226 | 239 |
| ENSG00000110092 | 27193 | 13341 | 15927 | 12479 | 13529 | 13853 |
| ENSG00000149716 | 939 | 444 | 544 | 402 | 468 | 521 |
| ENSG00000162344 | 0 | 0 | 0 | 0 | 0 | 0 |
| ENSG00000256441 | 0 | 0 | 0 | 0 | 0 | 0 |
| ENSG00000075388 | 0 | 0 | 0 | 0 | 0 | 0 |
| ENSG00000186895 | 0 | 0 | 0 | 0 | 0 | 0 |
| ENSG00000260348 | 0 | 0 | 0 | 0 | 0 | 0 |
| ENSG00000260679 | 0 | 0 | 0 | 0 | 0 | 0 |
| ENSG00000255191 | 0 | 0 | 0 | 0 | 0 | 0 |
| ENSG00000254605 | 0 | 0 | 0 | 0 | 0 | 0 |
| ENSG00000248844 | 0 | 0 | 0 | 0 | 0 | 0 |
| ENSG00000254417 | 0 | 0 | 0 | 0 | 0 | 0 |
| ENSG00000202070 | 0 | 0 | 0 | 0 | 0 | 0 |
| ENSG00000131620 | 4 | 1 | 4 | 3 | 1 | 5 |
| ENSG00000255143 | 0 | 0 | 0 | 0 | 0 | 0 |
| ENSG00000254902 | 0 | 0 | 0 | 0 | 0 | 0 |
| ENSG00000168040 | 408 | 232 | 273 | 289 | 294 | 308 |
| ENSG00000254721 | 0 | 0 | 0 | 0 | 0 | 0 |
| ENSG00000131626 | 2170 | 960 | 1402 | 858 | 1226 | 1311 |
| ENSG00000255329 | 0 | 0 | 0 | 2 | 0 | 0 |
| ENSG00000254604 | 9 | 1 | 4 | 4 | 3 | 4 |
| ENSG00000221333 | 0 | 0 | 0 | 0 | 0 | 0 |
| ENSG00000254484 | 0 | 0 | 0 | 0 | 2 | 0 |
| ENSG00000255539 | 0 | 0 | 0 | 0 | 0 | 0 |
| ENSG00000254495 | 0 | 2 | 2 | 0 | 0 | 0 |
| ENSG00000246889 | 38 | 23 | 22 | 34 | 32 | 32 |
| ENSG00000085733 | 14834 | 7096 | 9264 | 7793 | 9516 | 9415 |
| ENSG00000162105 | 1493 | 597 | 852 | 455 | 590 | 673 |
| ENSG00000227726 | 4 | 0 | 3 | 3 | 2 | 1 |
| ENSG00000279459 | 8 | 3 | 2 | 2 | 3 | 0 |
| ENSG00000226627 | 0 | 2 | 0 | 0 | 1 | 0 |
| ENSG00000236262 | 0 | 0 | 0 | 0 | 0 | 0 |
| ENSG00000207196 | 0 | 0 | 0 | 0 | 0 | 0 |
| ENSG00000280089 | 0 | 0 | 0 | 0 | 0 | 0 |
| ENSG00000171671 | 2 | 1 | 0 | 0 | 0 | 0 |
| ENSG00000263744 | 0 | 0 | 0 | 0 | 0 | 0 |
| ENSG00000172900 | 0 | 0 | 0 | 0 | 1 | 0 |
| ENSG00000172893 | 9010 | 4017 | 5215 | 5076 | 4444 | 4983 |
| ENSG00000254682 | 295 | 165 | 184 | 236 | 178 | 176 |
| ENSG00000172890 | 4452 | 2056 | 2888 | 2429 | 2735 | 2626 |
| ENSG00000278349 | 0 | 0 | 0 | 0 | 0 | 0 |
| ENSG00000254924 | 0 | 0 | 0 | 0 | 1 | 0 |
| ENSG00000244411 | 1 | 1 | 1 | 0 | 0 | 0 |
| ENSG00000241233 | 3 | 2 | 2 | 0 | 1 | 2 |

| | | | | | | |
|---|---|---|---|---|---|---|
| ENSG00000254997 | 9 | 5 | 8 | 4 | 3 | 4 |
| ENSG00000204572 | 27 | 4 | 9 | 11 | 10 | 6 |
| ENSG00000248903 | 0 | 0 | 1 | 0 | 0 | 0 |
| ENSG00000204571 | 0 | 0 | 3 | 0 | 0 | 0 |
| ENSG00000249387 | 0 | 0 | 0 | 0 | 0 | 0 |
| ENSG00000184055 | 0 | 0 | 0 | 0 | 0 | 0 |
| ENSG00000255562 | 0 | 0 | 0 | 0 | 0 | 0 |
| ENSG00000255261 | 0 | 0 | 0 | 0 | 0 | 0 |
| ENSG00000255415 | 0 | 0 | 0 | 0 | 0 | 0 |
| ENSG00000284625 | 0 | 0 | 0 | 0 | 0 | 0 |
| ENSG00000239481 | 0 | 0 | 0 | 0 | 0 | 0 |
| ENSG00000254805 | 0 | 0 | 0 | 0 | 0 | 0 |
| ENSG00000254972 | 0 | 0 | 0 | 0 | 0 | 0 |
| ENSG00000255319 | 0 | 0 | 0 | 0 | 1 | 1 |
| ENSG00000285933 | 4 | 2 | 3 | 5 | 10 | 2 |
| ENSG00000158483 | 248 | 142 | 185 | 183 | 132 | 190 |
| ENSG00000248671 | 17 | 18 | 17 | 10 | 6 | 17 |
| ENSG00000254978 | 12 | 7 | 6 | 9 | 8 | 14 |
| ENSG00000214534 | 1 | 0 | 1 | 1 | 5 | 4 |
| ENSG00000184276 | 0 | 0 | 0 | 0 | 0 | 0 |
| ENSG00000252902 | 0 | 0 | 0 | 0 | 0 | 0 |
| ENSG00000255157 | 0 | 0 | 0 | 0 | 0 | 0 |
| ENSG00000254469 | 49 | 31 | 41 | 15 | 19 | 21 |
| ENSG00000225805 | 3 | 2 | 3 | 0 | 0 | 0 |
| ENSG00000228915 | 6 | 6 | 6 | 2 | 6 | 5 |
| ENSG00000254593 | 6 | 5 | 5 | 2 | 4 | 0 |
| ENSG00000137522 | 1189 | 610 | 723 | 702 | 684 | 668 |
| ENSG00000255296 | 0 | 1 | 0 | 0 | 0 | 0 |
| ENSG00000137496 | 148 | 64 | 131 | 60 | 89 | 115 |
| ENSG00000137497 | 9690 | 3995 | 6211 | 4126 | 4464 | 4831 |
| ENSG00000251143 | 19 | 13 | 12 | 6 | 15 | 14 |
| ENSG00000207012 | 0 | 0 | 0 | 0 | 0 | 0 |
| ENSG00000263742 | 0 | 0 | 0 | 0 | 0 | 0 |
| ENSG00000184154 | 264 | 129 | 179 | 100 | 121 | 119 |
| ENSG00000284922 | 0 | 0 | 0 | 0 | 0 | 0 |
| ENSG00000149357 | 3565 | 1738 | 2045 | 2231 | 1842 | 1901 |
| ENSG00000284844 | 0 | 0 | 0 | 0 | 0 | 0 |
| ENSG00000110200 | 339 | 184 | 238 | 219 | 192 | 199 |
| ENSG00000110203 | 0 | 0 | 0 | 0 | 0 | 1 |
| ENSG00000257075 | 0 | 0 | 0 | 0 | 0 | 0 |
| ENSG00000256518 | 0 | 0 | 0 | 0 | 0 | 0 |
| ENSG00000204971 | 0 | 0 | 0 | 0 | 0 | 0 |
| ENSG00000255860 | 0 | 0 | 0 | 0 | 0 | 0 |
| ENSG00000110195 | 52 | 34 | 23 | 36 | 34 | 25 |
| ENSG00000165457 | 0 | 0 | 0 | 0 | 0 | 0 |
| ENSG00000165458 | 9089 | 3994 | 5415 | 4333 | 4732 | 4522 |
| ENSG00000165462 | 0 | 0 | 0 | 0 | 0 | 0 |
| ENSG00000228286 | 0 | 0 | 0 | 0 | 0 | 0 |
| ENSG00000285864 | 0 | 0 | 0 | 0 | 0 | 0 |
| ENSG00000213365 | 0 | 0 | 0 | 0 | 0 | 0 |

| | | | | | | |
|---|---|---|---|---|---|---|
| ENSG00000162129 | 610 | 289 | 389 | 372 | 347 | 401 |
| ENSG00000255843 | 2 | 0 | 1 | 1 | 2 | 3 |
| ENSG00000256739 | 0 | 0 | 0 | 0 | 0 | 0 |
| ENSG00000255672 | 0 | 0 | 0 | 0 | 0 | 0 |
| ENSG00000256403 | 0 | 0 | 0 | 0 | 0 | 0 |
| ENSG00000237408 | 0 | 0 | 0 | 0 | 0 | 0 |
| ENSG00000238048 | 0 | 0 | 0 | 0 | 0 | 0 |
| ENSG00000276109 | 0 | 0 | 0 | 0 | 0 | 0 |
| ENSG00000274664 | 0 | 0 | 0 | 0 | 0 | 0 |
| ENSG00000227467 | 0 | 0 | 0 | 0 | 0 | 0 |
| ENSG00000186642 | 0 | 2 | 0 | 1 | 2 | 3 |
| ENSG00000256633 | 0 | 0 | 0 | 0 | 0 | 0 |
| ENSG00000272036 | 0 | 0 | 0 | 0 | 0 | 0 |
| ENSG00000251919 | 0 | 0 | 0 | 0 | 0 | 0 |
| ENSG00000255808 | 0 | 0 | 0 | 0 | 0 | 0 |
| ENSG00000186635 | 1707 | 816 | 1040 | 857 | 816 | 913 |
| ENSG00000256007 | 0 | 0 | 0 | 0 | 0 | 0 |
| ENSG00000245148 | 0 | 0 | 1 | 0 | 1 | 0 |
| ENSG00000244134 | 0 | 1 | 0 | 0 | 0 | 0 |
| ENSG00000214530 | 15280 | 7258 | 8829 | 8884 | 6864 | 7009 |
| ENSG00000202522 | 17 | 21 | 5 | 3 | 5 | 5 |
| ENSG00000265064 | 0 | 1 | 1 | 0 | 2 | 0 |
| ENSG00000285693 | 220 | 105 | 140 | 140 | 114 | 113 |
| ENSG00000168010 | 912 | 384 | 652 | 380 | 488 | 544 |
| ENSG00000137478 | 664 | 333 | 395 | 277 | 359 | 365 |
| ENSG00000206638 | 0 | 0 | 0 | 0 | 0 | 0 |
| ENSG00000234751 | 0 | 0 | 0 | 0 | 0 | 0 |
| ENSG00000256568 | 0 | 0 | 0 | 0 | 0 | 3 |
| ENSG00000215841 | 8 | 6 | 16 | 5 | 7 | 7 |
| ENSG00000175591 | 1332 | 569 | 775 | 598 | 660 | 809 |
| ENSG00000260401 | 1080 | 550 | 693 | 697 | 732 | 736 |
| ENSG00000256347 | 2 | 1 | 0 | 4 | 5 | 0 |
| ENSG00000171631 | 595 | 247 | 321 | 325 | 316 | 359 |
| ENSG00000257038 | 14 | 3 | 9 | 8 | 6 | 6 |
| ENSG00000110237 | 1305 | 581 | 874 | 599 | 666 | 729 |
| ENSG00000054967 | 177 | 101 | 99 | 168 | 137 | 132 |
| ENSG00000256928 | 0 | 0 | 0 | 0 | 0 | 0 |
| ENSG00000054965 | 3571 | 1610 | 2221 | 1426 | 1694 | 1894 |
| ENSG00000256448 | 3 | 0 | 0 | 3 | 0 | 0 |
| ENSG00000255786 | 0 | 0 | 0 | 0 | 0 | 0 |
| ENSG00000256148 | 2 | 2 | 2 | 0 | 4 | 1 |
| ENSG00000231726 | 0 | 0 | 0 | 0 | 0 | 0 |
| ENSG00000270323 | 0 | 0 | 0 | 0 | 0 | 0 |
| ENSG00000021300 | 1051 | 481 | 636 | 480 | 407 | 477 |
| ENSG00000175582 | 5774 | 3069 | 3679 | 2415 | 3026 | 3008 |
| ENSG00000255928 | 0 | 1 | 0 | 0 | 0 | 1 |
| ENSG00000256034 | 0 | 0 | 0 | 0 | 0 | 0 |
| ENSG00000175581 | 831 | 381 | 481 | 463 | 468 | 442 |
| ENSG00000199975 | 0 | 0 | 0 | 0 | 0 | 0 |
| ENSG00000256681 | 1 | 0 | 0 | 0 | 0 | 0 |

| | | | | | | |
|---|---|---|---|---|---|---|
| ENSG00000181924 | 2106 | 1158 | 1310 | 1540 | 1382 | 1438 |
| ENSG00000175575 | 948 | 450 | 608 | 608 | 632 | 628 |
| ENSG00000256898 | 2 | 0 | 1 | 1 | 1 | 1 |
| ENSG00000187726 | 2 | 8 | 5 | 12 | 8 | 5 |
| ENSG00000255847 | 2 | 0 | 0 | 0 | 1 | 0 |
| ENSG00000175567 | 4375 | 2021 | 2639 | 1998 | 1654 | 2009 |
| ENSG00000256189 | 0 | 0 | 0 | 0 | 0 | 0 |
| ENSG00000256723 | 0 | 0 | 0 | 0 | 0 | 0 |
| ENSG00000175564 | 13 | 8 | 6 | 10 | 7 | 7 |
| ENSG00000168014 | 735 | 327 | 454 | 260 | 408 | 397 |
| ENSG00000256098 | 0 | 0 | 0 | 0 | 0 | 0 |
| ENSG00000214517 | 1496 | 664 | 822 | 892 | 793 | 831 |
| ENSG00000284736 | 0 | 0 | 0 | 0 | 0 | 0 |
| ENSG00000280269 | 4 | 0 | 0 | 1 | 0 | 0 |
| ENSG00000149380 | 14 | 4 | 5 | 2 | 5 | 2 |
| ENSG00000206913 | 0 | 1 | 0 | 1 | 0 | 1 |
| ENSG00000246211 | 0 | 0 | 0 | 0 | 0 | 0 |
| ENSG00000165434 | 506 | 306 | 197 | 131 | 373 | 203 |
| ENSG00000212961 | 1 | 0 | 0 | 0 | 0 | 0 |
| ENSG00000254631 | 0 | 2 | 1 | 0 | 0 | 0 |
| ENSG00000264402 | 0 | 0 | 0 | 0 | 0 | 0 |
| ENSG00000255440 | 0 | 1 | 0 | 0 | 0 | 0 |
| ENSG00000175538 | 0 | 0 | 0 | 0 | 0 | 0 |
| ENSG00000254928 | 0 | 0 | 0 | 0 | 0 | 0 |
| ENSG00000255444 | 0 | 0 | 0 | 0 | 0 | 0 |
| ENSG00000254974 | 0 | 0 | 0 | 0 | 0 | 0 |
| ENSG00000175536 | 82 | 48 | 49 | 30 | 30 | 49 |
| ENSG00000254837 | 143 | 62 | 90 | 55 | 69 | 75 |
| ENSG00000077514 | 655 | 331 | 355 | 247 | 353 | 365 |
| ENSG00000254500 | 0 | 0 | 0 | 0 | 0 | 0 |
| ENSG00000223202 | 0 | 0 | 0 | 0 | 0 | 0 |
| ENSG00000054938 | 0 | 0 | 1 | 0 | 0 | 0 |
| ENSG00000279353 | 0 | 0 | 0 | 1 | 0 | 0 |
| ENSG00000265902 | 0 | 0 | 0 | 0 | 0 | 0 |
| ENSG00000241097 | 0 | 0 | 0 | 0 | 0 | 0 |
| ENSG00000227615 | 52 | 23 | 24 | 26 | 21 | 21 |
| ENSG00000166439 | 478 | 224 | 295 | 164 | 298 | 288 |
| ENSG00000166435 | 196 | 122 | 122 | 74 | 113 | 105 |
| ENSG00000278879 | 2 | 1 | 2 | 0 | 4 | 1 |
| ENSG00000242999 | 2 | 0 | 0 | 0 | 0 | 2 |
| ENSG00000241170 | 18 | 10 | 12 | 5 | 7 | 8 |
| ENSG00000118363 | 1515 | 757 | 922 | 708 | 912 | 836 |
| ENSG00000200152 | 3 | 3 | 6 | 2 | 5 | 1 |
| ENSG00000162139 | 720 | 289 | 472 | 342 | 472 | 434 |
| ENSG00000284722 | 0 | 0 | 0 | 0 | 0 | 0 |
| ENSG00000234611 | 0 | 0 | 0 | 0 | 0 | 0 |
| ENSG00000213333 | 0 | 0 | 0 | 0 | 0 | 0 |
| ENSG00000171561 | 0 | 0 | 0 | 0 | 0 | 0 |
| ENSG00000255395 | 0 | 0 | 0 | 0 | 1 | 0 |
| ENSG00000137491 | 0 | 0 | 1 | 1 | 0 | 0 |

| | | | | | | |
|---|---|---|---|---|---|---|
| ENSG00000254475 | 0 | 0 | 0 | 0 | 0 | 0 |
| ENSG00000255136 | 1 | 0 | 0 | 0 | 0 | 0 |
| ENSG00000255460 | 0 | 0 | 1 | 0 | 0 | 0 |
| ENSG00000261594 | 0 | 1 | 1 | 6 | 2 | 1 |
| ENSG00000254429 | 0 | 0 | 0 | 0 | 0 | 0 |
| ENSG00000279117 | 905 | 505 | 544 | 480 | 631 | 520 |
| ENSG00000137486 | 1449 | 651 | 844 | 590 | 678 | 703 |
| ENSG00000254963 | 0 | 0 | 0 | 0 | 0 | 0 |
| ENSG00000199090 | 0 | 0 | 0 | 0 | 0 | 0 |
| ENSG00000149273 | 57798 | 29448 | 33942 | 44044 | 37238 | 35965 |
| ENSG00000206941 | 0 | 0 | 2 | 0 | 0 | 1 |
| ENSG00000207445 | 5 | 3 | 3 | 8 | 1 | 11 |
| ENSG00000149243 | 191 | 93 | 108 | 136 | 110 | 108 |
| ENSG00000158555 | 102 | 67 | 99 | 72 | 74 | 82 |
| ENSG00000254460 | 0 | 0 | 0 | 0 | 0 | 0 |
| ENSG00000149257 | 5223 | 2570 | 2867 | 2976 | 2690 | 2604 |
| ENSG00000255326 | 0 | 0 | 1 | 0 | 0 | 0 |
| ENSG00000171533 | 20 | 7 | 10 | 3 | 7 | 12 |
| ENSG00000255434 | 5 | 0 | 1 | 0 | 1 | 2 |
| ENSG00000254630 | 0 | 0 | 0 | 0 | 0 | 0 |
| ENSG00000255280 | 0 | 0 | 0 | 0 | 0 | 0 |
| ENSG00000166391 | 0 | 0 | 0 | 0 | 0 | 0 |
| ENSG00000263993 | 0 | 0 | 0 | 0 | 0 | 0 |
| ENSG00000247867 | 2 | 0 | 0 | 0 | 0 | 0 |
| ENSG00000062282 | 344 | 194 | 226 | 238 | 235 | 222 |
| ENSG00000254826 | 0 | 0 | 0 | 1 | 0 | 0 |
| ENSG00000254814 | 4 | 0 | 3 | 0 | 0 | 0 |
| ENSG00000255507 | 14 | 4 | 4 | 6 | 3 | 1 |
| ENSG00000198382 | 1311 | 583 | 681 | 457 | 687 | 577 |
| ENSG00000200256 | 0 | 2 | 0 | 0 | 0 | 0 |
| ENSG00000199362 | 0 | 0 | 0 | 0 | 0 | 0 |
| ENSG00000255255 | 0 | 0 | 1 | 0 | 0 | 0 |
| ENSG00000255081 | 0 | 0 | 0 | 0 | 0 | 0 |
| ENSG00000223013 | 0 | 0 | 0 | 0 | 0 | 0 |
| ENSG00000254915 | 0 | 0 | 0 | 0 | 0 | 0 |
| ENSG00000255421 | 3 | 1 | 1 | 1 | 1 | 1 |
| ENSG00000085741 | 19 | 23 | 9 | 8 | 2 | 3 |
| ENSG00000254933 | 4 | 0 | 1 | 0 | 0 | 0 |
| ENSG00000255362 | 0 | 1 | 0 | 0 | 0 | 0 |
| ENSG00000137492 | 1931 | 1006 | 1109 | 949 | 1155 | 1090 |
| ENSG00000179240 | 140 | 58 | 61 | 49 | 55 | 63 |
| ENSG00000272301 | 6 | 9 | 15 | 2 | 6 | 6 |
| ENSG00000271757 | 2 | 1 | 1 | 1 | 0 | 1 |
| ENSG00000201756 | 0 | 0 | 0 | 0 | 0 | 0 |
| ENSG00000255135 | 210 | 121 | 138 | 161 | 126 | 111 |
| ENSG00000158636 | 824 | 452 | 459 | 212 | 547 | 403 |
| ENSG00000254755 | 0 | 0 | 0 | 0 | 0 | 0 |
| ENSG00000255363 | 0 | 0 | 0 | 1 | 0 | 0 |
| ENSG00000255479 | 0 | 0 | 0 | 0 | 0 | 0 |
| ENSG00000254810 | 0 | 0 | 0 | 0 | 0 | 0 |

| | | | | | | |
|---|---|---|---|---|---|---|
| ENSG00000236304 | 0 | 0 | 0 | 0 | 0 | 0 |
| ENSG00000137507 | 0 | 0 | 1 | 0 | 1 | 0 |
| ENSG00000254975 | 0 | 0 | 0 | 0 | 0 | 0 |
| ENSG00000204529 | 0 | 1 | 1 | 1 | 1 | 5 |
| ENSG00000254632 | 9 | 5 | 19 | 10 | 27 | 21 |
| ENSG00000182704 | 3855 | 1845 | 2428 | 4501 | 4246 | 4924 |
| ENSG00000255100 | 23 | 9 | 19 | 18 | 25 | 17 |
| ENSG00000261578 | 374 | 214 | 231 | 378 | 523 | 509 |
| ENSG00000078124 | 777 | 420 | 416 | 393 | 629 | 539 |
| ENSG00000254988 | 0 | 1 | 0 | 0 | 0 | 0 |
| ENSG00000198488 | 0 | 0 | 0 | 0 | 0 | 0 |
| ENSG00000149260 | 1589 | 691 | 949 | 738 | 646 | 847 |
| ENSG00000254550 | 1 | 0 | 1 | 0 | 2 | 1 |
| ENSG00000137474 | 54 | 30 | 25 | 26 | 18 | 28 |
| ENSG00000178795 | 0 | 0 | 0 | 0 | 0 | 0 |
| ENSG00000224829 | 0 | 0 | 0 | 0 | 0 | 0 |
| ENSG00000149269 | 528 | 314 | 337 | 302 | 357 | 289 |
| ENSG00000268635 | 0 | 0 | 0 | 0 | 0 | 0 |
| ENSG00000074201 | 3147 | 1594 | 1947 | 1894 | 1978 | 2007 |
| ENSG00000238880 | 0 | 0 | 0 | 0 | 0 | 0 |
| ENSG00000178301 | 194 | 96 | 125 | 75 | 73 | 94 |
| ENSG00000048649 | 737 | 428 | 296 | 142 | 846 | 349 |
| ENSG00000206816 | 0 | 1 | 0 | 0 | 0 | 0 |
| ENSG00000254985 | 0 | 1 | 5 | 1 | 3 | 0 |
| ENSG00000227376 | 7 | 2 | 3 | 2 | 5 | 2 |
| ENSG00000255409 | 0 | 0 | 0 | 1 | 2 | 0 |
| ENSG00000219529 | 4 | 2 | 1 | 5 | 0 | 1 |
| ENSG00000087884 | 153 | 117 | 132 | 128 | 141 | 111 |
| ENSG00000254459 | 2 | 0 | 0 | 0 | 0 | 0 |
| ENSG00000254691 | 0 | 0 | 1 | 0 | 2 | 0 |
| ENSG00000255449 | 3 | 1 | 1 | 0 | 1 | 1 |
| ENSG00000241782 | 0 | 0 | 2 | 0 | 0 | 0 |
| ENSG00000149262 | 752 | 355 | 393 | 356 | 427 | 399 |
| ENSG00000255115 | 0 | 0 | 0 | 0 | 0 | 0 |
| ENSG00000254829 | 0 | 0 | 0 | 0 | 0 | 0 |
| ENSG00000151364 | 6 | 0 | 6 | 0 | 2 | 2 |
| ENSG00000259112 | 0 | 0 | 0 | 0 | 0 | 0 |
| ENSG00000254675 | 0 | 0 | 0 | 0 | 0 | 0 |
| ENSG00000151365 | 4 | 1 | 2 | 1 | 2 | 0 |
| ENSG00000151366 | 857 | 523 | 657 | 709 | 568 | 605 |
| ENSG00000159063 | 1802 | 900 | 961 | 862 | 1067 | 1038 |
| ENSG00000252494 | 1 | 0 | 0 | 1 | 1 | 0 |
| ENSG00000246174 | 56 | 28 | 28 | 19 | 29 | 34 |
| ENSG00000188997 | 685 | 336 | 338 | 310 | 357 | 311 |
| ENSG00000118369 | 535 | 249 | 306 | 217 | 286 | 284 |
| ENSG00000033327 | 613 | 262 | 352 | 315 | 367 | 400 |
| ENSG00000254420 | 0 | 1 | 0 | 0 | 0 | 0 |
| ENSG00000254649 | 0 | 0 | 0 | 0 | 0 | 0 |
| ENSG00000251323 | 2 | 2 | 3 | 1 | 6 | 4 |
| ENSG00000137513 | 711 | 358 | 451 | 400 | 490 | 457 |

| | | | | | | |
|---|---|---|---|---|---|---|
| ENSG00000255084 | 0 | 0 | 0 | 0 | 0 | 0 |
| ENSG00000252033 | 0 | 0 | 0 | 0 | 0 | 0 |
| ENSG00000254697 | 0 | 0 | 0 | 0 | 0 | 0 |
| ENSG00000149256 | 36 | 13 | 37 | 57 | 92 | 118 |
| ENSG00000254563 | 0 | 0 | 0 | 0 | 0 | 0 |
| ENSG00000255345 | 0 | 1 | 0 | 2 | 0 | 1 |
| ENSG00000254885 | 0 | 0 | 0 | 0 | 0 | 0 |
| ENSG00000211997 | 0 | 0 | 0 | 0 | 0 | 0 |
| ENSG00000266570 | 0 | 0 | 0 | 0 | 0 | 0 |
| ENSG00000255209 | 0 | 0 | 0 | 0 | 0 | 0 |
| ENSG00000279295 | 0 | 0 | 0 | 0 | 0 | 0 |
| ENSG00000254471 | 27 | 4 | 11 | 1 | 2 | 4 |
| ENSG00000254434 | 0 | 0 | 0 | 0 | 0 | 0 |
| ENSG00000200146 | 0 | 0 | 0 | 0 | 0 | 0 |
| ENSG00000255014 | 0 | 1 | 0 | 0 | 0 | 0 |
| ENSG00000255178 | 28 | 23 | 23 | 16 | 25 | 6 |
| ENSG00000254437 | 0 | 0 | 0 | 0 | 0 | 0 |
| ENSG00000285568 | 4 | 2 | 2 | 1 | 2 | 3 |
| ENSG00000254747 | 0 | 0 | 0 | 0 | 0 | 0 |
| ENSG00000255316 | 0 | 0 | 0 | 0 | 0 | 0 |
| ENSG00000270701 | 0 | 0 | 0 | 0 | 0 | 0 |
| ENSG00000245832 | 0 | 0 | 0 | 0 | 0 | 0 |
| ENSG00000264110 | 0 | 0 | 0 | 0 | 0 | 0 |
| ENSG00000254511 | 0 | 0 | 0 | 0 | 0 | 0 |
| ENSG00000255246 | 0 | 0 | 0 | 0 | 0 | 0 |
| ENSG00000227097 | 15 | 7 | 13 | 10 | 10 | 5 |
| ENSG00000182103 | 0 | 1 | 1 | 1 | 1 | 0 |
| ENSG00000255382 | 0 | 0 | 0 | 0 | 0 | 0 |
| ENSG00000255063 | 0 | 0 | 0 | 0 | 0 | 0 |
| ENSG00000137509 | 993 | 489 | 578 | 582 | 603 | 575 |
| ENSG00000254441 | 0 | 0 | 0 | 0 | 0 | 0 |
| ENSG00000165490 | 127 | 100 | 79 | 57 | 156 | 110 |
| ENSG00000242279 | 0 | 0 | 0 | 0 | 0 | 0 |
| ENSG00000254698 | 0 | 0 | 0 | 0 | 0 | 0 |
| ENSG00000137502 | 976 | 484 | 498 | 251 | 355 | 383 |
| ENSG00000254522 | 0 | 0 | 0 | 0 | 0 | 0 |
| ENSG00000207221 | 0 | 0 | 0 | 0 | 0 | 0 |
| ENSG00000246067 | 158 | 123 | 103 | 91 | 103 | 65 |
| ENSG00000255503 | 3 | 1 | 3 | 2 | 2 | 0 |
| ENSG00000279900 | 4 | 4 | 5 | 6 | 4 | 1 |
| ENSG00000255010 | 0 | 0 | 0 | 2 | 0 | 0 |
| ENSG00000254965 | 0 | 0 | 0 | 0 | 0 | 0 |
| ENSG00000255051 | 0 | 0 | 0 | 0 | 0 | 0 |
| ENSG00000165494 | 1930 | 994 | 1172 | 633 | 1310 | 951 |
| ENSG00000254676 | 28 | 10 | 11 | 10 | 8 | 8 |
| ENSG00000247137 | 56 | 30 | 40 | 21 | 38 | 31 |
| ENSG00000269939 | 0 | 0 | 0 | 0 | 0 | 0 |
| ENSG00000137494 | 697 | 323 | 415 | 264 | 346 | 370 |
| ENSG00000255541 | 0 | 0 | 0 | 0 | 0 | 0 |
| ENSG00000254551 | 0 | 0 | 0 | 0 | 0 | 0 |

| | | | | | | |
|---|---|---|---|---|---|---|
| ENSG00000241020 | 0 | 0 | 0 | 0 | 0 | 0 |
| ENSG00000137500 | 1495 | 675 | 779 | 677 | 941 | 799 |
| ENSG00000255234 | 4 | 2 | 2 | 1 | 1 | 0 |
| ENSG00000255003 | 0 | 0 | 0 | 0 | 0 | 0 |
| ENSG00000274004 | 0 | 0 | 0 | 0 | 0 | 0 |
| ENSG00000150672 | 30 | 17 | 33 | 13 | 25 | 18 |
| ENSG00000255311 | 0 | 0 | 0 | 0 | 0 | 0 |
| ENSG00000254629 | 0 | 0 | 0 | 0 | 0 | 0 |
| ENSG00000250257 | 0 | 0 | 0 | 0 | 0 | 0 |
| ENSG00000254713 | 0 | 0 | 0 | 0 | 0 | 0 |
| ENSG00000240174 | 0 | 0 | 0 | 0 | 0 | 0 |
| ENSG00000254787 | 0 | 0 | 1 | 0 | 0 | 0 |
| ENSG00000274006 | 0 | 0 | 0 | 0 | 0 | 0 |
| ENSG00000276179 | 0 | 0 | 0 | 0 | 0 | 0 |
| ENSG00000255555 | 0 | 0 | 0 | 0 | 0 | 0 |
| ENSG00000255396 | 0 | 0 | 0 | 0 | 0 | 0 |
| ENSG00000213305 | 0 | 0 | 0 | 0 | 0 | 0 |
| ENSG00000254684 | 0 | 0 | 0 | 0 | 0 | 0 |
| ENSG00000254897 | 0 | 0 | 0 | 0 | 0 | 0 |
| ENSG00000206583 | 0 | 0 | 0 | 0 | 0 | 0 |
| ENSG00000171204 | 1123 | 575 | 707 | 677 | 690 | 671 |
| ENSG00000171202 | 518 | 250 | 287 | 315 | 325 | 325 |
| ENSG00000137504 | 4345 | 2044 | 2783 | 1561 | 2306 | 2158 |
| ENSG00000179071 | 21 | 4 | 7 | 1 | 10 | 4 |
| ENSG00000137501 | 12799 | 7206 | 7628 | 2750 | 7291 | 5375 |
| ENSG00000279742 | 48 | 24 | 43 | 11 | 24 | 23 |
| ENSG00000150676 | 380 | 196 | 233 | 82 | 205 | 135 |
| ENSG00000255005 | 13 | 6 | 13 | 2 | 1 | 6 |
| ENSG00000254479 | 8 | 3 | 1 | 1 | 0 | 2 |
| ENSG00000073921 | 7827 | 3891 | 4854 | 2919 | 3905 | 3750 |
| ENSG00000200877 | 0 | 0 | 0 | 0 | 0 | 0 |
| ENSG00000254699 | 0 | 0 | 0 | 0 | 0 | 0 |
| ENSG00000254502 | 0 | 0 | 0 | 0 | 0 | 0 |
| ENSG00000276669 | 0 | 0 | 0 | 0 | 0 | 0 |
| ENSG00000074266 | 740 | 366 | 455 | 305 | 355 | 381 |
| ENSG00000283744 | 0 | 0 | 0 | 0 | 0 | 0 |
| ENSG00000254783 | 24 | 11 | 8 | 3 | 3 | 9 |
| ENSG00000255222 | 0 | 0 | 0 | 1 | 0 | 1 |
| ENSG00000149196 | 1175 | 559 | 667 | 730 | 703 | 579 |
| ENSG00000239856 | 0 | 0 | 0 | 0 | 0 | 0 |
| ENSG00000149201 | 13 | 7 | 10 | 3 | 6 | 10 |
| ENSG00000254733 | 0 | 0 | 0 | 0 | 0 | 0 |
| ENSG00000254727 | 0 | 0 | 0 | 0 | 0 | 0 |
| ENSG00000278989 | 0 | 1 | 0 | 0 | 0 | 0 |
| ENSG00000151376 | 0 | 0 | 0 | 3 | 1 | 0 |
| ENSG00000254731 | 0 | 0 | 0 | 0 | 0 | 0 |
| ENSG00000255250 | 0 | 0 | 0 | 0 | 0 | 0 |
| ENSG00000150687 | 4364 | 2279 | 2107 | 2236 | 1890 | 2186 |
| ENSG00000271318 | 0 | 0 | 0 | 0 | 0 | 1 |
| ENSG00000197882 | 3 | 0 | 0 | 0 | 0 | 0 |

| | | | | | | |
|---|---|---|---|---|---|---|
| ENSG00000269895 | 2 | 2 | 0 | 0 | 0 | 0 |
| ENSG00000255203 | 0 | 0 | 0 | 0 | 0 | 0 |
| ENSG00000255471 | 16 | 15 | 19 | 13 | 10 | 10 |
| ENSG00000280339 | 30 | 17 | 18 | 3 | 7 | 10 |
| ENSG00000174804 | 721 | 328 | 571 | 233 | 296 | 336 |
| ENSG00000246523 | 13 | 7 | 4 | 8 | 3 | 10 |
| ENSG00000254748 | 0 | 0 | 0 | 1 | 0 | 0 |
| ENSG00000166575 | 984 | 470 | 607 | 285 | 466 | 400 |
| ENSG00000254843 | 0 | 0 | 0 | 0 | 1 | 1 |
| ENSG00000279836 | 0 | 0 | 0 | 0 | 0 | 0 |
| ENSG00000213287 | 0 | 0 | 0 | 0 | 1 | 0 |
| ENSG00000270510 | 7 | 1 | 3 | 0 | 3 | 1 |
| ENSG00000254582 | 0 | 0 | 0 | 0 | 0 | 0 |
| ENSG00000255391 | 0 | 0 | 0 | 0 | 0 | 0 |
| ENSG00000279701 | 0 | 0 | 0 | 0 | 0 | 0 |
| ENSG00000278929 | 0 | 0 | 0 | 0 | 0 | 0 |
| ENSG00000223015 | 0 | 0 | 0 | 0 | 0 | 0 |
| ENSG00000283412 | 0 | 0 | 0 | 0 | 0 | 0 |
| ENSG00000255183 | 0 | 0 | 0 | 0 | 0 | 0 |
| ENSG00000279341 | 0 | 0 | 0 | 0 | 0 | 0 |
| ENSG00000254650 | 0 | 0 | 0 | 0 | 0 | 0 |
| ENSG00000255022 | 0 | 0 | 0 | 0 | 0 | 0 |
| ENSG00000279793 | 0 | 0 | 0 | 0 | 0 | 0 |
| ENSG00000285835 | 0 | 0 | 0 | 0 | 0 | 0 |
| ENSG00000280307 | 0 | 0 | 0 | 0 | 0 | 0 |
| ENSG00000254794 | 0 | 0 | 0 | 0 | 0 | 0 |
| ENSG00000255516 | 0 | 0 | 0 | 0 | 0 | 0 |
| ENSG00000255102 | 0 | 0 | 0 | 0 | 0 | 0 |
| ENSG00000255241 | 0 | 0 | 0 | 0 | 0 | 0 |
| ENSG00000206690 | 0 | 0 | 0 | 0 | 0 | 0 |
| ENSG00000123892 | 0 | 0 | 0 | 0 | 0 | 0 |
| ENSG00000279733 | 0 | 0 | 0 | 0 | 0 | 0 |
| ENSG00000266581 | 0 | 0 | 0 | 0 | 0 | 0 |
| ENSG00000109861 | 604 | 292 | 348 | 293 | 300 | 348 |
| ENSG00000249489 | 0 | 1 | 0 | 0 | 0 | 0 |
| ENSG00000168959 | 0 | 0 | 0 | 0 | 0 | 0 |
| ENSG00000255082 | 0 | 0 | 0 | 0 | 0 | 0 |
| ENSG00000207113 | 0 | 0 | 0 | 0 | 0 | 0 |
| ENSG00000077498 | 0 | 0 | 0 | 0 | 0 | 0 |
| ENSG00000197692 | 0 | 0 | 0 | 0 | 0 | 0 |
| ENSG00000279297 | 0 | 0 | 0 | 0 | 0 | 0 |
| ENSG00000279603 | 0 | 0 | 0 | 0 | 0 | 0 |
| ENSG00000086991 | 0 | 0 | 0 | 0 | 0 | 0 |
| ENSG00000255295 | 0 | 0 | 0 | 0 | 0 | 0 |
| ENSG00000279248 | 0 | 0 | 0 | 0 | 0 | 0 |
| ENSG00000279697 | 0 | 0 | 0 | 0 | 0 | 0 |
| ENSG00000255429 | 0 | 0 | 0 | 0 | 0 | 0 |
| ENSG00000279045 | 0 | 0 | 0 | 0 | 0 | 0 |
| ENSG00000279056 | 0 | 0 | 0 | 0 | 0 | 0 |
| ENSG00000280201 | 0 | 0 | 0 | 0 | 0 | 0 |

| | | | | | | |
|---|---|---|---|---|---|---|
| ENSG00000134612 | 0 | 0 | 0 | 0 | 1 | 0 |
| ENSG00000255385 | 0 | 0 | 0 | 0 | 0 | 0 |
| ENSG00000279163 | 0 | 0 | 0 | 0 | 0 | 0 |
| ENSG00000214414 | 0 | 0 | 0 | 0 | 0 | 0 |
| ENSG00000255486 | 0 | 0 | 0 | 0 | 0 | 0 |
| ENSG00000255540 | 0 | 0 | 0 | 0 | 0 | 0 |
| ENSG00000234376 | 0 | 0 | 0 | 0 | 0 | 0 |
| ENSG00000254971 | 0 | 0 | 0 | 0 | 0 | 0 |
| ENSG00000255170 | 0 | 0 | 0 | 0 | 0 | 0 |
| ENSG00000204456 | 0 | 0 | 0 | 0 | 0 | 0 |
| ENSG00000254751 | 0 | 0 | 0 | 0 | 0 | 0 |
| ENSG00000255162 | 0 | 0 | 0 | 0 | 0 | 0 |
| ENSG00000255285 | 0 | 0 | 0 | 0 | 0 | 0 |
| ENSG00000168930 | 0 | 0 | 0 | 0 | 0 | 0 |
| ENSG00000250946 | 0 | 0 | 0 | 0 | 0 | 0 |
| ENSG00000166013 | 0 | 0 | 0 | 0 | 0 | 0 |
| ENSG00000204455 | 0 | 0 | 0 | 0 | 0 | 0 |
| ENSG00000254558 | 0 | 0 | 0 | 0 | 0 | 0 |
| ENSG00000189253 | 0 | 0 | 0 | 0 | 0 | 0 |
| ENSG00000255235 | 0 | 0 | 0 | 0 | 0 | 0 |
| ENSG00000254617 | 0 | 0 | 0 | 0 | 0 | 0 |
| ENSG00000254785 | 0 | 0 | 0 | 0 | 0 | 0 |
| ENSG00000255055 | 0 | 0 | 0 | 0 | 0 | 0 |
| ENSG00000223417 | 0 | 0 | 0 | 0 | 0 | 0 |
| ENSG00000233802 | 0 | 0 | 0 | 0 | 0 | 0 |
| ENSG00000254818 | 0 | 0 | 0 | 0 | 0 | 0 |
| ENSG00000278837 | 0 | 0 | 0 | 0 | 0 | 0 |
| ENSG00000255360 | 0 | 0 | 0 | 0 | 0 | 0 |
| ENSG00000254655 | 0 | 0 | 0 | 0 | 0 | 0 |
| ENSG00000204450 | 0 | 0 | 0 | 0 | 0 | 0 |
| ENSG00000255305 | 0 | 0 | 0 | 0 | 0 | 0 |
| ENSG00000237706 | 0 | 0 | 0 | 0 | 0 | 0 |
| ENSG00000225581 | 0 | 0 | 0 | 0 | 0 | 0 |
| ENSG00000255184 | 0 | 0 | 0 | 0 | 0 | 0 |
| ENSG00000204449 | 0 | 0 | 0 | 0 | 0 | 0 |
| ENSG00000254803 | 0 | 0 | 0 | 0 | 0 | 0 |
| ENSG00000254916 | 0 | 0 | 0 | 0 | 0 | 0 |
| ENSG00000223709 | 1 | 0 | 0 | 0 | 0 | 0 |
| ENSG00000255011 | 0 | 0 | 0 | 0 | 0 | 0 |
| ENSG00000254888 | 0 | 0 | 0 | 0 | 0 | 0 |
| ENSG00000255009 | 0 | 0 | 0 | 0 | 0 | 0 |
| ENSG00000254436 | 0 | 0 | 0 | 0 | 0 | 0 |
| ENSG00000207299 | 0 | 0 | 0 | 0 | 0 | 0 |
| ENSG00000279299 | 0 | 0 | 0 | 0 | 0 | 0 |
| ENSG00000077616 | 6 | 0 | 2 | 0 | 3 | 0 |
| ENSG00000280385 | 10 | 5 | 6 | 1 | 9 | 6 |
| ENSG00000110172 | 545 | 333 | 308 | 224 | 396 | 290 |
| ENSG00000280367 | 16 | 8 | 8 | 0 | 1 | 3 |
| ENSG00000261645 | 1 | 0 | 1 | 0 | 0 | 0 |
| ENSG00000214391 | 6 | 1 | 4 | 6 | 3 | 4 |

| | | | | | | |
|---|---|---|---|---|---|---|
| ENSG00000266703 | 0 | 0 | 0 | 0 | 0 | 0 |
| ENSG00000280430 | 0 | 0 | 0 | 0 | 0 | 0 |
| ENSG00000221586 | 0 | 0 | 0 | 0 | 0 | 0 |
| ENSG00000280085 | 0 | 0 | 0 | 0 | 0 | 0 |
| ENSG00000279209 | 0 | 0 | 0 | 0 | 0 | 0 |
| ENSG00000279438 | 0 | 0 | 0 | 0 | 0 | 0 |
| ENSG00000254677 | 0 | 0 | 0 | 0 | 0 | 0 |
| ENSG00000255070 | 0 | 0 | 0 | 0 | 0 | 0 |
| ENSG00000255187 | 0 | 0 | 0 | 0 | 0 | 0 |
| ENSG00000280124 | 0 | 0 | 0 | 0 | 0 | 0 |
| ENSG00000278980 | 0 | 0 | 0 | 0 | 0 | 0 |
| ENSG00000280093 | 0 | 0 | 0 | 0 | 0 | 0 |
| ENSG00000255332 | 0 | 0 | 0 | 0 | 0 | 0 |
| ENSG00000254961 | 0 | 0 | 0 | 0 | 0 | 0 |
| ENSG00000278859 | 0 | 0 | 0 | 0 | 0 | 0 |
| ENSG00000242248 | 0 | 0 | 0 | 0 | 0 | 0 |
| ENSG00000235547 | 0 | 0 | 0 | 0 | 0 | 0 |
| ENSG00000165323 | 0 | 0 | 0 | 0 | 0 | 0 |
| ENSG00000249255 | 0 | 0 | 0 | 0 | 0 | 0 |
| ENSG00000254705 | 0 | 0 | 0 | 0 | 0 | 0 |
| ENSG00000244244 | 0 | 0 | 0 | 0 | 0 | 0 |
| ENSG00000255506 | 0 | 0 | 0 | 0 | 0 | 0 |
| ENSG00000282916 | 0 | 0 | 0 | 0 | 0 | 0 |
| ENSG00000255127 | 0 | 0 | 0 | 0 | 0 | 0 |
| ENSG00000255287 | 0 | 0 | 0 | 0 | 0 | 0 |
| ENSG00000254874 | 0 | 0 | 0 | 0 | 0 | 0 |
| ENSG00000134640 | 0 | 0 | 0 | 0 | 0 | 0 |
| ENSG00000242807 | 0 | 0 | 0 | 0 | 0 | 0 |
| ENSG00000278892 | 0 | 0 | 0 | 0 | 0 | 0 |
| ENSG00000279304 | 0 | 0 | 0 | 0 | 0 | 0 |
| ENSG00000180773 | 84 | 33 | 33 | 46 | 36 | 62 |
| ENSG00000255445 | 0 | 1 | 0 | 0 | 0 | 0 |
| ENSG00000280379 | 0 | 0 | 0 | 0 | 0 | 0 |
| ENSG00000279269 | 0 | 0 | 0 | 0 | 0 | 0 |
| ENSG00000233737 | 0 | 0 | 0 | 0 | 0 | 0 |
| ENSG00000255233 | 0 | 0 | 0 | 0 | 0 | 0 |
| ENSG00000279684 | 0 | 0 | 0 | 0 | 0 | 0 |
| ENSG00000165325 | 0 | 0 | 0 | 0 | 0 | 0 |
| ENSG00000166002 | 48 | 32 | 27 | 49 | 21 | 23 |
| ENSG00000234106 | 2 | 1 | 2 | 0 | 2 | 2 |
| ENSG00000240202 | 0 | 0 | 0 | 0 | 0 | 0 |
| ENSG00000279454 | 0 | 0 | 0 | 0 | 0 | 0 |
| ENSG00000278953 | 0 | 0 | 0 | 0 | 0 | 0 |
| ENSG00000255515 | 0 | 0 | 0 | 0 | 0 | 0 |
| ENSG00000166004 | 52 | 48 | 32 | 15 | 139 | 38 |
| ENSG00000199875 | 0 | 0 | 0 | 0 | 0 | 0 |
| ENSG00000254911 | 2 | 0 | 2 | 1 | 1 | 1 |
| ENSG00000275146 | 0 | 0 | 0 | 0 | 0 | 0 |
| ENSG00000273885 | 0 | 0 | 0 | 0 | 0 | 0 |
| ENSG00000279696 | 20 | 6 | 2 | 6 | 7 | 5 |

| | | | | | | |
|---|---|---|---|---|---|---|
| ENSG00000166012 | 1059 | 642 | 626 | 550 | 925 | 815 |
| ENSG00000207112 | 0 | 0 | 0 | 0 | 0 | 0 |
| ENSG00000206799 | 0 | 0 | 0 | 0 | 0 | 0 |
| ENSG00000202314 | 5 | 1 | 0 | 4 | 6 | 3 |
| ENSG00000206834 | 0 | 0 | 0 | 0 | 0 | 0 |
| ENSG00000207304 | 0 | 0 | 0 | 0 | 0 | 0 |
| ENSG00000239195 | 0 | 0 | 0 | 0 | 0 | 0 |
| ENSG00000207145 | 0 | 0 | 0 | 0 | 0 | 0 |
| ENSG00000284458 | 0 | 0 | 0 | 0 | 0 | 0 |
| ENSG00000210825 | 0 | 0 | 0 | 0 | 0 | 0 |
| ENSG00000182919 | 323 | 160 | 204 | 108 | 171 | 145 |
| ENSG00000284057 | 0 | 0 | 0 | 0 | 0 | 0 |
| ENSG00000042429 | 96 | 60 | 86 | 49 | 78 | 60 |
| ENSG00000214376 | 0 | 2 | 0 | 0 | 1 | 3 |
| ENSG00000206911 | 0 | 0 | 0 | 0 | 0 | 0 |
| ENSG00000255995 | 0 | 0 | 0 | 0 | 0 | 0 |
| ENSG00000256752 | 0 | 0 | 0 | 0 | 0 | 0 |
| ENSG00000181333 | 22 | 25 | 15 | 13 | 19 | 13 |
| ENSG00000227593 | 0 | 0 | 0 | 0 | 0 | 0 |
| ENSG00000110218 | 302 | 161 | 221 | 184 | 201 | 173 |
| ENSG00000256745 | 7 | 7 | 9 | 5 | 6 | 9 |
| ENSG00000250519 | 5 | 4 | 2 | 6 | 1 | 3 |
| ENSG00000183560 | 0 | 0 | 0 | 0 | 0 | 0 |
| ENSG00000123901 | 4 | 3 | 2 | 1 | 2 | 1 |
| ENSG00000020922 | 324 | 241 | 179 | 93 | 323 | 204 |
| ENSG00000221230 | 0 | 0 | 0 | 0 | 0 | 0 |
| ENSG00000255893 | 0 | 2 | 2 | 0 | 0 | 1 |
| ENSG00000168876 | 389 | 200 | 255 | 220 | 243 | 229 |
| ENSG00000257057 | 0 | 0 | 0 | 0 | 0 | 0 |
| ENSG00000134627 | 13 | 2 | 11 | 8 | 6 | 5 |
| ENSG00000196371 | 81 | 43 | 47 | 30 | 41 | 53 |
| ENSG00000255929 | 0 | 0 | 0 | 0 | 0 | 0 |
| ENSG00000280167 | 0 | 0 | 0 | 0 | 0 | 0 |
| ENSG00000233536 | 0 | 0 | 0 | 0 | 0 | 0 |
| ENSG00000255666 | 0 | 0 | 0 | 0 | 0 | 0 |
| ENSG00000166025 | 547 | 217 | 327 | 131 | 135 | 157 |
| ENSG00000256469 | 0 | 0 | 0 | 0 | 0 | 0 |
| ENSG00000213368 | 0 | 0 | 0 | 0 | 1 | 0 |
| ENSG00000150316 | 750 | 407 | 459 | 391 | 499 | 435 |
| ENSG00000186280 | 26 | 9 | 13 | 8 | 14 | 15 |
| ENSG00000255653 | 0 | 0 | 0 | 0 | 0 | 0 |
| ENSG00000235268 | 0 | 0 | 0 | 0 | 0 | 0 |
| ENSG00000274486 | 0 | 0 | 0 | 0 | 0 | 0 |
| ENSG00000256779 | 0 | 0 | 0 | 0 | 0 | 0 |
| ENSG00000257012 | 0 | 0 | 0 | 0 | 0 | 0 |
| ENSG00000255855 | 0 | 0 | 0 | 0 | 0 | 0 |
| ENSG00000277984 | 0 | 0 | 0 | 0 | 0 | 0 |
| ENSG00000263465 | 881 | 468 | 555 | 457 | 480 | 572 |
| ENSG00000149218 | 530 | 249 | 337 | 236 | 264 | 316 |
| ENSG00000270689 | 0 | 0 | 0 | 0 | 0 | 0 |

| | | | | | | |
|---|---|---|---|---|---|---|
| ENSG00000270578 | 0 | 0 | 0 | 0 | 0 | 0 |
| ENSG00000245552 | 12 | 3 | 10 | 1 | 3 | 3 |
| ENSG00000149212 | 312 | 179 | 184 | 64 | 176 | 131 |
| ENSG00000282834 | 0 | 0 | 0 | 0 | 0 | 0 |
| ENSG00000250390 | 0 | 0 | 0 | 0 | 0 | 0 |
| ENSG00000285842 | 0 | 0 | 0 | 0 | 0 | 0 |
| ENSG00000255605 | 0 | 0 | 0 | 0 | 0 | 0 |
| ENSG00000273600 | 0 | 0 | 0 | 0 | 0 | 0 |
| ENSG00000077458 | 311 | 130 | 146 | 70 | 126 | 105 |
| ENSG00000166037 | 461 | 253 | 234 | 136 | 369 | 220 |
| ENSG00000087053 | 490 | 270 | 312 | 254 | 419 | 316 |
| ENSG00000222578 | 0 | 0 | 0 | 0 | 0 | 0 |
| ENSG00000242993 | 0 | 0 | 0 | 0 | 0 | 0 |
| ENSG00000184384 | 54 | 31 | 36 | 9 | 20 | 32 |
| ENSG00000285921 | 0 | 0 | 1 | 0 | 0 | 0 |
| ENSG00000266192 | 0 | 0 | 0 | 0 | 0 | 0 |
| ENSG00000149231 | 32 | 12 | 16 | 11 | 39 | 16 |
| ENSG00000183340 | 208 | 91 | 129 | 72 | 131 | 87 |
| ENSG00000255679 | 0 | 0 | 1 | 0 | 0 | 0 |
| ENSG00000200411 | 0 | 0 | 0 | 0 | 0 | 0 |
| ENSG00000256684 | 0 | 0 | 0 | 0 | 0 | 0 |
| ENSG00000254587 | 0 | 0 | 0 | 0 | 0 | 0 |
| ENSG00000255376 | 0 | 0 | 0 | 0 | 0 | 0 |
| ENSG00000255205 | 0 | 0 | 0 | 0 | 0 | 0 |
| ENSG00000270753 | 0 | 0 | 0 | 0 | 0 | 0 |
| ENSG00000255039 | 0 | 0 | 0 | 0 | 0 | 0 |
| ENSG00000199315 | 0 | 0 | 0 | 0 | 0 | 0 |
| ENSG00000255502 | 0 | 0 | 0 | 0 | 0 | 0 |
| ENSG00000254555 | 0 | 0 | 0 | 0 | 0 | 0 |
| ENSG00000255380 | 0 | 0 | 0 | 0 | 0 | 0 |
| ENSG00000254939 | 0 | 0 | 0 | 0 | 0 | 0 |
| ENSG00000254599 | 0 | 0 | 0 | 0 | 0 | 0 |
| ENSG00000254830 | 0 | 0 | 0 | 0 | 0 | 0 |
| ENSG00000149972 | 0 | 0 | 0 | 0 | 0 | 0 |
| ENSG00000223269 | 0 | 0 | 0 | 0 | 0 | 0 |
| ENSG00000255547 | 0 | 0 | 0 | 0 | 0 | 0 |
| ENSG00000242165 | 0 | 0 | 0 | 0 | 0 | 0 |
| ENSG00000255059 | 0 | 0 | 0 | 0 | 0 | 0 |
| ENSG00000248027 | 8 | 3 | 4 | 2 | 5 | 5 |
| ENSG00000165895 | 142 | 69 | 63 | 28 | 85 | 73 |
| ENSG00000200047 | 0 | 0 | 0 | 0 | 0 | 0 |
| ENSG00000082175 | 140 | 68 | 50 | 48 | 94 | 52 |
| ENSG00000282728 | 8 | 3 | 5 | 3 | 7 | 1 |
| ENSG00000255504 | 0 | 0 | 0 | 0 | 0 | 0 |
| ENSG00000137672 | 0 | 1 | 0 | 0 | 0 | 1 |
| ENSG00000263885 | 0 | 0 | 0 | 0 | 0 | 0 |
| ENSG00000254506 | 0 | 0 | 0 | 0 | 1 | 0 |
| ENSG00000254534 | 0 | 0 | 0 | 0 | 0 | 0 |
| ENSG00000187151 | 0 | 0 | 1 | 2 | 0 | 0 |
| ENSG00000110318 | 74 | 39 | 37 | 7 | 50 | 23 |

| | | | | | | |
|---|---|---|---|---|---|---|
| ENSG00000137691 | 4 | 7 | 6 | 12 | 9 | 12 |
| ENSG00000274097 | 0 | 0 | 0 | 0 | 0 | 0 |
| ENSG00000260008 | 0 | 1 | 0 | 0 | 0 | 0 |
| ENSG00000277459 | 8 | 6 | 4 | 1 | 6 | 2 |
| ENSG00000137693 | 1685 | 753 | 954 | 558 | 781 | 804 |
| ENSG00000254422 | 0 | 0 | 0 | 0 | 0 | 0 |
| ENSG00000243777 | 0 | 0 | 0 | 0 | 0 | 0 |
| ENSG00000270868 | 0 | 0 | 0 | 0 | 0 | 0 |
| ENSG00000271600 | 0 | 0 | 0 | 0 | 0 | 0 |
| ENSG00000212466 | 0 | 0 | 0 | 0 | 0 | 0 |
| ENSG00000023445 | 54 | 26 | 29 | 20 | 49 | 29 |
| ENSG00000110330 | 655 | 368 | 344 | 191 | 497 | 363 |
| ENSG00000152558 | 2656 | 1348 | 1311 | 1050 | 1837 | 1493 |
| ENSG00000255337 | 5 | 6 | 3 | 4 | 4 | 9 |
| ENSG00000255482 | 0 | 0 | 0 | 0 | 0 | 0 |
| ENSG00000137673 | 0 | 0 | 0 | 0 | 0 | 0 |
| ENSG00000137674 | 0 | 0 | 0 | 0 | 0 | 0 |
| ENSG00000256916 | 0 | 0 | 0 | 0 | 0 | 0 |
| ENSG00000281655 | 3 | 0 | 2 | 0 | 0 | 0 |
| ENSG00000137675 | 0 | 0 | 0 | 0 | 0 | 0 |
| ENSG00000118113 | 0 | 0 | 0 | 0 | 0 | 0 |
| ENSG00000255282 | 2 | 1 | 1 | 3 | 3 | 3 |
| ENSG00000225678 | 4 | 0 | 3 | 0 | 0 | 3 |
| ENSG00000256254 | 0 | 0 | 0 | 0 | 0 | 0 |
| ENSG00000166670 | 0 | 0 | 0 | 1 | 0 | 0 |
| ENSG00000196611 | 1 | 1 | 0 | 0 | 1 | 1 |
| ENSG00000256035 | 0 | 0 | 0 | 0 | 1 | 0 |
| ENSG00000149968 | 0 | 0 | 1 | 0 | 1 | 0 |
| ENSG00000262406 | 0 | 0 | 0 | 0 | 0 | 0 |
| ENSG00000255469 | 0 | 0 | 0 | 0 | 0 | 0 |
| ENSG00000238562 | 0 | 0 | 0 | 0 | 0 | 0 |
| ENSG00000137745 | 6 | 5 | 4 | 10 | 10 | 5 |
| ENSG00000276521 | 0 | 0 | 0 | 0 | 0 | 0 |
| ENSG00000239861 | 0 | 0 | 0 | 0 | 0 | 0 |
| ENSG00000260966 | 16 | 9 | 3 | 5 | 16 | 16 |
| ENSG00000137692 | 873 | 418 | 552 | 413 | 461 | 461 |
| ENSG00000187240 | 72 | 32 | 26 | 18 | 77 | 49 |
| ENSG00000285878 | 0 | 0 | 0 | 0 | 0 | 0 |
| ENSG00000254895 | 0 | 0 | 0 | 0 | 0 | 0 |
| ENSG00000254931 | 0 | 0 | 0 | 0 | 0 | 0 |
| ENSG00000255417 | 0 | 0 | 0 | 0 | 0 | 0 |
| ENSG00000255350 | 0 | 0 | 0 | 0 | 0 | 0 |
| ENSG00000254565 | 0 | 0 | 0 | 0 | 0 | 0 |
| ENSG00000253245 | 0 | 0 | 0 | 0 | 0 | 0 |
| ENSG00000254824 | 0 | 0 | 0 | 0 | 0 | 0 |
| ENSG00000254987 | 0 | 0 | 0 | 0 | 0 | 0 |
| ENSG00000264200 | 0 | 0 | 0 | 0 | 0 | 0 |
| ENSG00000170962 | 0 | 0 | 0 | 0 | 0 | 0 |
| ENSG00000255548 | 0 | 0 | 0 | 0 | 0 | 0 |
| ENSG00000170967 | 0 | 0 | 0 | 0 | 0 | 0 |

| | | | | | | |
|---|---|---|---|---|---|---|
| ENSG00000280773 | 0 | 0 | 0 | 0 | 0 | 0 |
| ENSG00000256422 | 0 | 0 | 0 | 0 | 0 | 0 |
| ENSG00000270449 | 0 | 0 | 0 | 0 | 0 | 0 |
| ENSG00000254569 | 0 | 0 | 0 | 0 | 0 | 0 |
| ENSG00000204403 | 0 | 0 | 0 | 0 | 0 | 0 |
| ENSG00000235505 | 0 | 0 | 0 | 0 | 0 | 0 |
| ENSG00000196954 | 24 | 5 | 7 | 4 | 3 | 12 |
| ENSG00000284715 | 0 | 0 | 0 | 0 | 0 | 0 |
| ENSG00000137757 | 0 | 0 | 0 | 0 | 0 | 0 |
| ENSG00000137752 | 0 | 0 | 0 | 0 | 0 | 0 |
| ENSG00000204397 | 0 | 1 | 0 | 0 | 0 | 0 |
| ENSG00000254750 | 0 | 0 | 0 | 0 | 0 | 0 |
| ENSG00000255221 | 0 | 0 | 0 | 0 | 0 | 0 |
| ENSG00000255430 | 0 | 0 | 0 | 0 | 0 | 0 |
| ENSG00000255501 | 0 | 0 | 0 | 0 | 0 | 0 |
| ENSG00000254767 | 0 | 0 | 0 | 0 | 0 | 0 |
| ENSG00000255336 | 0 | 0 | 0 | 0 | 0 | 0 |
| ENSG00000278640 | 0 | 0 | 0 | 0 | 0 | 0 |
| ENSG00000152578 | 0 | 0 | 0 | 0 | 0 | 0 |
| ENSG00000255403 | 0 | 0 | 0 | 0 | 0 | 0 |
| ENSG00000222663 | 0 | 0 | 0 | 0 | 0 | 0 |
| ENSG00000252081 | 0 | 0 | 0 | 0 | 0 | 0 |
| ENSG00000285813 | 4 | 2 | 3 | 0 | 3 | 1 |
| ENSG00000170903 | 116 | 57 | 68 | 45 | 104 | 71 |
| ENSG00000182359 | 63 | 49 | 43 | 33 | 48 | 53 |
| ENSG00000149313 | 778 | 479 | 490 | 256 | 393 | 381 |
| ENSG00000254433 | 0 | 1 | 0 | 2 | 1 | 1 |
| ENSG00000254580 | 0 | 0 | 0 | 0 | 0 | 0 |
| ENSG00000254811 | 0 | 0 | 0 | 0 | 0 | 0 |
| ENSG00000255065 | 0 | 0 | 0 | 0 | 0 | 0 |
| ENSG00000152402 | 0 | 0 | 0 | 0 | 0 | 0 |
| ENSG00000274584 | 0 | 0 | 0 | 0 | 0 | 0 |
| ENSG00000213252 | 0 | 1 | 0 | 0 | 0 | 0 |
| ENSG00000255353 | 0 | 1 | 0 | 0 | 0 | 0 |
| ENSG00000261098 | 2 | 2 | 0 | 0 | 2 | 1 |
| ENSG00000152404 | 91 | 40 | 34 | 18 | 76 | 41 |
| ENSG00000225215 | 0 | 0 | 0 | 0 | 0 | 0 |
| ENSG00000137760 | 158 | 61 | 61 | 54 | 90 | 72 |
| ENSG00000270969 | 0 | 0 | 0 | 0 | 0 | 0 |
| ENSG00000110675 | 0 | 0 | 0 | 0 | 0 | 0 |
| ENSG00000255483 | 0 | 0 | 0 | 0 | 0 | 0 |
| ENSG00000254758 | 0 | 0 | 0 | 0 | 0 | 0 |
| ENSG00000170290 | 4 | 0 | 0 | 0 | 1 | 0 |
| ENSG00000254702 | 0 | 0 | 0 | 0 | 0 | 0 |
| ENSG00000110660 | 483 | 220 | 291 | 266 | 292 | 327 |
| ENSG00000240652 | 0 | 0 | 1 | 0 | 0 | 1 |
| ENSG00000179331 | 0 | 0 | 0 | 0 | 0 | 0 |
| ENSG00000166266 | 678 | 347 | 354 | 204 | 488 | 293 |
| ENSG00000200855 | 0 | 0 | 0 | 0 | 0 | 0 |
| ENSG00000255467 | 0 | 0 | 0 | 0 | 0 | 0 |

| | | | | | | |
|---|---|---|---|---|---|---|
| ENSG00000075239 | 755 | 506 | 557 | 457 | 553 | 524 |
| ENSG00000285696 | 0 | 0 | 0 | 0 | 0 | 2 |
| ENSG00000149308 | 82 | 78 | 48 | 18 | 111 | 61 |
| ENSG00000149311 | 296 | 160 | 106 | 51 | 245 | 146 |
| ENSG00000206967 | 0 | 0 | 0 | 0 | 0 | 0 |
| ENSG00000166323 | 15 | 16 | 18 | 5 | 13 | 11 |
| ENSG00000178202 | 526 | 224 | 310 | 96 | 226 | 181 |
| ENSG00000110723 | 53 | 43 | 32 | 10 | 32 | 36 |
| ENSG00000178105 | 200 | 141 | 141 | 105 | 195 | 167 |
| ENSG00000240698 | 0 | 0 | 0 | 0 | 0 | 0 |
| ENSG00000254730 | 0 | 0 | 0 | 0 | 0 | 0 |
| ENSG00000255379 | 0 | 0 | 0 | 0 | 0 | 0 |
| ENSG00000203334 | 0 | 0 | 0 | 0 | 0 | 0 |
| ENSG00000255528 | 0 | 0 | 0 | 0 | 0 | 0 |
| ENSG00000201243 | 0 | 0 | 0 | 0 | 0 | 0 |
| ENSG00000200613 | 0 | 0 | 0 | 0 | 0 | 0 |
| ENSG00000264997 | 0 | 0 | 0 | 0 | 0 | 0 |
| ENSG00000255028 | 0 | 0 | 0 | 0 | 0 | 0 |
| ENSG00000185742 | 0 | 0 | 0 | 0 | 0 | 0 |
| ENSG00000254890 | 0 | 0 | 0 | 0 | 0 | 0 |
| ENSG00000254482 | 0 | 0 | 0 | 0 | 0 | 0 |
| ENSG00000254659 | 0 | 0 | 0 | 0 | 0 | 0 |
| ENSG00000137710 | 5533 | 3109 | 3326 | 1226 | 3684 | 2934 |
| ENSG00000254798 | 0 | 0 | 0 | 0 | 0 | 0 |
| ENSG00000224776 | 0 | 0 | 0 | 0 | 0 | 0 |
| ENSG00000270423 | 0 | 0 | 0 | 0 | 0 | 0 |
| ENSG00000255210 | 0 | 0 | 0 | 0 | 0 | 0 |
| ENSG00000149289 | 63 | 34 | 31 | 18 | 53 | 42 |
| ENSG00000254416 | 162 | 76 | 153 | 79 | 78 | 145 |
| ENSG00000137714 | 804 | 399 | 586 | 434 | 468 | 559 |
| ENSG00000137727 | 2 | 1 | 1 | 0 | 0 | 0 |
| ENSG00000233497 | 0 | 0 | 0 | 0 | 0 | 0 |
| ENSG00000255149 | 0 | 0 | 0 | 0 | 0 | 0 |
| ENSG00000200168 | 0 | 0 | 0 | 0 | 0 | 0 |
| ENSG00000271390 | 0 | 0 | 0 | 0 | 0 | 0 |
| ENSG00000271584 | 0 | 0 | 0 | 0 | 0 | 0 |
| ENSG00000239354 | 0 | 0 | 0 | 0 | 0 | 0 |
| ENSG00000270202 | 0 | 0 | 0 | 0 | 0 | 0 |
| ENSG00000150750 | 0 | 0 | 0 | 0 | 0 | 0 |
| ENSG00000196167 | 1 | 3 | 2 | 0 | 0 | 1 |
| ENSG00000214290 | 15 | 1 | 4 | 1 | 1 | 1 |
| ENSG00000264032 | 0 | 0 | 0 | 0 | 0 | 0 |
| ENSG00000110777 | 0 | 1 | 0 | 0 | 0 | 0 |
| ENSG00000253099 | 0 | 0 | 0 | 0 | 0 | 0 |
| ENSG00000255428 | 0 | 0 | 0 | 0 | 0 | 0 |
| ENSG00000255093 | 0 | 0 | 0 | 0 | 0 | 0 |
| ENSG00000137707 | 0 | 0 | 0 | 0 | 1 | 0 |
| ENSG00000207811 | 0 | 0 | 0 | 0 | 0 | 0 |
| ENSG00000207562 | 0 | 0 | 0 | 0 | 0 | 0 |
| ENSG00000254980 | 0 | 0 | 0 | 0 | 0 | 0 |

| | | | | | | |
|---|---|---|---|---|---|---|
| ENSG00000183644 | 0 | 0 | 0 | 0 | 0 | 0 |
| ENSG00000204381 | 0 | 0 | 0 | 0 | 0 | 0 |
| ENSG00000170145 | 870 | 345 | 460 | 351 | 398 | 459 |
| ENSG00000235286 | 0 | 1 | 1 | 0 | 0 | 0 |
| ENSG00000223324 | 0 | 0 | 0 | 0 | 0 | 0 |
| ENSG00000137713 | 646 | 377 | 412 | 275 | 459 | 377 |
| ENSG00000254990 | 0 | 0 | 0 | 0 | 0 | 0 |
| ENSG00000086848 | 81 | 49 | 73 | 31 | 66 | 52 |
| ENSG00000258529 | 0 | 0 | 0 | 0 | 0 | 0 |
| ENSG00000254450 | 0 | 0 | 0 | 0 | 0 | 0 |
| ENSG00000254949 | 0 | 0 | 0 | 0 | 0 | 0 |
| ENSG00000255561 | 44 | 17 | 16 | 17 | 20 | 22 |
| ENSG00000137720 | 178 | 68 | 101 | 100 | 96 | 93 |
| ENSG00000176343 | 0 | 0 | 0 | 0 | 0 | 0 |
| ENSG00000109846 | 0 | 2 | 0 | 0 | 1 | 1 |
| ENSG00000170276 | 0 | 0 | 0 | 0 | 0 | 0 |
| ENSG00000254445 | 0 | 0 | 0 | 0 | 0 | 0 |
| ENSG00000149300 | 0 | 1 | 1 | 2 | 5 | 2 |
| ENSG00000150764 | 477 | 231 | 237 | 109 | 193 | 181 |
| ENSG00000238965 | 0 | 0 | 0 | 0 | 0 | 0 |
| ENSG00000255286 | 0 | 0 | 0 | 0 | 0 | 0 |
| ENSG00000255334 | 0 | 0 | 0 | 0 | 0 | 0 |
| ENSG00000150768 | 1373 | 667 | 771 | 611 | 786 | 781 |
| ENSG00000230911 | 1 | 0 | 0 | 2 | 2 | 1 |
| ENSG00000238444 | 0 | 0 | 0 | 0 | 0 | 0 |
| ENSG00000271025 | 0 | 0 | 0 | 0 | 0 | 0 |
| ENSG00000150773 | 19 | 13 | 8 | 4 | 9 | 3 |
| ENSG00000150776 | 831 | 401 | 559 | 390 | 496 | 469 |
| ENSG00000150779 | 900 | 434 | 538 | 608 | 545 | 541 |
| ENSG00000204370 | 1133 | 625 | 710 | 699 | 689 | 693 |
| ENSG00000255292 | 0 | 0 | 0 | 0 | 0 | 0 |
| ENSG00000150782 | 33 | 22 | 24 | 35 | 44 | 26 |
| ENSG00000254638 | 1 | 0 | 0 | 0 | 0 | 0 |
| ENSG00000150783 | 0 | 0 | 0 | 0 | 0 | 0 |
| ENSG00000197580 | 12 | 2 | 0 | 2 | 7 | 1 |
| ENSG00000214264 | 0 | 0 | 0 | 1 | 0 | 0 |
| ENSG00000255231 | 0 | 0 | 0 | 0 | 0 | 0 |
| ENSG00000243930 | 0 | 0 | 0 | 0 | 0 | 0 |
| ENSG00000150787 | 601 | 301 | 385 | 352 | 403 | 414 |
| ENSG00000243250 | 0 | 1 | 1 | 0 | 0 | 0 |
| ENSG00000188771 | 0 | 0 | 0 | 0 | 0 | 0 |
| ENSG00000268472 | 0 | 0 | 1 | 0 | 0 | 1 |
| ENSG00000213234 | 0 | 1 | 0 | 0 | 0 | 0 |
| ENSG00000250303 | 60 | 46 | 42 | 50 | 49 | 61 |
| ENSG00000206772 | 0 | 0 | 0 | 0 | 0 | 0 |
| ENSG00000254968 | 0 | 0 | 1 | 0 | 0 | 0 |
| ENSG00000213233 | 0 | 0 | 0 | 0 | 0 | 0 |
| ENSG00000255484 | 0 | 0 | 0 | 0 | 0 | 0 |
| ENSG00000212397 | 0 | 0 | 0 | 0 | 0 | 0 |
| ENSG00000285769 | 0 | 0 | 0 | 0 | 0 | 0 |

| | | | | | | |
|---|---|---|---|---|---|---|
| ENSG00000254626 | 0 | 0 | 0 | 0 | 0 | 0 |
| ENSG00000247416 | 0 | 0 | 0 | 0 | 0 | 0 |
| ENSG00000149294 | 0 | 0 | 0 | 2 | 0 | 1 |
| ENSG00000238998 | 0 | 0 | 0 | 0 | 0 | 0 |
| ENSG00000227487 | 0 | 0 | 0 | 0 | 0 | 0 |
| ENSG00000255129 | 0 | 0 | 0 | 0 | 0 | 0 |
| ENSG00000149292 | 325 | 165 | 211 | 168 | 179 | 154 |
| ENSG00000270179 | 9 | 4 | 11 | 4 | 8 | 3 |
| ENSG00000170209 | 0 | 0 | 1 | 0 | 0 | 2 |
| ENSG00000256757 | 0 | 0 | 0 | 0 | 0 | 0 |
| ENSG00000149295 | 0 | 0 | 0 | 0 | 0 | 0 |
| ENSG00000265140 | 0 | 0 | 0 | 0 | 0 | 0 |
| ENSG00000166682 | 10 | 1 | 2 | 4 | 2 | 0 |
| ENSG00000256091 | 0 | 0 | 0 | 0 | 0 | 0 |
| ENSG00000086827 | 514 | 263 | 288 | 231 | 293 | 263 |
| ENSG00000243353 | 1 | 0 | 0 | 0 | 2 | 0 |
| ENSG00000257070 | 2 | 0 | 0 | 0 | 0 | 0 |
| ENSG00000256603 | 0 | 4 | 0 | 0 | 0 | 1 |
| ENSG00000228607 | 0 | 0 | 0 | 0 | 0 | 0 |
| ENSG00000256167 | 0 | 0 | 0 | 0 | 0 | 0 |
| ENSG00000255870 | 0 | 0 | 0 | 0 | 0 | 0 |
| ENSG00000257101 | 0 | 0 | 0 | 0 | 0 | 0 |
| ENSG00000048028 | 437 | 221 | 268 | 127 | 216 | 222 |
| ENSG00000256452 | 0 | 0 | 1 | 0 | 2 | 0 |
| ENSG00000255710 | 0 | 1 | 1 | 1 | 0 | 0 |
| ENSG00000201687 | 0 | 0 | 0 | 0 | 0 | 0 |
| ENSG00000149305 | 0 | 0 | 0 | 1 | 0 | 0 |
| ENSG00000166736 | 0 | 0 | 0 | 0 | 0 | 0 |
| ENSG00000109906 | 0 | 0 | 0 | 0 | 0 | 0 |
| ENSG00000256947 | 0 | 0 | 0 | 0 | 0 | 0 |
| ENSG00000166741 | 0 | 0 | 0 | 0 | 0 | 0 |
| ENSG00000256195 | 0 | 0 | 0 | 0 | 0 | 0 |
| ENSG00000180425 | 209 | 88 | 113 | 83 | 122 | 122 |
| ENSG00000076053 | 228 | 132 | 154 | 106 | 146 | 97 |
| ENSG00000255663 | 0 | 0 | 0 | 0 | 1 | 0 |
| ENSG00000076043 | 523 | 286 | 365 | 392 | 376 | 388 |
| ENSG00000256533 | 0 | 0 | 0 | 0 | 0 | 0 |
| ENSG00000255982 | 0 | 0 | 0 | 0 | 0 | 0 |
| ENSG00000270204 | 0 | 0 | 0 | 0 | 0 | 0 |
| ENSG00000095110 | 0 | 0 | 0 | 0 | 0 | 0 |
| ENSG00000238724 | 0 | 0 | 0 | 0 | 0 | 0 |
| ENSG00000137634 | 0 | 0 | 0 | 0 | 0 | 0 |
| ENSG00000256674 | 0 | 0 | 0 | 0 | 0 | 0 |
| ENSG00000204361 | 0 | 0 | 0 | 0 | 0 | 0 |
| ENSG00000252870 | 0 | 0 | 0 | 0 | 0 | 0 |
| ENSG00000182985 | 894 | 418 | 488 | 325 | 402 | 380 |
| ENSG00000256972 | 0 | 0 | 0 | 0 | 0 | 0 |
| ENSG00000255580 | 0 | 0 | 0 | 0 | 0 | 0 |
| ENSG00000256315 | 0 | 0 | 0 | 0 | 0 | 0 |
| ENSG00000256281 | 0 | 0 | 0 | 0 | 0 | 0 |

| | | | | | | |
|---|---|---|---|---|---|---|
| ENSG00000257087 | 0 | 0 | 0 | 0 | 0 | 0 |
| ENSG00000255903 | 0 | 0 | 0 | 0 | 0 | 0 |
| ENSG00000255689 | 0 | 0 | 1 | 0 | 0 | 2 |
| ENSG00000239153 | 0 | 0 | 0 | 0 | 0 | 0 |
| ENSG00000255599 | 0 | 0 | 0 | 0 | 0 | 0 |
| ENSG00000260254 | 0 | 0 | 0 | 0 | 0 | 0 |
| ENSG00000279771 | 0 | 0 | 0 | 0 | 0 | 0 |
| ENSG00000256717 | 0 | 0 | 0 | 0 | 0 | 0 |
| ENSG00000279818 | 0 | 0 | 0 | 0 | 0 | 0 |
| ENSG00000246100 | 0 | 0 | 0 | 0 | 0 | 0 |
| ENSG00000257067 | 0 | 0 | 0 | 0 | 0 | 0 |
| ENSG00000244259 | 0 | 0 | 0 | 0 | 0 | 0 |
| ENSG00000239600 | 0 | 0 | 0 | 0 | 0 | 0 |
| ENSG00000236437 | 0 | 0 | 0 | 0 | 0 | 0 |
| ENSG00000278945 | 0 | 0 | 0 | 0 | 0 | 0 |
| ENSG00000237937 | 0 | 0 | 0 | 0 | 0 | 0 |
| ENSG00000137656 | 364 | 208 | 260 | 249 | 191 | 208 |
| ENSG00000226645 | 0 | 2 | 0 | 0 | 0 | 0 |
| ENSG00000109917 | 869 | 447 | 534 | 563 | 563 | 560 |
| ENSG00000110243 | 6 | 0 | 0 | 0 | 0 | 0 |
| ENSG00000236267 | 0 | 0 | 0 | 0 | 0 | 0 |
| ENSG00000285513 | 0 | 0 | 0 | 0 | 0 | 0 |
| ENSG00000110244 | 0 | 0 | 0 | 0 | 0 | 0 |
| ENSG00000110245 | 0 | 0 | 0 | 1 | 0 | 0 |
| ENSG00000118137 | 0 | 0 | 1 | 0 | 0 | 0 |
| ENSG00000235910 | 0 | 0 | 2 | 0 | 0 | 0 |
| ENSG00000160584 | 364 | 170 | 235 | 145 | 184 | 220 |
| ENSG00000200537 | 0 | 0 | 0 | 0 | 0 | 0 |
| ENSG00000234268 | 0 | 0 | 0 | 2 | 2 | 0 |
| ENSG00000224077 | 0 | 0 | 0 | 0 | 0 | 2 |
| ENSG00000225236 | 0 | 0 | 0 | 0 | 0 | 0 |
| ENSG00000254851 | 9 | 4 | 14 | 11 | 7 | 8 |
| ENSG00000254678 | 0 | 0 | 0 | 0 | 0 | 0 |
| ENSG00000168092 | 2516 | 1161 | 1567 | 1115 | 1366 | 1361 |
| ENSG00000149577 | 1054 | 441 | 686 | 520 | 481 | 547 |
| ENSG00000149591 | 55 | 21 | 46 | 34 | 32 | 33 |
| ENSG00000160613 | 601 | 233 | 326 | 289 | 328 | 316 |
| ENSG00000280143 | 2 | 2 | 1 | 0 | 3 | 3 |
| ENSG00000167257 | 382 | 171 | 187 | 136 | 195 | 169 |
| ENSG00000252992 | 0 | 0 | 0 | 0 | 0 | 0 |
| ENSG00000186318 | 1217 | 577 | 795 | 502 | 617 | 651 |
| ENSG00000278768 | 85 | 56 | 70 | 32 | 67 | 64 |
| ENSG00000276505 | 4 | 2 | 7 | 0 | 0 | 3 |
| ENSG00000110274 | 882 | 394 | 505 | 446 | 510 | 508 |
| ENSG00000250699 | 0 | 1 | 0 | 0 | 0 | 0 |
| ENSG00000271412 | 0 | 1 | 0 | 0 | 0 | 0 |
| ENSG00000177103 | 80 | 51 | 64 | 35 | 55 | 40 |
| ENSG00000279586 | 0 | 0 | 0 | 0 | 0 | 0 |
| ENSG00000270403 | 0 | 0 | 0 | 0 | 0 | 0 |
| ENSG00000137731 | 0 | 0 | 0 | 0 | 0 | 1 |

| | | | | | | |
|---|---|---|---|---|---|---|
| ENSG00000254844 | 0 | 0 | 0 | 0 | 0 | 0 |
| ENSG00000255245 | 0 | 0 | 0 | 0 | 0 | 0 |
| ENSG00000254528 | 0 | 2 | 3 | 2 | 0 | 1 |
| ENSG00000137726 | 4 | 2 | 12 | 1 | 4 | 3 |
| ENSG00000137747 | 375 | 172 | 242 | 265 | 303 | 231 |
| ENSG00000110324 | 0 | 2 | 5 | 1 | 0 | 0 |
| ENSG00000255274 | 0 | 0 | 0 | 1 | 0 | 1 |
| ENSG00000272075 | 0 | 0 | 0 | 0 | 0 | 0 |
| ENSG00000137648 | 185 | 140 | 169 | 86 | 74 | 95 |
| ENSG00000177098 | 0 | 1 | 1 | 0 | 0 | 0 |
| ENSG00000149575 | 0 | 0 | 0 | 0 | 0 | 0 |
| ENSG00000160593 | 0 | 0 | 2 | 0 | 1 | 0 |
| ENSG00000255163 | 0 | 0 | 0 | 0 | 0 | 0 |
| ENSG00000160588 | 1390 | 717 | 842 | 611 | 751 | 666 |
| ENSG00000149573 | 1350 | 714 | 935 | 677 | 908 | 975 |
| ENSG00000280032 | 4 | 6 | 1 | 2 | 1 | 3 |
| ENSG00000198851 | 0 | 0 | 0 | 1 | 0 | 0 |
| ENSG00000254992 | 0 | 0 | 0 | 0 | 0 | 0 |
| ENSG00000167286 | 0 | 0 | 0 | 0 | 0 | 0 |
| ENSG00000160654 | 0 | 0 | 0 | 0 | 0 | 0 |
| ENSG00000110344 | 1797 | 885 | 988 | 455 | 1085 | 863 |
| ENSG00000254873 | 0 | 1 | 1 | 0 | 1 | 1 |
| ENSG00000167283 | 1367 | 918 | 1204 | 1204 | 1080 | 1096 |
| ENSG00000285827 | 0 | 0 | 1 | 0 | 0 | 0 |
| ENSG00000269944 | 0 | 0 | 1 | 0 | 0 | 0 |
| ENSG00000255384 | 0 | 0 | 0 | 0 | 0 | 0 |
| ENSG00000118058 | 2393 | 1187 | 1096 | 439 | 1418 | 1047 |
| ENSG00000255435 | 11 | 3 | 8 | 9 | 11 | 14 |
| ENSG00000172425 | 18 | 9 | 17 | 15 | 13 | 13 |
| ENSG00000149582 | 524 | 218 | 330 | 242 | 217 | 251 |
| ENSG00000118096 | 1082 | 557 | 677 | 651 | 612 | 630 |
| ENSG00000243431 | 9 | 4 | 8 | 3 | 4 | 4 |
| ENSG00000095139 | 8136 | 3836 | 4863 | 3043 | 4386 | 4265 |
| ENSG00000207185 | 2 | 1 | 1 | 0 | 0 | 0 |
| ENSG00000019144 | 643 | 396 | 417 | 337 | 425 | 372 |
| ENSG00000255176 | 8 | 2 | 2 | 1 | 4 | 4 |
| ENSG00000284246 | 0 | 0 | 0 | 0 | 0 | 0 |
| ENSG00000118094 | 0 | 0 | 0 | 1 | 2 | 0 |
| ENSG00000255239 | 1 | 0 | 0 | 0 | 0 | 0 |
| ENSG00000255422 | 0 | 0 | 0 | 0 | 0 | 0 |
| ENSG00000207462 | 0 | 0 | 0 | 0 | 0 | 0 |
| ENSG00000110367 | 6844 | 2959 | 3865 | 2391 | 3554 | 3331 |
| ENSG00000278376 | 41 | 23 | 34 | 34 | 24 | 26 |
| ENSG00000254478 | 0 | 1 | 0 | 1 | 0 | 0 |
| ENSG00000254621 | 0 | 0 | 0 | 0 | 0 | 0 |
| ENSG00000201535 | 0 | 0 | 0 | 0 | 0 | 0 |
| ENSG00000160683 | 2 | 1 | 1 | 2 | 0 | 0 |
| ENSG00000245869 | 4 | 5 | 5 | 3 | 3 | 2 |
| ENSG00000186174 | 6777 | 2924 | 4385 | 3019 | 3144 | 3621 |
| ENSG00000264211 | 0 | 0 | 0 | 0 | 0 | 0 |

| | | | | | | |
|---|---|---|---|---|---|---|
| ENSG00000110375 | 434 | 269 | 327 | 272 | 230 | 200 |
| ENSG00000239726 | 9 | 14 | 5 | 1 | 4 | 1 |
| ENSG00000222529 | 0 | 0 | 0 | 0 | 0 | 0 |
| ENSG00000176302 | 2 | 2 | 3 | 0 | 3 | 4 |
| ENSG00000280502 | 5 | 1 | 8 | 6 | 7 | 16 |
| ENSG00000283691 | 1 | 1 | 0 | 0 | 0 | 1 |
| ENSG00000255121 | 56 | 30 | 55 | 41 | 26 | 56 |
| ENSG00000186166 | 605 | 282 | 460 | 264 | 366 | 348 |
| ENSG00000240970 | 19 | 15 | 36 | 3 | 28 | 23 |
| ENSG00000254428 | 51 | 22 | 47 | 21 | 50 | 24 |
| ENSG00000254909 | 26 | 15 | 20 | 9 | 21 | 13 |
| ENSG00000118181 | 15820 | 8379 | 9729 | 12062 | 10654 | 10291 |
| ENSG00000196655 | 1469 | 832 | 1043 | 891 | 992 | 911 |
| ENSG00000283704 | 0 | 0 | 0 | 0 | 0 | 0 |
| ENSG00000137700 | 1411 | 662 | 961 | 762 | 675 | 796 |
| ENSG00000255114 | 0 | 0 | 0 | 0 | 0 | 0 |
| ENSG00000149428 | 4608 | 2258 | 2872 | 2497 | 2594 | 2804 |
| ENSG00000271751 | 7 | 5 | 2 | 1 | 6 | 1 |
| ENSG00000272186 | 12 | 8 | 9 | 2 | 5 | 5 |
| ENSG00000160695 | 1974 | 925 | 1228 | 984 | 985 | 1097 |
| ENSG00000256269 | 1537 | 765 | 918 | 998 | 1059 | 1018 |
| ENSG00000188486 | 3394 | 1617 | 2016 | 2394 | 1861 | 2042 |
| ENSG00000172269 | 2148 | 1060 | 1263 | 1154 | 1190 | 1329 |
| ENSG00000172375 | 1022 | 437 | 584 | 527 | 453 | 542 |
| ENSG00000172273 | 911 | 416 | 574 | 502 | 492 | 518 |
| ENSG00000172350 | 4 | 4 | 1 | 1 | 0 | 0 |
| ENSG00000160703 | 227 | 125 | 143 | 164 | 143 | 149 |
| ENSG00000172367 | 6 | 1 | 2 | 2 | 1 | 1 |
| ENSG00000248712 | 131 | 57 | 80 | 77 | 57 | 62 |
| ENSG00000110395 | 1364 | 615 | 792 | 415 | 696 | 718 |
| ENSG00000222249 | 0 | 1 | 1 | 0 | 0 | 0 |
| ENSG00000076706 | 0 | 3 | 1 | 4 | 1 | 0 |
| ENSG00000284148 | 0 | 0 | 0 | 0 | 0 | 0 |
| ENSG00000252119 | 0 | 0 | 0 | 0 | 0 | 0 |
| ENSG00000173456 | 1157 | 588 | 745 | 864 | 711 | 797 |
| ENSG00000245385 | 0 | 0 | 0 | 0 | 0 | 0 |
| ENSG00000223953 | 0 | 0 | 0 | 0 | 0 | 0 |
| ENSG00000235718 | 0 | 0 | 0 | 0 | 0 | 0 |
| ENSG00000036672 | 110 | 37 | 87 | 63 | 57 | 56 |
| ENSG00000254740 | 0 | 0 | 0 | 0 | 0 | 0 |
| ENSG00000245248 | 3 | 7 | 2 | 9 | 3 | 5 |
| ENSG00000254590 | 0 | 1 | 0 | 0 | 0 | 0 |
| ENSG00000254892 | 0 | 0 | 0 | 0 | 0 | 0 |
| ENSG00000154096 | 0 | 0 | 0 | 0 | 0 | 0 |
| ENSG00000263873 | 0 | 0 | 0 | 0 | 0 | 0 |
| ENSG00000255377 | 0 | 0 | 0 | 0 | 0 | 0 |
| ENSG00000230716 | 3 | 1 | 1 | 0 | 0 | 1 |
| ENSG00000254561 | 0 | 0 | 0 | 0 | 0 | 0 |
| ENSG00000110400 | 918 | 386 | 492 | 497 | 432 | 467 |
| ENSG00000199217 | 0 | 0 | 0 | 0 | 0 | 0 |

| | | | | | | |
|---|---|---|---|---|---|---|
| ENSG00000255247 | 0 | 0 | 0 | 0 | 0 | 0 |
| ENSG00000254854 | 16 | 6 | 4 | 4 | 3 | 2 |
| ENSG00000254406 | 0 | 1 | 0 | 0 | 0 | 0 |
| ENSG00000254863 | 0 | 0 | 0 | 0 | 0 | 0 |
| ENSG00000255216 | 0 | 0 | 0 | 0 | 0 | 0 |
| ENSG00000137699 | 100 | 51 | 42 | 66 | 82 | 64 |
| ENSG00000176984 | 0 | 0 | 0 | 0 | 1 | 0 |
| ENSG00000184232 | 88 | 51 | 65 | 68 | 62 | 48 |
| ENSG00000137709 | 97 | 42 | 45 | 50 | 80 | 67 |
| ENSG00000259541 | 0 | 0 | 0 | 0 | 0 | 0 |
| ENSG00000181264 | 202 | 87 | 126 | 89 | 127 | 93 |
| ENSG00000196914 | 2566 | 1237 | 1370 | 639 | 1546 | 1229 |
| ENSG00000274303 | 0 | 0 | 0 | 0 | 0 | 1 |
| ENSG00000149403 | 4 | 1 | 3 | 3 | 1 | 1 |
| ENSG00000255006 | 0 | 0 | 0 | 0 | 0 | 0 |
| ENSG00000250493 | 0 | 0 | 0 | 0 | 0 | 0 |
| ENSG00000254735 | 0 | 0 | 0 | 0 | 0 | 0 |
| ENSG00000154114 | 486 | 213 | 286 | 198 | 262 | 221 |
| ENSG00000285509 | 4 | 1 | 4 | 2 | 4 | 1 |
| ENSG00000109927 | 6 | 2 | 3 | 3 | 4 | 7 |
| ENSG00000254515 | 0 | 0 | 0 | 0 | 0 | 0 |
| ENSG00000244400 | 0 | 0 | 0 | 0 | 0 | 0 |
| ENSG00000109929 | 1121 | 543 | 603 | 446 | 618 | 514 |
| ENSG00000232460 | 0 | 1 | 1 | 0 | 0 | 0 |
| ENSG00000271022 | 0 | 0 | 0 | 0 | 0 | 0 |
| ENSG00000246790 | 0 | 0 | 0 | 0 | 0 | 0 |
| ENSG00000137642 | 318 | 205 | 222 | 93 | 197 | 136 |
| ENSG00000252556 | 0 | 0 | 0 | 0 | 0 | 0 |
| ENSG00000255248 | 0 | 0 | 0 | 0 | 0 | 0 |
| ENSG00000207971 | 0 | 0 | 0 | 0 | 0 | 0 |
| ENSG00000259571 | 0 | 0 | 0 | 0 | 0 | 0 |
| ENSG00000198975 | 0 | 0 | 0 | 0 | 0 | 0 |
| ENSG00000207994 | 0 | 0 | 0 | 0 | 0 | 0 |
| ENSG00000255015 | 0 | 0 | 0 | 0 | 0 | 0 |
| ENSG00000255219 | 0 | 0 | 0 | 0 | 0 | 0 |
| ENSG00000252776 | 0 | 0 | 0 | 0 | 0 | 0 |
| ENSG00000253035 | 0 | 0 | 0 | 0 | 0 | 0 |
| ENSG00000223265 | 0 | 0 | 0 | 0 | 0 | 0 |
| ENSG00000254572 | 0 | 0 | 0 | 0 | 0 | 0 |
| ENSG00000254935 | 0 | 0 | 0 | 0 | 0 | 0 |
| ENSG00000154127 | 14 | 1 | 2 | 7 | 6 | 7 |
| ENSG00000239079 | 0 | 0 | 0 | 0 | 0 | 0 |
| ENSG00000285909 | 0 | 0 | 0 | 0 | 0 | 0 |
| ENSG00000109943 | 0 | 0 | 0 | 0 | 0 | 0 |
| ENSG00000109944 | 141 | 71 | 74 | 66 | 90 | 85 |
| ENSG00000199709 | 0 | 0 | 1 | 0 | 5 | 0 |
| ENSG00000254944 | 2 | 2 | 3 | 4 | 4 | 1 |
| ENSG00000188909 | 0 | 0 | 0 | 0 | 0 | 0 |
| ENSG00000213184 | 0 | 0 | 0 | 0 | 0 | 0 |
| ENSG00000243669 | 0 | 0 | 0 | 0 | 0 | 0 |

| | | | | | | |
|---|---|---|---|---|---|---|
| ENSG00000244641 | 0 | 0 | 0 | 0 | 0 | 0 |
| ENSG00000233541 | 0 | 0 | 0 | 0 | 0 | 0 |
| ENSG00000109971 | 32031 | 16392 | 18325 | 22196 | 21581 | 20352 |
| ENSG00000200879 | 1 | 4 | 3 | 2 | 4 | 1 |
| ENSG00000207118 | 0 | 0 | 0 | 0 | 0 | 0 |
| ENSG00000202252 | 0 | 0 | 0 | 0 | 0 | 0 |
| ENSG00000166250 | 10 | 7 | 12 | 17 | 16 | 12 |
| ENSG00000255342 | 0 | 0 | 0 | 0 | 0 | 0 |
| ENSG00000254710 | 0 | 0 | 0 | 0 | 0 | 0 |
| ENSG00000200496 | 0 | 0 | 0 | 0 | 0 | 0 |
| ENSG00000254823 | 0 | 0 | 0 | 0 | 0 | 0 |
| ENSG00000254407 | 0 | 0 | 0 | 0 | 0 | 0 |
| ENSG00000023171 | 14 | 2 | 1 | 4 | 9 | 5 |
| ENSG00000265357 | 0 | 0 | 0 | 0 | 0 | 0 |
| ENSG00000254449 | 0 | 0 | 0 | 0 | 0 | 0 |
| ENSG00000166257 | 0 | 0 | 0 | 0 | 0 | 0 |
| ENSG00000254467 | 0 | 0 | 0 | 0 | 0 | 0 |
| ENSG00000166261 | 293 | 139 | 233 | 145 | 211 | 230 |
| ENSG00000221931 | 0 | 0 | 0 | 0 | 0 | 0 |
| ENSG00000200197 | 0 | 0 | 0 | 0 | 0 | 0 |
| ENSG00000196099 | 0 | 0 | 0 | 0 | 0 | 0 |
| ENSG00000254613 | 0 | 0 | 0 | 0 | 0 | 0 |
| ENSG00000254430 | 0 | 0 | 0 | 0 | 0 | 0 |
| ENSG00000204300 | 0 | 0 | 0 | 0 | 0 | 0 |
| ENSG00000181518 | 0 | 0 | 0 | 0 | 0 | 0 |
| ENSG00000171014 | 0 | 0 | 0 | 0 | 0 | 0 |
| ENSG00000181499 | 0 | 0 | 0 | 0 | 0 | 0 |
| ENSG00000196248 | 0 | 0 | 0 | 0 | 0 | 0 |
| ENSG00000198674 | 0 | 0 | 0 | 0 | 0 | 0 |
| ENSG00000226461 | 0 | 0 | 0 | 0 | 0 | 0 |
| ENSG00000254737 | 0 | 0 | 0 | 0 | 0 | 0 |
| ENSG00000236981 | 0 | 0 | 0 | 0 | 0 | 0 |
| ENSG00000234560 | 0 | 0 | 0 | 0 | 0 | 0 |
| ENSG00000182634 | 2 | 0 | 0 | 0 | 0 | 0 |
| ENSG00000213182 | 0 | 0 | 0 | 0 | 0 | 0 |
| ENSG00000186268 | 0 | 0 | 0 | 0 | 0 | 0 |
| ENSG00000213181 | 0 | 0 | 0 | 0 | 0 | 0 |
| ENSG00000110002 | 6 | 3 | 2 | 7 | 3 | 4 |
| ENSG00000196403 | 0 | 0 | 1 | 0 | 0 | 0 |
| ENSG00000197309 | 2 | 0 | 0 | 0 | 0 | 0 |
| ENSG00000239426 | 0 | 0 | 0 | 0 | 0 | 1 |
| ENSG00000255425 | 2 | 0 | 1 | 1 | 2 | 1 |
| ENSG00000181214 | 0 | 0 | 0 | 0 | 0 | 0 |
| ENSG00000276316 | 0 | 0 | 0 | 0 | 0 | 0 |
| ENSG00000197849 | 0 | 0 | 1 | 0 | 0 | 0 |
| ENSG00000255298 | 4 | 2 | 4 | 3 | 3 | 4 |
| ENSG00000255116 | 0 | 0 | 0 | 0 | 0 | 0 |
| ENSG00000196341 | 4 | 6 | 6 | 1 | 8 | 0 |
| ENSG00000279116 | 0 | 0 | 0 | 0 | 1 | 0 |
| ENSG00000254976 | 0 | 0 | 0 | 0 | 0 | 0 |

| | | | | | | |
|---|---|---|---|---|---|---|
| ENSG00000255225 | 0 | 0 | 0 | 0 | 0 | 0 |
| ENSG00000255030 | 0 | 0 | 0 | 0 | 0 | 0 |
| ENSG00000227737 | 0 | 0 | 0 | 0 | 0 | 0 |
| ENSG00000229254 | 0 | 0 | 0 | 0 | 0 | 0 |
| ENSG00000284680 | 0 | 0 | 0 | 0 | 0 | 0 |
| ENSG00000284609 | 0 | 0 | 0 | 0 | 0 | 0 |
| ENSG00000255048 | 0 | 0 | 1 | 0 | 0 | 0 |
| ENSG00000285561 | 0 | 0 | 3 | 0 | 0 | 0 |
| ENSG00000280090 | 0 | 0 | 0 | 0 | 0 | 0 |
| ENSG00000197125 | 0 | 0 | 0 | 0 | 0 | 0 |
| ENSG00000255510 | 0 | 0 | 0 | 0 | 0 | 0 |
| ENSG00000229386 | 0 | 0 | 0 | 0 | 1 | 1 |
| ENSG00000254646 | 0 | 0 | 0 | 0 | 0 | 0 |
| ENSG00000255315 | 0 | 0 | 0 | 0 | 0 | 0 |
| ENSG00000254937 | 0 | 0 | 0 | 0 | 0 | 0 |
| ENSG00000170953 | 0 | 1 | 0 | 0 | 0 | 0 |
| ENSG00000196119 | 8 | 2 | 7 | 4 | 12 | 5 |
| ENSG00000255341 | 3 | 0 | 3 | 6 | 8 | 8 |
| ENSG00000154143 | 23 | 7 | 13 | 12 | 7 | 6 |
| ENSG00000154144 | 636 | 249 | 379 | 239 | 296 | 269 |
| ENSG00000110013 | 946 | 458 | 611 | 413 | 514 | 544 |
| ENSG00000200278 | 0 | 0 | 0 | 0 | 0 | 0 |
| ENSG00000064199 | 63 | 46 | 49 | 43 | 45 | 40 |
| ENSG00000154146 | 39 | 17 | 26 | 28 | 20 | 22 |
| ENSG00000255045 | 0 | 1 | 1 | 0 | 1 | 1 |
| ENSG00000019102 | 3 | 3 | 1 | 0 | 1 | 2 |
| ENSG00000149564 | 27 | 21 | 25 | 9 | 20 | 16 |
| ENSG00000250073 | 16 | 7 | 15 | 5 | 17 | 10 |
| ENSG00000120458 | 229 | 116 | 123 | 64 | 105 | 130 |
| ENSG00000279342 | 26 | 6 | 11 | 4 | 12 | 10 |
| ENSG00000254509 | 0 | 0 | 0 | 0 | 0 | 0 |
| ENSG00000245498 | 29 | 14 | 14 | 8 | 7 | 10 |
| ENSG00000254941 | 9 | 5 | 6 | 0 | 4 | 2 |
| ENSG00000154134 | 18 | 8 | 13 | 7 | 16 | 8 |
| ENSG00000254568 | 0 | 0 | 0 | 0 | 0 | 0 |
| ENSG00000154133 | 0 | 1 | 0 | 0 | 0 | 0 |
| ENSG00000254943 | 0 | 0 | 0 | 0 | 0 | 0 |
| ENSG00000165478 | 0 | 0 | 0 | 0 | 0 | 0 |
| ENSG00000221932 | 0 | 0 | 0 | 0 | 0 | 0 |
| ENSG00000285825 | 2 | 4 | 3 | 0 | 3 | 1 |
| ENSG00000149548 | 27 | 13 | 19 | 9 | 27 | 11 |
| ENSG00000134955 | 25 | 18 | 20 | 20 | 20 | 21 |
| ENSG00000150433 | 423 | 241 | 293 | 208 | 215 | 209 |
| ENSG00000187686 | 18 | 8 | 12 | 5 | 10 | 11 |
| ENSG00000255475 | 0 | 0 | 0 | 0 | 0 | 0 |
| ENSG00000254880 | 0 | 0 | 0 | 0 | 0 | 0 |
| ENSG00000165495 | 0 | 0 | 3 | 5 | 0 | 1 |
| ENSG00000259290 | 0 | 0 | 0 | 0 | 0 | 0 |
| ENSG00000254932 | 0 | 0 | 0 | 0 | 0 | 0 |
| ENSG00000222844 | 0 | 0 | 0 | 0 | 0 | 0 |

| | | | | | | |
|---|---|---|---|---|---|---|
| ENSG00000149557 | 348 | 170 | 230 | 188 | 169 | 184 |
| ENSG00000255537 | 0 | 1 | 1 | 0 | 0 | 0 |
| ENSG00000149547 | 3669 | 1702 | 2047 | 2063 | 2011 | 2118 |
| ENSG00000222297 | 0 | 0 | 0 | 0 | 0 | 0 |
| ENSG00000134910 | 3744 | 1797 | 2300 | 1975 | 2068 | 2129 |
| ENSG00000149554 | 368 | 205 | 198 | 154 | 204 | 226 |
| ENSG00000252255 | 0 | 0 | 0 | 0 | 0 | 0 |
| ENSG00000134940 | 7 | 3 | 3 | 6 | 3 | 1 |
| ENSG00000171053 | 0 | 0 | 0 | 0 | 0 | 0 |
| ENSG00000196844 | 1 | 1 | 1 | 0 | 1 | 1 |
| ENSG00000236027 | 2 | 0 | 0 | 0 | 0 | 0 |
| ENSG00000237353 | 0 | 2 | 0 | 1 | 0 | 0 |
| ENSG00000260209 | 0 | 2 | 1 | 0 | 0 | 0 |
| ENSG00000198331 | 258 | 141 | 155 | 170 | 198 | 151 |
| ENSG00000110060 | 172 | 111 | 88 | 90 | 99 | 109 |
| ENSG00000255027 | 0 | 1 | 0 | 0 | 0 | 0 |
| ENSG00000109832 | 2 | 1 | 2 | 0 | 0 | 1 |
| ENSG00000254790 | 0 | 0 | 0 | 0 | 0 | 0 |
| ENSG00000283703 | 0 | 0 | 0 | 0 | 0 | 0 |
| ENSG00000064309 | 133 | 53 | 76 | 44 | 56 | 53 |
| ENSG00000254759 | 2 | 3 | 2 | 1 | 0 | 0 |
| ENSG00000261257 | 0 | 0 | 0 | 0 | 0 | 0 |
| ENSG00000254833 | 0 | 0 | 0 | 0 | 0 | 0 |
| ENSG00000254771 | 0 | 0 | 0 | 0 | 0 | 0 |
| ENSG00000165526 | 561 | 269 | 289 | 370 | 380 | 347 |
| ENSG00000254694 | 11 | 5 | 8 | 1 | 1 | 3 |
| ENSG00000197798 | 203 | 140 | 127 | 137 | 146 | 154 |
| ENSG00000222067 | 0 | 0 | 0 | 0 | 0 | 0 |
| ENSG00000240098 | 0 | 0 | 0 | 0 | 0 | 0 |
| ENSG00000182934 | 3496 | 1571 | 1950 | 1678 | 1776 | 2029 |
| ENSG00000110074 | 770 | 370 | 455 | 418 | 393 | 414 |
| ENSG00000243607 | 0 | 0 | 0 | 0 | 0 | 0 |
| ENSG00000150455 | 162 | 94 | 105 | 88 | 99 | 100 |
| ENSG00000254905 | 0 | 0 | 0 | 0 | 0 | 0 |
| ENSG00000255062 | 25 | 13 | 19 | 11 | 25 | 27 |
| ENSG00000110063 | 393 | 243 | 238 | 282 | 212 | 244 |
| ENSG00000280832 | 324 | 148 | 183 | 185 | 201 | 186 |
| ENSG00000110080 | 3962 | 1824 | 2606 | 1713 | 1842 | 1874 |
| ENSG00000278215 | 0 | 2 | 0 | 0 | 0 | 0 |
| ENSG00000149571 | 14 | 9 | 10 | 7 | 7 | 7 |
| ENSG00000257271 | 0 | 0 | 0 | 0 | 0 | 0 |
| ENSG00000254607 | 0 | 0 | 0 | 0 | 0 | 0 |
| ENSG00000254938 | 0 | 0 | 0 | 0 | 0 | 0 |
| ENSG00000279358 | 0 | 0 | 0 | 0 | 0 | 0 |
| ENSG00000254960 | 0 | 0 | 0 | 0 | 0 | 0 |
| ENSG00000266215 | 0 | 0 | 0 | 0 | 0 | 0 |
| ENSG00000255317 | 0 | 0 | 0 | 0 | 0 | 0 |
| ENSG00000218109 | 0 | 0 | 0 | 0 | 0 | 0 |
| ENSG00000255087 | 0 | 0 | 0 | 0 | 0 | 0 |
| ENSG00000272642 | 0 | 0 | 0 | 0 | 0 | 0 |

| | | | | | | |
|---|---|---|---|---|---|---|
| ENSG00000273409 | 0 | 0 | 0 | 0 | 0 | 0 |
| ENSG00000222774 | 0 | 0 | 0 | 0 | 0 | 0 |
| ENSG00000223315 | 0 | 0 | 0 | 0 | 0 | 0 |
| ENSG00000254591 | 0 | 0 | 0 | 0 | 0 | 0 |
| ENSG00000254612 | 83 | 41 | 45 | 34 | 51 | 39 |
| ENSG00000273415 | 0 | 0 | 0 | 0 | 0 | 0 |
| ENSG00000272575 | 0 | 0 | 0 | 0 | 0 | 0 |
| ENSG00000134954 | 0 | 0 | 0 | 0 | 0 | 0 |
| ENSG00000276176 | 0 | 0 | 0 | 0 | 0 | 0 |
| ENSG00000254588 | 0 | 0 | 0 | 0 | 0 | 0 |
| ENSG00000245008 | 0 | 0 | 0 | 0 | 0 | 0 |
| ENSG00000151702 | 0 | 0 | 0 | 0 | 0 | 0 |
| ENSG00000254703 | 0 | 0 | 0 | 0 | 0 | 0 |
| ENSG00000151704 | 0 | 0 | 2 | 0 | 0 | 0 |
| ENSG00000255369 | 0 | 0 | 0 | 0 | 0 | 0 |
| ENSG00000120457 | 0 | 0 | 0 | 0 | 0 | 0 |
| ENSG00000174370 | 42 | 17 | 35 | 27 | 26 | 33 |
| ENSG00000120471 | 4 | 3 | 2 | 0 | 2 | 4 |
| ENSG00000134909 | 1018 | 512 | 630 | 247 | 460 | 428 |
| ENSG00000212597 | 0 | 0 | 0 | 0 | 0 | 0 |
| ENSG00000206847 | 0 | 0 | 0 | 0 | 0 | 0 |
| ENSG00000199260 | 0 | 0 | 0 | 0 | 0 | 0 |
| ENSG00000255259 | 0 | 0 | 0 | 0 | 0 | 0 |
| ENSG00000043039 | 100 | 82 | 96 | 67 | 73 | 80 |
| ENSG00000240121 | 0 | 0 | 0 | 0 | 0 | 0 |
| ENSG00000281386 | 0 | 1 | 1 | 0 | 0 | 0 |
| ENSG00000281097 | 4 | 1 | 1 | 0 | 2 | 2 |
| ENSG00000255188 | 0 | 0 | 0 | 0 | 0 | 0 |
| ENSG00000151715 | 724 | 467 | 472 | 904 | 1129 | 931 |
| ENSG00000170322 | 1333 | 641 | 797 | 888 | 1083 | 990 |
| ENSG00000170325 | 593 | 275 | 349 | 172 | 259 | 253 |
| ENSG00000233220 | 2 | 1 | 2 | 5 | 3 | 2 |
| ENSG00000255057 | 0 | 0 | 0 | 0 | 0 | 0 |
| ENSG00000255262 | 0 | 0 | 0 | 0 | 0 | 0 |
| ENSG00000255535 | 0 | 0 | 0 | 0 | 0 | 0 |
| ENSG00000084234 | 14453 | 6974 | 8724 | 6603 | 7917 | 7578 |
| ENSG00000244451 | 0 | 0 | 0 | 0 | 0 | 0 |
| ENSG00000274197 | 0 | 0 | 0 | 0 | 0 | 0 |
| ENSG00000149418 | 3120 | 1602 | 1975 | 2227 | 2133 | 1859 |
| ENSG00000196323 | 610 | 278 | 281 | 163 | 387 | 277 |
| ENSG00000255358 | 0 | 0 | 0 | 0 | 1 | 0 |
| ENSG00000255220 | 0 | 1 | 0 | 0 | 0 | 0 |
| ENSG00000272412 | 0 | 0 | 0 | 0 | 0 | 0 |
| ENSG00000175773 | 6 | 1 | 5 | 5 | 1 | 2 |
| ENSG00000134917 | 1 | 0 | 1 | 1 | 1 | 0 |
| ENSG00000166106 | 2 | 3 | 0 | 1 | 1 | 2 |
| ENSG00000255155 | 0 | 0 | 0 | 0 | 0 | 0 |
| ENSG00000236616 | 0 | 0 | 0 | 0 | 0 | 0 |
| ENSG00000274999 | 0 | 0 | 0 | 0 | 0 | 0 |
| ENSG00000175728 | 0 | 0 | 0 | 0 | 0 | 0 |

| | | | | | | |
|---|---|---|---|---|---|---|
| ENSG00000255352 | 0 | 0 | 0 | 0 | 0 | 0 |
| ENSG00000254842 | 0 | 0 | 0 | 0 | 0 | 0 |
| ENSG00000255455 | 60 | 19 | 32 | 12 | 39 | 20 |
| ENSG00000120451 | 1662 | 697 | 929 | 648 | 800 | 818 |
| ENSG00000242673 | 0 | 0 | 0 | 0 | 0 | 0 |
| ENSG00000236129 | 0 | 0 | 0 | 0 | 0 | 0 |
| ENSG00000227125 | 0 | 0 | 0 | 0 | 0 | 0 |
| ENSG00000237612 | 0 | 0 | 0 | 0 | 0 | 0 |
| ENSG00000231698 | 0 | 0 | 0 | 0 | 0 | 0 |
| ENSG00000182667 | 0 | 0 | 0 | 2 | 0 | 0 |
| ENSG00000252351 | 0 | 0 | 0 | 0 | 0 | 0 |
| ENSG00000237654 | 0 | 0 | 0 | 0 | 0 | 0 |
| ENSG00000285980 | 0 | 0 | 0 | 0 | 0 | 0 |
| ENSG00000224795 | 0 | 0 | 0 | 0 | 0 | 0 |
| ENSG00000271624 | 0 | 0 | 0 | 0 | 0 | 0 |
| ENSG00000238117 | 2 | 2 | 3 | 6 | 7 | 8 |
| ENSG00000224700 | 0 | 0 | 0 | 0 | 0 | 0 |
| ENSG00000238262 | 0 | 0 | 0 | 0 | 0 | 0 |
| ENSG00000279090 | 0 | 0 | 0 | 0 | 0 | 0 |
| ENSG00000183715 | 0 | 0 | 0 | 0 | 0 | 0 |
| ENSG00000283489 | 0 | 0 | 0 | 0 | 0 | 0 |
| ENSG00000279497 | 0 | 0 | 0 | 0 | 0 | 0 |
| ENSG00000255371 | 0 | 0 | 0 | 0 | 0 | 0 |
| ENSG00000279711 | 0 | 0 | 0 | 0 | 0 | 0 |
| ENSG00000279189 | 0 | 0 | 0 | 0 | 0 | 0 |
| ENSG00000254896 | 0 | 0 | 0 | 0 | 0 | 0 |
| ENSG00000255258 | 0 | 0 | 0 | 0 | 0 | 0 |
| ENSG00000166118 | 0 | 0 | 0 | 0 | 0 | 0 |
| ENSG00000213153 | 0 | 0 | 0 | 0 | 0 | 0 |
| ENSG00000280237 | 54 | 29 | 44 | 30 | 37 | 35 |
| ENSG00000284020 | 0 | 0 | 0 | 0 | 0 | 0 |
| ENSG00000080854 | 39 | 15 | 16 | 14 | 24 | 28 |
| ENSG00000255406 | 0 | 0 | 0 | 0 | 0 | 0 |
| ENSG00000204241 | 9 | 1 | 3 | 3 | 1 | 0 |
| ENSG00000254648 | 4 | 1 | 1 | 0 | 0 | 1 |
| ENSG00000166086 | 11 | 6 | 2 | 6 | 10 | 6 |
| ENSG00000254481 | 0 | 0 | 0 | 0 | 0 | 0 |
| ENSG00000151503 | 837 | 391 | 537 | 351 | 474 | 506 |
| ENSG00000254616 | 0 | 0 | 0 | 0 | 0 | 0 |
| ENSG00000255348 | 0 | 0 | 0 | 0 | 0 | 0 |
| ENSG00000151502 | 1693 | 793 | 1165 | 1059 | 902 | 977 |
| ENSG00000151500 | 433 | 210 | 348 | 229 | 255 | 255 |
| ENSG00000151498 | 774 | 349 | 516 | 358 | 378 | 460 |
| ENSG00000166105 | 0 | 0 | 0 | 0 | 0 | 0 |
| ENSG00000254442 | 0 | 0 | 0 | 0 | 0 | 0 |
| ENSG00000149328 | 248 | 138 | 172 | 244 | 179 | 241 |
| ENSG00000109956 | 2 | 0 | 0 | 0 | 0 | 2 |
| ENSG00000255545 | 0 | 0 | 0 | 0 | 0 | 0 |
| ENSG00000254573 | 0 | 0 | 0 | 0 | 0 | 0 |
| ENSG00000254989 | 0 | 0 | 0 | 0 | 0 | 0 |

| | | | | | | |
|---|---|---|---|---|---|---|
| ENSG00000285640 | 0 | 0 | 0 | 0 | 0 | 0 |
| ENSG00000255344 | 0 | 0 | 0 | 0 | 0 | 0 |
| ENSG00000251226 | 0 | 0 | 0 | 0 | 0 | 0 |
| ENSG00000255327 | 0 | 0 | 0 | 0 | 0 | 0 |
| ENSG00000279046 | 0 | 0 | 0 | 0 | 0 | 0 |
| ENSG00000285735 | 0 | 0 | 0 | 0 | 0 | 0 |
| ENSG00000255512 | 0 | 0 | 0 | 0 | 0 | 0 |
| ENSG00000260370 | 0 | 0 | 0 | 0 | 0 | 0 |
| ENSG00000237297 | 0 | 0 | 0 | 0 | 0 | 0 |
| ENSG00000261456 | 84 | 32 | 46 | 31 | 43 | 40 |
| ENSG00000232420 | 0 | 0 | 0 | 0 | 0 | 0 |
| ENSG00000015171 | 6250 | 2968 | 3544 | 2070 | 3241 | 2889 |
| ENSG00000276662 | 0 | 0 | 3 | 0 | 2 | 3 |
| ENSG00000212331 | 0 | 0 | 0 | 0 | 0 | 1 |
| ENSG00000151240 | 2554 | 1047 | 1476 | 924 | 1234 | 1346 |
| ENSG00000201861 | 0 | 0 | 1 | 0 | 0 | 0 |
| ENSG00000239822 | 0 | 0 | 0 | 0 | 0 | 0 |
| ENSG00000233021 | 0 | 0 | 1 | 0 | 0 | 0 |
| ENSG00000225140 | 0 | 1 | 0 | 0 | 0 | 0 |
| ENSG00000263511 | 0 | 0 | 0 | 0 | 0 | 0 |
| ENSG00000180525 | 6 | 2 | 2 | 1 | 1 | 0 |
| ENSG00000231601 | 0 | 0 | 0 | 0 | 0 | 0 |
| ENSG00000107929 | 5948 | 2514 | 3555 | 2231 | 2822 | 3085 |
| ENSG00000229869 | 1 | 0 | 5 | 0 | 1 | 5 |
| ENSG00000205740 | 49 | 32 | 18 | 16 | 15 | 14 |
| ENSG00000107937 | 3591 | 1838 | 2027 | 1798 | 2323 | 2238 |
| ENSG00000148377 | 0 | 0 | 1 | 0 | 0 | 1 |
| ENSG00000232656 | 4 | 1 | 1 | 1 | 4 | 0 |
| ENSG00000067064 | 4091 | 2087 | 2657 | 2280 | 2359 | 2294 |
| ENSG00000047056 | 1917 | 801 | 1105 | 688 | 922 | 1088 |
| ENSG00000229205 | 0 | 0 | 0 | 0 | 0 | 0 |
| ENSG00000185736 | 12 | 14 | 10 | 7 | 16 | 10 |
| ENSG00000282390 | 2 | 0 | 3 | 2 | 2 | 1 |
| ENSG00000233052 | 33 | 27 | 32 | 52 | 29 | 54 |
| ENSG00000205696 | 0 | 0 | 0 | 0 | 0 | 0 |
| ENSG00000234962 | 0 | 0 | 0 | 0 | 0 | 0 |
| ENSG00000226694 | 0 | 0 | 0 | 0 | 0 | 0 |
| ENSG00000278069 | 0 | 0 | 0 | 0 | 0 | 0 |
| ENSG00000252998 | 0 | 0 | 0 | 0 | 0 | 0 |
| ENSG00000227912 | 0 | 0 | 0 | 0 | 0 | 0 |
| ENSG00000231326 | 0 | 0 | 0 | 0 | 0 | 0 |
| ENSG00000212156 | 0 | 0 | 0 | 0 | 0 | 0 |
| ENSG00000234556 | 0 | 0 | 0 | 0 | 0 | 0 |
| ENSG00000234170 | 0 | 0 | 0 | 0 | 0 | 0 |
| ENSG00000237500 | 0 | 0 | 0 | 0 | 0 | 0 |
| ENSG00000235281 | 0 | 0 | 0 | 0 | 0 | 0 |
| ENSG00000231743 | 0 | 0 | 0 | 0 | 0 | 0 |
| ENSG00000234182 | 0 | 0 | 0 | 0 | 0 | 0 |
| ENSG00000273001 | 18 | 6 | 4 | 6 | 7 | 9 |
| ENSG00000067057 | 5198 | 2688 | 3376 | 4033 | 4061 | 4054 |

| | | | | | | |
|---|---|---|---|---|---|---|
| ENSG00000278419 | 0 | 0 | 0 | 0 | 0 | 0 |
| ENSG00000107959 | 4858 | 2329 | 3011 | 2397 | 2839 | 2957 |
| ENSG00000237399 | 21 | 15 | 30 | 10 | 16 | 27 |
| ENSG00000236968 | 0 | 0 | 0 | 0 | 0 | 0 |
| ENSG00000226762 | 0 | 0 | 0 | 0 | 0 | 0 |
| ENSG00000227338 | 0 | 0 | 0 | 0 | 0 | 0 |
| ENSG00000233321 | 0 | 0 | 0 | 0 | 0 | 0 |
| ENSG00000236892 | 0 | 0 | 0 | 0 | 0 | 0 |
| ENSG00000229672 | 0 | 0 | 0 | 0 | 0 | 0 |
| ENSG00000228353 | 0 | 0 | 0 | 0 | 0 | 0 |
| ENSG00000067082 | 2024 | 937 | 1128 | 808 | 891 | 1056 |
| ENSG00000230573 | 0 | 0 | 0 | 0 | 0 | 0 |
| ENSG00000226005 | 0 | 0 | 0 | 0 | 0 | 0 |
| ENSG00000274809 | 0 | 0 | 0 | 0 | 0 | 0 |
| ENSG00000227101 | 0 | 0 | 0 | 0 | 0 | 0 |
| ENSG00000236990 | 0 | 0 | 0 | 0 | 0 | 0 |
| ENSG00000233117 | 0 | 0 | 0 | 0 | 0 | 0 |
| ENSG00000224382 | 0 | 0 | 0 | 0 | 0 | 0 |
| ENSG00000230096 | 0 | 0 | 0 | 0 | 0 | 0 |
| ENSG00000207124 | 0 | 0 | 0 | 0 | 0 | 0 |
| ENSG00000231298 | 0 | 0 | 0 | 0 | 0 | 1 |
| ENSG00000225269 | 0 | 0 | 0 | 1 | 0 | 0 |
| ENSG00000165568 | 323 | 166 | 209 | 175 | 184 | 202 |
| ENSG00000151631 | 1 | 2 | 1 | 0 | 0 | 3 |
| ENSG00000238840 | 0 | 1 | 4 | 0 | 0 | 1 |
| ENSG00000187134 | 128 | 44 | 93 | 59 | 106 | 77 |
| ENSG00000151632 | 3068 | 1608 | 2176 | 2324 | 2457 | 2817 |
| ENSG00000224251 | 7 | 5 | 10 | 6 | 6 | 6 |
| ENSG00000239148 | 0 | 0 | 0 | 0 | 0 | 0 |
| ENSG00000196139 | 2465 | 1413 | 1847 | 1184 | 1246 | 1622 |
| ENSG00000239142 | 0 | 0 | 0 | 0 | 0 | 0 |
| ENSG00000225418 | 0 | 0 | 0 | 0 | 0 | 0 |
| ENSG00000264006 | 0 | 1 | 2 | 0 | 0 | 0 |
| ENSG00000198610 | 7 | 3 | 2 | 1 | 0 | 1 |
| ENSG00000228291 | 0 | 1 | 0 | 0 | 0 | 0 |
| ENSG00000231039 | 0 | 0 | 0 | 0 | 0 | 0 |
| ENSG00000224034 | 0 | 0 | 0 | 1 | 2 | 0 |
| ENSG00000215267 | 0 | 1 | 1 | 0 | 0 | 1 |
| ENSG00000223987 | 0 | 0 | 0 | 0 | 0 | 0 |
| ENSG00000280450 | 0 | 0 | 0 | 0 | 0 | 0 |
| ENSG00000178473 | 0 | 0 | 0 | 0 | 0 | 0 |
| ENSG00000178462 | 33 | 13 | 32 | 18 | 9 | 21 |
| ENSG00000173848 | 9614 | 4658 | 5886 | 3694 | 4740 | 4589 |
| ENSG00000178372 | 720 | 358 | 441 | 245 | 182 | 212 |
| ENSG00000205488 | 172 | 76 | 149 | 30 | 50 | 53 |
| ENSG00000178363 | 6 | 4 | 8 | 2 | 5 | 2 |
| ENSG00000256462 | 0 | 0 | 0 | 0 | 0 | 0 |
| ENSG00000242147 | 0 | 3 | 1 | 1 | 5 | 2 |
| ENSG00000231483 | 0 | 0 | 0 | 0 | 0 | 0 |
| ENSG00000228951 | 0 | 1 | 0 | 0 | 0 | 0 |

| | | | | | | |
|---|---|---|---|---|---|---|
| ENSG00000228685 | 0 | 0 | 0 | 0 | 0 | 0 |
| ENSG00000240577 | 0 | 0 | 0 | 0 | 0 | 0 |
| ENSG00000196372 | 1638 | 781 | 962 | 1016 | 972 | 1008 |
| ENSG00000108021 | 1766 | 1108 | 755 | 266 | 1478 | 760 |
| ENSG00000226647 | 0 | 1 | 3 | 0 | 0 | 0 |
| ENSG00000057608 | 11541 | 5849 | 7059 | 5980 | 7427 | 6849 |
| ENSG00000270427 | 2 | 1 | 7 | 1 | 6 | 3 |
| ENSG00000272764 | 20 | 8 | 11 | 7 | 13 | 10 |
| ENSG00000134461 | 498 | 230 | 294 | 272 | 221 | 297 |
| ENSG00000240180 | 0 | 0 | 0 | 0 | 0 | 0 |
| ENSG00000134452 | 3240 | 1510 | 1962 | 1606 | 1919 | 1874 |
| ENSG00000232807 | 26 | 17 | 26 | 13 | 15 | 7 |
| ENSG00000134470 | 50 | 20 | 29 | 31 | 35 | 15 |
| ENSG00000134460 | 0 | 0 | 0 | 0 | 0 | 0 |
| ENSG00000251922 | 0 | 0 | 0 | 0 | 0 | 0 |
| ENSG00000229664 | 0 | 0 | 0 | 0 | 0 | 0 |
| ENSG00000214015 | 0 | 0 | 0 | 0 | 0 | 0 |
| ENSG00000134453 | 3774 | 1933 | 2495 | 1907 | 2211 | 2283 |
| ENSG00000170525 | 6002 | 2983 | 3474 | 1963 | 2151 | 2414 |
| ENSG00000201581 | 0 | 0 | 0 | 0 | 1 | 1 |
| ENSG00000263628 | 0 | 0 | 0 | 0 | 0 | 0 |
| ENSG00000283634 | 0 | 0 | 0 | 0 | 0 | 0 |
| ENSG00000213994 | 5 | 4 | 3 | 1 | 0 | 1 |
| ENSG00000238366 | 0 | 0 | 0 | 0 | 0 | 0 |
| ENSG00000215244 | 43 | 20 | 29 | 6 | 16 | 12 |
| ENSG00000226705 | 0 | 0 | 0 | 0 | 0 | 0 |
| ENSG00000212743 | 7 | 15 | 22 | 8 | 3 | 17 |
| ENSG00000065675 | 0 | 1 | 0 | 0 | 0 | 1 |
| ENSG00000237943 | 0 | 0 | 0 | 0 | 0 | 0 |
| ENSG00000225948 | 0 | 0 | 0 | 0 | 0 | 0 |
| ENSG00000285743 | 0 | 0 | 0 | 0 | 0 | 0 |
| ENSG00000223784 | 0 | 0 | 0 | 0 | 0 | 0 |
| ENSG00000285988 | 0 | 0 | 0 | 0 | 0 | 0 |
| ENSG00000281186 | 0 | 0 | 0 | 0 | 0 | 0 |
| ENSG00000238266 | 0 | 0 | 0 | 0 | 0 | 0 |
| ENSG00000285845 | 0 | 0 | 0 | 0 | 0 | 0 |
| ENSG00000234248 | 0 | 0 | 0 | 0 | 0 | 0 |
| ENSG00000223581 | 0 | 0 | 0 | 0 | 0 | 0 |
| ENSG00000270811 | 0 | 0 | 0 | 0 | 0 | 0 |
| ENSG00000198879 | 235 | 92 | 110 | 65 | 99 | 104 |
| ENSG00000235141 | 0 | 0 | 0 | 0 | 0 | 0 |
| ENSG00000278982 | 0 | 1 | 1 | 1 | 2 | 0 |
| ENSG00000232591 | 0 | 0 | 0 | 0 | 0 | 0 |
| ENSG00000207453 | 0 | 0 | 0 | 0 | 0 | 0 |
| ENSG00000230121 | 0 | 0 | 0 | 0 | 0 | 0 |
| ENSG00000123243 | 0 | 0 | 0 | 0 | 0 | 0 |
| ENSG00000151655 | 10 | 2 | 8 | 3 | 2 | 5 |
| ENSG00000151657 | 443 | 320 | 320 | 201 | 373 | 262 |
| ENSG00000165629 | 9430 | 4653 | 5567 | 5300 | 5421 | 5257 |
| ENSG00000165632 | 820 | 409 | 427 | 306 | 446 | 405 |

| | | | | | | |
|---|---|---|---|---|---|---|
| ENSG00000233990 | 0 | 0 | 0 | 0 | 0 | 0 |
| ENSG00000107485 | 50842 | 22793 | 31224 | 21940 | 21439 | 25198 |
| ENSG00000197308 | 4887 | 2137 | 3115 | 2555 | 2312 | 2310 |
| ENSG00000232638 | 44 | 36 | 49 | 52 | 42 | 25 |
| ENSG00000225053 | 0 | 0 | 0 | 0 | 0 | 0 |
| ENSG00000232170 | 0 | 0 | 0 | 0 | 0 | 0 |
| ENSG00000226861 | 0 | 0 | 0 | 0 | 0 | 0 |
| ENSG00000226990 | 0 | 0 | 0 | 0 | 0 | 0 |
| ENSG00000271251 | 0 | 0 | 0 | 0 | 0 | 0 |
| ENSG00000213771 | 1 | 4 | 3 | 0 | 0 | 5 |
| ENSG00000236420 | 0 | 0 | 0 | 0 | 2 | 0 |
| ENSG00000212505 | 0 | 0 | 0 | 1 | 0 | 0 |
| ENSG00000270234 | 0 | 0 | 0 | 0 | 0 | 0 |
| ENSG00000234752 | 7 | 5 | 3 | 5 | 3 | 3 |
| ENSG00000223808 | 322 | 169 | 226 | 140 | 155 | 186 |
| ENSG00000230014 | 0 | 3 | 0 | 0 | 0 | 0 |
| ENSG00000235838 | 0 | 0 | 0 | 0 | 0 | 0 |
| ENSG00000228636 | 0 | 0 | 0 | 0 | 0 | 0 |
| ENSG00000224788 | 0 | 0 | 0 | 0 | 0 | 0 |
| ENSG00000229919 | 0 | 0 | 0 | 0 | 0 | 0 |
| ENSG00000200849 | 0 | 0 | 0 | 0 | 0 | 0 |
| ENSG00000272436 | 0 | 0 | 0 | 0 | 0 | 0 |
| ENSG00000234306 | 0 | 0 | 0 | 0 | 0 | 0 |
| ENSG00000279575 | 0 | 0 | 0 | 0 | 0 | 0 |
| ENSG00000270111 | 0 | 0 | 0 | 0 | 0 | 0 |
| ENSG00000228997 | 0 | 0 | 0 | 0 | 0 | 0 |
| ENSG00000225383 | 0 | 0 | 0 | 0 | 0 | 0 |
| ENSG00000048740 | 0 | 0 | 0 | 0 | 0 | 0 |
| ENSG00000229240 | 0 | 0 | 0 | 0 | 0 | 0 |
| ENSG00000231496 | 0 | 0 | 0 | 0 | 0 | 0 |
| ENSG00000228027 | 0 | 0 | 0 | 0 | 0 | 0 |
| ENSG00000229206 | 0 | 0 | 0 | 0 | 0 | 0 |
| ENSG00000237986 | 0 | 0 | 0 | 0 | 0 | 0 |
| ENSG00000230322 | 0 | 0 | 0 | 0 | 0 | 0 |
| ENSG00000181800 | 0 | 0 | 0 | 0 | 0 | 0 |
| ENSG00000280086 | 0 | 0 | 0 | 0 | 0 | 0 |
| ENSG00000148429 | 1485 | 742 | 883 | 535 | 831 | 897 |
| ENSG00000271360 | 0 | 0 | 0 | 0 | 0 | 0 |
| ENSG00000271046 | 0 | 0 | 0 | 0 | 0 | 0 |
| ENSG00000134463 | 181 | 92 | 108 | 84 | 116 | 92 |
| ENSG00000148426 | 576 | 291 | 370 | 340 | 347 | 297 |
| ENSG00000225778 | 7 | 6 | 7 | 2 | 0 | 1 |
| ENSG00000151461 | 813 | 521 | 335 | 176 | 766 | 384 |
| ENSG00000251957 | 2 | 1 | 0 | 0 | 0 | 1 |
| ENSG00000181192 | 3132 | 1574 | 2151 | 1089 | 1669 | 1675 |
| ENSG00000272507 | 2 | 2 | 4 | 2 | 2 | 0 |
| ENSG00000065665 | 329 | 206 | 228 | 164 | 203 | 191 |
| ENSG00000265653 | 0 | 0 | 0 | 0 | 0 | 0 |
| ENSG00000165609 | 1728 | 1038 | 1247 | 1082 | 1346 | 1260 |
| ENSG00000151465 | 1775 | 1028 | 1162 | 1147 | 1342 | 1287 |

| | | | | | | |
|---|---|---|---|---|---|---|
| ENSG00000228302 | 13 | 10 | 10 | 5 | 13 | 5 |
| ENSG00000264036 | 0 | 0 | 0 | 0 | 0 | 0 |
| ENSG00000183049 | 585 | 249 | 266 | 137 | 230 | 176 |
| ENSG00000252438 | 0 | 0 | 0 | 0 | 0 | 0 |
| ENSG00000252118 | 0 | 0 | 0 | 0 | 0 | 0 |
| ENSG00000285994 | 0 | 3 | 1 | 0 | 0 | 1 |
| ENSG00000263584 | 0 | 0 | 0 | 0 | 0 | 0 |
| ENSG00000283699 | 0 | 0 | 0 | 0 | 0 | 0 |
| ENSG00000221331 | 0 | 0 | 0 | 0 | 0 | 0 |
| ENSG00000285520 | 0 | 0 | 0 | 0 | 0 | 0 |
| ENSG00000151468 | 4 | 1 | 1 | 4 | 1 | 1 |
| ENSG00000212499 | 0 | 0 | 0 | 0 | 0 | 0 |
| ENSG00000225251 | 1 | 0 | 0 | 0 | 0 | 0 |
| ENSG00000123240 | 1239 | 704 | 783 | 553 | 795 | 669 |
| ENSG00000235763 | 6 | 0 | 3 | 2 | 1 | 2 |
| ENSG00000234175 | 1 | 1 | 1 | 0 | 2 | 0 |
| ENSG00000228330 | 0 | 0 | 0 | 0 | 0 | 1 |
| ENSG00000203414 | 74 | 30 | 42 | 30 | 31 | 39 |
| ENSG00000237540 | 1 | 0 | 0 | 0 | 0 | 0 |
| ENSG00000065328 | 382 | 183 | 240 | 131 | 229 | 207 |
| ENSG00000272055 | 4 | 2 | 2 | 3 | 3 | 1 |
| ENSG00000165623 | 0 | 0 | 0 | 0 | 0 | 0 |
| ENSG00000107537 | 834 | 440 | 518 | 455 | 578 | 503 |
| ENSG00000226209 | 0 | 0 | 0 | 0 | 0 | 0 |
| ENSG00000086475 | 4321 | 2052 | 2558 | 2449 | 2629 | 2599 |
| ENSG00000233256 | 0 | 0 | 0 | 0 | 0 | 0 |
| ENSG00000165626 | 621 | 350 | 439 | 283 | 413 | 383 |
| ENSG00000202508 | 0 | 0 | 1 | 0 | 0 | 0 |
| ENSG00000227175 | 0 | 0 | 0 | 0 | 0 | 0 |
| ENSG00000165630 | 181 | 153 | 145 | 126 | 240 | 149 |
| ENSG00000227474 | 0 | 0 | 0 | 0 | 0 | 0 |
| ENSG00000282246 | 0 | 0 | 1 | 0 | 0 | 0 |
| ENSG00000239665 | 133 | 93 | 91 | 40 | 81 | 79 |
| ENSG00000151474 | 237 | 130 | 133 | 120 | 166 | 164 |
| ENSG00000273474 | 4 | 0 | 1 | 0 | 1 | 2 |
| ENSG00000225112 | 2 | 0 | 0 | 0 | 0 | 0 |
| ENSG00000199407 | 0 | 0 | 0 | 0 | 0 | 0 |
| ENSG00000234091 | 0 | 0 | 0 | 0 | 0 | 0 |
| ENSG00000215227 | 0 | 0 | 0 | 0 | 0 | 0 |
| ENSG00000229751 | 0 | 0 | 0 | 0 | 0 | 0 |
| ENSG00000235410 | 0 | 0 | 0 | 0 | 0 | 0 |
| ENSG00000285671 | 0 | 0 | 1 | 0 | 0 | 0 |
| ENSG00000266321 | 0 | 0 | 0 | 0 | 0 | 0 |
| ENSG00000278100 | 0 | 0 | 0 | 0 | 0 | 0 |
| ENSG00000221371 | 0 | 0 | 0 | 0 | 0 | 0 |
| ENSG00000065809 | 2805 | 1239 | 1818 | 1120 | 1397 | 1556 |
| ENSG00000236495 | 2 | 0 | 0 | 0 | 0 | 1 |
| ENSG00000201766 | 0 | 0 | 0 | 0 | 0 | 0 |
| ENSG00000237032 | 0 | 0 | 0 | 0 | 0 | 0 |
| ENSG00000185267 | 52 | 36 | 57 | 23 | 28 | 40 |

| | | | | | | |
|---|---|---|---|---|---|---|
| ENSG00000284024 | 1079 | 499 | 616 | 527 | 616 | 581 |
| ENSG00000187522 | 442 | 281 | 334 | 207 | 324 | 311 |
| ENSG00000272853 | 27 | 16 | 15 | 18 | 21 | 18 |
| ENSG00000152455 | 371 | 174 | 208 | 142 | 267 | 202 |
| ENSG00000152457 | 261 | 133 | 156 | 127 | 177 | 147 |
| ENSG00000197889 | 20 | 9 | 12 | 22 | 12 | 13 |
| ENSG00000235623 | 0 | 0 | 0 | 0 | 0 | 0 |
| ENSG00000228547 | 0 | 1 | 0 | 0 | 0 | 1 |
| ENSG00000182531 | 0 | 0 | 0 | 0 | 0 | 0 |
| ENSG00000237470 | 0 | 0 | 0 | 0 | 0 | 0 |
| ENSG00000152463 | 3 | 4 | 0 | 0 | 0 | 1 |
| ENSG00000176244 | 210 | 97 | 135 | 104 | 112 | 108 |
| ENSG00000228312 | 0 | 0 | 0 | 0 | 0 | 0 |
| ENSG00000176236 | 9 | 5 | 2 | 2 | 4 | 3 |
| ENSG00000152464 | 399 | 188 | 254 | 253 | 251 | 313 |
| ENSG00000152465 | 427 | 205 | 243 | 152 | 279 | 250 |
| ENSG00000206448 | 2 | 1 | 0 | 0 | 0 | 0 |
| ENSG00000228181 | 0 | 0 | 0 | 0 | 0 | 0 |
| ENSG00000148468 | 48 | 24 | 20 | 18 | 29 | 21 |
| ENSG00000232739 | 0 | 0 | 0 | 0 | 1 | 0 |
| ENSG00000077943 | 8 | 3 | 5 | 1 | 3 | 4 |
| ENSG00000148481 | 652 | 325 | 427 | 278 | 401 | 372 |
| ENSG00000252537 | 0 | 0 | 0 | 0 | 0 | 0 |
| ENSG00000237913 | 0 | 0 | 0 | 0 | 0 | 0 |
| ENSG00000201260 | 0 | 0 | 0 | 0 | 0 | 0 |
| ENSG00000226140 | 0 | 0 | 0 | 0 | 0 | 0 |
| ENSG00000165983 | 362 | 196 | 198 | 128 | 253 | 200 |
| ENSG00000223156 | 0 | 0 | 0 | 0 | 0 | 0 |
| ENSG00000165985 | 2 | 4 | 1 | 0 | 1 | 5 |
| ENSG00000148484 | 2075 | 1125 | 1231 | 1030 | 1351 | 1196 |
| ENSG00000225213 | 1 | 2 | 1 | 1 | 4 | 2 |
| ENSG00000275001 | 0 | 0 | 0 | 0 | 0 | 0 |
| ENSG00000107611 | 0 | 0 | 0 | 0 | 0 | 0 |
| ENSG00000273153 | 0 | 0 | 0 | 0 | 0 | 0 |
| ENSG00000107614 | 88 | 34 | 52 | 21 | 45 | 46 |
| ENSG00000229124 | 4 | 1 | 2 | 2 | 1 | 0 |
| ENSG00000026025 | 7 | 9 | 2 | 2 | 2 | 3 |
| ENSG00000234961 | 0 | 0 | 0 | 0 | 0 | 0 |
| ENSG00000148488 | 171 | 96 | 114 | 60 | 92 | 108 |
| ENSG00000204832 | 9 | 13 | 6 | 4 | 2 | 4 |
| ENSG00000251803 | 0 | 0 | 0 | 0 | 0 | 0 |
| ENSG00000236582 | 0 | 0 | 0 | 0 | 0 | 0 |
| ENSG00000165996 | 384 | 226 | 242 | 230 | 224 | 211 |
| ENSG00000280126 | 0 | 0 | 0 | 0 | 0 | 0 |
| ENSG00000260589 | 20 | 8 | 17 | 6 | 11 | 6 |
| ENSG00000136738 | 856 | 450 | 510 | 366 | 513 | 520 |
| ENSG00000229190 | 0 | 0 | 0 | 0 | 1 | 1 |
| ENSG00000148483 | 0 | 1 | 4 | 0 | 2 | 1 |
| ENSG00000260314 | 0 | 0 | 0 | 0 | 0 | 0 |
| ENSG00000207938 | 0 | 0 | 0 | 0 | 0 | 0 |

| | | | | | | |
|---|---|---|---|---|---|---|
| ENSG00000235637 | 0 | 0 | 0 | 0 | 0 | 0 |
| ENSG00000148482 | 0 | 0 | 0 | 0 | 0 | 0 |
| ENSG00000226083 | 0 | 0 | 0 | 0 | 0 | 0 |
| ENSG00000165995 | 14 | 11 | 8 | 3 | 7 | 9 |
| ENSG00000235020 | 0 | 0 | 0 | 0 | 0 | 0 |
| ENSG00000240291 | 9 | 12 | 5 | 4 | 10 | 9 |
| ENSG00000225527 | 1 | 0 | 0 | 1 | 1 | 1 |
| ENSG00000241058 | 233 | 103 | 160 | 114 | 147 | 118 |
| ENSG00000165997 | 894 | 503 | 438 | 262 | 705 | 404 |
| ENSG00000225509 | 0 | 0 | 0 | 0 | 0 | 0 |
| ENSG00000234244 | 0 | 0 | 0 | 0 | 0 | 0 |
| ENSG00000234813 | 0 | 0 | 0 | 0 | 0 | 0 |
| ENSG00000204740 | 432 | 213 | 316 | 189 | 258 | 275 |
| ENSG00000226255 | 0 | 0 | 0 | 0 | 0 | 0 |
| ENSG00000227734 | 0 | 0 | 0 | 0 | 0 | 0 |
| ENSG00000239539 | 0 | 0 | 0 | 0 | 0 | 0 |
| ENSG00000236960 | 0 | 0 | 0 | 0 | 0 | 0 |
| ENSG00000200545 | 0 | 0 | 0 | 0 | 0 | 0 |
| ENSG00000252001 | 0 | 0 | 0 | 0 | 0 | 0 |
| ENSG00000233968 | 4 | 2 | 1 | 1 | 1 | 1 |
| ENSG00000283456 | 0 | 0 | 0 | 0 | 0 | 0 |
| ENSG00000230818 | 0 | 0 | 0 | 0 | 0 | 0 |
| ENSG00000252832 | 0 | 0 | 0 | 0 | 0 | 1 |
| ENSG00000285852 | 13 | 5 | 7 | 5 | 6 | 6 |
| ENSG00000120594 | 1361 | 666 | 757 | 356 | 691 | 498 |
| ENSG00000238246 | 0 | 0 | 0 | 0 | 1 | 0 |
| ENSG00000228339 | 0 | 3 | 1 | 0 | 0 | 0 |
| ENSG00000270727 | 0 | 0 | 0 | 0 | 0 | 0 |
| ENSG00000265372 | 0 | 0 | 0 | 0 | 0 | 0 |
| ENSG00000078114 | 2979 | 1619 | 1511 | 854 | 1938 | 1309 |
| ENSG00000235940 | 0 | 0 | 0 | 0 | 0 | 0 |
| ENSG00000230112 | 0 | 0 | 0 | 0 | 0 | 0 |
| ENSG00000228753 | 0 | 0 | 1 | 0 | 1 | 1 |
| ENSG00000228219 | 0 | 0 | 0 | 0 | 0 | 0 |
| ENSG00000204683 | 0 | 0 | 0 | 0 | 0 | 1 |
| ENSG00000231920 | 23 | 17 | 23 | 29 | 22 | 24 |
| ENSG00000228860 | 0 | 0 | 0 | 0 | 0 | 0 |
| ENSG00000232853 | 0 | 0 | 0 | 0 | 0 | 0 |
| ENSG00000207264 | 0 | 0 | 0 | 0 | 0 | 0 |
| ENSG00000230109 | 1 | 0 | 0 | 0 | 0 | 0 |
| ENSG00000231553 | 0 | 0 | 0 | 0 | 0 | 0 |
| ENSG00000278650 | 0 | 0 | 0 | 0 | 0 | 0 |
| ENSG00000273566 | 0 | 0 | 0 | 0 | 0 | 0 |
| ENSG00000199222 | 0 | 0 | 0 | 0 | 0 | 0 |
| ENSG00000204682 | 50 | 17 | 28 | 34 | 19 | 23 |
| ENSG00000222071 | 0 | 1 | 0 | 0 | 0 | 0 |
| ENSG00000180592 | 18 | 7 | 18 | 10 | 11 | 10 |
| ENSG00000078403 | 1045 | 575 | 678 | 436 | 615 | 554 |
| ENSG00000207347 | 0 | 0 | 0 | 0 | 0 | 0 |
| ENSG00000223984 | 0 | 0 | 0 | 0 | 0 | 3 |

| | | | | | | |
|---|---|---|---|---|---|---|
| ENSG00000201390 | 0 | 0 | 0 | 0 | 0 | 0 |
| ENSG00000136770 | 3086 | 1447 | 1813 | 1509 | 1561 | 1479 |
| ENSG00000252634 | 0 | 0 | 0 | 0 | 0 | 0 |
| ENSG00000279623 | 0 | 0 | 0 | 0 | 0 | 2 |
| ENSG00000201364 | 0 | 0 | 0 | 0 | 0 | 0 |
| ENSG00000232497 | 0 | 0 | 0 | 0 | 0 | 0 |
| ENSG00000236777 | 0 | 0 | 0 | 0 | 0 | 0 |
| ENSG00000223601 | 0 | 0 | 0 | 0 | 0 | 0 |
| ENSG00000260205 | 0 | 0 | 0 | 0 | 0 | 0 |
| ENSG00000227462 | 0 | 0 | 0 | 0 | 0 | 0 |
| ENSG00000279631 | 24 | 17 | 12 | 4 | 14 | 13 |
| ENSG00000261671 | 0 | 0 | 1 | 0 | 0 | 1 |
| ENSG00000233318 | 0 | 0 | 0 | 0 | 0 | 0 |
| ENSG00000148444 | 443 | 236 | 319 | 309 | 291 | 280 |
| ENSG00000269897 | 0 | 0 | 0 | 0 | 0 | 0 |
| ENSG00000168283 | 825 | 475 | 487 | 370 | 611 | 471 |
| ENSG00000272516 | 0 | 0 | 0 | 0 | 0 | 0 |
| ENSG00000271981 | 0 | 0 | 0 | 0 | 0 | 0 |
| ENSG00000272366 | 4 | 0 | 0 | 0 | 0 | 2 |
| ENSG00000077327 | 2 | 1 | 2 | 0 | 0 | 0 |
| ENSG00000233451 | 0 | 0 | 0 | 0 | 0 | 0 |
| ENSG00000285254 | 0 | 0 | 0 | 0 | 0 | 0 |
| ENSG00000150867 | 2896 | 1342 | 1809 | 1236 | 1427 | 1598 |
| ENSG00000279819 | 0 | 0 | 0 | 0 | 0 | 0 |
| ENSG00000206842 | 0 | 0 | 0 | 0 | 0 | 0 |
| ENSG00000165309 | 2 | 3 | 5 | 0 | 6 | 5 |
| ENSG00000223131 | 0 | 1 | 0 | 0 | 0 | 0 |
| ENSG00000148450 | 1237 | 565 | 720 | 717 | 634 | 594 |
| ENSG00000229932 | 13 | 5 | 8 | 3 | 5 | 3 |
| ENSG00000252049 | 0 | 0 | 0 | 0 | 0 | 0 |
| ENSG00000168267 | 4 | 0 | 0 | 0 | 0 | 0 |
| ENSG00000179133 | 5 | 0 | 0 | 1 | 4 | 2 |
| ENSG00000227537 | 0 | 0 | 0 | 0 | 0 | 0 |
| ENSG00000224215 | 0 | 0 | 0 | 0 | 0 | 0 |
| ENSG00000264574 | 0 | 0 | 0 | 0 | 0 | 0 |
| ENSG00000165312 | 491 | 235 | 296 | 247 | 282 | 274 |
| ENSG00000120549 | 544 | 299 | 341 | 219 | 374 | 360 |
| ENSG00000223928 | 0 | 0 | 0 | 0 | 0 | 0 |
| ENSG00000277563 | 0 | 0 | 0 | 0 | 0 | 0 |
| ENSG00000207930 | 0 | 0 | 0 | 0 | 0 | 0 |
| ENSG00000279029 | 0 | 0 | 0 | 0 | 0 | 0 |
| ENSG00000107863 | 734 | 372 | 329 | 189 | 663 | 387 |
| ENSG00000199535 | 0 | 0 | 0 | 0 | 0 | 0 |
| ENSG00000285750 | 0 | 0 | 0 | 0 | 0 | 0 |
| ENSG00000099256 | 33 | 18 | 13 | 24 | 43 | 25 |
| ENSG00000228667 | 0 | 0 | 0 | 0 | 0 | 0 |
| ENSG00000240294 | 0 | 0 | 0 | 0 | 0 | 0 |
| ENSG00000285859 | 1 | 0 | 0 | 0 | 0 | 0 |
| ENSG00000273107 | 0 | 0 | 0 | 0 | 0 | 2 |
| ENSG00000151023 | 1 | 0 | 0 | 0 | 3 | 0 |

| | | | | | | |
|---|---|---|---|---|---|---|
| ENSG00000185875 | 508 | 268 | 283 | 188 | 339 | 287 |
| ENSG00000231422 | 0 | 0 | 0 | 0 | 0 | 0 |
| ENSG00000233642 | 0 | 0 | 0 | 0 | 0 | 0 |
| ENSG00000151025 | 84 | 36 | 43 | 12 | 35 | 27 |
| ENSG00000274242 | 0 | 0 | 0 | 0 | 0 | 0 |
| ENSG00000222543 | 0 | 0 | 0 | 0 | 0 | 0 |
| ENSG00000227465 | 0 | 0 | 0 | 0 | 0 | 0 |
| ENSG00000280809 | 0 | 0 | 0 | 0 | 0 | 0 |
| ENSG00000271563 | 0 | 0 | 0 | 0 | 0 | 0 |
| ENSG00000223019 | 0 | 0 | 0 | 0 | 0 | 0 |
| ENSG00000251711 | 0 | 0 | 0 | 0 | 0 | 0 |
| ENSG00000226304 | 0 | 0 | 0 | 0 | 0 | 0 |
| ENSG00000095777 | 0 | 0 | 0 | 0 | 2 | 0 |
| ENSG00000136750 | 0 | 0 | 0 | 0 | 0 | 0 |
| ENSG00000284951 | 0 | 0 | 0 | 0 | 0 | 0 |
| ENSG00000077420 | 0 | 0 | 0 | 0 | 1 | 1 |
| ENSG00000199733 | 0 | 0 | 0 | 0 | 0 | 0 |
| ENSG00000236894 | 0 | 0 | 0 | 0 | 0 | 0 |
| ENSG00000233261 | 0 | 0 | 0 | 0 | 0 | 0 |
| ENSG00000234296 | 0 | 0 | 0 | 0 | 0 | 0 |
| ENSG00000231976 | 0 | 0 | 0 | 0 | 0 | 0 |
| ENSG00000227932 | 0 | 0 | 0 | 0 | 0 | 0 |
| ENSG00000279212 | 1 | 0 | 2 | 0 | 3 | 4 |
| ENSG00000148459 | 274 | 143 | 168 | 167 | 177 | 174 |
| ENSG00000234788 | 1 | 2 | 1 | 0 | 0 | 1 |
| ENSG00000235843 | 0 | 0 | 0 | 1 | 0 | 0 |
| ENSG00000136754 | 2547 | 1255 | 1524 | 1104 | 1440 | 1350 |
| ENSG00000223027 | 0 | 0 | 0 | 0 | 0 | 0 |
| ENSG00000206605 | 0 | 0 | 0 | 0 | 0 | 0 |
| ENSG00000283709 | 2 | 4 | 4 | 1 | 1 | 0 |
| ENSG00000107890 | 70 | 57 | 71 | 14 | 141 | 59 |
| ENSG00000199855 | 0 | 0 | 0 | 0 | 0 | 0 |
| ENSG00000136758 | 6070 | 3270 | 3433 | 1992 | 4046 | 3026 |
| ENSG00000120539 | 587 | 400 | 432 | 249 | 417 | 355 |
| ENSG00000252639 | 0 | 0 | 0 | 0 | 0 | 0 |
| ENSG00000107897 | 2774 | 1307 | 1660 | 777 | 1566 | 1494 |
| ENSG00000251839 | 0 | 0 | 0 | 0 | 0 | 0 |
| ENSG00000262412 | 34 | 2 | 5 | 9 | 7 | 5 |
| ENSG00000230445 | 58 | 23 | 36 | 8 | 44 | 23 |
| ENSG00000238021 | 7 | 1 | 2 | 1 | 0 | 4 |
| ENSG00000278830 | 0 | 0 | 0 | 0 | 0 | 0 |
| ENSG00000200369 | 0 | 1 | 3 | 0 | 0 | 0 |
| ENSG00000215409 | 4 | 1 | 0 | 0 | 1 | 1 |
| ENSG00000207135 | 0 | 0 | 0 | 0 | 0 | 0 |
| ENSG00000260151 | 0 | 0 | 0 | 0 | 0 | 0 |
| ENSG00000213770 | 0 | 0 | 0 | 0 | 0 | 0 |
| ENSG00000237956 | 0 | 0 | 0 | 0 | 0 | 0 |
| ENSG00000182077 | 0 | 0 | 0 | 0 | 1 | 0 |
| ENSG00000099246 | 3371 | 1727 | 1919 | 1301 | 1937 | 1640 |
| ENSG00000285737 | 1 | 1 | 0 | 0 | 0 | 0 |

| | | | | | | |
|---|---|---|---|---|---|---|
| ENSG00000150051 | 0 | 3 | 0 | 0 | 0 | 1 |
| ENSG00000230500 | 0 | 0 | 0 | 0 | 0 | 0 |
| ENSG00000251810 | 0 | 0 | 0 | 0 | 0 | 0 |
| ENSG00000169126 | 7 | 1 | 0 | 1 | 1 | 1 |
| ENSG00000226016 | 0 | 0 | 0 | 0 | 0 | 0 |
| ENSG00000233472 | 0 | 0 | 0 | 0 | 0 | 1 |
| ENSG00000226395 | 0 | 0 | 0 | 0 | 0 | 0 |
| ENSG00000150054 | 2248 | 1036 | 1207 | 680 | 1226 | 1041 |
| ENSG00000275036 | 0 | 0 | 0 | 0 | 0 | 0 |
| ENSG00000282996 | 4 | 3 | 0 | 0 | 3 | 1 |
| ENSG00000237746 | 0 | 0 | 0 | 0 | 0 | 0 |
| ENSG00000222666 | 0 | 0 | 0 | 0 | 0 | 0 |
| ENSG00000237128 | 1 | 1 | 0 | 0 | 0 | 1 |
| ENSG00000224439 | 0 | 0 | 0 | 0 | 0 | 0 |
| ENSG00000222705 | 0 | 0 | 0 | 0 | 0 | 0 |
| ENSG00000254635 | 1131 | 595 | 786 | 572 | 711 | 679 |
| ENSG00000095787 | 5557 | 2855 | 3010 | 1997 | 3186 | 2690 |
| ENSG00000252401 | 0 | 0 | 0 | 0 | 0 | 0 |
| ENSG00000231911 | 0 | 0 | 0 | 0 | 0 | 0 |
| ENSG00000207483 | 0 | 0 | 0 | 0 | 0 | 0 |
| ENSG00000095739 | 4658 | 2456 | 2779 | 2014 | 2072 | 2483 |
| ENSG00000232624 | 4 | 6 | 7 | 1 | 5 | 8 |
| ENSG00000236716 | 0 | 0 | 0 | 0 | 0 | 0 |
| ENSG00000201001 | 0 | 0 | 0 | 0 | 0 | 0 |
| ENSG00000235824 | 0 | 0 | 0 | 0 | 0 | 0 |
| ENSG00000271220 | 0 | 0 | 0 | 0 | 0 | 0 |
| ENSG00000199402 | 0 | 0 | 0 | 0 | 0 | 0 |
| ENSG00000229605 | 0 | 0 | 1 | 0 | 0 | 0 |
| ENSG00000120563 | 1 | 0 | 0 | 0 | 0 | 0 |
| ENSG00000224597 | 307 | 195 | 268 | 183 | 203 | 189 |
| ENSG00000274985 | 0 | 0 | 0 | 0 | 0 | 0 |
| ENSG00000197321 | 2022 | 902 | 1239 | 653 | 950 | 853 |
| ENSG00000207612 | 0 | 0 | 0 | 0 | 0 | 0 |
| ENSG00000212411 | 0 | 0 | 0 | 0 | 0 | 0 |
| ENSG00000216035 | 0 | 0 | 0 | 0 | 0 | 0 |
| ENSG00000225976 | 0 | 0 | 0 | 0 | 0 | 0 |
| ENSG00000222092 | 0 | 0 | 0 | 0 | 0 | 1 |
| ENSG00000237936 | 0 | 0 | 0 | 0 | 0 | 0 |
| ENSG00000165757 | 80 | 38 | 46 | 37 | 66 | 53 |
| ENSG00000200887 | 0 | 0 | 0 | 0 | 0 | 0 |
| ENSG00000230125 | 0 | 1 | 0 | 0 | 0 | 0 |
| ENSG00000259994 | 25 | 13 | 12 | 14 | 12 | 24 |
| ENSG00000107951 | 760 | 373 | 409 | 281 | 441 | 416 |
| ENSG00000285824 | 0 | 0 | 0 | 0 | 0 | 0 |
| ENSG00000250833 | 1 | 0 | 0 | 0 | 0 | 0 |
| ENSG00000241621 | 0 | 0 | 0 | 0 | 0 | 0 |
| ENSG00000239625 | 0 | 0 | 0 | 0 | 0 | 0 |
| ENSG00000284277 | 0 | 0 | 0 | 0 | 0 | 0 |
| ENSG00000214434 | 0 | 0 | 0 | 0 | 1 | 0 |
| ENSG00000235679 | 0 | 0 | 0 | 1 | 0 | 0 |

| | | | | | | |
|---|---|---|---|---|---|---|
| ENSG00000224473 | 5 | 1 | 1 | 0 | 1 | 2 |
| ENSG00000107968 | 317 | 143 | 236 | 153 | 197 | 202 |
| ENSG00000213778 | 2 | 0 | 0 | 0 | 0 | 0 |
| ENSG00000228800 | 0 | 0 | 0 | 0 | 0 | 0 |
| ENSG00000239744 | 0 | 0 | 0 | 0 | 0 | 0 |
| ENSG00000230298 | 0 | 0 | 0 | 0 | 0 | 0 |
| ENSG00000285630 | 2 | 0 | 0 | 0 | 0 | 0 |
| ENSG00000151033 | 10 | 2 | 2 | 2 | 1 | 2 |
| ENSG00000234814 | 4 | 7 | 2 | 6 | 16 | 4 |
| ENSG00000230720 | 0 | 0 | 0 | 0 | 0 | 0 |
| ENSG00000183621 | 145 | 80 | 85 | 79 | 91 | 77 |
| ENSG00000272914 | 2 | 0 | 1 | 0 | 1 | 2 |
| ENSG00000237135 | 2 | 0 | 0 | 2 | 0 | 0 |
| ENSG00000272381 | 0 | 0 | 0 | 0 | 0 | 0 |
| ENSG00000237036 | 152 | 72 | 90 | 66 | 109 | 67 |
| ENSG00000252479 | 0 | 0 | 0 | 0 | 0 | 0 |
| ENSG00000148516 | 0 | 1 | 0 | 0 | 0 | 3 |
| ENSG00000230397 | 2 | 0 | 2 | 1 | 0 | 0 |
| ENSG00000285781 | 0 | 0 | 0 | 0 | 0 | 0 |
| ENSG00000223834 | 1 | 1 | 3 | 3 | 4 | 0 |
| ENSG00000231855 | 0 | 0 | 0 | 0 | 0 | 0 |
| ENSG00000237797 | 0 | 0 | 0 | 0 | 0 | 0 |
| ENSG00000222412 | 0 | 0 | 0 | 0 | 0 | 0 |
| ENSG00000165322 | 1513 | 936 | 839 | 350 | 1333 | 787 |
| ENSG00000231148 | 0 | 0 | 0 | 0 | 0 | 0 |
| ENSG00000230240 | 0 | 0 | 0 | 0 | 0 | 0 |
| ENSG00000239731 | 0 | 0 | 0 | 0 | 1 | 0 |
| ENSG00000170759 | 3188 | 1932 | 1268 | 762 | 3395 | 1552 |
| ENSG00000206660 | 1 | 0 | 0 | 0 | 0 | 1 |
| ENSG00000234335 | 2 | 6 | 1 | 1 | 1 | 1 |
| ENSG00000235113 | 0 | 0 | 0 | 0 | 0 | 0 |
| ENSG00000234811 | 0 | 0 | 0 | 0 | 0 | 0 |
| ENSG00000252482 | 0 | 0 | 0 | 0 | 0 | 0 |
| ENSG00000217094 | 17 | 5 | 9 | 8 | 4 | 5 |
| ENSG00000233190 | 0 | 0 | 1 | 0 | 0 | 0 |
| ENSG00000227253 | 2 | 1 | 2 | 2 | 2 | 0 |
| ENSG00000120616 | 839 | 418 | 536 | 339 | 444 | 449 |
| ENSG00000226842 | 0 | 0 | 0 | 0 | 0 | 1 |
| ENSG00000233825 | 2 | 1 | 2 | 0 | 0 | 2 |
| ENSG00000229327 | 1 | 0 | 0 | 0 | 0 | 0 |
| ENSG00000200484 | 0 | 0 | 0 | 0 | 0 | 0 |
| ENSG00000216937 | 68 | 38 | 41 | 18 | 58 | 28 |
| ENSG00000231245 | 0 | 0 | 1 | 0 | 0 | 0 |
| ENSG00000273038 | 26 | 24 | 13 | 11 | 17 | 15 |
| ENSG00000150093 | 6497 | 3434 | 3900 | 2489 | 4463 | 3467 |
| ENSG00000228816 | 0 | 1 | 3 | 0 | 0 | 0 |
| ENSG00000229656 | 2 | 1 | 0 | 2 | 5 | 3 |
| ENSG00000234163 | 0 | 0 | 0 | 0 | 0 | 0 |
| ENSG00000265319 | 0 | 0 | 0 | 0 | 0 | 0 |
| ENSG00000229878 | 0 | 0 | 0 | 0 | 0 | 0 |

| | | | | | | |
|---|---|---|---|---|---|---|
| ENSG00000235962 | 0 | 0 | 0 | 0 | 0 | 1 |
| ENSG00000233387 | 0 | 0 | 0 | 0 | 0 | 0 |
| ENSG00000099250 | 3626 | 2054 | 2158 | 1512 | 2793 | 2380 |
| ENSG00000238258 | 1 | 0 | 0 | 0 | 0 | 0 |
| ENSG00000244356 | 0 | 0 | 0 | 0 | 0 | 0 |
| ENSG00000273012 | 2 | 1 | 0 | 2 | 2 | 2 |
| ENSG00000226412 | 0 | 0 | 0 | 0 | 0 | 0 |
| ENSG00000203565 | 0 | 0 | 0 | 0 | 0 | 0 |
| ENSG00000261683 | 0 | 0 | 0 | 0 | 0 | 0 |
| ENSG00000226407 | 0 | 0 | 0 | 0 | 0 | 0 |
| ENSG00000223470 | 0 | 0 | 0 | 0 | 0 | 0 |
| ENSG00000148498 | 2483 | 1202 | 1552 | 1209 | 1523 | 1389 |
| ENSG00000199200 | 0 | 0 | 0 | 0 | 0 | 0 |
| ENSG00000234167 | 0 | 0 | 0 | 0 | 0 | 0 |
| ENSG00000229235 | 0 | 0 | 0 | 0 | 0 | 0 |
| ENSG00000215184 | 0 | 0 | 0 | 0 | 0 | 1 |
| ENSG00000226386 | 9 | 11 | 6 | 7 | 4 | 5 |
| ENSG00000232525 | 0 | 0 | 0 | 0 | 0 | 0 |
| ENSG00000223047 | 0 | 0 | 0 | 0 | 0 | 0 |
| ENSG00000222909 | 0 | 0 | 0 | 0 | 0 | 0 |
| ENSG00000213724 | 0 | 0 | 0 | 0 | 0 | 0 |
| ENSG00000283000 | 0 | 3 | 7 | 8 | 6 | 8 |
| ENSG00000108094 | 1359 | 774 | 790 | 417 | 1027 | 700 |
| ENSG00000266228 | 0 | 0 | 0 | 0 | 0 | 0 |
| ENSG00000230534 | 1 | 1 | 3 | 1 | 0 | 0 |
| ENSG00000095794 | 453 | 249 | 232 | 212 | 261 | 213 |
| ENSG00000253054 | 0 | 0 | 0 | 0 | 0 | 0 |
| ENSG00000233343 | 0 | 0 | 0 | 0 | 0 | 0 |
| ENSG00000269952 | 4 | 1 | 3 | 3 | 1 | 2 |
| ENSG00000233200 | 0 | 0 | 0 | 0 | 0 | 1 |
| ENSG00000253070 | 0 | 0 | 0 | 0 | 0 | 0 |
| ENSG00000108100 | 2592 | 1175 | 1494 | 1349 | 1403 | 1497 |
| ENSG00000200097 | 0 | 0 | 0 | 0 | 0 | 0 |
| ENSG00000271335 | 45 | 35 | 45 | 29 | 30 | 36 |
| ENSG00000273312 | 2 | 0 | 0 | 1 | 2 | 2 |
| ENSG00000177291 | 2 | 0 | 0 | 1 | 1 | 0 |
| ENSG00000177283 | 1489 | 649 | 978 | 500 | 416 | 533 |
| ENSG00000284265 | 0 | 0 | 0 | 0 | 0 | 0 |
| ENSG00000226646 | 0 | 0 | 0 | 0 | 0 | 0 |
| ENSG00000280719 | 0 | 0 | 0 | 0 | 0 | 0 |
| ENSG00000227313 | 0 | 0 | 0 | 1 | 0 | 0 |
| ENSG00000227475 | 0 | 0 | 0 | 0 | 0 | 0 |
| ENSG00000236626 | 0 | 0 | 0 | 0 | 0 | 0 |
| ENSG00000228694 | 0 | 0 | 0 | 0 | 0 | 0 |
| ENSG00000237002 | 0 | 0 | 0 | 0 | 0 | 0 |
| ENSG00000274458 | 0 | 0 | 0 | 0 | 0 | 0 |
| ENSG00000229644 | 32 | 17 | 17 | 20 | 16 | 19 |
| ENSG00000207271 | 0 | 0 | 0 | 0 | 0 | 0 |
| ENSG00000238176 | 0 | 0 | 0 | 0 | 0 | 0 |
| ENSG00000233533 | 0 | 0 | 0 | 0 | 0 | 0 |

| | | | | | | |
|---|---|---|---|---|---|---|
| ENSG00000225355 | 2 | 0 | 0 | 0 | 0 | 1 |
| ENSG00000148513 | 0 | 1 | 1 | 3 | 2 | 0 |
| ENSG00000206796 | 0 | 0 | 0 | 0 | 0 | 0 |
| ENSG00000234918 | 0 | 0 | 0 | 0 | 0 | 0 |
| ENSG00000235687 | 0 | 0 | 1 | 0 | 0 | 0 |
| ENSG00000244402 | 0 | 0 | 0 | 0 | 0 | 0 |
| ENSG00000252621 | 0 | 0 | 0 | 0 | 0 | 0 |
| ENSG00000237970 | 0 | 0 | 0 | 0 | 0 | 0 |
| ENSG00000236313 | 0 | 0 | 0 | 0 | 0 | 0 |
| ENSG00000224920 | 0 | 0 | 0 | 0 | 0 | 0 |
| ENSG00000225276 | 0 | 0 | 0 | 0 | 0 | 0 |
| ENSG00000256892 | 0 | 0 | 0 | 0 | 0 | 0 |
| ENSG00000226578 | 2 | 1 | 1 | 1 | 1 | 0 |
| ENSG00000198105 | 605 | 315 | 391 | 248 | 418 | 354 |
| ENSG00000260137 | 0 | 0 | 0 | 0 | 0 | 0 |
| ENSG00000225192 | 0 | 0 | 0 | 0 | 0 | 0 |
| ENSG00000228828 | 0 | 0 | 0 | 0 | 0 | 0 |
| ENSG00000236514 | 11 | 5 | 12 | 5 | 6 | 8 |
| ENSG00000151963 | 0 | 0 | 1 | 1 | 0 | 1 |
| ENSG00000235197 | 0 | 0 | 0 | 0 | 0 | 0 |
| ENSG00000175395 | 207 | 107 | 110 | 66 | 148 | 105 |
| ENSG00000226447 | 0 | 0 | 0 | 0 | 0 | 0 |
| ENSG00000206840 | 0 | 0 | 0 | 0 | 0 | 0 |
| ENSG00000189180 | 840 | 474 | 412 | 207 | 723 | 406 |
| ENSG00000252132 | 0 | 1 | 1 | 0 | 0 | 0 |
| ENSG00000275675 | 0 | 0 | 0 | 0 | 0 | 0 |
| ENSG00000226086 | 0 | 0 | 2 | 0 | 0 | 0 |
| ENSG00000235964 | 0 | 0 | 0 | 0 | 0 | 0 |
| ENSG00000075407 | 468 | 280 | 198 | 84 | 415 | 220 |
| ENSG00000283930 | 32 | 12 | 24 | 10 | 21 | 26 |
| ENSG00000272983 | 10 | 6 | 4 | 2 | 4 | 1 |
| ENSG00000225965 | 0 | 0 | 0 | 0 | 0 | 0 |
| ENSG00000224761 | 0 | 0 | 0 | 0 | 0 | 0 |
| ENSG00000226113 | 0 | 0 | 0 | 0 | 0 | 0 |
| ENSG00000234941 | 0 | 0 | 0 | 0 | 0 | 0 |
| ENSG00000099251 | 88 | 50 | 62 | 37 | 58 | 49 |
| ENSG00000120555 | 10 | 3 | 10 | 4 | 3 | 8 |
| ENSG00000276805 | 242 | 104 | 160 | 101 | 141 | 151 |
| ENSG00000222314 | 0 | 0 | 0 | 0 | 0 | 0 |
| ENSG00000228882 | 0 | 0 | 0 | 2 | 0 | 1 |
| ENSG00000203496 | 0 | 0 | 2 | 0 | 0 | 0 |
| ENSG00000215151 | 0 | 0 | 0 | 0 | 0 | 0 |
| ENSG00000277133 | 0 | 0 | 0 | 0 | 0 | 0 |
| ENSG00000225638 | 0 | 0 | 0 | 0 | 0 | 0 |
| ENSG00000233867 | 0 | 0 | 0 | 0 | 0 | 0 |
| ENSG00000227264 | 0 | 0 | 0 | 0 | 0 | 0 |
| ENSG00000234541 | 0 | 0 | 0 | 0 | 0 | 0 |
| ENSG00000227679 | 0 | 0 | 0 | 0 | 0 | 0 |
| ENSG00000233832 | 0 | 0 | 0 | 0 | 0 | 0 |
| ENSG00000229485 | 2 | 1 | 0 | 0 | 0 | 0 |

| | | | | | | |
|---|---|---|---|---|---|---|
| ENSG00000237592 | 0 | 0 | 0 | 0 | 0 | 0 |
| ENSG00000271650 | 0 | 0 | 0 | 0 | 0 | 0 |
| ENSG00000274167 | 0 | 0 | 0 | 0 | 0 | 0 |
| ENSG00000228755 | 0 | 0 | 0 | 0 | 0 | 0 |
| ENSG00000215146 | 0 | 1 | 2 | 0 | 9 | 4 |
| ENSG00000182632 | 0 | 0 | 0 | 0 | 0 | 0 |
| ENSG00000279239 | 0 | 0 | 0 | 0 | 0 | 0 |
| ENSG00000185904 | 2 | 0 | 2 | 1 | 3 | 0 |
| ENSG00000234420 | 466 | 173 | 242 | 89 | 280 | 204 |
| ENSG00000270762 | 0 | 0 | 0 | 0 | 0 | 0 |
| ENSG00000233837 | 0 | 0 | 1 | 0 | 1 | 0 |
| ENSG00000196693 | 716 | 394 | 344 | 161 | 599 | 331 |
| ENSG00000277479 | 0 | 2 | 0 | 0 | 1 | 0 |
| ENSG00000272319 | 0 | 0 | 0 | 0 | 0 | 0 |
| ENSG00000272387 | 0 | 0 | 0 | 0 | 0 | 0 |
| ENSG00000251783 | 0 | 0 | 0 | 0 | 0 | 0 |
| ENSG00000285884 | 25 | 9 | 12 | 16 | 8 | 9 |
| ENSG00000232109 | 0 | 0 | 0 | 0 | 0 | 0 |
| ENSG00000233515 | 0 | 2 | 0 | 1 | 0 | 0 |
| ENSG00000231009 | 0 | 0 | 0 | 0 | 0 | 0 |
| ENSG00000234864 | 0 | 0 | 0 | 0 | 0 | 0 |
| ENSG00000230425 | 0 | 0 | 0 | 0 | 0 | 0 |
| ENSG00000270552 | 0 | 0 | 0 | 0 | 0 | 0 |
| ENSG00000259869 | 0 | 0 | 0 | 0 | 0 | 0 |
| ENSG00000165733 | 1857 | 929 | 1039 | 729 | 1261 | 1066 |
| ENSG00000252416 | 0 | 0 | 0 | 0 | 0 | 0 |
| ENSG00000234944 | 2 | 1 | 0 | 0 | 1 | 1 |
| ENSG00000229630 | 0 | 0 | 2 | 0 | 0 | 1 |
| ENSG00000263795 | 0 | 0 | 0 | 0 | 0 | 0 |
| ENSG00000165731 | 937 | 487 | 473 | 206 | 171 | 178 |
| ENSG00000273008 | 34 | 19 | 28 | 21 | 22 | 21 |
| ENSG00000169826 | 504 | 253 | 305 | 192 | 340 | 246 |
| ENSG00000198915 | 86 | 37 | 36 | 44 | 59 | 39 |
| ENSG00000224265 | 0 | 0 | 0 | 0 | 1 | 0 |
| ENSG00000285712 | 10 | 5 | 10 | 4 | 11 | 6 |
| ENSG00000221468 | 0 | 0 | 0 | 0 | 0 | 0 |
| ENSG00000150201 | 4 | 2 | 2 | 3 | 2 | 4 |
| ENSG00000169813 | 14223 | 6907 | 8482 | 6914 | 6973 | 7925 |
| ENSG00000221400 | 2 | 0 | 0 | 1 | 0 | 0 |
| ENSG00000230555 | 68 | 35 | 61 | 47 | 37 | 30 |
| ENSG00000243660 | 182 | 85 | 72 | 58 | 94 | 81 |
| ENSG00000252532 | 0 | 0 | 1 | 0 | 0 | 0 |
| ENSG00000236114 | 2 | 0 | 1 | 0 | 0 | 0 |
| ENSG00000196793 | 497 | 239 | 291 | 235 | 306 | 316 |
| ENSG00000232004 | 2 | 0 | 0 | 2 | 2 | 1 |
| ENSG00000198298 | 256 | 129 | 145 | 116 | 193 | 150 |
| ENSG00000223910 | 0 | 1 | 1 | 0 | 0 | 0 |
| ENSG00000237389 | 0 | 0 | 0 | 0 | 0 | 0 |
| ENSG00000238263 | 0 | 0 | 0 | 0 | 0 | 0 |
| ENSG00000169740 | 815 | 454 | 548 | 570 | 584 | 527 |

| | | | | | | |
|---|---|---|---|---|---|---|
| ENSG00000226245 | 7 | 6 | 4 | 9 | 7 | 5 |
| ENSG00000230565 | 37 | 20 | 48 | 26 | 31 | 43 |
| ENSG00000228426 | 0 | 0 | 0 | 0 | 0 | 0 |
| ENSG00000271173 | 0 | 0 | 0 | 0 | 0 | 0 |
| ENSG00000224162 | 0 | 0 | 0 | 0 | 0 | 0 |
| ENSG00000223746 | 0 | 0 | 0 | 0 | 0 | 0 |
| ENSG00000234973 | 0 | 0 | 0 | 0 | 0 | 0 |
| ENSG00000226790 | 0 | 0 | 0 | 0 | 0 | 0 |
| ENSG00000226808 | 0 | 0 | 0 | 2 | 1 | 0 |
| ENSG00000236769 | 0 | 0 | 0 | 0 | 0 | 0 |
| ENSG00000227029 | 0 | 0 | 0 | 0 | 0 | 0 |
| ENSG00000233395 | 0 | 0 | 0 | 0 | 0 | 0 |
| ENSG00000237590 | 0 | 0 | 0 | 0 | 0 | 0 |
| ENSG00000229116 | 0 | 0 | 0 | 0 | 0 | 0 |
| ENSG00000277288 | 0 | 2 | 0 | 0 | 1 | 0 |
| ENSG00000107562 | 334 | 166 | 115 | 149 | 145 | 180 |
| ENSG00000214089 | 1 | 0 | 0 | 0 | 0 | 0 |
| ENSG00000270767 | 0 | 0 | 0 | 0 | 0 | 0 |
| ENSG00000234580 | 0 | 0 | 0 | 0 | 0 | 0 |
| ENSG00000227395 | 0 | 0 | 0 | 0 | 0 | 0 |
| ENSG00000224812 | 2 | 1 | 0 | 0 | 0 | 3 |
| ENSG00000234504 | 0 | 1 | 0 | 0 | 1 | 0 |
| ENSG00000187783 | 0 | 0 | 0 | 0 | 0 | 0 |
| ENSG00000223462 | 0 | 0 | 2 | 1 | 2 | 0 |
| ENSG00000273363 | 6 | 0 | 1 | 1 | 0 | 1 |
| ENSG00000107551 | 133 | 51 | 79 | 34 | 19 | 26 |
| ENSG00000165507 | 166 | 77 | 106 | 29 | 41 | 37 |
| ENSG00000165511 | 5 | 3 | 12 | 2 | 1 | 4 |
| ENSG00000165512 | 326 | 177 | 183 | 173 | 175 | 200 |
| ENSG00000226937 | 6 | 0 | 2 | 1 | 0 | 2 |
| ENSG00000277757 | 0 | 0 | 0 | 0 | 0 | 0 |
| ENSG00000270258 | 0 | 0 | 0 | 0 | 0 | 0 |
| ENSG00000228462 | 0 | 0 | 0 | 0 | 0 | 0 |
| ENSG00000232554 | 0 | 0 | 0 | 0 | 0 | 0 |
| ENSG00000235690 | 0 | 0 | 0 | 0 | 0 | 0 |
| ENSG00000243349 | 0 | 0 | 0 | 0 | 0 | 0 |
| ENSG00000236442 | 0 | 0 | 0 | 0 | 0 | 0 |
| ENSG00000230501 | 0 | 0 | 0 | 0 | 0 | 0 |
| ENSG00000263476 | 0 | 0 | 0 | 0 | 0 | 0 |
| ENSG00000227683 | 0 | 0 | 0 | 0 | 0 | 0 |
| ENSG00000275312 | 0 | 0 | 0 | 0 | 0 | 0 |
| ENSG00000278531 | 0 | 0 | 0 | 0 | 0 | 0 |
| ENSG00000207071 | 0 | 0 | 0 | 0 | 0 | 0 |
| ENSG00000234993 | 0 | 0 | 0 | 0 | 0 | 0 |
| ENSG00000224402 | 0 | 0 | 0 | 0 | 0 | 0 |
| ENSG00000256574 | 0 | 0 | 0 | 0 | 0 | 0 |
| ENSG00000012779 | 14 | 5 | 9 | 6 | 10 | 8 |
| ENSG00000231964 | 0 | 0 | 1 | 0 | 0 | 1 |
| ENSG00000165406 | 592 | 254 | 360 | 241 | 297 | 315 |
| ENSG00000172671 | 56 | 28 | 38 | 24 | 52 | 52 |

| | | | | | | |
|---|---|---|---|---|---|---|
| ENSG00000230869 | 5 | 0 | 0 | 1 | 1 | 6 |
| ENSG00000237840 | 23 | 10 | 19 | 15 | 14 | 12 |
| ENSG00000228702 | 1 | 2 | 2 | 3 | 1 | 1 |
| ENSG00000172661 | 1629 | 699 | 898 | 581 | 766 | 681 |
| ENSG00000270434 | 0 | 0 | 0 | 0 | 0 | 0 |
| ENSG00000231122 | 0 | 0 | 0 | 0 | 0 | 0 |
| ENSG00000188234 | 119 | 56 | 83 | 52 | 52 | 65 |
| ENSG00000239883 | 103 | 51 | 66 | 23 | 36 | 52 |
| ENSG00000239152 | 0 | 0 | 0 | 0 | 0 | 0 |
| ENSG00000264217 | 0 | 0 | 0 | 0 | 0 | 0 |
| ENSG00000265354 | 2444 | 1295 | 1520 | 1484 | 1553 | 1625 |
| ENSG00000265733 | 2 | 0 | 0 | 0 | 0 | 0 |
| ENSG00000266412 | 6609 | 3163 | 3961 | 2824 | 3499 | 3342 |
| ENSG00000263639 | 42 | 35 | 31 | 17 | 38 | 30 |
| ENSG00000280012 | 0 | 0 | 0 | 0 | 0 | 0 |
| ENSG00000264204 | 62 | 28 | 40 | 20 | 34 | 40 |
| ENSG00000263482 | 0 | 0 | 0 | 0 | 0 | 0 |
| ENSG00000274209 | 0 | 0 | 1 | 0 | 0 | 0 |
| ENSG00000279058 | 14 | 2 | 5 | 4 | 4 | 3 |
| ENSG00000273225 | 0 | 0 | 0 | 0 | 0 | 0 |
| ENSG00000279458 | 0 | 0 | 0 | 0 | 0 | 0 |
| ENSG00000264230 | 0 | 1 | 1 | 0 | 1 | 0 |
| ENSG00000285402 | 0 | 0 | 0 | 0 | 0 | 0 |
| ENSG00000285294 | 0 | 0 | 0 | 1 | 0 | 0 |
| ENSG00000213412 | 0 | 0 | 0 | 0 | 0 | 0 |
| ENSG00000204174 | 0 | 0 | 0 | 0 | 0 | 0 |
| ENSG00000204175 | 4 | 0 | 1 | 0 | 0 | 0 |
| ENSG00000204176 | 26 | 15 | 28 | 14 | 19 | 20 |
| ENSG00000231187 | 9 | 3 | 6 | 1 | 3 | 8 |
| ENSG00000229227 | 0 | 1 | 0 | 0 | 0 | 0 |
| ENSG00000165874 | 1 | 0 | 0 | 0 | 3 | 0 |
| ENSG00000272430 | 0 | 0 | 3 | 2 | 0 | 0 |
| ENSG00000229927 | 0 | 0 | 0 | 0 | 0 | 0 |
| ENSG00000274186 | 0 | 0 | 0 | 0 | 0 | 0 |
| ENSG00000285662 | 0 | 0 | 0 | 0 | 0 | 0 |
| ENSG00000280913 | 0 | 0 | 0 | 2 | 0 | 0 |
| ENSG00000265366 | 8 | 4 | 11 | 6 | 11 | 2 |
| ENSG00000215097 | 10 | 2 | 0 | 0 | 7 | 3 |
| ENSG00000204177 | 97 | 47 | 49 | 25 | 34 | 53 |
| ENSG00000238405 | 4 | 1 | 1 | 1 | 0 | 0 |
| ENSG00000243289 | 3 | 1 | 1 | 2 | 5 | 1 |
| ENSG00000150175 | 0 | 0 | 0 | 0 | 0 | 0 |
| ENSG00000204179 | 0 | 0 | 0 | 0 | 0 | 0 |
| ENSG00000266524 | 0 | 0 | 0 | 0 | 0 | 0 |
| ENSG00000263761 | 0 | 0 | 0 | 0 | 0 | 0 |
| ENSG00000265203 | 0 | 0 | 0 | 0 | 0 | 0 |
| ENSG00000265763 | 40 | 27 | 24 | 39 | 33 | 29 |
| ENSG00000265190 | 0 | 0 | 0 | 0 | 1 | 0 |
| ENSG00000270369 | 0 | 0 | 0 | 0 | 0 | 1 |
| ENSG00000189090 | 0 | 0 | 0 | 0 | 0 | 0 |

| | | | | | | |
|---|---|---|---|---|---|---|
| ENSG00000224919 | 0 | 0 | 0 | 0 | 0 | 0 |
| ENSG00000204172 | 168 | 90 | 89 | 63 | 96 | 104 |
| ENSG00000254929 | 2 | 0 | 0 | 2 | 1 | 1 |
| ENSG00000252877 | 0 | 0 | 0 | 0 | 0 | 0 |
| ENSG00000251079 | 15 | 11 | 4 | 3 | 6 | 7 |
| ENSG00000265630 | 0 | 1 | 1 | 3 | 1 | 3 |
| ENSG00000215065 | 0 | 0 | 0 | 0 | 0 | 0 |
| ENSG00000266217 | 0 | 0 | 0 | 0 | 0 | 0 |
| ENSG00000264404 | 0 | 0 | 0 | 0 | 0 | 0 |
| ENSG00000189014 | 2 | 5 | 7 | 6 | 0 | 6 |
| ENSG00000276544 | 0 | 0 | 0 | 0 | 0 | 0 |
| ENSG00000226964 | 5 | 5 | 6 | 11 | 6 | 5 |
| ENSG00000277758 | 80 | 49 | 66 | 35 | 53 | 40 |
| ENSG00000276850 | 12 | 3 | 5 | 5 | 10 | 3 |
| ENSG00000264717 | 0 | 0 | 0 | 0 | 0 | 0 |
| ENSG00000273760 | 0 | 0 | 0 | 0 | 3 | 0 |
| ENSG00000276430 | 0 | 0 | 0 | 0 | 0 | 0 |
| ENSG00000265018 | 2 | 1 | 0 | 1 | 4 | 0 |
| ENSG00000252149 | 0 | 0 | 0 | 0 | 0 | 0 |
| ENSG00000270025 | 1 | 0 | 0 | 0 | 0 | 0 |
| ENSG00000278561 | 0 | 0 | 0 | 0 | 0 | 0 |
| ENSG00000170324 | 2 | 0 | 4 | 2 | 1 | 2 |
| ENSG00000285786 | 0 | 0 | 0 | 0 | 0 | 0 |
| ENSG00000225482 | 0 | 0 | 0 | 0 | 0 | 0 |
| ENSG00000107643 | 1281 | 579 | 785 | 472 | 684 | 630 |
| ENSG00000279822 | 0 | 0 | 0 | 1 | 0 | 0 |
| ENSG00000128805 | 6 | 3 | 10 | 11 | 11 | 8 |
| ENSG00000248682 | 0 | 0 | 0 | 0 | 0 | 0 |
| ENSG00000231906 | 0 | 0 | 1 | 0 | 0 | 0 |
| ENSG00000251413 | 0 | 0 | 0 | 0 | 0 | 0 |
| ENSG00000236800 | 0 | 0 | 0 | 0 | 0 | 0 |
| ENSG00000128815 | 32 | 13 | 40 | 35 | 21 | 46 |
| ENSG00000228754 | 0 | 0 | 0 | 0 | 0 | 0 |
| ENSG00000228403 | 0 | 0 | 0 | 0 | 0 | 0 |
| ENSG00000241577 | 0 | 0 | 0 | 0 | 0 | 0 |
| ENSG00000165383 | 0 | 0 | 0 | 0 | 0 | 0 |
| ENSG00000233665 | 0 | 0 | 0 | 0 | 0 | 0 |
| ENSG00000226576 | 0 | 0 | 0 | 0 | 0 | 0 |
| ENSG00000264800 | 0 | 0 | 0 | 0 | 0 | 0 |
| ENSG00000165633 | 0 | 0 | 0 | 0 | 0 | 0 |
| ENSG00000234736 | 0 | 0 | 0 | 0 | 0 | 0 |
| ENSG00000172538 | 0 | 0 | 0 | 0 | 0 | 0 |
| ENSG00000204161 | 0 | 0 | 0 | 0 | 0 | 0 |
| ENSG00000236208 | 0 | 0 | 0 | 0 | 0 | 0 |
| ENSG00000177354 | 0 | 0 | 0 | 0 | 0 | 0 |
| ENSG00000165606 | 0 | 0 | 0 | 0 | 0 | 0 |
| ENSG00000235939 | 1 | 8 | 6 | 4 | 2 | 0 |
| ENSG00000225830 | 1035 | 403 | 542 | 334 | 544 | 472 |
| ENSG00000271237 | 0 | 0 | 0 | 0 | 0 | 0 |
| ENSG00000070748 | 0 | 0 | 0 | 0 | 0 | 0 |

| | | | | | | |
|---|---|---|---|---|---|---|
| ENSG00000187714 | 0 | 0 | 0 | 0 | 0 | 0 |
| ENSG00000178645 | 0 | 0 | 0 | 0 | 0 | 0 |
| ENSG00000197444 | 135 | 54 | 83 | 65 | 85 | 66 |
| ENSG00000226389 | 0 | 0 | 0 | 0 | 0 | 0 |
| ENSG00000229870 | 2 | 0 | 1 | 1 | 0 | 0 |
| ENSG00000227345 | 1059 | 571 | 663 | 377 | 694 | 612 |
| ENSG00000230166 | 0 | 0 | 0 | 0 | 0 | 0 |
| ENSG00000204152 | 185 | 95 | 106 | 95 | 135 | 134 |
| ENSG00000223111 | 0 | 0 | 0 | 0 | 0 | 0 |
| ENSG00000178440 | 5 | 7 | 9 | 3 | 3 | 3 |
| ENSG00000222108 | 2 | 2 | 1 | 0 | 0 | 0 |
| ENSG00000285803 | 14 | 14 | 10 | 1 | 5 | 4 |
| ENSG00000204149 | 332 | 146 | 239 | 155 | 193 | 215 |
| ENSG00000235618 | 48 | 16 | 24 | 15 | 18 | 16 |
| ENSG00000226631 | 0 | 0 | 0 | 0 | 0 | 2 |
| ENSG00000099290 | 1968 | 968 | 1220 | 827 | 1115 | 1070 |
| ENSG00000233011 | 0 | 0 | 0 | 0 | 0 | 0 |
| ENSG00000188611 | 68 | 30 | 39 | 31 | 44 | 27 |
| ENSG00000225137 | 4 | 3 | 4 | 5 | 6 | 3 |
| ENSG00000198964 | 1560 | 651 | 925 | 593 | 836 | 797 |
| ENSG00000279863 | 42 | 18 | 22 | 10 | 11 | 20 |
| ENSG00000225303 | 0 | 0 | 0 | 0 | 0 | 0 |
| ENSG00000238523 | 0 | 0 | 0 | 0 | 0 | 0 |
| ENSG00000226200 | 296 | 94 | 175 | 86 | 106 | 127 |
| ENSG00000231588 | 0 | 0 | 0 | 0 | 0 | 0 |
| ENSG00000231345 | 220 | 80 | 114 | 72 | 91 | 106 |
| ENSG00000232706 | 2 | 1 | 0 | 0 | 0 | 0 |
| ENSG00000226168 | 0 | 0 | 0 | 0 | 0 | 0 |
| ENSG00000230011 | 0 | 0 | 0 | 0 | 0 | 0 |
| ENSG00000213667 | 2 | 2 | 2 | 1 | 2 | 1 |
| ENSG00000204147 | 336 | 156 | 197 | 136 | 187 | 185 |
| ENSG00000148584 | 1 | 0 | 0 | 0 | 0 | 0 |
| ENSG00000229711 | 0 | 0 | 0 | 0 | 0 | 0 |
| ENSG00000236944 | 0 | 0 | 0 | 0 | 0 | 0 |
| ENSG00000261368 | 0 | 0 | 0 | 0 | 0 | 0 |
| ENSG00000185532 | 40 | 14 | 21 | 4 | 12 | 18 |
| ENSG00000223502 | 0 | 1 | 1 | 0 | 1 | 0 |
| ENSG00000231132 | 0 | 0 | 0 | 0 | 0 | 0 |
| ENSG00000207813 | 0 | 0 | 0 | 0 | 1 | 0 |
| ENSG00000235279 | 0 | 0 | 0 | 0 | 0 | 0 |
| ENSG00000213659 | 0 | 0 | 0 | 0 | 0 | 0 |
| ENSG00000177613 | 1212 | 550 | 659 | 535 | 594 | 594 |
| ENSG00000236671 | 29 | 9 | 12 | 8 | 7 | 9 |
| ENSG00000107984 | 2199 | 1164 | 1619 | 683 | 659 | 839 |
| ENSG00000213657 | 2 | 0 | 0 | 2 | 0 | 0 |
| ENSG00000227972 | 0 | 0 | 3 | 0 | 1 | 0 |
| ENSG00000231131 | 1527 | 791 | 1005 | 586 | 827 | 711 |
| ENSG00000165471 | 2 | 0 | 0 | 0 | 0 | 1 |
| ENSG00000201196 | 0 | 0 | 0 | 0 | 0 | 0 |
| ENSG00000285958 | 0 | 2 | 1 | 0 | 0 | 1 |

| | | | | | | |
|---|---|---|---|---|---|---|
| ENSG00000227121 | 13 | 14 | 12 | 14 | 16 | 20 |
| ENSG00000231399 | 0 | 0 | 0 | 0 | 0 | 0 |
| ENSG00000226296 | 0 | 0 | 0 | 0 | 0 | 0 |
| ENSG00000252161 | 0 | 0 | 0 | 0 | 0 | 0 |
| ENSG00000233805 | 0 | 0 | 0 | 0 | 0 | 0 |
| ENSG00000150275 | 2 | 0 | 0 | 3 | 0 | 0 |
| ENSG00000252252 | 0 | 0 | 0 | 0 | 0 | 0 |
| ENSG00000234173 | 0 | 0 | 0 | 0 | 0 | 0 |
| ENSG00000224697 | 0 | 0 | 0 | 0 | 0 | 0 |
| ENSG00000221594 | 0 | 0 | 0 | 0 | 0 | 0 |
| ENSG00000236958 | 0 | 0 | 0 | 0 | 0 | 0 |
| ENSG00000236744 | 0 | 0 | 0 | 0 | 0 | 0 |
| ENSG00000223800 | 0 | 0 | 0 | 0 | 0 | 0 |
| ENSG00000228048 | 0 | 0 | 0 | 0 | 0 | 0 |
| ENSG00000249860 | 0 | 0 | 0 | 0 | 0 | 0 |
| ENSG00000236993 | 4 | 1 | 1 | 1 | 3 | 0 |
| ENSG00000270541 | 0 | 0 | 0 | 0 | 0 | 0 |
| ENSG00000122952 | 3032 | 1480 | 1916 | 1305 | 1495 | 1834 |
| ENSG00000238707 | 0 | 0 | 0 | 0 | 0 | 0 |
| ENSG00000264747 | 0 | 0 | 0 | 0 | 0 | 0 |
| ENSG00000235810 | 0 | 0 | 0 | 0 | 0 | 0 |
| ENSG00000223841 | 0 | 0 | 0 | 0 | 0 | 0 |
| ENSG00000270192 | 0 | 0 | 0 | 0 | 0 | 0 |
| ENSG00000232115 | 0 | 0 | 0 | 0 | 0 | 0 |
| ENSG00000151151 | 526 | 220 | 245 | 158 | 266 | 206 |
| ENSG00000180221 | 0 | 0 | 0 | 0 | 0 | 0 |
| ENSG00000122873 | 742 | 356 | 421 | 435 | 426 | 461 |
| ENSG00000228527 | 0 | 0 | 0 | 1 | 1 | 0 |
| ENSG00000261076 | 0 | 0 | 0 | 0 | 0 | 0 |
| ENSG00000072401 | 454 | 245 | 297 | 260 | 292 | 294 |
| ENSG00000108064 | 1432 | 813 | 937 | 694 | 955 | 888 |
| ENSG00000122870 | 0 | 0 | 0 | 0 | 0 | 0 |
| ENSG00000183055 | 0 | 1 | 0 | 0 | 0 | 5 |
| ENSG00000252076 | 0 | 0 | 0 | 0 | 0 | 0 |
| ENSG00000237949 | 0 | 0 | 0 | 0 | 0 | 0 |
| ENSG00000226888 | 0 | 0 | 0 | 0 | 0 | 0 |
| ENSG00000226557 | 0 | 0 | 0 | 0 | 0 | 0 |
| ENSG00000165443 | 0 | 0 | 0 | 0 | 0 | 0 |
| ENSG00000148541 | 0 | 0 | 0 | 0 | 0 | 0 |
| ENSG00000236556 | 0 | 0 | 0 | 0 | 0 | 0 |
| ENSG00000235469 | 0 | 0 | 0 | 0 | 0 | 0 |
| ENSG00000235140 | 0 | 0 | 0 | 0 | 0 | 0 |
| ENSG00000165449 | 203 | 130 | 143 | 90 | 94 | 108 |
| ENSG00000227877 | 2 | 0 | 1 | 0 | 0 | 0 |
| ENSG00000108091 | 5145 | 2619 | 2797 | 1580 | 2773 | 2174 |
| ENSG00000235931 | 0 | 0 | 0 | 0 | 0 | 0 |
| ENSG00000151150 | 2470 | 1457 | 1453 | 505 | 1499 | 986 |
| ENSG00000232682 | 13 | 3 | 6 | 4 | 3 | 0 |
| ENSG00000202190 | 0 | 1 | 0 | 0 | 0 | 0 |
| ENSG00000282121 | 1 | 1 | 0 | 0 | 0 | 0 |

| | | | | | | |
|---|---|---|---|---|---|---|
| ENSG00000122872 | 0 | 0 | 0 | 0 | 0 | 0 |
| ENSG00000254271 | 13 | 5 | 5 | 1 | 5 | 3 |
| ENSG00000170312 | 1039 | 593 | 589 | 466 | 849 | 645 |
| ENSG00000072422 | 350 | 157 | 174 | 177 | 235 | 209 |
| ENSG00000223107 | 0 | 0 | 0 | 0 | 0 | 0 |
| ENSG00000227244 | 0 | 0 | 0 | 0 | 1 | 0 |
| ENSG00000196932 | 53 | 31 | 39 | 22 | 37 | 36 |
| ENSG00000237233 | 1 | 0 | 0 | 0 | 1 | 0 |
| ENSG00000183346 | 7 | 4 | 2 | 1 | 4 | 1 |
| ENSG00000272592 | 0 | 0 | 0 | 0 | 0 | 0 |
| ENSG00000233643 | 0 | 0 | 1 | 0 | 1 | 1 |
| ENSG00000150347 | 842 | 496 | 461 | 215 | 507 | 364 |
| ENSG00000182010 | 116 | 82 | 68 | 29 | 93 | 67 |
| ENSG00000234756 | 0 | 0 | 0 | 0 | 0 | 0 |
| ENSG00000240940 | 0 | 0 | 0 | 0 | 0 | 0 |
| ENSG00000138311 | 72 | 28 | 44 | 32 | 52 | 52 |
| ENSG00000285837 | 0 | 0 | 0 | 0 | 0 | 0 |
| ENSG00000285551 | 0 | 0 | 0 | 0 | 0 | 0 |
| ENSG00000238280 | 3 | 0 | 2 | 1 | 0 | 2 |
| ENSG00000234489 | 0 | 0 | 0 | 0 | 0 | 0 |
| ENSG00000181915 | 1244 | 636 | 732 | 753 | 776 | 793 |
| ENSG00000122877 | 2 | 2 | 8 | 2 | 1 | 3 |
| ENSG00000199446 | 0 | 0 | 0 | 0 | 0 | 0 |
| ENSG00000224301 | 0 | 0 | 0 | 0 | 0 | 0 |
| ENSG00000148572 | 689 | 345 | 410 | 309 | 421 | 383 |
| ENSG00000171988 | 276 | 212 | 121 | 47 | 394 | 163 |
| ENSG00000234369 | 0 | 0 | 0 | 0 | 0 | 2 |
| ENSG00000270421 | 0 | 0 | 0 | 0 | 0 | 0 |
| ENSG00000221063 | 0 | 0 | 0 | 0 | 0 | 0 |
| ENSG00000235816 | 0 | 0 | 0 | 0 | 0 | 0 |
| ENSG00000272767 | 34 | 21 | 23 | 14 | 17 | 14 |
| ENSG00000224412 | 0 | 0 | 0 | 0 | 0 | 0 |
| ENSG00000165476 | 2108 | 974 | 1140 | 887 | 1127 | 1062 |
| ENSG00000232075 | 4 | 1 | 4 | 0 | 1 | 1 |
| ENSG00000270352 | 0 | 0 | 0 | 0 | 0 | 0 |
| ENSG00000228566 | 2 | 0 | 0 | 0 | 1 | 0 |
| ENSG00000213609 | 6 | 5 | 5 | 3 | 9 | 2 |
| ENSG00000226426 | 0 | 0 | 0 | 0 | 0 | 0 |
| ENSG00000235489 | 4 | 1 | 1 | 1 | 1 | 1 |
| ENSG00000233900 | 0 | 0 | 0 | 0 | 0 | 0 |
| ENSG00000270648 | 0 | 0 | 0 | 0 | 0 | 0 |
| ENSG00000216740 | 0 | 2 | 0 | 0 | 0 | 0 |
| ENSG00000224714 | 0 | 0 | 1 | 0 | 0 | 0 |
| ENSG00000270754 | 0 | 0 | 0 | 0 | 0 | 0 |
| ENSG00000282906 | 0 | 2 | 4 | 2 | 5 | 3 |
| ENSG00000227173 | 0 | 0 | 0 | 0 | 1 | 1 |
| ENSG00000233724 | 0 | 0 | 0 | 0 | 0 | 0 |
| ENSG00000235356 | 0 | 0 | 0 | 0 | 0 | 0 |
| ENSG00000228065 | 60 | 34 | 50 | 36 | 81 | 69 |
| ENSG00000183230 | 54 | 37 | 28 | 15 | 20 | 14 |

| | | | | | | |
|---|---|---|---|---|---|---|
| ENSG00000273360 | 0 | 0 | 0 | 0 | 0 | 0 |
| ENSG00000277056 | 0 | 0 | 0 | 0 | 0 | 0 |
| ENSG00000198739 | 2 | 1 | 0 | 2 | 3 | 3 |
| ENSG00000225299 | 0 | 0 | 0 | 0 | 0 | 0 |
| ENSG00000226448 | 0 | 0 | 0 | 0 | 0 | 0 |
| ENSG00000276866 | 0 | 0 | 0 | 0 | 0 | 0 |
| ENSG00000213606 | 0 | 0 | 0 | 0 | 0 | 0 |
| ENSG00000225922 | 2 | 0 | 0 | 0 | 0 | 0 |
| ENSG00000108176 | 253 | 114 | 176 | 115 | 151 | 136 |
| ENSG00000212520 | 0 | 0 | 0 | 0 | 0 | 0 |
| ENSG00000239942 | 0 | 0 | 0 | 1 | 0 | 0 |
| ENSG00000237141 | 0 | 0 | 0 | 0 | 0 | 0 |
| ENSG00000272892 | 17 | 7 | 7 | 6 | 10 | 7 |
| ENSG00000252113 | 0 | 0 | 0 | 0 | 0 | 0 |
| ENSG00000219932 | 14 | 5 | 2 | 5 | 5 | 10 |
| ENSG00000096717 | 975 | 446 | 530 | 322 | 475 | 457 |
| ENSG00000148634 | 1727 | 884 | 1042 | 587 | 1143 | 924 |
| ENSG00000226318 | 2 | 3 | 1 | 0 | 1 | 1 |
| ENSG00000266467 | 0 | 0 | 0 | 0 | 0 | 0 |
| ENSG00000236375 | 0 | 1 | 0 | 0 | 0 | 0 |
| ENSG00000138347 | 4 | 5 | 3 | 0 | 3 | 2 |
| ENSG00000222371 | 0 | 0 | 0 | 0 | 0 | 0 |
| ENSG00000179774 | 8 | 1 | 3 | 3 | 3 | 6 |
| ENSG00000233590 | 0 | 2 | 1 | 2 | 1 | 0 |
| ENSG00000234102 | 0 | 0 | 0 | 0 | 0 | 0 |
| ENSG00000108187 | 677 | 291 | 372 | 187 | 357 | 308 |
| ENSG00000096746 | 3379 | 1662 | 2101 | 1598 | 2357 | 2461 |
| ENSG00000204130 | 745 | 331 | 398 | 221 | 510 | 376 |
| ENSG00000138346 | 411 | 220 | 252 | 130 | 293 | 236 |
| ENSG00000234806 | 0 | 0 | 0 | 0 | 0 | 0 |
| ENSG00000199638 | 0 | 0 | 0 | 0 | 0 | 0 |
| ENSG00000122912 | 1000 | 457 | 632 | 383 | 492 | 474 |
| ENSG00000226493 | 1 | 1 | 2 | 1 | 3 | 1 |
| ENSG00000207438 | 0 | 0 | 0 | 1 | 0 | 0 |
| ENSG00000214881 | 2 | 0 | 1 | 0 | 1 | 2 |
| ENSG00000138336 | 54 | 29 | 19 | 14 | 38 | 28 |
| ENSG00000213025 | 95 | 58 | 60 | 79 | 80 | 65 |
| ENSG00000227583 | 0 | 0 | 0 | 0 | 0 | 0 |
| ENSG00000260400 | 8 | 13 | 7 | 7 | 5 | 5 |
| ENSG00000270494 | 0 | 0 | 0 | 0 | 0 | 0 |
| ENSG00000060339 | 1512 | 892 | 822 | 414 | 1365 | 778 |
| ENSG00000195401 | 0 | 0 | 0 | 1 | 0 | 0 |
| ENSG00000283551 | 1 | 0 | 0 | 0 | 0 | 0 |
| ENSG00000221184 | 7 | 14 | 17 | 4 | 9 | 10 |
| ENSG00000165730 | 129 | 82 | 66 | 39 | 76 | 67 |
| ENSG00000200218 | 0 | 0 | 0 | 0 | 0 | 0 |
| ENSG00000285871 | 0 | 2 | 3 | 1 | 2 | 1 |
| ENSG00000107625 | 1429 | 787 | 850 | 653 | 937 | 812 |
| ENSG00000206855 | 0 | 0 | 0 | 0 | 0 | 0 |
| ENSG00000165732 | 4293 | 2381 | 2486 | 1929 | 3518 | 3088 |

| | | | | | | |
|---|---|---|---|---|---|---|
| ENSG00000198954 | 2091 | 995 | 1280 | 923 | 1159 | 1031 |
| ENSG00000214875 | 0 | 2 | 1 | 2 | 1 | 0 |
| ENSG00000229001 | 0 | 0 | 0 | 0 | 1 | 0 |
| ENSG00000122862 | 0 | 0 | 0 | 0 | 0 | 0 |
| ENSG00000274008 | 0 | 0 | 0 | 0 | 0 | 0 |
| ENSG00000122958 | 2240 | 1135 | 1240 | 808 | 1502 | 1136 |
| ENSG00000233163 | 2 | 0 | 0 | 0 | 1 | 1 |
| ENSG00000156502 | 1127 | 458 | 622 | 503 | 609 | 655 |
| ENSG00000229261 | 0 | 1 | 0 | 0 | 0 | 0 |
| ENSG00000156510 | 0 | 1 | 0 | 0 | 0 | 0 |
| ENSG00000231748 | 0 | 0 | 0 | 0 | 0 | 0 |
| ENSG00000156515 | 14190 | 6571 | 8491 | 6304 | 6921 | 6872 |
| ENSG00000229197 | 0 | 0 | 0 | 0 | 0 | 0 |
| ENSG00000075073 | 23 | 10 | 13 | 9 | 13 | 15 |
| ENSG00000232734 | 0 | 0 | 0 | 0 | 0 | 0 |
| ENSG00000099282 | 10338 | 5061 | 6357 | 5557 | 5147 | 5633 |
| ENSG00000233197 | 0 | 0 | 2 | 0 | 0 | 0 |
| ENSG00000122859 | 0 | 0 | 0 | 3 | 0 | 1 |
| ENSG00000236154 | 0 | 0 | 0 | 0 | 0 | 0 |
| ENSG00000230755 | 0 | 0 | 0 | 0 | 0 | 0 |
| ENSG00000235803 | 0 | 0 | 0 | 0 | 0 | 0 |
| ENSG00000235524 | 0 | 0 | 0 | 0 | 0 | 0 |
| ENSG00000231152 | 0 | 0 | 0 | 0 | 0 | 0 |
| ENSG00000226794 | 0 | 0 | 0 | 0 | 0 | 0 |
| ENSG00000171224 | 617 | 253 | 343 | 369 | 313 | 324 |
| ENSG00000235645 | 0 | 0 | 0 | 0 | 0 | 0 |
| ENSG00000230469 | 0 | 0 | 0 | 0 | 0 | 0 |
| ENSG00000197467 | 1 | 4 | 1 | 0 | 3 | 0 |
| ENSG00000224222 | 0 | 0 | 0 | 0 | 0 | 0 |
| ENSG00000099284 | 1260 | 584 | 758 | 727 | 695 | 698 |
| ENSG00000042286 | 1370 | 693 | 893 | 834 | 951 | 954 |
| ENSG00000156521 | 2364 | 1096 | 1412 | 1631 | 1310 | 1398 |
| ENSG00000079332 | 3962 | 1877 | 2382 | 1892 | 2264 | 2089 |
| ENSG00000229097 | 18 | 3 | 2 | 5 | 8 | 9 |
| ENSG00000232142 | 0 | 0 | 0 | 0 | 0 | 0 |
| ENSG00000180817 | 6199 | 3254 | 3709 | 3268 | 4177 | 3840 |
| ENSG00000148734 | 0 | 1 | 2 | 0 | 1 | 0 |
| ENSG00000172731 | 878 | 363 | 490 | 495 | 387 | 497 |
| ENSG00000224725 | 0 | 1 | 0 | 0 | 0 | 0 |
| ENSG00000148730 | 5642 | 2563 | 3288 | 2277 | 2709 | 3017 |
| ENSG00000156574 | 0 | 3 | 2 | 0 | 1 | 4 |
| ENSG00000280401 | 1 | 0 | 0 | 3 | 0 | 1 |
| ENSG00000107719 | 12 | 8 | 7 | 9 | 2 | 5 |
| ENSG00000230661 | 0 | 0 | 0 | 0 | 0 | 0 |
| ENSG00000180644 | 0 | 0 | 0 | 0 | 0 | 0 |
| ENSG00000138316 | 8 | 3 | 2 | 1 | 3 | 6 |
| ENSG00000166220 | 0 | 0 | 1 | 0 | 0 | 1 |
| ENSG00000231366 | 0 | 0 | 0 | 0 | 0 | 0 |
| ENSG00000166224 | 4369 | 2101 | 2728 | 1807 | 2341 | 2341 |
| ENSG00000166228 | 2573 | 1394 | 1597 | 1684 | 1710 | 1550 |

| | | | | | | |
|---|---|---|---|---|---|---|
| ENSG00000285300 | 0 | 0 | 0 | 0 | 0 | 0 |
| ENSG00000271409 | 0 | 0 | 0 | 0 | 0 | 0 |
| ENSG00000259267 | 0 | 0 | 0 | 0 | 0 | 0 |
| ENSG00000107731 | 403 | 177 | 224 | 171 | 204 | 224 |
| ENSG00000237512 | 2 | 0 | 0 | 0 | 0 | 0 |
| ENSG00000198246 | 3012 | 1391 | 1541 | 1381 | 1306 | 1416 |
| ENSG00000279406 | 85 | 47 | 52 | 49 | 45 | 64 |
| ENSG00000107736 | 5 | 5 | 3 | 0 | 0 | 2 |
| ENSG00000223817 | 0 | 0 | 0 | 0 | 0 | 0 |
| ENSG00000214688 | 0 | 0 | 0 | 0 | 0 | 0 |
| ENSG00000107738 | 38 | 24 | 14 | 17 | 14 | 20 |
| ENSG00000274824 | 0 | 0 | 0 | 0 | 0 | 0 |
| ENSG00000197746 | 34864 | 16293 | 21126 | 18734 | 16774 | 18097 |
| ENSG00000238446 | 0 | 0 | 0 | 0 | 0 | 0 |
| ENSG00000230526 | 2 | 1 | 0 | 1 | 1 | 0 |
| ENSG00000122863 | 128 | 35 | 56 | 46 | 36 | 32 |
| ENSG00000272988 | 0 | 0 | 0 | 0 | 0 | 0 |
| ENSG00000107742 | 76 | 44 | 47 | 29 | 35 | 37 |
| ENSG00000138303 | 1441 | 762 | 998 | 754 | 972 | 922 |
| ENSG00000200294 | 0 | 0 | 0 | 0 | 0 | 0 |
| ENSG00000226701 | 11 | 0 | 0 | 4 | 2 | 1 |
| ENSG00000166295 | 4710 | 2333 | 2789 | 2534 | 2756 | 2674 |
| ENSG00000200170 | 3 | 2 | 6 | 5 | 4 | 6 |
| ENSG00000168209 | 2478 | 1374 | 1507 | 1995 | 1993 | 1857 |
| ENSG00000269926 | 0 | 0 | 0 | 0 | 0 | 0 |
| ENSG00000148719 | 2612 | 1074 | 1573 | 1334 | 1227 | 1356 |
| ENSG00000107745 | 1946 | 916 | 1219 | 972 | 1078 | 1087 |
| ENSG00000202513 | 0 | 0 | 1 | 0 | 1 | 0 |
| ENSG00000282915 | 0 | 0 | 0 | 0 | 0 | 0 |
| ENSG00000276531 | 3 | 4 | 2 | 0 | 0 | 1 |
| ENSG00000226163 | 0 | 0 | 0 | 0 | 0 | 0 |
| ENSG00000230609 | 0 | 0 | 0 | 0 | 0 | 0 |
| ENSG00000272536 | 0 | 0 | 0 | 0 | 0 | 0 |
| ENSG00000231471 | 0 | 0 | 0 | 0 | 0 | 0 |
| ENSG00000238215 | 0 | 0 | 1 | 0 | 0 | 0 |
| ENSG00000201881 | 0 | 0 | 0 | 0 | 0 | 0 |
| ENSG00000156026 | 1514 | 732 | 883 | 663 | 747 | 803 |
| ENSG00000266719 | 0 | 0 | 0 | 0 | 0 | 0 |
| ENSG00000280166 | 0 | 0 | 0 | 0 | 0 | 0 |
| ENSG00000279502 | 0 | 0 | 0 | 0 | 0 | 0 |
| ENSG00000272627 | 2 | 1 | 0 | 0 | 1 | 0 |
| ENSG00000138315 | 2 | 0 | 0 | 0 | 1 | 0 |
| ENSG00000215086 | 5 | 1 | 4 | 1 | 4 | 4 |
| ENSG00000138308 | 1 | 1 | 0 | 0 | 0 | 0 |
| ENSG00000213700 | 0 | 2 | 2 | 1 | 0 | 3 |
| ENSG00000122884 | 1264 | 674 | 803 | 551 | 724 | 672 |
| ENSG00000232646 | 0 | 0 | 0 | 0 | 0 | 0 |
| ENSG00000237768 | 0 | 0 | 0 | 0 | 0 | 0 |
| ENSG00000201047 | 0 | 0 | 0 | 0 | 0 | 0 |
| ENSG00000272630 | 10 | 6 | 3 | 0 | 5 | 7 |

| | | | | | | |
|---|---|---|---|---|---|---|
| ENSG00000166321 | 211 | 106 | 128 | 89 | 137 | 123 |
| ENSG00000272599 | 33 | 16 | 24 | 19 | 19 | 19 |
| ENSG00000221164 | 16 | 6 | 5 | 6 | 13 | 6 |
| ENSG00000122882 | 1404 | 779 | 903 | 591 | 955 | 822 |
| ENSG00000138286 | 984 | 522 | 697 | 479 | 596 | 546 |
| ENSG00000213551 | 697 | 356 | 527 | 422 | 485 | 469 |
| ENSG00000199201 | 0 | 0 | 0 | 0 | 0 | 0 |
| ENSG00000227382 | 7 | 1 | 2 | 3 | 2 | 1 |
| ENSG00000236756 | 0 | 0 | 0 | 0 | 0 | 1 |
| ENSG00000182180 | 3659 | 1922 | 2251 | 2295 | 2314 | 2454 |
| ENSG00000227540 | 56 | 43 | 48 | 40 | 54 | 48 |
| ENSG00000156042 | 41 | 28 | 36 | 17 | 17 | 31 |
| ENSG00000200356 | 0 | 0 | 2 | 0 | 1 | 0 |
| ENSG00000207473 | 0 | 0 | 1 | 0 | 0 | 0 |
| ENSG00000138279 | 5765 | 2958 | 3867 | 3061 | 3618 | 3512 |
| ENSG00000233144 | 0 | 0 | 3 | 0 | 1 | 0 |
| ENSG00000229659 | 0 | 2 | 1 | 1 | 0 | 1 |
| ENSG00000166343 | 35 | 8 | 21 | 11 | 19 | 21 |
| ENSG00000107758 | 2050 | 971 | 1310 | 1046 | 1161 | 1172 |
| ENSG00000221817 | 130 | 76 | 109 | 56 | 90 | 87 |
| ENSG00000166348 | 802 | 387 | 519 | 316 | 474 | 434 |
| ENSG00000207327 | 0 | 1 | 0 | 1 | 0 | 0 |
| ENSG00000237139 | 0 | 0 | 1 | 2 | 0 | 1 |
| ENSG00000268584 | 41 | 16 | 19 | 9 | 30 | 17 |
| ENSG00000272791 | 29 | 18 | 19 | 11 | 18 | 33 |
| ENSG00000177791 | 19 | 6 | 3 | 5 | 3 | 3 |
| ENSG00000166317 | 0 | 0 | 0 | 2 | 0 | 0 |
| ENSG00000271848 | 0 | 0 | 0 | 0 | 0 | 0 |
| ENSG00000172650 | 16 | 5 | 5 | 0 | 2 | 5 |
| ENSG00000242288 | 0 | 0 | 0 | 0 | 1 | 0 |
| ENSG00000271816 | 21 | 8 | 12 | 5 | 16 | 9 |
| ENSG00000272140 | 44 | 12 | 21 | 10 | 11 | 6 |
| ENSG00000252072 | 0 | 1 | 0 | 0 | 0 | 1 |
| ENSG00000242338 | 10 | 8 | 14 | 5 | 3 | 11 |
| ENSG00000250959 | 68 | 37 | 64 | 32 | 55 | 56 |
| ENSG00000235316 | 73 | 34 | 57 | 28 | 30 | 41 |
| ENSG00000270808 | 8 | 1 | 0 | 0 | 0 | 0 |
| ENSG00000279088 | 361 | 194 | 272 | 242 | 238 | 242 |
| ENSG00000176986 | 3885 | 1615 | 2339 | 1492 | 1886 | 1971 |
| ENSG00000279689 | 23 | 22 | 22 | 14 | 9 | 13 |
| ENSG00000196968 | 553 | 320 | 341 | 281 | 305 | 312 |
| ENSG00000172586 | 1814 | 930 | 1032 | 1137 | 1121 | 1023 |
| ENSG00000214655 | 2622 | 1216 | 1758 | 1173 | 1379 | 1421 |
| ENSG00000280238 | 1 | 0 | 0 | 0 | 0 | 0 |
| ENSG00000272589 | 0 | 0 | 0 | 0 | 0 | 1 |
| ENSG00000272916 | 0 | 0 | 2 | 1 | 2 | 3 |
| ENSG00000166507 | 21 | 13 | 19 | 7 | 17 | 25 |
| ENSG00000148660 | 2204 | 1072 | 1338 | 1086 | 1248 | 1184 |
| ENSG00000224195 | 11 | 7 | 5 | 3 | 7 | 2 |
| ENSG00000229990 | 0 | 0 | 0 | 1 | 1 | 1 |

| | | | | | | |
|---|---|---|---|---|---|---|
| ENSG00000122861 | 23 | 19 | 16 | 23 | 29 | 20 |
| ENSG00000222047 | 1 | 1 | 0 | 3 | 1 | 4 |
| ENSG00000035403 | 2491 | 1232 | 1531 | 974 | 1424 | 1293 |
| ENSG00000225761 | 0 | 0 | 0 | 0 | 0 | 0 |
| ENSG00000185009 | 2201 | 1225 | 1208 | 763 | 1469 | 1131 |
| ENSG00000156110 | 2744 | 1493 | 1705 | 1525 | 1815 | 1690 |
| ENSG00000236900 | 0 | 0 | 0 | 0 | 0 | 0 |
| ENSG00000214629 | 0 | 0 | 0 | 0 | 0 | 0 |
| ENSG00000213731 | 0 | 0 | 0 | 0 | 0 | 0 |
| ENSG00000232342 | 0 | 0 | 0 | 0 | 0 | 0 |
| ENSG00000226253 | 0 | 0 | 0 | 0 | 0 | 0 |
| ENSG00000206756 | 0 | 0 | 0 | 0 | 0 | 0 |
| ENSG00000271710 | 0 | 0 | 0 | 1 | 0 | 0 |
| ENSG00000214626 | 0 | 0 | 1 | 0 | 1 | 0 |
| ENSG00000213525 | 0 | 0 | 2 | 0 | 0 | 0 |
| ENSG00000272748 | 0 | 0 | 0 | 0 | 0 | 1 |
| ENSG00000156650 | 1343 | 640 | 689 | 265 | 616 | 467 |
| ENSG00000234149 | 0 | 0 | 0 | 0 | 0 | 0 |
| ENSG00000227186 | 0 | 0 | 0 | 0 | 0 | 0 |
| ENSG00000188716 | 0 | 0 | 2 | 0 | 0 | 0 |
| ENSG00000285810 | 0 | 0 | 0 | 1 | 1 | 0 |
| ENSG00000237882 | 0 | 0 | 0 | 0 | 0 | 1 |
| ENSG00000079393 | 53 | 20 | 34 | 34 | 40 | 38 |
| ENSG00000156671 | 777 | 350 | 498 | 189 | 379 | 332 |
| ENSG00000232437 | 0 | 0 | 0 | 0 | 0 | 0 |
| ENSG00000165637 | 4571 | 2295 | 2799 | 2615 | 2873 | 2874 |
| ENSG00000165644 | 1058 | 444 | 620 | 526 | 490 | 487 |
| ENSG00000227271 | 0 | 0 | 0 | 0 | 0 | 0 |
| ENSG00000226051 | 13 | 9 | 11 | 6 | 7 | 10 |
| ENSG00000233313 | 0 | 0 | 0 | 0 | 0 | 0 |
| ENSG00000270087 | 0 | 0 | 0 | 0 | 0 | 0 |
| ENSG00000223428 | 0 | 0 | 0 | 0 | 0 | 0 |
| ENSG00000236797 | 4 | 0 | 0 | 0 | 2 | 0 |
| ENSG00000165655 | 245 | 88 | 144 | 145 | 146 | 147 |
| ENSG00000237149 | 49 | 30 | 37 | 26 | 31 | 25 |
| ENSG00000273248 | 1 | 0 | 2 | 1 | 2 | 1 |
| ENSG00000272692 | 17 | 11 | 19 | 13 | 8 | 5 |
| ENSG00000236842 | 0 | 0 | 0 | 0 | 0 | 0 |
| ENSG00000148655 | 17 | 4 | 11 | 10 | 11 | 16 |
| ENSG00000279814 | 0 | 0 | 0 | 0 | 0 | 0 |
| ENSG00000207583 | 0 | 0 | 0 | 0 | 0 | 0 |
| ENSG00000268659 | 0 | 0 | 0 | 0 | 0 | 0 |
| ENSG00000228280 | 0 | 0 | 0 | 0 | 0 | 0 |
| ENSG00000265486 | 0 | 0 | 0 | 0 | 0 | 0 |
| ENSG00000201954 | 0 | 0 | 0 | 0 | 0 | 0 |
| ENSG00000278992 | 0 | 0 | 0 | 0 | 0 | 0 |
| ENSG00000230575 | 0 | 0 | 0 | 0 | 0 | 0 |
| ENSG00000269256 | 0 | 0 | 0 | 0 | 0 | 0 |
| ENSG00000252888 | 0 | 0 | 0 | 0 | 0 | 0 |
| ENSG00000224948 | 0 | 0 | 0 | 0 | 0 | 0 |

| | | | | | | |
|---|---|---|---|---|---|---|
| ENSG00000156113 | 66 | 26 | 40 | 14 | 23 | 34 |
| ENSG00000236467 | 0 | 0 | 0 | 0 | 0 | 0 |
| ENSG00000225497 | 0 | 0 | 0 | 0 | 0 | 0 |
| ENSG00000224500 | 0 | 0 | 0 | 0 | 0 | 0 |
| ENSG00000225652 | 2 | 0 | 1 | 0 | 0 | 0 |
| ENSG00000228092 | 0 | 0 | 0 | 0 | 0 | 0 |
| ENSG00000199592 | 0 | 0 | 0 | 0 | 0 | 0 |
| ENSG00000213514 | 4 | 4 | 5 | 2 | 2 | 5 |
| ENSG00000277981 | 0 | 1 | 0 | 0 | 0 | 0 |
| ENSG00000199664 | 1 | 3 | 1 | 0 | 2 | 2 |
| ENSG00000213513 | 31 | 16 | 27 | 15 | 12 | 23 |
| ENSG00000228748 | 18 | 10 | 19 | 14 | 14 | 25 |
| ENSG00000151208 | 4548 | 2004 | 3126 | 2129 | 2774 | 2887 |
| ENSG00000204049 | 0 | 1 | 2 | 1 | 4 | 6 |
| ENSG00000243446 | 0 | 0 | 0 | 0 | 0 | 0 |
| ENSG00000233871 | 39 | 21 | 31 | 26 | 23 | 33 |
| ENSG00000234612 | 0 | 0 | 0 | 0 | 0 | 0 |
| ENSG00000274686 | 0 | 0 | 0 | 1 | 1 | 0 |
| ENSG00000148606 | 1630 | 739 | 1004 | 803 | 896 | 943 |
| ENSG00000138326 | 28212 | 14624 | 17557 | 17776 | 18081 | 16831 |
| ENSG00000231957 | 0 | 0 | 0 | 0 | 0 | 0 |
| ENSG00000283096 | 0 | 0 | 0 | 0 | 0 | 0 |
| ENSG00000282863 | 0 | 0 | 0 | 0 | 0 | 0 |
| ENSG00000230417 | 4 | 7 | 7 | 5 | 3 | 7 |
| ENSG00000229543 | 0 | 0 | 0 | 0 | 0 | 2 |
| ENSG00000230229 | 0 | 0 | 0 | 0 | 0 | 0 |
| ENSG00000201393 | 0 | 0 | 0 | 0 | 0 | 0 |
| ENSG00000282952 | 0 | 0 | 0 | 0 | 0 | 0 |
| ENSG00000228683 | 0 | 0 | 0 | 0 | 0 | 0 |
| ENSG00000224596 | 20 | 9 | 8 | 4 | 9 | 9 |
| ENSG00000229569 | 0 | 0 | 0 | 0 | 0 | 0 |
| ENSG00000108175 | 8461 | 3596 | 5149 | 3030 | 3486 | 4309 |
| ENSG00000108179 | 2162 | 1094 | 1288 | 1504 | 1367 | 1360 |
| ENSG00000165424 | 257 | 133 | 162 | 127 | 142 | 128 |
| ENSG00000235426 | 0 | 0 | 1 | 0 | 0 | 0 |
| ENSG00000259021 | 0 | 0 | 0 | 0 | 0 | 0 |
| ENSG00000241317 | 0 | 0 | 0 | 0 | 0 | 0 |
| ENSG00000235705 | 0 | 0 | 0 | 0 | 0 | 0 |
| ENSG00000253626 | 60 | 22 | 22 | 37 | 36 | 34 |
| ENSG00000271985 | 0 | 0 | 0 | 0 | 0 | 0 |
| ENSG00000232965 | 0 | 0 | 0 | 0 | 0 | 0 |
| ENSG00000185303 | 10 | 9 | 2 | 1 | 10 | 1 |
| ENSG00000219430 | 0 | 0 | 0 | 0 | 0 | 0 |
| ENSG00000225827 | 0 | 0 | 0 | 0 | 0 | 0 |
| ENSG00000122852 | 0 | 0 | 0 | 0 | 0 | 0 |
| ENSG00000226676 | 0 | 0 | 0 | 0 | 0 | 0 |
| ENSG00000244733 | 16 | 6 | 4 | 3 | 9 | 4 |
| ENSG00000225484 | 539 | 286 | 312 | 281 | 341 | 376 |
| ENSG00000272489 | 0 | 0 | 0 | 0 | 0 | 0 |
| ENSG00000280355 | 25 | 12 | 14 | 11 | 5 | 10 |

| | | | | | | |
|---|---|---|---|---|---|---|
| ENSG00000278616 | 12 | 13 | 21 | 5 | 5 | 10 |
| ENSG00000224886 | 0 | 0 | 0 | 0 | 0 | 0 |
| ENSG00000188199 | 62 | 16 | 29 | 34 | 30 | 32 |
| ENSG00000272447 | 138 | 61 | 58 | 53 | 50 | 47 |
| ENSG00000228570 | 0 | 0 | 0 | 0 | 0 | 0 |
| ENSG00000214695 | 0 | 0 | 0 | 0 | 0 | 0 |
| ENSG00000204437 | 0 | 0 | 0 | 0 | 0 | 0 |
| ENSG00000214584 | 0 | 0 | 0 | 0 | 0 | 0 |
| ENSG00000283913 | 0 | 0 | 0 | 0 | 2 | 0 |
| ENSG00000283101 | 0 | 0 | 0 | 0 | 0 | 0 |
| ENSG00000242600 | 0 | 0 | 0 | 0 | 0 | 0 |
| ENSG00000133661 | 35 | 25 | 55 | 29 | 20 | 34 |
| ENSG00000273372 | 0 | 0 | 0 | 0 | 0 | 0 |
| ENSG00000280259 | 0 | 0 | 0 | 0 | 0 | 0 |
| ENSG00000235924 | 0 | 0 | 1 | 0 | 2 | 0 |
| ENSG00000225100 | 0 | 0 | 0 | 0 | 0 | 0 |
| ENSG00000229616 | 0 | 0 | 0 | 0 | 2 | 0 |
| ENSG00000224760 | 0 | 0 | 0 | 0 | 0 | 0 |
| ENSG00000236217 | 0 | 0 | 0 | 0 | 0 | 0 |
| ENSG00000227515 | 2 | 0 | 0 | 0 | 0 | 0 |
| ENSG00000230091 | 289 | 115 | 192 | 124 | 165 | 168 |
| ENSG00000282432 | 0 | 0 | 0 | 0 | 0 | 0 |
| ENSG00000133678 | 1263 | 715 | 929 | 724 | 707 | 806 |
| ENSG00000279399 | 0 | 0 | 0 | 0 | 0 | 0 |
| ENSG00000230171 | 0 | 0 | 0 | 0 | 0 | 0 |
| ENSG00000189129 | 12 | 6 | 3 | 1 | 4 | 3 |
| ENSG00000122359 | 5345 | 2674 | 3187 | 3234 | 2888 | 2882 |
| ENSG00000237523 | 7 | 6 | 6 | 9 | 4 | 6 |
| ENSG00000234319 | 0 | 1 | 0 | 0 | 0 | 0 |
| ENSG00000234382 | 0 | 0 | 0 | 0 | 0 | 0 |
| ENSG00000234743 | 0 | 0 | 0 | 1 | 1 | 1 |
| ENSG00000279359 | 0 | 0 | 0 | 0 | 0 | 0 |
| ENSG00000151224 | 4 | 1 | 1 | 2 | 0 | 0 |
| ENSG00000232950 | 0 | 0 | 0 | 0 | 0 | 0 |
| ENSG00000235858 | 0 | 0 | 0 | 0 | 0 | 0 |
| ENSG00000170788 | 0 | 0 | 0 | 0 | 0 | 0 |
| ENSG00000133665 | 0 | 0 | 0 | 0 | 0 | 0 |
| ENSG00000122378 | 5326 | 2703 | 3208 | 2339 | 3078 | 3116 |
| ENSG00000108219 | 4037 | 1924 | 2550 | 1890 | 2222 | 2200 |
| ENSG00000226659 | 1 | 7 | 5 | 2 | 5 | 2 |
| ENSG00000178217 | 9 | 2 | 2 | 0 | 0 | 1 |
| ENSG00000231082 | 0 | 0 | 0 | 0 | 0 | 0 |
| ENSG00000233353 | 0 | 0 | 0 | 0 | 0 | 0 |
| ENSG00000233703 | 0 | 0 | 0 | 0 | 0 | 0 |
| ENSG00000231060 | 0 | 0 | 0 | 0 | 0 | 0 |
| ENSG00000227209 | 4 | 5 | 1 | 0 | 1 | 2 |
| ENSG00000226466 | 0 | 0 | 0 | 0 | 0 | 0 |
| ENSG00000185737 | 80 | 38 | 36 | 22 | 30 | 24 |
| ENSG00000285739 | 0 | 0 | 0 | 0 | 0 | 0 |
| ENSG00000229458 | 0 | 0 | 0 | 0 | 0 | 0 |

| | | | | | | |
|---|---|---|---|---|---|---|
| ENSG00000225738 | 0 | 0 | 0 | 0 | 0 | 0 |
| ENSG00000207275 | 0 | 0 | 0 | 0 | 0 | 0 |
| ENSG00000200774 | 0 | 0 | 0 | 0 | 0 | 0 |
| ENSG00000234931 | 0 | 0 | 0 | 0 | 0 | 0 |
| ENSG00000233945 | 0 | 0 | 0 | 0 | 0 | 0 |
| ENSG00000223565 | 0 | 0 | 0 | 0 | 0 | 0 |
| ENSG00000212324 | 0 | 0 | 0 | 0 | 0 | 0 |
| ENSG00000233258 | 0 | 0 | 0 | 0 | 0 | 0 |
| ENSG00000200789 | 0 | 0 | 0 | 0 | 0 | 0 |
| ENSG00000212769 | 6 | 1 | 0 | 0 | 0 | 2 |
| ENSG00000271933 | 17 | 14 | 9 | 10 | 11 | 16 |
| ENSG00000165678 | 11413 | 6140 | 7217 | 6695 | 7247 | 7039 |
| ENSG00000285972 | 0 | 0 | 0 | 0 | 0 | 0 |
| ENSG00000188373 | 0 | 0 | 0 | 0 | 0 | 0 |
| ENSG00000148600 | 0 | 0 | 0 | 0 | 0 | 0 |
| ENSG00000204033 | 0 | 0 | 0 | 0 | 0 | 0 |
| ENSG00000148602 | 0 | 0 | 0 | 1 | 0 | 2 |
| ENSG00000148604 | 0 | 0 | 0 | 0 | 0 | 0 |
| ENSG00000229404 | 100 | 70 | 78 | 42 | 95 | 65 |
| ENSG00000107771 | 614 | 332 | 303 | 179 | 570 | 331 |
| ENSG00000233189 | 2 | 0 | 0 | 0 | 0 | 0 |
| ENSG00000225423 | 0 | 1 | 0 | 0 | 1 | 0 |
| ENSG00000225928 | 0 | 0 | 0 | 0 | 0 | 1 |
| ENSG00000178429 | 8 | 6 | 2 | 5 | 6 | 3 |
| ENSG00000237267 | 0 | 0 | 0 | 0 | 0 | 0 |
| ENSG00000223993 | 0 | 0 | 0 | 0 | 0 | 0 |
| ENSG00000224504 | 0 | 0 | 0 | 0 | 0 | 0 |
| ENSG00000230962 | 0 | 0 | 0 | 0 | 0 | 0 |
| ENSG00000279910 | 0 | 0 | 0 | 0 | 0 | 0 |
| ENSG00000223064 | 0 | 0 | 0 | 0 | 0 | 0 |
| ENSG00000280134 | 0 | 0 | 0 | 0 | 0 | 0 |
| ENSG00000234942 | 0 | 0 | 0 | 0 | 0 | 0 |
| ENSG00000182771 | 15 | 3 | 7 | 3 | 4 | 1 |
| ENSG00000278333 | 0 | 0 | 0 | 0 | 0 | 0 |
| ENSG00000270002 | 0 | 0 | 0 | 0 | 0 | 0 |
| ENSG00000252292 | 0 | 0 | 0 | 0 | 0 | 0 |
| ENSG00000236606 | 0 | 0 | 0 | 0 | 0 | 0 |
| ENSG00000200487 | 0 | 0 | 0 | 0 | 0 | 0 |
| ENSG00000199104 | 0 | 0 | 0 | 0 | 0 | 0 |
| ENSG00000062650 | 2262 | 1086 | 1307 | 741 | 1340 | 1142 |
| ENSG00000212332 | 0 | 0 | 2 | 0 | 0 | 0 |
| ENSG00000227896 | 124 | 53 | 40 | 33 | 39 | 31 |
| ENSG00000230507 | 0 | 0 | 0 | 0 | 0 | 0 |
| ENSG00000122375 | 0 | 0 | 0 | 0 | 0 | 0 |
| ENSG00000122367 | 49 | 39 | 33 | 21 | 31 | 34 |
| ENSG00000272631 | 112 | 43 | 77 | 42 | 64 | 50 |
| ENSG00000107779 | 1157 | 509 | 583 | 396 | 663 | 575 |
| ENSG00000270874 | 0 | 3 | 0 | 0 | 0 | 0 |
| ENSG00000200176 | 0 | 0 | 0 | 0 | 0 | 0 |
| ENSG00000173269 | 26 | 18 | 9 | 8 | 13 | 11 |

| | | | | | | |
|---|---|---|---|---|---|---|
| ENSG00000281974 | 2 | 0 | 0 | 0 | 0 | 0 |
| ENSG00000173267 | 78 | 39 | 40 | 91 | 82 | 75 |
| ENSG00000272734 | 270 | 161 | 197 | 134 | 175 | 127 |
| ENSG00000148671 | 0 | 0 | 0 | 0 | 0 | 0 |
| ENSG00000273413 | 0 | 0 | 0 | 0 | 0 | 0 |
| ENSG00000271880 | 2 | 7 | 5 | 4 | 3 | 4 |
| ENSG00000240089 | 0 | 0 | 0 | 0 | 0 | 0 |
| ENSG00000151303 | 3 | 0 | 3 | 2 | 0 | 3 |
| ENSG00000272508 | 6 | 2 | 2 | 0 | 0 | 0 |
| ENSG00000261011 | 0 | 0 | 0 | 0 | 0 | 0 |
| ENSG00000229969 | 0 | 0 | 0 | 0 | 0 | 0 |
| ENSG00000188100 | 0 | 5 | 2 | 3 | 4 | 1 |
| ENSG00000271573 | 0 | 1 | 0 | 0 | 0 | 0 |
| ENSG00000200253 | 1 | 0 | 1 | 0 | 1 | 1 |
| ENSG00000148672 | 5522 | 2685 | 3279 | 2584 | 2984 | 2879 |
| ENSG00000122376 | 872 | 409 | 467 | 322 | 719 | 436 |
| ENSG00000237280 | 0 | 1 | 0 | 0 | 0 | 0 |
| ENSG00000273946 | 0 | 0 | 1 | 0 | 0 | 0 |
| ENSG00000223482 | 434 | 205 | 253 | 218 | 306 | 251 |
| ENSG00000285257 | 11 | 9 | 10 | 3 | 14 | 8 |
| ENSG00000184923 | 29 | 7 | 9 | 9 | 14 | 10 |
| ENSG00000224914 | 215 | 102 | 147 | 112 | 127 | 119 |
| ENSG00000214562 | 49 | 21 | 28 | 23 | 25 | 27 |
| ENSG00000237740 | 0 | 0 | 1 | 0 | 0 | 0 |
| ENSG00000236417 | 0 | 0 | 0 | 0 | 0 | 0 |
| ENSG00000228055 | 0 | 0 | 0 | 0 | 0 | 0 |
| ENSG00000231569 | 0 | 0 | 0 | 0 | 0 | 0 |
| ENSG00000283672 | 2 | 0 | 1 | 0 | 0 | 0 |
| ENSG00000107789 | 1062 | 537 | 671 | 499 | 662 | 535 |
| ENSG00000223761 | 0 | 0 | 0 | 0 | 0 | 0 |
| ENSG00000225913 | 8 | 3 | 5 | 3 | 2 | 9 |
| ENSG00000196566 | 0 | 1 | 1 | 0 | 1 | 1 |
| ENSG00000234192 | 0 | 0 | 0 | 0 | 0 | 0 |
| ENSG00000198682 | 4866 | 2471 | 3560 | 2044 | 2525 | 2835 |
| ENSG00000138138 | 1619 | 873 | 1104 | 804 | 1116 | 1001 |
| ENSG00000223820 | 11 | 6 | 14 | 3 | 6 | 3 |
| ENSG00000243782 | 0 | 0 | 1 | 0 | 0 | 0 |
| ENSG00000227268 | 131 | 68 | 65 | 48 | 52 | 67 |
| ENSG00000171862 | 3781 | 1806 | 1915 | 1404 | 2244 | 1741 |
| ENSG00000224745 | 1 | 1 | 0 | 0 | 1 | 1 |
| ENSG00000213613 | 2 | 0 | 0 | 0 | 0 | 0 |
| ENSG00000200891 | 0 | 0 | 0 | 0 | 0 | 0 |
| ENSG00000227905 | 0 | 0 | 0 | 0 | 0 | 0 |
| ENSG00000275015 | 0 | 0 | 0 | 0 | 0 | 0 |
| ENSG00000184719 | 199 | 112 | 147 | 104 | 142 | 142 |
| ENSG00000201548 | 0 | 0 | 0 | 0 | 0 | 0 |
| ENSG00000204022 | 4 | 1 | 3 | 1 | 1 | 1 |
| ENSG00000237205 | 0 | 0 | 0 | 0 | 0 | 0 |
| ENSG00000182333 | 0 | 0 | 0 | 0 | 0 | 0 |
| ENSG00000271408 | 0 | 0 | 0 | 0 | 0 | 0 |

| | | | | | | |
|---|---|---|---|---|---|---|
| ENSG00000204021 | 0 | 0 | 1 | 2 | 0 | 0 |
| ENSG00000226358 | 0 | 0 | 0 | 0 | 0 | 0 |
| ENSG00000204020 | 0 | 0 | 1 | 0 | 0 | 0 |
| ENSG00000235150 | 0 | 0 | 0 | 0 | 0 | 0 |
| ENSG00000173239 | 0 | 2 | 1 | 1 | 2 | 1 |
| ENSG00000152766 | 18 | 9 | 14 | 11 | 12 | 18 |
| ENSG00000234256 | 0 | 0 | 0 | 0 | 0 | 1 |
| ENSG00000138134 | 14 | 8 | 5 | 4 | 6 | 4 |
| ENSG00000180139 | 0 | 0 | 0 | 0 | 0 | 0 |
| ENSG00000107796 | 11 | 13 | 17 | 21 | 12 | 9 |
| ENSG00000026103 | 34 | 28 | 20 | 15 | 32 | 18 |
| ENSG00000275693 | 0 | 0 | 0 | 0 | 0 | 0 |
| ENSG00000261438 | 0 | 1 | 0 | 0 | 0 | 0 |
| ENSG00000265375 | 0 | 0 | 0 | 0 | 0 | 0 |
| ENSG00000283664 | 0 | 0 | 0 | 0 | 0 | 0 |
| ENSG00000233292 | 0 | 0 | 0 | 0 | 0 | 0 |
| ENSG00000138135 | 2 | 0 | 0 | 0 | 0 | 0 |
| ENSG00000107798 | 3927 | 1798 | 2139 | 1860 | 2006 | 2070 |
| ENSG00000119922 | 8 | 12 | 15 | 12 | 8 | 7 |
| ENSG00000232110 | 0 | 0 | 0 | 0 | 0 | 0 |
| ENSG00000119917 | 26 | 11 | 5 | 4 | 1 | 3 |
| ENSG00000224289 | 0 | 0 | 0 | 0 | 0 | 0 |
| ENSG00000204010 | 0 | 0 | 0 | 0 | 0 | 0 |
| ENSG00000185745 | 60 | 42 | 54 | 45 | 24 | 28 |
| ENSG00000152778 | 312 | 166 | 190 | 134 | 200 | 206 |
| ENSG00000152779 | 0 | 0 | 1 | 0 | 1 | 0 |
| ENSG00000234452 | 0 | 0 | 0 | 0 | 0 | 0 |
| ENSG00000152782 | 675 | 366 | 371 | 307 | 409 | 387 |
| ENSG00000198997 | 0 | 0 | 0 | 0 | 0 | 0 |
| ENSG00000249962 | 0 | 0 | 0 | 0 | 0 | 0 |
| ENSG00000232936 | 0 | 0 | 0 | 2 | 2 | 0 |
| ENSG00000225836 | 0 | 0 | 0 | 0 | 0 | 1 |
| ENSG00000235100 | 7 | 0 | 4 | 1 | 3 | 6 |
| ENSG00000240996 | 3 | 1 | 5 | 1 | 7 | 2 |
| ENSG00000138182 | 46 | 61 | 59 | 19 | 119 | 78 |
| ENSG00000283578 | 0 | 0 | 0 | 0 | 0 | 0 |
| ENSG00000232229 | 19 | 13 | 27 | 12 | 15 | 23 |
| ENSG00000270670 | 0 | 0 | 0 | 0 | 0 | 0 |
| ENSG00000280560 | 0 | 0 | 0 | 0 | 0 | 0 |
| ENSG00000226159 | 0 | 0 | 0 | 0 | 0 | 0 |
| ENSG00000228938 | 0 | 0 | 0 | 0 | 1 | 0 |
| ENSG00000222451 | 0 | 0 | 0 | 0 | 0 | 0 |
| ENSG00000236373 | 0 | 0 | 0 | 0 | 0 | 0 |
| ENSG00000224750 | 0 | 0 | 0 | 0 | 0 | 0 |
| ENSG00000238291 | 0 | 0 | 0 | 0 | 0 | 0 |
| ENSG00000283053 | 0 | 0 | 0 | 0 | 0 | 0 |
| ENSG00000148680 | 0 | 0 | 3 | 1 | 0 | 0 |
| ENSG00000148688 | 884 | 382 | 468 | 408 | 466 | 482 |
| ENSG00000222305 | 0 | 0 | 0 | 0 | 0 | 0 |
| ENSG00000148677 | 6 | 6 | 6 | 13 | 15 | 12 |

| | | | | | | |
|---|---|---|---|---|---|---|
| ENSG00000201604 | 0 | 0 | 0 | 0 | 0 | 0 |
| ENSG00000225519 | 0 | 0 | 0 | 0 | 0 | 0 |
| ENSG00000273124 | 0 | 0 | 0 | 0 | 0 | 0 |
| ENSG00000224851 | 0 | 0 | 0 | 0 | 0 | 0 |
| ENSG00000220585 | 0 | 0 | 0 | 0 | 0 | 0 |
| ENSG00000234043 | 6 | 1 | 1 | 0 | 2 | 1 |
| ENSG00000180628 | 676 | 379 | 338 | 235 | 438 | 331 |
| ENSG00000165338 | 373 | 221 | 223 | 82 | 217 | 124 |
| ENSG00000271385 | 0 | 0 | 0 | 0 | 0 | 0 |
| ENSG00000223876 | 0 | 1 | 0 | 0 | 0 | 1 |
| ENSG00000119938 | 4314 | 1973 | 2459 | 1284 | 1425 | 1488 |
| ENSG00000213449 | 0 | 0 | 0 | 0 | 1 | 0 |
| ENSG00000228759 | 2 | 0 | 1 | 1 | 1 | 1 |
| ENSG00000228701 | 20 | 11 | 14 | 11 | 12 | 11 |
| ENSG00000107854 | 3113 | 1573 | 1721 | 915 | 1941 | 1358 |
| ENSG00000180581 | 9 | 5 | 6 | 4 | 5 | 10 |
| ENSG00000273497 | 0 | 0 | 0 | 0 | 0 | 0 |
| ENSG00000174721 | 100 | 77 | 99 | 86 | 89 | 70 |
| ENSG00000272817 | 6 | 3 | 0 | 9 | 4 | 7 |
| ENSG00000095564 | 1478 | 793 | 945 | 364 | 1019 | 818 |
| ENSG00000107864 | 183 | 75 | 116 | 46 | 67 | 85 |
| ENSG00000252993 | 0 | 0 | 0 | 0 | 0 | 0 |
| ENSG00000214549 | 0 | 0 | 0 | 1 | 0 | 0 |
| ENSG00000213484 | 2 | 0 | 0 | 0 | 0 | 0 |
| ENSG00000221042 | 0 | 0 | 0 | 0 | 0 | 0 |
| ENSG00000105988 | 1 | 3 | 2 | 1 | 1 | 1 |
| ENSG00000199881 | 0 | 0 | 0 | 0 | 0 | 0 |
| ENSG00000198060 | 2706 | 1387 | 1632 | 1317 | 1594 | 1449 |
| ENSG00000202297 | 0 | 0 | 0 | 0 | 0 | 0 |
| ENSG00000232709 | 0 | 0 | 2 | 1 | 1 | 2 |
| ENSG00000119912 | 1693 | 974 | 900 | 525 | 1324 | 921 |
| ENSG00000138160 | 677 | 430 | 333 | 180 | 656 | 409 |
| ENSG00000184111 | 0 | 0 | 0 | 0 | 0 | 1 |
| ENSG00000264313 | 0 | 0 | 0 | 0 | 0 | 0 |
| ENSG00000236493 | 0 | 1 | 2 | 1 | 0 | 1 |
| ENSG00000152804 | 687 | 260 | 271 | 212 | 251 | 216 |
| ENSG00000201412 | 0 | 0 | 0 | 0 | 0 | 0 |
| ENSG00000138190 | 746 | 437 | 446 | 260 | 552 | 355 |
| ENSG00000225118 | 0 | 0 | 0 | 0 | 0 | 0 |
| ENSG00000226425 | 0 | 0 | 0 | 0 | 0 | 0 |
| ENSG00000187553 | 1 | 0 | 0 | 0 | 1 | 1 |
| ENSG00000285846 | 0 | 0 | 0 | 0 | 0 | 0 |
| ENSG00000279607 | 2 | 0 | 3 | 2 | 1 | 1 |
| ENSG00000095596 | 22 | 8 | 11 | 9 | 4 | 10 |
| ENSG00000232805 | 0 | 0 | 0 | 0 | 0 | 0 |
| ENSG00000237417 | 0 | 0 | 0 | 0 | 0 | 0 |
| ENSG00000213432 | 0 | 0 | 1 | 0 | 0 | 2 |
| ENSG00000138119 | 17104 | 9511 | 10157 | 5136 | 10020 | 8566 |
| ENSG00000138180 | 619 | 368 | 362 | 265 | 651 | 447 |
| ENSG00000212396 | 1 | 1 | 0 | 3 | 0 | 0 |

| | | | | | | |
|---|---|---|---|---|---|---|
| ENSG00000186188 | 0 | 0 | 0 | 0 | 0 | 0 |
| ENSG00000138207 | 0 | 1 | 0 | 3 | 0 | 2 |
| ENSG00000095464 | 0 | 3 | 3 | 2 | 3 | 3 |
| ENSG00000148690 | 547 | 272 | 314 | 210 | 349 | 313 |
| ENSG00000108231 | 18 | 15 | 14 | 9 | 14 | 13 |
| ENSG00000280660 | 15 | 4 | 11 | 2 | 5 | 4 |
| ENSG00000280485 | 0 | 0 | 0 | 0 | 0 | 0 |
| ENSG00000227995 | 0 | 0 | 0 | 0 | 0 | 0 |
| ENSG00000176273 | 308 | 113 | 152 | 101 | 141 | 164 |
| ENSG00000180764 | 6 | 6 | 3 | 8 | 11 | 1 |
| ENSG00000138193 | 21 | 17 | 13 | 3 | 14 | 8 |
| ENSG00000232913 | 0 | 0 | 0 | 0 | 0 | 0 |
| ENSG00000231575 | 0 | 0 | 0 | 0 | 0 | 0 |
| ENSG00000234724 | 0 | 0 | 0 | 0 | 0 | 0 |
| ENSG00000206631 | 0 | 0 | 0 | 0 | 0 | 0 |
| ENSG00000228553 | 0 | 0 | 0 | 0 | 0 | 0 |
| ENSG00000268894 | 0 | 0 | 0 | 0 | 0 | 0 |
| ENSG00000273450 | 0 | 0 | 0 | 0 | 0 | 0 |
| ENSG00000173145 | 378 | 222 | 229 | 155 | 372 | 269 |
| ENSG00000108239 | 596 | 282 | 355 | 258 | 393 | 356 |
| ENSG00000119969 | 519 | 284 | 294 | 117 | 300 | 236 |
| ENSG00000244332 | 0 | 0 | 2 | 0 | 0 | 1 |
| ENSG00000235746 | 0 | 0 | 0 | 0 | 0 | 0 |
| ENSG00000108242 | 0 | 0 | 0 | 0 | 0 | 0 |
| ENSG00000276490 | 0 | 0 | 0 | 0 | 0 | 0 |
| ENSG00000165841 | 0 | 0 | 0 | 0 | 0 | 0 |
| ENSG00000230338 | 0 | 0 | 0 | 0 | 0 | 0 |
| ENSG00000235199 | 0 | 0 | 0 | 0 | 0 | 0 |
| ENSG00000225354 | 0 | 0 | 0 | 0 | 0 | 0 |
| ENSG00000138109 | 0 | 1 | 0 | 0 | 0 | 0 |
| ENSG00000233377 | 0 | 0 | 0 | 0 | 0 | 0 |
| ENSG00000228460 | 0 | 0 | 0 | 0 | 0 | 0 |
| ENSG00000260863 | 0 | 0 | 0 | 0 | 0 | 0 |
| ENSG00000237885 | 0 | 0 | 0 | 0 | 0 | 0 |
| ENSG00000138115 | 38 | 17 | 35 | 23 | 33 | 30 |
| ENSG00000231829 | 2 | 5 | 6 | 1 | 1 | 1 |
| ENSG00000274691 | 0 | 0 | 0 | 0 | 0 | 2 |
| ENSG00000225533 | 0 | 0 | 1 | 0 | 0 | 2 |
| ENSG00000173124 | 3 | 4 | 8 | 5 | 8 | 8 |
| ENSG00000234026 | 2 | 11 | 9 | 4 | 2 | 2 |
| ENSG00000107438 | 3799 | 2293 | 2241 | 2383 | 2600 | 2280 |
| ENSG00000095637 | 33 | 35 | 40 | 19 | 38 | 24 |
| ENSG00000214362 | 2 | 0 | 0 | 0 | 0 | 0 |
| ENSG00000059573 | 5413 | 2493 | 3184 | 2555 | 2602 | 2857 |
| ENSG00000119977 | 2085 | 1015 | 1269 | 1090 | 1201 | 1251 |
| ENSG00000138185 | 62 | 39 | 51 | 33 | 54 | 44 |
| ENSG00000226688 | 140 | 90 | 96 | 57 | 84 | 92 |
| ENSG00000240527 | 61 | 37 | 76 | 36 | 46 | 61 |
| ENSG00000270099 | 0 | 0 | 0 | 0 | 0 | 0 |
| ENSG00000188649 | 5 | 1 | 0 | 3 | 1 | 1 |

| | | | | | | |
|---|---|---|---|---|---|---|
| ENSG00000227805 | 0 | 0 | 0 | 0 | 0 | 0 |
| ENSG00000107443 | 736 | 330 | 422 | 294 | 530 | 416 |
| ENSG00000266407 | 0 | 0 | 0 | 0 | 0 | 0 |
| ENSG00000200728 | 0 | 0 | 0 | 0 | 0 | 0 |
| ENSG00000177853 | 200 | 134 | 108 | 40 | 233 | 121 |
| ENSG00000229417 | 2 | 0 | 1 | 2 | 1 | 2 |
| ENSG00000095585 | 147 | 73 | 115 | 57 | 85 | 83 |
| ENSG00000229418 | 0 | 0 | 0 | 0 | 0 | 0 |
| ENSG00000107447 | 0 | 0 | 0 | 2 | 0 | 0 |
| ENSG00000197430 | 0 | 0 | 0 | 0 | 0 | 0 |
| ENSG00000095587 | 12 | 5 | 6 | 6 | 2 | 3 |
| ENSG00000077147 | 8265 | 4167 | 4723 | 3203 | 5019 | 4339 |
| ENSG00000235677 | 2 | 3 | 1 | 1 | 2 | 1 |
| ENSG00000155629 | 64 | 15 | 30 | 5 | 10 | 19 |
| ENSG00000202047 | 0 | 0 | 0 | 0 | 0 | 0 |
| ENSG00000227032 | 0 | 0 | 0 | 0 | 0 | 0 |
| ENSG00000207287 | 0 | 0 | 0 | 0 | 0 | 0 |
| ENSG00000236552 | 184 | 127 | 108 | 127 | 90 | 265 |
| ENSG00000207976 | 0 | 0 | 0 | 0 | 0 | 0 |
| ENSG00000196233 | 1352 | 708 | 521 | 263 | 904 | 544 |
| ENSG00000232875 | 0 | 0 | 0 | 0 | 0 | 0 |
| ENSG00000187122 | 150 | 85 | 117 | 32 | 25 | 35 |
| ENSG00000234855 | 0 | 0 | 0 | 0 | 0 | 0 |
| ENSG00000269891 | 4 | 1 | 1 | 3 | 0 | 0 |
| ENSG00000213390 | 461 | 227 | 339 | 212 | 307 | 247 |
| ENSG00000231398 | 0 | 0 | 0 | 0 | 0 | 0 |
| ENSG00000277308 | 0 | 0 | 0 | 0 | 0 | 0 |
| ENSG00000237169 | 1 | 1 | 0 | 4 | 1 | 0 |
| ENSG00000165879 | 330 | 179 | 229 | 194 | 195 | 176 |
| ENSG00000181274 | 1158 | 671 | 713 | 743 | 698 | 714 |
| ENSG00000225850 | 2 | 3 | 3 | 4 | 8 | 3 |
| ENSG00000052749 | 1702 | 846 | 1050 | 1380 | 1107 | 1209 |
| ENSG00000231508 | 0 | 0 | 0 | 0 | 0 | 0 |
| ENSG00000231970 | 2 | 0 | 1 | 0 | 0 | 0 |
| ENSG00000200737 | 2 | 1 | 3 | 0 | 0 | 0 |
| ENSG00000171314 | 4013 | 1965 | 2475 | 2413 | 2297 | 2310 |
| ENSG00000171311 | 1209 | 562 | 661 | 647 | 722 | 636 |
| ENSG00000171307 | 2481 | 1258 | 1427 | 1231 | 1270 | 1148 |
| ENSG00000155229 | 2986 | 1366 | 1704 | 1726 | 1751 | 1677 |
| ENSG00000165886 | 466 | 286 | 364 | 392 | 367 | 334 |
| ENSG00000165887 | 6 | 5 | 1 | 4 | 4 | 7 |
| ENSG00000241935 | 13 | 7 | 8 | 18 | 2 | 10 |
| ENSG00000249967 | 0 | 0 | 0 | 0 | 0 | 0 |
| ENSG00000203942 | 2 | 0 | 0 | 1 | 0 | 0 |
| ENSG00000171160 | 599 | 291 | 315 | 309 | 346 | 334 |
| ENSG00000155252 | 841 | 393 | 508 | 585 | 518 | 515 |
| ENSG00000119986 | 672 | 301 | 464 | 561 | 389 | 441 |
| ENSG00000155254 | 1667 | 883 | 1103 | 1016 | 990 | 1027 |
| ENSG00000155256 | 1593 | 752 | 997 | 826 | 896 | 909 |
| ENSG00000120057 | 2 | 0 | 0 | 0 | 2 | 1 |

| | | | | | | |
|---|---|---|---|---|---|---|
| ENSG00000227356 | 9 | 5 | 2 | 1 | 3 | 2 |
| ENSG00000155265 | 283 | 170 | 221 | 186 | 216 | 204 |
| ENSG00000095713 | 0 | 0 | 1 | 0 | 0 | 0 |
| ENSG00000278752 | 0 | 0 | 0 | 0 | 0 | 0 |
| ENSG00000166024 | 399 | 247 | 241 | 133 | 349 | 232 |
| ENSG00000138131 | 5 | 3 | 1 | 1 | 1 | 2 |
| ENSG00000230928 | 1 | 0 | 1 | 0 | 1 | 0 |
| ENSG00000119943 | 993 | 435 | 589 | 453 | 429 | 490 |
| ENSG00000284415 | 0 | 0 | 0 | 0 | 0 | 0 |
| ENSG00000107521 | 1937 | 833 | 1089 | 1127 | 965 | 1042 |
| ENSG00000264610 | 1 | 0 | 0 | 0 | 0 | 1 |
| ENSG00000172987 | 3 | 2 | 3 | 0 | 0 | 3 |
| ENSG00000274056 | 0 | 0 | 0 | 0 | 0 | 0 |
| ENSG00000236818 | 0 | 2 | 0 | 0 | 0 | 0 |
| ENSG00000234109 | 0 | 0 | 0 | 0 | 0 | 0 |
| ENSG00000119946 | 456 | 219 | 282 | 175 | 181 | 224 |
| ENSG00000120053 | 1674 | 779 | 970 | 918 | 845 | 931 |
| ENSG00000224934 | 59 | 33 | 58 | 18 | 36 | 18 |
| ENSG00000257582 | 0 | 0 | 0 | 0 | 0 | 0 |
| ENSG00000228778 | 0 | 0 | 0 | 0 | 0 | 0 |
| ENSG00000119919 | 4 | 6 | 4 | 0 | 3 | 5 |
| ENSG00000155287 | 534 | 306 | 416 | 331 | 393 | 344 |
| ENSG00000260475 | 4 | 7 | 8 | 2 | 9 | 3 |
| ENSG00000285932 | 0 | 0 | 0 | 0 | 0 | 0 |
| ENSG00000229278 | 0 | 0 | 2 | 2 | 1 | 0 |
| ENSG00000198018 | 814 | 387 | 447 | 302 | 425 | 358 |
| ENSG00000233690 | 6 | 1 | 1 | 2 | 1 | 0 |
| ENSG00000119929 | 459 | 234 | 261 | 247 | 296 | 323 |
| ENSG00000014919 | 1403 | 661 | 916 | 680 | 760 | 826 |
| ENSG00000023839 | 49 | 38 | 51 | 23 | 60 | 70 |
| ENSG00000227351 | 0 | 0 | 0 | 0 | 0 | 0 |
| ENSG00000107554 | 1587 | 794 | 1107 | 1182 | 1547 | 1447 |
| ENSG00000227695 | 0 | 0 | 1 | 0 | 0 | 0 |
| ENSG00000120054 | 0 | 0 | 0 | 0 | 0 | 0 |
| ENSG00000283232 | 0 | 0 | 0 | 0 | 0 | 0 |
| ENSG00000232230 | 0 | 0 | 0 | 0 | 0 | 0 |
| ENSG00000270623 | 0 | 0 | 0 | 0 | 0 | 0 |
| ENSG00000107566 | 1705 | 769 | 957 | 772 | 811 | 964 |
| ENSG00000213341 | 1231 | 623 | 705 | 520 | 791 | 700 |
| ENSG00000236308 | 15 | 4 | 4 | 3 | 10 | 7 |
| ENSG00000227492 | 13 | 4 | 6 | 2 | 6 | 4 |
| ENSG00000095485 | 1126 | 610 | 742 | 761 | 809 | 751 |
| ENSG00000212464 | 2 | 5 | 1 | 0 | 0 | 1 |
| ENSG00000230224 | 18 | 12 | 18 | 12 | 18 | 8 |
| ENSG00000207362 | 2 | 1 | 0 | 0 | 0 | 0 |
| ENSG00000196072 | 1984 | 1022 | 1306 | 1081 | 1285 | 1214 |
| ENSG00000107593 | 4 | 0 | 2 | 0 | 0 | 0 |
| ENSG00000231188 | 0 | 1 | 1 | 3 | 1 | 1 |
| ENSG00000099194 | 141892 | 56147 | 72996 | 48292 | 52903 | 58442 |
| ENSG00000235823 | 1255 | 631 | 707 | 549 | 670 | 653 |

| | | | | | | |
|---|---|---|---|---|---|---|
| ENSG00000212325 | 0 | 0 | 0 | 0 | 0 | 0 |
| ENSG00000075290 | 11 | 7 | 7 | 10 | 12 | 10 |
| ENSG00000075826 | 75 | 37 | 49 | 34 | 30 | 27 |
| ENSG00000255339 | 0 | 1 | 0 | 0 | 0 | 0 |
| ENSG00000166136 | 246 | 192 | 230 | 266 | 228 | 243 |
| ENSG00000166135 | 4504 | 2112 | 2638 | 2053 | 2355 | 2393 |
| ENSG00000274660 | 0 | 0 | 0 | 0 | 0 | 0 |
| ENSG00000075891 | 0 | 1 | 1 | 1 | 1 | 0 |
| ENSG00000222072 | 1 | 0 | 0 | 0 | 0 | 0 |
| ENSG00000272572 | 32 | 9 | 15 | 12 | 6 | 17 |
| ENSG00000119906 | 1071 | 517 | 573 | 269 | 696 | 510 |
| ENSG00000273476 | 0 | 0 | 0 | 0 | 0 | 0 |
| ENSG00000055950 | 2828 | 1439 | 1758 | 1914 | 1547 | 1613 |
| ENSG00000095539 | 243 | 120 | 144 | 131 | 124 | 137 |
| ENSG00000207551 | 0 | 0 | 0 | 0 | 0 | 0 |
| ENSG00000236662 | 7 | 2 | 4 | 1 | 2 | 0 |
| ENSG00000107815 | 707 | 348 | 430 | 537 | 501 | 546 |
| ENSG00000107816 | 3031 | 1379 | 1914 | 1762 | 1656 | 1672 |
| ENSG00000186862 | 149 | 64 | 83 | 108 | 110 | 110 |
| ENSG00000107819 | 1268 | 589 | 800 | 767 | 911 | 778 |
| ENSG00000273162 | 69 | 31 | 61 | 29 | 33 | 34 |
| ENSG00000107821 | 1125 | 476 | 667 | 616 | 455 | 519 |
| ENSG00000236311 | 0 | 1 | 4 | 0 | 3 | 4 |
| ENSG00000107807 | 6 | 1 | 2 | 4 | 4 | 2 |
| ENSG00000237579 | 0 | 0 | 0 | 0 | 0 | 0 |
| ENSG00000270973 | 0 | 0 | 0 | 0 | 0 | 0 |
| ENSG00000138136 | 0 | 0 | 0 | 0 | 0 | 0 |
| ENSG00000227128 | 0 | 0 | 0 | 0 | 0 | 0 |
| ENSG00000229649 | 0 | 0 | 0 | 0 | 0 | 0 |
| ENSG00000222238 | 0 | 0 | 0 | 0 | 0 | 0 |
| ENSG00000225208 | 2 | 0 | 0 | 0 | 0 | 2 |
| ENSG00000230967 | 0 | 0 | 0 | 0 | 0 | 0 |
| ENSG00000166167 | 1421 | 613 | 800 | 521 | 696 | 697 |
| ENSG00000222414 | 0 | 0 | 0 | 0 | 0 | 0 |
| ENSG00000166171 | 584 | 290 | 327 | 416 | 388 | 385 |
| ENSG00000166169 | 1257 | 644 | 819 | 801 | 792 | 706 |
| ENSG00000283558 | 0 | 0 | 0 | 0 | 0 | 0 |
| ENSG00000263436 | 0 | 0 | 0 | 0 | 0 | 0 |
| ENSG00000107829 | 1752 | 860 | 1155 | 1088 | 1029 | 1050 |
| ENSG00000222051 | 0 | 0 | 0 | 0 | 0 | 0 |
| ENSG00000224817 | 0 | 0 | 0 | 0 | 0 | 0 |
| ENSG00000107831 | 0 | 0 | 4 | 1 | 1 | 1 |
| ENSG00000107833 | 798 | 436 | 445 | 661 | 547 | 545 |
| ENSG00000198408 | 5340 | 2601 | 3509 | 1746 | 3343 | 2903 |
| ENSG00000226009 | 2 | 0 | 0 | 0 | 0 | 1 |
| ENSG00000120049 | 27 | 13 | 19 | 4 | 12 | 11 |
| ENSG00000120029 | 1496 | 734 | 963 | 758 | 866 | 820 |
| ENSG00000274742 | 1 | 0 | 0 | 0 | 0 | 0 |
| ENSG00000222430 | 0 | 0 | 0 | 0 | 0 | 0 |
| ENSG00000166189 | 1270 | 583 | 759 | 883 | 717 | 626 |

| | | | | | | |
|---|---|---|---|---|---|---|
| ENSG00000224302 | 1 | 1 | 1 | 0 | 0 | 1 |
| ENSG00000229629 | 0 | 0 | 0 | 0 | 0 | 0 |
| ENSG00000198728 | 3219 | 1350 | 1920 | 1488 | 1552 | 1735 |
| ENSG00000148840 | 1285 | 577 | 779 | 781 | 738 | 892 |
| ENSG00000166197 | 3933 | 2033 | 2532 | 2153 | 2633 | 2904 |
| ENSG00000119915 | 60 | 51 | 75 | 30 | 29 | 63 |
| ENSG00000107859 | 16 | 7 | 12 | 14 | 12 | 14 |
| ENSG00000107862 | 4240 | 1832 | 2558 | 1713 | 2106 | 2148 |
| ENSG00000077150 | 1793 | 896 | 1242 | 1111 | 1173 | 1081 |
| ENSG00000059915 | 236 | 110 | 165 | 144 | 126 | 158 |
| ENSG00000107872 | 937 | 433 | 572 | 612 | 493 | 492 |
| ENSG00000107874 | 1442 | 687 | 940 | 1052 | 814 | 877 |
| ENSG00000202569 | 0 | 0 | 0 | 0 | 0 | 0 |
| ENSG00000269609 | 599 | 344 | 413 | 338 | 432 | 361 |
| ENSG00000120055 | 32 | 23 | 39 | 28 | 23 | 40 |
| ENSG00000138111 | 156 | 71 | 104 | 108 | 79 | 94 |
| ENSG00000138107 | 6443 | 3345 | 4102 | 4025 | 3759 | 3868 |
| ENSG00000273262 | 0 | 0 | 0 | 0 | 0 | 0 |
| ENSG00000107882 | 1263 | 562 | 690 | 568 | 592 | 615 |
| ENSG00000228657 | 0 | 2 | 1 | 0 | 0 | 0 |
| ENSG00000207029 | 0 | 0 | 0 | 0 | 1 | 0 |
| ENSG00000171206 | 3585 | 1725 | 2331 | 2076 | 1913 | 1984 |
| ENSG00000272933 | 0 | 0 | 0 | 0 | 0 | 0 |
| ENSG00000138175 | 1519 | 751 | 924 | 990 | 930 | 931 |
| ENSG00000156398 | 902 | 442 | 538 | 344 | 388 | 381 |
| ENSG00000166272 | 3771 | 1663 | 2169 | 1977 | 1970 | 2030 |
| ENSG00000252994 | 0 | 0 | 0 | 0 | 0 | 0 |
| ENSG00000148795 | 0 | 0 | 0 | 0 | 1 | 0 |
| ENSG00000213061 | 3 | 3 | 2 | 2 | 0 | 5 |
| ENSG00000236937 | 2 | 0 | 0 | 1 | 1 | 0 |
| ENSG00000282772 | 4 | 3 | 0 | 0 | 0 | 0 |
| ENSG00000166275 | 690 | 341 | 476 | 303 | 403 | 420 |
| ENSG00000270316 | 0 | 0 | 0 | 0 | 0 | 0 |
| ENSG00000283337 | 0 | 0 | 1 | 0 | 0 | 0 |
| ENSG00000214435 | 1 | 0 | 0 | 1 | 0 | 1 |
| ENSG00000235266 | 0 | 0 | 0 | 0 | 0 | 1 |
| ENSG00000272912 | 0 | 0 | 0 | 0 | 0 | 0 |
| ENSG00000148842 | 985 | 419 | 505 | 374 | 411 | 463 |
| ENSG00000278498 | 0 | 0 | 0 | 0 | 0 | 0 |
| ENSG00000076685 | 2736 | 1327 | 1868 | 881 | 1531 | 1426 |
| ENSG00000237827 | 0 | 0 | 1 | 0 | 1 | 1 |
| ENSG00000213277 | 12 | 3 | 2 | 1 | 7 | 1 |
| ENSG00000229256 | 0 | 0 | 0 | 0 | 0 | 0 |
| ENSG00000235376 | 2 | 0 | 0 | 0 | 0 | 0 |
| ENSG00000148798 | 0 | 0 | 0 | 0 | 0 | 0 |
| ENSG00000156374 | 501 | 230 | 321 | 221 | 278 | 333 |
| ENSG00000212413 | 0 | 0 | 0 | 0 | 0 | 0 |
| ENSG00000148835 | 187 | 97 | 138 | 126 | 136 | 117 |
| ENSG00000173915 | 2976 | 1512 | 1759 | 1989 | 1964 | 1846 |
| ENSG00000283867 | 0 | 0 | 1 | 0 | 1 | 0 |

| | | | | | | |
|---|---|---|---|---|---|---|
| ENSG00000148843 | 2358 | 1074 | 1317 | 1260 | 1447 | 1528 |
| ENSG00000138172 | 1650 | 895 | 1015 | 950 | 912 | 816 |
| ENSG00000273485 | 0 | 0 | 4 | 2 | 1 | 0 |
| ENSG00000234699 | 5 | 3 | 6 | 3 | 7 | 5 |
| ENSG00000185933 | 16 | 8 | 15 | 7 | 5 | 0 |
| ENSG00000183128 | 8 | 2 | 1 | 7 | 5 | 1 |
| ENSG00000235470 | 0 | 0 | 0 | 0 | 0 | 0 |
| ENSG00000107954 | 213 | 89 | 157 | 86 | 76 | 87 |
| ENSG00000226431 | 0 | 1 | 0 | 1 | 0 | 0 |
| ENSG00000107957 | 140 | 47 | 60 | 40 | 34 | 77 |
| ENSG00000273108 | 16 | 5 | 6 | 2 | 5 | 2 |
| ENSG00000222503 | 0 | 0 | 0 | 0 | 0 | 0 |
| ENSG00000280693 | 0 | 0 | 0 | 0 | 0 | 0 |
| ENSG00000260461 | 74 | 43 | 44 | 39 | 38 | 48 |
| ENSG00000107960 | 666 | 389 | 486 | 407 | 531 | 487 |
| ENSG00000065613 | 642 | 397 | 276 | 137 | 878 | 327 |
| ENSG00000266754 | 0 | 0 | 0 | 0 | 0 | 0 |
| ENSG00000065618 | 12 | 0 | 3 | 7 | 0 | 2 |
| ENSG00000283788 | 0 | 0 | 0 | 0 | 0 | 0 |
| ENSG00000156384 | 159 | 56 | 67 | 57 | 81 | 76 |
| ENSG00000197748 | 26 | 22 | 20 | 11 | 18 | 19 |
| ENSG00000208033 | 0 | 0 | 1 | 0 | 0 | 0 |
| ENSG00000148834 | 1500 | 823 | 902 | 1044 | 987 | 1010 |
| ENSG00000266852 | 5 | 3 | 4 | 1 | 2 | 2 |
| ENSG00000065621 | 1622 | 729 | 900 | 877 | 806 | 889 |
| ENSG00000148841 | 273 | 134 | 105 | 117 | 150 | 129 |
| ENSG00000270075 | 0 | 0 | 0 | 0 | 0 | 0 |
| ENSG00000228261 | 6 | 5 | 2 | 5 | 2 | 0 |
| ENSG00000231233 | 10 | 7 | 5 | 10 | 4 | 6 |
| ENSG00000120051 | 0 | 3 | 0 | 2 | 3 | 3 |
| ENSG00000225768 | 26 | 6 | 13 | 3 | 5 | 12 |
| ENSG00000237761 | 0 | 1 | 2 | 0 | 0 | 0 |
| ENSG00000156395 | 0 | 0 | 0 | 0 | 0 | 0 |
| ENSG00000226387 | 0 | 0 | 0 | 0 | 0 | 0 |
| ENSG00000207068 | 0 | 0 | 0 | 0 | 0 | 0 |
| ENSG00000233477 | 0 | 0 | 0 | 0 | 0 | 0 |
| ENSG00000213260 | 3 | 1 | 1 | 1 | 0 | 1 |
| ENSG00000229466 | 0 | 0 | 0 | 0 | 0 | 0 |
| ENSG00000229775 | 0 | 0 | 0 | 0 | 0 | 0 |
| ENSG00000215002 | 0 | 0 | 0 | 0 | 0 | 0 |
| ENSG00000108018 | 8 | 2 | 6 | 5 | 7 | 3 |
| ENSG00000200626 | 0 | 0 | 0 | 0 | 0 | 0 |
| ENSG00000200079 | 0 | 0 | 0 | 0 | 0 | 0 |
| ENSG00000229981 | 0 | 0 | 0 | 0 | 0 | 0 |
| ENSG00000234544 | 0 | 0 | 0 | 0 | 0 | 0 |
| ENSG00000203434 | 0 | 0 | 0 | 0 | 0 | 0 |
| ENSG00000223381 | 0 | 0 | 0 | 0 | 0 | 0 |
| ENSG00000224498 | 0 | 0 | 0 | 0 | 0 | 0 |
| ENSG00000222436 | 0 | 0 | 0 | 0 | 0 | 0 |
| ENSG00000252574 | 0 | 0 | 0 | 0 | 0 | 0 |

| | | | | | | |
|---|---|---|---|---|---|---|
| ENSG00000200217 | 0 | 0 | 0 | 0 | 0 | 0 |
| ENSG00000232728 | 0 | 0 | 0 | 0 | 0 | 0 |
| ENSG00000237242 | 0 | 0 | 0 | 0 | 0 | 0 |
| ENSG00000270333 | 0 | 0 | 0 | 0 | 0 | 0 |
| ENSG00000270294 | 0 | 0 | 0 | 0 | 0 | 0 |
| ENSG00000224907 | 0 | 0 | 0 | 0 | 0 | 0 |
| ENSG00000108039 | 1764 | 877 | 1176 | 931 | 1063 | 1123 |
| ENSG00000272160 | 0 | 0 | 0 | 0 | 0 | 0 |
| ENSG00000203876 | 2 | 3 | 5 | 0 | 0 | 0 |
| ENSG00000148700 | 613 | 307 | 338 | 205 | 371 | 310 |
| ENSG00000226734 | 0 | 0 | 0 | 0 | 0 | 0 |
| ENSG00000119950 | 1245 | 674 | 765 | 833 | 901 | 917 |
| ENSG00000228417 | 0 | 0 | 0 | 0 | 0 | 0 |
| ENSG00000214067 | 0 | 0 | 0 | 0 | 0 | 0 |
| ENSG00000119953 | 1063 | 551 | 696 | 557 | 621 | 619 |
| ENSG00000273143 | 4 | 2 | 2 | 1 | 2 | 3 |
| ENSG00000237642 | 0 | 0 | 0 | 0 | 0 | 0 |
| ENSG00000138166 | 182 | 81 | 91 | 112 | 129 | 87 |
| ENSG00000233518 | 0 | 0 | 0 | 0 | 0 | 0 |
| ENSG00000108055 | 377 | 264 | 194 | 67 | 605 | 259 |
| ENSG00000223302 | 0 | 0 | 0 | 0 | 0 | 0 |
| ENSG00000203867 | 12 | 4 | 10 | 6 | 3 | 5 |
| ENSG00000252036 | 0 | 0 | 0 | 0 | 0 | 0 |
| ENSG00000199364 | 0 | 0 | 0 | 0 | 0 | 0 |
| ENSG00000232470 | 14 | 2 | 6 | 2 | 8 | 7 |
| ENSG00000203497 | 96 | 67 | 73 | 77 | 75 | 72 |
| ENSG00000150593 | 2143 | 1126 | 1334 | 788 | 1344 | 1088 |
| ENSG00000284482 | 0 | 0 | 0 | 0 | 0 | 0 |
| ENSG00000214413 | 442 | 224 | 254 | 212 | 301 | 253 |
| ENSG00000278601 | 5 | 0 | 1 | 0 | 4 | 1 |
| ENSG00000270589 | 12 | 2 | 5 | 1 | 4 | 5 |
| ENSG00000108061 | 1146 | 595 | 547 | 390 | 768 | 615 |
| ENSG00000234118 | 5 | 0 | 2 | 0 | 0 | 1 |
| ENSG00000221214 | 0 | 0 | 0 | 0 | 0 | 0 |
| ENSG00000150594 | 2 | 1 | 1 | 1 | 1 | 0 |
| ENSG00000213247 | 0 | 0 | 0 | 0 | 0 | 0 |
| ENSG00000237618 | 0 | 0 | 0 | 0 | 0 | 0 |
| ENSG00000227851 | 0 | 0 | 0 | 0 | 0 | 0 |
| ENSG00000230809 | 0 | 0 | 0 | 0 | 0 | 0 |
| ENSG00000119927 | 784 | 372 | 421 | 252 | 458 | 371 |
| ENSG00000119913 | 0 | 0 | 0 | 0 | 0 | 0 |
| ENSG00000283688 | 0 | 0 | 0 | 0 | 0 | 0 |
| ENSG00000276857 | 0 | 0 | 0 | 0 | 0 | 0 |
| ENSG00000243316 | 0 | 0 | 0 | 0 | 0 | 0 |
| ENSG00000197142 | 1 | 3 | 2 | 2 | 1 | 1 |
| ENSG00000232934 | 0 | 0 | 0 | 0 | 1 | 1 |
| ENSG00000023041 | 1864 | 888 | 1130 | 823 | 1089 | 1069 |
| ENSG00000151532 | 1188 | 593 | 786 | 580 | 762 | 701 |
| ENSG00000234017 | 0 | 0 | 0 | 0 | 1 | 1 |
| ENSG00000264763 | 0 | 0 | 0 | 0 | 0 | 0 |

| | | | | | | |
|---|---|---|---|---|---|---|
| ENSG00000233340 | 0 | 0 | 0 | 0 | 0 | 0 |
| ENSG00000285676 | 11 | 6 | 13 | 2 | 10 | 9 |
| ENSG00000260917 | 276 | 99 | 158 | 101 | 129 | 140 |
| ENSG00000279315 | 0 | 0 | 0 | 0 | 1 | 1 |
| ENSG00000225292 | 2 | 0 | 0 | 0 | 0 | 0 |
| ENSG00000148737 | 800 | 305 | 408 | 287 | 334 | 412 |
| ENSG00000233547 | 0 | 2 | 4 | 0 | 1 | 1 |
| ENSG00000227560 | 0 | 0 | 1 | 0 | 0 | 2 |
| ENSG00000238380 | 0 | 0 | 1 | 0 | 0 | 0 |
| ENSG00000230018 | 0 | 0 | 1 | 0 | 0 | 0 |
| ENSG00000148702 | 2 | 2 | 4 | 1 | 4 | 3 |
| ENSG00000197893 | 0 | 0 | 0 | 0 | 0 | 2 |
| ENSG00000165806 | 1898 | 868 | 1184 | 800 | 1066 | 1074 |
| ENSG00000234393 | 6 | 1 | 3 | 1 | 2 | 2 |
| ENSG00000148735 | 619 | 324 | 298 | 160 | 304 | 286 |
| ENSG00000283656 | 0 | 0 | 0 | 0 | 0 | 0 |
| ENSG00000212589 | 0 | 0 | 0 | 0 | 0 | 0 |
| ENSG00000198924 | 443 | 249 | 253 | 153 | 308 | 222 |
| ENSG00000196865 | 926 | 457 | 483 | 223 | 568 | 435 |
| ENSG00000043591 | 32 | 12 | 9 | 10 | 16 | 10 |
| ENSG00000285540 | 0 | 0 | 0 | 0 | 0 | 0 |
| ENSG00000253066 | 0 | 0 | 0 | 0 | 0 | 0 |
| ENSG00000226582 | 0 | 0 | 0 | 0 | 0 | 0 |
| ENSG00000165813 | 388 | 228 | 178 | 66 | 410 | 167 |
| ENSG00000273904 | 0 | 0 | 0 | 0 | 0 | 0 |
| ENSG00000284442 | 0 | 0 | 0 | 0 | 0 | 0 |
| ENSG00000095627 | 389 | 197 | 229 | 135 | 286 | 184 |
| ENSG00000165816 | 9 | 1 | 1 | 5 | 5 | 1 |
| ENSG00000230165 | 0 | 0 | 0 | 0 | 0 | 0 |
| ENSG00000275024 | 0 | 1 | 0 | 0 | 0 | 0 |
| ENSG00000169129 | 422 | 256 | 284 | 193 | 272 | 231 |
| ENSG00000242912 | 0 | 0 | 0 | 0 | 0 | 0 |
| ENSG00000279796 | 0 | 0 | 0 | 0 | 0 | 0 |
| ENSG00000099204 | 957 | 436 | 626 | 329 | 508 | 464 |
| ENSG00000229668 | 0 | 0 | 0 | 0 | 0 | 0 |
| ENSG00000228169 | 0 | 0 | 0 | 0 | 0 | 0 |
| ENSG00000228484 | 2 | 2 | 1 | 2 | 3 | 1 |
| ENSG00000215760 | 1 | 0 | 1 | 0 | 0 | 0 |
| ENSG00000151553 | 1769 | 810 | 980 | 435 | 866 | 782 |
| ENSG00000226428 | 0 | 0 | 0 | 0 | 0 | 0 |
| ENSG00000165832 | 1212 | 651 | 778 | 510 | 690 | 754 |
| ENSG00000252611 | 1 | 0 | 0 | 0 | 0 | 1 |
| ENSG00000236799 | 0 | 1 | 0 | 0 | 1 | 0 |
| ENSG00000107518 | 146 | 65 | 72 | 67 | 85 | 77 |
| ENSG00000285582 | 0 | 0 | 0 | 0 | 0 | 0 |
| ENSG00000270965 | 0 | 0 | 0 | 0 | 0 | 0 |
| ENSG00000151892 | 20581 | 8619 | 12967 | 7037 | 6955 | 9090 |
| ENSG00000285587 | 134 | 48 | 74 | 63 | 40 | 65 |
| ENSG00000182645 | 0 | 0 | 0 | 0 | 0 | 0 |
| ENSG00000205433 | 0 | 0 | 0 | 0 | 0 | 0 |

| | | | | | | |
|---|---|---|---|---|---|---|
| ENSG00000203837 | 0 | 0 | 0 | 0 | 0 | 0 |
| ENSG00000227096 | 2 | 0 | 0 | 0 | 0 | 0 |
| ENSG00000200935 | 0 | 0 | 0 | 0 | 0 | 0 |
| ENSG00000175535 | 0 | 0 | 0 | 0 | 0 | 1 |
| ENSG00000232091 | 0 | 0 | 0 | 0 | 0 | 0 |
| ENSG00000187021 | 4 | 1 | 3 | 1 | 1 | 0 |
| ENSG00000266200 | 0 | 0 | 0 | 0 | 1 | 0 |
| ENSG00000165863 | 347 | 191 | 260 | 191 | 206 | 197 |
| ENSG00000232767 | 1 | 1 | 0 | 1 | 0 | 2 |
| ENSG00000165868 | 380 | 168 | 242 | 156 | 165 | 201 |
| ENSG00000228195 | 0 | 0 | 2 | 0 | 1 | 0 |
| ENSG00000188316 | 16 | 5 | 13 | 8 | 6 | 9 |
| ENSG00000187164 | 3273 | 1693 | 1901 | 1211 | 2016 | 1761 |
| ENSG00000235742 | 0 | 0 | 2 | 0 | 0 | 0 |
| ENSG00000148704 | 2 | 0 | 0 | 0 | 0 | 1 |
| ENSG00000234474 | 0 | 0 | 0 | 0 | 0 | 0 |
| ENSG00000266782 | 0 | 0 | 0 | 0 | 0 | 0 |
| ENSG00000236171 | 0 | 0 | 0 | 0 | 0 | 0 |
| ENSG00000186795 | 0 | 0 | 0 | 0 | 0 | 0 |
| ENSG00000225936 | 0 | 1 | 1 | 1 | 0 | 0 |
| ENSG00000165646 | 12 | 3 | 6 | 2 | 9 | 7 |
| ENSG00000277879 | 35 | 30 | 22 | 22 | 21 | 19 |
| ENSG00000165650 | 762 | 367 | 374 | 227 | 404 | 400 |
| ENSG00000258114 | 0 | 0 | 0 | 0 | 0 | 0 |
| ENSG00000229847 | 0 | 0 | 0 | 0 | 0 | 0 |
| ENSG00000170370 | 0 | 0 | 0 | 0 | 0 | 0 |
| ENSG00000235198 | 0 | 0 | 0 | 0 | 0 | 0 |
| ENSG00000234952 | 0 | 0 | 0 | 0 | 0 | 0 |
| ENSG00000263041 | 0 | 0 | 0 | 0 | 0 | 0 |
| ENSG00000107560 | 252 | 118 | 113 | 60 | 161 | 115 |
| ENSG00000231104 | 1 | 1 | 0 | 0 | 0 | 0 |
| ENSG00000177640 | 40 | 20 | 22 | 21 | 19 | 25 |
| ENSG00000238276 | 0 | 0 | 0 | 0 | 1 | 0 |
| ENSG00000165669 | 1113 | 515 | 619 | 477 | 605 | 596 |
| ENSG00000232139 | 0 | 1 | 0 | 1 | 0 | 2 |
| ENSG00000234309 | 0 | 0 | 0 | 0 | 0 | 0 |
| ENSG00000119973 | 0 | 0 | 0 | 0 | 0 | 1 |
| ENSG00000225155 | 0 | 0 | 0 | 0 | 0 | 0 |
| ENSG00000151893 | 3593 | 1764 | 2082 | 1342 | 2052 | 1918 |
| ENSG00000277687 | 6 | 4 | 9 | 4 | 5 | 4 |
| ENSG00000236058 | 0 | 1 | 0 | 1 | 0 | 1 |
| ENSG00000213574 | 0 | 1 | 1 | 0 | 1 | 1 |
| ENSG00000229272 | 0 | 0 | 1 | 0 | 0 | 1 |
| ENSG00000188613 | 207 | 115 | 95 | 60 | 90 | 82 |
| ENSG00000107581 | 7220 | 3992 | 3618 | 1906 | 5631 | 3392 |
| ENSG00000207468 | 0 | 0 | 0 | 0 | 0 | 0 |
| ENSG00000222588 | 0 | 0 | 0 | 0 | 1 | 1 |
| ENSG00000271343 | 0 | 0 | 0 | 0 | 0 | 0 |
| ENSG00000119979 | 1066 | 661 | 696 | 490 | 718 | 586 |
| ENSG00000183605 | 889 | 444 | 533 | 607 | 572 | 625 |

| | | | | | | |
|---|---|---|---|---|---|---|
| ENSG00000165672 | 6779 | 3352 | 4066 | 3122 | 3538 | 3463 |
| ENSG00000198873 | 70 | 36 | 30 | 46 | 52 | 35 |
| ENSG00000228485 | 0 | 1 | 0 | 0 | 0 | 1 |
| ENSG00000242853 | 0 | 0 | 0 | 0 | 0 | 1 |
| ENSG00000236426 | 2 | 1 | 0 | 0 | 0 | 0 |
| ENSG00000265719 | 0 | 0 | 0 | 0 | 0 | 0 |
| ENSG00000148908 | 827 | 474 | 568 | 569 | 507 | 463 |
| ENSG00000271353 | 0 | 0 | 0 | 0 | 0 | 0 |
| ENSG00000151923 | 4468 | 2118 | 2712 | 1887 | 2399 | 2435 |
| ENSG00000234569 | 0 | 0 | 0 | 0 | 0 | 0 |
| ENSG00000227437 | 0 | 0 | 0 | 0 | 0 | 0 |
| ENSG00000151929 | 10599 | 5074 | 6870 | 4316 | 4262 | 4819 |
| ENSG00000232823 | 0 | 2 | 0 | 0 | 0 | 0 |
| ENSG00000198825 | 512 | 282 | 369 | 153 | 366 | 280 |
| ENSG00000242818 | 0 | 0 | 0 | 0 | 0 | 0 |
| ENSG00000270300 | 6 | 1 | 3 | 1 | 4 | 2 |
| ENSG00000282413 | 0 | 0 | 0 | 0 | 0 | 0 |
| ENSG00000197771 | 2526 | 1344 | 1535 | 1199 | 1400 | 1594 |
| ENSG00000107651 | 2223 | 1190 | 1248 | 740 | 1387 | 1169 |
| ENSG00000265370 | 0 | 0 | 0 | 0 | 0 | 0 |
| ENSG00000180230 | 0 | 0 | 0 | 0 | 0 | 0 |
| ENSG00000223455 | 0 | 0 | 0 | 0 | 0 | 0 |
| ENSG00000220842 | 8 | 5 | 3 | 2 | 8 | 6 |
| ENSG00000234522 | 0 | 0 | 0 | 0 | 0 | 0 |
| ENSG00000203805 | 0 | 0 | 0 | 0 | 0 | 0 |
| ENSG00000177234 | 0 | 0 | 0 | 0 | 0 | 0 |
| ENSG00000227165 | 0 | 0 | 0 | 0 | 0 | 0 |
| ENSG00000274461 | 0 | 0 | 0 | 0 | 0 | 0 |
| ENSG00000120008 | 2045 | 973 | 1292 | 636 | 1246 | 1102 |
| ENSG00000271670 | 1 | 0 | 0 | 0 | 0 | 0 |
| ENSG00000227307 | 0 | 0 | 0 | 0 | 0 | 0 |
| ENSG00000278484 | 20 | 5 | 5 | 3 | 4 | 6 |
| ENSG00000236762 | 0 | 1 | 0 | 1 | 0 | 0 |
| ENSG00000227143 | 0 | 0 | 0 | 0 | 0 | 1 |
| ENSG00000222979 | 0 | 0 | 0 | 0 | 0 | 0 |
| ENSG00000066468 | 258 | 154 | 191 | 101 | 114 | 123 |
| ENSG00000223432 | 0 | 0 | 0 | 0 | 0 | 0 |
| ENSG00000223540 | 0 | 0 | 0 | 0 | 0 | 0 |
| ENSG00000224250 | 0 | 0 | 0 | 0 | 0 | 0 |
| ENSG00000107669 | 1041 | 485 | 584 | 338 | 581 | 542 |
| ENSG00000201884 | 0 | 0 | 0 | 0 | 0 | 0 |
| ENSG00000226864 | 17 | 12 | 5 | 3 | 16 | 3 |
| ENSG00000273767 | 0 | 0 | 0 | 0 | 0 | 0 |
| ENSG00000276742 | 4 | 1 | 1 | 3 | 4 | 1 |
| ENSG00000107672 | 1253 | 652 | 795 | 637 | 733 | 771 |
| ENSG00000273891 | 34 | 25 | 32 | 14 | 18 | 14 |
| ENSG00000138162 | 1506 | 746 | 979 | 843 | 824 | 932 |
| ENSG00000285715 | 0 | 0 | 0 | 0 | 0 | 0 |
| ENSG00000285973 | 0 | 1 | 0 | 0 | 0 | 0 |
| ENSG00000138152 | 0 | 0 | 0 | 0 | 2 | 0 |

| | | | | | | |
|---|---|---|---|---|---|---|
| ENSG00000202245 | 0 | 0 | 0 | 0 | 0 | 0 |
| ENSG00000107679 | 1105 | 491 | 547 | 342 | 537 | 497 |
| ENSG00000265442 | 0 | 2 | 0 | 0 | 0 | 0 |
| ENSG00000285955 | 0 | 0 | 3 | 0 | 0 | 2 |
| ENSG00000254636 | 0 | 1 | 0 | 0 | 0 | 0 |
| ENSG00000166033 | 544 | 316 | 294 | 301 | 297 | 242 |
| ENSG00000187908 | 0 | 0 | 0 | 0 | 0 | 0 |
| ENSG00000183559 | 0 | 0 | 0 | 0 | 0 | 0 |
| ENSG00000176584 | 0 | 0 | 0 | 0 | 0 | 0 |
| ENSG00000138161 | 82 | 43 | 44 | 33 | 51 | 36 |
| ENSG00000213185 | 45 | 11 | 32 | 32 | 32 | 34 |
| ENSG00000255624 | 42 | 29 | 29 | 21 | 25 | 50 |
| ENSG00000203795 | 0 | 0 | 0 | 0 | 0 | 0 |
| ENSG00000119965 | 349 | 208 | 244 | 219 | 244 | 254 |
| ENSG00000179988 | 171 | 88 | 117 | 95 | 82 | 129 |
| ENSG00000095574 | 505 | 264 | 284 | 191 | 307 | 252 |
| ENSG00000196177 | 1102 | 529 | 730 | 308 | 586 | 523 |
| ENSG00000188620 | 0 | 0 | 0 | 0 | 0 | 0 |
| ENSG00000188816 | 0 | 0 | 0 | 0 | 0 | 0 |
| ENSG00000154473 | 6302 | 3241 | 4159 | 3109 | 3754 | 3898 |
| ENSG00000227586 | 0 | 0 | 1 | 0 | 0 | 0 |
| ENSG00000270902 | 0 | 0 | 0 | 0 | 0 | 0 |
| ENSG00000230131 | 0 | 1 | 2 | 0 | 2 | 0 |
| ENSG00000225152 | 0 | 0 | 0 | 0 | 0 | 0 |
| ENSG00000234542 | 0 | 0 | 0 | 0 | 0 | 0 |
| ENSG00000154478 | 0 | 0 | 0 | 0 | 0 | 0 |
| ENSG00000121898 | 2 | 1 | 2 | 2 | 1 | 0 |
| ENSG00000231138 | 0 | 0 | 0 | 0 | 0 | 0 |
| ENSG00000234677 | 0 | 0 | 0 | 0 | 0 | 0 |
| ENSG00000213438 | 0 | 0 | 0 | 0 | 0 | 0 |
| ENSG00000232334 | 3 | 2 | 1 | 1 | 2 | 0 |
| ENSG00000182022 | 2974 | 1431 | 1964 | 1258 | 1456 | 1615 |
| ENSG00000065154 | 2125 | 1179 | 1364 | 1187 | 1356 | 1342 |
| ENSG00000229544 | 0 | 0 | 0 | 0 | 0 | 1 |
| ENSG00000271434 | 0 | 0 | 0 | 0 | 0 | 0 |
| ENSG00000107902 | 575 | 268 | 322 | 381 | 291 | 329 |
| ENSG00000229455 | 2 | 1 | 0 | 0 | 0 | 2 |
| ENSG00000258539 | 1 | 7 | 3 | 3 | 9 | 2 |
| ENSG00000189319 | 2931 | 1319 | 1668 | 1399 | 1332 | 1373 |
| ENSG00000278831 | 6 | 1 | 5 | 2 | 4 | 4 |
| ENSG00000233334 | 0 | 1 | 0 | 1 | 0 | 0 |
| ENSG00000203791 | 164 | 156 | 141 | 140 | 194 | 174 |
| ENSG00000199466 | 0 | 0 | 1 | 2 | 1 | 0 |
| ENSG00000165660 | 785 | 421 | 484 | 346 | 526 | 427 |
| ENSG00000207494 | 0 | 1 | 4 | 0 | 2 | 1 |
| ENSG00000214807 | 0 | 0 | 0 | 0 | 0 | 0 |
| ENSG00000249456 | 8 | 7 | 2 | 1 | 7 | 13 |
| ENSG00000019995 | 718 | 368 | 456 | 309 | 449 | 417 |
| ENSG00000226899 | 0 | 0 | 0 | 0 | 0 | 0 |
| ENSG00000175029 | 3224 | 1661 | 2067 | 1452 | 1597 | 1753 |

| | | | | | | |
|---|---|---|---|---|---|---|
| ENSG00000273599 | 132 | 58 | 94 | 37 | 56 | 55 |
| ENSG00000264572 | 0 | 1 | 0 | 0 | 0 | 0 |
| ENSG00000282787 | 13 | 11 | 7 | 4 | 4 | 7 |
| ENSG00000214298 | 0 | 0 | 1 | 0 | 1 | 0 |
| ENSG00000233135 | 0 | 0 | 0 | 0 | 0 | 0 |
| ENSG00000237675 | 0 | 0 | 0 | 0 | 0 | 0 |
| ENSG00000175018 | 0 | 0 | 0 | 0 | 1 | 0 |
| ENSG00000214297 | 2 | 0 | 0 | 0 | 1 | 0 |
| ENSG00000228021 | 6 | 0 | 2 | 2 | 2 | 0 |
| ENSG00000224023 | 21 | 9 | 8 | 10 | 11 | 5 |
| ENSG00000280561 | 0 | 0 | 0 | 0 | 0 | 0 |
| ENSG00000234134 | 10 | 7 | 7 | 3 | 1 | 6 |
| ENSG00000107938 | 773 | 395 | 467 | 222 | 493 | 388 |
| ENSG00000236991 | 5 | 13 | 12 | 6 | 5 | 7 |
| ENSG00000154485 | 3 | 0 | 2 | 0 | 1 | 0 |
| ENSG00000188690 | 1921 | 969 | 1187 | 1231 | 1207 | 1147 |
| ENSG00000265092 | 0 | 0 | 0 | 0 | 0 | 0 |
| ENSG00000107949 | 1624 | 977 | 1059 | 895 | 1165 | 1047 |
| ENSG00000089876 | 1689 | 876 | 1073 | 991 | 1228 | 1104 |
| ENSG00000277371 | 0 | 4 | 3 | 2 | 1 | 1 |
| ENSG00000222629 | 0 | 1 | 0 | 0 | 0 | 0 |
| ENSG00000203780 | 258 | 129 | 182 | 187 | 211 | 172 |
| ENSG00000234016 | 0 | 0 | 0 | 0 | 0 | 0 |
| ENSG00000233409 | 0 | 0 | 0 | 0 | 0 | 0 |
| ENSG00000148848 | 0 | 6 | 1 | 0 | 1 | 2 |
| ENSG00000222740 | 0 | 0 | 0 | 0 | 0 | 0 |
| ENSG00000213170 | 0 | 0 | 0 | 0 | 0 | 0 |
| ENSG00000235180 | 0 | 0 | 0 | 0 | 0 | 0 |
| ENSG00000154493 | 0 | 0 | 1 | 0 | 1 | 5 |
| ENSG00000199321 | 0 | 0 | 0 | 0 | 0 | 0 |
| ENSG00000150760 | 1015 | 454 | 600 | 283 | 441 | 479 |
| ENSG00000223528 | 1 | 0 | 0 | 0 | 0 | 0 |
| ENSG00000232935 | 0 | 0 | 0 | 0 | 0 | 0 |
| ENSG00000188916 | 26 | 15 | 8 | 6 | 7 | 10 |
| ENSG00000214285 | 0 | 0 | 0 | 0 | 0 | 0 |
| ENSG00000279725 | 0 | 0 | 0 | 0 | 0 | 0 |
| ENSG00000186766 | 0 | 0 | 0 | 0 | 0 | 0 |
| ENSG00000270778 | 0 | 0 | 0 | 0 | 0 | 0 |
| ENSG00000180745 | 6 | 1 | 3 | 5 | 1 | 4 |
| ENSG00000132334 | 635 | 320 | 404 | 274 | 346 | 383 |
| ENSG00000232259 | 10 | 11 | 7 | 3 | 6 | 3 |
| ENSG00000227076 | 9 | 7 | 13 | 7 | 9 | 8 |
| ENSG00000148773 | 982 | 728 | 551 | 285 | 1467 | 846 |
| ENSG00000280953 | 0 | 0 | 0 | 0 | 0 | 0 |
| ENSG00000234640 | 0 | 0 | 0 | 0 | 0 | 0 |
| ENSG00000224190 | 0 | 0 | 0 | 0 | 0 | 0 |
| ENSG00000232985 | 0 | 0 | 0 | 0 | 0 | 0 |
| ENSG00000225424 | 0 | 0 | 0 | 0 | 0 | 0 |
| ENSG00000285615 | 0 | 0 | 0 | 0 | 0 | 0 |
| ENSG00000170430 | 1576 | 769 | 902 | 1053 | 745 | 790 |

| | | | | | | |
|---|---|---|---|---|---|---|
| ENSG00000237224 | 0 | 0 | 0 | 0 | 0 | 0 |
| ENSG00000227374 | 0 | 0 | 0 | 0 | 0 | 1 |
| ENSG00000283141 | 3 | 4 | 0 | 0 | 2 | 1 |
| ENSG00000108001 | 10 | 7 | 4 | 0 | 15 | 6 |
| ENSG00000275327 | 0 | 0 | 0 | 0 | 0 | 0 |
| ENSG00000266676 | 0 | 0 | 0 | 0 | 0 | 0 |
| ENSG00000275005 | 2 | 1 | 0 | 0 | 1 | 0 |
| ENSG00000237489 | 167 | 75 | 99 | 89 | 92 | 82 |
| ENSG00000277218 | 19 | 7 | 6 | 9 | 7 | 2 |
| ENSG00000233122 | 46 | 15 | 36 | 21 | 17 | 14 |
| ENSG00000233319 | 0 | 0 | 0 | 0 | 0 | 0 |
| ENSG00000108010 | 3128 | 1507 | 1787 | 1678 | 1950 | 1803 |
| ENSG00000236303 | 1 | 3 | 2 | 4 | 1 | 1 |
| ENSG00000276375 | 0 | 0 | 0 | 0 | 1 | 0 |
| ENSG00000222997 | 0 | 0 | 0 | 0 | 0 | 0 |
| ENSG00000285529 | 0 | 0 | 0 | 0 | 0 | 0 |
| ENSG00000264803 | 0 | 0 | 0 | 0 | 0 | 0 |
| ENSG00000176769 | 0 | 0 | 0 | 0 | 0 | 0 |
| ENSG00000230098 | 0 | 1 | 0 | 1 | 4 | 1 |
| ENSG00000189275 | 0 | 0 | 2 | 3 | 1 | 1 |
| ENSG00000280446 | 0 | 0 | 0 | 0 | 0 | 0 |
| ENSG00000283095 | 0 | 0 | 0 | 0 | 0 | 0 |
| ENSG00000175470 | 2265 | 1088 | 1256 | 1143 | 1220 | 1177 |
| ENSG00000176171 | 2000 | 996 | 1185 | 1110 | 1214 | 1136 |
| ENSG00000277959 | 21 | 8 | 13 | 5 | 2 | 11 |
| ENSG00000273521 | 2 | 2 | 3 | 2 | 3 | 7 |
| ENSG00000279982 | 10 | 16 | 4 | 3 | 8 | 4 |
| ENSG00000188385 | 55 | 27 | 29 | 14 | 40 | 34 |
| ENSG00000235010 | 1 | 0 | 0 | 0 | 2 | 1 |
| ENSG00000151640 | 74 | 58 | 78 | 77 | 72 | 44 |
| ENSG00000165752 | 414 | 187 | 274 | 301 | 274 | 281 |
| ENSG00000148814 | 443 | 197 | 305 | 270 | 257 | 257 |
| ENSG00000171813 | 1898 | 879 | 1203 | 1072 | 967 | 964 |
| ENSG00000231705 | 4 | 1 | 4 | 0 | 1 | 2 |
| ENSG00000234311 | 2 | 1 | 0 | 2 | 2 | 5 |
| ENSG00000180066 | 8 | 1 | 2 | 4 | 3 | 3 |
| ENSG00000226900 | 4 | 4 | 6 | 1 | 2 | 1 |
| ENSG00000229081 | 2 | 8 | 10 | 1 | 8 | 3 |
| ENSG00000068383 | 1436 | 694 | 876 | 733 | 643 | 696 |
| ENSG00000148826 | 0 | 0 | 0 | 0 | 0 | 0 |
| ENSG00000171811 | 30 | 9 | 18 | 10 | 11 | 16 |
| ENSG00000232903 | 63 | 37 | 43 | 22 | 38 | 34 |
| ENSG00000224758 | 13 | 3 | 8 | 5 | 7 | 8 |
| ENSG00000240707 | 11 | 3 | 9 | 5 | 15 | 5 |
| ENSG00000197177 | 0 | 0 | 1 | 0 | 0 | 0 |
| ENSG00000256925 | 0 | 0 | 0 | 0 | 0 | 0 |
| ENSG00000235264 | 0 | 0 | 0 | 0 | 0 | 0 |
| ENSG00000171798 | 1009 | 443 | 617 | 491 | 531 | 516 |
| ENSG00000171794 | 0 | 0 | 0 | 0 | 0 | 0 |
| ENSG00000151650 | 2 | 1 | 0 | 0 | 0 | 0 |

| | | | | | | |
|---|---|---|---|---|---|---|
| ENSG00000166917 | 0 | 0 | 0 | 0 | 0 | 0 |
| ENSG00000284219 | 0 | 0 | 0 | 0 | 0 | 0 |
| ENSG00000273980 | 0 | 0 | 3 | 1 | 2 | 4 |
| ENSG00000151651 | 264 | 105 | 200 | 110 | 120 | 158 |
| ENSG00000130640 | 3782 | 1885 | 2354 | 2056 | 1929 | 2137 |
| ENSG00000235245 | 72 | 24 | 39 | 13 | 29 | 27 |
| ENSG00000198546 | 383 | 224 | 270 | 329 | 273 | 252 |
| ENSG00000283496 | 0 | 0 | 0 | 0 | 0 | 0 |
| ENSG00000130643 | 11 | 9 | 7 | 3 | 15 | 9 |
| ENSG00000230306 | 1 | 4 | 1 | 3 | 4 | 1 |
| ENSG00000226699 | 0 | 0 | 2 | 1 | 1 | 0 |
| ENSG00000165828 | 7 | 2 | 2 | 2 | 2 | 1 |
| ENSG00000148803 | 137 | 62 | 61 | 72 | 58 | 53 |
| ENSG00000127884 | 8470 | 4318 | 5142 | 5393 | 4708 | 4865 |
| ENSG00000265395 | 0 | 0 | 1 | 0 | 0 | 0 |
| ENSG00000274685 | 0 | 1 | 5 | 1 | 0 | 2 |
| ENSG00000148832 | 161 | 91 | 96 | 103 | 104 | 97 |
| ENSG00000254536 | 0 | 3 | 0 | 0 | 0 | 0 |
| ENSG00000148824 | 266 | 170 | 222 | 207 | 221 | 195 |
| ENSG00000203772 | 198 | 82 | 120 | 111 | 91 | 104 |
| ENSG00000273327 | 0 | 0 | 1 | 0 | 0 | 1 |
| ENSG00000214279 | 62 | 30 | 57 | 14 | 24 | 40 |
| ENSG00000273336 | 2 | 0 | 0 | 0 | 0 | 0 |
| ENSG00000130649 | 22 | 11 | 16 | 16 | 7 | 9 |
| ENSG00000278518 | 0 | 0 | 0 | 0 | 0 | 0 |
| ENSG00000171772 | 6 | 4 | 2 | 3 | 2 | 4 |
| ENSG00000233685 | 0 | 0 | 0 | 0 | 0 | 0 |
| ENSG00000225899 | 0 | 1 | 1 | 1 | 0 | 0 |
| ENSG00000213543 | 0 | 0 | 0 | 0 | 0 | 0 |
| ENSG00000233435 | 0 | 0 | 0 | 0 | 0 | 0 |
| ENSG00000230468 | 0 | 0 | 0 | 0 | 0 | 0 |
| ENSG00000279626 | 0 | 0 | 0 | 0 | 0 | 0 |
| ENSG00000280337 | 0 | 0 | 0 | 0 | 0 | 0 |
| ENSG00000274599 | 0 | 0 | 0 | 0 | 0 | 0 |
| ENSG00000276964 | 0 | 0 | 0 | 0 | 0 | 0 |
| ENSG00000277274 | 0 | 0 | 0 | 0 | 0 | 0 |
| ENSG00000275610 | 0 | 0 | 0 | 0 | 0 | 0 |
| ENSG00000276164 | 0 | 0 | 0 | 0 | 0 | 0 |
| ENSG00000237635 | 0 | 0 | 0 | 0 | 0 | 0 |
| ENSG00000278664 | 0 | 0 | 0 | 0 | 0 | 0 |
| ENSG00000278761 | 0 | 0 | 0 | 0 | 0 | 0 |
| ENSG00000278790 | 0 | 0 | 0 | 0 | 0 | 0 |
| ENSG00000276046 | 0 | 0 | 0 | 0 | 0 | 0 |
| ENSG00000276904 | 0 | 0 | 0 | 0 | 0 | 0 |
| ENSG00000278641 | 0 | 0 | 0 | 0 | 0 | 0 |
| ENSG00000213147 | 0 | 0 | 0 | 0 | 0 | 0 |
| ENSG00000282875 | 0 | 0 | 0 | 0 | 0 | 0 |
| ENSG00000256263 | 1 | 2 | 0 | 0 | 1 | 1 |
| ENSG00000226210 | 560 | 308 | 356 | 300 | 301 | 340 |
| ENSG00000249054 | 0 | 0 | 0 | 0 | 0 | 0 |

| | | | | | | |
|---|---|---|---|---|---|---|
| ENSG00000120645 | 0 | 0 | 0 | 0 | 1 | 0 |
| ENSG00000256948 | 0 | 0 | 0 | 0 | 0 | 0 |
| ENSG00000249695 | 0 | 0 | 0 | 0 | 0 | 0 |
| ENSG00000256694 | 0 | 0 | 0 | 0 | 0 | 0 |
| ENSG00000256540 | 0 | 0 | 0 | 0 | 0 | 0 |
| ENSG00000111181 | 6 | 2 | 2 | 3 | 3 | 0 |
| ENSG00000255671 | 0 | 0 | 0 | 0 | 0 | 0 |
| ENSG00000256577 | 2 | 1 | 0 | 0 | 0 | 0 |
| ENSG00000010379 | 0 | 1 | 0 | 1 | 0 | 0 |
| ENSG00000255746 | 0 | 0 | 0 | 0 | 0 | 0 |
| ENSG00000261799 | 51 | 36 | 35 | 9 | 46 | 33 |
| ENSG00000073614 | 1281 | 649 | 852 | 424 | 871 | 666 |
| ENSG00000120647 | 109 | 88 | 99 | 71 | 81 | 67 |
| ENSG00000139044 | 0 | 0 | 2 | 4 | 1 | 2 |
| ENSG00000171840 | 0 | 0 | 0 | 0 | 0 | 0 |
| ENSG00000256020 | 0 | 0 | 0 | 0 | 0 | 0 |
| ENSG00000238370 | 0 | 0 | 0 | 0 | 0 | 0 |
| ENSG00000255825 | 0 | 0 | 0 | 0 | 0 | 0 |
| ENSG00000177406 | 129 | 92 | 75 | 87 | 91 | 75 |
| ENSG00000256672 | 0 | 0 | 0 | 0 | 0 | 0 |
| ENSG00000060237 | 3891 | 1672 | 1864 | 953 | 2066 | 1904 |
| ENSG00000221439 | 0 | 0 | 0 | 0 | 0 | 0 |
| ENSG00000285704 | 0 | 0 | 0 | 0 | 0 | 0 |
| ENSG00000002016 | 273 | 132 | 179 | 123 | 120 | 128 |
| ENSG00000250132 | 92 | 37 | 66 | 48 | 64 | 39 |
| ENSG00000082805 | 847 | 365 | 471 | 229 | 475 | 414 |
| ENSG00000275811 | 0 | 0 | 0 | 0 | 0 | 0 |
| ENSG00000249028 | 0 | 2 | 0 | 0 | 1 | 0 |
| ENSG00000267285 | 0 | 2 | 0 | 0 | 0 | 0 |
| ENSG00000249628 | 4 | 1 | 1 | 2 | 0 | 0 |
| ENSG00000111186 | 12 | 12 | 11 | 5 | 2 | 2 |
| ENSG00000171823 | 374 | 192 | 217 | 181 | 207 | 193 |
| ENSG00000266043 | 0 | 0 | 0 | 0 | 0 | 0 |
| ENSG00000006831 | 8666 | 4327 | 5541 | 2695 | 3132 | 3268 |
| ENSG00000271500 | 0 | 2 | 4 | 0 | 1 | 1 |
| ENSG00000285627 | 0 | 3 | 1 | 1 | 2 | 0 |
| ENSG00000243663 | 0 | 1 | 0 | 0 | 1 | 1 |
| ENSG00000151062 | 11 | 9 | 9 | 4 | 2 | 8 |
| ENSG00000166159 | 0 | 0 | 0 | 0 | 1 | 1 |
| ENSG00000256706 | 0 | 0 | 0 | 0 | 0 | 0 |
| ENSG00000235049 | 0 | 1 | 0 | 0 | 0 | 0 |
| ENSG00000151065 | 418 | 182 | 220 | 165 | 240 | 198 |
| ENSG00000151067 | 5 | 3 | 2 | 0 | 0 | 4 |
| ENSG00000203593 | 1 | 0 | 0 | 1 | 3 | 1 |
| ENSG00000256837 | 0 | 0 | 0 | 0 | 0 | 0 |
| ENSG00000256257 | 0 | 0 | 0 | 0 | 0 | 0 |
| ENSG00000278255 | 0 | 0 | 0 | 0 | 0 | 0 |
| ENSG00000256025 | 0 | 0 | 0 | 0 | 0 | 0 |
| ENSG00000256721 | 0 | 0 | 0 | 0 | 0 | 0 |
| ENSG00000285555 | 0 | 0 | 0 | 0 | 0 | 0 |

| | | | | | | |
|---|---|---|---|---|---|---|
| ENSG00000285734 | 0 | 0 | 0 | 0 | 0 | 0 |
| ENSG00000256769 | 0 | 0 | 0 | 0 | 0 | 0 |
| ENSG00000203648 | 0 | 0 | 0 | 0 | 0 | 0 |
| ENSG00000256271 | 0 | 0 | 0 | 0 | 0 | 0 |
| ENSG00000246627 | 4 | 0 | 0 | 0 | 0 | 1 |
| ENSG00000256150 | 0 | 0 | 0 | 2 | 0 | 0 |
| ENSG00000283587 | 0 | 0 | 0 | 0 | 0 | 0 |
| ENSG00000274659 | 0 | 1 | 0 | 0 | 0 | 0 |
| ENSG00000256902 | 0 | 0 | 0 | 0 | 0 | 2 |
| ENSG00000255669 | 0 | 0 | 1 | 1 | 2 | 0 |
| ENSG00000224438 | 0 | 0 | 0 | 0 | 0 | 0 |
| ENSG00000256030 | 3 | 1 | 3 | 1 | 0 | 0 |
| ENSG00000004478 | 10342 | 5627 | 6128 | 7076 | 6404 | 6172 |
| ENSG00000258325 | 0 | 1 | 0 | 1 | 0 | 1 |
| ENSG00000258092 | 0 | 0 | 1 | 1 | 0 | 1 |
| ENSG00000111203 | 480 | 244 | 338 | 246 | 260 | 283 |
| ENSG00000053702 | 37 | 34 | 16 | 18 | 15 | 40 |
| ENSG00000283297 | 6 | 2 | 5 | 1 | 7 | 2 |
| ENSG00000111206 | 1088 | 534 | 709 | 736 | 661 | 729 |
| ENSG00000171792 | 905 | 455 | 535 | 449 | 490 | 570 |
| ENSG00000078246 | 951 | 430 | 549 | 363 | 468 | 496 |
| ENSG00000278356 | 11 | 4 | 7 | 3 | 9 | 7 |
| ENSG00000252996 | 0 | 0 | 1 | 0 | 0 | 0 |
| ENSG00000197905 | 278 | 153 | 170 | 227 | 210 | 185 |
| ENSG00000250899 | 82 | 33 | 34 | 46 | 51 | 57 |
| ENSG00000234589 | 0 | 1 | 0 | 0 | 0 | 1 |
| ENSG00000011105 | 1204 | 561 | 770 | 542 | 477 | 530 |
| ENSG00000278469 | 0 | 0 | 0 | 0 | 0 | 0 |
| ENSG00000256197 | 0 | 0 | 0 | 0 | 0 | 0 |
| ENSG00000275850 | 0 | 0 | 0 | 0 | 0 | 0 |
| ENSG00000227081 | 48 | 18 | 27 | 23 | 23 | 16 |
| ENSG00000250770 | 2 | 8 | 3 | 8 | 17 | 6 |
| ENSG00000236908 | 8 | 5 | 5 | 0 | 7 | 1 |
| ENSG00000257048 | 0 | 0 | 3 | 1 | 0 | 0 |
| ENSG00000111218 | 0 | 0 | 0 | 0 | 0 | 0 |
| ENSG00000256691 | 0 | 0 | 0 | 0 | 0 | 0 |
| ENSG00000243507 | 0 | 0 | 0 | 0 | 0 | 0 |
| ENSG00000130038 | 289 | 144 | 184 | 100 | 100 | 144 |
| ENSG00000280202 | 79 | 30 | 58 | 34 | 42 | 47 |
| ENSG00000222338 | 0 | 0 | 0 | 0 | 0 | 0 |
| ENSG00000283138 | 0 | 1 | 0 | 0 | 0 | 0 |
| ENSG00000111224 | 305 | 156 | 194 | 101 | 170 | 137 |
| ENSG00000118976 | 0 | 0 | 0 | 1 | 0 | 0 |
| ENSG00000256862 | 3 | 1 | 6 | 1 | 2 | 3 |
| ENSG00000256969 | 0 | 0 | 0 | 0 | 0 | 0 |
| ENSG00000256356 | 4 | 0 | 4 | 0 | 1 | 1 |
| ENSG00000242444 | 0 | 0 | 0 | 0 | 0 | 1 |
| ENSG00000255920 | 0 | 0 | 0 | 0 | 0 | 0 |
| ENSG00000285901 | 0 | 0 | 0 | 0 | 0 | 0 |
| ENSG00000118971 | 0 | 0 | 0 | 0 | 0 | 1 |

| | | | | | | |
|---|---|---|---|---|---|---|
| ENSG00000078237 | 18 | 19 | 15 | 17 | 21 | 22 |
| ENSG00000213970 | 0 | 5 | 1 | 1 | 0 | 0 |
| ENSG00000118972 | 0 | 0 | 0 | 0 | 0 | 0 |
| ENSG00000111241 | 0 | 0 | 0 | 0 | 0 | 0 |
| ENSG00000047621 | 197 | 122 | 114 | 59 | 132 | 112 |
| ENSG00000111247 | 146 | 89 | 91 | 62 | 97 | 93 |
| ENSG00000010219 | 139 | 83 | 96 | 85 | 77 | 93 |
| ENSG00000256748 | 0 | 0 | 0 | 0 | 0 | 0 |
| ENSG00000272921 | 0 | 0 | 0 | 0 | 0 | 0 |
| ENSG00000111254 | 18 | 5 | 9 | 3 | 4 | 8 |
| ENSG00000256799 | 2 | 0 | 0 | 0 | 0 | 2 |
| ENSG00000256381 | 0 | 1 | 1 | 0 | 0 | 0 |
| ENSG00000139180 | 257 | 186 | 252 | 262 | 227 | 222 |
| ENSG00000255639 | 0 | 0 | 0 | 0 | 0 | 0 |
| ENSG00000255474 | 0 | 0 | 0 | 0 | 0 | 0 |
| ENSG00000130035 | 0 | 0 | 0 | 0 | 0 | 0 |
| ENSG00000256988 | 0 | 0 | 0 | 0 | 0 | 0 |
| ENSG00000151079 | 0 | 0 | 0 | 0 | 0 | 0 |
| ENSG00000255608 | 0 | 0 | 0 | 0 | 0 | 0 |
| ENSG00000256654 | 0 | 0 | 0 | 0 | 0 | 0 |
| ENSG00000111262 | 0 | 0 | 0 | 0 | 0 | 0 |
| ENSG00000284513 | 0 | 0 | 0 | 0 | 0 | 0 |
| ENSG00000255996 | 0 | 0 | 0 | 0 | 0 | 0 |
| ENSG00000130037 | 0 | 0 | 0 | 0 | 0 | 0 |
| ENSG00000285523 | 0 | 0 | 0 | 0 | 2 | 0 |
| ENSG00000256115 | 17 | 7 | 7 | 1 | 3 | 5 |
| ENSG00000256417 | 132 | 55 | 75 | 40 | 59 | 45 |
| ENSG00000256146 | 0 | 1 | 0 | 0 | 0 | 0 |
| ENSG00000256218 | 0 | 0 | 0 | 0 | 0 | 0 |
| ENSG00000255983 | 60 | 15 | 22 | 12 | 12 | 18 |
| ENSG00000185652 | 0 | 0 | 0 | 0 | 0 | 1 |
| ENSG00000047617 | 0 | 2 | 2 | 1 | 1 | 0 |
| ENSG00000255973 | 0 | 0 | 0 | 0 | 0 | 0 |
| ENSG00000110799 | 16 | 3 | 3 | 1 | 3 | 1 |
| ENSG00000240533 | 0 | 0 | 0 | 0 | 0 | 0 |
| ENSG00000255775 | 0 | 0 | 0 | 0 | 0 | 0 |
| ENSG00000010278 | 15175 | 7264 | 8619 | 6196 | 6377 | 6279 |
| ENSG00000202318 | 0 | 2 | 3 | 2 | 2 | 4 |
| ENSG00000256103 | 0 | 0 | 0 | 1 | 0 | 0 |
| ENSG00000008323 | 756 | 352 | 453 | 496 | 457 | 406 |
| ENSG00000067182 | 6295 | 3116 | 3861 | 3979 | 3773 | 3672 |
| ENSG00000265388 | 0 | 0 | 0 | 0 | 0 | 0 |
| ENSG00000111319 | 4631 | 1915 | 2562 | 1318 | 1327 | 1717 |
| ENSG00000111321 | 8706 | 3985 | 5380 | 5015 | 4583 | 5126 |
| ENSG00000256433 | 35 | 17 | 37 | 12 | 28 | 32 |
| ENSG00000213942 | 2 | 3 | 3 | 1 | 1 | 2 |
| ENSG00000224906 | 0 | 0 | 0 | 0 | 1 | 0 |
| ENSG00000215039 | 689 | 342 | 537 | 335 | 393 | 434 |
| ENSG00000139193 | 16 | 6 | 8 | 2 | 8 | 2 |
| ENSG00000139192 | 496 | 219 | 340 | 244 | 239 | 250 |

| | | | | | | |
|---|---|---|---|---|---|---|
| ENSG00000139190 | 412 | 158 | 265 | 125 | 165 | 201 |
| ENSG00000276718 | 7 | 3 | 12 | 2 | 6 | 9 |
| ENSG00000256913 | 0 | 0 | 1 | 0 | 0 | 0 |
| ENSG00000111639 | 2968 | 1572 | 1846 | 2091 | 2031 | 1969 |
| ENSG00000010292 | 3654 | 1641 | 2100 | 1498 | 2245 | 1976 |
| ENSG00000239002 | 0 | 0 | 0 | 0 | 0 | 0 |
| ENSG00000276232 | 0 | 0 | 0 | 0 | 0 | 0 |
| ENSG00000255966 | 11 | 2 | 5 | 3 | 2 | 0 |
| ENSG00000111640 | 71064 | 38844 | 46598 | 50046 | 45494 | 44959 |
| ENSG00000269968 | 0 | 0 | 0 | 0 | 0 | 0 |
| ENSG00000010295 | 8 | 2 | 1 | 6 | 3 | 6 |
| ENSG00000245667 | 0 | 0 | 0 | 0 | 0 | 1 |
| ENSG00000111641 | 26 | 20 | 24 | 35 | 22 | 28 |
| ENSG00000285238 | 0 | 0 | 0 | 0 | 0 | 0 |
| ENSG00000111642 | 1124 | 627 | 675 | 676 | 685 | 647 |
| ENSG00000247853 | 12 | 5 | 11 | 4 | 7 | 9 |
| ENSG00000251898 | 2 | 0 | 1 | 0 | 0 | 1 |
| ENSG00000184574 | 22 | 18 | 22 | 17 | 20 | 14 |
| ENSG00000111644 | 14 | 7 | 8 | 4 | 11 | 6 |
| ENSG00000111653 | 972 | 513 | 647 | 528 | 490 | 520 |
| ENSG00000219410 | 10 | 3 | 2 | 3 | 7 | 5 |
| ENSG00000126746 | 866 | 353 | 533 | 358 | 432 | 484 |
| ENSG00000139200 | 20 | 9 | 16 | 9 | 15 | 15 |
| ENSG00000252186 | 0 | 0 | 0 | 0 | 0 | 0 |
| ENSG00000270068 | 2 | 1 | 0 | 0 | 3 | 0 |
| ENSG00000111652 | 1903 | 948 | 1221 | 1164 | 1025 | 1100 |
| ENSG00000269892 | 3 | 1 | 5 | 2 | 12 | 2 |
| ENSG00000089693 | 9379 | 4520 | 5811 | 5512 | 5147 | 5125 |
| ENSG00000159335 | 3944 | 1724 | 2397 | 2484 | 1825 | 2155 |
| ENSG00000089692 | 14 | 4 | 18 | 9 | 7 | 8 |
| ENSG00000244532 | 0 | 0 | 0 | 0 | 0 | 0 |
| ENSG00000010610 | 8 | 0 | 1 | 0 | 0 | 0 |
| ENSG00000250510 | 4 | 1 | 1 | 1 | 0 | 1 |
| ENSG00000110811 | 6 | 3 | 4 | 7 | 1 | 1 |
| ENSG00000111664 | 2 | 0 | 1 | 1 | 4 | 4 |
| ENSG00000111665 | 514 | 224 | 290 | 312 | 383 | 387 |
| ENSG00000111667 | 4936 | 2268 | 2947 | 3257 | 2613 | 2896 |
| ENSG00000111669 | 20149 | 10193 | 12054 | 13695 | 11521 | 11488 |
| ENSG00000111671 | 489 | 225 | 317 | 273 | 288 | 273 |
| ENSG00000240370 | 69 | 35 | 41 | 27 | 40 | 29 |
| ENSG00000010626 | 201 | 90 | 134 | 129 | 96 | 101 |
| ENSG00000248593 | 138 | 64 | 95 | 55 | 66 | 76 |
| ENSG00000275703 | 0 | 3 | 9 | 2 | 1 | 6 |
| ENSG00000111674 | 649 | 290 | 419 | 373 | 351 | 408 |
| ENSG00000111676 | 4323 | 1876 | 2555 | 2197 | 2104 | 2227 |
| ENSG00000111678 | 614 | 394 | 509 | 544 | 440 | 433 |
| ENSG00000272173 | 21 | 11 | 13 | 12 | 10 | 14 |
| ENSG00000238923 | 0 | 0 | 0 | 0 | 0 | 0 |
| ENSG00000111679 | 2143 | 924 | 1242 | 1151 | 1113 | 1157 |
| ENSG00000257084 | 196 | 95 | 147 | 115 | 110 | 97 |

| | | | | | | |
|---|---|---|---|---|---|---|
| ENSG00000207713 | 1 | 0 | 0 | 0 | 2 | 0 |
| ENSG00000207708 | 6 | 2 | 1 | 0 | 1 | 1 |
| ENSG00000271969 | 6 | 8 | 11 | 4 | 4 | 0 |
| ENSG00000215021 | 6045 | 2719 | 3544 | 3953 | 3200 | 3617 |
| ENSG00000238795 | 0 | 0 | 0 | 1 | 0 | 3 |
| ENSG00000126749 | 1003 | 548 | 592 | 644 | 627 | 655 |
| ENSG00000111684 | 537 | 290 | 349 | 253 | 283 | 297 |
| ENSG00000182326 | 18 | 20 | 20 | 8 | 6 | 10 |
| ENSG00000239701 | 0 | 0 | 0 | 0 | 1 | 0 |
| ENSG00000257078 | 0 | 0 | 0 | 0 | 0 | 0 |
| ENSG00000159403 | 29 | 12 | 16 | 15 | 13 | 15 |
| ENSG00000139178 | 6 | 3 | 6 | 7 | 9 | 4 |
| ENSG00000205885 | 0 | 1 | 3 | 1 | 1 | 1 |
| ENSG00000276144 | 0 | 0 | 0 | 0 | 0 | 0 |
| ENSG00000200345 | 0 | 0 | 0 | 0 | 0 | 0 |
| ENSG00000139194 | 4 | 3 | 4 | 3 | 7 | 9 |
| ENSG00000256967 | 3 | 2 | 1 | 8 | 3 | 3 |
| ENSG00000139182 | 1363 | 606 | 813 | 724 | 773 | 686 |
| ENSG00000285770 | 2 | 0 | 0 | 0 | 1 | 0 |
| ENSG00000255572 | 0 | 0 | 0 | 0 | 0 | 0 |
| ENSG00000139197 | 1458 | 627 | 847 | 737 | 717 | 819 |
| ENSG00000256480 | 0 | 0 | 0 | 0 | 0 | 0 |
| ENSG00000215009 | 2 | 0 | 1 | 0 | 6 | 0 |
| ENSG00000177675 | 52 | 33 | 31 | 16 | 29 | 22 |
| ENSG00000255836 | 127 | 64 | 68 | 36 | 68 | 49 |
| ENSG00000177575 | 0 | 0 | 0 | 0 | 0 | 0 |
| ENSG00000235868 | 0 | 0 | 0 | 0 | 0 | 0 |
| ENSG00000256331 | 0 | 0 | 0 | 0 | 0 | 0 |
| ENSG00000255977 | 0 | 0 | 0 | 0 | 0 | 0 |
| ENSG00000284327 | 0 | 0 | 0 | 0 | 0 | 0 |
| ENSG00000284591 | 0 | 0 | 0 | 0 | 0 | 0 |
| ENSG00000284522 | 0 | 0 | 0 | 0 | 0 | 0 |
| ENSG00000285142 | 0 | 0 | 0 | 0 | 0 | 0 |
| ENSG00000111701 | 0 | 0 | 0 | 0 | 0 | 0 |
| ENSG00000184344 | 0 | 0 | 0 | 0 | 0 | 0 |
| ENSG00000187569 | 0 | 0 | 0 | 0 | 0 | 0 |
| ENSG00000198178 | 0 | 0 | 0 | 0 | 0 | 0 |
| ENSG00000205857 | 0 | 0 | 0 | 0 | 0 | 0 |
| ENSG00000201549 | 0 | 0 | 0 | 0 | 0 | 0 |
| ENSG00000111704 | 0 | 0 | 0 | 0 | 1 | 0 |
| ENSG00000199912 | 0 | 0 | 0 | 0 | 0 | 0 |
| ENSG00000173262 | 0 | 0 | 0 | 0 | 0 | 3 |
| ENSG00000201663 | 0 | 0 | 0 | 0 | 0 | 0 |
| ENSG00000278459 | 0 | 0 | 0 | 0 | 0 | 0 |
| ENSG00000241828 | 0 | 0 | 0 | 0 | 0 | 0 |
| ENSG00000255885 | 0 | 0 | 0 | 0 | 0 | 0 |
| ENSG00000176654 | 0 | 0 | 0 | 0 | 0 | 0 |
| ENSG00000255356 | 0 | 0 | 0 | 0 | 0 | 0 |
| ENSG00000059804 | 0 | 0 | 1 | 0 | 1 | 0 |
| ENSG00000206636 | 0 | 0 | 0 | 0 | 0 | 0 |

| | | | | | | |
|---|---|---|---|---|---|---|
| ENSG00000240739 | 0 | 0 | 0 | 0 | 0 | 0 |
| ENSG00000282080 | 0 | 0 | 0 | 0 | 0 | 0 |
| ENSG00000255964 | 0 | 0 | 0 | 0 | 0 | 0 |
| ENSG00000255581 | 0 | 1 | 0 | 1 | 0 | 0 |
| ENSG00000065970 | 1095 | 425 | 587 | 482 | 520 | 527 |
| ENSG00000171860 | 0 | 0 | 0 | 0 | 1 | 0 |
| ENSG00000089818 | 1551 | 745 | 981 | 732 | 812 | 858 |
| ENSG00000279865 | 10 | 7 | 9 | 4 | 8 | 4 |
| ENSG00000284697 | 0 | 1 | 3 | 0 | 1 | 1 |
| ENSG00000111729 | 0 | 0 | 0 | 0 | 0 | 0 |
| ENSG00000235602 | 2 | 2 | 0 | 2 | 2 | 1 |
| ENSG00000284393 | 0 | 0 | 0 | 0 | 0 | 0 |
| ENSG00000256171 | 0 | 0 | 0 | 0 | 0 | 0 |
| ENSG00000196946 | 0 | 0 | 0 | 0 | 0 | 0 |
| ENSG00000226711 | 112 | 62 | 75 | 48 | 47 | 53 |
| ENSG00000244050 | 4 | 2 | 6 | 1 | 0 | 3 |
| ENSG00000276417 | 3 | 0 | 0 | 0 | 1 | 0 |
| ENSG00000275367 | 8 | 2 | 5 | 1 | 2 | 0 |
| ENSG00000171847 | 11 | 3 | 3 | 3 | 3 | 0 |
| ENSG00000254016 | 4 | 3 | 5 | 1 | 2 | 3 |
| ENSG00000164845 | 112 | 91 | 108 | 47 | 55 | 69 |
| ENSG00000215241 | 41 | 20 | 29 | 7 | 22 | 16 |
| ENSG00000256589 | 4 | 1 | 0 | 0 | 0 | 0 |
| ENSG00000226091 | 5 | 7 | 3 | 1 | 1 | 2 |
| ENSG00000256552 | 5 | 5 | 4 | 0 | 0 | 0 |
| ENSG00000256101 | 0 | 0 | 0 | 0 | 0 | 0 |
| ENSG00000277952 | 2 | 0 | 0 | 0 | 0 | 0 |
| ENSG00000242661 | 0 | 0 | 0 | 1 | 0 | 0 |
| ENSG00000284687 | 0 | 0 | 0 | 0 | 0 | 0 |
| ENSG00000284673 | 0 | 0 | 0 | 0 | 0 | 0 |
| ENSG00000256136 | 0 | 0 | 0 | 0 | 0 | 0 |
| ENSG00000238152 | 0 | 0 | 0 | 0 | 0 | 0 |
| ENSG00000214487 | 0 | 0 | 0 | 0 | 0 | 0 |
| ENSG00000177586 | 0 | 0 | 0 | 0 | 0 | 0 |
| ENSG00000205846 | 0 | 0 | 0 | 0 | 0 | 0 |
| ENSG00000243830 | 1 | 0 | 0 | 0 | 0 | 0 |
| ENSG00000201780 | 0 | 0 | 0 | 0 | 0 | 0 |
| ENSG00000166527 | 0 | 0 | 0 | 0 | 0 | 0 |
| ENSG00000246115 | 0 | 0 | 0 | 0 | 0 | 0 |
| ENSG00000166523 | 0 | 0 | 0 | 0 | 0 | 0 |
| ENSG00000255801 | 0 | 0 | 0 | 0 | 0 | 0 |
| ENSG00000238943 | 0 | 0 | 0 | 0 | 0 | 0 |
| ENSG00000252727 | 0 | 0 | 0 | 0 | 0 | 0 |
| ENSG00000111732 | 0 | 0 | 0 | 1 | 0 | 0 |
| ENSG00000280345 | 0 | 0 | 0 | 0 | 0 | 0 |
| ENSG00000255967 | 0 | 0 | 0 | 1 | 0 | 0 |
| ENSG00000197614 | 0 | 0 | 0 | 0 | 0 | 0 |
| ENSG00000255829 | 0 | 0 | 0 | 0 | 0 | 0 |
| ENSG00000256004 | 0 | 0 | 0 | 0 | 0 | 0 |
| ENSG00000166532 | 878 | 414 | 598 | 324 | 488 | 462 |

| | | | | | | |
|---|---|---|---|---|---|---|
| ENSG00000244131 | 0 | 0 | 0 | 0 | 0 | 0 |
| ENSG00000256661 | 0 | 0 | 0 | 0 | 0 | 0 |
| ENSG00000249790 | 0 | 0 | 3 | 0 | 1 | 0 |
| ENSG00000256904 | 0 | 0 | 0 | 0 | 0 | 0 |
| ENSG00000166535 | 30 | 14 | 24 | 6 | 16 | 7 |
| ENSG00000282022 | 0 | 1 | 1 | 1 | 0 | 0 |
| ENSG00000111752 | 500 | 224 | 361 | 211 | 241 | 297 |
| ENSG00000003056 | 2817 | 1443 | 1797 | 1434 | 1662 | 1780 |
| ENSG00000139187 | 2 | 4 | 0 | 2 | 2 | 2 |
| ENSG00000257105 | 0 | 0 | 0 | 0 | 0 | 0 |
| ENSG00000213049 | 0 | 0 | 0 | 0 | 0 | 0 |
| ENSG00000255776 | 0 | 0 | 0 | 0 | 0 | 0 |
| ENSG00000214851 | 10 | 3 | 11 | 5 | 12 | 10 |
| ENSG00000245105 | 36 | 21 | 19 | 14 | 12 | 23 |
| ENSG00000175899 | 2 | 6 | 1 | 5 | 8 | 1 |
| ENSG00000256937 | 0 | 0 | 1 | 1 | 0 | 0 |
| ENSG00000256720 | 0 | 0 | 0 | 0 | 0 | 0 |
| ENSG00000126838 | 6 | 2 | 3 | 4 | 1 | 1 |
| ENSG00000256817 | 0 | 0 | 0 | 0 | 0 | 0 |
| ENSG00000256069 | 13 | 24 | 17 | 2 | 11 | 14 |
| ENSG00000283429 | 4 | 6 | 3 | 3 | 2 | 4 |
| ENSG00000231503 | 24 | 9 | 19 | 15 | 15 | 22 |
| ENSG00000237248 | 1 | 5 | 2 | 0 | 4 | 3 |
| ENSG00000256427 | 0 | 2 | 2 | 0 | 0 | 0 |
| ENSG00000111788 | 115 | 76 | 103 | 68 | 71 | 99 |
| ENSG00000212440 | 0 | 1 | 0 | 0 | 0 | 1 |
| ENSG00000255753 | 0 | 0 | 0 | 0 | 1 | 0 |
| ENSG00000260423 | 5 | 0 | 6 | 2 | 1 | 3 |
| ENSG00000256673 | 3 | 3 | 5 | 3 | 9 | 9 |
| ENSG00000278635 | 1 | 4 | 3 | 1 | 4 | 0 |
| ENSG00000214826 | 104 | 69 | 92 | 65 | 80 | 85 |
| ENSG00000212432 | 0 | 0 | 0 | 0 | 0 | 1 |
| ENSG00000284634 | 16 | 8 | 14 | 4 | 11 | 13 |
| ENSG00000214776 | 0 | 1 | 0 | 0 | 1 | 0 |
| ENSG00000256975 | 0 | 0 | 0 | 0 | 0 | 0 |
| ENSG00000111796 | 2 | 0 | 0 | 0 | 0 | 0 |
| ENSG00000212345 | 0 | 0 | 0 | 0 | 0 | 0 |
| ENSG00000233719 | 0 | 0 | 2 | 1 | 0 | 0 |
| ENSG00000272917 | 0 | 1 | 0 | 0 | 0 | 0 |
| ENSG00000256442 | 2 | 3 | 1 | 0 | 0 | 1 |
| ENSG00000256594 | 145 | 91 | 140 | 77 | 105 | 96 |
| ENSG00000257027 | 353 | 170 | 203 | 142 | 158 | 180 |
| ENSG00000069493 | 628 | 308 | 331 | 119 | 273 | 271 |
| ENSG00000213443 | 0 | 0 | 0 | 0 | 0 | 0 |
| ENSG00000256582 | 3 | 0 | 1 | 0 | 0 | 0 |
| ENSG00000184293 | 0 | 0 | 0 | 0 | 0 | 0 |
| ENSG00000110848 | 0 | 0 | 0 | 0 | 0 | 1 |
| ENSG00000256912 | 0 | 0 | 0 | 0 | 0 | 0 |
| ENSG00000150045 | 0 | 0 | 0 | 0 | 0 | 0 |
| ENSG00000110852 | 0 | 0 | 0 | 0 | 0 | 0 |

| | | | | | | |
|---|---|---|---|---|---|---|
| ENSG00000255882 | 0 | 0 | 0 | 0 | 0 | 0 |
| ENSG00000256797 | 0 | 0 | 0 | 1 | 0 | 0 |
| ENSG00000188393 | 0 | 0 | 0 | 0 | 0 | 0 |
| ENSG00000225231 | 0 | 0 | 0 | 0 | 0 | 0 |
| ENSG00000231560 | 0 | 0 | 0 | 0 | 0 | 0 |
| ENSG00000172322 | 0 | 0 | 0 | 0 | 0 | 0 |
| ENSG00000165682 | 0 | 0 | 0 | 0 | 0 | 0 |
| ENSG00000256660 | 0 | 0 | 0 | 0 | 0 | 0 |
| ENSG00000256803 | 0 | 0 | 0 | 0 | 0 | 0 |
| ENSG00000197992 | 0 | 0 | 0 | 0 | 0 | 0 |
| ENSG00000256626 | 0 | 0 | 0 | 0 | 0 | 0 |
| ENSG00000150048 | 0 | 0 | 0 | 0 | 0 | 0 |
| ENSG00000223042 | 0 | 0 | 0 | 0 | 0 | 0 |
| ENSG00000255734 | 0 | 1 | 0 | 0 | 0 | 0 |
| ENSG00000172243 | 30 | 24 | 16 | 7 | 22 | 17 |
| ENSG00000173391 | 0 | 0 | 0 | 0 | 0 | 0 |
| ENSG00000165685 | 0 | 1 | 0 | 1 | 0 | 0 |
| ENSG00000139112 | 453 | 260 | 308 | 245 | 266 | 301 |
| ENSG00000255958 | 0 | 0 | 0 | 0 | 0 | 0 |
| ENSG00000134539 | 0 | 0 | 0 | 0 | 0 | 0 |
| ENSG00000256288 | 0 | 0 | 0 | 0 | 0 | 0 |
| ENSG00000256155 | 0 | 0 | 0 | 0 | 1 | 0 |
| ENSG00000245648 | 2 | 1 | 0 | 0 | 2 | 1 |
| ENSG00000255819 | 0 | 0 | 0 | 0 | 0 | 0 |
| ENSG00000213809 | 0 | 0 | 0 | 0 | 0 | 0 |
| ENSG00000183542 | 0 | 0 | 0 | 0 | 0 | 0 |
| ENSG00000255641 | 0 | 0 | 0 | 0 | 0 | 0 |
| ENSG00000205810 | 0 | 0 | 0 | 0 | 0 | 0 |
| ENSG00000205809 | 0 | 0 | 0 | 0 | 0 | 0 |
| ENSG00000134545 | 2 | 0 | 0 | 0 | 0 | 0 |
| ENSG00000180574 | 8 | 3 | 9 | 1 | 4 | 5 |
| ENSG00000257016 | 0 | 0 | 0 | 0 | 0 | 0 |
| ENSG00000256039 | 0 | 0 | 0 | 0 | 0 | 0 |
| ENSG00000256667 | 28 | 22 | 25 | 9 | 29 | 29 |
| ENSG00000111196 | 393 | 212 | 214 | 197 | 282 | 216 |
| ENSG00000060140 | 76 | 31 | 62 | 28 | 56 | 66 |
| ENSG00000060138 | 2693 | 1274 | 1643 | 1856 | 1595 | 1694 |
| ENSG00000256888 | 0 | 0 | 0 | 0 | 0 | 0 |
| ENSG00000235930 | 0 | 0 | 0 | 0 | 0 | 0 |
| ENSG00000121377 | 0 | 0 | 0 | 0 | 0 | 0 |
| ENSG00000121314 | 0 | 0 | 0 | 0 | 0 | 0 |
| ENSG00000121381 | 0 | 0 | 0 | 0 | 0 | 0 |
| ENSG00000231887 | 11 | 7 | 12 | 2 | 8 | 5 |
| ENSG00000121318 | 4 | 0 | 2 | 0 | 0 | 1 |
| ENSG00000256651 | 0 | 0 | 0 | 0 | 0 | 0 |
| ENSG00000111215 | 11 | 6 | 3 | 1 | 2 | 7 |
| ENSG00000275778 | 0 | 5 | 2 | 0 | 0 | 0 |
| ENSG00000284826 | 0 | 0 | 0 | 0 | 0 | 0 |
| ENSG00000256682 | 0 | 0 | 0 | 0 | 2 | 0 |
| ENSG00000212128 | 0 | 0 | 1 | 0 | 0 | 0 |

| | | | | | | |
|---|---|---|---|---|---|---|
| ENSG00000134551 | 2 | 1 | 0 | 0 | 0 | 0 |
| ENSG00000212127 | 14 | 3 | 11 | 5 | 7 | 7 |
| ENSG00000212125 | 9 | 6 | 2 | 1 | 8 | 5 |
| ENSG00000212126 | 0 | 0 | 0 | 0 | 0 | 0 |
| ENSG00000255837 | 0 | 0 | 0 | 0 | 0 | 1 |
| ENSG00000212124 | 0 | 2 | 0 | 2 | 2 | 0 |
| ENSG00000256436 | 8 | 2 | 0 | 1 | 2 | 1 |
| ENSG00000256019 | 2 | 0 | 0 | 0 | 0 | 0 |
| ENSG00000226761 | 0 | 0 | 0 | 0 | 0 | 0 |
| ENSG00000256274 | 0 | 1 | 0 | 0 | 0 | 0 |
| ENSG00000255374 | 0 | 0 | 1 | 0 | 0 | 0 |
| ENSG00000256400 | 0 | 0 | 0 | 0 | 0 | 0 |
| ENSG00000256188 | 0 | 0 | 0 | 1 | 0 | 0 |
| ENSG00000256981 | 2 | 0 | 0 | 0 | 0 | 0 |
| ENSG00000256712 | 0 | 0 | 0 | 0 | 0 | 0 |
| ENSG00000256537 | 1089 | 542 | 695 | 557 | 569 | 613 |
| ENSG00000186136 | 0 | 0 | 0 | 0 | 0 | 0 |
| ENSG00000256657 | 0 | 0 | 0 | 0 | 0 | 2 |
| ENSG00000275119 | 0 | 0 | 0 | 0 | 0 | 0 |
| ENSG00000197870 | 12 | 4 | 1 | 5 | 4 | 3 |
| ENSG00000230657 | 0 | 0 | 0 | 0 | 0 | 0 |
| ENSG00000251655 | 0 | 0 | 0 | 0 | 0 | 0 |
| ENSG00000121335 | 0 | 0 | 0 | 0 | 0 | 0 |
| ENSG00000255790 | 2 | 0 | 0 | 2 | 1 | 0 |
| ENSG00000237303 | 0 | 0 | 1 | 0 | 0 | 1 |
| ENSG00000278654 | 0 | 1 | 0 | 1 | 1 | 0 |
| ENSG00000277705 | 0 | 0 | 0 | 0 | 0 | 0 |
| ENSG00000270863 | 4 | 2 | 2 | 0 | 2 | 2 |
| ENSG00000274551 | 0 | 0 | 0 | 0 | 0 | 0 |
| ENSG00000256237 | 0 | 0 | 0 | 0 | 0 | 0 |
| ENSG00000251747 | 0 | 0 | 0 | 0 | 0 | 0 |
| ENSG00000247157 | 62 | 17 | 36 | 20 | 28 | 18 |
| ENSG00000139083 | 396 | 175 | 237 | 164 | 167 | 170 |
| ENSG00000121380 | 4 | 7 | 1 | 3 | 1 | 2 |
| ENSG00000198134 | 6 | 6 | 5 | 10 | 4 | 8 |
| ENSG00000283475 | 1 | 0 | 2 | 2 | 0 | 1 |
| ENSG00000070018 | 706 | 365 | 415 | 172 | 424 | 340 |
| ENSG00000242405 | 0 | 0 | 0 | 0 | 0 | 0 |
| ENSG00000199551 | 0 | 0 | 0 | 0 | 0 | 0 |
| ENSG00000270857 | 2 | 0 | 0 | 0 | 0 | 0 |
| ENSG00000207010 | 0 | 0 | 0 | 0 | 0 | 0 |
| ENSG00000111261 | 989 | 426 | 606 | 300 | 420 | 409 |
| ENSG00000243141 | 2 | 0 | 2 | 0 | 0 | 0 |
| ENSG00000205791 | 16 | 8 | 11 | 6 | 10 | 5 |
| ENSG00000165714 | 242 | 128 | 160 | 114 | 133 | 122 |
| ENSG00000271370 | 0 | 0 | 0 | 0 | 0 | 0 |
| ENSG00000111266 | 2077 | 898 | 1108 | 1046 | 1414 | 1342 |
| ENSG00000255670 | 2 | 2 | 1 | 0 | 0 | 0 |
| ENSG00000240424 | 0 | 1 | 0 | 0 | 0 | 0 |
| ENSG00000111269 | 1432 | 684 | 955 | 725 | 873 | 842 |

| | | | | | | |
|---|---|---|---|---|---|---|
| ENSG00000271215 | 0 | 0 | 0 | 0 | 0 | 0 |
| ENSG00000275963 | 2 | 0 | 0 | 0 | 2 | 0 |
| ENSG00000183150 | 15 | 12 | 14 | 15 | 16 | 19 |
| ENSG00000257004 | 0 | 0 | 0 | 0 | 0 | 0 |
| ENSG00000111276 | 3447 | 1568 | 2059 | 1751 | 1967 | 1992 |
| ENSG00000275560 | 38 | 15 | 29 | 12 | 29 | 20 |
| ENSG00000256658 | 0 | 0 | 0 | 0 | 0 | 1 |
| ENSG00000178878 | 161 | 68 | 130 | 78 | 112 | 118 |
| ENSG00000207983 | 0 | 0 | 0 | 0 | 0 | 1 |
| ENSG00000270296 | 0 | 0 | 0 | 0 | 0 | 0 |
| ENSG00000213782 | 26 | 23 | 31 | 21 | 21 | 28 |
| ENSG00000242329 | 0 | 0 | 0 | 0 | 0 | 0 |
| ENSG00000241352 | 3 | 1 | 1 | 2 | 3 | 1 |
| ENSG00000234498 | 5 | 4 | 7 | 1 | 3 | 4 |
| ENSG00000013588 | 10100 | 5246 | 5506 | 3544 | 4694 | 4188 |
| ENSG00000283759 | 0 | 0 | 0 | 0 | 0 | 0 |
| ENSG00000247498 | 107 | 94 | 99 | 64 | 65 | 76 |
| ENSG00000111291 | 14 | 5 | 3 | 3 | 6 | 2 |
| ENSG00000256011 | 0 | 0 | 0 | 0 | 0 | 0 |
| ENSG00000221611 | 0 | 1 | 0 | 2 | 0 | 0 |
| ENSG00000013583 | 1254 | 578 | 752 | 643 | 724 | 681 |
| ENSG00000183935 | 314 | 152 | 193 | 136 | 141 | 163 |
| ENSG00000255621 | 3 | 5 | 7 | 1 | 4 | 3 |
| ENSG00000243129 | 0 | 0 | 0 | 0 | 0 | 0 |
| ENSG00000084444 | 1006 | 484 | 570 | 291 | 373 | 376 |
| ENSG00000111305 | 3 | 0 | 0 | 0 | 1 | 0 |
| ENSG00000134531 | 10 | 6 | 3 | 2 | 5 | 6 |
| ENSG00000180861 | 0 | 0 | 0 | 0 | 0 | 0 |
| ENSG00000273079 | 0 | 0 | 0 | 0 | 0 | 0 |
| ENSG00000201476 | 0 | 0 | 0 | 0 | 0 | 0 |
| ENSG00000201909 | 0 | 0 | 0 | 0 | 0 | 0 |
| ENSG00000256306 | 0 | 0 | 0 | 0 | 0 | 0 |
| ENSG00000277935 | 0 | 0 | 0 | 0 | 0 | 0 |
| ENSG00000200475 | 0 | 0 | 0 | 0 | 0 | 0 |
| ENSG00000256346 | 0 | 0 | 0 | 0 | 0 | 0 |
| ENSG00000227760 | 0 | 0 | 0 | 0 | 0 | 0 |
| ENSG00000256084 | 0 | 0 | 0 | 0 | 1 | 0 |
| ENSG00000255763 | 0 | 0 | 0 | 0 | 0 | 0 |
| ENSG00000251908 | 0 | 0 | 0 | 0 | 0 | 0 |
| ENSG00000255749 | 0 | 0 | 0 | 0 | 1 | 0 |
| ENSG00000244573 | 4 | 0 | 0 | 0 | 0 | 0 |
| ENSG00000171681 | 1483 | 832 | 714 | 312 | 952 | 601 |
| ENSG00000121316 | 471 | 223 | 251 | 95 | 114 | 140 |
| ENSG00000255649 | 0 | 1 | 0 | 0 | 0 | 0 |
| ENSG00000200830 | 0 | 0 | 0 | 0 | 0 | 0 |
| ENSG00000256751 | 7 | 3 | 5 | 4 | 3 | 2 |
| ENSG00000070019 | 16 | 16 | 11 | 9 | 14 | 12 |
| ENSG00000214772 | 0 | 0 | 1 | 0 | 0 | 0 |
| ENSG00000261324 | 185 | 81 | 120 | 54 | 86 | 136 |
| ENSG00000197837 | 102 | 54 | 56 | 34 | 42 | 60 |

| | | | | | | |
|---|---|---|---|---|---|---|
| ENSG00000246705 | 5406 | 2785 | 3227 | 3461 | 3036 | 2691 |
| ENSG00000084463 | 2595 | 1371 | 1525 | 1412 | 1547 | 1568 |
| ENSG00000182993 | 30 | 27 | 14 | 27 | 37 | 22 |
| ENSG00000179256 | 2 | 5 | 0 | 2 | 4 | 0 |
| ENSG00000111339 | 4 | 2 | 1 | 2 | 5 | 0 |
| ENSG00000256339 | 0 | 0 | 0 | 0 | 0 | 0 |
| ENSG00000111341 | 84 | 38 | 23 | 45 | 48 | 33 |
| ENSG00000139055 | 52 | 38 | 28 | 11 | 29 | 15 |
| ENSG00000111348 | 24 | 23 | 18 | 28 | 36 | 22 |
| ENSG00000139053 | 0 | 0 | 0 | 1 | 0 | 1 |
| ENSG00000255727 | 0 | 0 | 0 | 0 | 0 | 0 |
| ENSG00000134533 | 419 | 149 | 240 | 243 | 195 | 303 |
| ENSG00000256650 | 0 | 0 | 1 | 0 | 0 | 3 |
| ENSG00000255660 | 0 | 0 | 0 | 0 | 0 | 0 |
| ENSG00000256283 | 0 | 0 | 0 | 0 | 0 | 0 |
| ENSG00000151490 | 0 | 0 | 1 | 0 | 0 | 0 |
| ENSG00000243916 | 0 | 0 | 0 | 0 | 0 | 0 |
| ENSG00000151491 | 1636 | 779 | 917 | 451 | 919 | 714 |
| ENSG00000200105 | 0 | 1 | 0 | 0 | 0 | 0 |
| ENSG00000255565 | 0 | 0 | 0 | 0 | 0 | 0 |
| ENSG00000256627 | 0 | 0 | 0 | 0 | 0 | 0 |
| ENSG00000023734 | 3596 | 1848 | 2146 | 1866 | 2286 | 2184 |
| ENSG00000023697 | 398 | 206 | 266 | 185 | 270 | 246 |
| ENSG00000256134 | 0 | 0 | 0 | 0 | 0 | 0 |
| ENSG00000188991 | 0 | 0 | 0 | 0 | 2 | 0 |
| ENSG00000008394 | 6398 | 3164 | 3759 | 3785 | 5104 | 4360 |
| ENSG00000256238 | 44 | 25 | 39 | 21 | 34 | 27 |
| ENSG00000256450 | 0 | 0 | 0 | 0 | 0 | 0 |
| ENSG00000048540 | 3 | 6 | 7 | 2 | 0 | 0 |
| ENSG00000256564 | 0 | 0 | 0 | 1 | 1 | 0 |
| ENSG00000283240 | 2 | 1 | 0 | 0 | 1 | 1 |
| ENSG00000255871 | 0 | 0 | 0 | 0 | 0 | 0 |
| ENSG00000256211 | 2 | 0 | 0 | 0 | 0 | 0 |
| ENSG00000213235 | 2 | 0 | 0 | 0 | 0 | 0 |
| ENSG00000212358 | 0 | 0 | 0 | 0 | 0 | 0 |
| ENSG00000285724 | 0 | 0 | 0 | 0 | 0 | 0 |
| ENSG00000242991 | 0 | 0 | 0 | 0 | 0 | 0 |
| ENSG00000256896 | 0 | 0 | 0 | 0 | 0 | 0 |
| ENSG00000256756 | 0 | 0 | 0 | 0 | 0 | 0 |
| ENSG00000256994 | 6 | 1 | 1 | 0 | 0 | 0 |
| ENSG00000256389 | 0 | 0 | 0 | 0 | 0 | 0 |
| ENSG00000283665 | 0 | 0 | 0 | 0 | 0 | 0 |
| ENSG00000206662 | 0 | 0 | 0 | 0 | 0 | 0 |
| ENSG00000111404 | 0 | 0 | 0 | 0 | 0 | 2 |
| ENSG00000139144 | 0 | 5 | 5 | 3 | 8 | 4 |
| ENSG00000255892 | 0 | 0 | 0 | 0 | 0 | 0 |
| ENSG00000255960 | 0 | 0 | 0 | 0 | 1 | 0 |
| ENSG00000256138 | 0 | 0 | 0 | 0 | 0 | 0 |
| ENSG00000139151 | 2 | 3 | 1 | 4 | 4 | 2 |
| ENSG00000255993 | 0 | 2 | 0 | 0 | 0 | 1 |

| | | | | | | |
|---|---|---|---|---|---|---|
| ENSG00000255648 | 0 | 0 | 0 | 0 | 0 | 0 |
| ENSG00000177938 | 0 | 0 | 0 | 0 | 0 | 0 |
| ENSG00000227525 | 0 | 0 | 1 | 0 | 0 | 0 |
| ENSG00000256844 | 0 | 0 | 0 | 0 | 0 | 0 |
| ENSG00000052126 | 430 | 311 | 291 | 120 | 348 | 227 |
| ENSG00000253284 | 0 | 1 | 5 | 1 | 5 | 2 |
| ENSG00000240993 | 0 | 0 | 0 | 0 | 0 | 0 |
| ENSG00000243854 | 0 | 0 | 0 | 0 | 0 | 0 |
| ENSG00000139154 | 668 | 294 | 334 | 230 | 336 | 316 |
| ENSG00000255909 | 0 | 0 | 2 | 2 | 1 | 2 |
| ENSG00000271462 | 0 | 0 | 0 | 0 | 0 | 0 |
| ENSG00000200247 | 0 | 0 | 0 | 0 | 0 | 0 |
| ENSG00000245205 | 17 | 6 | 5 | 3 | 8 | 5 |
| ENSG00000240328 | 0 | 1 | 0 | 0 | 0 | 0 |
| ENSG00000200885 | 0 | 0 | 0 | 0 | 0 | 0 |
| ENSG00000255910 | 0 | 0 | 0 | 0 | 0 | 0 |
| ENSG00000256849 | 0 | 0 | 0 | 0 | 0 | 0 |
| ENSG00000256287 | 0 | 0 | 0 | 0 | 0 | 0 |
| ENSG00000256499 | 0 | 0 | 0 | 0 | 0 | 0 |
| ENSG00000256879 | 0 | 0 | 0 | 0 | 0 | 0 |
| ENSG00000172572 | 0 | 0 | 0 | 0 | 0 | 0 |
| ENSG00000131982 | 0 | 0 | 0 | 0 | 0 | 0 |
| ENSG00000256663 | 47 | 24 | 47 | 20 | 24 | 28 |
| ENSG00000139155 | 0 | 0 | 0 | 0 | 0 | 0 |
| ENSG00000111700 | 0 | 0 | 0 | 0 | 0 | 0 |
| ENSG00000257046 | 0 | 0 | 0 | 0 | 0 | 0 |
| ENSG00000256157 | 0 | 0 | 0 | 0 | 0 | 0 |
| ENSG00000205754 | 0 | 0 | 0 | 0 | 0 | 0 |
| ENSG00000257062 | 0 | 0 | 0 | 0 | 0 | 0 |
| ENSG00000134538 | 0 | 0 | 1 | 0 | 0 | 0 |
| ENSG00000084453 | 0 | 1 | 4 | 1 | 1 | 0 |
| ENSG00000121351 | 2 | 1 | 3 | 0 | 0 | 2 |
| ENSG00000121350 | 107 | 61 | 79 | 40 | 102 | 76 |
| ENSG00000256021 | 0 | 0 | 0 | 0 | 0 | 1 |
| ENSG00000004700 | 236 | 180 | 151 | 65 | 295 | 161 |
| ENSG00000111711 | 907 | 427 | 456 | 347 | 558 | 428 |
| ENSG00000134548 | 2 | 0 | 1 | 0 | 0 | 0 |
| ENSG00000111713 | 0 | 0 | 0 | 0 | 0 | 0 |
| ENSG00000285854 | 0 | 0 | 0 | 0 | 0 | 0 |
| ENSG00000111716 | 0 | 0 | 1 | 0 | 0 | 0 |
| ENSG00000256615 | 0 | 0 | 0 | 0 | 0 | 0 |
| ENSG00000255644 | 1 | 2 | 2 | 1 | 1 | 0 |
| ENSG00000121361 | 1065 | 439 | 543 | 248 | 196 | 240 |
| ENSG00000069431 | 20 | 10 | 7 | 1 | 2 | 7 |
| ENSG00000257022 | 0 | 0 | 0 | 0 | 0 | 0 |
| ENSG00000275567 | 0 | 0 | 0 | 0 | 0 | 0 |
| ENSG00000255627 | 0 | 0 | 0 | 0 | 0 | 0 |
| ENSG00000111726 | 1973 | 1020 | 1136 | 953 | 1133 | 1163 |
| ENSG00000111728 | 7 | 10 | 11 | 6 | 13 | 4 |
| ENSG00000274624 | 0 | 0 | 0 | 0 | 0 | 0 |

| | | | | | | |
|---|---|---|---|---|---|---|
| ENSG00000283582 | 0 | 0 | 0 | 0 | 0 | 0 |
| ENSG00000212172 | 0 | 0 | 0 | 0 | 0 | 0 |
| ENSG00000256714 | 0 | 0 | 0 | 1 | 0 | 0 |
| ENSG00000111731 | 1019 | 574 | 670 | 311 | 637 | 524 |
| ENSG00000256973 | 0 | 1 | 0 | 1 | 3 | 0 |
| ENSG00000256371 | 0 | 0 | 0 | 0 | 0 | 0 |
| ENSG00000256774 | 0 | 0 | 0 | 0 | 0 | 0 |
| ENSG00000250166 | 0 | 0 | 0 | 0 | 0 | 0 |
| ENSG00000257023 | 0 | 2 | 0 | 0 | 2 | 0 |
| ENSG00000139163 | 817 | 412 | 404 | 263 | 545 | 405 |
| ENSG00000274885 | 0 | 0 | 0 | 0 | 0 | 0 |
| ENSG00000256995 | 0 | 1 | 1 | 1 | 3 | 1 |
| ENSG00000241451 | 0 | 0 | 0 | 0 | 0 | 0 |
| ENSG00000256923 | 0 | 0 | 0 | 0 | 0 | 0 |
| ENSG00000256321 | 0 | 0 | 0 | 0 | 0 | 1 |
| ENSG00000256353 | 0 | 0 | 0 | 0 | 0 | 0 |
| ENSG00000115934 | 0 | 0 | 0 | 0 | 0 | 0 |
| ENSG00000134532 | 0 | 0 | 0 | 0 | 0 | 0 |
| ENSG00000256473 | 0 | 0 | 0 | 0 | 0 | 0 |
| ENSG00000216192 | 0 | 0 | 0 | 0 | 0 | 0 |
| ENSG00000256120 | 0 | 0 | 0 | 0 | 0 | 0 |
| ENSG00000197503 | 0 | 0 | 0 | 0 | 0 | 0 |
| ENSG00000256852 | 0 | 0 | 0 | 0 | 0 | 0 |
| ENSG00000244662 | 0 | 0 | 0 | 0 | 0 | 0 |
| ENSG00000255745 | 0 | 0 | 0 | 0 | 0 | 0 |
| ENSG00000240481 | 0 | 0 | 0 | 0 | 0 | 0 |
| ENSG00000060982 | 2 | 2 | 5 | 4 | 5 | 3 |
| ENSG00000256482 | 0 | 0 | 0 | 0 | 0 | 0 |
| ENSG00000278969 | 0 | 0 | 0 | 0 | 0 | 0 |
| ENSG00000255921 | 0 | 0 | 0 | 0 | 0 | 0 |
| ENSG00000213736 | 0 | 0 | 0 | 0 | 0 | 0 |
| ENSG00000226397 | 0 | 0 | 0 | 0 | 0 | 0 |
| ENSG00000118308 | 2 | 0 | 0 | 0 | 0 | 0 |
| ENSG00000258546 | 0 | 0 | 0 | 0 | 0 | 0 |
| ENSG00000258449 | 0 | 0 | 0 | 0 | 0 | 0 |
| ENSG00000276842 | 0 | 0 | 0 | 0 | 0 | 0 |
| ENSG00000118307 | 6 | 3 | 8 | 2 | 5 | 4 |
| ENSG00000205707 | 228 | 126 | 152 | 136 | 149 | 138 |
| ENSG00000133703 | 1728 | 870 | 1026 | 456 | 959 | 822 |
| ENSG00000274987 | 0 | 0 | 0 | 1 | 1 | 0 |
| ENSG00000275197 | 0 | 0 | 0 | 0 | 0 | 0 |
| ENSG00000278743 | 2 | 1 | 3 | 0 | 5 | 3 |
| ENSG00000201439 | 0 | 0 | 0 | 0 | 0 | 0 |
| ENSG00000241846 | 0 | 0 | 0 | 0 | 0 | 0 |
| ENSG00000152936 | 4 | 3 | 3 | 1 | 2 | 3 |
| ENSG00000279173 | 0 | 0 | 0 | 0 | 0 | 0 |
| ENSG00000256079 | 0 | 0 | 0 | 0 | 0 | 0 |
| ENSG00000255988 | 0 | 0 | 0 | 0 | 0 | 0 |
| ENSG00000257009 | 0 | 0 | 0 | 0 | 0 | 0 |
| ENSG00000222950 | 0 | 0 | 0 | 0 | 0 | 0 |

| | | | | | | |
|---|---|---|---|---|---|---|
| ENSG00000256686 | 0 | 0 | 0 | 0 | 0 | 2 |
| ENSG00000255725 | 2 | 1 | 1 | 0 | 2 | 1 |
| ENSG00000256440 | 0 | 0 | 1 | 0 | 0 | 4 |
| ENSG00000283603 | 0 | 0 | 0 | 0 | 0 | 0 |
| ENSG00000256256 | 0 | 0 | 0 | 0 | 0 | 0 |
| ENSG00000246695 | 108 | 73 | 124 | 80 | 114 | 103 |
| ENSG00000123094 | 669 | 388 | 426 | 236 | 483 | 430 |
| ENSG00000123095 | 15 | 8 | 5 | 9 | 10 | 6 |
| ENSG00000123096 | 22 | 19 | 15 | 16 | 20 | 20 |
| ENSG00000256894 | 2 | 4 | 2 | 0 | 0 | 0 |
| ENSG00000256234 | 0 | 0 | 0 | 2 | 4 | 1 |
| ENSG00000255750 | 0 | 0 | 0 | 0 | 0 | 0 |
| ENSG00000278095 | 0 | 0 | 0 | 0 | 0 | 0 |
| ENSG00000255858 | 0 | 0 | 0 | 0 | 0 | 0 |
| ENSG00000255968 | 0 | 1 | 2 | 0 | 0 | 0 |
| ENSG00000256185 | 0 | 0 | 0 | 0 | 0 | 2 |
| ENSG00000123104 | 384 | 215 | 183 | 62 | 287 | 143 |
| ENSG00000212549 | 0 | 0 | 0 | 0 | 0 | 0 |
| ENSG00000234428 | 0 | 1 | 0 | 0 | 0 | 0 |
| ENSG00000256640 | 0 | 0 | 0 | 0 | 0 | 0 |
| ENSG00000064102 | 801 | 398 | 452 | 240 | 470 | 328 |
| ENSG00000111790 | 514 | 250 | 274 | 164 | 328 | 229 |
| ENSG00000247903 | 168 | 89 | 133 | 88 | 107 | 92 |
| ENSG00000064115 | 3257 | 1661 | 2119 | 1415 | 1548 | 1598 |
| ENSG00000152944 | 557 | 297 | 318 | 280 | 336 | 272 |
| ENSG00000275764 | 73 | 49 | 43 | 17 | 66 | 36 |
| ENSG00000214700 | 0 | 1 | 1 | 0 | 2 | 0 |
| ENSG00000271418 | 0 | 0 | 0 | 0 | 0 | 0 |
| ENSG00000256226 | 0 | 0 | 2 | 1 | 0 | 1 |
| ENSG00000256625 | 14 | 0 | 3 | 0 | 1 | 2 |
| ENSG00000211455 | 781 | 428 | 419 | 302 | 593 | 401 |
| ENSG00000029153 | 324 | 240 | 218 | 167 | 294 | 191 |
| ENSG00000245311 | 0 | 0 | 0 | 0 | 0 | 0 |
| ENSG00000165935 | 2 | 1 | 1 | 2 | 1 | 2 |
| ENSG00000257037 | 0 | 0 | 0 | 0 | 0 | 0 |
| ENSG00000110841 | 1141 | 597 | 615 | 331 | 650 | 526 |
| ENSG00000248100 | 0 | 0 | 0 | 0 | 0 | 0 |
| ENSG00000256056 | 0 | 0 | 0 | 0 | 0 | 0 |
| ENSG00000256557 | 0 | 0 | 0 | 0 | 0 | 0 |
| ENSG00000256377 | 3 | 0 | 1 | 0 | 0 | 2 |
| ENSG00000174236 | 1 | 0 | 0 | 1 | 2 | 0 |
| ENSG00000276261 | 0 | 0 | 0 | 3 | 2 | 2 |
| ENSG00000230519 | 0 | 0 | 0 | 0 | 0 | 0 |
| ENSG00000256504 | 0 | 0 | 0 | 0 | 0 | 0 |
| ENSG00000061794 | 2286 | 1054 | 1232 | 1056 | 1260 | 1239 |
| ENSG00000252585 | 0 | 0 | 0 | 0 | 0 | 3 |
| ENSG00000205693 | 0 | 0 | 1 | 1 | 0 | 0 |
| ENSG00000256747 | 0 | 0 | 0 | 0 | 1 | 1 |
| ENSG00000087448 | 1290 | 573 | 766 | 453 | 695 | 702 |
| ENSG00000256512 | 0 | 0 | 0 | 0 | 0 | 0 |

| | | | | | | |
|---|---|---|---|---|---|---|
| ENSG00000201612 | 0 | 0 | 0 | 0 | 0 | 0 |
| ENSG00000274075 | 0 | 0 | 0 | 0 | 0 | 0 |
| ENSG00000256378 | 0 | 0 | 0 | 0 | 0 | 0 |
| ENSG00000087494 | 0 | 4 | 1 | 6 | 3 | 2 |
| ENSG00000257042 | 0 | 0 | 0 | 0 | 0 | 0 |
| ENSG00000123106 | 370 | 219 | 218 | 139 | 268 | 210 |
| ENSG00000247934 | 53 | 20 | 34 | 10 | 14 | 12 |
| ENSG00000255951 | 0 | 0 | 0 | 0 | 0 | 0 |
| ENSG00000278733 | 4 | 2 | 2 | 1 | 0 | 1 |
| ENSG00000273989 | 0 | 0 | 0 | 0 | 0 | 0 |
| ENSG00000252237 | 0 | 0 | 0 | 0 | 0 | 0 |
| ENSG00000202187 | 0 | 0 | 0 | 0 | 0 | 0 |
| ENSG00000244712 | 0 | 0 | 0 | 1 | 1 | 0 |
| ENSG00000256513 | 0 | 0 | 0 | 0 | 0 | 0 |
| ENSG00000278687 | 0 | 0 | 0 | 0 | 0 | 0 |
| ENSG00000257258 | 0 | 0 | 0 | 0 | 0 | 0 |
| ENSG00000064763 | 47 | 19 | 27 | 14 | 17 | 33 |
| ENSG00000275278 | 0 | 0 | 0 | 0 | 0 | 0 |
| ENSG00000275476 | 0 | 0 | 0 | 0 | 0 | 0 |
| ENSG00000257176 | 16 | 1 | 6 | 1 | 5 | 5 |
| ENSG00000274315 | 0 | 1 | 0 | 0 | 1 | 0 |
| ENSG00000273680 | 0 | 7 | 4 | 0 | 5 | 3 |
| ENSG00000087502 | 797 | 503 | 430 | 309 | 681 | 411 |
| ENSG00000257599 | 0 | 3 | 0 | 0 | 1 | 1 |
| ENSG00000187950 | 0 | 0 | 2 | 0 | 0 | 0 |
| ENSG00000133687 | 182 | 80 | 130 | 55 | 108 | 80 |
| ENSG00000257456 | 1 | 0 | 0 | 0 | 0 | 0 |
| ENSG00000242986 | 0 | 0 | 0 | 0 | 0 | 0 |
| ENSG00000276431 | 0 | 0 | 0 | 0 | 0 | 0 |
| ENSG00000257262 | 0 | 0 | 0 | 0 | 0 | 0 |
| ENSG00000257756 | 0 | 0 | 0 | 0 | 0 | 0 |
| ENSG00000284682 | 0 | 0 | 0 | 0 | 0 | 0 |
| ENSG00000251781 | 0 | 0 | 0 | 0 | 0 | 0 |
| ENSG00000257932 | 0 | 0 | 0 | 0 | 0 | 0 |
| ENSG00000258202 | 0 | 0 | 0 | 0 | 0 | 0 |
| ENSG00000133704 | 1709 | 940 | 1028 | 609 | 1061 | 898 |
| ENSG00000285870 | 0 | 1 | 0 | 0 | 1 | 0 |
| ENSG00000110888 | 230 | 99 | 132 | 71 | 124 | 103 |
| ENSG00000246331 | 0 | 0 | 0 | 0 | 0 | 2 |
| ENSG00000235884 | 0 | 0 | 0 | 0 | 0 | 0 |
| ENSG00000285517 | 0 | 0 | 0 | 0 | 0 | 0 |
| ENSG00000258118 | 0 | 0 | 0 | 0 | 0 | 0 |
| ENSG00000110900 | 8 | 6 | 5 | 2 | 4 | 5 |
| ENSG00000226472 | 13 | 5 | 2 | 7 | 5 | 5 |
| ENSG00000256984 | 0 | 0 | 0 | 0 | 0 | 0 |
| ENSG00000245614 | 63 | 10 | 37 | 32 | 27 | 32 |
| ENSG00000013573 | 1291 | 604 | 768 | 698 | 634 | 721 |
| ENSG00000212533 | 0 | 0 | 0 | 0 | 0 | 0 |
| ENSG00000270926 | 0 | 0 | 1 | 0 | 0 | 0 |
| ENSG00000177359 | 4 | 3 | 1 | 0 | 0 | 5 |

| | | | | | | |
|---|---|---|---|---|---|---|
| ENSG00000270766 | 0 | 0 | 0 | 0 | 0 | 0 |
| ENSG00000243517 | 3 | 2 | 4 | 1 | 0 | 2 |
| ENSG00000270395 | 0 | 0 | 0 | 0 | 0 | 0 |
| ENSG00000275097 | 1 | 1 | 0 | 0 | 0 | 0 |
| ENSG00000139146 | 1393 | 706 | 902 | 652 | 871 | 901 |
| ENSG00000256176 | 0 | 1 | 0 | 0 | 0 | 0 |
| ENSG00000177340 | 6 | 0 | 9 | 2 | 6 | 5 |
| ENSG00000256232 | 0 | 0 | 0 | 0 | 0 | 0 |
| ENSG00000283256 | 0 | 0 | 0 | 0 | 0 | 0 |
| ENSG00000170456 | 260 | 122 | 131 | 80 | 109 | 92 |
| ENSG00000200388 | 0 | 0 | 0 | 0 | 0 | 0 |
| ENSG00000275769 | 0 | 0 | 1 | 0 | 0 | 0 |
| ENSG00000203437 | 0 | 1 | 1 | 0 | 0 | 0 |
| ENSG00000256159 | 0 | 0 | 0 | 0 | 0 | 0 |
| ENSG00000201228 | 0 | 0 | 0 | 0 | 2 | 0 |
| ENSG00000225349 | 0 | 0 | 2 | 0 | 0 | 0 |
| ENSG00000255867 | 3 | 2 | 0 | 1 | 0 | 1 |
| ENSG00000252390 | 0 | 0 | 0 | 0 | 0 | 0 |
| ENSG00000231788 | 0 | 0 | 0 | 0 | 0 | 0 |
| ENSG00000233381 | 1 | 0 | 3 | 0 | 0 | 0 |
| ENSG00000139160 | 62 | 17 | 37 | 20 | 33 | 28 |
| ENSG00000151743 | 134 | 42 | 61 | 68 | 82 | 56 |
| ENSG00000276058 | 6 | 1 | 0 | 1 | 2 | 2 |
| ENSG00000276900 | 61 | 40 | 48 | 24 | 26 | 46 |
| ENSG00000256843 | 0 | 0 | 0 | 0 | 0 | 0 |
| ENSG00000223722 | 0 | 0 | 0 | 0 | 0 | 1 |
| ENSG00000188375 | 2 | 2 | 6 | 1 | 4 | 4 |
| ENSG00000252421 | 0 | 0 | 0 | 0 | 0 | 0 |
| ENSG00000252204 | 0 | 0 | 0 | 0 | 0 | 0 |
| ENSG00000256372 | 0 | 0 | 0 | 0 | 0 | 0 |
| ENSG00000255760 | 0 | 0 | 0 | 0 | 0 | 0 |
| ENSG00000256465 | 0 | 0 | 0 | 0 | 0 | 0 |
| ENSG00000244436 | 0 | 0 | 0 | 0 | 0 | 0 |
| ENSG00000242314 | 0 | 0 | 0 | 0 | 0 | 0 |
| ENSG00000256987 | 0 | 0 | 0 | 0 | 0 | 0 |
| ENSG00000174718 | 82 | 67 | 53 | 16 | 130 | 71 |
| ENSG00000276136 | 0 | 0 | 0 | 3 | 2 | 1 |
| ENSG00000257094 | 0 | 0 | 0 | 0 | 0 | 0 |
| ENSG00000257852 | 0 | 0 | 0 | 0 | 0 | 0 |
| ENSG00000271357 | 0 | 0 | 0 | 0 | 0 | 0 |
| ENSG00000257530 | 0 | 0 | 0 | 0 | 0 | 0 |
| ENSG00000151746 | 279 | 173 | 213 | 109 | 267 | 231 |
| ENSG00000277342 | 2 | 0 | 0 | 4 | 1 | 3 |
| ENSG00000258134 | 0 | 0 | 0 | 0 | 0 | 0 |
| ENSG00000274964 | 22 | 11 | 11 | 16 | 15 | 21 |
| ENSG00000276115 | 2 | 2 | 1 | 2 | 6 | 3 |
| ENSG00000207176 | 0 | 0 | 0 | 0 | 0 | 0 |
| ENSG00000139132 | 352 | 186 | 201 | 68 | 273 | 136 |
| ENSG00000253063 | 0 | 0 | 0 | 0 | 0 | 0 |
| ENSG00000087470 | 1199 | 768 | 663 | 386 | 1225 | 840 |

| | | | | | | |
|---|---|---|---|---|---|---|
| ENSG00000257643 | 0 | 0 | 0 | 0 | 0 | 0 |
| ENSG00000201563 | 0 | 0 | 0 | 0 | 0 | 1 |
| ENSG00000276148 | 2 | 5 | 3 | 0 | 0 | 2 |
| ENSG00000257511 | 14 | 7 | 12 | 2 | 13 | 6 |
| ENSG00000139131 | 758 | 392 | 445 | 510 | 483 | 448 |
| ENSG00000274105 | 2 | 4 | 1 | 2 | 6 | 3 |
| ENSG00000275854 | 12 | 4 | 1 | 3 | 5 | 4 |
| ENSG00000057294 | 709 | 348 | 427 | 284 | 351 | 359 |
| ENSG00000274976 | 0 | 0 | 1 | 4 | 1 | 2 |
| ENSG00000244743 | 0 | 0 | 0 | 0 | 0 | 0 |
| ENSG00000257316 | 14 | 13 | 9 | 4 | 8 | 6 |
| ENSG00000257435 | 3 | 2 | 1 | 1 | 0 | 1 |
| ENSG00000110975 | 311 | 152 | 234 | 76 | 177 | 176 |
| ENSG00000259937 | 36 | 9 | 20 | 15 | 16 | 16 |
| ENSG00000273853 | 25 | 8 | 14 | 4 | 13 | 13 |
| ENSG00000256070 | 0 | 0 | 0 | 0 | 0 | 0 |
| ENSG00000212475 | 0 | 0 | 0 | 0 | 0 | 0 |
| ENSG00000270807 | 1 | 0 | 0 | 0 | 0 | 0 |
| ENSG00000207026 | 0 | 0 | 0 | 0 | 0 | 0 |
| ENSG00000201624 | 0 | 0 | 0 | 0 | 0 | 0 |
| ENSG00000245482 | 0 | 0 | 0 | 0 | 0 | 0 |
| ENSG00000139133 | 208 | 99 | 125 | 81 | 141 | 144 |
| ENSG00000256538 | 0 | 1 | 0 | 0 | 0 | 0 |
| ENSG00000255628 | 0 | 0 | 0 | 0 | 0 | 0 |
| ENSG00000256986 | 0 | 0 | 0 | 0 | 0 | 0 |
| ENSG00000250788 | 0 | 0 | 0 | 0 | 0 | 0 |
| ENSG00000255652 | 1 | 1 | 4 | 1 | 1 | 0 |
| ENSG00000272435 | 0 | 0 | 0 | 0 | 0 | 0 |
| ENSG00000258794 | 0 | 0 | 1 | 2 | 1 | 1 |
| ENSG00000256614 | 0 | 0 | 0 | 0 | 0 | 0 |
| ENSG00000258368 | 0 | 0 | 0 | 0 | 0 | 0 |
| ENSG00000257173 | 0 | 0 | 0 | 0 | 0 | 0 |
| ENSG00000258278 | 0 | 0 | 0 | 0 | 0 | 0 |
| ENSG00000274293 | 0 | 0 | 0 | 0 | 0 | 0 |
| ENSG00000199843 | 0 | 0 | 0 | 0 | 0 | 0 |
| ENSG00000201708 | 0 | 0 | 0 | 0 | 0 | 0 |
| ENSG00000254381 | 0 | 0 | 0 | 0 | 0 | 0 |
| ENSG00000257292 | 0 | 0 | 0 | 0 | 0 | 0 |
| ENSG00000258360 | 0 | 0 | 0 | 0 | 0 | 0 |
| ENSG00000253016 | 0 | 0 | 0 | 0 | 0 | 0 |
| ENSG00000175548 | 242 | 120 | 146 | 70 | 124 | 101 |
| ENSG00000257645 | 0 | 0 | 0 | 0 | 0 | 0 |
| ENSG00000258119 | 0 | 0 | 0 | 0 | 0 | 0 |
| ENSG00000139117 | 0 | 0 | 0 | 0 | 0 | 0 |
| ENSG00000257718 | 0 | 0 | 0 | 0 | 0 | 0 |
| ENSG00000257586 | 0 | 0 | 0 | 0 | 0 | 0 |
| ENSG00000257237 | 0 | 0 | 0 | 0 | 0 | 0 |
| ENSG00000258144 | 0 | 0 | 0 | 0 | 0 | 0 |
| ENSG00000270718 | 0 | 0 | 0 | 0 | 0 | 0 |
| ENSG00000139116 | 671 | 460 | 250 | 91 | 633 | 282 |

| | | | | | | |
|---|---|---|---|---|---|---|
| ENSG00000255991 | 0 | 0 | 0 | 0 | 1 | 1 |
| ENSG00000270273 | 0 | 0 | 0 | 0 | 0 | 0 |
| ENSG00000173208 | 0 | 0 | 0 | 0 | 0 | 0 |
| ENSG00000285732 | 0 | 0 | 0 | 0 | 0 | 0 |
| ENSG00000180116 | 0 | 0 | 0 | 0 | 0 | 0 |
| ENSG00000151229 | 207 | 98 | 102 | 46 | 78 | 92 |
| ENSG00000199571 | 0 | 1 | 0 | 0 | 0 | 0 |
| ENSG00000229014 | 0 | 0 | 0 | 0 | 0 | 0 |
| ENSG00000260943 | 0 | 0 | 0 | 0 | 0 | 0 |
| ENSG00000223914 | 0 | 0 | 0 | 0 | 0 | 0 |
| ENSG00000225342 | 0 | 0 | 0 | 0 | 0 | 0 |
| ENSG00000188906 | 2 | 0 | 1 | 0 | 3 | 0 |
| ENSG00000229899 | 0 | 0 | 0 | 0 | 0 | 1 |
| ENSG00000205592 | 2 | 0 | 2 | 1 | 0 | 1 |
| ENSG00000258167 | 0 | 0 | 0 | 0 | 0 | 0 |
| ENSG00000207492 | 0 | 0 | 0 | 0 | 0 | 0 |
| ENSG00000018236 | 0 | 1 | 0 | 0 | 2 | 0 |
| ENSG00000257680 | 0 | 0 | 0 | 0 | 0 | 0 |
| ENSG00000274682 | 0 | 0 | 0 | 0 | 0 | 0 |
| ENSG00000165966 | 0 | 0 | 0 | 0 | 0 | 0 |
| ENSG00000257228 | 0 | 0 | 0 | 0 | 0 | 0 |
| ENSG00000252364 | 0 | 0 | 0 | 0 | 0 | 0 |
| ENSG00000258239 | 0 | 0 | 0 | 0 | 0 | 0 |
| ENSG00000257858 | 0 | 0 | 0 | 0 | 0 | 0 |
| ENSG00000257784 | 0 | 0 | 0 | 0 | 0 | 0 |
| ENSG00000257239 | 0 | 0 | 0 | 0 | 0 | 0 |
| ENSG00000257128 | 0 | 0 | 0 | 0 | 0 | 0 |
| ENSG00000271379 | 4 | 0 | 0 | 0 | 0 | 0 |
| ENSG00000151233 | 626 | 357 | 262 | 147 | 471 | 278 |
| ENSG00000015153 | 318 | 185 | 211 | 157 | 195 | 174 |
| ENSG00000134283 | 1552 | 838 | 988 | 636 | 987 | 900 |
| ENSG00000257674 | 0 | 0 | 1 | 0 | 0 | 1 |
| ENSG00000139168 | 1635 | 794 | 880 | 790 | 1036 | 818 |
| ENSG00000278520 | 1 | 0 | 1 | 0 | 0 | 0 |
| ENSG00000252917 | 0 | 0 | 2 | 0 | 0 | 0 |
| ENSG00000274943 | 13 | 5 | 4 | 7 | 9 | 5 |
| ENSG00000199886 | 0 | 0 | 0 | 0 | 0 | 0 |
| ENSG00000257376 | 2 | 0 | 2 | 0 | 0 | 0 |
| ENSG00000207142 | 0 | 0 | 0 | 0 | 0 | 0 |
| ENSG00000139174 | 6 | 4 | 3 | 5 | 2 | 1 |
| ENSG00000257225 | 0 | 0 | 0 | 0 | 0 | 1 |
| ENSG00000258068 | 0 | 0 | 0 | 0 | 0 | 0 |
| ENSG00000257510 | 0 | 0 | 0 | 0 | 0 | 0 |
| ENSG00000257687 | 0 | 0 | 0 | 0 | 0 | 0 |
| ENSG00000257405 | 0 | 0 | 0 | 0 | 0 | 0 |
| ENSG00000257164 | 0 | 0 | 0 | 0 | 0 | 0 |
| ENSG00000240759 | 0 | 0 | 0 | 0 | 1 | 0 |
| ENSG00000257114 | 0 | 0 | 1 | 1 | 1 | 0 |
| ENSG00000271272 | 0 | 0 | 0 | 0 | 0 | 0 |
| ENSG00000257849 | 0 | 0 | 0 | 0 | 0 | 0 |

| | | | | | | |
|---|---|---|---|---|---|---|
| ENSG00000257373 | 0 | 0 | 0 | 0 | 0 | 0 |
| ENSG00000257927 | 0 | 0 | 0 | 0 | 0 | 0 |
| ENSG00000258331 | 0 | 0 | 0 | 0 | 0 | 0 |
| ENSG00000173157 | 36 | 16 | 10 | 5 | 22 | 20 |
| ENSG00000241251 | 0 | 0 | 0 | 0 | 0 | 0 |
| ENSG00000257813 | 0 | 0 | 0 | 0 | 0 | 0 |
| ENSG00000271508 | 0 | 0 | 0 | 0 | 0 | 0 |
| ENSG00000257907 | 2 | 1 | 0 | 1 | 0 | 0 |
| ENSG00000129317 | 374 | 269 | 203 | 112 | 344 | 198 |
| ENSG00000257896 | 0 | 0 | 3 | 1 | 2 | 1 |
| ENSG00000198001 | 881 | 410 | 482 | 281 | 506 | 432 |
| ENSG00000151239 | 3147 | 1733 | 1529 | 973 | 2199 | 1431 |
| ENSG00000139173 | 397 | 177 | 191 | 144 | 187 | 176 |
| ENSG00000258212 | 0 | 0 | 0 | 0 | 0 | 0 |
| ENSG00000257947 | 0 | 0 | 0 | 0 | 0 | 0 |
| ENSG00000257864 | 0 | 0 | 0 | 0 | 0 | 0 |
| ENSG00000275286 | 0 | 0 | 0 | 0 | 0 | 0 |
| ENSG00000184613 | 156 | 72 | 136 | 28 | 18 | 57 |
| ENSG00000201788 | 0 | 0 | 0 | 0 | 0 | 0 |
| ENSG00000185610 | 0 | 0 | 0 | 0 | 0 | 0 |
| ENSG00000257319 | 0 | 0 | 0 | 0 | 0 | 0 |
| ENSG00000257331 | 0 | 0 | 0 | 0 | 0 | 0 |
| ENSG00000257572 | 0 | 0 | 0 | 0 | 0 | 0 |
| ENSG00000199556 | 0 | 0 | 0 | 0 | 0 | 0 |
| ENSG00000134297 | 89 | 78 | 64 | 35 | 92 | 92 |
| ENSG00000257738 | 0 | 0 | 0 | 0 | 0 | 0 |
| ENSG00000177119 | 2327 | 1365 | 1549 | 727 | 1462 | 1340 |
| ENSG00000278351 | 0 | 0 | 1 | 0 | 0 | 0 |
| ENSG00000257368 | 0 | 0 | 0 | 0 | 0 | 0 |
| ENSG00000257657 | 0 | 0 | 0 | 0 | 0 | 0 |
| ENSG00000283545 | 0 | 0 | 0 | 0 | 0 | 0 |
| ENSG00000273015 | 909 | 432 | 568 | 374 | 411 | 434 |
| ENSG00000189079 | 1262 | 677 | 694 | 288 | 962 | 678 |
| ENSG00000278475 | 0 | 0 | 1 | 0 | 0 | 0 |
| ENSG00000265093 | 0 | 0 | 0 | 0 | 2 | 0 |
| ENSG00000270429 | 0 | 0 | 1 | 0 | 0 | 0 |
| ENSG00000139218 | 1703 | 982 | 781 | 333 | 1559 | 822 |
| ENSG00000239397 | 2 | 0 | 0 | 1 | 0 | 0 |
| ENSG00000111371 | 23969 | 12032 | 12993 | 5757 | 12683 | 10352 |
| ENSG00000274591 | 0 | 1 | 3 | 0 | 1 | 0 |
| ENSG00000134294 | 13902 | 6929 | 7428 | 4498 | 8727 | 6490 |
| ENSG00000258096 | 2 | 0 | 1 | 0 | 4 | 4 |
| ENSG00000257261 | 128 | 81 | 124 | 92 | 87 | 99 |
| ENSG00000257496 | 4 | 0 | 2 | 1 | 2 | 2 |
| ENSG00000275481 | 22 | 8 | 10 | 9 | 11 | 12 |
| ENSG00000278896 | 0 | 1 | 1 | 0 | 0 | 3 |
| ENSG00000272369 | 59 | 28 | 38 | 28 | 31 | 21 |
| ENSG00000272963 | 0 | 0 | 0 | 0 | 0 | 0 |
| ENSG00000271642 | 0 | 0 | 0 | 0 | 0 | 0 |
| ENSG00000139209 | 0 | 0 | 0 | 0 | 0 | 0 |

| | | | | | | |
|---|---|---|---|---|---|---|
| ENSG00000274723 | 0 | 1 | 3 | 0 | 4 | 1 |
| ENSG00000139211 | 1050 | 525 | 877 | 464 | 683 | 728 |
| ENSG00000179715 | 295 | 127 | 190 | 131 | 135 | 168 |
| ENSG00000258352 | 2 | 1 | 1 | 3 | 1 | 1 |
| ENSG00000263838 | 0 | 0 | 1 | 0 | 0 | 0 |
| ENSG00000247774 | 0 | 2 | 3 | 0 | 1 | 1 |
| ENSG00000257925 | 0 | 0 | 0 | 0 | 0 | 2 |
| ENSG00000258181 | 6 | 4 | 5 | 1 | 3 | 6 |
| ENSG00000276454 | 0 | 0 | 0 | 0 | 0 | 0 |
| ENSG00000258116 | 0 | 0 | 0 | 0 | 0 | 0 |
| ENSG00000199566 | 0 | 0 | 0 | 0 | 0 | 0 |
| ENSG00000257924 | 0 | 0 | 0 | 0 | 0 | 0 |
| ENSG00000264906 | 0 | 0 | 0 | 0 | 0 | 0 |
| ENSG00000257906 | 0 | 0 | 0 | 1 | 0 | 0 |
| ENSG00000258369 | 0 | 0 | 0 | 0 | 0 | 0 |
| ENSG00000257807 | 0 | 0 | 0 | 0 | 0 | 0 |
| ENSG00000257905 | 0 | 0 | 0 | 0 | 1 | 0 |
| ENSG00000005175 | 699 | 424 | 304 | 150 | 602 | 364 |
| ENSG00000276390 | 8 | 4 | 8 | 2 | 11 | 14 |
| ENSG00000257433 | 45 | 19 | 20 | 6 | 17 | 19 |
| ENSG00000111405 | 4 | 1 | 5 | 0 | 2 | 2 |
| ENSG00000280054 | 23 | 11 | 14 | 9 | 20 | 9 |
| ENSG00000274902 | 4 | 4 | 7 | 2 | 4 | 5 |
| ENSG00000079337 | 199 | 95 | 120 | 73 | 93 | 57 |
| ENSG00000211584 | 1211 | 577 | 737 | 702 | 765 | 692 |
| ENSG00000277173 | 0 | 0 | 0 | 1 | 0 | 0 |
| ENSG00000061273 | 2207 | 1017 | 1282 | 1133 | 1136 | 996 |
| ENSG00000268069 | 2 | 1 | 2 | 0 | 1 | 0 |
| ENSG00000276691 | 2 | 2 | 2 | 3 | 0 | 2 |
| ENSG00000274737 | 2 | 3 | 2 | 0 | 3 | 1 |
| ENSG00000257488 | 1 | 2 | 2 | 1 | 2 | 0 |
| ENSG00000111424 | 588 | 327 | 368 | 336 | 495 | 500 |
| ENSG00000205537 | 0 | 0 | 0 | 0 | 0 | 0 |
| ENSG00000278385 | 79 | 50 | 61 | 57 | 40 | 54 |
| ENSG00000279840 | 0 | 0 | 0 | 0 | 0 | 0 |
| ENSG00000134291 | 7926 | 4044 | 5164 | 3795 | 4079 | 3999 |
| ENSG00000139219 | 90 | 36 | 42 | 18 | 24 | 29 |
| ENSG00000258203 | 46 | 14 | 25 | 28 | 19 | 16 |
| ENSG00000257985 | 0 | 0 | 0 | 0 | 1 | 0 |
| ENSG00000257955 | 0 | 1 | 0 | 0 | 0 | 0 |
| ENSG00000276814 | 6 | 2 | 3 | 1 | 3 | 4 |
| ENSG00000079387 | 989 | 477 | 553 | 324 | 539 | 456 |
| ENSG00000240399 | 4 | 8 | 5 | 3 | 4 | 1 |
| ENSG00000222635 | 1 | 0 | 0 | 0 | 0 | 0 |
| ENSG00000226138 | 0 | 0 | 0 | 0 | 0 | 0 |
| ENSG00000152556 | 2587 | 1171 | 1593 | 1298 | 1360 | 1534 |
| ENSG00000275770 | 0 | 0 | 0 | 0 | 1 | 0 |
| ENSG00000177981 | 1328 | 628 | 747 | 501 | 681 | 582 |
| ENSG00000274124 | 2 | 0 | 3 | 1 | 6 | 5 |
| ENSG00000258051 | 0 | 0 | 0 | 0 | 0 | 0 |

| | | | | | | |
|---|---|---|---|---|---|---|
| ENSG00000257885 | 0 | 0 | 0 | 0 | 0 | 0 |
| ENSG00000257848 | 0 | 0 | 0 | 0 | 0 | 0 |
| ENSG00000177875 | 20 | 7 | 8 | 4 | 6 | 7 |
| ENSG00000279875 | 0 | 0 | 0 | 0 | 0 | 0 |
| ENSG00000269514 | 4 | 1 | 8 | 1 | 1 | 2 |
| ENSG00000172640 | 0 | 2 | 1 | 1 | 0 | 0 |
| ENSG00000258234 | 0 | 0 | 0 | 0 | 0 | 0 |
| ENSG00000275228 | 0 | 0 | 0 | 0 | 0 | 0 |
| ENSG00000187166 | 0 | 0 | 0 | 0 | 0 | 1 |
| ENSG00000258273 | 1 | 0 | 0 | 0 | 0 | 2 |
| ENSG00000167528 | 1344 | 633 | 878 | 560 | 660 | 617 |
| ENSG00000257735 | 2 | 1 | 1 | 0 | 0 | 2 |
| ENSG00000257763 | 2 | 1 | 1 | 1 | 1 | 1 |
| ENSG00000273765 | 2 | 1 | 1 | 0 | 0 | 0 |
| ENSG00000258024 | 0 | 0 | 0 | 0 | 0 | 0 |
| ENSG00000257792 | 0 | 0 | 0 | 0 | 0 | 0 |
| ENSG00000273434 | 0 | 0 | 0 | 0 | 0 | 0 |
| ENSG00000226413 | 0 | 0 | 0 | 0 | 1 | 1 |
| ENSG00000139223 | 0 | 0 | 0 | 0 | 0 | 1 |
| ENSG00000177627 | 6 | 2 | 0 | 1 | 1 | 1 |
| ENSG00000258121 | 0 | 0 | 0 | 0 | 1 | 0 |
| ENSG00000240443 | 0 | 0 | 0 | 0 | 0 | 0 |
| ENSG00000197376 | 0 | 0 | 0 | 0 | 0 | 1 |
| ENSG00000284723 | 0 | 0 | 0 | 0 | 0 | 0 |
| ENSG00000198678 | 0 | 0 | 0 | 0 | 0 | 0 |
| ENSG00000167531 | 0 | 0 | 0 | 0 | 0 | 0 |
| ENSG00000224709 | 0 | 0 | 0 | 0 | 0 | 0 |
| ENSG00000139620 | 1040 | 526 | 654 | 530 | 584 | 632 |
| ENSG00000221491 | 1 | 0 | 0 | 3 | 1 | 4 |
| ENSG00000281842 | 1 | 0 | 0 | 0 | 2 | 0 |
| ENSG00000206612 | 0 | 3 | 1 | 2 | 0 | 5 |
| ENSG00000207313 | 2 | 0 | 1 | 2 | 0 | 1 |
| ENSG00000129315 | 2079 | 1017 | 1238 | 753 | 1149 | 1054 |
| ENSG00000257987 | 1 | 0 | 0 | 0 | 0 | 0 |
| ENSG00000174233 | 3120 | 1304 | 1870 | 1356 | 1632 | 1546 |
| ENSG00000257653 | 0 | 0 | 0 | 0 | 0 | 0 |
| ENSG00000264201 | 5 | 1 | 4 | 0 | 2 | 5 |
| ENSG00000257660 | 2 | 3 | 0 | 1 | 2 | 4 |
| ENSG00000271547 | 11 | 2 | 10 | 7 | 8 | 6 |
| ENSG00000207467 | 4 | 2 | 0 | 0 | 1 | 1 |
| ENSG00000167535 | 2522 | 1116 | 1546 | 1387 | 1270 | 1352 |
| ENSG00000174243 | 6110 | 2922 | 3505 | 3201 | 3416 | 3530 |
| ENSG00000172602 | 239 | 130 | 164 | 372 | 463 | 384 |
| ENSG00000199394 | 0 | 0 | 0 | 0 | 0 | 1 |
| ENSG00000239617 | 0 | 1 | 0 | 2 | 0 | 0 |
| ENSG00000272822 | 1 | 0 | 0 | 3 | 0 | 2 |
| ENSG00000139537 | 59 | 46 | 23 | 29 | 32 | 39 |
| ENSG00000134285 | 456 | 216 | 295 | 277 | 290 | 327 |
| ENSG00000255863 | 0 | 0 | 0 | 0 | 0 | 0 |
| ENSG00000134287 | 14046 | 7370 | 8739 | 7741 | 8460 | 8276 |

| | | | | | | |
|---|---|---|---|---|---|---|
| ENSG00000169884 | 54 | 19 | 26 | 28 | 24 | 25 |
| ENSG00000200303 | 0 | 0 | 0 | 0 | 0 | 0 |
| ENSG00000125084 | 0 | 0 | 0 | 0 | 0 | 0 |
| ENSG00000181418 | 13 | 7 | 10 | 10 | 6 | 13 |
| ENSG00000258283 | 0 | 0 | 0 | 0 | 0 | 0 |
| ENSG00000257913 | 5 | 7 | 2 | 3 | 0 | 4 |
| ENSG00000181929 | 2860 | 1426 | 1731 | 1494 | 1616 | 1562 |
| ENSG00000167548 | 4608 | 1702 | 2929 | 1644 | 2167 | 2305 |
| ENSG00000167550 | 9 | 7 | 13 | 7 | 4 | 7 |
| ENSG00000277672 | 0 | 0 | 0 | 0 | 0 | 0 |
| ENSG00000139549 | 0 | 0 | 0 | 2 | 0 | 0 |
| ENSG00000257346 | 0 | 0 | 0 | 0 | 0 | 0 |
| ENSG00000139636 | 898 | 368 | 551 | 528 | 460 | 460 |
| ENSG00000258017 | 2 | 0 | 4 | 0 | 3 | 0 |
| ENSG00000123416 | 5215 | 3926 | 4265 | 5211 | 5871 | 5597 |
| ENSG00000200309 | 0 | 0 | 0 | 0 | 0 | 0 |
| ENSG00000167552 | 4165 | 2633 | 2542 | 3491 | 3881 | 3231 |
| ENSG00000167553 | 4489 | 3047 | 3475 | 4323 | 4817 | 4648 |
| ENSG00000274017 | 0 | 0 | 0 | 0 | 0 | 0 |
| ENSG00000258101 | 44 | 20 | 21 | 33 | 29 | 26 |
| ENSG00000258232 | 0 | 1 | 0 | 0 | 0 | 0 |
| ENSG00000258334 | 8 | 8 | 3 | 1 | 4 | 3 |
| ENSG00000135406 | 40 | 27 | 30 | 31 | 23 | 26 |
| ENSG00000135451 | 714 | 342 | 441 | 517 | 491 | 534 |
| ENSG00000186897 | 6 | 3 | 7 | 6 | 4 | 3 |
| ENSG00000178401 | 480 | 245 | 253 | 180 | 223 | 200 |
| ENSG00000123352 | 1787 | 826 | 1058 | 624 | 1078 | 899 |
| ENSG00000257954 | 2 | 0 | 2 | 1 | 3 | 1 |
| ENSG00000257464 | 0 | 0 | 0 | 0 | 1 | 1 |
| ENSG00000135519 | 79 | 40 | 71 | 47 | 37 | 40 |
| ENSG00000187778 | 3532 | 1712 | 2225 | 2294 | 1968 | 1944 |
| ENSG00000110844 | 248 | 109 | 163 | 119 | 120 | 118 |
| ENSG00000257964 | 0 | 2 | 1 | 1 | 0 | 1 |
| ENSG00000135436 | 24 | 8 | 22 | 8 | 7 | 17 |
| ENSG00000199237 | 3 | 0 | 1 | 0 | 2 | 0 |
| ENSG00000258284 | 0 | 3 | 4 | 0 | 2 | 1 |
| ENSG00000257243 | 3 | 2 | 3 | 0 | 3 | 0 |
| ENSG00000258104 | 0 | 0 | 0 | 0 | 0 | 0 |
| ENSG00000161791 | 225 | 119 | 141 | 112 | 117 | 104 |
| ENSG00000139644 | 57482 | 27421 | 33758 | 29457 | 32222 | 32549 |
| ENSG00000257570 | 0 | 0 | 0 | 0 | 0 | 0 |
| ENSG00000257730 | 0 | 0 | 0 | 0 | 0 | 0 |
| ENSG00000167566 | 1929 | 791 | 986 | 1005 | 917 | 936 |
| ENSG00000258057 | 39 | 14 | 35 | 15 | 43 | 18 |
| ENSG00000186666 | 307 | 176 | 206 | 184 | 217 | 194 |
| ENSG00000257253 | 0 | 0 | 0 | 0 | 0 | 0 |
| ENSG00000242041 | 0 | 0 | 0 | 0 | 0 | 0 |
| ENSG00000135472 | 0 | 4 | 1 | 1 | 4 | 4 |
| ENSG00000258135 | 0 | 0 | 3 | 0 | 0 | 0 |
| ENSG00000257771 | 2 | 3 | 0 | 0 | 2 | 1 |

| | | | | | | |
|---|---|---|---|---|---|---|
| ENSG00000167580 | 2 | 1 | 1 | 0 | 1 | 0 |
| ENSG00000257588 | 2 | 4 | 3 | 1 | 4 | 2 |
| ENSG00000257378 | 0 | 0 | 0 | 0 | 0 | 0 |
| ENSG00000161798 | 0 | 1 | 0 | 0 | 0 | 0 |
| ENSG00000086159 | 4 | 6 | 7 | 3 | 6 | 3 |
| ENSG00000161800 | 2467 | 1270 | 1484 | 1271 | 1673 | 1503 |
| ENSG00000110881 | 387 | 174 | 234 | 193 | 203 | 213 |
| ENSG00000066117 | 2851 | 1275 | 1629 | 1392 | 1352 | 1522 |
| ENSG00000167588 | 23 | 10 | 17 | 14 | 16 | 9 |
| ENSG00000178449 | 1462 | 870 | 1108 | 1276 | 972 | 1039 |
| ENSG00000272368 | 22 | 10 | 22 | 24 | 14 | 25 |
| ENSG00000139624 | 1412 | 717 | 880 | 887 | 912 | 890 |
| ENSG00000050405 | 2133 | 1082 | 1294 | 766 | 1219 | 1021 |
| ENSG00000257298 | 7 | 2 | 4 | 0 | 0 | 1 |
| ENSG00000257531 | 0 | 0 | 0 | 0 | 0 | 0 |
| ENSG00000257256 | 0 | 0 | 1 | 1 | 0 | 1 |
| ENSG00000221604 | 0 | 0 | 0 | 0 | 0 | 0 |
| ENSG00000212496 | 0 | 0 | 0 | 0 | 0 | 0 |
| ENSG00000278842 | 2 | 2 | 1 | 0 | 2 | 1 |
| ENSG00000185958 | 12 | 1 | 17 | 10 | 15 | 11 |
| ENSG00000244266 | 0 | 0 | 0 | 0 | 0 | 0 |
| ENSG00000161813 | 2532 | 1314 | 1301 | 649 | 1677 | 1158 |
| ENSG00000066084 | 3941 | 1706 | 2458 | 1268 | 2092 | 1966 |
| ENSG00000207136 | 0 | 0 | 0 | 0 | 0 | 0 |
| ENSG00000200183 | 0 | 1 | 0 | 0 | 0 | 0 |
| ENSG00000200428 | 0 | 1 | 0 | 0 | 2 | 1 |
| ENSG00000271018 | 2 | 1 | 2 | 0 | 1 | 1 |
| ENSG00000123268 | 941 | 461 | 452 | 329 | 479 | 383 |
| ENSG00000271490 | 0 | 0 | 0 | 0 | 0 | 0 |
| ENSG00000243075 | 0 | 0 | 0 | 0 | 0 | 0 |
| ENSG00000186452 | 0 | 1 | 0 | 1 | 0 | 0 |
| ENSG00000271621 | 0 | 0 | 0 | 0 | 0 | 0 |
| ENSG00000185432 | 330 | 137 | 164 | 85 | 69 | 102 |
| ENSG00000274156 | 0 | 0 | 0 | 0 | 0 | 0 |
| ENSG00000199740 | 0 | 0 | 0 | 1 | 1 | 0 |
| ENSG00000271596 | 0 | 0 | 0 | 0 | 0 | 0 |
| ENSG00000214511 | 2 | 0 | 0 | 0 | 0 | 0 |
| ENSG00000274797 | 0 | 0 | 0 | 0 | 0 | 0 |
| ENSG00000110911 | 2771 | 1258 | 1662 | 1057 | 1470 | 1425 |
| ENSG00000272028 | 0 | 0 | 0 | 0 | 0 | 0 |
| ENSG00000199903 | 0 | 0 | 0 | 0 | 0 | 0 |
| ENSG00000050426 | 3016 | 1425 | 2018 | 1813 | 2053 | 1830 |
| ENSG00000110925 | 1515 | 744 | 885 | 694 | 847 | 833 |
| ENSG00000135457 | 0 | 0 | 0 | 0 | 0 | 0 |
| ENSG00000244540 | 0 | 0 | 0 | 0 | 0 | 0 |
| ENSG00000257246 | 0 | 0 | 0 | 0 | 0 | 0 |
| ENSG00000200953 | 0 | 0 | 0 | 0 | 0 | 0 |
| ENSG00000199824 | 0 | 0 | 0 | 0 | 0 | 0 |
| ENSG00000184271 | 7 | 2 | 2 | 1 | 5 | 5 |
| ENSG00000278126 | 0 | 0 | 0 | 1 | 3 | 2 |

| | | | | | | |
|---|---|---|---|---|---|---|
| ENSG00000183283 | 13147 | 6279 | 8100 | 5600 | 6130 | 7003 |
| ENSG00000170545 | 739 | 389 | 441 | 401 | 356 | 373 |
| ENSG00000110934 | 22 | 8 | 5 | 1 | 5 | 2 |
| ENSG00000139610 | 0 | 0 | 0 | 0 | 0 | 0 |
| ENSG00000139629 | 1944 | 1030 | 1198 | 945 | 1324 | 1115 |
| ENSG00000050438 | 466 | 214 | 254 | 65 | 163 | 128 |
| ENSG00000271065 | 0 | 0 | 0 | 0 | 0 | 0 |
| ENSG00000196876 | 267 | 120 | 136 | 32 | 66 | 86 |
| ENSG00000257851 | 2 | 0 | 2 | 1 | 1 | 0 |
| ENSG00000260122 | 0 | 0 | 0 | 0 | 0 | 0 |
| ENSG00000284730 | 0 | 0 | 0 | 0 | 0 | 0 |
| ENSG00000260473 | 0 | 0 | 0 | 0 | 0 | 0 |
| ENSG00000261308 | 4 | 0 | 0 | 0 | 0 | 2 |
| ENSG00000261586 | 0 | 0 | 0 | 0 | 0 | 0 |
| ENSG00000259887 | 0 | 0 | 0 | 0 | 0 | 0 |
| ENSG00000167612 | 0 | 0 | 0 | 0 | 0 | 0 |
| ENSG00000258021 | 0 | 0 | 0 | 0 | 0 | 0 |
| ENSG00000139567 | 0 | 0 | 1 | 1 | 1 | 0 |
| ENSG00000135503 | 1562 | 772 | 1010 | 745 | 834 | 938 |
| ENSG00000206992 | 1 | 1 | 2 | 0 | 0 | 0 |
| ENSG00000161835 | 0 | 4 | 2 | 3 | 4 | 3 |
| ENSG00000123358 | 125 | 83 | 111 | 99 | 95 | 78 |
| ENSG00000259884 | 1 | 1 | 1 | 1 | 1 | 0 |
| ENSG00000123395 | 1052 | 544 | 701 | 798 | 695 | 669 |
| ENSG00000257663 | 0 | 0 | 0 | 0 | 0 | 0 |
| ENSG00000284791 | 0 | 0 | 0 | 0 | 0 | 0 |
| ENSG00000257542 | 0 | 0 | 0 | 0 | 0 | 0 |
| ENSG00000257989 | 0 | 1 | 0 | 0 | 0 | 0 |
| ENSG00000258279 | 44 | 39 | 34 | 39 | 62 | 28 |
| ENSG00000167767 | 34648 | 18967 | 18894 | 22715 | 25434 | 19681 |
| ENSG00000257137 | 43 | 24 | 14 | 18 | 29 | 18 |
| ENSG00000257649 | 0 | 0 | 1 | 0 | 1 | 0 |
| ENSG00000135480 | 245 | 158 | 160 | 251 | 264 | 192 |
| ENSG00000257671 | 31 | 13 | 14 | 12 | 20 | 16 |
| ENSG00000170442 | 353 | 249 | 218 | 405 | 403 | 243 |
| ENSG00000135477 | 20 | 16 | 17 | 14 | 21 | 17 |
| ENSG00000284883 | 2 | 2 | 4 | 1 | 1 | 0 |
| ENSG00000257830 | 2 | 2 | 4 | 3 | 5 | 4 |
| ENSG00000285102 | 2 | 0 | 1 | 0 | 1 | 0 |
| ENSG00000205426 | 1246 | 708 | 576 | 1447 | 1213 | 924 |
| ENSG00000257829 | 54 | 47 | 39 | 52 | 57 | 29 |
| ENSG00000170523 | 187 | 112 | 105 | 171 | 149 | 114 |
| ENSG00000274928 | 148 | 117 | 111 | 80 | 108 | 75 |
| ENSG00000135443 | 2 | 0 | 0 | 0 | 0 | 0 |
| ENSG00000161849 | 0 | 0 | 0 | 0 | 0 | 0 |
| ENSG00000258253 | 0 | 0 | 0 | 0 | 0 | 0 |
| ENSG00000161850 | 0 | 0 | 0 | 0 | 0 | 0 |
| ENSG00000257500 | 0 | 0 | 0 | 0 | 0 | 0 |
| ENSG00000257844 | 0 | 0 | 0 | 0 | 0 | 0 |
| ENSG00000170454 | 0 | 0 | 0 | 0 | 0 | 0 |

| | | | | | | |
|---|---|---|---|---|---|---|
| ENSG00000185479 | 2 | 0 | 1 | 1 | 0 | 0 |
| ENSG00000170465 | 0 | 0 | 0 | 1 | 0 | 0 |
| ENSG00000205420 | 4 | 0 | 0 | 0 | 1 | 0 |
| ENSG00000186081 | 0 | 0 | 0 | 0 | 0 | 1 |
| ENSG00000139648 | 0 | 0 | 0 | 0 | 0 | 0 |
| ENSG00000170484 | 2 | 0 | 0 | 0 | 0 | 0 |
| ENSG00000170486 | 0 | 0 | 0 | 0 | 0 | 0 |
| ENSG00000257495 | 0 | 0 | 0 | 0 | 0 | 0 |
| ENSG00000186049 | 0 | 0 | 0 | 0 | 0 | 0 |
| ENSG00000258120 | 0 | 0 | 0 | 0 | 0 | 0 |
| ENSG00000172867 | 0 | 0 | 0 | 0 | 0 | 0 |
| ENSG00000167768 | 4 | 0 | 0 | 0 | 0 | 0 |
| ENSG00000189182 | 0 | 0 | 0 | 0 | 0 | 0 |
| ENSG00000257700 | 0 | 0 | 0 | 0 | 0 | 0 |
| ENSG00000257402 | 0 | 0 | 0 | 0 | 0 | 0 |
| ENSG00000258036 | 0 | 0 | 0 | 0 | 0 | 0 |
| ENSG00000257616 | 1 | 0 | 0 | 0 | 0 | 0 |
| ENSG00000257343 | 0 | 0 | 0 | 0 | 0 | 0 |
| ENSG00000257675 | 0 | 0 | 0 | 0 | 0 | 0 |
| ENSG00000257125 | 0 | 0 | 0 | 0 | 0 | 0 |
| ENSG00000258071 | 0 | 0 | 0 | 0 | 0 | 0 |
| ENSG00000185069 | 0 | 0 | 0 | 0 | 0 | 0 |
| ENSG00000257404 | 0 | 0 | 0 | 0 | 0 | 0 |
| ENSG00000186442 | 12 | 6 | 0 | 0 | 1 | 2 |
| ENSG00000170477 | 10 | 5 | 6 | 1 | 4 | 4 |
| ENSG00000185640 | 0 | 0 | 0 | 0 | 0 | 0 |
| ENSG00000257389 | 0 | 0 | 0 | 0 | 0 | 0 |
| ENSG00000170423 | 0 | 0 | 0 | 0 | 0 | 0 |
| ENSG00000241146 | 0 | 0 | 2 | 0 | 0 | 1 |
| ENSG00000170421 | 546703 | 252028 | 322286 | 271067 | 201086 | 221019 |
| ENSG00000111057 | 178967 | 97725 | 115288 | 95412 | 78083 | 80157 |
| ENSG00000063046 | 31766 | 16468 | 20559 | 16492 | 19760 | 18323 |
| ENSG00000257475 | 1 | 1 | 1 | 0 | 0 | 2 |
| ENSG00000257337 | 467 | 281 | 390 | 223 | 280 | 246 |
| ENSG00000111077 | 282 | 141 | 196 | 127 | 109 | 126 |
| ENSG00000278108 | 0 | 0 | 1 | 0 | 0 | 0 |
| ENSG00000167778 | 3763 | 1640 | 2106 | 2082 | 1894 | 1934 |
| ENSG00000167779 | 26 | 7 | 14 | 24 | 11 | 11 |
| ENSG00000167780 | 0 | 0 | 1 | 0 | 2 | 0 |
| ENSG00000258074 | 0 | 0 | 0 | 0 | 0 | 0 |
| ENSG00000238669 | 0 | 0 | 0 | 0 | 0 | 0 |
| ENSG00000258016 | 0 | 1 | 0 | 2 | 0 | 1 |
| ENSG00000257790 | 0 | 2 | 0 | 0 | 1 | 1 |
| ENSG00000139631 | 1050 | 501 | 665 | 341 | 402 | 447 |
| ENSG00000257808 | 3 | 0 | 1 | 1 | 0 | 0 |
| ENSG00000139651 | 3182 | 1426 | 1943 | 1214 | 1512 | 1510 |
| ENSG00000139626 | 49 | 18 | 28 | 9 | 18 | 9 |
| ENSG00000172819 | 5455 | 2506 | 3096 | 2970 | 2686 | 2697 |
| ENSG00000283536 | 0 | 0 | 0 | 0 | 1 | 0 |
| ENSG00000182544 | 2274 | 1048 | 1153 | 1233 | 1113 | 1078 |

| | | | | | | |
|---|---|---|---|---|---|---|
| ENSG00000135476 | 1162 | 588 | 724 | 671 | 685 | 781 |
| ENSG00000123349 | 8479 | 4227 | 5098 | 5221 | 4958 | 4596 |
| ENSG00000257605 | 69 | 11 | 25 | 18 | 19 | 22 |
| ENSG00000139637 | 1519 | 844 | 980 | 1115 | 907 | 890 |
| ENSG00000094914 | 2131 | 962 | 1392 | 1211 | 1031 | 1131 |
| ENSG00000170374 | 2 | 0 | 0 | 0 | 1 | 6 |
| ENSG00000185591 | 6008 | 2648 | 3698 | 1781 | 2542 | 2654 |
| ENSG00000135409 | 4 | 7 | 4 | 0 | 5 | 2 |
| ENSG00000205352 | 2440 | 1397 | 1610 | 1621 | 1620 | 1610 |
| ENSG00000257379 | 0 | 0 | 4 | 2 | 2 | 0 |
| ENSG00000197111 | 8086 | 4508 | 5542 | 5305 | 4998 | 5076 |
| ENSG00000282977 | 9 | 1 | 6 | 3 | 1 | 3 |
| ENSG00000273658 | 0 | 0 | 0 | 0 | 0 | 0 |
| ENSG00000139625 | 589 | 222 | 386 | 218 | 268 | 263 |
| ENSG00000270175 | 40 | 13 | 22 | 15 | 10 | 12 |
| ENSG00000139546 | 1960 | 869 | 1093 | 1174 | 998 | 990 |
| ENSG00000267281 | 18 | 5 | 15 | 10 | 14 | 21 |
| ENSG00000139574 | 0 | 0 | 0 | 0 | 0 | 0 |
| ENSG00000170653 | 847 | 374 | 501 | 350 | 420 | 468 |
| ENSG00000257550 | 0 | 0 | 0 | 0 | 1 | 1 |
| ENSG00000285692 | 0 | 1 | 0 | 0 | 1 | 0 |
| ENSG00000275488 | 0 | 0 | 1 | 0 | 0 | 0 |
| ENSG00000135390 | 8295 | 4246 | 5047 | 5076 | 4569 | 4533 |
| ENSG00000213470 | 0 | 0 | 0 | 0 | 0 | 0 |
| ENSG00000012822 | 3252 | 1339 | 2176 | 1336 | 1460 | 1560 |
| ENSG00000260492 | 0 | 0 | 0 | 0 | 0 | 0 |
| ENSG00000260030 | 0 | 0 | 0 | 0 | 0 | 0 |
| ENSG00000253053 | 0 | 0 | 0 | 0 | 0 | 0 |
| ENSG00000249641 | 209 | 128 | 118 | 123 | 133 | 129 |
| ENSG00000123364 | 1307 | 697 | 751 | 733 | 711 | 721 |
| ENSG00000123407 | 7 | 4 | 8 | 3 | 4 | 3 |
| ENSG00000228630 | 269 | 141 | 162 | 132 | 204 | 162 |
| ENSG00000275589 | 0 | 0 | 0 | 0 | 0 | 0 |
| ENSG00000277994 | 0 | 0 | 0 | 0 | 0 | 0 |
| ENSG00000274817 | 2 | 1 | 1 | 3 | 3 | 3 |
| ENSG00000277129 | 5 | 1 | 4 | 4 | 3 | 0 |
| ENSG00000123388 | 373 | 141 | 198 | 209 | 225 | 198 |
| ENSG00000251151 | 2 | 3 | 4 | 5 | 8 | 2 |
| ENSG00000180818 | 74 | 26 | 33 | 40 | 34 | 26 |
| ENSG00000273049 | 0 | 0 | 0 | 0 | 0 | 0 |
| ENSG00000197757 | 263 | 124 | 199 | 168 | 157 | 149 |
| ENSG00000207924 | 0 | 0 | 0 | 0 | 0 | 0 |
| ENSG00000250133 | 1 | 1 | 0 | 1 | 2 | 4 |
| ENSG00000180806 | 251 | 134 | 134 | 126 | 140 | 149 |
| ENSG00000250451 | 49 | 23 | 41 | 38 | 24 | 27 |
| ENSG00000037965 | 12 | 10 | 10 | 17 | 17 | 10 |
| ENSG00000198353 | 71 | 33 | 44 | 24 | 36 | 26 |
| ENSG00000273046 | 0 | 0 | 0 | 0 | 0 | 0 |
| ENSG00000260597 | 48 | 18 | 25 | 12 | 23 | 15 |
| ENSG00000172789 | 2 | 2 | 0 | 3 | 1 | 0 |

| | | | | | | |
|---|---|---|---|---|---|---|
| ENSG00000207571 | 0 | 0 | 1 | 1 | 0 | 0 |
| ENSG00000248265 | 11 | 4 | 2 | 5 | 7 | 4 |
| ENSG00000250654 | 0 | 0 | 0 | 0 | 0 | 0 |
| ENSG00000249388 | 0 | 0 | 0 | 0 | 0 | 0 |
| ENSG00000250432 | 0 | 1 | 4 | 0 | 2 | 0 |
| ENSG00000202146 | 0 | 0 | 0 | 0 | 0 | 0 |
| ENSG00000123415 | 1824 | 907 | 1138 | 1057 | 972 | 1039 |
| ENSG00000250742 | 5 | 5 | 7 | 0 | 0 | 0 |
| ENSG00000283073 | 0 | 2 | 0 | 0 | 0 | 0 |
| ENSG00000260470 | 2 | 4 | 1 | 0 | 0 | 0 |
| ENSG00000257534 | 4 | 1 | 4 | 0 | 0 | 0 |
| ENSG00000248576 | 4 | 2 | 0 | 0 | 1 | 0 |
| ENSG00000094916 | 5954 | 2684 | 3441 | 1959 | 2744 | 3033 |
| ENSG00000283803 | 0 | 0 | 0 | 0 | 0 | 0 |
| ENSG00000241785 | 2 | 0 | 0 | 0 | 0 | 0 |
| ENSG00000257596 | 19 | 9 | 11 | 14 | 11 | 6 |
| ENSG00000258344 | 4 | 4 | 0 | 1 | 2 | 2 |
| ENSG00000135486 | 22357 | 11887 | 14262 | 10805 | 15292 | 14485 |
| ENSG00000123405 | 28 | 7 | 13 | 8 | 19 | 11 |
| ENSG00000111481 | 5982 | 2851 | 3547 | 3283 | 3194 | 3224 |
| ENSG00000207381 | 0 | 0 | 0 | 0 | 0 | 0 |
| ENSG00000199122 | 0 | 0 | 0 | 0 | 0 | 0 |
| ENSG00000264028 | 0 | 0 | 0 | 0 | 0 | 0 |
| ENSG00000258137 | 1 | 0 | 0 | 0 | 0 | 0 |
| ENSG00000258086 | 1 | 3 | 0 | 0 | 0 | 0 |
| ENSG00000139572 | 2 | 0 | 0 | 0 | 0 | 0 |
| ENSG00000161642 | 1740 | 885 | 969 | 975 | 835 | 747 |
| ENSG00000161638 | 736 | 314 | 338 | 428 | 437 | 355 |
| ENSG00000257477 | 0 | 0 | 0 | 0 | 0 | 0 |
| ENSG00000170627 | 0 | 0 | 0 | 0 | 0 | 0 |
| ENSG00000123338 | 0 | 2 | 3 | 0 | 0 | 1 |
| ENSG00000257824 | 0 | 0 | 0 | 0 | 0 | 0 |
| ENSG00000123360 | 2 | 2 | 2 | 0 | 2 | 2 |
| ENSG00000135447 | 2 | 0 | 2 | 0 | 2 | 2 |
| ENSG00000257780 | 0 | 0 | 0 | 0 | 0 | 0 |
| ENSG00000135413 | 0 | 0 | 0 | 0 | 0 | 0 |
| ENSG00000161634 | 0 | 0 | 0 | 0 | 0 | 0 |
| ENSG00000257634 | 0 | 0 | 0 | 0 | 0 | 0 |
| ENSG00000270858 | 0 | 0 | 1 | 0 | 3 | 0 |
| ENSG00000172551 | 257 | 111 | 141 | 156 | 92 | 102 |
| ENSG00000135426 | 0 | 0 | 0 | 0 | 0 | 0 |
| ENSG00000257870 | 0 | 0 | 0 | 0 | 0 | 0 |
| ENSG00000123307 | 0 | 0 | 0 | 0 | 0 | 0 |
| ENSG00000196534 | 0 | 0 | 0 | 0 | 0 | 0 |
| ENSG00000257350 | 0 | 0 | 0 | 0 | 0 | 0 |
| ENSG00000170605 | 0 | 0 | 0 | 0 | 0 | 0 |
| ENSG00000224622 | 0 | 0 | 0 | 0 | 0 | 0 |
| ENSG00000227423 | 0 | 0 | 0 | 0 | 0 | 0 |
| ENSG00000179919 | 0 | 0 | 0 | 0 | 0 | 0 |
| ENSG00000197706 | 0 | 0 | 0 | 0 | 0 | 0 |

| | | | | | | |
|---|---|---|---|---|---|---|
| ENSG00000213451 | 0 | 0 | 0 | 0 | 0 | 0 |
| ENSG00000205331 | 0 | 0 | 0 | 0 | 0 | 0 |
| ENSG00000188324 | 0 | 0 | 0 | 0 | 0 | 0 |
| ENSG00000230307 | 0 | 0 | 0 | 0 | 0 | 0 |
| ENSG00000205330 | 0 | 0 | 0 | 0 | 0 | 0 |
| ENSG00000205329 | 0 | 0 | 0 | 0 | 0 | 0 |
| ENSG00000257757 | 0 | 0 | 0 | 0 | 0 | 0 |
| ENSG00000187857 | 0 | 0 | 0 | 0 | 0 | 0 |
| ENSG00000203408 | 0 | 0 | 0 | 0 | 0 | 0 |
| ENSG00000233606 | 0 | 0 | 0 | 0 | 0 | 0 |
| ENSG00000257414 | 0 | 0 | 0 | 0 | 0 | 0 |
| ENSG00000205328 | 0 | 0 | 0 | 0 | 0 | 0 |
| ENSG00000179899 | 40 | 8 | 28 | 13 | 8 | 12 |
| ENSG00000185821 | 0 | 0 | 0 | 0 | 0 | 0 |
| ENSG00000258763 | 0 | 0 | 1 | 0 | 0 | 0 |
| ENSG00000179695 | 0 | 0 | 0 | 0 | 0 | 0 |
| ENSG00000184954 | 0 | 0 | 0 | 0 | 0 | 0 |
| ENSG00000205327 | 0 | 0 | 0 | 0 | 0 | 0 |
| ENSG00000234670 | 0 | 0 | 0 | 0 | 0 | 0 |
| ENSG00000179626 | 0 | 0 | 0 | 0 | 0 | 0 |
| ENSG00000179615 | 0 | 0 | 0 | 0 | 0 | 0 |
| ENSG00000272937 | 0 | 0 | 0 | 0 | 0 | 0 |
| ENSG00000175398 | 0 | 0 | 0 | 0 | 0 | 0 |
| ENSG00000258921 | 0 | 0 | 0 | 0 | 0 | 0 |
| ENSG00000272837 | 0 | 0 | 0 | 0 | 0 | 0 |
| ENSG00000258907 | 0 | 0 | 0 | 0 | 0 | 0 |
| ENSG00000270458 | 0 | 0 | 0 | 0 | 0 | 0 |
| ENSG00000170439 | 104 | 60 | 60 | 81 | 54 | 56 |
| ENSG00000135424 | 64 | 34 | 41 | 24 | 33 | 31 |
| ENSG00000258311 | 11 | 12 | 15 | 11 | 16 | 19 |
| ENSG00000135441 | 314 | 181 | 274 | 315 | 247 | 220 |
| ENSG00000135437 | 18 | 12 | 13 | 7 | 5 | 19 |
| ENSG00000135404 | 23618 | 11864 | 14151 | 15052 | 13556 | 12413 |
| ENSG00000258056 | 201 | 107 | 136 | 136 | 110 | 94 |
| ENSG00000258045 | 7 | 3 | 0 | 1 | 1 | 1 |
| ENSG00000257384 | 0 | 1 | 0 | 0 | 0 | 0 |
| ENSG00000135414 | 1153 | 561 | 676 | 695 | 776 | 749 |
| ENSG00000205323 | 5 | 13 | 31 | 18 | 16 | 16 |
| ENSG00000257390 | 2 | 1 | 3 | 1 | 0 | 2 |
| ENSG00000257509 | 15 | 3 | 6 | 4 | 4 | 8 |
| ENSG00000123353 | 798 | 459 | 495 | 486 | 489 | 522 |
| ENSG00000135392 | 978 | 488 | 620 | 500 | 595 | 651 |
| ENSG00000182796 | 798 | 289 | 524 | 292 | 371 | 414 |
| ENSG00000123342 | 181 | 79 | 151 | 79 | 99 | 95 |
| ENSG00000257966 | 63 | 16 | 16 | 9 | 13 | 20 |
| ENSG00000257569 | 0 | 0 | 1 | 0 | 0 | 1 |
| ENSG00000170473 | 1980 | 920 | 1200 | 1218 | 1039 | 1165 |
| ENSG00000065357 | 837 | 365 | 456 | 381 | 471 | 428 |
| ENSG00000273973 | 2 | 0 | 4 | 6 | 1 | 0 |
| ENSG00000185664 | 143 | 94 | 110 | 86 | 108 | 84 |

| | | | | | | |
|---|---|---|---|---|---|---|
| ENSG00000123374 | 1798 | 1000 | 1220 | 909 | 1079 | 1181 |
| ENSG00000258554 | 0 | 0 | 0 | 0 | 0 | 0 |
| ENSG00000111540 | 11010 | 5194 | 6491 | 4548 | 5371 | 5471 |
| ENSG00000237493 | 10 | 6 | 2 | 3 | 2 | 3 |
| ENSG00000139531 | 1670 | 795 | 920 | 739 | 718 | 811 |
| ENSG00000123411 | 376 | 150 | 270 | 159 | 167 | 182 |
| ENSG00000273890 | 13 | 0 | 2 | 1 | 0 | 0 |
| ENSG00000257449 | 1 | 0 | 0 | 0 | 1 | 0 |
| ENSG00000197728 | 4711 | 2255 | 2657 | 3175 | 2654 | 2709 |
| ENSG00000065361 | 16098 | 7218 | 9750 | 5094 | 7566 | 7225 |
| ENSG00000257411 | 0 | 0 | 0 | 0 | 0 | 0 |
| ENSG00000170515 | 5166 | 2860 | 3450 | 4049 | 3448 | 3929 |
| ENSG00000257553 | 0 | 0 | 0 | 0 | 0 | 0 |
| ENSG00000229117 | 17539 | 9244 | 10484 | 11770 | 10871 | 10600 |
| ENSG00000135482 | 340 | 148 | 193 | 185 | 190 | 191 |
| ENSG00000139641 | 6951 | 3349 | 4503 | 3852 | 3678 | 3896 |
| ENSG00000258345 | 0 | 0 | 2 | 0 | 0 | 0 |
| ENSG00000258317 | 4 | 2 | 7 | 9 | 2 | 5 |
| ENSG00000257809 | 8 | 0 | 4 | 1 | 2 | 2 |
| ENSG00000196465 | 877 | 439 | 611 | 676 | 546 | 615 |
| ENSG00000092841 | 21952 | 13661 | 16037 | 17000 | 14977 | 14250 |
| ENSG00000258199 | 48 | 20 | 37 | 37 | 42 | 22 |
| ENSG00000139613 | 3775 | 1740 | 2407 | 1582 | 1964 | 1895 |
| ENSG00000181852 | 1807 | 926 | 1141 | 1044 | 1059 | 1046 |
| ENSG00000139579 | 1248 | 634 | 831 | 860 | 887 | 830 |
| ENSG00000139540 | 10 | 1 | 3 | 2 | 2 | 3 |
| ENSG00000139645 | 7387 | 3161 | 4206 | 3541 | 3911 | 4073 |
| ENSG00000135469 | 133 | 66 | 86 | 116 | 107 | 87 |
| ENSG00000258260 | 0 | 0 | 0 | 0 | 3 | 0 |
| ENSG00000062485 | 7991 | 4110 | 4730 | 4102 | 4686 | 4873 |
| ENSG00000144785 | 0 | 1 | 0 | 0 | 0 | 0 |
| ENSG00000257303 | 14 | 10 | 18 | 9 | 10 | 7 |
| ENSG00000257740 | 16 | 6 | 7 | 5 | 9 | 9 |
| ENSG00000257727 | 309 | 272 | 312 | 385 | 374 | 356 |
| ENSG00000135473 | 2613 | 999 | 1769 | 955 | 1132 | 1463 |
| ENSG00000110944 | 12 | 5 | 11 | 17 | 6 | 17 |
| ENSG00000170581 | 2067 | 976 | 1296 | 782 | 986 | 974 |
| ENSG00000252206 | 0 | 0 | 3 | 0 | 0 | 0 |
| ENSG00000175336 | 0 | 1 | 4 | 4 | 2 | 1 |
| ENSG00000274823 | 0 | 0 | 0 | 0 | 0 | 0 |
| ENSG00000259099 | 0 | 0 | 0 | 0 | 0 | 0 |
| ENSG00000276272 | 8 | 1 | 4 | 7 | 4 | 4 |
| ENSG00000274569 | 3 | 2 | 0 | 0 | 0 | 0 |
| ENSG00000111602 | 2666 | 1298 | 1825 | 1368 | 1535 | 1705 |
| ENSG00000135517 | 8 | 4 | 3 | 0 | 1 | 3 |
| ENSG00000285528 | 0 | 0 | 0 | 0 | 0 | 0 |
| ENSG00000176422 | 967 | 574 | 602 | 570 | 614 | 601 |
| ENSG00000135423 | 12 | 3 | 8 | 7 | 3 | 13 |
| ENSG00000257576 | 16 | 2 | 4 | 3 | 1 | 2 |
| ENSG00000076067 | 423 | 237 | 258 | 182 | 255 | 219 |

| | | | | | | |
|---|---|---|---|---|---|---|
| ENSG00000201579 | 1 | 0 | 1 | 0 | 0 | 0 |
| ENSG00000076108 | 6632 | 2691 | 4045 | 2309 | 3058 | 3528 |
| ENSG00000110955 | 25285 | 12898 | 15243 | 15951 | 14575 | 14907 |
| ENSG00000207031 | 9 | 4 | 4 | 3 | 0 | 4 |
| ENSG00000110958 | 10827 | 5112 | 6191 | 5600 | 6301 | 6403 |
| ENSG00000241217 | 8 | 1 | 1 | 0 | 1 | 1 |
| ENSG00000196531 | 8710 | 5374 | 5593 | 5201 | 7094 | 5298 |
| ENSG00000285625 | 0 | 0 | 0 | 0 | 0 | 0 |
| ENSG00000198056 | 231 | 175 | 170 | 146 | 180 | 188 |
| ENSG00000025423 | 30 | 14 | 17 | 11 | 6 | 17 |
| ENSG00000258679 | 0 | 0 | 0 | 0 | 0 | 0 |
| ENSG00000273536 | 0 | 0 | 0 | 0 | 0 | 0 |
| ENSG00000258816 | 0 | 0 | 0 | 0 | 0 | 0 |
| ENSG00000212383 | 0 | 0 | 0 | 0 | 0 | 0 |
| ENSG00000170426 | 0 | 1 | 3 | 3 | 0 | 1 |
| ENSG00000278400 | 0 | 0 | 0 | 1 | 0 | 0 |
| ENSG00000139547 | 28 | 5 | 7 | 9 | 15 | 12 |
| ENSG00000278399 | 10 | 13 | 10 | 10 | 13 | 20 |
| ENSG00000166856 | 0 | 0 | 0 | 1 | 1 | 0 |
| ENSG00000166860 | 568 | 202 | 280 | 214 | 264 | 299 |
| ENSG00000166863 | 0 | 1 | 0 | 0 | 0 | 0 |
| ENSG00000166866 | 0 | 1 | 0 | 0 | 2 | 0 |
| ENSG00000166881 | 897 | 374 | 472 | 354 | 485 | 494 |
| ENSG00000166886 | 1028 | 453 | 658 | 633 | 554 | 576 |
| ENSG00000166888 | 177 | 86 | 188 | 128 | 126 | 170 |
| ENSG00000123384 | 524 | 171 | 282 | 202 | 213 | 250 |
| ENSG00000259125 | 0 | 0 | 0 | 0 | 0 | 0 |
| ENSG00000221365 | 0 | 0 | 0 | 0 | 0 | 0 |
| ENSG00000182379 | 51 | 19 | 25 | 40 | 22 | 30 |
| ENSG00000182199 | 4997 | 2514 | 2962 | 3602 | 2938 | 2975 |
| ENSG00000276727 | 0 | 0 | 0 | 1 | 0 | 0 |
| ENSG00000185633 | 4 | 10 | 7 | 18 | 13 | 9 |
| ENSG00000185482 | 173 | 65 | 102 | 87 | 72 | 86 |
| ENSG00000258830 | 0 | 1 | 1 | 4 | 3 | 1 |
| ENSG00000179912 | 1393 | 658 | 963 | 696 | 849 | 897 |
| ENSG00000201198 | 0 | 0 | 0 | 0 | 0 | 0 |
| ENSG00000258001 | 2 | 1 | 1 | 1 | 4 | 2 |
| ENSG00000175189 | 5 | 1 | 2 | 3 | 1 | 2 |
| ENSG00000139269 | 2 | 1 | 0 | 2 | 0 | 1 |
| ENSG00000111087 | 0 | 0 | 1 | 0 | 2 | 1 |
| ENSG00000123329 | 0 | 0 | 1 | 1 | 0 | 1 |
| ENSG00000166986 | 4861 | 2322 | 2867 | 2861 | 2963 | 2914 |
| ENSG00000284152 | 0 | 0 | 0 | 0 | 0 | 0 |
| ENSG00000175197 | 214 | 113 | 132 | 131 | 155 | 118 |
| ENSG00000285133 | 0 | 0 | 0 | 0 | 0 | 0 |
| ENSG00000208028 | 4 | 3 | 3 | 2 | 2 | 3 |
| ENSG00000166987 | 2507 | 1071 | 1442 | 1198 | 1113 | 1140 |
| ENSG00000175203 | 5723 | 2698 | 3325 | 3149 | 3046 | 3203 |
| ENSG00000155980 | 10 | 1 | 6 | 3 | 5 | 3 |
| ENSG00000166908 | 6056 | 2806 | 3465 | 2787 | 3341 | 3533 |

| | | | | | | |
|---|---|---|---|---|---|---|
| ENSG00000178498 | 1197 | 580 | 755 | 641 | 650 | 678 |
| ENSG00000240771 | 128 | 114 | 129 | 114 | 99 | 129 |
| ENSG00000224713 | 12 | 6 | 9 | 7 | 11 | 5 |
| ENSG00000135502 | 0 | 0 | 6 | 4 | 4 | 7 |
| ENSG00000135454 | 508 | 263 | 348 | 337 | 412 | 372 |
| ENSG00000242990 | 0 | 0 | 1 | 0 | 0 | 0 |
| ENSG00000135506 | 9561 | 4648 | 6331 | 5398 | 5171 | 5255 |
| ENSG00000257342 | 0 | 0 | 0 | 0 | 0 | 0 |
| ENSG00000135439 | 228 | 109 | 148 | 126 | 149 | 163 |
| ENSG00000255737 | 99 | 82 | 123 | 144 | 105 | 116 |
| ENSG00000135452 | 1682 | 883 | 1120 | 783 | 866 | 928 |
| ENSG00000135446 | 4133 | 2307 | 2718 | 3116 | 2630 | 2878 |
| ENSG00000283621 | 0 | 0 | 0 | 0 | 0 | 0 |
| ENSG00000139266 | 1539 | 774 | 876 | 901 | 754 | 835 |
| ENSG00000111012 | 179 | 54 | 92 | 87 | 68 | 79 |
| ENSG00000037897 | 398 | 244 | 333 | 352 | 294 | 298 |
| ENSG00000123427 | 315 | 162 | 177 | 166 | 202 | 207 |
| ENSG00000257921 | 0 | 0 | 0 | 1 | 0 | 0 |
| ENSG00000123297 | 1404 | 784 | 854 | 948 | 875 | 916 |
| ENSG00000135407 | 118 | 52 | 75 | 45 | 51 | 74 |
| ENSG00000270039 | 18 | 4 | 8 | 8 | 6 | 8 |
| ENSG00000206749 | 0 | 0 | 0 | 0 | 0 | 0 |
| ENSG00000269903 | 4 | 0 | 4 | 1 | 1 | 0 |
| ENSG00000175215 | 8662 | 4097 | 5314 | 3686 | 4127 | 4145 |
| ENSG00000207789 | 0 | 0 | 0 | 0 | 0 | 0 |
| ENSG00000257953 | 0 | 0 | 2 | 3 | 0 | 0 |
| ENSG00000245651 | 0 | 0 | 0 | 0 | 0 | 0 |
| ENSG00000273805 | 0 | 0 | 0 | 0 | 0 | 0 |
| ENSG00000166896 | 256 | 144 | 155 | 147 | 167 | 165 |
| ENSG00000257698 | 46 | 16 | 24 | 29 | 22 | 19 |
| ENSG00000257159 | 0 | 0 | 0 | 0 | 2 | 0 |
| ENSG00000222210 | 0 | 0 | 0 | 0 | 0 | 0 |
| ENSG00000257668 | 0 | 0 | 0 | 0 | 0 | 0 |
| ENSG00000257541 | 0 | 0 | 0 | 0 | 0 | 0 |
| ENSG00000279134 | 0 | 0 | 0 | 1 | 0 | 0 |
| ENSG00000279411 | 0 | 0 | 0 | 0 | 0 | 0 |
| ENSG00000241423 | 0 | 0 | 0 | 0 | 0 | 0 |
| ENSG00000258133 | 0 | 0 | 0 | 0 | 0 | 0 |
| ENSG00000258231 | 0 | 0 | 0 | 0 | 0 | 0 |
| ENSG00000257259 | 0 | 0 | 0 | 0 | 0 | 0 |
| ENSG00000139263 | 194 | 103 | 116 | 88 | 109 | 108 |
| ENSG00000257443 | 6 | 1 | 0 | 0 | 0 | 0 |
| ENSG00000244390 | 0 | 0 | 0 | 0 | 0 | 0 |
| ENSG00000257288 | 0 | 0 | 0 | 0 | 0 | 0 |
| ENSG00000258214 | 0 | 0 | 0 | 0 | 0 | 0 |
| ENSG00000212599 | 0 | 0 | 0 | 0 | 0 | 0 |
| ENSG00000251931 | 0 | 0 | 0 | 0 | 0 | 0 |
| ENSG00000257230 | 0 | 0 | 0 | 0 | 0 | 0 |
| ENSG00000202471 | 0 | 0 | 0 | 0 | 0 | 0 |
| ENSG00000118596 | 830 | 406 | 348 | 226 | 670 | 483 |

| | | | | | | |
|---|---|---|---|---|---|---|
| ENSG00000257146 | 0 | 0 | 0 | 0 | 0 | 1 |
| ENSG00000237176 | 0 | 0 | 0 | 0 | 0 | 0 |
| ENSG00000257294 | 0 | 0 | 0 | 0 | 0 | 0 |
| ENSG00000258184 | 0 | 0 | 0 | 0 | 0 | 0 |
| ENSG00000252607 | 0 | 0 | 0 | 0 | 0 | 0 |
| ENSG00000251822 | 0 | 0 | 0 | 0 | 0 | 0 |
| ENSG00000257865 | 0 | 0 | 0 | 0 | 0 | 0 |
| ENSG00000257308 | 0 | 1 | 0 | 0 | 0 | 0 |
| ENSG00000258336 | 0 | 0 | 0 | 0 | 0 | 0 |
| ENSG00000198673 | 10 | 5 | 6 | 5 | 8 | 4 |
| ENSG00000242922 | 0 | 0 | 0 | 0 | 0 | 0 |
| ENSG00000257528 | 0 | 0 | 0 | 0 | 0 | 0 |
| ENSG00000213363 | 0 | 0 | 0 | 1 | 0 | 2 |
| ENSG00000257880 | 0 | 0 | 0 | 0 | 0 | 0 |
| ENSG00000258316 | 0 | 0 | 0 | 0 | 0 | 0 |
| ENSG00000135655 | 1021 | 654 | 483 | 265 | 996 | 508 |
| ENSG00000284360 | 0 | 0 | 0 | 0 | 0 | 0 |
| ENSG00000200814 | 0 | 1 | 0 | 0 | 1 | 0 |
| ENSG00000275881 | 0 | 0 | 2 | 0 | 0 | 0 |
| ENSG00000061987 | 2474 | 1188 | 1376 | 612 | 1427 | 1101 |
| ENSG00000257568 | 0 | 0 | 0 | 0 | 0 | 0 |
| ENSG00000202034 | 1 | 0 | 0 | 0 | 0 | 0 |
| ENSG00000221949 | 17 | 7 | 10 | 18 | 5 | 7 |
| ENSG00000257354 | 56 | 24 | 37 | 20 | 30 | 27 |
| ENSG00000199179 | 0 | 1 | 0 | 0 | 0 | 0 |
| ENSG00000275180 | 28 | 23 | 22 | 13 | 25 | 8 |
| ENSG00000111110 | 2989 | 1358 | 1710 | 1166 | 1389 | 1478 |
| ENSG00000241941 | 0 | 0 | 0 | 0 | 0 | 0 |
| ENSG00000213352 | 0 | 0 | 0 | 0 | 0 | 1 |
| ENSG00000200296 | 0 | 0 | 0 | 0 | 0 | 0 |
| ENSG00000252660 | 0 | 0 | 0 | 0 | 0 | 0 |
| ENSG00000139239 | 40 | 16 | 19 | 29 | 23 | 28 |
| ENSG00000257779 | 0 | 0 | 0 | 0 | 0 | 0 |
| ENSG00000250517 | 0 | 0 | 0 | 0 | 0 | 0 |
| ENSG00000257235 | 0 | 0 | 0 | 0 | 0 | 0 |
| ENSG00000257664 | 0 | 0 | 0 | 0 | 0 | 0 |
| ENSG00000166148 | 0 | 0 | 0 | 0 | 0 | 0 |
| ENSG00000279444 | 0 | 1 | 0 | 0 | 0 | 0 |
| ENSG00000257332 | 0 | 0 | 0 | 0 | 0 | 0 |
| ENSG00000258117 | 0 | 0 | 0 | 0 | 0 | 0 |
| ENSG00000177990 | 8 | 2 | 1 | 1 | 4 | 3 |
| ENSG00000249753 | 0 | 0 | 0 | 1 | 0 | 1 |
| ENSG00000257005 | 0 | 0 | 0 | 0 | 0 | 0 |
| ENSG00000255583 | 0 | 0 | 0 | 0 | 0 | 0 |
| ENSG00000240027 | 0 | 0 | 0 | 0 | 0 | 0 |
| ENSG00000118600 | 532 | 279 | 319 | 295 | 358 | 359 |
| ENSG00000255850 | 0 | 0 | 0 | 0 | 0 | 0 |
| ENSG00000255642 | 6 | 3 | 3 | 2 | 3 | 4 |
| ENSG00000196935 | 990 | 535 | 662 | 301 | 614 | 498 |
| ENSG00000242087 | 0 | 0 | 0 | 0 | 0 | 0 |

| | | | | | | |
|---|---|---|---|---|---|---|
| ENSG00000256571 | 0 | 1 | 1 | 0 | 0 | 1 |
| ENSG00000256399 | 0 | 0 | 0 | 0 | 2 | 0 |
| ENSG00000255780 | 0 | 1 | 0 | 0 | 0 | 0 |
| ENSG00000255886 | 0 | 0 | 0 | 0 | 0 | 0 |
| ENSG00000255817 | 1 | 0 | 1 | 0 | 0 | 0 |
| ENSG00000255629 | 0 | 0 | 0 | 0 | 0 | 0 |
| ENSG00000174206 | 584 | 321 | 313 | 288 | 380 | 410 |
| ENSG00000212298 | 0 | 0 | 0 | 0 | 0 | 0 |
| ENSG00000243024 | 8 | 2 | 3 | 3 | 3 | 7 |
| ENSG00000251788 | 0 | 0 | 1 | 0 | 0 | 0 |
| ENSG00000185306 | 1 | 0 | 3 | 0 | 1 | 4 |
| ENSG00000256192 | 0 | 0 | 0 | 0 | 0 | 0 |
| ENSG00000256293 | 0 | 0 | 0 | 0 | 0 | 0 |
| ENSG00000238440 | 0 | 0 | 0 | 0 | 0 | 0 |
| ENSG00000184575 | 7633 | 3807 | 4564 | 3101 | 4494 | 4402 |
| ENSG00000255566 | 16 | 1 | 11 | 0 | 3 | 2 |
| ENSG00000277895 | 1 | 5 | 6 | 2 | 1 | 2 |
| ENSG00000183735 | 1432 | 877 | 750 | 545 | 1362 | 853 |
| ENSG00000153179 | 1297 | 628 | 795 | 624 | 787 | 870 |
| ENSG00000256199 | 1 | 0 | 0 | 0 | 0 | 0 |
| ENSG00000223294 | 0 | 0 | 0 | 0 | 0 | 0 |
| ENSG00000256670 | 0 | 0 | 0 | 1 | 0 | 0 |
| ENSG00000207546 | 0 | 0 | 0 | 0 | 0 | 0 |
| ENSG00000266016 | 0 | 0 | 0 | 0 | 0 | 0 |
| ENSG00000225195 | 0 | 0 | 0 | 0 | 0 | 0 |
| ENSG00000256314 | 0 | 0 | 0 | 0 | 0 | 0 |
| ENSG00000280320 | 0 | 0 | 0 | 0 | 0 | 0 |
| ENSG00000280181 | 0 | 2 | 0 | 0 | 0 | 0 |
| ENSG00000135677 | 11208 | 4911 | 7081 | 4226 | 5228 | 5548 |
| ENSG00000215159 | 2 | 0 | 0 | 3 | 0 | 0 |
| ENSG00000111490 | 1786 | 896 | 1228 | 850 | 978 | 1032 |
| ENSG00000243020 | 2 | 3 | 2 | 1 | 0 | 2 |
| ENSG00000255693 | 0 | 3 | 0 | 0 | 3 | 2 |
| ENSG00000248995 | 0 | 0 | 0 | 0 | 0 | 0 |
| ENSG00000221564 | 0 | 0 | 0 | 0 | 0 | 0 |
| ENSG00000198671 | 0 | 0 | 0 | 0 | 0 | 0 |
| ENSG00000156076 | 2 | 1 | 0 | 0 | 0 | 0 |
| ENSG00000207099 | 0 | 0 | 0 | 0 | 0 | 0 |
| ENSG00000255700 | 0 | 0 | 0 | 0 | 0 | 0 |
| ENSG00000174106 | 1422 | 638 | 852 | 553 | 776 | 717 |
| ENSG00000276853 | 6 | 0 | 10 | 4 | 0 | 7 |
| ENSG00000174099 | 14 | 6 | 2 | 1 | 4 | 0 |
| ENSG00000250280 | 0 | 0 | 0 | 0 | 0 | 0 |
| ENSG00000215208 | 0 | 0 | 1 | 0 | 1 | 0 |
| ENSG00000250748 | 0 | 0 | 0 | 0 | 0 | 0 |
| ENSG00000255866 | 0 | 0 | 0 | 0 | 0 | 0 |
| ENSG00000256268 | 2 | 0 | 2 | 0 | 5 | 0 |
| ENSG00000256915 | 0 | 0 | 0 | 0 | 0 | 0 |
| ENSG00000213344 | 0 | 0 | 0 | 1 | 0 | 0 |
| ENSG00000241749 | 0 | 0 | 0 | 0 | 0 | 0 |

| | | | | | | |
|---|---|---|---|---|---|---|
| ENSG00000149948 | 4 | 0 | 3 | 1 | 3 | 1 |
| ENSG00000277945 | 0 | 0 | 0 | 0 | 0 | 0 |
| ENSG00000197301 | 0 | 0 | 0 | 1 | 1 | 0 |
| ENSG00000256083 | 0 | 0 | 0 | 0 | 0 | 0 |
| ENSG00000276878 | 0 | 0 | 0 | 0 | 0 | 0 |
| ENSG00000256259 | 0 | 0 | 0 | 0 | 0 | 0 |
| ENSG00000213343 | 4 | 0 | 0 | 0 | 0 | 0 |
| ENSG00000251857 | 0 | 0 | 0 | 0 | 0 | 0 |
| ENSG00000139233 | 683 | 437 | 411 | 381 | 540 | 436 |
| ENSG00000228144 | 0 | 0 | 0 | 0 | 0 | 0 |
| ENSG00000239335 | 25 | 20 | 18 | 22 | 18 | 13 |
| ENSG00000155957 | 253 | 146 | 187 | 140 | 189 | 192 |
| ENSG00000090376 | 0 | 2 | 0 | 0 | 0 | 0 |
| ENSG00000225422 | 0 | 0 | 0 | 0 | 1 | 5 |
| ENSG00000284011 | 0 | 0 | 0 | 0 | 0 | 0 |
| ENSG00000256072 | 0 | 0 | 0 | 0 | 0 | 0 |
| ENSG00000256777 | 0 | 0 | 0 | 0 | 0 | 0 |
| ENSG00000222744 | 0 | 0 | 0 | 0 | 0 | 0 |
| ENSG00000127311 | 205 | 144 | 100 | 68 | 117 | 106 |
| ENSG00000155974 | 458 | 223 | 212 | 139 | 241 | 192 |
| ENSG00000256285 | 0 | 0 | 0 | 0 | 0 | 0 |
| ENSG00000256420 | 0 | 0 | 0 | 0 | 0 | 0 |
| ENSG00000257083 | 0 | 1 | 0 | 0 | 0 | 0 |
| ENSG00000279310 | 0 | 0 | 0 | 0 | 0 | 1 |
| ENSG00000255976 | 0 | 0 | 0 | 0 | 0 | 0 |
| ENSG00000237766 | 3 | 1 | 0 | 1 | 3 | 3 |
| ENSG00000111530 | 5531 | 3049 | 2940 | 1748 | 3996 | 2774 |
| ENSG00000256037 | 2 | 0 | 2 | 0 | 0 | 1 |
| ENSG00000256355 | 0 | 0 | 0 | 0 | 0 | 0 |
| ENSG00000256172 | 0 | 0 | 0 | 0 | 0 | 0 |
| ENSG00000203585 | 3 | 0 | 2 | 0 | 0 | 0 |
| ENSG00000256077 | 0 | 0 | 0 | 0 | 0 | 0 |
| ENSG00000127334 | 1930 | 1023 | 1180 | 675 | 1108 | 1025 |
| ENSG00000235872 | 0 | 1 | 0 | 3 | 0 | 5 |
| ENSG00000255970 | 0 | 1 | 2 | 0 | 1 | 0 |
| ENSG00000279505 | 0 | 0 | 0 | 0 | 0 | 0 |
| ENSG00000255772 | 0 | 0 | 0 | 0 | 0 | 0 |
| ENSG00000256835 | 0 | 0 | 0 | 0 | 0 | 0 |
| ENSG00000255733 | 0 | 0 | 0 | 0 | 0 | 0 |
| ENSG00000244432 | 0 | 0 | 0 | 0 | 0 | 0 |
| ENSG00000236946 | 0 | 0 | 0 | 0 | 0 | 0 |
| ENSG00000256708 | 0 | 0 | 0 | 0 | 0 | 0 |
| ENSG00000111537 | 0 | 0 | 0 | 0 | 0 | 0 |
| ENSG00000111536 | 0 | 0 | 0 | 0 | 0 | 0 |
| ENSG00000127318 | 0 | 0 | 0 | 0 | 2 | 0 |
| ENSG00000111554 | 303 | 159 | 156 | 127 | 192 | 164 |
| ENSG00000251301 | 0 | 1 | 1 | 0 | 0 | 3 |
| ENSG00000256273 | 0 | 0 | 0 | 0 | 0 | 0 |
| ENSG00000273824 | 0 | 0 | 0 | 0 | 0 | 0 |
| ENSG00000256075 | 0 | 0 | 0 | 0 | 0 | 0 |

| | | | | | | |
|---|---|---|---|---|---|---|
| ENSG00000256917 | 0 | 0 | 0 | 0 | 0 | 0 |
| ENSG00000240087 | 6 | 2 | 2 | 6 | 5 | 2 |
| ENSG00000274459 | 0 | 0 | 0 | 0 | 0 | 0 |
| ENSG00000127314 | 2408 | 1319 | 1335 | 995 | 1559 | 1466 |
| ENSG00000234040 | 4 | 1 | 0 | 0 | 0 | 0 |
| ENSG00000206650 | 0 | 0 | 0 | 0 | 0 | 0 |
| ENSG00000234925 | 6 | 1 | 5 | 0 | 1 | 1 |
| ENSG00000247363 | 12 | 6 | 8 | 9 | 9 | 6 |
| ENSG00000241825 | 2 | 0 | 1 | 0 | 0 | 0 |
| ENSG00000111581 | 1254 | 693 | 709 | 555 | 1028 | 862 |
| ENSG00000233560 | 6 | 5 | 7 | 2 | 1 | 3 |
| ENSG00000175782 | 1084 | 498 | 635 | 398 | 640 | 616 |
| ENSG00000256678 | 1 | 0 | 0 | 0 | 0 | 0 |
| ENSG00000135679 | 2760 | 1518 | 1557 | 933 | 1663 | 1485 |
| ENSG00000256325 | 4 | 1 | 0 | 0 | 2 | 0 |
| ENSG00000257181 | 0 | 0 | 0 | 0 | 0 | 0 |
| ENSG00000256664 | 0 | 0 | 0 | 0 | 0 | 0 |
| ENSG00000135678 | 62 | 46 | 42 | 39 | 57 | 33 |
| ENSG00000252770 | 0 | 0 | 0 | 0 | 0 | 0 |
| ENSG00000257289 | 0 | 0 | 0 | 0 | 0 | 0 |
| ENSG00000257336 | 0 | 0 | 0 | 0 | 0 | 0 |
| ENSG00000258140 | 0 | 0 | 0 | 0 | 0 | 0 |
| ENSG00000111605 | 2958 | 1487 | 1683 | 1257 | 1780 | 1682 |
| ENSG00000283476 | 0 | 0 | 0 | 0 | 0 | 0 |
| ENSG00000257818 | 2 | 5 | 3 | 0 | 2 | 0 |
| ENSG00000274979 | 0 | 1 | 0 | 0 | 0 | 0 |
| ENSG00000090382 | 2 | 1 | 2 | 1 | 1 | 6 |
| ENSG00000257764 | 0 | 0 | 0 | 0 | 0 | 0 |
| ENSG00000127337 | 613 | 328 | 418 | 393 | 364 | 379 |
| ENSG00000266185 | 0 | 0 | 0 | 0 | 0 | 0 |
| ENSG00000241765 | 0 | 0 | 0 | 0 | 2 | 0 |
| ENSG00000257507 | 0 | 1 | 0 | 0 | 1 | 0 |
| ENSG00000166225 | 1046 | 519 | 467 | 267 | 654 | 495 |
| ENSG00000264405 | 0 | 0 | 0 | 0 | 0 | 0 |
| ENSG00000283677 | 0 | 0 | 0 | 0 | 0 | 0 |
| ENSG00000166226 | 9690 | 5024 | 5804 | 5850 | 6617 | 6209 |
| ENSG00000198812 | 0 | 0 | 0 | 0 | 0 | 0 |
| ENSG00000127325 | 0 | 2 | 1 | 0 | 4 | 1 |
| ENSG00000279551 | 0 | 0 | 0 | 0 | 0 | 0 |
| ENSG00000247131 | 2 | 3 | 2 | 2 | 0 | 1 |
| ENSG00000127328 | 798 | 454 | 485 | 386 | 604 | 485 |
| ENSG00000258052 | 0 | 0 | 0 | 0 | 0 | 0 |
| ENSG00000166268 | 32 | 9 | 15 | 4 | 19 | 16 |
| ENSG00000257815 | 15 | 23 | 28 | 14 | 16 | 16 |
| ENSG00000257241 | 0 | 0 | 0 | 0 | 0 | 0 |
| ENSG00000257139 | 0 | 0 | 0 | 0 | 0 | 1 |
| ENSG00000257613 | 0 | 0 | 0 | 0 | 0 | 0 |
| ENSG00000111596 | 2738 | 1298 | 1493 | 1021 | 1568 | 1249 |
| ENSG00000279530 | 0 | 0 | 1 | 1 | 4 | 6 |
| ENSG00000135643 | 127 | 60 | 80 | 48 | 44 | 54 |

| | | | | | | |
|---|---|---|---|---|---|---|
| ENSG00000222405 | 0 | 0 | 0 | 0 | 0 | 0 |
| ENSG00000258168 | 0 | 0 | 0 | 2 | 1 | 1 |
| ENSG00000127329 | 2 | 0 | 0 | 0 | 1 | 0 |
| ENSG00000277247 | 0 | 0 | 0 | 0 | 0 | 0 |
| ENSG00000153233 | 0 | 0 | 0 | 0 | 1 | 0 |
| ENSG00000257302 | 0 | 0 | 0 | 0 | 0 | 0 |
| ENSG00000207387 | 0 | 0 | 0 | 0 | 0 | 0 |
| ENSG00000257265 | 0 | 0 | 0 | 0 | 0 | 0 |
| ENSG00000257454 | 0 | 0 | 0 | 0 | 0 | 0 |
| ENSG00000258053 | 0 | 0 | 0 | 0 | 0 | 0 |
| ENSG00000127324 | 0 | 0 | 0 | 0 | 0 | 0 |
| ENSG00000139292 | 0 | 0 | 2 | 0 | 0 | 0 |
| ENSG00000277223 | 0 | 0 | 0 | 0 | 0 | 0 |
| ENSG00000257761 | 0 | 0 | 0 | 0 | 0 | 0 |
| ENSG00000133858 | 1021 | 586 | 472 | 209 | 1040 | 414 |
| ENSG00000212461 | 0 | 0 | 0 | 0 | 0 | 0 |
| ENSG00000173451 | 126 | 80 | 72 | 80 | 98 | 71 |
| ENSG00000258064 | 0 | 0 | 0 | 0 | 0 | 0 |
| ENSG00000139291 | 796 | 355 | 444 | 384 | 405 | 423 |
| ENSG00000257515 | 0 | 0 | 0 | 0 | 0 | 0 |
| ENSG00000080371 | 2949 | 1401 | 1774 | 1190 | 1620 | 1512 |
| ENSG00000257410 | 0 | 0 | 0 | 0 | 0 | 0 |
| ENSG00000257991 | 0 | 0 | 0 | 0 | 0 | 0 |
| ENSG00000121749 | 601 | 337 | 343 | 175 | 469 | 281 |
| ENSG00000257802 | 0 | 1 | 0 | 0 | 0 | 0 |
| ENSG00000139287 | 0 | 0 | 0 | 0 | 0 | 0 |
| ENSG00000258115 | 0 | 0 | 0 | 0 | 0 | 0 |
| ENSG00000072657 | 0 | 0 | 0 | 0 | 0 | 0 |
| ENSG00000236333 | 0 | 0 | 0 | 0 | 0 | 0 |
| ENSG00000257287 | 0 | 0 | 0 | 0 | 0 | 0 |
| ENSG00000257723 | 0 | 0 | 0 | 0 | 0 | 0 |
| ENSG00000258235 | 0 | 0 | 0 | 0 | 0 | 0 |
| ENSG00000258294 | 0 | 0 | 0 | 0 | 0 | 0 |
| ENSG00000258123 | 0 | 0 | 0 | 0 | 0 | 0 |
| ENSG00000257682 | 0 | 0 | 0 | 0 | 0 | 0 |
| ENSG00000243164 | 0 | 0 | 0 | 0 | 0 | 0 |
| ENSG00000252415 | 0 | 0 | 0 | 0 | 0 | 0 |
| ENSG00000257750 | 0 | 0 | 0 | 0 | 0 | 0 |
| ENSG00000201809 | 0 | 0 | 0 | 0 | 0 | 0 |
| ENSG00000257587 | 1 | 0 | 0 | 0 | 0 | 0 |
| ENSG00000258332 | 0 | 0 | 0 | 0 | 0 | 0 |
| ENSG00000251138 | 9 | 7 | 17 | 3 | 16 | 11 |
| ENSG00000258320 | 0 | 0 | 0 | 0 | 0 | 0 |
| ENSG00000257183 | 0 | 0 | 1 | 0 | 0 | 0 |
| ENSG00000257113 | 0 | 0 | 0 | 0 | 0 | 0 |
| ENSG00000257364 | 0 | 0 | 1 | 1 | 0 | 0 |
| ENSG00000253719 | 8915 | 4145 | 5151 | 4100 | 4621 | 4724 |
| ENSG00000257386 | 0 | 0 | 0 | 0 | 0 | 0 |
| ENSG00000257323 | 0 | 0 | 0 | 0 | 0 | 0 |
| ENSG00000188646 | 0 | 0 | 0 | 0 | 0 | 0 |

| | | | | | | |
|---|---|---|---|---|---|---|
| ENSG00000257998 | 0 | 0 | 0 | 0 | 0 | 0 |
| ENSG00000257434 | 0 | 0 | 0 | 0 | 0 | 0 |
| ENSG00000166006 | 0 | 0 | 0 | 0 | 0 | 0 |
| ENSG00000254451 | 0 | 0 | 0 | 0 | 0 | 0 |
| ENSG00000258112 | 0 | 0 | 0 | 0 | 0 | 0 |
| ENSG00000180881 | 30 | 22 | 21 | 24 | 22 | 24 |
| ENSG00000257823 | 0 | 0 | 0 | 0 | 0 | 0 |
| ENSG00000273987 | 8 | 2 | 1 | 3 | 2 | 3 |
| ENSG00000173401 | 0 | 1 | 1 | 1 | 2 | 6 |
| ENSG00000180481 | 2 | 5 | 9 | 2 | 5 | 7 |
| ENSG00000139278 | 34 | 24 | 18 | 7 | 27 | 11 |
| ENSG00000257497 | 25 | 8 | 9 | 11 | 10 | 15 |
| ENSG00000111615 | 746 | 456 | 452 | 330 | 729 | 459 |
| ENSG00000258077 | 2 | 0 | 1 | 1 | 1 | 0 |
| ENSG00000257777 | 0 | 0 | 0 | 0 | 0 | 0 |
| ENSG00000258245 | 0 | 1 | 0 | 1 | 0 | 0 |
| ENSG00000251893 | 0 | 0 | 1 | 0 | 0 | 0 |
| ENSG00000258088 | 0 | 0 | 0 | 0 | 1 | 0 |
| ENSG00000258091 | 0 | 0 | 0 | 0 | 0 | 0 |
| ENSG00000243420 | 0 | 0 | 0 | 0 | 0 | 0 |
| ENSG00000257329 | 0 | 0 | 0 | 0 | 0 | 0 |
| ENSG00000198923 | 0 | 0 | 0 | 0 | 0 | 0 |
| ENSG00000139289 | 529 | 292 | 327 | 617 | 778 | 739 |
| ENSG00000257453 | 0 | 0 | 0 | 0 | 2 | 0 |
| ENSG00000257839 | 0 | 0 | 0 | 0 | 0 | 0 |
| ENSG00000187109 | 11076 | 5970 | 6395 | 5923 | 8475 | 6801 |
| ENSG00000257941 | 2 | 1 | 0 | 1 | 3 | 3 |
| ENSG00000258304 | 0 | 0 | 0 | 0 | 0 | 0 |
| ENSG00000257872 | 0 | 0 | 0 | 0 | 0 | 0 |
| ENSG00000252858 | 0 | 0 | 0 | 0 | 0 | 0 |
| ENSG00000257219 | 0 | 0 | 0 | 0 | 0 | 0 |
| ENSG00000223273 | 0 | 0 | 0 | 0 | 0 | 0 |
| ENSG00000179941 | 729 | 355 | 390 | 209 | 393 | 315 |
| ENSG00000091039 | 1218 | 651 | 491 | 241 | 1017 | 502 |
| ENSG00000257526 | 5 | 1 | 7 | 5 | 5 | 8 |
| ENSG00000213272 | 0 | 0 | 1 | 0 | 2 | 1 |
| ENSG00000271142 | 0 | 1 | 0 | 1 | 0 | 0 |
| ENSG00000243071 | 4 | 1 | 2 | 2 | 2 | 1 |
| ENSG00000241045 | 0 | 0 | 0 | 0 | 0 | 0 |
| ENSG00000258090 | 2 | 0 | 0 | 1 | 1 | 1 |
| ENSG00000186908 | 802 | 365 | 455 | 181 | 512 | 382 |
| ENSG00000175183 | 14 | 8 | 8 | 19 | 10 | 10 |
| ENSG00000257910 | 0 | 0 | 0 | 0 | 0 | 0 |
| ENSG00000257199 | 0 | 0 | 0 | 0 | 0 | 0 |
| ENSG00000165891 | 219 | 115 | 135 | 72 | 119 | 118 |
| ENSG00000257677 | 0 | 0 | 0 | 0 | 0 | 0 |
| ENSG00000067798 | 0 | 1 | 1 | 1 | 0 | 0 |
| ENSG00000257835 | 0 | 0 | 0 | 0 | 0 | 0 |
| ENSG00000258066 | 0 | 0 | 0 | 0 | 0 | 0 |
| ENSG00000258225 | 0 | 0 | 0 | 0 | 0 | 0 |

| | | | | | | |
|---|---|---|---|---|---|---|
| ENSG00000257220 | 0 | 0 | 0 | 0 | 0 | 0 |
| ENSG00000258220 | 0 | 0 | 0 | 0 | 0 | 0 |
| ENSG00000258084 | 0 | 0 | 0 | 0 | 0 | 0 |
| ENSG00000257879 | 0 | 0 | 0 | 0 | 0 | 0 |
| ENSG00000257165 | 0 | 0 | 0 | 0 | 0 | 0 |
| ENSG00000257564 | 0 | 0 | 0 | 0 | 0 | 0 |
| ENSG00000257948 | 0 | 0 | 0 | 0 | 0 | 0 |
| ENSG00000067715 | 1689 | 881 | 923 | 296 | 902 | 677 |
| ENSG00000257191 | 25 | 18 | 14 | 12 | 17 | 31 |
| ENSG00000257894 | 11 | 2 | 1 | 2 | 2 | 7 |
| ENSG00000221788 | 0 | 0 | 0 | 0 | 0 | 0 |
| ENSG00000257604 | 2 | 1 | 0 | 0 | 0 | 0 |
| ENSG00000257474 | 0 | 1 | 2 | 1 | 0 | 0 |
| ENSG00000257956 | 0 | 2 | 5 | 0 | 0 | 3 |
| ENSG00000177425 | 3482 | 1749 | 2040 | 1109 | 1913 | 1625 |
| ENSG00000258048 | 0 | 0 | 0 | 0 | 1 | 1 |
| ENSG00000058272 | 2092 | 1108 | 1031 | 544 | 1396 | 961 |
| ENSG00000277130 | 2 | 0 | 0 | 1 | 0 | 0 |
| ENSG00000258044 | 1 | 3 | 0 | 1 | 0 | 0 |
| ENSG00000201942 | 0 | 0 | 0 | 0 | 0 | 0 |
| ENSG00000257557 | 118 | 26 | 38 | 36 | 50 | 59 |
| ENSG00000271496 | 0 | 0 | 0 | 0 | 0 | 0 |
| ENSG00000239473 | 0 | 0 | 0 | 0 | 0 | 0 |
| ENSG00000238842 | 0 | 0 | 0 | 0 | 0 | 0 |
| ENSG00000165899 | 4 | 2 | 2 | 0 | 2 | 3 |
| ENSG00000230291 | 2 | 1 | 2 | 1 | 0 | 2 |
| ENSG00000222880 | 0 | 0 | 0 | 0 | 0 | 0 |
| ENSG00000139304 | 26 | 16 | 14 | 9 | 40 | 24 |
| ENSG00000257786 | 0 | 0 | 0 | 0 | 0 | 0 |
| ENSG00000257429 | 0 | 0 | 0 | 0 | 0 | 0 |
| ENSG00000111046 | 0 | 0 | 0 | 0 | 0 | 0 |
| ENSG00000111049 | 0 | 0 | 0 | 0 | 0 | 0 |
| ENSG00000257741 | 0 | 0 | 0 | 0 | 0 | 0 |
| ENSG00000111052 | 699 | 390 | 473 | 181 | 396 | 349 |
| ENSG00000207763 | 0 | 0 | 0 | 0 | 0 | 0 |
| ENSG00000258215 | 2 | 0 | 0 | 0 | 1 | 0 |
| ENSG00000208022 | 0 | 0 | 0 | 0 | 0 | 0 |
| ENSG00000111058 | 784 | 377 | 554 | 233 | 358 | 413 |
| ENSG00000258026 | 0 | 0 | 0 | 0 | 0 | 0 |
| ENSG00000265227 | 0 | 0 | 0 | 0 | 0 | 0 |
| ENSG00000139220 | 18 | 16 | 8 | 0 | 5 | 10 |
| ENSG00000257467 | 0 | 1 | 0 | 0 | 0 | 0 |
| ENSG00000258162 | 16 | 7 | 17 | 3 | 9 | 7 |
| ENSG00000258375 | 0 | 0 | 0 | 0 | 0 | 0 |
| ENSG00000257609 | 0 | 0 | 0 | 0 | 0 | 0 |
| ENSG00000257747 | 0 | 0 | 0 | 0 | 0 | 0 |
| ENSG00000258193 | 0 | 0 | 0 | 0 | 0 | 0 |
| ENSG00000133773 | 961 | 524 | 544 | 454 | 668 | 562 |
| ENSG00000127720 | 294 | 129 | 183 | 138 | 134 | 142 |
| ENSG00000257875 | 0 | 1 | 0 | 0 | 1 | 0 |

| | | | | | | |
|---|---|---|---|---|---|---|
| ENSG00000258170 | 0 | 0 | 0 | 0 | 0 | 0 |
| ENSG00000257837 | 0 | 0 | 0 | 0 | 0 | 0 |
| ENSG00000257431 | 0 | 0 | 0 | 0 | 0 | 0 |
| ENSG00000179104 | 937 | 430 | 569 | 461 | 751 | 710 |
| ENSG00000252220 | 0 | 0 | 0 | 0 | 0 | 0 |
| ENSG00000213270 | 0 | 1 | 0 | 0 | 0 | 0 |
| ENSG00000258338 | 0 | 0 | 0 | 0 | 0 | 0 |
| ENSG00000257124 | 0 | 0 | 0 | 0 | 0 | 0 |
| ENSG00000285191 | 0 | 0 | 0 | 0 | 0 | 0 |
| ENSG00000257729 | 0 | 0 | 0 | 0 | 0 | 0 |
| ENSG00000221148 | 0 | 0 | 0 | 0 | 0 | 0 |
| ENSG00000258358 | 0 | 0 | 0 | 0 | 0 | 0 |
| ENSG00000257296 | 0 | 0 | 0 | 0 | 0 | 0 |
| ENSG00000072041 | 0 | 1 | 0 | 0 | 0 | 0 |
| ENSG00000258073 | 0 | 0 | 0 | 0 | 0 | 0 |
| ENSG00000231738 | 0 | 0 | 0 | 0 | 0 | 0 |
| ENSG00000133640 | 2 | 0 | 0 | 1 | 0 | 0 |
| ENSG00000180318 | 0 | 0 | 0 | 0 | 0 | 0 |
| ENSG00000258815 | 7 | 4 | 2 | 2 | 9 | 0 |
| ENSG00000257855 | 0 | 0 | 0 | 0 | 0 | 0 |
| ENSG00000258206 | 0 | 0 | 0 | 0 | 0 | 0 |
| ENSG00000198774 | 0 | 0 | 0 | 0 | 0 | 0 |
| ENSG00000133636 | 0 | 0 | 0 | 1 | 0 | 0 |
| ENSG00000182050 | 0 | 0 | 0 | 0 | 0 | 0 |
| ENSG00000258178 | 0 | 0 | 0 | 0 | 0 | 0 |
| ENSG00000257897 | 0 | 0 | 0 | 0 | 0 | 0 |
| ENSG00000258185 | 0 | 0 | 0 | 0 | 0 | 0 |
| ENSG00000280049 | 0 | 0 | 0 | 0 | 0 | 0 |
| ENSG00000242850 | 0 | 0 | 0 | 0 | 0 | 0 |
| ENSG00000258205 | 0 | 0 | 0 | 0 | 0 | 0 |
| ENSG00000257648 | 0 | 0 | 0 | 0 | 0 | 0 |
| ENSG00000258173 | 0 | 0 | 0 | 0 | 0 | 0 |
| ENSG00000258128 | 0 | 0 | 0 | 0 | 0 | 1 |
| ENSG00000257912 | 0 | 0 | 0 | 0 | 0 | 0 |
| ENSG00000257940 | 0 | 0 | 0 | 0 | 0 | 0 |
| ENSG00000258179 | 0 | 0 | 0 | 0 | 0 | 0 |
| ENSG00000240721 | 0 | 0 | 0 | 0 | 0 | 0 |
| ENSG00000165805 | 0 | 0 | 0 | 0 | 0 | 0 |
| ENSG00000133641 | 404 | 234 | 267 | 218 | 283 | 221 |
| ENSG00000198707 | 68 | 56 | 59 | 33 | 93 | 87 |
| ENSG00000252747 | 2 | 0 | 0 | 0 | 0 | 0 |
| ENSG00000257752 | 10 | 3 | 0 | 1 | 2 | 1 |
| ENSG00000139324 | 972 | 561 | 408 | 168 | 710 | 368 |
| ENSG00000199245 | 0 | 0 | 0 | 0 | 0 | 0 |
| ENSG00000049130 | 1045 | 516 | 543 | 128 | 364 | 269 |
| ENSG00000281333 | 3 | 1 | 9 | 3 | 6 | 1 |
| ENSG00000252850 | 0 | 0 | 0 | 0 | 0 | 0 |
| ENSG00000278011 | 0 | 0 | 0 | 0 | 1 | 0 |
| ENSG00000246363 | 0 | 0 | 0 | 1 | 1 | 0 |
| ENSG00000257629 | 0 | 0 | 0 | 0 | 0 | 0 |

| | | | | | | |
|---|---|---|---|---|---|---|
| ENSG00000257156 | 0 | 0 | 0 | 0 | 0 | 0 |
| ENSG00000238302 | 0 | 0 | 0 | 0 | 0 | 0 |
| ENSG00000257820 | 0 | 0 | 0 | 0 | 0 | 0 |
| ENSG00000139318 | 44 | 29 | 24 | 27 | 18 | 30 |
| ENSG00000274021 | 0 | 0 | 0 | 0 | 0 | 0 |
| ENSG00000271327 | 17 | 3 | 11 | 6 | 11 | 7 |
| ENSG00000271259 | 4 | 7 | 0 | 5 | 1 | 4 |
| ENSG00000139323 | 745 | 425 | 545 | 439 | 491 | 534 |
| ENSG00000226982 | 0 | 0 | 0 | 0 | 0 | 0 |
| ENSG00000259075 | 1 | 0 | 5 | 3 | 5 | 8 |
| ENSG00000257594 | 17 | 5 | 4 | 3 | 8 | 5 |
| ENSG00000270344 | 131 | 80 | 73 | 63 | 55 | 75 |
| ENSG00000279939 | 0 | 0 | 0 | 0 | 0 | 0 |
| ENSG00000258302 | 5 | 3 | 6 | 2 | 7 | 10 |
| ENSG00000070961 | 1378 | 781 | 615 | 203 | 1096 | 537 |
| ENSG00000271614 | 92 | 54 | 76 | 45 | 38 | 42 |
| ENSG00000252207 | 0 | 1 | 0 | 0 | 0 | 0 |
| ENSG00000257480 | 0 | 0 | 0 | 0 | 0 | 0 |
| ENSG00000252823 | 0 | 0 | 0 | 0 | 0 | 0 |
| ENSG00000258216 | 5 | 4 | 4 | 8 | 1 | 3 |
| ENSG00000258290 | 0 | 1 | 1 | 1 | 1 | 1 |
| ENSG00000257725 | 0 | 0 | 0 | 0 | 0 | 0 |
| ENSG00000257194 | 8 | 5 | 7 | 1 | 6 | 4 |
| ENSG00000258183 | 0 | 0 | 0 | 0 | 0 | 0 |
| ENSG00000257995 | 0 | 0 | 0 | 0 | 0 | 0 |
| ENSG00000257787 | 0 | 0 | 0 | 0 | 0 | 0 |
| ENSG00000197651 | 0 | 0 | 0 | 0 | 0 | 0 |
| ENSG00000196243 | 0 | 0 | 0 | 0 | 0 | 0 |
| ENSG00000083782 | 0 | 0 | 0 | 0 | 0 | 0 |
| ENSG00000139330 | 0 | 0 | 0 | 0 | 0 | 0 |
| ENSG00000139329 | 0 | 0 | 0 | 4 | 5 | 3 |
| ENSG00000011465 | 0 | 0 | 3 | 0 | 1 | 0 |
| ENSG00000258148 | 0 | 0 | 0 | 0 | 0 | 0 |
| ENSG00000258100 | 0 | 0 | 0 | 0 | 0 | 0 |
| ENSG00000274647 | 0 | 0 | 0 | 0 | 1 | 0 |
| ENSG00000277738 | 4 | 1 | 0 | 2 | 8 | 2 |
| ENSG00000258125 | 0 | 0 | 0 | 0 | 0 | 0 |
| ENSG00000258224 | 0 | 0 | 0 | 0 | 0 | 0 |
| ENSG00000257893 | 0 | 0 | 0 | 0 | 0 | 0 |
| ENSG00000257863 | 0 | 0 | 0 | 0 | 0 | 0 |
| ENSG00000257242 | 4 | 4 | 5 | 1 | 4 | 3 |
| ENSG00000133639 | 3250 | 1611 | 2113 | 1171 | 1402 | 1463 |
| ENSG00000245904 | 34 | 10 | 17 | 8 | 7 | 7 |
| ENSG00000239201 | 0 | 0 | 0 | 0 | 0 | 0 |
| ENSG00000199895 | 0 | 0 | 0 | 0 | 0 | 0 |
| ENSG00000277851 | 0 | 0 | 0 | 0 | 0 | 0 |
| ENSG00000205057 | 0 | 0 | 0 | 0 | 0 | 0 |
| ENSG00000266923 | 0 | 0 | 0 | 0 | 0 | 0 |
| ENSG00000257127 | 0 | 0 | 0 | 0 | 0 | 0 |
| ENSG00000205056 | 2 | 0 | 0 | 0 | 0 | 0 |

| | | | | | | |
|---|---|---|---|---|---|---|
| ENSG00000258262 | 0 | 0 | 0 | 0 | 0 | 0 |
| ENSG00000187510 | 0 | 0 | 0 | 0 | 1 | 0 |
| ENSG00000102189 | 166 | 118 | 108 | 27 | 218 | 123 |
| ENSG00000258012 | 0 | 0 | 0 | 0 | 0 | 0 |
| ENSG00000257195 | 2 | 0 | 0 | 0 | 0 | 2 |
| ENSG00000202534 | 0 | 0 | 0 | 0 | 0 | 0 |
| ENSG00000257345 | 0 | 0 | 0 | 0 | 0 | 0 |
| ENSG00000257322 | 0 | 0 | 0 | 2 | 0 | 0 |
| ENSG00000207365 | 0 | 0 | 0 | 0 | 0 | 0 |
| ENSG00000270506 | 0 | 0 | 0 | 0 | 0 | 0 |
| ENSG00000256393 | 3 | 3 | 2 | 3 | 7 | 2 |
| ENSG00000257746 | 0 | 0 | 0 | 0 | 0 | 0 |
| ENSG00000257210 | 0 | 0 | 0 | 0 | 0 | 2 |
| ENSG00000258171 | 0 | 0 | 0 | 0 | 0 | 0 |
| ENSG00000201502 | 0 | 0 | 0 | 0 | 0 | 0 |
| ENSG00000257512 | 56 | 33 | 32 | 22 | 34 | 22 |
| ENSG00000257252 | 0 | 2 | 1 | 0 | 0 | 0 |
| ENSG00000173598 | 1160 | 597 | 680 | 419 | 602 | 629 |
| ENSG00000177889 | 3681 | 1886 | 2087 | 1861 | 2151 | 2026 |
| ENSG00000200011 | 0 | 0 | 0 | 0 | 0 | 0 |
| ENSG00000198015 | 1641 | 818 | 950 | 806 | 1136 | 988 |
| ENSG00000243015 | 0 | 0 | 0 | 0 | 0 | 0 |
| ENSG00000258271 | 0 | 0 | 0 | 0 | 0 | 1 |
| ENSG00000246985 | 349 | 135 | 230 | 159 | 156 | 167 |
| ENSG00000120833 | 1927 | 733 | 1114 | 845 | 786 | 1049 |
| ENSG00000186076 | 7 | 8 | 4 | 4 | 3 | 4 |
| ENSG00000169372 | 389 | 210 | 242 | 211 | 185 | 189 |
| ENSG00000258274 | 0 | 0 | 1 | 0 | 0 | 0 |
| ENSG00000264978 | 0 | 0 | 0 | 0 | 0 | 0 |
| ENSG00000258303 | 2 | 1 | 2 | 5 | 3 | 1 |
| ENSG00000257283 | 0 | 2 | 3 | 4 | 1 | 2 |
| ENSG00000271382 | 4 | 0 | 0 | 0 | 0 | 1 |
| ENSG00000223126 | 0 | 0 | 0 | 0 | 0 | 0 |
| ENSG00000257360 | 0 | 0 | 0 | 0 | 0 | 0 |
| ENSG00000136040 | 30 | 13 | 12 | 7 | 18 | 4 |
| ENSG00000258035 | 0 | 0 | 0 | 0 | 0 | 0 |
| ENSG00000258172 | 0 | 0 | 0 | 0 | 0 | 0 |
| ENSG00000258365 | 0 | 0 | 1 | 0 | 1 | 0 |
| ENSG00000173588 | 203 | 105 | 99 | 53 | 166 | 97 |
| ENSG00000244391 | 0 | 0 | 0 | 0 | 0 | 0 |
| ENSG00000213250 | 2 | 3 | 1 | 1 | 2 | 0 |
| ENSG00000278916 | 30 | 13 | 20 | 12 | 12 | 17 |
| ENSG00000257400 | 0 | 0 | 0 | 0 | 0 | 0 |
| ENSG00000244091 | 0 | 0 | 0 | 0 | 0 | 0 |
| ENSG00000258357 | 0 | 0 | 1 | 0 | 0 | 0 |
| ENSG00000236349 | 2 | 2 | 2 | 1 | 0 | 3 |
| ENSG00000266099 | 0 | 0 | 0 | 0 | 0 | 0 |
| ENSG00000057704 | 84 | 46 | 51 | 31 | 61 | 41 |
| ENSG00000283818 | 0 | 0 | 0 | 0 | 0 | 0 |
| ENSG00000257847 | 0 | 0 | 0 | 0 | 0 | 0 |

| | | | | | | |
|---|---|---|---|---|---|---|
| ENSG00000216306 | 0 | 0 | 1 | 0 | 1 | 0 |
| ENSG00000283998 | 0 | 0 | 0 | 0 | 0 | 0 |
| ENSG00000184752 | 1749 | 894 | 1018 | 1086 | 1171 | 1074 |
| ENSG00000120798 | 821 | 408 | 472 | 273 | 464 | 395 |
| ENSG00000180263 | 150 | 77 | 83 | 34 | 157 | 86 |
| ENSG00000257121 | 0 | 0 | 0 | 0 | 0 | 0 |
| ENSG00000237774 | 0 | 0 | 0 | 0 | 0 | 0 |
| ENSG00000028203 | 1683 | 904 | 956 | 536 | 1173 | 843 |
| ENSG00000202470 | 0 | 0 | 0 | 0 | 0 | 0 |
| ENSG00000257666 | 0 | 0 | 0 | 0 | 1 | 1 |
| ENSG00000212535 | 0 | 0 | 0 | 0 | 0 | 0 |
| ENSG00000199172 | 0 | 0 | 0 | 0 | 0 | 0 |
| ENSG00000265917 | 12 | 7 | 7 | 5 | 5 | 5 |
| ENSG00000257943 | 0 | 0 | 0 | 0 | 0 | 0 |
| ENSG00000258313 | 0 | 0 | 0 | 0 | 0 | 0 |
| ENSG00000222544 | 0 | 0 | 0 | 0 | 0 | 0 |
| ENSG00000241556 | 5 | 0 | 1 | 0 | 2 | 3 |
| ENSG00000111142 | 2273 | 1295 | 1324 | 891 | 1836 | 1290 |
| ENSG00000136014 | 0 | 1 | 0 | 0 | 0 | 0 |
| ENSG00000257150 | 12 | 3 | 5 | 3 | 1 | 2 |
| ENSG00000212448 | 0 | 0 | 0 | 0 | 0 | 0 |
| ENSG00000074527 | 1116 | 629 | 777 | 634 | 973 | 945 |
| ENSG00000199506 | 0 | 0 | 0 | 0 | 0 | 0 |
| ENSG00000258343 | 0 | 0 | 0 | 0 | 0 | 0 |
| ENSG00000258292 | 0 | 0 | 0 | 0 | 0 | 0 |
| ENSG00000139343 | 1854 | 920 | 1114 | 1278 | 1244 | 1135 |
| ENSG00000165972 | 0 | 2 | 0 | 2 | 1 | 0 |
| ENSG00000285030 | 0 | 0 | 0 | 0 | 0 | 0 |
| ENSG00000139344 | 6 | 1 | 6 | 2 | 5 | 4 |
| ENSG00000084110 | 10 | 8 | 3 | 3 | 2 | 10 |
| ENSG00000257878 | 0 | 1 | 1 | 0 | 0 | 0 |
| ENSG00000111144 | 3251 | 1760 | 2164 | 1644 | 2116 | 1846 |
| ENSG00000257715 | 0 | 0 | 0 | 0 | 0 | 0 |
| ENSG00000258251 | 0 | 0 | 0 | 0 | 0 | 0 |
| ENSG00000277062 | 0 | 0 | 0 | 0 | 0 | 0 |
| ENSG00000283422 | 0 | 0 | 0 | 0 | 0 | 0 |
| ENSG00000111145 | 446 | 196 | 221 | 174 | 264 | 260 |
| ENSG00000258177 | 0 | 0 | 0 | 3 | 0 | 0 |
| ENSG00000059758 | 851 | 428 | 516 | 257 | 423 | 371 |
| ENSG00000201435 | 0 | 0 | 0 | 0 | 0 | 0 |
| ENSG00000257169 | 0 | 0 | 0 | 0 | 2 | 0 |
| ENSG00000258272 | 0 | 0 | 0 | 0 | 0 | 0 |
| ENSG00000252827 | 0 | 0 | 0 | 0 | 0 | 0 |
| ENSG00000188596 | 0 | 0 | 2 | 0 | 2 | 0 |
| ENSG00000258846 | 0 | 0 | 1 | 0 | 2 | 0 |
| ENSG00000139350 | 932 | 498 | 443 | 317 | 721 | 496 |
| ENSG00000202368 | 0 | 0 | 0 | 0 | 0 | 0 |
| ENSG00000258131 | 2 | 0 | 0 | 0 | 0 | 0 |
| ENSG00000257470 | 0 | 0 | 0 | 0 | 0 | 0 |
| ENSG00000279190 | 0 | 0 | 0 | 0 | 0 | 0 |

| | | | | | | |
|---|---|---|---|---|---|---|
| ENSG00000257920 | 0 | 0 | 0 | 0 | 0 | 0 |
| ENSG00000255794 | 4 | 3 | 7 | 8 | 10 | 13 |
| ENSG00000221479 | 0 | 0 | 0 | 0 | 0 | 1 |
| ENSG00000273942 | 0 | 0 | 0 | 0 | 0 | 0 |
| ENSG00000274230 | 0 | 0 | 0 | 0 | 0 | 0 |
| ENSG00000275231 | 0 | 0 | 0 | 0 | 0 | 0 |
| ENSG00000273570 | 0 | 0 | 0 | 0 | 0 | 0 |
| ENSG00000275971 | 0 | 0 | 0 | 0 | 0 | 0 |
| ENSG00000274819 | 0 | 0 | 0 | 0 | 0 | 0 |
| ENSG00000251844 | 0 | 0 | 0 | 0 | 0 | 0 |
| ENSG00000278802 | 0 | 0 | 0 | 0 | 0 | 0 |
| ENSG00000276398 | 0 | 0 | 0 | 0 | 0 | 0 |
| ENSG00000279798 | 11 | 3 | 2 | 2 | 2 | 1 |
| ENSG00000207586 | 0 | 0 | 0 | 0 | 0 | 0 |
| ENSG00000277081 | 0 | 0 | 0 | 0 | 0 | 0 |
| ENSG00000257157 | 0 | 0 | 0 | 0 | 0 | 0 |
| ENSG00000257501 | 0 | 0 | 0 | 0 | 0 | 0 |
| ENSG00000206899 | 2 | 2 | 3 | 0 | 0 | 1 |
| ENSG00000265861 | 0 | 0 | 0 | 0 | 0 | 0 |
| ENSG00000263890 | 0 | 0 | 0 | 0 | 0 | 0 |
| ENSG00000258312 | 0 | 0 | 0 | 0 | 0 | 0 |
| ENSG00000257580 | 0 | 0 | 0 | 0 | 0 | 0 |
| ENSG00000201296 | 0 | 0 | 0 | 0 | 0 | 0 |
| ENSG00000227825 | 12 | 3 | 8 | 5 | 1 | 2 |
| ENSG00000245017 | 2 | 0 | 2 | 3 | 1 | 1 |
| ENSG00000257167 | 120 | 69 | 81 | 54 | 71 | 63 |
| ENSG00000120802 | 3579 | 1784 | 2188 | 1708 | 2016 | 2243 |
| ENSG00000225185 | 0 | 0 | 0 | 0 | 0 | 0 |
| ENSG00000075415 | 15577 | 7581 | 9024 | 8781 | 9507 | 9541 |
| ENSG00000212443 | 4 | 2 | 1 | 2 | 3 | 1 |
| ENSG00000166130 | 248 | 117 | 153 | 147 | 204 | 149 |
| ENSG00000120868 | 577 | 292 | 344 | 157 | 353 | 259 |
| ENSG00000185046 | 49 | 32 | 21 | 11 | 22 | 20 |
| ENSG00000258204 | 0 | 0 | 0 | 0 | 0 | 0 |
| ENSG00000257458 | 0 | 0 | 0 | 0 | 0 | 0 |
| ENSG00000202324 | 0 | 0 | 0 | 0 | 0 | 0 |
| ENSG00000258039 | 0 | 0 | 0 | 0 | 0 | 0 |
| ENSG00000180219 | 0 | 3 | 0 | 0 | 0 | 0 |
| ENSG00000214203 | 2 | 4 | 0 | 1 | 1 | 4 |
| ENSG00000279148 | 14 | 4 | 8 | 2 | 15 | 3 |
| ENSG00000111647 | 556 | 337 | 232 | 125 | 542 | 276 |
| ENSG00000280088 | 101 | 45 | 21 | 22 | 59 | 37 |
| ENSG00000257696 | 19 | 13 | 10 | 3 | 9 | 3 |
| ENSG00000238105 | 685 | 285 | 421 | 212 | 287 | 363 |
| ENSG00000266610 | 0 | 0 | 0 | 0 | 0 | 0 |
| ENSG00000257489 | 44 | 17 | 32 | 9 | 21 | 21 |
| ENSG00000221476 | 2 | 1 | 0 | 0 | 0 | 0 |
| ENSG00000075089 | 1036 | 540 | 668 | 516 | 692 | 561 |
| ENSG00000166153 | 28 | 31 | 26 | 20 | 24 | 22 |
| ENSG00000206790 | 0 | 0 | 0 | 0 | 0 | 0 |

| | | | | | | |
|---|---|---|---|---|---|---|
| ENSG00000136021 | 1759 | 1024 | 863 | 447 | 1559 | 852 |
| ENSG00000179520 | 0 | 2 | 0 | 0 | 0 | 0 |
| ENSG00000012504 | 0 | 0 | 0 | 0 | 0 | 0 |
| ENSG00000271177 | 0 | 0 | 0 | 0 | 0 | 0 |
| ENSG00000139354 | 254 | 130 | 124 | 78 | 221 | 158 |
| ENSG00000257803 | 0 | 0 | 2 | 1 | 0 | 1 |
| ENSG00000151572 | 0 | 2 | 0 | 0 | 1 | 1 |
| ENSG00000257325 | 0 | 0 | 0 | 0 | 0 | 0 |
| ENSG00000258007 | 0 | 0 | 0 | 0 | 0 | 0 |
| ENSG00000258033 | 0 | 0 | 2 | 0 | 1 | 0 |
| ENSG00000256870 | 0 | 0 | 0 | 0 | 0 | 0 |
| ENSG00000207414 | 0 | 0 | 0 | 0 | 0 | 0 |
| ENSG00000241744 | 0 | 0 | 0 | 0 | 0 | 0 |
| ENSG00000120800 | 341 | 242 | 167 | 95 | 446 | 200 |
| ENSG00000120805 | 3063 | 1461 | 1846 | 1519 | 1752 | 1636 |
| ENSG00000257543 | 5 | 2 | 8 | 0 | 10 | 2 |
| ENSG00000222890 | 0 | 0 | 0 | 0 | 0 | 0 |
| ENSG00000239413 | 0 | 0 | 0 | 0 | 0 | 0 |
| ENSG00000206929 | 0 | 0 | 0 | 0 | 0 | 0 |
| ENSG00000202249 | 0 | 0 | 0 | 0 | 0 | 0 |
| ENSG00000252366 | 0 | 0 | 0 | 0 | 0 | 0 |
| ENSG00000166211 | 0 | 0 | 0 | 0 | 0 | 0 |
| ENSG00000196091 | 1 | 0 | 1 | 2 | 2 | 1 |
| ENSG00000257514 | 0 | 0 | 0 | 0 | 0 | 0 |
| ENSG00000274560 | 2 | 1 | 7 | 1 | 3 | 0 |
| ENSG00000111666 | 1662 | 747 | 1014 | 874 | 876 | 962 |
| ENSG00000199933 | 2 | 1 | 0 | 0 | 0 | 0 |
| ENSG00000139351 | 11 | 6 | 2 | 1 | 4 | 2 |
| ENSG00000111670 | 2111 | 1132 | 1164 | 662 | 1421 | 1028 |
| ENSG00000222255 | 0 | 0 | 0 | 0 | 0 | 0 |
| ENSG00000258230 | 0 | 2 | 0 | 0 | 0 | 0 |
| ENSG00000201168 | 0 | 0 | 0 | 0 | 0 | 0 |
| ENSG00000222932 | 0 | 0 | 0 | 0 | 0 | 0 |
| ENSG00000223046 | 0 | 0 | 0 | 0 | 0 | 0 |
| ENSG00000252863 | 0 | 0 | 0 | 0 | 0 | 0 |
| ENSG00000258153 | 0 | 0 | 0 | 0 | 1 | 0 |
| ENSG00000136048 | 1885 | 1048 | 1195 | 699 | 1130 | 947 |
| ENSG00000257202 | 0 | 1 | 0 | 0 | 0 | 0 |
| ENSG00000241917 | 0 | 0 | 0 | 0 | 0 | 0 |
| ENSG00000258308 | 0 | 0 | 0 | 0 | 0 | 0 |
| ENSG00000258288 | 0 | 0 | 0 | 0 | 0 | 0 |
| ENSG00000120860 | 829 | 408 | 517 | 392 | 460 | 407 |
| ENSG00000257222 | 2 | 3 | 0 | 1 | 2 | 2 |
| ENSG00000075188 | 1016 | 532 | 611 | 557 | 614 | 605 |
| ENSG00000185480 | 304 | 183 | 183 | 159 | 249 | 199 |
| ENSG00000183395 | 0 | 0 | 0 | 0 | 0 | 0 |
| ENSG00000281344 | 5 | 3 | 3 | 0 | 2 | 6 |
| ENSG00000264554 | 0 | 0 | 0 | 0 | 0 | 0 |
| ENSG00000258142 | 0 | 0 | 0 | 0 | 0 | 0 |
| ENSG00000283052 | 0 | 0 | 0 | 0 | 0 | 0 |

| | | | | | | |
|---|---|---|---|---|---|---|
| ENSG00000017427 | 0 | 0 | 0 | 0 | 0 | 0 |
| ENSG00000257254 | 0 | 0 | 0 | 0 | 0 | 0 |
| ENSG00000258169 | 0 | 0 | 0 | 0 | 0 | 0 |
| ENSG00000171759 | 3 | 2 | 3 | 1 | 1 | 1 |
| ENSG00000282964 | 0 | 0 | 0 | 0 | 0 | 0 |
| ENSG00000139352 | 40 | 21 | 28 | 34 | 26 | 49 |
| ENSG00000257703 | 0 | 0 | 0 | 0 | 0 | 1 |
| ENSG00000257860 | 0 | 0 | 0 | 0 | 0 | 0 |
| ENSG00000179088 | 13 | 16 | 22 | 8 | 24 | 16 |
| ENSG00000257415 | 0 | 0 | 0 | 0 | 0 | 0 |
| ENSG00000257994 | 0 | 0 | 0 | 0 | 0 | 0 |
| ENSG00000257762 | 2 | 0 | 0 | 0 | 0 | 0 |
| ENSG00000136011 | 0 | 0 | 0 | 0 | 0 | 0 |
| ENSG00000212594 | 0 | 0 | 0 | 0 | 0 | 0 |
| ENSG00000257737 | 0 | 0 | 0 | 0 | 0 | 0 |
| ENSG00000257766 | 0 | 0 | 0 | 0 | 0 | 0 |
| ENSG00000257681 | 4 | 2 | 2 | 0 | 0 | 2 |
| ENSG00000111696 | 263 | 104 | 164 | 102 | 150 | 155 |
| ENSG00000257754 | 0 | 0 | 0 | 0 | 0 | 0 |
| ENSG00000257327 | 1 | 1 | 1 | 0 | 2 | 0 |
| ENSG00000214198 | 3 | 4 | 7 | 4 | 9 | 6 |
| ENSG00000166598 | 8710 | 5469 | 5242 | 3263 | 7333 | 4888 |
| ENSG00000284503 | 0 | 0 | 0 | 0 | 0 | 0 |
| ENSG00000204954 | 511 | 268 | 293 | 327 | 359 | 306 |
| ENSG00000139372 | 3013 | 1527 | 1902 | 1402 | 1800 | 1800 |
| ENSG00000120820 | 6 | 1 | 13 | 1 | 6 | 8 |
| ENSG00000216285 | 8 | 1 | 3 | 2 | 1 | 1 |
| ENSG00000111727 | 382 | 176 | 199 | 95 | 213 | 152 |
| ENSG00000120837 | 951 | 505 | 611 | 491 | 726 | 633 |
| ENSG00000199415 | 0 | 0 | 0 | 0 | 0 | 0 |
| ENSG00000257193 | 2 | 0 | 1 | 0 | 3 | 4 |
| ENSG00000198431 | 8327 | 4402 | 5585 | 6313 | 9367 | 8021 |
| ENSG00000257732 | 2 | 1 | 0 | 1 | 2 | 1 |
| ENSG00000213442 | 84 | 54 | 46 | 73 | 59 | 62 |
| ENSG00000255150 | 3 | 3 | 2 | 6 | 7 | 7 |
| ENSG00000171310 | 34 | 16 | 27 | 21 | 44 | 29 |
| ENSG00000258111 | 0 | 0 | 0 | 0 | 0 | 0 |
| ENSG00000279176 | 0 | 0 | 0 | 0 | 0 | 0 |
| ENSG00000264295 | 0 | 0 | 0 | 0 | 0 | 0 |
| ENSG00000136052 | 341 | 201 | 219 | 153 | 220 | 199 |
| ENSG00000222579 | 0 | 0 | 0 | 0 | 0 | 0 |
| ENSG00000257758 | 2 | 1 | 1 | 0 | 0 | 1 |
| ENSG00000151131 | 409 | 203 | 248 | 224 | 271 | 247 |
| ENSG00000136010 | 7 | 5 | 4 | 1 | 6 | 4 |
| ENSG00000257999 | 6 | 1 | 0 | 0 | 0 | 0 |
| ENSG00000136051 | 886 | 499 | 423 | 157 | 978 | 490 |
| ENSG00000136044 | 1866 | 928 | 1131 | 932 | 1271 | 1142 |
| ENSG00000235162 | 785 | 352 | 345 | 454 | 480 | 395 |
| ENSG00000257642 | 0 | 0 | 0 | 0 | 0 | 0 |
| ENSG00000257859 | 0 | 0 | 0 | 1 | 0 | 0 |

| | | | | | | |
|---|---|---|---|---|---|---|
| ENSG00000257773 | 0 | 0 | 1 | 0 | 1 | 0 |
| ENSG00000257890 | 0 | 2 | 2 | 0 | 0 | 0 |
| ENSG00000074590 | 817 | 404 | 478 | 313 | 408 | 397 |
| ENSG00000257438 | 2 | 1 | 0 | 0 | 0 | 0 |
| ENSG00000257611 | 0 | 0 | 1 | 0 | 0 | 0 |
| ENSG00000136026 | 3583 | 1680 | 1924 | 2099 | 1930 | 1931 |
| ENSG00000258355 | 0 | 0 | 0 | 0 | 0 | 1 |
| ENSG00000277715 | 28 | 10 | 16 | 10 | 15 | 8 |
| ENSG00000238609 | 0 | 0 | 0 | 0 | 0 | 0 |
| ENSG00000166046 | 337 | 186 | 209 | 109 | 221 | 190 |
| ENSG00000013503 | 477 | 229 | 300 | 233 | 311 | 258 |
| ENSG00000257545 | 0 | 0 | 3 | 0 | 0 | 0 |
| ENSG00000111783 | 0 | 0 | 0 | 0 | 1 | 0 |
| ENSG00000257711 | 0 | 0 | 0 | 0 | 3 | 0 |
| ENSG00000257918 | 2 | 2 | 0 | 0 | 0 | 0 |
| ENSG00000111785 | 432 | 220 | 255 | 167 | 250 | 216 |
| ENSG00000257119 | 0 | 0 | 0 | 0 | 0 | 0 |
| ENSG00000240631 | 0 | 0 | 0 | 0 | 0 | 0 |
| ENSG00000260329 | 26 | 18 | 35 | 10 | 20 | 19 |
| ENSG00000151135 | 1366 | 712 | 681 | 522 | 781 | 606 |
| ENSG00000120832 | 495 | 258 | 332 | 168 | 225 | 248 |
| ENSG00000008405 | 664 | 341 | 424 | 275 | 390 | 380 |
| ENSG00000257548 | 0 | 0 | 0 | 0 | 0 | 0 |
| ENSG00000257444 | 0 | 0 | 0 | 0 | 0 | 0 |
| ENSG00000151136 | 174 | 91 | 83 | 70 | 81 | 71 |
| ENSG00000200897 | 0 | 0 | 0 | 0 | 0 | 0 |
| ENSG00000222302 | 0 | 0 | 0 | 0 | 0 | 0 |
| ENSG00000257579 | 0 | 0 | 0 | 0 | 0 | 0 |
| ENSG00000222160 | 0 | 0 | 0 | 0 | 0 | 0 |
| ENSG00000136045 | 2129 | 1123 | 1332 | 1135 | 1457 | 1370 |
| ENSG00000240441 | 0 | 1 | 0 | 0 | 0 | 0 |
| ENSG00000110851 | 2154 | 1043 | 1258 | 999 | 1310 | 1159 |
| ENSG00000258136 | 14 | 12 | 12 | 0 | 11 | 6 |
| ENSG00000187855 | 0 | 0 | 0 | 0 | 0 | 0 |
| ENSG00000257141 | 0 | 0 | 0 | 0 | 0 | 0 |
| ENSG00000274395 | 0 | 0 | 0 | 0 | 0 | 0 |
| ENSG00000257398 | 0 | 0 | 0 | 0 | 0 | 0 |
| ENSG00000257951 | 29 | 15 | 19 | 31 | 27 | 19 |
| ENSG00000257392 | 0 | 0 | 0 | 0 | 0 | 0 |
| ENSG00000257426 | 2 | 2 | 0 | 0 | 2 | 0 |
| ENSG00000258072 | 41 | 20 | 18 | 23 | 31 | 25 |
| ENSG00000257129 | 0 | 1 | 0 | 0 | 0 | 0 |
| ENSG00000075035 | 0 | 1 | 0 | 0 | 0 | 0 |
| ENSG00000257281 | 0 | 0 | 0 | 0 | 0 | 0 |
| ENSG00000174600 | 0 | 0 | 0 | 0 | 0 | 0 |
| ENSG00000247213 | 0 | 0 | 0 | 0 | 0 | 0 |
| ENSG00000198855 | 267 | 132 | 178 | 141 | 167 | 132 |
| ENSG00000075856 | 2252 | 1032 | 1549 | 1028 | 1161 | 1280 |
| ENSG00000136003 | 2585 | 1252 | 1374 | 1287 | 1468 | 1303 |
| ENSG00000183160 | 0 | 0 | 0 | 0 | 0 | 0 |

| | | | | | | |
|---|---|---|---|---|---|---|
| ENSG00000110876 | 0 | 2 | 0 | 0 | 2 | 3 |
| ENSG00000257221 | 0 | 0 | 0 | 0 | 0 | 2 |
| ENSG00000284388 | 0 | 0 | 0 | 0 | 0 | 0 |
| ENSG00000110880 | 2291 | 1156 | 1374 | 1025 | 1472 | 1288 |
| ENSG00000238457 | 0 | 0 | 0 | 0 | 0 | 0 |
| ENSG00000200060 | 0 | 0 | 0 | 0 | 0 | 0 |
| ENSG00000273499 | 0 | 0 | 0 | 0 | 0 | 0 |
| ENSG00000084112 | 1987 | 860 | 1165 | 659 | 983 | 1086 |
| ENSG00000280110 | 0 | 0 | 0 | 0 | 0 | 0 |
| ENSG00000274598 | 0 | 2 | 0 | 0 | 0 | 1 |
| ENSG00000207622 | 0 | 1 | 0 | 0 | 0 | 0 |
| ENSG00000110887 | 0 | 0 | 0 | 0 | 1 | 0 |
| ENSG00000166111 | 0 | 0 | 0 | 0 | 1 | 0 |
| ENSG00000201324 | 0 | 0 | 0 | 0 | 0 | 0 |
| ENSG00000279630 | 0 | 0 | 0 | 0 | 0 | 0 |
| ENSG00000135093 | 940 | 442 | 579 | 398 | 505 | 490 |
| ENSG00000256262 | 0 | 0 | 0 | 0 | 0 | 1 |
| ENSG00000212138 | 0 | 0 | 0 | 0 | 0 | 0 |
| ENSG00000189046 | 507 | 233 | 326 | 363 | 355 | 369 |
| ENSG00000076248 | 2410 | 1181 | 1488 | 1324 | 1149 | 1349 |
| ENSG00000256139 | 0 | 0 | 0 | 0 | 0 | 0 |
| ENSG00000076555 | 201 | 84 | 143 | 58 | 89 | 99 |
| ENSG00000139445 | 0 | 0 | 0 | 0 | 0 | 1 |
| ENSG00000174527 | 7 | 5 | 9 | 3 | 3 | 1 |
| ENSG00000256560 | 0 | 0 | 0 | 0 | 0 | 0 |
| ENSG00000255655 | 0 | 0 | 0 | 0 | 0 | 0 |
| ENSG00000110906 | 3314 | 1548 | 1995 | 1443 | 1794 | 1838 |
| ENSG00000151148 | 2723 | 1274 | 1600 | 1112 | 1343 | 1385 |
| ENSG00000139428 | 1217 | 549 | 705 | 803 | 750 | 745 |
| ENSG00000200274 | 0 | 0 | 0 | 0 | 0 | 0 |
| ENSG00000110921 | 1024 | 462 | 569 | 652 | 487 | 548 |
| ENSG00000200794 | 0 | 0 | 0 | 0 | 0 | 0 |
| ENSG00000139438 | 284 | 152 | 190 | 165 | 151 | 177 |
| ENSG00000255650 | 0 | 1 | 0 | 4 | 0 | 1 |
| ENSG00000111199 | 177 | 70 | 122 | 116 | 109 | 88 |
| ENSG00000263510 | 0 | 0 | 0 | 0 | 0 | 0 |
| ENSG00000139433 | 2776 | 1447 | 1650 | 1131 | 1517 | 1348 |
| ENSG00000256351 | 1 | 0 | 1 | 1 | 0 | 0 |
| ENSG00000241413 | 0 | 1 | 0 | 0 | 0 | 0 |
| ENSG00000249094 | 0 | 0 | 0 | 1 | 0 | 0 |
| ENSG00000139437 | 711 | 319 | 476 | 288 | 372 | 386 |
| ENSG00000139436 | 599 | 293 | 374 | 215 | 373 | 335 |
| ENSG00000277299 | 2 | 1 | 0 | 0 | 2 | 0 |
| ENSG00000280426 | 2 | 1 | 3 | 1 | 0 | 2 |
| ENSG00000076513 | 1090 | 534 | 666 | 454 | 572 | 580 |
| ENSG00000174456 | 514 | 227 | 303 | 269 | 283 | 291 |
| ENSG00000277595 | 0 | 2 | 1 | 2 | 1 | 4 |
| ENSG00000279360 | 0 | 0 | 0 | 0 | 0 | 0 |
| ENSG00000122970 | 177 | 89 | 99 | 56 | 168 | 114 |
| ENSG00000174437 | 7955 | 3723 | 4934 | 3689 | 4563 | 4733 |

| | | | | | | |
|---|---|---|---|---|---|---|
| ENSG00000276956 | 0 | 0 | 0 | 0 | 0 | 0 |
| ENSG00000196510 | 1483 | 775 | 953 | 944 | 1010 | 1050 |
| ENSG00000258210 | 3 | 2 | 2 | 3 | 3 | 3 |
| ENSG00000258011 | 0 | 0 | 0 | 0 | 0 | 0 |
| ENSG00000111229 | 8526 | 4693 | 5412 | 5098 | 5419 | 5069 |
| ENSG00000111231 | 527 | 236 | 273 | 237 | 298 | 265 |
| ENSG00000241680 | 0 | 1 | 2 | 1 | 1 | 1 |
| ENSG00000204856 | 167 | 97 | 121 | 154 | 159 | 151 |
| ENSG00000111237 | 2124 | 1141 | 1389 | 1178 | 1290 | 1242 |
| ENSG00000202335 | 0 | 0 | 0 | 0 | 0 | 0 |
| ENSG00000278993 | 16 | 12 | 17 | 9 | 10 | 10 |
| ENSG00000151164 | 8 | 5 | 6 | 0 | 8 | 1 |
| ENSG00000196850 | 765 | 361 | 430 | 314 | 416 | 434 |
| ENSG00000204852 | 884 | 418 | 557 | 483 | 470 | 518 |
| ENSG00000263537 | 0 | 0 | 0 | 0 | 0 | 0 |
| ENSG00000122986 | 3 | 5 | 3 | 1 | 1 | 9 |
| ENSG00000186298 | 4926 | 2425 | 2918 | 2510 | 3005 | 2866 |
| ENSG00000279925 | 0 | 0 | 0 | 0 | 0 | 0 |
| ENSG00000257268 | 5 | 1 | 2 | 5 | 4 | 3 |
| ENSG00000241286 | 0 | 0 | 0 | 1 | 2 | 1 |
| ENSG00000173093 | 156 | 39 | 71 | 54 | 66 | 58 |
| ENSG00000111245 | 0 | 0 | 0 | 0 | 0 | 0 |
| ENSG00000185847 | 0 | 0 | 0 | 0 | 0 | 0 |
| ENSG00000258240 | 0 | 0 | 0 | 0 | 0 | 0 |
| ENSG00000111249 | 2 | 0 | 2 | 2 | 5 | 1 |
| ENSG00000273700 | 0 | 0 | 0 | 0 | 0 | 0 |
| ENSG00000253080 | 0 | 0 | 0 | 0 | 0 | 0 |
| ENSG00000198324 | 558 | 311 | 349 | 400 | 332 | 283 |
| ENSG00000257595 | 0 | 0 | 1 | 0 | 0 | 0 |
| ENSG00000257539 | 0 | 0 | 0 | 0 | 0 | 0 |
| ENSG00000111252 | 154 | 97 | 102 | 70 | 85 | 83 |
| ENSG00000204842 | 1680 | 903 | 1010 | 647 | 919 | 855 |
| ENSG00000272215 | 0 | 0 | 0 | 0 | 0 | 0 |
| ENSG00000238168 | 0 | 2 | 0 | 0 | 0 | 0 |
| ENSG00000258099 | 3 | 3 | 12 | 8 | 3 | 6 |
| ENSG00000089234 | 1119 | 538 | 654 | 513 | 711 | 707 |
| ENSG00000258359 | 14 | 11 | 6 | 3 | 4 | 4 |
| ENSG00000111271 | 1030 | 470 | 746 | 580 | 579 | 603 |
| ENSG00000257767 | 0 | 0 | 1 | 0 | 0 | 0 |
| ENSG00000111275 | 13 | 6 | 2 | 12 | 6 | 5 |
| ENSG00000274697 | 0 | 0 | 0 | 0 | 0 | 0 |
| ENSG00000234608 | 831 | 383 | 486 | 455 | 503 | 397 |
| ENSG00000089022 | 1384 | 694 | 892 | 700 | 882 | 843 |
| ENSG00000244060 | 2 | 0 | 0 | 0 | 0 | 0 |
| ENSG00000229186 | 0 | 0 | 0 | 0 | 0 | 0 |
| ENSG00000226469 | 6 | 3 | 5 | 2 | 4 | 5 |
| ENSG00000198270 | 576 | 305 | 366 | 294 | 286 | 287 |
| ENSG00000258373 | 0 | 0 | 0 | 0 | 0 | 1 |
| ENSG00000257624 | 7 | 0 | 1 | 0 | 1 | 0 |
| ENSG00000089248 | 7087 | 3495 | 4349 | 4040 | 3805 | 3559 |

| | | | | | | |
|---|---|---|---|---|---|---|
| ENSG00000274227 | 2 | 7 | 3 | 2 | 4 | 3 |
| ENSG00000111300 | 1682 | 892 | 1009 | 614 | 1099 | 934 |
| ENSG00000266370 | 0 | 0 | 0 | 0 | 0 | 0 |
| ENSG00000258323 | 0 | 0 | 0 | 0 | 0 | 0 |
| ENSG00000200688 | 0 | 0 | 0 | 0 | 0 | 0 |
| ENSG00000135148 | 8362 | 4162 | 4881 | 4665 | 5342 | 5410 |
| ENSG00000200135 | 3 | 0 | 0 | 0 | 0 | 2 |
| ENSG00000173064 | 2420 | 864 | 1431 | 721 | 1053 | 1137 |
| ENSG00000283793 | 0 | 0 | 0 | 0 | 0 | 0 |
| ENSG00000257494 | 0 | 0 | 0 | 0 | 0 | 0 |
| ENSG00000201428 | 0 | 0 | 0 | 0 | 0 | 0 |
| ENSG00000213152 | 2 | 3 | 2 | 2 | 2 | 2 |
| ENSG00000257658 | 0 | 0 | 0 | 0 | 0 | 0 |
| ENSG00000089009 | 34085 | 17900 | 20070 | 21153 | 21912 | 19754 |
| ENSG00000179295 | 5865 | 2660 | 3278 | 1933 | 3370 | 3057 |
| ENSG00000089169 | 0 | 0 | 0 | 0 | 0 | 0 |
| ENSG00000283507 | 0 | 0 | 0 | 0 | 0 | 0 |
| ENSG00000089127 | 125 | 56 | 97 | 85 | 82 | 75 |
| ENSG00000257452 | 0 | 0 | 0 | 0 | 0 | 0 |
| ENSG00000111331 | 776 | 344 | 464 | 295 | 376 | 399 |
| ENSG00000111335 | 24 | 7 | 20 | 20 | 3 | 9 |
| ENSG00000258159 | 0 | 0 | 0 | 0 | 0 | 0 |
| ENSG00000242461 | 0 | 0 | 0 | 0 | 0 | 0 |
| ENSG00000135144 | 134 | 61 | 66 | 40 | 72 | 70 |
| ENSG00000111344 | 1224 | 526 | 756 | 668 | 582 | 637 |
| ENSG00000186710 | 22 | 10 | 16 | 10 | 6 | 8 |
| ENSG00000199899 | 1 | 1 | 0 | 0 | 2 | 1 |
| ENSG00000123064 | 3765 | 1933 | 2305 | 2529 | 2596 | 2396 |
| ENSG00000276908 | 0 | 0 | 0 | 0 | 0 | 0 |
| ENSG00000201208 | 3 | 1 | 3 | 0 | 1 | 1 |
| ENSG00000139405 | 1706 | 889 | 1025 | 1281 | 1072 | 1146 |
| ENSG00000257286 | 0 | 0 | 0 | 0 | 0 | 0 |
| ENSG00000166578 | 227 | 84 | 133 | 123 | 186 | 143 |
| ENSG00000186815 | 2170 | 998 | 1354 | 1285 | 1544 | 1556 |
| ENSG00000277566 | 7 | 10 | 8 | 4 | 13 | 7 |
| ENSG00000274822 | 0 | 0 | 0 | 0 | 0 | 0 |
| ENSG00000089060 | 527 | 260 | 361 | 331 | 310 | 327 |
| ENSG00000151176 | 2910 | 1281 | 1730 | 1419 | 1482 | 1607 |
| ENSG00000135094 | 10 | 5 | 6 | 4 | 6 | 5 |
| ENSG00000139410 | 758 | 320 | 455 | 482 | 379 | 395 |
| ENSG00000089116 | 0 | 0 | 0 | 0 | 0 | 0 |
| ENSG00000257935 | 0 | 0 | 0 | 0 | 0 | 0 |
| ENSG00000249550 | 0 | 0 | 1 | 1 | 1 | 1 |
| ENSG00000257231 | 0 | 0 | 0 | 0 | 0 | 1 |
| ENSG00000122965 | 1038 | 504 | 643 | 566 | 603 | 534 |
| ENSG00000257359 | 0 | 0 | 0 | 0 | 0 | 0 |
| ENSG00000257603 | 0 | 0 | 0 | 0 | 0 | 0 |
| ENSG00000257997 | 0 | 0 | 0 | 0 | 0 | 0 |
| ENSG00000257300 | 0 | 0 | 0 | 0 | 0 | 0 |
| ENSG00000257915 | 0 | 0 | 0 | 0 | 0 | 0 |

| | | | | | | |
|---|---|---|---|---|---|---|
| ENSG00000257476 | 0 | 0 | 0 | 0 | 0 | 0 |
| ENSG00000089225 | 2 | 0 | 0 | 2 | 0 | 0 |
| ENSG00000255399 | 0 | 0 | 0 | 0 | 0 | 0 |
| ENSG00000222982 | 0 | 0 | 0 | 0 | 0 | 0 |
| ENSG00000255830 | 0 | 0 | 0 | 0 | 0 | 0 |
| ENSG00000258244 | 0 | 0 | 1 | 0 | 0 | 0 |
| ENSG00000258254 | 0 | 0 | 0 | 0 | 0 | 0 |
| ENSG00000199220 | 0 | 0 | 0 | 0 | 0 | 0 |
| ENSG00000135111 | 567 | 214 | 235 | 231 | 321 | 265 |
| ENSG00000257817 | 8 | 1 | 2 | 3 | 1 | 0 |
| ENSG00000252459 | 0 | 0 | 0 | 0 | 0 | 0 |
| ENSG00000257517 | 0 | 0 | 0 | 0 | 0 | 0 |
| ENSG00000283459 | 0 | 0 | 0 | 0 | 0 | 0 |
| ENSG00000274879 | 0 | 0 | 1 | 0 | 0 | 1 |
| ENSG00000257958 | 0 | 0 | 0 | 0 | 0 | 0 |
| ENSG00000257519 | 0 | 0 | 0 | 0 | 0 | 0 |
| ENSG00000271579 | 0 | 0 | 0 | 0 | 0 | 0 |
| ENSG00000276308 | 0 | 0 | 0 | 0 | 0 | 0 |
| ENSG00000276972 | 0 | 0 | 0 | 0 | 0 | 0 |
| ENSG00000257726 | 0 | 0 | 0 | 0 | 0 | 1 |
| ENSG00000258108 | 0 | 0 | 0 | 0 | 0 | 0 |
| ENSG00000258280 | 0 | 0 | 0 | 0 | 0 | 0 |
| ENSG00000257407 | 0 | 0 | 0 | 0 | 0 | 0 |
| ENSG00000241656 | 0 | 0 | 0 | 0 | 0 | 0 |
| ENSG00000240205 | 0 | 0 | 0 | 0 | 0 | 0 |
| ENSG00000257781 | 0 | 0 | 0 | 0 | 0 | 0 |
| ENSG00000257683 | 1 | 1 | 0 | 3 | 2 | 2 |
| ENSG00000200112 | 0 | 0 | 0 | 0 | 0 | 0 |
| ENSG00000123066 | 2990 | 1244 | 1680 | 888 | 1660 | 1578 |
| ENSG00000258034 | 0 | 0 | 0 | 1 | 2 | 0 |
| ENSG00000200665 | 0 | 0 | 0 | 0 | 0 | 0 |
| ENSG00000207967 | 0 | 0 | 0 | 0 | 0 | 0 |
| ENSG00000257979 | 0 | 0 | 0 | 0 | 0 | 0 |
| ENSG00000258337 | 0 | 0 | 0 | 0 | 0 | 1 |
| ENSG00000258346 | 0 | 0 | 0 | 0 | 0 | 0 |
| ENSG00000264037 | 0 | 0 | 0 | 0 | 0 | 0 |
| ENSG00000258018 | 0 | 0 | 0 | 0 | 0 | 0 |
| ENSG00000196668 | 7 | 3 | 7 | 3 | 8 | 8 |
| ENSG00000258102 | 16 | 11 | 16 | 22 | 14 | 20 |
| ENSG00000275898 | 0 | 0 | 0 | 0 | 0 | 0 |
| ENSG00000258249 | 0 | 0 | 0 | 0 | 0 | 0 |
| ENSG00000257883 | 2 | 0 | 0 | 0 | 0 | 0 |
| ENSG00000257654 | 1 | 0 | 0 | 0 | 0 | 0 |
| ENSG00000111412 | 4330 | 2055 | 2377 | 2056 | 2433 | 2447 |
| ENSG00000135119 | 120 | 79 | 80 | 83 | 66 | 84 |
| ENSG00000239374 | 0 | 1 | 0 | 1 | 0 | 1 |
| ENSG00000201382 | 0 | 0 | 0 | 0 | 0 | 0 |
| ENSG00000275467 | 2 | 0 | 1 | 3 | 1 | 0 |
| ENSG00000135116 | 26 | 10 | 5 | 8 | 8 | 11 |
| ENSG00000240100 | 0 | 0 | 0 | 0 | 0 | 0 |

| | | | | | | |
|---|---|---|---|---|---|---|
| ENSG00000174989 | 1122 | 585 | 734 | 528 | 633 | 637 |
| ENSG00000274554 | 2 | 3 | 6 | 1 | 2 | 2 |
| ENSG00000257279 | 4 | 2 | 1 | 1 | 0 | 1 |
| ENSG00000277840 | 4 | 1 | 2 | 1 | 2 | 0 |
| ENSG00000088992 | 38 | 21 | 38 | 38 | 28 | 46 |
| ENSG00000258285 | 0 | 0 | 0 | 0 | 0 | 0 |
| ENSG00000135108 | 3630 | 1602 | 2081 | 1423 | 1851 | 1881 |
| ENSG00000089250 | 0 | 0 | 0 | 0 | 0 | 2 |
| ENSG00000257465 | 0 | 0 | 0 | 0 | 0 | 0 |
| ENSG00000255686 | 0 | 0 | 0 | 0 | 0 | 0 |
| ENSG00000171435 | 348 | 131 | 197 | 168 | 208 | 261 |
| ENSG00000256071 | 0 | 0 | 0 | 0 | 0 | 0 |
| ENSG00000111445 | 1198 | 616 | 684 | 601 | 738 | 837 |
| ENSG00000277873 | 3 | 1 | 3 | 4 | 2 | 4 |
| ENSG00000176871 | 3858 | 1910 | 2294 | 1834 | 2042 | 2052 |
| ENSG00000276292 | 1 | 2 | 2 | 0 | 1 | 1 |
| ENSG00000176834 | 2256 | 1118 | 1272 | 954 | 1199 | 1169 |
| ENSG00000274859 | 0 | 0 | 0 | 0 | 0 | 0 |
| ENSG00000089220 | 14680 | 6980 | 8539 | 8192 | 7689 | 7839 |
| ENSG00000135090 | 2583 | 1129 | 1395 | 908 | 1303 | 1235 |
| ENSG00000240342 | 10620 | 5123 | 5961 | 7468 | 6341 | 6200 |
| ENSG00000270482 | 7 | 8 | 7 | 4 | 10 | 5 |
| ENSG00000111707 | 5335 | 2338 | 3026 | 2097 | 2689 | 2630 |
| ENSG00000275759 | 4 | 1 | 2 | 2 | 1 | 1 |
| ENSG00000275409 | 0 | 0 | 2 | 0 | 0 | 0 |
| ENSG00000255714 | 0 | 0 | 0 | 0 | 0 | 0 |
| ENSG00000241590 | 0 | 0 | 0 | 0 | 0 | 0 |
| ENSG00000256750 | 0 | 0 | 0 | 0 | 0 | 0 |
| ENSG00000255618 | 0 | 0 | 0 | 0 | 0 | 0 |
| ENSG00000255814 | 0 | 0 | 0 | 0 | 0 | 0 |
| ENSG00000199809 | 0 | 0 | 0 | 0 | 0 | 0 |
| ENSG00000256149 | 0 | 0 | 0 | 0 | 0 | 0 |
| ENSG00000201042 | 0 | 0 | 0 | 0 | 0 | 0 |
| ENSG00000139767 | 0 | 0 | 0 | 0 | 0 | 0 |
| ENSG00000256031 | 0 | 0 | 0 | 0 | 0 | 0 |
| ENSG00000244112 | 0 | 0 | 0 | 0 | 0 | 0 |
| ENSG00000257095 | 0 | 0 | 0 | 0 | 0 | 0 |
| ENSG00000256884 | 0 | 0 | 0 | 0 | 0 | 0 |
| ENSG00000152137 | 416 | 195 | 230 | 316 | 333 | 245 |
| ENSG00000256609 | 0 | 0 | 0 | 0 | 0 | 0 |
| ENSG00000213144 | 4 | 8 | 4 | 5 | 0 | 5 |
| ENSG00000256311 | 0 | 0 | 0 | 0 | 0 | 0 |
| ENSG00000281196 | 0 | 0 | 0 | 0 | 0 | 0 |
| ENSG00000183273 | 0 | 1 | 0 | 1 | 0 | 0 |
| ENSG00000248636 | 2 | 0 | 1 | 0 | 0 | 0 |
| ENSG00000224982 | 0 | 0 | 0 | 0 | 0 | 0 |
| ENSG00000252886 | 0 | 0 | 0 | 0 | 0 | 0 |
| ENSG00000111725 | 1857 | 948 | 1253 | 1014 | 988 | 1076 |
| ENSG00000122966 | 702 | 283 | 444 | 287 | 449 | 438 |
| ENSG00000255692 | 0 | 0 | 2 | 0 | 0 | 2 |

| | | | | | | |
|---|---|---|---|---|---|---|
| ENSG00000283768 | 0 | 0 | 0 | 0 | 0 | 0 |
| ENSG00000226729 | 0 | 0 | 0 | 0 | 0 | 0 |
| ENSG00000135127 | 1327 | 624 | 841 | 726 | 714 | 805 |
| ENSG00000278250 | 0 | 0 | 0 | 0 | 1 | 1 |
| ENSG00000255575 | 0 | 0 | 0 | 0 | 0 | 0 |
| ENSG00000111737 | 2909 | 1412 | 1744 | 1700 | 1663 | 1709 |
| ENSG00000277283 | 176 | 92 | 125 | 60 | 100 | 117 |
| ENSG00000089154 | 8156 | 3542 | 4621 | 3980 | 4425 | 4466 |
| ENSG00000266704 | 0 | 0 | 0 | 0 | 0 | 0 |
| ENSG00000089157 | 67883 | 34274 | 39585 | 49907 | 41038 | 39562 |
| ENSG00000255857 | 152 | 92 | 96 | 101 | 95 | 93 |
| ENSG00000089159 | 9162 | 5173 | 5997 | 4690 | 5115 | 4872 |
| ENSG00000275936 | 2 | 1 | 3 | 1 | 0 | 4 |
| ENSG00000240925 | 0 | 0 | 0 | 1 | 0 | 0 |
| ENSG00000123009 | 8 | 8 | 16 | 10 | 12 | 12 |
| ENSG00000202538 | 0 | 1 | 0 | 0 | 0 | 1 |
| ENSG00000200795 | 8 | 1 | 6 | 2 | 2 | 2 |
| ENSG00000089163 | 59 | 37 | 43 | 32 | 23 | 31 |
| ENSG00000252659 | 0 | 0 | 0 | 0 | 0 | 0 |
| ENSG00000170890 | 0 | 0 | 0 | 0 | 0 | 0 |
| ENSG00000135097 | 440 | 215 | 311 | 295 | 254 | 261 |
| ENSG00000239881 | 0 | 0 | 1 | 0 | 1 | 1 |
| ENSG00000256695 | 0 | 0 | 0 | 0 | 3 | 0 |
| ENSG00000111775 | 3221 | 1962 | 2448 | 2620 | 2470 | 2172 |
| ENSG00000111780 | 0 | 0 | 0 | 0 | 0 | 0 |
| ENSG00000170855 | 1369 | 720 | 820 | 848 | 818 | 763 |
| ENSG00000257218 | 1271 | 669 | 739 | 707 | 752 | 710 |
| ENSG00000219355 | 3 | 0 | 6 | 0 | 0 | 0 |
| ENSG00000111786 | 4285 | 2511 | 2994 | 2628 | 2736 | 2768 |
| ENSG00000088986 | 5510 | 2647 | 2843 | 3554 | 3013 | 2839 |
| ENSG00000248008 | 271 | 137 | 157 | 147 | 156 | 132 |
| ENSG00000278344 | 2 | 2 | 1 | 4 | 1 | 0 |
| ENSG00000110871 | 1328 | 616 | 826 | 606 | 690 | 613 |
| ENSG00000222601 | 2 | 0 | 2 | 0 | 0 | 0 |
| ENSG00000241030 | 5 | 3 | 2 | 0 | 2 | 1 |
| ENSG00000022840 | 4674 | 2094 | 2549 | 2463 | 2471 | 2460 |
| ENSG00000167272 | 1097 | 487 | 613 | 634 | 611 | 629 |
| ENSG00000242137 | 0 | 0 | 0 | 0 | 0 | 0 |
| ENSG00000256008 | 0 | 0 | 0 | 0 | 0 | 0 |
| ENSG00000157782 | 2 | 0 | 0 | 2 | 0 | 2 |
| ENSG00000110917 | 7601 | 3638 | 4867 | 3912 | 4499 | 4669 |
| ENSG00000256364 | 0 | 0 | 0 | 0 | 0 | 0 |
| ENSG00000277423 | 6 | 4 | 2 | 3 | 0 | 1 |
| ENSG00000276188 | 0 | 0 | 0 | 2 | 1 | 0 |
| ENSG00000175970 | 2518 | 1252 | 1603 | 1118 | 1339 | 1329 |
| ENSG00000256569 | 0 | 0 | 0 | 0 | 0 | 0 |
| ENSG00000284143 | 0 | 0 | 0 | 0 | 0 | 0 |
| ENSG00000122971 | 428 | 161 | 261 | 254 | 236 | 223 |
| ENSG00000255946 | 1 | 0 | 0 | 0 | 0 | 1 |
| ENSG00000157837 | 2027 | 970 | 1241 | 1066 | 1071 | 1172 |

| | | | | | | |
|---|---|---|---|---|---|---|
| ENSG00000213137 | 0 | 0 | 0 | 0 | 0 | 0 |
| ENSG00000231313 | 0 | 0 | 2 | 4 | 1 | 0 |
| ENSG00000244585 | 0 | 0 | 1 | 0 | 1 | 1 |
| ENSG00000241388 | 0 | 0 | 0 | 0 | 0 | 0 |
| ENSG00000135100 | 0 | 1 | 0 | 0 | 1 | 0 |
| ENSG00000157895 | 619 | 301 | 365 | 324 | 347 | 382 |
| ENSG00000135114 | 88 | 32 | 58 | 33 | 22 | 24 |
| ENSG00000283542 | 0 | 0 | 0 | 0 | 0 | 0 |
| ENSG00000256963 | 0 | 0 | 0 | 0 | 0 | 0 |
| ENSG00000279001 | 0 | 0 | 0 | 0 | 0 | 0 |
| ENSG00000201945 | 0 | 0 | 0 | 0 | 0 | 0 |
| ENSG00000223538 | 0 | 0 | 0 | 0 | 0 | 0 |
| ENSG00000089041 | 0 | 2 | 3 | 0 | 0 | 1 |
| ENSG00000257053 | 0 | 0 | 0 | 0 | 0 | 0 |
| ENSG00000274029 | 0 | 0 | 0 | 0 | 0 | 0 |
| ENSG00000135124 | 1391 | 701 | 863 | 631 | 607 | 604 |
| ENSG00000110931 | 2608 | 1190 | 1488 | 1156 | 1345 | 1370 |
| ENSG00000089053 | 6027 | 2982 | 3753 | 3277 | 3458 | 3550 |
| ENSG00000258435 | 0 | 0 | 0 | 0 | 1 | 0 |
| ENSG00000170633 | 1542 | 744 | 939 | 640 | 813 | 773 |
| ENSG00000089094 | 1245 | 590 | 865 | 661 | 742 | 787 |
| ENSG00000284402 | 0 | 0 | 0 | 1 | 0 | 2 |
| ENSG00000256742 | 2 | 2 | 6 | 4 | 2 | 3 |
| ENSG00000252393 | 0 | 0 | 0 | 0 | 0 | 0 |
| ENSG00000276045 | 1002 | 472 | 632 | 693 | 606 | 635 |
| ENSG00000139714 | 74 | 42 | 61 | 52 | 40 | 58 |
| ENSG00000188735 | 892 | 417 | 537 | 871 | 896 | 1074 |
| ENSG00000139725 | 189 | 95 | 143 | 166 | 155 | 138 |
| ENSG00000212694 | 446 | 185 | 326 | 222 | 238 | 265 |
| ENSG00000272849 | 54 | 26 | 47 | 35 | 35 | 46 |
| ENSG00000274292 | 28 | 13 | 21 | 11 | 4 | 24 |
| ENSG00000139718 | 2402 | 1013 | 1493 | 1137 | 1240 | 1334 |
| ENSG00000158104 | 13 | 20 | 17 | 20 | 17 | 18 |
| ENSG00000256811 | 0 | 0 | 1 | 0 | 0 | 3 |
| ENSG00000278084 | 0 | 0 | 1 | 0 | 1 | 0 |
| ENSG00000110801 | 318 | 295 | 351 | 330 | 310 | 322 |
| ENSG00000256950 | 0 | 0 | 2 | 0 | 2 | 3 |
| ENSG00000283389 | 0 | 0 | 0 | 0 | 0 | 0 |
| ENSG00000239082 | 0 | 0 | 3 | 1 | 0 | 1 |
| ENSG00000158023 | 198 | 89 | 128 | 90 | 131 | 111 |
| ENSG00000255856 | 2 | 0 | 0 | 0 | 1 | 1 |
| ENSG00000110987 | 1448 | 599 | 806 | 565 | 670 | 728 |
| ENSG00000256546 | 12 | 2 | 6 | 1 | 1 | 1 |
| ENSG00000175727 | 3724 | 1702 | 2210 | 1666 | 1944 | 1890 |
| ENSG00000158113 | 1 | 1 | 0 | 0 | 0 | 0 |
| ENSG00000204671 | 0 | 0 | 0 | 0 | 0 | 0 |
| ENSG00000176383 | 74 | 45 | 47 | 43 | 38 | 31 |
| ENSG00000284934 | 1 | 1 | 2 | 0 | 2 | 1 |
| ENSG00000184047 | 146 | 83 | 120 | 76 | 95 | 94 |
| ENSG00000256861 | 0 | 0 | 0 | 0 | 0 | 0 |

| | | | | | | |
|---|---|---|---|---|---|---|
| ENSG00000139719 | 823 | 432 | 503 | 455 | 443 | 429 |
| ENSG00000130779 | 2167 | 1291 | 1166 | 442 | 1788 | 960 |
| ENSG00000214760 | 0 | 0 | 0 | 0 | 0 | 0 |
| ENSG00000257097 | 1 | 0 | 3 | 0 | 1 | 0 |
| ENSG00000033030 | 1113 | 581 | 710 | 415 | 547 | 544 |
| ENSG00000252192 | 3 | 1 | 1 | 0 | 1 | 1 |
| ENSG00000275265 | 0 | 0 | 0 | 1 | 0 | 0 |
| ENSG00000111011 | 1745 | 921 | 878 | 499 | 1199 | 883 |
| ENSG00000184445 | 574 | 352 | 317 | 147 | 608 | 329 |
| ENSG00000255972 | 0 | 0 | 0 | 0 | 0 | 0 |
| ENSG00000274191 | 0 | 2 | 2 | 1 | 3 | 0 |
| ENSG00000256249 | 0 | 2 | 0 | 0 | 0 | 0 |
| ENSG00000182782 | 57 | 18 | 19 | 22 | 24 | 6 |
| ENSG00000255398 | 3 | 8 | 6 | 3 | 6 | 7 |
| ENSG00000196917 | 795 | 414 | 489 | 199 | 298 | 352 |
| ENSG00000256044 | 0 | 0 | 1 | 0 | 0 | 0 |
| ENSG00000139726 | 4128 | 1899 | 2281 | 1650 | 2283 | 2092 |
| ENSG00000206914 | 1 | 4 | 1 | 0 | 4 | 0 |
| ENSG00000130783 | 2 | 3 | 3 | 1 | 2 | 2 |
| ENSG00000130787 | 3829 | 1865 | 2301 | 2155 | 2171 | 2035 |
| ENSG00000139722 | 608 | 324 | 390 | 415 | 358 | 392 |
| ENSG00000256152 | 0 | 0 | 0 | 0 | 0 | 0 |
| ENSG00000280138 | 41 | 22 | 41 | 9 | 11 | 46 |
| ENSG00000150967 | 728 | 348 | 456 | 407 | 405 | 404 |
| ENSG00000111325 | 64 | 35 | 60 | 32 | 27 | 55 |
| ENSG00000256028 | 7 | 10 | 8 | 4 | 5 | 7 |
| ENSG00000182196 | 23 | 17 | 16 | 22 | 11 | 20 |
| ENSG00000090975 | 303 | 126 | 163 | 316 | 407 | 384 |
| ENSG00000265526 | 0 | 0 | 0 | 0 | 0 | 0 |
| ENSG00000251497 | 0 | 1 | 0 | 0 | 1 | 1 |
| ENSG00000280381 | 0 | 0 | 1 | 1 | 2 | 0 |
| ENSG00000051825 | 429 | 244 | 214 | 92 | 370 | 235 |
| ENSG00000280120 | 36 | 21 | 20 | 16 | 26 | 28 |
| ENSG00000130921 | 817 | 437 | 513 | 362 | 483 | 456 |
| ENSG00000235423 | 153 | 89 | 107 | 59 | 75 | 60 |
| ENSG00000111328 | 1413 | 840 | 936 | 898 | 903 | 874 |
| ENSG00000269980 | 2 | 1 | 0 | 1 | 0 | 0 |
| ENSG00000199845 | 0 | 0 | 0 | 0 | 0 | 0 |
| ENSG00000139697 | 2014 | 1155 | 1012 | 455 | 1630 | 1054 |
| ENSG00000207189 | 1 | 0 | 1 | 0 | 0 | 0 |
| ENSG00000256092 | 126 | 48 | 69 | 66 | 57 | 57 |
| ENSG00000284425 | 0 | 0 | 0 | 0 | 0 | 0 |
| ENSG00000183955 | 1827 | 835 | 1129 | 1058 | 1022 | 1164 |
| ENSG00000150977 | 819 | 407 | 464 | 396 | 406 | 441 |
| ENSG00000270445 | 0 | 0 | 3 | 0 | 0 | 2 |
| ENSG00000184209 | 912 | 419 | 540 | 486 | 471 | 487 |
| ENSG00000188026 | 303 | 188 | 238 | 169 | 192 | 172 |
| ENSG00000274427 | 5 | 1 | 1 | 2 | 2 | 3 |
| ENSG00000278112 | 10 | 4 | 10 | 3 | 8 | 11 |
| ENSG00000266655 | 0 | 0 | 0 | 0 | 0 | 0 |

| | | | | | | |
|---|---|---|---|---|---|---|
| ENSG00000247373 | 107 | 58 | 85 | 56 | 57 | 50 |
| ENSG00000086598 | 9263 | 4759 | 5851 | 4985 | 5633 | 5573 |
| ENSG00000111364 | 976 | 486 | 578 | 454 | 639 | 620 |
| ENSG00000206897 | 4 | 0 | 2 | 0 | 0 | 0 |
| ENSG00000111361 | 2174 | 1129 | 1288 | 1102 | 1340 | 1199 |
| ENSG00000111358 | 1542 | 727 | 923 | 548 | 920 | 936 |
| ENSG00000279953 | 9 | 6 | 5 | 7 | 5 | 4 |
| ENSG00000278861 | 5 | 5 | 5 | 3 | 8 | 7 |
| ENSG00000168778 | 275 | 122 | 170 | 92 | 141 | 150 |
| ENSG00000255839 | 0 | 0 | 0 | 0 | 0 | 0 |
| ENSG00000185344 | 848 | 428 | 554 | 458 | 512 | 556 |
| ENSG00000239215 | 0 | 0 | 0 | 0 | 0 | 0 |
| ENSG00000280300 | 9 | 3 | 8 | 2 | 12 | 4 |
| ENSG00000197653 | 90 | 37 | 60 | 27 | 46 | 47 |
| ENSG00000119242 | 966 | 507 | 575 | 552 | 562 | 561 |
| ENSG00000250091 | 276 | 117 | 189 | 123 | 145 | 139 |
| ENSG00000270130 | 2 | 1 | 4 | 0 | 3 | 0 |
| ENSG00000270048 | 0 | 2 | 0 | 0 | 2 | 1 |
| ENSG00000269997 | 3 | 3 | 3 | 4 | 1 | 3 |
| ENSG00000269938 | 10 | 8 | 4 | 2 | 5 | 5 |
| ENSG00000270061 | 0 | 2 | 2 | 2 | 2 | 4 |
| ENSG00000270095 | 2 | 2 | 0 | 2 | 0 | 1 |
| ENSG00000179195 | 8355 | 3994 | 5044 | 3103 | 4632 | 4265 |
| ENSG00000178882 | 2 | 0 | 0 | 1 | 0 | 2 |
| ENSG00000274874 | 0 | 0 | 0 | 0 | 0 | 0 |
| ENSG00000256827 | 0 | 1 | 0 | 0 | 0 | 0 |
| ENSG00000275389 | 14 | 5 | 2 | 4 | 3 | 2 |
| ENSG00000256596 | 1 | 0 | 1 | 0 | 3 | 1 |
| ENSG00000196498 | 5827 | 2539 | 3479 | 3284 | 3422 | 3437 |
| ENSG00000275967 | 0 | 1 | 2 | 0 | 0 | 0 |
| ENSG00000255965 | 0 | 0 | 0 | 0 | 0 | 0 |
| ENSG00000214650 | 0 | 0 | 3 | 0 | 3 | 1 |
| ENSG00000279527 | 2 | 1 | 2 | 1 | 0 | 0 |
| ENSG00000279071 | 0 | 0 | 1 | 1 | 0 | 2 |
| ENSG00000280354 | 0 | 0 | 1 | 0 | 0 | 0 |
| ENSG00000279121 | 0 | 0 | 1 | 0 | 0 | 3 |
| ENSG00000279334 | 0 | 0 | 0 | 0 | 0 | 0 |
| ENSG00000280097 | 0 | 0 | 0 | 0 | 0 | 0 |
| ENSG00000278973 | 0 | 0 | 0 | 0 | 1 | 0 |
| ENSG00000280444 | 0 | 0 | 1 | 0 | 1 | 0 |
| ENSG00000279087 | 0 | 0 | 0 | 1 | 0 | 0 |
| ENSG00000279931 | 0 | 0 | 0 | 0 | 0 | 0 |
| ENSG00000073060 | 3089 | 1574 | 1748 | 1894 | 1774 | 1643 |
| ENSG00000252008 | 0 | 0 | 0 | 0 | 0 | 0 |
| ENSG00000150991 | 18672 | 8754 | 11341 | 10885 | 10536 | 11085 |
| ENSG00000265345 | 2 | 0 | 1 | 0 | 0 | 0 |
| ENSG00000241129 | 0 | 0 | 0 | 0 | 0 | 0 |
| ENSG00000150990 | 1356 | 669 | 857 | 1044 | 852 | 1008 |
| ENSG00000184992 | 2572 | 1329 | 1590 | 1140 | 1576 | 1705 |
| ENSG00000280634 | 0 | 0 | 0 | 0 | 0 | 0 |

| | | | | | | |
|---|---|---|---|---|---|---|
| ENSG00000256093 | 0 | 0 | 0 | 0 | 0 | 0 |
| ENSG00000081760 | 3193 | 1460 | 1810 | 1881 | 1732 | 1792 |
| ENSG00000279233 | 99 | 45 | 75 | 46 | 34 | 49 |
| ENSG00000256814 | 0 | 0 | 0 | 0 | 0 | 0 |
| ENSG00000280117 | 0 | 0 | 0 | 0 | 0 | 0 |
| ENSG00000139364 | 42 | 9 | 8 | 11 | 15 | 10 |
| ENSG00000279462 | 0 | 0 | 0 | 0 | 0 | 0 |
| ENSG00000280125 | 0 | 0 | 0 | 0 | 0 | 0 |
| ENSG00000280415 | 0 | 0 | 0 | 0 | 0 | 0 |
| ENSG00000271184 | 0 | 0 | 0 | 0 | 0 | 0 |
| ENSG00000249267 | 2 | 3 | 3 | 1 | 3 | 3 |
| ENSG00000275212 | 0 | 0 | 0 | 0 | 3 | 0 |
| ENSG00000284848 | 0 | 0 | 0 | 0 | 0 | 0 |
| ENSG00000280364 | 0 | 0 | 0 | 0 | 0 | 0 |
| ENSG00000280398 | 0 | 0 | 0 | 0 | 0 | 0 |
| ENSG00000280272 | 0 | 0 | 0 | 0 | 0 | 0 |
| ENSG00000255595 | 0 | 0 | 0 | 0 | 0 | 0 |
| ENSG00000256494 | 0 | 0 | 0 | 0 | 0 | 0 |
| ENSG00000214043 | 0 | 0 | 0 | 0 | 0 | 0 |
| ENSG00000256125 | 0 | 0 | 0 | 0 | 0 | 0 |
| ENSG00000255853 | 0 | 0 | 0 | 0 | 0 | 0 |
| ENSG00000256310 | 0 | 0 | 0 | 0 | 0 | 0 |
| ENSG00000256732 | 0 | 0 | 0 | 0 | 0 | 0 |
| ENSG00000255900 | 0 | 0 | 0 | 0 | 0 | 0 |
| ENSG00000255944 | 0 | 0 | 0 | 0 | 0 | 0 |
| ENSG00000278866 | 0 | 0 | 0 | 0 | 0 | 0 |
| ENSG00000279171 | 0 | 0 | 0 | 0 | 0 | 0 |
| ENSG00000255998 | 0 | 0 | 0 | 0 | 0 | 0 |
| ENSG00000189238 | 0 | 0 | 0 | 0 | 0 | 0 |
| ENSG00000256128 | 0 | 0 | 0 | 0 | 1 | 0 |
| ENSG00000255923 | 0 | 0 | 0 | 0 | 0 | 0 |
| ENSG00000249873 | 0 | 0 | 0 | 0 | 0 | 0 |
| ENSG00000256286 | 0 | 0 | 0 | 0 | 0 | 0 |
| ENSG00000257021 | 0 | 0 | 0 | 0 | 0 | 0 |
| ENSG00000249345 | 0 | 0 | 0 | 0 | 0 | 0 |
| ENSG00000256001 | 0 | 0 | 0 | 0 | 0 | 0 |
| ENSG00000278266 | 0 | 0 | 0 | 0 | 0 | 0 |
| ENSG00000256292 | 0 | 0 | 0 | 0 | 0 | 0 |
| ENSG00000256496 | 0 | 0 | 0 | 0 | 0 | 0 |
| ENSG00000253089 | 0 | 0 | 0 | 0 | 0 | 0 |
| ENSG00000256362 | 0 | 0 | 0 | 0 | 0 | 0 |
| ENSG00000256502 | 0 | 0 | 0 | 0 | 0 | 0 |
| ENSG00000280024 | 0 | 0 | 0 | 0 | 0 | 0 |
| ENSG00000279952 | 0 | 0 | 0 | 0 | 0 | 0 |
| ENSG00000255945 | 0 | 0 | 0 | 0 | 0 | 0 |
| ENSG00000199584 | 0 | 0 | 0 | 0 | 0 | 0 |
| ENSG00000257035 | 0 | 0 | 0 | 0 | 0 | 0 |
| ENSG00000256022 | 0 | 0 | 0 | 0 | 0 | 0 |
| ENSG00000256922 | 0 | 0 | 0 | 0 | 0 | 0 |
| ENSG00000279475 | 0 | 0 | 0 | 0 | 0 | 0 |

| | | | | | | |
|---|---|---|---|---|---|---|
| ENSG00000255757 | 0 | 0 | 0 | 0 | 0 | 0 |
| ENSG00000256597 | 0 | 0 | 0 | 0 | 0 | 0 |
| ENSG00000256193 | 0 | 0 | 0 | 0 | 0 | 0 |
| ENSG00000256971 | 0 | 0 | 0 | 0 | 0 | 0 |
| ENSG00000256643 | 0 | 0 | 0 | 0 | 0 | 0 |
| ENSG00000256551 | 0 | 0 | 0 | 0 | 0 | 0 |
| ENSG00000256659 | 0 | 0 | 0 | 0 | 0 | 0 |
| ENSG00000276487 | 0 | 0 | 0 | 0 | 0 | 0 |
| ENSG00000265061 | 0 | 0 | 0 | 0 | 0 | 0 |
| ENSG00000181234 | 0 | 0 | 0 | 0 | 0 | 0 |
| ENSG00000279180 | 0 | 0 | 0 | 0 | 0 | 0 |
| ENSG00000265635 | 0 | 0 | 0 | 0 | 0 | 0 |
| ENSG00000257025 | 0 | 0 | 0 | 0 | 0 | 0 |
| ENSG00000256630 | 0 | 0 | 0 | 0 | 0 | 0 |
| ENSG00000139370 | 994 | 507 | 619 | 478 | 508 | 569 |
| ENSG00000279500 | 37 | 19 | 20 | 7 | 11 | 8 |
| ENSG00000274695 | 6 | 4 | 3 | 3 | 1 | 1 |
| ENSG00000151948 | 0 | 0 | 0 | 0 | 1 | 0 |
| ENSG00000255838 | 0 | 0 | 0 | 0 | 0 | 0 |
| ENSG00000256581 | 0 | 0 | 0 | 0 | 0 | 0 |
| ENSG00000151952 | 0 | 0 | 0 | 0 | 0 | 0 |
| ENSG00000249196 | 0 | 0 | 0 | 1 | 1 | 0 |
| ENSG00000277217 | 0 | 0 | 0 | 0 | 0 | 0 |
| ENSG00000256699 | 0 | 0 | 0 | 0 | 0 | 0 |
| ENSG00000280051 | 0 | 0 | 0 | 0 | 0 | 0 |
| ENSG00000256085 | 0 | 0 | 0 | 0 | 0 | 0 |
| ENSG00000256137 | 0 | 0 | 0 | 0 | 0 | 0 |
| ENSG00000280196 | 0 | 0 | 0 | 0 | 0 | 0 |
| ENSG00000251536 | 0 | 0 | 0 | 0 | 0 | 0 |
| ENSG00000261650 | 0 | 0 | 0 | 0 | 0 | 0 |
| ENSG00000214039 | 0 | 0 | 0 | 0 | 0 | 0 |
| ENSG00000256298 | 0 | 0 | 0 | 0 | 0 | 0 |
| ENSG00000256906 | 0 | 0 | 0 | 0 | 0 | 0 |
| ENSG00000279193 | 0 | 0 | 0 | 0 | 0 | 0 |
| ENSG00000276122 | 2 | 0 | 4 | 12 | 14 | 6 |
| ENSG00000250208 | 2 | 1 | 0 | 3 | 4 | 3 |
| ENSG00000280405 | 0 | 0 | 0 | 0 | 0 | 0 |
| ENSG00000111432 | 0 | 0 | 0 | 0 | 0 | 0 |
| ENSG00000242963 | 0 | 0 | 0 | 0 | 0 | 0 |
| ENSG00000259862 | 0 | 0 | 0 | 0 | 0 | 0 |
| ENSG00000125207 | 0 | 0 | 0 | 0 | 1 | 0 |
| ENSG00000279343 | 0 | 0 | 0 | 0 | 0 | 0 |
| ENSG00000279478 | 0 | 0 | 0 | 0 | 0 | 0 |
| ENSG00000060709 | 0 | 1 | 0 | 0 | 0 | 1 |
| ENSG00000256064 | 0 | 0 | 0 | 0 | 0 | 0 |
| ENSG00000279905 | 0 | 0 | 0 | 0 | 0 | 0 |
| ENSG00000279146 | 0 | 0 | 0 | 0 | 0 | 0 |
| ENSG00000256343 | 0 | 0 | 0 | 0 | 0 | 0 |
| ENSG00000279650 | 0 | 0 | 0 | 0 | 0 | 0 |
| ENSG00000111450 | 490 | 260 | 324 | 226 | 324 | 308 |

| | | | | | | |
|---|---|---|---|---|---|---|
| ENSG00000256250 | 3 | 2 | 0 | 0 | 0 | 1 |
| ENSG00000256299 | 1 | 1 | 1 | 0 | 0 | 1 |
| ENSG00000222438 | 0 | 0 | 0 | 0 | 0 | 0 |
| ENSG00000132341 | 14222 | 7466 | 8416 | 8604 | 9308 | 9359 |
| ENSG00000206850 | 0 | 0 | 0 | 0 | 0 | 0 |
| ENSG00000111452 | 68 | 29 | 44 | 51 | 31 | 37 |
| ENSG00000279940 | 0 | 0 | 0 | 0 | 0 | 0 |
| ENSG00000279976 | 0 | 0 | 0 | 0 | 0 | 0 |
| ENSG00000280200 | 0 | 0 | 0 | 0 | 0 | 0 |
| ENSG00000256204 | 0 | 0 | 0 | 0 | 0 | 0 |
| ENSG00000279581 | 0 | 0 | 0 | 0 | 0 | 0 |
| ENSG00000280002 | 0 | 0 | 0 | 1 | 1 | 0 |
| ENSG00000256151 | 0 | 0 | 1 | 0 | 0 | 0 |
| ENSG00000279128 | 0 | 0 | 0 | 0 | 0 | 0 |
| ENSG00000279993 | 0 | 0 | 1 | 0 | 0 | 0 |
| ENSG00000256810 | 0 | 0 | 0 | 0 | 0 | 0 |
| ENSG00000204603 | 0 | 0 | 0 | 0 | 0 | 0 |
| ENSG00000280217 | 160 | 56 | 93 | 78 | 69 | 123 |
| ENSG00000226356 | 0 | 0 | 0 | 0 | 0 | 0 |
| ENSG00000279746 | 0 | 0 | 0 | 0 | 0 | 0 |
| ENSG00000248703 | 0 | 0 | 0 | 0 | 0 | 0 |
| ENSG00000212251 | 0 | 0 | 0 | 0 | 0 | 0 |
| ENSG00000230294 | 0 | 0 | 0 | 0 | 0 | 0 |
| ENSG00000256484 | 0 | 0 | 0 | 0 | 0 | 0 |
| ENSG00000280319 | 13 | 0 | 4 | 3 | 1 | 1 |
| ENSG00000275232 | 1 | 0 | 3 | 1 | 0 | 1 |
| ENSG00000256209 | 2 | 1 | 5 | 1 | 2 | 2 |
| ENSG00000279388 | 0 | 0 | 0 | 1 | 0 | 0 |
| ENSG00000255704 | 0 | 0 | 0 | 0 | 0 | 0 |
| ENSG00000279455 | 0 | 0 | 0 | 0 | 0 | 0 |
| ENSG00000256258 | 0 | 0 | 0 | 0 | 0 | 0 |
| ENSG00000279153 | 0 | 0 | 0 | 0 | 0 | 0 |
| ENSG00000256424 | 0 | 0 | 0 | 0 | 0 | 0 |
| ENSG00000212154 | 0 | 0 | 0 | 0 | 0 | 0 |
| ENSG00000249926 | 0 | 0 | 0 | 0 | 0 | 0 |
| ENSG00000243499 | 0 | 0 | 0 | 0 | 0 | 0 |
| ENSG00000061936 | 1640 | 797 | 1075 | 825 | 894 | 876 |
| ENSG00000255933 | 0 | 1 | 3 | 1 | 1 | 0 |
| ENSG00000200516 | 0 | 0 | 0 | 0 | 0 | 0 |
| ENSG00000200483 | 0 | 0 | 0 | 0 | 0 | 1 |
| ENSG00000198598 | 834 | 408 | 488 | 624 | 550 | 558 |
| ENSG00000256955 | 26 | 11 | 24 | 18 | 20 | 17 |
| ENSG00000177169 | 6590 | 3107 | 3957 | 3520 | 3740 | 3583 |
| ENSG00000279283 | 7 | 17 | 10 | 2 | 4 | 4 |
| ENSG00000255992 | 20 | 7 | 7 | 9 | 11 | 4 |
| ENSG00000177192 | 892 | 479 | 517 | 659 | 596 | 654 |
| ENSG00000273568 | 44 | 25 | 27 | 22 | 20 | 28 |
| ENSG00000183495 | 3019 | 1317 | 1798 | 1196 | 1501 | 1734 |
| ENSG00000208892 | 0 | 0 | 0 | 0 | 0 | 0 |
| ENSG00000257000 | 0 | 0 | 0 | 0 | 0 | 0 |

| | | | | | | |
|---|---|---|---|---|---|---|
| ENSG00000274670 | 0 | 0 | 0 | 1 | 0 | 0 |
| ENSG00000185684 | 641 | 279 | 412 | 243 | 325 | 355 |
| ENSG00000279640 | 2 | 3 | 2 | 0 | 1 | 1 |
| ENSG00000256804 | 0 | 0 | 0 | 0 | 0 | 0 |
| ENSG00000185163 | 817 | 426 | 541 | 436 | 481 | 493 |
| ENSG00000184967 | 718 | 383 | 429 | 466 | 404 | 427 |
| ENSG00000276693 | 0 | 0 | 0 | 0 | 0 | 0 |
| ENSG00000256576 | 50 | 38 | 46 | 37 | 29 | 47 |
| ENSG00000256312 | 2 | 3 | 3 | 3 | 1 | 4 |
| ENSG00000182870 | 8 | 0 | 0 | 0 | 1 | 0 |
| ENSG00000279480 | 0 | 3 | 0 | 0 | 0 | 0 |
| ENSG00000256943 | 0 | 1 | 1 | 0 | 2 | 1 |
| ENSG00000256542 | 2 | 4 | 0 | 1 | 0 | 1 |
| ENSG00000278872 | 0 | 0 | 0 | 0 | 0 | 0 |
| ENSG00000277011 | 0 | 0 | 0 | 0 | 0 | 0 |
| ENSG00000255916 | 0 | 0 | 0 | 0 | 0 | 0 |
| ENSG00000279113 | 0 | 0 | 0 | 0 | 0 | 0 |
| ENSG00000279704 | 0 | 0 | 0 | 0 | 0 | 0 |
| ENSG00000279023 | 0 | 0 | 0 | 0 | 0 | 0 |
| ENSG00000280150 | 0 | 0 | 0 | 0 | 0 | 0 |
| ENSG00000274373 | 2 | 0 | 0 | 2 | 0 | 1 |
| ENSG00000256783 | 2 | 0 | 0 | 0 | 0 | 0 |
| ENSG00000256875 | 0 | 0 | 0 | 0 | 1 | 0 |
| ENSG00000112787 | 2595 | 1099 | 1505 | 1475 | 1369 | 1375 |
| ENSG00000280287 | 66 | 29 | 38 | 50 | 40 | 36 |
| ENSG00000279700 | 15 | 2 | 8 | 6 | 6 | 5 |
| ENSG00000277052 | 1 | 0 | 0 | 1 | 0 | 0 |
| ENSG00000277186 | 12 | 9 | 13 | 12 | 8 | 12 |
| ENSG00000204583 | 0 | 0 | 0 | 0 | 0 | 0 |
| ENSG00000187848 | 1623 | 542 | 920 | 602 | 506 | 572 |
| ENSG00000280311 | 2 | 3 | 1 | 1 | 2 | 0 |
| ENSG00000177084 | 2417 | 1065 | 1417 | 982 | 1181 | 1272 |
| ENSG00000176894 | 311 | 253 | 301 | 313 | 262 | 269 |
| ENSG00000256632 | 0 | 0 | 0 | 0 | 1 | 0 |
| ENSG00000247077 | 2129 | 1150 | 1314 | 1462 | 1500 | 1409 |
| ENSG00000201469 | 0 | 1 | 0 | 0 | 0 | 0 |
| ENSG00000176915 | 3316 | 1476 | 2078 | 1524 | 1842 | 1993 |
| ENSG00000256108 | 0 | 0 | 0 | 0 | 0 | 0 |
| ENSG00000090615 | 4497 | 2221 | 2767 | 2096 | 2386 | 2521 |
| ENSG00000072609 | 2139 | 902 | 1273 | 996 | 1176 | 1126 |
| ENSG00000232888 | 17 | 10 | 13 | 13 | 10 | 9 |
| ENSG00000252079 | 0 | 0 | 0 | 0 | 0 | 0 |
| ENSG00000278949 | 25 | 17 | 13 | 2 | 4 | 2 |
| ENSG00000236617 | 41 | 21 | 51 | 18 | 31 | 31 |
| ENSG00000255703 | 1 | 0 | 0 | 0 | 0 | 0 |
| ENSG00000250790 | 107 | 51 | 86 | 43 | 48 | 42 |
| ENSG00000196458 | 233 | 117 | 105 | 48 | 196 | 91 |
| ENSG00000256563 | 0 | 0 | 1 | 0 | 0 | 0 |
| ENSG00000198393 | 329 | 168 | 181 | 132 | 199 | 161 |
| ENSG00000252269 | 0 | 0 | 0 | 1 | 0 | 0 |

| | | | | | | |
|---|---|---|---|---|---|---|
| ENSG00000246394 | 0 | 0 | 0 | 0 | 1 | 0 |
| ENSG00000255807 | 2 | 0 | 0 | 0 | 0 | 0 |
| ENSG00000198040 | 669 | 371 | 371 | 142 | 525 | 302 |
| ENSG00000279466 | 2 | 0 | 0 | 0 | 0 | 0 |
| ENSG00000279817 | 0 | 1 | 0 | 0 | 0 | 2 |
| ENSG00000196387 | 690 | 315 | 438 | 268 | 365 | 343 |
| ENSG00000214029 | 197 | 137 | 119 | 73 | 111 | 101 |
| ENSG00000271963 | 0 | 0 | 3 | 0 | 0 | 0 |
| ENSG00000240991 | 4 | 1 | 1 | 1 | 1 | 3 |
| ENSG00000256223 | 122 | 63 | 65 | 21 | 92 | 66 |
| ENSG00000256825 | 0 | 0 | 0 | 0 | 0 | 0 |
| ENSG00000090612 | 290 | 188 | 145 | 49 | 274 | 139 |
| ENSG00000255919 | 0 | 0 | 0 | 0 | 0 | 0 |
| ENSG00000227059 | 0 | 0 | 0 | 0 | 0 | 0 |
| ENSG00000239190 | 0 | 0 | 0 | 0 | 0 | 0 |
| ENSG00000238443 | 0 | 0 | 0 | 0 | 0 | 0 |
| ENSG00000279924 | 0 | 0 | 0 | 0 | 0 | 0 |
| ENSG00000279516 | 0 | 0 | 0 | 0 | 0 | 0 |
| ENSG00000279081 | 0 | 0 | 0 | 0 | 0 | 0 |
| ENSG00000284679 | 0 | 0 | 0 | 0 | 0 | 0 |
| ENSG00000280208 | 0 | 0 | 2 | 3 | 2 | 1 |
| ENSG00000279124 | 0 | 0 | 0 | 0 | 0 | 0 |
| ENSG00000279730 | 2 | 4 | 1 | 2 | 2 | 1 |
| ENSG00000279231 | 0 | 0 | 0 | 0 | 0 | 0 |
| ENSG00000268486 | 0 | 0 | 0 | 0 | 0 | 0 |
| ENSG00000276183 | 0 | 0 | 0 | 0 | 0 | 0 |
| ENSG00000215604 | 0 | 0 | 0 | 0 | 0 | 0 |
| ENSG00000270496 | 0 | 0 | 0 | 0 | 0 | 0 |
| ENSG00000231238 | 0 | 0 | 0 | 0 | 0 | 0 |
| ENSG00000233905 | 0 | 0 | 0 | 0 | 0 | 0 |
| ENSG00000229788 | 0 | 0 | 0 | 0 | 0 | 0 |
| ENSG00000235876 | 0 | 0 | 0 | 0 | 0 | 0 |
| ENSG00000231914 | 0 | 0 | 0 | 0 | 0 | 0 |
| ENSG00000226361 | 0 | 0 | 0 | 0 | 0 | 0 |
| ENSG00000228797 | 21 | 8 | 12 | 7 | 13 | 9 |
| ENSG00000230931 | 0 | 0 | 0 | 0 | 0 | 0 |
| ENSG00000204718 | 4 | 0 | 0 | 0 | 0 | 0 |
| ENSG00000227784 | 0 | 0 | 0 | 0 | 0 | 0 |
| ENSG00000244155 | 0 | 0 | 0 | 0 | 0 | 0 |
| ENSG00000227568 | 0 | 0 | 0 | 0 | 0 | 0 |
| ENSG00000225016 | 0 | 0 | 1 | 0 | 0 | 0 |
| ENSG00000206192 | 0 | 0 | 0 | 0 | 0 | 0 |
| ENSG00000242756 | 0 | 0 | 0 | 0 | 0 | 0 |
| ENSG00000223024 | 0 | 0 | 0 | 0 | 0 | 0 |
| ENSG00000206787 | 0 | 0 | 0 | 0 | 0 | 0 |
| ENSG00000235500 | 0 | 0 | 0 | 0 | 0 | 1 |
| ENSG00000226250 | 0 | 0 | 1 | 0 | 0 | 0 |
| ENSG00000231358 | 0 | 0 | 1 | 0 | 0 | 0 |
| ENSG00000226057 | 1 | 2 | 9 | 0 | 0 | 2 |
| ENSG00000232685 | 0 | 0 | 0 | 0 | 0 | 0 |

| | | | | | | |
|---|---|---|---|---|---|---|
| ENSG00000229375 | 0 | 0 | 0 | 0 | 0 | 0 |
| ENSG00000187721 | 0 | 0 | 0 | 0 | 0 | 0 |
| ENSG00000226619 | 0 | 0 | 0 | 0 | 0 | 0 |
| ENSG00000252287 | 0 | 0 | 0 | 0 | 0 | 0 |
| ENSG00000224778 | 0 | 1 | 0 | 0 | 0 | 0 |
| ENSG00000234894 | 0 | 0 | 0 | 0 | 0 | 0 |
| ENSG00000229147 | 0 | 0 | 0 | 0 | 0 | 0 |
| ENSG00000278337 | 0 | 0 | 0 | 0 | 0 | 0 |
| ENSG00000198033 | 0 | 0 | 0 | 0 | 0 | 0 |
| ENSG00000121388 | 0 | 0 | 0 | 0 | 0 | 0 |
| ENSG00000227879 | 3 | 0 | 3 | 1 | 2 | 3 |
| ENSG00000232243 | 0 | 0 | 0 | 0 | 0 | 0 |
| ENSG00000237636 | 0 | 0 | 0 | 0 | 0 | 0 |
| ENSG00000215349 | 0 | 0 | 0 | 1 | 0 | 0 |
| ENSG00000236834 | 0 | 0 | 0 | 0 | 0 | 0 |
| ENSG00000224976 | 0 | 0 | 0 | 0 | 0 | 0 |
| ENSG00000231184 | 0 | 0 | 0 | 0 | 0 | 0 |
| ENSG00000229791 | 0 | 0 | 0 | 0 | 0 | 0 |
| ENSG00000237164 | 0 | 0 | 0 | 0 | 0 | 0 |
| ENSG00000132958 | 2 | 1 | 2 | 4 | 1 | 3 |
| ENSG00000223625 | 0 | 0 | 0 | 0 | 0 | 0 |
| ENSG00000232894 | 4 | 0 | 0 | 0 | 1 | 1 |
| ENSG00000215572 | 0 | 0 | 0 | 0 | 0 | 0 |
| ENSG00000225316 | 0 | 0 | 0 | 0 | 0 | 0 |
| ENSG00000238132 | 2 | 1 | 1 | 0 | 2 | 0 |
| ENSG00000196199 | 2598 | 1221 | 1434 | 845 | 1461 | 1251 |
| ENSG00000226352 | 12 | 11 | 17 | 5 | 7 | 11 |
| ENSG00000121390 | 2509 | 1307 | 1701 | 1362 | 1674 | 1713 |
| ENSG00000275014 | 0 | 1 | 0 | 0 | 0 | 0 |
| ENSG00000233469 | 3 | 2 | 2 | 0 | 4 | 0 |
| ENSG00000132950 | 809 | 450 | 560 | 275 | 474 | 423 |
| ENSG00000223576 | 0 | 5 | 2 | 3 | 2 | 3 |
| ENSG00000275964 | 516 | 208 | 287 | 261 | 230 | 228 |
| ENSG00000121741 | 3495 | 2029 | 1804 | 807 | 2755 | 1708 |
| ENSG00000237672 | 10 | 5 | 4 | 1 | 2 | 2 |
| ENSG00000279951 | 0 | 2 | 0 | 0 | 0 | 0 |
| ENSG00000236076 | 0 | 0 | 0 | 2 | 0 | 0 |
| ENSG00000121743 | 0 | 0 | 1 | 3 | 2 | 1 |
| ENSG00000226633 | 0 | 0 | 0 | 0 | 0 | 0 |
| ENSG00000260131 | 0 | 0 | 0 | 0 | 0 | 0 |
| ENSG00000165474 | 82 | 29 | 29 | 27 | 40 | 38 |
| ENSG00000279065 | 0 | 0 | 0 | 0 | 0 | 0 |
| ENSG00000121742 | 0 | 2 | 2 | 0 | 0 | 0 |
| ENSG00000165475 | 1723 | 834 | 1023 | 927 | 964 | 873 |
| ENSG00000263978 | 0 | 0 | 0 | 0 | 0 | 0 |
| ENSG00000277020 | 6 | 2 | 6 | 2 | 0 | 1 |
| ENSG00000032742 | 485 | 241 | 264 | 139 | 306 | 227 |
| ENSG00000238286 | 2 | 9 | 3 | 5 | 3 | 5 |
| ENSG00000222726 | 2 | 2 | 0 | 0 | 3 | 2 |
| ENSG00000278291 | 95 | 60 | 68 | 65 | 70 | 99 |

| | | | | | | |
|---|---|---|---|---|---|---|
| ENSG00000172458 | 122 | 37 | 57 | 69 | 77 | 57 |
| ENSG00000279135 | 0 | 0 | 2 | 0 | 0 | 1 |
| ENSG00000150456 | 403 | 223 | 272 | 232 | 266 | 228 |
| ENSG00000233524 | 0 | 0 | 0 | 0 | 0 | 0 |
| ENSG00000275485 | 9 | 6 | 4 | 3 | 1 | 7 |
| ENSG00000132953 | 1192 | 661 | 611 | 340 | 860 | 617 |
| ENSG00000236704 | 0 | 0 | 0 | 0 | 0 | 0 |
| ENSG00000233780 | 0 | 0 | 0 | 0 | 0 | 0 |
| ENSG00000234363 | 0 | 0 | 0 | 0 | 0 | 0 |
| ENSG00000213621 | 7 | 3 | 3 | 2 | 2 | 2 |
| ENSG00000150457 | 1249 | 597 | 822 | 513 | 590 | 633 |
| ENSG00000233851 | 0 | 0 | 0 | 0 | 0 | 0 |
| ENSG00000201821 | 0 | 0 | 0 | 0 | 0 | 0 |
| ENSG00000180172 | 0 | 0 | 0 | 0 | 0 | 0 |
| ENSG00000225585 | 0 | 0 | 0 | 0 | 0 | 0 |
| ENSG00000275689 | 0 | 0 | 0 | 0 | 0 | 0 |
| ENSG00000252128 | 0 | 0 | 0 | 0 | 0 | 0 |
| ENSG00000150459 | 6027 | 3183 | 3574 | 3395 | 3585 | 3406 |
| ENSG00000165480 | 344 | 231 | 230 | 227 | 261 | 284 |
| ENSG00000173141 | 2933 | 1416 | 1734 | 1731 | 1620 | 1587 |
| ENSG00000271250 | 0 | 0 | 0 | 0 | 0 | 0 |
| ENSG00000233405 | 0 | 0 | 0 | 0 | 0 | 0 |
| ENSG00000235438 | 2 | 1 | 0 | 1 | 2 | 1 |
| ENSG00000277077 | 0 | 0 | 0 | 0 | 0 | 0 |
| ENSG00000233325 | 35 | 22 | 17 | 18 | 35 | 26 |
| ENSG00000224429 | 11 | 10 | 14 | 4 | 7 | 4 |
| ENSG00000215571 | 2 | 1 | 1 | 0 | 1 | 0 |
| ENSG00000235076 | 1 | 1 | 2 | 1 | 1 | 5 |
| ENSG00000222747 | 0 | 0 | 0 | 0 | 0 | 1 |
| ENSG00000180776 | 6326 | 2811 | 3060 | 1826 | 3448 | 2787 |
| ENSG00000236953 | 0 | 2 | 0 | 0 | 1 | 0 |
| ENSG00000226908 | 0 | 0 | 1 | 0 | 0 | 1 |
| ENSG00000165487 | 1853 | 914 | 1084 | 652 | 1116 | 896 |
| ENSG00000230022 | 0 | 1 | 0 | 0 | 0 | 1 |
| ENSG00000207518 | 0 | 0 | 0 | 0 | 0 | 0 |
| ENSG00000226525 | 10 | 7 | 8 | 10 | 16 | 5 |
| ENSG00000102678 | 0 | 0 | 0 | 0 | 0 | 1 |
| ENSG00000244197 | 0 | 0 | 0 | 0 | 0 | 0 |
| ENSG00000226722 | 0 | 0 | 0 | 0 | 0 | 0 |
| ENSG00000276476 | 0 | 0 | 0 | 0 | 0 | 0 |
| ENSG00000237175 | 0 | 0 | 0 | 0 | 0 | 0 |
| ENSG00000226118 | 0 | 0 | 0 | 0 | 0 | 0 |
| ENSG00000232187 | 13 | 4 | 4 | 9 | 1 | 4 |
| ENSG00000225777 | 0 | 0 | 0 | 0 | 0 | 0 |
| ENSG00000253094 | 0 | 0 | 0 | 0 | 0 | 0 |
| ENSG00000237952 | 0 | 0 | 0 | 0 | 0 | 0 |
| ENSG00000226507 | 0 | 0 | 0 | 0 | 0 | 0 |
| ENSG00000231650 | 0 | 0 | 0 | 0 | 0 | 0 |
| ENSG00000262619 | 16 | 2 | 4 | 4 | 4 | 5 |
| ENSG00000230535 | 0 | 0 | 1 | 0 | 0 | 0 |

| | | | | | | |
|---|---|---|---|---|---|---|
| ENSG00000234685 | 12 | 14 | 23 | 22 | 21 | 23 |
| ENSG00000262198 | 8 | 5 | 21 | 9 | 6 | 15 |
| ENSG00000283075 | 31 | 15 | 43 | 13 | 5 | 4 |
| ENSG00000233440 | 0 | 0 | 0 | 0 | 0 | 0 |
| ENSG00000207157 | 0 | 0 | 0 | 0 | 0 | 0 |
| ENSG00000229483 | 2 | 0 | 1 | 0 | 0 | 0 |
| ENSG00000102683 | 853 | 374 | 665 | 330 | 322 | 434 |
| ENSG00000252952 | 0 | 2 | 0 | 0 | 0 | 0 |
| ENSG00000235205 | 49 | 20 | 44 | 10 | 24 | 20 |
| ENSG00000236803 | 2 | 1 | 1 | 0 | 1 | 0 |
| ENSG00000151835 | 72 | 48 | 39 | 26 | 81 | 68 |
| ENSG00000232163 | 0 | 0 | 0 | 0 | 0 | 0 |
| ENSG00000229558 | 0 | 0 | 0 | 0 | 0 | 0 |
| ENSG00000232977 | 3 | 0 | 0 | 0 | 0 | 0 |
| ENSG00000227893 | 0 | 0 | 0 | 0 | 0 | 0 |
| ENSG00000127863 | 4 | 1 | 0 | 1 | 1 | 1 |
| ENSG00000027001 | 698 | 300 | 423 | 335 | 376 | 377 |
| ENSG00000232102 | 0 | 0 | 0 | 0 | 0 | 0 |
| ENSG00000205861 | 0 | 1 | 1 | 0 | 0 | 1 |
| ENSG00000205863 | 4 | 0 | 0 | 2 | 0 | 1 |
| ENSG00000196593 | 0 | 0 | 0 | 2 | 0 | 0 |
| ENSG00000273167 | 1 | 0 | 0 | 0 | 0 | 1 |
| ENSG00000182957 | 2671 | 1123 | 1825 | 723 | 1062 | 1245 |
| ENSG00000228741 | 0 | 3 | 3 | 1 | 0 | 3 |
| ENSG00000225674 | 2 | 1 | 0 | 0 | 1 | 0 |
| ENSG00000252695 | 0 | 0 | 0 | 0 | 0 | 1 |
| ENSG00000227213 | 0 | 0 | 3 | 0 | 0 | 0 |
| ENSG00000240654 | 0 | 0 | 1 | 0 | 0 | 0 |
| ENSG00000240868 | 0 | 0 | 0 | 0 | 0 | 0 |
| ENSG00000277040 | 0 | 0 | 0 | 0 | 0 | 0 |
| ENSG00000234627 | 0 | 0 | 0 | 0 | 0 | 0 |
| ENSG00000261498 | 0 | 0 | 0 | 0 | 0 | 0 |
| ENSG00000233545 | 0 | 0 | 0 | 0 | 0 | 0 |
| ENSG00000205822 | 0 | 0 | 1 | 0 | 0 | 0 |
| ENSG00000102699 | 2040 | 1049 | 1249 | 892 | 1711 | 1354 |
| ENSG00000244615 | 0 | 0 | 0 | 0 | 0 | 0 |
| ENSG00000243008 | 0 | 1 | 0 | 0 | 0 | 0 |
| ENSG00000285228 | 0 | 0 | 0 | 0 | 0 | 0 |
| ENSG00000075673 | 2 | 0 | 5 | 1 | 0 | 2 |
| ENSG00000207474 | 0 | 0 | 0 | 0 | 0 | 0 |
| ENSG00000225404 | 1 | 0 | 0 | 0 | 0 | 0 |
| ENSG00000285806 | 0 | 0 | 0 | 0 | 0 | 0 |
| ENSG00000231417 | 0 | 0 | 0 | 0 | 0 | 0 |
| ENSG00000226695 | 1 | 0 | 1 | 0 | 1 | 0 |
| ENSG00000132972 | 6 | 2 | 5 | 1 | 5 | 1 |
| ENSG00000151849 | 202 | 121 | 112 | 79 | 238 | 126 |
| ENSG00000253771 | 10 | 0 | 7 | 3 | 2 | 2 |
| ENSG00000273628 | 0 | 0 | 3 | 0 | 0 | 0 |
| ENSG00000251299 | 0 | 0 | 0 | 0 | 0 | 0 |
| ENSG00000232858 | 0 | 0 | 0 | 0 | 2 | 2 |

| | | | | | | |
|---|---|---|---|---|---|---|
| ENSG00000269099 | 0 | 0 | 0 | 0 | 1 | 0 |
| ENSG00000276958 | 0 | 0 | 0 | 0 | 0 | 0 |
| ENSG00000151846 | 18 | 3 | 5 | 9 | 9 | 9 |
| ENSG00000279695 | 0 | 0 | 0 | 0 | 0 | 0 |
| ENSG00000165566 | 562 | 299 | 319 | 486 | 906 | 679 |
| ENSG00000234056 | 0 | 0 | 0 | 1 | 0 | 2 |
| ENSG00000238169 | 0 | 0 | 0 | 0 | 0 | 0 |
| ENSG00000279550 | 0 | 0 | 0 | 0 | 0 | 0 |
| ENSG00000225105 | 0 | 0 | 0 | 0 | 0 | 2 |
| ENSG00000235315 | 0 | 0 | 0 | 1 | 0 | 0 |
| ENSG00000139505 | 1283 | 650 | 759 | 763 | 1423 | 1187 |
| ENSG00000260509 | 8 | 2 | 1 | 0 | 3 | 1 |
| ENSG00000139496 | 1980 | 891 | 1085 | 740 | 1482 | 1218 |
| ENSG00000234152 | 0 | 0 | 0 | 0 | 0 | 0 |
| ENSG00000132932 | 108 | 41 | 72 | 49 | 40 | 76 |
| ENSG00000199767 | 0 | 0 | 0 | 0 | 0 | 0 |
| ENSG00000180803 | 1 | 0 | 0 | 1 | 0 | 0 |
| ENSG00000242381 | 0 | 0 | 0 | 0 | 0 | 0 |
| ENSG00000223158 | 0 | 0 | 0 | 0 | 0 | 0 |
| ENSG00000180730 | 427 | 239 | 321 | 219 | 285 | 372 |
| ENSG00000231983 | 0 | 0 | 0 | 0 | 0 | 0 |
| ENSG00000233963 | 0 | 0 | 0 | 0 | 0 | 0 |
| ENSG00000234706 | 0 | 0 | 0 | 0 | 0 | 0 |
| ENSG00000127870 | 784 | 489 | 419 | 210 | 820 | 430 |
| ENSG00000277368 | 3 | 0 | 2 | 3 | 1 | 4 |
| ENSG00000271559 | 0 | 0 | 0 | 0 | 0 | 0 |
| ENSG00000227882 | 0 | 0 | 0 | 0 | 0 | 0 |
| ENSG00000132964 | 906 | 405 | 497 | 393 | 546 | 461 |
| ENSG00000278413 | 0 | 0 | 0 | 0 | 0 | 0 |
| ENSG00000285929 | 0 | 1 | 1 | 0 | 1 | 1 |
| ENSG00000132970 | 161 | 96 | 116 | 108 | 120 | 127 |
| ENSG00000234031 | 0 | 0 | 1 | 0 | 0 | 0 |
| ENSG00000237001 | 0 | 0 | 0 | 0 | 0 | 0 |
| ENSG00000285935 | 0 | 1 | 0 | 1 | 0 | 0 |
| ENSG00000132975 | 0 | 0 | 0 | 0 | 0 | 0 |
| ENSG00000230256 | 0 | 0 | 0 | 0 | 0 | 0 |
| ENSG00000283579 | 0 | 0 | 0 | 0 | 0 | 0 |
| ENSG00000223782 | 0 | 0 | 0 | 0 | 0 | 0 |
| ENSG00000218198 | 0 | 0 | 0 | 0 | 0 | 0 |
| ENSG00000152484 | 1508 | 726 | 839 | 660 | 973 | 846 |
| ENSG00000278496 | 0 | 0 | 0 | 0 | 0 | 0 |
| ENSG00000232162 | 0 | 0 | 0 | 0 | 0 | 0 |
| ENSG00000230641 | 0 | 0 | 0 | 1 | 0 | 0 |
| ENSG00000275294 | 2 | 1 | 1 | 4 | 3 | 0 |
| ENSG00000234772 | 3 | 0 | 0 | 0 | 0 | 1 |
| ENSG00000122026 | 14852 | 7650 | 9023 | 10091 | 9838 | 8960 |
| ENSG00000207500 | 0 | 0 | 0 | 0 | 0 | 0 |
| ENSG00000207051 | 0 | 0 | 0 | 0 | 0 | 0 |
| ENSG00000122035 | 22 | 16 | 19 | 13 | 11 | 16 |
| ENSG00000252247 | 0 | 0 | 0 | 0 | 0 | 0 |

| | | | | | | |
|---|---|---|---|---|---|---|
| ENSG00000229609 | 0 | 0 | 0 | 0 | 0 | 0 |
| ENSG00000201242 | 0 | 0 | 0 | 0 | 0 | 0 |
| ENSG00000122034 | 3375 | 1729 | 1899 | 2432 | 2156 | 2347 |
| ENSG00000122033 | 974 | 550 | 549 | 481 | 685 | 470 |
| ENSG00000252499 | 0 | 0 | 0 | 0 | 0 | 0 |
| ENSG00000139517 | 1155 | 561 | 722 | 355 | 640 | 608 |
| ENSG00000186184 | 3369 | 1706 | 2116 | 2139 | 2096 | 2065 |
| ENSG00000273552 | 0 | 0 | 0 | 0 | 0 | 0 |
| ENSG00000226703 | 0 | 0 | 0 | 0 | 0 | 0 |
| ENSG00000169840 | 2 | 0 | 0 | 0 | 0 | 0 |
| ENSG00000247381 | 0 | 0 | 0 | 0 | 0 | 0 |
| ENSG00000207330 | 0 | 0 | 0 | 0 | 0 | 0 |
| ENSG00000139515 | 0 | 0 | 0 | 0 | 1 | 0 |
| ENSG00000180389 | 0 | 4 | 4 | 2 | 1 | 3 |
| ENSG00000260704 | 0 | 0 | 0 | 0 | 0 | 0 |
| ENSG00000165556 | 0 | 0 | 0 | 0 | 0 | 0 |
| ENSG00000183463 | 0 | 0 | 0 | 0 | 0 | 0 |
| ENSG00000242360 | 0 | 0 | 0 | 0 | 0 | 0 |
| ENSG00000122025 | 18 | 9 | 10 | 14 | 15 | 11 |
| ENSG00000271026 | 0 | 0 | 0 | 0 | 0 | 0 |
| ENSG00000235115 | 0 | 0 | 0 | 0 | 0 | 0 |
| ENSG00000261485 | 27 | 22 | 25 | 22 | 20 | 21 |
| ENSG00000152520 | 1476 | 667 | 864 | 421 | 839 | 663 |
| ENSG00000232587 | 0 | 0 | 0 | 1 | 0 | 0 |
| ENSG00000200840 | 0 | 0 | 0 | 0 | 0 | 0 |
| ENSG00000102755 | 10 | 5 | 4 | 4 | 3 | 1 |
| ENSG00000279393 | 0 | 0 | 0 | 0 | 0 | 0 |
| ENSG00000235472 | 0 | 1 | 3 | 1 | 0 | 0 |
| ENSG00000273964 | 0 | 0 | 0 | 0 | 0 | 0 |
| ENSG00000132963 | 2604 | 1359 | 1597 | 1529 | 1862 | 1630 |
| ENSG00000139508 | 780 | 363 | 503 | 314 | 380 | 371 |
| ENSG00000202237 | 0 | 0 | 0 | 0 | 0 | 0 |
| ENSG00000233588 | 4 | 2 | 2 | 0 | 2 | 2 |
| ENSG00000277227 | 0 | 0 | 0 | 0 | 0 | 0 |
| ENSG00000236339 | 0 | 0 | 0 | 0 | 0 | 0 |
| ENSG00000132938 | 2 | 0 | 2 | 0 | 0 | 2 |
| ENSG00000236758 | 0 | 0 | 0 | 0 | 0 | 0 |
| ENSG00000223460 | 0 | 0 | 0 | 0 | 1 | 0 |
| ENSG00000270680 | 0 | 0 | 0 | 0 | 0 | 0 |
| ENSG00000179141 | 0 | 1 | 0 | 0 | 0 | 1 |
| ENSG00000139514 | 2416 | 1274 | 1494 | 1314 | 1510 | 1871 |
| ENSG00000122042 | 2725 | 1249 | 1694 | 979 | 1328 | 1298 |
| ENSG00000278172 | 0 | 0 | 0 | 0 | 0 | 0 |
| ENSG00000236258 | 0 | 0 | 0 | 0 | 0 | 0 |
| ENSG00000224329 | 0 | 0 | 0 | 0 | 0 | 0 |
| ENSG00000224405 | 0 | 1 | 0 | 1 | 1 | 2 |
| ENSG00000122043 | 0 | 0 | 0 | 0 | 0 | 0 |
| ENSG00000266505 | 2 | 1 | 0 | 0 | 2 | 1 |
| ENSG00000224511 | 37 | 29 | 38 | 11 | 16 | 11 |
| ENSG00000232117 | 0 | 0 | 0 | 0 | 0 | 0 |

| | | | | | | |
|---|---|---|---|---|---|---|
| ENSG00000232643 | 0 | 0 | 0 | 0 | 0 | 0 |
| ENSG00000279499 | 0 | 0 | 0 | 0 | 0 | 0 |
| ENSG00000102781 | 184 | 112 | 95 | 76 | 123 | 120 |
| ENSG00000252928 | 0 | 0 | 0 | 0 | 0 | 0 |
| ENSG00000215049 | 1 | 0 | 0 | 0 | 0 | 0 |
| ENSG00000236463 | 2 | 0 | 0 | 0 | 0 | 0 |
| ENSG00000238121 | 0 | 1 | 1 | 0 | 0 | 0 |
| ENSG00000279149 | 4 | 0 | 1 | 2 | 0 | 0 |
| ENSG00000285773 | 0 | 0 | 0 | 0 | 0 | 0 |
| ENSG00000225039 | 0 | 1 | 1 | 0 | 1 | 0 |
| ENSG00000236444 | 0 | 1 | 3 | 1 | 2 | 0 |
| ENSG00000189403 | 11202 | 5878 | 6637 | 5662 | 7240 | 6773 |
| ENSG00000285840 | 0 | 0 | 0 | 0 | 0 | 0 |
| ENSG00000213411 | 0 | 1 | 0 | 0 | 1 | 1 |
| ENSG00000232489 | 0 | 0 | 0 | 0 | 0 | 1 |
| ENSG00000224117 | 0 | 0 | 0 | 0 | 0 | 0 |
| ENSG00000132952 | 715 | 302 | 323 | 167 | 468 | 339 |
| ENSG00000275788 | 0 | 0 | 0 | 0 | 0 | 0 |
| ENSG00000132965 | 0 | 0 | 0 | 0 | 1 | 2 |
| ENSG00000237879 | 0 | 0 | 0 | 0 | 0 | 0 |
| ENSG00000224743 | 0 | 0 | 0 | 0 | 0 | 0 |
| ENSG00000236094 | 0 | 0 | 0 | 0 | 0 | 0 |
| ENSG00000102802 | 0 | 0 | 0 | 0 | 0 | 0 |
| ENSG00000175664 | 0 | 0 | 0 | 0 | 0 | 0 |
| ENSG00000238185 | 0 | 0 | 0 | 0 | 0 | 0 |
| ENSG00000230403 | 0 | 0 | 0 | 0 | 0 | 0 |
| ENSG00000231894 | 0 | 0 | 0 | 0 | 0 | 0 |
| ENSG00000120694 | 2586 | 1608 | 1126 | 832 | 2761 | 1432 |
| ENSG00000187676 | 344 | 163 | 191 | 133 | 178 | 175 |
| ENSG00000229427 | 0 | 0 | 0 | 3 | 0 | 0 |
| ENSG00000278305 | 0 | 0 | 0 | 0 | 0 | 0 |
| ENSG00000277319 | 0 | 0 | 0 | 0 | 0 | 0 |
| ENSG00000133105 | 0 | 0 | 0 | 0 | 0 | 0 |
| ENSG00000073910 | 74 | 31 | 37 | 20 | 53 | 31 |
| ENSG00000229715 | 0 | 0 | 0 | 0 | 0 | 0 |
| ENSG00000279314 | 0 | 0 | 0 | 0 | 0 | 2 |
| ENSG00000278732 | 0 | 0 | 0 | 0 | 0 | 0 |
| ENSG00000237637 | 0 | 0 | 1 | 0 | 1 | 0 |
| ENSG00000189167 | 0 | 1 | 3 | 1 | 2 | 3 |
| ENSG00000139618 | 82 | 31 | 26 | 13 | 66 | 66 |
| ENSG00000215515 | 0 | 0 | 0 | 0 | 0 | 1 |
| ENSG00000139597 | 89 | 73 | 68 | 41 | 40 | 51 |
| ENSG00000212293 | 0 | 0 | 0 | 0 | 0 | 0 |
| ENSG00000270008 | 0 | 0 | 0 | 0 | 0 | 0 |
| ENSG00000244754 | 1797 | 859 | 1009 | 528 | 1150 | 871 |
| ENSG00000229800 | 0 | 3 | 1 | 1 | 1 | 2 |
| ENSG00000281026 | 20 | 7 | 5 | 7 | 15 | 9 |
| ENSG00000083642 | 845 | 510 | 378 | 156 | 702 | 345 |
| ENSG00000207325 | 0 | 0 | 1 | 0 | 0 | 0 |
| ENSG00000277151 | 0 | 2 | 1 | 0 | 1 | 2 |

| | | | | | | |
|---|---|---|---|---|---|---|
| ENSG00000226968 | 0 | 0 | 0 | 0 | 0 | 0 |
| ENSG00000227604 | 0 | 0 | 0 | 0 | 0 | 0 |
| ENSG00000133116 | 0 | 0 | 0 | 0 | 0 | 2 |
| ENSG00000133121 | 789 | 418 | 456 | 321 | 400 | 459 |
| ENSG00000230300 | 0 | 1 | 0 | 0 | 1 | 0 |
| ENSG00000236581 | 1 | 2 | 1 | 1 | 1 | 0 |
| ENSG00000227911 | 2 | 0 | 0 | 0 | 1 | 1 |
| ENSG00000285621 | 0 | 2 | 2 | 0 | 1 | 1 |
| ENSG00000277386 | 3 | 8 | 3 | 4 | 3 | 1 |
| ENSG00000230490 | 157 | 75 | 90 | 67 | 66 | 78 |
| ENSG00000234535 | 100 | 53 | 63 | 48 | 49 | 61 |
| ENSG00000232986 | 1 | 6 | 1 | 1 | 2 | 2 |
| ENSG00000133119 | 862 | 461 | 568 | 417 | 544 | 518 |
| ENSG00000252397 | 0 | 0 | 0 | 0 | 0 | 0 |
| ENSG00000276672 | 37 | 17 | 26 | 12 | 16 | 29 |
| ENSG00000280431 | 0 | 0 | 0 | 0 | 0 | 0 |
| ENSG00000227254 | 0 | 0 | 0 | 0 | 0 | 0 |
| ENSG00000199196 | 0 | 0 | 0 | 0 | 0 | 0 |
| ENSG00000278477 | 0 | 0 | 0 | 0 | 0 | 0 |
| ENSG00000271850 | 6 | 7 | 8 | 3 | 5 | 10 |
| ENSG00000225179 | 0 | 1 | 0 | 0 | 0 | 0 |
| ENSG00000271901 | 0 | 0 | 1 | 0 | 1 | 0 |
| ENSG00000231910 | 0 | 0 | 0 | 0 | 0 | 0 |
| ENSG00000273886 | 0 | 0 | 0 | 0 | 0 | 0 |
| ENSG00000261728 | 2 | 0 | 1 | 0 | 1 | 0 |
| ENSG00000172915 | 623 | 340 | 289 | 76 | 463 | 240 |
| ENSG00000232795 | 0 | 0 | 0 | 0 | 0 | 0 |
| ENSG00000234835 | 1 | 2 | 1 | 0 | 0 | 0 |
| ENSG00000180660 | 0 | 0 | 0 | 0 | 0 | 0 |
| ENSG00000236036 | 0 | 0 | 0 | 0 | 0 | 0 |
| ENSG00000133083 | 389 | 205 | 342 | 129 | 190 | 248 |
| ENSG00000120669 | 2 | 1 | 2 | 0 | 2 | 0 |
| ENSG00000250709 | 0 | 1 | 0 | 0 | 0 | 0 |
| ENSG00000242715 | 4 | 1 | 4 | 4 | 10 | 6 |
| ENSG00000207203 | 0 | 0 | 0 | 0 | 0 | 0 |
| ENSG00000133104 | 519 | 262 | 368 | 219 | 390 | 334 |
| ENSG00000120664 | 1 | 1 | 0 | 1 | 2 | 2 |
| ENSG00000133101 | 0 | 1 | 1 | 1 | 2 | 0 |
| ENSG00000223383 | 0 | 0 | 0 | 0 | 0 | 0 |
| ENSG00000180440 | 0 | 0 | 0 | 0 | 0 | 0 |
| ENSG00000235306 | 0 | 0 | 0 | 0 | 0 | 0 |
| ENSG00000223543 | 0 | 0 | 0 | 0 | 0 | 0 |
| ENSG00000215486 | 0 | 0 | 0 | 0 | 0 | 0 |
| ENSG00000271337 | 0 | 0 | 0 | 0 | 0 | 0 |
| ENSG00000133111 | 129 | 94 | 109 | 76 | 83 | 95 |
| ENSG00000120693 | 20 | 10 | 6 | 14 | 13 | 5 |
| ENSG00000236711 | 1 | 0 | 0 | 0 | 0 | 0 |
| ENSG00000228825 | 0 | 0 | 0 | 0 | 0 | 0 |
| ENSG00000232378 | 0 | 0 | 0 | 0 | 0 | 0 |
| ENSG00000226161 | 0 | 0 | 0 | 0 | 1 | 0 |

| | | | | | | |
|---|---|---|---|---|---|---|
| ENSG00000120697 | 876 | 477 | 544 | 488 | 563 | 551 |
| ENSG00000120699 | 694 | 337 | 507 | 304 | 428 | 379 |
| ENSG00000102710 | 1646 | 778 | 916 | 530 | 1000 | 811 |
| ENSG00000180138 | 0 | 0 | 0 | 0 | 0 | 0 |
| ENSG00000274549 | 0 | 0 | 0 | 0 | 0 | 0 |
| ENSG00000279511 | 0 | 0 | 0 | 0 | 0 | 0 |
| ENSG00000202395 | 0 | 0 | 0 | 0 | 0 | 0 |
| ENSG00000230283 | 0 | 0 | 0 | 0 | 0 | 0 |
| ENSG00000230390 | 0 | 0 | 0 | 0 | 0 | 0 |
| ENSG00000275226 | 2 | 0 | 0 | 0 | 0 | 0 |
| ENSG00000133110 | 0 | 0 | 0 | 0 | 0 | 1 |
| ENSG00000133107 | 0 | 0 | 0 | 0 | 0 | 0 |
| ENSG00000199276 | 0 | 0 | 0 | 0 | 0 | 0 |
| ENSG00000274317 | 0 | 1 | 2 | 0 | 2 | 1 |
| ENSG00000228036 | 0 | 0 | 0 | 0 | 0 | 0 |
| ENSG00000223685 | 1 | 4 | 1 | 1 | 2 | 0 |
| ENSG00000120686 | 1848 | 870 | 965 | 664 | 1169 | 877 |
| ENSG00000279025 | 0 | 0 | 0 | 0 | 0 | 0 |
| ENSG00000236354 | 0 | 0 | 3 | 0 | 1 | 0 |
| ENSG00000229437 | 0 | 0 | 0 | 0 | 0 | 0 |
| ENSG00000234701 | 0 | 0 | 0 | 0 | 1 | 0 |
| ENSG00000150893 | 1318 | 717 | 660 | 287 | 1207 | 769 |
| ENSG00000252795 | 0 | 0 | 0 | 0 | 0 | 1 |
| ENSG00000225350 | 0 | 0 | 0 | 0 | 0 | 0 |
| ENSG00000225221 | 0 | 0 | 0 | 0 | 0 | 0 |
| ENSG00000235550 | 0 | 0 | 0 | 0 | 0 | 0 |
| ENSG00000133115 | 0 | 0 | 0 | 0 | 0 | 0 |
| ENSG00000120685 | 2125 | 1016 | 1070 | 746 | 1211 | 1105 |
| ENSG00000188811 | 945 | 435 | 596 | 363 | 457 | 495 |
| ENSG00000273507 | 2 | 0 | 0 | 1 | 0 | 0 |
| ENSG00000214823 | 0 | 0 | 0 | 0 | 0 | 0 |
| ENSG00000183722 | 0 | 0 | 0 | 0 | 0 | 0 |
| ENSG00000133103 | 1286 | 597 | 705 | 368 | 801 | 623 |
| ENSG00000264171 | 0 | 0 | 0 | 0 | 0 | 2 |
| ENSG00000200526 | 0 | 0 | 0 | 0 | 0 | 0 |
| ENSG00000224351 | 0 | 0 | 0 | 0 | 0 | 0 |
| ENSG00000270444 | 0 | 0 | 0 | 0 | 0 | 0 |
| ENSG00000212553 | 0 | 0 | 0 | 0 | 0 | 0 |
| ENSG00000215483 | 0 | 0 | 1 | 0 | 0 | 4 |
| ENSG00000230710 | 0 | 0 | 0 | 0 | 0 | 0 |
| ENSG00000271429 | 0 | 0 | 0 | 0 | 0 | 0 |
| ENSG00000207458 | 0 | 0 | 0 | 0 | 0 | 0 |
| ENSG00000252812 | 0 | 0 | 0 | 0 | 0 | 0 |
| ENSG00000150907 | 224 | 134 | 157 | 95 | 154 | 133 |
| ENSG00000229456 | 0 | 0 | 0 | 0 | 0 | 0 |
| ENSG00000211491 | 0 | 0 | 0 | 0 | 0 | 0 |
| ENSG00000102738 | 530 | 230 | 296 | 207 | 307 | 299 |
| ENSG00000277794 | 0 | 0 | 0 | 0 | 1 | 0 |
| ENSG00000102743 | 617 | 309 | 367 | 260 | 320 | 426 |
| ENSG00000207652 | 89 | 61 | 74 | 83 | 66 | 83 |

| | | | | | | |
|---|---|---|---|---|---|---|
| ENSG00000168852 | 9 | 6 | 9 | 6 | 5 | 11 |
| ENSG00000176268 | 4 | 4 | 2 | 9 | 5 | 4 |
| ENSG00000239827 | 20 | 17 | 20 | 11 | 12 | 19 |
| ENSG00000120690 | 3198 | 1440 | 1873 | 1092 | 1609 | 1538 |
| ENSG00000229473 | 2 | 0 | 2 | 0 | 4 | 0 |
| ENSG00000120688 | 372 | 237 | 224 | 144 | 308 | 209 |
| ENSG00000264226 | 0 | 0 | 0 | 0 | 0 | 0 |
| ENSG00000244264 | 0 | 0 | 0 | 0 | 0 | 0 |
| ENSG00000165572 | 538 | 209 | 292 | 168 | 239 | 229 |
| ENSG00000278390 | 13 | 7 | 10 | 12 | 15 | 11 |
| ENSG00000215482 | 0 | 0 | 0 | 0 | 0 | 0 |
| ENSG00000120696 | 783 | 327 | 440 | 289 | 352 | 380 |
| ENSG00000225133 | 0 | 0 | 0 | 0 | 0 | 0 |
| ENSG00000120662 | 259 | 121 | 118 | 110 | 159 | 165 |
| ENSG00000277662 | 8 | 3 | 2 | 2 | 8 | 4 |
| ENSG00000231404 | 0 | 0 | 0 | 0 | 0 | 0 |
| ENSG00000172766 | 360 | 195 | 205 | 89 | 270 | 186 |
| ENSG00000223280 | 0 | 0 | 0 | 0 | 0 | 0 |
| ENSG00000214222 | 0 | 1 | 0 | 0 | 0 | 0 |
| ENSG00000205240 | 0 | 0 | 0 | 0 | 0 | 0 |
| ENSG00000234655 | 0 | 0 | 0 | 0 | 0 | 0 |
| ENSG00000215480 | 0 | 0 | 0 | 0 | 0 | 0 |
| ENSG00000102760 | 1 | 2 | 3 | 1 | 2 | 4 |
| ENSG00000102763 | 553 | 353 | 346 | 201 | 393 | 326 |
| ENSG00000284584 | 0 | 0 | 0 | 0 | 0 | 2 |
| ENSG00000241406 | 0 | 0 | 0 | 0 | 0 | 0 |
| ENSG00000206962 | 0 | 0 | 0 | 0 | 0 | 0 |
| ENSG00000229696 | 0 | 1 | 0 | 0 | 0 | 0 |
| ENSG00000278338 | 2 | 1 | 3 | 6 | 1 | 3 |
| ENSG00000226093 | 0 | 0 | 0 | 0 | 0 | 0 |
| ENSG00000102780 | 291 | 169 | 141 | 60 | 221 | 146 |
| ENSG00000231530 | 13 | 10 | 11 | 10 | 8 | 7 |
| ENSG00000237263 | 0 | 0 | 1 | 0 | 1 | 0 |
| ENSG00000270809 | 0 | 0 | 0 | 0 | 0 | 0 |
| ENSG00000234213 | 0 | 0 | 0 | 0 | 0 | 0 |
| ENSG00000023516 | 750 | 428 | 317 | 83 | 730 | 425 |
| ENSG00000283554 | 0 | 0 | 0 | 0 | 0 | 0 |
| ENSG00000233259 | 0 | 0 | 0 | 0 | 0 | 0 |
| ENSG00000120659 | 0 | 0 | 2 | 3 | 0 | 0 |
| ENSG00000179813 | 0 | 0 | 0 | 0 | 0 | 0 |
| ENSG00000271216 | 0 | 0 | 0 | 0 | 0 | 0 |
| ENSG00000229546 | 0 | 0 | 0 | 0 | 0 | 0 |
| ENSG00000133106 | 10 | 3 | 1 | 2 | 2 | 1 |
| ENSG00000238025 | 0 | 0 | 0 | 0 | 0 | 0 |
| ENSG00000120675 | 1623 | 861 | 886 | 727 | 827 | 724 |
| ENSG00000229928 | 2 | 0 | 0 | 0 | 0 | 0 |
| ENSG00000276569 | 0 | 0 | 0 | 0 | 0 | 0 |
| ENSG00000120658 | 10 | 9 | 8 | 5 | 6 | 3 |
| ENSG00000232540 | 0 | 0 | 0 | 0 | 0 | 0 |
| ENSG00000238189 | 0 | 0 | 0 | 0 | 0 | 0 |

| | | | | | | |
|---|---|---|---|---|---|---|
| ENSG00000233821 | 0 | 0 | 0 | 0 | 0 | 0 |
| ENSG00000270522 | 0 | 0 | 0 | 0 | 0 | 0 |
| ENSG00000151773 | 228 | 110 | 128 | 109 | 167 | 119 |
| ENSG00000274001 | 1 | 1 | 2 | 4 | 4 | 5 |
| ENSG00000179630 | 159 | 105 | 73 | 39 | 170 | 94 |
| ENSG00000281883 | 3 | 1 | 0 | 1 | 2 | 1 |
| ENSG00000280060 | 0 | 0 | 0 | 0 | 0 | 0 |
| ENSG00000233725 | 8 | 12 | 11 | 5 | 14 | 8 |
| ENSG00000179611 | 34 | 21 | 23 | 25 | 12 | 17 |
| ENSG00000226519 | 0 | 0 | 0 | 1 | 0 | 0 |
| ENSG00000227258 | 10 | 8 | 9 | 7 | 4 | 6 |
| ENSG00000139656 | 0 | 2 | 1 | 1 | 4 | 2 |
| ENSG00000235285 | 0 | 1 | 0 | 0 | 0 | 0 |
| ENSG00000276854 | 0 | 0 | 0 | 0 | 0 | 0 |
| ENSG00000276319 | 0 | 0 | 0 | 0 | 0 | 0 |
| ENSG00000230731 | 0 | 0 | 0 | 1 | 0 | 0 |
| ENSG00000277831 | 0 | 0 | 0 | 0 | 0 | 0 |
| ENSG00000151778 | 0 | 0 | 0 | 0 | 0 | 0 |
| ENSG00000237361 | 7 | 0 | 0 | 0 | 2 | 1 |
| ENSG00000102804 | 4520 | 2020 | 2624 | 1964 | 2488 | 2471 |
| ENSG00000280357 | 0 | 0 | 0 | 0 | 0 | 0 |
| ENSG00000225727 | 0 | 0 | 0 | 0 | 0 | 2 |
| ENSG00000278156 | 23 | 10 | 27 | 9 | 15 | 11 |
| ENSG00000230793 | 0 | 0 | 0 | 1 | 1 | 2 |
| ENSG00000237585 | 0 | 0 | 0 | 0 | 0 | 1 |
| ENSG00000276527 | 0 | 0 | 0 | 0 | 0 | 0 |
| ENSG00000235097 | 0 | 0 | 1 | 0 | 0 | 0 |
| ENSG00000083635 | 331 | 128 | 174 | 182 | 182 | 221 |
| ENSG00000223732 | 0 | 0 | 0 | 0 | 0 | 0 |
| ENSG00000133114 | 910 | 478 | 548 | 423 | 508 | 502 |
| ENSG00000265802 | 3 | 2 | 3 | 1 | 1 | 0 |
| ENSG00000223341 | 0 | 0 | 0 | 0 | 0 | 0 |
| ENSG00000188342 | 877 | 496 | 468 | 549 | 700 | 569 |
| ENSG00000271818 | 0 | 0 | 0 | 0 | 0 | 0 |
| ENSG00000277358 | 0 | 0 | 0 | 0 | 0 | 0 |
| ENSG00000180332 | 0 | 0 | 0 | 0 | 0 | 0 |
| ENSG00000270381 | 0 | 0 | 0 | 0 | 0 | 0 |
| ENSG00000133112 | 49135 | 26891 | 34574 | 34328 | 31381 | 31239 |
| ENSG00000253051 | 0 | 0 | 0 | 0 | 0 | 0 |
| ENSG00000199477 | 6 | 1 | 0 | 4 | 7 | 2 |
| ENSG00000273149 | 0 | 0 | 0 | 0 | 0 | 0 |
| ENSG00000170919 | 115 | 72 | 101 | 76 | 71 | 89 |
| ENSG00000228886 | 0 | 1 | 0 | 0 | 0 | 0 |
| ENSG00000214455 | 3 | 3 | 5 | 2 | 9 | 3 |
| ENSG00000174032 | 983 | 470 | 690 | 400 | 480 | 516 |
| ENSG00000251015 | 4 | 1 | 2 | 2 | 2 | 1 |
| ENSG00000229303 | 0 | 0 | 0 | 0 | 0 | 0 |
| ENSG00000136152 | 1627 | 837 | 1018 | 549 | 859 | 845 |
| ENSG00000165837 | 7 | 5 | 10 | 3 | 4 | 5 |
| ENSG00000200709 | 0 | 0 | 0 | 0 | 0 | 0 |

| | | | | | | |
|---|---|---|---|---|---|---|
| ENSG00000232786 | 0 | 0 | 0 | 0 | 0 | 0 |
| ENSG00000228869 | 4 | 1 | 3 | 1 | 2 | 0 |
| ENSG00000235366 | 0 | 0 | 0 | 0 | 1 | 1 |
| ENSG00000237876 | 0 | 0 | 0 | 0 | 0 | 0 |
| ENSG00000174015 | 23 | 12 | 12 | 14 | 9 | 11 |
| ENSG00000274357 | 0 | 0 | 0 | 0 | 0 | 0 |
| ENSG00000215475 | 2 | 0 | 0 | 0 | 0 | 0 |
| ENSG00000123200 | 541 | 329 | 213 | 102 | 658 | 311 |
| ENSG00000235903 | 0 | 1 | 3 | 2 | 3 | 2 |
| ENSG00000080618 | 0 | 0 | 0 | 0 | 0 | 0 |
| ENSG00000136167 | 148 | 82 | 114 | 81 | 63 | 94 |
| ENSG00000240767 | 0 | 0 | 0 | 0 | 0 | 0 |
| ENSG00000173988 | 0 | 0 | 0 | 0 | 0 | 0 |
| ENSG00000252854 | 0 | 0 | 0 | 0 | 0 | 0 |
| ENSG00000251106 | 0 | 0 | 0 | 0 | 0 | 0 |
| ENSG00000261097 | 0 | 0 | 0 | 0 | 0 | 0 |
| ENSG00000102445 | 29 | 9 | 16 | 26 | 20 | 17 |
| ENSG00000223336 | 0 | 0 | 0 | 0 | 0 | 0 |
| ENSG00000252385 | 0 | 0 | 0 | 0 | 0 | 0 |
| ENSG00000241353 | 0 | 0 | 0 | 0 | 0 | 0 |
| ENSG00000231817 | 0 | 0 | 0 | 0 | 0 | 0 |
| ENSG00000285576 | 0 | 0 | 0 | 0 | 0 | 0 |
| ENSG00000227019 | 0 | 0 | 0 | 0 | 0 | 0 |
| ENSG00000277228 | 0 | 0 | 0 | 0 | 0 | 0 |
| ENSG00000205105 | 0 | 0 | 1 | 1 | 1 | 1 |
| ENSG00000230243 | 0 | 0 | 0 | 0 | 0 | 0 |
| ENSG00000136141 | 1378 | 643 | 710 | 451 | 660 | 620 |
| ENSG00000278462 | 0 | 0 | 0 | 0 | 0 | 0 |
| ENSG00000139684 | 2269 | 1309 | 1423 | 1205 | 1552 | 1409 |
| ENSG00000102468 | 0 | 0 | 0 | 0 | 0 | 0 |
| ENSG00000224517 | 0 | 0 | 0 | 0 | 0 | 0 |
| ENSG00000234590 | 0 | 0 | 0 | 0 | 0 | 0 |
| ENSG00000244521 | 0 | 0 | 0 | 0 | 0 | 0 |
| ENSG00000136143 | 848 | 456 | 462 | 281 | 682 | 475 |
| ENSG00000234145 | 0 | 1 | 0 | 1 | 0 | 1 |
| ENSG00000229423 | 0 | 0 | 0 | 0 | 0 | 0 |
| ENSG00000228573 | 0 | 0 | 0 | 0 | 0 | 0 |
| ENSG00000260388 | 4 | 0 | 1 | 1 | 1 | 3 |
| ENSG00000227848 | 15 | 10 | 19 | 15 | 11 | 14 |
| ENSG00000136159 | 957 | 462 | 587 | 354 | 501 | 499 |
| ENSG00000276968 | 2 | 1 | 5 | 1 | 1 | 1 |
| ENSG00000136146 | 732 | 382 | 525 | 375 | 443 | 412 |
| ENSG00000229111 | 0 | 0 | 0 | 2 | 0 | 0 |
| ENSG00000224510 | 0 | 0 | 0 | 0 | 0 | 0 |
| ENSG00000136156 | 6482 | 3168 | 3830 | 2905 | 3331 | 3142 |
| ENSG00000231473 | 2 | 1 | 1 | 1 | 2 | 0 |
| ENSG00000139687 | 1496 | 810 | 753 | 440 | 874 | 706 |
| ENSG00000238086 | 0 | 0 | 0 | 0 | 0 | 0 |
| ENSG00000177197 | 0 | 1 | 0 | 1 | 0 | 0 |
| ENSG00000277684 | 0 | 0 | 0 | 0 | 0 | 0 |

| | | | | | | |
|---|---|---|---|---|---|---|
| ENSG00000139679 | 28 | 15 | 19 | 20 | 14 | 12 |
| ENSG00000277233 | 0 | 0 | 0 | 0 | 0 | 0 |
| ENSG00000136161 | 46 | 31 | 23 | 23 | 20 | 9 |
| ENSG00000274929 | 8 | 3 | 1 | 1 | 3 | 6 |
| ENSG00000233456 | 0 | 0 | 0 | 0 | 0 | 0 |
| ENSG00000233610 | 0 | 0 | 0 | 0 | 0 | 0 |
| ENSG00000152207 | 0 | 0 | 0 | 0 | 0 | 0 |
| ENSG00000225131 | 28 | 13 | 15 | 5 | 16 | 11 |
| ENSG00000275202 | 45 | 21 | 31 | 21 | 19 | 19 |
| ENSG00000102531 | 1034 | 509 | 530 | 221 | 761 | 478 |
| ENSG00000201662 | 0 | 0 | 0 | 0 | 0 | 0 |
| ENSG00000223724 | 0 | 0 | 0 | 0 | 0 | 0 |
| ENSG00000199788 | 0 | 0 | 0 | 0 | 0 | 0 |
| ENSG00000235957 | 2 | 2 | 1 | 2 | 2 | 0 |
| ENSG00000231665 | 0 | 1 | 0 | 1 | 0 | 0 |
| ENSG00000102539 | 0 | 0 | 0 | 2 | 1 | 1 |
| ENSG00000274351 | 0 | 0 | 0 | 0 | 0 | 0 |
| ENSG00000102543 | 174 | 94 | 94 | 90 | 93 | 107 |
| ENSG00000102547 | 887 | 430 | 482 | 321 | 550 | 474 |
| ENSG00000136169 | 500 | 227 | 272 | 122 | 245 | 200 |
| ENSG00000236577 | 2 | 1 | 1 | 0 | 7 | 0 |
| ENSG00000136147 | 661 | 325 | 386 | 338 | 404 | 394 |
| ENSG00000136144 | 1324 | 581 | 726 | 495 | 666 | 630 |
| ENSG00000176882 | 0 | 0 | 0 | 0 | 0 | 0 |
| ENSG00000152213 | 38 | 20 | 33 | 20 | 43 | 32 |
| ENSG00000123179 | 919 | 510 | 532 | 689 | 621 | 651 |
| ENSG00000102753 | 1261 | 730 | 713 | 523 | 955 | 780 |
| ENSG00000276436 | 5 | 0 | 0 | 0 | 0 | 0 |
| ENSG00000222148 | 0 | 0 | 0 | 0 | 0 | 0 |
| ENSG00000181358 | 0 | 0 | 0 | 1 | 1 | 0 |
| ENSG00000200064 | 0 | 0 | 0 | 0 | 0 | 0 |
| ENSG00000123178 | 467 | 234 | 310 | 254 | 286 | 289 |
| ENSG00000231607 | 42 | 17 | 30 | 23 | 27 | 29 |
| ENSG00000204977 | 488 | 250 | 317 | 212 | 296 | 279 |
| ENSG00000264864 | 2 | 0 | 0 | 0 | 1 | 0 |
| ENSG00000198553 | 3 | 4 | 2 | 3 | 3 | 2 |
| ENSG00000277913 | 0 | 0 | 0 | 0 | 0 | 0 |
| ENSG00000277066 | 0 | 0 | 0 | 0 | 0 | 0 |
| ENSG00000208006 | 0 | 0 | 0 | 0 | 0 | 0 |
| ENSG00000283785 | 0 | 0 | 0 | 0 | 0 | 0 |
| ENSG00000274878 | 0 | 0 | 0 | 0 | 0 | 0 |
| ENSG00000273612 | 0 | 0 | 0 | 0 | 0 | 0 |
| ENSG00000277752 | 0 | 0 | 0 | 0 | 0 | 0 |
| ENSG00000275559 | 0 | 0 | 0 | 0 | 0 | 0 |
| ENSG00000176124 | 81 | 62 | 74 | 54 | 74 | 75 |
| ENSG00000276089 | 0 | 0 | 0 | 0 | 0 | 0 |
| ENSG00000214359 | 2 | 0 | 1 | 0 | 1 | 0 |
| ENSG00000273541 | 0 | 0 | 0 | 0 | 0 | 0 |
| ENSG00000274270 | 36 | 12 | 16 | 14 | 17 | 21 |
| ENSG00000232150 | 0 | 0 | 3 | 2 | 3 | 1 |

| | | | | | | |
|---|---|---|---|---|---|---|
| ENSG00000223676 | 0 | 0 | 0 | 0 | 0 | 0 |
| ENSG00000229323 | 0 | 0 | 1 | 0 | 0 | 0 |
| ENSG00000186047 | 33 | 17 | 18 | 8 | 14 | 18 |
| ENSG00000237152 | 11 | 5 | 11 | 5 | 11 | 3 |
| ENSG00000200711 | 0 | 0 | 0 | 0 | 0 | 0 |
| ENSG00000233672 | 10 | 2 | 4 | 3 | 3 | 6 |
| ENSG00000136104 | 566 | 254 | 350 | 258 | 343 | 324 |
| ENSG00000123201 | 175 | 96 | 119 | 48 | 169 | 108 |
| ENSG00000222920 | 0 | 0 | 2 | 0 | 1 | 1 |
| ENSG00000226792 | 0 | 0 | 0 | 0 | 0 | 0 |
| ENSG00000214878 | 0 | 0 | 0 | 0 | 0 | 0 |
| ENSG00000274652 | 0 | 0 | 0 | 0 | 0 | 0 |
| ENSG00000271242 | 0 | 0 | 0 | 0 | 0 | 0 |
| ENSG00000150510 | 6 | 14 | 13 | 10 | 16 | 12 |
| ENSG00000253309 | 0 | 1 | 0 | 2 | 1 | 2 |
| ENSG00000266072 | 0 | 0 | 0 | 0 | 0 | 0 |
| ENSG00000102786 | 1435 | 694 | 745 | 491 | 815 | 723 |
| ENSG00000236778 | 39 | 18 | 26 | 13 | 22 | 19 |
| ENSG00000224892 | 8 | 3 | 10 | 5 | 5 | 8 |
| ENSG00000243900 | 0 | 0 | 0 | 0 | 0 | 0 |
| ENSG00000229906 | 0 | 0 | 0 | 0 | 0 | 0 |
| ENSG00000266611 | 0 | 0 | 0 | 0 | 0 | 0 |
| ENSG00000252141 | 0 | 0 | 0 | 0 | 0 | 0 |
| ENSG00000139668 | 377 | 159 | 228 | 189 | 204 | 198 |
| ENSG00000206920 | 0 | 0 | 0 | 0 | 0 | 0 |
| ENSG00000242893 | 0 | 0 | 0 | 0 | 0 | 0 |
| ENSG00000224451 | 0 | 0 | 0 | 0 | 0 | 0 |
| ENSG00000285444 | 0 | 0 | 0 | 0 | 0 | 0 |
| ENSG00000102796 | 144 | 104 | 113 | 120 | 121 | 97 |
| ENSG00000231856 | 88 | 30 | 60 | 40 | 55 | 39 |
| ENSG00000281106 | 2 | 0 | 0 | 0 | 0 | 1 |
| ENSG00000123171 | 0 | 0 | 0 | 0 | 0 | 0 |
| ENSG00000232872 | 2 | 1 | 1 | 2 | 4 | 2 |
| ENSG00000279677 | 0 | 1 | 0 | 0 | 0 | 0 |
| ENSG00000123191 | 905 | 378 | 521 | 293 | 371 | 381 |
| ENSG00000236044 | 0 | 0 | 1 | 0 | 0 | 0 |
| ENSG00000253710 | 298 | 131 | 152 | 108 | 160 | 124 |
| ENSG00000253797 | 235 | 141 | 135 | 60 | 154 | 94 |
| ENSG00000197168 | 42 | 22 | 21 | 8 | 14 | 18 |
| ENSG00000273523 | 2 | 0 | 4 | 1 | 0 | 1 |
| ENSG00000136098 | 236 | 111 | 141 | 80 | 123 | 95 |
| ENSG00000217576 | 0 | 0 | 0 | 0 | 0 | 0 |
| ENSG00000243406 | 14 | 9 | 11 | 2 | 15 | 11 |
| ENSG00000285160 | 0 | 0 | 0 | 0 | 0 | 0 |
| ENSG00000285323 | 0 | 0 | 0 | 0 | 0 | 0 |
| ENSG00000272281 | 23 | 4 | 9 | 5 | 3 | 5 |
| ENSG00000271564 | 0 | 0 | 0 | 0 | 0 | 0 |
| ENSG00000136114 | 7 | 6 | 6 | 3 | 6 | 4 |
| ENSG00000199605 | 0 | 0 | 0 | 0 | 0 | 0 |
| ENSG00000136100 | 1221 | 643 | 801 | 507 | 750 | 696 |

| | | | | | | |
|---|---|---|---|---|---|---|
| ENSG00000278238 | 56 | 22 | 31 | 38 | 24 | 25 |
| ENSG00000136108 | 1069 | 669 | 468 | 281 | 978 | 555 |
| ENSG00000235660 | 9 | 2 | 7 | 2 | 1 | 4 |
| ENSG00000244471 | 10 | 6 | 7 | 3 | 4 | 9 |
| ENSG00000280296 | 4 | 0 | 0 | 0 | 0 | 0 |
| ENSG00000273784 | 2 | 6 | 2 | 3 | 2 | 10 |
| ENSG00000250299 | 27 | 16 | 15 | 20 | 19 | 16 |
| ENSG00000139675 | 102 | 37 | 56 | 61 | 67 | 76 |
| ENSG00000273723 | 18 | 7 | 16 | 12 | 24 | 12 |
| ENSG00000165416 | 996 | 600 | 622 | 504 | 808 | 549 |
| ENSG00000136110 | 2 | 0 | 0 | 0 | 0 | 0 |
| ENSG00000211579 | 0 | 0 | 0 | 0 | 0 | 0 |
| ENSG00000150276 | 0 | 0 | 0 | 0 | 0 | 0 |
| ENSG00000136099 | 46 | 20 | 34 | 37 | 35 | 45 |
| ENSG00000102837 | 0 | 0 | 0 | 0 | 0 | 0 |
| ENSG00000237092 | 0 | 0 | 0 | 0 | 0 | 0 |
| ENSG00000279325 | 0 | 0 | 0 | 0 | 0 | 0 |
| ENSG00000225510 | 0 | 0 | 0 | 0 | 0 | 0 |
| ENSG00000241613 | 0 | 0 | 0 | 0 | 0 | 0 |
| ENSG00000273919 | 0 | 0 | 0 | 0 | 0 | 0 |
| ENSG00000274316 | 0 | 0 | 0 | 0 | 0 | 0 |
| ENSG00000261517 | 0 | 0 | 0 | 0 | 0 | 0 |
| ENSG00000234787 | 0 | 0 | 0 | 0 | 0 | 0 |
| ENSG00000271652 | 0 | 0 | 0 | 0 | 0 | 0 |
| ENSG00000283126 | 0 | 0 | 0 | 0 | 0 | 0 |
| ENSG00000136149 | 10 | 9 | 8 | 10 | 10 | 6 |
| ENSG00000277047 | 2 | 3 | 5 | 0 | 0 | 3 |
| ENSG00000278630 | 0 | 1 | 0 | 0 | 0 | 0 |
| ENSG00000275071 | 0 | 0 | 0 | 0 | 0 | 0 |
| ENSG00000264387 | 0 | 0 | 0 | 0 | 0 | 0 |
| ENSG00000274090 | 0 | 0 | 0 | 0 | 0 | 0 |
| ENSG00000228611 | 0 | 0 | 0 | 0 | 0 | 0 |
| ENSG00000228319 | 0 | 0 | 0 | 0 | 0 | 0 |
| ENSG00000201665 | 0 | 0 | 0 | 0 | 0 | 0 |
| ENSG00000204919 | 0 | 0 | 0 | 0 | 0 | 0 |
| ENSG00000229665 | 0 | 0 | 0 | 0 | 0 | 0 |
| ENSG00000204918 | 0 | 0 | 0 | 0 | 0 | 0 |
| ENSG00000227151 | 0 | 0 | 0 | 0 | 0 | 0 |
| ENSG00000234278 | 0 | 0 | 0 | 0 | 0 | 0 |
| ENSG00000271140 | 0 | 0 | 0 | 0 | 0 | 0 |
| ENSG00000230009 | 0 | 0 | 0 | 0 | 0 | 0 |
| ENSG00000234421 | 0 | 0 | 0 | 0 | 0 | 0 |
| ENSG00000225352 | 0 | 0 | 0 | 0 | 0 | 0 |
| ENSG00000118946 | 0 | 0 | 0 | 0 | 0 | 1 |
| ENSG00000278722 | 0 | 0 | 0 | 0 | 0 | 0 |
| ENSG00000202422 | 0 | 0 | 0 | 0 | 0 | 0 |
| ENSG00000277448 | 0 | 0 | 0 | 0 | 0 | 0 |
| ENSG00000232954 | 0 | 0 | 0 | 0 | 0 | 0 |
| ENSG00000276208 | 0 | 0 | 0 | 0 | 0 | 0 |
| ENSG00000222733 | 0 | 0 | 0 | 0 | 0 | 0 |

| | | | | | | |
|---|---|---|---|---|---|---|
| ENSG00000214335 | 0 | 0 | 0 | 0 | 0 | 0 |
| ENSG00000223717 | 0 | 0 | 0 | 0 | 0 | 0 |
| ENSG00000214281 | 0 | 0 | 0 | 0 | 0 | 0 |
| ENSG00000232647 | 0 | 0 | 0 | 0 | 0 | 0 |
| ENSG00000271190 | 0 | 0 | 0 | 0 | 0 | 0 |
| ENSG00000239003 | 0 | 0 | 0 | 0 | 0 | 0 |
| ENSG00000139734 | 334 | 206 | 211 | 155 | 255 | 186 |
| ENSG00000227528 | 0 | 0 | 0 | 0 | 0 | 0 |
| ENSG00000265745 | 0 | 0 | 0 | 0 | 0 | 0 |
| ENSG00000223815 | 0 | 0 | 0 | 0 | 0 | 0 |
| ENSG00000202151 | 0 | 0 | 0 | 0 | 0 | 0 |
| ENSG00000227336 | 0 | 0 | 0 | 0 | 0 | 0 |
| ENSG00000238213 | 0 | 0 | 0 | 0 | 1 | 0 |
| ENSG00000277549 | 0 | 0 | 0 | 0 | 0 | 0 |
| ENSG00000083544 | 380 | 165 | 200 | 136 | 212 | 174 |
| ENSG00000223076 | 0 | 0 | 0 | 0 | 0 | 0 |
| ENSG00000236608 | 0 | 0 | 1 | 0 | 0 | 0 |
| ENSG00000225249 | 0 | 0 | 0 | 0 | 0 | 0 |
| ENSG00000279657 | 0 | 0 | 0 | 0 | 0 | 0 |
| ENSG00000206854 | 0 | 0 | 0 | 0 | 0 | 0 |
| ENSG00000227510 | 0 | 0 | 0 | 0 | 0 | 0 |
| ENSG00000266663 | 0 | 0 | 0 | 0 | 0 | 0 |
| ENSG00000280165 | 0 | 0 | 0 | 0 | 0 | 0 |
| ENSG00000197991 | 0 | 0 | 0 | 0 | 0 | 0 |
| ENSG00000279797 | 0 | 0 | 0 | 0 | 0 | 0 |
| ENSG00000234033 | 0 | 0 | 0 | 0 | 0 | 0 |
| ENSG00000270856 | 0 | 0 | 0 | 0 | 0 | 0 |
| ENSG00000229578 | 0 | 0 | 0 | 0 | 0 | 0 |
| ENSG00000230142 | 0 | 0 | 0 | 0 | 0 | 0 |
| ENSG00000227611 | 0 | 0 | 0 | 0 | 0 | 0 |
| ENSG00000229307 | 0 | 0 | 0 | 0 | 0 | 0 |
| ENSG00000232105 | 0 | 0 | 0 | 0 | 0 | 0 |
| ENSG00000276586 | 0 | 0 | 0 | 0 | 0 | 0 |
| ENSG00000271465 | 0 | 0 | 0 | 0 | 0 | 0 |
| ENSG00000228669 | 0 | 0 | 0 | 0 | 0 | 0 |
| ENSG00000273550 | 0 | 0 | 0 | 0 | 0 | 0 |
| ENSG00000227564 | 0 | 0 | 0 | 0 | 0 | 0 |
| ENSG00000231061 | 0 | 0 | 0 | 0 | 0 | 0 |
| ENSG00000274052 | 0 | 0 | 0 | 0 | 0 | 0 |
| ENSG00000285764 | 0 | 0 | 0 | 0 | 0 | 0 |
| ENSG00000237378 | 0 | 0 | 0 | 0 | 0 | 0 |
| ENSG00000202478 | 0 | 0 | 0 | 0 | 0 | 0 |
| ENSG00000213232 | 0 | 0 | 0 | 0 | 0 | 0 |
| ENSG00000219926 | 0 | 0 | 0 | 0 | 0 | 0 |
| ENSG00000285566 | 0 | 0 | 0 | 0 | 0 | 0 |
| ENSG00000227674 | 0 | 0 | 0 | 0 | 0 | 0 |
| ENSG00000237849 | 0 | 0 | 0 | 0 | 0 | 0 |
| ENSG00000214269 | 8 | 5 | 2 | 3 | 2 | 6 |
| ENSG00000214266 | 0 | 0 | 0 | 0 | 0 | 0 |
| ENSG00000236565 | 5 | 2 | 4 | 1 | 3 | 3 |

| | | | | | | |
|---|---|---|---|---|---|---|
| ENSG00000234767 | 0 | 0 | 0 | 0 | 0 | 0 |
| ENSG00000263581 | 0 | 0 | 0 | 0 | 0 | 0 |
| ENSG00000263561 | 0 | 0 | 0 | 0 | 0 | 0 |
| ENSG00000235737 | 0 | 0 | 0 | 0 | 0 | 0 |
| ENSG00000184226 | 58 | 28 | 25 | 23 | 36 | 48 |
| ENSG00000234527 | 0 | 0 | 0 | 0 | 0 | 0 |
| ENSG00000238500 | 0 | 0 | 0 | 0 | 0 | 0 |
| ENSG00000228842 | 0 | 0 | 0 | 0 | 0 | 0 |
| ENSG00000225263 | 0 | 0 | 0 | 0 | 0 | 0 |
| ENSG00000233840 | 0 | 0 | 0 | 0 | 0 | 0 |
| ENSG00000285588 | 0 | 0 | 0 | 1 | 1 | 1 |
| ENSG00000214263 | 0 | 0 | 2 | 0 | 0 | 1 |
| ENSG00000230040 | 0 | 0 | 0 | 0 | 0 | 0 |
| ENSG00000271287 | 0 | 0 | 0 | 0 | 0 | 0 |
| ENSG00000227969 | 0 | 0 | 0 | 0 | 0 | 0 |
| ENSG00000231667 | 0 | 0 | 0 | 0 | 0 | 0 |
| ENSG00000226537 | 0 | 0 | 0 | 0 | 0 | 0 |
| ENSG00000234765 | 0 | 0 | 0 | 0 | 0 | 0 |
| ENSG00000229910 | 0 | 0 | 0 | 0 | 0 | 0 |
| ENSG00000236433 | 0 | 0 | 0 | 0 | 0 | 0 |
| ENSG00000230405 | 0 | 0 | 0 | 0 | 0 | 0 |
| ENSG00000229824 | 0 | 0 | 0 | 0 | 0 | 0 |
| ENSG00000243671 | 0 | 0 | 0 | 0 | 0 | 0 |
| ENSG00000281778 | 0 | 0 | 0 | 0 | 0 | 0 |
| ENSG00000280580 | 0 | 0 | 0 | 0 | 0 | 0 |
| ENSG00000232380 | 14 | 8 | 7 | 4 | 7 | 3 |
| ENSG00000225501 | 0 | 0 | 0 | 0 | 0 | 0 |
| ENSG00000235221 | 0 | 0 | 0 | 0 | 0 | 0 |
| ENSG00000224640 | 0 | 0 | 0 | 0 | 0 | 0 |
| ENSG00000150361 | 0 | 0 | 0 | 0 | 0 | 0 |
| ENSG00000207495 | 0 | 0 | 0 | 0 | 0 | 0 |
| ENSG00000232314 | 0 | 0 | 0 | 0 | 0 | 0 |
| ENSG00000230223 | 0 | 0 | 0 | 0 | 0 | 0 |
| ENSG00000202433 | 0 | 0 | 0 | 0 | 0 | 0 |
| ENSG00000276841 | 0 | 0 | 0 | 0 | 0 | 0 |
| ENSG00000226554 | 0 | 0 | 0 | 0 | 0 | 0 |
| ENSG00000226846 | 0 | 0 | 0 | 0 | 0 | 0 |
| ENSG00000225801 | 0 | 0 | 0 | 0 | 0 | 0 |
| ENSG00000276644 | 0 | 0 | 0 | 0 | 0 | 0 |
| ENSG00000236534 | 1 | 1 | 1 | 2 | 0 | 2 |
| ENSG00000237816 | 0 | 0 | 0 | 0 | 0 | 0 |
| ENSG00000233460 | 0 | 0 | 0 | 0 | 0 | 0 |
| ENSG00000232881 | 0 | 0 | 0 | 0 | 0 | 0 |
| ENSG00000212377 | 0 | 0 | 0 | 0 | 0 | 0 |
| ENSG00000227952 | 0 | 0 | 0 | 0 | 0 | 0 |
| ENSG00000251715 | 0 | 0 | 0 | 0 | 0 | 0 |
| ENSG00000278113 | 0 | 0 | 0 | 0 | 0 | 0 |
| ENSG00000199282 | 0 | 0 | 0 | 0 | 0 | 0 |
| ENSG00000226617 | 0 | 0 | 4 | 1 | 0 | 2 |
| ENSG00000206922 | 0 | 0 | 0 | 0 | 0 | 0 |

| | | | | | | |
|---|---|---|---|---|---|---|
| ENSG00000204899 | 444 | 196 | 227 | 224 | 269 | 236 |
| ENSG00000136122 | 126 | 51 | 64 | 50 | 91 | 89 |
| ENSG00000083520 | 455 | 287 | 261 | 126 | 468 | 256 |
| ENSG00000083535 | 53 | 37 | 23 | 11 | 73 | 32 |
| ENSG00000199381 | 0 | 0 | 0 | 0 | 0 | 0 |
| ENSG00000232041 | 0 | 0 | 0 | 0 | 0 | 0 |
| ENSG00000102554 | 663 | 327 | 365 | 206 | 306 | 254 |
| ENSG00000236972 | 0 | 0 | 0 | 0 | 0 | 0 |
| ENSG00000200267 | 0 | 0 | 0 | 0 | 0 | 0 |
| ENSG00000201253 | 0 | 0 | 0 | 0 | 0 | 0 |
| ENSG00000206697 | 0 | 0 | 0 | 0 | 0 | 0 |
| ENSG00000279754 | 0 | 0 | 0 | 0 | 0 | 0 |
| ENSG00000229708 | 0 | 0 | 0 | 0 | 0 | 0 |
| ENSG00000224853 | 0 | 0 | 0 | 0 | 0 | 0 |
| ENSG00000228295 | 0 | 0 | 0 | 0 | 0 | 0 |
| ENSG00000118922 | 131 | 75 | 51 | 37 | 112 | 62 |
| ENSG00000280348 | 2 | 0 | 0 | 0 | 0 | 0 |
| ENSG00000279463 | 0 | 0 | 0 | 0 | 0 | 0 |
| ENSG00000235532 | 0 | 0 | 0 | 0 | 0 | 1 |
| ENSG00000206617 | 0 | 0 | 0 | 0 | 0 | 0 |
| ENSG00000232701 | 0 | 0 | 0 | 0 | 0 | 0 |
| ENSG00000177596 | 0 | 0 | 0 | 0 | 0 | 0 |
| ENSG00000226240 | 0 | 0 | 1 | 0 | 0 | 0 |
| ENSG00000281167 | 0 | 0 | 0 | 0 | 0 | 0 |
| ENSG00000236678 | 0 | 0 | 0 | 0 | 0 | 0 |
| ENSG00000225944 | 0 | 0 | 0 | 0 | 0 | 0 |
| ENSG00000206812 | 0 | 0 | 0 | 0 | 0 | 0 |
| ENSG00000226665 | 0 | 0 | 0 | 0 | 0 | 0 |
| ENSG00000214249 | 6 | 4 | 2 | 0 | 5 | 2 |
| ENSG00000223880 | 0 | 0 | 0 | 0 | 0 | 0 |
| ENSG00000136111 | 515 | 204 | 216 | 132 | 250 | 202 |
| ENSG00000225203 | 0 | 0 | 0 | 0 | 0 | 0 |
| ENSG00000188243 | 670 | 346 | 428 | 459 | 463 | 435 |
| ENSG00000118939 | 42 | 56 | 70 | 55 | 70 | 57 |
| ENSG00000261553 | 0 | 1 | 0 | 0 | 1 | 0 |
| ENSG00000261105 | 0 | 0 | 0 | 0 | 0 | 0 |
| ENSG00000136153 | 430 | 215 | 213 | 167 | 348 | 256 |
| ENSG00000228444 | 2 | 2 | 1 | 1 | 1 | 0 |
| ENSG00000230902 | 0 | 0 | 0 | 0 | 0 | 0 |
| ENSG00000178734 | 0 | 1 | 0 | 0 | 0 | 0 |
| ENSG00000223458 | 0 | 0 | 0 | 0 | 0 | 0 |
| ENSG00000261206 | 0 | 0 | 0 | 0 | 0 | 0 |
| ENSG00000224933 | 0 | 0 | 0 | 0 | 0 | 0 |
| ENSG00000243274 | 0 | 0 | 0 | 0 | 0 | 0 |
| ENSG00000285572 | 0 | 0 | 0 | 0 | 0 | 0 |
| ENSG00000284809 | 0 | 0 | 0 | 0 | 0 | 0 |
| ENSG00000178695 | 10 | 5 | 7 | 4 | 4 | 5 |
| ENSG00000278727 | 0 | 0 | 0 | 0 | 0 | 0 |
| ENSG00000118903 | 0 | 0 | 0 | 0 | 0 | 0 |
| ENSG00000102794 | 0 | 0 | 0 | 0 | 0 | 0 |

| | | | | | | |
|---|---|---|---|---|---|---|
| ENSG00000233873 | 0 | 0 | 0 | 0 | 0 | 0 |
| ENSG00000228002 | 0 | 0 | 2 | 1 | 1 | 1 |
| ENSG00000102805 | 108 | 55 | 54 | 48 | 73 | 66 |
| ENSG00000283208 | 3 | 1 | 0 | 0 | 1 | 2 |
| ENSG00000005812 | 631 | 332 | 354 | 182 | 374 | 325 |
| ENSG00000236051 | 0 | 0 | 1 | 0 | 0 | 0 |
| ENSG00000274898 | 2 | 0 | 1 | 1 | 2 | 2 |
| ENSG00000005810 | 461 | 277 | 240 | 69 | 404 | 235 |
| ENSG00000229521 | 0 | 0 | 0 | 0 | 0 | 0 |
| ENSG00000285714 | 0 | 0 | 0 | 2 | 1 | 0 |
| ENSG00000136155 | 0 | 0 | 0 | 0 | 0 | 0 |
| ENSG00000224347 | 0 | 0 | 0 | 0 | 0 | 0 |
| ENSG00000206800 | 0 | 0 | 0 | 0 | 0 | 0 |
| ENSG00000227922 | 0 | 0 | 0 | 0 | 0 | 0 |
| ENSG00000285746 | 0 | 0 | 0 | 0 | 0 | 0 |
| ENSG00000139737 | 76 | 38 | 48 | 30 | 41 | 43 |
| ENSG00000266325 | 0 | 0 | 0 | 0 | 0 | 0 |
| ENSG00000225579 | 0 | 0 | 0 | 0 | 0 | 0 |
| ENSG00000136160 | 0 | 0 | 0 | 0 | 0 | 0 |
| ENSG00000234377 | 0 | 0 | 0 | 0 | 0 | 0 |
| ENSG00000233379 | 0 | 0 | 0 | 0 | 0 | 0 |
| ENSG00000243035 | 0 | 0 | 0 | 0 | 0 | 0 |
| ENSG00000236133 | 0 | 0 | 0 | 0 | 0 | 0 |
| ENSG00000229249 | 0 | 0 | 0 | 0 | 0 | 0 |
| ENSG00000235625 | 0 | 0 | 0 | 0 | 0 | 0 |
| ENSG00000271386 | 0 | 0 | 0 | 0 | 0 | 0 |
| ENSG00000200686 | 0 | 0 | 0 | 0 | 0 | 0 |
| ENSG00000229865 | 0 | 0 | 0 | 0 | 0 | 0 |
| ENSG00000224686 | 0 | 0 | 0 | 0 | 0 | 0 |
| ENSG00000271776 | 0 | 0 | 0 | 0 | 0 | 0 |
| ENSG00000152192 | 0 | 0 | 0 | 0 | 0 | 0 |
| ENSG00000152193 | 57 | 46 | 34 | 33 | 63 | 38 |
| ENSG00000237109 | 0 | 0 | 0 | 0 | 0 | 0 |
| ENSG00000225427 | 0 | 0 | 0 | 0 | 0 | 0 |
| ENSG00000215357 | 0 | 0 | 0 | 0 | 0 | 0 |
| ENSG00000230584 | 0 | 0 | 0 | 1 | 0 | 1 |
| ENSG00000236277 | 0 | 0 | 0 | 0 | 0 | 0 |
| ENSG00000226670 | 0 | 0 | 0 | 0 | 0 | 1 |
| ENSG00000139746 | 701 | 351 | 348 | 185 | 542 | 323 |
| ENSG00000252496 | 0 | 0 | 0 | 0 | 0 | 0 |
| ENSG00000227354 | 24 | 19 | 18 | 15 | 18 | 21 |
| ENSG00000232132 | 0 | 0 | 0 | 0 | 0 | 0 |
| ENSG00000102471 | 886 | 357 | 517 | 344 | 504 | 424 |
| ENSG00000227676 | 0 | 0 | 0 | 0 | 2 | 0 |
| ENSG00000285902 | 0 | 0 | 0 | 0 | 0 | 0 |
| ENSG00000229011 | 0 | 0 | 0 | 0 | 0 | 0 |
| ENSG00000229175 | 0 | 0 | 0 | 0 | 0 | 0 |
| ENSG00000284196 | 0 | 0 | 0 | 0 | 1 | 0 |
| ENSG00000281721 | 0 | 0 | 0 | 0 | 0 | 0 |
| ENSG00000285684 | 0 | 0 | 0 | 0 | 0 | 0 |

| | | | | | | |
|---|---|---|---|---|---|---|
| ENSG00000136158 | 30 | 23 | 26 | 20 | 30 | 25 |
| ENSG00000202398 | 0 | 0 | 0 | 0 | 0 | 0 |
| ENSG00000227862 | 0 | 0 | 0 | 0 | 0 | 0 |
| ENSG00000226683 | 0 | 0 | 0 | 0 | 0 | 0 |
| ENSG00000231222 | 0 | 0 | 0 | 0 | 0 | 0 |
| ENSG00000229246 | 0 | 0 | 0 | 0 | 0 | 0 |
| ENSG00000270415 | 0 | 0 | 0 | 0 | 0 | 0 |
| ENSG00000260094 | 0 | 0 | 0 | 0 | 0 | 0 |
| ENSG00000229309 | 0 | 0 | 0 | 0 | 0 | 0 |
| ENSG00000222486 | 0 | 0 | 0 | 0 | 0 | 0 |
| ENSG00000237148 | 0 | 0 | 0 | 0 | 0 | 0 |
| ENSG00000214182 | 38 | 24 | 25 | 18 | 32 | 25 |
| ENSG00000237099 | 0 | 0 | 0 | 0 | 0 | 0 |
| ENSG00000270400 | 0 | 0 | 0 | 0 | 0 | 0 |
| ENSG00000222791 | 0 | 0 | 0 | 0 | 0 | 0 |
| ENSG00000178235 | 4 | 3 | 7 | 1 | 0 | 1 |
| ENSG00000285680 | 0 | 0 | 0 | 0 | 0 | 0 |
| ENSG00000215037 | 0 | 0 | 0 | 0 | 0 | 0 |
| ENSG00000224188 | 0 | 0 | 0 | 0 | 0 | 0 |
| ENSG00000285742 | 0 | 0 | 0 | 0 | 0 | 0 |
| ENSG00000228597 | 0 | 0 | 0 | 0 | 0 | 0 |
| ENSG00000232447 | 0 | 0 | 0 | 0 | 0 | 0 |
| ENSG00000233349 | 0 | 0 | 0 | 0 | 0 | 0 |
| ENSG00000226370 | 0 | 0 | 0 | 0 | 0 | 0 |
| ENSG00000226317 | 0 | 0 | 0 | 0 | 0 | 0 |
| ENSG00000285834 | 0 | 0 | 0 | 0 | 0 | 0 |
| ENSG00000184564 | 417 | 291 | 217 | 84 | 457 | 244 |
| ENSG00000278580 | 0 | 0 | 0 | 0 | 0 | 0 |
| ENSG00000228241 | 0 | 0 | 0 | 0 | 0 | 0 |
| ENSG00000232252 | 0 | 0 | 0 | 0 | 0 | 0 |
| ENSG00000283371 | 0 | 0 | 0 | 0 | 0 | 0 |
| ENSG00000277914 | 0 | 0 | 0 | 0 | 0 | 0 |
| ENSG00000231879 | 0 | 0 | 0 | 0 | 0 | 0 |
| ENSG00000233528 | 0 | 0 | 0 | 0 | 0 | 0 |
| ENSG00000228074 | 0 | 0 | 0 | 0 | 0 | 0 |
| ENSG00000228473 | 0 | 0 | 0 | 0 | 0 | 0 |
| ENSG00000285699 | 0 | 0 | 0 | 0 | 0 | 0 |
| ENSG00000232636 | 0 | 0 | 0 | 0 | 0 | 0 |
| ENSG00000228824 | 0 | 0 | 0 | 0 | 1 | 0 |
| ENSG00000272515 | 0 | 0 | 0 | 0 | 0 | 0 |
| ENSG00000266052 | 0 | 0 | 0 | 0 | 0 | 0 |
| ENSG00000165300 | 2 | 0 | 1 | 0 | 0 | 1 |
| ENSG00000277640 | 0 | 0 | 0 | 0 | 0 | 0 |
| ENSG00000223404 | 0 | 0 | 1 | 0 | 0 | 0 |
| ENSG00000272046 | 0 | 0 | 0 | 0 | 0 | 0 |
| ENSG00000232204 | 0 | 0 | 0 | 0 | 0 | 0 |
| ENSG00000231019 | 0 | 0 | 0 | 0 | 0 | 0 |
| ENSG00000225289 | 0 | 0 | 0 | 0 | 0 | 0 |
| ENSG00000285811 | 0 | 0 | 0 | 0 | 0 | 0 |
| ENSG00000229443 | 0 | 0 | 0 | 0 | 0 | 0 |

| | | | | | | |
|---|---|---|---|---|---|---|
| ENSG00000283130 | 0 | 0 | 0 | 0 | 0 | 0 |
| ENSG00000261666 | 0 | 0 | 0 | 0 | 0 | 0 |
| ENSG00000233211 | 0 | 0 | 0 | 0 | 0 | 0 |
| ENSG00000232225 | 0 | 0 | 0 | 0 | 0 | 0 |
| ENSG00000227247 | 0 | 0 | 0 | 0 | 0 | 0 |
| ENSG00000234660 | 0 | 0 | 0 | 0 | 0 | 0 |
| ENSG00000225530 | 0 | 0 | 0 | 0 | 0 | 0 |
| ENSG00000226037 | 0 | 0 | 0 | 0 | 0 | 0 |
| ENSG00000236176 | 0 | 0 | 0 | 0 | 0 | 0 |
| ENSG00000271395 | 0 | 0 | 0 | 0 | 0 | 0 |
| ENSG00000234227 | 0 | 0 | 0 | 0 | 0 | 0 |
| ENSG00000200733 | 0 | 0 | 0 | 0 | 0 | 0 |
| ENSG00000215418 | 0 | 0 | 0 | 0 | 0 | 0 |
| ENSG00000261446 | 0 | 0 | 0 | 0 | 0 | 0 |
| ENSG00000252696 | 0 | 0 | 0 | 0 | 0 | 0 |
| ENSG00000235879 | 0 | 0 | 0 | 0 | 0 | 0 |
| ENSG00000224419 | 0 | 0 | 0 | 0 | 0 | 0 |
| ENSG00000283783 | 0 | 0 | 0 | 0 | 0 | 0 |
| ENSG00000234384 | 0 | 0 | 0 | 0 | 0 | 0 |
| ENSG00000251753 | 0 | 0 | 0 | 0 | 0 | 0 |
| ENSG00000231674 | 0 | 0 | 0 | 0 | 0 | 0 |
| ENSG00000227983 | 0 | 0 | 0 | 0 | 0 | 0 |
| ENSG00000234625 | 0 | 0 | 0 | 0 | 0 | 0 |
| ENSG00000229557 | 0 | 0 | 0 | 0 | 0 | 0 |
| ENSG00000214178 | 0 | 0 | 0 | 0 | 0 | 0 |
| ENSG00000215417 | 7 | 4 | 5 | 1 | 4 | 1 |
| ENSG00000284536 | 0 | 0 | 0 | 0 | 0 | 0 |
| ENSG00000283815 | 0 | 0 | 0 | 0 | 0 | 0 |
| ENSG00000284204 | 0 | 0 | 0 | 0 | 0 | 0 |
| ENSG00000283762 | 0 | 0 | 0 | 0 | 0 | 0 |
| ENSG00000284375 | 0 | 0 | 0 | 0 | 0 | 0 |
| ENSG00000283705 | 0 | 0 | 0 | 0 | 0 | 0 |
| ENSG00000179399 | 0 | 0 | 0 | 0 | 0 | 0 |
| ENSG00000282864 | 0 | 0 | 0 | 0 | 0 | 0 |
| ENSG00000282997 | 0 | 0 | 0 | 0 | 0 | 0 |
| ENSG00000252508 | 0 | 0 | 0 | 0 | 0 | 0 |
| ENSG00000229287 | 0 | 0 | 0 | 0 | 0 | 0 |
| ENSG00000282865 | 0 | 0 | 0 | 0 | 0 | 0 |
| ENSG00000232885 | 0 | 0 | 0 | 0 | 0 | 0 |
| ENSG00000283097 | 0 | 0 | 0 | 0 | 0 | 0 |
| ENSG00000236240 | 0 | 0 | 0 | 0 | 0 | 0 |
| ENSG00000263741 | 0 | 0 | 0 | 0 | 0 | 0 |
| ENSG00000282993 | 0 | 0 | 0 | 0 | 0 | 0 |
| ENSG00000235984 | 0 | 0 | 0 | 0 | 0 | 0 |
| ENSG00000232849 | 0 | 0 | 0 | 0 | 0 | 0 |
| ENSG00000278177 | 5 | 4 | 3 | 6 | 2 | 4 |
| ENSG00000183098 | 431 | 208 | 203 | 139 | 194 | 153 |
| ENSG00000235784 | 0 | 0 | 0 | 0 | 0 | 1 |
| ENSG00000224394 | 0 | 0 | 0 | 0 | 0 | 0 |
| ENSG00000212559 | 0 | 0 | 0 | 0 | 0 | 0 |

| | | | | | | |
|---|---|---|---|---|---|---|
| ENSG00000236520 | 0 | 0 | 0 | 0 | 0 | 0 |
| ENSG00000080166 | 2 | 0 | 0 | 1 | 3 | 0 |
| ENSG00000088451 | 239 | 104 | 129 | 87 | 159 | 147 |
| ENSG00000152749 | 198 | 129 | 120 | 58 | 130 | 114 |
| ENSG00000222129 | 0 | 0 | 0 | 0 | 0 | 0 |
| ENSG00000274168 | 0 | 0 | 0 | 0 | 0 | 0 |
| ENSG00000238230 | 0 | 0 | 0 | 0 | 0 | 0 |
| ENSG00000125285 | 0 | 0 | 0 | 0 | 0 | 1 |
| ENSG00000227640 | 0 | 0 | 0 | 0 | 0 | 1 |
| ENSG00000235768 | 0 | 0 | 0 | 0 | 0 | 0 |
| ENSG00000237924 | 0 | 0 | 0 | 0 | 0 | 0 |
| ENSG00000260962 | 0 | 0 | 0 | 0 | 0 | 0 |
| ENSG00000252335 | 0 | 0 | 0 | 0 | 0 | 0 |
| ENSG00000125257 | 443 | 241 | 258 | 156 | 260 | 216 |
| ENSG00000223298 | 2 | 0 | 2 | 1 | 1 | 5 |
| ENSG00000200211 | 0 | 0 | 0 | 0 | 0 | 0 |
| ENSG00000271492 | 0 | 0 | 0 | 0 | 0 | 0 |
| ENSG00000134873 | 0 | 0 | 0 | 0 | 0 | 0 |
| ENSG00000223392 | 0 | 0 | 0 | 0 | 0 | 0 |
| ENSG00000134874 | 2 | 0 | 0 | 0 | 0 | 0 |
| ENSG00000247400 | 81 | 78 | 78 | 68 | 95 | 78 |
| ENSG00000102580 | 1008 | 492 | 578 | 374 | 746 | 561 |
| ENSG00000229622 | 0 | 0 | 0 | 0 | 0 | 1 |
| ENSG00000227714 | 0 | 1 | 0 | 0 | 0 | 0 |
| ENSG00000232373 | 0 | 0 | 0 | 0 | 0 | 1 |
| ENSG00000276809 | 2 | 0 | 0 | 0 | 1 | 0 |
| ENSG00000102595 | 310 | 180 | 185 | 104 | 275 | 173 |
| ENSG00000226977 | 0 | 0 | 0 | 0 | 0 | 0 |
| ENSG00000185352 | 791 | 378 | 528 | 333 | 507 | 561 |
| ENSG00000242614 | 0 | 0 | 0 | 0 | 0 | 0 |
| ENSG00000283487 | 0 | 0 | 0 | 0 | 0 | 0 |
| ENSG00000233273 | 2 | 0 | 2 | 2 | 0 | 2 |
| ENSG00000235137 | 0 | 1 | 0 | 0 | 0 | 0 |
| ENSG00000234267 | 0 | 0 | 0 | 0 | 0 | 0 |
| ENSG00000272329 | 0 | 0 | 0 | 0 | 0 | 0 |
| ENSG00000222472 | 0 | 0 | 0 | 0 | 0 | 0 |
| ENSG00000165621 | 0 | 2 | 0 | 0 | 1 | 2 |
| ENSG00000233124 | 0 | 1 | 1 | 0 | 0 | 0 |
| ENSG00000139793 | 907 | 576 | 366 | 158 | 603 | 341 |
| ENSG00000202290 | 1 | 0 | 2 | 0 | 1 | 0 |
| ENSG00000125249 | 836 | 482 | 432 | 306 | 497 | 469 |
| ENSG00000276573 | 0 | 0 | 0 | 0 | 0 | 0 |
| ENSG00000276704 | 0 | 0 | 0 | 0 | 1 | 0 |
| ENSG00000271625 | 0 | 0 | 0 | 0 | 0 | 0 |
| ENSG00000226134 | 0 | 0 | 0 | 0 | 1 | 0 |
| ENSG00000233291 | 0 | 0 | 0 | 0 | 0 | 0 |
| ENSG00000270725 | 0 | 0 | 0 | 0 | 0 | 0 |
| ENSG00000065150 | 1951 | 1101 | 1044 | 814 | 1406 | 1049 |
| ENSG00000232217 | 0 | 0 | 0 | 0 | 0 | 0 |
| ENSG00000152767 | 716 | 382 | 420 | 373 | 423 | 394 |

| | | | | | | |
|---|---|---|---|---|---|---|
| ENSG00000139797 | 0 | 0 | 0 | 0 | 0 | 0 |
| ENSG00000222969 | 0 | 0 | 0 | 0 | 0 | 0 |
| ENSG00000263399 | 0 | 0 | 0 | 0 | 0 | 0 |
| ENSG00000269189 | 0 | 0 | 0 | 0 | 0 | 0 |
| ENSG00000269599 | 0 | 0 | 0 | 0 | 0 | 0 |
| ENSG00000231194 | 0 | 0 | 0 | 0 | 0 | 0 |
| ENSG00000102572 | 1328 | 617 | 785 | 686 | 758 | 739 |
| ENSG00000271437 | 0 | 0 | 0 | 0 | 0 | 0 |
| ENSG00000224418 | 7 | 6 | 6 | 3 | 3 | 8 |
| ENSG00000217686 | 0 | 0 | 0 | 0 | 0 | 0 |
| ENSG00000233549 | 0 | 0 | 0 | 0 | 0 | 0 |
| ENSG00000233662 | 0 | 0 | 0 | 0 | 0 | 0 |
| ENSG00000243366 | 0 | 0 | 0 | 0 | 0 | 0 |
| ENSG00000088386 | 0 | 0 | 0 | 0 | 0 | 0 |
| ENSG00000088387 | 591 | 264 | 365 | 177 | 322 | 288 |
| ENSG00000229918 | 0 | 0 | 0 | 0 | 1 | 0 |
| ENSG00000275919 | 0 | 0 | 0 | 0 | 0 | 0 |
| ENSG00000231981 | 0 | 0 | 0 | 0 | 0 | 0 |
| ENSG00000207298 | 0 | 0 | 0 | 0 | 0 | 0 |
| ENSG00000260992 | 28 | 9 | 10 | 22 | 15 | 15 |
| ENSG00000230871 | 0 | 0 | 0 | 0 | 0 | 0 |
| ENSG00000234005 | 0 | 3 | 0 | 2 | 0 | 0 |
| ENSG00000228889 | 52 | 25 | 36 | 29 | 46 | 17 |
| ENSG00000134882 | 1984 | 867 | 1046 | 1169 | 1108 | 1122 |
| ENSG00000201793 | 0 | 0 | 0 | 0 | 0 | 0 |
| ENSG00000218502 | 2 | 1 | 1 | 1 | 0 | 1 |
| ENSG00000125245 | 2 | 0 | 3 | 0 | 0 | 0 |
| ENSG00000169508 | 0 | 0 | 0 | 0 | 0 | 0 |
| ENSG00000207719 | 0 | 0 | 0 | 0 | 0 | 0 |
| ENSG00000228808 | 4 | 1 | 0 | 0 | 2 | 0 |
| ENSG00000238241 | 0 | 0 | 0 | 0 | 0 | 0 |
| ENSG00000285448 | 0 | 1 | 0 | 0 | 0 | 0 |
| ENSG00000125304 | 7420 | 3454 | 4383 | 2968 | 3759 | 3678 |
| ENSG00000280734 | 165 | 66 | 90 | 64 | 90 | 62 |
| ENSG00000280710 | 0 | 0 | 0 | 0 | 0 | 0 |
| ENSG00000203441 | 2 | 3 | 0 | 1 | 0 | 1 |
| ENSG00000234168 | 0 | 0 | 0 | 0 | 0 | 0 |
| ENSG00000234239 | 0 | 0 | 0 | 0 | 0 | 0 |
| ENSG00000125246 | 47 | 19 | 27 | 19 | 23 | 17 |
| ENSG00000263615 | 0 | 0 | 0 | 0 | 0 | 0 |
| ENSG00000227659 | 0 | 0 | 0 | 0 | 0 | 0 |
| ENSG00000234303 | 0 | 0 | 0 | 0 | 0 | 0 |
| ENSG00000201245 | 0 | 0 | 0 | 0 | 0 | 0 |
| ENSG00000139800 | 14 | 16 | 13 | 12 | 20 | 18 |
| ENSG00000043355 | 50 | 12 | 43 | 34 | 36 | 57 |
| ENSG00000260738 | 0 | 0 | 0 | 0 | 0 | 0 |
| ENSG00000225999 | 0 | 0 | 0 | 0 | 0 | 0 |
| ENSG00000236554 | 0 | 0 | 0 | 0 | 0 | 0 |
| ENSG00000274605 | 97 | 55 | 58 | 45 | 55 | 45 |
| ENSG00000175198 | 282 | 126 | 169 | 110 | 149 | 119 |

| | | | | | | |
|---|---|---|---|---|---|---|
| ENSG00000228501 | 2 | 0 | 2 | 0 | 0 | 0 |
| ENSG00000270632 | 0 | 0 | 0 | 0 | 0 | 0 |
| ENSG00000234650 | 0 | 0 | 2 | 0 | 0 | 0 |
| ENSG00000134864 | 152 | 60 | 84 | 68 | 67 | 71 |
| ENSG00000224356 | 6 | 1 | 1 | 2 | 3 | 0 |
| ENSG00000280169 | 0 | 0 | 0 | 0 | 0 | 2 |
| ENSG00000234354 | 2 | 0 | 0 | 0 | 0 | 0 |
| ENSG00000125247 | 447 | 238 | 307 | 221 | 257 | 231 |
| ENSG00000237082 | 1 | 0 | 5 | 0 | 0 | 0 |
| ENSG00000233009 | 0 | 0 | 0 | 0 | 0 | 0 |
| ENSG00000214051 | 0 | 0 | 0 | 0 | 0 | 0 |
| ENSG00000229599 | 0 | 0 | 0 | 0 | 0 | 0 |
| ENSG00000102452 | 0 | 0 | 0 | 0 | 0 | 0 |
| ENSG00000198542 | 0 | 0 | 0 | 0 | 0 | 0 |
| ENSG00000102466 | 0 | 0 | 0 | 0 | 0 | 0 |
| ENSG00000276012 | 0 | 0 | 0 | 0 | 0 | 0 |
| ENSG00000201155 | 0 | 0 | 0 | 0 | 0 | 0 |
| ENSG00000215399 | 0 | 0 | 0 | 0 | 0 | 0 |
| ENSG00000265164 | 0 | 0 | 0 | 0 | 0 | 0 |
| ENSG00000229975 | 0 | 0 | 0 | 0 | 0 | 0 |
| ENSG00000201690 | 0 | 0 | 0 | 0 | 0 | 0 |
| ENSG00000264482 | 0 | 0 | 0 | 0 | 0 | 0 |
| ENSG00000236860 | 0 | 0 | 0 | 0 | 0 | 0 |
| ENSG00000243319 | 0 | 0 | 0 | 0 | 0 | 0 |
| ENSG00000279702 | 0 | 0 | 0 | 0 | 0 | 0 |
| ENSG00000234445 | 0 | 0 | 0 | 0 | 0 | 0 |
| ENSG00000272143 | 0 | 0 | 0 | 0 | 0 | 0 |
| ENSG00000261057 | 0 | 0 | 0 | 0 | 0 | 0 |
| ENSG00000276957 | 2 | 0 | 1 | 2 | 0 | 0 |
| ENSG00000134900 | 730 | 408 | 445 | 215 | 526 | 334 |
| ENSG00000139780 | 0 | 0 | 0 | 0 | 0 | 0 |
| ENSG00000175820 | 0 | 0 | 0 | 0 | 0 | 0 |
| ENSG00000231633 | 0 | 0 | 0 | 0 | 0 | 0 |
| ENSG00000151287 | 187 | 96 | 104 | 64 | 103 | 74 |
| ENSG00000134901 | 205 | 89 | 125 | 125 | 103 | 121 |
| ENSG00000134897 | 256 | 111 | 156 | 102 | 169 | 128 |
| ENSG00000270181 | 0 | 0 | 0 | 0 | 0 | 0 |
| ENSG00000134899 | 38 | 30 | 32 | 18 | 34 | 20 |
| ENSG00000250878 | 0 | 0 | 1 | 0 | 0 | 0 |
| ENSG00000272542 | 0 | 0 | 0 | 0 | 0 | 0 |
| ENSG00000278445 | 0 | 0 | 0 | 0 | 0 | 0 |
| ENSG00000125255 | 0 | 0 | 0 | 0 | 0 | 0 |
| ENSG00000285789 | 0 | 0 | 0 | 0 | 0 | 0 |
| ENSG00000234551 | 0 | 0 | 0 | 0 | 0 | 0 |
| ENSG00000230474 | 0 | 0 | 0 | 0 | 0 | 0 |
| ENSG00000225823 | 0 | 0 | 0 | 0 | 0 | 0 |
| ENSG00000232307 | 0 | 0 | 0 | 0 | 0 | 0 |
| ENSG00000275968 | 0 | 0 | 0 | 0 | 0 | 0 |
| ENSG00000275502 | 0 | 0 | 0 | 0 | 0 | 0 |
| ENSG00000182346 | 0 | 0 | 0 | 0 | 0 | 0 |

| | | | | | | |
|---|---|---|---|---|---|---|
| ENSG00000280455 | 0 | 0 | 0 | 0 | 0 | 0 |
| ENSG00000226620 | 0 | 0 | 0 | 0 | 0 | 0 |
| ENSG00000252550 | 0 | 0 | 0 | 0 | 0 | 0 |
| ENSG00000222682 | 0 | 0 | 0 | 0 | 0 | 0 |
| ENSG00000233532 | 0 | 0 | 0 | 0 | 0 | 0 |
| ENSG00000174418 | 0 | 0 | 0 | 0 | 0 | 0 |
| ENSG00000125266 | 1052 | 535 | 822 | 404 | 492 | 639 |
| ENSG00000284966 | 4 | 6 | 2 | 0 | 2 | 2 |
| ENSG00000274204 | 0 | 0 | 2 | 0 | 1 | 1 |
| ENSG00000134884 | 3941 | 1789 | 2559 | 1283 | 2028 | 2004 |
| ENSG00000272274 | 0 | 1 | 1 | 1 | 0 | 0 |
| ENSG00000230156 | 0 | 0 | 0 | 0 | 0 | 0 |
| ENSG00000227590 | 0 | 0 | 0 | 0 | 0 | 0 |
| ENSG00000278698 | 2 | 0 | 1 | 0 | 0 | 0 |
| ENSG00000174408 | 0 | 0 | 0 | 0 | 0 | 0 |
| ENSG00000277863 | 0 | 0 | 0 | 0 | 0 | 0 |
| ENSG00000279157 | 0 | 0 | 0 | 0 | 0 | 0 |
| ENSG00000285643 | 0 | 0 | 0 | 0 | 0 | 0 |
| ENSG00000204442 | 55 | 17 | 28 | 12 | 13 | 11 |
| ENSG00000201847 | 0 | 0 | 0 | 0 | 0 | 0 |
| ENSG00000276740 | 0 | 1 | 0 | 0 | 1 | 0 |
| ENSG00000221650 | 0 | 0 | 0 | 0 | 0 | 0 |
| ENSG00000227248 | 0 | 0 | 0 | 0 | 0 | 0 |
| ENSG00000274718 | 2 | 0 | 0 | 0 | 4 | 0 |
| ENSG00000174405 | 240 | 125 | 113 | 51 | 204 | 148 |
| ENSG00000139826 | 392 | 176 | 194 | 163 | 222 | 205 |
| ENSG00000102524 | 7 | 4 | 2 | 1 | 7 | 3 |
| ENSG00000277246 | 2 | 1 | 0 | 0 | 0 | 0 |
| ENSG00000223177 | 0 | 0 | 0 | 0 | 0 | 0 |
| ENSG00000283384 | 0 | 0 | 0 | 0 | 0 | 0 |
| ENSG00000232087 | 0 | 0 | 0 | 0 | 0 | 0 |
| ENSG00000041515 | 18 | 14 | 18 | 29 | 39 | 49 |
| ENSG00000229938 | 0 | 0 | 0 | 0 | 0 | 0 |
| ENSG00000236242 | 0 | 0 | 0 | 0 | 0 | 0 |
| ENSG00000275216 | 0 | 0 | 0 | 0 | 1 | 0 |
| ENSG00000223617 | 0 | 0 | 0 | 0 | 0 | 0 |
| ENSG00000236053 | 0 | 0 | 0 | 0 | 0 | 0 |
| ENSG00000229792 | 0 | 0 | 0 | 0 | 0 | 0 |
| ENSG00000285534 | 0 | 0 | 0 | 0 | 0 | 0 |
| ENSG00000234854 | 0 | 0 | 0 | 0 | 0 | 0 |
| ENSG00000185950 | 219 | 102 | 150 | 109 | 132 | 152 |
| ENSG00000275741 | 0 | 0 | 0 | 0 | 0 | 0 |
| ENSG00000275248 | 0 | 0 | 1 | 0 | 0 | 0 |
| ENSG00000201161 | 0 | 0 | 0 | 0 | 0 | 0 |
| ENSG00000274331 | 0 | 0 | 0 | 0 | 0 | 0 |
| ENSG00000275830 | 0 | 0 | 0 | 0 | 0 | 0 |
| ENSG00000276269 | 0 | 0 | 0 | 0 | 0 | 0 |
| ENSG00000231428 | 0 | 0 | 0 | 0 | 0 | 0 |
| ENSG00000265885 | 0 | 0 | 0 | 0 | 0 | 0 |
| ENSG00000279965 | 0 | 0 | 0 | 0 | 0 | 0 |

| | | | | | | |
|---|---|---|---|---|---|---|
| ENSG00000187498 | 0 | 0 | 0 | 1 | 3 | 0 |
| ENSG00000279237 | 0 | 0 | 0 | 0 | 0 | 0 |
| ENSG00000134871 | 31 | 15 | 12 | 19 | 20 | 13 |
| ENSG00000276765 | 0 | 0 | 0 | 0 | 0 | 0 |
| ENSG00000224821 | 0 | 0 | 0 | 0 | 0 | 0 |
| ENSG00000232814 | 1 | 0 | 0 | 0 | 0 | 1 |
| ENSG00000139832 | 681 | 305 | 374 | 304 | 295 | 314 |
| ENSG00000275880 | 78 | 51 | 45 | 44 | 76 | 71 |
| ENSG00000213995 | 2237 | 1100 | 1337 | 1182 | 1093 | 1182 |
| ENSG00000134905 | 643 | 287 | 338 | 332 | 327 | 357 |
| ENSG00000153487 | 414 | 224 | 219 | 203 | 220 | 250 |
| ENSG00000225239 | 0 | 0 | 0 | 0 | 0 | 0 |
| ENSG00000259831 | 0 | 0 | 0 | 0 | 0 | 0 |
| ENSG00000255874 | 112 | 39 | 48 | 36 | 58 | 51 |
| ENSG00000088448 | 1009 | 502 | 580 | 424 | 618 | 495 |
| ENSG00000229152 | 55 | 28 | 35 | 16 | 39 | 21 |
| ENSG00000277767 | 0 | 1 | 5 | 2 | 0 | 0 |
| ENSG00000227105 | 0 | 0 | 0 | 0 | 0 | 0 |
| ENSG00000275989 | 0 | 0 | 0 | 0 | 0 | 0 |
| ENSG00000225760 | 0 | 0 | 0 | 0 | 0 | 0 |
| ENSG00000225870 | 0 | 0 | 0 | 0 | 0 | 0 |
| ENSG00000235875 | 0 | 0 | 0 | 0 | 0 | 0 |
| ENSG00000102606 | 1188 | 532 | 732 | 533 | 659 | 676 |
| ENSG00000233644 | 0 | 0 | 0 | 0 | 0 | 0 |
| ENSG00000227352 | 0 | 0 | 0 | 0 | 0 | 0 |
| ENSG00000285856 | 2 | 2 | 4 | 0 | 0 | 1 |
| ENSG00000274614 | 0 | 0 | 0 | 0 | 0 | 0 |
| ENSG00000153495 | 2 | 2 | 1 | 0 | 0 | 1 |
| ENSG00000204398 | 0 | 0 | 0 | 0 | 0 | 0 |
| ENSG00000215881 | 0 | 0 | 0 | 0 | 0 | 0 |
| ENSG00000226903 | 0 | 0 | 0 | 0 | 0 | 0 |
| ENSG00000200072 | 0 | 0 | 0 | 0 | 0 | 0 |
| ENSG00000182968 | 0 | 0 | 0 | 0 | 0 | 0 |
| ENSG00000260385 | 0 | 0 | 0 | 0 | 0 | 0 |
| ENSG00000229520 | 0 | 0 | 0 | 0 | 0 | 0 |
| ENSG00000224243 | 0 | 0 | 0 | 0 | 0 | 0 |
| ENSG00000260102 | 0 | 0 | 0 | 0 | 0 | 2 |
| ENSG00000260343 | 0 | 0 | 0 | 0 | 0 | 0 |
| ENSG00000223626 | 0 | 0 | 0 | 0 | 0 | 0 |
| ENSG00000153498 | 0 | 0 | 0 | 0 | 0 | 0 |
| ENSG00000126216 | 723 | 355 | 447 | 268 | 361 | 394 |
| ENSG00000277159 | 82 | 38 | 62 | 12 | 18 | 23 |
| ENSG00000197595 | 0 | 0 | 0 | 0 | 0 | 0 |
| ENSG00000274922 | 22 | 5 | 9 | 19 | 17 | 16 |
| ENSG00000068650 | 1295 | 564 | 836 | 529 | 633 | 748 |
| ENSG00000232684 | 0 | 0 | 0 | 0 | 0 | 0 |
| ENSG00000277545 | 0 | 0 | 0 | 0 | 0 | 0 |
| ENSG00000126217 | 2136 | 913 | 1261 | 883 | 920 | 953 |
| ENSG00000267868 | 54 | 25 | 49 | 38 | 30 | 35 |
| ENSG00000235280 | 34 | 12 | 20 | 13 | 21 | 33 |

| | | | | | | |
|---|---|---|---|---|---|---|
| ENSG00000269356 | 24 | 10 | 19 | 18 | 17 | 12 |
| ENSG00000269376 | 19 | 13 | 14 | 12 | 12 | 22 |
| ENSG00000057593 | 9 | 6 | 6 | 8 | 4 | 6 |
| ENSG00000126218 | 8 | 8 | 8 | 3 | 3 | 2 |
| ENSG00000231882 | 0 | 0 | 0 | 2 | 0 | 0 |
| ENSG00000230371 | 0 | 0 | 0 | 1 | 1 | 0 |
| ENSG00000283828 | 14 | 7 | 14 | 11 | 7 | 8 |
| ENSG00000126231 | 17 | 8 | 5 | 3 | 13 | 12 |
| ENSG00000269125 | 4 | 1 | 1 | 1 | 1 | 1 |
| ENSG00000126226 | 894 | 447 | 630 | 474 | 588 | 580 |
| ENSG00000139842 | 2151 | 1100 | 1410 | 997 | 1269 | 1206 |
| ENSG00000277942 | 0 | 0 | 0 | 0 | 0 | 0 |
| ENSG00000232662 | 0 | 0 | 0 | 1 | 0 | 0 |
| ENSG00000185896 | 5488 | 2739 | 3321 | 3117 | 3020 | 2939 |
| ENSG00000139835 | 887 | 390 | 520 | 386 | 399 | 456 |
| ENSG00000276434 | 9 | 10 | 4 | 1 | 5 | 1 |
| ENSG00000225083 | 4 | 4 | 1 | 1 | 2 | 1 |
| ENSG00000153531 | 527 | 272 | 363 | 272 | 296 | 286 |
| ENSG00000150401 | 404 | 194 | 277 | 137 | 216 | 253 |
| ENSG00000233613 | 1 | 2 | 4 | 0 | 0 | 1 |
| ENSG00000202347 | 0 | 1 | 0 | 1 | 1 | 0 |
| ENSG00000150403 | 2186 | 955 | 1296 | 1057 | 1172 | 1180 |
| ENSG00000276916 | 5 | 4 | 1 | 1 | 2 | 0 |
| ENSG00000276248 | 37 | 20 | 5 | 9 | 15 | 7 |
| ENSG00000198176 | 2480 | 1133 | 1456 | 1143 | 1245 | 1448 |
| ENSG00000186009 | 0 | 0 | 0 | 2 | 0 | 1 |
| ENSG00000185974 | 2 | 0 | 1 | 0 | 0 | 0 |
| ENSG00000279770 | 2 | 0 | 0 | 0 | 0 | 0 |
| ENSG00000184497 | 14 | 7 | 12 | 19 | 11 | 12 |
| ENSG00000233695 | 0 | 1 | 0 | 1 | 1 | 1 |
| ENSG00000183087 | 149 | 85 | 110 | 92 | 104 | 86 |
| ENSG00000272695 | 6 | 5 | 0 | 4 | 3 | 3 |
| ENSG00000237699 | 0 | 0 | 0 | 0 | 0 | 0 |
| ENSG00000226921 | 0 | 0 | 0 | 0 | 0 | 0 |
| ENSG00000274766 | 0 | 0 | 0 | 0 | 0 | 0 |
| ENSG00000229373 | 0 | 0 | 0 | 0 | 0 | 0 |
| ENSG00000260910 | 0 | 0 | 3 | 0 | 7 | 2 |
| ENSG00000283199 | 2 | 1 | 0 | 2 | 0 | 1 |
| ENSG00000185989 | 315 | 168 | 215 | 249 | 345 | 280 |
| ENSG00000232487 | 0 | 2 | 0 | 0 | 0 | 0 |
| ENSG00000283361 | 0 | 0 | 0 | 0 | 0 | 0 |
| ENSG00000283347 | 0 | 0 | 0 | 0 | 0 | 0 |
| ENSG00000130177 | 1398 | 660 | 841 | 611 | 831 | 773 |
| ENSG00000264539 | 1 | 0 | 0 | 0 | 0 | 0 |
| ENSG00000265450 | 0 | 0 | 0 | 0 | 0 | 0 |
| ENSG00000285672 | 1 | 1 | 2 | 2 | 2 | 0 |
| ENSG00000169062 | 974 | 475 | 680 | 415 | 588 | 558 |
| ENSG00000226158 | 0 | 0 | 0 | 0 | 0 | 0 |
| ENSG00000198824 | 493 | 247 | 282 | 183 | 245 | 264 |
| ENSG00000229723 | 0 | 0 | 1 | 0 | 0 | 0 |

| | | | | | | |
|---|---|---|---|---|---|---|
| ENSG00000260615 | 0 | 0 | 1 | 2 | 1 | 0 |
| ENSG00000283274 | 0 | 0 | 0 | 0 | 0 | 0 |
| ENSG00000215398 | 0 | 0 | 0 | 1 | 1 | 2 |
| ENSG00000206906 | 0 | 0 | 0 | 0 | 0 | 0 |
| ENSG00000257171 | 0 | 0 | 0 | 0 | 0 | 0 |
| ENSG00000257356 | 0 | 0 | 0 | 0 | 0 | 0 |
| ENSG00000257959 | 0 | 0 | 0 | 0 | 0 | 0 |
| ENSG00000257721 | 0 | 0 | 0 | 0 | 0 | 0 |
| ENSG00000258367 | 0 | 0 | 0 | 0 | 0 | 0 |
| ENSG00000257825 | 0 | 0 | 0 | 0 | 0 | 0 |
| ENSG00000257115 | 0 | 0 | 0 | 0 | 0 | 0 |
| ENSG00000257672 | 0 | 0 | 0 | 0 | 0 | 0 |
| ENSG00000258233 | 0 | 0 | 0 | 0 | 0 | 0 |
| ENSG00000257504 | 0 | 0 | 0 | 0 | 0 | 0 |
| ENSG00000257644 | 0 | 0 | 0 | 0 | 0 | 0 |
| ENSG00000257175 | 2 | 0 | 0 | 0 | 0 | 0 |
| ENSG00000258364 | 0 | 0 | 0 | 0 | 0 | 0 |
| ENSG00000257731 | 0 | 0 | 1 | 0 | 0 | 0 |
| ENSG00000258076 | 0 | 0 | 0 | 0 | 0 | 0 |
| ENSG00000257558 | 0 | 0 | 0 | 0 | 0 | 0 |
| ENSG00000239200 | 0 | 0 | 0 | 0 | 0 | 0 |
| ENSG00000257635 | 0 | 0 | 0 | 0 | 0 | 0 |
| ENSG00000257224 | 0 | 0 | 0 | 0 | 0 | 0 |
| ENSG00000277529 | 0 | 0 | 0 | 0 | 0 | 0 |
| ENSG00000277367 | 0 | 0 | 0 | 0 | 0 | 0 |
| ENSG00000278143 | 0 | 0 | 0 | 0 | 0 | 0 |
| ENSG00000257853 | 0 | 0 | 0 | 0 | 0 | 0 |
| ENSG00000257891 | 0 | 0 | 0 | 0 | 0 | 0 |
| ENSG00000258265 | 0 | 0 | 0 | 0 | 0 | 0 |
| ENSG00000222036 | 0 | 0 | 0 | 0 | 0 | 0 |
| ENSG00000275563 | 0 | 0 | 0 | 0 | 0 | 0 |
| ENSG00000212612 | 0 | 0 | 0 | 0 | 0 | 0 |
| ENSG00000258314 | 2 | 0 | 1 | 3 | 0 | 2 |
| ENSG00000274369 | 0 | 0 | 0 | 0 | 0 | 0 |
| ENSG00000225210 | 43 | 22 | 44 | 25 | 47 | 41 |
| ENSG00000277128 | 13 | 2 | 7 | 1 | 1 | 4 |
| ENSG00000277710 | 0 | 0 | 0 | 0 | 0 | 0 |
| ENSG00000278594 | 22 | 6 | 5 | 9 | 10 | 11 |
| ENSG00000196143 | 0 | 0 | 0 | 0 | 0 | 0 |
| ENSG00000271632 | 0 | 0 | 0 | 0 | 0 | 0 |
| ENSG00000277252 | 0 | 0 | 0 | 0 | 0 | 0 |
| ENSG00000258781 | 0 | 0 | 0 | 0 | 0 | 0 |
| ENSG00000285990 | 0 | 0 | 0 | 0 | 0 | 0 |
| ENSG00000257310 | 0 | 0 | 0 | 0 | 0 | 0 |
| ENSG00000277156 | 4 | 0 | 6 | 6 | 2 | 2 |
| ENSG00000244306 | 359 | 177 | 176 | 205 | 249 | 225 |
| ENSG00000228294 | 0 | 1 | 0 | 1 | 0 | 1 |
| ENSG00000274827 | 0 | 0 | 0 | 0 | 0 | 0 |
| ENSG00000278301 | 0 | 0 | 0 | 0 | 0 | 0 |
| ENSG00000187537 | 0 | 0 | 0 | 0 | 0 | 0 |

| | | | | | | |
|---|---|---|---|---|---|---|
| ENSG00000276888 | 0 | 0 | 0 | 0 | 0 | 0 |
| ENSG00000212516 | 0 | 0 | 0 | 0 | 0 | 0 |
| ENSG00000257162 | 0 | 0 | 0 | 0 | 0 | 0 |
| ENSG00000258027 | 0 | 0 | 0 | 0 | 0 | 0 |
| ENSG00000273639 | 0 | 0 | 0 | 0 | 0 | 0 |
| ENSG00000274649 | 0 | 0 | 0 | 0 | 0 | 0 |
| ENSG00000278184 | 0 | 0 | 0 | 0 | 0 | 0 |
| ENSG00000257432 | 0 | 0 | 0 | 0 | 0 | 0 |
| ENSG00000273558 | 0 | 0 | 0 | 0 | 0 | 0 |
| ENSG00000258324 | 0 | 0 | 0 | 0 | 0 | 0 |
| ENSG00000257493 | 0 | 0 | 0 | 0 | 0 | 0 |
| ENSG00000257751 | 0 | 0 | 0 | 0 | 0 | 0 |
| ENSG00000257846 | 2 | 0 | 0 | 0 | 0 | 0 |
| ENSG00000259069 | 0 | 0 | 0 | 0 | 0 | 0 |
| ENSG00000257884 | 0 | 0 | 0 | 0 | 0 | 0 |
| ENSG00000257395 | 0 | 0 | 0 | 0 | 0 | 0 |
| ENSG00000258080 | 0 | 0 | 0 | 0 | 0 | 0 |
| ENSG00000257142 | 0 | 0 | 0 | 0 | 0 | 0 |
| ENSG00000258453 | 0 | 0 | 0 | 0 | 0 | 0 |
| ENSG00000176294 | 0 | 0 | 0 | 0 | 0 | 0 |
| ENSG00000258438 | 0 | 0 | 0 | 0 | 0 | 0 |
| ENSG00000182652 | 0 | 0 | 0 | 0 | 0 | 0 |
| ENSG00000176312 | 0 | 0 | 0 | 0 | 0 | 0 |
| ENSG00000176299 | 0 | 0 | 0 | 0 | 0 | 0 |
| ENSG00000258564 | 0 | 0 | 0 | 0 | 0 | 0 |
| ENSG00000228304 | 0 | 0 | 0 | 0 | 0 | 0 |
| ENSG00000176290 | 0 | 0 | 0 | 0 | 0 | 0 |
| ENSG00000165762 | 0 | 0 | 0 | 0 | 0 | 0 |
| ENSG00000258541 | 0 | 0 | 0 | 0 | 0 | 0 |
| ENSG00000259016 | 0 | 0 | 0 | 0 | 0 | 0 |
| ENSG00000176281 | 0 | 0 | 0 | 0 | 0 | 0 |
| ENSG00000155249 | 0 | 0 | 0 | 0 | 0 | 0 |
| ENSG00000258822 | 0 | 0 | 0 | 0 | 0 | 0 |
| ENSG00000169488 | 0 | 0 | 0 | 0 | 0 | 0 |
| ENSG00000196383 | 0 | 0 | 0 | 0 | 0 | 0 |
| ENSG00000169484 | 0 | 0 | 0 | 0 | 0 | 0 |
| ENSG00000176253 | 0 | 0 | 0 | 0 | 0 | 0 |
| ENSG00000258899 | 0 | 0 | 0 | 0 | 0 | 0 |
| ENSG00000176246 | 0 | 0 | 0 | 0 | 0 | 0 |
| ENSG00000252999 | 0 | 0 | 0 | 0 | 0 | 0 |
| ENSG00000258641 | 0 | 0 | 0 | 0 | 0 | 0 |
| ENSG00000176230 | 0 | 0 | 0 | 0 | 0 | 0 |
| ENSG00000184394 | 0 | 0 | 0 | 0 | 0 | 0 |
| ENSG00000259074 | 0 | 0 | 0 | 0 | 0 | 0 |
| ENSG00000259095 | 0 | 0 | 0 | 0 | 0 | 0 |
| ENSG00000258468 | 0 | 0 | 0 | 0 | 0 | 0 |
| ENSG00000284742 | 0 | 0 | 0 | 0 | 0 | 0 |
| ENSG00000196832 | 0 | 0 | 0 | 0 | 0 | 0 |
| ENSG00000258625 | 0 | 0 | 0 | 0 | 0 | 0 |
| ENSG00000176219 | 0 | 0 | 0 | 0 | 0 | 0 |

| | | | | | | |
|---|---|---|---|---|---|---|
| ENSG00000258806 | 2 | 0 | 0 | 1 | 0 | 0 |
| ENSG00000176198 | 0 | 0 | 0 | 0 | 0 | 0 |
| ENSG00000136319 | 1116 | 573 | 638 | 574 | 622 | 574 |
| ENSG00000258768 | 40 | 9 | 9 | 4 | 12 | 11 |
| ENSG00000258459 | 0 | 0 | 0 | 0 | 0 | 0 |
| ENSG00000100814 | 1678 | 823 | 905 | 1101 | 1103 | 1119 |
| ENSG00000222489 | 2 | 2 | 2 | 0 | 1 | 0 |
| ENSG00000238344 | 0 | 0 | 0 | 0 | 0 | 0 |
| ENSG00000259001 | 0 | 0 | 0 | 0 | 0 | 0 |
| ENSG00000277209 | 0 | 0 | 0 | 0 | 0 | 0 |
| ENSG00000129484 | 911 | 507 | 592 | 416 | 586 | 514 |
| ENSG00000254846 | 0 | 1 | 0 | 0 | 1 | 0 |
| ENSG00000129566 | 1573 | 673 | 871 | 618 | 842 | 709 |
| ENSG00000200225 | 0 | 0 | 0 | 0 | 0 | 1 |
| ENSG00000185271 | 0 | 0 | 1 | 0 | 0 | 0 |
| ENSG00000259162 | 0 | 0 | 0 | 0 | 0 | 0 |
| ENSG00000092094 | 1064 | 463 | 738 | 459 | 502 | 560 |
| ENSG00000258515 | 32 | 9 | 19 | 13 | 12 | 10 |
| ENSG00000100823 | 9368 | 4849 | 5915 | 5780 | 5614 | 5673 |
| ENSG00000165782 | 1870 | 940 | 1273 | 1133 | 1139 | 1139 |
| ENSG00000198805 | 1326 | 872 | 882 | 1007 | 1069 | 1020 |
| ENSG00000258908 | 0 | 0 | 0 | 0 | 0 | 0 |
| ENSG00000182545 | 1 | 0 | 0 | 1 | 0 | 1 |
| ENSG00000258456 | 0 | 0 | 0 | 0 | 0 | 0 |
| ENSG00000258863 | 0 | 0 | 0 | 0 | 0 | 0 |
| ENSG00000188655 | 0 | 0 | 0 | 0 | 0 | 0 |
| ENSG00000259060 | 0 | 0 | 0 | 0 | 0 | 0 |
| ENSG00000173464 | 0 | 0 | 0 | 0 | 0 | 0 |
| ENSG00000258573 | 0 | 0 | 0 | 0 | 0 | 0 |
| ENSG00000258436 | 0 | 0 | 0 | 0 | 0 | 0 |
| ENSG00000239327 | 0 | 0 | 0 | 0 | 0 | 0 |
| ENSG00000181803 | 0 | 0 | 0 | 0 | 0 | 0 |
| ENSG00000199461 | 0 | 0 | 0 | 0 | 0 | 0 |
| ENSG00000258818 | 0 | 0 | 0 | 0 | 0 | 0 |
| ENSG00000214274 | 14 | 15 | 12 | 11 | 11 | 8 |
| ENSG00000259171 | 0 | 0 | 0 | 0 | 0 | 0 |
| ENSG00000258451 | 0 | 1 | 1 | 0 | 0 | 0 |
| ENSG00000259144 | 0 | 0 | 0 | 0 | 0 | 0 |
| ENSG00000258963 | 0 | 0 | 0 | 0 | 0 | 0 |
| ENSG00000181562 | 0 | 0 | 0 | 0 | 0 | 0 |
| ENSG00000181552 | 0 | 0 | 1 | 0 | 0 | 0 |
| ENSG00000169413 | 0 | 0 | 0 | 0 | 0 | 0 |
| ENSG00000258810 | 0 | 0 | 0 | 0 | 0 | 0 |
| ENSG00000129538 | 4 | 6 | 0 | 0 | 1 | 2 |
| ENSG00000259130 | 0 | 0 | 0 | 0 | 0 | 0 |
| ENSG00000258642 | 0 | 0 | 0 | 0 | 0 | 0 |
| ENSG00000169397 | 0 | 0 | 0 | 0 | 0 | 0 |
| ENSG00000258918 | 0 | 1 | 0 | 0 | 0 | 0 |
| ENSG00000258772 | 0 | 0 | 0 | 0 | 0 | 0 |
| ENSG00000136315 | 0 | 0 | 0 | 0 | 0 | 0 |

| | | | | | | |
|---|---|---|---|---|---|---|
| ENSG00000169385 | 0 | 0 | 0 | 0 | 0 | 0 |
| ENSG00000243817 | 0 | 0 | 0 | 0 | 0 | 0 |
| ENSG00000258599 | 1 | 1 | 0 | 0 | 0 | 0 |
| ENSG00000165792 | 1951 | 938 | 1313 | 1060 | 1113 | 1201 |
| ENSG00000258471 | 1 | 3 | 3 | 0 | 0 | 1 |
| ENSG00000165794 | 0 | 6 | 3 | 2 | 5 | 4 |
| ENSG00000165795 | 566 | 298 | 343 | 280 | 297 | 255 |
| ENSG00000284122 | 0 | 0 | 0 | 0 | 0 | 0 |
| ENSG00000179636 | 2 | 0 | 0 | 0 | 0 | 0 |
| ENSG00000258604 | 0 | 0 | 0 | 0 | 0 | 0 |
| ENSG00000255472 | 0 | 0 | 0 | 0 | 0 | 0 |
| ENSG00000206150 | 0 | 0 | 0 | 0 | 0 | 0 |
| ENSG00000165799 | 0 | 0 | 0 | 0 | 0 | 0 |
| ENSG00000173431 | 0 | 0 | 0 | 0 | 0 | 0 |
| ENSG00000165801 | 241 | 84 | 111 | 108 | 97 | 91 |
| ENSG00000165804 | 1261 | 561 | 835 | 846 | 757 | 789 |
| ENSG00000232070 | 4 | 0 | 2 | 3 | 0 | 4 |
| ENSG00000178107 | 0 | 0 | 0 | 0 | 0 | 1 |
| ENSG00000258722 | 0 | 0 | 1 | 1 | 0 | 0 |
| ENSG00000200102 | 0 | 0 | 0 | 0 | 0 | 0 |
| ENSG00000169327 | 0 | 0 | 0 | 0 | 0 | 0 |
| ENSG00000259102 | 4 | 0 | 1 | 0 | 0 | 0 |
| ENSG00000258906 | 0 | 0 | 0 | 0 | 0 | 0 |
| ENSG00000258441 | 168 | 83 | 110 | 27 | 109 | 109 |
| ENSG00000092199 | 25704 | 12567 | 15162 | 13037 | 14122 | 13819 |
| ENSG00000092200 | 54 | 27 | 28 | 19 | 32 | 17 |
| ENSG00000092201 | 6411 | 3696 | 4066 | 2466 | 4549 | 3751 |
| ENSG00000260830 | 12 | 3 | 4 | 0 | 2 | 5 |
| ENSG00000100888 | 4579 | 1945 | 2453 | 1419 | 2301 | 2263 |
| ENSG00000199436 | 2 | 3 | 0 | 1 | 0 | 3 |
| ENSG00000200785 | 0 | 0 | 0 | 0 | 0 | 0 |
| ENSG00000241035 | 0 | 0 | 0 | 0 | 0 | 0 |
| ENSG00000258870 | 0 | 3 | 2 | 0 | 0 | 0 |
| ENSG00000274475 | 0 | 0 | 0 | 0 | 0 | 0 |
| ENSG00000129472 | 927 | 429 | 541 | 417 | 484 | 508 |
| ENSG00000092203 | 5184 | 2451 | 3113 | 2320 | 2639 | 2726 |
| ENSG00000165819 | 1834 | 868 | 1213 | 880 | 1117 | 1131 |
| ENSG00000165821 | 414 | 192 | 211 | 152 | 175 | 185 |
| ENSG00000257096 | 0 | 0 | 0 | 0 | 0 | 0 |
| ENSG00000256737 | 0 | 0 | 0 | 0 | 1 | 0 |
| ENSG00000169208 | 0 | 0 | 0 | 0 | 0 | 0 |
| ENSG00000256081 | 0 | 0 | 1 | 0 | 0 | 0 |
| ENSG00000273586 | 0 | 0 | 0 | 0 | 0 | 0 |
| ENSG00000256385 | 0 | 0 | 0 | 1 | 0 | 0 |
| ENSG00000255569 | 0 | 0 | 0 | 0 | 0 | 0 |
| ENSG00000255582 | 0 | 0 | 0 | 0 | 0 | 0 |
| ENSG00000256553 | 0 | 0 | 0 | 0 | 0 | 0 |
| ENSG00000255664 | 0 | 0 | 0 | 0 | 0 | 0 |
| ENSG00000221977 | 0 | 0 | 0 | 0 | 0 | 0 |
| ENSG00000276240 | 0 | 0 | 0 | 0 | 0 | 0 |

| | | | | | | |
|---|---|---|---|---|---|---|
| ENSG00000256350 | 0 | 0 | 0 | 0 | 0 | 0 |
| ENSG00000211776 | 0 | 0 | 0 | 0 | 0 | 0 |
| ENSG00000248358 | 0 | 0 | 0 | 0 | 0 | 0 |
| ENSG00000211777 | 0 | 0 | 0 | 0 | 0 | 0 |
| ENSG00000211778 | 0 | 0 | 0 | 0 | 0 | 0 |
| ENSG00000211779 | 0 | 0 | 0 | 0 | 0 | 0 |
| ENSG00000240954 | 0 | 0 | 0 | 0 | 0 | 0 |
| ENSG00000211780 | 0 | 0 | 0 | 0 | 0 | 0 |
| ENSG00000211781 | 0 | 0 | 0 | 0 | 0 | 0 |
| ENSG00000211782 | 0 | 0 | 0 | 0 | 0 | 0 |
| ENSG00000211783 | 0 | 0 | 0 | 0 | 0 | 0 |
| ENSG00000211784 | 0 | 0 | 0 | 0 | 0 | 0 |
| ENSG00000256474 | 0 | 0 | 0 | 0 | 0 | 0 |
| ENSG00000211785 | 0 | 0 | 0 | 0 | 0 | 0 |
| ENSG00000211786 | 0 | 0 | 0 | 0 | 0 | 0 |
| ENSG00000211787 | 0 | 0 | 0 | 0 | 0 | 0 |
| ENSG00000211788 | 0 | 0 | 0 | 0 | 0 | 0 |
| ENSG00000211789 | 0 | 0 | 0 | 0 | 0 | 0 |
| ENSG00000211790 | 0 | 0 | 0 | 0 | 0 | 0 |
| ENSG00000256379 | 0 | 0 | 0 | 0 | 0 | 0 |
| ENSG00000211791 | 0 | 0 | 0 | 0 | 0 | 0 |
| ENSG00000211792 | 0 | 0 | 0 | 0 | 0 | 0 |
| ENSG00000211793 | 0 | 0 | 0 | 0 | 0 | 0 |
| ENSG00000258482 | 0 | 0 | 0 | 0 | 0 | 0 |
| ENSG00000258557 | 0 | 0 | 0 | 0 | 0 | 0 |
| ENSG00000211794 | 0 | 0 | 0 | 0 | 0 | 0 |
| ENSG00000211795 | 0 | 0 | 0 | 0 | 0 | 0 |
| ENSG00000211796 | 0 | 0 | 0 | 0 | 0 | 0 |
| ENSG00000211797 | 0 | 0 | 0 | 0 | 0 | 0 |
| ENSG00000211798 | 0 | 0 | 0 | 1 | 0 | 0 |
| ENSG00000211799 | 0 | 0 | 0 | 0 | 0 | 0 |
| ENSG00000258705 | 0 | 0 | 0 | 0 | 0 | 0 |
| ENSG00000211800 | 0 | 0 | 0 | 0 | 0 | 0 |
| ENSG00000211801 | 0 | 0 | 0 | 0 | 0 | 0 |
| ENSG00000276428 | 0 | 0 | 0 | 0 | 0 | 0 |
| ENSG00000211802 | 0 | 0 | 0 | 0 | 0 | 0 |
| ENSG00000211803 | 0 | 0 | 0 | 0 | 0 | 0 |
| ENSG00000211804 | 0 | 0 | 0 | 0 | 0 | 0 |
| ENSG00000211805 | 0 | 0 | 0 | 0 | 0 | 0 |
| ENSG00000211806 | 1 | 0 | 0 | 0 | 0 | 0 |
| ENSG00000211807 | 0 | 0 | 0 | 0 | 0 | 0 |
| ENSG00000211808 | 0 | 0 | 0 | 0 | 0 | 0 |
| ENSG00000211809 | 0 | 1 | 0 | 0 | 0 | 0 |
| ENSG00000258835 | 0 | 0 | 0 | 0 | 0 | 0 |
| ENSG00000211810 | 2 | 4 | 3 | 3 | 2 | 1 |
| ENSG00000259092 | 16 | 9 | 9 | 4 | 3 | 6 |
| ENSG00000249048 | 0 | 0 | 0 | 0 | 0 | 0 |
| ENSG00000258905 | 4 | 0 | 0 | 0 | 0 | 0 |
| ENSG00000258812 | 0 | 0 | 0 | 0 | 0 | 0 |
| ENSG00000211812 | 0 | 0 | 0 | 0 | 0 | 0 |

| | | | | | | |
|---|---|---|---|---|---|---|
| ENSG00000211813 | 2 | 0 | 2 | 0 | 1 | 0 |
| ENSG00000211814 | 0 | 0 | 0 | 2 | 0 | 0 |
| ENSG00000211815 | 6 | 1 | 2 | 1 | 2 | 4 |
| ENSG00000259068 | 0 | 0 | 1 | 0 | 0 | 0 |
| ENSG00000211816 | 10 | 3 | 3 | 1 | 1 | 1 |
| ENSG00000211817 | 0 | 2 | 1 | 0 | 0 | 0 |
| ENSG00000211818 | 8 | 7 | 27 | 4 | 2 | 9 |
| ENSG00000211819 | 0 | 0 | 0 | 0 | 0 | 0 |
| ENSG00000211820 | 5 | 1 | 0 | 1 | 1 | 2 |
| ENSG00000256221 | 0 | 0 | 0 | 0 | 0 | 0 |
| ENSG00000251002 | 0 | 0 | 0 | 0 | 0 | 0 |
| ENSG00000259042 | 0 | 0 | 0 | 0 | 0 | 0 |
| ENSG00000211821 | 0 | 0 | 0 | 0 | 0 | 0 |
| ENSG00000223997 | 0 | 0 | 0 | 0 | 0 | 0 |
| ENSG00000237235 | 0 | 0 | 0 | 0 | 0 | 0 |
| ENSG00000228985 | 0 | 0 | 0 | 0 | 0 | 0 |
| ENSG00000211825 | 0 | 0 | 0 | 0 | 0 | 0 |
| ENSG00000211826 | 0 | 0 | 0 | 0 | 0 | 0 |
| ENSG00000211827 | 0 | 0 | 0 | 0 | 0 | 0 |
| ENSG00000211828 | 0 | 0 | 0 | 0 | 0 | 0 |
| ENSG00000211829 | 0 | 0 | 0 | 0 | 0 | 0 |
| ENSG00000256590 | 0 | 0 | 0 | 0 | 0 | 0 |
| ENSG00000211831 | 0 | 0 | 0 | 0 | 0 | 0 |
| ENSG00000249446 | 0 | 0 | 0 | 0 | 0 | 0 |
| ENSG00000211832 | 0 | 0 | 0 | 0 | 0 | 0 |
| ENSG00000211833 | 0 | 0 | 0 | 0 | 0 | 0 |
| ENSG00000211834 | 0 | 0 | 0 | 0 | 0 | 0 |
| ENSG00000211835 | 0 | 0 | 0 | 0 | 0 | 0 |
| ENSG00000250688 | 0 | 0 | 0 | 0 | 0 | 0 |
| ENSG00000211836 | 0 | 0 | 0 | 0 | 0 | 0 |
| ENSG00000211837 | 0 | 0 | 0 | 0 | 0 | 0 |
| ENSG00000211838 | 0 | 0 | 0 | 0 | 0 | 0 |
| ENSG00000248366 | 0 | 0 | 0 | 0 | 0 | 0 |
| ENSG00000211839 | 0 | 0 | 0 | 0 | 0 | 0 |
| ENSG00000211840 | 0 | 0 | 0 | 0 | 0 | 0 |
| ENSG00000211841 | 0 | 0 | 0 | 0 | 0 | 0 |
| ENSG00000211842 | 0 | 0 | 0 | 0 | 0 | 0 |
| ENSG00000211843 | 0 | 0 | 0 | 0 | 0 | 0 |
| ENSG00000211844 | 0 | 0 | 0 | 0 | 0 | 0 |
| ENSG00000211845 | 0 | 0 | 0 | 0 | 0 | 0 |
| ENSG00000211846 | 0 | 0 | 0 | 0 | 0 | 0 |
| ENSG00000211847 | 0 | 1 | 0 | 0 | 0 | 0 |
| ENSG00000211848 | 0 | 0 | 0 | 0 | 0 | 0 |
| ENSG00000211849 | 0 | 0 | 0 | 0 | 0 | 0 |
| ENSG00000211850 | 0 | 0 | 0 | 0 | 0 | 0 |
| ENSG00000211851 | 0 | 0 | 0 | 0 | 0 | 0 |
| ENSG00000278661 | 0 | 0 | 0 | 0 | 0 | 0 |
| ENSG00000276699 | 0 | 0 | 0 | 0 | 0 | 0 |
| ENSG00000211854 | 0 | 0 | 0 | 0 | 0 | 0 |
| ENSG00000211855 | 0 | 0 | 0 | 0 | 0 | 0 |

| | | | | | | |
|---|---|---|---|---|---|---|
| ENSG00000211856 | 0 | 0 | 0 | 0 | 0 | 0 |
| ENSG00000211857 | 0 | 0 | 0 | 0 | 0 | 0 |
| ENSG00000211858 | 0 | 0 | 0 | 0 | 0 | 0 |
| ENSG00000211859 | 0 | 0 | 0 | 0 | 0 | 0 |
| ENSG00000211860 | 0 | 0 | 0 | 0 | 0 | 0 |
| ENSG00000211861 | 0 | 0 | 0 | 0 | 0 | 0 |
| ENSG00000211862 | 0 | 0 | 0 | 0 | 0 | 0 |
| ENSG00000211863 | 0 | 0 | 0 | 0 | 0 | 0 |
| ENSG00000211864 | 0 | 0 | 0 | 0 | 0 | 0 |
| ENSG00000211865 | 0 | 0 | 0 | 0 | 0 | 0 |
| ENSG00000211866 | 0 | 0 | 0 | 0 | 0 | 0 |
| ENSG00000211867 | 0 | 0 | 0 | 0 | 0 | 0 |
| ENSG00000211868 | 0 | 0 | 0 | 0 | 0 | 0 |
| ENSG00000211869 | 0 | 0 | 0 | 0 | 0 | 0 |
| ENSG00000211870 | 0 | 0 | 0 | 0 | 0 | 0 |
| ENSG00000211871 | 0 | 0 | 0 | 0 | 0 | 0 |
| ENSG00000211872 | 0 | 0 | 0 | 0 | 0 | 0 |
| ENSG00000211873 | 0 | 0 | 0 | 0 | 0 | 0 |
| ENSG00000211875 | 0 | 0 | 0 | 0 | 0 | 0 |
| ENSG00000211876 | 0 | 0 | 0 | 0 | 0 | 0 |
| ENSG00000211877 | 0 | 0 | 0 | 0 | 0 | 0 |
| ENSG00000211878 | 0 | 0 | 0 | 0 | 0 | 0 |
| ENSG00000211879 | 0 | 0 | 0 | 0 | 0 | 0 |
| ENSG00000211880 | 0 | 0 | 0 | 0 | 0 | 0 |
| ENSG00000211881 | 0 | 0 | 0 | 0 | 0 | 0 |
| ENSG00000211882 | 0 | 0 | 0 | 0 | 0 | 0 |
| ENSG00000211883 | 0 | 0 | 0 | 0 | 0 | 0 |
| ENSG00000211884 | 0 | 0 | 0 | 0 | 0 | 0 |
| ENSG00000211885 | 0 | 0 | 0 | 0 | 0 | 0 |
| ENSG00000211886 | 0 | 0 | 0 | 0 | 0 | 0 |
| ENSG00000211887 | 0 | 0 | 0 | 0 | 0 | 0 |
| ENSG00000211888 | 0 | 0 | 0 | 0 | 0 | 0 |
| ENSG00000277734 | 0 | 0 | 0 | 0 | 0 | 0 |
| ENSG00000259054 | 0 | 0 | 0 | 0 | 0 | 0 |
| ENSG00000259003 | 0 | 0 | 0 | 0 | 0 | 0 |
| ENSG00000129562 | 3665 | 1751 | 2069 | 2518 | 2117 | 2066 |
| ENSG00000275552 | 3 | 0 | 6 | 1 | 2 | 3 |
| ENSG00000100439 | 2635 | 1259 | 1612 | 1474 | 1584 | 1614 |
| ENSG00000255804 | 2 | 0 | 0 | 0 | 0 | 0 |
| ENSG00000242061 | 1 | 0 | 0 | 0 | 0 | 0 |
| ENSG00000258458 | 15 | 17 | 15 | 10 | 16 | 8 |
| ENSG00000235213 | 0 | 0 | 0 | 0 | 0 | 0 |
| ENSG00000258823 | 2 | 1 | 3 | 2 | 1 | 1 |
| ENSG00000212302 | 5 | 0 | 0 | 0 | 0 | 0 |
| ENSG00000155463 | 5215 | 2594 | 3107 | 2681 | 3144 | 2943 |
| ENSG00000155465 | 10 | 2 | 10 | 6 | 8 | 12 |
| ENSG00000172590 | 1648 | 782 | 923 | 1165 | 983 | 960 |
| ENSG00000157227 | 33 | 18 | 29 | 22 | 24 | 20 |
| ENSG00000212335 | 0 | 0 | 0 | 0 | 1 | 0 |
| ENSG00000199716 | 0 | 0 | 0 | 0 | 0 | 0 |

| | | | | | | |
|---|---|---|---|---|---|---|
| ENSG00000197324 | 16927 | 7208 | 9862 | 7913 | 7668 | 7810 |
| ENSG00000139890 | 50 | 25 | 42 | 40 | 28 | 41 |
| ENSG00000100461 | 3654 | 1749 | 2207 | 1821 | 1867 | 1939 |
| ENSG00000237054 | 8 | 3 | 6 | 3 | 9 | 4 |
| ENSG00000100462 | 2102 | 1375 | 1783 | 1818 | 1781 | 1974 |
| ENSG00000257285 | 14 | 6 | 21 | 6 | 12 | 11 |
| ENSG00000092036 | 573 | 340 | 485 | 501 | 434 | 468 |
| ENSG00000259132 | 0 | 0 | 0 | 0 | 0 | 0 |
| ENSG00000284118 | 0 | 0 | 0 | 0 | 0 | 0 |
| ENSG00000129474 | 1494 | 875 | 911 | 972 | 1223 | 955 |
| ENSG00000258457 | 36 | 31 | 27 | 25 | 27 | 31 |
| ENSG00000100802 | 567 | 334 | 423 | 373 | 383 | 341 |
| ENSG00000280129 | 29 | 16 | 20 | 10 | 11 | 13 |
| ENSG00000100804 | 10464 | 5158 | 6180 | 7334 | 6571 | 6302 |
| ENSG00000279656 | 20 | 9 | 4 | 9 | 14 | 17 |
| ENSG00000222028 | 0 | 1 | 0 | 0 | 0 | 0 |
| ENSG00000139880 | 1096 | 455 | 635 | 626 | 574 | 581 |
| ENSG00000100813 | 8656 | 3613 | 5156 | 3255 | 4461 | 4557 |
| ENSG00000179933 | 2682 | 1475 | 1711 | 1720 | 1767 | 1707 |
| ENSG00000283654 | 1 | 0 | 0 | 0 | 1 | 1 |
| ENSG00000092067 | 0 | 0 | 0 | 0 | 0 | 0 |
| ENSG00000092068 | 68 | 28 | 58 | 54 | 53 | 23 |
| ENSG00000202229 | 0 | 0 | 0 | 0 | 0 | 0 |
| ENSG00000215277 | 30 | 24 | 14 | 14 | 15 | 17 |
| ENSG00000240131 | 0 | 0 | 0 | 0 | 0 | 0 |
| ENSG00000252347 | 2 | 1 | 0 | 0 | 0 | 0 |
| ENSG00000215271 | 1581 | 767 | 921 | 728 | 768 | 831 |
| ENSG00000270433 | 8 | 0 | 4 | 1 | 4 | 0 |
| ENSG00000235194 | 479 | 197 | 356 | 207 | 262 | 285 |
| ENSG00000129473 | 1418 | 632 | 926 | 625 | 841 | 815 |
| ENSG00000258643 | 12 | 13 | 23 | 16 | 23 | 16 |
| ENSG00000100836 | 4169 | 1939 | 2719 | 2126 | 2186 | 2437 |
| ENSG00000092096 | 289 | 133 | 173 | 136 | 126 | 132 |
| ENSG00000100842 | 8 | 3 | 2 | 1 | 1 | 0 |
| ENSG00000259018 | 0 | 0 | 0 | 0 | 0 | 0 |
| ENSG00000166090 | 0 | 0 | 0 | 0 | 0 | 0 |
| ENSG00000166091 | 0 | 0 | 0 | 0 | 0 | 0 |
| ENSG00000197616 | 0 | 0 | 1 | 0 | 0 | 1 |
| ENSG00000199157 | 0 | 0 | 0 | 0 | 0 | 0 |
| ENSG00000092054 | 0 | 1 | 1 | 3 | 0 | 0 |
| ENSG00000258444 | 0 | 0 | 0 | 0 | 0 | 0 |
| ENSG00000215991 | 0 | 0 | 0 | 0 | 0 | 0 |
| ENSG00000129460 | 1124 | 556 | 682 | 693 | 647 | 630 |
| ENSG00000157306 | 14 | 9 | 19 | 9 | 17 | 16 |
| ENSG00000136367 | 216 | 89 | 108 | 72 | 86 | 82 |
| ENSG00000273618 | 2 | 1 | 1 | 2 | 0 | 1 |
| ENSG00000259431 | 184 | 96 | 132 | 136 | 108 | 121 |
| ENSG00000213983 | 2152 | 969 | 1498 | 1242 | 1236 | 1307 |
| ENSG00000258727 | 23 | 8 | 23 | 8 | 11 | 25 |
| ENSG00000092051 | 0 | 0 | 1 | 0 | 0 | 1 |

| | | | | | | |
|---|---|---|---|---|---|---|
| ENSG00000274002 | 2 | 2 | 2 | 1 | 1 | 2 |
| ENSG00000100867 | 367 | 225 | 223 | 237 | 197 | 227 |
| ENSG00000258464 | 0 | 0 | 0 | 0 | 0 | 0 |
| ENSG00000259043 | 0 | 0 | 0 | 0 | 0 | 0 |
| ENSG00000270618 | 0 | 0 | 0 | 0 | 0 | 0 |
| ENSG00000222931 | 0 | 0 | 0 | 0 | 0 | 0 |
| ENSG00000258887 | 0 | 0 | 0 | 0 | 0 | 1 |
| ENSG00000285727 | 4 | 1 | 0 | 0 | 0 | 0 |
| ENSG00000259334 | 0 | 0 | 0 | 0 | 0 | 0 |
| ENSG00000215256 | 1410 | 762 | 841 | 554 | 856 | 849 |
| ENSG00000157326 | 861 | 501 | 630 | 678 | 610 | 618 |
| ENSG00000187630 | 954 | 469 | 494 | 697 | 558 | 519 |
| ENSG00000285467 | 5 | 1 | 6 | 7 | 2 | 6 |
| ENSG00000225766 | 12 | 4 | 8 | 2 | 4 | 5 |
| ENSG00000186648 | 67 | 24 | 55 | 29 | 48 | 28 |
| ENSG00000100884 | 0 | 0 | 1 | 1 | 0 | 0 |
| ENSG00000129535 | 62 | 41 | 34 | 35 | 47 | 33 |
| ENSG00000100889 | 3059 | 1394 | 1862 | 1986 | 1585 | 1668 |
| ENSG00000100897 | 5381 | 2638 | 3545 | 2887 | 2831 | 2989 |
| ENSG00000259371 | 0 | 0 | 0 | 0 | 0 | 0 |
| ENSG00000139914 | 4 | 3 | 9 | 14 | 10 | 7 |
| ENSG00000092010 | 6216 | 3127 | 3805 | 3202 | 3189 | 3260 |
| ENSG00000100908 | 659 | 372 | 462 | 531 | 405 | 402 |
| ENSG00000259321 | 32 | 15 | 15 | 14 | 17 | 13 |
| ENSG00000100911 | 4216 | 2240 | 2844 | 2631 | 2555 | 2564 |
| ENSG00000283856 | 0 | 0 | 0 | 0 | 0 | 0 |
| ENSG00000092098 | 1039 | 431 | 573 | 498 | 531 | 561 |
| ENSG00000259529 | 15 | 13 | 21 | 17 | 10 | 21 |
| ENSG00000199804 | 5 | 1 | 7 | 1 | 3 | 4 |
| ENSG00000213928 | 84 | 25 | 46 | 37 | 53 | 41 |
| ENSG00000100918 | 71 | 31 | 45 | 37 | 53 | 34 |
| ENSG00000196497 | 41 | 34 | 42 | 35 | 48 | 49 |
| ENSG00000259522 | 1 | 3 | 0 | 1 | 0 | 0 |
| ENSG00000100926 | 326 | 205 | 266 | 219 | 242 | 253 |
| ENSG00000254692 | 0 | 0 | 0 | 0 | 0 | 0 |
| ENSG00000276698 | 44 | 17 | 50 | 44 | 24 | 23 |
| ENSG00000278784 | 119 | 59 | 101 | 54 | 59 | 91 |
| ENSG00000139908 | 66 | 53 | 72 | 43 | 39 | 43 |
| ENSG00000254505 | 26 | 28 | 31 | 19 | 38 | 29 |
| ENSG00000260669 | 12 | 15 | 19 | 16 | 17 | 13 |
| ENSG00000213920 | 9 | 2 | 7 | 2 | 2 | 5 |
| ENSG00000255526 | 0 | 1 | 2 | 0 | 4 | 3 |
| ENSG00000129559 | 2655 | 1835 | 2089 | 2416 | 2216 | 1957 |
| ENSG00000100938 | 4086 | 2004 | 2591 | 2341 | 2366 | 2251 |
| ENSG00000092330 | 2301 | 979 | 1326 | 1063 | 1124 | 1095 |
| ENSG00000092295 | 78 | 37 | 54 | 66 | 51 | 49 |
| ENSG00000100949 | 1585 | 752 | 949 | 1006 | 776 | 857 |
| ENSG00000157379 | 2026 | 913 | 1156 | 1123 | 1053 | 1045 |
| ENSG00000196943 | 1812 | 874 | 1202 | 1007 | 1061 | 1250 |
| ENSG00000136305 | 0 | 0 | 0 | 0 | 0 | 0 |

| | | | | | | |
|---|---|---|---|---|---|---|
| ENSG00000213906 | 143 | 75 | 111 | 50 | 84 | 95 |
| ENSG00000213903 | 916 | 385 | 536 | 391 | 411 | 478 |
| ENSG00000129467 | 7 | 2 | 6 | 3 | 1 | 1 |
| ENSG00000258973 | 0 | 0 | 0 | 0 | 0 | 0 |
| ENSG00000129465 | 0 | 1 | 0 | 0 | 0 | 0 |
| ENSG00000283290 | 0 | 0 | 0 | 0 | 0 | 0 |
| ENSG00000100968 | 161 | 75 | 103 | 90 | 74 | 85 |
| ENSG00000205978 | 349 | 174 | 242 | 143 | 132 | 176 |
| ENSG00000139899 | 250 | 116 | 169 | 113 | 120 | 137 |
| ENSG00000100441 | 4869 | 2230 | 3187 | 2271 | 2736 | 2778 |
| ENSG00000100445 | 223 | 104 | 167 | 132 | 111 | 137 |
| ENSG00000258744 | 17 | 6 | 17 | 3 | 13 | 12 |
| ENSG00000092009 | 0 | 3 | 0 | 0 | 0 | 0 |
| ENSG00000100448 | 0 | 0 | 0 | 0 | 0 | 0 |
| ENSG00000258657 | 0 | 0 | 0 | 0 | 0 | 1 |
| ENSG00000100450 | 0 | 0 | 0 | 0 | 0 | 0 |
| ENSG00000100453 | 4 | 0 | 4 | 0 | 0 | 0 |
| ENSG00000168952 | 4 | 4 | 3 | 6 | 2 | 2 |
| ENSG00000258098 | 0 | 0 | 0 | 0 | 0 | 0 |
| ENSG00000257179 | 1 | 3 | 1 | 0 | 0 | 0 |
| ENSG00000258275 | 0 | 0 | 0 | 0 | 0 | 0 |
| ENSG00000257976 | 0 | 0 | 0 | 0 | 0 | 0 |
| ENSG00000257986 | 0 | 0 | 0 | 0 | 0 | 0 |
| ENSG00000212270 | 0 | 0 | 0 | 0 | 0 | 0 |
| ENSG00000279521 | 0 | 0 | 0 | 0 | 0 | 0 |
| ENSG00000285633 | 0 | 0 | 0 | 0 | 0 | 0 |
| ENSG00000257503 | 0 | 0 | 0 | 0 | 0 | 0 |
| ENSG00000139910 | 158 | 59 | 74 | 71 | 89 | 89 |
| ENSG00000262119 | 0 | 1 | 0 | 0 | 0 | 0 |
| ENSG00000257842 | 65 | 40 | 64 | 49 | 49 | 45 |
| ENSG00000257845 | 0 | 1 | 0 | 0 | 3 | 0 |
| ENSG00000257185 | 7 | 3 | 2 | 0 | 6 | 4 |
| ENSG00000258081 | 0 | 0 | 0 | 0 | 0 | 0 |
| ENSG00000257612 | 0 | 0 | 0 | 0 | 0 | 0 |
| ENSG00000265165 | 0 | 0 | 0 | 0 | 0 | 0 |
| ENSG00000258932 | 0 | 0 | 1 | 0 | 0 | 1 |
| ENSG00000258548 | 0 | 0 | 0 | 0 | 0 | 0 |
| ENSG00000264657 | 0 | 0 | 0 | 0 | 0 | 0 |
| ENSG00000197358 | 2 | 2 | 4 | 1 | 0 | 1 |
| ENSG00000257869 | 0 | 0 | 1 | 0 | 3 | 0 |
| ENSG00000240179 | 0 | 0 | 0 | 0 | 0 | 0 |
| ENSG00000258175 | 0 | 0 | 0 | 0 | 0 | 1 |
| ENSG00000257662 | 0 | 0 | 0 | 0 | 0 | 0 |
| ENSG00000258282 | 0 | 0 | 0 | 1 | 0 | 0 |
| ENSG00000257126 | 0 | 0 | 0 | 0 | 0 | 0 |
| ENSG00000176165 | 0 | 0 | 0 | 0 | 0 | 0 |
| ENSG00000186960 | 0 | 0 | 0 | 0 | 0 | 0 |
| ENSG00000257056 | 0 | 0 | 0 | 0 | 0 | 0 |
| ENSG00000257748 | 0 | 0 | 0 | 0 | 0 | 0 |
| ENSG00000274762 | 0 | 0 | 0 | 0 | 0 | 0 |

| | | | | | | |
|---|---|---|---|---|---|---|
| ENSG00000258038 | 0 | 0 | 0 | 0 | 0 | 0 |
| ENSG00000258065 | 0 | 0 | 0 | 0 | 0 | 0 |
| ENSG00000257523 | 0 | 0 | 0 | 0 | 0 | 0 |
| ENSG00000207164 | 0 | 0 | 0 | 0 | 0 | 0 |
| ENSG00000258028 | 0 | 0 | 0 | 0 | 0 | 0 |
| ENSG00000252065 | 0 | 0 | 0 | 0 | 0 | 0 |
| ENSG00000257522 | 0 | 0 | 0 | 0 | 0 | 0 |
| ENSG00000258107 | 0 | 0 | 0 | 0 | 0 | 0 |
| ENSG00000212518 | 0 | 0 | 0 | 0 | 0 | 0 |
| ENSG00000184304 | 148 | 61 | 67 | 43 | 89 | 50 |
| ENSG00000257120 | 0 | 0 | 0 | 0 | 0 | 0 |
| ENSG00000252686 | 0 | 0 | 1 | 0 | 0 | 0 |
| ENSG00000257904 | 0 | 0 | 0 | 0 | 0 | 0 |
| ENSG00000248975 | 0 | 0 | 0 | 0 | 0 | 0 |
| ENSG00000240201 | 0 | 0 | 0 | 0 | 0 | 0 |
| ENSG00000279032 | 0 | 0 | 0 | 0 | 0 | 0 |
| ENSG00000257636 | 0 | 0 | 0 | 0 | 0 | 0 |
| ENSG00000278587 | 0 | 0 | 0 | 0 | 0 | 0 |
| ENSG00000258350 | 0 | 0 | 0 | 0 | 0 | 0 |
| ENSG00000092140 | 486 | 277 | 277 | 153 | 498 | 309 |
| ENSG00000092108 | 1299 | 803 | 755 | 529 | 1061 | 733 |
| ENSG00000258648 | 4 | 0 | 1 | 0 | 1 | 0 |
| ENSG00000258558 | 2 | 0 | 0 | 0 | 0 | 0 |
| ENSG00000243855 | 0 | 0 | 0 | 0 | 0 | 0 |
| ENSG00000240513 | 0 | 0 | 0 | 0 | 0 | 0 |
| ENSG00000213867 | 2 | 0 | 0 | 0 | 0 | 0 |
| ENSG00000100473 | 63 | 29 | 34 | 25 | 18 | 25 |
| ENSG00000258525 | 0 | 2 | 0 | 2 | 0 | 0 |
| ENSG00000196792 | 1301 | 635 | 773 | 357 | 765 | 538 |
| ENSG00000258886 | 0 | 0 | 0 | 0 | 0 | 0 |
| ENSG00000207952 | 0 | 0 | 0 | 0 | 0 | 0 |
| ENSG00000258968 | 0 | 0 | 1 | 0 | 0 | 0 |
| ENSG00000100478 | 470 | 259 | 295 | 246 | 284 | 275 |
| ENSG00000092148 | 4369 | 2323 | 1926 | 959 | 3462 | 1816 |
| ENSG00000258640 | 0 | 0 | 0 | 0 | 0 | 0 |
| ENSG00000207440 | 0 | 1 | 0 | 0 | 0 | 0 |
| ENSG00000202402 | 0 | 0 | 0 | 0 | 0 | 0 |
| ENSG00000257831 | 2 | 3 | 2 | 0 | 2 | 4 |
| ENSG00000199291 | 2 | 0 | 0 | 0 | 0 | 0 |
| ENSG00000129493 | 670 | 358 | 314 | 110 | 447 | 227 |
| ENSG00000283016 | 2 | 0 | 0 | 1 | 2 | 0 |
| ENSG00000203546 | 14 | 19 | 18 | 14 | 15 | 18 |
| ENSG00000250365 | 9 | 9 | 10 | 4 | 5 | 7 |
| ENSG00000129480 | 1092 | 515 | 585 | 483 | 574 | 546 |
| ENSG00000239323 | 0 | 0 | 0 | 0 | 0 | 0 |
| ENSG00000214943 | 0 | 0 | 0 | 0 | 0 | 0 |
| ENSG00000151413 | 512 | 239 | 297 | 215 | 319 | 288 |
| ENSG00000258196 | 0 | 0 | 0 | 0 | 0 | 0 |
| ENSG00000257155 | 0 | 0 | 0 | 0 | 0 | 0 |
| ENSG00000223087 | 0 | 0 | 0 | 0 | 0 | 0 |

| | | | | | | |
|---|---|---|---|---|---|---|
| ENSG00000207412 | 0 | 0 | 0 | 0 | 0 | 0 |
| ENSG00000257472 | 0 | 0 | 0 | 0 | 0 | 0 |
| ENSG00000258474 | 0 | 0 | 0 | 0 | 0 | 2 |
| ENSG00000275002 | 0 | 0 | 0 | 0 | 0 | 0 |
| ENSG00000258386 | 0 | 0 | 0 | 0 | 2 | 1 |
| ENSG00000258655 | 53 | 21 | 26 | 16 | 31 | 28 |
| ENSG00000100852 | 437 | 270 | 167 | 62 | 437 | 178 |
| ENSG00000285608 | 0 | 0 | 0 | 0 | 0 | 0 |
| ENSG00000201654 | 0 | 0 | 0 | 0 | 1 | 0 |
| ENSG00000202337 | 27 | 26 | 30 | 26 | 27 | 36 |
| ENSG00000258619 | 0 | 0 | 0 | 0 | 0 | 0 |
| ENSG00000259086 | 0 | 0 | 1 | 0 | 0 | 0 |
| ENSG00000151320 | 17 | 7 | 12 | 2 | 16 | 8 |
| ENSG00000258580 | 2 | 0 | 0 | 0 | 0 | 0 |
| ENSG00000278746 | 0 | 0 | 0 | 0 | 0 | 0 |
| ENSG00000259045 | 0 | 0 | 1 | 1 | 0 | 1 |
| ENSG00000280281 | 0 | 0 | 0 | 0 | 0 | 0 |
| ENSG00000151322 | 10 | 4 | 5 | 0 | 2 | 2 |
| ENSG00000129521 | 1088 | 583 | 610 | 545 | 517 | 503 |
| ENSG00000258897 | 0 | 0 | 0 | 0 | 0 | 0 |
| ENSG00000223593 | 2 | 0 | 0 | 0 | 0 | 1 |
| ENSG00000165389 | 1765 | 940 | 1115 | 1323 | 1740 | 1526 |
| ENSG00000237450 | 0 | 0 | 0 | 0 | 0 | 0 |
| ENSG00000129518 | 2002 | 1004 | 1233 | 973 | 1155 | 1024 |
| ENSG00000279423 | 1 | 0 | 0 | 0 | 0 | 0 |
| ENSG00000259002 | 0 | 0 | 0 | 0 | 0 | 0 |
| ENSG00000206596 | 0 | 0 | 0 | 0 | 0 | 0 |
| ENSG00000206588 | 0 | 0 | 0 | 0 | 0 | 0 |
| ENSG00000259135 | 0 | 0 | 0 | 0 | 0 | 0 |
| ENSG00000129515 | 2617 | 1363 | 1436 | 1219 | 1815 | 1349 |
| ENSG00000240463 | 5 | 1 | 4 | 1 | 4 | 7 |
| ENSG00000275978 | 0 | 0 | 0 | 0 | 0 | 0 |
| ENSG00000253003 | 0 | 0 | 0 | 0 | 0 | 0 |
| ENSG00000241582 | 0 | 0 | 0 | 0 | 1 | 0 |
| ENSG00000165410 | 1004 | 520 | 565 | 545 | 773 | 632 |
| ENSG00000243824 | 3 | 0 | 0 | 0 | 1 | 0 |
| ENSG00000198604 | 1564 | 945 | 785 | 430 | 1629 | 899 |
| ENSG00000251726 | 0 | 0 | 0 | 0 | 0 | 0 |
| ENSG00000258738 | 71 | 39 | 56 | 44 | 32 | 48 |
| ENSG00000258704 | 20 | 17 | 20 | 14 | 21 | 16 |
| ENSG00000258493 | 0 | 0 | 0 | 0 | 0 | 0 |
| ENSG00000226677 | 0 | 0 | 2 | 1 | 1 | 0 |
| ENSG00000100883 | 3276 | 1947 | 2152 | 1314 | 2253 | 1893 |
| ENSG00000151327 | 2469 | 1229 | 1549 | 1176 | 1549 | 1448 |
| ENSG00000092020 | 314 | 234 | 234 | 188 | 300 | 247 |
| ENSG00000279434 | 10 | 13 | 10 | 4 | 9 | 3 |
| ENSG00000100890 | 269 | 143 | 143 | 94 | 146 | 173 |
| ENSG00000258790 | 3 | 9 | 6 | 5 | 2 | 0 |
| ENSG00000241052 | 0 | 0 | 0 | 1 | 0 | 0 |
| ENSG00000259090 | 0 | 0 | 0 | 0 | 0 | 1 |

| | | | | | | |
|---|---|---|---|---|---|---|
| ENSG00000282625 | 0 | 0 | 0 | 0 | 0 | 0 |
| ENSG00000282308 | 0 | 1 | 0 | 0 | 0 | 1 |
| ENSG00000256243 | 0 | 1 | 0 | 0 | 0 | 0 |
| ENSG00000240392 | 0 | 0 | 0 | 0 | 0 | 0 |
| ENSG00000100902 | 589 | 525 | 681 | 575 | 719 | 577 |
| ENSG00000258860 | 0 | 0 | 0 | 1 | 0 | 0 |
| ENSG00000240023 | 5 | 4 | 2 | 7 | 3 | 3 |
| ENSG00000100906 | 9051 | 4073 | 4940 | 4811 | 4930 | 4786 |
| ENSG00000259022 | 0 | 2 | 2 | 3 | 0 | 3 |
| ENSG00000259017 | 0 | 0 | 0 | 0 | 0 | 0 |
| ENSG00000259105 | 2 | 0 | 0 | 0 | 0 | 0 |
| ENSG00000258783 | 0 | 0 | 0 | 0 | 1 | 0 |
| ENSG00000168348 | 0 | 0 | 1 | 0 | 1 | 1 |
| ENSG00000174373 | 886 | 476 | 412 | 188 | 692 | 435 |
| ENSG00000253059 | 0 | 1 | 0 | 0 | 0 | 0 |
| ENSG00000257957 | 0 | 0 | 0 | 0 | 0 | 0 |
| ENSG00000258938 | 46 | 16 | 41 | 12 | 17 | 15 |
| ENSG00000258300 | 0 | 0 | 0 | 0 | 0 | 0 |
| ENSG00000100916 | 559 | 339 | 352 | 236 | 376 | 356 |
| ENSG00000257272 | 0 | 0 | 0 | 0 | 0 | 1 |
| ENSG00000257307 | 0 | 1 | 0 | 0 | 0 | 0 |
| ENSG00000258342 | 1 | 0 | 0 | 0 | 0 | 0 |
| ENSG00000257614 | 2 | 0 | 0 | 0 | 0 | 0 |
| ENSG00000257720 | 0 | 0 | 0 | 0 | 0 | 0 |
| ENSG00000257826 | 0 | 0 | 0 | 0 | 0 | 0 |
| ENSG00000257585 | 0 | 0 | 0 | 0 | 0 | 0 |
| ENSG00000259104 | 0 | 0 | 0 | 0 | 0 | 0 |
| ENSG00000258844 | 0 | 0 | 0 | 0 | 0 | 0 |
| ENSG00000252312 | 0 | 0 | 0 | 0 | 0 | 0 |
| ENSG00000151332 | 485 | 245 | 259 | 219 | 317 | 297 |
| ENSG00000283098 | 0 | 1 | 1 | 0 | 2 | 0 |
| ENSG00000188831 | 0 | 0 | 0 | 0 | 0 | 0 |
| ENSG00000238540 | 0 | 0 | 0 | 0 | 0 | 0 |
| ENSG00000257520 | 0 | 0 | 0 | 0 | 0 | 0 |
| ENSG00000229415 | 0 | 0 | 0 | 0 | 0 | 0 |
| ENSG00000136352 | 0 | 0 | 0 | 0 | 0 | 0 |
| ENSG00000253563 | 0 | 0 | 0 | 0 | 0 | 0 |
| ENSG00000258467 | 0 | 0 | 0 | 0 | 0 | 0 |
| ENSG00000243730 | 0 | 0 | 0 | 0 | 0 | 0 |
| ENSG00000136327 | 10 | 9 | 10 | 6 | 11 | 7 |
| ENSG00000201395 | 0 | 0 | 0 | 0 | 0 | 0 |
| ENSG00000258661 | 2 | 0 | 0 | 0 | 0 | 0 |
| ENSG00000198807 | 349 | 170 | 199 | 153 | 169 | 178 |
| ENSG00000183032 | 32 | 17 | 25 | 9 | 18 | 15 |
| ENSG00000258690 | 0 | 0 | 0 | 0 | 0 | 0 |
| ENSG00000258601 | 0 | 0 | 0 | 0 | 0 | 0 |
| ENSG00000266327 | 0 | 0 | 0 | 0 | 0 | 0 |
| ENSG00000258827 | 0 | 0 | 0 | 0 | 0 | 0 |
| ENSG00000258708 | 79 | 37 | 60 | 44 | 50 | 38 |
| ENSG00000252746 | 0 | 0 | 2 | 0 | 0 | 0 |

| | | | | | | |
|---|---|---|---|---|---|---|
| ENSG00000151338 | 140 | 77 | 60 | 32 | 105 | 53 |
| ENSG00000207046 | 0 | 0 | 0 | 0 | 0 | 0 |
| ENSG00000259087 | 11 | 4 | 5 | 1 | 5 | 4 |
| ENSG00000258414 | 40 | 46 | 28 | 15 | 36 | 22 |
| ENSG00000129514 | 17451 | 7772 | 9722 | 7338 | 8103 | 9097 |
| ENSG00000139865 | 396 | 250 | 322 | 246 | 271 | 267 |
| ENSG00000200385 | 2 | 0 | 0 | 0 | 1 | 0 |
| ENSG00000201431 | 0 | 0 | 0 | 0 | 0 | 0 |
| ENSG00000259091 | 0 | 0 | 0 | 0 | 0 | 0 |
| ENSG00000258696 | 0 | 0 | 0 | 0 | 0 | 0 |
| ENSG00000259048 | 0 | 0 | 0 | 0 | 1 | 0 |
| ENSG00000258649 | 0 | 0 | 0 | 0 | 0 | 0 |
| ENSG00000139874 | 0 | 4 | 2 | 1 | 3 | 2 |
| ENSG00000176435 | 0 | 0 | 0 | 0 | 0 | 0 |
| ENSG00000259072 | 0 | 0 | 0 | 0 | 0 | 0 |
| ENSG00000271468 | 0 | 0 | 0 | 0 | 0 | 0 |
| ENSG00000258948 | 0 | 0 | 0 | 0 | 0 | 0 |
| ENSG00000259070 | 10 | 7 | 5 | 5 | 7 | 6 |
| ENSG00000258680 | 0 | 0 | 0 | 0 | 0 | 0 |
| ENSG00000270713 | 0 | 1 | 0 | 0 | 0 | 0 |
| ENSG00000199285 | 0 | 0 | 0 | 0 | 0 | 0 |
| ENSG00000100934 | 3330 | 1712 | 2043 | 1276 | 1974 | 1727 |
| ENSG00000259121 | 0 | 0 | 0 | 0 | 0 | 0 |
| ENSG00000258651 | 0 | 0 | 0 | 0 | 0 | 0 |
| ENSG00000258591 | 0 | 2 | 7 | 1 | 3 | 1 |
| ENSG00000092208 | 476 | 259 | 296 | 290 | 381 | 334 |
| ENSG00000182400 | 572 | 291 | 374 | 196 | 322 | 305 |
| ENSG00000285830 | 13 | 8 | 8 | 9 | 12 | 11 |
| ENSG00000241984 | 3 | 8 | 4 | 1 | 1 | 6 |
| ENSG00000259083 | 3 | 1 | 4 | 3 | 4 | 9 |
| ENSG00000100941 | 3022 | 2074 | 1791 | 798 | 3301 | 1865 |
| ENSG00000258671 | 0 | 0 | 0 | 0 | 0 | 0 |
| ENSG00000270503 | 2 | 3 | 1 | 2 | 1 | 0 |
| ENSG00000150527 | 1531 | 758 | 890 | 561 | 1034 | 862 |
| ENSG00000258940 | 38 | 28 | 35 | 33 | 20 | 28 |
| ENSG00000271424 | 0 | 0 | 1 | 2 | 6 | 3 |
| ENSG00000259100 | 4 | 0 | 0 | 1 | 1 | 2 |
| ENSG00000165355 | 1148 | 503 | 609 | 480 | 630 | 605 |
| ENSG00000258526 | 3 | 2 | 0 | 4 | 7 | 1 |
| ENSG00000258418 | 0 | 1 | 1 | 0 | 0 | 0 |
| ENSG00000258480 | 0 | 0 | 1 | 0 | 0 | 0 |
| ENSG00000251363 | 0 | 0 | 0 | 0 | 0 | 0 |
| ENSG00000258385 | 0 | 0 | 0 | 0 | 0 | 0 |
| ENSG00000279972 | 0 | 0 | 0 | 0 | 0 | 0 |
| ENSG00000258901 | 0 | 0 | 0 | 0 | 0 | 0 |
| ENSG00000258636 | 0 | 0 | 0 | 0 | 0 | 0 |
| ENSG00000251858 | 0 | 0 | 0 | 0 | 0 | 0 |
| ENSG00000165379 | 14 | 11 | 12 | 6 | 8 | 13 |
| ENSG00000258536 | 0 | 0 | 0 | 0 | 0 | 0 |
| ENSG00000259149 | 0 | 0 | 0 | 0 | 0 | 0 |

| | | | | | | |
|---|---|---|---|---|---|---|
| ENSG00000258394 | 0 | 0 | 0 | 0 | 0 | 0 |
| ENSG00000258850 | 0 | 0 | 0 | 0 | 0 | 0 |
| ENSG00000258505 | 0 | 0 | 0 | 0 | 0 | 0 |
| ENSG00000258622 | 0 | 0 | 0 | 0 | 0 | 0 |
| ENSG00000258934 | 0 | 0 | 0 | 0 | 0 | 0 |
| ENSG00000259047 | 0 | 0 | 0 | 0 | 0 | 0 |
| ENSG00000258406 | 0 | 0 | 0 | 0 | 0 | 0 |
| ENSG00000259051 | 0 | 4 | 0 | 1 | 1 | 1 |
| ENSG00000258828 | 0 | 0 | 2 | 1 | 0 | 1 |
| ENSG00000258387 | 1 | 0 | 3 | 1 | 3 | 1 |
| ENSG00000258894 | 0 | 0 | 0 | 0 | 0 | 1 |
| ENSG00000258969 | 2 | 0 | 0 | 0 | 0 | 0 |
| ENSG00000259019 | 0 | 0 | 0 | 0 | 0 | 0 |
| ENSG00000259137 | 0 | 0 | 0 | 0 | 1 | 0 |
| ENSG00000259148 | 0 | 0 | 0 | 0 | 0 | 0 |
| ENSG00000259126 | 1 | 0 | 1 | 0 | 0 | 0 |
| ENSG00000258487 | 0 | 0 | 0 | 0 | 0 | 0 |
| ENSG00000189139 | 4 | 2 | 1 | 2 | 0 | 0 |
| ENSG00000258747 | 2 | 2 | 1 | 1 | 0 | 0 |
| ENSG00000258998 | 3 | 1 | 1 | 0 | 0 | 0 |
| ENSG00000259000 | 0 | 0 | 0 | 0 | 0 | 0 |
| ENSG00000258633 | 1 | 2 | 1 | 0 | 1 | 2 |
| ENSG00000258509 | 0 | 0 | 0 | 0 | 0 | 1 |
| ENSG00000259046 | 0 | 0 | 0 | 0 | 1 | 0 |
| ENSG00000179476 | 328 | 141 | 164 | 134 | 177 | 167 |
| ENSG00000258949 | 4 | 6 | 3 | 2 | 5 | 8 |
| ENSG00000179454 | 569 | 287 | 229 | 92 | 340 | 244 |
| ENSG00000198718 | 809 | 473 | 391 | 196 | 500 | 365 |
| ENSG00000278147 | 2 | 1 | 1 | 0 | 0 | 1 |
| ENSG00000249163 | 0 | 0 | 0 | 0 | 1 | 0 |
| ENSG00000185246 | 1273 | 715 | 759 | 286 | 780 | 604 |
| ENSG00000212615 | 0 | 0 | 1 | 0 | 0 | 0 |
| ENSG00000239043 | 1 | 0 | 0 | 0 | 0 | 2 |
| ENSG00000100442 | 2370 | 1282 | 1457 | 1125 | 1655 | 1372 |
| ENSG00000187790 | 109 | 55 | 65 | 20 | 134 | 78 |
| ENSG00000129534 | 315 | 236 | 173 | 78 | 479 | 223 |
| ENSG00000199739 | 0 | 0 | 0 | 0 | 0 | 0 |
| ENSG00000258608 | 0 | 0 | 0 | 0 | 1 | 0 |
| ENSG00000260046 | 4 | 2 | 6 | 3 | 4 | 3 |
| ENSG00000257900 | 2 | 2 | 2 | 3 | 0 | 1 |
| ENSG00000258746 | 0 | 0 | 0 | 0 | 0 | 0 |
| ENSG00000258845 | 2 | 0 | 0 | 1 | 0 | 0 |
| ENSG00000258616 | 0 | 0 | 0 | 0 | 0 | 0 |
| ENSG00000258700 | 0 | 0 | 0 | 0 | 1 | 0 |
| ENSG00000165496 | 0 | 0 | 0 | 1 | 0 | 0 |
| ENSG00000139915 | 0 | 0 | 1 | 0 | 0 | 1 |
| ENSG00000251735 | 0 | 0 | 0 | 0 | 0 | 0 |
| ENSG00000258392 | 0 | 0 | 0 | 0 | 0 | 0 |
| ENSG00000244053 | 0 | 0 | 0 | 0 | 0 | 0 |
| ENSG00000214903 | 0 | 0 | 0 | 0 | 0 | 0 |

| | | | | | | |
|---|---|---|---|---|---|---|
| ENSG00000263945 | 0 | 0 | 0 | 0 | 0 | 0 |
| ENSG00000259129 | 6 | 2 | 3 | 1 | 1 | 0 |
| ENSG00000259117 | 0 | 0 | 0 | 0 | 0 | 0 |
| ENSG00000258639 | 0 | 0 | 0 | 0 | 0 | 0 |
| ENSG00000270359 | 0 | 0 | 0 | 0 | 0 | 0 |
| ENSG00000207366 | 0 | 0 | 0 | 0 | 0 | 0 |
| ENSG00000258751 | 0 | 0 | 0 | 0 | 0 | 0 |
| ENSG00000258750 | 0 | 0 | 0 | 0 | 0 | 0 |
| ENSG00000242417 | 0 | 0 | 0 | 0 | 0 | 0 |
| ENSG00000258381 | 0 | 0 | 0 | 0 | 0 | 0 |
| ENSG00000258868 | 0 | 0 | 0 | 0 | 1 | 0 |
| ENSG00000258401 | 0 | 0 | 0 | 0 | 0 | 0 |
| ENSG00000252424 | 0 | 0 | 0 | 0 | 0 | 0 |
| ENSG00000213741 | 9889 | 4989 | 5968 | 7340 | 6044 | 5895 |
| ENSG00000244270 | 0 | 0 | 0 | 0 | 0 | 0 |
| ENSG00000283029 | 0 | 0 | 0 | 0 | 0 | 0 |
| ENSG00000276168 | 0 | 0 | 0 | 0 | 0 | 0 |
| ENSG00000252919 | 0 | 0 | 0 | 0 | 0 | 0 |
| ENSG00000165501 | 400 | 238 | 280 | 239 | 242 | 310 |
| ENSG00000258568 | 18 | 12 | 17 | 16 | 12 | 18 |
| ENSG00000278009 | 22 | 0 | 5 | 8 | 4 | 5 |
| ENSG00000165502 | 3433 | 1969 | 2203 | 1994 | 2481 | 2108 |
| ENSG00000168282 | 40 | 33 | 30 | 48 | 35 | 41 |
| ENSG00000258377 | 0 | 0 | 0 | 0 | 0 | 0 |
| ENSG00000165506 | 928 | 450 | 550 | 417 | 567 | 541 |
| ENSG00000258450 | 11 | 11 | 8 | 5 | 7 | 8 |
| ENSG00000100479 | 170 | 101 | 125 | 73 | 127 | 111 |
| ENSG00000259155 | 0 | 1 | 1 | 0 | 0 | 0 |
| ENSG00000258462 | 0 | 0 | 0 | 0 | 0 | 0 |
| ENSG00000221114 | 0 | 0 | 0 | 0 | 0 | 0 |
| ENSG00000197776 | 46 | 19 | 27 | 8 | 11 | 10 |
| ENSG00000258400 | 0 | 0 | 0 | 0 | 0 | 0 |
| ENSG00000165516 | 2584 | 1287 | 1652 | 1085 | 1286 | 1316 |
| ENSG00000165525 | 300 | 191 | 139 | 63 | 452 | 203 |
| ENSG00000277634 | 0 | 0 | 0 | 0 | 0 | 0 |
| ENSG00000252474 | 0 | 0 | 0 | 0 | 0 | 0 |
| ENSG00000278771 | 51 | 136 | 18 | 16 | 11 | 18 |
| ENSG00000282885 | 7 | 2 | 2 | 4 | 4 | 3 |
| ENSG00000274012 | 327 | 1974 | 192 | 208 | 149 | 188 |
| ENSG00000278002 | 3 | 4 | 0 | 0 | 1 | 1 |
| ENSG00000165527 | 4533 | 2327 | 2625 | 2170 | 2717 | 2533 |
| ENSG00000251929 | 0 | 0 | 0 | 0 | 0 | 0 |
| ENSG00000274704 | 0 | 0 | 0 | 0 | 0 | 0 |
| ENSG00000214900 | 95 | 21 | 37 | 49 | 44 | 56 |
| ENSG00000278038 | 0 | 0 | 0 | 0 | 0 | 0 |
| ENSG00000270788 | 0 | 0 | 0 | 0 | 0 | 0 |
| ENSG00000243038 | 0 | 0 | 0 | 0 | 0 | 0 |
| ENSG00000201358 | 0 | 0 | 0 | 0 | 0 | 0 |
| ENSG00000251792 | 0 | 1 | 1 | 0 | 0 | 2 |
| ENSG00000100483 | 465 | 249 | 292 | 192 | 298 | 259 |

| | | | | | | |
|---|---|---|---|---|---|---|
| ENSG00000100485 | 1182 | 652 | 495 | 245 | 759 | 456 |
| ENSG00000087299 | 1159 | 515 | 703 | 401 | 723 | 680 |
| ENSG00000264712 | 0 | 0 | 2 | 0 | 0 | 0 |
| ENSG00000125375 | 1011 | 507 | 594 | 401 | 544 | 572 |
| ENSG00000259071 | 0 | 0 | 0 | 0 | 0 | 0 |
| ENSG00000100490 | 2 | 3 | 3 | 3 | 5 | 5 |
| ENSG00000258857 | 0 | 0 | 0 | 0 | 0 | 1 |
| ENSG00000012983 | 1146 | 686 | 727 | 356 | 868 | 599 |
| ENSG00000259113 | 0 | 0 | 0 | 0 | 0 | 0 |
| ENSG00000201573 | 0 | 0 | 0 | 0 | 1 | 0 |
| ENSG00000273675 | 0 | 0 | 0 | 0 | 0 | 0 |
| ENSG00000198513 | 39 | 16 | 16 | 22 | 10 | 18 |
| ENSG00000258577 | 0 | 0 | 0 | 0 | 0 | 0 |
| ENSG00000151748 | 572 | 252 | 305 | 256 | 356 | 325 |
| ENSG00000243103 | 0 | 0 | 0 | 0 | 0 | 0 |
| ENSG00000269906 | 14 | 9 | 7 | 13 | 15 | 4 |
| ENSG00000258491 | 0 | 0 | 0 | 0 | 0 | 0 |
| ENSG00000100503 | 961 | 597 | 539 | 351 | 1046 | 603 |
| ENSG00000270062 | 14 | 4 | 4 | 2 | 3 | 6 |
| ENSG00000258843 | 0 | 0 | 0 | 0 | 0 | 0 |
| ENSG00000285747 | 0 | 2 | 0 | 0 | 1 | 2 |
| ENSG00000252945 | 0 | 0 | 0 | 0 | 0 | 0 |
| ENSG00000258687 | 0 | 0 | 0 | 0 | 0 | 0 |
| ENSG00000100504 | 1620 | 982 | 1160 | 1475 | 2005 | 1862 |
| ENSG00000131969 | 6 | 0 | 0 | 0 | 4 | 7 |
| ENSG00000258745 | 0 | 1 | 0 | 1 | 1 | 0 |
| ENSG00000258711 | 2 | 1 | 0 | 2 | 3 | 5 |
| ENSG00000100505 | 8 | 5 | 10 | 4 | 4 | 11 |
| ENSG00000259055 | 0 | 0 | 0 | 0 | 0 | 0 |
| ENSG00000139921 | 2338 | 1047 | 1269 | 1090 | 1340 | 1257 |
| ENSG00000201376 | 0 | 1 | 0 | 0 | 0 | 0 |
| ENSG00000201820 | 2 | 0 | 1 | 1 | 6 | 3 |
| ENSG00000258955 | 0 | 0 | 0 | 0 | 0 | 0 |
| ENSG00000258479 | 0 | 0 | 0 | 1 | 0 | 1 |
| ENSG00000258942 | 0 | 0 | 0 | 0 | 0 | 1 |
| ENSG00000258808 | 0 | 0 | 0 | 0 | 0 | 0 |
| ENSG00000258893 | 0 | 1 | 0 | 0 | 0 | 2 |
| ENSG00000258537 | 0 | 0 | 0 | 0 | 0 | 0 |
| ENSG00000139926 | 1115 | 584 | 591 | 369 | 696 | 562 |
| ENSG00000252400 | 0 | 0 | 0 | 0 | 0 | 0 |
| ENSG00000277050 | 19 | 10 | 12 | 3 | 9 | 12 |
| ENSG00000273888 | 84 | 51 | 49 | 42 | 37 | 33 |
| ENSG00000251756 | 0 | 0 | 0 | 0 | 0 | 0 |
| ENSG00000207004 | 0 | 0 | 1 | 0 | 0 | 0 |
| ENSG00000259111 | 0 | 0 | 0 | 0 | 0 | 0 |
| ENSG00000258423 | 0 | 0 | 0 | 0 | 0 | 0 |
| ENSG00000258550 | 0 | 0 | 0 | 0 | 0 | 0 |
| ENSG00000258535 | 0 | 0 | 0 | 0 | 0 | 0 |
| ENSG00000258849 | 0 | 0 | 0 | 0 | 0 | 0 |
| ENSG00000259157 | 0 | 0 | 0 | 0 | 0 | 0 |

| | | | | | | |
|---|---|---|---|---|---|---|
| ENSG00000186469 | 0 | 0 | 0 | 0 | 0 | 0 |
| ENSG00000258928 | 0 | 0 | 0 | 0 | 0 | 0 |
| ENSG00000259007 | 0 | 0 | 0 | 0 | 0 | 0 |
| ENSG00000258854 | 0 | 0 | 0 | 0 | 0 | 0 |
| ENSG00000087302 | 3908 | 2043 | 2295 | 2246 | 2622 | 2328 |
| ENSG00000087303 | 4 | 0 | 2 | 4 | 0 | 3 |
| ENSG00000258694 | 0 | 0 | 0 | 0 | 0 | 0 |
| ENSG00000258981 | 0 | 0 | 0 | 0 | 0 | 0 |
| ENSG00000258440 | 0 | 0 | 0 | 0 | 0 | 0 |
| ENSG00000168229 | 0 | 0 | 0 | 0 | 0 | 0 |
| ENSG00000261208 | 0 | 0 | 0 | 0 | 0 | 0 |
| ENSG00000125384 | 36 | 12 | 10 | 15 | 22 | 11 |
| ENSG00000087301 | 472 | 252 | 284 | 111 | 351 | 221 |
| ENSG00000180998 | 257 | 119 | 152 | 78 | 154 | 114 |
| ENSG00000197930 | 2233 | 1091 | 1368 | 1032 | 1632 | 1439 |
| ENSG00000258757 | 0 | 0 | 0 | 0 | 0 | 0 |
| ENSG00000100519 | 3260 | 1776 | 1871 | 1278 | 2458 | 1680 |
| ENSG00000198252 | 1872 | 853 | 934 | 545 | 904 | 774 |
| ENSG00000100522 | 2359 | 1327 | 1635 | 1234 | 1434 | 1637 |
| ENSG00000259049 | 0 | 0 | 0 | 0 | 0 | 0 |
| ENSG00000285664 | 0 | 2 | 5 | 2 | 3 | 2 |
| ENSG00000258698 | 0 | 0 | 0 | 0 | 0 | 0 |
| ENSG00000258802 | 2 | 0 | 0 | 1 | 0 | 0 |
| ENSG00000073712 | 1349 | 697 | 797 | 489 | 836 | 708 |
| ENSG00000271075 | 2 | 1 | 2 | 1 | 0 | 1 |
| ENSG00000258503 | 0 | 0 | 0 | 0 | 0 | 0 |
| ENSG00000258993 | 0 | 0 | 0 | 0 | 0 | 0 |
| ENSG00000100523 | 1079 | 546 | 548 | 377 | 736 | 511 |
| ENSG00000258985 | 0 | 0 | 0 | 0 | 0 | 0 |
| ENSG00000258731 | 0 | 1 | 3 | 1 | 2 | 1 |
| ENSG00000237356 | 10 | 3 | 15 | 7 | 9 | 7 |
| ENSG00000225680 | 0 | 0 | 0 | 0 | 0 | 0 |
| ENSG00000226132 | 0 | 0 | 0 | 0 | 0 | 0 |
| ENSG00000235269 | 0 | 1 | 0 | 1 | 0 | 0 |
| ENSG00000231922 | 0 | 0 | 0 | 0 | 0 | 0 |
| ENSG00000266431 | 0 | 0 | 0 | 0 | 0 | 0 |
| ENSG00000125378 | 766 | 296 | 393 | 248 | 237 | 278 |
| ENSG00000224004 | 0 | 1 | 1 | 1 | 1 | 1 |
| ENSG00000100526 | 649 | 359 | 380 | 443 | 489 | 441 |
| ENSG00000100528 | 3280 | 1573 | 1931 | 1699 | 1964 | 1876 |
| ENSG00000258753 | 0 | 0 | 0 | 1 | 0 | 0 |
| ENSG00000197045 | 2188 | 1461 | 1097 | 854 | 2192 | 1234 |
| ENSG00000100532 | 457 | 200 | 250 | 233 | 280 | 255 |
| ENSG00000020577 | 626 | 342 | 375 | 347 | 530 | 469 |
| ENSG00000285774 | 0 | 0 | 0 | 0 | 0 | 0 |
| ENSG00000131979 | 1459 | 704 | 910 | 669 | 869 | 844 |
| ENSG00000252019 | 0 | 2 | 0 | 0 | 0 | 0 |
| ENSG00000265432 | 1 | 1 | 0 | 0 | 0 | 0 |
| ENSG00000258872 | 0 | 2 | 1 | 1 | 0 | 0 |
| ENSG00000198554 | 359 | 225 | 149 | 68 | 326 | 190 |

| | | | | | | |
|---|---|---|---|---|---|---|
| ENSG00000233924 | 0 | 0 | 0 | 0 | 0 | 0 |
| ENSG00000180008 | 960 | 536 | 432 | 233 | 696 | 474 |
| ENSG00000168175 | 3242 | 1522 | 1794 | 1309 | 1605 | 1675 |
| ENSG00000131981 | 16398 | 7738 | 10659 | 8527 | 8006 | 9049 |
| ENSG00000126787 | 458 | 315 | 278 | 202 | 626 | 388 |
| ENSG00000258050 | 0 | 1 | 1 | 0 | 0 | 0 |
| ENSG00000279392 | 2 | 2 | 0 | 3 | 1 | 1 |
| ENSG00000271180 | 0 | 0 | 0 | 0 | 0 | 0 |
| ENSG00000258413 | 1 | 0 | 0 | 0 | 3 | 3 |
| ENSG00000178974 | 1375 | 655 | 792 | 568 | 732 | 741 |
| ENSG00000239686 | 0 | 0 | 0 | 0 | 1 | 1 |
| ENSG00000278818 | 0 | 0 | 1 | 0 | 0 | 0 |
| ENSG00000258469 | 0 | 0 | 1 | 1 | 1 | 2 |
| ENSG00000258455 | 0 | 0 | 0 | 0 | 0 | 0 |
| ENSG00000126775 | 1119 | 613 | 766 | 539 | 729 | 624 |
| ENSG00000259318 | 6 | 6 | 7 | 1 | 3 | 3 |
| ENSG00000182521 | 0 | 0 | 0 | 0 | 0 | 0 |
| ENSG00000239199 | 0 | 0 | 0 | 0 | 0 | 0 |
| ENSG00000186615 | 80 | 46 | 53 | 60 | 82 | 70 |
| ENSG00000213641 | 0 | 0 | 0 | 0 | 1 | 0 |
| ENSG00000271484 | 0 | 0 | 0 | 0 | 0 | 0 |
| ENSG00000126777 | 783 | 527 | 411 | 134 | 1234 | 534 |
| ENSG00000200742 | 0 | 0 | 0 | 0 | 0 | 0 |
| ENSG00000258784 | 0 | 0 | 0 | 0 | 0 | 0 |
| ENSG00000177350 | 4 | 0 | 0 | 0 | 0 | 1 |
| ENSG00000258791 | 0 | 0 | 0 | 0 | 0 | 0 |
| ENSG00000259868 | 0 | 0 | 0 | 0 | 0 | 0 |
| ENSG00000277763 | 0 | 0 | 0 | 0 | 0 | 0 |
| ENSG00000139946 | 0 | 0 | 0 | 0 | 0 | 0 |
| ENSG00000259483 | 0 | 0 | 0 | 0 | 1 | 0 |
| ENSG00000275569 | 0 | 0 | 0 | 0 | 0 | 0 |
| ENSG00000259719 | 0 | 0 | 0 | 0 | 0 | 0 |
| ENSG00000070269 | 589 | 306 | 410 | 292 | 393 | 381 |
| ENSG00000258803 | 0 | 0 | 0 | 0 | 0 | 0 |
| ENSG00000258428 | 0 | 0 | 1 | 0 | 3 | 0 |
| ENSG00000259133 | 0 | 0 | 0 | 0 | 0 | 0 |
| ENSG00000242995 | 0 | 0 | 0 | 0 | 0 | 0 |
| ENSG00000258703 | 0 | 0 | 0 | 0 | 0 | 0 |
| ENSG00000165588 | 0 | 0 | 0 | 0 | 0 | 0 |
| ENSG00000248550 | 0 | 0 | 0 | 0 | 0 | 0 |
| ENSG00000258776 | 0 | 0 | 0 | 0 | 0 | 0 |
| ENSG00000243801 | 0 | 0 | 0 | 0 | 0 | 0 |
| ENSG00000200818 | 0 | 0 | 0 | 0 | 0 | 0 |
| ENSG00000243388 | 0 | 0 | 0 | 1 | 1 | 0 |
| ENSG00000258758 | 0 | 0 | 0 | 0 | 0 | 0 |
| ENSG00000259136 | 0 | 0 | 0 | 0 | 0 | 0 |
| ENSG00000258592 | 8 | 6 | 3 | 1 | 5 | 5 |
| ENSG00000070367 | 1487 | 812 | 619 | 367 | 1179 | 669 |
| ENSG00000053770 | 1012 | 498 | 507 | 349 | 740 | 535 |
| ENSG00000259008 | 0 | 0 | 0 | 0 | 0 | 0 |

| | | | | | | |
|---|---|---|---|---|---|---|
| ENSG00000139977 | 1730 | 806 | 938 | 646 | 981 | 898 |
| ENSG00000100557 | 0 | 0 | 0 | 1 | 0 | 0 |
| ENSG00000151812 | 0 | 0 | 0 | 0 | 0 | 0 |
| ENSG00000259039 | 0 | 0 | 1 | 0 | 0 | 2 |
| ENSG00000258856 | 0 | 0 | 0 | 0 | 0 | 0 |
| ENSG00000252837 | 0 | 0 | 0 | 0 | 0 | 0 |
| ENSG00000259969 | 0 | 1 | 0 | 0 | 0 | 0 |
| ENSG00000139971 | 18 | 3 | 3 | 1 | 1 | 3 |
| ENSG00000258836 | 0 | 0 | 0 | 0 | 0 | 0 |
| ENSG00000243023 | 0 | 0 | 0 | 0 | 0 | 0 |
| ENSG00000243700 | 0 | 0 | 0 | 0 | 0 | 0 |
| ENSG00000258519 | 0 | 0 | 0 | 0 | 0 | 0 |
| ENSG00000131966 | 2640 | 1302 | 1596 | 1145 | 1589 | 1417 |
| ENSG00000100567 | 4660 | 2538 | 2850 | 2702 | 3349 | 2778 |
| ENSG00000258682 | 16 | 12 | 16 | 6 | 24 | 8 |
| ENSG00000257621 | 1179 | 574 | 829 | 299 | 619 | 617 |
| ENSG00000180189 | 3 | 2 | 2 | 1 | 0 | 0 |
| ENSG00000279636 | 16 | 2 | 7 | 4 | 3 | 6 |
| ENSG00000252782 | 1 | 1 | 0 | 0 | 1 | 0 |
| ENSG00000032219 | 522 | 324 | 190 | 68 | 514 | 229 |
| ENSG00000258658 | 2 | 2 | 0 | 1 | 2 | 0 |
| ENSG00000196860 | 7 | 8 | 5 | 2 | 1 | 7 |
| ENSG00000258378 | 0 | 1 | 2 | 0 | 1 | 0 |
| ENSG00000100575 | 702 | 402 | 517 | 467 | 540 | 484 |
| ENSG00000100578 | 311 | 144 | 133 | 73 | 338 | 159 |
| ENSG00000252198 | 0 | 0 | 0 | 0 | 0 | 0 |
| ENSG00000258650 | 0 | 0 | 0 | 0 | 0 | 0 |
| ENSG00000258900 | 6 | 2 | 5 | 8 | 3 | 3 |
| ENSG00000239510 | 3 | 0 | 0 | 0 | 0 | 0 |
| ENSG00000165617 | 0 | 1 | 0 | 0 | 0 | 0 |
| ENSG00000258583 | 0 | 0 | 0 | 0 | 0 | 0 |
| ENSG00000213598 | 14 | 4 | 10 | 10 | 2 | 1 |
| ENSG00000258734 | 0 | 0 | 0 | 0 | 0 | 0 |
| ENSG00000258685 | 0 | 0 | 0 | 0 | 0 | 0 |
| ENSG00000258996 | 0 | 0 | 0 | 0 | 0 | 0 |
| ENSG00000100592 | 1872 | 1030 | 933 | 443 | 1270 | 764 |
| ENSG00000181619 | 71 | 29 | 31 | 23 | 32 | 41 |
| ENSG00000126790 | 235 | 98 | 169 | 140 | 180 | 194 |
| ENSG00000258782 | 0 | 0 | 0 | 0 | 0 | 0 |
| ENSG00000050130 | 1923 | 953 | 1143 | 873 | 1171 | 1051 |
| ENSG00000151838 | 0 | 0 | 0 | 0 | 0 | 0 |
| ENSG00000139970 | 2 | 7 | 3 | 0 | 0 | 0 |
| ENSG00000263629 | 0 | 0 | 0 | 0 | 0 | 0 |
| ENSG00000261120 | 0 | 0 | 0 | 0 | 0 | 0 |
| ENSG00000131951 | 0 | 0 | 0 | 0 | 0 | 0 |
| ENSG00000258553 | 0 | 0 | 0 | 0 | 1 | 0 |
| ENSG00000126773 | 1548 | 746 | 912 | 514 | 869 | 735 |
| ENSG00000100612 | 1691 | 986 | 1174 | 1176 | 1247 | 1173 |
| ENSG00000258896 | 2 | 0 | 1 | 0 | 1 | 1 |
| ENSG00000258563 | 0 | 0 | 0 | 0 | 0 | 0 |

| | | | | | | |
|---|---|---|---|---|---|---|
| ENSG00000242135 | 0 | 0 | 0 | 0 | 0 | 0 |
| ENSG00000254718 | 22 | 14 | 15 | 11 | 19 | 13 |
| ENSG00000100614 | 3188 | 1574 | 1992 | 1343 | 1897 | 1788 |
| ENSG00000258556 | 0 | 0 | 0 | 1 | 0 | 0 |
| ENSG00000179008 | 4 | 1 | 0 | 0 | 0 | 1 |
| ENSG00000258427 | 2 | 1 | 4 | 6 | 1 | 4 |
| ENSG00000259169 | 0 | 0 | 0 | 0 | 0 | 0 |
| ENSG00000258692 | 0 | 2 | 0 | 0 | 0 | 0 |
| ENSG00000184302 | 0 | 0 | 0 | 0 | 0 | 0 |
| ENSG00000258670 | 0 | 0 | 0 | 0 | 0 | 0 |
| ENSG00000258764 | 0 | 0 | 0 | 0 | 0 | 0 |
| ENSG00000244756 | 0 | 0 | 0 | 0 | 0 | 0 |
| ENSG00000241354 | 0 | 0 | 0 | 0 | 0 | 0 |
| ENSG00000258952 | 3 | 3 | 4 | 1 | 1 | 2 |
| ENSG00000126778 | 714 | 274 | 352 | 224 | 210 | 262 |
| ENSG00000100625 | 1734 | 789 | 991 | 419 | 640 | 585 |
| ENSG00000020426 | 1502 | 797 | 993 | 745 | 830 | 752 |
| ENSG00000258445 | 0 | 0 | 0 | 0 | 0 | 0 |
| ENSG00000258892 | 0 | 0 | 0 | 0 | 0 | 0 |
| ENSG00000258485 | 0 | 0 | 0 | 0 | 0 | 0 |
| ENSG00000126814 | 862 | 465 | 544 | 402 | 537 | 477 |
| ENSG00000258656 | 0 | 0 | 0 | 0 | 0 | 0 |
| ENSG00000206870 | 0 | 0 | 0 | 0 | 0 | 0 |
| ENSG00000139974 | 678 | 357 | 440 | 273 | 293 | 362 |
| ENSG00000027075 | 1304 | 669 | 868 | 585 | 791 | 745 |
| ENSG00000182107 | 3663 | 1665 | 2175 | 1509 | 1704 | 1951 |
| ENSG00000247287 | 43 | 16 | 27 | 27 | 27 | 25 |
| ENSG00000258989 | 0 | 0 | 0 | 0 | 0 | 0 |
| ENSG00000258926 | 3 | 0 | 0 | 0 | 2 | 0 |
| ENSG00000250548 | 6 | 0 | 0 | 1 | 2 | 0 |
| ENSG00000232774 | 15 | 2 | 3 | 2 | 5 | 5 |
| ENSG00000258777 | 0 | 0 | 0 | 0 | 0 | 0 |
| ENSG00000100644 | 2342 | 1536 | 1204 | 558 | 2349 | 1314 |
| ENSG00000258667 | 3 | 3 | 0 | 0 | 0 | 0 |
| ENSG00000258964 | 0 | 0 | 0 | 0 | 0 | 0 |
| ENSG00000023608 | 163 | 90 | 96 | 72 | 162 | 99 |
| ENSG00000279633 | 0 | 0 | 0 | 0 | 0 | 0 |
| ENSG00000139973 | 0 | 0 | 1 | 0 | 0 | 0 |
| ENSG00000258956 | 0 | 0 | 0 | 0 | 0 | 0 |
| ENSG00000258842 | 0 | 0 | 0 | 0 | 0 | 0 |
| ENSG00000258903 | 0 | 0 | 0 | 0 | 0 | 0 |
| ENSG00000186369 | 0 | 0 | 0 | 0 | 0 | 0 |
| ENSG00000259142 | 0 | 0 | 0 | 0 | 0 | 0 |
| ENSG00000270852 | 0 | 0 | 0 | 0 | 0 | 0 |
| ENSG00000258638 | 0 | 0 | 0 | 0 | 0 | 0 |
| ENSG00000258877 | 0 | 0 | 0 | 0 | 0 | 0 |
| ENSG00000140015 | 0 | 0 | 0 | 0 | 0 | 0 |
| ENSG00000258995 | 0 | 0 | 0 | 0 | 0 | 0 |
| ENSG00000259093 | 0 | 0 | 0 | 0 | 0 | 0 |
| ENSG00000126785 | 0 | 0 | 0 | 0 | 0 | 0 |

| | | | | | | |
|---|---|---|---|---|---|---|
| ENSG00000258943 | 0 | 0 | 0 | 0 | 0 | 0 |
| ENSG00000179600 | 0 | 0 | 0 | 0 | 0 | 0 |
| ENSG00000154001 | 2931 | 1437 | 1684 | 1141 | 1690 | 1532 |
| ENSG00000252800 | 0 | 0 | 0 | 0 | 0 | 0 |
| ENSG00000261242 | 5 | 6 | 6 | 0 | 11 | 3 |
| ENSG00000205767 | 0 | 0 | 0 | 0 | 0 | 0 |
| ENSG00000270878 | 10 | 4 | 8 | 5 | 4 | 1 |
| ENSG00000140006 | 419 | 228 | 242 | 212 | 322 | 258 |
| ENSG00000258800 | 0 | 0 | 0 | 0 | 0 | 0 |
| ENSG00000258645 | 0 | 1 | 0 | 1 | 1 | 0 |
| ENSG00000202490 | 0 | 0 | 0 | 0 | 0 | 0 |
| ENSG00000206822 | 0 | 0 | 0 | 0 | 0 | 0 |
| ENSG00000207172 | 0 | 0 | 0 | 0 | 0 | 0 |
| ENSG00000274015 | 14 | 1 | 2 | 3 | 3 | 4 |
| ENSG00000200693 | 6 | 5 | 4 | 3 | 5 | 3 |
| ENSG00000126821 | 1164 | 493 | 669 | 515 | 574 | 619 |
| ENSG00000258778 | 0 | 0 | 0 | 0 | 0 | 0 |
| ENSG00000252749 | 0 | 0 | 0 | 0 | 0 | 0 |
| ENSG00000258411 | 0 | 0 | 0 | 0 | 0 | 0 |
| ENSG00000270912 | 0 | 0 | 0 | 0 | 0 | 0 |
| ENSG00000202182 | 0 | 0 | 0 | 0 | 0 | 0 |
| ENSG00000054654 | 666 | 258 | 337 | 169 | 445 | 418 |
| ENSG00000140009 | 18 | 3 | 4 | 1 | 6 | 2 |
| ENSG00000221537 | 0 | 0 | 0 | 0 | 0 | 0 |
| ENSG00000214770 | 17 | 9 | 12 | 7 | 8 | 8 |
| ENSG00000234911 | 19 | 20 | 12 | 8 | 18 | 11 |
| ENSG00000284664 | 16 | 2 | 1 | 4 | 9 | 3 |
| ENSG00000271511 | 7 | 0 | 1 | 0 | 0 | 1 |
| ENSG00000100714 | 3652 | 1939 | 2563 | 1988 | 2439 | 2637 |
| ENSG00000258824 | 24 | 18 | 13 | 14 | 14 | 12 |
| ENSG00000272909 | 47 | 19 | 23 | 25 | 27 | 23 |
| ENSG00000089775 | 809 | 390 | 474 | 300 | 515 | 424 |
| ENSG00000179841 | 308 | 210 | 140 | 87 | 113 | 109 |
| ENSG00000126804 | 1787 | 876 | 932 | 480 | 962 | 824 |
| ENSG00000259116 | 20 | 5 | 7 | 5 | 2 | 6 |
| ENSG00000259160 | 0 | 0 | 0 | 0 | 0 | 0 |
| ENSG00000126803 | 440 | 199 | 249 | 165 | 170 | 136 |
| ENSG00000165807 | 26 | 15 | 8 | 7 | 18 | 8 |
| ENSG00000259076 | 3 | 2 | 1 | 4 | 0 | 1 |
| ENSG00000259010 | 10 | 2 | 1 | 5 | 1 | 3 |
| ENSG00000126822 | 1806 | 926 | 1174 | 1063 | 1085 | 1110 |
| ENSG00000070182 | 21 | 6 | 7 | 5 | 14 | 6 |
| ENSG00000284269 | 0 | 0 | 0 | 0 | 0 | 0 |
| ENSG00000252497 | 0 | 0 | 0 | 0 | 0 | 0 |
| ENSG00000258289 | 2704 | 1586 | 1397 | 815 | 1939 | 1269 |
| ENSG00000125954 | 0 | 1 | 1 | 0 | 0 | 0 |
| ENSG00000176153 | 438 | 221 | 241 | 247 | 257 | 301 |
| ENSG00000139998 | 6395 | 3334 | 3761 | 3899 | 4288 | 4017 |
| ENSG00000257365 | 412 | 235 | 282 | 248 | 234 | 247 |
| ENSG00000272158 | 0 | 0 | 2 | 0 | 2 | 0 |

| | | | | | | |
|---|---|---|---|---|---|---|
| ENSG00000125952 | 2004 | 975 | 1186 | 1115 | 1060 | 1130 |
| ENSG00000266531 | 0 | 0 | 0 | 0 | 0 | 0 |
| ENSG00000259118 | 9 | 2 | 5 | 1 | 3 | 1 |
| ENSG00000222985 | 0 | 0 | 0 | 1 | 0 | 0 |
| ENSG00000255002 | 0 | 2 | 3 | 1 | 1 | 0 |
| ENSG00000258878 | 1 | 2 | 1 | 0 | 4 | 2 |
| ENSG00000258760 | 2 | 0 | 9 | 3 | 4 | 4 |
| ENSG00000258613 | 1 | 0 | 0 | 1 | 0 | 0 |
| ENSG00000214759 | 0 | 0 | 0 | 0 | 0 | 0 |
| ENSG00000259078 | 2 | 0 | 0 | 1 | 2 | 1 |
| ENSG00000266740 | 0 | 0 | 0 | 0 | 0 | 0 |
| ENSG00000033170 | 4912 | 2503 | 3033 | 1984 | 2722 | 2731 |
| ENSG00000276116 | 157 | 92 | 125 | 106 | 102 | 103 |
| ENSG00000180662 | 0 | 1 | 0 | 0 | 0 | 0 |
| ENSG00000207781 | 0 | 0 | 0 | 0 | 0 | 0 |
| ENSG00000258980 | 0 | 0 | 0 | 0 | 0 | 0 |
| ENSG00000258629 | 0 | 0 | 0 | 0 | 0 | 0 |
| ENSG00000258960 | 0 | 0 | 0 | 0 | 0 | 0 |
| ENSG00000258847 | 0 | 0 | 0 | 0 | 0 | 0 |
| ENSG00000277651 | 0 | 0 | 0 | 0 | 0 | 0 |
| ENSG00000224861 | 10 | 11 | 5 | 8 | 12 | 3 |
| ENSG00000258502 | 0 | 0 | 0 | 0 | 0 | 2 |
| ENSG00000200860 | 0 | 0 | 0 | 0 | 0 | 0 |
| ENSG00000258561 | 5 | 3 | 1 | 2 | 6 | 8 |
| ENSG00000196553 | 0 | 0 | 0 | 0 | 0 | 1 |
| ENSG00000171723 | 412 | 236 | 254 | 217 | 273 | 241 |
| ENSG00000258796 | 0 | 0 | 0 | 0 | 0 | 0 |
| ENSG00000258490 | 0 | 0 | 0 | 0 | 0 | 1 |
| ENSG00000172717 | 4 | 2 | 2 | 0 | 0 | 0 |
| ENSG00000258988 | 2 | 0 | 0 | 0 | 0 | 0 |
| ENSG00000072415 | 1301 | 704 | 736 | 394 | 859 | 610 |
| ENSG00000100554 | 2057 | 1129 | 1260 | 1022 | 1327 | 973 |
| ENSG00000199755 | 0 | 0 | 0 | 0 | 0 | 0 |
| ENSG00000134001 | 4591 | 2345 | 2742 | 2337 | 2845 | 2672 |
| ENSG00000100558 | 159 | 71 | 92 | 68 | 114 | 81 |
| ENSG00000265993 | 0 | 0 | 0 | 0 | 0 | 0 |
| ENSG00000198133 | 551 | 264 | 335 | 355 | 278 | 346 |
| ENSG00000054690 | 1222 | 515 | 827 | 489 | 738 | 790 |
| ENSG00000100564 | 506 | 273 | 325 | 335 | 285 | 353 |
| ENSG00000259502 | 4 | 3 | 9 | 2 | 5 | 6 |
| ENSG00000259648 | 6 | 3 | 6 | 0 | 2 | 1 |
| ENSG00000281228 | 0 | 0 | 0 | 0 | 0 | 0 |
| ENSG00000201529 | 0 | 0 | 1 | 0 | 0 | 0 |
| ENSG00000081181 | 21 | 14 | 24 | 11 | 17 | 15 |
| ENSG00000100568 | 2982 | 1784 | 2059 | 1820 | 2283 | 2129 |
| ENSG00000258626 | 0 | 0 | 2 | 0 | 0 | 0 |
| ENSG00000272253 | 0 | 3 | 0 | 0 | 0 | 0 |
| ENSG00000258466 | 0 | 1 | 1 | 1 | 0 | 0 |
| ENSG00000072042 | 2770 | 1338 | 1795 | 1495 | 1826 | 1849 |
| ENSG00000139988 | 6 | 0 | 1 | 0 | 2 | 1 |

| | | | | | | |
|---|---|---|---|---|---|---|
| ENSG00000241542 | 0 | 0 | 0 | 0 | 0 | 0 |
| ENSG00000258618 | 0 | 0 | 0 | 0 | 0 | 0 |
| ENSG00000072121 | 1747 | 709 | 945 | 680 | 922 | 860 |
| ENSG00000271697 | 0 | 0 | 0 | 0 | 0 | 0 |
| ENSG00000239820 | 0 | 0 | 0 | 0 | 2 | 0 |
| ENSG00000258759 | 2 | 1 | 0 | 0 | 0 | 1 |
| ENSG00000182185 | 279 | 115 | 140 | 101 | 129 | 129 |
| ENSG00000274666 | 0 | 0 | 0 | 0 | 0 | 0 |
| ENSG00000258837 | 1 | 3 | 5 | 3 | 5 | 5 |
| ENSG00000244677 | 0 | 0 | 0 | 0 | 0 | 0 |
| ENSG00000243546 | 0 | 0 | 0 | 0 | 0 | 0 |
| ENSG00000240210 | 0 | 0 | 1 | 0 | 1 | 0 |
| ENSG00000258477 | 0 | 1 | 1 | 0 | 1 | 0 |
| ENSG00000259038 | 10 | 6 | 3 | 8 | 8 | 7 |
| ENSG00000258623 | 2 | 0 | 0 | 0 | 3 | 1 |
| ENSG00000242941 | 0 | 1 | 0 | 0 | 1 | 0 |
| ENSG00000207089 | 0 | 0 | 0 | 0 | 0 | 0 |
| ENSG00000185650 | 5475 | 2759 | 3232 | 2568 | 2804 | 3203 |
| ENSG00000270975 | 0 | 0 | 0 | 0 | 0 | 1 |
| ENSG00000258565 | 0 | 0 | 0 | 0 | 0 | 0 |
| ENSG00000072110 | 6253 | 3031 | 3715 | 3297 | 3763 | 3552 |
| ENSG00000258967 | 0 | 0 | 0 | 0 | 0 | 0 |
| ENSG00000258531 | 1 | 0 | 0 | 0 | 0 | 1 |
| ENSG00000259062 | 4 | 0 | 2 | 1 | 1 | 0 |
| ENSG00000242293 | 0 | 0 | 0 | 0 | 0 | 0 |
| ENSG00000139990 | 3845 | 1683 | 2290 | 1303 | 1886 | 1839 |
| ENSG00000259165 | 2 | 0 | 0 | 0 | 1 | 0 |
| ENSG00000206768 | 0 | 0 | 0 | 0 | 0 | 0 |
| ENSG00000276063 | 1 | 0 | 2 | 0 | 1 | 0 |
| ENSG00000258957 | 5 | 7 | 6 | 6 | 1 | 5 |
| ENSG00000081177 | 1948 | 826 | 1144 | 843 | 959 | 1023 |
| ENSG00000258520 | 2 | 2 | 5 | 2 | 0 | 3 |
| ENSG00000100626 | 50 | 25 | 32 | 8 | 19 | 15 |
| ENSG00000280073 | 12 | 4 | 6 | 7 | 1 | 5 |
| ENSG00000100632 | 5485 | 2980 | 3310 | 3357 | 3452 | 3288 |
| ENSG00000029364 | 4204 | 1907 | 2500 | 1694 | 2150 | 2028 |
| ENSG00000175985 | 93 | 36 | 46 | 19 | 23 | 31 |
| ENSG00000244237 | 0 | 0 | 0 | 0 | 0 | 0 |
| ENSG00000267909 | 0 | 0 | 0 | 0 | 0 | 0 |
| ENSG00000100647 | 2158 | 1020 | 1344 | 841 | 947 | 1082 |
| ENSG00000273797 | 0 | 4 | 0 | 0 | 0 | 0 |
| ENSG00000100650 | 11038 | 5427 | 7952 | 4445 | 6114 | 5860 |
| ENSG00000100652 | 0 | 8 | 2 | 1 | 6 | 1 |
| ENSG00000221060 | 0 | 0 | 0 | 0 | 0 | 0 |
| ENSG00000198732 | 4 | 4 | 1 | 1 | 1 | 2 |
| ENSG00000242071 | 50 | 30 | 30 | 31 | 40 | 39 |
| ENSG00000100678 | 0 | 1 | 3 | 0 | 0 | 1 |
| ENSG00000258422 | 0 | 0 | 0 | 0 | 0 | 0 |
| ENSG00000274330 | 0 | 0 | 0 | 0 | 0 | 0 |
| ENSG00000270610 | 0 | 0 | 0 | 0 | 0 | 0 |

| | | | | | | |
|---|---|---|---|---|---|---|
| ENSG00000235812 | 0 | 1 | 0 | 0 | 0 | 0 |
| ENSG00000259033 | 0 | 0 | 1 | 1 | 0 | 0 |
| ENSG00000133983 | 82 | 38 | 52 | 52 | 51 | 72 |
| ENSG00000258644 | 0 | 0 | 2 | 2 | 4 | 1 |
| ENSG00000222640 | 1 | 2 | 2 | 0 | 0 | 0 |
| ENSG00000213463 | 3101 | 1773 | 1752 | 610 | 1072 | 1023 |
| ENSG00000257759 | 0 | 1 | 1 | 0 | 0 | 0 |
| ENSG00000139985 | 6 | 0 | 4 | 0 | 0 | 1 |
| ENSG00000259158 | 19 | 11 | 12 | 6 | 10 | 5 |
| ENSG00000252263 | 0 | 0 | 0 | 0 | 0 | 0 |
| ENSG00000134007 | 2 | 0 | 0 | 1 | 2 | 0 |
| ENSG00000274397 | 0 | 0 | 0 | 0 | 0 | 2 |
| ENSG00000133997 | 758 | 377 | 460 | 325 | 502 | 392 |
| ENSG00000259115 | 0 | 0 | 0 | 0 | 0 | 0 |
| ENSG00000258951 | 8 | 2 | 2 | 1 | 5 | 1 |
| ENSG00000245466 | 0 | 1 | 0 | 1 | 0 | 2 |
| ENSG00000240837 | 0 | 0 | 0 | 0 | 0 | 0 |
| ENSG00000133985 | 1546 | 774 | 1091 | 810 | 942 | 949 |
| ENSG00000258689 | 10 | 10 | 31 | 13 | 10 | 13 |
| ENSG00000006432 | 1965 | 923 | 1093 | 770 | 988 | 1025 |
| ENSG00000259153 | 313 | 158 | 173 | 163 | 207 | 185 |
| ENSG00000275630 | 0 | 0 | 0 | 0 | 1 | 0 |
| ENSG00000258431 | 0 | 0 | 0 | 0 | 0 | 0 |
| ENSG00000274818 | 8 | 1 | 2 | 6 | 5 | 2 |
| ENSG00000100731 | 2206 | 982 | 1152 | 630 | 1219 | 990 |
| ENSG00000258930 | 0 | 0 | 0 | 0 | 0 | 0 |
| ENSG00000258966 | 0 | 0 | 0 | 0 | 0 | 0 |
| ENSG00000258571 | 3 | 0 | 0 | 2 | 1 | 0 |
| ENSG00000269927 | 0 | 0 | 0 | 0 | 0 | 0 |
| ENSG00000259907 | 0 | 0 | 1 | 0 | 0 | 0 |
| ENSG00000258517 | 0 | 0 | 0 | 0 | 0 | 0 |
| ENSG00000269945 | 0 | 0 | 0 | 0 | 0 | 0 |
| ENSG00000259146 | 56 | 37 | 50 | 50 | 66 | 50 |
| ENSG00000197555 | 2214 | 1116 | 1458 | 1104 | 1444 | 1532 |
| ENSG00000259079 | 0 | 0 | 2 | 0 | 0 | 0 |
| ENSG00000207444 | 0 | 0 | 0 | 0 | 0 | 0 |
| ENSG00000242330 | 0 | 0 | 0 | 0 | 0 | 0 |
| ENSG00000285612 | 2 | 0 | 0 | 0 | 0 | 1 |
| ENSG00000285518 | 4 | 3 | 6 | 5 | 3 | 3 |
| ENSG00000266869 | 97 | 54 | 59 | 50 | 85 | 62 |
| ENSG00000182732 | 259 | 137 | 180 | 104 | 139 | 146 |
| ENSG00000200298 | 0 | 0 | 0 | 0 | 0 | 0 |
| ENSG00000273830 | 0 | 0 | 0 | 0 | 0 | 0 |
| ENSG00000258871 | 5 | 1 | 1 | 1 | 4 | 0 |
| ENSG00000205683 | 36 | 10 | 10 | 10 | 9 | 11 |
| ENSG00000285936 | 2 | 0 | 0 | 0 | 0 | 0 |
| ENSG00000206751 | 0 | 0 | 0 | 0 | 0 | 0 |
| ENSG00000119599 | 1571 | 791 | 1137 | 764 | 801 | 934 |
| ENSG00000176043 | 0 | 0 | 1 | 1 | 1 | 2 |
| ENSG00000259015 | 57 | 18 | 38 | 22 | 23 | 35 |

| | | | | | | |
|---|---|---|---|---|---|---|
| ENSG00000239544 | 2 | 0 | 0 | 0 | 1 | 1 |
| ENSG00000165861 | 1695 | 756 | 957 | 759 | 814 | 822 |
| ENSG00000241487 | 0 | 0 | 0 | 0 | 0 | 0 |
| ENSG00000258813 | 2 | 3 | 6 | 2 | 7 | 4 |
| ENSG00000119707 | 2399 | 1576 | 1301 | 641 | 1949 | 1281 |
| ENSG00000258448 | 0 | 0 | 0 | 0 | 0 | 1 |
| ENSG00000080815 | 3914 | 1874 | 2387 | 1614 | 2216 | 2113 |
| ENSG00000259109 | 0 | 0 | 1 | 0 | 0 | 0 |
| ENSG00000100767 | 41 | 22 | 29 | 26 | 22 | 34 |
| ENSG00000258376 | 14 | 5 | 7 | 9 | 4 | 6 |
| ENSG00000252839 | 0 | 0 | 0 | 0 | 0 | 0 |
| ENSG00000258944 | 3 | 3 | 0 | 2 | 3 | 5 |
| ENSG00000133961 | 2867 | 1394 | 1740 | 1104 | 1653 | 1477 |
| ENSG00000285006 | 0 | 0 | 0 | 0 | 0 | 0 |
| ENSG00000284930 | 0 | 0 | 0 | 0 | 0 | 0 |
| ENSG00000251393 | 0 | 0 | 0 | 0 | 0 | 0 |
| ENSG00000187105 | 15 | 10 | 12 | 15 | 14 | 14 |
| ENSG00000170468 | 740 | 389 | 508 | 547 | 489 | 554 |
| ENSG00000258443 | 2 | 0 | 1 | 2 | 0 | 2 |
| ENSG00000258695 | 1 | 0 | 1 | 0 | 0 | 1 |
| ENSG00000184227 | 139 | 91 | 80 | 70 | 66 | 70 |
| ENSG00000258408 | 0 | 0 | 0 | 0 | 0 | 0 |
| ENSG00000119673 | 885 | 408 | 538 | 411 | 419 | 403 |
| ENSG00000258524 | 2 | 1 | 3 | 0 | 3 | 1 |
| ENSG00000279026 | 31 | 9 | 24 | 12 | 18 | 8 |
| ENSG00000177465 | 609 | 280 | 363 | 336 | 285 | 316 |
| ENSG00000205669 | 4 | 3 | 5 | 2 | 1 | 3 |
| ENSG00000258603 | 3 | 0 | 1 | 0 | 2 | 3 |
| ENSG00000258972 | 0 | 0 | 0 | 0 | 0 | 0 |
| ENSG00000119661 | 820 | 423 | 405 | 357 | 465 | 423 |
| ENSG00000251907 | 0 | 0 | 0 | 0 | 0 | 0 |
| ENSG00000258660 | 0 | 0 | 0 | 0 | 0 | 0 |
| ENSG00000176903 | 3184 | 1573 | 1765 | 1712 | 1736 | 1746 |
| ENSG00000156030 | 5359 | 2370 | 2994 | 1802 | 2430 | 2592 |
| ENSG00000264741 | 0 | 0 | 1 | 0 | 0 | 0 |
| ENSG00000259065 | 70 | 40 | 36 | 36 | 68 | 46 |
| ENSG00000258586 | 0 | 1 | 2 | 0 | 0 | 0 |
| ENSG00000270576 | 0 | 0 | 0 | 0 | 0 | 0 |
| ENSG00000140043 | 187 | 138 | 188 | 91 | 144 | 155 |
| ENSG00000258653 | 2 | 0 | 2 | 0 | 0 | 0 |
| ENSG00000223300 | 3 | 0 | 2 | 2 | 2 | 2 |
| ENSG00000273711 | 3 | 4 | 4 | 1 | 3 | 4 |
| ENSG00000119725 | 15 | 13 | 26 | 14 | 17 | 25 |
| ENSG00000258891 | 6 | 1 | 0 | 0 | 0 | 1 |
| ENSG00000270140 | 2 | 0 | 3 | 0 | 0 | 1 |
| ENSG00000199472 | 2 | 1 | 2 | 0 | 0 | 1 |
| ENSG00000156050 | 771 | 301 | 407 | 144 | 213 | 210 |
| ENSG00000119723 | 518 | 368 | 476 | 342 | 367 | 389 |
| ENSG00000187097 | 2344 | 988 | 1501 | 632 | 880 | 937 |
| ENSG00000279461 | 5 | 0 | 1 | 5 | 1 | 3 |

| | | | | | | |
|---|---|---|---|---|---|---|
| ENSG00000119636 | 219 | 120 | 152 | 84 | 132 | 86 |
| ENSG00000119711 | 1504 | 688 | 985 | 514 | 711 | 728 |
| ENSG00000205659 | 426 | 158 | 259 | 136 | 206 | 210 |
| ENSG00000239910 | 0 | 0 | 0 | 0 | 0 | 0 |
| ENSG00000258513 | 0 | 0 | 0 | 0 | 0 | 0 |
| ENSG00000119614 | 0 | 0 | 0 | 0 | 0 | 0 |
| ENSG00000119688 | 1125 | 571 | 721 | 622 | 616 | 655 |
| ENSG00000258559 | 42 | 12 | 12 | 14 | 15 | 10 |
| ENSG00000133980 | 0 | 0 | 0 | 1 | 0 | 0 |
| ENSG00000258737 | 0 | 0 | 0 | 0 | 0 | 0 |
| ENSG00000177776 | 0 | 0 | 0 | 0 | 0 | 0 |
| ENSG00000183379 | 0 | 0 | 0 | 0 | 0 | 0 |
| ENSG00000270000 | 0 | 0 | 0 | 0 | 0 | 0 |
| ENSG00000259005 | 0 | 0 | 0 | 0 | 0 | 0 |
| ENSG00000119655 | 2566 | 1307 | 1788 | 1705 | 1629 | 1562 |
| ENSG00000283944 | 0 | 0 | 0 | 0 | 0 | 0 |
| ENSG00000258769 | 13 | 13 | 19 | 6 | 10 | 12 |
| ENSG00000165898 | 994 | 528 | 602 | 663 | 539 | 601 |
| ENSG00000119681 | 305 | 120 | 158 | 107 | 120 | 102 |
| ENSG00000258425 | 0 | 0 | 0 | 0 | 0 | 0 |
| ENSG00000258976 | 0 | 0 | 0 | 0 | 0 | 0 |
| ENSG00000119682 | 3650 | 1642 | 2107 | 1614 | 1952 | 2131 |
| ENSG00000258439 | 0 | 1 | 2 | 0 | 5 | 2 |
| ENSG00000222604 | 0 | 0 | 0 | 0 | 0 | 0 |
| ENSG00000119616 | 2312 | 1118 | 1228 | 1125 | 1273 | 1320 |
| ENSG00000119596 | 2905 | 1287 | 1815 | 1108 | 1482 | 1656 |
| ENSG00000119608 | 6 | 0 | 2 | 0 | 4 | 3 |
| ENSG00000258587 | 0 | 0 | 0 | 0 | 0 | 0 |
| ENSG00000119689 | 5498 | 2694 | 3392 | 2943 | 3143 | 3349 |
| ENSG00000198208 | 772 | 321 | 439 | 461 | 463 | 495 |
| ENSG00000119630 | 4 | 5 | 4 | 2 | 6 | 8 |
| ENSG00000119718 | 1940 | 1094 | 1250 | 1298 | 1266 | 1207 |
| ENSG00000258646 | 12 | 7 | 7 | 1 | 4 | 6 |
| ENSG00000279594 | 0 | 3 | 1 | 2 | 2 | 1 |
| ENSG00000119684 | 574 | 333 | 289 | 183 | 536 | 303 |
| ENSG00000206924 | 0 | 1 | 0 | 0 | 0 | 1 |
| ENSG00000119640 | 192 | 112 | 127 | 130 | 150 | 124 |
| ENSG00000119703 | 45 | 25 | 21 | 18 | 19 | 23 |
| ENSG00000119638 | 601 | 272 | 377 | 299 | 324 | 383 |
| ENSG00000258978 | 0 | 0 | 0 | 0 | 0 | 0 |
| ENSG00000259138 | 1 | 0 | 0 | 0 | 0 | 0 |
| ENSG00000170348 | 13345 | 6558 | 7915 | 6010 | 7103 | 6761 |
| ENSG00000273565 | 8 | 2 | 6 | 4 | 1 | 4 |
| ENSG00000252013 | 0 | 0 | 0 | 0 | 0 | 0 |
| ENSG00000258740 | 0 | 0 | 0 | 0 | 0 | 1 |
| ENSG00000258820 | 2 | 0 | 0 | 0 | 0 | 0 |
| ENSG00000283519 | 0 | 0 | 0 | 0 | 0 | 0 |
| ENSG00000170345 | 132 | 46 | 97 | 85 | 42 | 126 |
| ENSG00000119660 | 0 | 0 | 0 | 0 | 0 | 0 |
| ENSG00000259687 | 10 | 4 | 2 | 5 | 3 | 2 |

| | | | | | | |
|---|---|---|---|---|---|---|
| ENSG00000259319 | 14 | 11 | 8 | 5 | 9 | 12 |
| ENSG00000140044 | 194 | 69 | 93 | 104 | 89 | 86 |
| ENSG00000156127 | 1054 | 554 | 629 | 847 | 804 | 697 |
| ENSG00000224721 | 0 | 1 | 3 | 0 | 0 | 0 |
| ENSG00000119686 | 1023 | 570 | 696 | 802 | 858 | 868 |
| ENSG00000242951 | 2 | 0 | 2 | 2 | 3 | 0 |
| ENSG00000201096 | 1 | 0 | 0 | 0 | 0 | 0 |
| ENSG00000280494 | 0 | 0 | 0 | 0 | 0 | 0 |
| ENSG00000119685 | 1294 | 641 | 762 | 576 | 775 | 745 |
| ENSG00000133935 | 2980 | 1454 | 1719 | 1773 | 1776 | 1691 |
| ENSG00000259103 | 0 | 0 | 0 | 0 | 1 | 0 |
| ENSG00000242488 | 0 | 0 | 1 | 0 | 0 | 1 |
| ENSG00000119650 | 1520 | 686 | 950 | 860 | 815 | 793 |
| ENSG00000119699 | 1131 | 490 | 829 | 302 | 303 | 447 |
| ENSG00000258876 | 6 | 0 | 1 | 0 | 0 | 1 |
| ENSG00000258578 | 0 | 0 | 0 | 0 | 0 | 0 |
| ENSG00000283629 | 0 | 0 | 0 | 0 | 0 | 0 |
| ENSG00000089916 | 1497 | 737 | 784 | 498 | 1070 | 785 |
| ENSG00000258454 | 4 | 2 | 1 | 0 | 0 | 0 |
| ENSG00000280078 | 0 | 0 | 0 | 0 | 0 | 0 |
| ENSG00000258402 | 0 | 0 | 0 | 0 | 0 | 0 |
| ENSG00000119715 | 0 | 0 | 0 | 2 | 1 | 0 |
| ENSG00000259124 | 0 | 0 | 0 | 0 | 0 | 0 |
| ENSG00000271423 | 0 | 0 | 0 | 0 | 0 | 0 |
| ENSG00000242952 | 0 | 1 | 2 | 0 | 1 | 0 |
| ENSG00000258566 | 0 | 0 | 0 | 0 | 0 | 0 |
| ENSG00000258569 | 0 | 0 | 0 | 0 | 0 | 0 |
| ENSG00000071246 | 407 | 208 | 230 | 232 | 243 | 234 |
| ENSG00000259081 | 5 | 6 | 9 | 2 | 5 | 5 |
| ENSG00000258610 | 0 | 0 | 0 | 0 | 0 | 0 |
| ENSG00000258301 | 4 | 9 | 19 | 14 | 12 | 8 |
| ENSG00000013523 | 2562 | 1165 | 1442 | 1385 | 1403 | 1374 |
| ENSG00000100565 | 0 | 0 | 0 | 0 | 0 | 0 |
| ENSG00000241081 | 5 | 7 | 4 | 7 | 9 | 10 |
| ENSG00000223174 | 0 | 1 | 0 | 0 | 0 | 1 |
| ENSG00000259058 | 0 | 0 | 0 | 0 | 0 | 0 |
| ENSG00000285966 | 0 | 3 | 1 | 0 | 3 | 0 |
| ENSG00000258602 | 0 | 0 | 0 | 2 | 3 | 0 |
| ENSG00000266553 | 0 | 0 | 0 | 0 | 0 | 0 |
| ENSG00000243225 | 0 | 0 | 0 | 1 | 0 | 0 |
| ENSG00000119669 | 3658 | 1619 | 2133 | 2043 | 1971 | 2282 |
| ENSG00000273729 | 43 | 11 | 47 | 18 | 13 | 25 |
| ENSG00000246548 | 18 | 10 | 15 | 13 | 4 | 11 |
| ENSG00000258473 | 0 | 0 | 0 | 0 | 0 | 1 |
| ENSG00000258819 | 14 | 1 | 4 | 3 | 2 | 5 |
| ENSG00000198894 | 1337 | 609 | 682 | 424 | 755 | 569 |
| ENSG00000259164 | 0 | 0 | 0 | 0 | 0 | 0 |
| ENSG00000165548 | 276 | 140 | 135 | 148 | 179 | 127 |
| ENSG00000177108 | 6 | 3 | 6 | 5 | 3 | 3 |
| ENSG00000269883 | 0 | 0 | 0 | 0 | 0 | 0 |

| | | | | | | |
|---|---|---|---|---|---|---|
| ENSG00000258552 | 0 | 0 | 1 | 0 | 0 | 0 |
| ENSG00000201384 | 0 | 1 | 0 | 0 | 0 | 0 |
| ENSG00000165553 | 0 | 0 | 0 | 0 | 0 | 1 |
| ENSG00000221754 | 0 | 0 | 0 | 0 | 0 | 0 |
| ENSG00000009830 | 1555 | 687 | 805 | 812 | 838 | 880 |
| ENSG00000100577 | 2595 | 1338 | 1558 | 1574 | 1391 | 1367 |
| ENSG00000100580 | 2116 | 988 | 1238 | 645 | 1031 | 999 |
| ENSG00000258510 | 0 | 1 | 1 | 0 | 0 | 1 |
| ENSG00000100583 | 102 | 73 | 93 | 63 | 48 | 47 |
| ENSG00000165555 | 64 | 18 | 38 | 16 | 40 | 27 |
| ENSG00000151445 | 1592 | 760 | 848 | 757 | 807 | 816 |
| ENSG00000100591 | 5471 | 2878 | 3067 | 3836 | 4018 | 3762 |
| ENSG00000212371 | 7 | 3 | 1 | 1 | 2 | 3 |
| ENSG00000100593 | 6 | 0 | 1 | 2 | 0 | 2 |
| ENSG00000100596 | 4049 | 1906 | 2313 | 1329 | 2351 | 1864 |
| ENSG00000240233 | 0 | 1 | 0 | 0 | 0 | 0 |
| ENSG00000258668 | 0 | 0 | 0 | 0 | 0 | 0 |
| ENSG00000100601 | 674 | 363 | 412 | 364 | 421 | 367 |
| ENSG00000239272 | 4 | 2 | 2 | 1 | 4 | 2 |
| ENSG00000258917 | 0 | 2 | 0 | 0 | 0 | 0 |
| ENSG00000119705 | 1646 | 841 | 947 | 1158 | 1005 | 1084 |
| ENSG00000100603 | 2439 | 1411 | 1788 | 1257 | 1691 | 1440 |
| ENSG00000197734 | 2 | 1 | 8 | 3 | 3 | 1 |
| ENSG00000063761 | 540 | 221 | 256 | 303 | 256 | 297 |
| ENSG00000199440 | 0 | 0 | 0 | 0 | 0 | 0 |
| ENSG00000258421 | 0 | 0 | 0 | 0 | 0 | 0 |
| ENSG00000021645 | 0 | 0 | 0 | 0 | 0 | 0 |
| ENSG00000199454 | 0 | 0 | 0 | 0 | 0 | 0 |
| ENSG00000239438 | 0 | 0 | 0 | 0 | 0 | 0 |
| ENSG00000258723 | 0 | 0 | 0 | 0 | 0 | 0 |
| ENSG00000258719 | 0 | 0 | 0 | 0 | 0 | 0 |
| ENSG00000258874 | 0 | 0 | 0 | 0 | 0 | 0 |
| ENSG00000258478 | 0 | 0 | 0 | 0 | 0 | 0 |
| ENSG00000258829 | 0 | 0 | 0 | 0 | 0 | 0 |
| ENSG00000258419 | 0 | 0 | 0 | 0 | 0 | 0 |
| ENSG00000258662 | 0 | 0 | 0 | 0 | 0 | 0 |
| ENSG00000259106 | 0 | 0 | 0 | 0 | 0 | 0 |
| ENSG00000258637 | 0 | 0 | 0 | 0 | 0 | 0 |
| ENSG00000258416 | 0 | 0 | 0 | 0 | 0 | 0 |
| ENSG00000211448 | 243 | 122 | 51 | 66 | 129 | 45 |
| ENSG00000258766 | 0 | 1 | 2 | 1 | 0 | 0 |
| ENSG00000100629 | 40 | 32 | 19 | 17 | 38 | 34 |
| ENSG00000258452 | 0 | 0 | 0 | 0 | 0 | 0 |
| ENSG00000165409 | 2 | 0 | 1 | 0 | 0 | 0 |
| ENSG00000258915 | 0 | 0 | 0 | 0 | 0 | 0 |
| ENSG00000271705 | 0 | 0 | 0 | 0 | 0 | 0 |
| ENSG00000271656 | 0 | 0 | 0 | 0 | 0 | 0 |
| ENSG00000284959 | 12 | 2 | 6 | 5 | 7 | 7 |
| ENSG00000259167 | 0 | 0 | 0 | 0 | 0 | 0 |
| ENSG00000165417 | 1579 | 729 | 836 | 432 | 724 | 829 |

| | | | | | | |
|---|---|---|---|---|---|---|
| ENSG00000221303 | 0 | 0 | 0 | 0 | 0 | 0 |
| ENSG00000273783 | 113 | 58 | 52 | 38 | 50 | 67 |
| ENSG00000205579 | 0 | 0 | 0 | 0 | 1 | 0 |
| ENSG00000270343 | 0 | 0 | 0 | 1 | 0 | 0 |
| ENSG00000140022 | 139 | 96 | 50 | 30 | 61 | 38 |
| ENSG00000242163 | 0 | 0 | 0 | 0 | 0 | 0 |
| ENSG00000258605 | 0 | 0 | 0 | 0 | 0 | 0 |
| ENSG00000258675 | 0 | 0 | 0 | 0 | 0 | 0 |
| ENSG00000071537 | 4011 | 2017 | 2404 | 1067 | 2355 | 1845 |
| ENSG00000258977 | 0 | 0 | 0 | 0 | 0 | 0 |
| ENSG00000258841 | 0 | 0 | 0 | 0 | 0 | 0 |
| ENSG00000258718 | 0 | 0 | 0 | 0 | 0 | 0 |
| ENSG00000259035 | 0 | 0 | 0 | 0 | 0 | 0 |
| ENSG00000241891 | 0 | 0 | 0 | 0 | 0 | 0 |
| ENSG00000258501 | 0 | 0 | 0 | 0 | 0 | 0 |
| ENSG00000276782 | 0 | 0 | 0 | 0 | 0 | 0 |
| ENSG00000258743 | 0 | 0 | 0 | 0 | 0 | 0 |
| ENSG00000259032 | 6 | 7 | 10 | 8 | 6 | 7 |
| ENSG00000258683 | 0 | 0 | 0 | 0 | 0 | 0 |
| ENSG00000258600 | 0 | 0 | 0 | 0 | 0 | 0 |
| ENSG00000258426 | 0 | 0 | 0 | 0 | 0 | 0 |
| ENSG00000252369 | 0 | 0 | 0 | 0 | 0 | 0 |
| ENSG00000238561 | 0 | 0 | 0 | 0 | 0 | 0 |
| ENSG00000258549 | 0 | 0 | 0 | 0 | 0 | 0 |
| ENSG00000258532 | 0 | 0 | 0 | 0 | 0 | 0 |
| ENSG00000285826 | 0 | 0 | 0 | 0 | 0 | 0 |
| ENSG00000258762 | 0 | 0 | 0 | 0 | 0 | 0 |
| ENSG00000258923 | 0 | 0 | 0 | 0 | 0 | 0 |
| ENSG00000259012 | 0 | 0 | 0 | 0 | 0 | 0 |
| ENSG00000251895 | 0 | 0 | 0 | 0 | 0 | 0 |
| ENSG00000258506 | 0 | 0 | 0 | 0 | 0 | 0 |
| ENSG00000271712 | 0 | 0 | 0 | 0 | 0 | 0 |
| ENSG00000259044 | 0 | 0 | 0 | 0 | 0 | 0 |
| ENSG00000258902 | 0 | 0 | 0 | 0 | 0 | 0 |
| ENSG00000258814 | 0 | 0 | 0 | 0 | 0 | 0 |
| ENSG00000259107 | 0 | 0 | 0 | 0 | 0 | 0 |
| ENSG00000205562 | 0 | 0 | 0 | 0 | 0 | 0 |
| ENSG00000258945 | 0 | 0 | 0 | 0 | 0 | 0 |
| ENSG00000185070 | 0 | 0 | 0 | 2 | 0 | 0 |
| ENSG00000258733 | 0 | 0 | 0 | 0 | 0 | 0 |
| ENSG00000258912 | 0 | 0 | 0 | 0 | 0 | 0 |
| ENSG00000258994 | 0 | 0 | 0 | 0 | 0 | 0 |
| ENSG00000258804 | 0 | 0 | 0 | 0 | 0 | 0 |
| ENSG00000259061 | 0 | 0 | 0 | 0 | 0 | 0 |
| ENSG00000258496 | 0 | 0 | 0 | 0 | 0 | 0 |
| ENSG00000258958 | 0 | 0 | 0 | 0 | 0 | 0 |
| ENSG00000258859 | 0 | 0 | 0 | 0 | 0 | 0 |
| ENSG00000259028 | 0 | 0 | 0 | 0 | 0 | 0 |
| ENSG00000258770 | 0 | 0 | 0 | 0 | 0 | 0 |
| ENSG00000258807 | 0 | 0 | 0 | 0 | 0 | 0 |

| | | | | | | |
|---|---|---|---|---|---|---|
| ENSG00000259077 | 0 | 0 | 0 | 0 | 0 | 0 |
| ENSG00000054983 | 1348 | 671 | 899 | 489 | 649 | 632 |
| ENSG00000259096 | 0 | 0 | 0 | 0 | 0 | 0 |
| ENSG00000252783 | 0 | 0 | 0 | 0 | 0 | 0 |
| ENSG00000140030 | 2 | 1 | 5 | 1 | 0 | 0 |
| ENSG00000258407 | 0 | 0 | 0 | 0 | 0 | 0 |
| ENSG00000258826 | 0 | 0 | 0 | 0 | 0 | 0 |
| ENSG00000258867 | 0 | 0 | 0 | 0 | 0 | 0 |
| ENSG00000258975 | 0 | 0 | 0 | 0 | 0 | 0 |
| ENSG00000259998 | 0 | 0 | 0 | 0 | 0 | 0 |
| ENSG00000258538 | 0 | 0 | 0 | 0 | 0 | 0 |
| ENSG00000100433 | 0 | 0 | 0 | 0 | 0 | 0 |
| ENSG00000042317 | 365 | 175 | 196 | 142 | 196 | 178 |
| ENSG00000070778 | 910 | 422 | 541 | 403 | 487 | 449 |
| ENSG00000258983 | 0 | 2 | 1 | 0 | 0 | 3 |
| ENSG00000222990 | 0 | 0 | 2 | 0 | 0 | 0 |
| ENSG00000258789 | 0 | 0 | 0 | 0 | 0 | 0 |
| ENSG00000100722 | 2220 | 1291 | 1379 | 786 | 1447 | 1169 |
| ENSG00000278576 | 16 | 5 | 6 | 3 | 8 | 2 |
| ENSG00000165521 | 166 | 63 | 59 | 31 | 74 | 46 |
| ENSG00000274492 | 8 | 4 | 3 | 3 | 2 | 1 |
| ENSG00000200653 | 0 | 0 | 0 | 0 | 0 | 0 |
| ENSG00000165533 | 1931 | 808 | 1126 | 734 | 966 | 1023 |
| ENSG00000207342 | 2 | 2 | 4 | 0 | 1 | 0 |
| ENSG00000283627 | 0 | 1 | 1 | 0 | 1 | 0 |
| ENSG00000258990 | 0 | 0 | 0 | 0 | 0 | 0 |
| ENSG00000053254 | 1006 | 459 | 565 | 371 | 538 | 472 |
| ENSG00000277801 | 0 | 0 | 0 | 0 | 0 | 0 |
| ENSG00000259151 | 0 | 1 | 0 | 0 | 1 | 1 |
| ENSG00000258752 | 0 | 0 | 0 | 0 | 0 | 1 |
| ENSG00000258699 | 0 | 0 | 0 | 0 | 0 | 0 |
| ENSG00000258380 | 0 | 0 | 0 | 0 | 0 | 0 |
| ENSG00000259053 | 0 | 0 | 0 | 0 | 0 | 0 |
| ENSG00000258920 | 30 | 11 | 11 | 19 | 8 | 19 |
| ENSG00000259073 | 0 | 0 | 0 | 0 | 0 | 0 |
| ENSG00000201027 | 0 | 0 | 0 | 0 | 0 | 0 |
| ENSG00000258792 | 0 | 0 | 0 | 0 | 0 | 0 |
| ENSG00000252655 | 0 | 0 | 0 | 0 | 0 | 0 |
| ENSG00000200312 | 0 | 0 | 0 | 0 | 0 | 0 |
| ENSG00000259050 | 0 | 0 | 0 | 0 | 0 | 0 |
| ENSG00000140025 | 397 | 194 | 255 | 247 | 274 | 344 |
| ENSG00000230997 | 0 | 0 | 0 | 0 | 0 | 0 |
| ENSG00000042088 | 1194 | 589 | 727 | 560 | 689 | 752 |
| ENSG00000152315 | 6 | 4 | 1 | 3 | 3 | 3 |
| ENSG00000258544 | 0 | 0 | 0 | 0 | 0 | 0 |
| ENSG00000201351 | 0 | 0 | 0 | 0 | 0 | 0 |
| ENSG00000100764 | 2343 | 1282 | 1581 | 1281 | 1563 | 1383 |
| ENSG00000119720 | 1080 | 519 | 724 | 449 | 525 | 548 |
| ENSG00000242571 | 2 | 0 | 0 | 0 | 3 | 0 |
| ENSG00000275198 | 86 | 46 | 74 | 43 | 58 | 52 |

| | | | | | | |
|---|---|---|---|---|---|---|
| ENSG00000198668 | 21716 | 10600 | 13243 | 10365 | 13146 | 12265 |
| ENSG00000258424 | 39 | 19 | 20 | 14 | 9 | 9 |
| ENSG00000258678 | 0 | 4 | 1 | 0 | 1 | 2 |
| ENSG00000233208 | 2 | 0 | 1 | 5 | 1 | 0 |
| ENSG00000259789 | 0 | 0 | 0 | 0 | 0 | 0 |
| ENSG00000165914 | 1003 | 458 | 587 | 495 | 627 | 576 |
| ENSG00000258935 | 4 | 0 | 0 | 0 | 0 | 0 |
| ENSG00000259163 | 1 | 1 | 1 | 2 | 3 | 0 |
| ENSG00000258437 | 0 | 0 | 0 | 0 | 0 | 0 |
| ENSG00000258716 | 0 | 1 | 0 | 0 | 0 | 0 |
| ENSG00000213315 | 4 | 2 | 0 | 1 | 2 | 1 |
| ENSG00000258884 | 30 | 16 | 27 | 32 | 36 | 36 |
| ENSG00000100784 | 534 | 296 | 300 | 124 | 319 | 253 |
| ENSG00000133943 | 121 | 65 | 74 | 54 | 68 | 81 |
| ENSG00000252644 | 0 | 0 | 0 | 0 | 0 | 0 |
| ENSG00000221102 | 0 | 0 | 2 | 2 | 0 | 0 |
| ENSG00000119714 | 331 | 197 | 211 | 170 | 153 | 181 |
| ENSG00000258875 | 14 | 8 | 11 | 2 | 3 | 4 |
| ENSG00000260810 | 65 | 31 | 33 | 18 | 28 | 16 |
| ENSG00000015133 | 1924 | 831 | 1168 | 689 | 781 | 907 |
| ENSG00000258798 | 3 | 3 | 5 | 5 | 1 | 4 |
| ENSG00000258446 | 5 | 4 | 3 | 4 | 1 | 2 |
| ENSG00000100796 | 3814 | 1852 | 2032 | 1383 | 2145 | 1861 |
| ENSG00000278342 | 0 | 0 | 0 | 0 | 0 | 0 |
| ENSG00000279286 | 8 | 3 | 4 | 0 | 1 | 0 |
| ENSG00000133962 | 31 | 25 | 19 | 9 | 51 | 28 |
| ENSG00000268295 | 0 | 1 | 0 | 1 | 1 | 0 |
| ENSG00000258774 | 6 | 1 | 0 | 0 | 2 | 2 |
| ENSG00000258962 | 0 | 1 | 0 | 0 | 0 | 0 |
| ENSG00000275154 | 0 | 0 | 1 | 0 | 0 | 0 |
| ENSG00000260711 | 98 | 45 | 40 | 24 | 56 | 47 |
| ENSG00000165929 | 3296 | 1718 | 1968 | 1066 | 1904 | 1684 |
| ENSG00000140092 | 2 | 1 | 0 | 0 | 1 | 4 |
| ENSG00000100815 | 451 | 210 | 214 | 85 | 461 | 282 |
| ENSG00000259020 | 0 | 0 | 0 | 0 | 4 | 0 |
| ENSG00000066427 | 589 | 380 | 316 | 230 | 459 | 288 |
| ENSG00000183648 | 2596 | 1284 | 1451 | 1365 | 1235 | 1341 |
| ENSG00000165934 | 2821 | 1462 | 1578 | 1012 | 1882 | 1409 |
| ENSG00000258481 | 0 | 0 | 0 | 0 | 0 | 0 |
| ENSG00000201097 | 0 | 0 | 0 | 0 | 0 | 0 |
| ENSG00000140090 | 0 | 0 | 0 | 0 | 0 | 0 |
| ENSG00000100599 | 120 | 63 | 108 | 138 | 102 | 101 |
| ENSG00000100600 | 1394 | 667 | 805 | 784 | 803 | 771 |
| ENSG00000066455 | 2240 | 1177 | 1370 | 880 | 1529 | 1268 |
| ENSG00000258742 | 0 | 0 | 0 | 0 | 0 | 0 |
| ENSG00000258499 | 0 | 0 | 0 | 0 | 0 | 0 |
| ENSG00000100604 | 2 | 0 | 2 | 0 | 0 | 4 |
| ENSG00000100605 | 4915 | 2469 | 3022 | 2763 | 2511 | 2587 |
| ENSG00000258730 | 6 | 5 | 1 | 0 | 1 | 2 |
| ENSG00000258595 | 0 | 0 | 0 | 0 | 0 | 0 |

| | | | | | | |
|---|---|---|---|---|---|---|
| ENSG00000165943 | 2546 | 1224 | 1691 | 1149 | 1485 | 1522 |
| ENSG00000153485 | 513 | 337 | 388 | 298 | 371 | 370 |
| ENSG00000259066 | 3 | 1 | 3 | 3 | 4 | 1 |
| ENSG00000170270 | 725 | 299 | 421 | 398 | 435 | 426 |
| ENSG00000012963 | 2858 | 1383 | 1693 | 1025 | 1359 | 1418 |
| ENSG00000240096 | 4 | 1 | 4 | 0 | 3 | 3 |
| ENSG00000011114 | 2447 | 1173 | 1463 | 622 | 1165 | 1078 |
| ENSG00000241499 | 0 | 1 | 0 | 0 | 0 | 0 |
| ENSG00000133958 | 16 | 8 | 21 | 12 | 12 | 13 |
| ENSG00000278396 | 10 | 7 | 10 | 4 | 8 | 9 |
| ENSG00000187581 | 0 | 1 | 1 | 0 | 2 | 0 |
| ENSG00000202306 | 0 | 0 | 0 | 0 | 0 | 0 |
| ENSG00000252720 | 0 | 0 | 0 | 0 | 0 | 0 |
| ENSG00000277826 | 0 | 0 | 0 | 0 | 0 | 0 |
| ENSG00000175785 | 6 | 11 | 1 | 4 | 1 | 8 |
| ENSG00000279593 | 0 | 0 | 0 | 0 | 0 | 0 |
| ENSG00000258584 | 0 | 0 | 0 | 0 | 0 | 0 |
| ENSG00000140067 | 0 | 0 | 0 | 0 | 0 | 0 |
| ENSG00000100628 | 0 | 0 | 1 | 0 | 0 | 0 |
| ENSG00000258987 | 0 | 0 | 0 | 0 | 0 | 0 |
| ENSG00000265768 | 0 | 0 | 0 | 0 | 0 | 0 |
| ENSG00000175699 | 0 | 0 | 0 | 0 | 0 | 0 |
| ENSG00000089723 | 303 | 145 | 162 | 154 | 183 | 146 |
| ENSG00000089737 | 5583 | 2737 | 3393 | 2802 | 3019 | 3193 |
| ENSG00000165948 | 175 | 89 | 125 | 127 | 130 | 140 |
| ENSG00000165949 | 99 | 53 | 55 | 58 | 56 | 50 |
| ENSG00000119632 | 138 | 77 | 97 | 84 | 81 | 84 |
| ENSG00000119698 | 6 | 4 | 2 | 0 | 7 | 4 |
| ENSG00000140093 | 0 | 0 | 0 | 0 | 0 | 0 |
| ENSG00000170099 | 0 | 0 | 0 | 0 | 0 | 0 |
| ENSG00000258597 | 0 | 0 | 0 | 0 | 0 | 0 |
| ENSG00000197249 | 77 | 30 | 45 | 22 | 15 | 23 |
| ENSG00000256357 | 0 | 0 | 0 | 0 | 0 | 0 |
| ENSG00000186910 | 4 | 6 | 4 | 5 | 2 | 4 |
| ENSG00000170054 | 1 | 0 | 3 | 1 | 1 | 0 |
| ENSG00000165953 | 5 | 2 | 3 | 1 | 0 | 1 |
| ENSG00000100665 | 6 | 1 | 1 | 0 | 2 | 2 |
| ENSG00000188488 | 401 | 229 | 299 | 157 | 115 | 218 |
| ENSG00000273259 | 6 | 3 | 1 | 0 | 2 | 6 |
| ENSG00000196136 | 49 | 26 | 34 | 9 | 10 | 14 |
| ENSG00000258805 | 0 | 0 | 0 | 0 | 0 | 0 |
| ENSG00000187483 | 0 | 0 | 0 | 0 | 1 | 0 |
| ENSG00000259140 | 0 | 0 | 0 | 0 | 0 | 0 |
| ENSG00000239269 | 0 | 0 | 0 | 1 | 1 | 0 |
| ENSG00000133937 | 108 | 33 | 59 | 60 | 43 | 38 |
| ENSG00000285205 | 7 | 4 | 16 | 5 | 6 | 8 |
| ENSG00000240914 | 0 | 2 | 0 | 1 | 1 | 1 |
| ENSG00000274379 | 0 | 0 | 0 | 0 | 0 | 0 |
| ENSG00000258933 | 0 | 0 | 0 | 0 | 0 | 0 |
| ENSG00000258866 | 0 | 0 | 0 | 0 | 0 | 0 |

| | | | | | | |
|---|---|---|---|---|---|---|
| ENSG00000100697 | 2028 | 1113 | 990 | 425 | 1570 | 905 |
| ENSG00000264607 | 0 | 0 | 0 | 0 | 0 | 0 |
| ENSG00000235706 | 243 | 98 | 164 | 137 | 168 | 159 |
| ENSG00000165959 | 2049 | 1018 | 1345 | 791 | 1053 | 1255 |
| ENSG00000259143 | 0 | 0 | 0 | 0 | 0 | 0 |
| ENSG00000258615 | 2 | 0 | 0 | 0 | 0 | 0 |
| ENSG00000258630 | 0 | 2 | 3 | 3 | 0 | 0 |
| ENSG00000176438 | 36 | 23 | 17 | 12 | 18 | 10 |
| ENSG00000258572 | 0 | 0 | 1 | 1 | 0 | 0 |
| ENSG00000247092 | 335 | 166 | 253 | 219 | 225 | 233 |
| ENSG00000252481 | 7 | 15 | 14 | 6 | 13 | 12 |
| ENSG00000182512 | 2263 | 1213 | 1295 | 1453 | 1402 | 1340 |
| ENSG00000258390 | 0 | 0 | 0 | 3 | 0 | 0 |
| ENSG00000258927 | 0 | 0 | 0 | 0 | 0 | 0 |
| ENSG00000270038 | 0 | 0 | 0 | 0 | 0 | 0 |
| ENSG00000187621 | 0 | 0 | 1 | 0 | 0 | 0 |
| ENSG00000275033 | 0 | 0 | 0 | 0 | 0 | 0 |
| ENSG00000259084 | 0 | 0 | 0 | 0 | 0 | 0 |
| ENSG00000277468 | 0 | 0 | 0 | 0 | 0 | 0 |
| ENSG00000213231 | 0 | 0 | 0 | 0 | 0 | 0 |
| ENSG00000100721 | 0 | 0 | 0 | 0 | 0 | 0 |
| ENSG00000257275 | 0 | 0 | 0 | 0 | 0 | 0 |
| ENSG00000258516 | 0 | 0 | 0 | 0 | 0 | 0 |
| ENSG00000250366 | 0 | 0 | 0 | 0 | 0 | 0 |
| ENSG00000271358 | 0 | 0 | 2 | 1 | 0 | 0 |
| ENSG00000227051 | 7582 | 3380 | 3582 | 2360 | 4156 | 2843 |
| ENSG00000168398 | 559 | 292 | 326 | 105 | 136 | 161 |
| ENSG00000258691 | 0 | 0 | 0 | 0 | 0 | 0 |
| ENSG00000258412 | 0 | 2 | 1 | 0 | 0 | 0 |
| ENSG00000258788 | 4 | 3 | 4 | 3 | 4 | 1 |
| ENSG00000259036 | 0 | 0 | 1 | 0 | 0 | 0 |
| ENSG00000100739 | 11 | 2 | 2 | 1 | 1 | 0 |
| ENSG00000258793 | 0 | 0 | 0 | 0 | 0 | 0 |
| ENSG00000258729 | 0 | 0 | 1 | 0 | 0 | 0 |
| ENSG00000066739 | 1624 | 842 | 708 | 357 | 1004 | 715 |
| ENSG00000100744 | 1288 | 728 | 648 | 472 | 853 | 642 |
| ENSG00000222276 | 0 | 1 | 1 | 0 | 0 | 0 |
| ENSG00000140057 | 801 | 372 | 471 | 275 | 431 | 383 |
| ENSG00000259059 | 0 | 0 | 0 | 0 | 0 | 0 |
| ENSG00000240051 | 3 | 3 | 0 | 1 | 1 | 0 |
| ENSG00000260806 | 114 | 52 | 84 | 69 | 64 | 52 |
| ENSG00000090060 | 6461 | 3674 | 3347 | 2280 | 4699 | 3499 |
| ENSG00000276182 | 31 | 17 | 12 | 16 | 17 | 16 |
| ENSG00000258702 | 0 | 1 | 1 | 0 | 0 | 1 |
| ENSG00000223299 | 0 | 0 | 0 | 0 | 0 | 0 |
| ENSG00000258979 | 0 | 0 | 0 | 0 | 0 | 0 |
| ENSG00000100749 | 642 | 323 | 394 | 361 | 426 | 375 |
| ENSG00000225163 | 6 | 0 | 2 | 2 | 3 | 2 |
| ENSG00000280070 | 0 | 0 | 0 | 0 | 0 | 0 |
| ENSG00000259026 | 0 | 0 | 0 | 0 | 0 | 0 |

| | | | | | | |
|---|---|---|---|---|---|---|
| ENSG00000258393 | 0 | 0 | 0 | 0 | 0 | 0 |
| ENSG00000259110 | 0 | 0 | 0 | 0 | 0 | 0 |
| ENSG00000285584 | 0 | 0 | 0 | 0 | 0 | 0 |
| ENSG00000246084 | 0 | 0 | 0 | 0 | 0 | 0 |
| ENSG00000197176 | 0 | 0 | 0 | 0 | 0 | 0 |
| ENSG00000258869 | 0 | 0 | 0 | 0 | 0 | 0 |
| ENSG00000258379 | 0 | 0 | 0 | 0 | 0 | 0 |
| ENSG00000246223 | 0 | 0 | 0 | 0 | 0 | 0 |
| ENSG00000259097 | 0 | 0 | 0 | 0 | 0 | 0 |
| ENSG00000258511 | 0 | 0 | 0 | 0 | 0 | 0 |
| ENSG00000241757 | 0 | 0 | 0 | 0 | 0 | 0 |
| ENSG00000258795 | 0 | 0 | 0 | 0 | 0 | 0 |
| ENSG00000176605 | 0 | 0 | 0 | 0 | 0 | 0 |
| ENSG00000237342 | 0 | 0 | 0 | 0 | 0 | 0 |
| ENSG00000278990 | 0 | 0 | 0 | 0 | 0 | 0 |
| ENSG00000259119 | 0 | 0 | 0 | 0 | 0 | 0 |
| ENSG00000232573 | 36 | 35 | 30 | 27 | 38 | 34 |
| ENSG00000234648 | 6 | 3 | 10 | 3 | 4 | 5 |
| ENSG00000229402 | 0 | 0 | 0 | 0 | 0 | 0 |
| ENSG00000127152 | 110 | 56 | 58 | 65 | 45 | 55 |
| ENSG00000235785 | 0 | 0 | 0 | 0 | 0 | 0 |
| ENSG00000258383 | 0 | 0 | 0 | 0 | 0 | 0 |
| ENSG00000183576 | 3301 | 1571 | 2017 | 1615 | 2043 | 1968 |
| ENSG00000272439 | 0 | 0 | 0 | 0 | 0 | 0 |
| ENSG00000090061 | 596 | 271 | 371 | 262 | 313 | 366 |
| ENSG00000205476 | 2094 | 1029 | 1379 | 1286 | 1195 | 1261 |
| ENSG00000258749 | 0 | 0 | 0 | 0 | 0 | 0 |
| ENSG00000200506 | 0 | 0 | 0 | 0 | 0 | 0 |
| ENSG00000247970 | 5 | 1 | 8 | 4 | 5 | 1 |
| ENSG00000182218 | 24 | 6 | 7 | 3 | 3 | 5 |
| ENSG00000036530 | 56 | 31 | 36 | 26 | 33 | 32 |
| ENSG00000258672 | 0 | 0 | 0 | 0 | 2 | 0 |
| ENSG00000066629 | 1472 | 773 | 875 | 545 | 715 | 686 |
| ENSG00000199836 | 0 | 0 | 0 | 0 | 0 | 0 |
| ENSG00000271198 | 2 | 0 | 1 | 0 | 0 | 0 |
| ENSG00000196405 | 8053 | 3081 | 6619 | 2813 | 2687 | 3722 |
| ENSG00000242888 | 0 | 0 | 4 | 2 | 0 | 1 |
| ENSG00000258693 | 0 | 0 | 0 | 0 | 0 | 1 |
| ENSG00000258560 | 0 | 0 | 0 | 0 | 0 | 0 |
| ENSG00000265154 | 2 | 0 | 0 | 0 | 0 | 1 |
| ENSG00000199082 | 0 | 0 | 0 | 0 | 0 | 0 |
| ENSG00000168350 | 1006 | 377 | 663 | 435 | 263 | 363 |
| ENSG00000258982 | 13 | 10 | 27 | 7 | 9 | 21 |
| ENSG00000100811 | 5313 | 2400 | 2981 | 2344 | 2927 | 2914 |
| ENSG00000258904 | 5 | 2 | 5 | 2 | 5 | 4 |
| ENSG00000283840 | 0 | 0 | 0 | 0 | 0 | 0 |
| ENSG00000259052 | 0 | 1 | 1 | 1 | 1 | 3 |
| ENSG00000197119 | 3740 | 1827 | 2496 | 1784 | 1866 | 1956 |
| ENSG00000258504 | 0 | 0 | 0 | 0 | 0 | 0 |
| ENSG00000198984 | 0 | 0 | 1 | 0 | 0 | 0 |

| | | | | | | |
|---|---|---|---|---|---|---|
| ENSG00000243976 | 0 | 0 | 0 | 0 | 0 | 0 |
| ENSG00000140107 | 0 | 3 | 1 | 1 | 0 | 1 |
| ENSG00000140105 | 3960 | 1950 | 2516 | 2339 | 2180 | 2449 |
| ENSG00000258666 | 0 | 0 | 0 | 0 | 0 | 0 |
| ENSG00000258521 | 2 | 2 | 3 | 1 | 0 | 0 |
| ENSG00000258581 | 3 | 2 | 0 | 0 | 1 | 1 |
| ENSG00000176473 | 420 | 212 | 268 | 252 | 210 | 207 |
| ENSG00000258620 | 0 | 1 | 0 | 0 | 0 | 0 |
| ENSG00000258500 | 0 | 1 | 0 | 0 | 0 | 0 |
| ENSG00000253075 | 0 | 0 | 0 | 0 | 0 | 0 |
| ENSG00000183092 | 45 | 24 | 38 | 34 | 47 | 43 |
| ENSG00000259031 | 9 | 2 | 1 | 2 | 0 | 3 |
| ENSG00000258576 | 0 | 0 | 0 | 0 | 0 | 0 |
| ENSG00000196273 | 0 | 0 | 0 | 0 | 0 | 0 |
| ENSG00000258717 | 0 | 0 | 0 | 0 | 0 | 0 |
| ENSG00000185559 | 0 | 1 | 1 | 0 | 0 | 0 |
| ENSG00000214548 | 0 | 0 | 0 | 0 | 0 | 0 |
| ENSG00000266461 | 0 | 0 | 0 | 0 | 0 | 0 |
| ENSG00000275161 | 0 | 0 | 0 | 0 | 0 | 0 |
| ENSG00000274884 | 0 | 0 | 0 | 0 | 0 | 0 |
| ENSG00000276919 | 0 | 0 | 0 | 0 | 0 | 0 |
| ENSG00000258663 | 0 | 0 | 0 | 0 | 0 | 0 |
| ENSG00000258399 | 0 | 0 | 0 | 0 | 0 | 0 |
| ENSG00000211574 | 0 | 0 | 0 | 0 | 0 | 0 |
| ENSG00000207989 | 0 | 0 | 0 | 0 | 0 | 0 |
| ENSG00000199151 | 0 | 0 | 0 | 0 | 0 | 0 |
| ENSG00000283159 | 0 | 0 | 0 | 0 | 0 | 0 |
| ENSG00000254656 | 0 | 0 | 0 | 0 | 0 | 1 |
| ENSG00000208001 | 0 | 0 | 0 | 0 | 0 | 0 |
| ENSG00000207569 | 0 | 0 | 0 | 0 | 0 | 0 |
| ENSG00000207608 | 0 | 0 | 0 | 0 | 0 | 0 |
| ENSG00000272458 | 0 | 0 | 0 | 0 | 0 | 0 |
| ENSG00000207942 | 0 | 0 | 0 | 0 | 0 | 0 |
| ENSG00000225746 | 0 | 0 | 0 | 1 | 0 | 0 |
| ENSG00000276225 | 0 | 0 | 0 | 0 | 0 | 0 |
| ENSG00000275662 | 0 | 0 | 0 | 0 | 0 | 0 |
| ENSG00000275134 | 0 | 0 | 0 | 0 | 0 | 0 |
| ENSG00000199005 | 0 | 0 | 0 | 0 | 0 | 0 |
| ENSG00000202191 | 0 | 0 | 0 | 0 | 0 | 0 |
| ENSG00000212384 | 0 | 0 | 0 | 0 | 0 | 0 |
| ENSG00000201700 | 0 | 0 | 0 | 0 | 0 | 0 |
| ENSG00000201672 | 0 | 0 | 0 | 0 | 0 | 0 |
| ENSG00000272474 | 0 | 0 | 0 | 0 | 0 | 0 |
| ENSG00000200215 | 0 | 0 | 0 | 0 | 0 | 0 |
| ENSG00000200632 | 0 | 0 | 0 | 0 | 0 | 0 |
| ENSG00000200367 | 0 | 0 | 0 | 0 | 0 | 0 |
| ENSG00000201950 | 0 | 0 | 0 | 0 | 0 | 0 |
| ENSG00000199575 | 0 | 0 | 0 | 0 | 0 | 0 |
| ENSG00000200823 | 0 | 0 | 0 | 0 | 0 | 0 |
| ENSG00000201839 | 0 | 0 | 0 | 0 | 0 | 0 |

| | | | | | | |
|---|---|---|---|---|---|---|
| ENSG00000200832 | 0 | 0 | 0 | 0 | 0 | 0 |
| ENSG00000199798 | 0 | 0 | 0 | 0 | 0 | 0 |
| ENSG00000200150 | 0 | 0 | 0 | 0 | 0 | 0 |
| ENSG00000201263 | 0 | 0 | 0 | 0 | 0 | 0 |
| ENSG00000199390 | 0 | 0 | 0 | 0 | 0 | 0 |
| ENSG00000201240 | 0 | 0 | 0 | 0 | 0 | 0 |
| ENSG00000200279 | 0 | 0 | 0 | 0 | 0 | 0 |
| ENSG00000200608 | 0 | 0 | 0 | 0 | 0 | 0 |
| ENSG00000202270 | 0 | 0 | 0 | 0 | 0 | 0 |
| ENSG00000201247 | 0 | 0 | 0 | 0 | 0 | 0 |
| ENSG00000199593 | 0 | 0 | 0 | 0 | 0 | 0 |
| ENSG00000201557 | 0 | 0 | 0 | 0 | 0 | 0 |
| ENSG00000199914 | 0 | 0 | 0 | 0 | 0 | 0 |
| ENSG00000201569 | 0 | 0 | 0 | 0 | 0 | 0 |
| ENSG00000202142 | 0 | 0 | 0 | 0 | 0 | 0 |
| ENSG00000199942 | 0 | 0 | 0 | 0 | 0 | 0 |
| ENSG00000201500 | 0 | 0 | 0 | 0 | 0 | 0 |
| ENSG00000201036 | 0 | 0 | 0 | 0 | 0 | 0 |
| ENSG00000201710 | 0 | 0 | 0 | 0 | 0 | 0 |
| ENSG00000202048 | 0 | 0 | 0 | 0 | 0 | 0 |
| ENSG00000272344 | 0 | 0 | 0 | 0 | 0 | 0 |
| ENSG00000202293 | 0 | 0 | 0 | 0 | 0 | 0 |
| ENSG00000200406 | 0 | 0 | 0 | 0 | 0 | 0 |
| ENSG00000201899 | 0 | 0 | 0 | 0 | 0 | 0 |
| ENSG00000200612 | 0 | 0 | 0 | 0 | 0 | 0 |
| ENSG00000200413 | 0 | 0 | 0 | 0 | 0 | 0 |
| ENSG00000200636 | 0 | 0 | 0 | 0 | 0 | 0 |
| ENSG00000200480 | 0 | 0 | 0 | 0 | 0 | 0 |
| ENSG00000201689 | 0 | 0 | 0 | 0 | 0 | 0 |
| ENSG00000201318 | 0 | 0 | 0 | 0 | 0 | 0 |
| ENSG00000200089 | 0 | 0 | 0 | 0 | 0 | 0 |
| ENSG00000222095 | 0 | 0 | 0 | 0 | 0 | 0 |
| ENSG00000222185 | 0 | 0 | 0 | 0 | 0 | 0 |
| ENSG00000199088 | 0 | 0 | 0 | 0 | 0 | 0 |
| ENSG00000199109 | 0 | 0 | 0 | 0 | 0 | 0 |
| ENSG00000207749 | 0 | 0 | 0 | 0 | 0 | 0 |
| ENSG00000198982 | 0 | 0 | 0 | 0 | 0 | 0 |
| ENSG00000221745 | 0 | 0 | 0 | 0 | 0 | 0 |
| ENSG00000199069 | 0 | 0 | 0 | 0 | 0 | 0 |
| ENSG00000211582 | 0 | 0 | 0 | 0 | 0 | 0 |
| ENSG00000207761 | 0 | 0 | 0 | 0 | 0 | 0 |
| ENSG00000207762 | 0 | 0 | 0 | 0 | 0 | 0 |
| ENSG00000194717 | 0 | 0 | 0 | 0 | 0 | 0 |
| ENSG00000221036 | 0 | 0 | 0 | 0 | 0 | 0 |
| ENSG00000212040 | 0 | 0 | 0 | 0 | 0 | 0 |
| ENSG00000207743 | 0 | 0 | 0 | 0 | 0 | 0 |
| ENSG00000283279 | 0 | 0 | 0 | 0 | 0 | 0 |
| ENSG00000283561 | 0 | 0 | 0 | 0 | 0 | 0 |
| ENSG00000207934 | 0 | 0 | 0 | 0 | 0 | 0 |
| ENSG00000283556 | 0 | 0 | 0 | 0 | 0 | 0 |

| | | | | | | |
|---|---|---|---|---|---|---|
| ENSG00000283588 | 0 | 0 | 0 | 0 | 0 | 0 |
| ENSG00000215957 | 0 | 0 | 0 | 0 | 0 | 0 |
| ENSG00000221525 | 0 | 0 | 0 | 0 | 0 | 0 |
| ENSG00000221614 | 0 | 0 | 0 | 0 | 0 | 0 |
| ENSG00000258861 | 0 | 0 | 0 | 0 | 0 | 0 |
| ENSG00000199020 | 0 | 0 | 0 | 0 | 0 | 0 |
| ENSG00000207754 | 0 | 0 | 0 | 0 | 0 | 0 |
| ENSG00000202560 | 0 | 0 | 0 | 0 | 0 | 0 |
| ENSG00000216099 | 0 | 0 | 0 | 0 | 0 | 0 |
| ENSG00000207587 | 0 | 0 | 0 | 0 | 0 | 0 |
| ENSG00000207646 | 0 | 0 | 0 | 0 | 0 | 0 |
| ENSG00000207742 | 0 | 0 | 0 | 0 | 0 | 0 |
| ENSG00000283170 | 0 | 0 | 0 | 0 | 0 | 0 |
| ENSG00000207993 | 0 | 0 | 0 | 0 | 0 | 0 |
| ENSG00000276352 | 0 | 0 | 0 | 0 | 0 | 0 |
| ENSG00000208027 | 0 | 0 | 0 | 0 | 0 | 0 |
| ENSG00000208004 | 0 | 0 | 0 | 0 | 0 | 0 |
| ENSG00000207978 | 0 | 0 | 0 | 0 | 0 | 0 |
| ENSG00000207961 | 0 | 0 | 0 | 0 | 0 | 0 |
| ENSG00000199015 | 0 | 0 | 0 | 0 | 0 | 0 |
| ENSG00000223403 | 2 | 2 | 4 | 1 | 2 | 0 |
| ENSG00000216179 | 0 | 0 | 0 | 0 | 0 | 0 |
| ENSG00000199107 | 0 | 0 | 0 | 0 | 0 | 0 |
| ENSG00000199012 | 0 | 0 | 0 | 0 | 0 | 0 |
| ENSG00000199025 | 0 | 0 | 0 | 0 | 0 | 0 |
| ENSG00000199092 | 0 | 0 | 0 | 0 | 0 | 0 |
| ENSG00000207959 | 0 | 0 | 0 | 0 | 0 | 0 |
| ENSG00000230805 | 0 | 0 | 0 | 0 | 0 | 0 |
| ENSG00000259004 | 0 | 0 | 0 | 0 | 0 | 0 |
| ENSG00000277754 | 0 | 0 | 0 | 0 | 0 | 0 |
| ENSG00000258681 | 0 | 0 | 0 | 0 | 0 | 0 |
| ENSG00000258497 | 0 | 0 | 0 | 0 | 0 | 0 |
| ENSG00000258460 | 0 | 0 | 0 | 0 | 0 | 0 |
| ENSG00000259023 | 0 | 0 | 1 | 1 | 0 | 2 |
| ENSG00000259082 | 0 | 0 | 0 | 0 | 0 | 0 |
| ENSG00000259166 | 0 | 0 | 0 | 0 | 0 | 0 |
| ENSG00000258498 | 6 | 6 | 7 | 1 | 4 | 6 |
| ENSG00000283857 | 0 | 0 | 0 | 0 | 0 | 0 |
| ENSG00000197406 | 16 | 12 | 12 | 9 | 7 | 12 |
| ENSG00000258919 | 1 | 0 | 0 | 0 | 0 | 0 |
| ENSG00000258404 | 0 | 0 | 0 | 0 | 0 | 0 |
| ENSG00000241494 | 0 | 1 | 0 | 0 | 2 | 1 |
| ENSG00000258512 | 0 | 0 | 0 | 1 | 1 | 1 |
| ENSG00000078304 | 5186 | 2624 | 2953 | 2431 | 3251 | 2984 |
| ENSG00000259088 | 12 | 14 | 18 | 14 | 11 | 13 |
| ENSG00000207208 | 0 | 0 | 0 | 0 | 0 | 0 |
| ENSG00000256705 | 0 | 1 | 0 | 0 | 2 | 1 |
| ENSG00000259161 | 0 | 0 | 0 | 0 | 0 | 0 |
| ENSG00000271780 | 127 | 61 | 74 | 80 | 77 | 81 |
| ENSG00000272444 | 25 | 11 | 21 | 19 | 13 | 14 |

| | | | | | | |
|---|---|---|---|---|---|---|
| ENSG00000197102 | 14751 | 6473 | 8714 | 5124 | 8131 | 7518 |
| ENSG00000259013 | 0 | 2 | 0 | 0 | 1 | 0 |
| ENSG00000258959 | 0 | 0 | 0 | 0 | 0 | 0 |
| ENSG00000240457 | 0 | 1 | 0 | 0 | 1 | 0 |
| ENSG00000080824 | 18734 | 13645 | 9681 | 7414 | 27799 | 13543 |
| ENSG00000140153 | 1400 | 636 | 761 | 595 | 686 | 749 |
| ENSG00000080823 | 353 | 157 | 237 | 171 | 206 | 175 |
| ENSG00000022976 | 549 | 266 | 367 | 247 | 283 | 318 |
| ENSG00000100865 | 1186 | 609 | 730 | 777 | 697 | 738 |
| ENSG00000196663 | 1550 | 717 | 909 | 748 | 811 | 814 |
| ENSG00000212330 | 0 | 0 | 0 | 0 | 0 | 0 |
| ENSG00000156381 | 2158 | 1024 | 1238 | 1393 | 1290 | 1265 |
| ENSG00000266015 | 0 | 0 | 0 | 0 | 0 | 0 |
| ENSG00000259230 | 0 | 0 | 0 | 0 | 0 | 0 |
| ENSG00000239224 | 0 | 0 | 0 | 0 | 0 | 0 |
| ENSG00000089902 | 2535 | 1149 | 1457 | 919 | 1218 | 1382 |
| ENSG00000230988 | 0 | 0 | 1 | 0 | 0 | 0 |
| ENSG00000202459 | 0 | 0 | 0 | 0 | 1 | 0 |
| ENSG00000259508 | 0 | 0 | 0 | 0 | 0 | 0 |
| ENSG00000131323 | 1481 | 701 | 881 | 631 | 854 | 868 |
| ENSG00000206969 | 0 | 0 | 0 | 0 | 0 | 0 |
| ENSG00000166126 | 11 | 7 | 5 | 3 | 4 | 7 |
| ENSG00000198752 | 5212 | 2411 | 3360 | 2337 | 3003 | 3187 |
| ENSG00000259515 | 8 | 6 | 12 | 3 | 6 | 3 |
| ENSG00000213176 | 0 | 0 | 0 | 0 | 0 | 0 |
| ENSG00000283071 | 2 | 0 | 0 | 0 | 0 | 0 |
| ENSG00000259444 | 0 | 0 | 0 | 0 | 0 | 0 |
| ENSG00000205436 | 37 | 24 | 23 | 17 | 20 | 18 |
| ENSG00000259717 | 0 | 0 | 0 | 0 | 0 | 0 |
| ENSG00000185215 | 648 | 403 | 451 | 275 | 343 | 372 |
| ENSG00000259374 | 0 | 0 | 0 | 0 | 0 | 0 |
| ENSG00000259535 | 0 | 0 | 0 | 0 | 0 | 0 |
| ENSG00000259525 | 0 | 0 | 0 | 0 | 0 | 0 |
| ENSG00000251533 | 0 | 0 | 0 | 0 | 0 | 0 |
| ENSG00000270705 | 0 | 0 | 0 | 0 | 0 | 0 |
| ENSG00000278071 | 10 | 5 | 7 | 2 | 6 | 4 |
| ENSG00000242837 | 0 | 0 | 0 | 0 | 0 | 0 |
| ENSG00000270938 | 0 | 0 | 0 | 0 | 0 | 0 |
| ENSG00000240624 | 0 | 0 | 0 | 0 | 0 | 0 |
| ENSG00000259775 | 6 | 8 | 13 | 10 | 10 | 15 |
| ENSG00000100664 | 9092 | 4531 | 5262 | 3490 | 5318 | 4933 |
| ENSG00000272533 | 1 | 1 | 2 | 0 | 1 | 1 |
| ENSG00000259428 | 0 | 0 | 0 | 0 | 0 | 0 |
| ENSG00000243904 | 0 | 1 | 1 | 0 | 0 | 0 |
| ENSG00000075413 | 3687 | 1739 | 2186 | 1570 | 2056 | 2085 |
| ENSG00000244691 | 2 | 0 | 0 | 0 | 0 | 2 |
| ENSG00000166165 | 1486 | 711 | 1033 | 860 | 608 | 660 |
| ENSG00000260285 | 898 | 294 | 566 | 266 | 289 | 342 |
| ENSG00000166166 | 1328 | 646 | 771 | 888 | 684 | 745 |
| ENSG00000252469 | 0 | 1 | 0 | 1 | 0 | 0 |

| | | | | | | |
|---|---|---|---|---|---|---|
| ENSG00000258851 | 10 | 0 | 2 | 0 | 1 | 1 |
| ENSG00000166170 | 2159 | 1041 | 1243 | 922 | 1161 | 1201 |
| ENSG00000126214 | 1672 | 859 | 1151 | 858 | 1031 | 1107 |
| ENSG00000256500 | 0 | 0 | 1 | 0 | 0 | 0 |
| ENSG00000256053 | 636 | 430 | 436 | 437 | 428 | 422 |
| ENSG00000201184 | 0 | 0 | 0 | 0 | 0 | 0 |
| ENSG00000246451 | 182 | 93 | 106 | 57 | 100 | 81 |
| ENSG00000270108 | 11 | 8 | 5 | 5 | 3 | 6 |
| ENSG00000269910 | 11 | 18 | 19 | 8 | 23 | 17 |
| ENSG00000269940 | 38 | 26 | 48 | 39 | 44 | 51 |
| ENSG00000269958 | 472 | 210 | 269 | 235 | 288 | 250 |
| ENSG00000126215 | 668 | 298 | 393 | 411 | 398 | 376 |
| ENSG00000100711 | 2338 | 1145 | 1470 | 1233 | 1274 | 1292 |
| ENSG00000253096 | 0 | 0 | 0 | 0 | 0 | 0 |
| ENSG00000088808 | 1950 | 854 | 1157 | 817 | 957 | 904 |
| ENSG00000270591 | 3 | 2 | 1 | 0 | 1 | 0 |
| ENSG00000202275 | 0 | 0 | 0 | 0 | 0 | 0 |
| ENSG00000258735 | 15 | 2 | 1 | 4 | 3 | 7 |
| ENSG00000258534 | 2 | 0 | 4 | 2 | 1 | 2 |
| ENSG00000258914 | 0 | 0 | 0 | 1 | 0 | 2 |
| ENSG00000156411 | 4370 | 2247 | 2574 | 2737 | 2565 | 2426 |
| ENSG00000156414 | 0 | 0 | 0 | 0 | 0 | 0 |
| ENSG00000227729 | 0 | 0 | 0 | 0 | 0 | 0 |
| ENSG00000242894 | 0 | 0 | 0 | 0 | 0 | 0 |
| ENSG00000166183 | 15 | 21 | 6 | 4 | 5 | 11 |
| ENSG00000207568 | 0 | 0 | 0 | 0 | 0 | 0 |
| ENSG00000283165 | 0 | 0 | 0 | 0 | 0 | 0 |
| ENSG00000258748 | 18 | 6 | 10 | 10 | 11 | 2 |
| ENSG00000066735 | 491 | 189 | 238 | 241 | 212 | 224 |
| ENSG00000258913 | 0 | 0 | 0 | 0 | 0 | 0 |
| ENSG00000258598 | 0 | 0 | 0 | 0 | 0 | 0 |
| ENSG00000222761 | 0 | 0 | 0 | 0 | 0 | 0 |
| ENSG00000213159 | 0 | 0 | 0 | 0 | 0 | 0 |
| ENSG00000258986 | 2 | 0 | 1 | 0 | 0 | 1 |
| ENSG00000184601 | 0 | 0 | 0 | 0 | 0 | 0 |
| ENSG00000259037 | 0 | 0 | 0 | 0 | 0 | 0 |
| ENSG00000258736 | 2 | 1 | 0 | 0 | 0 | 0 |
| ENSG00000260792 | 0 | 0 | 0 | 0 | 0 | 0 |
| ENSG00000265291 | 0 | 0 | 0 | 0 | 0 | 0 |
| ENSG00000256050 | 0 | 2 | 0 | 1 | 7 | 3 |
| ENSG00000203485 | 5093 | 2547 | 3003 | 3400 | 3186 | 2865 |
| ENSG00000258858 | 0 | 1 | 0 | 2 | 0 | 0 |
| ENSG00000185100 | 327 | 184 | 250 | 222 | 186 | 220 |
| ENSG00000184990 | 2635 | 1420 | 1538 | 1767 | 1416 | 1393 |
| ENSG00000142208 | 11739 | 6189 | 7824 | 6878 | 6538 | 6457 |
| ENSG00000258430 | 0 | 0 | 0 | 0 | 0 | 0 |
| ENSG00000179627 | 2020 | 946 | 1252 | 1023 | 854 | 921 |
| ENSG00000279140 | 54 | 23 | 28 | 23 | 32 | 31 |
| ENSG00000258701 | 174 | 76 | 100 | 90 | 80 | 75 |
| ENSG00000239365 | 18 | 8 | 8 | 6 | 9 | 10 |

| | | | | | | |
|---|---|---|---|---|---|---|
| ENSG00000196183 | 15 | 11 | 12 | 2 | 3 | 5 |
| ENSG00000258811 | 23 | 31 | 35 | 15 | 32 | 16 |
| ENSG00000258593 | 15 | 7 | 14 | 8 | 7 | 3 |
| ENSG00000099814 | 4580 | 1945 | 2854 | 2535 | 2492 | 2517 |
| ENSG00000259067 | 0 | 0 | 0 | 0 | 0 | 0 |
| ENSG00000166428 | 10 | 2 | 4 | 3 | 0 | 2 |
| ENSG00000185567 | 170 | 46 | 88 | 42 | 66 | 49 |
| ENSG00000140104 | 918 | 447 | 539 | 509 | 484 | 563 |
| ENSG00000170779 | 994 | 476 | 648 | 541 | 566 | 666 |
| ENSG00000183484 | 56 | 41 | 58 | 54 | 80 | 63 |
| ENSG00000257556 | 0 | 1 | 3 | 0 | 0 | 0 |
| ENSG00000280244 | 0 | 0 | 0 | 0 | 0 | 0 |
| ENSG00000258154 | 0 | 0 | 0 | 0 | 0 | 0 |
| ENSG00000184916 | 1764 | 799 | 1062 | 897 | 863 | 908 |
| ENSG00000273778 | 0 | 0 | 0 | 0 | 0 | 0 |
| ENSG00000257622 | 0 | 0 | 0 | 0 | 0 | 1 |
| ENSG00000183828 | 776 | 372 | 488 | 543 | 393 | 438 |
| ENSG00000185024 | 2401 | 1077 | 1453 | 1364 | 1135 | 1254 |
| ENSG00000184887 | 1821 | 961 | 1133 | 1287 | 1001 | 1073 |
| ENSG00000179364 | 3873 | 1706 | 2306 | 2230 | 2076 | 2142 |
| ENSG00000279495 | 77 | 46 | 43 | 26 | 46 | 35 |
| ENSG00000242085 | 22 | 8 | 7 | 2 | 11 | 5 |
| ENSG00000226174 | 89 | 38 | 48 | 39 | 41 | 22 |
| ENSG00000251602 | 24 | 24 | 18 | 19 | 18 | 20 |
| ENSG00000182979 | 4059 | 1839 | 2509 | 2279 | 2239 | 2107 |
| ENSG00000257270 | 119 | 57 | 98 | 64 | 66 | 56 |
| ENSG00000182809 | 10658 | 4563 | 6166 | 6096 | 4563 | 4692 |
| ENSG00000213145 | 71 | 20 | 37 | 35 | 32 | 30 |
| ENSG00000257341 | 0 | 0 | 2 | 0 | 0 | 1 |
| ENSG00000185347 | 998 | 499 | 704 | 659 | 525 | 570 |
| ENSG00000184986 | 288 | 142 | 170 | 155 | 125 | 132 |
| ENSG00000230157 | 0 | 0 | 0 | 1 | 1 | 0 |
| ENSG00000234402 | 0 | 0 | 0 | 0 | 0 | 0 |
| ENSG00000211890 | 0 | 1 | 1 | 0 | 0 | 0 |
| ENSG00000211891 | 104 | 40 | 49 | 37 | 26 | 33 |
| ENSG00000211892 | 2 | 0 | 0 | 0 | 1 | 1 |
| ENSG00000274172 | 0 | 0 | 0 | 0 | 0 | 0 |
| ENSG00000211893 | 1 | 0 | 0 | 0 | 0 | 0 |
| ENSG00000277030 | 0 | 0 | 0 | 0 | 0 | 0 |
| ENSG00000253364 | 0 | 0 | 0 | 0 | 0 | 0 |
| ENSG00000253755 | 0 | 0 | 0 | 0 | 0 | 0 |
| ENSG00000213140 | 0 | 0 | 0 | 0 | 0 | 0 |
| ENSG00000211895 | 0 | 0 | 4 | 2 | 2 | 0 |
| ENSG00000253692 | 2 | 1 | 1 | 0 | 0 | 3 |
| ENSG00000211896 | 0 | 0 | 0 | 0 | 0 | 0 |
| ENSG00000211897 | 0 | 0 | 0 | 0 | 2 | 0 |
| ENSG00000270371 | 0 | 0 | 0 | 0 | 0 | 0 |
| ENSG00000211898 | 0 | 0 | 0 | 0 | 0 | 0 |
| ENSG00000211899 | 2 | 0 | 1 | 0 | 1 | 0 |
| ENSG00000265612 | 0 | 0 | 0 | 0 | 0 | 0 |

| | | | | | | |
|---|---|---|---|---|---|---|
| ENSG00000264024 | 0 | 0 | 0 | 0 | 0 | 0 |
| ENSG00000263413 | 0 | 0 | 0 | 0 | 0 | 0 |
| ENSG00000264781 | 0 | 0 | 0 | 0 | 0 | 0 |
| ENSG00000211900 | 0 | 0 | 0 | 0 | 0 | 0 |
| ENSG00000237111 | 0 | 0 | 0 | 0 | 0 | 0 |
| ENSG00000242472 | 0 | 0 | 0 | 0 | 0 | 0 |
| ENSG00000240041 | 0 | 0 | 0 | 0 | 0 | 0 |
| ENSG00000242887 | 0 | 0 | 0 | 0 | 0 | 0 |
| ENSG00000237547 | 0 | 0 | 0 | 0 | 0 | 0 |
| ENSG00000211904 | 0 | 0 | 0 | 0 | 0 | 0 |
| ENSG00000211905 | 0 | 0 | 0 | 0 | 0 | 0 |
| ENSG00000236597 | 0 | 0 | 0 | 0 | 0 | 0 |
| ENSG00000227335 | 0 | 0 | 0 | 0 | 0 | 0 |
| ENSG00000211907 | 0 | 0 | 0 | 0 | 0 | 0 |
| ENSG00000225825 | 0 | 0 | 0 | 0 | 0 | 0 |
| ENSG00000211909 | 0 | 0 | 0 | 0 | 0 | 0 |
| ENSG00000227196 | 0 | 0 | 0 | 0 | 0 | 0 |
| ENSG00000211911 | 0 | 0 | 0 | 0 | 0 | 0 |
| ENSG00000211912 | 0 | 0 | 0 | 0 | 0 | 0 |
| ENSG00000244620 | 0 | 0 | 0 | 1 | 0 | 0 |
| ENSG00000237020 | 0 | 0 | 0 | 0 | 0 | 0 |
| ENSG00000211914 | 0 | 0 | 0 | 0 | 0 | 0 |
| ENSG00000211915 | 0 | 0 | 0 | 0 | 0 | 0 |
| ENSG00000227800 | 0 | 0 | 0 | 0 | 0 | 0 |
| ENSG00000211917 | 0 | 0 | 0 | 0 | 0 | 0 |
| ENSG00000211918 | 0 | 0 | 0 | 0 | 0 | 0 |
| ENSG00000227108 | 0 | 0 | 0 | 0 | 0 | 0 |
| ENSG00000211920 | 0 | 0 | 0 | 0 | 0 | 0 |
| ENSG00000211921 | 0 | 0 | 0 | 0 | 0 | 0 |
| ENSG00000232543 | 0 | 0 | 0 | 0 | 0 | 0 |
| ENSG00000211923 | 0 | 0 | 0 | 0 | 0 | 0 |
| ENSG00000211924 | 0 | 0 | 0 | 0 | 0 | 0 |
| ENSG00000211925 | 0 | 0 | 0 | 0 | 0 | 0 |
| ENSG00000237197 | 0 | 0 | 0 | 0 | 0 | 0 |
| ENSG00000228131 | 0 | 0 | 0 | 0 | 0 | 0 |
| ENSG00000211928 | 0 | 0 | 0 | 0 | 0 | 0 |
| ENSG00000233655 | 0 | 0 | 0 | 0 | 0 | 0 |
| ENSG00000211930 | 0 | 0 | 0 | 0 | 0 | 0 |
| ENSG00000211931 | 0 | 0 | 0 | 0 | 0 | 0 |
| ENSG00000226777 | 1 | 2 | 2 | 0 | 1 | 2 |
| ENSG00000236170 | 0 | 0 | 0 | 0 | 0 | 0 |
| ENSG00000211933 | 0 | 0 | 0 | 0 | 0 | 0 |
| ENSG00000254279 | 0 | 0 | 0 | 0 | 0 | 0 |
| ENSG00000271968 | 0 | 0 | 0 | 0 | 0 | 0 |
| ENSG00000225200 | 10 | 2 | 3 | 8 | 3 | 6 |
| ENSG00000211934 | 0 | 0 | 0 | 0 | 0 | 0 |
| ENSG00000253780 | 0 | 0 | 0 | 0 | 0 | 0 |
| ENSG00000211935 | 0 | 0 | 0 | 0 | 0 | 0 |
| ENSG00000276775 | 0 | 0 | 0 | 0 | 0 | 0 |
| ENSG00000282122 | 0 | 0 | 0 | 0 | 0 | 0 |

| | | | | | | |
|---|---|---|---|---|---|---|
| ENSG00000211937 | 0 | 0 | 0 | 0 | 0 | 0 |
| ENSG00000253387 | 0 | 0 | 0 | 0 | 0 | 0 |
| ENSG00000254053 | 0 | 0 | 0 | 0 | 0 | 0 |
| ENSG00000253763 | 0 | 0 | 0 | 0 | 0 | 0 |
| ENSG00000211938 | 0 | 0 | 0 | 0 | 0 | 0 |
| ENSG00000282639 | 0 | 0 | 0 | 0 | 0 | 0 |
| ENSG00000282651 | 0 | 0 | 0 | 0 | 0 | 0 |
| ENSG00000211941 | 0 | 0 | 0 | 0 | 0 | 0 |
| ENSG00000254215 | 0 | 0 | 0 | 0 | 0 | 0 |
| ENSG00000254174 | 0 | 0 | 0 | 0 | 0 | 0 |
| ENSG00000211942 | 0 | 0 | 0 | 0 | 0 | 0 |
| ENSG00000253412 | 0 | 0 | 0 | 0 | 0 | 0 |
| ENSG00000253709 | 0 | 0 | 0 | 0 | 0 | 0 |
| ENSG00000211943 | 0 | 0 | 0 | 0 | 0 | 0 |
| ENSG00000253458 | 0 | 0 | 0 | 0 | 0 | 0 |
| ENSG00000211944 | 0 | 0 | 0 | 0 | 0 | 0 |
| ENSG00000283948 | 0 | 0 | 0 | 0 | 0 | 0 |
| ENSG00000254046 | 0 | 0 | 0 | 0 | 0 | 0 |
| ENSG00000273894 | 0 | 0 | 0 | 0 | 0 | 0 |
| ENSG00000211945 | 0 | 0 | 0 | 0 | 0 | 0 |
| ENSG00000253883 | 0 | 0 | 0 | 0 | 0 | 0 |
| ENSG00000271201 | 0 | 0 | 0 | 0 | 0 | 0 |
| ENSG00000211946 | 0 | 0 | 0 | 0 | 0 | 0 |
| ENSG00000211947 | 0 | 0 | 0 | 0 | 0 | 0 |
| ENSG00000253957 | 0 | 0 | 0 | 0 | 0 | 0 |
| ENSG00000253345 | 0 | 0 | 0 | 0 | 0 | 0 |
| ENSG00000254045 | 0 | 0 | 0 | 0 | 0 | 0 |
| ENSG00000211949 | 0 | 0 | 0 | 0 | 0 | 0 |
| ENSG00000211950 | 0 | 0 | 0 | 0 | 0 | 0 |
| ENSG00000238275 | 0 | 0 | 0 | 0 | 0 | 0 |
| ENSG00000276210 | 0 | 0 | 0 | 0 | 0 | 0 |
| ENSG00000253441 | 0 | 0 | 0 | 0 | 0 | 0 |
| ENSG00000253367 | 0 | 0 | 0 | 0 | 0 | 0 |
| ENSG00000211951 | 0 | 0 | 0 | 0 | 0 | 0 |
| ENSG00000253462 | 0 | 0 | 0 | 0 | 0 | 0 |
| ENSG00000253482 | 0 | 0 | 0 | 0 | 0 | 0 |
| ENSG00000254326 | 0 | 0 | 0 | 0 | 0 | 0 |
| ENSG00000211952 | 0 | 0 | 0 | 0 | 0 | 0 |
| ENSG00000253149 | 0 | 0 | 0 | 0 | 0 | 0 |
| ENSG00000254289 | 0 | 0 | 0 | 0 | 0 | 0 |
| ENSG00000270550 | 0 | 0 | 0 | 0 | 0 | 0 |
| ENSG00000253491 | 0 | 0 | 0 | 0 | 0 | 0 |
| ENSG00000253587 | 0 | 0 | 0 | 0 | 0 | 0 |
| ENSG00000231475 | 0 | 0 | 1 | 0 | 0 | 0 |
| ENSG00000283607 | 0 | 0 | 0 | 0 | 0 | 0 |
| ENSG00000270474 | 0 | 0 | 0 | 0 | 0 | 0 |
| ENSG00000211955 | 0 | 0 | 0 | 0 | 0 | 0 |
| ENSG00000254203 | 0 | 0 | 0 | 0 | 0 | 0 |
| ENSG00000253440 | 0 | 0 | 0 | 0 | 0 | 0 |
| ENSG00000211956 | 0 | 0 | 0 | 0 | 0 | 0 |

| | | | | | | |
|---|---|---|---|---|---|---|
| ENSG00000253325 | 0 | 0 | 0 | 0 | 0 | 0 |
| ENSG00000211957 | 0 | 0 | 0 | 0 | 0 | 0 |
| ENSG00000253240 | 0 | 0 | 0 | 0 | 0 | 0 |
| ENSG00000253359 | 0 | 0 | 0 | 0 | 0 | 0 |
| ENSG00000211958 | 0 | 0 | 0 | 0 | 0 | 0 |
| ENSG00000253989 | 0 | 0 | 0 | 0 | 0 | 0 |
| ENSG00000211959 | 0 | 0 | 0 | 0 | 0 | 0 |
| ENSG00000253467 | 0 | 0 | 0 | 0 | 0 | 0 |
| ENSG00000228757 | 0 | 0 | 0 | 0 | 0 | 0 |
| ENSG00000253294 | 0 | 0 | 0 | 0 | 0 | 0 |
| ENSG00000278473 | 0 | 0 | 0 | 0 | 0 | 0 |
| ENSG00000228966 | 0 | 0 | 0 | 0 | 1 | 0 |
| ENSG00000254228 | 0 | 0 | 0 | 0 | 0 | 0 |
| ENSG00000232216 | 0 | 0 | 0 | 0 | 0 | 0 |
| ENSG00000283195 | 0 | 0 | 0 | 0 | 0 | 0 |
| ENSG00000283562 | 0 | 0 | 0 | 0 | 0 | 0 |
| ENSG00000283464 | 0 | 0 | 0 | 0 | 0 | 0 |
| ENSG00000270816 | 0 | 0 | 0 | 0 | 0 | 0 |
| ENSG00000253465 | 0 | 0 | 0 | 0 | 0 | 0 |
| ENSG00000253895 | 0 | 0 | 0 | 0 | 0 | 0 |
| ENSG00000211961 | 0 | 0 | 0 | 0 | 0 | 0 |
| ENSG00000211962 | 0 | 0 | 0 | 0 | 0 | 0 |
| ENSG00000253808 | 0 | 0 | 0 | 0 | 0 | 0 |
| ENSG00000229092 | 0 | 0 | 0 | 0 | 0 | 0 |
| ENSG00000253862 | 0 | 0 | 0 | 0 | 0 | 0 |
| ENSG00000211964 | 0 | 0 | 0 | 0 | 0 | 0 |
| ENSG00000211965 | 0 | 0 | 0 | 0 | 0 | 0 |
| ENSG00000253386 | 0 | 0 | 0 | 0 | 0 | 0 |
| ENSG00000253241 | 0 | 0 | 0 | 0 | 0 | 0 |
| ENSG00000211966 | 0 | 0 | 0 | 0 | 0 | 0 |
| ENSG00000254167 | 0 | 0 | 0 | 0 | 0 | 0 |
| ENSG00000253941 | 0 | 0 | 0 | 0 | 0 | 0 |
| ENSG00000253545 | 0 | 0 | 0 | 0 | 0 | 0 |
| ENSG00000211967 | 0 | 0 | 0 | 0 | 0 | 0 |
| ENSG00000253714 | 0 | 0 | 0 | 0 | 0 | 0 |
| ENSG00000275475 | 0 | 0 | 0 | 0 | 0 | 0 |
| ENSG00000254395 | 0 | 0 | 0 | 0 | 0 | 0 |
| ENSG00000253131 | 0 | 0 | 0 | 0 | 0 | 0 |
| ENSG00000253759 | 0 | 0 | 0 | 0 | 0 | 0 |
| ENSG00000211968 | 0 | 0 | 0 | 0 | 0 | 0 |
| ENSG00000224373 | 0 | 0 | 0 | 0 | 0 | 0 |
| ENSG00000253742 | 0 | 0 | 0 | 0 | 0 | 0 |
| ENSG00000199480 | 0 | 0 | 0 | 0 | 0 | 0 |
| ENSG00000254329 | 0 | 0 | 0 | 0 | 0 | 0 |
| ENSG00000211970 | 0 | 0 | 0 | 0 | 0 | 0 |
| ENSG00000253132 | 0 | 0 | 0 | 0 | 0 | 0 |
| ENSG00000253747 | 0 | 0 | 0 | 0 | 0 | 0 |
| ENSG00000253936 | 0 | 0 | 0 | 0 | 0 | 0 |
| ENSG00000223648 | 0 | 0 | 0 | 0 | 0 | 0 |
| ENSG00000253209 | 0 | 0 | 0 | 0 | 0 | 0 |

| | | | | | | |
|---|---|---|---|---|---|---|
| ENSG00000253169 | 0 | 0 | 0 | 0 | 0 | 0 |
| ENSG00000211972 | 0 | 0 | 0 | 0 | 0 | 0 |
| ENSG00000253274 | 0 | 0 | 0 | 0 | 0 | 0 |
| ENSG00000223506 | 0 | 0 | 0 | 0 | 0 | 0 |
| ENSG00000253820 | 0 | 0 | 0 | 0 | 0 | 0 |
| ENSG00000254036 | 0 | 0 | 0 | 0 | 0 | 0 |
| ENSG00000253635 | 0 | 0 | 0 | 0 | 0 | 0 |
| ENSG00000254052 | 0 | 0 | 0 | 0 | 0 | 0 |
| ENSG00000253703 | 0 | 0 | 0 | 0 | 0 | 0 |
| ENSG00000211973 | 0 | 0 | 0 | 0 | 0 | 0 |
| ENSG00000211974 | 0 | 0 | 0 | 0 | 0 | 0 |
| ENSG00000282600 | 0 | 0 | 0 | 0 | 0 | 0 |
| ENSG00000281990 | 0 | 0 | 0 | 0 | 0 | 0 |
| ENSG00000283571 | 0 | 0 | 0 | 0 | 0 | 0 |
| ENSG00000280411 | 0 | 0 | 0 | 0 | 0 | 0 |
| ENSG00000274576 | 0 | 0 | 0 | 0 | 0 | 0 |
| ENSG00000254056 | 0 | 0 | 0 | 0 | 0 | 0 |
| ENSG00000225698 | 0 | 0 | 0 | 0 | 0 | 0 |
| ENSG00000211976 | 0 | 0 | 0 | 0 | 0 | 0 |
| ENSG00000224650 | 0 | 0 | 1 | 0 | 3 | 0 |
| ENSG00000254134 | 0 | 0 | 0 | 0 | 0 | 0 |
| ENSG00000254176 | 0 | 0 | 0 | 0 | 0 | 0 |
| ENSG00000253247 | 0 | 0 | 0 | 0 | 0 | 0 |
| ENSG00000253310 | 0 | 0 | 0 | 0 | 0 | 0 |
| ENSG00000265929 | 0 | 0 | 0 | 0 | 0 | 0 |
| ENSG00000211978 | 0 | 0 | 0 | 0 | 1 | 0 |
| ENSG00000253674 | 0 | 0 | 0 | 0 | 0 | 0 |
| ENSG00000275747 | 0 | 0 | 1 | 0 | 0 | 0 |
| ENSG00000283906 | 0 | 0 | 0 | 0 | 0 | 0 |
| ENSG00000211979 | 0 | 0 | 0 | 0 | 0 | 0 |
| ENSG00000253303 | 0 | 0 | 0 | 0 | 0 | 0 |
| ENSG00000215567 | 0 | 0 | 0 | 0 | 0 | 0 |
| ENSG00000201241 | 0 | 0 | 0 | 0 | 0 | 0 |
| ENSG00000258463 | 0 | 0 | 0 | 0 | 0 | 0 |
| ENSG00000274347 | 0 | 0 | 0 | 0 | 0 | 0 |
| ENSG00000188403 | 0 | 0 | 0 | 0 | 0 | 0 |
| ENSG00000258706 | 0 | 0 | 0 | 0 | 0 | 0 |
| ENSG00000259337 | 0 | 0 | 0 | 0 | 0 | 0 |
| ENSG00000259490 | 0 | 0 | 0 | 0 | 0 | 0 |
| ENSG00000270961 | 0 | 0 | 0 | 0 | 0 | 0 |
| ENSG00000271317 | 0 | 0 | 0 | 0 | 0 | 0 |
| ENSG00000282520 | 0 | 0 | 0 | 0 | 0 | 0 |
| ENSG00000282599 | 0 | 0 | 0 | 0 | 0 | 0 |
| ENSG00000271336 | 0 | 0 | 0 | 0 | 0 | 0 |
| ENSG00000277988 | 0 | 0 | 0 | 0 | 0 | 0 |
| ENSG00000259769 | 0 | 0 | 0 | 0 | 0 | 0 |
| ENSG00000258397 | 0 | 0 | 0 | 0 | 0 | 0 |
| ENSG00000239471 | 0 | 0 | 0 | 0 | 0 | 0 |
| ENSG00000258410 | 0 | 0 | 0 | 0 | 0 | 0 |
| ENSG00000278935 | 0 | 0 | 0 | 0 | 0 | 0 |

| | | | | | | |
|---|---|---|---|---|---|---|
| ENSG00000258780 | 0 | 0 | 0 | 0 | 0 | 0 |
| ENSG00000258628 | 0 | 0 | 0 | 0 | 0 | 0 |
| ENSG00000259156 | 0 | 0 | 0 | 0 | 0 | 0 |
| ENSG00000258654 | 0 | 0 | 0 | 0 | 0 | 0 |
| ENSG00000180229 | 24 | 10 | 18 | 7 | 17 | 13 |
| ENSG00000258488 | 0 | 0 | 0 | 0 | 0 | 0 |
| ENSG00000277322 | 0 | 0 | 0 | 0 | 0 | 0 |
| ENSG00000258797 | 0 | 0 | 0 | 0 | 0 | 0 |
| ENSG00000181984 | 0 | 0 | 0 | 0 | 0 | 0 |
| ENSG00000278497 | 0 | 0 | 0 | 0 | 0 | 0 |
| ENSG00000258916 | 0 | 0 | 0 | 0 | 0 | 0 |
| ENSG00000258707 | 0 | 0 | 0 | 0 | 0 | 0 |
| ENSG00000270685 | 0 | 0 | 0 | 0 | 0 | 0 |
| ENSG00000278626 | 2 | 0 | 0 | 0 | 0 | 0 |
| ENSG00000258590 | 3 | 0 | 5 | 1 | 1 | 0 |
| ENSG00000279628 | 0 | 0 | 0 | 0 | 0 | 0 |
| ENSG00000260409 | 6 | 3 | 0 | 0 | 0 | 0 |
| ENSG00000238478 | 0 | 0 | 0 | 0 | 0 | 0 |
| ENSG00000259383 | 0 | 0 | 0 | 0 | 0 | 0 |
| ENSG00000259443 | 0 | 0 | 0 | 0 | 0 | 0 |
| ENSG00000258420 | 0 | 0 | 0 | 0 | 0 | 0 |
| ENSG00000274499 | 0 | 0 | 0 | 0 | 0 | 0 |
| ENSG00000259089 | 0 | 0 | 0 | 0 | 0 | 0 |
| ENSG00000259214 | 0 | 0 | 0 | 0 | 0 | 0 |
| ENSG00000265322 | 0 | 0 | 0 | 0 | 0 | 0 |
| ENSG00000230031 | 0 | 0 | 0 | 0 | 0 | 0 |
| ENSG00000238489 | 0 | 0 | 0 | 0 | 0 | 0 |
| ENSG00000261048 | 0 | 0 | 0 | 0 | 0 | 0 |
| ENSG00000261554 | 0 | 0 | 0 | 0 | 0 | 0 |
| ENSG00000258799 | 0 | 0 | 0 | 0 | 0 | 0 |
| ENSG00000270831 | 0 | 0 | 0 | 0 | 0 | 0 |
| ENSG00000266545 | 0 | 0 | 0 | 0 | 0 | 0 |
| ENSG00000258710 | 0 | 0 | 0 | 0 | 0 | 0 |
| ENSG00000258855 | 0 | 0 | 0 | 0 | 0 | 0 |
| ENSG00000258494 | 0 | 0 | 0 | 0 | 0 | 0 |
| ENSG00000270824 | 0 | 0 | 0 | 0 | 0 | 0 |
| ENSG00000270451 | 0 | 0 | 0 | 0 | 0 | 0 |
| ENSG00000282089 | 0 | 0 | 0 | 0 | 0 | 0 |
| ENSG00000282268 | 0 | 0 | 0 | 0 | 0 | 0 |
| ENSG00000270185 | 0 | 0 | 0 | 0 | 0 | 0 |
| ENSG00000259698 | 0 | 0 | 0 | 0 | 0 | 0 |
| ENSG00000259025 | 0 | 0 | 0 | 0 | 0 | 0 |
| ENSG00000258771 | 0 | 0 | 0 | 0 | 0 | 0 |
| ENSG00000243059 | 0 | 0 | 0 | 0 | 0 | 0 |
| ENSG00000260986 | 0 | 0 | 0 | 0 | 0 | 0 |
| ENSG00000279595 | 0 | 0 | 0 | 0 | 0 | 0 |
| ENSG00000258684 | 0 | 0 | 0 | 0 | 0 | 0 |
| ENSG00000258883 | 0 | 0 | 0 | 0 | 0 | 0 |
| ENSG00000237161 | 0 | 0 | 0 | 0 | 0 | 0 |
| ENSG00000247765 | 0 | 0 | 0 | 0 | 0 | 0 |

| | | | | | | |
|---|---|---|---|---|---|---|
| ENSG00000207289 | 0 | 0 | 0 | 0 | 0 | 0 |
| ENSG00000280881 | 0 | 0 | 0 | 0 | 0 | 0 |
| ENSG00000280703 | 0 | 0 | 0 | 0 | 0 | 0 |
| ENSG00000258415 | 0 | 0 | 0 | 0 | 0 | 0 |
| ENSG00000268531 | 0 | 0 | 0 | 0 | 0 | 0 |
| ENSG00000258712 | 0 | 0 | 0 | 0 | 0 | 0 |
| ENSG00000258767 | 0 | 0 | 0 | 0 | 0 | 0 |
| ENSG00000258848 | 0 | 0 | 0 | 0 | 0 | 0 |
| ENSG00000277243 | 0 | 0 | 0 | 0 | 0 | 0 |
| ENSG00000278522 | 0 | 0 | 0 | 0 | 0 | 0 |
| ENSG00000275174 | 0 | 0 | 0 | 0 | 0 | 0 |
| ENSG00000281347 | 0 | 0 | 0 | 0 | 0 | 0 |
| ENSG00000277841 | 0 | 0 | 0 | 0 | 0 | 0 |
| ENSG00000284834 | 0 | 0 | 0 | 0 | 0 | 0 |
| ENSG00000284788 | 0 | 0 | 0 | 0 | 0 | 0 |
| ENSG00000280709 | 0 | 0 | 0 | 0 | 0 | 0 |
| ENSG00000284988 | 0 | 0 | 0 | 0 | 0 | 0 |
| ENSG00000279639 | 0 | 0 | 0 | 0 | 0 | 0 |
| ENSG00000259324 | 0 | 0 | 0 | 0 | 0 | 0 |
| ENSG00000285491 | 0 | 0 | 0 | 0 | 0 | 0 |
| ENSG00000285116 | 0 | 0 | 0 | 0 | 0 | 0 |
| ENSG00000182974 | 0 | 0 | 0 | 0 | 0 | 0 |
| ENSG00000279408 | 0 | 0 | 0 | 0 | 0 | 0 |
| ENSG00000279445 | 0 | 0 | 0 | 0 | 0 | 0 |
| ENSG00000281113 | 0 | 0 | 0 | 0 | 0 | 0 |
| ENSG00000281087 | 27 | 0 | 0 | 0 | 0 | 0 |
| ENSG00000281179 | 0 | 0 | 0 | 0 | 0 | 0 |
| ENSG00000281550 | 0 | 0 | 0 | 0 | 0 | 0 |
| ENSG00000278263 | 0 | 0 | 0 | 0 | 0 | 0 |
| ENSG00000280655 | 0 | 0 | 0 | 0 | 0 | 0 |
| ENSG00000265793 | 0 | 0 | 0 | 0 | 0 | 0 |
| ENSG00000233917 | 0 | 0 | 0 | 0 | 0 | 0 |
| ENSG00000200790 | 0 | 0 | 0 | 0 | 0 | 0 |
| ENSG00000258585 | 0 | 0 | 0 | 0 | 0 | 0 |
| ENSG00000258997 | 0 | 0 | 0 | 0 | 0 | 0 |
| ENSG00000264902 | 0 | 0 | 0 | 0 | 0 | 0 |
| ENSG00000283524 | 0 | 0 | 0 | 0 | 0 | 0 |
| ENSG00000285405 | 0 | 0 | 0 | 0 | 0 | 0 |
| ENSG00000284500 | 0 | 0 | 0 | 0 | 0 | 0 |
| ENSG00000258721 | 0 | 0 | 0 | 0 | 0 | 0 |
| ENSG00000285472 | 0 | 0 | 0 | 0 | 0 | 0 |
| ENSG00000283680 | 0 | 0 | 0 | 0 | 0 | 0 |
| ENSG00000285135 | 0 | 0 | 0 | 0 | 0 | 0 |
| ENSG00000284892 | 0 | 0 | 0 | 0 | 0 | 0 |
| ENSG00000274102 | 0 | 0 | 0 | 0 | 0 | 0 |
| ENSG00000183706 | 0 | 0 | 0 | 0 | 0 | 0 |
| ENSG00000259435 | 0 | 0 | 0 | 0 | 0 | 0 |
| ENSG00000260739 | 0 | 0 | 0 | 0 | 0 | 0 |
| ENSG00000223877 | 27 | 0 | 0 | 0 | 0 | 0 |
| ENSG00000270505 | 0 | 0 | 0 | 0 | 0 | 0 |

| | | | | | | |
|---|---|---|---|---|---|---|
| ENSG00000271288 | 0 | 0 | 0 | 0 | 0 | 0 |
| ENSG00000259261 | 0 | 0 | 0 | 0 | 0 | 0 |
| ENSG00000270356 | 0 | 0 | 0 | 0 | 0 | 0 |
| ENSG00000271507 | 0 | 0 | 0 | 0 | 0 | 0 |
| ENSG00000221641 | 0 | 0 | 0 | 0 | 0 | 0 |
| ENSG00000259098 | 0 | 0 | 0 | 0 | 0 | 0 |
| ENSG00000258732 | 0 | 0 | 0 | 0 | 0 | 0 |
| ENSG00000274835 | 0 | 0 | 0 | 0 | 0 | 1 |
| ENSG00000273679 | 0 | 0 | 0 | 0 | 0 | 0 |
| ENSG00000274279 | 0 | 0 | 0 | 0 | 0 | 0 |
| ENSG00000185182 | 0 | 0 | 0 | 0 | 0 | 0 |
| ENSG00000274568 | 0 | 0 | 0 | 0 | 0 | 0 |
| ENSG00000275363 | 0 | 0 | 0 | 0 | 0 | 0 |
| ENSG00000277865 | 0 | 0 | 0 | 0 | 0 | 0 |
| ENSG00000281909 | 4 | 0 | 0 | 0 | 0 | 0 |
| ENSG00000276550 | 1183 | 486 | 739 | 294 | 561 | 571 |
| ENSG00000277561 | 2 | 0 | 0 | 0 | 0 | 0 |
| ENSG00000277515 | 0 | 0 | 0 | 0 | 0 | 0 |
| ENSG00000276141 | 18 | 6 | 15 | 4 | 12 | 8 |
| ENSG00000277435 | 0 | 0 | 1 | 1 | 0 | 0 |
| ENSG00000283273 | 0 | 0 | 0 | 0 | 0 | 0 |
| ENSG00000277867 | 0 | 0 | 0 | 0 | 0 | 0 |
| ENSG00000275325 | 0 | 0 | 0 | 0 | 0 | 0 |
| ENSG00000277755 | 0 | 0 | 0 | 0 | 0 | 0 |
| ENSG00000274253 | 4 | 7 | 6 | 5 | 2 | 6 |
| ENSG00000259425 | 2 | 2 | 1 | 0 | 0 | 0 |
| ENSG00000170113 | 1609 | 766 | 940 | 596 | 854 | 847 |
| ENSG00000140157 | 1543 | 802 | 967 | 777 | 931 | 967 |
| ENSG00000273749 | 4767 | 2333 | 2871 | 2325 | 2609 | 2625 |
| ENSG00000275835 | 749 | 345 | 379 | 237 | 400 | 322 |
| ENSG00000276172 | 0 | 0 | 0 | 0 | 0 | 0 |
| ENSG00000273976 | 0 | 0 | 0 | 0 | 0 | 0 |
| ENSG00000277505 | 0 | 0 | 0 | 0 | 0 | 0 |
| ENSG00000175676 | 0 | 0 | 0 | 0 | 0 | 0 |
| ENSG00000273981 | 0 | 0 | 0 | 0 | 0 | 0 |
| ENSG00000274099 | 0 | 0 | 0 | 0 | 0 | 0 |
| ENSG00000278541 | 0 | 0 | 0 | 0 | 0 | 0 |
| ENSG00000276941 | 0 | 0 | 0 | 0 | 0 | 0 |
| ENSG00000278552 | 0 | 0 | 0 | 0 | 0 | 0 |
| ENSG00000259414 | 0 | 3 | 0 | 2 | 0 | 0 |
| ENSG00000274471 | 9 | 2 | 12 | 2 | 11 | 10 |
| ENSG00000273756 | 0 | 0 | 0 | 0 | 0 | 0 |
| ENSG00000260399 | 0 | 0 | 0 | 0 | 0 | 0 |
| ENSG00000261739 | 0 | 0 | 0 | 0 | 0 | 0 |
| ENSG00000278222 | 0 | 0 | 0 | 0 | 0 | 0 |
| ENSG00000261418 | 0 | 0 | 0 | 0 | 0 | 0 |
| ENSG00000174450 | 0 | 0 | 0 | 0 | 0 | 0 |
| ENSG00000265673 | 0 | 0 | 0 | 0 | 0 | 0 |
| ENSG00000179455 | 4 | 0 | 0 | 0 | 1 | 1 |
| ENSG00000260978 | 0 | 1 | 0 | 0 | 0 | 0 |

| | | | | | | |
|---|---|---|---|---|---|---|
| ENSG00000254585 | 0 | 0 | 0 | 0 | 0 | 0 |
| ENSG00000182636 | 0 | 0 | 0 | 0 | 0 | 0 |
| ENSG00000206616 | 0 | 0 | 0 | 0 | 0 | 0 |
| ENSG00000261622 | 0 | 0 | 0 | 0 | 0 | 0 |
| ENSG00000260232 | 0 | 0 | 0 | 0 | 0 | 0 |
| ENSG00000260551 | 0 | 0 | 0 | 0 | 0 | 0 |
| ENSG00000260780 | 0 | 0 | 0 | 0 | 0 | 0 |
| ENSG00000261621 | 0 | 0 | 0 | 0 | 0 | 0 |
| ENSG00000260959 | 0 | 0 | 0 | 0 | 0 | 0 |
| ENSG00000261598 | 0 | 0 | 0 | 0 | 0 | 0 |
| ENSG00000260760 | 0 | 0 | 0 | 0 | 0 | 0 |
| ENSG00000259905 | 0 | 0 | 0 | 0 | 0 | 0 |
| ENSG00000185823 | 0 | 0 | 0 | 0 | 0 | 0 |
| ENSG00000128739 | 1 | 0 | 0 | 0 | 0 | 0 |
| ENSG00000280118 | 0 | 0 | 0 | 0 | 0 | 0 |
| ENSG00000241061 | 0 | 0 | 0 | 4 | 1 | 2 |
| ENSG00000242088 | 4 | 0 | 4 | 1 | 4 | 2 |
| ENSG00000273173 | 0 | 0 | 0 | 0 | 0 | 0 |
| ENSG00000214265 | 0 | 0 | 0 | 0 | 0 | 0 |
| ENSG00000224078 | 2 | 1 | 1 | 0 | 3 | 1 |
| ENSG00000276314 | 0 | 0 | 0 | 0 | 0 | 0 |
| ENSG00000270704 | 0 | 0 | 0 | 0 | 0 | 0 |
| ENSG00000279192 | 0 | 0 | 0 | 0 | 0 | 0 |
| ENSG00000276610 | 0 | 0 | 0 | 0 | 0 | 0 |
| ENSG00000239014 | 0 | 0 | 0 | 0 | 0 | 0 |
| ENSG00000271347 | 0 | 0 | 0 | 0 | 0 | 0 |
| ENSG00000257647 | 0 | 0 | 0 | 0 | 0 | 0 |
| ENSG00000257151 | 0 | 0 | 0 | 0 | 0 | 0 |
| ENSG00000274640 | 0 | 0 | 0 | 0 | 0 | 0 |
| ENSG00000207063 | 0 | 0 | 0 | 0 | 0 | 0 |
| ENSG00000207001 | 0 | 0 | 0 | 0 | 0 | 0 |
| ENSG00000207014 | 0 | 0 | 0 | 0 | 0 | 0 |
| ENSG00000275529 | 0 | 0 | 0 | 0 | 0 | 0 |
| ENSG00000207191 | 0 | 0 | 0 | 0 | 0 | 0 |
| ENSG00000207442 | 0 | 0 | 0 | 0 | 0 | 0 |
| ENSG00000207133 | 0 | 0 | 0 | 0 | 0 | 0 |
| ENSG00000207093 | 0 | 0 | 0 | 0 | 0 | 0 |
| ENSG00000206727 | 0 | 0 | 0 | 0 | 0 | 0 |
| ENSG00000200661 | 0 | 0 | 0 | 0 | 0 | 0 |
| ENSG00000206609 | 0 | 0 | 0 | 0 | 0 | 0 |
| ENSG00000207197 | 0 | 0 | 0 | 0 | 0 | 0 |
| ENSG00000207137 | 0 | 0 | 0 | 0 | 0 | 0 |
| ENSG00000206621 | 0 | 0 | 0 | 0 | 0 | 0 |
| ENSG00000207174 | 0 | 0 | 0 | 0 | 0 | 0 |
| ENSG00000207263 | 0 | 0 | 0 | 0 | 0 | 0 |
| ENSG00000206656 | 0 | 0 | 0 | 0 | 0 | 0 |
| ENSG00000206688 | 0 | 0 | 0 | 0 | 0 | 0 |
| ENSG00000207460 | 0 | 0 | 0 | 0 | 0 | 0 |
| ENSG00000261069 | 0 | 0 | 0 | 0 | 0 | 0 |
| ENSG00000278715 | 0 | 0 | 0 | 0 | 0 | 0 |

| | | | | | | |
|---|---|---|---|---|---|---|
| ENSG00000277785 | 0 | 0 | 0 | 0 | 0 | 0 |
| ENSG00000275127 | 0 | 0 | 0 | 0 | 0 | 0 |
| ENSG00000207375 | 0 | 0 | 0 | 0 | 0 | 0 |
| ENSG00000279735 | 0 | 0 | 0 | 0 | 0 | 0 |
| ENSG00000207279 | 0 | 0 | 0 | 0 | 0 | 0 |
| ENSG00000252326 | 0 | 0 | 0 | 0 | 0 | 0 |
| ENSG00000251815 | 0 | 0 | 0 | 0 | 0 | 0 |
| ENSG00000251896 | 0 | 0 | 0 | 0 | 0 | 0 |
| ENSG00000278123 | 0 | 0 | 0 | 0 | 0 | 0 |
| ENSG00000207245 | 0 | 0 | 0 | 0 | 0 | 0 |
| ENSG00000252277 | 0 | 0 | 0 | 0 | 0 | 0 |
| ENSG00000270523 | 0 | 0 | 0 | 0 | 0 | 0 |
| ENSG00000279050 | 0 | 0 | 0 | 0 | 0 | 0 |
| ENSG00000212428 | 0 | 0 | 0 | 0 | 0 | 0 |
| ENSG00000270814 | 0 | 0 | 0 | 0 | 0 | 0 |
| ENSG00000201831 | 0 | 0 | 0 | 0 | 0 | 0 |
| ENSG00000199712 | 0 | 0 | 0 | 0 | 0 | 0 |
| ENSG00000199970 | 0 | 0 | 0 | 0 | 0 | 0 |
| ENSG00000200680 | 0 | 0 | 0 | 0 | 0 | 0 |
| ENSG00000200503 | 0 | 0 | 0 | 0 | 0 | 0 |
| ENSG00000200812 | 0 | 0 | 0 | 0 | 0 | 0 |
| ENSG00000278089 | 0 | 0 | 0 | 0 | 0 | 0 |
| ENSG00000200726 | 0 | 0 | 0 | 0 | 0 | 0 |
| ENSG00000199782 | 0 | 0 | 0 | 0 | 0 | 0 |
| ENSG00000201943 | 0 | 0 | 0 | 0 | 0 | 0 |
| ENSG00000200486 | 0 | 0 | 0 | 0 | 0 | 0 |
| ENSG00000199453 | 0 | 0 | 0 | 0 | 0 | 0 |
| ENSG00000273835 | 0 | 0 | 0 | 0 | 0 | 0 |
| ENSG00000199960 | 0 | 0 | 0 | 0 | 0 | 0 |
| ENSG00000201679 | 0 | 0 | 0 | 0 | 0 | 0 |
| ENSG00000200757 | 0 | 0 | 0 | 0 | 0 | 0 |
| ENSG00000201482 | 0 | 0 | 0 | 0 | 0 | 0 |
| ENSG00000200163 | 0 | 0 | 0 | 0 | 0 | 0 |
| ENSG00000199968 | 0 | 0 | 0 | 0 | 0 | 0 |
| ENSG00000201969 | 0 | 0 | 0 | 0 | 0 | 0 |
| ENSG00000199833 | 0 | 0 | 0 | 0 | 0 | 0 |
| ENSG00000201326 | 0 | 0 | 0 | 0 | 0 | 0 |
| ENSG00000201331 | 0 | 0 | 0 | 0 | 0 | 0 |
| ENSG00000200398 | 0 | 0 | 0 | 0 | 0 | 0 |
| ENSG00000199489 | 0 | 0 | 0 | 0 | 0 | 0 |
| ENSG00000275524 | 0 | 0 | 0 | 0 | 0 | 0 |
| ENSG00000201300 | 0 | 0 | 0 | 0 | 0 | 0 |
| ENSG00000200801 | 0 | 0 | 0 | 0 | 0 | 0 |
| ENSG00000199704 | 0 | 0 | 0 | 0 | 0 | 0 |
| ENSG00000200987 | 0 | 0 | 0 | 0 | 0 | 0 |
| ENSG00000202188 | 0 | 0 | 0 | 0 | 0 | 0 |
| ENSG00000200949 | 0 | 0 | 0 | 0 | 0 | 0 |
| ENSG00000200593 | 0 | 0 | 0 | 0 | 0 | 0 |
| ENSG00000199311 | 0 | 0 | 0 | 0 | 0 | 0 |
| ENSG00000201992 | 0 | 0 | 0 | 0 | 0 | 0 |

| | | | | | | |
|---|---|---|---|---|---|---|
| ENSG00000202499 | 0 | 0 | 0 | 0 | 0 | 0 |
| ENSG00000200638 | 0 | 0 | 0 | 0 | 0 | 0 |
| ENSG00000201907 | 0 | 0 | 0 | 0 | 0 | 0 |
| ENSG00000200564 | 0 | 0 | 0 | 0 | 0 | 0 |
| ENSG00000272460 | 0 | 0 | 0 | 0 | 0 | 0 |
| ENSG00000200478 | 0 | 0 | 0 | 0 | 0 | 0 |
| ENSG00000201143 | 0 | 0 | 0 | 0 | 0 | 0 |
| ENSG00000202373 | 0 | 0 | 0 | 0 | 0 | 0 |
| ENSG00000202261 | 0 | 0 | 0 | 0 | 0 | 0 |
| ENSG00000280234 | 0 | 0 | 0 | 0 | 0 | 0 |
| ENSG00000212380 | 0 | 0 | 0 | 0 | 0 | 0 |
| ENSG00000276844 | 0 | 0 | 0 | 0 | 0 | 0 |
| ENSG00000212528 | 0 | 0 | 0 | 0 | 0 | 0 |
| ENSG00000201634 | 0 | 0 | 0 | 0 | 0 | 0 |
| ENSG00000239169 | 0 | 0 | 0 | 0 | 0 | 0 |
| ENSG00000114062 | 2111 | 1088 | 1302 | 757 | 1282 | 1001 |
| ENSG00000261529 | 0 | 1 | 0 | 0 | 1 | 0 |
| ENSG00000235731 | 0 | 0 | 0 | 0 | 0 | 0 |
| ENSG00000223628 | 0 | 0 | 0 | 0 | 0 | 0 |
| ENSG00000206190 | 0 | 0 | 0 | 0 | 0 | 0 |
| ENSG00000274307 | 0 | 0 | 0 | 0 | 0 | 0 |
| ENSG00000222123 | 0 | 0 | 0 | 0 | 0 | 0 |
| ENSG00000266517 | 0 | 0 | 0 | 0 | 0 | 0 |
| ENSG00000259011 | 0 | 0 | 0 | 0 | 0 | 0 |
| ENSG00000206187 | 0 | 0 | 0 | 0 | 0 | 0 |
| ENSG00000258965 | 0 | 0 | 0 | 0 | 1 | 0 |
| ENSG00000212604 | 0 | 0 | 0 | 0 | 0 | 0 |
| ENSG00000258931 | 0 | 0 | 0 | 0 | 0 | 0 |
| ENSG00000259150 | 0 | 0 | 0 | 0 | 0 | 0 |
| ENSG00000235160 | 0 | 0 | 0 | 0 | 0 | 0 |
| ENSG00000166206 | 0 | 0 | 0 | 0 | 0 | 0 |
| ENSG00000278840 | 0 | 0 | 0 | 0 | 0 | 0 |
| ENSG00000235518 | 0 | 0 | 0 | 0 | 0 | 0 |
| ENSG00000242793 | 0 | 0 | 0 | 0 | 0 | 0 |
| ENSG00000186297 | 0 | 0 | 0 | 0 | 0 | 0 |
| ENSG00000259122 | 0 | 0 | 0 | 0 | 0 | 0 |
| ENSG00000182256 | 0 | 0 | 0 | 0 | 0 | 0 |
| ENSG00000228740 | 0 | 0 | 0 | 0 | 0 | 0 |
| ENSG00000214254 | 0 | 0 | 0 | 0 | 0 | 0 |
| ENSG00000200326 | 0 | 0 | 0 | 0 | 0 | 0 |
| ENSG00000258624 | 0 | 0 | 0 | 0 | 0 | 0 |
| ENSG00000261426 | 0 | 0 | 0 | 0 | 0 | 0 |
| ENSG00000258970 | 0 | 0 | 0 | 0 | 0 | 0 |
| ENSG00000259152 | 0 | 0 | 0 | 0 | 0 | 0 |
| ENSG00000259168 | 0 | 0 | 0 | 0 | 0 | 0 |
| ENSG00000258594 | 0 | 0 | 0 | 0 | 0 | 0 |
| ENSG00000258853 | 0 | 0 | 0 | 0 | 0 | 0 |
| ENSG00000104044 | 0 | 0 | 0 | 0 | 0 | 0 |
| ENSG00000232394 | 0 | 0 | 0 | 0 | 0 | 0 |
| ENSG00000228992 | 0 | 0 | 0 | 0 | 0 | 0 |

| | | | | | | |
|---|---|---|---|---|---|---|
| ENSG00000128731 | 2952 | 1176 | 1850 | 1010 | 1329 | 1529 |
| ENSG00000256338 | 12 | 4 | 7 | 7 | 5 | 4 |
| ENSG00000261401 | 0 | 0 | 0 | 0 | 0 | 0 |
| ENSG00000261561 | 0 | 0 | 0 | 0 | 0 | 0 |
| ENSG00000237850 | 0 | 0 | 0 | 0 | 0 | 0 |
| ENSG00000261497 | 0 | 0 | 0 | 0 | 0 | 0 |
| ENSG00000153684 | 0 | 0 | 0 | 0 | 0 | 0 |
| ENSG00000276891 | 0 | 0 | 0 | 0 | 0 | 0 |
| ENSG00000261524 | 0 | 0 | 0 | 0 | 0 | 0 |
| ENSG00000276928 | 0 | 0 | 0 | 0 | 0 | 0 |
| ENSG00000274532 | 0 | 0 | 0 | 0 | 0 | 0 |
| ENSG00000266039 | 0 | 0 | 0 | 0 | 0 | 0 |
| ENSG00000284628 | 0 | 0 | 0 | 0 | 0 | 0 |
| ENSG00000260159 | 0 | 0 | 0 | 0 | 0 | 0 |
| ENSG00000261365 | 0 | 0 | 0 | 0 | 0 | 0 |
| ENSG00000263964 | 0 | 0 | 0 | 0 | 0 | 0 |
| ENSG00000274814 | 0 | 0 | 0 | 0 | 0 | 0 |
| ENSG00000260053 | 0 | 0 | 0 | 0 | 0 | 0 |
| ENSG00000183629 | 0 | 0 | 0 | 0 | 0 | 0 |
| ENSG00000276955 | 0 | 0 | 0 | 0 | 0 | 0 |
| ENSG00000261041 | 0 | 0 | 0 | 0 | 0 | 0 |
| ENSG00000227717 | 0 | 0 | 0 | 0 | 0 | 0 |
| ENSG00000260444 | 0 | 0 | 0 | 0 | 0 | 0 |
| ENSG00000261581 | 2 | 0 | 1 | 0 | 0 | 0 |
| ENSG00000206149 | 362 | 166 | 291 | 128 | 194 | 226 |
| ENSG00000254398 | 0 | 0 | 0 | 1 | 1 | 0 |
| ENSG00000188626 | 0 | 0 | 1 | 0 | 0 | 1 |
| ENSG00000274632 | 0 | 0 | 0 | 0 | 0 | 0 |
| ENSG00000261480 | 0 | 0 | 0 | 0 | 0 | 0 |
| ENSG00000248334 | 22 | 8 | 10 | 9 | 16 | 14 |
| ENSG00000260844 | 0 | 0 | 0 | 0 | 0 | 0 |
| ENSG00000232431 | 0 | 0 | 0 | 0 | 0 | 0 |
| ENSG00000256951 | 0 | 0 | 0 | 0 | 0 | 0 |
| ENSG00000261377 | 0 | 0 | 0 | 0 | 0 | 0 |
| ENSG00000271616 | 0 | 0 | 0 | 0 | 0 | 0 |
| ENSG00000261649 | 2 | 0 | 0 | 0 | 0 | 0 |
| ENSG00000270301 | 0 | 0 | 0 | 0 | 0 | 0 |
| ENSG00000034053 | 97 | 58 | 56 | 70 | 87 | 52 |
| ENSG00000259277 | 0 | 0 | 0 | 0 | 0 | 0 |
| ENSG00000104059 | 0 | 1 | 4 | 2 | 2 | 4 |
| ENSG00000259299 | 0 | 0 | 0 | 0 | 0 | 0 |
| ENSG00000252868 | 0 | 0 | 0 | 0 | 0 | 0 |
| ENSG00000185115 | 544 | 271 | 313 | 322 | 322 | 319 |
| ENSG00000259690 | 0 | 0 | 0 | 0 | 0 | 0 |
| ENSG00000259814 | 0 | 0 | 0 | 0 | 0 | 0 |
| ENSG00000259300 | 0 | 0 | 1 | 0 | 0 | 0 |
| ENSG00000256802 | 6 | 4 | 6 | 3 | 9 | 2 |
| ENSG00000104067 | 2272 | 1240 | 1138 | 468 | 1505 | 1080 |
| ENSG00000259644 | 2 | 2 | 2 | 0 | 1 | 3 |
| ENSG00000234664 | 283 | 137 | 152 | 133 | 123 | 123 |

| | | | | | | |
|---|---|---|---|---|---|---|
| ENSG00000259523 | 13 | 7 | 7 | 6 | 8 | 10 |
| ENSG00000259749 | 0 | 0 | 0 | 1 | 0 | 0 |
| ENSG00000259647 | 0 | 0 | 0 | 0 | 0 | 0 |
| ENSG00000260174 | 2 | 0 | 0 | 1 | 1 | 0 |
| ENSG00000229389 | 0 | 0 | 0 | 0 | 0 | 0 |
| ENSG00000179938 | 0 | 0 | 0 | 0 | 0 | 0 |
| ENSG00000273818 | 0 | 0 | 0 | 0 | 0 | 0 |
| ENSG00000261792 | 0 | 0 | 0 | 0 | 0 | 0 |
| ENSG00000178081 | 0 | 0 | 0 | 0 | 1 | 0 |
| ENSG00000207432 | 0 | 0 | 0 | 0 | 0 | 0 |
| ENSG00000261247 | 0 | 0 | 0 | 0 | 1 | 0 |
| ENSG00000277637 | 0 | 0 | 0 | 0 | 0 | 0 |
| ENSG00000260897 | 0 | 0 | 0 | 0 | 0 | 0 |
| ENSG00000261747 | 0 | 0 | 0 | 0 | 0 | 0 |
| ENSG00000259906 | 0 | 0 | 0 | 0 | 0 | 0 |
| ENSG00000225930 | 0 | 0 | 0 | 0 | 0 | 0 |
| ENSG00000259993 | 0 | 0 | 0 | 0 | 1 | 0 |
| ENSG00000206972 | 0 | 0 | 0 | 0 | 0 | 0 |
| ENSG00000270955 | 0 | 0 | 0 | 0 | 0 | 0 |
| ENSG00000279417 | 0 | 0 | 1 | 0 | 0 | 0 |
| ENSG00000238783 | 0 | 0 | 0 | 0 | 0 | 0 |
| ENSG00000166664 | 0 | 0 | 0 | 0 | 0 | 0 |
| ENSG00000263070 | 0 | 0 | 0 | 0 | 0 | 0 |
| ENSG00000238519 | 0 | 0 | 0 | 0 | 0 | 0 |
| ENSG00000274966 | 0 | 0 | 0 | 0 | 0 | 0 |
| ENSG00000186399 | 8 | 7 | 8 | 2 | 4 | 5 |
| ENSG00000274424 | 0 | 0 | 0 | 0 | 0 | 0 |
| ENSG00000271333 | 0 | 0 | 0 | 0 | 0 | 0 |
| ENSG00000215302 | 24 | 9 | 4 | 3 | 11 | 13 |
| ENSG00000270055 | 60 | 23 | 29 | 23 | 35 | 40 |
| ENSG00000283345 | 0 | 0 | 0 | 0 | 0 | 0 |
| ENSG00000260784 | 0 | 0 | 0 | 0 | 0 | 0 |
| ENSG00000260693 | 0 | 0 | 1 | 2 | 0 | 1 |
| ENSG00000178115 | 4 | 0 | 1 | 0 | 0 | 0 |
| ENSG00000277031 | 0 | 0 | 0 | 0 | 0 | 0 |
| ENSG00000259890 | 0 | 0 | 0 | 0 | 0 | 0 |
| ENSG00000260128 | 4 | 1 | 0 | 0 | 0 | 0 |
| ENSG00000207430 | 0 | 0 | 0 | 0 | 0 | 0 |
| ENSG00000261794 | 2 | 6 | 3 | 2 | 2 | 1 |
| ENSG00000270016 | 6 | 0 | 3 | 2 | 1 | 1 |
| ENSG00000277467 | 0 | 2 | 0 | 0 | 0 | 0 |
| ENSG00000269930 | 0 | 0 | 0 | 1 | 0 | 0 |
| ENSG00000247728 | 0 | 1 | 6 | 3 | 1 | 1 |
| ENSG00000285077 | 0 | 1 | 1 | 1 | 2 | 3 |
| ENSG00000284906 | 24 | 14 | 17 | 10 | 14 | 24 |
| ENSG00000285035 | 0 | 0 | 1 | 0 | 1 | 0 |
| ENSG00000269974 | 20 | 8 | 10 | 5 | 6 | 13 |
| ENSG00000187951 | 125 | 53 | 82 | 53 | 93 | 97 |
| ENSG00000206846 | 0 | 0 | 0 | 0 | 1 | 0 |
| ENSG00000284783 | 0 | 0 | 0 | 0 | 1 | 0 |

| | | | | | | |
|---|---|---|---|---|---|---|
| ENSG00000103832 | 0 | 0 | 0 | 0 | 0 | 1 |
| ENSG00000278696 | 0 | 0 | 0 | 0 | 0 | 0 |
| ENSG00000259845 | 4 | 0 | 2 | 0 | 2 | 0 |
| ENSG00000260382 | 6 | 0 | 0 | 1 | 0 | 0 |
| ENSG00000261628 | 0 | 0 | 0 | 1 | 0 | 0 |
| ENSG00000198690 | 440 | 234 | 296 | 175 | 245 | 302 |
| ENSG00000270015 | 75 | 39 | 82 | 32 | 40 | 48 |
| ENSG00000166912 | 469 | 202 | 303 | 130 | 254 | 214 |
| ENSG00000212526 | 0 | 0 | 0 | 0 | 0 | 0 |
| ENSG00000134160 | 0 | 0 | 0 | 0 | 0 | 0 |
| ENSG00000259720 | 0 | 0 | 0 | 0 | 0 | 0 |
| ENSG00000207702 | 0 | 0 | 0 | 0 | 0 | 0 |
| ENSG00000259448 | 0 | 2 | 4 | 0 | 3 | 1 |
| ENSG00000259772 | 2 | 0 | 0 | 1 | 0 | 0 |
| ENSG00000169926 | 852 | 438 | 506 | 319 | 405 | 366 |
| ENSG00000259179 | 0 | 0 | 0 | 0 | 0 | 0 |
| ENSG00000259696 | 0 | 0 | 0 | 0 | 0 | 0 |
| ENSG00000169918 | 45 | 26 | 32 | 25 | 18 | 23 |
| ENSG00000259358 | 0 | 0 | 0 | 0 | 0 | 0 |
| ENSG00000277500 | 0 | 0 | 0 | 0 | 0 | 0 |
| ENSG00000175344 | 0 | 1 | 0 | 0 | 0 | 0 |
| ENSG00000206849 | 0 | 0 | 0 | 0 | 0 | 0 |
| ENSG00000201084 | 0 | 0 | 0 | 0 | 0 | 0 |
| ENSG00000279912 | 2 | 0 | 0 | 1 | 1 | 1 |
| ENSG00000271078 | 0 | 0 | 0 | 0 | 0 | 0 |
| ENSG00000207257 | 0 | 0 | 0 | 0 | 0 | 0 |
| ENSG00000260211 | 0 | 0 | 0 | 0 | 0 | 0 |
| ENSG00000260642 | 0 | 0 | 0 | 0 | 0 | 0 |
| ENSG00000261491 | 0 | 0 | 0 | 0 | 0 | 0 |
| ENSG00000249931 | 4 | 0 | 1 | 0 | 0 | 0 |
| ENSG00000275776 | 0 | 0 | 0 | 0 | 0 | 0 |
| ENSG00000215304 | 0 | 0 | 0 | 0 | 0 | 0 |
| ENSG00000261279 | 0 | 0 | 0 | 0 | 0 | 0 |
| ENSG00000206987 | 0 | 0 | 0 | 0 | 0 | 0 |
| ENSG00000261708 | 0 | 0 | 0 | 0 | 0 | 0 |
| ENSG00000206127 | 0 | 0 | 1 | 0 | 1 | 0 |
| ENSG00000274076 | 0 | 0 | 0 | 0 | 0 | 0 |
| ENSG00000270789 | 0 | 0 | 0 | 0 | 0 | 0 |
| ENSG00000261375 | 0 | 0 | 1 | 0 | 0 | 0 |
| ENSG00000254912 | 0 | 0 | 0 | 0 | 0 | 0 |
| ENSG00000223509 | 46 | 30 | 52 | 16 | 28 | 44 |
| ENSG00000261064 | 7 | 7 | 7 | 3 | 4 | 15 |
| ENSG00000276724 | 2 | 0 | 0 | 0 | 0 | 0 |
| ENSG00000262728 | 4 | 5 | 3 | 4 | 3 | 5 |
| ENSG00000232653 | 48 | 13 | 14 | 13 | 33 | 16 |
| ENSG00000277464 | 0 | 0 | 0 | 0 | 0 | 0 |
| ENSG00000244952 | 3 | 2 | 0 | 0 | 1 | 1 |
| ENSG00000198826 | 431 | 232 | 222 | 152 | 406 | 251 |
| ENSG00000285948 | 0 | 0 | 0 | 0 | 0 | 1 |
| ENSG00000166922 | 6 | 0 | 3 | 1 | 1 | 4 |

| | | | | | | |
|---|---|---|---|---|---|---|
| ENSG00000241818 | 0 | 0 | 0 | 0 | 0 | 0 |
| ENSG00000259721 | 0 | 0 | 0 | 0 | 0 | 0 |
| ENSG00000166923 | 0 | 0 | 0 | 0 | 1 | 3 |
| ENSG00000248905 | 352 | 176 | 199 | 90 | 241 | 214 |
| ENSG00000259392 | 0 | 0 | 0 | 1 | 0 | 1 |
| ENSG00000212415 | 0 | 0 | 0 | 0 | 0 | 0 |
| ENSG00000259686 | 2 | 0 | 0 | 0 | 0 | 0 |
| ENSG00000279958 | 0 | 0 | 0 | 0 | 0 | 0 |
| ENSG00000215296 | 0 | 0 | 0 | 0 | 0 | 0 |
| ENSG00000259446 | 0 | 0 | 0 | 0 | 0 | 0 |
| ENSG00000198838 | 11 | 19 | 20 | 3 | 13 | 14 |
| ENSG00000200008 | 0 | 0 | 0 | 0 | 0 | 0 |
| ENSG00000276702 | 0 | 0 | 0 | 0 | 0 | 0 |
| ENSG00000259408 | 2 | 3 | 3 | 0 | 2 | 1 |
| ENSG00000259287 | 7 | 3 | 4 | 5 | 1 | 2 |
| ENSG00000169857 | 367 | 193 | 199 | 239 | 242 | 232 |
| ENSG00000184984 | 4 | 3 | 1 | 1 | 0 | 2 |
| ENSG00000278472 | 6 | 4 | 0 | 1 | 2 | 0 |
| ENSG00000259505 | 0 | 0 | 0 | 0 | 0 | 0 |
| ENSG00000134153 | 1870 | 968 | 1100 | 1057 | 1069 | 1132 |
| ENSG00000243094 | 0 | 1 | 3 | 1 | 1 | 0 |
| ENSG00000182405 | 124 | 66 | 81 | 46 | 58 | 50 |
| ENSG00000134152 | 769 | 334 | 412 | 356 | 415 | 408 |
| ENSG00000128463 | 2176 | 1181 | 1366 | 1395 | 1369 | 1417 |
| ENSG00000140199 | 560 | 275 | 350 | 171 | 311 | 286 |
| ENSG00000259468 | 0 | 0 | 0 | 0 | 0 | 0 |
| ENSG00000207091 | 0 | 0 | 0 | 0 | 0 | 0 |
| ENSG00000182117 | 2322 | 1103 | 1407 | 1533 | 1390 | 1347 |
| ENSG00000184507 | 0 | 0 | 2 | 0 | 1 | 0 |
| ENSG00000176454 | 992 | 543 | 661 | 575 | 590 | 567 |
| ENSG00000259904 | 6 | 0 | 1 | 0 | 1 | 3 |
| ENSG00000175265 | 850 | 339 | 592 | 246 | 440 | 470 |
| ENSG00000221649 | 0 | 0 | 0 | 0 | 0 | 0 |
| ENSG00000260639 | 0 | 0 | 0 | 0 | 0 | 0 |
| ENSG00000261081 | 0 | 0 | 0 | 0 | 0 | 0 |
| ENSG00000279092 | 0 | 0 | 0 | 0 | 0 | 0 |
| ENSG00000259917 | 6 | 5 | 2 | 3 | 2 | 4 |
| ENSG00000261559 | 0 | 0 | 0 | 0 | 0 | 0 |
| ENSG00000278582 | 0 | 0 | 0 | 0 | 0 | 0 |
| ENSG00000215252 | 280 | 101 | 235 | 62 | 134 | 164 |
| ENSG00000221065 | 0 | 0 | 0 | 0 | 0 | 0 |
| ENSG00000259892 | 0 | 0 | 0 | 0 | 0 | 0 |
| ENSG00000252425 | 0 | 0 | 0 | 0 | 0 | 0 |
| ENSG00000261304 | 0 | 0 | 0 | 0 | 0 | 0 |
| ENSG00000159248 | 0 | 0 | 0 | 0 | 0 | 0 |
| ENSG00000250007 | 0 | 0 | 0 | 0 | 0 | 0 |
| ENSG00000159251 | 0 | 0 | 0 | 0 | 0 | 0 |
| ENSG00000259751 | 0 | 0 | 1 | 1 | 1 | 0 |
| ENSG00000021776 | 1524 | 851 | 794 | 421 | 1038 | 731 |
| ENSG00000231409 | 0 | 2 | 1 | 1 | 2 | 1 |

| | | | | | | |
|---|---|---|---|---|---|---|
| ENSG00000198146 | 717 | 353 | 297 | 137 | 418 | 281 |
| ENSG00000279364 | 0 | 0 | 0 | 0 | 0 | 0 |
| ENSG00000259181 | 0 | 0 | 0 | 0 | 0 | 0 |
| ENSG00000280396 | 0 | 0 | 0 | 0 | 0 | 0 |
| ENSG00000259587 | 0 | 0 | 0 | 0 | 0 | 0 |
| ENSG00000259665 | 0 | 1 | 0 | 1 | 1 | 0 |
| ENSG00000255192 | 0 | 0 | 0 | 0 | 0 | 0 |
| ENSG00000259397 | 2 | 2 | 0 | 1 | 0 | 3 |
| ENSG00000259336 | 1 | 0 | 0 | 1 | 2 | 1 |
| ENSG00000259592 | 0 | 1 | 2 | 1 | 0 | 0 |
| ENSG00000259516 | 2 | 0 | 0 | 0 | 1 | 1 |
| ENSG00000134146 | 164 | 118 | 155 | 127 | 121 | 136 |
| ENSG00000259710 | 0 | 0 | 1 | 0 | 0 | 1 |
| ENSG00000259585 | 0 | 0 | 0 | 0 | 0 | 1 |
| ENSG00000265102 | 1 | 2 | 1 | 1 | 2 | 3 |
| ENSG00000259688 | 1 | 0 | 0 | 0 | 0 | 0 |
| ENSG00000259411 | 0 | 0 | 0 | 0 | 0 | 0 |
| ENSG00000248079 | 4 | 2 | 1 | 1 | 1 | 1 |
| ENSG00000259217 | 0 | 0 | 0 | 0 | 0 | 0 |
| ENSG00000265098 | 0 | 0 | 0 | 0 | 0 | 0 |
| ENSG00000259639 | 0 | 0 | 0 | 0 | 1 | 1 |
| ENSG00000259245 | 2 | 0 | 0 | 0 | 0 | 0 |
| ENSG00000259395 | 0 | 0 | 0 | 0 | 0 | 0 |
| ENSG00000259737 | 0 | 0 | 0 | 0 | 0 | 0 |
| ENSG00000270500 | 0 | 0 | 0 | 0 | 0 | 0 |
| ENSG00000186073 | 247 | 146 | 143 | 127 | 180 | 152 |
| ENSG00000261635 | 2 | 0 | 0 | 0 | 0 | 0 |
| ENSG00000261315 | 0 | 1 | 0 | 0 | 0 | 0 |
| ENSG00000260582 | 0 | 0 | 0 | 0 | 0 | 1 |
| ENSG00000261191 | 0 | 0 | 0 | 0 | 0 | 0 |
| ENSG00000223518 | 0 | 0 | 0 | 0 | 0 | 1 |
| ENSG00000212511 | 0 | 0 | 0 | 0 | 0 | 0 |
| ENSG00000259280 | 0 | 0 | 0 | 0 | 0 | 0 |
| ENSG00000134138 | 266 | 120 | 145 | 66 | 101 | 100 |
| ENSG00000277202 | 1 | 0 | 0 | 0 | 0 | 0 |
| ENSG00000206676 | 0 | 0 | 0 | 0 | 0 | 0 |
| ENSG00000259460 | 0 | 0 | 0 | 0 | 0 | 0 |
| ENSG00000274444 | 6 | 3 | 0 | 1 | 1 | 0 |
| ENSG00000274216 | 0 | 0 | 0 | 0 | 0 | 0 |
| ENSG00000270304 | 0 | 0 | 0 | 0 | 0 | 0 |
| ENSG00000259434 | 0 | 0 | 0 | 0 | 0 | 0 |
| ENSG00000243122 | 0 | 0 | 0 | 0 | 0 | 0 |
| ENSG00000166069 | 0 | 0 | 0 | 0 | 0 | 0 |
| ENSG00000259225 | 0 | 0 | 0 | 0 | 0 | 0 |
| ENSG00000236914 | 2 | 1 | 0 | 1 | 1 | 1 |
| ENSG00000259380 | 0 | 0 | 0 | 0 | 0 | 0 |
| ENSG00000166068 | 158 | 78 | 86 | 77 | 94 | 91 |
| ENSG00000171262 | 1116 | 543 | 565 | 388 | 635 | 581 |
| ENSG00000259192 | 0 | 0 | 0 | 0 | 0 | 0 |
| ENSG00000172575 | 474 | 294 | 204 | 199 | 395 | 231 |

| | | | | | | |
|---|---|---|---|---|---|---|
| ENSG00000259326 | 8 | 1 | 0 | 1 | 0 | 1 |
| ENSG00000259598 | 0 | 0 | 1 | 0 | 0 | 0 |
| ENSG00000175779 | 0 | 0 | 0 | 0 | 1 | 0 |
| ENSG00000259731 | 0 | 0 | 0 | 0 | 0 | 0 |
| ENSG00000274281 | 0 | 0 | 0 | 0 | 0 | 0 |
| ENSG00000259345 | 31 | 13 | 18 | 7 | 16 | 15 |
| ENSG00000259278 | 0 | 0 | 0 | 0 | 0 | 0 |
| ENSG00000259423 | 0 | 0 | 0 | 0 | 0 | 0 |
| ENSG00000175746 | 0 | 0 | 1 | 0 | 0 | 0 |
| ENSG00000259447 | 0 | 0 | 0 | 0 | 0 | 0 |
| ENSG00000259269 | 0 | 0 | 0 | 0 | 0 | 0 |
| ENSG00000259390 | 0 | 0 | 0 | 0 | 0 | 0 |
| ENSG00000137801 | 8021 | 3834 | 5268 | 2583 | 3173 | 3586 |
| ENSG00000276107 | 2 | 6 | 8 | 3 | 1 | 2 |
| ENSG00000278621 | 0 | 0 | 0 | 0 | 0 | 0 |
| ENSG00000150667 | 31 | 21 | 26 | 19 | 12 | 23 |
| ENSG00000259580 | 0 | 0 | 0 | 0 | 0 | 0 |
| ENSG00000261136 | 52 | 32 | 47 | 15 | 34 | 29 |
| ENSG00000166073 | 268 | 117 | 154 | 120 | 166 | 164 |
| ENSG00000259432 | 0 | 0 | 0 | 0 | 0 | 0 |
| ENSG00000246863 | 2 | 4 | 1 | 1 | 0 | 7 |
| ENSG00000200305 | 0 | 0 | 0 | 0 | 0 | 0 |
| ENSG00000128829 | 1198 | 529 | 699 | 438 | 695 | 559 |
| ENSG00000259389 | 0 | 1 | 0 | 0 | 0 | 0 |
| ENSG00000140319 | 6851 | 3580 | 4197 | 4196 | 4488 | 3955 |
| ENSG00000248508 | 173 | 86 | 105 | 126 | 110 | 115 |
| ENSG00000259584 | 0 | 1 | 3 | 0 | 2 | 0 |
| ENSG00000275636 | 1 | 0 | 0 | 0 | 0 | 0 |
| ENSG00000104081 | 1920 | 1047 | 1256 | 1857 | 2525 | 1894 |
| ENSG00000259409 | 0 | 0 | 0 | 0 | 0 | 0 |
| ENSG00000259239 | 0 | 0 | 0 | 0 | 0 | 0 |
| ENSG00000156970 | 411 | 292 | 309 | 252 | 490 | 365 |
| ENSG00000259288 | 0 | 0 | 0 | 0 | 0 | 0 |
| ENSG00000273786 | 0 | 0 | 0 | 0 | 0 | 0 |
| ENSG00000260648 | 0 | 0 | 0 | 0 | 0 | 1 |
| ENSG00000137843 | 61 | 18 | 36 | 16 | 21 | 17 |
| ENSG00000279537 | 0 | 0 | 0 | 0 | 0 | 0 |
| ENSG00000176753 | 6 | 1 | 1 | 4 | 4 | 0 |
| ENSG00000279409 | 2 | 1 | 0 | 0 | 0 | 0 |
| ENSG00000137841 | 16 | 15 | 10 | 4 | 1 | 8 |
| ENSG00000230778 | 5 | 0 | 0 | 0 | 2 | 2 |
| ENSG00000273855 | 0 | 0 | 0 | 0 | 0 | 0 |
| ENSG00000259307 | 0 | 0 | 0 | 0 | 0 | 0 |
| ENSG00000244705 | 0 | 0 | 0 | 0 | 0 | 0 |
| ENSG00000259198 | 0 | 1 | 0 | 0 | 0 | 0 |
| ENSG00000259330 | 276 | 143 | 166 | 133 | 150 | 126 |
| ENSG00000188549 | 133 | 42 | 62 | 23 | 25 | 29 |
| ENSG00000252714 | 0 | 0 | 0 | 0 | 0 | 0 |
| ENSG00000233041 | 0 | 0 | 1 | 0 | 0 | 0 |
| ENSG00000140323 | 75 | 57 | 38 | 48 | 34 | 36 |

| | | | | | | |
|---|---|---|---|---|---|---|
| ENSG00000259368 | 0 | 0 | 0 | 0 | 0 | 0 |
| ENSG00000128944 | 627 | 351 | 382 | 398 | 477 | 460 |
| ENSG00000128928 | 3284 | 1655 | 2122 | 1498 | 1784 | 1737 |
| ENSG00000140320 | 1518 | 739 | 1020 | 897 | 858 | 819 |
| ENSG00000259364 | 0 | 3 | 1 | 1 | 0 | 1 |
| ENSG00000259211 | 3 | 2 | 1 | 1 | 1 | 2 |
| ENSG00000169105 | 672 | 328 | 404 | 409 | 309 | 353 |
| ENSG00000244251 | 0 | 0 | 0 | 0 | 0 | 0 |
| ENSG00000259536 | 3 | 3 | 2 | 3 | 0 | 1 |
| ENSG00000128891 | 408 | 278 | 315 | 251 | 301 | 271 |
| ENSG00000223313 | 5 | 8 | 5 | 9 | 6 | 5 |
| ENSG00000166133 | 371 | 166 | 189 | 243 | 216 | 234 |
| ENSG00000203573 | 0 | 0 | 0 | 0 | 0 | 0 |
| ENSG00000137812 | 101 | 65 | 52 | 23 | 158 | 69 |
| ENSG00000265814 | 0 | 0 | 0 | 0 | 0 | 0 |
| ENSG00000259254 | 2 | 1 | 0 | 0 | 0 | 1 |
| ENSG00000245849 | 319 | 164 | 270 | 114 | 182 | 184 |
| ENSG00000051180 | 11 | 15 | 20 | 13 | 8 | 12 |
| ENSG00000285396 | 0 | 0 | 0 | 0 | 0 | 0 |
| ENSG00000137824 | 3016 | 1369 | 1795 | 1456 | 1332 | 1467 |
| ENSG00000271236 | 0 | 0 | 0 | 0 | 0 | 0 |
| ENSG00000137880 | 52 | 24 | 46 | 36 | 40 | 49 |
| ENSG00000104129 | 585 | 276 | 365 | 290 | 269 | 296 |
| ENSG00000188277 | 45 | 36 | 47 | 28 | 36 | 27 |
| ENSG00000166140 | 1526 | 660 | 903 | 721 | 634 | 673 |
| ENSG00000166143 | 4 | 2 | 8 | 1 | 3 | 3 |
| ENSG00000261183 | 377 | 238 | 296 | 280 | 262 | 255 |
| ENSG00000166145 | 7184 | 3141 | 4198 | 3459 | 3045 | 3111 |
| ENSG00000104140 | 830 | 375 | 465 | 547 | 487 | 490 |
| ENSG00000274798 | 4 | 1 | 2 | 0 | 4 | 1 |
| ENSG00000104142 | 1347 | 642 | 843 | 817 | 796 | 765 |
| ENSG00000251161 | 11 | 14 | 13 | 12 | 10 | 17 |
| ENSG00000128917 | 52 | 21 | 26 | 30 | 23 | 23 |
| ENSG00000259405 | 2 | 3 | 7 | 10 | 4 | 5 |
| ENSG00000128965 | 78 | 39 | 50 | 52 | 41 | 38 |
| ENSG00000128908 | 1517 | 661 | 983 | 567 | 821 | 793 |
| ENSG00000259617 | 0 | 1 | 0 | 0 | 0 | 1 |
| ENSG00000259463 | 0 | 1 | 0 | 0 | 0 | 0 |
| ENSG00000259521 | 0 | 0 | 1 | 0 | 1 | 1 |
| ENSG00000280036 | 0 | 0 | 0 | 2 | 0 | 0 |
| ENSG00000259547 | 0 | 0 | 0 | 0 | 0 | 0 |
| ENSG00000283133 | 0 | 0 | 0 | 0 | 0 | 0 |
| ENSG00000157021 | 4 | 1 | 3 | 1 | 5 | 3 |
| ENSG00000178997 | 2 | 1 | 2 | 5 | 2 | 1 |
| ENSG00000240847 | 0 | 0 | 0 | 0 | 0 | 0 |
| ENSG00000187446 | 2787 | 1285 | 1552 | 1190 | 1629 | 1409 |
| ENSG00000247556 | 2756 | 1508 | 1570 | 1022 | 1537 | 1331 |
| ENSG00000285920 | 0 | 0 | 0 | 0 | 0 | 0 |
| ENSG00000104147 | 138 | 73 | 75 | 95 | 105 | 120 |
| ENSG00000137804 | 109 | 63 | 126 | 71 | 101 | 110 |

| | | | | | | |
|---|---|---|---|---|---|---|
| ENSG00000259396 | 0 | 0 | 0 | 0 | 0 | 0 |
| ENSG00000137806 | 388 | 170 | 238 | 179 | 209 | 212 |
| ENSG00000137815 | 2551 | 1282 | 1606 | 825 | 1445 | 1322 |
| ENSG00000200407 | 0 | 0 | 0 | 0 | 0 | 0 |
| ENSG00000137825 | 131 | 78 | 113 | 74 | 80 | 58 |
| ENSG00000062524 | 4 | 2 | 5 | 4 | 2 | 1 |
| ENSG00000103932 | 653 | 317 | 450 | 385 | 417 | 358 |
| ENSG00000259838 | 0 | 2 | 0 | 0 | 0 | 1 |
| ENSG00000092445 | 425 | 191 | 184 | 245 | 203 | 168 |
| ENSG00000260926 | 0 | 0 | 0 | 0 | 0 | 0 |
| ENSG00000174197 | 1060 | 595 | 489 | 232 | 759 | 557 |
| ENSG00000207766 | 0 | 0 | 0 | 0 | 0 | 0 |
| ENSG00000270702 | 0 | 0 | 0 | 0 | 0 | 0 |
| ENSG00000260814 | 0 | 0 | 0 | 0 | 0 | 0 |
| ENSG00000137802 | 351 | 145 | 218 | 144 | 203 | 219 |
| ENSG00000250379 | 0 | 0 | 0 | 0 | 0 | 0 |
| ENSG00000243789 | 5 | 6 | 6 | 3 | 4 | 4 |
| ENSG00000168970 | 71 | 49 | 82 | 38 | 36 | 52 |
| ENSG00000243708 | 7 | 1 | 4 | 2 | 5 | 5 |
| ENSG00000137877 | 21 | 12 | 13 | 13 | 8 | 8 |
| ENSG00000200293 | 0 | 0 | 0 | 0 | 0 | 0 |
| ENSG00000264850 | 0 | 0 | 0 | 0 | 0 | 0 |
| ENSG00000174171 | 0 | 0 | 0 | 0 | 0 | 0 |
| ENSG00000103966 | 1544 | 721 | 950 | 742 | 760 | 872 |
| ENSG00000278493 | 4 | 4 | 2 | 4 | 0 | 2 |
| ENSG00000259883 | 0 | 2 | 1 | 3 | 1 | 0 |
| ENSG00000246740 | 0 | 0 | 1 | 2 | 5 | 1 |
| ENSG00000188089 | 1 | 0 | 1 | 0 | 1 | 1 |
| ENSG00000257797 | 0 | 0 | 0 | 0 | 0 | 0 |
| ENSG00000159337 | 0 | 0 | 0 | 0 | 0 | 0 |
| ENSG00000168907 | 321 | 109 | 196 | 133 | 120 | 171 |
| ENSG00000166887 | 1709 | 776 | 1115 | 716 | 919 | 902 |
| ENSG00000207712 | 0 | 0 | 0 | 0 | 0 | 0 |
| ENSG00000261002 | 2 | 0 | 0 | 0 | 0 | 3 |
| ENSG00000103978 | 1354 | 832 | 1100 | 680 | 872 | 885 |
| ENSG00000214013 | 109 | 69 | 74 | 57 | 53 | 58 |
| ENSG00000260571 | 0 | 0 | 0 | 0 | 0 | 1 |
| ENSG00000258461 | 0 | 0 | 2 | 0 | 0 | 0 |
| ENSG00000092529 | 0 | 0 | 0 | 0 | 0 | 0 |
| ENSG00000103994 | 918 | 467 | 507 | 263 | 559 | 430 |
| ENSG00000201077 | 0 | 0 | 0 | 0 | 0 | 0 |
| ENSG00000092531 | 315 | 187 | 255 | 162 | 205 | 184 |
| ENSG00000261684 | 0 | 0 | 0 | 0 | 1 | 0 |
| ENSG00000285942 | 0 | 0 | 0 | 0 | 0 | 0 |
| ENSG00000180979 | 275 | 142 | 184 | 120 | 176 | 133 |
| ENSG00000137814 | 417 | 222 | 258 | 181 | 276 | 217 |
| ENSG00000261822 | 0 | 0 | 0 | 0 | 0 | 0 |
| ENSG00000260490 | 0 | 0 | 0 | 1 | 2 | 1 |
| ENSG00000159433 | 14 | 5 | 15 | 3 | 7 | 7 |
| ENSG00000260282 | 0 | 0 | 0 | 0 | 0 | 0 |

| | | | | | | |
|---|---|---|---|---|---|---|
| ENSG00000140326 | 429 | 225 | 226 | 197 | 255 | 235 |
| ENSG00000278769 | 0 | 0 | 0 | 0 | 0 | 0 |
| ENSG00000274403 | 4 | 0 | 0 | 2 | 4 | 1 |
| ENSG00000128881 | 167 | 64 | 90 | 27 | 84 | 55 |
| ENSG00000246283 | 0 | 0 | 0 | 0 | 0 | 0 |
| ENSG00000260799 | 0 | 0 | 0 | 0 | 0 | 0 |
| ENSG00000254793 | 1 | 0 | 1 | 1 | 0 | 0 |
| ENSG00000159459 | 531 | 276 | 235 | 116 | 388 | 252 |
| ENSG00000285080 | 0 | 0 | 0 | 0 | 0 | 0 |
| ENSG00000281269 | 0 | 0 | 0 | 0 | 0 | 0 |
| ENSG00000205871 | 1 | 3 | 2 | 0 | 1 | 2 |
| ENSG00000137842 | 255 | 139 | 190 | 151 | 173 | 176 |
| ENSG00000260928 | 0 | 0 | 0 | 0 | 0 | 0 |
| ENSG00000285117 | 1 | 0 | 0 | 0 | 1 | 0 |
| ENSG00000261687 | 0 | 0 | 0 | 0 | 0 | 0 |
| ENSG00000166946 | 1236 | 590 | 722 | 599 | 625 | 557 |
| ENSG00000166947 | 0 | 0 | 0 | 0 | 0 | 0 |
| ENSG00000104055 | 2 | 0 | 0 | 0 | 0 | 0 |
| ENSG00000202211 | 0 | 0 | 0 | 0 | 0 | 0 |
| ENSG00000261679 | 0 | 0 | 0 | 0 | 0 | 0 |
| ENSG00000159495 | 0 | 0 | 0 | 0 | 0 | 0 |
| ENSG00000260406 | 0 | 0 | 0 | 0 | 0 | 0 |
| ENSG00000168806 | 548 | 213 | 267 | 257 | 236 | 283 |
| ENSG00000168803 | 246 | 87 | 117 | 59 | 108 | 91 |
| ENSG00000140265 | 660 | 286 | 415 | 226 | 402 | 350 |
| ENSG00000137822 | 340 | 155 | 175 | 141 | 177 | 148 |
| ENSG00000243871 | 0 | 0 | 0 | 0 | 0 | 0 |
| ENSG00000067369 | 1226 | 638 | 770 | 395 | 772 | 598 |
| ENSG00000166963 | 10 | 9 | 14 | 5 | 7 | 19 |
| ENSG00000168781 | 477 | 177 | 288 | 206 | 227 | 195 |
| ENSG00000237289 | 76 | 39 | 63 | 60 | 55 | 48 |
| ENSG00000242866 | 0 | 0 | 1 | 0 | 0 | 1 |
| ENSG00000222398 | 0 | 0 | 0 | 0 | 0 | 0 |
| ENSG00000284772 | 0 | 0 | 0 | 0 | 0 | 0 |
| ENSG00000166762 | 58 | 12 | 62 | 19 | 38 | 22 |
| ENSG00000206991 | 4 | 1 | 1 | 0 | 0 | 2 |
| ENSG00000275601 | 11 | 10 | 10 | 7 | 6 | 3 |
| ENSG00000224677 | 8 | 2 | 2 | 2 | 0 | 1 |
| ENSG00000249839 | 11 | 1 | 1 | 1 | 1 | 1 |
| ENSG00000223572 | 91 | 49 | 49 | 50 | 41 | 55 |
| ENSG00000166763 | 4 | 1 | 1 | 0 | 0 | 0 |
| ENSG00000201136 | 0 | 0 | 0 | 0 | 0 | 0 |
| ENSG00000205771 | 6 | 4 | 4 | 0 | 5 | 7 |
| ENSG00000206589 | 0 | 0 | 0 | 0 | 0 | 0 |
| ENSG00000167004 | 5357 | 2826 | 3526 | 2605 | 2967 | 3037 |
| ENSG00000128886 | 13 | 11 | 25 | 18 | 23 | 20 |
| ENSG00000262560 | 0 | 0 | 0 | 0 | 0 | 0 |
| ENSG00000140264 | 6184 | 4110 | 4916 | 5190 | 4319 | 4190 |
| ENSG00000221792 | 0 | 0 | 0 | 0 | 0 | 0 |
| ENSG00000184716 | 0 | 0 | 0 | 0 | 0 | 1 |

| | | | | | | |
|---|---|---|---|---|---|---|
| ENSG00000242028 | 54 | 27 | 26 | 15 | 28 | 16 |
| ENSG00000140259 | 667 | 366 | 376 | 262 | 454 | 360 |
| ENSG00000092470 | 176 | 116 | 81 | 69 | 100 | 73 |
| ENSG00000200391 | 0 | 0 | 0 | 0 | 0 | 0 |
| ENSG00000171877 | 74 | 50 | 38 | 31 | 41 | 47 |
| ENSG00000227973 | 2 | 0 | 1 | 0 | 0 | 0 |
| ENSG00000185607 | 0 | 0 | 0 | 1 | 1 | 0 |
| ENSG00000224055 | 2 | 0 | 0 | 0 | 0 | 0 |
| ENSG00000166734 | 1734 | 945 | 1145 | 706 | 1183 | 1074 |
| ENSG00000250130 | 0 | 0 | 0 | 0 | 0 | 0 |
| ENSG00000259499 | 0 | 0 | 0 | 0 | 0 | 0 |
| ENSG00000259595 | 0 | 2 | 1 | 1 | 0 | 2 |
| ENSG00000137770 | 530 | 295 | 293 | 152 | 345 | 273 |
| ENSG00000259335 | 0 | 0 | 0 | 0 | 0 | 0 |
| ENSG00000259563 | 2 | 1 | 0 | 0 | 2 | 2 |
| ENSG00000179523 | 183 | 99 | 116 | 71 | 127 | 99 |
| ENSG00000104131 | 1459 | 817 | 957 | 725 | 1103 | 945 |
| ENSG00000259659 | 0 | 2 | 6 | 4 | 4 | 1 |
| ENSG00000104133 | 1146 | 601 | 598 | 277 | 712 | 474 |
| ENSG00000229474 | 10 | 6 | 7 | 2 | 6 | 5 |
| ENSG00000206957 | 0 | 0 | 0 | 0 | 0 | 0 |
| ENSG00000166710 | 5786 | 2957 | 3474 | 2929 | 3003 | 3123 |
| ENSG00000185880 | 164 | 69 | 91 | 116 | 108 | 86 |
| ENSG00000276121 | 0 | 0 | 0 | 0 | 0 | 0 |
| ENSG00000259479 | 33 | 19 | 20 | 16 | 17 | 19 |
| ENSG00000259187 | 0 | 0 | 0 | 0 | 0 | 0 |
| ENSG00000238845 | 0 | 0 | 0 | 0 | 0 | 0 |
| ENSG00000212424 | 0 | 0 | 0 | 0 | 0 | 0 |
| ENSG00000259742 | 0 | 0 | 0 | 0 | 0 | 0 |
| ENSG00000259231 | 0 | 0 | 0 | 0 | 0 | 0 |
| ENSG00000252117 | 0 | 0 | 0 | 0 | 0 | 0 |
| ENSG00000167014 | 0 | 0 | 0 | 0 | 0 | 0 |
| ENSG00000252475 | 0 | 0 | 0 | 0 | 0 | 0 |
| ENSG00000259304 | 0 | 0 | 0 | 0 | 0 | 0 |
| ENSG00000252288 | 0 | 0 | 0 | 0 | 0 | 0 |
| ENSG00000212550 | 0 | 0 | 0 | 0 | 0 | 0 |
| ENSG00000140263 | 976 | 544 | 615 | 610 | 562 | 671 |
| ENSG00000259418 | 0 | 0 | 0 | 0 | 0 | 0 |
| ENSG00000252171 | 0 | 0 | 0 | 0 | 0 | 0 |
| ENSG00000259352 | 0 | 0 | 0 | 0 | 0 | 0 |
| ENSG00000140279 | 2 | 1 | 1 | 4 | 2 | 3 |
| ENSG00000140274 | 0 | 2 | 1 | 0 | 0 | 0 |
| ENSG00000140254 | 1 | 2 | 6 | 2 | 3 | 4 |
| ENSG00000137857 | 28 | 20 | 22 | 14 | 16 | 25 |
| ENSG00000259539 | 0 | 0 | 0 | 0 | 0 | 0 |
| ENSG00000138606 | 43 | 21 | 16 | 34 | 27 | 16 |
| ENSG00000259519 | 0 | 0 | 0 | 0 | 0 | 0 |
| ENSG00000259932 | 0 | 0 | 1 | 0 | 0 | 0 |
| ENSG00000260035 | 2 | 0 | 0 | 0 | 0 | 0 |
| ENSG00000279014 | 0 | 0 | 0 | 0 | 0 | 0 |

| | | | | | | |
|---|---|---|---|---|---|---|
| ENSG00000261709 | 0 | 0 | 0 | 0 | 0 | 0 |
| ENSG00000259338 | 0 | 0 | 0 | 0 | 0 | 0 |
| ENSG00000259520 | 2 | 0 | 2 | 1 | 2 | 1 |
| ENSG00000137860 | 1 | 1 | 2 | 2 | 1 | 0 |
| ENSG00000285597 | 2 | 0 | 1 | 0 | 0 | 1 |
| ENSG00000235390 | 0 | 0 | 0 | 0 | 0 | 0 |
| ENSG00000259433 | 0 | 0 | 0 | 0 | 0 | 0 |
| ENSG00000238941 | 0 | 0 | 0 | 0 | 0 | 0 |
| ENSG00000171766 | 4 | 2 | 6 | 8 | 1 | 5 |
| ENSG00000275672 | 0 | 0 | 0 | 0 | 2 | 0 |
| ENSG00000171763 | 244 | 137 | 141 | 169 | 174 | 184 |
| ENSG00000166920 | 52 | 57 | 71 | 51 | 66 | 59 |
| ENSG00000259342 | 0 | 0 | 1 | 0 | 0 | 0 |
| ENSG00000284386 | 0 | 0 | 0 | 0 | 0 | 0 |
| ENSG00000259354 | 3 | 1 | 2 | 0 | 2 | 4 |
| ENSG00000104154 | 107 | 64 | 79 | 52 | 63 | 67 |
| ENSG00000179362 | 9 | 4 | 5 | 5 | 3 | 5 |
| ENSG00000207516 | 0 | 0 | 0 | 0 | 0 | 0 |
| ENSG00000259681 | 0 | 0 | 0 | 0 | 0 | 0 |
| ENSG00000259427 | 0 | 0 | 0 | 0 | 0 | 0 |
| ENSG00000275709 | 2 | 0 | 2 | 1 | 0 | 1 |
| ENSG00000104164 | 684 | 344 | 370 | 220 | 393 | 388 |
| ENSG00000260170 | 0 | 0 | 0 | 0 | 0 | 0 |
| ENSG00000200419 | 0 | 0 | 0 | 0 | 0 | 0 |
| ENSG00000137767 | 335 | 181 | 215 | 196 | 223 | 206 |
| ENSG00000273972 | 0 | 0 | 0 | 1 | 0 | 1 |
| ENSG00000259200 | 0 | 0 | 0 | 0 | 0 | 0 |
| ENSG00000259743 | 0 | 0 | 0 | 0 | 0 | 0 |
| ENSG00000259593 | 0 | 0 | 0 | 0 | 0 | 0 |
| ENSG00000238819 | 0 | 0 | 0 | 0 | 0 | 0 |
| ENSG00000285934 | 0 | 0 | 0 | 0 | 0 | 0 |
| ENSG00000238456 | 0 | 0 | 0 | 0 | 0 | 0 |
| ENSG00000259255 | 0 | 0 | 0 | 0 | 0 | 0 |
| ENSG00000223308 | 0 | 0 | 0 | 0 | 0 | 0 |
| ENSG00000259509 | 0 | 0 | 0 | 0 | 0 | 0 |
| ENSG00000284540 | 0 | 0 | 0 | 0 | 0 | 0 |
| ENSG00000137872 | 2 | 0 | 5 | 0 | 1 | 6 |
| ENSG00000222343 | 0 | 0 | 0 | 0 | 0 | 0 |
| ENSG00000259554 | 0 | 0 | 0 | 0 | 0 | 0 |
| ENSG00000259221 | 0 | 0 | 0 | 0 | 0 | 0 |
| ENSG00000259360 | 0 | 0 | 0 | 0 | 0 | 0 |
| ENSG00000259740 | 0 | 0 | 0 | 0 | 0 | 0 |
| ENSG00000259588 | 0 | 0 | 0 | 0 | 0 | 0 |
| ENSG00000213862 | 20 | 5 | 5 | 4 | 7 | 10 |
| ENSG00000259572 | 0 | 0 | 0 | 0 | 0 | 0 |
| ENSG00000259754 | 0 | 0 | 0 | 0 | 0 | 0 |
| ENSG00000259730 | 0 | 0 | 0 | 0 | 0 | 0 |
| ENSG00000188467 | 0 | 0 | 1 | 0 | 0 | 0 |
| ENSG00000104177 | 1035 | 579 | 722 | 375 | 734 | 617 |
| ENSG00000233932 | 0 | 0 | 0 | 0 | 0 | 0 |

| | | | | | | |
|---|---|---|---|---|---|---|
| ENSG00000074803 | 7 | 10 | 16 | 4 | 7 | 6 |
| ENSG00000259235 | 0 | 0 | 0 | 0 | 0 | 0 |
| ENSG00000259385 | 1 | 0 | 0 | 0 | 0 | 0 |
| ENSG00000259488 | 13 | 6 | 15 | 4 | 4 | 7 |
| ENSG00000202542 | 0 | 0 | 0 | 0 | 0 | 0 |
| ENSG00000128951 | 702 | 330 | 434 | 368 | 361 | 415 |
| ENSG00000166147 | 38 | 27 | 29 | 15 | 34 | 20 |
| ENSG00000274654 | 0 | 0 | 0 | 0 | 0 | 0 |
| ENSG00000259705 | 0 | 0 | 0 | 0 | 1 | 2 |
| ENSG00000273925 | 0 | 0 | 0 | 0 | 0 | 0 |
| ENSG00000103995 | 30 | 31 | 16 | 13 | 61 | 36 |
| ENSG00000259216 | 0 | 0 | 2 | 0 | 0 | 0 |
| ENSG00000259469 | 0 | 0 | 0 | 0 | 0 | 0 |
| ENSG00000259670 | 0 | 0 | 0 | 0 | 0 | 0 |
| ENSG00000259700 | 0 | 0 | 0 | 0 | 0 | 0 |
| ENSG00000185634 | 30 | 11 | 13 | 17 | 12 | 10 |
| ENSG00000255302 | 4062 | 1820 | 2397 | 1650 | 2125 | 2072 |
| ENSG00000259602 | 0 | 0 | 0 | 0 | 1 | 0 |
| ENSG00000259500 | 0 | 0 | 1 | 0 | 0 | 1 |
| ENSG00000138593 | 1054 | 492 | 569 | 196 | 534 | 366 |
| ENSG00000207105 | 0 | 0 | 0 | 0 | 0 | 0 |
| ENSG00000265942 | 0 | 0 | 0 | 0 | 0 | 0 |
| ENSG00000259656 | 0 | 0 | 0 | 0 | 0 | 0 |
| ENSG00000166200 | 964 | 650 | 487 | 336 | 945 | 485 |
| ENSG00000200120 | 3 | 0 | 0 | 1 | 0 | 0 |
| ENSG00000156958 | 490 | 229 | 299 | 194 | 279 | 262 |
| ENSG00000259467 | 0 | 0 | 1 | 0 | 1 | 0 |
| ENSG00000264210 | 0 | 0 | 0 | 0 | 0 | 0 |
| ENSG00000259545 | 0 | 0 | 0 | 0 | 3 | 0 |
| ENSG00000243338 | 0 | 0 | 1 | 0 | 0 | 0 |
| ENSG00000166262 | 180 | 85 | 91 | 85 | 107 | 91 |
| ENSG00000275580 | 2 | 6 | 1 | 0 | 2 | 3 |
| ENSG00000276593 | 0 | 1 | 0 | 0 | 1 | 0 |
| ENSG00000259531 | 0 | 1 | 1 | 0 | 3 | 2 |
| ENSG00000140285 | 0 | 0 | 3 | 0 | 1 | 1 |
| ENSG00000104047 | 845 | 383 | 445 | 373 | 480 | 439 |
| ENSG00000259388 | 0 | 0 | 0 | 0 | 0 | 0 |
| ENSG00000104043 | 2 | 3 | 1 | 0 | 1 | 2 |
| ENSG00000259188 | 0 | 1 | 1 | 1 | 0 | 0 |
| ENSG00000201086 | 0 | 0 | 0 | 0 | 0 | 0 |
| ENSG00000140284 | 399 | 251 | 257 | 343 | 392 | 346 |
| ENSG00000200605 | 0 | 0 | 0 | 0 | 0 | 0 |
| ENSG00000140287 | 0 | 0 | 0 | 1 | 0 | 1 |
| ENSG00000241175 | 0 | 0 | 0 | 0 | 0 | 0 |
| ENSG00000104064 | 725 | 452 | 462 | 403 | 488 | 436 |
| ENSG00000271819 | 0 | 0 | 2 | 1 | 3 | 0 |
| ENSG00000285410 | 1329 | 572 | 810 | 668 | 708 | 756 |
| ENSG00000244879 | 2235 | 1276 | 1378 | 832 | 1559 | 1471 |
| ENSG00000259715 | 0 | 0 | 0 | 0 | 0 | 0 |
| ENSG00000284284 | 0 | 0 | 0 | 0 | 0 | 0 |

| | | | | | | |
|---|---|---|---|---|---|---|
| ENSG00000259465 | 0 | 0 | 0 | 0 | 0 | 0 |
| ENSG00000138592 | 2683 | 1633 | 1479 | 711 | 2346 | 1334 |
| ENSG00000252376 | 0 | 0 | 0 | 0 | 3 | 0 |
| ENSG00000242737 | 0 | 2 | 1 | 0 | 1 | 0 |
| ENSG00000259618 | 1 | 2 | 6 | 0 | 1 | 6 |
| ENSG00000259298 | 5 | 13 | 16 | 6 | 6 | 7 |
| ENSG00000170236 | 0 | 0 | 0 | 0 | 0 | 0 |
| ENSG00000092439 | 2766 | 1683 | 1239 | 635 | 2595 | 1419 |
| ENSG00000259684 | 0 | 0 | 0 | 0 | 0 | 0 |
| ENSG00000243027 | 0 | 0 | 0 | 0 | 0 | 0 |
| ENSG00000138600 | 5746 | 2860 | 3494 | 2372 | 3486 | 3270 |
| ENSG00000279665 | 0 | 0 | 0 | 0 | 0 | 0 |
| ENSG00000259773 | 0 | 0 | 0 | 0 | 0 | 0 |
| ENSG00000241130 | 0 | 0 | 0 | 0 | 0 | 0 |
| ENSG00000273674 | 9 | 6 | 9 | 10 | 8 | 10 |
| ENSG00000081014 | 1346 | 667 | 677 | 295 | 875 | 583 |
| ENSG00000259378 | 4 | 2 | 1 | 1 | 6 | 3 |
| ENSG00000259240 | 47 | 40 | 33 | 29 | 39 | 35 |
| ENSG00000183578 | 26 | 16 | 7 | 8 | 7 | 7 |
| ENSG00000259204 | 0 | 0 | 1 | 0 | 0 | 0 |
| ENSG00000137869 | 29 | 23 | 36 | 12 | 22 | 24 |
| ENSG00000266593 | 0 | 0 | 0 | 0 | 0 | 0 |
| ENSG00000259701 | 7 | 2 | 3 | 0 | 3 | 3 |
| ENSG00000283614 | 0 | 0 | 0 | 0 | 0 | 0 |
| ENSG00000278098 | 0 | 0 | 0 | 0 | 0 | 0 |
| ENSG00000259306 | 56 | 33 | 34 | 25 | 33 | 41 |
| ENSG00000186417 | 382 | 172 | 279 | 320 | 473 | 448 |
| ENSG00000270378 | 0 | 0 | 0 | 0 | 0 | 0 |
| ENSG00000259194 | 0 | 3 | 0 | 1 | 0 | 2 |
| ENSG00000259678 | 1 | 0 | 0 | 0 | 0 | 0 |
| ENSG00000104093 | 773 | 519 | 400 | 115 | 1008 | 410 |
| ENSG00000259668 | 7 | 8 | 8 | 4 | 8 | 5 |
| ENSG00000104112 | 2 | 0 | 0 | 0 | 0 | 0 |
| ENSG00000259241 | 0 | 0 | 0 | 0 | 0 | 0 |
| ENSG00000259296 | 0 | 0 | 0 | 0 | 0 | 0 |
| ENSG00000140280 | 157 | 67 | 82 | 90 | 89 | 76 |
| ENSG00000128872 | 224 | 137 | 101 | 57 | 179 | 101 |
| ENSG00000259377 | 0 | 0 | 0 | 0 | 0 | 0 |
| ENSG00000138594 | 3923 | 2020 | 2481 | 1162 | 2530 | 1964 |
| ENSG00000274954 | 0 | 0 | 0 | 0 | 0 | 0 |
| ENSG00000277677 | 0 | 0 | 0 | 0 | 0 | 0 |
| ENSG00000259632 | 0 | 1 | 0 | 3 | 0 | 0 |
| ENSG00000207484 | 0 | 1 | 0 | 0 | 0 | 0 |
| ENSG00000259201 | 0 | 0 | 1 | 0 | 0 | 0 |
| ENSG00000272337 | 2 | 0 | 0 | 0 | 0 | 0 |
| ENSG00000259185 | 0 | 0 | 0 | 0 | 0 | 0 |
| ENSG00000259556 | 13 | 6 | 16 | 3 | 4 | 13 |
| ENSG00000166477 | 1193 | 810 | 658 | 383 | 1218 | 858 |
| ENSG00000069956 | 3508 | 1879 | 1847 | 1263 | 2631 | 2105 |
| ENSG00000259438 | 3 | 4 | 5 | 5 | 1 | 11 |

| | | | | | | |
|---|---|---|---|---|---|---|
| ENSG00000274528 | 35 | 5 | 9 | 13 | 18 | 24 |
| ENSG00000259398 | 0 | 0 | 0 | 1 | 0 | 0 |
| ENSG00000259712 | 2 | 2 | 3 | 1 | 4 | 1 |
| ENSG00000259178 | 0 | 0 | 0 | 0 | 0 | 0 |
| ENSG00000242327 | 0 | 0 | 1 | 0 | 0 | 0 |
| ENSG00000137875 | 4 | 3 | 1 | 3 | 1 | 1 |
| ENSG00000069966 | 847 | 472 | 559 | 477 | 580 | 514 |
| ENSG00000259327 | 0 | 0 | 0 | 0 | 0 | 0 |
| ENSG00000259709 | 0 | 1 | 1 | 0 | 0 | 0 |
| ENSG00000206995 | 0 | 0 | 0 | 0 | 0 | 0 |
| ENSG00000259577 | 16 | 9 | 7 | 12 | 6 | 11 |
| ENSG00000128833 | 4101 | 1886 | 2282 | 1176 | 2372 | 1947 |
| ENSG00000221052 | 0 | 0 | 1 | 0 | 0 | 0 |
| ENSG00000197535 | 808 | 451 | 389 | 180 | 672 | 380 |
| ENSG00000231169 | 0 | 1 | 0 | 0 | 0 | 0 |
| ENSG00000128989 | 10330 | 4978 | 5803 | 3806 | 5909 | 5252 |
| ENSG00000260618 | 21 | 15 | 13 | 8 | 21 | 18 |
| ENSG00000047346 | 2652 | 1393 | 1400 | 607 | 1665 | 1107 |
| ENSG00000259935 | 0 | 0 | 1 | 0 | 0 | 1 |
| ENSG00000169856 | 18 | 11 | 5 | 8 | 9 | 5 |
| ENSG00000259203 | 0 | 0 | 0 | 0 | 0 | 0 |
| ENSG00000214896 | 0 | 0 | 1 | 0 | 1 | 0 |
| ENSG00000259612 | 8 | 2 | 6 | 5 | 5 | 3 |
| ENSG00000259237 | 0 | 0 | 0 | 0 | 0 | 0 |
| ENSG00000166415 | 0 | 0 | 0 | 0 | 0 | 0 |
| ENSG00000252066 | 0 | 0 | 0 | 0 | 0 | 0 |
| ENSG00000206641 | 0 | 0 | 0 | 0 | 0 | 0 |
| ENSG00000259301 | 0 | 0 | 0 | 0 | 0 | 0 |
| ENSG00000280362 | 0 | 0 | 0 | 0 | 0 | 0 |
| ENSG00000259669 | 0 | 0 | 0 | 0 | 0 | 0 |
| ENSG00000137766 | 0 | 0 | 0 | 0 | 0 | 0 |
| ENSG00000259619 | 0 | 0 | 0 | 0 | 0 | 0 |
| ENSG00000259550 | 0 | 0 | 0 | 0 | 0 | 0 |
| ENSG00000260937 | 0 | 0 | 0 | 0 | 0 | 0 |
| ENSG00000279145 | 0 | 0 | 0 | 0 | 1 | 2 |
| ENSG00000137876 | 1837 | 1008 | 1165 | 1136 | 1312 | 1131 |
| ENSG00000069974 | 321 | 137 | 168 | 98 | 200 | 141 |
| ENSG00000276533 | 4 | 1 | 0 | 0 | 1 | 1 |
| ENSG00000225973 | 141 | 72 | 85 | 85 | 85 | 84 |
| ENSG00000069943 | 289 | 185 | 195 | 140 | 184 | 150 |
| ENSG00000260916 | 123 | 113 | 58 | 44 | 196 | 80 |
| ENSG00000274765 | 0 | 1 | 0 | 0 | 1 | 0 |
| ENSG00000277548 | 8 | 1 | 4 | 2 | 0 | 1 |
| ENSG00000261771 | 0 | 0 | 0 | 0 | 0 | 0 |
| ENSG00000283891 | 0 | 0 | 0 | 0 | 0 | 0 |
| ENSG00000261652 | 12 | 19 | 9 | 8 | 19 | 15 |
| ENSG00000256061 | 4 | 1 | 7 | 2 | 2 | 2 |
| ENSG00000260036 | 0 | 0 | 0 | 0 | 0 | 0 |
| ENSG00000171016 | 0 | 3 | 1 | 1 | 4 | 4 |
| ENSG00000285805 | 0 | 0 | 0 | 0 | 0 | 0 |

| | | | | | | |
|---|---|---|---|---|---|---|
| ENSG00000166450 | 90 | 51 | 59 | 26 | 98 | 78 |
| ENSG00000259180 | 0 | 0 | 0 | 0 | 0 | 0 |
| ENSG00000259369 | 0 | 0 | 0 | 0 | 0 | 0 |
| ENSG00000069869 | 545 | 375 | 296 | 201 | 623 | 339 |
| ENSG00000230183 | 0 | 0 | 1 | 0 | 0 | 0 |
| ENSG00000239703 | 0 | 0 | 0 | 0 | 0 | 0 |
| ENSG00000261333 | 0 | 0 | 0 | 0 | 0 | 0 |
| ENSG00000181827 | 460 | 215 | 184 | 85 | 318 | 201 |
| ENSG00000261072 | 0 | 0 | 0 | 0 | 0 | 0 |
| ENSG00000151575 | 25 | 21 | 17 | 13 | 36 | 15 |
| ENSG00000275417 | 0 | 2 | 0 | 0 | 0 | 0 |
| ENSG00000199784 | 0 | 0 | 0 | 0 | 0 | 0 |
| ENSG00000261174 | 0 | 0 | 0 | 0 | 0 | 0 |
| ENSG00000259941 | 0 | 0 | 0 | 0 | 0 | 0 |
| ENSG00000261823 | 0 | 0 | 0 | 0 | 0 | 0 |
| ENSG00000138587 | 6 | 2 | 1 | 0 | 4 | 7 |
| ENSG00000277245 | 0 | 0 | 0 | 0 | 1 | 0 |
| ENSG00000260392 | 0 | 0 | 0 | 0 | 0 | 0 |
| ENSG00000137871 | 171 | 133 | 96 | 82 | 142 | 104 |
| ENSG00000276696 | 0 | 0 | 0 | 0 | 0 | 0 |
| ENSG00000285253 | 0 | 1 | 0 | 0 | 0 | 0 |
| ENSG00000274667 | 12 | 7 | 10 | 7 | 8 | 5 |
| ENSG00000279683 | 0 | 0 | 0 | 0 | 0 | 0 |
| ENSG00000285331 | 14 | 12 | 19 | 18 | 16 | 15 |
| ENSG00000225625 | 0 | 0 | 0 | 0 | 0 | 0 |
| ENSG00000276524 | 0 | 0 | 0 | 0 | 0 | 1 |
| ENSG00000275343 | 0 | 0 | 2 | 0 | 3 | 0 |
| ENSG00000140262 | 675 | 343 | 314 | 187 | 416 | 294 |
| ENSG00000260689 | 0 | 0 | 1 | 0 | 0 | 0 |
| ENSG00000239035 | 0 | 0 | 0 | 0 | 0 | 0 |
| ENSG00000247982 | 10 | 3 | 1 | 4 | 2 | 6 |
| ENSG00000260172 | 0 | 0 | 0 | 0 | 0 | 0 |
| ENSG00000260870 | 0 | 0 | 0 | 0 | 0 | 0 |
| ENSG00000128849 | 149 | 71 | 66 | 34 | 57 | 47 |
| ENSG00000252984 | 0 | 0 | 0 | 0 | 0 | 0 |
| ENSG00000261489 | 0 | 0 | 0 | 0 | 0 | 0 |
| ENSG00000261265 | 0 | 0 | 0 | 0 | 0 | 0 |
| ENSG00000276772 | 4 | 0 | 0 | 0 | 0 | 0 |
| ENSG00000240874 | 0 | 0 | 0 | 0 | 1 | 0 |
| ENSG00000137878 | 91 | 53 | 46 | 28 | 36 | 27 |
| ENSG00000263155 | 50 | 29 | 16 | 7 | 20 | 10 |
| ENSG00000255529 | 370 | 211 | 177 | 151 | 238 | 193 |
| ENSG00000261219 | 0 | 0 | 0 | 0 | 0 | 0 |
| ENSG00000128918 | 27 | 22 | 15 | 9 | 4 | 15 |
| ENSG00000259477 | 0 | 0 | 0 | 0 | 0 | 0 |
| ENSG00000274719 | 0 | 0 | 0 | 0 | 0 | 0 |
| ENSG00000259285 | 0 | 0 | 0 | 0 | 0 | 0 |
| ENSG00000103569 | 0 | 0 | 0 | 0 | 0 | 0 |
| ENSG00000259651 | 0 | 0 | 0 | 0 | 0 | 0 |
| ENSG00000259626 | 0 | 0 | 0 | 0 | 0 | 0 |

| | | | | | | |
|---|---|---|---|---|---|---|
| ENSG00000259600 | 0 | 0 | 0 | 0 | 0 | 0 |
| ENSG00000259379 | 0 | 0 | 0 | 0 | 0 | 0 |
| ENSG00000259671 | 0 | 0 | 0 | 0 | 0 | 0 |
| ENSG00000240934 | 0 | 0 | 0 | 0 | 0 | 0 |
| ENSG00000166035 | 2 | 1 | 0 | 0 | 0 | 0 |
| ENSG00000259293 | 0 | 0 | 0 | 0 | 0 | 0 |
| ENSG00000277229 | 0 | 0 | 0 | 0 | 0 | 0 |
| ENSG00000277790 | 0 | 0 | 0 | 0 | 0 | 0 |
| ENSG00000259476 | 0 | 0 | 0 | 0 | 0 | 0 |
| ENSG00000259250 | 9 | 7 | 10 | 4 | 12 | 4 |
| ENSG00000137845 | 2226 | 1173 | 1238 | 442 | 1278 | 936 |
| ENSG00000270986 | 0 | 0 | 0 | 0 | 0 | 0 |
| ENSG00000182625 | 0 | 0 | 0 | 0 | 0 | 0 |
| ENSG00000282100 | 2 | 0 | 0 | 1 | 1 | 2 |
| ENSG00000199730 | 0 | 0 | 0 | 0 | 0 | 0 |
| ENSG00000278513 | 0 | 0 | 0 | 0 | 0 | 0 |
| ENSG00000259173 | 0 | 0 | 0 | 0 | 0 | 0 |
| ENSG00000200318 | 0 | 0 | 0 | 0 | 0 | 0 |
| ENSG00000245975 | 7 | 6 | 12 | 7 | 4 | 4 |
| ENSG00000128923 | 258 | 111 | 160 | 54 | 121 | 78 |
| ENSG00000259402 | 2 | 1 | 0 | 0 | 0 | 0 |
| ENSG00000259353 | 0 | 1 | 0 | 0 | 0 | 0 |
| ENSG00000259738 | 0 | 0 | 0 | 0 | 0 | 0 |
| ENSG00000157450 | 582 | 296 | 304 | 195 | 300 | 250 |
| ENSG00000242747 | 0 | 0 | 0 | 0 | 0 | 0 |
| ENSG00000137776 | 1065 | 660 | 489 | 254 | 1087 | 567 |
| ENSG00000225798 | 0 | 0 | 0 | 0 | 0 | 0 |
| ENSG00000259393 | 0 | 0 | 0 | 0 | 0 | 0 |
| ENSG00000241640 | 0 | 0 | 0 | 0 | 0 | 0 |
| ENSG00000157456 | 676 | 371 | 370 | 433 | 532 | 515 |
| ENSG00000277144 | 0 | 0 | 3 | 0 | 0 | 1 |
| ENSG00000259732 | 0 | 0 | 1 | 0 | 2 | 0 |
| ENSG00000157483 | 757 | 373 | 469 | 350 | 488 | 415 |
| ENSG00000253030 | 0 | 0 | 0 | 0 | 1 | 1 |
| ENSG00000171989 | 5 | 1 | 2 | 0 | 1 | 0 |
| ENSG00000200070 | 0 | 1 | 0 | 0 | 0 | 0 |
| ENSG00000259771 | 4 | 0 | 0 | 2 | 2 | 2 |
| ENSG00000199512 | 2 | 0 | 0 | 0 | 0 | 0 |
| ENSG00000259735 | 0 | 1 | 1 | 0 | 1 | 0 |
| ENSG00000157470 | 33 | 12 | 21 | 18 | 25 | 16 |
| ENSG00000259750 | 0 | 0 | 0 | 0 | 0 | 0 |
| ENSG00000259674 | 0 | 0 | 0 | 0 | 0 | 0 |
| ENSG00000200252 | 0 | 0 | 0 | 0 | 0 | 0 |
| ENSG00000243780 | 0 | 0 | 0 | 0 | 0 | 0 |
| ENSG00000240238 | 0 | 0 | 0 | 0 | 0 | 0 |
| ENSG00000201704 | 0 | 0 | 0 | 0 | 0 | 0 |
| ENSG00000140297 | 0 | 0 | 1 | 3 | 2 | 1 |
| ENSG00000140307 | 924 | 427 | 521 | 552 | 490 | 505 |
| ENSG00000227161 | 0 | 0 | 0 | 0 | 0 | 0 |
| ENSG00000140299 | 856 | 437 | 481 | 324 | 609 | 433 |

| | | | | | | |
|---|---|---|---|---|---|---|
| ENSG00000259657 | 7 | 4 | 2 | 1 | 2 | 1 |
| ENSG00000259238 | 0 | 2 | 0 | 0 | 0 | 0 |
| ENSG00000234797 | 254 | 128 | 150 | 143 | 158 | 141 |
| ENSG00000259601 | 0 | 0 | 0 | 0 | 0 | 0 |
| ENSG00000259573 | 0 | 0 | 0 | 0 | 0 | 0 |
| ENSG00000171956 | 0 | 0 | 0 | 1 | 0 | 0 |
| ENSG00000259558 | 0 | 0 | 0 | 0 | 0 | 0 |
| ENSG00000259223 | 0 | 0 | 0 | 0 | 0 | 0 |
| ENSG00000182718 | 15190 | 8553 | 9191 | 9085 | 10052 | 8635 |
| ENSG00000240163 | 0 | 1 | 0 | 0 | 0 | 0 |
| ENSG00000242667 | 0 | 1 | 2 | 0 | 1 | 0 |
| ENSG00000273721 | 0 | 0 | 0 | 0 | 0 | 0 |
| ENSG00000128915 | 352 | 241 | 191 | 129 | 332 | 190 |
| ENSG00000245534 | 15 | 8 | 5 | 1 | 9 | 9 |
| ENSG00000069667 | 30 | 7 | 16 | 13 | 26 | 12 |
| ENSG00000259274 | 0 | 0 | 0 | 0 | 0 | 1 |
| ENSG00000259513 | 0 | 0 | 0 | 0 | 0 | 0 |
| ENSG00000273958 | 0 | 0 | 0 | 0 | 0 | 1 |
| ENSG00000277413 | 0 | 0 | 0 | 0 | 0 | 0 |
| ENSG00000259482 | 0 | 0 | 0 | 0 | 0 | 0 |
| ENSG00000212625 | 0 | 0 | 0 | 0 | 0 | 0 |
| ENSG00000259591 | 0 | 0 | 0 | 0 | 0 | 0 |
| ENSG00000273792 | 0 | 0 | 0 | 0 | 0 | 0 |
| ENSG00000259481 | 0 | 0 | 0 | 0 | 0 | 0 |
| ENSG00000259575 | 0 | 0 | 0 | 0 | 0 | 0 |
| ENSG00000276473 | 0 | 0 | 0 | 0 | 0 | 0 |
| ENSG00000259616 | 0 | 0 | 0 | 0 | 0 | 0 |
| ENSG00000259530 | 0 | 0 | 0 | 0 | 0 | 0 |
| ENSG00000259675 | 0 | 0 | 0 | 0 | 0 | 0 |
| ENSG00000271195 | 0 | 0 | 0 | 0 | 0 | 0 |
| ENSG00000259284 | 0 | 0 | 0 | 0 | 0 | 0 |
| ENSG00000259564 | 0 | 0 | 1 | 0 | 0 | 0 |
| ENSG00000129003 | 287 | 172 | 134 | 51 | 345 | 187 |
| ENSG00000285863 | 3 | 2 | 3 | 0 | 3 | 0 |
| ENSG00000259251 | 0 | 0 | 1 | 0 | 0 | 3 |
| ENSG00000198535 | 4 | 3 | 3 | 8 | 3 | 2 |
| ENSG00000259466 | 0 | 0 | 0 | 0 | 0 | 0 |
| ENSG00000205502 | 0 | 0 | 0 | 0 | 0 | 0 |
| ENSG00000259697 | 0 | 0 | 0 | 0 | 0 | 0 |
| ENSG00000166104 | 0 | 0 | 0 | 0 | 0 | 0 |
| ENSG00000259878 | 0 | 0 | 0 | 0 | 0 | 0 |
| ENSG00000276713 | 0 | 0 | 0 | 0 | 0 | 0 |
| ENSG00000260062 | 0 | 0 | 0 | 0 | 0 | 0 |
| ENSG00000260375 | 0 | 0 | 0 | 0 | 0 | 0 |
| ENSG00000261296 | 0 | 0 | 0 | 0 | 0 | 0 |
| ENSG00000260957 | 0 | 0 | 0 | 0 | 0 | 0 |
| ENSG00000277919 | 0 | 0 | 0 | 0 | 0 | 0 |
| ENSG00000275451 | 0 | 0 | 0 | 0 | 0 | 0 |
| ENSG00000259557 | 0 | 0 | 0 | 0 | 0 | 0 |
| ENSG00000274340 | 2 | 0 | 0 | 0 | 0 | 3 |

| | | | | | | |
|---|---|---|---|---|---|---|
| ENSG00000171914 | 480 | 210 | 282 | 234 | 250 | 292 |
| ENSG00000259458 | 0 | 0 | 0 | 0 | 0 | 0 |
| ENSG00000259756 | 0 | 0 | 0 | 0 | 0 | 0 |
| ENSG00000211137 | 0 | 0 | 0 | 0 | 0 | 0 |
| ENSG00000259370 | 4 | 5 | 5 | 9 | 9 | 7 |
| ENSG00000259727 | 4 | 0 | 3 | 1 | 2 | 6 |
| ENSG00000259394 | 0 | 0 | 0 | 0 | 0 | 0 |
| ENSG00000140416 | 3858 | 2430 | 2269 | 2473 | 3459 | 2557 |
| ENSG00000259498 | 2 | 5 | 2 | 1 | 3 | 3 |
| ENSG00000259627 | 0 | 0 | 0 | 0 | 0 | 0 |
| ENSG00000259672 | 0 | 0 | 1 | 0 | 0 | 0 |
| ENSG00000103642 | 2192 | 1195 | 1088 | 1363 | 1832 | 1494 |
| ENSG00000185088 | 3029 | 1605 | 1635 | 1873 | 2190 | 1786 |
| ENSG00000166128 | 589 | 311 | 330 | 232 | 453 | 313 |
| ENSG00000138613 | 821 | 428 | 439 | 421 | 503 | 456 |
| ENSG00000074410 | 5730 | 2894 | 3172 | 3391 | 3932 | 4236 |
| ENSG00000259459 | 50 | 20 | 19 | 18 | 25 | 33 |
| ENSG00000259268 | 2 | 0 | 1 | 1 | 1 | 4 |
| ENSG00000276651 | 6 | 2 | 1 | 4 | 1 | 5 |
| ENSG00000140455 | 1446 | 838 | 976 | 989 | 1303 | 1178 |
| ENSG00000259248 | 29 | 15 | 21 | 11 | 27 | 23 |
| ENSG00000197361 | 84 | 45 | 58 | 56 | 71 | 56 |
| ENSG00000103657 | 1495 | 850 | 887 | 352 | 976 | 795 |
| ENSG00000259589 | 0 | 0 | 0 | 0 | 0 | 1 |
| ENSG00000199156 | 0 | 0 | 0 | 0 | 0 | 0 |
| ENSG00000275785 | 0 | 0 | 1 | 0 | 0 | 0 |
| ENSG00000035664 | 221 | 129 | 138 | 88 | 97 | 80 |
| ENSG00000259351 | 1 | 4 | 2 | 0 | 0 | 0 |
| ENSG00000166797 | 1690 | 843 | 1004 | 953 | 960 | 997 |
| ENSG00000028528 | 4982 | 2314 | 2812 | 1998 | 2565 | 2507 |
| ENSG00000201071 | 0 | 1 | 0 | 0 | 0 | 0 |
| ENSG00000157734 | 49 | 21 | 34 | 21 | 34 | 19 |
| ENSG00000166794 | 3356 | 2065 | 2339 | 2479 | 2328 | 2235 |
| ENSG00000169118 | 1343 | 561 | 762 | 435 | 656 | 621 |
| ENSG00000259316 | 0 | 0 | 1 | 0 | 1 | 0 |
| ENSG00000252774 | 0 | 0 | 0 | 0 | 0 | 0 |
| ENSG00000166803 | 164 | 91 | 118 | 83 | 94 | 115 |
| ENSG00000103671 | 1184 | 643 | 729 | 563 | 740 | 691 |
| ENSG00000206781 | 0 | 0 | 0 | 0 | 0 | 0 |
| ENSG00000180357 | 3058 | 1307 | 1728 | 996 | 1416 | 1549 |
| ENSG00000259682 | 0 | 0 | 0 | 0 | 0 | 0 |
| ENSG00000248415 | 0 | 2 | 0 | 0 | 0 | 0 |
| ENSG00000243403 | 0 | 0 | 0 | 0 | 1 | 0 |
| ENSG00000239465 | 0 | 1 | 0 | 0 | 0 | 0 |
| ENSG00000207223 | 0 | 0 | 0 | 0 | 0 | 0 |
| ENSG00000207162 | 0 | 0 | 0 | 0 | 0 | 0 |
| ENSG00000180304 | 3006 | 1421 | 1673 | 1603 | 1456 | 1635 |
| ENSG00000265967 | 0 | 0 | 0 | 0 | 0 | 0 |
| ENSG00000264937 | 4 | 3 | 4 | 0 | 0 | 3 |
| ENSG00000259635 | 5 | 3 | 4 | 0 | 2 | 3 |

| | | | | | | |
|---|---|---|---|---|---|---|
| ENSG00000166831 | 60 | 37 | 39 | 37 | 37 | 20 |
| ENSG00000221033 | 0 | 0 | 0 | 0 | 0 | 0 |
| ENSG00000140451 | 119 | 55 | 81 | 94 | 115 | 87 |
| ENSG00000249240 | 0 | 0 | 0 | 0 | 0 | 0 |
| ENSG00000241839 | 152 | 73 | 94 | 108 | 94 | 87 |
| ENSG00000166839 | 42 | 17 | 25 | 13 | 33 | 17 |
| ENSG00000275332 | 0 | 0 | 1 | 0 | 0 | 1 |
| ENSG00000274383 | 0 | 2 | 0 | 0 | 5 | 1 |
| ENSG00000090487 | 4260 | 2098 | 2434 | 2437 | 2476 | 2412 |
| ENSG00000103707 | 456 | 239 | 262 | 274 | 292 | 278 |
| ENSG00000270218 | 4 | 0 | 0 | 0 | 1 | 1 |
| ENSG00000186198 | 0 | 0 | 0 | 0 | 0 | 0 |
| ENSG00000277351 | 0 | 0 | 0 | 0 | 0 | 0 |
| ENSG00000103710 | 0 | 0 | 0 | 0 | 0 | 1 |
| ENSG00000234438 | 0 | 0 | 0 | 0 | 0 | 0 |
| ENSG00000278737 | 0 | 1 | 0 | 0 | 0 | 0 |
| ENSG00000246922 | 45 | 40 | 51 | 33 | 40 | 42 |
| ENSG00000259258 | 0 | 0 | 1 | 0 | 0 | 0 |
| ENSG00000090470 | 1092 | 477 | 582 | 440 | 631 | 585 |
| ENSG00000166855 | 2049 | 1079 | 1264 | 814 | 1206 | 1174 |
| ENSG00000138615 | 12 | 4 | 4 | 5 | 7 | 4 |
| ENSG00000279268 | 0 | 0 | 0 | 0 | 0 | 0 |
| ENSG00000201709 | 0 | 0 | 0 | 0 | 0 | 0 |
| ENSG00000138617 | 571 | 247 | 317 | 260 | 288 | 322 |
| ENSG00000206903 | 0 | 0 | 0 | 0 | 1 | 0 |
| ENSG00000199568 | 4 | 0 | 1 | 4 | 1 | 3 |
| ENSG00000259346 | 0 | 0 | 1 | 0 | 1 | 0 |
| ENSG00000200156 | 0 | 0 | 1 | 0 | 0 | 0 |
| ENSG00000174498 | 989 | 451 | 600 | 409 | 456 | 480 |
| ENSG00000103742 | 10 | 9 | 6 | 1 | 4 | 4 |
| ENSG00000074603 | 1598 | 797 | 889 | 484 | 984 | 766 |
| ENSG00000270690 | 0 | 3 | 1 | 0 | 0 | 3 |
| ENSG00000259232 | 1 | 2 | 0 | 0 | 0 | 1 |
| ENSG00000252217 | 0 | 0 | 0 | 0 | 0 | 0 |
| ENSG00000074696 | 9251 | 4252 | 5250 | 3449 | 4610 | 4374 |
| ENSG00000207449 | 0 | 1 | 0 | 0 | 0 | 0 |
| ENSG00000138614 | 2061 | 953 | 1376 | 1053 | 1170 | 1186 |
| ENSG00000074621 | 682 | 452 | 473 | 260 | 326 | 397 |
| ENSG00000275638 | 0 | 0 | 0 | 0 | 0 | 0 |
| ENSG00000174485 | 535 | 343 | 258 | 125 | 469 | 245 |
| ENSG00000238311 | 0 | 0 | 0 | 0 | 0 | 0 |
| ENSG00000264737 | 0 | 0 | 0 | 0 | 0 | 0 |
| ENSG00000103769 | 9655 | 4350 | 5477 | 4222 | 4993 | 4896 |
| ENSG00000261544 | 0 | 0 | 1 | 0 | 1 | 0 |
| ENSG00000259924 | 0 | 0 | 2 | 0 | 0 | 0 |
| ENSG00000249271 | 2 | 0 | 0 | 0 | 0 | 0 |
| ENSG00000283249 | 0 | 0 | 0 | 0 | 0 | 0 |
| ENSG00000157890 | 10 | 6 | 5 | 3 | 4 | 4 |
| ENSG00000263512 | 0 | 0 | 0 | 0 | 0 | 0 |
| ENSG00000261318 | 2 | 0 | 0 | 0 | 0 | 1 |

| | | | | | | |
|---|---|---|---|---|---|---|
| ENSG00000166938 | 845 | 384 | 522 | 282 | 429 | 436 |
| ENSG00000260773 | 5 | 3 | 1 | 3 | 3 | 2 |
| ENSG00000075131 | 201 | 108 | 117 | 119 | 123 | 126 |
| ENSG00000252712 | 0 | 0 | 1 | 0 | 0 | 0 |
| ENSG00000223271 | 0 | 0 | 0 | 0 | 0 | 0 |
| ENSG00000240821 | 0 | 0 | 0 | 0 | 0 | 0 |
| ENSG00000169032 | 2151 | 1104 | 1335 | 1064 | 1053 | 1078 |
| ENSG00000261102 | 0 | 0 | 0 | 0 | 0 | 0 |
| ENSG00000243404 | 0 | 1 | 0 | 1 | 0 | 3 |
| ENSG00000261351 | 38 | 26 | 33 | 28 | 25 | 24 |
| ENSG00000174446 | 586 | 318 | 368 | 355 | 388 | 388 |
| ENSG00000266589 | 1 | 1 | 1 | 1 | 0 | 1 |
| ENSG00000174444 | 58676 | 29048 | 34167 | 38393 | 36257 | 35064 |
| ENSG00000199574 | 0 | 0 | 0 | 0 | 0 | 0 |
| ENSG00000280554 | 0 | 0 | 0 | 0 | 0 | 0 |
| ENSG00000202529 | 0 | 0 | 0 | 0 | 0 | 0 |
| ENSG00000199673 | 0 | 0 | 0 | 0 | 0 | 0 |
| ENSG00000200623 | 0 | 0 | 0 | 0 | 0 | 0 |
| ENSG00000174442 | 664 | 374 | 419 | 297 | 474 | 401 |
| ENSG00000188501 | 8 | 4 | 11 | 4 | 3 | 1 |
| ENSG00000259471 | 0 | 1 | 1 | 0 | 0 | 0 |
| ENSG00000137834 | 919 | 367 | 445 | 430 | 490 | 468 |
| ENSG00000274995 | 0 | 0 | 0 | 1 | 0 | 0 |
| ENSG00000270490 | 4 | 1 | 0 | 1 | 2 | 3 |
| ENSG00000277152 | 6 | 3 | 1 | 2 | 3 | 0 |
| ENSG00000259437 | 0 | 2 | 0 | 0 | 0 | 1 |
| ENSG00000259289 | 0 | 2 | 0 | 0 | 1 | 0 |
| ENSG00000259347 | 19 | 12 | 8 | 12 | 6 | 14 |
| ENSG00000259246 | 0 | 0 | 0 | 0 | 0 | 0 |
| ENSG00000166949 | 5322 | 2728 | 3094 | 2365 | 2933 | 2815 |
| ENSG00000259202 | 0 | 1 | 0 | 0 | 0 | 1 |
| ENSG00000103591 | 2924 | 1606 | 1894 | 1486 | 2096 | 1846 |
| ENSG00000242634 | 0 | 0 | 0 | 0 | 0 | 0 |
| ENSG00000103599 | 118 | 57 | 85 | 53 | 85 | 67 |
| ENSG00000259673 | 280 | 159 | 177 | 137 | 171 | 176 |
| ENSG00000189227 | 186 | 116 | 146 | 125 | 137 | 134 |
| ENSG00000270964 | 80 | 46 | 50 | 36 | 31 | 43 |
| ENSG00000137764 | 668 | 314 | 352 | 369 | 357 | 348 |
| ENSG00000259512 | 0 | 0 | 0 | 0 | 0 | 0 |
| ENSG00000188779 | 11 | 2 | 5 | 0 | 7 | 3 |
| ENSG00000259410 | 0 | 0 | 0 | 0 | 0 | 0 |
| ENSG00000245719 | 0 | 0 | 0 | 0 | 0 | 0 |
| ENSG00000206625 | 2 | 4 | 2 | 5 | 0 | 1 |
| ENSG00000261702 | 0 | 0 | 0 | 0 | 0 | 0 |
| ENSG00000260109 | 0 | 0 | 0 | 0 | 0 | 0 |
| ENSG00000033800 | 812 | 554 | 526 | 272 | 583 | 464 |
| ENSG00000260007 | 45 | 31 | 36 | 19 | 43 | 45 |
| ENSG00000129007 | 9 | 7 | 18 | 3 | 12 | 8 |
| ENSG00000128973 | 858 | 427 | 535 | 565 | 476 | 597 |
| ENSG00000260602 | 0 | 0 | 0 | 0 | 1 | 0 |

| | | | | | | |
|---|---|---|---|---|---|---|
| ENSG00000260657 | 0 | 2 | 0 | 0 | 2 | 1 |
| ENSG00000169018 | 2510 | 1218 | 1340 | 991 | 1619 | 1309 |
| ENSG00000137809 | 4 | 2 | 1 | 1 | 2 | 1 |
| ENSG00000103647 | 40 | 15 | 18 | 10 | 7 | 11 |
| ENSG00000271232 | 0 | 0 | 0 | 0 | 0 | 0 |
| ENSG00000140350 | 4252 | 2238 | 2459 | 2233 | 2827 | 2841 |
| ENSG00000265195 | 0 | 2 | 0 | 0 | 0 | 1 |
| ENSG00000258484 | 100 | 47 | 51 | 43 | 67 | 55 |
| ENSG00000259286 | 6 | 3 | 1 | 2 | 3 | 1 |
| ENSG00000285919 | 0 | 0 | 0 | 0 | 0 | 0 |
| ENSG00000137808 | 0 | 0 | 0 | 0 | 0 | 0 |
| ENSG00000255346 | 129 | 49 | 90 | 33 | 49 | 41 |
| ENSG00000273851 | 0 | 0 | 0 | 0 | 0 | 2 |
| ENSG00000259265 | 0 | 0 | 0 | 0 | 0 | 0 |
| ENSG00000212766 | 6 | 5 | 9 | 0 | 3 | 3 |
| ENSG00000259222 | 0 | 0 | 0 | 0 | 0 | 0 |
| ENSG00000138604 | 1242 | 542 | 655 | 375 | 538 | 441 |
| ENSG00000261634 | 1 | 1 | 2 | 0 | 1 | 0 |
| ENSG00000259504 | 0 | 1 | 1 | 0 | 0 | 0 |
| ENSG00000137819 | 143 | 69 | 85 | 49 | 39 | 52 |
| ENSG00000259645 | 3 | 1 | 2 | 0 | 2 | 0 |
| ENSG00000259426 | 18 | 14 | 12 | 13 | 8 | 9 |
| ENSG00000207395 | 0 | 0 | 0 | 0 | 0 | 0 |
| ENSG00000137807 | 1037 | 635 | 634 | 415 | 919 | 641 |
| ENSG00000259191 | 0 | 0 | 0 | 0 | 1 | 0 |
| ENSG00000137818 | 24033 | 12362 | 13550 | 17296 | 14338 | 13557 |
| ENSG00000207119 | 2 | 0 | 0 | 0 | 1 | 1 |
| ENSG00000259215 | 4 | 4 | 0 | 2 | 0 | 0 |
| ENSG00000245750 | 29 | 17 | 17 | 5 | 12 | 10 |
| ENSG00000259457 | 0 | 1 | 1 | 1 | 0 | 0 |
| ENSG00000259496 | 0 | 0 | 0 | 0 | 0 | 0 |
| ENSG00000259309 | 0 | 0 | 0 | 0 | 0 | 0 |
| ENSG00000140332 | 2290 | 1042 | 1244 | 906 | 936 | 974 |
| ENSG00000207965 | 0 | 0 | 0 | 0 | 0 | 0 |
| ENSG00000200216 | 0 | 0 | 0 | 0 | 0 | 0 |
| ENSG00000259252 | 0 | 0 | 0 | 0 | 0 | 0 |
| ENSG00000259503 | 0 | 0 | 0 | 0 | 0 | 0 |
| ENSG00000259473 | 0 | 0 | 0 | 0 | 0 | 0 |
| ENSG00000280639 | 0 | 0 | 1 | 0 | 1 | 0 |
| ENSG00000259227 | 0 | 0 | 0 | 0 | 0 | 0 |
| ENSG00000281264 | 0 | 0 | 0 | 0 | 0 | 0 |
| ENSG00000280515 | 0 | 1 | 0 | 0 | 0 | 0 |
| ENSG00000137831 | 282 | 168 | 126 | 51 | 380 | 139 |
| ENSG00000259532 | 0 | 0 | 0 | 0 | 0 | 3 |
| ENSG00000274297 | 2 | 0 | 0 | 1 | 1 | 1 |
| ENSG00000259624 | 0 | 1 | 0 | 0 | 0 | 0 |
| ENSG00000259452 | 0 | 0 | 0 | 0 | 0 | 0 |
| ENSG00000235420 | 0 | 0 | 0 | 0 | 0 | 0 |
| ENSG00000166173 | 179 | 91 | 105 | 79 | 83 | 82 |
| ENSG00000259744 | 0 | 0 | 1 | 0 | 2 | 0 |

| | | | | | | |
|---|---|---|---|---|---|---|
| ENSG00000137821 | 23 | 8 | 10 | 14 | 15 | 15 |
| ENSG00000259470 | 6 | 2 | 1 | 1 | 1 | 0 |
| ENSG00000129028 | 275 | 153 | 150 | 108 | 140 | 137 |
| ENSG00000242276 | 2 | 0 | 1 | 1 | 0 | 0 |
| ENSG00000187720 | 1304 | 548 | 770 | 537 | 603 | 654 |
| ENSG00000225362 | 17 | 9 | 13 | 8 | 11 | 14 |
| ENSG00000259964 | 0 | 0 | 0 | 0 | 0 | 0 |
| ENSG00000259781 | 260 | 144 | 171 | 138 | 182 | 181 |
| ENSG00000260586 | 0 | 0 | 0 | 0 | 1 | 2 |
| ENSG00000212664 | 0 | 2 | 7 | 2 | 1 | 1 |
| ENSG00000278408 | 0 | 0 | 0 | 0 | 0 | 0 |
| ENSG00000278570 | 12 | 4 | 8 | 8 | 10 | 12 |
| ENSG00000260037 | 1 | 0 | 0 | 1 | 5 | 2 |
| ENSG00000066933 | 680 | 314 | 300 | 135 | 561 | 335 |
| ENSG00000200873 | 0 | 0 | 0 | 0 | 0 | 0 |
| ENSG00000261632 | 0 | 0 | 0 | 0 | 0 | 0 |
| ENSG00000260576 | 2 | 0 | 0 | 0 | 1 | 0 |
| ENSG00000222094 | 0 | 0 | 0 | 0 | 0 | 0 |
| ENSG00000166192 | 197 | 108 | 117 | 67 | 106 | 96 |
| ENSG00000243568 | 0 | 0 | 0 | 0 | 1 | 0 |
| ENSG00000260173 | 0 | 1 | 1 | 0 | 0 | 1 |
| ENSG00000175318 | 58 | 28 | 37 | 24 | 19 | 24 |
| ENSG00000067225 | 47295 | 22903 | 26494 | 33750 | 29334 | 28578 |
| ENSG00000137817 | 2619 | 1160 | 1783 | 1309 | 1586 | 1598 |
| ENSG00000273025 | 0 | 0 | 0 | 0 | 0 | 0 |
| ENSG00000261460 | 0 | 3 | 6 | 3 | 2 | 4 |
| ENSG00000140488 | 0 | 0 | 0 | 0 | 0 | 0 |
| ENSG00000260729 | 6 | 1 | 1 | 2 | 2 | 0 |
| ENSG00000213614 | 2171 | 1590 | 1944 | 1674 | 1733 | 1707 |
| ENSG00000260339 | 52 | 17 | 26 | 15 | 30 | 16 |
| ENSG00000243007 | 2 | 4 | 3 | 2 | 0 | 0 |
| ENSG00000187806 | 0 | 0 | 0 | 0 | 0 | 0 |
| ENSG00000261423 | 35 | 23 | 15 | 23 | 19 | 26 |
| ENSG00000271072 | 0 | 0 | 0 | 0 | 1 | 0 |
| ENSG00000261187 | 11 | 6 | 7 | 3 | 7 | 9 |
| ENSG00000166233 | 3540 | 1748 | 1974 | 1341 | 2091 | 1688 |
| ENSG00000283798 | 0 | 0 | 0 | 0 | 0 | 0 |
| ENSG00000260534 | 0 | 0 | 0 | 0 | 0 | 0 |
| ENSG00000278313 | 0 | 0 | 2 | 0 | 0 | 0 |
| ENSG00000260144 | 7 | 0 | 1 | 1 | 0 | 1 |
| ENSG00000259783 | 0 | 1 | 0 | 0 | 0 | 0 |
| ENSG00000260672 | 2 | 1 | 1 | 3 | 1 | 1 |
| ENSG00000259909 | 1 | 0 | 0 | 0 | 0 | 0 |
| ENSG00000215186 | 2 | 1 | 0 | 1 | 0 | 0 |
| ENSG00000274350 | 0 | 0 | 0 | 0 | 0 | 0 |
| ENSG00000175202 | 0 | 1 | 2 | 4 | 1 | 1 |
| ENSG00000261281 | 0 | 0 | 0 | 0 | 0 | 0 |
| ENSG00000140463 | 1261 | 577 | 759 | 551 | 618 | 646 |
| ENSG00000159322 | 2096 | 1110 | 1442 | 1166 | 1366 | 1356 |
| ENSG00000260898 | 4 | 1 | 3 | 0 | 2 | 0 |

| | | | | | | |
|---|---|---|---|---|---|---|
| ENSG00000285729 | 0 | 0 | 0 | 0 | 0 | 0 |
| ENSG00000259259 | 0 | 0 | 0 | 0 | 0 | 0 |
| ENSG00000067141 | 3128 | 1267 | 1865 | 1137 | 1561 | 1561 |
| ENSG00000259691 | 0 | 0 | 0 | 0 | 0 | 0 |
| ENSG00000259440 | 0 | 0 | 3 | 0 | 0 | 1 |
| ENSG00000259528 | 0 | 1 | 0 | 0 | 0 | 0 |
| ENSG00000138622 | 10 | 6 | 14 | 13 | 2 | 4 |
| ENSG00000259650 | 0 | 0 | 0 | 0 | 0 | 0 |
| ENSG00000183324 | 6 | 1 | 0 | 0 | 0 | 0 |
| ENSG00000259186 | 0 | 0 | 0 | 0 | 0 | 0 |
| ENSG00000156642 | 4286 | 1963 | 2554 | 2029 | 2367 | 2272 |
| ENSG00000281183 | 100 | 37 | 77 | 22 | 46 | 38 |
| ENSG00000103855 | 5805 | 2738 | 3263 | 3438 | 2889 | 3068 |
| ENSG00000276807 | 0 | 0 | 0 | 0 | 0 | 0 |
| ENSG00000205363 | 112 | 54 | 43 | 78 | 51 | 48 |
| ENSG00000260469 | 8 | 3 | 4 | 1 | 3 | 5 |
| ENSG00000260624 | 0 | 0 | 0 | 0 | 0 | 0 |
| ENSG00000167139 | 0 | 0 | 0 | 0 | 0 | 0 |
| ENSG00000261801 | 1080 | 561 | 685 | 636 | 711 | 695 |
| ENSG00000129038 | 1363 | 739 | 753 | 1255 | 1071 | 1099 |
| ENSG00000067221 | 335 | 185 | 220 | 273 | 245 | 194 |
| ENSG00000140464 | 957 | 404 | 555 | 558 | 497 | 507 |
| ENSG00000260096 | 0 | 0 | 0 | 0 | 0 | 0 |
| ENSG00000159289 | 4 | 0 | 1 | 1 | 0 | 0 |
| ENSG00000278604 | 0 | 0 | 0 | 0 | 0 | 0 |
| ENSG00000214702 | 4 | 0 | 0 | 0 | 0 | 0 |
| ENSG00000167178 | 6 | 0 | 2 | 3 | 0 | 5 |
| ENSG00000248540 | 0 | 0 | 0 | 0 | 0 | 4 |
| ENSG00000261543 | 0 | 0 | 0 | 0 | 0 | 0 |
| ENSG00000278348 | 0 | 0 | 0 | 0 | 0 | 0 |
| ENSG00000129009 | 0 | 0 | 0 | 2 | 1 | 8 |
| ENSG00000137868 | 329 | 219 | 205 | 355 | 401 | 456 |
| ENSG00000140481 | 71 | 40 | 44 | 84 | 113 | 103 |
| ENSG00000280309 | 0 | 2 | 0 | 0 | 0 | 0 |
| ENSG00000261384 | 0 | 0 | 0 | 0 | 0 | 0 |
| ENSG00000277749 | 0 | 0 | 0 | 0 | 0 | 0 |
| ENSG00000140459 | 0 | 0 | 0 | 0 | 0 | 0 |
| ENSG00000260266 | 0 | 0 | 1 | 0 | 0 | 0 |
| ENSG00000261821 | 0 | 0 | 0 | 0 | 1 | 0 |
| ENSG00000274937 | 4 | 0 | 0 | 0 | 0 | 0 |
| ENSG00000277718 | 0 | 0 | 0 | 0 | 0 | 0 |
| ENSG00000138623 | 121 | 61 | 54 | 97 | 98 | 74 |
| ENSG00000277391 | 0 | 0 | 0 | 0 | 0 | 0 |
| ENSG00000138629 | 1958 | 817 | 1052 | 1174 | 886 | 980 |
| ENSG00000247240 | 160 | 103 | 109 | 98 | 117 | 143 |
| ENSG00000260103 | 17 | 3 | 10 | 4 | 5 | 2 |
| ENSG00000261775 | 2 | 1 | 4 | 1 | 6 | 1 |
| ENSG00000212279 | 0 | 0 | 0 | 0 | 0 | 0 |
| ENSG00000179361 | 314 | 137 | 210 | 174 | 197 | 195 |
| ENSG00000179335 | 3753 | 1761 | 2282 | 2459 | 2674 | 2322 |

| | | | | | | |
|---|---|---|---|---|---|---|
| ENSG00000275527 | 0 | 0 | 0 | 0 | 0 | 0 |
| ENSG00000260919 | 2 | 4 | 1 | 1 | 4 | 2 |
| ENSG00000179151 | 2503 | 1273 | 1666 | 2619 | 2844 | 2635 |
| ENSG00000140465 | 562 | 347 | 490 | 23793 | 30805 | 25007 |
| ENSG00000140505 | 0 | 0 | 0 | 7 | 6 | 10 |
| ENSG00000103653 | 4214 | 2158 | 2601 | 3184 | 3010 | 2933 |
| ENSG00000264386 | 4 | 2 | 0 | 1 | 0 | 0 |
| ENSG00000140506 | 0 | 0 | 1 | 0 | 0 | 0 |
| ENSG00000261606 | 5 | 0 | 1 | 3 | 2 | 0 |
| ENSG00000213578 | 0 | 0 | 0 | 0 | 0 | 0 |
| ENSG00000140474 | 2415 | 1072 | 1564 | 1410 | 1359 | 1546 |
| ENSG00000275411 | 0 | 0 | 0 | 0 | 0 | 0 |
| ENSG00000140497 | 6358 | 2904 | 3654 | 3512 | 3008 | 3409 |
| ENSG00000178802 | 1853 | 902 | 1090 | 1088 | 885 | 936 |
| ENSG00000178761 | 694 | 311 | 403 | 285 | 389 | 400 |
| ENSG00000178741 | 3158 | 1527 | 1788 | 2024 | 2039 | 1804 |
| ENSG00000178718 | 458 | 203 | 241 | 375 | 267 | 288 |
| ENSG00000198794 | 1716 | 821 | 1045 | 971 | 907 | 1011 |
| ENSG00000261813 | 0 | 0 | 0 | 0 | 0 | 0 |
| ENSG00000199580 | 0 | 0 | 0 | 0 | 0 | 0 |
| ENSG00000260483 | 0 | 0 | 0 | 0 | 0 | 0 |
| ENSG00000138621 | 767 | 414 | 463 | 449 | 391 | 436 |
| ENSG00000260152 | 0 | 0 | 0 | 0 | 0 | 0 |
| ENSG00000203411 | 0 | 0 | 6 | 1 | 0 | 1 |
| ENSG00000252722 | 0 | 0 | 0 | 0 | 0 | 0 |
| ENSG00000240366 | 0 | 0 | 0 | 0 | 0 | 0 |
| ENSG00000167173 | 976 | 433 | 533 | 612 | 515 | 525 |
| ENSG00000261779 | 0 | 0 | 0 | 0 | 0 | 0 |
| ENSG00000260660 | 1 | 0 | 0 | 1 | 0 | 0 |
| ENSG00000260104 | 0 | 0 | 0 | 0 | 0 | 0 |
| ENSG00000261205 | 0 | 0 | 0 | 0 | 0 | 0 |
| ENSG00000167195 | 0 | 0 | 0 | 0 | 0 | 0 |
| ENSG00000277295 | 0 | 0 | 0 | 0 | 0 | 0 |
| ENSG00000140478 | 0 | 0 | 0 | 0 | 0 | 0 |
| ENSG00000275992 | 0 | 0 | 0 | 0 | 0 | 0 |
| ENSG00000260357 | 0 | 0 | 0 | 0 | 0 | 0 |
| ENSG00000259790 | 2 | 3 | 0 | 1 | 0 | 0 |
| ENSG00000140365 | 2475 | 1300 | 1503 | 1599 | 1405 | 1368 |
| ENSG00000275645 | 2 | 3 | 5 | 7 | 10 | 6 |
| ENSG00000140398 | 287 | 131 | 177 | 116 | 107 | 123 |
| ENSG00000284343 | 0 | 0 | 0 | 0 | 0 | 0 |
| ENSG00000140400 | 1912 | 861 | 1391 | 991 | 1026 | 1162 |
| ENSG00000260274 | 133 | 60 | 93 | 65 | 82 | 71 |
| ENSG00000169375 | 3445 | 1488 | 1893 | 1198 | 1765 | 1698 |
| ENSG00000241890 | 2 | 1 | 1 | 2 | 0 | 0 |
| ENSG00000261714 | 1 | 0 | 0 | 0 | 0 | 3 |
| ENSG00000276744 | 6 | 9 | 9 | 6 | 5 | 7 |
| ENSG00000169410 | 1819 | 887 | 1069 | 894 | 973 | 1068 |
| ENSG00000260165 | 0 | 1 | 0 | 1 | 0 | 0 |
| ENSG00000259931 | 0 | 0 | 0 | 0 | 0 | 0 |

| | | | | | | |
|---|---|---|---|---|---|---|
| ENSG00000260269 | 1 | 0 | 0 | 1 | 0 | 0 |
| ENSG00000169371 | 110 | 77 | 82 | 60 | 87 | 95 |
| ENSG00000260235 | 0 | 0 | 0 | 0 | 0 | 0 |
| ENSG00000260206 | 3 | 0 | 0 | 1 | 0 | 1 |
| ENSG00000177971 | 1243 | 742 | 879 | 1033 | 783 | 840 |
| ENSG00000275454 | 10 | 10 | 12 | 7 | 6 | 5 |
| ENSG00000274515 | 6 | 5 | 6 | 0 | 1 | 6 |
| ENSG00000173548 | 1707 | 672 | 957 | 633 | 621 | 629 |
| ENSG00000173546 | 78 | 54 | 52 | 21 | 36 | 32 |
| ENSG00000260892 | 0 | 0 | 0 | 0 | 0 | 0 |
| ENSG00000203392 | 65 | 31 | 49 | 22 | 26 | 29 |
| ENSG00000182950 | 0 | 0 | 0 | 0 | 0 | 0 |
| ENSG00000246877 | 16 | 9 | 10 | 6 | 12 | 7 |
| ENSG00000260288 | 2 | 3 | 2 | 3 | 4 | 4 |
| ENSG00000260815 | 4 | 1 | 1 | 0 | 2 | 0 |
| ENSG00000261043 | 2 | 0 | 0 | 0 | 0 | 0 |
| ENSG00000284385 | 0 | 0 | 0 | 0 | 0 | 0 |
| ENSG00000187812 | 0 | 0 | 0 | 0 | 0 | 0 |
| ENSG00000241807 | 0 | 0 | 0 | 0 | 0 | 0 |
| ENSG00000261820 | 0 | 0 | 0 | 0 | 0 | 0 |
| ENSG00000140367 | 1201 | 607 | 633 | 425 | 758 | 563 |
| ENSG00000266308 | 0 | 1 | 0 | 0 | 0 | 0 |
| ENSG00000167196 | 2098 | 1093 | 1219 | 1126 | 1160 | 1208 |
| ENSG00000169752 | 7 | 3 | 1 | 0 | 5 | 4 |
| ENSG00000260685 | 0 | 0 | 0 | 0 | 0 | 1 |
| ENSG00000169758 | 53 | 26 | 31 | 19 | 20 | 23 |
| ENSG00000259422 | 8 | 6 | 2 | 5 | 3 | 9 |
| ENSG00000140374 | 9451 | 4737 | 5270 | 4279 | 5284 | 4697 |
| ENSG00000196274 | 4 | 0 | 3 | 1 | 0 | 0 |
| ENSG00000259581 | 6 | 1 | 4 | 1 | 2 | 1 |
| ENSG00000159556 | 23 | 16 | 24 | 22 | 18 | 21 |
| ENSG00000259514 | 0 | 1 | 5 | 4 | 1 | 3 |
| ENSG00000270036 | 2 | 0 | 0 | 0 | 1 | 0 |
| ENSG00000140386 | 205 | 132 | 113 | 54 | 202 | 100 |
| ENSG00000261232 | 0 | 0 | 0 | 0 | 0 | 0 |
| ENSG00000266449 | 0 | 0 | 0 | 0 | 0 | 0 |
| ENSG00000201510 | 0 | 0 | 0 | 0 | 0 | 0 |
| ENSG00000117906 | 3494 | 1633 | 2136 | 1678 | 1989 | 1973 |
| ENSG00000243365 | 0 | 0 | 0 | 0 | 0 | 0 |
| ENSG00000259565 | 0 | 0 | 0 | 1 | 0 | 0 |
| ENSG00000140368 | 1 | 1 | 2 | 0 | 0 | 0 |
| ENSG00000140391 | 1669 | 807 | 1024 | 949 | 1015 | 1012 |
| ENSG00000259652 | 0 | 0 | 0 | 0 | 0 | 0 |
| ENSG00000278991 | 2 | 1 | 0 | 0 | 3 | 1 |
| ENSG00000269951 | 2 | 0 | 3 | 1 | 4 | 2 |
| ENSG00000173517 | 728 | 452 | 378 | 174 | 437 | 286 |
| ENSG00000259722 | 0 | 0 | 0 | 0 | 0 | 0 |
| ENSG00000230459 | 0 | 0 | 0 | 0 | 0 | 0 |
| ENSG00000140382 | 1121 | 522 | 654 | 352 | 595 | 504 |
| ENSG00000279033 | 0 | 0 | 0 | 0 | 0 | 0 |

| | | | | | | |
|---|---|---|---|---|---|---|
| ENSG00000259362 | 4 | 4 | 7 | 5 | 4 | 3 |
| ENSG00000259420 | 4 | 0 | 0 | 0 | 0 | 0 |
| ENSG00000169783 | 224 | 123 | 107 | 109 | 82 | 69 |
| ENSG00000259666 | 0 | 1 | 0 | 0 | 0 | 0 |
| ENSG00000259281 | 0 | 0 | 0 | 0 | 0 | 0 |
| ENSG00000280359 | 0 | 0 | 0 | 0 | 0 | 0 |
| ENSG00000259213 | 0 | 0 | 0 | 0 | 0 | 0 |
| ENSG00000260139 | 0 | 0 | 0 | 0 | 0 | 0 |
| ENSG00000260776 | 0 | 0 | 1 | 0 | 0 | 0 |
| ENSG00000261244 | 0 | 0 | 0 | 0 | 0 | 0 |
| ENSG00000242066 | 0 | 0 | 0 | 0 | 0 | 0 |
| ENSG00000214646 | 2 | 0 | 0 | 1 | 0 | 0 |
| ENSG00000261403 | 0 | 0 | 0 | 0 | 0 | 0 |
| ENSG00000261143 | 1 | 1 | 1 | 1 | 4 | 1 |
| ENSG00000167202 | 2168 | 1056 | 1311 | 856 | 1206 | 1068 |
| ENSG00000259792 | 6 | 0 | 1 | 5 | 2 | 3 |
| ENSG00000183476 | 0 | 1 | 1 | 0 | 3 | 0 |
| ENSG00000199633 | 0 | 0 | 0 | 0 | 0 | 0 |
| ENSG00000136425 | 4 | 3 | 4 | 6 | 4 | 2 |
| ENSG00000166411 | 1178 | 626 | 765 | 575 | 642 | 704 |
| ENSG00000260988 | 2 | 1 | 5 | 2 | 4 | 3 |
| ENSG00000103740 | 10 | 5 | 4 | 1 | 11 | 5 |
| ENSG00000259708 | 0 | 0 | 0 | 0 | 0 | 0 |
| ENSG00000140403 | 1984 | 942 | 1289 | 938 | 1073 | 1100 |
| ENSG00000140395 | 3796 | 1940 | 2432 | 2048 | 2238 | 2083 |
| ENSG00000272418 | 0 | 0 | 0 | 0 | 0 | 0 |
| ENSG00000259562 | 55 | 35 | 38 | 22 | 31 | 30 |
| ENSG00000259322 | 7 | 7 | 6 | 2 | 1 | 2 |
| ENSG00000277482 | 0 | 0 | 0 | 0 | 0 | 0 |
| ENSG00000279694 | 0 | 0 | 2 | 1 | 0 | 0 |
| ENSG00000166426 | 120 | 65 | 85 | 98 | 70 | 63 |
| ENSG00000259413 | 0 | 0 | 0 | 0 | 0 | 0 |
| ENSG00000136381 | 4366 | 2132 | 2094 | 1034 | 2853 | 1847 |
| ENSG00000259474 | 2 | 2 | 1 | 0 | 0 | 0 |
| ENSG00000188266 | 288 | 137 | 171 | 104 | 143 | 114 |
| ENSG00000279373 | 0 | 2 | 2 | 0 | 2 | 0 |
| ENSG00000041357 | 9029 | 4885 | 5582 | 5448 | 6003 | 5368 |
| ENSG00000169684 | 184 | 105 | 116 | 132 | 150 | 155 |
| ENSG00000261762 | 105 | 57 | 56 | 50 | 72 | 87 |
| ENSG00000080644 | 53 | 21 | 21 | 17 | 24 | 24 |
| ENSG00000279421 | 0 | 0 | 0 | 2 | 1 | 1 |
| ENSG00000117971 | 119 | 62 | 81 | 69 | 64 | 62 |
| ENSG00000259555 | 2 | 0 | 0 | 0 | 1 | 0 |
| ENSG00000261303 | 9 | 1 | 5 | 0 | 1 | 4 |
| ENSG00000213307 | 5 | 2 | 2 | 0 | 4 | 3 |
| ENSG00000273512 | 0 | 0 | 0 | 0 | 0 | 0 |
| ENSG00000238166 | 0 | 0 | 0 | 0 | 0 | 0 |
| ENSG00000136378 | 74 | 45 | 32 | 43 | 36 | 21 |
| ENSG00000185787 | 13570 | 6631 | 7992 | 6500 | 7489 | 7172 |
| ENSG00000243711 | 0 | 0 | 0 | 0 | 0 | 0 |

| | | | | | | |
|---|---|---|---|---|---|---|
| ENSG00000252061 | 0 | 2 | 1 | 2 | 3 | 5 |
| ENSG00000279235 | 0 | 2 | 0 | 0 | 1 | 1 |
| ENSG00000103811 | 4121 | 2080 | 2535 | 1720 | 1528 | 1739 |
| ENSG00000058335 | 2 | 0 | 1 | 0 | 0 | 0 |
| ENSG00000177699 | 0 | 0 | 0 | 0 | 0 | 0 |
| ENSG00000259234 | 0 | 0 | 1 | 0 | 0 | 0 |
| ENSG00000207695 | 0 | 0 | 0 | 0 | 0 | 0 |
| ENSG00000259419 | 0 | 0 | 0 | 1 | 0 | 0 |
| ENSG00000235711 | 0 | 0 | 0 | 0 | 0 | 0 |
| ENSG00000166557 | 6574 | 3090 | 3639 | 4286 | 3588 | 3419 |
| ENSG00000169330 | 9 | 10 | 4 | 5 | 9 | 14 |
| ENSG00000259493 | 0 | 0 | 0 | 0 | 0 | 0 |
| ENSG00000259208 | 0 | 0 | 0 | 0 | 0 | 0 |
| ENSG00000260674 | 0 | 0 | 0 | 0 | 0 | 0 |
| ENSG00000213527 | 0 | 0 | 0 | 0 | 0 | 0 |
| ENSG00000259229 | 0 | 0 | 0 | 0 | 0 | 0 |
| ENSG00000252995 | 0 | 0 | 0 | 0 | 0 | 0 |
| ENSG00000259548 | 0 | 0 | 0 | 0 | 0 | 0 |
| ENSG00000136371 | 84 | 83 | 97 | 96 | 117 | 105 |
| ENSG00000261229 | 2 | 6 | 4 | 2 | 3 | 2 |
| ENSG00000259332 | 0 | 1 | 0 | 0 | 1 | 0 |
| ENSG00000279394 | 0 | 0 | 0 | 0 | 0 | 0 |
| ENSG00000180953 | 236 | 165 | 155 | 139 | 173 | 143 |
| ENSG00000278600 | 232 | 77 | 176 | 70 | 121 | 94 |
| ENSG00000259642 | 283 | 109 | 159 | 144 | 147 | 146 |
| ENSG00000259770 | 0 | 0 | 0 | 0 | 0 | 0 |
| ENSG00000140379 | 2 | 0 | 1 | 0 | 0 | 0 |
| ENSG00000271364 | 0 | 0 | 0 | 0 | 0 | 0 |
| ENSG00000279719 | 0 | 0 | 0 | 0 | 0 | 0 |
| ENSG00000279705 | 0 | 0 | 1 | 0 | 0 | 0 |
| ENSG00000275460 | 0 | 0 | 0 | 0 | 0 | 0 |
| ENSG00000086666 | 3193 | 1574 | 1866 | 1559 | 1820 | 1620 |
| ENSG00000103876 | 763 | 352 | 431 | 610 | 612 | 552 |
| ENSG00000259967 | 0 | 0 | 3 | 0 | 2 | 0 |
| ENSG00000259417 | 2 | 0 | 0 | 1 | 3 | 2 |
| ENSG00000259361 | 0 | 0 | 0 | 0 | 0 | 0 |
| ENSG00000259495 | 43 | 17 | 36 | 19 | 29 | 10 |
| ENSG00000172379 | 1982 | 790 | 1286 | 471 | 508 | 662 |
| ENSG00000258010 | 0 | 1 | 0 | 0 | 0 | 0 |
| ENSG00000259175 | 2 | 0 | 0 | 0 | 0 | 0 |
| ENSG00000270919 | 0 | 0 | 2 | 0 | 0 | 0 |
| ENSG00000284059 | 0 | 1 | 0 | 0 | 0 | 0 |
| ENSG00000222139 | 0 | 0 | 0 | 0 | 0 | 0 |
| ENSG00000136379 | 1131 | 525 | 671 | 652 | 673 | 665 |
| ENSG00000271983 | 7 | 3 | 2 | 1 | 4 | 2 |
| ENSG00000103888 | 89 | 45 | 60 | 50 | 52 | 49 |
| ENSG00000208003 | 0 | 0 | 0 | 0 | 0 | 0 |
| ENSG00000259546 | 0 | 0 | 0 | 0 | 0 | 0 |
| ENSG00000259649 | 0 | 2 | 1 | 0 | 2 | 0 |
| ENSG00000117899 | 3376 | 1749 | 2050 | 1660 | 1840 | 1871 |

| | | | | | | |
|---|---|---|---|---|---|---|
| ENSG00000260619 | 16 | 7 | 20 | 9 | 12 | 1 |
| ENSG00000265855 | 1 | 0 | 1 | 0 | 0 | 0 |
| ENSG00000277782 | 94 | 49 | 56 | 43 | 63 | 49 |
| ENSG00000140406 | 1076 | 594 | 705 | 747 | 761 | 713 |
| ENSG00000156206 | 8 | 10 | 6 | 3 | 5 | 8 |
| ENSG00000279834 | 2 | 0 | 1 | 0 | 1 | 0 |
| ENSG00000259461 | 0 | 0 | 0 | 0 | 0 | 0 |
| ENSG00000172349 | 4 | 6 | 5 | 3 | 7 | 5 |
| ENSG00000271725 | 0 | 0 | 0 | 0 | 0 | 0 |
| ENSG00000172345 | 146 | 73 | 95 | 59 | 56 | 85 |
| ENSG00000259343 | 14 | 6 | 4 | 4 | 9 | 8 |
| ENSG00000188869 | 3 | 1 | 0 | 0 | 2 | 1 |
| ENSG00000273920 | 4 | 2 | 0 | 1 | 0 | 0 |
| ENSG00000284709 | 2 | 0 | 0 | 0 | 0 | 0 |
| ENSG00000275995 | 0 | 0 | 0 | 0 | 0 | 0 |
| ENSG00000259543 | 0 | 1 | 0 | 0 | 2 | 1 |
| ENSG00000259594 | 0 | 3 | 0 | 0 | 4 | 2 |
| ENSG00000259610 | 27 | 19 | 18 | 8 | 9 | 14 |
| ENSG00000259692 | 41 | 20 | 26 | 31 | 20 | 14 |
| ENSG00000259445 | 0 | 0 | 0 | 0 | 0 | 0 |
| ENSG00000259622 | 0 | 0 | 0 | 0 | 0 | 0 |
| ENSG00000183496 | 496 | 209 | 241 | 194 | 220 | 193 |
| ENSG00000259518 | 0 | 0 | 0 | 0 | 0 | 0 |
| ENSG00000140598 | 1539 | 749 | 907 | 608 | 983 | 802 |
| ENSG00000212170 | 0 | 1 | 0 | 0 | 0 | 0 |
| ENSG00000285974 | 0 | 0 | 0 | 0 | 0 | 0 |
| ENSG00000188659 | 59 | 26 | 17 | 21 | 15 | 13 |
| ENSG00000274376 | 0 | 0 | 0 | 0 | 0 | 0 |
| ENSG00000278662 | 48 | 21 | 40 | 13 | 39 | 43 |
| ENSG00000259429 | 97 | 59 | 91 | 32 | 59 | 64 |
| ENSG00000237550 | 3503 | 1757 | 2071 | 2380 | 1983 | 2071 |
| ENSG00000278202 | 0 | 5 | 2 | 1 | 4 | 1 |
| ENSG00000197978 | 193 | 83 | 166 | 81 | 133 | 140 |
| ENSG00000275695 | 0 | 0 | 0 | 0 | 0 | 0 |
| ENSG00000276710 | 103 | 43 | 68 | 39 | 57 | 37 |
| ENSG00000278603 | 0 | 0 | 0 | 0 | 0 | 0 |
| ENSG00000255769 | 199 | 91 | 175 | 134 | 161 | 122 |
| ENSG00000277741 | 6 | 0 | 1 | 0 | 0 | 0 |
| ENSG00000259637 | 0 | 0 | 0 | 0 | 0 | 0 |
| ENSG00000284803 | 0 | 0 | 0 | 0 | 0 | 0 |
| ENSG00000182774 | 526 | 356 | 442 | 425 | 496 | 325 |
| ENSG00000260836 | 0 | 0 | 0 | 0 | 0 | 0 |
| ENSG00000278013 | 0 | 0 | 0 | 0 | 0 | 0 |
| ENSG00000214575 | 12 | 3 | 8 | 4 | 2 | 1 |
| ENSG00000259462 | 0 | 0 | 0 | 1 | 0 | 0 |
| ENSG00000103723 | 118 | 60 | 66 | 80 | 84 | 66 |
| ENSG00000259707 | 1 | 0 | 0 | 1 | 1 | 0 |
| ENSG00000228141 | 0 | 0 | 0 | 0 | 0 | 0 |
| ENSG00000259315 | 6 | 5 | 8 | 4 | 7 | 7 |
| ENSG00000279758 | 3 | 3 | 3 | 1 | 0 | 2 |

| | | | | | | |
|---|---|---|---|---|---|---|
| ENSG00000259442 | 11 | 7 | 3 | 3 | 7 | 7 |
| ENSG00000250988 | 34 | 25 | 36 | 33 | 40 | 24 |
| ENSG00000252690 | 21 | 25 | 29 | 14 | 24 | 14 |
| ENSG00000186628 | 2 | 2 | 3 | 0 | 2 | 0 |
| ENSG00000277864 | 0 | 0 | 0 | 0 | 0 | 0 |
| ENSG00000156232 | 811 | 336 | 421 | 318 | 429 | 385 |
| ENSG00000103942 | 1694 | 773 | 954 | 710 | 856 | 789 |
| ENSG00000259805 | 0 | 0 | 0 | 0 | 3 | 0 |
| ENSG00000169612 | 472 | 355 | 414 | 371 | 436 | 402 |
| ENSG00000169609 | 1087 | 593 | 670 | 521 | 701 | 610 |
| ENSG00000260608 | 4 | 9 | 3 | 3 | 3 | 3 |
| ENSG00000064726 | 3139 | 1539 | 1838 | 1670 | 1854 | 1750 |
| ENSG00000273747 | 14 | 6 | 5 | 4 | 5 | 6 |
| ENSG00000259767 | 5 | 1 | 0 | 1 | 2 | 2 |
| ENSG00000260351 | 5 | 1 | 4 | 1 | 1 | 1 |
| ENSG00000259308 | 2 | 0 | 1 | 0 | 0 | 0 |
| ENSG00000263643 | 0 | 0 | 0 | 0 | 0 | 0 |
| ENSG00000136404 | 3 | 2 | 6 | 1 | 4 | 4 |
| ENSG00000166503 | 5462 | 2709 | 3302 | 2078 | 2807 | 2671 |
| ENSG00000260579 | 0 | 0 | 0 | 0 | 0 | 0 |
| ENSG00000259986 | 0 | 1 | 2 | 0 | 0 | 0 |
| ENSG00000169594 | 14 | 15 | 15 | 6 | 8 | 6 |
| ENSG00000140600 | 14 | 15 | 7 | 9 | 10 | 12 |
| ENSG00000156218 | 99 | 60 | 36 | 34 | 30 | 44 |
| ENSG00000212374 | 0 | 0 | 0 | 0 | 0 | 0 |
| ENSG00000261665 | 0 | 0 | 0 | 0 | 0 | 0 |
| ENSG00000200444 | 0 | 0 | 0 | 0 | 0 | 0 |
| ENSG00000274128 | 0 | 1 | 0 | 0 | 0 | 0 |
| ENSG00000259609 | 0 | 0 | 0 | 0 | 0 | 0 |
| ENSG00000259404 | 0 | 0 | 0 | 1 | 0 | 0 |
| ENSG00000280038 | 0 | 0 | 0 | 0 | 0 | 0 |
| ENSG00000259694 | 0 | 0 | 0 | 0 | 0 | 0 |
| ENSG00000259511 | 0 | 1 | 0 | 3 | 1 | 0 |
| ENSG00000259726 | 6 | 1 | 1 | 1 | 0 | 1 |
| ENSG00000225151 | 62 | 44 | 67 | 50 | 40 | 35 |
| ENSG00000278422 | 0 | 0 | 0 | 0 | 0 | 0 |
| ENSG00000184206 | 0 | 1 | 0 | 1 | 2 | 0 |
| ENSG00000259633 | 0 | 0 | 0 | 0 | 0 | 0 |
| ENSG00000259683 | 2 | 1 | 0 | 0 | 0 | 0 |
| ENSG00000244056 | 0 | 0 | 0 | 0 | 0 | 0 |
| ENSG00000259570 | 5 | 1 | 0 | 0 | 2 | 1 |
| ENSG00000235370 | 3 | 0 | 0 | 1 | 1 | 0 |
| ENSG00000272887 | 0 | 0 | 0 | 0 | 0 | 0 |
| ENSG00000259538 | 0 | 0 | 0 | 0 | 0 | 0 |
| ENSG00000259774 | 2 | 2 | 1 | 0 | 0 | 0 |
| ENSG00000278785 | 0 | 0 | 0 | 0 | 0 | 0 |
| ENSG00000197627 | 0 | 0 | 0 | 0 | 0 | 0 |
| ENSG00000259551 | 0 | 0 | 0 | 0 | 0 | 0 |
| ENSG00000259302 | 0 | 0 | 0 | 0 | 0 | 0 |
| ENSG00000230373 | 7 | 3 | 4 | 2 | 3 | 0 |

| | | | | | | |
|---|---|---|---|---|---|---|
| ENSG00000259244 | 1 | 3 | 1 | 1 | 1 | 3 |
| ENSG00000189136 | 33 | 15 | 12 | 6 | 27 | 11 |
| ENSG00000259728 | 11 | 11 | 7 | 8 | 12 | 10 |
| ENSG00000275120 | 0 | 2 | 5 | 4 | 2 | 4 |
| ENSG00000176371 | 380 | 211 | 282 | 262 | 295 | 236 |
| ENSG00000256278 | 6 | 2 | 3 | 0 | 2 | 5 |
| ENSG00000276278 | 0 | 0 | 0 | 0 | 0 | 0 |
| ENSG00000176700 | 68 | 32 | 55 | 36 | 36 | 50 |
| ENSG00000254779 | 0 | 0 | 0 | 0 | 0 | 0 |
| ENSG00000177082 | 504 | 258 | 342 | 201 | 295 | 285 |
| ENSG00000277578 | 0 | 0 | 0 | 0 | 0 | 0 |
| ENSG00000197696 | 279 | 180 | 186 | 212 | 187 | 156 |
| ENSG00000140612 | 8019 | 4142 | 4677 | 4972 | 4915 | 4773 |
| ENSG00000259654 | 0 | 0 | 0 | 0 | 1 | 0 |
| ENSG00000285667 | 43 | 14 | 31 | 12 | 18 | 18 |
| ENSG00000166716 | 2959 | 1201 | 1620 | 1326 | 1293 | 1494 |
| ENSG00000259415 | 0 | 2 | 0 | 0 | 0 | 0 |
| ENSG00000136383 | 135 | 55 | 78 | 65 | 80 | 65 |
| ENSG00000200991 | 0 | 0 | 0 | 1 | 1 | 0 |
| ENSG00000156222 | 10 | 7 | 5 | 10 | 18 | 10 |
| ENSG00000207037 | 2 | 1 | 1 | 0 | 1 | 1 |
| ENSG00000212388 | 0 | 0 | 0 | 1 | 0 | 0 |
| ENSG00000271044 | 0 | 0 | 0 | 0 | 0 | 0 |
| ENSG00000073417 | 3319 | 1445 | 1966 | 1170 | 1576 | 1617 |
| ENSG00000259270 | 0 | 0 | 0 | 0 | 0 | 0 |
| ENSG00000229212 | 12 | 22 | 23 | 13 | 12 | 20 |
| ENSG00000259449 | 0 | 0 | 0 | 0 | 1 | 0 |
| ENSG00000259295 | 57 | 26 | 26 | 16 | 26 | 26 |
| ENSG00000277582 | 0 | 0 | 0 | 0 | 0 | 0 |
| ENSG00000188388 | 0 | 0 | 0 | 0 | 0 | 1 |
| ENSG00000218052 | 0 | 0 | 0 | 0 | 0 | 0 |
| ENSG00000259317 | 0 | 0 | 0 | 0 | 0 | 0 |
| ENSG00000170776 | 2118 | 930 | 1131 | 613 | 1133 | 954 |
| ENSG00000284160 | 0 | 0 | 0 | 0 | 0 | 0 |
| ENSG00000251891 | 0 | 0 | 0 | 0 | 0 | 0 |
| ENSG00000259630 | 2 | 0 | 0 | 1 | 3 | 5 |
| ENSG00000259375 | 0 | 0 | 0 | 0 | 0 | 0 |
| ENSG00000259367 | 0 | 1 | 0 | 0 | 0 | 0 |
| ENSG00000259276 | 0 | 0 | 0 | 0 | 0 | 0 |
| ENSG00000202081 | 0 | 0 | 2 | 0 | 0 | 0 |
| ENSG00000259453 | 0 | 0 | 0 | 0 | 0 | 0 |
| ENSG00000259544 | 0 | 0 | 0 | 0 | 0 | 0 |
| ENSG00000259407 | 2 | 7 | 3 | 3 | 7 | 3 |
| ENSG00000259762 | 0 | 4 | 0 | 4 | 1 | 0 |
| ENSG00000259416 | 2 | 5 | 5 | 0 | 0 | 1 |
| ENSG00000183655 | 869 | 403 | 460 | 468 | 418 | 393 |
| ENSG00000221634 | 0 | 0 | 0 | 0 | 0 | 1 |
| ENSG00000259195 | 0 | 0 | 1 | 0 | 0 | 0 |
| ENSG00000252931 | 0 | 0 | 0 | 0 | 0 | 0 |
| ENSG00000264406 | 0 | 0 | 0 | 0 | 0 | 0 |

| | | | | | | |
|---|---|---|---|---|---|---|
| ENSG00000259608 | 0 | 0 | 0 | 0 | 0 | 0 |
| ENSG00000260477 | 0 | 0 | 0 | 0 | 0 | 0 |
| ENSG00000273540 | 2 | 1 | 0 | 0 | 0 | 2 |
| ENSG00000252964 | 0 | 0 | 0 | 0 | 0 | 0 |
| ENSG00000261077 | 0 | 0 | 0 | 0 | 0 | 0 |
| ENSG00000200206 | 0 | 0 | 0 | 0 | 0 | 0 |
| ENSG00000260125 | 0 | 0 | 0 | 0 | 0 | 0 |
| ENSG00000259734 | 0 | 0 | 0 | 0 | 0 | 0 |
| ENSG00000259636 | 1 | 0 | 0 | 0 | 0 | 0 |
| ENSG00000259620 | 6 | 3 | 2 | 3 | 0 | 3 |
| ENSG00000259761 | 7 | 7 | 17 | 4 | 3 | 6 |
| ENSG00000259560 | 2 | 1 | 1 | 0 | 0 | 2 |
| ENSG00000207150 | 0 | 0 | 0 | 0 | 0 | 0 |
| ENSG00000259527 | 1910 | 893 | 1408 | 632 | 767 | 1090 |
| ENSG00000140538 | 16 | 5 | 5 | 1 | 3 | 2 |
| ENSG00000259183 | 0 | 0 | 0 | 0 | 0 | 0 |
| ENSG00000260305 | 0 | 0 | 0 | 0 | 0 | 0 |
| ENSG00000259494 | 210 | 138 | 197 | 159 | 147 | 143 |
| ENSG00000173867 | 1 | 0 | 0 | 0 | 1 | 3 |
| ENSG00000181991 | 1329 | 650 | 800 | 902 | 855 | 753 |
| ENSG00000279945 | 0 | 0 | 0 | 0 | 0 | 0 |
| ENSG00000140543 | 53 | 23 | 45 | 12 | 31 | 24 |
| ENSG00000259569 | 0 | 0 | 0 | 0 | 0 | 0 |
| ENSG00000249487 | 0 | 0 | 0 | 0 | 0 | 0 |
| ENSG00000221630 | 0 | 0 | 0 | 0 | 0 | 0 |
| ENSG00000207703 | 0 | 0 | 0 | 0 | 0 | 0 |
| ENSG00000283484 | 0 | 0 | 0 | 0 | 0 | 0 |
| ENSG00000181026 | 1300 | 682 | 775 | 1001 | 932 | 884 |
| ENSG00000172183 | 358 | 182 | 282 | 311 | 249 | 265 |
| ENSG00000259676 | 0 | 0 | 0 | 0 | 0 | 0 |
| ENSG00000157766 | 2 | 0 | 2 | 5 | 7 | 4 |
| ENSG00000140511 | 57 | 23 | 25 | 32 | 34 | 17 |
| ENSG00000140545 | 560 | 261 | 362 | 306 | 292 | 296 |
| ENSG00000259350 | 0 | 0 | 0 | 0 | 0 | 0 |
| ENSG00000259489 | 0 | 0 | 0 | 0 | 0 | 0 |
| ENSG00000261549 | 0 | 0 | 0 | 0 | 0 | 0 |
| ENSG00000260275 | 0 | 0 | 0 | 0 | 0 | 0 |
| ENSG00000260123 | 0 | 0 | 0 | 0 | 0 | 0 |
| ENSG00000261407 | 0 | 0 | 2 | 0 | 0 | 0 |
| ENSG00000140526 | 3703 | 1673 | 2131 | 949 | 1577 | 1504 |
| ENSG00000239151 | 0 | 0 | 0 | 0 | 0 | 0 |
| ENSG00000259699 | 0 | 0 | 1 | 0 | 3 | 1 |
| ENSG00000259948 | 0 | 0 | 0 | 0 | 0 | 0 |
| ENSG00000140522 | 0 | 0 | 1 | 0 | 0 | 0 |
| ENSG00000140525 | 1470 | 939 | 1056 | 549 | 1216 | 926 |
| ENSG00000140521 | 2010 | 833 | 1206 | 937 | 994 | 1042 |
| ENSG00000275101 | 0 | 0 | 0 | 0 | 0 | 0 |
| ENSG00000261441 | 2 | 3 | 7 | 3 | 2 | 3 |
| ENSG00000279708 | 0 | 0 | 0 | 0 | 1 | 0 |
| ENSG00000255571 | 52 | 43 | 71 | 56 | 57 | 44 |

| | | | | | | |
|---|---|---|---|---|---|---|
| ENSG00000284329 | 0 | 0 | 0 | 0 | 0 | 0 |
| ENSG00000140519 | 36 | 11 | 13 | 29 | 20 | 29 |
| ENSG00000259218 | 0 | 0 | 0 | 0 | 2 | 0 |
| ENSG00000261478 | 0 | 1 | 0 | 0 | 0 | 0 |
| ENSG00000259713 | 0 | 1 | 0 | 0 | 0 | 0 |
| ENSG00000140534 | 390 | 175 | 249 | 201 | 250 | 245 |
| ENSG00000259615 | 0 | 0 | 0 | 0 | 0 | 0 |
| ENSG00000166813 | 51 | 40 | 37 | 46 | 40 | 42 |
| ENSG00000166819 | 38 | 10 | 17 | 8 | 16 | 8 |
| ENSG00000166821 | 1276 | 656 | 763 | 597 | 644 | 625 |
| ENSG00000241362 | 18 | 5 | 6 | 2 | 3 | 9 |
| ENSG00000140527 | 12 | 12 | 23 | 15 | 17 | 19 |
| ENSG00000259561 | 0 | 1 | 0 | 1 | 0 | 0 |
| ENSG00000206590 | 0 | 0 | 0 | 0 | 0 | 0 |
| ENSG00000166823 | 150 | 67 | 122 | 88 | 87 | 121 |
| ENSG00000259441 | 0 | 1 | 1 | 1 | 0 | 2 |
| ENSG00000188095 | 11 | 12 | 10 | 14 | 3 | 16 |
| ENSG00000166825 | 4 | 4 | 9 | 4 | 3 | 4 |
| ENSG00000157823 | 1216 | 564 | 733 | 599 | 662 | 730 |
| ENSG00000250021 | 0 | 0 | 0 | 0 | 0 | 0 |
| ENSG00000264966 | 0 | 0 | 0 | 0 | 1 | 0 |
| ENSG00000259677 | 3 | 6 | 4 | 0 | 4 | 3 |
| ENSG00000264796 | 0 | 0 | 0 | 0 | 0 | 0 |
| ENSG00000212420 | 0 | 0 | 0 | 0 | 0 | 0 |
| ENSG00000242498 | 526 | 328 | 365 | 162 | 173 | 210 |
| ENSG00000252645 | 0 | 0 | 0 | 0 | 0 | 0 |
| ENSG00000259273 | 0 | 0 | 0 | 0 | 0 | 0 |
| ENSG00000280377 | 0 | 0 | 0 | 0 | 0 | 0 |
| ENSG00000140548 | 803 | 412 | 532 | 337 | 352 | 369 |
| ENSG00000265871 | 0 | 1 | 0 | 0 | 0 | 0 |
| ENSG00000273691 | 0 | 1 | 0 | 2 | 0 | 0 |
| ENSG00000259291 | 301 | 146 | 195 | 123 | 137 | 165 |
| ENSG00000276376 | 0 | 0 | 0 | 0 | 0 | 0 |
| ENSG00000182054 | 12635 | 6142 | 7811 | 6555 | 5915 | 5861 |
| ENSG00000259685 | 2 | 0 | 0 | 1 | 3 | 2 |
| ENSG00000185033 | 3245 | 1503 | 2004 | 1564 | 1574 | 1587 |
| ENSG00000240470 | 0 | 0 | 0 | 0 | 0 | 0 |
| ENSG00000225193 | 7 | 4 | 3 | 1 | 2 | 2 |
| ENSG00000185043 | 4732 | 2436 | 3112 | 2862 | 2561 | 2522 |
| ENSG00000183208 | 103 | 64 | 66 | 67 | 71 | 82 |
| ENSG00000284626 | 0 | 0 | 0 | 0 | 0 | 1 |
| ENSG00000213471 | 3 | 5 | 0 | 1 | 0 | 2 |
| ENSG00000261147 | 0 | 0 | 1 | 0 | 0 | 0 |
| ENSG00000182768 | 1940 | 1114 | 1257 | 1235 | 1166 | 1362 |
| ENSG00000275674 | 0 | 1 | 0 | 0 | 0 | 1 |
| ENSG00000228998 | 0 | 0 | 0 | 0 | 0 | 0 |
| ENSG00000270322 | 0 | 0 | 0 | 0 | 0 | 0 |
| ENSG00000275345 | 1 | 0 | 0 | 0 | 1 | 0 |
| ENSG00000271396 | 5 | 2 | 4 | 2 | 4 | 5 |
| ENSG00000214433 | 3 | 1 | 1 | 1 | 0 | 1 |

| | | | | | | |
|---|---|---|---|---|---|---|
| ENSG00000275803 | 0 | 0 | 1 | 0 | 1 | 0 |
| ENSG00000279980 | 0 | 0 | 0 | 0 | 0 | 0 |
| ENSG00000238244 | 0 | 0 | 0 | 0 | 0 | 0 |
| ENSG00000196391 | 165 | 95 | 89 | 84 | 110 | 96 |
| ENSG00000259262 | 0 | 0 | 0 | 1 | 1 | 0 |
| ENSG00000140575 | 7075 | 4104 | 3267 | 1742 | 5623 | 3327 |
| ENSG00000259177 | 0 | 2 | 0 | 2 | 0 | 0 |
| ENSG00000140577 | 1092 | 534 | 664 | 551 | 588 | 607 |
| ENSG00000259212 | 4 | 1 | 1 | 1 | 1 | 0 |
| ENSG00000259314 | 1 | 0 | 4 | 1 | 0 | 0 |
| ENSG00000259736 | 6 | 12 | 17 | 22 | 30 | 19 |
| ENSG00000285560 | 0 | 1 | 0 | 3 | 5 | 3 |
| ENSG00000259746 | 0 | 2 | 1 | 1 | 2 | 0 |
| ENSG00000278370 | 0 | 0 | 0 | 0 | 0 | 1 |
| ENSG00000245479 | 8 | 6 | 5 | 1 | 4 | 3 |
| ENSG00000197299 | 214 | 136 | 114 | 79 | 189 | 148 |
| ENSG00000200677 | 0 | 0 | 0 | 0 | 0 | 3 |
| ENSG00000259704 | 0 | 0 | 0 | 0 | 0 | 0 |
| ENSG00000241869 | 0 | 0 | 0 | 0 | 0 | 0 |
| ENSG00000140564 | 2645 | 1247 | 1359 | 1294 | 1325 | 1060 |
| ENSG00000182511 | 44 | 35 | 32 | 24 | 26 | 25 |
| ENSG00000196547 | 3117 | 1300 | 1833 | 1294 | 1538 | 1621 |
| ENSG00000259661 | 0 | 0 | 0 | 0 | 0 | 0 |
| ENSG00000184508 | 549 | 258 | 378 | 394 | 282 | 346 |
| ENSG00000140553 | 6438 | 3016 | 3930 | 4130 | 3729 | 3581 |
| ENSG00000258384 | 10 | 7 | 16 | 10 | 11 | 11 |
| ENSG00000166965 | 1074 | 509 | 626 | 586 | 594 | 660 |
| ENSG00000198901 | 46 | 40 | 51 | 45 | 56 | 68 |
| ENSG00000284946 | 196 | 101 | 124 | 56 | 88 | 100 |
| ENSG00000258725 | 7 | 4 | 10 | 7 | 2 | 11 |
| ENSG00000200142 | 6 | 0 | 0 | 0 | 0 | 0 |
| ENSG00000184056 | 190 | 112 | 128 | 133 | 159 | 140 |
| ENSG00000214432 | 22 | 14 | 12 | 12 | 18 | 15 |
| ENSG00000278514 | 3 | 4 | 7 | 2 | 2 | 1 |
| ENSG00000258542 | 0 | 0 | 0 | 0 | 0 | 0 |
| ENSG00000275929 | 0 | 0 | 2 | 1 | 0 | 1 |
| ENSG00000258665 | 0 | 0 | 0 | 0 | 0 | 0 |
| ENSG00000285661 | 0 | 0 | 0 | 0 | 0 | 0 |
| ENSG00000185518 | 18 | 17 | 14 | 5 | 10 | 11 |
| ENSG00000285883 | 0 | 0 | 1 | 0 | 0 | 0 |
| ENSG00000258551 | 20 | 13 | 16 | 14 | 19 | 12 |
| ENSG00000258954 | 0 | 0 | 0 | 0 | 0 | 0 |
| ENSG00000258765 | 0 | 1 | 0 | 0 | 0 | 0 |
| ENSG00000258540 | 2 | 0 | 0 | 0 | 0 | 0 |
| ENSG00000259057 | 0 | 0 | 0 | 0 | 1 | 0 |
| ENSG00000176463 | 338 | 196 | 243 | 166 | 146 | 176 |
| ENSG00000260661 | 1 | 0 | 0 | 0 | 0 | 0 |
| ENSG00000258761 | 0 | 0 | 0 | 0 | 0 | 0 |
| ENSG00000258726 | 0 | 0 | 0 | 0 | 0 | 0 |
| ENSG00000258925 | 0 | 0 | 0 | 0 | 0 | 1 |

| | | | | | | |
|---|---|---|---|---|---|---|
| ENSG00000140557 | 0 | 0 | 0 | 0 | 0 | 0 |
| ENSG00000239197 | 0 | 0 | 0 | 0 | 0 | 0 |
| ENSG00000258739 | 0 | 0 | 0 | 0 | 0 | 0 |
| ENSG00000259170 | 0 | 0 | 0 | 0 | 0 | 0 |
| ENSG00000183643 | 0 | 0 | 0 | 0 | 0 | 0 |
| ENSG00000258647 | 23 | 8 | 20 | 26 | 32 | 46 |
| ENSG00000258676 | 39 | 13 | 15 | 8 | 23 | 35 |
| ENSG00000260337 | 70 | 29 | 61 | 33 | 75 | 107 |
| ENSG00000271763 | 7 | 4 | 1 | 4 | 3 | 6 |
| ENSG00000185442 | 4548 | 2261 | 3039 | 3362 | 3144 | 3827 |
| ENSG00000259064 | 3 | 2 | 1 | 0 | 4 | 4 |
| ENSG00000275965 | 2 | 2 | 3 | 1 | 2 | 2 |
| ENSG00000272298 | 0 | 0 | 0 | 0 | 0 | 0 |
| ENSG00000253954 | 116 | 50 | 47 | 61 | 81 | 47 |
| ENSG00000258741 | 8 | 0 | 4 | 4 | 4 | 2 |
| ENSG00000240484 | 0 | 0 | 0 | 0 | 0 | 0 |
| ENSG00000258562 | 0 | 0 | 0 | 0 | 0 | 0 |
| ENSG00000260361 | 14 | 12 | 7 | 10 | 11 | 17 |
| ENSG00000258527 | 2 | 2 | 2 | 1 | 0 | 0 |
| ENSG00000258922 | 3 | 0 | 3 | 0 | 0 | 0 |
| ENSG00000274215 | 0 | 1 | 2 | 4 | 2 | 3 |
| ENSG00000275175 | 0 | 0 | 1 | 1 | 0 | 0 |
| ENSG00000272888 | 1493 | 820 | 1053 | 799 | 732 | 977 |
| ENSG00000279765 | 4 | 2 | 5 | 5 | 3 | 5 |
| ENSG00000173575 | 3151 | 1601 | 1874 | 886 | 1915 | 1496 |
| ENSG00000284324 | 0 | 0 | 0 | 0 | 0 | 0 |
| ENSG00000182175 | 4 | 2 | 4 | 2 | 4 | 6 |
| ENSG00000258611 | 0 | 0 | 0 | 0 | 0 | 0 |
| ENSG00000285663 | 0 | 0 | 0 | 0 | 0 | 0 |
| ENSG00000280076 | 0 | 0 | 0 | 0 | 0 | 0 |
| ENSG00000258888 | 0 | 0 | 0 | 0 | 0 | 0 |
| ENSG00000258971 | 0 | 0 | 0 | 0 | 0 | 0 |
| ENSG00000257060 | 0 | 0 | 0 | 0 | 0 | 0 |
| ENSG00000258909 | 0 | 0 | 0 | 0 | 0 | 0 |
| ENSG00000258787 | 0 | 0 | 0 | 0 | 0 | 0 |
| ENSG00000258631 | 0 | 0 | 0 | 0 | 0 | 0 |
| ENSG00000258754 | 0 | 0 | 0 | 0 | 0 | 0 |
| ENSG00000258627 | 0 | 0 | 0 | 0 | 0 | 0 |
| ENSG00000259123 | 0 | 0 | 0 | 0 | 0 | 0 |
| ENSG00000258476 | 0 | 0 | 0 | 0 | 0 | 0 |
| ENSG00000258831 | 0 | 0 | 0 | 0 | 0 | 0 |
| ENSG00000258785 | 0 | 0 | 0 | 0 | 0 | 0 |
| ENSG00000259724 | 0 | 0 | 0 | 0 | 0 | 0 |
| ENSG00000258895 | 0 | 0 | 1 | 1 | 0 | 0 |
| ENSG00000258570 | 0 | 0 | 0 | 0 | 0 | 0 |
| ENSG00000140563 | 2 | 1 | 0 | 0 | 0 | 0 |
| ENSG00000258632 | 0 | 0 | 0 | 0 | 0 | 0 |
| ENSG00000285866 | 0 | 0 | 0 | 0 | 0 | 0 |
| ENSG00000259331 | 0 | 0 | 0 | 0 | 0 | 0 |
| ENSG00000277654 | 9 | 4 | 4 | 1 | 1 | 2 |

| | | | | | | |
|---|---|---|---|---|---|---|
| ENSG00000270017 | 0 | 0 | 0 | 0 | 0 | 0 |
| ENSG00000258433 | 0 | 0 | 0 | 0 | 0 | 0 |
| ENSG00000258773 | 0 | 0 | 0 | 0 | 0 | 0 |
| ENSG00000248441 | 0 | 1 | 0 | 0 | 0 | 0 |
| ENSG00000273923 | 0 | 0 | 0 | 0 | 0 | 0 |
| ENSG00000280304 | 0 | 0 | 0 | 0 | 0 | 0 |
| ENSG00000285700 | 0 | 0 | 0 | 0 | 0 | 0 |
| ENSG00000259134 | 0 | 0 | 0 | 0 | 0 | 0 |
| ENSG00000258489 | 0 | 0 | 0 | 0 | 0 | 0 |
| ENSG00000275016 | 0 | 0 | 0 | 0 | 0 | 0 |
| ENSG00000222076 | 0 | 0 | 0 | 0 | 0 | 0 |
| ENSG00000275443 | 2 | 0 | 0 | 1 | 0 | 0 |
| ENSG00000259702 | 9 | 2 | 1 | 1 | 3 | 0 |
| ENSG00000273771 | 2 | 0 | 1 | 0 | 1 | 1 |
| ENSG00000247809 | 179 | 96 | 116 | 49 | 61 | 68 |
| ENSG00000280048 | 0 | 0 | 0 | 0 | 0 | 0 |
| ENSG00000277135 | 0 | 0 | 0 | 0 | 1 | 0 |
| ENSG00000259359 | 0 | 0 | 0 | 0 | 0 | 0 |
| ENSG00000278146 | 1 | 1 | 0 | 0 | 0 | 0 |
| ENSG00000259763 | 7 | 3 | 1 | 0 | 0 | 2 |
| ENSG00000279645 | 6 | 0 | 5 | 0 | 0 | 3 |
| ENSG00000280291 | 0 | 1 | 0 | 0 | 0 | 0 |
| ENSG00000185551 | 15522 | 6915 | 8619 | 5490 | 6803 | 7457 |
| ENSG00000283888 | 0 | 0 | 0 | 0 | 0 | 0 |
| ENSG00000275322 | 91 | 43 | 71 | 38 | 31 | 51 |
| ENSG00000259275 | 290 | 115 | 204 | 70 | 85 | 145 |
| ENSG00000259614 | 10 | 2 | 6 | 3 | 1 | 1 |
| ENSG00000259542 | 9 | 9 | 6 | 2 | 6 | 3 |
| ENSG00000259312 | 0 | 0 | 0 | 0 | 0 | 0 |
| ENSG00000250825 | 0 | 0 | 1 | 0 | 0 | 0 |
| ENSG00000242295 | 0 | 0 | 0 | 0 | 0 | 0 |
| ENSG00000259534 | 2 | 0 | 0 | 0 | 0 | 0 |
| ENSG00000222617 | 0 | 0 | 0 | 0 | 0 | 0 |
| ENSG00000274079 | 0 | 0 | 0 | 0 | 0 | 0 |
| ENSG00000259340 | 0 | 0 | 0 | 0 | 0 | 0 |
| ENSG00000214424 | 0 | 0 | 0 | 0 | 0 | 0 |
| ENSG00000259282 | 0 | 0 | 0 | 0 | 0 | 0 |
| ENSG00000185594 | 0 | 0 | 0 | 0 | 0 | 0 |
| ENSG00000223120 | 0 | 0 | 0 | 0 | 0 | 0 |
| ENSG00000259403 | 0 | 0 | 0 | 0 | 0 | 0 |
| ENSG00000259485 | 64 | 36 | 42 | 31 | 31 | 37 |
| ENSG00000242863 | 0 | 0 | 0 | 0 | 0 | 0 |
| ENSG00000259664 | 0 | 3 | 0 | 1 | 0 | 0 |
| ENSG00000251729 | 0 | 0 | 0 | 0 | 0 | 0 |
| ENSG00000279091 | 2 | 0 | 0 | 0 | 0 | 0 |
| ENSG00000259582 | 0 | 0 | 0 | 0 | 0 | 0 |
| ENSG00000251209 | 0 | 0 | 0 | 0 | 0 | 0 |
| ENSG00000259870 | 0 | 0 | 0 | 0 | 0 | 0 |
| ENSG00000140450 | 921 | 496 | 550 | 470 | 635 | 509 |
| ENSG00000259478 | 0 | 0 | 0 | 0 | 0 | 0 |

| | | | | | | |
|---|---|---|---|---|---|---|
| ENSG00000258483 | 2 | 0 | 0 | 0 | 0 | 0 |
| ENSG00000283372 | 0 | 0 | 0 | 0 | 0 | 0 |
| ENSG00000259199 | 0 | 0 | 0 | 0 | 0 | 0 |
| ENSG00000222361 | 0 | 0 | 0 | 0 | 0 | 0 |
| ENSG00000259611 | 0 | 0 | 0 | 0 | 0 | 0 |
| ENSG00000259266 | 0 | 0 | 0 | 0 | 0 | 0 |
| ENSG00000259320 | 0 | 0 | 0 | 0 | 0 | 0 |
| ENSG00000259041 | 0 | 0 | 0 | 0 | 0 | 0 |
| ENSG00000274312 | 0 | 0 | 0 | 0 | 0 | 0 |
| ENSG00000277987 | 0 | 0 | 0 | 0 | 0 | 0 |
| ENSG00000283597 | 0 | 0 | 0 | 0 | 0 | 0 |
| ENSG00000259507 | 0 | 0 | 0 | 0 | 0 | 0 |
| ENSG00000259424 | 101 | 62 | 88 | 140 | 162 | 171 |
| ENSG00000140443 | 12058 | 5202 | 8152 | 7461 | 9042 | 10534 |
| ENSG00000278022 | 8 | 3 | 4 | 6 | 9 | 11 |
| ENSG00000264480 | 0 | 0 | 0 | 0 | 0 | 0 |
| ENSG00000259621 | 0 | 0 | 0 | 0 | 0 | 1 |
| ENSG00000259475 | 8 | 7 | 7 | 2 | 9 | 15 |
| ENSG00000183571 | 0 | 2 | 0 | 2 | 3 | 1 |
| ENSG00000278090 | 0 | 0 | 0 | 1 | 0 | 0 |
| ENSG00000182253 | 211 | 94 | 131 | 102 | 137 | 110 |
| ENSG00000261054 | 2 | 1 | 2 | 0 | 2 | 3 |
| ENSG00000103852 | 470 | 223 | 293 | 238 | 275 | 279 |
| ENSG00000261616 | 14 | 5 | 8 | 2 | 9 | 7 |
| ENSG00000259921 | 2 | 2 | 2 | 0 | 0 | 0 |
| ENSG00000168904 | 743 | 366 | 420 | 333 | 439 | 402 |
| ENSG00000259706 | 0 | 4 | 2 | 3 | 0 | 4 |
| ENSG00000259228 | 0 | 0 | 0 | 0 | 0 | 0 |
| ENSG00000259257 | 3 | 2 | 2 | 1 | 1 | 0 |
| ENSG00000259590 | 2 | 0 | 0 | 0 | 0 | 1 |
| ENSG00000259760 | 0 | 0 | 1 | 0 | 0 | 0 |
| ENSG00000274294 | 6 | 1 | 0 | 1 | 3 | 0 |
| ENSG00000259341 | 0 | 0 | 0 | 0 | 0 | 0 |
| ENSG00000279846 | 0 | 0 | 0 | 0 | 0 | 0 |
| ENSG00000241562 | 0 | 0 | 0 | 0 | 0 | 0 |
| ENSG00000068305 | 1273 | 602 | 707 | 371 | 669 | 552 |
| ENSG00000201724 | 0 | 0 | 0 | 0 | 0 | 1 |
| ENSG00000183060 | 374 | 179 | 247 | 240 | 267 | 225 |
| ENSG00000182397 | 0 | 0 | 0 | 0 | 0 | 0 |
| ENSG00000259655 | 0 | 0 | 0 | 0 | 0 | 0 |
| ENSG00000241588 | 0 | 0 | 0 | 0 | 0 | 0 |
| ENSG00000259363 | 2 | 2 | 5 | 1 | 4 | 3 |
| ENSG00000254744 | 0 | 0 | 0 | 0 | 0 | 2 |
| ENSG00000140470 | 171 | 75 | 100 | 71 | 97 | 105 |
| ENSG00000259219 | 2 | 2 | 2 | 0 | 1 | 4 |
| ENSG00000252957 | 0 | 0 | 0 | 0 | 0 | 0 |
| ENSG00000259356 | 0 | 0 | 0 | 0 | 0 | 0 |
| ENSG00000189419 | 30 | 11 | 12 | 13 | 24 | 19 |
| ENSG00000259430 | 0 | 0 | 0 | 0 | 0 | 0 |
| ENSG00000154227 | 0 | 0 | 0 | 0 | 0 | 0 |

| | | | | | | |
|---|---|---|---|---|---|---|
| ENSG00000270127 | 30 | 16 | 17 | 11 | 20 | 16 |
| ENSG00000251819 | 3 | 3 | 8 | 0 | 2 | 3 |
| ENSG00000259205 | 10 | 4 | 10 | 6 | 16 | 8 |
| ENSG00000259540 | 2 | 2 | 0 | 2 | 0 | 1 |
| ENSG00000140471 | 793 | 343 | 398 | 312 | 418 | 381 |
| ENSG00000200095 | 0 | 0 | 1 | 1 | 2 | 3 |
| ENSG00000183475 | 1240 | 591 | 691 | 564 | 653 | 711 |
| ENSG00000212306 | 0 | 0 | 0 | 0 | 0 | 0 |
| ENSG00000259381 | 7 | 5 | 7 | 0 | 1 | 3 |
| ENSG00000259579 | 0 | 1 | 0 | 2 | 3 | 2 |
| ENSG00000279588 | 2 | 1 | 0 | 2 | 7 | 3 |
| ENSG00000259586 | 0 | 0 | 0 | 0 | 0 | 0 |
| ENSG00000232386 | 71 | 45 | 49 | 101 | 137 | 110 |
| ENSG00000272808 | 161 | 91 | 66 | 331 | 461 | 369 |
| ENSG00000259604 | 0 | 0 | 0 | 0 | 0 | 0 |
| ENSG00000272639 | 0 | 0 | 0 | 0 | 0 | 0 |
| ENSG00000184254 | 157 | 145 | 97 | 448 | 684 | 592 |
| ENSG00000278456 | 0 | 0 | 0 | 2 | 3 | 1 |
| ENSG00000259583 | 12 | 6 | 10 | 23 | 40 | 35 |
| ENSG00000154237 | 119 | 61 | 70 | 105 | 132 | 109 |
| ENSG00000259376 | 2 | 0 | 2 | 2 | 0 | 0 |
| ENSG00000259755 | 0 | 0 | 0 | 0 | 0 | 2 |
| ENSG00000259365 | 0 | 0 | 0 | 0 | 0 | 0 |
| ENSG00000259182 | 5 | 1 | 0 | 5 | 0 | 0 |
| ENSG00000131873 | 1617 | 849 | 1084 | 787 | 933 | 1174 |
| ENSG00000131871 | 2419 | 1085 | 1433 | 1300 | 1372 | 1445 |
| ENSG00000131876 | 3303 | 1714 | 1955 | 1932 | 2422 | 2120 |
| ENSG00000259172 | 50 | 34 | 46 | 19 | 19 | 20 |
| ENSG00000140479 | 910 | 461 | 520 | 292 | 352 | 347 |
| ENSG00000279970 | 60 | 33 | 30 | 20 | 24 | 19 |
| ENSG00000259764 | 0 | 1 | 0 | 0 | 0 | 0 |
| ENSG00000274139 | 0 | 0 | 0 | 1 | 1 | 1 |
| ENSG00000277210 | 0 | 0 | 0 | 0 | 0 | 0 |
| ENSG00000282793 | 0 | 0 | 0 | 0 | 0 | 0 |
| ENSG00000259464 | 0 | 0 | 0 | 0 | 0 | 0 |
| ENSG00000259695 | 0 | 0 | 0 | 0 | 0 | 0 |
| ENSG00000273585 | 0 | 0 | 0 | 0 | 0 | 0 |
| ENSG00000184277 | 1440 | 624 | 780 | 694 | 856 | 825 |
| ENSG00000252614 | 4 | 2 | 0 | 0 | 1 | 3 |
| ENSG00000185418 | 422 | 219 | 246 | 211 | 235 | 251 |
| ENSG00000259658 | 2 | 0 | 2 | 0 | 2 | 1 |
| ENSG00000259660 | 0 | 0 | 0 | 0 | 0 | 0 |
| ENSG00000259236 | 0 | 0 | 0 | 0 | 0 | 0 |
| ENSG00000240927 | 0 | 0 | 0 | 0 | 0 | 0 |
| ENSG00000184140 | 0 | 0 | 0 | 0 | 0 | 0 |
| ENSG00000182854 | 0 | 0 | 1 | 0 | 0 | 0 |
| ENSG00000232459 | 0 | 0 | 0 | 0 | 0 | 0 |
| ENSG00000214344 | 0 | 0 | 0 | 0 | 0 | 0 |
| ENSG00000257109 | 0 | 0 | 0 | 0 | 0 | 0 |
| ENSG00000259454 | 0 | 0 | 0 | 0 | 0 | 0 |

| | | | | | | |
|---|---|---|---|---|---|---|
| ENSG00000259646 | 0 | 0 | 0 | 0 | 0 | 0 |
| ENSG00000177693 | 0 | 0 | 0 | 0 | 0 | 0 |
| ENSG00000183909 | 0 | 0 | 0 | 0 | 0 | 0 |
| ENSG00000275771 | 0 | 0 | 0 | 0 | 0 | 0 |
| ENSG00000248893 | 0 | 0 | 0 | 0 | 0 | 0 |
| ENSG00000259553 | 0 | 0 | 0 | 0 | 0 | 0 |
| ENSG00000284557 | 0 | 0 | 0 | 0 | 0 | 0 |
| ENSG00000185596 | 259 | 139 | 202 | 168 | 147 | 162 |
| ENSG00000275449 | 0 | 0 | 0 | 0 | 0 | 0 |
| ENSG00000248472 | 4 | 0 | 1 | 3 | 4 | 3 |
| ENSG00000233614 | 24 | 10 | 22 | 18 | 19 | 15 |
| ENSG00000234769 | 8 | 8 | 12 | 11 | 9 | 8 |
| ENSG00000278739 | 0 | 0 | 1 | 0 | 0 | 0 |
| ENSG00000260803 | 5 | 3 | 0 | 0 | 4 | 0 |
| ENSG00000231439 | 82 | 35 | 52 | 33 | 34 | 58 |
| ENSG00000226942 | 33 | 15 | 26 | 14 | 7 | 17 |
| ENSG00000161980 | 813 | 445 | 558 | 593 | 538 | 556 |
| ENSG00000161981 | 1415 | 805 | 892 | 1047 | 804 | 866 |
| ENSG00000007384 | 3673 | 1698 | 2225 | 1681 | 1583 | 1700 |
| ENSG00000278995 | 25 | 12 | 23 | 19 | 9 | 23 |
| ENSG00000103152 | 1583 | 845 | 1051 | 1201 | 1000 | 961 |
| ENSG00000103148 | 1162 | 527 | 699 | 723 | 577 | 705 |
| ENSG00000269482 | 0 | 0 | 0 | 1 | 0 | 1 |
| ENSG00000228779 | 2 | 3 | 1 | 0 | 1 | 0 |
| ENSG00000130656 | 0 | 0 | 0 | 2 | 0 | 1 |
| ENSG00000206177 | 0 | 0 | 0 | 0 | 0 | 0 |
| ENSG00000206178 | 0 | 0 | 2 | 0 | 1 | 0 |
| ENSG00000225323 | 0 | 0 | 0 | 0 | 0 | 0 |
| ENSG00000188536 | 1 | 0 | 0 | 0 | 0 | 0 |
| ENSG00000206172 | 3 | 2 | 0 | 2 | 0 | 2 |
| ENSG00000207243 | 0 | 0 | 0 | 0 | 0 | 0 |
| ENSG00000086506 | 2 | 2 | 4 | 2 | 3 | 0 |
| ENSG00000268836 | 6 | 0 | 0 | 1 | 0 | 1 |
| ENSG00000007392 | 2040 | 967 | 1282 | 957 | 1036 | 1176 |
| ENSG00000206168 | 2 | 4 | 15 | 7 | 1 | 5 |
| ENSG00000167930 | 1490 | 777 | 949 | 1094 | 868 | 840 |
| ENSG00000269881 | 15 | 2 | 5 | 3 | 3 | 4 |
| ENSG00000076344 | 109 | 66 | 83 | 48 | 28 | 31 |
| ENSG00000242173 | 148 | 70 | 103 | 106 | 88 | 92 |
| ENSG00000185615 | 84 | 29 | 27 | 33 | 35 | 46 |
| ENSG00000103126 | 1497 | 666 | 838 | 904 | 757 | 852 |
| ENSG00000086504 | 4177 | 1898 | 2359 | 2549 | 2134 | 2011 |
| ENSG00000129925 | 3079 | 1439 | 1940 | 1844 | 1689 | 1603 |
| ENSG00000236829 | 89 | 36 | 51 | 33 | 60 | 46 |
| ENSG00000103202 | 3088 | 1422 | 1805 | 1834 | 1492 | 1723 |
| ENSG00000242612 | 510 | 285 | 335 | 361 | 301 | 303 |
| ENSG00000276984 | 2 | 0 | 2 | 5 | 2 | 1 |
| ENSG00000090565 | 2001 | 949 | 1354 | 1041 | 1046 | 1124 |
| ENSG00000201034 | 0 | 0 | 0 | 0 | 0 | 0 |
| ENSG00000230428 | 0 | 1 | 0 | 0 | 0 | 0 |

| | | | | | | |
|---|---|---|---|---|---|---|
| ENSG00000277142 | 77 | 42 | 60 | 47 | 53 | 46 |
| ENSG00000103326 | 1431 | 727 | 999 | 976 | 819 | 893 |
| ENSG00000266124 | 3 | 2 | 4 | 3 | 2 | 2 |
| ENSG00000266235 | 3 | 2 | 3 | 0 | 1 | 2 |
| ENSG00000261691 | 0 | 0 | 0 | 0 | 0 | 0 |
| ENSG00000161992 | 0 | 0 | 0 | 0 | 0 | 0 |
| ENSG00000007541 | 1274 | 613 | 713 | 771 | 636 | 616 |
| ENSG00000257108 | 0 | 0 | 3 | 4 | 0 | 2 |
| ENSG00000282907 | 0 | 0 | 0 | 0 | 0 | 0 |
| ENSG00000197562 | 1277 | 561 | 749 | 739 | 664 | 636 |
| ENSG00000127578 | 46 | 20 | 15 | 35 | 34 | 25 |
| ENSG00000130731 | 1190 | 567 | 738 | 959 | 683 | 728 |
| ENSG00000172366 | 1056 | 490 | 586 | 746 | 678 | 623 |
| ENSG00000228201 | 1 | 0 | 0 | 0 | 0 | 0 |
| ENSG00000161996 | 564 | 247 | 360 | 262 | 310 | 330 |
| ENSG00000262528 | 0 | 0 | 0 | 0 | 0 | 0 |
| ENSG00000140983 | 3212 | 1335 | 2030 | 1807 | 1656 | 1673 |
| ENSG00000103269 | 154 | 72 | 57 | 74 | 73 | 60 |
| ENSG00000260394 | 3 | 2 | 0 | 2 | 2 | 2 |
| ENSG00000279441 | 0 | 0 | 0 | 0 | 0 | 0 |
| ENSG00000103266 | 1399 | 840 | 1011 | 1160 | 975 | 980 |
| ENSG00000161999 | 552 | 277 | 367 | 375 | 301 | 351 |
| ENSG00000127580 | 366 | 150 | 214 | 243 | 193 | 208 |
| ENSG00000261659 | 2 | 18 | 9 | 11 | 7 | 22 |
| ENSG00000127585 | 1222 | 557 | 717 | 661 | 659 | 636 |
| ENSG00000279255 | 12 | 2 | 10 | 11 | 2 | 6 |
| ENSG00000259840 | 6 | 2 | 5 | 7 | 5 | 5 |
| ENSG00000103260 | 3131 | 1391 | 1762 | 1970 | 1487 | 1552 |
| ENSG00000103254 | 631 | 261 | 382 | 426 | 290 | 353 |
| ENSG00000162004 | 510 | 189 | 347 | 312 | 239 | 294 |
| ENSG00000103253 | 1260 | 626 | 753 | 808 | 612 | 722 |
| ENSG00000103245 | 1021 | 530 | 662 | 623 | 554 | 523 |
| ENSG00000102854 | 0 | 2 | 2 | 0 | 0 | 0 |
| ENSG00000279136 | 0 | 0 | 0 | 0 | 0 | 0 |
| ENSG00000162006 | 0 | 0 | 0 | 2 | 2 | 0 |
| ENSG00000207579 | 0 | 0 | 0 | 0 | 0 | 0 |
| ENSG00000007376 | 1295 | 583 | 766 | 995 | 770 | 801 |
| ENSG00000127586 | 569 | 289 | 337 | 313 | 329 | 305 |
| ENSG00000127588 | 14 | 10 | 9 | 5 | 5 | 7 |
| ENSG00000275438 | 4 | 2 | 7 | 5 | 4 | 7 |
| ENSG00000103227 | 458 | 251 | 297 | 257 | 240 | 273 |
| ENSG00000260022 | 2 | 7 | 7 | 5 | 1 | 4 |
| ENSG00000260316 | 2 | 0 | 3 | 0 | 0 | 1 |
| ENSG00000260439 | 0 | 0 | 4 | 1 | 1 | 1 |
| ENSG00000276931 | 2 | 6 | 2 | 2 | 1 | 5 |
| ENSG00000260807 | 16 | 7 | 8 | 17 | 14 | 9 |
| ENSG00000005513 | 12 | 8 | 8 | 6 | 5 | 9 |
| ENSG00000260496 | 0 | 0 | 0 | 0 | 0 | 0 |
| ENSG00000261713 | 17 | 14 | 16 | 7 | 8 | 2 |
| ENSG00000261720 | 0 | 0 | 0 | 0 | 0 | 0 |

| | | | | | | |
|---|---|---|---|---|---|---|
| ENSG00000162009 | 20 | 6 | 9 | 9 | 8 | 9 |
| ENSG00000184471 | 0 | 0 | 0 | 0 | 0 | 1 |
| ENSG00000260702 | 0 | 0 | 0 | 0 | 0 | 0 |
| ENSG00000260532 | 4 | 0 | 0 | 0 | 0 | 2 |
| ENSG00000273551 | 0 | 2 | 0 | 2 | 1 | 0 |
| ENSG00000196557 | 2931 | 1308 | 1752 | 1569 | 1629 | 1526 |
| ENSG00000260403 | 0 | 0 | 2 | 0 | 0 | 0 |
| ENSG00000259910 | 2 | 0 | 0 | 0 | 0 | 0 |
| ENSG00000261294 | 0 | 0 | 0 | 0 | 0 | 0 |
| ENSG00000116176 | 15 | 1 | 8 | 9 | 10 | 10 |
| ENSG00000277010 | 6 | 0 | 3 | 3 | 4 | 3 |
| ENSG00000197253 | 0 | 1 | 1 | 0 | 1 | 0 |
| ENSG00000172236 | 0 | 0 | 0 | 0 | 0 | 2 |
| ENSG00000095917 | 0 | 0 | 0 | 0 | 0 | 0 |
| ENSG00000260182 | 0 | 0 | 0 | 0 | 0 | 0 |
| ENSG00000196364 | 7 | 9 | 9 | 9 | 8 | 5 |
| ENSG00000261564 | 0 | 0 | 0 | 0 | 0 | 0 |
| ENSG00000260991 | 0 | 0 | 0 | 0 | 0 | 0 |
| ENSG00000260284 | 12 | 2 | 24 | 1 | 3 | 6 |
| ENSG00000260710 | 9 | 14 | 14 | 5 | 3 | 6 |
| ENSG00000274751 | 0 | 1 | 6 | 2 | 1 | 0 |
| ENSG00000103275 | 3124 | 1739 | 2244 | 2025 | 1941 | 1986 |
| ENSG00000261505 | 136 | 68 | 82 | 52 | 67 | 76 |
| ENSG00000219274 | 2 | 0 | 2 | 0 | 0 | 0 |
| ENSG00000007516 | 632 | 271 | 396 | 274 | 227 | 262 |
| ENSG00000007520 | 1117 | 631 | 758 | 858 | 705 | 641 |
| ENSG00000090581 | 920 | 406 | 623 | 513 | 488 | 533 |
| ENSG00000260425 | 7 | 4 | 8 | 2 | 3 | 6 |
| ENSG00000059145 | 709 | 375 | 435 | 390 | 472 | 455 |
| ENSG00000260132 | 0 | 0 | 2 | 0 | 2 | 0 |
| ENSG00000174109 | 567 | 305 | 347 | 339 | 319 | 337 |
| ENSG00000284395 | 2 | 1 | 2 | 0 | 2 | 0 |
| ENSG00000197599 | 29 | 6 | 19 | 12 | 2 | 4 |
| ENSG00000279930 | 4 | 9 | 14 | 3 | 0 | 3 |
| ENSG00000103249 | 1325 | 580 | 863 | 715 | 610 | 693 |
| ENSG00000261641 | 0 | 0 | 0 | 0 | 0 | 0 |
| ENSG00000260051 | 7 | 0 | 5 | 3 | 8 | 4 |
| ENSG00000261430 | 4 | 0 | 0 | 1 | 2 | 0 |
| ENSG00000219027 | 0 | 2 | 1 | 1 | 0 | 0 |
| ENSG00000251692 | 10 | 4 | 6 | 3 | 1 | 3 |
| ENSG00000100726 | 1902 | 935 | 1156 | 1143 | 930 | 941 |
| ENSG00000187535 | 1342 | 571 | 836 | 576 | 481 | 554 |
| ENSG00000260646 | 0 | 7 | 1 | 1 | 1 | 0 |
| ENSG00000131634 | 7 | 4 | 3 | 1 | 1 | 5 |
| ENSG00000280062 | 4 | 1 | 1 | 0 | 0 | 2 |
| ENSG00000280231 | 1 | 2 | 0 | 1 | 0 | 1 |
| ENSG00000280327 | 27 | 10 | 15 | 8 | 17 | 15 |
| ENSG00000260954 | 12 | 4 | 6 | 1 | 3 | 2 |
| ENSG00000260989 | 1 | 0 | 3 | 1 | 0 | 7 |
| ENSG00000007545 | 656 | 325 | 455 | 327 | 338 | 395 |

| | | | | | | |
|---|---|---|---|---|---|---|
| ENSG00000226890 | 0 | 0 | 0 | 0 | 2 | 0 |
| ENSG00000261732 | 8 | 7 | 7 | 1 | 6 | 4 |
| ENSG00000206053 | 8145 | 3905 | 4958 | 3581 | 3944 | 4201 |
| ENSG00000278987 | 15 | 8 | 11 | 0 | 11 | 8 |
| ENSG00000138834 | 2150 | 931 | 1367 | 828 | 1069 | 929 |
| ENSG00000275092 | 0 | 1 | 1 | 0 | 1 | 1 |
| ENSG00000261399 | 0 | 0 | 0 | 0 | 0 | 0 |
| ENSG00000265820 | 0 | 0 | 0 | 0 | 0 | 0 |
| ENSG00000261207 | 4 | 4 | 2 | 3 | 2 | 2 |
| ENSG00000103024 | 1401 | 658 | 932 | 845 | 715 | 747 |
| ENSG00000074071 | 3637 | 1791 | 2173 | 2664 | 2162 | 2208 |
| ENSG00000197774 | 1048 | 440 | 623 | 534 | 544 | 573 |
| ENSG00000162032 | 12 | 1 | 9 | 10 | 8 | 9 |
| ENSG00000095906 | 1477 | 780 | 898 | 1116 | 865 | 776 |
| ENSG00000099769 | 43 | 8 | 21 | 10 | 8 | 19 |
| ENSG00000063854 | 2049 | 989 | 1265 | 1228 | 1036 | 941 |
| ENSG00000180185 | 1161 | 614 | 699 | 598 | 672 | 613 |
| ENSG00000162039 | 0 | 3 | 5 | 6 | 1 | 4 |
| ENSG00000260541 | 8 | 2 | 3 | 8 | 5 | 2 |
| ENSG00000281219 | 0 | 0 | 0 | 0 | 0 | 0 |
| ENSG00000259947 | 0 | 0 | 0 | 0 | 0 | 0 |
| ENSG00000162040 | 0 | 0 | 5 | 1 | 0 | 0 |
| ENSG00000198736 | 1691 | 849 | 990 | 1101 | 1033 | 977 |
| ENSG00000140986 | 6 | 6 | 5 | 9 | 5 | 3 |
| ENSG00000140990 | 2659 | 1318 | 1581 | 1590 | 1490 | 1384 |
| ENSG00000140988 | 47998 | 23469 | 26498 | 35501 | 29469 | 28417 |
| ENSG00000206811 | 0 | 0 | 0 | 0 | 0 | 0 |
| ENSG00000207405 | 0 | 0 | 0 | 0 | 0 | 0 |
| ENSG00000255513 | 0 | 0 | 0 | 0 | 0 | 0 |
| ENSG00000255198 | 116 | 67 | 71 | 83 | 53 | 74 |
| ENSG00000273587 | 0 | 0 | 0 | 0 | 0 | 0 |
| ENSG00000179580 | 11 | 0 | 0 | 0 | 1 | 0 |
| ENSG00000277602 | 0 | 0 | 1 | 1 | 1 | 0 |
| ENSG00000183751 | 1282 | 600 | 750 | 910 | 689 | 745 |
| ENSG00000196408 | 1 | 0 | 1 | 0 | 0 | 0 |
| ENSG00000127554 | 694 | 375 | 512 | 474 | 408 | 427 |
| ENSG00000261790 | 0 | 0 | 0 | 0 | 0 | 1 |
| ENSG00000127561 | 565 | 284 | 317 | 334 | 304 | 305 |
| ENSG00000260107 | 0 | 0 | 0 | 0 | 0 | 0 |
| ENSG00000167962 | 1323 | 642 | 802 | 875 | 846 | 943 |
| ENSG00000183971 | 38 | 9 | 25 | 21 | 17 | 21 |
| ENSG00000265386 | 2 | 4 | 6 | 7 | 4 | 4 |
| ENSG00000065054 | 1353 | 626 | 791 | 1115 | 1023 | 831 |
| ENSG00000065057 | 486 | 269 | 332 | 481 | 331 | 330 |
| ENSG00000103197 | 2828 | 1199 | 1581 | 1421 | 1378 | 1418 |
| ENSG00000008710 | 1301 | 544 | 853 | 657 | 712 | 745 |
| ENSG00000221656 | 0 | 0 | 0 | 0 | 0 | 0 |
| ENSG00000259933 | 3 | 3 | 7 | 0 | 2 | 1 |
| ENSG00000261123 | 5 | 1 | 1 | 0 | 1 | 6 |
| ENSG00000284008 | 0 | 0 | 0 | 0 | 0 | 0 |

| | | | | | | |
|---|---|---|---|---|---|---|
| ENSG00000261240 | 0 | 0 | 0 | 0 | 0 | 0 |
| ENSG00000260447 | 2 | 0 | 1 | 2 | 0 | 0 |
| ENSG00000200059 | 0 | 0 | 2 | 0 | 0 | 0 |
| ENSG00000265867 | 0 | 0 | 0 | 1 | 0 | 0 |
| ENSG00000264397 | 0 | 0 | 0 | 0 | 0 | 0 |
| ENSG00000167964 | 1272 | 576 | 887 | 444 | 401 | 471 |
| ENSG00000260260 | 645 | 355 | 380 | 461 | 385 | 342 |
| ENSG00000206630 | 0 | 0 | 0 | 0 | 0 | 0 |
| ENSG00000281010 | 0 | 0 | 0 | 0 | 0 | 0 |
| ENSG00000131653 | 3674 | 1750 | 2387 | 2386 | 2286 | 2152 |
| ENSG00000167971 | 348 | 159 | 197 | 178 | 160 | 173 |
| ENSG00000167965 | 921 | 481 | 565 | 611 | 479 | 499 |
| ENSG00000182685 | 206 | 125 | 159 | 132 | 128 | 129 |
| ENSG00000261532 | 0 | 0 | 0 | 0 | 0 | 0 |
| ENSG00000184207 | 1125 | 535 | 657 | 780 | 653 | 682 |
| ENSG00000167967 | 1126 | 493 | 638 | 642 | 536 | 546 |
| ENSG00000259780 | 0 | 0 | 0 | 1 | 0 | 1 |
| ENSG00000167968 | 140 | 59 | 83 | 85 | 44 | 52 |
| ENSG00000167969 | 2081 | 961 | 1119 | 1245 | 1052 | 962 |
| ENSG00000261663 | 41 | 24 | 24 | 23 | 26 | 39 |
| ENSG00000205937 | 3959 | 1954 | 2366 | 2339 | 2240 | 2303 |
| ENSG00000260778 | 59 | 21 | 31 | 30 | 27 | 22 |
| ENSG00000266643 | 1 | 7 | 5 | 6 | 3 | 5 |
| ENSG00000284346 | 0 | 0 | 0 | 0 | 0 | 0 |
| ENSG00000264004 | 0 | 0 | 0 | 0 | 0 | 0 |
| ENSG00000167972 | 2460 | 1098 | 1370 | 1286 | 1113 | 1415 |
| ENSG00000243445 | 2 | 1 | 1 | 0 | 0 | 1 |
| ENSG00000238098 | 32 | 21 | 15 | 20 | 20 | 27 |
| ENSG00000162063 | 562 | 341 | 382 | 393 | 327 | 377 |
| ENSG00000285970 | 0 | 0 | 0 | 0 | 0 | 1 |
| ENSG00000283180 | 0 | 0 | 0 | 0 | 0 | 0 |
| ENSG00000260095 | 19 | 3 | 5 | 2 | 2 | 4 |
| ENSG00000260874 | 5 | 2 | 4 | 4 | 3 | 1 |
| ENSG00000162062 | 230 | 105 | 122 | 143 | 149 | 136 |
| ENSG00000274805 | 0 | 0 | 0 | 0 | 0 | 0 |
| ENSG00000259895 | 0 | 3 | 2 | 4 | 2 | 2 |
| ENSG00000162068 | 2 | 1 | 3 | 0 | 0 | 2 |
| ENSG00000162065 | 702 | 289 | 391 | 312 | 355 | 361 |
| ENSG00000260293 | 10 | 3 | 6 | 6 | 4 | 5 |
| ENSG00000260272 | 0 | 0 | 0 | 0 | 0 | 0 |
| ENSG00000185883 | 9 | 52 | 67 | 80 | 88 | 81 |
| ENSG00000259784 | 0 | 0 | 0 | 0 | 0 | 0 |
| ENSG00000279520 | 19 | 13 | 26 | 13 | 18 | 21 |
| ENSG00000162066 | 582 | 244 | 386 | 313 | 234 | 327 |
| ENSG00000205923 | 0 | 0 | 0 | 0 | 0 | 0 |
| ENSG00000266232 | 4 | 0 | 1 | 0 | 0 | 1 |
| ENSG00000140992 | 1844 | 845 | 1131 | 794 | 936 | 1095 |
| ENSG00000261613 | 0 | 0 | 0 | 0 | 0 | 0 |
| ENSG00000261288 | 0 | 0 | 0 | 0 | 0 | 0 |
| ENSG00000269937 | 2 | 2 | 4 | 4 | 2 | 3 |

| | | | | | | |
|---|---|---|---|---|---|---|
| ENSG00000261140 | 0 | 0 | 0 | 1 | 1 | 0 |
| ENSG00000260436 | 0 | 0 | 0 | 0 | 0 | 0 |
| ENSG00000279568 | 4 | 0 | 0 | 0 | 1 | 5 |
| ENSG00000280402 | 0 | 1 | 1 | 0 | 1 | 0 |
| ENSG00000279162 | 0 | 0 | 5 | 0 | 2 | 0 |
| ENSG00000261093 | 0 | 0 | 0 | 0 | 0 | 0 |
| ENSG00000215154 | 63 | 37 | 38 | 18 | 31 | 42 |
| ENSG00000205918 | 28 | 17 | 16 | 10 | 20 | 16 |
| ENSG00000279057 | 1 | 2 | 2 | 0 | 2 | 1 |
| ENSG00000260176 | 5 | 0 | 2 | 6 | 4 | 3 |
| ENSG00000260565 | 508 | 194 | 277 | 102 | 198 | 218 |
| ENSG00000167977 | 2532 | 1171 | 1490 | 1337 | 1276 | 1399 |
| ENSG00000279901 | 1 | 0 | 1 | 2 | 3 | 0 |
| ENSG00000172382 | 157 | 60 | 74 | 44 | 36 | 52 |
| ENSG00000205913 | 99 | 55 | 86 | 42 | 47 | 45 |
| ENSG00000167978 | 14017 | 5575 | 8235 | 5143 | 6654 | 7354 |
| ENSG00000103363 | 3602 | 1845 | 2125 | 2498 | 2077 | 1803 |
| ENSG00000276791 | 209 | 105 | 120 | 73 | 115 | 71 |
| ENSG00000103355 | 0 | 0 | 0 | 0 | 0 | 0 |
| ENSG00000221719 | 0 | 0 | 0 | 0 | 0 | 0 |
| ENSG00000215148 | 0 | 0 | 0 | 0 | 1 | 0 |
| ENSG00000007038 | 2 | 0 | 0 | 0 | 0 | 1 |
| ENSG00000262154 | 1 | 0 | 0 | 0 | 0 | 0 |
| ENSG00000162078 | 948 | 422 | 604 | 454 | 380 | 396 |
| ENSG00000172460 | 10 | 4 | 12 | 1 | 1 | 6 |
| ENSG00000005001 | 952 | 375 | 525 | 405 | 409 | 337 |
| ENSG00000263325 | 6 | 7 | 8 | 2 | 3 | 1 |
| ENSG00000263280 | 15 | 6 | 7 | 3 | 8 | 2 |
| ENSG00000162076 | 1368 | 710 | 773 | 774 | 639 | 633 |
| ENSG00000059122 | 1429 | 661 | 817 | 815 | 792 | 750 |
| ENSG00000262959 | 2 | 0 | 0 | 0 | 0 | 1 |
| ENSG00000262482 | 0 | 1 | 2 | 1 | 3 | 2 |
| ENSG00000131650 | 115 | 46 | 65 | 95 | 70 | 59 |
| ENSG00000127564 | 428 | 292 | 342 | 284 | 312 | 349 |
| ENSG00000162073 | 394 | 225 | 284 | 276 | 242 | 282 |
| ENSG00000274367 | 15 | 5 | 3 | 3 | 5 | 6 |
| ENSG00000270168 | 92 | 58 | 78 | 83 | 88 | 96 |
| ENSG00000262152 | 39 | 27 | 41 | 13 | 22 | 12 |
| ENSG00000262362 | 24 | 10 | 18 | 10 | 10 | 13 |
| ENSG00000272079 | 508 | 221 | 272 | 258 | 257 | 266 |
| ENSG00000213937 | 1052 | 495 | 653 | 585 | 449 | 503 |
| ENSG00000184697 | 10 | 6 | 17 | 1 | 1 | 5 |
| ENSG00000006327 | 4974 | 2597 | 3091 | 3619 | 3133 | 2997 |
| ENSG00000103145 | 1024 | 439 | 709 | 511 | 357 | 426 |
| ENSG00000131652 | 655 | 332 | 425 | 394 | 306 | 383 |
| ENSG00000162069 | 1583 | 644 | 862 | 693 | 772 | 803 |
| ENSG00000205890 | 119 | 45 | 89 | 39 | 67 | 62 |
| ENSG00000261971 | 186 | 79 | 115 | 79 | 100 | 114 |
| ENSG00000008516 | 49 | 5 | 26 | 11 | 12 | 26 |
| ENSG00000008517 | 37 | 13 | 15 | 27 | 12 | 21 |

| | | | | | | |
|---|---|---|---|---|---|---|
| ENSG00000252561 | 0 | 0 | 0 | 0 | 0 | 0 |
| ENSG00000262370 | 0 | 0 | 0 | 0 | 0 | 0 |
| ENSG00000200204 | 0 | 0 | 0 | 0 | 0 | 0 |
| ENSG00000130182 | 0 | 1 | 0 | 1 | 0 | 0 |
| ENSG00000263011 | 7 | 0 | 3 | 5 | 4 | 5 |
| ENSG00000263072 | 374 | 178 | 212 | 178 | 160 | 158 |
| ENSG00000122386 | 287 | 153 | 207 | 213 | 188 | 174 |
| ENSG00000085644 | 520 | 266 | 382 | 314 | 299 | 291 |
| ENSG00000228146 | 37 | 24 | 20 | 18 | 19 | 16 |
| ENSG00000261889 | 23 | 14 | 23 | 9 | 11 | 12 |
| ENSG00000262521 | 1 | 0 | 1 | 1 | 0 | 2 |
| ENSG00000262668 | 1 | 0 | 0 | 0 | 0 | 0 |
| ENSG00000168124 | 4 | 0 | 1 | 1 | 0 | 0 |
| ENSG00000203581 | 0 | 0 | 2 | 1 | 0 | 0 |
| ENSG00000010539 | 439 | 216 | 247 | 165 | 214 | 212 |
| ENSG00000103313 | 14 | 10 | 3 | 3 | 1 | 0 |
| ENSG00000281005 | 29 | 13 | 17 | 13 | 17 | 23 |
| ENSG00000006194 | 1113 | 547 | 685 | 664 | 652 | 605 |
| ENSG00000279330 | 2 | 1 | 9 | 1 | 7 | 1 |
| ENSG00000279031 | 2 | 0 | 3 | 1 | 4 | 3 |
| ENSG00000140993 | 28 | 20 | 25 | 10 | 24 | 18 |
| ENSG00000162086 | 598 | 336 | 360 | 248 | 358 | 327 |
| ENSG00000262899 | 0 | 0 | 0 | 0 | 0 | 0 |
| ENSG00000262554 | 0 | 0 | 0 | 0 | 0 | 0 |
| ENSG00000262316 | 0 | 0 | 0 | 0 | 0 | 0 |
| ENSG00000168158 | 0 | 0 | 0 | 0 | 0 | 0 |
| ENSG00000285329 | 0 | 1 | 1 | 0 | 2 | 2 |
| ENSG00000262118 | 5 | 6 | 2 | 1 | 2 | 2 |
| ENSG00000263177 | 5 | 4 | 0 | 2 | 0 | 0 |
| ENSG00000232196 | 8 | 2 | 3 | 0 | 1 | 0 |
| ENSG00000140987 | 41 | 23 | 26 | 21 | 41 | 32 |
| ENSG00000262621 | 0 | 0 | 0 | 0 | 0 | 1 |
| ENSG00000103343 | 502 | 236 | 336 | 288 | 312 | 339 |
| ENSG00000167981 | 355 | 154 | 184 | 118 | 208 | 146 |
| ENSG00000122390 | 2223 | 1046 | 1412 | 1379 | 1184 | 1304 |
| ENSG00000263212 | 0 | 0 | 0 | 0 | 0 | 0 |
| ENSG00000273776 | 0 | 0 | 0 | 0 | 0 | 0 |
| ENSG00000215131 | 0 | 1 | 0 | 2 | 2 | 0 |
| ENSG00000240311 | 2 | 0 | 1 | 2 | 1 | 0 |
| ENSG00000103351 | 514 | 252 | 286 | 177 | 255 | 278 |
| ENSG00000278942 | 0 | 0 | 0 | 0 | 0 | 0 |
| ENSG00000167984 | 20 | 12 | 19 | 15 | 18 | 15 |
| ENSG00000262312 | 6 | 0 | 1 | 0 | 1 | 0 |
| ENSG00000188827 | 494 | 225 | 276 | 221 | 316 | 260 |
| ENSG00000261938 | 0 | 0 | 0 | 0 | 0 | 0 |
| ENSG00000213918 | 1668 | 840 | 1105 | 713 | 1034 | 916 |
| ENSG00000263235 | 30 | 15 | 15 | 14 | 20 | 16 |
| ENSG00000126602 | 2203 | 1269 | 1498 | 1868 | 1626 | 1696 |
| ENSG00000276754 | 0 | 2 | 2 | 0 | 0 | 0 |
| ENSG00000005339 | 3478 | 1400 | 1995 | 1169 | 1606 | 1656 |

| | | | | | | |
|---|---|---|---|---|---|---|
| ENSG00000262516 | 0 | 0 | 0 | 0 | 0 | 1 |
| ENSG00000262185 | 18 | 16 | 14 | 4 | 7 | 12 |
| ENSG00000162104 | 1785 | 676 | 1054 | 655 | 716 | 816 |
| ENSG00000263159 | 0 | 0 | 0 | 0 | 0 | 0 |
| ENSG00000262471 | 0 | 0 | 0 | 0 | 0 | 0 |
| ENSG00000263105 | 16 | 5 | 13 | 9 | 12 | 17 |
| ENSG00000185739 | 7 | 4 | 5 | 3 | 3 | 3 |
| ENSG00000262468 | 397 | 194 | 250 | 195 | 185 | 202 |
| ENSG00000090447 | 409 | 189 | 231 | 254 | 238 | 283 |
| ENSG00000126603 | 885 | 405 | 490 | 544 | 447 | 415 |
| ENSG00000262686 | 2 | 5 | 2 | 1 | 4 | 1 |
| ENSG00000217930 | 80 | 70 | 100 | 43 | 90 | 65 |
| ENSG00000262712 | 21 | 7 | 13 | 19 | 16 | 13 |
| ENSG00000103426 | 0 | 0 | 0 | 0 | 0 | 0 |
| ENSG00000280063 | 40 | 19 | 46 | 11 | 19 | 15 |
| ENSG00000262246 | 221 | 120 | 181 | 180 | 136 | 146 |
| ENSG00000168140 | 509 | 192 | 311 | 319 | 297 | 233 |
| ENSG00000103423 | 2736 | 1338 | 1670 | 1574 | 1548 | 1665 |
| ENSG00000277170 | 0 | 2 | 2 | 0 | 0 | 3 |
| ENSG00000277440 | 0 | 0 | 1 | 2 | 0 | 0 |
| ENSG00000275515 | 1 | 0 | 0 | 0 | 0 | 0 |
| ENSG00000153406 | 1651 | 767 | 1011 | 1083 | 965 | 936 |
| ENSG00000103415 | 1558 | 740 | 847 | 1036 | 967 | 943 |
| ENSG00000089486 | 40 | 20 | 25 | 16 | 24 | 29 |
| ENSG00000278434 | 1 | 0 | 0 | 0 | 0 | 0 |
| ENSG00000205832 | 0 | 0 | 1 | 0 | 0 | 1 |
| ENSG00000261789 | 0 | 0 | 0 | 0 | 0 | 0 |
| ENSG00000261612 | 0 | 0 | 0 | 0 | 0 | 1 |
| ENSG00000153443 | 663 | 296 | 420 | 485 | 354 | 374 |
| ENSG00000102858 | 2160 | 1023 | 1367 | 1191 | 1057 | 1171 |
| ENSG00000263955 | 0 | 0 | 0 | 0 | 0 | 0 |
| ENSG00000261442 | 0 | 3 | 2 | 0 | 0 | 1 |
| ENSG00000278700 | 0 | 0 | 0 | 1 | 0 | 0 |
| ENSG00000276641 | 0 | 0 | 0 | 0 | 1 | 3 |
| ENSG00000168101 | 1536 | 789 | 910 | 922 | 765 | 777 |
| ENSG00000168096 | 399 | 218 | 242 | 235 | 242 | 219 |
| ENSG00000266994 | 0 | 0 | 1 | 0 | 1 | 1 |
| ENSG00000166246 | 36 | 18 | 21 | 15 | 18 | 13 |
| ENSG00000103199 | 439 | 209 | 242 | 210 | 202 | 217 |
| ENSG00000276096 | 4 | 2 | 1 | 0 | 1 | 1 |
| ENSG00000140623 | 0 | 1 | 3 | 1 | 0 | 2 |
| ENSG00000285952 | 2 | 0 | 0 | 1 | 1 | 4 |
| ENSG00000267795 | 213 | 110 | 175 | 166 | 152 | 160 |
| ENSG00000267077 | 3 | 0 | 6 | 0 | 3 | 4 |
| ENSG00000067836 | 908 | 443 | 600 | 511 | 450 | 480 |
| ENSG00000140632 | 5010 | 2322 | 2878 | 2209 | 2582 | 2685 |
| ENSG00000275056 | 14 | 3 | 3 | 3 | 4 | 5 |
| ENSG00000118900 | 3351 | 1363 | 1961 | 1445 | 1708 | 1809 |
| ENSG00000118898 | 3821 | 1746 | 2239 | 1568 | 1867 | 1712 |
| ENSG00000103184 | 9 | 9 | 9 | 10 | 18 | 3 |

| | | | | | | |
|---|---|---|---|---|---|---|
| ENSG00000267072 | 2 | 0 | 1 | 0 | 1 | 1 |
| ENSG00000103174 | 320 | 218 | 221 | 220 | 200 | 224 |
| ENSG00000033011 | 544 | 285 | 358 | 362 | 351 | 293 |
| ENSG00000153446 | 4 | 1 | 0 | 0 | 0 | 1 |
| ENSG00000260882 | 0 | 1 | 0 | 1 | 1 | 0 |
| ENSG00000118894 | 275 | 187 | 184 | 219 | 202 | 221 |
| ENSG00000267070 | 0 | 0 | 0 | 0 | 0 | 0 |
| ENSG00000285567 | 0 | 0 | 0 | 0 | 0 | 0 |
| ENSG00000273963 | 0 | 0 | 0 | 0 | 0 | 1 |
| ENSG00000260431 | 0 | 0 | 0 | 0 | 0 | 0 |
| ENSG00000261811 | 0 | 0 | 0 | 0 | 0 | 0 |
| ENSG00000078328 | 0 | 0 | 1 | 0 | 0 | 1 |
| ENSG00000260817 | 0 | 0 | 0 | 0 | 0 | 0 |
| ENSG00000242278 | 0 | 0 | 0 | 0 | 0 | 0 |
| ENSG00000267182 | 0 | 0 | 0 | 0 | 0 | 0 |
| ENSG00000261007 | 0 | 0 | 0 | 0 | 0 | 0 |
| ENSG00000260338 | 0 | 0 | 0 | 0 | 0 | 0 |
| ENSG00000273648 | 0 | 0 | 0 | 0 | 0 | 0 |
| ENSG00000270440 | 0 | 0 | 0 | 0 | 0 | 0 |
| ENSG00000280000 | 0 | 0 | 0 | 0 | 0 | 0 |
| ENSG00000257180 | 0 | 0 | 0 | 0 | 0 | 0 |
| ENSG00000260835 | 0 | 0 | 0 | 0 | 0 | 0 |
| ENSG00000278975 | 0 | 0 | 0 | 0 | 0 | 0 |
| ENSG00000259963 | 0 | 0 | 0 | 0 | 0 | 0 |
| ENSG00000252080 | 0 | 0 | 0 | 0 | 0 | 0 |
| ENSG00000200869 | 0 | 0 | 0 | 0 | 0 | 0 |
| ENSG00000212189 | 0 | 0 | 0 | 0 | 0 | 0 |
| ENSG00000240419 | 0 | 0 | 0 | 0 | 0 | 0 |
| ENSG00000283584 | 0 | 0 | 0 | 0 | 0 | 0 |
| ENSG00000261198 | 0 | 0 | 0 | 0 | 0 | 0 |
| ENSG00000252138 | 0 | 0 | 0 | 0 | 0 | 0 |
| ENSG00000260289 | 0 | 0 | 0 | 0 | 0 | 0 |
| ENSG00000260003 | 0 | 0 | 0 | 0 | 0 | 0 |
| ENSG00000261319 | 0 | 0 | 0 | 0 | 0 | 0 |
| ENSG00000260058 | 0 | 0 | 0 | 0 | 0 | 0 |
| ENSG00000259819 | 0 | 0 | 0 | 0 | 0 | 0 |
| ENSG00000260379 | 0 | 0 | 0 | 0 | 0 | 0 |
| ENSG00000261066 | 0 | 0 | 0 | 0 | 0 | 0 |
| ENSG00000232258 | 0 | 0 | 0 | 0 | 0 | 0 |
| ENSG00000067365 | 666 | 313 | 378 | 336 | 340 | 336 |
| ENSG00000279357 | 1 | 0 | 0 | 0 | 0 | 0 |
| ENSG00000183044 | 3187 | 1575 | 2011 | 1417 | 1770 | 1842 |
| ENSG00000243954 | 0 | 0 | 0 | 0 | 0 | 0 |
| ENSG00000238417 | 0 | 0 | 0 | 0 | 0 | 0 |
| ENSG00000260764 | 0 | 0 | 0 | 0 | 0 | 0 |
| ENSG00000184857 | 372 | 192 | 244 | 304 | 300 | 288 |
| ENSG00000140650 | 785 | 413 | 517 | 478 | 573 | 607 |
| ENSG00000260350 | 1 | 1 | 0 | 0 | 0 | 0 |
| ENSG00000260276 | 39 | 22 | 39 | 16 | 30 | 22 |
| ENSG00000153048 | 1575 | 883 | 1096 | 1078 | 1127 | 937 |

| | | | | | | |
|---|---|---|---|---|---|---|
| ENSG00000259939 | 0 | 0 | 0 | 0 | 0 | 0 |
| ENSG00000261481 | 2 | 0 | 0 | 1 | 1 | 1 |
| ENSG00000283516 | 0 | 0 | 1 | 0 | 0 | 0 |
| ENSG00000187555 | 5355 | 2535 | 3269 | 2350 | 3068 | 2962 |
| ENSG00000260979 | 0 | 0 | 3 | 1 | 4 | 1 |
| ENSG00000261392 | 0 | 0 | 0 | 0 | 0 | 0 |
| ENSG00000182831 | 2252 | 1182 | 1231 | 970 | 1301 | 1128 |
| ENSG00000263244 | 0 | 0 | 0 | 0 | 0 | 0 |
| ENSG00000260349 | 0 | 0 | 0 | 0 | 0 | 0 |
| ENSG00000220793 | 2 | 1 | 1 | 2 | 2 | 1 |
| ENSG00000261617 | 0 | 0 | 0 | 0 | 0 | 0 |
| ENSG00000261397 | 0 | 0 | 0 | 0 | 0 | 0 |
| ENSG00000261075 | 0 | 0 | 0 | 0 | 0 | 0 |
| ENSG00000260071 | 0 | 0 | 0 | 1 | 0 | 0 |
| ENSG00000283003 | 0 | 0 | 0 | 0 | 0 | 0 |
| ENSG00000200468 | 0 | 0 | 0 | 0 | 0 | 0 |
| ENSG00000252927 | 0 | 0 | 0 | 0 | 0 | 0 |
| ENSG00000260362 | 0 | 0 | 0 | 0 | 0 | 0 |
| ENSG00000183454 | 0 | 0 | 0 | 0 | 1 | 0 |
| ENSG00000283221 | 0 | 0 | 0 | 0 | 0 | 0 |
| ENSG00000261810 | 0 | 0 | 0 | 0 | 0 | 0 |
| ENSG00000261647 | 0 | 0 | 0 | 0 | 0 | 0 |
| ENSG00000285543 | 0 | 0 | 0 | 0 | 0 | 0 |
| ENSG00000266439 | 0 | 0 | 0 | 0 | 0 | 0 |
| ENSG00000166669 | 398 | 203 | 255 | 163 | 259 | 232 |
| ENSG00000260756 | 0 | 0 | 0 | 0 | 0 | 0 |
| ENSG00000279662 | 194 | 58 | 121 | 43 | 64 | 115 |
| ENSG00000199482 | 0 | 2 | 1 | 0 | 0 | 0 |
| ENSG00000260468 | 11 | 6 | 16 | 6 | 12 | 5 |
| ENSG00000213853 | 9928 | 5986 | 6932 | 4569 | 5289 | 5850 |
| ENSG00000256013 | 0 | 0 | 0 | 0 | 0 | 0 |
| ENSG00000260310 | 3 | 3 | 3 | 1 | 2 | 1 |
| ENSG00000153060 | 31 | 11 | 18 | 15 | 14 | 17 |
| ENSG00000263279 | 0 | 0 | 0 | 1 | 0 | 0 |
| ENSG00000263241 | 0 | 1 | 0 | 0 | 0 | 0 |
| ENSG00000262107 | 0 | 0 | 0 | 0 | 0 | 0 |
| ENSG00000261904 | 0 | 0 | 2 | 0 | 0 | 0 |
| ENSG00000262322 | 0 | 0 | 0 | 0 | 0 | 0 |
| ENSG00000262259 | 0 | 0 | 0 | 0 | 0 | 0 |
| ENSG00000263196 | 0 | 0 | 0 | 0 | 0 | 0 |
| ENSG00000262158 | 0 | 0 | 0 | 0 | 0 | 0 |
| ENSG00000262381 | 0 | 0 | 1 | 0 | 0 | 0 |
| ENSG00000103274 | 667 | 355 | 454 | 471 | 410 | 429 |
| ENSG00000166676 | 10 | 12 | 22 | 3 | 19 | 14 |
| ENSG00000262488 | 0 | 1 | 1 | 0 | 1 | 0 |
| ENSG00000262151 | 0 | 0 | 0 | 0 | 0 | 0 |
| ENSG00000179583 | 61 | 36 | 60 | 34 | 42 | 25 |
| ENSG00000182108 | 182 | 83 | 114 | 76 | 97 | 88 |
| ENSG00000280153 | 0 | 0 | 0 | 0 | 0 | 0 |
| ENSG00000038532 | 1221 | 506 | 775 | 554 | 647 | 754 |

| | | | | | | |
|---|---|---|---|---|---|---|
| ENSG00000229917 | 0 | 0 | 0 | 0 | 2 | 0 |
| ENSG00000274038 | 10 | 2 | 1 | 1 | 0 | 2 |
| ENSG00000262020 | 0 | 0 | 0 | 0 | 0 | 0 |
| ENSG00000263033 | 4 | 1 | 8 | 0 | 5 | 2 |
| ENSG00000263179 | 0 | 2 | 1 | 1 | 0 | 1 |
| ENSG00000175643 | 910 | 493 | 564 | 404 | 426 | 480 |
| ENSG00000185338 | 50 | 33 | 29 | 37 | 17 | 29 |
| ENSG00000178279 | 2 | 0 | 0 | 0 | 0 | 0 |
| ENSG00000178257 | 0 | 0 | 0 | 0 | 0 | 0 |
| ENSG00000212228 | 0 | 0 | 0 | 0 | 0 | 0 |
| ENSG00000122304 | 0 | 0 | 0 | 0 | 0 | 0 |
| ENSG00000175646 | 0 | 0 | 0 | 0 | 0 | 0 |
| ENSG00000221801 | 0 | 0 | 0 | 0 | 0 | 0 |
| ENSG00000199668 | 0 | 0 | 0 | 0 | 0 | 0 |
| ENSG00000263080 | 10 | 6 | 2 | 2 | 0 | 1 |
| ENSG00000262703 | 10 | 4 | 9 | 2 | 9 | 8 |
| ENSG00000188897 | 51 | 18 | 25 | 16 | 21 | 14 |
| ENSG00000262636 | 0 | 0 | 0 | 0 | 0 | 0 |
| ENSG00000262999 | 2 | 0 | 0 | 0 | 0 | 0 |
| ENSG00000189067 | 9512 | 4899 | 5579 | 4826 | 5550 | 5525 |
| ENSG00000184602 | 1516 | 677 | 863 | 730 | 866 | 876 |
| ENSG00000153066 | 1622 | 874 | 1104 | 898 | 984 | 991 |
| ENSG00000262420 | 6 | 3 | 5 | 5 | 6 | 8 |
| ENSG00000122299 | 2150 | 1100 | 1275 | 825 | 1272 | 1140 |
| ENSG00000274505 | 0 | 0 | 1 | 1 | 0 | 0 |
| ENSG00000277369 | 26 | 6 | 10 | 16 | 6 | 8 |
| ENSG00000262117 | 2 | 4 | 1 | 1 | 0 | 2 |
| ENSG00000171490 | 4220 | 2089 | 2133 | 1747 | 3043 | 2327 |
| ENSG00000263307 | 5 | 5 | 5 | 3 | 9 | 8 |
| ENSG00000103342 | 7444 | 3859 | 4332 | 3355 | 4667 | 4245 |
| ENSG00000261560 | 33 | 17 | 12 | 10 | 16 | 13 |
| ENSG00000260318 | 2 | 4 | 2 | 1 | 1 | 3 |
| ENSG00000261216 | 3 | 0 | 1 | 1 | 1 | 3 |
| ENSG00000277458 | 0 | 0 | 0 | 0 | 0 | 0 |
| ENSG00000234719 | 101 | 57 | 64 | 67 | 53 | 57 |
| ENSG00000048462 | 0 | 0 | 1 | 0 | 0 | 0 |
| ENSG00000260224 | 0 | 0 | 0 | 0 | 0 | 0 |
| ENSG00000048471 | 1073 | 439 | 576 | 378 | 535 | 522 |
| ENSG00000260488 | 0 | 0 | 0 | 0 | 0 | 0 |
| ENSG00000241641 | 0 | 0 | 0 | 0 | 0 | 0 |
| ENSG00000175604 | 0 | 0 | 0 | 0 | 0 | 0 |
| ENSG00000261293 | 0 | 0 | 1 | 0 | 1 | 0 |
| ENSG00000238685 | 0 | 0 | 0 | 0 | 0 | 0 |
| ENSG00000261394 | 0 | 1 | 0 | 0 | 0 | 0 |
| ENSG00000260723 | 0 | 1 | 1 | 0 | 0 | 0 |
| ENSG00000278979 | 0 | 0 | 1 | 0 | 0 | 1 |
| ENSG00000260601 | 1 | 0 | 2 | 0 | 0 | 1 |
| ENSG00000259876 | 0 | 0 | 2 | 0 | 0 | 3 |
| ENSG00000259899 | 0 | 3 | 4 | 1 | 2 | 0 |
| ENSG00000274834 | 0 | 0 | 1 | 0 | 0 | 0 |

| | | | | | | |
|---|---|---|---|---|---|---|
| ENSG00000103381 | 1031 | 519 | 658 | 479 | 456 | 473 |
| ENSG00000264733 | 0 | 0 | 0 | 0 | 0 | 0 |
| ENSG00000260378 | 0 | 0 | 0 | 0 | 0 | 0 |
| ENSG00000261158 | 0 | 2 | 0 | 0 | 0 | 0 |
| ENSG00000199474 | 0 | 0 | 0 | 0 | 0 | 0 |
| ENSG00000241932 | 0 | 0 | 0 | 0 | 0 | 0 |
| ENSG00000237515 | 390 | 174 | 327 | 106 | 162 | 156 |
| ENSG00000262116 | 2 | 0 | 0 | 0 | 0 | 0 |
| ENSG00000262801 | 5 | 7 | 5 | 4 | 2 | 1 |
| ENSG00000262011 | 0 | 0 | 0 | 0 | 0 | 0 |
| ENSG00000262235 | 0 | 0 | 0 | 0 | 0 | 0 |
| ENSG00000262267 | 0 | 0 | 0 | 0 | 0 | 0 |
| ENSG00000175595 | 303 | 150 | 185 | 109 | 232 | 174 |
| ENSG00000242307 | 2 | 0 | 0 | 0 | 0 | 0 |
| ENSG00000262732 | 2 | 3 | 0 | 2 | 1 | 2 |
| ENSG00000263234 | 0 | 0 | 0 | 0 | 0 | 0 |
| ENSG00000262097 | 0 | 0 | 0 | 0 | 0 | 0 |
| ENSG00000261856 | 0 | 0 | 0 | 0 | 0 | 0 |
| ENSG00000186260 | 2046 | 1013 | 1070 | 413 | 1149 | 819 |
| ENSG00000276564 | 20 | 7 | 13 | 2 | 11 | 10 |
| ENSG00000262529 | 0 | 0 | 0 | 0 | 0 | 0 |
| ENSG00000262470 | 0 | 0 | 0 | 0 | 0 | 0 |
| ENSG00000201075 | 0 | 0 | 0 | 0 | 0 | 0 |
| ENSG00000262454 | 296 | 176 | 207 | 257 | 251 | 258 |
| ENSG00000207639 | 0 | 0 | 0 | 0 | 0 | 1 |
| ENSG00000199130 | 0 | 1 | 0 | 0 | 0 | 0 |
| ENSG00000260009 | 18 | 6 | 4 | 4 | 14 | 5 |
| ENSG00000263257 | 2 | 1 | 1 | 0 | 0 | 0 |
| ENSG00000140694 | 1759 | 844 | 1152 | 786 | 1136 | 997 |
| ENSG00000242430 | 0 | 0 | 0 | 0 | 0 | 0 |
| ENSG00000103429 | 2662 | 1277 | 1456 | 1215 | 1611 | 1394 |
| ENSG00000238964 | 0 | 0 | 1 | 0 | 0 | 0 |
| ENSG00000069764 | 68 | 57 | 54 | 29 | 16 | 30 |
| ENSG00000261523 | 0 | 0 | 0 | 0 | 0 | 0 |
| ENSG00000224712 | 0 | 2 | 2 | 1 | 1 | 0 |
| ENSG00000254609 | 8 | 2 | 3 | 0 | 1 | 1 |
| ENSG00000261695 | 0 | 0 | 0 | 0 | 0 | 0 |
| ENSG00000254852 | 0 | 1 | 0 | 1 | 1 | 0 |
| ENSG00000255037 | 0 | 0 | 0 | 0 | 0 | 0 |
| ENSG00000255277 | 2 | 1 | 2 | 1 | 1 | 1 |
| ENSG00000103512 | 3794 | 1790 | 2280 | 1826 | 2032 | 2056 |
| ENSG00000270734 | 0 | 0 | 0 | 0 | 0 | 0 |
| ENSG00000257264 | 0 | 0 | 0 | 0 | 0 | 0 |
| ENSG00000284305 | 0 | 0 | 0 | 0 | 0 | 0 |
| ENSG00000263918 | 0 | 0 | 0 | 0 | 0 | 0 |
| ENSG00000258354 | 8 | 5 | 1 | 2 | 5 | 2 |
| ENSG00000265537 | 0 | 0 | 0 | 0 | 0 | 0 |
| ENSG00000183458 | 202 | 59 | 128 | 56 | 71 | 113 |
| ENSG00000207425 | 27 | 23 | 31 | 8 | 22 | 13 |
| ENSG00000183426 | 153 | 75 | 116 | 85 | 90 | 108 |

| | | | | | | |
|---|---|---|---|---|---|---|
| ENSG00000275259 | 0 | 0 | 0 | 0 | 0 | 0 |
| ENSG00000278265 | 0 | 0 | 0 | 0 | 0 | 0 |
| ENSG00000273804 | 0 | 0 | 0 | 0 | 0 | 0 |
| ENSG00000179889 | 7499 | 3590 | 4630 | 3110 | 3974 | 3900 |
| ENSG00000261819 | 0 | 2 | 3 | 2 | 5 | 8 |
| ENSG00000238728 | 0 | 0 | 0 | 0 | 0 | 2 |
| ENSG00000275910 | 4 | 7 | 8 | 6 | 6 | 8 |
| ENSG00000157045 | 844 | 444 | 457 | 525 | 511 | 435 |
| ENSG00000085721 | 2160 | 1104 | 1215 | 1021 | 1333 | 1197 |
| ENSG00000260735 | 6 | 6 | 9 | 5 | 4 | 9 |
| ENSG00000188599 | 50 | 17 | 30 | 29 | 23 | 42 |
| ENSG00000270580 | 10 | 7 | 14 | 15 | 17 | 16 |
| ENSG00000250251 | 119 | 59 | 111 | 65 | 86 | 83 |
| ENSG00000283346 | 0 | 0 | 0 | 0 | 0 | 0 |
| ENSG00000207294 | 0 | 0 | 0 | 0 | 0 | 0 |
| ENSG00000264115 | 0 | 0 | 0 | 0 | 0 | 0 |
| ENSG00000257391 | 0 | 0 | 0 | 0 | 0 | 0 |
| ENSG00000277770 | 0 | 0 | 0 | 0 | 0 | 0 |
| ENSG00000183793 | 0 | 3 | 1 | 0 | 0 | 1 |
| ENSG00000156968 | 0 | 0 | 0 | 0 | 0 | 0 |
| ENSG00000261130 | 0 | 0 | 0 | 0 | 0 | 0 |
| ENSG00000166780 | 466 | 217 | 318 | 155 | 192 | 189 |
| ENSG00000166783 | 1025 | 542 | 676 | 308 | 674 | 542 |
| ENSG00000257769 | 0 | 0 | 0 | 0 | 0 | 1 |
| ENSG00000284184 | 0 | 0 | 0 | 0 | 0 | 0 |
| ENSG00000072864 | 443 | 291 | 286 | 236 | 309 | 317 |
| ENSG00000283736 | 0 | 0 | 0 | 0 | 0 | 0 |
| ENSG00000262380 | 0 | 0 | 0 | 0 | 0 | 0 |
| ENSG00000280206 | 95 | 60 | 70 | 61 | 69 | 63 |
| ENSG00000133392 | 15 | 16 | 7 | 2 | 8 | 4 |
| ENSG00000263335 | 14 | 10 | 10 | 7 | 13 | 5 |
| ENSG00000263065 | 0 | 0 | 0 | 0 | 0 | 0 |
| ENSG00000280429 | 0 | 0 | 0 | 0 | 0 | 0 |
| ENSG00000133393 | 1300 | 615 | 786 | 608 | 842 | 771 |
| ENSG00000206778 | 0 | 0 | 0 | 0 | 0 | 0 |
| ENSG00000262171 | 0 | 2 | 3 | 0 | 1 | 2 |
| ENSG00000215003 | 0 | 0 | 0 | 0 | 0 | 1 |
| ENSG00000103222 | 2473 | 1092 | 1479 | 1056 | 1288 | 1473 |
| ENSG00000241067 | 0 | 0 | 0 | 0 | 0 | 0 |
| ENSG00000091262 | 20 | 7 | 13 | 8 | 10 | 9 |
| ENSG00000262332 | 0 | 0 | 0 | 1 | 0 | 0 |
| ENSG00000103226 | 485 | 194 | 316 | 235 | 265 | 242 |
| ENSG00000263029 | 0 | 0 | 0 | 0 | 0 | 0 |
| ENSG00000262848 | 0 | 0 | 0 | 0 | 0 | 0 |
| ENSG00000257381 | 0 | 0 | 0 | 0 | 0 | 0 |
| ENSG00000264722 | 0 | 0 | 0 | 0 | 0 | 0 |
| ENSG00000257366 | 0 | 0 | 0 | 0 | 0 | 0 |
| ENSG00000265373 | 0 | 0 | 0 | 0 | 0 | 0 |
| ENSG00000244257 | 0 | 0 | 1 | 0 | 0 | 1 |
| ENSG00000206706 | 0 | 0 | 0 | 0 | 0 | 0 |

| | | | | | | |
|---|---|---|---|---|---|---|
| ENSG00000183889 | 19 | 7 | 14 | 15 | 13 | 19 |
| ENSG00000278221 | 0 | 0 | 0 | 0 | 0 | 0 |
| ENSG00000277698 | 0 | 0 | 0 | 0 | 0 | 0 |
| ENSG00000277920 | 0 | 0 | 0 | 0 | 0 | 0 |
| ENSG00000227827 | 15 | 1 | 2 | 0 | 1 | 2 |
| ENSG00000276311 | 0 | 0 | 0 | 0 | 0 | 0 |
| ENSG00000214967 | 1 | 1 | 0 | 0 | 0 | 0 |
| ENSG00000261181 | 0 | 0 | 0 | 0 | 0 | 0 |
| ENSG00000261362 | 0 | 0 | 0 | 0 | 0 | 0 |
| ENSG00000260126 | 0 | 0 | 0 | 0 | 0 | 0 |
| ENSG00000279729 | 0 | 0 | 0 | 0 | 0 | 0 |
| ENSG00000261748 | 0 | 0 | 0 | 0 | 0 | 0 |
| ENSG00000279490 | 0 | 0 | 0 | 0 | 0 | 0 |
| ENSG00000260675 | 0 | 0 | 0 | 0 | 0 | 0 |
| ENSG00000260405 | 0 | 0 | 0 | 0 | 0 | 0 |
| ENSG00000103489 | 139 | 67 | 89 | 47 | 59 | 42 |
| ENSG00000261448 | 0 | 0 | 0 | 0 | 0 | 0 |
| ENSG00000279410 | 0 | 0 | 0 | 0 | 0 | 0 |
| ENSG00000279415 | 2 | 2 | 4 | 2 | 1 | 1 |
| ENSG00000279620 | 0 | 1 | 0 | 0 | 0 | 0 |
| ENSG00000275927 | 0 | 0 | 0 | 0 | 0 | 1 |
| ENSG00000285850 | 0 | 0 | 0 | 0 | 0 | 0 |
| ENSG00000260945 | 0 | 0 | 0 | 0 | 0 | 0 |
| ENSG00000261555 | 0 | 0 | 0 | 0 | 0 | 0 |
| ENSG00000228071 | 0 | 0 | 0 | 1 | 0 | 1 |
| ENSG00000259929 | 0 | 0 | 0 | 0 | 0 | 0 |
| ENSG00000280283 | 0 | 0 | 0 | 0 | 0 | 0 |
| ENSG00000260242 | 0 | 0 | 0 | 0 | 0 | 0 |
| ENSG00000260933 | 0 | 0 | 0 | 0 | 0 | 0 |
| ENSG00000278994 | 0 | 0 | 0 | 0 | 0 | 0 |
| ENSG00000276484 | 0 | 0 | 0 | 0 | 0 | 0 |
| ENSG00000214940 | 0 | 0 | 0 | 0 | 0 | 0 |
| ENSG00000285628 | 0 | 0 | 0 | 0 | 0 | 0 |
| ENSG00000205746 | 57 | 21 | 36 | 28 | 32 | 24 |
| ENSG00000273613 | 0 | 0 | 0 | 0 | 0 | 0 |
| ENSG00000277647 | 0 | 0 | 0 | 0 | 0 | 0 |
| ENSG00000233024 | 2 | 1 | 0 | 0 | 0 | 0 |
| ENSG00000254681 | 48 | 20 | 30 | 34 | 33 | 41 |
| ENSG00000275391 | 0 | 0 | 0 | 0 | 0 | 0 |
| ENSG00000206927 | 0 | 0 | 0 | 0 | 0 | 0 |
| ENSG00000257563 | 0 | 0 | 0 | 0 | 0 | 0 |
| ENSG00000266291 | 0 | 0 | 0 | 0 | 0 | 0 |
| ENSG00000265776 | 0 | 0 | 0 | 0 | 0 | 0 |
| ENSG00000257527 | 1 | 0 | 0 | 0 | 0 | 0 |
| ENSG00000266454 | 0 | 0 | 0 | 0 | 0 | 0 |
| ENSG00000185164 | 1209 | 566 | 723 | 583 | 594 | 656 |
| ENSG00000274025 | 0 | 0 | 0 | 0 | 0 | 0 |
| ENSG00000277014 | 0 | 0 | 0 | 0 | 0 | 0 |
| ENSG00000262974 | 0 | 0 | 0 | 0 | 0 | 0 |
| ENSG00000260550 | 0 | 0 | 0 | 0 | 0 | 0 |

| | | | | | | |
|---|---|---|---|---|---|---|
| ENSG00000256340 | 2 | 0 | 0 | 0 | 0 | 0 |
| ENSG00000134419 | 7273 | 5659 | 6588 | 7851 | 6637 | 6626 |
| ENSG00000260342 | 0 | 0 | 0 | 0 | 0 | 0 |
| ENSG00000170540 | 6310 | 3322 | 3666 | 3324 | 4463 | 3729 |
| ENSG00000260017 | 1 | 0 | 1 | 1 | 0 | 0 |
| ENSG00000157106 | 1707 | 1059 | 844 | 346 | 1750 | 933 |
| ENSG00000260352 | 15 | 5 | 5 | 13 | 4 | 10 |
| ENSG00000170537 | 122 | 71 | 66 | 58 | 116 | 71 |
| ENSG00000207167 | 0 | 0 | 0 | 0 | 0 | 0 |
| ENSG00000261465 | 0 | 0 | 1 | 0 | 0 | 1 |
| ENSG00000261357 | 1 | 1 | 0 | 2 | 1 | 2 |
| ENSG00000167186 | 776 | 399 | 497 | 420 | 387 | 500 |
| ENSG00000260430 | 11 | 8 | 6 | 13 | 3 | 10 |
| ENSG00000278389 | 0 | 0 | 0 | 1 | 0 | 0 |
| ENSG00000205730 | 114 | 81 | 91 | 42 | 49 | 72 |
| ENSG00000261427 | 0 | 0 | 0 | 0 | 0 | 0 |
| ENSG00000261759 | 0 | 0 | 0 | 0 | 0 | 0 |
| ENSG00000103528 | 46 | 27 | 31 | 22 | 37 | 32 |
| ENSG00000261210 | 0 | 0 | 0 | 0 | 0 | 0 |
| ENSG00000259871 | 0 | 0 | 0 | 0 | 0 | 0 |
| ENSG00000285869 | 0 | 0 | 0 | 0 | 0 | 0 |
| ENSG00000259925 | 0 | 2 | 0 | 1 | 0 | 0 |
| ENSG00000280265 | 0 | 0 | 0 | 0 | 0 | 0 |
| ENSG00000103534 | 42 | 18 | 21 | 13 | 29 | 28 |
| ENSG00000260681 | 0 | 0 | 0 | 0 | 0 | 0 |
| ENSG00000260592 | 0 | 0 | 0 | 0 | 0 | 0 |
| ENSG00000222750 | 0 | 0 | 0 | 0 | 0 | 0 |
| ENSG00000006007 | 4782 | 2244 | 2844 | 2345 | 2632 | 2650 |
| ENSG00000260934 | 0 | 0 | 0 | 0 | 0 | 0 |
| ENSG00000279747 | 0 | 0 | 0 | 0 | 0 | 0 |
| ENSG00000103540 | 415 | 225 | 197 | 69 | 337 | 155 |
| ENSG00000103544 | 1612 | 772 | 970 | 683 | 836 | 796 |
| ENSG00000276571 | 4 | 8 | 9 | 1 | 2 | 4 |
| ENSG00000103550 | 691 | 432 | 485 | 476 | 521 | 528 |
| ENSG00000261312 | 0 | 0 | 0 | 0 | 0 | 0 |
| ENSG00000174628 | 198 | 88 | 151 | 76 | 130 | 118 |
| ENSG00000261195 | 2 | 0 | 1 | 0 | 1 | 0 |
| ENSG00000261939 | 0 | 0 | 0 | 0 | 0 | 0 |
| ENSG00000167191 | 101 | 48 | 57 | 29 | 35 | 44 |
| ENSG00000180269 | 0 | 0 | 0 | 0 | 0 | 0 |
| ENSG00000277211 | 0 | 0 | 0 | 0 | 0 | 0 |
| ENSG00000263253 | 0 | 0 | 0 | 0 | 0 | 0 |
| ENSG00000227624 | 0 | 0 | 0 | 0 | 0 | 0 |
| ENSG00000169347 | 106 | 51 | 55 | 30 | 45 | 46 |
| ENSG00000169344 | 2 | 0 | 0 | 0 | 0 | 0 |
| ENSG00000169340 | 0 | 0 | 0 | 0 | 0 | 0 |
| ENSG00000263237 | 0 | 0 | 0 | 0 | 0 | 0 |
| ENSG00000183549 | 0 | 0 | 0 | 0 | 0 | 0 |
| ENSG00000262995 | 0 | 0 | 0 | 0 | 0 | 0 |
| ENSG00000183747 | 0 | 0 | 0 | 0 | 0 | 0 |

| | | | | | | |
|---|---|---|---|---|---|---|
| ENSG00000280306 | 0 | 0 | 0 | 0 | 0 | 0 |
| ENSG00000066813 | 0 | 0 | 0 | 0 | 0 | 0 |
| ENSG00000260201 | 0 | 0 | 0 | 0 | 0 | 0 |
| ENSG00000260762 | 0 | 0 | 0 | 0 | 0 | 0 |
| ENSG00000005187 | 33 | 13 | 18 | 14 | 17 | 25 |
| ENSG00000275315 | 0 | 0 | 0 | 0 | 0 | 0 |
| ENSG00000274730 | 0 | 0 | 0 | 0 | 0 | 0 |
| ENSG00000166743 | 6 | 2 | 2 | 5 | 0 | 0 |
| ENSG00000066654 | 1781 | 888 | 932 | 467 | 1120 | 806 |
| ENSG00000283399 | 0 | 2 | 0 | 0 | 1 | 1 |
| ENSG00000285303 | 0 | 0 | 0 | 0 | 0 | 0 |
| ENSG00000260510 | 0 | 0 | 0 | 0 | 0 | 0 |
| ENSG00000196678 | 628 | 353 | 456 | 238 | 477 | 350 |
| ENSG00000200893 | 0 | 0 | 0 | 0 | 0 | 0 |
| ENSG00000005189 | 309 | 159 | 211 | 134 | 163 | 168 |
| ENSG00000275908 | 0 | 0 | 0 | 0 | 0 | 0 |
| ENSG00000188215 | 466 | 208 | 258 | 214 | 246 | 216 |
| ENSG00000102897 | 592 | 320 | 426 | 362 | 410 | 363 |
| ENSG00000158486 | 261 | 130 | 195 | 97 | 127 | 112 |
| ENSG00000263331 | 0 | 0 | 0 | 0 | 0 | 0 |
| ENSG00000011638 | 1580 | 878 | 995 | 886 | 986 | 892 |
| ENSG00000262983 | 0 | 0 | 0 | 0 | 0 | 0 |
| ENSG00000103310 | 0 | 0 | 0 | 0 | 0 | 1 |
| ENSG00000175311 | 0 | 0 | 0 | 0 | 0 | 1 |
| ENSG00000103316 | 1 | 3 | 1 | 2 | 1 | 0 |
| ENSG00000283421 | 0 | 0 | 0 | 0 | 0 | 0 |
| ENSG00000189149 | 8 | 17 | 12 | 6 | 10 | 14 |
| ENSG00000257403 | 2 | 1 | 1 | 1 | 1 | 0 |
| ENSG00000158482 | 1 | 1 | 2 | 0 | 2 | 0 |
| ENSG00000276548 | 1 | 0 | 0 | 0 | 0 | 0 |
| ENSG00000169246 | 11 | 3 | 11 | 2 | 4 | 5 |
| ENSG00000180747 | 42 | 27 | 35 | 18 | 24 | 42 |
| ENSG00000265462 | 0 | 0 | 3 | 0 | 1 | 0 |
| ENSG00000258186 | 0 | 0 | 0 | 0 | 0 | 0 |
| ENSG00000273956 | 0 | 0 | 0 | 0 | 0 | 0 |
| ENSG00000257639 | 0 | 0 | 0 | 0 | 0 | 0 |
| ENSG00000271609 | 0 | 0 | 0 | 0 | 0 | 0 |
| ENSG00000252798 | 0 | 0 | 0 | 0 | 0 | 0 |
| ENSG00000197006 | 4743 | 2407 | 2712 | 2583 | 2865 | 2572 |
| ENSG00000261596 | 7 | 7 | 6 | 2 | 3 | 6 |
| ENSG00000140749 | 4 | 8 | 9 | 4 | 4 | 0 |
| ENSG00000207248 | 0 | 1 | 0 | 0 | 0 | 0 |
| ENSG00000207042 | 0 | 0 | 0 | 0 | 0 | 0 |
| ENSG00000155719 | 0 | 0 | 2 | 0 | 0 | 2 |
| ENSG00000260306 | 0 | 0 | 0 | 0 | 0 | 0 |
| ENSG00000248124 | 0 | 0 | 1 | 2 | 0 | 1 |
| ENSG00000276721 | 0 | 0 | 0 | 0 | 1 | 0 |
| ENSG00000185864 | 40 | 24 | 35 | 18 | 36 | 47 |
| ENSG00000185710 | 29 | 21 | 19 | 15 | 27 | 27 |
| ENSG00000257408 | 2 | 0 | 0 | 0 | 0 | 0 |

| | | | | | | |
|---|---|---|---|---|---|---|
| ENSG00000274460 | 33 | 18 | 19 | 10 | 33 | 13 |
| ENSG00000140740 | 5473 | 2830 | 3312 | 2599 | 3283 | 2967 |
| ENSG00000155714 | 0 | 0 | 0 | 0 | 1 | 0 |
| ENSG00000260395 | 0 | 0 | 0 | 0 | 0 | 0 |
| ENSG00000275445 | 1 | 0 | 0 | 1 | 1 | 2 |
| ENSG00000185716 | 457 | 201 | 253 | 184 | 235 | 221 |
| ENSG00000260277 | 7 | 3 | 3 | 0 | 5 | 0 |
| ENSG00000175267 | 2 | 0 | 1 | 6 | 2 | 1 |
| ENSG00000183921 | 1 | 0 | 1 | 5 | 3 | 0 |
| ENSG00000103319 | 1233 | 671 | 808 | 714 | 738 | 797 |
| ENSG00000261113 | 0 | 0 | 0 | 0 | 0 | 0 |
| ENSG00000058600 | 1776 | 855 | 1018 | 944 | 1056 | 1051 |
| ENSG00000140743 | 503 | 269 | 337 | 250 | 338 | 315 |
| ENSG00000260635 | 1 | 0 | 1 | 0 | 3 | 1 |
| ENSG00000260790 | 0 | 0 | 0 | 1 | 1 | 1 |
| ENSG00000230872 | 4 | 3 | 1 | 3 | 1 | 3 |
| ENSG00000257122 | 117 | 48 | 84 | 70 | 57 | 74 |
| ENSG00000237296 | 15 | 6 | 5 | 4 | 11 | 5 |
| ENSG00000243716 | 12 | 8 | 12 | 9 | 10 | 14 |
| ENSG00000277041 | 0 | 1 | 0 | 0 | 0 | 0 |
| ENSG00000257838 | 4 | 0 | 0 | 0 | 1 | 1 |
| ENSG00000260973 | 0 | 0 | 0 | 0 | 0 | 0 |
| ENSG00000260905 | 0 | 0 | 0 | 0 | 0 | 0 |
| ENSG00000283213 | 0 | 0 | 0 | 0 | 0 | 0 |
| ENSG00000122254 | 0 | 0 | 0 | 0 | 0 | 0 |
| ENSG00000261090 | 0 | 0 | 0 | 0 | 0 | 0 |
| ENSG00000103404 | 583 | 273 | 346 | 213 | 291 | 329 |
| ENSG00000260566 | 0 | 0 | 0 | 0 | 0 | 0 |
| ENSG00000280400 | 0 | 0 | 0 | 0 | 0 | 0 |
| ENSG00000285613 | 0 | 0 | 0 | 0 | 0 | 0 |
| ENSG00000166828 | 2 | 3 | 0 | 1 | 3 | 1 |
| ENSG00000260741 | 0 | 0 | 0 | 0 | 0 | 0 |
| ENSG00000168447 | 18 | 5 | 4 | 17 | 13 | 7 |
| ENSG00000168434 | 1679 | 853 | 1127 | 772 | 979 | 970 |
| ENSG00000280039 | 10 | 6 | 9 | 4 | 7 | 9 |
| ENSG00000212593 | 0 | 0 | 0 | 0 | 0 | 0 |
| ENSG00000259782 | 0 | 1 | 1 | 1 | 0 | 0 |
| ENSG00000260136 | 37 | 22 | 37 | 27 | 28 | 38 |
| ENSG00000103365 | 4184 | 1820 | 2586 | 1840 | 2136 | 2149 |
| ENSG00000261528 | 6 | 1 | 0 | 0 | 1 | 0 |
| ENSG00000103356 | 1967 | 951 | 1250 | 1147 | 1147 | 1167 |
| ENSG00000260247 | 0 | 0 | 0 | 0 | 0 | 0 |
| ENSG00000103353 | 7425 | 3296 | 4229 | 3378 | 3910 | 3931 |
| ENSG00000260751 | 30 | 22 | 32 | 13 | 13 | 32 |
| ENSG00000004779 | 4417 | 2157 | 2566 | 2862 | 2679 | 2628 |
| ENSG00000083093 | 619 | 320 | 327 | 225 | 456 | 376 |
| ENSG00000261723 | 7 | 2 | 1 | 1 | 2 | 0 |
| ENSG00000166847 | 6466 | 2950 | 3756 | 2828 | 3446 | 3358 |
| ENSG00000260482 | 0 | 0 | 0 | 0 | 0 | 0 |
| ENSG00000166851 | 1074 | 482 | 748 | 973 | 812 | 956 |

| | | | | | | |
|---|---|---|---|---|---|---|
| ENSG00000261266 | 0 | 1 | 0 | 1 | 0 | 0 |
| ENSG00000134398 | 0 | 1 | 0 | 0 | 0 | 0 |
| ENSG00000278912 | 0 | 0 | 0 | 0 | 0 | 0 |
| ENSG00000261583 | 0 | 0 | 0 | 0 | 0 | 0 |
| ENSG00000279756 | 0 | 0 | 0 | 0 | 0 | 0 |
| ENSG00000166869 | 0 | 0 | 0 | 0 | 0 | 0 |
| ENSG00000166501 | 0 | 0 | 0 | 0 | 0 | 0 |
| ENSG00000279276 | 0 | 0 | 0 | 0 | 0 | 0 |
| ENSG00000279202 | 0 | 0 | 0 | 0 | 0 | 0 |
| ENSG00000274466 | 0 | 0 | 0 | 0 | 0 | 0 |
| ENSG00000279554 | 0 | 0 | 0 | 0 | 0 | 0 |
| ENSG00000260264 | 0 | 0 | 0 | 0 | 0 | 0 |
| ENSG00000006116 | 0 | 0 | 0 | 0 | 0 | 0 |
| ENSG00000201541 | 0 | 0 | 0 | 0 | 0 | 0 |
| ENSG00000239081 | 0 | 0 | 0 | 0 | 0 | 0 |
| ENSG00000260072 | 0 | 1 | 0 | 0 | 0 | 0 |
| ENSG00000122257 | 1457 | 762 | 739 | 529 | 1097 | 793 |
| ENSG00000090905 | 2584 | 1145 | 1412 | 629 | 1320 | 1095 |
| ENSG00000224310 | 0 | 0 | 0 | 0 | 0 | 0 |
| ENSG00000261669 | 1 | 2 | 2 | 0 | 1 | 1 |
| ENSG00000158865 | 10 | 6 | 3 | 3 | 4 | 1 |
| ENSG00000140750 | 1085 | 585 | 712 | 583 | 629 | 631 |
| ENSG00000260714 | 0 | 0 | 0 | 0 | 0 | 0 |
| ENSG00000262587 | 113 | 58 | 59 | 38 | 40 | 63 |
| ENSG00000262155 | 38 | 16 | 16 | 9 | 13 | 16 |
| ENSG00000271105 | 25 | 3 | 4 | 1 | 7 | 5 |
| ENSG00000263326 | 27 | 10 | 13 | 4 | 10 | 5 |
| ENSG00000260448 | 15 | 8 | 7 | 11 | 11 | 9 |
| ENSG00000262700 | 2 | 2 | 1 | 0 | 1 | 4 |
| ENSG00000275494 | 38 | 30 | 42 | 59 | 36 | 38 |
| ENSG00000205629 | 1687 | 916 | 1124 | 1037 | 1031 | 1054 |
| ENSG00000260034 | 13 | 8 | 12 | 2 | 3 | 10 |
| ENSG00000266166 | 0 | 0 | 0 | 0 | 0 | 0 |
| ENSG00000103375 | 2 | 1 | 0 | 0 | 0 | 1 |
| ENSG00000155592 | 275 | 140 | 193 | 123 | 147 | 167 |
| ENSG00000259955 | 0 | 0 | 0 | 0 | 0 | 0 |
| ENSG00000274925 | 138 | 68 | 81 | 53 | 65 | 48 |
| ENSG00000261636 | 0 | 0 | 0 | 0 | 0 | 0 |
| ENSG00000260690 | 0 | 0 | 0 | 0 | 0 | 0 |
| ENSG00000182601 | 0 | 0 | 0 | 0 | 0 | 0 |
| ENSG00000285882 | 0 | 0 | 0 | 0 | 0 | 0 |
| ENSG00000260255 | 0 | 0 | 0 | 0 | 0 | 0 |
| ENSG00000265005 | 0 | 0 | 0 | 0 | 0 | 0 |
| ENSG00000199929 | 0 | 0 | 0 | 0 | 0 | 0 |
| ENSG00000230330 | 44 | 33 | 38 | 25 | 30 | 25 |
| ENSG00000237140 | 0 | 0 | 0 | 0 | 0 | 0 |
| ENSG00000231876 | 0 | 0 | 0 | 0 | 0 | 0 |
| ENSG00000228480 | 0 | 0 | 0 | 0 | 0 | 0 |
| ENSG00000236481 | 0 | 0 | 0 | 0 | 0 | 0 |
| ENSG00000260580 | 0 | 0 | 0 | 0 | 0 | 0 |

| | | | | | | |
|---|---|---|---|---|---|---|
| ENSG00000234186 | 0 | 0 | 0 | 0 | 0 | 0 |
| ENSG00000261482 | 0 | 0 | 0 | 0 | 0 | 0 |
| ENSG00000261557 | 1 | 2 | 4 | 2 | 3 | 1 |
| ENSG00000261045 | 0 | 0 | 0 | 0 | 0 | 0 |
| ENSG00000155666 | 148 | 72 | 111 | 89 | 69 | 78 |
| ENSG00000259940 | 27 | 7 | 9 | 9 | 8 | 9 |
| ENSG00000169189 | 1208 | 608 | 702 | 756 | 622 | 639 |
| ENSG00000245888 | 44 | 30 | 37 | 20 | 32 | 26 |
| ENSG00000274092 | 0 | 0 | 0 | 0 | 3 | 1 |
| ENSG00000077238 | 868 | 404 | 475 | 365 | 404 | 353 |
| ENSG00000103522 | 7 | 2 | 5 | 2 | 5 | 2 |
| ENSG00000259954 | 4 | 0 | 3 | 3 | 2 | 1 |
| ENSG00000077235 | 8863 | 3615 | 5428 | 3150 | 3207 | 3587 |
| ENSG00000047578 | 2222 | 814 | 1350 | 747 | 960 | 1032 |
| ENSG00000261736 | 0 | 0 | 0 | 0 | 0 | 0 |
| ENSG00000261329 | 5 | 1 | 0 | 0 | 1 | 2 |
| ENSG00000273553 | 0 | 0 | 0 | 0 | 0 | 0 |
| ENSG00000201371 | 0 | 0 | 0 | 0 | 0 | 0 |
| ENSG00000169181 | 2 | 2 | 0 | 1 | 3 | 1 |
| ENSG00000212382 | 0 | 0 | 0 | 0 | 0 | 0 |
| ENSG00000207266 | 0 | 0 | 0 | 0 | 0 | 0 |
| ENSG00000169180 | 4449 | 2067 | 2671 | 2248 | 2571 | 2610 |
| ENSG00000260452 | 0 | 0 | 1 | 0 | 0 | 0 |
| ENSG00000200652 | 0 | 0 | 0 | 0 | 0 | 0 |
| ENSG00000200138 | 0 | 0 | 1 | 0 | 0 | 0 |
| ENSG00000250982 | 2 | 1 | 2 | 2 | 3 | 2 |
| ENSG00000283662 | 2 | 1 | 1 | 6 | 5 | 1 |
| ENSG00000246465 | 48 | 27 | 32 | 63 | 54 | 50 |
| ENSG00000188322 | 1210 | 583 | 791 | 662 | 692 | 681 |
| ENSG00000277191 | 0 | 0 | 0 | 0 | 0 | 0 |
| ENSG00000198156 | 9 | 11 | 2 | 5 | 8 | 4 |
| ENSG00000271623 | 0 | 0 | 1 | 0 | 0 | 0 |
| ENSG00000205609 | 48 | 24 | 35 | 30 | 43 | 29 |
| ENSG00000275429 | 0 | 0 | 0 | 0 | 0 | 0 |
| ENSG00000261089 | 5 | 2 | 1 | 0 | 4 | 0 |
| ENSG00000275654 | 0 | 0 | 0 | 0 | 0 | 0 |
| ENSG00000270118 | 0 | 0 | 0 | 0 | 0 | 0 |
| ENSG00000261832 | 2 | 1 | 2 | 0 | 0 | 1 |
| ENSG00000233232 | 0 | 0 | 0 | 0 | 0 | 0 |
| ENSG00000188603 | 54 | 37 | 51 | 26 | 32 | 45 |
| ENSG00000184730 | 92 | 24 | 50 | 44 | 36 | 42 |
| ENSG00000197272 | 2 | 4 | 2 | 1 | 1 | 3 |
| ENSG00000176046 | 16 | 8 | 5 | 13 | 7 | 5 |
| ENSG00000271495 | 0 | 0 | 0 | 0 | 0 | 0 |
| ENSG00000275441 | 11 | 7 | 11 | 3 | 11 | 7 |
| ENSG00000176476 | 361 | 213 | 248 | 238 | 182 | 194 |
| ENSG00000197165 | 125 | 24 | 35 | 29 | 20 | 30 |
| ENSG00000278665 | 1 | 0 | 1 | 0 | 1 | 0 |
| ENSG00000278725 | 0 | 0 | 0 | 0 | 1 | 0 |
| ENSG00000196502 | 1598 | 694 | 1096 | 738 | 685 | 781 |

| | | | | | | |
|---|---|---|---|---|---|---|
| ENSG00000285020 | 0 | 1 | 1 | 0 | 1 | 1 |
| ENSG00000255524 | 0 | 0 | 0 | 0 | 0 | 0 |
| ENSG00000261419 | 6 | 5 | 0 | 1 | 2 | 2 |
| ENSG00000184110 | 606 | 296 | 391 | 328 | 377 | 394 |
| ENSG00000259982 | 2 | 1 | 3 | 1 | 0 | 1 |
| ENSG00000278340 | 0 | 0 | 0 | 0 | 0 | 0 |
| ENSG00000270424 | 0 | 2 | 1 | 1 | 1 | 0 |
| ENSG00000196993 | 13 | 0 | 1 | 0 | 5 | 4 |
| ENSG00000278528 | 0 | 0 | 0 | 0 | 0 | 0 |
| ENSG00000251417 | 39 | 14 | 26 | 13 | 21 | 20 |
| ENSG00000240634 | 2 | 4 | 3 | 2 | 3 | 4 |
| ENSG00000260796 | 4 | 2 | 3 | 0 | 6 | 2 |
| ENSG00000275807 | 51 | 13 | 17 | 24 | 19 | 13 |
| ENSG00000168488 | 3762 | 1762 | 2312 | 2192 | 2022 | 2179 |
| ENSG00000260570 | 0 | 0 | 0 | 0 | 0 | 0 |
| ENSG00000178952 | 10048 | 5041 | 6250 | 7703 | 6098 | 6422 |
| ENSG00000283845 | 0 | 0 | 0 | 0 | 0 | 0 |
| ENSG00000178188 | 2333 | 950 | 1478 | 1192 | 1150 | 1197 |
| ENSG00000261766 | 12 | 6 | 14 | 4 | 7 | 8 |
| ENSG00000196296 | 41 | 33 | 32 | 21 | 19 | 25 |
| ENSG00000260442 | 14 | 5 | 7 | 20 | 7 | 12 |
| ENSG00000252461 | 0 | 0 | 0 | 1 | 2 | 0 |
| ENSG00000177548 | 1306 | 656 | 891 | 781 | 799 | 651 |
| ENSG00000177455 | 0 | 0 | 0 | 2 | 0 | 0 |
| ENSG00000176953 | 2314 | 1139 | 1557 | 953 | 1222 | 1263 |
| ENSG00000260853 | 76 | 64 | 76 | 39 | 50 | 46 |
| ENSG00000284541 | 0 | 0 | 0 | 0 | 0 | 0 |
| ENSG00000260367 | 13 | 4 | 8 | 6 | 8 | 5 |
| ENSG00000169682 | 15 | 11 | 16 | 2 | 7 | 7 |
| ENSG00000261067 | 4 | 9 | 13 | 13 | 10 | 7 |
| ENSG00000213658 | 2 | 4 | 5 | 3 | 3 | 1 |
| ENSG00000261552 | 0 | 0 | 0 | 0 | 0 | 0 |
| ENSG00000196796 | 1 | 0 | 2 | 0 | 1 | 0 |
| ENSG00000260908 | 9 | 8 | 15 | 7 | 8 | 3 |
| ENSG00000103472 | 0 | 0 | 0 | 0 | 0 | 2 |
| ENSG00000284671 | 0 | 0 | 0 | 0 | 0 | 0 |
| ENSG00000284649 | 0 | 1 | 0 | 0 | 0 | 0 |
| ENSG00000284685 | 0 | 0 | 0 | 2 | 0 | 0 |
| ENSG00000260517 | 0 | 0 | 3 | 0 | 0 | 0 |
| ENSG00000279106 | 0 | 0 | 0 | 0 | 0 | 0 |
| ENSG00000259807 | 0 | 0 | 0 | 0 | 0 | 0 |
| ENSG00000273582 | 0 | 0 | 0 | 0 | 0 | 1 |
| ENSG00000260953 | 0 | 0 | 1 | 0 | 0 | 0 |
| ENSG00000277999 | 0 | 0 | 3 | 0 | 0 | 0 |
| ENSG00000198106 | 0 | 0 | 0 | 0 | 0 | 4 |
| ENSG00000260413 | 0 | 0 | 0 | 0 | 0 | 0 |
| ENSG00000271699 | 0 | 0 | 0 | 0 | 0 | 0 |
| ENSG00000278078 | 0 | 0 | 0 | 0 | 0 | 0 |
| ENSG00000254206 | 8 | 0 | 17 | 6 | 6 | 5 |
| ENSG00000254634 | 2 | 6 | 4 | 3 | 2 | 6 |

| | | | | | | |
|---|---|---|---|---|---|---|
| ENSG00000183336 | 0 | 0 | 0 | 1 | 0 | 0 |
| ENSG00000261740 | 14 | 12 | 11 | 12 | 18 | 11 |
| ENSG00000260335 | 1 | 0 | 0 | 0 | 0 | 0 |
| ENSG00000181625 | 0 | 0 | 0 | 3 | 1 | 0 |
| ENSG00000260280 | 0 | 0 | 0 | 0 | 0 | 0 |
| ENSG00000213648 | 0 | 0 | 0 | 0 | 0 | 0 |
| ENSG00000258150 | 0 | 0 | 0 | 0 | 0 | 0 |
| ENSG00000279228 | 0 | 0 | 0 | 0 | 0 | 0 |
| ENSG00000257691 | 0 | 0 | 0 | 0 | 0 | 0 |
| ENSG00000257506 | 0 | 0 | 0 | 0 | 0 | 0 |
| ENSG00000169203 | 12 | 13 | 8 | 2 | 3 | 14 |
| ENSG00000205534 | 67 | 34 | 36 | 20 | 29 | 46 |
| ENSG00000279583 | 4 | 4 | 2 | 7 | 3 | 6 |
| ENSG00000266758 | 0 | 1 | 0 | 0 | 0 | 0 |
| ENSG00000260727 | 8 | 3 | 5 | 9 | 4 | 6 |
| ENSG00000260133 | 0 | 0 | 0 | 0 | 0 | 0 |
| ENSG00000260514 | 0 | 0 | 0 | 0 | 0 | 0 |
| ENSG00000197471 | 0 | 1 | 0 | 0 | 0 | 1 |
| ENSG00000277669 | 0 | 0 | 0 | 0 | 0 | 0 |
| ENSG00000103485 | 3314 | 1730 | 2198 | 1686 | 1598 | 1705 |
| ENSG00000222375 | 0 | 0 | 0 | 0 | 0 | 0 |
| ENSG00000185905 | 0 | 0 | 0 | 0 | 1 | 3 |
| ENSG00000260719 | 0 | 0 | 1 | 0 | 0 | 0 |
| ENSG00000174992 | 6 | 3 | 15 | 1 | 4 | 6 |
| ENSG00000079616 | 3298 | 1661 | 2326 | 2172 | 1976 | 2018 |
| ENSG00000275857 | 0 | 0 | 0 | 0 | 0 | 0 |
| ENSG00000103495 | 2221 | 1573 | 1998 | 1935 | 1681 | 1730 |
| ENSG00000259952 | 0 | 0 | 0 | 0 | 0 | 0 |
| ENSG00000238045 | 332 | 152 | 167 | 156 | 163 | 188 |
| ENSG00000280607 | 0 | 0 | 0 | 0 | 0 | 0 |
| ENSG00000167371 | 229 | 102 | 154 | 68 | 75 | 74 |
| ENSG00000280893 | 0 | 0 | 0 | 0 | 0 | 0 |
| ENSG00000280789 | 327 | 185 | 263 | 251 | 205 | 209 |
| ENSG00000281348 | 0 | 2 | 1 | 3 | 2 | 0 |
| ENSG00000013364 | 2705 | 1302 | 1806 | 1648 | 1459 | 1517 |
| ENSG00000103502 | 5928 | 2898 | 3491 | 3606 | 3120 | 3133 |
| ENSG00000278713 | 0 | 0 | 0 | 0 | 0 | 0 |
| ENSG00000214725 | 0 | 1 | 2 | 0 | 0 | 1 |
| ENSG00000174938 | 3649 | 1545 | 2214 | 1994 | 1522 | 1648 |
| ENSG00000174939 | 180 | 87 | 120 | 170 | 127 | 150 |
| ENSG00000174943 | 1670 | 867 | 1029 | 1011 | 973 | 939 |
| ENSG00000279789 | 199 | 106 | 128 | 54 | 113 | 92 |
| ENSG00000247735 | 134 | 80 | 91 | 57 | 98 | 63 |
| ENSG00000149932 | 4070 | 1871 | 2522 | 2707 | 2225 | 2156 |
| ENSG00000149930 | 7649 | 3301 | 4533 | 3829 | 3696 | 3632 |
| ENSG00000149929 | 663 | 391 | 472 | 424 | 358 | 445 |
| ENSG00000169592 | 2442 | 1193 | 1683 | 1680 | 1509 | 1448 |
| ENSG00000149927 | 491 | 189 | 283 | 244 | 245 | 284 |
| ENSG00000167194 | 2 | 1 | 0 | 0 | 2 | 1 |
| ENSG00000149926 | 65 | 33 | 52 | 26 | 35 | 32 |

| | | | | | | |
|---|---|---|---|---|---|---|
| ENSG00000285043 | 71 | 44 | 66 | 43 | 33 | 34 |
| ENSG00000149925 | 2591 | 1949 | 2689 | 2559 | 1995 | 2445 |
| ENSG00000274904 | 0 | 0 | 0 | 0 | 0 | 0 |
| ENSG00000149923 | 12393 | 5756 | 7240 | 7087 | 6513 | 5988 |
| ENSG00000149922 | 210 | 61 | 121 | 92 | 91 | 83 |
| ENSG00000090238 | 5804 | 2528 | 3623 | 2482 | 2332 | 2417 |
| ENSG00000250616 | 78 | 34 | 76 | 45 | 42 | 45 |
| ENSG00000102886 | 1349 | 815 | 1131 | 669 | 562 | 619 |
| ENSG00000261367 | 9 | 5 | 7 | 5 | 3 | 1 |
| ENSG00000275371 | 24 | 15 | 16 | 8 | 8 | 2 |
| ENSG00000102882 | 12484 | 5945 | 7780 | 6409 | 5689 | 5895 |
| ENSG00000102879 | 974 | 465 | 583 | 578 | 414 | 481 |
| ENSG00000261416 | 2 | 0 | 2 | 0 | 2 | 1 |
| ENSG00000169627 | 42 | 25 | 25 | 29 | 33 | 33 |
| ENSG00000132207 | 6 | 1 | 4 | 0 | 3 | 2 |
| ENSG00000213599 | 0 | 1 | 1 | 1 | 1 | 1 |
| ENSG00000261052 | 14 | 10 | 15 | 16 | 17 | 15 |
| ENSG00000258130 | 13 | 10 | 14 | 10 | 4 | 15 |
| ENSG00000278887 | 0 | 0 | 0 | 0 | 0 | 0 |
| ENSG00000261444 | 0 | 0 | 0 | 1 | 0 | 0 |
| ENSG00000261203 | 0 | 0 | 0 | 0 | 0 | 0 |
| ENSG00000198064 | 1 | 1 | 2 | 0 | 1 | 1 |
| ENSG00000183604 | 64 | 32 | 38 | 17 | 28 | 29 |
| ENSG00000273724 | 6 | 11 | 6 | 3 | 7 | 6 |
| ENSG00000169217 | 6026 | 2861 | 3601 | 3820 | 3412 | 3371 |
| ENSG00000260219 | 137 | 70 | 101 | 79 | 58 | 102 |
| ENSG00000169221 | 2854 | 1312 | 1662 | 1741 | 1521 | 1607 |
| ENSG00000274653 | 13 | 9 | 19 | 14 | 9 | 14 |
| ENSG00000180209 | 7 | 1 | 4 | 7 | 6 | 0 |
| ENSG00000180035 | 1132 | 488 | 699 | 756 | 603 | 663 |
| ENSG00000180096 | 73 | 34 | 47 | 47 | 42 | 31 |
| ENSG00000280137 | 0 | 0 | 0 | 0 | 0 | 0 |
| ENSG00000179965 | 310 | 126 | 180 | 221 | 158 | 181 |
| ENSG00000199787 | 0 | 0 | 0 | 0 | 1 | 0 |
| ENSG00000179958 | 2982 | 1622 | 1994 | 2458 | 1820 | 2023 |
| ENSG00000179918 | 10719 | 5122 | 6277 | 6005 | 6044 | 6402 |
| ENSG00000202476 | 0 | 0 | 0 | 0 | 0 | 0 |
| ENSG00000222701 | 0 | 0 | 0 | 0 | 0 | 0 |
| ENSG00000005844 | 29 | 17 | 19 | 31 | 21 | 26 |
| ENSG00000261346 | 0 | 0 | 0 | 0 | 0 | 0 |
| ENSG00000260487 | 0 | 0 | 0 | 0 | 0 | 1 |
| ENSG00000251706 | 0 | 0 | 0 | 0 | 0 | 0 |
| ENSG00000261332 | 0 | 0 | 0 | 0 | 0 | 0 |
| ENSG00000266305 | 0 | 0 | 0 | 0 | 0 | 0 |
| ENSG00000169957 | 2871 | 1322 | 1684 | 1799 | 1560 | 1607 |
| ENSG00000261459 | 0 | 0 | 0 | 0 | 0 | 0 |
| ENSG00000278922 | 189 | 65 | 95 | 78 | 91 | 87 |
| ENSG00000169955 | 156 | 74 | 96 | 89 | 89 | 84 |
| ENSG00000260869 | 0 | 0 | 0 | 0 | 0 | 0 |
| ENSG00000235560 | 99 | 43 | 62 | 55 | 72 | 79 |

| | | | | | | |
|---|---|---|---|---|---|---|
| ENSG00000260494 | 3 | 0 | 1 | 0 | 0 | 0 |
| ENSG00000169951 | 616 | 277 | 378 | 338 | 248 | 344 |
| ENSG00000229809 | 661 | 288 | 373 | 401 | 269 | 337 |
| ENSG00000239791 | 0 | 2 | 4 | 1 | 1 | 2 |
| ENSG00000197162 | 524 | 325 | 337 | 295 | 287 | 319 |
| ENSG00000260167 | 13 | 12 | 4 | 12 | 6 | 7 |
| ENSG00000261588 | 11 | 2 | 3 | 3 | 1 | 4 |
| ENSG00000156853 | 630 | 265 | 360 | 309 | 295 | 341 |
| ENSG00000260678 | 2 | 0 | 4 | 0 | 0 | 0 |
| ENSG00000261680 | 0 | 0 | 0 | 0 | 0 | 0 |
| ENSG00000260113 | 0 | 0 | 0 | 0 | 1 | 0 |
| ENSG00000156858 | 1589 | 743 | 1024 | 1029 | 912 | 850 |
| ENSG00000156860 | 4109 | 1825 | 2366 | 2246 | 2093 | 2118 |
| ENSG00000252074 | 0 | 0 | 0 | 0 | 0 | 0 |
| ENSG00000261840 | 47 | 28 | 45 | 30 | 20 | 21 |
| ENSG00000080603 | 194 | 113 | 175 | 90 | 113 | 131 |
| ENSG00000222287 | 0 | 1 | 0 | 0 | 0 | 0 |
| ENSG00000282034 | 75 | 33 | 41 | 27 | 52 | 57 |
| ENSG00000206755 | 0 | 0 | 0 | 0 | 0 | 0 |
| ENSG00000281991 | 0 | 0 | 0 | 0 | 0 | 0 |
| ENSG00000260899 | 0 | 0 | 3 | 3 | 1 | 4 |
| ENSG00000156873 | 1755 | 862 | 1145 | 1079 | 1018 | 987 |
| ENSG00000196118 | 80 | 28 | 45 | 30 | 27 | 23 |
| ENSG00000103549 | 2992 | 1306 | 1807 | 1579 | 1554 | 1639 |
| ENSG00000280211 | 17 | 8 | 4 | 5 | 5 | 6 |
| ENSG00000102870 | 2177 | 884 | 1167 | 1164 | 1112 | 1127 |
| ENSG00000274848 | 0 | 0 | 1 | 0 | 0 | 0 |
| ENSG00000260082 | 2 | 0 | 0 | 0 | 2 | 1 |
| ENSG00000274678 | 0 | 1 | 2 | 2 | 0 | 0 |
| ENSG00000099385 | 854 | 604 | 775 | 796 | 654 | 653 |
| ENSG00000260083 | 24 | 19 | 21 | 18 | 21 | 18 |
| ENSG00000265991 | 0 | 0 | 0 | 0 | 0 | 0 |
| ENSG00000262721 | 0 | 0 | 0 | 0 | 0 | 0 |
| ENSG00000211591 | 0 | 0 | 0 | 0 | 0 | 0 |
| ENSG00000150281 | 13 | 4 | 3 | 8 | 1 | 11 |
| ENSG00000230447 | 0 | 0 | 0 | 0 | 0 | 0 |
| ENSG00000260852 | 262 | 140 | 213 | 142 | 171 | 169 |
| ENSG00000099364 | 2104 | 1015 | 1229 | 1327 | 1294 | 1205 |
| ENSG00000261487 | 0 | 0 | 0 | 0 | 1 | 0 |
| ENSG00000175938 | 560 | 282 | 373 | 305 | 248 | 311 |
| ENSG00000275263 | 3 | 0 | 1 | 3 | 3 | 0 |
| ENSG00000099381 | 1434 | 666 | 883 | 799 | 686 | 864 |
| ENSG00000279196 | 20 | 18 | 7 | 14 | 21 | 13 |
| ENSG00000099377 | 475 | 232 | 312 | 359 | 254 | 257 |
| ENSG00000099365 | 53 | 11 | 21 | 8 | 12 | 23 |
| ENSG00000103496 | 2171 | 1003 | 1238 | 1180 | 1228 | 1200 |
| ENSG00000260911 | 5 | 3 | 8 | 6 | 6 | 5 |
| ENSG00000232748 | 6 | 3 | 10 | 9 | 15 | 14 |
| ENSG00000167394 | 246 | 136 | 141 | 134 | 121 | 147 |
| ENSG00000261124 | 0 | 1 | 1 | 1 | 1 | 0 |

| | | | | | | |
|---|---|---|---|---|---|---|
| ENSG00000167395 | 1284 | 574 | 821 | 738 | 728 | 727 |
| ENSG00000151006 | 44 | 18 | 16 | 13 | 28 | 25 |
| ENSG00000255439 | 0 | 1 | 0 | 2 | 0 | 0 |
| ENSG00000167397 | 279 | 166 | 222 | 271 | 222 | 232 |
| ENSG00000280160 | 2 | 4 | 6 | 4 | 1 | 3 |
| ENSG00000103507 | 2168 | 960 | 1276 | 1298 | 1038 | 1088 |
| ENSG00000103510 | 1242 | 602 | 869 | 652 | 650 | 697 |
| ENSG00000262766 | 18 | 9 | 20 | 8 | 15 | 14 |
| ENSG00000278133 | 258 | 137 | 200 | 116 | 138 | 160 |
| ENSG00000261385 | 0 | 0 | 0 | 0 | 0 | 0 |
| ENSG00000052344 | 5094 | 2508 | 3444 | 2941 | 2446 | 2671 |
| ENSG00000178226 | 22 | 10 | 25 | 12 | 13 | 16 |
| ENSG00000263343 | 0 | 0 | 0 | 0 | 0 | 0 |
| ENSG00000262366 | 0 | 0 | 0 | 0 | 0 | 0 |
| ENSG00000089280 | 7524 | 3370 | 5216 | 3817 | 4049 | 5127 |
| ENSG00000260304 | 0 | 0 | 0 | 0 | 0 | 0 |
| ENSG00000260060 | 2 | 1 | 4 | 0 | 0 | 0 |
| ENSG00000103490 | 459 | 272 | 315 | 355 | 316 | 267 |
| ENSG00000261359 | 0 | 1 | 1 | 1 | 0 | 0 |
| ENSG00000177238 | 0 | 2 | 4 | 0 | 0 | 1 |
| ENSG00000169900 | 0 | 1 | 2 | 0 | 1 | 0 |
| ENSG00000169896 | 10 | 6 | 7 | 9 | 24 | 14 |
| ENSG00000140678 | 0 | 1 | 1 | 0 | 0 | 0 |
| ENSG00000261245 | 0 | 0 | 0 | 0 | 0 | 0 |
| ENSG00000156886 | 0 | 0 | 0 | 0 | 0 | 0 |
| ENSG00000260757 | 0 | 0 | 0 | 0 | 0 | 1 |
| ENSG00000156885 | 0 | 0 | 0 | 0 | 0 | 0 |
| ENSG00000277543 | 0 | 0 | 0 | 0 | 0 | 0 |
| ENSG00000176723 | 16 | 9 | 16 | 13 | 19 | 16 |
| ENSG00000261474 | 47 | 25 | 33 | 46 | 49 | 42 |
| ENSG00000260267 | 161 | 114 | 133 | 85 | 89 | 109 |
| ENSG00000140691 | 389 | 152 | 208 | 199 | 154 | 184 |
| ENSG00000280132 | 12 | 2 | 2 | 3 | 4 | 6 |
| ENSG00000140682 | 22 | 5 | 10 | 3 | 13 | 6 |
| ENSG00000140675 | 2 | 0 | 1 | 4 | 3 | 2 |
| ENSG00000260740 | 1 | 0 | 1 | 0 | 3 | 0 |
| ENSG00000140688 | 1523 | 748 | 996 | 902 | 950 | 863 |
| ENSG00000260625 | 21 | 13 | 11 | 13 | 6 | 14 |
| ENSG00000169877 | 0 | 1 | 1 | 0 | 0 | 1 |
| ENSG00000261648 | 0 | 0 | 0 | 0 | 0 | 0 |
| ENSG00000213547 | 0 | 0 | 0 | 0 | 0 | 0 |
| ENSG00000261475 | 0 | 0 | 0 | 0 | 0 | 0 |
| ENSG00000260883 | 0 | 0 | 0 | 0 | 0 | 0 |
| ENSG00000261741 | 0 | 0 | 0 | 0 | 0 | 0 |
| ENSG00000278885 | 0 | 0 | 0 | 0 | 0 | 0 |
| ENSG00000261614 | 0 | 0 | 0 | 0 | 0 | 1 |
| ENSG00000260010 | 0 | 0 | 0 | 0 | 0 | 1 |
| ENSG00000237185 | 0 | 0 | 0 | 0 | 0 | 0 |
| ENSG00000260631 | 0 | 0 | 0 | 0 | 0 | 0 |
| ENSG00000260472 | 0 | 0 | 0 | 0 | 0 | 0 |

| | | | | | | |
|---|---|---|---|---|---|---|
| ENSG00000260568 | 0 | 0 | 0 | 0 | 0 | 0 |
| ENSG00000131797 | 846 | 354 | 523 | 332 | 480 | 434 |
| ENSG00000276867 | 4 | 1 | 3 | 3 | 5 | 3 |
| ENSG00000261731 | 0 | 0 | 0 | 0 | 0 | 0 |
| ENSG00000197302 | 690 | 364 | 388 | 255 | 543 | 368 |
| ENSG00000259810 | 4 | 1 | 2 | 0 | 0 | 1 |
| ENSG00000260722 | 0 | 0 | 0 | 0 | 0 | 0 |
| ENSG00000261457 | 0 | 0 | 0 | 0 | 0 | 0 |
| ENSG00000180663 | 0 | 0 | 0 | 0 | 0 | 0 |
| ENSG00000261284 | 0 | 0 | 0 | 0 | 0 | 0 |
| ENSG00000259874 | 0 | 0 | 0 | 0 | 0 | 0 |
| ENSG00000185947 | 88 | 93 | 55 | 42 | 199 | 70 |
| ENSG00000259950 | 0 | 0 | 0 | 0 | 0 | 0 |
| ENSG00000261289 | 0 | 0 | 0 | 0 | 1 | 0 |
| ENSG00000197476 | 0 | 0 | 0 | 0 | 0 | 0 |
| ENSG00000260218 | 0 | 0 | 0 | 0 | 0 | 0 |
| ENSG00000260628 | 0 | 0 | 0 | 0 | 0 | 0 |
| ENSG00000261727 | 0 | 0 | 0 | 0 | 0 | 0 |
| ENSG00000261704 | 0 | 0 | 0 | 0 | 0 | 0 |
| ENSG00000223931 | 0 | 0 | 0 | 0 | 0 | 0 |
| ENSG00000260048 | 0 | 0 | 0 | 0 | 0 | 0 |
| ENSG00000270472 | 0 | 0 | 0 | 0 | 0 | 0 |
| ENSG00000230267 | 0 | 0 | 0 | 1 | 0 | 0 |
| ENSG00000260584 | 0 | 0 | 0 | 0 | 0 | 0 |
| ENSG00000279997 | 0 | 0 | 0 | 0 | 0 | 0 |
| ENSG00000260847 | 0 | 0 | 0 | 0 | 0 | 0 |
| ENSG00000260649 | 0 | 0 | 0 | 0 | 0 | 0 |
| ENSG00000260540 | 0 | 0 | 0 | 0 | 0 | 0 |
| ENSG00000260344 | 0 | 0 | 0 | 0 | 0 | 0 |
| ENSG00000260575 | 0 | 0 | 0 | 0 | 0 | 0 |
| ENSG00000205456 | 0 | 0 | 0 | 0 | 0 | 0 |
| ENSG00000260402 | 0 | 0 | 0 | 0 | 0 | 0 |
| ENSG00000259822 | 0 | 0 | 0 | 0 | 0 | 0 |
| ENSG00000261127 | 0 | 0 | 0 | 0 | 0 | 0 |
| ENSG00000260516 | 0 | 0 | 0 | 0 | 0 | 0 |
| ENSG00000260327 | 0 | 0 | 0 | 0 | 0 | 0 |
| ENSG00000260866 | 0 | 0 | 0 | 0 | 0 | 0 |
| ENSG00000260307 | 0 | 0 | 0 | 0 | 0 | 0 |
| ENSG00000261541 | 0 | 0 | 0 | 0 | 0 | 0 |
| ENSG00000260662 | 0 | 0 | 0 | 0 | 0 | 0 |
| ENSG00000261233 | 0 | 0 | 0 | 0 | 0 | 0 |
| ENSG00000259966 | 0 | 0 | 0 | 0 | 0 | 0 |
| ENSG00000262187 | 0 | 0 | 0 | 0 | 0 | 0 |
| ENSG00000261111 | 0 | 0 | 0 | 0 | 0 | 0 |
| ENSG00000261569 | 0 | 0 | 0 | 0 | 0 | 0 |
| ENSG00000261263 | 0 | 0 | 0 | 0 | 0 | 0 |
| ENSG00000260311 | 0 | 0 | 0 | 0 | 0 | 0 |
| ENSG00000183632 | 0 | 0 | 0 | 0 | 0 | 0 |
| ENSG00000260974 | 0 | 0 | 0 | 0 | 0 | 0 |
| ENSG00000261391 | 0 | 0 | 0 | 0 | 0 | 0 |

| | | | | | | |
|---|---|---|---|---|---|---|
| ENSG00000260864 | 0 | 0 | 0 | 0 | 0 | 0 |
| ENSG00000260845 | 0 | 0 | 0 | 0 | 0 | 0 |
| ENSG00000261108 | 0 | 0 | 0 | 0 | 0 | 0 |
| ENSG00000260644 | 0 | 0 | 0 | 0 | 0 | 0 |
| ENSG00000278950 | 0 | 0 | 0 | 0 | 0 | 0 |
| ENSG00000279780 | 0 | 0 | 0 | 0 | 0 | 0 |
| ENSG00000279795 | 0 | 0 | 0 | 0 | 0 | 0 |
| ENSG00000260158 | 0 | 0 | 0 | 0 | 0 | 0 |
| ENSG00000261719 | 0 | 0 | 0 | 0 | 0 | 0 |
| ENSG00000259934 | 0 | 0 | 0 | 0 | 0 | 0 |
| ENSG00000259303 | 0 | 0 | 0 | 0 | 0 | 0 |
| ENSG00000260047 | 0 | 0 | 1 | 0 | 1 | 0 |
| ENSG00000214617 | 0 | 1 | 0 | 0 | 0 | 0 |
| ENSG00000214614 | 0 | 0 | 0 | 0 | 0 | 0 |
| ENSG00000261834 | 0 | 0 | 0 | 0 | 0 | 0 |
| ENSG00000270864 | 0 | 0 | 0 | 0 | 0 | 0 |
| ENSG00000277304 | 0 | 0 | 0 | 0 | 0 | 0 |
| ENSG00000260610 | 0 | 0 | 0 | 0 | 0 | 0 |
| ENSG00000261259 | 0 | 0 | 0 | 0 | 0 | 0 |
| ENSG00000259852 | 0 | 0 | 0 | 0 | 0 | 0 |
| ENSG00000233732 | 0 | 0 | 0 | 0 | 0 | 0 |
| ENSG00000260921 | 0 | 0 | 0 | 0 | 0 | 0 |
| ENSG00000259997 | 0 | 0 | 0 | 0 | 0 | 0 |
| ENSG00000271130 | 0 | 0 | 0 | 0 | 0 | 0 |
| ENSG00000260141 | 0 | 0 | 0 | 0 | 0 | 0 |
| ENSG00000260900 | 0 | 0 | 0 | 0 | 0 | 0 |
| ENSG00000261599 | 0 | 0 | 0 | 0 | 0 | 0 |
| ENSG00000261682 | 0 | 0 | 0 | 0 | 0 | 0 |
| ENSG00000259800 | 0 | 0 | 0 | 0 | 0 | 0 |
| ENSG00000261536 | 0 | 0 | 0 | 0 | 0 | 0 |
| ENSG00000260414 | 0 | 0 | 0 | 0 | 0 | 0 |
| ENSG00000260419 | 0 | 0 | 0 | 0 | 0 | 0 |
| ENSG00000205457 | 0 | 0 | 0 | 0 | 0 | 0 |
| ENSG00000260827 | 0 | 0 | 0 | 0 | 0 | 0 |
| ENSG00000261507 | 0 | 0 | 0 | 0 | 0 | 0 |
| ENSG00000275034 | 0 | 0 | 0 | 0 | 0 | 0 |
| ENSG00000282973 | 0 | 0 | 0 | 0 | 0 | 0 |
| ENSG00000261509 | 0 | 0 | 0 | 0 | 0 | 0 |
| ENSG00000261009 | 0 | 0 | 0 | 0 | 0 | 0 |
| ENSG00000283110 | 0 | 0 | 0 | 0 | 0 | 0 |
| ENSG00000278848 | 0 | 0 | 0 | 0 | 0 | 0 |
| ENSG00000282927 | 0 | 0 | 0 | 0 | 0 | 0 |
| ENSG00000262090 | 0 | 0 | 0 | 0 | 0 | 0 |
| ENSG00000260626 | 0 | 0 | 0 | 0 | 0 | 0 |
| ENSG00000261200 | 0 | 0 | 0 | 0 | 0 | 0 |
| ENSG00000261466 | 0 | 0 | 0 | 0 | 0 | 0 |
| ENSG00000261405 | 3 | 1 | 0 | 0 | 0 | 0 |
| ENSG00000259882 | 9 | 4 | 9 | 3 | 6 | 2 |
| ENSG00000260518 | 0 | 2 | 2 | 0 | 1 | 0 |
| ENSG00000282924 | 0 | 0 | 0 | 0 | 0 | 0 |

| | | | | | | |
|---|---|---|---|---|---|---|
| ENSG00000261580 | 0 | 0 | 0 | 0 | 0 | 0 |
| ENSG00000260308 | 0 | 0 | 0 | 0 | 0 | 0 |
| ENSG00000270467 | 0 | 0 | 0 | 0 | 0 | 0 |
| ENSG00000261153 | 0 | 0 | 0 | 0 | 0 | 0 |
| ENSG00000271178 | 0 | 0 | 0 | 0 | 0 | 0 |
| ENSG00000259680 | 0 | 0 | 0 | 0 | 0 | 0 |
| ENSG00000270924 | 0 | 0 | 0 | 0 | 0 | 0 |
| ENSG00000271691 | 0 | 0 | 0 | 0 | 0 | 0 |
| ENSG00000260312 | 0 | 0 | 0 | 0 | 0 | 0 |
| ENSG00000270318 | 0 | 0 | 0 | 0 | 0 | 0 |
| ENSG00000260525 | 0 | 0 | 0 | 0 | 0 | 0 |
| ENSG00000260781 | 0 | 0 | 0 | 0 | 0 | 0 |
| ENSG00000271620 | 0 | 0 | 0 | 0 | 0 | 0 |
| ENSG00000261607 | 0 | 0 | 0 | 0 | 0 | 0 |
| ENSG00000259842 | 0 | 0 | 0 | 0 | 0 | 0 |
| ENSG00000205452 | 0 | 0 | 0 | 0 | 0 | 0 |
| ENSG00000270401 | 0 | 0 | 0 | 0 | 0 | 0 |
| ENSG00000198555 | 0 | 0 | 0 | 0 | 0 | 0 |
| ENSG00000261217 | 0 | 0 | 0 | 0 | 0 | 0 |
| ENSG00000259990 | 0 | 0 | 0 | 0 | 0 | 0 |
| ENSG00000283065 | 0 | 0 | 0 | 0 | 0 | 0 |
| ENSG00000261197 | 0 | 0 | 0 | 0 | 0 | 0 |
| ENSG00000262561 | 0 | 0 | 0 | 0 | 0 | 0 |
| ENSG00000261440 | 0 | 0 | 0 | 0 | 0 | 0 |
| ENSG00000260087 | 0 | 0 | 0 | 0 | 0 | 0 |
| ENSG00000259987 | 0 | 0 | 0 | 0 | 0 | 0 |
| ENSG00000260207 | 0 | 0 | 0 | 0 | 0 | 0 |
| ENSG00000256642 | 0 | 0 | 0 | 0 | 0 | 0 |
| ENSG00000200434 | 0 | 0 | 0 | 0 | 0 | 0 |
| ENSG00000279800 | 0 | 0 | 2 | 0 | 3 | 0 |
| ENSG00000284209 | 0 | 0 | 0 | 0 | 0 | 0 |
| ENSG00000279165 | 0 | 0 | 0 | 0 | 0 | 0 |
| ENSG00000260854 | 0 | 0 | 0 | 0 | 0 | 0 |
| ENSG00000179755 | 0 | 0 | 0 | 0 | 0 | 0 |
| ENSG00000261566 | 0 | 0 | 0 | 0 | 0 | 0 |
| ENSG00000262885 | 0 | 0 | 0 | 0 | 0 | 0 |
| ENSG00000216671 | 0 | 0 | 0 | 0 | 0 | 0 |
| ENSG00000278138 | 0 | 0 | 0 | 0 | 0 | 0 |
| ENSG00000261053 | 0 | 0 | 0 | 0 | 0 | 0 |
| ENSG00000260809 | 0 | 1 | 0 | 1 | 0 | 0 |
| ENSG00000259979 | 0 | 0 | 0 | 0 | 0 | 0 |
| ENSG00000260724 | 0 | 0 | 0 | 0 | 0 | 0 |
| ENSG00000261800 | 0 | 0 | 0 | 0 | 0 | 0 |
| ENSG00000280180 | 0 | 0 | 0 | 0 | 0 | 0 |
| ENSG00000261461 | 12 | 0 | 1 | 6 | 1 | 4 |
| ENSG00000261608 | 0 | 0 | 0 | 0 | 0 | 0 |
| ENSG00000260073 | 0 | 0 | 0 | 0 | 0 | 0 |
| ENSG00000260590 | 0 | 0 | 0 | 0 | 0 | 0 |
| ENSG00000261398 | 0 | 0 | 0 | 0 | 0 | 0 |
| ENSG00000261274 | 0 | 0 | 0 | 0 | 0 | 0 |

| | | | | | | |
|---|---|---|---|---|---|---|
| ENSG00000260958 | 0 | 0 | 0 | 0 | 0 | 0 |
| ENSG00000260967 | 0 | 0 | 0 | 0 | 0 | 0 |
| ENSG00000260984 | 0 | 0 | 0 | 0 | 0 | 0 |
| ENSG00000259897 | 0 | 0 | 0 | 0 | 0 | 0 |
| ENSG00000261010 | 0 | 0 | 0 | 0 | 0 | 0 |
| ENSG00000261752 | 0 | 0 | 0 | 0 | 0 | 0 |
| ENSG00000260846 | 0 | 0 | 0 | 0 | 0 | 0 |
| ENSG00000261486 | 0 | 0 | 0 | 0 | 0 | 0 |
| ENSG00000260427 | 0 | 0 | 0 | 0 | 0 | 0 |
| ENSG00000260480 | 0 | 0 | 0 | 0 | 0 | 0 |
| ENSG00000259841 | 0 | 0 | 0 | 0 | 0 | 0 |
| ENSG00000260291 | 0 | 0 | 0 | 0 | 0 | 0 |
| ENSG00000261689 | 0 | 0 | 0 | 0 | 0 | 0 |
| ENSG00000260598 | 0 | 0 | 0 | 0 | 0 | 0 |
| ENSG00000260812 | 0 | 0 | 0 | 0 | 0 | 0 |
| ENSG00000260680 | 0 | 0 | 0 | 0 | 0 | 0 |
| ENSG00000261299 | 0 | 0 | 0 | 0 | 0 | 0 |
| ENSG00000278681 | 0 | 0 | 0 | 0 | 0 | 0 |
| ENSG00000214581 | 0 | 0 | 1 | 0 | 0 | 0 |
| ENSG00000261711 | 0 | 0 | 0 | 0 | 0 | 0 |
| ENSG00000260153 | 0 | 0 | 0 | 0 | 0 | 0 |
| ENSG00000259836 | 0 | 0 | 0 | 0 | 0 | 0 |
| ENSG00000261350 | 0 | 1 | 0 | 0 | 0 | 0 |
| ENSG00000260341 | 0 | 0 | 0 | 0 | 0 | 0 |
| ENSG00000261445 | 0 | 0 | 0 | 0 | 0 | 0 |
| ENSG00000260857 | 0 | 0 | 0 | 0 | 0 | 0 |
| ENSG00000269622 | 0 | 0 | 0 | 0 | 0 | 0 |
| ENSG00000282946 | 0 | 2 | 0 | 0 | 1 | 1 |
| ENSG00000259996 | 0 | 0 | 0 | 0 | 0 | 0 |
| ENSG00000260994 | 0 | 0 | 0 | 0 | 0 | 0 |
| ENSG00000261046 | 0 | 0 | 0 | 0 | 0 | 0 |
| ENSG00000260134 | 0 | 0 | 0 | 0 | 0 | 0 |
| ENSG00000260746 | 0 | 0 | 0 | 0 | 0 | 0 |
| ENSG00000259791 | 0 | 0 | 0 | 0 | 0 | 0 |
| ENSG00000260522 | 0 | 0 | 0 | 0 | 0 | 0 |
| ENSG00000251915 | 0 | 0 | 0 | 0 | 0 | 0 |
| ENSG00000222207 | 0 | 0 | 0 | 0 | 0 | 0 |
| ENSG00000251924 | 0 | 0 | 0 | 0 | 0 | 0 |
| ENSG00000252624 | 0 | 0 | 0 | 0 | 0 | 0 |
| ENSG00000261122 | 0 | 0 | 0 | 0 | 0 | 0 |
| ENSG00000252094 | 0 | 0 | 0 | 0 | 0 | 0 |
| ENSG00000276599 | 0 | 0 | 0 | 0 | 0 | 0 |
| ENSG00000277004 | 0 | 0 | 0 | 0 | 0 | 0 |
| ENSG00000252182 | 0 | 0 | 0 | 0 | 0 | 0 |
| ENSG00000274228 | 0 | 0 | 0 | 0 | 0 | 0 |
| ENSG00000253055 | 0 | 0 | 0 | 0 | 0 | 0 |
| ENSG00000252647 | 0 | 0 | 0 | 0 | 0 | 0 |
| ENSG00000252268 | 0 | 0 | 0 | 0 | 0 | 0 |
| ENSG00000251760 | 0 | 0 | 0 | 0 | 0 | 0 |
| ENSG00000251717 | 0 | 0 | 0 | 0 | 0 | 0 |

| | | | | | | |
|---|---|---|---|---|---|---|
| ENSG00000252806 | 0 | 0 | 0 | 0 | 0 | 0 |
| ENSG00000252673 | 0 | 0 | 0 | 0 | 0 | 0 |
| ENSG00000252587 | 0 | 0 | 0 | 0 | 0 | 0 |
| ENSG00000252936 | 0 | 0 | 0 | 0 | 0 | 0 |
| ENSG00000260611 | 0 | 0 | 0 | 0 | 0 | 0 |
| ENSG00000261620 | 2 | 4 | 8 | 3 | 1 | 11 |
| ENSG00000261515 | 0 | 0 | 0 | 0 | 0 | 0 |
| ENSG00000261782 | 0 | 0 | 0 | 0 | 0 | 0 |
| ENSG00000279666 | 0 | 0 | 0 | 0 | 0 | 0 |
| ENSG00000261084 | 0 | 0 | 0 | 0 | 0 | 0 |
| ENSG00000261239 | 0 | 0 | 0 | 0 | 0 | 0 |
| ENSG00000199448 | 0 | 0 | 0 | 0 | 0 | 0 |
| ENSG00000261356 | 0 | 0 | 0 | 0 | 0 | 0 |
| ENSG00000260251 | 0 | 0 | 0 | 0 | 0 | 0 |
| ENSG00000171241 | 282 | 130 | 153 | 102 | 213 | 171 |
| ENSG00000261512 | 133 | 61 | 62 | 64 | 69 | 56 |
| ENSG00000259856 | 2 | 0 | 3 | 0 | 2 | 2 |
| ENSG00000260909 | 0 | 0 | 0 | 0 | 0 | 0 |
| ENSG00000069329 | 3124 | 1684 | 1711 | 1208 | 1995 | 1591 |
| ENSG00000261131 | 0 | 1 | 1 | 1 | 0 | 0 |
| ENSG00000280376 | 0 | 0 | 0 | 0 | 0 | 0 |
| ENSG00000091651 | 212 | 106 | 142 | 96 | 140 | 164 |
| ENSG00000140795 | 8 | 3 | 4 | 2 | 2 | 3 |
| ENSG00000260782 | 0 | 0 | 1 | 2 | 2 | 0 |
| ENSG00000155330 | 284 | 129 | 153 | 107 | 158 | 147 |
| ENSG00000260889 | 0 | 0 | 0 | 0 | 0 | 0 |
| ENSG00000261725 | 0 | 0 | 0 | 0 | 0 | 0 |
| ENSG00000166123 | 951 | 442 | 518 | 557 | 498 | 527 |
| ENSG00000069345 | 2653 | 1216 | 1515 | 1206 | 1385 | 1350 |
| ENSG00000261173 | 0 | 4 | 2 | 0 | 4 | 4 |
| ENSG00000259821 | 0 | 0 | 0 | 0 | 0 | 0 |
| ENSG00000259983 | 0 | 0 | 0 | 0 | 0 | 0 |
| ENSG00000171208 | 480 | 294 | 319 | 278 | 335 | 314 |
| ENSG00000260281 | 5 | 1 | 5 | 0 | 4 | 2 |
| ENSG00000239840 | 0 | 1 | 0 | 0 | 0 | 0 |
| ENSG00000129636 | 1270 | 676 | 797 | 528 | 842 | 701 |
| ENSG00000260744 | 0 | 1 | 0 | 0 | 0 | 0 |
| ENSG00000279509 | 0 | 0 | 0 | 0 | 0 | 0 |
| ENSG00000260012 | 0 | 0 | 0 | 0 | 0 | 0 |
| ENSG00000261369 | 0 | 0 | 0 | 0 | 0 | 1 |
| ENSG00000222767 | 0 | 0 | 0 | 0 | 0 | 0 |
| ENSG00000102893 | 1154 | 556 | 624 | 306 | 649 | 561 |
| ENSG00000222268 | 0 | 0 | 0 | 0 | 0 | 0 |
| ENSG00000262038 | 0 | 0 | 0 | 0 | 0 | 0 |
| ENSG00000259866 | 0 | 0 | 0 | 0 | 0 | 0 |
| ENSG00000261336 | 2 | 1 | 0 | 2 | 1 | 0 |
| ENSG00000259918 | 0 | 0 | 0 | 0 | 0 | 0 |
| ENSG00000261231 | 0 | 0 | 0 | 0 | 0 | 0 |
| ENSG00000260497 | 0 | 0 | 0 | 0 | 0 | 0 |
| ENSG00000261325 | 0 | 0 | 0 | 0 | 0 | 0 |

| | | | | | | |
|---|---|---|---|---|---|---|
| ENSG00000260450 | 0 | 0 | 0 | 0 | 0 | 0 |
| ENSG00000261017 | 0 | 0 | 0 | 0 | 0 | 0 |
| ENSG00000140798 | 0 | 0 | 0 | 0 | 0 | 0 |
| ENSG00000261538 | 0 | 0 | 0 | 0 | 0 | 0 |
| ENSG00000121270 | 12 | 4 | 3 | 3 | 4 | 3 |
| ENSG00000102910 | 2700 | 1149 | 1474 | 816 | 1208 | 1253 |
| ENSG00000240793 | 4 | 2 | 0 | 0 | 0 | 1 |
| ENSG00000280067 | 8 | 7 | 5 | 4 | 7 | 4 |
| ENSG00000196470 | 156 | 89 | 117 | 110 | 98 | 121 |
| ENSG00000275909 | 0 | 0 | 0 | 0 | 0 | 0 |
| ENSG00000261802 | 0 | 0 | 0 | 0 | 0 | 0 |
| ENSG00000260688 | 0 | 0 | 0 | 0 | 0 | 0 |
| ENSG00000260347 | 0 | 0 | 0 | 0 | 0 | 0 |
| ENSG00000102921 | 828 | 383 | 514 | 337 | 423 | 478 |
| ENSG00000261267 | 0 | 1 | 2 | 0 | 1 | 0 |
| ENSG00000243609 | 0 | 0 | 0 | 0 | 0 | 0 |
| ENSG00000260033 | 0 | 0 | 0 | 0 | 0 | 0 |
| ENSG00000259912 | 0 | 0 | 0 | 0 | 0 | 0 |
| ENSG00000260086 | 2 | 1 | 1 | 1 | 2 | 1 |
| ENSG00000260052 | 0 | 0 | 0 | 1 | 0 | 0 |
| ENSG00000272545 | 0 | 0 | 0 | 0 | 0 | 0 |
| ENSG00000260726 | 0 | 0 | 0 | 0 | 0 | 0 |
| ENSG00000222170 | 0 | 0 | 0 | 0 | 0 | 0 |
| ENSG00000260184 | 0 | 0 | 0 | 0 | 0 | 0 |
| ENSG00000260067 | 0 | 0 | 0 | 0 | 0 | 0 |
| ENSG00000261421 | 0 | 0 | 0 | 0 | 0 | 0 |
| ENSG00000261815 | 0 | 0 | 0 | 0 | 0 | 0 |
| ENSG00000102924 | 0 | 0 | 0 | 0 | 1 | 0 |
| ENSG00000279249 | 0 | 0 | 0 | 0 | 0 | 0 |
| ENSG00000280189 | 0 | 0 | 0 | 0 | 0 | 0 |
| ENSG00000166152 | 0 | 0 | 0 | 0 | 0 | 0 |
| ENSG00000262950 | 0 | 0 | 0 | 0 | 0 | 0 |
| ENSG00000263110 | 0 | 0 | 0 | 0 | 0 | 0 |
| ENSG00000261623 | 0 | 0 | 0 | 0 | 0 | 0 |
| ENSG00000102935 | 66 | 26 | 35 | 25 | 18 | 26 |
| ENSG00000260089 | 0 | 0 | 0 | 0 | 0 | 0 |
| ENSG00000260024 | 0 | 0 | 0 | 0 | 0 | 0 |
| ENSG00000280131 | 0 | 0 | 0 | 0 | 0 | 1 |
| ENSG00000259978 | 0 | 0 | 0 | 0 | 0 | 0 |
| ENSG00000275155 | 8 | 0 | 4 | 1 | 1 | 2 |
| ENSG00000279842 | 0 | 0 | 0 | 0 | 0 | 0 |
| ENSG00000261751 | 0 | 0 | 0 | 0 | 0 | 0 |
| ENSG00000244551 | 0 | 0 | 0 | 0 | 0 | 0 |
| ENSG00000283365 | 0 | 0 | 0 | 0 | 0 | 0 |
| ENSG00000205423 | 264 | 115 | 138 | 103 | 147 | 122 |
| ENSG00000259843 | 0 | 2 | 0 | 0 | 0 | 0 |
| ENSG00000261835 | 0 | 1 | 0 | 0 | 0 | 0 |
| ENSG00000273829 | 0 | 0 | 0 | 0 | 0 | 0 |
| ENSG00000260298 | 0 | 0 | 0 | 0 | 0 | 0 |
| ENSG00000155393 | 506 | 292 | 294 | 308 | 307 | 332 |

| | | | | | | |
|---|---|---|---|---|---|---|
| ENSG00000279356 | 3 | 4 | 2 | 4 | 3 | 5 |
| ENSG00000202124 | 0 | 1 | 0 | 0 | 0 | 0 |
| ENSG00000260381 | 3 | 1 | 0 | 1 | 0 | 1 |
| ENSG00000260031 | 0 | 0 | 0 | 0 | 0 | 0 |
| ENSG00000252526 | 0 | 0 | 0 | 0 | 0 | 0 |
| ENSG00000121274 | 599 | 292 | 312 | 167 | 295 | 268 |
| ENSG00000121281 | 26 | 21 | 14 | 12 | 25 | 30 |
| ENSG00000274969 | 0 | 0 | 0 | 0 | 0 | 0 |
| ENSG00000166164 | 1366 | 753 | 871 | 787 | 936 | 879 |
| ENSG00000261393 | 0 | 0 | 0 | 0 | 0 | 0 |
| ENSG00000261092 | 0 | 0 | 0 | 1 | 1 | 0 |
| ENSG00000260573 | 0 | 1 | 0 | 0 | 0 | 0 |
| ENSG00000140807 | 10 | 9 | 5 | 8 | 17 | 8 |
| ENSG00000260029 | 0 | 0 | 0 | 0 | 0 | 0 |
| ENSG00000278909 | 0 | 0 | 0 | 0 | 0 | 0 |
| ENSG00000205414 | 0 | 0 | 2 | 0 | 0 | 0 |
| ENSG00000260249 | 0 | 0 | 0 | 0 | 0 | 0 |
| ENSG00000167208 | 0 | 0 | 0 | 0 | 1 | 0 |
| ENSG00000167207 | 6 | 9 | 3 | 4 | 2 | 3 |
| ENSG00000270120 | 0 | 0 | 0 | 0 | 0 | 0 |
| ENSG00000261644 | 0 | 0 | 0 | 1 | 0 | 1 |
| ENSG00000083799 | 391 | 170 | 198 | 147 | 173 | 165 |
| ENSG00000264947 | 0 | 0 | 0 | 0 | 0 | 0 |
| ENSG00000260616 | 0 | 0 | 0 | 0 | 0 | 0 |
| ENSG00000261703 | 0 | 0 | 0 | 0 | 0 | 0 |
| ENSG00000261241 | 4 | 0 | 0 | 0 | 0 | 0 |
| ENSG00000260042 | 0 | 0 | 0 | 0 | 0 | 0 |
| ENSG00000199771 | 0 | 0 | 0 | 0 | 0 | 0 |
| ENSG00000261637 | 0 | 0 | 0 | 0 | 0 | 0 |
| ENSG00000260605 | 0 | 0 | 0 | 0 | 0 | 0 |
| ENSG00000260620 | 0 | 0 | 0 | 0 | 0 | 0 |
| ENSG00000261282 | 0 | 0 | 0 | 0 | 0 | 0 |
| ENSG00000103449 | 0 | 0 | 0 | 0 | 0 | 0 |
| ENSG00000261238 | 0 | 0 | 0 | 0 | 0 | 0 |
| ENSG00000285367 | 0 | 0 | 0 | 0 | 0 | 0 |
| ENSG00000280344 | 0 | 0 | 0 | 0 | 0 | 0 |
| ENSG00000261047 | 0 | 0 | 0 | 0 | 0 | 0 |
| ENSG00000260818 | 0 | 0 | 0 | 0 | 0 | 0 |
| ENSG00000260850 | 0 | 0 | 0 | 0 | 0 | 0 |
| ENSG00000224578 | 102 | 51 | 56 | 48 | 67 | 74 |
| ENSG00000223168 | 0 | 0 | 0 | 0 | 0 | 0 |
| ENSG00000260057 | 0 | 0 | 0 | 0 | 0 | 0 |
| ENSG00000261190 | 0 | 0 | 0 | 0 | 0 | 0 |
| ENSG00000261470 | 0 | 0 | 0 | 0 | 0 | 0 |
| ENSG00000279779 | 0 | 0 | 0 | 0 | 0 | 0 |
| ENSG00000260268 | 0 | 0 | 0 | 0 | 0 | 0 |
| ENSG00000261749 | 0 | 0 | 0 | 0 | 0 | 0 |
| ENSG00000260975 | 0 | 0 | 0 | 0 | 0 | 0 |
| ENSG00000260478 | 0 | 0 | 0 | 0 | 0 | 0 |
| ENSG00000260887 | 0 | 0 | 0 | 0 | 0 | 0 |

| | | | | | | |
|---|---|---|---|---|---|---|
| ENSG00000279427 | 0 | 0 | 0 | 0 | 0 | 0 |
| ENSG00000103460 | 0 | 0 | 0 | 0 | 0 | 0 |
| ENSG00000249231 | 0 | 0 | 0 | 0 | 0 | 0 |
| ENSG00000285800 | 0 | 0 | 0 | 0 | 0 | 0 |
| ENSG00000260963 | 0 | 0 | 0 | 0 | 0 | 0 |
| ENSG00000260393 | 0 | 0 | 0 | 0 | 0 | 0 |
| ENSG00000261261 | 0 | 0 | 0 | 0 | 0 | 0 |
| ENSG00000234337 | 6 | 2 | 4 | 1 | 3 | 1 |
| ENSG00000260326 | 0 | 0 | 1 | 0 | 1 | 1 |
| ENSG00000260258 | 0 | 0 | 0 | 0 | 2 | 0 |
| ENSG00000260939 | 0 | 0 | 0 | 0 | 0 | 0 |
| ENSG00000277639 | 13 | 5 | 8 | 10 | 20 | 6 |
| ENSG00000177200 | 242 | 160 | 145 | 47 | 291 | 158 |
| ENSG00000261291 | 2 | 0 | 0 | 1 | 0 | 0 |
| ENSG00000280227 | 0 | 0 | 0 | 0 | 0 | 0 |
| ENSG00000261056 | 0 | 1 | 1 | 0 | 0 | 0 |
| ENSG00000279589 | 4 | 2 | 0 | 0 | 0 | 0 |
| ENSG00000238645 | 0 | 0 | 0 | 0 | 0 | 0 |
| ENSG00000202193 | 0 | 0 | 0 | 0 | 0 | 0 |
| ENSG00000259962 | 0 | 0 | 0 | 0 | 0 | 0 |
| ENSG00000260078 | 4 | 1 | 1 | 1 | 3 | 1 |
| ENSG00000261804 | 58 | 31 | 32 | 17 | 24 | 36 |
| ENSG00000262714 | 42 | 27 | 30 | 16 | 18 | 20 |
| ENSG00000279741 | 13 | 10 | 9 | 5 | 12 | 7 |
| ENSG00000280027 | 20 | 17 | 21 | 11 | 16 | 12 |
| ENSG00000103479 | 3616 | 1632 | 2386 | 890 | 1614 | 1449 |
| ENSG00000252569 | 2 | 1 | 0 | 0 | 1 | 4 |
| ENSG00000259926 | 4 | 2 | 0 | 0 | 0 | 0 |
| ENSG00000279344 | 0 | 2 | 3 | 0 | 3 | 0 |
| ENSG00000279722 | 16 | 9 | 11 | 1 | 8 | 4 |
| ENSG00000166971 | 1141 | 560 | 715 | 418 | 595 | 515 |
| ENSG00000103494 | 133 | 62 | 77 | 54 | 103 | 51 |
| ENSG00000275191 | 0 | 0 | 0 | 2 | 1 | 0 |
| ENSG00000140718 | 593 | 257 | 382 | 200 | 329 | 338 |
| ENSG00000261630 | 0 | 0 | 0 | 0 | 0 | 0 |
| ENSG00000280392 | 0 | 0 | 1 | 0 | 0 | 0 |
| ENSG00000260194 | 0 | 0 | 0 | 0 | 0 | 0 |
| ENSG00000261049 | 0 | 0 | 0 | 0 | 0 | 0 |
| ENSG00000280454 | 0 | 0 | 0 | 0 | 0 | 0 |
| ENSG00000259759 | 0 | 0 | 0 | 0 | 0 | 0 |
| ENSG00000259517 | 0 | 0 | 0 | 0 | 0 | 1 |
| ENSG00000177508 | 263 | 139 | 160 | 154 | 137 | 131 |
| ENSG00000283689 | 16 | 6 | 8 | 8 | 6 | 11 |
| ENSG00000277559 | 48 | 18 | 34 | 21 | 8 | 25 |
| ENSG00000261803 | 0 | 0 | 0 | 0 | 0 | 0 |
| ENSG00000283304 | 0 | 0 | 0 | 0 | 0 | 0 |
| ENSG00000283258 | 0 | 0 | 0 | 0 | 0 | 0 |
| ENSG00000245694 | 467 | 242 | 281 | 215 | 237 | 184 |
| ENSG00000176842 | 631 | 292 | 292 | 315 | 261 | 286 |
| ENSG00000259711 | 8 | 4 | 6 | 5 | 2 | 4 |

| | | | | | | |
|---|---|---|---|---|---|---|
| ENSG00000259725 | 2 | 0 | 0 | 0 | 0 | 0 |
| ENSG00000260147 | 0 | 0 | 0 | 0 | 0 | 0 |
| ENSG00000285775 | 0 | 0 | 0 | 0 | 0 | 0 |
| ENSG00000259283 | 0 | 0 | 0 | 0 | 0 | 0 |
| ENSG00000239555 | 0 | 0 | 0 | 0 | 0 | 0 |
| ENSG00000283423 | 0 | 1 | 0 | 0 | 0 | 2 |
| ENSG00000159387 | 6 | 2 | 1 | 3 | 2 | 0 |
| ENSG00000263207 | 0 | 0 | 0 | 0 | 0 | 0 |
| ENSG00000240760 | 0 | 0 | 0 | 0 | 0 | 0 |
| ENSG00000087245 | 0 | 0 | 0 | 1 | 0 | 0 |
| ENSG00000274508 | 0 | 0 | 0 | 0 | 0 | 0 |
| ENSG00000260135 | 0 | 0 | 0 | 0 | 0 | 0 |
| ENSG00000279030 | 0 | 0 | 0 | 0 | 0 | 0 |
| ENSG00000087253 | 192 | 78 | 119 | 76 | 119 | 102 |
| ENSG00000261997 | 6 | 1 | 1 | 1 | 1 | 1 |
| ENSG00000256812 | 0 | 0 | 0 | 0 | 0 | 0 |
| ENSG00000103546 | 0 | 0 | 0 | 0 | 0 | 0 |
| ENSG00000278928 | 0 | 0 | 0 | 0 | 0 | 0 |
| ENSG00000260765 | 0 | 0 | 0 | 0 | 0 | 0 |
| ENSG00000228695 | 0 | 0 | 0 | 0 | 0 | 0 |
| ENSG00000198848 | 0 | 0 | 0 | 0 | 0 | 0 |
| ENSG00000159398 | 0 | 0 | 0 | 0 | 0 | 0 |
| ENSG00000246379 | 0 | 0 | 0 | 0 | 0 | 0 |
| ENSG00000279523 | 0 | 0 | 0 | 0 | 0 | 0 |
| ENSG00000260041 | 0 | 0 | 0 | 0 | 0 | 0 |
| ENSG00000087258 | 2 | 5 | 2 | 4 | 4 | 8 |
| ENSG00000261439 | 0 | 0 | 0 | 0 | 0 | 0 |
| ENSG00000265281 | 0 | 0 | 0 | 0 | 0 | 0 |
| ENSG00000260198 | 0 | 0 | 0 | 0 | 0 | 0 |
| ENSG00000279764 | 0 | 0 | 0 | 0 | 0 | 0 |
| ENSG00000272372 | 0 | 1 | 0 | 0 | 0 | 0 |
| ENSG00000159461 | 2190 | 1078 | 1371 | 1088 | 1087 | 1176 |
| ENSG00000260621 | 0 | 0 | 0 | 0 | 0 | 0 |
| ENSG00000167005 | 2366 | 1178 | 1382 | 889 | 1329 | 1206 |
| ENSG00000087263 | 709 | 429 | 499 | 381 | 526 | 450 |
| ENSG00000274031 | 4 | 0 | 1 | 0 | 0 | 3 |
| ENSG00000125124 | 1092 | 528 | 727 | 489 | 604 | 606 |
| ENSG00000102891 | 0 | 0 | 0 | 0 | 0 | 0 |
| ENSG00000087250 | 1 | 0 | 0 | 0 | 0 | 0 |
| ENSG00000125148 | 825 | 336 | 417 | 362 | 286 | 318 |
| ENSG00000260823 | 0 | 2 | 0 | 0 | 0 | 0 |
| ENSG00000260549 | 0 | 0 | 0 | 0 | 0 | 0 |
| ENSG00000169715 | 0 | 0 | 0 | 0 | 0 | 0 |
| ENSG00000205364 | 0 | 0 | 0 | 0 | 0 | 0 |
| ENSG00000255986 | 0 | 0 | 0 | 0 | 0 | 0 |
| ENSG00000205362 | 0 | 0 | 0 | 0 | 0 | 0 |
| ENSG00000205361 | 0 | 0 | 0 | 0 | 0 | 0 |
| ENSG00000205360 | 0 | 0 | 0 | 0 | 0 | 0 |
| ENSG00000259923 | 0 | 0 | 0 | 0 | 0 | 0 |
| ENSG00000169688 | 0 | 0 | 1 | 0 | 0 | 0 |

| | | | | | | |
|---|---|---|---|---|---|---|
| ENSG00000198417 | 74 | 45 | 51 | 55 | 40 | 34 |
| ENSG00000125144 | 0 | 0 | 0 | 0 | 0 | 0 |
| ENSG00000205358 | 0 | 0 | 0 | 0 | 0 | 0 |
| ENSG00000275691 | 0 | 0 | 0 | 0 | 0 | 0 |
| ENSG00000187193 | 68 | 45 | 57 | 46 | 31 | 42 |
| ENSG00000259827 | 0 | 0 | 0 | 0 | 0 | 0 |
| ENSG00000261429 | 0 | 0 | 0 | 0 | 0 | 0 |
| ENSG00000261302 | 347 | 215 | 188 | 143 | 211 | 182 |
| ENSG00000102900 | 1947 | 1008 | 1087 | 1011 | 1091 | 1089 |
| ENSG00000207649 | 0 | 0 | 0 | 0 | 0 | 0 |
| ENSG00000070915 | 6 | 0 | 1 | 0 | 0 | 0 |
| ENSG00000283471 | 0 | 0 | 0 | 0 | 0 | 0 |
| ENSG00000243988 | 0 | 0 | 0 | 0 | 1 | 1 |
| ENSG00000051108 | 1827 | 840 | 1047 | 711 | 783 | 754 |
| ENSG00000261270 | 14 | 12 | 6 | 5 | 11 | 7 |
| ENSG00000261114 | 15 | 2 | 6 | 3 | 5 | 6 |
| ENSG00000087237 | 0 | 0 | 0 | 0 | 0 | 0 |
| ENSG00000140853 | 20 | 5 | 8 | 2 | 1 | 7 |
| ENSG00000259922 | 0 | 0 | 0 | 0 | 0 | 0 |
| ENSG00000260145 | 0 | 0 | 0 | 0 | 0 | 0 |
| ENSG00000260148 | 0 | 0 | 0 | 0 | 0 | 0 |
| ENSG00000140848 | 576 | 240 | 336 | 401 | 318 | 359 |
| ENSG00000279019 | 10 | 1 | 5 | 2 | 4 | 1 |
| ENSG00000279803 | 0 | 0 | 1 | 3 | 0 | 0 |
| ENSG00000172775 | 2715 | 1256 | 1575 | 1354 | 1431 | 1496 |
| ENSG00000285979 | 98 | 44 | 53 | 22 | 47 | 30 |
| ENSG00000159579 | 1042 | 461 | 530 | 341 | 554 | 462 |
| ENSG00000102931 | 63 | 56 | 51 | 60 | 57 | 48 |
| ENSG00000276663 | 2 | 1 | 5 | 1 | 3 | 2 |
| ENSG00000260038 | 7 | 1 | 2 | 3 | 3 | 1 |
| ENSG00000102934 | 503 | 249 | 291 | 195 | 163 | 205 |
| ENSG00000260965 | 0 | 0 | 0 | 0 | 0 | 0 |
| ENSG00000271456 | 0 | 0 | 0 | 0 | 0 | 0 |
| ENSG00000102962 | 86 | 43 | 74 | 64 | 44 | 76 |
| ENSG00000006210 | 2 | 0 | 1 | 0 | 2 | 4 |
| ENSG00000102970 | 0 | 4 | 1 | 5 | 0 | 0 |
| ENSG00000005194 | 2096 | 951 | 1179 | 1197 | 1144 | 1232 |
| ENSG00000088682 | 1252 | 613 | 764 | 688 | 733 | 691 |
| ENSG00000260345 | 0 | 0 | 0 | 0 | 0 | 0 |
| ENSG00000102978 | 2682 | 1363 | 1534 | 1478 | 1392 | 1359 |
| ENSG00000125170 | 924 | 450 | 506 | 450 | 445 | 398 |
| ENSG00000135736 | 166 | 94 | 87 | 92 | 106 | 67 |
| ENSG00000159618 | 27 | 25 | 15 | 24 | 16 | 19 |
| ENSG00000260828 | 0 | 0 | 2 | 0 | 0 | 0 |
| ENSG00000205336 | 6274 | 2984 | 3358 | 3010 | 3206 | 2868 |
| ENSG00000261633 | 0 | 3 | 1 | 2 | 2 | 1 |
| ENSG00000182885 | 0 | 0 | 0 | 0 | 0 | 0 |
| ENSG00000260467 | 0 | 0 | 0 | 0 | 0 | 0 |
| ENSG00000159625 | 2 | 0 | 1 | 0 | 1 | 2 |
| ENSG00000260438 | 0 | 0 | 0 | 0 | 0 | 0 |

| | | | | | | |
|---|---|---|---|---|---|---|
| ENSG00000140854 | 793 | 435 | 542 | 534 | 515 | 560 |
| ENSG00000140859 | 758 | 366 | 532 | 354 | 358 | 309 |
| ENSG00000276166 | 3 | 6 | 2 | 0 | 3 | 7 |
| ENSG00000274816 | 2 | 1 | 0 | 0 | 1 | 0 |
| ENSG00000187185 | 3 | 2 | 6 | 1 | 1 | 7 |
| ENSG00000206833 | 0 | 0 | 0 | 0 | 0 | 0 |
| ENSG00000070729 | 2 | 1 | 0 | 0 | 1 | 0 |
| ENSG00000159648 | 0 | 0 | 0 | 0 | 0 | 0 |
| ENSG00000166188 | 376 | 168 | 209 | 205 | 202 | 201 |
| ENSG00000103005 | 1355 | 648 | 740 | 719 | 761 | 692 |
| ENSG00000102996 | 2257 | 1054 | 1329 | 1452 | 1315 | 1307 |
| ENSG00000207173 | 0 | 0 | 0 | 0 | 0 | 0 |
| ENSG00000261151 | 0 | 0 | 0 | 0 | 0 | 0 |
| ENSG00000070761 | 854 | 459 | 563 | 574 | 566 | 524 |
| ENSG00000260545 | 0 | 2 | 1 | 0 | 0 | 0 |
| ENSG00000260927 | 4 | 2 | 2 | 2 | 4 | 1 |
| ENSG00000070770 | 1958 | 1003 | 1151 | 1051 | 1237 | 1176 |
| ENSG00000260867 | 0 | 0 | 1 | 0 | 0 | 0 |
| ENSG00000240863 | 0 | 0 | 0 | 0 | 0 | 0 |
| ENSG00000103021 | 185 | 87 | 105 | 98 | 134 | 114 |
| ENSG00000103023 | 0 | 0 | 0 | 0 | 0 | 0 |
| ENSG00000181938 | 254 | 166 | 206 | 148 | 155 | 181 |
| ENSG00000212379 | 0 | 0 | 0 | 0 | 0 | 0 |
| ENSG00000261078 | 0 | 0 | 0 | 0 | 0 | 0 |
| ENSG00000252271 | 0 | 0 | 0 | 0 | 0 | 0 |
| ENSG00000276131 | 0 | 0 | 0 | 0 | 0 | 0 |
| ENSG00000260186 | 0 | 0 | 0 | 0 | 0 | 0 |
| ENSG00000103034 | 853 | 426 | 509 | 619 | 654 | 554 |
| ENSG00000200556 | 0 | 0 | 0 | 0 | 0 | 0 |
| ENSG00000103037 | 622 | 282 | 396 | 348 | 411 | 362 |
| ENSG00000125107 | 4056 | 2252 | 2253 | 1143 | 2975 | 2027 |
| ENSG00000276259 | 11 | 4 | 3 | 0 | 8 | 1 |
| ENSG00000207493 | 2 | 1 | 0 | 0 | 0 | 1 |
| ENSG00000206952 | 0 | 0 | 0 | 0 | 0 | 0 |
| ENSG00000281910 | 0 | 0 | 0 | 0 | 0 | 0 |
| ENSG00000260865 | 0 | 0 | 0 | 1 | 0 | 0 |
| ENSG00000103042 | 918 | 414 | 526 | 443 | 468 | 470 |
| ENSG00000279889 | 0 | 0 | 0 | 0 | 0 | 0 |
| ENSG00000125166 | 5035 | 2285 | 2754 | 2566 | 2636 | 2626 |
| ENSG00000245768 | 2 | 0 | 1 | 0 | 1 | 1 |
| ENSG00000200424 | 0 | 0 | 0 | 0 | 0 | 0 |
| ENSG00000244003 | 0 | 0 | 0 | 0 | 0 | 0 |
| ENSG00000260451 | 0 | 0 | 1 | 0 | 0 | 0 |
| ENSG00000242423 | 0 | 0 | 0 | 0 | 0 | 0 |
| ENSG00000261638 | 0 | 0 | 0 | 0 | 0 | 0 |
| ENSG00000275808 | 0 | 0 | 0 | 0 | 0 | 0 |
| ENSG00000244043 | 0 | 0 | 0 | 0 | 0 | 0 |
| ENSG00000261144 | 0 | 0 | 0 | 0 | 0 | 0 |
| ENSG00000200062 | 0 | 0 | 0 | 0 | 0 | 0 |
| ENSG00000270222 | 0 | 0 | 0 | 0 | 0 | 0 |

| | | | | | | |
|---|---|---|---|---|---|---|
| ENSG00000260240 | 0 | 0 | 0 | 0 | 0 | 0 |
| ENSG00000261807 | 0 | 0 | 0 | 0 | 0 | 0 |
| ENSG00000261278 | 0 | 0 | 0 | 0 | 0 | 0 |
| ENSG00000261590 | 0 | 0 | 0 | 0 | 0 | 0 |
| ENSG00000278499 | 0 | 0 | 0 | 0 | 0 | 0 |
| ENSG00000261436 | 0 | 0 | 0 | 0 | 0 | 0 |
| ENSG00000261310 | 0 | 0 | 0 | 0 | 0 | 0 |
| ENSG00000259844 | 0 | 0 | 1 | 0 | 0 | 1 |
| ENSG00000261178 | 0 | 0 | 0 | 0 | 0 | 0 |
| ENSG00000224631 | 191 | 166 | 94 | 152 | 287 | 222 |
| ENSG00000150394 | 7 | 5 | 4 | 4 | 9 | 12 |
| ENSG00000260600 | 0 | 0 | 0 | 0 | 0 | 0 |
| ENSG00000201289 | 0 | 0 | 0 | 0 | 0 | 0 |
| ENSG00000260115 | 6 | 5 | 7 | 4 | 8 | 7 |
| ENSG00000207441 | 0 | 0 | 0 | 0 | 0 | 0 |
| ENSG00000280261 | 0 | 0 | 0 | 0 | 0 | 0 |
| ENSG00000237901 | 0 | 0 | 0 | 0 | 0 | 0 |
| ENSG00000231589 | 0 | 0 | 0 | 0 | 0 | 0 |
| ENSG00000260332 | 0 | 0 | 0 | 0 | 0 | 0 |
| ENSG00000261502 | 9 | 7 | 10 | 7 | 10 | 4 |
| ENSG00000269986 | 0 | 2 | 0 | 2 | 0 | 0 |
| ENSG00000261199 | 0 | 1 | 0 | 0 | 0 | 0 |
| ENSG00000260658 | 13 | 2 | 6 | 2 | 2 | 6 |
| ENSG00000241808 | 0 | 0 | 0 | 0 | 0 | 0 |
| ENSG00000261743 | 0 | 0 | 0 | 0 | 0 | 0 |
| ENSG00000279225 | 2 | 0 | 0 | 0 | 0 | 0 |
| ENSG00000259864 | 0 | 0 | 0 | 0 | 0 | 0 |
| ENSG00000261014 | 0 | 0 | 0 | 1 | 0 | 0 |
| ENSG00000261653 | 0 | 0 | 0 | 0 | 0 | 0 |
| ENSG00000260715 | 0 | 0 | 0 | 0 | 0 | 0 |
| ENSG00000261028 | 0 | 0 | 0 | 0 | 0 | 0 |
| ENSG00000260668 | 0 | 0 | 0 | 0 | 0 | 0 |
| ENSG00000259846 | 0 | 0 | 0 | 0 | 0 | 0 |
| ENSG00000260187 | 0 | 0 | 0 | 0 | 0 | 0 |
| ENSG00000260068 | 0 | 0 | 0 | 0 | 0 | 0 |
| ENSG00000140937 | 40 | 13 | 13 | 16 | 15 | 16 |
| ENSG00000278893 | 0 | 0 | 0 | 0 | 0 | 0 |
| ENSG00000259847 | 0 | 0 | 0 | 0 | 0 | 0 |
| ENSG00000260834 | 0 | 0 | 0 | 0 | 0 | 0 |
| ENSG00000260364 | 0 | 0 | 0 | 0 | 0 | 0 |
| ENSG00000261742 | 0 | 0 | 1 | 0 | 0 | 0 |
| ENSG00000261818 | 0 | 0 | 0 | 0 | 0 | 0 |
| ENSG00000260004 | 0 | 0 | 0 | 0 | 0 | 0 |
| ENSG00000260695 | 0 | 0 | 0 | 0 | 0 | 0 |
| ENSG00000261234 | 0 | 0 | 0 | 0 | 0 | 0 |
| ENSG00000201999 | 0 | 0 | 0 | 0 | 0 | 0 |
| ENSG00000179776 | 30 | 13 | 9 | 2 | 3 | 12 |
| ENSG00000246898 | 2 | 0 | 1 | 0 | 0 | 0 |
| ENSG00000166546 | 4 | 2 | 1 | 1 | 4 | 5 |
| ENSG00000260146 | 0 | 0 | 0 | 0 | 0 | 0 |

| | | | | | | |
|---|---|---|---|---|---|---|
| ENSG00000261656 | 0 | 0 | 0 | 0 | 0 | 0 |
| ENSG00000260851 | 0 | 0 | 0 | 0 | 0 | 0 |
| ENSG00000166548 | 190 | 87 | 111 | 112 | 102 | 121 |
| ENSG00000260755 | 0 | 0 | 0 | 0 | 0 | 0 |
| ENSG00000261519 | 0 | 0 | 0 | 0 | 0 | 0 |
| ENSG00000275745 | 0 | 0 | 0 | 0 | 0 | 0 |
| ENSG00000217555 | 4 | 13 | 17 | 23 | 18 | 21 |
| ENSG00000254788 | 4 | 4 | 5 | 3 | 4 | 7 |
| ENSG00000277978 | 4 | 1 | 5 | 3 | 4 | 9 |
| ENSG00000089505 | 12 | 9 | 18 | 7 | 20 | 7 |
| ENSG00000140932 | 3 | 1 | 0 | 3 | 1 | 0 |
| ENSG00000260650 | 0 | 0 | 3 | 1 | 0 | 0 |
| ENSG00000140931 | 76 | 55 | 30 | 63 | 49 | 65 |
| ENSG00000183723 | 6140 | 2717 | 3476 | 2257 | 2995 | 2977 |
| ENSG00000135720 | 4676 | 2205 | 2717 | 1536 | 2454 | 2308 |
| ENSG00000260465 | 0 | 0 | 0 | 0 | 0 | 0 |
| ENSG00000260558 | 2 | 0 | 0 | 1 | 1 | 0 |
| ENSG00000246777 | 10 | 1 | 2 | 3 | 1 | 6 |
| ENSG00000249961 | 2 | 0 | 2 | 0 | 2 | 0 |
| ENSG00000159593 | 1044 | 604 | 614 | 468 | 799 | 639 |
| ENSG00000258122 | 0 | 0 | 0 | 0 | 0 | 0 |
| ENSG00000168748 | 0 | 0 | 0 | 0 | 0 | 1 |
| ENSG00000172840 | 695 | 359 | 412 | 325 | 375 | 341 |
| ENSG00000166589 | 0 | 0 | 0 | 0 | 0 | 0 |
| ENSG00000166592 | 0 | 0 | 0 | 1 | 1 | 1 |
| ENSG00000166595 | 1173 | 648 | 666 | 803 | 717 | 703 |
| ENSG00000172831 | 1200 | 582 | 768 | 723 | 653 | 647 |
| ENSG00000265408 | 0 | 0 | 0 | 0 | 0 | 0 |
| ENSG00000280416 | 0 | 0 | 0 | 0 | 0 | 0 |
| ENSG00000172828 | 2 | 4 | 0 | 2 | 4 | 6 |
| ENSG00000172824 | 241 | 100 | 179 | 89 | 119 | 113 |
| ENSG00000280334 | 29 | 11 | 9 | 3 | 11 | 7 |
| ENSG00000265918 | 0 | 0 | 0 | 0 | 0 | 0 |
| ENSG00000067955 | 1418 | 598 | 808 | 560 | 764 | 760 |
| ENSG00000125149 | 987 | 434 | 526 | 433 | 503 | 522 |
| ENSG00000237172 | 187 | 92 | 80 | 98 | 119 | 89 |
| ENSG00000102871 | 1053 | 500 | 678 | 742 | 595 | 565 |
| ENSG00000135722 | 239 | 126 | 204 | 163 | 153 | 178 |
| ENSG00000265690 | 0 | 1 | 0 | 0 | 1 | 2 |
| ENSG00000102878 | 488 | 174 | 293 | 189 | 226 | 228 |
| ENSG00000140939 | 1408 | 648 | 962 | 860 | 736 | 814 |
| ENSG00000196123 | 728 | 324 | 435 | 302 | 336 | 388 |
| ENSG00000179044 | 70 | 31 | 34 | 48 | 36 | 40 |
| ENSG00000205250 | 2566 | 1182 | 1505 | 1539 | 1575 | 1495 |
| ENSG00000102890 | 1036 | 496 | 609 | 622 | 535 | 469 |
| ENSG00000207948 | 0 | 0 | 0 | 0 | 0 | 1 |
| ENSG00000280163 | 3 | 3 | 0 | 1 | 1 | 2 |
| ENSG00000125122 | 61 | 29 | 29 | 40 | 27 | 27 |
| ENSG00000168701 | 1516 | 749 | 951 | 1066 | 919 | 874 |
| ENSG00000135723 | 698 | 372 | 467 | 550 | 544 | 461 |

| | | | | | | |
|---|---|---|---|---|---|---|
| ENSG00000135740 | 56 | 28 | 40 | 37 | 53 | 37 |
| ENSG00000262691 | 23 | 17 | 15 | 21 | 16 | 16 |
| ENSG00000196155 | 245 | 82 | 183 | 109 | 77 | 139 |
| ENSG00000168676 | 0 | 0 | 0 | 1 | 0 | 0 |
| ENSG00000201201 | 0 | 0 | 0 | 0 | 0 | 0 |
| ENSG00000159708 | 0 | 0 | 0 | 0 | 0 | 0 |
| ENSG00000159713 | 14 | 15 | 10 | 18 | 8 | 9 |
| ENSG00000239194 | 0 | 0 | 0 | 0 | 0 | 0 |
| ENSG00000159714 | 72 | 38 | 52 | 61 | 51 | 41 |
| ENSG00000176387 | 67 | 37 | 36 | 36 | 18 | 23 |
| ENSG00000261320 | 0 | 0 | 0 | 0 | 0 | 0 |
| ENSG00000159720 | 5507 | 2475 | 3068 | 3211 | 2608 | 2854 |
| ENSG00000270049 | 93 | 28 | 39 | 30 | 34 | 28 |
| ENSG00000159723 | 2 | 0 | 0 | 0 | 2 | 0 |
| ENSG00000276075 | 35 | 11 | 16 | 19 | 13 | 13 |
| ENSG00000039523 | 1695 | 798 | 1063 | 948 | 862 | 880 |
| ENSG00000261396 | 2 | 0 | 0 | 1 | 0 | 0 |
| ENSG00000259945 | 0 | 0 | 0 | 0 | 0 | 0 |
| ENSG00000260894 | 0 | 0 | 0 | 0 | 0 | 0 |
| ENSG00000261386 | 6 | 4 | 9 | 2 | 4 | 4 |
| ENSG00000280214 | 12 | 2 | 4 | 3 | 4 | 5 |
| ENSG00000259804 | 12 | 3 | 5 | 2 | 4 | 2 |
| ENSG00000102974 | 3663 | 1710 | 2256 | 1457 | 1836 | 1983 |
| ENSG00000237718 | 0 | 0 | 0 | 0 | 0 | 0 |
| ENSG00000159753 | 25 | 19 | 15 | 24 | 25 | 15 |
| ENSG00000102977 | 791 | 346 | 411 | 479 | 411 | 425 |
| ENSG00000102981 | 230 | 133 | 127 | 141 | 133 | 92 |
| ENSG00000124074 | 544 | 231 | 315 | 336 | 239 | 218 |
| ENSG00000159761 | 14 | 7 | 19 | 13 | 9 | 17 |
| ENSG00000141098 | 1378 | 702 | 806 | 753 | 811 | 757 |
| ENSG00000141084 | 1975 | 891 | 1153 | 870 | 976 | 1082 |
| ENSG00000270165 | 11 | 3 | 2 | 1 | 8 | 9 |
| ENSG00000102904 | 45 | 20 | 33 | 26 | 22 | 21 |
| ENSG00000102901 | 1175 | 583 | 748 | 664 | 749 | 710 |
| ENSG00000168286 | 1243 | 690 | 781 | 805 | 716 | 705 |
| ENSG00000102898 | 4228 | 2335 | 2533 | 2467 | 2724 | 2465 |
| ENSG00000262141 | 0 | 0 | 0 | 0 | 0 | 0 |
| ENSG00000038358 | 2215 | 998 | 1347 | 1100 | 1134 | 1159 |
| ENSG00000263126 | 15 | 15 | 14 | 8 | 11 | 13 |
| ENSG00000188038 | 0 | 0 | 0 | 0 | 0 | 0 |
| ENSG00000159792 | 1768 | 785 | 1194 | 1088 | 1046 | 1011 |
| ENSG00000141086 | 0 | 0 | 0 | 0 | 0 | 0 |
| ENSG00000261884 | 36 | 19 | 22 | 12 | 23 | 25 |
| ENSG00000205220 | 231 | 136 | 175 | 184 | 130 | 157 |
| ENSG00000213398 | 80 | 46 | 44 | 31 | 33 | 33 |
| ENSG00000124067 | 434 | 188 | 265 | 240 | 236 | 196 |
| ENSG00000141096 | 0 | 0 | 0 | 0 | 0 | 0 |
| ENSG00000167261 | 0 | 3 | 1 | 0 | 0 | 0 |
| ENSG00000167264 | 787 | 384 | 476 | 573 | 549 | 507 |
| ENSG00000262962 | 0 | 0 | 0 | 0 | 0 | 0 |

| | | | | | | |
|---|---|---|---|---|---|---|
| ENSG00000263201 | 0 | 0 | 0 | 0 | 0 | 0 |
| ENSG00000202336 | 0 | 0 | 0 | 0 | 0 | 0 |
| ENSG00000182810 | 505 | 231 | 287 | 328 | 263 | 300 |
| ENSG00000261864 | 1 | 1 | 3 | 1 | 4 | 0 |
| ENSG00000072736 | 887 | 402 | 492 | 363 | 448 | 438 |
| ENSG00000201850 | 0 | 1 | 0 | 0 | 0 | 0 |
| ENSG00000212445 | 0 | 0 | 0 | 0 | 0 | 0 |
| ENSG00000262514 | 0 | 0 | 0 | 0 | 0 | 0 |
| ENSG00000279621 | 0 | 0 | 0 | 0 | 0 | 0 |
| ENSG00000240435 | 2 | 2 | 0 | 0 | 0 | 0 |
| ENSG00000260891 | 3 | 2 | 1 | 5 | 5 | 4 |
| ENSG00000263276 | 10 | 3 | 4 | 2 | 2 | 3 |
| ENSG00000262160 | 0 | 0 | 0 | 0 | 0 | 0 |
| ENSG00000103067 | 5853 | 2716 | 3374 | 2637 | 2796 | 2775 |
| ENSG00000284259 | 0 | 0 | 0 | 0 | 0 | 0 |
| ENSG00000261469 | 21 | 13 | 28 | 10 | 10 | 10 |
| ENSG00000103066 | 978 | 476 | 658 | 651 | 572 | 536 |
| ENSG00000260441 | 5 | 1 | 5 | 2 | 5 | 2 |
| ENSG00000103064 | 626 | 352 | 399 | 285 | 388 | 402 |
| ENSG00000252026 | 4 | 3 | 3 | 2 | 2 | 1 |
| ENSG00000103061 | 263 | 111 | 152 | 106 | 140 | 135 |
| ENSG00000279649 | 22 | 12 | 17 | 6 | 11 | 16 |
| ENSG00000132600 | 1028 | 512 | 695 | 671 | 649 | 668 |
| ENSG00000259797 | 2 | 3 | 2 | 0 | 2 | 1 |
| ENSG00000222177 | 0 | 0 | 0 | 0 | 0 | 0 |
| ENSG00000103056 | 50 | 29 | 28 | 31 | 24 | 27 |
| ENSG00000279693 | 0 | 0 | 0 | 0 | 0 | 0 |
| ENSG00000274698 | 17 | 12 | 9 | 9 | 14 | 9 |
| ENSG00000199263 | 0 | 0 | 0 | 1 | 2 | 0 |
| ENSG00000241334 | 0 | 0 | 0 | 0 | 0 | 0 |
| ENSG00000184939 | 1787 | 906 | 950 | 628 | 992 | 748 |
| ENSG00000201164 | 2 | 1 | 0 | 0 | 0 | 0 |
| ENSG00000260084 | 0 | 0 | 0 | 1 | 2 | 0 |
| ENSG00000275383 | 3 | 0 | 1 | 0 | 3 | 1 |
| ENSG00000252040 | 0 | 0 | 0 | 0 | 0 | 0 |
| ENSG00000252640 | 0 | 0 | 0 | 0 | 0 | 0 |
| ENSG00000062038 | 5140 | 2449 | 2895 | 2327 | 2662 | 2464 |
| ENSG00000260577 | 11 | 11 | 10 | 6 | 14 | 4 |
| ENSG00000260860 | 2 | 0 | 0 | 0 | 0 | 0 |
| ENSG00000261395 | 0 | 0 | 0 | 0 | 2 | 0 |
| ENSG00000039068 | 29473 | 14955 | 18862 | 10752 | 13840 | 13998 |
| ENSG00000200558 | 0 | 0 | 0 | 0 | 0 | 0 |
| ENSG00000260798 | 0 | 0 | 0 | 0 | 0 | 0 |
| ENSG00000260459 | 42 | 20 | 30 | 10 | 26 | 18 |
| ENSG00000103047 | 235 | 110 | 103 | 103 | 140 | 130 |
| ENSG00000199979 | 0 | 0 | 0 | 0 | 0 | 0 |
| ENSG00000275504 | 0 | 0 | 0 | 0 | 0 | 0 |
| ENSG00000260999 | 0 | 0 | 0 | 0 | 0 | 0 |
| ENSG00000266801 | 0 | 1 | 0 | 2 | 0 | 1 |
| ENSG00000202497 | 0 | 0 | 0 | 0 | 0 | 0 |

| | | | | | | |
|---|---|---|---|---|---|---|
| ENSG00000271129 | 0 | 0 | 0 | 0 | 0 | 0 |
| ENSG00000244378 | 0 | 0 | 0 | 0 | 1 | 0 |
| ENSG00000103044 | 26 | 22 | 21 | 7 | 19 | 8 |
| ENSG00000168802 | 4463 | 2324 | 2610 | 2610 | 2575 | 2461 |
| ENSG00000141076 | 1866 | 989 | 1119 | 1197 | 1239 | 1241 |
| ENSG00000207083 | 0 | 0 | 0 | 0 | 0 | 0 |
| ENSG00000168807 | 1762 | 881 | 877 | 495 | 785 | 621 |
| ENSG00000200164 | 0 | 0 | 0 | 1 | 0 | 0 |
| ENSG00000265113 | 0 | 0 | 0 | 0 | 0 | 0 |
| ENSG00000260914 | 0 | 0 | 1 | 0 | 0 | 1 |
| ENSG00000132612 | 2268 | 1229 | 1270 | 1488 | 1350 | 1362 |
| ENSG00000213380 | 782 | 562 | 587 | 521 | 491 | 569 |
| ENSG00000258429 | 98 | 104 | 92 | 74 | 75 | 91 |
| ENSG00000272617 | 0 | 0 | 0 | 0 | 0 | 0 |
| ENSG00000260371 | 0 | 0 | 0 | 0 | 0 | 0 |
| ENSG00000259900 | 0 | 0 | 0 | 0 | 0 | 0 |
| ENSG00000132603 | 1293 | 608 | 707 | 606 | 810 | 813 |
| ENSG00000157315 | 5 | 5 | 4 | 5 | 7 | 9 |
| ENSG00000132604 | 962 | 535 | 645 | 492 | 552 | 513 |
| ENSG00000103018 | 3851 | 1747 | 2118 | 1779 | 2025 | 2118 |
| ENSG00000261527 | 0 | 3 | 0 | 0 | 0 | 0 |
| ENSG00000260108 | 0 | 0 | 0 | 0 | 0 | 0 |
| ENSG00000102908 | 1138 | 581 | 556 | 142 | 721 | 489 |
| ENSG00000223109 | 0 | 0 | 0 | 0 | 0 | 0 |
| ENSG00000274093 | 0 | 1 | 0 | 0 | 0 | 2 |
| ENSG00000260772 | 0 | 0 | 0 | 0 | 0 | 0 |
| ENSG00000181019 | 50899 | 26487 | 33411 | 32751 | 34163 | 35636 |
| ENSG00000261602 | 0 | 0 | 0 | 0 | 0 | 0 |
| ENSG00000262136 | 46 | 25 | 23 | 25 | 24 | 26 |
| ENSG00000141101 | 2153 | 1244 | 1386 | 1637 | 1453 | 1423 |
| ENSG00000260290 | 9 | 9 | 5 | 1 | 2 | 6 |
| ENSG00000198373 | 3132 | 1399 | 1898 | 1449 | 1480 | 1705 |
| ENSG00000252443 | 0 | 0 | 0 | 0 | 0 | 0 |
| ENSG00000208017 | 0 | 0 | 0 | 0 | 1 | 0 |
| ENSG00000157322 | 8 | 7 | 4 | 1 | 4 | 3 |
| ENSG00000277003 | 0 | 0 | 0 | 0 | 0 | 0 |
| ENSG00000196696 | 324 | 150 | 256 | 134 | 167 | 218 |
| ENSG00000226232 | 0 | 0 | 0 | 0 | 0 | 0 |
| ENSG00000255185 | 2 | 2 | 1 | 4 | 2 | 3 |
| ENSG00000239118 | 0 | 1 | 0 | 0 | 1 | 0 |
| ENSG00000241183 | 0 | 0 | 0 | 0 | 0 | 0 |
| ENSG00000090857 | 1881 | 959 | 1316 | 717 | 977 | 1015 |
| ENSG00000247228 | 0 | 0 | 1 | 0 | 0 | 0 |
| ENSG00000157335 | 3 | 1 | 1 | 0 | 0 | 0 |
| ENSG00000261145 | 0 | 0 | 0 | 0 | 0 | 0 |
| ENSG00000261556 | 112 | 38 | 60 | 28 | 55 | 67 |
| ENSG00000223496 | 1991 | 977 | 1164 | 1011 | 1051 | 1026 |
| ENSG00000090861 | 8322 | 4002 | 5025 | 4546 | 4555 | 4555 |
| ENSG00000265648 | 0 | 0 | 0 | 0 | 0 | 0 |
| ENSG00000157349 | 583 | 322 | 394 | 361 | 355 | 383 |

| | | | | | | |
|---|---|---|---|---|---|---|
| ENSG00000260537 | 0 | 1 | 1 | 0 | 2 | 5 |
| ENSG00000261777 | 31 | 14 | 18 | 17 | 24 | 21 |
| ENSG00000168872 | 538 | 309 | 353 | 289 | 365 | 315 |
| ENSG00000285710 | 2 | 0 | 0 | 0 | 1 | 0 |
| ENSG00000157350 | 279 | 102 | 138 | 106 | 114 | 108 |
| ENSG00000260111 | 0 | 0 | 0 | 0 | 0 | 0 |
| ENSG00000240043 | 0 | 0 | 0 | 0 | 0 | 0 |
| ENSG00000200153 | 0 | 0 | 0 | 0 | 0 | 0 |
| ENSG00000157353 | 1316 | 579 | 765 | 774 | 648 | 627 |
| ENSG00000103051 | 3807 | 1772 | 2230 | 1924 | 1815 | 1857 |
| ENSG00000189091 | 13713 | 6520 | 8299 | 6828 | 7452 | 8129 |
| ENSG00000221514 | 0 | 0 | 0 | 0 | 0 | 0 |
| ENSG00000221066 | 0 | 0 | 1 | 0 | 0 | 0 |
| ENSG00000157368 | 0 | 0 | 0 | 0 | 0 | 0 |
| ENSG00000132613 | 1197 | 659 | 851 | 698 | 766 | 770 |
| ENSG00000279569 | 14 | 5 | 5 | 2 | 3 | 7 |
| ENSG00000103043 | 1675 | 774 | 1049 | 1015 | 972 | 934 |
| ENSG00000279412 | 12 | 9 | 11 | 1 | 6 | 0 |
| ENSG00000260156 | 0 | 2 | 1 | 1 | 1 | 4 |
| ENSG00000214353 | 3 | 8 | 3 | 0 | 1 | 2 |
| ENSG00000279815 | 0 | 0 | 0 | 0 | 0 | 0 |
| ENSG00000279122 | 0 | 0 | 0 | 0 | 0 | 0 |
| ENSG00000279123 | 8 | 1 | 4 | 0 | 1 | 5 |
| ENSG00000262890 | 0 | 0 | 0 | 0 | 0 | 0 |
| ENSG00000157423 | 0 | 2 | 0 | 0 | 0 | 0 |
| ENSG00000221725 | 0 | 0 | 0 | 0 | 0 | 0 |
| ENSG00000259833 | 0 | 0 | 0 | 0 | 0 | 0 |
| ENSG00000259798 | 0 | 0 | 0 | 0 | 0 | 0 |
| ENSG00000180917 | 1149 | 623 | 592 | 367 | 754 | 579 |
| ENSG00000261260 | 0 | 0 | 0 | 0 | 0 | 0 |
| ENSG00000172137 | 4 | 2 | 1 | 1 | 0 | 0 |
| ENSG00000261348 | 0 | 0 | 0 | 0 | 0 | 0 |
| ENSG00000262120 | 0 | 0 | 0 | 0 | 0 | 0 |
| ENSG00000260734 | 0 | 0 | 0 | 1 | 0 | 0 |
| ENSG00000262008 | 0 | 0 | 0 | 0 | 0 | 0 |
| ENSG00000167377 | 34 | 21 | 27 | 19 | 16 | 24 |
| ENSG00000261611 | 4 | 1 | 0 | 0 | 1 | 0 |
| ENSG00000247324 | 0 | 0 | 2 | 1 | 0 | 1 |
| ENSG00000157429 | 128 | 59 | 107 | 44 | 69 | 60 |
| ENSG00000260520 | 0 | 0 | 0 | 0 | 0 | 1 |
| ENSG00000140835 | 2 | 1 | 4 | 2 | 1 | 1 |
| ENSG00000260015 | 0 | 0 | 1 | 0 | 1 | 0 |
| ENSG00000252339 | 0 | 1 | 0 | 0 | 0 | 0 |
| ENSG00000198650 | 2 | 3 | 2 | 3 | 4 | 2 |
| ENSG00000260886 | 7 | 3 | 3 | 1 | 1 | 0 |
| ENSG00000260593 | 0 | 2 | 0 | 2 | 3 | 0 |
| ENSG00000140832 | 1213 | 574 | 665 | 557 | 708 | 641 |
| ENSG00000040199 | 1610 | 701 | 845 | 534 | 850 | 719 |
| ENSG00000260185 | 2 | 3 | 1 | 0 | 1 | 1 |
| ENSG00000199301 | 2 | 1 | 1 | 0 | 0 | 0 |

| | | | | | | |
|---|---|---|---|---|---|---|
| ENSG00000263666 | 0 | 0 | 0 | 1 | 1 | 0 |
| ENSG00000261513 | 3 | 4 | 4 | 4 | 5 | 1 |
| ENSG00000260612 | 0 | 1 | 0 | 0 | 1 | 0 |
| ENSG00000166747 | 6974 | 3258 | 3789 | 2040 | 3836 | 3117 |
| ENSG00000223224 | 0 | 0 | 1 | 0 | 0 | 0 |
| ENSG00000283057 | 0 | 1 | 0 | 0 | 1 | 0 |
| ENSG00000262140 | 0 | 0 | 0 | 0 | 0 | 0 |
| ENSG00000224470 | 3331 | 1358 | 1897 | 1185 | 1703 | 1732 |
| ENSG00000182149 | 8464 | 3946 | 4753 | 3680 | 4184 | 3869 |
| ENSG00000279592 | 0 | 0 | 0 | 0 | 0 | 0 |
| ENSG00000102984 | 151 | 102 | 143 | 114 | 116 | 106 |
| ENSG00000261765 | 0 | 3 | 1 | 0 | 1 | 0 |
| ENSG00000277481 | 6 | 3 | 6 | 0 | 2 | 2 |
| ENSG00000242176 | 0 | 0 | 0 | 0 | 0 | 0 |
| ENSG00000263311 | 0 | 0 | 0 | 0 | 0 | 0 |
| ENSG00000263232 | 0 | 2 | 0 | 1 | 0 | 0 |
| ENSG00000102967 | 492 | 294 | 321 | 354 | 342 | 386 |
| ENSG00000140830 | 905 | 514 | 577 | 539 | 593 | 567 |
| ENSG00000257017 | 0 | 1 | 0 | 1 | 0 | 0 |
| ENSG00000261701 | 0 | 1 | 0 | 0 | 0 | 0 |
| ENSG00000260252 | 9 | 8 | 1 | 5 | 3 | 5 |
| ENSG00000140829 | 3229 | 1465 | 1961 | 1577 | 1849 | 1866 |
| ENSG00000118557 | 27 | 4 | 5 | 2 | 11 | 3 |
| ENSG00000261673 | 0 | 0 | 0 | 0 | 0 | 0 |
| ENSG00000261774 | 0 | 0 | 0 | 0 | 0 | 0 |
| ENSG00000261008 | 43 | 24 | 39 | 28 | 30 | 36 |
| ENSG00000283564 | 0 | 0 | 0 | 0 | 0 | 0 |
| ENSG00000260664 | 0 | 0 | 0 | 0 | 0 | 0 |
| ENSG00000259768 | 174 | 96 | 97 | 71 | 105 | 107 |
| ENSG00000238731 | 0 | 0 | 0 | 0 | 0 | 0 |
| ENSG00000261764 | 2 | 2 | 0 | 1 | 0 | 1 |
| ENSG00000279206 | 0 | 0 | 0 | 0 | 2 | 1 |
| ENSG00000140836 | 2211 | 994 | 1279 | 681 | 1033 | 1179 |
| ENSG00000259209 | 10 | 4 | 9 | 1 | 5 | 5 |
| ENSG00000251868 | 0 | 0 | 0 | 0 | 0 | 0 |
| ENSG00000279757 | 0 | 0 | 2 | 0 | 1 | 0 |
| ENSG00000280092 | 0 | 0 | 0 | 0 | 0 | 3 |
| ENSG00000272330 | 0 | 0 | 0 | 0 | 0 | 1 |
| ENSG00000279591 | 0 | 1 | 1 | 3 | 1 | 2 |
| ENSG00000271009 | 34 | 22 | 29 | 17 | 25 | 12 |
| ENSG00000260880 | 2 | 0 | 1 | 2 | 4 | 0 |
| ENSG00000284675 | 0 | 1 | 0 | 0 | 0 | 0 |
| ENSG00000261227 | 0 | 0 | 0 | 0 | 0 | 0 |
| ENSG00000279120 | 0 | 0 | 0 | 0 | 0 | 0 |
| ENSG00000258779 | 0 | 0 | 0 | 0 | 0 | 0 |
| ENSG00000260848 | 0 | 0 | 0 | 0 | 0 | 0 |
| ENSG00000283457 | 0 | 0 | 0 | 0 | 0 | 0 |
| ENSG00000259817 | 0 | 0 | 0 | 0 | 0 | 0 |
| ENSG00000284670 | 0 | 0 | 0 | 0 | 0 | 0 |
| ENSG00000261404 | 6 | 1 | 3 | 6 | 5 | 5 |

| | | | | | | |
|---|---|---|---|---|---|---|
| ENSG00000279420 | 0 | 0 | 0 | 0 | 0 | 0 |
| ENSG00000279100 | 0 | 0 | 0 | 0 | 0 | 0 |
| ENSG00000260471 | 0 | 0 | 0 | 0 | 0 | 0 |
| ENSG00000261248 | 0 | 2 | 1 | 0 | 0 | 0 |
| ENSG00000275236 | 10 | 4 | 13 | 6 | 2 | 5 |
| ENSG00000103035 | 3877 | 2336 | 2830 | 2442 | 2799 | 2715 |
| ENSG00000259972 | 188 | 61 | 122 | 65 | 83 | 93 |
| ENSG00000239763 | 21 | 16 | 17 | 13 | 25 | 8 |
| ENSG00000260884 | 22 | 18 | 11 | 5 | 13 | 16 |
| ENSG00000214331 | 141 | 82 | 82 | 33 | 61 | 78 |
| ENSG00000261079 | 4 | 5 | 17 | 2 | 5 | 5 |
| ENSG00000196436 | 141 | 56 | 109 | 57 | 55 | 77 |
| ENSG00000276822 | 0 | 0 | 0 | 0 | 0 | 0 |
| ENSG00000140839 | 8 | 2 | 11 | 6 | 2 | 7 |
| ENSG00000261170 | 0 | 2 | 0 | 0 | 0 | 0 |
| ENSG00000090863 | 7752 | 3432 | 4462 | 2671 | 3521 | 3461 |
| ENSG00000251794 | 0 | 1 | 0 | 0 | 4 | 0 |
| ENSG00000207525 | 0 | 0 | 4 | 0 | 3 | 1 |
| ENSG00000240861 | 0 | 0 | 0 | 0 | 0 | 0 |
| ENSG00000270945 | 0 | 0 | 0 | 0 | 0 | 0 |
| ENSG00000168411 | 2522 | 1309 | 1607 | 1059 | 1335 | 1372 |
| ENSG00000262904 | 2 | 9 | 6 | 6 | 9 | 8 |
| ENSG00000168404 | 110 | 67 | 90 | 79 | 80 | 87 |
| ENSG00000103089 | 385 | 198 | 227 | 211 | 165 | 161 |
| ENSG00000261458 | 0 | 0 | 0 | 0 | 0 | 0 |
| ENSG00000103091 | 2781 | 1328 | 1747 | 1475 | 1689 | 1676 |
| ENSG00000186187 | 3228 | 1696 | 1926 | 2081 | 1792 | 1854 |
| ENSG00000247033 | 0 | 0 | 0 | 0 | 0 | 0 |
| ENSG00000166816 | 340 | 148 | 228 | 124 | 112 | 146 |
| ENSG00000261058 | 0 | 0 | 0 | 1 | 1 | 1 |
| ENSG00000184517 | 657 | 340 | 386 | 203 | 533 | 337 |
| ENSG00000213318 | 0 | 0 | 0 | 0 | 0 | 0 |
| ENSG00000168928 | 13 | 13 | 14 | 5 | 9 | 6 |
| ENSG00000168925 | 29 | 14 | 9 | 3 | 6 | 2 |
| ENSG00000240338 | 6 | 7 | 6 | 2 | 9 | 7 |
| ENSG00000050820 | 4337 | 2074 | 2536 | 2876 | 2568 | 2340 |
| ENSG00000280152 | 60 | 24 | 33 | 30 | 34 | 24 |
| ENSG00000252101 | 0 | 0 | 1 | 0 | 0 | 0 |
| ENSG00000153774 | 1517 | 831 | 944 | 826 | 1002 | 837 |
| ENSG00000259999 | 2 | 0 | 0 | 0 | 0 | 0 |
| ENSG00000261783 | 22 | 8 | 11 | 9 | 8 | 13 |
| ENSG00000252122 | 0 | 0 | 1 | 1 | 0 | 0 |
| ENSG00000261717 | 0 | 0 | 0 | 0 | 0 | 0 |
| ENSG00000274220 | 97 | 72 | 77 | 41 | 71 | 68 |
| ENSG00000166822 | 1355 | 550 | 757 | 580 | 686 | 673 |
| ENSG00000261476 | 3 | 8 | 5 | 1 | 1 | 4 |
| ENSG00000183196 | 4 | 0 | 3 | 1 | 2 | 2 |
| ENSG00000203472 | 0 | 0 | 0 | 0 | 0 | 0 |
| ENSG00000273971 | 0 | 1 | 1 | 0 | 1 | 2 |
| ENSG00000262583 | 20 | 10 | 7 | 12 | 8 | 14 |

| | | | | | | |
|---|---|---|---|---|---|---|
| ENSG00000260092 | 0 | 0 | 0 | 0 | 0 | 0 |
| ENSG00000135702 | 1 | 2 | 2 | 0 | 1 | 1 |
| ENSG00000205084 | 429 | 238 | 264 | 270 | 260 | 301 |
| ENSG00000259992 | 0 | 0 | 0 | 0 | 0 | 0 |
| ENSG00000276408 | 4 | 8 | 1 | 7 | 4 | 6 |
| ENSG00000034713 | 5456 | 2755 | 3340 | 2420 | 2871 | 2631 |
| ENSG00000276523 | 17 | 16 | 24 | 2 | 9 | 10 |
| ENSG00000065457 | 1949 | 946 | 1246 | 792 | 1167 | 1170 |
| ENSG00000065427 | 9342 | 4764 | 5487 | 5725 | 6093 | 6019 |
| ENSG00000166848 | 4954 | 2478 | 2991 | 2495 | 2601 | 2520 |
| ENSG00000284484 | 0 | 1 | 6 | 1 | 2 | 0 |
| ENSG00000282757 | 4 | 1 | 3 | 0 | 1 | 2 |
| ENSG00000240199 | 0 | 0 | 0 | 0 | 0 | 0 |
| ENSG00000261424 | 0 | 0 | 0 | 0 | 0 | 0 |
| ENSG00000283755 | 0 | 0 | 0 | 0 | 0 | 0 |
| ENSG00000261313 | 0 | 0 | 0 | 0 | 0 | 0 |
| ENSG00000199299 | 0 | 0 | 0 | 0 | 0 | 0 |
| ENSG00000260223 | 0 | 0 | 0 | 0 | 0 | 0 |
| ENSG00000260983 | 0 | 0 | 0 | 0 | 0 | 0 |
| ENSG00000244485 | 0 | 1 | 0 | 0 | 0 | 0 |
| ENSG00000152910 | 0 | 0 | 0 | 0 | 0 | 0 |
| ENSG00000252814 | 0 | 0 | 0 | 0 | 0 | 0 |
| ENSG00000252022 | 0 | 0 | 0 | 0 | 0 | 0 |
| ENSG00000261833 | 0 | 0 | 0 | 0 | 0 | 0 |
| ENSG00000273659 | 0 | 0 | 0 | 0 | 0 | 0 |
| ENSG00000250514 | 0 | 0 | 0 | 0 | 0 | 0 |
| ENSG00000259995 | 0 | 0 | 0 | 0 | 0 | 0 |
| ENSG00000266426 | 0 | 0 | 0 | 0 | 0 | 0 |
| ENSG00000260797 | 0 | 0 | 0 | 0 | 0 | 0 |
| ENSG00000279856 | 0 | 0 | 0 | 0 | 0 | 0 |
| ENSG00000103111 | 3184 | 1413 | 1822 | 1565 | 1726 | 1668 |
| ENSG00000205078 | 33 | 25 | 24 | 22 | 16 | 8 |
| ENSG00000261063 | 0 | 0 | 0 | 0 | 0 | 0 |
| ENSG00000226348 | 0 | 0 | 0 | 0 | 0 | 0 |
| ENSG00000260922 | 0 | 0 | 0 | 0 | 0 | 1 |
| ENSG00000140873 | 12 | 4 | 4 | 3 | 3 | 1 |
| ENSG00000260701 | 0 | 0 | 0 | 0 | 0 | 0 |
| ENSG00000261154 | 2 | 0 | 1 | 1 | 1 | 0 |
| ENSG00000278862 | 4 | 3 | 4 | 1 | 0 | 1 |
| ENSG00000140876 | 166 | 109 | 126 | 88 | 80 | 84 |
| ENSG00000261707 | 0 | 5 | 10 | 1 | 4 | 3 |
| ENSG00000285538 | 0 | 0 | 0 | 0 | 0 | 0 |
| ENSG00000171724 | 207 | 106 | 122 | 46 | 53 | 66 |
| ENSG00000278926 | 0 | 0 | 0 | 0 | 0 | 0 |
| ENSG00000261330 | 0 | 2 | 1 | 0 | 0 | 0 |
| ENSG00000261540 | 20 | 13 | 16 | 0 | 3 | 4 |
| ENSG00000166509 | 2424 | 1466 | 2136 | 371 | 535 | 768 |
| ENSG00000260731 | 13 | 6 | 10 | 3 | 2 | 7 |
| ENSG00000261776 | 56 | 46 | 46 | 2 | 7 | 15 |
| ENSG00000186153 | 2274 | 1072 | 1403 | 953 | 878 | 1036 |

| | | | | | | |
|---|---|---|---|---|---|---|
| ENSG00000276007 | 127 | 45 | 88 | 54 | 46 | 69 |
| ENSG00000279991 | 0 | 1 | 0 | 0 | 2 | 0 |
| ENSG00000260969 | 2 | 1 | 0 | 0 | 2 | 1 |
| ENSG00000260014 | 0 | 0 | 0 | 0 | 0 | 0 |
| ENSG00000260733 | 2 | 2 | 0 | 0 | 0 | 0 |
| ENSG00000279887 | 0 | 0 | 2 | 0 | 0 | 1 |
| ENSG00000261837 | 0 | 0 | 4 | 0 | 0 | 1 |
| ENSG00000279129 | 2 | 1 | 3 | 0 | 3 | 3 |
| ENSG00000243101 | 0 | 0 | 0 | 0 | 0 | 1 |
| ENSG00000280115 | 0 | 0 | 0 | 0 | 0 | 0 |
| ENSG00000280190 | 0 | 0 | 0 | 0 | 0 | 2 |
| ENSG00000260816 | 8 | 1 | 7 | 1 | 0 | 0 |
| ENSG00000279732 | 2 | 1 | 1 | 0 | 1 | 1 |
| ENSG00000261537 | 0 | 2 | 3 | 1 | 0 | 0 |
| ENSG00000280274 | 2 | 0 | 1 | 1 | 0 | 0 |
| ENSG00000260694 | 0 | 0 | 0 | 0 | 0 | 0 |
| ENSG00000260479 | 0 | 0 | 0 | 0 | 0 | 0 |
| ENSG00000260511 | 0 | 0 | 0 | 0 | 0 | 0 |
| ENSG00000277954 | 4 | 0 | 0 | 0 | 1 | 2 |
| ENSG00000285918 | 2 | 1 | 1 | 0 | 0 | 0 |
| ENSG00000261722 | 0 | 0 | 0 | 0 | 0 | 0 |
| ENSG00000222244 | 0 | 0 | 0 | 0 | 0 | 0 |
| ENSG00000261472 | 0 | 0 | 0 | 0 | 0 | 0 |
| ENSG00000178573 | 68 | 21 | 18 | 16 | 41 | 24 |
| ENSG00000277214 | 0 | 0 | 0 | 0 | 0 | 0 |
| ENSG00000278058 | 0 | 0 | 0 | 0 | 2 | 1 |
| ENSG00000275040 | 0 | 0 | 0 | 0 | 0 | 0 |
| ENSG00000279378 | 0 | 0 | 0 | 0 | 0 | 0 |
| ENSG00000260876 | 4 | 2 | 0 | 1 | 1 | 1 |
| ENSG00000261390 | 2 | 3 | 2 | 6 | 6 | 2 |
| ENSG00000261082 | 0 | 0 | 0 | 0 | 0 | 0 |
| ENSG00000260237 | 0 | 0 | 0 | 0 | 0 | 0 |
| ENSG00000260706 | 0 | 0 | 0 | 0 | 0 | 1 |
| ENSG00000259867 | 3 | 2 | 2 | 0 | 2 | 2 |
| ENSG00000261376 | 0 | 0 | 0 | 0 | 0 | 0 |
| ENSG00000168589 | 33 | 18 | 24 | 25 | 17 | 18 |
| ENSG00000260183 | 4 | 1 | 1 | 0 | 1 | 0 |
| ENSG00000260737 | 0 | 1 | 0 | 0 | 0 | 0 |
| ENSG00000166446 | 2016 | 1012 | 1332 | 642 | 997 | 997 |
| ENSG00000260064 | 2 | 0 | 0 | 0 | 1 | 0 |
| ENSG00000260594 | 0 | 0 | 0 | 0 | 0 | 0 |
| ENSG00000260896 | 3 | 7 | 7 | 2 | 5 | 9 |
| ENSG00000103121 | 1497 | 833 | 933 | 1048 | 1020 | 1083 |
| ENSG00000278985 | 2 | 4 | 3 | 5 | 5 | 2 |
| ENSG00000166451 | 773 | 474 | 593 | 486 | 584 | 573 |
| ENSG00000260213 | 1 | 1 | 4 | 5 | 6 | 3 |
| ENSG00000284512 | 7 | 4 | 3 | 0 | 1 | 7 |
| ENSG00000261061 | 194 | 124 | 155 | 152 | 179 | 164 |
| ENSG00000166454 | 5672 | 2805 | 3296 | 2273 | 3310 | 2808 |
| ENSG00000166455 | 2 | 2 | 1 | 4 | 0 | 2 |

| | | | | | | |
|---|---|---|---|---|---|---|
| ENSG00000260643 | 42 | 42 | 34 | 22 | 36 | 26 |
| ENSG00000261141 | 17 | 7 | 9 | 6 | 7 | 19 |
| ENSG00000261838 | 2 | 3 | 1 | 1 | 0 | 0 |
| ENSG00000245059 | 7 | 3 | 3 | 3 | 0 | 2 |
| ENSG00000140905 | 85 | 50 | 66 | 52 | 54 | 59 |
| ENSG00000166473 | 257 | 104 | 166 | 85 | 137 | 110 |
| ENSG00000252608 | 0 | 0 | 1 | 0 | 1 | 2 |
| ENSG00000135697 | 95 | 49 | 62 | 63 | 62 | 57 |
| ENSG00000280182 | 0 | 3 | 2 | 0 | 3 | 1 |
| ENSG00000260495 | 39 | 30 | 34 | 26 | 23 | 16 |
| ENSG00000261609 | 2951 | 1695 | 1568 | 1037 | 1440 | 1403 |
| ENSG00000284473 | 0 | 0 | 0 | 0 | 0 | 0 |
| ENSG00000272923 | 0 | 0 | 0 | 0 | 0 | 0 |
| ENSG00000153815 | 1619 | 838 | 912 | 809 | 742 | 867 |
| ENSG00000279021 | 0 | 1 | 0 | 0 | 0 | 0 |
| ENSG00000260229 | 0 | 3 | 0 | 0 | 0 | 1 |
| ENSG00000279476 | 0 | 1 | 1 | 0 | 0 | 1 |
| ENSG00000277255 | 0 | 0 | 0 | 0 | 0 | 0 |
| ENSG00000275109 | 0 | 0 | 0 | 0 | 0 | 0 |
| ENSG00000279681 | 0 | 0 | 0 | 0 | 0 | 0 |
| ENSG00000279294 | 2 | 0 | 0 | 0 | 0 | 1 |
| ENSG00000279841 | 0 | 0 | 0 | 3 | 1 | 5 |
| ENSG00000261218 | 0 | 0 | 0 | 0 | 0 | 0 |
| ENSG00000197943 | 379 | 169 | 212 | 201 | 228 | 206 |
| ENSG00000260682 | 0 | 0 | 0 | 0 | 0 | 0 |
| ENSG00000276519 | 0 | 0 | 0 | 0 | 1 | 1 |
| ENSG00000184860 | 908 | 447 | 591 | 473 | 538 | 579 |
| ENSG00000086696 | 0 | 0 | 3 | 1 | 0 | 3 |
| ENSG00000261235 | 77 | 38 | 67 | 25 | 27 | 36 |
| ENSG00000260161 | 0 | 0 | 0 | 0 | 0 | 0 |
| ENSG00000135698 | 460 | 266 | 309 | 238 | 299 | 265 |
| ENSG00000261029 | 0 | 0 | 0 | 0 | 0 | 0 |
| ENSG00000251888 | 0 | 0 | 0 | 0 | 0 | 0 |
| ENSG00000259873 | 0 | 0 | 0 | 0 | 0 | 0 |
| ENSG00000261285 | 0 | 0 | 0 | 0 | 0 | 0 |
| ENSG00000140945 | 146 | 60 | 67 | 39 | 63 | 56 |
| ENSG00000277387 | 0 | 0 | 0 | 0 | 0 | 0 |
| ENSG00000280044 | 0 | 0 | 0 | 0 | 0 | 0 |
| ENSG00000260862 | 0 | 0 | 0 | 0 | 0 | 0 |
| ENSG00000263360 | 0 | 0 | 0 | 0 | 0 | 0 |
| ENSG00000260832 | 0 | 0 | 0 | 0 | 0 | 0 |
| ENSG00000261410 | 0 | 0 | 0 | 0 | 0 | 0 |
| ENSG00000263785 | 0 | 0 | 0 | 0 | 0 | 0 |
| ENSG00000261103 | 0 | 0 | 0 | 0 | 0 | 0 |
| ENSG00000260788 | 0 | 0 | 0 | 0 | 0 | 0 |
| ENSG00000259914 | 0 | 0 | 0 | 0 | 0 | 0 |
| ENSG00000260228 | 119 | 86 | 96 | 52 | 61 | 60 |
| ENSG00000230989 | 10338 | 4985 | 6006 | 5902 | 5878 | 5541 |
| ENSG00000103150 | 909 | 469 | 540 | 555 | 556 | 597 |
| ENSG00000260300 | 21 | 16 | 17 | 9 | 18 | 10 |

| | | | | | | |
|---|---|---|---|---|---|---|
| ENSG00000260932 | 2 | 2 | 3 | 3 | 1 | 0 |
| ENSG00000140961 | 1251 | 591 | 733 | 1178 | 808 | 953 |
| ENSG00000103154 | 57 | 26 | 33 | 27 | 22 | 45 |
| ENSG00000166558 | 0 | 1 | 0 | 0 | 0 | 0 |
| ENSG00000199350 | 0 | 0 | 0 | 0 | 0 | 0 |
| ENSG00000285792 | 2 | 1 | 0 | 1 | 0 | 0 |
| ENSG00000140943 | 10680 | 4950 | 6720 | 5290 | 6227 | 6147 |
| ENSG00000274677 | 17 | 3 | 4 | 5 | 2 | 5 |
| ENSG00000260018 | 249 | 102 | 154 | 166 | 144 | 119 |
| ENSG00000260410 | 0 | 3 | 2 | 3 | 7 | 5 |
| ENSG00000103160 | 3039 | 1512 | 1750 | 1202 | 1905 | 1538 |
| ENSG00000154099 | 156 | 92 | 137 | 124 | 95 | 107 |
| ENSG00000103168 | 2495 | 1074 | 1619 | 1288 | 1376 | 1425 |
| ENSG00000140955 | 9 | 3 | 2 | 7 | 9 | 5 |
| ENSG00000250685 | 0 | 0 | 0 | 1 | 0 | 0 |
| ENSG00000168418 | 0 | 0 | 1 | 0 | 0 | 0 |
| ENSG00000260858 | 0 | 0 | 0 | 0 | 0 | 0 |
| ENSG00000201361 | 0 | 0 | 0 | 0 | 0 | 0 |
| ENSG00000103175 | 0 | 0 | 0 | 0 | 0 | 1 |
| ENSG00000260530 | 0 | 0 | 0 | 0 | 0 | 0 |
| ENSG00000260504 | 0 | 0 | 0 | 0 | 0 | 0 |
| ENSG00000064270 | 2023 | 942 | 1282 | 764 | 788 | 908 |
| ENSG00000261286 | 42 | 17 | 26 | 13 | 17 | 16 |
| ENSG00000140950 | 1659 | 964 | 1125 | 986 | 1099 | 968 |
| ENSG00000261243 | 5 | 3 | 2 | 3 | 3 | 1 |
| ENSG00000103187 | 728 | 334 | 311 | 495 | 451 | 343 |
| ENSG00000261471 | 0 | 2 | 0 | 1 | 0 | 0 |
| ENSG00000135686 | 5948 | 2635 | 3692 | 2574 | 3356 | 3379 |
| ENSG00000103194 | 4544 | 2195 | 2690 | 2096 | 2867 | 2525 |
| ENSG00000285848 | 9 | 9 | 11 | 7 | 6 | 9 |
| ENSG00000103196 | 966 | 468 | 615 | 507 | 667 | 782 |
| ENSG00000260859 | 0 | 0 | 2 | 2 | 6 | 2 |
| ENSG00000279622 | 2 | 0 | 0 | 1 | 1 | 1 |
| ENSG00000260340 | 0 | 0 | 0 | 0 | 0 | 0 |
| ENSG00000153786 | 9894 | 4767 | 6212 | 4147 | 4784 | 5116 |
| ENSG00000135709 | 3344 | 1550 | 2113 | 1234 | 1537 | 1413 |
| ENSG00000153789 | 0 | 0 | 1 | 0 | 0 | 0 |
| ENSG00000278214 | 0 | 0 | 1 | 0 | 0 | 0 |
| ENSG00000275088 | 4 | 3 | 3 | 5 | 5 | 5 |
| ENSG00000131149 | 15697 | 6847 | 9507 | 6220 | 7821 | 8052 |
| ENSG00000270313 | 0 | 0 | 0 | 0 | 0 | 0 |
| ENSG00000179219 | 0 | 2 | 1 | 3 | 1 | 3 |
| ENSG00000266307 | 1 | 0 | 0 | 0 | 0 | 0 |
| ENSG00000278716 | 10 | 1 | 2 | 7 | 4 | 3 |
| ENSG00000270124 | 3 | 1 | 2 | 1 | 1 | 2 |
| ENSG00000260417 | 2 | 3 | 5 | 0 | 1 | 0 |
| ENSG00000263968 | 34 | 19 | 11 | 6 | 18 | 11 |
| ENSG00000131153 | 964 | 562 | 680 | 565 | 492 | 584 |
| ENSG00000154102 | 561 | 332 | 305 | 376 | 338 | 349 |
| ENSG00000275393 | 13 | 12 | 11 | 14 | 11 | 8 |

| | | | | | | |
|---|---|---|---|---|---|---|
| ENSG00000283416 | 0 | 0 | 0 | 0 | 0 | 0 |
| ENSG00000269898 | 0 | 0 | 2 | 0 | 0 | 0 |
| ENSG00000131148 | 1793 | 869 | 1049 | 1205 | 1140 | 1084 |
| ENSG00000252311 | 15 | 4 | 6 | 9 | 4 | 3 |
| ENSG00000270184 | 16 | 10 | 18 | 5 | 7 | 13 |
| ENSG00000270159 | 2 | 0 | 1 | 0 | 0 | 1 |
| ENSG00000131143 | 16149 | 7986 | 9650 | 10809 | 9521 | 9354 |
| ENSG00000270698 | 0 | 0 | 0 | 0 | 0 | 0 |
| ENSG00000214259 | 0 | 0 | 0 | 0 | 0 | 0 |
| ENSG00000279907 | 0 | 0 | 1 | 0 | 1 | 0 |
| ENSG00000140968 | 0 | 0 | 0 | 1 | 0 | 2 |
| ENSG00000274134 | 0 | 0 | 0 | 0 | 0 | 0 |
| ENSG00000285163 | 0 | 0 | 0 | 0 | 0 | 0 |
| ENSG00000268804 | 0 | 0 | 0 | 0 | 0 | 0 |
| ENSG00000285040 | 0 | 0 | 0 | 0 | 0 | 0 |
| ENSG00000269667 | 0 | 0 | 0 | 0 | 0 | 0 |
| ENSG00000285012 | 0 | 0 | 0 | 0 | 0 | 0 |
| ENSG00000279604 | 0 | 0 | 0 | 0 | 0 | 0 |
| ENSG00000261177 | 0 | 0 | 0 | 0 | 0 | 0 |
| ENSG00000268078 | 0 | 0 | 0 | 0 | 0 | 0 |
| ENSG00000268505 | 0 | 0 | 0 | 0 | 0 | 0 |
| ENSG00000269186 | 0 | 0 | 0 | 0 | 0 | 0 |
| ENSG00000268754 | 0 | 0 | 0 | 0 | 0 | 0 |
| ENSG00000199949 | 0 | 0 | 0 | 0 | 0 | 0 |
| ENSG00000268532 | 0 | 0 | 0 | 0 | 0 | 0 |
| ENSG00000168367 | 0 | 0 | 0 | 0 | 0 | 0 |
| ENSG00000269826 | 0 | 0 | 0 | 0 | 0 | 0 |
| ENSG00000268473 | 0 | 0 | 0 | 0 | 0 | 0 |
| ENSG00000268388 | 0 | 0 | 0 | 0 | 0 | 0 |
| ENSG00000103241 | 4 | 2 | 1 | 2 | 0 | 1 |
| ENSG00000270020 | 0 | 2 | 4 | 2 | 1 | 3 |
| ENSG00000243697 | 0 | 0 | 2 | 0 | 1 | 2 |
| ENSG00000103248 | 1147 | 534 | 678 | 573 | 551 | 597 |
| ENSG00000280278 | 4 | 1 | 2 | 4 | 5 | 4 |
| ENSG00000260944 | 0 | 0 | 0 | 0 | 0 | 0 |
| ENSG00000176692 | 0 | 2 | 0 | 1 | 0 | 0 |
| ENSG00000176678 | 0 | 0 | 0 | 0 | 0 | 1 |
| ENSG00000279142 | 0 | 0 | 0 | 0 | 0 | 0 |
| ENSG00000260387 | 0 | 0 | 0 | 0 | 0 | 0 |
| ENSG00000261161 | 0 | 0 | 0 | 0 | 0 | 0 |
| ENSG00000274478 | 0 | 0 | 0 | 0 | 0 | 0 |
| ENSG00000261175 | 0 | 0 | 0 | 0 | 0 | 0 |
| ENSG00000260026 | 0 | 0 | 0 | 0 | 0 | 0 |
| ENSG00000260795 | 0 | 0 | 0 | 0 | 0 | 0 |
| ENSG00000232190 | 0 | 0 | 0 | 0 | 0 | 0 |
| ENSG00000261363 | 0 | 0 | 0 | 0 | 0 | 0 |
| ENSG00000260456 | 25 | 19 | 21 | 14 | 14 | 19 |
| ENSG00000285527 | 0 | 0 | 0 | 0 | 0 | 0 |
| ENSG00000261095 | 0 | 0 | 0 | 0 | 0 | 0 |
| ENSG00000261651 | 0 | 0 | 0 | 0 | 0 | 0 |

| | | | | | | |
|---|---|---|---|---|---|---|
| ENSG00000261697 | 0 | 0 | 0 | 0 | 0 | 0 |
| ENSG00000243705 | 0 | 0 | 0 | 0 | 0 | 0 |
| ENSG00000280419 | 0 | 0 | 0 | 0 | 0 | 0 |
| ENSG00000131152 | 0 | 1 | 1 | 0 | 0 | 0 |
| ENSG00000270006 | 79 | 27 | 24 | 50 | 47 | 23 |
| ENSG00000270082 | 0 | 0 | 0 | 0 | 0 | 0 |
| ENSG00000103264 | 1473 | 690 | 857 | 793 | 807 | 794 |
| ENSG00000269901 | 4 | 7 | 4 | 1 | 5 | 3 |
| ENSG00000140941 | 4332 | 2128 | 2680 | 2286 | 2540 | 2392 |
| ENSG00000261592 | 6 | 0 | 3 | 0 | 4 | 4 |
| ENSG00000140948 | 2046 | 904 | 1244 | 843 | 899 | 1045 |
| ENSG00000276393 | 0 | 0 | 0 | 0 | 0 | 0 |
| ENSG00000276337 | 8 | 2 | 3 | 1 | 0 | 0 |
| ENSG00000260750 | 8 | 2 | 7 | 4 | 0 | 6 |
| ENSG00000269935 | 0 | 0 | 7 | 1 | 2 | 1 |
| ENSG00000154118 | 314 | 147 | 173 | 208 | 169 | 170 |
| ENSG00000277504 | 14 | 1 | 1 | 3 | 5 | 6 |
| ENSG00000226180 | 46 | 20 | 31 | 11 | 12 | 18 |
| ENSG00000104731 | 2205 | 1043 | 1312 | 1326 | 1264 | 1356 |
| ENSG00000260671 | 4 | 2 | 0 | 0 | 0 | 0 |
| ENSG00000260498 | 0 | 0 | 0 | 0 | 0 | 0 |
| ENSG00000260177 | 0 | 1 | 0 | 0 | 0 | 0 |
| ENSG00000103257 | 1710 | 920 | 1016 | 7792 | 7694 | 6691 |
| ENSG00000278598 | 0 | 0 | 1 | 2 | 0 | 1 |
| ENSG00000260466 | 0 | 0 | 0 | 4 | 1 | 1 |
| ENSG00000174990 | 0 | 1 | 0 | 1 | 0 | 0 |
| ENSG00000260603 | 0 | 0 | 0 | 0 | 0 | 0 |
| ENSG00000270385 | 0 | 0 | 0 | 0 | 0 | 0 |
| ENSG00000260443 | 18 | 12 | 13 | 7 | 9 | 17 |
| ENSG00000172530 | 920 | 361 | 417 | 390 | 406 | 386 |
| ENSG00000261193 | 2 | 0 | 0 | 0 | 0 | 0 |
| ENSG00000260166 | 0 | 0 | 0 | 0 | 0 | 0 |
| ENSG00000205037 | 8 | 1 | 3 | 0 | 1 | 3 |
| ENSG00000260162 | 0 | 0 | 0 | 0 | 0 | 0 |
| ENSG00000261327 | 0 | 0 | 0 | 0 | 0 | 0 |
| ENSG00000261816 | 0 | 0 | 0 | 0 | 0 | 1 |
| ENSG00000260420 | 0 | 0 | 0 | 0 | 0 | 0 |
| ENSG00000261273 | 0 | 0 | 0 | 0 | 0 | 0 |
| ENSG00000225614 | 578 | 256 | 353 | 183 | 249 | 225 |
| ENSG00000179588 | 548 | 281 | 333 | 301 | 251 | 307 |
| ENSG00000263456 | 0 | 0 | 0 | 0 | 0 | 0 |
| ENSG00000261744 | 12 | 3 | 11 | 4 | 10 | 7 |
| ENSG00000158545 | 3134 | 1309 | 1745 | 1604 | 1636 | 1681 |
| ENSG00000124391 | 12 | 6 | 9 | 5 | 8 | 9 |
| ENSG00000051523 | 9628 | 4473 | 5371 | 6573 | 4728 | 5088 |
| ENSG00000167508 | 3311 | 1519 | 1993 | 2476 | 1900 | 1820 |
| ENSG00000260630 | 122 | 49 | 59 | 53 | 45 | 54 |
| ENSG00000185669 | 69 | 35 | 41 | 28 | 43 | 37 |
| ENSG00000158717 | 935 | 500 | 661 | 651 | 607 | 599 |
| ENSG00000259813 | 0 | 0 | 0 | 0 | 0 | 0 |

| | | | | | | |
|---|---|---|---|---|---|---|
| ENSG00000174177 | 873 | 410 | 586 | 792 | 573 | 580 |
| ENSG00000278341 | 44 | 27 | 22 | 23 | 31 | 32 |
| ENSG00000103335 | 16240 | 7569 | 9965 | 7569 | 8273 | 8022 |
| ENSG00000284585 | 0 | 0 | 0 | 0 | 0 | 0 |
| ENSG00000260121 | 131 | 96 | 138 | 56 | 93 | 99 |
| ENSG00000224888 | 11 | 10 | 9 | 2 | 9 | 4 |
| ENSG00000260617 | 0 | 0 | 0 | 0 | 0 | 0 |
| ENSG00000182376 | 0 | 1 | 1 | 1 | 2 | 2 |
| ENSG00000167513 | 1256 | 653 | 758 | 756 | 599 | 693 |
| ENSG00000198931 | 7721 | 4086 | 4460 | 5888 | 4438 | 4449 |
| ENSG00000141012 | 2023 | 904 | 1232 | 1092 | 1031 | 1086 |
| ENSG00000167515 | 3284 | 1652 | 1963 | 2060 | 1762 | 1713 |
| ENSG00000205022 | 13 | 6 | 4 | 1 | 5 | 8 |
| ENSG00000129993 | 1819 | 1000 | 1149 | 822 | 827 | 848 |
| ENSG00000259881 | 65 | 28 | 48 | 25 | 36 | 23 |
| ENSG00000261226 | 22 | 12 | 17 | 18 | 9 | 10 |
| ENSG00000205018 | 2 | 5 | 7 | 4 | 4 | 6 |
| ENSG00000259989 | 0 | 0 | 0 | 0 | 0 | 0 |
| ENSG00000256982 | 13 | 6 | 18 | 14 | 17 | 11 |
| ENSG00000176715 | 1784 | 889 | 1059 | 1004 | 898 | 914 |
| ENSG00000261546 | 3 | 1 | 5 | 0 | 5 | 3 |
| ENSG00000180422 | 14 | 5 | 11 | 7 | 5 | 1 |
| ENSG00000205015 | 2 | 0 | 2 | 0 | 0 | 1 |
| ENSG00000129910 | 7 | 10 | 1 | 5 | 4 | 5 |
| ENSG00000259803 | 16 | 18 | 9 | 14 | 7 | 15 |
| ENSG00000259877 | 357 | 155 | 210 | 148 | 167 | 216 |
| ENSG00000170100 | 903 | 467 | 554 | 358 | 503 | 488 |
| ENSG00000260659 | 0 | 0 | 0 | 0 | 0 | 0 |
| ENSG00000167522 | 4185 | 1882 | 2602 | 1846 | 2307 | 2372 |
| ENSG00000268218 | 48 | 29 | 49 | 26 | 29 | 31 |
| ENSG00000260279 | 147 | 76 | 92 | 43 | 79 | 64 |
| ENSG00000261253 | 0 | 1 | 2 | 0 | 1 | 1 |
| ENSG00000261692 | 0 | 0 | 1 | 0 | 0 | 1 |
| ENSG00000252887 | 0 | 1 | 0 | 0 | 0 | 0 |
| ENSG00000270832 | 0 | 0 | 1 | 0 | 0 | 0 |
| ENSG00000197912 | 5439 | 2420 | 3277 | 2592 | 2767 | 2878 |
| ENSG00000261118 | 78 | 41 | 42 | 31 | 30 | 24 |
| ENSG00000167526 | 69357 | 33208 | 39831 | 54454 | 40715 | 40847 |
| ENSG00000200084 | 0 | 0 | 0 | 0 | 0 | 0 |
| ENSG00000178773 | 27 | 6 | 13 | 24 | 12 | 18 |
| ENSG00000015413 | 8 | 7 | 4 | 1 | 4 | 0 |
| ENSG00000131165 | 9270 | 4583 | 5448 | 5457 | 4915 | 4841 |
| ENSG00000167523 | 429 | 188 | 259 | 285 | 218 | 238 |
| ENSG00000275734 | 31 | 26 | 34 | 24 | 32 | 20 |
| ENSG00000185324 | 2969 | 1384 | 1854 | 1520 | 1551 | 1599 |
| ENSG00000260259 | 14 | 15 | 15 | 4 | 17 | 11 |
| ENSG00000158792 | 1840 | 804 | 982 | 1211 | 878 | 909 |
| ENSG00000075399 | 1061 | 527 | 688 | 609 | 471 | 485 |
| ENSG00000261373 | 201 | 121 | 161 | 208 | 168 | 215 |
| ENSG00000158805 | 871 | 393 | 515 | 442 | 509 | 476 |

| | | | | | | |
|---|---|---|---|---|---|---|
| ENSG00000187741 | 904 | 484 | 609 | 375 | 476 | 560 |
| ENSG00000204991 | 1073 | 530 | 677 | 512 | 584 | 557 |
| ENSG00000141002 | 7386 | 3712 | 4871 | 4204 | 4222 | 4012 |
| ENSG00000267048 | 3 | 0 | 4 | 4 | 0 | 0 |
| ENSG00000258839 | 398 | 237 | 319 | 246 | 196 | 252 |
| ENSG00000198211 | 0 | 0 | 0 | 0 | 0 | 0 |
| ENSG00000259006 | 1 | 2 | 0 | 2 | 0 | 1 |
| ENSG00000258947 | 96 | 61 | 83 | 125 | 156 | 117 |
| ENSG00000140995 | 3296 | 1697 | 1983 | 1977 | 1908 | 1834 |
| ENSG00000177946 | 1036 | 515 | 597 | 535 | 597 | 580 |
| ENSG00000223959 | 917 | 406 | 591 | 425 | 493 | 529 |
| ENSG00000003249 | 6135 | 2814 | 4094 | 3842 | 3105 | 3318 |
| ENSG00000141013 | 1231 | 705 | 975 | 705 | 740 | 776 |
| ENSG00000221819 | 62 | 22 | 36 | 28 | 23 | 40 |
| ENSG00000222019 | 71 | 57 | 62 | 29 | 62 | 50 |
| ENSG00000126856 | 0 | 4 | 5 | 2 | 1 | 0 |
| ENSG00000261812 | 11 | 11 | 11 | 7 | 7 | 9 |
| ENSG00000260528 | 20 | 8 | 11 | 8 | 8 | 6 |
| ENSG00000260507 | 0 | 0 | 0 | 0 | 0 | 0 |
| ENSG00000261172 | 0 | 0 | 1 | 0 | 1 | 0 |
| ENSG00000222359 | 0 | 0 | 0 | 0 | 0 | 0 |
| ENSG00000260923 | 12 | 4 | 7 | 7 | 4 | 6 |
| ENSG00000277349 | 0 | 0 | 0 | 0 | 0 | 0 |
| ENSG00000280279 | 8 | 3 | 5 | 3 | 4 | 4 |
| ENSG00000280136 | 0 | 0 | 0 | 0 | 0 | 0 |
| ENSG00000277613 | 0 | 0 | 0 | 0 | 0 | 0 |
| ENSG00000268320 | 0 | 0 | 0 | 0 | 0 | 0 |
| ENSG00000272636 | 0 | 1 | 0 | 0 | 1 | 1 |
| ENSG00000273172 | 0 | 1 | 0 | 0 | 0 | 0 |
| ENSG00000181031 | 244 | 147 | 168 | 212 | 196 | 230 |
| ENSG00000262920 | 0 | 0 | 0 | 0 | 0 | 0 |
| ENSG00000262061 | 0 | 0 | 1 | 2 | 0 | 0 |
| ENSG00000262294 | 2 | 1 | 0 | 2 | 4 | 2 |
| ENSG00000262558 | 1 | 2 | 1 | 0 | 0 | 2 |
| ENSG00000241525 | 0 | 0 | 0 | 0 | 0 | 0 |
| ENSG00000187624 | 39 | 45 | 56 | 52 | 49 | 44 |
| ENSG00000183688 | 38 | 23 | 20 | 25 | 25 | 12 |
| ENSG00000263015 | 0 | 0 | 0 | 0 | 0 | 1 |
| ENSG00000141252 | 1356 | 718 | 897 | 596 | 761 | 736 |
| ENSG00000263300 | 0 | 0 | 1 | 0 | 0 | 0 |
| ENSG00000262905 | 0 | 2 | 1 | 0 | 0 | 0 |
| ENSG00000262708 | 0 | 0 | 0 | 0 | 0 | 0 |
| ENSG00000244097 | 6 | 0 | 1 | 5 | 1 | 3 |
| ENSG00000167695 | 356 | 211 | 258 | 242 | 248 | 281 |
| ENSG00000179409 | 965 | 474 | 604 | 595 | 551 | 702 |
| ENSG00000231784 | 18 | 10 | 7 | 12 | 8 | 10 |
| ENSG00000167699 | 2029 | 1020 | 1353 | 1030 | 1204 | 1252 |
| ENSG00000262228 | 0 | 2 | 3 | 1 | 4 | 3 |
| ENSG00000171861 | 445 | 236 | 297 | 314 | 271 | 308 |
| ENSG00000262434 | 3 | 1 | 7 | 3 | 6 | 4 |

| | | | | | | |
|---|---|---|---|---|---|---|
| ENSG00000277491 | 0 | 0 | 2 | 0 | 1 | 0 |
| ENSG00000167693 | 150 | 83 | 82 | 108 | 88 | 90 |
| ENSG00000262133 | 0 | 0 | 0 | 0 | 0 | 0 |
| ENSG00000262003 | 0 | 0 | 0 | 0 | 0 | 0 |
| ENSG00000263017 | 0 | 0 | 0 | 0 | 0 | 0 |
| ENSG00000177370 | 736 | 358 | 394 | 441 | 429 | 407 |
| ENSG00000159842 | 5365 | 2334 | 3328 | 2477 | 2548 | 2791 |
| ENSG00000264429 | 0 | 1 | 0 | 0 | 0 | 0 |
| ENSG00000278794 | 5 | 1 | 0 | 0 | 0 | 1 |
| ENSG00000275997 | 0 | 0 | 0 | 1 | 0 | 0 |
| ENSG00000235361 | 0 | 0 | 0 | 0 | 0 | 0 |
| ENSG00000262213 | 0 | 1 | 0 | 2 | 2 | 1 |
| ENSG00000205899 | 0 | 0 | 0 | 0 | 0 | 0 |
| ENSG00000184811 | 0 | 0 | 0 | 0 | 0 | 0 |
| ENSG00000108953 | 17428 | 8236 | 10080 | 8116 | 9388 | 9482 |
| ENSG00000280408 | 0 | 1 | 0 | 0 | 1 | 2 |
| ENSG00000167193 | 3318 | 1504 | 1881 | 1194 | 1816 | 1667 |
| ENSG00000262777 | 0 | 4 | 6 | 3 | 0 | 1 |
| ENSG00000197879 | 8543 | 4016 | 5151 | 4412 | 4316 | 4396 |
| ENSG00000132376 | 1180 | 548 | 740 | 665 | 565 | 627 |
| ENSG00000236618 | 56 | 50 | 66 | 49 | 61 | 43 |
| ENSG00000174238 | 2850 | 1476 | 1846 | 1388 | 1495 | 1646 |
| ENSG00000167703 | 1634 | 731 | 1010 | 755 | 590 | 755 |
| ENSG00000243704 | 0 | 0 | 0 | 0 | 0 | 0 |
| ENSG00000074660 | 30 | 12 | 32 | 11 | 9 | 12 |
| ENSG00000167705 | 121 | 54 | 83 | 46 | 60 | 50 |
| ENSG00000174231 | 13410 | 5364 | 7720 | 4909 | 6013 | 6960 |
| ENSG00000277597 | 0 | 3 | 1 | 4 | 1 | 0 |
| ENSG00000185561 | 344 | 208 | 200 | 173 | 225 | 194 |
| ENSG00000186594 | 549 | 263 | 341 | 234 | 302 | 320 |
| ENSG00000283824 | 0 | 0 | 0 | 0 | 0 | 0 |
| ENSG00000167716 | 1134 | 493 | 729 | 609 | 547 | 566 |
| ENSG00000262791 | 0 | 0 | 0 | 0 | 0 | 0 |
| ENSG00000167711 | 8 | 5 | 11 | 4 | 2 | 4 |
| ENSG00000132386 | 54 | 23 | 41 | 26 | 20 | 33 |
| ENSG00000186532 | 475 | 289 | 308 | 213 | 261 | 295 |
| ENSG00000132383 | 3456 | 1746 | 2330 | 1548 | 1892 | 1955 |
| ENSG00000108958 | 2 | 6 | 6 | 3 | 5 | 2 |
| ENSG00000185924 | 865 | 361 | 437 | 504 | 464 | 482 |
| ENSG00000262445 | 0 | 3 | 2 | 0 | 1 | 0 |
| ENSG00000228133 | 0 | 0 | 0 | 0 | 0 | 2 |
| ENSG00000262869 | 0 | 0 | 0 | 0 | 0 | 0 |
| ENSG00000108963 | 623 | 294 | 413 | 390 | 306 | 384 |
| ENSG00000262664 | 0 | 0 | 0 | 0 | 0 | 0 |
| ENSG00000262533 | 0 | 0 | 0 | 0 | 0 | 0 |
| ENSG00000263050 | 17 | 10 | 21 | 15 | 4 | 17 |
| ENSG00000267200 | 0 | 0 | 0 | 0 | 0 | 0 |
| ENSG00000267195 | 0 | 0 | 0 | 0 | 0 | 1 |
| ENSG00000177374 | 21 | 16 | 10 | 14 | 22 | 12 |
| ENSG00000070366 | 829 | 346 | 484 | 352 | 391 | 431 |

| | | | | | | |
|---|---|---|---|---|---|---|
| ENSG00000280333 | 0 | 2 | 0 | 0 | 1 | 0 |
| ENSG00000262810 | 0 | 1 | 0 | 0 | 0 | 0 |
| ENSG00000279040 | 0 | 0 | 0 | 0 | 0 | 2 |
| ENSG00000279562 | 0 | 0 | 0 | 0 | 0 | 0 |
| ENSG00000279340 | 0 | 0 | 0 | 0 | 0 | 0 |
| ENSG00000262402 | 0 | 0 | 0 | 0 | 0 | 0 |
| ENSG00000265777 | 0 | 0 | 0 | 0 | 0 | 0 |
| ENSG00000236838 | 0 | 0 | 0 | 0 | 0 | 0 |
| ENSG00000236457 | 0 | 0 | 0 | 0 | 0 | 0 |
| ENSG00000225084 | 0 | 0 | 0 | 0 | 0 | 0 |
| ENSG00000278941 | 0 | 0 | 0 | 0 | 0 | 0 |
| ENSG00000280242 | 0 | 0 | 0 | 0 | 0 | 0 |
| ENSG00000167720 | 270 | 156 | 218 | 132 | 185 | 164 |
| ENSG00000262333 | 57 | 24 | 32 | 11 | 19 | 32 |
| ENSG00000167721 | 1807 | 933 | 1246 | 862 | 1181 | 1259 |
| ENSG00000275084 | 6 | 2 | 0 | 2 | 0 | 1 |
| ENSG00000212163 | 0 | 1 | 0 | 0 | 0 | 0 |
| ENSG00000141258 | 4518 | 2119 | 3110 | 2105 | 2428 | 2484 |
| ENSG00000274758 | 0 | 0 | 0 | 0 | 0 | 0 |
| ENSG00000263345 | 108 | 35 | 63 | 30 | 35 | 44 |
| ENSG00000070444 | 810 | 322 | 505 | 321 | 358 | 454 |
| ENSG00000262456 | 0 | 0 | 0 | 0 | 0 | 0 |
| ENSG00000127804 | 995 | 564 | 590 | 491 | 598 | 556 |
| ENSG00000279872 | 0 | 0 | 1 | 0 | 0 | 0 |
| ENSG00000280280 | 0 | 0 | 0 | 0 | 0 | 0 |
| ENSG00000243049 | 0 | 0 | 0 | 0 | 0 | 0 |
| ENSG00000262953 | 0 | 1 | 0 | 0 | 0 | 1 |
| ENSG00000007168 | 3204 | 1577 | 1882 | 1325 | 1930 | 1717 |
| ENSG00000262480 | 0 | 0 | 0 | 0 | 0 | 0 |
| ENSG00000279432 | 2 | 5 | 1 | 2 | 0 | 0 |
| ENSG00000239884 | 4 | 4 | 3 | 0 | 1 | 0 |
| ENSG00000272770 | 0 | 0 | 0 | 0 | 0 | 0 |
| ENSG00000272911 | 2 | 1 | 1 | 0 | 0 | 0 |
| ENSG00000132361 | 4446 | 1874 | 2528 | 2809 | 2380 | 2641 |
| ENSG00000273606 | 0 | 1 | 0 | 0 | 0 | 0 |
| ENSG00000262050 | 4 | 4 | 4 | 5 | 1 | 4 |
| ENSG00000277200 | 4 | 1 | 5 | 1 | 2 | 2 |
| ENSG00000261963 | 4 | 0 | 0 | 0 | 3 | 0 |
| ENSG00000221200 | 0 | 0 | 0 | 0 | 0 | 0 |
| ENSG00000272920 | 0 | 0 | 0 | 0 | 0 | 0 |
| ENSG00000132359 | 1130 | 562 | 783 | 420 | 497 | 554 |
| ENSG00000262884 | 47 | 30 | 32 | 23 | 22 | 18 |
| ENSG00000262628 | 0 | 0 | 0 | 0 | 0 | 0 |
| ENSG00000184166 | 0 | 0 | 0 | 0 | 0 | 0 |
| ENSG00000142163 | 0 | 0 | 0 | 0 | 0 | 0 |
| ENSG00000183024 | 0 | 0 | 0 | 0 | 0 | 0 |
| ENSG00000261848 | 0 | 0 | 0 | 0 | 0 | 0 |
| ENSG00000262085 | 0 | 0 | 0 | 0 | 0 | 0 |
| ENSG00000172150 | 0 | 0 | 0 | 0 | 0 | 0 |
| ENSG00000172146 | 0 | 0 | 0 | 0 | 0 | 0 |

| | | | | | | |
|---|---|---|---|---|---|---|
| ENSG00000255095 | 0 | 0 | 0 | 0 | 0 | 0 |
| ENSG00000262106 | 0 | 0 | 0 | 0 | 0 | 0 |
| ENSG00000221882 | 0 | 0 | 0 | 0 | 0 | 0 |
| ENSG00000180090 | 0 | 0 | 0 | 0 | 0 | 0 |
| ENSG00000180068 | 0 | 0 | 0 | 0 | 0 | 0 |
| ENSG00000285760 | 0 | 0 | 0 | 0 | 0 | 0 |
| ENSG00000267129 | 0 | 0 | 0 | 0 | 0 | 0 |
| ENSG00000180042 | 0 | 0 | 0 | 0 | 0 | 0 |
| ENSG00000180016 | 0 | 0 | 0 | 0 | 0 | 0 |
| ENSG00000159961 | 0 | 0 | 0 | 0 | 0 | 0 |
| ENSG00000127780 | 0 | 0 | 0 | 0 | 0 | 0 |
| ENSG00000280268 | 0 | 0 | 0 | 0 | 0 | 0 |
| ENSG00000141255 | 0 | 0 | 0 | 0 | 0 | 0 |
| ENSG00000108381 | 0 | 0 | 0 | 0 | 0 | 0 |
| ENSG00000167723 | 40 | 16 | 27 | 18 | 25 | 19 |
| ENSG00000196689 | 202 | 109 | 153 | 93 | 79 | 105 |
| ENSG00000263338 | 0 | 0 | 2 | 0 | 1 | 1 |
| ENSG00000262304 | 3 | 0 | 0 | 0 | 0 | 0 |
| ENSG00000261916 | 1 | 2 | 4 | 1 | 2 | 1 |
| ENSG00000197417 | 164 | 119 | 152 | 122 | 105 | 106 |
| ENSG00000262248 | 2 | 4 | 5 | 2 | 2 | 6 |
| ENSG00000040531 | 682 | 335 | 409 | 373 | 376 | 388 |
| ENSG00000262903 | 17 | 16 | 23 | 15 | 17 | 11 |
| ENSG00000213977 | 205 | 162 | 167 | 181 | 153 | 154 |
| ENSG00000257950 | 45 | 16 | 19 | 21 | 15 | 18 |
| ENSG00000240898 | 12 | 7 | 11 | 2 | 8 | 9 |
| ENSG00000127774 | 51 | 43 | 49 | 53 | 43 | 44 |
| ENSG00000083454 | 3 | 7 | 2 | 7 | 7 | 3 |
| ENSG00000083457 | 558 | 302 | 341 | 351 | 359 | 311 |
| ENSG00000262692 | 15 | 17 | 15 | 11 | 17 | 5 |
| ENSG00000177602 | 77 | 63 | 53 | 54 | 62 | 81 |
| ENSG00000262358 | 0 | 0 | 0 | 3 | 1 | 1 |
| ENSG00000074356 | 2016 | 807 | 1133 | 704 | 965 | 1036 |
| ENSG00000004660 | 99 | 57 | 75 | 94 | 64 | 76 |
| ENSG00000108405 | 2 | 0 | 0 | 0 | 0 | 0 |
| ENSG00000074370 | 12197 | 5455 | 7846 | 5480 | 5268 | 5461 |
| ENSG00000263312 | 0 | 0 | 0 | 0 | 0 | 0 |
| ENSG00000074755 | 2867 | 1219 | 1757 | 1074 | 1378 | 1504 |
| ENSG00000252456 | 0 | 1 | 0 | 0 | 1 | 0 |
| ENSG00000167740 | 1087 | 544 | 719 | 510 | 500 | 618 |
| ENSG00000185722 | 2402 | 1042 | 1443 | 937 | 1090 | 1418 |
| ENSG00000263165 | 0 | 0 | 0 | 0 | 0 | 0 |
| ENSG00000251812 | 0 | 0 | 0 | 0 | 0 | 0 |
| ENSG00000263219 | 16 | 6 | 3 | 1 | 3 | 3 |
| ENSG00000274363 | 6 | 1 | 3 | 2 | 1 | 5 |
| ENSG00000132388 | 1969 | 902 | 1252 | 912 | 1159 | 1142 |
| ENSG00000261868 | 0 | 1 | 0 | 0 | 0 | 0 |
| ENSG00000263882 | 0 | 0 | 0 | 0 | 0 | 0 |
| ENSG00000182557 | 0 | 0 | 0 | 0 | 0 | 0 |
| ENSG00000262823 | 0 | 0 | 0 | 1 | 0 | 0 |

| | | | | | | |
|---|---|---|---|---|---|---|
| ENSG00000229782 | 1 | 0 | 0 | 0 | 0 | 0 |
| ENSG00000183018 | 120 | 57 | 94 | 102 | 95 | 80 |
| ENSG00000132382 | 1481 | 665 | 1003 | 1082 | 892 | 1147 |
| ENSG00000167741 | 4 | 3 | 1 | 1 | 2 | 0 |
| ENSG00000262921 | 0 | 0 | 0 | 0 | 1 | 0 |
| ENSG00000262519 | 4 | 0 | 3 | 2 | 0 | 2 |
| ENSG00000188176 | 30 | 18 | 6 | 25 | 25 | 23 |
| ENSG00000261863 | 0 | 0 | 0 | 0 | 0 | 0 |
| ENSG00000222429 | 0 | 0 | 0 | 0 | 0 | 0 |
| ENSG00000161905 | 706 | 388 | 488 | 376 | 367 | 435 |
| ENSG00000141456 | 1461 | 705 | 924 | 901 | 767 | 910 |
| ENSG00000235085 | 0 | 0 | 0 | 0 | 1 | 0 |
| ENSG00000244184 | 19 | 9 | 16 | 13 | 6 | 13 |
| ENSG00000213939 | 12 | 4 | 7 | 12 | 5 | 7 |
| ENSG00000141480 | 788 | 343 | 477 | 527 | 423 | 470 |
| ENSG00000161920 | 482 | 307 | 389 | 360 | 336 | 305 |
| ENSG00000261898 | 0 | 2 | 1 | 1 | 0 | 0 |
| ENSG00000161921 | 1314 | 692 | 800 | 748 | 717 | 845 |
| ENSG00000141497 | 26 | 24 | 25 | 17 | 16 | 20 |
| ENSG00000142484 | 0 | 0 | 0 | 1 | 3 | 1 |
| ENSG00000182853 | 76 | 22 | 36 | 42 | 22 | 39 |
| ENSG00000280254 | 309 | 159 | 244 | 181 | 189 | 176 |
| ENSG00000182327 | 84 | 55 | 67 | 59 | 59 | 59 |
| ENSG00000142507 | 4415 | 2225 | 2649 | 2850 | 2604 | 2513 |
| ENSG00000262165 | 18 | 4 | 16 | 10 | 15 | 9 |
| ENSG00000129219 | 931 | 474 | 606 | 555 | 538 | 499 |
| ENSG00000141503 | 4264 | 1865 | 2643 | 2324 | 2328 | 2517 |
| ENSG00000230201 | 0 | 1 | 0 | 0 | 0 | 0 |
| ENSG00000264113 | 0 | 0 | 0 | 0 | 0 | 0 |
| ENSG00000108556 | 6 | 1 | 1 | 3 | 1 | 6 |
| ENSG00000205710 | 21 | 10 | 20 | 17 | 12 | 11 |
| ENSG00000185245 | 49 | 35 | 37 | 28 | 23 | 21 |
| ENSG00000108528 | 2370 | 1146 | 1489 | 1427 | 1196 | 1245 |
| ENSG00000108523 | 2738 | 1264 | 1650 | 1639 | 1378 | 1564 |
| ENSG00000108518 | 18853 | 10781 | 12645 | 13688 | 11533 | 11934 |
| ENSG00000108515 | 264 | 142 | 205 | 147 | 121 | 186 |
| ENSG00000091640 | 2204 | 1010 | 1396 | 1179 | 1086 | 1103 |
| ENSG00000108509 | 1375 | 638 | 889 | 707 | 709 | 773 |
| ENSG00000262429 | 0 | 0 | 0 | 0 | 0 | 0 |
| ENSG00000274300 | 1 | 0 | 1 | 0 | 1 | 0 |
| ENSG00000278517 | 1 | 0 | 0 | 0 | 0 | 0 |
| ENSG00000234203 | 0 | 0 | 0 | 0 | 1 | 2 |
| ENSG00000262678 | 2 | 4 | 7 | 3 | 6 | 2 |
| ENSG00000262227 | 10 | 1 | 2 | 0 | 4 | 2 |
| ENSG00000196388 | 69 | 42 | 69 | 48 | 63 | 52 |
| ENSG00000129250 | 5103 | 2710 | 3381 | 3506 | 3445 | 2989 |
| ENSG00000227495 | 2 | 0 | 5 | 2 | 1 | 2 |
| ENSG00000132517 | 4 | 5 | 5 | 4 | 2 | 4 |
| ENSG00000262495 | 0 | 0 | 0 | 0 | 0 | 0 |
| ENSG00000262693 | 10 | 4 | 8 | 1 | 7 | 9 |

| | | | | | | |
|---|---|---|---|---|---|---|
| ENSG00000180787 | 508 | 223 | 283 | 154 | 262 | 256 |
| ENSG00000167840 | 200 | 95 | 111 | 115 | 130 | 118 |
| ENSG00000234327 | 48 | 33 | 34 | 52 | 39 | 53 |
| ENSG00000129204 | 18 | 16 | 25 | 12 | 17 | 22 |
| ENSG00000262855 | 0 | 0 | 0 | 0 | 0 | 0 |
| ENSG00000180626 | 325 | 138 | 183 | 75 | 200 | 114 |
| ENSG00000261879 | 23 | 17 | 16 | 19 | 13 | 13 |
| ENSG00000161929 | 0 | 4 | 5 | 1 | 2 | 5 |
| ENSG00000263164 | 4 | 0 | 1 | 2 | 0 | 1 |
| ENSG00000029725 | 1689 | 982 | 898 | 490 | 1169 | 784 |
| ENSG00000108559 | 1805 | 941 | 1105 | 929 | 1233 | 1123 |
| ENSG00000263220 | 3 | 0 | 0 | 0 | 0 | 0 |
| ENSG00000129197 | 1080 | 633 | 727 | 665 | 753 | 762 |
| ENSG00000263272 | 140 | 65 | 85 | 41 | 93 | 57 |
| ENSG00000108561 | 4905 | 2548 | 2955 | 3502 | 3347 | 3586 |
| ENSG00000005100 | 1396 | 649 | 945 | 623 | 921 | 948 |
| ENSG00000262099 | 0 | 0 | 1 | 0 | 0 | 0 |
| ENSG00000072849 | 1856 | 921 | 1050 | 891 | 1053 | 1014 |
| ENSG00000167842 | 1149 | 504 | 668 | 425 | 666 | 561 |
| ENSG00000091592 | 22 | 10 | 11 | 15 | 8 | 15 |
| ENSG00000282980 | 0 | 0 | 0 | 0 | 0 | 0 |
| ENSG00000262961 | 0 | 0 | 0 | 0 | 0 | 0 |
| ENSG00000284837 | 0 | 0 | 0 | 0 | 0 | 0 |
| ENSG00000285471 | 0 | 0 | 0 | 0 | 0 | 0 |
| ENSG00000179314 | 55 | 25 | 22 | 95 | 81 | 78 |
| ENSG00000262231 | 0 | 0 | 0 | 0 | 0 | 0 |
| ENSG00000206618 | 0 | 0 | 0 | 0 | 0 | 0 |
| ENSG00000262609 | 0 | 0 | 0 | 0 | 0 | 0 |
| ENSG00000285593 | 0 | 2 | 1 | 0 | 4 | 2 |
| ENSG00000129221 | 2 | 1 | 1 | 0 | 0 | 2 |
| ENSG00000262818 | 0 | 0 | 0 | 0 | 0 | 0 |
| ENSG00000129195 | 359 | 199 | 264 | 319 | 269 | 330 |
| ENSG00000091622 | 97 | 32 | 52 | 28 | 28 | 40 |
| ENSG00000274860 | 0 | 0 | 0 | 0 | 0 | 0 |
| ENSG00000198920 | 556 | 293 | 331 | 192 | 322 | 332 |
| ENSG00000200914 | 0 | 0 | 0 | 0 | 0 | 0 |
| ENSG00000282936 | 22 | 15 | 15 | 2 | 16 | 21 |
| ENSG00000129235 | 1184 | 649 | 728 | 780 | 751 | 695 |
| ENSG00000108590 | 124 | 77 | 103 | 71 | 88 | 98 |
| ENSG00000256806 | 147 | 57 | 86 | 66 | 69 | 70 |
| ENSG00000261996 | 0 | 3 | 3 | 3 | 1 | 8 |
| ENSG00000264468 | 0 | 0 | 1 | 0 | 0 | 0 |
| ENSG00000283301 | 0 | 0 | 0 | 0 | 0 | 0 |
| ENSG00000274114 | 6 | 6 | 3 | 5 | 3 | 2 |
| ENSG00000141485 | 1 | 0 | 0 | 0 | 0 | 0 |
| ENSG00000262990 | 0 | 0 | 0 | 0 | 0 | 0 |
| ENSG00000244283 | 0 | 0 | 0 | 0 | 0 | 0 |
| ENSG00000132530 | 0 | 0 | 1 | 0 | 0 | 0 |
| ENSG00000279296 | 0 | 0 | 0 | 0 | 0 | 0 |
| ENSG00000177294 | 0 | 0 | 0 | 0 | 0 | 0 |

| | | | | | | |
|---|---|---|---|---|---|---|
| ENSG00000167858 | 0 | 0 | 0 | 0 | 0 | 0 |
| ENSG00000262943 | 395 | 180 | 215 | 137 | 215 | 194 |
| ENSG00000215067 | 98 | 67 | 75 | 38 | 62 | 72 |
| ENSG00000262503 | 15 | 7 | 8 | 0 | 9 | 7 |
| ENSG00000263316 | 31 | 9 | 17 | 7 | 13 | 22 |
| ENSG00000267047 | 0 | 0 | 0 | 0 | 0 | 0 |
| ENSG00000262089 | 20 | 17 | 17 | 11 | 21 | 13 |
| ENSG00000108839 | 53 | 29 | 35 | 22 | 28 | 26 |
| ENSG00000219200 | 69 | 33 | 60 | 54 | 57 | 81 |
| ENSG00000161939 | 0 | 1 | 0 | 3 | 0 | 0 |
| ENSG00000258315 | 17 | 11 | 9 | 6 | 15 | 21 |
| ENSG00000267532 | 0 | 5 | 6 | 3 | 1 | 5 |
| ENSG00000284112 | 0 | 0 | 0 | 0 | 0 | 0 |
| ENSG00000284027 | 0 | 0 | 0 | 0 | 0 | 0 |
| ENSG00000161940 | 4 | 0 | 0 | 0 | 1 | 1 |
| ENSG00000174327 | 353 | 195 | 245 | 226 | 219 | 224 |
| ENSG00000174326 | 3 | 5 | 9 | 4 | 9 | 3 |
| ENSG00000262067 | 0 | 0 | 0 | 0 | 0 | 0 |
| ENSG00000132514 | 0 | 0 | 0 | 0 | 0 | 0 |
| ENSG00000161944 | 0 | 0 | 0 | 0 | 0 | 0 |
| ENSG00000213876 | 0 | 0 | 1 | 0 | 0 | 0 |
| ENSG00000141505 | 2 | 1 | 5 | 0 | 2 | 4 |
| ENSG00000132535 | 223 | 67 | 149 | 79 | 85 | 96 |
| ENSG00000279812 | 2 | 0 | 0 | 0 | 0 | 0 |
| ENSG00000072778 | 10146 | 4232 | 6302 | 5743 | 5098 | 5618 |
| ENSG00000199053 | 11 | 0 | 2 | 2 | 1 | 0 |
| ENSG00000004975 | 1602 | 688 | 907 | 782 | 799 | 880 |
| ENSG00000040633 | 2064 | 1059 | 1244 | 1160 | 1181 | 1130 |
| ENSG00000170296 | 23 | 160 | 215 | 168 | 174 | 234 |
| ENSG00000262526 | 0 | 0 | 0 | 0 | 0 | 0 |
| ENSG00000279641 | 4 | 3 | 4 | 2 | 5 | 9 |
| ENSG00000175826 | 1388 | 879 | 1193 | 1168 | 970 | 1124 |
| ENSG00000262302 | 0 | 0 | 0 | 0 | 0 | 0 |
| ENSG00000170291 | 845 | 403 | 531 | 467 | 451 | 510 |
| ENSG00000181885 | 4942 | 2574 | 3675 | 3241 | 2803 | 3118 |
| ENSG00000202251 | 0 | 0 | 0 | 0 | 0 | 0 |
| ENSG00000181856 | 46 | 23 | 43 | 22 | 27 | 27 |
| ENSG00000263342 | 0 | 0 | 0 | 0 | 0 | 0 |
| ENSG00000006047 | 764 | 341 | 380 | 338 | 291 | 337 |
| ENSG00000132507 | 7533 | 4289 | 5125 | 6459 | 5493 | 5712 |
| ENSG00000132522 | 28 | 16 | 35 | 20 | 25 | 32 |
| ENSG00000261915 | 0 | 0 | 0 | 0 | 0 | 1 |
| ENSG00000215041 | 814 | 362 | 578 | 390 | 478 | 473 |
| ENSG00000224647 | 4 | 0 | 0 | 1 | 3 | 0 |
| ENSG00000072818 | 88 | 48 | 54 | 47 | 36 | 40 |
| ENSG00000213859 | 724 | 401 | 482 | 485 | 403 | 396 |
| ENSG00000263171 | 0 | 0 | 0 | 0 | 0 | 0 |
| ENSG00000182896 | 0 | 0 | 1 | 0 | 0 | 0 |
| ENSG00000174292 | 860 | 391 | 466 | 536 | 444 | 498 |
| ENSG00000187838 | 9 | 9 | 20 | 5 | 11 | 3 |

| | | | | | | |
|---|---|---|---|---|---|---|
| ENSG00000262481 | 0 | 0 | 1 | 2 | 0 | 1 |
| ENSG00000205544 | 246 | 170 | 197 | 218 | 170 | 183 |
| ENSG00000169992 | 1555 | 650 | 955 | 782 | 728 | 877 |
| ENSG00000262880 | 0 | 0 | 0 | 0 | 0 | 0 |
| ENSG00000181323 | 2 | 0 | 0 | 0 | 0 | 1 |
| ENSG00000184560 | 4 | 2 | 0 | 0 | 0 | 2 |
| ENSG00000283439 | 0 | 0 | 0 | 2 | 0 | 1 |
| ENSG00000181284 | 540 | 284 | 369 | 381 | 334 | 356 |
| ENSG00000262624 | 0 | 0 | 0 | 0 | 0 | 0 |
| ENSG00000161958 | 48 | 24 | 41 | 28 | 42 | 41 |
| ENSG00000263301 | 0 | 0 | 0 | 0 | 0 | 0 |
| ENSG00000272884 | 0 | 0 | 0 | 0 | 0 | 0 |
| ENSG00000170175 | 208 | 118 | 141 | 125 | 136 | 140 |
| ENSG00000174282 | 5279 | 2451 | 3148 | 3033 | 2984 | 3054 |
| ENSG00000259224 | 7 | 1 | 2 | 2 | 2 | 2 |
| ENSG00000181222 | 7401 | 3254 | 4560 | 3993 | 3958 | 4331 |
| ENSG00000201489 | 0 | 1 | 0 | 0 | 0 | 1 |
| ENSG00000239697 | 2 | 0 | 0 | 0 | 0 | 1 |
| ENSG00000248871 | 0 | 0 | 0 | 0 | 0 | 0 |
| ENSG00000276384 | 12 | 1 | 0 | 1 | 2 | 1 |
| ENSG00000161955 | 33 | 17 | 25 | 17 | 18 | 16 |
| ENSG00000161956 | 684 | 349 | 435 | 512 | 455 | 534 |
| ENSG00000277957 | 4 | 1 | 0 | 1 | 2 | 1 |
| ENSG00000161960 | 109 | 51 | 100 | 73 | 78 | 101 |
| ENSG00000264772 | 4 | 0 | 2 | 3 | 4 | 0 |
| ENSG00000209582 | 0 | 0 | 0 | 0 | 0 | 0 |
| ENSG00000238917 | 0 | 0 | 0 | 0 | 0 | 0 |
| ENSG00000277985 | 0 | 0 | 0 | 0 | 0 | 0 |
| ENSG00000129226 | 4 | 0 | 1 | 0 | 2 | 0 |
| ENSG00000233223 | 115 | 47 | 102 | 81 | 71 | 73 |
| ENSG00000129255 | 2070 | 996 | 1326 | 1294 | 1076 | 1104 |
| ENSG00000129194 | 32 | 35 | 32 | 28 | 15 | 23 |
| ENSG00000129245 | 2085 | 934 | 1216 | 1204 | 1159 | 1168 |
| ENSG00000129214 | 8 | 7 | 4 | 6 | 2 | 3 |
| ENSG00000141504 | 256 | 139 | 207 | 174 | 170 | 140 |
| ENSG00000129244 | 2 | 3 | 0 | 0 | 1 | 1 |
| ENSG00000141510 | 2904 | 1598 | 1902 | 1821 | 1845 | 1828 |
| ENSG00000141499 | 500 | 230 | 268 | 331 | 277 | 261 |
| ENSG00000108947 | 482 | 217 | 290 | 247 | 240 | 245 |
| ENSG00000183914 | 4 | 5 | 6 | 1 | 2 | 3 |
| ENSG00000240480 | 0 | 0 | 0 | 0 | 0 | 0 |
| ENSG00000132510 | 2228 | 970 | 1360 | 983 | 980 | 1110 |
| ENSG00000167874 | 17 | 15 | 29 | 13 | 9 | 10 |
| ENSG00000183011 | 1228 | 644 | 806 | 829 | 644 | 695 |
| ENSG00000182224 | 13 | 12 | 18 | 12 | 19 | 19 |
| ENSG00000280046 | 42 | 24 | 35 | 8 | 17 | 18 |
| ENSG00000170004 | 7702 | 3130 | 4622 | 2900 | 3658 | 3593 |
| ENSG00000252835 | 0 | 0 | 3 | 0 | 0 | 0 |
| ENSG00000179859 | 60 | 35 | 54 | 31 | 40 | 35 |
| ENSG00000170049 | 43 | 16 | 37 | 25 | 27 | 25 |

| | | | | | | |
|---|---|---|---|---|---|---|
| ENSG00000262730 | 0 | 0 | 0 | 0 | 0 | 0 |
| ENSG00000170043 | 1770 | 877 | 1195 | 1234 | 1055 | 1096 |
| ENSG00000170037 | 963 | 473 | 629 | 550 | 571 | 536 |
| ENSG00000262492 | 0 | 1 | 0 | 0 | 0 | 0 |
| ENSG00000132518 | 0 | 0 | 0 | 2 | 0 | 1 |
| ENSG00000279174 | 0 | 0 | 0 | 0 | 0 | 0 |
| ENSG00000179593 | 0 | 0 | 1 | 0 | 0 | 0 |
| ENSG00000263427 | 0 | 1 | 0 | 0 | 1 | 0 |
| ENSG00000275011 | 0 | 0 | 0 | 0 | 0 | 1 |
| ENSG00000179477 | 2 | 1 | 0 | 3 | 0 | 0 |
| ENSG00000214999 | 2 | 0 | 1 | 0 | 1 | 2 |
| ENSG00000264005 | 0 | 0 | 0 | 0 | 0 | 0 |
| ENSG00000179148 | 56 | 31 | 42 | 60 | 89 | 66 |
| ENSG00000179111 | 2 | 0 | 1 | 2 | 3 | 2 |
| ENSG00000179094 | 548 | 234 | 330 | 290 | 246 | 292 |
| ENSG00000284117 | 0 | 0 | 0 | 0 | 0 | 0 |
| ENSG00000263620 | 0 | 0 | 0 | 0 | 0 | 0 |
| ENSG00000220205 | 1021 | 433 | 757 | 435 | 488 | 545 |
| ENSG00000179029 | 358 | 207 | 247 | 193 | 212 | 227 |
| ENSG00000200463 | 0 | 0 | 0 | 0 | 0 | 0 |
| ENSG00000266824 | 4 | 1 | 1 | 2 | 0 | 2 |
| ENSG00000283160 | 0 | 0 | 0 | 0 | 0 | 0 |
| ENSG00000196544 | 364 | 202 | 217 | 276 | 197 | 223 |
| ENSG00000279152 | 0 | 0 | 0 | 0 | 0 | 0 |
| ENSG00000271002 | 5 | 2 | 1 | 1 | 1 | 1 |
| ENSG00000178999 | 422 | 233 | 328 | 365 | 336 | 381 |
| ENSG00000178977 | 15 | 8 | 11 | 11 | 10 | 3 |
| ENSG00000178971 | 1366 | 584 | 874 | 580 | 626 | 722 |
| ENSG00000178921 | 874 | 408 | 523 | 503 | 496 | 598 |
| ENSG00000269947 | 10 | 0 | 4 | 2 | 7 | 6 |
| ENSG00000125434 | 146 | 77 | 154 | 77 | 76 | 101 |
| ENSG00000108961 | 10 | 2 | 6 | 2 | 6 | 6 |
| ENSG00000271029 | 0 | 1 | 2 | 0 | 1 | 1 |
| ENSG00000198844 | 0 | 0 | 0 | 0 | 0 | 0 |
| ENSG00000226871 | 0 | 0 | 0 | 0 | 0 | 0 |
| ENSG00000212206 | 0 | 0 | 0 | 0 | 0 | 0 |
| ENSG00000266279 | 0 | 0 | 0 | 0 | 0 | 0 |
| ENSG00000184650 | 0 | 0 | 0 | 0 | 0 | 0 |
| ENSG00000184619 | 17 | 11 | 11 | 4 | 12 | 12 |
| ENSG00000265749 | 12 | 18 | 19 | 14 | 16 | 18 |
| ENSG00000263809 | 0 | 0 | 0 | 0 | 0 | 0 |
| ENSG00000161970 | 2875 | 2556 | 3113 | 3460 | 3058 | 3223 |
| ENSG00000189051 | 12 | 1 | 3 | 0 | 1 | 1 |
| ENSG00000166579 | 1176 | 573 | 756 | 579 | 690 | 649 |
| ENSG00000133026 | 2137 | 1089 | 1400 | 720 | 1257 | 1136 |
| ENSG00000252363 | 0 | 1 | 0 | 0 | 0 | 0 |
| ENSG00000270606 | 0 | 0 | 0 | 1 | 0 | 0 |
| ENSG00000244604 | 0 | 0 | 0 | 0 | 1 | 1 |
| ENSG00000161973 | 0 | 0 | 0 | 0 | 0 | 0 |
| ENSG00000183318 | 0 | 0 | 0 | 0 | 0 | 1 |

| | | | | | | |
|---|---|---|---|---|---|---|
| ENSG00000185156 | 0 | 2 | 0 | 0 | 0 | 0 |
| ENSG00000276231 | 0 | 0 | 0 | 0 | 0 | 0 |
| ENSG00000141506 | 14 | 11 | 8 | 12 | 15 | 13 |
| ENSG00000266389 | 6 | 7 | 8 | 6 | 7 | 3 |
| ENSG00000265975 | 16 | 11 | 10 | 8 | 14 | 9 |
| ENSG00000065320 | 480 | 237 | 325 | 238 | 259 | 284 |
| ENSG00000262966 | 2 | 0 | 0 | 0 | 0 | 0 |
| ENSG00000273816 | 1 | 0 | 2 | 3 | 2 | 4 |
| ENSG00000170310 | 392 | 201 | 271 | 247 | 279 | 221 |
| ENSG00000263708 | 0 | 0 | 0 | 0 | 0 | 1 |
| ENSG00000225751 | 0 | 0 | 0 | 2 | 0 | 0 |
| ENSG00000262815 | 0 | 0 | 0 | 0 | 0 | 0 |
| ENSG00000263051 | 0 | 0 | 0 | 0 | 0 | 0 |
| ENSG00000271851 | 34 | 15 | 22 | 24 | 29 | 24 |
| ENSG00000166596 | 16 | 13 | 13 | 12 | 11 | 16 |
| ENSG00000241654 | 0 | 0 | 0 | 0 | 0 | 0 |
| ENSG00000265349 | 1 | 2 | 2 | 0 | 0 | 2 |
| ENSG00000154914 | 472 | 191 | 253 | 182 | 237 | 215 |
| ENSG00000262296 | 0 | 0 | 0 | 0 | 0 | 0 |
| ENSG00000282882 | 0 | 0 | 0 | 0 | 0 | 2 |
| ENSG00000184544 | 0 | 0 | 0 | 0 | 0 | 0 |
| ENSG00000278944 | 0 | 0 | 0 | 0 | 0 | 0 |
| ENSG00000214978 | 0 | 1 | 0 | 0 | 0 | 0 |
| ENSG00000283025 | 0 | 0 | 0 | 0 | 0 | 0 |
| ENSG00000065325 | 28 | 17 | 12 | 31 | 32 | 18 |
| ENSG00000265593 | 0 | 0 | 0 | 0 | 0 | 0 |
| ENSG00000109047 | 0 | 0 | 0 | 0 | 0 | 0 |
| ENSG00000007237 | 4 | 0 | 0 | 0 | 0 | 0 |
| ENSG00000242692 | 0 | 0 | 1 | 2 | 0 | 1 |
| ENSG00000273388 | 0 | 0 | 0 | 0 | 0 | 0 |
| ENSG00000006788 | 0 | 0 | 0 | 0 | 0 | 0 |
| ENSG00000264067 | 0 | 0 | 0 | 0 | 0 | 0 |
| ENSG00000272736 | 0 | 0 | 0 | 0 | 0 | 0 |
| ENSG00000272975 | 0 | 0 | 0 | 0 | 0 | 0 |
| ENSG00000133020 | 0 | 1 | 0 | 0 | 0 | 0 |
| ENSG00000264424 | 4 | 1 | 2 | 2 | 0 | 0 |
| ENSG00000109061 | 0 | 2 | 0 | 0 | 0 | 1 |
| ENSG00000125414 | 2 | 0 | 0 | 0 | 0 | 1 |
| ENSG00000214970 | 0 | 0 | 0 | 0 | 0 | 0 |
| ENSG00000109063 | 12 | 5 | 6 | 3 | 4 | 11 |
| ENSG00000261433 | 0 | 0 | 0 | 0 | 1 | 1 |
| ENSG00000133028 | 1410 | 672 | 857 | 737 | 729 | 880 |
| ENSG00000263388 | 0 | 0 | 1 | 0 | 0 | 0 |
| ENSG00000170222 | 137 | 71 | 84 | 88 | 124 | 74 |
| ENSG00000187824 | 2 | 2 | 0 | 0 | 2 | 3 |
| ENSG00000264176 | 2 | 0 | 0 | 0 | 0 | 0 |
| ENSG00000263400 | 25 | 11 | 8 | 9 | 8 | 16 |
| ENSG00000285220 | 0 | 0 | 0 | 0 | 0 | 0 |
| ENSG00000264016 | 0 | 0 | 0 | 0 | 0 | 0 |
| ENSG00000240813 | 0 | 0 | 0 | 0 | 1 | 0 |

| | | | | | | |
|---|---|---|---|---|---|---|
| ENSG00000266114 | 0 | 0 | 0 | 0 | 0 | 0 |
| ENSG00000284876 | 0 | 0 | 0 | 0 | 0 | 0 |
| ENSG00000263429 | 0 | 1 | 0 | 0 | 0 | 5 |
| ENSG00000279542 | 0 | 0 | 0 | 0 | 0 | 0 |
| ENSG00000233670 | 2 | 2 | 3 | 0 | 1 | 0 |
| ENSG00000263508 | 4 | 3 | 2 | 2 | 3 | 2 |
| ENSG00000207242 | 1 | 0 | 2 | 0 | 2 | 1 |
| ENSG00000244753 | 0 | 2 | 1 | 0 | 1 | 0 |
| ENSG00000243346 | 0 | 0 | 0 | 0 | 0 | 0 |
| ENSG00000285541 | 0 | 0 | 0 | 0 | 0 | 0 |
| ENSG00000188803 | 0 | 0 | 0 | 0 | 0 | 0 |
| ENSG00000264727 | 0 | 0 | 0 | 0 | 0 | 0 |
| ENSG00000007174 | 0 | 1 | 1 | 0 | 0 | 2 |
| ENSG00000263684 | 0 | 0 | 0 | 0 | 0 | 0 |
| ENSG00000266368 | 0 | 0 | 0 | 2 | 2 | 0 |
| ENSG00000154957 | 234 | 140 | 167 | 126 | 154 | 137 |
| ENSG00000241185 | 0 | 0 | 1 | 0 | 1 | 0 |
| ENSG00000065559 | 1021 | 554 | 617 | 408 | 694 | 592 |
| ENSG00000252707 | 0 | 0 | 0 | 0 | 0 | 0 |
| ENSG00000266297 | 0 | 0 | 0 | 0 | 0 | 0 |
| ENSG00000263648 | 0 | 3 | 0 | 0 | 0 | 0 |
| ENSG00000265400 | 0 | 0 | 0 | 0 | 0 | 0 |
| ENSG00000179136 | 0 | 0 | 0 | 0 | 0 | 0 |
| ENSG00000141052 | 0 | 1 | 0 | 0 | 0 | 0 |
| ENSG00000227274 | 0 | 0 | 0 | 0 | 0 | 0 |
| ENSG00000265489 | 0 | 1 | 1 | 1 | 1 | 1 |
| ENSG00000006740 | 656 | 336 | 509 | 312 | 374 | 470 |
| ENSG00000265503 | 0 | 1 | 0 | 0 | 0 | 0 |
| ENSG00000280168 | 9 | 5 | 5 | 2 | 5 | 0 |
| ENSG00000277621 | 1 | 0 | 2 | 1 | 0 | 1 |
| ENSG00000263707 | 0 | 0 | 0 | 0 | 0 | 0 |
| ENSG00000006744 | 3061 | 1490 | 1848 | 1962 | 1857 | 1914 |
| ENSG00000273098 | 0 | 0 | 0 | 0 | 0 | 0 |
| ENSG00000234634 | 0 | 0 | 0 | 0 | 0 | 0 |
| ENSG00000153976 | 67 | 23 | 63 | 32 | 39 | 45 |
| ENSG00000221698 | 0 | 0 | 0 | 0 | 0 | 0 |
| ENSG00000236088 | 153 | 67 | 88 | 80 | 95 | 97 |
| ENSG00000265494 | 0 | 0 | 0 | 0 | 0 | 0 |
| ENSG00000233852 | 0 | 0 | 0 | 0 | 0 | 0 |
| ENSG00000266744 | 0 | 0 | 1 | 0 | 0 | 0 |
| ENSG00000141028 | 16 | 9 | 7 | 5 | 13 | 9 |
| ENSG00000006695 | 436 | 233 | 276 | 285 | 253 | 281 |
| ENSG00000252305 | 0 | 0 | 0 | 0 | 0 | 0 |
| ENSG00000231595 | 11 | 10 | 8 | 2 | 8 | 8 |
| ENSG00000223510 | 1 | 0 | 0 | 0 | 2 | 2 |
| ENSG00000262786 | 0 | 0 | 0 | 1 | 0 | 1 |
| ENSG00000125430 | 54 | 31 | 27 | 28 | 29 | 40 |
| ENSG00000266709 | 14 | 3 | 12 | 3 | 6 | 1 |
| ENSG00000266378 | 2 | 0 | 0 | 0 | 0 | 0 |
| ENSG00000230647 | 2 | 0 | 0 | 0 | 0 | 0 |

| | | | | | | |
|---|---|---|---|---|---|---|
| ENSG00000230897 | 2 | 7 | 3 | 5 | 4 | 4 |
| ENSG00000226130 | 0 | 0 | 0 | 0 | 0 | 0 |
| ENSG00000205325 | 0 | 0 | 0 | 0 | 0 | 0 |
| ENSG00000238212 | 0 | 0 | 0 | 0 | 0 | 0 |
| ENSG00000226156 | 0 | 0 | 0 | 0 | 0 | 0 |
| ENSG00000232058 | 0 | 0 | 0 | 0 | 0 | 0 |
| ENSG00000259944 | 0 | 0 | 0 | 0 | 0 | 0 |
| ENSG00000265163 | 0 | 0 | 0 | 0 | 0 | 0 |
| ENSG00000109099 | 2537 | 1250 | 1477 | 673 | 727 | 758 |
| ENSG00000265110 | 0 | 1 | 0 | 0 | 0 | 0 |
| ENSG00000279660 | 17 | 8 | 5 | 5 | 7 | 2 |
| ENSG00000265445 | 2 | 2 | 1 | 0 | 0 | 2 |
| ENSG00000230971 | 4 | 0 | 2 | 1 | 0 | 0 |
| ENSG00000237377 | 0 | 0 | 0 | 0 | 0 | 0 |
| ENSG00000125409 | 8 | 4 | 6 | 1 | 2 | 2 |
| ENSG00000200437 | 0 | 0 | 0 | 0 | 0 | 0 |
| ENSG00000279567 | 0 | 0 | 0 | 1 | 0 | 0 |
| ENSG00000266667 | 0 | 0 | 0 | 0 | 0 | 0 |
| ENSG00000239888 | 0 | 0 | 0 | 0 | 0 | 0 |
| ENSG00000239704 | 7 | 1 | 4 | 1 | 4 | 3 |
| ENSG00000259024 | 0 | 3 | 1 | 2 | 1 | 1 |
| ENSG00000234324 | 0 | 0 | 1 | 1 | 2 | 0 |
| ENSG00000175106 | 56 | 26 | 30 | 22 | 30 | 21 |
| ENSG00000223878 | 1 | 6 | 5 | 0 | 3 | 3 |
| ENSG00000266538 | 0 | 1 | 1 | 0 | 0 | 0 |
| ENSG00000241322 | 8 | 12 | 5 | 10 | 11 | 10 |
| ENSG00000251537 | 2 | 3 | 5 | 4 | 5 | 4 |
| ENSG00000252129 | 0 | 0 | 0 | 0 | 0 | 0 |
| ENSG00000223544 | 0 | 0 | 0 | 0 | 0 | 0 |
| ENSG00000221926 | 1773 | 1074 | 1095 | 1469 | 1577 | 1558 |
| ENSG00000266261 | 0 | 0 | 1 | 1 | 1 | 4 |
| ENSG00000108452 | 0 | 2 | 0 | 2 | 0 | 1 |
| ENSG00000187607 | 296 | 155 | 168 | 113 | 225 | 178 |
| ENSG00000255104 | 0 | 1 | 0 | 0 | 0 | 0 |
| ENSG00000233966 | 10 | 8 | 4 | 2 | 4 | 7 |
| ENSG00000214946 | 4 | 0 | 1 | 2 | 0 | 2 |
| ENSG00000233002 | 0 | 0 | 0 | 0 | 0 | 0 |
| ENSG00000267227 | 0 | 0 | 0 | 0 | 0 | 0 |
| ENSG00000233193 | 0 | 0 | 0 | 0 | 0 | 0 |
| ENSG00000227255 | 0 | 0 | 0 | 0 | 0 | 0 |
| ENSG00000235430 | 0 | 0 | 0 | 0 | 0 | 0 |
| ENSG00000251829 | 0 | 0 | 0 | 0 | 0 | 0 |
| ENSG00000227018 | 0 | 0 | 0 | 0 | 0 | 0 |
| ENSG00000179277 | 268 | 135 | 164 | 123 | 159 | 159 |
| ENSG00000265519 | 124 | 76 | 92 | 61 | 101 | 106 |
| ENSG00000276855 | 211 | 110 | 121 | 135 | 145 | 137 |
| ENSG00000233090 | 0 | 0 | 0 | 0 | 0 | 1 |
| ENSG00000237057 | 0 | 0 | 0 | 0 | 0 | 0 |
| ENSG00000227790 | 0 | 0 | 0 | 0 | 0 | 0 |
| ENSG00000170425 | 23 | 21 | 25 | 32 | 25 | 22 |

| | | | | | | |
|---|---|---|---|---|---|---|
| ENSG00000214941 | 466 | 265 | 302 | 299 | 312 | 332 |
| ENSG00000011295 | 1949 | 998 | 1278 | 941 | 1188 | 1220 |
| ENSG00000275413 | 0 | 1 | 0 | 1 | 3 | 0 |
| ENSG00000141027 | 2585 | 1342 | 1499 | 740 | 1486 | 1324 |
| ENSG00000227782 | 9 | 3 | 8 | 2 | 6 | 4 |
| ENSG00000243686 | 0 | 0 | 0 | 0 | 0 | 0 |
| ENSG00000233979 | 0 | 0 | 0 | 0 | 0 | 0 |
| ENSG00000252383 | 0 | 0 | 0 | 0 | 0 | 0 |
| ENSG00000199674 | 0 | 0 | 0 | 0 | 0 | 0 |
| ENSG00000239221 | 0 | 1 | 0 | 0 | 0 | 0 |
| ENSG00000108474 | 288 | 157 | 228 | 132 | 189 | 179 |
| ENSG00000221355 | 0 | 0 | 0 | 0 | 0 | 1 |
| ENSG00000166582 | 91 | 53 | 68 | 50 | 83 | 58 |
| ENSG00000170315 | 6591 | 3932 | 4670 | 4315 | 4686 | 4349 |
| ENSG00000265401 | 0 | 0 | 0 | 0 | 0 | 0 |
| ENSG00000265095 | 3 | 1 | 3 | 7 | 1 | 3 |
| ENSG00000264739 | 0 | 0 | 0 | 0 | 0 | 0 |
| ENSG00000187688 | 2 | 0 | 1 | 1 | 0 | 3 |
| ENSG00000239203 | 0 | 0 | 0 | 0 | 0 | 0 |
| ENSG00000175061 | 8288 | 4698 | 5658 | 6837 | 6299 | 6737 |
| ENSG00000277108 | 0 | 0 | 0 | 0 | 0 | 0 |
| ENSG00000277370 | 0 | 0 | 0 | 0 | 0 | 0 |
| ENSG00000266651 | 7 | 15 | 10 | 9 | 9 | 9 |
| ENSG00000277512 | 0 | 0 | 0 | 0 | 0 | 0 |
| ENSG00000181350 | 94 | 49 | 70 | 47 | 75 | 61 |
| ENSG00000141040 | 316 | 217 | 220 | 148 | 187 | 186 |
| ENSG00000266803 | 0 | 0 | 2 | 1 | 1 | 0 |
| ENSG00000235554 | 0 | 1 | 3 | 2 | 3 | 0 |
| ENSG00000197566 | 154 | 94 | 85 | 38 | 123 | 89 |
| ENSG00000264765 | 0 | 0 | 0 | 0 | 0 | 0 |
| ENSG00000264359 | 0 | 0 | 0 | 0 | 0 | 0 |
| ENSG00000231458 | 2 | 0 | 0 | 1 | 0 | 3 |
| ENSG00000170160 | 0 | 0 | 0 | 0 | 0 | 0 |
| ENSG00000266302 | 0 | 0 | 0 | 0 | 0 | 0 |
| ENSG00000252446 | 0 | 0 | 0 | 0 | 0 | 0 |
| ENSG00000276088 | 0 | 0 | 0 | 0 | 0 | 0 |
| ENSG00000226478 | 6 | 0 | 2 | 0 | 0 | 1 |
| ENSG00000188933 | 0 | 0 | 1 | 1 | 0 | 2 |
| ENSG00000266129 | 0 | 0 | 0 | 0 | 0 | 0 |
| ENSG00000272815 | 0 | 0 | 0 | 0 | 0 | 0 |
| ENSG00000264673 | 0 | 0 | 0 | 0 | 0 | 0 |
| ENSG00000264729 | 0 | 0 | 0 | 0 | 0 | 0 |
| ENSG00000264892 | 0 | 0 | 0 | 0 | 0 | 0 |
| ENSG00000226145 | 0 | 0 | 0 | 0 | 0 | 0 |
| ENSG00000227300 | 0 | 0 | 0 | 0 | 0 | 0 |
| ENSG00000131885 | 0 | 0 | 0 | 0 | 0 | 0 |
| ENSG00000205312 | 0 | 0 | 0 | 0 | 0 | 0 |
| ENSG00000229749 | 0 | 0 | 0 | 0 | 0 | 0 |
| ENSG00000283033 | 0 | 0 | 0 | 0 | 0 | 0 |
| ENSG00000265853 | 0 | 0 | 0 | 0 | 0 | 0 |

| | | | | | | |
|---|---|---|---|---|---|---|
| ENSG00000128438 | 6 | 0 | 0 | 0 | 0 | 0 |
| ENSG00000240505 | 0 | 0 | 0 | 0 | 0 | 0 |
| ENSG00000230709 | 2 | 1 | 0 | 1 | 1 | 0 |
| ENSG00000230969 | 0 | 0 | 0 | 0 | 0 | 0 |
| ENSG00000260328 | 0 | 0 | 0 | 0 | 0 | 0 |
| ENSG00000133030 | 2996 | 1438 | 1905 | 1307 | 1794 | 1879 |
| ENSG00000225442 | 0 | 0 | 1 | 1 | 1 | 0 |
| ENSG00000263624 | 0 | 1 | 1 | 0 | 0 | 1 |
| ENSG00000206859 | 0 | 0 | 0 | 0 | 0 | 0 |
| ENSG00000243370 | 0 | 0 | 1 | 0 | 0 | 2 |
| ENSG00000278864 | 114 | 53 | 87 | 43 | 53 | 55 |
| ENSG00000179598 | 148 | 62 | 122 | 124 | 138 | 109 |
| ENSG00000264187 | 0 | 1 | 2 | 0 | 1 | 1 |
| ENSG00000154803 | 732 | 406 | 532 | 351 | 501 | 440 |
| ENSG00000266498 | 0 | 0 | 0 | 0 | 1 | 1 |
| ENSG00000226359 | 2 | 1 | 0 | 0 | 2 | 1 |
| ENSG00000141030 | 1855 | 973 | 1097 | 1023 | 1132 | 1049 |
| ENSG00000205309 | 50 | 24 | 36 | 26 | 26 | 34 |
| ENSG00000215030 | 4622 | 2170 | 2474 | 3497 | 2745 | 2548 |
| ENSG00000227158 | 0 | 0 | 1 | 0 | 0 | 0 |
| ENSG00000230865 | 0 | 0 | 0 | 0 | 0 | 0 |
| ENSG00000201741 | 0 | 0 | 0 | 0 | 0 | 0 |
| ENSG00000141026 | 458 | 207 | 242 | 266 | 190 | 254 |
| ENSG00000108551 | 143 | 72 | 128 | 183 | 167 | 195 |
| ENSG00000133027 | 507 | 279 | 307 | 431 | 330 | 363 |
| ENSG00000265511 | 5 | 6 | 20 | 11 | 4 | 11 |
| ENSG00000206730 | 0 | 0 | 0 | 0 | 0 | 0 |
| ENSG00000264666 | 0 | 0 | 0 | 0 | 0 | 2 |
| ENSG00000279200 | 0 | 0 | 0 | 1 | 0 | 0 |
| ENSG00000223979 | 0 | 1 | 0 | 1 | 2 | 2 |
| ENSG00000273948 | 0 | 0 | 0 | 0 | 0 | 0 |
| ENSG00000274836 | 0 | 0 | 0 | 0 | 0 | 0 |
| ENSG00000108557 | 4461 | 1985 | 2881 | 2036 | 2194 | 2498 |
| ENSG00000237328 | 0 | 0 | 0 | 1 | 0 | 2 |
| ENSG00000226746 | 3 | 0 | 0 | 0 | 2 | 0 |
| ENSG00000264167 | 0 | 0 | 0 | 0 | 0 | 0 |
| ENSG00000072310 | 16803 | 6776 | 10099 | 7160 | 5931 | 7292 |
| ENSG00000274111 | 0 | 0 | 0 | 0 | 0 | 0 |
| ENSG00000207839 | 1 | 0 | 2 | 1 | 2 | 1 |
| ENSG00000175662 | 5160 | 2340 | 3315 | 1965 | 2012 | 2171 |
| ENSG00000197815 | 74 | 41 | 44 | 12 | 11 | 28 |
| ENSG00000171962 | 64 | 33 | 75 | 37 | 30 | 46 |
| ENSG00000171953 | 607 | 324 | 414 | 434 | 382 | 389 |
| ENSG00000230273 | 0 | 0 | 0 | 0 | 0 | 0 |
| ENSG00000232344 | 7 | 3 | 1 | 1 | 1 | 2 |
| ENSG00000280198 | 4 | 0 | 5 | 0 | 4 | 7 |
| ENSG00000141034 | 405 | 189 | 269 | 198 | 224 | 213 |
| ENSG00000108591 | 1315 | 633 | 768 | 830 | 684 | 713 |
| ENSG00000279428 | 0 | 0 | 1 | 0 | 0 | 0 |
| ENSG00000091536 | 14 | 11 | 14 | 7 | 11 | 10 |

| | | | | | | |
|---|---|---|---|---|---|---|
| ENSG00000266677 | 5 | 0 | 1 | 1 | 0 | 3 |
| ENSG00000091542 | 2969 | 1360 | 1748 | 1611 | 1630 | 1578 |
| ENSG00000131899 | 1750 | 724 | 981 | 879 | 846 | 861 |
| ENSG00000177731 | 7160 | 3222 | 4171 | 4176 | 4077 | 4007 |
| ENSG00000177427 | 833 | 382 | 527 | 469 | 517 | 464 |
| ENSG00000260647 | 3 | 2 | 6 | 6 | 4 | 6 |
| ENSG00000177302 | 1160 | 565 | 667 | 558 | 601 | 625 |
| ENSG00000228000 | 0 | 2 | 1 | 0 | 0 | 0 |
| ENSG00000237382 | 0 | 0 | 0 | 0 | 0 | 0 |
| ENSG00000176994 | 1537 | 636 | 951 | 553 | 742 | 812 |
| ENSG00000176974 | 2803 | 1357 | 1645 | 1373 | 1413 | 1630 |
| ENSG00000283772 | 0 | 0 | 0 | 0 | 0 | 0 |
| ENSG00000214860 | 15 | 7 | 14 | 8 | 8 | 6 |
| ENSG00000264177 | 16 | 5 | 3 | 2 | 3 | 1 |
| ENSG00000267441 | 0 | 0 | 0 | 0 | 0 | 0 |
| ENSG00000275206 | 0 | 0 | 0 | 0 | 0 | 0 |
| ENSG00000204471 | 0 | 0 | 0 | 0 | 0 | 0 |
| ENSG00000220161 | 3 | 0 | 2 | 1 | 1 | 0 |
| ENSG00000205266 | 0 | 0 | 0 | 0 | 0 | 0 |
| ENSG00000266162 | 0 | 0 | 0 | 0 | 0 | 0 |
| ENSG00000186831 | 0 | 0 | 0 | 0 | 0 | 0 |
| ENSG00000214856 | 0 | 0 | 0 | 0 | 0 | 0 |
| ENSG00000229526 | 0 | 0 | 0 | 0 | 0 | 0 |
| ENSG00000227919 | 0 | 0 | 0 | 0 | 0 | 0 |
| ENSG00000267492 | 0 | 0 | 0 | 0 | 0 | 0 |
| ENSG00000230662 | 0 | 0 | 1 | 0 | 0 | 1 |
| ENSG00000171916 | 0 | 1 | 0 | 0 | 0 | 0 |
| ENSG00000167494 | 0 | 0 | 0 | 0 | 0 | 0 |
| ENSG00000273018 | 31 | 9 | 14 | 3 | 17 | 16 |
| ENSG00000233327 | 0 | 3 | 6 | 1 | 3 | 1 |
| ENSG00000227689 | 0 | 0 | 1 | 0 | 1 | 0 |
| ENSG00000154874 | 39 | 13 | 13 | 1 | 13 | 13 |
| ENSG00000227077 | 4 | 0 | 1 | 0 | 1 | 0 |
| ENSG00000276532 | 0 | 0 | 0 | 0 | 0 | 0 |
| ENSG00000189375 | 2 | 0 | 0 | 0 | 0 | 2 |
| ENSG00000262262 | 0 | 0 | 0 | 0 | 0 | 0 |
| ENSG00000174977 | 62 | 71 | 85 | 71 | 77 | 87 |
| ENSG00000249459 | 86 | 59 | 34 | 43 | 80 | 48 |
| ENSG00000264885 | 38 | 34 | 23 | 32 | 39 | 45 |
| ENSG00000240445 | 70 | 17 | 17 | 35 | 26 | 20 |
| ENSG00000224126 | 0 | 0 | 0 | 2 | 0 | 0 |
| ENSG00000108448 | 1165 | 665 | 822 | 1275 | 1352 | 1447 |
| ENSG00000265693 | 0 | 0 | 0 | 0 | 0 | 0 |
| ENSG00000171931 | 3 | 2 | 3 | 2 | 8 | 1 |
| ENSG00000171928 | 606 | 327 | 355 | 267 | 442 | 303 |
| ENSG00000265478 | 0 | 3 | 1 | 0 | 0 | 1 |
| ENSG00000250111 | 0 | 0 | 0 | 0 | 0 | 0 |
| ENSG00000141127 | 974 | 482 | 637 | 521 | 605 | 613 |
| ENSG00000274747 | 0 | 0 | 0 | 0 | 0 | 0 |
| ENSG00000264273 | 0 | 0 | 0 | 0 | 0 | 2 |

| | | | | | | |
|---|---|---|---|---|---|---|
| ENSG00000235672 | 4 | 0 | 1 | 0 | 0 | 0 |
| ENSG00000212618 | 0 | 0 | 0 | 0 | 0 | 0 |
| ENSG00000154025 | 9 | 3 | 4 | 1 | 1 | 2 |
| ENSG00000196893 | 0 | 0 | 0 | 0 | 0 | 0 |
| ENSG00000188522 | 1272 | 573 | 852 | 879 | 748 | 810 |
| ENSG00000263045 | 0 | 0 | 0 | 0 | 0 | 0 |
| ENSG00000154016 | 0 | 0 | 0 | 0 | 0 | 0 |
| ENSG00000279825 | 0 | 0 | 0 | 0 | 0 | 0 |
| ENSG00000265185 | 2 | 1 | 0 | 2 | 0 | 0 |
| ENSG00000262074 | 0 | 1 | 0 | 1 | 2 | 0 |
| ENSG00000230197 | 0 | 0 | 0 | 0 | 0 | 0 |
| ENSG00000265746 | 0 | 0 | 0 | 0 | 0 | 0 |
| ENSG00000263394 | 0 | 0 | 0 | 0 | 0 | 0 |
| ENSG00000228157 | 0 | 0 | 0 | 0 | 0 | 0 |
| ENSG00000262202 | 4 | 0 | 0 | 0 | 1 | 0 |
| ENSG00000277947 | 0 | 0 | 0 | 0 | 0 | 0 |
| ENSG00000281000 | 0 | 0 | 0 | 0 | 0 | 0 |
| ENSG00000189152 | 0 | 0 | 0 | 0 | 0 | 0 |
| ENSG00000262319 | 0 | 0 | 0 | 0 | 0 | 0 |
| ENSG00000266664 | 0 | 0 | 0 | 0 | 0 | 0 |
| ENSG00000197665 | 0 | 0 | 0 | 0 | 0 | 0 |
| ENSG00000262292 | 0 | 0 | 0 | 0 | 0 | 0 |
| ENSG00000266311 | 0 | 0 | 0 | 0 | 0 | 0 |
| ENSG00000261987 | 0 | 0 | 0 | 0 | 0 | 0 |
| ENSG00000263934 | 6 | 18 | 5 | 3 | 3 | 5 |
| ENSG00000264940 | 0 | 0 | 0 | 0 | 0 | 0 |
| ENSG00000236022 | 0 | 0 | 0 | 0 | 0 | 0 |
| ENSG00000263206 | 0 | 0 | 0 | 0 | 0 | 0 |
| ENSG00000072134 | 898 | 460 | 596 | 423 | 468 | 502 |
| ENSG00000281856 | 0 | 0 | 0 | 0 | 0 | 0 |
| ENSG00000235397 | 2 | 3 | 1 | 3 | 1 | 2 |
| ENSG00000265263 | 0 | 0 | 0 | 0 | 0 | 0 |
| ENSG00000108641 | 383 | 144 | 203 | 191 | 214 | 214 |
| ENSG00000221540 | 2 | 0 | 1 | 0 | 0 | 1 |
| ENSG00000166484 | 299 | 158 | 205 | 211 | 184 | 190 |
| ENSG00000166482 | 2 | 0 | 0 | 2 | 1 | 1 |
| ENSG00000128482 | 0 | 0 | 0 | 1 | 1 | 2 |
| ENSG00000265126 | 0 | 0 | 0 | 0 | 0 | 0 |
| ENSG00000235979 | 0 | 0 | 0 | 0 | 0 | 0 |
| ENSG00000227078 | 0 | 0 | 0 | 0 | 0 | 0 |
| ENSG00000283675 | 0 | 0 | 0 | 0 | 0 | 0 |
| ENSG00000189343 | 156 | 60 | 77 | 87 | 74 | 65 |
| ENSG00000231477 | 0 | 0 | 0 | 0 | 0 | 0 |
| ENSG00000142494 | 2 | 4 | 6 | 14 | 6 | 11 |
| ENSG00000228331 | 2 | 0 | 0 | 1 | 1 | 0 |
| ENSG00000266079 | 0 | 0 | 0 | 0 | 0 | 0 |
| ENSG00000262769 | 0 | 0 | 0 | 0 | 0 | 0 |
| ENSG00000228983 | 0 | 0 | 0 | 0 | 0 | 0 |
| ENSG00000263107 | 0 | 0 | 0 | 0 | 0 | 0 |
| ENSG00000266179 | 0 | 0 | 0 | 0 | 0 | 0 |

| | | | | | | |
|---|---|---|---|---|---|---|
| ENSG00000271900 | 0 | 0 | 0 | 0 | 0 | 0 |
| ENSG00000227225 | 0 | 0 | 0 | 0 | 0 | 0 |
| ENSG00000242752 | 0 | 0 | 0 | 0 | 0 | 0 |
| ENSG00000228725 | 0 | 1 | 0 | 0 | 0 | 0 |
| ENSG00000224254 | 0 | 0 | 0 | 0 | 0 | 0 |
| ENSG00000231625 | 2 | 0 | 0 | 1 | 0 | 1 |
| ENSG00000072210 | 1703 | 839 | 1090 | 791 | 1038 | 973 |
| ENSG00000264932 | 1 | 0 | 0 | 1 | 0 | 1 |
| ENSG00000275982 | 0 | 0 | 0 | 0 | 0 | 0 |
| ENSG00000252349 | 0 | 0 | 0 | 0 | 0 | 1 |
| ENSG00000180638 | 0 | 2 | 4 | 2 | 0 | 1 |
| ENSG00000262681 | 2 | 0 | 0 | 0 | 0 | 0 |
| ENSG00000264785 | 0 | 0 | 0 | 0 | 0 | 0 |
| ENSG00000108602 | 30 | 16 | 9 | 72 | 92 | 46 |
| ENSG00000230607 | 0 | 0 | 0 | 0 | 0 | 0 |
| ENSG00000083290 | 897 | 433 | 640 | 344 | 463 | 502 |
| ENSG00000231096 | 0 | 0 | 1 | 0 | 0 | 0 |
| ENSG00000279251 | 0 | 0 | 0 | 0 | 0 | 0 |
| ENSG00000276406 | 2 | 1 | 0 | 0 | 0 | 2 |
| ENSG00000270091 | 17 | 4 | 11 | 8 | 7 | 4 |
| ENSG00000108599 | 1121 | 595 | 736 | 457 | 621 | 585 |
| ENSG00000266126 | 4 | 0 | 0 | 0 | 0 | 0 |
| ENSG00000261033 | 8 | 3 | 0 | 8 | 9 | 4 |
| ENSG00000128487 | 2087 | 1172 | 1182 | 793 | 1124 | 984 |
| ENSG00000225681 | 0 | 0 | 0 | 0 | 0 | 0 |
| ENSG00000225014 | 0 | 0 | 0 | 0 | 0 | 0 |
| ENSG00000212186 | 0 | 0 | 0 | 0 | 0 | 0 |
| ENSG00000263494 | 0 | 1 | 0 | 0 | 0 | 0 |
| ENSG00000252971 | 0 | 1 | 0 | 0 | 0 | 0 |
| ENSG00000154898 | 10 | 1 | 1 | 0 | 2 | 2 |
| ENSG00000276577 | 0 | 0 | 0 | 0 | 0 | 0 |
| ENSG00000266839 | 0 | 0 | 0 | 0 | 0 | 0 |
| ENSG00000278701 | 0 | 0 | 0 | 0 | 0 | 0 |
| ENSG00000214832 | 4 | 0 | 0 | 0 | 0 | 0 |
| ENSG00000189423 | 20 | 5 | 11 | 1 | 3 | 7 |
| ENSG00000237911 | 0 | 0 | 0 | 0 | 0 | 0 |
| ENSG00000265556 | 0 | 0 | 0 | 0 | 0 | 0 |
| ENSG00000230528 | 0 | 0 | 0 | 0 | 2 | 0 |
| ENSG00000170298 | 0 | 0 | 0 | 0 | 0 | 0 |
| ENSG00000229586 | 2 | 0 | 0 | 0 | 0 | 0 |
| ENSG00000263976 | 0 | 0 | 0 | 0 | 0 | 0 |
| ENSG00000230088 | 0 | 0 | 0 | 0 | 0 | 0 |
| ENSG00000214822 | 0 | 0 | 0 | 0 | 0 | 0 |
| ENSG00000266925 | 0 | 0 | 0 | 0 | 0 | 0 |
| ENSG00000231645 | 0 | 0 | 0 | 0 | 0 | 0 |
| ENSG00000205215 | 0 | 0 | 0 | 0 | 0 | 0 |
| ENSG00000266042 | 0 | 0 | 0 | 0 | 0 | 0 |
| ENSG00000260907 | 0 | 0 | 0 | 0 | 0 | 0 |
| ENSG00000265916 | 2 | 1 | 2 | 0 | 0 | 0 |
| ENSG00000264981 | 0 | 0 | 0 | 0 | 0 | 0 |

| | | | | | | |
|---|---|---|---|---|---|---|
| ENSG00000235203 | 0 | 0 | 0 | 0 | 0 | 0 |
| ENSG00000267075 | 0 | 0 | 0 | 0 | 0 | 0 |
| ENSG00000230339 | 0 | 0 | 0 | 0 | 0 | 0 |
| ENSG00000263946 | 15 | 7 | 8 | 8 | 7 | 3 |
| ENSG00000235323 | 0 | 0 | 0 | 0 | 0 | 0 |
| ENSG00000264422 | 0 | 0 | 0 | 0 | 0 | 0 |
| ENSG00000214819 | 0 | 0 | 0 | 0 | 0 | 0 |
| ENSG00000231258 | 0 | 0 | 0 | 0 | 0 | 1 |
| ENSG00000188013 | 20 | 19 | 22 | 11 | 24 | 19 |
| ENSG00000232889 | 0 | 0 | 0 | 0 | 0 | 0 |
| ENSG00000226096 | 0 | 0 | 0 | 0 | 0 | 0 |
| ENSG00000235546 | 0 | 0 | 0 | 0 | 0 | 0 |
| ENSG00000227685 | 0 | 0 | 0 | 0 | 0 | 0 |
| ENSG00000266364 | 0 | 0 | 0 | 0 | 0 | 0 |
| ENSG00000236504 | 0 | 0 | 0 | 0 | 0 | 0 |
| ENSG00000264874 | 0 | 0 | 0 | 0 | 0 | 0 |
| ENSG00000226521 | 4 | 1 | 3 | 1 | 0 | 0 |
| ENSG00000187870 | 48 | 8 | 16 | 15 | 8 | 15 |
| ENSG00000218454 | 0 | 0 | 0 | 0 | 0 | 0 |
| ENSG00000213671 | 0 | 0 | 0 | 0 | 0 | 0 |
| ENSG00000226549 | 8 | 4 | 6 | 0 | 3 | 1 |
| ENSG00000264215 | 0 | 1 | 1 | 0 | 1 | 0 |
| ENSG00000264662 | 0 | 0 | 0 | 0 | 0 | 0 |
| ENSG00000226981 | 0 | 0 | 0 | 0 | 1 | 0 |
| ENSG00000205212 | 0 | 0 | 0 | 0 | 0 | 0 |
| ENSG00000266369 | 0 | 0 | 0 | 0 | 0 | 0 |
| ENSG00000233098 | 0 | 1 | 2 | 0 | 1 | 0 |
| ENSG00000252483 | 0 | 0 | 0 | 0 | 0 | 0 |
| ENSG00000265790 | 0 | 0 | 0 | 0 | 0 | 0 |
| ENSG00000265099 | 0 | 0 | 0 | 0 | 0 | 0 |
| ENSG00000244456 | 0 | 0 | 0 | 0 | 0 | 0 |
| ENSG00000264660 | 0 | 0 | 0 | 0 | 0 | 0 |
| ENSG00000266009 | 0 | 0 | 0 | 0 | 0 | 0 |
| ENSG00000263729 | 0 | 0 | 0 | 0 | 0 | 0 |
| ENSG00000124422 | 7031 | 3227 | 4562 | 3156 | 3717 | 4193 |
| ENSG00000263986 | 0 | 0 | 0 | 0 | 0 | 0 |
| ENSG00000236819 | 0 | 0 | 0 | 0 | 0 | 0 |
| ENSG00000109016 | 843 | 434 | 557 | 476 | 410 | 449 |
| ENSG00000178307 | 1240 | 528 | 656 | 694 | 683 | 651 |
| ENSG00000235530 | 15 | 3 | 7 | 8 | 7 | 8 |
| ENSG00000263815 | 1 | 1 | 1 | 0 | 0 | 0 |
| ENSG00000274180 | 760 | 383 | 521 | 323 | 308 | 346 |
| ENSG00000266563 | 0 | 0 | 0 | 0 | 0 | 0 |
| ENSG00000034152 | 1466 | 782 | 947 | 974 | 842 | 889 |
| ENSG00000184185 | 6 | 4 | 1 | 2 | 3 | 1 |
| ENSG00000212719 | 0 | 0 | 0 | 0 | 0 | 0 |
| ENSG00000265881 | 0 | 0 | 0 | 0 | 0 | 0 |
| ENSG00000265043 | 0 | 0 | 0 | 0 | 0 | 0 |
| ENSG00000264023 | 0 | 0 | 0 | 0 | 0 | 0 |
| ENSG00000244582 | 0 | 0 | 0 | 0 | 0 | 2 |

| | | | | | | |
|---|---|---|---|---|---|---|
| ENSG00000265265 | 0 | 0 | 0 | 0 | 0 | 0 |
| ENSG00000266466 | 0 | 0 | 0 | 0 | 0 | 0 |
| ENSG00000266673 | 0 | 0 | 0 | 0 | 0 | 0 |
| ENSG00000266050 | 0 | 0 | 0 | 0 | 0 | 0 |
| ENSG00000264245 | 0 | 0 | 0 | 0 | 0 | 0 |
| ENSG00000263375 | 0 | 0 | 0 | 0 | 0 | 0 |
| ENSG00000260157 | 0 | 0 | 0 | 0 | 0 | 0 |
| ENSG00000264617 | 0 | 0 | 0 | 0 | 0 | 0 |
| ENSG00000274748 | 0 | 0 | 0 | 0 | 0 | 0 |
| ENSG00000260458 | 0 | 0 | 0 | 0 | 0 | 0 |
| ENSG00000283566 | 0 | 0 | 0 | 0 | 1 | 0 |
| ENSG00000265233 | 0 | 0 | 0 | 0 | 0 | 0 |
| ENSG00000265019 | 0 | 0 | 0 | 0 | 0 | 0 |
| ENSG00000263563 | 38 | 9 | 10 | 3 | 17 | 18 |
| ENSG00000263725 | 0 | 0 | 0 | 0 | 0 | 0 |
| ENSG00000263609 | 0 | 0 | 0 | 0 | 0 | 0 |
| ENSG00000264956 | 0 | 0 | 0 | 0 | 0 | 0 |
| ENSG00000266691 | 0 | 0 | 0 | 0 | 0 | 0 |
| ENSG00000178130 | 0 | 0 | 0 | 0 | 0 | 0 |
| ENSG00000264811 | 0 | 0 | 0 | 0 | 0 | 0 |
| ENSG00000265567 | 0 | 0 | 0 | 0 | 0 | 0 |
| ENSG00000264562 | 0 | 0 | 0 | 0 | 0 | 0 |
| ENSG00000266806 | 0 | 0 | 0 | 0 | 0 | 0 |
| ENSG00000276399 | 0 | 0 | 0 | 0 | 0 | 0 |
| ENSG00000266746 | 0 | 0 | 0 | 0 | 0 | 0 |
| ENSG00000261020 | 0 | 0 | 0 | 0 | 0 | 0 |
| ENSG00000266885 | 0 | 0 | 0 | 0 | 0 | 0 |
| ENSG00000283879 | 0 | 0 | 0 | 0 | 0 | 0 |
| ENSG00000266529 | 0 | 0 | 0 | 0 | 0 | 0 |
| ENSG00000256618 | 4 | 0 | 0 | 3 | 3 | 3 |
| ENSG00000243655 | 0 | 0 | 0 | 0 | 0 | 0 |
| ENSG00000264168 | 0 | 0 | 0 | 0 | 0 | 0 |
| ENSG00000263914 | 0 | 0 | 0 | 0 | 0 | 0 |
| ENSG00000240103 | 0 | 0 | 0 | 0 | 0 | 0 |
| ENSG00000241225 | 0 | 0 | 0 | 0 | 0 | 0 |
| ENSG00000263483 | 0 | 0 | 0 | 0 | 0 | 0 |
| ENSG00000264316 | 0 | 0 | 0 | 0 | 0 | 0 |
| ENSG00000284300 | 0 | 0 | 0 | 0 | 0 | 0 |
| ENSG00000284328 | 0 | 0 | 0 | 0 | 0 | 0 |
| ENSG00000284263 | 0 | 0 | 0 | 0 | 0 | 0 |
| ENSG00000275480 | 0 | 0 | 0 | 0 | 0 | 0 |
| ENSG00000264930 | 0 | 0 | 0 | 0 | 0 | 0 |
| ENSG00000264970 | 0 | 0 | 0 | 0 | 0 | 0 |
| ENSG00000264262 | 0 | 0 | 0 | 0 | 0 | 0 |
| ENSG00000264431 | 0 | 0 | 0 | 0 | 0 | 0 |
| ENSG00000263433 | 0 | 0 | 0 | 0 | 0 | 0 |
| ENSG00000264734 | 0 | 0 | 0 | 0 | 0 | 0 |
| ENSG00000264172 | 0 | 0 | 0 | 0 | 0 | 0 |
| ENSG00000266736 | 0 | 0 | 0 | 0 | 0 | 0 |
| ENSG00000264837 | 0 | 0 | 0 | 0 | 0 | 0 |

| | | | | | | |
|---|---|---|---|---|---|---|
| ENSG00000265646 | 0 | 1 | 1 | 1 | 0 | 0 |
| ENSG00000285822 | 0 | 0 | 0 | 0 | 0 | 0 |
| ENSG00000265246 | 324 | 164 | 229 | 169 | 160 | 147 |
| ENSG00000239223 | 0 | 0 | 0 | 0 | 0 | 0 |
| ENSG00000213664 | 0 | 0 | 0 | 0 | 0 | 0 |
| ENSG00000265453 | 0 | 0 | 0 | 0 | 0 | 0 |
| ENSG00000263583 | 2 | 0 | 0 | 0 | 0 | 0 |
| ENSG00000109046 | 2862 | 1195 | 1780 | 846 | 1343 | 1357 |
| ENSG00000266313 | 4 | 4 | 6 | 4 | 3 | 8 |
| ENSG00000265683 | 32 | 16 | 14 | 8 | 24 | 7 |
| ENSG00000266356 | 0 | 0 | 0 | 0 | 0 | 0 |
| ENSG00000266433 | 0 | 0 | 0 | 0 | 1 | 0 |
| ENSG00000141068 | 196 | 107 | 176 | 129 | 200 | 207 |
| ENSG00000265801 | 0 | 0 | 0 | 0 | 0 | 0 |
| ENSG00000263368 | 2 | 5 | 2 | 1 | 0 | 0 |
| ENSG00000266853 | 0 | 0 | 0 | 0 | 0 | 0 |
| ENSG00000266728 | 0 | 0 | 0 | 0 | 0 | 0 |
| ENSG00000266872 | 0 | 0 | 0 | 0 | 0 | 0 |
| ENSG00000168961 | 12 | 5 | 7 | 6 | 4 | 7 |
| ENSG00000265788 | 0 | 1 | 0 | 0 | 0 | 0 |
| ENSG00000265041 | 0 | 0 | 0 | 0 | 0 | 0 |
| ENSG00000266771 | 0 | 0 | 0 | 0 | 0 | 0 |
| ENSG00000264240 | 0 | 0 | 0 | 0 | 0 | 0 |
| ENSG00000264689 | 0 | 0 | 0 | 0 | 0 | 0 |
| ENSG00000263604 | 0 | 0 | 0 | 0 | 0 | 0 |
| ENSG00000265833 | 0 | 0 | 0 | 0 | 0 | 0 |
| ENSG00000266416 | 0 | 0 | 0 | 0 | 0 | 0 |
| ENSG00000266786 | 0 | 0 | 0 | 0 | 0 | 0 |
| ENSG00000007171 | 0 | 0 | 1 | 0 | 1 | 0 |
| ENSG00000266202 | 0 | 0 | 0 | 0 | 0 | 0 |
| ENSG00000266527 | 0 | 1 | 0 | 0 | 2 | 0 |
| ENSG00000232859 | 236 | 91 | 143 | 104 | 133 | 116 |
| ENSG00000260019 | 0 | 0 | 0 | 0 | 0 | 0 |
| ENSG00000251818 | 0 | 0 | 0 | 0 | 0 | 0 |
| ENSG00000266190 | 0 | 0 | 0 | 0 | 0 | 0 |
| ENSG00000202389 | 0 | 0 | 0 | 0 | 0 | 0 |
| ENSG00000087095 | 902 | 406 | 575 | 347 | 473 | 500 |
| ENSG00000252283 | 0 | 0 | 0 | 0 | 0 | 0 |
| ENSG00000240873 | 0 | 0 | 0 | 0 | 0 | 0 |
| ENSG00000267644 | 6 | 0 | 0 | 1 | 1 | 1 |
| ENSG00000237575 | 9 | 11 | 9 | 12 | 10 | 8 |
| ENSG00000267259 | 89 | 56 | 49 | 24 | 48 | 48 |
| ENSG00000266830 | 0 | 0 | 0 | 0 | 0 | 0 |
| ENSG00000265060 | 0 | 0 | 0 | 0 | 0 | 0 |
| ENSG00000263788 | 0 | 0 | 0 | 0 | 0 | 0 |
| ENSG00000264486 | 0 | 0 | 0 | 0 | 0 | 0 |
| ENSG00000260777 | 2 | 3 | 3 | 0 | 0 | 0 |
| ENSG00000266306 | 0 | 2 | 0 | 0 | 0 | 0 |
| ENSG00000264029 | 0 | 0 | 0 | 0 | 0 | 0 |
| ENSG00000265480 | 0 | 2 | 2 | 1 | 3 | 1 |

| | | | | | | |
|---|---|---|---|---|---|---|
| ENSG00000109084 | 2037 | 1109 | 1031 | 898 | 1099 | 947 |
| ENSG00000109083 | 626 | 271 | 432 | 302 | 367 | 332 |
| ENSG00000109079 | 3169 | 1381 | 1797 | 1486 | 1680 | 1713 |
| ENSG00000004142 | 3701 | 1820 | 2182 | 2378 | 2022 | 2252 |
| ENSG00000244045 | 258 | 188 | 233 | 219 | 217 | 189 |
| ENSG00000258924 | 0 | 0 | 0 | 0 | 0 | 0 |
| ENSG00000284532 | 0 | 0 | 0 | 0 | 0 | 0 |
| ENSG00000265618 | 0 | 0 | 0 | 0 | 0 | 0 |
| ENSG00000274529 | 0 | 0 | 0 | 0 | 0 | 0 |
| ENSG00000273171 | 0 | 0 | 0 | 0 | 0 | 0 |
| ENSG00000004139 | 118 | 49 | 107 | 61 | 74 | 94 |
| ENSG00000109072 | 14 | 6 | 7 | 11 | 7 | 6 |
| ENSG00000277450 | 0 | 0 | 0 | 0 | 0 | 0 |
| ENSG00000279532 | 0 | 2 | 1 | 0 | 2 | 0 |
| ENSG00000076351 | 194 | 110 | 109 | 122 | 113 | 145 |
| ENSG00000265254 | 0 | 0 | 0 | 0 | 0 | 0 |
| ENSG00000265541 | 0 | 0 | 0 | 0 | 0 | 0 |
| ENSG00000263870 | 0 | 0 | 0 | 0 | 0 | 0 |
| ENSG00000258472 | 72 | 36 | 37 | 30 | 36 | 51 |
| ENSG00000263266 | 148 | 57 | 56 | 71 | 58 | 74 |
| ENSG00000007216 | 0 | 0 | 0 | 0 | 0 | 1 |
| ENSG00000109101 | 4 | 10 | 17 | 8 | 7 | 4 |
| ENSG00000109103 | 1277 | 556 | 679 | 735 | 729 | 723 |
| ENSG00000087111 | 3470 | 1643 | 2173 | 1684 | 1514 | 1811 |
| ENSG00000109107 | 150 | 126 | 175 | 145 | 118 | 142 |
| ENSG00000265168 | 0 | 0 | 0 | 0 | 0 | 0 |
| ENSG00000076382 | 729 | 363 | 508 | 409 | 585 | 520 |
| ENSG00000227543 | 32 | 21 | 23 | 11 | 10 | 14 |
| ENSG00000264608 | 12 | 11 | 19 | 7 | 6 | 15 |
| ENSG00000265287 | 0 | 0 | 0 | 0 | 0 | 0 |
| ENSG00000167524 | 80 | 49 | 68 | 47 | 43 | 62 |
| ENSG00000007202 | 6633 | 3080 | 4472 | 3358 | 3833 | 4108 |
| ENSG00000264044 | 0 | 0 | 0 | 1 | 1 | 1 |
| ENSG00000132581 | 1372 | 694 | 863 | 844 | 851 | 831 |
| ENSG00000240494 | 0 | 0 | 0 | 0 | 0 | 0 |
| ENSG00000109111 | 5002 | 2296 | 3031 | 2066 | 2873 | 2512 |
| ENSG00000265205 | 12 | 5 | 14 | 3 | 8 | 6 |
| ENSG00000167525 | 220 | 114 | 151 | 112 | 115 | 106 |
| ENSG00000109113 | 0 | 1 | 3 | 3 | 1 | 2 |
| ENSG00000198242 | 3756 | 2506 | 3001 | 3158 | 3363 | 2939 |
| ENSG00000238423 | 0 | 0 | 1 | 0 | 0 | 0 |
| ENSG00000264577 | 62 | 23 | 41 | 46 | 39 | 68 |
| ENSG00000238578 | 0 | 0 | 0 | 0 | 0 | 0 |
| ENSG00000238649 | 0 | 0 | 0 | 0 | 0 | 0 |
| ENSG00000238597 | 0 | 0 | 0 | 0 | 0 | 0 |
| ENSG00000160606 | 365 | 223 | 258 | 236 | 211 | 253 |
| ENSG00000160602 | 273 | 153 | 193 | 142 | 170 | 161 |
| ENSG00000267729 | 0 | 3 | 2 | 1 | 2 | 1 |
| ENSG00000265073 | 0 | 0 | 0 | 0 | 0 | 1 |
| ENSG00000076604 | 4406 | 2212 | 2978 | 2824 | 2517 | 2699 |

| | | | | | | |
|---|---|---|---|---|---|---|
| ENSG00000265474 | 20 | 10 | 7 | 1 | 11 | 10 |
| ENSG00000265840 | 0 | 0 | 0 | 0 | 0 | 0 |
| ENSG00000173065 | 664 | 319 | 485 | 451 | 369 | 400 |
| ENSG00000243053 | 0 | 0 | 0 | 0 | 0 | 0 |
| ENSG00000238516 | 0 | 1 | 0 | 0 | 0 | 0 |
| ENSG00000264138 | 0 | 0 | 0 | 1 | 0 | 0 |
| ENSG00000132591 | 1618 | 809 | 974 | 991 | 887 | 951 |
| ENSG00000264066 | 0 | 0 | 0 | 0 | 0 | 0 |
| ENSG00000284565 | 0 | 0 | 0 | 0 | 0 | 0 |
| ENSG00000283364 | 0 | 0 | 0 | 0 | 0 | 0 |
| ENSG00000283819 | 0 | 0 | 0 | 0 | 0 | 0 |
| ENSG00000284085 | 0 | 0 | 0 | 0 | 0 | 0 |
| ENSG00000132589 | 3032 | 1425 | 1880 | 1764 | 1477 | 1716 |
| ENSG00000264304 | 9 | 1 | 2 | 0 | 4 | 1 |
| ENSG00000266642 | 0 | 0 | 0 | 1 | 0 | 0 |
| ENSG00000167536 | 175 | 87 | 90 | 79 | 99 | 78 |
| ENSG00000109118 | 1815 | 799 | 1068 | 699 | 811 | 944 |
| ENSG00000265845 | 1 | 2 | 0 | 0 | 0 | 0 |
| ENSG00000265908 | 5 | 6 | 3 | 0 | 0 | 0 |
| ENSG00000179761 | 15 | 7 | 11 | 4 | 7 | 10 |
| ENSG00000063015 | 5 | 0 | 1 | 0 | 2 | 0 |
| ENSG00000263613 | 0 | 0 | 0 | 0 | 0 | 0 |
| ENSG00000238007 | 0 | 0 | 0 | 0 | 0 | 0 |
| ENSG00000263781 | 2 | 0 | 2 | 0 | 0 | 0 |
| ENSG00000196535 | 1938 | 830 | 1180 | 844 | 909 | 1034 |
| ENSG00000221995 | 52 | 24 | 38 | 13 | 19 | 22 |
| ENSG00000263709 | 0 | 0 | 0 | 0 | 0 | 0 |
| ENSG00000264050 | 0 | 0 | 0 | 0 | 0 | 0 |
| ENSG00000178082 | 2 | 3 | 1 | 1 | 2 | 4 |
| ENSG00000108255 | 0 | 0 | 1 | 1 | 0 | 0 |
| ENSG00000108256 | 3469 | 1704 | 1804 | 1008 | 2371 | 1748 |
| ENSG00000239256 | 0 | 0 | 0 | 0 | 0 | 0 |
| ENSG00000266111 | 0 | 0 | 0 | 0 | 2 | 0 |
| ENSG00000264808 | 1 | 0 | 2 | 2 | 2 | 1 |
| ENSG00000222363 | 0 | 0 | 0 | 0 | 0 | 0 |
| ENSG00000160551 | 1703 | 943 | 767 | 302 | 1362 | 816 |
| ENSG00000284162 | 0 | 0 | 0 | 0 | 0 | 0 |
| ENSG00000253064 | 0 | 0 | 0 | 0 | 0 | 0 |
| ENSG00000202205 | 0 | 0 | 0 | 0 | 0 | 0 |
| ENSG00000168792 | 812 | 442 | 532 | 487 | 502 | 537 |
| ENSG00000264031 | 0 | 0 | 0 | 0 | 0 | 0 |
| ENSG00000167543 | 1382 | 659 | 871 | 845 | 772 | 754 |
| ENSG00000264290 | 9 | 6 | 3 | 5 | 2 | 7 |
| ENSG00000108262 | 2215 | 1008 | 1262 | 1423 | 1282 | 1344 |
| ENSG00000198720 | 281 | 109 | 143 | 177 | 118 | 127 |
| ENSG00000264647 | 1 | 1 | 0 | 1 | 0 | 1 |
| ENSG00000167549 | 12 | 4 | 18 | 3 | 6 | 2 |
| ENSG00000264007 | 0 | 0 | 0 | 0 | 0 | 0 |
| ENSG00000141298 | 587 | 237 | 285 | 165 | 268 | 278 |
| ENSG00000263370 | 2 | 0 | 0 | 0 | 3 | 0 |

| | | | | | | |
|---|---|---|---|---|---|---|
| ENSG00000222858 | 0 | 0 | 0 | 0 | 0 | 0 |
| ENSG00000265625 | 0 | 0 | 0 | 0 | 0 | 0 |
| ENSG00000240531 | 0 | 0 | 0 | 0 | 0 | 0 |
| ENSG00000263657 | 0 | 0 | 1 | 0 | 1 | 1 |
| ENSG00000265713 | 0 | 4 | 1 | 0 | 1 | 0 |
| ENSG00000252657 | 0 | 0 | 0 | 0 | 0 | 0 |
| ENSG00000240074 | 0 | 0 | 1 | 0 | 0 | 0 |
| ENSG00000263477 | 0 | 0 | 0 | 0 | 0 | 0 |
| ENSG00000239129 | 0 | 0 | 0 | 0 | 0 | 0 |
| ENSG00000176927 | 1 | 2 | 4 | 1 | 6 | 0 |
| ENSG00000264435 | 0 | 0 | 0 | 0 | 0 | 0 |
| ENSG00000207011 | 0 | 0 | 0 | 0 | 0 | 0 |
| ENSG00000267482 | 0 | 0 | 0 | 0 | 0 | 0 |
| ENSG00000265289 | 0 | 0 | 0 | 0 | 0 | 0 |
| ENSG00000266876 | 0 | 0 | 0 | 0 | 0 | 0 |
| ENSG00000265394 | 2 | 0 | 0 | 0 | 1 | 0 |
| ENSG00000126653 | 172 | 99 | 66 | 44 | 126 | 93 |
| ENSG00000283935 | 0 | 0 | 0 | 0 | 0 | 0 |
| ENSG00000266919 | 0 | 0 | 0 | 0 | 0 | 0 |
| ENSG00000284399 | 0 | 0 | 0 | 0 | 0 | 0 |
| ENSG00000265739 | 0 | 0 | 0 | 0 | 0 | 0 |
| ENSG00000266987 | 0 | 0 | 0 | 0 | 0 | 0 |
| ENSG00000108576 | 0 | 0 | 0 | 0 | 0 | 0 |
| ENSG00000264125 | 0 | 0 | 0 | 0 | 2 | 0 |
| ENSG00000266120 | 0 | 0 | 0 | 0 | 0 | 0 |
| ENSG00000252112 | 0 | 0 | 0 | 0 | 0 | 0 |
| ENSG00000108578 | 547 | 260 | 318 | 250 | 313 | 320 |
| ENSG00000222679 | 0 | 0 | 0 | 0 | 0 | 0 |
| ENSG00000182271 | 2 | 0 | 0 | 0 | 0 | 0 |
| ENSG00000207132 | 0 | 0 | 0 | 0 | 0 | 0 |
| ENSG00000201255 | 0 | 0 | 0 | 0 | 0 | 0 |
| ENSG00000108582 | 1207 | 587 | 592 | 311 | 638 | 435 |
| ENSG00000108587 | 1013 | 421 | 558 | 380 | 473 | 508 |
| ENSG00000264958 | 1 | 0 | 0 | 0 | 0 | 0 |
| ENSG00000263860 | 0 | 0 | 0 | 0 | 0 | 0 |
| ENSG00000279431 | 0 | 0 | 0 | 0 | 0 | 1 |
| ENSG00000266733 | 0 | 0 | 0 | 0 | 0 | 0 |
| ENSG00000266775 | 0 | 0 | 0 | 0 | 0 | 0 |
| ENSG00000259928 | 0 | 0 | 0 | 0 | 0 | 0 |
| ENSG00000231870 | 0 | 0 | 0 | 0 | 0 | 0 |
| ENSG00000231421 | 0 | 0 | 0 | 0 | 0 | 0 |
| ENSG00000214719 | 11 | 7 | 15 | 11 | 8 | 16 |
| ENSG00000248121 | 11 | 10 | 5 | 0 | 2 | 5 |
| ENSG00000273609 | 0 | 0 | 0 | 0 | 0 | 0 |
| ENSG00000264943 | 4 | 2 | 1 | 1 | 1 | 0 |
| ENSG00000250462 | 122 | 56 | 60 | 42 | 78 | 55 |
| ENSG00000241631 | 0 | 0 | 1 | 0 | 2 | 4 |
| ENSG00000264538 | 237 | 85 | 131 | 98 | 108 | 118 |
| ENSG00000279337 | 0 | 0 | 0 | 0 | 0 | 0 |
| ENSG00000265443 | 4 | 1 | 1 | 0 | 1 | 1 |

| | | | | | | |
|---|---|---|---|---|---|---|
| ENSG00000263603 | 0 | 2 | 4 | 0 | 0 | 1 |
| ENSG00000280069 | 0 | 1 | 0 | 1 | 2 | 2 |
| ENSG00000176390 | 270 | 107 | 154 | 105 | 146 | 142 |
| ENSG00000265791 | 5 | 0 | 3 | 1 | 4 | 1 |
| ENSG00000266490 | 6 | 11 | 6 | 5 | 10 | 6 |
| ENSG00000276250 | 2 | 0 | 3 | 5 | 4 | 1 |
| ENSG00000242439 | 0 | 0 | 0 | 0 | 0 | 0 |
| ENSG00000176208 | 40 | 38 | 30 | 10 | 52 | 35 |
| ENSG00000265334 | 0 | 0 | 0 | 0 | 0 | 0 |
| ENSG00000212190 | 0 | 0 | 0 | 0 | 0 | 0 |
| ENSG00000263531 | 3 | 0 | 1 | 0 | 0 | 2 |
| ENSG00000172171 | 116 | 74 | 73 | 79 | 98 | 96 |
| ENSG00000275185 | 0 | 0 | 0 | 0 | 0 | 0 |
| ENSG00000184060 | 77 | 29 | 36 | 38 | 41 | 38 |
| ENSG00000230113 | 0 | 0 | 0 | 0 | 0 | 0 |
| ENSG00000266274 | 0 | 2 | 0 | 0 | 0 | 0 |
| ENSG00000265743 | 2 | 0 | 0 | 0 | 0 | 3 |
| ENSG00000181481 | 647 | 370 | 449 | 466 | 421 | 426 |
| ENSG00000264456 | 29 | 13 | 25 | 13 | 9 | 14 |
| ENSG00000264743 | 2 | 0 | 2 | 4 | 1 | 0 |
| ENSG00000266340 | 2 | 0 | 3 | 0 | 0 | 1 |
| ENSG00000264148 | 0 | 1 | 6 | 1 | 1 | 0 |
| ENSG00000266865 | 23 | 14 | 28 | 13 | 27 | 25 |
| ENSG00000265798 | 3 | 1 | 2 | 2 | 0 | 1 |
| ENSG00000264107 | 6 | 4 | 3 | 2 | 1 | 4 |
| ENSG00000265444 | 0 | 0 | 0 | 0 | 0 | 0 |
| ENSG00000196712 | 1445 | 810 | 702 | 286 | 1039 | 732 |
| ENSG00000266371 | 0 | 0 | 1 | 0 | 0 | 0 |
| ENSG00000126861 | 0 | 2 | 1 | 1 | 0 | 0 |
| ENSG00000185862 | 0 | 0 | 0 | 0 | 0 | 0 |
| ENSG00000265118 | 0 | 0 | 0 | 0 | 0 | 0 |
| ENSG00000126860 | 2 | 0 | 0 | 1 | 1 | 2 |
| ENSG00000263535 | 0 | 1 | 0 | 1 | 0 | 0 |
| ENSG00000131242 | 2421 | 1143 | 1619 | 1083 | 1182 | 1195 |
| ENSG00000239595 | 0 | 0 | 0 | 0 | 0 | 1 |
| ENSG00000264862 | 0 | 1 | 0 | 0 | 0 | 0 |
| ENSG00000278977 | 0 | 0 | 0 | 0 | 0 | 0 |
| ENSG00000284424 | 0 | 0 | 0 | 0 | 0 | 0 |
| ENSG00000207614 | 0 | 0 | 0 | 4 | 0 | 1 |
| ENSG00000278546 | 2 | 1 | 1 | 2 | 2 | 1 |
| ENSG00000221038 | 0 | 0 | 0 | 0 | 0 | 0 |
| ENSG00000228768 | 0 | 0 | 0 | 0 | 1 | 0 |
| ENSG00000265976 | 0 | 0 | 0 | 0 | 0 | 0 |
| ENSG00000283978 | 0 | 0 | 0 | 0 | 0 | 0 |
| ENSG00000266877 | 0 | 0 | 3 | 0 | 0 | 1 |
| ENSG00000266448 | 0 | 0 | 0 | 0 | 0 | 0 |
| ENSG00000202026 | 0 | 0 | 0 | 0 | 0 | 0 |
| ENSG00000265863 | 0 | 0 | 0 | 0 | 0 | 0 |
| ENSG00000263567 | 0 | 0 | 0 | 0 | 0 | 0 |
| ENSG00000265046 | 0 | 0 | 0 | 0 | 0 | 0 |

| | | | | | | |
|---|---|---|---|---|---|---|
| ENSG00000172301 | 708 | 378 | 452 | 514 | 423 | 437 |
| ENSG00000108651 | 819 | 387 | 517 | 436 | 502 | 498 |
| ENSG00000263990 | 3 | 3 | 6 | 4 | 4 | 5 |
| ENSG00000178691 | 701 | 365 | 361 | 175 | 476 | 330 |
| ENSG00000253058 | 0 | 0 | 0 | 0 | 0 | 0 |
| ENSG00000264300 | 0 | 0 | 0 | 0 | 0 | 0 |
| ENSG00000185158 | 111 | 63 | 86 | 62 | 66 | 82 |
| ENSG00000264497 | 0 | 0 | 0 | 0 | 0 | 0 |
| ENSG00000266777 | 14 | 5 | 11 | 4 | 5 | 10 |
| ENSG00000278867 | 0 | 6 | 4 | 6 | 9 | 5 |
| ENSG00000264164 | 0 | 0 | 0 | 0 | 0 | 0 |
| ENSG00000266379 | 0 | 0 | 0 | 0 | 0 | 0 |
| ENSG00000263369 | 0 | 0 | 0 | 0 | 0 | 0 |
| ENSG00000265574 | 18 | 12 | 23 | 9 | 9 | 15 |
| ENSG00000277511 | 2 | 2 | 1 | 0 | 1 | 1 |
| ENSG00000214708 | 7 | 3 | 7 | 3 | 2 | 7 |
| ENSG00000126858 | 735 | 388 | 457 | 297 | 475 | 411 |
| ENSG00000265293 | 0 | 1 | 0 | 0 | 0 | 1 |
| ENSG00000280033 | 3 | 1 | 3 | 2 | 1 | 1 |
| ENSG00000280020 | 3 | 0 | 1 | 0 | 1 | 1 |
| ENSG00000266282 | 0 | 0 | 0 | 0 | 0 | 0 |
| ENSG00000141314 | 16 | 9 | 9 | 0 | 3 | 1 |
| ENSG00000263674 | 0 | 0 | 0 | 0 | 0 | 0 |
| ENSG00000265794 | 0 | 0 | 0 | 0 | 0 | 0 |
| ENSG00000108666 | 347 | 205 | 235 | 183 | 209 | 244 |
| ENSG00000265139 | 0 | 0 | 0 | 0 | 0 | 0 |
| ENSG00000263749 | 0 | 0 | 0 | 0 | 0 | 0 |
| ENSG00000283774 | 0 | 0 | 0 | 0 | 0 | 0 |
| ENSG00000010244 | 3140 | 1698 | 1997 | 1577 | 1962 | 1944 |
| ENSG00000274341 | 0 | 0 | 0 | 0 | 0 | 0 |
| ENSG00000278668 | 2 | 0 | 0 | 0 | 1 | 1 |
| ENSG00000266385 | 0 | 1 | 0 | 0 | 1 | 0 |
| ENSG00000279762 | 0 | 1 | 1 | 0 | 0 | 0 |
| ENSG00000264083 | 0 | 0 | 0 | 0 | 0 | 3 |
| ENSG00000264373 | 0 | 0 | 1 | 0 | 0 | 0 |
| ENSG00000108671 | 3514 | 1646 | 2030 | 1684 | 1996 | 2078 |
| ENSG00000202100 | 0 | 0 | 1 | 0 | 0 | 0 |
| ENSG00000176749 | 118 | 58 | 58 | 64 | 53 | 60 |
| ENSG00000176658 | 2766 | 1424 | 1791 | 1468 | 1712 | 1620 |
| ENSG00000266718 | 0 | 0 | 4 | 0 | 4 | 4 |
| ENSG00000280245 | 0 | 0 | 0 | 0 | 0 | 0 |
| ENSG00000265222 | 2 | 0 | 3 | 0 | 0 | 1 |
| ENSG00000263717 | 0 | 0 | 1 | 0 | 0 | 0 |
| ENSG00000279781 | 2 | 0 | 0 | 1 | 0 | 0 |
| ENSG00000266599 | 0 | 0 | 0 | 0 | 0 | 0 |
| ENSG00000265337 | 0 | 0 | 0 | 2 | 0 | 0 |
| ENSG00000264458 | 0 | 0 | 0 | 0 | 0 | 0 |
| ENSG00000201178 | 0 | 0 | 0 | 0 | 0 | 0 |
| ENSG00000226377 | 14 | 4 | 2 | 4 | 3 | 2 |
| ENSG00000236377 | 2 | 1 | 1 | 1 | 1 | 1 |

| | | | | | | |
|---|---|---|---|---|---|---|
| ENSG00000006042 | 2 | 1 | 2 | 1 | 2 | 0 |
| ENSG00000141316 | 0 | 0 | 0 | 0 | 0 | 0 |
| ENSG00000108684 | 0 | 0 | 0 | 0 | 0 | 0 |
| ENSG00000264598 | 0 | 0 | 0 | 0 | 0 | 0 |
| ENSG00000280349 | 0 | 0 | 0 | 0 | 0 | 0 |
| ENSG00000266535 | 0 | 0 | 0 | 0 | 0 | 0 |
| ENSG00000265544 | 0 | 0 | 0 | 0 | 0 | 0 |
| ENSG00000265697 | 0 | 0 | 0 | 0 | 0 | 0 |
| ENSG00000265125 | 0 | 0 | 0 | 0 | 0 | 0 |
| ENSG00000265115 | 0 | 0 | 0 | 0 | 0 | 0 |
| ENSG00000264643 | 0 | 0 | 0 | 0 | 0 | 0 |
| ENSG00000265689 | 0 | 0 | 0 | 0 | 0 | 0 |
| ENSG00000283381 | 0 | 0 | 0 | 0 | 0 | 0 |
| ENSG00000283417 | 0 | 0 | 0 | 0 | 0 | 0 |
| ENSG00000279668 | 0 | 0 | 0 | 0 | 0 | 0 |
| ENSG00000265356 | 0 | 0 | 0 | 0 | 0 | 0 |
| ENSG00000263435 | 0 | 0 | 0 | 0 | 0 | 0 |
| ENSG00000279674 | 0 | 0 | 0 | 0 | 0 | 0 |
| ENSG00000226049 | 10 | 11 | 10 | 8 | 7 | 8 |
| ENSG00000263485 | 0 | 0 | 0 | 0 | 0 | 0 |
| ENSG00000265775 | 0 | 0 | 0 | 0 | 0 | 0 |
| ENSG00000252370 | 0 | 0 | 0 | 0 | 0 | 0 |
| ENSG00000264791 | 0 | 0 | 0 | 0 | 0 | 0 |
| ENSG00000263571 | 0 | 0 | 0 | 0 | 0 | 0 |
| ENSG00000261156 | 0 | 0 | 0 | 0 | 0 | 0 |
| ENSG00000264990 | 0 | 0 | 0 | 0 | 0 | 0 |
| ENSG00000108691 | 0 | 0 | 2 | 2 | 2 | 3 |
| ENSG00000108688 | 0 | 0 | 0 | 0 | 0 | 0 |
| ENSG00000172156 | 0 | 0 | 0 | 0 | 0 | 0 |
| ENSG00000108700 | 0 | 0 | 0 | 0 | 0 | 0 |
| ENSG00000265614 | 0 | 0 | 0 | 0 | 0 | 0 |
| ENSG00000181374 | 0 | 0 | 0 | 0 | 0 | 0 |
| ENSG00000108702 | 0 | 0 | 0 | 0 | 0 | 0 |
| ENSG00000225582 | 0 | 0 | 0 | 0 | 0 | 0 |
| ENSG00000197322 | 0 | 0 | 0 | 0 | 0 | 0 |
| ENSG00000181291 | 2 | 2 | 0 | 0 | 0 | 0 |
| ENSG00000278860 | 0 | 0 | 0 | 0 | 0 | 0 |
| ENSG00000264174 | 0 | 0 | 0 | 0 | 0 | 0 |
| ENSG00000264622 | 0 | 0 | 0 | 0 | 0 | 0 |
| ENSG00000132141 | 74 | 46 | 39 | 26 | 33 | 25 |
| ENSG00000285559 | 0 | 0 | 0 | 0 | 0 | 0 |
| ENSG00000198783 | 273 | 141 | 169 | 132 | 160 | 125 |
| ENSG00000005156 | 826 | 411 | 522 | 396 | 480 | 534 |
| ENSG00000092871 | 316 | 161 | 212 | 169 | 260 | 230 |
| ENSG00000267618 | 0 | 0 | 0 | 0 | 0 | 0 |
| ENSG00000273687 | 0 | 3 | 1 | 0 | 1 | 0 |
| ENSG00000267457 | 0 | 2 | 0 | 1 | 0 | 0 |
| ENSG00000243423 | 0 | 0 | 0 | 0 | 0 | 0 |
| ENSG00000185379 | 379 | 201 | 226 | 199 | 189 | 218 |
| ENSG00000253084 | 0 | 0 | 0 | 0 | 0 | 0 |

| | | | | | | |
|---|---|---|---|---|---|---|
| ENSG00000252328 | 0 | 0 | 0 | 0 | 0 | 0 |
| ENSG00000073598 | 0 | 0 | 0 | 1 | 0 | 1 |
| ENSG00000073536 | 490 | 259 | 304 | 424 | 373 | 362 |
| ENSG00000141161 | 0 | 0 | 0 | 0 | 0 | 0 |
| ENSG00000267782 | 0 | 0 | 0 | 0 | 0 | 0 |
| ENSG00000266981 | 0 | 0 | 0 | 0 | 0 | 0 |
| ENSG00000267744 | 0 | 0 | 0 | 0 | 0 | 0 |
| ENSG00000164729 | 0 | 0 | 0 | 0 | 0 | 0 |
| ENSG00000266947 | 99 | 29 | 55 | 36 | 36 | 46 |
| ENSG00000166750 | 1396 | 688 | 693 | 338 | 831 | 629 |
| ENSG00000267364 | 0 | 2 | 0 | 0 | 1 | 1 |
| ENSG00000267271 | 0 | 0 | 0 | 0 | 0 | 0 |
| ENSG00000172716 | 270 | 145 | 184 | 71 | 102 | 131 |
| ENSG00000267315 | 0 | 0 | 0 | 0 | 0 | 0 |
| ENSG00000267711 | 0 | 0 | 0 | 0 | 0 | 0 |
| ENSG00000267547 | 0 | 0 | 0 | 0 | 0 | 0 |
| ENSG00000267745 | 0 | 0 | 0 | 0 | 0 | 0 |
| ENSG00000267648 | 0 | 0 | 0 | 0 | 0 | 0 |
| ENSG00000267102 | 0 | 0 | 0 | 0 | 0 | 0 |
| ENSG00000172123 | 0 | 0 | 0 | 0 | 0 | 0 |
| ENSG00000154760 | 8 | 3 | 5 | 1 | 2 | 4 |
| ENSG00000267312 | 0 | 0 | 0 | 0 | 0 | 0 |
| ENSG00000205045 | 0 | 0 | 0 | 0 | 0 | 0 |
| ENSG00000267554 | 0 | 0 | 0 | 0 | 0 | 0 |
| ENSG00000267349 | 0 | 0 | 0 | 0 | 0 | 0 |
| ENSG00000267046 | 0 | 0 | 0 | 0 | 0 | 0 |
| ENSG00000267369 | 0 | 0 | 0 | 0 | 0 | 0 |
| ENSG00000267074 | 0 | 0 | 0 | 0 | 0 | 0 |
| ENSG00000255987 | 0 | 0 | 0 | 0 | 0 | 0 |
| ENSG00000267035 | 0 | 0 | 0 | 0 | 0 | 0 |
| ENSG00000236320 | 0 | 0 | 0 | 0 | 0 | 0 |
| ENSG00000267359 | 0 | 0 | 0 | 0 | 0 | 0 |
| ENSG00000267625 | 0 | 0 | 0 | 0 | 0 | 0 |
| ENSG00000242660 | 0 | 0 | 1 | 0 | 1 | 0 |
| ENSG00000267321 | 226 | 107 | 132 | 124 | 158 | 132 |
| ENSG00000207297 | 0 | 0 | 0 | 0 | 0 | 0 |
| ENSG00000108733 | 125 | 76 | 84 | 61 | 66 | 60 |
| ENSG00000006125 | 5290 | 2493 | 3104 | 2197 | 2931 | 2872 |
| ENSG00000267592 | 0 | 0 | 0 | 0 | 0 | 0 |
| ENSG00000270647 | 2811 | 1393 | 1701 | 1132 | 1488 | 1617 |
| ENSG00000270885 | 42 | 22 | 17 | 9 | 19 | 5 |
| ENSG00000270765 | 0 | 2 | 0 | 1 | 4 | 0 |
| ENSG00000271447 | 21 | 11 | 8 | 6 | 8 | 9 |
| ENSG00000271392 | 1 | 0 | 0 | 2 | 1 | 0 |
| ENSG00000270806 | 6 | 0 | 4 | 1 | 0 | 1 |
| ENSG00000270829 | 0 | 0 | 0 | 0 | 0 | 0 |
| ENSG00000270871 | 5 | 2 | 13 | 2 | 5 | 6 |
| ENSG00000270894 | 2 | 0 | 0 | 1 | 0 | 1 |
| ENSG00000270379 | 0 | 0 | 2 | 0 | 0 | 0 |
| ENSG00000271268 | 0 | 0 | 2 | 0 | 0 | 1 |

| | | | | | | |
|---|---|---|---|---|---|---|
| ENSG00000270240 | 0 | 0 | 0 | 0 | 1 | 0 |
| ENSG00000271503 | 12 | 7 | 9 | 5 | 2 | 7 |
| ENSG00000271394 | 0 | 0 | 0 | 0 | 0 | 0 |
| ENSG00000270977 | 16 | 3 | 3 | 3 | 7 | 13 |
| ENSG00000271013 | 2 | 5 | 3 | 1 | 4 | 3 |
| ENSG00000278023 | 39 | 17 | 32 | 26 | 27 | 33 |
| ENSG00000275722 | 0 | 0 | 0 | 0 | 0 | 0 |
| ENSG00000273736 | 0 | 0 | 0 | 0 | 0 | 0 |
| ENSG00000275152 | 0 | 0 | 0 | 0 | 0 | 0 |
| ENSG00000276409 | 0 | 0 | 0 | 0 | 0 | 0 |
| ENSG00000275688 | 0 | 0 | 0 | 0 | 0 | 0 |
| ENSG00000275431 | 0 | 0 | 0 | 0 | 0 | 0 |
| ENSG00000275718 | 0 | 0 | 0 | 0 | 0 | 0 |
| ENSG00000275944 | 0 | 0 | 0 | 0 | 0 | 0 |
| ENSG00000278690 | 0 | 0 | 0 | 0 | 0 | 0 |
| ENSG00000274736 | 0 | 0 | 0 | 0 | 0 | 0 |
| ENSG00000275385 | 0 | 0 | 0 | 0 | 0 | 0 |
| ENSG00000277089 | 0 | 0 | 0 | 0 | 0 | 0 |
| ENSG00000277632 | 0 | 0 | 0 | 0 | 0 | 0 |
| ENSG00000275302 | 0 | 0 | 0 | 0 | 1 | 0 |
| ENSG00000276014 | 0 | 0 | 0 | 0 | 0 | 0 |
| ENSG00000276241 | 0 | 0 | 0 | 0 | 0 | 0 |
| ENSG00000282738 | 0 | 0 | 0 | 0 | 0 | 0 |
| ENSG00000274808 | 0 | 0 | 0 | 0 | 1 | 0 |
| ENSG00000274767 | 29 | 4 | 11 | 4 | 9 | 3 |
| ENSG00000276085 | 0 | 1 | 1 | 1 | 0 | 1 |
| ENSG00000276070 | 0 | 0 | 0 | 1 | 0 | 0 |
| ENSG00000278097 | 0 | 0 | 0 | 0 | 0 | 0 |
| ENSG00000276755 | 0 | 0 | 1 | 0 | 0 | 0 |
| ENSG00000278395 | 0 | 0 | 0 | 0 | 0 | 0 |
| ENSG00000274933 | 0 | 0 | 0 | 0 | 0 | 0 |
| ENSG00000261499 | 6 | 3 | 7 | 1 | 4 | 4 |
| ENSG00000265889 | 0 | 0 | 0 | 0 | 0 | 0 |
| ENSG00000260287 | 0 | 0 | 0 | 0 | 1 | 0 |
| ENSG00000274226 | 0 | 0 | 0 | 0 | 0 | 0 |
| ENSG00000277029 | 0 | 0 | 0 | 0 | 0 | 0 |
| ENSG00000275954 | 0 | 0 | 0 | 0 | 0 | 0 |
| ENSG00000274658 | 0 | 0 | 0 | 0 | 0 | 0 |
| ENSG00000273611 | 1071 | 510 | 682 | 646 | 624 | 554 |
| ENSG00000274164 | 0 | 0 | 0 | 0 | 0 | 0 |
| ENSG00000278259 | 1342 | 570 | 894 | 630 | 738 | 861 |
| ENSG00000277161 | 311 | 170 | 216 | 154 | 193 | 245 |
| ENSG00000278311 | 1301 | 660 | 766 | 421 | 804 | 705 |
| ENSG00000274756 | 15 | 3 | 3 | 2 | 5 | 5 |
| ENSG00000278535 | 425 | 164 | 205 | 215 | 217 | 192 |
| ENSG00000278619 | 189 | 94 | 123 | 164 | 120 | 162 |
| ENSG00000275720 | 0 | 1 | 1 | 0 | 2 | 1 |
| ENSG00000276508 | 0 | 0 | 0 | 0 | 0 | 0 |
| ENSG00000275613 | 0 | 0 | 0 | 0 | 0 | 0 |
| ENSG00000277268 | 62 | 32 | 68 | 38 | 35 | 42 |

| | | | | | | |
|---|---|---|---|---|---|---|
| ENSG00000273706 | 352 | 149 | 235 | 135 | 173 | 223 |
| ENSG00000276707 | 11 | 4 | 5 | 1 | 4 | 3 |
| ENSG00000275700 | 1462 | 699 | 912 | 885 | 883 | 888 |
| ENSG00000277911 | 0 | 0 | 0 | 0 | 0 | 0 |
| ENSG00000277589 | 2 | 2 | 0 | 0 | 0 | 0 |
| ENSG00000276326 | 0 | 0 | 0 | 0 | 0 | 0 |
| ENSG00000277579 | 0 | 0 | 0 | 1 | 0 | 0 |
| ENSG00000278638 | 2 | 0 | 0 | 1 | 1 | 1 |
| ENSG00000278540 | 3898 | 1548 | 2245 | 1223 | 1722 | 1926 |
| ENSG00000274308 | 0 | 0 | 0 | 0 | 1 | 0 |
| ENSG00000275029 | 0 | 0 | 2 | 0 | 0 | 0 |
| ENSG00000276810 | 0 | 1 | 0 | 0 | 0 | 0 |
| ENSG00000278505 | 0 | 1 | 2 | 0 | 0 | 0 |
| ENSG00000276054 | 1 | 0 | 2 | 0 | 0 | 1 |
| ENSG00000276234 | 307 | 167 | 206 | 159 | 186 | 186 |
| ENSG00000273965 | 1 | 0 | 1 | 0 | 0 | 0 |
| ENSG00000275839 | 0 | 1 | 0 | 0 | 0 | 2 |
| ENSG00000276023 | 614 | 299 | 338 | 380 | 390 | 371 |
| ENSG00000275066 | 1225 | 586 | 652 | 378 | 593 | 596 |
| ENSG00000278053 | 572 | 296 | 306 | 231 | 338 | 267 |
| ENSG00000277688 | 0 | 0 | 0 | 0 | 0 | 0 |
| ENSG00000274620 | 0 | 0 | 0 | 0 | 0 | 0 |
| ENSG00000277501 | 0 | 0 | 0 | 0 | 0 | 0 |
| ENSG00000274244 | 0 | 0 | 0 | 0 | 0 | 0 |
| ENSG00000275410 | 0 | 0 | 0 | 1 | 0 | 0 |
| ENSG00000275173 | 0 | 0 | 0 | 0 | 0 | 0 |
| ENSG00000276715 | 0 | 0 | 0 | 0 | 0 | 0 |
| ENSG00000277341 | 0 | 0 | 0 | 0 | 0 | 0 |
| ENSG00000273513 | 0 | 0 | 0 | 2 | 0 | 0 |
| ENSG00000277488 | 0 | 0 | 0 | 0 | 0 | 0 |
| ENSG00000274512 | 94 | 25 | 50 | 47 | 55 | 52 |
| ENSG00000274419 | 0 | 0 | 0 | 0 | 0 | 0 |
| ENSG00000278299 | 0 | 0 | 0 | 0 | 0 | 0 |
| ENSG00000274615 | 0 | 4 | 0 | 0 | 0 | 1 |
| ENSG00000275006 | 0 | 0 | 0 | 0 | 0 | 0 |
| ENSG00000278599 | 1 | 0 | 1 | 0 | 1 | 0 |
| ENSG00000274663 | 0 | 0 | 0 | 0 | 0 | 0 |
| ENSG00000274611 | 0 | 0 | 0 | 0 | 0 | 0 |
| ENSG00000274487 | 49 | 27 | 35 | 24 | 36 | 23 |
| ENSG00000274284 | 0 | 0 | 0 | 0 | 0 | 0 |
| ENSG00000278845 | 1298 | 574 | 725 | 696 | 710 | 675 |
| ENSG00000277399 | 8 | 6 | 6 | 6 | 3 | 6 |
| ENSG00000274211 | 929 | 422 | 615 | 352 | 474 | 502 |
| ENSG00000275832 | 2 | 0 | 0 | 1 | 0 | 2 |
| ENSG00000276170 | 0 | 0 | 0 | 0 | 0 | 0 |
| ENSG00000277363 | 386 | 183 | 186 | 154 | 172 | 196 |
| ENSG00000274996 | 0 | 0 | 0 | 0 | 1 | 0 |
| ENSG00000273604 | 250 | 128 | 132 | 200 | 178 | 204 |
| ENSG00000275238 | 0 | 0 | 0 | 0 | 0 | 0 |
| ENSG00000277969 | 61 | 41 | 58 | 38 | 47 | 47 |

| | | | | | | |
|---|---|---|---|---|---|---|
| ENSG00000275532 | 1 | 0 | 3 | 1 | 3 | 1 |
| ENSG00000275023 | 2037 | 914 | 1326 | 945 | 1092 | 1251 |
| ENSG00000275665 | 0 | 0 | 0 | 0 | 0 | 0 |
| ENSG00000273627 | 0 | 0 | 0 | 0 | 0 | 0 |
| ENSG00000279119 | 0 | 0 | 0 | 0 | 0 | 0 |
| ENSG00000277972 | 552 | 225 | 314 | 372 | 277 | 309 |
| ENSG00000276036 | 0 | 0 | 0 | 0 | 4 | 0 |
| ENSG00000277258 | 1527 | 622 | 927 | 750 | 697 | 734 |
| ENSG00000277182 | 53 | 26 | 28 | 14 | 20 | 26 |
| ENSG00000277791 | 2872 | 1470 | 1699 | 2084 | 1734 | 1655 |
| ENSG00000274607 | 0 | 0 | 0 | 0 | 0 | 0 |
| ENSG00000276293 | 1797 | 744 | 1068 | 771 | 854 | 1032 |
| ENSG00000273559 | 439 | 177 | 260 | 209 | 254 | 213 |
| ENSG00000274054 | 0 | 0 | 0 | 0 | 0 | 0 |
| ENSG00000275489 | 0 | 0 | 0 | 0 | 0 | 0 |
| ENSG00000125691 | 12684 | 6287 | 7341 | 8971 | 7804 | 7605 |
| ENSG00000252699 | 0 | 0 | 0 | 0 | 0 | 0 |
| ENSG00000199293 | 0 | 0 | 0 | 0 | 0 | 0 |
| ENSG00000244086 | 0 | 0 | 0 | 0 | 0 | 0 |
| ENSG00000002834 | 12193 | 6196 | 7573 | 6351 | 6299 | 6495 |
| ENSG00000263466 | 0 | 1 | 1 | 1 | 0 | 0 |
| ENSG00000277057 | 0 | 0 | 0 | 0 | 0 | 0 |
| ENSG00000265784 | 0 | 0 | 0 | 0 | 0 | 0 |
| ENSG00000263874 | 105 | 38 | 53 | 20 | 44 | 40 |
| ENSG00000204952 | 0 | 1 | 0 | 0 | 0 | 1 |
| ENSG00000265428 | 0 | 0 | 0 | 0 | 0 | 0 |
| ENSG00000266588 | 0 | 0 | 0 | 0 | 0 | 0 |
| ENSG00000277426 | 0 | 0 | 0 | 0 | 0 | 0 |
| ENSG00000266013 | 0 | 0 | 0 | 0 | 0 | 0 |
| ENSG00000214553 | 0 | 1 | 0 | 0 | 1 | 0 |
| ENSG00000266048 | 0 | 0 | 0 | 0 | 0 | 0 |
| ENSG00000263818 | 29 | 13 | 12 | 9 | 14 | 17 |
| ENSG00000161381 | 1 | 10 | 4 | 3 | 10 | 8 |
| ENSG00000141748 | 2 | 0 | 0 | 0 | 0 | 0 |
| ENSG00000280177 | 0 | 0 | 0 | 0 | 0 | 0 |
| ENSG00000266101 | 0 | 0 | 0 | 0 | 0 | 0 |
| ENSG00000067191 | 110 | 67 | 57 | 81 | 92 | 94 |
| ENSG00000108298 | 17842 | 9205 | 10592 | 12081 | 10825 | 10729 |
| ENSG00000141750 | 0 | 0 | 0 | 0 | 0 | 0 |
| ENSG00000265460 | 0 | 0 | 1 | 0 | 0 | 0 |
| ENSG00000108306 | 559 | 292 | 364 | 194 | 282 | 253 |
| ENSG00000266469 | 242 | 122 | 162 | 98 | 159 | 127 |
| ENSG00000125686 | 1087 | 511 | 666 | 344 | 573 | 594 |
| ENSG00000167258 | 1566 | 745 | 899 | 605 | 865 | 833 |
| ENSG00000222777 | 0 | 0 | 0 | 0 | 0 | 0 |
| ENSG00000273576 | 63 | 26 | 38 | 24 | 28 | 26 |
| ENSG00000171532 | 0 | 1 | 0 | 0 | 1 | 0 |
| ENSG00000214546 | 0 | 0 | 0 | 0 | 0 | 0 |
| ENSG00000131771 | 0 | 0 | 0 | 0 | 1 | 1 |
| ENSG00000131748 | 1039 | 503 | 602 | 611 | 558 | 490 |

| | | | | | | |
|---|---|---|---|---|---|---|
| ENSG00000173991 | 24 | 12 | 10 | 23 | 20 | 16 |
| ENSG00000141744 | 0 | 0 | 0 | 0 | 0 | 0 |
| ENSG00000161395 | 564 | 300 | 391 | 309 | 287 | 285 |
| ENSG00000141736 | 6000 | 2782 | 3913 | 2965 | 3163 | 3083 |
| ENSG00000265178 | 1 | 0 | 1 | 0 | 0 | 0 |
| ENSG00000141741 | 791 | 432 | 544 | 539 | 452 | 488 |
| ENSG00000141738 | 545 | 277 | 371 | 288 | 263 | 304 |
| ENSG00000161405 | 2 | 4 | 2 | 11 | 7 | 10 |
| ENSG00000264663 | 0 | 0 | 0 | 0 | 0 | 0 |
| ENSG00000226117 | 0 | 0 | 0 | 0 | 0 | 0 |
| ENSG00000186075 | 0 | 0 | 0 | 0 | 0 | 0 |
| ENSG00000073605 | 192 | 100 | 108 | 86 | 119 | 79 |
| ENSG00000172057 | 1673 | 744 | 982 | 869 | 837 | 879 |
| ENSG00000264968 | 0 | 0 | 2 | 1 | 0 | 0 |
| ENSG00000204913 | 0 | 0 | 0 | 0 | 0 | 0 |
| ENSG00000167914 | 2 | 0 | 2 | 0 | 0 | 0 |
| ENSG00000108344 | 3563 | 1790 | 2172 | 2232 | 2161 | 2210 |
| ENSG00000265799 | 0 | 0 | 0 | 0 | 0 | 0 |
| ENSG00000108342 | 0 | 0 | 0 | 0 | 0 | 0 |
| ENSG00000008838 | 1318 | 553 | 728 | 747 | 663 | 749 |
| ENSG00000283721 | 0 | 0 | 0 | 0 | 0 | 0 |
| ENSG00000238793 | 4 | 0 | 0 | 0 | 1 | 0 |
| ENSG00000126351 | 1066 | 486 | 591 | 468 | 463 | 473 |
| ENSG00000126368 | 94 | 36 | 70 | 54 | 58 | 51 |
| ENSG00000279199 | 6 | 5 | 4 | 1 | 5 | 5 |
| ENSG00000188895 | 3462 | 1557 | 1920 | 1465 | 1618 | 1808 |
| ENSG00000108349 | 1879 | 788 | 1105 | 875 | 961 | 1044 |
| ENSG00000284144 | 0 | 0 | 0 | 0 | 0 | 0 |
| ENSG00000108352 | 552 | 241 | 282 | 376 | 282 | 308 |
| ENSG00000274621 | 0 | 0 | 4 | 0 | 0 | 0 |
| ENSG00000171475 | 1701 | 796 | 1039 | 693 | 807 | 960 |
| ENSG00000094804 | 481 | 292 | 330 | 217 | 270 | 349 |
| ENSG00000131759 | 2732 | 1258 | 1645 | 1222 | 976 | 1074 |
| ENSG00000265666 | 4 | 1 | 5 | 3 | 1 | 4 |
| ENSG00000278918 | 27 | 8 | 18 | 5 | 11 | 15 |
| ENSG00000266208 | 764 | 443 | 520 | 442 | 478 | 434 |
| ENSG00000183153 | 8 | 4 | 4 | 9 | 7 | 5 |
| ENSG00000264655 | 10 | 12 | 15 | 6 | 8 | 8 |
| ENSG00000252618 | 0 | 0 | 5 | 1 | 0 | 0 |
| ENSG00000278346 | 0 | 0 | 0 | 0 | 0 | 0 |
| ENSG00000131747 | 1365 | 966 | 742 | 337 | 2076 | 959 |
| ENSG00000222881 | 0 | 3 | 1 | 0 | 0 | 0 |
| ENSG00000240125 | 0 | 0 | 0 | 0 | 0 | 0 |
| ENSG00000141753 | 4229 | 1967 | 2274 | 2194 | 1558 | 1877 |
| ENSG00000279806 | 4 | 0 | 0 | 0 | 4 | 0 |
| ENSG00000131746 | 73 | 43 | 49 | 30 | 34 | 23 |
| ENSG00000266088 | 1 | 0 | 1 | 1 | 0 | 1 |
| ENSG00000126353 | 31 | 17 | 17 | 12 | 15 | 11 |
| ENSG00000279775 | 0 | 0 | 0 | 0 | 0 | 0 |
| ENSG00000073584 | 314 | 171 | 191 | 103 | 140 | 142 |

| | | | | | | |
|---|---|---|---|---|---|---|
| ENSG00000264058 | 0 | 0 | 0 | 0 | 0 | 0 |
| ENSG00000278834 | 13 | 11 | 24 | 23 | 16 | 29 |
| ENSG00000213424 | 0 | 2 | 2 | 0 | 0 | 1 |
| ENSG00000224244 | 5 | 8 | 6 | 2 | 1 | 3 |
| ENSG00000167916 | 24 | 5 | 7 | 6 | 1 | 2 |
| ENSG00000229028 | 2 | 4 | 3 | 1 | 0 | 4 |
| ENSG00000204897 | 0 | 0 | 0 | 0 | 0 | 0 |
| ENSG00000186393 | 0 | 0 | 0 | 0 | 0 | 0 |
| ENSG00000171446 | 0 | 0 | 0 | 0 | 0 | 0 |
| ENSG00000173908 | 0 | 0 | 0 | 0 | 0 | 0 |
| ENSG00000264488 | 0 | 0 | 0 | 0 | 0 | 0 |
| ENSG00000186395 | 517 | 239 | 305 | 302 | 311 | 286 |
| ENSG00000167920 | 367 | 170 | 248 | 194 | 181 | 167 |
| ENSG00000265359 | 1 | 0 | 0 | 0 | 0 | 0 |
| ENSG00000187242 | 2 | 6 | 0 | 0 | 1 | 0 |
| ENSG00000171431 | 13 | 5 | 7 | 5 | 3 | 3 |
| ENSG00000234477 | 3 | 5 | 6 | 0 | 2 | 3 |
| ENSG00000108244 | 7 | 2 | 9 | 0 | 0 | 1 |
| ENSG00000196859 | 0 | 2 | 0 | 0 | 0 | 0 |
| ENSG00000204889 | 4 | 1 | 4 | 2 | 0 | 1 |
| ENSG00000212899 | 0 | 0 | 0 | 0 | 0 | 0 |
| ENSG00000212900 | 2 | 0 | 0 | 0 | 0 | 0 |
| ENSG00000226776 | 0 | 0 | 0 | 0 | 0 | 0 |
| ENSG00000212901 | 0 | 3 | 3 | 0 | 0 | 2 |
| ENSG00000221852 | 0 | 0 | 0 | 0 | 0 | 0 |
| ENSG00000204887 | 0 | 0 | 0 | 0 | 0 | 0 |
| ENSG00000221880 | 0 | 0 | 0 | 1 | 0 | 0 |
| ENSG00000188581 | 0 | 0 | 0 | 0 | 0 | 0 |
| ENSG00000212725 | 0 | 0 | 0 | 0 | 0 | 0 |
| ENSG00000214518 | 0 | 0 | 0 | 0 | 0 | 0 |
| ENSG00000212724 | 0 | 0 | 0 | 0 | 0 | 0 |
| ENSG00000213417 | 0 | 0 | 0 | 0 | 0 | 0 |
| ENSG00000237230 | 0 | 0 | 0 | 0 | 0 | 0 |
| ENSG00000240871 | 0 | 0 | 0 | 0 | 0 | 0 |
| ENSG00000204880 | 0 | 0 | 0 | 0 | 0 | 0 |
| ENSG00000241241 | 0 | 0 | 0 | 0 | 0 | 0 |
| ENSG00000212722 | 0 | 0 | 0 | 0 | 0 | 0 |
| ENSG00000212721 | 0 | 0 | 0 | 0 | 0 | 0 |
| ENSG00000213416 | 0 | 0 | 0 | 0 | 0 | 0 |
| ENSG00000198090 | 0 | 0 | 0 | 0 | 0 | 0 |
| ENSG00000198271 | 0 | 0 | 0 | 0 | 0 | 0 |
| ENSG00000171396 | 0 | 0 | 0 | 0 | 0 | 0 |
| ENSG00000196156 | 0 | 0 | 0 | 0 | 0 | 0 |
| ENSG00000244537 | 0 | 0 | 0 | 0 | 0 | 0 |
| ENSG00000198443 | 28 | 12 | 21 | 17 | 7 | 12 |
| ENSG00000251439 | 0 | 0 | 0 | 0 | 1 | 0 |
| ENSG00000240542 | 0 | 0 | 1 | 0 | 0 | 0 |
| ENSG00000248807 | 0 | 0 | 0 | 0 | 0 | 0 |
| ENSG00000239886 | 0 | 0 | 0 | 0 | 0 | 0 |
| ENSG00000204873 | 0 | 0 | 0 | 0 | 0 | 0 |

| | | | | | | |
|---|---|---|---|---|---|---|
| ENSG00000187272 | 0 | 0 | 0 | 0 | 0 | 0 |
| ENSG00000241595 | 0 | 0 | 0 | 0 | 0 | 0 |
| ENSG00000198083 | 0 | 0 | 0 | 0 | 0 | 0 |
| ENSG00000212659 | 0 | 0 | 0 | 0 | 0 | 0 |
| ENSG00000229351 | 0 | 0 | 0 | 0 | 0 | 0 |
| ENSG00000180386 | 0 | 0 | 0 | 0 | 0 | 0 |
| ENSG00000237183 | 0 | 0 | 0 | 0 | 0 | 0 |
| ENSG00000212658 | 0 | 0 | 0 | 0 | 0 | 0 |
| ENSG00000212657 | 0 | 0 | 0 | 0 | 0 | 0 |
| ENSG00000186860 | 0 | 0 | 0 | 0 | 0 | 0 |
| ENSG00000233014 | 0 | 0 | 0 | 0 | 1 | 1 |
| ENSG00000108516 | 0 | 0 | 0 | 0 | 0 | 0 |
| ENSG00000006059 | 0 | 0 | 0 | 0 | 0 | 0 |
| ENSG00000131738 | 0 | 0 | 0 | 0 | 0 | 0 |
| ENSG00000131737 | 0 | 0 | 0 | 0 | 0 | 0 |
| ENSG00000094796 | 13 | 0 | 0 | 0 | 0 | 2 |
| ENSG00000234859 | 1 | 2 | 0 | 0 | 0 | 0 |
| ENSG00000225438 | 0 | 0 | 3 | 1 | 1 | 1 |
| ENSG00000108417 | 0 | 5 | 6 | 13 | 7 | 8 |
| ENSG00000171360 | 0 | 0 | 1 | 1 | 1 | 0 |
| ENSG00000236473 | 0 | 0 | 0 | 0 | 1 | 0 |
| ENSG00000108759 | 4 | 1 | 0 | 2 | 1 | 5 |
| ENSG00000223125 | 0 | 0 | 0 | 0 | 0 | 0 |
| ENSG00000197079 | 0 | 1 | 0 | 2 | 2 | 0 |
| ENSG00000126337 | 0 | 0 | 0 | 0 | 0 | 0 |
| ENSG00000171401 | 7 | 1 | 2 | 5 | 0 | 1 |
| ENSG00000229732 | 0 | 0 | 0 | 0 | 0 | 0 |
| ENSG00000171346 | 269 | 101 | 128 | 66 | 57 | 100 |
| ENSG00000283796 | 0 | 0 | 0 | 0 | 0 | 0 |
| ENSG00000171345 | 105006 | 51987 | 62718 | 53185 | 42110 | 42272 |
| ENSG00000226629 | 2 | 0 | 0 | 0 | 0 | 0 |
| ENSG00000171403 | 0 | 0 | 1 | 0 | 1 | 0 |
| ENSG00000186847 | 1 | 1 | 0 | 1 | 0 | 0 |
| ENSG00000186832 | 20 | 3 | 3 | 7 | 4 | 3 |
| ENSG00000128422 | 13 | 3 | 0 | 2 | 2 | 0 |
| ENSG00000214514 | 0 | 0 | 0 | 0 | 0 | 0 |
| ENSG00000278954 | 0 | 0 | 0 | 0 | 0 | 0 |
| ENSG00000173812 | 9213 | 4745 | 5733 | 6506 | 5997 | 5516 |
| ENSG00000184502 | 1 | 0 | 0 | 0 | 0 | 0 |
| ENSG00000173805 | 88 | 38 | 43 | 47 | 59 | 49 |
| ENSG00000201920 | 0 | 0 | 0 | 0 | 0 | 0 |
| ENSG00000239542 | 2 | 0 | 0 | 2 | 0 | 0 |
| ENSG00000173801 | 42606 | 20042 | 24047 | 30067 | 26666 | 25691 |
| ENSG00000141696 | 1127 | 562 | 718 | 783 | 755 | 689 |
| ENSG00000141756 | 119 | 62 | 70 | 75 | 62 | 89 |
| ENSG00000141698 | 1124 | 533 | 572 | 674 | 705 | 606 |
| ENSG00000161594 | 0 | 4 | 4 | 1 | 1 | 1 |
| ENSG00000259623 | 277 | 162 | 155 | 142 | 187 | 134 |
| ENSG00000178502 | 131 | 65 | 71 | 24 | 51 | 67 |
| ENSG00000131473 | 9528 | 4612 | 5834 | 4835 | 5370 | 5709 |

| | | | | | | |
|---|---|---|---|---|---|---|
| ENSG00000274630 | 0 | 0 | 0 | 0 | 0 | 0 |
| ENSG00000204815 | 40 | 30 | 39 | 35 | 23 | 20 |
| ENSG00000173786 | 1515 | 715 | 897 | 716 | 649 | 677 |
| ENSG00000168259 | 1799 | 951 | 1097 | 968 | 1166 | 1066 |
| ENSG00000168256 | 1842 | 822 | 1128 | 1034 | 959 | 978 |
| ENSG00000187595 | 46 | 15 | 35 | 31 | 20 | 20 |
| ENSG00000267221 | 118 | 44 | 78 | 58 | 63 | 62 |
| ENSG00000108771 | 4 | 3 | 4 | 0 | 2 | 0 |
| ENSG00000108773 | 1202 | 590 | 685 | 818 | 735 | 822 |
| ENSG00000267261 | 0 | 0 | 0 | 0 | 0 | 0 |
| ENSG00000260325 | 12 | 3 | 2 | 9 | 7 | 5 |
| ENSG00000108774 | 1981 | 1269 | 1498 | 1590 | 1373 | 1377 |
| ENSG00000267658 | 2 | 0 | 1 | 0 | 0 | 0 |
| ENSG00000089558 | 0 | 0 | 3 | 0 | 0 | 0 |
| ENSG00000161610 | 0 | 0 | 0 | 0 | 0 | 0 |
| ENSG00000167925 | 614 | 250 | 335 | 330 | 267 | 294 |
| ENSG00000173757 | 1304 | 512 | 682 | 637 | 678 | 676 |
| ENSG00000280295 | 0 | 0 | 0 | 1 | 1 | 0 |
| ENSG00000267758 | 0 | 0 | 0 | 0 | 0 | 2 |
| ENSG00000236194 | 0 | 1 | 0 | 0 | 0 | 0 |
| ENSG00000278829 | 0 | 1 | 0 | 1 | 0 | 0 |
| ENSG00000126561 | 42 | 30 | 31 | 24 | 28 | 32 |
| ENSG00000168610 | 3653 | 1546 | 1997 | 1268 | 1725 | 1630 |
| ENSG00000177469 | 54 | 44 | 20 | 20 | 35 | 19 |
| ENSG00000238704 | 0 | 0 | 0 | 0 | 0 | 0 |
| ENSG00000033627 | 1694 | 826 | 1151 | 828 | 975 | 937 |
| ENSG00000264314 | 1 | 0 | 0 | 0 | 0 | 0 |
| ENSG00000267222 | 4 | 0 | 1 | 0 | 0 | 0 |
| ENSG00000267632 | 11 | 4 | 7 | 3 | 8 | 4 |
| ENSG00000283929 | 7 | 3 | 4 | 1 | 2 | 2 |
| ENSG00000267185 | 5 | 1 | 2 | 2 | 5 | 3 |
| ENSG00000108784 | 1224 | 584 | 820 | 738 | 674 | 696 |
| ENSG00000266929 | 0 | 0 | 0 | 0 | 0 | 0 |
| ENSG00000108785 | 35 | 23 | 32 | 34 | 43 | 28 |
| ENSG00000108786 | 74 | 45 | 67 | 60 | 69 | 58 |
| ENSG00000266962 | 14 | 18 | 26 | 20 | 10 | 17 |
| ENSG00000068120 | 2240 | 1042 | 1272 | 1374 | 1270 | 1173 |
| ENSG00000108788 | 2395 | 1205 | 1426 | 1356 | 1381 | 1311 |
| ENSG00000131470 | 106 | 46 | 71 | 59 | 60 | 70 |
| ENSG00000141699 | 2099 | 932 | 1230 | 876 | 1006 | 1064 |
| ENSG00000280183 | 6 | 0 | 3 | 0 | 1 | 0 |
| ENSG00000131462 | 1649 | 929 | 1135 | 1278 | 1157 | 1228 |
| ENSG00000267189 | 0 | 0 | 0 | 0 | 0 | 0 |
| ENSG00000267132 | 0 | 0 | 0 | 0 | 0 | 0 |
| ENSG00000037042 | 530 | 239 | 305 | 236 | 266 | 255 |
| ENSG00000068137 | 1693 | 711 | 965 | 922 | 769 | 817 |
| ENSG00000184451 | 9 | 3 | 6 | 1 | 6 | 7 |
| ENSG00000267042 | 1 | 0 | 1 | 0 | 0 | 0 |
| ENSG00000108797 | 75 | 28 | 42 | 28 | 25 | 34 |
| ENSG00000267765 | 1 | 1 | 0 | 0 | 0 | 0 |

| | | | | | | |
|---|---|---|---|---|---|---|
| ENSG00000108799 | 1286 | 504 | 864 | 476 | 633 | 599 |
| ENSG00000275273 | 0 | 0 | 0 | 0 | 0 | 0 |
| ENSG00000239671 | 0 | 0 | 0 | 0 | 0 | 0 |
| ENSG00000273650 | 0 | 0 | 1 | 1 | 0 | 0 |
| ENSG00000214578 | 1 | 0 | 1 | 1 | 0 | 0 |
| ENSG00000197291 | 0 | 2 | 4 | 1 | 0 | 0 |
| ENSG00000131477 | 0 | 0 | 0 | 2 | 8 | 2 |
| ENSG00000131475 | 2860 | 1533 | 1741 | 1964 | 1493 | 1532 |
| ENSG00000126562 | 18 | 12 | 17 | 7 | 10 | 9 |
| ENSG00000183978 | 1178 | 599 | 762 | 794 | 603 | 619 |
| ENSG00000176563 | 22 | 15 | 18 | 9 | 18 | 9 |
| ENSG00000126581 | 1975 | 1025 | 1291 | 899 | 1146 | 1197 |
| ENSG00000278447 | 0 | 0 | 0 | 1 | 0 | 0 |
| ENSG00000131467 | 4699 | 2326 | 2896 | 2630 | 2746 | 3210 |
| ENSG00000131480 | 26 | 16 | 17 | 15 | 16 | 12 |
| ENSG00000131471 | 2 | 2 | 10 | 3 | 2 | 8 |
| ENSG00000260105 | 0 | 0 | 0 | 0 | 0 | 0 |
| ENSG00000213373 | 0 | 0 | 1 | 0 | 0 | 0 |
| ENSG00000252039 | 0 | 0 | 0 | 0 | 0 | 0 |
| ENSG00000131482 | 0 | 0 | 0 | 0 | 0 | 0 |
| ENSG00000212149 | 0 | 0 | 0 | 0 | 0 | 0 |
| ENSG00000266967 | 31 | 19 | 28 | 19 | 17 | 26 |
| ENSG00000108825 | 0 | 0 | 1 | 0 | 0 | 1 |
| ENSG00000267060 | 8 | 10 | 12 | 2 | 3 | 13 |
| ENSG00000198863 | 1218 | 606 | 749 | 572 | 602 | 647 |
| ENSG00000131469 | 15157 | 7657 | 8825 | 10608 | 9115 | 8798 |
| ENSG00000068079 | 199 | 101 | 121 | 107 | 108 | 101 |
| ENSG00000108828 | 7434 | 3656 | 4659 | 6600 | 5865 | 5775 |
| ENSG00000108830 | 66 | 30 | 31 | 17 | 16 | 34 |
| ENSG00000012048 | 301 | 178 | 138 | 67 | 234 | 136 |
| ENSG00000240828 | 0 | 0 | 0 | 0 | 0 | 1 |
| ENSG00000198496 | 206 | 122 | 150 | 119 | 163 | 137 |
| ENSG00000285057 | 0 | 0 | 0 | 0 | 0 | 0 |
| ENSG00000267681 | 7 | 8 | 7 | 1 | 5 | 3 |
| ENSG00000267002 | 340 | 181 | 213 | 129 | 206 | 153 |
| ENSG00000267595 | 2 | 6 | 3 | 0 | 3 | 4 |
| ENSG00000267340 | 5 | 2 | 8 | 0 | 3 | 2 |
| ENSG00000188554 | 5233 | 2394 | 3261 | 1912 | 2698 | 2526 |
| ENSG00000184988 | 23 | 12 | 13 | 18 | 11 | 16 |
| ENSG00000236383 | 5 | 6 | 7 | 3 | 6 | 2 |
| ENSG00000274585 | 0 | 0 | 0 | 0 | 0 | 0 |
| ENSG00000275616 | 0 | 0 | 0 | 0 | 0 | 0 |
| ENSG00000273709 | 0 | 0 | 0 | 0 | 0 | 0 |
| ENSG00000274432 | 0 | 0 | 0 | 0 | 0 | 0 |
| ENSG00000278048 | 0 | 0 | 0 | 0 | 0 | 0 |
| ENSG00000276596 | 0 | 0 | 0 | 0 | 0 | 0 |
| ENSG00000277903 | 0 | 0 | 0 | 0 | 0 | 0 |
| ENSG00000278774 | 0 | 0 | 0 | 0 | 0 | 0 |
| ENSG00000274862 | 0 | 0 | 0 | 0 | 0 | 0 |
| ENSG00000274062 | 0 | 0 | 0 | 0 | 0 | 0 |

| | | | | | | |
|---|---|---|---|---|---|---|
| ENSG00000274452 | 0 | 0 | 0 | 0 | 0 | 0 |
| ENSG00000275219 | 0 | 0 | 0 | 0 | 0 | 0 |
| ENSG00000278591 | 0 | 0 | 0 | 0 | 0 | 0 |
| ENSG00000279602 | 6 | 10 | 8 | 8 | 6 | 7 |
| ENSG00000188825 | 172 | 113 | 130 | 78 | 105 | 89 |
| ENSG00000277084 | 0 | 0 | 0 | 0 | 0 | 0 |
| ENSG00000252882 | 0 | 0 | 0 | 0 | 0 | 0 |
| ENSG00000175906 | 30 | 7 | 11 | 4 | 7 | 11 |
| ENSG00000251763 | 0 | 0 | 0 | 0 | 0 | 0 |
| ENSG00000267151 | 1 | 0 | 0 | 0 | 0 | 0 |
| ENSG00000284344 | 0 | 0 | 0 | 0 | 0 | 0 |
| ENSG00000252279 | 0 | 0 | 0 | 0 | 0 | 0 |
| ENSG00000252729 | 0 | 0 | 0 | 0 | 0 | 0 |
| ENSG00000237888 | 0 | 0 | 0 | 0 | 0 | 2 |
| ENSG00000067596 | 2093 | 1001 | 1235 | 975 | 1154 | 1110 |
| ENSG00000175832 | 39 | 13 | 10 | 23 | 15 | 14 |
| ENSG00000278109 | 0 | 0 | 0 | 0 | 0 | 0 |
| ENSG00000005102 | 0 | 0 | 0 | 0 | 0 | 0 |
| ENSG00000267440 | 0 | 0 | 0 | 0 | 0 | 0 |
| ENSG00000267253 | 0 | 0 | 0 | 0 | 0 | 0 |
| ENSG00000167941 | 0 | 0 | 0 | 0 | 0 | 2 |
| ENSG00000108861 | 3938 | 1897 | 2181 | 1799 | 2193 | 2017 |
| ENSG00000231256 | 0 | 0 | 2 | 1 | 2 | 2 |
| ENSG00000267604 | 0 | 0 | 0 | 0 | 0 | 0 |
| ENSG00000161647 | 312 | 126 | 159 | 127 | 179 | 182 |
| ENSG00000161649 | 0 | 0 | 0 | 0 | 0 | 0 |
| ENSG00000108852 | 284 | 108 | 193 | 141 | 117 | 129 |
| ENSG00000282199 | 0 | 0 | 0 | 0 | 0 | 0 |
| ENSG00000267496 | 0 | 0 | 0 | 0 | 0 | 0 |
| ENSG00000267166 | 0 | 0 | 0 | 0 | 0 | 0 |
| ENSG00000267420 | 0 | 0 | 0 | 0 | 0 | 0 |
| ENSG00000261514 | 0 | 0 | 0 | 0 | 0 | 0 |
| ENSG00000108849 | 0 | 0 | 0 | 0 | 0 | 0 |
| ENSG00000131096 | 2 | 0 | 0 | 0 | 0 | 2 |
| ENSG00000161653 | 45 | 18 | 27 | 19 | 28 | 16 |
| ENSG00000091947 | 1404 | 679 | 813 | 957 | 782 | 791 |
| ENSG00000221044 | 1 | 2 | 3 | 0 | 3 | 3 |
| ENSG00000161654 | 668 | 352 | 418 | 334 | 372 | 403 |
| ENSG00000141349 | 2017 | 1012 | 1201 | 1366 | 1126 | 1154 |
| ENSG00000108840 | 976 | 461 | 554 | 464 | 402 | 392 |
| ENSG00000267638 | 0 | 0 | 0 | 0 | 0 | 0 |
| ENSG00000212446 | 0 | 0 | 0 | 0 | 0 | 0 |
| ENSG00000125319 | 136 | 50 | 79 | 66 | 69 | 79 |
| ENSG00000161664 | 61 | 40 | 40 | 25 | 34 | 35 |
| ENSG00000267080 | 353 | 188 | 221 | 171 | 227 | 235 |
| ENSG00000168591 | 1847 | 733 | 1022 | 904 | 904 | 895 |
| ENSG00000087152 | 2749 | 1190 | 1613 | 1288 | 1315 | 1338 |
| ENSG00000267394 | 32 | 8 | 13 | 14 | 17 | 14 |
| ENSG00000108312 | 4274 | 1992 | 2601 | 2278 | 2098 | 2363 |
| ENSG00000275107 | 0 | 0 | 0 | 0 | 0 | 0 |

| | | | | | | |
|---|---|---|---|---|---|---|
| ENSG00000260793 | 536 | 166 | 326 | 204 | 184 | 258 |
| ENSG00000267691 | 33 | 19 | 22 | 14 | 14 | 9 |
| ENSG00000004939 | 2 | 1 | 1 | 1 | 1 | 0 |
| ENSG00000285486 | 0 | 2 | 1 | 1 | 1 | 1 |
| ENSG00000239702 | 0 | 1 | 0 | 0 | 0 | 0 |
| ENSG00000267750 | 569 | 335 | 363 | 232 | 261 | 275 |
| ENSG00000221496 | 0 | 0 | 0 | 0 | 0 | 0 |
| ENSG00000252628 | 2 | 0 | 0 | 2 | 1 | 0 |
| ENSG00000108309 | 72 | 23 | 40 | 33 | 34 | 24 |
| ENSG00000013306 | 8925 | 4088 | 4938 | 6324 | 5000 | 5273 |
| ENSG00000267211 | 0 | 0 | 0 | 0 | 0 | 1 |
| ENSG00000030582 | 17534 | 7834 | 10145 | 9544 | 7664 | 7799 |
| ENSG00000161682 | 568 | 229 | 364 | 331 | 306 | 300 |
| ENSG00000267634 | 0 | 0 | 0 | 0 | 0 | 0 |
| ENSG00000005961 | 25 | 13 | 16 | 4 | 14 | 7 |
| ENSG00000186566 | 1541 | 679 | 867 | 484 | 812 | 742 |
| ENSG00000240589 | 0 | 1 | 0 | 0 | 0 | 0 |
| ENSG00000283045 | 0 | 1 | 0 | 0 | 0 | 0 |
| ENSG00000180340 | 408 | 193 | 234 | 261 | 192 | 195 |
| ENSG00000271222 | 0 | 0 | 0 | 0 | 0 | 0 |
| ENSG00000267384 | 0 | 0 | 0 | 0 | 0 | 0 |
| ENSG00000266979 | 0 | 0 | 0 | 0 | 0 | 0 |
| ENSG00000180336 | 6 | 4 | 4 | 2 | 6 | 3 |
| ENSG00000180329 | 861 | 510 | 603 | 479 | 618 | 592 |
| ENSG00000267160 | 1 | 0 | 0 | 0 | 1 | 0 |
| ENSG00000279879 | 18 | 8 | 13 | 3 | 2 | 6 |
| ENSG00000161692 | 216 | 111 | 131 | 158 | 163 | 153 |
| ENSG00000264251 | 0 | 0 | 0 | 0 | 0 | 0 |
| ENSG00000240305 | 1 | 2 | 0 | 1 | 0 | 1 |
| ENSG00000073670 | 83 | 47 | 58 | 53 | 40 | 57 |
| ENSG00000267505 | 0 | 0 | 0 | 0 | 0 | 0 |
| ENSG00000267405 | 0 | 0 | 0 | 0 | 1 | 0 |
| ENSG00000182963 | 0 | 0 | 1 | 1 | 1 | 0 |
| ENSG00000131097 | 3 | 3 | 0 | 1 | 3 | 2 |
| ENSG00000108883 | 4641 | 2238 | 2851 | 2529 | 2574 | 2891 |
| ENSG00000264540 | 0 | 0 | 0 | 0 | 0 | 0 |
| ENSG00000167131 | 0 | 0 | 0 | 1 | 0 | 0 |
| ENSG00000214447 | 0 | 0 | 0 | 0 | 0 | 0 |
| ENSG00000131095 | 0 | 0 | 0 | 4 | 1 | 2 |
| ENSG00000186185 | 299 | 182 | 208 | 206 | 234 | 263 |
| ENSG00000278223 | 0 | 0 | 0 | 0 | 0 | 0 |
| ENSG00000267334 | 0 | 0 | 0 | 0 | 0 | 0 |
| ENSG00000131094 | 67 | 31 | 39 | 40 | 24 | 23 |
| ENSG00000267788 | 0 | 0 | 0 | 0 | 0 | 0 |
| ENSG00000172992 | 1434 | 634 | 853 | 723 | 710 | 776 |
| ENSG00000136448 | 4063 | 1903 | 2385 | 1890 | 2025 | 2219 |
| ENSG00000200888 | 0 | 0 | 0 | 0 | 0 | 1 |
| ENSG00000161714 | 3680 | 1737 | 2150 | 1784 | 1681 | 1676 |
| ENSG00000277249 | 4 | 1 | 1 | 2 | 2 | 0 |
| ENSG00000181513 | 394 | 207 | 350 | 232 | 219 | 221 |

| | | | | | | |
|---|---|---|---|---|---|---|
| ENSG00000276728 | 84 | 48 | 69 | 49 | 32 | 62 |
| ENSG00000186834 | 3765 | 1637 | 1997 | 1627 | 1712 | 1785 |
| ENSG00000224505 | 10 | 12 | 5 | 4 | 4 | 4 |
| ENSG00000168517 | 85 | 67 | 81 | 51 | 56 | 53 |
| ENSG00000267288 | 47 | 27 | 36 | 34 | 25 | 27 |
| ENSG00000267121 | 35 | 7 | 13 | 9 | 10 | 10 |
| ENSG00000184922 | 308 | 141 | 232 | 140 | 159 | 148 |
| ENSG00000233175 | 30 | 6 | 18 | 14 | 18 | 18 |
| ENSG00000233483 | 32 | 10 | 17 | 10 | 15 | 17 |
| ENSG00000267278 | 229 | 153 | 171 | 147 | 203 | 171 |
| ENSG00000184361 | 14 | 8 | 20 | 2 | 10 | 8 |
| ENSG00000006062 | 282 | 156 | 214 | 182 | 199 | 164 |
| ENSG00000285757 | 0 | 0 | 0 | 0 | 0 | 0 |
| ENSG00000199953 | 0 | 0 | 0 | 0 | 0 | 0 |
| ENSG00000267446 | 0 | 1 | 2 | 2 | 3 | 0 |
| ENSG00000159314 | 1270 | 660 | 768 | 802 | 863 | 814 |
| ENSG00000267344 | 9 | 7 | 11 | 2 | 7 | 8 |
| ENSG00000225190 | 1142 | 513 | 732 | 604 | 639 | 551 |
| ENSG00000236234 | 1 | 2 | 4 | 1 | 1 | 1 |
| ENSG00000284055 | 0 | 0 | 0 | 0 | 0 | 0 |
| ENSG00000214425 | 10 | 6 | 4 | 1 | 1 | 2 |
| ENSG00000278770 | 0 | 0 | 0 | 0 | 0 | 0 |
| ENSG00000266918 | 0 | 0 | 0 | 1 | 2 | 1 |
| ENSG00000131484 | 0 | 0 | 0 | 0 | 0 | 1 |
| ENSG00000267198 | 1 | 3 | 1 | 0 | 0 | 0 |
| ENSG00000267246 | 1 | 0 | 2 | 1 | 0 | 0 |
| ENSG00000266504 | 0 | 0 | 0 | 0 | 0 | 0 |
| ENSG00000264070 | 0 | 0 | 0 | 0 | 0 | 0 |
| ENSG00000285668 | 0 | 0 | 0 | 0 | 0 | 0 |
| ENSG00000280022 | 2 | 2 | 0 | 0 | 2 | 0 |
| ENSG00000263503 | 0 | 0 | 1 | 0 | 0 | 0 |
| ENSG00000204652 | 0 | 0 | 0 | 0 | 1 | 0 |
| ENSG00000204650 | 632 | 288 | 380 | 325 | 358 | 312 |
| ENSG00000263715 | 0 | 0 | 0 | 0 | 0 | 0 |
| ENSG00000265547 | 0 | 0 | 0 | 0 | 0 | 0 |
| ENSG00000265964 | 0 | 0 | 0 | 0 | 0 | 0 |
| ENSG00000120088 | 0 | 0 | 0 | 0 | 0 | 0 |
| ENSG00000264589 | 0 | 7 | 0 | 3 | 5 | 4 |
| ENSG00000185294 | 0 | 0 | 0 | 0 | 0 | 0 |
| ENSG00000186868 | 1743 | 705 | 1004 | 751 | 807 | 906 |
| ENSG00000279685 | 0 | 1 | 0 | 0 | 1 | 1 |
| ENSG00000262881 | 0 | 0 | 0 | 0 | 0 | 0 |
| ENSG00000274883 | 0 | 0 | 0 | 0 | 0 | 0 |
| ENSG00000256762 | 4 | 1 | 1 | 0 | 0 | 0 |
| ENSG00000120071 | 1429 | 572 | 903 | 476 | 674 | 657 |
| ENSG00000262372 | 0 | 0 | 0 | 0 | 0 | 0 |
| ENSG00000214401 | 45 | 11 | 26 | 24 | 16 | 22 |
| ENSG00000238723 | 0 | 1 | 0 | 0 | 2 | 0 |
| ENSG00000262500 | 2 | 0 | 0 | 0 | 0 | 0 |
| ENSG00000262539 | 0 | 0 | 0 | 1 | 1 | 0 |

| | | | | | | |
|---|---|---|---|---|---|---|
| ENSG00000261575 | 17 | 8 | 9 | 6 | 5 | 14 |
| ENSG00000228696 | 5 | 0 | 4 | 0 | 4 | 2 |
| ENSG00000176681 | 18 | 9 | 12 | 11 | 6 | 3 |
| ENSG00000265411 | 0 | 0 | 0 | 0 | 0 | 0 |
| ENSG00000260075 | 3 | 0 | 0 | 0 | 0 | 1 |
| ENSG00000266497 | 0 | 0 | 0 | 0 | 0 | 0 |
| ENSG00000238083 | 97 | 24 | 50 | 28 | 36 | 48 |
| ENSG00000185829 | 47 | 25 | 37 | 6 | 26 | 23 |
| ENSG00000265315 | 0 | 0 | 0 | 0 | 0 | 0 |
| ENSG00000232300 | 0 | 2 | 0 | 0 | 1 | 4 |
| ENSG00000073969 | 1892 | 956 | 1137 | 736 | 980 | 918 |
| ENSG00000213326 | 18 | 9 | 8 | 7 | 12 | 10 |
| ENSG00000108379 | 112 | 56 | 45 | 37 | 44 | 49 |
| ENSG00000158955 | 0 | 0 | 0 | 0 | 0 | 0 |
| ENSG00000262031 | 0 | 0 | 0 | 0 | 0 | 0 |
| ENSG00000261886 | 0 | 1 | 2 | 1 | 1 | 4 |
| ENSG00000210709 | 0 | 1 | 0 | 0 | 0 | 0 |
| ENSG00000108433 | 1742 | 774 | 1011 | 857 | 956 | 892 |
| ENSG00000262633 | 0 | 0 | 0 | 0 | 0 | 0 |
| ENSG00000264999 | 0 | 0 | 0 | 0 | 0 | 0 |
| ENSG00000179673 | 1 | 0 | 0 | 0 | 0 | 0 |
| ENSG00000263142 | 119 | 95 | 85 | 42 | 59 | 71 |
| ENSG00000262879 | 384 | 162 | 234 | 190 | 201 | 227 |
| ENSG00000277198 | 0 | 2 | 0 | 0 | 0 | 0 |
| ENSG00000004897 | 2137 | 1084 | 1100 | 664 | 1402 | 1033 |
| ENSG00000239291 | 0 | 1 | 0 | 0 | 0 | 0 |
| ENSG00000262265 | 0 | 1 | 1 | 1 | 0 | 0 |
| ENSG00000261872 | 0 | 1 | 0 | 0 | 0 | 1 |
| ENSG00000198336 | 0 | 0 | 0 | 0 | 0 | 0 |
| ENSG00000259207 | 5 | 3 | 5 | 1 | 4 | 2 |
| ENSG00000259753 | 0 | 0 | 0 | 0 | 0 | 0 |
| ENSG00000238419 | 0 | 0 | 0 | 0 | 0 | 0 |
| ENSG00000276790 | 0 | 0 | 0 | 0 | 0 | 0 |
| ENSG00000263293 | 3 | 0 | 2 | 0 | 1 | 0 |
| ENSG00000178852 | 28 | 14 | 19 | 7 | 32 | 23 |
| ENSG00000253347 | 2 | 1 | 0 | 1 | 2 | 2 |
| ENSG00000228782 | 180 | 101 | 99 | 119 | 104 | 116 |
| ENSG00000141279 | 4479 | 2482 | 3088 | 2667 | 3661 | 3451 |
| ENSG00000206734 | 0 | 0 | 0 | 0 | 0 | 0 |
| ENSG00000263766 | 18 | 10 | 14 | 9 | 6 | 12 |
| ENSG00000108424 | 8895 | 4577 | 5597 | 3680 | 5501 | 5138 |
| ENSG00000264558 | 0 | 0 | 0 | 0 | 0 | 0 |
| ENSG00000198933 | 610 | 273 | 318 | 305 | 361 | 312 |
| ENSG00000073861 | 0 | 0 | 0 | 1 | 0 | 0 |
| ENSG00000006025 | 1558 | 619 | 986 | 778 | 818 | 851 |
| ENSG00000159111 | 3263 | 1536 | 1996 | 1788 | 1601 | 1824 |
| ENSG00000141294 | 85 | 43 | 46 | 41 | 34 | 43 |
| ENSG00000141295 | 1462 | 718 | 985 | 764 | 685 | 802 |
| ENSG00000189120 | 662 | 331 | 403 | 296 | 308 | 290 |
| ENSG00000264243 | 0 | 0 | 0 | 0 | 0 | 0 |

| | | | | | | |
|---|---|---|---|---|---|---|
| ENSG00000266821 | 12 | 1 | 6 | 0 | 7 | 8 |
| ENSG00000264920 | 120 | 69 | 108 | 78 | 77 | 87 |
| ENSG00000167182 | 1226 | 501 | 790 | 517 | 525 | 662 |
| ENSG00000234494 | 135 | 74 | 85 | 55 | 65 | 78 |
| ENSG00000266601 | 18 | 14 | 12 | 13 | 4 | 20 |
| ENSG00000108439 | 1221 | 784 | 951 | 811 | 952 | 982 |
| ENSG00000263798 | 0 | 0 | 0 | 0 | 0 | 0 |
| ENSG00000264019 | 0 | 0 | 0 | 0 | 0 | 0 |
| ENSG00000167183 | 3126 | 1488 | 1882 | 1533 | 1464 | 1561 |
| ENSG00000108465 | 1253 | 647 | 1178 | 711 | 827 | 980 |
| ENSG00000264701 | 0 | 0 | 0 | 0 | 0 | 0 |
| ENSG00000005243 | 249 | 134 | 113 | 107 | 95 | 66 |
| ENSG00000207947 | 0 | 0 | 0 | 0 | 0 | 0 |
| ENSG00000263412 | 76 | 50 | 55 | 28 | 49 | 26 |
| ENSG00000082641 | 10682 | 5213 | 6673 | 4988 | 5516 | 5845 |
| ENSG00000266341 | 0 | 0 | 0 | 0 | 0 | 0 |
| ENSG00000278765 | 6 | 8 | 11 | 7 | 12 | 18 |
| ENSG00000108468 | 6156 | 3111 | 3718 | 2868 | 3830 | 3917 |
| ENSG00000206954 | 0 | 0 | 2 | 0 | 0 | 0 |
| ENSG00000002919 | 1438 | 657 | 837 | 857 | 844 | 842 |
| ENSG00000141293 | 52 | 28 | 30 | 37 | 33 | 23 |
| ENSG00000221739 | 0 | 0 | 0 | 0 | 0 | 0 |
| ENSG00000263787 | 0 | 0 | 0 | 0 | 0 | 0 |
| ENSG00000207306 | 0 | 0 | 0 | 0 | 0 | 0 |
| ENSG00000235300 | 0 | 0 | 0 | 0 | 0 | 0 |
| ENSG00000266222 | 0 | 0 | 0 | 0 | 0 | 0 |
| ENSG00000200538 | 0 | 0 | 0 | 0 | 0 | 0 |
| ENSG00000264451 | 0 | 0 | 0 | 0 | 0 | 0 |
| ENSG00000206805 | 0 | 0 | 0 | 0 | 0 | 0 |
| ENSG00000120094 | 8 | 0 | 4 | 1 | 0 | 0 |
| ENSG00000173917 | 86 | 34 | 34 | 21 | 17 | 14 |
| ENSG00000230148 | 9 | 4 | 20 | 5 | 4 | 5 |
| ENSG00000120093 | 60 | 30 | 54 | 25 | 29 | 25 |
| ENSG00000233101 | 2 | 0 | 1 | 1 | 0 | 2 |
| ENSG00000239552 | 0 | 0 | 0 | 0 | 0 | 0 |
| ENSG00000182742 | 13 | 2 | 3 | 0 | 0 | 1 |
| ENSG00000257178 | 0 | 0 | 0 | 0 | 0 | 0 |
| ENSG00000284038 | 0 | 0 | 0 | 0 | 0 | 0 |
| ENSG00000120075 | 0 | 1 | 0 | 0 | 0 | 3 |
| ENSG00000108511 | 24 | 14 | 19 | 13 | 6 | 8 |
| ENSG00000260027 | 458 | 223 | 254 | 255 | 239 | 246 |
| ENSG00000120068 | 8 | 0 | 0 | 1 | 2 | 1 |
| ENSG00000170689 | 26 | 23 | 5 | 9 | 16 | 12 |
| ENSG00000242207 | 0 | 0 | 0 | 0 | 0 | 0 |
| ENSG00000210741 | 0 | 0 | 0 | 0 | 0 | 0 |
| ENSG00000272763 | 0 | 0 | 0 | 0 | 1 | 0 |
| ENSG00000244649 | 0 | 0 | 0 | 0 | 0 | 0 |
| ENSG00000244559 | 0 | 0 | 0 | 0 | 0 | 0 |
| ENSG00000242067 | 0 | 0 | 0 | 0 | 0 | 0 |
| ENSG00000241788 | 0 | 0 | 0 | 0 | 0 | 0 |

| | | | | | | |
|---|---|---|---|---|---|---|
| ENSG00000159182 | 0 | 0 | 0 | 0 | 0 | 0 |
| ENSG00000229637 | 0 | 0 | 0 | 0 | 0 | 0 |
| ENSG00000273750 | 0 | 0 | 0 | 0 | 0 | 0 |
| ENSG00000277542 | 0 | 0 | 0 | 0 | 0 | 0 |
| ENSG00000263602 | 0 | 0 | 0 | 0 | 0 | 0 |
| ENSG00000159184 | 22 | 12 | 9 | 4 | 6 | 7 |
| ENSG00000242407 | 0 | 0 | 0 | 0 | 0 | 0 |
| ENSG00000170703 | 6 | 1 | 2 | 0 | 5 | 0 |
| ENSG00000244514 | 0 | 0 | 0 | 0 | 0 | 0 |
| ENSG00000136436 | 3074 | 1619 | 1882 | 1157 | 1692 | 1575 |
| ENSG00000248278 | 62 | 26 | 44 | 28 | 34 | 33 |
| ENSG00000212565 | 0 | 0 | 0 | 0 | 0 | 0 |
| ENSG00000248895 | 0 | 0 | 0 | 0 | 0 | 0 |
| ENSG00000159199 | 2855 | 1366 | 1711 | 2248 | 2008 | 1916 |
| ENSG00000270781 | 1 | 0 | 3 | 0 | 2 | 2 |
| ENSG00000159202 | 10860 | 5050 | 6268 | 5267 | 5731 | 6070 |
| ENSG00000159210 | 1835 | 1145 | 1423 | 1378 | 1540 | 1397 |
| ENSG00000230532 | 0 | 0 | 0 | 0 | 0 | 0 |
| ENSG00000200903 | 0 | 0 | 0 | 0 | 0 | 0 |
| ENSG00000238804 | 0 | 0 | 0 | 0 | 0 | 0 |
| ENSG00000159224 | 0 | 0 | 0 | 0 | 0 | 0 |
| ENSG00000250838 | 0 | 0 | 0 | 0 | 0 | 0 |
| ENSG00000159217 | 32 | 8 | 14 | 13 | 9 | 11 |
| ENSG00000251461 | 0 | 0 | 0 | 0 | 0 | 0 |
| ENSG00000255851 | 0 | 0 | 0 | 0 | 0 | 0 |
| ENSG00000252636 | 0 | 0 | 0 | 0 | 0 | 0 |
| ENSG00000251550 | 0 | 0 | 0 | 0 | 0 | 0 |
| ENSG00000248883 | 0 | 0 | 0 | 0 | 0 | 0 |
| ENSG00000167080 | 6 | 7 | 6 | 1 | 10 | 7 |
| ENSG00000240995 | 0 | 1 | 0 | 0 | 0 | 0 |
| ENSG00000167083 | 0 | 0 | 0 | 0 | 0 | 0 |
| ENSG00000108798 | 0 | 0 | 4 | 1 | 0 | 0 |
| ENSG00000173868 | 0 | 0 | 1 | 1 | 0 | 0 |
| ENSG00000177369 | 0 | 0 | 0 | 0 | 0 | 0 |
| ENSG00000273500 | 0 | 0 | 0 | 0 | 2 | 0 |
| ENSG00000198740 | 1791 | 999 | 924 | 597 | 977 | 926 |
| ENSG00000248714 | 0 | 0 | 0 | 1 | 1 | 1 |
| ENSG00000262039 | 2 | 1 | 2 | 2 | 2 | 0 |
| ENSG00000250948 | 2 | 0 | 0 | 0 | 0 | 0 |
| ENSG00000186244 | 0 | 0 | 0 | 0 | 0 | 0 |
| ENSG00000167085 | 5026 | 2847 | 3407 | 4086 | 3708 | 3604 |
| ENSG00000250186 | 0 | 0 | 0 | 0 | 0 | 0 |
| ENSG00000207127 | 0 | 1 | 1 | 0 | 0 | 0 |
| ENSG00000229944 | 8 | 9 | 16 | 3 | 10 | 9 |
| ENSG00000249497 | 0 | 0 | 0 | 0 | 0 | 0 |
| ENSG00000279036 | 0 | 0 | 0 | 0 | 0 | 0 |
| ENSG00000064300 | 12 | 13 | 9 | 6 | 5 | 8 |
| ENSG00000249906 | 0 | 0 | 0 | 0 | 0 | 0 |
| ENSG00000277478 | 0 | 0 | 0 | 0 | 0 | 0 |
| ENSG00000250310 | 0 | 0 | 0 | 0 | 0 | 0 |

| | | | | | | |
|---|---|---|---|---|---|---|
| ENSG00000182575 | 34 | 30 | 30 | 19 | 25 | 16 |
| ENSG00000121067 | 2659 | 1295 | 1668 | 1099 | 1243 | 1299 |
| ENSG00000121073 | 3172 | 1574 | 2021 | 1932 | 1833 | 1874 |
| ENSG00000250751 | 0 | 1 | 1 | 0 | 0 | 1 |
| ENSG00000121104 | 368 | 189 | 231 | 199 | 212 | 176 |
| ENSG00000136504 | 2182 | 1172 | 1334 | 873 | 1029 | 1160 |
| ENSG00000235183 | 0 | 2 | 1 | 0 | 0 | 0 |
| ENSG00000262837 | 0 | 0 | 1 | 0 | 1 | 0 |
| ENSG00000176358 | 1 | 2 | 0 | 0 | 2 | 1 |
| ENSG00000204584 | 6 | 2 | 3 | 1 | 2 | 4 |
| ENSG00000248954 | 9 | 8 | 2 | 1 | 3 | 1 |
| ENSG00000254039 | 0 | 1 | 2 | 2 | 1 | 3 |
| ENSG00000249176 | 0 | 0 | 1 | 0 | 0 | 1 |
| ENSG00000199492 | 0 | 0 | 0 | 0 | 0 | 0 |
| ENSG00000108813 | 151 | 74 | 93 | 91 | 100 | 90 |
| ENSG00000064195 | 376 | 160 | 237 | 234 | 258 | 220 |
| ENSG00000248172 | 2 | 0 | 1 | 0 | 0 | 1 |
| ENSG00000246640 | 10 | 7 | 6 | 5 | 1 | 9 |
| ENSG00000005884 | 4703 | 1952 | 2778 | 2548 | 2463 | 2437 |
| ENSG00000276851 | 2 | 0 | 2 | 0 | 1 | 1 |
| ENSG00000005882 | 823 | 353 | 565 | 403 | 391 | 429 |
| ENSG00000275025 | 0 | 0 | 0 | 0 | 0 | 0 |
| ENSG00000250282 | 0 | 1 | 1 | 0 | 1 | 1 |
| ENSG00000167100 | 2 | 4 | 7 | 2 | 7 | 0 |
| ENSG00000108819 | 2887 | 1416 | 1752 | 1787 | 1708 | 1609 |
| ENSG00000236472 | 0 | 1 | 0 | 1 | 0 | 0 |
| ENSG00000253730 | 2 | 0 | 4 | 0 | 1 | 1 |
| ENSG00000261959 | 0 | 0 | 0 | 0 | 0 | 0 |
| ENSG00000108823 | 0 | 0 | 0 | 0 | 0 | 0 |
| ENSG00000188662 | 0 | 0 | 0 | 0 | 0 | 0 |
| ENSG00000108821 | 402 | 155 | 242 | 155 | 165 | 194 |
| ENSG00000249406 | 0 | 0 | 0 | 0 | 0 | 0 |
| ENSG00000263176 | 0 | 0 | 0 | 0 | 0 | 0 |
| ENSG00000249838 | 0 | 0 | 0 | 0 | 0 | 0 |
| ENSG00000167105 | 2 | 0 | 0 | 1 | 1 | 1 |
| ENSG00000251179 | 0 | 0 | 1 | 0 | 0 | 0 |
| ENSG00000015532 | 2258 | 1022 | 1442 | 1220 | 1252 | 1231 |
| ENSG00000279792 | 6 | 3 | 3 | 6 | 2 | 2 |
| ENSG00000108826 | 1829 | 864 | 1077 | 1239 | 992 | 992 |
| ENSG00000154920 | 154 | 90 | 121 | 91 | 110 | 118 |
| ENSG00000108829 | 5297 | 2867 | 3392 | 3266 | 3411 | 3381 |
| ENSG00000206824 | 2 | 0 | 0 | 0 | 0 | 0 |
| ENSG00000253102 | 1 | 1 | 0 | 0 | 0 | 0 |
| ENSG00000167107 | 1233 | 521 | 713 | 569 | 572 | 529 |
| ENSG00000262759 | 0 | 0 | 0 | 0 | 0 | 0 |
| ENSG00000136457 | 14 | 3 | 10 | 4 | 7 | 13 |
| ENSG00000275897 | 1 | 1 | 5 | 5 | 8 | 3 |
| ENSG00000136444 | 2034 | 972 | 1292 | 1183 | 1099 | 1274 |
| ENSG00000249451 | 0 | 0 | 1 | 0 | 0 | 0 |
| ENSG00000136449 | 0 | 2 | 5 | 1 | 4 | 0 |

| | | | | | | |
|---|---|---|---|---|---|---|
| ENSG00000049283 | 3232 | 1397 | 1793 | 1698 | 1481 | 1536 |
| ENSG00000250286 | 14 | 6 | 5 | 7 | 3 | 1 |
| ENSG00000006282 | 4864 | 2204 | 2994 | 2790 | 2506 | 2582 |
| ENSG00000250107 | 1 | 2 | 2 | 1 | 1 | 3 |
| ENSG00000006283 | 19 | 8 | 4 | 5 | 12 | 4 |
| ENSG00000250976 | 0 | 0 | 0 | 0 | 0 | 0 |
| ENSG00000251239 | 0 | 0 | 0 | 0 | 0 | 0 |
| ENSG00000108846 | 5503 | 2633 | 2979 | 3268 | 3892 | 3408 |
| ENSG00000154945 | 1728 | 869 | 1050 | 761 | 981 | 917 |
| ENSG00000262967 | 0 | 0 | 0 | 0 | 0 | 0 |
| ENSG00000201340 | 0 | 1 | 2 | 0 | 0 | 0 |
| ENSG00000108848 | 3375 | 2008 | 1987 | 906 | 2840 | 1815 |
| ENSG00000278963 | 4 | 4 | 4 | 0 | 0 | 1 |
| ENSG00000167117 | 4 | 0 | 2 | 0 | 2 | 0 |
| ENSG00000278672 | 0 | 0 | 0 | 0 | 0 | 0 |
| ENSG00000285829 | 0 | 0 | 0 | 0 | 0 | 0 |
| ENSG00000248702 | 0 | 0 | 0 | 0 | 0 | 0 |
| ENSG00000173714 | 0 | 0 | 0 | 0 | 0 | 0 |
| ENSG00000261976 | 0 | 0 | 0 | 0 | 0 | 0 |
| ENSG00000243521 | 0 | 0 | 0 | 0 | 0 | 0 |
| ENSG00000141232 | 5303 | 2226 | 2998 | 1736 | 1884 | 2319 |
| ENSG00000229980 | 97 | 93 | 82 | 64 | 81 | 60 |
| ENSG00000262006 | 4 | 1 | 2 | 1 | 2 | 2 |
| ENSG00000247011 | 4 | 4 | 6 | 3 | 2 | 0 |
| ENSG00000251665 | 0 | 1 | 2 | 0 | 0 | 0 |
| ENSG00000008294 | 3588 | 1751 | 2067 | 1224 | 2152 | 1762 |
| ENSG00000249870 | 0 | 0 | 0 | 0 | 0 | 0 |
| ENSG00000239672 | 1730 | 1195 | 1262 | 1978 | 1618 | 1723 |
| ENSG00000011052 | 0 | 3 | 0 | 5 | 8 | 5 |
| ENSG00000243678 | 83 | 58 | 76 | 79 | 90 | 81 |
| ENSG00000011258 | 1993 | 964 | 1150 | 529 | 1077 | 980 |
| ENSG00000279089 | 67 | 33 | 27 | 17 | 30 | 36 |
| ENSG00000264895 | 4 | 5 | 0 | 0 | 0 | 3 |
| ENSG00000011260 | 2290 | 1189 | 1513 | 1292 | 1443 | 1494 |
| ENSG00000249383 | 0 | 0 | 0 | 0 | 0 | 0 |
| ENSG00000249982 | 0 | 0 | 0 | 0 | 0 | 0 |
| ENSG00000249658 | 0 | 0 | 0 | 0 | 0 | 2 |
| ENSG00000267452 | 0 | 0 | 0 | 0 | 0 | 0 |
| ENSG00000187504 | 0 | 0 | 0 | 0 | 0 | 0 |
| ENSG00000154975 | 0 | 0 | 0 | 0 | 0 | 0 |
| ENSG00000279743 | 0 | 0 | 0 | 0 | 0 | 0 |
| ENSG00000263317 | 0 | 0 | 0 | 0 | 0 | 0 |
| ENSG00000226364 | 0 | 0 | 0 | 0 | 0 | 0 |
| ENSG00000227011 | 0 | 0 | 0 | 0 | 0 | 0 |
| ENSG00000262902 | 10 | 3 | 16 | 8 | 12 | 11 |
| ENSG00000285939 | 0 | 1 | 0 | 0 | 0 | 0 |
| ENSG00000263252 | 0 | 0 | 0 | 0 | 0 | 0 |
| ENSG00000262271 | 0 | 0 | 0 | 0 | 0 | 0 |
| ENSG00000262518 | 0 | 0 | 0 | 0 | 0 | 0 |
| ENSG00000262384 | 0 | 0 | 0 | 0 | 0 | 0 |

| | | | | | | |
|---|---|---|---|---|---|---|
| ENSG00000233380 | 0 | 0 | 0 | 1 | 0 | 0 |
| ENSG00000141200 | 0 | 0 | 0 | 0 | 0 | 0 |
| ENSG00000263189 | 0 | 0 | 0 | 0 | 0 | 0 |
| ENSG00000261965 | 0 | 0 | 0 | 0 | 0 | 0 |
| ENSG00000263199 | 0 | 0 | 0 | 0 | 0 | 0 |
| ENSG00000263125 | 0 | 0 | 0 | 0 | 0 | 0 |
| ENSG00000251809 | 0 | 0 | 0 | 0 | 0 | 0 |
| ENSG00000141198 | 1884 | 1137 | 1211 | 679 | 1329 | 1006 |
| ENSG00000166260 | 1878 | 921 | 1131 | 881 | 1033 | 1063 |
| ENSG00000279059 | 232 | 118 | 142 | 82 | 100 | 104 |
| ENSG00000275710 | 1 | 2 | 1 | 0 | 1 | 0 |
| ENSG00000166263 | 426 | 215 | 202 | 96 | 289 | 204 |
| ENSG00000108924 | 38 | 16 | 20 | 19 | 16 | 22 |
| ENSG00000263096 | 0 | 0 | 0 | 0 | 0 | 0 |
| ENSG00000262079 | 0 | 0 | 0 | 0 | 0 | 0 |
| ENSG00000108960 | 285 | 127 | 135 | 129 | 164 | 152 |
| ENSG00000261873 | 2 | 0 | 2 | 2 | 1 | 0 |
| ENSG00000236319 | 0 | 0 | 0 | 0 | 0 | 1 |
| ENSG00000262135 | 0 | 0 | 0 | 0 | 0 | 0 |
| ENSG00000200107 | 0 | 0 | 0 | 0 | 0 | 0 |
| ENSG00000166292 | 0 | 0 | 0 | 0 | 0 | 0 |
| ENSG00000141179 | 1562 | 755 | 938 | 699 | 832 | 847 |
| ENSG00000262052 | 0 | 0 | 0 | 0 | 0 | 0 |
| ENSG00000153930 | 0 | 0 | 2 | 0 | 0 | 0 |
| ENSG00000261882 | 0 | 0 | 0 | 0 | 0 | 0 |
| ENSG00000279606 | 0 | 0 | 0 | 0 | 0 | 0 |
| ENSG00000243058 | 0 | 0 | 0 | 0 | 0 | 0 |
| ENSG00000183691 | 4 | 0 | 0 | 4 | 2 | 4 |
| ENSG00000214226 | 101 | 66 | 63 | 56 | 75 | 77 |
| ENSG00000283209 | 7 | 0 | 2 | 0 | 3 | 0 |
| ENSG00000153933 | 991 | 485 | 615 | 274 | 622 | 452 |
| ENSG00000121060 | 7080 | 3313 | 4403 | 3314 | 3764 | 4208 |
| ENSG00000212469 | 0 | 0 | 0 | 0 | 0 | 0 |
| ENSG00000262408 | 0 | 0 | 0 | 0 | 1 | 0 |
| ENSG00000262112 | 0 | 0 | 0 | 0 | 0 | 0 |
| ENSG00000284542 | 0 | 0 | 0 | 0 | 0 | 0 |
| ENSG00000274213 | 74 | 41 | 35 | 46 | 53 | 38 |
| ENSG00000121058 | 1875 | 955 | 1199 | 771 | 1094 | 983 |
| ENSG00000262298 | 3 | 5 | 1 | 0 | 0 | 0 |
| ENSG00000262623 | 0 | 0 | 1 | 0 | 0 | 0 |
| ENSG00000121064 | 1932 | 886 | 1102 | 875 | 999 | 1021 |
| ENSG00000263120 | 0 | 2 | 5 | 1 | 0 | 0 |
| ENSG00000261192 | 0 | 0 | 0 | 0 | 0 | 0 |
| ENSG00000263004 | 238 | 134 | 180 | 140 | 177 | 162 |
| ENSG00000121057 | 3452 | 1713 | 2059 | 1863 | 2016 | 2314 |
| ENSG00000263089 | 12 | 4 | 5 | 2 | 3 | 4 |
| ENSG00000285785 | 0 | 0 | 0 | 0 | 0 | 0 |
| ENSG00000153944 | 5063 | 2184 | 3156 | 1854 | 2596 | 2882 |
| ENSG00000279555 | 4 | 2 | 3 | 1 | 1 | 5 |
| ENSG00000279281 | 0 | 1 | 1 | 0 | 0 | 0 |

| | | | | | | |
|---|---|---|---|---|---|---|
| ENSG00000266100 | 0 | 0 | 0 | 0 | 0 | 0 |
| ENSG00000280028 | 0 | 0 | 0 | 0 | 0 | 0 |
| ENSG00000263499 | 0 | 0 | 0 | 0 | 1 | 0 |
| ENSG00000242889 | 2 | 0 | 0 | 0 | 0 | 0 |
| ENSG00000238447 | 0 | 0 | 0 | 0 | 0 | 0 |
| ENSG00000166329 | 0 | 0 | 0 | 0 | 0 | 0 |
| ENSG00000265542 | 6 | 1 | 0 | 3 | 1 | 0 |
| ENSG00000222976 | 0 | 0 | 0 | 0 | 0 | 0 |
| ENSG00000181610 | 3746 | 2021 | 2326 | 2547 | 2427 | 2485 |
| ENSG00000180891 | 2696 | 1284 | 1650 | 1786 | 1604 | 1618 |
| ENSG00000264914 | 0 | 3 | 0 | 0 | 0 | 2 |
| ENSG00000278642 | 2 | 2 | 1 | 1 | 0 | 1 |
| ENSG00000136451 | 3232 | 1561 | 1791 | 1194 | 1809 | 1664 |
| ENSG00000280233 | 7 | 6 | 1 | 0 | 2 | 1 |
| ENSG00000266086 | 0 | 1 | 0 | 0 | 0 | 0 |
| ENSG00000264112 | 294 | 117 | 134 | 52 | 123 | 153 |
| ENSG00000136450 | 2526 | 1554 | 1792 | 1424 | 1543 | 1604 |
| ENSG00000279069 | 84 | 65 | 55 | 51 | 61 | 34 |
| ENSG00000285930 | 0 | 0 | 0 | 0 | 0 | 0 |
| ENSG00000266290 | 2 | 0 | 0 | 1 | 1 | 0 |
| ENSG00000264364 | 12439 | 6026 | 7591 | 6245 | 6715 | 7333 |
| ENSG00000279207 | 1325 | 576 | 854 | 405 | 549 | 597 |
| ENSG00000141194 | 0 | 0 | 0 | 0 | 0 | 1 |
| ENSG00000229590 | 0 | 0 | 0 | 0 | 0 | 0 |
| ENSG00000255713 | 0 | 0 | 0 | 0 | 0 | 0 |
| ENSG00000121053 | 8 | 5 | 5 | 2 | 10 | 4 |
| ENSG00000278627 | 4 | 3 | 6 | 2 | 5 | 2 |
| ENSG00000011143 | 1352 | 622 | 849 | 639 | 693 | 688 |
| ENSG00000167419 | 2 | 0 | 1 | 2 | 1 | 2 |
| ENSG00000005381 | 0 | 0 | 0 | 0 | 0 | 0 |
| ENSG00000005379 | 2 | 0 | 0 | 0 | 0 | 1 |
| ENSG00000265148 | 2 | 7 | 9 | 1 | 7 | 4 |
| ENSG00000265206 | 0 | 0 | 0 | 0 | 0 | 0 |
| ENSG00000284353 | 0 | 0 | 0 | 0 | 0 | 0 |
| ENSG00000264399 | 0 | 0 | 0 | 0 | 0 | 0 |
| ENSG00000213246 | 0 | 77 | 71 | 62 | 74 | 68 |
| ENSG00000285897 | 12 | 6 | 5 | 3 | 3 | 0 |
| ENSG00000108375 | 568 | 300 | 359 | 265 | 256 | 355 |
| ENSG00000176160 | 6 | 0 | 0 | 0 | 0 | 0 |
| ENSG00000266648 | 9 | 0 | 2 | 2 | 0 | 1 |
| ENSG00000108389 | 4597 | 1919 | 2842 | 1768 | 2203 | 2545 |
| ENSG00000264672 | 18 | 11 | 7 | 9 | 10 | 10 |
| ENSG00000108387 | 22 | 13 | 11 | 3 | 9 | 11 |
| ENSG00000181013 | 4 | 3 | 4 | 4 | 2 | 1 |
| ENSG00000121101 | 67 | 43 | 50 | 16 | 30 | 32 |
| ENSG00000212195 | 4 | 1 | 4 | 1 | 1 | 4 |
| ENSG00000266826 | 0 | 3 | 0 | 0 | 0 | 0 |
| ENSG00000199426 | 0 | 0 | 0 | 0 | 0 | 0 |
| ENSG00000266561 | 0 | 0 | 0 | 0 | 0 | 0 |
| ENSG00000206917 | 0 | 0 | 0 | 0 | 0 | 0 |

| | | | | | | |
|---|---|---|---|---|---|---|
| ENSG00000200997 | 0 | 0 | 0 | 0 | 0 | 0 |
| ENSG00000108384 | 3405 | 1879 | 1997 | 1990 | 2204 | 2077 |
| ENSG00000274198 | 0 | 0 | 0 | 0 | 0 | 0 |
| ENSG00000175175 | 339 | 151 | 186 | 92 | 144 | 159 |
| ENSG00000276601 | 0 | 0 | 0 | 0 | 0 | 0 |
| ENSG00000251865 | 0 | 0 | 0 | 0 | 0 | 0 |
| ENSG00000241462 | 0 | 0 | 0 | 0 | 0 | 0 |
| ENSG00000275680 | 0 | 0 | 0 | 0 | 0 | 0 |
| ENSG00000263938 | 4 | 2 | 1 | 2 | 2 | 1 |
| ENSG00000108395 | 36813 | 19532 | 21710 | 12869 | 23771 | 18992 |
| ENSG00000263558 | 0 | 2 | 1 | 0 | 0 | 0 |
| ENSG00000224738 | 1230 | 707 | 794 | 836 | 1076 | 975 |
| ENSG00000182628 | 4112 | 2127 | 2646 | 1665 | 2491 | 2305 |
| ENSG00000252212 | 0 | 0 | 1 | 0 | 0 | 0 |
| ENSG00000211514 | 0 | 0 | 0 | 0 | 0 | 0 |
| ENSG00000207996 | 0 | 0 | 0 | 0 | 0 | 0 |
| ENSG00000068489 | 1443 | 807 | 983 | 789 | 1384 | 1167 |
| ENSG00000266537 | 4 | 0 | 0 | 0 | 0 | 0 |
| ENSG00000265303 | 0 | 0 | 0 | 0 | 0 | 0 |
| ENSG00000265415 | 167 | 99 | 127 | 111 | 180 | 142 |
| ENSG00000167447 | 3057 | 1339 | 1783 | 1358 | 1678 | 1763 |
| ENSG00000153982 | 259 | 117 | 161 | 129 | 161 | 162 |
| ENSG00000207235 | 1 | 1 | 0 | 0 | 1 | 0 |
| ENSG00000271063 | 0 | 0 | 2 | 0 | 0 | 1 |
| ENSG00000175155 | 3078 | 1372 | 1706 | 805 | 1103 | 1072 |
| ENSG00000278863 | 0 | 2 | 0 | 0 | 0 | 0 |
| ENSG00000263857 | 0 | 0 | 1 | 0 | 0 | 0 |
| ENSG00000266002 | 0 | 0 | 0 | 0 | 0 | 0 |
| ENSG00000265313 | 0 | 0 | 0 | 0 | 0 | 0 |
| ENSG00000200889 | 0 | 0 | 0 | 0 | 0 | 0 |
| ENSG00000108406 | 6336 | 3113 | 3946 | 2618 | 3833 | 3482 |
| ENSG00000273702 | 348 | 177 | 190 | 166 | 166 | 182 |
| ENSG00000141367 | 39614 | 21281 | 21831 | 13716 | 27758 | 20602 |
| ENSG00000141378 | 1756 | 1047 | 1077 | 1079 | 1273 | 1286 |
| ENSG00000062716 | 43534 | 23377 | 29037 | 20214 | 29340 | 24135 |
| ENSG00000267637 | 0 | 0 | 0 | 0 | 0 | 0 |
| ENSG00000284190 | 0 | 0 | 0 | 0 | 0 | 0 |
| ENSG00000201524 | 2 | 2 | 2 | 0 | 4 | 2 |
| ENSG00000108423 | 3824 | 1886 | 1951 | 1291 | 2604 | 1868 |
| ENSG00000270264 | 51 | 41 | 40 | 30 | 40 | 36 |
| ENSG00000108443 | 22010 | 11830 | 12621 | 6726 | 14362 | 11392 |
| ENSG00000241913 | 4 | 5 | 0 | 7 | 5 | 7 |
| ENSG00000267318 | 0 | 1 | 0 | 0 | 0 | 0 |
| ENSG00000189050 | 1148 | 600 | 757 | 483 | 633 | 669 |
| ENSG00000267104 | 0 | 1 | 0 | 0 | 0 | 0 |
| ENSG00000267302 | 12 | 13 | 6 | 3 | 5 | 6 |
| ENSG00000266992 | 1 | 1 | 1 | 0 | 1 | 0 |
| ENSG00000238283 | 2 | 1 | 4 | 0 | 2 | 1 |
| ENSG00000275860 | 0 | 0 | 0 | 0 | 0 | 0 |
| ENSG00000274378 | 0 | 1 | 3 | 0 | 0 | 1 |

| | | | | | | |
|---|---|---|---|---|---|---|
| ENSG00000264049 | 85 | 13 | 28 | 14 | 28 | 23 |
| ENSG00000068097 | 10792 | 4790 | 6594 | 4395 | 5689 | 5967 |
| ENSG00000267526 | 2 | 3 | 3 | 0 | 2 | 4 |
| ENSG00000267416 | 35 | 25 | 35 | 16 | 23 | 19 |
| ENSG00000261040 | 28 | 23 | 28 | 19 | 16 | 19 |
| ENSG00000280852 | 2 | 4 | 0 | 0 | 1 | 0 |
| ENSG00000267295 | 0 | 0 | 0 | 1 | 0 | 0 |
| ENSG00000277532 | 0 | 1 | 0 | 0 | 0 | 0 |
| ENSG00000267248 | 247 | 124 | 125 | 105 | 127 | 133 |
| ENSG00000267095 | 8 | 3 | 5 | 4 | 3 | 4 |
| ENSG00000167434 | 645 | 315 | 524 | 273 | 319 | 400 |
| ENSG00000170832 | 18069 | 8553 | 10415 | 5474 | 10152 | 8510 |
| ENSG00000252577 | 7 | 4 | 5 | 0 | 2 | 5 |
| ENSG00000241157 | 6 | 4 | 3 | 2 | 4 | 4 |
| ENSG00000275562 | 0 | 1 | 1 | 1 | 1 | 0 |
| ENSG00000271626 | 0 | 2 | 0 | 0 | 0 | 0 |
| ENSG00000141371 | 3 | 1 | 0 | 5 | 3 | 3 |
| ENSG00000213228 | 4 | 1 | 1 | 3 | 5 | 4 |
| ENSG00000062725 | 29491 | 13991 | 16311 | 10148 | 16242 | 13647 |
| ENSG00000270553 | 5 | 2 | 1 | 0 | 2 | 0 |
| ENSG00000259349 | 255 | 122 | 169 | 125 | 156 | 115 |
| ENSG00000267678 | 3 | 0 | 0 | 0 | 1 | 2 |
| ENSG00000267772 | 0 | 1 | 1 | 1 | 0 | 1 |
| ENSG00000266943 | 0 | 0 | 0 | 0 | 0 | 0 |
| ENSG00000170836 | 8478 | 3989 | 4899 | 3463 | 4366 | 4258 |
| ENSG00000200013 | 7 | 1 | 4 | 6 | 4 | 5 |
| ENSG00000141376 | 46927 | 23193 | 28208 | 26709 | 23244 | 24393 |
| ENSG00000239932 | 2 | 1 | 1 | 0 | 2 | 0 |
| ENSG00000224011 | 0 | 0 | 4 | 0 | 1 | 0 |
| ENSG00000252534 | 3 | 2 | 0 | 0 | 2 | 0 |
| ENSG00000267083 | 2 | 4 | 0 | 2 | 1 | 1 |
| ENSG00000254114 | 0 | 0 | 0 | 0 | 0 | 0 |
| ENSG00000264322 | 0 | 0 | 0 | 0 | 0 | 0 |
| ENSG00000267207 | 0 | 0 | 0 | 0 | 0 | 0 |
| ENSG00000267449 | 0 | 0 | 0 | 0 | 0 | 0 |
| ENSG00000227694 | 0 | 0 | 0 | 0 | 1 | 1 |
| ENSG00000267667 | 0 | 1 | 2 | 0 | 1 | 0 |
| ENSG00000273982 | 80 | 42 | 65 | 37 | 35 | 41 |
| ENSG00000267131 | 284 | 205 | 253 | 154 | 117 | 157 |
| ENSG00000266934 | 274 | 132 | 210 | 63 | 124 | 138 |
| ENSG00000267280 | 2777 | 1221 | 1848 | 1809 | 1547 | 1547 |
| ENSG00000121068 | 10890 | 4734 | 6791 | 5571 | 5273 | 5581 |
| ENSG00000187013 | 374 | 167 | 227 | 186 | 216 | 191 |
| ENSG00000121075 | 144 | 65 | 102 | 55 | 67 | 69 |
| ENSG00000267137 | 4 | 0 | 2 | 2 | 4 | 1 |
| ENSG00000253506 | 4 | 8 | 4 | 3 | 8 | 6 |
| ENSG00000136492 | 1826 | 1129 | 1093 | 430 | 1325 | 977 |
| ENSG00000108506 | 2466 | 1176 | 1314 | 637 | 1561 | 1112 |
| ENSG00000202361 | 0 | 0 | 0 | 0 | 0 | 0 |
| ENSG00000108510 | 4654 | 2723 | 2261 | 889 | 4196 | 2433 |

| | | | | | | |
|---|---|---|---|---|---|---|
| ENSG00000279133 | 29 | 27 | 26 | 21 | 22 | 32 |
| ENSG00000242398 | 0 | 0 | 0 | 0 | 0 | 0 |
| ENSG00000285879 | 2 | 0 | 0 | 0 | 0 | 0 |
| ENSG00000200842 | 0 | 0 | 0 | 0 | 0 | 0 |
| ENSG00000207123 | 0 | 0 | 0 | 0 | 0 | 0 |
| ENSG00000266066 | 44 | 19 | 31 | 27 | 27 | 35 |
| ENSG00000266365 | 6 | 2 | 5 | 3 | 3 | 6 |
| ENSG00000266647 | 0 | 0 | 0 | 0 | 0 | 0 |
| ENSG00000188755 | 0 | 0 | 0 | 0 | 0 | 0 |
| ENSG00000263887 | 2 | 1 | 1 | 0 | 0 | 0 |
| ENSG00000242123 | 0 | 0 | 0 | 0 | 0 | 0 |
| ENSG00000172421 | 0 | 1 | 0 | 0 | 0 | 0 |
| ENSG00000239809 | 0 | 0 | 0 | 0 | 0 | 0 |
| ENSG00000251981 | 0 | 0 | 0 | 0 | 0 | 0 |
| ENSG00000239527 | 0 | 0 | 0 | 0 | 0 | 0 |
| ENSG00000087995 | 2157 | 1124 | 1381 | 1256 | 1410 | 1506 |
| ENSG00000146872 | 4831 | 2302 | 3036 | 2221 | 2785 | 2799 |
| ENSG00000239246 | 7 | 2 | 0 | 3 | 2 | 2 |
| ENSG00000264546 | 4 | 1 | 0 | 0 | 1 | 0 |
| ENSG00000277463 | 43 | 31 | 25 | 33 | 20 | 32 |
| ENSG00000274565 | 11 | 17 | 15 | 7 | 12 | 15 |
| ENSG00000011028 | 826 | 431 | 470 | 476 | 493 | 481 |
| ENSG00000207382 | 0 | 0 | 0 | 0 | 0 | 0 |
| ENSG00000279713 | 2 | 0 | 1 | 0 | 4 | 1 |
| ENSG00000199697 | 0 | 1 | 0 | 0 | 0 | 0 |
| ENSG00000265702 | 56 | 51 | 29 | 52 | 60 | 45 |
| ENSG00000173838 | 107 | 65 | 101 | 79 | 93 | 106 |
| ENSG00000265000 | 4 | 5 | 7 | 0 | 0 | 1 |
| ENSG00000264907 | 2 | 0 | 1 | 0 | 0 | 1 |
| ENSG00000283538 | 64 | 36 | 51 | 27 | 34 | 31 |
| ENSG00000207552 | 0 | 0 | 0 | 0 | 0 | 0 |
| ENSG00000265262 | 0 | 0 | 1 | 1 | 0 | 2 |
| ENSG00000270324 | 0 | 0 | 0 | 0 | 0 | 0 |
| ENSG00000272164 | 0 | 0 | 0 | 0 | 0 | 0 |
| ENSG00000170921 | 7571 | 3325 | 4729 | 2043 | 3355 | 3856 |
| ENSG00000272383 | 6 | 10 | 5 | 4 | 7 | 1 |
| ENSG00000264513 | 3 | 0 | 0 | 0 | 0 | 0 |
| ENSG00000264529 | 0 | 0 | 3 | 0 | 0 | 0 |
| ENSG00000233635 | 0 | 2 | 0 | 0 | 1 | 0 |
| ENSG00000263501 | 0 | 1 | 0 | 0 | 1 | 0 |
| ENSG00000271919 | 0 | 1 | 1 | 0 | 0 | 0 |
| ENSG00000226797 | 2 | 1 | 1 | 0 | 2 | 0 |
| ENSG00000263644 | 0 | 2 | 0 | 0 | 0 | 0 |
| ENSG00000271749 | 3 | 1 | 2 | 1 | 6 | 4 |
| ENSG00000265282 | 12 | 2 | 8 | 3 | 7 | 8 |
| ENSG00000008283 | 16817 | 7956 | 10506 | 9385 | 9344 | 9619 |
| ENSG00000265971 | 0 | 1 | 0 | 0 | 1 | 0 |
| ENSG00000228979 | 0 | 0 | 0 | 0 | 0 | 0 |
| ENSG00000159640 | 8 | 9 | 5 | 16 | 6 | 4 |
| ENSG00000264813 | 0 | 0 | 0 | 0 | 0 | 0 |

| | | | | | | |
|---|---|---|---|---|---|---|
| ENSG00000224353 | 0 | 0 | 0 | 0 | 0 | 0 |
| ENSG00000173826 | 3 | 0 | 0 | 0 | 0 | 1 |
| ENSG00000136485 | 13378 | 6020 | 8364 | 5635 | 6961 | 7635 |
| ENSG00000200560 | 4 | 0 | 0 | 0 | 0 | 1 |
| ENSG00000136463 | 2531 | 1243 | 1648 | 1630 | 1481 | 1484 |
| ENSG00000270587 | 20 | 11 | 17 | 13 | 17 | 13 |
| ENSG00000198909 | 2971 | 1372 | 1873 | 1395 | 1479 | 1594 |
| ENSG00000263883 | 1 | 1 | 0 | 0 | 2 | 0 |
| ENSG00000266173 | 301 | 179 | 230 | 142 | 173 | 198 |
| ENSG00000136490 | 217 | 112 | 97 | 99 | 93 | 77 |
| ENSG00000279369 | 1 | 4 | 4 | 2 | 4 | 4 |
| ENSG00000125695 | 0 | 0 | 0 | 0 | 0 | 0 |
| ENSG00000108588 | 5640 | 3224 | 3853 | 2384 | 4413 | 3834 |
| ENSG00000198231 | 7692 | 3721 | 5242 | 2848 | 4204 | 4331 |
| ENSG00000108592 | 7129 | 3410 | 4219 | 3566 | 3702 | 4005 |
| ENSG00000087191 | 10271 | 5413 | 6868 | 5889 | 6323 | 6336 |
| ENSG00000108604 | 20008 | 9602 | 12069 | 11864 | 10324 | 11044 |
| ENSG00000240280 | 0 | 1 | 0 | 0 | 0 | 1 |
| ENSG00000213218 | 0 | 0 | 0 | 0 | 0 | 0 |
| ENSG00000136487 | 0 | 0 | 0 | 0 | 0 | 0 |
| ENSG00000259533 | 0 | 0 | 0 | 0 | 0 | 0 |
| ENSG00000136488 | 0 | 0 | 0 | 0 | 0 | 0 |
| ENSG00000259333 | 0 | 0 | 0 | 0 | 0 | 0 |
| ENSG00000204414 | 0 | 0 | 0 | 0 | 0 | 0 |
| ENSG00000259384 | 0 | 0 | 0 | 0 | 0 | 0 |
| ENSG00000285947 | 0 | 0 | 0 | 0 | 0 | 0 |
| ENSG00000007312 | 0 | 0 | 0 | 0 | 0 | 0 |
| ENSG00000007314 | 0 | 0 | 0 | 0 | 0 | 0 |
| ENSG00000263489 | 0 | 0 | 0 | 0 | 0 | 0 |
| ENSG00000264954 | 0 | 0 | 0 | 0 | 2 | 0 |
| ENSG00000224383 | 10 | 8 | 9 | 7 | 9 | 5 |
| ENSG00000108622 | 0 | 0 | 0 | 1 | 1 | 0 |
| ENSG00000178607 | 886 | 338 | 497 | 333 | 430 | 459 |
| ENSG00000266402 | 38 | 9 | 10 | 18 | 13 | 5 |
| ENSG00000199753 | 121 | 48 | 59 | 89 | 93 | 79 |
| ENSG00000281311 | 0 | 0 | 0 | 0 | 0 | 0 |
| ENSG00000277887 | 0 | 0 | 0 | 0 | 0 | 0 |
| ENSG00000136478 | 2765 | 1323 | 1671 | 1247 | 1619 | 1729 |
| ENSG00000256358 | 0 | 0 | 0 | 0 | 0 | 0 |
| ENSG00000261371 | 2 | 1 | 0 | 0 | 2 | 0 |
| ENSG00000239823 | 0 | 0 | 0 | 0 | 0 | 0 |
| ENSG00000265031 | 0 | 0 | 0 | 0 | 0 | 0 |
| ENSG00000271605 | 0 | 0 | 0 | 1 | 1 | 0 |
| ENSG00000256525 | 1103 | 559 | 682 | 452 | 583 | 571 |
| ENSG00000256826 | 0 | 0 | 0 | 0 | 0 | 0 |
| ENSG00000108654 | 48949 | 25654 | 37708 | 20082 | 31062 | 30971 |
| ENSG00000284564 | 0 | 0 | 0 | 0 | 0 | 0 |
| ENSG00000284368 | 0 | 0 | 0 | 0 | 0 | 0 |
| ENSG00000258890 | 1586 | 742 | 1037 | 568 | 859 | 921 |
| ENSG00000108854 | 1654 | 747 | 928 | 497 | 888 | 773 |

| | | | | | | |
|---|---|---|---|---|---|---|
| ENSG00000271779 | 2 | 0 | 0 | 0 | 1 | 0 |
| ENSG00000270714 | 0 | 2 | 1 | 0 | 0 | 4 |
| ENSG00000215769 | 92 | 89 | 141 | 76 | 106 | 104 |
| ENSG00000266820 | 0 | 0 | 0 | 0 | 0 | 0 |
| ENSG00000265298 | 311 | 188 | 204 | 149 | 198 | 192 |
| ENSG00000263647 | 0 | 0 | 0 | 0 | 0 | 0 |
| ENSG00000265982 | 63 | 21 | 42 | 23 | 23 | 28 |
| ENSG00000214176 | 1332 | 511 | 784 | 424 | 629 | 695 |
| ENSG00000278581 | 0 | 0 | 0 | 0 | 0 | 0 |
| ENSG00000284172 | 0 | 0 | 0 | 0 | 0 | 0 |
| ENSG00000252242 | 0 | 0 | 0 | 0 | 0 | 1 |
| ENSG00000272400 | 0 | 0 | 0 | 0 | 0 | 0 |
| ENSG00000176809 | 630 | 260 | 360 | 223 | 259 | 270 |
| ENSG00000264840 | 0 | 0 | 0 | 0 | 0 | 0 |
| ENSG00000272134 | 0 | 0 | 1 | 0 | 0 | 0 |
| ENSG00000272220 | 0 | 0 | 0 | 0 | 0 | 0 |
| ENSG00000265218 | 8 | 3 | 8 | 3 | 5 | 6 |
| ENSG00000266644 | 8 | 2 | 1 | 2 | 5 | 3 |
| ENSG00000271974 | 0 | 0 | 0 | 0 | 0 | 0 |
| ENSG00000264057 | 5 | 7 | 7 | 1 | 8 | 7 |
| ENSG00000266155 | 2 | 0 | 0 | 0 | 0 | 1 |
| ENSG00000280299 | 0 | 0 | 0 | 0 | 0 | 1 |
| ENSG00000272346 | 0 | 0 | 0 | 0 | 0 | 0 |
| ENSG00000265912 | 9 | 6 | 1 | 2 | 3 | 7 |
| ENSG00000272006 | 0 | 1 | 2 | 3 | 1 | 5 |
| ENSG00000232457 | 3 | 0 | 7 | 2 | 1 | 6 |
| ENSG00000214174 | 475 | 196 | 309 | 210 | 211 | 232 |
| ENSG00000120063 | 3257 | 1810 | 1659 | 946 | 2287 | 1647 |
| ENSG00000266598 | 0 | 0 | 0 | 0 | 0 | 0 |
| ENSG00000108370 | 59 | 21 | 24 | 30 | 48 | 25 |
| ENSG00000265883 | 0 | 0 | 0 | 0 | 0 | 0 |
| ENSG00000263520 | 0 | 0 | 0 | 0 | 0 | 0 |
| ENSG00000168646 | 105 | 54 | 42 | 26 | 58 | 22 |
| ENSG00000266076 | 0 | 0 | 0 | 0 | 0 | 0 |
| ENSG00000154240 | 35 | 22 | 28 | 8 | 39 | 23 |
| ENSG00000271587 | 0 | 0 | 0 | 0 | 0 | 0 |
| ENSG00000270870 | 0 | 0 | 0 | 0 | 0 | 0 |
| ENSG00000266707 | 0 | 0 | 0 | 1 | 0 | 0 |
| ENSG00000091583 | 0 | 0 | 0 | 0 | 0 | 0 |
| ENSG00000252653 | 0 | 0 | 0 | 0 | 0 | 0 |
| ENSG00000284242 | 0 | 0 | 0 | 0 | 0 | 0 |
| ENSG00000154229 | 366 | 188 | 258 | 178 | 271 | 224 |
| ENSG00000244044 | 0 | 0 | 0 | 0 | 0 | 0 |
| ENSG00000264630 | 0 | 0 | 1 | 0 | 0 | 0 |
| ENSG00000252685 | 0 | 0 | 0 | 0 | 0 | 0 |
| ENSG00000251698 | 0 | 0 | 0 | 0 | 0 | 0 |
| ENSG00000229330 | 0 | 0 | 0 | 0 | 0 | 0 |
| ENSG00000207943 | 0 | 0 | 0 | 0 | 0 | 0 |
| ENSG00000075429 | 0 | 0 | 0 | 0 | 0 | 0 |
| ENSG00000251764 | 0 | 0 | 0 | 0 | 0 | 0 |

| | | | | | | |
|---|---|---|---|---|---|---|
| ENSG00000075461 | 11498 | 5578 | 7264 | 6559 | 5935 | 6335 |
| ENSG00000264491 | 49 | 24 | 24 | 9 | 18 | 25 |
| ENSG00000265664 | 0 | 0 | 0 | 0 | 0 | 0 |
| ENSG00000108878 | 21 | 9 | 7 | 8 | 6 | 7 |
| ENSG00000198265 | 1967 | 1201 | 877 | 368 | 1564 | 935 |
| ENSG00000264421 | 0 | 1 | 0 | 0 | 0 | 0 |
| ENSG00000213180 | 0 | 0 | 0 | 0 | 0 | 0 |
| ENSG00000285877 | 56 | 28 | 25 | 19 | 17 | 22 |
| ENSG00000266473 | 16 | 9 | 15 | 11 | 14 | 15 |
| ENSG00000200619 | 0 | 0 | 0 | 0 | 0 | 0 |
| ENSG00000207410 | 0 | 0 | 0 | 0 | 0 | 0 |
| ENSG00000264093 | 0 | 0 | 0 | 0 | 0 | 0 |
| ENSG00000197170 | 5859 | 3025 | 3501 | 2587 | 3821 | 3509 |
| ENSG00000154217 | 1010 | 575 | 653 | 671 | 634 | 644 |
| ENSG00000201547 | 0 | 0 | 0 | 0 | 0 | 0 |
| ENSG00000244610 | 0 | 0 | 0 | 0 | 0 | 0 |
| ENSG00000263690 | 0 | 0 | 0 | 0 | 0 | 0 |
| ENSG00000207688 | 0 | 0 | 0 | 0 | 0 | 0 |
| ENSG00000264754 | 0 | 0 | 0 | 0 | 0 | 0 |
| ENSG00000279917 | 2 | 0 | 1 | 0 | 0 | 1 |
| ENSG00000279361 | 0 | 0 | 0 | 3 | 0 | 0 |
| ENSG00000130935 | 3712 | 2011 | 2273 | 1767 | 2663 | 2322 |
| ENSG00000200394 | 0 | 0 | 0 | 0 | 0 | 2 |
| ENSG00000266858 | 0 | 0 | 0 | 0 | 0 | 0 |
| ENSG00000213179 | 0 | 0 | 0 | 0 | 0 | 0 |
| ENSG00000171634 | 3100 | 1665 | 1466 | 659 | 2575 | 1741 |
| ENSG00000266717 | 0 | 2 | 0 | 0 | 1 | 0 |
| ENSG00000279573 | 0 | 0 | 3 | 0 | 0 | 3 |
| ENSG00000279880 | 7 | 3 | 8 | 1 | 2 | 4 |
| ENSG00000265052 | 0 | 0 | 1 | 0 | 0 | 0 |
| ENSG00000186665 | 1120 | 529 | 725 | 634 | 588 | 652 |
| ENSG00000182481 | 6208 | 3384 | 3682 | 3805 | 5332 | 4611 |
| ENSG00000266117 | 3 | 0 | 0 | 0 | 0 | 0 |
| ENSG00000265055 | 20 | 17 | 7 | 11 | 15 | 18 |
| ENSG00000237854 | 311 | 217 | 275 | 145 | 217 | 242 |
| ENSG00000267023 | 486 | 366 | 499 | 275 | 389 | 394 |
| ENSG00000278730 | 1742 | 870 | 1168 | 879 | 1040 | 1025 |
| ENSG00000274561 | 115 | 50 | 80 | 62 | 62 | 66 |
| ENSG00000277476 | 400 | 190 | 257 | 153 | 223 | 203 |
| ENSG00000267352 | 0 | 0 | 0 | 0 | 0 | 0 |
| ENSG00000267708 | 0 | 1 | 0 | 0 | 3 | 1 |
| ENSG00000278740 | 3 | 3 | 0 | 1 | 5 | 2 |
| ENSG00000267731 | 6 | 1 | 4 | 0 | 1 | 0 |
| ENSG00000267472 | 7 | 5 | 1 | 3 | 5 | 2 |
| ENSG00000274712 | 352 | 200 | 225 | 207 | 206 | 192 |
| ENSG00000265100 | 17 | 4 | 6 | 3 | 15 | 11 |
| ENSG00000196704 | 4473 | 2182 | 2830 | 2511 | 2555 | 2628 |
| ENSG00000141337 | 716 | 367 | 491 | 296 | 354 | 365 |
| ENSG00000108932 | 664 | 395 | 533 | 261 | 392 | 373 |
| ENSG00000267009 | 9 | 13 | 16 | 6 | 14 | 11 |

| | | | | | | |
|---|---|---|---|---|---|---|
| ENSG00000070540 | 1569 | 788 | 1001 | 895 | 956 | 1001 |
| ENSG00000207561 | 34 | 25 | 26 | 23 | 24 | 26 |
| ENSG00000108946 | 18438 | 8657 | 11816 | 6644 | 8959 | 9458 |
| ENSG00000108950 | 10 | 7 | 9 | 2 | 10 | 6 |
| ENSG00000267461 | 0 | 1 | 0 | 0 | 0 | 0 |
| ENSG00000267659 | 0 | 0 | 0 | 0 | 0 | 0 |
| ENSG00000283376 | 0 | 0 | 0 | 0 | 0 | 0 |
| ENSG00000267250 | 0 | 0 | 0 | 0 | 0 | 0 |
| ENSG00000141338 | 2 | 0 | 0 | 0 | 0 | 0 |
| ENSG00000231749 | 0 | 0 | 0 | 0 | 0 | 0 |
| ENSG00000154258 | 4 | 0 | 2 | 0 | 4 | 0 |
| ENSG00000154262 | 4 | 0 | 4 | 2 | 0 | 0 |
| ENSG00000267361 | 0 | 0 | 0 | 0 | 0 | 0 |
| ENSG00000265331 | 0 | 0 | 0 | 0 | 0 | 0 |
| ENSG00000283385 | 0 | 0 | 0 | 0 | 0 | 0 |
| ENSG00000154263 | 40 | 17 | 23 | 9 | 17 | 13 |
| ENSG00000278873 | 0 | 0 | 0 | 0 | 0 | 0 |
| ENSG00000285931 | 0 | 1 | 0 | 0 | 0 | 0 |
| ENSG00000154265 | 1083 | 510 | 620 | 207 | 608 | 469 |
| ENSG00000267203 | 13 | 8 | 6 | 0 | 4 | 4 |
| ENSG00000252352 | 0 | 0 | 0 | 0 | 0 | 0 |
| ENSG00000108984 | 28 | 14 | 13 | 13 | 7 | 35 |
| ENSG00000278972 | 0 | 0 | 0 | 0 | 0 | 0 |
| ENSG00000267653 | 0 | 0 | 0 | 0 | 0 | 0 |
| ENSG00000227517 | 0 | 0 | 0 | 0 | 0 | 0 |
| ENSG00000267184 | 0 | 0 | 0 | 0 | 0 | 0 |
| ENSG00000237560 | 0 | 0 | 0 | 0 | 0 | 0 |
| ENSG00000199801 | 0 | 0 | 0 | 0 | 0 | 0 |
| ENSG00000230258 | 0 | 0 | 0 | 0 | 0 | 0 |
| ENSG00000267603 | 0 | 0 | 0 | 0 | 0 | 0 |
| ENSG00000153822 | 0 | 0 | 0 | 0 | 0 | 0 |
| ENSG00000267365 | 0 | 0 | 0 | 0 | 0 | 0 |
| ENSG00000123700 | 0 | 0 | 0 | 0 | 0 | 0 |
| ENSG00000267170 | 0 | 0 | 0 | 0 | 0 | 0 |
| ENSG00000267109 | 0 | 0 | 0 | 0 | 0 | 0 |
| ENSG00000267471 | 0 | 0 | 0 | 0 | 0 | 0 |
| ENSG00000271239 | 0 | 0 | 0 | 0 | 0 | 0 |
| ENSG00000271101 | 0 | 0 | 0 | 0 | 0 | 0 |
| ENSG00000260785 | 0 | 0 | 0 | 0 | 0 | 0 |
| ENSG00000238759 | 0 | 0 | 0 | 0 | 0 | 0 |
| ENSG00000225818 | 0 | 0 | 0 | 0 | 0 | 0 |
| ENSG00000228118 | 0 | 1 | 0 | 1 | 0 | 0 |
| ENSG00000283517 | 0 | 0 | 0 | 0 | 0 | 0 |
| ENSG00000228639 | 0 | 0 | 0 | 0 | 0 | 0 |
| ENSG00000256124 | 0 | 0 | 0 | 0 | 0 | 0 |
| ENSG00000234899 | 0 | 1 | 0 | 0 | 0 | 1 |
| ENSG00000226101 | 0 | 0 | 0 | 0 | 0 | 0 |
| ENSG00000125398 | 488 | 269 | 311 | 142 | 119 | 225 |
| ENSG00000227036 | 169 | 72 | 101 | 116 | 135 | 130 |
| ENSG00000264026 | 0 | 0 | 0 | 0 | 0 | 0 |

| | | | | | | |
|---|---|---|---|---|---|---|
| ENSG00000263680 | 0 | 0 | 0 | 0 | 0 | 0 |
| ENSG00000243514 | 0 | 0 | 0 | 0 | 0 | 0 |
| ENSG00000263893 | 5 | 3 | 7 | 4 | 3 | 2 |
| ENSG00000133195 | 2300 | 1019 | 1402 | 1204 | 1192 | 1242 |
| ENSG00000200783 | 0 | 0 | 0 | 0 | 1 | 0 |
| ENSG00000252274 | 0 | 0 | 0 | 0 | 1 | 0 |
| ENSG00000264196 | 2 | 0 | 0 | 0 | 0 | 0 |
| ENSG00000266501 | 0 | 1 | 2 | 1 | 0 | 1 |
| ENSG00000265469 | 2 | 0 | 0 | 0 | 0 | 0 |
| ENSG00000264860 | 0 | 2 | 0 | 0 | 0 | 2 |
| ENSG00000180616 | 8 | 2 | 4 | 2 | 5 | 3 |
| ENSG00000166685 | 1378 | 637 | 848 | 715 | 709 | 760 |
| ENSG00000277728 | 6 | 4 | 8 | 3 | 5 | 3 |
| ENSG00000133193 | 965 | 540 | 714 | 508 | 524 | 628 |
| ENSG00000141219 | 876 | 460 | 526 | 316 | 518 | 481 |
| ENSG00000265010 | 1 | 1 | 1 | 1 | 2 | 1 |
| ENSG00000187959 | 2 | 1 | 7 | 2 | 1 | 1 |
| ENSG00000179604 | 2257 | 987 | 1445 | 1124 | 1056 | 1122 |
| ENSG00000069188 | 10 | 13 | 7 | 14 | 18 | 12 |
| ENSG00000264985 | 0 | 0 | 0 | 0 | 0 | 0 |
| ENSG00000264750 | 0 | 0 | 0 | 0 | 0 | 0 |
| ENSG00000265935 | 0 | 0 | 0 | 0 | 0 | 0 |
| ENSG00000263574 | 0 | 0 | 0 | 0 | 0 | 0 |
| ENSG00000177338 | 0 | 0 | 0 | 1 | 0 | 0 |
| ENSG00000234721 | 0 | 0 | 0 | 1 | 1 | 0 |
| ENSG00000265927 | 0 | 0 | 0 | 0 | 0 | 0 |
| ENSG00000266765 | 0 | 0 | 0 | 0 | 0 | 0 |
| ENSG00000285648 | 0 | 0 | 0 | 0 | 0 | 0 |
| ENSG00000266357 | 0 | 0 | 0 | 0 | 0 | 0 |
| ENSG00000172809 | 8680 | 4052 | 4917 | 5702 | 5311 | 4854 |
| ENSG00000246731 | 2 | 0 | 5 | 0 | 0 | 2 |
| ENSG00000141540 | 30 | 16 | 23 | 15 | 22 | 21 |
| ENSG00000264272 | 0 | 0 | 0 | 0 | 0 | 0 |
| ENSG00000171595 | 0 | 0 | 0 | 0 | 0 | 0 |
| ENSG00000266106 | 0 | 0 | 0 | 0 | 0 | 0 |
| ENSG00000196169 | 0 | 0 | 0 | 0 | 0 | 0 |
| ENSG00000204347 | 0 | 0 | 0 | 0 | 0 | 0 |
| ENSG00000257008 | 0 | 0 | 0 | 0 | 0 | 0 |
| ENSG00000200021 | 0 | 0 | 0 | 0 | 0 | 0 |
| ENSG00000170412 | 3621 | 1634 | 2149 | 2759 | 2493 | 2206 |
| ENSG00000167851 | 0 | 0 | 0 | 0 | 0 | 0 |
| ENSG00000178789 | 0 | 0 | 0 | 0 | 0 | 0 |
| ENSG00000167850 | 0 | 0 | 1 | 3 | 2 | 0 |
| ENSG00000236770 | 0 | 0 | 0 | 0 | 0 | 0 |
| ENSG00000284690 | 0 | 0 | 0 | 0 | 0 | 0 |
| ENSG00000204345 | 0 | 0 | 0 | 0 | 0 | 0 |
| ENSG00000182352 | 0 | 0 | 0 | 0 | 0 | 0 |
| ENSG00000261222 | 0 | 0 | 0 | 0 | 0 | 0 |
| ENSG00000186407 | 0 | 0 | 0 | 0 | 0 | 0 |
| ENSG00000172794 | 164 | 66 | 113 | 64 | 70 | 80 |

| | | | | | | |
|---|---|---|---|---|---|---|
| ENSG00000264659 | 0 | 0 | 0 | 0 | 0 | 0 |
| ENSG00000186074 | 0 | 0 | 1 | 0 | 1 | 0 |
| ENSG00000266036 | 8 | 9 | 14 | 7 | 8 | 4 |
| ENSG00000284186 | 0 | 0 | 0 | 0 | 0 | 0 |
| ENSG00000109062 | 20098 | 9793 | 11713 | 10430 | 8027 | 8942 |
| ENSG00000109065 | 1049 | 530 | 629 | 757 | 668 | 671 |
| ENSG00000109066 | 793 | 347 | 416 | 505 | 437 | 436 |
| ENSG00000161509 | 74 | 36 | 69 | 44 | 37 | 55 |
| ENSG00000161513 | 961 | 408 | 498 | 554 | 498 | 414 |
| ENSG00000172782 | 0 | 0 | 0 | 0 | 0 | 0 |
| ENSG00000182040 | 10 | 2 | 13 | 4 | 4 | 5 |
| ENSG00000183034 | 0 | 0 | 0 | 0 | 0 | 0 |
| ENSG00000182938 | 0 | 2 | 0 | 0 | 0 | 0 |
| ENSG00000167861 | 10268 | 4436 | 6100 | 5015 | 4749 | 4677 |
| ENSG00000263586 | 0 | 2 | 3 | 1 | 1 | 2 |
| ENSG00000109089 | 900 | 452 | 551 | 606 | 515 | 503 |
| ENSG00000167862 | 1461 | 698 | 879 | 762 | 836 | 755 |
| ENSG00000252545 | 0 | 0 | 0 | 0 | 0 | 0 |
| ENSG00000180901 | 2178 | 964 | 1210 | 1063 | 1080 | 1135 |
| ENSG00000167863 | 3300 | 2168 | 2424 | 2512 | 2491 | 2300 |
| ENSG00000239607 | 0 | 0 | 0 | 0 | 0 | 0 |
| ENSG00000270993 | 2 | 0 | 1 | 0 | 4 | 4 |
| ENSG00000232724 | 0 | 0 | 0 | 1 | 0 | 0 |
| ENSG00000170190 | 4 | 3 | 0 | 2 | 3 | 2 |
| ENSG00000206713 | 0 | 0 | 0 | 0 | 0 | 0 |
| ENSG00000125449 | 590 | 249 | 274 | 316 | 275 | 270 |
| ENSG00000125458 | 1040 | 621 | 825 | 760 | 651 | 693 |
| ENSG00000279035 | 0 | 0 | 2 | 4 | 1 | 4 |
| ENSG00000189159 | 4467 | 1943 | 2429 | 3427 | 2773 | 2991 |
| ENSG00000265800 | 19 | 10 | 6 | 9 | 8 | 9 |
| ENSG00000263786 | 12 | 12 | 12 | 7 | 6 | 8 |
| ENSG00000252042 | 0 | 0 | 0 | 0 | 0 | 0 |
| ENSG00000188612 | 8226 | 4415 | 5457 | 4286 | 4665 | 4878 |
| ENSG00000125450 | 2263 | 1063 | 1296 | 1234 | 1312 | 1317 |
| ENSG00000125447 | 1722 | 726 | 1085 | 832 | 924 | 967 |
| ENSG00000125445 | 3081 | 1654 | 1922 | 1943 | 1898 | 1767 |
| ENSG00000125457 | 1092 | 583 | 708 | 670 | 574 | 643 |
| ENSG00000263843 | 111 | 43 | 65 | 68 | 64 | 77 |
| ENSG00000125454 | 222 | 99 | 136 | 150 | 143 | 156 |
| ENSG00000177885 | 13802 | 6439 | 8226 | 6465 | 7014 | 7509 |
| ENSG00000228007 | 4 | 0 | 0 | 2 | 0 | 0 |
| ENSG00000265987 | 0 | 0 | 0 | 0 | 0 | 0 |
| ENSG00000264853 | 0 | 1 | 0 | 0 | 0 | 3 |
| ENSG00000265342 | 0 | 1 | 2 | 4 | 4 | 2 |
| ENSG00000264511 | 0 | 0 | 0 | 2 | 0 | 0 |
| ENSG00000223217 | 0 | 0 | 0 | 0 | 0 | 0 |
| ENSG00000202332 | 0 | 0 | 0 | 0 | 0 | 0 |
| ENSG00000177728 | 1155 | 471 | 757 | 487 | 485 | 644 |
| ENSG00000284595 | 0 | 0 | 0 | 0 | 0 | 0 |
| ENSG00000177303 | 2420 | 1016 | 1465 | 1362 | 1292 | 1264 |

| | | | | | | |
|---|---|---|---|---|---|---|
| ENSG00000182173 | 4200 | 2036 | 2691 | 2587 | 2253 | 2309 |
| ENSG00000073350 | 19669 | 8927 | 12492 | 9389 | 9810 | 9496 |
| ENSG00000266714 | 1037 | 458 | 859 | 346 | 388 | 469 |
| ENSG00000108469 | 2519 | 1033 | 1445 | 1374 | 1361 | 1491 |
| ENSG00000204323 | 268 | 141 | 185 | 265 | 293 | 301 |
| ENSG00000259120 | 16 | 1 | 3 | 1 | 10 | 7 |
| ENSG00000161526 | 4203 | 1971 | 2567 | 2513 | 2452 | 2470 |
| ENSG00000264829 | 0 | 1 | 1 | 2 | 0 | 0 |
| ENSG00000264270 | 0 | 0 | 1 | 0 | 1 | 1 |
| ENSG00000132470 | 16044 | 7062 | 10137 | 7886 | 7366 | 7655 |
| ENSG00000108479 | 2078 | 1062 | 1295 | 1610 | 1217 | 1260 |
| ENSG00000132475 | 32715 | 16347 | 20882 | 20476 | 20453 | 22212 |
| ENSG00000283853 | 0 | 0 | 0 | 0 | 0 | 0 |
| ENSG00000132478 | 1767 | 928 | 1236 | 957 | 1044 | 1044 |
| ENSG00000266980 | 6 | 2 | 2 | 0 | 1 | 0 |
| ENSG00000092929 | 3490 | 1597 | 1908 | 1769 | 1574 | 1705 |
| ENSG00000132471 | 5403 | 2349 | 3341 | 3107 | 2648 | 2757 |
| ENSG00000132481 | 161 | 95 | 110 | 101 | 92 | 97 |
| ENSG00000267801 | 6 | 2 | 2 | 7 | 1 | 3 |
| ENSG00000141569 | 1424 | 781 | 951 | 850 | 805 | 856 |
| ENSG00000267342 | 19 | 8 | 3 | 12 | 12 | 10 |
| ENSG00000204316 | 122 | 43 | 72 | 59 | 59 | 50 |
| ENSG00000267426 | 1 | 0 | 1 | 1 | 1 | 2 |
| ENSG00000188878 | 173 | 84 | 120 | 96 | 108 | 120 |
| ENSG00000161533 | 2618 | 1322 | 1544 | 992 | 1305 | 1128 |
| ENSG00000267615 | 0 | 0 | 0 | 0 | 0 | 0 |
| ENSG00000257949 | 0 | 2 | 1 | 2 | 0 | 1 |
| ENSG00000261408 | 31 | 18 | 27 | 21 | 21 | 21 |
| ENSG00000250506 | 13 | 9 | 17 | 11 | 13 | 11 |
| ENSG00000167880 | 3168 | 1362 | 1847 | 1592 | 1419 | 1463 |
| ENSG00000167881 | 3566 | 1642 | 2004 | 1510 | 1926 | 1788 |
| ENSG00000186919 | 10 | 4 | 6 | 5 | 8 | 7 |
| ENSG00000182687 | 5 | 6 | 1 | 9 | 1 | 6 |
| ENSG00000182473 | 3151 | 1424 | 1971 | 1426 | 1764 | 1652 |
| ENSG00000279382 | 199 | 87 | 148 | 74 | 119 | 108 |
| ENSG00000275360 | 0 | 0 | 0 | 0 | 0 | 0 |
| ENSG00000129654 | 0 | 1 | 1 | 3 | 0 | 0 |
| ENSG00000267128 | 8 | 0 | 4 | 0 | 3 | 7 |
| ENSG00000141576 | 393 | 231 | 271 | 234 | 291 | 319 |
| ENSG00000267376 | 0 | 0 | 1 | 0 | 0 | 0 |
| ENSG00000228218 | 3 | 0 | 0 | 4 | 1 | 2 |
| ENSG00000185262 | 724 | 315 | 464 | 562 | 486 | 441 |
| ENSG00000129646 | 94 | 28 | 48 | 30 | 37 | 29 |
| ENSG00000161542 | 2139 | 1102 | 1297 | 1026 | 1170 | 1132 |
| ENSG00000251779 | 0 | 0 | 0 | 0 | 0 | 0 |
| ENSG00000207169 | 0 | 0 | 0 | 0 | 0 | 0 |
| ENSG00000267518 | 0 | 0 | 0 | 0 | 0 | 0 |
| ENSG00000176170 | 516 | 269 | 322 | 442 | 389 | 336 |
| ENSG00000175931 | 1858 | 832 | 1087 | 1083 | 1105 | 1142 |
| ENSG00000242931 | 0 | 0 | 0 | 0 | 0 | 0 |

| | | | | | | |
|---|---|---|---|---|---|---|
| ENSG00000129673 | 7 | 3 | 6 | 3 | 1 | 7 |
| ENSG00000129667 | 1008 | 440 | 572 | 630 | 565 | 526 |
| ENSG00000161544 | 3 | 4 | 5 | 2 | 5 | 2 |
| ENSG00000214140 | 20 | 5 | 16 | 5 | 10 | 6 |
| ENSG00000267546 | 21 | 11 | 23 | 12 | 10 | 9 |
| ENSG00000267543 | 4 | 0 | 3 | 3 | 3 | 7 |
| ENSG00000272386 | 6 | 1 | 7 | 3 | 3 | 2 |
| ENSG00000163597 | 1950 | 1094 | 1203 | 1080 | 1289 | 1171 |
| ENSG00000274091 | 0 | 0 | 0 | 0 | 0 | 0 |
| ENSG00000199961 | 6 | 1 | 1 | 0 | 1 | 5 |
| ENSG00000278261 | 0 | 0 | 0 | 0 | 0 | 0 |
| ENSG00000284526 | 120 | 82 | 98 | 39 | 59 | 51 |
| ENSG00000070731 | 1109 | 546 | 776 | 502 | 489 | 532 |
| ENSG00000267078 | 9 | 2 | 15 | 4 | 10 | 9 |
| ENSG00000267084 | 0 | 0 | 3 | 0 | 0 | 0 |
| ENSG00000070526 | 18 | 9 | 20 | 2 | 5 | 6 |
| ENSG00000199735 | 0 | 0 | 0 | 0 | 0 | 0 |
| ENSG00000261335 | 9 | 3 | 5 | 9 | 3 | 11 |
| ENSG00000182534 | 270 | 130 | 141 | 128 | 109 | 124 |
| ENSG00000212418 | 0 | 0 | 0 | 0 | 0 | 0 |
| ENSG00000277382 | 0 | 0 | 2 | 0 | 1 | 0 |
| ENSG00000070495 | 713 | 349 | 385 | 410 | 427 | 392 |
| ENSG00000274051 | 0 | 1 | 2 | 3 | 2 | 3 |
| ENSG00000181038 | 721 | 380 | 428 | 461 | 466 | 469 |
| ENSG00000267168 | 0 | 0 | 0 | 0 | 0 | 0 |
| ENSG00000161547 | 4743 | 2579 | 3289 | 2570 | 2858 | 3091 |
| ENSG00000092931 | 1405 | 671 | 850 | 666 | 744 | 773 |
| ENSG00000283805 | 0 | 0 | 0 | 0 | 0 | 0 |
| ENSG00000200257 | 0 | 3 | 0 | 0 | 1 | 0 |
| ENSG00000267065 | 2 | 0 | 0 | 0 | 0 | 0 |
| ENSG00000267535 | 0 | 0 | 0 | 0 | 0 | 0 |
| ENSG00000167889 | 58 | 50 | 53 | 46 | 68 | 45 |
| ENSG00000267568 | 0 | 0 | 2 | 0 | 0 | 0 |
| ENSG00000267521 | 0 | 0 | 0 | 0 | 0 | 0 |
| ENSG00000280122 | 0 | 0 | 0 | 0 | 0 | 0 |
| ENSG00000276202 | 0 | 0 | 0 | 0 | 0 | 0 |
| ENSG00000234912 | 266 | 153 | 176 | 183 | 160 | 212 |
| ENSG00000129657 | 2397 | 1006 | 1302 | 907 | 1283 | 1164 |
| ENSG00000275143 | 0 | 0 | 0 | 0 | 0 | 0 |
| ENSG00000284250 | 0 | 0 | 0 | 0 | 0 | 0 |
| ENSG00000222808 | 4 | 3 | 2 | 0 | 6 | 1 |
| ENSG00000262870 | 1 | 1 | 0 | 0 | 0 | 0 |
| ENSG00000263718 | 16 | 7 | 11 | 8 | 7 | 12 |
| ENSG00000265121 | 0 | 0 | 2 | 0 | 3 | 0 |
| ENSG00000184640 | 9363 | 4754 | 5609 | 5493 | 5211 | 5007 |
| ENSG00000266998 | 0 | 0 | 0 | 0 | 0 | 2 |
| ENSG00000264060 | 0 | 0 | 0 | 0 | 0 | 0 |
| ENSG00000200651 | 2 | 0 | 1 | 1 | 0 | 0 |
| ENSG00000279801 | 8 | 2 | 5 | 2 | 2 | 3 |
| ENSG00000267263 | 0 | 0 | 0 | 0 | 0 | 0 |

| | | | | | | |
|---|---|---|---|---|---|---|
| ENSG00000267016 | 0 | 2 | 0 | 0 | 0 | 0 |
| ENSG00000267665 | 0 | 0 | 0 | 0 | 0 | 0 |
| ENSG00000267506 | 0 | 0 | 0 | 0 | 0 | 0 |
| ENSG00000267466 | 0 | 0 | 0 | 0 | 0 | 0 |
| ENSG00000279572 | 0 | 0 | 0 | 0 | 0 | 0 |
| ENSG00000285535 | 0 | 1 | 0 | 0 | 0 | 0 |
| ENSG00000267790 | 0 | 0 | 0 | 0 | 0 | 0 |
| ENSG00000204283 | 0 | 0 | 0 | 0 | 0 | 0 |
| ENSG00000238898 | 0 | 0 | 0 | 0 | 0 | 0 |
| ENSG00000078687 | 619 | 280 | 354 | 189 | 287 | 287 |
| ENSG00000279600 | 0 | 0 | 0 | 0 | 0 | 0 |
| ENSG00000267463 | 0 | 0 | 0 | 0 | 0 | 0 |
| ENSG00000238658 | 0 | 0 | 0 | 0 | 0 | 0 |
| ENSG00000204282 | 189 | 102 | 121 | 81 | 81 | 80 |
| ENSG00000141524 | 2322 | 1072 | 1262 | 1140 | 1070 | 961 |
| ENSG00000167895 | 58 | 31 | 21 | 24 | 23 | 29 |
| ENSG00000187997 | 0 | 0 | 0 | 0 | 0 | 0 |
| ENSG00000267679 | 0 | 0 | 0 | 0 | 0 | 0 |
| ENSG00000108639 | 9352 | 4253 | 5497 | 5386 | 4414 | 4525 |
| ENSG00000167900 | 2139 | 1147 | 1337 | 1227 | 983 | 1155 |
| ENSG00000183077 | 623 | 366 | 414 | 380 | 436 | 421 |
| ENSG00000267624 | 0 | 0 | 0 | 0 | 0 | 0 |
| ENSG00000089685 | 1431 | 758 | 942 | 1126 | 989 | 1054 |
| ENSG00000204278 | 0 | 0 | 0 | 0 | 0 | 0 |
| ENSG00000283914 | 0 | 0 | 2 | 0 | 0 | 0 |
| ENSG00000267676 | 5 | 2 | 0 | 6 | 0 | 5 |
| ENSG00000279886 | 0 | 0 | 0 | 0 | 0 | 0 |
| ENSG00000204277 | 0 | 0 | 0 | 0 | 0 | 0 |
| ENSG00000267737 | 4 | 0 | 3 | 1 | 4 | 1 |
| ENSG00000184557 | 264 | 152 | 171 | 165 | 155 | 182 |
| ENSG00000266970 | 56 | 31 | 47 | 37 | 37 | 47 |
| ENSG00000243870 | 1 | 1 | 4 | 3 | 0 | 0 |
| ENSG00000087157 | 1323 | 672 | 821 | 724 | 699 | 731 |
| ENSG00000200063 | 4 | 0 | 0 | 0 | 0 | 0 |
| ENSG00000187775 | 98 | 67 | 80 | 58 | 64 | 52 |
| ENSG00000267432 | 2 | 0 | 1 | 0 | 2 | 2 |
| ENSG00000267770 | 0 | 0 | 0 | 0 | 0 | 0 |
| ENSG00000243426 | 0 | 0 | 0 | 0 | 0 | 0 |
| ENSG00000267123 | 0 | 0 | 0 | 0 | 0 | 0 |
| ENSG00000108669 | 1831 | 849 | 1123 | 823 | 985 | 996 |
| ENSG00000279570 | 0 | 2 | 0 | 0 | 0 | 0 |
| ENSG00000252391 | 0 | 2 | 1 | 0 | 1 | 0 |
| ENSG00000055483 | 1600 | 765 | 972 | 944 | 909 | 1072 |
| ENSG00000267355 | 19 | 14 | 13 | 11 | 13 | 11 |
| ENSG00000035862 | 1416 | 734 | 953 | 946 | 878 | 957 |
| ENSG00000267601 | 0 | 0 | 0 | 0 | 0 | 0 |
| ENSG00000178404 | 2 | 2 | 2 | 0 | 0 | 0 |
| ENSG00000279339 | 0 | 0 | 0 | 0 | 0 | 0 |
| ENSG00000267491 | 0 | 0 | 0 | 0 | 0 | 0 |
| ENSG00000108679 | 1565 | 601 | 1008 | 779 | 620 | 794 |

| | | | | | | |
|---|---|---|---|---|---|---|
| ENSG00000171302 | 5828 | 2513 | 3365 | 2454 | 2403 | 2741 |
| ENSG00000265096 | 0 | 0 | 0 | 0 | 0 | 0 |
| ENSG00000173918 | 4 | 1 | 3 | 2 | 0 | 3 |
| ENSG00000167280 | 658 | 280 | 368 | 332 | 311 | 295 |
| ENSG00000167281 | 5 | 2 | 0 | 1 | 2 | 3 |
| ENSG00000266711 | 0 | 0 | 0 | 0 | 0 | 0 |
| ENSG00000263278 | 0 | 0 | 0 | 0 | 0 | 0 |
| ENSG00000266665 | 0 | 0 | 0 | 0 | 0 | 0 |
| ENSG00000267719 | 1 | 0 | 0 | 0 | 0 | 0 |
| ENSG00000182156 | 0 | 0 | 0 | 1 | 0 | 0 |
| ENSG00000173894 | 2622 | 1215 | 1517 | 1513 | 1418 | 1545 |
| ENSG00000141570 | 1128 | 470 | 586 | 604 | 566 | 498 |
| ENSG00000262768 | 9 | 4 | 5 | 5 | 11 | 2 |
| ENSG00000262772 | 79 | 36 | 64 | 31 | 31 | 39 |
| ENSG00000141582 | 3352 | 1467 | 1908 | 1602 | 1531 | 1597 |
| ENSG00000262343 | 0 | 0 | 0 | 0 | 1 | 1 |
| ENSG00000262585 | 59 | 19 | 21 | 16 | 15 | 19 |
| ENSG00000262188 | 20 | 9 | 20 | 13 | 13 | 6 |
| ENSG00000167291 | 2475 | 1132 | 1453 | 1392 | 1648 | 1563 |
| ENSG00000275516 | 0 | 0 | 0 | 0 | 2 | 0 |
| ENSG00000262172 | 2 | 2 | 4 | 0 | 3 | 2 |
| ENSG00000261978 | 0 | 1 | 0 | 0 | 0 | 0 |
| ENSG00000141519 | 94 | 50 | 68 | 46 | 68 | 63 |
| ENSG00000265561 | 0 | 0 | 0 | 0 | 0 | 0 |
| ENSG00000171298 | 6742 | 2734 | 4190 | 3794 | 2917 | 3336 |
| ENSG00000141543 | 4624 | 2311 | 2876 | 2592 | 2719 | 2710 |
| ENSG00000279259 | 38 | 20 | 35 | 17 | 30 | 12 |
| ENSG00000275479 | 6 | 3 | 6 | 1 | 2 | 0 |
| ENSG00000141527 | 235 | 111 | 167 | 130 | 161 | 150 |
| ENSG00000262580 | 172 | 83 | 139 | 82 | 95 | 83 |
| ENSG00000181523 | 1561 | 766 | 1002 | 800 | 909 | 844 |
| ENSG00000181045 | 1249 | 587 | 831 | 626 | 647 | 663 |
| ENSG00000173821 | 6817 | 2559 | 4097 | 1908 | 2790 | 3254 |
| ENSG00000262979 | 8 | 5 | 8 | 8 | 8 | 11 |
| ENSG00000276863 | 1 | 4 | 1 | 1 | 0 | 0 |
| ENSG00000280248 | 0 | 0 | 2 | 0 | 2 | 0 |
| ENSG00000263069 | 35 | 13 | 21 | 11 | 9 | 11 |
| ENSG00000173818 | 1174 | 607 | 667 | 738 | 655 | 634 |
| ENSG00000264961 | 0 | 1 | 1 | 0 | 0 | 2 |
| ENSG00000260369 | 2 | 0 | 1 | 1 | 0 | 1 |
| ENSG00000279273 | 0 | 0 | 0 | 0 | 0 | 0 |
| ENSG00000171246 | 8 | 3 | 6 | 2 | 3 | 3 |
| ENSG00000262000 | 0 | 0 | 0 | 0 | 0 | 0 |
| ENSG00000271340 | 0 | 0 | 0 | 0 | 0 | 0 |
| ENSG00000213128 | 0 | 0 | 0 | 0 | 0 | 0 |
| ENSG00000141564 | 1225 | 544 | 770 | 635 | 605 | 700 |
| ENSG00000232083 | 0 | 0 | 0 | 1 | 0 | 0 |
| ENSG00000262833 | 0 | 1 | 0 | 0 | 0 | 0 |
| ENSG00000262313 | 0 | 0 | 0 | 0 | 0 | 0 |
| ENSG00000262662 | 1 | 0 | 0 | 0 | 0 | 0 |

| | | | | | | |
|---|---|---|---|---|---|---|
| ENSG00000261924 | 14 | 4 | 5 | 2 | 4 | 9 |
| ENSG00000240270 | 2 | 1 | 0 | 0 | 1 | 0 |
| ENSG00000176108 | 642 | 317 | 377 | 389 | 264 | 338 |
| ENSG00000263218 | 0 | 0 | 0 | 0 | 0 | 0 |
| ENSG00000280351 | 69 | 36 | 38 | 44 | 45 | 34 |
| ENSG00000262873 | 2 | 2 | 1 | 0 | 2 | 1 |
| ENSG00000262098 | 0 | 2 | 0 | 1 | 2 | 1 |
| ENSG00000226137 | 1116 | 541 | 656 | 482 | 581 | 574 |
| ENSG00000175866 | 1099 | 586 | 747 | 727 | 646 | 687 |
| ENSG00000181409 | 18 | 7 | 14 | 8 | 8 | 10 |
| ENSG00000207736 | 0 | 0 | 0 | 0 | 0 | 0 |
| ENSG00000211563 | 0 | 0 | 0 | 0 | 0 | 0 |
| ENSG00000283604 | 0 | 0 | 0 | 0 | 0 | 0 |
| ENSG00000221025 | 0 | 0 | 0 | 0 | 0 | 0 |
| ENSG00000263167 | 0 | 0 | 2 | 1 | 0 | 0 |
| ENSG00000225180 | 5 | 2 | 1 | 1 | 2 | 3 |
| ENSG00000141577 | 709 | 386 | 438 | 496 | 431 | 459 |
| ENSG00000262115 | 0 | 1 | 3 | 1 | 2 | 1 |
| ENSG00000167302 | 333 | 154 | 223 | 208 | 191 | 168 |
| ENSG00000260005 | 16 | 14 | 8 | 5 | 9 | 7 |
| ENSG00000224877 | 504 | 281 | 336 | 392 | 358 | 326 |
| ENSG00000157637 | 4868 | 2144 | 2979 | 2897 | 2592 | 2621 |
| ENSG00000276101 | 0 | 0 | 0 | 0 | 0 | 0 |
| ENSG00000279187 | 6 | 4 | 5 | 0 | 3 | 10 |
| ENSG00000185168 | 140 | 75 | 110 | 67 | 70 | 56 |
| ENSG00000185332 | 170 | 89 | 122 | 73 | 85 | 69 |
| ENSG00000275542 | 60 | 19 | 19 | 20 | 15 | 12 |
| ENSG00000275966 | 11 | 8 | 23 | 17 | 22 | 10 |
| ENSG00000263154 | 2 | 2 | 3 | 0 | 1 | 1 |
| ENSG00000262223 | 54 | 17 | 41 | 19 | 19 | 38 |
| ENSG00000274833 | 5 | 0 | 1 | 2 | 2 | 0 |
| ENSG00000262877 | 248 | 105 | 169 | 86 | 89 | 131 |
| ENSG00000266074 | 699 | 264 | 442 | 291 | 366 | 345 |
| ENSG00000279692 | 123 | 63 | 79 | 60 | 56 | 73 |
| ENSG00000266392 | 2 | 0 | 0 | 0 | 0 | 0 |
| ENSG00000266189 | 2 | 0 | 0 | 3 | 2 | 3 |
| ENSG00000263271 | 0 | 0 | 0 | 0 | 0 | 0 |
| ENSG00000278200 | 0 | 0 | 0 | 1 | 0 | 0 |
| ENSG00000184009 | 84680 | 45692 | 55234 | 61234 | 56781 | 50783 |
| ENSG00000229848 | 57 | 28 | 46 | 25 | 23 | 33 |
| ENSG00000186765 | 121 | 52 | 72 | 35 | 58 | 60 |
| ENSG00000185504 | 1542 | 691 | 1012 | 926 | 762 | 830 |
| ENSG00000182446 | 7193 | 3272 | 4324 | 3651 | 4077 | 3999 |
| ENSG00000207021 | 0 | 1 | 0 | 0 | 1 | 0 |
| ENSG00000182612 | 66 | 33 | 22 | 22 | 23 | 27 |
| ENSG00000185527 | 9 | 4 | 3 | 12 | 6 | 5 |
| ENSG00000204237 | 1123 | 613 | 721 | 814 | 729 | 667 |
| ENSG00000185298 | 1640 | 814 | 1033 | 1056 | 908 | 1053 |
| ENSG00000214087 | 400 | 208 | 246 | 226 | 273 | 207 |
| ENSG00000185359 | 3285 | 1533 | 2029 | 1821 | 1813 | 1771 |

| | | | | | | |
|---|---|---|---|---|---|---|
| ENSG00000277784 | 0 | 0 | 1 | 0 | 0 | 0 |
| ENSG00000275902 | 0 | 0 | 0 | 0 | 0 | 0 |
| ENSG00000262049 | 182 | 110 | 152 | 113 | 90 | 126 |
| ENSG00000262814 | 586 | 361 | 422 | 683 | 436 | 514 |
| ENSG00000262660 | 0 | 0 | 0 | 0 | 0 | 0 |
| ENSG00000183048 | 751 | 403 | 532 | 565 | 392 | 442 |
| ENSG00000215644 | 6 | 1 | 6 | 1 | 0 | 3 |
| ENSG00000225663 | 1461 | 753 | 811 | 1054 | 878 | 798 |
| ENSG00000182676 | 2 | 3 | 3 | 1 | 0 | 1 |
| ENSG00000185624 | 23909 | 12183 | 16267 | 14893 | 13007 | 13689 |
| ENSG00000262831 | 0 | 0 | 0 | 0 | 0 | 0 |
| ENSG00000278876 | 5 | 9 | 7 | 3 | 8 | 8 |
| ENSG00000141522 | 4288 | 2157 | 2813 | 3004 | 2389 | 2579 |
| ENSG00000262413 | 0 | 0 | 0 | 0 | 0 | 0 |
| ENSG00000263731 | 19 | 7 | 31 | 28 | 16 | 24 |
| ENSG00000263859 | 0 | 0 | 0 | 0 | 0 | 0 |
| ENSG00000183684 | 2303 | 1274 | 1402 | 1640 | 1399 | 1422 |
| ENSG00000141552 | 2174 | 1087 | 1259 | 1521 | 1287 | 1148 |
| ENSG00000183979 | 0 | 0 | 0 | 0 | 0 | 1 |
| ENSG00000185813 | 1542 | 745 | 994 | 1010 | 840 | 881 |
| ENSG00000187531 | 1729 | 762 | 1009 | 1095 | 1030 | 996 |
| ENSG00000197063 | 1305 | 592 | 751 | 860 | 894 | 866 |
| ENSG00000264769 | 7 | 2 | 2 | 2 | 2 | 6 |
| ENSG00000265688 | 38 | 23 | 26 | 32 | 21 | 27 |
| ENSG00000183010 | 1440 | 807 | 962 | 1364 | 1056 | 1101 |
| ENSG00000263585 | 0 | 0 | 0 | 0 | 0 | 0 |
| ENSG00000185105 | 5 | 5 | 1 | 4 | 2 | 3 |
| ENSG00000235296 | 1 | 0 | 2 | 1 | 1 | 1 |
| ENSG00000185269 | 4 | 1 | 2 | 2 | 1 | 0 |
| ENSG00000264735 | 0 | 0 | 0 | 0 | 0 | 0 |
| ENSG00000169696 | 1253 | 552 | 772 | 856 | 609 | 705 |
| ENSG00000169689 | 3494 | 1761 | 2094 | 2277 | 2046 | 1908 |
| ENSG00000169683 | 1502 | 631 | 863 | 916 | 741 | 814 |
| ENSG00000169750 | 771 | 453 | 488 | 584 | 450 | 500 |
| ENSG00000169738 | 3054 | 1442 | 1774 | 1955 | 1509 | 1465 |
| ENSG00000279372 | 1 | 1 | 1 | 3 | 0 | 2 |
| ENSG00000264569 | 0 | 6 | 3 | 2 | 6 | 2 |
| ENSG00000169733 | 1262 | 562 | 712 | 736 | 629 | 626 |
| ENSG00000169727 | 4909 | 2263 | 2912 | 3272 | 2547 | 2590 |
| ENSG00000169718 | 3833 | 1976 | 2283 | 2805 | 2595 | 2376 |
| ENSG00000169710 | 53609 | 19929 | 32070 | 25982 | 21053 | 25314 |
| ENSG00000176155 | 1600 | 717 | 976 | 794 | 726 | 848 |
| ENSG00000265678 | 18 | 13 | 17 | 10 | 11 | 15 |
| ENSG00000266654 | 0 | 0 | 0 | 0 | 0 | 0 |
| ENSG00000264548 | 43 | 12 | 16 | 9 | 13 | 19 |
| ENSG00000141526 | 957 | 432 | 648 | 549 | 467 | 509 |
| ENSG00000284574 | 0 | 0 | 0 | 0 | 0 | 0 |
| ENSG00000141551 | 6873 | 3365 | 4534 | 3526 | 3673 | 4040 |
| ENSG00000275888 | 27 | 27 | 22 | 20 | 11 | 12 |
| ENSG00000280407 | 40 | 19 | 13 | 7 | 14 | 16 |

| | | | | | | |
|---|---|---|---|---|---|---|
| ENSG00000265692 | 6 | 7 | 6 | 5 | 7 | 5 |
| ENSG00000260563 | 105 | 54 | 62 | 38 | 70 | 51 |
| ENSG00000173762 | 2 | 0 | 0 | 1 | 1 | 0 |
| ENSG00000141574 | 92 | 30 | 62 | 109 | 85 | 117 |
| ENSG00000182459 | 25 | 4 | 13 | 16 | 13 | 24 |
| ENSG00000278964 | 0 | 0 | 0 | 0 | 0 | 0 |
| ENSG00000181408 | 0 | 0 | 0 | 0 | 0 | 1 |
| ENSG00000260011 | 5 | 2 | 3 | 0 | 1 | 1 |
| ENSG00000181396 | 1371 | 570 | 848 | 794 | 743 | 772 |
| ENSG00000264812 | 5 | 2 | 1 | 1 | 2 | 1 |
| ENSG00000201239 | 0 | 0 | 0 | 0 | 0 | 0 |
| ENSG00000169660 | 822 | 343 | 456 | 483 | 422 | 436 |
| ENSG00000279066 | 16 | 7 | 2 | 3 | 1 | 4 |
| ENSG00000178927 | 2535 | 1152 | 1509 | 1540 | 1308 | 1411 |
| ENSG00000265458 | 45 | 19 | 36 | 31 | 30 | 29 |
| ENSG00000141562 | 1622 | 803 | 965 | 958 | 975 | 968 |
| ENSG00000279744 | 11 | 19 | 13 | 4 | 12 | 8 |
| ENSG00000266445 | 12 | 2 | 6 | 3 | 8 | 3 |
| ENSG00000266236 | 4 | 3 | 9 | 1 | 4 | 3 |
| ENSG00000141568 | 2890 | 1477 | 1929 | 1683 | 1698 | 1800 |
| ENSG00000265136 | 8 | 4 | 6 | 4 | 6 | 5 |
| ENSG00000262652 | 3 | 7 | 4 | 1 | 7 | 3 |
| ENSG00000262147 | 4 | 5 | 8 | 4 | 2 | 4 |
| ENSG00000261845 | 0 | 0 | 0 | 0 | 0 | 0 |
| ENSG00000141580 | 4413 | 2152 | 2740 | 2268 | 2631 | 2726 |
| ENSG00000279613 | 2 | 0 | 8 | 0 | 0 | 0 |
| ENSG00000141542 | 501 | 261 | 355 | 251 | 300 | 275 |
| ENSG00000266107 | 0 | 0 | 0 | 0 | 0 | 0 |
| ENSG00000263063 | 15 | 2 | 1 | 2 | 3 | 4 |
| ENSG00000141560 | 1067 | 506 | 595 | 535 | 588 | 549 |
| ENSG00000263321 | 0 | 3 | 3 | 1 | 1 | 3 |
| ENSG00000167363 | 270 | 132 | 186 | 171 | 135 | 142 |
| ENSG00000262410 | 7 | 5 | 4 | 1 | 5 | 8 |
| ENSG00000141556 | 2871 | 1319 | 1682 | 1702 | 1489 | 1746 |
| ENSG00000263098 | 0 | 0 | 1 | 0 | 0 | 0 |
| ENSG00000141579 | 116 | 43 | 91 | 35 | 61 | 77 |
| ENSG00000262663 | 4 | 2 | 3 | 0 | 1 | 5 |
| ENSG00000175711 | 198 | 96 | 157 | 117 | 106 | 129 |
| ENSG00000262339 | 0 | 1 | 3 | 0 | 3 | 0 |
| ENSG00000176845 | 4712 | 2190 | 2671 | 2883 | 2438 | 2503 |
| ENSG00000274370 | 63 | 14 | 55 | 21 | 35 | 24 |
| ENSG00000261888 | 460 | 208 | 277 | 216 | 188 | 205 |
| ENSG00000279143 | 0 | 0 | 2 | 0 | 2 | 0 |
| ENSG00000262898 | 0 | 1 | 0 | 0 | 0 | 0 |
| ENSG00000262952 | 0 | 1 | 1 | 1 | 0 | 1 |
| ENSG00000262094 | 69 | 36 | 37 | 30 | 40 | 43 |
| ENSG00000232938 | 53 | 19 | 45 | 19 | 31 | 38 |
| ENSG00000262352 | 0 | 0 | 0 | 0 | 0 | 0 |
| ENSG00000263305 | 0 | 0 | 0 | 0 | 0 | 0 |
| ENSG00000262181 | 0 | 0 | 0 | 0 | 0 | 0 |

| | | | | | | |
|---|---|---|---|---|---|---|
| ENSG00000173213 | 16 | 4 | 6 | 5 | 11 | 5 |
| ENSG00000282965 | 0 | 1 | 2 | 0 | 0 | 0 |
| ENSG00000262081 | 0 | 0 | 0 | 0 | 0 | 0 |
| ENSG00000274446 | 0 | 0 | 0 | 0 | 0 | 0 |
| ENSG00000263006 | 46 | 19 | 17 | 9 | 26 | 25 |
| ENSG00000277521 | 0 | 2 | 1 | 0 | 3 | 0 |
| ENSG00000101557 | 2968 | 1482 | 1863 | 1345 | 1812 | 1716 |
| ENSG00000079134 | 562 | 292 | 308 | 196 | 388 | 295 |
| ENSG00000263884 | 2 | 3 | 13 | 2 | 8 | 4 |
| ENSG00000264514 | 0 | 0 | 0 | 0 | 0 | 0 |
| ENSG00000158270 | 8 | 0 | 3 | 1 | 3 | 5 |
| ENSG00000265477 | 0 | 0 | 0 | 0 | 0 | 0 |
| ENSG00000264265 | 0 | 0 | 0 | 0 | 0 | 0 |
| ENSG00000177143 | 0 | 0 | 0 | 0 | 0 | 0 |
| ENSG00000222174 | 0 | 0 | 0 | 0 | 0 | 0 |
| ENSG00000079101 | 5 | 5 | 2 | 0 | 2 | 2 |
| ENSG00000176912 | 24 | 21 | 19 | 31 | 21 | 18 |
| ENSG00000266456 | 0 | 3 | 4 | 0 | 5 | 1 |
| ENSG00000263727 | 2 | 2 | 0 | 0 | 3 | 1 |
| ENSG00000176890 | 262 | 175 | 243 | 178 | 190 | 240 |
| ENSG00000132199 | 1296 | 638 | 982 | 550 | 714 | 738 |
| ENSG00000265490 | 3 | 1 | 2 | 0 | 2 | 0 |
| ENSG00000176105 | 1651 | 765 | 792 | 458 | 1052 | 723 |
| ENSG00000266171 | 4 | 1 | 3 | 1 | 3 | 4 |
| ENSG00000264339 | 0 | 0 | 0 | 0 | 0 | 0 |
| ENSG00000264635 | 10 | 4 | 6 | 2 | 1 | 4 |
| ENSG00000206687 | 0 | 0 | 0 | 0 | 1 | 0 |
| ENSG00000273355 | 26 | 19 | 31 | 28 | 20 | 14 |
| ENSG00000266575 | 0 | 0 | 0 | 0 | 0 | 0 |
| ENSG00000264179 | 0 | 0 | 0 | 0 | 0 | 0 |
| ENSG00000265671 | 0 | 0 | 0 | 0 | 0 | 0 |
| ENSG00000265179 | 0 | 0 | 0 | 0 | 0 | 0 |
| ENSG00000141433 | 0 | 0 | 0 | 0 | 0 | 0 |
| ENSG00000259837 | 0 | 0 | 0 | 0 | 0 | 0 |
| ENSG00000272461 | 0 | 0 | 0 | 0 | 0 | 0 |
| ENSG00000263551 | 0 | 0 | 0 | 0 | 0 | 1 |
| ENSG00000263505 | 0 | 0 | 0 | 0 | 0 | 0 |
| ENSG00000266251 | 0 | 0 | 0 | 0 | 0 | 0 |
| ENSG00000132204 | 4 | 3 | 6 | 3 | 1 | 2 |
| ENSG00000199197 | 0 | 0 | 0 | 0 | 0 | 0 |
| ENSG00000263396 | 0 | 0 | 0 | 0 | 0 | 0 |
| ENSG00000265417 | 0 | 1 | 1 | 1 | 0 | 0 |
| ENSG00000266602 | 0 | 0 | 0 | 0 | 0 | 1 |
| ENSG00000266450 | 0 | 0 | 0 | 0 | 0 | 0 |
| ENSG00000264080 | 0 | 0 | 0 | 0 | 0 | 0 |
| ENSG00000263745 | 2 | 0 | 3 | 1 | 0 | 1 |
| ENSG00000264189 | 0 | 0 | 0 | 0 | 0 | 0 |
| ENSG00000101574 | 244 | 149 | 168 | 110 | 183 | 139 |
| ENSG00000238575 | 1 | 0 | 0 | 0 | 0 | 0 |
| ENSG00000266783 | 16 | 8 | 6 | 6 | 6 | 7 |

| | | | | | | |
|---|---|---|---|---|---|---|
| ENSG00000264150 | 0 | 0 | 0 | 0 | 0 | 0 |
| ENSG00000080986 | 210 | 144 | 119 | 91 | 245 | 177 |
| ENSG00000235422 | 0 | 0 | 1 | 0 | 0 | 0 |
| ENSG00000202224 | 0 | 0 | 0 | 0 | 0 | 0 |
| ENSG00000200875 | 0 | 0 | 0 | 0 | 0 | 0 |
| ENSG00000266405 | 17 | 4 | 7 | 5 | 3 | 8 |
| ENSG00000101596 | 525 | 352 | 257 | 108 | 577 | 271 |
| ENSG00000266049 | 2 | 0 | 0 | 0 | 0 | 1 |
| ENSG00000207034 | 0 | 0 | 0 | 0 | 0 | 0 |
| ENSG00000132205 | 8 | 2 | 0 | 11 | 1 | 5 |
| ENSG00000101577 | 1160 | 534 | 716 | 428 | 594 | 600 |
| ENSG00000272625 | 0 | 0 | 0 | 0 | 0 | 0 |
| ENSG00000263606 | 2 | 0 | 0 | 0 | 0 | 0 |
| ENSG00000265907 | 0 | 0 | 0 | 0 | 0 | 0 |
| ENSG00000266397 | 0 | 1 | 0 | 0 | 0 | 0 |
| ENSG00000264186 | 0 | 0 | 0 | 0 | 0 | 0 |
| ENSG00000277447 | 0 | 0 | 0 | 0 | 3 | 0 |
| ENSG00000252258 | 0 | 0 | 0 | 0 | 0 | 0 |
| ENSG00000101605 | 35 | 22 | 27 | 8 | 16 | 14 |
| ENSG00000252353 | 0 | 0 | 0 | 0 | 0 | 0 |
| ENSG00000265399 | 46 | 27 | 43 | 26 | 43 | 35 |
| ENSG00000272688 | 0 | 0 | 0 | 0 | 0 | 0 |
| ENSG00000101608 | 11076 | 7056 | 8156 | 7677 | 8557 | 7467 |
| ENSG00000264235 | 10 | 6 | 4 | 3 | 7 | 8 |
| ENSG00000118680 | 26919 | 14119 | 16107 | 14026 | 15152 | 13675 |
| ENSG00000266578 | 1 | 0 | 0 | 0 | 1 | 2 |
| ENSG00000259256 | 0 | 0 | 0 | 0 | 0 | 0 |
| ENSG00000240364 | 0 | 0 | 0 | 0 | 0 | 0 |
| ENSG00000265243 | 0 | 0 | 0 | 2 | 0 | 0 |
| ENSG00000243720 | 0 | 0 | 0 | 0 | 0 | 0 |
| ENSG00000177426 | 2230 | 1016 | 1308 | 1351 | 1483 | 1400 |
| ENSG00000263460 | 0 | 0 | 0 | 0 | 0 | 0 |
| ENSG00000266835 | 0 | 4 | 0 | 0 | 0 | 0 |
| ENSG00000170579 | 0 | 1 | 1 | 1 | 0 | 0 |
| ENSG00000241223 | 0 | 0 | 0 | 0 | 0 | 0 |
| ENSG00000266373 | 0 | 0 | 0 | 0 | 0 | 1 |
| ENSG00000177337 | 126 | 63 | 92 | 64 | 71 | 70 |
| ENSG00000262001 | 90 | 33 | 56 | 43 | 44 | 35 |
| ENSG00000266401 | 0 | 0 | 0 | 0 | 2 | 0 |
| ENSG00000266513 | 0 | 0 | 0 | 0 | 0 | 0 |
| ENSG00000263724 | 0 | 0 | 0 | 0 | 0 | 0 |
| ENSG00000275518 | 0 | 0 | 0 | 0 | 0 | 0 |
| ENSG00000207278 | 0 | 0 | 0 | 0 | 0 | 0 |
| ENSG00000263878 | 0 | 0 | 0 | 0 | 0 | 0 |
| ENSG00000250933 | 0 | 0 | 0 | 0 | 0 | 0 |
| ENSG00000261520 | 0 | 0 | 0 | 0 | 0 | 0 |
| ENSG00000264246 | 0 | 0 | 0 | 0 | 0 | 0 |
| ENSG00000265545 | 0 | 0 | 0 | 0 | 0 | 0 |
| ENSG00000266268 | 0 | 0 | 0 | 0 | 0 | 0 |
| ENSG00000264775 | 0 | 0 | 0 | 1 | 0 | 0 |

| | | | | | | |
|---|---|---|---|---|---|---|
| ENSG00000265514 | 0 | 0 | 0 | 0 | 0 | 0 |
| ENSG00000215368 | 0 | 0 | 0 | 0 | 0 | 0 |
| ENSG00000231824 | 0 | 0 | 0 | 0 | 0 | 0 |
| ENSG00000266153 | 0 | 0 | 0 | 0 | 0 | 0 |
| ENSG00000285575 | 0 | 0 | 0 | 0 | 0 | 0 |
| ENSG00000265091 | 0 | 0 | 0 | 0 | 0 | 0 |
| ENSG00000264575 | 21 | 4 | 12 | 5 | 6 | 3 |
| ENSG00000263753 | 49 | 44 | 40 | 32 | 34 | 46 |
| ENSG00000265188 | 0 | 0 | 0 | 0 | 0 | 0 |
| ENSG00000198081 | 668 | 314 | 405 | 293 | 315 | 376 |
| ENSG00000264254 | 0 | 0 | 0 | 0 | 0 | 0 |
| ENSG00000266288 | 0 | 0 | 0 | 0 | 0 | 0 |
| ENSG00000082397 | 0 | 0 | 0 | 0 | 0 | 0 |
| ENSG00000265316 | 0 | 0 | 0 | 0 | 0 | 0 |
| ENSG00000264000 | 0 | 0 | 0 | 0 | 0 | 0 |
| ENSG00000261738 | 0 | 0 | 0 | 0 | 0 | 0 |
| ENSG00000272380 | 0 | 0 | 0 | 0 | 0 | 0 |
| ENSG00000206432 | 0 | 0 | 0 | 0 | 0 | 0 |
| ENSG00000264449 | 0 | 0 | 0 | 0 | 0 | 0 |
| ENSG00000154655 | 0 | 0 | 0 | 0 | 0 | 0 |
| ENSG00000265487 | 0 | 0 | 0 | 0 | 0 | 0 |
| ENSG00000200637 | 0 | 0 | 0 | 0 | 0 | 0 |
| ENSG00000266846 | 0 | 0 | 0 | 0 | 0 | 0 |
| ENSG00000264707 | 0 | 0 | 0 | 0 | 0 | 0 |
| ENSG00000283532 | 0 | 0 | 0 | 0 | 0 | 0 |
| ENSG00000279885 | 0 | 0 | 0 | 0 | 0 | 0 |
| ENSG00000199272 | 0 | 0 | 0 | 0 | 0 | 0 |
| ENSG00000235552 | 92 | 71 | 65 | 71 | 92 | 69 |
| ENSG00000266065 | 0 | 0 | 0 | 0 | 0 | 0 |
| ENSG00000265944 | 0 | 0 | 0 | 0 | 0 | 0 |
| ENSG00000265174 | 0 | 0 | 0 | 0 | 0 | 0 |
| ENSG00000243591 | 0 | 0 | 0 | 0 | 0 | 0 |
| ENSG00000261627 | 0 | 0 | 0 | 0 | 0 | 0 |
| ENSG00000279479 | 0 | 0 | 0 | 0 | 0 | 0 |
| ENSG00000266441 | 0 | 0 | 0 | 0 | 0 | 0 |
| ENSG00000088756 | 0 | 0 | 0 | 0 | 0 | 0 |
| ENSG00000263797 | 0 | 0 | 0 | 0 | 0 | 0 |
| ENSG00000265933 | 0 | 0 | 0 | 0 | 0 | 0 |
| ENSG00000264127 | 0 | 0 | 0 | 0 | 0 | 0 |
| ENSG00000251897 | 0 | 0 | 0 | 0 | 0 | 0 |
| ENSG00000101680 | 0 | 0 | 0 | 0 | 0 | 0 |
| ENSG00000265069 | 0 | 0 | 0 | 0 | 0 | 0 |
| ENSG00000264475 | 0 | 0 | 0 | 0 | 0 | 0 |
| ENSG00000263716 | 0 | 0 | 0 | 0 | 0 | 0 |
| ENSG00000206422 | 0 | 0 | 0 | 0 | 0 | 0 |
| ENSG00000212626 | 0 | 0 | 0 | 0 | 0 | 0 |
| ENSG00000173482 | 0 | 0 | 0 | 1 | 0 | 0 |
| ENSG00000266767 | 0 | 0 | 0 | 0 | 0 | 0 |
| ENSG00000264596 | 0 | 0 | 0 | 0 | 0 | 0 |
| ENSG00000199856 | 0 | 0 | 0 | 0 | 0 | 0 |

| | | | | | | |
|---|---|---|---|---|---|---|
| ENSG00000273368 | 0 | 0 | 0 | 0 | 0 | 0 |
| ENSG00000266149 | 0 | 0 | 0 | 0 | 0 | 0 |
| ENSG00000265413 | 0 | 0 | 0 | 0 | 0 | 0 |
| ENSG00000263970 | 0 | 0 | 0 | 0 | 0 | 0 |
| ENSG00000266613 | 0 | 0 | 0 | 0 | 0 | 0 |
| ENSG00000242985 | 0 | 0 | 0 | 0 | 0 | 0 |
| ENSG00000229442 | 0 | 0 | 0 | 0 | 0 | 0 |
| ENSG00000225212 | 0 | 0 | 0 | 0 | 0 | 0 |
| ENSG00000206418 | 1125 | 693 | 798 | 600 | 700 | 790 |
| ENSG00000266708 | 0 | 1 | 5 | 1 | 1 | 5 |
| ENSG00000263935 | 0 | 0 | 0 | 0 | 0 | 0 |
| ENSG00000265962 | 2 | 0 | 0 | 2 | 0 | 0 |
| ENSG00000168502 | 262 | 122 | 160 | 128 | 167 | 120 |
| ENSG00000252319 | 0 | 0 | 0 | 0 | 0 | 0 |
| ENSG00000272788 | 0 | 0 | 2 | 0 | 0 | 2 |
| ENSG00000242058 | 0 | 0 | 0 | 0 | 0 | 0 |
| ENSG00000178127 | 39 | 24 | 40 | 4 | 25 | 22 |
| ENSG00000265257 | 0 | 0 | 0 | 0 | 0 | 0 |
| ENSG00000263847 | 3 | 5 | 6 | 1 | 3 | 8 |
| ENSG00000266053 | 25 | 22 | 23 | 15 | 33 | 24 |
| ENSG00000101745 | 102 | 68 | 56 | 17 | 122 | 86 |
| ENSG00000200883 | 0 | 0 | 0 | 0 | 0 | 0 |
| ENSG00000207092 | 0 | 0 | 0 | 0 | 0 | 0 |
| ENSG00000273284 | 0 | 0 | 0 | 0 | 1 | 0 |
| ENSG00000264964 | 36 | 20 | 25 | 20 | 25 | 14 |
| ENSG00000128791 | 665 | 270 | 316 | 201 | 263 | 235 |
| ENSG00000273335 | 0 | 1 | 0 | 1 | 3 | 2 |
| ENSG00000017797 | 2669 | 1242 | 1519 | 1143 | 1430 | 1480 |
| ENSG00000266805 | 2 | 0 | 1 | 1 | 1 | 0 |
| ENSG00000251994 | 0 | 0 | 0 | 0 | 0 | 0 |
| ENSG00000154845 | 1666 | 843 | 984 | 756 | 846 | 807 |
| ENSG00000266541 | 0 | 0 | 1 | 0 | 0 | 0 |
| ENSG00000263627 | 4 | 3 | 0 | 2 | 1 | 0 |
| ENSG00000212572 | 1 | 0 | 0 | 0 | 0 | 0 |
| ENSG00000285594 | 0 | 0 | 0 | 0 | 0 | 0 |
| ENSG00000264644 | 2 | 0 | 1 | 1 | 1 | 0 |
| ENSG00000168461 | 794 | 360 | 489 | 448 | 490 | 476 |
| ENSG00000266891 | 0 | 0 | 0 | 0 | 0 | 0 |
| ENSG00000266127 | 0 | 0 | 0 | 0 | 0 | 0 |
| ENSG00000242651 | 0 | 0 | 0 | 0 | 0 | 0 |
| ENSG00000285653 | 0 | 0 | 1 | 0 | 0 | 0 |
| ENSG00000252680 | 0 | 0 | 0 | 0 | 0 | 1 |
| ENSG00000265564 | 0 | 0 | 0 | 0 | 0 | 0 |
| ENSG00000168454 | 0 | 0 | 0 | 1 | 0 | 0 |
| ENSG00000272799 | 0 | 2 | 0 | 0 | 3 | 1 |
| ENSG00000101558 | 3679 | 1803 | 2110 | 1854 | 2164 | 1928 |
| ENSG00000223138 | 0 | 0 | 0 | 0 | 2 | 0 |
| ENSG00000265554 | 0 | 0 | 0 | 0 | 0 | 0 |
| ENSG00000264876 | 0 | 0 | 0 | 0 | 0 | 0 |
| ENSG00000266102 | 0 | 0 | 0 | 0 | 0 | 0 |

| | | | | | | |
|---|---|---|---|---|---|---|
| ENSG00000260913 | 0 | 0 | 0 | 0 | 0 | 0 |
| ENSG00000154856 | 54 | 19 | 20 | 27 | 14 | 19 |
| ENSG00000134265 | 1672 | 826 | 1051 | 667 | 859 | 919 |
| ENSG00000265728 | 2 | 3 | 1 | 0 | 0 | 0 |
| ENSG00000266604 | 0 | 0 | 0 | 0 | 0 | 0 |
| ENSG00000264503 | 0 | 0 | 0 | 0 | 0 | 0 |
| ENSG00000264311 | 0 | 0 | 0 | 0 | 0 | 0 |
| ENSG00000264072 | 0 | 0 | 0 | 0 | 0 | 0 |
| ENSG00000266470 | 0 | 0 | 0 | 0 | 0 | 0 |
| ENSG00000265038 | 0 | 0 | 0 | 0 | 0 | 0 |
| ENSG00000260779 | 0 | 0 | 0 | 0 | 0 | 0 |
| ENSG00000154864 | 0 | 3 | 5 | 0 | 2 | 0 |
| ENSG00000264843 | 0 | 0 | 0 | 0 | 0 | 0 |
| ENSG00000264714 | 8 | 10 | 6 | 1 | 6 | 6 |
| ENSG00000274838 | 0 | 0 | 0 | 0 | 0 | 0 |
| ENSG00000263952 | 0 | 0 | 0 | 1 | 0 | 0 |
| ENSG00000263682 | 0 | 0 | 0 | 0 | 0 | 0 |
| ENSG00000267051 | 0 | 0 | 0 | 0 | 0 | 0 |
| ENSG00000273119 | 0 | 0 | 0 | 0 | 0 | 0 |
| ENSG00000267564 | 0 | 0 | 0 | 0 | 0 | 0 |
| ENSG00000267425 | 0 | 0 | 0 | 0 | 0 | 0 |
| ENSG00000267252 | 0 | 0 | 0 | 0 | 0 | 0 |
| ENSG00000267371 | 0 | 0 | 0 | 0 | 0 | 0 |
| ENSG00000236396 | 0 | 0 | 0 | 0 | 0 | 0 |
| ENSG00000267794 | 0 | 0 | 0 | 0 | 0 | 0 |
| ENSG00000257513 | 0 | 0 | 0 | 0 | 0 | 1 |
| ENSG00000267143 | 1 | 0 | 0 | 0 | 0 | 1 |
| ENSG00000267773 | 0 | 0 | 0 | 0 | 0 | 0 |
| ENSG00000283492 | 0 | 0 | 0 | 0 | 0 | 0 |
| ENSG00000260759 | 0 | 0 | 0 | 0 | 0 | 0 |
| ENSG00000141404 | 10 | 16 | 17 | 12 | 15 | 14 |
| ENSG00000267333 | 0 | 0 | 0 | 0 | 0 | 0 |
| ENSG00000267455 | 0 | 0 | 0 | 0 | 0 | 0 |
| ENSG00000255112 | 1802 | 1021 | 1286 | 895 | 1101 | 1097 |
| ENSG00000267165 | 0 | 0 | 0 | 0 | 0 | 0 |
| ENSG00000272703 | 0 | 0 | 0 | 0 | 0 | 0 |
| ENSG00000154889 | 452 | 211 | 342 | 206 | 239 | 305 |
| ENSG00000273141 | 0 | 0 | 0 | 0 | 0 | 0 |
| ENSG00000267079 | 0 | 0 | 0 | 0 | 0 | 0 |
| ENSG00000267661 | 0 | 0 | 0 | 0 | 0 | 0 |
| ENSG00000141401 | 440 | 208 | 264 | 338 | 252 | 280 |
| ENSG00000266955 | 2 | 0 | 0 | 1 | 0 | 0 |
| ENSG00000267480 | 0 | 0 | 2 | 4 | 2 | 3 |
| ENSG00000266995 | 0 | 0 | 0 | 0 | 0 | 0 |
| ENSG00000267292 | 0 | 0 | 0 | 0 | 0 | 0 |
| ENSG00000212712 | 0 | 0 | 0 | 0 | 0 | 0 |
| ENSG00000267533 | 0 | 1 | 1 | 1 | 1 | 2 |
| ENSG00000256616 | 0 | 0 | 0 | 0 | 0 | 0 |
| ENSG00000267722 | 1 | 0 | 0 | 0 | 0 | 0 |
| ENSG00000267478 | 0 | 0 | 0 | 0 | 0 | 0 |

| | | | | | | |
|---|---|---|---|---|---|---|
| ENSG00000181626 | 2 | 0 | 0 | 0 | 1 | 1 |
| ENSG00000200827 | 0 | 0 | 0 | 0 | 0 | 0 |
| ENSG00000267162 | 0 | 0 | 0 | 0 | 0 | 0 |
| ENSG00000267116 | 2 | 0 | 0 | 0 | 0 | 0 |
| ENSG00000279285 | 2 | 0 | 1 | 0 | 0 | 1 |
| ENSG00000267643 | 0 | 0 | 2 | 0 | 0 | 1 |
| ENSG00000267733 | 21 | 11 | 15 | 9 | 19 | 16 |
| ENSG00000267076 | 0 | 0 | 0 | 0 | 0 | 0 |
| ENSG00000267247 | 0 | 0 | 0 | 0 | 0 | 0 |
| ENSG00000267663 | 0 | 0 | 0 | 0 | 0 | 0 |
| ENSG00000176194 | 0 | 0 | 0 | 0 | 0 | 0 |
| ENSG00000280302 | 0 | 0 | 0 | 0 | 0 | 0 |
| ENSG00000267069 | 0 | 0 | 0 | 0 | 0 | 0 |
| ENSG00000199702 | 0 | 0 | 0 | 0 | 0 | 0 |
| ENSG00000176014 | 679 | 338 | 403 | 514 | 465 | 416 |
| ENSG00000141385 | 1927 | 981 | 1259 | 1035 | 1092 | 1250 |
| ENSG00000251937 | 0 | 0 | 0 | 0 | 0 | 0 |
| ENSG00000141391 | 145 | 51 | 81 | 71 | 76 | 71 |
| ENSG00000267108 | 2 | 0 | 1 | 0 | 0 | 2 |
| ENSG00000267199 | 32 | 6 | 7 | 6 | 9 | 4 |
| ENSG00000134278 | 1387 | 645 | 839 | 581 | 736 | 748 |
| ENSG00000267515 | 6 | 0 | 1 | 0 | 0 | 0 |
| ENSG00000128789 | 1781 | 864 | 1121 | 955 | 1178 | 1030 |
| ENSG00000101624 | 245 | 136 | 182 | 125 | 192 | 191 |
| ENSG00000267249 | 16 | 9 | 7 | 9 | 8 | 7 |
| ENSG00000266961 | 0 | 0 | 0 | 0 | 0 | 0 |
| ENSG00000260302 | 0 | 0 | 0 | 0 | 0 | 0 |
| ENSG00000201466 | 0 | 0 | 0 | 0 | 0 | 0 |
| ENSG00000175354 | 889 | 437 | 487 | 398 | 618 | 437 |
| ENSG00000267570 | 0 | 0 | 0 | 0 | 0 | 0 |
| ENSG00000267336 | 0 | 0 | 0 | 0 | 0 | 0 |
| ENSG00000085415 | 1002 | 565 | 626 | 548 | 648 | 679 |
| ENSG00000266969 | 0 | 1 | 0 | 1 | 0 | 0 |
| ENSG00000101639 | 702 | 422 | 407 | 170 | 476 | 367 |
| ENSG00000267656 | 0 | 0 | 0 | 0 | 0 | 0 |
| ENSG00000267032 | 0 | 0 | 0 | 0 | 0 | 0 |
| ENSG00000267286 | 0 | 0 | 0 | 0 | 0 | 0 |
| ENSG00000267239 | 7 | 3 | 3 | 4 | 5 | 2 |
| ENSG00000168675 | 1999 | 920 | 1285 | 561 | 693 | 642 |
| ENSG00000267177 | 0 | 0 | 0 | 0 | 0 | 0 |
| ENSG00000279020 | 0 | 0 | 0 | 0 | 0 | 0 |
| ENSG00000279275 | 0 | 0 | 0 | 0 | 0 | 1 |
| ENSG00000267393 | 0 | 0 | 0 | 0 | 0 | 0 |
| ENSG00000267690 | 0 | 2 | 1 | 0 | 2 | 2 |
| ENSG00000266146 | 0 | 0 | 0 | 0 | 0 | 0 |
| ENSG00000267694 | 0 | 0 | 0 | 0 | 0 | 0 |
| ENSG00000267529 | 0 | 0 | 0 | 0 | 0 | 0 |
| ENSG00000267702 | 32 | 11 | 11 | 4 | 10 | 14 |
| ENSG00000267366 | 12 | 3 | 6 | 4 | 5 | 3 |
| ENSG00000267136 | 12 | 8 | 10 | 2 | 2 | 1 |

| | | | | | | |
|---|---|---|---|---|---|---|
| ENSG00000272746 | 6 | 10 | 7 | 3 | 0 | 2 |
| ENSG00000267503 | 6 | 2 | 2 | 0 | 1 | 1 |
| ENSG00000263527 | 0 | 0 | 2 | 0 | 0 | 0 |
| ENSG00000266954 | 1 | 0 | 1 | 0 | 0 | 1 |
| ENSG00000177150 | 520 | 231 | 279 | 244 | 275 | 299 |
| ENSG00000279962 | 0 | 0 | 0 | 0 | 0 | 0 |
| ENSG00000242707 | 0 | 0 | 0 | 0 | 0 | 0 |
| ENSG00000101654 | 1009 | 527 | 489 | 244 | 632 | 482 |
| ENSG00000176136 | 8 | 4 | 11 | 3 | 1 | 2 |
| ENSG00000185231 | 0 | 0 | 0 | 1 | 0 | 0 |
| ENSG00000243779 | 0 | 0 | 0 | 0 | 0 | 0 |
| ENSG00000279352 | 2 | 1 | 0 | 2 | 0 | 0 |
| ENSG00000270469 | 2 | 0 | 0 | 0 | 1 | 0 |
| ENSG00000175322 | 98 | 34 | 34 | 33 | 68 | 47 |
| ENSG00000267756 | 0 | 0 | 0 | 0 | 0 | 0 |
| ENSG00000267150 | 0 | 1 | 1 | 0 | 2 | 0 |
| ENSG00000267356 | 0 | 0 | 0 | 0 | 0 | 0 |
| ENSG00000267704 | 0 | 0 | 0 | 0 | 0 | 0 |
| ENSG00000267324 | 0 | 0 | 0 | 0 | 0 | 0 |
| ENSG00000175319 | 0 | 0 | 0 | 0 | 0 | 0 |
| ENSG00000186481 | 73 | 27 | 50 | 19 | 47 | 53 |
| ENSG00000212329 | 0 | 1 | 0 | 1 | 2 | 0 |
| ENSG00000266145 | 7 | 2 | 4 | 0 | 3 | 3 |
| ENSG00000283294 | 9 | 2 | 4 | 0 | 3 | 2 |
| ENSG00000274214 | 12 | 0 | 1 | 0 | 1 | 2 |
| ENSG00000265787 | 7 | 3 | 1 | 0 | 1 | 7 |
| ENSG00000277743 | 0 | 0 | 0 | 0 | 0 | 0 |
| ENSG00000265437 | 0 | 0 | 0 | 0 | 0 | 0 |
| ENSG00000187589 | 0 | 0 | 0 | 0 | 0 | 0 |
| ENSG00000265296 | 0 | 0 | 2 | 1 | 3 | 0 |
| ENSG00000266605 | 0 | 1 | 1 | 2 | 0 | 0 |
| ENSG00000265766 | 2 | 0 | 0 | 0 | 0 | 0 |
| ENSG00000266242 | 2 | 1 | 0 | 1 | 0 | 0 |
| ENSG00000183206 | 0 | 0 | 0 | 0 | 0 | 0 |
| ENSG00000200132 | 0 | 0 | 0 | 0 | 0 | 0 |
| ENSG00000265340 | 0 | 0 | 0 | 0 | 0 | 0 |
| ENSG00000266693 | 0 | 0 | 0 | 0 | 0 | 0 |
| ENSG00000264570 | 0 | 1 | 3 | 1 | 1 | 0 |
| ENSG00000273732 | 0 | 0 | 0 | 0 | 0 | 0 |
| ENSG00000276318 | 0 | 0 | 0 | 0 | 0 | 0 |
| ENSG00000180777 | 15 | 6 | 3 | 2 | 4 | 7 |
| ENSG00000200645 | 0 | 0 | 0 | 0 | 0 | 0 |
| ENSG00000265737 | 0 | 0 | 0 | 0 | 0 | 1 |
| ENSG00000265499 | 0 | 0 | 0 | 0 | 0 | 0 |
| ENSG00000265786 | 0 | 0 | 0 | 0 | 0 | 0 |
| ENSG00000264061 | 0 | 0 | 0 | 0 | 0 | 1 |
| ENSG00000263618 | 0 | 0 | 0 | 0 | 0 | 2 |
| ENSG00000266554 | 0 | 0 | 0 | 0 | 0 | 0 |
| ENSG00000264301 | 0 | 0 | 2 | 0 | 2 | 0 |
| ENSG00000263821 | 0 | 0 | 0 | 0 | 0 | 0 |

| | | | | | | |
|---|---|---|---|---|---|---|
| ENSG00000279042 | 18 | 2 | 5 | 4 | 6 | 11 |
| ENSG00000266522 | 0 | 0 | 0 | 0 | 0 | 1 |
| ENSG00000278923 | 0 | 0 | 0 | 0 | 0 | 0 |
| ENSG00000263635 | 0 | 0 | 0 | 0 | 0 | 0 |
| ENSG00000264880 | 0 | 0 | 0 | 0 | 0 | 0 |
| ENSG00000265015 | 0 | 0 | 0 | 0 | 0 | 0 |
| ENSG00000265631 | 0 | 0 | 0 | 0 | 0 | 0 |
| ENSG00000266818 | 0 | 0 | 0 | 0 | 0 | 0 |
| ENSG00000215512 | 0 | 0 | 0 | 0 | 0 | 0 |
| ENSG00000222087 | 0 | 0 | 0 | 0 | 0 | 0 |
| ENSG00000067900 | 794 | 536 | 413 | 156 | 1015 | 439 |
| ENSG00000244527 | 0 | 0 | 0 | 0 | 0 | 0 |
| ENSG00000251886 | 0 | 0 | 0 | 0 | 0 | 0 |
| ENSG00000263748 | 0 | 0 | 0 | 0 | 0 | 0 |
| ENSG00000265984 | 2 | 0 | 0 | 0 | 0 | 0 |
| ENSG00000141449 | 450 | 208 | 215 | 116 | 171 | 126 |
| ENSG00000265948 | 0 | 0 | 0 | 0 | 0 | 0 |
| ENSG00000265751 | 0 | 0 | 0 | 0 | 0 | 0 |
| ENSG00000221139 | 0 | 0 | 0 | 0 | 0 | 0 |
| ENSG00000141446 | 217 | 198 | 122 | 58 | 380 | 171 |
| ENSG00000167088 | 1380 | 809 | 913 | 881 | 1046 | 988 |
| ENSG00000158201 | 433 | 198 | 231 | 207 | 242 | 230 |
| ENSG00000264188 | 0 | 1 | 3 | 0 | 0 | 0 |
| ENSG00000265656 | 2 | 0 | 0 | 2 | 0 | 0 |
| ENSG00000221493 | 0 | 0 | 0 | 0 | 0 | 0 |
| ENSG00000101752 | 1565 | 744 | 653 | 352 | 977 | 644 |
| ENSG00000252677 | 0 | 0 | 0 | 0 | 0 | 0 |
| ENSG00000200750 | 0 | 0 | 0 | 0 | 0 | 0 |
| ENSG00000242971 | 0 | 0 | 0 | 0 | 0 | 0 |
| ENSG00000199977 | 0 | 1 | 0 | 0 | 0 | 0 |
| ENSG00000265142 | 0 | 0 | 1 | 0 | 0 | 0 |
| ENSG00000283927 | 0 | 0 | 0 | 0 | 0 | 0 |
| ENSG00000284453 | 0 | 0 | 0 | 0 | 0 | 0 |
| ENSG00000243303 | 0 | 0 | 0 | 0 | 0 | 0 |
| ENSG00000264149 | 0 | 0 | 0 | 0 | 0 | 0 |
| ENSG00000252325 | 0 | 0 | 0 | 0 | 0 | 0 |
| ENSG00000264289 | 0 | 1 | 0 | 0 | 0 | 0 |
| ENSG00000222520 | 0 | 0 | 0 | 0 | 0 | 0 |
| ENSG00000265758 | 0 | 0 | 0 | 0 | 0 | 0 |
| ENSG00000263588 | 0 | 0 | 0 | 0 | 0 | 0 |
| ENSG00000221363 | 0 | 0 | 0 | 0 | 0 | 0 |
| ENSG00000266010 | 21 | 5 | 7 | 10 | 7 | 4 |
| ENSG00000141448 | 384 | 199 | 225 | 156 | 154 | 174 |
| ENSG00000264012 | 3 | 0 | 2 | 0 | 0 | 0 |
| ENSG00000201852 | 0 | 0 | 0 | 0 | 0 | 0 |
| ENSG00000266283 | 2 | 2 | 1 | 0 | 0 | 0 |
| ENSG00000264825 | 0 | 1 | 1 | 0 | 0 | 0 |
| ENSG00000273348 | 0 | 0 | 0 | 0 | 0 | 0 |
| ENSG00000212710 | 0 | 0 | 0 | 0 | 0 | 0 |
| ENSG00000270855 | 0 | 0 | 0 | 0 | 0 | 0 |

| | | | | | | |
|---|---|---|---|---|---|---|
| ENSG00000239490 | 0 | 0 | 1 | 0 | 0 | 0 |
| ENSG00000206827 | 0 | 0 | 0 | 0 | 0 | 0 |
| ENSG00000265943 | 14 | 4 | 9 | 3 | 3 | 7 |
| ENSG00000266850 | 2 | 2 | 1 | 1 | 1 | 2 |
| ENSG00000101773 | 319 | 199 | 209 | 125 | 383 | 202 |
| ENSG00000273232 | 0 | 1 | 1 | 0 | 0 | 1 |
| ENSG00000265939 | 2 | 0 | 1 | 0 | 2 | 2 |
| ENSG00000284331 | 0 | 0 | 0 | 0 | 0 | 0 |
| ENSG00000242182 | 0 | 0 | 0 | 0 | 0 | 0 |
| ENSG00000223023 | 0 | 0 | 0 | 0 | 0 | 0 |
| ENSG00000134508 | 1553 | 818 | 1009 | 1189 | 1395 | 1406 |
| ENSG00000134490 | 783 | 348 | 476 | 379 | 404 | 391 |
| ENSG00000266495 | 1 | 0 | 0 | 0 | 0 | 1 |
| ENSG00000101782 | 1423 | 739 | 809 | 393 | 805 | 552 |
| ENSG00000199357 | 0 | 0 | 0 | 0 | 0 | 0 |
| ENSG00000283084 | 0 | 0 | 0 | 0 | 0 | 0 |
| ENSG00000141452 | 581 | 290 | 403 | 381 | 434 | 399 |
| ENSG00000141458 | 2035 | 940 | 1283 | 1040 | 1348 | 1326 |
| ENSG00000199366 | 0 | 0 | 0 | 0 | 0 | 0 |
| ENSG00000154065 | 23 | 8 | 16 | 68 | 109 | 99 |
| ENSG00000267752 | 0 | 0 | 0 | 0 | 0 | 0 |
| ENSG00000053747 | 342 | 183 | 249 | 376 | 775 | 735 |
| ENSG00000267301 | 0 | 0 | 0 | 0 | 1 | 1 |
| ENSG00000265752 | 0 | 0 | 0 | 0 | 1 | 0 |
| ENSG00000168234 | 319 | 173 | 202 | 200 | 219 | 248 |
| ENSG00000264745 | 0 | 0 | 0 | 0 | 3 | 3 |
| ENSG00000265204 | 3 | 0 | 1 | 0 | 0 | 0 |
| ENSG00000264924 | 2 | 0 | 0 | 1 | 0 | 0 |
| ENSG00000206863 | 0 | 0 | 0 | 0 | 0 | 0 |
| ENSG00000279332 | 7 | 1 | 1 | 3 | 3 | 6 |
| ENSG00000154040 | 66 | 41 | 56 | 42 | 76 | 45 |
| ENSG00000265750 | 1 | 2 | 3 | 1 | 3 | 1 |
| ENSG00000141447 | 674 | 370 | 396 | 250 | 353 | 300 |
| ENSG00000199874 | 0 | 0 | 0 | 0 | 0 | 0 |
| ENSG00000273524 | 0 | 0 | 0 | 0 | 0 | 0 |
| ENSG00000206766 | 0 | 0 | 0 | 0 | 0 | 0 |
| ENSG00000212051 | 0 | 0 | 0 | 0 | 0 | 0 |
| ENSG00000264365 | 1 | 1 | 0 | 0 | 0 | 4 |
| ENSG00000273321 | 0 | 0 | 0 | 0 | 0 | 3 |
| ENSG00000154059 | 1076 | 551 | 610 | 346 | 607 | 578 |
| ENSG00000199636 | 0 | 0 | 0 | 0 | 0 | 0 |
| ENSG00000265094 | 0 | 2 | 0 | 0 | 1 | 0 |
| ENSG00000134489 | 0 | 0 | 0 | 0 | 0 | 0 |
| ENSG00000264695 | 0 | 0 | 0 | 0 | 0 | 0 |
| ENSG00000266489 | 0 | 0 | 0 | 0 | 0 | 0 |
| ENSG00000266181 | 0 | 0 | 0 | 0 | 0 | 0 |
| ENSG00000265485 | 0 | 0 | 0 | 0 | 0 | 0 |
| ENSG00000266520 | 0 | 0 | 0 | 0 | 0 | 0 |
| ENSG00000266809 | 0 | 0 | 0 | 0 | 0 | 0 |
| ENSG00000266573 | 0 | 0 | 0 | 0 | 0 | 0 |

| | | | | | | |
|---|---|---|---|---|---|---|
| ENSG00000264345 | 0 | 0 | 0 | 0 | 0 | 0 |
| ENSG00000229248 | 0 | 0 | 0 | 0 | 0 | 0 |
| ENSG00000265552 | 0 | 0 | 0 | 0 | 0 | 0 |
| ENSG00000198795 | 0 | 2 | 0 | 0 | 0 | 0 |
| ENSG00000285595 | 0 | 0 | 0 | 0 | 0 | 0 |
| ENSG00000279439 | 0 | 0 | 0 | 0 | 0 | 0 |
| ENSG00000264434 | 0 | 0 | 0 | 0 | 0 | 0 |
| ENSG00000242216 | 0 | 0 | 0 | 0 | 0 | 0 |
| ENSG00000264193 | 0 | 0 | 0 | 0 | 1 | 0 |
| ENSG00000141380 | 3396 | 1688 | 2410 | 1339 | 2016 | 1980 |
| ENSG00000243925 | 0 | 0 | 0 | 0 | 0 | 0 |
| ENSG00000154611 | 0 | 0 | 0 | 0 | 0 | 0 |
| ENSG00000188985 | 4 | 5 | 2 | 0 | 4 | 3 |
| ENSG00000266227 | 0 | 0 | 0 | 0 | 0 | 0 |
| ENSG00000141384 | 112 | 84 | 89 | 45 | 85 | 69 |
| ENSG00000263829 | 0 | 2 | 2 | 0 | 1 | 1 |
| ENSG00000266237 | 2 | 0 | 0 | 0 | 0 | 0 |
| ENSG00000264911 | 0 | 0 | 0 | 0 | 0 | 0 |
| ENSG00000207160 | 0 | 0 | 0 | 0 | 0 | 0 |
| ENSG00000263862 | 0 | 0 | 0 | 0 | 0 | 0 |
| ENSG00000134504 | 566 | 264 | 309 | 222 | 303 | 250 |
| ENSG00000277534 | 236 | 112 | 117 | 113 | 127 | 108 |
| ENSG00000275805 | 2 | 0 | 0 | 0 | 0 | 0 |
| ENSG00000276221 | 0 | 0 | 0 | 0 | 0 | 0 |
| ENSG00000263846 | 0 | 0 | 1 | 0 | 0 | 1 |
| ENSG00000260372 | 0 | 0 | 2 | 0 | 0 | 0 |
| ENSG00000265369 | 0 | 0 | 0 | 0 | 0 | 0 |
| ENSG00000263677 | 0 | 0 | 0 | 0 | 0 | 0 |
| ENSG00000275900 | 0 | 0 | 0 | 0 | 0 | 0 |
| ENSG00000263382 | 0 | 0 | 0 | 0 | 0 | 0 |
| ENSG00000266184 | 0 | 0 | 0 | 0 | 0 | 0 |
| ENSG00000171885 | 0 | 0 | 1 | 0 | 0 | 0 |
| ENSG00000154080 | 0 | 0 | 0 | 0 | 0 | 0 |
| ENSG00000266549 | 0 | 0 | 0 | 0 | 0 | 0 |
| ENSG00000265374 | 0 | 0 | 0 | 0 | 0 | 0 |
| ENSG00000264151 | 0 | 0 | 0 | 0 | 0 | 0 |
| ENSG00000264790 | 0 | 0 | 0 | 0 | 0 | 0 |
| ENSG00000244329 | 0 | 0 | 0 | 0 | 0 | 0 |
| ENSG00000263611 | 0 | 0 | 0 | 0 | 0 | 0 |
| ENSG00000265097 | 0 | 0 | 0 | 0 | 0 | 0 |
| ENSG00000263405 | 0 | 0 | 0 | 0 | 0 | 0 |
| ENSG00000170558 | 0 | 0 | 0 | 0 | 0 | 0 |
| ENSG00000227279 | 0 | 0 | 0 | 0 | 0 | 0 |
| ENSG00000274578 | 0 | 0 | 0 | 0 | 0 | 0 |
| ENSG00000265994 | 0 | 0 | 0 | 0 | 0 | 0 |
| ENSG00000266296 | 0 | 0 | 0 | 0 | 0 | 0 |
| ENSG00000271238 | 0 | 0 | 0 | 0 | 0 | 0 |
| ENSG00000251719 | 0 | 0 | 0 | 0 | 0 | 0 |
| ENSG00000266479 | 0 | 0 | 0 | 0 | 0 | 0 |
| ENSG00000265980 | 0 | 0 | 0 | 0 | 0 | 0 |

| | | | | | | |
|---|---|---|---|---|---|---|
| ENSG00000264587 | 0 | 0 | 0 | 0 | 0 | 0 |
| ENSG00000283218 | 0 | 0 | 0 | 0 | 0 | 0 |
| ENSG00000266196 | 0 | 0 | 0 | 0 | 0 | 0 |
| ENSG00000279734 | 0 | 0 | 0 | 0 | 0 | 0 |
| ENSG00000265670 | 0 | 0 | 0 | 0 | 0 | 0 |
| ENSG00000251702 | 0 | 0 | 0 | 0 | 0 | 0 |
| ENSG00000134762 | 0 | 0 | 0 | 0 | 0 | 0 |
| ENSG00000134755 | 214 | 120 | 156 | 98 | 173 | 125 |
| ENSG00000265888 | 0 | 0 | 1 | 0 | 0 | 0 |
| ENSG00000134765 | 3 | 0 | 0 | 0 | 0 | 1 |
| ENSG00000263698 | 0 | 0 | 0 | 0 | 0 | 0 |
| ENSG00000134760 | 0 | 0 | 0 | 0 | 0 | 0 |
| ENSG00000266729 | 0 | 0 | 0 | 0 | 0 | 0 |
| ENSG00000207019 | 0 | 0 | 0 | 0 | 0 | 0 |
| ENSG00000175065 | 0 | 0 | 0 | 0 | 0 | 0 |
| ENSG00000207240 | 0 | 0 | 0 | 0 | 0 | 0 |
| ENSG00000134757 | 0 | 0 | 0 | 0 | 1 | 0 |
| ENSG00000266521 | 14 | 2 | 1 | 2 | 1 | 4 |
| ENSG00000046604 | 1334 | 684 | 667 | 286 | 896 | 514 |
| ENSG00000264859 | 2 | 0 | 1 | 0 | 0 | 0 |
| ENSG00000118271 | 6 | 5 | 2 | 5 | 2 | 3 |
| ENSG00000118276 | 141 | 62 | 81 | 60 | 62 | 70 |
| ENSG00000259985 | 14 | 2 | 1 | 9 | 10 | 2 |
| ENSG00000200472 | 0 | 0 | 0 | 0 | 0 | 0 |
| ENSG00000265158 | 0 | 0 | 0 | 0 | 0 | 0 |
| ENSG00000263772 | 0 | 0 | 0 | 0 | 0 | 0 |
| ENSG00000141437 | 0 | 0 | 0 | 0 | 0 | 2 |
| ENSG00000153339 | 865 | 431 | 383 | 193 | 635 | 419 |
| ENSG00000263924 | 0 | 0 | 0 | 0 | 0 | 0 |
| ENSG00000276486 | 0 | 0 | 0 | 0 | 0 | 0 |
| ENSG00000276282 | 0 | 0 | 0 | 0 | 1 | 1 |
| ENSG00000222320 | 0 | 0 | 0 | 0 | 0 | 0 |
| ENSG00000263823 | 8 | 4 | 7 | 5 | 5 | 3 |
| ENSG00000265273 | 0 | 0 | 2 | 0 | 3 | 0 |
| ENSG00000266105 | 0 | 0 | 0 | 0 | 0 | 0 |
| ENSG00000280380 | 0 | 0 | 0 | 0 | 0 | 0 |
| ENSG00000101695 | 66 | 52 | 58 | 33 | 33 | 45 |
| ENSG00000263917 | 0 | 0 | 0 | 0 | 0 | 0 |
| ENSG00000274275 | 0 | 0 | 0 | 0 | 1 | 0 |
| ENSG00000214917 | 0 | 0 | 0 | 0 | 0 | 0 |
| ENSG00000265008 | 3 | 6 | 3 | 0 | 3 | 3 |
| ENSG00000134758 | 296 | 188 | 185 | 154 | 215 | 207 |
| ENSG00000263393 | 0 | 3 | 3 | 0 | 2 | 0 |
| ENSG00000141441 | 421 | 197 | 229 | 121 | 208 | 171 |
| ENSG00000199373 | 0 | 0 | 0 | 0 | 0 | 0 |
| ENSG00000141434 | 4 | 0 | 1 | 0 | 1 | 0 |
| ENSG00000265279 | 0 | 0 | 0 | 0 | 0 | 0 |
| ENSG00000263904 | 0 | 0 | 0 | 0 | 0 | 0 |
| ENSG00000222395 | 0 | 0 | 0 | 0 | 0 | 0 |
| ENSG00000264982 | 0 | 0 | 0 | 0 | 0 | 0 |

| | | | | | | |
|---|---|---|---|---|---|---|
| ENSG00000262477 | 0 | 0 | 0 | 1 | 0 | 0 |
| ENSG00000215492 | 1 | 4 | 3 | 3 | 9 | 7 |
| ENSG00000285095 | 0 | 0 | 0 | 1 | 2 | 0 |
| ENSG00000260389 | 0 | 0 | 0 | 0 | 0 | 0 |
| ENSG00000263688 | 0 | 0 | 0 | 0 | 0 | 0 |
| ENSG00000197705 | 0 | 0 | 0 | 0 | 0 | 1 |
| ENSG00000228835 | 0 | 0 | 0 | 2 | 0 | 0 |
| ENSG00000243355 | 0 | 0 | 0 | 0 | 0 | 0 |
| ENSG00000263765 | 1 | 0 | 1 | 0 | 1 | 0 |
| ENSG00000166960 | 0 | 0 | 0 | 0 | 0 | 3 |
| ENSG00000263450 | 0 | 0 | 0 | 0 | 0 | 0 |
| ENSG00000267642 | 0 | 0 | 0 | 0 | 0 | 0 |
| ENSG00000141431 | 0 | 1 | 0 | 0 | 0 | 0 |
| ENSG00000101746 | 0 | 0 | 0 | 0 | 0 | 0 |
| ENSG00000199204 | 0 | 0 | 0 | 0 | 0 | 0 |
| ENSG00000280406 | 0 | 0 | 0 | 0 | 0 | 0 |
| ENSG00000267746 | 0 | 0 | 0 | 0 | 0 | 0 |
| ENSG00000278052 | 0 | 0 | 0 | 0 | 0 | 0 |
| ENSG00000134769 | 2310 | 1192 | 1475 | 970 | 1511 | 1392 |
| ENSG00000267585 | 0 | 0 | 0 | 0 | 0 | 0 |
| ENSG00000268873 | 0 | 0 | 0 | 0 | 1 | 0 |
| ENSG00000278464 | 0 | 0 | 0 | 0 | 0 | 0 |
| ENSG00000166974 | 146 | 81 | 90 | 126 | 154 | 172 |
| ENSG00000280096 | 0 | 0 | 0 | 0 | 0 | 0 |
| ENSG00000274400 | 0 | 0 | 0 | 0 | 0 | 0 |
| ENSG00000186812 | 990 | 517 | 591 | 341 | 499 | 436 |
| ENSG00000186814 | 393 | 143 | 177 | 108 | 158 | 151 |
| ENSG00000279989 | 2 | 2 | 2 | 2 | 7 | 2 |
| ENSG00000274184 | 2 | 0 | 0 | 0 | 0 | 0 |
| ENSG00000268573 | 56 | 24 | 28 | 25 | 30 | 39 |
| ENSG00000257267 | 505 | 291 | 266 | 186 | 375 | 239 |
| ENSG00000274918 | 0 | 2 | 1 | 1 | 3 | 0 |
| ENSG00000172466 | 4160 | 1905 | 2287 | 1347 | 2067 | 1769 |
| ENSG00000186496 | 173 | 109 | 95 | 49 | 92 | 40 |
| ENSG00000239683 | 0 | 0 | 0 | 0 | 0 | 0 |
| ENSG00000275512 | 1 | 1 | 0 | 1 | 0 | 0 |
| ENSG00000267583 | 0 | 0 | 1 | 0 | 1 | 0 |
| ENSG00000267140 | 0 | 0 | 0 | 0 | 0 | 0 |
| ENSG00000153391 | 457 | 227 | 285 | 230 | 224 | 202 |
| ENSG00000279074 | 0 | 0 | 3 | 0 | 0 | 0 |
| ENSG00000141429 | 2126 | 1160 | 1153 | 660 | 1243 | 948 |
| ENSG00000283514 | 0 | 0 | 0 | 0 | 0 | 0 |
| ENSG00000266965 | 0 | 0 | 0 | 0 | 0 | 0 |
| ENSG00000267397 | 0 | 0 | 0 | 0 | 0 | 0 |
| ENSG00000267797 | 0 | 0 | 0 | 0 | 0 | 0 |
| ENSG00000207797 | 0 | 0 | 0 | 0 | 0 | 0 |
| ENSG00000265641 | 0 | 0 | 0 | 0 | 1 | 0 |
| ENSG00000267627 | 2 | 0 | 0 | 1 | 0 | 0 |
| ENSG00000141428 | 365 | 199 | 199 | 187 | 222 | 196 |
| ENSG00000278986 | 6 | 1 | 8 | 0 | 5 | 14 |

| | | | | | | |
|---|---|---|---|---|---|---|
| ENSG00000141425 | 2389 | 1190 | 1459 | 887 | 1333 | 1248 |
| ENSG00000141424 | 62943 | 26724 | 41955 | 20375 | 23313 | 26759 |
| ENSG00000134759 | 9844 | 4320 | 6724 | 3453 | 4025 | 4533 |
| ENSG00000260552 | 34 | 10 | 13 | 14 | 22 | 23 |
| ENSG00000075643 | 393 | 235 | 264 | 247 | 237 | 273 |
| ENSG00000274849 | 15 | 17 | 14 | 6 | 13 | 14 |
| ENSG00000267404 | 2 | 0 | 0 | 0 | 0 | 0 |
| ENSG00000267456 | 0 | 0 | 0 | 0 | 0 | 0 |
| ENSG00000134775 | 6 | 4 | 11 | 12 | 23 | 12 |
| ENSG00000134779 | 2070 | 944 | 1233 | 1062 | 1150 | 1105 |
| ENSG00000150477 | 98 | 58 | 67 | 39 | 42 | 49 |
| ENSG00000267039 | 0 | 0 | 0 | 0 | 0 | 0 |
| ENSG00000267651 | 0 | 1 | 0 | 0 | 0 | 0 |
| ENSG00000101489 | 4 | 4 | 6 | 4 | 5 | 1 |
| ENSG00000267707 | 6 | 0 | 2 | 1 | 1 | 2 |
| ENSG00000267202 | 0 | 0 | 0 | 0 | 0 | 0 |
| ENSG00000279678 | 0 | 0 | 0 | 0 | 0 | 0 |
| ENSG00000279543 | 0 | 0 | 0 | 0 | 0 | 0 |
| ENSG00000285940 | 0 | 0 | 0 | 0 | 0 | 0 |
| ENSG00000266530 | 0 | 0 | 0 | 0 | 0 | 0 |
| ENSG00000243516 | 0 | 0 | 0 | 0 | 0 | 0 |
| ENSG00000268566 | 0 | 0 | 0 | 0 | 0 | 0 |
| ENSG00000222704 | 0 | 0 | 0 | 0 | 0 | 0 |
| ENSG00000253038 | 0 | 0 | 0 | 0 | 0 | 0 |
| ENSG00000267374 | 24 | 16 | 13 | 11 | 23 | 24 |
| ENSG00000283458 | 0 | 0 | 0 | 0 | 0 | 0 |
| ENSG00000175886 | 2 | 2 | 4 | 6 | 1 | 2 |
| ENSG00000283615 | 0 | 0 | 0 | 0 | 0 | 0 |
| ENSG00000283645 | 0 | 0 | 0 | 0 | 0 | 0 |
| ENSG00000266148 | 0 | 0 | 0 | 0 | 0 | 0 |
| ENSG00000212354 | 0 | 0 | 0 | 0 | 0 | 0 |
| ENSG00000267413 | 0 | 0 | 0 | 0 | 0 | 0 |
| ENSG00000283503 | 0 | 0 | 0 | 0 | 0 | 0 |
| ENSG00000279034 | 0 | 0 | 0 | 0 | 0 | 0 |
| ENSG00000261715 | 0 | 0 | 0 | 0 | 0 | 0 |
| ENSG00000267664 | 0 | 0 | 0 | 0 | 0 | 0 |
| ENSG00000267313 | 0 | 0 | 0 | 0 | 0 | 0 |
| ENSG00000267652 | 0 | 0 | 0 | 0 | 0 | 0 |
| ENSG00000225253 | 0 | 0 | 0 | 0 | 0 | 0 |
| ENSG00000078142 | 350 | 198 | 215 | 160 | 270 | 198 |
| ENSG00000276174 | 0 | 0 | 0 | 0 | 0 | 0 |
| ENSG00000267088 | 0 | 0 | 0 | 0 | 0 | 0 |
| ENSG00000267586 | 0 | 0 | 0 | 0 | 1 | 0 |
| ENSG00000253040 | 0 | 0 | 0 | 0 | 0 | 0 |
| ENSG00000152214 | 0 | 0 | 0 | 0 | 0 | 0 |
| ENSG00000132872 | 0 | 0 | 0 | 0 | 0 | 0 |
| ENSG00000278933 | 0 | 0 | 0 | 0 | 0 | 0 |
| ENSG00000267716 | 0 | 0 | 0 | 0 | 0 | 0 |
| ENSG00000202060 | 0 | 0 | 0 | 0 | 0 | 0 |
| ENSG00000285550 | 0 | 0 | 0 | 0 | 0 | 0 |

| | | | | | | |
|---|---|---|---|---|---|---|
| ENSG00000267573 | 4 | 3 | 3 | 0 | 2 | 0 |
| ENSG00000267337 | 0 | 0 | 0 | 0 | 0 | 0 |
| ENSG00000277899 | 0 | 0 | 0 | 0 | 0 | 0 |
| ENSG00000267414 | 8 | 5 | 2 | 2 | 3 | 2 |
| ENSG00000152217 | 61 | 20 | 29 | 26 | 34 | 20 |
| ENSG00000280365 | 0 | 0 | 0 | 0 | 0 | 0 |
| ENSG00000265957 | 0 | 0 | 0 | 0 | 0 | 0 |
| ENSG00000279319 | 0 | 0 | 0 | 0 | 0 | 0 |
| ENSG00000267101 | 0 | 1 | 0 | 0 | 0 | 0 |
| ENSG00000132874 | 0 | 1 | 3 | 0 | 2 | 0 |
| ENSG00000267097 | 0 | 0 | 0 | 0 | 0 | 0 |
| ENSG00000266988 | 0 | 0 | 0 | 0 | 0 | 0 |
| ENSG00000285790 | 0 | 0 | 0 | 0 | 0 | 0 |
| ENSG00000267354 | 0 | 0 | 0 | 0 | 0 | 0 |
| ENSG00000266968 | 0 | 0 | 0 | 0 | 0 | 0 |
| ENSG00000267193 | 2 | 4 | 1 | 4 | 0 | 1 |
| ENSG00000141469 | 145 | 69 | 73 | 35 | 42 | 79 |
| ENSG00000267558 | 0 | 1 | 0 | 0 | 0 | 0 |
| ENSG00000197046 | 282 | 173 | 203 | 114 | 132 | 208 |
| ENSG00000152223 | 820 | 402 | 452 | 207 | 501 | 398 |
| ENSG00000152229 | 190 | 88 | 93 | 54 | 91 | 68 |
| ENSG00000222179 | 0 | 0 | 0 | 0 | 0 | 0 |
| ENSG00000267293 | 4 | 1 | 0 | 0 | 0 | 0 |
| ENSG00000252034 | 0 | 0 | 0 | 0 | 0 | 3 |
| ENSG00000152234 | 12470 | 6385 | 7700 | 6630 | 6912 | 6756 |
| ENSG00000152240 | 508 | 259 | 324 | 254 | 277 | 296 |
| ENSG00000251939 | 0 | 0 | 0 | 0 | 0 | 0 |
| ENSG00000152242 | 1168 | 503 | 652 | 355 | 533 | 481 |
| ENSG00000141622 | 24 | 6 | 11 | 11 | 10 | 11 |
| ENSG00000279829 | 0 | 0 | 0 | 0 | 0 | 0 |
| ENSG00000285993 | 0 | 0 | 0 | 0 | 0 | 0 |
| ENSG00000167210 | 0 | 0 | 0 | 0 | 0 | 0 |
| ENSG00000101638 | 0 | 1 | 1 | 0 | 0 | 2 |
| ENSG00000240983 | 0 | 0 | 0 | 0 | 0 | 0 |
| ENSG00000270112 | 0 | 0 | 0 | 0 | 0 | 0 |
| ENSG00000274776 | 8 | 0 | 0 | 0 | 0 | 0 |
| ENSG00000078043 | 417 | 223 | 191 | 162 | 224 | 200 |
| ENSG00000267587 | 0 | 0 | 0 | 0 | 0 | 0 |
| ENSG00000279354 | 0 | 0 | 0 | 0 | 0 | 0 |
| ENSG00000167216 | 59 | 36 | 51 | 31 | 56 | 46 |
| ENSG00000274744 | 0 | 0 | 0 | 0 | 0 | 0 |
| ENSG00000275553 | 0 | 0 | 0 | 0 | 0 | 0 |
| ENSG00000278674 | 0 | 0 | 0 | 0 | 0 | 0 |
| ENSG00000183791 | 0 | 0 | 0 | 0 | 0 | 0 |
| ENSG00000206181 | 0 | 0 | 0 | 0 | 0 | 0 |
| ENSG00000280212 | 0 | 0 | 0 | 0 | 3 | 2 |
| ENSG00000266957 | 0 | 1 | 1 | 0 | 0 | 0 |
| ENSG00000267724 | 4 | 0 | 0 | 0 | 0 | 0 |
| ENSG00000167220 | 18 | 21 | 44 | 43 | 41 | 35 |
| ENSG00000267228 | 0 | 0 | 0 | 0 | 0 | 0 |

| | | | | | | |
|---|---|---|---|---|---|---|
| ENSG00000267800 | 7 | 1 | 3 | 2 | 0 | 1 |
| ENSG00000134049 | 517 | 273 | 336 | 257 | 331 | 246 |
| ENSG00000215474 | 0 | 0 | 0 | 0 | 0 | 0 |
| ENSG00000201825 | 0 | 0 | 0 | 0 | 0 | 0 |
| ENSG00000267761 | 0 | 0 | 0 | 0 | 0 | 0 |
| ENSG00000266582 | 0 | 0 | 0 | 0 | 0 | 0 |
| ENSG00000261307 | 0 | 0 | 0 | 0 | 0 | 0 |
| ENSG00000279367 | 0 | 0 | 0 | 0 | 0 | 0 |
| ENSG00000267156 | 0 | 2 | 0 | 0 | 0 | 1 |
| ENSG00000267187 | 0 | 0 | 0 | 0 | 0 | 0 |
| ENSG00000175387 | 1465 | 714 | 719 | 649 | 961 | 787 |
| ENSG00000267541 | 4 | 5 | 2 | 2 | 4 | 1 |
| ENSG00000279707 | 0 | 0 | 0 | 0 | 0 | 1 |
| ENSG00000269365 | 0 | 0 | 0 | 0 | 1 | 0 |
| ENSG00000279366 | 0 | 0 | 0 | 0 | 0 | 0 |
| ENSG00000184828 | 0 | 3 | 2 | 0 | 0 | 3 |
| ENSG00000278891 | 0 | 0 | 0 | 0 | 0 | 0 |
| ENSG00000206944 | 0 | 0 | 0 | 0 | 0 | 0 |
| ENSG00000280197 | 0 | 0 | 0 | 0 | 0 | 0 |
| ENSG00000200872 | 0 | 0 | 0 | 0 | 0 | 0 |
| ENSG00000277907 | 0 | 0 | 0 | 0 | 0 | 0 |
| ENSG00000266984 | 0 | 0 | 0 | 0 | 0 | 0 |
| ENSG00000134030 | 530 | 238 | 312 | 305 | 298 | 291 |
| ENSG00000278983 | 6 | 1 | 0 | 0 | 0 | 0 |
| ENSG00000279629 | 0 | 0 | 0 | 0 | 0 | 0 |
| ENSG00000266276 | 0 | 0 | 0 | 0 | 0 | 0 |
| ENSG00000267762 | 0 | 0 | 0 | 0 | 0 | 0 |
| ENSG00000279250 | 0 | 0 | 0 | 0 | 0 | 0 |
| ENSG00000267764 | 2 | 0 | 0 | 0 | 0 | 0 |
| ENSG00000101665 | 482 | 187 | 216 | 211 | 233 | 213 |
| ENSG00000266905 | 2 | 0 | 0 | 0 | 0 | 0 |
| ENSG00000280335 | 4 | 4 | 2 | 0 | 3 | 0 |
| ENSG00000264269 | 6 | 4 | 3 | 6 | 6 | 6 |
| ENSG00000141627 | 955 | 454 | 559 | 511 | 611 | 538 |
| ENSG00000263849 | 0 | 0 | 0 | 0 | 0 | 0 |
| ENSG00000265128 | 0 | 0 | 0 | 0 | 0 | 0 |
| ENSG00000266696 | 0 | 0 | 0 | 0 | 0 | 0 |
| ENSG00000263895 | 0 | 0 | 0 | 0 | 0 | 0 |
| ENSG00000264971 | 0 | 0 | 0 | 0 | 0 | 0 |
| ENSG00000177576 | 75 | 62 | 63 | 48 | 76 | 56 |
| ENSG00000215472 | 0 | 1 | 3 | 5 | 2 | 1 |
| ENSG00000263916 | 6 | 1 | 2 | 8 | 1 | 3 |
| ENSG00000265496 | 0 | 0 | 6 | 2 | 0 | 1 |
| ENSG00000284239 | 0 | 0 | 0 | 0 | 0 | 0 |
| ENSG00000265681 | 217 | 682 | 809 | 953 | 937 | 904 |
| ENSG00000202093 | 0 | 0 | 0 | 0 | 0 | 0 |
| ENSG00000206602 | 0 | 0 | 0 | 0 | 0 | 0 |
| ENSG00000271982 | 0 | 0 | 0 | 0 | 0 | 0 |
| ENSG00000265182 | 0 | 0 | 0 | 0 | 0 | 0 |
| ENSG00000264464 | 0 | 0 | 0 | 0 | 0 | 0 |

| | | | | | | |
|---|---|---|---|---|---|---|
| ENSG00000101670 | 2 | 3 | 4 | 2 | 3 | 2 |
| ENSG00000267444 | 0 | 0 | 0 | 0 | 0 | 0 |
| ENSG00000267674 | 0 | 0 | 0 | 0 | 0 | 0 |
| ENSG00000167315 | 1844 | 1041 | 1153 | 664 | 772 | 738 |
| ENSG00000266997 | 0 | 0 | 0 | 0 | 0 | 0 |
| ENSG00000267322 | 11 | 4 | 4 | 0 | 3 | 4 |
| ENSG00000251992 | 0 | 0 | 0 | 0 | 0 | 0 |
| ENSG00000252139 | 0 | 0 | 0 | 0 | 0 | 0 |
| ENSG00000167306 | 1865 | 896 | 1114 | 796 | 1090 | 1030 |
| ENSG00000201668 | 0 | 0 | 0 | 0 | 0 | 0 |
| ENSG00000200527 | 0 | 0 | 0 | 0 | 0 | 0 |
| ENSG00000267196 | 0 | 1 | 0 | 0 | 0 | 0 |
| ENSG00000267511 | 0 | 0 | 0 | 0 | 0 | 0 |
| ENSG00000267591 | 0 | 0 | 0 | 0 | 0 | 0 |
| ENSG00000283343 | 0 | 0 | 0 | 0 | 0 | 0 |
| ENSG00000240663 | 0 | 0 | 0 | 0 | 0 | 0 |
| ENSG00000172361 | 70 | 37 | 35 | 28 | 25 | 33 |
| ENSG00000277310 | 5 | 1 | 4 | 2 | 2 | 1 |
| ENSG00000141644 | 1571 | 664 | 878 | 713 | 721 | 801 |
| ENSG00000154832 | 1025 | 463 | 578 | 612 | 501 | 545 |
| ENSG00000267695 | 0 | 0 | 0 | 0 | 0 | 0 |
| ENSG00000274930 | 0 | 0 | 0 | 0 | 0 | 0 |
| ENSG00000251997 | 0 | 0 | 0 | 0 | 0 | 0 |
| ENSG00000154839 | 176 | 84 | 98 | 94 | 129 | 146 |
| ENSG00000239780 | 0 | 0 | 0 | 0 | 0 | 1 |
| ENSG00000141639 | 0 | 1 | 11 | 10 | 8 | 11 |
| ENSG00000267713 | 0 | 0 | 0 | 0 | 0 | 0 |
| ENSG00000134042 | 0 | 0 | 0 | 0 | 0 | 0 |
| ENSG00000267233 | 0 | 0 | 0 | 0 | 0 | 0 |
| ENSG00000239689 | 0 | 0 | 0 | 0 | 0 | 0 |
| ENSG00000082212 | 605 | 351 | 372 | 356 | 420 | 391 |
| ENSG00000200752 | 0 | 0 | 0 | 0 | 0 | 0 |
| ENSG00000141642 | 68 | 41 | 55 | 54 | 69 | 83 |
| ENSG00000267699 | 0 | 0 | 0 | 0 | 0 | 0 |
| ENSG00000141646 | 703 | 391 | 424 | 327 | 432 | 364 |
| ENSG00000267717 | 0 | 0 | 0 | 0 | 0 | 0 |
| ENSG00000176624 | 1200 | 597 | 708 | 536 | 643 | 652 |
| ENSG00000207154 | 0 | 0 | 0 | 0 | 0 | 0 |
| ENSG00000227115 | 0 | 0 | 0 | 0 | 0 | 0 |
| ENSG00000266201 | 0 | 0 | 0 | 0 | 0 | 0 |
| ENSG00000266003 | 0 | 0 | 0 | 0 | 0 | 0 |
| ENSG00000265402 | 0 | 0 | 0 | 0 | 0 | 0 |
| ENSG00000265712 | 0 | 0 | 0 | 0 | 0 | 0 |
| ENSG00000215457 | 0 | 0 | 0 | 0 | 0 | 0 |
| ENSG00000285695 | 0 | 0 | 0 | 0 | 0 | 0 |
| ENSG00000266335 | 0 | 0 | 0 | 0 | 0 | 0 |
| ENSG00000187323 | 0 | 0 | 0 | 0 | 0 | 0 |
| ENSG00000280098 | 0 | 0 | 0 | 0 | 0 | 0 |
| ENSG00000273762 | 0 | 0 | 0 | 0 | 0 | 0 |
| ENSG00000263872 | 0 | 0 | 0 | 0 | 0 | 0 |

| | | | | | | |
|---|---|---|---|---|---|---|
| ENSG00000260433 | 0 | 0 | 0 | 0 | 0 | 0 |
| ENSG00000263438 | 0 | 0 | 0 | 0 | 0 | 0 |
| ENSG00000242945 | 0 | 0 | 0 | 0 | 0 | 0 |
| ENSG00000264296 | 0 | 0 | 1 | 0 | 0 | 1 |
| ENSG00000134046 | 1545 | 825 | 1035 | 718 | 803 | 894 |
| ENSG00000207233 | 1 | 1 | 0 | 0 | 0 | 0 |
| ENSG00000277324 | 10 | 8 | 6 | 5 | 2 | 8 |
| ENSG00000101751 | 731 | 382 | 442 | 171 | 469 | 371 |
| ENSG00000174448 | 0 | 0 | 0 | 0 | 0 | 0 |
| ENSG00000166845 | 84 | 48 | 22 | 22 | 59 | 51 |
| ENSG00000264350 | 129 | 69 | 70 | 91 | 81 | 70 |
| ENSG00000283613 | 0 | 0 | 0 | 0 | 0 | 0 |
| ENSG00000264564 | 0 | 0 | 0 | 0 | 0 | 0 |
| ENSG00000267428 | 0 | 0 | 0 | 0 | 0 | 0 |
| ENSG00000178690 | 8 | 4 | 2 | 0 | 3 | 3 |
| ENSG00000041353 | 1264 | 767 | 733 | 248 | 819 | 489 |
| ENSG00000267524 | 0 | 0 | 0 | 0 | 0 | 0 |
| ENSG00000267311 | 0 | 1 | 0 | 0 | 0 | 0 |
| ENSG00000267112 | 1 | 0 | 1 | 0 | 0 | 0 |
| ENSG00000166510 | 218 | 124 | 146 | 64 | 134 | 126 |
| ENSG00000267647 | 0 | 0 | 0 | 0 | 0 | 0 |
| ENSG00000267013 | 0 | 0 | 0 | 0 | 0 | 0 |
| ENSG00000252437 | 0 | 0 | 0 | 0 | 0 | 0 |
| ENSG00000196628 | 0 | 0 | 0 | 0 | 0 | 0 |
| ENSG00000267028 | 0 | 0 | 0 | 0 | 0 | 0 |
| ENSG00000264571 | 0 | 0 | 0 | 0 | 0 | 0 |
| ENSG00000267402 | 0 | 0 | 0 | 0 | 0 | 0 |
| ENSG00000242320 | 0 | 0 | 0 | 0 | 0 | 0 |
| ENSG00000267284 | 0 | 0 | 0 | 0 | 0 | 0 |
| ENSG00000267325 | 0 | 0 | 0 | 0 | 0 | 0 |
| ENSG00000267172 | 0 | 0 | 0 | 0 | 0 | 0 |
| ENSG00000260930 | 0 | 0 | 0 | 0 | 0 | 0 |
| ENSG00000267327 | 0 | 0 | 0 | 0 | 0 | 0 |
| ENSG00000206129 | 0 | 0 | 0 | 0 | 0 | 0 |
| ENSG00000267057 | 0 | 0 | 0 | 0 | 0 | 0 |
| ENSG00000267399 | 0 | 0 | 0 | 0 | 0 | 0 |
| ENSG00000201816 | 0 | 0 | 0 | 0 | 0 | 0 |
| ENSG00000267712 | 0 | 0 | 0 | 0 | 0 | 0 |
| ENSG00000267732 | 0 | 0 | 0 | 0 | 0 | 0 |
| ENSG00000091164 | 2191 | 1037 | 1294 | 1070 | 1183 | 1299 |
| ENSG00000091157 | 353 | 191 | 200 | 104 | 191 | 175 |
| ENSG00000206865 | 0 | 0 | 0 | 0 | 0 | 0 |
| ENSG00000267146 | 0 | 0 | 0 | 0 | 0 | 0 |
| ENSG00000212539 | 0 | 0 | 0 | 0 | 0 | 0 |
| ENSG00000267225 | 0 | 0 | 0 | 0 | 0 | 0 |
| ENSG00000258609 | 0 | 0 | 0 | 0 | 0 | 0 |
| ENSG00000267789 | 0 | 0 | 0 | 0 | 0 | 0 |
| ENSG00000228075 | 0 | 0 | 0 | 0 | 1 | 0 |
| ENSG00000267726 | 0 | 0 | 0 | 0 | 0 | 0 |
| ENSG00000202240 | 0 | 0 | 0 | 0 | 0 | 0 |

| | | | | | | |
|---|---|---|---|---|---|---|
| ENSG00000177511 | 0 | 0 | 0 | 0 | 0 | 0 |
| ENSG00000277837 | 0 | 0 | 0 | 0 | 0 | 0 |
| ENSG00000119547 | 158 | 67 | 63 | 38 | 91 | 73 |
| ENSG00000066926 | 2081 | 1068 | 1403 | 1316 | 1507 | 1641 |
| ENSG00000278703 | 2 | 0 | 0 | 0 | 1 | 0 |
| ENSG00000134440 | 3440 | 1900 | 2256 | 1592 | 2286 | 1923 |
| ENSG00000267040 | 31 | 8 | 14 | 6 | 10 | 20 |
| ENSG00000267787 | 6 | 3 | 4 | 4 | 2 | 7 |
| ENSG00000081923 | 2633 | 1232 | 1572 | 796 | 1519 | 1335 |
| ENSG00000202159 | 0 | 0 | 0 | 0 | 0 | 0 |
| ENSG00000231544 | 0 | 0 | 0 | 0 | 0 | 0 |
| ENSG00000267209 | 0 | 0 | 0 | 0 | 0 | 0 |
| ENSG00000235028 | 4 | 0 | 0 | 0 | 0 | 0 |
| ENSG00000049759 | 1475 | 659 | 883 | 489 | 630 | 670 |
| ENSG00000267743 | 0 | 0 | 0 | 0 | 0 | 0 |
| ENSG00000267504 | 0 | 1 | 0 | 0 | 0 | 1 |
| ENSG00000267396 | 0 | 0 | 0 | 0 | 0 | 0 |
| ENSG00000267675 | 0 | 0 | 0 | 0 | 0 | 0 |
| ENSG00000267391 | 0 | 0 | 0 | 0 | 0 | 0 |
| ENSG00000284440 | 0 | 0 | 0 | 0 | 0 | 0 |
| ENSG00000207778 | 0 | 0 | 0 | 0 | 0 | 0 |
| ENSG00000276403 | 0 | 0 | 0 | 0 | 0 | 0 |
| ENSG00000198796 | 0 | 0 | 0 | 0 | 0 | 0 |
| ENSG00000274354 | 0 | 0 | 0 | 0 | 0 | 0 |
| ENSG00000267257 | 0 | 0 | 0 | 0 | 0 | 0 |
| ENSG00000267579 | 0 | 0 | 0 | 0 | 0 | 0 |
| ENSG00000252284 | 0 | 0 | 0 | 0 | 0 | 0 |
| ENSG00000239659 | 0 | 0 | 0 | 0 | 0 | 0 |
| ENSG00000275418 | 0 | 0 | 0 | 0 | 0 | 0 |
| ENSG00000267226 | 2 | 0 | 0 | 0 | 0 | 1 |
| ENSG00000172175 | 201 | 116 | 120 | 78 | 141 | 105 |
| ENSG00000267476 | 0 | 0 | 0 | 0 | 0 | 0 |
| ENSG00000266946 | 0 | 0 | 0 | 0 | 0 | 0 |
| ENSG00000267705 | 0 | 0 | 0 | 0 | 0 | 0 |
| ENSG00000267501 | 0 | 0 | 0 | 0 | 0 | 0 |
| ENSG00000201598 | 0 | 0 | 0 | 0 | 0 | 0 |
| ENSG00000267593 | 0 | 0 | 0 | 0 | 0 | 0 |
| ENSG00000199713 | 0 | 0 | 0 | 0 | 0 | 0 |
| ENSG00000074657 | 68 | 35 | 31 | 17 | 48 | 36 |
| ENSG00000251870 | 0 | 0 | 0 | 0 | 0 | 0 |
| ENSG00000224367 | 0 | 0 | 0 | 0 | 0 | 0 |
| ENSG00000267215 | 0 | 0 | 0 | 0 | 0 | 0 |
| ENSG00000166562 | 709 | 352 | 398 | 449 | 412 | 387 |
| ENSG00000134443 | 0 | 0 | 0 | 0 | 0 | 0 |
| ENSG00000134438 | 0 | 0 | 0 | 0 | 0 | 0 |
| ENSG00000166569 | 0 | 0 | 0 | 0 | 0 | 0 |
| ENSG00000074695 | 1756 | 945 | 919 | 475 | 1143 | 808 |
| ENSG00000267677 | 0 | 2 | 0 | 1 | 0 | 1 |
| ENSG00000183287 | 0 | 0 | 0 | 0 | 1 | 0 |
| ENSG00000267000 | 0 | 0 | 0 | 0 | 0 | 0 |

| | | | | | | |
|---|---|---|---|---|---|---|
| ENSG00000267061 | 0 | 0 | 0 | 0 | 0 | 0 |
| ENSG00000267269 | 0 | 0 | 0 | 0 | 0 | 0 |
| ENSG00000267774 | 0 | 0 | 0 | 0 | 0 | 0 |
| ENSG00000273664 | 0 | 0 | 0 | 0 | 0 | 0 |
| ENSG00000224161 | 0 | 0 | 0 | 0 | 0 | 0 |
| ENSG00000267486 | 0 | 0 | 0 | 0 | 0 | 0 |
| ENSG00000267669 | 0 | 0 | 0 | 0 | 0 | 0 |
| ENSG00000141682 | 90 | 40 | 43 | 38 | 26 | 72 |
| ENSG00000267066 | 0 | 0 | 0 | 0 | 0 | 0 |
| ENSG00000267497 | 0 | 0 | 0 | 0 | 0 | 0 |
| ENSG00000239398 | 0 | 0 | 0 | 0 | 0 | 0 |
| ENSG00000256704 | 0 | 0 | 0 | 0 | 0 | 0 |
| ENSG00000267462 | 0 | 0 | 0 | 0 | 0 | 0 |
| ENSG00000267742 | 2 | 0 | 0 | 0 | 1 | 0 |
| ENSG00000252555 | 0 | 0 | 0 | 0 | 0 | 0 |
| ENSG00000267686 | 0 | 0 | 0 | 0 | 0 | 0 |
| ENSG00000242060 | 1 | 0 | 1 | 1 | 0 | 1 |
| ENSG00000202468 | 0 | 0 | 0 | 0 | 0 | 0 |
| ENSG00000267620 | 0 | 0 | 0 | 0 | 0 | 0 |
| ENSG00000267401 | 0 | 0 | 0 | 0 | 0 | 0 |
| ENSG00000285681 | 0 | 0 | 0 | 0 | 0 | 0 |
| ENSG00000166603 | 0 | 0 | 1 | 0 | 0 | 0 |
| ENSG00000267400 | 0 | 0 | 0 | 0 | 0 | 0 |
| ENSG00000266915 | 0 | 0 | 0 | 0 | 0 | 0 |
| ENSG00000267687 | 0 | 0 | 0 | 0 | 0 | 0 |
| ENSG00000267382 | 0 | 0 | 0 | 0 | 0 | 0 |
| ENSG00000267098 | 0 | 0 | 0 | 0 | 0 | 0 |
| ENSG00000267153 | 0 | 0 | 0 | 0 | 0 | 0 |
| ENSG00000267513 | 0 | 0 | 0 | 0 | 0 | 0 |
| ENSG00000101542 | 0 | 0 | 0 | 0 | 1 | 0 |
| ENSG00000206769 | 0 | 0 | 0 | 0 | 0 | 0 |
| ENSG00000243256 | 0 | 0 | 0 | 0 | 0 | 0 |
| ENSG00000267316 | 0 | 0 | 0 | 0 | 0 | 0 |
| ENSG00000267279 | 2 | 0 | 0 | 0 | 0 | 0 |
| ENSG00000267175 | 0 | 0 | 2 | 0 | 0 | 0 |
| ENSG00000267038 | 0 | 0 | 0 | 0 | 0 | 0 |
| ENSG00000260440 | 0 | 0 | 0 | 0 | 0 | 0 |
| ENSG00000176641 | 303 | 155 | 179 | 98 | 155 | 126 |
| ENSG00000197563 | 594 | 293 | 330 | 217 | 326 | 268 |
| ENSG00000267597 | 0 | 0 | 0 | 0 | 0 | 0 |
| ENSG00000134444 | 851 | 450 | 493 | 278 | 516 | 409 |
| ENSG00000266900 | 0 | 0 | 0 | 1 | 0 | 0 |
| ENSG00000267560 | 2 | 1 | 1 | 1 | 3 | 1 |
| ENSG00000141655 | 134 | 66 | 98 | 83 | 96 | 118 |
| ENSG00000199867 | 0 | 0 | 0 | 0 | 0 | 0 |
| ENSG00000267341 | 0 | 0 | 2 | 1 | 0 | 0 |
| ENSG00000241088 | 0 | 0 | 0 | 0 | 0 | 0 |
| ENSG00000267487 | 0 | 0 | 0 | 0 | 0 | 0 |
| ENSG00000266920 | 0 | 0 | 0 | 0 | 0 | 0 |
| ENSG00000141664 | 628 | 246 | 391 | 221 | 327 | 342 |

| | | | | | | |
|---|---|---|---|---|---|---|
| ENSG00000278017 | 0 | 1 | 1 | 1 | 0 | 2 |
| ENSG00000279236 | 0 | 0 | 0 | 0 | 0 | 0 |
| ENSG00000243549 | 0 | 0 | 0 | 0 | 0 | 0 |
| ENSG00000081913 | 420 | 202 | 267 | 185 | 239 | 223 |
| ENSG00000206746 | 0 | 0 | 0 | 0 | 0 | 0 |
| ENSG00000171791 | 52 | 12 | 24 | 22 | 23 | 31 |
| ENSG00000267766 | 0 | 0 | 0 | 0 | 0 | 0 |
| ENSG00000283125 | 0 | 0 | 0 | 0 | 0 | 0 |
| ENSG00000119537 | 634 | 246 | 390 | 273 | 327 | 340 |
| ENSG00000267390 | 10 | 3 | 5 | 4 | 6 | 6 |
| ENSG00000119541 | 1502 | 663 | 894 | 458 | 815 | 730 |
| ENSG00000267430 | 2 | 0 | 0 | 0 | 0 | 0 |
| ENSG00000206075 | 12 | 7 | 8 | 2 | 7 | 7 |
| ENSG00000269989 | 0 | 0 | 0 | 0 | 0 | 0 |
| ENSG00000214318 | 0 | 0 | 0 | 0 | 0 | 0 |
| ENSG00000166634 | 0 | 0 | 0 | 0 | 0 | 0 |
| ENSG00000197641 | 0 | 0 | 0 | 0 | 0 | 0 |
| ENSG00000206073 | 0 | 0 | 0 | 0 | 0 | 0 |
| ENSG00000206072 | 0 | 0 | 0 | 0 | 0 | 0 |
| ENSG00000057149 | 0 | 0 | 0 | 0 | 0 | 1 |
| ENSG00000166396 | 0 | 0 | 0 | 0 | 0 | 0 |
| ENSG00000197632 | 0 | 0 | 0 | 0 | 0 | 0 |
| ENSG00000242550 | 0 | 0 | 0 | 0 | 0 | 0 |
| ENSG00000221887 | 0 | 0 | 0 | 0 | 0 | 1 |
| ENSG00000283667 | 14 | 3 | 5 | 3 | 5 | 4 |
| ENSG00000166401 | 34 | 25 | 28 | 29 | 35 | 46 |
| ENSG00000267134 | 0 | 0 | 0 | 0 | 0 | 0 |
| ENSG00000237314 | 0 | 0 | 0 | 0 | 0 | 0 |
| ENSG00000179676 | 0 | 0 | 0 | 0 | 0 | 0 |
| ENSG00000266952 | 0 | 0 | 0 | 0 | 0 | 0 |
| ENSG00000275448 | 0 | 0 | 0 | 0 | 0 | 0 |
| ENSG00000263863 | 0 | 0 | 0 | 0 | 0 | 0 |
| ENSG00000264131 | 0 | 0 | 0 | 0 | 0 | 0 |
| ENSG00000264232 | 0 | 0 | 0 | 0 | 0 | 0 |
| ENSG00000265217 | 0 | 0 | 0 | 0 | 0 | 0 |
| ENSG00000081138 | 10 | 0 | 1 | 1 | 2 | 2 |
| ENSG00000263720 | 0 | 0 | 0 | 0 | 0 | 0 |
| ENSG00000263622 | 0 | 0 | 0 | 0 | 0 | 0 |
| ENSG00000264685 | 0 | 0 | 0 | 0 | 0 | 0 |
| ENSG00000071991 | 0 | 0 | 0 | 0 | 0 | 0 |
| ENSG00000238680 | 0 | 0 | 0 | 0 | 0 | 0 |
| ENSG00000263354 | 0 | 0 | 0 | 0 | 0 | 0 |
| ENSG00000264054 | 0 | 0 | 0 | 0 | 0 | 0 |
| ENSG00000265936 | 0 | 0 | 0 | 0 | 1 | 1 |
| ENSG00000260578 | 0 | 0 | 1 | 0 | 0 | 0 |
| ENSG00000171451 | 2 | 0 | 3 | 2 | 4 | 6 |
| ENSG00000263424 | 0 | 0 | 0 | 0 | 0 | 0 |
| ENSG00000265533 | 0 | 0 | 0 | 0 | 0 | 0 |
| ENSG00000264040 | 0 | 0 | 0 | 0 | 0 | 0 |
| ENSG00000263594 | 0 | 0 | 0 | 0 | 0 | 0 |

| | | | | | | |
|---|---|---|---|---|---|---|
| ENSG00000265472 | 0 | 0 | 0 | 0 | 0 | 0 |
| ENSG00000264104 | 0 | 0 | 0 | 0 | 0 | 0 |
| ENSG00000265555 | 0 | 0 | 0 | 0 | 0 | 0 |
| ENSG00000265240 | 0 | 0 | 0 | 0 | 0 | 0 |
| ENSG00000264634 | 0 | 0 | 0 | 0 | 0 | 0 |
| ENSG00000264869 | 0 | 0 | 0 | 0 | 0 | 0 |
| ENSG00000264699 | 0 | 0 | 0 | 0 | 0 | 0 |
| ENSG00000260747 | 12 | 19 | 14 | 15 | 26 | 7 |
| ENSG00000265425 | 0 | 0 | 0 | 0 | 0 | 0 |
| ENSG00000263355 | 0 | 0 | 0 | 0 | 0 | 0 |
| ENSG00000264697 | 0 | 0 | 0 | 0 | 0 | 0 |
| ENSG00000166479 | 499 | 297 | 278 | 161 | 394 | 343 |
| ENSG00000150636 | 2 | 1 | 3 | 1 | 2 | 2 |
| ENSG00000276092 | 0 | 0 | 2 | 0 | 0 | 0 |
| ENSG00000263637 | 0 | 0 | 0 | 0 | 0 | 0 |
| ENSG00000207072 | 0 | 0 | 0 | 0 | 2 | 0 |
| ENSG00000264444 | 0 | 0 | 0 | 0 | 0 | 0 |
| ENSG00000264472 | 0 | 0 | 0 | 0 | 0 | 0 |
| ENSG00000273584 | 0 | 0 | 0 | 0 | 0 | 0 |
| ENSG00000264236 | 0 | 0 | 0 | 0 | 0 | 0 |
| ENSG00000264705 | 0 | 0 | 0 | 0 | 0 | 0 |
| ENSG00000285748 | 0 | 0 | 0 | 0 | 0 | 0 |
| ENSG00000278532 | 0 | 0 | 0 | 0 | 0 | 0 |
| ENSG00000206052 | 0 | 0 | 0 | 0 | 0 | 0 |
| ENSG00000264845 | 0 | 0 | 0 | 0 | 0 | 0 |
| ENSG00000265643 | 0 | 0 | 0 | 0 | 0 | 0 |
| ENSG00000279221 | 0 | 0 | 0 | 0 | 0 | 0 |
| ENSG00000266840 | 0 | 0 | 0 | 0 | 0 | 0 |
| ENSG00000150637 | 0 | 0 | 1 | 0 | 0 | 0 |
| ENSG00000176225 | 137 | 78 | 81 | 43 | 85 | 69 |
| ENSG00000170677 | 394 | 215 | 237 | 119 | 250 | 198 |
| ENSG00000266258 | 7 | 0 | 0 | 4 | 2 | 3 |
| ENSG00000266304 | 0 | 1 | 1 | 1 | 2 | 2 |
| ENSG00000266278 | 0 | 0 | 0 | 0 | 0 | 0 |
| ENSG00000265698 | 0 | 0 | 0 | 0 | 0 | 0 |
| ENSG00000214255 | 2 | 0 | 0 | 0 | 0 | 0 |
| ENSG00000278971 | 2 | 0 | 0 | 0 | 0 | 0 |
| ENSG00000239748 | 0 | 0 | 0 | 0 | 0 | 0 |
| ENSG00000263417 | 0 | 0 | 0 | 0 | 0 | 0 |
| ENSG00000265781 | 0 | 0 | 0 | 0 | 0 | 0 |
| ENSG00000265639 | 0 | 0 | 0 | 0 | 0 | 0 |
| ENSG00000265484 | 0 | 0 | 0 | 0 | 0 | 0 |
| ENSG00000263733 | 0 | 0 | 0 | 0 | 0 | 0 |
| ENSG00000260676 | 0 | 0 | 0 | 0 | 0 | 0 |
| ENSG00000264388 | 0 | 0 | 0 | 0 | 0 | 0 |
| ENSG00000265946 | 0 | 0 | 0 | 0 | 0 | 0 |
| ENSG00000265352 | 0 | 0 | 0 | 0 | 0 | 0 |
| ENSG00000260457 | 0 | 0 | 0 | 0 | 0 | 0 |
| ENSG00000265995 | 0 | 0 | 0 | 0 | 0 | 0 |
| ENSG00000271313 | 0 | 0 | 0 | 0 | 0 | 0 |

| | | | | | | |
|---|---|---|---|---|---|---|
| ENSG00000141668 | 0 | 0 | 0 | 1 | 0 | 0 |
| ENSG00000264315 | 0 | 0 | 0 | 0 | 0 | 0 |
| ENSG00000166342 | 0 | 0 | 0 | 0 | 0 | 0 |
| ENSG00000265579 | 0 | 0 | 0 | 0 | 0 | 0 |
| ENSG00000252598 | 0 | 0 | 0 | 0 | 0 | 0 |
| ENSG00000263750 | 0 | 0 | 0 | 0 | 0 | 0 |
| ENSG00000263958 | 0 | 0 | 0 | 0 | 0 | 0 |
| ENSG00000263711 | 0 | 0 | 1 | 0 | 0 | 0 |
| ENSG00000261780 | 0 | 0 | 0 | 0 | 0 | 0 |
| ENSG00000265380 | 0 | 0 | 0 | 0 | 0 | 0 |
| ENSG00000241866 | 0 | 0 | 0 | 0 | 0 | 0 |
| ENSG00000263655 | 0 | 0 | 0 | 0 | 0 | 0 |
| ENSG00000278902 | 0 | 0 | 0 | 0 | 0 | 0 |
| ENSG00000141665 | 83 | 26 | 56 | 45 | 30 | 34 |
| ENSG00000075336 | 406 | 190 | 224 | 187 | 238 | 233 |
| ENSG00000260569 | 0 | 0 | 0 | 0 | 0 | 0 |
| ENSG00000243856 | 0 | 0 | 0 | 0 | 0 | 0 |
| ENSG00000166347 | 1569 | 760 | 993 | 982 | 957 | 927 |
| ENSG00000206043 | 0 | 0 | 0 | 0 | 0 | 0 |
| ENSG00000235297 | 4 | 0 | 2 | 2 | 0 | 3 |
| ENSG00000259779 | 0 | 0 | 0 | 0 | 0 | 0 |
| ENSG00000187773 | 14 | 9 | 7 | 6 | 1 | 7 |
| ENSG00000133313 | 4901 | 2343 | 3249 | 2466 | 2830 | 3065 |
| ENSG00000276934 | 6 | 1 | 1 | 1 | 1 | 2 |
| ENSG00000150656 | 22 | 11 | 24 | 14 | 16 | 28 |
| ENSG00000264340 | 0 | 0 | 0 | 0 | 0 | 0 |
| ENSG00000264247 | 206 | 104 | 142 | 88 | 154 | 136 |
| ENSG00000215421 | 280 | 154 | 181 | 85 | 178 | 147 |
| ENSG00000278607 | 0 | 0 | 0 | 0 | 1 | 1 |
| ENSG00000273669 | 0 | 0 | 1 | 1 | 3 | 2 |
| ENSG00000283452 | 0 | 0 | 0 | 0 | 0 | 0 |
| ENSG00000278911 | 0 | 0 | 0 | 0 | 0 | 0 |
| ENSG00000285915 | 0 | 0 | 0 | 0 | 0 | 0 |
| ENSG00000180011 | 1128 | 464 | 693 | 551 | 659 | 647 |
| ENSG00000179981 | 408 | 178 | 226 | 174 | 197 | 223 |
| ENSG00000276993 | 0 | 0 | 0 | 0 | 0 | 2 |
| ENSG00000264116 | 0 | 0 | 0 | 0 | 0 | 0 |
| ENSG00000206026 | 0 | 0 | 0 | 0 | 0 | 0 |
| ENSG00000266774 | 0 | 0 | 0 | 0 | 0 | 0 |
| ENSG00000261194 | 0 | 0 | 0 | 0 | 0 | 0 |
| ENSG00000266460 | 0 | 0 | 0 | 0 | 0 | 0 |
| ENSG00000275827 | 0 | 0 | 0 | 0 | 0 | 0 |
| ENSG00000264108 | 0 | 0 | 0 | 0 | 0 | 0 |
| ENSG00000263547 | 0 | 0 | 0 | 0 | 0 | 0 |
| ENSG00000266743 | 0 | 0 | 0 | 0 | 0 | 0 |
| ENSG00000264260 | 0 | 0 | 0 | 0 | 0 | 0 |
| ENSG00000264595 | 0 | 0 | 0 | 0 | 0 | 0 |
| ENSG00000266014 | 0 | 0 | 0 | 0 | 0 | 0 |
| ENSG00000265717 | 0 | 0 | 0 | 0 | 0 | 0 |
| ENSG00000264212 | 0 | 0 | 0 | 0 | 0 | 0 |

| | | | | | | |
|---|---|---|---|---|---|---|
| ENSG00000266614 | 0 | 0 | 0 | 0 | 0 | 0 |
| ENSG00000101493 | 1209 | 461 | 754 | 542 | 585 | 668 |
| ENSG00000263982 | 0 | 7 | 3 | 0 | 5 | 2 |
| ENSG00000280205 | 2 | 0 | 0 | 0 | 0 | 0 |
| ENSG00000265778 | 61 | 30 | 33 | 40 | 38 | 22 |
| ENSG00000275763 | 24 | 9 | 20 | 15 | 21 | 10 |
| ENSG00000263812 | 0 | 1 | 1 | 1 | 0 | 1 |
| ENSG00000266256 | 9 | 8 | 14 | 9 | 7 | 9 |
| ENSG00000266312 | 0 | 1 | 0 | 0 | 2 | 1 |
| ENSG00000276397 | 0 | 1 | 1 | 0 | 1 | 3 |
| ENSG00000265934 | 0 | 0 | 0 | 0 | 0 | 0 |
| ENSG00000265836 | 0 | 0 | 0 | 0 | 0 | 0 |
| ENSG00000265626 | 0 | 0 | 0 | 0 | 0 | 0 |
| ENSG00000264278 | 35 | 19 | 27 | 24 | 17 | 15 |
| ENSG00000252097 | 0 | 0 | 0 | 0 | 1 | 0 |
| ENSG00000278107 | 17 | 6 | 17 | 19 | 5 | 6 |
| ENSG00000130856 | 442 | 176 | 260 | 113 | 195 | 181 |
| ENSG00000244229 | 0 | 0 | 0 | 0 | 0 | 0 |
| ENSG00000275355 | 0 | 0 | 0 | 0 | 0 | 0 |
| ENSG00000265261 | 0 | 0 | 1 | 0 | 0 | 0 |
| ENSG00000266844 | 2 | 0 | 0 | 0 | 0 | 0 |
| ENSG00000197971 | 372 | 179 | 235 | 128 | 216 | 182 |
| ENSG00000279811 | 0 | 0 | 0 | 0 | 0 | 0 |
| ENSG00000275178 | 0 | 0 | 0 | 0 | 0 | 1 |
| ENSG00000279433 | 0 | 0 | 0 | 0 | 0 | 0 |
| ENSG00000278330 | 0 | 0 | 3 | 0 | 3 | 0 |
| ENSG00000265844 | 0 | 0 | 0 | 0 | 0 | 0 |
| ENSG00000166573 | 0 | 0 | 0 | 0 | 0 | 0 |
| ENSG00000264015 | 0 | 0 | 0 | 0 | 0 | 0 |
| ENSG00000199392 | 0 | 0 | 0 | 0 | 0 | 0 |
| ENSG00000264736 | 0 | 0 | 0 | 0 | 0 | 0 |
| ENSG00000264693 | 0 | 0 | 0 | 0 | 0 | 0 |
| ENSG00000271702 | 0 | 0 | 0 | 0 | 0 | 0 |
| ENSG00000252260 | 0 | 0 | 0 | 0 | 0 | 0 |
| ENSG00000265843 | 0 | 0 | 0 | 0 | 0 | 0 |
| ENSG00000264334 | 0 | 0 | 0 | 0 | 0 | 0 |
| ENSG00000199728 | 0 | 0 | 0 | 0 | 0 | 0 |
| ENSG00000266213 | 0 | 0 | 0 | 0 | 0 | 0 |
| ENSG00000265644 | 0 | 0 | 0 | 0 | 0 | 0 |
| ENSG00000265378 | 0 | 0 | 0 | 0 | 0 | 0 |
| ENSG00000201723 | 0 | 0 | 0 | 0 | 0 | 0 |
| ENSG00000266273 | 0 | 0 | 0 | 0 | 0 | 0 |
| ENSG00000266586 | 0 | 0 | 0 | 0 | 0 | 0 |
| ENSG00000265101 | 0 | 0 | 0 | 0 | 0 | 0 |
| ENSG00000263146 | 0 | 0 | 0 | 0 | 0 | 0 |
| ENSG00000256463 | 2 | 0 | 1 | 0 | 0 | 0 |
| ENSG00000166377 | 326 | 163 | 172 | 121 | 171 | 159 |
| ENSG00000279416 | 0 | 0 | 0 | 0 | 0 | 0 |
| ENSG00000267655 | 11 | 3 | 7 | 1 | 5 | 1 |
| ENSG00000279474 | 0 | 0 | 0 | 1 | 0 | 0 |

| | | | | | | |
|---|---|---|---|---|---|---|
| ENSG00000267628 | 0 | 0 | 0 | 0 | 0 | 0 |
| ENSG00000267015 | 0 | 0 | 0 | 1 | 0 | 0 |
| ENSG00000279077 | 2 | 0 | 0 | 0 | 0 | 0 |
| ENSG00000131196 | 328 | 136 | 156 | 164 | 160 | 148 |
| ENSG00000279637 | 0 | 1 | 1 | 0 | 0 | 0 |
| ENSG00000275186 | 1 | 0 | 0 | 0 | 0 | 0 |
| ENSG00000279981 | 0 | 0 | 0 | 0 | 1 | 0 |
| ENSG00000279397 | 0 | 0 | 0 | 0 | 0 | 0 |
| ENSG00000267287 | 0 | 0 | 2 | 0 | 0 | 1 |
| ENSG00000267409 | 0 | 0 | 0 | 0 | 0 | 0 |
| ENSG00000178412 | 0 | 0 | 0 | 0 | 1 | 0 |
| ENSG00000274828 | 182 | 87 | 126 | 93 | 90 | 90 |
| ENSG00000060069 | 675 | 303 | 382 | 382 | 376 | 396 |
| ENSG00000266901 | 0 | 0 | 0 | 0 | 0 | 0 |
| ENSG00000266924 | 0 | 0 | 0 | 0 | 0 | 0 |
| ENSG00000267780 | 0 | 0 | 0 | 0 | 0 | 0 |
| ENSG00000178342 | 0 | 0 | 2 | 0 | 0 | 4 |
| ENSG00000278075 | 1 | 0 | 0 | 1 | 0 | 0 |
| ENSG00000122490 | 505 | 268 | 290 | 332 | 284 | 287 |
| ENSG00000226742 | 378 | 209 | 238 | 225 | 279 | 231 |
| ENSG00000141759 | 1210 | 646 | 701 | 832 | 785 | 754 |
| ENSG00000267127 | 4 | 3 | 7 | 2 | 2 | 3 |
| ENSG00000101546 | 310 | 168 | 207 | 241 | 216 | 183 |
| ENSG00000261126 | 34 | 15 | 19 | 6 | 13 | 15 |
| ENSG00000266909 | 0 | 0 | 1 | 0 | 0 | 0 |
| ENSG00000101544 | 1143 | 529 | 636 | 513 | 575 | 573 |
| ENSG00000267270 | 15 | 1 | 9 | 3 | 4 | 3 |
| ENSG00000178184 | 190 | 83 | 106 | 97 | 112 | 118 |
| ENSG00000278000 | 4 | 3 | 3 | 3 | 4 | 9 |
| ENSG00000267251 | 4 | 2 | 4 | 3 | 1 | 0 |
| ENSG00000178591 | 0 | 0 | 0 | 0 | 0 | 0 |
| ENSG00000125788 | 0 | 0 | 0 | 0 | 0 | 0 |
| ENSG00000088782 | 0 | 0 | 0 | 0 | 0 | 0 |
| ENSG00000185982 | 0 | 0 | 0 | 0 | 0 | 0 |
| ENSG00000125903 | 0 | 0 | 0 | 0 | 0 | 0 |
| ENSG00000186458 | 0 | 0 | 0 | 0 | 0 | 0 |
| ENSG00000272874 | 0 | 0 | 0 | 0 | 0 | 0 |
| ENSG00000196476 | 222 | 72 | 117 | 95 | 118 | 107 |
| ENSG00000247315 | 570 | 279 | 338 | 304 | 300 | 300 |
| ENSG00000225377 | 128 | 84 | 67 | 52 | 60 | 48 |
| ENSG00000177732 | 1701 | 798 | 1025 | 1033 | 928 | 867 |
| ENSG00000125841 | 1261 | 563 | 698 | 706 | 582 | 598 |
| ENSG00000101255 | 1939 | 929 | 1145 | 1249 | 1052 | 1025 |
| ENSG00000125826 | 2273 | 1089 | 1424 | 1312 | 1130 | 1221 |
| ENSG00000125875 | 1321 | 577 | 764 | 582 | 630 | 766 |
| ENSG00000206797 | 0 | 0 | 0 | 0 | 0 | 0 |
| ENSG00000101266 | 3071 | 1494 | 1890 | 1370 | 1775 | 1808 |
| ENSG00000125878 | 0 | 0 | 0 | 0 | 0 | 0 |
| ENSG00000271303 | 12 | 26 | 28 | 24 | 29 | 24 |
| ENSG00000270299 | 1 | 0 | 2 | 1 | 1 | 2 |

| | | | | | | |
|---|---|---|---|---|---|---|
| ENSG00000215397 | 0 | 0 | 0 | 0 | 0 | 0 |
| ENSG00000101276 | 78 | 32 | 44 | 32 | 49 | 39 |
| ENSG00000125898 | 1039 | 420 | 640 | 427 | 381 | 445 |
| ENSG00000101278 | 21 | 17 | 24 | 16 | 11 | 11 |
| ENSG00000101280 | 8 | 1 | 6 | 2 | 3 | 3 |
| ENSG00000101282 | 0 | 1 | 0 | 0 | 0 | 0 |
| ENSG00000125818 | 2286 | 999 | 1360 | 1389 | 1115 | 1365 |
| ENSG00000276026 | 1 | 1 | 2 | 4 | 2 | 3 |
| ENSG00000215388 | 4 | 0 | 2 | 2 | 2 | 0 |
| ENSG00000125895 | 6 | 3 | 1 | 0 | 2 | 2 |
| ENSG00000274269 | 0 | 0 | 0 | 0 | 0 | 0 |
| ENSG00000215595 | 2 | 0 | 0 | 0 | 0 | 0 |
| ENSG00000244588 | 0 | 0 | 0 | 0 | 0 | 1 |
| ENSG00000101298 | 0 | 0 | 3 | 0 | 0 | 3 |
| ENSG00000125775 | 54 | 43 | 25 | 39 | 64 | 37 |
| ENSG00000274322 | 0 | 1 | 0 | 1 | 1 | 0 |
| ENSG00000280162 | 0 | 0 | 0 | 0 | 1 | 0 |
| ENSG00000234684 | 65 | 36 | 65 | 35 | 49 | 54 |
| ENSG00000229728 | 14 | 11 | 8 | 10 | 10 | 9 |
| ENSG00000088832 | 4035 | 2188 | 2475 | 2427 | 2435 | 2449 |
| ENSG00000284436 | 0 | 0 | 0 | 0 | 0 | 0 |
| ENSG00000088833 | 1850 | 846 | 1077 | 1106 | 1011 | 1093 |
| ENSG00000196209 | 0 | 1 | 0 | 0 | 0 | 0 |
| ENSG00000242324 | 0 | 0 | 0 | 0 | 0 | 0 |
| ENSG00000252103 | 0 | 0 | 0 | 0 | 0 | 0 |
| ENSG00000125900 | 0 | 0 | 0 | 0 | 0 | 0 |
| ENSG00000260861 | 0 | 0 | 0 | 0 | 0 | 0 |
| ENSG00000101307 | 0 | 3 | 2 | 2 | 1 | 1 |
| ENSG00000271523 | 0 | 0 | 0 | 0 | 0 | 0 |
| ENSG00000089012 | 0 | 0 | 0 | 0 | 0 | 0 |
| ENSG00000237914 | 0 | 0 | 0 | 0 | 0 | 0 |
| ENSG00000223750 | 0 | 0 | 0 | 0 | 0 | 0 |
| ENSG00000242348 | 0 | 0 | 0 | 0 | 0 | 0 |
| ENSG00000236550 | 0 | 0 | 0 | 0 | 0 | 0 |
| ENSG00000234282 | 0 | 0 | 0 | 0 | 0 | 0 |
| ENSG00000271461 | 0 | 0 | 0 | 0 | 0 | 0 |
| ENSG00000232528 | 0 | 0 | 0 | 0 | 0 | 0 |
| ENSG00000230839 | 0 | 0 | 0 | 0 | 0 | 0 |
| ENSG00000276649 | 0 | 0 | 0 | 0 | 0 | 0 |
| ENSG00000198053 | 51 | 17 | 23 | 31 | 45 | 22 |
| ENSG00000233896 | 0 | 0 | 0 | 0 | 0 | 0 |
| ENSG00000101327 | 0 | 0 | 0 | 0 | 0 | 0 |
| ENSG00000227663 | 0 | 0 | 0 | 0 | 0 | 0 |
| ENSG00000125834 | 1250 | 595 | 735 | 587 | 610 | 688 |
| ENSG00000226644 | 22 | 8 | 11 | 25 | 21 | 26 |
| ENSG00000125780 | 0 | 1 | 5 | 0 | 0 | 0 |
| ENSG00000166948 | 0 | 0 | 0 | 0 | 0 | 0 |
| ENSG00000207305 | 0 | 0 | 0 | 0 | 0 | 0 |
| ENSG00000125835 | 3905 | 2038 | 2556 | 3142 | 2575 | 2615 |
| ENSG00000256566 | 0 | 0 | 0 | 0 | 1 | 0 |

| | | | | | | |
|---|---|---|---|---|---|---|
| ENSG00000088876 | 169 | 68 | 116 | 72 | 100 | 107 |
| ENSG00000149488 | 6 | 0 | 0 | 1 | 0 | 1 |
| ENSG00000101361 | 2187 | 1169 | 1467 | 1430 | 1392 | 1585 |
| ENSG00000284481 | 0 | 0 | 0 | 0 | 0 | 0 |
| ENSG00000221116 | 0 | 0 | 0 | 0 | 0 | 0 |
| ENSG00000271798 | 0 | 0 | 0 | 0 | 0 | 1 |
| ENSG00000212498 | 0 | 0 | 0 | 0 | 0 | 0 |
| ENSG00000229686 | 0 | 0 | 0 | 0 | 0 | 0 |
| ENSG00000226572 | 0 | 0 | 0 | 0 | 0 | 0 |
| ENSG00000101365 | 1887 | 899 | 1107 | 1109 | 928 | 962 |
| ENSG00000228293 | 0 | 0 | 3 | 1 | 1 | 0 |
| ENSG00000088881 | 79 | 43 | 59 | 37 | 36 | 40 |
| ENSG00000226217 | 0 | 0 | 0 | 0 | 0 | 0 |
| ENSG00000088882 | 0 | 0 | 0 | 0 | 0 | 1 |
| ENSG00000241690 | 0 | 0 | 0 | 0 | 0 | 0 |
| ENSG00000258713 | 0 | 0 | 0 | 0 | 0 | 0 |
| ENSG00000198326 | 0 | 0 | 0 | 0 | 0 | 0 |
| ENSG00000132635 | 713 | 275 | 416 | 392 | 338 | 393 |
| ENSG00000215305 | 816 | 450 | 590 | 516 | 470 | 486 |
| ENSG00000132670 | 1776 | 866 | 1220 | 807 | 1008 | 966 |
| ENSG00000125787 | 0 | 2 | 3 | 1 | 2 | 1 |
| ENSG00000125901 | 1097 | 581 | 759 | 861 | 667 | 780 |
| ENSG00000101405 | 0 | 0 | 1 | 0 | 0 | 0 |
| ENSG00000101200 | 0 | 0 | 0 | 0 | 0 | 0 |
| ENSG00000263905 | 0 | 0 | 0 | 0 | 0 | 0 |
| ENSG00000235958 | 7 | 3 | 2 | 2 | 1 | 1 |
| ENSG00000185019 | 242 | 150 | 168 | 161 | 124 | 160 |
| ENSG00000215251 | 489 | 263 | 353 | 288 | 315 | 365 |
| ENSG00000088899 | 142 | 63 | 77 | 74 | 90 | 95 |
| ENSG00000198171 | 1576 | 851 | 1039 | 1041 | 869 | 992 |
| ENSG00000125877 | 1044 | 574 | 664 | 854 | 682 | 766 |
| ENSG00000088836 | 146 | 74 | 101 | 105 | 85 | 100 |
| ENSG00000277287 | 20 | 7 | 10 | 5 | 10 | 7 |
| ENSG00000088854 | 358 | 154 | 266 | 156 | 222 | 194 |
| ENSG00000201294 | 0 | 0 | 0 | 0 | 0 | 0 |
| ENSG00000226632 | 2 | 0 | 2 | 0 | 0 | 0 |
| ENSG00000201346 | 0 | 0 | 0 | 0 | 0 | 0 |
| ENSG00000225116 | 0 | 0 | 0 | 0 | 0 | 0 |
| ENSG00000088812 | 1873 | 843 | 1208 | 634 | 931 | 941 |
| ENSG00000230568 | 0 | 0 | 0 | 0 | 0 | 0 |
| ENSG00000125861 | 0 | 0 | 0 | 0 | 0 | 0 |
| ENSG00000149451 | 0 | 0 | 0 | 0 | 0 | 0 |
| ENSG00000088827 | 0 | 0 | 0 | 0 | 0 | 0 |
| ENSG00000132622 | 43 | 15 | 20 | 19 | 24 | 28 |
| ENSG00000101220 | 1470 | 831 | 987 | 1406 | 959 | 1098 |
| ENSG00000101222 | 10 | 9 | 13 | 5 | 11 | 9 |
| ENSG00000125817 | 4806 | 2389 | 3184 | 3687 | 2915 | 3104 |
| ENSG00000101224 | 3462 | 1581 | 2273 | 2966 | 2798 | 2847 |
| ENSG00000275491 | 5 | 4 | 1 | 5 | 7 | 5 |
| ENSG00000274949 | 2 | 1 | 5 | 6 | 3 | 3 |

| | | | | | | |
|---|---|---|---|---|---|---|
| ENSG00000125843 | 903 | 446 | 582 | 541 | 505 | 580 |
| ENSG00000088888 | 8667 | 4718 | 5408 | 3613 | 4412 | 4696 |
| ENSG00000229539 | 21 | 8 | 11 | 10 | 9 | 7 |
| ENSG00000125779 | 570 | 370 | 390 | 306 | 332 | 334 |
| ENSG00000199024 | 0 | 0 | 1 | 0 | 0 | 0 |
| ENSG00000283320 | 0 | 0 | 0 | 0 | 0 | 0 |
| ENSG00000275582 | 22 | 4 | 16 | 13 | 8 | 7 |
| ENSG00000101236 | 794 | 365 | 503 | 341 | 480 | 476 |
| ENSG00000226608 | 8 | 3 | 0 | 3 | 3 | 2 |
| ENSG00000215063 | 0 | 0 | 0 | 0 | 1 | 0 |
| ENSG00000205300 | 0 | 0 | 0 | 0 | 0 | 0 |
| ENSG00000088826 | 76 | 31 | 36 | 63 | 59 | 37 |
| ENSG00000230176 | 0 | 0 | 0 | 0 | 1 | 0 |
| ENSG00000171873 | 0 | 0 | 0 | 0 | 0 | 0 |
| ENSG00000236387 | 0 | 0 | 0 | 0 | 0 | 0 |
| ENSG00000238282 | 0 | 0 | 0 | 0 | 0 | 0 |
| ENSG00000228210 | 0 | 0 | 0 | 0 | 0 | 0 |
| ENSG00000226948 | 0 | 1 | 0 | 0 | 1 | 0 |
| ENSG00000171867 | 512 | 282 | 364 | 347 | 403 | 374 |
| ENSG00000171864 | 0 | 0 | 0 | 0 | 0 | 0 |
| ENSG00000180259 | 0 | 0 | 0 | 0 | 0 | 0 |
| ENSG00000231071 | 0 | 0 | 0 | 0 | 0 | 0 |
| ENSG00000278595 | 0 | 1 | 0 | 0 | 0 | 0 |
| ENSG00000101265 | 42 | 38 | 39 | 9 | 30 | 18 |
| ENSG00000089057 | 836 | 440 | 574 | 318 | 384 | 523 |
| ENSG00000252096 | 0 | 0 | 0 | 0 | 0 | 0 |
| ENSG00000234573 | 0 | 0 | 0 | 0 | 0 | 0 |
| ENSG00000277087 | 0 | 0 | 0 | 0 | 0 | 0 |
| ENSG00000276768 | 0 | 0 | 0 | 0 | 0 | 2 |
| ENSG00000278816 | 2 | 0 | 0 | 0 | 0 | 0 |
| ENSG00000277425 | 0 | 0 | 0 | 0 | 0 | 0 |
| ENSG00000089063 | 1683 | 850 | 1072 | 824 | 981 | 987 |
| ENSG00000212536 | 0 | 0 | 0 | 2 | 0 | 2 |
| ENSG00000252367 | 0 | 0 | 0 | 0 | 0 | 2 |
| ENSG00000132646 | 1777 | 1041 | 1134 | 830 | 940 | 1029 |
| ENSG00000212517 | 1 | 1 | 0 | 0 | 0 | 0 |
| ENSG00000101290 | 1626 | 721 | 941 | 692 | 817 | 867 |
| ENSG00000225022 | 0 | 2 | 1 | 0 | 3 | 1 |
| ENSG00000101292 | 0 | 0 | 0 | 0 | 0 | 0 |
| ENSG00000200313 | 0 | 0 | 0 | 0 | 0 | 0 |
| ENSG00000276688 | 0 | 0 | 0 | 0 | 0 | 0 |
| ENSG00000226995 | 0 | 0 | 0 | 0 | 0 | 0 |
| ENSG00000230563 | 0 | 0 | 0 | 0 | 0 | 0 |
| ENSG00000205181 | 0 | 0 | 0 | 0 | 0 | 0 |
| ENSG00000266908 | 0 | 0 | 0 | 0 | 0 | 0 |
| ENSG00000225640 | 0 | 0 | 0 | 0 | 0 | 0 |
| ENSG00000225831 | 0 | 0 | 0 | 0 | 0 | 0 |
| ENSG00000235820 | 0 | 0 | 0 | 0 | 0 | 0 |
| ENSG00000125772 | 185 | 131 | 160 | 71 | 100 | 117 |
| ENSG00000233578 | 2 | 1 | 0 | 1 | 0 | 1 |

| | | | | | | |
|---|---|---|---|---|---|---|
| ENSG00000171984 | 70 | 25 | 34 | 52 | 36 | 50 |
| ENSG00000202380 | 0 | 0 | 0 | 0 | 0 | 0 |
| ENSG00000089199 | 1 | 0 | 0 | 0 | 1 | 0 |
| ENSG00000215589 | 0 | 0 | 0 | 0 | 0 | 0 |
| ENSG00000089195 | 387 | 206 | 251 | 187 | 214 | 247 |
| ENSG00000125885 | 301 | 149 | 156 | 89 | 175 | 136 |
| ENSG00000207380 | 0 | 0 | 0 | 0 | 0 | 0 |
| ENSG00000231961 | 0 | 0 | 0 | 0 | 0 | 0 |
| ENSG00000278719 | 2 | 0 | 0 | 0 | 0 | 0 |
| ENSG00000275632 | 3 | 2 | 2 | 2 | 6 | 2 |
| ENSG00000263755 | 0 | 0 | 0 | 0 | 0 | 0 |
| ENSG00000088766 | 535 | 244 | 344 | 330 | 348 | 340 |
| ENSG00000125872 | 0 | 0 | 2 | 0 | 0 | 0 |
| ENSG00000278192 | 0 | 1 | 0 | 0 | 1 | 0 |
| ENSG00000101311 | 24 | 19 | 16 | 14 | 24 | 16 |
| ENSG00000233402 | 0 | 0 | 0 | 0 | 0 | 0 |
| ENSG00000234241 | 10 | 1 | 2 | 1 | 2 | 2 |
| ENSG00000271565 | 0 | 0 | 0 | 0 | 0 | 0 |
| ENSG00000276915 | 0 | 0 | 0 | 0 | 0 | 0 |
| ENSG00000229876 | 0 | 0 | 0 | 0 | 0 | 0 |
| ENSG00000228482 | 0 | 0 | 0 | 0 | 0 | 0 |
| ENSG00000125845 | 0 | 0 | 0 | 2 | 0 | 0 |
| ENSG00000232271 | 0 | 0 | 0 | 0 | 0 | 0 |
| ENSG00000228888 | 0 | 0 | 0 | 0 | 0 | 0 |
| ENSG00000238102 | 0 | 0 | 0 | 0 | 0 | 0 |
| ENSG00000261249 | 0 | 0 | 0 | 0 | 0 | 0 |
| ENSG00000224711 | 0 | 0 | 0 | 0 | 0 | 0 |
| ENSG00000278159 | 0 | 0 | 0 | 0 | 0 | 0 |
| ENSG00000240584 | 0 | 0 | 0 | 0 | 0 | 0 |
| ENSG00000215586 | 0 | 0 | 0 | 0 | 0 | 0 |
| ENSG00000277315 | 0 | 0 | 0 | 0 | 0 | 0 |
| ENSG00000101323 | 0 | 0 | 0 | 0 | 0 | 0 |
| ENSG00000125827 | 1422 | 763 | 929 | 584 | 746 | 756 |
| ENSG00000229766 | 1 | 2 | 2 | 1 | 2 | 1 |
| ENSG00000182621 | 641 | 411 | 314 | 92 | 431 | 277 |
| ENSG00000226262 | 0 | 0 | 0 | 0 | 0 | 0 |
| ENSG00000225479 | 0 | 0 | 0 | 0 | 0 | 0 |
| ENSG00000201348 | 0 | 0 | 0 | 0 | 0 | 0 |
| ENSG00000228438 | 0 | 0 | 0 | 0 | 0 | 0 |
| ENSG00000276385 | 0 | 0 | 0 | 0 | 0 | 0 |
| ENSG00000275222 | 0 | 0 | 0 | 0 | 0 | 0 |
| ENSG00000101333 | 262 | 149 | 99 | 68 | 208 | 127 |
| ENSG00000225988 | 0 | 0 | 0 | 0 | 0 | 0 |
| ENSG00000125869 | 0 | 0 | 0 | 0 | 0 | 0 |
| ENSG00000101349 | 0 | 2 | 0 | 0 | 0 | 0 |
| ENSG00000232738 | 0 | 0 | 0 | 0 | 0 | 0 |
| ENSG00000243961 | 0 | 0 | 0 | 0 | 0 | 0 |
| ENSG00000132623 | 781 | 376 | 468 | 309 | 368 | 385 |
| ENSG00000227906 | 0 | 0 | 0 | 0 | 0 | 1 |
| ENSG00000276525 | 0 | 0 | 0 | 0 | 0 | 0 |

| | | | | | | |
|---|---|---|---|---|---|---|
| ENSG00000230506 | 0 | 0 | 0 | 0 | 0 | 0 |
| ENSG00000232448 | 0 | 0 | 0 | 0 | 0 | 0 |
| ENSG00000132639 | 0 | 0 | 1 | 3 | 2 | 4 |
| ENSG00000237005 | 0 | 0 | 0 | 0 | 0 | 0 |
| ENSG00000214835 | 0 | 0 | 0 | 0 | 0 | 0 |
| ENSG00000230750 | 0 | 0 | 0 | 0 | 0 | 0 |
| ENSG00000125863 | 745 | 376 | 453 | 375 | 477 | 425 |
| ENSG00000285508 | 0 | 0 | 0 | 0 | 0 | 0 |
| ENSG00000285723 | 0 | 0 | 0 | 0 | 0 | 0 |
| ENSG00000149346 | 557 | 223 | 320 | 187 | 287 | 246 |
| ENSG00000235036 | 8 | 2 | 1 | 1 | 0 | 3 |
| ENSG00000101384 | 243 | 128 | 174 | 62 | 67 | 91 |
| ENSG00000273745 | 0 | 0 | 0 | 0 | 0 | 0 |
| ENSG00000270792 | 0 | 0 | 0 | 0 | 0 | 0 |
| ENSG00000224961 | 0 | 0 | 0 | 0 | 0 | 0 |
| ENSG00000234900 | 0 | 0 | 0 | 0 | 0 | 0 |
| ENSG00000232900 | 0 | 0 | 0 | 0 | 0 | 0 |
| ENSG00000217809 | 0 | 0 | 0 | 0 | 0 | 0 |
| ENSG00000270777 | 0 | 0 | 0 | 0 | 0 | 0 |
| ENSG00000125899 | 0 | 0 | 0 | 0 | 0 | 0 |
| ENSG00000230990 | 0 | 0 | 0 | 0 | 0 | 0 |
| ENSG00000235544 | 0 | 0 | 0 | 0 | 0 | 0 |
| ENSG00000282478 | 0 | 0 | 0 | 0 | 0 | 0 |
| ENSG00000271397 | 0 | 0 | 0 | 0 | 0 | 0 |
| ENSG00000237256 | 0 | 0 | 0 | 0 | 0 | 0 |
| ENSG00000261411 | 0 | 0 | 0 | 0 | 0 | 0 |
| ENSG00000228422 | 0 | 0 | 0 | 0 | 0 | 0 |
| ENSG00000222281 | 0 | 0 | 0 | 0 | 0 | 0 |
| ENSG00000271298 | 0 | 0 | 0 | 0 | 0 | 0 |
| ENSG00000132640 | 771 | 388 | 461 | 363 | 443 | 412 |
| ENSG00000236526 | 3 | 0 | 0 | 0 | 3 | 1 |
| ENSG00000235292 | 0 | 0 | 0 | 0 | 0 | 0 |
| ENSG00000230437 | 0 | 0 | 0 | 0 | 0 | 0 |
| ENSG00000235698 | 2 | 0 | 0 | 1 | 0 | 0 |
| ENSG00000233048 | 0 | 1 | 0 | 1 | 3 | 1 |
| ENSG00000269931 | 0 | 0 | 0 | 0 | 0 | 0 |
| ENSG00000234265 | 0 | 0 | 0 | 0 | 0 | 0 |
| ENSG00000172296 | 12 | 16 | 10 | 3 | 6 | 10 |
| ENSG00000271426 | 0 | 0 | 0 | 0 | 0 | 0 |
| ENSG00000101230 | 16 | 9 | 1 | 1 | 5 | 3 |
| ENSG00000226263 | 0 | 1 | 1 | 0 | 3 | 1 |
| ENSG00000225956 | 0 | 1 | 0 | 0 | 0 | 0 |
| ENSG00000278153 | 2 | 1 | 0 | 0 | 0 | 1 |
| ENSG00000089123 | 484 | 293 | 354 | 226 | 330 | 282 |
| ENSG00000236811 | 6 | 0 | 0 | 0 | 0 | 0 |
| ENSG00000089048 | 140 | 80 | 65 | 21 | 191 | 86 |
| ENSG00000101247 | 348 | 180 | 201 | 216 | 210 | 239 |
| ENSG00000101251 | 0 | 0 | 0 | 0 | 1 | 0 |
| ENSG00000212247 | 0 | 0 | 0 | 0 | 0 | 0 |
| ENSG00000172264 | 299 | 138 | 193 | 94 | 201 | 183 |

| | | | | | | |
|---|---|---|---|---|---|---|
| ENSG00000230935 | 0 | 0 | 0 | 0 | 1 | 0 |
| ENSG00000234187 | 0 | 0 | 2 | 0 | 0 | 0 |
| ENSG00000239923 | 0 | 0 | 0 | 0 | 0 | 0 |
| ENSG00000125848 | 232 | 133 | 196 | 141 | 185 | 254 |
| ENSG00000199570 | 1 | 0 | 0 | 0 | 1 | 0 |
| ENSG00000227563 | 0 | 0 | 0 | 0 | 0 | 0 |
| ENSG00000227927 | 0 | 0 | 1 | 0 | 0 | 0 |
| ENSG00000213950 | 4 | 2 | 2 | 2 | 2 | 3 |
| ENSG00000235914 | 2 | 1 | 1 | 0 | 1 | 0 |
| ENSG00000237832 | 0 | 0 | 0 | 0 | 0 | 0 |
| ENSG00000199803 | 0 | 0 | 0 | 0 | 0 | 0 |
| ENSG00000201180 | 0 | 0 | 0 | 0 | 0 | 0 |
| ENSG00000222500 | 0 | 0 | 0 | 0 | 0 | 0 |
| ENSG00000225181 | 0 | 0 | 0 | 0 | 0 | 0 |
| ENSG00000224274 | 0 | 0 | 0 | 0 | 0 | 0 |
| ENSG00000225622 | 0 | 0 | 0 | 0 | 0 | 0 |
| ENSG00000089177 | 803 | 445 | 426 | 228 | 550 | 409 |
| ENSG00000273998 | 1 | 1 | 1 | 2 | 1 | 2 |
| ENSG00000275728 | 0 | 0 | 0 | 0 | 2 | 0 |
| ENSG00000226229 | 0 | 0 | 0 | 0 | 0 | 0 |
| ENSG00000200494 | 0 | 0 | 0 | 0 | 0 | 0 |
| ENSG00000235208 | 0 | 0 | 0 | 0 | 0 | 0 |
| ENSG00000228809 | 0 | 4 | 2 | 0 | 0 | 2 |
| ENSG00000125870 | 3246 | 1823 | 2042 | 1917 | 2114 | 1969 |
| ENSG00000125879 | 2 | 0 | 2 | 1 | 0 | 5 |
| ENSG00000236985 | 0 | 0 | 0 | 0 | 0 | 0 |
| ENSG00000212165 | 0 | 0 | 0 | 0 | 0 | 0 |
| ENSG00000251955 | 0 | 0 | 0 | 0 | 0 | 0 |
| ENSG00000220540 | 0 | 0 | 0 | 0 | 0 | 0 |
| ENSG00000238833 | 0 | 0 | 0 | 0 | 0 | 0 |
| ENSG00000125851 | 358 | 127 | 305 | 91 | 64 | 134 |
| ENSG00000232241 | 0 | 0 | 0 | 0 | 0 | 0 |
| ENSG00000125864 | 165 | 84 | 92 | 141 | 186 | 132 |
| ENSG00000232333 | 2 | 0 | 5 | 1 | 0 | 0 |
| ENSG00000277373 | 14 | 3 | 4 | 4 | 3 | 1 |
| ENSG00000125868 | 17921 | 9068 | 10488 | 9400 | 11117 | 9710 |
| ENSG00000202260 | 0 | 3 | 1 | 0 | 0 | 0 |
| ENSG00000230495 | 5 | 0 | 1 | 0 | 0 | 0 |
| ENSG00000125844 | 4463 | 2045 | 2667 | 2392 | 2608 | 2517 |
| ENSG00000125888 | 0 | 0 | 0 | 0 | 0 | 0 |
| ENSG00000233625 | 0 | 0 | 0 | 0 | 0 | 0 |
| ENSG00000212555 | 0 | 0 | 0 | 0 | 0 | 0 |
| ENSG00000089006 | 4399 | 2338 | 2643 | 2148 | 2951 | 2478 |
| ENSG00000125850 | 137 | 52 | 60 | 53 | 63 | 38 |
| ENSG00000212232 | 5 | 13 | 9 | 4 | 7 | 13 |
| ENSG00000125871 | 1219 | 658 | 793 | 580 | 754 | 755 |
| ENSG00000218902 | 1 | 0 | 1 | 0 | 0 | 0 |
| ENSG00000229262 | 0 | 0 | 0 | 0 | 0 | 0 |
| ENSG00000227343 | 0 | 0 | 0 | 0 | 0 | 0 |
| ENSG00000252597 | 0 | 0 | 0 | 0 | 0 | 0 |

| | | | | | | |
|---|---|---|---|---|---|---|
| ENSG00000207282 | 0 | 0 | 0 | 0 | 0 | 0 |
| ENSG00000232838 | 46 | 38 | 42 | 59 | 69 | 61 |
| ENSG00000149474 | 877 | 432 | 616 | 457 | 503 | 524 |
| ENSG00000232943 | 0 | 0 | 0 | 0 | 0 | 0 |
| ENSG00000266720 | 0 | 0 | 0 | 0 | 0 | 0 |
| ENSG00000252635 | 0 | 0 | 0 | 0 | 0 | 0 |
| ENSG00000125846 | 426 | 216 | 233 | 224 | 339 | 258 |
| ENSG00000199719 | 0 | 0 | 0 | 0 | 1 | 1 |
| ENSG00000237259 | 0 | 0 | 0 | 1 | 0 | 0 |
| ENSG00000230010 | 0 | 1 | 0 | 0 | 0 | 1 |
| ENSG00000237282 | 0 | 0 | 0 | 0 | 0 | 0 |
| ENSG00000089091 | 212 | 96 | 125 | 59 | 107 | 89 |
| ENSG00000236474 | 4 | 2 | 0 | 0 | 1 | 1 |
| ENSG00000252422 | 0 | 0 | 0 | 0 | 0 | 0 |
| ENSG00000132664 | 727 | 389 | 489 | 372 | 523 | 398 |
| ENSG00000265137 | 1 | 1 | 0 | 1 | 0 | 0 |
| ENSG00000225419 | 3 | 0 | 2 | 2 | 2 | 4 |
| ENSG00000089050 | 1472 | 687 | 853 | 631 | 731 | 820 |
| ENSG00000214612 | 2 | 0 | 1 | 0 | 0 | 0 |
| ENSG00000101310 | 5936 | 2664 | 3473 | 2613 | 2935 | 3098 |
| ENSG00000232388 | 1004 | 619 | 781 | 778 | 748 | 715 |
| ENSG00000284776 | 0 | 1 | 0 | 0 | 0 | 1 |
| ENSG00000125821 | 1035 | 535 | 662 | 618 | 608 | 634 |
| ENSG00000243702 | 0 | 0 | 1 | 0 | 0 | 0 |
| ENSG00000221806 | 0 | 0 | 0 | 0 | 0 | 0 |
| ENSG00000232136 | 0 | 2 | 0 | 0 | 0 | 0 |
| ENSG00000233993 | 1 | 0 | 0 | 1 | 0 | 0 |
| ENSG00000233324 | 0 | 0 | 0 | 0 | 0 | 0 |
| ENSG00000179935 | 12 | 5 | 9 | 2 | 13 | 5 |
| ENSG00000273148 | 100 | 60 | 70 | 38 | 59 | 84 |
| ENSG00000149443 | 0 | 0 | 0 | 0 | 0 | 0 |
| ENSG00000132631 | 0 | 0 | 0 | 0 | 0 | 0 |
| ENSG00000274863 | 0 | 0 | 0 | 0 | 0 | 0 |
| ENSG00000185052 | 1543 | 690 | 880 | 544 | 489 | 614 |
| ENSG00000235996 | 1 | 0 | 0 | 0 | 0 | 1 |
| ENSG00000179447 | 0 | 1 | 0 | 0 | 2 | 4 |
| ENSG00000232675 | 0 | 0 | 0 | 0 | 0 | 0 |
| ENSG00000268628 | 38 | 26 | 34 | 31 | 22 | 44 |
| ENSG00000132669 | 758 | 380 | 483 | 336 | 397 | 348 |
| ENSG00000236992 | 6 | 1 | 2 | 1 | 1 | 5 |
| ENSG00000173418 | 3366 | 1852 | 2143 | 2191 | 2450 | 2403 |
| ENSG00000101343 | 1474 | 847 | 842 | 566 | 983 | 768 |
| ENSG00000089101 | 7 | 2 | 7 | 1 | 3 | 4 |
| ENSG00000241054 | 0 | 0 | 0 | 0 | 0 | 0 |
| ENSG00000225417 | 0 | 0 | 0 | 0 | 0 | 0 |
| ENSG00000229500 | 0 | 0 | 0 | 0 | 0 | 0 |
| ENSG00000243810 | 0 | 0 | 0 | 0 | 0 | 0 |
| ENSG00000264879 | 0 | 0 | 0 | 0 | 0 | 0 |
| ENSG00000280229 | 0 | 2 | 0 | 0 | 0 | 0 |
| ENSG00000173404 | 2 | 1 | 0 | 1 | 1 | 0 |

| | | | | | | |
|---|---|---|---|---|---|---|
| ENSG00000188559 | 3481 | 1484 | 1989 | 1125 | 1566 | 1626 |
| ENSG00000213384 | 0 | 0 | 0 | 0 | 0 | 0 |
| ENSG00000270793 | 0 | 0 | 0 | 0 | 0 | 0 |
| ENSG00000278525 | 0 | 0 | 0 | 0 | 0 | 0 |
| ENSG00000225803 | 0 | 0 | 0 | 0 | 0 | 0 |
| ENSG00000234104 | 0 | 0 | 0 | 0 | 0 | 0 |
| ENSG00000225127 | 0 | 0 | 0 | 0 | 0 | 0 |
| ENSG00000283405 | 0 | 0 | 0 | 0 | 0 | 0 |
| ENSG00000235065 | 26 | 8 | 22 | 19 | 15 | 16 |
| ENSG00000088970 | 211 | 145 | 156 | 86 | 176 | 129 |
| ENSG00000199509 | 1 | 0 | 1 | 0 | 1 | 2 |
| ENSG00000228604 | 0 | 1 | 0 | 0 | 2 | 1 |
| ENSG00000232712 | 1 | 1 | 5 | 0 | 3 | 2 |
| ENSG00000214535 | 8 | 3 | 3 | 5 | 7 | 5 |
| ENSG00000226875 | 0 | 0 | 0 | 0 | 0 | 0 |
| ENSG00000275457 | 20 | 15 | 8 | 17 | 10 | 19 |
| ENSG00000088930 | 7149 | 3553 | 4437 | 3018 | 4053 | 3980 |
| ENSG00000125816 | 0 | 0 | 0 | 0 | 0 | 0 |
| ENSG00000225280 | 0 | 0 | 0 | 0 | 0 | 0 |
| ENSG00000223128 | 0 | 0 | 0 | 0 | 0 | 0 |
| ENSG00000226405 | 0 | 0 | 0 | 0 | 0 | 0 |
| ENSG00000278041 | 0 | 0 | 1 | 0 | 2 | 0 |
| ENSG00000227693 | 0 | 0 | 0 | 0 | 0 | 0 |
| ENSG00000125820 | 4 | 1 | 0 | 0 | 1 | 1 |
| ENSG00000258197 | 0 | 0 | 0 | 0 | 0 | 0 |
| ENSG00000233492 | 0 | 0 | 0 | 0 | 0 | 0 |
| ENSG00000279082 | 0 | 0 | 0 | 0 | 0 | 0 |
| ENSG00000280350 | 0 | 0 | 0 | 0 | 0 | 0 |
| ENSG00000278410 | 0 | 0 | 0 | 0 | 0 | 0 |
| ENSG00000125813 | 0 | 0 | 0 | 0 | 0 | 0 |
| ENSG00000233015 | 0 | 0 | 0 | 0 | 1 | 0 |
| ENSG00000227063 | 384 | 434 | 439 | 593 | 416 | 469 |
| ENSG00000238034 | 0 | 0 | 0 | 0 | 0 | 0 |
| ENSG00000234435 | 0 | 0 | 0 | 0 | 0 | 0 |
| ENSG00000234862 | 0 | 0 | 0 | 0 | 0 | 0 |
| ENSG00000225321 | 0 | 0 | 0 | 0 | 0 | 0 |
| ENSG00000225170 | 0 | 0 | 0 | 0 | 0 | 0 |
| ENSG00000270838 | 0 | 0 | 0 | 0 | 0 | 0 |
| ENSG00000204684 | 0 | 0 | 0 | 0 | 0 | 0 |
| ENSG00000259974 | 0 | 0 | 0 | 0 | 0 | 0 |
| ENSG00000283932 | 0 | 0 | 0 | 0 | 0 | 0 |
| ENSG00000125798 | 0 | 0 | 0 | 0 | 0 | 0 |
| ENSG00000237396 | 0 | 0 | 0 | 0 | 0 | 0 |
| ENSG00000230400 | 0 | 0 | 0 | 0 | 1 | 0 |
| ENSG00000283072 | 0 | 0 | 0 | 0 | 1 | 1 |
| ENSG00000215553 | 2 | 0 | 0 | 0 | 1 | 0 |
| ENSG00000213882 | 0 | 0 | 0 | 0 | 0 | 0 |
| ENSG00000228935 | 4 | 10 | 0 | 9 | 6 | 2 |
| ENSG00000230492 | 0 | 0 | 0 | 0 | 0 | 0 |
| ENSG00000234646 | 0 | 0 | 0 | 0 | 0 | 0 |

| | | | | | | |
|---|---|---|---|---|---|---|
| ENSG00000132671 | 0 | 0 | 0 | 0 | 0 | 0 |
| ENSG00000178726 | 940 | 503 | 404 | 890 | 1163 | 925 |
| ENSG00000279447 | 0 | 0 | 0 | 0 | 0 | 0 |
| ENSG00000125810 | 0 | 0 | 0 | 0 | 0 | 0 |
| ENSG00000233746 | 0 | 0 | 0 | 0 | 0 | 0 |
| ENSG00000279164 | 0 | 0 | 0 | 0 | 0 | 0 |
| ENSG00000201527 | 0 | 0 | 0 | 0 | 0 | 0 |
| ENSG00000275179 | 0 | 0 | 1 | 0 | 1 | 0 |
| ENSG00000230387 | 46 | 27 | 29 | 27 | 55 | 23 |
| ENSG00000277901 | 4 | 0 | 1 | 0 | 0 | 2 |
| ENSG00000234832 | 2 | 1 | 2 | 0 | 2 | 3 |
| ENSG00000132661 | 624 | 312 | 367 | 469 | 393 | 413 |
| ENSG00000232645 | 53 | 19 | 38 | 22 | 18 | 30 |
| ENSG00000125812 | 1211 | 585 | 720 | 649 | 661 | 643 |
| ENSG00000125814 | 787 | 360 | 528 | 349 | 447 | 471 |
| ENSG00000201728 | 0 | 0 | 0 | 0 | 1 | 0 |
| ENSG00000125823 | 5 | 4 | 10 | 2 | 11 | 2 |
| ENSG00000125831 | 0 | 0 | 1 | 0 | 1 | 0 |
| ENSG00000200208 | 0 | 0 | 0 | 0 | 0 | 0 |
| ENSG00000273838 | 0 | 1 | 2 | 0 | 2 | 2 |
| ENSG00000226344 | 0 | 0 | 0 | 0 | 0 | 0 |
| ENSG00000260202 | 0 | 0 | 0 | 0 | 0 | 0 |
| ENSG00000276638 | 0 | 0 | 0 | 0 | 0 | 0 |
| ENSG00000125815 | 0 | 0 | 0 | 0 | 0 | 0 |
| ENSG00000204663 | 0 | 0 | 0 | 0 | 0 | 0 |
| ENSG00000204662 | 0 | 0 | 0 | 0 | 0 | 0 |
| ENSG00000101435 | 0 | 0 | 0 | 0 | 0 | 0 |
| ENSG00000204658 | 0 | 0 | 0 | 0 | 0 | 0 |
| ENSG00000173335 | 0 | 0 | 3 | 0 | 1 | 0 |
| ENSG00000101439 | 7807 | 3544 | 4472 | 4677 | 3901 | 4091 |
| ENSG00000270001 | 7 | 3 | 2 | 5 | 2 | 1 |
| ENSG00000225056 | 0 | 0 | 0 | 0 | 0 | 0 |
| ENSG00000101441 | 0 | 0 | 0 | 2 | 0 | 1 |
| ENSG00000232595 | 0 | 0 | 0 | 0 | 0 | 0 |
| ENSG00000170373 | 4 | 0 | 3 | 2 | 0 | 0 |
| ENSG00000277975 | 0 | 0 | 0 | 0 | 0 | 0 |
| ENSG00000230908 | 0 | 0 | 0 | 0 | 0 | 0 |
| ENSG00000170369 | 10 | 7 | 13 | 9 | 6 | 3 |
| ENSG00000170367 | 8 | 3 | 2 | 1 | 2 | 2 |
| ENSG00000228476 | 0 | 0 | 0 | 2 | 2 | 4 |
| ENSG00000236168 | 0 | 0 | 0 | 0 | 0 | 0 |
| ENSG00000149435 | 0 | 0 | 0 | 0 | 1 | 0 |
| ENSG00000167390 | 0 | 0 | 0 | 0 | 0 | 0 |
| ENSG00000236151 | 0 | 0 | 0 | 0 | 0 | 0 |
| ENSG00000230133 | 0 | 0 | 0 | 0 | 0 | 0 |
| ENSG00000200231 | 0 | 0 | 0 | 0 | 0 | 0 |
| ENSG00000232200 | 0 | 0 | 0 | 0 | 0 | 0 |
| ENSG00000234069 | 0 | 0 | 0 | 0 | 0 | 0 |
| ENSG00000101463 | 25 | 14 | 18 | 9 | 13 | 10 |
| ENSG00000228539 | 0 | 0 | 0 | 0 | 0 | 0 |

| | | | | | | |
|---|---|---|---|---|---|---|
| ENSG00000231015 | 0 | 0 | 0 | 0 | 0 | 0 |
| ENSG00000274173 | 2133 | 1154 | 1522 | 1483 | 1489 | 1538 |
| ENSG00000077984 | 3 | 1 | 4 | 0 | 1 | 1 |
| ENSG00000101474 | 10738 | 5405 | 6190 | 5195 | 5082 | 5185 |
| ENSG00000207355 | 0 | 0 | 0 | 0 | 0 | 0 |
| ENSG00000154930 | 728 | 356 | 509 | 155 | 184 | 212 |
| ENSG00000275358 | 0 | 0 | 0 | 0 | 0 | 0 |
| ENSG00000100987 | 2 | 0 | 4 | 0 | 3 | 2 |
| ENSG00000225069 | 0 | 0 | 0 | 0 | 0 | 0 |
| ENSG00000230725 | 0 | 0 | 0 | 0 | 0 | 0 |
| ENSG00000225344 | 0 | 0 | 0 | 0 | 0 | 0 |
| ENSG00000197586 | 3291 | 1580 | 2016 | 1984 | 1883 | 1851 |
| ENSG00000202414 | 0 | 0 | 0 | 0 | 0 | 0 |
| ENSG00000279322 | 0 | 0 | 0 | 0 | 0 | 0 |
| ENSG00000277938 | 23 | 12 | 16 | 13 | 9 | 21 |
| ENSG00000274414 | 32 | 19 | 23 | 14 | 21 | 13 |
| ENSG00000100994 | 8040 | 3578 | 5005 | 4918 | 4352 | 4382 |
| ENSG00000274507 | 0 | 0 | 0 | 0 | 0 | 1 |
| ENSG00000276952 | 25 | 11 | 10 | 7 | 17 | 7 |
| ENSG00000100997 | 4017 | 1790 | 2321 | 2015 | 1939 | 1961 |
| ENSG00000227379 | 2 | 0 | 2 | 0 | 0 | 0 |
| ENSG00000101003 | 1167 | 665 | 764 | 546 | 704 | 769 |
| ENSG00000101004 | 908 | 397 | 599 | 396 | 521 | 481 |
| ENSG00000251883 | 0 | 0 | 0 | 0 | 1 | 0 |
| ENSG00000170191 | 572 | 215 | 267 | 190 | 297 | 342 |
| ENSG00000213742 | 128 | 78 | 98 | 42 | 66 | 52 |
| ENSG00000242236 | 0 | 0 | 0 | 0 | 1 | 0 |
| ENSG00000236542 | 0 | 1 | 0 | 1 | 0 | 0 |
| ENSG00000130684 | 111 | 38 | 66 | 40 | 62 | 67 |
| ENSG00000278383 | 21 | 10 | 5 | 7 | 9 | 3 |
| ENSG00000226465 | 98 | 45 | 42 | 42 | 46 | 48 |
| ENSG00000230772 | 0 | 0 | 1 | 0 | 0 | 2 |
| ENSG00000175170 | 31 | 25 | 25 | 15 | 21 | 14 |
| ENSG00000275048 | 0 | 0 | 0 | 0 | 0 | 0 |
| ENSG00000250090 | 2 | 0 | 0 | 0 | 0 | 0 |
| ENSG00000274973 | 1 | 0 | 1 | 0 | 1 | 0 |
| ENSG00000228593 | 0 | 0 | 0 | 0 | 0 | 0 |
| ENSG00000226203 | 0 | 0 | 0 | 0 | 0 | 0 |
| ENSG00000231081 | 7 | 1 | 6 | 0 | 2 | 2 |
| ENSG00000204556 | 42 | 28 | 41 | 32 | 47 | 54 |
| ENSG00000125804 | 0 | 0 | 0 | 0 | 0 | 0 |
| ENSG00000277508 | 0 | 0 | 0 | 0 | 0 | 0 |
| ENSG00000240108 | 0 | 0 | 0 | 0 | 0 | 0 |
| ENSG00000230440 | 0 | 0 | 0 | 0 | 0 | 0 |
| ENSG00000227195 | 0 | 0 | 0 | 0 | 0 | 0 |
| ENSG00000238129 | 0 | 0 | 0 | 0 | 0 | 0 |
| ENSG00000284419 | 0 | 0 | 0 | 0 | 0 | 0 |
| ENSG00000282826 | 29 | 14 | 18 | 8 | 14 | 13 |
| ENSG00000282870 | 29 | 22 | 39 | 5 | 22 | 18 |
| ENSG00000283568 | 0 | 0 | 0 | 0 | 0 | 0 |

| | | | | | | |
|---|---|---|---|---|---|---|
| ENSG00000283069 | 0 | 0 | 0 | 0 | 0 | 0 |
| ENSG00000282949 | 0 | 0 | 0 | 0 | 0 | 0 |
| ENSG00000282894 | 0 | 0 | 0 | 0 | 0 | 0 |
| ENSG00000282935 | 0 | 0 | 0 | 0 | 0 | 1 |
| ENSG00000282842 | 0 | 0 | 0 | 0 | 0 | 0 |
| ENSG00000283123 | 0 | 0 | 0 | 0 | 0 | 0 |
| ENSG00000282968 | 0 | 0 | 0 | 0 | 0 | 0 |
| ENSG00000282911 | 0 | 0 | 0 | 0 | 0 | 0 |
| ENSG00000282995 | 14 | 13 | 15 | 11 | 5 | 16 |
| ENSG00000283592 | 0 | 0 | 0 | 0 | 0 | 0 |
| ENSG00000282921 | 0 | 0 | 0 | 0 | 0 | 0 |
| ENSG00000283291 | 0 | 0 | 0 | 0 | 0 | 0 |
| ENSG00000283005 | 0 | 0 | 1 | 0 | 0 | 0 |
| ENSG00000283433 | 2 | 0 | 0 | 0 | 0 | 0 |
| ENSG00000283020 | 8 | 2 | 5 | 2 | 11 | 2 |
| ENSG00000204555 | 0 | 0 | 0 | 0 | 0 | 0 |
| ENSG00000205611 | 0 | 0 | 0 | 0 | 0 | 0 |
| ENSG00000200325 | 0 | 0 | 0 | 0 | 0 | 0 |
| ENSG00000231934 | 0 | 0 | 0 | 0 | 0 | 0 |
| ENSG00000149531 | 406 | 212 | 316 | 225 | 274 | 243 |
| ENSG00000238151 | 2 | 4 | 3 | 1 | 3 | 0 |
| ENSG00000275877 | 0 | 0 | 0 | 0 | 0 | 0 |
| ENSG00000283443 | 0 | 0 | 0 | 0 | 0 | 2 |
| ENSG00000277112 | 6 | 0 | 1 | 0 | 0 | 1 |
| ENSG00000274385 | 0 | 0 | 0 | 0 | 0 | 0 |
| ENSG00000275305 | 0 | 0 | 0 | 0 | 0 | 0 |
| ENSG00000215547 | 0 | 0 | 0 | 0 | 0 | 0 |
| ENSG00000215546 | 0 | 0 | 0 | 0 | 0 | 0 |
| ENSG00000224628 | 4 | 0 | 0 | 1 | 0 | 0 |
| ENSG00000215545 | 0 | 0 | 0 | 0 | 0 | 0 |
| ENSG00000237929 | 0 | 0 | 0 | 0 | 0 | 0 |
| ENSG00000236121 | 0 | 0 | 0 | 0 | 0 | 0 |
| ENSG00000252649 | 0 | 0 | 0 | 0 | 0 | 0 |
| ENSG00000212717 | 0 | 0 | 0 | 0 | 0 | 0 |
| ENSG00000131068 | 0 | 0 | 0 | 0 | 0 | 0 |
| ENSG00000212134 | 0 | 0 | 0 | 0 | 0 | 0 |
| ENSG00000180483 | 0 | 0 | 0 | 0 | 0 | 0 |
| ENSG00000204548 | 0 | 0 | 0 | 0 | 0 | 0 |
| ENSG00000204547 | 0 | 0 | 0 | 0 | 0 | 0 |
| ENSG00000180424 | 0 | 0 | 0 | 0 | 0 | 0 |
| ENSG00000180383 | 0 | 0 | 0 | 0 | 0 | 0 |
| ENSG00000088320 | 0 | 0 | 0 | 0 | 0 | 0 |
| ENSG00000233354 | 0 | 0 | 0 | 0 | 0 | 0 |
| ENSG00000101294 | 6973 | 3448 | 4182 | 4191 | 3813 | 3956 |
| ENSG00000274364 | 2 | 1 | 1 | 2 | 1 | 0 |
| ENSG00000101898 | 0 | 0 | 0 | 0 | 0 | 0 |
| ENSG00000235313 | 4 | 7 | 10 | 7 | 5 | 5 |
| ENSG00000230613 | 4 | 6 | 2 | 4 | 1 | 2 |
| ENSG00000278276 | 1 | 2 | 2 | 1 | 3 | 1 |
| ENSG00000201770 | 0 | 0 | 0 | 0 | 0 | 0 |

| | | | | | | |
|---|---|---|---|---|---|---|
| ENSG00000125968 | 4385 | 1930 | 2024 | 2391 | 2068 | 2128 |
| ENSG00000264395 | 4 | 0 | 1 | 3 | 0 | 0 |
| ENSG00000131055 | 0 | 1 | 0 | 0 | 1 | 0 |
| ENSG00000171552 | 3626 | 1755 | 2078 | 1740 | 1522 | 1611 |
| ENSG00000236559 | 3 | 2 | 0 | 0 | 3 | 0 |
| ENSG00000281376 | 26 | 11 | 17 | 17 | 15 | 19 |
| ENSG00000088325 | 2754 | 1333 | 1695 | 1672 | 1959 | 1842 |
| ENSG00000230801 | 0 | 0 | 0 | 0 | 0 | 1 |
| ENSG00000101306 | 4 | 3 | 5 | 4 | 3 | 5 |
| ENSG00000179772 | 0 | 0 | 0 | 0 | 0 | 0 |
| ENSG00000149599 | 29 | 13 | 24 | 23 | 24 | 26 |
| ENSG00000131044 | 38 | 20 | 34 | 28 | 28 | 32 |
| ENSG00000199497 | 0 | 0 | 0 | 0 | 0 | 0 |
| ENSG00000088356 | 1686 | 821 | 974 | 1069 | 995 | 957 |
| ENSG00000260903 | 10 | 6 | 6 | 8 | 6 | 8 |
| ENSG00000278012 | 0 | 0 | 0 | 1 | 1 | 1 |
| ENSG00000252623 | 0 | 0 | 0 | 0 | 0 | 0 |
| ENSG00000101331 | 25 | 24 | 22 | 20 | 28 | 25 |
| ENSG00000226239 | 2 | 2 | 4 | 0 | 3 | 0 |
| ENSG00000212571 | 0 | 0 | 0 | 0 | 0 | 0 |
| ENSG00000101336 | 37 | 25 | 20 | 25 | 14 | 16 |
| ENSG00000101337 | 4962 | 2182 | 2913 | 2024 | 2294 | 2633 |
| ENSG00000275576 | 6 | 1 | 3 | 0 | 0 | 3 |
| ENSG00000224452 | 0 | 0 | 0 | 4 | 0 | 1 |
| ENSG00000235217 | 89 | 49 | 63 | 38 | 47 | 43 |
| ENSG00000126003 | 1737 | 717 | 954 | 666 | 790 | 863 |
| ENSG00000101346 | 2331 | 1067 | 1377 | 1066 | 1158 | 1345 |
| ENSG00000284125 | 0 | 0 | 0 | 0 | 0 | 0 |
| ENSG00000101350 | 2753 | 1197 | 1641 | 1021 | 1610 | 1345 |
| ENSG00000277692 | 0 | 3 | 1 | 1 | 0 | 0 |
| ENSG00000171456 | 1801 | 705 | 1036 | 746 | 950 | 1013 |
| ENSG00000197183 | 1546 | 770 | 997 | 967 | 1086 | 1212 |
| ENSG00000236772 | 7 | 3 | 8 | 3 | 10 | 6 |
| ENSG00000277301 | 13 | 2 | 7 | 12 | 23 | 22 |
| ENSG00000233293 | 0 | 0 | 0 | 0 | 0 | 0 |
| ENSG00000204393 | 0 | 0 | 2 | 0 | 1 | 0 |
| ENSG00000228156 | 2 | 1 | 3 | 1 | 0 | 0 |
| ENSG00000198547 | 4 | 0 | 0 | 0 | 0 | 0 |
| ENSG00000285382 | 0 | 0 | 0 | 0 | 0 | 0 |
| ENSG00000175730 | 6 | 1 | 5 | 4 | 2 | 4 |
| ENSG00000149600 | 996 | 530 | 659 | 690 | 611 | 598 |
| ENSG00000088305 | 683 | 321 | 403 | 286 | 408 | 353 |
| ENSG00000101367 | 4428 | 2348 | 2724 | 2412 | 2856 | 2766 |
| ENSG00000260536 | 0 | 0 | 0 | 0 | 1 | 0 |
| ENSG00000260257 | 30 | 8 | 21 | 11 | 18 | 11 |
| ENSG00000215529 | 0 | 1 | 1 | 1 | 0 | 1 |
| ENSG00000167098 | 0 | 0 | 0 | 0 | 0 | 0 |
| ENSG00000078898 | 0 | 0 | 0 | 1 | 1 | 0 |
| ENSG00000167104 | 0 | 0 | 0 | 0 | 0 | 0 |
| ENSG00000214245 | 0 | 0 | 0 | 0 | 0 | 0 |

| | | | | | | |
|---|---|---|---|---|---|---|
| ENSG00000186190 | 0 | 0 | 0 | 0 | 0 | 0 |
| ENSG00000186191 | 0 | 0 | 0 | 0 | 1 | 0 |
| ENSG00000131050 | 2 | 0 | 0 | 2 | 2 | 3 |
| ENSG00000237662 | 0 | 0 | 5 | 0 | 0 | 1 |
| ENSG00000183566 | 2 | 4 | 9 | 7 | 13 | 15 |
| ENSG00000224876 | 0 | 0 | 0 | 0 | 0 | 0 |
| ENSG00000131059 | 0 | 0 | 0 | 0 | 0 | 0 |
| ENSG00000198183 | 4 | 1 | 0 | 0 | 5 | 4 |
| ENSG00000233106 | 0 | 0 | 0 | 0 | 0 | 0 |
| ENSG00000125999 | 83 | 30 | 57 | 55 | 43 | 98 |
| ENSG00000233146 | 0 | 0 | 0 | 0 | 0 | 0 |
| ENSG00000125997 | 0 | 1 | 1 | 1 | 0 | 0 |
| ENSG00000101391 | 878 | 455 | 514 | 518 | 488 | 474 |
| ENSG00000101400 | 217 | 107 | 121 | 131 | 102 | 97 |
| ENSG00000078699 | 943 | 446 | 565 | 400 | 495 | 460 |
| ENSG00000275223 | 21 | 8 | 14 | 5 | 9 | 14 |
| ENSG00000125967 | 862 | 436 | 540 | 521 | 480 | 472 |
| ENSG00000149609 | 13 | 4 | 5 | 9 | 8 | 8 |
| ENSG00000182584 | 126 | 56 | 78 | 85 | 73 | 87 |
| ENSG00000271803 | 0 | 1 | 1 | 0 | 0 | 1 |
| ENSG00000101412 | 690 | 363 | 458 | 407 | 320 | 417 |
| ENSG00000101417 | 842 | 428 | 528 | 336 | 397 | 445 |
| ENSG00000229188 | 0 | 0 | 0 | 0 | 0 | 0 |
| ENSG00000131061 | 281 | 117 | 166 | 127 | 124 | 115 |
| ENSG00000230837 | 0 | 0 | 0 | 1 | 0 | 1 |
| ENSG00000230753 | 0 | 2 | 2 | 1 | 3 | 1 |
| ENSG00000101421 | 3936 | 1865 | 2278 | 2046 | 2176 | 2192 |
| ENSG00000214200 | 0 | 0 | 0 | 0 | 0 | 0 |
| ENSG00000228958 | 0 | 0 | 0 | 0 | 0 | 0 |
| ENSG00000285230 | 197 | 134 | 137 | 101 | 127 | 131 |
| ENSG00000125970 | 4323 | 2113 | 2547 | 2852 | 2619 | 2498 |
| ENSG00000228386 | 0 | 0 | 1 | 0 | 0 | 0 |
| ENSG00000264616 | 0 | 0 | 0 | 0 | 0 | 0 |
| ENSG00000125977 | 2832 | 1643 | 1559 | 1355 | 2033 | 1693 |
| ENSG00000225246 | 0 | 0 | 0 | 0 | 0 | 0 |
| ENSG00000101440 | 0 | 1 | 1 | 0 | 2 | 0 |
| ENSG00000214185 | 4 | 2 | 3 | 2 | 1 | 3 |
| ENSG00000250917 | 1 | 0 | 0 | 0 | 1 | 1 |
| ENSG00000236456 | 0 | 0 | 3 | 0 | 1 | 1 |
| ENSG00000101444 | 6370 | 3017 | 4009 | 4713 | 3870 | 4106 |
| ENSG00000272945 | 8 | 3 | 9 | 4 | 2 | 5 |
| ENSG00000234087 | 0 | 0 | 0 | 0 | 0 | 0 |
| ENSG00000078747 | 1763 | 893 | 989 | 594 | 1067 | 887 |
| ENSG00000201498 | 0 | 0 | 0 | 0 | 0 | 0 |
| ENSG00000234933 | 0 | 0 | 0 | 1 | 0 | 0 |
| ENSG00000236388 | 0 | 0 | 0 | 0 | 0 | 0 |
| ENSG00000231795 | 0 | 0 | 0 | 0 | 0 | 0 |
| ENSG00000207997 | 0 | 0 | 0 | 0 | 0 | 0 |
| ENSG00000181741 | 0 | 0 | 1 | 1 | 0 | 0 |
| ENSG00000277558 | 0 | 0 | 0 | 0 | 0 | 0 |

| | | | | | | |
|---|---|---|---|---|---|---|
| ENSG00000125971 | 2476 | 1206 | 1455 | 1445 | 1353 | 1267 |
| ENSG00000206582 | 1 | 1 | 0 | 0 | 0 | 0 |
| ENSG00000101460 | 1 | 0 | 1 | 1 | 0 | 2 |
| ENSG00000101464 | 783 | 443 | 486 | 507 | 481 | 502 |
| ENSG00000231034 | 0 | 0 | 0 | 0 | 0 | 0 |
| ENSG00000198646 | 1698 | 718 | 1025 | 627 | 888 | 907 |
| ENSG00000078804 | 625 | 320 | 341 | 209 | 274 | 222 |
| ENSG00000225336 | 0 | 0 | 0 | 0 | 0 | 0 |
| ENSG00000131067 | 399 | 148 | 260 | 163 | 195 | 191 |
| ENSG00000131069 | 1045 | 567 | 679 | 642 | 561 | 628 |
| ENSG00000100983 | 2354 | 1251 | 1419 | 1462 | 1350 | 1257 |
| ENSG00000078814 | 17 | 10 | 22 | 4 | 6 | 17 |
| ENSG00000207635 | 0 | 0 | 0 | 0 | 0 | 0 |
| ENSG00000283441 | 0 | 0 | 0 | 0 | 0 | 0 |
| ENSG00000100991 | 3006 | 1330 | 1925 | 1540 | 1588 | 1663 |
| ENSG00000202150 | 0 | 0 | 0 | 0 | 1 | 0 |
| ENSG00000088298 | 972 | 508 | 589 | 617 | 594 | 581 |
| ENSG00000201151 | 0 | 0 | 0 | 2 | 1 | 0 |
| ENSG00000101000 | 35 | 21 | 23 | 51 | 44 | 39 |
| ENSG00000278367 | 0 | 0 | 0 | 0 | 0 | 0 |
| ENSG00000200301 | 0 | 0 | 0 | 0 | 0 | 0 |
| ENSG00000126005 | 540 | 285 | 332 | 339 | 293 | 278 |
| ENSG00000229230 | 0 | 0 | 0 | 0 | 0 | 0 |
| ENSG00000125966 | 18 | 12 | 13 | 5 | 16 | 7 |
| ENSG00000279253 | 30 | 16 | 27 | 15 | 27 | 25 |
| ENSG00000261582 | 8 | 3 | 9 | 2 | 4 | 2 |
| ENSG00000242372 | 1906 | 1179 | 1395 | 1751 | 1331 | 1410 |
| ENSG00000235214 | 2 | 0 | 0 | 1 | 0 | 0 |
| ENSG00000125998 | 0 | 0 | 1 | 0 | 0 | 0 |
| ENSG00000101019 | 1501 | 699 | 866 | 690 | 823 | 731 |
| ENSG00000204183 | 0 | 0 | 0 | 0 | 0 | 0 |
| ENSG00000125965 | 0 | 3 | 0 | 1 | 3 | 0 |
| ENSG00000221763 | 0 | 0 | 1 | 0 | 0 | 0 |
| ENSG00000126001 | 1173 | 436 | 691 | 529 | 600 | 599 |
| ENSG00000230155 | 0 | 0 | 0 | 0 | 1 | 1 |
| ENSG00000242507 | 0 | 0 | 0 | 0 | 0 | 0 |
| ENSG00000125975 | 0 | 0 | 0 | 0 | 0 | 0 |
| ENSG00000125991 | 9452 | 4840 | 5778 | 5667 | 5162 | 5081 |
| ENSG00000224497 | 0 | 0 | 0 | 0 | 0 | 0 |
| ENSG00000088340 | 1733 | 737 | 1224 | 837 | 1072 | 1196 |
| ENSG00000227401 | 2 | 0 | 0 | 3 | 0 | 4 |
| ENSG00000061656 | 232 | 118 | 159 | 126 | 131 | 140 |
| ENSG00000214078 | 2308 | 1073 | 1372 | 1298 | 1254 | 1180 |
| ENSG00000272897 | 0 | 0 | 0 | 0 | 0 | 0 |
| ENSG00000222460 | 0 | 0 | 0 | 0 | 1 | 0 |
| ENSG00000252549 | 0 | 0 | 0 | 0 | 0 | 0 |
| ENSG00000244462 | 2757 | 1384 | 1691 | 1300 | 1603 | 1662 |
| ENSG00000244005 | 1285 | 662 | 781 | 796 | 798 | 804 |
| ENSG00000125995 | 2006 | 965 | 1197 | 1350 | 1117 | 999 |
| ENSG00000131051 | 6776 | 3381 | 4006 | 2317 | 3666 | 3232 |

| | | | | | | |
|---|---|---|---|---|---|---|
| ENSG00000225357 | 0 | 1 | 0 | 0 | 0 | 0 |
| ENSG00000283527 | 0 | 0 | 0 | 0 | 0 | 0 |
| ENSG00000025293 | 1050 | 543 | 631 | 345 | 601 | 537 |
| ENSG00000234274 | 0 | 0 | 0 | 0 | 0 | 2 |
| ENSG00000199676 | 0 | 0 | 0 | 0 | 0 | 0 |
| ENSG00000219608 | 0 | 0 | 0 | 0 | 1 | 0 |
| ENSG00000201221 | 0 | 0 | 0 | 0 | 0 | 0 |
| ENSG00000207053 | 0 | 0 | 0 | 0 | 0 | 0 |
| ENSG00000171222 | 1689 | 768 | 920 | 1138 | 808 | 800 |
| ENSG00000149646 | 4 | 4 | 2 | 3 | 0 | 1 |
| ENSG00000260032 | 9353 | 4054 | 5048 | 2619 | 5370 | 4520 |
| ENSG00000277235 | 4 | 6 | 6 | 2 | 4 | 5 |
| ENSG00000234139 | 1 | 2 | 5 | 1 | 2 | 1 |
| ENSG00000231470 | 0 | 0 | 0 | 0 | 1 | 0 |
| ENSG00000237063 | 5 | 2 | 2 | 1 | 3 | 2 |
| ENSG00000088367 | 2860 | 1192 | 1551 | 1318 | 1484 | 1507 |
| ENSG00000232406 | 6 | 3 | 5 | 4 | 1 | 2 |
| ENSG00000278035 | 5 | 1 | 5 | 3 | 1 | 2 |
| ENSG00000131043 | 1350 | 604 | 823 | 879 | 739 | 742 |
| ENSG00000080845 | 1492 | 799 | 989 | 928 | 898 | 931 |
| ENSG00000232907 | 3 | 2 | 0 | 4 | 0 | 3 |
| ENSG00000101335 | 0 | 0 | 0 | 0 | 0 | 0 |
| ENSG00000273986 | 0 | 0 | 0 | 0 | 0 | 0 |
| ENSG00000118707 | 1480 | 763 | 894 | 860 | 929 | 952 |
| ENSG00000259399 | 0 | 0 | 1 | 0 | 0 | 2 |
| ENSG00000101084 | 358 | 251 | 260 | 336 | 281 | 324 |
| ENSG00000101082 | 2 | 1 | 0 | 1 | 2 | 0 |
| ENSG00000227688 | 2 | 0 | 0 | 0 | 0 | 0 |
| ENSG00000101079 | 1592 | 791 | 938 | 728 | 872 | 813 |
| ENSG00000202255 | 0 | 0 | 0 | 0 | 0 | 0 |
| ENSG00000149636 | 446 | 228 | 301 | 152 | 238 | 219 |
| ENSG00000149639 | 2921 | 1303 | 1750 | 1134 | 1374 | 1603 |
| ENSG00000242509 | 0 | 0 | 0 | 0 | 0 | 0 |
| ENSG00000101342 | 3 | 0 | 0 | 0 | 1 | 0 |
| ENSG00000101347 | 883 | 340 | 440 | 261 | 359 | 339 |
| ENSG00000080839 | 170 | 109 | 112 | 56 | 117 | 88 |
| ENSG00000231241 | 0 | 0 | 1 | 0 | 2 | 1 |
| ENSG00000226144 | 0 | 0 | 0 | 0 | 0 | 0 |
| ENSG00000269846 | 0 | 0 | 1 | 1 | 0 | 1 |
| ENSG00000101353 | 20 | 9 | 15 | 10 | 9 | 7 |
| ENSG00000236159 | 0 | 0 | 0 | 0 | 0 | 0 |
| ENSG00000118705 | 10412 | 5387 | 6217 | 5043 | 6162 | 5590 |
| ENSG00000118702 | 0 | 0 | 0 | 0 | 2 | 0 |
| ENSG00000101363 | 1194 | 577 | 729 | 748 | 658 | 698 |
| ENSG00000197122 | 1644 | 770 | 983 | 951 | 966 | 916 |
| ENSG00000213979 | 2 | 0 | 0 | 0 | 0 | 0 |
| ENSG00000166619 | 1847 | 937 | 1094 | 1018 | 1052 | 967 |
| ENSG00000053438 | 12 | 9 | 13 | 4 | 10 | 8 |
| ENSG00000276603 | 5 | 2 | 4 | 4 | 6 | 4 |
| ENSG00000235044 | 1 | 0 | 2 | 1 | 1 | 2 |

| | | | | | | |
|---|---|---|---|---|---|---|
| ENSG00000274469 | 0 | 0 | 0 | 0 | 0 | 0 |
| ENSG00000228234 | 0 | 0 | 0 | 0 | 0 | 0 |
| ENSG00000225759 | 0 | 0 | 0 | 0 | 0 | 0 |
| ENSG00000204117 | 0 | 0 | 0 | 0 | 0 | 0 |
| ENSG00000132792 | 1372 | 704 | 842 | 742 | 712 | 730 |
| ENSG00000132821 | 2 | 1 | 1 | 1 | 0 | 0 |
| ENSG00000199683 | 0 | 1 | 0 | 0 | 0 | 0 |
| ENSG00000101407 | 1023 | 463 | 532 | 422 | 469 | 502 |
| ENSG00000101413 | 2034 | 907 | 1191 | 708 | 957 | 1030 |
| ENSG00000264767 | 0 | 0 | 0 | 0 | 0 | 0 |
| ENSG00000277550 | 0 | 0 | 0 | 2 | 0 | 0 |
| ENSG00000198959 | 168 | 160 | 81 | 137 | 171 | 129 |
| ENSG00000149633 | 0 | 0 | 0 | 0 | 0 | 0 |
| ENSG00000277829 | 0 | 0 | 0 | 0 | 0 | 0 |
| ENSG00000285144 | 0 | 0 | 0 | 0 | 0 | 0 |
| ENSG00000285347 | 0 | 0 | 0 | 0 | 0 | 0 |
| ENSG00000285100 | 0 | 0 | 0 | 0 | 0 | 0 |
| ENSG00000206249 | 0 | 0 | 1 | 0 | 0 | 0 |
| ENSG00000101425 | 1 | 0 | 0 | 0 | 0 | 0 |
| ENSG00000129988 | 4 | 6 | 2 | 8 | 9 | 4 |
| ENSG00000275285 | 0 | 0 | 0 | 0 | 0 | 0 |
| ENSG00000224635 | 10 | 5 | 4 | 3 | 9 | 10 |
| ENSG00000275401 | 2 | 0 | 2 | 1 | 2 | 1 |
| ENSG00000196756 | 375 | 171 | 242 | 257 | 204 | 265 |
| ENSG00000277034 | 0 | 0 | 0 | 0 | 0 | 0 |
| ENSG00000235408 | 5 | 4 | 4 | 1 | 10 | 3 |
| ENSG00000225091 | 0 | 2 | 0 | 2 | 1 | 1 |
| ENSG00000201512 | 2 | 3 | 0 | 2 | 5 | 4 |
| ENSG00000200354 | 0 | 1 | 0 | 1 | 0 | 0 |
| ENSG00000201811 | 0 | 0 | 0 | 0 | 0 | 0 |
| ENSG00000174365 | 409 | 191 | 253 | 249 | 217 | 235 |
| ENSG00000274309 | 0 | 0 | 0 | 0 | 0 | 0 |
| ENSG00000199266 | 10 | 7 | 7 | 4 | 3 | 2 |
| ENSG00000170471 | 2264 | 1076 | 1216 | 584 | 1148 | 971 |
| ENSG00000275453 | 0 | 0 | 0 | 1 | 0 | 0 |
| ENSG00000231307 | 2 | 0 | 1 | 0 | 0 | 0 |
| ENSG00000182035 | 0 | 0 | 0 | 0 | 0 | 0 |
| ENSG00000124143 | 2 | 1 | 0 | 0 | 0 | 0 |
| ENSG00000213959 | 0 | 0 | 0 | 0 | 0 | 0 |
| ENSG00000277965 | 0 | 0 | 0 | 0 | 0 | 0 |
| ENSG00000101438 | 0 | 0 | 0 | 0 | 0 | 0 |
| ENSG00000101442 | 367 | 137 | 221 | 206 | 218 | 219 |
| ENSG00000199691 | 5 | 2 | 5 | 1 | 3 | 5 |
| ENSG00000101445 | 0 | 0 | 0 | 0 | 0 | 0 |
| ENSG00000240474 | 0 | 0 | 0 | 0 | 0 | 0 |
| ENSG00000101447 | 495 | 232 | 285 | 390 | 359 | 378 |
| ENSG00000274825 | 0 | 4 | 0 | 2 | 2 | 1 |
| ENSG00000261431 | 2 | 1 | 2 | 1 | 2 | 1 |
| ENSG00000101452 | 357 | 133 | 209 | 140 | 185 | 174 |
| ENSG00000277581 | 2 | 0 | 0 | 0 | 0 | 0 |

| | | | | | | |
|---|---|---|---|---|---|---|
| ENSG00000223864 | 0 | 1 | 0 | 1 | 1 | 0 |
| ENSG00000230324 | 0 | 0 | 0 | 0 | 0 | 0 |
| ENSG00000224357 | 0 | 0 | 0 | 0 | 0 | 0 |
| ENSG00000275337 | 0 | 0 | 0 | 0 | 0 | 0 |
| ENSG00000229976 | 0 | 0 | 0 | 0 | 0 | 0 |
| ENSG00000234878 | 0 | 0 | 0 | 0 | 0 | 0 |
| ENSG00000237767 | 0 | 0 | 0 | 0 | 0 | 0 |
| ENSG00000233415 | 0 | 0 | 0 | 0 | 0 | 0 |
| ENSG00000223410 | 0 | 0 | 0 | 0 | 0 | 0 |
| ENSG00000204103 | 638 | 243 | 318 | 184 | 246 | 259 |
| ENSG00000229771 | 0 | 1 | 0 | 0 | 1 | 0 |
| ENSG00000238908 | 0 | 0 | 0 | 0 | 0 | 0 |
| ENSG00000222612 | 33 | 9 | 17 | 13 | 10 | 25 |
| ENSG00000198900 | 7429 | 3715 | 4242 | 2965 | 4651 | 3955 |
| ENSG00000226648 | 6 | 3 | 4 | 2 | 7 | 4 |
| ENSG00000124181 | 7923 | 3338 | 4924 | 3851 | 4587 | 4592 |
| ENSG00000276169 | 2 | 0 | 0 | 0 | 0 | 0 |
| ENSG00000174306 | 3364 | 1399 | 1956 | 1307 | 1648 | 1716 |
| ENSG00000272407 | 0 | 0 | 0 | 0 | 0 | 0 |
| ENSG00000242748 | 1 | 1 | 0 | 0 | 1 | 0 |
| ENSG00000263989 | 0 | 0 | 0 | 0 | 0 | 0 |
| ENSG00000235102 | 0 | 0 | 0 | 0 | 0 | 0 |
| ENSG00000132793 | 1696 | 716 | 1069 | 667 | 877 | 911 |
| ENSG00000183798 | 6 | 6 | 2 | 3 | 2 | 6 |
| ENSG00000229398 | 0 | 0 | 0 | 0 | 0 | 0 |
| ENSG00000124177 | 5847 | 2690 | 3193 | 1326 | 2942 | 2577 |
| ENSG00000235088 | 3 | 0 | 0 | 0 | 0 | 0 |
| ENSG00000273951 | 10 | 6 | 1 | 2 | 2 | 8 |
| ENSG00000232493 | 0 | 1 | 0 | 0 | 0 | 2 |
| ENSG00000252386 | 0 | 0 | 0 | 0 | 0 | 0 |
| ENSG00000212224 | 0 | 0 | 0 | 0 | 0 | 0 |
| ENSG00000225458 | 0 | 0 | 0 | 0 | 0 | 0 |
| ENSG00000223651 | 0 | 0 | 0 | 0 | 0 | 0 |
| ENSG00000228959 | 0 | 0 | 0 | 0 | 0 | 0 |
| ENSG00000196090 | 20 | 8 | 14 | 7 | 9 | 7 |
| ENSG00000229892 | 0 | 0 | 0 | 0 | 0 | 0 |
| ENSG00000229042 | 0 | 0 | 0 | 0 | 0 | 0 |
| ENSG00000227599 | 0 | 0 | 0 | 0 | 0 | 0 |
| ENSG00000223190 | 0 | 0 | 0 | 0 | 0 | 0 |
| ENSG00000233508 | 0 | 0 | 0 | 0 | 0 | 0 |
| ENSG00000240639 | 0 | 0 | 0 | 0 | 0 | 0 |
| ENSG00000226038 | 0 | 0 | 0 | 0 | 0 | 1 |
| ENSG00000252193 | 0 | 0 | 0 | 0 | 0 | 0 |
| ENSG00000237555 | 0 | 0 | 1 | 0 | 1 | 0 |
| ENSG00000201021 | 0 | 0 | 0 | 0 | 0 | 0 |
| ENSG00000233711 | 0 | 0 | 0 | 0 | 0 | 0 |
| ENSG00000206998 | 0 | 0 | 0 | 0 | 0 | 0 |
| ENSG00000124193 | 7566 | 3451 | 4851 | 3537 | 3786 | 4201 |
| ENSG00000201372 | 0 | 0 | 0 | 0 | 0 | 0 |
| ENSG00000185513 | 117 | 48 | 84 | 45 | 78 | 57 |

| | | | | | | |
|---|---|---|---|---|---|---|
| ENSG00000234271 | 0 | 0 | 0 | 0 | 0 | 0 |
| ENSG00000277611 | 0 | 0 | 0 | 0 | 0 | 0 |
| ENSG00000226143 | 0 | 0 | 0 | 0 | 0 | 0 |
| ENSG00000101049 | 13 | 1 | 7 | 7 | 2 | 6 |
| ENSG00000282876 | 4 | 4 | 4 | 4 | 0 | 5 |
| ENSG00000101052 | 826 | 414 | 531 | 470 | 557 | 487 |
| ENSG00000215467 | 0 | 0 | 0 | 0 | 0 | 0 |
| ENSG00000101057 | 2375 | 1193 | 1418 | 1606 | 1287 | 1463 |
| ENSG00000124196 | 0 | 0 | 0 | 0 | 0 | 0 |
| ENSG00000206801 | 0 | 0 | 0 | 0 | 0 | 0 |
| ENSG00000233277 | 0 | 0 | 0 | 0 | 0 | 0 |
| ENSG00000124191 | 50 | 27 | 25 | 31 | 24 | 20 |
| ENSG00000241229 | 0 | 0 | 0 | 0 | 0 | 0 |
| ENSG00000149596 | 4 | 0 | 2 | 1 | 0 | 1 |
| ENSG00000132823 | 987 | 469 | 643 | 471 | 522 | 541 |
| ENSG00000223891 | 162 | 82 | 112 | 92 | 115 | 80 |
| ENSG00000124194 | 1 | 3 | 1 | 0 | 0 | 1 |
| ENSG00000197296 | 841 | 454 | 568 | 427 | 566 | 545 |
| ENSG00000101074 | 0 | 0 | 0 | 0 | 0 | 0 |
| ENSG00000201363 | 0 | 0 | 0 | 0 | 0 | 0 |
| ENSG00000226812 | 0 | 0 | 0 | 0 | 0 | 0 |
| ENSG00000101076 | 0 | 0 | 1 | 0 | 1 | 0 |
| ENSG00000229005 | 0 | 0 | 0 | 0 | 0 | 0 |
| ENSG00000233376 | 0 | 0 | 0 | 0 | 0 | 0 |
| ENSG00000266151 | 0 | 0 | 0 | 0 | 0 | 0 |
| ENSG00000168746 | 0 | 0 | 0 | 0 | 0 | 0 |
| ENSG00000226243 | 6 | 2 | 1 | 1 | 0 | 1 |
| ENSG00000124120 | 1330 | 603 | 727 | 470 | 647 | 640 |
| ENSG00000132824 | 6336 | 2921 | 3774 | 2259 | 3429 | 3248 |
| ENSG00000168734 | 466 | 210 | 286 | 328 | 245 | 303 |
| ENSG00000196839 | 102 | 47 | 42 | 98 | 67 | 71 |
| ENSG00000283440 | 0 | 0 | 0 | 0 | 0 | 0 |
| ENSG00000132832 | 0 | 1 | 0 | 0 | 1 | 1 |
| ENSG00000244558 | 27 | 24 | 20 | 35 | 18 | 22 |
| ENSG00000064205 | 931 | 490 | 532 | 1044 | 871 | 853 |
| ENSG00000124249 | 244 | 169 | 183 | 267 | 191 | 227 |
| ENSG00000276223 | 23 | 11 | 12 | 24 | 17 | 12 |
| ENSG00000101098 | 232 | 119 | 184 | 167 | 170 | 135 |
| ENSG00000277708 | 0 | 0 | 0 | 0 | 0 | 0 |
| ENSG00000166913 | 11004 | 5528 | 6273 | 4758 | 6574 | 5707 |
| ENSG00000101104 | 928 | 400 | 606 | 343 | 460 | 555 |
| ENSG00000025772 | 1164 | 651 | 714 | 733 | 778 | 862 |
| ENSG00000227477 | 40 | 6 | 18 | 12 | 7 | 18 |
| ENSG00000101109 | 1612 | 774 | 954 | 646 | 865 | 875 |
| ENSG00000252021 | 0 | 0 | 0 | 0 | 0 | 0 |
| ENSG00000124134 | 0 | 0 | 0 | 0 | 0 | 0 |
| ENSG00000175121 | 0 | 0 | 0 | 0 | 0 | 0 |
| ENSG00000168703 | 0 | 1 | 0 | 0 | 0 | 0 |
| ENSG00000283142 | 0 | 0 | 0 | 0 | 1 | 0 |
| ENSG00000124102 | 0 | 0 | 0 | 1 | 2 | 0 |

| | | | | | | |
|---|---|---|---|---|---|---|
| ENSG00000124233 | 0 | 0 | 0 | 0 | 0 | 0 |
| ENSG00000124157 | 0 | 0 | 0 | 0 | 0 | 0 |
| ENSG00000233352 | 0 | 0 | 0 | 0 | 0 | 0 |
| ENSG00000243995 | 0 | 0 | 0 | 0 | 0 | 0 |
| ENSG00000124107 | 6 | 6 | 8 | 1 | 1 | 5 |
| ENSG00000124159 | 6 | 1 | 3 | 1 | 3 | 1 |
| ENSG00000124232 | 0 | 2 | 0 | 0 | 1 | 0 |
| ENSG00000124145 | 7975 | 3800 | 5013 | 3705 | 3991 | 4226 |
| ENSG00000275894 | 0 | 2 | 0 | 1 | 0 | 1 |
| ENSG00000204070 | 1122 | 522 | 651 | 545 | 560 | 557 |
| ENSG00000254806 | 0 | 0 | 2 | 2 | 0 | 1 |
| ENSG00000124251 | 28 | 16 | 17 | 5 | 11 | 19 |
| ENSG00000232880 | 0 | 0 | 0 | 0 | 0 | 1 |
| ENSG00000244274 | 18 | 16 | 9 | 13 | 10 | 17 |
| ENSG00000124155 | 4929 | 2428 | 3106 | 2556 | 2399 | 2599 |
| ENSG00000273555 | 0 | 1 | 1 | 1 | 0 | 0 |
| ENSG00000277022 | 6 | 1 | 7 | 8 | 5 | 2 |
| ENSG00000101443 | 19 | 6 | 14 | 7 | 7 | 9 |
| ENSG00000237464 | 0 | 0 | 0 | 0 | 0 | 0 |
| ENSG00000237068 | 0 | 0 | 0 | 0 | 0 | 0 |
| ENSG00000101446 | 0 | 0 | 0 | 0 | 0 | 0 |
| ENSG00000243543 | 0 | 0 | 0 | 0 | 0 | 0 |
| ENSG00000232135 | 0 | 0 | 0 | 0 | 0 | 0 |
| ENSG00000249139 | 0 | 0 | 0 | 0 | 0 | 0 |
| ENSG00000229900 | 0 | 0 | 0 | 0 | 0 | 0 |
| ENSG00000101448 | 0 | 0 | 0 | 0 | 0 | 0 |
| ENSG00000158901 | 0 | 0 | 0 | 0 | 0 | 0 |
| ENSG00000201923 | 0 | 0 | 0 | 0 | 0 | 0 |
| ENSG00000230860 | 0 | 0 | 0 | 0 | 0 | 0 |
| ENSG00000180205 | 0 | 0 | 0 | 0 | 0 | 0 |
| ENSG00000180305 | 0 | 0 | 0 | 0 | 0 | 0 |
| ENSG00000235508 | 8 | 9 | 3 | 14 | 6 | 8 |
| ENSG00000180083 | 0 | 0 | 0 | 0 | 0 | 0 |
| ENSG00000182931 | 0 | 0 | 1 | 0 | 0 | 0 |
| ENSG00000168634 | 10 | 1 | 4 | 3 | 2 | 2 |
| ENSG00000266071 | 0 | 0 | 0 | 0 | 0 | 0 |
| ENSG00000149651 | 0 | 0 | 0 | 0 | 0 | 0 |
| ENSG00000226188 | 0 | 0 | 0 | 0 | 0 | 0 |
| ENSG00000124116 | 0 | 2 | 5 | 2 | 1 | 1 |
| ENSG00000168630 | 0 | 0 | 0 | 0 | 0 | 0 |
| ENSG00000221046 | 0 | 0 | 0 | 0 | 0 | 0 |
| ENSG00000101457 | 1090 | 516 | 604 | 643 | 595 | 582 |
| ENSG00000175063 | 878 | 442 | 572 | 806 | 819 | 710 |
| ENSG00000101470 | 9 | 1 | 11 | 4 | 1 | 2 |
| ENSG00000124104 | 589 | 326 | 437 | 320 | 286 | 332 |
| ENSG00000101473 | 876 | 476 | 523 | 505 | 465 | 416 |
| ENSG00000277051 | 2 | 1 | 0 | 0 | 0 | 0 |
| ENSG00000132801 | 236 | 119 | 159 | 121 | 123 | 143 |
| ENSG00000168612 | 564 | 286 | 325 | 282 | 284 | 302 |
| ENSG00000149634 | 35 | 35 | 32 | 9 | 19 | 20 |

| | | | | | | |
|---|---|---|---|---|---|---|
| ENSG00000124257 | 8 | 8 | 9 | 5 | 8 | 7 |
| ENSG00000064601 | 4673 | 2638 | 3344 | 2792 | 2602 | 2605 |
| ENSG00000271984 | 0 | 0 | 0 | 0 | 0 | 0 |
| ENSG00000100979 | 2 | 4 | 7 | 3 | 9 | 2 |
| ENSG00000285796 | 418 | 197 | 240 | 179 | 176 | 176 |
| ENSG00000100982 | 1104 | 536 | 700 | 569 | 545 | 591 |
| ENSG00000198026 | 1317 | 588 | 812 | 599 | 630 | 625 |
| ENSG00000235525 | 0 | 0 | 0 | 0 | 0 | 0 |
| ENSG00000100985 | 30 | 39 | 24 | 27 | 25 | 18 |
| ENSG00000204044 | 42 | 35 | 48 | 33 | 28 | 29 |
| ENSG00000124140 | 428 | 185 | 255 | 146 | 190 | 189 |
| ENSG00000124160 | 1998 | 941 | 1097 | 1017 | 1031 | 1110 |
| ENSG00000213820 | 0 | 0 | 0 | 0 | 0 | 0 |
| ENSG00000101017 | 0 | 0 | 0 | 0 | 0 | 1 |
| ENSG00000229957 | 0 | 0 | 0 | 0 | 0 | 0 |
| ENSG00000149654 | 0 | 2 | 0 | 0 | 0 | 0 |
| ENSG00000080189 | 3172 | 1484 | 1970 | 1819 | 1617 | 1726 |
| ENSG00000273828 | 5 | 5 | 1 | 0 | 2 | 1 |
| ENSG00000062598 | 2196 | 946 | 1331 | 1000 | 1223 | 1193 |
| ENSG00000215452 | 0 | 0 | 0 | 3 | 0 | 0 |
| ENSG00000283757 | 0 | 0 | 0 | 0 | 0 | 0 |
| ENSG00000225849 | 0 | 0 | 0 | 0 | 0 | 0 |
| ENSG00000184617 | 0 | 0 | 0 | 0 | 0 | 0 |
| ENSG00000198185 | 0 | 1 | 1 | 0 | 0 | 1 |
| ENSG00000149635 | 0 | 0 | 0 | 0 | 0 | 0 |
| ENSG00000158296 | 69 | 36 | 57 | 52 | 40 | 67 |
| ENSG00000273893 | 21 | 17 | 14 | 30 | 15 | 31 |
| ENSG00000172315 | 1197 | 583 | 743 | 648 | 684 | 693 |
| ENSG00000271784 | 2 | 0 | 1 | 1 | 5 | 3 |
| ENSG00000197496 | 3456 | 1508 | 2106 | 1255 | 1504 | 1691 |
| ENSG00000222874 | 0 | 0 | 0 | 0 | 0 | 0 |
| ENSG00000235922 | 0 | 0 | 0 | 0 | 0 | 0 |
| ENSG00000064655 | 131 | 111 | 69 | 64 | 83 | 48 |
| ENSG00000236028 | 0 | 0 | 0 | 0 | 0 | 0 |
| ENSG00000226769 | 0 | 0 | 0 | 0 | 0 | 0 |
| ENSG00000239783 | 0 | 0 | 0 | 0 | 0 | 0 |
| ENSG00000215450 | 1 | 0 | 0 | 0 | 0 | 0 |
| ENSG00000264901 | 0 | 0 | 0 | 0 | 0 | 0 |
| ENSG00000101040 | 4239 | 2030 | 2445 | 1147 | 1952 | 1935 |
| ENSG00000267882 | 10 | 2 | 2 | 1 | 5 | 7 |
| ENSG00000273451 | 18 | 12 | 13 | 2 | 5 | 7 |
| ENSG00000231119 | 23 | 9 | 17 | 3 | 8 | 13 |
| ENSG00000224565 | 4 | 6 | 2 | 6 | 3 | 9 |
| ENSG00000234075 | 0 | 0 | 0 | 0 | 1 | 0 |
| ENSG00000202186 | 0 | 0 | 0 | 0 | 1 | 0 |
| ENSG00000201742 | 0 | 0 | 0 | 0 | 0 | 0 |
| ENSG00000124151 | 52026 | 23943 | 29757 | 11796 | 23464 | 20376 |
| ENSG00000229853 | 8 | 10 | 4 | 3 | 4 | 5 |
| ENSG00000196562 | 24412 | 12034 | 15586 | 13885 | 15046 | 15000 |
| ENSG00000255438 | 37 | 22 | 27 | 15 | 28 | 21 |

| | | | | | | |
|---|---|---|---|---|---|---|
| ENSG00000238950 | 0 | 0 | 0 | 0 | 0 | 0 |
| ENSG00000215448 | 0 | 0 | 0 | 0 | 0 | 0 |
| ENSG00000251770 | 0 | 0 | 0 | 0 | 0 | 0 |
| ENSG00000238785 | 0 | 0 | 0 | 0 | 0 | 0 |
| ENSG00000234967 | 48 | 49 | 59 | 31 | 44 | 45 |
| ENSG00000228503 | 35 | 29 | 39 | 12 | 23 | 36 |
| ENSG00000237423 | 11 | 2 | 7 | 4 | 8 | 5 |
| ENSG00000229522 | 1 | 0 | 0 | 0 | 0 | 1 |
| ENSG00000276923 | 0 | 0 | 0 | 0 | 0 | 0 |
| ENSG00000235621 | 34 | 10 | 22 | 11 | 23 | 18 |
| ENSG00000236874 | 0 | 0 | 0 | 0 | 0 | 0 |
| ENSG00000274261 | 0 | 0 | 0 | 0 | 0 | 0 |
| ENSG00000281365 | 0 | 0 | 0 | 0 | 0 | 0 |
| ENSG00000281460 | 0 | 0 | 0 | 0 | 0 | 0 |
| ENSG00000238452 | 0 | 0 | 0 | 0 | 0 | 0 |
| ENSG00000124126 | 21143 | 8762 | 11848 | 7508 | 7321 | 7531 |
| ENSG00000278231 | 44 | 26 | 30 | 24 | 14 | 29 |
| ENSG00000124198 | 3834 | 1739 | 2093 | 1030 | 2110 | 1774 |
| ENSG00000230758 | 0 | 1 | 5 | 1 | 1 | 4 |
| ENSG00000227431 | 4 | 0 | 4 | 2 | 6 | 1 |
| ENSG00000124207 | 4209 | 2513 | 2180 | 1620 | 3794 | 2583 |
| ENSG00000124214 | 8600 | 4129 | 5276 | 4296 | 5173 | 5217 |
| ENSG00000226284 | 5 | 1 | 3 | 1 | 1 | 3 |
| ENSG00000124228 | 3315 | 1790 | 2156 | 1966 | 2166 | 2197 |
| ENSG00000124201 | 4189 | 1889 | 2492 | 1394 | 2076 | 1935 |
| ENSG00000177410 | 1703 | 1493 | 1964 | 2176 | 2087 | 2068 |
| ENSG00000277830 | 0 | 0 | 0 | 0 | 0 | 0 |
| ENSG00000209042 | 0 | 1 | 0 | 0 | 0 | 3 |
| ENSG00000274760 | 0 | 0 | 0 | 0 | 0 | 0 |
| ENSG00000222365 | 7 | 7 | 3 | 4 | 0 | 6 |
| ENSG00000277967 | 0 | 0 | 0 | 0 | 0 | 0 |
| ENSG00000212304 | 0 | 0 | 0 | 0 | 0 | 0 |
| ENSG00000158445 | 11 | 13 | 10 | 6 | 10 | 8 |
| ENSG00000283529 | 2 | 0 | 0 | 0 | 0 | 0 |
| ENSG00000124212 | 0 | 1 | 0 | 0 | 0 | 0 |
| ENSG00000158470 | 1988 | 967 | 1140 | 878 | 1071 | 1117 |
| ENSG00000276031 | 0 | 0 | 0 | 0 | 0 | 0 |
| ENSG00000231878 | 0 | 2 | 0 | 0 | 2 | 0 |
| ENSG00000252540 | 0 | 0 | 0 | 0 | 0 | 0 |
| ENSG00000197818 | 1097 | 454 | 686 | 513 | 543 | 610 |
| ENSG00000237788 | 2 | 0 | 4 | 3 | 0 | 2 |
| ENSG00000158480 | 1342 | 640 | 819 | 846 | 742 | 816 |
| ENSG00000199490 | 0 | 1 | 2 | 0 | 0 | 0 |
| ENSG00000269103 | 0 | 0 | 0 | 0 | 0 | 0 |
| ENSG00000124226 | 7399 | 3389 | 4465 | 3742 | 4047 | 4471 |
| ENSG00000229222 | 14 | 5 | 6 | 4 | 3 | 6 |
| ENSG00000206710 | 0 | 0 | 0 | 0 | 0 | 0 |
| ENSG00000124216 | 10 | 5 | 12 | 8 | 18 | 17 |
| ENSG00000231265 | 0 | 0 | 0 | 0 | 0 | 0 |
| ENSG00000124208 | 0 | 0 | 0 | 0 | 0 | 0 |

| | | | | | | |
|---|---|---|---|---|---|---|
| ENSG00000244687 | 389 | 194 | 318 | 181 | 251 | 253 |
| ENSG00000240849 | 5036 | 2583 | 3124 | 3260 | 3024 | 3130 |
| ENSG00000237595 | 2 | 0 | 1 | 0 | 3 | 0 |
| ENSG00000234698 | 0 | 0 | 0 | 0 | 0 | 3 |
| ENSG00000231742 | 0 | 0 | 1 | 0 | 0 | 0 |
| ENSG00000277449 | 98 | 57 | 64 | 25 | 50 | 50 |
| ENSG00000172216 | 1412 | 704 | 878 | 984 | 914 | 897 |
| ENSG00000224397 | 2 | 0 | 0 | 0 | 0 | 0 |
| ENSG00000203999 | 14 | 7 | 5 | 5 | 8 | 9 |
| ENSG00000233077 | 2 | 0 | 1 | 0 | 1 | 1 |
| ENSG00000244376 | 0 | 0 | 0 | 0 | 0 | 0 |
| ENSG00000231715 | 0 | 0 | 0 | 0 | 1 | 0 |
| ENSG00000196396 | 5711 | 2740 | 3603 | 2853 | 3089 | 3295 |
| ENSG00000239742 | 0 | 0 | 1 | 0 | 0 | 0 |
| ENSG00000201501 | 0 | 0 | 0 | 0 | 0 | 0 |
| ENSG00000232043 | 11 | 6 | 14 | 1 | 6 | 7 |
| ENSG00000208018 | 0 | 0 | 0 | 0 | 0 | 1 |
| ENSG00000042062 | 150 | 106 | 116 | 58 | 121 | 67 |
| ENSG00000221091 | 0 | 0 | 0 | 0 | 0 | 0 |
| ENSG00000231302 | 0 | 0 | 0 | 0 | 0 | 0 |
| ENSG00000234693 | 1 | 1 | 2 | 1 | 3 | 1 |
| ENSG00000124171 | 5092 | 2193 | 2959 | 1579 | 2494 | 2126 |
| ENSG00000124243 | 3023 | 1419 | 1813 | 1746 | 1516 | 1611 |
| ENSG00000230043 | 0 | 0 | 0 | 0 | 0 | 0 |
| ENSG00000101126 | 6535 | 3020 | 3875 | 2391 | 3337 | 3422 |
| ENSG00000228663 | 0 | 1 | 1 | 0 | 1 | 4 |
| ENSG00000259456 | 41 | 26 | 27 | 16 | 23 | 13 |
| ENSG00000000419 | 3016 | 1607 | 1928 | 1695 | 1885 | 1784 |
| ENSG00000124217 | 962 | 523 | 645 | 572 | 597 | 653 |
| ENSG00000232358 | 0 | 0 | 0 | 0 | 0 | 0 |
| ENSG00000026559 | 22 | 14 | 15 | 19 | 26 | 21 |
| ENSG00000279835 | 0 | 0 | 0 | 0 | 0 | 0 |
| ENSG00000228820 | 0 | 0 | 0 | 0 | 0 | 0 |
| ENSG00000101096 | 414 | 159 | 263 | 146 | 203 | 226 |
| ENSG00000266761 | 0 | 0 | 0 | 0 | 0 | 0 |
| ENSG00000054793 | 14961 | 6603 | 9516 | 6376 | 8882 | 9453 |
| ENSG00000215444 | 1 | 0 | 4 | 0 | 2 | 3 |
| ENSG00000101115 | 236 | 114 | 148 | 99 | 166 | 141 |
| ENSG00000227964 | 0 | 0 | 0 | 0 | 1 | 2 |
| ENSG00000252467 | 0 | 0 | 0 | 0 | 0 | 0 |
| ENSG00000252654 | 0 | 0 | 0 | 0 | 0 | 0 |
| ENSG00000020256 | 898 | 444 | 527 | 499 | 495 | 509 |
| ENSG00000233347 | 2 | 1 | 0 | 0 | 0 | 0 |
| ENSG00000234948 | 0 | 0 | 3 | 0 | 0 | 1 |
| ENSG00000225785 | 0 | 0 | 0 | 2 | 0 | 0 |
| ENSG00000275958 | 0 | 0 | 0 | 0 | 0 | 0 |
| ENSG00000232465 | 0 | 0 | 0 | 0 | 0 | 0 |
| ENSG00000234821 | 0 | 0 | 0 | 0 | 0 | 0 |
| ENSG00000232294 | 0 | 0 | 0 | 0 | 0 | 0 |
| ENSG00000232286 | 0 | 0 | 0 | 0 | 0 | 0 |

| | | | | | | |
|---|---|---|---|---|---|---|
| ENSG00000230636 | 0 | 0 | 0 | 0 | 0 | 0 |
| ENSG00000182463 | 20 | 27 | 18 | 11 | 17 | 17 |
| ENSG00000199218 | 0 | 0 | 0 | 0 | 0 | 0 |
| ENSG00000223492 | 0 | 0 | 0 | 0 | 0 | 0 |
| ENSG00000227705 | 0 | 0 | 0 | 0 | 0 | 0 |
| ENSG00000271774 | 0 | 0 | 0 | 0 | 0 | 0 |
| ENSG00000231703 | 18 | 10 | 12 | 10 | 13 | 24 |
| ENSG00000224530 | 4 | 0 | 0 | 0 | 0 | 0 |
| ENSG00000259723 | 53 | 38 | 46 | 25 | 55 | 41 |
| ENSG00000206959 | 0 | 0 | 0 | 0 | 0 | 0 |
| ENSG00000197670 | 255 | 182 | 229 | 50 | 137 | 122 |
| ENSG00000171940 | 22523 | 10356 | 13618 | 5899 | 9808 | 9451 |
| ENSG00000235415 | 8 | 9 | 14 | 9 | 6 | 8 |
| ENSG00000238468 | 0 | 0 | 0 | 0 | 0 | 0 |
| ENSG00000225563 | 1 | 3 | 2 | 1 | 0 | 4 |
| ENSG00000241721 | 0 | 2 | 0 | 0 | 1 | 0 |
| ENSG00000064787 | 4836 | 2430 | 2956 | 1720 | 2851 | 2417 |
| ENSG00000236352 | 0 | 0 | 0 | 0 | 1 | 0 |
| ENSG00000265595 | 4 | 2 | 0 | 0 | 2 | 0 |
| ENSG00000019186 | 33 | 13 | 8 | 4 | 8 | 19 |
| ENSG00000101132 | 9450 | 4418 | 4956 | 5129 | 5890 | 5528 |
| ENSG00000101134 | 0 | 0 | 0 | 0 | 0 | 0 |
| ENSG00000252089 | 0 | 0 | 0 | 0 | 0 | 0 |
| ENSG00000185834 | 58 | 32 | 35 | 35 | 31 | 23 |
| ENSG00000235166 | 1 | 0 | 2 | 1 | 5 | 5 |
| ENSG00000224008 | 112 | 32 | 50 | 36 | 39 | 48 |
| ENSG00000275923 | 0 | 0 | 0 | 0 | 0 | 0 |
| ENSG00000054803 | 56 | 13 | 35 | 15 | 25 | 14 |
| ENSG00000222118 | 0 | 0 | 0 | 0 | 0 | 0 |
| ENSG00000124089 | 0 | 0 | 0 | 0 | 0 | 0 |
| ENSG00000235594 | 0 | 0 | 0 | 0 | 0 | 0 |
| ENSG00000230843 | 0 | 0 | 0 | 0 | 0 | 0 |
| ENSG00000124098 | 3243 | 1590 | 2010 | 1373 | 1900 | 1779 |
| ENSG00000087586 | 2655 | 1330 | 1571 | 1643 | 2088 | 1882 |
| ENSG00000101138 | 2642 | 1354 | 1537 | 1292 | 1475 | 1453 |
| ENSG00000087589 | 23 | 16 | 14 | 34 | 30 | 34 |
| ENSG00000228601 | 0 | 0 | 0 | 0 | 0 | 0 |
| ENSG00000022277 | 5285 | 3065 | 3539 | 3434 | 3385 | 3463 |
| ENSG00000124091 | 12 | 10 | 19 | 1 | 6 | 5 |
| ENSG00000124103 | 3 | 0 | 5 | 2 | 3 | 2 |
| ENSG00000280387 | 2 | 1 | 1 | 0 | 5 | 4 |
| ENSG00000213714 | 1 | 0 | 2 | 3 | 3 | 2 |
| ENSG00000225657 | 0 | 0 | 0 | 0 | 1 | 0 |
| ENSG00000214541 | 0 | 2 | 0 | 1 | 0 | 1 |
| ENSG00000087510 | 4552 | 2218 | 2879 | 2085 | 2342 | 2518 |
| ENSG00000252091 | 0 | 1 | 0 | 0 | 0 | 0 |
| ENSG00000241821 | 0 | 0 | 0 | 0 | 0 | 0 |
| ENSG00000237000 | 0 | 0 | 0 | 0 | 0 | 0 |
| ENSG00000231604 | 0 | 0 | 0 | 0 | 0 | 0 |
| ENSG00000207158 | 0 | 0 | 0 | 0 | 0 | 0 |

| | | | | | | |
|---|---|---|---|---|---|---|
| ENSG00000231078 | 0 | 0 | 0 | 0 | 0 | 0 |
| ENSG00000203266 | 9 | 6 | 7 | 2 | 2 | 4 |
| ENSG00000280220 | 0 | 1 | 2 | 0 | 0 | 1 |
| ENSG00000101144 | 6760 | 3238 | 4053 | 2685 | 2608 | 2271 |
| ENSG00000235032 | 0 | 1 | 0 | 0 | 0 | 0 |
| ENSG00000226308 | 6 | 8 | 5 | 2 | 1 | 1 |
| ENSG00000252064 | 0 | 0 | 0 | 0 | 0 | 0 |
| ENSG00000266666 | 0 | 0 | 0 | 0 | 0 | 0 |
| ENSG00000231005 | 0 | 0 | 0 | 0 | 0 | 0 |
| ENSG00000054796 | 0 | 0 | 1 | 0 | 0 | 0 |
| ENSG00000101146 | 6219 | 3163 | 3895 | 3526 | 3669 | 3774 |
| ENSG00000228995 | 2 | 1 | 0 | 1 | 0 | 0 |
| ENSG00000256222 | 2 | 0 | 3 | 0 | 2 | 1 |
| ENSG00000218018 | 7 | 2 | 4 | 9 | 5 | 3 |
| ENSG00000132819 | 2477 | 1240 | 1417 | 1725 | 1461 | 1564 |
| ENSG00000124097 | 36 | 27 | 40 | 16 | 35 | 28 |
| ENSG00000124092 | 0 | 0 | 0 | 0 | 0 | 0 |
| ENSG00000124253 | 0 | 0 | 0 | 0 | 0 | 0 |
| ENSG00000227297 | 0 | 0 | 0 | 0 | 0 | 0 |
| ENSG00000124256 | 0 | 0 | 1 | 1 | 1 | 0 |
| ENSG00000124225 | 4069 | 2076 | 2093 | 2535 | 2786 | 2142 |
| ENSG00000278709 | 376 | 219 | 257 | 254 | 243 | 234 |
| ENSG00000263453 | 0 | 0 | 0 | 0 | 0 | 0 |
| ENSG00000203971 | 0 | 0 | 0 | 0 | 0 | 0 |
| ENSG00000275852 | 2 | 0 | 2 | 0 | 0 | 0 |
| ENSG00000225708 | 0 | 0 | 0 | 0 | 0 | 0 |
| ENSG00000270411 | 2 | 0 | 4 | 1 | 6 | 5 |
| ENSG00000124237 | 0 | 0 | 0 | 0 | 0 | 0 |
| ENSG00000124227 | 0 | 0 | 0 | 0 | 0 | 0 |
| ENSG00000124224 | 1084 | 406 | 585 | 304 | 476 | 456 |
| ENSG00000124209 | 5418 | 2981 | 3430 | 2537 | 3477 | 3155 |
| ENSG00000124164 | 8389 | 3897 | 5125 | 3799 | 3984 | 4504 |
| ENSG00000198768 | 4 | 0 | 0 | 0 | 1 | 0 |
| ENSG00000231290 | 6 | 1 | 3 | 2 | 0 | 3 |
| ENSG00000254620 | 2 | 0 | 0 | 0 | 0 | 1 |
| ENSG00000268941 | 2 | 0 | 0 | 1 | 0 | 2 |
| ENSG00000124222 | 8763 | 3708 | 5415 | 3084 | 4873 | 4758 |
| ENSG00000254995 | 49 | 23 | 58 | 25 | 20 | 37 |
| ENSG00000215440 | 1687 | 853 | 1282 | 1086 | 961 | 1077 |
| ENSG00000254419 | 58 | 16 | 43 | 25 | 30 | 31 |
| ENSG00000237121 | 0 | 3 | 1 | 0 | 0 | 0 |
| ENSG00000270279 | 0 | 0 | 0 | 0 | 0 | 0 |
| ENSG00000268649 | 34 | 10 | 28 | 18 | 19 | 13 |
| ENSG00000284040 | 0 | 0 | 0 | 0 | 0 | 0 |
| ENSG00000216031 | 6 | 0 | 0 | 1 | 0 | 0 |
| ENSG00000235590 | 35 | 36 | 36 | 25 | 37 | 35 |
| ENSG00000268333 | 9 | 10 | 9 | 8 | 15 | 16 |
| ENSG00000087460 | 59330 | 29152 | 36322 | 32409 | 30891 | 30950 |
| ENSG00000276859 | 0 | 0 | 0 | 0 | 0 | 0 |
| ENSG00000276722 | 0 | 0 | 0 | 0 | 0 | 0 |

| | | | | | | |
|---|---|---|---|---|---|---|
| ENSG00000274491 | 0 | 0 | 0 | 0 | 0 | 0 |
| ENSG00000275069 | 0 | 0 | 0 | 0 | 0 | 0 |
| ENSG00000225806 | 10 | 7 | 10 | 6 | 6 | 4 |
| ENSG00000270951 | 6 | 1 | 6 | 1 | 1 | 0 |
| ENSG00000285091 | 25 | 19 | 24 | 12 | 22 | 14 |
| ENSG00000284985 | 0 | 0 | 0 | 0 | 0 | 0 |
| ENSG00000101158 | 9218 | 4078 | 5779 | 5037 | 5090 | 5918 |
| ENSG00000101160 | 67 | 28 | 53 | 29 | 36 | 38 |
| ENSG00000101162 | 14 | 7 | 10 | 2 | 5 | 6 |
| ENSG00000124172 | 14233 | 6831 | 8145 | 8484 | 8185 | 7555 |
| ENSG00000101166 | 6157 | 3168 | 3636 | 2910 | 4219 | 3715 |
| ENSG00000232925 | 0 | 0 | 0 | 0 | 0 | 0 |
| ENSG00000124203 | 0 | 0 | 0 | 3 | 0 | 0 |
| ENSG00000124205 | 0 | 0 | 0 | 0 | 0 | 0 |
| ENSG00000225129 | 0 | 0 | 0 | 0 | 0 | 0 |
| ENSG00000230352 | 0 | 0 | 0 | 0 | 0 | 0 |
| ENSG00000233686 | 0 | 0 | 0 | 0 | 0 | 0 |
| ENSG00000227282 | 0 | 0 | 0 | 0 | 0 | 0 |
| ENSG00000087495 | 3 | 0 | 0 | 0 | 1 | 0 |
| ENSG00000238194 | 0 | 0 | 0 | 0 | 0 | 0 |
| ENSG00000238777 | 0 | 0 | 0 | 0 | 1 | 0 |
| ENSG00000196074 | 396 | 215 | 240 | 88 | 356 | 245 |
| ENSG00000196227 | 1312 | 558 | 740 | 409 | 739 | 642 |
| ENSG00000132825 | 1448 | 667 | 937 | 703 | 769 | 812 |
| ENSG00000124215 | 60 | 42 | 29 | 65 | 72 | 51 |
| ENSG00000176659 | 0 | 1 | 2 | 1 | 1 | 2 |
| ENSG00000276093 | 0 | 0 | 0 | 0 | 0 | 0 |
| ENSG00000228340 | 2 | 0 | 1 | 0 | 0 | 0 |
| ENSG00000207802 | 0 | 0 | 0 | 0 | 0 | 0 |
| ENSG00000229690 | 0 | 0 | 0 | 0 | 0 | 0 |
| ENSG00000266140 | 0 | 0 | 0 | 0 | 0 | 0 |
| ENSG00000265617 | 0 | 0 | 0 | 0 | 0 | 0 |
| ENSG00000276627 | 0 | 0 | 0 | 0 | 0 | 0 |
| ENSG00000261035 | 0 | 0 | 0 | 0 | 0 | 0 |
| ENSG00000229755 | 0 | 0 | 0 | 0 | 0 | 0 |
| ENSG00000229606 | 0 | 0 | 0 | 0 | 0 | 0 |
| ENSG00000179242 | 40 | 18 | 32 | 9 | 13 | 23 |
| ENSG00000276786 | 0 | 0 | 0 | 0 | 0 | 0 |
| ENSG00000277270 | 0 | 0 | 0 | 0 | 0 | 1 |
| ENSG00000179253 | 0 | 0 | 0 | 0 | 0 | 0 |
| ENSG00000225106 | 0 | 0 | 0 | 0 | 0 | 0 |
| ENSG00000130699 | 1407 | 711 | 867 | 591 | 806 | 792 |
| ENSG00000284193 | 0 | 0 | 0 | 0 | 0 | 0 |
| ENSG00000235018 | 0 | 0 | 1 | 0 | 0 | 0 |
| ENSG00000265306 | 0 | 0 | 0 | 0 | 0 | 0 |
| ENSG00000283078 | 70 | 27 | 32 | 32 | 42 | 26 |
| ENSG00000149657 | 3933 | 1835 | 2526 | 1992 | 2167 | 2450 |
| ENSG00000101182 | 12775 | 6317 | 7578 | 8062 | 7756 | 7698 |
| ENSG00000184402 | 1909 | 830 | 1239 | 752 | 1117 | 1010 |
| ENSG00000101181 | 1908 | 938 | 1269 | 1024 | 1077 | 1052 |

| | | | | | | |
|---|---|---|---|---|---|---|
| ENSG00000101180 | 2 | 0 | 4 | 1 | 4 | 4 |
| ENSG00000130703 | 4830 | 2275 | 3022 | 2209 | 2618 | 2777 |
| ENSG00000130706 | 5228 | 3317 | 4190 | 4426 | 3783 | 3965 |
| ENSG00000226332 | 143 | 67 | 101 | 91 | 70 | 80 |
| ENSG00000130702 | 10262 | 3818 | 5986 | 4307 | 4323 | 4872 |
| ENSG00000284074 | 0 | 0 | 0 | 0 | 0 | 0 |
| ENSG00000228812 | 2 | 0 | 1 | 0 | 1 | 1 |
| ENSG00000273619 | 267 | 140 | 127 | 159 | 121 | 121 |
| ENSG00000171858 | 16252 | 8141 | 9764 | 12268 | 10184 | 10012 |
| ENSG00000149679 | 1867 | 928 | 1202 | 906 | 1046 | 999 |
| ENSG00000275437 | 98 | 42 | 84 | 46 | 55 | 57 |
| ENSG00000130701 | 885 | 380 | 515 | 471 | 446 | 383 |
| ENSG00000233017 | 24 | 14 | 14 | 10 | 7 | 17 |
| ENSG00000130700 | 2 | 0 | 2 | 2 | 1 | 2 |
| ENSG00000272050 | 0 | 0 | 0 | 0 | 0 | 0 |
| ENSG00000260542 | 0 | 0 | 0 | 0 | 0 | 0 |
| ENSG00000174403 | 0 | 0 | 0 | 0 | 0 | 0 |
| ENSG00000174407 | 0 | 0 | 0 | 0 | 0 | 0 |
| ENSG00000199017 | 0 | 0 | 0 | 0 | 0 | 0 |
| ENSG00000284508 | 0 | 0 | 0 | 0 | 0 | 0 |
| ENSG00000226094 | 0 | 0 | 0 | 0 | 0 | 0 |
| ENSG00000264490 | 0 | 0 | 0 | 0 | 0 | 0 |
| ENSG00000273812 | 0 | 0 | 0 | 0 | 0 | 0 |
| ENSG00000232121 | 0 | 0 | 0 | 0 | 0 | 0 |
| ENSG00000229882 | 0 | 1 | 0 | 0 | 0 | 0 |
| ENSG00000167046 | 19 | 5 | 6 | 4 | 3 | 8 |
| ENSG00000101187 | 892 | 417 | 531 | 705 | 674 | 649 |
| ENSG00000277496 | 42 | 25 | 22 | 22 | 18 | 13 |
| ENSG00000276317 | 9 | 5 | 5 | 5 | 4 | 8 |
| ENSG00000232803 | 3 | 0 | 7 | 6 | 4 | 9 |
| ENSG00000237687 | 2 | 0 | 0 | 0 | 0 | 0 |
| ENSG00000101188 | 49 | 26 | 46 | 38 | 31 | 29 |
| ENSG00000223669 | 8 | 1 | 7 | 1 | 1 | 4 |
| ENSG00000228705 | 45 | 22 | 35 | 36 | 23 | 23 |
| ENSG00000101189 | 2513 | 1206 | 1505 | 1547 | 1382 | 1473 |
| ENSG00000229873 | 4 | 5 | 7 | 1 | 3 | 5 |
| ENSG00000060491 | 3641 | 1820 | 2413 | 2306 | 1989 | 2020 |
| ENSG00000092758 | 25 | 25 | 34 | 18 | 31 | 27 |
| ENSG00000101190 | 1728 | 891 | 1082 | 974 | 1015 | 1045 |
| ENSG00000233838 | 0 | 0 | 0 | 0 | 0 | 0 |
| ENSG00000236527 | 5 | 2 | 3 | 2 | 0 | 5 |
| ENSG00000101191 | 5432 | 2405 | 3547 | 2085 | 2678 | 2866 |
| ENSG00000273759 | 22 | 11 | 14 | 15 | 26 | 19 |
| ENSG00000101193 | 8227 | 4058 | 5205 | 4088 | 4335 | 4778 |
| ENSG00000101194 | 0 | 0 | 0 | 1 | 2 | 0 |
| ENSG00000125533 | 0 | 0 | 0 | 0 | 0 | 0 |
| ENSG00000272259 | 0 | 1 | 0 | 0 | 1 | 2 |
| ENSG00000125514 | 0 | 0 | 0 | 0 | 0 | 0 |
| ENSG00000237119 | 1 | 0 | 0 | 0 | 0 | 0 |
| ENSG00000231133 | 51 | 29 | 38 | 69 | 78 | 61 |

| | | | | | | |
|---|---|---|---|---|---|---|
| ENSG00000274915 | 0 | 0 | 0 | 0 | 0 | 0 |
| ENSG00000225978 | 6 | 3 | 8 | 23 | 12 | 22 |
| ENSG00000260416 | 0 | 0 | 0 | 0 | 0 | 0 |
| ENSG00000231977 | 0 | 1 | 0 | 0 | 0 | 1 |
| ENSG00000207598 | 0 | 0 | 0 | 0 | 0 | 0 |
| ENSG00000149658 | 4438 | 2163 | 2753 | 2340 | 2327 | 2711 |
| ENSG00000273821 | 30 | 11 | 15 | 22 | 15 | 15 |
| ENSG00000101197 | 5 | 2 | 6 | 2 | 1 | 7 |
| ENSG00000266463 | 0 | 0 | 0 | 0 | 0 | 0 |
| ENSG00000101198 | 5 | 3 | 3 | 1 | 2 | 0 |
| ENSG00000275620 | 0 | 1 | 0 | 0 | 0 | 0 |
| ENSG00000101199 | 4400 | 2011 | 2652 | 2568 | 2560 | 2329 |
| ENSG00000266104 | 1 | 0 | 1 | 0 | 0 | 0 |
| ENSG00000101203 | 11 | 3 | 8 | 2 | 6 | 3 |
| ENSG00000222439 | 0 | 0 | 0 | 0 | 0 | 0 |
| ENSG00000101204 | 0 | 0 | 0 | 0 | 0 | 1 |
| ENSG00000203900 | 0 | 0 | 0 | 0 | 0 | 0 |
| ENSG00000075043 | 9 | 6 | 14 | 10 | 12 | 14 |
| ENSG00000280936 | 0 | 0 | 0 | 0 | 0 | 0 |
| ENSG00000226390 | 0 | 0 | 0 | 0 | 0 | 1 |
| ENSG00000101210 | 30407 | 15393 | 18291 | 23778 | 20291 | 17708 |
| ENSG00000230226 | 2 | 2 | 0 | 1 | 1 | 2 |
| ENSG00000125534 | 12515 | 5319 | 7860 | 7922 | 5396 | 6189 |
| ENSG00000101213 | 1722 | 821 | 1077 | 842 | 878 | 812 |
| ENSG00000125508 | 96 | 41 | 67 | 58 | 45 | 67 |
| ENSG00000275812 | 2 | 5 | 3 | 5 | 9 | 6 |
| ENSG00000125531 | 2 | 2 | 1 | 2 | 1 | 0 |
| ENSG00000130589 | 2819 | 1040 | 1799 | 1450 | 1421 | 1554 |
| ENSG00000101216 | 1518 | 736 | 916 | 805 | 844 | 760 |
| ENSG00000232442 | 380 | 181 | 295 | 244 | 251 | 279 |
| ENSG00000197457 | 1010 | 495 | 693 | 653 | 573 | 593 |
| ENSG00000258366 | 50 | 23 | 36 | 28 | 35 | 50 |
| ENSG00000026036 | 146 | 65 | 109 | 61 | 99 | 80 |
| ENSG00000243509 | 0 | 0 | 0 | 0 | 0 | 0 |
| ENSG00000101246 | 1806 | 845 | 1265 | 1023 | 821 | 937 |
| ENSG00000197114 | 283 | 141 | 224 | 205 | 192 | 222 |
| ENSG00000273154 | 28 | 18 | 29 | 42 | 41 | 46 |
| ENSG00000274501 | 2 | 0 | 0 | 0 | 1 | 1 |
| ENSG00000203896 | 215 | 106 | 136 | 123 | 150 | 127 |
| ENSG00000273047 | 0 | 0 | 0 | 0 | 0 | 0 |
| ENSG00000125520 | 5299 | 2529 | 3289 | 3378 | 2761 | 2757 |
| ENSG00000130584 | 1392 | 656 | 794 | 692 | 724 | 727 |
| ENSG00000229299 | 0 | 0 | 0 | 0 | 0 | 0 |
| ENSG00000231208 | 0 | 0 | 0 | 0 | 0 | 0 |
| ENSG00000279610 | 0 | 0 | 0 | 0 | 1 | 0 |
| ENSG00000268858 | 587 | 245 | 363 | 235 | 259 | 289 |
| ENSG00000183260 | 0 | 0 | 0 | 0 | 0 | 0 |
| ENSG00000101150 | 10588 | 4979 | 6406 | 6195 | 5695 | 6393 |
| ENSG00000101152 | 7861 | 3894 | 5169 | 4275 | 4289 | 4483 |
| ENSG00000199805 | 0 | 1 | 2 | 1 | 0 | 0 |

| | | | | | | |
|---|---|---|---|---|---|---|
| ENSG00000283206 | 0 | 2 | 0 | 0 | 0 | 0 |
| ENSG00000216141 | 0 | 0 | 0 | 0 | 0 | 0 |
| ENSG00000283513 | 0 | 0 | 0 | 0 | 0 | 0 |
| ENSG00000216195 | 0 | 0 | 0 | 0 | 0 | 0 |
| ENSG00000275842 | 0 | 0 | 0 | 0 | 0 | 0 |
| ENSG00000198276 | 1887 | 789 | 1169 | 1075 | 937 | 1065 |
| ENSG00000284433 | 0 | 0 | 0 | 0 | 0 | 0 |
| ENSG00000207554 | 12 | 9 | 18 | 9 | 15 | 14 |
| ENSG00000280213 | 49 | 24 | 32 | 13 | 26 | 26 |
| ENSG00000196700 | 5922 | 2649 | 3783 | 3016 | 3157 | 3317 |
| ENSG00000130590 | 693 | 305 | 424 | 406 | 390 | 380 |
| ENSG00000101161 | 9175 | 4357 | 5516 | 5450 | 4654 | 5069 |
| ENSG00000196421 | 95 | 33 | 61 | 32 | 33 | 34 |
| ENSG00000203883 | 52 | 15 | 17 | 19 | 25 | 17 |
| ENSG00000171703 | 2218 | 1067 | 1435 | 1397 | 1212 | 1230 |
| ENSG00000237371 | 0 | 2 | 2 | 3 | 1 | 0 |
| ENSG00000171700 | 1212 | 533 | 639 | 727 | 582 | 624 |
| ENSG00000274034 | 2 | 0 | 1 | 0 | 0 | 0 |
| ENSG00000125510 | 97 | 41 | 56 | 45 | 44 | 41 |
| ENSG00000171695 | 6 | 1 | 5 | 0 | 0 | 4 |
| ENSG00000196132 | 6 | 2 | 12 | 14 | 7 | 8 |
| ENSG00000125522 | 0 | 0 | 0 | 0 | 0 | 0 |
| ENSG00000203880 | 3430 | 1485 | 2138 | 1018 | 1606 | 1702 |
| ENSG00000278171 | 2 | 0 | 0 | 0 | 0 | 0 |
| ENSG00000232852 | 4 | 2 | 0 | 0 | 1 | 1 |
| ENSG00000149656 | 2 | 1 | 1 | 0 | 0 | 0 |
| ENSG00000274727 | 0 | 0 | 0 | 0 | 0 | 0 |
| ENSG00000282458 | 1 | 3 | 2 | 3 | 3 | 2 |
| ENSG00000282542 | 0 | 0 | 0 | 0 | 0 | 0 |
| ENSG00000282798 | 0 | 0 | 0 | 0 | 0 | 0 |
| ENSG00000282807 | 0 | 0 | 0 | 0 | 0 | 0 |
| ENSG00000283801 | 0 | 0 | 0 | 0 | 0 | 0 |
| ENSG00000282591 | 0 | 0 | 0 | 0 | 0 | 0 |
| ENSG00000282137 | 0 | 0 | 0 | 0 | 0 | 0 |
| ENSG00000267310 | 0 | 0 | 0 | 0 | 0 | 0 |
| ENSG00000176695 | 0 | 0 | 0 | 0 | 0 | 0 |
| ENSG00000267792 | 0 | 0 | 0 | 0 | 0 | 0 |
| ENSG00000266971 | 0 | 0 | 0 | 0 | 0 | 0 |
| ENSG00000282535 | 0 | 0 | 0 | 0 | 0 | 0 |
| ENSG00000281379 | 0 | 0 | 0 | 0 | 0 | 0 |
| ENSG00000282059 | 0 | 0 | 0 | 0 | 0 | 0 |
| ENSG00000282416 | 0 | 0 | 0 | 0 | 0 | 0 |
| ENSG00000282051 | 0 | 4 | 1 | 1 | 0 | 0 |
| ENSG00000282508 | 67 | 20 | 42 | 19 | 23 | 34 |
| ENSG00000282339 | 0 | 0 | 0 | 0 | 0 | 0 |
| ENSG00000222329 | 0 | 0 | 0 | 0 | 0 | 0 |
| ENSG00000282393 | 2 | 1 | 2 | 1 | 0 | 3 |
| ENSG00000141934 | 1774 | 939 | 1190 | 1136 | 1045 | 1045 |
| ENSG00000105556 | 544 | 234 | 319 | 340 | 286 | 316 |
| ENSG00000267124 | 0 | 0 | 0 | 0 | 0 | 0 |

| | | | | | | |
|---|---|---|---|---|---|---|
| ENSG00000105549 | 8 | 13 | 7 | 5 | 6 | 8 |
| ENSG00000267443 | 0 | 0 | 0 | 0 | 0 | 0 |
| ENSG00000183186 | 48 | 30 | 22 | 18 | 18 | 18 |
| ENSG00000129946 | 192 | 113 | 120 | 138 | 107 | 105 |
| ENSG00000252539 | 0 | 0 | 0 | 0 | 0 | 0 |
| ENSG00000181781 | 2 | 1 | 0 | 0 | 1 | 1 |
| ENSG00000099866 | 22 | 11 | 21 | 12 | 13 | 8 |
| ENSG00000266933 | 3 | 3 | 7 | 4 | 2 | 6 |
| ENSG00000141933 | 162 | 93 | 90 | 100 | 86 | 68 |
| ENSG00000099804 | 1840 | 912 | 1008 | 1135 | 1013 | 885 |
| ENSG00000197540 | 2 | 4 | 1 | 0 | 2 | 0 |
| ENSG00000267751 | 71 | 60 | 91 | 46 | 75 | 77 |
| ENSG00000172270 | 25409 | 11313 | 14907 | 15736 | 11597 | 13119 |
| ENSG00000099822 | 1189 | 686 | 765 | 809 | 716 | 694 |
| ENSG00000267036 | 1 | 0 | 0 | 0 | 0 | 0 |
| ENSG00000099821 | 1744 | 846 | 1079 | 1136 | 1000 | 1018 |
| ENSG00000261204 | 11 | 6 | 1 | 7 | 1 | 4 |
| ENSG00000070388 | 12 | 10 | 8 | 7 | 15 | 3 |
| ENSG00000070423 | 857 | 495 | 580 | 686 | 622 | 597 |
| ENSG00000267666 | 60 | 18 | 27 | 23 | 21 | 25 |
| ENSG00000070404 | 746 | 367 | 466 | 581 | 511 | 405 |
| ENSG00000185198 | 2 | 0 | 0 | 0 | 0 | 0 |
| ENSG00000213726 | 0 | 0 | 0 | 0 | 0 | 0 |
| ENSG00000099864 | 1474 | 627 | 834 | 846 | 715 | 649 |
| ENSG00000099812 | 4627 | 2022 | 2701 | 2943 | 2109 | 2173 |
| ENSG00000272473 | 12 | 5 | 4 | 5 | 4 | 6 |
| ENSG00000267530 | 43 | 22 | 27 | 16 | 24 | 20 |
| ENSG00000011304 | 11947 | 5661 | 7030 | 6318 | 6416 | 5947 |
| ENSG00000284361 | 0 | 0 | 0 | 0 | 0 | 0 |
| ENSG00000129951 | 124 | 40 | 64 | 101 | 61 | 49 |
| ENSG00000263414 | 0 | 0 | 0 | 0 | 0 | 0 |
| ENSG00000172232 | 16 | 9 | 6 | 2 | 4 | 2 |
| ENSG00000196415 | 2 | 0 | 2 | 0 | 0 | 1 |
| ENSG00000197561 | 0 | 0 | 1 | 0 | 1 | 0 |
| ENSG00000197766 | 136 | 59 | 116 | 60 | 64 | 46 |
| ENSG00000175221 | 1452 | 732 | 972 | 883 | 703 | 705 |
| ENSG00000207507 | 6 | 1 | 4 | 2 | 1 | 1 |
| ENSG00000198858 | 2461 | 1139 | 1364 | 1428 | 1248 | 1169 |
| ENSG00000116014 | 0 | 0 | 0 | 0 | 0 | 0 |
| ENSG00000116017 | 1550 | 739 | 961 | 708 | 853 | 757 |
| ENSG00000267159 | 0 | 0 | 0 | 0 | 0 | 0 |
| ENSG00000275162 | 2 | 2 | 2 | 0 | 3 | 3 |
| ENSG00000065268 | 963 | 454 | 542 | 703 | 546 | 525 |
| ENSG00000266990 | 0 | 1 | 0 | 0 | 3 | 0 |
| ENSG00000116032 | 124 | 60 | 85 | 64 | 56 | 50 |
| ENSG00000182087 | 2138 | 1132 | 1533 | 1362 | 1466 | 1377 |
| ENSG00000274177 | 0 | 0 | 0 | 0 | 0 | 0 |
| ENSG00000207357 | 2 | 2 | 4 | 5 | 4 | 3 |
| ENSG00000064666 | 727 | 414 | 450 | 691 | 666 | 584 |
| ENSG00000279753 | 0 | 0 | 0 | 0 | 0 | 0 |

| | | | | | | |
|---|---|---|---|---|---|---|
| ENSG00000064687 | 570 | 226 | 293 | 212 | 270 | 311 |
| ENSG00000180448 | 638 | 293 | 390 | 386 | 323 | 317 |
| ENSG00000099817 | 3132 | 1567 | 1800 | 2142 | 1801 | 1672 |
| ENSG00000167468 | 5991 | 2744 | 3457 | 4185 | 2979 | 2989 |
| ENSG00000064932 | 1913 | 915 | 1261 | 1104 | 1065 | 1063 |
| ENSG00000118046 | 2272 | 1025 | 1307 | 1281 | 1247 | 1159 |
| ENSG00000267736 | 0 | 0 | 6 | 0 | 5 | 0 |
| ENSG00000099625 | 145 | 64 | 78 | 106 | 91 | 102 |
| ENSG00000267778 | 2 | 5 | 2 | 1 | 3 | 0 |
| ENSG00000099624 | 2196 | 1037 | 1269 | 1542 | 1163 | 1160 |
| ENSG00000167470 | 4515 | 2056 | 2582 | 2372 | 2146 | 2379 |
| ENSG00000099622 | 7597 | 3359 | 5059 | 3536 | 3307 | 3764 |
| ENSG00000267493 | 55 | 21 | 33 | 11 | 13 | 29 |
| ENSG00000228300 | 491 | 195 | 295 | 418 | 261 | 279 |
| ENSG00000099617 | 32 | 9 | 11 | 8 | 9 | 4 |
| ENSG00000240846 | 0 | 0 | 0 | 0 | 0 | 0 |
| ENSG00000267372 | 0 | 0 | 0 | 0 | 0 | 0 |
| ENSG00000160953 | 1023 | 497 | 629 | 566 | 498 | 523 |
| ENSG00000279488 | 13 | 7 | 3 | 2 | 1 | 3 |
| ENSG00000267755 | 0 | 0 | 0 | 0 | 0 | 0 |
| ENSG00000115286 | 379 | 336 | 417 | 487 | 350 | 330 |
| ENSG00000280486 | 5 | 2 | 3 | 0 | 2 | 3 |
| ENSG00000248015 | 4 | 3 | 6 | 2 | 3 | 5 |
| ENSG00000130005 | 390 | 137 | 185 | 151 | 107 | 126 |
| ENSG00000071626 | 4148 | 2027 | 2545 | 2495 | 2495 | 2394 |
| ENSG00000115268 | 11849 | 5851 | 6770 | 8042 | 6622 | 6318 |
| ENSG00000268798 | 9 | 1 | 5 | 3 | 4 | 3 |
| ENSG00000115266 | 163 | 94 | 133 | 115 | 107 | 105 |
| ENSG00000267317 | 124 | 86 | 84 | 131 | 83 | 77 |
| ENSG00000119559 | 567 | 296 | 449 | 431 | 334 | 365 |
| ENSG00000115257 | 77 | 36 | 66 | 53 | 40 | 42 |
| ENSG00000115255 | 748 | 318 | 460 | 452 | 348 | 345 |
| ENSG00000185761 | 195 | 70 | 99 | 79 | 88 | 68 |
| ENSG00000267092 | 1 | 2 | 2 | 1 | 2 | 9 |
| ENSG00000185988 | 16 | 3 | 6 | 1 | 2 | 12 |
| ENSG00000181588 | 1062 | 456 | 663 | 741 | 681 | 658 |
| ENSG00000239367 | 4 | 0 | 0 | 1 | 0 | 0 |
| ENSG00000071655 | 1143 | 606 | 786 | 897 | 738 | 759 |
| ENSG00000279009 | 0 | 0 | 0 | 0 | 0 | 0 |
| ENSG00000267059 | 0 | 0 | 0 | 0 | 0 | 0 |
| ENSG00000127540 | 617 | 398 | 500 | 514 | 405 | 446 |
| ENSG00000071564 | 3098 | 1383 | 1699 | 1737 | 1722 | 1620 |
| ENSG00000252933 | 3 | 1 | 2 | 4 | 1 | 0 |
| ENSG00000275642 | 0 | 0 | 0 | 0 | 0 | 0 |
| ENSG00000267073 | 2 | 0 | 0 | 0 | 0 | 0 |
| ENSG00000205922 | 0 | 0 | 0 | 0 | 0 | 0 |
| ENSG00000130270 | 20 | 11 | 12 | 16 | 8 | 9 |
| ENSG00000079313 | 569 | 290 | 361 | 383 | 332 | 333 |
| ENSG00000267007 | 0 | 0 | 0 | 0 | 0 | 0 |
| ENSG00000284216 | 0 | 0 | 0 | 0 | 0 | 0 |

| | | | | | | |
|---|---|---|---|---|---|---|
| ENSG00000267244 | 41 | 13 | 10 | 6 | 8 | 13 |
| ENSG00000267125 | 0 | 0 | 0 | 0 | 0 | 0 |
| ENSG00000129911 | 499 | 236 | 301 | 347 | 311 | 313 |
| ENSG00000267141 | 2 | 0 | 1 | 1 | 1 | 0 |
| ENSG00000261526 | 55 | 53 | 41 | 33 | 29 | 37 |
| ENSG00000267232 | 0 | 0 | 0 | 0 | 0 | 0 |
| ENSG00000129968 | 1612 | 646 | 864 | 947 | 823 | 876 |
| ENSG00000227500 | 1822 | 829 | 1152 | 989 | 798 | 930 |
| ENSG00000213638 | 107 | 54 | 65 | 81 | 56 | 57 |
| ENSG00000133275 | 2112 | 1002 | 1302 | 1289 | 1226 | 1147 |
| ENSG00000180846 | 5 | 0 | 1 | 2 | 0 | 1 |
| ENSG00000133243 | 2625 | 1208 | 1586 | 1748 | 1292 | 1443 |
| ENSG00000267283 | 1 | 1 | 0 | 1 | 0 | 1 |
| ENSG00000279653 | 0 | 0 | 0 | 0 | 0 | 1 |
| ENSG00000275468 | 2 | 5 | 0 | 3 | 1 | 0 |
| ENSG00000267180 | 0 | 0 | 0 | 0 | 0 | 0 |
| ENSG00000099875 | 5073 | 2120 | 2718 | 2508 | 2596 | 2495 |
| ENSG00000172081 | 846 | 399 | 537 | 471 | 377 | 475 |
| ENSG00000099840 | 108 | 33 | 41 | 37 | 34 | 26 |
| ENSG00000065000 | 4931 | 2241 | 2821 | 2408 | 2522 | 2558 |
| ENSG00000104885 | 1110 | 501 | 700 | 601 | 614 | 660 |
| ENSG00000267122 | 0 | 0 | 0 | 0 | 0 | 0 |
| ENSG00000104886 | 1476 | 802 | 952 | 1074 | 839 | 816 |
| ENSG00000221411 | 0 | 0 | 0 | 0 | 0 | 0 |
| ENSG00000284129 | 0 | 0 | 0 | 0 | 0 | 0 |
| ENSG00000104897 | 998 | 430 | 619 | 608 | 553 | 514 |
| ENSG00000104899 | 372 | 149 | 239 | 211 | 205 | 203 |
| ENSG00000284272 | 0 | 0 | 0 | 0 | 0 | 0 |
| ENSG00000167476 | 8 | 8 | 1 | 0 | 4 | 5 |
| ENSG00000104904 | 12036 | 6511 | 7638 | 8284 | 6826 | 6630 |
| ENSG00000273734 | 0 | 0 | 0 | 0 | 0 | 0 |
| ENSG00000188305 | 1 | 1 | 1 | 2 | 1 | 0 |
| ENSG00000220008 | 6 | 6 | 3 | 6 | 5 | 12 |
| ENSG00000130332 | 1222 | 615 | 721 | 853 | 764 | 684 |
| ENSG00000005206 | 1231 | 618 | 773 | 607 | 625 | 619 |
| ENSG00000178297 | 28 | 14 | 24 | 23 | 19 | 17 |
| ENSG00000099800 | 531 | 283 | 321 | 318 | 278 | 315 |
| ENSG00000176619 | 3798 | 1684 | 2043 | 1855 | 1980 | 1889 |
| ENSG00000283728 | 0 | 0 | 0 | 0 | 0 | 0 |
| ENSG00000267201 | 1 | 0 | 1 | 0 | 1 | 0 |
| ENSG00000099860 | 919 | 440 | 537 | 552 | 490 | 392 |
| ENSG00000252962 | 0 | 0 | 0 | 0 | 0 | 0 |
| ENSG00000176533 | 122 | 58 | 62 | 50 | 36 | 56 |
| ENSG00000105694 | 0 | 0 | 2 | 0 | 0 | 1 |
| ENSG00000263974 | 0 | 0 | 0 | 0 | 0 | 0 |
| ENSG00000267412 | 0 | 0 | 0 | 0 | 0 | 2 |
| ENSG00000277904 | 0 | 0 | 0 | 0 | 0 | 0 |
| ENSG00000267214 | 0 | 0 | 0 | 0 | 0 | 0 |
| ENSG00000267749 | 0 | 0 | 0 | 1 | 0 | 0 |
| ENSG00000276097 | 0 | 0 | 0 | 0 | 0 | 1 |

| | | | | | | |
|---|---|---|---|---|---|---|
| ENSG00000176490 | 139 | 57 | 77 | 105 | 73 | 63 |
| ENSG00000267001 | 0 | 0 | 0 | 0 | 0 | 0 |
| ENSG00000261342 | 13 | 22 | 17 | 23 | 18 | 13 |
| ENSG00000141873 | 359 | 172 | 188 | 263 | 181 | 226 |
| ENSG00000104969 | 1753 | 754 | 1071 | 1224 | 912 | 970 |
| ENSG00000172009 | 1116 | 562 | 686 | 843 | 695 | 681 |
| ENSG00000172006 | 166 | 74 | 112 | 78 | 90 | 94 |
| ENSG00000186300 | 90 | 51 | 65 | 30 | 67 | 60 |
| ENSG00000267063 | 0 | 1 | 0 | 1 | 0 | 0 |
| ENSG00000172000 | 0 | 1 | 0 | 0 | 0 | 0 |
| ENSG00000267398 | 0 | 0 | 0 | 0 | 0 | 0 |
| ENSG00000266938 | 0 | 1 | 0 | 0 | 0 | 0 |
| ENSG00000171970 | 102 | 48 | 51 | 47 | 73 | 62 |
| ENSG00000253392 | 0 | 0 | 3 | 1 | 1 | 1 |
| ENSG00000175691 | 101 | 58 | 60 | 58 | 56 | 65 |
| ENSG00000279452 | 18 | 11 | 13 | 16 | 9 | 12 |
| ENSG00000104953 | 100 | 33 | 59 | 32 | 35 | 39 |
| ENSG00000065717 | 771 | 314 | 446 | 360 | 314 | 321 |
| ENSG00000104964 | 3454 | 1902 | 2477 | 2512 | 2054 | 2104 |
| ENSG00000267469 | 0 | 0 | 0 | 0 | 0 | 0 |
| ENSG00000088256 | 1909 | 1087 | 1272 | 1272 | 1206 | 1223 |
| ENSG00000267139 | 0 | 0 | 0 | 0 | 0 | 0 |
| ENSG00000267688 | 0 | 0 | 0 | 0 | 0 | 0 |
| ENSG00000266944 | 0 | 0 | 2 | 0 | 0 | 0 |
| ENSG00000060558 | 0 | 0 | 0 | 0 | 0 | 0 |
| ENSG00000267551 | 0 | 0 | 0 | 0 | 0 | 0 |
| ENSG00000125910 | 0 | 0 | 0 | 0 | 0 | 1 |
| ENSG00000125912 | 1898 | 906 | 1086 | 1422 | 1225 | 1133 |
| ENSG00000161082 | 45 | 38 | 62 | 38 | 51 | 53 |
| ENSG00000267448 | 0 | 0 | 0 | 0 | 0 | 0 |
| ENSG00000141905 | 1667 | 720 | 964 | 769 | 705 | 803 |
| ENSG00000280103 | 2 | 0 | 1 | 1 | 0 | 0 |
| ENSG00000279599 | 0 | 0 | 0 | 0 | 0 | 0 |
| ENSG00000095932 | 0 | 0 | 1 | 0 | 0 | 1 |
| ENSG00000284638 | 0 | 0 | 0 | 0 | 0 | 0 |
| ENSG00000129932 | 298 | 133 | 173 | 190 | 158 | 183 |
| ENSG00000105325 | 1301 | 614 | 656 | 750 | 690 | 603 |
| ENSG00000252408 | 0 | 0 | 0 | 0 | 0 | 0 |
| ENSG00000161091 | 726 | 399 | 481 | 538 | 534 | 432 |
| ENSG00000183397 | 40 | 28 | 38 | 18 | 35 | 24 |
| ENSG00000267436 | 0 | 0 | 0 | 0 | 0 | 0 |
| ENSG00000267231 | 0 | 0 | 1 | 0 | 0 | 0 |
| ENSG00000279590 | 0 | 0 | 2 | 0 | 0 | 0 |
| ENSG00000266959 | 0 | 0 | 1 | 1 | 0 | 0 |
| ENSG00000064961 | 3892 | 1964 | 2281 | 2394 | 2024 | 2033 |
| ENSG00000179855 | 0 | 0 | 0 | 0 | 0 | 0 |
| ENSG00000006638 | 0 | 1 | 4 | 3 | 1 | 0 |
| ENSG00000226800 | 6 | 4 | 4 | 1 | 3 | 3 |
| ENSG00000105298 | 476 | 226 | 275 | 285 | 270 | 271 |
| ENSG00000186111 | 1429 | 607 | 888 | 800 | 643 | 790 |

| | | | | | | |
|---|---|---|---|---|---|---|
| ENSG00000267304 | 0 | 0 | 0 | 0 | 0 | 0 |
| ENSG00000105289 | 2649 | 1162 | 1531 | 1298 | 1225 | 1197 |
| ENSG00000011132 | 331 | 193 | 193 | 220 | 167 | 203 |
| ENSG00000267205 | 0 | 0 | 0 | 0 | 0 | 0 |
| ENSG00000267138 | 0 | 0 | 1 | 0 | 0 | 1 |
| ENSG00000183617 | 608 | 297 | 366 | 444 | 286 | 323 |
| ENSG00000173976 | 0 | 0 | 0 | 0 | 0 | 0 |
| ENSG00000267148 | 0 | 0 | 0 | 0 | 0 | 0 |
| ENSG00000007264 | 230 | 121 | 141 | 281 | 170 | 185 |
| ENSG00000105278 | 0 | 0 | 0 | 0 | 0 | 0 |
| ENSG00000266989 | 0 | 0 | 0 | 0 | 0 | 0 |
| ENSG00000167654 | 0 | 0 | 0 | 1 | 0 | 1 |
| ENSG00000266627 | 0 | 0 | 0 | 0 | 0 | 0 |
| ENSG00000077009 | 1 | 1 | 3 | 0 | 1 | 0 |
| ENSG00000167657 | 1499 | 729 | 854 | 901 | 882 | 780 |
| ENSG00000283928 | 0 | 0 | 0 | 0 | 0 | 0 |
| ENSG00000167658 | 106743 | 48074 | 63240 | 71336 | 59834 | 62046 |
| ENSG00000206775 | 2 | 0 | 0 | 0 | 0 | 0 |
| ENSG00000105229 | 1456 | 539 | 720 | 735 | 640 | 652 |
| ENSG00000178951 | 2973 | 1276 | 1742 | 1506 | 1351 | 1418 |
| ENSG00000268670 | 11 | 3 | 7 | 4 | 4 | 5 |
| ENSG00000126934 | 3866 | 1895 | 2361 | 2664 | 2319 | 2088 |
| ENSG00000278007 | 0 | 0 | 0 | 0 | 0 | 0 |
| ENSG00000060566 | 14 | 7 | 4 | 5 | 9 | 6 |
| ENSG00000077463 | 800 | 339 | 440 | 512 | 407 | 429 |
| ENSG00000089847 | 75 | 31 | 46 | 36 | 32 | 39 |
| ENSG00000105246 | 4 | 0 | 0 | 3 | 0 | 1 |
| ENSG00000280247 | 2 | 0 | 3 | 3 | 3 | 0 |
| ENSG00000105248 | 532 | 295 | 325 | 327 | 302 | 353 |
| ENSG00000105251 | 2 | 0 | 0 | 0 | 0 | 1 |
| ENSG00000167664 | 0 | 0 | 0 | 0 | 0 | 0 |
| ENSG00000105255 | 292 | 104 | 149 | 178 | 170 | 175 |
| ENSG00000178078 | 1939 | 1007 | 1162 | 1154 | 1186 | 1026 |
| ENSG00000269425 | 6 | 1 | 2 | 3 | 1 | 7 |
| ENSG00000008382 | 344 | 234 | 281 | 270 | 255 | 248 |
| ENSG00000269807 | 3 | 0 | 1 | 1 | 0 | 1 |
| ENSG00000267938 | 9 | 5 | 8 | 6 | 1 | 5 |
| ENSG00000269318 | 24 | 23 | 13 | 5 | 8 | 10 |
| ENSG00000141985 | 3040 | 1571 | 1855 | 1816 | 1685 | 1466 |
| ENSG00000267980 | 25 | 16 | 30 | 19 | 18 | 24 |
| ENSG00000167670 | 470 | 221 | 286 | 210 | 246 | 257 |
| ENSG00000267255 | 0 | 0 | 1 | 0 | 0 | 0 |
| ENSG00000167671 | 2204 | 1130 | 1331 | 1430 | 1171 | 1158 |
| ENSG00000266437 | 0 | 0 | 0 | 0 | 0 | 0 |
| ENSG00000267030 | 0 | 0 | 0 | 0 | 0 | 0 |
| ENSG00000280239 | 14 | 7 | 20 | 18 | 19 | 15 |
| ENSG00000267769 | 2 | 4 | 6 | 3 | 4 | 5 |
| ENSG00000267011 | 6 | 1 | 3 | 4 | 2 | 2 |
| ENSG00000167674 | 2254 | 1093 | 1323 | 1363 | 1216 | 1222 |
| ENSG00000167676 | 94 | 40 | 47 | 34 | 37 | 69 |

| | | | | | | |
|---|---|---|---|---|---|---|
| ENSG00000214456 | 33 | 16 | 15 | 15 | 19 | 9 |
| ENSG00000267385 | 2 | 0 | 0 | 0 | 0 | 0 |
| ENSG00000171236 | 69 | 29 | 20 | 20 | 28 | 31 |
| ENSG00000167680 | 803 | 346 | 496 | 420 | 345 | 333 |
| ENSG00000267408 | 0 | 0 | 0 | 0 | 0 | 0 |
| ENSG00000185361 | 135 | 54 | 64 | 98 | 91 | 82 |
| ENSG00000074842 | 1606 | 758 | 992 | 1060 | 955 | 905 |
| ENSG00000268565 | 0 | 0 | 0 | 0 | 0 | 0 |
| ENSG00000142002 | 1498 | 757 | 975 | 900 | 926 | 1033 |
| ENSG00000205790 | 3 | 3 | 5 | 1 | 1 | 2 |
| ENSG00000176840 | 0 | 0 | 0 | 0 | 1 | 0 |
| ENSG00000207630 | 0 | 0 | 0 | 0 | 0 | 0 |
| ENSG00000268536 | 0 | 0 | 1 | 1 | 0 | 1 |
| ENSG00000141965 | 85 | 39 | 41 | 44 | 64 | 66 |
| ENSG00000269604 | 0 | 0 | 0 | 0 | 0 | 0 |
| ENSG00000127666 | 306 | 141 | 198 | 198 | 173 | 173 |
| ENSG00000267484 | 0 | 0 | 1 | 0 | 0 | 0 |
| ENSG00000105355 | 2068 | 1002 | 1218 | 1221 | 1204 | 1081 |
| ENSG00000205784 | 0 | 0 | 0 | 0 | 0 | 0 |
| ENSG00000276043 | 952 | 393 | 588 | 368 | 368 | 506 |
| ENSG00000263409 | 0 | 0 | 0 | 0 | 0 | 0 |
| ENSG00000127663 | 1182 | 548 | 712 | 545 | 660 | 622 |
| ENSG00000105426 | 2727 | 1083 | 1773 | 1183 | 1575 | 1529 |
| ENSG00000267550 | 0 | 0 | 0 | 0 | 0 | 0 |
| ENSG00000239524 | 0 | 0 | 0 | 0 | 1 | 0 |
| ENSG00000274447 | 9 | 3 | 13 | 8 | 7 | 10 |
| ENSG00000105428 | 0 | 0 | 0 | 0 | 0 | 0 |
| ENSG00000223573 | 1467 | 694 | 882 | 1042 | 1191 | 940 |
| ENSG00000233270 | 20 | 9 | 11 | 16 | 7 | 12 |
| ENSG00000130254 | 1885 | 824 | 1149 | 891 | 1051 | 1091 |
| ENSG00000160633 | 2864 | 1353 | 1702 | 1328 | 1679 | 1643 |
| ENSG00000130255 | 7391 | 3684 | 4468 | 5253 | 4337 | 4239 |
| ENSG00000174917 | 606 | 276 | 333 | 399 | 315 | 324 |
| ENSG00000167733 | 76 | 37 | 44 | 37 | 36 | 33 |
| ENSG00000196365 | 1728 | 833 | 1104 | 1208 | 1014 | 1004 |
| ENSG00000174898 | 2 | 2 | 2 | 0 | 1 | 0 |
| ENSG00000212123 | 49 | 18 | 35 | 34 | 19 | 21 |
| ENSG00000267157 | 0 | 0 | 0 | 0 | 1 | 0 |
| ENSG00000141994 | 304 | 168 | 178 | 261 | 202 | 211 |
| ENSG00000171119 | 42 | 19 | 21 | 26 | 22 | 18 |
| ENSG00000156413 | 0 | 0 | 0 | 0 | 0 | 0 |
| ENSG00000171124 | 2 | 0 | 0 | 0 | 1 | 0 |
| ENSG00000267709 | 0 | 0 | 0 | 0 | 0 | 0 |
| ENSG00000130383 | 2 | 0 | 0 | 0 | 0 | 0 |
| ENSG00000267740 | 0 | 0 | 0 | 1 | 0 | 0 |
| ENSG00000174886 | 248 | 170 | 213 | 277 | 188 | 214 |
| ENSG00000266941 | 0 | 0 | 0 | 0 | 1 | 0 |
| ENSG00000267314 | 0 | 0 | 0 | 0 | 0 | 0 |
| ENSG00000187650 | 176 | 104 | 112 | 67 | 90 | 81 |
| ENSG00000267571 | 13 | 8 | 6 | 3 | 7 | 6 |

| | | | | | | |
|---|---|---|---|---|---|---|
| ENSG00000105519 | 686 | 397 | 482 | 435 | 410 | 375 |
| ENSG00000031823 | 1070 | 506 | 618 | 595 | 596 | 604 |
| ENSG00000266983 | 2 | 5 | 7 | 3 | 2 | 5 |
| ENSG00000087903 | 182 | 73 | 103 | 87 | 76 | 92 |
| ENSG00000267262 | 0 | 0 | 0 | 0 | 0 | 0 |
| ENSG00000267589 | 0 | 0 | 0 | 0 | 0 | 0 |
| ENSG00000267299 | 0 | 0 | 3 | 0 | 1 | 0 |
| ENSG00000130377 | 9 | 6 | 2 | 3 | 3 | 4 |
| ENSG00000267415 | 0 | 0 | 0 | 0 | 0 | 0 |
| ENSG00000284490 | 0 | 0 | 0 | 0 | 0 | 0 |
| ENSG00000130382 | 2151 | 956 | 1288 | 1237 | 1187 | 1118 |
| ENSG00000267563 | 0 | 0 | 0 | 0 | 0 | 0 |
| ENSG00000280394 | 0 | 0 | 0 | 0 | 0 | 0 |
| ENSG00000167769 | 0 | 1 | 1 | 0 | 0 | 0 |
| ENSG00000269802 | 0 | 0 | 0 | 0 | 0 | 0 |
| ENSG00000125656 | 1104 | 436 | 569 | 803 | 539 | 616 |
| ENSG00000125652 | 691 | 279 | 399 | 419 | 303 | 305 |
| ENSG00000125650 | 69 | 45 | 48 | 45 | 53 | 51 |
| ENSG00000125651 | 2598 | 1210 | 1452 | 1480 | 1488 | 1450 |
| ENSG00000269444 | 0 | 1 | 0 | 0 | 0 | 1 |
| ENSG00000274390 | 0 | 0 | 0 | 0 | 0 | 0 |
| ENSG00000273758 | 0 | 0 | 0 | 0 | 0 | 0 |
| ENSG00000214347 | 0 | 0 | 0 | 0 | 0 | 0 |
| ENSG00000088247 | 4460 | 2114 | 2864 | 2682 | 2426 | 2715 |
| ENSG00000284159 | 0 | 0 | 0 | 0 | 0 | 0 |
| ENSG00000181240 | 11 | 1 | 4 | 6 | 4 | 4 |
| ENSG00000125648 | 1502 | 682 | 943 | 724 | 659 | 742 |
| ENSG00000130545 | 742 | 293 | 406 | 348 | 354 | 346 |
| ENSG00000205744 | 605 | 297 | 406 | 305 | 360 | 387 |
| ENSG00000275234 | 285 | 165 | 205 | 194 | 201 | 179 |
| ENSG00000104833 | 10 | 7 | 8 | 4 | 7 | 4 |
| ENSG00000268191 | 0 | 0 | 0 | 0 | 0 | 0 |
| ENSG00000268203 | 0 | 0 | 0 | 0 | 0 | 0 |
| ENSG00000125657 | 38 | 12 | 12 | 16 | 22 | 31 |
| ENSG00000268742 | 0 | 0 | 0 | 0 | 0 | 0 |
| ENSG00000269546 | 0 | 0 | 0 | 0 | 0 | 0 |
| ENSG00000199223 | 0 | 0 | 0 | 0 | 0 | 0 |
| ENSG00000125726 | 19 | 2 | 7 | 9 | 7 | 3 |
| ENSG00000268785 | 6 | 3 | 8 | 4 | 5 | 6 |
| ENSG00000125735 | 0 | 0 | 0 | 0 | 0 | 1 |
| ENSG00000125730 | 299 | 198 | 199 | 156 | 257 | 182 |
| ENSG00000276980 | 0 | 0 | 0 | 0 | 0 | 0 |
| ENSG00000125734 | 1731 | 802 | 1049 | 953 | 895 | 813 |
| ENSG00000283950 | 0 | 0 | 0 | 0 | 0 | 0 |
| ENSG00000125733 | 1166 | 551 | 548 | 534 | 578 | 496 |
| ENSG00000269680 | 11 | 10 | 14 | 7 | 4 | 9 |
| ENSG00000125731 | 547 | 298 | 344 | 353 | 291 | 272 |
| ENSG00000141968 | 4 | 3 | 6 | 7 | 3 | 2 |
| ENSG00000174837 | 6 | 0 | 1 | 2 | 1 | 4 |
| ENSG00000280091 | 0 | 0 | 0 | 0 | 0 | 0 |

| | | | | | | |
|---|---|---|---|---|---|---|
| ENSG00000269385 | 0 | 0 | 0 | 0 | 0 | 0 |
| ENSG00000276568 | 0 | 0 | 0 | 0 | 0 | 0 |
| ENSG00000268758 | 0 | 0 | 0 | 0 | 0 | 0 |
| ENSG00000268845 | 0 | 0 | 0 | 0 | 0 | 0 |
| ENSG00000269761 | 0 | 0 | 0 | 0 | 0 | 0 |
| ENSG00000283679 | 0 | 0 | 0 | 0 | 0 | 0 |
| ENSG00000196589 | 0 | 0 | 0 | 0 | 0 | 0 |
| ENSG00000237247 | 0 | 0 | 0 | 0 | 0 | 0 |
| ENSG00000205718 | 0 | 0 | 0 | 0 | 0 | 0 |
| ENSG00000230522 | 0 | 0 | 0 | 0 | 0 | 0 |
| ENSG00000182315 | 0 | 0 | 0 | 0 | 0 | 0 |
| ENSG00000130544 | 231 | 107 | 152 | 103 | 136 | 130 |
| ENSG00000171105 | 723 | 333 | 478 | 223 | 304 | 323 |
| ENSG00000263264 | 4 | 4 | 3 | 2 | 5 | 4 |
| ENSG00000268861 | 0 | 0 | 0 | 0 | 0 | 0 |
| ENSG00000267852 | 0 | 2 | 3 | 0 | 0 | 0 |
| ENSG00000104880 | 60 | 39 | 49 | 43 | 38 | 55 |
| ENSG00000104883 | 141 | 69 | 85 | 64 | 58 | 66 |
| ENSG00000198723 | 17 | 12 | 15 | 9 | 16 | 12 |
| ENSG00000267952 | 0 | 0 | 0 | 1 | 0 | 0 |
| ENSG00000198816 | 402 | 226 | 278 | 385 | 231 | 244 |
| ENSG00000269371 | 0 | 0 | 0 | 0 | 0 | 0 |
| ENSG00000090674 | 370 | 143 | 203 | 183 | 202 | 165 |
| ENSG00000268614 | 0 | 0 | 0 | 0 | 0 | 0 |
| ENSG00000032444 | 1620 | 686 | 883 | 915 | 737 | 749 |
| ENSG00000076826 | 1749 | 725 | 887 | 946 | 854 | 857 |
| ENSG00000273657 | 0 | 0 | 0 | 0 | 0 | 0 |
| ENSG00000076924 | 1209 | 621 | 726 | 712 | 609 | 665 |
| ENSG00000229833 | 142 | 134 | 178 | 153 | 168 | 109 |
| ENSG00000268400 | 3 | 0 | 1 | 1 | 1 | 0 |
| ENSG00000174788 | 44 | 21 | 52 | 21 | 18 | 31 |
| ENSG00000268204 | 35 | 12 | 22 | 14 | 10 | 30 |
| ENSG00000076944 | 1085 | 567 | 717 | 800 | 696 | 620 |
| ENSG00000269228 | 0 | 0 | 0 | 0 | 0 | 0 |
| ENSG00000104918 | 0 | 0 | 0 | 0 | 1 | 1 |
| ENSG00000183019 | 0 | 0 | 0 | 0 | 1 | 0 |
| ENSG00000269711 | 0 | 0 | 0 | 0 | 0 | 0 |
| ENSG00000181029 | 300 | 131 | 157 | 182 | 194 | 163 |
| ENSG00000104921 | 0 | 0 | 0 | 0 | 0 | 0 |
| ENSG00000182566 | 0 | 0 | 0 | 0 | 0 | 0 |
| ENSG00000090659 | 0 | 0 | 0 | 0 | 0 | 0 |
| ENSG00000242636 | 1 | 0 | 0 | 0 | 0 | 0 |
| ENSG00000104938 | 2 | 0 | 0 | 0 | 2 | 0 |
| ENSG00000268297 | 0 | 2 | 3 | 0 | 0 | 0 |
| ENSG00000269763 | 0 | 0 | 0 | 0 | 0 | 0 |
| ENSG00000142459 | 957 | 464 | 567 | 567 | 523 | 432 |
| ENSG00000183248 | 410 | 200 | 278 | 257 | 228 | 248 |
| ENSG00000268120 | 0 | 0 | 0 | 0 | 0 | 0 |
| ENSG00000269153 | 0 | 0 | 1 | 1 | 0 | 2 |
| ENSG00000199900 | 0 | 0 | 0 | 0 | 0 | 0 |

| | | | | | | |
|---|---|---|---|---|---|---|
| ENSG00000285674 | 0 | 0 | 0 | 0 | 0 | 0 |
| ENSG00000171017 | 44 | 33 | 41 | 36 | 33 | 37 |
| ENSG00000214248 | 11 | 9 | 7 | 3 | 8 | 3 |
| ENSG00000076984 | 1092 | 453 | 638 | 579 | 572 | 561 |
| ENSG00000268149 | 0 | 0 | 0 | 0 | 0 | 0 |
| ENSG00000260001 | 35 | 7 | 32 | 35 | 21 | 19 |
| ENSG00000260500 | 7 | 0 | 0 | 5 | 1 | 4 |
| ENSG00000104976 | 444 | 180 | 207 | 322 | 211 | 224 |
| ENSG00000178531 | 571 | 283 | 288 | 425 | 283 | 327 |
| ENSG00000269139 | 2 | 1 | 2 | 3 | 1 | 0 |
| ENSG00000104980 | 695 | 433 | 483 | 498 | 451 | 448 |
| ENSG00000066044 | 2620 | 1499 | 1684 | 1394 | 1577 | 1569 |
| ENSG00000269813 | 0 | 0 | 0 | 0 | 0 | 0 |
| ENSG00000267939 | 22 | 7 | 7 | 7 | 4 | 7 |
| ENSG00000131142 | 4 | 1 | 2 | 2 | 1 | 3 |
| ENSG00000142449 | 0 | 0 | 0 | 2 | 0 | 0 |
| ENSG00000269651 | 0 | 0 | 0 | 0 | 0 | 0 |
| ENSG00000090661 | 1663 | 713 | 998 | 791 | 731 | 643 |
| ENSG00000271717 | 0 | 0 | 0 | 0 | 0 | 1 |
| ENSG00000167775 | 492 | 223 | 279 | 391 | 270 | 304 |
| ENSG00000167774 | 0 | 4 | 2 | 0 | 1 | 1 |
| ENSG00000267855 | 6 | 5 | 9 | 8 | 11 | 7 |
| ENSG00000233927 | 4849 | 2503 | 2860 | 3470 | 2905 | 2632 |
| ENSG00000186994 | 13 | 7 | 10 | 17 | 7 | 11 |
| ENSG00000268884 | 0 | 0 | 1 | 1 | 0 | 0 |
| ENSG00000167772 | 296 | 132 | 127 | 147 | 151 | 132 |
| ENSG00000269386 | 75 | 48 | 69 | 52 | 63 | 38 |
| ENSG00000265390 | 0 | 0 | 2 | 0 | 0 | 0 |
| ENSG00000185236 | 1328 | 678 | 830 | 820 | 689 | 673 |
| ENSG00000099785 | 655 | 307 | 342 | 344 | 306 | 292 |
| ENSG00000268931 | 0 | 0 | 3 | 0 | 0 | 0 |
| ENSG00000279827 | 0 | 1 | 0 | 0 | 0 | 0 |
| ENSG00000099783 | 7500 | 3651 | 4380 | 3486 | 4188 | 4193 |
| ENSG00000133246 | 9 | 5 | 6 | 10 | 2 | 5 |
| ENSG00000133250 | 199 | 80 | 117 | 111 | 91 | 90 |
| ENSG00000142347 | 79 | 43 | 71 | 56 | 57 | 67 |
| ENSG00000268618 | 0 | 0 | 0 | 0 | 0 | 0 |
| ENSG00000142303 | 0 | 1 | 2 | 1 | 2 | 0 |
| ENSG00000267986 | 0 | 0 | 0 | 0 | 0 | 0 |
| ENSG00000268480 | 0 | 0 | 0 | 0 | 0 | 0 |
| ENSG00000261866 | 0 | 0 | 0 | 0 | 0 | 0 |
| ENSG00000279426 | 0 | 0 | 0 | 0 | 0 | 0 |
| ENSG00000181786 | 0 | 0 | 0 | 0 | 0 | 0 |
| ENSG00000181733 | 0 | 0 | 0 | 0 | 0 | 0 |
| ENSG00000269244 | 0 | 0 | 0 | 0 | 0 | 0 |
| ENSG00000268922 | 0 | 0 | 0 | 0 | 0 | 0 |
| ENSG00000167785 | 316 | 181 | 245 | 146 | 224 | 200 |
| ENSG00000269300 | 0 | 1 | 1 | 0 | 2 | 2 |
| ENSG00000170948 | 0 | 0 | 0 | 0 | 0 | 0 |
| ENSG00000269400 | 0 | 0 | 0 | 0 | 0 | 0 |

| | | | | | | |
|---|---|---|---|---|---|---|
| ENSG00000181143 | 4 | 1 | 11 | 2 | 2 | 1 |
| ENSG00000267332 | 0 | 0 | 0 | 0 | 0 | 0 |
| ENSG00000267154 | 0 | 0 | 0 | 0 | 0 | 0 |
| ENSG00000170929 | 0 | 0 | 0 | 0 | 0 | 0 |
| ENSG00000170923 | 0 | 0 | 0 | 0 | 0 | 0 |
| ENSG00000161807 | 0 | 0 | 0 | 0 | 0 | 0 |
| ENSG00000275787 | 0 | 0 | 0 | 0 | 0 | 0 |
| ENSG00000170920 | 0 | 0 | 0 | 0 | 0 | 0 |
| ENSG00000130803 | 855 | 375 | 515 | 430 | 480 | 461 |
| ENSG00000188000 | 0 | 0 | 0 | 0 | 1 | 0 |
| ENSG00000267418 | 0 | 0 | 0 | 0 | 0 | 0 |
| ENSG00000267775 | 0 | 0 | 0 | 0 | 0 | 0 |
| ENSG00000187847 | 0 | 0 | 0 | 0 | 0 | 0 |
| ENSG00000174667 | 0 | 0 | 0 | 0 | 0 | 0 |
| ENSG00000188314 | 0 | 0 | 0 | 0 | 0 | 0 |
| ENSG00000237521 | 0 | 0 | 0 | 0 | 0 | 0 |
| ENSG00000180926 | 0 | 0 | 0 | 0 | 0 | 0 |
| ENSG00000225980 | 0 | 0 | 0 | 0 | 0 | 0 |
| ENSG00000226480 | 0 | 0 | 0 | 0 | 0 | 0 |
| ENSG00000267510 | 4 | 3 | 3 | 5 | 0 | 1 |
| ENSG00000196110 | 39 | 16 | 21 | 4 | 19 | 13 |
| ENSG00000188321 | 102 | 42 | 58 | 23 | 44 | 30 |
| ENSG00000270011 | 0 | 0 | 0 | 0 | 0 | 0 |
| ENSG00000270614 | 0 | 0 | 0 | 0 | 0 | 0 |
| ENSG00000188629 | 0 | 0 | 0 | 0 | 0 | 0 |
| ENSG00000283108 | 0 | 0 | 0 | 0 | 0 | 0 |
| ENSG00000174652 | 344 | 127 | 194 | 137 | 173 | 155 |
| ENSG00000198028 | 0 | 0 | 0 | 0 | 0 | 0 |
| ENSG00000267273 | 0 | 0 | 0 | 0 | 0 | 0 |
| ENSG00000260112 | 0 | 0 | 0 | 0 | 0 | 0 |
| ENSG00000130818 | 0 | 0 | 1 | 1 | 1 | 1 |
| ENSG00000278611 | 0 | 0 | 0 | 0 | 0 | 0 |
| ENSG00000197961 | 414 | 194 | 210 | 180 | 253 | 228 |
| ENSG00000171469 | 830 | 464 | 489 | 282 | 429 | 385 |
| ENSG00000267106 | 147 | 88 | 112 | 73 | 89 | 95 |
| ENSG00000171466 | 752 | 374 | 416 | 313 | 431 | 379 |
| ENSG00000239830 | 6 | 1 | 6 | 5 | 4 | 2 |
| ENSG00000224689 | 0 | 0 | 0 | 0 | 0 | 0 |
| ENSG00000277587 | 0 | 0 | 0 | 0 | 0 | 0 |
| ENSG00000267490 | 0 | 0 | 0 | 0 | 0 | 0 |
| ENSG00000267612 | 0 | 0 | 1 | 0 | 0 | 0 |
| ENSG00000196605 | 123 | 59 | 88 | 52 | 64 | 50 |
| ENSG00000267370 | 14 | 4 | 13 | 2 | 13 | 3 |
| ENSG00000267741 | 2 | 1 | 0 | 1 | 2 | 0 |
| ENSG00000266950 | 0 | 1 | 0 | 1 | 1 | 1 |
| ENSG00000127452 | 480 | 217 | 290 | 280 | 263 | 270 |
| ENSG00000200237 | 3 | 0 | 0 | 0 | 1 | 0 |
| ENSG00000267119 | 2 | 2 | 1 | 0 | 1 | 1 |
| ENSG00000198258 | 2149 | 1133 | 1260 | 1391 | 1308 | 1213 |
| ENSG00000267289 | 0 | 0 | 0 | 0 | 0 | 0 |

| | | | | | | |
|---|---|---|---|---|---|---|
| ENSG00000127445 | 1370 | 580 | 745 | 840 | 700 | 679 |
| ENSG00000105088 | 336 | 158 | 221 | 196 | 194 | 202 |
| ENSG00000080573 | 37 | 22 | 49 | 16 | 30 | 33 |
| ENSG00000267650 | 0 | 0 | 0 | 2 | 1 | 0 |
| ENSG00000080511 | 0 | 0 | 0 | 1 | 0 | 0 |
| ENSG00000167798 | 0 | 0 | 0 | 0 | 0 | 0 |
| ENSG00000266204 | 0 | 0 | 0 | 0 | 0 | 0 |
| ENSG00000130813 | 100 | 59 | 73 | 74 | 56 | 58 |
| ENSG00000267387 | 1 | 0 | 0 | 0 | 1 | 0 |
| ENSG00000130812 | 7 | 5 | 10 | 7 | 8 | 6 |
| ENSG00000243207 | 0 | 0 | 0 | 0 | 0 | 1 |
| ENSG00000130810 | 112 | 58 | 75 | 94 | 66 | 90 |
| ENSG00000209645 | 0 | 0 | 0 | 1 | 0 | 0 |
| ENSG00000238531 | 0 | 0 | 0 | 0 | 0 | 0 |
| ENSG00000244165 | 101 | 91 | 115 | 122 | 114 | 140 |
| ENSG00000130811 | 4074 | 2107 | 2834 | 3008 | 2491 | 2718 |
| ENSG00000130816 | 4172 | 1955 | 2508 | 1784 | 2230 | 2302 |
| ENSG00000267534 | 169 | 106 | 128 | 118 | 158 | 134 |
| ENSG00000264266 | 0 | 0 | 0 | 0 | 0 | 0 |
| ENSG00000105364 | 1623 | 824 | 1053 | 1323 | 1012 | 1084 |
| ENSG00000266978 | 0 | 3 | 0 | 2 | 0 | 0 |
| ENSG00000267105 | 0 | 0 | 0 | 0 | 0 | 0 |
| ENSG00000090339 | 294 | 136 | 181 | 164 | 184 | 224 |
| ENSG00000267607 | 0 | 0 | 3 | 2 | 0 | 1 |
| ENSG00000105371 | 0 | 0 | 0 | 0 | 0 | 0 |
| ENSG00000105376 | 81 | 30 | 31 | 41 | 44 | 51 |
| ENSG00000220201 | 44 | 15 | 15 | 31 | 23 | 25 |
| ENSG00000167807 | 4 | 2 | 2 | 2 | 4 | 6 |
| ENSG00000267673 | 5 | 1 | 3 | 4 | 7 | 3 |
| ENSG00000267303 | 0 | 0 | 1 | 0 | 0 | 0 |
| ENSG00000161847 | 884 | 424 | 629 | 592 | 439 | 522 |
| ENSG00000274425 | 6 | 2 | 7 | 10 | 9 | 3 |
| ENSG00000076662 | 0 | 1 | 0 | 0 | 1 | 0 |
| ENSG00000105397 | 2336 | 969 | 1382 | 1151 | 1157 | 1218 |
| ENSG00000105401 | 6470 | 3405 | 4241 | 4319 | 4143 | 4102 |
| ENSG00000284268 | 0 | 0 | 0 | 0 | 0 | 0 |
| ENSG00000065989 | 28 | 19 | 11 | 15 | 13 | 10 |
| ENSG00000079999 | 3268 | 1594 | 1982 | 1871 | 1659 | 1741 |
| ENSG00000267197 | 22 | 5 | 11 | 6 | 11 | 10 |
| ENSG00000180739 | 123 | 51 | 64 | 57 | 42 | 35 |
| ENSG00000130734 | 1076 | 529 | 678 | 680 | 578 | 539 |
| ENSG00000238364 | 0 | 0 | 0 | 1 | 1 | 0 |
| ENSG00000221410 | 0 | 0 | 0 | 0 | 0 | 0 |
| ENSG00000129347 | 903 | 471 | 560 | 503 | 574 | 508 |
| ENSG00000129355 | 235 | 102 | 142 | 124 | 121 | 140 |
| ENSG00000129354 | 7539 | 3734 | 4490 | 4531 | 4095 | 4192 |
| ENSG00000271010 | 0 | 0 | 0 | 0 | 0 | 0 |
| ENSG00000129353 | 14473 | 6877 | 8411 | 6947 | 6854 | 7091 |
| ENSG00000267100 | 622 | 354 | 440 | 342 | 441 | 394 |
| ENSG00000129351 | 10508 | 4857 | 6359 | 5446 | 5887 | 6265 |

| | | | | | | |
|---|---|---|---|---|---|---|
| ENSG00000213339 | 1019 | 418 | 551 | 550 | 505 | 535 |
| ENSG00000079805 | 11365 | 5214 | 6575 | 4877 | 5030 | 4839 |
| ENSG00000207972 | 0 | 2 | 0 | 0 | 0 | 0 |
| ENSG00000265879 | 0 | 1 | 0 | 0 | 0 | 0 |
| ENSG00000207752 | 3 | 3 | 2 | 4 | 0 | 2 |
| ENSG00000275640 | 0 | 0 | 0 | 0 | 2 | 0 |
| ENSG00000099203 | 303 | 184 | 246 | 216 | 208 | 200 |
| ENSG00000214212 | 60 | 20 | 18 | 29 | 19 | 14 |
| ENSG00000267544 | 2 | 1 | 0 | 2 | 1 | 0 |
| ENSG00000142453 | 2999 | 1536 | 1857 | 1664 | 1563 | 1554 |
| ENSG00000130733 | 2039 | 1052 | 1219 | 1167 | 1060 | 996 |
| ENSG00000142444 | 915 | 508 | 562 | 579 | 541 | 551 |
| ENSG00000127616 | 5994 | 2745 | 3598 | 3182 | 3130 | 3389 |
| ENSG00000266936 | 1 | 1 | 0 | 0 | 0 | 0 |
| ENSG00000276757 | 2 | 0 | 0 | 0 | 0 | 2 |
| ENSG00000278643 | 0 | 0 | 0 | 0 | 0 | 0 |
| ENSG00000130164 | 3474 | 1632 | 2088 | 1635 | 1856 | 1982 |
| ENSG00000284553 | 0 | 0 | 0 | 0 | 0 | 0 |
| ENSG00000161888 | 462 | 240 | 299 | 298 | 320 | 256 |
| ENSG00000197256 | 3594 | 1779 | 2050 | 1484 | 1775 | 1647 |
| ENSG00000130158 | 2186 | 1040 | 1509 | 1176 | 1230 | 1270 |
| ENSG00000267082 | 14 | 12 | 12 | 9 | 10 | 11 |
| ENSG00000279807 | 5 | 3 | 1 | 1 | 0 | 2 |
| ENSG00000273733 | 8 | 2 | 1 | 0 | 3 | 4 |
| ENSG00000130173 | 68 | 46 | 60 | 58 | 65 | 42 |
| ENSG00000130167 | 2 | 4 | 5 | 1 | 2 | 0 |
| ENSG00000267174 | 16 | 8 | 13 | 6 | 8 | 8 |
| ENSG00000105514 | 2677 | 1406 | 1955 | 871 | 969 | 1105 |
| ENSG00000267576 | 0 | 0 | 0 | 0 | 0 | 0 |
| ENSG00000105518 | 1544 | 988 | 1433 | 1129 | 822 | 977 |
| ENSG00000183401 | 397 | 146 | 273 | 150 | 125 | 151 |
| ENSG00000105520 | 2237 | 936 | 1430 | 1053 | 872 | 941 |
| ENSG00000267277 | 22 | 9 | 24 | 13 | 11 | 10 |
| ENSG00000173928 | 245 | 133 | 167 | 125 | 153 | 145 |
| ENSG00000187266 | 271 | 124 | 187 | 87 | 88 | 108 |
| ENSG00000205517 | 4 | 1 | 2 | 0 | 0 | 1 |
| ENSG00000202357 | 0 | 0 | 0 | 0 | 0 | 0 |
| ENSG00000198003 | 36 | 12 | 21 | 18 | 26 | 8 |
| ENSG00000130175 | 7991 | 3685 | 5307 | 5494 | 4269 | 4793 |
| ENSG00000196361 | 4 | 1 | 0 | 2 | 1 | 1 |
| ENSG00000267477 | 0 | 0 | 0 | 0 | 0 | 0 |
| ENSG00000161914 | 144 | 75 | 106 | 92 | 87 | 81 |
| ENSG00000274713 | 0 | 0 | 0 | 0 | 0 | 0 |
| ENSG00000130159 | 485 | 260 | 326 | 394 | 340 | 298 |
| ENSG00000244080 | 0 | 2 | 0 | 1 | 0 | 0 |
| ENSG00000213304 | 32 | 23 | 22 | 20 | 18 | 26 |
| ENSG00000213303 | 9 | 1 | 1 | 2 | 6 | 4 |
| ENSG00000130176 | 0 | 1 | 0 | 0 | 0 | 0 |
| ENSG00000130165 | 2112 | 1035 | 1254 | 1441 | 1182 | 1181 |
| ENSG00000198551 | 677 | 352 | 397 | 268 | 402 | 300 |

| | | | | | | |
|---|---|---|---|---|---|---|
| ENSG00000102575 | 4 | 4 | 9 | 3 | 1 | 7 |
| ENSG00000249840 | 0 | 0 | 0 | 0 | 1 | 0 |
| ENSG00000197332 | 21 | 8 | 7 | 6 | 6 | 5 |
| ENSG00000267500 | 14 | 18 | 5 | 9 | 8 | 9 |
| ENSG00000214223 | 6 | 9 | 4 | 5 | 7 | 6 |
| ENSG00000267343 | 0 | 0 | 0 | 0 | 0 | 0 |
| ENSG00000197933 | 254 | 119 | 154 | 103 | 175 | 131 |
| ENSG00000266922 | 0 | 0 | 0 | 0 | 1 | 5 |
| ENSG00000220949 | 0 | 0 | 0 | 0 | 0 | 0 |
| ENSG00000197044 | 0 | 0 | 0 | 0 | 0 | 0 |
| ENSG00000267646 | 0 | 0 | 0 | 0 | 0 | 0 |
| ENSG00000177599 | 24 | 13 | 7 | 9 | 14 | 12 |
| ENSG00000171295 | 221 | 124 | 107 | 79 | 172 | 104 |
| ENSG00000267089 | 1 | 0 | 0 | 0 | 0 | 0 |
| ENSG00000171291 | 2 | 0 | 1 | 1 | 0 | 1 |
| ENSG00000278897 | 0 | 0 | 0 | 0 | 0 | 0 |
| ENSG00000198429 | 18 | 15 | 12 | 8 | 19 | 11 |
| ENSG00000196757 | 228 | 130 | 191 | 57 | 178 | 144 |
| ENSG00000267179 | 1 | 0 | 2 | 0 | 0 | 1 |
| ENSG00000267274 | 1 | 2 | 4 | 1 | 3 | 1 |
| ENSG00000197054 | 12 | 2 | 3 | 3 | 2 | 5 |
| ENSG00000219665 | 96 | 43 | 67 | 42 | 87 | 62 |
| ENSG00000252060 | 0 | 2 | 0 | 0 | 0 | 0 |
| ENSG00000197647 | 240 | 138 | 149 | 92 | 125 | 114 |
| ENSG00000257355 | 0 | 0 | 0 | 0 | 0 | 0 |
| ENSG00000212497 | 0 | 0 | 1 | 0 | 2 | 1 |
| ENSG00000257446 | 11 | 5 | 3 | 11 | 8 | 4 |
| ENSG00000223547 | 3 | 0 | 2 | 0 | 3 | 0 |
| ENSG00000252546 | 0 | 0 | 0 | 0 | 0 | 0 |
| ENSG00000199444 | 0 | 0 | 0 | 0 | 0 | 0 |
| ENSG00000214189 | 0 | 2 | 0 | 0 | 0 | 0 |
| ENSG00000132010 | 8 | 2 | 6 | 7 | 7 | 1 |
| ENSG00000237273 | 0 | 0 | 0 | 0 | 0 | 0 |
| ENSG00000212576 | 0 | 0 | 0 | 0 | 0 | 0 |
| ENSG00000213297 | 0 | 0 | 0 | 1 | 0 | 0 |
| ENSG00000242615 | 38 | 17 | 12 | 15 | 22 | 15 |
| ENSG00000257591 | 5 | 11 | 1 | 2 | 1 | 4 |
| ENSG00000196646 | 123 | 72 | 82 | 17 | 65 | 28 |
| ENSG00000234773 | 2 | 7 | 1 | 0 | 2 | 0 |
| ENSG00000197857 | 202 | 89 | 131 | 53 | 115 | 77 |
| ENSG00000213293 | 0 | 0 | 0 | 0 | 1 | 0 |
| ENSG00000234750 | 6 | 4 | 2 | 0 | 2 | 2 |
| ENSG00000188868 | 54 | 41 | 30 | 30 | 42 | 27 |
| ENSG00000198342 | 54 | 35 | 27 | 29 | 45 | 30 |
| ENSG00000231361 | 0 | 0 | 0 | 0 | 0 | 0 |
| ENSG00000268744 | 6 | 1 | 0 | 1 | 0 | 0 |
| ENSG00000248406 | 0 | 0 | 0 | 0 | 0 | 0 |
| ENSG00000196466 | 164 | 84 | 95 | 52 | 117 | 63 |
| ENSG00000268870 | 0 | 0 | 0 | 0 | 0 | 0 |
| ENSG00000180855 | 201 | 125 | 128 | 79 | 119 | 108 |

| | | | | | | |
|---|---|---|---|---|---|---|
| ENSG00000269755 | 0 | 0 | 0 | 0 | 0 | 0 |
| ENSG00000234848 | 0 | 0 | 0 | 1 | 2 | 0 |
| ENSG00000242852 | 11 | 7 | 5 | 3 | 7 | 7 |
| ENSG00000196826 | 0 | 0 | 0 | 0 | 0 | 1 |
| ENSG00000236483 | 0 | 4 | 5 | 1 | 1 | 6 |
| ENSG00000232579 | 2 | 0 | 1 | 0 | 2 | 1 |
| ENSG00000230321 | 1 | 2 | 0 | 1 | 0 | 1 |
| ENSG00000234003 | 0 | 2 | 1 | 0 | 0 | 5 |
| ENSG00000235686 | 1 | 0 | 0 | 0 | 0 | 1 |
| ENSG00000249709 | 10 | 4 | 6 | 3 | 3 | 10 |
| ENSG00000269693 | 36 | 16 | 15 | 12 | 19 | 14 |
| ENSG00000269560 | 4 | 0 | 0 | 0 | 0 | 0 |
| ENSG00000230310 | 9 | 2 | 1 | 1 | 0 | 1 |
| ENSG00000213290 | 1 | 2 | 1 | 1 | 2 | 1 |
| ENSG00000188033 | 63 | 36 | 31 | 13 | 21 | 35 |
| ENSG00000173875 | 765 | 372 | 407 | 215 | 451 | 309 |
| ENSG00000178464 | 61 | 61 | 68 | 59 | 35 | 46 |
| ENSG00000269242 | 0 | 1 | 0 | 0 | 1 | 0 |
| ENSG00000104774 | 1802 | 994 | 1207 | 1034 | 1225 | 1058 |
| ENSG00000269590 | 0 | 0 | 0 | 0 | 0 | 0 |
| ENSG00000123154 | 125 | 91 | 113 | 131 | 85 | 120 |
| ENSG00000105583 | 0 | 0 | 0 | 1 | 1 | 0 |
| ENSG00000285589 | 57 | 30 | 37 | 16 | 42 | 40 |
| ENSG00000285583 | 0 | 0 | 0 | 0 | 0 | 0 |
| ENSG00000095059 | 85 | 35 | 49 | 50 | 54 | 63 |
| ENSG00000268945 | 6 | 1 | 4 | 1 | 4 | 5 |
| ENSG00000283980 | 0 | 0 | 0 | 0 | 0 | 0 |
| ENSG00000132004 | 589 | 293 | 370 | 369 | 312 | 350 |
| ENSG00000267791 | 0 | 0 | 0 | 0 | 0 | 0 |
| ENSG00000105576 | 3701 | 1612 | 2299 | 1924 | 1935 | 2172 |
| ENSG00000209702 | 0 | 0 | 0 | 0 | 0 | 0 |
| ENSG00000267004 | 0 | 0 | 1 | 0 | 0 | 2 |
| ENSG00000123144 | 5370 | 2517 | 3077 | 3555 | 3121 | 3179 |
| ENSG00000198356 | 3249 | 1553 | 1913 | 2186 | 1896 | 1875 |
| ENSG00000039987 | 0 | 1 | 1 | 0 | 0 | 2 |
| ENSG00000095066 | 2474 | 1109 | 1658 | 1300 | 1321 | 1296 |
| ENSG00000267212 | 0 | 4 | 5 | 1 | 1 | 1 |
| ENSG00000263800 | 2 | 0 | 0 | 0 | 0 | 2 |
| ENSG00000171223 | 4670 | 2476 | 2811 | 3022 | 2643 | 2449 |
| ENSG00000167815 | 5673 | 2928 | 4005 | 4097 | 2912 | 3615 |
| ENSG00000267062 | 0 | 0 | 0 | 0 | 0 | 0 |
| ENSG00000284491 | 3 | 2 | 0 | 2 | 3 | 1 |
| ENSG00000104889 | 7 | 9 | 9 | 9 | 10 | 10 |
| ENSG00000132026 | 0 | 1 | 6 | 1 | 4 | 3 |
| ENSG00000267424 | 0 | 0 | 0 | 0 | 0 | 0 |
| ENSG00000105613 | 112 | 47 | 61 | 52 | 62 | 49 |
| ENSG00000276181 | 0 | 0 | 0 | 0 | 0 | 0 |
| ENSG00000105612 | 2177 | 1200 | 1424 | 1253 | 1258 | 1217 |
| ENSG00000267735 | 0 | 1 | 0 | 0 | 0 | 0 |
| ENSG00000105610 | 0 | 2 | 5 | 1 | 0 | 2 |

| | | | | | | |
|---|---|---|---|---|---|---|
| ENSG00000105607 | 518 | 241 | 283 | 289 | 276 | 283 |
| ENSG00000240616 | 0 | 5 | 2 | 3 | 7 | 3 |
| ENSG00000161860 | 16 | 9 | 12 | 7 | 8 | 12 |
| ENSG00000266721 | 0 | 1 | 0 | 0 | 0 | 0 |
| ENSG00000179115 | 2797 | 1453 | 1835 | 2019 | 1686 | 1806 |
| ENSG00000266975 | 0 | 1 | 1 | 0 | 1 | 0 |
| ENSG00000179218 | 16854 | 8458 | 10275 | 9162 | 8957 | 8676 |
| ENSG00000274417 | 0 | 0 | 0 | 0 | 0 | 0 |
| ENSG00000267458 | 0 | 0 | 0 | 0 | 0 | 0 |
| ENSG00000179262 | 2160 | 962 | 1337 | 1470 | 1203 | 1342 |
| ENSG00000179271 | 1397 | 661 | 785 | 974 | 797 | 770 |
| ENSG00000179284 | 0 | 1 | 1 | 1 | 1 | 2 |
| ENSG00000008441 | 128 | 68 | 80 | 65 | 58 | 48 |
| ENSG00000267417 | 0 | 0 | 0 | 0 | 0 | 0 |
| ENSG00000267610 | 0 | 0 | 0 | 0 | 0 | 0 |
| ENSG00000279044 | 2 | 0 | 1 | 0 | 0 | 0 |
| ENSG00000104903 | 0 | 0 | 0 | 0 | 0 | 0 |
| ENSG00000104907 | 955 | 430 | 510 | 607 | 558 | 581 |
| ENSG00000160877 | 5399 | 2648 | 3326 | 3287 | 3203 | 3137 |
| ENSG00000266897 | 12 | 3 | 4 | 2 | 6 | 3 |
| ENSG00000267512 | 0 | 0 | 0 | 0 | 0 | 0 |
| ENSG00000104915 | 1280 | 644 | 793 | 792 | 759 | 752 |
| ENSG00000160888 | 1682 | 904 | 1220 | 1146 | 1095 | 1051 |
| ENSG00000267598 | 0 | 0 | 0 | 0 | 0 | 0 |
| ENSG00000213253 | 7 | 4 | 3 | 3 | 2 | 1 |
| ENSG00000141837 | 4 | 3 | 0 | 1 | 1 | 0 |
| ENSG00000104957 | 670 | 301 | 439 | 388 | 324 | 370 |
| ENSG00000037757 | 742 | 438 | 492 | 448 | 547 | 486 |
| ENSG00000267633 | 0 | 0 | 3 | 0 | 0 | 0 |
| ENSG00000104979 | 1979 | 943 | 1170 | 1352 | 1112 | 1102 |
| ENSG00000132003 | 162 | 84 | 107 | 79 | 110 | 91 |
| ENSG00000267582 | 0 | 0 | 0 | 1 | 0 | 0 |
| ENSG00000266770 | 0 | 2 | 5 | 2 | 3 | 2 |
| ENSG00000267519 | 376 | 285 | 257 | 185 | 262 | 180 |
| ENSG00000284387 | 0 | 0 | 0 | 0 | 0 | 0 |
| ENSG00000207808 | 0 | 0 | 0 | 0 | 0 | 0 |
| ENSG00000207980 | 0 | 0 | 0 | 0 | 0 | 0 |
| ENSG00000187556 | 0 | 3 | 1 | 2 | 3 | 2 |
| ENSG00000207613 | 0 | 0 | 0 | 0 | 0 | 0 |
| ENSG00000207585 | 1 | 0 | 0 | 0 | 0 | 0 |
| ENSG00000132016 | 66 | 35 | 64 | 44 | 39 | 27 |
| ENSG00000132024 | 2713 | 1286 | 1594 | 1523 | 1404 | 1584 |
| ENSG00000132000 | 119 | 53 | 79 | 88 | 75 | 53 |
| ENSG00000132017 | 1251 | 577 | 793 | 722 | 718 | 683 |
| ENSG00000132005 | 608 | 259 | 380 | 335 | 325 | 289 |
| ENSG00000275091 | 4 | 1 | 2 | 5 | 1 | 0 |
| ENSG00000171136 | 0 | 0 | 0 | 0 | 0 | 0 |
| ENSG00000267670 | 0 | 0 | 0 | 0 | 0 | 0 |
| ENSG00000104998 | 237 | 117 | 128 | 153 | 124 | 110 |
| ENSG00000187867 | 574 | 281 | 321 | 307 | 323 | 311 |

| | | | | | | |
|---|---|---|---|---|---|---|
| ENSG00000228887 | 28 | 14 | 13 | 7 | 13 | 12 |
| ENSG00000141854 | 1545 | 665 | 943 | 794 | 743 | 722 |
| ENSG00000284081 | 0 | 0 | 0 | 0 | 0 | 0 |
| ENSG00000188032 | 8 | 6 | 3 | 1 | 3 | 3 |
| ENSG00000141858 | 2074 | 1028 | 1197 | 1106 | 953 | 1014 |
| ENSG00000072062 | 3624 | 1665 | 2181 | 1716 | 1623 | 1783 |
| ENSG00000267783 | 0 | 0 | 1 | 2 | 1 | 0 |
| ENSG00000105011 | 1032 | 548 | 644 | 585 | 506 | 635 |
| ENSG00000267169 | 55 | 26 | 38 | 31 | 32 | 41 |
| ENSG00000072071 | 2365 | 1103 | 1411 | 1141 | 1138 | 1221 |
| ENSG00000240803 | 0 | 0 | 0 | 0 | 0 | 0 |
| ENSG00000267723 | 0 | 0 | 1 | 0 | 0 | 0 |
| ENSG00000267406 | 0 | 0 | 0 | 0 | 0 | 0 |
| ENSG00000266913 | 0 | 0 | 0 | 0 | 0 | 2 |
| ENSG00000267147 | 20 | 12 | 11 | 14 | 14 | 12 |
| ENSG00000123146 | 625 | 330 | 384 | 322 | 353 | 300 |
| ENSG00000267379 | 0 | 0 | 0 | 0 | 0 | 0 |
| ENSG00000123136 | 2228 | 1093 | 1369 | 1474 | 1344 | 1348 |
| ENSG00000123143 | 4319 | 2104 | 2694 | 2996 | 2546 | 2595 |
| ENSG00000267474 | 6 | 4 | 1 | 0 | 2 | 0 |
| ENSG00000160951 | 15 | 5 | 3 | 4 | 0 | 7 |
| ENSG00000123159 | 10191 | 4917 | 5952 | 6299 | 5618 | 5256 |
| ENSG00000224543 | 60 | 50 | 60 | 24 | 29 | 33 |
| ENSG00000132002 | 6799 | 3345 | 3857 | 3575 | 3504 | 3391 |
| ENSG00000099797 | 3440 | 1444 | 2014 | 1925 | 1372 | 1760 |
| ENSG00000283822 | 0 | 0 | 0 | 0 | 0 | 0 |
| ENSG00000099795 | 2434 | 1244 | 1364 | 1806 | 1219 | 1307 |
| ENSG00000187912 | 0 | 0 | 0 | 0 | 0 | 0 |
| ENSG00000243488 | 0 | 0 | 0 | 0 | 0 | 0 |
| ENSG00000243066 | 0 | 0 | 0 | 0 | 0 | 0 |
| ENSG00000131355 | 0 | 0 | 0 | 1 | 0 | 0 |
| ENSG00000269378 | 40 | 20 | 21 | 13 | 31 | 26 |
| ENSG00000275432 | 0 | 0 | 0 | 0 | 0 | 0 |
| ENSG00000160961 | 197 | 101 | 114 | 50 | 121 | 121 |
| ENSG00000127507 | 8 | 4 | 4 | 10 | 11 | 11 |
| ENSG00000127530 | 2 | 2 | 2 | 0 | 2 | 1 |
| ENSG00000256210 | 0 | 0 | 0 | 0 | 0 | 0 |
| ENSG00000188269 | 0 | 0 | 0 | 0 | 0 | 0 |
| ENSG00000127515 | 0 | 0 | 0 | 0 | 0 | 0 |
| ENSG00000269552 | 0 | 0 | 0 | 0 | 0 | 0 |
| ENSG00000172148 | 0 | 0 | 0 | 0 | 0 | 0 |
| ENSG00000185385 | 0 | 0 | 0 | 0 | 0 | 0 |
| ENSG00000269678 | 0 | 0 | 0 | 0 | 0 | 0 |
| ENSG00000268834 | 0 | 0 | 0 | 0 | 0 | 0 |
| ENSG00000269787 | 0 | 0 | 0 | 0 | 0 | 0 |
| ENSG00000198367 | 0 | 0 | 0 | 0 | 0 | 0 |
| ENSG00000176923 | 0 | 0 | 0 | 0 | 0 | 0 |
| ENSG00000127529 | 0 | 0 | 0 | 0 | 0 | 0 |
| ENSG00000105143 | 0 | 1 | 0 | 0 | 0 | 0 |
| ENSG00000160994 | 0 | 0 | 0 | 0 | 0 | 0 |

| | | | | | | |
|---|---|---|---|---|---|---|
| ENSG00000105141 | 0 | 0 | 1 | 1 | 1 | 0 |
| ENSG00000094661 | 0 | 0 | 0 | 0 | 0 | 0 |
| ENSG00000105137 | 32 | 21 | 31 | 22 | 22 | 31 |
| ENSG00000105135 | 2291 | 1069 | 1526 | 1665 | 1373 | 1368 |
| ENSG00000268564 | 2 | 1 | 2 | 2 | 4 | 0 |
| ENSG00000267961 | 0 | 0 | 0 | 0 | 0 | 0 |
| ENSG00000252661 | 0 | 0 | 0 | 0 | 0 | 0 |
| ENSG00000074181 | 10451 | 4191 | 6100 | 4791 | 4716 | 5171 |
| ENSG00000275711 | 0 | 0 | 0 | 0 | 0 | 0 |
| ENSG00000269635 | 0 | 0 | 0 | 0 | 0 | 0 |
| ENSG00000105131 | 8 | 4 | 4 | 3 | 2 | 7 |
| ENSG00000141867 | 1566 | 658 | 1005 | 823 | 785 | 834 |
| ENSG00000270802 | 2 | 2 | 0 | 0 | 0 | 0 |
| ENSG00000273218 | 17 | 9 | 8 | 3 | 5 | 10 |
| ENSG00000105127 | 1582 | 748 | 971 | 755 | 832 | 926 |
| ENSG00000268189 | 8 | 2 | 10 | 4 | 8 | 8 |
| ENSG00000011243 | 1758 | 844 | 1112 | 1011 | 1047 | 1050 |
| ENSG00000279203 | 2 | 4 | 9 | 2 | 2 | 2 |
| ENSG00000279716 | 24 | 20 | 33 | 16 | 24 | 21 |
| ENSG00000011451 | 2502 | 1141 | 1483 | 1325 | 1380 | 1366 |
| ENSG00000269782 | 0 | 0 | 0 | 0 | 0 | 0 |
| ENSG00000105122 | 4 | 3 | 4 | 3 | 6 | 6 |
| ENSG00000161031 | 0 | 2 | 1 | 3 | 3 | 9 |
| ENSG00000171954 | 4 | 2 | 1 | 2 | 0 | 3 |
| ENSG00000269765 | 0 | 0 | 0 | 0 | 0 | 0 |
| ENSG00000269516 | 103 | 42 | 87 | 45 | 44 | 65 |
| ENSG00000267135 | 0 | 0 | 0 | 0 | 0 | 0 |
| ENSG00000225067 | 3 | 2 | 0 | 2 | 0 | 2 |
| ENSG00000266951 | 0 | 0 | 0 | 0 | 0 | 0 |
| ENSG00000186526 | 6 | 2 | 4 | 1 | 3 | 6 |
| ENSG00000186529 | 0 | 2 | 4 | 1 | 1 | 2 |
| ENSG00000267241 | 0 | 0 | 0 | 0 | 0 | 0 |
| ENSG00000186204 | 29 | 9 | 13 | 9 | 6 | 8 |
| ENSG00000171942 | 0 | 0 | 0 | 0 | 0 | 0 |
| ENSG00000171936 | 0 | 0 | 0 | 0 | 0 | 0 |
| ENSG00000267594 | 0 | 0 | 0 | 0 | 0 | 0 |
| ENSG00000268673 | 0 | 0 | 0 | 0 | 0 | 0 |
| ENSG00000172519 | 0 | 0 | 0 | 0 | 0 | 0 |
| ENSG00000186723 | 0 | 1 | 1 | 0 | 2 | 0 |
| ENSG00000267235 | 0 | 0 | 1 | 0 | 0 | 0 |
| ENSG00000267308 | 0 | 0 | 0 | 3 | 0 | 2 |
| ENSG00000214049 | 16 | 11 | 15 | 8 | 14 | 9 |
| ENSG00000273782 | 0 | 0 | 0 | 0 | 0 | 0 |
| ENSG00000267453 | 0 | 0 | 0 | 0 | 0 | 0 |
| ENSG00000273330 | 0 | 0 | 0 | 0 | 0 | 0 |
| ENSG00000186115 | 0 | 0 | 0 | 1 | 0 | 0 |
| ENSG00000267218 | 0 | 0 | 0 | 1 | 0 | 1 |
| ENSG00000279717 | 0 | 0 | 1 | 0 | 0 | 0 |
| ENSG00000267056 | 38 | 12 | 14 | 13 | 15 | 20 |
| ENSG00000171903 | 325 | 147 | 228 | 159 | 231 | 241 |

| | | | | | | |
|---|---|---|---|---|---|---|
| ENSG00000176231 | 0 | 0 | 0 | 0 | 0 | 0 |
| ENSG00000267392 | 0 | 0 | 0 | 0 | 0 | 0 |
| ENSG00000205396 | 0 | 0 | 0 | 0 | 0 | 0 |
| ENSG00000280412 | 0 | 0 | 0 | 0 | 0 | 0 |
| ENSG00000167459 | 0 | 0 | 1 | 0 | 0 | 0 |
| ENSG00000267609 | 0 | 0 | 0 | 0 | 0 | 0 |
| ENSG00000277766 | 0 | 0 | 0 | 0 | 0 | 0 |
| ENSG00000267450 | 0 | 0 | 0 | 0 | 0 | 0 |
| ENSG00000267517 | 0 | 0 | 0 | 0 | 0 | 0 |
| ENSG00000167460 | 32717 | 17551 | 20115 | 16241 | 22899 | 20397 |
| ENSG00000279198 | 24 | 16 | 14 | 11 | 16 | 13 |
| ENSG00000267373 | 0 | 0 | 0 | 0 | 0 | 0 |
| ENSG00000167461 | 1266 | 785 | 1088 | 718 | 901 | 964 |
| ENSG00000269243 | 29 | 11 | 13 | 3 | 4 | 12 |
| ENSG00000196684 | 1474 | 685 | 1118 | 524 | 479 | 586 |
| ENSG00000141977 | 42 | 11 | 40 | 14 | 14 | 14 |
| ENSG00000105058 | 3800 | 1801 | 2216 | 1998 | 1972 | 2094 |
| ENSG00000267033 | 2 | 6 | 3 | 0 | 7 | 1 |
| ENSG00000275901 | 0 | 2 | 0 | 0 | 1 | 0 |
| ENSG00000072958 | 1607 | 634 | 876 | 875 | 792 | 825 |
| ENSG00000267275 | 0 | 0 | 0 | 0 | 0 | 0 |
| ENSG00000127528 | 104 | 57 | 40 | 66 | 53 | 64 |
| ENSG00000277825 | 1 | 0 | 0 | 0 | 0 | 0 |
| ENSG00000127527 | 3159 | 1529 | 1951 | 1774 | 1758 | 1863 |
| ENSG00000280332 | 59 | 29 | 48 | 35 | 48 | 48 |
| ENSG00000240418 | 2 | 0 | 1 | 0 | 0 | 1 |
| ENSG00000267703 | 0 | 0 | 0 | 0 | 0 | 1 |
| ENSG00000269058 | 2 | 0 | 0 | 0 | 0 | 0 |
| ENSG00000141979 | 0 | 1 | 0 | 0 | 0 | 0 |
| ENSG00000105072 | 84 | 59 | 74 | 79 | 86 | 72 |
| ENSG00000085872 | 1200 | 572 | 758 | 701 | 691 | 795 |
| ENSG00000240106 | 0 | 1 | 1 | 0 | 1 | 0 |
| ENSG00000269399 | 74 | 41 | 78 | 33 | 50 | 24 |
| ENSG00000127526 | 3782 | 2214 | 2745 | 2001 | 2328 | 2425 |
| ENSG00000268309 | 0 | 0 | 0 | 0 | 0 | 0 |
| ENSG00000269085 | 0 | 0 | 0 | 0 | 0 | 0 |
| ENSG00000279529 | 57 | 16 | 34 | 20 | 22 | 15 |
| ENSG00000105085 | 248 | 110 | 173 | 158 | 167 | 135 |
| ENSG00000268790 | 0 | 3 | 1 | 0 | 0 | 0 |
| ENSG00000279748 | 47 | 8 | 18 | 11 | 18 | 18 |
| ENSG00000269578 | 0 | 0 | 0 | 0 | 0 | 0 |
| ENSG00000279977 | 2 | 1 | 3 | 1 | 0 | 0 |
| ENSG00000268087 | 22 | 26 | 20 | 15 | 27 | 23 |
| ENSG00000269427 | 27 | 28 | 15 | 7 | 12 | 17 |
| ENSG00000214046 | 1430 | 884 | 919 | 766 | 732 | 731 |
| ENSG00000269044 | 140 | 63 | 82 | 67 | 70 | 57 |
| ENSG00000267904 | 32 | 44 | 24 | 17 | 16 | 14 |
| ENSG00000072954 | 442 | 200 | 263 | 240 | 251 | 214 |
| ENSG00000269419 | 0 | 0 | 0 | 0 | 1 | 0 |
| ENSG00000188039 | 0 | 1 | 1 | 0 | 1 | 1 |

| | | | | | | |
|---|---|---|---|---|---|---|
| ENSG00000269799 | 0 | 0 | 0 | 0 | 0 | 1 |
| ENSG00000127511 | 1564 | 714 | 957 | 786 | 970 | 970 |
| ENSG00000268743 | 0 | 2 | 1 | 0 | 0 | 0 |
| ENSG00000127533 | 0 | 1 | 1 | 0 | 0 | 0 |
| ENSG00000160111 | 366 | 203 | 268 | 154 | 208 | 197 |
| ENSG00000264250 | 0 | 1 | 0 | 0 | 0 | 0 |
| ENSG00000263595 | 0 | 0 | 1 | 0 | 0 | 0 |
| ENSG00000268985 | 0 | 0 | 0 | 0 | 0 | 0 |
| ENSG00000131351 | 420 | 179 | 254 | 222 | 194 | 275 |
| ENSG00000268289 | 0 | 0 | 0 | 0 | 0 | 0 |
| ENSG00000099331 | 1581 | 685 | 927 | 795 | 862 | 876 |
| ENSG00000269066 | 0 | 1 | 0 | 1 | 0 | 0 |
| ENSG00000268056 | 0 | 0 | 0 | 0 | 0 | 0 |
| ENSG00000269836 | 0 | 1 | 3 | 0 | 0 | 0 |
| ENSG00000269480 | 0 | 0 | 0 | 0 | 0 | 0 |
| ENSG00000053501 | 169 | 108 | 120 | 133 | 93 | 111 |
| ENSG00000099330 | 373 | 188 | 223 | 216 | 183 | 199 |
| ENSG00000160113 | 2022 | 1093 | 1213 | 1479 | 1223 | 1176 |
| ENSG00000269095 | 0 | 0 | 0 | 0 | 0 | 0 |
| ENSG00000130307 | 9 | 4 | 2 | 4 | 3 | 8 |
| ENSG00000105393 | 518 | 375 | 429 | 483 | 435 | 467 |
| ENSG00000269307 | 0 | 0 | 0 | 0 | 0 | 0 |
| ENSG00000160117 | 1 | 3 | 7 | 7 | 5 | 5 |
| ENSG00000127220 | 258 | 120 | 146 | 154 | 157 | 172 |
| ENSG00000130312 | 1049 | 499 | 607 | 640 | 591 | 579 |
| ENSG00000269815 | 6 | 4 | 6 | 5 | 8 | 2 |
| ENSG00000130311 | 1382 | 685 | 908 | 878 | 829 | 792 |
| ENSG00000074855 | 791 | 325 | 482 | 347 | 439 | 424 |
| ENSG00000130299 | 592 | 295 | 368 | 318 | 346 | 342 |
| ENSG00000130300 | 0 | 0 | 0 | 0 | 0 | 0 |
| ENSG00000269350 | 0 | 0 | 0 | 0 | 0 | 0 |
| ENSG00000269720 | 0 | 0 | 0 | 0 | 0 | 0 |
| ENSG00000130303 | 28 | 9 | 15 | 14 | 13 | 12 |
| ENSG00000282851 | 4 | 1 | 12 | 7 | 9 | 5 |
| ENSG00000141971 | 905 | 410 | 606 | 593 | 444 | 448 |
| ENSG00000269481 | 11 | 5 | 6 | 2 | 3 | 4 |
| ENSG00000254503 | 0 | 2 | 1 | 0 | 0 | 0 |
| ENSG00000269053 | 0 | 0 | 0 | 0 | 0 | 0 |
| ENSG00000188051 | 19 | 14 | 11 | 15 | 15 | 11 |
| ENSG00000269035 | 0 | 0 | 0 | 0 | 0 | 0 |
| ENSG00000269736 | 0 | 0 | 0 | 0 | 0 | 0 |
| ENSG00000171773 | 0 | 0 | 0 | 0 | 0 | 0 |
| ENSG00000130304 | 416 | 196 | 254 | 223 | 217 | 224 |
| ENSG00000269439 | 24 | 7 | 8 | 9 | 4 | 8 |
| ENSG00000130313 | 1765 | 911 | 1104 | 1318 | 966 | 1071 |
| ENSG00000269161 | 2 | 5 | 4 | 2 | 7 | 2 |
| ENSG00000167483 | 10 | 6 | 0 | 7 | 3 | 6 |
| ENSG00000130309 | 1373 | 841 | 887 | 894 | 898 | 933 |
| ENSG00000243771 | 0 | 0 | 0 | 0 | 1 | 0 |
| ENSG00000130477 | 6 | 3 | 6 | 1 | 2 | 2 |

| | | | | | | |
|---|---|---|---|---|---|---|
| ENSG00000269752 | 0 | 0 | 0 | 0 | 0 | 0 |
| ENSG00000279300 | 0 | 0 | 0 | 0 | 0 | 0 |
| ENSG00000130479 | 447 | 233 | 278 | 379 | 269 | 247 |
| ENSG00000268112 | 0 | 0 | 0 | 0 | 0 | 0 |
| ENSG00000130475 | 530 | 252 | 355 | 360 | 338 | 332 |
| ENSG00000179913 | 259 | 136 | 142 | 185 | 197 | 153 |
| ENSG00000248099 | 4 | 5 | 6 | 11 | 6 | 7 |
| ENSG00000105639 | 49 | 35 | 41 | 44 | 37 | 25 |
| ENSG00000105640 | 17270 | 8626 | 10216 | 13497 | 10274 | 10377 |
| ENSG00000207166 | 0 | 0 | 0 | 0 | 0 | 0 |
| ENSG00000279617 | 0 | 1 | 0 | 3 | 0 | 0 |
| ENSG00000105641 | 18 | 18 | 14 | 12 | 20 | 11 |
| ENSG00000007080 | 1414 | 732 | 872 | 1052 | 875 | 817 |
| ENSG00000105642 | 2 | 2 | 2 | 3 | 3 | 3 |
| ENSG00000252591 | 0 | 0 | 0 | 0 | 0 | 0 |
| ENSG00000105643 | 839 | 403 | 578 | 551 | 492 | 482 |
| ENSG00000279172 | 0 | 0 | 0 | 0 | 0 | 0 |
| ENSG00000273654 | 6 | 3 | 1 | 5 | 6 | 4 |
| ENSG00000268623 | 0 | 0 | 0 | 0 | 0 | 1 |
| ENSG00000243455 | 2 | 1 | 3 | 0 | 2 | 0 |
| ENSG00000096996 | 0 | 0 | 3 | 1 | 0 | 1 |
| ENSG00000099308 | 750 | 369 | 500 | 355 | 361 | 401 |
| ENSG00000269145 | 0 | 0 | 0 | 0 | 0 | 0 |
| ENSG00000105647 | 0 | 3 | 3 | 2 | 3 | 5 |
| ENSG00000268173 | 2 | 1 | 1 | 4 | 3 | 2 |
| ENSG00000216490 | 3 | 3 | 2 | 0 | 2 | 2 |
| ENSG00000254858 | 687 | 325 | 380 | 428 | 369 | 352 |
| ENSG00000105649 | 179 | 82 | 112 | 116 | 96 | 99 |
| ENSG00000268650 | 0 | 5 | 2 | 1 | 1 | 0 |
| ENSG00000285188 | 1 | 1 | 1 | 0 | 0 | 1 |
| ENSG00000105650 | 1 | 1 | 4 | 1 | 1 | 1 |
| ENSG00000284797 | 2 | 1 | 1 | 2 | 0 | 0 |
| ENSG00000130518 | 1085 | 370 | 716 | 484 | 416 | 619 |
| ENSG00000130522 | 5567 | 2422 | 3162 | 3759 | 2930 | 3244 |
| ENSG00000267959 | 0 | 0 | 0 | 0 | 0 | 0 |
| ENSG00000241464 | 2 | 0 | 2 | 0 | 0 | 1 |
| ENSG00000130520 | 3793 | 1943 | 2259 | 2667 | 2296 | 2345 |
| ENSG00000239821 | 0 | 1 | 0 | 1 | 1 | 3 |
| ENSG00000130517 | 2334 | 1230 | 1450 | 1285 | 1458 | 1491 |
| ENSG00000130513 | 2002 | 932 | 1462 | 3114 | 2742 | 3023 |
| ENSG00000264175 | 1 | 7 | 9 | 6 | 11 | 8 |
| ENSG00000175489 | 1 | 0 | 0 | 0 | 0 | 0 |
| ENSG00000130511 | 1566 | 735 | 1006 | 1075 | 930 | 1017 |
| ENSG00000105655 | 1398 | 581 | 937 | 852 | 588 | 756 |
| ENSG00000268199 | 4 | 1 | 2 | 0 | 1 | 4 |
| ENSG00000105656 | 697 | 294 | 423 | 393 | 361 | 352 |
| ENSG00000280121 | 0 | 0 | 0 | 0 | 0 | 0 |
| ENSG00000279262 | 0 | 0 | 0 | 0 | 0 | 0 |
| ENSG00000268938 | 9 | 4 | 5 | 3 | 4 | 4 |
| ENSG00000105701 | 10228 | 4864 | 6229 | 6622 | 4882 | 5042 |

| | | | | | | |
|---|---|---|---|---|---|---|
| ENSG00000269191 | 10 | 2 | 3 | 5 | 1 | 2 |
| ENSG00000105700 | 1437 | 859 | 1077 | 1065 | 1046 | 1043 |
| ENSG00000268030 | 51 | 22 | 29 | 34 | 35 | 34 |
| ENSG00000268983 | 0 | 0 | 0 | 0 | 0 | 0 |
| ENSG00000221983 | 11787 | 6025 | 6945 | 8041 | 7328 | 6730 |
| ENSG00000006016 | 299 | 112 | 157 | 129 | 122 | 112 |
| ENSG00000006015 | 887 | 429 | 585 | 649 | 518 | 544 |
| ENSG00000105696 | 50 | 31 | 51 | 39 | 49 | 46 |
| ENSG00000263490 | 0 | 0 | 0 | 0 | 0 | 0 |
| ENSG00000167487 | 349 | 208 | 217 | 210 | 194 | 199 |
| ENSG00000105662 | 919 | 381 | 592 | 478 | 474 | 465 |
| ENSG00000105664 | 4 | 1 | 0 | 0 | 1 | 0 |
| ENSG00000005007 | 6528 | 2964 | 3792 | 3234 | 3256 | 3567 |
| ENSG00000223802 | 85 | 40 | 69 | 75 | 68 | 41 |
| ENSG00000130283 | 0 | 0 | 0 | 0 | 0 | 0 |
| ENSG00000269694 | 1 | 1 | 1 | 1 | 3 | 3 |
| ENSG00000105669 | 3977 | 2023 | 2864 | 3380 | 2344 | 2665 |
| ENSG00000268193 | 0 | 0 | 0 | 0 | 0 | 0 |
| ENSG00000105671 | 2190 | 943 | 1337 | 1585 | 1179 | 1289 |
| ENSG00000051128 | 408 | 248 | 263 | 399 | 357 | 284 |
| ENSG00000269019 | 2 | 1 | 0 | 1 | 0 | 1 |
| ENSG00000241172 | 0 | 0 | 0 | 0 | 0 | 0 |
| ENSG00000064607 | 2593 | 1058 | 1670 | 1084 | 1479 | 1437 |
| ENSG00000105676 | 1130 | 593 | 770 | 891 | 685 | 837 |
| ENSG00000181035 | 603 | 278 | 313 | 262 | 255 | 231 |
| ENSG00000064545 | 1021 | 453 | 597 | 711 | 628 | 684 |
| ENSG00000064489 | 5 | 3 | 1 | 3 | 0 | 4 |
| ENSG00000213999 | 5 | 2 | 5 | 3 | 3 | 4 |
| ENSG00000254901 | 495 | 233 | 342 | 240 | 267 | 270 |
| ENSG00000064490 | 1460 | 728 | 844 | 1001 | 825 | 865 |
| ENSG00000184162 | 630 | 367 | 493 | 519 | 397 | 485 |
| ENSG00000130287 | 0 | 0 | 0 | 0 | 0 | 0 |
| ENSG00000252973 | 0 | 0 | 0 | 0 | 0 | 0 |
| ENSG00000267629 | 0 | 0 | 0 | 0 | 0 | 0 |
| ENSG00000280282 | 0 | 0 | 0 | 0 | 0 | 0 |
| ENSG00000187664 | 0 | 0 | 0 | 0 | 0 | 0 |
| ENSG00000213996 | 0 | 0 | 0 | 0 | 1 | 1 |
| ENSG00000105705 | 1170 | 509 | 677 | 667 | 536 | 594 |
| ENSG00000129933 | 2473 | 1167 | 1625 | 1032 | 1256 | 1311 |
| ENSG00000167491 | 2837 | 1506 | 1847 | 1713 | 1761 | 1877 |
| ENSG00000267234 | 0 | 0 | 0 | 0 | 0 | 0 |
| ENSG00000207821 | 0 | 0 | 0 | 0 | 0 | 0 |
| ENSG00000178093 | 107 | 51 | 65 | 76 | 62 | 50 |
| ENSG00000186010 | 68 | 36 | 33 | 42 | 51 | 39 |
| ENSG00000258674 | 15 | 5 | 9 | 4 | 14 | 8 |
| ENSG00000250067 | 107 | 22 | 52 | 37 | 35 | 48 |
| ENSG00000160161 | 86 | 45 | 52 | 56 | 53 | 50 |
| ENSG00000105717 | 77 | 30 | 51 | 60 | 47 | 51 |
| ENSG00000267178 | 0 | 0 | 0 | 0 | 0 | 0 |
| ENSG00000259242 | 0 | 0 | 0 | 0 | 0 | 1 |

| | | | | | | |
|---|---|---|---|---|---|---|
| ENSG00000064547 | 1130 | 549 | 699 | 823 | 621 | 680 |
| ENSG00000089639 | 1272 | 617 | 702 | 785 | 804 | 703 |
| ENSG00000105726 | 2650 | 1089 | 1480 | 1624 | 1312 | 1419 |
| ENSG00000181896 | 462 | 241 | 241 | 203 | 228 | 220 |
| ENSG00000271283 | 0 | 0 | 1 | 0 | 0 | 0 |
| ENSG00000270402 | 6 | 1 | 1 | 1 | 2 | 0 |
| ENSG00000105708 | 136 | 91 | 66 | 25 | 110 | 50 |
| ENSG00000275540 | 4 | 2 | 4 | 1 | 1 | 0 |
| ENSG00000271499 | 0 | 1 | 0 | 0 | 0 | 0 |
| ENSG00000266904 | 58 | 23 | 30 | 37 | 34 | 27 |
| ENSG00000267581 | 1 | 0 | 1 | 1 | 0 | 1 |
| ENSG00000267693 | 2 | 0 | 1 | 0 | 0 | 0 |
| ENSG00000267419 | 33 | 17 | 18 | 18 | 19 | 18 |
| ENSG00000081665 | 152 | 113 | 93 | 59 | 109 | 76 |
| ENSG00000267481 | 26 | 23 | 14 | 14 | 12 | 12 |
| ENSG00000270325 | 0 | 0 | 0 | 0 | 0 | 0 |
| ENSG00000256771 | 240 | 125 | 94 | 70 | 177 | 135 |
| ENSG00000270270 | 2 | 0 | 0 | 0 | 0 | 1 |
| ENSG00000267565 | 21 | 17 | 11 | 7 | 26 | 18 |
| ENSG00000184635 | 39 | 34 | 30 | 22 | 68 | 26 |
| ENSG00000273791 | 0 | 0 | 0 | 0 | 0 | 0 |
| ENSG00000274800 | 0 | 0 | 0 | 0 | 0 | 0 |
| ENSG00000197124 | 94 | 45 | 58 | 25 | 55 | 30 |
| ENSG00000268070 | 3 | 2 | 2 | 0 | 1 | 1 |
| ENSG00000274033 | 0 | 0 | 0 | 0 | 0 | 0 |
| ENSG00000269154 | 0 | 0 | 0 | 0 | 0 | 0 |
| ENSG00000213988 | 2 | 2 | 3 | 0 | 6 | 4 |
| ENSG00000240673 | 0 | 0 | 0 | 0 | 0 | 0 |
| ENSG00000274612 | 0 | 0 | 0 | 0 | 0 | 0 |
| ENSG00000224864 | 0 | 0 | 0 | 0 | 0 | 0 |
| ENSG00000267383 | 12 | 4 | 5 | 3 | 2 | 2 |
| ENSG00000267220 | 0 | 0 | 0 | 0 | 0 | 0 |
| ENSG00000270716 | 0 | 0 | 0 | 0 | 0 | 0 |
| ENSG00000256229 | 43 | 17 | 11 | 7 | 14 | 9 |
| ENSG00000267641 | 0 | 0 | 0 | 0 | 0 | 0 |
| ENSG00000267771 | 0 | 0 | 0 | 0 | 0 | 0 |
| ENSG00000279039 | 4 | 2 | 1 | 2 | 1 | 0 |
| ENSG00000280079 | 8 | 9 | 4 | 0 | 5 | 4 |
| ENSG00000271524 | 0 | 1 | 1 | 0 | 1 | 1 |
| ENSG00000268620 | 0 | 0 | 0 | 0 | 0 | 0 |
| ENSG00000213985 | 6 | 2 | 1 | 0 | 2 | 1 |
| ENSG00000269055 | 0 | 0 | 0 | 0 | 0 | 0 |
| ENSG00000269274 | 0 | 0 | 0 | 0 | 0 | 0 |
| ENSG00000271638 | 0 | 0 | 0 | 0 | 0 | 0 |
| ENSG00000276294 | 0 | 0 | 0 | 0 | 0 | 0 |
| ENSG00000231205 | 0 | 0 | 0 | 0 | 0 | 0 |
| ENSG00000271704 | 0 | 0 | 0 | 0 | 0 | 0 |
| ENSG00000268461 | 0 | 0 | 0 | 0 | 0 | 0 |
| ENSG00000238517 | 0 | 0 | 0 | 0 | 0 | 0 |
| ENSG00000268174 | 0 | 0 | 0 | 0 | 0 | 0 |

| | | | | | | |
|---|---|---|---|---|---|---|
| ENSG00000269043 | 0 | 0 | 0 | 0 | 0 | 0 |
| ENSG00000268335 | 0 | 0 | 0 | 0 | 0 | 0 |
| ENSG00000237440 | 153 | 86 | 82 | 47 | 119 | 73 |
| ENSG00000269110 | 0 | 0 | 0 | 0 | 0 | 0 |
| ENSG00000268272 | 0 | 0 | 0 | 0 | 0 | 0 |
| ENSG00000268560 | 0 | 0 | 0 | 0 | 0 | 0 |
| ENSG00000188171 | 2 | 0 | 0 | 0 | 0 | 0 |
| ENSG00000280171 | 0 | 0 | 0 | 0 | 0 | 0 |
| ENSG00000269779 | 0 | 0 | 0 | 0 | 0 | 0 |
| ENSG00000268416 | 6 | 7 | 8 | 6 | 6 | 7 |
| ENSG00000269885 | 0 | 0 | 0 | 0 | 0 | 0 |
| ENSG00000160229 | 10 | 13 | 19 | 7 | 17 | 15 |
| ENSG00000269040 | 0 | 2 | 0 | 0 | 0 | 0 |
| ENSG00000269373 | 2 | 1 | 0 | 1 | 1 | 0 |
| ENSG00000105750 | 109 | 80 | 64 | 37 | 67 | 49 |
| ENSG00000279404 | 0 | 0 | 1 | 0 | 0 | 0 |
| ENSG00000269012 | 2 | 0 | 0 | 0 | 0 | 0 |
| ENSG00000268326 | 0 | 0 | 0 | 0 | 0 | 0 |
| ENSG00000118620 | 165 | 63 | 57 | 24 | 89 | 51 |
| ENSG00000271598 | 0 | 0 | 0 | 0 | 0 | 0 |
| ENSG00000268598 | 0 | 0 | 0 | 0 | 0 | 0 |
| ENSG00000160352 | 380 | 196 | 212 | 94 | 203 | 185 |
| ENSG00000201035 | 0 | 0 | 1 | 0 | 0 | 0 |
| ENSG00000268357 | 25 | 7 | 8 | 12 | 15 | 8 |
| ENSG00000196705 | 364 | 182 | 190 | 112 | 251 | 161 |
| ENSG00000240522 | 9 | 12 | 11 | 2 | 9 | 3 |
| ENSG00000268995 | 0 | 0 | 0 | 0 | 0 | 0 |
| ENSG00000268892 | 0 | 0 | 0 | 0 | 0 | 0 |
| ENSG00000269839 | 0 | 0 | 0 | 0 | 0 | 0 |
| ENSG00000242291 | 0 | 0 | 0 | 0 | 1 | 0 |
| ENSG00000268747 | 1 | 0 | 0 | 0 | 0 | 2 |
| ENSG00000268521 | 7 | 6 | 4 | 2 | 6 | 5 |
| ENSG00000182141 | 154 | 65 | 86 | 34 | 93 | 67 |
| ENSG00000268322 | 4 | 2 | 0 | 0 | 0 | 1 |
| ENSG00000172687 | 227 | 100 | 134 | 77 | 125 | 103 |
| ENSG00000268731 | 0 | 0 | 0 | 0 | 1 | 0 |
| ENSG00000213976 | 4 | 5 | 6 | 3 | 9 | 10 |
| ENSG00000196268 | 47 | 42 | 35 | 21 | 66 | 42 |
| ENSG00000269237 | 0 | 0 | 0 | 0 | 0 | 0 |
| ENSG00000268119 | 6 | 4 | 3 | 3 | 2 | 2 |
| ENSG00000268658 | 0 | 4 | 6 | 0 | 1 | 0 |
| ENSG00000197013 | 245 | 144 | 135 | 50 | 168 | 119 |
| ENSG00000268705 | 2 | 4 | 0 | 0 | 1 | 1 |
| ENSG00000268081 | 0 | 0 | 3 | 4 | 2 | 1 |
| ENSG00000268555 | 2 | 0 | 0 | 0 | 0 | 0 |
| ENSG00000268240 | 0 | 0 | 0 | 0 | 0 | 0 |
| ENSG00000274503 | 0 | 0 | 0 | 0 | 0 | 0 |
| ENSG00000271182 | 0 | 0 | 0 | 0 | 0 | 0 |
| ENSG00000268278 | 0 | 0 | 0 | 0 | 0 | 0 |
| ENSG00000268433 | 0 | 0 | 0 | 0 | 0 | 0 |

| | | | | | | |
|---|---|---|---|---|---|---|
| ENSG00000268535 | 0 | 0 | 0 | 0 | 0 | 0 |
| ENSG00000268117 | 1 | 2 | 0 | 0 | 0 | 0 |
| ENSG00000197020 | 191 | 102 | 82 | 54 | 142 | 82 |
| ENSG00000269845 | 8 | 1 | 7 | 4 | 3 | 6 |
| ENSG00000268184 | 0 | 0 | 1 | 0 | 0 | 0 |
| ENSG00000269136 | 0 | 0 | 0 | 0 | 0 | 0 |
| ENSG00000198521 | 246 | 157 | 116 | 96 | 214 | 127 |
| ENSG00000268438 | 2 | 2 | 0 | 0 | 0 | 0 |
| ENSG00000160321 | 0 | 1 | 0 | 0 | 0 | 1 |
| ENSG00000269615 | 0 | 0 | 0 | 0 | 0 | 0 |
| ENSG00000279377 | 0 | 0 | 0 | 0 | 0 | 0 |
| ENSG00000271095 | 0 | 0 | 0 | 0 | 0 | 0 |
| ENSG00000269504 | 0 | 0 | 0 | 0 | 0 | 0 |
| ENSG00000269199 | 0 | 0 | 0 | 0 | 0 | 0 |
| ENSG00000197134 | 25 | 12 | 15 | 16 | 23 | 11 |
| ENSG00000269742 | 2 | 0 | 0 | 0 | 0 | 0 |
| ENSG00000269505 | 0 | 0 | 0 | 0 | 0 | 0 |
| ENSG00000268116 | 0 | 0 | 0 | 0 | 0 | 0 |
| ENSG00000268140 | 12 | 2 | 5 | 1 | 3 | 7 |
| ENSG00000269575 | 0 | 0 | 0 | 0 | 0 | 0 |
| ENSG00000269345 | 0 | 0 | 0 | 0 | 0 | 0 |
| ENSG00000196109 | 0 | 0 | 0 | 0 | 0 | 0 |
| ENSG00000268079 | 0 | 0 | 0 | 0 | 0 | 0 |
| ENSG00000282732 | 0 | 0 | 0 | 0 | 0 | 0 |
| ENSG00000267872 | 0 | 0 | 0 | 0 | 0 | 0 |
| ENSG00000268597 | 0 | 0 | 0 | 0 | 0 | 0 |
| ENSG00000196350 | 0 | 0 | 0 | 0 | 0 | 0 |
| ENSG00000270191 | 0 | 0 | 0 | 0 | 0 | 0 |
| ENSG00000268685 | 0 | 0 | 0 | 0 | 0 | 0 |
| ENSG00000236312 | 0 | 0 | 0 | 0 | 0 | 0 |
| ENSG00000197360 | 6 | 0 | 0 | 0 | 0 | 2 |
| ENSG00000269796 | 0 | 0 | 0 | 0 | 0 | 0 |
| ENSG00000268830 | 0 | 0 | 0 | 0 | 0 | 0 |
| ENSG00000270947 | 0 | 0 | 0 | 0 | 0 | 0 |
| ENSG00000268717 | 0 | 0 | 0 | 0 | 0 | 0 |
| ENSG00000269138 | 0 | 0 | 0 | 0 | 0 | 0 |
| ENSG00000270317 | 0 | 0 | 0 | 0 | 0 | 0 |
| ENSG00000260599 | 0 | 0 | 0 | 0 | 0 | 0 |
| ENSG00000269364 | 0 | 0 | 0 | 0 | 0 | 0 |
| ENSG00000207397 | 0 | 0 | 0 | 0 | 0 | 0 |
| ENSG00000271149 | 0 | 0 | 0 | 0 | 0 | 0 |
| ENSG00000269332 | 0 | 0 | 0 | 0 | 0 | 0 |
| ENSG00000240713 | 0 | 0 | 0 | 0 | 0 | 0 |
| ENSG00000268823 | 0 | 0 | 0 | 0 | 0 | 0 |
| ENSG00000269025 | 0 | 0 | 0 | 0 | 0 | 0 |
| ENSG00000269316 | 0 | 0 | 0 | 0 | 0 | 0 |
| ENSG00000268038 | 0 | 0 | 0 | 0 | 0 | 0 |
| ENSG00000268589 | 0 | 0 | 0 | 0 | 0 | 0 |
| ENSG00000229676 | 0 | 1 | 3 | 2 | 3 | 2 |
| ENSG00000268981 | 0 | 0 | 0 | 0 | 0 | 0 |

| | | | | | | |
|---|---|---|---|---|---|---|
| ENSG00000198153 | 0 | 0 | 0 | 0 | 0 | 0 |
| ENSG00000241282 | 0 | 1 | 0 | 0 | 0 | 0 |
| ENSG00000270475 | 0 | 0 | 0 | 0 | 0 | 0 |
| ENSG00000268789 | 0 | 0 | 0 | 0 | 0 | 0 |
| ENSG00000213973 | 1 | 2 | 0 | 0 | 1 | 0 |
| ENSG00000279966 | 0 | 0 | 0 | 0 | 0 | 0 |
| ENSG00000270518 | 0 | 0 | 0 | 0 | 0 | 0 |
| ENSG00000269509 | 0 | 1 | 0 | 0 | 0 | 0 |
| ENSG00000268696 | 0 | 0 | 0 | 0 | 0 | 1 |
| ENSG00000271655 | 0 | 0 | 0 | 0 | 0 | 0 |
| ENSG00000268541 | 0 | 0 | 0 | 0 | 0 | 0 |
| ENSG00000268296 | 0 | 0 | 0 | 0 | 0 | 0 |
| ENSG00000269458 | 0 | 0 | 0 | 0 | 0 | 0 |
| ENSG00000269067 | 0 | 0 | 0 | 0 | 0 | 0 |
| ENSG00000271661 | 0 | 0 | 0 | 0 | 0 | 0 |
| ENSG00000261558 | 0 | 0 | 0 | 0 | 0 | 0 |
| ENSG00000261615 | 0 | 0 | 0 | 0 | 0 | 0 |
| ENSG00000269543 | 0 | 0 | 0 | 0 | 0 | 0 |
| ENSG00000267886 | 19 | 13 | 22 | 7 | 17 | 14 |
| ENSG00000183850 | 16 | 10 | 12 | 3 | 10 | 9 |
| ENSG00000270479 | 0 | 0 | 0 | 0 | 0 | 0 |
| ENSG00000268105 | 0 | 0 | 0 | 0 | 0 | 0 |
| ENSG00000267920 | 0 | 0 | 0 | 0 | 0 | 0 |
| ENSG00000269471 | 0 | 0 | 0 | 0 | 0 | 0 |
| ENSG00000196081 | 16 | 22 | 9 | 11 | 19 | 17 |
| ENSG00000283201 | 0 | 0 | 0 | 0 | 0 | 0 |
| ENSG00000268469 | 0 | 0 | 0 | 0 | 1 | 0 |
| ENSG00000269837 | 401 | 224 | 228 | 216 | 268 | 220 |
| ENSG00000269107 | 0 | 0 | 0 | 0 | 0 | 0 |
| ENSG00000284428 | 125 | 67 | 86 | 53 | 70 | 72 |
| ENSG00000268794 | 0 | 0 | 0 | 0 | 0 | 0 |
| ENSG00000268148 | 0 | 0 | 0 | 0 | 0 | 0 |
| ENSG00000167232 | 221 | 136 | 102 | 49 | 198 | 98 |
| ENSG00000267934 | 4 | 5 | 2 | 0 | 7 | 6 |
| ENSG00000269431 | 0 | 1 | 0 | 0 | 1 | 0 |
| ENSG00000268992 | 0 | 0 | 2 | 0 | 0 | 0 |
| ENSG00000269416 | 0 | 0 | 1 | 0 | 0 | 0 |
| ENSG00000268908 | 0 | 0 | 0 | 0 | 0 | 0 |
| ENSG00000275201 | 0 | 0 | 0 | 0 | 0 | 1 |
| ENSG00000268100 | 5 | 1 | 1 | 0 | 0 | 1 |
| ENSG00000197372 | 246 | 117 | 125 | 59 | 133 | 97 |
| ENSG00000268455 | 0 | 0 | 0 | 0 | 0 | 0 |
| ENSG00000240231 | 0 | 1 | 0 | 1 | 0 | 1 |
| ENSG00000269320 | 0 | 0 | 0 | 0 | 0 | 1 |
| ENSG00000196172 | 99 | 70 | 37 | 22 | 74 | 36 |
| ENSG00000233836 | 41 | 29 | 29 | 20 | 37 | 15 |
| ENSG00000269662 | 2 | 0 | 0 | 1 | 1 | 1 |
| ENSG00000267924 | 1 | 2 | 3 | 0 | 0 | 1 |
| ENSG00000225178 | 2560 | 1371 | 1627 | 2030 | 1752 | 1583 |
| ENSG00000277493 | 0 | 0 | 0 | 0 | 0 | 1 |

| | | | | | | |
|---|---|---|---|---|---|---|
| ENSG00000213967 | 105 | 52 | 58 | 47 | 75 | 64 |
| ENSG00000269289 | 2 | 0 | 2 | 0 | 0 | 0 |
| ENSG00000269397 | 22 | 7 | 8 | 13 | 13 | 5 |
| ENSG00000269421 | 0 | 0 | 1 | 0 | 0 | 0 |
| ENSG00000201966 | 0 | 0 | 0 | 0 | 0 | 0 |
| ENSG00000213096 | 155 | 95 | 87 | 48 | 201 | 92 |
| ENSG00000268362 | 400 | 178 | 243 | 176 | 215 | 217 |
| ENSG00000279425 | 40 | 11 | 23 | 11 | 12 | 23 |
| ENSG00000268058 | 1 | 4 | 0 | 0 | 0 | 1 |
| ENSG00000276030 | 5 | 2 | 2 | 1 | 1 | 0 |
| ENSG00000268442 | 4 | 0 | 1 | 2 | 6 | 0 |
| ENSG00000267696 | 0 | 1 | 0 | 0 | 0 | 0 |
| ENSG00000279882 | 0 | 0 | 0 | 0 | 0 | 0 |
| ENSG00000261824 | 140 | 61 | 88 | 92 | 89 | 93 |
| ENSG00000261770 | 6 | 1 | 2 | 1 | 2 | 5 |
| ENSG00000267575 | 84 | 75 | 62 | 62 | 62 | 43 |
| ENSG00000272635 | 0 | 0 | 1 | 0 | 0 | 0 |
| ENSG00000281468 | 19 | 24 | 13 | 20 | 31 | 22 |
| ENSG00000267264 | 4 | 0 | 2 | 0 | 0 | 0 |
| ENSG00000267323 | 2 | 0 | 0 | 0 | 0 | 0 |
| ENSG00000266977 | 0 | 0 | 0 | 0 | 0 | 0 |
| ENSG00000267389 | 0 | 0 | 0 | 0 | 0 | 0 |
| ENSG00000267623 | 0 | 0 | 0 | 0 | 0 | 0 |
| ENSG00000267768 | 0 | 0 | 0 | 0 | 0 | 0 |
| ENSG00000267630 | 0 | 0 | 0 | 0 | 0 | 0 |
| ENSG00000266906 | 0 | 0 | 0 | 0 | 0 | 0 |
| ENSG00000267509 | 0 | 0 | 0 | 0 | 0 | 0 |
| ENSG00000266928 | 0 | 0 | 0 | 0 | 0 | 0 |
| ENSG00000267320 | 0 | 0 | 0 | 0 | 0 | 0 |
| ENSG00000267238 | 0 | 0 | 0 | 0 | 0 | 0 |
| ENSG00000260725 | 0 | 0 | 0 | 0 | 0 | 0 |
| ENSG00000283403 | 0 | 0 | 0 | 0 | 0 | 0 |
| ENSG00000267243 | 0 | 0 | 0 | 0 | 0 | 0 |
| ENSG00000266893 | 0 | 0 | 0 | 0 | 0 | 0 |
| ENSG00000267423 | 0 | 0 | 0 | 0 | 0 | 0 |
| ENSG00000267537 | 0 | 0 | 0 | 0 | 0 | 0 |
| ENSG00000266976 | 0 | 0 | 0 | 0 | 0 | 0 |
| ENSG00000283269 | 0 | 0 | 0 | 0 | 0 | 0 |
| ENSG00000267799 | 0 | 0 | 0 | 0 | 0 | 0 |
| ENSG00000267617 | 0 | 0 | 0 | 0 | 0 | 0 |
| ENSG00000267528 | 0 | 0 | 0 | 0 | 0 | 0 |
| ENSG00000267683 | 0 | 0 | 0 | 0 | 0 | 0 |
| ENSG00000267027 | 0 | 0 | 0 | 0 | 0 | 0 |
| ENSG00000267240 | 0 | 0 | 0 | 0 | 0 | 0 |
| ENSG00000267339 | 0 | 0 | 0 | 0 | 0 | 0 |
| ENSG00000252272 | 0 | 0 | 0 | 0 | 0 | 0 |
| ENSG00000267014 | 0 | 1 | 0 | 0 | 0 | 0 |
| ENSG00000169021 | 2982 | 1423 | 1814 | 1953 | 1777 | 1808 |
| ENSG00000267498 | 3 | 1 | 3 | 1 | 0 | 3 |
| ENSG00000276251 | 0 | 0 | 0 | 0 | 0 | 0 |

| | | | | | | |
|---|---|---|---|---|---|---|
| ENSG00000276631 | 0 | 0 | 0 | 0 | 0 | 0 |
| ENSG00000241604 | 0 | 0 | 0 | 0 | 0 | 0 |
| ENSG00000264515 | 0 | 0 | 0 | 0 | 0 | 3 |
| ENSG00000267574 | 0 | 0 | 0 | 0 | 0 | 0 |
| ENSG00000266248 | 0 | 0 | 0 | 0 | 0 | 0 |
| ENSG00000279387 | 0 | 0 | 0 | 0 | 0 | 0 |
| ENSG00000187135 | 0 | 0 | 0 | 0 | 0 | 0 |
| ENSG00000105171 | 1515 | 672 | 950 | 884 | 903 | 903 |
| ENSG00000166289 | 84 | 61 | 71 | 131 | 87 | 86 |
| ENSG00000131943 | 768 | 363 | 497 | 468 | 501 | 500 |
| ENSG00000105173 | 420 | 227 | 258 | 239 | 247 | 261 |
| ENSG00000267433 | 0 | 0 | 0 | 0 | 0 | 0 |
| ENSG00000278895 | 2 | 1 | 0 | 0 | 0 | 0 |
| ENSG00000267258 | 0 | 0 | 0 | 0 | 0 | 0 |
| ENSG00000105176 | 2413 | 1261 | 1421 | 1149 | 1402 | 1321 |
| ENSG00000242100 | 4 | 0 | 0 | 1 | 1 | 0 |
| ENSG00000267006 | 0 | 0 | 0 | 0 | 0 | 0 |
| ENSG00000267692 | 1 | 2 | 0 | 2 | 0 | 0 |
| ENSG00000266910 | 4 | 0 | 0 | 2 | 2 | 3 |
| ENSG00000267223 | 0 | 0 | 0 | 0 | 0 | 0 |
| ENSG00000198597 | 0 | 0 | 0 | 1 | 0 | 0 |
| ENSG00000280061 | 0 | 0 | 0 | 0 | 0 | 0 |
| ENSG00000267760 | 0 | 0 | 0 | 0 | 0 | 0 |
| ENSG00000267081 | 0 | 0 | 0 | 0 | 0 | 0 |
| ENSG00000261316 | 0 | 0 | 0 | 0 | 0 | 0 |
| ENSG00000267378 | 0 | 0 | 0 | 0 | 0 | 0 |
| ENSG00000284430 | 0 | 0 | 0 | 0 | 0 | 0 |
| ENSG00000267777 | 0 | 0 | 0 | 0 | 0 | 0 |
| ENSG00000121297 | 26 | 19 | 24 | 13 | 24 | 14 |
| ENSG00000267662 | 2 | 0 | 0 | 0 | 0 | 0 |
| ENSG00000278875 | 0 | 4 | 1 | 0 | 0 | 0 |
| ENSG00000267636 | 0 | 0 | 0 | 0 | 0 | 0 |
| ENSG00000267599 | 0 | 0 | 0 | 0 | 0 | 0 |
| ENSG00000267407 | 0 | 0 | 0 | 0 | 0 | 0 |
| ENSG00000261400 | 0 | 0 | 0 | 0 | 0 | 0 |
| ENSG00000267465 | 0 | 0 | 0 | 0 | 0 | 0 |
| ENSG00000252780 | 0 | 0 | 0 | 0 | 0 | 0 |
| ENSG00000201894 | 0 | 0 | 0 | 0 | 0 | 0 |
| ENSG00000267489 | 0 | 0 | 0 | 0 | 0 | 0 |
| ENSG00000267012 | 0 | 0 | 0 | 0 | 0 | 0 |
| ENSG00000267779 | 0 | 0 | 0 | 0 | 0 | 0 |
| ENSG00000267204 | 0 | 0 | 0 | 0 | 0 | 0 |
| ENSG00000267133 | 0 | 0 | 0 | 0 | 0 | 0 |
| ENSG00000200738 | 0 | 0 | 0 | 0 | 0 | 0 |
| ENSG00000168813 | 625 | 330 | 316 | 118 | 431 | 294 |
| ENSG00000267213 | 34 | 18 | 28 | 30 | 23 | 19 |
| ENSG00000178904 | 1052 | 579 | 549 | 310 | 726 | 498 |
| ENSG00000105185 | 1311 | 750 | 836 | 856 | 898 | 866 |
| ENSG00000105186 | 2351 | 1135 | 1366 | 957 | 1085 | 1128 |
| ENSG00000201388 | 3 | 2 | 1 | 1 | 0 | 0 |

| | | | | | | |
|---|---|---|---|---|---|---|
| ENSG00000242325 | 0 | 1 | 0 | 0 | 1 | 1 |
| ENSG00000242436 | 0 | 0 | 0 | 0 | 0 | 0 |
| ENSG00000267557 | 0 | 0 | 0 | 0 | 1 | 0 |
| ENSG00000186326 | 34 | 32 | 44 | 23 | 18 | 21 |
| ENSG00000267475 | 52 | 23 | 43 | 12 | 13 | 13 |
| ENSG00000213965 | 1119 | 536 | 752 | 390 | 485 | 548 |
| ENSG00000267567 | 5 | 3 | 7 | 5 | 0 | 3 |
| ENSG00000173809 | 19 | 11 | 4 | 1 | 6 | 7 |
| ENSG00000267245 | 0 | 0 | 1 | 0 | 0 | 0 |
| ENSG00000021488 | 2 | 2 | 2 | 1 | 0 | 0 |
| ENSG00000267555 | 0 | 0 | 0 | 0 | 0 | 0 |
| ENSG00000201967 | 0 | 0 | 0 | 0 | 0 | 0 |
| ENSG00000121289 | 1186 | 686 | 792 | 612 | 664 | 626 |
| ENSG00000239988 | 0 | 1 | 0 | 0 | 0 | 0 |
| ENSG00000131944 | 95 | 82 | 65 | 68 | 75 | 60 |
| ENSG00000131941 | 1781 | 857 | 1172 | 656 | 755 | 916 |
| ENSG00000267611 | 0 | 0 | 0 | 0 | 0 | 0 |
| ENSG00000267725 | 0 | 0 | 0 | 0 | 0 | 0 |
| ENSG00000076650 | 511 | 269 | 278 | 204 | 296 | 287 |
| ENSG00000166359 | 57 | 53 | 71 | 26 | 45 | 57 |
| ENSG00000130881 | 1348 | 675 | 866 | 959 | 732 | 862 |
| ENSG00000267054 | 0 | 0 | 0 | 0 | 0 | 0 |
| ENSG00000273420 | 0 | 0 | 0 | 0 | 0 | 0 |
| ENSG00000130876 | 0 | 0 | 0 | 2 | 0 | 0 |
| ENSG00000267714 | 0 | 0 | 0 | 0 | 0 | 0 |
| ENSG00000270212 | 0 | 0 | 0 | 0 | 0 | 0 |
| ENSG00000267580 | 0 | 0 | 0 | 0 | 0 | 0 |
| ENSG00000245848 | 240 | 147 | 198 | 193 | 182 | 182 |
| ENSG00000267727 | 0 | 0 | 0 | 0 | 0 | 0 |
| ENSG00000267296 | 145 | 64 | 84 | 76 | 58 | 53 |
| ENSG00000267130 | 0 | 1 | 1 | 3 | 4 | 4 |
| ENSG00000243864 | 0 | 0 | 0 | 0 | 0 | 0 |
| ENSG00000266899 | 0 | 0 | 0 | 0 | 0 | 0 |
| ENSG00000153879 | 918 | 491 | 594 | 517 | 567 | 528 |
| ENSG00000124299 | 755 | 391 | 482 | 484 | 446 | 411 |
| ENSG00000266985 | 0 | 0 | 0 | 0 | 0 | 0 |
| ENSG00000232429 | 0 | 0 | 0 | 0 | 0 | 0 |
| ENSG00000124302 | 17 | 6 | 8 | 14 | 5 | 10 |
| ENSG00000153885 | 1598 | 705 | 1057 | 828 | 887 | 916 |
| ENSG00000277013 | 85 | 40 | 56 | 43 | 50 | 45 |
| ENSG00000240626 | 0 | 0 | 0 | 0 | 0 | 0 |
| ENSG00000239253 | 0 | 0 | 0 | 0 | 0 | 0 |
| ENSG00000249987 | 7 | 0 | 0 | 0 | 1 | 0 |
| ENSG00000186008 | 7 | 0 | 1 | 0 | 0 | 0 |
| ENSG00000270849 | 1 | 0 | 0 | 0 | 0 | 0 |
| ENSG00000257103 | 4034 | 1925 | 2524 | 1595 | 2466 | 2275 |
| ENSG00000166398 | 586 | 235 | 388 | 209 | 270 | 261 |
| ENSG00000257074 | 0 | 0 | 0 | 0 | 0 | 0 |
| ENSG00000267219 | 9 | 4 | 3 | 10 | 6 | 2 |
| ENSG00000105220 | 10779 | 5123 | 6491 | 6205 | 5469 | 6104 |

| | | | | | | |
|---|---|---|---|---|---|---|
| ENSG00000266953 | 0 | 0 | 0 | 0 | 0 | 0 |
| ENSG00000126249 | 183 | 81 | 128 | 153 | 153 | 161 |
| ENSG00000264317 | 0 | 0 | 0 | 0 | 3 | 0 |
| ENSG00000267024 | 3 | 0 | 8 | 1 | 0 | 3 |
| ENSG00000126261 | 2822 | 1455 | 1626 | 1283 | 2008 | 1674 |
| ENSG00000142279 | 156 | 68 | 70 | 57 | 45 | 53 |
| ENSG00000239210 | 0 | 0 | 0 | 0 | 0 | 0 |
| ENSG00000248830 | 8 | 0 | 2 | 1 | 1 | 0 |
| ENSG00000268751 | 0 | 3 | 2 | 2 | 6 | 4 |
| ENSG00000205209 | 1 | 1 | 5 | 3 | 4 | 2 |
| ENSG00000271049 | 0 | 0 | 1 | 0 | 2 | 1 |
| ENSG00000252230 | 0 | 0 | 0 | 0 | 0 | 0 |
| ENSG00000268683 | 0 | 0 | 0 | 0 | 0 | 0 |
| ENSG00000279971 | 0 | 0 | 0 | 0 | 0 | 0 |
| ENSG00000269811 | 0 | 0 | 0 | 0 | 0 | 0 |
| ENSG00000279329 | 34 | 11 | 21 | 15 | 10 | 17 |
| ENSG00000089335 | 563 | 279 | 299 | 156 | 389 | 247 |
| ENSG00000274104 | 22 | 13 | 20 | 12 | 12 | 12 |
| ENSG00000197841 | 311 | 116 | 120 | 72 | 177 | 116 |
| ENSG00000153896 | 199 | 87 | 140 | 86 | 104 | 87 |
| ENSG00000270281 | 0 | 0 | 0 | 0 | 0 | 1 |
| ENSG00000267767 | 39 | 23 | 49 | 11 | 13 | 15 |
| ENSG00000261754 | 122 | 66 | 87 | 26 | 45 | 41 |
| ENSG00000271666 | 1 | 0 | 0 | 0 | 0 | 0 |
| ENSG00000269086 | 17 | 5 | 6 | 2 | 7 | 4 |
| ENSG00000271109 | 78 | 30 | 41 | 27 | 31 | 29 |
| ENSG00000270624 | 3 | 3 | 2 | 3 | 3 | 4 |
| ENSG00000271649 | 0 | 0 | 0 | 0 | 0 | 0 |
| ENSG00000270544 | 0 | 0 | 0 | 0 | 0 | 0 |
| ENSG00000271171 | 0 | 0 | 0 | 0 | 0 | 0 |
| ENSG00000269037 | 0 | 0 | 0 | 0 | 0 | 0 |
| ENSG00000270876 | 3 | 1 | 0 | 0 | 0 | 2 |
| ENSG00000168661 | 164 | 82 | 100 | 61 | 128 | 88 |
| ENSG00000280106 | 36 | 17 | 24 | 26 | 13 | 18 |
| ENSG00000180884 | 260 | 135 | 165 | 121 | 121 | 141 |
| ENSG00000089351 | 2339 | 1090 | 1439 | 1176 | 1286 | 1065 |
| ENSG00000269303 | 0 | 2 | 0 | 0 | 0 | 0 |
| ENSG00000271032 | 11 | 7 | 8 | 8 | 3 | 5 |
| ENSG00000105711 | 1142 | 567 | 651 | 505 | 514 | 435 |
| ENSG00000105707 | 32 | 14 | 22 | 18 | 14 | 8 |
| ENSG00000227392 | 0 | 0 | 0 | 0 | 0 | 0 |
| ENSG00000179066 | 0 | 0 | 0 | 0 | 0 | 1 |
| ENSG00000285526 | 0 | 0 | 0 | 0 | 0 | 0 |
| ENSG00000089356 | 266 | 126 | 239 | 196 | 225 | 191 |
| ENSG00000284065 | 0 | 0 | 0 | 0 | 0 | 0 |
| ENSG00000153902 | 0 | 0 | 0 | 0 | 3 | 0 |
| ENSG00000279619 | 0 | 0 | 0 | 0 | 0 | 0 |
| ENSG00000266964 | 0 | 0 | 0 | 0 | 0 | 0 |
| ENSG00000221857 | 0 | 0 | 0 | 0 | 0 | 0 |
| ENSG00000221946 | 0 | 0 | 0 | 0 | 0 | 0 |

| | | | | | | |
|---|---|---|---|---|---|---|
| ENSG00000089327 | 4 | 6 | 2 | 10 | 8 | 6 |
| ENSG00000177558 | 0 | 0 | 0 | 0 | 0 | 0 |
| ENSG00000262497 | 0 | 2 | 5 | 0 | 2 | 0 |
| ENSG00000105699 | 5122 | 2544 | 3118 | 3498 | 2980 | 2711 |
| ENSG00000271366 | 5 | 10 | 14 | 5 | 5 | 2 |
| ENSG00000268947 | 22 | 18 | 22 | 12 | 17 | 16 |
| ENSG00000105698 | 2864 | 1239 | 1628 | 1675 | 1398 | 1454 |
| ENSG00000105697 | 0 | 0 | 1 | 0 | 0 | 0 |
| ENSG00000105695 | 12 | 3 | 3 | 7 | 2 | 4 |
| ENSG00000012124 | 78 | 20 | 32 | 79 | 63 | 48 |
| ENSG00000269553 | 0 | 1 | 3 | 0 | 1 | 0 |
| ENSG00000284034 | 0 | 0 | 0 | 0 | 0 | 0 |
| ENSG00000126266 | 0 | 0 | 2 | 0 | 0 | 1 |
| ENSG00000185897 | 0 | 0 | 0 | 0 | 0 | 0 |
| ENSG00000126251 | 0 | 0 | 0 | 0 | 0 | 0 |
| ENSG00000268222 | 2 | 0 | 2 | 2 | 2 | 1 |
| ENSG00000205786 | 0 | 0 | 0 | 0 | 0 | 0 |
| ENSG00000283479 | 0 | 0 | 0 | 0 | 0 | 0 |
| ENSG00000264400 | 0 | 0 | 0 | 0 | 0 | 0 |
| ENSG00000233214 | 0 | 0 | 0 | 0 | 0 | 1 |
| ENSG00000232680 | 0 | 0 | 0 | 0 | 0 | 0 |
| ENSG00000126262 | 34 | 12 | 29 | 10 | 8 | 10 |
| ENSG00000188508 | 4 | 0 | 4 | 0 | 2 | 1 |
| ENSG00000161249 | 1251 | 584 | 722 | 800 | 738 | 765 |
| ENSG00000189001 | 6 | 4 | 7 | 5 | 5 | 2 |
| ENSG00000105679 | 4 | 0 | 3 | 1 | 1 | 2 |
| ENSG00000236144 | 242 | 141 | 172 | 155 | 156 | 180 |
| ENSG00000105677 | 2213 | 1236 | 1471 | 1767 | 1479 | 1505 |
| ENSG00000105675 | 2 | 0 | 1 | 0 | 0 | 0 |
| ENSG00000283907 | 0 | 0 | 0 | 0 | 0 | 0 |
| ENSG00000283758 | 0 | 0 | 0 | 0 | 0 | 0 |
| ENSG00000224910 | 0 | 0 | 0 | 0 | 1 | 0 |
| ENSG00000267626 | 0 | 0 | 0 | 0 | 0 | 0 |
| ENSG00000249115 | 503 | 243 | 345 | 258 | 309 | 295 |
| ENSG00000126254 | 2820 | 1342 | 1608 | 1960 | 1459 | 1490 |
| ENSG00000105672 | 51 | 15 | 19 | 22 | 13 | 17 |
| ENSG00000126267 | 4856 | 2402 | 2771 | 3096 | 2795 | 2597 |
| ENSG00000105668 | 33 | 26 | 23 | 19 | 21 | 9 |
| ENSG00000226510 | 0 | 0 | 1 | 0 | 2 | 1 |
| ENSG00000267190 | 0 | 0 | 0 | 0 | 0 | 0 |
| ENSG00000011590 | 12 | 16 | 22 | 12 | 15 | 24 |
| ENSG00000272333 | 2185 | 953 | 1329 | 1117 | 1106 | 1181 |
| ENSG00000126246 | 5 | 10 | 9 | 11 | 5 | 14 |
| ENSG00000267120 | 0 | 0 | 0 | 0 | 0 | 2 |
| ENSG00000161265 | 104 | 79 | 97 | 61 | 59 | 68 |
| ENSG00000205155 | 525 | 405 | 466 | 512 | 460 | 403 |
| ENSG00000188223 | 0 | 0 | 0 | 0 | 0 | 1 |
| ENSG00000267439 | 1 | 0 | 0 | 3 | 2 | 1 |
| ENSG00000267796 | 88 | 45 | 58 | 67 | 64 | 72 |
| ENSG00000267328 | 0 | 0 | 0 | 0 | 0 | 0 |

| | | | | | | |
|---|---|---|---|---|---|---|
| ENSG00000004776 | 3 | 0 | 0 | 1 | 0 | 1 |
| ENSG00000167595 | 442 | 233 | 318 | 206 | 230 | 253 |
| ENSG00000267049 | 6 | 9 | 11 | 14 | 16 | 10 |
| ENSG00000004777 | 235 | 108 | 171 | 125 | 157 | 187 |
| ENSG00000225872 | 9 | 1 | 3 | 3 | 10 | 3 |
| ENSG00000250799 | 0 | 0 | 0 | 0 | 0 | 0 |
| ENSG00000161270 | 8 | 1 | 3 | 5 | 8 | 2 |
| ENSG00000126259 | 0 | 0 | 0 | 0 | 0 | 0 |
| ENSG00000105290 | 326 | 177 | 193 | 189 | 144 | 138 |
| ENSG00000243968 | 0 | 0 | 0 | 0 | 0 | 1 |
| ENSG00000167604 | 177 | 67 | 115 | 96 | 99 | 113 |
| ENSG00000280194 | 7 | 2 | 5 | 4 | 8 | 10 |
| ENSG00000126264 | 1 | 1 | 1 | 0 | 0 | 1 |
| ENSG00000011600 | 0 | 0 | 0 | 0 | 0 | 0 |
| ENSG00000126243 | 391 | 143 | 199 | 241 | 179 | 166 |
| ENSG00000267786 | 2 | 4 | 2 | 1 | 1 | 1 |
| ENSG00000267113 | 0 | 0 | 0 | 1 | 0 | 0 |
| ENSG00000267388 | 0 | 0 | 1 | 0 | 0 | 0 |
| ENSG00000205138 | 722 | 346 | 364 | 451 | 376 | 384 |
| ENSG00000181392 | 1121 | 515 | 710 | 714 | 569 | 601 |
| ENSG00000248101 | 0 | 0 | 0 | 0 | 0 | 0 |
| ENSG00000239382 | 97 | 62 | 77 | 79 | 78 | 66 |
| ENSG00000267698 | 9 | 9 | 14 | 20 | 13 | 10 |
| ENSG00000105270 | 125 | 53 | 70 | 59 | 74 | 58 |
| ENSG00000161277 | 239 | 129 | 183 | 156 | 126 | 146 |
| ENSG00000075702 | 514 | 199 | 314 | 285 | 297 | 294 |
| ENSG00000280023 | 0 | 0 | 1 | 0 | 0 | 0 |
| ENSG00000105261 | 0 | 0 | 1 | 1 | 0 | 4 |
| ENSG00000105258 | 161 | 130 | 140 | 189 | 138 | 172 |
| ENSG00000105254 | 3016 | 1521 | 1867 | 2006 | 1719 | 1728 |
| ENSG00000279504 | 0 | 0 | 0 | 0 | 0 | 0 |
| ENSG00000126247 | 14450 | 7051 | 8751 | 8293 | 6944 | 7294 |
| ENSG00000270760 | 5 | 3 | 5 | 0 | 1 | 2 |
| ENSG00000161281 | 3 | 3 | 4 | 3 | 4 | 1 |
| ENSG00000267005 | 0 | 3 | 2 | 0 | 0 | 0 |
| ENSG00000196357 | 84 | 54 | 44 | 25 | 57 | 48 |
| ENSG00000200179 | 0 | 0 | 1 | 0 | 0 | 0 |
| ENSG00000167635 | 1088 | 770 | 559 | 380 | 1341 | 764 |
| ENSG00000267053 | 0 | 0 | 0 | 0 | 0 | 0 |
| ENSG00000232677 | 41 | 22 | 39 | 26 | 35 | 33 |
| ENSG00000142065 | 392 | 176 | 240 | 118 | 214 | 174 |
| ENSG00000266973 | 0 | 0 | 0 | 0 | 0 | 0 |
| ENSG00000267363 | 0 | 0 | 0 | 0 | 0 | 0 |
| ENSG00000181007 | 0 | 0 | 0 | 0 | 0 | 0 |
| ENSG00000267142 | 0 | 0 | 0 | 0 | 0 | 0 |
| ENSG00000186017 | 216 | 103 | 98 | 69 | 118 | 79 |
| ENSG00000267309 | 42 | 7 | 21 | 21 | 18 | 13 |
| ENSG00000267091 | 4 | 0 | 0 | 0 | 2 | 0 |
| ENSG00000254004 | 449 | 247 | 184 | 95 | 361 | 198 |
| ENSG00000228629 | 0 | 0 | 1 | 0 | 3 | 0 |

| | | | | | | |
|---|---|---|---|---|---|---|
| ENSG00000186020 | 193 | 79 | 99 | 77 | 147 | 111 |
| ENSG00000233527 | 87 | 59 | 60 | 48 | 59 | 45 |
| ENSG00000161298 | 4 | 5 | 4 | 1 | 6 | 3 |
| ENSG00000197808 | 133 | 75 | 97 | 50 | 62 | 78 |
| ENSG00000276071 | 8 | 9 | 7 | 2 | 6 | 2 |
| ENSG00000225975 | 50 | 18 | 26 | 16 | 34 | 34 |
| ENSG00000189042 | 166 | 85 | 74 | 54 | 117 | 80 |
| ENSG00000267041 | 78 | 38 | 68 | 34 | 32 | 43 |
| ENSG00000267260 | 0 | 1 | 2 | 0 | 0 | 1 |
| ENSG00000267353 | 0 | 0 | 0 | 0 | 0 | 0 |
| ENSG00000267254 | 69 | 55 | 68 | 71 | 76 | 42 |
| ENSG00000197863 | 64 | 46 | 53 | 20 | 47 | 39 |
| ENSG00000251247 | 111 | 52 | 50 | 29 | 57 | 38 |
| ENSG00000206669 | 0 | 0 | 0 | 0 | 0 | 0 |
| ENSG00000185869 | 97 | 55 | 47 | 25 | 56 | 35 |
| ENSG00000243297 | 0 | 0 | 0 | 0 | 0 | 0 |
| ENSG00000198453 | 44 | 38 | 42 | 21 | 48 | 31 |
| ENSG00000197050 | 53 | 52 | 30 | 17 | 73 | 43 |
| ENSG00000275547 | 0 | 0 | 0 | 0 | 0 | 0 |
| ENSG00000267345 | 0 | 0 | 0 | 0 | 2 | 0 |
| ENSG00000267672 | 4 | 0 | 0 | 0 | 0 | 0 |
| ENSG00000267360 | 0 | 0 | 0 | 0 | 0 | 0 |
| ENSG00000196967 | 108 | 61 | 52 | 14 | 103 | 50 |
| ENSG00000267437 | 0 | 0 | 0 | 0 | 0 | 0 |
| ENSG00000245680 | 262 | 148 | 132 | 88 | 175 | 104 |
| ENSG00000188283 | 186 | 90 | 100 | 79 | 110 | 96 |
| ENSG00000226686 | 7 | 7 | 5 | 6 | 8 | 4 |
| ENSG00000267605 | 2 | 2 | 0 | 1 | 2 | 0 |
| ENSG00000181666 | 995 | 480 | 596 | 491 | 561 | 563 |
| ENSG00000276846 | 40 | 33 | 37 | 23 | 27 | 31 |
| ENSG00000267682 | 6 | 2 | 4 | 1 | 1 | 5 |
| ENSG00000189164 | 129 | 52 | 49 | 36 | 70 | 34 |
| ENSG00000279936 | 0 | 0 | 0 | 0 | 0 | 0 |
| ENSG00000196437 | 60 | 46 | 37 | 20 | 38 | 27 |
| ENSG00000171827 | 163 | 67 | 83 | 41 | 106 | 65 |
| ENSG00000266916 | 32 | 21 | 18 | 13 | 19 | 11 |
| ENSG00000267115 | 0 | 0 | 0 | 0 | 2 | 0 |
| ENSG00000188227 | 289 | 200 | 164 | 110 | 205 | 142 |
| ENSG00000180458 | 0 | 0 | 0 | 0 | 0 | 0 |
| ENSG00000267470 | 5 | 6 | 4 | 3 | 6 | 1 |
| ENSG00000171817 | 6 | 5 | 2 | 2 | 5 | 1 |
| ENSG00000180479 | 80 | 37 | 47 | 28 | 54 | 49 |
| ENSG00000120784 | 257 | 136 | 172 | 88 | 162 | 96 |
| ENSG00000196381 | 0 | 0 | 0 | 0 | 0 | 0 |
| ENSG00000267552 | 0 | 0 | 0 | 0 | 0 | 0 |
| ENSG00000267319 | 3 | 0 | 0 | 3 | 0 | 0 |
| ENSG00000198182 | 356 | 161 | 204 | 123 | 158 | 147 |
| ENSG00000267152 | 23 | 14 | 10 | 7 | 17 | 12 |
| ENSG00000189144 | 63 | 33 | 33 | 10 | 29 | 23 |
| ENSG00000267422 | 8 | 0 | 3 | 0 | 1 | 2 |

| | | | | | | |
|---|---|---|---|---|---|---|
| ENSG00000267640 | 0 | 0 | 0 | 0 | 0 | 0 |
| ENSG00000225868 | 0 | 0 | 0 | 0 | 0 | 0 |
| ENSG00000171804 | 0 | 4 | 0 | 0 | 3 | 3 |
| ENSG00000105738 | 1417 | 669 | 904 | 636 | 725 | 855 |
| ENSG00000268051 | 0 | 0 | 0 | 0 | 0 | 0 |
| ENSG00000268764 | 1 | 2 | 1 | 0 | 0 | 1 |
| ENSG00000269653 | 0 | 1 | 1 | 1 | 0 | 0 |
| ENSG00000268499 | 0 | 1 | 0 | 1 | 0 | 2 |
| ENSG00000275132 | 0 | 0 | 0 | 1 | 0 | 0 |
| ENSG00000011332 | 87 | 39 | 62 | 45 | 55 | 44 |
| ENSG00000167642 | 16851 | 9865 | 12315 | 10971 | 10357 | 10071 |
| ENSG00000167641 | 2 | 0 | 0 | 0 | 0 | 0 |
| ENSG00000267748 | 0 | 0 | 0 | 0 | 0 | 1 |
| ENSG00000200209 | 5 | 1 | 4 | 0 | 0 | 0 |
| ENSG00000167644 | 46 | 40 | 39 | 59 | 65 | 52 |
| ENSG00000167645 | 722 | 392 | 467 | 515 | 394 | 383 |
| ENSG00000099337 | 1850 | 1012 | 1167 | 948 | 1110 | 1066 |
| ENSG00000099338 | 194 | 97 | 124 | 54 | 109 | 94 |
| ENSG00000266963 | 1 | 0 | 3 | 1 | 0 | 2 |
| ENSG00000099341 | 5673 | 2831 | 3528 | 3503 | 3255 | 3289 |
| ENSG00000179168 | 27 | 14 | 16 | 9 | 10 | 8 |
| ENSG00000267090 | 0 | 0 | 0 | 0 | 0 | 0 |
| ENSG00000188766 | 51 | 29 | 47 | 34 | 32 | 25 |
| ENSG00000130244 | 1137 | 545 | 676 | 623 | 546 | 619 |
| ENSG00000171777 | 3 | 4 | 2 | 1 | 3 | 0 |
| ENSG00000196218 | 24 | 12 | 12 | 10 | 3 | 14 |
| ENSG00000269445 | 0 | 0 | 0 | 0 | 0 | 0 |
| ENSG00000268055 | 0 | 0 | 0 | 0 | 0 | 0 |
| ENSG00000104814 | 41 | 8 | 25 | 17 | 11 | 8 |
| ENSG00000267291 | 0 | 0 | 0 | 0 | 0 | 0 |
| ENSG00000178982 | 3806 | 1910 | 2326 | 2657 | 2420 | 2367 |
| ENSG00000130402 | 16184 | 7588 | 9665 | 8129 | 7906 | 8120 |
| ENSG00000267375 | 0 | 2 | 0 | 0 | 0 | 0 |
| ENSG00000182472 | 45 | 15 | 22 | 17 | 22 | 23 |
| ENSG00000267892 | 7 | 7 | 6 | 5 | 15 | 12 |
| ENSG00000205076 | 0 | 0 | 0 | 0 | 0 | 0 |
| ENSG00000178934 | 1 | 0 | 1 | 0 | 0 | 0 |
| ENSG00000207296 | 0 | 0 | 0 | 0 | 0 | 0 |
| ENSG00000171747 | 6 | 4 | 4 | 2 | 1 | 0 |
| ENSG00000104823 | 3190 | 1806 | 2111 | 2113 | 1579 | 1734 |
| ENSG00000268083 | 0 | 0 | 0 | 0 | 0 | 0 |
| ENSG00000268756 | 3 | 6 | 6 | 1 | 3 | 3 |
| ENSG00000104824 | 4746 | 2359 | 3087 | 2361 | 2646 | 2860 |
| ENSG00000269688 | 34 | 10 | 15 | 14 | 9 | 17 |
| ENSG00000187994 | 282 | 126 | 180 | 178 | 173 | 183 |
| ENSG00000269050 | 0 | 0 | 0 | 0 | 0 | 0 |
| ENSG00000068903 | 1531 | 621 | 885 | 785 | 762 | 714 |
| ENSG00000104825 | 593 | 317 | 420 | 431 | 387 | 372 |
| ENSG00000262484 | 6 | 4 | 4 | 2 | 4 | 9 |
| ENSG00000104835 | 50 | 59 | 80 | 54 | 59 | 79 |

| | | | | | | |
|---|---|---|---|---|---|---|
| ENSG00000269547 | 0 | 0 | 0 | 0 | 0 | 0 |
| ENSG00000128626 | 1754 | 879 | 1100 | 1522 | 1139 | 1162 |
| ENSG00000269486 | 16 | 4 | 11 | 9 | 3 | 6 |
| ENSG00000269190 | 208 | 112 | 145 | 132 | 108 | 120 |
| ENSG00000161243 | 250 | 114 | 141 | 130 | 135 | 150 |
| ENSG00000267992 | 0 | 0 | 0 | 0 | 0 | 0 |
| ENSG00000183760 | 18 | 6 | 8 | 8 | 4 | 3 |
| ENSG00000130669 | 3934 | 1833 | 2517 | 2332 | 2067 | 1907 |
| ENSG00000269172 | 0 | 1 | 0 | 0 | 0 | 0 |
| ENSG00000188505 | 13 | 6 | 8 | 17 | 14 | 17 |
| ENSG00000179751 | 0 | 0 | 0 | 0 | 0 | 0 |
| ENSG00000268510 | 6 | 2 | 1 | 0 | 0 | 0 |
| ENSG00000197110 | 0 | 0 | 0 | 0 | 0 | 1 |
| ENSG00000272395 | 0 | 0 | 0 | 0 | 0 | 0 |
| ENSG00000213926 | 0 | 0 | 0 | 0 | 0 | 0 |
| ENSG00000272311 | 0 | 0 | 0 | 0 | 0 | 0 |
| ENSG00000183709 | 0 | 0 | 0 | 2 | 0 | 0 |
| ENSG00000182393 | 4 | 0 | 1 | 0 | 1 | 0 |
| ENSG00000128011 | 173 | 63 | 78 | 132 | 87 | 125 |
| ENSG00000268262 | 0 | 0 | 5 | 4 | 7 | 5 |
| ENSG00000130755 | 2 | 2 | 5 | 0 | 0 | 2 |
| ENSG00000269246 | 4 | 0 | 0 | 4 | 4 | 4 |
| ENSG00000179134 | 3319 | 1423 | 1962 | 1637 | 1742 | 1871 |
| ENSG00000241983 | 2 | 3 | 2 | 0 | 1 | 0 |
| ENSG00000006712 | 3031 | 1394 | 1793 | 1502 | 1513 | 1616 |
| ENSG00000063322 | 2624 | 1237 | 1573 | 1134 | 1342 | 1394 |
| ENSG00000128016 | 1231 | 580 | 766 | 710 | 703 | 735 |
| ENSG00000266559 | 0 | 0 | 0 | 0 | 0 | 0 |
| ENSG00000090924 | 1140 | 492 | 656 | 463 | 496 | 533 |
| ENSG00000105193 | 14797 | 7258 | 8520 | 11489 | 9437 | 8882 |
| ENSG00000196235 | 4628 | 2068 | 2709 | 2605 | 2488 | 2600 |
| ENSG00000105197 | 1813 | 963 | 1185 | 1315 | 1213 | 1260 |
| ENSG00000090932 | 0 | 1 | 0 | 0 | 0 | 0 |
| ENSG00000186838 | 0 | 0 | 0 | 1 | 0 | 0 |
| ENSG00000176401 | 77 | 42 | 66 | 48 | 59 | 55 |
| ENSG00000269792 | 0 | 0 | 0 | 0 | 0 | 0 |
| ENSG00000269584 | 0 | 0 | 0 | 0 | 0 | 2 |
| ENSG00000176396 | 452 | 212 | 271 | 223 | 245 | 258 |
| ENSG00000269588 | 0 | 0 | 0 | 0 | 0 | 0 |
| ENSG00000269188 | 0 | 0 | 0 | 0 | 0 | 0 |
| ENSG00000244253 | 0 | 0 | 0 | 0 | 0 | 0 |
| ENSG00000243581 | 0 | 0 | 0 | 0 | 0 | 0 |
| ENSG00000239320 | 0 | 0 | 0 | 0 | 0 | 0 |
| ENSG00000105198 | 0 | 0 | 0 | 0 | 0 | 0 |
| ENSG00000269519 | 0 | 0 | 0 | 0 | 0 | 0 |
| ENSG00000268153 | 0 | 0 | 0 | 0 | 0 | 0 |
| ENSG00000268243 | 0 | 0 | 0 | 0 | 0 | 0 |
| ENSG00000249861 | 0 | 0 | 0 | 0 | 0 | 0 |
| ENSG00000226025 | 0 | 0 | 0 | 0 | 1 | 0 |
| ENSG00000269460 | 0 | 0 | 0 | 0 | 0 | 0 |

| | | | | | | |
|---|---|---|---|---|---|---|
| ENSG00000268088 | 0 | 0 | 2 | 0 | 0 | 0 |
| ENSG00000240320 | 0 | 0 | 0 | 0 | 0 | 0 |
| ENSG00000006659 | 0 | 0 | 0 | 0 | 0 | 0 |
| ENSG00000278813 | 0 | 0 | 0 | 0 | 0 | 0 |
| ENSG00000105205 | 0 | 0 | 0 | 0 | 0 | 0 |
| ENSG00000213921 | 0 | 0 | 0 | 0 | 0 | 0 |
| ENSG00000276445 | 10 | 1 | 2 | 5 | 4 | 5 |
| ENSG00000105204 | 1094 | 489 | 631 | 586 | 460 | 525 |
| ENSG00000277759 | 1 | 1 | 1 | 1 | 0 | 1 |
| ENSG00000105202 | 3578 | 1871 | 2137 | 2708 | 2217 | 2476 |
| ENSG00000275395 | 50 | 24 | 24 | 39 | 27 | 30 |
| ENSG00000187534 | 46 | 20 | 19 | 25 | 30 | 35 |
| ENSG00000013275 | 5001 | 2570 | 3051 | 2981 | 2892 | 2977 |
| ENSG00000187187 | 54 | 19 | 24 | 11 | 30 | 18 |
| ENSG00000269069 | 23 | 8 | 11 | 2 | 23 | 11 |
| ENSG00000128000 | 192 | 154 | 127 | 51 | 142 | 110 |
| ENSG00000269749 | 0 | 1 | 1 | 0 | 1 | 0 |
| ENSG00000197782 | 379 | 244 | 191 | 115 | 303 | 185 |
| ENSG00000269296 | 2 | 0 | 1 | 0 | 0 | 0 |
| ENSG00000230086 | 0 | 0 | 0 | 0 | 0 | 0 |
| ENSG00000130758 | 559 | 273 | 280 | 345 | 331 | 308 |
| ENSG00000174521 | 8 | 2 | 8 | 2 | 6 | 2 |
| ENSG00000105219 | 22 | 16 | 15 | 18 | 18 | 17 |
| ENSG00000105221 | 5398 | 2547 | 3384 | 3027 | 2801 | 3072 |
| ENSG00000205041 | 7 | 3 | 4 | 4 | 2 | 4 |
| ENSG00000207631 | 3 | 1 | 0 | 1 | 0 | 0 |
| ENSG00000206674 | 0 | 1 | 0 | 0 | 0 | 0 |
| ENSG00000160392 | 500 | 224 | 302 | 263 | 219 | 256 |
| ENSG00000207281 | 0 | 0 | 0 | 0 | 0 | 0 |
| ENSG00000206925 | 0 | 0 | 0 | 0 | 0 | 0 |
| ENSG00000105223 | 4263 | 1808 | 2827 | 2708 | 2143 | 2413 |
| ENSG00000279759 | 101 | 60 | 58 | 12 | 47 | 47 |
| ENSG00000275652 | 0 | 0 | 0 | 0 | 0 | 0 |
| ENSG00000160396 | 16 | 8 | 12 | 10 | 7 | 11 |
| ENSG00000105227 | 278 | 142 | 163 | 178 | 156 | 132 |
| ENSG00000197019 | 726 | 334 | 409 | 540 | 445 | 390 |
| ENSG00000277453 | 2 | 1 | 0 | 3 | 2 | 1 |
| ENSG00000167565 | 2369 | 1032 | 1386 | 1191 | 1143 | 1166 |
| ENSG00000268366 | 14 | 10 | 12 | 11 | 7 | 15 |
| ENSG00000090013 | 3494 | 1715 | 2102 | 2231 | 2059 | 1776 |
| ENSG00000277510 | 0 | 0 | 0 | 0 | 0 | 0 |
| ENSG00000160460 | 39 | 13 | 27 | 28 | 20 | 17 |
| ENSG00000160410 | 2213 | 1053 | 1377 | 1443 | 1311 | 1353 |
| ENSG00000090006 | 1483 | 651 | 886 | 803 | 706 | 765 |
| ENSG00000105245 | 622 | 271 | 362 | 351 | 338 | 397 |
| ENSG00000123815 | 828 | 403 | 568 | 532 | 400 | 436 |
| ENSG00000223284 | 1 | 0 | 0 | 0 | 1 | 0 |
| ENSG00000086544 | 1225 | 634 | 832 | 781 | 817 | 718 |
| ENSG00000188493 | 471 | 184 | 282 | 229 | 191 | 243 |
| ENSG00000077312 | 1816 | 887 | 1204 | 1339 | 1094 | 1309 |

| | | | | | | |
|---|---|---|---|---|---|---|
| ENSG00000268975 | 0 | 0 | 0 | 0 | 0 | 0 |
| ENSG00000261857 | 0 | 2 | 0 | 0 | 0 | 0 |
| ENSG00000167578 | 31 | 11 | 18 | 11 | 9 | 13 |
| ENSG00000171570 | 0 | 0 | 1 | 0 | 0 | 0 |
| ENSG00000282951 | 7 | 3 | 2 | 4 | 3 | 1 |
| ENSG00000269858 | 171 | 84 | 115 | 79 | 98 | 85 |
| ENSG00000268797 | 0 | 0 | 0 | 0 | 0 | 0 |
| ENSG00000233622 | 62 | 33 | 49 | 21 | 33 | 45 |
| ENSG00000237118 | 25 | 19 | 27 | 4 | 11 | 14 |
| ENSG00000269843 | 33 | 36 | 27 | 15 | 29 | 20 |
| ENSG00000279108 | 56 | 30 | 38 | 21 | 20 | 21 |
| ENSG00000255974 | 12 | 6 | 4 | 5 | 11 | 3 |
| ENSG00000198077 | 0 | 1 | 0 | 0 | 0 | 0 |
| ENSG00000130612 | 2 | 3 | 6 | 0 | 1 | 3 |
| ENSG00000198251 | 0 | 0 | 0 | 0 | 0 | 0 |
| ENSG00000256612 | 2 | 3 | 3 | 2 | 9 | 3 |
| ENSG00000269353 | 0 | 0 | 0 | 0 | 0 | 0 |
| ENSG00000197408 | 0 | 0 | 0 | 0 | 0 | 0 |
| ENSG00000213908 | 0 | 0 | 0 | 0 | 0 | 0 |
| ENSG00000268101 | 0 | 0 | 0 | 0 | 0 | 0 |
| ENSG00000271299 | 0 | 0 | 0 | 0 | 0 | 0 |
| ENSG00000197838 | 0 | 0 | 0 | 0 | 0 | 0 |
| ENSG00000197446 | 0 | 0 | 1 | 0 | 0 | 2 |
| ENSG00000268529 | 0 | 0 | 0 | 0 | 0 | 0 |
| ENSG00000224476 | 0 | 0 | 0 | 0 | 0 | 0 |
| ENSG00000264423 | 0 | 0 | 0 | 0 | 0 | 0 |
| ENSG00000167600 | 4 | 5 | 1 | 5 | 2 | 6 |
| ENSG00000167601 | 0 | 0 | 0 | 0 | 3 | 0 |
| ENSG00000269652 | 0 | 0 | 0 | 0 | 0 | 0 |
| ENSG00000105323 | 10048 | 4730 | 6325 | 5145 | 5203 | 5658 |
| ENSG00000239868 | 0 | 0 | 0 | 0 | 1 | 0 |
| ENSG00000105329 | 672 | 335 | 474 | 441 | 317 | 403 |
| ENSG00000142039 | 1272 | 558 | 753 | 760 | 650 | 717 |
| ENSG00000255730 | 0 | 0 | 0 | 0 | 0 | 0 |
| ENSG00000142046 | 72 | 34 | 53 | 33 | 44 | 54 |
| ENSG00000123810 | 137 | 67 | 92 | 101 | 69 | 78 |
| ENSG00000277744 | 19 | 12 | 6 | 14 | 9 | 14 |
| ENSG00000077348 | 551 | 315 | 360 | 518 | 354 | 436 |
| ENSG00000248098 | 44 | 26 | 49 | 33 | 39 | 35 |
| ENSG00000268987 | 0 | 0 | 0 | 0 | 0 | 0 |
| ENSG00000177191 | 17 | 7 | 8 | 7 | 8 | 7 |
| ENSG00000268475 | 0 | 0 | 0 | 0 | 0 | 0 |
| ENSG00000105341 | 1593 | 745 | 947 | 948 | 854 | 869 |
| ENSG00000277136 | 0 | 0 | 0 | 0 | 0 | 0 |
| ENSG00000204978 | 0 | 0 | 0 | 0 | 0 | 0 |
| ENSG00000267107 | 0 | 1 | 0 | 0 | 1 | 0 |
| ENSG00000268034 | 4 | 1 | 0 | 2 | 1 | 2 |
| ENSG00000269800 | 0 | 0 | 0 | 1 | 0 | 0 |
| ENSG00000268355 | 0 | 0 | 1 | 0 | 1 | 0 |
| ENSG00000270164 | 0 | 0 | 0 | 0 | 1 | 1 |

| | | | | | | |
|---|---|---|---|---|---|---|
| ENSG00000268027 | 0 | 0 | 0 | 0 | 0 | 0 |
| ENSG00000007129 | 0 | 2 | 1 | 0 | 0 | 0 |
| ENSG00000280014 | 0 | 0 | 0 | 0 | 0 | 0 |
| ENSG00000239736 | 0 | 0 | 0 | 0 | 0 | 0 |
| ENSG00000269266 | 0 | 0 | 0 | 0 | 0 | 0 |
| ENSG00000105352 | 0 | 0 | 0 | 0 | 0 | 0 |
| ENSG00000268997 | 0 | 0 | 0 | 0 | 0 | 0 |
| ENSG00000269354 | 0 | 0 | 0 | 0 | 0 | 0 |
| ENSG00000268582 | 0 | 0 | 0 | 0 | 0 | 0 |
| ENSG00000007306 | 2 | 1 | 3 | 1 | 0 | 0 |
| ENSG00000105388 | 352 | 178 | 178 | 166 | 250 | 205 |
| ENSG00000267881 | 0 | 0 | 0 | 0 | 0 | 0 |
| ENSG00000086548 | 722 | 466 | 301 | 386 | 515 | 343 |
| ENSG00000268833 | 0 | 0 | 0 | 0 | 0 | 0 |
| ENSG00000170956 | 0 | 0 | 0 | 0 | 0 | 0 |
| ENSG00000269119 | 0 | 0 | 0 | 0 | 0 | 0 |
| ENSG00000278239 | 0 | 0 | 0 | 0 | 0 | 0 |
| ENSG00000273111 | 0 | 0 | 0 | 0 | 0 | 0 |
| ENSG00000142025 | 0 | 0 | 0 | 0 | 0 | 0 |
| ENSG00000105372 | 20901 | 10934 | 12507 | 15988 | 13258 | 12462 |
| ENSG00000276926 | 1 | 5 | 0 | 2 | 1 | 3 |
| ENSG00000105369 | 2 | 4 | 5 | 5 | 4 | 1 |
| ENSG00000076928 | 2197 | 948 | 1291 | 1338 | 1253 | 1319 |
| ENSG00000268041 | 36 | 3 | 18 | 9 | 10 | 8 |
| ENSG00000105404 | 353 | 238 | 306 | 354 | 307 | 273 |
| ENSG00000285505 | 0 | 0 | 0 | 0 | 0 | 0 |
| ENSG00000105409 | 5 | 1 | 4 | 10 | 5 | 2 |
| ENSG00000105737 | 8 | 10 | 9 | 4 | 5 | 9 |
| ENSG00000105732 | 884 | 423 | 487 | 531 | 432 | 515 |
| ENSG00000028277 | 10 | 7 | 17 | 10 | 9 | 9 |
| ENSG00000254887 | 0 | 0 | 0 | 0 | 0 | 1 |
| ENSG00000266226 | 0 | 0 | 0 | 0 | 0 | 0 |
| ENSG00000259436 | 0 | 1 | 0 | 0 | 0 | 0 |
| ENSG00000160570 | 689 | 341 | 467 | 422 | 398 | 384 |
| ENSG00000167625 | 418 | 196 | 241 | 239 | 264 | 178 |
| ENSG00000105723 | 1668 | 781 | 985 | 1112 | 928 | 1014 |
| ENSG00000268643 | 0 | 0 | 0 | 0 | 0 | 0 |
| ENSG00000279539 | 22 | 8 | 18 | 12 | 17 | 13 |
| ENSG00000105722 | 2844 | 1347 | 1641 | 1746 | 1458 | 1462 |
| ENSG00000079432 | 2369 | 959 | 1476 | 1197 | 1085 | 1295 |
| ENSG00000079462 | 1187 | 661 | 784 | 889 | 676 | 684 |
| ENSG00000188368 | 73 | 27 | 50 | 50 | 30 | 27 |
| ENSG00000167619 | 8 | 0 | 3 | 1 | 2 | 1 |
| ENSG00000105429 | 1032 | 391 | 558 | 428 | 457 | 474 |
| ENSG00000277533 | 0 | 0 | 0 | 0 | 0 | 0 |
| ENSG00000105427 | 62 | 23 | 31 | 43 | 29 | 28 |
| ENSG00000213904 | 15 | 7 | 12 | 15 | 13 | 13 |
| ENSG00000079435 | 14 | 6 | 11 | 7 | 6 | 5 |
| ENSG00000268605 | 0 | 0 | 0 | 0 | 0 | 0 |
| ENSG00000189377 | 4 | 1 | 0 | 0 | 2 | 0 |

| | | | | | | |
|---|---|---|---|---|---|---|
| ENSG00000253048 | 0 | 0 | 0 | 0 | 0 | 0 |
| ENSG00000269374 | 9 | 4 | 2 | 2 | 6 | 6 |
| ENSG00000079385 | 5 | 11 | 6 | 2 | 5 | 13 |
| ENSG00000230558 | 0 | 0 | 0 | 0 | 0 | 0 |
| ENSG00000124469 | 0 | 0 | 0 | 0 | 0 | 0 |
| ENSG00000233681 | 0 | 1 | 0 | 0 | 0 | 0 |
| ENSG00000231561 | 0 | 0 | 0 | 1 | 0 | 0 |
| ENSG00000243167 | 0 | 0 | 0 | 0 | 0 | 0 |
| ENSG00000221826 | 0 | 0 | 0 | 0 | 1 | 0 |
| ENSG00000124467 | 8 | 3 | 7 | 7 | 15 | 3 |
| ENSG00000238092 | 0 | 0 | 0 | 0 | 0 | 0 |
| ENSG00000225877 | 0 | 0 | 0 | 0 | 0 | 0 |
| ENSG00000248257 | 0 | 0 | 0 | 0 | 0 | 1 |
| ENSG00000231924 | 0 | 0 | 1 | 3 | 1 | 3 |
| ENSG00000170848 | 0 | 0 | 0 | 0 | 0 | 0 |
| ENSG00000221878 | 0 | 0 | 0 | 0 | 0 | 0 |
| ENSG00000227349 | 0 | 0 | 0 | 0 | 0 | 0 |
| ENSG00000282943 | 0 | 0 | 2 | 0 | 0 | 1 |
| ENSG00000243130 | 0 | 0 | 0 | 0 | 0 | 0 |
| ENSG00000236932 | 0 | 0 | 0 | 0 | 0 | 0 |
| ENSG00000242221 | 0 | 0 | 0 | 0 | 0 | 0 |
| ENSG00000230529 | 0 | 0 | 0 | 0 | 0 | 0 |
| ENSG00000227606 | 0 | 0 | 0 | 0 | 0 | 0 |
| ENSG00000204941 | 10 | 5 | 11 | 11 | 6 | 9 |
| ENSG00000243137 | 15 | 14 | 14 | 23 | 26 | 32 |
| ENSG00000183668 | 108 | 60 | 85 | 98 | 74 | 109 |
| ENSG00000241104 | 1 | 1 | 6 | 2 | 2 | 1 |
| ENSG00000236123 | 0 | 0 | 2 | 1 | 5 | 2 |
| ENSG00000230681 | 0 | 0 | 0 | 0 | 0 | 0 |
| ENSG00000231412 | 0 | 0 | 0 | 0 | 0 | 0 |
| ENSG00000204936 | 4 | 0 | 1 | 4 | 3 | 3 |
| ENSG00000272396 | 0 | 0 | 0 | 0 | 0 | 0 |
| ENSG00000204933 | 0 | 0 | 0 | 0 | 0 | 0 |
| ENSG00000131126 | 1 | 2 | 0 | 0 | 0 | 0 |
| ENSG00000124466 | 4384 | 2109 | 2637 | 2392 | 2086 | 2151 |
| ENSG00000176531 | 637 | 306 | 436 | 356 | 315 | 307 |
| ENSG00000105755 | 295 | 199 | 223 | 271 | 180 | 217 |
| ENSG00000176472 | 22 | 10 | 4 | 8 | 8 | 15 |
| ENSG00000073050 | 1540 | 855 | 997 | 832 | 849 | 898 |
| ENSG00000269177 | 0 | 0 | 0 | 0 | 1 | 0 |
| ENSG00000268361 | 0 | 0 | 0 | 0 | 0 | 0 |
| ENSG00000269583 | 0 | 0 | 0 | 0 | 0 | 0 |
| ENSG00000234465 | 8 | 9 | 4 | 2 | 4 | 11 |
| ENSG00000167378 | 1638 | 806 | 1047 | 829 | 967 | 978 |
| ENSG00000124444 | 227 | 121 | 143 | 115 | 121 | 119 |
| ENSG00000226763 | 41 | 18 | 23 | 5 | 15 | 16 |
| ENSG00000131116 | 928 | 514 | 646 | 678 | 602 | 605 |
| ENSG00000105767 | 559 | 264 | 312 | 328 | 302 | 297 |
| ENSG00000011422 | 50 | 18 | 33 | 40 | 28 | 34 |
| ENSG00000239948 | 0 | 0 | 0 | 0 | 0 | 0 |

| | | | | | | |
|---|---|---|---|---|---|---|
| ENSG00000283525 | 0 | 0 | 0 | 1 | 1 | 0 |
| ENSG00000124449 | 0 | 1 | 0 | 0 | 0 | 0 |
| ENSG00000105771 | 1473 | 659 | 903 | 853 | 729 | 795 |
| ENSG00000104783 | 1007 | 460 | 572 | 602 | 586 | 503 |
| ENSG00000268601 | 0 | 0 | 0 | 0 | 0 | 0 |
| ENSG00000159871 | 0 | 1 | 0 | 0 | 0 | 0 |
| ENSG00000167637 | 191 | 99 | 121 | 74 | 138 | 82 |
| ENSG00000176222 | 48 | 23 | 27 | 14 | 26 | 28 |
| ENSG00000278917 | 12 | 4 | 7 | 8 | 4 | 4 |
| ENSG00000267058 | 11 | 6 | 4 | 2 | 5 | 9 |
| ENSG00000267191 | 2 | 1 | 0 | 0 | 2 | 0 |
| ENSG00000124459 | 333 | 184 | 202 | 125 | 166 | 171 |
| ENSG00000159905 | 3 | 4 | 2 | 2 | 5 | 3 |
| ENSG00000204920 | 117 | 76 | 88 | 45 | 64 | 46 |
| ENSG00000277806 | 2 | 0 | 0 | 0 | 0 | 0 |
| ENSG00000278492 | 0 | 0 | 0 | 0 | 0 | 0 |
| ENSG00000266921 | 23 | 5 | 14 | 10 | 13 | 6 |
| ENSG00000159882 | 114 | 60 | 95 | 56 | 70 | 63 |
| ENSG00000267144 | 0 | 0 | 0 | 0 | 0 | 0 |
| ENSG00000159885 | 149 | 84 | 109 | 58 | 84 | 62 |
| ENSG00000267022 | 0 | 0 | 0 | 0 | 0 | 0 |
| ENSG00000178386 | 24 | 16 | 26 | 10 | 23 | 17 |
| ENSG00000186026 | 66 | 20 | 23 | 15 | 23 | 27 |
| ENSG00000253021 | 0 | 0 | 0 | 0 | 0 | 0 |
| ENSG00000267680 | 321 | 144 | 190 | 70 | 144 | 121 |
| ENSG00000186019 | 3 | 4 | 2 | 2 | 1 | 1 |
| ENSG00000256294 | 136 | 74 | 71 | 42 | 76 | 51 |
| ENSG00000263002 | 282 | 146 | 151 | 96 | 192 | 134 |
| ENSG00000167380 | 642 | 335 | 382 | 188 | 320 | 243 |
| ENSG00000131115 | 220 | 133 | 148 | 67 | 139 | 78 |
| ENSG00000159917 | 96 | 43 | 48 | 26 | 53 | 63 |
| ENSG00000279103 | 2 | 0 | 1 | 0 | 1 | 1 |
| ENSG00000159915 | 113 | 46 | 63 | 27 | 61 | 34 |
| ENSG00000270679 | 0 | 0 | 0 | 0 | 1 | 0 |
| ENSG00000267590 | 2 | 2 | 1 | 0 | 1 | 0 |
| ENSG00000062370 | 60 | 41 | 31 | 18 | 63 | 31 |
| ENSG00000267173 | 0 | 0 | 0 | 0 | 0 | 0 |
| ENSG00000267508 | 35 | 6 | 20 | 13 | 26 | 21 |
| ENSG00000267188 | 0 | 0 | 0 | 0 | 0 | 0 |
| ENSG00000278318 | 304 | 112 | 161 | 79 | 125 | 95 |
| ENSG00000176761 | 0 | 1 | 0 | 0 | 1 | 1 |
| ENSG00000167384 | 247 | 146 | 130 | 91 | 144 | 127 |
| ENSG00000273777 | 0 | 1 | 0 | 1 | 0 | 0 |
| ENSG00000267614 | 0 | 0 | 0 | 0 | 0 | 0 |
| ENSG00000230666 | 0 | 0 | 0 | 0 | 1 | 0 |
| ENSG00000224366 | 0 | 0 | 0 | 0 | 0 | 0 |
| ENSG00000216588 | 0 | 2 | 1 | 0 | 0 | 0 |
| ENSG00000266903 | 38 | 35 | 35 | 30 | 16 | 23 |
| ENSG00000073008 | 886 | 423 | 539 | 534 | 528 | 578 |
| ENSG00000283673 | 0 | 0 | 0 | 0 | 0 | 0 |

| | | | | | | |
|---|---|---|---|---|---|---|
| ENSG00000186567 | 101 | 56 | 55 | 74 | 85 | 52 |
| ENSG00000279095 | 141 | 50 | 55 | 61 | 64 | 78 |
| ENSG00000213892 | 1 | 1 | 0 | 1 | 1 | 0 |
| ENSG00000252200 | 0 | 1 | 1 | 1 | 1 | 1 |
| ENSG00000069399 | 1413 | 635 | 641 | 819 | 762 | 692 |
| ENSG00000284258 | 0 | 0 | 0 | 0 | 0 | 0 |
| ENSG00000253027 | 0 | 0 | 0 | 0 | 0 | 0 |
| ENSG00000236807 | 0 | 0 | 0 | 0 | 0 | 0 |
| ENSG00000142273 | 0 | 2 | 0 | 0 | 2 | 2 |
| ENSG00000285128 | 0 | 0 | 0 | 1 | 0 | 0 |
| ENSG00000187244 | 5869 | 2344 | 3687 | 3070 | 2089 | 2694 |
| ENSG00000130202 | 8503 | 3801 | 5619 | 4551 | 4050 | 4349 |
| ENSG00000267282 | 4 | 2 | 2 | 1 | 3 | 0 |
| ENSG00000130204 | 2868 | 1432 | 1753 | 2068 | 1653 | 1783 |
| ENSG00000130203 | 28 | 7 | 17 | 18 | 22 | 16 |
| ENSG00000280087 | 0 | 1 | 1 | 0 | 1 | 0 |
| ENSG00000130208 | 13 | 5 | 11 | 12 | 9 | 8 |
| ENSG00000214855 | 4 | 0 | 0 | 0 | 0 | 0 |
| ENSG00000267467 | 0 | 0 | 0 | 0 | 0 | 0 |
| ENSG00000224916 | 0 | 1 | 0 | 0 | 0 | 0 |
| ENSG00000234906 | 0 | 0 | 0 | 0 | 0 | 0 |
| ENSG00000267114 | 7 | 9 | 6 | 4 | 5 | 1 |
| ENSG00000104853 | 6510 | 2736 | 3728 | 3844 | 3182 | 3548 |
| ENSG00000104856 | 154 | 95 | 141 | 146 | 167 | 164 |
| ENSG00000104859 | 865 | 379 | 562 | 467 | 423 | 457 |
| ENSG00000207003 | 0 | 1 | 1 | 0 | 0 | 0 |
| ENSG00000170684 | 221 | 109 | 170 | 133 | 126 | 146 |
| ENSG00000267348 | 37 | 26 | 33 | 19 | 18 | 27 |
| ENSG00000142252 | 545 | 328 | 393 | 406 | 365 | 387 |
| ENSG00000007047 | 1160 | 529 | 713 | 645 | 648 | 633 |
| ENSG00000104866 | 3888 | 1881 | 2538 | 2227 | 2053 | 1948 |
| ENSG00000267346 | 2 | 1 | 3 | 0 | 1 | 3 |
| ENSG00000267044 | 0 | 0 | 1 | 2 | 1 | 1 |
| ENSG00000179846 | 59 | 21 | 20 | 19 | 16 | 22 |
| ENSG00000007255 | 1337 | 612 | 781 | 937 | 771 | 759 |
| ENSG00000189114 | 673 | 313 | 426 | 393 | 330 | 379 |
| ENSG00000267545 | 0 | 0 | 0 | 0 | 0 | 1 |
| ENSG00000266958 | 3 | 1 | 2 | 1 | 1 | 1 |
| ENSG00000283632 | 6 | 0 | 7 | 1 | 7 | 4 |
| ENSG00000104879 | 12 | 5 | 1 | 2 | 0 | 5 |
| ENSG00000242675 | 0 | 0 | 0 | 0 | 0 | 0 |
| ENSG00000104892 | 172 | 79 | 88 | 101 | 89 | 78 |
| ENSG00000104884 | 1598 | 699 | 968 | 902 | 890 | 891 |
| ENSG00000104881 | 1223 | 593 | 768 | 636 | 658 | 601 |
| ENSG00000117877 | 204 | 118 | 112 | 145 | 142 | 173 |
| ENSG00000012061 | 1270 | 699 | 934 | 749 | 703 | 660 |
| ENSG00000275726 | 0 | 0 | 0 | 0 | 0 | 0 |
| ENSG00000125740 | 42 | 13 | 36 | 28 | 16 | 36 |
| ENSG00000125744 | 303 | 180 | 226 | 176 | 189 | 168 |
| ENSG00000213889 | 63 | 49 | 51 | 47 | 35 | 36 |

| | | | | | | |
|---|---|---|---|---|---|---|
| ENSG00000125753 | 5333 | 2338 | 2881 | 2506 | 2341 | 2241 |
| ENSG00000125741 | 1282 | 662 | 780 | 617 | 689 | 715 |
| ENSG00000177464 | 0 | 1 | 0 | 2 | 0 | 0 |
| ENSG00000125746 | 1663 | 777 | 1092 | 920 | 951 | 902 |
| ENSG00000284544 | 0 | 0 | 0 | 0 | 0 | 0 |
| ENSG00000267757 | 0 | 0 | 1 | 0 | 1 | 0 |
| ENSG00000241226 | 0 | 0 | 0 | 0 | 0 | 0 |
| ENSG00000010310 | 54 | 15 | 32 | 25 | 22 | 18 |
| ENSG00000207773 | 0 | 0 | 0 | 0 | 0 | 0 |
| ENSG00000283212 | 0 | 0 | 0 | 0 | 0 | 0 |
| ENSG00000125743 | 3788 | 1946 | 2306 | 2716 | 2404 | 2292 |
| ENSG00000011478 | 420 | 241 | 352 | 340 | 303 | 297 |
| ENSG00000177051 | 616 | 291 | 391 | 408 | 341 | 323 |
| ENSG00000279407 | 46 | 25 | 26 | 19 | 36 | 25 |
| ENSG00000237452 | 0 | 1 | 0 | 1 | 0 | 0 |
| ENSG00000259605 | 29 | 8 | 7 | 12 | 12 | 11 |
| ENSG00000177045 | 281 | 134 | 217 | 202 | 161 | 200 |
| ENSG00000267395 | 21 | 5 | 10 | 10 | 10 | 13 |
| ENSG00000104936 | 1151 | 632 | 852 | 837 | 867 | 753 |
| ENSG00000268434 | 0 | 0 | 0 | 0 | 0 | 0 |
| ENSG00000185800 | 865 | 418 | 593 | 544 | 520 | 545 |
| ENSG00000104941 | 0 | 1 | 0 | 1 | 0 | 1 |
| ENSG00000125755 | 2756 | 1195 | 1604 | 1454 | 1392 | 1353 |
| ENSG00000269148 | 8 | 1 | 6 | 1 | 8 | 5 |
| ENSG00000170608 | 2 | 0 | 3 | 0 | 0 | 4 |
| ENSG00000170604 | 1036 | 482 | 570 | 692 | 554 | 530 |
| ENSG00000176182 | 301 | 184 | 171 | 213 | 181 | 179 |
| ENSG00000188425 | 0 | 0 | 1 | 0 | 0 | 0 |
| ENSG00000104967 | 10 | 5 | 8 | 6 | 4 | 4 |
| ENSG00000104983 | 291 | 123 | 158 | 165 | 160 | 160 |
| ENSG00000211580 | 0 | 0 | 0 | 0 | 0 | 0 |
| ENSG00000008438 | 0 | 0 | 0 | 0 | 0 | 0 |
| ENSG00000268401 | 1 | 1 | 0 | 0 | 4 | 1 |
| ENSG00000204869 | 1 | 2 | 0 | 0 | 0 | 1 |
| ENSG00000267922 | 0 | 0 | 0 | 0 | 0 | 0 |
| ENSG00000268053 | 0 | 0 | 0 | 0 | 0 | 0 |
| ENSG00000188624 | 0 | 0 | 0 | 1 | 0 | 0 |
| ENSG00000259339 | 0 | 0 | 0 | 0 | 0 | 0 |
| ENSG00000204866 | 0 | 0 | 1 | 0 | 0 | 0 |
| ENSG00000269729 | 0 | 0 | 0 | 0 | 0 | 0 |
| ENSG00000268621 | 3 | 0 | 0 | 1 | 2 | 3 |
| ENSG00000268879 | 0 | 0 | 0 | 0 | 0 | 0 |
| ENSG00000268460 | 0 | 2 | 0 | 0 | 0 | 0 |
| ENSG00000269637 | 0 | 0 | 0 | 0 | 0 | 0 |
| ENSG00000269745 | 0 | 0 | 0 | 0 | 0 | 0 |
| ENSG00000188293 | 5 | 0 | 3 | 0 | 3 | 0 |
| ENSG00000277748 | 0 | 0 | 0 | 0 | 0 | 0 |
| ENSG00000268238 | 0 | 0 | 0 | 0 | 0 | 0 |
| ENSG00000124440 | 0 | 0 | 0 | 1 | 2 | 0 |
| ENSG00000199796 | 0 | 0 | 0 | 0 | 0 | 0 |

| | | | | | | |
|---|---|---|---|---|---|---|
| ENSG00000269124 | 0 | 0 | 0 | 0 | 0 | 0 |
| ENSG00000011485 | 2272 | 1207 | 1370 | 1295 | 1357 | 1331 |
| ENSG00000268810 | 0 | 0 | 0 | 0 | 0 | 0 |
| ENSG00000269151 | 0 | 0 | 0 | 0 | 0 | 0 |
| ENSG00000169515 | 0 | 0 | 0 | 0 | 1 | 0 |
| ENSG00000277531 | 0 | 0 | 0 | 0 | 0 | 0 |
| ENSG00000182013 | 0 | 0 | 0 | 0 | 0 | 0 |
| ENSG00000230510 | 23 | 16 | 10 | 10 | 13 | 3 |
| ENSG00000204851 | 0 | 0 | 0 | 0 | 0 | 0 |
| ENSG00000204850 | 0 | 0 | 0 | 0 | 0 | 0 |
| ENSG00000278797 | 0 | 0 | 0 | 0 | 0 | 0 |
| ENSG00000268423 | 6 | 3 | 5 | 2 | 4 | 4 |
| ENSG00000249210 | 4 | 1 | 3 | 1 | 4 | 4 |
| ENSG00000160014 | 6617 | 3444 | 4103 | 3954 | 3489 | 3823 |
| ENSG00000269292 | 0 | 0 | 0 | 0 | 0 | 0 |
| ENSG00000160013 | 0 | 0 | 0 | 0 | 0 | 0 |
| ENSG00000167414 | 0 | 0 | 2 | 0 | 0 | 0 |
| ENSG00000279161 | 0 | 1 | 8 | 0 | 1 | 1 |
| ENSG00000197380 | 10 | 9 | 12 | 4 | 10 | 9 |
| ENSG00000245598 | 2 | 3 | 1 | 1 | 1 | 0 |
| ENSG00000105287 | 2069 | 947 | 1249 | 1187 | 1142 | 1107 |
| ENSG00000243560 | 0 | 0 | 0 | 0 | 1 | 1 |
| ENSG00000211513 | 0 | 3 | 0 | 2 | 0 | 1 |
| ENSG00000090372 | 3782 | 1758 | 2437 | 2336 | 2074 | 2202 |
| ENSG00000269487 | 0 | 0 | 2 | 1 | 0 | 3 |
| ENSG00000222614 | 0 | 0 | 0 | 0 | 0 | 0 |
| ENSG00000181027 | 428 | 198 | 250 | 204 | 232 | 225 |
| ENSG00000105281 | 7494 | 3945 | 4702 | 5712 | 4811 | 4736 |
| ENSG00000275719 | 14 | 9 | 1 | 2 | 6 | 7 |
| ENSG00000275862 | 0 | 0 | 0 | 0 | 0 | 0 |
| ENSG00000269576 | 0 | 0 | 0 | 0 | 0 | 0 |
| ENSG00000269014 | 0 | 0 | 0 | 0 | 0 | 0 |
| ENSG00000042753 | 3399 | 1725 | 2018 | 2383 | 2058 | 1946 |
| ENSG00000160007 | 5709 | 2408 | 3176 | 2195 | 3094 | 2965 |
| ENSG00000279948 | 10 | 9 | 7 | 6 | 5 | 13 |
| ENSG00000130751 | 70 | 28 | 30 | 38 | 25 | 29 |
| ENSG00000130748 | 560 | 264 | 298 | 413 | 345 | 380 |
| ENSG00000130749 | 2097 | 936 | 1229 | 1023 | 945 | 1118 |
| ENSG00000142230 | 5423 | 2734 | 3271 | 3005 | 3107 | 3224 |
| ENSG00000105327 | 744 | 332 | 432 | 407 | 414 | 365 |
| ENSG00000265134 | 0 | 0 | 0 | 0 | 0 | 0 |
| ENSG00000284411 | 0 | 0 | 0 | 0 | 0 | 0 |
| ENSG00000105321 | 586 | 219 | 399 | 309 | 317 | 308 |
| ENSG00000257704 | 219 | 106 | 115 | 135 | 96 | 97 |
| ENSG00000197405 | 111 | 57 | 61 | 33 | 45 | 52 |
| ENSG00000134830 | 203 | 107 | 127 | 198 | 170 | 164 |
| ENSG00000134815 | 602 | 287 | 404 | 419 | 358 | 444 |
| ENSG00000105419 | 1209 | 535 | 646 | 631 | 603 | 577 |
| ENSG00000118160 | 2 | 1 | 0 | 0 | 0 | 1 |
| ENSG00000118162 | 377 | 185 | 232 | 226 | 213 | 219 |

| | | | | | | |
|---|---|---|---|---|---|---|
| ENSG00000268061 | 24 | 12 | 14 | 17 | 14 | 19 |
| ENSG00000105402 | 1642 | 896 | 1211 | 1018 | 921 | 950 |
| ENSG00000279861 | 42 | 13 | 16 | 10 | 13 | 18 |
| ENSG00000118156 | 30 | 6 | 17 | 11 | 13 | 18 |
| ENSG00000276588 | 0 | 0 | 0 | 0 | 0 | 0 |
| ENSG00000268746 | 0 | 0 | 0 | 0 | 0 | 0 |
| ENSG00000277383 | 38 | 28 | 35 | 19 | 27 | 13 |
| ENSG00000063169 | 1031 | 422 | 521 | 337 | 352 | 416 |
| ENSG00000269806 | 2 | 0 | 0 | 0 | 0 | 2 |
| ENSG00000278999 | 0 | 1 | 0 | 0 | 0 | 1 |
| ENSG00000024422 | 152 | 63 | 92 | 92 | 74 | 94 |
| ENSG00000105373 | 7308 | 3231 | 4521 | 5215 | 3962 | 4112 |
| ENSG00000221803 | 0 | 0 | 0 | 0 | 0 | 0 |
| ENSG00000269656 | 4 | 3 | 9 | 7 | 9 | 8 |
| ENSG00000178980 | 2886 | 1417 | 1533 | 1888 | 1698 | 1507 |
| ENSG00000213035 | 0 | 0 | 0 | 0 | 0 | 0 |
| ENSG00000271150 | 0 | 0 | 0 | 0 | 0 | 0 |
| ENSG00000178928 | 0 | 0 | 0 | 0 | 0 | 2 |
| ENSG00000105392 | 0 | 0 | 0 | 0 | 0 | 0 |
| ENSG00000259009 | 0 | 0 | 0 | 0 | 0 | 0 |
| ENSG00000259108 | 0 | 0 | 0 | 0 | 0 | 0 |
| ENSG00000105398 | 0 | 0 | 0 | 0 | 0 | 0 |
| ENSG00000188334 | 0 | 0 | 0 | 0 | 0 | 0 |
| ENSG00000169393 | 0 | 2 | 0 | 0 | 0 | 0 |
| ENSG00000105507 | 0 | 0 | 0 | 0 | 0 | 0 |
| ENSG00000105499 | 126 | 59 | 88 | 60 | 68 | 62 |
| ENSG00000269420 | 0 | 0 | 2 | 0 | 0 | 2 |
| ENSG00000105486 | 1077 | 622 | 810 | 569 | 651 | 656 |
| ENSG00000269534 | 1 | 2 | 1 | 3 | 1 | 2 |
| ENSG00000269321 | 0 | 2 | 1 | 0 | 3 | 0 |
| ENSG00000185453 | 248 | 112 | 134 | 133 | 159 | 142 |
| ENSG00000105483 | 495 | 242 | 347 | 191 | 291 | 275 |
| ENSG00000280353 | 2 | 0 | 1 | 1 | 1 | 0 |
| ENSG00000268583 | 3 | 1 | 2 | 2 | 1 | 0 |
| ENSG00000268001 | 41 | 16 | 18 | 18 | 29 | 19 |
| ENSG00000268186 | 1 | 1 | 6 | 0 | 2 | 0 |
| ENSG00000178150 | 117 | 76 | 72 | 52 | 73 | 79 |
| ENSG00000105479 | 41 | 28 | 34 | 34 | 19 | 30 |
| ENSG00000142227 | 114 | 59 | 115 | 68 | 62 | 69 |
| ENSG00000161558 | 257 | 129 | 181 | 139 | 98 | 165 |
| ENSG00000105467 | 4 | 4 | 3 | 7 | 1 | 2 |
| ENSG00000105438 | 8772 | 4197 | 5348 | 5030 | 4497 | 4673 |
| ENSG00000105464 | 20 | 10 | 9 | 13 | 7 | 10 |
| ENSG00000105447 | 1811 | 898 | 995 | 1225 | 982 | 1111 |
| ENSG00000182324 | 34 | 12 | 23 | 22 | 14 | 34 |
| ENSG00000268530 | 0 | 0 | 0 | 0 | 0 | 0 |
| ENSG00000268465 | 0 | 0 | 0 | 0 | 0 | 0 |
| ENSG00000105443 | 3159 | 1448 | 1945 | 1560 | 1662 | 1726 |
| ENSG00000142235 | 499 | 225 | 337 | 317 | 236 | 238 |
| ENSG00000269814 | 4 | 5 | 7 | 12 | 9 | 10 |

| | | | | | | |
|---|---|---|---|---|---|---|
| ENSG00000088002 | 1274 | 596 | 760 | 586 | 527 | 520 |
| ENSG00000105523 | 100 | 43 | 84 | 40 | 42 | 49 |
| ENSG00000177202 | 14 | 6 | 6 | 6 | 7 | 2 |
| ENSG00000063177 | 23620 | 11523 | 14472 | 16515 | 13293 | 13386 |
| ENSG00000268093 | 1 | 5 | 3 | 2 | 2 | 1 |
| ENSG00000063176 | 516 | 222 | 352 | 279 | 231 | 307 |
| ENSG00000105516 | 676 | 241 | 375 | 302 | 215 | 230 |
| ENSG00000063180 | 233 | 102 | 135 | 101 | 77 | 87 |
| ENSG00000232871 | 10 | 5 | 21 | 14 | 14 | 13 |
| ENSG00000142233 | 57 | 23 | 57 | 42 | 32 | 38 |
| ENSG00000176920 | 48 | 31 | 28 | 13 | 14 | 14 |
| ENSG00000176909 | 21 | 15 | 15 | 12 | 3 | 11 |
| ENSG00000105538 | 61 | 19 | 47 | 37 | 30 | 30 |
| ENSG00000182264 | 29 | 18 | 34 | 18 | 14 | 15 |
| ENSG00000174951 | 356 | 181 | 249 | 157 | 209 | 193 |
| ENSG00000105550 | 2 | 0 | 0 | 0 | 0 | 0 |
| ENSG00000206758 | 0 | 0 | 0 | 0 | 0 | 0 |
| ENSG00000105552 | 1956 | 977 | 1271 | 1190 | 1030 | 1154 |
| ENSG00000087076 | 18 | 4 | 10 | 4 | 2 | 6 |
| ENSG00000105559 | 34 | 15 | 23 | 9 | 15 | 16 |
| ENSG00000087074 | 698 | 347 | 448 | 443 | 355 | 433 |
| ENSG00000104804 | 2 | 0 | 0 | 1 | 0 | 1 |
| ENSG00000104805 | 4363 | 1894 | 2844 | 2609 | 2096 | 2405 |
| ENSG00000235191 | 0 | 0 | 1 | 0 | 0 | 0 |
| ENSG00000260366 | 0 | 0 | 0 | 0 | 0 | 0 |
| ENSG00000104808 | 2 | 0 | 5 | 2 | 3 | 2 |
| ENSG00000087088 | 2267 | 1185 | 1400 | 1723 | 1432 | 1437 |
| ENSG00000267898 | 2 | 7 | 5 | 4 | 1 | 2 |
| ENSG00000087086 | 10616 | 5204 | 7086 | 8107 | 6443 | 7436 |
| ENSG00000104812 | 2753 | 1272 | 1589 | 1489 | 1507 | 1391 |
| ENSG00000183207 | 3293 | 1612 | 1925 | 2411 | 1883 | 2107 |
| ENSG00000273898 | 0 | 0 | 0 | 0 | 0 | 0 |
| ENSG00000104826 | 0 | 0 | 1 | 0 | 0 | 0 |
| ENSG00000268655 | 0 | 0 | 0 | 3 | 3 | 2 |
| ENSG00000104827 | 2 | 0 | 0 | 0 | 0 | 0 |
| ENSG00000267335 | 0 | 0 | 0 | 0 | 0 | 0 |
| ENSG00000267019 | 0 | 0 | 0 | 0 | 0 | 0 |
| ENSG00000267099 | 0 | 0 | 0 | 0 | 0 | 0 |
| ENSG00000104818 | 0 | 0 | 0 | 0 | 0 | 0 |
| ENSG00000267631 | 0 | 0 | 0 | 0 | 0 | 0 |
| ENSG00000267685 | 0 | 0 | 0 | 0 | 0 | 0 |
| ENSG00000268669 | 0 | 0 | 0 | 0 | 0 | 0 |
| ENSG00000189052 | 0 | 0 | 0 | 0 | 0 | 0 |
| ENSG00000213030 | 0 | 0 | 0 | 0 | 0 | 0 |
| ENSG00000269706 | 0 | 0 | 0 | 0 | 0 | 0 |
| ENSG00000196337 | 1 | 0 | 0 | 0 | 1 | 2 |
| ENSG00000283663 | 0 | 0 | 0 | 0 | 0 | 0 |
| ENSG00000283251 | 0 | 0 | 0 | 0 | 0 | 0 |
| ENSG00000268108 | 0 | 0 | 0 | 0 | 0 | 0 |
| ENSG00000225950 | 7 | 1 | 3 | 8 | 1 | 2 |

| | | | | | | |
|---|---|---|---|---|---|---|
| ENSG00000268287 | 1 | 0 | 0 | 0 | 0 | 0 |
| ENSG00000104848 | 0 | 3 | 2 | 0 | 0 | 1 |
| ENSG00000265675 | 0 | 0 | 0 | 0 | 0 | 0 |
| ENSG00000104852 | 5320 | 2448 | 3298 | 3309 | 3255 | 2983 |
| ENSG00000104863 | 231 | 114 | 156 | 141 | 131 | 132 |
| ENSG00000221916 | 36 | 21 | 24 | 29 | 24 | 24 |
| ENSG00000177380 | 1073 | 519 | 647 | 617 | 534 | 541 |
| ENSG00000130528 | 8 | 2 | 7 | 2 | 2 | 0 |
| ENSG00000130529 | 2006 | 841 | 1209 | 1037 | 935 | 998 |
| ENSG00000269009 | 0 | 0 | 0 | 0 | 0 | 0 |
| ENSG00000235555 | 0 | 0 | 0 | 0 | 0 | 0 |
| ENSG00000063127 | 57 | 18 | 12 | 17 | 32 | 27 |
| ENSG00000265407 | 3 | 0 | 2 | 1 | 0 | 0 |
| ENSG00000197813 | 32 | 15 | 13 | 12 | 13 | 15 |
| ENSG00000104894 | 80 | 36 | 44 | 16 | 29 | 41 |
| ENSG00000074219 | 1497 | 752 | 1063 | 909 | 784 | 883 |
| ENSG00000268157 | 4 | 8 | 2 | 1 | 1 | 0 |
| ENSG00000104901 | 19 | 12 | 11 | 8 | 7 | 8 |
| ENSG00000268686 | 2 | 2 | 7 | 0 | 1 | 2 |
| ENSG00000161609 | 0 | 0 | 0 | 0 | 0 | 0 |
| ENSG00000142538 | 0 | 1 | 1 | 0 | 0 | 2 |
| ENSG00000261949 | 1 | 0 | 0 | 0 | 0 | 0 |
| ENSG00000104888 | 10 | 4 | 5 | 6 | 2 | 5 |
| ENSG00000104872 | 1438 | 695 | 943 | 894 | 816 | 834 |
| ENSG00000161618 | 1268 | 701 | 828 | 845 | 747 | 742 |
| ENSG00000269469 | 0 | 0 | 0 | 0 | 0 | 0 |
| ENSG00000090554 | 3 | 2 | 0 | 5 | 0 | 0 |
| ENSG00000273189 | 0 | 0 | 0 | 0 | 1 | 1 |
| ENSG00000142541 | 58849 | 30702 | 35869 | 47209 | 38200 | 36848 |
| ENSG00000201675 | 0 | 0 | 0 | 0 | 0 | 0 |
| ENSG00000199631 | 0 | 0 | 0 | 0 | 0 | 0 |
| ENSG00000202503 | 0 | 0 | 0 | 0 | 0 | 0 |
| ENSG00000200259 | 0 | 0 | 0 | 0 | 0 | 0 |
| ENSG00000142534 | 25159 | 12609 | 15188 | 18072 | 14804 | 14470 |
| ENSG00000200530 | 0 | 0 | 0 | 0 | 0 | 0 |
| ENSG00000207782 | 0 | 0 | 0 | 0 | 0 | 0 |
| ENSG00000268681 | 0 | 0 | 0 | 0 | 0 | 0 |
| ENSG00000104870 | 22 | 12 | 23 | 6 | 3 | 1 |
| ENSG00000142552 | 2 | 0 | 2 | 3 | 1 | 3 |
| ENSG00000142546 | 1682 | 828 | 999 | 1217 | 1040 | 935 |
| ENSG00000207073 | 0 | 0 | 0 | 0 | 0 | 0 |
| ENSG00000126460 | 141 | 62 | 96 | 70 | 90 | 73 |
| ENSG00000126464 | 1336 | 656 | 872 | 755 | 617 | 748 |
| ENSG00000268636 | 0 | 0 | 0 | 0 | 0 | 0 |
| ENSG00000126458 | 714 | 372 | 443 | 501 | 438 | 365 |
| ENSG00000126461 | 1829 | 790 | 1062 | 1185 | 925 | 1006 |
| ENSG00000126456 | 2682 | 1306 | 1697 | 1973 | 1783 | 1766 |
| ENSG00000126453 | 763 | 352 | 433 | 556 | 477 | 503 |
| ENSG00000126457 | 6723 | 3466 | 4040 | 4665 | 3918 | 4390 |
| ENSG00000274306 | 0 | 0 | 0 | 0 | 0 | 0 |

| | | | | | | |
|---|---|---|---|---|---|---|
| ENSG00000224420 | 56 | 30 | 45 | 65 | 42 | 75 |
| ENSG00000268677 | 0 | 0 | 0 | 0 | 0 | 0 |
| ENSG00000169169 | 33 | 16 | 30 | 17 | 24 | 18 |
| ENSG00000243829 | 4 | 2 | 2 | 1 | 0 | 1 |
| ENSG00000126467 | 0 | 0 | 0 | 0 | 0 | 0 |
| ENSG00000206599 | 0 | 0 | 0 | 0 | 0 | 1 |
| ENSG00000196961 | 4533 | 2233 | 2875 | 2865 | 2647 | 2629 |
| ENSG00000276270 | 0 | 0 | 0 | 0 | 0 | 0 |
| ENSG00000010361 | 331 | 178 | 232 | 211 | 148 | 184 |
| ENSG00000269194 | 0 | 2 | 3 | 1 | 0 | 1 |
| ENSG00000104973 | 1180 | 574 | 765 | 654 | 599 | 617 |
| ENSG00000284114 | 0 | 0 | 0 | 0 | 0 | 0 |
| ENSG00000268006 | 74 | 43 | 30 | 22 | 33 | 39 |
| ENSG00000104960 | 1517 | 1014 | 1312 | 1377 | 1211 | 1129 |
| ENSG00000268047 | 9 | 0 | 4 | 0 | 2 | 2 |
| ENSG00000265954 | 0 | 0 | 0 | 0 | 0 | 0 |
| ENSG00000269352 | 57 | 24 | 38 | 37 | 35 | 20 |
| ENSG00000039650 | 1026 | 460 | 605 | 493 | 513 | 485 |
| ENSG00000204673 | 3365 | 1595 | 1962 | 2047 | 1681 | 1856 |
| ENSG00000104946 | 1151 | 561 | 749 | 673 | 630 | 637 |
| ENSG00000284394 | 0 | 0 | 0 | 0 | 0 | 0 |
| ENSG00000104951 | 4 | 3 | 6 | 2 | 4 | 4 |
| ENSG00000213024 | 2107 | 1130 | 1351 | 1029 | 1170 | 1218 |
| ENSG00000269179 | 0 | 0 | 0 | 0 | 0 | 0 |
| ENSG00000169136 | 987 | 510 | 653 | 655 | 535 | 650 |
| ENSG00000283842 | 0 | 0 | 0 | 0 | 0 | 0 |
| ENSG00000161640 | 0 | 0 | 1 | 0 | 0 | 0 |
| ENSG00000221125 | 0 | 0 | 0 | 0 | 0 | 0 |
| ENSG00000161643 | 4 | 0 | 2 | 3 | 2 | 1 |
| ENSG00000105053 | 1030 | 477 | 555 | 616 | 528 | 511 |
| ENSG00000276729 | 0 | 0 | 0 | 0 | 0 | 0 |
| ENSG00000142528 | 510 | 217 | 340 | 257 | 325 | 298 |
| ENSG00000269091 | 3 | 1 | 2 | 1 | 2 | 0 |
| ENSG00000204666 | 2 | 1 | 4 | 3 | 3 | 7 |
| ENSG00000269540 | 0 | 0 | 1 | 0 | 0 | 0 |
| ENSG00000267890 | 8 | 2 | 4 | 3 | 0 | 1 |
| ENSG00000161652 | 1 | 0 | 0 | 0 | 1 | 1 |
| ENSG00000105357 | 7372 | 3273 | 4749 | 4358 | 4144 | 4199 |
| ENSG00000276002 | 0 | 0 | 0 | 0 | 0 | 1 |
| ENSG00000267815 | 0 | 0 | 0 | 0 | 0 | 0 |
| ENSG00000131398 | 415 | 156 | 269 | 105 | 138 | 137 |
| ENSG00000131408 | 2465 | 1122 | 1422 | 1530 | 1325 | 1221 |
| ENSG00000131401 | 1 | 1 | 4 | 2 | 3 | 5 |
| ENSG00000131400 | 148 | 63 | 82 | 83 | 73 | 69 |
| ENSG00000269392 | 20 | 7 | 6 | 7 | 3 | 5 |
| ENSG00000062822 | 889 | 427 | 589 | 543 | 503 | 523 |
| ENSG00000142539 | 0 | 0 | 0 | 0 | 0 | 0 |
| ENSG00000269404 | 2 | 2 | 1 | 0 | 0 | 1 |
| ENSG00000086967 | 0 | 1 | 1 | 4 | 1 | 3 |
| ENSG00000142530 | 60 | 43 | 36 | 57 | 52 | 58 |

| | | | | | | |
|---|---|---|---|---|---|---|
| ENSG00000161671 | 3408 | 2046 | 2388 | 2364 | 2245 | 2070 |
| ENSG00000268854 | 0 | 0 | 0 | 0 | 0 | 0 |
| ENSG00000268518 | 0 | 0 | 0 | 0 | 0 | 0 |
| ENSG00000161677 | 1087 | 547 | 699 | 884 | 696 | 632 |
| ENSG00000204653 | 27 | 2 | 7 | 6 | 8 | 12 |
| ENSG00000131409 | 32 | 9 | 10 | 15 | 21 | 18 |
| ENSG00000268231 | 0 | 0 | 0 | 0 | 0 | 0 |
| ENSG00000213023 | 97 | 48 | 66 | 40 | 65 | 66 |
| ENSG00000235034 | 105 | 33 | 57 | 61 | 48 | 50 |
| ENSG00000161681 | 4 | 3 | 4 | 3 | 0 | 4 |
| ENSG00000105472 | 208 | 110 | 127 | 145 | 116 | 104 |
| ENSG00000269565 | 0 | 0 | 0 | 0 | 0 | 0 |
| ENSG00000142511 | 0 | 0 | 0 | 0 | 0 | 0 |
| ENSG00000261341 | 1 | 2 | 0 | 3 | 0 | 1 |
| ENSG00000142513 | 2 | 2 | 1 | 3 | 3 | 1 |
| ENSG00000268375 | 0 | 0 | 0 | 0 | 0 | 0 |
| ENSG00000167747 | 2599 | 1358 | 1520 | 2164 | 1717 | 1711 |
| ENSG00000221381 | 0 | 0 | 0 | 0 | 0 | 0 |
| ENSG00000221241 | 0 | 0 | 0 | 1 | 2 | 2 |
| ENSG00000220988 | 0 | 0 | 0 | 0 | 0 | 0 |
| ENSG00000284711 | 0 | 2 | 0 | 1 | 1 | 0 |
| ENSG00000180279 | 16 | 2 | 9 | 5 | 4 | 4 |
| ENSG00000167748 | 0 | 0 | 3 | 0 | 0 | 0 |
| ENSG00000174562 | 0 | 0 | 0 | 0 | 0 | 0 |
| ENSG00000267968 | 0 | 0 | 0 | 0 | 0 | 0 |
| ENSG00000142515 | 0 | 0 | 0 | 0 | 0 | 0 |
| ENSG00000167751 | 0 | 0 | 0 | 0 | 0 | 0 |
| ENSG00000197588 | 0 | 0 | 0 | 1 | 0 | 0 |
| ENSG00000167749 | 0 | 0 | 1 | 0 | 0 | 0 |
| ENSG00000228674 | 0 | 0 | 0 | 1 | 0 | 0 |
| ENSG00000167754 | 4 | 4 | 8 | 4 | 1 | 7 |
| ENSG00000269495 | 0 | 0 | 0 | 2 | 0 | 0 |
| ENSG00000167755 | 251 | 162 | 160 | 127 | 140 | 132 |
| ENSG00000279676 | 0 | 0 | 0 | 0 | 0 | 0 |
| ENSG00000267879 | 0 | 0 | 1 | 0 | 0 | 1 |
| ENSG00000169035 | 2 | 0 | 0 | 1 | 0 | 0 |
| ENSG00000129455 | 2 | 0 | 0 | 0 | 0 | 1 |
| ENSG00000269741 | 0 | 0 | 0 | 1 | 0 | 0 |
| ENSG00000213022 | 0 | 0 | 0 | 0 | 0 | 0 |
| ENSG00000129451 | 1 | 0 | 2 | 3 | 0 | 3 |
| ENSG00000268739 | 0 | 0 | 0 | 0 | 0 | 0 |
| ENSG00000167757 | 32 | 10 | 10 | 13 | 14 | 9 |
| ENSG00000186474 | 2 | 0 | 1 | 0 | 0 | 0 |
| ENSG00000268906 | 0 | 0 | 0 | 0 | 0 | 0 |
| ENSG00000268390 | 0 | 0 | 0 | 0 | 0 | 1 |
| ENSG00000167759 | 0 | 0 | 0 | 0 | 0 | 1 |
| ENSG00000129437 | 15 | 4 | 10 | 11 | 10 | 8 |
| ENSG00000142544 | 171 | 77 | 97 | 139 | 122 | 101 |
| ENSG00000268581 | 0 | 0 | 0 | 0 | 0 | 0 |
| ENSG00000129450 | 0 | 0 | 0 | 0 | 0 | 0 |

| | | | | | | |
|---|---|---|---|---|---|---|
| ENSG00000267990 | 0 | 0 | 0 | 0 | 0 | 0 |
| ENSG00000168995 | 0 | 0 | 0 | 0 | 0 | 0 |
| ENSG00000267895 | 0 | 0 | 0 | 0 | 0 | 0 |
| ENSG00000269072 | 0 | 0 | 0 | 0 | 0 | 0 |
| ENSG00000269681 | 0 | 0 | 0 | 0 | 0 | 0 |
| ENSG00000171101 | 0 | 0 | 0 | 0 | 0 | 0 |
| ENSG00000268336 | 0 | 0 | 0 | 0 | 0 | 0 |
| ENSG00000268652 | 0 | 0 | 0 | 0 | 0 | 0 |
| ENSG00000269253 | 0 | 0 | 0 | 0 | 0 | 0 |
| ENSG00000278617 | 0 | 0 | 0 | 0 | 0 | 0 |
| ENSG00000268849 | 0 | 0 | 0 | 0 | 0 | 0 |
| ENSG00000105383 | 2 | 0 | 0 | 0 | 0 | 0 |
| ENSG00000179213 | 0 | 0 | 0 | 0 | 0 | 0 |
| ENSG00000268595 | 0 | 0 | 0 | 0 | 0 | 0 |
| ENSG00000268318 | 0 | 0 | 0 | 0 | 0 | 0 |
| ENSG00000269021 | 0 | 0 | 0 | 0 | 0 | 0 |
| ENSG00000142549 | 6 | 1 | 2 | 0 | 2 | 5 |
| ENSG00000186806 | 222 | 88 | 106 | 80 | 87 | 88 |
| ENSG00000267905 | 0 | 0 | 2 | 1 | 0 | 0 |
| ENSG00000267984 | 0 | 0 | 0 | 0 | 0 | 0 |
| ENSG00000105379 | 947 | 705 | 786 | 862 | 724 | 703 |
| ENSG00000269403 | 0 | 0 | 1 | 0 | 0 | 0 |
| ENSG00000268520 | 4 | 5 | 5 | 3 | 1 | 5 |
| ENSG00000160318 | 30 | 8 | 17 | 12 | 16 | 17 |
| ENSG00000105374 | 0 | 3 | 4 | 1 | 0 | 3 |
| ENSG00000105370 | 0 | 0 | 0 | 0 | 0 | 0 |
| ENSG00000262874 | 0 | 0 | 0 | 0 | 0 | 0 |
| ENSG00000268889 | 0 | 0 | 1 | 1 | 2 | 0 |
| ENSG00000268144 | 1 | 0 | 0 | 0 | 0 | 0 |
| ENSG00000142512 | 0 | 0 | 2 | 0 | 4 | 1 |
| ENSG00000254760 | 0 | 0 | 0 | 0 | 0 | 0 |
| ENSG00000255441 | 0 | 0 | 1 | 1 | 0 | 1 |
| ENSG00000268711 | 0 | 0 | 0 | 0 | 0 | 0 |
| ENSG00000269181 | 0 | 0 | 0 | 0 | 0 | 0 |
| ENSG00000105366 | 0 | 0 | 0 | 0 | 0 | 0 |
| ENSG00000268957 | 0 | 0 | 0 | 0 | 0 | 0 |
| ENSG00000213822 | 0 | 0 | 0 | 0 | 0 | 0 |
| ENSG00000254521 | 0 | 0 | 0 | 0 | 0 | 0 |
| ENSG00000268839 | 0 | 0 | 0 | 0 | 0 | 0 |
| ENSG00000269580 | 0 | 0 | 0 | 0 | 0 | 0 |
| ENSG00000105492 | 0 | 0 | 0 | 0 | 0 | 0 |
| ENSG00000268777 | 0 | 0 | 0 | 0 | 0 | 0 |
| ENSG00000269130 | 0 | 0 | 0 | 0 | 0 | 0 |
| ENSG00000268267 | 0 | 0 | 0 | 0 | 0 | 0 |
| ENSG00000105497 | 4 | 5 | 0 | 5 | 9 | 3 |
| ENSG00000269388 | 12 | 19 | 8 | 17 | 20 | 21 |
| ENSG00000267808 | 0 | 0 | 0 | 0 | 1 | 0 |
| ENSG00000105501 | 0 | 0 | 0 | 0 | 0 | 0 |
| ENSG00000268500 | 0 | 0 | 0 | 0 | 0 | 0 |
| ENSG00000244071 | 0 | 0 | 0 | 0 | 0 | 0 |

| | | | | | | |
|---|---|---|---|---|---|---|
| ENSG00000273837 | 0 | 0 | 0 | 0 | 0 | 0 |
| ENSG00000254415 | 0 | 0 | 0 | 0 | 0 | 0 |
| ENSG00000243469 | 0 | 0 | 0 | 0 | 0 | 0 |
| ENSG00000269959 | 53 | 17 | 25 | 13 | 17 | 6 |
| ENSG00000182310 | 223 | 106 | 154 | 72 | 97 | 108 |
| ENSG00000207550 | 0 | 0 | 0 | 0 | 0 | 0 |
| ENSG00000198972 | 0 | 0 | 0 | 0 | 0 | 0 |
| ENSG00000208008 | 0 | 0 | 0 | 0 | 0 | 0 |
| ENSG00000105509 | 3 | 0 | 2 | 1 | 2 | 1 |
| ENSG00000171051 | 0 | 0 | 0 | 0 | 0 | 0 |
| ENSG00000171049 | 0 | 0 | 0 | 0 | 0 | 0 |
| ENSG00000187474 | 2 | 0 | 0 | 1 | 0 | 1 |
| ENSG00000161551 | 0 | 0 | 0 | 0 | 0 | 0 |
| ENSG00000268316 | 0 | 0 | 0 | 0 | 0 | 0 |
| ENSG00000269483 | 0 | 0 | 0 | 0 | 0 | 0 |
| ENSG00000283236 | 0 | 0 | 0 | 0 | 0 | 0 |
| ENSG00000268095 | 0 | 0 | 0 | 0 | 0 | 0 |
| ENSG00000198093 | 0 | 0 | 0 | 0 | 0 | 0 |
| ENSG00000283285 | 0 | 0 | 0 | 0 | 0 | 0 |
| ENSG00000176024 | 65 | 40 | 42 | 24 | 50 | 32 |
| ENSG00000269235 | 0 | 1 | 1 | 0 | 2 | 0 |
| ENSG00000256683 | 160 | 105 | 107 | 55 | 152 | 82 |
| ENSG00000197619 | 104 | 60 | 46 | 27 | 81 | 40 |
| ENSG00000142556 | 182 | 101 | 87 | 51 | 129 | 85 |
| ENSG00000256087 | 180 | 109 | 82 | 53 | 142 | 85 |
| ENSG00000275055 | 3 | 2 | 2 | 6 | 3 | 2 |
| ENSG00000260160 | 7 | 8 | 9 | 6 | 6 | 8 |
| ENSG00000197608 | 304 | 156 | 175 | 73 | 129 | 114 |
| ENSG00000270248 | 0 | 1 | 0 | 0 | 3 | 0 |
| ENSG00000204611 | 265 | 149 | 114 | 76 | 229 | 134 |
| ENSG00000267827 | 0 | 0 | 0 | 0 | 0 | 0 |
| ENSG00000268458 | 0 | 0 | 0 | 0 | 0 | 0 |
| ENSG00000243680 | 2 | 1 | 1 | 2 | 3 | 0 |
| ENSG00000196267 | 97 | 78 | 93 | 52 | 98 | 69 |
| ENSG00000267927 | 0 | 0 | 0 | 1 | 2 | 2 |
| ENSG00000105568 | 10495 | 5025 | 6254 | 6391 | 5558 | 5504 |
| ENSG00000274380 | 0 | 2 | 0 | 1 | 1 | 0 |
| ENSG00000268015 | 0 | 0 | 0 | 0 | 0 | 0 |
| ENSG00000277562 | 0 | 0 | 0 | 0 | 0 | 0 |
| ENSG00000207265 | 0 | 0 | 0 | 0 | 0 | 0 |
| ENSG00000196214 | 483 | 263 | 299 | 125 | 305 | 221 |
| ENSG00000208002 | 0 | 0 | 1 | 0 | 0 | 0 |
| ENSG00000269102 | 1 | 1 | 1 | 1 | 2 | 1 |
| ENSG00000198464 | 300 | 189 | 149 | 73 | 359 | 149 |
| ENSG00000269535 | 2 | 3 | 0 | 0 | 1 | 0 |
| ENSG00000167554 | 135 | 46 | 68 | 31 | 70 | 46 |
| ENSG00000221923 | 91 | 50 | 59 | 25 | 56 | 26 |
| ENSG00000269834 | 32 | 14 | 7 | 8 | 18 | 8 |
| ENSG00000277977 | 2 | 0 | 0 | 1 | 1 | 0 |
| ENSG00000167555 | 129 | 58 | 87 | 26 | 73 | 51 |

| | | | | | | |
|---|---|---|---|---|---|---|
| ENSG00000269776 | 0 | 0 | 0 | 0 | 0 | 0 |
| ENSG00000198633 | 0 | 0 | 2 | 0 | 1 | 0 |
| ENSG00000258405 | 0 | 0 | 0 | 0 | 0 | 0 |
| ENSG00000278543 | 0 | 0 | 0 | 0 | 0 | 0 |
| ENSG00000269349 | 0 | 0 | 0 | 0 | 0 | 0 |
| ENSG00000198482 | 96 | 74 | 55 | 27 | 77 | 35 |
| ENSG00000167562 | 284 | 116 | 139 | 70 | 138 | 109 |
| ENSG00000242255 | 0 | 0 | 0 | 0 | 0 | 0 |
| ENSG00000268886 | 0 | 0 | 0 | 0 | 0 | 0 |
| ENSG00000123870 | 62 | 33 | 47 | 32 | 55 | 30 |
| ENSG00000167766 | 905 | 461 | 544 | 166 | 589 | 372 |
| ENSG00000268970 | 30 | 12 | 16 | 8 | 35 | 17 |
| ENSG00000269825 | 78 | 38 | 30 | 3 | 22 | 18 |
| ENSG00000213020 | 179 | 122 | 91 | 55 | 106 | 78 |
| ENSG00000189190 | 190 | 116 | 103 | 43 | 163 | 102 |
| ENSG00000198538 | 409 | 234 | 122 | 95 | 330 | 141 |
| ENSG00000255875 | 0 | 1 | 0 | 0 | 0 | 1 |
| ENSG00000204604 | 605 | 321 | 236 | 155 | 381 | 266 |
| ENSG00000268225 | 0 | 0 | 0 | 1 | 0 | 0 |
| ENSG00000182986 | 0 | 0 | 0 | 0 | 0 | 0 |
| ENSG00000283695 | 0 | 0 | 0 | 0 | 0 | 0 |
| ENSG00000283088 | 0 | 0 | 0 | 0 | 0 | 0 |
| ENSG00000213793 | 44 | 28 | 29 | 13 | 55 | 32 |
| ENSG00000269646 | 7 | 0 | 2 | 0 | 0 | 0 |
| ENSG00000213801 | 31 | 14 | 18 | 17 | 16 | 11 |
| ENSG00000221874 | 0 | 0 | 0 | 1 | 0 | 0 |
| ENSG00000180257 | 157 | 74 | 59 | 38 | 93 | 62 |
| ENSG00000269082 | 0 | 0 | 0 | 0 | 0 | 0 |
| ENSG00000242779 | 0 | 0 | 0 | 0 | 0 | 0 |
| ENSG00000268282 | 0 | 1 | 2 | 0 | 1 | 0 |
| ENSG00000267943 | 0 | 0 | 0 | 0 | 0 | 0 |
| ENSG00000269526 | 0 | 0 | 0 | 0 | 0 | 0 |
| ENSG00000268964 | 0 | 0 | 0 | 0 | 0 | 0 |
| ENSG00000170949 | 410 | 245 | 244 | 93 | 275 | 181 |
| ENSG00000170954 | 0 | 0 | 1 | 0 | 2 | 0 |
| ENSG00000197937 | 72 | 43 | 43 | 20 | 48 | 23 |
| ENSG00000197497 | 31 | 8 | 6 | 9 | 14 | 11 |
| ENSG00000268842 | 0 | 0 | 0 | 0 | 0 | 0 |
| ENSG00000269001 | 502 | 285 | 235 | 118 | 326 | 217 |
| ENSG00000267809 | 50 | 25 | 38 | 19 | 27 | 31 |
| ENSG00000197928 | 0 | 0 | 0 | 0 | 0 | 0 |
| ENSG00000269288 | 0 | 0 | 0 | 0 | 0 | 0 |
| ENSG00000196131 | 0 | 0 | 0 | 0 | 0 | 0 |
| ENSG00000228567 | 0 | 0 | 0 | 0 | 0 | 0 |
| ENSG00000269758 | 0 | 0 | 0 | 0 | 0 | 0 |
| ENSG00000189348 | 0 | 0 | 0 | 0 | 0 | 0 |
| ENSG00000163098 | 0 | 0 | 0 | 0 | 0 | 0 |
| ENSG00000269118 | 0 | 0 | 0 | 0 | 0 | 0 |
| ENSG00000174677 | 0 | 0 | 0 | 0 | 0 | 0 |
| ENSG00000213799 | 103 | 66 | 39 | 28 | 107 | 68 |

| | | | | | | |
|---|---|---|---|---|---|---|
| ENSG00000203326 | 95 | 64 | 40 | 31 | 107 | 55 |
| ENSG00000241069 | 0 | 0 | 0 | 0 | 0 | 0 |
| ENSG00000196417 | 132 | 88 | 75 | 37 | 151 | 50 |
| ENSG00000239912 | 3 | 3 | 0 | 3 | 3 | 3 |
| ENSG00000249435 | 0 | 0 | 0 | 0 | 0 | 0 |
| ENSG00000241015 | 224 | 85 | 93 | 66 | 128 | 95 |
| ENSG00000160336 | 305 | 201 | 156 | 59 | 307 | 164 |
| ENSG00000198346 | 126 | 97 | 72 | 38 | 106 | 73 |
| ENSG00000250731 | 11 | 5 | 4 | 6 | 5 | 7 |
| ENSG00000241434 | 0 | 0 | 0 | 0 | 0 | 0 |
| ENSG00000213777 | 1 | 2 | 0 | 0 | 3 | 1 |
| ENSG00000274892 | 0 | 3 | 0 | 0 | 0 | 0 |
| ENSG00000130844 | 755 | 387 | 502 | 283 | 363 | 404 |
| ENSG00000268864 | 89 | 66 | 73 | 27 | 46 | 41 |
| ENSG00000269608 | 0 | 0 | 0 | 0 | 0 | 0 |
| ENSG00000242668 | 0 | 0 | 0 | 0 | 0 | 0 |
| ENSG00000204595 | 0 | 3 | 1 | 0 | 0 | 0 |
| ENSG00000200393 | 0 | 0 | 0 | 0 | 0 | 0 |
| ENSG00000270347 | 0 | 0 | 0 | 0 | 0 | 0 |
| ENSG00000207645 | 0 | 0 | 0 | 0 | 0 | 0 |
| ENSG00000207644 | 0 | 0 | 0 | 0 | 0 | 0 |
| ENSG00000221017 | 0 | 0 | 0 | 0 | 0 | 0 |
| ENSG00000207869 | 0 | 0 | 0 | 0 | 0 | 0 |
| ENSG00000207599 | 0 | 0 | 0 | 0 | 0 | 0 |
| ENSG00000207616 | 0 | 0 | 0 | 0 | 0 | 0 |
| ENSG00000207810 | 0 | 0 | 0 | 0 | 0 | 0 |
| ENSG00000283540 | 0 | 0 | 0 | 0 | 0 | 0 |
| ENSG00000207615 | 0 | 0 | 0 | 0 | 0 | 0 |
| ENSG00000207788 | 0 | 0 | 0 | 0 | 0 | 0 |
| ENSG00000221421 | 0 | 0 | 0 | 0 | 0 | 0 |
| ENSG00000207594 | 0 | 0 | 0 | 0 | 0 | 0 |
| ENSG00000207580 | 0 | 0 | 0 | 0 | 0 | 0 |
| ENSG00000207825 | 0 | 0 | 0 | 0 | 0 | 0 |
| ENSG00000207711 | 0 | 0 | 0 | 0 | 0 | 0 |
| ENSG00000283455 | 0 | 0 | 0 | 0 | 0 | 0 |
| ENSG00000207706 | 0 | 0 | 0 | 0 | 0 | 0 |
| ENSG00000207722 | 0 | 0 | 0 | 0 | 0 | 0 |
| ENSG00000207862 | 0 | 0 | 0 | 0 | 0 | 0 |
| ENSG00000207629 | 0 | 0 | 0 | 0 | 0 | 0 |
| ENSG00000207738 | 0 | 0 | 0 | 0 | 0 | 0 |
| ENSG00000283490 | 0 | 0 | 0 | 0 | 0 | 0 |
| ENSG00000283289 | 0 | 0 | 0 | 0 | 0 | 0 |
| ENSG00000207734 | 0 | 0 | 0 | 0 | 0 | 0 |
| ENSG00000207981 | 0 | 0 | 0 | 0 | 0 | 0 |
| ENSG00000207549 | 0 | 0 | 0 | 0 | 0 | 0 |
| ENSG00000251843 | 0 | 0 | 0 | 0 | 0 | 0 |
| ENSG00000207735 | 0 | 0 | 0 | 0 | 0 | 0 |
| ENSG00000207837 | 0 | 0 | 0 | 0 | 0 | 0 |
| ENSG00000207799 | 0 | 0 | 0 | 0 | 0 | 0 |
| ENSG00000207925 | 0 | 0 | 0 | 0 | 0 | 0 |

| | | | | | | |
|---|---|---|---|---|---|---|
| ENSG00000211532 | 0 | 0 | 0 | 0 | 0 | 0 |
| ENSG00000207987 | 0 | 0 | 0 | 0 | 0 | 0 |
| ENSG00000207803 | 0 | 0 | 0 | 0 | 0 | 0 |
| ENSG00000252734 | 0 | 0 | 0 | 0 | 0 | 0 |
| ENSG00000283330 | 0 | 0 | 0 | 0 | 0 | 0 |
| ENSG00000207946 | 0 | 0 | 0 | 0 | 0 | 0 |
| ENSG00000207699 | 0 | 0 | 0 | 0 | 0 | 0 |
| ENSG00000207838 | 0 | 0 | 0 | 0 | 0 | 0 |
| ENSG00000207861 | 0 | 0 | 0 | 0 | 0 | 0 |
| ENSG00000252025 | 0 | 0 | 0 | 0 | 0 | 0 |
| ENSG00000207634 | 0 | 0 | 0 | 0 | 0 | 0 |
| ENSG00000252191 | 0 | 0 | 0 | 0 | 0 | 0 |
| ENSG00000283685 | 0 | 0 | 0 | 0 | 0 | 0 |
| ENSG00000207992 | 0 | 0 | 0 | 0 | 0 | 0 |
| ENSG00000207979 | 0 | 0 | 0 | 0 | 0 | 0 |
| ENSG00000207767 | 0 | 0 | 0 | 0 | 0 | 0 |
| ENSG00000221548 | 0 | 0 | 0 | 0 | 0 | 0 |
| ENSG00000252063 | 0 | 0 | 0 | 0 | 0 | 0 |
| ENSG00000207620 | 0 | 0 | 0 | 0 | 0 | 0 |
| ENSG00000269842 | 0 | 0 | 0 | 0 | 0 | 0 |
| ENSG00000284362 | 0 | 0 | 0 | 0 | 0 | 0 |
| ENSG00000252979 | 0 | 0 | 0 | 0 | 0 | 0 |
| ENSG00000237589 | 0 | 0 | 0 | 0 | 0 | 0 |
| ENSG00000229000 | 0 | 0 | 0 | 0 | 0 | 0 |
| ENSG00000238150 | 0 | 0 | 0 | 0 | 0 | 0 |
| ENSG00000269877 | 0 | 0 | 0 | 0 | 0 | 0 |
| ENSG00000199031 | 0 | 0 | 0 | 0 | 0 | 0 |
| ENSG00000284568 | 0 | 0 | 0 | 0 | 0 | 0 |
| ENSG00000199095 | 0 | 0 | 0 | 0 | 0 | 0 |
| ENSG00000199143 | 0 | 0 | 0 | 0 | 0 | 0 |
| ENSG00000269564 | 0 | 0 | 0 | 0 | 0 | 0 |
| ENSG00000142405 | 0 | 0 | 3 | 0 | 0 | 0 |
| ENSG00000228323 | 12 | 8 | 12 | 7 | 3 | 13 |
| ENSG00000232324 | 2 | 4 | 1 | 1 | 2 | 1 |
| ENSG00000179820 | 1113 | 717 | 769 | 611 | 660 | 642 |
| ENSG00000232220 | 0 | 0 | 0 | 0 | 0 | 0 |
| ENSG00000126583 | 6 | 0 | 3 | 2 | 5 | 1 |
| ENSG00000105605 | 0 | 0 | 0 | 0 | 0 | 0 |
| ENSG00000142408 | 10 | 11 | 11 | 5 | 14 | 12 |
| ENSG00000284583 | 0 | 0 | 0 | 0 | 0 | 0 |
| ENSG00000130433 | 63 | 36 | 31 | 48 | 30 | 15 |
| ENSG00000276206 | 0 | 0 | 0 | 0 | 0 | 0 |
| ENSG00000189068 | 0 | 0 | 0 | 0 | 0 | 0 |
| ENSG00000248385 | 0 | 0 | 0 | 0 | 0 | 0 |
| ENSG00000170909 | 31 | 9 | 25 | 16 | 10 | 18 |
| ENSG00000170906 | 1434 | 694 | 861 | 862 | 766 | 768 |
| ENSG00000105619 | 1064 | 607 | 691 | 795 | 716 | 632 |
| ENSG00000105618 | 1308 | 752 | 911 | 898 | 814 | 877 |
| ENSG00000237017 | 19 | 12 | 10 | 8 | 14 | 9 |
| ENSG00000088038 | 1993 | 921 | 1234 | 1063 | 1058 | 1136 |

| | | | | | | |
|---|---|---|---|---|---|---|
| ENSG00000105617 | 429 | 166 | 287 | 228 | 223 | 215 |
| ENSG00000167608 | 2591 | 1149 | 1757 | 1124 | 960 | 1202 |
| ENSG00000125505 | 7580 | 3679 | 4505 | 4840 | 4465 | 4175 |
| ENSG00000170892 | 3472 | 1668 | 2026 | 2274 | 2034 | 2023 |
| ENSG00000223660 | 0 | 0 | 0 | 0 | 0 | 0 |
| ENSG00000170889 | 17624 | 8729 | 10372 | 13382 | 10389 | 9881 |
| ENSG00000224579 | 0 | 0 | 0 | 0 | 0 | 0 |
| ENSG00000204577 | 2 | 4 | 2 | 3 | 0 | 2 |
| ENSG00000269524 | 0 | 0 | 0 | 0 | 0 | 0 |
| ENSG00000235081 | 2 | 0 | 0 | 0 | 3 | 0 |
| ENSG00000244482 | 1 | 1 | 1 | 0 | 0 | 0 |
| ENSG00000249189 | 0 | 0 | 0 | 0 | 0 | 0 |
| ENSG00000105609 | 0 | 0 | 0 | 0 | 0 | 0 |
| ENSG00000212314 | 0 | 0 | 0 | 0 | 0 | 0 |
| ENSG00000240296 | 0 | 0 | 0 | 0 | 0 | 0 |
| ENSG00000131042 | 0 | 0 | 0 | 0 | 0 | 0 |
| ENSG00000264703 | 0 | 0 | 0 | 0 | 0 | 0 |
| ENSG00000240197 | 0 | 0 | 0 | 0 | 0 | 0 |
| ENSG00000276758 | 0 | 0 | 0 | 0 | 0 | 0 |
| ENSG00000251431 | 0 | 0 | 0 | 0 | 0 | 0 |
| ENSG00000234436 | 0 | 0 | 0 | 0 | 0 | 0 |
| ENSG00000187116 | 0 | 0 | 0 | 0 | 0 | 0 |
| ENSG00000275210 | 0 | 0 | 0 | 0 | 0 | 0 |
| ENSG00000249654 | 0 | 0 | 0 | 0 | 0 | 0 |
| ENSG00000250507 | 0 | 0 | 0 | 0 | 0 | 0 |
| ENSG00000239961 | 0 | 0 | 0 | 0 | 0 | 0 |
| ENSG00000167613 | 0 | 0 | 0 | 0 | 0 | 0 |
| ENSG00000237955 | 0 | 0 | 0 | 0 | 0 | 0 |
| ENSG00000167614 | 27 | 10 | 11 | 9 | 11 | 5 |
| ENSG00000227407 | 0 | 0 | 0 | 0 | 1 | 0 |
| ENSG00000268496 | 1 | 0 | 0 | 0 | 2 | 0 |
| ENSG00000269873 | 0 | 1 | 1 | 0 | 0 | 0 |
| ENSG00000267838 | 4 | 6 | 4 | 5 | 4 | 7 |
| ENSG00000226696 | 71 | 32 | 53 | 20 | 35 | 32 |
| ENSG00000235681 | 5 | 10 | 8 | 4 | 6 | 6 |
| ENSG00000167615 | 3471 | 1329 | 2211 | 1341 | 1651 | 1845 |
| ENSG00000275183 | 330 | 177 | 173 | 208 | 192 | 196 |
| ENSG00000167617 | 14 | 5 | 11 | 12 | 15 | 5 |
| ENSG00000167618 | 0 | 0 | 0 | 0 | 0 | 0 |
| ENSG00000213016 | 0 | 0 | 0 | 0 | 0 | 0 |
| ENSG00000104970 | 0 | 0 | 0 | 0 | 0 | 0 |
| ENSG00000235181 | 0 | 0 | 0 | 0 | 0 | 0 |
| ENSG00000239998 | 0 | 0 | 0 | 0 | 0 | 0 |
| ENSG00000269271 | 0 | 0 | 0 | 0 | 0 | 0 |
| ENSG00000104974 | 0 | 0 | 0 | 0 | 0 | 0 |
| ENSG00000238094 | 0 | 0 | 0 | 0 | 0 | 0 |
| ENSG00000104972 | 0 | 0 | 0 | 0 | 0 | 0 |
| ENSG00000224730 | 0 | 0 | 0 | 0 | 0 | 0 |
| ENSG00000186818 | 0 | 0 | 0 | 0 | 0 | 0 |
| ENSG00000277636 | 0 | 0 | 0 | 0 | 0 | 0 |

| | | | | | | |
|---|---|---|---|---|---|---|
| ENSG00000242856 | 0 | 0 | 0 | 0 | 0 | 0 |
| ENSG00000237323 | 0 | 0 | 0 | 0 | 0 | 0 |
| ENSG00000225370 | 0 | 0 | 0 | 0 | 0 | 0 |
| ENSG00000186152 | 0 | 0 | 0 | 0 | 0 | 0 |
| ENSG00000228790 | 0 | 0 | 0 | 0 | 0 | 0 |
| ENSG00000170858 | 0 | 0 | 0 | 0 | 0 | 0 |
| ENSG00000242019 | 0 | 0 | 0 | 0 | 0 | 0 |
| ENSG00000215765 | 0 | 0 | 0 | 0 | 0 | 0 |
| ENSG00000243772 | 0 | 0 | 0 | 0 | 0 | 0 |
| ENSG00000242473 | 0 | 0 | 0 | 0 | 0 | 0 |
| ENSG00000125498 | 0 | 0 | 0 | 0 | 0 | 0 |
| ENSG00000264257 | 0 | 0 | 0 | 0 | 0 | 0 |
| ENSG00000189013 | 0 | 0 | 0 | 0 | 0 | 0 |
| ENSG00000167633 | 0 | 0 | 0 | 0 | 0 | 0 |
| ENSG00000221957 | 0 | 0 | 0 | 0 | 0 | 0 |
| ENSG00000240403 | 0 | 0 | 0 | 0 | 0 | 0 |
| ENSG00000199308 | 0 | 0 | 0 | 0 | 0 | 0 |
| ENSG00000186431 | 0 | 0 | 0 | 0 | 0 | 0 |
| ENSG00000268734 | 0 | 0 | 0 | 0 | 0 | 0 |
| ENSG00000189430 | 0 | 0 | 0 | 0 | 0 | 0 |
| ENSG00000167634 | 0 | 0 | 0 | 0 | 0 | 0 |
| ENSG00000275012 | 0 | 0 | 0 | 0 | 0 | 0 |
| ENSG00000022556 | 0 | 0 | 0 | 0 | 0 | 0 |
| ENSG00000243494 | 0 | 0 | 0 | 0 | 0 | 0 |
| ENSG00000267265 | 30 | 35 | 37 | 23 | 50 | 40 |
| ENSG00000088053 | 0 | 2 | 3 | 5 | 2 | 2 |
| ENSG00000267149 | 2 | 0 | 1 | 0 | 1 | 0 |
| ENSG00000160439 | 843 | 480 | 596 | 426 | 438 | 490 |
| ENSG00000131037 | 1359 | 572 | 823 | 615 | 541 | 512 |
| ENSG00000125503 | 2333 | 1023 | 1348 | 1431 | 1338 | 1365 |
| ENSG00000267507 | 0 | 0 | 2 | 0 | 2 | 1 |
| ENSG00000275863 | 0 | 0 | 0 | 0 | 0 | 1 |
| ENSG00000105048 | 1289 | 594 | 819 | 853 | 666 | 715 |
| ENSG00000129991 | 2 | 0 | 3 | 1 | 0 | 0 |
| ENSG00000267110 | 0 | 0 | 0 | 0 | 0 | 0 |
| ENSG00000167646 | 81 | 45 | 67 | 70 | 87 | 65 |
| ENSG00000267577 | 0 | 0 | 0 | 0 | 0 | 0 |
| ENSG00000239137 | 0 | 0 | 0 | 0 | 0 | 0 |
| ENSG00000129990 | 4 | 2 | 3 | 4 | 3 | 1 |
| ENSG00000080031 | 12 | 9 | 8 | 4 | 4 | 10 |
| ENSG00000267649 | 16 | 3 | 4 | 2 | 12 | 5 |
| ENSG00000180089 | 44 | 15 | 21 | 26 | 19 | 26 |
| ENSG00000276570 | 24 | 12 | 12 | 12 | 12 | 19 |
| ENSG00000105063 | 3200 | 1496 | 1934 | 1975 | 1969 | 1834 |
| ENSG00000275519 | 0 | 0 | 0 | 1 | 0 | 0 |
| ENSG00000278328 | 0 | 0 | 0 | 0 | 0 | 0 |
| ENSG00000278264 | 0 | 0 | 0 | 0 | 0 | 0 |
| ENSG00000133265 | 1102 | 504 | 588 | 868 | 697 | 623 |
| ENSG00000160469 | 64 | 30 | 40 | 38 | 28 | 30 |
| ENSG00000268729 | 0 | 0 | 0 | 0 | 0 | 0 |

| | | | | | | |
|---|---|---|---|---|---|---|
| ENSG00000180061 | 0 | 0 | 0 | 0 | 0 | 0 |
| ENSG00000276831 | 1 | 0 | 1 | 0 | 1 | 0 |
| ENSG00000133247 | 701 | 312 | 405 | 401 | 366 | 341 |
| ENSG00000160471 | 2 | 0 | 1 | 2 | 0 | 3 |
| ENSG00000267706 | 0 | 0 | 0 | 0 | 0 | 0 |
| ENSG00000180043 | 0 | 0 | 0 | 0 | 0 | 0 |
| ENSG00000095752 | 93 | 34 | 56 | 43 | 54 | 29 |
| ENSG00000160472 | 10 | 12 | 7 | 10 | 9 | 5 |
| ENSG00000269275 | 0 | 0 | 1 | 0 | 0 | 0 |
| ENSG00000233493 | 178 | 77 | 114 | 100 | 76 | 94 |
| ENSG00000108107 | 12212 | 5586 | 6593 | 8394 | 6835 | 6915 |
| ENSG00000284029 | 0 | 0 | 0 | 0 | 0 | 0 |
| ENSG00000108106 | 2133 | 897 | 1216 | 1927 | 1412 | 1515 |
| ENSG00000187902 | 4 | 0 | 0 | 0 | 0 | 1 |
| ENSG00000063241 | 1891 | 902 | 1107 | 1459 | 1050 | 1129 |
| ENSG00000283567 | 0 | 0 | 1 | 0 | 1 | 0 |
| ENSG00000279063 | 0 | 0 | 0 | 0 | 0 | 0 |
| ENSG00000269859 | 2 | 0 | 0 | 0 | 0 | 0 |
| ENSG00000197483 | 187 | 64 | 101 | 140 | 103 | 107 |
| ENSG00000090971 | 986 | 463 | 584 | 590 | 500 | 543 |
| ENSG00000179954 | 4 | 2 | 3 | 1 | 3 | 4 |
| ENSG00000187550 | 4 | 4 | 0 | 4 | 2 | 1 |
| ENSG00000231274 | 0 | 0 | 0 | 0 | 0 | 0 |
| ENSG00000218891 | 521 | 188 | 290 | 334 | 269 | 304 |
| ENSG00000179943 | 488 | 266 | 347 | 305 | 259 | 253 |
| ENSG00000171443 | 378 | 232 | 264 | 254 | 199 | 208 |
| ENSG00000261221 | 658 | 315 | 393 | 410 | 299 | 369 |
| ENSG00000276488 | 34 | 25 | 23 | 12 | 10 | 16 |
| ENSG00000179922 | 197 | 85 | 104 | 103 | 89 | 104 |
| ENSG00000213015 | 618 | 280 | 381 | 397 | 286 | 307 |
| ENSG00000171425 | 882 | 422 | 542 | 677 | 543 | 560 |
| ENSG00000173581 | 466 | 154 | 242 | 252 | 170 | 222 |
| ENSG00000063244 | 7976 | 3844 | 5054 | 5032 | 4468 | 5008 |
| ENSG00000267523 | 36 | 5 | 13 | 11 | 18 | 12 |
| ENSG00000267096 | 0 | 4 | 1 | 0 | 1 | 0 |
| ENSG00000063245 | 4165 | 1777 | 2343 | 2685 | 2189 | 2107 |
| ENSG00000267117 | 0 | 0 | 3 | 1 | 0 | 0 |
| ENSG00000267689 | 0 | 0 | 0 | 0 | 0 | 0 |
| ENSG00000268593 | 0 | 0 | 0 | 0 | 0 | 0 |
| ENSG00000185792 | 0 | 5 | 1 | 1 | 0 | 0 |
| ENSG00000222524 | 0 | 0 | 0 | 0 | 0 | 0 |
| ENSG00000223638 | 0 | 0 | 0 | 0 | 0 | 0 |
| ENSG00000229292 | 0 | 0 | 0 | 0 | 0 | 0 |
| ENSG00000255238 | 0 | 0 | 0 | 0 | 0 | 0 |
| ENSG00000179873 | 0 | 0 | 0 | 0 | 0 | 0 |
| ENSG00000160505 | 0 | 0 | 0 | 0 | 0 | 0 |
| ENSG00000173572 | 0 | 0 | 0 | 0 | 0 | 0 |
| ENSG00000179709 | 0 | 0 | 0 | 0 | 0 | 0 |
| ENSG00000171487 | 0 | 0 | 0 | 0 | 0 | 0 |
| ENSG00000267522 | 0 | 0 | 0 | 1 | 1 | 1 |

| | | | | | | |
|---|---|---|---|---|---|---|
| ENSG00000142409 | 877 | 385 | 469 | 606 | 497 | 489 |
| ENSG00000223060 | 1 | 2 | 1 | 2 | 0 | 0 |
| ENSG00000279959 | 0 | 0 | 0 | 0 | 0 | 0 |
| ENSG00000167685 | 1869 | 877 | 1126 | 1016 | 875 | 878 |
| ENSG00000279794 | 33 | 17 | 19 | 17 | 14 | 13 |
| ENSG00000197487 | 0 | 1 | 0 | 0 | 0 | 0 |
| ENSG00000197213 | 0 | 0 | 0 | 1 | 0 | 0 |
| ENSG00000204532 | 2 | 1 | 0 | 0 | 1 | 1 |
| ENSG00000242411 | 0 | 2 | 0 | 0 | 0 | 0 |
| ENSG00000131848 | 104 | 58 | 89 | 60 | 62 | 84 |
| ENSG00000267908 | 0 | 0 | 1 | 0 | 0 | 1 |
| ENSG00000267710 | 3 | 5 | 5 | 0 | 5 | 1 |
| ENSG00000267298 | 0 | 2 | 0 | 0 | 1 | 0 |
| ENSG00000267549 | 3 | 2 | 4 | 4 | 7 | 3 |
| ENSG00000200646 | 0 | 0 | 0 | 0 | 0 | 0 |
| ENSG00000213014 | 5 | 0 | 0 | 0 | 1 | 0 |
| ENSG00000266907 | 0 | 0 | 0 | 0 | 0 | 0 |
| ENSG00000267145 | 0 | 0 | 0 | 0 | 0 | 0 |
| ENSG00000266932 | 0 | 1 | 0 | 0 | 0 | 0 |
| ENSG00000267429 | 0 | 0 | 0 | 0 | 0 | 0 |
| ENSG00000267606 | 0 | 0 | 0 | 0 | 0 | 0 |
| ENSG00000240225 | 0 | 0 | 0 | 0 | 0 | 0 |
| ENSG00000018869 | 0 | 0 | 0 | 0 | 0 | 0 |
| ENSG00000267776 | 0 | 0 | 0 | 0 | 0 | 0 |
| ENSG00000267459 | 0 | 0 | 0 | 0 | 0 | 0 |
| ENSG00000267781 | 0 | 0 | 0 | 0 | 0 | 0 |
| ENSG00000267192 | 0 | 0 | 0 | 0 | 0 | 0 |
| ENSG00000267454 | 2 | 0 | 0 | 1 | 0 | 0 |
| ENSG00000198440 | 26 | 17 | 17 | 4 | 15 | 13 |
| ENSG00000198046 | 0 | 0 | 0 | 0 | 0 | 0 |
| ENSG00000166770 | 0 | 0 | 0 | 0 | 0 | 0 |
| ENSG00000279405 | 0 | 0 | 0 | 0 | 0 | 0 |
| ENSG00000276578 | 0 | 0 | 0 | 0 | 0 | 0 |
| ENSG00000196263 | 0 | 0 | 0 | 0 | 0 | 0 |
| ENSG00000267224 | 0 | 0 | 0 | 0 | 0 | 0 |
| ENSG00000267421 | 0 | 0 | 0 | 0 | 0 | 0 |
| ENSG00000196867 | 0 | 0 | 0 | 0 | 0 | 0 |
| ENSG00000269696 | 0 | 0 | 0 | 0 | 0 | 0 |
| ENSG00000197016 | 0 | 0 | 0 | 0 | 0 | 0 |
| ENSG00000268847 | 0 | 0 | 0 | 0 | 0 | 0 |
| ENSG00000197951 | 0 | 0 | 0 | 0 | 0 | 1 |
| ENSG00000268182 | 0 | 0 | 0 | 0 | 0 | 0 |
| ENSG00000127903 | 0 | 0 | 0 | 0 | 0 | 0 |
| ENSG00000268352 | 0 | 0 | 0 | 0 | 0 | 0 |
| ENSG00000268568 | 38 | 20 | 19 | 22 | 24 | 21 |
| ENSG00000268067 | 0 | 0 | 0 | 0 | 0 | 0 |
| ENSG00000284729 | 0 | 0 | 0 | 0 | 0 | 0 |
| ENSG00000269793 | 0 | 0 | 0 | 0 | 0 | 0 |
| ENSG00000269699 | 0 | 0 | 0 | 0 | 0 | 0 |
| ENSG00000268407 | 0 | 0 | 0 | 0 | 0 | 0 |

| | | | | | | |
|---|---|---|---|---|---|---|
| ENSG00000198300 | 0 | 0 | 0 | 0 | 0 | 0 |
| ENSG00000268654 | 0 | 0 | 0 | 0 | 0 | 0 |
| ENSG00000274777 | 0 | 0 | 0 | 0 | 0 | 0 |
| ENSG00000269786 | 0 | 0 | 0 | 0 | 0 | 0 |
| ENSG00000268483 | 0 | 0 | 0 | 0 | 0 | 0 |
| ENSG00000274219 | 0 | 0 | 0 | 0 | 0 | 0 |
| ENSG00000131864 | 0 | 0 | 0 | 0 | 0 | 0 |
| ENSG00000141946 | 0 | 0 | 0 | 0 | 0 | 0 |
| ENSG00000270293 | 0 | 0 | 0 | 0 | 0 | 0 |
| ENSG00000270790 | 0 | 0 | 0 | 0 | 0 | 0 |
| ENSG00000258873 | 0 | 0 | 0 | 0 | 0 | 0 |
| ENSG00000268379 | 0 | 0 | 0 | 0 | 0 | 0 |
| ENSG00000083844 | 30 | 15 | 23 | 10 | 16 | 19 |
| ENSG00000105146 | 0 | 1 | 0 | 0 | 0 | 0 |
| ENSG00000204524 | 51 | 25 | 23 | 11 | 26 | 21 |
| ENSG00000268713 | 44 | 15 | 12 | 15 | 23 | 12 |
| ENSG00000267871 | 10 | 6 | 1 | 7 | 9 | 3 |
| ENSG00000197714 | 51 | 50 | 41 | 8 | 24 | 24 |
| ENSG00000279541 | 7 | 11 | 5 | 2 | 2 | 4 |
| ENSG00000268205 | 333 | 149 | 135 | 139 | 164 | 148 |
| ENSG00000268723 | 0 | 0 | 0 | 0 | 0 | 0 |
| ENSG00000178229 | 67 | 34 | 42 | 23 | 30 | 40 |
| ENSG00000268678 | 1 | 1 | 0 | 0 | 1 | 1 |
| ENSG00000131845 | 260 | 111 | 188 | 89 | 152 | 135 |
| ENSG00000152433 | 30 | 9 | 20 | 9 | 9 | 8 |
| ENSG00000256060 | 83 | 50 | 71 | 58 | 66 | 29 |
| ENSG00000268133 | 0 | 0 | 0 | 0 | 0 | 0 |
| ENSG00000188785 | 246 | 111 | 172 | 79 | 135 | 117 |
| ENSG00000269533 | 2 | 0 | 2 | 1 | 2 | 2 |
| ENSG00000268533 | 0 | 0 | 0 | 0 | 0 | 0 |
| ENSG00000186272 | 104 | 62 | 64 | 47 | 53 | 56 |
| ENSG00000186230 | 128 | 94 | 92 | 68 | 86 | 91 |
| ENSG00000268163 | 0 | 0 | 0 | 0 | 0 | 1 |
| ENSG00000276449 | 4 | 2 | 5 | 6 | 1 | 0 |
| ENSG00000178201 | 6 | 7 | 6 | 3 | 6 | 5 |
| ENSG00000268545 | 0 | 1 | 0 | 0 | 0 | 0 |
| ENSG00000197128 | 306 | 154 | 221 | 83 | 161 | 151 |
| ENSG00000268266 | 0 | 0 | 0 | 0 | 0 | 0 |
| ENSG00000105136 | 205 | 102 | 136 | 54 | 98 | 82 |
| ENSG00000268107 | 0 | 0 | 0 | 0 | 0 | 0 |
| ENSG00000152439 | 66 | 36 | 25 | 18 | 36 | 26 |
| ENSG00000121406 | 121 | 52 | 76 | 17 | 53 | 46 |
| ENSG00000251369 | 250 | 126 | 152 | 53 | 127 | 96 |
| ENSG00000083817 | 175 | 78 | 75 | 67 | 80 | 72 |
| ENSG00000171649 | 0 | 0 | 0 | 0 | 0 | 0 |
| ENSG00000183647 | 91 | 32 | 57 | 25 | 37 | 29 |
| ENSG00000213762 | 684 | 310 | 374 | 194 | 272 | 264 |
| ENSG00000268392 | 0 | 1 | 0 | 0 | 0 | 0 |
| ENSG00000121417 | 344 | 150 | 220 | 132 | 184 | 158 |
| ENSG00000269097 | 0 | 0 | 0 | 0 | 0 | 0 |

| | | | | | | |
|---|---|---|---|---|---|---|
| ENSG00000180532 | 0 | 0 | 0 | 0 | 0 | 0 |
| ENSG00000204519 | 158 | 90 | 119 | 52 | 87 | 65 |
| ENSG00000269026 | 0 | 0 | 0 | 0 | 0 | 0 |
| ENSG00000179909 | 12 | 10 | 2 | 5 | 3 | 4 |
| ENSG00000083814 | 2 | 1 | 3 | 0 | 0 | 3 |
| ENSG00000152443 | 433 | 230 | 292 | 126 | 194 | 209 |
| ENSG00000083828 | 234 | 107 | 123 | 105 | 85 | 120 |
| ENSG00000178935 | 263 | 164 | 153 | 123 | 163 | 146 |
| ENSG00000268750 | 0 | 1 | 0 | 0 | 0 | 0 |
| ENSG00000269343 | 303 | 164 | 258 | 175 | 200 | 233 |
| ENSG00000269304 | 0 | 0 | 0 | 0 | 0 | 1 |
| ENSG00000204514 | 133 | 53 | 100 | 58 | 92 | 76 |
| ENSG00000198466 | 568 | 333 | 358 | 162 | 320 | 322 |
| ENSG00000269867 | 52 | 25 | 40 | 34 | 21 | 26 |
| ENSG00000270804 | 44 | 24 | 16 | 16 | 22 | 32 |
| ENSG00000269476 | 0 | 0 | 0 | 0 | 0 | 1 |
| ENSG00000173480 | 202 | 114 | 131 | 66 | 81 | 86 |
| ENSG00000196724 | 182 | 76 | 108 | 38 | 73 | 82 |
| ENSG00000152454 | 113 | 60 | 58 | 26 | 61 | 39 |
| ENSG00000243234 | 0 | 0 | 0 | 0 | 0 | 0 |
| ENSG00000177025 | 8 | 0 | 6 | 1 | 2 | 1 |
| ENSG00000166704 | 351 | 188 | 225 | 93 | 226 | 156 |
| ENSG00000176593 | 148 | 79 | 81 | 44 | 70 | 46 |
| ENSG00000235974 | 0 | 0 | 0 | 0 | 0 | 0 |
| ENSG00000152467 | 3 | 2 | 3 | 3 | 2 | 0 |
| ENSG00000268660 | 0 | 0 | 0 | 0 | 0 | 0 |
| ENSG00000270837 | 0 | 0 | 0 | 0 | 0 | 0 |
| ENSG00000176293 | 0 | 0 | 0 | 1 | 0 | 0 |
| ENSG00000243642 | 0 | 0 | 0 | 0 | 0 | 0 |
| ENSG00000121413 | 233 | 113 | 138 | 102 | 119 | 90 |
| ENSG00000181894 | 426 | 236 | 263 | 184 | 190 | 177 |
| ENSG00000268293 | 1 | 0 | 0 | 0 | 0 | 0 |
| ENSG00000269236 | 0 | 0 | 0 | 0 | 0 | 0 |
| ENSG00000171606 | 743 | 322 | 407 | 290 | 435 | 377 |
| ENSG00000213013 | 6 | 1 | 0 | 1 | 5 | 1 |
| ENSG00000268866 | 0 | 0 | 0 | 0 | 2 | 0 |
| ENSG00000198131 | 495 | 265 | 259 | 202 | 322 | 218 |
| ENSG00000283515 | 0 | 0 | 0 | 0 | 0 | 0 |
| ENSG00000268516 | 61 | 61 | 49 | 36 | 51 | 54 |
| ENSG00000268201 | 8 | 8 | 5 | 3 | 6 | 0 |
| ENSG00000278129 | 370 | 177 | 180 | 155 | 143 | 187 |
| ENSG00000267216 | 11 | 3 | 8 | 1 | 7 | 7 |
| ENSG00000142396 | 281 | 149 | 238 | 107 | 138 | 144 |
| ENSG00000283103 | 78 | 47 | 59 | 59 | 71 | 51 |
| ENSG00000269794 | 4 | 0 | 2 | 0 | 3 | 0 |
| ENSG00000182318 | 221 | 122 | 118 | 107 | 133 | 125 |
| ENSG00000277588 | 0 | 0 | 0 | 0 | 0 | 0 |
| ENSG00000121410 | 13 | 7 | 15 | 5 | 5 | 6 |
| ENSG00000268230 | 46 | 32 | 27 | 14 | 17 | 21 |
| ENSG00000268895 | 84 | 60 | 78 | 65 | 76 | 56 |

| | | | | | | |
|---|---|---|---|---|---|---|
| ENSG00000279611 | 23 | 19 | 14 | 8 | 17 | 7 |
| ENSG00000174586 | 135 | 66 | 99 | 64 | 79 | 94 |
| ENSG00000268049 | 56 | 17 | 44 | 23 | 22 | 28 |
| ENSG00000269054 | 6 | 8 | 9 | 4 | 8 | 7 |
| ENSG00000252211 | 0 | 2 | 4 | 0 | 0 | 0 |
| ENSG00000152475 | 135 | 60 | 56 | 80 | 83 | 89 |
| ENSG00000083845 | 16907 | 8260 | 9875 | 13038 | 10233 | 9717 |
| ENSG00000266640 | 4 | 0 | 1 | 0 | 0 | 0 |
| ENSG00000269855 | 0 | 0 | 4 | 0 | 4 | 1 |
| ENSG00000268307 | 0 | 3 | 4 | 4 | 2 | 3 |
| ENSG00000171574 | 257 | 146 | 181 | 136 | 140 | 153 |
| ENSG00000232098 | 369 | 187 | 233 | 197 | 207 | 254 |
| ENSG00000268543 | 18 | 12 | 4 | 1 | 4 | 10 |
| ENSG00000268912 | 45 | 10 | 20 | 6 | 8 | 15 |
| ENSG00000131849 | 66 | 33 | 45 | 14 | 41 | 45 |
| ENSG00000269473 | 58 | 21 | 19 | 16 | 17 | 16 |
| ENSG00000249471 | 181 | 87 | 95 | 95 | 78 | 116 |
| ENSG00000083812 | 438 | 226 | 265 | 285 | 231 | 247 |
| ENSG00000083838 | 279 | 177 | 235 | 223 | 175 | 196 |
| ENSG00000269106 | 0 | 0 | 0 | 0 | 0 | 0 |
| ENSG00000083807 | 218 | 116 | 138 | 175 | 134 | 151 |
| ENSG00000252334 | 0 | 1 | 2 | 1 | 0 | 0 |
| ENSG00000265272 | 0 | 0 | 0 | 0 | 0 | 0 |
| ENSG00000273901 | 7 | 9 | 17 | 8 | 6 | 9 |
| ENSG00000119574 | 617 | 302 | 377 | 401 | 366 | 358 |
| ENSG00000264910 | 0 | 0 | 0 | 0 | 0 | 0 |
| ENSG00000130726 | 13123 | 6204 | 8012 | 9073 | 7243 | 7540 |
| ENSG00000275924 | 0 | 0 | 0 | 0 | 0 | 0 |
| ENSG00000130724 | 2432 | 1265 | 1623 | 1376 | 1276 | 1269 |
| ENSG00000130725 | 2157 | 995 | 1308 | 1414 | 1230 | 1200 |
| ENSG00000267858 | 131 | 55 | 96 | 67 | 71 | 63 |
| ENSG00000099326 | 698 | 301 | 455 | 353 | 289 | 378 |
| ENSG00000213753 | 869 | 418 | 550 | 543 | 553 | 539 |
| ENSG00000285138 | 0 | 0 | 0 | 0 | 0 | 0 |
| ENSG00000269032 | 0 | 0 | 0 | 0 | 0 | 0 |
| ENSG00000269600 | 0 | 0 | 0 | 0 | 1 | 0 |
| ENSG00000251841 | 0 | 0 | 0 | 0 | 0 | 0 |
| ENSG00000184895 | 0 | 0 | 0 | 0 | 0 | 0 |
| ENSG00000237659 | 0 | 0 | 0 | 0 | 0 | 0 |
| ENSG00000232195 | 0 | 0 | 0 | 0 | 0 | 0 |
| ENSG00000129824 | 0 | 0 | 0 | 0 | 0 | 0 |
| ENSG00000227289 | 0 | 0 | 0 | 0 | 0 | 0 |
| ENSG00000229163 | 0 | 0 | 0 | 0 | 0 | 0 |
| ENSG00000278847 | 0 | 0 | 0 | 0 | 0 | 0 |
| ENSG00000067646 | 0 | 0 | 0 | 0 | 0 | 0 |
| ENSG00000233070 | 0 | 0 | 0 | 0 | 0 | 0 |
| ENSG00000223600 | 0 | 0 | 0 | 0 | 0 | 0 |
| ENSG00000231535 | 0 | 0 | 0 | 0 | 0 | 0 |
| ENSG00000234652 | 0 | 0 | 0 | 0 | 0 | 0 |
| ENSG00000216777 | 0 | 0 | 0 | 0 | 0 | 0 |

| | | | | | | |
|---|---|---|---|---|---|---|
| ENSG00000176679 | 0 | 0 | 0 | 0 | 0 | 0 |
| ENSG00000232927 | 0 | 0 | 0 | 0 | 1 | 0 |
| ENSG00000225653 | 0 | 0 | 0 | 0 | 0 | 0 |
| ENSG00000218410 | 0 | 0 | 0 | 0 | 0 | 0 |
| ENSG00000229518 | 0 | 0 | 0 | 0 | 0 | 0 |
| ENSG00000229308 | 0 | 0 | 0 | 0 | 0 | 0 |
| ENSG00000252900 | 0 | 0 | 0 | 0 | 0 | 0 |
| ENSG00000225878 | 0 | 0 | 0 | 0 | 0 | 0 |
| ENSG00000277930 | 0 | 0 | 0 | 0 | 0 | 0 |
| ENSG00000099715 | 0 | 0 | 0 | 0 | 0 | 0 |
| ENSG00000252468 | 0 | 0 | 0 | 0 | 0 | 0 |
| ENSG00000231341 | 2 | 0 | 1 | 0 | 0 | 0 |
| ENSG00000235001 | 8 | 3 | 4 | 5 | 4 | 4 |
| ENSG00000214207 | 28 | 11 | 11 | 8 | 7 | 5 |
| ENSG00000278585 | 0 | 0 | 0 | 0 | 0 | 0 |
| ENSG00000235175 | 0 | 0 | 0 | 0 | 0 | 0 |
| ENSG00000285470 | 0 | 0 | 0 | 0 | 0 | 0 |
| ENSG00000237195 | 0 | 0 | 0 | 0 | 0 | 0 |
| ENSG00000250204 | 0 | 0 | 0 | 0 | 0 | 0 |
| ENSG00000168757 | 0 | 0 | 0 | 0 | 0 | 0 |
| ENSG00000234830 | 0 | 0 | 0 | 0 | 0 | 0 |
| ENSG00000238235 | 0 | 0 | 0 | 0 | 0 | 0 |
| ENSG00000275352 | 0 | 0 | 0 | 0 | 0 | 0 |
| ENSG00000234583 | 0 | 0 | 0 | 0 | 0 | 0 |
| ENSG00000235094 | 0 | 0 | 0 | 0 | 0 | 0 |
| ENSG00000242879 | 0 | 0 | 0 | 0 | 0 | 0 |
| ENSG00000223744 | 0 | 0 | 0 | 0 | 0 | 0 |
| ENSG00000229643 | 0 | 0 | 0 | 0 | 0 | 0 |
| ENSG00000129816 | 0 | 0 | 0 | 0 | 0 | 0 |
| ENSG00000212856 | 0 | 0 | 0 | 0 | 0 | 0 |
| ENSG00000237563 | 0 | 0 | 0 | 0 | 0 | 0 |
| ENSG00000147753 | 0 | 0 | 0 | 0 | 0 | 0 |
| ENSG00000185700 | 0 | 0 | 0 | 0 | 0 | 0 |
| ENSG00000235895 | 0 | 0 | 0 | 0 | 0 | 0 |
| ENSG00000232620 | 0 | 0 | 0 | 0 | 0 | 0 |
| ENSG00000224953 | 0 | 0 | 0 | 0 | 0 | 0 |
| ENSG00000274365 | 0 | 0 | 0 | 0 | 0 | 0 |
| ENSG00000233634 | 0 | 0 | 0 | 0 | 0 | 0 |
| ENSG00000099721 | 0 | 0 | 0 | 0 | 0 | 0 |
| ENSG00000237701 | 0 | 0 | 0 | 0 | 0 | 0 |
| ENSG00000092377 | 0 | 0 | 0 | 0 | 0 | 0 |
| ENSG00000273906 | 0 | 0 | 0 | 0 | 0 | 0 |
| ENSG00000232583 | 0 | 0 | 0 | 0 | 0 | 0 |
| ENSG00000099725 | 2 | 0 | 0 | 0 | 0 | 0 |
| ENSG00000252633 | 0 | 0 | 0 | 0 | 0 | 0 |
| ENSG00000251996 | 0 | 0 | 0 | 0 | 0 | 0 |
| ENSG00000252155 | 0 | 0 | 0 | 0 | 0 | 0 |
| ENSG00000252472 | 0 | 0 | 0 | 0 | 0 | 0 |
| ENSG00000275280 | 0 | 0 | 0 | 0 | 0 | 0 |
| ENSG00000238073 | 0 | 0 | 0 | 0 | 0 | 0 |

| | | | | | | |
|---|---|---|---|---|---|---|
| ENSG00000223422 | 0 | 0 | 0 | 0 | 0 | 0 |
| ENSG00000236435 | 0 | 0 | 0 | 0 | 0 | 0 |
| ENSG00000236690 | 0 | 0 | 0 | 0 | 0 | 0 |
| ENSG00000225520 | 0 | 0 | 0 | 0 | 0 | 0 |
| ENSG00000234795 | 0 | 0 | 0 | 0 | 0 | 0 |
| ENSG00000237048 | 0 | 0 | 0 | 0 | 0 | 0 |
| ENSG00000278854 | 0 | 0 | 0 | 0 | 0 | 0 |
| ENSG00000215603 | 0 | 0 | 0 | 0 | 0 | 0 |
| ENSG00000273863 | 0 | 0 | 0 | 0 | 0 | 0 |
| ENSG00000244246 | 0 | 0 | 0 | 0 | 0 | 0 |
| ENSG00000229745 | 0 | 0 | 0 | 0 | 0 | 0 |
| ENSG00000241200 | 0 | 0 | 0 | 0 | 0 | 0 |
| ENSG00000239893 | 0 | 0 | 0 | 0 | 0 | 0 |
| ENSG00000227204 | 0 | 0 | 0 | 0 | 0 | 0 |
| ENSG00000225809 | 0 | 0 | 0 | 0 | 0 | 0 |
| ENSG00000215601 | 0 | 0 | 0 | 0 | 0 | 0 |
| ENSG00000224634 | 0 | 0 | 0 | 0 | 0 | 0 |
| ENSG00000223955 | 0 | 0 | 0 | 0 | 0 | 0 |
| ENSG00000234179 | 0 | 0 | 0 | 0 | 0 | 0 |
| ENSG00000226529 | 0 | 0 | 0 | 0 | 0 | 0 |
| ENSG00000234850 | 0 | 0 | 0 | 0 | 0 | 0 |
| ENSG00000227830 | 0 | 0 | 0 | 0 | 0 | 0 |
| ENSG00000216824 | 0 | 0 | 0 | 0 | 0 | 0 |
| ENSG00000275866 | 0 | 0 | 0 | 0 | 0 | 0 |
| ENSG00000232348 | 0 | 0 | 0 | 0 | 0 | 0 |
| ENSG00000233699 | 0 | 0 | 0 | 0 | 0 | 0 |
| ENSG00000232419 | 0 | 0 | 0 | 0 | 0 | 0 |
| ENSG00000180910 | 0 | 0 | 0 | 0 | 0 | 0 |
| ENSG00000228207 | 0 | 0 | 0 | 0 | 0 | 0 |
| ENSG00000173357 | 0 | 0 | 0 | 0 | 0 | 0 |
| ENSG00000273731 | 0 | 0 | 0 | 0 | 0 | 0 |
| ENSG00000230025 | 0 | 0 | 0 | 0 | 0 | 0 |
| ENSG00000231988 | 0 | 0 | 0 | 0 | 0 | 0 |
| ENSG00000232235 | 0 | 0 | 0 | 0 | 0 | 0 |
| ENSG00000224408 | 0 | 0 | 0 | 0 | 0 | 0 |
| ENSG00000238154 | 0 | 0 | 0 | 0 | 0 | 0 |
| ENSG00000232924 | 0 | 0 | 0 | 0 | 0 | 0 |
| ENSG00000197038 | 0 | 0 | 0 | 0 | 0 | 0 |
| ENSG00000275310 | 0 | 0 | 0 | 0 | 0 | 0 |
| ENSG00000232808 | 0 | 0 | 0 | 0 | 0 | 0 |
| ENSG00000233803 | 0 | 0 | 0 | 0 | 0 | 0 |
| ENSG00000225560 | 0 | 0 | 0 | 0 | 0 | 0 |
| ENSG00000229549 | 0 | 0 | 0 | 0 | 0 | 0 |
| ENSG00000228383 | 0 | 0 | 0 | 0 | 0 | 0 |
| ENSG00000188656 | 0 | 0 | 0 | 0 | 0 | 0 |
| ENSG00000237802 | 0 | 0 | 0 | 0 | 0 | 0 |
| ENSG00000228927 | 0 | 0 | 0 | 0 | 0 | 0 |
| ENSG00000225516 | 0 | 0 | 0 | 0 | 0 | 0 |
| ENSG00000258992 | 0 | 0 | 0 | 0 | 0 | 0 |
| ENSG00000274196 | 0 | 0 | 0 | 0 | 0 | 0 |

| | | | | | | |
|---|---|---|---|---|---|---|
| ENSG00000238074 | 0 | 0 | 0 | 0 | 0 | 0 |
| ENSG00000230066 | 0 | 0 | 0 | 0 | 0 | 0 |
| ENSG00000270073 | 0 | 0 | 0 | 0 | 0 | 0 |
| ENSG00000234803 | 0 | 0 | 0 | 0 | 0 | 0 |
| ENSG00000236424 | 0 | 0 | 0 | 0 | 0 | 0 |
| ENSG00000224336 | 0 | 0 | 0 | 0 | 0 | 0 |
| ENSG00000236786 | 0 | 0 | 0 | 0 | 0 | 0 |
| ENSG00000231436 | 0 | 0 | 0 | 0 | 0 | 0 |
| ENSG00000234110 | 0 | 0 | 0 | 0 | 0 | 0 |
| ENSG00000226223 | 0 | 0 | 0 | 0 | 0 | 0 |
| ENSG00000232617 | 0 | 0 | 0 | 0 | 0 | 0 |
| ENSG00000183385 | 0 | 0 | 0 | 0 | 0 | 0 |
| ENSG00000147761 | 0 | 0 | 0 | 0 | 0 | 0 |
| ENSG00000228890 | 0 | 0 | 0 | 0 | 0 | 0 |
| ENSG00000212855 | 0 | 0 | 0 | 0 | 0 | 0 |
| ENSG00000129845 | 0 | 0 | 0 | 0 | 0 | 0 |
| ENSG00000224075 | 0 | 0 | 0 | 0 | 0 | 0 |
| ENSG00000228379 | 0 | 0 | 0 | 0 | 0 | 0 |
| ENSG00000229208 | 0 | 0 | 0 | 0 | 0 | 0 |
| ENSG00000274445 | 0 | 0 | 0 | 0 | 0 | 0 |
| ENSG00000231874 | 0 | 0 | 0 | 0 | 0 | 0 |
| ENSG00000187657 | 0 | 0 | 0 | 0 | 0 | 0 |
| ENSG00000239225 | 0 | 0 | 0 | 0 | 0 | 0 |
| ENSG00000234950 | 0 | 0 | 0 | 0 | 0 | 0 |
| ENSG00000236718 | 0 | 0 | 0 | 0 | 0 | 0 |
| ENSG00000243643 | 0 | 0 | 0 | 0 | 0 | 0 |
| ENSG00000225685 | 0 | 0 | 0 | 0 | 0 | 0 |
| ENSG00000278478 | 0 | 0 | 0 | 0 | 0 | 0 |
| ENSG00000235193 | 0 | 0 | 0 | 0 | 0 | 0 |
| ENSG00000251766 | 0 | 0 | 0 | 0 | 0 | 0 |
| ENSG00000235691 | 0 | 0 | 0 | 0 | 0 | 0 |
| ENSG00000252289 | 0 | 0 | 0 | 0 | 0 | 0 |
| ENSG00000226975 | 0 | 0 | 0 | 0 | 0 | 0 |
| ENSG00000231411 | 0 | 0 | 0 | 0 | 0 | 0 |
| ENSG00000226061 | 0 | 0 | 0 | 0 | 0 | 0 |
| ENSG00000237447 | 0 | 0 | 0 | 0 | 0 | 0 |
| ENSG00000225840 | 0 | 0 | 0 | 0 | 0 | 0 |
| ENSG00000251705 | 0 | 0 | 0 | 0 | 0 | 0 |
| ENSG00000273858 | 0 | 0 | 0 | 0 | 0 | 0 |
| ENSG00000273693 | 0 | 0 | 0 | 0 | 0 | 0 |
| ENSG00000271365 | 0 | 0 | 0 | 0 | 0 | 0 |
| ENSG00000271309 | 0 | 0 | 0 | 0 | 0 | 0 |
| ENSG00000270570 | 0 | 0 | 0 | 0 | 0 | 0 |
| ENSG00000274231 | 0 | 0 | 0 | 0 | 0 | 0 |
| ENSG00000278212 | 0 | 0 | 0 | 0 | 3 | 0 |
| ENSG00000274502 | 0 | 0 | 0 | 0 | 0 | 0 |
| ENSG00000274162 | 0 | 0 | 0 | 0 | 0 | 0 |
| ENSG00000258567 | 0 | 0 | 0 | 0 | 0 | 0 |
| ENSG00000259154 | 0 | 0 | 0 | 0 | 0 | 0 |
| ENSG00000259029 | 0 | 0 | 0 | 0 | 0 | 0 |

| | | | | | | |
|---|---|---|---|---|---|---|
| ENSG00000258991 | 0 | 0 | 0 | 0 | 1 | 0 |
| ENSG00000270455 | 0 | 0 | 0 | 0 | 0 | 0 |
| ENSG00000183704 | 0 | 0 | 0 | 0 | 0 | 0 |
| ENSG00000270242 | 0 | 0 | 0 | 0 | 0 | 0 |
| ENSG00000275828 | 0 | 0 | 0 | 0 | 0 | 0 |
| ENSG00000225433 | 0 | 0 | 0 | 0 | 0 | 0 |
| ENSG00000224567 | 0 | 0 | 0 | 0 | 0 | 0 |
| ENSG00000271375 | 0 | 0 | 0 | 0 | 0 | 0 |
| ENSG00000234385 | 0 | 0 | 0 | 0 | 0 | 0 |
| ENSG00000215583 | 0 | 0 | 0 | 0 | 0 | 0 |
| ENSG00000235649 | 0 | 0 | 0 | 0 | 0 | 0 |
| ENSG00000236477 | 0 | 0 | 0 | 0 | 0 | 0 |
| ENSG00000232226 | 0 | 0 | 0 | 0 | 0 | 0 |
| ENSG00000243980 | 0 | 0 | 0 | 0 | 0 | 0 |
| ENSG00000232730 | 0 | 0 | 0 | 0 | 0 | 0 |
| ENSG00000224060 | 0 | 0 | 0 | 0 | 0 | 0 |
| ENSG00000225117 | 0 | 0 | 0 | 0 | 0 | 0 |
| ENSG00000206159 | 0 | 0 | 0 | 0 | 0 | 0 |
| ENSG00000227447 | 0 | 0 | 0 | 0 | 0 | 0 |
| ENSG00000226863 | 0 | 0 | 0 | 0 | 0 | 0 |
| ENSG00000233774 | 0 | 0 | 0 | 0 | 0 | 0 |
| ENSG00000233864 | 0 | 0 | 0 | 0 | 0 | 0 |
| ENSG00000114374 | 0 | 0 | 0 | 0 | 0 | 1 |
| ENSG00000228411 | 0 | 0 | 0 | 0 | 0 | 0 |
| ENSG00000067048 | 0 | 0 | 0 | 0 | 0 | 0 |
| ENSG00000234059 | 0 | 0 | 0 | 0 | 0 | 0 |
| ENSG00000224035 | 0 | 0 | 0 | 0 | 0 | 0 |
| ENSG00000235462 | 0 | 0 | 0 | 0 | 0 | 0 |
| ENSG00000223915 | 0 | 0 | 0 | 0 | 0 | 0 |
| ENSG00000183878 | 0 | 0 | 0 | 0 | 0 | 0 |
| ENSG00000215414 | 16 | 7 | 6 | 5 | 4 | 3 |
| ENSG00000154620 | 0 | 0 | 0 | 0 | 0 | 0 |
| ENSG00000241859 | 0 | 0 | 0 | 0 | 0 | 0 |
| ENSG00000129864 | 0 | 0 | 0 | 0 | 0 | 0 |
| ENSG00000129862 | 0 | 0 | 0 | 0 | 0 | 0 |
| ENSG00000235451 | 0 | 0 | 0 | 0 | 0 | 0 |
| ENSG00000223517 | 0 | 0 | 0 | 0 | 0 | 0 |
| ENSG00000165246 | 0 | 0 | 0 | 0 | 0 | 0 |
| ENSG00000226555 | 0 | 0 | 0 | 0 | 0 | 0 |
| ENSG00000228787 | 0 | 0 | 0 | 0 | 0 | 0 |
| ENSG00000236131 | 0 | 0 | 0 | 0 | 0 | 0 |
| ENSG00000227949 | 0 | 0 | 0 | 0 | 0 | 0 |
| ENSG00000224518 | 0 | 0 | 0 | 0 | 0 | 0 |
| ENSG00000234620 | 0 | 0 | 0 | 0 | 0 | 0 |
| ENSG00000227166 | 0 | 0 | 0 | 0 | 0 | 0 |
| ENSG00000252689 | 0 | 0 | 0 | 0 | 0 | 0 |
| ENSG00000252173 | 0 | 0 | 0 | 0 | 0 | 0 |
| ENSG00000252323 | 0 | 0 | 0 | 0 | 0 | 0 |
| ENSG00000228518 | 0 | 0 | 0 | 0 | 0 | 0 |
| ENSG00000271595 | 0 | 0 | 0 | 0 | 0 | 0 |

| | | | | | | |
|---|---|---|---|---|---|---|
| ENSG00000224989 | 0 | 0 | 0 | 0 | 0 | 0 |
| ENSG00000283076 | 0 | 0 | 0 | 0 | 0 | 0 |
| ENSG00000228578 | 0 | 0 | 0 | 0 | 0 | 0 |
| ENSG00000230663 | 0 | 0 | 0 | 0 | 0 | 0 |
| ENSG00000252315 | 0 | 0 | 0 | 0 | 0 | 0 |
| ENSG00000252012 | 0 | 0 | 0 | 0 | 0 | 0 |
| ENSG00000273966 | 0 | 0 | 0 | 0 | 0 | 0 |
| ENSG00000238191 | 0 | 0 | 0 | 0 | 0 | 0 |
| ENSG00000226353 | 0 | 0 | 0 | 0 | 0 | 0 |
| ENSG00000275578 | 0 | 0 | 0 | 0 | 0 | 0 |
| ENSG00000226449 | 0 | 0 | 0 | 0 | 0 | 0 |
| ENSG00000225896 | 0 | 0 | 0 | 0 | 0 | 0 |
| ENSG00000224166 | 0 | 0 | 0 | 0 | 0 | 0 |
| ENSG00000229129 | 2 | 0 | 0 | 1 | 0 | 0 |
| ENSG00000250868 | 0 | 0 | 0 | 0 | 0 | 0 |
| ENSG00000223555 | 0 | 0 | 0 | 0 | 0 | 0 |
| ENSG00000235521 | 0 | 0 | 0 | 0 | 0 | 0 |
| ENSG00000252513 | 0 | 0 | 0 | 0 | 0 | 0 |
| ENSG00000224964 | 0 | 0 | 0 | 0 | 0 | 0 |
| ENSG00000226011 | 0 | 0 | 0 | 0 | 0 | 0 |
| ENSG00000225740 | 0 | 0 | 0 | 0 | 0 | 0 |
| ENSG00000129873 | 0 | 0 | 0 | 0 | 0 | 0 |
| ENSG00000229138 | 0 | 0 | 0 | 0 | 0 | 0 |
| ENSG00000224485 | 0 | 0 | 0 | 0 | 0 | 0 |
| ENSG00000226116 | 0 | 0 | 0 | 0 | 0 | 0 |
| ENSG00000237558 | 0 | 0 | 0 | 0 | 0 | 0 |
| ENSG00000233378 | 0 | 0 | 0 | 0 | 0 | 0 |
| ENSG00000237616 | 0 | 0 | 0 | 0 | 0 | 0 |
| ENSG00000224033 | 0 | 0 | 0 | 0 | 0 | 0 |
| ENSG00000182415 | 0 | 0 | 0 | 0 | 0 | 0 |
| ENSG00000230412 | 0 | 0 | 0 | 0 | 0 | 0 |
| ENSG00000226611 | 0 | 0 | 0 | 0 | 0 | 0 |
| ENSG00000225895 | 0 | 0 | 0 | 0 | 0 | 0 |
| ENSG00000233120 | 0 | 0 | 0 | 0 | 0 | 0 |
| ENSG00000252166 | 0 | 0 | 0 | 0 | 0 | 0 |
| ENSG00000232744 | 0 | 0 | 0 | 0 | 0 | 0 |
| ENSG00000244646 | 0 | 0 | 0 | 0 | 0 | 0 |
| ENSG00000229465 | 0 | 0 | 0 | 0 | 0 | 0 |
| ENSG00000231311 | 0 | 0 | 0 | 0 | 0 | 0 |
| ENSG00000236599 | 0 | 0 | 0 | 0 | 0 | 0 |
| ENSG00000232899 | 0 | 0 | 0 | 0 | 0 | 0 |
| ENSG00000229302 | 0 | 0 | 0 | 0 | 0 | 0 |
| ENSG00000228945 | 0 | 0 | 0 | 0 | 0 | 0 |
| ENSG00000233522 | 0 | 0 | 0 | 0 | 0 | 0 |
| ENSG00000278391 | 0 | 0 | 0 | 0 | 0 | 0 |
| ENSG00000251953 | 0 | 0 | 0 | 0 | 0 | 0 |
| ENSG00000252667 | 0 | 0 | 0 | 0 | 0 | 0 |
| ENSG00000249726 | 0 | 0 | 0 | 0 | 0 | 0 |
| ENSG00000282909 | 0 | 0 | 0 | 0 | 0 | 0 |
| ENSG00000226362 | 0 | 0 | 0 | 0 | 0 | 0 |

| | | | | | | |
|---|---|---|---|---|---|---|
| ENSG00000225716 | 0 | 0 | 0 | 0 | 0 | 0 |
| ENSG00000229406 | 0 | 0 | 0 | 0 | 0 | 0 |
| ENSG00000227494 | 0 | 0 | 0 | 0 | 0 | 0 |
| ENSG00000252209 | 0 | 0 | 0 | 0 | 0 | 0 |
| ENSG00000249634 | 0 | 0 | 0 | 0 | 0 | 0 |
| ENSG00000251510 | 0 | 0 | 0 | 0 | 0 | 0 |
| ENSG00000238067 | 0 | 0 | 0 | 0 | 0 | 0 |
| ENSG00000233546 | 0 | 0 | 0 | 0 | 0 | 0 |
| ENSG00000230377 | 0 | 0 | 0 | 0 | 0 | 0 |
| ENSG00000250951 | 0 | 0 | 0 | 0 | 0 | 0 |
| ENSG00000172468 | 0 | 0 | 0 | 0 | 0 | 0 |
| ENSG00000230458 | 0 | 0 | 0 | 0 | 0 | 0 |
| ENSG00000131007 | 0 | 0 | 0 | 0 | 0 | 0 |
| ENSG00000240438 | 0 | 0 | 0 | 0 | 0 | 0 |
| ENSG00000235479 | 0 | 0 | 0 | 0 | 0 | 0 |
| ENSG00000249606 | 0 | 0 | 0 | 0 | 0 | 0 |
| ENSG00000242153 | 0 | 0 | 0 | 0 | 0 | 0 |
| ENSG00000169953 | 0 | 0 | 0 | 0 | 0 | 0 |
| ENSG00000236429 | 0 | 0 | 0 | 0 | 0 | 0 |
| ENSG00000249501 | 0 | 0 | 0 | 0 | 0 | 0 |
| ENSG00000228193 | 0 | 0 | 0 | 0 | 0 | 0 |
| ENSG00000248573 | 0 | 0 | 0 | 0 | 0 | 0 |
| ENSG00000230904 | 0 | 0 | 0 | 0 | 0 | 0 |
| ENSG00000238135 | 0 | 0 | 0 | 0 | 0 | 0 |
| ENSG00000224151 | 0 | 0 | 0 | 0 | 0 | 0 |
| ENSG00000251970 | 0 | 0 | 0 | 0 | 0 | 0 |
| ENSG00000238088 | 0 | 0 | 0 | 0 | 0 | 0 |
| ENSG00000217179 | 0 | 0 | 0 | 0 | 0 | 0 |
| ENSG00000176728 | 0 | 0 | 0 | 0 | 0 | 0 |
| ENSG00000217896 | 0 | 0 | 0 | 0 | 0 | 0 |
| ENSG00000185275 | 3 | 2 | 6 | 3 | 2 | 0 |
| ENSG00000252766 | 0 | 0 | 0 | 0 | 0 | 0 |
| ENSG00000274282 | 0 | 0 | 0 | 0 | 0 | 0 |
| ENSG00000234529 | 0 | 0 | 0 | 0 | 0 | 0 |
| ENSG00000215580 | 0 | 0 | 0 | 0 | 0 | 0 |
| ENSG00000131002 | 0 | 1 | 0 | 0 | 0 | 0 |
| ENSG00000267793 | 0 | 0 | 0 | 0 | 0 | 0 |
| ENSG00000260197 | 0 | 0 | 0 | 0 | 0 | 0 |
| ENSG00000012817 | 0 | 0 | 0 | 0 | 0 | 0 |
| ENSG00000274837 | 0 | 0 | 0 | 0 | 0 | 0 |
| ENSG00000237997 | 0 | 0 | 0 | 0 | 0 | 0 |
| ENSG00000232522 | 0 | 0 | 0 | 0 | 0 | 0 |
| ENSG00000228764 | 0 | 0 | 0 | 0 | 0 | 0 |
| ENSG00000229236 | 0 | 0 | 0 | 0 | 0 | 0 |
| ENSG00000277438 | 0 | 0 | 0 | 0 | 0 | 0 |
| ENSG00000198692 | 0 | 0 | 0 | 0 | 0 | 0 |
| ENSG00000225624 | 0 | 0 | 0 | 0 | 0 | 0 |
| ENSG00000280969 | 0 | 0 | 0 | 0 | 0 | 0 |
| ENSG00000224482 | 0 | 0 | 0 | 0 | 0 | 0 |
| ENSG00000216844 | 0 | 0 | 0 | 0 | 0 | 0 |

| | | | | | | |
|---|---|---|---|---|---|---|
| ENSG00000229551 | 0 | 0 | 0 | 0 | 0 | 0 |
| ENSG00000226504 | 0 | 0 | 0 | 0 | 0 | 0 |
| ENSG00000254488 | 0 | 0 | 0 | 0 | 0 | 0 |
| ENSG00000237427 | 0 | 0 | 0 | 0 | 0 | 0 |
| ENSG00000232634 | 0 | 0 | 0 | 0 | 0 | 0 |
| ENSG00000226918 | 0 | 0 | 0 | 0 | 0 | 0 |
| ENSG00000183146 | 0 | 0 | 0 | 0 | 0 | 0 |
| ENSG00000223637 | 0 | 0 | 0 | 0 | 0 | 0 |
| ENSG00000236615 | 0 | 0 | 0 | 0 | 0 | 0 |
| ENSG00000237269 | 0 | 0 | 0 | 0 | 0 | 0 |
| ENSG00000235719 | 0 | 0 | 0 | 0 | 0 | 0 |
| ENSG00000169849 | 0 | 0 | 0 | 0 | 0 | 0 |
| ENSG00000169811 | 0 | 0 | 0 | 0 | 0 | 0 |
| ENSG00000242393 | 0 | 0 | 0 | 0 | 0 | 0 |
| ENSG00000242875 | 0 | 0 | 0 | 0 | 0 | 0 |
| ENSG00000234414 | 0 | 0 | 0 | 0 | 0 | 0 |
| ENSG00000228257 | 0 | 0 | 0 | 0 | 0 | 0 |
| ENSG00000227989 | 0 | 0 | 0 | 0 | 0 | 0 |
| ENSG00000184991 | 0 | 0 | 0 | 0 | 0 | 0 |
| ENSG00000271123 | 0 | 0 | 0 | 0 | 0 | 0 |
| ENSG00000226042 | 0 | 0 | 0 | 0 | 0 | 0 |
| ENSG00000237023 | 0 | 0 | 0 | 0 | 0 | 0 |
| ENSG00000229416 | 0 | 0 | 0 | 0 | 0 | 0 |
| ENSG00000230029 | 0 | 0 | 0 | 0 | 0 | 0 |
| ENSG00000277519 | 0 | 0 | 0 | 0 | 0 | 0 |
| ENSG00000229725 | 0 | 0 | 0 | 0 | 0 | 0 |
| ENSG00000244395 | 0 | 0 | 0 | 0 | 0 | 0 |
| ENSG00000251618 | 0 | 0 | 0 | 0 | 0 | 0 |
| ENSG00000242389 | 0 | 0 | 0 | 0 | 0 | 0 |
| ENSG00000227444 | 0 | 0 | 0 | 0 | 0 | 0 |
| ENSG00000226092 | 0 | 0 | 0 | 0 | 0 | 0 |
| ENSG00000237968 | 0 | 0 | 0 | 0 | 0 | 0 |
| ENSG00000231159 | 0 | 0 | 0 | 0 | 0 | 0 |
| ENSG00000232475 | 0 | 0 | 0 | 0 | 0 | 0 |
| ENSG00000229553 | 0 | 0 | 0 | 0 | 0 | 0 |
| ENSG00000228850 | 0 | 0 | 0 | 0 | 0 | 0 |
| ENSG00000232029 | 0 | 0 | 0 | 0 | 0 | 0 |
| ENSG00000169807 | 0 | 0 | 0 | 0 | 0 | 0 |
| ENSG00000236951 | 0 | 0 | 0 | 0 | 0 | 0 |
| ENSG00000131548 | 0 | 0 | 0 | 0 | 0 | 0 |
| ENSG00000169800 | 0 | 0 | 0 | 0 | 0 | 0 |
| ENSG00000229159 | 0 | 0 | 0 | 0 | 0 | 0 |
| ENSG00000225615 | 0 | 0 | 0 | 0 | 0 | 0 |
| ENSG00000229234 | 0 | 0 | 0 | 0 | 0 | 0 |
| ENSG00000215560 | 0 | 0 | 0 | 0 | 0 | 0 |
| ENSG00000229940 | 0 | 0 | 0 | 0 | 0 | 0 |
| ENSG00000243040 | 0 | 0 | 0 | 0 | 0 | 0 |
| ENSG00000259247 | 0 | 0 | 0 | 0 | 0 | 0 |
| ENSG00000237902 | 0 | 0 | 0 | 0 | 0 | 0 |
| ENSG00000226941 | 0 | 0 | 0 | 0 | 0 | 0 |

| | | | | | | |
|---|---|---|---|---|---|---|
| ENSG00000235059 | 0 | 0 | 0 | 0 | 0 | 0 |
| ENSG00000131538 | 0 | 0 | 0 | 0 | 0 | 0 |
| ENSG00000169789 | 0 | 0 | 0 | 0 | 0 | 0 |
| ENSG00000234131 | 0 | 0 | 0 | 0 | 0 | 0 |
| ENSG00000224873 | 0 | 0 | 0 | 0 | 0 | 0 |
| ENSG00000230814 | 0 | 0 | 0 | 0 | 0 | 0 |
| ENSG00000228571 | 0 | 0 | 0 | 0 | 0 | 0 |
| ENSG00000230476 | 0 | 0 | 0 | 0 | 0 | 0 |
| ENSG00000273589 | 0 | 0 | 0 | 0 | 0 | 0 |
| ENSG00000224657 | 0 | 0 | 0 | 0 | 0 | 0 |
| ENSG00000224917 | 0 | 0 | 0 | 0 | 0 | 0 |
| ENSG00000230727 | 0 | 0 | 0 | 0 | 0 | 0 |
| ENSG00000228240 | 0 | 0 | 0 | 0 | 0 | 0 |
| ENSG00000276829 | 0 | 0 | 0 | 0 | 0 | 0 |
| ENSG00000215537 | 0 | 0 | 0 | 0 | 0 | 0 |
| ENSG00000226906 | 0 | 0 | 0 | 0 | 0 | 0 |
| ENSG00000183753 | 0 | 0 | 0 | 0 | 0 | 0 |
| ENSG00000244000 | 0 | 0 | 0 | 0 | 0 | 0 |
| ENSG00000232764 | 0 | 0 | 0 | 0 | 0 | 0 |
| ENSG00000233126 | 0 | 0 | 0 | 0 | 0 | 0 |
| ENSG00000227251 | 0 | 0 | 0 | 0 | 0 | 0 |
| ENSG00000188120 | 0 | 0 | 0 | 0 | 0 | 0 |
| ENSG00000205944 | 0 | 0 | 0 | 0 | 0 | 0 |
| ENSG00000205936 | 0 | 0 | 0 | 0 | 0 | 0 |
| ENSG00000251925 | 0 | 0 | 0 | 0 | 0 | 0 |
| ENSG00000225189 | 0 | 0 | 0 | 0 | 0 | 0 |
| ENSG00000230073 | 0 | 0 | 0 | 0 | 0 | 0 |
| ENSG00000215540 | 0 | 0 | 0 | 0 | 0 | 0 |
| ENSG00000236647 | 0 | 0 | 0 | 0 | 0 | 0 |
| ENSG00000225466 | 0 | 0 | 0 | 0 | 0 | 0 |
| ENSG00000224169 | 0 | 0 | 0 | 0 | 0 | 0 |
| ENSG00000223407 | 0 | 0 | 0 | 0 | 0 | 0 |
| ENSG00000226873 | 0 | 0 | 0 | 0 | 0 | 0 |
| ENSG00000227915 | 0 | 0 | 0 | 0 | 0 | 0 |
| ENSG00000169763 | 0 | 0 | 0 | 0 | 0 | 0 |
| ENSG00000236620 | 0 | 0 | 0 | 0 | 0 | 0 |
| ENSG00000224866 | 0 | 0 | 0 | 0 | 0 | 0 |
| ENSG00000232424 | 0 | 0 | 0 | 0 | 0 | 0 |
| ENSG00000252681 | 0 | 0 | 0 | 0 | 0 | 0 |
| ENSG00000232910 | 0 | 0 | 0 | 0 | 0 | 0 |
| ENSG00000232845 | 0 | 0 | 0 | 0 | 0 | 0 |
| ENSG00000237302 | 0 | 0 | 0 | 0 | 0 | 0 |
| ENSG00000231540 | 0 | 0 | 0 | 0 | 0 | 0 |
| ENSG00000223362 | 0 | 0 | 0 | 0 | 0 | 0 |
| ENSG00000231026 | 0 | 0 | 0 | 0 | 0 | 0 |
| ENSG00000224571 | 0 | 0 | 0 | 0 | 0 | 0 |
| ENSG00000226369 | 0 | 0 | 0 | 0 | 0 | 0 |
| ENSG00000280961 | 0 | 0 | 0 | 0 | 0 | 0 |
| ENSG00000227871 | 0 | 0 | 0 | 0 | 0 | 0 |
| ENSG00000251917 | 0 | 0 | 0 | 0 | 0 | 0 |

| | | | | | | |
|---|---|---|---|---|---|---|
| ENSG00000231423 | 0 | 0 | 0 | 0 | 0 | 0 |
| ENSG00000228465 | 0 | 0 | 0 | 0 | 0 | 0 |
| ENSG00000232585 | 0 | 0 | 0 | 0 | 0 | 0 |
| ENSG00000234081 | 0 | 0 | 0 | 0 | 0 | 0 |
| ENSG00000172352 | 0 | 0 | 0 | 0 | 0 | 0 |
| ENSG00000231375 | 0 | 0 | 0 | 0 | 0 | 0 |
| ENSG00000237467 | 0 | 0 | 0 | 0 | 0 | 0 |
| ENSG00000229250 | 0 | 0 | 0 | 0 | 0 | 0 |
| ENSG00000232205 | 0 | 0 | 0 | 0 | 0 | 0 |
| ENSG00000235981 | 0 | 0 | 0 | 0 | 0 | 0 |
| ENSG00000223698 | 0 | 0 | 0 | 0 | 0 | 0 |
| ENSG00000242854 | 0 | 0 | 0 | 0 | 0 | 0 |
| ENSG00000230977 | 0 | 0 | 0 | 0 | 0 | 0 |
| ENSG00000244231 | 0 | 0 | 0 | 0 | 0 | 0 |
| ENSG00000172342 | 0 | 0 | 0 | 0 | 0 | 0 |
| ENSG00000284071 | 0 | 0 | 0 | 0 | 0 | 0 |
| ENSG00000239533 | 0 | 0 | 0 | 0 | 0 | 0 |
| ENSG00000274234 | 0 | 0 | 0 | 0 | 0 | 0 |
| ENSG00000225491 | 0 | 0 | 0 | 0 | 0 | 0 |
| ENSG00000224827 | 0 | 0 | 0 | 0 | 0 | 0 |
| ENSG00000233740 | 0 | 0 | 0 | 0 | 0 | 0 |
| ENSG00000248792 | 0 | 0 | 0 | 0 | 0 | 0 |
| ENSG00000279274 | 0 | 0 | 0 | 0 | 0 | 0 |
| ENSG00000234399 | 0 | 0 | 0 | 0 | 0 | 0 |
| ENSG00000227439 | 0 | 0 | 0 | 0 | 0 | 0 |
| ENSG00000224210 | 0 | 0 | 0 | 0 | 0 | 0 |
| ENSG00000232003 | 0 | 0 | 0 | 0 | 0 | 0 |
| ENSG00000235412 | 0 | 0 | 0 | 0 | 0 | 0 |
| ENSG00000183795 | 0 | 0 | 0 | 0 | 0 | 0 |
| ENSG00000240566 | 0 | 0 | 0 | 0 | 0 | 0 |
| ENSG00000232976 | 0 | 0 | 0 | 0 | 0 | 0 |
| ENSG00000226270 | 0 | 0 | 0 | 0 | 0 | 0 |
| ENSG00000227837 | 0 | 0 | 0 | 0 | 0 | 0 |
| ENSG00000187191 | 0 | 0 | 0 | 0 | 0 | 0 |
| ENSG00000205916 | 0 | 0 | 0 | 0 | 0 | 0 |
| ENSG00000278602 | 0 | 0 | 0 | 0 | 0 | 0 |
| ENSG00000223978 | 0 | 0 | 0 | 0 | 0 | 0 |
| ENSG00000278197 | 0 | 0 | 0 | 0 | 0 | 0 |
| ENSG00000230819 | 0 | 0 | 0 | 0 | 0 | 0 |
| ENSG00000185894 | 0 | 0 | 0 | 0 | 0 | 0 |
| ENSG00000228296 | 0 | 0 | 0 | 0 | 0 | 0 |
| ENSG00000236379 | 0 | 0 | 0 | 0 | 0 | 0 |
| ENSG00000274899 | 0 | 0 | 0 | 0 | 0 | 0 |
| ENSG00000223641 | 0 | 0 | 0 | 0 | 0 | 0 |
| ENSG00000227633 | 0 | 0 | 0 | 0 | 0 | 0 |
| ENSG00000279115 | 0 | 0 | 0 | 0 | 0 | 0 |
| ENSG00000228786 | 0 | 0 | 0 | 0 | 0 | 0 |
| ENSG00000233652 | 0 | 0 | 0 | 0 | 0 | 0 |
| ENSG00000233944 | 0 | 0 | 0 | 0 | 0 | 0 |
| ENSG00000223636 | 0 | 0 | 0 | 0 | 0 | 0 |

| | | | | | | |
|---|---|---|---|---|---|---|
| ENSG00000172297 | 0 | 0 | 0 | 0 | 0 | 0 |
| ENSG00000278242 | 0 | 0 | 0 | 0 | 0 | 0 |
| ENSG00000284380 | 0 | 0 | 0 | 0 | 0 | 0 |
| ENSG00000172294 | 0 | 0 | 0 | 0 | 0 | 0 |
| ENSG00000240450 | 0 | 0 | 0 | 0 | 0 | 0 |
| ENSG00000239304 | 0 | 0 | 0 | 0 | 0 | 0 |
| ENSG00000233619 | 0 | 0 | 0 | 0 | 0 | 0 |
| ENSG00000197092 | 0 | 0 | 0 | 0 | 0 | 0 |
| ENSG00000235583 | 2 | 0 | 0 | 0 | 0 | 0 |
| ENSG00000237823 | 0 | 0 | 0 | 0 | 0 | 0 |
| ENSG00000229709 | 0 | 0 | 0 | 0 | 0 | 0 |
| ENSG00000232064 | 0 | 0 | 0 | 0 | 0 | 0 |
| ENSG00000225609 | 0 | 0 | 0 | 0 | 0 | 0 |
| ENSG00000172288 | 0 | 0 | 0 | 0 | 0 | 0 |
| ENSG00000270535 | 0 | 0 | 0 | 0 | 0 | 0 |
| ENSG00000225287 | 0 | 0 | 0 | 0 | 0 | 0 |
| ENSG00000225256 | 0 | 0 | 0 | 0 | 0 | 0 |
| ENSG00000223856 | 0 | 0 | 0 | 0 | 0 | 0 |
| ENSG00000235004 | 0 | 0 | 0 | 0 | 0 | 0 |
| ENSG00000252426 | 0 | 0 | 0 | 0 | 0 | 0 |
| ENSG00000231141 | 0 | 0 | 0 | 0 | 0 | 0 |
| ENSG00000232614 | 0 | 0 | 0 | 0 | 0 | 0 |
| ENSG00000225326 | 0 | 0 | 0 | 0 | 0 | 0 |
| ENSG00000223406 | 0 | 0 | 0 | 0 | 0 | 0 |
| ENSG00000229343 | 0 | 0 | 0 | 0 | 0 | 0 |
| ENSG00000232695 | 0 | 0 | 0 | 0 | 0 | 0 |
| ENSG00000235511 | 0 | 0 | 0 | 0 | 0 | 0 |
| ENSG00000232914 | 0 | 0 | 0 | 0 | 0 | 0 |
| ENSG00000223655 | 0 | 0 | 0 | 0 | 0 | 0 |
| ENSG00000230854 | 0 | 0 | 0 | 0 | 0 | 0 |
| ENSG00000252625 | 0 | 0 | 0 | 0 | 0 | 0 |
| ENSG00000227635 | 0 | 0 | 0 | 0 | 0 | 0 |
| ENSG00000237546 | 0 | 0 | 0 | 0 | 0 | 0 |
| ENSG00000172283 | 0 | 0 | 0 | 0 | 0 | 0 |
| ENSG00000227867 | 0 | 0 | 0 | 0 | 0 | 0 |
| ENSG00000231716 | 0 | 0 | 0 | 0 | 0 | 0 |
| ENSG00000234744 | 0 | 0 | 0 | 0 | 0 | 0 |
| ENSG00000233156 | 0 | 0 | 0 | 0 | 0 | 0 |
| ENSG00000234888 | 0 | 0 | 0 | 0 | 0 | 0 |
| ENSG00000215507 | 0 | 0 | 0 | 0 | 0 | 0 |
| ENSG00000235014 | 0 | 0 | 0 | 0 | 0 | 0 |
| ENSG00000225876 | 0 | 0 | 0 | 0 | 0 | 0 |
| ENSG00000251796 | 0 | 0 | 0 | 0 | 0 | 0 |
| ENSG00000229238 | 2 | 0 | 0 | 0 | 0 | 0 |
| ENSG00000252948 | 0 | 0 | 0 | 0 | 0 | 0 |
| ENSG00000233843 | 0 | 0 | 0 | 0 | 0 | 0 |
| ENSG00000188399 | 0 | 0 | 0 | 0 | 0 | 0 |
| ENSG00000277146 | 0 | 0 | 0 | 0 | 0 | 0 |
| ENSG00000215506 | 0 | 0 | 0 | 0 | 0 | 0 |
| ENSG00000224240 | 0 | 0 | 0 | 0 | 0 | 0 |

| | | | | | | |
|---|---|---|---|---|---|---|
| ENSG00000227629 | 0 | 0 | 0 | 0 | 0 | 0 |
| ENSG00000237917 | 2 | 1 | 1 | 0 | 2 | 0 |
| ENSG00000231514 | 0 | 0 | 0 | 0 | 0 | 0 |
| ENSG00000235857 | 0 | 0 | 0 | 0 | 0 | 0 |
| ENSG00000277248 | 0 | 0 | 0 | 0 | 0 | 0 |
| ENSG00000283047 | 6 | 6 | 6 | 3 | 7 | 2 |
| ENSG00000279973 | 1 | 0 | 0 | 0 | 0 | 0 |
| ENSG00000226444 | 0 | 0 | 0 | 0 | 0 | 0 |
| ENSG00000276871 | 0 | 0 | 0 | 0 | 0 | 0 |
| ENSG00000283023 | 22 | 14 | 16 | 8 | 5 | 11 |
| ENSG00000276138 | 0 | 0 | 0 | 0 | 0 | 0 |
| ENSG00000280341 | 0 | 0 | 0 | 0 | 0 | 0 |
| ENSG00000236235 | 0 | 0 | 0 | 0 | 0 | 0 |
| ENSG00000279442 | 0 | 0 | 0 | 0 | 0 | 0 |
| ENSG00000184624 | 0 | 0 | 0 | 0 | 0 | 0 |
| ENSG00000236097 | 0 | 0 | 0 | 0 | 0 | 0 |
| ENSG00000236831 | 0 | 0 | 0 | 0 | 0 | 0 |
| ENSG00000130538 | 0 | 0 | 0 | 0 | 0 | 0 |
| ENSG00000237299 | 0 | 0 | 0 | 0 | 0 | 0 |
| ENSG00000235759 | 0 | 0 | 0 | 0 | 0 | 0 |
| ENSG00000233408 | 0 | 0 | 0 | 0 | 0 | 0 |
| ENSG00000215268 | 0 | 0 | 0 | 0 | 0 | 0 |
| ENSG00000213727 | 0 | 0 | 0 | 0 | 0 | 0 |
| ENSG00000230643 | 0 | 0 | 0 | 0 | 0 | 0 |
| ENSG00000230471 | 0 | 0 | 0 | 0 | 0 | 0 |
| ENSG00000231565 | 0 | 0 | 0 | 0 | 0 | 0 |
| ENSG00000226474 | 0 | 0 | 0 | 0 | 0 | 0 |
| ENSG00000224435 | 0 | 0 | 0 | 0 | 0 | 0 |
| ENSG00000234381 | 0 | 0 | 0 | 0 | 0 | 0 |
| ENSG00000198062 | 0 | 0 | 0 | 0 | 0 | 0 |
| ENSG00000236666 | 0 | 0 | 0 | 0 | 0 | 0 |
| ENSG00000212216 | 0 | 0 | 0 | 0 | 0 | 0 |
| ENSG00000241838 | 0 | 0 | 0 | 0 | 0 | 0 |
| ENSG00000225255 | 2 | 0 | 1 | 0 | 0 | 0 |
| ENSG00000235992 | 0 | 0 | 0 | 0 | 0 | 0 |
| ENSG00000206195 | 428 | 238 | 297 | 176 | 260 | 243 |
| ENSG00000271127 | 2 | 6 | 2 | 2 | 1 | 6 |
| ENSG00000232775 | 16 | 14 | 13 | 8 | 8 | 8 |
| ENSG00000272872 | 72 | 33 | 37 | 47 | 40 | 27 |
| ENSG00000271672 | 0 | 0 | 0 | 0 | 0 | 0 |
| ENSG00000223875 | 0 | 0 | 0 | 0 | 0 | 0 |
| ENSG00000215270 | 4 | 4 | 7 | 2 | 7 | 0 |
| ENSG00000229286 | 0 | 0 | 0 | 0 | 0 | 0 |
| ENSG00000233866 | 0 | 0 | 0 | 0 | 0 | 0 |
| ENSG00000278188 | 0 | 0 | 0 | 0 | 0 | 0 |
| ENSG00000225293 | 0 | 0 | 0 | 0 | 0 | 0 |
| ENSG00000226160 | 0 | 0 | 0 | 0 | 0 | 0 |
| ENSG00000229658 | 0 | 0 | 0 | 0 | 0 | 0 |
| ENSG00000227367 | 2 | 0 | 0 | 0 | 0 | 1 |
| ENSG00000280263 | 0 | 0 | 0 | 0 | 0 | 0 |

| | | | | | | |
|---|---|---|---|---|---|---|
| ENSG00000235969 | 0 | 0 | 0 | 0 | 0 | 0 |
| ENSG00000273362 | 0 | 0 | 0 | 0 | 0 | 0 |
| ENSG00000233995 | 0 | 0 | 0 | 0 | 0 | 0 |
| ENSG00000239435 | 0 | 0 | 1 | 0 | 0 | 0 |
| ENSG00000198445 | 0 | 0 | 0 | 0 | 1 | 0 |
| ENSG00000240122 | 0 | 0 | 1 | 0 | 0 | 0 |
| ENSG00000100181 | 20 | 9 | 12 | 1 | 4 | 4 |
| ENSG00000283633 | 0 | 0 | 0 | 0 | 0 | 1 |
| ENSG00000267338 | 0 | 0 | 0 | 0 | 0 | 0 |
| ENSG00000249288 | 1 | 0 | 1 | 1 | 1 | 2 |
| ENSG00000249263 | 0 | 3 | 0 | 0 | 0 | 1 |
| ENSG00000189295 | 2 | 3 | 4 | 1 | 3 | 2 |
| ENSG00000259271 | 8 | 1 | 6 | 1 | 4 | 4 |
| ENSG00000273244 | 0 | 0 | 0 | 0 | 0 | 0 |
| ENSG00000241717 | 8 | 4 | 6 | 1 | 1 | 0 |
| ENSG00000235343 | 10 | 6 | 5 | 1 | 0 | 3 |
| ENSG00000172967 | 0 | 1 | 0 | 0 | 0 | 0 |
| ENSG00000229027 | 0 | 0 | 0 | 0 | 0 | 0 |
| ENSG00000234121 | 0 | 0 | 0 | 0 | 0 | 0 |
| ENSG00000283535 | 0 | 0 | 0 | 0 | 0 | 0 |
| ENSG00000237835 | 0 | 0 | 0 | 0 | 0 | 0 |
| ENSG00000220248 | 0 | 0 | 0 | 0 | 0 | 0 |
| ENSG00000237689 | 0 | 0 | 0 | 0 | 0 | 0 |
| ENSG00000228122 | 0 | 0 | 0 | 0 | 0 | 0 |
| ENSG00000237358 | 0 | 0 | 0 | 0 | 0 | 0 |
| ENSG00000225480 | 0 | 0 | 0 | 0 | 0 | 0 |
| ENSG00000230481 | 2 | 0 | 0 | 0 | 0 | 0 |
| ENSG00000253691 | 0 | 0 | 0 | 0 | 0 | 0 |
| ENSG00000253460 | 0 | 0 | 0 | 0 | 0 | 0 |
| ENSG00000254264 | 0 | 0 | 0 | 0 | 0 | 0 |
| ENSG00000253481 | 0 | 0 | 0 | 0 | 0 | 0 |
| ENSG00000285045 | 0 | 0 | 0 | 0 | 0 | 0 |
| ENSG00000215568 | 0 | 0 | 0 | 0 | 0 | 0 |
| ENSG00000270226 | 0 | 0 | 0 | 0 | 0 | 0 |
| ENSG00000276400 | 0 | 0 | 0 | 0 | 0 | 1 |
| ENSG00000237438 | 2 | 0 | 0 | 1 | 0 | 0 |
| ENSG00000273203 | 0 | 0 | 0 | 0 | 0 | 0 |
| ENSG00000273442 | 0 | 0 | 0 | 0 | 0 | 0 |
| ENSG00000177663 | 1203 | 572 | 678 | 640 | 631 | 614 |
| ENSG00000235328 | 2 | 1 | 2 | 0 | 0 | 1 |
| ENSG00000183307 | 4 | 3 | 7 | 9 | 4 | 4 |
| ENSG00000235478 | 2 | 0 | 1 | 0 | 0 | 0 |
| ENSG00000069998 | 1653 | 744 | 983 | 1087 | 938 | 1070 |
| ENSG00000185837 | 13 | 4 | 15 | 12 | 11 | 5 |
| ENSG00000093072 | 17 | 20 | 10 | 1 | 12 | 12 |
| ENSG00000236325 | 0 | 2 | 0 | 2 | 2 | 3 |
| ENSG00000203416 | 2 | 2 | 2 | 1 | 5 | 1 |
| ENSG00000241832 | 0 | 0 | 0 | 0 | 0 | 0 |
| ENSG00000231004 | 0 | 0 | 0 | 0 | 0 | 0 |
| ENSG00000239547 | 0 | 0 | 0 | 0 | 0 | 0 |

| | | | | | | |
|---|---|---|---|---|---|---|
| ENSG00000099954 | 18 | 18 | 14 | 10 | 10 | 10 |
| ENSG00000229492 | 0 | 0 | 0 | 0 | 0 | 0 |
| ENSG00000232305 | 0 | 0 | 0 | 0 | 0 | 0 |
| ENSG00000228045 | 0 | 0 | 0 | 0 | 0 | 0 |
| ENSG00000182902 | 35 | 24 | 23 | 20 | 21 | 16 |
| ENSG00000236754 | 0 | 2 | 0 | 1 | 1 | 1 |
| ENSG00000131100 | 4540 | 2243 | 2630 | 2171 | 2572 | 2308 |
| ENSG00000099968 | 2629 | 1296 | 1659 | 1218 | 1468 | 1572 |
| ENSG00000015475 | 1920 | 1060 | 1182 | 1347 | 1289 | 1262 |
| ENSG00000264757 | 0 | 0 | 1 | 0 | 0 | 0 |
| ENSG00000269220 | 0 | 0 | 1 | 2 | 1 | 0 |
| ENSG00000243156 | 480 | 205 | 304 | 192 | 267 | 272 |
| ENSG00000093100 | 0 | 0 | 2 | 2 | 0 | 1 |
| ENSG00000207780 | 0 | 0 | 0 | 0 | 0 | 0 |
| ENSG00000235445 | 0 | 0 | 0 | 0 | 0 | 0 |
| ENSG00000234913 | 0 | 0 | 0 | 0 | 0 | 0 |
| ENSG00000237446 | 0 | 0 | 0 | 0 | 0 | 1 |
| ENSG00000235295 | 0 | 0 | 0 | 0 | 0 | 0 |
| ENSG00000225335 | 63 | 30 | 35 | 33 | 22 | 22 |
| ENSG00000215193 | 1888 | 1001 | 1144 | 1036 | 1062 | 1115 |
| ENSG00000236890 | 2 | 0 | 1 | 1 | 3 | 0 |
| ENSG00000235617 | 0 | 0 | 0 | 0 | 0 | 0 |
| ENSG00000225225 | 0 | 0 | 0 | 0 | 0 | 0 |
| ENSG00000183785 | 4 | 4 | 3 | 2 | 1 | 0 |
| ENSG00000280007 | 18 | 7 | 13 | 3 | 3 | 9 |
| ENSG00000184979 | 86 | 44 | 53 | 25 | 43 | 36 |
| ENSG00000187979 | 0 | 0 | 0 | 0 | 0 | 0 |
| ENSG00000276160 | 0 | 0 | 0 | 0 | 0 | 0 |
| ENSG00000274044 | 0 | 0 | 0 | 0 | 0 | 0 |
| ENSG00000276874 | 0 | 0 | 0 | 0 | 0 | 0 |
| ENSG00000275060 | 0 | 0 | 0 | 0 | 0 | 0 |
| ENSG00000277870 | 0 | 0 | 0 | 0 | 0 | 0 |
| ENSG00000277017 | 0 | 0 | 0 | 0 | 0 | 0 |
| ENSG00000278044 | 0 | 0 | 0 | 0 | 0 | 0 |
| ENSG00000274252 | 1 | 0 | 0 | 0 | 0 | 0 |
| ENSG00000275319 | 0 | 0 | 0 | 0 | 0 | 0 |
| ENSG00000277690 | 0 | 1 | 0 | 0 | 0 | 0 |
| ENSG00000278558 | 110 | 48 | 62 | 48 | 38 | 39 |
| ENSG00000274602 | 236 | 105 | 148 | 95 | 118 | 116 |
| ENSG00000274625 | 0 | 0 | 0 | 0 | 0 | 0 |
| ENSG00000222630 | 0 | 0 | 0 | 0 | 0 | 0 |
| ENSG00000275793 | 68 | 22 | 44 | 19 | 15 | 19 |
| ENSG00000275362 | 0 | 0 | 0 | 0 | 0 | 0 |
| ENSG00000278110 | 0 | 0 | 0 | 0 | 0 | 0 |
| ENSG00000273907 | 0 | 0 | 0 | 0 | 0 | 0 |
| ENSG00000273643 | 0 | 0 | 0 | 0 | 0 | 0 |
| ENSG00000278008 | 0 | 2 | 0 | 0 | 0 | 0 |
| ENSG00000182824 | 0 | 0 | 0 | 0 | 0 | 0 |
| ENSG00000197421 | 1 | 0 | 0 | 0 | 0 | 0 |
| ENSG00000239989 | 0 | 0 | 0 | 0 | 0 | 0 |

| | | | | | | |
|---|---|---|---|---|---|---|
| ENSG00000234764 | 0 | 0 | 0 | 0 | 0 | 0 |
| ENSG00000161103 | 0 | 0 | 0 | 1 | 0 | 0 |
| ENSG00000215544 | 0 | 0 | 0 | 0 | 1 | 0 |
| ENSG00000276095 | 0 | 0 | 0 | 0 | 0 | 0 |
| ENSG00000279987 | 0 | 0 | 0 | 0 | 0 | 0 |
| ENSG00000183628 | 10 | 1 | 1 | 0 | 2 | 4 |
| ENSG00000283809 | 0 | 0 | 0 | 0 | 0 | 0 |
| ENSG00000100033 | 0 | 0 | 2 | 0 | 0 | 0 |
| ENSG00000284294 | 0 | 0 | 0 | 0 | 0 | 0 |
| ENSG00000280418 | 0 | 0 | 0 | 0 | 0 | 0 |
| ENSG00000237517 | 23 | 23 | 16 | 11 | 17 | 12 |
| ENSG00000273032 | 54 | 33 | 38 | 24 | 32 | 34 |
| ENSG00000283583 | 0 | 0 | 0 | 0 | 0 | 0 |
| ENSG00000271275 | 0 | 0 | 0 | 0 | 0 | 0 |
| ENSG00000283366 | 0 | 0 | 0 | 0 | 0 | 0 |
| ENSG00000270393 | 0 | 0 | 0 | 0 | 0 | 0 |
| ENSG00000273164 | 5 | 3 | 4 | 3 | 3 | 0 |
| ENSG00000241527 | 8 | 13 | 7 | 2 | 1 | 2 |
| ENSG00000070413 | 3619 | 1659 | 2326 | 1984 | 1729 | 2023 |
| ENSG00000206739 | 2 | 2 | 1 | 0 | 3 | 0 |
| ENSG00000273311 | 78 | 37 | 53 | 31 | 35 | 40 |
| ENSG00000237347 | 2 | 1 | 0 | 1 | 0 | 0 |
| ENSG00000281530 | 5 | 2 | 7 | 2 | 6 | 3 |
| ENSG00000223461 | 3 | 4 | 2 | 2 | 3 | 8 |
| ENSG00000272682 | 0 | 2 | 2 | 0 | 2 | 1 |
| ENSG00000231086 | 0 | 0 | 0 | 0 | 0 | 0 |
| ENSG00000100056 | 821 | 346 | 461 | 529 | 461 | 427 |
| ENSG00000206203 | 0 | 0 | 0 | 0 | 0 | 0 |
| ENSG00000063515 | 0 | 0 | 1 | 0 | 0 | 0 |
| ENSG00000260924 | 60 | 26 | 37 | 33 | 42 | 31 |
| ENSG00000100075 | 5697 | 2897 | 3320 | 3483 | 2885 | 2650 |
| ENSG00000070371 | 397 | 170 | 220 | 121 | 155 | 113 |
| ENSG00000230194 | 0 | 0 | 0 | 0 | 0 | 0 |
| ENSG00000251940 | 0 | 0 | 0 | 0 | 0 | 0 |
| ENSG00000233471 | 0 | 0 | 0 | 1 | 0 | 0 |
| ENSG00000100084 | 306 | 218 | 228 | 228 | 195 | 241 |
| ENSG00000242259 | 825 | 403 | 555 | 467 | 523 | 501 |
| ENSG00000244296 | 0 | 0 | 0 | 0 | 0 | 0 |
| ENSG00000185608 | 817 | 389 | 558 | 569 | 475 | 458 |
| ENSG00000185065 | 28 | 15 | 29 | 21 | 14 | 21 |
| ENSG00000070010 | 2277 | 1279 | 1593 | 1433 | 1499 | 1468 |
| ENSG00000273300 | 0 | 0 | 0 | 0 | 0 | 0 |
| ENSG00000273212 | 7 | 0 | 2 | 2 | 3 | 3 |
| ENSG00000093009 | 414 | 215 | 351 | 202 | 208 | 265 |
| ENSG00000184113 | 0 | 0 | 2 | 2 | 2 | 2 |
| ENSG00000281548 | 0 | 0 | 1 | 0 | 0 | 0 |
| ENSG00000230485 | 0 | 0 | 0 | 0 | 0 | 0 |
| ENSG00000225007 | 0 | 0 | 0 | 0 | 0 | 0 |
| ENSG00000184702 | 7 | 1 | 16 | 10 | 6 | 7 |
| ENSG00000284874 | 3 | 2 | 0 | 4 | 1 | 3 |

| | | | | | | |
|---|---|---|---|---|---|---|
| ENSG00000203618 | 0 | 0 | 0 | 0 | 0 | 0 |
| ENSG00000184058 | 8 | 11 | 5 | 5 | 4 | 7 |
| ENSG00000185838 | 202 | 108 | 102 | 150 | 105 | 120 |
| ENSG00000235776 | 5 | 2 | 2 | 3 | 5 | 7 |
| ENSG00000215012 | 1079 | 481 | 566 | 520 | 573 | 567 |
| ENSG00000184470 | 1108 | 516 | 710 | 729 | 543 | 547 |
| ENSG00000232926 | 14 | 6 | 9 | 3 | 11 | 10 |
| ENSG00000093010 | 5022 | 2727 | 3233 | 2664 | 2404 | 2563 |
| ENSG00000284031 | 0 | 1 | 0 | 1 | 0 | 0 |
| ENSG00000099889 | 751 | 335 | 426 | 431 | 384 | 374 |
| ENSG00000183597 | 695 | 460 | 508 | 491 | 495 | 490 |
| ENSG00000208023 | 0 | 0 | 0 | 0 | 0 | 0 |
| ENSG00000236540 | 79 | 47 | 77 | 61 | 64 | 72 |
| ENSG00000268292 | 0 | 0 | 0 | 0 | 0 | 0 |
| ENSG00000277950 | 0 | 0 | 0 | 0 | 0 | 0 |
| ENSG00000128191 | 1392 | 622 | 859 | 659 | 737 | 835 |
| ENSG00000284140 | 0 | 0 | 0 | 0 | 0 | 0 |
| ENSG00000284464 | 0 | 0 | 0 | 0 | 0 | 0 |
| ENSG00000243762 | 0 | 0 | 0 | 0 | 0 | 0 |
| ENSG00000099899 | 1336 | 583 | 767 | 828 | 687 | 707 |
| ENSG00000278278 | 0 | 0 | 0 | 0 | 0 | 0 |
| ENSG00000099901 | 2328 | 1264 | 1434 | 1598 | 1514 | 1500 |
| ENSG00000264346 | 0 | 0 | 0 | 0 | 0 | 0 |
| ENSG00000099904 | 1032 | 470 | 645 | 580 | 610 | 589 |
| ENSG00000234409 | 5 | 1 | 7 | 7 | 3 | 3 |
| ENSG00000249923 | 0 | 0 | 0 | 0 | 0 | 0 |
| ENSG00000236499 | 0 | 0 | 1 | 0 | 0 | 0 |
| ENSG00000040608 | 133 | 44 | 81 | 112 | 68 | 69 |
| ENSG00000221039 | 0 | 0 | 0 | 0 | 0 | 0 |
| ENSG00000161132 | 5 | 2 | 4 | 3 | 8 | 3 |
| ENSG00000128185 | 1341 | 693 | 833 | 947 | 693 | 747 |
| ENSG00000273343 | 25 | 8 | 18 | 8 | 12 | 14 |
| ENSG00000273139 | 12 | 5 | 7 | 4 | 6 | 2 |
| ENSG00000188280 | 0 | 0 | 0 | 0 | 0 | 0 |
| ENSG00000235578 | 0 | 0 | 0 | 0 | 0 | 0 |
| ENSG00000161133 | 0 | 0 | 0 | 0 | 0 | 0 |
| ENSG00000185252 | 674 | 317 | 367 | 400 | 407 | 410 |
| ENSG00000207343 | 0 | 0 | 0 | 0 | 0 | 0 |
| ENSG00000244486 | 67 | 19 | 27 | 44 | 25 | 28 |
| ENSG00000277971 | 0 | 0 | 0 | 0 | 0 | 0 |
| ENSG00000099910 | 795 | 429 | 576 | 509 | 461 | 512 |
| ENSG00000215493 | 41 | 15 | 13 | 4 | 17 | 13 |
| ENSG00000255156 | 1 | 0 | 1 | 0 | 0 | 0 |
| ENSG00000201916 | 0 | 0 | 1 | 0 | 0 | 1 |
| ENSG00000242876 | 0 | 1 | 1 | 0 | 1 | 0 |
| ENSG00000236670 | 6 | 1 | 1 | 1 | 3 | 1 |
| ENSG00000099917 | 1909 | 941 | 1141 | 1051 | 1123 | 1117 |
| ENSG00000236003 | 0 | 0 | 0 | 0 | 0 | 0 |
| ENSG00000250261 | 0 | 0 | 0 | 0 | 0 | 0 |
| ENSG00000223553 | 0 | 0 | 0 | 0 | 0 | 0 |

| | | | | | | |
|---|---|---|---|---|---|---|
| ENSG00000276427 | 0 | 0 | 0 | 0 | 0 | 0 |
| ENSG00000238125 | 0 | 0 | 0 | 0 | 0 | 0 |
| ENSG00000229107 | 0 | 0 | 0 | 0 | 0 | 0 |
| ENSG00000217261 | 1 | 1 | 0 | 0 | 1 | 0 |
| ENSG00000235062 | 2 | 2 | 2 | 0 | 1 | 1 |
| ENSG00000226287 | 368 | 180 | 215 | 275 | 212 | 210 |
| ENSG00000241973 | 3981 | 1719 | 2422 | 1608 | 1912 | 2051 |
| ENSG00000099937 | 0 | 6 | 1 | 0 | 0 | 1 |
| ENSG00000099940 | 1670 | 862 | 1091 | 700 | 938 | 840 |
| ENSG00000272600 | 0 | 0 | 0 | 0 | 0 | 0 |
| ENSG00000099942 | 3192 | 1399 | 2056 | 1442 | 1802 | 1831 |
| ENSG00000237476 | 27 | 24 | 18 | 20 | 19 | 17 |
| ENSG00000183773 | 80 | 54 | 63 | 50 | 49 | 53 |
| ENSG00000285314 | 0 | 0 | 0 | 0 | 0 | 0 |
| ENSG00000272829 | 15 | 8 | 10 | 7 | 17 | 5 |
| ENSG00000099949 | 373 | 218 | 239 | 266 | 288 | 245 |
| ENSG00000184436 | 853 | 408 | 471 | 533 | 423 | 427 |
| ENSG00000230513 | 154 | 66 | 105 | 117 | 85 | 99 |
| ENSG00000284060 | 0 | 0 | 0 | 0 | 0 | 0 |
| ENSG00000161149 | 0 | 0 | 1 | 0 | 0 | 0 |
| ENSG00000099957 | 6 | 1 | 9 | 2 | 6 | 8 |
| ENSG00000284130 | 0 | 0 | 0 | 0 | 0 | 0 |
| ENSG00000099960 | 0 | 0 | 1 | 1 | 0 | 0 |
| ENSG00000226872 | 0 | 0 | 0 | 0 | 0 | 0 |
| ENSG00000207575 | 0 | 0 | 0 | 0 | 0 | 0 |
| ENSG00000206145 | 0 | 0 | 0 | 0 | 0 | 0 |
| ENSG00000187905 | 0 | 1 | 0 | 0 | 1 | 0 |
| ENSG00000249680 | 0 | 0 | 0 | 0 | 0 | 0 |
| ENSG00000284665 | 0 | 0 | 0 | 0 | 0 | 0 |
| ENSG00000169668 | 2 | 0 | 0 | 3 | 0 | 0 |
| ENSG00000197210 | 0 | 0 | 0 | 0 | 0 | 0 |
| ENSG00000239511 | 0 | 0 | 0 | 0 | 0 | 0 |
| ENSG00000224688 | 0 | 0 | 0 | 0 | 0 | 0 |
| ENSG00000215498 | 0 | 0 | 0 | 0 | 0 | 0 |
| ENSG00000234503 | 0 | 0 | 0 | 0 | 0 | 0 |
| ENSG00000283972 | 0 | 0 | 0 | 0 | 0 | 0 |
| ENSG00000133475 | 0 | 0 | 0 | 0 | 0 | 0 |
| ENSG00000283145 | 0 | 0 | 0 | 0 | 0 | 0 |
| ENSG00000226885 | 0 | 0 | 0 | 0 | 0 | 0 |
| ENSG00000229266 | 0 | 0 | 0 | 0 | 0 | 0 |
| ENSG00000169662 | 1 | 0 | 0 | 0 | 0 | 0 |
| ENSG00000206142 | 0 | 0 | 0 | 0 | 0 | 0 |
| ENSG00000237407 | 0 | 0 | 0 | 0 | 0 | 0 |
| ENSG00000232771 | 0 | 0 | 0 | 0 | 0 | 0 |
| ENSG00000279920 | 0 | 0 | 0 | 0 | 0 | 0 |
| ENSG00000226534 | 0 | 0 | 0 | 0 | 0 | 0 |
| ENSG00000273846 | 0 | 0 | 0 | 0 | 0 | 0 |
| ENSG00000274600 | 2 | 1 | 1 | 1 | 3 | 6 |
| ENSG00000200057 | 0 | 0 | 0 | 0 | 0 | 0 |
| ENSG00000252020 | 0 | 0 | 0 | 0 | 0 | 0 |

| | | | | | | |
|---|---|---|---|---|---|---|
| ENSG00000275840 | 0 | 0 | 0 | 0 | 0 | 0 |
| ENSG00000169635 | 256 | 109 | 137 | 100 | 112 | 112 |
| ENSG00000276800 | 0 | 0 | 0 | 0 | 0 | 0 |
| ENSG00000206140 | 36 | 22 | 19 | 25 | 14 | 17 |
| ENSG00000183506 | 154 | 88 | 128 | 79 | 84 | 96 |
| ENSG00000252143 | 0 | 0 | 0 | 0 | 0 | 0 |
| ENSG00000222352 | 0 | 0 | 0 | 0 | 0 | 0 |
| ENSG00000183246 | 0 | 0 | 0 | 1 | 0 | 0 |
| ENSG00000185651 | 4013 | 1939 | 2242 | 2087 | 2120 | 2217 |
| ENSG00000275293 | 0 | 0 | 0 | 0 | 0 | 0 |
| ENSG00000161179 | 1217 | 565 | 661 | 890 | 670 | 807 |
| ENSG00000161180 | 6 | 2 | 5 | 0 | 0 | 2 |
| ENSG00000273342 | 0 | 0 | 0 | 0 | 1 | 1 |
| ENSG00000128228 | 617 | 300 | 329 | 467 | 368 | 389 |
| ENSG00000207751 | 6 | 8 | 6 | 4 | 3 | 3 |
| ENSG00000212102 | 0 | 0 | 0 | 0 | 0 | 0 |
| ENSG00000283871 | 0 | 0 | 0 | 0 | 0 | 0 |
| ENSG00000284630 | 3 | 5 | 15 | 3 | 1 | 4 |
| ENSG00000284651 | 3 | 0 | 1 | 1 | 4 | 4 |
| ENSG00000284654 | 9 | 4 | 3 | 3 | 3 | 1 |
| ENSG00000284621 | 4 | 2 | 4 | 2 | 3 | 2 |
| ENSG00000272954 | 7 | 2 | 12 | 5 | 9 | 7 |
| ENSG00000100023 | 2071 | 958 | 1345 | 947 | 1058 | 1123 |
| ENSG00000100027 | 48 | 25 | 42 | 19 | 35 | 29 |
| ENSG00000244671 | 2 | 0 | 0 | 0 | 0 | 0 |
| ENSG00000100030 | 3843 | 1906 | 2204 | 1548 | 2167 | 1896 |
| ENSG00000200985 | 0 | 0 | 0 | 0 | 0 | 0 |
| ENSG00000225544 | 2 | 0 | 0 | 0 | 1 | 0 |
| ENSG00000100034 | 1153 | 432 | 683 | 610 | 573 | 624 |
| ENSG00000224086 | 130 | 83 | 89 | 63 | 72 | 55 |
| ENSG00000100038 | 11 | 0 | 3 | 4 | 3 | 7 |
| ENSG00000197549 | 0 | 0 | 0 | 0 | 0 | 0 |
| ENSG00000253239 | 0 | 0 | 0 | 0 | 0 | 0 |
| ENSG00000211637 | 0 | 0 | 0 | 0 | 0 | 0 |
| ENSG00000254355 | 0 | 0 | 0 | 0 | 0 | 0 |
| ENSG00000278657 | 0 | 0 | 0 | 0 | 0 | 0 |
| ENSG00000211642 | 0 | 0 | 0 | 0 | 0 | 0 |
| ENSG00000249333 | 0 | 0 | 0 | 0 | 0 | 0 |
| ENSG00000253794 | 0 | 0 | 0 | 0 | 0 | 0 |
| ENSG00000228161 | 0 | 0 | 0 | 0 | 0 | 0 |
| ENSG00000253874 | 0 | 0 | 0 | 0 | 0 | 0 |
| ENSG00000254075 | 0 | 0 | 0 | 0 | 0 | 0 |
| ENSG00000254161 | 0 | 0 | 0 | 0 | 0 | 0 |
| ENSG00000253242 | 0 | 0 | 0 | 0 | 0 | 0 |
| ENSG00000253752 | 0 | 0 | 0 | 0 | 0 | 0 |
| ENSG00000253823 | 0 | 0 | 0 | 0 | 0 | 0 |
| ENSG00000275361 | 0 | 0 | 0 | 0 | 0 | 0 |
| ENSG00000211638 | 0 | 0 | 0 | 0 | 0 | 0 |
| ENSG00000225741 | 0 | 0 | 0 | 0 | 0 | 0 |
| ENSG00000263366 | 0 | 0 | 0 | 0 | 0 | 0 |

| | | | | | | |
|---|---|---|---|---|---|---|
| ENSG00000236118 | 0 | 0 | 0 | 0 | 0 | 0 |
| ENSG00000233720 | 0 | 0 | 0 | 0 | 0 | 0 |
| ENSG00000211639 | 0 | 0 | 0 | 0 | 0 | 0 |
| ENSG00000227710 | 0 | 0 | 0 | 0 | 0 | 0 |
| ENSG00000224465 | 0 | 0 | 0 | 0 | 0 | 0 |
| ENSG00000254308 | 0 | 0 | 0 | 0 | 0 | 0 |
| ENSG00000253637 | 0 | 0 | 0 | 0 | 0 | 0 |
| ENSG00000236323 | 0 | 0 | 0 | 0 | 0 | 0 |
| ENSG00000211640 | 0 | 0 | 0 | 0 | 0 | 0 |
| ENSG00000253126 | 0 | 0 | 0 | 0 | 0 | 0 |
| ENSG00000211641 | 0 | 0 | 0 | 0 | 0 | 0 |
| ENSG00000253935 | 0 | 0 | 0 | 0 | 0 | 0 |
| ENSG00000228050 | 1 | 0 | 0 | 0 | 0 | 1 |
| ENSG00000234726 | 0 | 0 | 0 | 0 | 0 | 0 |
| ENSG00000169575 | 0 | 0 | 0 | 0 | 0 | 0 |
| ENSG00000279278 | 27 | 13 | 18 | 4 | 23 | 17 |
| ENSG00000274422 | 2 | 2 | 0 | 0 | 2 | 3 |
| ENSG00000234630 | 8 | 4 | 2 | 2 | 6 | 5 |
| ENSG00000272779 | 438 | 254 | 273 | 215 | 292 | 280 |
| ENSG00000211643 | 31 | 24 | 28 | 11 | 34 | 32 |
| ENSG00000211644 | 15 | 4 | 8 | 2 | 7 | 1 |
| ENSG00000211645 | 0 | 1 | 0 | 0 | 0 | 0 |
| ENSG00000225724 | 0 | 0 | 0 | 0 | 0 | 0 |
| ENSG00000223350 | 0 | 0 | 0 | 0 | 0 | 0 |
| ENSG00000232515 | 0 | 0 | 0 | 0 | 0 | 0 |
| ENSG00000211647 | 0 | 0 | 0 | 0 | 0 | 0 |
| ENSG00000211648 | 0 | 0 | 0 | 0 | 0 | 0 |
| ENSG00000237428 | 0 | 0 | 0 | 0 | 0 | 0 |
| ENSG00000211649 | 0 | 0 | 0 | 0 | 0 | 0 |
| ENSG00000232603 | 0 | 0 | 0 | 0 | 0 | 0 |
| ENSG00000211650 | 0 | 0 | 0 | 0 | 0 | 0 |
| ENSG00000211651 | 0 | 0 | 0 | 0 | 0 | 0 |
| ENSG00000211652 | 0 | 0 | 0 | 0 | 0 | 0 |
| ENSG00000254175 | 0 | 0 | 0 | 0 | 0 | 0 |
| ENSG00000254073 | 0 | 0 | 0 | 0 | 0 | 0 |
| ENSG00000253818 | 0 | 0 | 0 | 0 | 0 | 0 |
| ENSG00000211653 | 0 | 0 | 0 | 0 | 0 | 0 |
| ENSG00000224144 | 0 | 0 | 0 | 0 | 0 | 0 |
| ENSG00000253889 | 0 | 0 | 0 | 0 | 0 | 0 |
| ENSG00000211654 | 0 | 0 | 0 | 0 | 0 | 0 |
| ENSG00000211655 | 0 | 0 | 0 | 0 | 0 | 0 |
| ENSG00000230821 | 0 | 0 | 0 | 0 | 0 | 0 |
| ENSG00000253631 | 0 | 0 | 0 | 0 | 0 | 0 |
| ENSG00000226595 | 0 | 0 | 0 | 0 | 0 | 0 |
| ENSG00000275004 | 4 | 2 | 3 | 1 | 0 | 1 |
| ENSG00000169548 | 2 | 4 | 1 | 5 | 1 | 5 |
| ENSG00000185686 | 4 | 1 | 0 | 0 | 2 | 1 |
| ENSG00000220891 | 0 | 0 | 0 | 0 | 0 | 0 |
| ENSG00000253120 | 0 | 0 | 0 | 0 | 0 | 0 |
| ENSG00000211656 | 0 | 0 | 0 | 0 | 0 | 0 |

| | | | | | | |
|---|---|---|---|---|---|---|
| ENSG00000211657 | 0 | 0 | 0 | 0 | 0 | 0 |
| ENSG00000253920 | 0 | 0 | 0 | 0 | 0 | 0 |
| ENSG00000253329 | 0 | 0 | 0 | 0 | 0 | 0 |
| ENSG00000215456 | 0 | 3 | 1 | 0 | 0 | 1 |
| ENSG00000183169 | 0 | 0 | 0 | 0 | 0 | 0 |
| ENSG00000100121 | 0 | 0 | 0 | 0 | 0 | 0 |
| ENSG00000253338 | 0 | 0 | 0 | 0 | 0 | 0 |
| ENSG00000253451 | 0 | 0 | 0 | 0 | 0 | 0 |
| ENSG00000211658 | 0 | 0 | 0 | 0 | 0 | 0 |
| ENSG00000253913 | 0 | 0 | 0 | 0 | 0 | 0 |
| ENSG00000253779 | 0 | 0 | 0 | 0 | 0 | 0 |
| ENSG00000211659 | 0 | 0 | 0 | 0 | 0 | 0 |
| ENSG00000232350 | 0 | 0 | 0 | 0 | 0 | 0 |
| ENSG00000223999 | 0 | 0 | 0 | 0 | 0 | 0 |
| ENSG00000253822 | 0 | 0 | 0 | 0 | 0 | 0 |
| ENSG00000211660 | 0 | 0 | 0 | 0 | 0 | 0 |
| ENSG00000253546 | 0 | 0 | 0 | 0 | 0 | 0 |
| ENSG00000211661 | 0 | 0 | 0 | 0 | 0 | 0 |
| ENSG00000211662 | 0 | 0 | 0 | 0 | 0 | 0 |
| ENSG00000254240 | 0 | 0 | 0 | 0 | 0 | 0 |
| ENSG00000211663 | 0 | 0 | 0 | 0 | 0 | 0 |
| ENSG00000278667 | 0 | 0 | 0 | 0 | 0 | 0 |
| ENSG00000211664 | 0 | 0 | 0 | 0 | 0 | 0 |
| ENSG00000253152 | 0 | 0 | 0 | 0 | 0 | 0 |
| ENSG00000211665 | 0 | 0 | 0 | 0 | 0 | 0 |
| ENSG00000253786 | 0 | 0 | 0 | 0 | 0 | 0 |
| ENSG00000211666 | 0 | 0 | 0 | 0 | 0 | 0 |
| ENSG00000253590 | 0 | 0 | 0 | 0 | 0 | 0 |
| ENSG00000211667 | 0 | 0 | 0 | 0 | 0 | 0 |
| ENSG00000231392 | 0 | 0 | 0 | 0 | 0 | 0 |
| ENSG00000275772 | 0 | 0 | 0 | 0 | 0 | 0 |
| ENSG00000211668 | 0 | 0 | 0 | 0 | 0 | 0 |
| ENSG00000274381 | 0 | 0 | 0 | 0 | 0 | 0 |
| ENSG00000211669 | 0 | 0 | 0 | 0 | 0 | 0 |
| ENSG00000226477 | 0 | 0 | 0 | 0 | 0 | 0 |
| ENSG00000211670 | 0 | 0 | 0 | 0 | 0 | 0 |
| ENSG00000278196 | 0 | 0 | 0 | 0 | 0 | 0 |
| ENSG00000284049 | 0 | 0 | 0 | 0 | 0 | 0 |
| ENSG00000237127 | 0 | 0 | 0 | 0 | 0 | 0 |
| ENSG00000254077 | 0 | 0 | 0 | 0 | 0 | 0 |
| ENSG00000253448 | 0 | 0 | 0 | 0 | 0 | 0 |
| ENSG00000253234 | 0 | 0 | 0 | 0 | 0 | 0 |
| ENSG00000226420 | 0 | 0 | 0 | 0 | 0 | 0 |
| ENSG00000211672 | 0 | 0 | 0 | 0 | 0 | 0 |
| ENSG00000253963 | 0 | 0 | 0 | 0 | 0 | 0 |
| ENSG00000211673 | 0 | 0 | 0 | 0 | 0 | 0 |
| ENSG00000264824 | 0 | 0 | 0 | 0 | 0 | 0 |
| ENSG00000254709 | 0 | 0 | 0 | 0 | 0 | 0 |
| ENSG00000211674 | 0 | 0 | 0 | 0 | 0 | 0 |
| ENSG00000211675 | 0 | 0 | 0 | 0 | 0 | 0 |

| | | | | | | |
|---|---|---|---|---|---|---|
| ENSG00000211676 | 0 | 0 | 0 | 0 | 0 | 0 |
| ENSG00000211677 | 0 | 0 | 0 | 0 | 0 | 0 |
| ENSG00000211678 | 0 | 0 | 0 | 0 | 0 | 0 |
| ENSG00000211679 | 0 | 0 | 0 | 0 | 0 | 0 |
| ENSG00000211680 | 0 | 0 | 0 | 0 | 0 | 0 |
| ENSG00000254029 | 0 | 0 | 0 | 0 | 0 | 0 |
| ENSG00000211681 | 0 | 0 | 0 | 0 | 0 | 0 |
| ENSG00000254030 | 0 | 0 | 0 | 0 | 0 | 0 |
| ENSG00000211682 | 0 | 0 | 0 | 0 | 0 | 0 |
| ENSG00000222037 | 0 | 0 | 0 | 0 | 0 | 0 |
| ENSG00000211684 | 0 | 0 | 0 | 0 | 0 | 0 |
| ENSG00000211685 | 0 | 0 | 0 | 0 | 0 | 0 |
| ENSG00000237472 | 0 | 0 | 0 | 0 | 0 | 0 |
| ENSG00000223679 | 0 | 0 | 0 | 0 | 0 | 0 |
| ENSG00000100218 | 9 | 3 | 3 | 3 | 1 | 3 |
| ENSG00000128266 | 222 | 129 | 143 | 123 | 134 | 119 |
| ENSG00000283438 | 0 | 0 | 0 | 0 | 0 | 0 |
| ENSG00000275635 | 0 | 0 | 0 | 0 | 0 | 0 |
| ENSG00000100228 | 300 | 135 | 203 | 150 | 154 | 142 |
| ENSG00000186716 | 2044 | 916 | 1296 | 1176 | 1032 | 1050 |
| ENSG00000236794 | 6 | 1 | 0 | 0 | 0 | 0 |
| ENSG00000240160 | 0 | 0 | 2 | 0 | 0 | 1 |
| ENSG00000230701 | 6 | 8 | 6 | 10 | 5 | 5 |
| ENSG00000214526 | 0 | 0 | 0 | 0 | 0 | 1 |
| ENSG00000236611 | 0 | 0 | 0 | 0 | 0 | 0 |
| ENSG00000215478 | 13 | 3 | 5 | 3 | 0 | 7 |
| ENSG00000133519 | 251 | 69 | 132 | 114 | 117 | 125 |
| ENSG00000248409 | 42 | 12 | 24 | 20 | 16 | 15 |
| ENSG00000234928 | 11 | 8 | 8 | 3 | 8 | 9 |
| ENSG00000227755 | 15 | 4 | 8 | 4 | 5 | 3 |
| ENSG00000178248 | 4 | 1 | 2 | 4 | 2 | 2 |
| ENSG00000225413 | 0 | 0 | 0 | 0 | 0 | 0 |
| ENSG00000280623 | 0 | 0 | 0 | 0 | 0 | 0 |
| ENSG00000224277 | 0 | 0 | 0 | 0 | 0 | 0 |
| ENSG00000128322 | 0 | 0 | 0 | 0 | 0 | 1 |
| ENSG00000272733 | 7 | 3 | 5 | 3 | 6 | 3 |
| ENSG00000278655 | 0 | 0 | 1 | 0 | 0 | 0 |
| ENSG00000189269 | 10 | 7 | 12 | 7 | 2 | 6 |
| ENSG00000275609 | 0 | 0 | 0 | 0 | 1 | 0 |
| ENSG00000211683 | 1 | 2 | 2 | 1 | 2 | 0 |
| ENSG00000228315 | 1 | 2 | 4 | 4 | 7 | 5 |
| ENSG00000234353 | 9 | 12 | 13 | 4 | 8 | 4 |
| ENSG00000244723 | 11 | 4 | 9 | 5 | 5 | 4 |
| ENSG00000273000 | 2 | 3 | 2 | 0 | 3 | 0 |
| ENSG00000272578 | 27 | 23 | 20 | 7 | 15 | 18 |
| ENSG00000159496 | 33 | 15 | 11 | 10 | 25 | 15 |
| ENSG00000187792 | 177 | 93 | 96 | 75 | 62 | 75 |
| ENSG00000128218 | 11 | 3 | 8 | 8 | 9 | 12 |
| ENSG00000169314 | 6 | 1 | 2 | 1 | 3 | 3 |
| ENSG00000250479 | 348 | 153 | 161 | 219 | 145 | 154 |

| | | | | | | |
|---|---|---|---|---|---|---|
| ENSG00000099953 | 130 | 61 | 68 | 53 | 46 | 44 |
| ENSG00000099956 | 3313 | 1637 | 1939 | 2008 | 1714 | 1887 |
| ENSG00000099958 | 0 | 0 | 0 | 0 | 0 | 0 |
| ENSG00000272973 | 5 | 4 | 0 | 1 | 2 | 2 |
| ENSG00000133460 | 582 | 293 | 481 | 396 | 542 | 555 |
| ENSG00000251357 | 0 | 0 | 0 | 0 | 0 | 0 |
| ENSG00000279467 | 13 | 2 | 5 | 6 | 13 | 10 |
| ENSG00000244425 | 4 | 3 | 12 | 10 | 16 | 24 |
| ENSG00000240972 | 5 | 10 | 11 | 17 | 14 | 22 |
| ENSG00000218537 | 28 | 10 | 27 | 15 | 15 | 30 |
| ENSG00000273295 | 47 | 21 | 26 | 12 | 26 | 20 |
| ENSG00000225282 | 8 | 5 | 2 | 3 | 2 | 2 |
| ENSG00000206090 | 0 | 0 | 0 | 0 | 0 | 0 |
| ENSG00000231271 | 0 | 0 | 0 | 0 | 1 | 0 |
| ENSG00000133433 | 4 | 0 | 1 | 1 | 0 | 0 |
| ENSG00000228039 | 0 | 0 | 2 | 0 | 1 | 0 |
| ENSG00000099974 | 72 | 25 | 40 | 35 | 36 | 29 |
| ENSG00000272787 | 18 | 6 | 11 | 8 | 14 | 10 |
| ENSG00000099977 | 2 | 3 | 8 | 11 | 1 | 4 |
| ENSG00000285762 | 0 | 0 | 0 | 0 | 0 | 0 |
| ENSG00000224205 | 0 | 0 | 0 | 1 | 0 | 0 |
| ENSG00000250470 | 0 | 0 | 0 | 0 | 0 | 0 |
| ENSG00000099984 | 0 | 0 | 0 | 0 | 0 | 0 |
| ENSG00000276950 | 0 | 0 | 0 | 0 | 0 | 0 |
| ENSG00000235689 | 1 | 0 | 0 | 1 | 1 | 0 |
| ENSG00000099991 | 1951 | 826 | 1159 | 1013 | 1000 | 1096 |
| ENSG00000232545 | 8 | 5 | 8 | 2 | 7 | 6 |
| ENSG00000099994 | 175 | 99 | 116 | 61 | 47 | 55 |
| ENSG00000099998 | 4 | 4 | 4 | 2 | 1 | 2 |
| ENSG00000230712 | 0 | 0 | 0 | 0 | 0 | 2 |
| ENSG00000128262 | 2 | 0 | 0 | 0 | 0 | 0 |
| ENSG00000225098 | 0 | 0 | 0 | 0 | 0 | 0 |
| ENSG00000215464 | 0 | 0 | 0 | 1 | 0 | 0 |
| ENSG00000100014 | 778 | 412 | 480 | 399 | 439 | 482 |
| ENSG00000258555 | 0 | 1 | 0 | 1 | 0 | 0 |
| ENSG00000128271 | 1 | 2 | 3 | 0 | 0 | 2 |
| ENSG00000178803 | 14 | 5 | 5 | 1 | 4 | 4 |
| ENSG00000100024 | 2 | 0 | 0 | 0 | 1 | 0 |
| ENSG00000228923 | 0 | 0 | 0 | 0 | 0 | 0 |
| ENSG00000138867 | 2590 | 1247 | 1557 | 1370 | 1335 | 1368 |
| ENSG00000100028 | 2615 | 1444 | 1644 | 1366 | 1666 | 1656 |
| ENSG00000100031 | 14 | 11 | 21 | 19 | 6 | 11 |
| ENSG00000178026 | 167 | 70 | 85 | 70 | 77 | 60 |
| ENSG00000230947 | 0 | 0 | 0 | 0 | 0 | 0 |
| ENSG00000215481 | 0 | 1 | 0 | 0 | 0 | 0 |
| ENSG00000284233 | 0 | 0 | 0 | 0 | 0 | 0 |
| ENSG00000284128 | 0 | 0 | 0 | 1 | 1 | 1 |
| ENSG00000284070 | 10 | 5 | 6 | 11 | 7 | 6 |
| ENSG00000224124 | 0 | 0 | 0 | 0 | 0 | 0 |
| ENSG00000224806 | 0 | 0 | 0 | 0 | 0 | 0 |

| | | | | | | |
|---|---|---|---|---|---|---|
| ENSG00000224334 | 0 | 0 | 0 | 0 | 0 | 1 |
| ENSG00000282012 | 0 | 0 | 0 | 0 | 0 | 0 |
| ENSG00000237601 | 0 | 0 | 0 | 0 | 0 | 0 |
| ENSG00000182351 | 0 | 0 | 0 | 0 | 0 | 0 |
| ENSG00000184571 | 0 | 1 | 0 | 0 | 0 | 1 |
| ENSG00000167037 | 64 | 35 | 23 | 11 | 16 | 12 |
| ENSG00000199783 | 0 | 0 | 0 | 0 | 0 | 0 |
| ENSG00000206069 | 0 | 0 | 0 | 0 | 0 | 0 |
| ENSG00000197077 | 2223 | 892 | 1301 | 754 | 1012 | 1160 |
| ENSG00000279085 | 0 | 0 | 0 | 0 | 0 | 0 |
| ENSG00000279548 | 0 | 0 | 1 | 0 | 1 | 1 |
| ENSG00000279006 | 0 | 1 | 0 | 0 | 1 | 0 |
| ENSG00000279110 | 8 | 10 | 4 | 3 | 2 | 5 |
| ENSG00000203280 | 1 | 2 | 1 | 1 | 6 | 2 |
| ENSG00000271181 | 0 | 1 | 0 | 0 | 0 | 0 |
| ENSG00000100053 | 3 | 2 | 8 | 2 | 6 | 7 |
| ENSG00000236641 | 0 | 2 | 3 | 0 | 1 | 1 |
| ENSG00000244752 | 4 | 1 | 2 | 1 | 3 | 2 |
| ENSG00000279298 | 0 | 0 | 0 | 0 | 0 | 0 |
| ENSG00000233577 | 0 | 1 | 0 | 1 | 1 | 1 |
| ENSG00000285968 | 0 | 0 | 0 | 0 | 0 | 0 |
| ENSG00000285690 | 0 | 0 | 0 | 0 | 0 | 0 |
| ENSG00000206066 | 0 | 0 | 0 | 0 | 0 | 0 |
| ENSG00000230637 | 0 | 5 | 0 | 1 | 3 | 2 |
| ENSG00000231466 | 11 | 5 | 2 | 1 | 3 | 4 |
| ENSG00000100068 | 100 | 38 | 76 | 58 | 62 | 67 |
| ENSG00000272942 | 0 | 0 | 0 | 0 | 0 | 0 |
| ENSG00000272798 | 0 | 0 | 0 | 1 | 0 | 0 |
| ENSG00000271138 | 0 | 0 | 0 | 0 | 0 | 0 |
| ENSG00000100058 | 296 | 122 | 205 | 133 | 176 | 144 |
| ENSG00000277872 | 0 | 0 | 0 | 0 | 0 | 0 |
| ENSG00000272977 | 116 | 30 | 50 | 40 | 48 | 54 |
| ENSG00000234884 | 0 | 0 | 0 | 0 | 0 | 0 |
| ENSG00000100077 | 340 | 154 | 193 | 137 | 162 | 161 |
| ENSG00000237387 | 0 | 0 | 0 | 0 | 0 | 0 |
| ENSG00000224003 | 0 | 1 | 0 | 0 | 0 | 0 |
| ENSG00000222585 | 0 | 2 | 0 | 0 | 0 | 0 |
| ENSG00000133454 | 0 | 0 | 0 | 0 | 0 | 0 |
| ENSG00000278960 | 0 | 0 | 0 | 0 | 0 | 0 |
| ENSG00000231933 | 0 | 0 | 0 | 0 | 0 | 0 |
| ENSG00000232464 | 0 | 0 | 0 | 0 | 0 | 0 |
| ENSG00000223056 | 0 | 0 | 0 | 0 | 0 | 0 |
| ENSG00000232050 | 0 | 0 | 0 | 0 | 0 | 0 |
| ENSG00000236464 | 0 | 0 | 0 | 0 | 0 | 0 |
| ENSG00000229770 | 0 | 0 | 0 | 0 | 0 | 0 |
| ENSG00000100095 | 0 | 0 | 0 | 0 | 0 | 0 |
| ENSG00000224192 | 0 | 0 | 0 | 0 | 0 | 0 |
| ENSG00000252267 | 0 | 0 | 0 | 0 | 0 | 0 |
| ENSG00000230203 | 0 | 0 | 0 | 0 | 0 | 0 |
| ENSG00000128203 | 30 | 16 | 28 | 19 | 36 | 27 |

| | | | | | | |
|---|---|---|---|---|---|---|
| ENSG00000100099 | 1046 | 433 | 632 | 446 | 501 | 590 |
| ENSG00000100104 | 373 | 210 | 231 | 213 | 252 | 250 |
| ENSG00000100109 | 1020 | 507 | 619 | 511 | 580 | 581 |
| ENSG00000261188 | 116 | 97 | 116 | 66 | 122 | 104 |
| ENSG00000128294 | 253 | 158 | 179 | 174 | 202 | 180 |
| ENSG00000221760 | 0 | 0 | 0 | 0 | 0 | 0 |
| ENSG00000213707 | 31 | 30 | 26 | 31 | 41 | 36 |
| ENSG00000100122 | 0 | 0 | 0 | 0 | 0 | 0 |
| ENSG00000196431 | 0 | 0 | 0 | 0 | 1 | 0 |
| ENSG00000226912 | 2 | 0 | 2 | 0 | 3 | 2 |
| ENSG00000225783 | 16 | 4 | 16 | 1 | 6 | 4 |
| ENSG00000278404 | 0 | 0 | 0 | 0 | 0 | 0 |
| ENSG00000206028 | 0 | 0 | 0 | 0 | 0 | 0 |
| ENSG00000277941 | 0 | 0 | 0 | 0 | 0 | 0 |
| ENSG00000275942 | 0 | 0 | 0 | 0 | 0 | 0 |
| ENSG00000244625 | 49 | 42 | 59 | 54 | 48 | 54 |
| ENSG00000276991 | 0 | 0 | 0 | 0 | 0 | 0 |
| ENSG00000223704 | 0 | 0 | 1 | 0 | 0 | 0 |
| ENSG00000231369 | 0 | 0 | 1 | 1 | 0 | 1 |
| ENSG00000235271 | 0 | 0 | 0 | 0 | 0 | 0 |
| ENSG00000279440 | 0 | 0 | 0 | 0 | 0 | 0 |
| ENSG00000200443 | 0 | 0 | 0 | 0 | 0 | 0 |
| ENSG00000279914 | 0 | 0 | 0 | 0 | 0 | 0 |
| ENSG00000279848 | 0 | 0 | 0 | 0 | 0 | 0 |
| ENSG00000236858 | 0 | 0 | 0 | 0 | 0 | 0 |
| ENSG00000280025 | 0 | 0 | 0 | 0 | 0 | 0 |
| ENSG00000231405 | 0 | 0 | 0 | 0 | 0 | 0 |
| ENSG00000238195 | 0 | 0 | 0 | 0 | 0 | 0 |
| ENSG00000233521 | 0 | 0 | 0 | 0 | 0 | 0 |
| ENSG00000233574 | 0 | 0 | 0 | 0 | 0 | 0 |
| ENSG00000223726 | 0 | 0 | 0 | 0 | 0 | 0 |
| ENSG00000226741 | 0 | 0 | 0 | 0 | 0 | 0 |
| ENSG00000230051 | 0 | 0 | 0 | 0 | 0 | 0 |
| ENSG00000280445 | 0 | 0 | 0 | 0 | 0 | 0 |
| ENSG00000227880 | 0 | 0 | 0 | 0 | 0 | 0 |
| ENSG00000224027 | 0 | 0 | 0 | 0 | 0 | 0 |
| ENSG00000227838 | 0 | 0 | 0 | 0 | 0 | 0 |
| ENSG00000231495 | 0 | 0 | 0 | 0 | 0 | 0 |
| ENSG00000226169 | 0 | 0 | 0 | 0 | 0 | 0 |
| ENSG00000169184 | 8 | 1 | 4 | 1 | 3 | 1 |
| ENSG00000180957 | 2147 | 1124 | 1410 | 944 | 1326 | 1367 |
| ENSG00000235954 | 198 | 116 | 141 | 102 | 130 | 126 |
| ENSG00000277460 | 0 | 0 | 0 | 0 | 0 | 0 |
| ENSG00000264073 | 0 | 0 | 0 | 0 | 0 | 0 |
| ENSG00000283225 | 0 | 0 | 0 | 0 | 0 | 0 |
| ENSG00000276150 | 0 | 0 | 0 | 0 | 0 | 0 |
| ENSG00000100154 | 257 | 95 | 163 | 78 | 111 | 128 |
| ENSG00000274342 | 0 | 0 | 0 | 0 | 0 | 0 |
| ENSG00000276675 | 0 | 0 | 0 | 0 | 0 | 0 |
| ENSG00000239249 | 0 | 0 | 0 | 0 | 0 | 0 |

| | | | | | | |
|---|---|---|---|---|---|---|
| ENSG00000201209 | 0 | 0 | 0 | 0 | 0 | 0 |
| ENSG00000199797 | 0 | 0 | 0 | 0 | 0 | 0 |
| ENSG00000276162 | 0 | 0 | 0 | 0 | 0 | 0 |
| ENSG00000266503 | 0 | 0 | 0 | 0 | 0 | 0 |
| ENSG00000183765 | 1001 | 466 | 582 | 514 | 506 | 500 |
| ENSG00000100209 | 504 | 294 | 327 | 286 | 314 | 313 |
| ENSG00000159873 | 2573 | 1119 | 1397 | 865 | 1090 | 1215 |
| ENSG00000100219 | 28142 | 12895 | 15184 | 12753 | 11297 | 12926 |
| ENSG00000226471 | 47 | 35 | 36 | 28 | 28 | 21 |
| ENSG00000272858 | 20 | 9 | 13 | 3 | 4 | 5 |
| ENSG00000224256 | 0 | 0 | 0 | 0 | 0 | 0 |
| ENSG00000183579 | 399 | 158 | 211 | 136 | 179 | 211 |
| ENSG00000235786 | 0 | 0 | 0 | 0 | 0 | 0 |
| ENSG00000177993 | 0 | 0 | 0 | 0 | 1 | 0 |
| ENSG00000100249 | 0 | 0 | 0 | 0 | 0 | 0 |
| ENSG00000183762 | 1012 | 488 | 576 | 442 | 523 | 518 |
| ENSG00000226772 | 0 | 1 | 1 | 0 | 0 | 0 |
| ENSG00000200871 | 0 | 0 | 0 | 0 | 0 | 0 |
| ENSG00000251952 | 0 | 0 | 0 | 0 | 0 | 0 |
| ENSG00000186998 | 364 | 162 | 260 | 148 | 123 | 157 |
| ENSG00000100263 | 1112 | 553 | 762 | 610 | 610 | 600 |
| ENSG00000237015 | 24 | 11 | 17 | 12 | 11 | 14 |
| ENSG00000182944 | 11030 | 5222 | 6769 | 5437 | 5511 | 6068 |
| ENSG00000185340 | 1068 | 424 | 610 | 548 | 492 | 459 |
| ENSG00000100276 | 97 | 39 | 56 | 51 | 44 | 22 |
| ENSG00000273216 | 3 | 3 | 5 | 1 | 2 | 3 |
| ENSG00000100280 | 6164 | 2899 | 3464 | 3552 | 3326 | 3079 |
| ENSG00000280968 | 0 | 3 | 1 | 0 | 2 | 0 |
| ENSG00000239127 | 0 | 0 | 0 | 0 | 0 | 0 |
| ENSG00000285697 | 0 | 0 | 1 | 0 | 1 | 0 |
| ENSG00000242156 | 0 | 0 | 0 | 0 | 0 | 0 |
| ENSG00000278292 | 0 | 0 | 0 | 0 | 0 | 0 |
| ENSG00000225465 | 2 | 0 | 0 | 2 | 2 | 0 |
| ENSG00000128250 | 0 | 0 | 0 | 0 | 0 | 0 |
| ENSG00000282816 | 0 | 0 | 0 | 0 | 0 | 0 |
| ENSG00000100285 | 0 | 2 | 2 | 0 | 1 | 4 |
| ENSG00000100296 | 1295 | 702 | 847 | 891 | 919 | 887 |
| ENSG00000234208 | 0 | 0 | 0 | 0 | 1 | 0 |
| ENSG00000184117 | 5361 | 2849 | 3498 | 3301 | 3084 | 3455 |
| ENSG00000186575 | 1879 | 802 | 1153 | 991 | 969 | 1072 |
| ENSG00000233010 | 0 | 0 | 0 | 0 | 0 | 0 |
| ENSG00000232396 | 0 | 0 | 0 | 0 | 0 | 0 |
| ENSG00000239446 | 0 | 0 | 0 | 0 | 0 | 0 |
| ENSG00000100314 | 3 | 3 | 1 | 8 | 10 | 9 |
| ENSG00000100319 | 218 | 136 | 165 | 199 | 150 | 162 |
| ENSG00000200976 | 3 | 0 | 0 | 0 | 2 | 0 |
| ENSG00000184076 | 2788 | 1466 | 1615 | 2031 | 1818 | 1669 |
| ENSG00000100325 | 2115 | 995 | 1254 | 1218 | 1284 | 1180 |
| ENSG00000100330 | 277 | 138 | 198 | 135 | 168 | 179 |
| ENSG00000207455 | 0 | 0 | 0 | 0 | 0 | 0 |

| | | | | | | |
|---|---|---|---|---|---|---|
| ENSG00000279159 | 79 | 35 | 47 | 23 | 41 | 38 |
| ENSG00000275818 | 0 | 0 | 0 | 0 | 0 | 0 |
| ENSG00000227117 | 0 | 0 | 0 | 0 | 0 | 0 |
| ENSG00000227201 | 0 | 1 | 1 | 1 | 0 | 0 |
| ENSG00000176635 | 0 | 0 | 0 | 0 | 0 | 0 |
| ENSG00000226573 | 0 | 0 | 0 | 0 | 0 | 0 |
| ENSG00000225971 | 16 | 11 | 3 | 4 | 13 | 11 |
| ENSG00000225676 | 0 | 0 | 0 | 0 | 0 | 0 |
| ENSG00000232530 | 0 | 0 | 0 | 0 | 0 | 0 |
| ENSG00000128342 | 37 | 23 | 20 | 40 | 61 | 40 |
| ENSG00000268812 | 3 | 2 | 0 | 1 | 5 | 2 |
| ENSG00000099985 | 2 | 0 | 1 | 1 | 0 | 1 |
| ENSG00000279699 | 5 | 5 | 5 | 3 | 2 | 0 |
| ENSG00000239282 | 20 | 17 | 26 | 12 | 21 | 12 |
| ENSG00000248751 | 0 | 0 | 0 | 2 | 0 | 1 |
| ENSG00000099992 | 691 | 381 | 437 | 386 | 358 | 321 |
| ENSG00000099995 | 2698 | 1222 | 1627 | 1471 | 1304 | 1464 |
| ENSG00000187860 | 63 | 37 | 46 | 48 | 39 | 44 |
| ENSG00000099999 | 532 | 230 | 337 | 248 | 271 | 252 |
| ENSG00000100003 | 53 | 30 | 42 | 34 | 33 | 32 |
| ENSG00000249590 | 0 | 0 | 0 | 0 | 0 | 0 |
| ENSG00000273350 | 0 | 0 | 0 | 2 | 0 | 0 |
| ENSG00000222915 | 1 | 0 | 0 | 0 | 0 | 1 |
| ENSG00000272689 | 1 | 3 | 1 | 0 | 0 | 0 |
| ENSG00000242114 | 60 | 62 | 45 | 80 | 59 | 61 |
| ENSG00000273428 | 0 | 0 | 0 | 0 | 0 | 0 |
| ENSG00000230137 | 0 | 0 | 0 | 0 | 0 | 0 |
| ENSG00000100012 | 0 | 0 | 0 | 0 | 0 | 0 |
| ENSG00000181123 | 0 | 0 | 0 | 0 | 0 | 0 |
| ENSG00000214161 | 0 | 0 | 0 | 0 | 0 | 0 |
| ENSG00000133488 | 0 | 2 | 0 | 1 | 2 | 1 |
| ENSG00000214491 | 0 | 0 | 0 | 0 | 0 | 0 |
| ENSG00000225774 | 0 | 0 | 0 | 2 | 1 | 1 |
| ENSG00000128242 | 0 | 1 | 0 | 0 | 0 | 1 |
| ENSG00000100029 | 2516 | 1304 | 1464 | 1772 | 1476 | 1562 |
| ENSG00000223831 | 0 | 0 | 0 | 0 | 0 | 0 |
| ENSG00000185339 | 10 | 2 | 3 | 0 | 8 | 7 |
| ENSG00000100036 | 126 | 50 | 68 | 48 | 57 | 68 |
| ENSG00000167065 | 118 | 68 | 72 | 65 | 75 | 57 |
| ENSG00000250318 | 17 | 11 | 16 | 9 | 22 | 18 |
| ENSG00000184792 | 599 | 252 | 404 | 280 | 266 | 300 |
| ENSG00000264661 | 0 | 0 | 0 | 0 | 0 | 0 |
| ENSG00000237977 | 30 | 10 | 8 | 7 | 9 | 9 |
| ENSG00000235989 | 2 | 0 | 3 | 1 | 1 | 0 |
| ENSG00000133422 | 1937 | 969 | 1229 | 976 | 1193 | 1100 |
| ENSG00000253352 | 7138 | 4064 | 4465 | 2387 | 4993 | 3914 |
| ENSG00000276664 | 0 | 0 | 0 | 0 | 0 | 0 |
| ENSG00000276057 | 0 | 0 | 0 | 0 | 0 | 0 |
| ENSG00000276965 | 0 | 0 | 0 | 0 | 0 | 0 |
| ENSG00000275307 | 0 | 0 | 0 | 0 | 0 | 0 |

| | | | | | | |
|---|---|---|---|---|---|---|
| ENSG00000269987 | 0 | 0 | 0 | 0 | 0 | 0 |
| ENSG00000269972 | 0 | 0 | 0 | 0 | 0 | 0 |
| ENSG00000278920 | 1 | 0 | 1 | 1 | 0 | 0 |
| ENSG00000240186 | 0 | 0 | 0 | 0 | 0 | 0 |
| ENSG00000183963 | 917 | 410 | 532 | 606 | 640 | 456 |
| ENSG00000273387 | 7 | 3 | 8 | 2 | 5 | 6 |
| ENSG00000198832 | 140 | 68 | 113 | 63 | 64 | 56 |
| ENSG00000235573 | 0 | 0 | 0 | 0 | 0 | 0 |
| ENSG00000185133 | 556 | 210 | 336 | 166 | 129 | 170 |
| ENSG00000100078 | 50 | 26 | 32 | 18 | 19 | 8 |
| ENSG00000214125 | 0 | 1 | 0 | 0 | 1 | 1 |
| ENSG00000264141 | 0 | 0 | 0 | 0 | 0 | 0 |
| ENSG00000138942 | 1318 | 599 | 762 | 650 | 674 | 709 |
| ENSG00000254835 | 0 | 0 | 0 | 0 | 0 | 0 |
| ENSG00000182541 | 1263 | 623 | 768 | 607 | 626 | 615 |
| ENSG00000199695 | 0 | 0 | 0 | 0 | 0 | 0 |
| ENSG00000202019 | 0 | 0 | 0 | 0 | 0 | 0 |
| ENSG00000100100 | 1147 | 445 | 678 | 442 | 422 | 452 |
| ENSG00000228839 | 13 | 4 | 10 | 4 | 12 | 3 |
| ENSG00000212542 | 1 | 2 | 1 | 0 | 1 | 0 |
| ENSG00000100105 | 3489 | 1484 | 2072 | 1863 | 1727 | 1889 |
| ENSG00000213888 | 264 | 124 | 196 | 132 | 141 | 197 |
| ENSG00000206615 | 0 | 0 | 0 | 0 | 0 | 0 |
| ENSG00000185721 | 1651 | 832 | 876 | 966 | 951 | 954 |
| ENSG00000184708 | 793 | 396 | 529 | 412 | 455 | 482 |
| ENSG00000240591 | 2 | 2 | 1 | 1 | 0 | 2 |
| ENSG00000199248 | 0 | 0 | 1 | 0 | 0 | 0 |
| ENSG00000198089 | 978 | 453 | 629 | 449 | 468 | 451 |
| ENSG00000233733 | 0 | 1 | 1 | 0 | 0 | 0 |
| ENSG00000229169 | 0 | 0 | 0 | 0 | 0 | 0 |
| ENSG00000214093 | 0 | 0 | 3 | 0 | 1 | 0 |
| ENSG00000241878 | 2112 | 1079 | 1436 | 1130 | 1002 | 1248 |
| ENSG00000284082 | 0 | 0 | 0 | 0 | 0 | 0 |
| ENSG00000183530 | 1084 | 556 | 531 | 253 | 942 | 646 |
| ENSG00000100150 | 344 | 159 | 213 | 147 | 176 | 191 |
| ENSG00000285404 | 0 | 0 | 0 | 0 | 0 | 0 |
| ENSG00000242251 | 0 | 0 | 0 | 0 | 0 | 0 |
| ENSG00000236132 | 0 | 0 | 0 | 0 | 3 | 0 |
| ENSG00000252909 | 0 | 0 | 0 | 0 | 0 | 0 |
| ENSG00000227813 | 0 | 0 | 0 | 0 | 0 | 0 |
| ENSG00000128254 | 4 | 0 | 4 | 2 | 0 | 2 |
| ENSG00000128245 | 2743 | 1430 | 1744 | 1380 | 1442 | 1584 |
| ENSG00000227519 | 0 | 0 | 1 | 0 | 0 | 0 |
| ENSG00000230866 | 0 | 0 | 0 | 0 | 0 | 0 |
| ENSG00000240647 | 0 | 0 | 0 | 0 | 0 | 0 |
| ENSG00000232346 | 204 | 104 | 86 | 121 | 125 | 101 |
| ENSG00000100170 | 0 | 1 | 0 | 0 | 0 | 0 |
| ENSG00000234479 | 0 | 0 | 0 | 0 | 0 | 0 |
| ENSG00000237439 | 0 | 0 | 0 | 0 | 0 | 0 |
| ENSG00000232707 | 0 | 0 | 0 | 0 | 0 | 0 |

| | | | | | | |
|---|---|---|---|---|---|---|
| ENSG00000239674 | 0 | 0 | 0 | 0 | 0 | 0 |
| ENSG00000205856 | 0 | 0 | 0 | 0 | 0 | 0 |
| ENSG00000230120 | 0 | 0 | 0 | 0 | 0 | 0 |
| ENSG00000275656 | 0 | 0 | 0 | 0 | 0 | 0 |
| ENSG00000128253 | 4 | 2 | 5 | 2 | 0 | 3 |
| ENSG00000254127 | 2 | 0 | 0 | 0 | 0 | 0 |
| ENSG00000242082 | 0 | 0 | 0 | 1 | 0 | 0 |
| ENSG00000100191 | 2 | 1 | 7 | 0 | 4 | 1 |
| ENSG00000214076 | 2 | 1 | 3 | 3 | 3 | 5 |
| ENSG00000224050 | 4 | 4 | 1 | 0 | 0 | 0 |
| ENSG00000273325 | 0 | 0 | 0 | 0 | 0 | 0 |
| ENSG00000234626 | 0 | 0 | 0 | 0 | 1 | 0 |
| ENSG00000128276 | 0 | 0 | 0 | 0 | 0 | 0 |
| ENSG00000271093 | 0 | 0 | 0 | 0 | 0 | 0 |
| ENSG00000205853 | 4 | 2 | 2 | 2 | 0 | 0 |
| ENSG00000274617 | 0 | 0 | 0 | 0 | 0 | 0 |
| ENSG00000274170 | 0 | 0 | 0 | 0 | 0 | 0 |
| ENSG00000230736 | 1 | 0 | 3 | 3 | 7 | 1 |
| ENSG00000243519 | 0 | 0 | 0 | 0 | 0 | 1 |
| ENSG00000241954 | 1 | 0 | 0 | 0 | 1 | 0 |
| ENSG00000232218 | 1 | 1 | 0 | 0 | 0 | 3 |
| ENSG00000100220 | 3150 | 1536 | 1980 | 1871 | 1895 | 1872 |
| ENSG00000184459 | 0 | 0 | 0 | 0 | 0 | 0 |
| ENSG00000100225 | 3611 | 1763 | 2145 | 1871 | 2073 | 2046 |
| ENSG00000185666 | 8 | 6 | 11 | 1 | 2 | 5 |
| ENSG00000236054 | 10 | 10 | 6 | 12 | 5 | 7 |
| ENSG00000251890 | 0 | 0 | 0 | 0 | 0 | 0 |
| ENSG00000229673 | 0 | 0 | 0 | 0 | 0 | 0 |
| ENSG00000100234 | 1900 | 885 | 1103 | 994 | 1099 | 1231 |
| ENSG00000228194 | 0 | 0 | 0 | 0 | 0 | 0 |
| ENSG00000228622 | 0 | 0 | 0 | 0 | 0 | 0 |
| ENSG00000231253 | 0 | 0 | 0 | 0 | 0 | 0 |
| ENSG00000233632 | 0 | 0 | 0 | 0 | 0 | 0 |
| ENSG00000133424 | 2110 | 981 | 1241 | 843 | 967 | 1000 |
| ENSG00000232073 | 0 | 0 | 0 | 0 | 0 | 0 |
| ENSG00000230741 | 0 | 1 | 0 | 0 | 0 | 0 |
| ENSG00000266012 | 0 | 0 | 0 | 0 | 0 | 0 |
| ENSG00000253007 | 0 | 0 | 0 | 0 | 0 | 0 |
| ENSG00000281295 | 0 | 0 | 0 | 0 | 0 | 0 |
| ENSG00000232081 | 0 | 0 | 0 | 0 | 0 | 0 |
| ENSG00000224973 | 0 | 4 | 0 | 1 | 0 | 1 |
| ENSG00000273082 | 1 | 0 | 0 | 0 | 1 | 0 |
| ENSG00000236052 | 0 | 0 | 0 | 0 | 0 | 0 |
| ENSG00000224404 | 0 | 0 | 0 | 0 | 0 | 0 |
| ENSG00000235485 | 0 | 0 | 0 | 0 | 0 | 0 |
| ENSG00000280356 | 0 | 2 | 0 | 0 | 3 | 0 |
| ENSG00000234891 | 0 | 0 | 0 | 0 | 0 | 0 |
| ENSG00000237316 | 0 | 0 | 0 | 0 | 0 | 0 |
| ENSG00000227895 | 0 | 0 | 0 | 0 | 0 | 0 |
| ENSG00000175329 | 0 | 0 | 0 | 0 | 0 | 0 |

| | | | | | | |
|---|---|---|---|---|---|---|
| ENSG00000233080 | 0 | 0 | 1 | 0 | 0 | 0 |
| ENSG00000273068 | 0 | 0 | 0 | 0 | 0 | 0 |
| ENSG00000238153 | 0 | 0 | 0 | 0 | 0 | 0 |
| ENSG00000238584 | 0 | 0 | 0 | 0 | 0 | 0 |
| ENSG00000100281 | 1114 | 488 | 699 | 355 | 470 | 550 |
| ENSG00000273176 | 0 | 0 | 1 | 2 | 0 | 0 |
| ENSG00000100284 | 968 | 474 | 562 | 522 | 444 | 450 |
| ENSG00000266320 | 0 | 0 | 0 | 0 | 0 | 0 |
| ENSG00000274280 | 0 | 0 | 0 | 0 | 0 | 0 |
| ENSG00000282602 | 0 | 0 | 0 | 0 | 0 | 0 |
| ENSG00000282041 | 0 | 0 | 0 | 0 | 0 | 0 |
| ENSG00000100292 | 178 | 78 | 99 | 115 | 116 | 101 |
| ENSG00000100297 | 1368 | 738 | 897 | 793 | 622 | 869 |
| ENSG00000233388 | 0 | 0 | 0 | 0 | 0 | 0 |
| ENSG00000273044 | 0 | 0 | 0 | 0 | 0 | 0 |
| ENSG00000100302 | 16 | 9 | 0 | 6 | 12 | 6 |
| ENSG00000198125 | 1097 | 616 | 740 | 473 | 438 | 490 |
| ENSG00000268818 | 0 | 0 | 1 | 0 | 0 | 0 |
| ENSG00000221963 | 12 | 1 | 6 | 3 | 3 | 5 |
| ENSG00000228587 | 0 | 0 | 0 | 0 | 0 | 0 |
| ENSG00000226336 | 0 | 0 | 0 | 0 | 0 | 0 |
| ENSG00000128313 | 0 | 0 | 0 | 0 | 0 | 0 |
| ENSG00000100320 | 1311 | 537 | 673 | 474 | 625 | 677 |
| ENSG00000237406 | 2 | 0 | 0 | 1 | 1 | 0 |
| ENSG00000279652 | 6 | 3 | 2 | 0 | 1 | 1 |
| ENSG00000279927 | 0 | 0 | 0 | 0 | 0 | 0 |
| ENSG00000279217 | 0 | 0 | 0 | 0 | 1 | 1 |
| ENSG00000279714 | 0 | 0 | 0 | 0 | 0 | 0 |
| ENSG00000279805 | 0 | 0 | 0 | 0 | 0 | 0 |
| ENSG00000128284 | 14 | 14 | 17 | 3 | 8 | 10 |
| ENSG00000233764 | 0 | 0 | 0 | 0 | 0 | 0 |
| ENSG00000231576 | 0 | 0 | 0 | 0 | 0 | 0 |
| ENSG00000237948 | 0 | 0 | 0 | 0 | 0 | 0 |
| ENSG00000225557 | 0 | 0 | 0 | 0 | 0 | 0 |
| ENSG00000237129 | 0 | 0 | 0 | 0 | 0 | 0 |
| ENSG00000229088 | 0 | 0 | 0 | 0 | 0 | 0 |
| ENSG00000100336 | 0 | 0 | 1 | 2 | 0 | 1 |
| ENSG00000128335 | 457 | 233 | 328 | 177 | 271 | 204 |
| ENSG00000100342 | 6 | 10 | 14 | 12 | 12 | 7 |
| ENSG00000100345 | 20715 | 9173 | 12311 | 11165 | 12259 | 12270 |
| ENSG00000278420 | 1 | 0 | 0 | 0 | 0 | 0 |
| ENSG00000223695 | 0 | 0 | 3 | 0 | 2 | 1 |
| ENSG00000237668 | 3 | 1 | 1 | 3 | 1 | 1 |
| ENSG00000252225 | 0 | 0 | 0 | 0 | 0 | 1 |
| ENSG00000228719 | 6 | 3 | 2 | 6 | 10 | 4 |
| ENSG00000100348 | 2388 | 1144 | 1367 | 1539 | 1236 | 1229 |
| ENSG00000100350 | 715 | 395 | 505 | 388 | 362 | 510 |
| ENSG00000100353 | 4068 | 2097 | 2586 | 2712 | 2497 | 2530 |
| ENSG00000188078 | 0 | 0 | 0 | 0 | 0 | 0 |
| ENSG00000229971 | 0 | 0 | 0 | 0 | 0 | 0 |

| | | | | | | |
|---|---|---|---|---|---|---|
| ENSG00000261675 | 0 | 0 | 0 | 0 | 0 | 0 |
| ENSG00000166862 | 0 | 0 | 0 | 0 | 0 | 0 |
| ENSG00000234688 | 0 | 0 | 0 | 0 | 0 | 0 |
| ENSG00000100360 | 829 | 329 | 538 | 391 | 421 | 353 |
| ENSG00000100362 | 0 | 2 | 0 | 0 | 0 | 0 |
| ENSG00000234979 | 0 | 0 | 0 | 0 | 0 | 0 |
| ENSG00000183822 | 2 | 0 | 0 | 0 | 1 | 0 |
| ENSG00000100365 | 4 | 1 | 0 | 0 | 3 | 5 |
| ENSG00000100368 | 0 | 0 | 0 | 0 | 0 | 0 |
| ENSG00000232254 | 0 | 0 | 0 | 0 | 0 | 0 |
| ENSG00000215403 | 0 | 0 | 0 | 0 | 0 | 0 |
| ENSG00000185264 | 0 | 0 | 0 | 0 | 0 | 0 |
| ENSG00000128311 | 2143 | 1028 | 1395 | 1580 | 1263 | 1219 |
| ENSG00000252412 | 6 | 5 | 1 | 4 | 4 | 4 |
| ENSG00000128309 | 1583 | 757 | 1005 | 1236 | 911 | 967 |
| ENSG00000100379 | 382 | 213 | 236 | 277 | 197 | 250 |
| ENSG00000201078 | 0 | 0 | 0 | 0 | 0 | 0 |
| ENSG00000187045 | 90 | 49 | 76 | 35 | 37 | 44 |
| ENSG00000231467 | 3 | 1 | 2 | 0 | 0 | 1 |
| ENSG00000100385 | 0 | 1 | 2 | 0 | 1 | 1 |
| ENSG00000235237 | 0 | 0 | 0 | 0 | 0 | 0 |
| ENSG00000133466 | 1037 | 464 | 719 | 498 | 495 | 523 |
| ENSG00000278195 | 0 | 0 | 0 | 0 | 0 | 0 |
| ENSG00000226233 | 0 | 0 | 0 | 0 | 0 | 0 |
| ENSG00000128340 | 0 | 0 | 0 | 0 | 0 | 0 |
| ENSG00000100055 | 0 | 1 | 3 | 0 | 1 | 1 |
| ENSG00000243902 | 0 | 1 | 0 | 0 | 0 | 0 |
| ENSG00000237862 | 0 | 1 | 0 | 0 | 0 | 0 |
| ENSG00000166897 | 55 | 27 | 29 | 22 | 33 | 21 |
| ENSG00000272694 | 0 | 0 | 0 | 0 | 0 | 0 |
| ENSG00000100060 | 14 | 3 | 4 | 3 | 3 | 8 |
| ENSG00000100065 | 280 | 158 | 210 | 211 | 196 | 222 |
| ENSG00000225867 | 0 | 0 | 0 | 0 | 0 | 0 |
| ENSG00000128283 | 240 | 121 | 151 | 184 | 151 | 165 |
| ENSG00000100079 | 0 | 2 | 4 | 2 | 3 | 1 |
| ENSG00000273866 | 0 | 0 | 1 | 0 | 1 | 0 |
| ENSG00000100083 | 1777 | 750 | 1047 | 892 | 960 | 867 |
| ENSG00000100092 | 81 | 39 | 36 | 63 | 42 | 44 |
| ENSG00000285304 | 0 | 0 | 0 | 0 | 0 | 0 |
| ENSG00000233360 | 5 | 2 | 2 | 3 | 5 | 4 |
| ENSG00000241360 | 21 | 15 | 15 | 27 | 14 | 20 |
| ENSG00000241333 | 0 | 1 | 0 | 0 | 0 | 0 |
| ENSG00000100097 | 4592 | 2250 | 2268 | 2733 | 2268 | 2064 |
| ENSG00000273899 | 91 | 40 | 45 | 30 | 35 | 34 |
| ENSG00000100101 | 0 | 0 | 0 | 0 | 0 | 0 |
| ENSG00000100106 | 1602 | 815 | 1016 | 861 | 843 | 842 |
| ENSG00000189060 | 5492 | 2529 | 3446 | 2743 | 2932 | 3089 |
| ENSG00000100116 | 352 | 161 | 231 | 233 | 142 | 195 |
| ENSG00000128310 | 1 | 0 | 0 | 1 | 0 | 0 |
| ENSG00000100124 | 790 | 426 | 436 | 453 | 407 | 426 |

| | | | | | | |
|---|---|---|---|---|---|---|
| ENSG00000284197 | 0 | 0 | 0 | 0 | 0 | 0 |
| ENSG00000207696 | 1 | 2 | 4 | 0 | 6 | 2 |
| ENSG00000100129 | 3480 | 2009 | 2251 | 1923 | 2321 | 2091 |
| ENSG00000279738 | 28 | 18 | 16 | 13 | 14 | 20 |
| ENSG00000207227 | 0 | 0 | 0 | 0 | 0 | 0 |
| ENSG00000100139 | 855 | 384 | 462 | 429 | 436 | 422 |
| ENSG00000278948 | 10 | 12 | 16 | 18 | 14 | 12 |
| ENSG00000128346 | 22 | 25 | 14 | 5 | 16 | 21 |
| ENSG00000272582 | 0 | 2 | 2 | 1 | 1 | 1 |
| ENSG00000100142 | 810 | 408 | 543 | 579 | 516 | 508 |
| ENSG00000284289 | 0 | 0 | 0 | 0 | 0 | 0 |
| ENSG00000100146 | 0 | 0 | 0 | 0 | 0 | 1 |
| ENSG00000284266 | 0 | 0 | 0 | 0 | 0 | 0 |
| ENSG00000222044 | 0 | 2 | 0 | 0 | 0 | 0 |
| ENSG00000273214 | 0 | 0 | 0 | 0 | 0 | 0 |
| ENSG00000100151 | 656 | 306 | 419 | 451 | 383 | 408 |
| ENSG00000233739 | 0 | 0 | 0 | 0 | 0 | 0 |
| ENSG00000100156 | 30 | 19 | 27 | 7 | 16 | 19 |
| ENSG00000128298 | 28 | 9 | 13 | 5 | 7 | 13 |
| ENSG00000272720 | 30 | 7 | 17 | 10 | 11 | 17 |
| ENSG00000184381 | 281 | 130 | 171 | 112 | 142 | 138 |
| ENSG00000279080 | 4 | 0 | 1 | 0 | 0 | 0 |
| ENSG00000185022 | 53 | 38 | 42 | 57 | 52 | 42 |
| ENSG00000198792 | 2714 | 1219 | 1604 | 1495 | 1532 | 1574 |
| ENSG00000235246 | 0 | 0 | 1 | 0 | 2 | 0 |
| ENSG00000241693 | 3 | 1 | 0 | 0 | 0 | 0 |
| ENSG00000277385 | 0 | 0 | 0 | 0 | 1 | 0 |
| ENSG00000213923 | 1936 | 1045 | 1192 | 1112 | 1274 | 1147 |
| ENSG00000283900 | 0 | 0 | 0 | 0 | 0 | 0 |
| ENSG00000244627 | 51 | 27 | 16 | 24 | 42 | 17 |
| ENSG00000229955 | 2 | 2 | 0 | 0 | 0 | 0 |
| ENSG00000228620 | 1 | 0 | 2 | 0 | 1 | 1 |
| ENSG00000168135 | 15 | 2 | 7 | 12 | 10 | 9 |
| ENSG00000100196 | 545 | 241 | 286 | 262 | 268 | 290 |
| ENSG00000100201 | 15194 | 6310 | 9348 | 4378 | 6857 | 6650 |
| ENSG00000100206 | 32 | 15 | 12 | 8 | 13 | 10 |
| ENSG00000224279 | 0 | 0 | 0 | 0 | 0 | 0 |
| ENSG00000184949 | 130 | 56 | 92 | 35 | 84 | 69 |
| ENSG00000100211 | 378 | 219 | 264 | 268 | 281 | 245 |
| ENSG00000230912 | 4 | 3 | 2 | 0 | 2 | 2 |
| ENSG00000228274 | 18 | 10 | 14 | 5 | 6 | 11 |
| ENSG00000100216 | 1767 | 859 | 1024 | 1089 | 1053 | 1068 |
| ENSG00000100221 | 1437 | 710 | 840 | 730 | 793 | 873 |
| ENSG00000100226 | 725 | 345 | 441 | 322 | 381 | 359 |
| ENSG00000229598 | 4 | 3 | 2 | 1 | 3 | 2 |
| ENSG00000244491 | 0 | 0 | 0 | 0 | 0 | 0 |
| ENSG00000100242 | 865 | 420 | 604 | 494 | 521 | 531 |
| ENSG00000230149 | 2 | 4 | 3 | 4 | 2 | 4 |
| ENSG00000273096 | 1 | 0 | 1 | 2 | 1 | 1 |
| ENSG00000225450 | 0 | 1 | 4 | 1 | 2 | 0 |

| | | | | | | |
|---|---|---|---|---|---|---|
| ENSG00000272669 | 15 | 12 | 9 | 7 | 20 | 14 |
| ENSG00000273076 | 2 | 0 | 2 | 2 | 0 | 1 |
| ENSG00000100246 | 586 | 279 | 359 | 343 | 319 | 285 |
| ENSG00000221890 | 729 | 391 | 519 | 369 | 339 | 380 |
| ENSG00000183741 | 7168 | 3155 | 4511 | 3457 | 3231 | 3711 |
| ENSG00000235209 | 10 | 0 | 5 | 2 | 1 | 3 |
| ENSG00000128383 | 1 | 1 | 0 | 0 | 0 | 4 |
| ENSG00000179750 | 125 | 77 | 130 | 69 | 78 | 83 |
| ENSG00000249310 | 0 | 0 | 0 | 0 | 0 | 0 |
| ENSG00000244509 | 4 | 0 | 2 | 0 | 1 | 1 |
| ENSG00000284554 | 0 | 0 | 0 | 0 | 0 | 0 |
| ENSG00000243811 | 0 | 0 | 0 | 0 | 0 | 0 |
| ENSG00000128394 | 0 | 0 | 1 | 0 | 0 | 0 |
| ENSG00000259421 | 0 | 0 | 0 | 0 | 0 | 0 |
| ENSG00000239713 | 0 | 0 | 0 | 0 | 0 | 0 |
| ENSG00000225720 | 0 | 0 | 0 | 0 | 0 | 0 |
| ENSG00000100298 | 12 | 7 | 13 | 10 | 6 | 8 |
| ENSG00000100307 | 406 | 198 | 263 | 232 | 214 | 222 |
| ENSG00000226024 | 0 | 0 | 0 | 0 | 0 | 0 |
| ENSG00000279833 | 24 | 13 | 20 | 15 | 16 | 21 |
| ENSG00000227009 | 0 | 0 | 0 | 0 | 0 | 0 |
| ENSG00000100311 | 1357 | 611 | 818 | 684 | 671 | 679 |
| ENSG00000284633 | 43 | 13 | 22 | 22 | 16 | 9 |
| ENSG00000224794 | 0 | 1 | 6 | 4 | 0 | 1 |
| ENSG00000100316 | 45247 | 22964 | 26438 | 33683 | 28470 | 27779 |
| ENSG00000209480 | 0 | 0 | 0 | 0 | 0 | 0 |
| ENSG00000209482 | 10 | 1 | 3 | 5 | 4 | 5 |
| ENSG00000263764 | 0 | 0 | 0 | 0 | 0 | 0 |
| ENSG00000100321 | 1024 | 521 | 642 | 445 | 415 | 455 |
| ENSG00000280216 | 3 | 4 | 5 | 1 | 5 | 7 |
| ENSG00000100324 | 826 | 356 | 502 | 459 | 403 | 400 |
| ENSG00000128268 | 4 | 3 | 5 | 1 | 6 | 2 |
| ENSG00000227188 | 0 | 0 | 0 | 0 | 0 | 0 |
| ENSG00000100335 | 990 | 469 | 657 | 519 | 638 | 594 |
| ENSG00000285025 | 0 | 0 | 0 | 0 | 0 | 0 |
| ENSG00000128272 | 7834 | 3702 | 4749 | 4763 | 4662 | 4546 |
| ENSG00000187051 | 1556 | 738 | 923 | 1182 | 883 | 850 |
| ENSG00000100346 | 4 | 6 | 5 | 9 | 10 | 9 |
| ENSG00000176177 | 0 | 0 | 0 | 0 | 0 | 0 |
| ENSG00000235647 | 0 | 0 | 0 | 0 | 0 | 0 |
| ENSG00000238875 | 0 | 0 | 0 | 0 | 0 | 0 |
| ENSG00000226085 | 44 | 24 | 22 | 26 | 30 | 20 |
| ENSG00000100351 | 0 | 0 | 0 | 0 | 0 | 0 |
| ENSG00000225528 | 0 | 0 | 0 | 0 | 0 | 0 |
| ENSG00000133477 | 214 | 127 | 127 | 92 | 110 | 96 |
| ENSG00000261202 | 0 | 1 | 2 | 0 | 0 | 2 |
| ENSG00000100354 | 806 | 389 | 422 | 200 | 360 | 326 |
| ENSG00000228599 | 0 | 0 | 0 | 0 | 0 | 0 |
| ENSG00000239900 | 131 | 95 | 93 | 95 | 96 | 101 |
| ENSG00000284431 | 0 | 6 | 2 | 0 | 1 | 3 |

| | | | | | | |
|---|---|---|---|---|---|---|
| ENSG00000100359 | 43 | 14 | 28 | 22 | 29 | 13 |
| ENSG00000229999 | 0 | 2 | 1 | 0 | 0 | 0 |
| ENSG00000196588 | 803 | 377 | 475 | 502 | 432 | 410 |
| ENSG00000272834 | 0 | 2 | 1 | 0 | 4 | 0 |
| ENSG00000227413 | 0 | 0 | 1 | 0 | 2 | 0 |
| ENSG00000232564 | 0 | 0 | 0 | 1 | 1 | 0 |
| ENSG00000172912 | 0 | 0 | 0 | 0 | 0 | 0 |
| ENSG00000230071 | 7 | 3 | 5 | 4 | 3 | 3 |
| ENSG00000231907 | 0 | 0 | 0 | 0 | 0 | 0 |
| ENSG00000128285 | 0 | 1 | 1 | 0 | 0 | 2 |
| ENSG00000100372 | 899 | 465 | 538 | 494 | 515 | 493 |
| ENSG00000230769 | 2 | 1 | 0 | 0 | 0 | 1 |
| ENSG00000233603 | 0 | 0 | 0 | 0 | 0 | 0 |
| ENSG00000266594 | 0 | 0 | 1 | 0 | 1 | 1 |
| ENSG00000100380 | 5625 | 2974 | 3447 | 2851 | 3448 | 3216 |
| ENSG00000196236 | 732 | 372 | 350 | 296 | 373 | 329 |
| ENSG00000172404 | 9 | 1 | 1 | 0 | 0 | 1 |
| ENSG00000273979 | 0 | 0 | 0 | 0 | 0 | 0 |
| ENSG00000200683 | 0 | 0 | 0 | 0 | 0 | 0 |
| ENSG00000100387 | 1118 | 490 | 599 | 607 | 601 | 623 |
| ENSG00000222698 | 0 | 0 | 0 | 0 | 0 | 0 |
| ENSG00000237214 | 6 | 5 | 5 | 4 | 3 | 3 |
| ENSG00000199515 | 0 | 0 | 0 | 0 | 0 | 0 |
| ENSG00000213857 | 2 | 1 | 1 | 0 | 0 | 0 |
| ENSG00000100393 | 2600 | 962 | 1336 | 653 | 1044 | 1158 |
| ENSG00000284015 | 0 | 0 | 0 | 0 | 0 | 0 |
| ENSG00000252859 | 0 | 0 | 0 | 0 | 0 | 0 |
| ENSG00000232754 | 0 | 0 | 0 | 0 | 0 | 0 |
| ENSG00000231993 | 13 | 6 | 2 | 1 | 8 | 6 |
| ENSG00000215771 | 0 | 0 | 0 | 0 | 0 | 0 |
| ENSG00000100395 | 826 | 439 | 566 | 506 | 452 | 481 |
| ENSG00000235513 | 0 | 4 | 4 | 0 | 2 | 0 |
| ENSG00000100399 | 31 | 10 | 15 | 21 | 5 | 17 |
| ENSG00000100401 | 3325 | 1710 | 2114 | 2356 | 2095 | 2080 |
| ENSG00000274552 | 0 | 0 | 0 | 0 | 0 | 0 |
| ENSG00000100403 | 3134 | 1403 | 1938 | 1376 | 1696 | 1658 |
| ENSG00000167074 | 1326 | 544 | 815 | 603 | 590 | 568 |
| ENSG00000199865 | 0 | 0 | 0 | 0 | 0 | 1 |
| ENSG00000273424 | 4 | 1 | 1 | 3 | 2 | 0 |
| ENSG00000183864 | 4446 | 1990 | 2427 | 1928 | 2050 | 2142 |
| ENSG00000100410 | 1048 | 530 | 685 | 672 | 599 | 640 |
| ENSG00000100412 | 3338 | 1674 | 2145 | 2059 | 1949 | 1928 |
| ENSG00000100413 | 1484 | 759 | 955 | 1021 | 867 | 1112 |
| ENSG00000172346 | 2 | 0 | 0 | 0 | 0 | 0 |
| ENSG00000100417 | 514 | 309 | 404 | 382 | 272 | 395 |
| ENSG00000100418 | 1017 | 431 | 550 | 572 | 542 | 607 |
| ENSG00000196419 | 10441 | 5100 | 6140 | 6032 | 5897 | 5956 |
| ENSG00000200332 | 0 | 0 | 0 | 0 | 0 | 0 |
| ENSG00000100138 | 2261 | 1114 | 1341 | 1476 | 1359 | 1398 |
| ENSG00000285531 | 8 | 8 | 4 | 5 | 8 | 11 |

| | | | | | | |
|---|---|---|---|---|---|---|
| ENSG00000207457 | 1 | 0 | 1 | 0 | 0 | 0 |
| ENSG00000184208 | 579 | 229 | 289 | 193 | 324 | 244 |
| ENSG00000167077 | 0 | 3 | 4 | 0 | 1 | 1 |
| ENSG00000231261 | 0 | 0 | 0 | 0 | 0 | 0 |
| ENSG00000252603 | 0 | 0 | 0 | 0 | 0 | 0 |
| ENSG00000206813 | 0 | 0 | 0 | 0 | 0 | 0 |
| ENSG00000100147 | 72 | 45 | 65 | 44 | 48 | 59 |
| ENSG00000184068 | 76 | 42 | 36 | 46 | 51 | 39 |
| ENSG00000198911 | 4986 | 2243 | 2995 | 2565 | 2473 | 2766 |
| ENSG00000207932 | 0 | 0 | 0 | 0 | 0 | 1 |
| ENSG00000234965 | 22 | 8 | 7 | 5 | 8 | 8 |
| ENSG00000159958 | 0 | 1 | 10 | 6 | 0 | 4 |
| ENSG00000283829 | 0 | 0 | 0 | 0 | 0 | 0 |
| ENSG00000100162 | 251 | 113 | 154 | 173 | 150 | 145 |
| ENSG00000205704 | 20 | 6 | 5 | 11 | 5 | 4 |
| ENSG00000100167 | 142 | 59 | 78 | 39 | 62 | 62 |
| ENSG00000224883 | 0 | 0 | 0 | 0 | 0 | 0 |
| ENSG00000183066 | 17 | 6 | 9 | 7 | 15 | 5 |
| ENSG00000215347 | 0 | 0 | 1 | 1 | 2 | 0 |
| ENSG00000198951 | 820 | 364 | 529 | 395 | 360 | 440 |
| ENSG00000177096 | 30 | 23 | 34 | 23 | 29 | 27 |
| ENSG00000238498 | 0 | 0 | 0 | 0 | 0 | 0 |
| ENSG00000183172 | 915 | 404 | 569 | 561 | 469 | 452 |
| ENSG00000184983 | 1586 | 739 | 920 | 931 | 833 | 825 |
| ENSG00000270083 | 9 | 4 | 9 | 1 | 1 | 1 |
| ENSG00000237037 | 39 | 14 | 15 | 12 | 12 | 19 |
| ENSG00000213790 | 2 | 1 | 0 | 2 | 1 | 3 |
| ENSG00000100197 | 8 | 4 | 9 | 4 | 4 | 1 |
| ENSG00000227370 | 2 | 1 | 2 | 3 | 1 | 0 |
| ENSG00000232710 | 2 | 1 | 2 | 0 | 4 | 1 |
| ENSG00000281538 | 21 | 5 | 4 | 8 | 4 | 6 |
| ENSG00000205702 | 26 | 5 | 13 | 9 | 12 | 17 |
| ENSG00000226450 | 6 | 3 | 3 | 2 | 2 | 3 |
| ENSG00000100207 | 1248 | 480 | 701 | 481 | 591 | 641 |
| ENSG00000182057 | 12 | 7 | 9 | 10 | 9 | 10 |
| ENSG00000233903 | 11 | 13 | 5 | 10 | 10 | 16 |
| ENSG00000273366 | 11 | 2 | 7 | 3 | 7 | 9 |
| ENSG00000229891 | 14 | 11 | 4 | 10 | 13 | 8 |
| ENSG00000235568 | 4 | 0 | 1 | 0 | 1 | 1 |
| ENSG00000230107 | 0 | 0 | 0 | 0 | 0 | 0 |
| ENSG00000172250 | 29 | 22 | 27 | 24 | 27 | 25 |
| ENSG00000189306 | 1032 | 559 | 702 | 704 | 586 | 627 |
| ENSG00000183569 | 178 | 108 | 175 | 119 | 94 | 142 |
| ENSG00000182841 | 122 | 49 | 63 | 52 | 50 | 63 |
| ENSG00000202058 | 2 | 2 | 2 | 1 | 3 | 4 |
| ENSG00000251913 | 0 | 0 | 0 | 0 | 0 | 0 |
| ENSG00000100227 | 2533 | 1117 | 1416 | 1250 | 1094 | 1132 |
| ENSG00000270022 | 11 | 4 | 5 | 7 | 6 | 7 |
| ENSG00000276027 | 0 | 0 | 0 | 0 | 0 | 0 |
| ENSG00000100243 | 3402 | 1563 | 2099 | 1929 | 1637 | 1848 |

| | | | | | | |
|---|---|---|---|---|---|---|
| ENSG00000249222 | 2 | 0 | 0 | 0 | 0 | 0 |
| ENSG00000128274 | 631 | 285 | 341 | 480 | 352 | 350 |
| ENSG00000200448 | 0 | 0 | 0 | 0 | 0 | 0 |
| ENSG00000234009 | 180 | 92 | 130 | 115 | 123 | 98 |
| ENSG00000229608 | 0 | 0 | 0 | 0 | 1 | 0 |
| ENSG00000274717 | 1 | 0 | 2 | 0 | 0 | 0 |
| ENSG00000242247 | 963 | 501 | 667 | 371 | 499 | 540 |
| ENSG00000100266 | 1722 | 787 | 1023 | 741 | 894 | 908 |
| ENSG00000226989 | 0 | 0 | 0 | 0 | 0 | 0 |
| ENSG00000230319 | 1 | 0 | 1 | 2 | 2 | 1 |
| ENSG00000100271 | 153 | 98 | 122 | 81 | 92 | 93 |
| ENSG00000234735 | 0 | 1 | 0 | 0 | 0 | 0 |
| ENSG00000100290 | 536 | 230 | 322 | 214 | 194 | 201 |
| ENSG00000100294 | 499 | 267 | 306 | 309 | 253 | 258 |
| ENSG00000100300 | 862 | 380 | 449 | 579 | 445 | 426 |
| ENSG00000100304 | 1593 | 711 | 953 | 978 | 812 | 905 |
| ENSG00000159307 | 2 | 2 | 3 | 4 | 1 | 1 |
| ENSG00000236272 | 0 | 0 | 0 | 0 | 0 | 0 |
| ENSG00000234892 | 0 | 0 | 0 | 0 | 0 | 0 |
| ENSG00000203527 | 0 | 0 | 0 | 0 | 0 | 0 |
| ENSG00000236117 | 0 | 0 | 0 | 0 | 0 | 0 |
| ENSG00000186732 | 0 | 0 | 0 | 0 | 0 | 0 |
| ENSG00000223843 | 2 | 1 | 0 | 0 | 0 | 0 |
| ENSG00000186976 | 49 | 19 | 27 | 16 | 32 | 24 |
| ENSG00000233157 | 0 | 0 | 0 | 0 | 0 | 0 |
| ENSG00000180150 | 0 | 0 | 0 | 0 | 0 | 0 |
| ENSG00000261251 | 26 | 3 | 7 | 3 | 6 | 10 |
| ENSG00000130540 | 2 | 0 | 0 | 0 | 0 | 0 |
| ENSG00000100341 | 0 | 0 | 0 | 0 | 0 | 0 |
| ENSG00000100344 | 6 | 7 | 5 | 4 | 6 | 0 |
| ENSG00000100347 | 1164 | 621 | 749 | 762 | 659 | 731 |
| ENSG00000224856 | 0 | 3 | 0 | 0 | 0 | 0 |
| ENSG00000188677 | 7 | 9 | 4 | 9 | 12 | 10 |
| ENSG00000279175 | 0 | 0 | 0 | 0 | 0 | 1 |
| ENSG00000280434 | 0 | 0 | 0 | 0 | 0 | 3 |
| ENSG00000138964 | 0 | 0 | 0 | 0 | 0 | 0 |
| ENSG00000279933 | 0 | 0 | 0 | 0 | 0 | 0 |
| ENSG00000280011 | 0 | 0 | 0 | 0 | 0 | 0 |
| ENSG00000138944 | 0 | 0 | 0 | 0 | 0 | 0 |
| ENSG00000220702 | 0 | 0 | 0 | 0 | 0 | 0 |
| ENSG00000236079 | 0 | 0 | 0 | 0 | 0 | 0 |
| ENSG00000232655 | 0 | 0 | 0 | 0 | 0 | 0 |
| ENSG00000228807 | 0 | 0 | 0 | 0 | 0 | 0 |
| ENSG00000279642 | 0 | 0 | 0 | 0 | 0 | 0 |
| ENSG00000188636 | 1498 | 672 | 833 | 711 | 794 | 918 |
| ENSG00000215333 | 2 | 1 | 4 | 2 | 0 | 1 |
| ENSG00000187012 | 0 | 0 | 0 | 0 | 0 | 0 |
| ENSG00000234300 | 0 | 0 | 0 | 0 | 0 | 0 |
| ENSG00000231801 | 0 | 0 | 0 | 0 | 0 | 0 |
| ENSG00000186654 | 134 | 99 | 99 | 111 | 101 | 91 |

| | | | | | | |
|---|---|---|---|---|---|---|
| ENSG00000248405 | 54 | 27 | 37 | 11 | 22 | 24 |
| ENSG00000241484 | 155 | 81 | 105 | 38 | 61 | 69 |
| ENSG00000056487 | 31 | 12 | 22 | 16 | 15 | 15 |
| ENSG00000223730 | 0 | 0 | 0 | 0 | 0 | 0 |
| ENSG00000230922 | 3 | 1 | 0 | 2 | 2 | 2 |
| ENSG00000226328 | 162 | 76 | 120 | 100 | 115 | 107 |
| ENSG00000273243 | 8 | 5 | 2 | 0 | 6 | 2 |
| ENSG00000093000 | 1531 | 681 | 796 | 516 | 858 | 706 |
| ENSG00000273353 | 2 | 2 | 1 | 0 | 0 | 0 |
| ENSG00000100364 | 1640 | 765 | 948 | 766 | 765 | 774 |
| ENSG00000221598 | 0 | 1 | 1 | 0 | 0 | 0 |
| ENSG00000273287 | 2 | 0 | 1 | 3 | 1 | 6 |
| ENSG00000213752 | 0 | 0 | 0 | 0 | 0 | 0 |
| ENSG00000100373 | 150 | 90 | 97 | 30 | 32 | 66 |
| ENSG00000100376 | 679 | 404 | 499 | 338 | 380 | 396 |
| ENSG00000077935 | 10 | 3 | 8 | 4 | 8 | 2 |
| ENSG00000128408 | 62 | 31 | 40 | 23 | 22 | 54 |
| ENSG00000232363 | 3 | 0 | 2 | 1 | 1 | 1 |
| ENSG00000077942 | 104 | 38 | 43 | 44 | 38 | 53 |
| ENSG00000238120 | 0 | 0 | 0 | 0 | 0 | 0 |
| ENSG00000251985 | 0 | 0 | 0 | 0 | 0 | 0 |
| ENSG00000280383 | 2 | 10 | 2 | 2 | 1 | 4 |
| ENSG00000130638 | 1831 | 970 | 1152 | 852 | 1087 | 1030 |
| ENSG00000264160 | 0 | 1 | 0 | 0 | 0 | 0 |
| ENSG00000235347 | 0 | 0 | 0 | 0 | 0 | 0 |
| ENSG00000279954 | 4 | 1 | 2 | 0 | 0 | 2 |
| ENSG00000235091 | 1 | 0 | 2 | 0 | 0 | 0 |
| ENSG00000188064 | 1202 | 574 | 814 | 780 | 672 | 749 |
| ENSG00000280424 | 12 | 6 | 5 | 5 | 6 | 4 |
| ENSG00000273145 | 56 | 21 | 31 | 28 | 36 | 27 |
| ENSG00000231711 | 4 | 0 | 3 | 1 | 3 | 5 |
| ENSG00000182257 | 4 | 3 | 6 | 3 | 6 | 9 |
| ENSG00000273289 | 0 | 0 | 0 | 0 | 0 | 0 |
| ENSG00000241990 | 63 | 41 | 34 | 47 | 41 | 39 |
| ENSG00000197182 | 43 | 37 | 32 | 25 | 28 | 33 |
| ENSG00000231010 | 2 | 0 | 4 | 4 | 2 | 0 |
| ENSG00000235159 | 17 | 12 | 8 | 6 | 6 | 7 |
| ENSG00000266533 | 0 | 0 | 0 | 0 | 0 | 0 |
| ENSG00000283990 | 0 | 0 | 0 | 0 | 0 | 0 |
| ENSG00000284175 | 0 | 0 | 0 | 0 | 0 | 0 |
| ENSG00000284520 | 0 | 0 | 0 | 0 | 0 | 0 |
| ENSG00000186951 | 605 | 258 | 319 | 157 | 264 | 272 |
| ENSG00000280384 | 0 | 0 | 0 | 0 | 4 | 1 |
| ENSG00000205643 | 182 | 79 | 119 | 105 | 109 | 109 |
| ENSG00000130943 | 18 | 9 | 7 | 9 | 10 | 15 |
| ENSG00000075234 | 529 | 213 | 277 | 269 | 211 | 253 |
| ENSG00000277232 | 48 | 15 | 15 | 23 | 21 | 9 |
| ENSG00000075218 | 425 | 215 | 264 | 205 | 265 | 283 |
| ENSG00000100416 | 710 | 321 | 434 | 467 | 432 | 443 |
| ENSG00000075275 | 4229 | 1738 | 2680 | 1748 | 2017 | 2167 |

| | | | | | | |
|---|---|---|---|---|---|---|
| ENSG00000230634 | 0 | 0 | 0 | 0 | 0 | 0 |
| ENSG00000234869 | 166 | 74 | 93 | 56 | 62 | 63 |
| ENSG00000075240 | 1564 | 762 | 1016 | 775 | 859 | 836 |
| ENSG00000100422 | 450 | 239 | 308 | 188 | 248 | 237 |
| ENSG00000260708 | 66 | 38 | 53 | 39 | 61 | 53 |
| ENSG00000054611 | 793 | 354 | 493 | 426 | 362 | 385 |
| ENSG00000280080 | 2 | 0 | 0 | 0 | 0 | 0 |
| ENSG00000278881 | 4 | 0 | 1 | 2 | 0 | 0 |
| ENSG00000280312 | 0 | 0 | 0 | 0 | 0 | 0 |
| ENSG00000224715 | 0 | 0 | 0 | 0 | 0 | 0 |
| ENSG00000218357 | 0 | 0 | 0 | 0 | 0 | 0 |
| ENSG00000205634 | 6 | 3 | 8 | 3 | 2 | 3 |
| ENSG00000224271 | 37 | 15 | 27 | 11 | 14 | 25 |
| ENSG00000232897 | 0 | 0 | 0 | 0 | 0 | 0 |
| ENSG00000230345 | 0 | 1 | 0 | 1 | 2 | 1 |
| ENSG00000279712 | 2 | 1 | 4 | 0 | 1 | 4 |
| ENSG00000235154 | 0 | 0 | 0 | 0 | 0 | 0 |
| ENSG00000278890 | 0 | 0 | 0 | 0 | 0 | 0 |
| ENSG00000238192 | 0 | 0 | 0 | 0 | 0 | 0 |
| ENSG00000229275 | 0 | 0 | 0 | 0 | 0 | 0 |
| ENSG00000285707 | 0 | 0 | 0 | 0 | 0 | 0 |
| ENSG00000266508 | 0 | 0 | 0 | 0 | 0 | 0 |
| ENSG00000285855 | 0 | 0 | 0 | 0 | 0 | 0 |
| ENSG00000233179 | 0 | 0 | 0 | 0 | 0 | 0 |
| ENSG00000219438 | 242 | 103 | 129 | 142 | 146 | 120 |
| ENSG00000281732 | 0 | 0 | 0 | 1 | 0 | 0 |
| ENSG00000266887 | 0 | 0 | 0 | 0 | 0 | 0 |
| ENSG00000205632 | 4 | 0 | 1 | 0 | 0 | 1 |
| ENSG00000285623 | 0 | 0 | 0 | 0 | 0 | 0 |
| ENSG00000226142 | 0 | 0 | 0 | 0 | 0 | 0 |
| ENSG00000280199 | 0 | 0 | 0 | 0 | 0 | 0 |
| ENSG00000285722 | 5 | 2 | 5 | 2 | 4 | 8 |
| ENSG00000188511 | 0 | 0 | 0 | 0 | 0 | 0 |
| ENSG00000280224 | 0 | 0 | 0 | 0 | 0 | 0 |
| ENSG00000279597 | 0 | 0 | 0 | 0 | 0 | 0 |
| ENSG00000264139 | 0 | 0 | 0 | 0 | 0 | 0 |
| ENSG00000212939 | 2 | 1 | 1 | 0 | 2 | 0 |
| ENSG00000213279 | 0 | 0 | 0 | 0 | 0 | 0 |
| ENSG00000235111 | 0 | 0 | 0 | 0 | 0 | 0 |
| ENSG00000226954 | 0 | 0 | 0 | 1 | 1 | 0 |
| ENSG00000223142 | 0 | 0 | 0 | 0 | 0 | 0 |
| ENSG00000214727 | 0 | 0 | 0 | 0 | 0 | 0 |
| ENSG00000100425 | 1037 | 492 | 667 | 551 | 627 | 602 |
| ENSG00000279345 | 3 | 2 | 3 | 5 | 6 | 1 |
| ENSG00000236867 | 0 | 2 | 3 | 0 | 2 | 0 |
| ENSG00000260613 | 0 | 0 | 1 | 0 | 0 | 0 |
| ENSG00000229409 | 0 | 0 | 0 | 0 | 0 | 1 |
| ENSG00000277771 | 0 | 0 | 0 | 0 | 0 | 0 |
| ENSG00000279494 | 2 | 0 | 0 | 0 | 1 | 0 |
| ENSG00000100426 | 760 | 342 | 454 | 363 | 463 | 492 |

| | | | | | | |
|---|---|---|---|---|---|---|
| ENSG00000182858 | 391 | 177 | 255 | 205 | 184 | 201 |
| ENSG00000273192 | 2 | 2 | 6 | 0 | 1 | 1 |
| ENSG00000184164 | 848 | 381 | 553 | 593 | 530 | 490 |
| ENSG00000278869 | 12 | 7 | 7 | 7 | 7 | 6 |
| ENSG00000198355 | 2497 | 1120 | 1590 | 1790 | 1565 | 1763 |
| ENSG00000276753 | 0 | 0 | 0 | 0 | 0 | 0 |
| ENSG00000188263 | 43 | 26 | 25 | 16 | 47 | 47 |
| ENSG00000138892 | 0 | 1 | 0 | 0 | 0 | 0 |
| ENSG00000100427 | 0 | 0 | 0 | 0 | 0 | 0 |
| ENSG00000073146 | 0 | 2 | 0 | 4 | 0 | 1 |
| ENSG00000280395 | 0 | 0 | 0 | 0 | 0 | 0 |
| ENSG00000073150 | 702 | 316 | 402 | 512 | 505 | 445 |
| ENSG00000170638 | 924 | 462 | 660 | 667 | 653 | 641 |
| ENSG00000273188 | 12 | 1 | 1 | 3 | 11 | 4 |
| ENSG00000273253 | 37 | 16 | 19 | 16 | 20 | 20 |
| ENSG00000073169 | 770 | 367 | 476 | 543 | 520 | 516 |
| ENSG00000272836 | 27 | 8 | 7 | 5 | 4 | 6 |
| ENSG00000273137 | 19 | 17 | 24 | 15 | 15 | 29 |
| ENSG00000128159 | 906 | 351 | 539 | 362 | 401 | 426 |
| ENSG00000100429 | 31 | 21 | 38 | 34 | 27 | 22 |
| ENSG00000188130 | 368 | 189 | 260 | 215 | 201 | 180 |
| ENSG00000185386 | 461 | 254 | 316 | 248 | 209 | 205 |
| ENSG00000196576 | 11809 | 4942 | 7187 | 5367 | 5230 | 5412 |
| ENSG00000205593 | 188 | 93 | 123 | 115 | 88 | 76 |
| ENSG00000227484 | 0 | 0 | 1 | 2 | 0 | 0 |
| ENSG00000279182 | 0 | 0 | 0 | 0 | 0 | 0 |
| ENSG00000100239 | 2055 | 924 | 1123 | 1050 | 1113 | 1010 |
| ENSG00000100241 | 2236 | 969 | 1368 | 1320 | 1320 | 1272 |
| ENSG00000128165 | 90 | 27 | 45 | 36 | 40 | 42 |
| ENSG00000100253 | 0 | 2 | 1 | 1 | 0 | 0 |
| ENSG00000100258 | 1133 | 515 | 701 | 674 | 590 | 549 |
| ENSG00000025770 | 743 | 322 | 461 | 421 | 366 | 375 |
| ENSG00000284194 | 0 | 0 | 0 | 0 | 0 | 0 |
| ENSG00000130489 | 12 | 9 | 6 | 21 | 8 | 8 |
| ENSG00000272821 | 15 | 9 | 19 | 20 | 15 | 20 |
| ENSG00000025708 | 42 | 17 | 24 | 24 | 12 | 16 |
| ENSG00000177989 | 57 | 21 | 30 | 25 | 20 | 23 |
| ENSG00000272666 | 0 | 2 | 3 | 0 | 0 | 1 |
| ENSG00000273272 | 0 | 0 | 0 | 0 | 0 | 0 |
| ENSG00000130487 | 8 | 7 | 4 | 9 | 10 | 6 |
| ENSG00000217442 | 16 | 7 | 17 | 8 | 12 | 8 |
| ENSG00000205560 | 13 | 13 | 14 | 5 | 13 | 12 |
| ENSG00000254413 | 8 | 10 | 17 | 12 | 5 | 7 |
| ENSG00000100288 | 10 | 11 | 17 | 7 | 9 | 20 |
| ENSG00000205559 | 17 | 12 | 8 | 10 | 13 | 14 |
| ENSG00000272940 | 0 | 2 | 2 | 2 | 7 | 3 |
| ENSG00000008735 | 158 | 90 | 113 | 92 | 75 | 88 |
| ENSG00000100299 | 459 | 200 | 290 | 250 | 240 | 203 |
| ENSG00000212569 | 0 | 0 | 1 | 0 | 0 | 0 |
| ENSG00000251322 | 238 | 90 | 136 | 157 | 125 | 123 |

| | | | | | | |
|---|---|---|---|---|---|---|
| ENSG00000206841 | 0 | 0 | 0 | 0 | 0 | 0 |
| ENSG00000225929 | 0 | 0 | 0 | 0 | 0 | 0 |
| ENSG00000100312 | 5 | 1 | 0 | 2 | 0 | 0 |
| ENSG00000254499 | 0 | 0 | 0 | 0 | 0 | 0 |
| ENSG00000213683 | 2 | 2 | 1 | 2 | 1 | 2 |
| ENSG00000184319 | 106 | 49 | 79 | 89 | 113 | 86 |
| ENSG00000079974 | 356 | 181 | 250 | 201 | 220 | 208 |
| ENSG00000279493 | 0 | 0 | 0 | 0 | 0 | 0 |
| ENSG00000277117 | 61 | 35 | 46 | 27 | 33 | 30 |
| ENSG00000279687 | 0 | 0 | 0 | 0 | 0 | 0 |
| ENSG00000280071 | 310 | 141 | 199 | 209 | 159 | 199 |
| ENSG00000276612 | 0 | 0 | 0 | 0 | 0 | 0 |
| ENSG00000275464 | 356 | 174 | 208 | 238 | 209 | 225 |
| ENSG00000280433 | 128 | 49 | 72 | 62 | 63 | 68 |
| ENSG00000279669 | 0 | 0 | 0 | 0 | 0 | 0 |
| ENSG00000279094 | 0 | 2 | 0 | 1 | 0 | 1 |
| ENSG00000274333 | 0 | 0 | 0 | 0 | 0 | 0 |
| ENSG00000277777 | 0 | 0 | 0 | 0 | 0 | 0 |
| ENSG00000279186 | 0 | 0 | 0 | 0 | 0 | 0 |
| ENSG00000279784 | 0 | 0 | 0 | 0 | 0 | 0 |
| ENSG00000279064 | 0 | 0 | 0 | 0 | 0 | 0 |
| ENSG00000274559 | 0 | 0 | 0 | 0 | 0 | 0 |
| ENSG00000280013 | 0 | 0 | 0 | 0 | 0 | 0 |
| ENSG00000279769 | 0 | 0 | 0 | 0 | 0 | 0 |
| ENSG00000280191 | 0 | 0 | 0 | 0 | 0 | 0 |
| ENSG00000280095 | 0 | 0 | 0 | 0 | 0 | 0 |
| ENSG00000280179 | 0 | 0 | 0 | 0 | 1 | 0 |
| ENSG00000275993 | 314 | 134 | 210 | 160 | 174 | 174 |
| ENSG00000274790 | 0 | 0 | 0 | 0 | 0 | 0 |
| ENSG00000275496 | 0 | 0 | 0 | 0 | 0 | 0 |
| ENSG00000280019 | 0 | 0 | 0 | 0 | 0 | 0 |
| ENSG00000279709 | 0 | 0 | 0 | 0 | 0 | 0 |
| ENSG00000278903 | 0 | 0 | 0 | 0 | 0 | 0 |
| ENSG00000276902 | 0 | 0 | 0 | 0 | 0 | 0 |
| ENSG00000274276 | 6 | 4 | 6 | 1 | 2 | 2 |
| ENSG00000275895 | 4 | 0 | 5 | 0 | 2 | 1 |
| ENSG00000280346 | 0 | 0 | 0 | 0 | 0 | 0 |
| ENSG00000279226 | 0 | 0 | 0 | 0 | 0 | 0 |
| ENSG00000278927 | 0 | 0 | 0 | 0 | 0 | 0 |
| ENSG00000276076 | 0 | 0 | 0 | 0 | 0 | 0 |
| ENSG00000280145 | 0 | 1 | 0 | 0 | 0 | 0 |
| ENSG00000278878 | 0 | 0 | 0 | 0 | 0 | 0 |
| ENSG00000279788 | 0 | 0 | 0 | 0 | 0 | 0 |
| ENSG00000279728 | 0 | 0 | 0 | 0 | 0 | 0 |
| ENSG00000280164 | 2 | 2 | 0 | 0 | 0 | 0 |
| ENSG00000279998 | 0 | 0 | 0 | 0 | 0 | 0 |
| ENSG00000278955 | 2 | 0 | 1 | 0 | 3 | 1 |
| ENSG00000284550 | 0 | 0 | 0 | 0 | 0 | 0 |
| ENSG00000280172 | 8 | 1 | 4 | 0 | 1 | 1 |
| ENSG00000279751 | 0 | 0 | 0 | 0 | 0 | 0 |

| | | | | | | |
|---|---|---|---|---|---|---|
| ENSG00000280018 | 5 | 1 | 6 | 2 | 7 | 4 |
| ENSG00000279313 | 0 | 0 | 0 | 0 | 0 | 0 |
| ENSG00000279477 | 0 | 0 | 0 | 0 | 0 | 0 |
| ENSG00000278884 | 0 | 0 | 0 | 0 | 0 | 0 |
| ENSG00000277067 | 2 | 0 | 0 | 0 | 0 | 0 |
| ENSG00000274046 | 0 | 0 | 0 | 0 | 0 | 0 |
| ENSG00000280330 | 0 | 0 | 0 | 0 | 0 | 0 |
| ENSG00000279303 | 0 | 0 | 0 | 0 | 0 | 0 |
| ENSG00000279647 | 0 | 0 | 0 | 0 | 0 | 0 |
| ENSG00000279895 | 0 | 0 | 0 | 0 | 0 | 0 |
| ENSG00000276077 | 2 | 0 | 0 | 0 | 0 | 0 |
| ENSG00000274484 | 0 | 0 | 0 | 0 | 0 | 0 |
| ENSG00000279177 | 0 | 0 | 0 | 0 | 0 | 0 |
| ENSG00000276546 | 0 | 0 | 0 | 0 | 0 | 0 |
| ENSG00000277991 | 0 | 0 | 0 | 0 | 0 | 0 |
| ENSG00000273590 | 0 | 0 | 0 | 0 | 0 | 0 |
| ENSG00000277277 | 0 | 0 | 0 | 0 | 0 | 0 |
| ENSG00000278961 | 0 | 0 | 0 | 0 | 0 | 0 |
| ENSG00000276289 | 0 | 0 | 0 | 0 | 1 | 0 |
| ENSG00000273614 | 0 | 0 | 0 | 0 | 0 | 0 |
| ENSG00000279967 | 0 | 0 | 0 | 0 | 0 | 0 |
| ENSG00000278996 | 0 | 0 | 0 | 0 | 0 | 0 |
| ENSG00000275950 | 0 | 0 | 0 | 0 | 0 | 0 |
| ENSG00000275664 | 0 | 0 | 0 | 0 | 0 | 0 |
| ENSG00000275708 | 0 | 0 | 0 | 0 | 0 | 0 |
| ENSG00000277437 | 0 | 0 | 0 | 0 | 0 | 0 |
| ENSG00000280800 | 0 | 0 | 0 | 0 | 0 | 0 |
| ENSG00000278233 | 0 | 0 | 0 | 0 | 0 | 0 |
| ENSG00000274060 | 0 | 0 | 0 | 0 | 0 | 0 |
| ENSG00000277671 | 0 | 0 | 0 | 0 | 0 | 0 |
| ENSG00000281383 | 88 | 88 | 106 | 91 | 76 | 82 |
| ENSG00000277739 | 0 | 0 | 0 | 0 | 0 | 0 |
| ENSG00000280441 | 0 | 0 | 0 | 0 | 0 | 0 |
| ENSG00000277379 | 0 | 0 | 0 | 0 | 0 | 0 |
| ENSG00000274868 | 0 | 0 | 0 | 0 | 0 | 0 |
| ENSG00000280614 | 0 | 0 | 0 | 0 | 0 | 0 |
| ENSG00000275215 | 0 | 0 | 0 | 0 | 0 | 0 |
| ENSG00000275692 | 0 | 0 | 0 | 0 | 0 | 0 |
| ENSG00000278775 | 0 | 0 | 0 | 0 | 0 | 0 |
| ENSG00000281181 | 0 | 0 | 0 | 0 | 0 | 0 |
| ENSG00000278189 | 0 | 0 | 0 | 0 | 0 | 0 |
| ENSG00000279990 | 0 | 1 | 0 | 0 | 0 | 0 |
| ENSG00000279615 | 0 | 0 | 0 | 0 | 0 | 0 |
| ENSG00000279167 | 0 | 0 | 0 | 0 | 0 | 0 |
| ENSG00000279579 | 0 | 0 | 3 | 0 | 0 | 0 |
| ENSG00000279213 | 0 | 0 | 0 | 0 | 0 | 0 |
| ENSG00000280243 | 0 | 0 | 0 | 0 | 0 | 0 |
| ENSG00000278931 | 0 | 0 | 1 | 0 | 0 | 0 |
| ENSG00000264462 | 0 | 0 | 0 | 0 | 0 | 0 |
| ENSG00000264063 | 0 | 0 | 0 | 0 | 0 | 0 |

| | | | | | | |
|---|---|---|---|---|---|---|
| ENSG00000279501 | 0 | 0 | 0 | 0 | 0 | 0 |
| ENSG00000279414 | 0 | 0 | 0 | 0 | 0 | 0 |
| ENSG00000280436 | 0 | 0 | 0 | 0 | 0 | 0 |
| ENSG00000188681 | 14 | 7 | 6 | 8 | 8 | 6 |
| ENSG00000279720 | 0 | 0 | 0 | 0 | 1 | 0 |
| ENSG00000279381 | 0 | 0 | 0 | 0 | 0 | 0 |
| ENSG00000278932 | 18 | 5 | 7 | 2 | 5 | 8 |
| ENSG00000279718 | 1 | 0 | 0 | 0 | 0 | 0 |
| ENSG00000279208 | 0 | 0 | 1 | 0 | 0 | 0 |
| ENSG00000279864 | 0 | 0 | 0 | 0 | 0 | 0 |
| ENSG00000279211 | 0 | 0 | 0 | 0 | 0 | 0 |
| ENSG00000275631 | 0 | 0 | 0 | 0 | 0 | 0 |
| ENSG00000280081 | 0 | 0 | 0 | 0 | 0 | 0 |
| ENSG00000280075 | 0 | 0 | 0 | 0 | 0 | 0 |
| ENSG00000279783 | 0 | 0 | 0 | 0 | 0 | 0 |
| ENSG00000264002 | 0 | 0 | 0 | 0 | 0 | 0 |
| ENSG00000252199 | 0 | 0 | 0 | 0 | 0 | 0 |
| ENSG00000279321 | 0 | 0 | 0 | 0 | 0 | 0 |
| ENSG00000270533 | 8 | 3 | 5 | 5 | 4 | 5 |
| ENSG00000280372 | 0 | 0 | 0 | 0 | 0 | 0 |
| ENSG00000279851 | 2 | 0 | 0 | 0 | 0 | 0 |
| ENSG00000279062 | 0 | 0 | 0 | 0 | 0 | 0 |
| ENSG00000277693 | 0 | 0 | 0 | 0 | 1 | 0 |
| ENSG00000275945 | 0 | 0 | 0 | 0 | 0 | 0 |
| ENSG00000275592 | 0 | 0 | 0 | 0 | 0 | 0 |
| ENSG00000278106 | 0 | 1 | 0 | 0 | 0 | 0 |
| ENSG00000187172 | 151 | 46 | 73 | 39 | 53 | 51 |
| ENSG00000273840 | 0 | 2 | 2 | 2 | 5 | 3 |
| ENSG00000274391 | 16 | 5 | 9 | 1 | 4 | 7 |
| ENSG00000278678 | 0 | 0 | 0 | 0 | 0 | 0 |
| ENSG00000273872 | 2 | 3 | 2 | 1 | 3 | 5 |
| ENSG00000276556 | 0 | 0 | 0 | 0 | 0 | 0 |
| ENSG00000277282 | 0 | 0 | 0 | 0 | 0 | 0 |
| ENSG00000229306 | 0 | 0 | 0 | 0 | 0 | 0 |
| ENSG00000224309 | 0 | 0 | 0 | 0 | 0 | 0 |
| ENSG00000207097 | 0 | 0 | 0 | 0 | 0 | 0 |
| ENSG00000168122 | 0 | 0 | 0 | 0 | 0 | 0 |
| ENSG00000237202 | 0 | 0 | 0 | 0 | 0 | 0 |
| ENSG00000280108 | 0 | 0 | 0 | 0 | 0 | 0 |
| ENSG00000185390 | 0 | 0 | 0 | 0 | 0 | 0 |
| ENSG00000175302 | 0 | 0 | 0 | 0 | 0 | 0 |
| ENSG00000266211 | 0 | 0 | 0 | 0 | 0 | 0 |
| ENSG00000226930 | 0 | 0 | 0 | 0 | 0 | 0 |
| ENSG00000219280 | 0 | 0 | 0 | 0 | 0 | 0 |
| ENSG00000228184 | 0 | 0 | 0 | 0 | 0 | 0 |
| ENSG00000179381 | 0 | 0 | 0 | 0 | 0 | 0 |
| ENSG00000228159 | 0 | 0 | 0 | 0 | 0 | 0 |
| ENSG00000218549 | 0 | 0 | 0 | 0 | 0 | 0 |
| ENSG00000166351 | 0 | 0 | 0 | 0 | 0 | 0 |
| ENSG00000207476 | 0 | 0 | 0 | 0 | 0 | 0 |

| | | | | | | |
|---|---|---|---|---|---|---|
| ENSG00000266299 | 0 | 0 | 0 | 0 | 0 | 0 |
| ENSG00000278381 | 0 | 0 | 0 | 0 | 0 | 0 |
| ENSG00000227874 | 0 | 0 | 0 | 0 | 0 | 0 |
| ENSG00000214319 | 0 | 0 | 0 | 0 | 0 | 0 |
| ENSG00000279773 | 0 | 0 | 0 | 0 | 0 | 0 |
| ENSG00000273692 | 0 | 0 | 0 | 0 | 0 | 0 |
| ENSG00000278618 | 0 | 0 | 0 | 0 | 0 | 0 |
| ENSG00000229231 | 0 | 0 | 0 | 0 | 0 | 0 |
| ENSG00000224922 | 0 | 0 | 0 | 0 | 0 | 0 |
| ENSG00000173231 | 0 | 0 | 0 | 0 | 0 | 1 |
| ENSG00000232797 | 0 | 0 | 0 | 0 | 0 | 0 |
| ENSG00000224860 | 0 | 0 | 0 | 0 | 0 | 0 |
| ENSG00000215562 | 0 | 0 | 0 | 0 | 0 | 0 |
| ENSG00000234538 | 0 | 0 | 0 | 0 | 0 | 0 |
| ENSG00000228314 | 26 | 13 | 23 | 18 | 12 | 14 |
| ENSG00000230965 | 4 | 4 | 2 | 1 | 5 | 3 |
| ENSG00000215559 | 12 | 6 | 13 | 5 | 12 | 14 |
| ENSG00000203616 | 0 | 0 | 0 | 1 | 4 | 4 |
| ENSG00000223287 | 0 | 0 | 0 | 0 | 0 | 0 |
| ENSG00000183249 | 0 | 0 | 0 | 0 | 0 | 0 |
| ENSG00000233442 | 0 | 0 | 0 | 0 | 0 | 0 |
| ENSG00000275170 | 0 | 0 | 0 | 0 | 0 | 0 |
| ENSG00000224905 | 0 | 0 | 0 | 0 | 0 | 0 |
| ENSG00000249493 | 5 | 1 | 1 | 0 | 0 | 3 |
| ENSG00000201812 | 0 | 0 | 0 | 0 | 0 | 0 |
| ENSG00000277572 | 0 | 0 | 0 | 0 | 0 | 0 |
| ENSG00000188992 | 0 | 1 | 0 | 0 | 1 | 0 |
| ENSG00000240755 | 0 | 0 | 0 | 0 | 0 | 0 |
| ENSG00000185272 | 14 | 6 | 4 | 2 | 8 | 4 |
| ENSG00000243064 | 2 | 0 | 1 | 0 | 1 | 5 |
| ENSG00000155304 | 615 | 345 | 353 | 264 | 459 | 312 |
| ENSG00000155307 | 0 | 0 | 0 | 0 | 0 | 0 |
| ENSG00000223662 | 0 | 0 | 0 | 0 | 0 | 1 |
| ENSG00000232884 | 0 | 0 | 0 | 0 | 0 | 0 |
| ENSG00000228600 | 0 | 0 | 0 | 0 | 0 | 0 |
| ENSG00000226751 | 0 | 0 | 0 | 0 | 0 | 0 |
| ENSG00000225502 | 0 | 0 | 0 | 0 | 0 | 0 |
| ENSG00000235609 | 66 | 43 | 26 | 23 | 59 | 38 |
| ENSG00000281903 | 5 | 7 | 4 | 1 | 2 | 5 |
| ENSG00000226406 | 0 | 0 | 0 | 0 | 0 | 0 |
| ENSG00000235277 | 191 | 102 | 103 | 127 | 141 | 135 |
| ENSG00000180530 | 1375 | 761 | 576 | 281 | 1404 | 814 |
| ENSG00000229047 | 0 | 0 | 0 | 0 | 0 | 0 |
| ENSG00000231201 | 0 | 0 | 0 | 0 | 0 | 4 |
| ENSG00000279390 | 0 | 1 | 0 | 0 | 1 | 2 |
| ENSG00000236471 | 1 | 2 | 3 | 1 | 2 | 1 |
| ENSG00000280082 | 0 | 0 | 0 | 0 | 0 | 0 |
| ENSG00000229425 | 0 | 0 | 0 | 0 | 1 | 0 |
| ENSG00000224524 | 0 | 0 | 0 | 0 | 0 | 0 |
| ENSG00000224247 | 0 | 0 | 0 | 0 | 0 | 0 |

| | | | | | | |
|---|---|---|---|---|---|---|
| ENSG00000212564 | 0 | 0 | 0 | 0 | 0 | 0 |
| ENSG00000226298 | 0 | 0 | 0 | 0 | 0 | 0 |
| ENSG00000155313 | 744 | 401 | 353 | 209 | 522 | 344 |
| ENSG00000234159 | 0 | 0 | 0 | 0 | 0 | 0 |
| ENSG00000215386 | 136 | 55 | 56 | 42 | 34 | 31 |
| ENSG00000252273 | 0 | 0 | 0 | 0 | 0 | 0 |
| ENSG00000214976 | 0 | 0 | 0 | 0 | 0 | 0 |
| ENSG00000224602 | 0 | 0 | 0 | 0 | 0 | 0 |
| ENSG00000201025 | 0 | 0 | 0 | 0 | 0 | 0 |
| ENSG00000270071 | 0 | 0 | 0 | 0 | 0 | 0 |
| ENSG00000270139 | 0 | 1 | 0 | 0 | 0 | 0 |
| ENSG00000280432 | 0 | 0 | 0 | 0 | 0 | 0 |
| ENSG00000207638 | 0 | 0 | 0 | 0 | 0 | 0 |
| ENSG00000199030 | 0 | 0 | 0 | 0 | 0 | 0 |
| ENSG00000269950 | 0 | 0 | 0 | 0 | 0 | 0 |
| ENSG00000207863 | 0 | 0 | 0 | 0 | 0 | 0 |
| ENSG00000239023 | 0 | 0 | 0 | 0 | 0 | 0 |
| ENSG00000237735 | 0 | 0 | 0 | 0 | 0 | 0 |
| ENSG00000232886 | 0 | 0 | 0 | 0 | 0 | 0 |
| ENSG00000225735 | 0 | 0 | 0 | 0 | 0 | 0 |
| ENSG00000278181 | 0 | 0 | 0 | 0 | 0 | 0 |
| ENSG00000252462 | 0 | 0 | 0 | 0 | 0 | 0 |
| ENSG00000232560 | 0 | 0 | 0 | 0 | 0 | 0 |
| ENSG00000226580 | 0 | 0 | 0 | 0 | 0 | 0 |
| ENSG00000266195 | 0 | 0 | 0 | 0 | 0 | 0 |
| ENSG00000154639 | 24 | 6 | 9 | 4 | 10 | 3 |
| ENSG00000232260 | 0 | 0 | 0 | 0 | 0 | 0 |
| ENSG00000200754 | 0 | 0 | 0 | 0 | 0 | 0 |
| ENSG00000199962 | 0 | 0 | 0 | 0 | 0 | 0 |
| ENSG00000154640 | 41 | 15 | 22 | 18 | 19 | 22 |
| ENSG00000280594 | 0 | 0 | 0 | 0 | 0 | 0 |
| ENSG00000226956 | 0 | 0 | 0 | 0 | 0 | 0 |
| ENSG00000279648 | 0 | 0 | 0 | 0 | 0 | 0 |
| ENSG00000240770 | 0 | 0 | 0 | 0 | 0 | 0 |
| ENSG00000154642 | 162 | 97 | 78 | 64 | 139 | 91 |
| ENSG00000244676 | 0 | 0 | 0 | 0 | 2 | 0 |
| ENSG00000231755 | 0 | 0 | 0 | 0 | 0 | 0 |
| ENSG00000215369 | 0 | 0 | 0 | 0 | 0 | 0 |
| ENSG00000154645 | 0 | 1 | 0 | 0 | 1 | 0 |
| ENSG00000227330 | 0 | 0 | 0 | 0 | 0 | 0 |
| ENSG00000154646 | 0 | 1 | 0 | 0 | 0 | 0 |
| ENSG00000228708 | 0 | 0 | 0 | 0 | 0 | 0 |
| ENSG00000224141 | 8 | 7 | 14 | 6 | 5 | 7 |
| ENSG00000230233 | 0 | 0 | 0 | 0 | 0 | 0 |
| ENSG00000265841 | 0 | 0 | 0 | 0 | 0 | 0 |
| ENSG00000198618 | 2010 | 952 | 1121 | 1156 | 1182 | 1098 |
| ENSG00000279156 | 0 | 0 | 0 | 0 | 0 | 0 |
| ENSG00000232193 | 0 | 0 | 0 | 0 | 0 | 0 |
| ENSG00000226204 | 0 | 0 | 0 | 0 | 0 | 0 |
| ENSG00000235965 | 0 | 0 | 0 | 0 | 0 | 0 |

| | | | | | | |
|---|---|---|---|---|---|---|
| ENSG00000229289 | 0 | 0 | 0 | 0 | 0 | 0 |
| ENSG00000226818 | 0 | 0 | 0 | 0 | 0 | 0 |
| ENSG00000231620 | 0 | 0 | 0 | 0 | 0 | 0 |
| ENSG00000212609 | 0 | 0 | 0 | 0 | 0 | 0 |
| ENSG00000230198 | 0 | 0 | 0 | 0 | 0 | 0 |
| ENSG00000271486 | 0 | 0 | 0 | 0 | 0 | 0 |
| ENSG00000229336 | 0 | 0 | 1 | 0 | 0 | 0 |
| ENSG00000215353 | 0 | 0 | 0 | 0 | 0 | 0 |
| ENSG00000233480 | 0 | 0 | 0 | 0 | 0 | 0 |
| ENSG00000233236 | 0 | 0 | 0 | 0 | 0 | 0 |
| ENSG00000251851 | 0 | 0 | 0 | 0 | 0 | 0 |
| ENSG00000252963 | 0 | 0 | 0 | 0 | 0 | 0 |
| ENSG00000233676 | 0 | 0 | 0 | 0 | 0 | 0 |
| ENSG00000234439 | 2 | 0 | 1 | 0 | 0 | 0 |
| ENSG00000233206 | 0 | 0 | 0 | 0 | 0 | 0 |
| ENSG00000276738 | 0 | 0 | 0 | 0 | 0 | 0 |
| ENSG00000224924 | 0 | 0 | 0 | 0 | 0 | 0 |
| ENSG00000215351 | 0 | 0 | 0 | 0 | 0 | 0 |
| ENSG00000154654 | 2709 | 1382 | 1562 | 472 | 931 | 935 |
| ENSG00000226771 | 0 | 1 | 0 | 0 | 0 | 0 |
| ENSG00000234730 | 0 | 0 | 0 | 0 | 0 | 0 |
| ENSG00000237527 | 32 | 30 | 27 | 19 | 25 | 19 |
| ENSG00000238265 | 0 | 0 | 0 | 0 | 0 | 0 |
| ENSG00000233997 | 75 | 34 | 67 | 80 | 70 | 73 |
| ENSG00000227075 | 0 | 0 | 0 | 0 | 0 | 0 |
| ENSG00000233215 | 0 | 0 | 0 | 0 | 0 | 0 |
| ENSG00000184856 | 0 | 0 | 0 | 0 | 0 | 0 |
| ENSG00000234340 | 0 | 0 | 0 | 0 | 0 | 0 |
| ENSG00000231136 | 0 | 0 | 0 | 0 | 0 | 0 |
| ENSG00000202339 | 0 | 0 | 0 | 0 | 0 | 0 |
| ENSG00000226043 | 0 | 0 | 0 | 0 | 0 | 0 |
| ENSG00000223488 | 0 | 0 | 0 | 0 | 0 | 0 |
| ENSG00000234034 | 0 | 0 | 0 | 0 | 0 | 0 |
| ENSG00000225906 | 0 | 0 | 0 | 0 | 0 | 0 |
| ENSG00000230972 | 0 | 0 | 0 | 0 | 0 | 0 |
| ENSG00000275469 | 0 | 0 | 0 | 0 | 0 | 0 |
| ENSG00000219368 | 0 | 0 | 0 | 0 | 0 | 0 |
| ENSG00000231058 | 0 | 0 | 0 | 0 | 0 | 0 |
| ENSG00000223078 | 0 | 0 | 0 | 0 | 0 | 0 |
| ENSG00000228592 | 0 | 0 | 0 | 0 | 0 | 0 |
| ENSG00000223822 | 1 | 0 | 0 | 0 | 0 | 1 |
| ENSG00000237569 | 0 | 0 | 0 | 0 | 0 | 0 |
| ENSG00000199698 | 0 | 0 | 0 | 0 | 0 | 0 |
| ENSG00000227716 | 0 | 0 | 0 | 0 | 0 | 0 |
| ENSG00000231986 | 0 | 0 | 0 | 0 | 0 | 0 |
| ENSG00000235564 | 0 | 0 | 0 | 0 | 0 | 0 |
| ENSG00000226996 | 0 | 0 | 0 | 0 | 0 | 0 |
| ENSG00000224018 | 0 | 0 | 0 | 0 | 0 | 0 |
| ENSG00000224832 | 0 | 0 | 0 | 0 | 0 | 0 |
| ENSG00000237484 | 0 | 0 | 0 | 0 | 0 | 0 |

| | | | | | | |
|---|---|---|---|---|---|---|
| ENSG00000226983 | 0 | 0 | 0 | 0 | 0 | 0 |
| ENSG00000227090 | 0 | 0 | 0 | 0 | 0 | 0 |
| ENSG00000230379 | 0 | 0 | 0 | 0 | 0 | 0 |
| ENSG00000237444 | 0 | 0 | 0 | 0 | 0 | 0 |
| ENSG00000244278 | 0 | 0 | 0 | 0 | 0 | 0 |
| ENSG00000232512 | 0 | 0 | 0 | 0 | 0 | 0 |
| ENSG00000274662 | 0 | 0 | 0 | 0 | 0 | 0 |
| ENSG00000223870 | 0 | 0 | 0 | 0 | 0 | 0 |
| ENSG00000222042 | 0 | 0 | 0 | 0 | 0 | 0 |
| ENSG00000238627 | 0 | 0 | 0 | 0 | 0 | 0 |
| ENSG00000213885 | 2 | 5 | 0 | 1 | 0 | 0 |
| ENSG00000279534 | 0 | 0 | 0 | 0 | 0 | 0 |
| ENSG00000185433 | 0 | 0 | 0 | 0 | 0 | 0 |
| ENSG00000229962 | 0 | 0 | 0 | 0 | 0 | 0 |
| ENSG00000234883 | 0 | 0 | 0 | 0 | 0 | 0 |
| ENSG00000283904 | 0 | 0 | 0 | 0 | 0 | 0 |
| ENSG00000260583 | 0 | 0 | 0 | 0 | 2 | 0 |
| ENSG00000154719 | 666 | 357 | 416 | 368 | 438 | 408 |
| ENSG00000154721 | 8 | 10 | 13 | 8 | 7 | 7 |
| ENSG00000237731 | 0 | 0 | 0 | 0 | 0 | 0 |
| ENSG00000227054 | 0 | 0 | 0 | 0 | 0 | 0 |
| ENSG00000154723 | 1764 | 1084 | 1159 | 1123 | 1212 | 1204 |
| ENSG00000154727 | 561 | 283 | 264 | 128 | 380 | 322 |
| ENSG00000235514 | 0 | 0 | 0 | 0 | 0 | 0 |
| ENSG00000142192 | 12449 | 6052 | 7733 | 5000 | 5797 | 5631 |
| ENSG00000233783 | 0 | 1 | 0 | 0 | 0 | 0 |
| ENSG00000251972 | 0 | 0 | 0 | 0 | 0 | 0 |
| ENSG00000224541 | 0 | 0 | 0 | 0 | 0 | 0 |
| ENSG00000273492 | 0 | 2 | 2 | 0 | 0 | 1 |
| ENSG00000206802 | 0 | 0 | 0 | 0 | 0 | 0 |
| ENSG00000229025 | 0 | 0 | 0 | 0 | 0 | 0 |
| ENSG00000232692 | 0 | 0 | 0 | 0 | 0 | 0 |
| ENSG00000197934 | 0 | 0 | 0 | 0 | 0 | 0 |
| ENSG00000273115 | 0 | 0 | 0 | 0 | 0 | 0 |
| ENSG00000166265 | 0 | 0 | 0 | 0 | 0 | 0 |
| ENSG00000154734 | 0 | 0 | 0 | 0 | 0 | 0 |
| ENSG00000223563 | 0 | 0 | 0 | 0 | 0 | 0 |
| ENSG00000154736 | 0 | 0 | 0 | 0 | 0 | 0 |
| ENSG00000266133 | 0 | 0 | 0 | 0 | 0 | 0 |
| ENSG00000215326 | 0 | 0 | 0 | 0 | 0 | 0 |
| ENSG00000231236 | 0 | 0 | 0 | 0 | 0 | 0 |
| ENSG00000236332 | 0 | 0 | 0 | 0 | 0 | 0 |
| ENSG00000233300 | 0 | 0 | 0 | 0 | 0 | 0 |
| ENSG00000217026 | 0 | 0 | 0 | 0 | 0 | 0 |
| ENSG00000219592 | 0 | 0 | 0 | 0 | 0 | 0 |
| ENSG00000234052 | 0 | 0 | 0 | 0 | 0 | 0 |
| ENSG00000225298 | 0 | 0 | 0 | 0 | 0 | 0 |
| ENSG00000234083 | 0 | 0 | 0 | 0 | 0 | 0 |
| ENSG00000178457 | 0 | 0 | 0 | 0 | 0 | 0 |
| ENSG00000232079 | 0 | 0 | 0 | 0 | 0 | 0 |

| | | | | | | |
|---|---|---|---|---|---|---|
| ENSG00000236532 | 0 | 0 | 0 | 0 | 0 | 0 |
| ENSG00000232855 | 0 | 1 | 2 | 1 | 0 | 1 |
| ENSG00000226935 | 0 | 0 | 0 | 1 | 0 | 0 |
| ENSG00000283300 | 0 | 0 | 0 | 0 | 0 | 0 |
| ENSG00000156239 | 471 | 271 | 308 | 241 | 295 | 225 |
| ENSG00000215005 | 0 | 0 | 0 | 0 | 0 | 0 |
| ENSG00000215317 | 0 | 0 | 0 | 0 | 0 | 0 |
| ENSG00000198862 | 432 | 298 | 194 | 79 | 630 | 219 |
| ENSG00000176054 | 7 | 4 | 2 | 1 | 9 | 4 |
| ENSG00000156253 | 1227 | 577 | 763 | 603 | 683 | 785 |
| ENSG00000273254 | 3 | 0 | 3 | 5 | 4 | 3 |
| ENSG00000156256 | 588 | 392 | 315 | 170 | 570 | 314 |
| ENSG00000156261 | 5164 | 2949 | 3081 | 2684 | 3907 | 3299 |
| ENSG00000231125 | 8 | 1 | 4 | 1 | 6 | 4 |
| ENSG00000156265 | 0 | 0 | 3 | 1 | 0 | 2 |
| ENSG00000232687 | 0 | 0 | 0 | 0 | 0 | 0 |
| ENSG00000201984 | 0 | 0 | 0 | 0 | 0 | 0 |
| ENSG00000212479 | 0 | 0 | 0 | 0 | 0 | 0 |
| ENSG00000224649 | 0 | 0 | 0 | 1 | 0 | 0 |
| ENSG00000215533 | 0 | 0 | 0 | 0 | 0 | 0 |
| ENSG00000156273 | 325 | 187 | 163 | 61 | 238 | 178 |
| ENSG00000236056 | 2 | 0 | 0 | 0 | 0 | 0 |
| ENSG00000248476 | 4 | 0 | 5 | 1 | 2 | 1 |
| ENSG00000273017 | 1 | 0 | 1 | 0 | 1 | 0 |
| ENSG00000232118 | 23 | 7 | 13 | 10 | 4 | 16 |
| ENSG00000228817 | 2 | 0 | 1 | 1 | 1 | 1 |
| ENSG00000234293 | 0 | 0 | 0 | 0 | 0 | 0 |
| ENSG00000171189 | 5 | 0 | 1 | 0 | 0 | 1 |
| ENSG00000273464 | 0 | 0 | 0 | 0 | 0 | 0 |
| ENSG00000174680 | 0 | 0 | 0 | 0 | 0 | 0 |
| ENSG00000259981 | 0 | 0 | 0 | 0 | 0 | 0 |
| ENSG00000156282 | 0 | 0 | 0 | 0 | 0 | 0 |
| ENSG00000227342 | 0 | 0 | 0 | 0 | 0 | 0 |
| ENSG00000156284 | 0 | 0 | 0 | 0 | 0 | 0 |
| ENSG00000225267 | 0 | 0 | 0 | 0 | 0 | 0 |
| ENSG00000188694 | 0 | 0 | 0 | 0 | 0 | 0 |
| ENSG00000232263 | 0 | 0 | 0 | 0 | 0 | 0 |
| ENSG00000197683 | 0 | 0 | 0 | 0 | 0 | 0 |
| ENSG00000206107 | 0 | 0 | 0 | 0 | 0 | 0 |
| ENSG00000186980 | 0 | 0 | 0 | 0 | 0 | 0 |
| ENSG00000250973 | 0 | 0 | 0 | 0 | 0 | 0 |
| ENSG00000182816 | 0 | 0 | 0 | 0 | 0 | 0 |
| ENSG00000265007 | 0 | 0 | 0 | 0 | 0 | 0 |
| ENSG00000198390 | 0 | 0 | 0 | 0 | 0 | 0 |
| ENSG00000240432 | 0 | 0 | 0 | 0 | 0 | 0 |
| ENSG00000186971 | 0 | 0 | 0 | 0 | 0 | 0 |
| ENSG00000233640 | 0 | 0 | 0 | 0 | 0 | 0 |
| ENSG00000186970 | 0 | 0 | 0 | 0 | 0 | 0 |
| ENSG00000184351 | 0 | 0 | 0 | 0 | 0 | 0 |
| ENSG00000186965 | 0 | 0 | 0 | 0 | 0 | 0 |

| | | | | | | |
|---|---|---|---|---|---|---|
| ENSG00000244025 | 0 | 0 | 0 | 0 | 0 | 0 |
| ENSG00000186967 | 0 | 0 | 0 | 0 | 0 | 0 |
| ENSG00000186977 | 0 | 0 | 0 | 0 | 0 | 0 |
| ENSG00000237841 | 0 | 0 | 0 | 0 | 0 | 0 |
| ENSG00000235312 | 0 | 0 | 0 | 0 | 0 | 0 |
| ENSG00000218125 | 0 | 0 | 0 | 0 | 0 | 0 |
| ENSG00000236612 | 0 | 0 | 0 | 0 | 0 | 0 |
| ENSG00000186925 | 0 | 0 | 0 | 0 | 0 | 0 |
| ENSG00000244362 | 0 | 0 | 0 | 0 | 0 | 0 |
| ENSG00000206106 | 0 | 0 | 0 | 0 | 0 | 0 |
| ENSG00000212938 | 0 | 0 | 0 | 0 | 0 | 0 |
| ENSG00000186930 | 0 | 0 | 0 | 0 | 0 | 0 |
| ENSG00000186924 | 0 | 0 | 0 | 0 | 0 | 0 |
| ENSG00000184724 | 0 | 0 | 0 | 0 | 0 | 0 |
| ENSG00000244624 | 0 | 0 | 0 | 0 | 0 | 0 |
| ENSG00000206105 | 0 | 0 | 0 | 0 | 0 | 0 |
| ENSG00000184032 | 0 | 0 | 0 | 0 | 0 | 0 |
| ENSG00000206104 | 0 | 0 | 0 | 0 | 0 | 0 |
| ENSG00000231068 | 0 | 0 | 0 | 0 | 0 | 0 |
| ENSG00000236400 | 0 | 0 | 0 | 0 | 0 | 0 |
| ENSG00000187026 | 0 | 0 | 0 | 0 | 0 | 0 |
| ENSG00000187005 | 0 | 0 | 0 | 0 | 0 | 0 |
| ENSG00000238257 | 0 | 0 | 0 | 0 | 0 | 0 |
| ENSG00000233036 | 0 | 0 | 0 | 0 | 0 | 0 |
| ENSG00000227840 | 0 | 0 | 0 | 0 | 0 | 0 |
| ENSG00000183640 | 0 | 0 | 0 | 0 | 0 | 0 |
| ENSG00000274749 | 0 | 0 | 0 | 0 | 0 | 0 |
| ENSG00000182591 | 0 | 0 | 0 | 0 | 0 | 0 |
| ENSG00000206102 | 0 | 0 | 0 | 0 | 0 | 0 |
| ENSG00000228941 | 0 | 0 | 0 | 0 | 0 | 0 |
| ENSG00000156299 | 345 | 173 | 164 | 141 | 191 | 233 |
| ENSG00000237325 | 0 | 0 | 0 | 0 | 0 | 0 |
| ENSG00000237594 | 38 | 20 | 17 | 17 | 15 | 19 |
| ENSG00000230870 | 10 | 14 | 5 | 5 | 9 | 9 |
| ENSG00000234509 | 4 | 2 | 6 | 2 | 5 | 2 |
| ENSG00000142168 | 7077 | 3767 | 4206 | 4284 | 4279 | 4031 |
| ENSG00000238390 | 1 | 3 | 3 | 2 | 3 | 6 |
| ENSG00000273271 | 22 | 14 | 13 | 10 | 4 | 5 |
| ENSG00000156304 | 1075 | 479 | 628 | 477 | 625 | 624 |
| ENSG00000229046 | 0 | 0 | 0 | 0 | 0 | 0 |
| ENSG00000273091 | 11 | 7 | 5 | 1 | 7 | 6 |
| ENSG00000234107 | 0 | 0 | 0 | 0 | 0 | 0 |
| ENSG00000142149 | 199 | 87 | 136 | 54 | 51 | 88 |
| ENSG00000237138 | 0 | 0 | 0 | 0 | 0 | 0 |
| ENSG00000230323 | 0 | 0 | 0 | 0 | 0 | 0 |
| ENSG00000261610 | 0 | 0 | 0 | 0 | 0 | 1 |
| ENSG00000159055 | 523 | 262 | 298 | 258 | 334 | 349 |
| ENSG00000227256 | 10 | 10 | 12 | 4 | 2 | 5 |
| ENSG00000170262 | 0 | 0 | 0 | 0 | 0 | 0 |
| ENSG00000232623 | 2 | 2 | 1 | 1 | 0 | 1 |

| | | | | | | |
|---|---|---|---|---|---|---|
| ENSG00000142207 | 1309 | 601 | 796 | 587 | 808 | 853 |
| ENSG00000200792 | 0 | 0 | 0 | 0 | 0 | 0 |
| ENSG00000256073 | 338 | 172 | 191 | 187 | 215 | 179 |
| ENSG00000166979 | 14 | 11 | 21 | 11 | 14 | 16 |
| ENSG00000229007 | 3 | 0 | 1 | 0 | 0 | 0 |
| ENSG00000252045 | 0 | 0 | 0 | 0 | 0 | 0 |
| ENSG00000252950 | 0 | 0 | 0 | 0 | 2 | 0 |
| ENSG00000186842 | 0 | 0 | 0 | 0 | 0 | 0 |
| ENSG00000242220 | 0 | 2 | 2 | 2 | 0 | 2 |
| ENSG00000265590 | 0 | 0 | 0 | 0 | 0 | 0 |
| ENSG00000159079 | 438 | 278 | 332 | 294 | 313 | 307 |
| ENSG00000228433 | 0 | 0 | 0 | 0 | 0 | 0 |
| ENSG00000238220 | 0 | 0 | 0 | 0 | 0 | 0 |
| ENSG00000159082 | 336 | 170 | 166 | 74 | 188 | 132 |
| ENSG00000238197 | 28 | 17 | 32 | 19 | 20 | 32 |
| ENSG00000159086 | 850 | 531 | 458 | 268 | 729 | 479 |
| ENSG00000205930 | 29 | 12 | 8 | 5 | 18 | 18 |
| ENSG00000279690 | 0 | 1 | 0 | 0 | 0 | 0 |
| ENSG00000205929 | 1 | 1 | 2 | 0 | 0 | 1 |
| ENSG00000224427 | 2 | 0 | 1 | 1 | 0 | 2 |
| ENSG00000207098 | 0 | 0 | 0 | 0 | 0 | 0 |
| ENSG00000232360 | 0 | 0 | 0 | 0 | 0 | 0 |
| ENSG00000227757 | 0 | 0 | 0 | 0 | 0 | 0 |
| ENSG00000228961 | 0 | 0 | 0 | 0 | 0 | 0 |
| ENSG00000205927 | 2 | 0 | 0 | 0 | 0 | 0 |
| ENSG00000232539 | 0 | 0 | 0 | 0 | 0 | 0 |
| ENSG00000184221 | 0 | 0 | 0 | 0 | 0 | 0 |
| ENSG00000226527 | 0 | 0 | 0 | 0 | 0 | 0 |
| ENSG00000226433 | 0 | 0 | 0 | 0 | 0 | 0 |
| ENSG00000229086 | 0 | 0 | 0 | 0 | 0 | 0 |
| ENSG00000159110 | 17 | 22 | 12 | 17 | 15 | 10 |
| ENSG00000249624 | 0 | 0 | 0 | 0 | 0 | 0 |
| ENSG00000223799 | 4 | 5 | 4 | 1 | 1 | 5 |
| ENSG00000243646 | 24 | 20 | 21 | 10 | 26 | 16 |
| ENSG00000142166 | 1603 | 879 | 948 | 594 | 991 | 898 |
| ENSG00000226501 | 0 | 0 | 0 | 0 | 0 | 1 |
| ENSG00000159128 | 680 | 358 | 437 | 452 | 448 | 489 |
| ENSG00000142188 | 1389 | 629 | 829 | 620 | 731 | 658 |
| ENSG00000215088 | 0 | 0 | 0 | 0 | 0 | 0 |
| ENSG00000231355 | 0 | 0 | 0 | 0 | 4 | 0 |
| ENSG00000177692 | 85 | 30 | 42 | 25 | 39 | 37 |
| ENSG00000159131 | 2242 | 998 | 1314 | 1143 | 1329 | 1338 |
| ENSG00000233956 | 0 | 0 | 0 | 0 | 0 | 0 |
| ENSG00000159140 | 6374 | 3193 | 3951 | 2264 | 3743 | 3597 |
| ENSG00000284448 | 0 | 0 | 0 | 0 | 0 | 0 |
| ENSG00000159147 | 344 | 184 | 198 | 139 | 234 | 188 |
| ENSG00000249209 | 0 | 0 | 0 | 0 | 0 | 0 |
| ENSG00000205758 | 426 | 211 | 297 | 193 | 272 | 247 |
| ENSG00000205726 | 670 | 313 | 388 | 241 | 366 | 389 |
| ENSG00000241837 | 784 | 446 | 594 | 575 | 539 | 580 |

| | | | | | | |
|---|---|---|---|---|---|---|
| ENSG00000237945 | 24 | 19 | 9 | 11 | 6 | 4 |
| ENSG00000244294 | 0 | 0 | 0 | 0 | 0 | 0 |
| ENSG00000273102 | 0 | 0 | 0 | 0 | 0 | 0 |
| ENSG00000243927 | 600 | 308 | 357 | 374 | 376 | 356 |
| ENSG00000198743 | 296 | 161 | 101 | 62 | 227 | 149 |
| ENSG00000272657 | 0 | 0 | 0 | 0 | 1 | 0 |
| ENSG00000224598 | 0 | 0 | 0 | 0 | 0 | 0 |
| ENSG00000227456 | 2 | 1 | 0 | 2 | 1 | 1 |
| ENSG00000214955 | 1 | 2 | 0 | 0 | 0 | 0 |
| ENSG00000159197 | 9 | 6 | 2 | 3 | 1 | 1 |
| ENSG00000225555 | 4 | 1 | 0 | 0 | 1 | 1 |
| ENSG00000205670 | 0 | 0 | 0 | 0 | 0 | 0 |
| ENSG00000222018 | 0 | 0 | 0 | 0 | 0 | 0 |
| ENSG00000273104 | 0 | 0 | 0 | 0 | 0 | 0 |
| ENSG00000243627 | 0 | 0 | 0 | 0 | 0 | 0 |
| ENSG00000272958 | 0 | 0 | 0 | 0 | 0 | 0 |
| ENSG00000180509 | 0 | 0 | 0 | 0 | 0 | 0 |
| ENSG00000221398 | 0 | 0 | 0 | 0 | 0 | 0 |
| ENSG00000159200 | 367 | 161 | 164 | 161 | 203 | 186 |
| ENSG00000159212 | 6 | 1 | 1 | 2 | 4 | 3 |
| ENSG00000230978 | 0 | 2 | 3 | 2 | 3 | 2 |
| ENSG00000234380 | 0 | 0 | 0 | 0 | 0 | 0 |
| ENSG00000159216 | 1112 | 487 | 767 | 717 | 854 | 1038 |
| ENSG00000234703 | 0 | 0 | 0 | 0 | 0 | 0 |
| ENSG00000223671 | 0 | 0 | 0 | 0 | 0 | 0 |
| ENSG00000231300 | 2 | 0 | 0 | 0 | 0 | 0 |
| ENSG00000230794 | 0 | 0 | 0 | 0 | 0 | 0 |
| ENSG00000211590 | 0 | 0 | 0 | 0 | 0 | 0 |
| ENSG00000229761 | 0 | 0 | 0 | 0 | 0 | 0 |
| ENSG00000234008 | 0 | 0 | 0 | 0 | 0 | 0 |
| ENSG00000231106 | 0 | 0 | 0 | 0 | 0 | 0 |
| ENSG00000214914 | 0 | 1 | 0 | 0 | 0 | 0 |
| ENSG00000185917 | 664 | 272 | 427 | 277 | 368 | 380 |
| ENSG00000189089 | 2 | 1 | 1 | 1 | 0 | 1 |
| ENSG00000236677 | 0 | 0 | 0 | 0 | 0 | 0 |
| ENSG00000200213 | 0 | 0 | 0 | 0 | 0 | 0 |
| ENSG00000230212 | 17 | 9 | 15 | 7 | 9 | 10 |
| ENSG00000159228 | 1397 | 674 | 831 | 840 | 772 | 811 |
| ENSG00000236119 | 0 | 0 | 0 | 0 | 0 | 1 |
| ENSG00000233393 | 2 | 1 | 3 | 0 | 2 | 1 |
| ENSG00000226054 | 26 | 22 | 28 | 8 | 12 | 20 |
| ENSG00000236830 | 441 | 258 | 308 | 177 | 195 | 239 |
| ENSG00000214889 | 0 | 0 | 0 | 0 | 0 | 0 |
| ENSG00000159231 | 96 | 70 | 87 | 87 | 83 | 84 |
| ENSG00000142197 | 3783 | 1696 | 2129 | 968 | 1556 | 1433 |
| ENSG00000228149 | 0 | 0 | 0 | 0 | 0 | 2 |
| ENSG00000214867 | 12 | 9 | 6 | 5 | 9 | 7 |
| ENSG00000273199 | 0 | 1 | 2 | 7 | 7 | 1 |
| ENSG00000159256 | 545 | 296 | 284 | 102 | 403 | 226 |
| ENSG00000228107 | 0 | 2 | 3 | 0 | 1 | 0 |

| | | | | | | |
|---|---|---|---|---|---|---|
| ENSG00000159259 | 484 | 218 | 321 | 202 | 248 | 292 |
| ENSG00000224421 | 0 | 0 | 0 | 0 | 0 | 0 |
| ENSG00000230479 | 0 | 1 | 6 | 2 | 1 | 2 |
| ENSG00000233818 | 1 | 5 | 0 | 0 | 3 | 2 |
| ENSG00000159261 | 0 | 0 | 0 | 0 | 0 | 0 |
| ENSG00000279365 | 0 | 0 | 1 | 0 | 0 | 0 |
| ENSG00000223741 | 0 | 0 | 0 | 0 | 0 | 0 |
| ENSG00000231324 | 8 | 0 | 3 | 2 | 0 | 3 |
| ENSG00000224269 | 0 | 0 | 0 | 0 | 0 | 0 |
| ENSG00000159263 | 174 | 95 | 123 | 93 | 90 | 129 |
| ENSG00000159267 | 1208 | 476 | 696 | 519 | 664 | 658 |
| ENSG00000237646 | 0 | 0 | 0 | 0 | 0 | 0 |
| ENSG00000199806 | 0 | 0 | 0 | 0 | 0 | 0 |
| ENSG00000270652 | 0 | 0 | 1 | 0 | 0 | 0 |
| ENSG00000224790 | 8 | 7 | 5 | 3 | 7 | 5 |
| ENSG00000207416 | 0 | 0 | 0 | 0 | 0 | 0 |
| ENSG00000215734 | 0 | 0 | 0 | 0 | 0 | 0 |
| ENSG00000183145 | 331 | 132 | 195 | 147 | 179 | 175 |
| ENSG00000212136 | 0 | 0 | 0 | 1 | 0 | 0 |
| ENSG00000185808 | 713 | 316 | 431 | 319 | 361 | 362 |
| ENSG00000182670 | 3186 | 1926 | 1378 | 602 | 3215 | 1573 |
| ENSG00000270116 | 8 | 0 | 2 | 1 | 0 | 4 |
| ENSG00000228677 | 2 | 1 | 1 | 0 | 0 | 0 |
| ENSG00000230366 | 11 | 12 | 4 | 6 | 9 | 3 |
| ENSG00000263969 | 0 | 0 | 0 | 0 | 0 | 0 |
| ENSG00000242553 | 30 | 23 | 20 | 8 | 23 | 18 |
| ENSG00000157538 | 1718 | 758 | 983 | 725 | 944 | 924 |
| ENSG00000272948 | 20 | 4 | 12 | 5 | 7 | 8 |
| ENSG00000273210 | 0 | 1 | 0 | 0 | 3 | 0 |
| ENSG00000157540 | 1547 | 744 | 979 | 548 | 840 | 789 |
| ENSG00000157542 | 0 | 0 | 0 | 0 | 2 | 0 |
| ENSG00000233213 | 0 | 0 | 0 | 0 | 0 | 0 |
| ENSG00000184029 | 0 | 0 | 0 | 0 | 0 | 0 |
| ENSG00000223608 | 0 | 0 | 0 | 0 | 0 | 0 |
| ENSG00000198054 | 1 | 2 | 2 | 1 | 3 | 0 |
| ENSG00000157551 | 2 | 0 | 2 | 0 | 1 | 0 |
| ENSG00000270835 | 0 | 0 | 0 | 0 | 0 | 0 |
| ENSG00000233316 | 0 | 0 | 0 | 0 | 0 | 0 |
| ENSG00000226012 | 0 | 0 | 0 | 0 | 0 | 0 |
| ENSG00000231123 | 0 | 0 | 0 | 0 | 0 | 0 |
| ENSG00000231231 | 0 | 0 | 0 | 0 | 0 | 0 |
| ENSG00000157554 | 14 | 7 | 8 | 7 | 5 | 4 |
| ENSG00000231480 | 0 | 0 | 0 | 0 | 0 | 0 |
| ENSG00000223806 | 0 | 0 | 0 | 0 | 0 | 0 |
| ENSG00000157557 | 757 | 340 | 466 | 399 | 500 | 474 |
| ENSG00000229986 | 0 | 0 | 0 | 0 | 0 | 0 |
| ENSG00000205622 | 0 | 0 | 0 | 0 | 0 | 0 |
| ENSG00000229925 | 0 | 0 | 0 | 0 | 0 | 0 |
| ENSG00000272015 | 0 | 0 | 0 | 0 | 0 | 0 |
| ENSG00000227721 | 0 | 0 | 0 | 0 | 0 | 0 |

| | | | | | | |
|---|---|---|---|---|---|---|
| ENSG00000232837 | 0 | 0 | 0 | 0 | 0 | 0 |
| ENSG00000235888 | 0 | 0 | 0 | 0 | 0 | 0 |
| ENSG00000237721 | 0 | 0 | 0 | 1 | 0 | 0 |
| ENSG00000237609 | 0 | 0 | 0 | 0 | 0 | 0 |
| ENSG00000228861 | 0 | 0 | 0 | 0 | 0 | 0 |
| ENSG00000235701 | 2 | 1 | 0 | 0 | 1 | 0 |
| ENSG00000276873 | 0 | 0 | 0 | 0 | 0 | 0 |
| ENSG00000183527 | 1270 | 718 | 870 | 787 | 796 | 887 |
| ENSG00000185658 | 1921 | 1179 | 1078 | 599 | 2006 | 1219 |
| ENSG00000272991 | 0 | 0 | 0 | 0 | 0 | 0 |
| ENSG00000232608 | 0 | 1 | 0 | 1 | 2 | 1 |
| ENSG00000237373 | 0 | 0 | 0 | 0 | 1 | 0 |
| ENSG00000229623 | 0 | 1 | 0 | 0 | 0 | 0 |
| ENSG00000255568 | 96 | 41 | 52 | 47 | 45 | 46 |
| ENSG00000238141 | 10 | 2 | 6 | 1 | 4 | 5 |
| ENSG00000205581 | 21508 | 10548 | 12500 | 11173 | 12907 | 11888 |
| ENSG00000252915 | 13 | 7 | 12 | 4 | 6 | 3 |
| ENSG00000227406 | 0 | 0 | 1 | 0 | 0 | 1 |
| ENSG00000182093 | 1178 | 630 | 713 | 634 | 719 | 622 |
| ENSG00000285815 | 0 | 0 | 0 | 0 | 0 | 1 |
| ENSG00000157578 | 251 | 118 | 120 | 88 | 110 | 108 |
| ENSG00000185437 | 14 | 6 | 18 | 13 | 13 | 13 |
| ENSG00000275523 | 0 | 0 | 0 | 0 | 0 | 0 |
| ENSG00000235808 | 1 | 0 | 1 | 0 | 0 | 0 |
| ENSG00000228349 | 0 | 0 | 0 | 0 | 0 | 0 |
| ENSG00000235012 | 8 | 0 | 3 | 1 | 1 | 3 |
| ENSG00000183778 | 525 | 165 | 212 | 52 | 70 | 51 |
| ENSG00000184809 | 19 | 11 | 5 | 2 | 1 | 5 |
| ENSG00000225330 | 1 | 2 | 0 | 0 | 5 | 1 |
| ENSG00000231713 | 6 | 3 | 6 | 1 | 0 | 0 |
| ENSG00000183067 | 29 | 35 | 42 | 21 | 32 | 30 |
| ENSG00000183036 | 129 | 68 | 69 | 78 | 92 | 75 |
| ENSG00000171587 | 4690 | 2042 | 2754 | 1998 | 2338 | 2589 |
| ENSG00000263973 | 0 | 0 | 0 | 0 | 0 | 0 |
| ENSG00000235123 | 33885 | 16914 | 20020 | 16587 | 20488 | 17262 |
| ENSG00000207147 | 0 | 0 | 0 | 0 | 0 | 0 |
| ENSG00000233756 | 0 | 0 | 0 | 0 | 0 | 0 |
| ENSG00000230859 | 0 | 0 | 0 | 0 | 0 | 0 |
| ENSG00000226496 | 0 | 0 | 0 | 0 | 0 | 0 |
| ENSG00000263681 | 1 | 0 | 0 | 0 | 0 | 0 |
| ENSG00000182240 | 1071 | 516 | 570 | 544 | 485 | 529 |
| ENSG00000280109 | 6 | 4 | 1 | 0 | 2 | 1 |
| ENSG00000224388 | 0 | 0 | 0 | 0 | 0 | 0 |
| ENSG00000183844 | 4 | 1 | 3 | 0 | 0 | 1 |
| ENSG00000183486 | 8 | 4 | 5 | 5 | 3 | 5 |
| ENSG00000157601 | 38 | 25 | 34 | 13 | 23 | 22 |
| ENSG00000228318 | 0 | 0 | 0 | 0 | 0 | 0 |
| ENSG00000184012 | 2300 | 1052 | 1364 | 884 | 1082 | 1017 |
| ENSG00000207503 | 0 | 0 | 0 | 2 | 0 | 1 |
| ENSG00000232806 | 0 | 0 | 0 | 0 | 0 | 0 |

| | | | | | | |
|---|---|---|---|---|---|---|
| ENSG00000223400 | 0 | 0 | 0 | 0 | 0 | 0 |
| ENSG00000227702 | 1 | 1 | 1 | 0 | 2 | 0 |
| ENSG00000236384 | 6 | 1 | 2 | 0 | 0 | 0 |
| ENSG00000232401 | 0 | 0 | 0 | 0 | 0 | 0 |
| ENSG00000183421 | 422 | 187 | 275 | 221 | 246 | 217 |
| ENSG00000236883 | 0 | 0 | 0 | 0 | 0 | 0 |
| ENSG00000275166 | 0 | 0 | 0 | 1 | 0 | 0 |
| ENSG00000141956 | 805 | 376 | 499 | 355 | 430 | 458 |
| ENSG00000236545 | 2 | 1 | 0 | 0 | 0 | 4 |
| ENSG00000227698 | 3 | 3 | 4 | 3 | 4 | 6 |
| ENSG00000251778 | 0 | 0 | 0 | 0 | 0 | 0 |
| ENSG00000157617 | 805 | 377 | 489 | 346 | 375 | 409 |
| ENSG00000173276 | 591 | 334 | 326 | 147 | 442 | 327 |
| ENSG00000237232 | 7 | 1 | 4 | 4 | 3 | 5 |
| ENSG00000177398 | 10 | 13 | 16 | 12 | 13 | 13 |
| ENSG00000184385 | 0 | 0 | 0 | 0 | 0 | 0 |
| ENSG00000160179 | 3800 | 1470 | 2031 | 1180 | 1282 | 1318 |
| ENSG00000223262 | 0 | 0 | 0 | 0 | 0 | 0 |
| ENSG00000160180 | 128 | 51 | 86 | 104 | 75 | 80 |
| ENSG00000160181 | 0 | 0 | 0 | 0 | 0 | 0 |
| ENSG00000160182 | 6536 | 3517 | 3833 | 6926 | 6644 | 5294 |
| ENSG00000160183 | 41 | 10 | 27 | 12 | 22 | 22 |
| ENSG00000160185 | 0 | 0 | 1 | 0 | 1 | 0 |
| ENSG00000252619 | 0 | 0 | 0 | 0 | 0 | 0 |
| ENSG00000160188 | 189 | 108 | 147 | 103 | 117 | 129 |
| ENSG00000160190 | 3137 | 1537 | 1925 | 1648 | 1840 | 1957 |
| ENSG00000239930 | 0 | 0 | 2 | 1 | 8 | 0 |
| ENSG00000231867 | 0 | 0 | 0 | 0 | 0 | 0 |
| ENSG00000235772 | 20 | 9 | 12 | 7 | 10 | 11 |
| ENSG00000225431 | 3 | 0 | 0 | 0 | 0 | 0 |
| ENSG00000235023 | 2 | 2 | 1 | 0 | 0 | 0 |
| ENSG00000160191 | 354 | 208 | 204 | 201 | 219 | 196 |
| ENSG00000225731 | 0 | 1 | 0 | 0 | 0 | 0 |
| ENSG00000283051 | 0 | 0 | 0 | 0 | 1 | 0 |
| ENSG00000233754 | 0 | 0 | 0 | 0 | 0 | 0 |
| ENSG00000225218 | 12 | 4 | 3 | 7 | 8 | 8 |
| ENSG00000160193 | 469 | 256 | 276 | 292 | 304 | 312 |
| ENSG00000160194 | 1864 | 968 | 1084 | 1095 | 1089 | 1093 |
| ENSG00000233056 | 4 | 6 | 2 | 0 | 2 | 4 |
| ENSG00000264580 | 0 | 0 | 1 | 0 | 2 | 0 |
| ENSG00000224100 | 2 | 2 | 3 | 1 | 0 | 3 |
| ENSG00000160199 | 792 | 334 | 446 | 345 | 399 | 380 |
| ENSG00000160200 | 172 | 99 | 137 | 137 | 141 | 131 |
| ENSG00000160201 | 115 | 49 | 61 | 25 | 64 | 68 |
| ENSG00000232777 | 0 | 0 | 0 | 0 | 0 | 0 |
| ENSG00000236663 | 0 | 0 | 0 | 0 | 0 | 0 |
| ENSG00000228120 | 0 | 0 | 0 | 0 | 0 | 0 |
| ENSG00000160202 | 0 | 0 | 0 | 0 | 0 | 0 |
| ENSG00000237864 | 9 | 3 | 5 | 1 | 3 | 2 |
| ENSG00000237989 | 0 | 1 | 0 | 0 | 0 | 0 |

| | | | | | | |
|---|---|---|---|---|---|---|
| ENSG00000225637 | 0 | 0 | 0 | 0 | 0 | 0 |
| ENSG00000142178 | 4 | 4 | 0 | 0 | 0 | 2 |
| ENSG00000188660 | 0 | 0 | 0 | 0 | 0 | 0 |
| ENSG00000185186 | 0 | 0 | 1 | 0 | 0 | 0 |
| ENSG00000223975 | 0 | 0 | 0 | 0 | 0 | 0 |
| ENSG00000160207 | 32 | 17 | 23 | 25 | 24 | 22 |
| ENSG00000214326 | 0 | 0 | 0 | 0 | 0 | 0 |
| ENSG00000234289 | 3 | 0 | 0 | 0 | 0 | 0 |
| ENSG00000278433 | 0 | 0 | 0 | 0 | 0 | 1 |
| ENSG00000160208 | 2181 | 1020 | 1288 | 993 | 1179 | 1353 |
| ENSG00000160209 | 6576 | 3284 | 3962 | 3625 | 3853 | 3621 |
| ENSG00000281420 | 4 | 3 | 1 | 4 | 1 | 1 |
| ENSG00000160213 | 4916 | 2405 | 2789 | 2884 | 2734 | 2659 |
| ENSG00000234030 | 0 | 0 | 0 | 0 | 0 | 0 |
| ENSG00000160214 | 1737 | 923 | 1072 | 1352 | 1168 | 1160 |
| ENSG00000215458 | 2 | 5 | 9 | 3 | 8 | 3 |
| ENSG00000226543 | 0 | 0 | 0 | 1 | 0 | 0 |
| ENSG00000160216 | 5308 | 2496 | 3355 | 2622 | 2565 | 3111 |
| ENSG00000199598 | 0 | 0 | 0 | 0 | 0 | 0 |
| ENSG00000252606 | 0 | 0 | 0 | 0 | 0 | 0 |
| ENSG00000160218 | 1617 | 723 | 1002 | 624 | 929 | 949 |
| ENSG00000213440 | 0 | 0 | 1 | 0 | 0 | 1 |
| ENSG00000241945 | 22 | 9 | 11 | 13 | 14 | 8 |
| ENSG00000160221 | 6 | 4 | 6 | 3 | 7 | 5 |
| ENSG00000225331 | 0 | 0 | 0 | 0 | 0 | 0 |
| ENSG00000285413 | 0 | 0 | 0 | 0 | 0 | 0 |
| ENSG00000237604 | 0 | 0 | 2 | 1 | 1 | 0 |
| ENSG00000232124 | 0 | 2 | 0 | 1 | 4 | 1 |
| ENSG00000160223 | 0 | 0 | 6 | 0 | 1 | 0 |
| ENSG00000278158 | 0 | 0 | 0 | 0 | 0 | 0 |
| ENSG00000275799 | 0 | 0 | 0 | 0 | 0 | 0 |
| ENSG00000142182 | 0 | 0 | 0 | 1 | 0 | 0 |
| ENSG00000232010 | 0 | 0 | 0 | 0 | 0 | 0 |
| ENSG00000160224 | 0 | 0 | 1 | 1 | 0 | 1 |
| ENSG00000141959 | 9521 | 4420 | 5973 | 6029 | 5375 | 5539 |
| ENSG00000160226 | 648 | 393 | 537 | 477 | 492 | 454 |
| ENSG00000241728 | 0 | 0 | 0 | 0 | 0 | 0 |
| ENSG00000184441 | 126 | 85 | 101 | 110 | 97 | 117 |
| ENSG00000232969 | 3 | 2 | 2 | 4 | 1 | 1 |
| ENSG00000260256 | 0 | 0 | 0 | 0 | 0 | 0 |
| ENSG00000142185 | 0 | 0 | 0 | 0 | 3 | 0 |
| ENSG00000230061 | 0 | 0 | 0 | 0 | 0 | 0 |
| ENSG00000266692 | 0 | 0 | 0 | 0 | 0 | 0 |
| ENSG00000264452 | 0 | 0 | 0 | 0 | 0 | 0 |
| ENSG00000229356 | 0 | 0 | 0 | 1 | 0 | 2 |
| ENSG00000160233 | 264 | 118 | 140 | 169 | 137 | 154 |
| ENSG00000224747 | 4 | 3 | 2 | 3 | 0 | 0 |
| ENSG00000223431 | 1 | 3 | 0 | 1 | 0 | 4 |
| ENSG00000227999 | 4 | 1 | 2 | 2 | 4 | 0 |
| ENSG00000274225 | 22 | 8 | 8 | 7 | 7 | 11 |

| | | | | | | |
|---|---|---|---|---|---|---|
| ENSG00000228709 | 0 | 0 | 1 | 0 | 0 | 0 |
| ENSG00000277352 | 0 | 0 | 0 | 0 | 0 | 0 |
| ENSG00000175894 | 0 | 2 | 0 | 2 | 1 | 2 |
| ENSG00000235890 | 0 | 0 | 1 | 1 | 0 | 0 |
| ENSG00000182912 | 0 | 0 | 2 | 0 | 2 | 0 |
| ENSG00000215455 | 0 | 0 | 0 | 0 | 0 | 0 |
| ENSG00000205445 | 0 | 0 | 0 | 0 | 0 | 0 |
| ENSG00000212935 | 0 | 0 | 0 | 0 | 0 | 0 |
| ENSG00000215454 | 0 | 0 | 0 | 0 | 0 | 0 |
| ENSG00000241123 | 0 | 0 | 0 | 0 | 0 | 0 |
| ENSG00000188155 | 0 | 0 | 0 | 0 | 0 | 0 |
| ENSG00000272804 | 0 | 0 | 0 | 0 | 0 | 0 |
| ENSG00000187766 | 0 | 0 | 0 | 0 | 0 | 0 |
| ENSG00000221837 | 0 | 0 | 0 | 0 | 0 | 0 |
| ENSG00000221859 | 0 | 0 | 0 | 0 | 0 | 0 |
| ENSG00000243489 | 0 | 0 | 0 | 0 | 0 | 0 |
| ENSG00000212933 | 0 | 0 | 0 | 0 | 0 | 0 |
| ENSG00000205439 | 0 | 0 | 0 | 0 | 0 | 0 |
| ENSG00000221864 | 0 | 0 | 0 | 0 | 0 | 0 |
| ENSG00000229880 | 0 | 0 | 0 | 0 | 0 | 0 |
| ENSG00000187175 | 0 | 0 | 0 | 0 | 0 | 0 |
| ENSG00000276647 | 0 | 0 | 0 | 0 | 0 | 0 |
| ENSG00000189169 | 0 | 0 | 0 | 0 | 0 | 0 |
| ENSG00000236382 | 0 | 0 | 0 | 0 | 0 | 0 |
| ENSG00000184787 | 4109 | 1933 | 2553 | 1827 | 2339 | 2502 |
| ENSG00000236519 | 13 | 4 | 5 | 6 | 4 | 12 |
| ENSG00000184900 | 9453 | 4424 | 5618 | 4750 | 4718 | 4975 |
| ENSG00000183255 | 12908 | 6338 | 7837 | 7240 | 7107 | 7432 |
| ENSG00000160255 | 78 | 45 | 67 | 33 | 28 | 30 |
| ENSG00000227039 | 14 | 12 | 8 | 16 | 10 | 14 |
| ENSG00000273027 | 29 | 6 | 27 | 23 | 17 | 16 |
| ENSG00000183250 | 368 | 131 | 202 | 138 | 152 | 162 |
| ENSG00000272825 | 19 | 12 | 11 | 9 | 5 | 7 |
| ENSG00000160256 | 875 | 440 | 522 | 706 | 546 | 574 |
| ENSG00000276529 | 50 | 27 | 34 | 55 | 45 | 41 |
| ENSG00000234880 | 0 | 0 | 0 | 0 | 0 | 0 |
| ENSG00000261706 | 0 | 0 | 0 | 0 | 0 | 0 |
| ENSG00000275874 | 2 | 0 | 1 | 1 | 0 | 0 |
| ENSG00000235374 | 158 | 60 | 93 | 89 | 67 | 76 |
| ENSG00000197381 | 1368 | 584 | 814 | 454 | 646 | 609 |
| ENSG00000267857 | 0 | 0 | 0 | 0 | 0 | 0 |
| ENSG00000182586 | 5 | 5 | 2 | 1 | 2 | 4 |
| ENSG00000186866 | 1062 | 512 | 725 | 515 | 574 | 538 |
| ENSG00000223768 | 2793 | 1171 | 1627 | 1258 | 1472 | 1504 |
| ENSG00000184274 | 7 | 3 | 3 | 1 | 1 | 0 |
| ENSG00000229382 | 0 | 2 | 1 | 0 | 1 | 0 |
| ENSG00000237664 | 1 | 2 | 4 | 2 | 0 | 0 |
| ENSG00000228930 | 0 | 0 | 0 | 0 | 0 | 0 |
| ENSG00000228355 | 0 | 0 | 0 | 0 | 0 | 0 |
| ENSG00000273796 | 56 | 23 | 27 | 22 | 23 | 24 |

| | | | | | | |
|---|---|---|---|---|---|---|
| ENSG00000182871 | 6065 | 2885 | 4538 | 3395 | 2929 | 3353 |
| ENSG00000224574 | 0 | 3 | 0 | 0 | 4 | 0 |
| ENSG00000183535 | 6 | 6 | 3 | 0 | 4 | 0 |
| ENSG00000275167 | 0 | 0 | 0 | 0 | 0 | 0 |
| ENSG00000173638 | 1753 | 829 | 1029 | 1408 | 1135 | 1270 |
| ENSG00000233922 | 2 | 1 | 2 | 0 | 0 | 0 |
| ENSG00000183570 | 341 | 175 | 212 | 240 | 230 | 243 |
| ENSG00000275139 | 4 | 1 | 0 | 0 | 2 | 0 |
| ENSG00000205424 | 0 | 0 | 2 | 0 | 0 | 0 |
| ENSG00000276633 | 0 | 0 | 0 | 1 | 0 | 1 |
| ENSG00000280604 | 74 | 39 | 45 | 28 | 45 | 35 |
| ENSG00000274248 | 0 | 0 | 0 | 0 | 0 | 0 |
| ENSG00000142156 | 318 | 127 | 187 | 157 | 110 | 127 |
| ENSG00000224413 | 0 | 0 | 0 | 0 | 0 | 0 |
| ENSG00000228235 | 0 | 0 | 0 | 0 | 0 | 0 |
| ENSG00000226115 | 0 | 0 | 0 | 0 | 0 | 0 |
| ENSG00000233767 | 0 | 0 | 0 | 0 | 0 | 0 |
| ENSG00000227438 | 0 | 0 | 0 | 0 | 0 | 0 |
| ENSG00000142173 | 546 | 233 | 339 | 330 | 257 | 250 |
| ENSG00000160282 | 88 | 26 | 76 | 41 | 24 | 49 |
| ENSG00000237338 | 2 | 0 | 1 | 0 | 0 | 1 |
| ENSG00000160284 | 370 | 144 | 187 | 231 | 168 | 202 |
| ENSG00000228404 | 5 | 1 | 3 | 2 | 1 | 0 |
| ENSG00000160285 | 9684 | 4241 | 6108 | 5113 | 4581 | 4957 |
| ENSG00000223901 | 18 | 21 | 21 | 11 | 16 | 18 |
| ENSG00000215424 | 411 | 203 | 223 | 175 | 231 | 213 |
| ENSG00000160294 | 3651 | 1694 | 2311 | 1435 | 1661 | 1970 |
| ENSG00000228137 | 24 | 23 | 23 | 14 | 16 | 17 |
| ENSG00000239415 | 142 | 85 | 100 | 71 | 98 | 106 |
| ENSG00000182362 | 438 | 236 | 288 | 224 | 271 | 208 |
| ENSG00000160298 | 766 | 385 | 490 | 363 | 439 | 420 |
| ENSG00000160299 | 3019 | 1243 | 1886 | 1026 | 1361 | 1475 |
| ENSG00000225043 | 3 | 0 | 6 | 4 | 0 | 0 |
| ENSG00000160305 | 2762 | 1181 | 1658 | 924 | 1396 | 1500 |
| ENSG00000223692 | 0 | 0 | 3 | 0 | 0 | 2 |
| ENSG00000202239 | 0 | 0 | 0 | 0 | 0 | 0 |
| ENSG00000160307 | 0 | 0 | 1 | 1 | 2 | 0 |
| ENSG00000160310 | 3923 | 1850 | 2272 | 2176 | 2346 | 2180 |
| ENSG00000230982 | 34 | 18 | 17 | 17 | 28 | 14 |
| ENSG00000212932 | 0 | 0 | 0 | 0 | 0 | 1 |
| ENSG00000210049 | 0 | 0 | 0 | 0 | 0 | 0 |
| ENSG00000211459 | 0 | 0 | 0 | 0 | 0 | 0 |
| ENSG00000210077 | 0 | 0 | 0 | 0 | 0 | 0 |
| ENSG00000210082 | 0 | 0 | 0 | 0 | 0 | 0 |
| ENSG00000209082 | 0 | 0 | 0 | 0 | 0 | 0 |
| ENSG00000198888 | 0 | 0 | 0 | 0 | 0 | 0 |
| ENSG00000210100 | 0 | 0 | 0 | 0 | 0 | 0 |
| ENSG00000210107 | 0 | 0 | 0 | 0 | 0 | 0 |
| ENSG00000210112 | 0 | 0 | 0 | 0 | 0 | 0 |
| ENSG00000198763 | 0 | 0 | 0 | 0 | 0 | 0 |

| | | | | | | |
|---|---|---|---|---|---|---|
| ENSG00000210117 | 0 | 0 | 0 | 0 | 0 | 0 |
| ENSG00000210127 | 0 | 0 | 0 | 0 | 0 | 0 |
| ENSG00000210135 | 0 | 0 | 0 | 0 | 0 | 0 |
| ENSG00000210140 | 0 | 0 | 0 | 0 | 0 | 0 |
| ENSG00000210144 | 0 | 0 | 0 | 0 | 0 | 0 |
| ENSG00000198804 | 0 | 0 | 0 | 0 | 0 | 0 |
| ENSG00000210151 | 0 | 0 | 0 | 0 | 0 | 0 |
| ENSG00000210154 | 0 | 0 | 0 | 0 | 0 | 0 |
| ENSG00000198712 | 0 | 0 | 0 | 0 | 0 | 0 |
| ENSG00000210156 | 0 | 0 | 0 | 0 | 0 | 0 |
| ENSG00000228253 | 0 | 0 | 0 | 0 | 0 | 0 |
| ENSG00000198899 | 0 | 0 | 0 | 0 | 0 | 0 |
| ENSG00000198938 | 0 | 0 | 0 | 0 | 0 | 0 |
| ENSG00000210164 | 0 | 0 | 0 | 0 | 0 | 0 |
| ENSG00000198840 | 0 | 0 | 0 | 0 | 0 | 0 |
| ENSG00000210174 | 0 | 0 | 0 | 0 | 0 | 0 |
| ENSG00000212907 | 0 | 0 | 0 | 0 | 0 | 0 |
| ENSG00000198886 | 0 | 0 | 0 | 0 | 0 | 0 |
| ENSG00000210176 | 0 | 0 | 0 | 0 | 0 | 0 |
| ENSG00000210184 | 0 | 0 | 0 | 0 | 0 | 0 |
| ENSG00000210191 | 0 | 0 | 0 | 0 | 0 | 0 |
| ENSG00000198786 | 0 | 0 | 0 | 0 | 0 | 0 |
| ENSG00000198695 | 0 | 0 | 0 | 0 | 0 | 0 |
| ENSG00000210194 | 0 | 0 | 0 | 0 | 0 | 0 |
| ENSG00000198727 | 0 | 0 | 0 | 0 | 0 | 0 |
| ENSG00000210195 | 0 | 0 | 0 | 0 | 0 | 0 |
| ENSG00000210196 | 0 | 0 | 0 | 0 | 0 | 0 |
| ENSG00000274175 | 0 | 0 | 0 | 0 | 0 | 0 |
| ENSG00000273843 | 0 | 0 | 0 | 0 | 0 | 0 |
| ENSG00000275869 | 0 | 0 | 0 | 0 | 0 | 0 |
| ENSG00000278510 | 0 | 0 | 0 | 0 | 0 | 0 |
| ENSG00000273554 | 0 | 0 | 0 | 0 | 0 | 0 |
| ENSG00000278782 | 0 | 0 | 0 | 0 | 0 | 0 |
| ENSG00000277761 | 0 | 0 | 0 | 0 | 0 | 0 |
| ENSG00000277836 | 0 | 0 | 0 | 0 | 0 | 0 |
| ENSG00000278673 | 8 | 1 | 2 | 1 | 1 | 1 |
| ENSG00000277890 | 0 | 0 | 0 | 0 | 0 | 0 |
| ENSG00000277666 | 0 | 0 | 0 | 0 | 0 | 0 |
| ENSG00000276760 | 0 | 0 | 0 | 0 | 0 | 0 |
| ENSG00000277927 | 0 | 0 | 0 | 0 | 0 | 0 |
| ENSG00000275249 | 0 | 0 | 0 | 0 | 0 | 0 |
| ENSG00000274792 | 0 | 0 | 0 | 0 | 0 | 0 |
| ENSG00000276872 | 0 | 0 | 0 | 0 | 0 | 0 |
| ENSG00000278294 | 0 | 0 | 0 | 0 | 0 | 0 |
| ENSG00000278457 | 0 | 0 | 0 | 0 | 1 | 1 |
| ENSG00000276711 | 0 | 0 | 0 | 0 | 0 | 0 |
| ENSG00000278704 | 6 | 7 | 8 | 9 | 8 | 5 |
| ENSG00000277400 | 17 | 5 | 8 | 9 | 9 | 8 |
| ENSG00000274847 | 0 | 0 | 0 | 0 | 0 | 0 |
| ENSG00000276932 | 0 | 0 | 0 | 0 | 0 | 0 |

| | | | | | | |
|---|---|---|---|---|---|---|
| ENSG00000273496 | 0 | 0 | 0 | 0 | 0 | 0 |
| ENSG00000277428 | 0 | 0 | 0 | 0 | 0 | 0 |
| ENSG00000276256 | 16 | 2 | 12 | 20 | 8 | 15 |
| ENSG00000278198 | 0 | 0 | 0 | 0 | 0 | 0 |
| ENSG00000273739 | 0 | 0 | 0 | 0 | 0 | 0 |
| ENSG00000276700 | 3 | 4 | 1 | 3 | 0 | 5 |
| ENSG00000276312 | 0 | 0 | 0 | 0 | 0 | 0 |
| ENSG00000275757 | 0 | 0 | 0 | 0 | 0 | 0 |
| ENSG00000273748 | 50 | 44 | 66 | 34 | 47 | 53 |
| ENSG00000274202 | 0 | 0 | 0 | 0 | 0 | 0 |
| ENSG00000278625 | 1 | 0 | 0 | 0 | 0 | 0 |
| ENSG00000278573 | 0 | 0 | 0 | 0 | 0 | 0 |
| ENSG00000276017 | 0 | 0 | 0 | 0 | 0 | 0 |
| ENSG00000278817 | 28 | 4 | 21 | 19 | 17 | 20 |
| ENSG00000277196 | 14 | 21 | 16 | 27 | 11 | 5 |
| ENSG00000277630 | 0 | 0 | 0 | 0 | 0 | 0 |
| ENSG00000278793 | 0 | 0 | 0 | 0 | 0 | 0 |
| ENSG00000276197 | 0 | 0 | 0 | 0 | 0 | 0 |
| ENSG00000274917 | 0 | 0 | 0 | 0 | 0 | 0 |
| ENSG00000273937 | 0 | 0 | 0 | 0 | 0 | 0 |
| ENSG00000273730 | 0 | 0 | 0 | 0 | 0 | 0 |
| ENSG00000278384 | 53 | 18 | 38 | 23 | 28 | 43 |
| ENSG00000278633 | 0 | 0 | 0 | 0 | 0 | 0 |
| ENSG00000278066 | 0 | 0 | 0 | 0 | 0 | 0 |
| ENSG00000277374 | 0 | 0 | 0 | 0 | 0 | 0 |
| ENSG00000276345 | 1 | 0 | 0 | 1 | 0 | 0 |
| ENSG00000273532 | 0 | 0 | 0 | 0 | 0 | 0 |
| ENSG00000276351 | 0 | 0 | 0 | 0 | 0 | 0 |
| ENSG00000275661 | 0 | 0 | 0 | 0 | 0 | 0 |
| ENSG00000277856 | 0 | 0 | 0 | 0 | 0 | 0 |
| ENSG00000275063 | 0 | 0 | 0 | 0 | 0 | 0 |
| ENSG00000271254 | 1493 | 683 | 929 | 864 | 1369 | 1288 |
| ENSG00000275405 | 0 | 1 | 0 | 0 | 1 | 1 |
| ENSG00000275987 | 0 | 0 | 0 | 0 | 0 | 0 |
| ENSG00000277475 | 2 | 1 | 1 | 2 | 0 | 0 |
| ENSG00000268674 | 0 | 0 | 0 | 0 | 0 | 0 |
