## Supplementary material for "Global transcriptome analysis reveals partial estrogen-like effects of karanjin in MCF-7 breast cancer cells": All supplementary material: Supplementary data 7.pdf

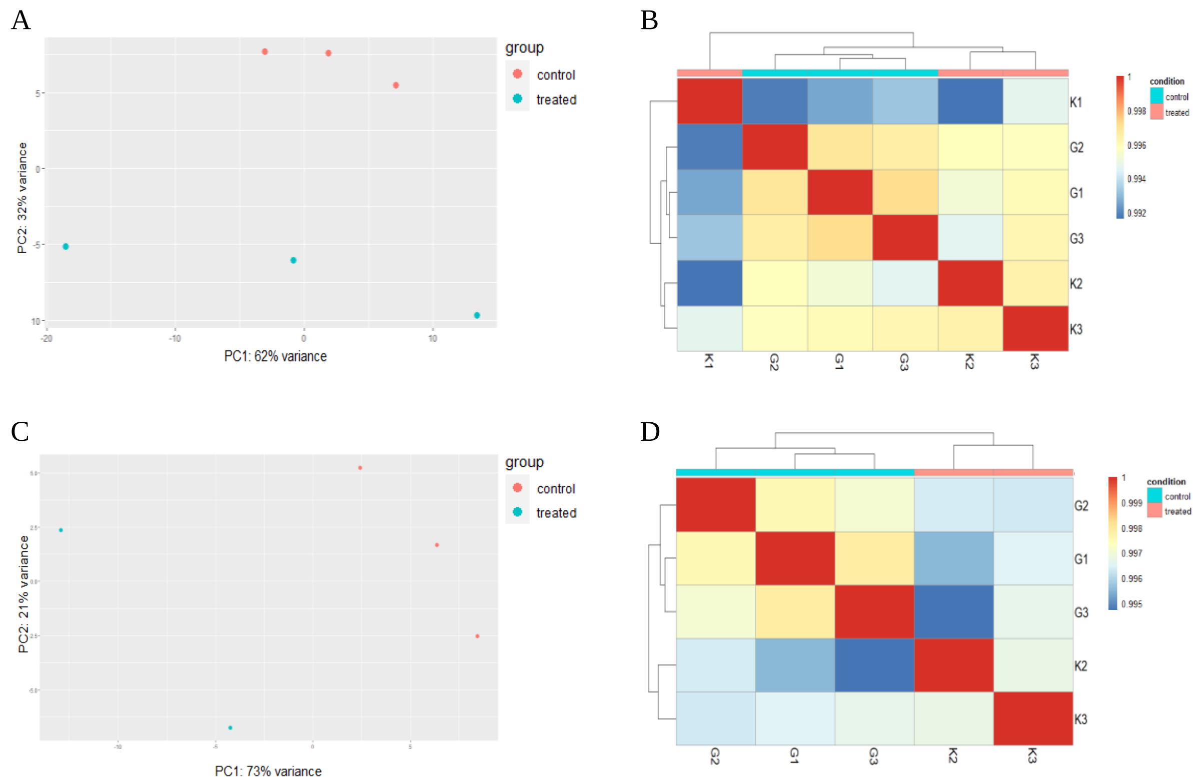


**Supplementary data 7**. **Quality assessment of the read count data**. Read count data were analysed using DESeq2 package in R. Data were assessed using PCA plot (panels A, C), and correlation heatmap (panels B, D). **A, B**. Quality assessment for three control (G1, G2, G3) and three karanjin treated samples (K1, K2, K3). PCA reveals that control and treated groups separate across PC1 with one treated sample (K1) separating farther from other treated samples (panel A). The correlation heatmap also shows that K1 is separated from the other two karanjin treated samples, which is positioned distant from K2 and K3 (panel B). This observation, combined with interpretation from the PCA plot indicates that K1 is an outlier. **C, D**. Quality assessment results for re-analysis of the raw counts after omission of sample K1. The PCA plot after removal of K1 shows better separation between the control and treated groups (panel C). Correlation heatmap also shows better hierarchical clustering, with the two experimental groups separating distinctly (panel D).
