## Supplementary material for "Global transcriptome analysis reveals partial estrogen-like effects of karanjin in MCF-7 breast cancer cells": All supplementary material: Supplementary data 8.pdf

| ensgene | log2FoldChange | lfcSE | pvalue | padj | symbol |
|---|---|---|---|---|---|
| ENSG00000140465 | 6.177330103 | 0.140890312 | 0 | 0 | CYP1A1 |
| ENSG00000103257 | 2.882405822 | 0.089815159 | 5.5844E-226 | 3.9342E-222 | SLC7A5 |
| ENSG00000138061 | 2.723353677 | 0.116423682 | 5.1852E-121 | 2.4353E-117 | CYP1B1 |
| ENSG00000113739 | 1.107099037 | 0.075441753 | 9.33739E-49 | 3.2891E-45 | STC2 |
| ENSG00000053747 | 1.840310185 | 0.132547121 | 7.89819E-44 | 2.22571E-40 | LAMA3 |
| ENSG00000163659 | 2.058824044 | 0.152633236 | 1.82284E-41 | 4.28065E-38 | TIPARP |
| ENSG00000168003 | 0.97688135 | 0.079730521 | 1.63321E-34 | 3.28742E-31 | SLC3A2 |
| ENSG00000272808 | 2.248500986 | 0.203913947 | 2.84052E-28 | 5.00286E-25 | AC015712.6 |
| ENSG00000139289 | 1.299553933 | 0.118750209 | 7.13165E-28 | 1.1165E-24 | PHLDA1 |
| ENSG00000182704 | 1.103894549 | 0.102526314 | 4.93125E-27 | 6.94813E-24 | TSKU |
| ENSG00000184254 | 2.466328562 | 0.229743624 | 6.9618E-27 | 8.91743E-24 | ALDH1A3 |
| ENSG00000130513 | 1.315919326 | 0.1232221 | 1.27299E-26 | 1.4947E-23 | GDF15 |
| ENSG00000071282 | 1.311817458 | 0.123008798 | 1.49328E-26 | 1.61849E-23 | LMCD1 |
| ENSG00000179151 | 0.903527975 | 0.086343978 | 1.26025E-25 | 1.26835E-22 | EDC3 |
| ENSG00000219481 | 0.927719222 | 0.089481937 | 3.47903E-25 | 3.26797E-22 | NBPF1 |
| ENSG00000118777 | 1.226270914 | 0.12031267 | 2.14477E-24 | 1.79683E-21 | ABCG2 |
| ENSG00000172602 | 1.560014279 | 0.153072791 | 2.16793E-24 | 1.79683E-21 | RND1 |
| ENSG00000075426 | 0.904466907 | 0.090347151 | 1.36343E-23 | 1.06726E-20 | FOSL2 |
| ENSG00000109099 | -0.922794736 | 0.094043684 | 9.95666E-23 | 7.38365E-20 | PMP22 |
| ENSG00000165731 | -1.487156862 | 0.152009229 | 1.32765E-22 | 9.19232E-20 | RET |
| ENSG00000124813 | 1.436230302 | 0.146851525 | 1.37004E-22 | 9.19232E-20 | RUNX2 |
| ENSG00000154065 | 3.122873431 | 0.330874072 | 3.79044E-21 | 2.4276E-18 | ANKRD29 |
| ENSG00000115594 | -1.169660153 | 0.126796996 | 2.84442E-20 | 1.74251E-17 | IL1R1 |
| ENSG00000175874 | 2.5829749 | 0.280670451 | 3.485E-20 | 2.04599E-17 | CREG2 |
| ENSG00000165566 | 1.338992411 | 0.145826576 | 4.22813E-20 | 2.38298E-17 | AMER2 |
| ENSG00000108448 | 0.974444784 | 0.106362615 | 5.11644E-20 | 2.77272E-17 | TRIM16L |
| ENSG00000143412 | -0.831699692 | 0.091004614 | 6.29779E-20 | 3.28651E-17 | ANXA9 |
| ENSG00000145808 | -1.113268405 | 0.122321888 | 8.93901E-20 | 4.49824E-17 | ADAMTS19 |
| ENSG00000181830 | -1.161193635 | 0.128358235 | 1.47615E-19 | 7.17206E-17 | SLC35C1 |
| ENSG00000137331 | 0.774851493 | 0.08591497 | 1.90129E-19 | 8.92974E-17 | IER3 |
| ENSG00000286169 | 0.997267037 | 0.111255444 | 3.13793E-19 | 1.38708E-16 | AHRR |
| ENSG00000261578 | 1.214655664 | 0.135513906 | 3.15022E-19 | 1.38708E-16 | AP003119.3 |
| ENSG00000249846 | -1.65804719 | 0.185574579 | 4.08404E-19 | 1.74376E-16 | LINC02021 |
| ENSG00000151012 | 2.437851795 | 0.274616706 | 6.85098E-19 | 2.83913E-16 | SLC7A11 |
| ENSG00000198431 | 0.828475474 | 0.093437807 | 7.54145E-19 | 3.03597E-16 | TXNRD1 |
| ENSG00000090975 | 1.347564568 | 0.15287647 | 1.19976E-18 | 4.69571E-16 | PITPNM2 |
| ENSG00000151715 | 1.185055103 | 0.1395611 | 2.04339E-17 | 7.78146E-15 | TMEM45B |
| ENSG00000254231 | -1.169776011 | 0.138942729 | 3.79311E-17 | 1.33612E-14 | AC103760.1 |
| ENSG00000167653 | -0.985226333 | 0.116984434 | 3.70559E-17 | 1.33612E-14 | PSCA |
| ENSG00000109321 | 1.487118745 | 0.176588675 | 3.72104E-17 | 1.33612E-14 | AREG |
| ENSG00000139505 | 0.880014084 | 0.106150297 | 1.12997E-16 | 3.88322E-14 | MTMR6 |
| ENSG00000166509 | -1.401684318 | 0.170106911 | 1.72182E-16 | 5.7763E-14 | CLEC3A |
| ENSG00000121671 | -0.85709644 | 0.104638788 | 2.59062E-16 | 8.4888E-14 | CRY2 |
| ENSG00000240875 | 1.19268675 | 0.14634348 | 3.64184E-16 | 1.16622E-13 | LINC00886 |
| ENSG00000197191 | 0.965349399 | 0.119397706 | 6.20833E-16 | 1.9439E-13 | CYSRT1 |
| ENSG00000168874 | 1.42633265 | 0.176493324 | 6.39694E-16 | 1.95941E-13 | ATOH8 |

| | | | | | |
|---|---|---|---|---|---|
| ENSG00000008513 | 0.856684851 | 0.106494659 | 8.66742E-16 | 2.59838E-13 | ST3GAL1 |
| ENSG00000183778 | -1.897183833 | 0.240361546 | 2.94908E-15 | 8.65677E-13 | B3GALT5 |
| ENSG00000104081 | 0.945723932 | 0.120659004 | 4.57819E-15 | 1.31646E-12 | BMF |
| ENSG00000197043 | -0.793630817 | 0.101530976 | 5.42531E-15 | 1.52885E-12 | ANXA6 |
| ENSG00000100504 | 0.870774336 | 0.111917842 | 7.22509E-15 | 1.99611E-12 | PYGL |
| ENSG00000164850 | -1.102710484 | 0.144407101 | 2.23859E-14 | 6.06573E-12 | GPER1 |
| ENSG00000026508 | 0.882847926 | 0.115793936 | 2.45359E-14 | 6.52283E-12 | CD44 |
| ENSG00000188735 | 0.987441767 | 0.129728356 | 2.70701E-14 | 7.06328E-12 | TMEM120B |
| ENSG00000121361 | -1.264497746 | 0.166277409 | 2.8546E-14 | 7.31297E-12 | KCNJ8 |
| ENSG00000064651 | -0.649463337 | 0.086071053 | 4.49985E-14 | 1.1322E-11 | SLC12A2 |
| ENSG00000168743 | -0.666528644 | 0.088638575 | 5.49345E-14 | 1.35794E-11 | NPNT |
| ENSG00000091831 | -0.798391743 | 0.10743012 | 1.07186E-13 | 2.60387E-11 | ESR1 |
| ENSG00000006831 | -0.615200932 | 0.082938835 | 1.19329E-13 | 2.84973E-11 | ADIPOR2 |
| ENSG00000144674 | 1.214692922 | 0.163834216 | 1.22399E-13 | 2.87434E-11 | GOLGA4 |
| ENSG00000154930 | -1.112544026 | 0.150561216 | 1.47589E-13 | 3.40906E-11 | ACSS1 |
| ENSG00000072952 | 2.763353459 | 0.376932418 | 2.28162E-13 | 5.18516E-11 | MRVI1 |
| ENSG00000119938 | -0.636754592 | 0.086971418 | 2.45354E-13 | 5.48736E-11 | PPP1R3C |
| ENSG00000154654 | -0.675334386 | 0.093146985 | 4.16153E-13 | 9.16187E-11 | NCAM2 |
| ENSG00000186417 | 1.094229364 | 0.15274734 | 7.85535E-13 | 1.7028E-10 | GLDN |
| ENSG00000178372 | -1.023916831 | 0.143341706 | 9.11904E-13 | 1.94678E-10 | CALML5 |
| ENSG00000104419 | -1.003416663 | 0.142096525 | 1.64698E-12 | 3.41263E-10 | NDRG1 |
| ENSG00000101144 | -0.602075127 | 0.085252521 | 1.63821E-12 | 3.41263E-10 | BMP7 |
| ENSG00000168675 | -0.72902516 | 0.103532606 | 1.90177E-12 | 3.88347E-10 | LDLRAD4 |
| ENSG00000271254 | 0.688690719 | 0.098721627 | 3.03513E-12 | 6.10929E-10 | AC240274.1 |
| ENSG00000213463 | -0.764962169 | 0.110317401 | 4.08546E-12 | 8.10763E-10 | SYNJ2BP |
| ENSG00000165389 | 0.681402037 | 0.098381832 | 4.32612E-12 | 8.46598E-10 | SPTSSA |
| ENSG00000134508 | 0.609233344 | 0.091447044 | 2.69862E-11 | 5.20871E-09 | CABLES1 |
| ENSG00000135083 | 1.052186776 | 0.158268694 | 2.96892E-11 | 5.65298E-09 | CCNJL |
| ENSG00000162337 | 0.527736752 | 0.07997928 | 4.15567E-11 | 7.80712E-09 | LRP5 |
| ENSG00000107554 | 0.664312833 | 0.100931698 | 4.64769E-11 | 8.61656E-09 | DNMBP |
| ENSG00000111885 | -0.582396911 | 0.088521252 | 4.73062E-11 | 8.65642E-09 | MAN1A1 |
| ENSG00000064205 | 0.724032373 | 0.110285485 | 5.2007E-11 | 9.3946E-09 | WISP2 |
| ENSG00000124496 | -0.859648455 | 0.131771526 | 6.85572E-11 | 1.22275E-08 | TRERF1 |
| ENSG00000178726 | 1.101115441 | 0.168865135 | 6.99895E-11 | 1.23269E-08 | THBD |
| ENSG00000108828 | 0.484073787 | 0.074320613 | 7.35087E-11 | 1.27869E-08 | VAT1 |
| ENSG00000160182 | 0.696876832 | 0.107980462 | 1.0913E-10 | 1.87517E-08 | TFF1 |
| ENSG00000171345 | -0.448477024 | 0.069947682 | 1.43994E-10 | 2.44443E-08 | KRT19 |
| ENSG00000149256 | 2.261598789 | 0.353618248 | 1.59924E-10 | 2.68254E-08 | TENM4 |
| ENSG00000168398 | -1.034762446 | 0.163152132 | 2.26336E-10 | 3.75185E-08 | BDKRB2 |
| ENSG00000167553 | 0.580709555 | 0.091647001 | 2.35237E-10 | 3.85406E-08 | TUBA1C |
| ENSG00000143418 | -0.486874662 | 0.077135904 | 2.75618E-10 | 4.43072E-08 | CERS2 |
| ENSG00000134258 | 1.691280411 | 0.267977086 | 2.76724E-10 | 4.43072E-08 | VTCN1 |
| ENSG00000218416 | -0.912307909 | 0.144680895 | 2.86976E-10 | 4.54326E-08 | PP14571 |
| ENSG00000157514 | -0.491632197 | 0.078048683 | 2.99484E-10 | 4.63707E-08 | TSC22D3 |
| ENSG00000188886 | 1.421012456 | 0.225565315 | 2.98063E-10 | 4.63707E-08 | ASTL |
| ENSG00000105514 | -0.69953631 | 0.111086796 | 3.03061E-10 | 4.64144E-08 | RAB3D |
| ENSG00000117724 | 1.81302524 | 0.288547497 | 3.31499E-10 | 5.02239E-08 | CENPF |

| | | | | | |
|---|---|---|---|---|---|
| ENSG00000115414 | -0.733887915 | 0.116998644 | 3.55025E-10 | 5.26558E-08 | FN1 |
| ENSG00000115216 | -0.477946211 | 0.076190114 | 3.53989E-10 | 5.26558E-08 | NRBP1 |
| ENSG00000090539 | -1.144091324 | 0.185225374 | 6.54335E-10 | 9.60372E-08 | CHRD |
| ENSG00000175155 | -0.560849848 | 0.091234676 | 7.87971E-10 | 1.1414E-07 | YPEL2 |
| ENSG00000137868 | 1.06361885 | 0.173054567 | 7.93876E-10 | 1.1414E-07 | STRA6 |
| ENSG00000107984 | -0.862231662 | 0.141529646 | 1.11346E-09 | 1.58471E-07 | DKK1 |
| ENSG00000066583 | -0.591344436 | 0.097408096 | 1.2728E-09 | 1.78549E-07 | ISOC1 |
| ENSG00000132170 | 0.779603571 | 0.128437564 | 1.27987E-09 | 1.78549E-07 | PPARG |
| ENSG00000158258 | 0.669791663 | 0.11063237 | 1.41108E-09 | 1.94923E-07 | CLSTN2 |
| ENSG00000188042 | 1.020286785 | 0.168664308 | 1.45553E-09 | 1.9911E-07 | ARL4C |
| ENSG00000040933 | 0.552043917 | 0.091458568 | 1.57981E-09 | 2.14034E-07 | INPP4A |
| ENSG00000204323 | 0.905155291 | 0.152619245 | 3.01449E-09 | 4.04516E-07 | SMIM5 |
| ENSG00000102034 | 0.475651801 | 0.080426646 | 3.33681E-09 | 4.43543E-07 | ELF4 |
| ENSG00000100625 | -0.58179383 | 0.098555252 | 3.56464E-09 | 4.69399E-07 | SIX4 |
| ENSG00000124126 | -0.538475314 | 0.091314284 | 3.70293E-09 | 4.83095E-07 | PREX1 |
| ENSG00000198797 | -2.3965739 | 0.406627871 | 3.77467E-09 | 4.87937E-07 | BRINP2 |
| ENSG00000138759 | -1.001384959 | 0.170282741 | 4.08488E-09 | 5.23236E-07 | FRAS1 |
| ENSG00000124664 | -0.520617183 | 0.088666176 | 4.31466E-09 | 5.4769E-07 | SPDEF |
| ENSG00000144476 | -0.624834817 | 0.106550332 | 4.51243E-09 | 5.6768E-07 | ACKR3 |
| ENSG00000187122 | -1.65670714 | 0.284473917 | 5.7539E-09 | 7.17455E-07 | SLIT1 |
| ENSG00000150556 | 0.676869132 | 0.117140134 | 7.54657E-09 | 9.3273E-07 | LYPD6B |
| ENSG00000111057 | -0.429809161 | 0.074550113 | 8.14777E-09 | 9.9828E-07 | KRT18 |
| ENSG00000101224 | 0.553287896 | 0.096279031 | 9.09953E-09 | 1.10528E-06 | CDC25B |
| ENSG00000175063 | 0.618160135 | 0.10836196 | 1.16626E-08 | 1.4045E-06 | UBE2C |
| ENSG00000147224 | 0.588942824 | 0.103569344 | 1.29701E-08 | 1.5357E-06 | PRPS1 |
| ENSG00000159167 | 0.596706417 | 0.104926859 | 1.29382E-08 | 1.5357E-06 | STC1 |
| ENSG00000232973 | 1.403669869 | 0.247205565 | 1.36161E-08 | 1.59875E-06 | CYP1B1-AS1 |
| ENSG00000170525 | -0.526307812 | 0.093547111 | 1.84302E-08 | 2.14613E-06 | PFKFB3 |
| ENSG00000143341 | -0.465178594 | 0.08287212 | 1.98608E-08 | 2.29375E-06 | HMCN1 |
| ENSG00000116299 | -0.686857798 | 0.122668985 | 2.15246E-08 | 2.4657E-06 | KIAA1324 |
| ENSG00000173230 | 0.901652025 | 0.161597261 | 2.41038E-08 | 2.7389E-06 | GOLGB1 |
| ENSG00000076864 | 0.63542571 | 0.114045637 | 2.52296E-08 | 2.84388E-06 | RAP1GAP |
| ENSG00000115461 | -1.010679917 | 0.181801443 | 2.70934E-08 | 3.02973E-06 | IGFBP5 |
| ENSG00000117472 | -0.713278223 | 0.12850473 | 2.84692E-08 | 3.15852E-06 | TSPAN1 |
| ENSG00000170421 | -0.468687891 | 0.084514442 | 2.92857E-08 | 3.22371E-06 | KRT8 |
| ENSG00000177283 | -0.800453362 | 0.144846503 | 3.27207E-08 | 3.57392E-06 | FZD8 |
| ENSG00000167996 | 0.462224755 | 0.083847851 | 3.53449E-08 | 3.83085E-06 | FTH1 |
| ENSG00000137203 | 0.664246728 | 0.120845626 | 3.87063E-08 | 4.16314E-06 | TFAP2A |
| ENSG00000172379 | -0.844846283 | 0.154392104 | 4.44778E-08 | 4.74767E-06 | ARNT2 |
| ENSG00000092969 | -0.926479818 | 0.17008755 | 5.12046E-08 | 5.34424E-06 | TGFB2 |
| ENSG00000163931 | 0.428199886 | 0.078607816 | 5.11419E-08 | 5.34424E-06 | TKT |
| ENSG00000233198 | 1.113299702 | 0.204293604 | 5.05093E-08 | 5.34424E-06 | RNF224 |
| ENSG00000167315 | -0.55897537 | 0.102810605 | 5.42026E-08 | 5.61554E-06 | ACAA2 |
| ENSG00000138650 | -0.705190775 | 0.129916756 | 5.69826E-08 | 5.86047E-06 | PCDH10 |
| ENSG00000136997 | 1.002471024 | 0.184903068 | 5.9067E-08 | 6.03082E-06 | MYC |
| ENSG00000179314 | 1.682420806 | 0.31063889 | 6.09465E-08 | 6.17795E-06 | WSCD1 |
| ENSG00000156050 | -0.830184028 | 0.153611121 | 6.50066E-08 | 6.54245E-06 | FAM161B |

| | | | | | |
|---|---|---|---|---|---|
| ENSG00000079257 | 0.70516236 | 0.130957551 | 7.25803E-08 | 7.25288E-06 | LXN |
| ENSG00000008130 | 0.41873594 | 0.077836948 | 7.46286E-08 | 7.40505E-06 | NADK |
| ENSG00000121966 | -1.50612122 | 0.282427459 | 9.67248E-08 | 9.53043E-06 | CXCR4 |
| ENSG00000259424 | 1.187592372 | 0.22399273 | 1.14589E-07 | 1.12122E-05 | AC118658.1 |
| ENSG00000131759 | -0.529939216 | 0.100137203 | 1.20899E-07 | 1.17481E-05 | RARA |
| ENSG00000166025 | -0.920056514 | 0.173906584 | 1.21968E-07 | 1.17708E-05 | AMOTL1 |
| ENSG00000164938 | -0.454594654 | 0.086048533 | 1.27083E-07 | 1.2181E-05 | TP53INP1 |
| ENSG00000138778 | 1.969792845 | 0.374526244 | 1.44506E-07 | 1.37574E-05 | CENPE |
| ENSG00000116667 | -0.53868756 | 0.102538424 | 1.4922E-07 | 1.41108E-05 | C1orf21 |
| ENSG00000067057 | 0.402368777 | 0.077004567 | 1.73912E-07 | 1.63361E-05 | PFKP |
| ENSG00000112773 | -0.503609133 | 0.096578914 | 1.8433E-07 | 1.72001E-05 | FAM46A |
| ENSG00000170322 | 0.492557459 | 0.094647196 | 1.94895E-07 | 1.80662E-05 | NFRKB |
| ENSG00000173175 | -0.636886196 | 0.122697949 | 2.09524E-07 | 1.92954E-05 | ADCY5 |
| ENSG00000049130 | -0.770833404 | 0.149286389 | 2.42434E-07 | 2.21812E-05 | KITLG |
| ENSG00000182481 | 0.49407922 | 0.095734724 | 2.4574E-07 | 2.23385E-05 | KPNA2 |
| ENSG00000121316 | -0.970280654 | 0.188586473 | 2.67496E-07 | 2.41604E-05 | PLBD1 |
| ENSG00000087586 | 0.441092063 | 0.086015605 | 2.92763E-07 | 2.62741E-05 | AURKA |
| ENSG00000143416 | -0.537617617 | 0.104872848 | 2.95373E-07 | 2.63405E-05 | SELENBP1 |
| ENSG00000223523 | -0.937863609 | 0.18319517 | 3.06383E-07 | 2.71505E-05 | AC092598.1 |
| ENSG00000102886 | -0.655047762 | 0.128525769 | 3.45761E-07 | 3.02594E-05 | GDPD3 |
| ENSG00000138411 | 0.695140764 | 0.136379702 | 3.44899E-07 | 3.02594E-05 | HECW2 |
| ENSG00000103811 | -0.496358989 | 0.097606721 | 3.67055E-07 | 3.19247E-05 | CTSH |
| ENSG00000165507 | -1.271522282 | 0.250425184 | 3.82527E-07 | 3.30663E-05 | C10orf10 |
| ENSG00000198569 | 1.038684494 | 0.204765832 | 3.92524E-07 | 3.37236E-05 | SLC34A3 |
| ENSG00000168765 | 0.500996413 | 0.098870663 | 4.03732E-07 | 3.44762E-05 | GSTM4 |
| ENSG00000240204 | 1.052476589 | 0.207769157 | 4.07104E-07 | 3.45548E-05 | SMKR1 |
| ENSG00000130653 | -0.833623725 | 0.164797185 | 4.22605E-07 | 3.56557E-05 | PNPLA7 |
| ENSG00000134057 | 0.406704906 | 0.080715524 | 4.68595E-07 | 3.93006E-05 | CCNB1 |
| ENSG00000109436 | -0.587334663 | 0.116657867 | 4.78656E-07 | 3.99068E-05 | TBC1D9 |
| ENSG00000158050 | 0.601248374 | 0.119517659 | 4.88887E-07 | 4.05201E-05 | DUSP2 |
| ENSG00000188229 | 0.460557767 | 0.091610376 | 4.97294E-07 | 4.09759E-05 | TUBB4B |
| ENSG00000102316 | -0.383367264 | 0.076276502 | 5.00762E-07 | 4.10217E-05 | MAGED2 |
| ENSG00000182836 | 1.222982914 | 0.244066919 | 5.41899E-07 | 4.4135E-05 | PLCXD3 |
| ENSG00000258647 | 1.578261749 | 0.316611398 | 6.20085E-07 | 5.02126E-05 | LINC00930 |
| ENSG00000179104 | 0.531536929 | 0.106717722 | 6.33302E-07 | 5.09899E-05 | TMTC2 |
| ENSG00000224984 | -0.644970426 | 0.129583133 | 6.44869E-07 | 5.16261E-05 | AL512363.1 |
| ENSG00000135318 | -1.140286318 | 0.229594443 | 6.81636E-07 | 5.42613E-05 | NT5E |
| ENSG00000145147 | -1.143928643 | 0.230651773 | 7.06572E-07 | 5.59304E-05 | SLIT2 |
| ENSG00000164976 | -0.446508709 | 0.090124675 | 7.25744E-07 | 5.7127E-05 | KIAA1161 |
| ENSG00000167964 | -0.708497107 | 0.143096643 | 7.37652E-07 | 5.74227E-05 | RAB26 |
| ENSG00000106789 | -0.474002686 | 0.095725194 | 7.3567E-07 | 5.74227E-05 | CORO2A |
| ENSG00000010278 | -0.354163806 | 0.071631791 | 7.64467E-07 | 5.91832E-05 | CD9 |
| ENSG00000008394 | 0.428478143 | 0.086690518 | 7.70803E-07 | 5.93476E-05 | MGST1 |
| ENSG00000123975 | 0.498205838 | 0.101142387 | 8.40216E-07 | 6.43404E-05 | CKS2 |
| ENSG00000134333 | 0.411520251 | 0.083617726 | 8.59064E-07 | 6.54282E-05 | LDHA |
| ENSG00000233016 | 0.38730808 | 0.078818954 | 8.92845E-07 | 6.76354E-05 | SNHG7 |
| ENSG00000104267 | -0.690208348 | 0.140833068 | 9.54001E-07 | 7.18817E-05 | CA2 |

| | | | | | |
|---|---|---|---|---|---|
| ENSG00000172478 | -0.902853136 | 0.184435407 | 9.81925E-07 | 7.35921E-05 | C2orf54 |
| ENSG00000185133 | -0.926173141 | 0.189923246 | 1.07949E-06 | 8.01926E-05 | INPP5J |
| ENSG00000115241 | -0.372951837 | 0.076483789 | 1.08138E-06 | 8.01926E-05 | PPM1G |
| ENSG00000253540 | -0.992685133 | 0.203969941 | 1.13408E-06 | 8.32248E-05 | FAM86HP |
| ENSG00000113580 | 0.565543081 | 0.116198924 | 1.13291E-06 | 8.32248E-05 | NR3C1 |
| ENSG00000111319 | -0.625616091 | 0.129397355 | 1.3325E-06 | 9.72792E-05 | SCNN1A |
| ENSG00000111961 | -0.489978571 | 0.101739933 | 1.46471E-06 | 0.00010638 | SASH1 |
| ENSG00000205488 | -1.023599884 | 0.212592458 | 1.47314E-06 | 0.000106444 | CALML3-AS1 |
| ENSG00000112964 | -1.570889615 | 0.326442765 | 1.4932E-06 | 0.000106551 | GHR |
| ENSG00000175895 | -0.399333603 | 0.082994039 | 1.49731E-06 | 0.000106551 | PLEKHF2 |
| ENSG00000140481 | 1.265440105 | 0.262993808 | 1.49672E-06 | 0.000106551 | CCDC33 |
| ENSG00000066279 | 1.552671972 | 0.323793599 | 1.6247E-06 | 0.000115035 | ASPM |
| ENSG00000108602 | 2.135667051 | 0.44590996 | 1.67232E-06 | 0.000117815 | ALDH3A1 |
| ENSG00000158715 | 0.792568809 | 0.165876603 | 1.76992E-06 | 0.000124071 | SLC45A3 |
| ENSG00000072195 | -0.670223214 | 0.140385589 | 1.80459E-06 | 0.000125875 | SPEG |
| ENSG00000085224 | 0.945891602 | 0.198175537 | 1.81503E-06 | 0.000125979 | ATRX |
| ENSG00000165802 | 0.390687235 | 0.081899177 | 1.83912E-06 | 0.000127025 | NSMF |
| ENSG00000084444 | -0.53200462 | 0.111707344 | 1.91223E-06 | 0.000131248 | FAM234B |
| ENSG00000068489 | 0.52062148 | 0.109333261 | 1.91888E-06 | 0.000131248 | PRR11 |
| ENSG00000065833 | 0.470895335 | 0.09908594 | 2.01023E-06 | 0.000136832 | ME1 |
| ENSG00000153294 | 0.983667363 | 0.20764181 | 2.16555E-06 | 0.000146695 | ADGRF4 |
| ENSG00000049089 | -0.79366042 | 0.167622404 | 2.19259E-06 | 0.000147816 | COL9A2 |
| ENSG00000099219 | -0.33159349 | 0.070219993 | 2.33297E-06 | 0.000156531 | ERMP1 |
| ENSG00000221926 | 0.519419539 | 0.110056785 | 2.36356E-06 | 0.000157832 | TRIM16 |
| ENSG00000087085 | -0.851783966 | 0.180628225 | 2.40911E-06 | 0.000160115 | ACHE |
| ENSG00000106853 | 0.490262672 | 0.104162641 | 2.51755E-06 | 0.000166536 | PTGR1 |
| ENSG00000106003 | -0.592743752 | 0.12640268 | 2.74102E-06 | 0.000180472 | LFNG |
| ENSG00000141279 | 0.374101167 | 0.079896098 | 2.8361E-06 | 0.000185864 | NPEPPS |
| ENSG00000196684 | -0.71529613 | 0.152876741 | 2.88407E-06 | 0.000188132 | HSH2D |
| ENSG00000235584 | -2.581650792 | 0.55390618 | 3.14969E-06 | 0.000204512 | AC008268.1 |
| ENSG00000140443 | 0.557802568 | 0.119782858 | 3.21191E-06 | 0.000207596 | IGF1R |
| ENSG00000160179 | -0.515809905 | 0.110977558 | 3.35371E-06 | 0.00021577 | ABCG1 |
| ENSG00000259583 | 2.228644611 | 0.479608528 | 3.37125E-06 | 0.000215913 | AC015712.2 |
| ENSG00000047410 | 0.842476587 | 0.181822117 | 3.59498E-06 | 0.000229201 | TPR |
| ENSG00000153093 | 1.239374444 | 0.267784676 | 3.68769E-06 | 0.000234052 | ACOXL |
| ENSG00000050438 | -0.764281616 | 0.1652202 | 3.7309E-06 | 0.000235732 | SLC4A8 |
| ENSG00000232386 | 1.279873479 | 0.277268072 | 3.91179E-06 | 0.000246059 | AC015712.1 |
| ENSG00000221869 | 0.742932084 | 0.161154814 | 4.02568E-06 | 0.000252097 | CEBPD |
| ENSG00000107551 | -1.481323148 | 0.321561207 | 4.09188E-06 | 0.000255109 | RASSF4 |
| ENSG00000142197 | -0.395984627 | 0.086085123 | 4.22654E-06 | 0.000262344 | DOPEY2 |
| ENSG00000123595 | 0.416977205 | 0.090822642 | 4.40884E-06 | 0.000272459 | RAB9A |
| ENSG00000168453 | 0.621603947 | 0.135943837 | 4.81926E-06 | 0.000296522 | HR |
| ENSG00000244468 | -1.637249971 | 0.358139532 | 4.84148E-06 | 0.000296593 | AC093001.1 |
| ENSG00000180611 | 0.475123409 | 0.103980441 | 4.89231E-06 | 0.000298409 | MB21D2 |
| ENSG00000175264 | -1.813847622 | 0.397500959 | 5.03971E-06 | 0.000306076 | CHST1 |
| ENSG00000224738 | 0.473045837 | 0.104192232 | 5.62207E-06 | 0.000339978 | AC099850.1 |
| ENSG00000137801 | -0.44265697 | 0.097616942 | 5.77038E-06 | 0.000347456 | THBS1 |

| | | | | | |
|---|---|---|---|---|---|
| ENSG00000103876 | 0.516562188 | 0.113948364 | 5.80691E-06 | 0.000348167 | FAH |
| ENSG00000171940 | -0.342774277 | 0.07588725 | 6.27548E-06 | 0.000374668 | ZNF217 |
| ENSG00000172197 | -0.622484336 | 0.138001115 | 6.46081E-06 | 0.000384105 | MBOAT1 |
| ENSG00000213694 | 0.467184953 | 0.103972321 | 7.01085E-06 | 0.000415054 | S1PR3 |
| ENSG00000185989 | 0.713340985 | 0.158953437 | 7.19842E-06 | 0.000424376 | RASA3 |
| ENSG00000169583 | -0.446736053 | 0.099734428 | 7.49036E-06 | 0.000438431 | CLIC3 |
| ENSG00000242125 | 0.418258129 | 0.093381867 | 7.49907E-06 | 0.000438431 | SNHG3 |
| ENSG00000137166 | -0.42057312 | 0.093973956 | 7.62605E-06 | 0.000444013 | FOXP4 |
| ENSG00000171522 | 0.622755345 | 0.139243967 | 7.73449E-06 | 0.000448473 | PTGER4 |
| ENSG00000058091 | -0.82380919 | 0.184471837 | 7.97805E-06 | 0.0004607 | CDK14 |
| ENSG00000147852 | -0.655096753 | 0.146800292 | 8.10078E-06 | 0.000461887 | VLDLR |
| ENSG00000140479 | -0.528412164 | 0.118431897 | 8.12974E-06 | 0.000461887 | PCSK6 |
| ENSG00000159348 | -0.357167482 | 0.080015442 | 8.05443E-06 | 0.000461887 | CYB5R1 |
| ENSG00000166974 | 0.904043703 | 0.202601987 | 8.11353E-06 | 0.000461887 | MAPRE2 |
| ENSG00000123124 | -0.430337132 | 0.096639944 | 8.4681E-06 | 0.000479179 | WWP1 |
| ENSG00000261776 | -2.02451966 | 0.455181061 | 8.6785E-06 | 0.000485238 | AC079414.2 |
| ENSG00000131389 | 0.344900189 | 0.077543023 | 8.67317E-06 | 0.000485238 | SLC6A6 |
| ENSG00000074410 | 0.404887271 | 0.091003311 | 8.62113E-06 | 0.000485238 | CA12 |
| ENSG00000147041 | -0.501041366 | 0.112703825 | 8.76279E-06 | 0.000487592 | SYTL5 |
| ENSG00000175662 | -0.438568015 | 0.098665803 | 8.7898E-06 | 0.000487592 | TOM1L2 |
| ENSG00000146038 | -0.498633796 | 0.112324538 | 9.02838E-06 | 0.000498862 | DCDC2 |
| ENSG00000119686 | 0.459988234 | 0.103701774 | 9.17807E-06 | 0.000505152 | FLVCR2 |
| ENSG00000163162 | 0.405532027 | 0.09165209 | 9.65812E-06 | 0.000529505 | RNF149 |
| ENSG00000196230 | 0.397040483 | 0.090114904 | 1.05328E-05 | 0.000575222 | TUBB |
| ENSG00000074527 | 0.497170171 | 0.113177339 | 1.11878E-05 | 0.000608636 | NTN4 |
| ENSG00000119699 | -0.828196096 | 0.189276029 | 1.21099E-05 | 0.000654236 | TGFB3 |
| ENSG00000242498 | -0.761553918 | 0.174052068 | 1.21189E-05 | 0.000654236 | ARPIN |
| ENSG00000147100 | -0.57822413 | 0.132266375 | 1.23303E-05 | 0.000663108 | SLC16A2 |
| ENSG00000249242 | -0.454988301 | 0.104302744 | 1.28767E-05 | 0.000689861 | TMEM150C |
| ENSG00000144381 | 0.41979744 | 0.096262421 | 1.29488E-05 | 0.000691091 | HSPD1 |
| ENSG00000137393 | -0.590481433 | 0.135521993 | 1.31803E-05 | 0.000700797 | RNF144B |
| ENSG00000123416 | 0.463529264 | 0.106621943 | 1.37755E-05 | 0.000729686 | TUBA1B |
| ENSG00000109062 | -0.379647975 | 0.087416234 | 1.40555E-05 | 0.000741733 | SLC9A3R1 |
| ENSG00000180694 | -0.363974156 | 0.084085832 | 1.50058E-05 | 0.000788926 | TMEM64 |
| ENSG00000159199 | 0.349834435 | 0.08100675 | 1.57034E-05 | 0.000822529 | ATP5G1 |
| ENSG00000169862 | -0.518443379 | 0.120242021 | 1.62029E-05 | 0.000845553 | CTNND2 |
| ENSG00000234741 | 0.323994897 | 0.075220891 | 1.653E-05 | 0.000859436 | GAS5 |
| ENSG00000157110 | -0.509832294 | 0.118503503 | 1.69069E-05 | 0.0008758 | RBPMS |
| ENSG00000173559 | 0.703374379 | 0.163532499 | 1.6993E-05 | 0.000877038 | NABP1 |
| ENSG00000124151 | -0.347059901 | 0.08105159 | 1.85252E-05 | 0.000952627 | NCOA3 |
| ENSG00000239672 | 0.480225323 | 0.112214148 | 1.87277E-05 | 0.00095954 | NME1 |
| ENSG00000159216 | 0.622291467 | 0.145884233 | 1.99319E-05 | 0.001017536 | RUNX1 |
| ENSG00000153395 | 0.320075493 | 0.075051536 | 2.00133E-05 | 0.001018003 | LPCAT1 |
| ENSG00000064666 | 0.528153393 | 0.124006595 | 2.05275E-05 | 0.001040402 | CNN2 |
| ENSG00000162415 | -0.495821062 | 0.11664759 | 2.13208E-05 | 0.001075102 | ZSWIM5 |
| ENSG00000118855 | 0.3599268 | 0.084686989 | 2.13691E-05 | 0.001075102 | MFSD1 |
| ENSG00000115163 | 0.696428625 | 0.16389136 | 2.1441E-05 | 0.001075102 | CENPA |

| | | | | | |
|---|---|---|---|---|---|
| ENSG00000165732 | 0.439639355 | 0.103558222 | 2.18267E-05 | 0.001090561 | DDX21 |
| ENSG00000173281 | -0.484954892 | 0.114327805 | 2.21741E-05 | 0.001104004 | PPP1R3B |
| ENSG00000185630 | -0.399139333 | 0.094179802 | 2.25463E-05 | 0.001118582 | PBX1 |
| ENSG00000130956 | 0.491740476 | 0.116184303 | 2.31193E-05 | 0.001142984 | HABP4 |
| ENSG00000185052 | -0.566009909 | 0.133759324 | 2.32081E-05 | 0.001143366 | SLC24A3 |
| ENSG00000287900 | -0.672025276 | 0.159217594 | 2.43439E-05 | 0.001195142 | |
| ENSG00000129038 | 0.494769821 | 0.117339532 | 2.48051E-05 | 0.001213556 | LOXL1 |
| ENSG00000287566 | -1.58370586 | 0.375871543 | 2.51529E-05 | 0.001222085 | |
| ENSG00000140526 | -0.349808608 | 0.083016694 | 2.51205E-05 | 0.001222085 | ABHD2 |
| ENSG00000099994 | -1.02173679 | 0.24272089 | 2.55921E-05 | 0.001234909 | SUSD2 |
| ENSG00000169083 | -0.483796911 | 0.11492666 | 2.55807E-05 | 0.001234909 | AR |
| ENSG00000184216 | 0.332233552 | 0.079032076 | 2.62493E-05 | 0.001262294 | IRAK1 |
| ENSG00000147883 | -0.42067955 | 0.100132608 | 2.65475E-05 | 0.001272295 | CDKN2B |
| ENSG00000162989 | -0.472974162 | 0.112603011 | 2.66481E-05 | 0.001272784 | KCNJ3 |
| ENSG00000092978 | -0.414813768 | 0.098808021 | 2.6907E-05 | 0.001280385 | GPATCH2 |
| ENSG00000187017 | 0.445682134 | 0.106178248 | 2.6989E-05 | 0.001280385 | ESPN |
| ENSG00000231133 | 1.24838705 | 0.297712748 | 2.74974E-05 | 0.001300127 | HAR1B |
| ENSG00000153898 | 1.227130518 | 0.293335085 | 2.87214E-05 | 0.001353458 | MCOLN2 |
| ENSG00000140455 | 0.433713084 | 0.103711981 | 2.89084E-05 | 0.00135773 | USP3 |
| ENSG00000186815 | 0.391556736 | 0.093685306 | 2.92164E-05 | 0.00136764 | TPCN1 |
| ENSG00000171724 | -0.952794983 | 0.228253789 | 2.98931E-05 | 0.001394683 | VAT1L |
| ENSG00000116991 | 0.491616722 | 0.117823621 | 3.01302E-05 | 0.001401103 | SIPA1L2 |
| ENSG00000269893 | 0.365797768 | 0.087823214 | 3.11134E-05 | 0.001442064 | SNHG8 |
| ENSG00000143515 | -0.503946255 | 0.121056397 | 3.14224E-05 | 0.001447608 | ATP8B2 |
| ENSG00000111424 | 0.518354681 | 0.124521025 | 3.14385E-05 | 0.001447608 | VDR |
| ENSG00000182606 | -0.408539695 | 0.098166701 | 3.15897E-05 | 0.001449831 | TRAK1 |
| ENSG00000108551 | 0.993374945 | 0.239153605 | 3.27126E-05 | 0.001496497 | RASD1 |
| ENSG00000162670 | -0.610624542 | 0.147091212 | 3.30545E-05 | 0.001507242 | BRINP3 |
| ENSG00000187097 | -0.469602249 | 0.11332796 | 3.41679E-05 | 0.001552984 | ENTPD5 |
| ENSG00000108106 | 0.418517295 | 0.101227612 | 3.55855E-05 | 0.001612217 | UBE2S |
| ENSG00000111859 | 0.543568085 | 0.13157365 | 3.60743E-05 | 0.001629124 | NEDD9 |
| ENSG00000150753 | 0.352784252 | 0.085769476 | 3.90266E-05 | 0.001756822 | CCT5 |
| ENSG00000145386 | 0.494496342 | 0.12029382 | 3.94397E-05 | 0.001769763 | CCNA2 |
| ENSG00000159388 | -0.461060942 | 0.112254006 | 4.0031E-05 | 0.001790594 | BTG2 |
| ENSG00000155366 | -0.370483848 | 0.09032445 | 4.10125E-05 | 0.001828689 | RHOC |
| ENSG00000079805 | -0.30173634 | 0.074052025 | 4.60829E-05 | 0.002044406 | DNM2 |
| ENSG00000205426 | 0.718964741 | 0.176460677 | 4.61406E-05 | 0.002044406 | KRT81 |
| ENSG00000143641 | 0.416665828 | 0.102436194 | 4.75071E-05 | 0.002098353 | GALNT2 |
| ENSG00000185432 | -0.926742352 | 0.228003968 | 4.81172E-05 | 0.002118662 | METTL7A |
| ENSG00000268104 | -0.335727328 | 0.082637337 | 4.85169E-05 | 0.002129603 | SLC6A14 |
| ENSG00000080224 | -0.777964013 | 0.191617186 | 4.90746E-05 | 0.002147397 | EPHA6 |
| ENSG00000149809 | -0.36331701 | 0.089665262 | 5.07977E-05 | 0.002215913 | TM7SF2 |
| ENSG00000020577 | 0.500186505 | 0.123469939 | 5.09819E-05 | 0.002217085 | SAMD4A |
| ENSG00000101384 | -0.92466031 | 0.228386834 | 5.1512E-05 | 0.002233245 | JAG1 |
| ENSG00000152518 | -0.612099056 | 0.151796868 | 5.52204E-05 | 0.002379374 | ZFP36L2 |
| ENSG00000088325 | 0.333142938 | 0.082604454 | 5.50716E-05 | 0.002379374 | TPX2 |
| ENSG00000145912 | 0.408655764 | 0.101577532 | 5.7439E-05 | 0.002467427 | NHP2 |

| | | | | | |
|---|---|---|---|---|---|
| ENSG00000114790 | -0.597041457 | 0.148495872 | 5.80518E-05 | 0.002479646 | ARHGEF26 |
| ENSG00000100591 | 0.361264579 | 0.0898557 | 5.80755E-05 | 0.002479646 | AHSA1 |
| ENSG00000163938 | 0.445381488 | 0.111019361 | 6.02713E-05 | 0.002565628 | GNL3 |
| ENSG00000168350 | -0.769163881 | 0.191822748 | 6.07796E-05 | 0.002566076 | DEGS2 |
| ENSG00000077235 | -0.44079224 | 0.10993491 | 6.08282E-05 | 0.002566076 | GTF3C1 |
| ENSG00000103653 | 0.320121961 | 0.07983845 | 6.08162E-05 | 0.002566076 | CSK |
| ENSG00000125148 | -0.595222568 | 0.148739743 | 6.28698E-05 | 0.002628592 | MT2A |
| ENSG00000111665 | 0.520493564 | 0.130034371 | 6.26135E-05 | 0.002628592 | CDCA3 |
| ENSG00000144355 | 0.733832682 | 0.183357203 | 6.27555E-05 | 0.002628592 | DLX1 |
| ENSG00000176399 | -0.609040445 | 0.152307306 | 6.36751E-05 | 0.002654384 | DMRTA1 |
| ENSG00000173457 | 0.350824062 | 0.087855179 | 6.51852E-05 | 0.002709318 | PPP1R14B |
| ENSG00000189058 | -0.466329425 | 0.116808564 | 6.54483E-05 | 0.002712254 | APOD |
| ENSG00000198719 | 0.440296273 | 0.110507428 | 6.76754E-05 | 0.002796323 | DLL1 |
| ENSG00000138496 | -0.540168264 | 0.13568156 | 6.85835E-05 | 0.002825558 | PARP9 |
| ENSG00000090238 | -0.384572628 | 0.096754993 | 7.04663E-05 | 0.002894666 | YPEL3 |
| ENSG00000131238 | -0.306442727 | 0.077139883 | 7.11046E-05 | 0.002912397 | PPT1 |
| ENSG00000101447 | 0.517457199 | 0.13030292 | 7.15156E-05 | 0.002920737 | FAM83D |
| ENSG00000156103 | -0.870371609 | 0.219246987 | 7.19262E-05 | 0.002929018 | MMP16 |
| ENSG00000133639 | -0.364171689 | 0.091874713 | 7.37704E-05 | 0.002995462 | BTG1 |
| ENSG00000115884 | -0.378925765 | 0.095781222 | 7.61644E-05 | 0.003083783 | SDC1 |
| ENSG00000126787 | 0.798685307 | 0.202486411 | 8.00035E-05 | 0.00322994 | DLGAP5 |
| ENSG00000039068 | -0.279215344 | 0.070812804 | 8.04665E-05 | 0.003239351 | CDH1 |
| ENSG00000041515 | 1.67174174 | 0.424179522 | 8.11026E-05 | 0.003255656 | MYO16 |
| ENSG00000120708 | 0.494534115 | 0.125532301 | 8.16526E-05 | 0.003268426 | TGFBI |
| ENSG00000211452 | -1.401821296 | 0.355983306 | 8.2203E-05 | 0.003281134 | DIO1 |
| ENSG00000179148 | 1.137338739 | 0.289092245 | 8.34838E-05 | 0.003322845 | ALOXE3 |
| ENSG00000107099 | -0.40466021 | 0.103019182 | 8.56522E-05 | 0.00339 | DOCK8 |
| ENSG00000154237 | 0.879666537 | 0.22393929 | 8.56027E-05 | 0.00339 | LRRK1 |
| ENSG00000120725 | -0.347676203 | 0.088620951 | 8.73868E-05 | 0.003448962 | SIL1 |
| ENSG00000162783 | 0.398985757 | 0.101768863 | 8.83621E-05 | 0.003477715 | IER5 |
| ENSG00000165175 | -0.34749315 | 0.088714605 | 8.9666E-05 | 0.003509426 | MID1IP1 |
| ENSG00000229951 | 1.148806459 | 0.293277808 | 8.96105E-05 | 0.003509426 | FLJ31356 |
| ENSG00000171067 | 0.441176615 | 0.112893096 | 9.30961E-05 | 0.003633584 | C11orf24 |
| ENSG00000111266 | 0.394519057 | 0.101006411 | 9.38826E-05 | 0.003654159 | DUSP16 |
| ENSG00000131398 | -0.723115881 | 0.185228115 | 9.46468E-05 | 0.003673755 | KCNC3 |
| ENSG00000173467 | -0.587296513 | 0.150549425 | 9.57877E-05 | 0.003707827 | AGR3 |
| ENSG00000115310 | 0.297164835 | 0.076300617 | 9.83372E-05 | 0.003785715 | RTN4 |
| ENSG00000272405 | 0.706594876 | 0.181410975 | 9.82007E-05 | 0.003785715 | AL365181.3 |
| ENSG00000103479 | -0.401116301 | 0.103023588 | 9.88321E-05 | 0.003794398 | RBL2 |
| ENSG00000135245 | -0.382016665 | 0.098245038 | 0.000100904 | 0.003863433 | HILPDA |
| ENSG00000171552 | -0.342093233 | 0.088196284 | 0.000104985 | 0.004008789 | BCL2L1 |
| ENSG00000157456 | 0.481129893 | 0.124070353 | 0.000105371 | 0.004012635 | CCNB2 |
| ENSG00000151929 | -0.376800447 | 0.097224137 | 0.000106369 | 0.004039715 | BAG3 |
| ENSG00000178538 | 0.807357757 | 0.208412924 | 0.000107135 | 0.00405788 | CA8 |
| ENSG00000151632 | 0.473374301 | 0.122401847 | 0.000110012 | 0.004155663 | AKR1C2 |
| ENSG00000082482 | 0.934333502 | 0.241759762 | 0.000111217 | 0.004189976 | KCNK2 |
| ENSG00000120885 | -0.505380701 | 0.131198163 | 0.000117137 | 0.004374601 | CLU |

| | | | | | |
|---|---|---|---|---|---|
| ENSG00000141424 | -0.450941275 | 0.117049854 | 0.000116891 | 0.004374601 | SLC39A6 |
| ENSG00000131037 | -0.438608245 | 0.113877584 | 0.00011736 | 0.004374601 | EPS8L1 |
| ENSG00000123213 | 0.410708076 | 0.106621277 | 0.000117144 | 0.004374601 | NLN |
| ENSG00000166851 | 0.52835444 | 0.137216791 | 0.000117873 | 0.00438213 | PLK1 |
| ENSG00000186468 | 0.306661389 | 0.079701147 | 0.000119261 | 0.004410456 | RPS23 |
| ENSG00000072571 | 0.971191086 | 0.252379189 | 0.000119017 | 0.004410456 | HMMR |
| ENSG00000196208 | 0.495150849 | 0.128789774 | 0.000120728 | 0.004453012 | GREB1 |
| ENSG00000247809 | -0.87121907 | 0.226706697 | 0.000121572 | 0.004472444 | NR2F2-AS1 |
| ENSG00000272068 | 0.610322123 | 0.158880676 | 0.000122341 | 0.004489029 | AL365181.2 |
| ENSG00000150995 | 0.869168754 | 0.226362879 | 0.000123178 | 0.004507981 | ITPR1 |
| ENSG00000175061 | 0.320794266 | 0.083830339 | 0.000129868 | 0.004718645 | LRRC75A-AS1 |
| ENSG00000254632 | 1.601013341 | 0.418300744 | 0.000129493 | 0.004718645 | AP003119.1 |
| ENSG00000271857 | 1.974358414 | 0.515959677 | 0.000129939 | 0.004718645 | AL096865.1 |
| ENSG00000198125 | -0.490838606 | 0.12837461 | 0.000131578 | 0.004765887 | MB |
| ENSG00000164970 | 0.425918086 | 0.111536755 | 0.000134192 | 0.004848134 | FAM219A |
| ENSG00000173548 | -0.443906227 | 0.116382592 | 0.000136626 | 0.004923428 | SNX33 |
| ENSG00000129187 | 0.397082376 | 0.104327768 | 0.000141173 | 0.005074298 | DCTD |
| ENSG00000110315 | 0.455610228 | 0.119826664 | 0.000143391 | 0.005140926 | RNF141 |
| ENSG00000162949 | -0.360313257 | 0.094789504 | 0.000144001 | 0.00514967 | CAPN13 |
| ENSG00000140284 | 0.589523489 | 0.155118874 | 0.000144426 | 0.00515181 | SLC27A2 |
| ENSG00000164611 | 0.417332324 | 0.109876982 | 0.000145764 | 0.00518639 | PTTG1 |
| ENSG00000168056 | -0.4597947 | 0.121864121 | 0.000161289 | 0.005724338 | LTBP3 |
| ENSG00000095380 | -0.358943691 | 0.095201481 | 0.000163013 | 0.005771001 | NANS |
| ENSG00000143367 | 0.350925759 | 0.093162816 | 0.000165353 | 0.005839154 | TUFT1 |
| ENSG00000169679 | 0.586918212 | 0.156165739 | 0.000171069 | 0.006025921 | BUB1 |
| ENSG00000223573 | 0.420348269 | 0.111973234 | 0.00017403 | 0.006110451 | TINCR |
| ENSG00000189007 | 0.763568292 | 0.20342476 | 0.000174337 | 0.006110451 | ADAT2 |
| ENSG00000184809 | -2.057477209 | 0.548394021 | 0.000175554 | 0.006137839 | B3GALT5-AS1 |
| ENSG00000185442 | 0.430974772 | 0.115037726 | 0.000179406 | 0.00625702 | FAM174B |
| ENSG00000173227 | 0.462567067 | 0.123714462 | 0.000184761 | 0.006427875 | SYT12 |
| ENSG00000006652 | 0.35656998 | 0.095470958 | 0.000187825 | 0.006518359 | IFRD1 |
| ENSG00000106683 | 0.361057445 | 0.096797016 | 0.000191444 | 0.006627626 | LIMK1 |
| ENSG00000095015 | -0.408139509 | 0.109480894 | 0.000193042 | 0.00666658 | MAP3K1 |
| ENSG00000188994 | 1.050943337 | 0.281995857 | 0.000193923 | 0.006680624 | ZNF292 |
| ENSG00000167371 | -0.844397412 | 0.226615691 | 0.000194449 | 0.006682418 | PRRT2 |
| ENSG00000151458 | -0.413663072 | 0.111123162 | 0.00019721 | 0.006760799 | ANKRD50 |
| ENSG00000196876 | -0.814573694 | 0.219122656 | 0.000201258 | 0.006882818 | SCN8A |
| ENSG00000168758 | -0.430805071 | 0.116086025 | 0.000206373 | 0.007040668 | SEMA4C |
| ENSG00000187848 | -0.605425946 | 0.163526504 | 0.000213645 | 0.007236186 | P2RX2 |
| ENSG00000102699 | 0.410610266 | 0.110903336 | 0.000213556 | 0.007236186 | PARP4 |
| ENSG00000167552 | 0.498355036 | 0.134587109 | 0.000213196 | 0.007236186 | TUBA1A |
| ENSG00000141574 | 1.140128423 | 0.308333461 | 0.000217551 | 0.007350833 | SECTM1 |
| ENSG00000066926 | 0.380593398 | 0.103030291 | 0.000220758 | 0.00744134 | FECH |
| ENSG00000055044 | 0.555228922 | 0.15045639 | 0.000223991 | 0.007532314 | NOP58 |
| ENSG00000101187 | 0.423221669 | 0.114720593 | 0.000225003 | 0.007548317 | SLCO4A1 |
| ENSG00000139631 | -0.49623318 | 0.134540501 | 0.000225708 | 0.007553989 | CSAD |
| ENSG00000164920 | -0.447054194 | 0.121495486 | 0.000233605 | 0.007799736 | OSR2 |

| | | | | | |
|---|---|---|---|---|---|
| ENSG00000132554 | 1.460793961 | 0.397248873 | 0.000235737 | 0.007852341 | RGS22 |
| ENSG00000108561 | 0.324045276 | 0.088333461 | 0.000244043 | 0.008109817 | C1QBP |
| ENSG00000108846 | 0.327436804 | 0.08928006 | 0.00024491 | 0.00811949 | ABCC3 |
| ENSG00000114686 | 0.325022308 | 0.088648053 | 0.000245956 | 0.00813501 | MRPL3 |
| ENSG00000133460 | 0.553572356 | 0.151191774 | 0.000250849 | 0.008277415 | SLC2A11 |
| ENSG00000112379 | -0.385348701 | 0.105368501 | 0.000255032 | 0.00839581 | ARFGEF3 |
| ENSG00000137312 | -0.357348066 | 0.097872789 | 0.00026107 | 0.008574551 | FLOT1 |
| ENSG00000129682 | -0.612457277 | 0.167873307 | 0.00026395 | 0.00864897 | FGF13 |
| ENSG00000051108 | -0.34023175 | 0.093291546 | 0.000265348 | 0.008674595 | HERPUD1 |
| ENSG00000175792 | 0.335467426 | 0.092072036 | 0.000268922 | 0.00877109 | RUVBL1 |
| ENSG00000064270 | -0.416803873 | 0.114529025 | 0.000273395 | 0.008896389 | ATP2C2 |
| ENSG00000166801 | -0.405702932 | 0.111613758 | 0.000278107 | 0.009028865 | FAM111A |
| ENSG00000146386 | 0.376356377 | 0.103607851 | 0.00028068 | 0.009091437 | ABRACL |
| ENSG00000168079 | 1.483527092 | 0.409110622 | 0.000287595 | 0.009294067 | SCARA5 |
| ENSG00000125378 | -0.557013548 | 0.153890204 | 0.000295114 | 0.009515227 | BMP4 |
| ENSG00000245694 | -0.52013447 | 0.143748888 | 0.000296482 | 0.009537509 | CRNDE |
| ENSG00000164125 | 0.953930333 | 0.263844862 | 0.000299772 | 0.009621378 | FAM198B |
| ENSG00000243566 | -0.396004094 | 0.109558364 | 0.000300871 | 0.0096347 | UPK3B |
| ENSG00000184203 | 0.354148981 | 0.098089132 | 0.000305629 | 0.009764891 | PPP1R2 |
| ENSG00000166689 | -0.360609014 | 0.100044077 | 0.000312753 | 0.009969879 | PLEKHA7 |
| ENSG00000267131 | -0.722953999 | 0.200674453 | 0.000315025 | 0.009986639 | AC005746.2 |
| ENSG00000117298 | -0.449908426 | 0.124882969 | 0.000315001 | 0.009986639 | ECE1 |
| ENSG00000175356 | 0.574977037 | 0.159613532 | 0.000315405 | 0.009986639 | SCUBE2 |
| ENSG00000142192 | -0.264570988 | 0.073493848 | 0.000318332 | 0.010056707 | APP |
| ENSG00000103642 | 0.497801983 | 0.138342878 | 0.000320279 | 0.010095597 | LACTB |
| ENSG00000163870 | 0.370420309 | 0.103200011 | 0.000331512 | 0.010426333 | TPRA1 |
| ENSG00000179082 | -0.810758787 | 0.225979299 | 0.000333535 | 0.010443353 | C9orf106 |
| ENSG00000129993 | -0.353420524 | 0.098492484 | 0.000332842 | 0.010443353 | CBFA2T3 |
| ENSG00000135709 | -0.322428151 | 0.090131587 | 0.000347155 | 0.010838882 | KIAA0513 |
| ENSG00000196205 | 0.315285187 | 0.08814505 | 0.000347706 | 0.010838882 | EEF1A1P5 |
| ENSG00000174473 | -1.042214515 | 0.291574156 | 0.000350977 | 0.010916715 | GALNTL6 |
| ENSG00000142973 | -0.830973778 | 0.232758479 | 0.000356829 | 0.011063179 | CYP4B1 |
| ENSG00000126803 | -0.587284747 | 0.164514842 | 0.000357257 | 0.011063179 | HSPA2 |
| ENSG00000277159 | -1.277204194 | 0.358157183 | 0.000362411 | 0.011198178 | AL139384.2 |
| ENSG00000050344 | 0.489181315 | 0.137236434 | 0.00036453 | 0.011239002 | NFE2L3 |
| ENSG00000133706 | 0.347155028 | 0.097495184 | 0.000369811 | 0.011376929 | LARS |
| ENSG00000058085 | 0.626290092 | 0.175999796 | 0.000373021 | 0.011450676 | LAMC2 |
| ENSG00000186153 | -0.395734358 | 0.111284862 | 0.000376475 | 0.011531584 | WWOX |
| ENSG00000187535 | -0.456019558 | 0.128285337 | 0.000378369 | 0.011547902 | IFT140 |
| ENSG00000125864 | 0.78771811 | 0.221609263 | 0.000378647 | 0.011547902 | BFSP1 |
| ENSG00000277443 | -0.38211709 | 0.107815239 | 0.00039383 | 0.01198501 | MARCKS |
| ENSG00000119508 | -1.436836709 | 0.406008619 | 0.00040175 | 0.012199693 | NR4A3 |
| ENSG00000169851 | -1.00301381 | 0.283593555 | 0.000405006 | 0.012272104 | PCDH7 |
| ENSG00000088002 | -0.408070733 | 0.115461809 | 0.000408936 | 0.012364618 | SULT2B1 |
| ENSG00000136235 | -0.893277332 | 0.253059749 | 0.000415706 | 0.012542395 | GPNMB |
| ENSG00000150938 | -0.436922504 | 0.123933395 | 0.000422744 | 0.012700361 | CRIM1 |
| ENSG00000139496 | 0.375072794 | 0.10637714 | 0.000422082 | 0.012700361 | NUP58 |

| | | | | | |
|---|---|---|---|---|---|
| ENSG00000079819 | -0.411446601 | 0.116744368 | 0.000424542 | 0.012727241 | EPB41L2 |
| ENSG00000096746 | 0.303707451 | 0.086262871 | 0.000430376 | 0.012874735 | HNRNPH3 |
| ENSG00000183111 | -0.643148206 | 0.182957183 | 0.000439268 | 0.013112898 | ARHGEF37 |
| ENSG00000100815 | 0.718544132 | 0.204581385 | 0.000444304 | 0.013235186 | TRIP11 |
| ENSG00000078124 | 0.468837858 | 0.133565956 | 0.000447837 | 0.013312296 | ACER3 |
| ENSG00000104755 | -1.682714938 | 0.48193693 | 0.000480197 | 0.014187509 | ADAM2 |
| ENSG00000196141 | 0.404520724 | 0.115858436 | 0.000480301 | 0.014187509 | SPATS2L |
| ENSG00000103196 | 0.430584769 | 0.12329021 | 0.000478613 | 0.014187509 | CRISPLD2 |
| ENSG00000107815 | 0.412009779 | 0.118095949 | 0.000485246 | 0.014303585 | TWNK |
| ENSG00000076344 | -1.148498347 | 0.329399558 | 0.000489126 | 0.014387848 | RGS11 |
| ENSG00000145919 | 0.315118087 | 0.090461635 | 0.000495006 | 0.01450027 | BOD1 |
| ENSG00000134690 | 0.424687669 | 0.121907229 | 0.000494542 | 0.01450027 | CDCA8 |
| ENSG00000145012 | 0.55810132 | 0.160383853 | 0.000501816 | 0.014669279 | LPP |
| ENSG00000100321 | -0.392152043 | 0.112790446 | 0.000507399 | 0.014801762 | SYNGR1 |
| ENSG00000162078 | -0.426799034 | 0.122881394 | 0.000514177 | 0.014968498 | ZG16B |
| ENSG00000075399 | -0.395003006 | 0.113959282 | 0.000527916 | 0.015336782 | VPS9D1 |
| ENSG00000160796 | -0.324672507 | 0.093691174 | 0.000529546 | 0.015352478 | NBEAL2 |
| ENSG00000126878 | -0.312509306 | 0.09022343 | 0.000532747 | 0.015413576 | AIF1L |
| ENSG00000145220 | 0.595100527 | 0.17186387 | 0.000534929 | 0.015444973 | LYAR |
| ENSG00000166165 | -0.434044878 | 0.125425284 | 0.000539003 | 0.015530773 | CKB |
| ENSG00000197183 | 0.368010842 | 0.106421645 | 0.00054411 | 0.01562092 | NOL4L |
| ENSG00000115946 | 0.369697985 | 0.10691318 | 0.000544349 | 0.01562092 | PNO1 |
| ENSG00000072274 | 0.299578317 | 0.086682711 | 0.000548188 | 0.015687773 | TFRC |
| ENSG00000189159 | 0.326214551 | 0.094399505 | 0.000548905 | 0.015687773 | JPT1 |
| ENSG00000135899 | -0.624020528 | 0.180846358 | 0.000559433 | 0.015956286 | SP110 |
| ENSG00000184613 | -1.407638307 | 0.40824184 | 0.000564649 | 0.016072544 | NELL2 |
| ENSG00000152284 | -0.705065453 | 0.204890057 | 0.000579163 | 0.016446801 | TCF7L1 |
| ENSG00000124507 | -0.427209288 | 0.124162144 | 0.000580132 | 0.016446801 | PACSIN1 |
| ENSG00000115541 | 0.40782037 | 0.118588663 | 0.000583977 | 0.016522565 | HSPE1 |
| ENSG00000226950 | 0.360032158 | 0.104715101 | 0.000585581 | 0.016534732 | DANCR |
| ENSG00000116337 | 0.350666588 | 0.102100854 | 0.000593622 | 0.016728274 | AMPD2 |
| ENSG00000115306 | 0.311141674 | 0.09062118 | 0.000595991 | 0.016761513 | SPTBN1 |
| ENSG00000166145 | -0.297536698 | 0.086693371 | 0.000599019 | 0.0168131 | SPINT1 |
| ENSG00000135333 | -0.478974912 | 0.139808948 | 0.00061271 | 0.017129149 | EPHA7 |
| ENSG00000109790 | 0.539349293 | 0.157428769 | 0.000612564 | 0.017129149 | KLHL5 |
| ENSG00000108840 | -0.386594169 | 0.112886581 | 0.000615649 | 0.017177226 | HDAC5 |
| ENSG00000147255 | 0.502136703 | 0.146857143 | 0.000628012 | 0.017487529 | IGSF1 |
| ENSG00000159423 | -0.330886453 | 0.096824156 | 0.000632233 | 0.017570347 | ALDH4A1 |
| ENSG00000254726 | 0.413935972 | 0.121145843 | 0.000633525 | 0.017571583 | MEX3A |
| ENSG00000136928 | -1.088510934 | 0.318714448 | 0.000637078 | 0.017591601 | GABBR2 |
| ENSG00000124615 | -0.517234253 | 0.151462774 | 0.000637992 | 0.017591601 | MOCS1 |
| ENSG00000110492 | -0.472688373 | 0.138376099 | 0.000635556 | 0.017591601 | MDK |
| ENSG00000128342 | 1.248989416 | 0.365932253 | 0.000642116 | 0.017670731 | LIF |
| ENSG00000180758 | 0.335053101 | 0.09821293 | 0.000646073 | 0.017744954 | GPR157 |
| ENSG00000067334 | 0.894871294 | 0.262546263 | 0.000653372 | 0.017910521 | DNTTIP2 |
| ENSG00000105520 | -0.402412977 | 0.118131111 | 0.000658029 | 0.018003157 | PLPPR2 |
| ENSG00000105516 | -0.567676888 | 0.16687342 | 0.000669335 | 0.018276996 | DBP |

| | | | | | |
|---|---|---|---|---|---|
| ENSG00000176058 | 0.30097036 | 0.088668505 | 0.000687962 | 0.01874929 | TPRN |
| ENSG00000106484 | 0.308309634 | 0.090900602 | 0.000694547 | 0.018892215 | MEST |
| ENSG00000112742 | 0.905833389 | 0.267144836 | 0.000696902 | 0.018919746 | TTK |
| ENSG00000081377 | 0.510550659 | 0.150717711 | 0.000705423 | 0.019114237 | CDC14B |
| ENSG00000158286 | -0.532076329 | 0.157136612 | 0.000709 | 0.019158829 | RNF207 |
| ENSG00000135124 | -0.350917325 | 0.103644753 | 0.000709788 | 0.019158829 | P2RX4 |
| ENSG00000151892 | -0.447547431 | 0.132441815 | 0.00072697 | 0.0195851 | GFRA1 |
| ENSG00000100814 | 0.334745788 | 0.099086252 | 0.000729282 | 0.0196099 | CCNB1IP1 |
| ENSG00000137502 | -0.450782529 | 0.133490171 | 0.000733086 | 0.01967462 | RAB30 |
| ENSG00000011105 | -0.399387227 | 0.118385404 | 0.00074187 | 0.01987253 | TSPAN9 |
| ENSG00000165029 | -0.42763348 | 0.126824535 | 0.000746647 | 0.019962546 | ABCA1 |
| ENSG00000103145 | -0.541097663 | 0.160602025 | 0.000753913 | 0.020004953 | HCFC1R1 |
| ENSG00000182054 | -0.245435081 | 0.07284538 | 0.000753701 | 0.020004953 | IDH2 |
| ENSG00000131876 | 0.312162113 | 0.092625292 | 0.000751244 | 0.020004953 | SNRPA1 |
| ENSG00000164163 | 0.493803603 | 0.146563263 | 0.000753817 | 0.020004953 | ABCE1 |
| ENSG00000111799 | -1.104368575 | 0.328017648 | 0.000760469 | 0.020128207 | COL12A1 |
| ENSG00000134202 | 0.264287192 | 0.078506122 | 0.000761415 | 0.020128207 | GSTM3 |
| ENSG00000196878 | 0.686183564 | 0.203903857 | 0.000764796 | 0.020179736 | LAMB3 |
| ENSG00000138180 | 0.569056914 | 0.169406525 | 0.00078191 | 0.02059273 | CEP55 |
| ENSG00000188643 | -0.279969055 | 0.083370616 | 0.000784728 | 0.020628379 | S100A16 |
| ENSG00000074370 | -0.313421305 | 0.093458294 | 0.000797689 | 0.020930054 | ATP2A3 |
| ENSG00000102934 | -0.573383501 | 0.171263181 | 0.000814067 | 0.021320072 | PLLP |
| ENSG00000140332 | -0.324823258 | 0.097070046 | 0.000819045 | 0.021410669 | TLE3 |
| ENSG00000156049 | -0.537601511 | 0.160687191 | 0.000820929 | 0.021420159 | GNA14 |
| ENSG00000113396 | -0.684545482 | 0.204772037 | 0.000828887 | 0.021587845 | SLC27A6 |
| ENSG00000188549 | -1.11391118 | 0.334244144 | 0.000860301 | 0.022364647 | C15orf52 |
| ENSG00000213762 | -0.414491278 | 0.124454544 | 0.000867016 | 0.022497716 | ZNF134 |
| ENSG00000187840 | -0.341534867 | 0.102567805 | 0.000868944 | 0.022506294 | EIF4EBP1 |
| ENSG00000164741 | -0.745474607 | 0.224110188 | 0.000879834 | 0.022746524 | DLC1 |
| ENSG00000196917 | -0.455975497 | 0.137206978 | 0.000889695 | 0.022959355 | HCAR1 |
| ENSG00000131171 | -0.361742822 | 0.108897232 | 0.000894152 | 0.023032188 | SH3BGRL |
| ENSG00000144579 | -0.313227648 | 0.094353013 | 0.000901007 | 0.023110478 | CTDSP1 |
| ENSG00000173418 | 0.300703334 | 0.090573213 | 0.000900165 | 0.023110478 | NAA20 |
| ENSG00000111581 | 0.406309658 | 0.122404563 | 0.000902112 | 0.023110478 | NUP107 |
| ENSG00000157388 | 0.896578304 | 0.270276657 | 0.000909045 | 0.023245826 | CACNA1D |
| ENSG00000116574 | -0.40738617 | 0.122883929 | 0.000915739 | 0.023374578 | RHOU |
| ENSG00000173638 | 0.327645267 | 0.098903599 | 0.000923756 | 0.023536568 | SLC19A1 |
| ENSG00000197930 | 0.298075295 | 0.090022551 | 0.00092924 | 0.023633558 | ERO1A |
| ENSG00000065054 | 0.367806128 | 0.111158453 | 0.000936816 | 0.023783304 | SLC9A3R2 |
| ENSG00000168542 | -0.716166545 | 0.216612901 | 0.00094569 | 0.023922403 | COL3A1 |
| ENSG00000156127 | 0.357223551 | 0.108042056 | 0.00094524 | 0.023922403 | BATF |
| ENSG00000177000 | -0.464523812 | 0.140821297 | 0.00097142 | 0.024529228 | MTHFR |
| ENSG00000173917 | -1.327520166 | 0.402675112 | 0.000978097 | 0.024638241 | HOXB2 |
| ENSG00000100290 | -0.503644851 | 0.152808363 | 0.000980983 | 0.024638241 | BIK |
| ENSG00000165105 | -0.329550257 | 0.099986421 | 0.000980894 | 0.024638241 | RASEF |
| ENSG00000093010 | -0.278444824 | 0.084495504 | 0.000982886 | 0.024642094 | COMT |
| ENSG00000076716 | -0.487774569 | 0.148048845 | 0.000985314 | 0.024659113 | GPC4 |

| | | | | | |
|---|---|---|---|---|---|
| ENSG00000130826 | 0.333425896 | 0.101218951 | 0.000987355 | 0.024666359 | DKC1 |
| ENSG00000030582 | -0.252873789 | 0.076790498 | 0.000991122 | 0.024716649 | GRN |
| ENSG00000109971 | 0.251224344 | 0.076328708 | 0.000997085 | 0.024821418 | HSPA8 |
| ENSG00000166780 | -0.507413939 | 0.154217603 | 0.001000996 | 0.024835507 | C16orf45 |
| ENSG00000274180 | -0.454614071 | 0.138193123 | 0.001002939 | 0.024835507 | NATD1 |
| ENSG00000175567 | -0.361803683 | 0.109968611 | 0.001001649 | 0.024835507 | UCP2 |
| ENSG00000214530 | -0.244564736 | 0.074418404 | 0.001014957 | 0.025045092 | STARD10 |
| ENSG00000140961 | 0.389037051 | 0.118371805 | 0.001014161 | 0.025045092 | OSGIN1 |
| ENSG00000009413 | 0.801944951 | 0.244372885 | 0.001032036 | 0.025422007 | REV3L |
| ENSG00000105711 | -0.380233887 | 0.115943271 | 0.001039978 | 0.025484024 | SCN1B |
| ENSG00000156970 | 0.575438739 | 0.175438642 | 0.001038071 | 0.025484024 | BUB1B |
| ENSG00000114812 | 0.630090567 | 0.192131145 | 0.00103998 | 0.025484024 | VIPR1 |
| ENSG00000109805 | 0.851669558 | 0.259982635 | 0.001053367 | 0.025767256 | NCAPG |
| ENSG00000168209 | 0.32507288 | 0.099258396 | 0.001056558 | 0.025800529 | DDIT4 |
| ENSG00000165272 | 0.325152136 | 0.099341381 | 0.001063829 | 0.025933142 | AQP3 |
| ENSG00000023909 | 0.435089286 | 0.132972516 | 0.001067804 | 0.025985087 | GCLM |
| ENSG00000143061 | -0.245759607 | 0.075142442 | 0.00107326 | 0.026027934 | IGSF3 |
| ENSG00000118193 | 1.170057827 | 0.357740404 | 0.001072856 | 0.026027934 | KIF14 |
| ENSG00000181026 | 0.325267092 | 0.099646664 | 0.001097719 | 0.026575359 | AEN |
| ENSG00000164237 | -0.337757221 | 0.10357452 | 0.001110173 | 0.02683078 | CMBL |
| ENSG00000084731 | 0.374024915 | 0.11475168 | 0.001116369 | 0.026934302 | KIF3C |
| ENSG00000106077 | -0.305691819 | 0.093970484 | 0.001141686 | 0.027498032 | ABHD11 |
| ENSG00000070731 | -0.429259747 | 0.132096364 | 0.001155693 | 0.027655076 | ST6GALNAC2 |
| ENSG00000185591 | -0.300042883 | 0.092308063 | 0.001152219 | 0.027655076 | SP1 |
| ENSG00000096384 | 0.226454053 | 0.069701722 | 0.001158515 | 0.027655076 | HSP90AB1 |
| ENSG00000163811 | 0.307505908 | 0.094659678 | 0.001159982 | 0.027655076 | WDR43 |
| ENSG00000281398 | 0.486877712 | 0.149826754 | 0.001155656 | 0.027655076 | SNHG4 |
| ENSG00000118503 | 0.653889171 | 0.201282474 | 0.001159685 | 0.027655076 | TNFAIP3 |
| ENSG00000163683 | -0.235859867 | 0.072699848 | 0.001177411 | 0.028023173 | SMIM14 |
| ENSG00000142945 | 0.377241238 | 0.116336226 | 0.001184108 | 0.02813001 | KIF2C |
| ENSG00000198301 | 0.588439664 | 0.181491052 | 0.001185893 | 0.02813001 | SDAD1 |
| ENSG00000265415 | 0.619296566 | 0.191049222 | 0.001188795 | 0.02815146 | AC099850.3 |
| ENSG00000237330 | 0.367287308 | 0.113406815 | 0.001200882 | 0.028389981 | RNF223 |
| ENSG00000179051 | 0.28815275 | 0.089090262 | 0.001219024 | 0.028770602 | RCC2 |
| ENSG00000196924 | 0.234817849 | 0.072650942 | 0.001228678 | 0.028949962 | FLNA |
| ENSG00000090661 | -0.356103709 | 0.110238892 | 0.001236642 | 0.02908897 | CERS4 |
| ENSG00000170412 | 0.279443816 | 0.086550983 | 0.001243711 | 0.029206471 | GPRC5C |
| ENSG00000141068 | 0.655790414 | 0.203426008 | 0.001265329 | 0.029664697 | KSR1 |
| ENSG00000119950 | 0.337423431 | 0.104734609 | 0.001274327 | 0.029826035 | MXI1 |
| ENSG00000136810 | 0.256098899 | 0.079505325 | 0.001276755 | 0.029833298 | TXN |
| ENSG00000172878 | 0.653679533 | 0.203183979 | 0.001294571 | 0.030199519 | METAP1D |
| ENSG00000142961 | 0.450686918 | 0.14013669 | 0.001299672 | 0.030239667 | MOB3C |
| ENSG00000054654 | 0.453580078 | 0.141045129 | 0.001300585 | 0.030239667 | SYNE2 |
| ENSG00000114405 | -0.248838609 | 0.077531852 | 0.001329653 | 0.030833972 | C3orf14 |
| ENSG00000145734 | 0.7890618 | 0.245866205 | 0.001330522 | 0.030833972 | BDP1 |
| ENSG00000168143 | -0.436448082 | 0.136052272 | 0.001336875 | 0.030930324 | FAM83B |
| ENSG00000167291 | 0.274640912 | 0.08574755 | 0.001360508 | 0.031425493 | TBC1D16 |

| | | | | | |
|---|---|---|---|---|---|
| ENSG00000179627 | -0.317902285 | 0.099404711 | 0.001383552 | 0.031896101 | ZBTB42 |
| ENSG00000126705 | 0.338316074 | 0.105816267 | 0.001387673 | 0.031896101 | AHDC1 |
| ENSG00000115844 | 0.858247161 | 0.268403639 | 0.001385769 | 0.031896101 | DLX2 |
| ENSG00000134909 | -0.346198675 | 0.108426879 | 0.001408405 | 0.032319913 | ARHGAP32 |
| ENSG00000164659 | -0.351069402 | 0.109984552 | 0.001412971 | 0.03237196 | KIAA1324L |
| ENSG00000053372 | 0.342428359 | 0.107352703 | 0.001423956 | 0.032570677 | MRTO4 |
| ENSG00000160211 | 0.295075467 | 0.092523743 | 0.001426738 | 0.032581429 | G6PD |
| ENSG00000159166 | 0.324452317 | 0.101819548 | 0.001439843 | 0.032827497 | LAD1 |
| ENSG00000230836 | -0.689538445 | 0.216494826 | 0.001447484 | 0.032948385 | LINC01293 |
| ENSG00000072310 | -0.391263905 | 0.122868403 | 0.001450472 | 0.032963152 | SREBF1 |
| ENSG00000073150 | 0.384807016 | 0.120927369 | 0.001461945 | 0.033170373 | PANX2 |
| ENSG00000080561 | -0.33842741 | 0.106451494 | 0.001476974 | 0.033457491 | MID2 |
| ENSG00000102882 | -0.251106167 | 0.078999956 | 0.001480082 | 0.033474076 | MAPK3 |
| ENSG00000130940 | -0.481549407 | 0.151552332 | 0.001485782 | 0.033549142 | CASZ1 |
| ENSG00000156398 | -0.359985031 | 0.113467127 | 0.001510841 | 0.034060395 | SFXN2 |
| ENSG00000173267 | 0.858636589 | 0.270815105 | 0.001521434 | 0.034244412 | SNCG |
| ENSG00000197063 | 0.319735982 | 0.100968564 | 0.001541854 | 0.034648686 | MAFG |
| ENSG00000105971 | 0.395423994 | 0.124945926 | 0.001552104 | 0.034823476 | CAV2 |
| ENSG00000169607 | 0.712451983 | 0.225206497 | 0.001558576 | 0.034913104 | CKAP2L |
| ENSG00000162545 | -0.312210455 | 0.098819409 | 0.001580921 | 0.035357417 | CAMK2N1 |
| ENSG00000197969 | 0.784206754 | 0.248252083 | 0.001583587 | 0.035360922 | VPS13A |
| ENSG00000168807 | -0.416284197 | 0.131814007 | 0.001587912 | 0.035376477 | SNTB2 |
| ENSG00000156795 | 0.331199998 | 0.104896358 | 0.001591816 | 0.035376477 | WDYHV1 |
| ENSG00000184857 | 0.447271478 | 0.141650575 | 0.001590905 | 0.035376477 | TMEM186 |
| ENSG00000272150 | 0.57417586 | 0.182116736 | 0.00161718 | 0.035883566 | NBPF25P |
| ENSG00000166197 | 0.299794727 | 0.095220684 | 0.001641555 | 0.036367159 | NOLC1 |
| ENSG00000162804 | -0.465834813 | 0.147998643 | 0.001646385 | 0.036416906 | SNED1 |
| ENSG00000138413 | -0.269040169 | 0.085499659 | 0.001651349 | 0.03646945 | IDH1 |
| ENSG00000140859 | -0.43395051 | 0.138014497 | 0.001665196 | 0.036603139 | KIFC3 |
| ENSG00000167608 | -0.420466524 | 0.133707924 | 0.001662776 | 0.036603139 | TMC4 |
| ENSG00000141753 | -0.350774146 | 0.111551629 | 0.001663707 | 0.036603139 | IGFBP4 |
| ENSG00000186998 | -0.583672347 | 0.185797173 | 0.001681141 | 0.036896064 | EMID1 |
| ENSG00000168301 | -0.868189192 | 0.276827854 | 0.001711489 | 0.037450255 | KCTD6 |
| ENSG00000126778 | -0.526179333 | 0.167777759 | 0.001711708 | 0.037450255 | SIX1 |
| ENSG00000063180 | -0.665879114 | 0.212425349 | 0.001720593 | 0.037528099 | CA11 |
| ENSG00000087470 | 0.549950473 | 0.17542754 | 0.001719038 | 0.037528099 | DNM1L |
| ENSG00000179335 | 0.270796678 | 0.086450004 | 0.001733788 | 0.037757455 | CLK3 |
| ENSG00000013810 | 0.318839241 | 0.101862647 | 0.001747528 | 0.037997951 | TACC3 |
| ENSG00000137193 | 0.498345884 | 0.159322496 | 0.001760561 | 0.038222354 | PIM1 |
| ENSG00000096696 | 0.378178906 | 0.12094081 | 0.001766149 | 0.038284666 | DSP |
| ENSG00000165209 | 0.269541434 | 0.08622032 | 0.001770848 | 0.038327567 | STRBP |
| ENSG00000286836 | -1.446383202 | 0.462911714 | 0.001780872 | 0.038485405 | |
| ENSG00000013588 | -0.299441504 | 0.095927557 | 0.001799091 | 0.038819587 | GPRC5A |
| ENSG00000211689 | -0.847648648 | 0.271638853 | 0.001805452 | 0.038897273 | TRGC1 |
| ENSG00000250241 | -0.504617256 | 0.162021522 | 0.001842522 | 0.039635328 | AC105383.1 |
| ENSG00000185386 | -0.459887658 | 0.147798773 | 0.001860893 | 0.03991436 | MAPK11 |
| ENSG00000116977 | -0.288304535 | 0.09265662 | 0.001861159 | 0.03991436 | LGALS8 |

| | | | | | |
|---|---|---|---|---|---|
| ENSG00000188846 | 0.26447943 | 0.085016827 | 0.001865138 | 0.039938888 | RPL14 |
| ENSG00000165704 | 0.33405824 | 0.107458989 | 0.001879096 | 0.040176721 | HPRT1 |
| ENSG00000137955 | 0.364478566 | 0.117278281 | 0.001884795 | 0.040237508 | RABGGTB |
| ENSG00000136068 | -0.316304606 | 0.101959967 | 0.001920602 | 0.040939904 | FLNB |
| ENSG00000027697 | 0.361134614 | 0.11649636 | 0.001935437 | 0.04113855 | IFNGR1 |
| ENSG00000249364 | 0.731388096 | 0.235938092 | 0.00193576 | 0.04113855 | AC112206.2 |
| ENSG00000197670 | -0.62390328 | 0.201409015 | 0.001950334 | 0.041363476 | AL157838.1 |
| ENSG00000141232 | -0.370505254 | 0.119617871 | 0.001952215 | 0.041363476 | TOB1 |
| ENSG00000132746 | 0.465707913 | 0.150395899 | 0.001957891 | 0.041421441 | ALDH3B2 |
| ENSG00000085465 | 0.838018295 | 0.270808758 | 0.001971435 | 0.041645451 | OVGP1 |
| ENSG00000124766 | 0.228944078 | 0.074051249 | 0.001990154 | 0.041977946 | SOX4 |
| ENSG00000256269 | 0.285174584 | 0.092263074 | 0.001995607 | 0.042030041 | HMBS |
| ENSG00000125398 | -0.720388359 | 0.233184577 | 0.002005962 | 0.042185074 | SOX9 |
| ENSG00000258947 | 0.777200269 | 0.25167433 | 0.002014278 | 0.042296841 | TUBB3 |
| ENSG00000144452 | -0.674107308 | 0.218325835 | 0.00201766 | 0.042304806 | ABCA12 |
| ENSG00000135862 | -0.240687089 | 0.078046554 | 0.002043122 | 0.042606958 | LAMC1 |
| ENSG00000116260 | 0.244555485 | 0.079304851 | 0.002044166 | 0.042606958 | QSOX1 |
| ENSG00000179041 | 0.352336114 | 0.11423789 | 0.002040786 | 0.042606958 | RRS1 |
| ENSG00000164683 | 0.84998221 | 0.27552789 | 0.002036051 | 0.042606958 | HEY1 |
| ENSG00000069764 | -1.266591769 | 0.411752513 | 0.002097277 | 0.043649389 | PLA2G10 |
| ENSG00000260853 | -0.647275524 | 0.210480751 | 0.002103443 | 0.04371314 | AC109460.2 |
| ENSG00000116120 | 0.328790723 | 0.106969469 | 0.002114308 | 0.043745364 | FARSB |
| ENSG00000249859 | 0.407959454 | 0.132691201 | 0.002108538 | 0.043745364 | PVT1 |
| ENSG00000228065 | 0.677728045 | 0.220463426 | 0.002111341 | 0.043745364 | LINC01515 |
| ENSG00000168078 | 0.401468987 | 0.130855331 | 0.0021547 | 0.044515721 | PBK |
| ENSG00000164855 | -0.274380955 | 0.08944934 | 0.00215897 | 0.044538632 | TMEM184A |
| ENSG00000143179 | 0.331019531 | 0.107937984 | 0.002163949 | 0.044557058 | UCK2 |
| ENSG00000166012 | 0.415236528 | 0.135432151 | 0.00216935 | 0.044557058 | TAF1D |
| ENSG00000130768 | 0.889329509 | 0.290046236 | 0.002168239 | 0.044557058 | SMPDL3B |
| ENSG00000187266 | -0.626036889 | 0.204293677 | 0.002181094 | 0.044733067 | EPOR |
| ENSG00000176920 | -1.363363726 | 0.445233086 | 0.002197645 | 0.044960334 | FUT2 |
| ENSG00000107130 | 0.287636459 | 0.093937123 | 0.002198557 | 0.044960334 | NCS1 |
| ENSG00000222041 | -0.502004656 | 0.164073164 | 0.002216039 | 0.045252159 | CYTOR |
| ENSG00000137648 | -0.781537732 | 0.255548184 | 0.00222612 | 0.045261218 | TMPRSS4 |
| ENSG00000078804 | -0.448892112 | 0.146766316 | 0.00222411 | 0.045261218 | TP53INP2 |
| ENSG00000100526 | 0.385050903 | 0.125886594 | 0.002222903 | 0.045261218 | CDKN3 |
| ENSG00000272602 | 0.680824577 | 0.222671788 | 0.00223173 | 0.045309901 | ZNF595 |
| ENSG00000171603 | -0.266472264 | 0.087188578 | 0.002241057 | 0.045409148 | CLSTN1 |
| ENSG00000167165 | 0.728154756 | 0.238270026 | 0.002243064 | 0.045409148 | UGT1A6 |
| ENSG00000261754 | -0.803664566 | 0.263106975 | 0.002254237 | 0.045569871 | AC008555.1 |
| ENSG00000198707 | 0.885090135 | 0.289849691 | 0.002261 | 0.045641111 | CEP290 |
| ENSG00000070371 | -0.584434206 | 0.191506833 | 0.002274987 | 0.045857755 | CLTCL1 |
| ENSG00000173801 | 0.210670413 | 0.069058579 | 0.002283812 | 0.045969869 | JUP |
| ENSG00000100100 | -0.40348335 | 0.132307709 | 0.002291592 | 0.046060682 | PIK3IP1 |
| ENSG00000112246 | -1.445615859 | 0.474450016 | 0.002311917 | 0.046097656 | SIM1 |
| ENSG00000249661 | -1.010760586 | 0.331717102 | 0.002310946 | 0.046097656 | TNRC18P1 |
| ENSG00000105518 | -0.398087055 | 0.130658245 | 0.002313062 | 0.046097656 | TMEM205 |

| | | | | | |
|---|---|---|---|---|---|
| ENSG00000126522 | -0.336952678 | 0.110559527 | 0.002305972 | 0.046097656 | ASL |
| ENSG00000164574 | 0.307414393 | 0.100890566 | 0.002311313 | 0.046097656 | GALNT10 |
| ENSG00000101745 | 0.804724209 | 0.264058191 | 0.002307335 | 0.046097656 | ANKRD12 |
| ENSG00000141449 | -0.577179824 | 0.189497826 | 0.002320322 | 0.046177036 | GREB1L |
| ENSG00000267475 | -1.343989582 | 0.44176526 | 0.002347652 | 0.046655028 | AC008736.1 |
| ENSG00000130005 | -0.640542145 | 0.210695743 | 0.002364772 | 0.046929063 | GAMT |
| ENSG00000095383 | -0.332604863 | 0.109428386 | 0.002369907 | 0.04696483 | TBC1D2 |
| ENSG00000114850 | 0.269902157 | 0.088877755 | 0.002391202 | 0.047320271 | SSR3 |
| ENSG00000148341 | -0.258134203 | 0.085138351 | 0.002429891 | 0.047951209 | SH3GLB2 |
| ENSG00000158122 | 0.410853378 | 0.135492966 | 0.002427094 | 0.047951209 | AAED1 |
| ENSG00000135148 | 0.228548799 | 0.075446469 | 0.002451339 | 0.048306811 | TRAFD1 |
| ENSG00000117593 | 0.283556533 | 0.0936683 | 0.00246797 | 0.048542234 | DARS2 |
| ENSG00000169991 | 0.395277545 | 0.130585175 | 0.002470176 | 0.048542234 | IFFO2 |
| ENSG00000197451 | 0.263421417 | 0.087069433 | 0.002482898 | 0.048724287 | HNRNPAB |
| ENSG00000265972 | -0.305667777 | 0.101077716 | 0.002493849 | 0.048871124 | TXNIP |
| ENSG00000020181 | -1.225310206 | 0.405295875 | 0.002500767 | 0.048914996 | ADGRA2 |
| ENSG00000254285 | -0.380371865 | 0.125827008 | 0.002503031 | 0.048914996 | KRT8P3 |
| ENSG00000161714 | -0.24345936 | 0.080603594 | 0.002523969 | 0.049255858 | PLCD3 |
| ENSG00000103034 | 0.352855784 | 0.116877082 | 0.002535828 | 0.049418831 | NDRG4 |
| ENSG00000121552 | -1.710717068 | 0.567018702 | 0.002552577 | 0.049471534 | CSTA |
| ENSG00000138356 | -0.820822099 | 0.27200187 | 0.002546946 | 0.049471534 | AOX1 |
| ENSG00000286208 | -0.53586814 | 0.177598714 | 0.002550395 | 0.049471534 | |
| ENSG00000143409 | -0.270646876 | 0.089682038 | 0.002545755 | 0.049471534 | MINDY1 |
| ENSG00000180104 | 0.250437182 | 0.083019475 | 0.002556204 | 0.049473791 | EXOC3 |
| ENSG00000168077 | -0.408412781 | 0.135428818 | 0.002563862 | 0.049506164 | SCARA3 |
| ENSG00000175274 | -0.35578854 | 0.117983552 | 0.002564904 | 0.049506164 | TP53I11 |
| ENSG00000174791 | 0.711856428 | 0.236120198 | 0.002571443 | 0.049564468 | RIN1 |
| ENSG00000171793 | 0.337561384 | 0.11199945 | 0.002578654 | 0.049635572 | CTPS1 |
| ENSG00000214194 | 0.411283822 | 0.136480654 | 0.00258257 | 0.049643118 | SMIM30 |
| ENSG00000105976 | -0.355804121 | 0.118106314 | 0.002590415 | 0.049726083 | MET |
| ENSG00000158158 | -0.326598533 | 0.10843703 | 0.002596414 | 0.049773441 | CNNM4 |
| ENSG00000172508 | -0.901595104 | 0.299524017 | 0.002611678 | 0.049998026 | CARNS1 |
