## Supplementary material for "Global transcriptome analysis reveals partial estrogen-like effects of karanjin in MCF-7 breast cancer cells": All supplementary material: Supplementary data 9.pdf

**Table 1:** List of common genes regulated by tamoxifen and karanjin. Red cells represent negative $log_2FC$ values implying downregulation of the gene. Green cells represent positive $log_2FC$ implying upregulation of the gene.

| Gene | Tamoxifen | Karanjin |
|---|---|---|
| GREB1 | -2.55735 | 0.49515 |
| FOSL2 | 0.93781 | 0.90447 |
| IER3 | 0.84203 | 0.77485 |
| TMEM120B | 0.92590 | 0.98744 |
| ESR1 | -0.47007 | -0.79839 |
| RET | 0.78069 | -1.48716 |
| PHLDA1 | 0.83261 | 1.29955 |
| ANXA9 | 0.72957 | -0.83170 |
| SLC9A3R1 | 0.42184 | -0.37965 |
| THBS1 | -0.42953 | -0.44266 |
| SLC7A5 | 0.52527 | 2.88241 |
| IGFBP4 | -0.56627 | -0.35077 |
| DCDC2 | 0.55065 | -0.49863 |
| STARD10 | 0.39895 | -0.24456 |
| IGFBP5 | 0.61801 | -1.01068 |
| TMEM64 | 0.70083 | -0.36397 |
| TFF1 | -0.41660 | 0.69688 |
| KRT18 | -0.22320 | -0.42981 |
| IGF1R | 0.27579 | 0.55780 |
| WISP2 | 0.41743 | 0.72403 |
| ACSS1 | 0.84822 | -1.11254 |
| ABCG1 | -0.84740 | -0.51581 |
| SLC39A6 | -0.21267 | -0.45094 |
| SYTL5 | 0.47828 | -0.50104 |
| HR | 0.70226 | 0.62160 |
| TXNIP | 0.35485 | -0.30567 |
| CLIC3 | 0.58364 | -0.44674 |
| PREX1 | 0.22564 | -0.53848 |
| PLCD3 | -0.51035 | -0.24346 |
| LOXL1 | 0.46926 | 0.49477 |
| SLC24A3 | -0.56067 | -0.56601 |
| KRT8 | -0.16156 | -0.46869 |
| STC1 | 0.41809 | 0.59671 |
| TBC1D9 | -0.22039 | -0.58733 |
| PCDH7 | -0.46997 | -1.00301 |
| SP1 | 0.27299 | -0.30004 |
| TXNRD1 | 0.33188 | 0.82848 |
| SCNN1A | 0.49363 | -0.62562 |
| SPTBN1 | -0.26363 | 0.31114 |
| FLNB | 0.23159 | -0.31630 |

| | | |
|---|---|---|
| LTBP3 | -0.29467 | -0.45979 |
| CERS2 | 0.16688 | -0.48687 |
| SH3BGRL | 0.26594 | -0.36174 |
| SEMA4C | -0.20818 | -0.43081 |
| PITPNM2 | 0.73282 | 1.34756 |
| KRT19 | 0.14678 | -0.44848 |
| DDIT4 | 0.34856 | 0.32507 |
| ST3GAL1 | 0.23705 | 0.85668 |
| GPER1 | -0.40173 | -1.10271 |
| AGR3 | 1.42952 | -0.58730 |
| GFRA1 | 0.14759 | -0.44755 |
| PCSK6 | -0.31629 | -0.52841 |
| SOX4 | -0.21216 | 0.22894 |
| RUNX1 | -0.33836 | 0.62229 |
| AR | 0.34493 | -0.48380 |
| CHRD | -0.74749 | -1.14409 |
| DSP | -0.14198 | 0.37818 |
| GPRC5A | -0.22756 | -0.29944 |
| SYNE2 | -0.28570 | 0.45358 |
| GREB1L | 0.55469 | -0.57718 |
| MT2A | -0.64036 | -0.59522 |
| ALDH3B2 | -0.43826 | 0.46571 |
| CYP1B1 | -0.23035 | 2.72335 |
| ABHD2 | -0.23062 | -0.34981 |
| SPEG | 0.63428 | -0.67022 |
| CA12 | 0.17531 | 0.40489 |
| CLSTN2 | -0.63517 | 0.66979 |
| KITLG | -0.54133 | -0.77083 |
| TPCN1 | 0.28573 | 0.39156 |
| IDH1 | 0.25064 | -0.26904 |
| MAGED2 | 0.17328 | -0.38337 |
| LMCD1 | 0.44012 | 1.31182 |
| ADCY5 | 0.35298 | -0.63689 |
| SIM1 | 1.26923 | -1.44562 |
| CAMK2N1 | -0.33133 | -0.31221 |
| DEGS2 | 0.28426 | -0.76916 |
| LAMC2 | 0.70334 | 0.62629 |
| BAG3 | 0.15985 | -0.37680 |
| TUBB3 | -0.17962 | 0.77720 |
| PMP22 | -0.23010 | -0.92279 |
| KCNC3 | -0.36726 | -0.72312 |
| CD9 | -0.15108 | -0.35416 |
| CA2 | 0.26254 | -0.69021 |
| ACHE | -0.34861 | -0.85178 |
| TINCR | 0.25000 | 0.42035 |
| NT5E | -0.16356 | -1.14029 |

| | | |
|---|---|---|
| LACTB | -0.30758 | 0.49780 |
| RBL2 | 0.21375 | -0.40112 |
| TMEM150C | 0.20269 | -0.45499 |
| GDPD3 | -0.36800 | -0.65505 |
| DIO1 | 1.98040 | -1.40182 |
| HNRNPAB | -0.12155 | 0.26342 |
| EPS8L1 | -0.25663 | -0.43861 |
| LDHA | 0.15873 | 0.41152 |

**Table 2:** List of common genes regulated by estrogen and karanjin. Red cells represent negative $log_2FC$ values implying downregulation of the gene. Green cells represent positive $log_2FC$ implying upregulation of the gene.

| Gene | Estrogen | Karanjin |
|---|---|---|
| CYP1A1 | -1.75158 | 6.17733 |
| SLC7A5 | 1.688633 | 2.88241 |
| STC2 | 0.760613 | 1.1071 |
| LAMA3 | 1.378145 | 1.84031 |
| TIPARP | 1.127496 | 2.05882 |
| SLC3A2 | 0.572105 | 0.97688 |
| PHLDA1 | 0.434351 | 1.29955 |
| TSKU | 0.45057 | 1.10389 |
| ALDH1A3 | -1.50793 | 2.46633 |
| GDF15 | -0.60991 | 1.31592 |
| LMCD1 | -0.69094 | 1.31182 |
| RND1 | -0.86829 | 1.56001 |
| FOSL2 | 0.741212 | 0.90447 |
| PMP22 | -1.06477 | -0.9228 |
| RUNX2 | 0.477098 | 1.43623 |
| RET | 1.430755 | -1.4872 |
| IL1R1 | -2.17744 | -1.1697 |
| ANXA9 | -0.39428 | -0.8317 |
| ADAMTS19 | -0.62557 | -1.1133 |
| SLC35C1 | -0.79732 | -1.1612 |
| SLC7A11 | 0.983892 | 2.43785 |
| TXNRD1 | 0.277647 | 0.82848 |
| TMEM45B | -1.0134 | 1.18506 |
| AREG | 3.124479 | 1.48712 |
| PSCA | -2.22258 | -0.9852 |
| CRY2 | -0.81375 | -0.8571 |
| CYSRT1 | -0.80867 | 0.96535 |
| ATOH8 | -2.11823 | 1.42633 |
| ST3GAL1 | -0.58697 | 0.85668 |
| BMF | -2.44599 | 0.94572 |
| GPER1 | -0.97146 | -1.1027 |

| | | |
|---|---|---|
| CD44 | 1.116591 | 0.88285 |
| TMEM120B | 0.873404 | 0.98744 |
| KCNJ8 | -1.05771 | -1.2645 |
| SLC12A2 | -0.6015 | -0.6495 |
| NPNT | -0.79012 | -0.6665 |
| ESR1 | -0.47548 | -0.7984 |
| ACSS1 | -0.88194 | -1.1125 |
| PPP1R3C | -0.91188 | -0.6368 |
| NCAM2 | -0.45638 | -0.6753 |
| NDRG1 | -0.59345 | -1.0034 |
| BMP7 | -0.76204 | -0.6021 |
| LDLRAD4 | -0.76891 | -0.729 |
| SYNJ2BP | -0.44679 | -0.765 |
| SPTSSA | 0.352374 | 0.6814 |
| CABLES1 | -0.27484 | 0.60923 |
| LRP5 | -0.47271 | 0.52774 |
| DNMBP | 0.727415 | 0.66431 |
| WISP2 | 1.077031 | 0.72403 |
| TRERF1 | -0.95702 | -0.8596 |
| TFF1 | 2.298919 | 0.69688 |
| TUBA1C | 0.421166 | 0.58071 |
| VTCN1 | -0.79609 | 1.69128 |
| CERS2 | -0.51071 | -0.4869 |
| PP14571 | -0.95377 | -0.9123 |
| TSC22D3 | -0.98198 | -0.4916 |
| RAB3D | -1.00866 | -0.6995 |
| NRBP1 | -0.54008 | -0.4779 |
| FN1 | 0.46273 | -0.7339 |
| CHRD | -2.97177 | -1.1441 |
| YPEL2 | -1.31399 | -0.5608 |
| DKK1 | -0.69777 | -0.8622 |
| ISOC1 | 0.34044 | -0.5913 |
| INPP4A | -0.47411 | 0.55204 |
| SMIM5 | -0.93201 | 0.90516 |
| SIX4 | -0.52164 | -0.5818 |
| PREX1 | -0.17437 | -0.5385 |
| SPDEF | -0.54518 | -0.5206 |
| ACKR3 | -0.45018 | -0.6248 |
| KRT18 | -0.36464 | -0.4298 |
| UBE2C | 0.487766 | 0.61816 |
| PRPS1 | 0.510245 | 0.58894 |
| STC1 | 0.452835 | 0.59671 |
| PFKFB3 | -0.31107 | -0.5263 |
| HMCN1 | -0.41572 | -0.4652 |

| | | |
|---|---|---|
| KIAA1324 | 0.366045 | -0.6869 |
| GOLGB1 | -0.38362 | 0.90165 |
| RAP1GAP | -0.96451 | 0.63543 |
| IGFBP5 | 0.583799 | -1.0107 |
| TSPAN1 | -1.07738 | -0.7133 |
| KRT8 | -0.28676 | -0.4687 |
| FZD8 | -1.31306 | -0.8005 |
| ARNT2 | -0.9015 | -0.8448 |
| TGFB2 | -1.23017 | -0.9265 |
| RNF224 | -1.3355 | 1.1133 |
| PCDH10 | -0.98857 | -0.7052 |
| MYC | 0.66237 | 1.00247 |
| FAM161B | -0.64168 | -0.8302 |
| LXN | -0.93553 | 0.70516 |
| CXCR4 | -0.48187 | -1.5061 |
| RARA | 0.509096 | -0.5299 |
| TP53INP1 | -1.81276 | -0.4546 |
| CENPE | 0.288627 | 1.96979 |
| C1orf21 | -0.43756 | -0.5387 |
| PFKP | 0.422214 | 0.40237 |
| NFRKB | -0.38334 | 0.49256 |
| ADCY5 | -0.39958 | -0.6369 |
| KITLG | -1.08387 | -0.7708 |
| KPNA2 | 0.497823 | 0.49408 |
| AURKA | 0.479986 | 0.44109 |
| SELENBP1 | -1.10957 | -0.5376 |
| GDPD3 | -0.86798 | -0.655 |
| CTSH | -0.75853 | -0.4964 |
| SLC34A3 | -1.01086 | 1.03868 |
| PNPLA7 | -1.05749 | -0.8336 |
| CCNB1 | 0.457928 | 0.4067 |
| TBC1D9 | -0.72279 | -0.5873 |
| TUBB4B | 0.386359 | 0.46056 |
| TMTC2 | -1.02662 | 0.53154 |
| NT5E | -1.66106 | -1.1403 |
| SLIT2 | -1.18843 | -1.1439 |
| KIAA1161 | -0.42748 | -0.4465 |
| CORO2A | -0.87547 | -0.474 |
| RAB26 | -1.20989 | -0.7085 |
| CD9 | -0.25947 | -0.3542 |
| CKS2 | 0.448754 | 0.49821 |
| LDHA | 0.910461 | 0.41152 |
| SNHG7 | -0.25145 | 0.38731 |
| CA2 | 0.701731 | -0.6902 |

| | | |
|---|---|---|
| C2orf54 | -2.45978 | -0.9029 |
| INPP5J | -1.4691 | -0.9262 |
| NR3C1 | -0.47164 | 0.56554 |
| SCNN1A | -1.36197 | -0.6256 |
| SASH1 | -0.79742 | -0.49 |
| CALML3-AS1 | -0.91918 | -1.0236 |
| GHR | -1.40799 | -1.5709 |
| PLEKHF2 | -0.49242 | -0.3993 |
| SPEG | -0.51744 | -0.6702 |
| ATRX | -0.25797 | 0.94589 |
| PRR11 | 0.244951 | 0.52062 |
| ME1 | 0.463535 | 0.4709 |
| COL9A2 | -1.27725 | -0.7937 |
| ERMP1 | -0.57793 | -0.3316 |
| TRIM16 | -0.5793 | 0.51942 |
| ACHE | -3.43826 | -0.8518 |
| LFNG | -0.54835 | -0.5927 |
| HSH2D | -0.31324 | -0.7153 |
| IGF1R | 0.217154 | 0.5578 |
| ABCG1 | -1.06329 | -0.5158 |
| DOPEY2 | -0.37147 | -0.396 |
| RAB9A | -0.41178 | 0.41698 |
| HR | 0.724799 | 0.6216 |
| CHST1 | -1.33087 | -1.8138 |
| ZNF217 | -1.11263 | -0.3428 |
| S1PR3 | -0.82007 | 0.46718 |
| SNHG3 | 0.494196 | 0.41826 |
| FOXP4 | -0.33415 | -0.4206 |
| CYB5R1 | -0.31418 | -0.3572 |
| VLDLR | -0.51264 | -0.6551 |
| PCSK6 | -0.93851 | -0.5284 |
| WWP1 | -0.2666 | -0.4303 |
| CA12 | 0.692127 | 0.40489 |
| SYTL5 | 1.778343 | -0.501 |
| TOM1L2 | -1.02068 | -0.4386 |
| DCDC2 | -0.48971 | -0.4986 |
| TUBB | 0.56829 | 0.39704 |
| NTN4 | -1.11339 | 0.49717 |
| TGFB3 | -1.71313 | -0.8282 |
| ARPIN | -1.50722 | -0.7616 |
| TMEM150C | -0.65778 | -0.455 |
| HSPD1 | 0.590409 | 0.4198 |
| RNF144B | -1.42335 | -0.5905 |
| TUBA1B | 0.730043 | 0.46353 |

| | | |
|---|---|---|
| SLC9A3R1 | 0.861735 | -0.3796 |
| TMEM64 | 0.956628 | -0.364 |
| ATP5G1 | 0.664534 | 0.34983 |
| CTNND2 | -0.96865 | -0.5184 |
| NCOA3 | -0.56461 | -0.3471 |
| NME1 | 0.861508 | 0.48023 |
| RUNX1 | 0.424036 | 0.62229 |
| LPCAT1 | 0.387774 | 0.32008 |
| CENPA | 0.432966 | 0.69643 |
| DDX21 | 0.584913 | 0.43964 |
| ABHD2 | 0.691698 | -0.3498 |
| SUSD2 | -0.55383 | -1.0217 |
| IRAK1 | 0.369123 | 0.33223 |
| CDKN2B | -1.59539 | -0.4207 |
| KCNJ3 | -0.91568 | -0.473 |
| HAR1B | -0.80863 | 1.24839 |
| VAT1L | -1.87241 | -0.9528 |
| SIPA1L2 | 0.279529 | 0.49162 |
| ATP8B2 | -0.96893 | -0.5039 |
| TRAK1 | -0.52093 | -0.4085 |
| RASD1 | -0.78809 | 0.99337 |
| UBE2S | 0.568225 | 0.41852 |
| NEDD9 | -0.67199 | 0.54357 |
| CCT5 | 0.549373 | 0.35278 |
| CCNA2 | 0.438331 | 0.4945 |
| BTG2 | -1.52902 | -0.4611 |
| RHOC | -0.39684 | -0.3705 |
| KRT81 | -0.94708 | 0.71896 |
| TM7SF2 | -0.473 | -0.3633 |
| SAMD4A | -0.47248 | 0.50019 |
| JAG1 | -0.40645 | -0.9247 |
| ZFP36L2 | -0.19128 | -0.6121 |
| TPX2 | 0.392047 | 0.33314 |
| NHP2 | 0.283017 | 0.40866 |
| AHSA1 | 0.642493 | 0.36126 |
| GNL3 | 0.423313 | 0.44538 |
| CDCA3 | 0.573291 | 0.52049 |
| MT2A | 0.481078 | -0.5952 |
| DMRTA1 | -1.66979 | -0.609 |
| APOD | -0.6323 | -0.4663 |
| DLL1 | -0.51114 | 0.4403 |
| PARP9 | -0.48827 | -0.5402 |
| YPEL3 | -0.91846 | -0.3846 |
| PPT1 | -0.33853 | -0.3064 |

| | | |
|---|---|---|
| FAM83D | 0.368989 | 0.51746 |
| MMP16 | -1.59719 | -0.8704 |
| BTG1 | -0.55682 | -0.3642 |
| DLGAP5 | 0.360403 | 0.79869 |
| CDH1 | 0.378598 | -0.2792 |
| TGFBI | -0.46841 | 0.49453 |
| ALOXE3 | 0.667374 | 1.13734 |
| DOCK8 | -0.87052 | -0.4047 |
| C11orf24 | 0.598047 | 0.44118 |
| KCNC3 | -0.89787 | -0.7231 |
| AGR3 | 1.785798 | -0.5873 |
| RTN4 | 0.289675 | 0.29716 |
| RBL2 | -0.53701 | -0.4011 |
| HILPDA | -0.64705 | -0.382 |
| BCL2L1 | -0.47191 | -0.3421 |
| CCNB2 | 0.36822 | 0.48113 |
| BAG3 | -0.26922 | -0.3768 |
| CA8 | 1.706083 | 0.80736 |
| AKR1C2 | 0.495678 | 0.47337 |
| NLN | 0.609003 | 0.41071 |
| CLU | -0.60212 | -0.5054 |
| SLC39A6 | 0.319382 | -0.4509 |
| EPS8L1 | -0.39634 | -0.4386 |
| PLK1 | 0.353433 | 0.52835 |
| HMMR | 0.269244 | 0.97119 |
| GREB1 | 2.916463 | 0.49515 |
| LRRC75A-AS1 | 0.288952 | 0.32079 |
| MB | -1.19225 | -0.4908 |
| SNX33 | -0.5394 | -0.4439 |
| CAPN13 | -1.34403 | -0.3603 |
| SLC27A2 | 0.578858 | 0.58952 |
| PTTG1 | 0.48712 | 0.41733 |
| LTBP3 | -0.89805 | -0.4598 |
| TUFT1 | -0.52707 | 0.35093 |
| BUB1 | 0.365493 | 0.58692 |
| ADAT2 | 1.068878 | 0.76357 |
| TINCR | 0.356238 | 0.42035 |
| FAM174B | -0.70405 | 0.43097 |
| IFRD1 | 0.47622 | 0.35657 |
| PRRT2 | -0.51352 | -0.8444 |
| ANKRD50 | -0.7715 | -0.4137 |
| SEMA4C | -0.52421 | -0.4308 |
| P2RX2 | -2.15942 | -0.6054 |
| SECTM1 | -1.37958 | 1.14013 |

| | | |
|---|---|---|
| NOP58 | 0.489763 | 0.55523 |
| OSR2 | -0.37233 | -0.4471 |
| RGS22 | 1.910792 | 1.46079 |
| C1QBP | 0.445925 | 0.32405 |
| ABCC3 | -0.83471 | 0.32744 |
| MRPL3 | 0.339822 | 0.32502 |
| SLC2A11 | -0.46149 | 0.55357 |
| FLOT1 | -0.25824 | -0.3573 |
| FGF13 | -1.08767 | -0.6125 |
| RUVBL1 | 0.578492 | 0.33547 |
| ATP2C2 | -0.69536 | -0.4168 |
| BMP4 | -0.80989 | -0.557 |
| FAM198B | 0.859587 | 0.95393 |
| PLEKHA7 | -0.45517 | -0.3606 |
| ECE1 | 0.873938 | -0.4499 |
| SCUBE2 | -0.26608 | 0.57498 |
| APP | -0.22446 | -0.2646 |
| KIAA0513 | -1.1095 | -0.3224 |
| LARS | 0.299862 | 0.34716 |
| LAMC2 | 0.941714 | 0.62629 |
| WWOX | -0.42486 | -0.3957 |
| IFT140 | -0.56394 | -0.456 |
| MARCKS | -0.55908 | -0.3821 |
| PCDH7 | -0.77329 | -1.003 |
| CRIM1 | -0.51498 | -0.4369 |
| EPB41L2 | -0.53443 | -0.4114 |
| HNRNPH3 | 0.417431 | 0.30371 |
| ARHGEF37 | -1.06151 | -0.6431 |
| SPATS2L | 0.273344 | 0.40452 |
| CRISPLD2 | -0.52641 | 0.43058 |
| CDCA8 | 0.521254 | 0.42469 |
| BOD1 | 0.277616 | 0.31512 |
| ZG16B | -0.69532 | -0.4268 |
| NBEAL2 | -0.44377 | -0.3247 |
| AIF1L | -0.30348 | -0.3125 |
| LYAR | 0.873111 | 0.5951 |
| CKB | -0.62952 | -0.434 |
| PNO1 | 0.457455 | 0.3697 |
| NOL4L | -0.32054 | 0.36801 |
| TFRC | 0.434736 | 0.29958 |
| NELL2 | -2.06886 | -1.4076 |
| TCF7L1 | -1.28343 | -0.7051 |
| PACSIN1 | -0.68216 | -0.4272 |
| HSPE1 | 0.578903 | 0.40782 |

| | | |
|---|---|---|
| SPTBN1 | 0.29014 | 0.31114 |
| SPINT1 | -0.41111 | -0.2975 |
| KLHL5 | -0.50183 | 0.53935 |
| EPHA7 | -0.57402 | -0.479 |
| HDAC5 | -0.8136 | -0.3866 |
| IGSF1 | 3.329681 | 0.50214 |
| ALDH4A1 | -0.81534 | -0.3309 |
| MEX3A | -0.36707 | 0.41394 |
| GABBR2 | -1.82556 | -1.0885 |
| LIF | 0.702269 | 1.24899 |
| GPR157 | -0.65938 | 0.33505 |
| DNTTIP2 | 0.357915 | 0.89487 |
| DBP | -0.95665 | -0.5677 |
| MEST | 0.392134 | 0.30831 |
| TTK | 0.342633 | 0.90583 |
| RNF207 | -1.41223 | -0.5321 |
| GFRA1 | 0.422923 | -0.4475 |
| RAB30 | -0.43571 | -0.4508 |
| TSPAN9 | -0.57789 | -0.3994 |
| ABCA1 | -0.89092 | -0.4276 |
| ABCE1 | 0.668304 | 0.4938 |
| GSTM3 | -0.28895 | 0.26429 |
| COL12A1 | 0.599568 | -1.1044 |
| CEP55 | 0.608627 | 0.56906 |
| S100A16 | -0.50954 | -0.28 |
| ATP2A3 | -0.88584 | -0.3134 |
| PLLP | -1.05811 | -0.5734 |
| TLE3 | 0.36105 | -0.3248 |
| DLC1 | 0.471079 | -0.7455 |
| HCAR1 | -1.54447 | -0.456 |
| SH3BGRL | -0.51166 | -0.3617 |
| CTDSP1 | -0.37386 | -0.3132 |
| NUP107 | 0.524756 | 0.40631 |
| NAA20 | 0.239287 | 0.3007 |
| SLC9A3R2 | -0.49305 | 0.36781 |
| MTHFR | -0.91668 | -0.4645 |
| RASEF | -0.51392 | -0.3296 |
| BIK | -1.02725 | -0.5036 |
| COMT | -0.59362 | -0.2784 |
| GPC4 | -0.35121 | -0.4878 |
| DKC1 | 0.668535 | 0.33343 |
| GRN | -0.4755 | -0.2529 |
| HSPA8 | 0.551927 | 0.25122 |
| NATD1 | -1.12318 | -0.4546 |

| | | |
|---|---|---|
| OSGIN1 | 0.810572 | 0.38904 |
| BUB1B | 0.528073 | 0.57544 |
| SCN1B | -0.36402 | -0.3802 |
| NCAPG | 0.460454 | 0.85167 |
| DDIT4 | -1.08202 | 0.32507 |
| AQP3 | -0.61538 | 0.32515 |
| GCLM | 0.599053 | 0.43509 |
| KIF14 | 0.262833 | 1.17006 |
| AEN | 0.742537 | 0.32527 |
| ABHD11 | -0.40919 | -0.3057 |
| WDR43 | 0.545842 | 0.30751 |
| HSP90AB1 | 0.467331 | 0.22645 |
| KIF2C | 0.376307 | 0.37724 |
| RNF223 | 0.745848 | 0.36729 |
| CERS4 | -0.43523 | -0.3561 |
| GPRC5C | -0.60541 | 0.27944 |
| KSR1 | 0.597607 | 0.65579 |
| TXN | 0.483713 | 0.2561 |
| FAM83B | -0.70966 | -0.4364 |
| ZBTB42 | -0.5043 | -0.3179 |
| ARHGAP32 | -0.41477 | -0.3462 |
| MRTO4 | 0.611522 | 0.34243 |
| G6PD | 0.266311 | 0.29508 |
| SREBF1 | -0.2273 | -0.3913 |
| MID2 | -0.51096 | -0.3384 |
| MAPK3 | -0.28646 | -0.2511 |
| SFXN2 | 2.033102 | -0.36 |
| CAV2 | 0.441529 | 0.39542 |
| CAMK2N1 | -1.07375 | -0.3122 |
| WDYHV1 | 0.386264 | 0.3312 |
| NOLC1 | 0.591656 | 0.29979 |
| SNED1 | -0.41338 | -0.4658 |
| IDH1 | -0.28869 | -0.269 |
| KIFC3 | -1.406 | -0.434 |
| IGFBP4 | 1.494262 | -0.3508 |
| TMC4 | -0.93905 | -0.4205 |
| SIX1 | -0.60364 | -0.5262 |
| TACC3 | 0.445793 | 0.31884 |
| PIM1 | 0.687684 | 0.49835 |
| STRBP | 0.281979 | 0.26954 |
| HPRT1 | 0.871493 | 0.33406 |
| RABGGTB | 0.500559 | 0.36448 |
| FLNB | 0.582385 | -0.3163 |
| ALDH3B2 | -0.36838 | 0.46571 |

| | | |
|---|---|---|
| SOX4 | -0.60067 | 0.22894 |
| HMBS | 0.514845 | 0.28517 |
| ABCA12 | 0.70634 | -0.6741 |
| RRS1 | 0.521749 | 0.35234 |
| FARSB | 0.620703 | 0.32879 |
| PVT1 | 0.38793 | 0.40796 |
| PBK | 0.515338 | 0.40147 |
| TMEM184A | -0.4161 | -0.2744 |
| SMPDL3B | 0.919566 | 0.88933 |
| UCK2 | 0.57607 | 0.33102 |
| EPOR | -0.96045 | -0.626 |
| NCS1 | 0.387845 | 0.28764 |
| CDKN3 | 0.507873 | 0.38505 |
| TP53INP2 | -1.20973 | -0.4489 |
| CLSTN1 | -0.31378 | -0.2665 |
| UGT1A6 | -1.49938 | 0.72815 |
| JUP | -0.2376 | 0.21067 |
| PIK3IP1 | -1.44754 | -0.4035 |
| GALNT10 | -0.68066 | 0.30741 |
| SIM1 | -1.78121 | -1.4456 |
| TMEM205 | -0.43402 | -0.3981 |
| TBC1D2 | -0.77788 | -0.3326 |
| SSR3 | 0.296486 | 0.2699 |
| SH3GLB2 | -0.39772 | -0.2581 |
| TRAFD1 | -0.57143 | 0.22855 |
| IFFO2 | -0.45036 | 0.39528 |
| DARS2 | 0.603367 | 0.28356 |
| HNRNPAB | 0.517167 | 0.26342 |
| TXNIP | -0.7109 | -0.3057 |
| PLCD3 | 0.312313 | -0.2435 |
| NDRG4 | -0.44602 | 0.35286 |
| AOX1 | -1.28534 | -0.8208 |
| CSTA | -2.58213 | -1.7107 |
| TP53I11 | -0.44489 | -0.3558 |
| CTPS1 | 0.878573 | 0.33756 |
| AHRR | -1.00846 | 0.99727 |
